# A *Trypanosoma cruzi* Zinc Finger protein that controls expression of epimastigote specific genes and affects metacyclogenesis

**DOI:** 10.1101/2020.07.07.189688

**Authors:** Thais Silva Tavares, Fernanda Lins Brandão Mügge, Viviane Grazielle-Silva, Bruna Mattioly Valente, Wanessa Moreira Goes, Antonio Edson Rocha Oliveira, Ashton Trey Belew, Alessandra Aparecida Guarneri, Fabiano Sviatopolk-Mirsky Pais, Najib M. El-Sayed, Santuza Maria Ribeiro Teixeira

## Abstract

*Trypanosoma cruzi* has three biochemically and morphologically distinct developmental stages that are programed to rapidly respond to environmental changes the parasite faces during its life cycle. Unlike other eukaryotes, Trypanosomatid genomes contain protein coding genes that are transcribed into polycistronic pre-mRNAs and control of gene expression relies on mechanisms acting at the post-transcriptional level. Transcriptome analyses comparing epimastigote, trypomastigote and intracellular amastigote stages revealed changes in gene expression that reflect the parasite adaptation to distinct environments. Several genes encoding RNA binding proteins (RBP), known to act as key post-transcriptional regulatory factors, were also differentially expressed. We characterized one *T. cruzi* RBP (TcZH3H12) that contains a zinc finger domain, and whose transcripts are upregulated in epimastigotes compared to trypomastigotes and amastigotes. TcZC3H12 knockout epimastigotes showed decreased growth rates and increased capacity to differentiate into metacyclic trypomastigotes. Comparative transcriptome analysis revealed a TcZC3H12-dependent expression of epimastigote specific genes encoding amino acid transporters and proteins associated with differentiation (PAD), among others. RNA immunoprecipitation assays showed that transcripts from the PAD family interact with TcZC3H12. Taken together, these findings suggest that TcZC3H12 positively regulates the expression of genes involved in epimastigote proliferation and also acts as a negative regulator of metacyclogenesis.

## Introduction

*Trypanosoma cruzi* the causative agent of Chagas disease, affects approximately 6-8 million people worldwide. It is endemic in Latin America, where it is a major public health problem and causes over 10,000 deaths annually (WHO., 2020). There is no vaccine and the two drugs currently available, in addition to having unwanted side effects, are only effective in the acute phase of the infection (DNDi., 2020). Natural *T. cruzi* transmission to humans occurs from domiciliated triatomine insect vectors, mainly *Triatoma infestans* and *Rhodnius prolixus.* Although recent vector control programs have succeeded in reducing infection rates in endemic areas, non-vectorial transmission routes, including transmission via contaminated food, blood transfusions, organ donation and congenital transmission, frequently occurs (WHO., 2020).

When taking a blood meal on an infected mammal, the insect vector ingests circulating trypomastigotes, which, once inside the insect posterior midgut, differentiate into replicative epimastigotes. In the posterior end of the digestive tract, epimastigotes differentiate into infective, non-dividing metacyclic trypomastigotes, which are eliminated with the insect’s urine and feces during a blood meal. If inoculated through ruptures in the skin of a new mammalian host, trypomastigotes can enter the bloodstream and infect different cell types. Once in the host cell, trypomastigotes differentiate into amastigotes that replicate in the cytoplasm for 3 to 5 days before differentiating again into highly motile trypomastigotes. Highly motile trypomastigotes cause lysis of the infected cells and reach the circulatory system, propagating the infection (Brener, 1973).

The existence of various morphologically and biochemically distinct life cycle stages that alternate between invertebrate and vertebrate hosts requires a finely regulated developmental program. Similar to other members of the Trypanosomatid family, all *T. cruzi* protein coding genes are organized into long polycistronic transcription units that are transcribed into polycistronic pre-mRNAs, which are subsequently processed into mature, monocistronic mRNAs through coupled trans-splicing and poly-adenylation reactions (El-Sayed et al., 2005). Because of polycistronic transcription, control of gene expression in *T. cruzi* must rely on post-transcriptional mechanisms, which occur mainly through a fine tuning of steady state levels and translation efficiency of the mRNA populations (Clayton, 2019, Araújo and Teixeira, 2011).

Through direct or indirect RNA-protein interactions, RNA Binding Proteins (RBPs) are key elements that regulate the expression Trypanosomatid genes (Clayton, 2019). A large variety of these proteins are encoded in the *T. cruzi* genome, the majority of those having RNA recognition motifs (RRM) (De Gaudenzi et al., 2005), but also zinc finger domains (Kramer et al., 2010), Pumilio (Caro et al., 2006), among others. Studies on several RPBs have shown that they can bind to regulatory sequences present in the 3’ untranslated regions (UTR) of mRNAs and associate with additional cellular machinery to control target mRNA degradation (Dallagiovanna et al., 2005, Dallagiovanna et al., 2008, Pérez-Díaz et al., 2012, Pérez-Díaz et al., 2013) or to promote mRNA stabilization (Sabalette et al., 2019, Noé et al., 2008).

Although RBPs containing a CCCH-type zinc finger domain are known to play an important role in regulating gene expression in other eukaryotes, they are relatively less well characterized in Trypanosomatids. Previous *in silico* analysis have shown that the *T. cruzi* genome encodes approximately 50 different proteins with CCCH motif and studies with *Trypanosoma brucei* and different *Leishmania* species have demonstrated that these proteins can exert important functions in controlling stage-specific gene expression and parasite growth and differentiation (Kramer et al., 2010). Two small proteins named TcZFP1 and TcZFP2, whose homologues in *T. brucei* have been implicated in regulating parasite differentiation, have also been characterized in *T. cruzi* as zinc finger RBPs that interact with each other and are differentially expressed throughout the life cycle (Caro et al., 2005). While expression of TcZFP1 is increased in metacyclic trypomastigotes (Mörking et al., 2004, Espinosa et al., 2003), TcZFP2 levels are reduced in this stage (Mörking et al., 2012). In contrast, the zinc finger protein TcZC3H39 is expressed constitutively among all life cycle stages, but changes in its subcellular location within RNA granules and its association with distinct mRNA targets also indicate a role in regulating gene expression during *T. cruzi* differentiation (Alves et al., 2014). As the only *T. cruzi* RBP that has been functionally characterized using reverse genetics, TcZC3H31 was identified as a positive regulator of metacyclogenesis since TcZC3H31 gene knockout resulted in inhibited metacyclogenesis and an arrest of epimastigotes into an intermediate state (Alcantara et al., 2018).

Differentiation between life cycle stages that occurs into the insect and mammalian hosts is accompanied by changes in the expression of a large number of *T. cruzi* genes. By performing global gene expression analysis, we identified all RBPs with differential expression between epimastigotes, trypomastigotes and amastigotes. The gene TcCLB.506739.99 encoding a zinc finger protein with orthologs in other Trypanosomatids (Ouna et al., 2012) but not characterized in *T. cruzi,* showed a 10-fold higher expression in epimastigotes when compared to the mammalian stages, suggesting a role for TcZC3H12 as key regulator of parasite differentiation. Through phenotypical characterization of TcZC3H12 knockout cell lines, we present evidence indicating that this zinc finger protein controls the levels of epimastigote-specific transcripts, acting as a positive regulator of epimastigote growth and a negative regulator of metacyclogenesis.

## Results

### Identification of RNA binding proteins genes upregulated in epimastigotes

We have previously reported transcriptome profiling analyses comparing trypomastigotes and intracellular amastigotes from *T. cruzi*, revealing distinct virulence phenotypes and essential factors involved in the parasite’s adaptation to the mammalian host (Belew et al., 2017). Extending our RNA-seq data analyses to include the epimastigote stage (**Supplementary Table 1**), we compared here the transcriptome profiles obtained from tissue culture-derived trypomastigotes and intracellular amastigotes obtained 60 hours post-infection (hpi) with the transcriptome obtained from *in vitro* cultured epimastigotes of the CL Brener cloned strain. Mapped sequencing data derived from the epimastigote samples were analyzed using principal component analysis (PCA) to inspect relationships between samples. The resulting PCA plot (embedded in **Supplementary Table 2**), after normalization, showed the expected clustering between biological replicates.Differential expression (DE) analyses (**Supplementary Table 2**) revealed a total of 2489 transcripts with increased expression levels in epimastigotes (log2 fold change > 1, adj. *P* value <0.05) and 624 transcripts with reduced expression levels compared to amastigotes. Using similar criteria to identify statistically significant DE genes, we compared the epimastigote and trypomastigote global transcriptomes. A total of 2404 transcripts were identified as significantly upregulated in epimastigotes whereas 3525 transcripts showed reduced expression levels. Among a total of 3780 transcripts that showed increased levels in epimastigotes compared to the other forms (log2FC>1 and adj. P value<0.05), gene ontology (GO) analyses revealed that a significant group encodes proteins involved with oxidation-reduction process (GO:0055114), ion transport (GO:0006811), amino acid metabolism (GO:0006520), translation (GO:0006520) and ATP metabolic process (GO:0046034). Interestingly, members of the ß amastin sub family, whose epimastigote-specific expression have been previously described by our group (Kangussu-Marcolino et al., 2013), as well as members of the family of proteins associated with differentiation (PADs), which have been characterized in *Trypanosoma brucei* (Dean et al., 2009) but not in *T. cruzi,* were also found among the genes that are upregulated in epimastigotes (**Supplementary Table 3**).

Because post-transcriptional mechanisms mediated by RBPs play a major role in controlling gene expression in *T. cruzi,* we searched for RBPs that may act as regulators of epimastigote genes by comparing transcript levels of all RBP genes between epimastigotes and the two other forms of the parasite. Although 253 *T. cruzi* sequences encoding RBPs were previously identified in the CL Brener genome (Belew et al., 2017), homology searches based on protein domains that included additional motifs known to be present in RBPs resulted in the identification of 297 sequences that correspond to 175 RBP genes (**Supplementary Table 4**). As indicated in our previous work, because the CL Brener strain has a hybrid genome (El-Sayed et al., 2005), for most genes, we identified two distinct alleles, corresponding to the Esmeraldo and the non-Esmeraldo haplotypes. Differential expression analysis highlighted over 20 RBPs that are developmentally regulated in *T. cruzi*, with a higher amount being identified when epimastigote and trypomastigote stages were compared (**Supplementary Table 5**). Among all RBP genes whose expression is upregulated in epimastigotes, we identified the gene TcCLB.506739.99 as the one with higher differential expression when epimastigotes were compared to trypomastigotes. Based on the RNA-seq data, its transcript levels presented a 10-fold increase in epimastigotes compared to trypomastigotes and a 7-fold increase compared to amastigotes (**Fig. 1A-B**, respectively). Similar differences in expression levels were found when the non-esmeraldo allele TcCLB.510819.119 for the same gene was analyzed (4-fold higher expression in epimastigotes compared to trypomastigotes and 3-fold increase in epimastigotes compared to amastigotes). Quantitative PCR analysis performed with RNA obtained from cultured epimastigotes and tissue culture derived trypomastigotes confirmed that the TcCLB.506739.99 gene is upregulated in the insect stage of the parasite (**Fig. 1C**). Our data is also in accordance with previous RNA-Seq studies that showed higher expression of this gene in other *T.* cruzi strains, namely Y strain (Li et al., 2016), and Dm28c strain (Smircich et al., 2015). In the database TriTrypDB, TcCLB.506739.99 is annotated as a gene encoding a 19.5 kDa RBP that contains a zinc finger motif and presents orthologs in the genomes of several other members of Trypanosomatid family including *T. brucei.* Interestingly, no similar sequences were found in the genomes of other eukaryotes. The *T. brucei* ortholog, Tb927.5.15.70, encodes TbZC3H12, an RBP that is upregulated in the procyclic forms of the parasite, the stage found in the tsetse fly (Ouna et al., 2012, Fernández-Moya et al., 2014). Multiple sequence alignment revealed a high degree of conservation between orthologs at the N-terminal region (**Fig. 2A**), where the zinc finger domain is localized (C-X8-C-X5-C-X3-H), as well as in the C-terminal region where a HNPY motif is located. In *T. brucei*, the HNPY motif has been characterized as a MKT1-PBP1 binding motif (Singh et al., 2014), potentially forming a translation regulatory complex originally characterized in yeast (Tadauchi et al., 2004). As shown in **Fig. 2B**, a phylogenetic tree built with sequences derived from various members of the Trypanosomatid family showed that TcZC3H12 homologues can be clustered in 2 sub-families: a sub-family comprising *Trypanosoma* sequences and a sub-family that includes sequences derived from *Leishmania, Leptomonas* and *Crithidia* spp.

**Figure 1.**
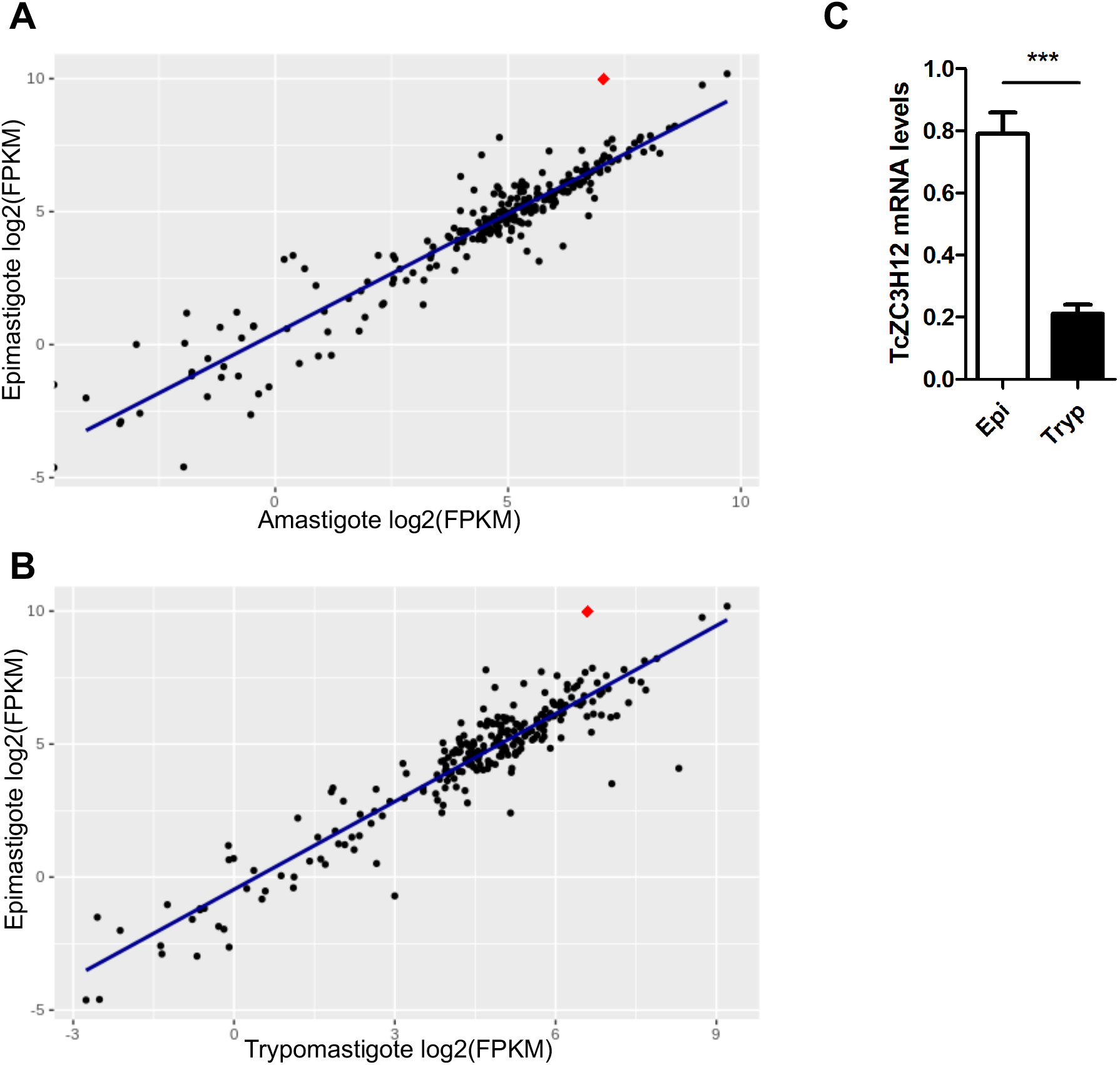
TcZC3H12 is an RNA binding protein that is upregulated in epimastigotes. RNA-Seq data published by Belew and cols (2017) were used to identify 175 *T. cruzi* genes coding for RBPs. (A) Scatterplot comparing mRNA levels of RBPs in epimastigote and amastigote or (B) epimastigote and trypomastigote. Each black dot corresponds to a different RBP transcript. Red dot in each plot corresponds to TcZC3H12 (TcCLB.506739.99). (C) Real time PCR data showing that TcZC3H12 is upregulated in epimastigotes compared with tissue culture derived trypomastigotes obtained after infection of LLC-MK2 cell monolayers (N=3) (***p < 0.001).

**Figure 2.**
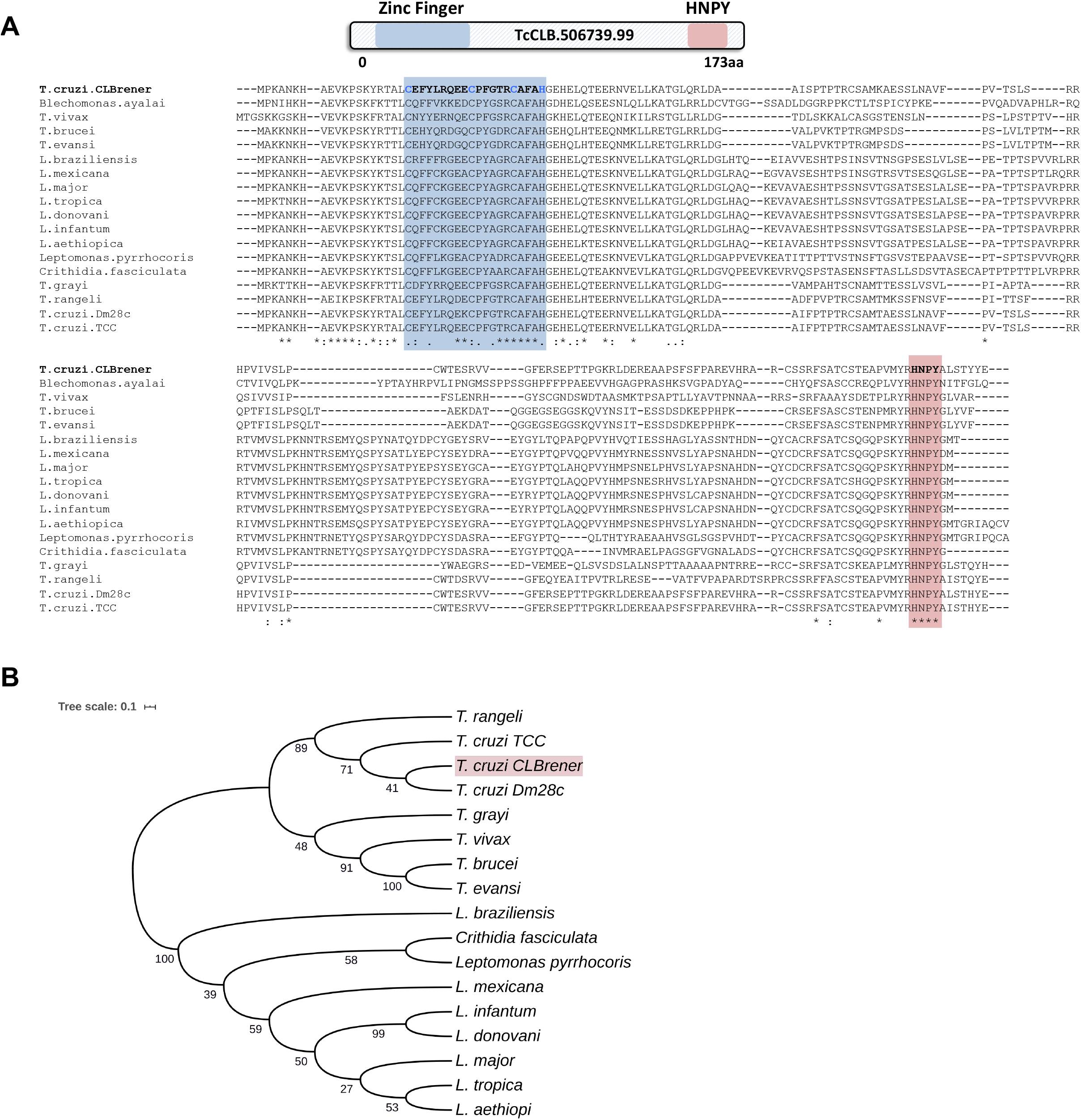
TcZC3H12 orthologs are conserved in Kinetoplastids. (A) Schematic representation of predicted relevant domains of TcZC3H12. Orthologs sequences were extracted from TriTrypDB data bank and used to generate a multiple sequence alignment and (B) a phylogenetic tree. Some ortholog genes were represented by the abbreviated genus name, where “L.” is *Leishmania* and “T.” is *Trypanosoma.* TritrypDB identifiers used were: TcCLB.506739.99 *(T. cruzi* Esmeraldo), TvY486_0501060 *(T. vivax),* TevSTIB805.5.1760 *(T. evansi),* Tb927.5.1570 *(T. brucei),* LbrM.15.0140 (*L. braziliensis),* LmxM.15.0140 (*L. mexicana),* LINF_150006300 (*L. infantum),* LdCL_150006300 (*L. danovani),* LMJLV39_150006500 (*L. major),* CFAC1_060021400 *(C. fasciculata),* DQ04_00711020 *(T. grayi),* TRSC58_03397 *(T. rangeli),* LpyrH10_38_0260 (*L. pyrrhocoris),* C4B63_83g24 (*T. cruzi* Dm28), C3747_176g18 *(T. cruzi* TCC), LAEL147_000208800 (*L. aethiopica),* LTRL590_150006300 (*L. tropica).*

### TcZC3H12 has a cytoplasmic localization and decreased protein levels after metacyclogenesis

To investigate the role of TcZC3H12 in *T. cruzi,* we generated epimastigote cell lines expressing an HA-tagged version of this protein. The coding region of TcCLB.506739.99 was amplified from DNA extracted from the CL Brener clone using a set of primers to add a C-terminal HA tag. The PCR product was then used to transfect CL Brener epimastigotes to allow homologous recombination with the endogenous gene. Using an anti-HA antibody, we determine the cellular localization of the protein by performing immunofluorescence assays of log-phase and stationary phase epimastigotes cultivated in LIT medium. As described by (Figueiredo et al., 2000), parasites that were maintained in LIT medium for more than 7 days without changing medium are submitted to a culture condition that induces epimastigote differentiation into metacyclic trypomastigotes due to nutritional starvation. As shown in the upper panels of **Fig. 3A**, TcZC3H12 has a cytoplasmic localization in log-phase, replicative epimastigotes, where it also accumulates in several foci located in the posterior portion of the parasite body. In metacyclic trypomastigotes (lower panels), a much weaker fluorescence signal was observed and no accumulation in specific foci within the cytoplasm was observed. Quantification of the fluorescence intensity comparing log-phase epimastigotes and metacyclic trypomastigotes shown in **Fig. 3B** indicates that the TcZC3H12 protein content decreases by 80% upon metacyclogenesis.

**Figure 3.**
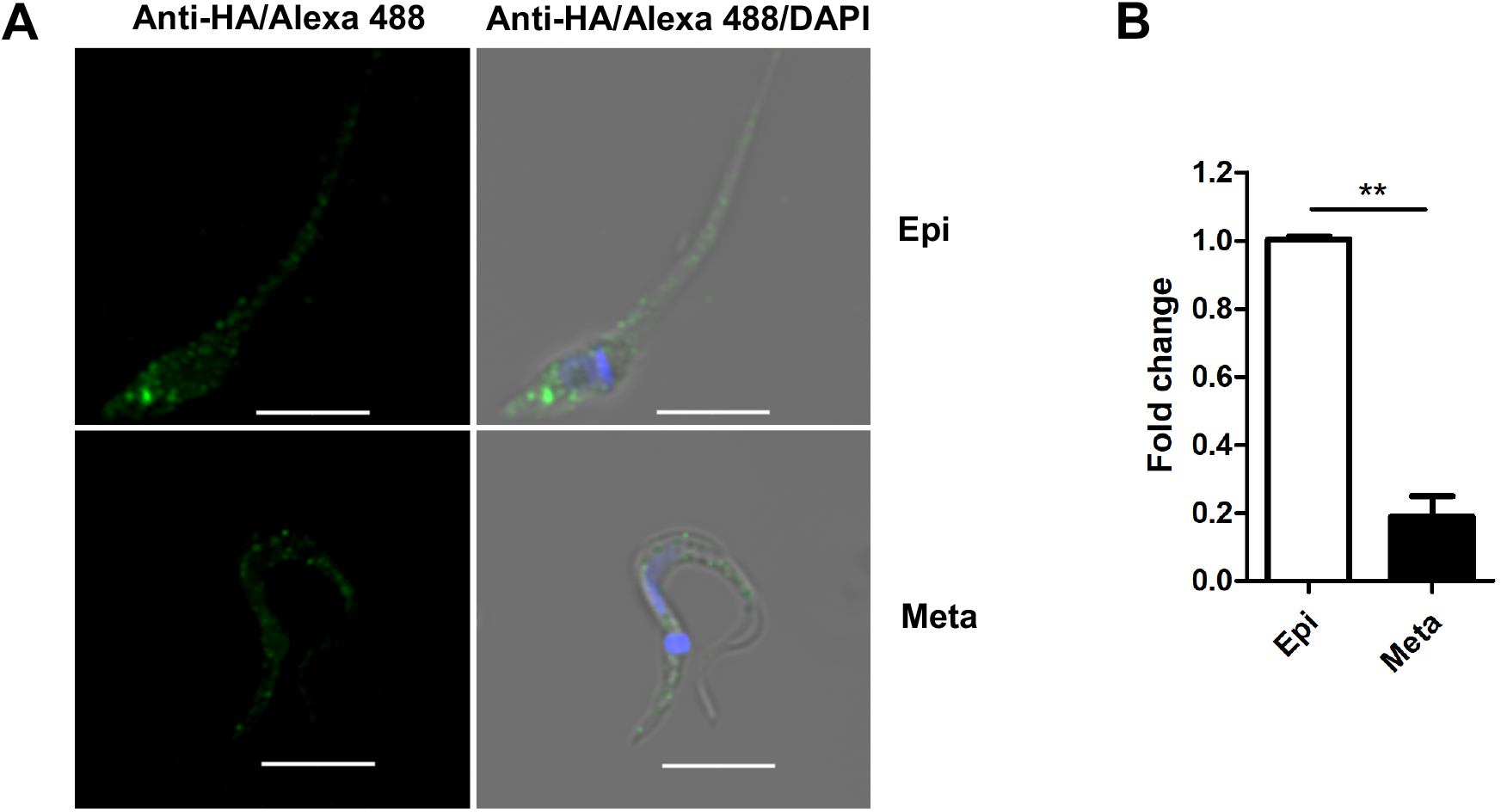
Expression levels and cellular localization of TcZC3H12 in epimastigotes and metacyclic trypomastigotes. (A) Epimastigotes expressing TcZC3H12 fused to an HA epitope were diluted every two days in LIT medium to maintain log-phase confluency. These parasites were fixed in 4% paraformaldehyde and incubated with primary mouse anti-HA antibody and secondary anti-mouse antibody conjugated to Alexa-488 (upper panels). To induce metacyclogenesis, epimastigotes were kept in LIT medium without dilution for nine days and parasites were fixed and stained using the same protocol (lower panels). Nuclei and kinetoplast were stained with DAPI to allow the identification of differentiated metacyclic trypomastigotes. (B) Fluorescence intensity of log-phase epimastigotes and metacyclic trypomastigotes were measured using ImageJ software and arbitrary unit values of intensity levels were used to calculate the fold change between epimastigotes and metacyclic trypomastigotes. Epi = epimastigote. Meta = metacyclic trypomastigote. ** p < 0.01.

### TcZC3H12 knockout impairs epimastigote growth and increases metacyclogenesis

To direct assess the role of TcZC3H12, we generated knockout cell lines by disrupting both alleles through homologous recombination with neomycin-resistance and hygromycin-resistance cassettes followed by selection in LIT medium containing G418 and hygromycin. To improve the efficiency of the process, the second allele was disrupted using the CRISPR/Cas9 technology by transfecting parasites with recombinant Cas9 and a small guide RNA targeting this allele (this strategy is illustrated in **Fig. 4A**). DNA extracted from two clones, selected from the G418 and hygromycin resistant population, were tested by PCR using different combinations of primers. The results of PCR amplification using primers P1 and P3 (generating a 1653 bp product) as well as P2 and P4 (generating a 1582 bp product) confirmed the correct insertion of the neomycin resistance cassette into the first allele, generating an heterozygous mutant. PCR products using primers P5 and P2 (generating a 1586 bp product) confirmed the insertion of the hygromycin resistance cassette into the second allele. As expected, PCR products generated with primers P1 and P2 amplified a fragment corresponding to the TcZC3H12 coding region and part of the UTR region (1274pb) only with DNA extracted from WT parasites but not with DNA from the clones 1 and 2, confirming the disruption of both alleles by the drug resistance cassettes (**Fig. 4B**). Furthermore, as shown in **Fig. 4C**, quantitative PCR (qPCR) using total RNA and primers targeting the TcZC3H12 mRNA resulted in PCR products only with RNA from wild type parasites but no PCR product were obtained with RNA from both KO cell lines. To test the effect of TcZC3H12 KO on growth of epimastigotes, we inoculated the two KO cell lines as well as WT epimastigotes in LIT medium and determined the cell densities for up to seven days. As shown in **Fig 5D**, the KO parasites presented lower growth rate compared to WT, with a difference that is highly evident on day 7. This observation indicates that expression of TcZC3H12 is required for epimastigote proliferation and its absence may cause the parasite to enter early in the stationary growth phase (**Fig. 4D**). As indicated before, metacyclogenesis can be reproduced *in vitro* under starving conditions that can be mimicked by culturing epimastigotes in aged LIT medium (Figueiredo et al., 2000). Comparing to wild type epimastigotes, which, upon entering the stationary phase of the growth curve, have about 10% of its parasite population differentiated into metacyclic trypomastigotes, the two TcZC3H12 KO cell lines presented a percentage of metacyclic trypomastigotes that is 20-30% higher after 9 days in culture in the LIT medium (**Fig. 4E**). Such differences in the growth and metacyclogenesis rates between WT parasites and TcZC3H12 KO mutants indicate that the TcZC3H12 RBP may act a regulatory factor that controls genes involved with epimastigote proliferation and differentiation in the insect vector.

**Figure 4.**
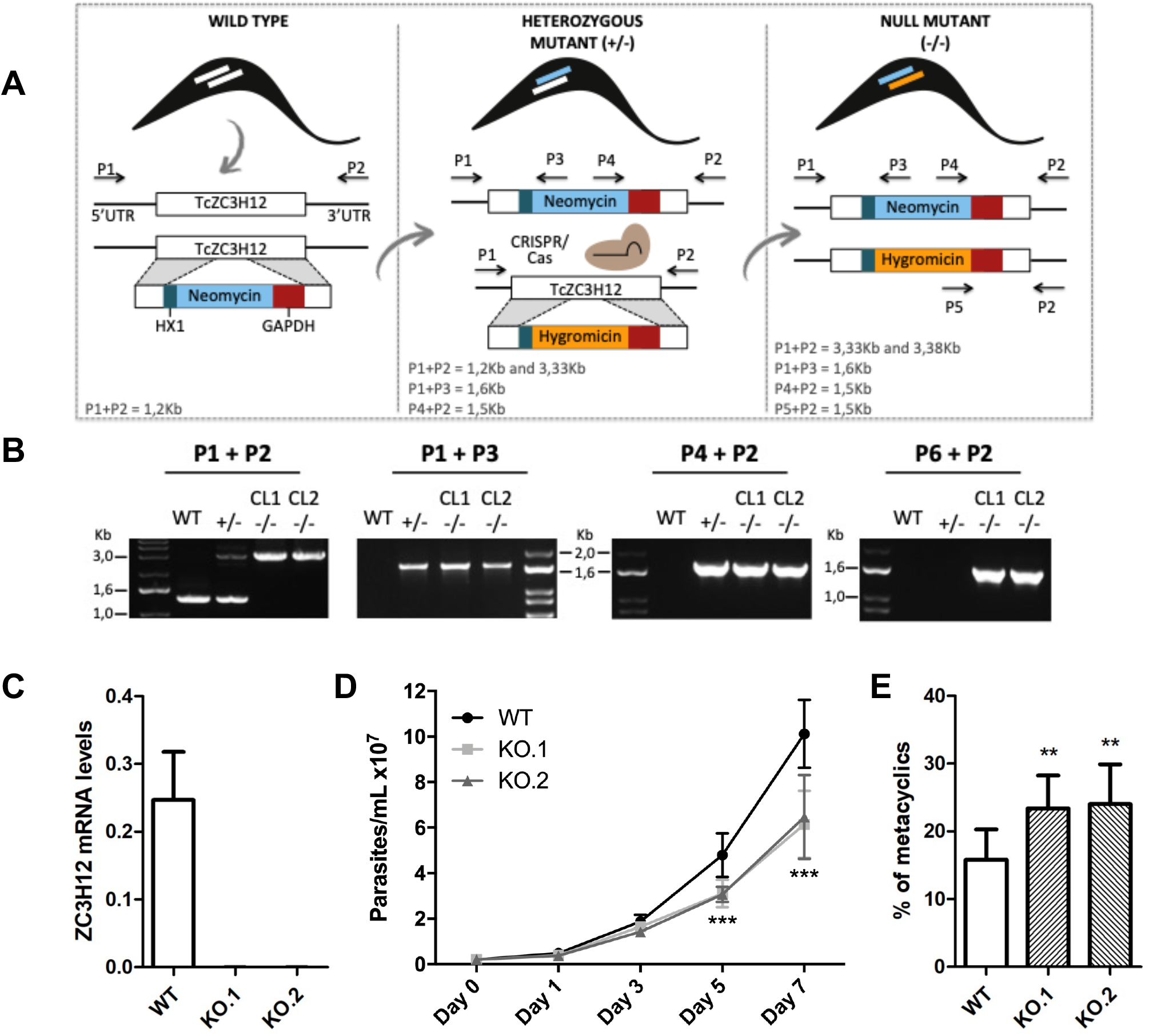
TcZC3H12 knockout parsites have altered growth rates and metacyclogenesis *in vitro*. (A) Schematic representation of the generation of knockout (KO) cell lines. The first allele was interrupted by a sequence containing the gene corresponding to neomycin resistance flanked by TcZC3H12 sequences that integrated into the genome by homologous recombination (+/-). To delete the second allele, one cloned cell line resistant to neomycin was transfected with a second DNA construct containing hygromycin resistance gene flanked by the same TcZC3H12 sequence plus the CRISPR/Cas system. (B) Agarose gel showing PCR products from different primer combinations to verify the correct integration of the DNA constructs. P1+P2 in WT parasites amplified the TcZC3H12 coding region and part of the UTR region (1274pb). In the KO cell lines the primers align outside to the inserted constructs and amplified their sequence plus some nucleotides of the UTR region (3,334bp - interruption by neomycin sequence and 3,382bp - interruption by hygromycin sequence). Neomycin resistance gene integration in TcZC3H12 locus was verified by amplification with primers P1+P3 (1653bp) and P2+P4 (1582bp). P5+P2 (1586bp) amplified hygromycin resistance gene integrated in TcZC3H12 locus. (C) Relative expression levels of TcZC3H12 quantified by RT-qPCR using RNA extracted from wild type (WT) and two knockout cell lines (KO.1 and KO.2). (D) Growth curves and (E) *in vitro* metacyclogenesis assays to compare the percent of metacyclic trypomastigotes of WT, KO.1 and KO.2 parasites. N=3 for (C), (D) and (E). ***p < 0.001 and **p<0.01.

To verify whether the differences in growth and differentiation between WT and TcZC3H12 KO parasites also occur *in vivo*, *Rhodnius prolixus* insects were infected with two KO cell lines or with WT epimastigotes. Fourth instar nymphs were fed with citrated heat-inactivated blood containing epimastigotes from each cell line. As previously described by Ferreira et al. (2016), parasites from CL strain migrate and colonize the insect digestive tract, differentiating into metacyclic trypomastigotes within 30 to 45 days (Ferreira et al., 2016). A second blood meal was offered 30 days after the infection and the urine eliminated by the insects immediately after the feeding was collected. Ten days after the second feeding, the bugs were dissected, and the rectum was analyzed under the microscope after maceration in PBS. Total parasite numbers and the percentage of metacyclic trypomastigotes were determined in the urine and rectum samples. As shown in **Fig. 5A**, no significant differences in total parasite numbers obtained from the digestive tract were observed, although a tendency towards higher numbers of WT parasites compared to KO cell lines can be noted. In urine samples, a natural source of infections by *T. cruzi*, the percentages of infectious metacyclic trypomastigotes were about 2-fold higher in insects infected with KO cell lines compared to insects infected with WT parasites **(Fig. 5B)**. Although less evident compared to the results observed in urine samples, the percentage of metacyclic trypomastigotes found in the rectum was also higher in insects infected with KO parasites than with WT (**Fig. 5C**). Thus, the presence of higher proportion of metacyclic trypomastigotes in the rectum and in the urine of infected triatomines corroborates the results obtained with *in vitro* experiments which showed that, in the absence of the TcZC3H12 gene, *T. cruzi* epimastigotes undergo metacyclogenesis at higher rates.

**Figure 5.**
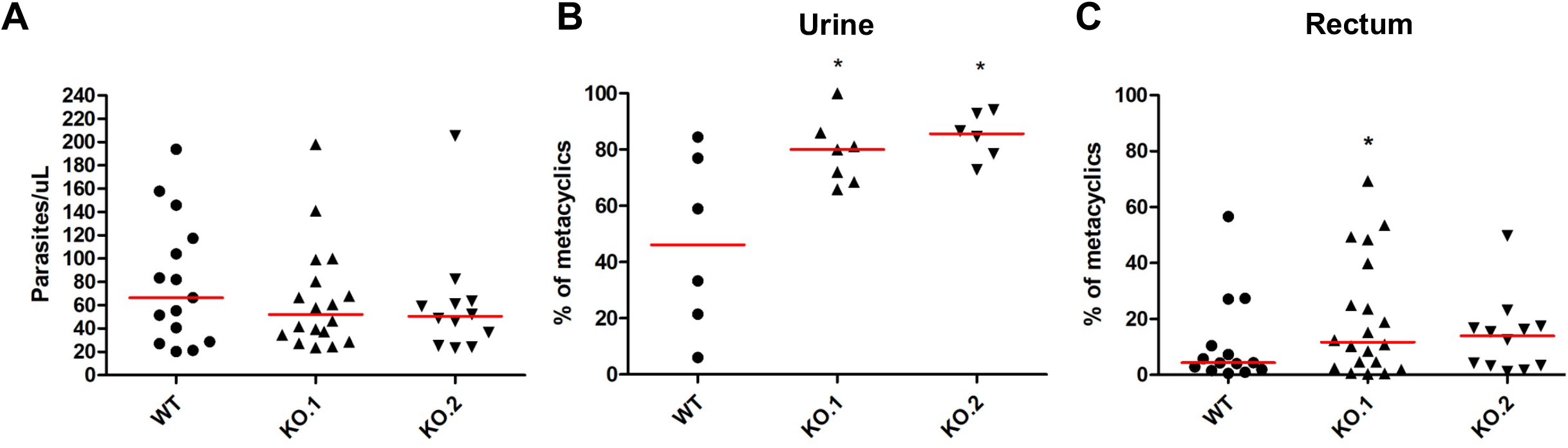
Increased numbers of metacyclic trypomastigotes in the excreta of triatomines infected with TcZC3H12 KO parasites. *Rhodnius prolixus* bugs were infected with WT and two KO cell lines (KO.1 and KO.2). Each point represents urine or rectum samples from different infected insects. Median values are displayed as red lines. (A) Total parasite numbers (epimastigotes + metacyclic trypomastigotes) inside the rectum were counted after maceration in PBS and without fixation (WT n=15; KO.1 n=18; KO.2 n=12). Percentage of metacyclic trypomastigotes found in samples of (B) urine (WT n=6; KO.1 n=7; KO.2 n=6) and (C) rectum (WT n=23; KO. 1 n=19; KO.2 n=24) were calculated after sample fixation and Giemsa staining (number of metacyclic trypomastigotes/number of total parasites). *p < 0.05 in Mann-Whitney’s test.

To further test the role of this RBP, we transfected one of the TcZC3H12 KO cell lines with the pROCK expression vector (DaRocha et al., 2004) containing the complete TcZC3H12 coding region fused to an HA-tag and the puromycin-resistance gene. After selection in LIT medium containing puromycin, we generated two addback cloned cell lines re-expressing TcZC3H12, as shown by western blots with an anti-HA antibody and qPCR using primers designed for TcZC3H12 mRNA amplification (**Fig. 6A-C**). However, as shown in **Fig. 6B**, qPCR assays demonstrated that the two TcZC3H12 addback cell lines have around 10-fold increase in transcripts levels compared to WT epimastigotes. It is thus expected that these addback cell lines will behave more similarly to an TcZC3H12 overexpressor population than with WT epimastigotes. Over-expression of TcZC3H12 was not surprising since we have used the pROCK expression vector, which contains the TcZC3H12 coding sequence driven by the ribosomal RNA promoter and integrated within the tubulin locus. In addition of being over-expressed the re-transfected cell lines present constitutive expression of the TcZC3H12 gene due to the processing signal sequences present in the 5’ and 3’ UTRs. When growth curves and metacyclogenesis rates of two TcZC3H12 re-expressor cell lines were compared to WT parasites, a partial reversion of the growth defect phenotype was observed since only a small difference in cell density was observed on day 7 (**Fig. 6C**). Although there is a tendency towards a reduction in the percentage of metacyclic trypomastigotes, no statistically significant differences in metacyclogenesis rates between WT and addback clones were detected (**Fig. 6D**). These results are matched by the results observed when we analyzed cell lines over-expressing the TcZC3H12 gene generated after transfecting WT epimastigotes with the pROCK-TcZC3H12::HA construct (**Supplementary Fig.1**). Similar to the qPCR data comparing WT and addback clones, transfection of pROCK-TcZC3H12::HA results in parasites expressing around 15-fold more TcZC3H12 transcripts than WT cells (**Supplementary Fig. 1B**). Also similar to the results obtained with the addback clones, a tendency to a reduced percentage of metacyclic trypomastigotes were observed with TcZC3H12 overexpressors compared to WT parasites and no differences in the growth curve were observed between WT parasites and over-expressing parasites as well as with parasites transfected with the empty pROCK vector (**Supplementary Fig. 1C-D**). Taken together, these data indicate that, in contrast to TcZC3H12 KO parasites, which have increased metacyclogenesis rates, over-expression of this RBP has little impact on growth or metacyclogenesis rates.

**Figure 6.**
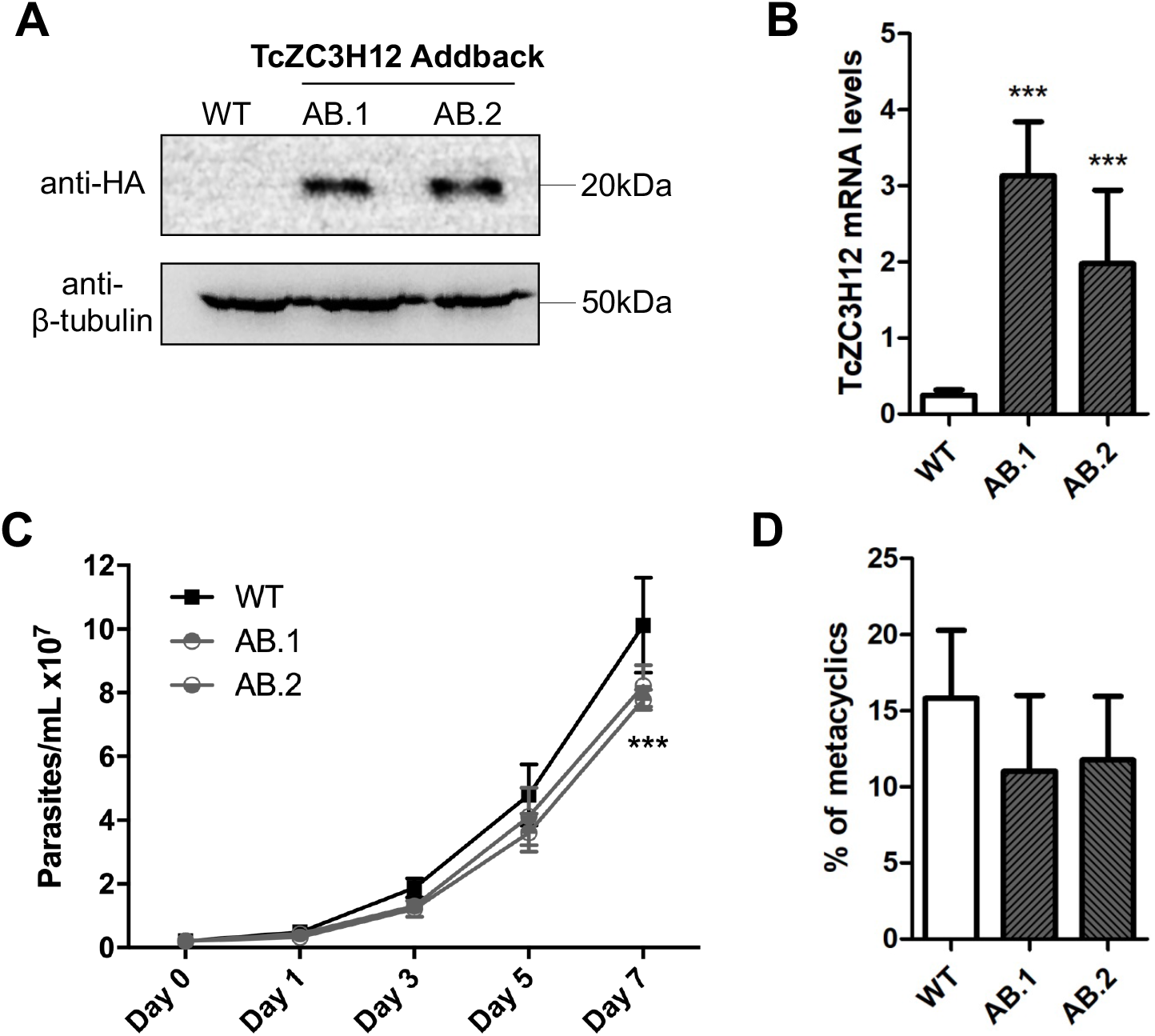
Re expression of TcZC3H12 in KO mutants partially restores growth and differentiation phenotypes. (A) Western blot with total parasite protein extract using anti-HA primary antibody. β-tubulin was used as loading control. (B) qPCR using specific primers for TcZC3H12 of wild type (WT) and two Addback clones TcZC3H12-HA (AB.1, AB.2). (C) Growth curve and (D) percent of metacyclic trypomastigotes were evaluated *in vitro* assay as previously described. (***p < 0.001)

### Epimastigote specific transcripts are targets of TcZC3H12

Based on the TcZC3H12 KO phenotype, observed both *in vitro* and *in vivo* we hypothesized that TcZC3H12 has a role in controlling parasite adaptation to nutritional restrictions and differentiation in the insect vector. Previous work based on proteomic and transcriptome analyses have shown that the parasite faces changes in gene expression when entering the stationary growth phase, which corresponds to the initial steps of metacyclogenesis (Avila et al., 2018, Berná et al., 2017, de Godoy et al., 2012, Santos et al., 2018). To investigate whether the absence of TcZC3H12 has an impact on global epimastigote gene expression, mRNA populations derived from biological triplicates of WT epimastigote cultures and duplicates of each TcZC3H12 KO cell line were sequenced using the MiSeq Illumina platform. After mapping the total number of reads to the *T. cruzi* CL Brener reference genome, mapped sequencing data derived from all seven libraries were analyzed using principal component analysis (PCA). As expected, the global transcriptome profiles of WT epimastigotes significantly differ from the profiles of both KO mutants (**Supplementary Fig.2**). From this data, it can be concluded that disruption of the TcZC3H12 gene significantly affects gene expression in epimastigotes. To identify the genes that have their expression affected, we performed differential gene expression analyses using an DESeq2 package. Because the *T. cruzi* genome contains large gene families, various of them with more than 1,000 copies (El-Sayed et al., 2005), for this differential expression analyses, all DE genes that belong to any of the six largest multigenic families (MASP, mucin, DGF-1, Trans-sialidase, RHS and GP63) were removed to avoid giving undue weight to those very repetitive transcripts. DE analyses identified 20 genes that are down-regulated in TcZC3H12 KO mutants compared to WT and 54 genes that are upregulated in KO (top 12 DE genes are shown in **Table 1**; scores for all DE genes are shown in **Supplementary Table 6**). Among the down-regulated genes, several members of the gene family encoding amino acid transporters were identified. Another gene family, encoding proteins associated with differentiation (PAD), has more than one member downregulated in KO parasites compared to WT. Although their role in *T. cruzi* has not yet been investigated, PAD genes have been characterized in *T. brucei* as a gene family encoding carboxylate-transporters that convey differentiation signals in this parasite (Dean et al., 2009). As shown in the heatmaps obtained from RNA-seq data comparing epimastigotes, trypomastigotes and amastigotes, PAD genes are upregulated only in epimastigote (**Fig.7 A)**. Amino acid transporters, on the other hand, are highly expressed in both replicative forms, epimastigotes and amastigotes (**Fig.7 C)** (for more detail on DE data see **Supplementary Table 2**). Similar results showing higher expression of PAD and amino acid transporter genes in epimastigotes were obtained from the RNA-seq data previously described for the Dm28c (Smircich et al., 2015) and Y strains (Li et al., 2016).

**Figure 7.**
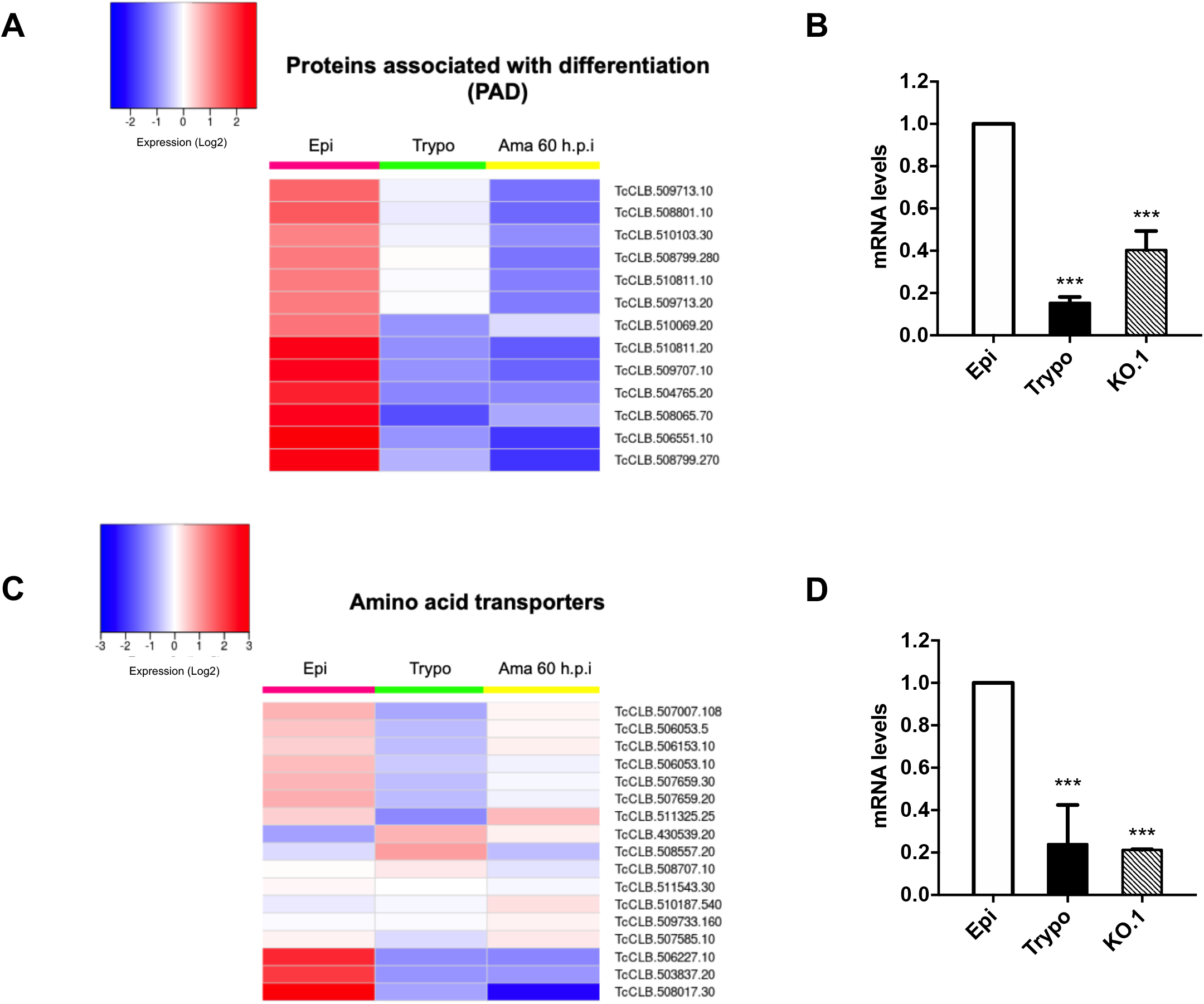
Potential targets of the TcZC3H12 among transcripts that are downregulated in KO parasites. Comparative RNA-Seq analyses from epimastigotes, tissue culture-derived trypomastigotes, and intracellular amastigotes 60 hours post infection (hpi) were used to determine the global expression of *T. cruzi* CL Brener genes. Heatmaps representing differential expression of mRNAs encoding the families: (A) proteins associated with differentiation (PAD) and (C) amino acid transporters in WT parasites. (B,D) RT-qPCRs to quantify PAD and amino acid transporters mRNA levels in WT epimastigotes (epi), WT trypomastigotes (tryp) and KO TcZC3H12 (KO.1) cell line using specific primers for one gene of the PAD family (TcCLB.506551.10) and one gene of the amino acid transporters (TcCLB.506153.10) family. ***p < 0.001.

**Table 1:**

| Gene ID | Gene annotation | log2FC | adjPvalue |
| --- | --- | --- | --- |
| Upregulated |  |  |  |
| TcCLB.510131.40 | haloacid dehalogenase-like hydrolase, putative | 1,446 | 9,35E-13 |
| TcCLB.507851.40 | hypothetical protein, conserved | 1,319 | 2,199E-08 |
| TcCLB.511037.20 | L-2-hydroxyglutarate dehydrogenase, mitochondrial, putative | 1,189 | 6,718E-20 |
| TcCLB.506247.300 | hypothetical protein, conserved | 1,058 | 1,797E-11 |
| TcCLB.504171.40 | Mechanosensitive ion channel MscS, putative | 1,048 | 6,948E-09 |
| Downregulated |  |  |  |
| TcCLB.506053.10 | amino acid transporter, putative | -1,882 | 2,121E-10 |
| TcCLB.503733.80 | hypothetical protein, conserved | -1,863 | 5,902E-20 |
| TcCLB.504173.40 | hypothetical protein | -1,848 | 2,136E-06 |
| TcCLB.506153.10 | amino acid transporter, putative | -1,785 | 3,603E-08 |
| TcCLB.504173.60 | RING-variant domain containing protein, putative | -1,508 | 1,894E-08 |
| TcCLB.506551.10 | protein associated with differentiation 8, putative | -1,141 | 3,707E-12 |
| TcCLB.508799.270 | protein associated with differentiation 8, putative | -1,048 | 4,872E-08 |

We validated our transcriptome analysis by quantitative PCR (qPCR) using primers designed to amplify one member of the amino acid transporter gene family (TcCLB.506153.10) and one member of the PAD (TcCLB.506551.10) family. qPCR amplification of RNA extracted from WT epimastigotes and tissue culture trypomastigotes as well as from TcZC3H12 KO epimastigotes confirmed that transcripts levels for PAD and amino acid transporter genes are upregulated in epimastigotes compared to trypomastigotes. Importantly, expression levels of both genes are significantly lower in TcZC3H12 KO epimastigotes than in WT epimastigotes (**Fig. 7 B** and **D**).

Due to the presence of the conserved HNPY sequence in the TcZC3H12 zinc finger protein, it can be predicted that this RBP interacts with the MKT1-PBP1 complex, for which the interaction with poly-A binding protein results in increased reporter mRNA abundance and translation rates in *T. brucei* (Singh et al., 2014). It is therefore less likely that genes that were found to be upregulated in TcZC3H12 KO mutants may constitute direct targets of this RBP. Based on this assumption, we hypothesized that the mRNAs that are downregulated in TcZC3H12 KO cell lines constitute targets of TcZC3H12 and that this RBP directly interacts with these mRNAs in epimastigotes. To test this hypothesis, WT epimastigotes and epimastigotes expressing an HA-tagged TcZC3H12 (inserted in the endogenous TcZC3H12 locus) were used in RNA immunoprecipitation assays. After UV crosslinking and cell lysis, sepharose beads conjugated to anti-HA antibody were used to immunoprecipitated the RNA-protein complexes. Total RNA purified from the immunoprecipitated beads as well as from input cell extracts prepared from both WT and TcZC3H12-HA transfected epimastigotes were used in RT-PCR assays. **Fig. 8A** shows the results of western blot analysis performed with total protein extracts (input) as well as with immunoprecipitated fractions (eluate) and non-immunoprecipitated protein (unbound) prepared from WT and TcZC3H12-HA epimastigotes. A 20 kDa band corresponding to the tagged protein was observed only in the total cell extract and in the eluate fraction of transfected parasites. Using specific primers for the TcCLB.506551.10 gene, which encodes one member of the PAD family and for the GAPDH gene (TcCLB.506943.50), used as an endogenous control, we performed RT-PCR with the RNA extracted from total cell extracts and from immunoprecipitated (eluate) fractions. Gel images of the PCR products shown **Fig. 8B** confirmed that PAD transcripts are present only in the immunoprecipitated fractions of TcZC3H12-HA transfected epimastigotes, demonstrating that TcZC3H12 binds to at least one member of the PAD family. qPCR shown in **Fig. 8C**, revealed a 20% enrichment of PAD transcripts in the eluate fraction derived from TcZC3H12-HA transfected epimastigotes compared to WT parasites, whereas GAPDH transcripts were found in the eluate fractions derived from both parasite cultures with roughly the same abundance. Taken together, the data presented here strongly indicates that TcZC3H12 is a trans-acting factor that is specifically and abundantly expressed in epimastigotes and controls the expression of genes that are upregulated in the insect stage of the *T. cruzi* life cycle. For at least one of these genes, we have shown that TcZC3H12 binds to its mRNA, possibly stabilizing it through the formation of complexes involving the MKT1 protein.

**Figure 8.**
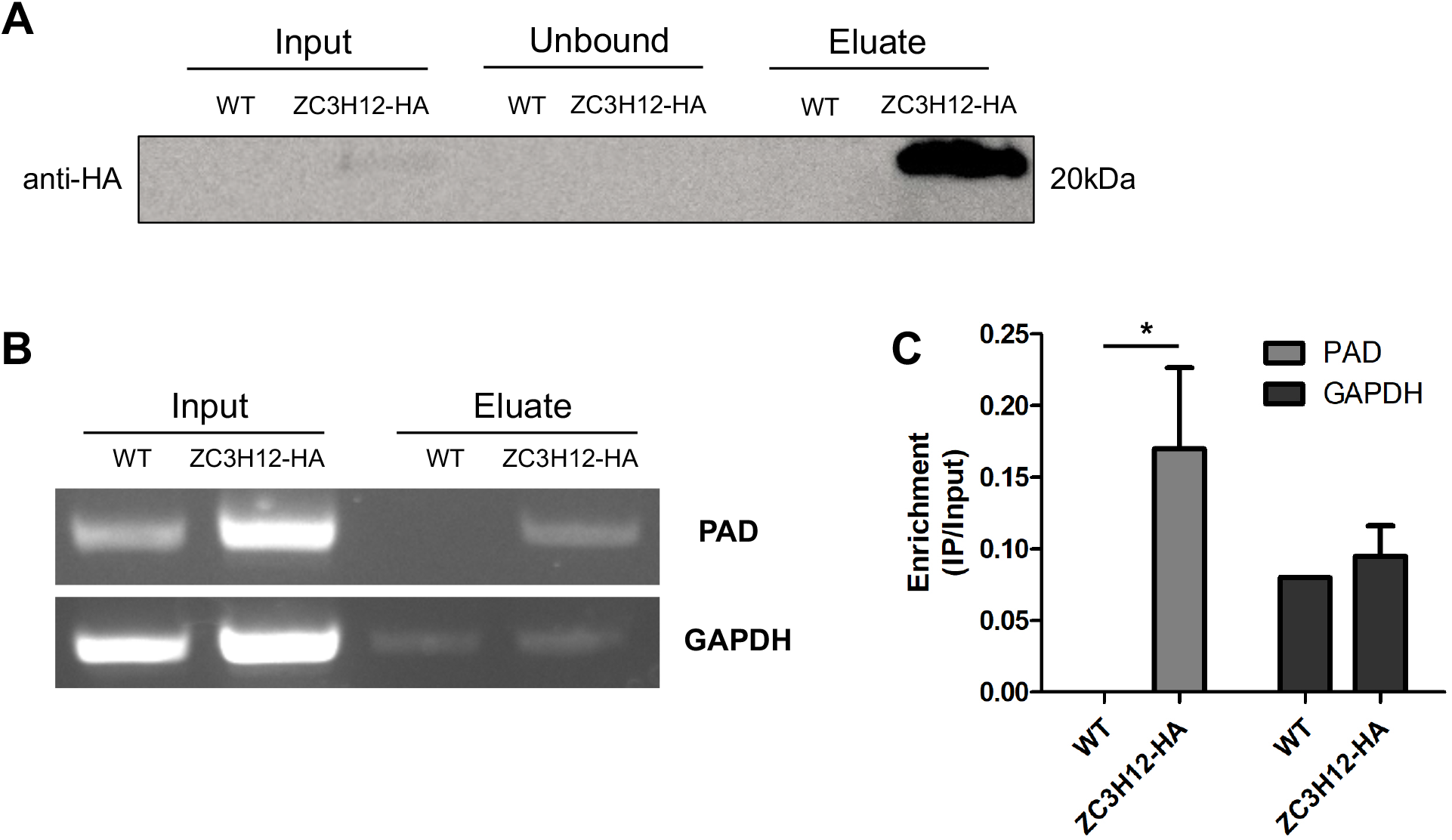
TcZC3H12 binds to PAD transcript. Parasites expressing endogenous TcZC3H12 with HA tag were used in immunoprecipitation assay with beads conjugated to anti-HA antibodies. (A) Western blot with total protein extract of input, unbound and eluate fractions incubated with anti HA antibodies. Total RNA was extracted from input and immunoprecipitated fractions, and one-step RT-PCR was done using specific primers to the coding region of one member of the PAD family (TcCLB.506551.10) and GAPDH (B) Samples were run in an agarose gel and (C) intensity of the bands was quantified. Fold change of the intensity of immunoprecipitated versus input samples was calculate and plotted. * p <a 0.05.

## Discussion

*T. cruzi* is a member of an early divergent group of eukaryotic organisms that is characterized by unique features with respect to genome organization and gene expression. Unlike most eukaryotes, transcription in Trypanosomatids is polycistronic and the processing of all mRNAs is dependent on coupled trans-splicing and polyadenylation reactions. Because of polycistronic transcription and the lack of individual promoters for every gene, regulation of gene expression must rely on RNA binding proteins, which are key elements involved in posttranscriptional control mechanisms. Previous works studying mechanisms controlling gene expression, mainly using *T. brucei* as a model organism, revealed the importance of stage-specific RBPs as trans-acting factors that modulate mRNA levels throughout the cell cycle, during different growth conditions and differentiation (de Pablos et al., 2019, Clayton, 2019). Our transcriptome analysis based on RNA-seq data obtained from the three stages of the *T. cruzi* life cycle revealed that among 175 genes encoding *T. cruzi* RBPs, transcript levels of 29 RBP genes are upregulated in the epimastigote insect stage compared to the mammalian stages (amastigotes and trypomastigotes). Among those, the gene encoding the zinc finger TcZC3H12 presented a 10-fold increase in transcript levels in epimastigotes compared to the other forms of the parasite. The same gene was also found to be highly upregulated in epimastigotes of the Dm28c strain but has decreased levels in metacyclic trypomastigotes (Smircich et al., 2015). This corroborates the results obtained with immunofluorescence assays of parasites that were HA-tagged at the TcZC3H12 locus, which showed decreased levels of the HA labelling in metacyclic forms. Similarly, RNA-Seq data obtained from the Y strain of *T. cruzi* showed that TcZC3H12 has increased levels in epimastigotes compared to the other forms (Li et al., 2016). The ortholog of the TcZC3H12 gene in *T. brucei,* TbZC3H12, also encodes a CCCH-type zinc finger whose mRNA levels are increased in the late logarithmic growth phase of procyclic parasites (Fernández-Moya et al., 2014). Interestingly, both *T. cruzi* and *T. brucei* genes are syntenic but the *T. brucei* genome possess a second gene encoding a zinc finger protein, named TbZC3H13 that is absent in the *T. cruzi* genome (Ouna et al., 2012). The fact that this RBP is exclusive of members of the Trypanosomatid family, and its expression is associated with differentiation between life cycle stages present in the mammalian and insect hosts points towards a role as regulator of cell division and differentiation. Studies on this type of RBP are highly relevant since these organisms belong to a group of early diverging eukaryotes, for which little is known about the mechanism involved in the regulation of gene expression during cell differentiation and development.

The two *T. brucei* orthologs, TbZC3H12 and TbZC3H13 encode cytosolic proteins that have similar expression levels in bloodstream and procyclic forms and are phosphorylated (Ouna et al., 2012). TbZC3H12 knockout mutants were generated in bloodstream parasites and this gene deletion had no effects in parasite growth. RNA interference targeting TbZC3H12 and TbZC3H13 separately or in double knockdown also did not result in any significant phenotypical changes either in bloodstream or in procyclic forms. The authors speculated that, most likely, these proteins have a role that can only be observed in the life cycle stages found in natural hosts (Ouna et al., 2012). In contrast, our findings clearly showed that TcZC3H12 has a regulatory role involved with epimastigote growth and differentiation in *T. cruzi,* since knockout parasites have decreased growth rate and increased differentiation capacity, observed both *in vitro* and *in vivo*. Thus, in the absence of TcZC3H12, epimastigotes undergo growth arrest and enter the stationary phase prematurely, which in turn, initiates metacyclogenesis. Based on these results, we hypothesized that TcZC3H12 positively regulates transcripts that are required for epimastigote proliferation at the same time that negatively regulates transcripts that are required for differentiation into metacyclic trypomastigotes. The decreased levels of the tagged protein observed in metacyclic trypomastigotes corroborate this hypothesis.

The cytoplasmic localization of TcZC3H12 is also in accordance with its role as a protein that binds mRNAs and controls their steady state levels. The accumulation of this protein in granular structures, as shown by confocal microscopy, suggests it might be part of RNPs involved with mRNA storage. Experiments comparing the dynamics of association mRNAs to RNP granules during epimastigote growth and differentiation would provide further evidence of the role of TcZC3H12. Although the mechanism behind the formation of these cytoplasmic ribonucleoprotein granules as well as their function are not yet fully understood, studies with *T. brucei* and *T. cruzi* showed that they are formed in response to nutritional stress and contain orthologous proteins to those present in P bodies and stress granules from metazoan organisms (Cassola et al., 2007).

The presence of the conserved HNPY motif suggesting an interaction between TcZC3H12 and the MKT1-PBP1 complex also points towards a role of this zinc finger as a factor involved with mRNA stabilization in *T. cruzi* epimastigotes. MKT1 has been shown to interact with multiple RBPs and other proteins involved in RNA regulation, acting as a master regulator of mRNA expression in *T. brucei* (Singh et al., 2014). A putative interaction between MKT1 and TbZC3H12 in *T. brucei* has been described by Lueong at al. (2016), who also showed that the TcZC3H12 ortholog acts as a positive regulator of gene expression in procyclic parasites.

Aiming at identifying targets of the TcZC3H12 RBP, we analyzed changes that occurred in the transcriptome of TcZC3H12 KO mutants. We identified 74 genes with altered expression, 20 of them showing down-regulation in the KO mutants compared to WT epimastigotes. For one member of the PAD gene family and one member of the amino acid transporter, the decreased levels of mRNA in KO epimastigotes were confirmed through quantitative RT-qPCR. When comparing epimastigotes in different growth phases, changes in gene expression were described as clusters of up and downregulated genes (Santos et al., 2018), as well as in differences in the proteome (de Godoy et al., 2012, Avila et al., 2018) and metabolome (Barisón et al., 2017). During the initial stages of metacyclogenesis, significant morphological changes of epimastigotes, which include the position of the flagellar basal body (Gonçalves et al., 2018, Avila et al., 2018) are easily observed under the microscope. Also, major changes in the metabolism of epimastigotes are triggered by culture conditions that mimics the conditions found in the insect hindgut (Shaw et al., 2016), which include changes in glucose levels and amino acid composition (De Lima et al., 2008). The fact that we identified amino acid transporters as part of the group of epimastigote-specific genes that were down-regulated in TcZC3H12 KO parasites is in agreement with data from the literature that showed that amino acids are the main source of energy for epimastigotes and, as a consequence, expression of these transporters are essential for epimastigote proliferation (Silber et al., 2005). Although we were not able to demonstrate the presence of transcripts encoding members of the amino acid transporter gene family in the immunoprecipitated RNA samples does not exclude the possibility that TcZC3H12 binds either directly or indirectly to these transcripts. High-throughput sequencing of all co-immunoprecipitated mRNAs using RIP-Seq protocol, which is underway, is a key experiment that will allow the identification of all mRNA targets of TcZC3H12.

For at least one member of the PAD gene family, RNA co-immunoprecipitation assay allowed us to show that TcZC3H12 interacts with transcripts that are upregulated in epimastigotes. Orthologs of PAD genes have been characterized in *T. brucei* as transmembrane proteins encoding carboxylate-transporters that are required for the perception of the differentiation signal mediated by citrate or cis-aconitate. TbPAD1 is expressed only by stumpy trypomastigotes, which can survive and differentiate inside the tsetse fly after a blood meal. TbPAD2, on the other hand, is highly expressed in procyclics and both proteins are not expressed by slender trypomastigotes that replicate in the mammalian bloodstream (Dean et al., 2009). Members of this gene family are thermoregulated in *T. brucei* (Dean et al., 2009) and in *Leishmania major* (Rastrojo et al., 2019). In *Trypanosoma congolense,* PAD expression was upregulated in parasites located in the cardia (procyclics) of infected *Glosinia morsitans* and downregulated in to parasites located in the proboscis (metacyclic trypomastigotes) (Awuoche et al., 2018). These data corroborate the role of PAD proteins as sensors of the concentrations of citrate/cis aconitate that allow a successful transition between insect and mammalian hosts. Although two recently described zinc finger proteins (TbZC3H20 and TbZC3H21) were identified as regulatory factors that bind hundreds of mRNAs in *T. brucei* procyclic forms, and although the depletion of TbZC3H20 causes a decrease in PAD1 expression, in contrast to the results shown here, none of these *T. brucei* proteins were shown to bind PAD transcripts (Liu et al., 2020). Considering the importance of this gene family to convey differentiation signal in trypanosomes, it is highly relevant to further investigate the role TcZC3H12 as an RBP that controls PAD expression in *T. cruzi*.

The evidence presented here suggest that TcZC3H12 is a regulatory RBP with a stage specific function most likely related to stabilization of transcripts that are upregulated in *T. cruzi* epimastigotes. Besides sequencing the RNA population that is present in immunoprecipitation complexes containing this RBP under different growth conditions, mass spectrometry analyses will be performed to identify parasite proteins that interact with TcZC3H12. These studies may help elucidate part of the molecular mechanisms that control epimastigote growth and differentiation, a key aspect of the biology of a protozoan parasite that still poses a significant threat to public health worldwide.

## Experimental procedures

### Parasite cultures

Epimastigotes forms of *T. cruzi* CL Brener clone were maintained at 28°C in liver infusion tryptose medium (LIT) supplemented with 10% of fetal bovine serum and Penicillin/Streptomycin as previously described (CAMARGO, 1964). Cultures were maintained in the exponential growth phase by doing 2 to 3 dilutions per week in fresh LIT medium.

### Differential expression analyses of total T. cruzi genes

Differential gene expression analyses were performed between epimastigotes samples and tissue cultured trypomastigotes or intracellular amastigotes samples from the RNA-Seq data published by Belew et al. (2017). A total of 22014 genes passed the low counts filter (applied to the raw counts of all 25099 genes deposited in CL Brener genome v.38) and the differential expression analysis was performed using the DESeq2 package (Love et al., 2014). Genes were considered differently expressed (DEG) when they presented an adjusted p-value (padj) < 0.05 and an absolute value of foldchange in log base 2 (ļlog2FCļ) ≥ 1. For gene ontology (GO) enrichment analysis the upregulated DEG in epimastigotes samples were analyzed using the goseq package (Young et al., 2010) in two approaches, including and excluding genes related to the six largest gene families in *T. cruzi* genome (“trans-sialidase”, “MASP”, “mucin”, “RHS”, “DGF-1”, “GP63”). The terms presenting an overrepresented p-value < 0.05 were considered enriched.

### In silico analyses of T. cruzi RBPs

To obtain the genes encoding for RBPs in the *T. cruzi* genome, Pfam and Interpro databases were used to obtain sequences that determine RNA binding domains (RRM, PABP, PUF, Alba KH (type I e II), Zinc Finger (CCHC, CCCH), S1, PAZ, PIWI, TRAP e SAM). Tool “Protein features and properties - Interpro Domain” from TritrypDB database (release 46) was used to identify the genes encoding RBPs based on the presence of the domain of interest in the analyzed gene sequence. All *T. cruzi* genes containing RNA binding domains identified in **Supplementary Table 4** were analyzed in the three main life cycle stages of this parasite using data from differential expression analysis. Amino acid sequences corresponding to orthologous of RBPs in different Trypanosomatids were extracted from TritrypDB database and used to perform multiple sequence alignments using Muscle algorithm (http://www.ebi.ac.uk/Tools/msa/muscle). Phylogenetic analysis was built using method “Neighbor joining” in the MEGA software.

### Expression of HA tagged TcZC3H12

Genomic DNA from CL Brener strain was extracted using Illustra blood genomic Prep Mini Spin Kit (GE). 200 ng of DNA were used to amplify the complete coding sequence of TcCLB.506739.99 by PCR, using forward For_TcZC3H12_XbaI and reverse Rev_TcZC3H12-HA_XhoI primers containing XbaI and XhoI restriction sites, respectively (**Supplementary Table 7**). Reverse primer also contains the sequence corresponding to HA epitope, to be inserted in the C-terminal region of the TcZC3H12 sequence. Amplicons were purified and inserted in pROCKGFPNeo (DaRocha et al., 2004) expressing vector, previously digested with XbaI and XhoI, generating pROCK.ZC3H12-HA plasmid. This plasmid was digested with XbaI and NheI enzymes to release TcZC3H12-HA sequence followed by the Neomycin expression cassette and treated with One Taq DNA polymerase (NEB) to allow for cloning in pCR2.1-TOPO plasmid (Invitrogen). Each construct was confirmed by PCR using specific primers. For transfection, (1) 5 μg of TOPO-TcZC3H12-HA were digested with AflII; and (2) 20 μg of pROCKGFPNeo, were digested with NotI. In each case, 2×10^7^ epimastigotes were resuspended in 100μl of Tb-BSF buffer (Schumann Burkard et al., 2011) and submitted to the program U033 in AMAXA Nucleofector (Lonza). Twenty-four hours after transfection, parasites were selected with G418 (200μg/ml) antibiotic. For pROCKGFPNeo, clones were obtained after serial dilutions to 0.5 parasites/well. Expression of TcZC3H12 with HA tag was confirmed by Western Blot assay. Briefly, epimastigotes were lysed in 2x SDS loading buffer containing protease inhibitor cocktail (Merck), and a volume corresponding to 2×10^6^ cells/μL was loaded into 12% SDS-polyacrylamide gel. After transferring to nitrocellulose membrane, incubation was done with 1:2000 primary anti-HA (Sigma) and 1:3000 secondary HRP anti-IgG (Sigma) antibodies.

### Fluorescence microscopy

For cellular localization, log-phase cultures or cultures aged in LIT medium from epimastigotes expressing endogenous TcZC3H12 with HA tag were centrifuged and washed with PBS and later fixed with 4% paraformaldehyde for 10 min. Following steps were done with parasites in suspension and washing steps consisted in resuspension followed by centrifugation at 400 x g. After washing with PBS, parasites were blocked and simultaneously permeabilized with blocking solution (PBS-1% BSA-0.2% Triton X-100) for 20 min at room temperature. Then, parasites were incubated with 1:250 anti-HA antibody (Millipore) in PBS-BSA 0.2% overnight at 4°C. After washing with PBS-BSA 0.2%, parasites were incubated with secondary anti-mouse IgG conjugated to Alexa Fluor 488 (Invitrogen) for 30 min in the dark. After washing, 1 μg/mL of DAPI (Molecular Probes/ Life Technologies) was used for nuclei staining. Parasites were resuspended in Prolong Gold anti-fade (Molecular Probes/Life Technologies) and mounted in glass slides topped with glass coverslips, sealed with nail polish. Images were captured on a Zeiss LSM 980 Airyscan 2 laser scanning confocal microscope using a 60x oil immersion objective, located in the Imaging Core of the Department of Cell Biology and Molecular Genetics of University of Maryland. Image processing was done in the Zen Black software and fluorescence quantification using ImageJ.

### Generation of TcZC3H12 knockout cell lines

DNA constructs were generated to disrupt both TcZC3H12 alleles. The first allele was disrupted by homologous recombination. For this, PCR amplicons corresponding to 5’ (from 150 bp before the first ATG until 250 bp after that) and 3’ (from 200 bp before stop codon until 200 bp after stop codon) sequences of the gene were obtained. Additionally, restriction sites for HindIII/SacI and XhoI/XbaI enzymes were added in 5’ and 3’, respectively (for specific primers sequences see **Supplementary Table 7**). PCR product corresponding to the 5’ was first cloned upstream of pTopo_HX1_Neo_GAPDH (Grazielle-Silva et al., 2015) and then the 3’ fragment was cloned downstream of GAPDH region, generating the plasmid TcZC3H12_Neo. To generate single KO parasites, this plasmid was used as PCR template using the primers For5’KO_TcZC3H12_HindIII and Rev3’KO_TcZC3H12_XbaI. These PCR products were used to transfect WT parasites. Twenty-four hours post transfection single KO parasites were selected with G418 (200 μg/ml) antibiotic. To delete the second allele, the plasmid TcZC3H12_Neo was digested with SpeI and NotI restriction enzymes and cloned into the plasmid Topo_HX1_Hygro_GAPDH, previously digested with the same enzymes. After that we obtained the plasmid TcZC3H12_Hygro. PCR products generated by amplification with primers For5’KO_TcZC3H12_HindIII and Rev3’KO_TcZC3H12_XbaI, were used as donor sequences to disrupt the second allele using sgRNAs (for specificities see **Supplementary Table 7**) and recombinant Cas9 derived from *Staphylococcus aureus* (rSaCas9) as previously described by (Soares Medeiros et al., 2017) and (Burle-Caldas et al., 2018). After twenty-four hours, hygromycin (200 μg/ml) was added to the medium already containing G418 for resistant parasites selection and clones were obtained by limiting dilution after platting 0.5 parasites/well. Addback parasites were generated from a KO clone transfected with pROCK.ZC3H12-HA plasmid linearized with NotI in which the Neomycin resistance gene was replaced by the Puromycin resistance sequence.

### In vitro metacyclogenesis

Metacyclogenesis was induced by starvation in aged LIT medium, as described by (Shaw et al., 2016). For this, CL Brener epimastigotes that were kept in exponential-growth phase were transferred to new culture flasks containing fresh LIT supplemented with fetal bovine serum and penicillin/streptomycin to a density of 2×10^6^/mL. The cell suspensions were left undisturbed for the parasites to attach to the bottom of the bottles for a period of 8 and 11 days. After these specific times, aliquots of each flask were collected, and parasites were fixed on glass slides and stained with Giemsa. Briefly, parasite smears were air dried, fixed with methanol 100% for 1 min and stained with 3% Giemsa stain (3% Giemsa stock in 0.95% m/v sodium phosphate dibasic, 0,9% m/v potassium phosphate monobasic, pH 7.0) for 30 minutes. After staining, the smears were washed in running water and air dried. This staining allows the identification of epimastigotes and metacyclic trypomastigotes based on the location of the kinetoplast in relation to the nucleus. Cells were counted on a light microscope using the 100x objective with immersion oil and the percentage of metacyclic trypomastigotes was calculated in relation to a total of 500 parasites per slide.

### Infection of Rhodnius prolixus bugs to assess in vivo metacyclogenesis

*Rhodnius prolixus* bugs used in this study were obtained from the Vector Behavior and Pathogen Interaction Group from René Rachou Institute (IRR, Fiocruz, MG, Brazil). Triatomines were maintained under a controlled environment with relative humidity of 26 ± 1 °C, 50 ± 5% and natural illumination. WT or TcZC3H12 KO epimastigotes were added to citrated heat-inactivated (56°C/30min) rabbit blood at a concentration of 1×10^7^ epimastigotes/mL. Rabbit blood was provided by CECAL (Centro de Criação de Animais de Laboratório, Fiocruz, Rio de Janeiro). Fourth instar nymphs were fed with these parasite suspensions using an artificial feeding apparatus (Guarneri, 2020). Insects from each group that did not feed were removed from the containers. After 15 days bugs were expected to molt. After additional 20 days, bugs were fed again, this time on chickens anesthetized with intraperitoneal injections of ketamine (20 mg/kg; Cristália, Brazil) and detomidine (0.3 mg/kg; Syntec, Brazil). The use of chickens followed established procedures of Fiocruz and was approved by the Ethics Committee on Animal Use (CEUA-FIOCRUZ) under the license number LW-8/17. After feeding, each bug was transferred to a 1.5 mL microcentrifuge tube for urine collection. Smears were prepared from the urine and afterwards stained for parasite counting. The bugs were returned to containers and kept for more 10 days in the same conditions. After this period the bugs were dissected and the intestine, separated into three portions (anterior midgut, posterior midgut and rectum), was macerated in 50 μL of PBS and analyzed under the microscope. As no parasites were found in the anterior midgut and forms in the posterior midgut were rarely observed, only those parasites of the rectum were counted. Urine smear staining was done with Giemsa as previously described. Differential counting of epimastigotes and metacyclic trypomastigotes was done under a 100x objective with immersion oil and the percentage of metacyclic trypomastigotes for each bug was calculated.

### RNA extraction and quantitative PCR analyses

Total RNA from epimastigote cultures was extracted using TRIzol reagent (Invitrogen) following manufacturer’s instructions. After DNase treatment (Invitrogen), first strand of cDNA was obtained from 200 ng of total RNA using SuperScript II Reverse Transcriptase (Invitrogen) and OligodT. Specific primers (**Supplementary Table 7**) and SSO SYBR Green Supermix (Bio-Rad) were used in qPCR reactions following manufacturer’s instructions. Applied Biosystems 7900HT Fast Real-Time PCR System (Life Technologies) platform was used to run the reactions. Relative mRNA levels were normalized to Ct values for the gene encoding for constitutive 60S ribosomal protein L9 (TcCLB.504181.10), following the 2^-△△ct^ method (Livak and Schmittgen, 2001).

### RNA sequencing and bioinformatics analysis

Total RNA was extracted from 10^8^ epimastigotes from exponentially growing cultures using Illustra RNAspin Mini kit (GE Healthcare). cDNA synthesis was done using SuperScript II Reverse Transcriptase kit first-strand synthesis (Invitrogen) and Oligo(dT)_18_, according to the manufacturer’s instructions. The quality of RNA was determined by Agilent 2100 bioanalyzer and quantified by qPCR using a KAPA Biosystems library quantification kit. RNA coming from duplicates of two clones TcZC3H12 null-mutants and triplicates from WT *T. cruzi* CL Brener clone were submitted to Illumina^®^ TruSeq^®^ RNA Sample Preparation Kit v2 to generate RNA-Seq libraries. Illumina next-generation sequencing was done using MiSeq platform. Sequence quality metrics were assessed using FastQC (Andrews 2010). Trimmomatic (Bolger et al., 2014) was used to remove any remaining Illumina adapter sequences from reads and to trim bases off the start or the end of a read when the quality score fell below a threshold of 25. The remaining reads were aligned against the concatenated *Esmeraldo, Non-Esmeraldo,* and *Unassigned* haplotypes of *Trypanosoma cruzi* using hisat2 (Kim et al., 2019). Reads from multigene families (trans-sialidase, mucin, MASP, RHS, GP63, DGF) were removed from the list with the aim to avoid interference in downstream analyses. The hisat2 derived accepted hits were sorted and indexed via samtools (Li et al., 2009) and passed to htseq (Anders et al., 2015) to generate count tables. The resulting data were passed to DESeq2 (Love et al., 2014) and a *de-novo* basic analysis which served as a ‘negative control’ for the statistical models employed. Pairwise contrasts were performed using the experimental condition and batch, or after applying sva (Leek et al., 2012) to the data. The pairwise results were compared across methods, and the interesting contrasts were extracted. Genes with differential expression were defined as genes with log2FC > 1 or log2FC < 1 and adjp-value < 0.05, between WT and null mutants.

### RNA immunoprecipitation

WT and endogenous TcZC3H12-HA expressing epimastigotes were incubated with 10 μg/mL proteasome inhibitor (MG-132 - Calbiochem) for one hour at 28° C. Following the incubation period, cells were centrifuged 3,000 x *g* for 10 minutes at 4° C, resuspended in PBS and UV crosslinked two times at 240 mJ/cm^2^ on ice (UV Crosslinker, Spectroline). After centrifugation at 3,000 x *g* for 10 minutes at 4° C and supernatant removal, cell pellets were snap frozen in liquid nitrogen. Pellets were then resuspended in RIPA buffer (Tris-HCl pH 8.0 5mM; NaCl 15 mM; NP-40 0.1%, 0.05% sodium deoxycholate, 0.01% SDS) and incubated at 4° C for 10 minutes. Shearing was done 10 times using 21 G needle and 5 times using a 30 G syringe. Lysates were centrifuged 8,200 x *g* for 10 minutes and the supernatant transferred to a new tube. 10% of cell lysate (input fraction) were transferred to a new tube and saved for further analysis. Remaining lysate was incubated with EZview Red Anti-HA Affinity Gel (Sigma - Aldrich) for 2 hours at 4° C under soft agitation. Beads were then centrifuged 8,000 x *g* for 5 minutes and unbound proteins, present in the supernatant, transferred to new tube. Beads were washed three times with lysis buffer and divided in two tubes: (1) for protein elution and (2) for RNA extraction. (1) Bound proteins were eluted in elution buffer (62.5 mM Tris-HCl pH 6.8; 10% glycerol; 2% SDS; 5% β-mercaptoethanol; 0.002% bromophenol blue). (2) Beads were resuspended in 1% SDS and treated with proteinase K (Invitrogen) for 30 minutes at 37° C. After proteinase K inactivation, RNA extraction was done using TRIzol reagent. The same was done for input fractions, following manufacturer’s instructions. SuperScript IV One-step RT-PCR kit (ThermoFisher) was used for detection of transcripts, using specific primers for PAD and GAPDH, following specifications (**Supplementary Table 7**). After RT-PCR, products were run on 0.5% agarose gel (Sigma), visualized and captured in an iBright equipment (ThermoFisher). Densitometry analysis were done using ImageJ.

### Statistical analysis

Two-tailed t-test was done for experiments in which variables had a normal distribution. Each experiment had at least 3 biological replicates. For variables that did not show a normal distribution Mann-Whitney non-parametric test was used. For all tests accepted significance level was p < 0.05.

## Acknowledgements

This work was supported by Fundação de Apoio a Pesquisa do Estado de Minas Gerais (FAPEMIG), Conselho Nacional de Desenvolvimento Científico e Tecnológico (CNPq) and the Instituto Nacional de Ciência e Tecnologia de Vacinas (INCTV). The research leading to these results has, in part, received funding from UK Research and Innovation via the Global Challenges Research Fund under grant agreement ‘A Global Network for Neglected Tropical Diseases’ (grant number MR/P027989/1). Purchase of the Zeiss LSM 980 Airyscan 2 (CBMG, University of Maryland, USA) was supported by Award Number 1S10OD025223-01A1 from the National Institute of Health. VGS is a postdoctoral fellow from PNPD/CAPES, Brazil (grant number 1785670). FLBM was a recipient of a postdoctoral fellowship from CNPq, Brazil (153195/2018-5) and is currently a DAAD-PRIME fellow, a program of the German Academic Exchange Service (DAAD) that is supported by funds from the German Federal Ministry of Education and Research (BMBF).

## Authors contributions

TST, FLBM and VGS did all the experiments. WMG, AERO, TAB did bioinformatics analysis. BMV and FMSP prepared the samples, libraries and ran RNA-Seq experiments from KO parasites. AAG conceptualized the *in vivo* differentiation experiments. NMES provided resources for microscopy, immunoprecipitation experiments and supervised bioinformatics analysis and quality control. TST, FLBM, VGS and SMRT wrote and revised the manuscript.

The authors declare there are no conflicts of interest. The data that supports the findings of this study are available in the supplementary material of this article

## Supporting Information

**Supplementary Figure 1. Overexpression of TcZC3H12 in epimastigotes.**

(A) Plasmid used to transfect WT parasites to over-express TcZC3H12. The expression vector pROCK_Neo contains neomycin (NEO) resistance gene and TcZC3H12 coding region fused to HA tag in C-terminus (TcZC3H12-HA). (B) Western blot using anti-HA primary antibody, of total protein extract from wild type (WT), two clones overexpressing TcZC3H12-HA (OE. 1 and OE.2) and parasites expressing the pROCK_Neo empty vector (EV). Anti-β-tubulin antibody was used as loading control. Detected proteins and molecular weight are indicated next to the panel. (C) RT-qPCR using primers specific for TcZC3H12 showing higher expression levels in OE.1 and OE.2 compared to WT (N=3; ***p < 0.001). (D) Growth curves of WT, OE.1, OE.2 and EV to evaluate the proliferation of parasites. Parasites were diluted to 5.0×10^6^ cells/mL in supplemented LIT medium and cell density estimated with a haemocytometer for up to 7 days. (E) Metacyclogenesis assays with WT, EV, OE.1 and OE.2 cell lines, kept in aged LIT medium. Percent of metacyclic trypomastigotes were evaluated in cell lines in the day 8. N=3 for (C) and (D). ns = not statistically different.

**Supplementary Figure 2. Global statistical assessment of biological replicates**

Principal Component Analysis (PCA) plots corresponding to 3 replicates of wild type parasites and 4 replicates, 2 for each cloned cell line, of TcZC3H12 KO mutants.

**Supplementary Table 1. Summary of samples collected, mapping statistics and experimental metadata.**

**Supplementary Table 2. Differently expressed (DE) genes in CL Brener epimastigotes, tissue culture derived trypomastigotes and intracellular amastigotes.**

**Supplementary Table 3. Gene ontology (GO) Enrichment Analysis of DE genes.**

GO enriched based on DE genes in CL Brener epimastigotes compared to trypomastigotes (1) and amastigotes (2).

**Supplementary Table 4. Annotated genes corresponding to RNA Binding Proteins identified in the CL Brener genome.**

**Supplementary Table 5. Differentially expressed (DE) RBP genes in CL Brener epimastigotes compared to tissue culture derived trypomastigotes and amastigotes**

**Supplementary Table 6. Complete list of DE genes in TcZC3H12 KO cell lines compared to wild type parasites.**

**Supplementary Table 7. Primer sequences**

