## Supplementary Figures and Tables for "A *Trypanosoma cruzi* Zinc Finger protein that controls expression of epimastigote specific genes and affects metacyclogenesis"

### Supp. Fig.1

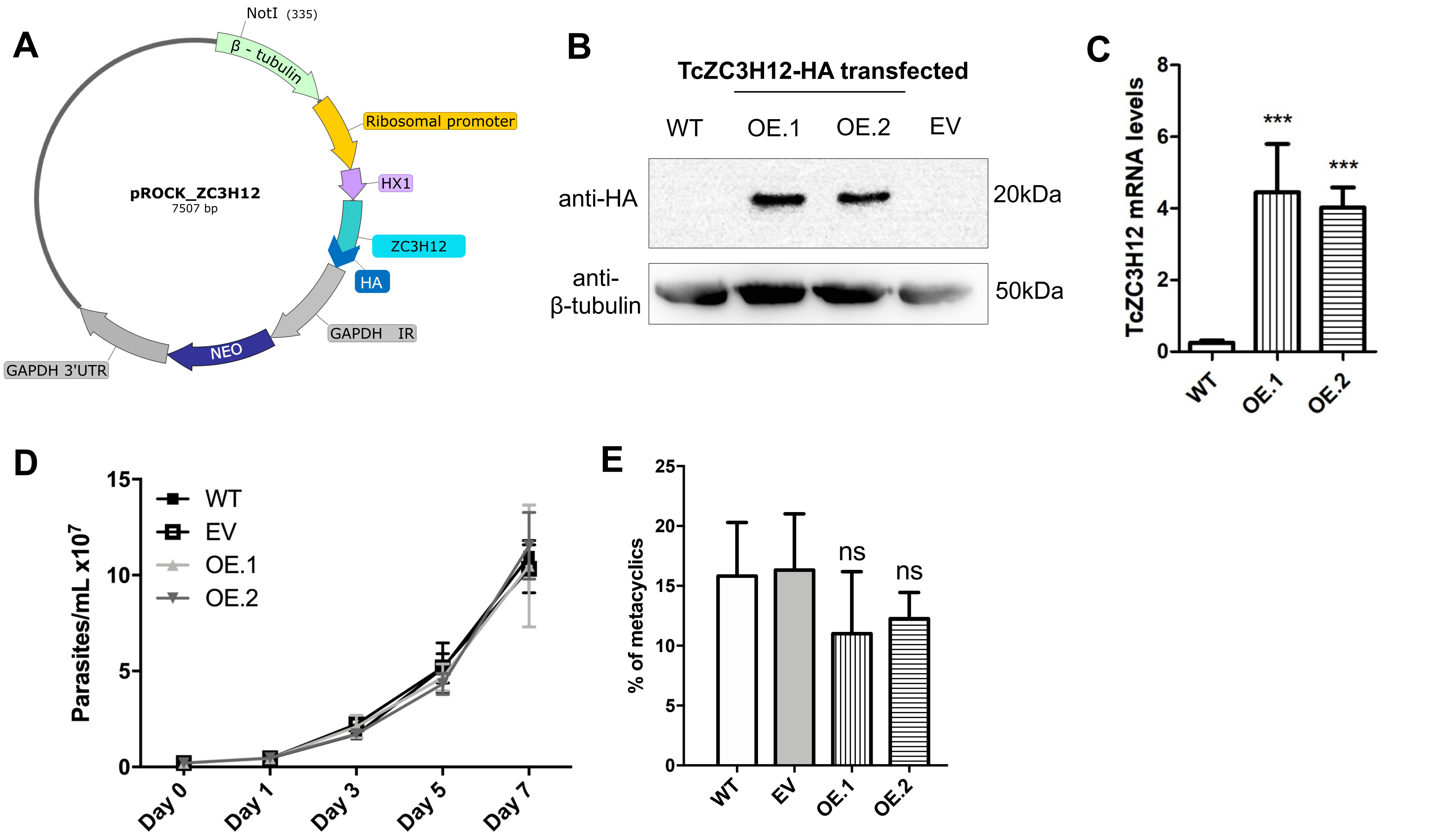

A
NotI (335)
β - tubulin
Ribosomal promoter
HX1
ZC3H12
HA
GAPDH IR
NEO
GAPDH 3'UTR
pROCK_ZC3H12
7507 bp
B
TcZC3H12-HA transfected
WT
OE.1
OE.2
EV
anti-HA
20kDa
anti-β-tubulin
50kDa
C
TcZC3H12 mRNA levels
***
***
WT
OE.1
OE.2
D
Parasites/mL x10^7
WT
EV
OE.1
OE.2
Day 0
Day 1
Day 3
Day 5
Day 7
E
% of metacyclics
ns
ns
WT
EV
OE.1
OE.2

**Supp. Fig.2**

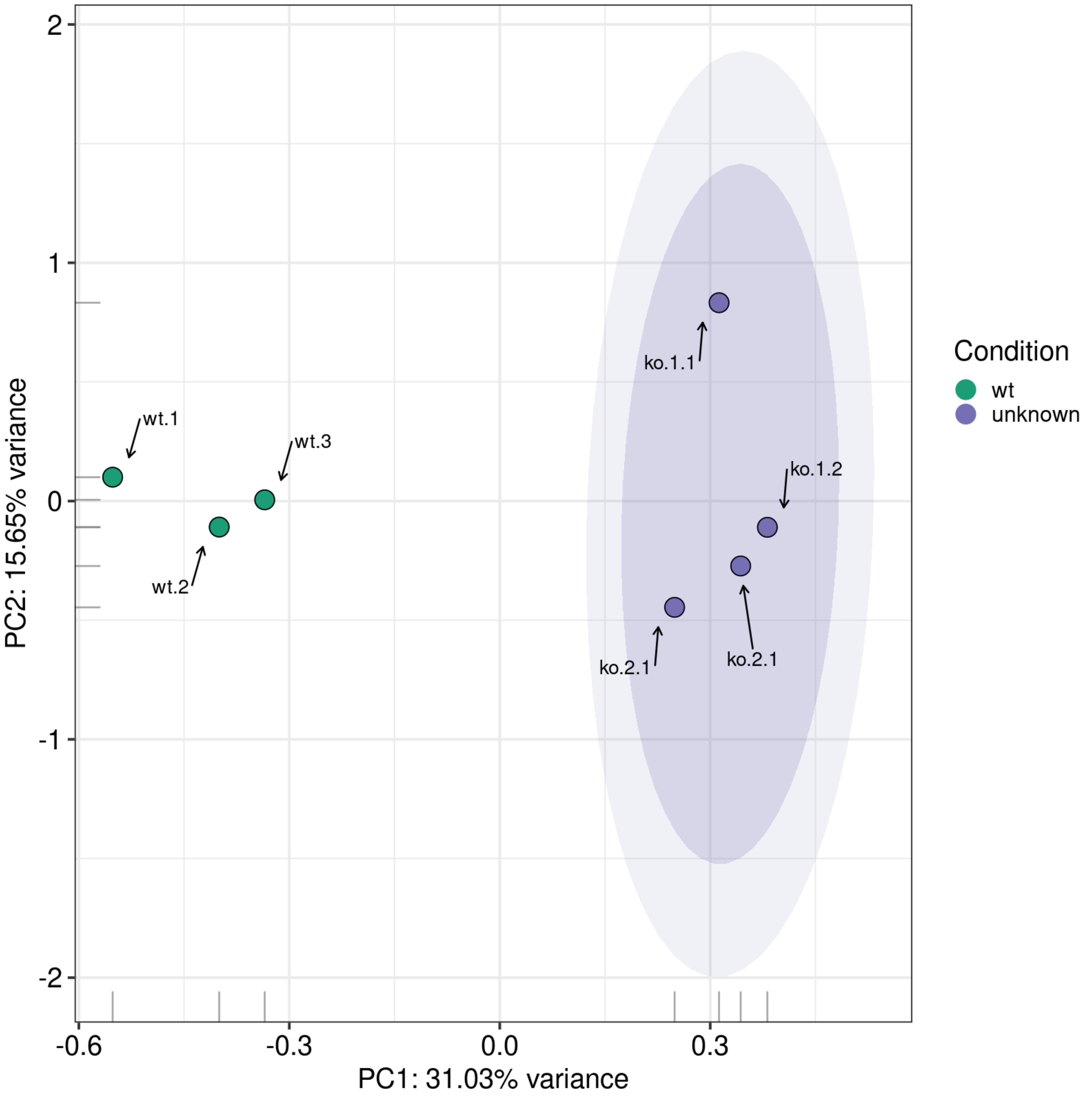

2
1
0
-1
-2
PC2: 15.65% variance
-0.6
-0.3
0.0
0.3
PC1: 31.03% variance
wt.1
wt.2
wt.3
ko.1.1
ko.1.2
ko.2.1
ko.2.1
Condition
wt
unknown

**Supplementary Table 1. Summary of samples collected, mapping statistics and experimental metadata for RNA-Seq analysis of epimastigote, trypomastigote and amastigote.**

**Table S1. Summary of samples collected and mapping statistics**

| Lab sample ID | Accession Number | Alias used in some figures | Developmental stage | Clone | Infected | Batch | Number of reads that pass Illumina filter | Total number of reads mapped | % of total reads mapped | Reads mapped to *T. cruzi* CL Brener 8.1 | Reads mapped to *H. sapiens* hg38 | % of mapped reads belonging to *T. cruzi* CL Brener 8.1 | % of mapped reads belonging to *H. sapiens* hg38 |
|---|---|---|---|---|---|---|---|---|---|---|---|---|---|
| HPGL0475 | SAMN07291202 | CLBr.A60.1 | Amastigote 60 hpi | CL Brener | Y | 20121216 | 122.215.032 | 112.803.140 | 92,30% | 14.141.199 | 98.661.941 | 12,54% | 87% |
| HPGL0476 | SAMN07291204 | CLBr.A60.3 | | | Y | 20121216 | 100.238.716 | 92.420.686 | 92,20% | 8.493.782 | 83.926.904 | 9,19% | 91% |
| HPGL0482 | SAMN07291203 | CLBr.A60.2 | | | Y | 20121216 | 94.665.058 | 87.115.475 | 92,02% | 8.947.962 | 78.167.513 | 10,27% | 90% |
| HPGL0481 | SAMN07290964 | CLBr.Tryp.3 | Trypomastigote | CL Brener | N | 20140601 | 102.571.998 | 92.052.266 | 89,74% | 92.052.266 | NA | 100,00% | NA |
| HPGL0490 | SAMN07290962 | CLBr.Tryp.1 | | | N | 19700101 | 93.463.730 | 82.016.187 | 87,75% | 82.016.187 | NA | 100,00% | NA |
| HPGL0488 | SAMN07290963 | CLBr.Tryp.2 | | | N | 20140601 | 109.810.272 | 98.242.222 | 89,47% | 98.242.222 | NA | 100,00% | NA |
| HPGL0483 | *Pending: PRJNA389926* | CLBr.Epi.1 | Epimastigote | CL Brener | N | 20140601 | 33.680.621 | 24.932.294 | 74,03% | 24.932.294 | NA | 100,00% | NA |
| HPGL0486 | *Pending: PRJNA389926* | CLBr.Epi.2 | | | N | 20140508 | 49.547.924 | 36.541.063 | 73,75% | 36.541.063 | NA | 100,00% | NA |
| HPGL0487 | *Pending: PRJNA389926* | CLBr.Epi.3 | | | N | 20140508 | 40.660.345 | 29.823.376 | 73,35% | 29.823.376 | NA | 100,00% | NA |
| Total | | | | | | | 746.853.696 | 655.946.709 | 87,83% | 395.190.351 | 260.756.358 | 60,25% | 40% |

| Alias used in some figures | Total number of reads mapped | Reads mapped to *T. cruzi* CL Brener 8.1 | Percent mapping |
|---|---|---|---|
| CLB_ama60.1 | 112.803.140 | 14.141.199 | 12,54% |
| CLB_ama60.2 | 92.420.686 | 8.493.782 | 9,19% |
| CLB_ama60.3 | 87.115.475 | 8.947.962 | 10,27% |
| **CLB_ama60** | 97446433,67 | 10527647,67 | 10,67% |
| CL14_ama60.1 | 58.860.711 | 4.205.223 | 7,14% |
| CL14_ama60.2 | 105.055.679 | 6.774.939 | 6,45% |
| CL14_ama60.3 | 60.855.520 | 3.909.457 | 6,42% |
| **CL14_ama60** | 74923970 | 4963206,333 | 6,67% |
| CLB_trypo.1 | 92.052.266 | 92.052.266 | 100,00% |
| CLB_trypo.2 | 82.016.187 | 82.016.187 | 100,00% |
| CLB_trypo.3 | 98.242.222 | 98.242.222 | 100,00% |
| **CLB_tryp** | 90770225 | 90770225 | 100,00% |
| CL14_trypo.1 | 66.239.477 | 66.239.477 | 100,00% |
| CL14_trypo.2 | 61.965.400 | 61.965.400 | 100,00% |
| CL14_trypo.3 | 74.965.544 | 74.965.544 | 100,00% |
| **CL14_tryp** | 67723473,67 | 67723473,67 | 100,00% |
| CLB_epi.1 | 58.695.502 | 58.695.502 | 100,00% |
| CLB_epi.2 | 86.709.132 | 86.709.132 | 100,00% |
| CLB_epi.3 | 70.999.582 | 70.999.582 | 100,00% |
| | 72134738,67 | 72134738,67 | 100,00% |
| CL14_epi.1 | 53.562.767 | 53.562.767 | 100,00% |
| CL14_epi.2 | 69.654.101 | 69.654.101 | 100,00% |
| CL14_epi.3 | 70.371.255 | 70.371.255 | 100,00% |
| | 64529374,33 | 64529374,33 | 100,00% |
| 24 | 2.048.069.040 | 1.086.849.589 | 53,07% |

**Supplementary Table 2. Differently expressed (DE) genes in CL Brener epimastigotes, tissue culture derived trypomastigotes and intracellular amastigotes.**

epi_over_ama

| geneid | genedescription | deseq_logfc | deseq_adjp |
|---|---|---|---|
| TcCLB.508741.80 | mucin TcSMUGS, putative | 7,293 | 5,33E-09 |
| TcCLB.509601.140 | hypothetical protein | 6,868 | 2,264E-260 |
| TcCLB.510103.24 | hypothetical protein, conserved | 6,775 | 0 |
| TcCLB.506357.50 | alcohol dehydrogenase, putative | 6,692 | 4,579E-295 |
| TcCLB.509713.30 | hypothetical protein, conserved | 6,64 | 0 |
| TcCLB.508741.60 | mucin TcSMUGS, putative | 6,125 | 1,052E-29 |
| TcCLB.508017.30 | amino acid transporter, putative (fragment) | 5,89 | 0 |
| TcCLB.511277.60 | alcohol dehydrogenase, putative | 5,877 | 1,114E-264 |
| TcCLB.511807.220 | hypothetical protein | 5,787 | 3,725E-283 |
| TcCLB.508741.70 | mucin TcSMUGS, putative | 5,715 | 4,931E-26 |
| TcCLB.508741.50 | mucin TcSMUGS, putative | 5,626 | 0 |
| TcCLB.508741.40 | mucin TcSMUGS, putative | 5,62 | 9,907E-203 |
| TcCLB.510257.80 | cytosolic nonspecific dipeptidase, putative | 5,544 | 0 |
| TcCLB.511685.20 | mucin TcSMUGS, putative | 5,522 | 1,844E-115 |
| TcCLB.511391.120 | glutamamyl carboxypeptidase, putative | 5,284 | 0 |
| TcCLB.509213.120 | cytosolic nonspecific dipeptidase, putative | 5,279 | 0 |
| TcCLB.511003.160 | hypothetical protein, conserved | 5,277 | 4,819E-184 |
| TcCLB.503411.30 | hypothetical protein, conserved | 5,272 | 2,245E-19 |
| TcCLB.509767.140 | hypothetical protein | 5,23 | 1,579E-136 |
| TcCLB.510679.100 | lipase, putative | 5,107 | 1,923E-113 |
| TcCLB.507047.150 | D-arabinono-1,4-lactone oxidase, putative (ALO) | 5,071 | 2,426E-98 |
| TcCLB.510357.20 | hypothetical protein, conserved | 5,054 | 9,182E-46 |
| TcCLB.506401.380 | surface protease GP63, putative | 4,954 | 9,832E-167 |
| TcCLB.511685.10 | mucin TcSMUGS, putative | 4,943 | 1,171E-88 |
| TcCLB.509041.10 | tyrosine aminotransferase, putative | 4,931 | 0,0001792 |
| TcCLB.509217.40 | trans-sialidase, Group V, putative | 4,827 | 0,00001215 |
| TcCLB.511685.30 | mucin TcSMUGS, putative | 4,811 | 6,586E-23 |
| TcCLB.511679.10 | mucin TcSMUGS, putative | 4,769 | 1,288E-38 |
| TcCLB.506551.10 | protein associated with differentiation 4, putative | 4,76 | 7,121E-239 |
| TcCLB.506563.79 | calpain-like cysteine peptidase (pseudogene), putative | 4,755 | 1,109E-116 |
| TcCLB.508799.270 | protein associated with differentiation 4, putative | 4,703 | 3,93E-255 |
| TcCLB.509003.30 | calpain-like cysteine peptidase, putative | 4,701 | 1,542E-115 |
| TcCLB.509965.390 | amastin, putative | 4,681 | 3,181E-87 |
| TcCLB.511201.49 | hypothetical protein, conserved | 4,647 | 2,102E-266 |
| TcCLB.506863.30 | hypothetical protein, conserved | 4,642 | 1,38E-23 |
| TcCLB.508545.40 | surface protease GP63, putative | 4,601 | 2,132E-230 |
| TcCLB.510187.50 | tyrosine aminotransferase | 4,54 | 0 |
| TcCLB.510795.10 | tyrosine aminotransferase (2.6.1.5) | 4,488 | 0 |
| TcCLB.511497.19 | amastin, putative | 4,476 | 2,32E-72 |
| TcCLB.510535.100 | cysteine peptidase C (CPC), putative (TCCB) | 4,472 | 0 |
| TcCLB.506753.130 | hypothetical protein, conserved | 4,408 | 8,41E-79 |
| TcCLB.506789.340 | hypothetical protein, conserved | 4,395 | 1,376E-88 |
| TcCLB.511779.60 | trans-sialidase, Group III, putative | 4,367 | 0,0007992 |

| | | | |
|---|---|---|---|
| TcCLB.510187.40 | tyrosine aminotransferase | 4,359 | 6,23E-299 |
| TcCLB.507689.40 | glutamamyl carboxypeptidase, putative | 4,349 | 0 |
| TcCLB.511859.200 | hypothetical protein, conserved | 4,303 | 1,274E-258 |
| TcCLB.506789.352 | hypothetical protein, conserved | 4,284 | 2,195E-22 |
| TcCLB.506885.310 | meiotic recombination protein DMC1, putative (dmc1) | 4,248 | 3,469E-91 |
| TcCLB.509051.20 | amastin, putative | 4,206 | 1,911E-79 |
| TcCLB.503975.100 | Present in the outer mitochondrial membrane proteome 2 | 4,178 | 4,847E-155 |
| TcCLB.508535.50 | tyrosine aminotransferase | 4,173 | 4,355E-97 |
| TcCLB.509965.394 | amastin, putative | 4,156 | 5,22E-46 |
| TcCLB.508263.5 | hypothetical protein, conserved (fragment) | 4,151 | 6,838E-53 |
| TcCLB.510837.9 | acetylornithine deacetylase, putative (fragment) | 4,149 | 6,447E-36 |
| TcCLB.509205.120 | hypothetical protein, conserved | 4,143 | 1,15E-289 |
| TcCLB.509023.80 | hypothetical protein, conserved | 4,137 | 2,13E-157 |
| TcCLB.510565.11 | tyrosine aminotransferase, putative (fragment) | 4,108 | 0 |
| TcCLB.509707.10 | protein associated with differentiation 8, putative | 4,084 | 4,845E-39 |
| TcCLB.504149.175 | hypothetical protein, conserved | 4,071 | 2,938E-186 |
| TcCLB.511779.10 | trans-sialidase, putative | 4,041 | 0,002643 |
| TcCLB.508799.290 | hypothetical protein, conserved (fragment) | 4,034 | 6,854E-222 |
| TcCLB.509769.40 | hypothetical protein, conserved | 4,034 | 1,448E-54 |
| TcCLB.510811.20 | protein associated with differentiation 4, putative | 4,027 | 1,62E-108 |
| TcCLB.511825.240 | hypothetical protein, conserved | 4,027 | 1,499E-38 |
| TcCLB.507689.50 | metallo-peptidase, clan MH, family M18, putative (fragme | 4,005 | 0 |
| TcCLB.503413.4 | Present in the outer mitochondrial membrane proteome 2 | 4,002 | 3,077E-82 |
| TcCLB.509197.50 | hypothetical protein, conserved | 3,979 | 1,438E-56 |
| TcCLB.509043.10 | tyrosine aminotransferase (pseudogene), putative | 3,971 | 2,694E-132 |
| TcCLB.508175.360 | cystathionine beta-synthase (CBS) | 3,956 | 0,0005739 |
| TcCLB.507519.150 | Adenine phosphoribosyltransferase, putative | 3,935 | 1,901E-85 |
| TcCLB.508999.260 | calpain-like cysteine peptidase (pseudogene), putative | 3,895 | 3,409E-193 |
| TcCLB.506773.50 | nucleoside transporter-like, putative | 3,877 | 8,176E-232 |
| TcCLB.510187.70 | tyrosine aminotransferase, putative | 3,871 | 6,107E-158 |
| TcCLB.511827.100 | cysteine peptidase C (CPC), putative (TCCB) | 3,86 | 2,898E-211 |
| TcCLB.503925.6 | hypothetical protein, conserved | 3,851 | 4,454E-37 |
| TcCLB.511437.30 | lipase, putative | 3,841 | 2,699E-46 |
| TcCLB.507689.30 | glutamamyl carboxypeptidase, putative | 3,824 | 9,359E-177 |
| TcCLB.505789.20 | hypothetical protein, conserved | 3,801 | 3,441E-91 |
| TcCLB.510101.80 | hypothetical protein, conserved | 3,797 | 4,032E-52 |
| TcCLB.506563.170 | calpain-like cysteine peptidase (pseudogene), putative | 3,769 | 1,265E-162 |
| TcCLB.503747.10 | glutamine aminotransferase, putative (fragment) (GlnAT) | 3,753 | 2,732E-82 |
| TcCLB.504533.50 | hypothetical protein, conserved | 3,749 | 6,26E-86 |
| TcCLB.510357.30 | hypothetical protein, conserved | 3,747 | 1,935E-64 |
| TcCLB.508601.141 | dipeptidyl-peptidase, putative (fragment) | 3,74 | 1,693E-176 |
| TcCLB.507657.20 | glutamamyl carboxypeptidase, putative | 3,734 | 0 |
| TcCLB.510187.30 | tyrosine aminotransferase | 3,683 | 5,263E-23 |
| TcCLB.507657.10 | glutamamyl carboxypeptidase, putative | 3,678 | 0 |
| TcCLB.507949.250 | hypothetical protein, conserved | 3,67 | 0 |

| | | | |
|---|---|---|---|
| TcCLB.508699.120 | dipeptidyl-peptidase, putative | 3,664 | 2,17E-232 |
| TcCLB.508173.160 | hypothetical protein, conserved | 3,648 | 1,805E-153 |
| TcCLB.509717.176 | retrotransposon hot spot (RHS) protein, putative (fragmen | 3,646 | 1,875E-09 |
| TcCLB.504045.60 | thermostable carboxypeptidase 2, putative | 3,638 | 2E-266 |
| TcCLB.509463.10 | transmembrane amino acid transporter, putative | 3,628 | 1,817E-138 |
| TcCLB.511367.240 | argonaute-like protein, putative | 3,617 | 4,378E-122 |
| TcCLB.507359.10 | hypothetical protein, conserved | 3,591 | 4,468E-25 |
| TcCLB.507143.20 | hypothetical protein, conserved | 3,589 | 4,044E-92 |
| TcCLB.506177.100 | actyltransferase-like, putative | 3,57 | 1,491E-45 |
| TcCLB.509179.100 | L-galactonolactone oxidase (ALO) | 3,558 | 3,094E-95 |
| TcCLB.507819.30 | trans-sialidase, Group II, putative | 3,554 | 1,031E-54 |
| TcCLB.509683.90 | hypothetical protein | 3,552 | 9,022E-53 |
| TcCLB.508033.39 | mucin-associated surface protein (MASP, pseudogene), pu | 3,545 | 2,871E-34 |
| TcCLB.504765.20 | protein associated with differentiation 8, putative | 3,545 | 2,562E-13 |
| TcCLB.507187.9 | dispersed gene family protein 1 (DGF-1), putative | 3,523 | 4,946E-100 |
| TcCLB.510381.10 | cystathionine beta-synthase, putative | 3,516 | 7,488E-09 |
| TcCLB.506247.220 | histidine ammonia-lyase, putative | 3,514 | 1,55E-260 |
| TcCLB.506227.10 | amino acid transporter, putative | 3,475 | 9,074E-109 |
| TcCLB.508647.180 | hypothetical protein, conserved | 3,47 | 3,039E-87 |
| TcCLB.508863.10 | hypothetical protein, conserved | 3,469 | 2,965E-77 |
| TcCLB.504421.19 | hypothetical protein, conserved | 3,466 | 1,922E-90 |
| TcCLB.508065.70 | protein associated with differentiation 8, putative | 3,461 | 1,336E-65 |
| TcCLB.511041.40 | hexose transporter, putative | 3,453 | 4,683E-36 |
| TcCLB.510977.9 | hypothetical protein, conserved (fragment) | 3,451 | 3,043E-140 |
| TcCLB.506001.90 | retrotransposon hot spot (RHS) protein, putative | 3,442 | 3,585E-114 |
| TcCLB.504003.30 | hypothetical protein | 3,43 | 9,329E-108 |
| TcCLB.510855.10 | peptide methionine sulfoxide reductase, putative | 3,423 | 1,589E-153 |
| TcCLB.506213.50 | prostaglandin F synthase (pseudogene), putative (PGFS) | 3,403 | 6,405E-153 |
| TcCLB.506513.110 | peptidase t, putative | 3,398 | 2,783E-119 |
| TcCLB.510605.10 | hypothetical protein, conserved | 3,388 | 1,212E-56 |
| TcCLB.401961.10 | trans-sialidase, putative (fragment) | 3,368 | 0,01501 |
| TcCLB.503837.20 | amino acid transporter, putative | 3,366 | 3,449E-114 |
| TcCLB.506753.140 | hypothetical protein, conserved | 3,364 | 1,279E-72 |
| TcCLB.510877.170 | hypothetical protein, conserved | 3,362 | 1,424E-134 |
| TcCLB.503697.70 | hypothetical protein, conserved | 3,36 | 2,186E-79 |
| TcCLB.506773.10 | hypothetical protein, conserved | 3,346 | 0,01548 |
| TcCLB.510837.20 | glutamamyl carboxypeptidase, putative | 3,336 | 3,181E-24 |
| TcCLB.506325.10 | retrotransposon hot spot (RHS) protein, putative | 3,334 | 6,319E-40 |
| TcCLB.509569.100 | nucleoside phosphorylase, putative | 3,332 | 3,896E-198 |
| TcCLB.509331.120 | hypothetical protein, conserved | 3,319 | 1,987E-66 |
| TcCLB.506879.10 | mucin-associated surface protein (MASP), putative | 3,315 | 0,0004173 |
| TcCLB.509153.20 | hypothetical protein, conserved | 3,313 | 5,417E-42 |
| TcCLB.507019.20 | nitrate reductase, putative | 3,309 | 1,377E-112 |
| TcCLB.510905.10 | dispersed gene family protein 1 (DGF-1), putative | 3,302 | 7,271E-06 |
| TcCLB.511691.10 | cystathionine beta-synthase (CBS) | 3,301 | 6,734E-20 |

| | | | |
|---|---|---|---|
| TcCLB.510613.40 | hypothetical protein, conserved (fragment) | 3,299 | 1,466E-21 |
| TcCLB.508045.131 | retrotransposon hot spot protein (RHS, pseudogene), putat | 3,289 | 1,32E-09 |
| TcCLB.504425.60 | NADP-dependent alcohol hydrogenase, putative | 3,286 | 2,304E-115 |
| TcCLB.509805.190 | hypothetical protein, conserved | 3,285 | 4,561E-67 |
| TcCLB.509331.74 | hypothetical protein, conserved | 3,245 | 8,7E-74 |
| TcCLB.504393.20 | hypothetical protein | 3,239 | 0,02463 |
| TcCLB.504105.220 | hypothetical protein, conserved | 3,232 | 1,248E-69 |
| TcCLB.507603.220 | hypothetical protein | 3,228 | 3,877E-51 |
| TcCLB.511439.40 | hypothetical protein | 3,222 | 4,195E-167 |
| TcCLB.504033.21 | UDP-Gal or UDP-GlcNAc-dependent glycosyltransferase, pu | 3,22 | 1,968E-07 |
| TcCLB.508917.49 | peptidase t, putative | 3,219 | 2,345E-37 |
| TcCLB.508439.40 | hypothetical protein | 3,214 | 0,0002256 |
| TcCLB.508039.70 | hypothetical protein, conserved | 3,213 | 6,752E-140 |
| TcCLB.511691.39 | hypothetical protein, conserved | 3,209 | 2,399E-136 |
| TcCLB.511207.10 | hypothetical protein, conserved | 3,207 | 4,041E-79 |
| TcCLB.503445.9 | retrotransposon hot spot (RHS) protein, putative | 3,205 | 1,117E-55 |
| TcCLB.508805.80 | short chain dehydrogenase, putative | 3,202 | 1,033E-49 |
| TcCLB.507247.14 | dispersed gene family protein 1 (DGF-1), putative (fragme | 3,188 | 8,983E-53 |
| TcCLB.508241.140 | cystathionine beta-synthase, putative (CBS) | 3,175 | 2,31E-26 |
| TcCLB.510643.40 | trans-sialidase, Group VIII, putative | 3,173 | 2,23E-12 |
| TcCLB.510679.40 | hypothetical protein | 3,171 | 9,155E-145 |
| TcCLB.508823.20 | hypothetical protein, conserved | 3,17 | 2,662E-157 |
| TcCLB.510065.5 | hypothetical protein, conserved (fragment) | 3,17 | 2,839E-42 |
| TcCLB.507891.29 | flagellar calcium-binding protein, putative | 3,169 | 0,0001865 |
| TcCLB.506011.10 | hypothetical protein, conserved | 3,159 | 2,996E-105 |
| TcCLB.506559.410 | hypothetical protein, conserved | 3,152 | 2,213E-142 |
| TcCLB.508919.10 | peptidase T (pseudogene), putative | 3,151 | 2,615E-38 |
| TcCLB.505615.20 | lipase (pseudogene), putative | 3,15 | 0,00001984 |
| TcCLB.511245.180 | guanin nucleotide-binding protein, putative | 3,141 | 2,133E-77 |
| TcCLB.462761.16 | hypothetical protein, conserved (fragment) | 3,14 | 8,77E-30 |
| TcCLB.511047.70 | hypothetical protein, conserved | 3,135 | 1,504E-171 |
| TcCLB.510729.110 | meiotic recombination protein DMC1, putative (dmc1) | 3,135 | 7,276E-100 |
| TcCLB.507117.9 | dispersed gene family protein 1 (DGF-1), putative | 3,129 | 2,106E-37 |
| TcCLB.510483.280 | 90 kDa surface protein, putative | 3,129 | 1,572E-18 |
| TcCLB.503419.30 | thiol-dependent reductase 1, putative (TcAc2) | 3,128 | 5,708E-204 |
| TcCLB.509107.9 | hypothetical protein, conserved (fragment) | 3,127 | 2,785E-25 |
| TcCLB.505047.19 | trans-sialidase (pseudogene), putative | 3,126 | 0,001112 |
| TcCLB.510063.30 | hypothetical protein, conserved | 3,118 | 6,072E-108 |
| TcCLB.503975.40 | hypothetical protein, conserved | 3,111 | 4,871E-146 |
| TcCLB.426675.9 | trans-sialidase, putative | 3,104 | 0,0001024 |
| TcCLB.429257.20 | cytochrome b5-dependent oleate desaturase | 3,103 | 5,711E-114 |
| TcCLB.511209.69 | membrane-bound acid phosphatase, putative (fragment) | 3,088 | 5,885E-70 |
| TcCLB.508827.50 | DNA repair and recombination helicase protein PIF3, putat | 3,087 | 2,283E-102 |
| TcCLB.508735.100 | hypothetical protein, conserved | 3,072 | 9,044E-107 |
| TcCLB.508999.250 | calpain-like cysteine peptidase (pseudogene), putative | 3,068 | 2,283E-129 |

| | | | |
|---|---|---|---|
| TcCLB.507473.40 | retrotransposon hot spot protein (RHS, pseudogene), putat | 3,068 | 2,656E-52 |
| TcCLB.506905.50 | cystathionine beta-synthase (CBS) | 3,066 | 0 |
| TcCLB.511051.10 | hypothetical protein | 3,064 | 1,879E-29 |
| TcCLB.506779.140 | hypothetical protein, conserved | 3,055 | 1,232E-101 |
| TcCLB.508003.10 | retrotransposon hot spot (RHS) protein, putative | 3,054 | 9,579E-50 |
| TcCLB.510347.29 | hypothetical protein, conserved | 3,053 | 7,191E-46 |
| TcCLB.508689.40 | hypothetical protein, conserved | 3,043 | 7,719E-114 |
| TcCLB.511575.130 | folate/pteridine transporter, putative | 3,04 | 6,218E-200 |
| TcCLB.504339.19 | NADP-dependent alcohol hydrogenase, putative | 3,036 | 0,00004197 |
| TcCLB.509705.10 | cellulosomal scaffoldin anchoring protein, putative | 3,026 | 1,477E-52 |
| TcCLB.506947.10 | hypothetical protein, conserved | 3,021 | 1,426E-72 |
| TcCLB.503459.10 | hypothetical protein, conserved | 3,021 | 3,806E-36 |
| TcCLB.511309.60 | hypothetical protein, conserved | 3,017 | 7,692E-34 |
| TcCLB.505971.10 | UDP-Gal or UDP-GlcNAc-dependent glycosyltransferase, pu | 3,01 | 3,753E-11 |
| TcCLB.507473.30 | trans-sialidase, Group II, putative | 3,007 | 7,002E-28 |
| TcCLB.507115.20 | dispersed gene family protein 1 (DGF-1), putative | 3,001 | 3,376E-31 |
| TcCLB.508177.110 | cystathionine beta-synthase | 2,998 | 6,313E-168 |
| TcCLB.508177.120 | cystathionine beta-synthase (CBS) | 2,997 | 7,574E-260 |
| TcCLB.503713.30 | 6-phosphogluconolactonase, putative | 2,997 | 9,363E-68 |
| TcCLB.505193.100 | Present in the outer mitochondrial membrane proteome 1 | 2,996 | 5,036E-60 |
| TcCLB.506909.60 | mucin TcMUCI, putative | 2,981 | 0,02959 |
| TcCLB.506931.4 | hypothetical protein, conserved | 2,98 | 7,302E-103 |
| TcCLB.510517.40 | hypothetical protein | 2,974 | 1,661E-74 |
| TcCLB.475133.10 | trans-sialidase, putative | 2,973 | 0,0002826 |
| TcCLB.508055.30 | trans-sialidase, Group II, putative | 2,972 | 6,063E-25 |
| TcCLB.511777.10 | trans-sialidase, putative (fragment) | 2,97 | 0,002146 |
| TcCLB.509367.30 | dispersed gene family protein 1 (DGF-1, pseudogene), puta | 2,966 | 5,342E-22 |
| TcCLB.510187.20 | tyrosine aminotransferase, putative | 2,964 | 3,637E-142 |
| TcCLB.509457.50 | retrotransposon hot spot (RHS) protein, putative | 2,962 | 3,037E-22 |
| TcCLB.404455.10 | trans-sialidase, putative (fragment) | 2,956 | 1,966E-18 |
| TcCLB.506563.120 | hypothetical protein, conserved | 2,955 | 6,854E-63 |
| TcCLB.506825.40 | hypothetical protein, conserved | 2,953 | 5,645E-23 |
| TcCLB.503683.11 | RNA-binding protein, putative | 2,952 | 3,458E-13 |
| TcCLB.510065.10 | hypothetical protein, conserved | 2,95 | 1,627E-153 |
| TcCLB.505609.40 | trans-sialidase, Group V, putative | 2,947 | 0,00199 |
| TcCLB.413409.10 | trans-sialidase, putative (fragment) | 2,943 | 4,01E-35 |
| TcCLB.508177.129 | cystathionine beta-synthase, putative | 2,939 | 1,087E-224 |
| TcCLB.509997.10 | hypothetical protein, conserved | 2,939 | 5,698E-20 |
| TcCLB.506195.120 | lysosomal alpha-mannosidase precursor, putative | 2,936 | 1,422E-88 |
| TcCLB.507831.20 | hypothetical protein, conserved | 2,932 | 0,002866 |
| TcCLB.510667.120 | galactokinase, putative | 2,93 | 2,755E-126 |
| TcCLB.507085.140 | mucin TcMUCII, putative | 2,928 | 0,0002868 |
| TcCLB.508827.10 | membrane-bound acid phosphatase, putative | 2,922 | 1,161E-106 |
| TcCLB.508801.10 | protein associated with differentiation 4, putative | 2,92 | 8,734E-118 |
| TcCLB.506739.99 | zinc finger (CCCH type) protein, putative | 2,915 | 6,318E-232 |

| | | | |
|---|---|---|---|
| TcCLB.508897.119 | FKBP-type peptidylprolyl cis-trans isomerase, putative | 2,905 | 1,436E-57 |
| TcCLB.511421.185 | hypothetical protein, conserved | 2,903 | 3,616E-27 |
| TcCLB.506937.10 | cytosolic malate dehydrogenase, putative | 2,9 | 2,747E-78 |
| TcCLB.511149.210 | mucin TcMUCII, putative | 2,872 | 0,00002839 |
| TcCLB.511911.90 | hypothetical protein, conserved | 2,868 | 6,46E-96 |
| TcCLB.503939.20 | hypothetical protein, conserved | 2,863 | 7,562E-78 |
| TcCLB.506825.200 | hypothetical protein | 2,857 | 3,403E-177 |
| TcCLB.509457.60 | retrotransposon hot spot (RHS) protein, putative | 2,852 | 2,461E-17 |
| TcCLB.510853.10 | dispersed gene family protein 1 (DGF-1, pseudogene), puta | 2,844 | 1,258E-31 |
| TcCLB.508503.20 | cytochrome c oxidase subunit V, putative | 2,841 | 5,923E-206 |
| TcCLB.510301.30 | hypothetical protein, conserved | 2,839 | 2,295E-40 |
| TcCLB.504137.100 | hypothetical protein, conserved | 2,837 | 9,559E-78 |
| TcCLB.511639.20 | hypothetical protein, conserved | 2,835 | 2,522E-144 |
| TcCLB.509713.10 | protein associated with differentiation 4, putative | 2,833 | 0 |
| TcCLB.510039.40 | nitrilase, putative | 2,83 | 5,695E-163 |
| TcCLB.509545.70 | mucin TcMUCII, putative | 2,824 | 0,01803 |
| TcCLB.511287.49 | aldo-keto reductase (AKR) | 2,823 | 1,717E-95 |
| TcCLB.508241.10 | glycerol-3-phosphate dehydrogenase, putative | 2,82 | 5,575E-81 |
| TcCLB.511923.10 | mucin-associated surface protein (MASP, pseudogene), pu | 2,816 | 0,05152 |
| TcCLB.511639.9 | hypothetical protein, conserved | 2,811 | 5,248E-42 |
| TcCLB.511817.40 | NADH-cytochrome b5 reductase, putative | 2,808 | 4,851E-103 |
| TcCLB.511793.30 | mucin-associated surface protein (MASP), putative | 2,807 | 0,04639 |
| TcCLB.507923.10 | L-threonine 3-dehydrogenase, putative | 2,806 | 5,315E-234 |
| TcCLB.506181.104 | actyltransferase-like protein | 2,806 | 6,133E-52 |
| TcCLB.403481.10 | trans-sialidase, putative (fragment) | 2,803 | 0,01839 |
| TcCLB.511613.189 | mucin TcMUCII, putative | 2,802 | 0,00003532 |
| TcCLB.511277.140 | serine/threonine protein kinase, putative | 2,801 | 6,346E-56 |
| TcCLB.510187.100 | glucokinase | 2,798 | 1,013E-100 |
| TcCLB.509331.210 | NADP-dependent alcohol hydrogenase, putative | 2,795 | 2,891E-40 |
| TcCLB.504005.54 | hypothetical protein, conserved | 2,79 | 8,646E-76 |
| TcCLB.508809.40 | hypothetical protein, conserved | 2,789 | 6,723E-38 |
| TcCLB.506763.30 | mucin-associated surface protein (MASP), putative | 2,784 | 0,04219 |
| TcCLB.511753.120 | L-threonine 3-dehydrogenase, putative | 2,783 | 5,371E-267 |
| TcCLB.508303.10 | alanine racemase 2, putative | 2,782 | 1,87E-71 |
| TcCLB.507255.20 | trans-sialidase, Group II, putative | 2,769 | 4,152E-51 |
| TcCLB.508501.110 | mucin TcMUCII, putative | 2,767 | 7,469E-06 |
| TcCLB.507009.40 | galactokinase, putative | 2,754 | 5,35E-89 |
| TcCLB.509999.140 | RNA-binding protein, putative | 2,747 | 6,087E-12 |
| TcCLB.508595.40 | cysteine peptidase, putative | 2,744 | 1,052E-109 |
| TcCLB.506257.98 | hypothetical protein, conserved (pseudogene) | 2,744 | 1,847E-64 |
| TcCLB.506025.4 | membrane-bound acid phosphatase, putative (fragment) | 2,743 | 7,433E-91 |
| TcCLB.504153.110 | hypothetical protein, conserved | 2,739 | 1,353E-72 |
| TcCLB.506147.9 | prostaglandin F2alpha synthase (TcPGFS) | 2,734 | 5,931E-182 |
| TcCLB.508139.130 | trans-sialidase, Group II, putative | 2,733 | 8,875E-81 |
| TcCLB.506319.60 | hypothetical protein, conserved | 2,732 | 1,398E-40 |

| | | | |
|---|---|---|---|
| TcCLB.508535.10 | NADH-dependent fumarate reductase, putative | 2,729 | 8,797E-234 |
| TcCLB.509641.79 | hypothetical protein, conserved (fragment) | 2,724 | 2,322E-37 |
| TcCLB.508923.10 | amino acid permease, putative | 2,722 | 4,793E-68 |
| TcCLB.506485.10 | hypothetical protein, conserved | 2,721 | 3,676E-37 |
| TcCLB.504557.10 | UDP-Gal or UDP-GlcNAc-dependent glycosyltransferase, pu | 2,719 | 1,487E-23 |
| TcCLB.508405.70 | protein kinase, putative | 2,716 | 7,073E-72 |
| TcCLB.510345.9 | protein kinase, putative (fragment) | 2,715 | 3,852E-12 |
| TcCLB.511295.10 | FKBP-type peptidylprolyl cis-trans isomerase, putative | 2,712 | 1,585E-54 |
| TcCLB.508677.80 | NADP-dependent alcohol hydrogenase, putative | 2,712 | 2,149E-42 |
| TcCLB.506563.180 | calpain-like cysteine peptidase (pseudogene), putative | 2,71 | 1,406E-82 |
| TcCLB.508761.260 | mucin TcMUCII, putative | 2,71 | 3,621E-14 |
| TcCLB.511303.34 | Macro domain containing protein, putative | 2,708 | 1,757E-57 |
| TcCLB.506779.200 | glycerol-3-phosphate dehydrogenase (pseudogene), putati | 2,707 | 6,657E-112 |
| TcCLB.507167.240 | retrotransposon hot spot (RHS) protein, putative | 2,699 | 7,508E-43 |
| TcCLB.505373.10 | retrotransposon hot spot (RHS) protein, putative | 2,697 | 8,42E-63 |
| TcCLB.506125.20 | hypothetical protein, conserved | 2,694 | 3,569E-80 |
| TcCLB.510131.44 | hypothetical protein, conserved | 2,679 | 6,665E-67 |
| TcCLB.427013.10 | trans-sialidase, putative (fragment) | 2,675 | 2,964E-20 |
| TcCLB.507153.30 | dispersed gene family protein 1 (DGF-1), putative | 2,671 | 1,01E-58 |
| TcCLB.509287.230 | trans-sialidase, Group II, putative | 2,666 | 2,8E-76 |
| TcCLB.506993.110 | mucin TcMUCII, putative | 2,663 | 0,00001867 |
| TcCLB.508257.30 | hypothetical protein, conserved | 2,662 | 1,164E-43 |
| TcCLB.511479.39 | hypothetical protein, conserved | 2,659 | 3,265E-63 |
| TcCLB.508081.30 | mucin-associated surface protein (MASP), putative | 2,654 | 0,06565 |
| TcCLB.504045.55 | hypothetical protein, conserved | 2,646 | 3,542E-115 |
| TcCLB.508831.110 | serine/threonine kinase, putative | 2,646 | 3,058E-46 |
| TcCLB.510561.35 | hypothetical protein | 2,646 | 0,00771 |
| TcCLB.508425.10 | dispersed gene family protein 1 (DGF-1), putative | 2,638 | 7,267E-08 |
| TcCLB.503841.80 | NADH-dependent fumarate reductase, putative | 2,634 | 4,815E-21 |
| TcCLB.510213.110 | NADH-dependent fumarate reductase, putative | 2,633 | 5,463E-32 |
| TcCLB.504127.20 | hypothetical protein | 2,632 | 2,423E-29 |
| TcCLB.504045.110 | urocanate hydratase, putative | 2,631 | 7,386E-218 |
| TcCLB.504223.10 | amino acid permease, putative | 2,623 | 3,208E-82 |
| TcCLB.510655.120 | aminopeptidase P, putative | 2,618 | 6,064E-121 |
| TcCLB.508611.30 | surface protease GP63, putative | 2,618 | 3,861E-32 |
| TcCLB.506789.160 | hypothetical protein, conserved | 2,617 | 2,945E-90 |
| TcCLB.506593.60 | nicotinamidase, putative (PNC1) | 2,615 | 6,441E-94 |
| TcCLB.506549.9 | dispersed gene family protein 1 (DGF-1), putative (fragme | 2,614 | 1,448E-75 |
| TcCLB.503969.20 | hypothetical protein, conserved | 2,613 | 0,003078 |
| TcCLB.506399.80 | hypothetical protein, conserved | 2,611 | 2,008E-49 |
| TcCLB.509601.59 | protoheme IX farnesyltransferase, putative | 2,61 | 2,58E-110 |
| TcCLB.510751.4 | aminopeptidase P, putative | 2,608 | 3,545E-100 |
| TcCLB.511325.40 | amino acid permease, putative | 2,608 | 6,012E-77 |
| TcCLB.438313.9 | dispersed gene family protein 1 (DGF-1), putative | 2,606 | 2,038E-49 |
| TcCLB.511277.500 | hypothetical protein, conserved | 2,601 | 2,894E-41 |

| | | | |
|---|---|---|---|
| TcCLB.506355.10 | glucose transporter | 2,597 | 8,236E-85 |
| TcCLB.509229.60 | hypothetical protein, conserved | 2,596 | 6,593E-36 |
| TcCLB.504223.30 | hypothetical protein, conserved (fragment) | 2,595 | 1,529E-08 |
| TcCLB.509713.20 | protein associated with differentiation 4, putative | 2,59 | 0 |
| TcCLB.506629.200 | 1,2-Dihydroxy-3-keto-5-methylthiopentene dioxygenase, pu | 2,589 | 6,845E-146 |
| TcCLB.509153.10 | hypothetical protein, conserved | 2,588 | 3,113E-133 |
| TcCLB.510299.20 | hypothetical protein, conserved | 2,588 | 5,075E-59 |
| TcCLB.508609.10 | surface protease GP63, putative | 2,588 | 5,121E-37 |
| TcCLB.464807.10 | serine/threonine protein kinase, putative | 2,587 | 3,523E-35 |
| TcCLB.503943.14 | hypothetical protein, conserved | 2,583 | 2,765E-70 |
| TcCLB.510249.30 | hypothetical protein, conserved | 2,583 | 3,023E-13 |
| TcCLB.509177.50 | hypothetical protein, conserved | 2,582 | 1,426E-55 |
| TcCLB.504933.10 | calpain-like cysteine peptidase, putative | 2,582 | 3,331E-37 |
| TcCLB.506775.180 | hypothetical protein, conserved | 2,581 | 1,876E-139 |
| TcCLB.509463.41 | retrotransposon hot spot (RHS) protein, putative (fragmen | 2,577 | 3,117E-93 |
| TcCLB.511545.80 | L-Lysine transport protein | 2,575 | 2,851E-106 |
| TcCLB.508799.280 | protein associated with differentiation 4, putative | 2,573 | 1,012E-89 |
| TcCLB.508551.39 | hexose transporter, putative | 2,565 | 5,623E-51 |
| TcCLB.507237.80 | mucin-associated surface protein (MASP), putative | 2,56 | 0,009281 |
| TcCLB.440219.10 | trans-sialidase, putative (fragment) | 2,559 | 2,217E-48 |
| TcCLB.402919.10 | trans-sialidase, putative | 2,559 | 7,621E-12 |
| TcCLB.508109.20 | mucin TcMUCII, putative | 2,558 | 0,00001731 |
| TcCLB.503837.10 | hypothetical protein, conserved | 2,552 | 4,464E-114 |
| TcCLB.509157.220 | hypothetical protein, conserved | 2,551 | 1,349E-66 |
| TcCLB.508399.10 | NADH-ubiquinone oxidoreductase, mitochondrial, putative | 2,55 | 6,567E-89 |
| TcCLB.506789.280 | methylthioadenosine phosphorylase, putative | 2,549 | 2,944E-89 |
| TcCLB.507101.10 | amino acid permease, putative | 2,541 | 9,575E-84 |
| TcCLB.508461.80 | prostaglandin F2alpha synthase (TcPGFS) | 2,537 | 2,047E-263 |
| TcCLB.510739.19 | cystathione gamma lyase, putative (fragment) | 2,535 | 8,261E-33 |
| TcCLB.509459.79 | actin interacting protein-like protein, putative | 2,531 | 1,227E-45 |
| TcCLB.507919.10 | surface protease GP63, putative | 2,529 | 8,185E-48 |
| TcCLB.508651.20 | cytochrome c oxidase subunit 10, putative | 2,528 | 2,399E-67 |
| TcCLB.510667.140 | ribulose-phosphate 3-epimerase, putative | 2,525 | 3,059E-27 |
| TcCLB.506777.64 | hypothetical protein, conserved | 2,52 | 3,462E-26 |
| TcCLB.506195.290 | hypothetical protein, conserved | 2,518 | 3,361E-82 |
| TcCLB.508501.250 | UMP-CMP kinase, mitochondrial, putative | 2,516 | 5,152E-60 |
| TcCLB.509319.90 | trypanothione synthetase (TRYS) | 2,514 | 1,239E-70 |
| TcCLB.508451.60 | trans-sialidase, Group II, putative | 2,511 | 0,0001621 |
| TcCLB.507125.10 | trans-sialidase, Group II, putative | 2,506 | 1,201E-16 |
| TcCLB.511129.60 | hypothetical protein, conserved | 2,505 | 6,047E-57 |
| TcCLB.510989.30 | guanin nucleotide-binding protein, putative | 2,5 | 8,219E-40 |
| TcCLB.506355.100 | hexose transporter (pseudogene), putative | 2,498 | 7,255E-84 |
| TcCLB.508837.189 | endoplasmic reticulum oxidoreductin, putative | 2,497 | 2,503E-88 |
| TcCLB.510603.169 | hypothetical protein, conserved | 2,496 | 8,53E-14 |
| TcCLB.511051.59 | hypothetical protein, conserved | 2,491 | 2,732E-112 |

| | | | |
|---|---|---|---|
| TcCLB.506323.30 | retrotransposon hot spot (RHS) protein, putative | 2,491 | 1,188E-80 |
| TcCLB.508177.90 | hypothetical protein, conserved | 2,491 | 2,276E-76 |
| TcCLB.507837.20 | protein kinase, putative | 2,485 | 3,901E-54 |
| TcCLB.507809.39 | hypothetical protein, conserved | 2,483 | 1,132E-28 |
| TcCLB.508495.170 | mucin TcMUCII, putative | 2,483 | 1,73E-08 |
| TcCLB.509615.50 | hypothetical protein | 2,478 | 0,003645 |
| TcCLB.511687.19 | voltage-dependent anion-selective channel, putative (fragr | 2,476 | 4,587E-09 |
| TcCLB.510155.110 | hypothetical protein, conserved | 2,475 | 1,86E-73 |
| TcCLB.504983.10 | hypothetical protein | 2,475 | 4,846E-45 |
| TcCLB.509237.151 | calpain-like cysteine peptidase, putative (fragment) | 2,47 | 5,872E-53 |
| TcCLB.504065.10 | retrotransposon hot spot protein (RHS, pseudogene), putat | 2,47 | 3,527E-43 |
| TcCLB.511909.30 | hypothetical protein, conserved | 2,467 | 1,345E-43 |
| TcCLB.511289.90 | hypothetical protein, conserved | 2,462 | 3,909E-99 |
| TcCLB.510605.19 | hypothetical protein, conserved | 2,457 | 8,711E-103 |
| TcCLB.506355.5 | hexose transporter, putative | 2,456 | 2,814E-27 |
| TcCLB.506275.121 | hypothetical protein | 2,456 | 0,004325 |
| TcCLB.511527.70 | Unconventional myosin | 2,455 | 7,905E-91 |
| TcCLB.506625.230 | hypothetical protein, conserved | 2,454 | 4,377E-24 |
| TcCLB.511073.10 | fatty acid desaturase, putative | 2,451 | 6,293E-91 |
| TcCLB.510749.99 | aminopeptidase P, putative | 2,45 | 2,431E-19 |
| TcCLB.510741.10 | cystathione gamma lyase, putative | 2,447 | 9,678E-44 |
| TcCLB.507167.70 | retrotransposon hot spot (RHS) protein, putative | 2,446 | 3,601E-42 |
| TcCLB.510811.10 | protein associated with differentiation 4, putative | 2,446 | 1,842E-20 |
| TcCLB.510101.330 | hypothetical protein, conserved | 2,443 | 1,399E-98 |
| TcCLB.509617.80 | ABC transporter, putative | 2,441 | 1,021E-49 |
| TcCLB.506993.20 | mucin TcMUCI, putative | 2,44 | 0,01394 |
| TcCLB.420293.20 | trans-sialidase, Group II, putative | 2,437 | 4,043E-29 |
| TcCLB.509703.10 | cellulosomal scaffoldin anchoring protein, putative | 2,436 | 2,099E-40 |
| TcCLB.511361.4 | hypothetical protein, conserved | 2,426 | 0,0001757 |
| TcCLB.509963.40 | dispersed gene family protein 1 (DGF-1), putative | 2,425 | 2,971E-46 |
| TcCLB.511165.109 | hypothetical protein, conserved | 2,423 | 2,839E-18 |
| TcCLB.472357.10 | trans-sialidase, Group V, putative | 2,415 | 1,442E-22 |
| TcCLB.511759.40 | hypothetical protein, conserved | 2,412 | 6,812E-17 |
| TcCLB.402375.10 | retrotransposon hot spot protein (RHS, pseudogene), putat | 2,411 | 6,333E-88 |
| TcCLB.506779.80 | pantothenate kinase subunit, putative | 2,408 | 3,446E-100 |
| TcCLB.463279.20 | trans-sialidase, Group II, putative | 2,408 | 8,884E-45 |
| TcCLB.407477.50 | cytochrome c oxidase VIII (COX VIII), putative | 2,407 | 2,649E-119 |
| TcCLB.506263.10 | D-isomer specific 2-hydroxyacid dehydrogenase-protein, pu | 2,404 | 5,972E-183 |
| TcCLB.509961.9 | cytochrome c oxidase subunit 10, putative | 2,403 | 5,233E-67 |
| TcCLB.510343.15 | protein kinase, putative | 2,402 | 3,367E-10 |
| TcCLB.511329.10 | Calpain-like protein 2 (CALP2) | 2,4 | 6,447E-72 |
| TcCLB.503959.78 | hypothetical protein, conserved | 2,395 | 1,625E-71 |
| TcCLB.510699.50 | mucin TcMUCII, putative | 2,395 | 5,219E-12 |
| TcCLB.468803.10 | surface protease GP63, putative | 2,392 | 0,0001478 |
| TcCLB.510407.30 | hypothetical protein, conserved | 2,391 | 4,351E-43 |

| | | | |
|---|---|---|---|
| TcCLB.507617.9 | prostaglandin F2alpha synthase (TcPGFS) | 2,386 | 6,616E-198 |
| TcCLB.505057.20 | hypothetical protein | 2,386 | 1,862E-36 |
| TcCLB.507951.299 | 5-histidylcysteine sulfoxide synthase | 2,385 | 1,022E-114 |
| TcCLB.426799.9 | hypothetical protein, conserved (fragment) | 2,385 | 1,267E-10 |
| TcCLB.508999.200 | cysteine peptidase, Clan CA, family C2, putative | 2,382 | 1,827E-85 |
| TcCLB.508551.30 | hexose transporter, putative | 2,382 | 7,074E-73 |
| TcCLB.509739.10 | trans-sialidase, Group II, putative | 2,382 | 3,499E-30 |
| TcCLB.416605.20 | mucin TcMUCII (pseudogene), putative | 2,375 | 0,00001375 |
| TcCLB.508823.70 | P27 protein, putative (P27) | 2,374 | 2,559E-226 |
| TcCLB.510581.20 | receptor-type adenylate cyclase, putative | 2,372 | 6,333E-26 |
| TcCLB.506529.360 | cytochrome oxidase subunit IV, putative (COXIV) | 2,369 | 2,228E-172 |
| TcCLB.510359.570 | mucin-associated surface protein (MASP), putative | 2,368 | 1,854E-28 |
| TcCLB.511907.89 | hypothetical protein, conserved | 2,368 | 8,036E-19 |
| TcCLB.511335.9 | calpain-like cysteine peptidase, putative (fragment) | 2,367 | 3,945E-11 |
| TcCLB.507125.30 | retrotransposon hot spot protein (RHS, pseudogene), putat | 2,366 | 6,119E-08 |
| TcCLB.508621.10 | protein kinase, putative | 2,363 | 2,03E-56 |
| TcCLB.509937.200 | hypothetical protein, conserved | 2,363 | 4,424E-46 |
| TcCLB.507149.20 | retrotransposon hot spot protein (RHS, pseudogene), putat | 2,361 | 9,815E-07 |
| TcCLB.511151.80 | hypothetical protein, conserved | 2,36 | 3,877E-68 |
| TcCLB.511255.604 | mucin TcMUCII, putative | 2,358 | 0,0006141 |
| TcCLB.510623.39 | mucin-associated surface protein (MASP, pseudogene), pu | 2,358 | 0,05597 |
| TcCLB.508207.70 | Adenine phosphoribosyltransferase, putative | 2,357 | 1,027E-51 |
| TcCLB.511075.9 | fatty acid desaturase, putative (fragment) | 2,354 | 6,439E-63 |
| TcCLB.507515.90 | hypothetical protein, conserved | 2,354 | 5,495E-55 |
| TcCLB.508023.100 | trans-sialidase, Group IV, putative | 2,354 | 2,092E-09 |
| TcCLB.506731.10 | hypothetical protein | 2,35 | 6,924E-13 |
| TcCLB.510339.80 | hypothetical protein, conserved | 2,349 | 1,961E-143 |
| TcCLB.511575.30 | hypothetical protein, conserved | 2,344 | 1,415E-45 |
| TcCLB.511671.80 | serine/threonine kinase, putative | 2,342 | 1,365E-34 |
| TcCLB.508669.10 | hypothetical protein, conserved | 2,338 | 1,522E-57 |
| TcCLB.511053.10 | hypothetical protein, conserved | 2,338 | 5,306E-31 |
| TcCLB.510659.240 | lactoylglutathione lyase-like protein, putative | 2,336 | 8,819E-59 |
| TcCLB.506717.130 | trans-sialidase (pseudogene), putative | 2,335 | 3,812E-34 |
| TcCLB.506211.140 | hypothetical protein, conserved | 2,334 | 2,289E-107 |
| TcCLB.511389.110 | cytochrome c oxidase VIII (COX VIII), putative | 2,333 | 1,316E-122 |
| TcCLB.508193.60 | UDP-Gal or UDP-GlcNAc-dependent glycosyltransferase, pu | 2,333 | 6,527E-23 |
| TcCLB.508493.110 | mucin-associated surface protein (MASP), putative | 2,331 | 3,998E-11 |
| TcCLB.511817.20 | hypothetical protein, conserved | 2,33 | 4,144E-73 |
| TcCLB.507561.10 | trans-sialidase, putative | 2,33 | 0,01825 |
| TcCLB.504131.90 | hypothetical protein, conserved | 2,327 | 6,211E-69 |
| TcCLB.509461.10 | actin interacting protein-like protein, putative | 2,327 | 7,824E-56 |
| TcCLB.504197.5 | mucin-associated surface protein (MASP), putative (fragm | 2,327 | 0,1053 |
| TcCLB.401183.10 | trans-sialidase, putative (fragment) | 2,325 | 9,797E-09 |
| TcCLB.510791.50 | mucin TcMUCII, putative | 2,325 | 0,0000185 |
| TcCLB.507085.150 | mucin-associated surface protein (MASP), putative | 2,324 | 9,267E-15 |

| | | | |
|---|---|---|---|
| TcCLB.511127.80 | hypothetical protein, conserved | 2,323 | 3,962E-47 |
| TcCLB.511039.30 | hypothetical protein, conserved | 2,322 | 9,614E-39 |
| TcCLB.508677.134 | hypothetical protein, conserved | 2,321 | 4,105E-39 |
| TcCLB.507869.10 | NADH-ubiquinone oxidoreductase, mitochondrial, putative | 2,318 | 6,72E-77 |
| TcCLB.508849.11 | trans-sialidase, putative (fragment) | 2,318 | 0,005908 |
| TcCLB.508741.229 | Mitochondrial outer membrane protein porin, putative (VD | 2,314 | 2,52E-144 |
| TcCLB.509747.80 | hypothetical protein, conserved | 2,314 | 1,759E-25 |
| TcCLB.511613.180 | mucin-associated surface protein (MASP), putative | 2,313 | 0,00004936 |
| TcCLB.509059.50 | pyridoxal kinase, putative | 2,311 | 3,675E-79 |
| TcCLB.504069.120 | amino acid permease 24, putative (AAP24) | 2,308 | 1,053E-43 |
| TcCLB.510747.10 | hypothetical protein, conserved | 2,307 | 6,61E-31 |
| TcCLB.424195.5 | hypothetical protein, conserved | 2,307 | 0,002473 |
| TcCLB.506459.219 | mucin TcMUCII, putative | 2,303 | 0,00002699 |
| TcCLB.508195.10 | UDP-Gal or UDP-GlcNAc-dependent glycosyltransferase, pu | 2,3 | 2,75E-22 |
| TcCLB.506759.140 | 90 kDa surface protein, putative | 2,3 | 0,05631 |
| TcCLB.509003.60 | hypothetical protein, conserved | 2,298 | 1,927E-52 |
| TcCLB.505945.20 | ribonuclease mar1, putative | 2,297 | 5,107E-57 |
| TcCLB.503943.20 | hypothetical protein, conserved | 2,294 | 1,383E-113 |
| TcCLB.510103.30 | protein associated with differentiation 4, putative | 2,292 | 3,27E-88 |
| TcCLB.503501.14 | mucin TcMUCII, putative | 2,289 | 0,005912 |
| TcCLB.510005.20 | trans-sialidase, Group II, putative | 2,288 | 4,079E-20 |
| TcCLB.508761.80 | mucin TcMUCII, putative | 2,288 | 0,001924 |
| TcCLB.510155.160 | protein kinase, putative | 2,286 | 2,059E-90 |
| TcCLB.510305.70 | hypothetical protein, conserved | 2,286 | 5,809E-39 |
| TcCLB.508141.90 | hypothetical protein, conserved | 2,283 | 1,093E-39 |
| TcCLB.509753.90 | trans-sialidase (pseudogene), putative | 2,283 | 8,761E-17 |
| TcCLB.508737.70 | cytochrome b5-dependent oleate desaturase | 2,28 | 2,7E-107 |
| TcCLB.511301.90 | protein kinase, putative | 2,28 | 6,406E-51 |
| TcCLB.504557.20 | UDP-Gal or UDP-GlcNAc-dependent glycosyltransferase, pu | 2,28 | 9,274E-22 |
| TcCLB.507723.100 | cytochrome oxidase subunit IX, putative (COXIX) | 2,276 | 3,158E-80 |
| TcCLB.509463.20 | hypothetical protein, conserved | 2,276 | 1,245E-43 |
| TcCLB.507317.34 | hypothetical protein, conserved | 2,272 | 1,01E-40 |
| TcCLB.507889.20 | hypothetical protein, conserved | 2,269 | 3,628E-21 |
| TcCLB.504587.40 | mucin TcMUCII, putative | 2,267 | 0,00004719 |
| TcCLB.504867.70 | cytochrome c oxidase assembly protein, putative | 2,265 | 1,877E-63 |
| TcCLB.510725.40 | hypothetical protein, conserved | 2,265 | 9,064E-58 |
| TcCLB.505945.90 | folate/pteridine transporter, putative | 2,264 | 7,66E-92 |
| TcCLB.506563.210 | cysteine peptidase, Clan CA, family C2, putative | 2,259 | 3,296E-157 |
| TcCLB.506405.43 | hypothetical protein, conserved | 2,259 | 2,296E-25 |
| TcCLB.506797.110 | hypothetical protein, conserved | 2,259 | 7,105E-20 |
| TcCLB.508799.150 | nucleoside transporter-like, putative | 2,258 | 2,717E-67 |
| TcCLB.506801.30 | hypothetical protein | 2,257 | 4,145E-15 |
| TcCLB.506879.40 | mucin-associated surface protein (MASP, pseudogene), pu | 2,256 | 0,0001499 |
| TcCLB.508055.40 | surface protease GP63 (pseudogene), putative | 2,254 | 0,06875 |
| TcCLB.509895.30 | hypothetical protein, conserved | 2,253 | 7,358E-146 |

| | | | |
|---|---|---|---|
| TcCLB.511807.130 | hypothetical protein, conserved | 2,253 | 3,561E-08 |
| TcCLB.507233.110 | mismatch repair protein MSH4, putative | 2,248 | 1,368E-87 |
| TcCLB.508495.80 | mucin-associated surface protein (MASP, pseudogene), pu | 2,248 | 0,02624 |
| TcCLB.511859.150 | mucin-like glycoprotein, putative | 2,247 | 1,257E-30 |
| TcCLB.508985.21 | mucin TcMUCII, putative (fragment) | 2,247 | 0,1324 |
| TcCLB.510507.40 | transmembrane amino acid transporter, putative | 2,246 | 1,832E-46 |
| TcCLB.511325.50 | amino acid permease, putative | 2,245 | 4,384E-19 |
| TcCLB.504669.50 | mucin TcMUCI, putative | 2,244 | 1,241E-07 |
| TcCLB.508231.9 | hexose transporter, putative | 2,243 | 9,965E-55 |
| TcCLB.508737.10 | surface antigen TASV-C2, putative (TASV-C2) | 2,242 | 8,197E-56 |
| TcCLB.507143.80 | hypothetical protein, conserved | 2,242 | 2,106E-29 |
| TcCLB.509197.20 | cation transporter, putative | 2,241 | 1,637E-06 |
| TcCLB.509669.30 | ABC transporter, putative | 2,24 | 6,136E-46 |
| TcCLB.510785.10 | hypothetical protein | 2,24 | 2,151E-38 |
| TcCLB.510561.30 | mucin-associated surface protein (MASP), putative | 2,239 | 0,1242 |
| TcCLB.511859.130 | mucin-like glycoprotein, putative | 2,237 | 2,669E-27 |
| TcCLB.503501.7 | mucin-associated surface protein (MASP), putative (fragm | 2,236 | 0,1228 |
| TcCLB.510661.250 | cystathione gamma lyase, putative | 2,234 | 5,303E-147 |
| TcCLB.503879.110 | hypothetical protein, conserved | 2,233 | 9,845E-73 |
| TcCLB.509333.10 | trans-sialidase, Group III, putative | 2,232 | 1,441E-64 |
| TcCLB.511657.40 | hypothetical protein, conserved | 2,232 | 1,605E-40 |
| TcCLB.507479.29 | retrotransposon hot spot protein (RHS, pseudogene), putat | 2,229 | 1,705E-16 |
| TcCLB.509267.30 | hypothetical protein, conserved | 2,228 | 5,833E-24 |
| TcCLB.505683.10 | sterol 24-c-methyltransferase, putative | 2,225 | 4,488E-78 |
| TcCLB.506753.150 | hypothetical protein, conserved | 2,225 | 2,57E-22 |
| TcCLB.510669.5 | ribulose-phosphate 3-epimerase, putative (fragment) | 2,225 | 0,00002284 |
| TcCLB.510843.10 | beta galactofuranosyl glycosyltransferase (pseudogene), pı | 2,224 | 1,21E-41 |
| TcCLB.509329.31 | trans-sialidase, putative | 2,224 | 0,008186 |
| TcCLB.510863.9 | hypothetical protein | 2,222 | 5,115E-12 |
| TcCLB.507641.64 | hypothetical protein, conserved | 2,221 | 1,917E-84 |
| TcCLB.506853.50 | 2-oxoisovalerate dehydrogenase alpha subunit, putative | 2,221 | 7,69E-76 |
| TcCLB.503791.30 | hypothetical protein, conserved | 2,22 | 2,735E-119 |
| TcCLB.506735.30 | hypothetical protein, conserved | 2,22 | 1,469E-45 |
| TcCLB.506123.70 | hypothetical protein, conserved | 2,218 | 3,665E-12 |
| TcCLB.508119.70 | mucin TcMUCII, putative | 2,218 | 1,717E-11 |
| TcCLB.509441.10 | hypothetical protein, conserved | 2,215 | 4,005E-55 |
| TcCLB.474079.10 | trans-sialidase, putative | 2,213 | 9,804E-26 |
| TcCLB.507969.40 | hypothetical protein, conserved | 2,209 | 9,058E-31 |
| TcCLB.506529.550 | cysteine peptidase, putative | 2,208 | 8,474E-61 |
| TcCLB.509053.30 | hypothetical protein, conserved | 2,207 | 3,297E-106 |
| TcCLB.509257.10 | trans-sialidase, Group II, putative | 2,206 | 2,221E-56 |
| TcCLB.410241.10 | retrotransposon hot spot protein (RHS, pseudogene), putat | 2,204 | 0,004482 |
| TcCLB.511099.160 | mucin TcMUCII, putative | 2,197 | 0,0003316 |
| TcCLB.508821.40 | hypothetical protein, conserved | 2,196 | 7,234E-106 |
| TcCLB.507211.20 | protein kinase, putative | 2,195 | 3,845E-07 |

| | | | |
|---|---|---|---|
| TcCLB.506203.60 | hypothetical protein, conserved | 2,189 | 2,742E-53 |
| TcCLB.504163.60 | hypothetical protein, conserved | 2,189 | 2,107E-35 |
| TcCLB.511591.10 | mucin-associated surface protein (MASP), putative | 2,189 | 0,03014 |
| TcCLB.504033.80 | hypothetical protein, conserved | 2,188 | 7,823E-118 |
| TcCLB.509207.90 | pyrroline-5-carboxylate reductase, putative (P5CR) | 2,183 | 3,866E-94 |
| TcCLB.507219.20 | retrotransposon hot spot protein (RHS, pseudogene), putat | 2,183 | 4,667E-76 |
| TcCLB.506931.10 | haloacid dehalogenase-like hydrolase, putative | 2,183 | 1,007E-38 |
| TcCLB.507491.39 | hypothetical protein, conserved | 2,183 | 2,16E-21 |
| TcCLB.510329.150 | hypothetical protein, conserved | 2,181 | 1,273E-74 |
| TcCLB.511361.10 | kinesin-like protein (pseudogene), putative | 2,181 | 3,606E-41 |
| TcCLB.509859.10 | hypothetical protein, conserved | 2,177 | 1,399E-98 |
| TcCLB.511037.20 | L-2-hydroxyglutarate dehydrogenase, mitochondrial, putati | 2,177 | 3,621E-59 |
| TcCLB.509853.10 | mannosyl-oligosaccharide 1,2-alpha-mannosidase IB, puta | 2,177 | 1,184E-51 |
| TcCLB.506179.5 | ferric reductase transmembrane protein, putative (fragme | 2,177 | 0,0004931 |
| TcCLB.507773.10 | trans-sialidase, Group IV, putative | 2,173 | 2,109E-24 |
| TcCLB.507523.75 | mucin TcMUCII (pseudogene), putative | 2,172 | 0,0001958 |
| TcCLB.509109.14 | hypothetical protein, conserved (fragment) | 2,171 | 1,014E-25 |
| TcCLB.507771.70 | hypothetical protein, conserved | 2,163 | 2,399E-79 |
| TcCLB.510565.30 | cytochrome c oxidase subunit V, putative | 2,161 | 5,077E-127 |
| TcCLB.511693.110 | hypothetical protein, conserved | 2,159 | 4,081E-36 |
| TcCLB.509329.9 | retrotransposon hot spot (RHS) protein, putative | 2,158 | 5,7E-22 |
| TcCLB.503731.9 | hypothetical protein (fragment) | 2,157 | 0,00001068 |
| TcCLB.505621.10 | mucin-associated surface protein (MASP, pseudogene), pu | 2,154 | 1,564E-25 |
| TcCLB.510591.30 | retrotransposon hot spot protein (RHS, pseudogene), putat | 2,152 | 9,441E-09 |
| TcCLB.511661.80 | hypothetical protein, conserved | 2,151 | 3,846E-31 |
| TcCLB.508211.60 | hypothetical protein, conserved | 2,149 | 1,469E-51 |
| TcCLB.504153.30 | hypothetical protein, conserved | 2,147 | 8,042E-51 |
| TcCLB.511613.120 | mucin TcMUCII, putative | 2,146 | 0,00629 |
| TcCLB.507611.280 | cytochrome oxidase subunit IX, putative (COXIX) | 2,145 | 5,514E-78 |
| TcCLB.508465.40 | hypothetical protein, conserved | 2,143 | 1,035E-51 |
| TcCLB.508121.120 | mucin TcMUCII, putative | 2,143 | 0,0004603 |
| TcCLB.509799.140 | L-2-hydroxyglutarate dehydrogenase, mitochondrial, putati | 2,142 | 8,535E-56 |
| TcCLB.508873.430 | trans-sialidase (pseudogene), putative | 2,142 | 2,884E-18 |
| TcCLB.509109.160 | hypothetical protein, conserved | 2,141 | 2,785E-15 |
| TcCLB.466593.10 | hypothetical protein, conserved | 2,14 | 1,646E-23 |
| TcCLB.503461.20 | per-hexamer repeat protein 5, putative | 2,139 | 7,173E-34 |
| TcCLB.508163.310 | mucin TcMUCII, putative | 2,138 | 2,293E-08 |
| TcCLB.507719.9 | dispersed gene family protein 1 (DGF-1), putative | 2,137 | 5,785E-52 |
| TcCLB.504143.10 | hypothetical protein, conserved | 2,136 | 1,103E-26 |
| TcCLB.506543.40 | hypothetical protein, conserved | 2,134 | 6,267E-22 |
| TcCLB.509429.329 | cruzipain precursor, putative (fragment) | 2,133 | 1,469E-213 |
| TcCLB.511151.90 | glycerol-3-phosphate dehydrogenase, putative | 2,133 | 3,448E-132 |
| TcCLB.506417.5 | endoribonuclease L-PSP (pb5), putative | 2,133 | 4,72E-63 |
| TcCLB.508701.20 | cation transporter, putative | 2,133 | 5,021E-06 |
| TcCLB.507129.5 | hypothetical protein, conserved (fragment) | 2,131 | 4,324E-09 |

| | | | |
|---|---|---|---|
| TcCLB.504081.500 | mucin TcMUCII, putative | 2,131 | 0,01658 |
| TcCLB.506243.10 | retrotransposon hot spot protein (RHS, pseudogene), putat | 2,129 | 3,22E-55 |
| TcCLB.509429.280 | hypothetical protein | 2,127 | 1,113E-50 |
| TcCLB.508173.140 | hypothetical protein | 2,126 | 6,448E-53 |
| TcCLB.507669.220 | hypothetical protein, conserved | 2,124 | 1,602E-57 |
| TcCLB.510245.10 | amino acid permease, putative | 2,122 | 5,177E-86 |
| TcCLB.508495.150 | mucin-associated surface protein (MASP, pseudogene), pu | 2,122 | 0,00000488 |
| TcCLB.506001.30 | hypothetical protein | 2,119 | 0,01619 |
| TcCLB.510101.50 | lanosterol 14-alpha-demethylase, putative | 2,117 | 1,118E-189 |
| TcCLB.508737.210 | Mitochondrial outer membrane protein porin, putative (VD | 2,117 | 4,214E-102 |
| TcCLB.510581.9 | receptor-type adenylate cyclase, putative (fragment) | 2,117 | 5,29E-27 |
| TcCLB.511439.30 | lipase, putative | 2,117 | 5,588E-16 |
| TcCLB.508257.220 | expression site-associated gene (ESAG-like) protein, putati | 2,116 | 1,459E-68 |
| TcCLB.510033.14 | mucin-associated surface protein (MASP, pseudogene), pu | 2,112 | 4,931E-27 |
| TcCLB.509001.30 | hypothetical protein, conserved | 2,109 | 2,933E-29 |
| TcCLB.482369.29 | arginine kinase (fragment) | 2,109 | 8,8E-18 |
| TcCLB.508241.20 | hypothetical protein, conserved | 2,108 | 3,936E-69 |
| TcCLB.510945.70 | aspartate aminotransferase, mitochondrial, putative | 2,107 | 3,823E-56 |
| TcCLB.426897.10 | hypothetical protein | 2,105 | 2,372E-40 |
| TcCLB.506679.270 | hypothetical protein, conserved | 2,102 | 9,456E-51 |
| TcCLB.510357.40 | hypothetical protein, conserved | 2,101 | 5,841E-24 |
| TcCLB.509437.160 | trans-sialidase, Group IV, putative | 2,099 | 5,64E-25 |
| TcCLB.508555.50 | calpain-like cysteine peptidase, putative | 2,097 | 2,403E-36 |
| TcCLB.509105.70 | thiol-dependent reductase 1, putative (TcAc2) | 2,095 | 2,147E-108 |
| TcCLB.508267.50 | hypothetical protein, conserved | 2,09 | 2,863E-39 |
| TcCLB.504069.20 | cytochrome c oxidase subunit VI, putative | 2,089 | 1,82E-64 |
| TcCLB.510093.10 | ATPase, putative | 2,087 | 0,03825 |
| TcCLB.508267.30 | hypothetical protein, conserved | 2,085 | 3,796E-57 |
| TcCLB.506767.310 | trans-sialidase (pseudogene), putative | 2,085 | 7,061E-10 |
| TcCLB.511605.55 | hypothetical protein | 2,083 | 0,001055 |
| TcCLB.511545.90 | hypothetical protein | 2,081 | 1,497E-44 |
| TcCLB.508567.70 | peptidyl-prolyl cis-trans isomerase (PIN1) | 2,081 | 9,3E-38 |
| TcCLB.510609.50 | nuclear receptor binding factor, putative | 2,08 | 2,477E-34 |
| TcCLB.511775.21 | mitochondrial RNA editing ligase 1 (pseudogene), putative | 2,079 | 0,158 |
| TcCLB.506635.60 | pyroglutamyl-peptidase I, putative (PPI) | 2,077 | 8,267E-47 |
| TcCLB.510943.50 | delta-1-pyrroline-5-carboxylate dehydrogenase, putative | 2,076 | 7,668E-85 |
| TcCLB.511817.210 | protein kinase, putative | 2,076 | 2,249E-29 |
| TcCLB.510483.210 | trans-sialidase (pseudogene), putative | 2,076 | 2,063E-22 |
| TcCLB.509239.10 | fatty acid desaturase, putative | 2,075 | 3,767E-92 |
| TcCLB.509365.20 | dispersed gene family protein 1 (DGF-1, pseudogene), puta | 2,073 | 1,656E-23 |
| TcCLB.510491.60 | trans-sialidase, Group IV, putative | 2,072 | 4,668E-37 |
| TcCLB.504801.20 | surface protease GP63, putative | 2,072 | 0,006731 |
| TcCLB.507359.20 | surface antigen TASV, putative | 2,072 | 0,01345 |
| TcCLB.505683.5 | hypothetical protein | 2,071 | 9,178E-67 |
| TcCLB.509693.209 | mismatch repair protein, putative | 2,071 | 7,812E-33 |

| | | | |
|---|---|---|---|
| TcCLB.511361.20 | hypothetical protein, conserved | 2,071 | 6,965E-10 |
| TcCLB.506297.230 | hypothetical protein, conserved | 2,07 | 1,199E-31 |
| TcCLB.506965.90 | mucin TcMUCII, putative | 2,069 | 0,0002161 |
| TcCLB.510185.10 | sterol 24-c-methyltransferase, putative | 2,065 | 7,307E-69 |
| TcCLB.510673.10 | dispersed gene family protein 1 (DGF-1, pseudogene), puta | 2,063 | 5,349E-27 |
| TcCLB.506893.100 | UMP-CMP kinase, mitochondrial, putative | 2,062 | 1,587E-37 |
| TcCLB.507393.9 | dispersed gene family protein 1 (DGF-1), putative | 2,061 | 5,066E-32 |
| TcCLB.503975.50 | hypothetical protein, conserved | 2,06 | 1,02E-83 |
| TcCLB.508739.21 | mucin TcSMUGS (pseudogene), putative | 2,06 | 5,087E-37 |
| TcCLB.504393.10 | hypothetical protein | 2,06 | 0,002387 |
| TcCLB.508823.10 | hypothetical protein, conserved | 2,059 | 1,476E-52 |
| TcCLB.508201.40 | mucin TcMUCII, putative | 2,059 | 7,99E-10 |
| TcCLB.507631.20 | hypothetical protein, conserved | 2,057 | 1,668E-26 |
| TcCLB.510829.10 | hypothetical protein, conserved | 2,053 | 1,354E-39 |
| TcCLB.508781.110 | 2-oxoisovalerate dehydrogenase alpha subunit, putative | 2,048 | 8,714E-51 |
| TcCLB.511393.10 | nonspecific lipid-transfer protein, putative | 2,048 | 1,503E-31 |
| TcCLB.508677.160 | hypothetical protein, conserved | 2,046 | 1,597E-38 |
| TcCLB.503943.10 | serine/threonine protein kinase, putative (fragment) | 2,045 | 8,852E-61 |
| TcCLB.510445.10 | casein kinase 1 isoform 2, putative | 2,044 | 4,98E-17 |
| TcCLB.507275.40 | hypothetical protein, conserved | 2,043 | 2,848E-18 |
| TcCLB.507429.10 | mucin TcMUCII (pseudogene), putative | 2,043 | 0,00007385 |
| TcCLB.511809.30 | hypothetical protein, conserved | 2,041 | 1,764E-53 |
| TcCLB.509449.10 | receptor-type adenylate cyclase, putative | 2,041 | 6,654E-24 |
| TcCLB.445783.10 | mucin-associated surface protein (MASP, pseudogene), pu | 2,041 | 0,001427 |
| TcCLB.507443.30 | UDP-Gal or UDP-GlcNAc-dependent glycosyltransferase, pu | 2,04 | 3,349E-09 |
| TcCLB.505945.60 | hypothetical protein, conserved | 2,039 | 2,177E-46 |
| TcCLB.511237.30 | proline oxidase (pseudogene), putative | 2,038 | 1,417E-48 |
| TcCLB.509327.10 | retrotransposon hot spot protein (RHS, pseudogene), putat | 2,038 | 5,805E-26 |
| TcCLB.506821.210 | cytochrome-b5 reductase, putative | 2,037 | 1,218E-63 |
| TcCLB.511843.10 | mucin-associated surface protein (MASP), putative | 2,037 | 0,00001027 |
| TcCLB.509397.9 | dispersed gene family protein 1 (DGF-1), putative (fragme | 2,036 | 9,873E-13 |
| TcCLB.511839.10 | mucin TcMUCII, putative | 2,036 | 0,001949 |
| TcCLB.508569.10 | hypothetical protein, conserved | 2,032 | 0,00002306 |
| TcCLB.506537.80 | trans-sialidase, Group II, putative | 2,031 | 5,064E-07 |
| TcCLB.510889.50 | cytochrome oxidase subunit IV, putative (COXIV) | 2,03 | 1,006E-89 |
| TcCLB.507483.39 | calmodulin (CALA2) | 2,03 | 6,005E-07 |
| TcCLB.511333.10 | calpain-like cysteine peptidase, putative | 2,024 | 2,203E-09 |
| TcCLB.506819.30 | superoxide dismutase, putative | 2,019 | 2,113E-38 |
| TcCLB.506693.20 | hypothetical protein, conserved | 2,019 | 2,364E-29 |
| TcCLB.511117.30 | trans-sialidase, Group II, putative | 2,018 | 1,485E-06 |
| TcCLB.507645.50 | hypothetical protein, conserved | 2,013 | 5,082E-30 |
| TcCLB.507967.20 | hypothetical protein, conserved | 2,012 | 5,806E-28 |
| TcCLB.503779.20 | hypothetical protein, conserved | 2,011 | 3,087E-42 |
| TcCLB.510039.80 | hypothetical protein, conserved | 2,011 | 1,131E-35 |
| TcCLB.509217.30 | retrotransposon hot spot protein (RHS, pseudogene), putat | 2,01 | 1,725E-06 |

| | | | |
|---|---|---|---|
| TcCLB.507357.130 | mucin TcMUCII, putative | 2,01 | 0,005202 |
| TcCLB.511895.20 | hypothetical protein | 2,008 | 2,394E-37 |
| TcCLB.510233.39 | trans-sialidase, putative (fragment) | 2,007 | 0,04311 |
| TcCLB.508013.60 | hypothetical protein | 2,005 | 0,00535 |
| TcCLB.508747.40 | mucin TcMUCII, putative | 2,004 | 7,283E-11 |
| TcCLB.506799.40 | hypothetical protein | 2,004 | 1,048E-10 |
| TcCLB.506717.140 | retrotransposon hot spot protein (RHS, pseudogene), putat | 2,003 | 0,00000144 |
| TcCLB.509141.40 | Mitochondrial outer membrane protein porin, putative (VD | 2,002 | 2,972E-51 |
| TcCLB.509205.60 | hypothetical protein, conserved | 2,001 | 1,867E-62 |
| TcCLB.426435.9 | fatty acid desaturase, putative | 2 | 6,427E-76 |
| TcCLB.506377.10 | trans-sialidase, putative | 2 | 1,653E-35 |
| TcCLB.508589.5 | trans-sialidase, putative (fragment) | 1,999 | 0,0003509 |
| TcCLB.510677.9 | dispersed gene family protein 1 (DGF-1), putative (fragme | 1,995 | 3,122E-32 |
| TcCLB.430539.10 | hypothetical protein, conserved (fragment) | 1,993 | 8,636E-08 |
| TcCLB.510139.10 | mucin-associated surface protein (MASP, pseudogene), pu | 1,992 | 8,846E-07 |
| TcCLB.508297.50 | short chain dehydrogenase, putative | 1,991 | 3,322E-34 |
| TcCLB.506683.110 | trans-sialidase, Group VIII, putative | 1,989 | 1,758E-30 |
| TcCLB.506417.10 | multidrug resistance protein A (pseudogene), putative | 1,988 | 6,739E-190 |
| TcCLB.506475.100 | conserved protein, unknown function | 1,987 | 3,602E-31 |
| TcCLB.507625.60 | dispersed gene family protein 1 (DGF-1, pseudogene), puta | 1,987 | 2,231E-27 |
| TcCLB.511817.60 | hypothetical protein, conserved | 1,987 | 4,652E-23 |
| TcCLB.509197.10 | cation transporter, putative | 1,987 | 0,00001192 |
| TcCLB.509453.79 | hypothetical protein, conserved | 1,984 | 4,099E-74 |
| TcCLB.506743.135 | trans-sialidase (pseudogene), putative | 1,984 | 1,282E-08 |
| TcCLB.509217.90 | mucin-associated surface protein (MASP), putative | 1,979 | 0,0003101 |
| TcCLB.511627.130 | hypothetical protein, conserved | 1,978 | 1,121E-48 |
| TcCLB.508023.40 | retrotransposon hot spot (RHS) protein, putative | 1,978 | 4,751E-25 |
| TcCLB.509265.10 | retrotransposon hot spot (RHS) protein, putative | 1,976 | 9,804E-50 |
| TcCLB.510843.19 | trans-sialidase, putative (fragment) | 1,974 | 6,71E-09 |
| TcCLB.508025.10 | retrotransposon hot spot (RHS) protein, putative (fragmen | 1,973 | 2,139E-08 |
| TcCLB.510191.20 | mucin TcMUCII, putative | 1,971 | 0,02365 |
| TcCLB.398477.10 | trans-sialidase, Group II, putative | 1,97 | 0,0001414 |
| TcCLB.511687.10 | Mitochondrial outer membrane protein porin, putative (VD | 1,969 | 4,737E-54 |
| TcCLB.509331.70 | 3-oxo-5-alpha-steroid 4-dehydrogenase, putative | 1,968 | 7,477E-43 |
| TcCLB.506199.20 | hypothetical protein, conserved | 1,968 | 1,998E-18 |
| TcCLB.511383.80 | hypothetical protein, conserved | 1,967 | 7,378E-20 |
| TcCLB.511255.700 | trans-sialidase, Group V, putative | 1,966 | 0,00000268 |
| TcCLB.509281.20 | trans-sialidase, Group II, putative | 1,965 | 0,00009676 |
| TcCLB.510659.20 | hypothetical protein, conserved | 1,964 | 2,386E-35 |
| TcCLB.508501.70 | mucin TcMUCII, putative | 1,964 | 1,503E-07 |
| TcCLB.509667.10 | dispersed gene family protein 1 (DGF-1), putative | 1,962 | 5,368E-18 |
| TcCLB.510039.30 | mucin-associated surface protein (MASP), putative | 1,961 | 9,536E-11 |
| TcCLB.507559.80 | deoxyribose-phosphate aldolase, putative | 1,959 | 2,28E-90 |
| TcCLB.503887.40 | hypothetical protein, conserved | 1,959 | 1,675E-76 |
| TcCLB.508387.20 | methylthioadenosine phosphorylase, putative | 1,958 | 4,214E-53 |

| | | | |
|---|---|---|---|
| TcCLB.503393.10 | amino acid permease, putative (fragment) | 1,958 | 1,807E-13 |
| TcCLB.510165.70 | retrotransposon hot spot protein (RHS, pseudogene), putat | 1,958 | 1,072E-07 |
| TcCLB.508495.50 | mucin TcMUCII, putative | 1,957 | 2,307E-09 |
| TcCLB.511573.30 | hypothetical protein, conserved | 1,949 | 8,238E-27 |
| TcCLB.508109.30 | mucin-associated surface protein (MASP), putative | 1,949 | 0,02538 |
| TcCLB.510669.49 | Gim5A protein, putative | 1,948 | 6,619E-58 |
| TcCLB.508677.60 | hypothetical protein | 1,948 | 5,082E-09 |
| TcCLB.506615.100 | mucin-associated surface protein (MASP), putative | 1,947 | 8,842E-15 |
| TcCLB.511693.10 | hypothetical protein | 1,946 | 1,214E-83 |
| TcCLB.508799.230 | hypothetical protein, conserved | 1,946 | 1,442E-44 |
| TcCLB.508811.40 | hypothetical protein | 1,945 | 3,168E-40 |
| TcCLB.508571.10 | hypothetical protein | 1,945 | 9,846E-16 |
| TcCLB.503975.70 | hypothetical protein, conserved | 1,941 | 5,106E-53 |
| TcCLB.503577.9 | delta-1-pyrroline-5-carboxylate dehydrogenase (pseudoger | 1,941 | 8,064E-30 |
| TcCLB.508971.40 | acetylornithine deacetylase-like, putative | 1,94 | 6,246E-114 |
| TcCLB.509493.20 | mucin TcMUCII, putative | 1,94 | 3,369E-06 |
| TcCLB.508557.29 | hypothetical protein, conserved | 1,937 | 2,984E-62 |
| TcCLB.506145.70 | hypothetical protein, conserved | 1,936 | 7,293E-26 |
| TcCLB.511821.20 | hypothetical protein, conserved | 1,936 | 6,48E-23 |
| TcCLB.508153.700 | hypothetical protein, conserved | 1,935 | 6,962E-29 |
| TcCLB.506417.83 | multidrug resistance-associated protein (pseudogene), put | 1,934 | 2,86E-34 |
| TcCLB.509493.80 | mucin TcMUCII, putative | 1,934 | 6,536E-12 |
| TcCLB.507709.90 | sterol C-24 reductase, putative | 1,933 | 9,532E-42 |
| TcCLB.509129.10 | Mitochondrial outer membrane protein porin, putative (VD | 1,932 | 1,178E-90 |
| TcCLB.504205.20 | trans-sialidase (pseudogene), putative | 1,932 | 1,728E-07 |
| TcCLB.506999.60 | hypothetical protein, conserved | 1,929 | 9,464E-46 |
| TcCLB.508293.44 | mucin-associated surface protein (MASP, pseudogene), pu | 1,928 | 0,06372 |
| TcCLB.503855.40 | calpain-like cysteine peptidase, putative | 1,927 | 3,462E-50 |
| TcCLB.503415.40 | hypothetical protein | 1,927 | 6,22E-28 |
| TcCLB.507611.170 | trans-sialidase, Group II, putative | 1,926 | 0,0003882 |
| TcCLB.509601.5 | mannosyl-oligosaccharide 1,2-alpha-mannosidase IB, puta | 1,925 | 3,454E-07 |
| TcCLB.508495.140 | mucin TcMUCII, putative | 1,924 | 3,367E-06 |
| TcCLB.508257.70 | phosphomannomutase-like protein, putative | 1,922 | 1,048E-57 |
| TcCLB.503745.25 | hypothetical protein, conserved | 1,922 | 3,267E-18 |
| TcCLB.509331.160 | hypothetical protein, conserved | 1,921 | 5,544E-29 |
| TcCLB.504085.60 | CMP-sialic acid transporter, putative | 1,921 | 2,558E-28 |
| TcCLB.510331.10 | trans-sialidase, Group III, putative | 1,921 | 1,316E-18 |
| TcCLB.504423.10 | hypothetical protein, conserved | 1,92 | 2,764E-19 |
| TcCLB.416619.10 | trans-sialidase, putative (fragment) | 1,92 | 0,01842 |
| TcCLB.508023.80 | retrotransposon hot spot protein (RHS, pseudogene), putat | 1,919 | 1,215E-21 |
| TcCLB.511487.90 | trans-sialidase, Group V, putative | 1,919 | 0,00000212 |
| TcCLB.505037.30 | hypothetical protein, conserved | 1,918 | 1,067E-12 |
| TcCLB.507963.20 | arginase, putative | 1,914 | 5,685E-47 |
| TcCLB.511815.100 | hypothetical protein, conserved | 1,913 | 7,639E-35 |
| TcCLB.507527.50 | cation transporter, putative | 1,913 | 0,0006014 |

| | | | |
|---|---|---|---|
| TcCLB.506407.10 | lysosomal alpha-mannosidase precursor, putative | 1,912 | 2,279E-70 |
| TcCLB.506759.190 | surface protease GP63 (pseudogene), putative | 1,911 | 0,06014 |
| TcCLB.506825.10 | RNA-binding protein, putative | 1,91 | 0,006898 |
| TcCLB.509967.20 | mismatch repair protein MSH4, putative | 1,908 | 3,571E-40 |
| TcCLB.511817.134 | hypothetical protein, conserved | 1,908 | 2,693E-26 |
| TcCLB.508501.40 | mucin TcMUCII, putative | 1,907 | 1,615E-10 |
| TcCLB.506529.430 | alanine aminotransferase, putative | 1,907 | 3,184E-07 |
| TcCLB.509699.150 | mucin TcMUCII, putative | 1,906 | 0,06774 |
| TcCLB.510625.79 | mucin TcMUCII (pseudogene), putative | 1,905 | 0,03027 |
| TcCLB.511585.120 | hypothetical protein, conserved | 1,904 | 4,697E-49 |
| TcCLB.509429.320 | cysteine peptidase, putative | 1,903 | 4,903E-191 |
| TcCLB.511553.90 | mucin TcMUCII, putative | 1,902 | 0,0001489 |
| TcCLB.510621.55 | mucin TcMUCII (pseudogene), putative | 1,901 | 0,009107 |
| TcCLB.508999.190 | calpain-like cysteine peptidase, putative | 1,9 | 2,205E-139 |
| TcCLB.509011.10 | UDP-Gal or UDP-GlcNAc-dependent glycosyltransferase (p | 1,898 | 3,122E-09 |
| TcCLB.509825.40 | hypothetical protein, conserved | 1,894 | 5,511E-48 |
| TcCLB.506989.140 | Elongation factor Ts, mitochondrial, putative (EF-Ts) | 1,893 | 8,861E-46 |
| TcCLB.508461.280 | protein kinase A catalytic subunit isoform 2, putative | 1,892 | 1,353E-32 |
| TcCLB.510129.11 | surface glycoprotein, putative | 1,891 | 3,038E-25 |
| TcCLB.510603.10 | hypothetical protein, conserved | 1,891 | 7,36E-14 |
| TcCLB.506219.20 | hypothetical protein | 1,891 | 1,575E-13 |
| TcCLB.511571.20 | hypothetical protein, conserved | 1,889 | 6,219E-50 |
| TcCLB.509419.70 | trans-sialidase, Group V, putative | 1,887 | 0,00004123 |
| TcCLB.506499.149 | mucin TcMUCII, putative | 1,887 | 0,0008859 |
| TcCLB.505201.9 | retrotransposon hot spot (RHS) protein, putative | 1,886 | 5,492E-24 |
| TcCLB.508043.10 | trans-sialidase, putative | 1,886 | 7,179E-08 |
| TcCLB.511761.60 | hypothetical protein, conserved | 1,885 | 5,629E-19 |
| TcCLB.507821.130 | trans-sialidase, Group II, putative | 1,885 | 1,665E-09 |
| TcCLB.510101.380 | hypothetical protein, conserved | 1,883 | 3,506E-59 |
| TcCLB.509695.100 | hypothetical protein, conserved | 1,883 | 1,658E-27 |
| TcCLB.508931.40 | hypothetical protein, conserved | 1,883 | 1,683E-15 |
| TcCLB.510005.41 | retrotransposon hot spot (RHS) protein, putative | 1,881 | 1,822E-52 |
| TcCLB.508207.90 | hypothetical protein, conserved | 1,879 | 2,818E-30 |
| TcCLB.509597.20 | carbonic anhydrase-like protein, putative | 1,878 | 7,88E-73 |
| TcCLB.509387.19 | trans-sialidase, putative | 1,878 | 1,898E-24 |
| TcCLB.508013.90 | hypothetical protein | 1,878 | 0,005692 |
| TcCLB.506945.190 | cytochrome p450-like protein, putative | 1,877 | 1,187E-78 |
| TcCLB.509157.90 | hypothetical protein, conserved | 1,874 | 2,002E-85 |
| TcCLB.511213.50 | mucin TcMUCII, putative | 1,873 | 9,702E-08 |
| TcCLB.503581.30 | hypothetical protein, conserved | 1,872 | 1,617E-29 |
| TcCLB.504203.40 | hypothetical protein, conserved | 1,87 | 2,84E-79 |
| TcCLB.508799.260 | hypothetical protein, conserved | 1,87 | 3,328E-66 |
| TcCLB.508023.60 | trans-sialidase (pseudogene), putative | 1,87 | 6,999E-22 |
| TcCLB.510017.40 | mucin TcMUCII, putative | 1,87 | 0,03101 |
| TcCLB.510583.100 | mucin TcMUCII, putative | 1,87 | 0,06515 |

| | | | |
|---|---|---|---|
| TcCLB.507241.30 | arginine kinase | 1,869 | 7,609E-18 |
| TcCLB.510621.70 | mucin TcMUCII, putative | 1,869 | 0,03052 |
| TcCLB.509601.150 | hypothetical protein, conserved | 1,865 | 1,029E-39 |
| TcCLB.506459.149 | mucin-associated surface protein (MASP), putative | 1,865 | 1,285E-21 |
| TcCLB.506189.20 | hypothetical protein, conserved | 1,864 | 2,104E-22 |
| TcCLB.505647.40 | endoribonuclease L-PSP (pb5), putative | 1,863 | 2,362E-119 |
| TcCLB.508699.70 | hypothetical protein, conserved | 1,862 | 7,382E-46 |
| TcCLB.507085.70 | mucin-associated surface protein (MASP, pseudogene), pu | 1,862 | 0,00004125 |
| TcCLB.511743.10 | hypothetical protein, conserved | 1,861 | 3,164E-31 |
| TcCLB.504055.130 | trans-sialidase (pseudogene), putative | 1,861 | 9,074E-22 |
| TcCLB.506779.120 | hypothetical protein, conserved | 1,86 | 3,748E-44 |
| TcCLB.511531.20 | hypothetical protein, conserved | 1,86 | 6,118E-23 |
| TcCLB.506383.10 | mannosyl-oligosaccharide 1,2-alpha-mannosidase IB, puta | 1,859 | 1,654E-26 |
| TcCLB.507479.51 | beta galactofuranosyl glycosyltransferase, putative | 1,856 | 0,0001129 |
| TcCLB.503411.4 | hypothetical protein, conserved | 1,854 | 5,773E-07 |
| TcCLB.509787.10 | retrotransposon hot spot (RHS) protein, putative | 1,853 | 8,999E-72 |
| TcCLB.509159.10 | ATP-dependent DNA DEAD/DEAH box helicase, putative (fr | 1,852 | 9,253E-39 |
| TcCLB.507711.200 | hypothetical protein, conserved | 1,85 | 4,902E-171 |
| TcCLB.506469.80 | protein kinase, putative | 1,85 | 4,953E-16 |
| TcCLB.509007.30 | hypothetical protein, conserved | 1,849 | 6,292E-23 |
| TcCLB.511583.40 | tricarboxylate carrier, putative | 1,847 | 3,139E-34 |
| TcCLB.415273.9 | mucin TcMUCII (pseudogene), putative | 1,847 | 0,07261 |
| TcCLB.509719.40 | cytochrome P450, putative | 1,846 | 6,135E-41 |
| TcCLB.511625.130 | trans-sialidase, Group V, putative | 1,846 | 4,285E-15 |
| TcCLB.511813.20 | ubiquitin-conjugating enzyme E2, putative | 1,845 | 4,039E-57 |
| TcCLB.505853.10 | trans-sialidase, putative (fragment) | 1,845 | 0,0002165 |
| TcCLB.508677.50 | beta galactofuranosyl glycosyltransferase, putative | 1,843 | 5,604E-29 |
| TcCLB.506873.4 | ATP-dependent DNA DEAD/DEAH box helicase, putative (fr | 1,842 | 2,518E-13 |
| TcCLB.506843.100 | retrotransposon hot spot (RHS) protein, putative | 1,841 | 2,373E-73 |
| TcCLB.506529.420 | alanine aminotransferase, putative | 1,841 | 1,788E-66 |
| TcCLB.505573.10 | trans-sialidase, putative (fragment) | 1,841 | 6,922E-07 |
| TcCLB.510887.50 | hypothetical protein, conserved | 1,836 | 6,038E-56 |
| TcCLB.508739.90 | hypothetical protein, conserved | 1,836 | 2,295E-20 |
| TcCLB.508547.160 | hypothetical protein, conserved | 1,835 | 1,128E-55 |
| TcCLB.511303.70 | hypothetical protein, conserved | 1,834 | 4,135E-31 |
| TcCLB.506169.21 | retrotransposon hot spot (RHS) protein, putative (fragmen | 1,833 | 1,299E-10 |
| TcCLB.511153.70 | hypothetical protein, conserved | 1,832 | 8,15E-30 |
| TcCLB.507479.20 | trans-sialidase, Group II, putative | 1,83 | 0,004172 |
| TcCLB.508461.84 | hypothetical protein | 1,829 | 3,609E-38 |
| TcCLB.467373.10 | trans-sialidase, putative (fragment) | 1,829 | 8,983E-29 |
| TcCLB.509351.10 | delta-1-pyrroline-5-carboxylate dehydrogenase, putative | 1,828 | 7,489E-30 |
| TcCLB.511843.30 | trans-sialidase (pseudogene), putative | 1,827 | 0,005942 |
| TcCLB.504151.60 | actin interacting protein-like protein, putative | 1,825 | 5,005E-51 |
| TcCLB.504777.40 | mucin TcMUCII, putative | 1,825 | 2,803E-07 |
| TcCLB.509151.10 | hypothetical protein, conserved | 1,824 | 3,543E-50 |

| | | | |
|---|---|---|---|
| TcCLB.508163.40 | mucin TcMUCII (pseudogene), putative | 1,824 | 0,006777 |
| TcCLB.509791.140 | hypothetical protein, conserved | 1,823 | 4,419E-27 |
| TcCLB.508419.20 | dispersed gene family protein 1 (DGF-1, pseudogene), puta | 1,823 | 2,955E-21 |
| TcCLB.509377.20 | trans-sialidase, Group VII, putative | 1,823 | 0,1787 |
| TcCLB.511173.496 | mucin-associated surface protein (MASP), putative (fragm | 1,822 | 0,02637 |
| TcCLB.506781.39 | mucin-associated surface protein (MASP), putative | 1,822 | 0,07343 |
| TcCLB.508163.50 | mucin-associated surface protein (MASP), putative | 1,821 | 3,164E-06 |
| TcCLB.511015.30 | Trichohyalin, putative (fragment) | 1,818 | 6,744E-103 |
| TcCLB.504655.9 | mucin-associated surface protein (MASP), putative | 1,818 | 1,504E-77 |
| TcCLB.510069.20 | protein associated with differentiation 4, putative | 1,818 | 1,581E-62 |
| TcCLB.508643.20 | hypothetical protein, conserved | 1,818 | 4,123E-48 |
| TcCLB.509295.80 | retrotransposon hot spot (RHS) protein, putative | 1,817 | 1,647E-43 |
| TcCLB.506795.80 | proline racemase | 1,817 | 3,443E-33 |
| TcCLB.503471.10 | protein tyrosine phosphatase, putative | 1,817 | 4,118E-31 |
| TcCLB.503827.30 | mucin TcMUCII, putative | 1,817 | 0,1451 |
| TcCLB.505647.60 | hypothetical protein, conserved | 1,816 | 1,111E-37 |
| TcCLB.503893.80 | NADH dehydrogenase [ubiquinone] iron-sulfur protein 7, m | 1,816 | 4,839E-28 |
| TcCLB.507695.14 | dispersed gene family protein 1 (DGF-1), putative | 1,816 | 0,001932 |
| TcCLB.506129.60 | beta galactofuranosyl glycosyltransferase, putative | 1,815 | 2,824E-06 |
| TcCLB.505919.70 | glycine dehydrogenase [decarboxylating] (pseudogene), pu | 1,814 | 1,471E-11 |
| TcCLB.510149.60 | hypothetical protein, conserved | 1,813 | 3,386E-27 |
| TcCLB.509029.80 | hypothetical protein | 1,812 | 1,761E-30 |
| TcCLB.510095.10 | hypothetical protein | 1,811 | 0,002041 |
| TcCLB.510515.30 | hypothetical protein, conserved | 1,81 | 3,092E-50 |
| TcCLB.510035.10 | trans-sialidase (pseudogene), putative | 1,81 | 1,143E-49 |
| TcCLB.507049.60 | NAD(P)-dependent oxidoreductase, putative | 1,808 | 8,388E-53 |
| TcCLB.511807.60 | hypothetical protein, conserved | 1,806 | 1,943E-49 |
| TcCLB.506041.20 | hypothetical protein, conserved | 1,805 | 7,987E-33 |
| TcCLB.509975.30 | mucin-associated surface protein (MASP), putative | 1,805 | 1,886E-17 |
| TcCLB.511471.120 | retrotransposon hot spot protein (RHS, pseudogene), putat | 1,804 | 4,298E-57 |
| TcCLB.506819.40 | hypothetical protein, conserved | 1,801 | 8,151E-26 |
| TcCLB.503615.10 | UDP-Gal or UDP-GlcNAc-dependent glycosyltransferase, pu | 1,801 | 2,481E-12 |
| TcCLB.508163.240 | hypothetical protein | 1,801 | 0,01466 |
| TcCLB.508493.130 | mucin TcMUCII, putative | 1,798 | 1,037E-06 |
| TcCLB.504105.250 | hypothetical protein | 1,795 | 6,538E-21 |
| TcCLB.508495.110 | hypothetical protein | 1,795 | 0,009086 |
| TcCLB.508023.150 | trans-sialidase (pseudogene), putative | 1,794 | 5,769E-26 |
| TcCLB.504539.10 | mucin TcSMUGL, putative | 1,794 | 0,00004215 |
| TcCLB.508109.10 | mucin-associated surface protein (MASP), putative | 1,794 | 0,03914 |
| TcCLB.503955.55 | hypothetical protein, conserved | 1,793 | 6,836E-20 |
| TcCLB.506171.10 | retrotransposon hot spot (RHS) protein, putative | 1,793 | 3,185E-12 |
| TcCLB.508013.110 | mucin-associated surface protein (MASP), putative | 1,793 | 0,01048 |
| TcCLB.508931.10 | hypothetical protein | 1,792 | 6,607E-24 |
| TcCLB.506373.39 | trans-sialidase, putative (fragment) | 1,792 | 1,332E-10 |
| TcCLB.510437.40 | cytosolic malate dehydrogenase, putative | 1,791 | 1,893E-53 |

| | | | |
|---|---|---|---|
| TcCLB.424937.10 | glucose transporter, putative | 1,79 | 7,503E-27 |
| TcCLB.509765.129 | trans-sialidase, putative (fragment) | 1,788 | 5,596E-26 |
| TcCLB.511513.14 | hypothetical protein, conserved | 1,788 | 2,678E-08 |
| TcCLB.509611.180 | peptide methionine sulfoxide reductase, putative | 1,785 | 6,261E-34 |
| TcCLB.511693.130 | hypothetical protein | 1,785 | 2,911E-31 |
| TcCLB.510153.19 | tricarboxylate carrier, putative | 1,785 | 7,819E-18 |
| TcCLB.506475.105 | hypothetical protein, conserved | 1,784 | 2,437E-25 |
| TcCLB.504427.20 | hypothetical protein, conserved | 1,78 | 1,994E-19 |
| TcCLB.507297.10 | cruzipain precursor, putative | 1,779 | 1,146E-122 |
| TcCLB.508285.50 | retrotransposon hot spot protein (RHS, pseudogene), putat | 1,779 | 2,181E-33 |
| TcCLB.511063.180 | trans-sialidase (pseudogene), putative | 1,779 | 0,00002494 |
| TcCLB.504769.170 | retrotransposon hot spot protein (RHS, pseudogene), putat | 1,778 | 5,958E-35 |
| TcCLB.508261.60 | mucin TcMUCII, putative | 1,776 | 0,02175 |
| TcCLB.509767.59 | protoheme IX farnesyltransferase, putative | 1,775 | 1,293E-48 |
| TcCLB.504393.30 | trans-sialidase (pseudogene), putative | 1,773 | 0,00003356 |
| TcCLB.503851.32 | trans-sialidase (pseudogene), putative | 1,772 | 2,096E-07 |
| TcCLB.503769.40 | cytochrome oxidase subunit VII (COXVII) | 1,771 | 2,183E-73 |
| TcCLB.503845.30 | retrotransposon hot spot protein (RHS, pseudogene), putat | 1,771 | 1,967E-50 |
| TcCLB.511295.60 | hypothetical protein, conserved (pseudogene) | 1,771 | 1,578E-37 |
| TcCLB.511179.150 | alanine racemase, putative | 1,771 | 2,541E-22 |
| TcCLB.506139.120 | trans-sialidase, Group V, putative | 1,771 | 2,037E-09 |
| TcCLB.506755.20 | paraflagellar rod component, putative | 1,769 | 1,604E-35 |
| TcCLB.507009.30 | ribulose-phosphate 3-epimerase, putative | 1,769 | 1,771E-21 |
| TcCLB.507465.10 | receptor-type adenylate cyclase, putative (fragment) | 1,769 | 6,483E-07 |
| TcCLB.511911.30 | mismatch repair protein MSH4 (pseudogene), putative | 1,768 | 6,035E-34 |
| TcCLB.508837.90 | hypothetical protein, conserved | 1,768 | 2,742E-30 |
| TcCLB.505193.110 | serine/threonine protein kinase, putative | 1,767 | 3,475E-83 |
| TcCLB.506975.40 | amidohydrolase, putative | 1,767 | 1,313E-61 |
| TcCLB.508041.40 | hypothetical protein | 1,766 | 2,077E-52 |
| TcCLB.449247.10 | hypothetical protein, conserved | 1,765 | 2,22E-15 |
| TcCLB.511821.160 | microtubial binding protein, putative | 1,765 | 1,372E-10 |
| TcCLB.509601.90 | alanine racemase, putative | 1,763 | 9,505E-48 |
| TcCLB.463451.10 | hypothetical protein, conserved | 1,758 | 2,031E-33 |
| TcCLB.506823.50 | retrotransposon hot spot protein (RHS, pseudogene), putat | 1,756 | 2,943E-45 |
| TcCLB.504177.30 | hypothetical protein, conserved | 1,754 | 1,092E-35 |
| TcCLB.506467.50 | serine/threonine protein kinase, putative | 1,753 | 6,097E-57 |
| TcCLB.508677.40 | trans-sialidase, Group VIII, putative | 1,75 | 3,545E-17 |
| TcCLB.509123.10 | surface antigen TASV, putative | 1,749 | 4,608E-24 |
| TcCLB.509337.5 | hypothetical protein, conserved (pseudogene) | 1,746 | 3,593E-27 |
| TcCLB.507515.4 | trypanothione/tryparedoxin dependent peroxidase 2, putat | 1,744 | 1,247E-58 |
| TcCLB.510349.50 | CDP-diacylglycerol--inositol 3-phosphatidyltransferase, put | 1,743 | 3,853E-21 |
| TcCLB.507537.20 | cysteine peptidase, putative | 1,742 | 1,963E-89 |
| TcCLB.509563.10 | hypothetical protein, conserved | 1,741 | 5,412E-33 |
| TcCLB.507009.50 | hypothetical protein, conserved | 1,741 | 6,964E-21 |
| TcCLB.508119.200 | hypothetical protein | 1,739 | 1,534E-58 |

| | | | |
|---|---|---|---|
| TcCLB.510977.20 | hypothetical protein, conserved (pseudogene) | 1,739 | 2,188E-23 |
| TcCLB.503401.29 | hypothetical protein, conserved | 1,739 | 7,281E-20 |
| TcCLB.506321.110 | mucin TcMUCII, putative | 1,739 | 1,394E-07 |
| TcCLB.511573.10 | leucine-rich repeat protein, putative | 1,737 | 7,204E-10 |
| TcCLB.506309.70 | mucin-associated surface protein (MASP), putative | 1,737 | 0,08318 |
| TcCLB.506633.10 | protein kinase, putative | 1,736 | 1,556E-17 |
| TcCLB.509391.20 | flagellar calcium-binding protein, putative | 1,736 | 0,06779 |
| TcCLB.511589.70 | ADP-ribosylation factor GTPase activating protein, putative | 1,735 | 1,143E-30 |
| TcCLB.511829.20 | retrotransposon hot spot protein (RHS, pseudogene), putat | 1,735 | 2,416E-10 |
| TcCLB.509427.50 | retrotransposon hot spot protein (RHS, pseudogene), putat | 1,733 | 1,909E-32 |
| TcCLB.511449.110 | surface protease GP63, putative | 1,733 | 0,0003215 |
| TcCLB.507357.60 | mucin TcMUCII, putative | 1,733 | 0,01601 |
| TcCLB.507091.50 | aspartate carbamoyltransferase, putative | 1,731 | 7,132E-27 |
| TcCLB.509721.60 | poly(ADP-ribose) polymerase, putative | 1,73 | 7,121E-47 |
| TcCLB.508493.40 | mucin TcMUCII, putative | 1,73 | 9,778E-09 |
| TcCLB.507615.130 | hypothetical protein, conserved | 1,728 | 1,391E-20 |
| TcCLB.504039.99 | mucin TcMUCII (pseudogene), putative | 1,728 | 0,003646 |
| TcCLB.505789.10 | hypothetical protein, conserved | 1,726 | 2,842E-20 |
| TcCLB.508147.140 | mucin TcMUCII, putative | 1,725 | 0,02921 |
| TcCLB.507487.30 | mucin TcMUCII, putative | 1,724 | 0,05143 |
| TcCLB.510675.9 | dispersed gene family protein 1 (DGF-1), putative (fragme | 1,723 | 0,00000491 |
| TcCLB.504055.100 | retrotransposon hot spot protein (RHS, pseudogene), putat | 1,723 | 0,0003316 |
| TcCLB.508165.270 | mucin TcMUCII, putative | 1,722 | 0,1679 |
| TcCLB.511597.40 | mucin-associated surface protein (MASP), putative | 1,721 | 0,03062 |
| TcCLB.507675.10 | hypothetical protein, conserved | 1,72 | 3,935E-70 |
| TcCLB.508885.5 | mucin-associated surface protein (MASP), putative | 1,717 | 0,257 |
| TcCLB.509223.20 | retrotransposon hot spot (RHS) protein, putative | 1,716 | 4,502E-38 |
| TcCLB.504199.4 | hypothetical protein, conserved | 1,716 | 1,055E-06 |
| TcCLB.504427.230 | trans-sialidase, Group VIII, putative | 1,716 | 0,1697 |
| TcCLB.511827.80 | hypothetical protein, conserved | 1,715 | 7,429E-20 |
| TcCLB.505945.30 | ribonuclease mar1, putative | 1,714 | 1,821E-32 |
| TcCLB.504111.40 | hypothetical protein, conserved | 1,713 | 1,136E-19 |
| TcCLB.508387.140 | hypothetical protein, conserved | 1,712 | 2,403E-37 |
| TcCLB.509147.50 | mucin TcSMUGL, putative | 1,712 | 1,328E-34 |
| TcCLB.483389.10 | retrotransposon hot spot protein (RHS, pseudogene), putat | 1,711 | 1,06E-39 |
| TcCLB.506783.39 | mucin-associated surface protein (MASP), putative | 1,71 | 0,04819 |
| TcCLB.422189.20 | leucine-rich repeat protein, putative | 1,709 | 6,12E-14 |
| TcCLB.509155.39 | surface protease GP63, putative | 1,709 | 0,1075 |
| TcCLB.505975.20 | trans-sialidase, Group IV, putative | 1,708 | 5,249E-16 |
| TcCLB.503897.60 | hypothetical protein, conserved | 1,706 | 5,658E-26 |
| TcCLB.511649.14 | hypothetical protein, conserved (fragment) | 1,706 | 1,259E-14 |
| TcCLB.510659.270 | hypothetical protein, conserved | 1,705 | 1,439E-29 |
| TcCLB.506563.130 | calpain-like cysteine peptidase (pseudogene), putative | 1,702 | 5,042E-24 |
| TcCLB.507801.110 | hypothetical protein, conserved | 1,701 | 5,114E-36 |
| TcCLB.507699.130 | mucin-associated surface protein (MASP), putative | 1,699 | 0,00000251 |

| | | | |
|---|---|---|---|
| TcCLB.510583.200 | mucin TcMUCII, putative | 1,699 | 0,03032 |
| TcCLB.506595.149 | retrotransposon hot spot (RHS) protein, putative | 1,698 | 8,757E-79 |
| TcCLB.509289.39 | hypothetical protein, conserved (pseudogene) | 1,698 | 3,514E-20 |
| TcCLB.511145.10 | cytochrome c oxidase subunit VI, putative | 1,697 | 1,077E-35 |
| TcCLB.506627.20 | hypothetical protein, conserved | 1,696 | 7,263E-69 |
| TcCLB.509097.20 | trans-sialidase, Group V, putative | 1,696 | 0,00006403 |
| TcCLB.509911.20 | hypothetical protein | 1,695 | 2,418E-10 |
| TcCLB.511553.209 | mucin TcMUC (pseudogene), putative | 1,695 | 0,008891 |
| TcCLB.507537.10 | cruzipain precursor (pseudogene), putative | 1,694 | 3,368E-54 |
| TcCLB.509815.20 | trans-sialidase (pseudogene), putative | 1,694 | 4,346E-18 |
| TcCLB.506057.15 | hypothetical protein | 1,694 | 0,0005483 |
| TcCLB.511259.9 | mucin TcMUCI, putative | 1,694 | 0,1084 |
| TcCLB.509677.10 | hypothetical protein, conserved | 1,693 | 8,867E-27 |
| TcCLB.510507.29 | hypothetical protein, conserved | 1,692 | 9,34E-21 |
| TcCLB.510095.5 | retrotransposon hot spot protein (RHS, pseudogene), putat | 1,692 | 0,004597 |
| TcCLB.506999.191 | hypothetical protein, conserved (fragment) | 1,691 | 0,0000515 |
| TcCLB.508555.20 | hypothetical protein, conserved | 1,69 | 5,541E-19 |
| TcCLB.511675.3 | surface antigen TASV, putative | 1,69 | 2,76E-18 |
| TcCLB.508001.70 | retrotransposon hot spot (RHS) protein, putative | 1,689 | 8,094E-35 |
| TcCLB.508569.140 | hypothetical protein, conserved | 1,689 | 2,023E-19 |
| TcCLB.509225.21 | hypothetical protein | 1,689 | 0,00001886 |
| TcCLB.507555.80 | retrotransposon hot spot (RHS) protein, putative | 1,688 | 1,261E-77 |
| TcCLB.503405.10 | glutamine synthetase, putative | 1,686 | 1,826E-46 |
| TcCLB.508151.10 | mucin TcMUCII, putative | 1,686 | 0,001004 |
| TcCLB.507067.40 | mucin TcMUCII, putative | 1,686 | 0,005503 |
| TcCLB.506611.70 | surface protease GP63, putative | 1,685 | 0,004464 |
| TcCLB.506923.21 | mucin TcMUCII, putative | 1,684 | 0,00001867 |
| TcCLB.503811.60 | Elongation factor Ts, mitochondrial, putative (EF-Ts) | 1,683 | 7,906E-42 |
| TcCLB.511573.40 | Mitogen-activated protein kinase 10, putative (MPK10) | 1,683 | 5,204E-37 |
| TcCLB.509791.100 | hypothetical protein, conserved | 1,683 | 4,598E-29 |
| TcCLB.504083.20 | retrotransposon hot spot protein (RHS, pseudogene), putat | 1,682 | 0,06636 |
| TcCLB.507225.19 | hypothetical protein | 1,681 | 1,992E-33 |
| TcCLB.507669.139 | hypothetical protein, conserved | 1,681 | 2,841E-24 |
| TcCLB.476833.10 | trans-sialidase, putative | 1,681 | 0,03878 |
| TcCLB.506573.5 | surface antigen TASV-A8, putative (TASV-A8) | 1,68 | 2,23E-46 |
| TcCLB.510479.20 | dispersed gene family protein 1 (DGF-1, pseudogene), puta | 1,68 | 3,065E-13 |
| TcCLB.506243.77 | retrotransposon hot spot protein (RHS, pseudogene), putat | 1,679 | 2,489E-07 |
| TcCLB.510741.170 | hypothetical protein | 1,678 | 2,066E-32 |
| TcCLB.509815.90 | retrotransposon hot spot protein (RHS, pseudogene), putat | 1,678 | 2,44E-23 |
| TcCLB.509901.160 | hypothetical protein, conserved | 1,678 | 1,036E-17 |
| TcCLB.508221.820 | mucin-associated surface protein (MASP), putative | 1,678 | 0,00001221 |
| TcCLB.506665.30 | trans-sialidase, putative (fragment) | 1,678 | 0,000644 |
| TcCLB.510693.220 | mucin TcMUCII, putative | 1,678 | 0,005071 |
| TcCLB.511575.10 | hypothetical protein, conserved | 1,677 | 6,007E-20 |
| TcCLB.508153.605 | hypothetical protein, conserved | 1,676 | 1,223E-27 |

| | | | |
|---|---|---|---|
| TcCLB.506989.10 | hypothetical protein, conserved | 1,676 | 4,213E-23 |
| TcCLB.509387.5 | trans-sialidase, putative | 1,676 | 3,788E-11 |
| TcCLB.506983.48 | calpain-like cysteine peptidase, putative | 1,675 | 1,615E-30 |
| TcCLB.506149.10 | folate/pteridine transporter, putative | 1,674 | 1,416E-29 |
| TcCLB.506713.80 | trans-sialidase, putative (fragment) | 1,674 | 1,614E-17 |
| TcCLB.506353.10 | trans-sialidase, Group II, putative | 1,674 | 1,321E-06 |
| TcCLB.506485.100 | hypothetical protein, conserved | 1,672 | 1,545E-25 |
| TcCLB.506633.90 | folate/pteridine transporter, putative | 1,67 | 1,655E-67 |
| TcCLB.506835.159 | casein kinase 1 isoform 2, putative (fragment) | 1,669 | 5,014E-08 |
| TcCLB.510957.9 | calpain-like cysteine peptidase, putative | 1,664 | 3,42E-86 |
| TcCLB.508173.150 | protein disulfide isomerase, putative | 1,664 | 1,195E-41 |
| TcCLB.506775.140 | hypothetical protein, conserved | 1,664 | 1,014E-17 |
| TcCLB.509785.50 | trans-sialidase, Group V, putative | 1,664 | 1,461E-09 |
| TcCLB.511277.390 | hypothetical protein, conserved | 1,663 | 4,397E-42 |
| TcCLB.508945.10 | hypothetical protein, conserved | 1,663 | 5,038E-29 |
| TcCLB.508893.10 | hypothetical protein, conserved | 1,662 | 4,423E-31 |
| TcCLB.506597.20 | mucin-associated surface protein (MASP, pseudogene), pu | 1,662 | 0,03304 |
| TcCLB.510025.70 | 90 kDa surface protein, putative | 1,661 | 0,009499 |
| TcCLB.510847.20 | beta galactofuranosyl glycosyltransferase, putative | 1,66 | 1,29E-13 |
| TcCLB.509495.49 | mucin TcMUCII, putative | 1,66 | 4,46E-07 |
| TcCLB.510743.70 | lactoylglutathione lyase-like protein, putative | 1,659 | 1,036E-50 |
| TcCLB.510381.30 | hypothetical protein, conserved | 1,658 | 3,825E-37 |
| TcCLB.509207.80 | glycogen synthase kinase-3 alpha, putative | 1,658 | 2,776E-35 |
| TcCLB.507925.40 | pyridoxal kinase, putative | 1,658 | 3,345E-25 |
| TcCLB.507357.135 | hypothetical protein | 1,658 | 0,01791 |
| TcCLB.507723.50 | trans-sialidase, Group IV, putative | 1,657 | 1,397E-06 |
| TcCLB.511099.230 | mucin TcMUCII, putative | 1,657 | 0,04013 |
| TcCLB.511821.150 | hypothetical protein, conserved | 1,656 | 7,082E-20 |
| TcCLB.507569.4 | electon transport protein SCO1/SCO2, putative | 1,654 | 4,001E-21 |
| TcCLB.507699.180 | mucin-associated surface protein (MASP, pseudogene), pu | 1,654 | 0,0001088 |
| TcCLB.506571.20 | retrotransposon hot spot protein (RHS, pseudogene), putat | 1,653 | 2,243E-18 |
| TcCLB.507625.140 | membrane transporter protein, putative | 1,652 | 7,051E-36 |
| TcCLB.511693.20 | hypothetical protein, conserved | 1,652 | 7,903E-19 |
| TcCLB.507157.10 | mucin TcSMUGL, putative | 1,652 | 3,845E-09 |
| TcCLB.511063.30 | trans-sialidase (pseudogene), putative | 1,651 | 0,00005022 |
| TcCLB.510861.140 | serine/threonine protein kinase, putative | 1,65 | 3,439E-68 |
| TcCLB.506773.44 | cytochrome b5, putative | 1,65 | 4,776E-41 |
| TcCLB.509259.50 | retrotransposon hot spot protein (RHS, pseudogene), putat | 1,65 | 2,407E-27 |
| TcCLB.509335.10 | retrotransposon hot spot (RHS) protein, putative | 1,649 | 3,505E-68 |
| TcCLB.446325.9 | retrotransposon hot spot (RHS) protein, putative (fragmen | 1,649 | 1,927E-24 |
| TcCLB.510207.21 | mucin-associated surface protein (MASP, pseudogene), pu | 1,648 | 0,002629 |
| TcCLB.510101.140 | pyruvate phosphate dikinase, putative | 1,647 | 2,089E-52 |
| TcCLB.503411.10 | hypothetical protein | 1,647 | 9,158E-35 |
| TcCLB.506015.40 | mucin TcMUCI, putative | 1,646 | 0,00008154 |
| TcCLB.507993.270 | hypothetical protein, conserved | 1,642 | 5,587E-29 |

| | | | |
|---|---|---|---|
| TcCLB.508549.20 | hypothetical protein, conserved | 1,641 | 1,53E-51 |
| TcCLB.509959.90 | retrotransposon hot spot protein (RHS, pseudogene), putat | 1,64 | 9,677E-21 |
| TcCLB.510693.40 | 90 kDa surface protein, putative | 1,64 | 0,007388 |
| TcCLB.509601.20 | hypothetical protein, conserved | 1,639 | 9,055E-12 |
| TcCLB.506565.20 | hypothetical protein, conserved | 1,638 | 3,128E-15 |
| TcCLB.506063.50 | retrotransposon hot spot protein (RHS, pseudogene), putat | 1,638 | 2,093E-12 |
| TcCLB.487455.10 | trans-sialidase, putative (fragment) | 1,638 | 0,02993 |
| TcCLB.511807.190 | hypothetical protein, conserved | 1,636 | 3,702E-57 |
| TcCLB.506277.10 | mucin-associated surface protein (MASP), putative | 1,636 | 0,0000129 |
| TcCLB.503973.250 | mucin TcMUCII (pseudogene), putative | 1,636 | 0,00007883 |
| TcCLB.506773.90 | amino acid permease/transporter, putative | 1,635 | 7,002E-63 |
| TcCLB.511351.30 | protein kinase, putative | 1,635 | 6,876E-60 |
| TcCLB.511367.210 | ubiquitin-conjugating enzyme variant Kua homologue, puta | 1,635 | 3,486E-23 |
| TcCLB.506721.50 | hypothetical protein | 1,635 | 1,735E-07 |
| TcCLB.507071.170 | mucin TcMUCII, putative | 1,634 | 5,614E-06 |
| TcCLB.504797.20 | hypothetical protein, conserved | 1,632 | 3,289E-31 |
| TcCLB.506399.30 | hypothetical protein, conserved | 1,632 | 1,497E-25 |
| TcCLB.510233.20 | hypothetical protein | 1,632 | 0,003017 |
| TcCLB.511057.20 | mucin TcMUCII, putative | 1,631 | 0,004444 |
| TcCLB.509849.30 | hypothetical protein | 1,63 | 1,92E-68 |
| TcCLB.510377.270 | retrotransposon hot spot protein (RHS, pseudogene), putat | 1,63 | 0,0000189 |
| TcCLB.507031.90 | arginase, putative | 1,629 | 3,408E-42 |
| TcCLB.507669.20 | hypothetical protein, conserved | 1,628 | 2,296E-38 |
| TcCLB.511323.50 | NADH-ubiquinone oxidoreductase, mitochondrial, putative | 1,628 | 1,708E-18 |
| TcCLB.508231.40 | hypothetical protein, conserved | 1,627 | 8,718E-58 |
| TcCLB.506381.10 | retrotransposon hot spot protein (RHS, pseudogene), putat | 1,627 | 3,491E-15 |
| TcCLB.509637.36 | procyclic form surface glycoprotein, putative (fragment) | 1,625 | 1,737E-20 |
| TcCLB.505919.31 | hypothetical protein, conserved (pseudogene) | 1,625 | 1,455E-13 |
| TcCLB.510791.20 | mucin TcMUCII, putative | 1,625 | 9,512E-07 |
| TcCLB.506399.70 | hypothetical protein, conserved | 1,624 | 3,855E-19 |
| TcCLB.509977.80 | mucin-associated surface protein (MASP), putative | 1,624 | 3,994E-09 |
| TcCLB.506131.199 | mucin TcMUCII, putative | 1,624 | 0,02862 |
| TcCLB.506195.160 | hypothetical protein, conserved | 1,623 | 9,246E-37 |
| TcCLB.510359.430 | trans-sialidase (pseudogene), putative | 1,623 | 9,582E-24 |
| TcCLB.510101.450 | hypothetical protein, conserved | 1,621 | 4,994E-49 |
| TcCLB.511821.170 | hypothetical protein, conserved | 1,62 | 2,338E-15 |
| TcCLB.439245.10 | mucin TcMUCII, putative (fragment) | 1,62 | 0,02145 |
| TcCLB.511599.80 | trans-sialidase, Group V, putative | 1,619 | 4,076E-13 |
| TcCLB.508887.20 | mucin TcMUCII, putative | 1,619 | 0,003079 |
| TcCLB.509329.20 | retrotransposon hot spot protein (RHS, pseudogene), putat | 1,617 | 1,771E-45 |
| TcCLB.504427.75 | hypothetical protein, conserved (pseudogene) | 1,617 | 1,672E-14 |
| TcCLB.509099.50 | trypanothione synthetase (TRYS) | 1,616 | 1,522E-72 |
| TcCLB.506835.70 | PAS-domain containing phosphoglycerate kinase, putative | 1,616 | 1,625E-50 |
| TcCLB.511821.179 | hypothetical protein, conserved | 1,616 | 5,521E-49 |
| TcCLB.509799.50 | hypothetical protein, conserved | 1,616 | 1,957E-38 |

| | | | |
|---|---|---|---|
| TcCLB.508789.204 | mucin-associated surface protein (MASP, pseudogene), pu | 1,615 | 0,007234 |
| TcCLB.507467.10 | receptor-type adenylate cyclase, putative (fragment) | 1,614 | 4,587E-09 |
| TcCLB.510503.70 | trans-sialidase (pseudogene), putative | 1,614 | 0,00002429 |
| TcCLB.507603.270 | cruzipain precursor, putative | 1,613 | 1,444E-34 |
| TcCLB.510283.70 | glycosyl hydrolase-like protein, putative | 1,612 | 1,859E-39 |
| TcCLB.510889.140 | alanine aminotransferase, putative | 1,611 | 1,853E-42 |
| TcCLB.511693.70 | hypothetical protein, conserved | 1,611 | 7,932E-33 |
| TcCLB.508159.60 | mucin TcMUCII, putative | 1,611 | 0,003259 |
| TcCLB.508045.14 | hypothetical protein, conserved | 1,609 | 1,661E-23 |
| TcCLB.511259.189 | mucin TcMUCII, putative | 1,609 | 0,04248 |
| TcCLB.506769.80 | mucin-associated surface protein (MASP), putative | 1,606 | 0,000641 |
| TcCLB.508651.10 | dispersed gene family protein 1 (DGF-1, pseudogene), puta | 1,606 | 0,003092 |
| TcCLB.506533.106 | mucin TcSMUGL, putative | 1,605 | 2,11E-34 |
| TcCLB.511701.19 | surface protease GP63, putative (fragment) | 1,604 | 0,1442 |
| TcCLB.510857.4 | hypothetical protein, conserved | 1,603 | 1,617E-29 |
| TcCLB.508071.90 | retrotransposon hot spot (RHS) protein, putative | 1,601 | 1,434E-44 |
| TcCLB.509493.100 | mucin-associated surface protein (MASP, pseudogene), pu | 1,601 | 1,646E-12 |
| TcCLB.509875.240 | hypothetical protein, conserved | 1,599 | 1,772E-42 |
| TcCLB.506675.110 | retrotransposon hot spot (RHS) protein, putative | 1,598 | 5,215E-27 |
| TcCLB.508493.20 | mucin-associated surface protein (MASP, pseudogene), pu | 1,598 | 4,54E-11 |
| TcCLB.509493.39 | mucin TcMUCII, putative | 1,595 | 2,098E-07 |
| TcCLB.508741.170 | glycerate kinase, putative | 1,594 | 7,331E-133 |
| TcCLB.506529.300 | hypothetical protein, conserved | 1,593 | 9,285E-50 |
| TcCLB.511003.110 | hypothetical protein, conserved | 1,593 | 1,058E-24 |
| TcCLB.508221.509 | mucin-associated surface protein (MASP, pseudogene), pu | 1,593 | 0,02976 |
| TcCLB.507499.30 | retrotransposon hot spot protein (RHS, pseudogene), putat | 1,592 | 3,935E-16 |
| TcCLB.511827.115 | hypothetical protein | 1,592 | 5,766E-09 |
| TcCLB.509153.100 | arginine N-methyltransferase, type III, putative (PRMT7) | 1,59 | 1,842E-59 |
| TcCLB.506941.180 | hypothetical protein, conserved | 1,589 | 1,194E-26 |
| TcCLB.508503.10 | fatty acid desaturase, putative | 1,589 | 3,096E-22 |
| TcCLB.508119.20 | mucin-associated surface protein (MASP), putative | 1,589 | 1,705E-10 |
| TcCLB.506989.110 | NADH-ubiquinone oxidoreductase complex I subunit, putati | 1,588 | 1,448E-26 |
| TcCLB.504239.240 | mucin-associated surface protein (MASP, pseudogene), pu | 1,588 | 0,2144 |
| TcCLB.508989.9 | nucleoside phosphorylase, putative | 1,586 | 2,141E-55 |
| TcCLB.505401.31 | trans-sialidase (pseudogene), putative | 1,586 | 4,025E-17 |
| TcCLB.508027.110 | pyroglutamyl-peptidase I, putative (PPI) | 1,585 | 1,821E-32 |
| TcCLB.511149.10 | mucin TcMUCII, putative | 1,585 | 0,05267 |
| TcCLB.510863.30 | hypothetical protein, conserved | 1,584 | 3,84E-22 |
| TcCLB.511303.150 | hypothetical protein, conserved | 1,584 | 3,02E-18 |
| TcCLB.510555.10 | mucin-associated surface protein (MASP, pseudogene), pu | 1,584 | 0,001669 |
| TcCLB.506763.190 | mucin TcMUCII, putative | 1,584 | 0,009992 |
| TcCLB.505609.10 | trans-sialidase, Group II, putative | 1,584 | 0,3048 |
| TcCLB.506763.80 | mucin TcMUCII, putative | 1,583 | 0,01459 |
| TcCLB.509601.120 | hypothetical protein, conserved | 1,581 | 2,571E-50 |
| TcCLB.510155.50 | hypothetical protein, conserved | 1,581 | 7,006E-29 |

| | | | |
|---|---|---|---|
| TcCLB.504081.270 | mucin TcMUCII, putative | 1,58 | 0,01503 |
| TcCLB.508873.289 | 90 kDa surface protein, putative | 1,578 | 0,02301 |
| TcCLB.509961.70 | dispersed gene family protein 1 (DGF-1, pseudogene), puta | 1,575 | 2,021E-22 |
| TcCLB.506047.20 | retrotransposon hot spot (RHS) protein, putative | 1,574 | 8,712E-38 |
| TcCLB.508989.120 | hypothetical protein, conserved | 1,574 | 7,901E-17 |
| TcCLB.509027.20 | ankyrin repeat protein, putative | 1,574 | 8,776E-13 |
| TcCLB.508493.10 | mucin TcMUCII, putative | 1,574 | 0,00001393 |
| TcCLB.506537.200 | trans-sialidase, Group VIII, putative | 1,573 | 2,761E-17 |
| TcCLB.507739.40 | uracil phosphoribosyltransferase, putative | 1,572 | 1,408E-43 |
| TcCLB.510497.90 | mucin TcMUCII, putative | 1,572 | 0,002791 |
| TcCLB.510237.60 | mucin TcMUCII, putative | 1,572 | 0,02768 |
| TcCLB.509427.30 | dispersed gene family protein 1 (DGF-1, pseudogene), puta | 1,57 | 0,00006112 |
| TcCLB.510431.20 | hypothetical protein, conserved | 1,569 | 1,982E-17 |
| TcCLB.509151.29 | hypothetical protein, conserved (pseudogene) | 1,569 | 1,202E-06 |
| TcCLB.508173.20 | retrotransposon hot spot protein (RHS, pseudogene), putat | 1,568 | 1,126E-103 |
| TcCLB.503859.40 | mucin-associated surface protein (MASP), putative | 1,568 | 0,002016 |
| TcCLB.509281.10 | hypothetical protein | 1,568 | 0,01863 |
| TcCLB.507811.100 | amino acid permease, putative | 1,567 | 3,776E-10 |
| TcCLB.503973.70 | serine-alanine-and proline-rich protein, putative | 1,567 | 0,0009151 |
| TcCLB.507959.90 | mucin TcMUCII, putative | 1,567 | 0,09488 |
| TcCLB.440101.10 | fatty acid desaturase, putative (fragment) | 1,566 | 1,033E-13 |
| TcCLB.508595.50 | cysteine peptidase, putative | 1,565 | 1,206E-61 |
| TcCLB.507065.89 | mucin TcMUC (pseudogene), putative | 1,565 | 0,003342 |
| TcCLB.509243.30 | hypothetical protein, conserved | 1,564 | 7,495E-23 |
| TcCLB.511385.30 | Pyrophosphate-energized vacuolar membrane proton pum | 1,563 | 4,21E-78 |
| TcCLB.511751.205 | hypothetical protein | 1,563 | 6,723E-32 |
| TcCLB.507487.20 | mucin-associated surface protein (MASP), putative | 1,563 | 0,09006 |
| TcCLB.510943.130 | cytochrome c oxidase assembly protein, putative | 1,562 | 7,294E-33 |
| TcCLB.510181.20 | delta-4 fatty acid desaturase, putative | 1,562 | 1,457E-24 |
| TcCLB.506411.30 | proline oxidase, putative | 1,56 | 1,41E-27 |
| TcCLB.507429.3 | mucin-associated surface protein (MASP), putative | 1,56 | 0,006765 |
| TcCLB.506401.140 | hypothetical protein, conserved | 1,557 | 3,803E-07 |
| TcCLB.510773.20 | Pyrophosphate-energized vacuolar membrane proton pum | 1,556 | 4,471E-71 |
| TcCLB.510889.120 | alanine aminotransferase, putative | 1,556 | 5,031E-49 |
| TcCLB.507063.10 | hypothetical protein | 1,555 | 5,547E-23 |
| TcCLB.509401.30 | cysteine peptidase, putative | 1,552 | 6,074E-94 |
| TcCLB.509683.124 | hypothetical protein, conserved | 1,552 | 2,168E-53 |
| TcCLB.510557.50 | mucin TcMUCII, putative | 1,552 | 0,0006094 |
| TcCLB.507963.10 | hypothetical protein, conserved | 1,551 | 1,886E-21 |
| TcCLB.511217.130 | hypothetical protein, conserved | 1,549 | 3,313E-51 |
| TcCLB.510669.40 | hypothetical protein | 1,549 | 7,511E-45 |
| TcCLB.508569.20 | ubiquitin-conjugating enzyme E2, putative | 1,549 | 6,129E-36 |
| TcCLB.511837.90 | protein kinase, putative | 1,549 | 5,957E-35 |
| TcCLB.428999.30 | hypothetical protein | 1,549 | 9,68E-24 |
| TcCLB.506773.100 | protein kinase, putative | 1,548 | 2,644E-33 |

| | | | |
|---|---|---|---|
| TcCLB.509413.10 | hypothetical protein, conserved | 1,547 | 5,731E-51 |
| TcCLB.509163.90 | retrotransposon hot spot protein (RHS, pseudogene), putat | 1,547 | 8,802E-38 |
| TcCLB.506145.24 | hypothetical protein, conserved | 1,547 | 1,664E-19 |
| TcCLB.504199.20 | hypothetical protein, conserved | 1,546 | 4,746E-39 |
| TcCLB.506129.10 | dispersed gene family protein 1 (DGF-1, pseudogene), puta | 1,546 | 3,872E-07 |
| TcCLB.439307.9 | hypothetical protein, conserved | 1,545 | 0,235 |
| TcCLB.504149.100 | hypothetical protein, conserved | 1,544 | 2,394E-58 |
| TcCLB.509647.13 | hypothetical protein, conserved (fragment) | 1,544 | 1,271E-57 |
| TcCLB.508965.14 | multidrug resistance protein E, putative (MRPE) | 1,544 | 3,878E-40 |
| TcCLB.510847.40 | retrotransposon hot spot protein (RHS, pseudogene), putat | 1,544 | 2,234E-22 |
| TcCLB.511109.130 | heterogeneous nuclear ribonucleoprotein H/F, putative | 1,543 | 2,06E-30 |
| TcCLB.507917.10 | surface protease GP63, putative | 1,542 | 6,185E-12 |
| TcCLB.506183.90 | trans-sialidase (pseudogene), putative | 1,54 | 2,987E-08 |
| TcCLB.506055.14 | trans-sialidase (pseudogene), putative | 1,54 | 0,00003001 |
| TcCLB.507625.10 | retrotransposon hot spot (RHS) protein, putative | 1,538 | 2,734E-44 |
| TcCLB.511823.14 | hypothetical protein, conserved | 1,538 | 2,109E-08 |
| TcCLB.510359.350 | hypothetical protein, conserved | 1,537 | 7,537E-45 |
| TcCLB.510941.3 | protein kinase, putative (fragment) | 1,537 | 1,532E-33 |
| TcCLB.506773.104 | hypothetical protein, conserved | 1,537 | 9,884E-24 |
| TcCLB.506563.220 | calpain-like cysteine peptidase, putative | 1,536 | 1,132E-47 |
| TcCLB.457101.30 | multidrug resistance protein E, putative | 1,536 | 7,052E-19 |
| TcCLB.507035.11 | trans-sialidase, putative (fragment) | 1,536 | 1,323E-06 |
| TcCLB.508219.60 | mucin TcMUCII (pseudogene), putative | 1,536 | 0,02552 |
| TcCLB.504769.20 | mucin-associated surface protein (MASP), putative | 1,535 | 2,775E-09 |
| TcCLB.505373.30 | retrotransposon hot spot protein (RHS, pseudogene), putat | 1,534 | 2,526E-11 |
| TcCLB.507221.60 | oxidoreductase-like protein (pseudogene), putative | 1,533 | 1,158E-20 |
| TcCLB.511245.120 | hypothetical protein, conserved | 1,533 | 9,539E-20 |
| TcCLB.509489.20 | hypothetical protein, conserved | 1,532 | 7,812E-22 |
| TcCLB.508119.120 | GDP-mannose 4,6 dehydratase, putative | 1,531 | 2,557E-38 |
| TcCLB.510697.40 | mucin-associated surface protein (MASP), putative | 1,531 | 0,002815 |
| TcCLB.511149.30 | mucin TcMUCII (pseudogene), putative | 1,527 | 0,0167 |
| TcCLB.509419.58 | mucin TcMUCII (pseudogene), putative | 1,526 | 0,06904 |
| TcCLB.509601.80 | hypothetical protein, conserved | 1,525 | 5,444E-25 |
| TcCLB.509977.90 | hypothetical protein | 1,524 | 5,155E-13 |
| TcCLB.507711.90 | procyclic form surface phosphoprotein, putative | 1,522 | 3,034E-29 |
| TcCLB.508307.125 | hypothetical protein, conserved | 1,522 | 1,573E-15 |
| TcCLB.505047.9 | dispersed gene family protein 1 (DGF-1), putative | 1,521 | 0,0001417 |
| TcCLB.507609.40 | delta-4 fatty acid desaturase, putative | 1,52 | 5,525E-21 |
| TcCLB.506529.610 | hypothetical protein | 1,519 | 4,082E-06 |
| TcCLB.504047.5 | urocanate hydratase, putative (fragment) | 1,518 | 0,00117 |
| TcCLB.511259.180 | mucin-associated surface protein (MASP), putative | 1,518 | 0,09026 |
| TcCLB.509339.10 | hypothetical protein, conserved | 1,516 | 7,603E-18 |
| TcCLB.503617.31 | nuclear receptor binding factor, putative | 1,515 | 1,599E-31 |
| TcCLB.504219.40 | trans-sialidase (pseudogene), putative | 1,514 | 0,00001966 |
| TcCLB.504269.10 | protein kinase (pseudogene), putative | 1,513 | 1,098E-28 |

| | | | |
|---|---|---|---|
| TcCLB.511807.40 | hypothetical protein, conserved | 1,513 | 1,331E-24 |
| TcCLB.504587.30 | mucin-associated surface protein (MASP), putative | 1,513 | 7,566E-09 |
| TcCLB.504239.280 | 90 kDa surface protein, putative | 1,512 | 0,01036 |
| TcCLB.511393.70 | hypothetical protein, conserved | 1,511 | 1,575E-36 |
| TcCLB.503597.10 | homoserine kinase, putative | 1,511 | 3,723E-31 |
| TcCLB.511909.50 | hypothetical protein, conserved | 1,511 | 1,171E-26 |
| TcCLB.507065.150 | mucin TcMUCII, putative | 1,511 | 0,001238 |
| TcCLB.504051.20 | ATP-dependent DNA DEAD/DEAH box helicase, putative | 1,51 | 4,276E-31 |
| TcCLB.508567.40 | hypothetical protein, conserved | 1,509 | 1,098E-28 |
| TcCLB.511545.120 | superoxide dismutase, putative | 1,509 | 2,429E-25 |
| TcCLB.508641.160 | glycogenin glucosyltransferase, putative | 1,509 | 1,693E-21 |
| TcCLB.506595.140 | hypothetical protein | 1,508 | 2,762E-11 |
| TcCLB.509147.40 | surface antigen TASV, putative | 1,507 | 6,542E-15 |
| TcCLB.507085.10 | mucin TcMUCII, putative | 1,507 | 2,457E-06 |
| TcCLB.508435.70 | mucin-associated surface protein (MASP), putative | 1,507 | 0,01994 |
| TcCLB.506533.142 | mucin TcSMUGL, putative | 1,506 | 9,836E-14 |
| TcCLB.504081.470 | mucin TcMUCII, putative | 1,505 | 0,09897 |
| TcCLB.508973.9 | hypothetical protein, conserved (fragment) | 1,504 | 3,25E-14 |
| TcCLB.506855.180 | adenylate kinase, putative | 1,503 | 1,003E-22 |
| TcCLB.509157.120 | hypothetical protein, conserved | 1,503 | 3,839E-19 |
| TcCLB.506471.120 | trans-sialidase, Group II, putative | 1,503 | 1,089E-18 |
| TcCLB.506945.20 | PAS-domain containing phosphoglycerate kinase, putative | 1,502 | 3,659E-45 |
| TcCLB.506829.80 | nicotinamidase, putative (PNC1) | 1,502 | 7,911E-22 |
| TcCLB.508113.14 | mucin TcMUC (pseudogene), putative | 1,502 | 1,397E-06 |
| TcCLB.511827.90 | hypothetical protein, conserved | 1,501 | 2,203E-06 |
| TcCLB.506173.270 | mucin TcMUCII, putative | 1,5 | 0,009803 |
| TcCLB.503705.19 | trans-sialidase, putative | 1,498 | 0,2538 |
| TcCLB.509693.180 | hypothetical protein, conserved | 1,497 | 1,318E-12 |
| TcCLB.509849.40 | mannosyl-oligosaccharide 1,2-alpha-mannosidase IB, puta | 1,497 | 1,807E-08 |
| TcCLB.510853.50 | beta galactofuranosyl glycosyltransferase, putative | 1,497 | 0,005593 |
| TcCLB.503527.60 | hypothetical protein, conserved | 1,496 | 3,425E-19 |
| TcCLB.508501.280 | hypothetical protein, conserved | 1,496 | 1,956E-16 |
| TcCLB.503971.50 | hypothetical protein, conserved | 1,495 | 2,771E-13 |
| TcCLB.509525.400 | mucin TcMUCII, putative | 1,495 | 0,007868 |
| TcCLB.510591.10 | trans-sialidase (pseudogene), putative | 1,495 | 0,04344 |
| TcCLB.509997.40 | hypothetical protein, conserved | 1,494 | 5,443E-26 |
| TcCLB.507357.80 | mucin TcMUCII, putative | 1,494 | 0,02722 |
| TcCLB.510101.320 | hypothetical protein | 1,493 | 3,163E-40 |
| TcCLB.506681.40 | hypothetical protein, conserved | 1,493 | 1,799E-06 |
| TcCLB.510583.114 | mucin-associated surface protein (MASP, pseudogene), pu | 1,493 | 0,2741 |
| TcCLB.505919.80 | retrotransposon hot spot (RHS) protein, putative | 1,491 | 6,267E-32 |
| TcCLB.510579.50 | hypothetical protein, conserved | 1,491 | 1,713E-28 |
| TcCLB.506593.10 | hypothetical protein, conserved | 1,491 | 1,2E-09 |
| TcCLB.506687.10 | retrotransposon hot spot protein (RHS, pseudogene), putat | 1,49 | 1,926E-26 |
| TcCLB.506321.40 | mucin TcMUCII, putative | 1,49 | 3,74E-11 |

| | | | |
|---|---|---|---|
| TcCLB.510861.110 | hypothetical protein | 1,489 | 6,187E-33 |
| TcCLB.509055.44 | hypothetical protein, conserved | 1,489 | 9,907E-21 |
| TcCLB.510359.660 | mucin TcMUCII, putative | 1,489 | 0,009727 |
| TcCLB.508507.59 | NADH dehydrogenase [ubiquinone] iron-sulfur protein 7, m | 1,488 | 2,243E-28 |
| TcCLB.511545.160 | hypothetical protein, conserved | 1,487 | 2,388E-14 |
| TcCLB.507429.49 | mucin TcMUCII, putative | 1,487 | 0,0005156 |
| TcCLB.510853.73 | retrotransposon hot spot protein (pseudogene), putative | 1,486 | 3,595E-59 |
| TcCLB.507089.110 | hypothetical protein, conserved | 1,486 | 9,283E-32 |
| TcCLB.508601.50 | NADH dehydrogenase subunit NB6M, putative | 1,486 | 4,638E-17 |
| TcCLB.503409.20 | hypothetical protein, conserved | 1,486 | 7,531E-15 |
| TcCLB.509369.50 | surface protease GP63, putative | 1,486 | 0,01817 |
| TcCLB.509807.11 | hypothetical protein, conserved (pseudogene) | 1,485 | 4,05E-12 |
| TcCLB.509449.5 | receptor-type adenylate cyclase, putative (fragment) | 1,485 | 0,03042 |
| TcCLB.507081.110 | aminopeptidase P1, putative | 1,484 | 3,306E-33 |
| TcCLB.505193.74 | hypothetical protein, conserved | 1,484 | 2,156E-26 |
| TcCLB.503487.50 | UDP-Gal or UDP-GlcNAc-dependent glycosyltransferase, pu | 1,483 | 5,619E-18 |
| TcCLB.504081.440 | 90 kDa surface protein, putative | 1,483 | 0,00008699 |
| TcCLB.510795.4 | CLC-type chloride channel protein (pseudogene), putative | 1,483 | 0,004355 |
| TcCLB.508851.189 | phosphoenolpyruvate mutase, putative | 1,482 | 1,2E-44 |
| TcCLB.507809.150 | hypothetical protein, conserved | 1,482 | 2,206E-21 |
| TcCLB.503653.60 | cytochrome b-domain protein, putative | 1,482 | 3,935E-20 |
| TcCLB.507715.10 | cysteine peptidase, Clan CA, family C51, putative | 1,481 | 2,763E-37 |
| TcCLB.504427.64 | hypothetical protein, conserved | 1,481 | 5,872E-22 |
| TcCLB.511621.60 | Asparagine-linked glycosylation protein 12 (ALG12) | 1,481 | 0,00004853 |
| TcCLB.506133.140 | mucin-associated surface protein (MASP), putative | 1,481 | 0,001748 |
| TcCLB.503981.30 | hypothetical protein, conserved | 1,48 | 6,928E-26 |
| TcCLB.505975.10 | hypothetical protein | 1,48 | 0,07993 |
| TcCLB.510359.20 | paraflagellar rod component, putative | 1,479 | 1,022E-07 |
| TcCLB.508495.130 | hypothetical protein | 1,479 | 0,01334 |
| TcCLB.506921.20 | hypothetical protein, conserved | 1,478 | 1,462E-26 |
| TcCLB.507699.110 | mucin-associated surface protein (MASP, pseudogene), pu | 1,477 | 0,001908 |
| TcCLB.511211.70 | cytochrome c oxidase assembly protein (COX15) | 1,476 | 6,421E-46 |
| TcCLB.505193.90 | hypothetical protein, conserved | 1,476 | 2,442E-28 |
| TcCLB.508461.310 | protein kinase A catalytic subunit isoform 1, putative | 1,476 | 1,048E-24 |
| TcCLB.511763.19 | hypothetical protein, conserved | 1,476 | 4,352E-14 |
| TcCLB.507787.50 | membrane transporter protein (pseudogene), putative | 1,476 | 1,827E-13 |
| TcCLB.507167.200 | beta galactofuranosyl glycosyltransferase, putative | 1,476 | 0,00005386 |
| TcCLB.511853.10 | mucin TcMUCI, putative | 1,475 | 0,04178 |
| TcCLB.511025.90 | hypothetical protein, conserved | 1,474 | 2,111E-16 |
| TcCLB.510025.30 | mucin-associated surface protein (MASP, pseudogene), pu | 1,474 | 0,02427 |
| TcCLB.509401.20 | cysteine peptidase (pseudogene), putative | 1,473 | 1,256E-72 |
| TcCLB.510073.17 | cysteine peptidase (C-terminal), putative | 1,473 | 6,559E-49 |
| TcCLB.509965.330 | hypothetical protein, conserved | 1,472 | 9,844E-69 |
| TcCLB.506755.110 | hypothetical protein, conserved | 1,471 | 8,573E-36 |
| TcCLB.511011.10 | retrotransposon hot spot (RHS) protein, putative | 1,471 | 8,587E-28 |

| | | | |
|---|---|---|---|
| TcCLB.508307.70 | hypothetical protein, conserved | 1,471 | 1,322E-25 |
| TcCLB.506951.80 | trans-sialidase, Group V, putative | 1,471 | 1,966E-18 |
| TcCLB.508879.20 | hypothetical protein | 1,471 | 1,082E-15 |
| TcCLB.508221.104 | mucin-associated surface protein (MASP), putative | 1,469 | 0,277 |
| TcCLB.511647.10 | hypothetical protein, conserved | 1,468 | 8,285E-51 |
| TcCLB.506297.4 | hypothetical protein, conserved (pseudogene) | 1,468 | 7,62E-15 |
| TcCLB.510743.30 | hypothetical protein, conserved | 1,467 | 1,914E-25 |
| TcCLB.510719.100 | hypothetical protein, conserved | 1,467 | 8,107E-12 |
| TcCLB.508761.160 | mucin TcMUCII, putative | 1,467 | 0,001792 |
| TcCLB.509973.80 | hypothetical protein | 1,466 | 0,00004708 |
| TcCLB.510503.100 | surface protease GP63, putative | 1,466 | 0,001201 |
| TcCLB.507699.120 | mucin TcMUCII, putative | 1,466 | 0,009131 |
| TcCLB.506955.184 | mucin-associated surface protein (MASP, pseudogene), pu | 1,465 | 0,003252 |
| TcCLB.511817.180 | histone H2A, putative | 1,464 | 0,003397 |
| TcCLB.506499.70 | mucin TcMUCII, putative | 1,464 | 0,00797 |
| TcCLB.410049.20 | hypothetical protein, conserved | 1,463 | 1,035E-09 |
| TcCLB.505807.180 | 2-hydroxy-3-oxopropionate reductase, putative | 1,462 | 3,109E-49 |
| TcCLB.506407.50 | hypothetical protein, conserved | 1,462 | 1,092E-30 |
| TcCLB.506779.110 | phopshatase, putative | 1,461 | 6,792E-40 |
| TcCLB.503781.80 | universal minicircle sequence binding protein 1 (UMSBP1) | 1,461 | 5,44E-34 |
| TcCLB.470827.10 | retrotransposon hot spot protein (RHS, pseudogene), putat | 1,461 | 1,393E-11 |
| TcCLB.511871.90 | hypothetical protein, conserved | 1,46 | 1,805E-34 |
| TcCLB.509769.33 | palmitoyl acyltransferase 12, putative | 1,46 | 1,149E-21 |
| TcCLB.510461.30 | mucin-associated surface protein (MASP), putative (fragm | 1,459 | 0,3438 |
| TcCLB.444777.10 | drug resistance protein, putative | 1,458 | 8,85E-07 |
| TcCLB.509233.150 | cytochrome oxidase subunit VII (COXVII) | 1,457 | 1,224E-53 |
| TcCLB.506325.40 | retrotransposon hot spot protein (RHS, pseudogene), putat | 1,456 | 1,164E-41 |
| TcCLB.508451.30 | retrotransposon hot spot protein (RHS, pseudogene), putat | 1,456 | 2,839E-07 |
| TcCLB.510349.30 | phosphatidylinositol-4-phosphate 5-kinase, putative | 1,455 | 2,798E-25 |
| TcCLB.510967.20 | hypothetical protein, conserved | 1,455 | 8,607E-11 |
| TcCLB.510759.140 | hypothetical protein, conserved | 1,454 | 1,816E-34 |
| TcCLB.508671.10 | serine carboxypeptidase (CBP1), putative | 1,454 | 1,375E-29 |
| TcCLB.508207.74 | Adenine phosphoribosyltransferase, putative | 1,454 | 3,072E-20 |
| TcCLB.504139.10 | retrotransposon hot spot protein (RHS, pseudogene), putat | 1,452 | 5,162E-11 |
| TcCLB.511819.40 | hypothetical protein, conserved | 1,451 | 5,627E-30 |
| TcCLB.510203.34 | mucin-associated surface protein (MASP, pseudogene), pu | 1,451 | 0,0001735 |
| TcCLB.508817.130 | carbonic anhydrase-like protein, putative | 1,45 | 7,712E-59 |
| TcCLB.506281.134 | mucin-associated surface protein (MASP), putative | 1,45 | 0,1049 |
| TcCLB.507477.5 | trans-sialidase, putative | 1,45 | 0,3372 |
| TcCLB.506513.190 | NADH-ubiquinone oxidoreductase, mitochondrial, putative | 1,449 | 1,236E-20 |
| TcCLB.511859.120 | mucin-like glycoprotein, putative | 1,449 | 8,941E-12 |
| TcCLB.510693.19 | mucin-associated surface protein (MASP, pseudogene), pu | 1,449 | 0,00003952 |
| TcCLB.511725.170 | hypothetical protein, conserved | 1,448 | 2,64E-29 |
| TcCLB.508417.29 | dispersed gene family protein 1 (DGF-1), putative (fragme | 1,447 | 9,15E-18 |
| TcCLB.511613.80 | mucin-associated surface protein (MASP), putative | 1,447 | 0,002219 |

| | | | |
|---|---|---|---|
| TcCLB.508909.90 | Protein kinase domain, putative | 1,446 | 1,112E-35 |
| TcCLB.506741.110 | mucin TcMUCII, putative | 1,446 | 2,778E-10 |
| TcCLB.506221.70 | Paraflagellar Rod Proteome Component 9, putative (PFC9) | 1,445 | 9,185E-14 |
| TcCLB.508389.138 | mucin TcMUCII, putative | 1,445 | 0,07303 |
| TcCLB.508401.20 | retrotransposon hot spot (RHS) protein, putative | 1,444 | 2,462E-39 |
| TcCLB.506297.260 | lanosterol 14-alpha-demethylase, putative (CYP51) | 1,443 | 2,295E-63 |
| TcCLB.511827.64 | hypothetical protein, conserved | 1,443 | 3,743E-32 |
| TcCLB.510345.40 | hypothetical protein, conserved | 1,443 | 1,177E-29 |
| TcCLB.507067.60 | mucin-associated surface protein (MASP), putative | 1,443 | 0,01155 |
| TcCLB.508125.100 | mucin TcMUCII, putative | 1,443 | 0,3475 |
| TcCLB.510149.150 | NADH-ubiquinone oxidoreductase complex I subunit, putati | 1,44 | 5,404E-18 |
| TcCLB.509973.10 | mucin-associated surface protein (MASP), putative | 1,44 | 8,255E-16 |
| TcCLB.506775.190 | serine/threonine protein kinase, putative | 1,439 | 4,205E-15 |
| TcCLB.508495.210 | mucin-associated surface protein (MASP, pseudogene), pu | 1,439 | 0,002989 |
| TcCLB.503753.29 | NADPH--cytochrome P450 reductase, putative | 1,438 | 2,198E-28 |
| TcCLB.507225.10 | retrotransposon hot spot (RHS) protein, putative | 1,437 | 7,943E-43 |
| TcCLB.509849.10 | ammonium transporter, putative | 1,437 | 1,435E-19 |
| TcCLB.509977.40 | hypothetical protein | 1,437 | 4,482E-07 |
| TcCLB.504039.180 | mucin-associated surface protein (MASP), putative | 1,437 | 0,00000578 |
| TcCLB.509725.80 | retrotransposon hot spot protein (RHS, pseudogene), putat | 1,436 | 1,064E-52 |
| TcCLB.508495.70 | mucin TcMUCII, putative | 1,436 | 2,245E-07 |
| TcCLB.510359.540 | hypothetical protein | 1,436 | 0,06193 |
| TcCLB.508113.7 | trans-sialidase (pseudogene), putative | 1,435 | 1,303E-07 |
| TcCLB.483623.10 | hypothetical protein, conserved | 1,434 | 1,11E-11 |
| TcCLB.508693.199 | hypothetical protein, conserved | 1,433 | 3,525E-20 |
| TcCLB.507639.30 | universal minicircle sequence binding protein 1 (pdz5) | 1,432 | 1,147E-18 |
| TcCLB.507023.70 | hypothetical protein, conserved | 1,432 | 4,556E-11 |
| TcCLB.510789.10 | mucin TcMUCII, putative | 1,432 | 0,00004288 |
| TcCLB.506909.20 | mucin TcMUCI, putative | 1,432 | 0,001744 |
| TcCLB.510279.140 | mucin-associated surface protein (MASP), putative | 1,432 | 0,004541 |
| TcCLB.506401.110 | protein kinase, putative | 1,431 | 4,109E-68 |
| TcCLB.506993.86 | mucin-associated surface protein (MASP, pseudogene), pu | 1,431 | 0,001095 |
| TcCLB.506337.160 | NADH-ubiquinone oxidoreductase complex I subunit, putati | 1,43 | 6,185E-28 |
| TcCLB.508023.50 | hypothetical protein | 1,43 | 1,426E-06 |
| TcCLB.510013.270 | trans-sialidase (pseudogene), putative | 1,43 | 7,662E-06 |
| TcCLB.510305.79 | hypothetical protein, conserved | 1,429 | 4,197E-59 |
| TcCLB.508671.20 | serine carboxypeptidase (CBP1), putative | 1,429 | 5,826E-34 |
| TcCLB.503521.50 | UTP-glucose-1-phosphate uridylyltransferase 2, putative | 1,428 | 6,286E-23 |
| TcCLB.507475.30 | retrotransposon hot spot protein (RHS, pseudogene), putat | 1,428 | 8,922E-17 |
| TcCLB.503653.20 | hypothetical protein, conserved | 1,428 | 1,236E-12 |
| TcCLB.509961.51 | tryptophanyl-tRNA synthetase (pseudogene), putative | 1,428 | 0,00008804 |
| TcCLB.511585.260 | dispersed gene family protein 1 (DGF-1, pseudogene), puta | 1,427 | 5,879E-13 |
| TcCLB.509699.30 | mucin TcMUCII (pseudogene), putative | 1,427 | 0,03861 |
| TcCLB.507739.59 | hypothetical protein, conserved | 1,426 | 3,463E-10 |
| TcCLB.507693.10 | dispersed gene family protein 1 (DGF-1), putative | 1,426 | 4,217E-07 |

| | | | |
|---|---|---|---|
| TcCLB.507163.60 | hypothetical protein | 1,426 | 0,002984 |
| TcCLB.510101.350 | hypothetical protein, conserved | 1,425 | 4,614E-73 |
| TcCLB.510535.90 | hypothetical protein, conserved | 1,425 | 0,0001701 |
| TcCLB.509463.30 | 3-ketoacyl-CoA thiolase, putative | 1,424 | 9,753E-68 |
| TcCLB.509921.20 | trans-sialidase (pseudogene), putative | 1,424 | 7,974E-08 |
| TcCLB.511815.170 | hypothetical protein, conserved | 1,423 | 8,607E-34 |
| TcCLB.511395.20 | phosphatidylglycerolphosphate synthase, mitochondrial, pu | 1,423 | 3E-30 |
| TcCLB.503643.30 | hypothetical protein, conserved | 1,423 | 1,299E-21 |
| TcCLB.510343.5 | hypothetical protein, conserved (fragment) | 1,423 | 1,153E-14 |
| TcCLB.510513.10 | mannosyl-oligosaccharide 1,2-alpha-mannosidase IB, puta | 1,422 | 7,298E-25 |
| TcCLB.508973.69 | hypothetical protein, conserved | 1,422 | 1,491E-14 |
| TcCLB.506183.9 | pyruvate dehydrogenase (lipoamide) kinase, putative (frag | 1,422 | 2,056E-13 |
| TcCLB.510795.30 | fatty acid desaturase (pseudogene), putative | 1,421 | 8,066E-27 |
| TcCLB.506501.190 | mucin TcMUCII, putative | 1,421 | 0,03885 |
| TcCLB.506775.100 | DNA repair and recombination helicase protein PIF2, putat | 1,42 | 1,254E-22 |
| TcCLB.507875.110 | retrotransposon hot spot protein (RHS, pseudogene), putat | 1,42 | 3,294E-20 |
| TcCLB.509843.20 | trans-sialidase, Group VII, putative | 1,419 | 3,191E-43 |
| TcCLB.511129.40 | trans-sialidase, Group III, putative | 1,419 | 0,2118 |
| TcCLB.509179.10 | choline/carnitine O-acyltransferase (pseudogene), putative | 1,418 | 1,978E-09 |
| TcCLB.509331.189 | hypothetical protein, conserved | 1,417 | 1,834E-21 |
| TcCLB.511911.60 | trans-sialidase, Group III, putative | 1,417 | 0,1034 |
| TcCLB.511817.120 | hypothetical protein, conserved | 1,416 | 2,31E-35 |
| TcCLB.510201.5 | mucin-associated surface protein (MASP), putative | 1,416 | 0,2782 |
| TcCLB.509331.200 | ABC transporter, putative | 1,415 | 5,044E-21 |
| TcCLB.508355.127 | hypothetical protein, conserved | 1,414 | 7,679E-28 |
| TcCLB.506271.30 | retrotransposon hot spot (RHS) protein, putative (fragmen | 1,414 | 2,851E-06 |
| TcCLB.510099.120 | D-isomer specific 2-hydroxyacid dehydrogenase-protein, pu | 1,413 | 1,316E-122 |
| TcCLB.508741.160 | hypothetical protein, conserved | 1,413 | 6,683E-38 |
| TcCLB.506299.9 | dispersed gene family protein 1 (DGF-1), putative (fragme | 1,413 | 3,143E-11 |
| TcCLB.510377.210 | trans-sialidase (pseudogene), putative | 1,413 | 0,008183 |
| TcCLB.507237.20 | mucin-associated surface protein (MASP), putative | 1,413 | 0,04037 |
| TcCLB.503865.60 | hypothetical protein, conserved | 1,412 | 7,272E-34 |
| TcCLB.454787.10 | mucin TcMUC, putative | 1,412 | 0,3926 |
| TcCLB.511599.30 | 90 kDa surface protein, putative | 1,411 | 0,2018 |
| TcCLB.508499.10 | mucin TcMUCII, putative | 1,41 | 0,00003349 |
| TcCLB.503987.30 | Enriched in surface-labeled proteome protein 16 (ESP16) | 1,409 | 5,64E-25 |
| TcCLB.510993.10 | (H )-ATPase G subunit | 1,408 | 5,359E-31 |
| TcCLB.511573.49 | hypothetical protein | 1,407 | 1,449E-23 |
| TcCLB.510079.10 | retrotransposon hot spot protein (RHS, pseudogene), putat | 1,407 | 8,999E-23 |
| TcCLB.509331.180 | aldose 1-epimerase-like protein, putative | 1,407 | 2,894E-18 |
| TcCLB.511825.30 | hypothetical protein, conserved | 1,407 | 5,619E-18 |
| TcCLB.511781.31 | trans-sialidase (pseudogene), putative | 1,407 | 0,3917 |
| TcCLB.503519.30 | mucin-associated surface protein (MASP), putative | 1,406 | 0,0001456 |
| TcCLB.506859.240 | hypothetical protein, conserved | 1,406 | 0,000266 |
| TcCLB.510423.50 | hypothetical protein, conserved | 1,405 | 2,315E-28 |

| | | | |
|---|---|---|---|
| TcCLB.509213.100 | 3-ketoacyl-CoA reductase, putative | 1,404 | 2,817E-35 |
| TcCLB.510265.4 | hypothetical protein, conserved | 1,404 | 4,295E-07 |
| TcCLB.508501.140 | mucin TcMUCII, putative | 1,404 | 0,00001219 |
| TcCLB.507737.40 | hypothetical protein, conserved (fragment) | 1,404 | 0,01083 |
| TcCLB.511331.10 | calpain cysteine peptidase, putative (fragment) | 1,403 | 0,01736 |
| TcCLB.511843.80 | retrotransposon hot spot (RHS) protein, putative | 1,402 | 2,5E-13 |
| TcCLB.506457.140 | hypothetical protein, conserved | 1,401 | 2,554E-09 |
| TcCLB.509331.30 | acetyl-CoA synthetase, putative | 1,4 | 2,407E-27 |
| TcCLB.507645.11 | hypothetical protein, conserved (fragment) | 1,4 | 1,159E-09 |
| TcCLB.508493.100 | mucin TcMUCII, putative | 1,4 | 7,139E-06 |
| TcCLB.506331.130 | trans-sialidase, Group II, putative | 1,4 | 0,0004584 |
| TcCLB.506285.50 | mucin TcMUCII, putative | 1,4 | 0,001009 |
| TcCLB.510377.190 | mucin TcMUCII, putative | 1,4 | 0,003857 |
| TcCLB.511121.30 | hypothetical protein | 1,4 | 0,05877 |
| TcCLB.507847.30 | mucin TcMUCII (pseudogene), putative | 1,397 | 0,006915 |
| TcCLB.506675.30 | retrotransposon hot spot protein (RHS, pseudogene), putat | 1,396 | 7,02E-30 |
| TcCLB.506961.130 | retrotransposon hot spot protein (RHS, pseudogene), putat | 1,396 | 4,026E-28 |
| TcCLB.508043.20 | metallo-beta-lactamase-like protein, putative | 1,395 | 2,802E-26 |
| TcCLB.511669.10 | trans-sialidase, Group VIII, putative | 1,394 | 8,579E-15 |
| TcCLB.509973.25 | mucin TcMUC (pseudogene), putative | 1,394 | 0,00001552 |
| TcCLB.508017.10 | hypothetical protein, conserved | 1,394 | 0,002514 |
| TcCLB.508777.110 | hypothetical protein, conserved | 1,393 | 2,045E-35 |
| TcCLB.510855.20 | hypothetical protein, conserved | 1,393 | 3,913E-20 |
| TcCLB.510131.40 | haloacid dehalogenase-like hydrolase, putative | 1,393 | 2,027E-17 |
| TcCLB.503779.109 | hypothetical protein, conserved (fragment) | 1,393 | 7,14E-16 |
| TcCLB.509535.9 | electon transport protein SCO1/SCO2, putative | 1,393 | 2,024E-12 |
| TcCLB.506219.40 | hypothetical protein, conserved | 1,392 | 1,589E-59 |
| TcCLB.508677.150 | glycosyl transferase, putative | 1,392 | 2,295E-52 |
| TcCLB.511817.260 | hypothetical protein, conserved | 1,392 | 9,233E-48 |
| TcCLB.509961.20 | dispersed gene family protein 1 (DGF-1), putative | 1,392 | 0,007968 |
| TcCLB.508225.4 | trans-sialidase, putative | 1,391 | 0,004822 |
| TcCLB.511809.120 | hypothetical protein, conserved | 1,388 | 6,725E-43 |
| TcCLB.503607.4 | retrotransposon hot spot (RHS) protein, putative | 1,388 | 6,999E-21 |
| TcCLB.509967.140 | hypothetical protein, conserved | 1,388 | 1,334E-14 |
| TcCLB.507699.64 | mucin TcMUCII, putative | 1,387 | 0,02247 |
| TcCLB.503823.40 | glycosyl hydrolase-like protein, putative | 1,385 | 3,452E-41 |
| TcCLB.510819.119 | zinc finger (CCCH type) protein, putative | 1,385 | 8,737E-29 |
| TcCLB.509797.30 | hypothetical protein, conserved | 1,385 | 2,329E-24 |
| TcCLB.506577.120 | sterol C-24 reductase, putative | 1,385 | 2,176E-16 |
| TcCLB.460127.30 | amino acid permease, putative | 1,385 | 4,781E-07 |
| TcCLB.504031.11 | hypothetical protein | 1,385 | 0,0008403 |
| TcCLB.509765.20 | retrotransposon hot spot protein (RHS, pseudogene), putat | 1,384 | 0,0001364 |
| TcCLB.510021.140 | mucin TcMUCII (pseudogene), putative | 1,384 | 0,001726 |
| TcCLB.506823.20 | retrotransposon hot spot protein (RHS, pseudogene), putat | 1,382 | 6,081E-10 |
| TcCLB.508799.240 | aminopeptidase, putative | 1,381 | 3,104E-54 |

| | | | |
|---|---|---|---|
| TcCLB.511861.120 | surface antigen TASV, putative | 1,381 | 5,544E-11 |
| TcCLB.508741.140 | imidazolonepropionase, putative | 1,38 | 9,248E-44 |
| TcCLB.511809.40 | hypothetical protein, conserved | 1,38 | 2,77E-18 |
| TcCLB.510861.50 | hypothetical protein, conserved | 1,38 | 0,0001774 |
| TcCLB.511391.160 | cytochrome c1, heme protein, mitochondrial, putative (CY( | 1,379 | 2,569E-68 |
| TcCLB.510759.40 | Protein kinase domain, putative | 1,379 | 5,569E-31 |
| TcCLB.506931.80 | p450 reductase, putative | 1,379 | 7,643E-29 |
| TcCLB.510883.10 | retrotransposon hot spot (RHS) protein, putative | 1,379 | 3,106E-17 |
| TcCLB.511821.40 | hypothetical protein, conserved | 1,379 | 6,259E-13 |
| TcCLB.509025.20 | hypothetical protein, conserved | 1,379 | 7,655E-09 |
| TcCLB.506777.110 | hypothetical protein, conserved | 1,377 | 2,54E-33 |
| TcCLB.504107.10 | calpain-like cysteine peptidase, putative (fragment) | 1,376 | 0,03135 |
| TcCLB.506005.10 | dispersed gene family protein 1 (DGF-1, pseudogene), puta | 1,375 | 5,193E-20 |
| TcCLB.506435.424 | hypothetical protein, conserved | 1,375 | 7,278E-11 |
| TcCLB.508495.200 | mucin TcMUCII, putative | 1,375 | 1,57E-10 |
| TcCLB.401569.10 | trans-sialidase, putative | 1,375 | 0,0006766 |
| TcCLB.506059.10 | trans-sialidase, putative | 1,375 | 0,0007609 |
| TcCLB.506823.80 | retrotransposon hot spot protein (RHS, pseudogene), putat | 1,374 | 2,853E-16 |
| TcCLB.505919.11 | mucin-associated surface protein (MASP, pseudogene), pu | 1,373 | 3,137E-23 |
| TcCLB.510279.180 | mucin TcMUCII (pseudogene), putative | 1,373 | 0,04372 |
| TcCLB.506501.80 | mucin-associated surface protein (MASP, pseudogene), pu | 1,372 | 0,000097 |
| TcCLB.507757.30 | hypothetical protein, conserved | 1,371 | 2,678E-14 |
| TcCLB.510279.120 | mucin-associated surface protein (MASP), putative | 1,371 | 0,03408 |
| TcCLB.506263.20 | D-isomer specific 2-hydroxyacid dehydrogenase-protein, pu | 1,37 | 1,01E-45 |
| TcCLB.510349.10 | hypothetical protein, conserved | 1,37 | 7,251E-19 |
| TcCLB.509545.79 | mucin-associated surface protein (MASP), putative | 1,37 | 0,02527 |
| TcCLB.511375.30 | hypothetical protein (fragment) | 1,369 | 0,02835 |
| TcCLB.399997.10 | retrotransposon hot spot (RHS) protein, putative | 1,368 | 2,676E-23 |
| TcCLB.508499.20 | mucin-associated surface protein (MASP), putative | 1,368 | 3,417E-06 |
| TcCLB.506157.30 | retrotransposon hot spot protein (RHS, pseudogene), putat | 1,365 | 2,754E-15 |
| TcCLB.508069.10 | retrotransposon hot spot protein (RHS, pseudogene), putat | 1,365 | 1,672E-10 |
| TcCLB.511607.60 | trans-sialidase (pseudogene), putative | 1,365 | 6,256E-09 |
| TcCLB.508493.70 | hypothetical protein | 1,365 | 0,00432 |
| TcCLB.510961.40 | mucin TcMUCII, putative | 1,365 | 0,02808 |
| TcCLB.511607.30 | mucin TcMUCII, putative | 1,365 | 0,07614 |
| TcCLB.508221.790 | trans-sialidase, Group V, putative | 1,364 | 0,006021 |
| TcCLB.511909.60 | hypothetical protein, conserved | 1,363 | 1,054E-16 |
| TcCLB.452475.10 | trans-sialidase, Group V, putative | 1,363 | 0,001126 |
| TcCLB.508443.30 | hypothetical protein, conserved | 1,362 | 1,959E-44 |
| TcCLB.506477.10 | cytochrome c oxidase assembly factor, putative | 1,361 | 3,029E-19 |
| TcCLB.504071.110 | ama1 protein, putative | 1,361 | 1,403E-13 |
| TcCLB.510265.10 | quinone oxidoreductase, putative | 1,36 | 1,76E-23 |
| TcCLB.511727.220 | hypothetical protein, conserved | 1,36 | 1,281E-19 |
| TcCLB.509387.10 | trans-sialidase, Group VIII, putative | 1,36 | 6,418E-18 |
| TcCLB.509965.144 | hypothetical protein, conserved | 1,359 | 4,077E-16 |

| | | | |
|---|---|---|---|
| TcCLB.510377.10 | trans-sialidase, Group V, putative | 1,359 | 0,0009629 |
| TcCLB.508677.30 | trans-sialidase (pseudogene), putative | 1,358 | 4,324E-11 |
| TcCLB.509281.30 | hypothetical protein | 1,358 | 1,358E-09 |
| TcCLB.508799.160 | cytochrome b5-like protein, putative | 1,357 | 2,415E-42 |
| TcCLB.511079.71 | mucin TcMUC (pseudogene), putative | 1,356 | 0,004355 |
| TcCLB.508119.144 | ribosomal protein L38, putative | 1,355 | 4,816E-32 |
| TcCLB.506585.70 | p450 reductase, putative | 1,354 | 1,245E-56 |
| TcCLB.506521.9 | dispersed gene family protein 1 (DGF-1), putative | 1,354 | 4,655E-16 |
| TcCLB.510025.50 | trans-sialidase, Group V, putative | 1,353 | 3,514E-10 |
| TcCLB.508903.100 | ATPase, putative | 1,353 | 3,794E-07 |
| TcCLB.511797.130 | mucin TcMUCII (pseudogene), putative | 1,353 | 4,903E-06 |
| TcCLB.403031.10 | trans-sialidase, putative (fragment) | 1,353 | 0,3862 |
| TcCLB.506865.2 | nucleoside phosphorylase, putative | 1,352 | 8,833E-15 |
| TcCLB.506187.20 | trans-sialidase, Group V, putative | 1,351 | 2,12E-10 |
| TcCLB.507427.20 | dispersed gene family protein 1 (DGF-1, pseudogene), puta | 1,351 | 0,00000292 |
| TcCLB.505843.49 | ERV/ALR sulfhydryl oxidase domain-containing protein (PC | 1,35 | 3,142E-06 |
| TcCLB.507699.90 | mucin TcMUCII, putative | 1,35 | 0,0002429 |
| TcCLB.503519.20 | mucin TcMUCII, putative | 1,35 | 0,009631 |
| TcCLB.507179.7 | trans-sialidase (pseudogene), putative | 1,349 | 3,069E-11 |
| TcCLB.510583.40 | mucin TcMUCII, putative | 1,349 | 0,06354 |
| TcCLB.510879.170 | hypothetical protein, conserved | 1,348 | 1,262E-14 |
| TcCLB.510115.70 | trans-sialidase, putative | 1,348 | 0,001291 |
| TcCLB.508449.31 | trans-sialidase, putative | 1,348 | 0,02539 |
| TcCLB.507085.30 | trans-sialidase, Group I, putative | 1,347 | 3,626E-16 |
| TcCLB.509665.10 | hypothetical protein | 1,347 | 0,06049 |
| TcCLB.506767.130 | mucin TcMUCII, putative | 1,347 | 0,296 |
| TcCLB.509477.10 | mucin-associated surface protein (MASP), putative | 1,347 | 0,3921 |
| TcCLB.504769.140 | trans-sialidase, Group VIII, putative | 1,346 | 0,0001969 |
| TcCLB.504093.10 | hypothetical protein, conserved (pseudogene) | 1,345 | 1,179E-25 |
| TcCLB.511511.40 | hypothetical protein, conserved | 1,345 | 4,294E-25 |
| TcCLB.510151.19 | fatty acyl CoA syntetase 1, putative | 1,345 | 1,16E-14 |
| TcCLB.511127.270 | ankyrin repeat protein, putative | 1,345 | 5,494E-10 |
| TcCLB.509079.10 | surface protease GP63 (pseudogene), putative | 1,344 | 0,05261 |
| TcCLB.507981.40 | hypothetical protein | 1,344 | 0,3487 |
| TcCLB.441241.10 | microtubule-associated protein Gb4, putative (fragment) | 1,343 | 2,021E-22 |
| TcCLB.506369.20 | transporter, putative | 1,343 | 2,494E-10 |
| TcCLB.506631.30 | hypothetical protein, conserved | 1,342 | 3,606E-23 |
| TcCLB.511019.84 | GDP-mannose 4,6 dehydratase, putative | 1,342 | 2,144E-17 |
| TcCLB.508873.25 | hypothetical protein | 1,342 | 0,004605 |
| TcCLB.509151.70 | hypothetical protein, conserved | 1,34 | 5,969E-39 |
| TcCLB.506615.29 | mucin-associated surface protein (MASP, pseudogene), pu | 1,34 | 9,012E-14 |
| TcCLB.509915.50 | trans-sialidase (pseudogene), putative | 1,34 | 0,03926 |
| TcCLB.511751.150 | hypothetical protein, conserved | 1,339 | 2,367E-11 |
| TcCLB.511203.5 | hypothetical protein, conserved | 1,339 | 0,002559 |
| TcCLB.510013.260 | mucin-associated surface protein (MASP), putative | 1,339 | 0,02865 |

| | | | |
|---|---|---|---|
| TcCLB.511655.40 | hypothetical protein, conserved | 1,338 | 5,419E-21 |
| TcCLB.506989.220 | hypothetical protein, conserved | 1,337 | 4,428E-22 |
| TcCLB.507021.110 | phosphopantothenoylcysteine decarboxylase, putative (PPC | 1,337 | 9,836E-22 |
| TcCLB.510671.10 | dispersed gene family protein 1 (DGF-1), putative | 1,336 | 1,922E-11 |
| TcCLB.503633.30 | surface protease GP63, putative | 1,336 | 0,01197 |
| TcCLB.508071.20 | trans-sialidase (pseudogene), putative | 1,336 | 0,04692 |
| TcCLB.506999.180 | hypothetical protein | 1,334 | 1,356E-37 |
| TcCLB.510523.20 | hypothetical protein, conserved | 1,334 | 6,653E-13 |
| TcCLB.508269.40 | hypothetical protein, conserved | 1,334 | 8,118E-12 |
| TcCLB.506063.20 | retrotransposon hot spot protein (RHS, pseudogene), putat | 1,334 | 2,051E-10 |
| TcCLB.459199.10 | retrotransposon hot spot (RHS) protein, putative | 1,333 | 4,47E-63 |
| TcCLB.510715.120 | mucin TcMUCII, putative | 1,333 | 0,00281 |
| TcCLB.506593.50 | hypothetical protein, conserved | 1,332 | 4,746E-20 |
| TcCLB.508163.234 | trans-sialidase, putative | 1,331 | 0,0021 |
| TcCLB.508267.20 | hypothetical protein, conserved | 1,33 | 5,913E-34 |
| TcCLB.508687.20 | retrotransposon hot spot protein (RHS, pseudogene), putat | 1,33 | 2,697E-14 |
| TcCLB.508593.130 | hypothetical protein, conserved | 1,329 | 5,854E-31 |
| TcCLB.506583.70 | cytosolic factor SEC14, putative | 1,329 | 2,148E-15 |
| TcCLB.510933.120 | surface protease GP63, putative | 1,329 | 8,866E-07 |
| TcCLB.511117.60 | mucin-associated surface protein (MASP, pseudogene), pu | 1,329 | 0,004757 |
| TcCLB.508079.10 | conserved protein | 1,328 | 2,536E-50 |
| TcCLB.508173.10 | retrotransposon hot spot protein (RHS, pseudogene), putat | 1,328 | 1,009E-48 |
| TcCLB.510769.60 | hypothetical protein, conserved (pseudogene) | 1,328 | 4,591E-25 |
| TcCLB.509869.10 | mucin TcMUCI, putative | 1,328 | 0,01037 |
| TcCLB.506777.40 | hypothetical protein, conserved | 1,327 | 2,126E-28 |
| TcCLB.504479.10 | trans-sialidase, Group II, putative | 1,327 | 5,171E-15 |
| TcCLB.506761.54 | mucin-associated surface protein (MASP, pseudogene), pu | 1,327 | 0,01203 |
| TcCLB.506287.100 | hypothetical protein, conserved | 1,326 | 1,835E-16 |
| TcCLB.424123.20 | paraflagellar rod component, putative (PFC16) | 1,326 | 1,357E-11 |
| TcCLB.508229.30 | mucin-associated surface protein (MASP), putative | 1,326 | 0,004976 |
| TcCLB.508389.50 | mucin-associated surface protein (MASP), putative | 1,326 | 0,008592 |
| TcCLB.508515.90 | hypothetical protein, conserved | 1,325 | 3,473E-10 |
| TcCLB.510979.60 | mucin TcMUCII, putative | 1,325 | 0,01329 |
| TcCLB.506459.60 | retrotransposon hot spot protein (RHS, pseudogene), putat | 1,323 | 1,182E-18 |
| TcCLB.503477.10 | hypothetical protein, conserved | 1,323 | 3,127E-14 |
| TcCLB.506469.30 | protein kinase, putative | 1,323 | 1,525E-09 |
| TcCLB.509725.113 | ATP-dependent DEAD/H RNA helicase (pseudogene), putat | 1,323 | 0,001162 |
| TcCLB.506423.20 | mucin TcMUCII, putative | 1,323 | 0,2226 |
| TcCLB.504533.40 | trans-sialidase, Group IV, putative | 1,322 | 1,793E-29 |
| TcCLB.510021.150 | hypothetical protein, conserved (pseudogene) | 1,322 | 0,0001239 |
| TcCLB.511613.50 | mucin TcMUCII, putative | 1,322 | 0,006495 |
| TcCLB.506015.50 | hypothetical protein (pseudogene) | 1,322 | 0,01342 |
| TcCLB.507031.100 | hypothetical protein, conserved | 1,321 | 1,551E-16 |
| TcCLB.509533.10 | trans-sialidase, Group II, putative | 1,321 | 1,096E-11 |
| TcCLB.504149.160 | phosphoinositide-specific phospholipase C, putative | 1,32 | 7,627E-45 |

| | | | |
|---|---|---|---|
| TcCLB.508173.40 | mercaptopyruvate sulfurtransferase, putative (MST) | 1,319 | 1,862E-27 |
| TcCLB.503411.20 | hypothetical protein, conserved | 1,319 | 1,137E-18 |
| TcCLB.510089.60 | hypothetical protein, conserved | 1,319 | 1,669E-12 |
| TcCLB.508851.120 | ADP-ribosylation factor GTPase activating protein, putative | 1,317 | 1,609E-20 |
| TcCLB.507089.210 | hypothetical protein, conserved | 1,316 | 3,193E-26 |
| TcCLB.509495.30 | trans-sialidase, Group I, putative | 1,316 | 1,102E-14 |
| TcCLB.506327.99 | serine/threonine protein kinase, putative (fragment) | 1,315 | 3,175E-20 |
| TcCLB.509109.120 | protein tyrosine phosphatase, putative | 1,314 | 8,935E-20 |
| TcCLB.507765.149 | hypothetical protein, conserved | 1,314 | 1,838E-16 |
| TcCLB.511257.100 | surface protease GP63, putative | 1,313 | 1,89E-13 |
| TcCLB.503991.20 | Xaa-Pro aminopeptidase, putative | 1,312 | 3,86E-29 |
| TcCLB.507953.180 | mucin TcMUCII, putative | 1,312 | 0,01154 |
| TcCLB.507587.50 | hypothetical protein | 1,312 | 0,06891 |
| TcCLB.506779.100 | RNA-binding protein, putative | 1,311 | 2,499E-14 |
| TcCLB.504039.230 | mucin-associated surface protein (MASP), putative | 1,311 | 0,0008225 |
| TcCLB.506857.20 | pyrroline-5-carboxylate reductase, putative (P5CR) | 1,31 | 2,672E-25 |
| TcCLB.509259.180 | retrotransposon hot spot (RHS) protein, putative | 1,31 | 4,075E-23 |
| TcCLB.510525.139 | protein kinase, putative | 1,31 | 8,335E-09 |
| TcCLB.506529.70 | hypothetical protein, conserved | 1,309 | 1,009E-19 |
| TcCLB.510325.40 | acyltransferase, putative | 1,309 | 7,708E-18 |
| TcCLB.506331.90 | trans-sialidase, Group II, putative | 1,309 | 0,00001329 |
| TcCLB.503889.10 | mucin TcMUCII, putative | 1,309 | 0,000023 |
| TcCLB.503553.10 | trans-sialidase (pseudogene), putative | 1,309 | 0,02092 |
| TcCLB.506779.160 | palmitoyl acyltransferase 9, putative | 1,308 | 1,246E-23 |
| TcCLB.509967.170 | dynein arm light chain, putative | 1,308 | 1,026E-22 |
| TcCLB.509295.50 | beta galactofuranosyl glycosyltransferase, putative | 1,308 | 4,625E-06 |
| TcCLB.509493.110 | mucin TcMUCII, putative | 1,308 | 0,0003966 |
| TcCLB.511865.10 | hypothetical protein, conserved | 1,307 | 1,804E-39 |
| TcCLB.506649.50 | cytochrome c oxidase assembly factor, putative | 1,307 | 1,472E-20 |
| TcCLB.511643.73 | mucin-associated surface protein (MASP, pseudogene), pu | 1,307 | 0,02286 |
| TcCLB.506401.320 | 60S ribosomal protein L7a, putative | 1,306 | 3,082E-64 |
| TcCLB.511817.100 | hypothetical protein, conserved | 1,306 | 1,01E-33 |
| TcCLB.503527.40 | GPI alpha-mannosyltransferase III (GPI10) | 1,306 | 2,261E-17 |
| TcCLB.508005.10 | hypothetical protein | 1,306 | 0,0752 |
| TcCLB.507711.20 | paraflagellar rod component, putative (PFC16) | 1,305 | 1,619E-14 |
| TcCLB.510093.20 | trans-sialidase, Group VIII, putative | 1,305 | 4,384E-07 |
| TcCLB.508009.10 | surface protease GP63 (pseudogene), putative | 1,305 | 0,003598 |
| TcCLB.507471.10 | surface protease GP63 (pseudogene), putative | 1,305 | 0,247 |
| TcCLB.509777.140 | hypothetical protein, conserved | 1,304 | 6,751E-17 |
| TcCLB.509259.40 | surface protease GP63, putative | 1,304 | 0,01857 |
| TcCLB.506465.14 | hypothetical protein, conserved | 1,302 | 1,092E-18 |
| TcCLB.511309.40 | hydrolase-like protein, putative | 1,302 | 7,685E-09 |
| TcCLB.503503.30 | mucin TcMUCII, putative | 1,302 | 0,0003353 |
| TcCLB.506325.36 | N-acetyltransferase complex ARD1 subunit (pseudogene), | 1,302 | 0,007342 |
| TcCLB.509973.40 | hypothetical protein | 1,302 | 0,01617 |

| | | | |
|---|---|---|---|
| TcCLB.508511.49 | trans-sialidase, putative | 1,301 | 0,0006983 |
| TcCLB.507391.20 | hypothetical protein, conserved (pseudogene) | 1,3 | 2,66E-19 |
| TcCLB.511885.20 | trans-sialidase, Group II, putative | 1,3 | 9,178E-11 |
| TcCLB.506217.40 | trans-sialidase, Group VIII, putative | 1,3 | 0,00004384 |
| TcCLB.509857.50 | bilobe protein, conserved | 1,299 | 5,667E-33 |
| TcCLB.506367.30 | 1,2-Dihydroxy-3-keto-5-methylthiopentene dioxygenase, pu | 1,298 | 1,448E-55 |
| TcCLB.510155.115 | hypothetical protein, conserved | 1,297 | 1,316E-18 |
| TcCLB.509759.10 | L-ribulokinase, putative | 1,297 | 1,047E-06 |
| TcCLB.511461.60 | NADH-dependent fumarate reductase (pseudogene), putat | 1,296 | 3,182E-45 |
| TcCLB.511807.284 | carnitine O-palmitoyltransferase II, putative | 1,296 | 1,245E-33 |
| TcCLB.508521.40 | hypothetical protein | 1,296 | 0,04737 |
| TcCLB.510101.250 | hypothetical protein, conserved | 1,295 | 3,784E-30 |
| TcCLB.510407.90 | serine hydroxymethyltransferase, putative | 1,295 | 4,027E-24 |
| TcCLB.506425.90 | hypothetical protein, conserved | 1,295 | 3,935E-20 |
| TcCLB.511057.30 | mucin-associated surface protein (MASP), putative | 1,295 | 0,1006 |
| TcCLB.506843.5 | trans-sialidase, putative (fragment) | 1,295 | 0,319 |
| TcCLB.509437.60 | hypothetical protein, conserved | 1,294 | 4,84E-29 |
| TcCLB.445635.10 | receptor-type adenylate cyclase (pseudogene), putative | 1,294 | 1,925E-17 |
| TcCLB.509331.100 | hypothetical protein, conserved | 1,294 | 1,642E-11 |
| TcCLB.509963.30 | tryptophanyl-tRNA synthetase (pseudogene), putative | 1,294 | 0,04191 |
| TcCLB.509149.80 | hypothetical protein, conserved | 1,293 | 1,445E-27 |
| TcCLB.508045.120 | trans-sialidase, Group III, putative | 1,293 | 4,115E-22 |
| TcCLB.504423.15 | hypothetical protein, conserved | 1,293 | 4,215E-14 |
| TcCLB.511233.160 | hypothetical protein | 1,293 | 0,001879 |
| TcCLB.511771.40 | trans-sialidase (pseudogene), putative | 1,292 | 0,0452 |
| TcCLB.506001.24 | hypothetical protein (pseudogene) | 1,292 | 0,1813 |
| TcCLB.507677.160 | hypothetical protein, conserved | 1,291 | 3,372E-37 |
| TcCLB.509977.30 | mucin-associated surface protein (MASP), putative | 1,291 | 4,543E-08 |
| TcCLB.503579.119 | hypothetical protein, conserved | 1,291 | 8,631E-07 |
| TcCLB.510483.300 | mucin TcMUCII, putative | 1,291 | 0,001175 |
| TcCLB.504269.20 | GPR1/FUN34/yaaH family, putative | 1,29 | 3,481E-31 |
| TcCLB.506467.29 | methyltransferase domain containing protein, putative (PC | 1,29 | 4,447E-31 |
| TcCLB.510667.110 | hypothetical protein, conserved | 1,29 | 1,015E-14 |
| TcCLB.506487.37 | hypothetical protein | 1,29 | 0,0008752 |
| TcCLB.509877.60 | hypothetical protein, conserved | 1,289 | 3,717E-23 |
| TcCLB.506241.160 | hypothetical protein, conserved | 1,288 | 5,703E-31 |
| TcCLB.506129.30 | trans-sialidase, Group II, putative | 1,288 | 4,887E-16 |
| TcCLB.506615.120 | trans-sialidase (pseudogene), putative | 1,288 | 2,775E-11 |
| TcCLB.511173.89 | mucin TcMUCII (pseudogene), putative | 1,287 | 0,00005026 |
| TcCLB.510495.11 | surface protease GP63 (pseudogene), putative | 1,287 | 0,002544 |
| TcCLB.511255.640 | mucin TcMUCII, putative | 1,287 | 0,0473 |
| TcCLB.510289.54 | hypothetical protein, conserved | 1,286 | 5,824E-09 |
| TcCLB.506129.20 | dispersed gene family protein 1 (DGF-1, pseudogene), puta | 1,286 | 0,003285 |
| TcCLB.505955.4 | mucin TcMUCII, putative (fragment) | 1,286 | 0,009403 |
| TcCLB.510593.20 | hypothetical protein, conserved | 1,285 | 1,145E-24 |

| | | | |
|---|---|---|---|
| TcCLB.510611.50 | hypothetical protein, conserved | 1,285 | 2,388E-20 |
| TcCLB.503531.20 | hypothetical protein, conserved | 1,285 | 1,173E-19 |
| TcCLB.506715.44 | hypothetical protein, conserved | 1,285 | 1,514E-09 |
| TcCLB.509461.50 | hypothetical protein, conserved | 1,284 | 2,587E-16 |
| TcCLB.508117.20 | mucin-associated surface protein (MASP, pseudogene), pu | 1,284 | 0,0001348 |
| TcCLB.511597.10 | trans-sialidase, putative | 1,284 | 0,2301 |
| TcCLB.511821.60 | protein kinase, putative | 1,283 | 1,719E-18 |
| TcCLB.510053.10 | hypothetical protein, conserved | 1,283 | 3,374E-08 |
| TcCLB.511827.125 | trans-sialidase (pseudogene), putative | 1,283 | 0,0001436 |
| TcCLB.506133.75 | mucin TcMUCII (pseudogene), putative | 1,283 | 0,0001848 |
| TcCLB.506599.350 | mucin TcMUCII, putative | 1,283 | 0,3337 |
| TcCLB.507707.20 | hypothetical protein, conserved | 1,282 | 4,518E-25 |
| TcCLB.504669.40 | mucin-associated surface protein (MASP, pseudogene), pu | 1,281 | 1,976E-17 |
| TcCLB.504015.30 | surface protease GP63, putative | 1,281 | 2,843E-07 |
| TcCLB.510849.11 | retrotransposon hot spot protein (RHS, pseudogene), putat | 1,281 | 0,00006471 |
| TcCLB.508641.220 | UDP-Gal or UDP-GlcNAc-dependent glycosyltransferase, pu | 1,28 | 3,269E-10 |
| TcCLB.510041.30 | hypothetical protein, conserved (fragment) | 1,279 | 9,104E-06 |
| TcCLB.508799.220 | hypothetical protein, conserved (pseudogene) | 1,279 | 0,009751 |
| TcCLB.511771.30 | beta galactofuranosyl glycosyltransferase (pseudogene), p | 1,278 | 0,00001174 |
| TcCLB.506995.40 | mucin TcMUCII, putative | 1,278 | 0,002859 |
| TcCLB.477983.10 | calpain cysteine peptidase, putative | 1,278 | 0,03956 |
| TcCLB.506145.30 | O-6 methyl-guanine alkyl transferase, putative | 1,277 | 2,511E-08 |
| TcCLB.509217.60 | trans-sialidase (pseudogene), putative | 1,277 | 0,003167 |
| TcCLB.511025.80 | hypothetical protein, conserved | 1,276 | 3,808E-08 |
| TcCLB.510595.30 | hypothetical protein | 1,276 | 3,203E-07 |
| TcCLB.506843.40 | retrotransposon hot spot (RHS) protein, putative | 1,275 | 7,301E-54 |
| TcCLB.509459.60 | Na/H antiporter-like protein, putative | 1,275 | 2,429E-39 |
| TcCLB.503469.9 | dispersed gene family protein 1 (DGF-1), putative (fragme | 1,275 | 9,542E-13 |
| TcCLB.510095.20 | trans-sialidase, Group II, putative | 1,275 | 0,08359 |
| TcCLB.507105.100 | retrotransposon hot spot protein (RHS, pseudogene), putat | 1,273 | 1,387E-41 |
| TcCLB.508995.60 | electron transport protein SCO1/2, putative | 1,273 | 7,364E-14 |
| TcCLB.415803.10 | hypothetical protein, conserved | 1,273 | 9,319E-07 |
| TcCLB.507163.20 | hypothetical protein | 1,273 | 0,0228 |
| TcCLB.507547.90 | Phosphoenolpyruvate carboxykinase [ATP], glycosomal | 1,272 | 7,495E-56 |
| TcCLB.508039.30 | retrotransposon hot spot (RHS) protein, putative | 1,272 | 1,481E-25 |
| TcCLB.511829.5 | trans-sialidase (pseudogene), putative | 1,272 | 3,564E-10 |
| TcCLB.508055.20 | retrotransposon hot spot protein (RHS, pseudogene), putat | 1,271 | 3,512E-07 |
| TcCLB.506401.170 | plasma membrane Ca2 ATPase | 1,27 | 4,217E-55 |
| TcCLB.504177.10 | fatty acyl CoA synthetase 2, putative | 1,27 | 1,307E-12 |
| TcCLB.507959.200 | mucin TcMUCII, putative | 1,27 | 0,03236 |
| TcCLB.506279.90 | mucin TcMUCII, putative | 1,27 | 0,06124 |
| TcCLB.509153.130 | hypothetical protein | 1,268 | 3,815E-21 |
| TcCLB.510303.200 | glucosamine-fructose-6-phosphate aminotransferase, puta | 1,266 | 2,828E-23 |
| TcCLB.507167.80 | trans-sialidase, Group II, putative | 1,266 | 0,00001489 |
| TcCLB.508157.40 | hypothetical protein | 1,266 | 0,005465 |

| | | | |
|---|---|---|---|
| TcCLB.507071.164 | mucin TcMUCII (pseudogene), putative | 1,266 | 0,1019 |
| TcCLB.510087.90 | hypothetical protein, conserved | 1,265 | 3,439E-12 |
| TcCLB.511807.22 | retrotransposon hot spot protein (RHS, pseudogene), putat | 1,264 | 3,01E-22 |
| TcCLB.507991.129 | cyclophilin, putative | 1,264 | 1,22E-12 |
| TcCLB.507925.60 | calcium-binding protein, putative | 1,263 | 1,246E-14 |
| TcCLB.509859.80 | coronin, putative (CRN12) | 1,262 | 4,436E-43 |
| TcCLB.503775.50 | mucin TcMUCII (pseudogene), putative | 1,261 | 0,02666 |
| TcCLB.506779.180 | surface protease GP63, putative | 1,26 | 4,579E-10 |
| TcCLB.511531.30 | hypothetical protein, conserved | 1,26 | 1,362E-06 |
| TcCLB.507895.80 | hypothetical protein, conserved | 1,259 | 2,027E-22 |
| TcCLB.510921.20 | mucin TcMUCII, putative | 1,259 | 0,001033 |
| TcCLB.510359.120 | hypothetical protein, conserved (pseudogene) | 1,258 | 7,782E-19 |
| TcCLB.511005.10 | retrotransposon hot spot (RHS) protein, putative (fragmen | 1,258 | 5,001E-16 |
| TcCLB.490989.10 | trans-sialidase, putative | 1,258 | 0,001253 |
| TcCLB.508441.20 | Phosphoenolpyruvate carboxykinase [ATP], glycosomal | 1,257 | 4,516E-63 |
| TcCLB.509137.30 | imidazolonepropionase, putative | 1,257 | 1,556E-34 |
| TcCLB.511065.23 | hypothetical protein, conserved | 1,257 | 1,068E-13 |
| TcCLB.511245.10 | hypothetical protein, conserved | 1,257 | 4,047E-12 |
| TcCLB.511593.20 | mucin TcMUCII, putative | 1,257 | 0,0347 |
| TcCLB.511517.44 | chaperone DNAJ protein, putative | 1,256 | 6,425E-22 |
| TcCLB.506811.20 | hypothetical protein, conserved | 1,255 | 1,417E-18 |
| TcCLB.505183.14 | hypothetical protein, conserved | 1,255 | 3,131E-12 |
| TcCLB.510187.439 | hypothetical protein | 1,254 | 1,419E-42 |
| TcCLB.511337.4 | hypothetical protein, conserved | 1,254 | 1,663E-06 |
| TcCLB.510603.160 | hypothetical protein, conserved | 1,253 | 2,06E-20 |
| TcCLB.508261.80 | trans-sialidase (pseudogene), putative | 1,253 | 0,001432 |
| TcCLB.508975.30 | UDP-Gal or UDP-GlcNAc-dependent glycosyltransferase, pu | 1,253 | 0,07595 |
| TcCLB.508433.170 | mucin TcMUCII, putative | 1,253 | 0,1859 |
| TcCLB.508045.90 | oxidoreductase, putative | 1,252 | 4,328E-19 |
| TcCLB.504669.30 | surface protease GP63, putative | 1,252 | 3,07E-15 |
| TcCLB.511773.110 | retrotransposon hot spot (RHS) protein, putative | 1,251 | 1,801E-37 |
| TcCLB.510479.11 | retrotransposon hot spot (RHS) protein, putative (fragmen | 1,251 | 2,494E-21 |
| TcCLB.458015.4 | hypothetical protein, conserved | 1,251 | 1,719E-18 |
| TcCLB.508501.190 | hypothetical protein, conserved | 1,251 | 4,466E-09 |
| TcCLB.509769.30 | hypothetical protein, conserved | 1,251 | 2,582E-06 |
| TcCLB.509373.30 | mucin-associated surface protein (MASP, pseudogene), pu | 1,251 | 0,005835 |
| TcCLB.508743.10 | surface antigen TASV, putative | 1,25 | 4,963E-25 |
| TcCLB.509217.180 | trans-sialidase (pseudogene), putative | 1,249 | 0,1088 |
| TcCLB.418221.30 | hypothetical protein, conserved | 1,248 | 9,615E-51 |
| TcCLB.508077.9 | conserved protein (fragment) | 1,248 | 2,458E-39 |
| TcCLB.506941.40 | hypothetical protein, conserved | 1,248 | 2,813E-16 |
| TcCLB.510361.160 | retrotransposon hot spot protein (RHS, pseudogene), putat | 1,248 | 1,366E-06 |
| TcCLB.507801.200 | dispersed gene family protein 1 (DGF-1), putative | 1,248 | 0,04702 |
| TcCLB.507357.90 | mucin-associated surface protein (MASP), putative | 1,248 | 0,09597 |
| TcCLB.506775.130 | hypothetical protein, conserved | 1,247 | 1,611E-31 |

| | | | |
|---|---|---|---|
| TcCLB.509199.24 | ribose 5-phosphate isomerase type B | 1,247 | 6,184E-22 |
| TcCLB.504105.70 | hypothetical protein, conserved | 1,247 | 1,918E-21 |
| TcCLB.506015.30 | mucin-associated surface protein (MASP, pseudogene), pu | 1,247 | 3,29E-12 |
| TcCLB.509237.90 | hypothetical protein, conserved | 1,247 | 1,161E-09 |
| TcCLB.507133.10 | trans-sialidase, putative | 1,247 | 0,00001371 |
| TcCLB.508115.10 | hypothetical protein | 1,247 | 0,000348 |
| TcCLB.494675.10 | fatty acyl CoA synthetase 2, putative | 1,246 | 5,264E-22 |
| TcCLB.503521.30 | Dynein light chain roadblock-type protein, putative | 1,246 | 4,589E-17 |
| TcCLB.508325.270 | retrotransposon hot spot protein (RHS, pseudogene), putat | 1,245 | 6,195E-15 |
| TcCLB.503891.80 | hypothetical protein, conserved | 1,245 | 1,078E-14 |
| TcCLB.509279.10 | hypothetical protein, conserved | 1,245 | 1,226E-08 |
| TcCLB.508873.30 | mucin-associated surface protein (MASP), putative | 1,245 | 2,432E-08 |
| TcCLB.510851.21 | retrotransposon hot spot protein (RHS, pseudogene), putat | 1,244 | 1,098E-27 |
| TcCLB.437121.9 | hypothetical protein, conserved (fragment) | 1,244 | 4,529E-20 |
| TcCLB.507485.100 | hypothetical protein, conserved | 1,244 | 3,015E-18 |
| TcCLB.511233.80 | calmodulin, putative | 1,244 | 1,486E-16 |
| TcCLB.511645.21 | mucin TcMUCII, putative | 1,244 | 0,0001993 |
| TcCLB.506799.120 | hypothetical protein | 1,244 | 0,000332 |
| TcCLB.503507.20 | mucin TcMUCII, putative (fragment) | 1,244 | 0,1028 |
| TcCLB.503849.60 | NADH-dependent fumarate reductase, putative | 1,243 | 4,347E-29 |
| TcCLB.508179.60 | protein kinase, putative | 1,243 | 1,064E-25 |
| TcCLB.506615.50 | mucin TcMUCII, putative | 1,243 | 3,298E-08 |
| TcCLB.510025.80 | hypothetical protein, conserved (pseudogene) | 1,243 | 0,00001025 |
| TcCLB.506599.250 | surface protease GP63 (pseudogene), putative | 1,243 | 0,09386 |
| TcCLB.510859.17 | nucleolar RNA-binding protein, putative | 1,242 | 6,03E-65 |
| TcCLB.508443.60 | hypothetical protein, conserved | 1,242 | 7,682E-19 |
| TcCLB.510861.30 | hypothetical protein, conserved | 1,241 | 4,707E-21 |
| TcCLB.505807.10 | cyclophilin, putative | 1,241 | 4,075E-11 |
| TcCLB.511797.260 | mucin-associated surface protein (MASP, pseudogene), pu | 1,241 | 0,05821 |
| TcCLB.510959.15 | elongation factor 1-gamma (EF-1-gamma, pseudogene), p | 1,241 | 0,08962 |
| TcCLB.505757.20 | transcriptional regulator, putative | 1,239 | 1,78E-13 |
| TcCLB.504797.100 | 35 kDa cyclophilin, putative (CYP35) | 1,238 | 1,521E-14 |
| TcCLB.509013.10 | calpain-like cysteine peptidase, putative | 1,238 | 3,025E-09 |
| TcCLB.511825.180 | ATP-dependent DEAD/H RNA helicase, putative | 1,237 | 4,805E-38 |
| TcCLB.509979.150 | nitrilase, putative | 1,237 | 4,523E-25 |
| TcCLB.510519.14 | hypothetical protein, conserved | 1,237 | 1,659E-07 |
| TcCLB.452283.10 | amino acid permease (pseudogene), putative | 1,236 | 8,552E-08 |
| TcCLB.511731.74 | hypothetical protein, conserved | 1,235 | 7,393E-13 |
| TcCLB.510037.10 | mucin-associated surface protein (MASP), putative | 1,235 | 7,907E-09 |
| TcCLB.508461.580 | hypothetical protein | 1,234 | 6,831E-37 |
| TcCLB.509721.40 | MIX protein, putative | 1,234 | 7,221E-15 |
| TcCLB.506567.70 | NAD(P)-dependent oxidoreductase, putative | 1,233 | 4,207E-26 |
| TcCLB.511393.49 | hypothetical protein, conserved | 1,233 | 1,095E-18 |
| TcCLB.507645.20 | hypothetical protein, conserved | 1,233 | 4,109E-16 |
| TcCLB.509767.230 | hypothetical protein, conserved | 1,233 | 1,167E-14 |

| | | | |
|---|---|---|---|
| TcCLB.507023.80 | hypothetical protein, conserved | 1,233 | 3,346E-08 |
| TcCLB.508167.20 | mucin TcMUCII, putative | 1,233 | 0,007308 |
| TcCLB.507957.43 | hypothetical protein, conserved (pseudogene) | 1,233 | 0,2 |
| TcCLB.506825.100 | hypothetical protein, conserved | 1,232 | 3,416E-23 |
| TcCLB.506261.10 | fatty acyl CoA synthetase 2, putative | 1,231 | 2,239E-24 |
| TcCLB.507787.100 | gamma-glutamylcysteine synthetase, putative | 1,23 | 1,076E-22 |
| TcCLB.509597.29 | hypothetical protein, conserved | 1,229 | 9,216E-12 |
| TcCLB.508501.10 | hypothetical protein | 1,228 | 0,002212 |
| TcCLB.507709.60 | hypothetical protein, conserved | 1,227 | 9,579E-35 |
| TcCLB.511589.74 | hypothetical protein | 1,227 | 1,13E-17 |
| TcCLB.506843.20 | retrotransposon hot spot protein (RHS, pseudogene), putat | 1,227 | 3,118E-11 |
| TcCLB.508143.50 | trans-sialidase (pseudogene), putative | 1,227 | 1,773E-10 |
| TcCLB.508343.10 | trans-sialidase, putative | 1,227 | 0,3665 |
| TcCLB.504071.30 | hypothetical protein, conserved | 1,226 | 9,661E-50 |
| TcCLB.509431.10 | hypothetical protein, conserved (pseudogene) | 1,226 | 6,708E-15 |
| TcCLB.510889.261 | cruzipain precursor, putative | 1,226 | 1,815E-08 |
| TcCLB.508541.160 | hypothetical protein, conserved | 1,226 | 3,442E-07 |
| TcCLB.510137.10 | trans-sialidase (pseudogene), putative | 1,226 | 0,00003394 |
| TcCLB.506327.90 | serine/threonine protein kinase, putative | 1,225 | 2,968E-35 |
| TcCLB.507011.240 | hypothetical protein, conserved | 1,224 | 1,433E-09 |
| TcCLB.510111.11 | trans-sialidase (pseudogene), putative | 1,224 | 0,00001071 |
| TcCLB.506055.30 | retrotransposon hot spot protein (RHS, pseudogene), putat | 1,224 | 0,00007329 |
| TcCLB.506779.90 | hypothetical protein, conserved | 1,223 | 2,622E-38 |
| TcCLB.506943.40 | hypothetical protein, conserved | 1,223 | 6,589E-13 |
| TcCLB.508297.41 | hypothetical protein, conserved | 1,223 | 7,648E-08 |
| TcCLB.509115.50 | mucin-like glycoprotein, putative | 1,223 | 0,00006225 |
| TcCLB.511671.20 | trans-sialidase (pseudogene), putative | 1,223 | 0,0003099 |
| TcCLB.511395.80 | hypothetical protein | 1,222 | 1,314E-20 |
| TcCLB.508831.10 | hypothetical protein, conserved | 1,222 | 1,145E-17 |
| TcCLB.507747.180 | trans-sialidase, Group VI, putative | 1,222 | 2,139E-12 |
| TcCLB.510165.90 | trans-sialidase (pseudogene), putative | 1,222 | 0,00003542 |
| TcCLB.508885.10 | surface protease GP63 (pseudogene), putative | 1,222 | 0,06277 |
| TcCLB.506775.30 | tubulin-tyrosine ligase-like protein, putative | 1,221 | 7,193E-37 |
| TcCLB.508547.150 | hypothetical protein, conserved | 1,221 | 9,555E-37 |
| TcCLB.510661.190 | protein kinase, putative | 1,221 | 0,0007532 |
| TcCLB.508081.50 | mucin-associated surface protein (MASP), putative | 1,221 | 0,001475 |
| TcCLB.511181.40 | hypothetical protein | 1,22 | 2,591E-10 |
| TcCLB.503993.20 | hypothetical protein | 1,22 | 0,04997 |
| TcCLB.510039.70 | hypothetical protein, conserved | 1,218 | 4,618E-16 |
| TcCLB.506349.47 | retrotransposon hot spot protein (RHS, pseudogene), putat | 1,218 | 0,0004415 |
| TcCLB.503613.60 | acidocalcisomal pyrophosphatase, putative | 1,217 | 3,026E-29 |
| TcCLB.506831.63 | hypothetical protein, conserved | 1,217 | 3,109E-13 |
| TcCLB.506501.70 | mucin TcMUCII, putative | 1,217 | 0,03847 |
| TcCLB.511805.20 | Trichohyalin, putative | 1,216 | 1,713E-08 |
| TcCLB.509331.150 | permease-like protein, putative | 1,216 | 5,252E-08 |

| | | | |
|---|---|---|---|
| TcCLB.509925.20 | mucin TcMUCII, putative | 1,216 | 2,252E-07 |
| TcCLB.506329.50 | mucin TcMUCII, putative | 1,216 | 0,03921 |
| TcCLB.510101.90 | hypothetical protein, conserved | 1,215 | 3,923E-44 |
| TcCLB.503925.40 | hypothetical protein | 1,215 | 6,022E-12 |
| TcCLB.503697.40 | hypothetical protein | 1,215 | 2,819E-09 |
| TcCLB.506153.3 | hypothetical protein, conserved | 1,214 | 4,469E-17 |
| TcCLB.503739.20 | trans-sialidase (pseudogene), putative | 1,214 | 1,464E-08 |
| TcCLB.507779.20 | trans-sialidase (pseudogene), putative | 1,214 | 1,917E-06 |
| TcCLB.511843.60 | retrotransposon hot spot protein (RHS, pseudogene), putat | 1,214 | 0,00003003 |
| TcCLB.508221.860 | mucin-associated surface protein (MASP, pseudogene), pu | 1,214 | 0,01745 |
| TcCLB.511773.90 | retrotransposon hot spot protein (RHS, pseudogene), putat | 1,213 | 7,222E-20 |
| TcCLB.441401.10 | imidazolonepropienase, putative | 1,213 | 1,247E-17 |
| TcCLB.508695.10 | hypothetical protein, conserved | 1,213 | 0,00001238 |
| TcCLB.508761.150 | mucin-associated surface protein (MASP, pseudogene), pu | 1,213 | 0,03543 |
| TcCLB.510101.430 | 40S ribosomal protein S21, putative | 1,212 | 4,732E-25 |
| TcCLB.506375.90 | hypothetical protein, conserved | 1,212 | 8,844E-16 |
| TcCLB.509695.20 | hypothetical protein, conserved | 1,212 | 3,085E-13 |
| TcCLB.509885.80 | hypothetical protein, conserved | 1,212 | 0,00002323 |
| TcCLB.504039.120 | mucin TcMUCII, putative | 1,212 | 0,0663 |
| TcCLB.418185.50 | retrotransposon hot spot protein (RHS, pseudogene), putat | 1,21 | 2,792E-17 |
| TcCLB.506305.29 | hypothetical protein, conserved | 1,209 | 5,625E-25 |
| TcCLB.508177.10 | hypothetical protein, conserved | 1,209 | 4,104E-18 |
| TcCLB.511373.20 | mucin TcMUCII, putative | 1,209 | 0,0005525 |
| TcCLB.509857.40 | hypothetical protein, conserved | 1,208 | 2,133E-36 |
| TcCLB.511595.50 | mucin-associated surface protein (MASP, pseudogene), pu | 1,208 | 0,0001022 |
| TcCLB.510715.64 | mucin-associated surface protein (MASP), putative | 1,208 | 0,1392 |
| TcCLB.504131.180 | ATPase subunit 9, putative | 1,207 | 2,978E-30 |
| TcCLB.511645.10 | mucin-associated surface protein (MASP), putative | 1,207 | 0,00009239 |
| TcCLB.508165.400 | mucin TcMUCII, putative | 1,207 | 0,006041 |
| TcCLB.510947.30 | hypothetical protein, conserved | 1,206 | 1,362E-18 |
| TcCLB.508507.40 | hypothetical protein, conserved | 1,206 | 4,157E-07 |
| TcCLB.508139.110 | hypothetical protein | 1,206 | 0,01237 |
| TcCLB.510279.40 | 90 kDa surface protein, putative | 1,206 | 0,03755 |
| TcCLB.509875.170 | pyruvate dehydrogenase (lipoamide) kinase, putative | 1,205 | 1,706E-23 |
| TcCLB.507669.10 | fumarate hydratase, putative | 1,204 | 1,44E-23 |
| TcCLB.506791.50 | Present in the outer mitochondrial membrane proteome 3 | 1,204 | 1,687E-19 |
| TcCLB.442383.19 | hypothetical protein | 1,204 | 6,293E-16 |
| TcCLB.503645.40 | mucin-associated surface protein (MASP), putative | 1,204 | 0,05722 |
| TcCLB.510267.20 | 60S acidic ribosomal protein P2, putative | 1,203 | 3,766E-20 |
| TcCLB.511255.30 | mucin-associated surface protein (MASP), putative | 1,203 | 0,001235 |
| TcCLB.506281.117 | mucin TcMUCII (pseudogene), putative | 1,203 | 0,01347 |
| TcCLB.508161.26 | mucin TcMUCII (pseudogene), putative | 1,203 | 0,04364 |
| TcCLB.508503.90 | hypothetical protein, conserved | 1,202 | 1,1E-16 |
| TcCLB.506941.220 | Present in the outer mitochondrial membrane proteome 3 | 1,202 | 2,858E-12 |
| TcCLB.508241.50 | surface protease GP63, putative | 1,201 | 1,693E-22 |

| | | | |
|---|---|---|---|
| TcCLB.511465.10 | proteasome activator protein pa26, putative | 1,201 | 2,755E-19 |
| TcCLB.503741.40 | mucin TcMUCII, putative | 1,201 | 0,06557 |
| TcCLB.506401.310 | hypothetical protein, conserved | 1,2 | 8,359E-14 |
| TcCLB.507519.120 | hypothetical protein, conserved | 1,2 | 3,589E-13 |
| TcCLB.509725.100 | hypothetical protein, conserved (pseudogene) | 1,2 | 3,234E-12 |
| TcCLB.406377.10 | hypothetical protein, conserved (pseudogene) | 1,2 | 0,00002983 |
| TcCLB.510553.30 | mucin-associated surface protein (MASP), putative | 1,199 | 0,0005997 |
| TcCLB.509331.221 | hypothetical protein, conserved | 1,199 | 0,008663 |
| TcCLB.508097.60 | mucin-associated surface protein (MASP), putative | 1,199 | 0,08759 |
| TcCLB.508831.220 | trans-sialidase (pseudogene), putative | 1,199 | 0,1503 |
| TcCLB.508587.10 | retrotransposon hot spot protein (RHS, pseudogene), putat | 1,197 | 1,411E-14 |
| TcCLB.508257.10 | serine-palmitoyl-CoA transferase, putative | 1,197 | 3,603E-09 |
| TcCLB.508273.120 | mucin TcMUCI, putative | 1,197 | 0,0008508 |
| TcCLB.508221.87 | retrotransposon hot spot protein (RHS, pseudogene), putat | 1,196 | 0,0005159 |
| TcCLB.504769.100 | trans-sialidase, Group II, putative | 1,195 | 3,008E-08 |
| TcCLB.511183.90 | mucin TcMUCII, putative | 1,195 | 0,05169 |
| TcCLB.506401.304 | mitochondrial carrier protein, putative | 1,194 | 5,725E-22 |
| TcCLB.509195.40 | mucin TcMUCII, putative | 1,194 | 0,003848 |
| TcCLB.507547.18 | mucin TcMUCI, putative | 1,194 | 0,07797 |
| TcCLB.506893.10 | mucin TcMUCII, putative | 1,194 | 0,1427 |
| TcCLB.421231.10 | hypothetical protein, conserved | 1,193 | 1,552E-17 |
| TcCLB.510359.650 | mucin-associated surface protein (MASP), putative | 1,193 | 0,1061 |
| TcCLB.510533.170 | hypothetical protein, conserved | 1,192 | 8,93E-10 |
| TcCLB.510561.20 | mucin TcMUCII, putative | 1,192 | 0,01308 |
| TcCLB.506773.80 | hypothetical protein, conserved | 1,191 | 1,059E-26 |
| TcCLB.506347.5 | trans-sialidase, putative (fragment) | 1,191 | 0,004524 |
| TcCLB.506283.20 | trans-sialidase, putative | 1,191 | 0,1189 |
| TcCLB.509765.50 | trans-sialidase, Group VIII, putative | 1,189 | 0,1554 |
| TcCLB.511929.30 | retrotransposon hot spot (RHS) protein, putative | 1,188 | 1,635E-13 |
| TcCLB.508975.10 | acyl-CoA dehydrogenase, mitochondrial precursor, putative | 1,187 | 6,897E-33 |
| TcCLB.511589.110 | hypothetical protein, conserved | 1,187 | 1,301E-28 |
| TcCLB.509855.20 | Probable methyltransferase BTM2 homolog | 1,187 | 6,538E-21 |
| TcCLB.507317.30 | hypothetical protein, conserved | 1,187 | 4,188E-11 |
| TcCLB.506825.70 | serine-threonine dehydratase, putative | 1,187 | 1,373E-08 |
| TcCLB.510003.41 | clathrin coat assembly protein ap19, putative | 1,186 | 9,71E-94 |
| TcCLB.510145.10 | hypothetical protein, conserved | 1,186 | 7,425E-08 |
| TcCLB.404431.10 | trans-sialidase, putative | 1,186 | 0,0001694 |
| TcCLB.506145.80 | surface protease GP63, putative | 1,185 | 7,387E-17 |
| TcCLB.504427.70 | 3-oxo-5-alpha-steroid 4-dehydrogenase, putative | 1,185 | 1,241E-13 |
| TcCLB.508119.140 | retrotransposon hot spot (RHS) protein, putative | 1,185 | 6,456E-08 |
| TcCLB.503965.20 | hypothetical protein, conserved | 1,184 | 1,262E-22 |
| TcCLB.509161.10 | ubiquitin-conjugating enzyme E2, putative | 1,184 | 1,253E-21 |
| TcCLB.511807.230 | hypothetical protein | 1,183 | 2,855E-34 |
| TcCLB.508267.40 | hypothetical protein, conserved | 1,183 | 1,172E-23 |
| TcCLB.511875.70 | mucin TcMUCII (pseudogene), putative | 1,183 | 2,214E-07 |

| | | | |
|---|---|---|---|
| TcCLB.509599.130 | serine/threonine protein kinase, putative | 1,182 | 2,765E-25 |
| TcCLB.510847.10 | trans-sialidase, Group VIII, putative | 1,182 | 2,665E-15 |
| TcCLB.508781.120 | hypothetical protein, conserved | 1,182 | 2,341E-12 |
| TcCLB.508751.20 | trans-sialidase (pseudogene), putative | 1,182 | 0,007131 |
| TcCLB.507221.40 | hypothetical protein | 1,181 | 3,296E-13 |
| TcCLB.506369.60 | hypothetical protein, conserved | 1,181 | 2,114E-11 |
| TcCLB.506995.60 | surface protease GP63, putative | 1,181 | 0,01077 |
| TcCLB.506741.70 | surface protease GP63 (pseudogene), putative | 1,181 | 0,134 |
| TcCLB.511827.30 | hypothetical protein | 1,18 | 3,785E-18 |
| TcCLB.508045.4 | hypothetical protein, conserved | 1,18 | 1,806E-17 |
| TcCLB.506239.40 | hypothetical protein, conserved | 1,18 | 5,14E-13 |
| TcCLB.509769.80 | hypothetical protein, conserved | 1,179 | 1,283E-54 |
| TcCLB.503861.10 | retrotransposon hot spot (RHS) protein, putative | 1,179 | 2,957E-40 |
| TcCLB.508965.80 | hypothetical protein, conserved | 1,179 | 9,948E-15 |
| TcCLB.508119.110 | kynureninase, putative | 1,179 | 4,084E-13 |
| TcCLB.511219.40 | trans-sialidase, Group II, putative | 1,179 | 4,729E-07 |
| TcCLB.509219.20 | retrotransposon hot spot (RHS) protein, putative | 1,178 | 2,194E-36 |
| TcCLB.506505.30 | hypothetical protein, conserved | 1,177 | 3,621E-23 |
| TcCLB.504239.340 | mucin TcMUCII, putative | 1,177 | 0,1065 |
| TcCLB.508713.10 | hypothetical protein, conserved | 1,176 | 2,955E-34 |
| TcCLB.511827.50 | hypothetical protein | 1,176 | 7,662E-10 |
| TcCLB.506435.50 | hypothetical protein, conserved | 1,176 | 7,782E-09 |
| TcCLB.506133.61 | mucin-associated surface protein (MASP), putative | 1,175 | 0,09389 |
| TcCLB.503479.30 | adenylate kinase 1 | 1,174 | 6,797E-20 |
| TcCLB.508241.70 | hypothetical protein, conserved | 1,174 | 1,913E-15 |
| TcCLB.511807.36 | hypothetical protein | 1,174 | 7,493E-15 |
| TcCLB.509651.10 | hypothetical protein, conserved | 1,174 | 4,836E-14 |
| TcCLB.510095.70 | retrotransposon hot spot protein (RHS, pseudogene), putat | 1,174 | 6,729E-13 |
| TcCLB.503949.40 | UDP-Gal or UDP-GlcNAc-dependent glycosyltransferase, pu | 1,174 | 2,504E-07 |
| TcCLB.505975.30 | beta galactofuranosyl glycosyltransferase (pseudogene), pu | 1,174 | 6,883E-06 |
| TcCLB.509623.19 | clathrin coat assembly protein ap19, putative (ClaATc) | 1,173 | 2,255E-92 |
| TcCLB.508173.60 | cis-prenyltransferase, putative | 1,173 | 1,702E-18 |
| TcCLB.504769.10 | hypothetical protein | 1,173 | 3,558E-08 |
| TcCLB.508851.90 | hypothetical protein, conserved | 1,172 | 1,585E-12 |
| TcCLB.509863.10 | UDP-Gal or UDP-GlcNAc-dependent glycosyltransferase, pu | 1,172 | 1,143E-11 |
| TcCLB.506843.137 | retrotransposon hot spot protein (RHS, pseudogene), putat | 1,172 | 0,01839 |
| TcCLB.508247.70 | mucin TcMUCII, putative | 1,172 | 0,1477 |
| TcCLB.505647.10 | hypothetical protein, conserved | 1,171 | 2,078E-41 |
| TcCLB.506595.100 | 60S acidic ribosomal protein P2, putative | 1,171 | 3,483E-27 |
| TcCLB.510155.100 | hypothetical protein | 1,171 | 7,755E-27 |
| TcCLB.509905.20 | retrotransposon hot spot protein (RHS, pseudogene), putat | 1,171 | 1,046E-06 |
| TcCLB.510329.10 | hypothetical protein, conserved | 1,17 | 7,773E-34 |
| TcCLB.511741.60 | hypothetical protein, conserved | 1,17 | 1,843E-18 |
| TcCLB.510369.10 | mucin-associated surface protein (MASP), putative | 1,17 | 0,01867 |
| TcCLB.511377.20 | trans-sialidase (pseudogene), putative | 1,17 | 0,04773 |

| | | | |
|---|---|---|---|
| TcCLB.508461.570 | Gim5A protein, putative | 1,169 | 2,384E-48 |
| TcCLB.511817.200 | Present in the outer mitochondrial membrane proteome 3: | 1,169 | 2,687E-17 |
| TcCLB.504907.9 | dispersed gene family protein 1 (DGF-1), putative (fragme | 1,169 | 3,406E-07 |
| TcCLB.507153.10 | hypothetical protein, conserved | 1,169 | 0,01114 |
| TcCLB.510345.50 | hypothetical protein, conserved | 1,168 | 2,937E-20 |
| TcCLB.511825.100 | hypothetical protein, conserved | 1,168 | 6,118E-16 |
| TcCLB.510661.174 | riboflavin kinase, putative | 1,168 | 1,32E-09 |
| TcCLB.511819.50 | hypothetical protein, conserved | 1,167 | 2,121E-35 |
| TcCLB.504109.200 | retrotransposon hot spot (RHS) protein, putative | 1,167 | 1,975E-33 |
| TcCLB.504065.50 | trans-sialidase (pseudogene), putative | 1,167 | 4,377E-22 |
| TcCLB.507255.30 | retrotransposon hot spot protein (RHS, pseudogene), putat | 1,167 | 1,815E-11 |
| TcCLB.506973.120 | trans-sialidase, Group V, putative | 1,167 | 0,0001333 |
| TcCLB.508165.210 | mucin-associated surface protein (MASP), putative | 1,167 | 0,007444 |
| TcCLB.510025.165 | hypothetical protein, conserved (pseudogene) | 1,167 | 0,09575 |
| TcCLB.510101.40 | 60S ribosomal protein L28, putative | 1,166 | 6,212E-49 |
| TcCLB.507569.10 | hypothetical protein, conserved | 1,166 | 7,378E-24 |
| TcCLB.505163.70 | Selenoprotein T, putative (SelT) | 1,166 | 2,494E-22 |
| TcCLB.481211.10 | mucin TcMUCII, putative | 1,166 | 0,000366 |
| TcCLB.511613.100 | elongation factor 1-gamma (EF-1-gamma), putative | 1,166 | 0,06912 |
| TcCLB.506775.150 | hypothetical protein, conserved | 1,165 | 5,736E-17 |
| TcCLB.507853.10 | lathosterol oxidase, putative | 1,165 | 3,59E-16 |
| TcCLB.511361.50 | hypothetical protein, conserved | 1,165 | 7,97E-09 |
| TcCLB.508267.60 | hypothetical protein, conserved (fragment) | 1,165 | 9,533E-09 |
| TcCLB.510763.30 | hypothetical protein, conserved | 1,165 | 0,0000175 |
| TcCLB.508159.13 | trans-sialidase (pseudogene), putative | 1,165 | 0,00006006 |
| TcCLB.507063.60 | hypothetical protein, conserved | 1,165 | 0,001729 |
| TcCLB.504081.330 | mucin TcMUCII, putative | 1,165 | 0,1195 |
| TcCLB.508331.90 | retrotransposon hot spot protein (RHS, pseudogene), putat | 1,164 | 1,019E-12 |
| TcCLB.503723.20 | hypothetical protein, conserved | 1,164 | 2,32E-11 |
| TcCLB.507659.20 | amino acid transporter, putative | 1,162 | 4,778E-66 |
| TcCLB.510309.20 | MFS transporter, putative | 1,162 | 4,066E-26 |
| TcCLB.506921.10 | surface protease GP63, putative | 1,162 | 8,798E-20 |
| TcCLB.508863.20 | hypothetical protein, conserved | 1,162 | 5E-10 |
| TcCLB.509875.165 | trans-sialidase (pseudogene), putative | 1,162 | 0,00002164 |
| TcCLB.510101.360 | hypothetical protein, conserved | 1,161 | 4,453E-33 |
| TcCLB.503873.10 | cytochrome-b5 reductase, putative | 1,161 | 1,675E-29 |
| TcCLB.508241.60 | phosphoadenosine phosphosulfate reductase-like protein, | 1,161 | 9,488E-24 |
| TcCLB.506885.90 | hypothetical protein, conserved | 1,161 | 2,47E-21 |
| TcCLB.511643.30 | aspartate carbamoyltransferase, putative | 1,161 | 1,553E-19 |
| TcCLB.506175.25 | hypothetical protein | 1,161 | 2,57E-18 |
| TcCLB.507245.20 | trans-sialidase (pseudogene), putative | 1,161 | 3,636E-09 |
| TcCLB.503417.30 | mucin TcMUCII, putative | 1,161 | 0,07755 |
| TcCLB.511815.50 | hypothetical protein, conserved | 1,16 | 2,07E-33 |
| TcCLB.507477.10 | dispersed gene family protein 1 (DGF-1), putative | 1,16 | 3,295E-13 |
| TcCLB.508033.69 | mucin-associated surface protein (MASP, pseudogene), pu | 1,16 | 0,001991 |

| | | | |
|---|---|---|---|
| TcCLB.508501.80 | mucin-associated surface protein (MASP), putative | 1,159 | 0,0000393 |
| TcCLB.506405.50 | serine-palmitoyl-CoA transferase, putative | 1,158 | 9,76E-38 |
| TcCLB.511395.30 | protein phosphatase 2A, regulatory subunit B, putative | 1,158 | 2,503E-35 |
| TcCLB.509973.70 | mucin-associated surface protein (MASP), putative | 1,158 | 0,00000341 |
| TcCLB.506877.60 | mucin-associated surface protein (MASP), putative | 1,158 | 0,009094 |
| TcCLB.510117.10 | retrotransposon hot spot protein (RHS, pseudogene), putat | 1,157 | 1,23E-12 |
| TcCLB.507011.10 | hypothetical protein, conserved | 1,156 | 1,06E-19 |
| TcCLB.509537.60 | meiosis recombination protein SPO11, putative | 1,156 | 1,633E-18 |
| TcCLB.506179.10 | ferric reductase transmembrane protein, putative (FR1) | 1,155 | 7,61E-45 |
| TcCLB.506775.14 | hypothetical protein, conserved | 1,155 | 5,375E-20 |
| TcCLB.509541.4 | paraflagellar rod component par4, putative | 1,155 | 1,168E-07 |
| TcCLB.509153.50 | hypothetical protein, conserved | 1,154 | 1,694E-13 |
| TcCLB.504797.90 | hypothetical protein, conserved | 1,154 | 3,387E-10 |
| TcCLB.508983.10 | mucin-associated surface protein (MASP, pseudogene), pu | 1,154 | 0,007514 |
| TcCLB.508495.30 | hypothetical protein | 1,154 | 0,01598 |
| TcCLB.509237.120 | calpain-like cysteine peptidase, putative | 1,153 | 1,103E-35 |
| TcCLB.503771.74 | mucin TcMUCII (pseudogene), putative | 1,153 | 0,07706 |
| TcCLB.510101.420 | 40S ribosomal protein S21, putative | 1,152 | 1,279E-59 |
| TcCLB.510901.230 | hypothetical protein, conserved | 1,152 | 1,368E-17 |
| TcCLB.511815.80 | 3'a2rel-related protein, putative | 1,152 | 1,6E-09 |
| TcCLB.511223.20 | trans-sialidase, putative | 1,152 | 0,143 |
| TcCLB.506925.110 | hypothetical protein, conserved | 1,151 | 7,651E-40 |
| TcCLB.509859.50 | hypothetical protein, conserved | 1,151 | 3,605E-11 |
| TcCLB.504343.10 | trans-sialidase, Group II, putative | 1,15 | 5,284E-08 |
| TcCLB.511077.90 | mucin TcMUCII, putative | 1,15 | 0,01925 |
| TcCLB.504421.10 | serine/threonine protein kinase, putative | 1,149 | 1,089E-51 |
| TcCLB.509151.100 | hypothetical protein, conserved | 1,149 | 1,495E-15 |
| TcCLB.506713.24 | hypothetical protein, conserved | 1,149 | 9,237E-06 |
| TcCLB.508119.30 | mucin TcMUCII, putative | 1,149 | 0,00005137 |
| TcCLB.508433.220 | mucin-associated surface protein (MASP), putative | 1,149 | 0,06486 |
| TcCLB.509185.30 | trans-sialidase, Group II, putative | 1,149 | 0,4735 |
| TcCLB.510039.90 | 60S ribosomal protein L35, putative | 1,148 | 1,947E-54 |
| TcCLB.508729.18 | hypothetical protein, conserved | 1,148 | 1,151E-13 |
| TcCLB.510173.130 | hypothetical protein, conserved | 1,148 | 8,466E-12 |
| TcCLB.504341.20 | trans-sialidase, putative | 1,148 | 0,4688 |
| TcCLB.507429.20 | mucin-associated surface protein (MASP), putative | 1,147 | 0,0000398 |
| TcCLB.508873.514 | mucin TcMUC (pseudogene), putative | 1,147 | 0,04765 |
| TcCLB.511153.120 | pantothenate kinase subunit, putative | 1,146 | 2,658E-29 |
| TcCLB.510307.230 | trans-sialidase, Group II, putative | 1,146 | 0,00002498 |
| TcCLB.510039.36 | hypothetical protein | 1,146 | 0,0001562 |
| TcCLB.476421.9 | protein kinase, putative | 1,145 | 5,398E-15 |
| TcCLB.511321.60 | hypothetical protein, conserved | 1,145 | 0,0004598 |
| TcCLB.508631.20 | retrotransposon hot spot protein (RHS, pseudogene), putat | 1,144 | 1,207E-30 |
| TcCLB.509865.9 | retrotransposon hot spot (RHS) protein, putative | 1,144 | 2,557E-22 |
| TcCLB.510039.50 | hypothetical protein, conserved | 1,144 | 1,227E-17 |

| | | | |
|---|---|---|---|
| TcCLB.506925.20 | hypothetical protein, conserved | 1,144 | 3,656E-17 |
| TcCLB.507769.20 | hypothetical protein, conserved | 1,143 | 2,494E-08 |
| TcCLB.503899.119 | trypanothione/tryparedoxin dependent peroxidase 2, putat | 1,142 | 2,626E-29 |
| TcCLB.508569.80 | phosphoacetylglucosamine mutase, putative | 1,142 | 2,743E-20 |
| TcCLB.506339.10 | trans-sialidase, Group V, putative | 1,142 | 1,104E-06 |
| TcCLB.509427.80 | hypothetical protein | 1,142 | 0,07225 |
| TcCLB.509891.40 | hypothetical protein, conserved | 1,141 | 6,241E-12 |
| TcCLB.510783.49 | cytochrome c oxidase subunit 10, putative | 1,141 | 0,00002761 |
| TcCLB.507047.40 | trans-sialidase, Group II, putative | 1,14 | 1,374E-11 |
| TcCLB.506825.180 | protein kinase, putative | 1,139 | 1,745E-41 |
| TcCLB.507063.170 | hypothetical protein | 1,139 | 0,00001642 |
| TcCLB.509081.100 | mucin TcMUCII, putative | 1,139 | 0,002474 |
| TcCLB.455721.9 | cytochrome c oxidase subunit 10, putative | 1,138 | 4,202E-14 |
| TcCLB.508901.10 | hypothetical protein, conserved | 1,138 | 6,227E-10 |
| TcCLB.509437.103 | dispersed gene family protein 1 (DGF-1, pseudogene), puta | 1,138 | 0,02116 |
| TcCLB.508221.714 | retrotransposon hot spot protein (RHS, pseudogene), putat | 1,138 | 0,04549 |
| TcCLB.506751.50 | trans-sialidase, Group II, putative | 1,137 | 0,0009444 |
| TcCLB.511595.39 | mucin TcMUCII, putative | 1,137 | 0,04913 |
| TcCLB.510173.90 | poly(ADP-ribose) polymerase, putative | 1,136 | 2,076E-21 |
| TcCLB.511183.450 | trans-sialidase (pseudogene), putative | 1,136 | 0,0000829 |
| TcCLB.506321.20 | mucin-associated surface protein (MASP), putative | 1,136 | 0,0006423 |
| TcCLB.510311.100 | hypothetical protein, conserved | 1,135 | 7,166E-10 |
| TcCLB.511819.20 | hypothetical protein, conserved | 1,134 | 9,607E-25 |
| TcCLB.509601.40 | hypothetical protein, conserved | 1,133 | 1,422E-23 |
| TcCLB.506715.30 | hypothetical protein | 1,133 | 2,599E-14 |
| TcCLB.511649.100 | amino acid permease, putative | 1,133 | 7,113E-11 |
| TcCLB.511263.40 | hypothetical protein, conserved | 1,133 | 0,07461 |
| TcCLB.510403.20 | retrotransposon hot spot protein (RHS, pseudogene), putat | 1,132 | 1,523E-18 |
| TcCLB.511603.470 | 90 kDa surface protein, putative | 1,132 | 0,0007626 |
| TcCLB.510101.240 | 60S ribosomal protein L24, putative | 1,131 | 1,872E-17 |
| TcCLB.511587.60 | retrotransposon hot spot protein (RHS, pseudogene), putat | 1,131 | 7,005E-14 |
| TcCLB.509461.40 | hypothetical protein, conserved | 1,131 | 3,637E-10 |
| TcCLB.510037.5 | elongation factor 1-gamma (EF-1-gamma), putative | 1,131 | 3,309E-07 |
| TcCLB.508495.100 | trans-sialidase, Group V, putative | 1,131 | 0,0007988 |
| TcCLB.508761.20 | mucin TcMUCII, putative | 1,131 | 0,004034 |
| TcCLB.506009.10 | hypothetical protein, conserved | 1,131 | 0,07059 |
| TcCLB.506775.110 | hypothetical protein, conserved | 1,13 | 3,756E-24 |
| TcCLB.506219.10 | hypothetical protein, conserved | 1,13 | 1,204E-18 |
| TcCLB.505097.35 | retrotransposon hot spot protein (RHS, pseudogene), putat | 1,13 | 1,063E-10 |
| TcCLB.509153.120 | acyl-CoA dehydrogenase, putative | 1,129 | 2,736E-26 |
| TcCLB.509915.60 | retrotransposon hot spot (RHS) protein, putative | 1,129 | 1,144E-10 |
| TcCLB.508159.50 | mucin-associated surface protein (MASP, pseudogene), pu | 1,129 | 0,002152 |
| TcCLB.506173.310 | mucin TcMUCII, putative | 1,129 | 0,006075 |
| TcCLB.510371.30 | hypothetical protein, conserved (pseudogene) | 1,129 | 0,1267 |
| TcCLB.509003.70 | beta tubulin (pseudogene), putative | 1,128 | 2,449E-36 |

| | | | |
|---|---|---|---|
| TcCLB.511815.180 | hypothetical protein, conserved | 1,128 | 1,64E-20 |
| TcCLB.507851.60 | hypothetical protein, conserved | 1,128 | 4,801E-16 |
| TcCLB.511923.110 | aspartate carbamoyltransferase, putative | 1,127 | 1,112E-14 |
| TcCLB.506661.40 | hypothetical protein, conserved | 1,127 | 1,188E-08 |
| TcCLB.503683.30 | RNA-binding protein, putative | 1,127 | 0,00003895 |
| TcCLB.508047.60 | trans-sialidase, Group V, putative | 1,127 | 0,0001365 |
| TcCLB.511149.70 | mucin TcMUCII, putative | 1,127 | 0,02046 |
| TcCLB.508951.20 | hexokinase, putative | 1,126 | 4,509E-35 |
| TcCLB.503595.10 | hypothetical protein, conserved | 1,126 | 8,677E-28 |
| TcCLB.510349.60 | hypothetical protein, conserved | 1,126 | 8,977E-06 |
| TcCLB.510693.190 | mucin-associated surface protein (MASP), putative | 1,126 | 0,01924 |
| TcCLB.508119.100 | hypothetical protein, conserved | 1,125 | 8,04E-27 |
| TcCLB.506003.39 | trans-sialidase, putative (fragment) | 1,125 | 0,3952 |
| TcCLB.506123.24 | dynein light chain, putative | 1,124 | 6,891E-08 |
| TcCLB.510005.30 | retrotransposon hot spot protein (RHS, pseudogene), putat | 1,124 | 1,422E-07 |
| TcCLB.508143.90 | trans-sialidase (pseudogene), putative | 1,124 | 7,838E-07 |
| TcCLB.510411.5 | retrotransposon hot spot protein (RHS, pseudogene), putat | 1,124 | 0,02521 |
| TcCLB.509237.70 | UDP-Gal or UDP-GlcNAc-dependent glycosyltransferase, pu | 1,123 | 0,00000736 |
| TcCLB.506825.150 | hypothetical protein, conserved | 1,122 | 1,373E-36 |
| TcCLB.508601.119 | ribose 5-phosphate isomerase, putative | 1,122 | 1,674E-14 |
| TcCLB.505939.80 | hypothetical protein, conserved | 1,122 | 9,772E-12 |
| TcCLB.507745.10 | mucin-associated surface protein (MASP, pseudogene), pu | 1,122 | 0,00393 |
| TcCLB.506387.9 | chaperone DNAJ protein, putative | 1,121 | 0,0000267 |
| TcCLB.506919.120 | retrotransposon hot spot protein (RHS, pseudogene), putat | 1,121 | 0,00004709 |
| TcCLB.506309.34 | mucin TcMUCII, putative | 1,121 | 0,4171 |
| TcCLB.506281.70 | mucin TcMUCI, putative | 1,12 | 0,02591 |
| TcCLB.507849.50 | cysteine proteinase, putative | 1,119 | 3,843E-34 |
| TcCLB.449247.20 | hypothetical protein, conserved | 1,119 | 6,647E-07 |
| TcCLB.506741.10 | trans-sialidase (pseudogene), putative | 1,119 | 0,0001763 |
| TcCLB.509207.100 | hypothetical protein, conserved | 1,118 | 1,42E-19 |
| TcCLB.510603.60 | protein tyrosine phosphatase-like protein, putative | 1,118 | 3,431E-08 |
| TcCLB.506599.50 | mucin-associated surface protein (MASP), putative | 1,118 | 0,007216 |
| TcCLB.508621.51 | peptidylprolyl isomerase-like protein (pseudogene), putativ | 1,118 | 0,1176 |
| TcCLB.508501.290 | Kinesin-13 1, putative (KIN13-1) | 1,117 | 1,469E-35 |
| TcCLB.506775.170 | NADH-cytochrome b5 reductase, putative | 1,117 | 2,149E-18 |
| TcCLB.509023.70 | hypothetical protein, conserved | 1,117 | 0,00002463 |
| TcCLB.506923.30 | hypothetical protein (pseudogene) | 1,117 | 0,005818 |
| TcCLB.510279.190 | mucin-associated surface protein (MASP), putative | 1,117 | 0,007471 |
| TcCLB.507509.20 | hypothetical protein, conserved | 1,115 | 9,556E-20 |
| TcCLB.506401.330 | hypothetical protein, conserved | 1,115 | 6,384E-13 |
| TcCLB.508375.80 | hypothetical protein, conserved | 1,115 | 3,416E-10 |
| TcCLB.507133.20 | trans-sialidase, putative (fragment) | 1,115 | 7,509E-06 |
| TcCLB.508501.90 | hypothetical protein | 1,115 | 0,01535 |
| TcCLB.511077.60 | mucin-associated surface protein (MASP), putative | 1,115 | 0,08742 |
| TcCLB.508675.29 | calpain-like cysteine peptidase, putative | 1,114 | 9,319E-25 |

| | | | |
|---|---|---|---|
| TcCLB.508729.9 | hypothetical protein, conserved | 1,114 | 2,327E-08 |
| TcCLB.510387.40 | mucin-associated surface protein (MASP), putative | 1,114 | 0,06471 |
| TcCLB.510101.390 | hypothetical protein, conserved | 1,113 | 2,176E-16 |
| TcCLB.510531.100 | hypothetical protein, conserved | 1,112 | 2,895E-31 |
| TcCLB.507683.40 | protein kinase, putative | 1,112 | 4,991E-20 |
| TcCLB.440379.10 | trans-sialidase, putative (fragment) | 1,112 | 0,002836 |
| TcCLB.510205.70 | surface protease GP63, putative | 1,112 | 0,02268 |
| TcCLB.510583.10 | mucin TcMUCII, putative | 1,111 | 0,04835 |
| TcCLB.504049.40 | retrotransposon hot spot protein (RHS, pseudogene), putat | 1,11 | 7,984E-15 |
| TcCLB.503767.10 | trans-sialidase, Group II, putative | 1,11 | 0,007351 |
| TcCLB.506959.70 | serine palmitoyltransferase, putative | 1,109 | 8,345E-30 |
| TcCLB.511827.120 | retrotransposon hot spot protein (RHS, pseudogene), putat | 1,109 | 1,457E-21 |
| TcCLB.506633.74 | hypothetical protein, conserved (pseudogene) | 1,109 | 6,604E-10 |
| TcCLB.510773.120 | hypothetical protein | 1,109 | 0,08502 |
| TcCLB.510101.170 | 40S ribosomal protein S5, putative | 1,108 | 4,53E-38 |
| TcCLB.506825.24 | hypothetical protein, conserved | 1,108 | 1,58E-17 |
| TcCLB.506401.210 | hypothetical protein, conserved | 1,108 | 4,096E-14 |
| TcCLB.509559.20 | retrotransposon hot spot protein (RHS, pseudogene), putat | 1,108 | 3,332E-13 |
| TcCLB.506649.20 | P-type H -ATPase, putative | 1,107 | 4,649E-30 |
| TcCLB.510039.64 | hypothetical protein, conserved | 1,107 | 7,819E-15 |
| TcCLB.509429.260 | hypothetical protein, conserved | 1,107 | 2,858E-14 |
| TcCLB.511217.10 | hypothetical protein | 1,107 | 6,818E-11 |
| TcCLB.506957.100 | hypothetical protein, conserved | 1,107 | 4,284E-09 |
| TcCLB.504011.80 | hypothetical protein, conserved | 1,107 | 8,412E-08 |
| TcCLB.510521.10 | NADH-cytochrome b5 reductase (pseudogene), putative | 1,106 | 3,612E-18 |
| TcCLB.507491.30 | hypothetical protein, conserved | 1,106 | 3,23E-16 |
| TcCLB.503559.90 | hypothetical protein, conserved | 1,106 | 9,354E-15 |
| TcCLB.506945.320 | hypothetical protein, conserved | 1,105 | 5,051E-28 |
| TcCLB.506295.160 | 2-oxoisovalerate dehydrogenase beta subunit, mitochondri | 1,105 | 2,148E-20 |
| TcCLB.507783.10 | ribosomal P protein AGP2beta-1, putative | 1,105 | 0,00002319 |
| TcCLB.507479.70 | trans-sialidase, Group IV, putative | 1,105 | 0,00003114 |
| TcCLB.460197.10 | cysteine peptidase (pseudogene), putative | 1,105 | 0,002299 |
| TcCLB.508523.30 | trans-sialidase, Group II, putative | 1,105 | 0,01574 |
| TcCLB.508125.30 | mucin TcMUCII, putative | 1,105 | 0,1717 |
| TcCLB.509617.20 | paraflagellar rod protein 3, putative (PAR3) | 1,104 | 6,497E-26 |
| TcCLB.508039.10 | retrotransposon hot spot protein (RHS, pseudogene), putat | 1,104 | 2,585E-19 |
| TcCLB.510421.110 | hypothetical protein | 1,104 | 3,839E-14 |
| TcCLB.510237.110 | mucin TcMUCII, putative | 1,104 | 0,02411 |
| TcCLB.508637.90 | phosphoglucomutase | 1,103 | 3,296E-19 |
| TcCLB.507167.110 | retrotransposon hot spot protein (RHS, pseudogene), putat | 1,103 | 2,464E-14 |
| TcCLB.503955.80 | hypothetical protein | 1,103 | 2,995E-12 |
| TcCLB.510463.200 | mucin-associated surface protein (MASP), putative | 1,103 | 0,03101 |
| TcCLB.509001.20 | hypothetical protein, conserved | 1,102 | 2,414E-20 |
| TcCLB.506623.40 | retrotransposon hot spot protein (RHS, pseudogene), putat | 1,102 | 5,463E-14 |
| TcCLB.503841.10 | proteasome activator protein pa26, putative | 1,102 | 7,159E-14 |

| | | | |
|---|---|---|---|
| TcCLB.503605.20 | retrotransposon hot spot (RHS) protein, putative | 1,102 | 0,00002532 |
| TcCLB.510557.70 | mucin TcMUCII, putative | 1,102 | 0,0008045 |
| TcCLB.510121.90 | hypothetical protein, conserved | 1,101 | 3,741E-21 |
| TcCLB.503991.39 | 2,4-dihydroxyhept-2-ene-1,7-dioic acid aldolase, putative | 1,1 | 2,329E-12 |
| TcCLB.509785.70 | retrotransposon hot spot (RHS) protein, putative | 1,1 | 4,118E-08 |
| TcCLB.510543.70 | trans-sialidase, Group II, putative | 1,1 | 0,0009103 |
| TcCLB.506715.59 | Monooxygenase, putative | 1,1 | 0,002511 |
| TcCLB.508873.240 | mucin TcMUCII, putative | 1,1 | 0,01974 |
| TcCLB.503421.50 | retrotransposon hot spot protein (RHS, pseudogene), putat | 1,099 | 2,58E-22 |
| TcCLB.510723.29 | 1-acyl-sn-glycerol-3-phosphate acyltransferase, putative (f | 1,099 | 2,149E-13 |
| TcCLB.511553.10 | mucin-associated surface protein (MASP), putative | 1,099 | 0,00868 |
| TcCLB.511173.110 | mucin TcMUCII, putative | 1,099 | 0,01939 |
| TcCLB.507615.100 | Present in the outer mitochondrial membrane proteome 2 | 1,098 | 2,572E-34 |
| TcCLB.508547.170 | hypothetical protein, conserved | 1,098 | 6,723E-33 |
| TcCLB.506825.130 | hypothetical protein, conserved | 1,098 | 2,345E-20 |
| TcCLB.506563.50 | calpain-like cysteine peptidase (pseudogene), putative | 1,098 | 9,952E-11 |
| TcCLB.503677.20 | DNA repair and recombination helicase protein PIF4, putat | 1,098 | 3,931E-09 |
| TcCLB.509753.150 | mucin TcMUCII, putative | 1,098 | 0,1934 |
| TcCLB.511889.11 | trans-sialidase (pseudogene), putative | 1,097 | 1,573E-16 |
| TcCLB.508041.20 | hypothetical protein, conserved | 1,097 | 3,058E-14 |
| TcCLB.506895.80 | trans-sialidase, putative | 1,097 | 0,00002227 |
| TcCLB.511827.139 | retrotransposon hot spot protein (RHS, pseudogene), putat | 1,097 | 0,001952 |
| TcCLB.511393.60 | protein kinase, putative | 1,096 | 6,406E-23 |
| TcCLB.508741.440 | surface antigen TASV, putative | 1,096 | 2,53E-21 |
| TcCLB.507221.20 | poly(A)-specific ribonuclease PARN, putative | 1,096 | 1,528E-16 |
| TcCLB.509259.160 | retrotransposon hot spot (RHS) protein, putative | 1,096 | 1,636E-16 |
| TcCLB.508569.100 | hypothetical protein, conserved | 1,096 | 1,169E-11 |
| TcCLB.511755.80 | formin, putative | 1,096 | 9,044E-06 |
| TcCLB.509149.50 | hypothetical protein, conserved | 1,095 | 5,751E-28 |
| TcCLB.509601.10 | neutral sphingomyelinase activation associated factor-like | 1,095 | 7,918E-13 |
| TcCLB.507603.190 | hypothetical protein, conserved | 1,094 | 1,237E-13 |
| TcCLB.510925.9 | dispersed gene family protein 1 (DGF-1), putative (fragme | 1,094 | 0,03056 |
| TcCLB.503525.4 | hypothetical protein, conserved | 1,093 | 4,673E-19 |
| TcCLB.511065.10 | STOP axonemal protein, putative | 1,093 | 9,044E-19 |
| TcCLB.511667.20 | retrotransposon hot spot (RHS) protein, putative | 1,093 | 3,22E-10 |
| TcCLB.506053.50 | trans-sialidase, Group IV, putative | 1,093 | 0,00202 |
| TcCLB.507411.20 | retrotransposon hot spot (RHS) protein, putative | 1,092 | 0,0004637 |
| TcCLB.507533.21 | clathrin assembly protein (pseudogene), putative | 1,091 | 5,773E-38 |
| TcCLB.508527.39 | retrotransposon hot spot (RHS) protein, putative, (fragmer | 1,091 | 5,921E-13 |
| TcCLB.504045.80 | hypothetical protein, conserved | 1,091 | 6,394E-10 |
| TcCLB.506183.40 | retrotransposon hot spot protein (RHS, pseudogene), putat | 1,091 | 3,436E-07 |
| TcCLB.506737.128 | mucin TcMUCI, putative | 1,091 | 0,0003405 |
| TcCLB.507067.20 | surface protease GP63, putative | 1,091 | 0,001952 |
| TcCLB.506801.110 | hypothetical protein, conserved | 1,09 | 5,812E-10 |
| TcCLB.504605.10 | retrotransposon hot spot (RHS) protein, putative (fragmen | 1,09 | 9,441E-09 |

| | | | |
|---|---|---|---|
| TcCLB.511223.10 | hypothetical protein | 1,09 | 0,02317 |
| TcCLB.504187.10 | retrotransposon hot spot protein (RHS, pseudogene), putat | 1,09 | 0,03178 |
| TcCLB.510699.80 | mucin-associated surface protein (MASP), putative | 1,09 | 0,09162 |
| TcCLB.509139.10 | glycerate kinase, putative | 1,089 | 1,64E-42 |
| TcCLB.506825.210 | hypothetical protein, conserved | 1,089 | 2,041E-31 |
| TcCLB.511825.20 | hypothetical protein, conserved | 1,089 | 2,282E-11 |
| TcCLB.509661.21 | ferric reductase (pseudogene), putative | 1,089 | 1,037E-07 |
| TcCLB.504769.5 | hypothetical protein | 1,089 | 1,269E-07 |
| TcCLB.507091.160 | trans-sialidase (pseudogene), putative | 1,089 | 4,382E-07 |
| TcCLB.511001.240 | proteasome activator protein pa26, putative | 1,089 | 3,458E-06 |
| TcCLB.506871.190 | hypothetical protein | 1,089 | 4,327E-06 |
| TcCLB.510101.180 | 40S ribosomal protein S5, putative | 1,089 | 0,0005418 |
| TcCLB.509187.20 | retrotransposon hot spot protein (RHS, pseudogene), putat | 1,089 | 0,03729 |
| TcCLB.511613.151 | syntaxin binding protein (pseudogene), putative | 1,089 | 0,08542 |
| TcCLB.508221.910 | mucin TcMUCII, putative | 1,088 | 0,09511 |
| TcCLB.511823.70 | farnesyl pyrophosphate synthase | 1,087 | 1,679E-18 |
| TcCLB.510689.14 | NADH-ubiquinone oxidoreductase complex I subunit, putati | 1,087 | 3,309E-13 |
| TcCLB.508201.30 | mucin-associated surface protein (MASP), putative | 1,087 | 7,582E-06 |
| TcCLB.510377.200 | mucin-associated surface protein (MASP), putative | 1,087 | 0,02026 |
| TcCLB.506925.530 | glycosomal transporter (GAT3), putative | 1,086 | 4,33E-41 |
| TcCLB.504797.14 | hypothetical protein, conserved | 1,085 | 4,83E-10 |
| TcCLB.463155.20 | retrotransposon hot spot (RHS) protein, putative | 1,084 | 3,592E-43 |
| TcCLB.508857.30 | trans-sialidase, Group I, putative | 1,084 | 4,119E-12 |
| TcCLB.508023.140 | dispersed gene family protein 1 (DGF-1), putative | 1,083 | 6,947E-09 |
| TcCLB.508511.30 | mucin TcMUCII, putative | 1,083 | 0,05086 |
| TcCLB.508221.550 | mucin-associated surface protein (MASP), putative | 1,083 | 0,1507 |
| TcCLB.506773.160 | hypothetical protein, conserved | 1,082 | 8,358E-21 |
| TcCLB.506325.30 | hypothetical protein, conserved (pseudogene) | 1,082 | 1,257E-10 |
| TcCLB.456139.10 | hypothetical protein, conserved | 1,082 | 0,0001814 |
| TcCLB.510687.60 | hypothetical protein, conserved | 1,081 | 9,051E-22 |
| TcCLB.509561.30 | citrate transporter, putative | 1,081 | 3,999E-11 |
| TcCLB.509667.20 | tryptophanyl-tRNA synthetase (pseudogene), putative | 1,081 | 7,028E-06 |
| TcCLB.509871.70 | retrotransposon hot spot protein (RHS, pseudogene), putat | 1,08 | 8,337E-10 |
| TcCLB.511007.20 | retrotransposon hot spot (RHS) protein, putative | 1,08 | 0,00000165 |
| TcCLB.506183.70 | beta galactofuranosyl glycosyltransferase, putative | 1,078 | 9,341E-13 |
| TcCLB.506433.10 | retrotransposon hot spot protein (RHS, pseudogene), putat | 1,078 | 2,333E-09 |
| TcCLB.508465.100 | hypothetical protein | 1,078 | 8,064E-08 |
| TcCLB.506763.260 | mucin-associated surface protein (MASP), putative | 1,078 | 0,0009327 |
| TcCLB.507163.30 | 90 kDa surface protein, putative | 1,076 | 0,06654 |
| TcCLB.510859.30 | hypothetical protein, conserved | 1,075 | 6,491E-38 |
| TcCLB.509979.30 | hypothetical protein, conserved | 1,075 | 6,457E-26 |
| TcCLB.507891.47 | flagellar calcium-binding protein, putative | 1,075 | 2,223E-24 |
| TcCLB.507021.40 | hypothetical protein, conserved | 1,075 | 1,096E-09 |
| TcCLB.509979.10 | hypothetical protein, conserved | 1,074 | 1,545E-33 |
| TcCLB.508663.10 | mucin TcMUC (pseudogene), putative | 1,074 | 2,947E-19 |

| | | | |
|---|---|---|---|
| TcCLB.507801.210 | retrotransposon hot spot protein (RHS, pseudogene), putat | 1,074 | 0,00004087 |
| TcCLB.508001.50 | trans-sialidase (pseudogene), putative | 1,074 | 0,03016 |
| TcCLB.482097.20 | hypothetical protein, conserved | 1,074 | 0,03147 |
| TcCLB.510483.310 | mucin-associated surface protein (MASP), putative | 1,074 | 0,04361 |
| TcCLB.507637.100 | mucin TcMUCII, putative | 1,074 | 0,05984 |
| TcCLB.506773.140 | DNA ligase, putative | 1,073 | 1,346E-17 |
| TcCLB.509317.50 | Asparagine-linked glycosylation protein 12 (ALG12) | 1,073 | 9,006E-16 |
| TcCLB.509065.50 | hypothetical protein, conserved | 1,073 | 1,501E-12 |
| TcCLB.506669.4 | MFS transporter, putative | 1,073 | 0,0009286 |
| TcCLB.507069.60 | mucin TcMUCII, putative | 1,073 | 0,05326 |
| TcCLB.511395.40 | hypothetical protein | 1,072 | 1,711E-29 |
| TcCLB.508065.50 | phosphatidylinositol-4-phosphate 5-kinase-like protein, put | 1,072 | 8,679E-23 |
| TcCLB.506513.7 | hypothetical protein, conserved | 1,072 | 1,189E-13 |
| TcCLB.508541.20 | mucin-associated surface protein (MASP), putative | 1,072 | 0,02145 |
| TcCLB.503843.20 | hypothetical protein, conserved | 1,071 | 1,236E-22 |
| TcCLB.509019.21 | retrotransposon hot spot protein (RHS, pseudogene), putat | 1,071 | 1,528E-10 |
| TcCLB.510801.10 | calpain-like cysteine peptidase, putative (fragment) | 1,071 | 8,777E-08 |
| TcCLB.503613.70 | hypothetical protein, conserved | 1,071 | 2,787E-06 |
| TcCLB.511373.10 | mucin-associated surface protein (MASP, pseudogene), pu | 1,071 | 0,2217 |
| TcCLB.511175.10 | mucin-associated surface protein (MASP), putative | 1,071 | 0,3842 |
| TcCLB.508851.160 | hypothetical protein, conserved | 1,07 | 8,374E-24 |
| TcCLB.504153.180 | UDP-N-acetylglucosamine pyrophosphorylase, putative | 1,07 | 6,244E-18 |
| TcCLB.433559.10 | trans-sialidase, putative (fragment) | 1,07 | 0,1952 |
| TcCLB.510347.50 | hypothetical protein, conserved | 1,069 | 5,375E-20 |
| TcCLB.511825.4 | hypothetical protein, conserved (fragment) | 1,069 | 3,695E-14 |
| TcCLB.506571.10 | dispersed gene family protein 1 (DGF-1), putative | 1,069 | 1,702E-07 |
| TcCLB.510101.260 | hypothetical protein, conserved | 1,068 | 1,623E-19 |
| TcCLB.510861.40 | hypothetical protein, conserved | 1,068 | 6,751E-17 |
| TcCLB.506359.70 | hypothetical protein, conserved | 1,068 | 7,365E-08 |
| TcCLB.506563.40 | beta tubulin, putative | 1,067 | 9,768E-17 |
| TcCLB.506933.43 | hypothetical protein, conserved | 1,067 | 5,23E-14 |
| TcCLB.508313.40 | mucin TcMUCII, putative | 1,066 | 0,0228 |
| TcCLB.509149.40 | 60S ribosomal protein L19, putative | 1,065 | 9,458E-72 |
| TcCLB.504797.110 | hypothetical protein, conserved | 1,065 | 4,051E-22 |
| TcCLB.507673.70 | hypothetical protein, conserved | 1,065 | 1,597E-13 |
| TcCLB.506283.10 | retrotransposon hot spot protein (RHS, pseudogene), putat | 1,065 | 8,66E-11 |
| TcCLB.506951.103 | hypothetical protein | 1,065 | 0,1243 |
| TcCLB.509419.94 | mucin TcMUCII (pseudogene), putative | 1,065 | 0,1866 |
| TcCLB.510585.10 | mucin-associated surface protein (MASP, pseudogene), pu | 1,065 | 0,2032 |
| TcCLB.506599.91 | surface protease GP63 (pseudogene), putative | 1,065 | 0,2118 |
| TcCLB.508359.50 | mucin TcMUCII, putative | 1,064 | 0,1228 |
| TcCLB.504127.30 | trans-sialidase (pseudogene), putative | 1,063 | 3,115E-20 |
| TcCLB.506983.39 | calpain-like cysteine peptidase, putative | 1,063 | 2,731E-17 |
| TcCLB.510139.50 | retrotransposon hot spot protein (RHS, pseudogene), putat | 1,063 | 1,288E-07 |
| TcCLB.510889.231 | ATP synthase subunit, putative | 1,063 | 0,00001006 |

| | | | |
|---|---|---|---|
| TcCLB.509207.70 | hypothetical protein, conserved | 1,062 | 1,561E-17 |
| TcCLB.510411.30 | retrotransposon hot spot protein (RHS, pseudogene), putat | 1,062 | 0,0001119 |
| TcCLB.508885.20 | mucin TcMUCI, putative | 1,062 | 0,1268 |
| TcCLB.508883.10 | mucin-associated surface protein (MASP, pseudogene), pu | 1,062 | 0,2833 |
| TcCLB.511825.200 | 60S ribosomal protein L14, putative | 1,061 | 2,409E-39 |
| TcCLB.507659.30 | amino acid transporter, putative (fragment) | 1,061 | 2,319E-36 |
| TcCLB.507611.10 | retrotransposon hot spot (RHS) protein, putative | 1,061 | 4,243E-35 |
| TcCLB.506777.50 | cell cycle sequence binding phosphoprotein (RBP45), putat | 1,061 | 1,666E-18 |
| TcCLB.508045.100 | mitochondrial DNA primase, putative (PRI1) | 1,06 | 9,273E-23 |
| TcCLB.504231.30 | hypothetical protein, conserved | 1,06 | 8,012E-21 |
| TcCLB.503477.20 | transaldolase, putative | 1,06 | 9,114E-18 |
| TcCLB.507515.10 | glutathione peroxidase-like protein, putative | 1,06 | 3,999E-15 |
| TcCLB.506183.15 | trans-sialidase (pseudogene), putative | 1,06 | 0,01338 |
| TcCLB.509773.20 | protein kinase, putative | 1,06 | 0,01736 |
| TcCLB.504087.20 | hypothetical protein | 1,058 | 6,686E-36 |
| TcCLB.506051.10 | retrotransposon hot spot protein (RHS, pseudogene), putat | 1,058 | 1,529E-07 |
| TcCLB.507875.170 | ribosomal P protein AGP2beta-1, putative | 1,058 | 8,308E-06 |
| TcCLB.506801.60 | mucin-like glycoprotein, putative | 1,058 | 0,001677 |
| TcCLB.508495.20 | trans-sialidase (pseudogene), putative | 1,057 | 3,981E-11 |
| TcCLB.507163.50 | mucin TcMUCII, putative | 1,057 | 0,04773 |
| TcCLB.508041.50 | hypothetical protein, conserved | 1,056 | 1,011E-23 |
| TcCLB.507601.70 | dihydrolipoamide branched chain transacylase, putative | 1,056 | 4,274E-17 |
| TcCLB.507165.30 | hypothetical protein, conserved | 1,056 | 1,197E-13 |
| TcCLB.510861.125 | hypothetical protein, conserved | 1,055 | 2,057E-13 |
| TcCLB.508029.10 | Hydroxymethylglutaryl-CoA lyase, mitochondrial, putative | 1,055 | 5,679E-13 |
| TcCLB.433237.10 | hypothetical protein | 1,055 | 0,231 |
| TcCLB.509901.100 | hypothetical protein, conserved | 1,054 | 6,2E-31 |
| TcCLB.509331.20 | hypothetical protein, conserved | 1,054 | 3,242E-14 |
| TcCLB.511827.70 | hypothetical protein, conserved | 1,054 | 3,837E-12 |
| TcCLB.507633.120 | mucin TcMUCII, putative | 1,054 | 0,04575 |
| TcCLB.509155.10 | hypothetical protein, conserved | 1,053 | 2,994E-34 |
| TcCLB.506975.80 | trans-sialidase, Group I, putative | 1,053 | 1,702E-07 |
| TcCLB.510349.80 | protein kinase, putative | 1,052 | 6,208E-27 |
| TcCLB.508037.10 | retrotransposon hot spot (RHS) protein, putative | 1,052 | 5,522E-17 |
| TcCLB.509081.130 | surface protease GP63 (pseudogene), putative | 1,052 | 0,2274 |
| TcCLB.504177.20 | fatty acyl CoA synthetase 2, putative | 1,051 | 3,419E-22 |
| TcCLB.511291.40 | protein kinase, putative | 1,051 | 8,479E-20 |
| TcCLB.509271.10 | trans-sialidase, putative | 1,051 | 3,001E-14 |
| TcCLB.508239.49 | dynein regulatory complex subunit, putative | 1,051 | 1,22E-09 |
| TcCLB.507871.9 | dispersed gene family protein 1 (DGF-1), putative | 1,051 | 0,04034 |
| TcCLB.504033.40 | hypothetical protein | 1,05 | 1,947E-20 |
| TcCLB.511081.30 | mucin TcMUCII, putative | 1,05 | 0,0414 |
| TcCLB.504769.150 | hypothetical protein | 1,05 | 0,1104 |
| TcCLB.504191.10 | sterol 24-c-methyltransferase, putative | 1,049 | 3,218E-38 |
| TcCLB.511825.230 | hypothetical protein | 1,049 | 8,077E-13 |

| | | | |
|---|---|---|---|
| TcCLB.511211.90 | surface protease GP63, putative | 1,049 | 1,958E-06 |
| TcCLB.506245.220 | mucin TcMUCII, putative | 1,049 | 0,1621 |
| TcCLB.430605.40 | ERV/ALR sulfhydryl oxidase domain-containing protein (PC | 1,048 | 2,748E-14 |
| TcCLB.503445.20 | trans-sialidase (pseudogene), putative | 1,048 | 3,473E-10 |
| TcCLB.418185.21 | retrotransposon hot spot protein (RHS, pseudogene), putat | 1,048 | 0,0003742 |
| TcCLB.508289.9 | mucin-associated surface protein (MASP), putative | 1,048 | 0,2625 |
| TcCLB.511019.40 | retrotransposon hot spot protein (RHS, pseudogene), putat | 1,047 | 2,253E-47 |
| TcCLB.507001.110 | galactokinase-like protein, putative | 1,047 | 1,253E-16 |
| TcCLB.510759.10 | hypothetical protein, conserved | 1,047 | 1,381E-13 |
| TcCLB.503465.40 | hypothetical protein | 1,047 | 5,188E-08 |
| TcCLB.511287.70 | hypothetical protein, conserved | 1,047 | 1,179E-06 |
| TcCLB.424499.10 | mucin-associated surface protein (MASP), putative | 1,047 | 0,06249 |
| TcCLB.509733.199 | Extradiol ring-cleavage dioxygenase, putative | 1,046 | 3,165E-14 |
| TcCLB.511711.110 | TLR4 regulator and MIR-interacting MSAP, putative (ESP16 | 1,046 | 2,455E-12 |
| TcCLB.510287.50 | hypothetical protein, conserved | 1,046 | 6,049E-09 |
| TcCLB.510155.10 | hypothetical protein, conserved | 1,045 | 1,223E-47 |
| TcCLB.506537.299 | retrotransposon hot spot (RHS) protein, putative | 1,045 | 9,621E-24 |
| TcCLB.404843.20 | hypothetical protein, conserved | 1,045 | 1,04E-19 |
| TcCLB.506615.10 | trans-sialidase (pseudogene), putative | 1,045 | 1,564E-10 |
| TcCLB.504157.10 | heterogeneous nuclear ribonucleoprotein H/F, putative | 1,044 | 1,253E-15 |
| TcCLB.508627.40 | trans-sialidase, Group VIII, putative | 1,044 | 4,733E-11 |
| TcCLB.507275.24 | ubiquitin fold modifier protein, putative (UFM1) | 1,044 | 1,321E-07 |
| TcCLB.504579.10 | retrotransposon hot spot protein (RHS, pseudogene), putat | 1,044 | 1,386E-07 |
| TcCLB.506825.140 | hypothetical protein, conserved | 1,043 | 6,874E-13 |
| TcCLB.503761.30 | mucin TcMUCII, putative | 1,043 | 0,0694 |
| TcCLB.509461.70 | hypothetical protein, conserved | 1,042 | 7,619E-19 |
| TcCLB.506773.110 | hypothetical protein, conserved | 1,04 | 1,265E-16 |
| TcCLB.509363.20 | retrotransposon hot spot protein (RHS, pseudogene), putat | 1,04 | 0,06265 |
| TcCLB.510859.40 | hypothetical protein, conserved | 1,039 | 3,147E-13 |
| TcCLB.510377.300 | trans-sialidase (pseudogene), putative | 1,039 | 0,01961 |
| TcCLB.507603.260 | cysteine peptidase, putative | 1,039 | 0,06318 |
| TcCLB.511149.130 | mucin TcMUCI, putative | 1,039 | 0,307 |
| TcCLB.506577.90 | retrotransposon hot spot protein (RHS, pseudogene), putat | 1,038 | 4,225E-15 |
| TcCLB.507691.40 | trans-sialidase (pseudogene), putative | 1,038 | 1,146E-07 |
| TcCLB.511127.74 | hypothetical protein, conserved | 1,038 | 3,203E-07 |
| TcCLB.510845.10 | retrotransposon hot spot (RHS) protein, putative (fragmen | 1,038 | 0,00003138 |
| TcCLB.511081.10 | mucin-associated surface protein (MASP), putative | 1,038 | 0,006627 |
| TcCLB.509153.70 | exosome-associated protein 3, putative (EAP3) | 1,037 | 5,523E-14 |
| TcCLB.511431.60 | hypothetical protein, conserved | 1,037 | 2,569E-07 |
| TcCLB.508633.130 | 90 kDa surface protein, putative | 1,037 | 0,006326 |
| TcCLB.506767.170 | trans-sialidase (pseudogene), putative | 1,037 | 0,008499 |
| TcCLB.505699.10 | trans-sialidase, putative | 1,036 | 2,116E-06 |
| TcCLB.508521.50 | surface protease GP63 (pseudogene), putative | 1,036 | 0,0003582 |
| TcCLB.509617.10 | paraflagellar rod protein 3, putative (pseudogene) (PAR3) | 1,035 | 1,089E-18 |
| TcCLB.507821.70 | dispersed gene family protein 1 (DGF-1, pseudogene), puta | 1,035 | 6,663E-07 |

| | | | |
|---|---|---|---|
| TcCLB.510669.20 | Gim5A protein, putative | 1,034 | 4,078E-25 |
| TcCLB.510225.59 | transporter, putative | 1,034 | 0,00004122 |
| TcCLB.510951.10 | hypothetical protein, conserved | 1,034 | 0,001416 |
| TcCLB.506501.27 | syntaxin binding protein (pseudogene), putative | 1,034 | 0,1923 |
| TcCLB.508569.120 | hypothetical protein, conserved | 1,033 | 7,345E-19 |
| TcCLB.510511.9 | Na/H antiporter-like protein, putative | 1,033 | 2,332E-07 |
| TcCLB.505937.80 | retrotransposon hot spot protein (RHS, pseudogene), putat | 1,033 | 0,00001886 |
| TcCLB.505925.10 | trans-sialidase, putative (fragment) | 1,033 | 0,00003709 |
| TcCLB.508873.70 | mucin-associated surface protein (MASP, pseudogene), pu | 1,033 | 0,0396 |
| TcCLB.507937.14 | mucin TcMUCII (pseudogene), putative | 1,033 | 0,07871 |
| TcCLB.507665.30 | ferric reductase transmembrane protein, putative (FR1) | 1,032 | 1,466E-25 |
| TcCLB.503453.100 | serine palmitoyltransferase, putative | 1,032 | 2,047E-20 |
| TcCLB.507559.70 | hypothetical protein, conserved | 1,032 | 5,708E-11 |
| TcCLB.508119.180 | hypothetical protein, conserved | 1,032 | 5,726E-07 |
| TcCLB.503715.30 | ras-related protein RAB1A, putative (Rab1A) | 1,031 | 2,15E-35 |
| TcCLB.506163.50 | hypothetical protein, conserved | 1,031 | 4,506E-18 |
| TcCLB.507473.20 | hypothetical protein | 1,031 | 3,482E-10 |
| TcCLB.506595.40 | trans-sialidase, Group II, putative | 1,031 | 3,524E-08 |
| TcCLB.509233.20 | retrotransposon hot spot protein (RHS, pseudogene), putat | 1,031 | 1,102E-06 |
| TcCLB.509779.10 | DNA repair protein, putative | 1,03 | 1,036E-19 |
| TcCLB.510373.30 | 90 kDa surface protein, putative | 1,03 | 0,08441 |
| TcCLB.507643.60 | trans-sialidase (pseudogene), putative | 1,029 | 4,912E-06 |
| TcCLB.507213.40 | trans-sialidase, Group II, putative | 1,029 | 0,0005977 |
| TcCLB.510101.270 | hypothetical protein, conserved | 1,027 | 9,669E-15 |
| TcCLB.509395.100 | cytochrome b-domain protein, putative | 1,027 | 6,554E-10 |
| TcCLB.503929.5 | hypothetical protein, conserved | 1,027 | 4,398E-07 |
| TcCLB.511173.400 | mucin-associated surface protein (MASP), putative | 1,027 | 0,1417 |
| TcCLB.503401.40 | glycosyl transferase, putative | 1,026 | 4,469E-29 |
| TcCLB.510861.94 | hypothetical protein, conserved | 1,026 | 1,426E-25 |
| TcCLB.511821.130 | hypothetical protein | 1,026 | 1,808E-21 |
| TcCLB.511305.10 | 5'-3' exoribonuclease B, putative | 1,026 | 6,491E-14 |
| TcCLB.505193.70 | hypothetical protein, conserved | 1,026 | 2,013E-13 |
| TcCLB.506401.190 | hypothetical protein, conserved | 1,026 | 1,164E-12 |
| TcCLB.504787.99 | mucin TcMUCII, putative | 1,026 | 0,04021 |
| TcCLB.509769.60 | succinate dehydrogenase iron-sulfur subunit (SDH2N) | 1,025 | 1,771E-27 |
| TcCLB.511807.253 | hypothetical protein, conserved | 1,025 | 6,37E-18 |
| TcCLB.505171.90 | hypothetical protein | 1,025 | 1,659E-11 |
| TcCLB.507221.70 | hypothetical protein, conserved | 1,025 | 1,055E-09 |
| TcCLB.503929.79 | serine-palmitoyl-CoA transferase, putative | 1,025 | 9,226E-08 |
| TcCLB.508501.240 | hypothetical protein, conserved | 1,025 | 1,939E-06 |
| TcCLB.511603.420 | syntaxin binding protein (pseudogene), putative | 1,025 | 0,09752 |
| TcCLB.509481.10 | trans-sialidase, putative | 1,025 | 0,1238 |
| TcCLB.506905.30 | hypothetical protein, conserved | 1,024 | 2,213E-12 |
| TcCLB.510879.180 | hypothetical protein | 1,024 | 1,013E-11 |
| TcCLB.506595.60 | beta galactofuranosyl glycosyltransferase (pseudogene), p | 1,024 | 2,859E-09 |

| | | | |
|---|---|---|---|
| TcCLB.504033.130 | spermidine synthase, putative | 1,024 | 0,0003186 |
| TcCLB.511787.10 | mucin-associated surface protein (MASP), putative | 1,024 | 0,2041 |
| TcCLB.509875.50 | beta galactofuranosyl glycosyltransferase, putative | 1,023 | 0,0006008 |
| TcCLB.510465.60 | hypothetical protein, conserved (pseudogene) | 1,023 | 0,006915 |
| TcCLB.511517.30 | hypothetical protein, conserved | 1,021 | 1,045E-14 |
| TcCLB.509491.20 | hypothetical protein, conserved (pseudogene) | 1,021 | 4,629E-14 |
| TcCLB.507747.230 | acid phosphatase type 5, putative | 1,021 | 1,798E-10 |
| TcCLB.511261.20 | chaperone DNAJ protein, putative | 1,02 | 4,908E-16 |
| TcCLB.507867.60 | protein phosphatase 1, regulatory subunit, putative | 1,02 | 8,356E-16 |
| TcCLB.506467.40 | serine/threonine protein kinase, putative | 1,019 | 3,879E-19 |
| TcCLB.511585.110 | hypothetical protein | 1,019 | 2,624E-18 |
| TcCLB.509151.60 | hypothetical protein, conserved | 1,019 | 3,335E-17 |
| TcCLB.476125.9 | dispersed gene family protein 1 (DGF-1), putative | 1,019 | 0,0002091 |
| TcCLB.509009.10 | dynein heavy chain, cytosolic, putative | 1,018 | 7,73E-11 |
| TcCLB.511117.10 | retrotransposon hot spot protein (RHS, pseudogene), putat | 1,018 | 6,137E-10 |
| TcCLB.507559.110 | surface protease GP63, putative | 1,018 | 3,966E-07 |
| TcCLB.509757.49 | L-ribulokinase, putative | 1,018 | 0,006249 |
| TcCLB.507559.60 | hypothetical protein, conserved | 1,017 | 3,785E-18 |
| TcCLB.506595.30 | retrotransposon hot spot protein (RHS, pseudogene), putat | 1,017 | 5,294E-12 |
| TcCLB.511545.130 | hypothetical protein, conserved | 1,017 | 2,603E-11 |
| TcCLB.511825.250 | hypothetical protein, conserved | 1,017 | 9,586E-09 |
| TcCLB.504827.159 | hypothetical protein, conserved | 1,017 | 2,444E-07 |
| TcCLB.509725.30 | hypothetical protein, conserved (pseudogene) | 1,017 | 4,617E-06 |
| TcCLB.508125.50 | mucin-associated surface protein (MASP, pseudogene), pu | 1,017 | 0,04565 |
| TcCLB.507637.130 | mucin TcMUCII, putative | 1,017 | 0,08822 |
| TcCLB.510627.70 | hypothetical protein, conserved (pseudogene) | 1,017 | 0,08948 |
| TcCLB.506771.9 | hypothetical protein, conserved | 1,016 | 3,227E-14 |
| TcCLB.503739.10 | beta galactofuranosyl glycosyltransferase (pseudogene), p | 1,016 | 0,01884 |
| TcCLB.506537.180 | retrotransposon hot spot protein (RHS, pseudogene), putat | 1,015 | 7,537E-22 |
| TcCLB.507615.76 | hypothetical protein, conserved | 1,015 | 4,629E-12 |
| TcCLB.511753.40 | hypothetical protein, conserved | 1,015 | 0,00003437 |
| TcCLB.511693.50 | hypothetical protein, conserved | 1,014 | 2,53E-18 |
| TcCLB.504229.110 | amino acid permease 24, putative (AAP24) | 1,014 | 6,333E-09 |
| TcCLB.509295.30 | trans-sialidase (pseudogene), putative | 1,014 | 0,000268 |
| TcCLB.509753.240 | mucin TcMUCII, putative | 1,014 | 0,3519 |
| TcCLB.506579.10 | ABC transporter, putative | 1,013 | 1,269E-34 |
| TcCLB.510101.59 | palmitoyl acyltransferase 7, putative | 1,013 | 1,415E-17 |
| TcCLB.506617.90 | ATP-dependent Clp protease subunit, heat shock protein 1( | 1,013 | 3,74E-10 |
| TcCLB.508297.61 | choline ethanolamine kinase, putative | 1,013 | 0,0007211 |
| TcCLB.507429.40 | mucin-associated surface protein (MASP, pseudogene), pu | 1,013 | 0,002406 |
| TcCLB.511123.11 | trans-sialidase, putative (fragment) | 1,013 | 0,004336 |
| TcCLB.507059.10 | mucin-like glycoprotein, putative | 1,013 | 0,01299 |
| TcCLB.504155.70 | mucin TcMUCII, putative | 1,013 | 0,1662 |
| TcCLB.506779.20 | hypothetical protein, conserved | 1,012 | 3,741E-22 |
| TcCLB.506775.22 | hypothetical protein, conserved | 1,012 | 1,383E-11 |

| | | | |
|---|---|---|---|
| TcCLB.447483.10 | hypothetical protein, conserved | 1,012 | 3,243E-11 |
| TcCLB.507721.20 | clathrin assembly protein (pseudogene), putative | 1,011 | 4,434E-27 |
| TcCLB.506959.40 | enoyl-CoA hydratase/isomerase family protein, putative | 1,011 | 1,841E-16 |
| TcCLB.506177.30 | hypothetical protein, conserved | 1,011 | 5,293E-12 |
| TcCLB.408437.20 | hypothetical protein, conserved | 1,011 | 4,943E-08 |
| TcCLB.511319.30 | hypothetical protein | 1,011 | 1,253E-07 |
| TcCLB.503865.70 | hypothetical protein, conserved | 1,01 | 1,184E-20 |
| TcCLB.508973.30 | NUDIX family hydrolase, putative | 1,01 | 1,082E-10 |
| TcCLB.509037.20 | surface protease GP63, putative | 1,01 | 0,001954 |
| TcCLB.506855.220 | surface protease GP63, putative | 1,01 | 0,005035 |
| TcCLB.510655.100 | hypothetical protein | 1,008 | 1,991E-12 |
| TcCLB.506537.10 | retrotransposon hot spot protein (RHS, pseudogene), putat | 1,007 | 1,031E-12 |
| TcCLB.506113.10 | retrotransposon hot spot protein (RHS, pseudogene), putat | 1,007 | 8,99E-12 |
| TcCLB.506285.120 | mucin TcMUCII, putative | 1,007 | 0,03566 |
| TcCLB.511825.220 | alanyl-tRNA synthetase, putative | 1,006 | 4,471E-41 |
| TcCLB.510155.140 | Meckel syndrome type 6 protein, putative (mks6) | 1,006 | 1,063E-32 |
| TcCLB.506945.110 | casein kinase 1 isoform 2, putative | 1,006 | 7,547E-20 |
| TcCLB.506793.10 | hypothetical protein, conserved (pseudogene) | 1,005 | 1,274E-10 |
| TcCLB.507517.104 | hypothetical protein, conserved | 1,005 | 6,253E-07 |
| TcCLB.511569.30 | trans-sialidase (pseudogene), putative | 1,005 | 0,0009643 |
| TcCLB.508221.210 | mucin-associated surface protein (MASP), putative | 1,005 | 0,1361 |
| TcCLB.510303.220 | hypothetical protein, conserved | 1,004 | 6,946E-23 |
| TcCLB.508153.520 | hypothetical protein, conserved (pseudogene) | 1,004 | 3,493E-20 |
| TcCLB.403985.10 | retrotransposon hot spot (RHS) protein, putative, (fragmer | 1,004 | 2,916E-11 |
| TcCLB.511117.50 | surface protease GP63, putative | 1,004 | 0,003046 |
| TcCLB.503741.10 | mucin TcMUCII, putative | 1,004 | 0,02917 |
| TcCLB.506053.10 | amino acid transporter, putative | 1,003 | 8,202E-42 |
| TcCLB.510565.20 | fatty acid desaturase, putative | 1,003 | 1,183E-19 |
| TcCLB.509935.29 | proline racemase (fragment) | 1,003 | 3,141E-10 |
| TcCLB.506163.30 | hypothetical protein, conserved | 1,003 | 1,28E-08 |
| TcCLB.507083.40 | hypothetical protein, conserved | 1,003 | 0,00001984 |
| TcCLB.510131.30 | hypothetical protein, conserved | 1,002 | 2,939E-19 |
| TcCLB.508317.50 | ammonium transporter, putative | 1,002 | 1,134E-18 |
| TcCLB.506289.40 | hypothetical protein, conserved | 1,002 | 8,321E-11 |
| TcCLB.504151.80 | Na/H antiporter-like protein, putative | 1,002 | 9,322E-10 |
| TcCLB.508413.5 | hypothetical protein, conserved | 1,002 | 0,0002889 |
| TcCLB.510479.30 | hypothetical protein | 1,002 | 0,00121 |
| TcCLB.509493.60 | hypothetical protein | 1,002 | 0,002063 |
| TcCLB.509513.10 | trans-sialidase, Group II, putative | 1,002 | 0,002861 |
| TcCLB.506597.9 | trans-sialidase, putative (fragment) | 1,002 | 0,5585 |
| TcCLB.511245.24 | hypothetical protein, conserved | 1,001 | 2,416E-22 |
| TcCLB.506201.70 | serine/threonine-protein phosphatase PP1 alpha | 1,001 | 0,0001411 |
| TcCLB.509699.220 | trans-sialidase (pseudogene), putative | 1,001 | 0,001155 |
| TcCLB.445281.9 | calpain-like cysteine peptidase, putative | 1,001 | 0,004896 |
| TcCLB.410275.9 | mucin-associated surface protein (MASP), putative | 1,001 | 0,4769 |

| | | | |
|---|---|---|---|
| TcCLB.509695.210 | serine carboxypeptidase (CBP1), putative | 0,9998 | 4,379E-20 |
| TcCLB.508989.100 | hypothetical protein, conserved | 0,9998 | 1,371E-12 |
| TcCLB.511173.483 | mucin TcMUCII (pseudogene), putative | 0,9998 | 0,04067 |
| TcCLB.509033.30 | cyclophilin, putative | 0,9994 | 4,498E-08 |
| TcCLB.507873.60 | 1,2-Dihydroxy-3-keto-5-methylthiopentene dioxygenase, pu | 0,9988 | 0,00000016 |
| TcCLB.509601.160 | hypothetical protein, conserved | 0,9987 | 0,00001351 |
| TcCLB.509149.30 | hypothetical protein, conserved | 0,9986 | 1,623E-26 |
| TcCLB.507851.50 | hypothetical protein, conserved | 0,9985 | 6,02E-20 |
| TcCLB.510889.221 | glucose-6-phosphate isomerase, glycosomal, putative | 0,9983 | 4,5E-20 |
| TcCLB.510433.10 | hypothetical protein | 0,9978 | 2,138E-17 |
| TcCLB.508859.118 | trans-sialidase, putative | 0,9977 | 3,797E-06 |
| TcCLB.511573.20 | hypothetical protein | 0,9973 | 9,534E-29 |
| TcCLB.511321.23 | cytidine deaminase-like protein | 0,9968 | 1,729E-09 |
| TcCLB.511171.40 | trans-sialidase (pseudogene), putative | 0,9964 | 0,000113 |
| TcCLB.506615.111 | elongation factor 1-gamma (EF-1-gamma, pseudogene), p | 0,9961 | 0,00008605 |
| TcCLB.509259.126 | hypothetical protein | 0,9956 | 0,0003585 |
| TcCLB.511823.30 | hypothetical protein, conserved | 0,9954 | 6,964E-16 |
| TcCLB.510347.20 | hypothetical protein, conserved | 0,9953 | 3,918E-17 |
| TcCLB.510625.54 | mucin-associated surface protein (MASP), putative | 0,9953 | 0,1206 |
| TcCLB.510307.60 | retrotransposon hot spot protein (RHS, pseudogene), putat | 0,9952 | 1,104E-23 |
| TcCLB.506285.30 | trans-sialidase (pseudogene), putative | 0,9952 | 0,0008558 |
| TcCLB.508811.30 | hypothetical protein, conserved | 0,9948 | 9,027E-21 |
| TcCLB.505647.20 | hypothetical protein, conserved | 0,9945 | 3,008E-27 |
| TcCLB.507427.30 | retrotransposon hot spot protein (RHS, pseudogene), putat | 0,9945 | 5,67E-07 |
| TcCLB.503599.60 | hypothetical protein, conserved | 0,9937 | 1,62E-09 |
| TcCLB.506345.40 | trans-sialidase, Group II, putative | 0,9936 | 5,082E-09 |
| TcCLB.510465.125 | trans-sialidase (pseudogene), putative | 0,9934 | 0,3826 |
| TcCLB.509217.70 | hypothetical protein, conserved (pseudogene) | 0,9931 | 0,00001003 |
| TcCLB.504787.20 | mucin TcMUCII, putative | 0,9931 | 0,01541 |
| TcCLB.506279.74 | mucin-associated surface protein (MASP, pseudogene), pu | 0,9925 | 0,05326 |
| TcCLB.508567.20 | hypothetical protein, conserved | 0,9922 | 6,293E-09 |
| TcCLB.509755.89 | hypothetical protein | 0,9921 | 0,02427 |
| TcCLB.510055.50 | hypothetical protein, conserved | 0,9918 | 2,7E-22 |
| TcCLB.511279.40 | hypothetical protein, conserved | 0,9918 | 5,036E-06 |
| TcCLB.510301.70 | hypothetical protein, conserved | 0,9917 | 3,471E-15 |
| TcCLB.427247.4 | hypothetical protein, conserved (fragment) | 0,9916 | 3,059E-09 |
| TcCLB.504155.20 | trans-sialidase, Group V, putative | 0,9912 | 0,003908 |
| TcCLB.471799.10 | trans-sialidase, putative (fragment) | 0,9912 | 0,01976 |
| TcCLB.511173.420 | mucin TcMUCII, putative | 0,991 | 0,08185 |
| TcCLB.508221.170 | mucin-associated surface protein (MASP), putative | 0,991 | 0,2936 |
| TcCLB.507017.40 | 2-hydroxy-3-oxopropionate reductase, putative | 0,9909 | 1,436E-21 |
| TcCLB.506871.70 | hypothetical protein, conserved | 0,9902 | 1,789E-14 |
| TcCLB.509179.150 | otubain cysteine peptidase, Clan CA, family C65, putative | 0,99 | 7,219E-15 |
| TcCLB.510575.5 | proton motive ATPase, putative | 0,9896 | 5,012E-15 |
| TcCLB.509631.30 | mucin TcMUCII (pseudogene), putative | 0,9896 | 0,05797 |

| | | | |
|---|---|---|---|
| TcCLB.509007.99 | P-glycoprotein (pseudogene) (tcpgp1) | 0,9894 | 9,079E-21 |
| TcCLB.509975.41 | mucin TcMUC, putative | 0,9894 | 0,009195 |
| TcCLB.508119.190 | hypothetical protein, conserved | 0,989 | 1,47E-23 |
| TcCLB.507883.40 | hypothetical protein, conserved | 0,989 | 2,831E-08 |
| TcCLB.508873.150 | mucin TcMUCII, putative | 0,9885 | 0,1337 |
| TcCLB.511255.169 | mucin-associated surface protein (MASP), putative | 0,9884 | 0,1144 |
| TcCLB.508905.30 | hypothetical protein, conserved (pseudogene) | 0,9883 | 3,912E-10 |
| TcCLB.511511.150 | sphingosine 1-phosphate lyase, putative (SPL) | 0,9879 | 5,852E-14 |
| TcCLB.507031.140 | hypothetical protein, conserved (pseudogene) | 0,9879 | 6,921E-11 |
| TcCLB.506801.80 | cysteine peptidase inhibitor (CHA) | 0,9878 | 1,515E-14 |
| TcCLB.509599.150 | serine/threonine protein kinase, putative | 0,9874 | 1,515E-08 |
| TcCLB.510643.100 | trans-sialidase (pseudogene), putative | 0,9873 | 0,3995 |
| TcCLB.507167.20 | retrotransposon hot spot (RHS) protein, putative | 0,9865 | 1,586E-37 |
| TcCLB.506409.170 | trans-sialidase, Group VI, putative | 0,9863 | 4,489E-08 |
| TcCLB.506829.110 | fatty acyl CoA synthetase 2, putative | 0,9862 | 2,706E-18 |
| TcCLB.510285.80 | hypothetical protein, conserved | 0,986 | 3,191E-10 |
| TcCLB.503559.109 | electron-transfer-flavoprotein, alpha polypeptide, putative | 0,9859 | 5,228E-19 |
| TcCLB.507955.20 | mucin-associated surface protein (MASP), putative | 0,9858 | 0,2 |
| TcCLB.506781.10 | mucin TcMUCII, putative | 0,9851 | 0,1124 |
| TcCLB.510679.70 | lipase, putative | 0,9845 | 0,006506 |
| TcCLB.506559.190 | dual specificity protein phosphatase, putative | 0,9842 | 3,12E-17 |
| TcCLB.511443.10 | calpain-like cysteine peptidase (pseudogene), putative | 0,9839 | 7,707E-06 |
| TcCLB.510663.70 | 6-phosphogluconate dehydrogenase, decarboxylating, puta | 0,9837 | 2,573E-13 |
| TcCLB.504333.20 | trans-sialidase (pseudogene), putative | 0,9835 | 9,258E-13 |
| TcCLB.511613.90 | mucin-associated surface protein (MASP), putative | 0,9834 | 0,007711 |
| TcCLB.509631.90 | mucin TcMUCII, putative | 0,9834 | 0,04159 |
| TcCLB.506083.20 | trans-sialidase (pseudogene), putative | 0,9832 | 1,617E-09 |
| TcCLB.511859.80 | hypothetical protein | 0,9831 | 2,583E-13 |
| TcCLB.503437.10 | retrotransposon hot spot protein (RHS, pseudogene), putat | 0,983 | 1,269E-07 |
| TcCLB.506173.240 | mucin TcMUCII, putative | 0,9829 | 0,03846 |
| TcCLB.508163.10 | mucin-associated surface protein (MASP, pseudogene), pu | 0,9829 | 0,3191 |
| TcCLB.510759.114 | hypothetical protein, conserved | 0,9826 | 2,577E-09 |
| TcCLB.506951.50 | mucin TcMUCII (pseudogene), putative | 0,9825 | 4,868E-06 |
| TcCLB.506715.10 | protein kinase, putative | 0,9819 | 3,257E-24 |
| TcCLB.504153.240 | hypothetical protein, conserved | 0,9816 | 1,312E-15 |
| TcCLB.507791.10 | retrotransposon hot spot (RHS) protein, putative (fragmen | 0,9816 | 0,05992 |
| TcCLB.507773.60 | phosphoribosylpyrophosphate synthetase, putative | 0,9812 | 2,448E-18 |
| TcCLB.511839.20 | mucin-associated surface protein (MASP), putative | 0,9808 | 0,052 |
| TcCLB.509317.20 | esterase, putative | 0,9807 | 4,483E-08 |
| TcCLB.506001.14 | mucin-associated surface protein (MASP), putative | 0,9806 | 0,09031 |
| TcCLB.506247.390 | hypothetical protein, conserved | 0,9804 | 3,381E-16 |
| TcCLB.510787.10 | trans-sialidase, Group I, putative | 0,9804 | 7,896E-08 |
| TcCLB.503613.40 | pterin-4-alpha-carbinolamine dehydratase, putative | 0,9803 | 1,968E-12 |
| TcCLB.435601.10 | trans-sialidase, putative (fragment) | 0,9802 | 0,02102 |
| TcCLB.511907.140 | hypothetical protein | 0,9801 | 0,0002178 |

| | | | |
|---|---|---|---|
| TcCLB.506355.140 | hypothetical protein, conserved | 0,9798 | 4,383E-19 |
| TcCLB.508121.30 | trans-sialidase (pseudogene), putative | 0,9793 | 2,795E-06 |
| TcCLB.508503.40 | hypothetical protein, conserved | 0,9791 | 2,596E-15 |
| TcCLB.506779.190 | Unconventional myosin | 0,979 | 2,927E-17 |
| TcCLB.506967.70 | mucin-associated surface protein (MASP), putative | 0,978 | 0,01404 |
| TcCLB.507491.20 | surface antigen TASV, putative | 0,9775 | 1,785E-22 |
| TcCLB.504769.60 | hypothetical protein, conserved | 0,9775 | 6,543E-07 |
| TcCLB.510979.30 | mucin TcMUCII (pseudogene), putative | 0,9775 | 0,001183 |
| TcCLB.508125.40 | hypothetical protein | 0,9771 | 0,07853 |
| TcCLB.506113.60 | retrotransposon hot spot (RHS) protein, putative | 0,9769 | 3,412E-37 |
| TcCLB.510557.40 | hypothetical protein | 0,9766 | 0,0001778 |
| TcCLB.509857.60 | hypothetical protein, conserved | 0,9765 | 1,284E-27 |
| TcCLB.504797.144 | hypothetical protein, conserved | 0,9762 | 1,451E-15 |
| TcCLB.508817.119 | hypothetical protein, conserved | 0,9762 | 5,882E-08 |
| TcCLB.509167.150 | homoserine kinase, putative | 0,9754 | 1,837E-15 |
| TcCLB.507153.55 | hypothetical protein, conserved (pseudogene) | 0,9754 | 0,00001567 |
| TcCLB.503975.60 | hypothetical protein | 0,9751 | 1,44E-15 |
| TcCLB.508841.10 | surface antigen TASV-A4 (TASV-A4) | 0,9751 | 2,337E-06 |
| TcCLB.511585.170 | protein kinase, putative | 0,9749 | 2,499E-17 |
| TcCLB.510359.340 | phospholipid:diacylglycerol acyltransferase, putative | 0,9746 | 2,504E-21 |
| TcCLB.506247.410 | protein-l-isoaspartate o-methyltransferase, putative | 0,9743 | 1,625E-13 |
| TcCLB.506769.50 | mucin TcMUCII, putative | 0,9737 | 0,1406 |
| TcCLB.511837.120 | hypothetical protein, conserved | 0,9733 | 0,0002174 |
| TcCLB.509151.140 | 60S ribosomal protein L23a, putative | 0,973 | 5,121E-33 |
| TcCLB.504081.220 | retrotransposon hot spot protein (RHS, pseudogene), putat | 0,9726 | 6,675E-13 |
| TcCLB.511653.69 | DNA repair protein, putative (fragment) | 0,972 | 3,45E-22 |
| TcCLB.503861.40 | trans-sialidase, Group VIII, putative | 0,972 | 2,753E-07 |
| TcCLB.503939.110 | hypothetical protein, conserved | 0,9719 | 1,597E-14 |
| TcCLB.506715.40 | hypothetical protein, conserved | 0,9714 | 2,143E-19 |
| TcCLB.511817.250 | hypothetical protein, conserved | 0,9712 | 1,797E-15 |
| TcCLB.511751.210 | phosphopantothenoylcysteine decarboxylase, putative (PPC | 0,971 | 3,175E-15 |
| TcCLB.510049.10 | trans-sialidase, putative | 0,9701 | 7,224E-07 |
| TcCLB.511903.180 | hypothetical protein, conserved | 0,97 | 6,44E-10 |
| TcCLB.508221.260 | 90 kDa surface protein, putative | 0,9698 | 0,1057 |
| TcCLB.511817.270 | hypothetical protein, conserved | 0,9694 | 1,652E-16 |
| TcCLB.506785.14 | mucin-associated surface protein (MASP), putative | 0,9692 | 0,2037 |
| TcCLB.505943.110 | retrotransposon hot spot protein (RHS, pseudogene), putat | 0,9686 | 2,973E-09 |
| TcCLB.509341.20 | hypothetical protein, conserved | 0,9684 | 5,738E-24 |
| TcCLB.506295.130 | prohibitin, putative | 0,9681 | 1,657E-15 |
| TcCLB.510279.210 | mucin-associated surface protein (MASP), putative | 0,968 | 0,09475 |
| TcCLB.510239.80 | trans-sialidase, Group V, putative | 0,9679 | 0,004352 |
| TcCLB.510623.10 | mucin TcMUCII, putative | 0,9677 | 0,1171 |
| TcCLB.508319.20 | Present in the outer mitochondrial membrane proteome 1[illegible] | 0,9673 | 1,428E-14 |
| TcCLB.507633.40 | mucin TcMUCII, putative | 0,9671 | 0,0005516 |
| TcCLB.507043.21 | mucin TcMUCI, putative (fragment) | 0,9671 | 0,02223 |

| | | | |
|---|---|---|---|
| TcCLB.510023.30 | mucin-associated surface protein (MASP), putative (fragm | 0,9668 | 0,07806 |
| TcCLB.511201.40 | hypothetical protein, conserved | 0,9666 | 1,334E-08 |
| TcCLB.507937.3 | trans-sialidase, putative | 0,9666 | 0,4926 |
| TcCLB.506401.130 | hypothetical protein, conserved | 0,9664 | 3,61E-09 |
| TcCLB.509903.5 | retrotransposon hot spot (RHS) protein, putative (fragmen | 0,9658 | 8,343E-21 |
| TcCLB.508071.40 | hypothetical protein | 0,9657 | 0,04287 |
| TcCLB.504123.30 | map kinase, putative | 0,9656 | 1,642E-14 |
| TcCLB.503453.10 | hypothetical protein, conserved | 0,9656 | 6,326E-12 |
| TcCLB.454403.30 | retrotransposon hot spot protein (RHS, pseudogene), putat | 0,9656 | 1,115E-08 |
| TcCLB.511149.100 | mucin TcMUCII, putative | 0,9652 | 0,00187 |
| TcCLB.506041.10 | hypothetical protein, conserved (fragment) | 0,9649 | 3,598E-07 |
| TcCLB.506519.51 | hypothetical protein, conserved | 0,9647 | 6,743E-13 |
| TcCLB.511077.100 | mucin TcMUCII, putative | 0,9643 | 0,2526 |
| TcCLB.508501.300 | hypothetical protein, conserved | 0,964 | 5,846E-12 |
| TcCLB.506409.190 | mucin-associated surface protein (MASP, pseudogene), pu | 0,9639 | 0,005942 |
| TcCLB.511589.40 | hypothetical protein, conserved | 0,9632 | 7,738E-10 |
| TcCLB.504257.60 | S-adenosylmethionine decarboxylase proenzyme, putative | 0,9632 | 7,662E-06 |
| TcCLB.509599.170 | hypothetical protein, conserved | 0,9629 | 1,529E-09 |
| TcCLB.510007.30 | RNA-binding protein, putative | 0,9627 | 2,857E-17 |
| TcCLB.508953.10 | hypothetical protein, conserved | 0,9627 | 4,142E-06 |
| TcCLB.509815.10 | retrotransposon hot spot (RHS) protein, putative | 0,9626 | 8,955E-08 |
| TcCLB.509805.90 | hypothetical protein, conserved | 0,9624 | 2,901E-16 |
| TcCLB.506359.80 | Dynein light chain roadblock-type protein, putative | 0,9624 | 6,012E-09 |
| TcCLB.510465.74 | mucin-associated surface protein (MASP), putative | 0,9624 | 0,1758 |
| TcCLB.510371.130 | mucin TcMUCII, putative | 0,9617 | 0,2056 |
| TcCLB.507491.162 | flagellar calcium-binding protein, putative (fragment) | 0,9616 | 3,069E-18 |
| TcCLB.506443.150 | retrotransposon hot spot (RHS) protein, putative | 0,9612 | 1,671E-39 |
| TcCLB.511809.10 | hypothetical protein | 0,961 | 7,122E-12 |
| TcCLB.506661.170 | hypothetical protein, conserved | 0,961 | 5,104E-07 |
| TcCLB.511385.60 | hypothetical protein | 0,961 | 8,906E-07 |
| TcCLB.510173.70 | MIX protein, putative | 0,9609 | 7,234E-08 |
| TcCLB.508163.230 | mucin-associated surface protein (MASP), putative | 0,9608 | 0,1535 |
| TcCLB.506471.100 | trans-sialidase, Group II, putative | 0,9605 | 0,00003161 |
| TcCLB.511173.180 | mucin-associated surface protein (MASP), putative | 0,9604 | 0,08039 |
| TcCLB.510643.10 | dispersed gene family protein 1 (DGF-1), putative | 0,9603 | 5,531E-07 |
| TcCLB.511771.80 | retrotransposon hot spot protein (RHS, pseudogene), putat | 0,9601 | 1,463E-09 |
| TcCLB.509589.20 | squalene monooxygenase, putative | 0,96 | 8,609E-22 |
| TcCLB.510555.30 | mucin TcMUCII, putative | 0,9597 | 0,0704 |
| TcCLB.504081.340 | hypothetical protein (pseudogene) | 0,9597 | 0,2295 |
| TcCLB.511891.10 | trans-sialidase, Group II, putative | 0,9592 | 7,592E-08 |
| TcCLB.508387.160 | MORN repeat-containing protein | 0,9591 | 1,502E-11 |
| TcCLB.511817.300 | hypothetical protein, conserved | 0,9586 | 3,734E-12 |
| TcCLB.510901.180 | hypothetical protein, conserved | 0,9586 | 9,421E-09 |
| TcCLB.507485.40 | hypothetical protein, conserved | 0,9584 | 2,229E-16 |
| TcCLB.452283.4 | hypothetical protein, conserved | 0,9584 | 0,01354 |

| | | | |
|---|---|---|---|
| TcCLB.508205.4 | hypothetical protein, conserved | 0,9582 | 0,0003256 |
| TcCLB.511655.30 | hypothetical protein | 0,9579 | 1,103E-22 |
| TcCLB.506501.240 | mucin-associated surface protein (MASP), putative | 0,9577 | 0,2588 |
| TcCLB.509419.80 | mucin-associated surface protein (MASP), putative | 0,9568 | 0,4026 |
| TcCLB.511815.90 | hypothetical protein | 0,9559 | 6,712E-14 |
| TcCLB.509259.120 | retrotransposon hot spot (RHS) protein, putative | 0,9553 | 1,577E-09 |
| TcCLB.509459.50 | hypothetical protein, conserved | 0,9552 | 2,456E-15 |
| TcCLB.509153.110 | hypothetical protein | 0,9545 | 1,395E-15 |
| TcCLB.511605.80 | mucin-associated surface protein (MASP), putative | 0,9543 | 0,2015 |
| TcCLB.506741.50 | mucin TcMUCII, putative | 0,9538 | 0,001519 |
| TcCLB.509153.90 | dihydrofolate reductase-thymidylate synthase | 0,9537 | 4,509E-28 |
| TcCLB.504769.70 | hypothetical protein, conserved | 0,9536 | 1,005E-27 |
| TcCLB.506607.10 | hypothetical protein, conserved | 0,9535 | 4,929E-06 |
| TcCLB.506667.70 | mucin TcMUCII, putative | 0,9532 | 0,05594 |
| TcCLB.505193.10 | serine/threonine protein phosphatase-like protein, putative | 0,953 | 2,099E-09 |
| TcCLB.508501.200 | hypothetical protein, conserved | 0,953 | 0,001738 |
| TcCLB.511259.50 | trans-sialidase (pseudogene), putative | 0,9526 | 0,002969 |
| TcCLB.506281.20 | mucin TcMUCI, putative | 0,9524 | 0,1446 |
| TcCLB.507529.20 | hypothetical protein, conserved | 0,9523 | 9,21E-23 |
| TcCLB.509217.150 | retrotransposon hot spot protein (RHS, pseudogene), putat | 0,9523 | 0,0001514 |
| TcCLB.508979.50 | trans-sialidase, Group VI, putative | 0,9522 | 0,003003 |
| TcCLB.509977.100 | hypothetical protein, conserved | 0,9517 | 2,111E-23 |
| TcCLB.508745.60 | mucin TcMUCII, putative | 0,9517 | 0,03065 |
| TcCLB.509215.30 | hypothetical protein, conserved | 0,9516 | 6,824E-06 |
| TcCLB.479847.20 | retrotransposon hot spot (RHS) protein, putative | 0,9511 | 0,008296 |
| TcCLB.509645.5 | hypothetical protein, conserved (pseudogene) | 0,951 | 9,393E-09 |
| TcCLB.508961.10 | hypothetical protein, conserved | 0,9508 | 1,735E-14 |
| TcCLB.509601.100 | hypothetical protein, conserved | 0,9507 | 1,732E-16 |
| TcCLB.511047.40 | NADH-cytochrome b5 reductase, putative | 0,9505 | 5,219E-21 |
| TcCLB.506599.224 | mucin TcMUCII, putative | 0,9504 | 0,1569 |
| TcCLB.507007.108 | amino acid transporter, putative | 0,9501 | 8,179E-10 |
| TcCLB.508563.30 | retrotransposon hot spot protein (RHS, pseudogene), putat | 0,9501 | 0,1306 |
| TcCLB.511261.71 | hypothetical protein, conserved (pseudogene) | 0,9499 | 0,00000129 |
| TcCLB.509417.9 | trans-sialidase, putative | 0,9496 | 0,0001509 |
| TcCLB.508387.90 | paraflagellar rod protein, putative | 0,9494 | 5,379E-17 |
| TcCLB.505207.30 | retrotransposon hot spot (RHS) protein, putative | 0,9494 | 1,333E-13 |
| TcCLB.510611.79 | Present in the outer mitochondrial membrane proteome 3 | 0,9494 | 6,256E-09 |
| TcCLB.511227.20 | retrotransposon hot spot protein (RHS, pseudogene), putat | 0,9493 | 1,427E-18 |
| TcCLB.507019.90 | hypothetical protein, conserved | 0,9493 | 0,00000153 |
| TcCLB.506683.100 | retrotransposon hot spot protein (RHS, pseudogene), putat | 0,9493 | 0,00007418 |
| TcCLB.503543.20 | divalent cation transporter, putative (fragment) | 0,9488 | 3,12E-15 |
| TcCLB.504125.40 | ATPase subunit 9, putative | 0,9483 | 1,214E-14 |
| TcCLB.510381.50 | hypothetical protein, conserved | 0,9475 | 6,151E-08 |
| TcCLB.506499.90 | 90 kDa surface protein, putative | 0,9475 | 0,2154 |
| TcCLB.452699.5 | trans-sialidase (pseudogene), putative | 0,9473 | 0,08877 |

| | | | |
|---|---|---|---|
| TcCLB.510943.33 | fatty acyl CoA synthetase, putative | 0,9472 | 2,119E-14 |
| TcCLB.510083.10 | retrotransposon hot spot protein (RHS, pseudogene), putat | 0,9472 | 4,999E-10 |
| TcCLB.510557.60 | mucin-associated surface protein (MASP), putative | 0,9471 | 0,2581 |
| TcCLB.505647.50 | hypothetical protein, conserved | 0,946 | 0,00002944 |
| TcCLB.506673.30 | hypothetical protein, conserved | 0,9459 | 0,00000102 |
| TcCLB.506245.190 | trans-sialidase, Group V, putative | 0,9459 | 0,01691 |
| TcCLB.510935.20 | surface protease GP63, putative | 0,9458 | 0,001785 |
| TcCLB.503431.10 | caltractin, putative | 0,9457 | 0,0003976 |
| TcCLB.503505.20 | MFS transporter, putative | 0,9452 | 1,553E-10 |
| TcCLB.511825.60 | aquaporin, putative | 0,9451 | 1,756E-13 |
| TcCLB.504637.21 | trans-sialidase (pseudogene), putative | 0,9451 | 0,00001449 |
| TcCLB.506813.120 | retrotransposon hot spot protein (RHS, pseudogene), putat | 0,9449 | 0,0007692 |
| TcCLB.508759.60 | mucin-associated surface protein (MASP), putative | 0,9445 | 0,2732 |
| TcCLB.506751.148 | mucin TcMUCII (pseudogene), putative | 0,9443 | 0,2114 |
| TcCLB.508241.104 | hypothetical protein, conserved | 0,9441 | 2,749E-20 |
| TcCLB.509869.30 | mucin TcMUCII, putative | 0,9441 | 0,1878 |
| TcCLB.504003.70 | S-adenosyl-methyltransferase mraW-like protein, putative | 0,944 | 3,954E-16 |
| TcCLB.503735.20 | hypothetical protein | 0,944 | 1,438E-09 |
| TcCLB.508873.454 | mucin TcMUC (pseudogene), putative | 0,944 | 0,1672 |
| TcCLB.510155.70 | heat shock protein 70 (hsp70), putative | 0,9435 | 2,403E-23 |
| TcCLB.510543.60 | retrotransposon hot spot protein (RHS, pseudogene), putat | 0,9434 | 6,37E-10 |
| TcCLB.508177.81 | surface protease GP63 (pseudogene), putative | 0,9433 | 0,00002859 |
| TcCLB.511577.180 | retrotransposon hot spot (RHS) protein, putative | 0,9431 | 1,418E-10 |
| TcCLB.511907.130 | mucin-like glycoprotein, putative | 0,9431 | 0,000242 |
| TcCLB.506183.80 | hypothetical protein | 0,9429 | 0,0769 |
| TcCLB.510095.50 | dispersed gene family protein 1 (DGF-1), putative | 0,9428 | 0,00001126 |
| TcCLB.506295.110 | hypothetical protein, conserved | 0,9426 | 0,00000996 |
| TcCLB.507085.170 | mucin TcMUCII, putative | 0,9425 | 0,008247 |
| TcCLB.511737.6 | hypothetical protein, conserved | 0,9421 | 0,00002612 |
| TcCLB.504689.10 | hypothetical protein | 0,9419 | 0,06484 |
| TcCLB.505779.20 | trans-sialidase, Group II, putative | 0,9414 | 2,721E-06 |
| TcCLB.510101.400 | hypothetical protein, conserved | 0,941 | 4,036E-20 |
| TcCLB.507275.61 | surface antigen TASV-A1 (TASV-A1) | 0,9405 | 0,0002094 |
| TcCLB.506131.84 | mucin-associated surface protein (MASP), putative | 0,9403 | 0,0612 |
| TcCLB.506777.90 | hypothetical protein, conserved | 0,9398 | 4,016E-13 |
| TcCLB.506909.160 | trans-sialidase, Group V, putative | 0,9397 | 0,0006286 |
| TcCLB.508529.10 | retrotransposon hot spot (RHS) protein, putative, (fragmer | 0,9395 | 1,405E-07 |
| TcCLB.506825.230 | hypothetical protein, conserved | 0,9394 | 3,805E-26 |
| TcCLB.506441.20 | hypothetical protein, conserved | 0,9394 | 1,747E-22 |
| TcCLB.503723.50 | hypothetical protein, conserved | 0,9393 | 2,98E-18 |
| TcCLB.507507.10 | hypothetical protein, conserved | 0,9392 | 6,257E-12 |
| TcCLB.511593.30 | mucin-associated surface protein (MASP), putative | 0,9391 | 0,006758 |
| TcCLB.511759.30 | hypothetical protein, conserved | 0,9388 | 5,666E-14 |
| TcCLB.503661.20 | retrotransposon hot spot (RHS) protein, putative | 0,9388 | 0,0302 |
| TcCLB.506927.20 | hypothetical protein, conserved | 0,9387 | 8,348E-11 |

| | | | |
|---|---|---|---|
| TcCLB.506825.20 | hypothetical protein, conserved | 0,9365 | 3,168E-23 |
| TcCLB.506401.90 | centrin, putative | 0,9363 | 1,39E-11 |
| TcCLB.511441.20 | calpain-like cysteine peptidase, putative (fragment) | 0,9363 | 0,1282 |
| TcCLB.508205.48 | hypothetical protein, conserved | 0,9359 | 0,001177 |
| TcCLB.506327.80 | serine/threonine protein kinase, putative | 0,9358 | 1,182E-10 |
| TcCLB.510881.9 | trans-sialidase, putative | 0,9354 | 0,09719 |
| TcCLB.503433.30 | hypothetical protein, conserved | 0,9345 | 1,126E-12 |
| TcCLB.511603.60 | mucin TcMUCII, putative | 0,9344 | 0,01511 |
| TcCLB.506763.140 | hypothetical protein | 0,9342 | 0,2901 |
| TcCLB.418185.40 | hypothetical protein, conserved (pseudogene) | 0,934 | 9,29E-09 |
| TcCLB.509331.60 | hypothetical protein, conserved | 0,934 | 1,724E-07 |
| TcCLB.503723.60 | tartrate-resistant acid phosphatase type 5 precursor, putat | 0,9335 | 1,28E-09 |
| TcCLB.509779.4 | hypothetical protein, conserved | 0,9335 | 1,801E-07 |
| TcCLB.511607.40 | mucin-associated surface protein (MASP), putative | 0,9333 | 0,006832 |
| TcCLB.508521.81 | elongation factor 1-gamma (EF-1-gamma), putative | 0,9333 | 0,03124 |
| TcCLB.508973.60 | hypothetical protein, conserved | 0,933 | 6,12E-14 |
| TcCLB.506791.79 | hypothetical protein, conserved | 0,933 | 8,846E-07 |
| TcCLB.510915.5 | dispersed gene family protein 1 (DGF-1), putative | 0,9329 | 0,01256 |
| TcCLB.511077.20 | beta galactofuranosyl glycosyltransferase (pseudogene), p | 0,9329 | 0,04938 |
| TcCLB.510719.90 | hypothetical protein, conserved | 0,9328 | 2,139E-08 |
| TcCLB.506763.4 | surface protease GP63 (pseudogene), putative | 0,9328 | 0,005071 |
| TcCLB.510087.80 | hypothetical protein, conserved | 0,9323 | 4,881E-16 |
| TcCLB.510347.40 | hypothetical protein, conserved | 0,9322 | 9,642E-18 |
| TcCLB.509207.40 | hypothetical protein, conserved | 0,9318 | 1,357E-35 |
| TcCLB.510155.90 | hypothetical protein, conserved | 0,9318 | 2,788E-28 |
| TcCLB.508961.4 | ADP-ribosylation factor, putative | 0,9315 | 2,243E-06 |
| TcCLB.508429.20 | mucin-like glycoprotein, putative | 0,9315 | 0,1206 |
| TcCLB.477935.9 | calpain-like cysteine peptidase (pseudogene), putative | 0,9314 | 6,542E-06 |
| TcCLB.511921.29 | mucin-associated surface protein (MASP, pseudogene), pu | 0,9311 | 0,1379 |
| TcCLB.508435.20 | mucin TcMUCII, putative | 0,9309 | 0,1365 |
| TcCLB.508899.54 | hypothetical protein, conserved | 0,9307 | 2,804E-13 |
| TcCLB.504249.65 | mucin TcMUCII, putative | 0,9307 | 0,03401 |
| TcCLB.506763.90 | mucin-associated surface protein (MASP, pseudogene), pu | 0,9302 | 0,2697 |
| TcCLB.505647.30 | hypothetical protein, conserved | 0,9295 | 3,309E-13 |
| TcCLB.507645.30 | hypothetical protein, conserved | 0,929 | 6,507E-10 |
| TcCLB.510095.80 | hypothetical protein | 0,929 | 0,0313 |
| TcCLB.511003.60 | 3-ketoacyl-CoA thiolase, putative (ACAT) | 0,9283 | 5,218E-14 |
| TcCLB.511411.16 | hypothetical protein, conserved | 0,9279 | 2,175E-15 |
| TcCLB.508461.240 | chaperone protein DNAJ, putatative | 0,9279 | 2,028E-11 |
| TcCLB.508641.110 | hypothetical protein, conserved | 0,9279 | 1,746E-06 |
| TcCLB.506683.210 | retrotransposon hot spot protein (RHS, pseudogene), putat | 0,9278 | 3,529E-06 |
| TcCLB.511825.190 | hypothetical protein, conserved | 0,9276 | 5,815E-19 |
| TcCLB.511153.19 | surface protease GP63, putative | 0,9268 | 2,42E-12 |
| TcCLB.506941.150 | sphingosine 1-phosphate lyase, putative (SPL) | 0,9265 | 8,934E-11 |
| TcCLB.509683.10 | hypothetical protein | 0,9263 | 1,483E-31 |

| | | | |
|---|---|---|---|
| TcCLB.511431.90 | conserved protein, unknown function | 0,9263 | 0,0178 |
| TcCLB.413293.9 | trans-sialidase, putative (fragment) | 0,9263 | 0,03387 |
| TcCLB.511911.130 | phosphoglucomutase | 0,9259 | 9,986E-13 |
| TcCLB.510025.150 | mucin TcMUCII, putative | 0,9259 | 0,1278 |
| TcCLB.503579.70 | ATPase subunit 9, putative | 0,9256 | 3,859E-13 |
| TcCLB.510187.530 | hypothetical protein, conserved | 0,9255 | 1,489E-16 |
| TcCLB.507491.151 | flagellar calcium-binding protein, putative | 0,925 | 4,997E-18 |
| TcCLB.506067.280 | mucin TcMUCII, putative | 0,925 | 0,01379 |
| TcCLB.503657.70 | tyrosine phosphatase, putative | 0,9248 | 1,044E-09 |
| TcCLB.506593.30 | ribosomal protein S7, putative | 0,9246 | 3,412E-37 |
| TcCLB.507485.20 | hypothetical protein, conserved | 0,9235 | 3,193E-08 |
| TcCLB.510155.220 | trans-sialidase (pseudogene), putative | 0,9234 | 4,765E-06 |
| TcCLB.507615.40 | hypothetical protein, conserved | 0,9229 | 3,398E-22 |
| TcCLB.509287.130 | surface protease GP63 (pseudogene), putative | 0,9229 | 4,186E-06 |
| TcCLB.511259.230 | trans-sialidase (pseudogene), putative | 0,9228 | 0,01611 |
| TcCLB.506337.80 | surface antigen TASV, putative | 0,9225 | 5,211E-17 |
| TcCLB.510155.94 | hypothetical protein, conserved | 0,9222 | 5,662E-11 |
| TcCLB.507747.114 | mucin-associated surface protein (MASP, pseudogene), pu | 0,9222 | 0,08552 |
| TcCLB.506475.90 | hypothetical protein, conserved | 0,9221 | 4,647E-13 |
| TcCLB.507747.195 | glyceraldehyde-3-phosphate dehydrogenase (pseudogene), | 0,9217 | 0,003228 |
| TcCLB.507145.60 | trans-sialidase, Group V, putative | 0,9214 | 0,007394 |
| TcCLB.509065.36 | hypothetical protein, conserved | 0,9213 | 2,319E-06 |
| TcCLB.503975.20 | hypothetical protein | 0,9211 | 9,752E-07 |
| TcCLB.510353.30 | paraflagellar rod component par4, putative | 0,921 | 1,787E-20 |
| TcCLB.506825.170 | lanosterol synthase, putative | 0,9205 | 3,135E-20 |
| TcCLB.505997.80 | trans-sialidase, Group VIII, putative | 0,9201 | 0,01662 |
| TcCLB.509461.80 | hypothetical protein, conserved | 0,919 | 1,017E-11 |
| TcCLB.508317.120 | mannosyl-oligosaccharide 1,2-alpha-mannosidase IB, puta | 0,9189 | 2,462E-14 |
| TcCLB.511571.40 | hypothetical protein, conserved | 0,9187 | 1,346E-15 |
| TcCLB.511647.20 | hypothetical protein, conserved | 0,9187 | 1,638E-06 |
| TcCLB.510643.50 | retrotransposon hot spot protein (RHS, pseudogene), putat | 0,9177 | 2,47E-10 |
| TcCLB.511643.130 | mucin-associated surface protein (MASP, pseudogene), pu | 0,9176 | 0,0001621 |
| TcCLB.509427.10 | trans-sialidase, Group II, putative | 0,9173 | 0,02504 |
| TcCLB.506759.40 | mucin-associated surface protein (MASP), putative | 0,9171 | 0,02235 |
| TcCLB.509525.250 | mucin TcMUCII, putative | 0,9171 | 0,2327 |
| TcCLB.511517.120 | hypothetical protein, conserved | 0,917 | 1,731E-18 |
| TcCLB.511233.59 | hypothetical protein, conserved | 0,9168 | 6,629E-11 |
| TcCLB.511817.50 | hypothetical protein, conserved | 0,9166 | 4,356E-14 |
| TcCLB.508241.80 | hypothetical protein, conserved | 0,9163 | 3,072E-08 |
| TcCLB.508857.160 | threonine synthase, putative | 0,9159 | 1,901E-14 |
| TcCLB.511807.64 | DNA repair protein, putative | 0,9155 | 2,937E-12 |
| TcCLB.507653.20 | trans-sialidase, Group II, putative | 0,9155 | 0,001024 |
| TcCLB.510377.250 | mucin TcMUCII (pseudogene), putative | 0,9154 | 0,1678 |
| TcCLB.511507.70 | calpain-like cysteine peptidase, putative | 0,9153 | 5,736E-10 |
| TcCLB.506837.10 | mucin TcMUCI, putative | 0,9153 | 0,00002292 |

| | | | |
|---|---|---|---|
| TcCLB.508045.20 | hypothetical protein, conserved | 0,9152 | 2,741E-10 |
| TcCLB.508871.10 | trans-sialidase, Group V, putative | 0,9146 | 0,00906 |
| TcCLB.506153.20 | amino acid transporter, putative | 0,9145 | 1,062E-24 |
| TcCLB.511333.4 | Calpain-like protein 1 (CAP5.5) | 0,9144 | 1,047E-11 |
| TcCLB.508435.30 | mucin-associated surface protein (MASP, pseudogene), pu | 0,914 | 0,1334 |
| TcCLB.511851.50 | mucin TcMUCI, putative | 0,9134 | 0,05569 |
| TcCLB.510377.430 | mucin-associated surface protein (MASP), putative | 0,9133 | 0,05643 |
| TcCLB.510823.30 | Isoprenylcysteine alpha-carbonyl methylesterase, putative | 0,9127 | 2,799E-17 |
| TcCLB.507645.40 | hypothetical protein, conserved | 0,9126 | 4,618E-18 |
| TcCLB.505065.30 | carnitine O-palmitoyltransferase (pseudogene), putative | 0,9126 | 0,07443 |
| TcCLB.510503.20 | mucin TcMUCII, putative | 0,9123 | 0,04354 |
| TcCLB.509119.30 | hypothetical protein, conserved | 0,9122 | 3,409E-12 |
| TcCLB.509109.130 | protein tyrosine phosphatase-like protein, putative | 0,9122 | 3,175E-08 |
| TcCLB.506853.40 | hypothetical protein, conserved | 0,9121 | 2,562E-08 |
| TcCLB.507953.210 | hypothetical protein | 0,9121 | 0,1152 |
| TcCLB.511445.10 | calpain-like cysteine peptidase, putative | 0,9116 | 0,0003075 |
| TcCLB.509975.15 | surface protease GP63 (pseudogene), putative | 0,9113 | 0,09413 |
| TcCLB.506195.10 | glucose-6-phosphate dehydrogenase (pseudogene), putativ | 0,9109 | 0,00002014 |
| TcCLB.506289.160 | hypothetical protein | 0,9107 | 0,003796 |
| TcCLB.509737.19 | retrotransposon hot spot protein (RHS, pseudogene), putat | 0,9106 | 0,005512 |
| TcCLB.508175.100 | hypothetical protein | 0,9104 | 6,237E-20 |
| TcCLB.508787.14 | hypothetical protein, conserved | 0,9104 | 0,0004156 |
| TcCLB.507699.149 | mucin TcMUCII, putative | 0,9104 | 0,08584 |
| TcCLB.511047.20 | mucin-like glycoprotein, putative | 0,9098 | 6,12E-14 |
| TcCLB.508567.60 | hypothetical protein | 0,9097 | 5,839E-15 |
| TcCLB.510101.150 | adaptin-related protein-like, putative | 0,9096 | 1,354E-18 |
| TcCLB.507691.20 | dispersed gene family protein 1 (DGF-1), putative | 0,9092 | 1,903E-12 |
| TcCLB.506529.595 | hypothetical protein | 0,9091 | 2,123E-10 |
| TcCLB.510351.90 | dihydrolipoamide branched chain transacylase, putative | 0,9089 | 1,615E-16 |
| TcCLB.511587.30 | retrotransposon hot spot protein (RHS, pseudogene), putat | 0,9087 | 2,082E-24 |
| TcCLB.510889.290 | hypothetical protein | 0,9087 | 0,0002671 |
| TcCLB.507543.30 | retrotransposon hot spot protein (RHS, pseudogene), putat | 0,9081 | 1,444E-20 |
| TcCLB.506885.416 | regulator of chromosome condensation (RCC1), putative | 0,9081 | 3,118E-11 |
| TcCLB.508149.70 | mucin TcMUCII, putative | 0,9079 | 0,1023 |
| TcCLB.510545.40 | mucin TcMUCII, putative | 0,9076 | 0,039 |
| TcCLB.506337.90 | surface antigen TASV, putative (TASV) | 0,9074 | 7,067E-22 |
| TcCLB.508721.4 | hypothetical protein, conserved | 0,9074 | 0,0000204 |
| TcCLB.506499.120 | hypothetical protein | 0,907 | 0,2226 |
| TcCLB.511245.130 | fatty acid elongase, putative | 0,9064 | 6,677E-21 |
| TcCLB.503505.10 | 60S acidic ribosomal protein P2, putative | 0,9064 | 2,674E-12 |
| TcCLB.511395.10 | hypothetical protein, conserved | 0,9061 | 2,337E-09 |
| TcCLB.505949.10 | mucin TcMUCII, putative | 0,9059 | 0,06304 |
| TcCLB.508541.100 | surface protease GP63, putative | 0,9056 | 0,06715 |
| TcCLB.504081.110 | mucin TcMUCII (pseudogene), putative | 0,9052 | 0,1533 |
| TcCLB.509747.90 | hypothetical protein, conserved | 0,9049 | 3,37E-08 |

| | | | |
|---|---|---|---|
| TcCLB.507645.70 | Meckel syndrome type 1 protein, putative (fragment) (mks | 0,9047 | 1,976E-14 |
| TcCLB.427789.20 | Paraflagellar Rod Proteome Component 9, putative (PFC9) | 0,9047 | 5,885E-08 |
| TcCLB.511127.330 | kinetoplastid kinetochore protein 19, putative (kkt19) | 0,9047 | 0,0005923 |
| TcCLB.510483.200 | hypothetical protein | 0,9047 | 0,1635 |
| TcCLB.506017.51 | retrotransposon hot spot (RHS) protein, putative (fragmen | 0,9046 | 3,92E-26 |
| TcCLB.508221.560 | mucin TcMUCII, putative | 0,9043 | 0,3027 |
| TcCLB.507781.10 | hypothetical protein, conserved (pseudogene) | 0,9042 | 0,0001273 |
| TcCLB.511661.90 | ribose-phosphate pyrophosphokinase, putative | 0,9037 | 1,043E-23 |
| TcCLB.510789.20 | mucin TcMUC (pseudogene), putative | 0,9035 | 0,0006265 |
| TcCLB.509429.160 | hypothetical protein | 0,9034 | 5,068E-15 |
| TcCLB.504797.140 | CLC-type chloride channel, putative | 0,9033 | 2,859E-21 |
| TcCLB.508323.50 | hypothetical protein, conserved | 0,9032 | 1,405E-06 |
| TcCLB.508763.20 | mucin TcMUCII, putative | 0,903 | 0,01618 |
| TcCLB.511621.9 | hypothetical protein, conserved | 0,9029 | 4,483E-08 |
| TcCLB.508117.10 | surface protease GP63 (pseudogene), putative | 0,9027 | 0,0004729 |
| TcCLB.509155.30 | hypothetical protein, conserved | 0,9024 | 4,986E-17 |
| TcCLB.508465.110 | acyl-CoA dehydrogenase, putative | 0,9024 | 8,963E-13 |
| TcCLB.509149.60 | 60S ribosomal protein L19, putative | 0,9022 | 5,825E-11 |
| TcCLB.507665.10 | hypothetical protein, conserved | 0,9022 | 6,124E-07 |
| TcCLB.506001.120 | hypothetical protein, conserved (pseudogene) | 0,9021 | 0,02531 |
| TcCLB.507167.150 | retrotransposon hot spot protein (RHS, pseudogene), putat | 0,9019 | 1,194E-07 |
| TcCLB.510733.5 | hypothetical protein, conserved (fragment) | 0,9019 | 0,00004658 |
| TcCLB.506137.10 | surface protease GP63, putative | 0,9015 | 0,01143 |
| TcCLB.508851.130 | hypothetical protein | 0,9012 | 7,664E-18 |
| TcCLB.505779.10 | trans-sialidase, Group II, putative | 0,9011 | 4,378E-06 |
| TcCLB.509539.67 | paraflagellar rod component par4, putative (fragment) | 0,901 | 0,001586 |
| TcCLB.445777.10 | retrotransposon hot spot (RHS) protein, putative, (fragmer | 0,9004 | 6,89E-11 |
| TcCLB.506325.33 | ATP-dependent DEAD/H RNA helicase (pseudogene), putat | 0,9004 | 0,04926 |
| TcCLB.509811.18 | mucin-associated surface protein (MASP, pseudogene), pu | 0,9003 | 0,1052 |
| TcCLB.508027.129 | Hydroxymethylglutaryl-CoA lyase, mitochondrial, putative | 0,9001 | 3,478E-06 |
| TcCLB.506779.150 | histone H2B variant, putative | 0,9 | 2,551E-16 |
| TcCLB.507357.100 | trans-sialidase, Group V, putative | 0,9 | 0,00002614 |
| TcCLB.506909.90 | mucin-associated surface protein (MASP, pseudogene), pu | 0,8998 | 0,1628 |
| TcCLB.508859.74 | SNF2 DNA repair protein, putative | 0,8997 | 3,622E-23 |
| TcCLB.509815.70 | retrotransposon hot spot protein (RHS, pseudogene), putat | 0,8993 | 6,344E-08 |
| TcCLB.506409.10 | hypothetical protein | 0,8992 | 0,1035 |
| TcCLB.508479.80 | retrotransposon hot spot (RHS) protein, putative | 0,8991 | 2,988E-08 |
| TcCLB.508503.60 | protein kinase, putative | 0,8989 | 3,631E-28 |
| TcCLB.508239.40 | palmitoyl acyltransferase 8, putative | 0,8989 | 1,593E-06 |
| TcCLB.508509.51 | mucin-associated surface protein (MASP, pseudogene), pu | 0,8985 | 0,1103 |
| TcCLB.508045.30 | hypothetical protein, conserved | 0,8984 | 3,718E-15 |
| TcCLB.503881.10 | kynureninase, putative | 0,8984 | 4,471E-08 |
| TcCLB.510101.34 | hypothetical protein, conserved | 0,8982 | 1,555E-10 |
| TcCLB.504081.250 | 90 kDa surface protein, putative | 0,8982 | 0,02843 |
| TcCLB.509571.60 | hypothetical protein, conserved | 0,8979 | 1,192E-08 |

| | | | |
|---|---|---|---|
| TcCLB.507317.60 | protein kinase-like protein, putative | 0,8978 | 0,00001731 |
| TcCLB.507807.30 | hypothetical protein | 0,8978 | 0,00153 |
| TcCLB.504741.140 | esterase, putative | 0,8975 | 1,972E-07 |
| TcCLB.509153.30 | hypothetical protein, conserved | 0,8972 | 1,082E-25 |
| TcCLB.510755.89 | kinetoplastid membrane protein KMP-11, putative | 0,8972 | 8,362E-17 |
| TcCLB.510847.70 | dispersed gene family protein 1 (DGF-1), putative | 0,8969 | 0,00009482 |
| TcCLB.511017.25 | hypothetical protein | 0,8967 | 0,001317 |
| TcCLB.509169.30 | hypothetical protein, conserved | 0,8966 | 3,798E-09 |
| TcCLB.508099.20 | serine-alanine-and proline-rich protein, putative | 0,8965 | 0,01113 |
| TcCLB.510781.10 | dispersed gene family protein 1 (DGF-1, pseudogene), puta | 0,8961 | 5,495E-06 |
| TcCLB.511367.150 | hypothetical protein, conserved | 0,896 | 2,917E-11 |
| TcCLB.508677.90 | hypothetical protein, conserved | 0,8953 | 0,0001927 |
| TcCLB.510343.10 | hypothetical protein, conserved | 0,8947 | 0,00002431 |
| TcCLB.430895.4 | hypothetical protein, conserved | 0,8943 | 0,1382 |
| TcCLB.507699.10 | mucin TcMUCII, putative | 0,8942 | 0,01752 |
| TcCLB.506613.9 | surface protease GP63, putative (fragment) | 0,8938 | 0,1004 |
| TcCLB.506969.6 | trans-sialidase (pseudogene), putative | 0,8936 | 0,4427 |
| TcCLB.507589.30 | Checkpoint protein HUS1, putative | 0,8935 | 1,994E-14 |
| TcCLB.510187.150 | guanylate kinase, putative | 0,8935 | 6,354E-13 |
| TcCLB.511729.70 | cationic amino acid transporter, putative | 0,8931 | 0,0001737 |
| TcCLB.504655.30 | protein kinase, putative | 0,8925 | 3,815E-16 |
| TcCLB.507063.290 | hypothetical protein | 0,8924 | 3,224E-13 |
| TcCLB.509599.164 | engulfment and cell motility domain 2, putative | 0,8923 | 7,335E-14 |
| TcCLB.507623.120 | hypothetical protein, conserved | 0,8921 | 1,365E-06 |
| TcCLB.507681.170 | hypothetical protein, conserved | 0,892 | 8,343E-09 |
| TcCLB.506683.190 | trans-sialidase, Group II, putative | 0,892 | 0,00005259 |
| TcCLB.506435.134 | Enriched in surface-labeled proteome protein 11, putative | 0,8919 | 2,257E-08 |
| TcCLB.509887.19 | dispersed gene family protein 1 (DGF-1), putative | 0,8915 | 0,00008872 |
| TcCLB.508247.130 | mucin-associated surface protein (MASP), putative | 0,8914 | 0,1954 |
| TcCLB.507809.160 | hypothetical protein, conserved | 0,891 | 9,082E-11 |
| TcCLB.506339.81 | elongation factor 1-gamma (EF-1-gamma, pseudogene), p | 0,8903 | 0,03417 |
| TcCLB.508411.70 | hypothetical protein, conserved | 0,8901 | 1,535E-10 |
| TcCLB.509391.30 | flagellar calcium-binding protein, putative | 0,89 | 0,227 |
| TcCLB.510947.50 | cyclophilin, putative (CYP) | 0,8899 | 1,42E-09 |
| TcCLB.508433.145 | mucin TcMUCII (pseudogene), putative | 0,8897 | 0,3109 |
| TcCLB.509767.150 | hypothetical protein, conserved | 0,8896 | 9,848E-12 |
| TcCLB.504173.40 | hypothetical protein | 0,8894 | 3,267E-06 |
| TcCLB.507873.40 | hypothetical protein, conserved | 0,8893 | 0,00002022 |
| TcCLB.503827.87 | hypothetical protein, conserved (pseudogene) | 0,8892 | 0,1959 |
| TcCLB.510679.60 | lipase, putative | 0,8889 | 0,00321 |
| TcCLB.506779.70 | hypothetical protein, conserved | 0,8888 | 2,131E-07 |
| TcCLB.511213.30 | surface protease GP63 (pseudogene), putative | 0,8886 | 0,0007345 |
| TcCLB.509777.110 | hypothetical protein, conserved | 0,8884 | 9,706E-06 |
| TcCLB.508777.50 | hypothetical protein | 0,8882 | 4,986E-12 |
| TcCLB.508543.180 | hypothetical protein, conserved | 0,8881 | 1,36E-16 |

| | | | |
|---|---|---|---|
| TcCLB.511827.40 | hypothetical protein, conserved | 0,8879 | 4,221E-11 |
| TcCLB.511433.10 | hypothetical protein, conserved | 0,8879 | 7,521E-07 |
| TcCLB.509607.20 | hypothetical protein, conserved | 0,8875 | 0,07779 |
| TcCLB.508501.260 | hypothetical protein, conserved | 0,8869 | 5,227E-10 |
| TcCLB.510849.20 | trans-sialidase (pseudogene), putative | 0,8869 | 0,00001859 |
| TcCLB.506773.20 | hypothetical protein, conserved | 0,8865 | 3,924E-16 |
| TcCLB.511469.50 | hypothetical protein, conserved | 0,8859 | 7,439E-06 |
| TcCLB.506769.70 | mucin TcMUCII, putative | 0,8858 | 0,1354 |
| TcCLB.509859.70 | hypothetical protein, conserved | 0,8857 | 6,586E-09 |
| TcCLB.504253.50 | hypothetical protein, conserved | 0,8853 | 3,541E-10 |
| TcCLB.508153.740 | hypothetical protein | 0,8846 | 2,815E-15 |
| TcCLB.510759.54 | hypothetical protein | 0,8845 | 0,0001998 |
| TcCLB.508061.20 | trans-sialidase, Group II, putative | 0,8843 | 0,0003101 |
| TcCLB.507849.10 | trans-sialidase (pseudogene), putative | 0,8842 | 0,2201 |
| TcCLB.510105.240 | short chain 3-hydroxyacyl-CoA dehydrogenase, putative | 0,8839 | 8,173E-17 |
| TcCLB.511817.80 | protein kinase, putative | 0,8835 | 2,628E-11 |
| TcCLB.505365.20 | retrotransposon hot spot protein (RHS, pseudogene), putat | 0,8833 | 1,593E-08 |
| TcCLB.508053.10 | trans-sialidase, putative | 0,8826 | 0,2502 |
| TcCLB.510799.60 | mucin TcMUCII, putative | 0,8825 | 0,0097 |
| TcCLB.506795.20 | hypothetical protein, conserved | 0,8823 | 6,992E-11 |
| TcCLB.409117.10 | hypothetical protein, conserved | 0,8823 | 0,00002509 |
| TcCLB.506401.250 | hypothetical protein, conserved | 0,8822 | 3,102E-23 |
| TcCLB.508711.30 | NADH dehydrogenase subunit NI8M, putative | 0,8822 | 5,409E-09 |
| TcCLB.510763.34 | hypothetical protein, conserved | 0,882 | 3,174E-07 |
| TcCLB.506499.130 | hypothetical protein | 0,8819 | 0,2699 |
| TcCLB.508273.100 | mucin TcMUCI, putative | 0,8816 | 0,02342 |
| TcCLB.508737.60 | hypothetical protein, conserved | 0,8814 | 1,493E-07 |
| TcCLB.511405.10 | mucin TcMUCI, putative | 0,8812 | 0,1599 |
| TcCLB.505807.130 | hypothetical protein, conserved | 0,8809 | 0,000242 |
| TcCLB.511335.19 | hypothetical protein, conserved (pseudogene) | 0,8808 | 0,0000123 |
| TcCLB.506931.60 | hypothetical protein | 0,8807 | 6,251E-07 |
| TcCLB.511517.50 | hypothetical protein, conserved | 0,8794 | 4,563E-10 |
| TcCLB.506797.20 | hypothetical protein | 0,8791 | 1,719E-21 |
| TcCLB.506775.60 | hypothetical protein, conserved | 0,8791 | 2,94E-15 |
| TcCLB.509683.80 | adenylosuccinate synthetase, putative | 0,8789 | 4,898E-10 |
| TcCLB.508887.24 | mucin-associated surface protein (MASP, pseudogene), pu | 0,8787 | 0,2179 |
| TcCLB.511603.390 | mucin TcMUCII, putative | 0,8785 | 0,3771 |
| TcCLB.510557.80 | mucin TcMUCII, putative | 0,8784 | 0,09607 |
| TcCLB.511925.20 | hypothetical protein, conserved | 0,8783 | 1,086E-09 |
| TcCLB.509419.100 | mucin-associated surface protein (MASP, pseudogene), pu | 0,8781 | 0,2087 |
| TcCLB.507083.100 | hypothetical protein, conserved | 0,878 | 7,649E-22 |
| TcCLB.503447.40 | retrotransposon hot spot (RHS) protein, putative | 0,878 | 0,0004347 |
| TcCLB.507091.40 | dihydroorotate dehydrogenase, putative (DHODH) | 0,8771 | 0,0001684 |
| TcCLB.506813.59 | mucin TcMUC (pseudogene), putative | 0,8771 | 0,09798 |
| TcCLB.509711.20 | retrotransposon hot spot protein (RHS, pseudogene), putat | 0,8764 | 0,06565 |

| | | | |
|---|---|---|---|
| TcCLB.505997.10 | retrotransposon hot spot (RHS) protein, putative | 0,8762 | 1,238E-06 |
| TcCLB.506201.50 | hypothetical protein, conserved | 0,8761 | 0,0001372 |
| TcCLB.506661.30 | fatty acid elongase, putative | 0,876 | 3,037E-25 |
| TcCLB.507429.30 | mucin TcMUCII, putative | 0,8755 | 0,001873 |
| TcCLB.509065.9 | aldehyde dehydrogenase, putative (fragment) | 0,8754 | 1,398E-17 |
| TcCLB.511151.110 | surface protease GP63, putative (fragment) | 0,8754 | 7,207E-14 |
| TcCLB.505915.10 | retrotransposon hot spot protein (RHS, pseudogene), putat | 0,8752 | 0,00000181 |
| TcCLB.511823.50 | palmitoyl acyltransferase 11, putative | 0,8748 | 3,588E-09 |
| TcCLB.505201.50 | trans-sialidase (pseudogene), putative | 0,8747 | 0,0001586 |
| TcCLB.507559.20 | hypothetical protein, conserved | 0,8745 | 2,537E-10 |
| TcCLB.506493.9 | hypothetical protein, conserved | 0,8745 | 3,273E-09 |
| TcCLB.508179.70 | GPR1/FUN34/yaaH family, putative | 0,8741 | 1,933E-09 |
| TcCLB.511153.40 | surface protease GP63, putative | 0,874 | 6,752E-16 |
| TcCLB.506175.120 | hypothetical protein, conserved | 0,8737 | 0,1296 |
| TcCLB.503699.19 | mucin-associated surface protein (MASP), putative | 0,8737 | 0,1359 |
| TcCLB.507653.50 | retrotransposon hot spot protein (RHS, pseudogene), putat | 0,8736 | 0,00002346 |
| TcCLB.508777.10 | hypothetical protein, conserved | 0,8728 | 1,582E-12 |
| TcCLB.511693.30 | hypothetical protein, conserved | 0,8725 | 1,136E-19 |
| TcCLB.511555.80 | ribonucleoside-diphosphate reductase small chain, putativ | 0,8725 | 5,275E-09 |
| TcCLB.510543.90 | retrotransposon hot spot (RHS) protein, putative | 0,8725 | 0,0007949 |
| TcCLB.506751.90 | trans-sialidase (pseudogene), putative | 0,872 | 0,1174 |
| TcCLB.510137.40 | retrotransposon hot spot protein (RHS, pseudogene), putat | 0,8719 | 1,881E-15 |
| TcCLB.506887.69 | dual specificity protein phosphatase, putative | 0,8718 | 3,258E-09 |
| TcCLB.510693.163 | mucin-associated surface protein (MASP), putative | 0,8713 | 0,0003345 |
| TcCLB.507233.70 | trans-sialidase (pseudogene), putative | 0,871 | 1,663E-10 |
| TcCLB.510377.220 | mucin-associated surface protein (MASP, pseudogene), pu | 0,8707 | 0,1063 |
| TcCLB.508693.190 | hypothetical protein, conserved | 0,8706 | 3,506E-21 |
| TcCLB.508285.10 | retrotransposon hot spot (RHS) protein, putative | 0,8706 | 0,00008605 |
| TcCLB.511827.20 | hypothetical protein, conserved | 0,8703 | 1,396E-21 |
| TcCLB.511219.60 | retrotransposon hot spot protein (RHS, pseudogene), putat | 0,8702 | 2,445E-06 |
| TcCLB.510371.40 | hypothetical protein | 0,8701 | 0,1284 |
| TcCLB.504491.10 | hypothetical protein | 0,8699 | 0,005655 |
| TcCLB.506499.190 | 90 kDa surface protein, putative | 0,8699 | 0,06365 |
| TcCLB.443403.9 | retrotransposon hot spot (RHS) protein, putative | 0,8695 | 3,743E-08 |
| TcCLB.507643.15 | beta galactofuranosyl glycosyltransferase, putative (fragm | 0,8695 | 0,007298 |
| TcCLB.509361.10 | surface protease GP63 (pseudogene), putative | 0,8692 | 0,2539 |
| TcCLB.510309.10 | hypothetical protein, conserved | 0,8689 | 4,702E-09 |
| TcCLB.509341.10 | hypothetical protein, conserved | 0,8686 | 2,002E-13 |
| TcCLB.508781.60 | hypothetical protein, conserved | 0,8685 | 8,641E-07 |
| TcCLB.503609.100 | mucin TcMUCII, putative | 0,8685 | 0,3444 |
| TcCLB.508593.150 | puromycin-sensitive aminopeptidase-like protein, putative | 0,8683 | 1,234E-15 |
| TcCLB.503579.20 | hypothetical protein, conserved | 0,8682 | 1,109E-10 |
| TcCLB.506659.35 | hypothetical protein, conserved | 0,8682 | 0,0001054 |
| TcCLB.508177.70 | hypothetical protein | 0,8674 | 4,027E-16 |
| TcCLB.511019.13 | retrotransposon hot spot (RHS) protein, putative (fragmen | 0,8673 | 1,117E-11 |

| | | | |
|---|---|---|---|
| TcCLB.504097.20 | hypothetical protein, conserved | 0,8672 | 1,231E-07 |
| TcCLB.507429.70 | mucin TcMUCII, putative | 0,8668 | 0,000116 |
| TcCLB.509733.185 | hypothetical protein, conserved (pseudogene) | 0,8662 | 7,864E-09 |
| TcCLB.447899.10 | trans-sialidase, putative | 0,8662 | 0,2548 |
| TcCLB.504125.120 | hypothetical protein, conserved | 0,8661 | 0,0000025 |
| TcCLB.510101.30 | 60S ribosomal protein L28, putative | 0,8657 | 1,847E-21 |
| TcCLB.505993.30 | trans-sialidase, Group VIII, putative | 0,8656 | 0,00008078 |
| TcCLB.509139.20 | hypothetical protein, conserved | 0,8653 | 1,546E-09 |
| TcCLB.511411.24 | hypothetical protein, conserved | 0,8649 | 1,034E-06 |
| TcCLB.510857.60 | cation-transporting ATPase, putative | 0,8648 | 3,01E-12 |
| TcCLB.507063.340 | mucin-associated surface protein (MASP), putative | 0,8647 | 0,02274 |
| TcCLB.508719.80 | hypothetical protein, conserved | 0,8646 | 7,845E-08 |
| TcCLB.506867.30 | hypothetical protein, conserved | 0,8645 | 0,00008674 |
| TcCLB.510483.290 | hypothetical protein | 0,8645 | 0,1477 |
| TcCLB.509179.20 | Macro domain containing protein, putative | 0,8644 | 2,601E-12 |
| TcCLB.506777.10 | ATP-dependent DEAD/H RNA helicase, putative | 0,8642 | 3,571E-10 |
| TcCLB.504797.30 | pseudouridylate synthase-like protein, putative | 0,8641 | 4,134E-11 |
| TcCLB.506145.20 | hypothetical protein, conserved | 0,864 | 1,925E-18 |
| TcCLB.511249.20 | ADP/ATP translocase, putative | 0,864 | 2,908E-14 |
| TcCLB.509331.90 | AMPK subunit beta, putative (AMPKB) | 0,864 | 2,645E-11 |
| TcCLB.510101.300 | hypothetical protein, conserved | 0,8637 | 6,408E-28 |
| TcCLB.507971.20 | syntaxin binding protein (pseudogene), putative | 0,8631 | 8,252E-16 |
| TcCLB.511033.29 | hypothetical protein | 0,863 | 3,698E-09 |
| TcCLB.511545.199 | AMP deaminase, putative | 0,8622 | 1,235E-07 |
| TcCLB.503781.90 | universal minicircle sequence binding protein 2, putative (U | 0,8621 | 4,735E-15 |
| TcCLB.510341.4 | Monooxygenase, putative (fragment) | 0,862 | 0,00001607 |
| TcCLB.509367.15 | tryptophanyl-tRNA synthetase (pseudogene), putative | 0,8617 | 0,01335 |
| TcCLB.506285.110 | mucin-associated surface protein (MASP, pseudogene), pu | 0,8616 | 0,1263 |
| TcCLB.508011.99 | mucin TcMUCII (pseudogene), putative | 0,8616 | 0,2005 |
| TcCLB.505699.20 | retrotransposon hot spot protein (RHS, pseudogene), putat | 0,8614 | 0,00007053 |
| TcCLB.503891.120 | dynein light chain, putative | 0,8613 | 2,161E-06 |
| TcCLB.510197.229 | trans-sialidase, putative | 0,8613 | 0,008964 |
| TcCLB.508745.20 | mucin-associated surface protein (MASP), putative | 0,8613 | 0,01678 |
| TcCLB.507691.61 | trans-sialidase (pseudogene), putative | 0,8612 | 0,0004143 |
| TcCLB.507663.20 | hypothetical protein, conserved | 0,8611 | 1,379E-10 |
| TcCLB.504769.160 | dispersed gene family protein 1 (DGF-1), putative | 0,8607 | 0,00002842 |
| TcCLB.507965.10 | receptor-type adenylate cyclase, putative | 0,8604 | 1,147E-09 |
| TcCLB.506885.80 | hypothetical protein | 0,8603 | 6,944E-11 |
| TcCLB.509695.220 | serine carboxypeptidase (CBP1), putative | 0,8601 | 1,189E-13 |
| TcCLB.511585.230 | trans-sialidase, Group II, putative | 0,8601 | 4,636E-07 |
| TcCLB.510971.5 | retrotransposon hot spot protein (RHS, pseudogene), putat | 0,8596 | 1,863E-07 |
| TcCLB.508995.50 | hypothetical protein, conserved | 0,8595 | 9,321E-06 |
| TcCLB.511907.310 | hypothetical protein, conserved | 0,859 | 4,821E-17 |
| TcCLB.510859.24 | hypothetical protein, conserved | 0,859 | 8,091E-14 |
| TcCLB.480561.10 | trans-sialidase, putative | 0,8589 | 0,0002398 |

| | | | |
|---|---|---|---|
| TcCLB.508789.60 | mucin TcMUCII, putative | 0,8586 | 0,008673 |
| TcCLB.509203.40 | glycosomal membrane protein, putative | 0,8585 | 3,131E-26 |
| TcCLB.510155.150 | hypothetical protein, conserved | 0,8585 | 1,054E-11 |
| TcCLB.510033.22 | surface protease GP63 (pseudogene), putative | 0,8585 | 1,493E-06 |
| TcCLB.503771.90 | mucin-associated surface protein (MASP), putative | 0,8585 | 0,1533 |
| TcCLB.507081.40 | hypothetical protein, conserved | 0,8579 | 9,249E-09 |
| TcCLB.511405.40 | mucin TcMUCII (pseudogene), putative | 0,8579 | 0,1261 |
| TcCLB.506799.30 | mucin-like glycoprotein, putative | 0,8576 | 0,003058 |
| TcCLB.506955.250 | mucin TcMUCII, putative | 0,8576 | 0,07348 |
| TcCLB.510103.17 | hypothetical protein, conserved | 0,8573 | 2,373E-11 |
| TcCLB.503977.15 | surface protease GP63 (pseudogene), putative | 0,8573 | 0,1859 |
| TcCLB.510583.220 | mucin TcMUCII, putative | 0,8572 | 0,3359 |
| TcCLB.507615.73 | hypothetical protein, conserved | 0,8571 | 2,639E-16 |
| TcCLB.507589.14 | tryptophanyl-tRNA synthetase, putative | 0,8569 | 6,792E-34 |
| TcCLB.509897.230 | trans-sialidase (pseudogene), putative | 0,8564 | 0,00001762 |
| TcCLB.504883.60 | hypothetical protein, conserved | 0,8562 | 2,299E-21 |
| TcCLB.507643.90 | beta galactofuranosyl glycosyltransferase (pseudogene), p | 0,8561 | 3,975E-08 |
| TcCLB.508999.220 | calpain cysteine peptidase, putative | 0,856 | 6,055E-10 |
| TcCLB.503495.20 | retrotransposon hot spot protein (RHS, pseudogene), putat | 0,8559 | 0,002057 |
| TcCLB.507981.51 | trans-sialidase (pseudogene), putative | 0,8554 | 0,0648 |
| TcCLB.506245.250 | trans-sialidase, Group V, putative | 0,8551 | 0,02318 |
| TcCLB.510741.140 | hypothetical protein, conserved | 0,8545 | 3,552E-11 |
| TcCLB.509493.120 | hypothetical protein | 0,8545 | 0,0005254 |
| TcCLB.504013.60 | regulatory subunit of protein kinase a-like protein, putative | 0,8544 | 1,682E-13 |
| TcCLB.506725.70 | hypothetical protein, conserved | 0,854 | 1,177E-12 |
| TcCLB.511135.30 | hypothetical protein, conserved | 0,8539 | 2,02E-08 |
| TcCLB.507957.240 | mucin-associated surface protein (MASP), putative | 0,8539 | 0,2186 |
| TcCLB.506775.40 | hypothetical protein, conserved | 0,8537 | 1,147E-11 |
| TcCLB.507891.56 | flagellar calcium-binding protein, putative (fragment) | 0,853 | 6,153E-11 |
| TcCLB.509747.104 | hypothetical protein, conserved | 0,8529 | 1,921E-08 |
| TcCLB.506967.40 | mucin TcMUCII, putative | 0,8528 | 0,1583 |
| TcCLB.511245.150 | fatty acid elongase, putative | 0,8524 | 3,324E-10 |
| TcCLB.511431.54 | hypothetical protein | 0,8523 | 0,006642 |
| TcCLB.504003.40 | hypothetical protein, conserved | 0,8522 | 1,585E-14 |
| TcCLB.509019.10 | trans-sialidase (pseudogene), putative | 0,8518 | 2,662E-09 |
| TcCLB.507737.20 | hypothetical protein, conserved | 0,8518 | 1,884E-08 |
| TcCLB.508173.110 | cysteine protease, putative | 0,8516 | 3,092E-12 |
| TcCLB.508045.40 | hypothetical protein, conserved | 0,8514 | 3,707E-12 |
| TcCLB.507119.20 | hypothetical protein, conserved (pseudogene) | 0,8513 | 2,676E-12 |
| TcCLB.510139.20 | surface protease GP63, putative | 0,851 | 0,0054 |
| TcCLB.511903.190 | STOP axonemal protein, putative | 0,8508 | 1,312E-10 |
| TcCLB.509439.10 | retrotransposon hot spot protein (RHS, pseudogene), putat | 0,8508 | 0,01004 |
| TcCLB.510175.90 | retrotransposon hot spot (RHS) protein, putative | 0,85 | 1,229E-07 |
| TcCLB.506401.120 | 40S ribosomal protein S9, putative | 0,8499 | 3,387E-27 |
| TcCLB.510755.70 | hypothetical protein, conserved | 0,8497 | 2,596E-08 |

| | | | |
|---|---|---|---|
| TcCLB.507501.20 | hypothetical protein, conserved | 0,8496 | 0,07445 |
| TcCLB.508567.30 | hypothetical protein | 0,8492 | 0,00004457 |
| TcCLB.507975.70 | retrotransposon hot spot protein (RHS, pseudogene), putat | 0,8489 | 8,817E-14 |
| TcCLB.504149.220 | retrotransposon hot spot protein (RHS, pseudogene), putat | 0,8488 | 1,334E-08 |
| TcCLB.508219.140 | retrotransposon hot spot protein (RHS, pseudogene), putat | 0,8487 | 0,008961 |
| TcCLB.507993.110 | hypothetical protein, conserved | 0,8486 | 3,377E-09 |
| TcCLB.508173.170 | protein kinase, putative | 0,8484 | 2,421E-16 |
| TcCLB.508973.20 | hypothetical protein, conserved | 0,848 | 1,04E-10 |
| TcCLB.510855.40 | proteasome regulatory non-ATPase subunit, putative | 0,8479 | 2,527E-13 |
| TcCLB.506797.50 | hypothetical protein, conserved | 0,8479 | 6,464E-13 |
| TcCLB.507073.49 | hypothetical protein, conserved | 0,8477 | 9,356E-08 |
| TcCLB.508115.20 | trans-sialidase, putative | 0,8476 | 0,2275 |
| TcCLB.507517.60 | ATPase subunit 9, putative | 0,8473 | 2,579E-12 |
| TcCLB.506227.20 | hypothetical protein, conserved | 0,8473 | 0,00001279 |
| TcCLB.506321.130 | mucin TcMUCII, putative | 0,8472 | 0,09631 |
| TcCLB.507765.40 | hypothetical protein, conserved | 0,8464 | 3,307E-11 |
| TcCLB.511815.160 | hypothetical protein, conserved | 0,846 | 1,232E-17 |
| TcCLB.506247.470 | hypothetical protein, conserved | 0,8456 | 8,831E-19 |
| TcCLB.504797.50 | hypothetical protein, conserved | 0,8456 | 4,159E-13 |
| TcCLB.430895.16 | hypothetical protein, conserved (fragment) | 0,8456 | 0,0007385 |
| TcCLB.447483.4 | Meckel syndrome type 1 protein, putative (fragment) (mks | 0,8456 | 0,002002 |
| TcCLB.503611.40 | ribosomal protein L24, putative | 0,8443 | 0,02552 |
| TcCLB.508079.20 | fatty acyl CoA syntetase 1 (pseudogene), putative | 0,8438 | 1,204E-12 |
| TcCLB.509331.50 | ribosomal RNA methyltransferase, putative | 0,8437 | 3,28E-08 |
| TcCLB.510237.120 | mucin-associated surface protein (MASP), putative | 0,8437 | 0,01314 |
| TcCLB.506951.90 | trans-sialidase, Group IV, putative | 0,8435 | 0,0007741 |
| TcCLB.506559.350 | asparaginyl-tRNA synthetase, putative | 0,8434 | 4,282E-34 |
| TcCLB.509295.10 | dispersed gene family protein 1 (DGF-1), putative | 0,8434 | 0,0006666 |
| TcCLB.505401.19 | dispersed gene family protein 1 (DGF-1), putative (fragme | 0,8428 | 0,00004167 |
| TcCLB.508891.30 | hypothetical protein, conserved | 0,8425 | 2,032E-18 |
| TcCLB.511467.10 | hypothetical protein, conserved | 0,8421 | 7,615E-10 |
| TcCLB.509765.60 | retrotransposon hot spot protein (RHS, pseudogene), putat | 0,8419 | 3,503E-09 |
| TcCLB.510513.70 | hypothetical protein, conserved | 0,8418 | 1,723E-09 |
| TcCLB.508231.174 | hypothetical protein, conserved | 0,8417 | 0,00004484 |
| TcCLB.508443.20 | heat shock protein-like protein, putative | 0,8416 | 9,499E-08 |
| TcCLB.506649.70 | fructose-1,6-bisphosphatase, cytosolic, putative | 0,8411 | 1,445E-13 |
| TcCLB.506741.40 | mucin TcMUCII (pseudogene), putative | 0,8411 | 0,2302 |
| TcCLB.504153.200 | hypothetical protein, conserved | 0,8409 | 1,23E-08 |
| TcCLB.504147.340 | hypothetical protein, conserved | 0,8405 | 8,057E-17 |
| TcCLB.511235.31 | surface protease GP63, putative | 0,8405 | 0,2782 |
| TcCLB.510735.90 | hypothetical protein, conserved | 0,8403 | 1,639E-09 |
| TcCLB.506595.110 | AGP2beta-2, putative | 0,8403 | 5,573E-08 |
| TcCLB.506657.20 | hypothetical protein, conserved | 0,8402 | 1,377E-09 |
| TcCLB.507849.80 | hypothetical protein, conserved | 0,8402 | 1,22E-07 |
| TcCLB.511183.350 | surface protease GP63 (pseudogene), putative | 0,8401 | 0,08141 |

| | | | |
|---|---|---|---|
| TcCLB.509207.150 | kinesin, putative | 0,8398 | 8,218E-23 |
| TcCLB.509857.30 | hypothetical protein, conserved | 0,8394 | 9,627E-08 |
| TcCLB.509407.10 | hypothetical protein | 0,8388 | 0,004402 |
| TcCLB.511825.210 | dephospho-CoA kinase, putative | 0,8382 | 4,528E-08 |
| TcCLB.511839.30 | mucin-associated surface protein (MASP), putative | 0,8381 | 0,09624 |
| TcCLB.511213.89 | dispersed gene family protein 1 (DGF-1), putative (fragme | 0,8379 | 0,00795 |
| TcCLB.507509.80 | hypothetical protein, conserved | 0,8377 | 1,427E-12 |
| TcCLB.509671.189 | hypothetical protein, conserved | 0,8376 | 5,432E-07 |
| TcCLB.508707.10 | amino acid transporter, putative | 0,8374 | 0,003238 |
| TcCLB.511395.14 | hypothetical protein, conserved | 0,8372 | 0,0001878 |
| TcCLB.508505.4 | hypothetical protein, conserved | 0,8371 | 0,0001002 |
| TcCLB.509811.9 | surface protease GP63, putative | 0,8367 | 0,00834 |
| TcCLB.504153.140 | metallo-peptidase, Clan MF, Family M17 | 0,8366 | 1,82E-19 |
| TcCLB.509647.130 | hypothetical protein, conserved | 0,8366 | 5,322E-10 |
| TcCLB.509175.9 | hypothetical protein, conserved | 0,8366 | 0,02212 |
| TcCLB.511817.90 | hypothetical protein, conserved | 0,8365 | 6,344E-08 |
| TcCLB.503617.10 | protein kinase, putative | 0,8361 | 1,877E-16 |
| TcCLB.507889.10 | transaldolase, putative | 0,8358 | 6,309E-12 |
| TcCLB.509211.30 | serine/threonine protein phosphatase, putative | 0,8356 | 1,511E-08 |
| TcCLB.509695.130 | dynein arm light chain, putative | 0,8355 | 6,077E-08 |
| TcCLB.507223.10 | dynein heavy chain, cytosolic, putative | 0,8352 | 0,002648 |
| TcCLB.511871.110 | hypothetical protein, conserved | 0,835 | 5,919E-11 |
| TcCLB.507959.280 | mucin-associated surface protein (MASP), putative | 0,8348 | 0,003894 |
| TcCLB.508555.34 | hypothetical protein, conserved | 0,8347 | 0,0003295 |
| TcCLB.509707.29 | phosphatidylinositol-4-phosphate 5-kinase-like protein | 0,8345 | 0,001269 |
| TcCLB.509649.40 | hypothetical protein | 0,8344 | 3,485E-13 |
| TcCLB.508887.5 | trans-sialidase, putative | 0,8344 | 0,2599 |
| TcCLB.507711.130 | hypothetical protein, conserved | 0,8343 | 0,0001159 |
| TcCLB.511183.270 | mucin TcMUCII (pseudogene), putative | 0,834 | 0,1635 |
| TcCLB.508163.380 | trans-sialidase (pseudogene), putative | 0,8338 | 0,1291 |
| TcCLB.508625.160 | pumilio/PUF RNA binding protein 1, putative (PUF1) | 0,8335 | 1,862E-14 |
| TcCLB.509165.34 | ribosomal P protein AGP2beta-1, putative | 0,8335 | 3,774E-08 |
| TcCLB.510715.140 | mucin-associated surface protein (MASP, pseudogene), pu | 0,8332 | 0,04864 |
| TcCLB.506287.9 | hypothetical protein, conserved | 0,8331 | 1,248E-07 |
| TcCLB.508397.70 | transporter, putative | 0,833 | 8,309E-08 |
| TcCLB.408445.10 | trans-sialidase, putative (fragment) | 0,8329 | 0,09555 |
| TcCLB.509151.130 | deoxyuridine triphosphatase, putative | 0,8328 | 3,474E-17 |
| TcCLB.508465.90 | galactokinase, putative | 0,8326 | 2,876E-13 |
| TcCLB.510603.80 | protein tyrosine phosphatase-like protein, putative | 0,8324 | 1,195E-08 |
| TcCLB.507611.110 | retrotransposon hot spot (RHS) protein, putative | 0,8324 | 5,586E-07 |
| TcCLB.505193.80 | hypothetical protein, conserved | 0,8315 | 1,826E-09 |
| TcCLB.511817.220 | kinesin, putative | 0,8314 | 1,354E-16 |
| TcCLB.507089.240 | hypothetical protein, conserved | 0,831 | 7,953E-07 |
| TcCLB.424795.10 | hypothetical protein, conserved | 0,8306 | 2,234E-07 |
| TcCLB.508277.40 | divalent cation transporter, putative | 0,8305 | 5,808E-28 |

| | | | |
|---|---|---|---|
| TcCLB.510021.200 | mucin TcMUCII, putative | 0,8299 | 0,3201 |
| TcCLB.508253.10 | mucin-associated surface protein (MASP), putative | 0,8298 | 0,0001687 |
| TcCLB.507051.11 | trans-sialidase, putative (fragment) | 0,8296 | 0,102 |
| TcCLB.510357.140 | hypothetical protein, conserved | 0,8294 | 0,00005621 |
| TcCLB.510491.70 | hypothetical protein | 0,8289 | 0,0005951 |
| TcCLB.506181.10 | hypothetical protein, conserved | 0,8288 | 3,707E-08 |
| TcCLB.506595.90 | damage-specific DNA binding protein (pseudogene), putati | 0,8287 | 1,12E-12 |
| TcCLB.506173.80 | hypothetical protein | 0,8287 | 0,1935 |
| TcCLB.511395.120 | UDP-Gal or UDP-GlcNAc-dependent glycosyltransferase, pu | 0,8286 | 0,001568 |
| TcCLB.506435.420 | hypothetical protein, conserved | 0,8283 | 0,00001316 |
| TcCLB.510025.210 | mucin TcMUCII, putative | 0,828 | 0,194 |
| TcCLB.506759.80 | mucin-associated surface protein (MASP), putative | 0,8276 | 0,1401 |
| TcCLB.506271.20 | N-acetyltransferase complex ARD1 subunit (pseudogene), | 0,8276 | 0,1587 |
| TcCLB.509365.11 | tryptophanyl-tRNA synthetase (pseudogene), putative | 0,8275 | 0,001877 |
| TcCLB.506173.200 | retrotransposon hot spot protein (RHS, pseudogene), putat | 0,8273 | 0,0001704 |
| TcCLB.508153.1000 | hypothetical protein, conserved | 0,8271 | 1,993E-06 |
| TcCLB.509753.300 | mucin TcMUCII, putative | 0,8271 | 0,3374 |
| TcCLB.506511.40 | hypothetical protein, conserved | 0,8269 | 2,594E-09 |
| TcCLB.511285.30 | hypothetical protein, conserved | 0,8268 | 2,105E-09 |
| TcCLB.506593.19 | ribosomal protein S7, putative | 0,8266 | 6,103E-29 |
| TcCLB.508093.29 | hypothetical protein, conserved (pseudogene) | 0,8263 | 0,04831 |
| TcCLB.511523.20 | hypothetical protein, conserved | 0,8255 | 7,313E-08 |
| TcCLB.506589.80 | surface protease GP63 (pseudogene), putative | 0,8253 | 0,133 |
| TcCLB.511525.20 | hypothetical protein, conserved | 0,825 | 3,192E-15 |
| TcCLB.507603.90 | hypothetical protein | 0,8249 | 6,356E-11 |
| TcCLB.454787.20 | elongation factor 1-gamma (EF-1-gamma, pseudogene), p | 0,8249 | 0,129 |
| TcCLB.510469.10 | mucin-associated surface protein (MASP), putative | 0,8248 | 0,3163 |
| TcCLB.511809.110 | hypothetical protein, conserved | 0,8247 | 1,739E-14 |
| TcCLB.509905.200 | trans-sialidase, Group II, putative | 0,8247 | 1,154E-13 |
| TcCLB.506201.80 | serine/threonine-protein phosphatase PP1, putative | 0,8246 | 0,006001 |
| TcCLB.510717.10 | surface antigen TASV, putative | 0,8242 | 5,455E-11 |
| TcCLB.506427.10 | trans-sialidase, Group VIII, putative | 0,8235 | 0,01949 |
| TcCLB.509979.20 | hypothetical protein, conserved | 0,8231 | 4,235E-10 |
| TcCLB.510311.90 | quinone oxidoreductase, putative | 0,823 | 2,465E-19 |
| TcCLB.510279.130 | mucin TcMUCII, putative | 0,823 | 0,1803 |
| TcCLB.511847.30 | hypothetical protein, conserved | 0,8225 | 2,42E-08 |
| TcCLB.504787.40 | trans-sialidase, Group V, putative | 0,8225 | 0,04661 |
| TcCLB.506737.161 | mucin TcMUCI, putative | 0,8222 | 0,002443 |
| TcCLB.508221.284 | retrotransposon hot spot protein (RHS, pseudogene), putat | 0,8222 | 0,03921 |
| TcCLB.506667.30 | 90 kDa surface protein, putative | 0,822 | 0,009023 |
| TcCLB.510439.50 | hypothetical protein, conserved | 0,8219 | 4,526E-11 |
| TcCLB.510947.40 | agmatinase, putative | 0,8218 | 3,09E-08 |
| TcCLB.505207.20 | beta galactofuranosyl glycosyltransferase, putative | 0,8218 | 0,002014 |
| TcCLB.509419.190 | mucin-associated surface protein (MASP, pseudogene), pu | 0,8218 | 0,3504 |
| TcCLB.510719.60 | Biotin--acetyl-CoA-carboxylase ligase, putative | 0,8216 | 1,131E-09 |

| | | | |
|---|---|---|---|
| TcCLB.504013.40 | serine acetyltransferase, putative (SAT) | 0,8212 | 8,646E-11 |
| TcCLB.506767.40 | mucin-associated surface protein (MASP), putative | 0,8211 | 0,04238 |
| TcCLB.506757.20 | hypothetical protein, conserved | 0,8207 | 4,006E-09 |
| TcCLB.510101.190 | hypothetical protein, conserved | 0,8206 | 1,078E-08 |
| TcCLB.510731.59 | dual specificity protein phosphatase, putative | 0,8206 | 2,089E-07 |
| TcCLB.510609.70 | protein kinase, putative | 0,8205 | 2,866E-16 |
| TcCLB.511819.30 | hypothetical protein, conserved | 0,8199 | 1,72E-07 |
| TcCLB.510945.50 | hypothetical protein, conserved | 0,8198 | 9,282E-18 |
| TcCLB.509225.30 | trans-sialidase (pseudogene), putative | 0,8198 | 2,123E-08 |
| TcCLB.511807.33 | expression site-associated gene (ESAG-like) protein, putati | 0,8195 | 2,924E-10 |
| TcCLB.510359.420 | mucin-associated surface protein (MASP, pseudogene), pu | 0,8193 | 0,001298 |
| TcCLB.508473.10 | retrotransposon hot spot (RHS) protein, putative | 0,8192 | 2,539E-16 |
| TcCLB.511817.30 | hypothetical protein, conserved | 0,8189 | 0,0000147 |
| TcCLB.509207.120 | hypothetical protein, conserved | 0,8187 | 7,787E-19 |
| TcCLB.509801.20 | hypothetical protein, conserved | 0,8186 | 3,251E-09 |
| TcCLB.506451.50 | trans-sialidase (pseudogene), putative | 0,8183 | 0,06876 |
| TcCLB.504233.4 | trifunctional enzyme alpha subunit, mitochondrial precurso | 0,8182 | 0,08134 |
| TcCLB.509331.80 | hypothetical protein, conserved | 0,8181 | 8,953E-08 |
| TcCLB.506951.30 | trans-sialidase (pseudogene), putative | 0,8181 | 0,01976 |
| TcCLB.507603.140 | leucine-rich repeat protein (LRRP), putative | 0,8179 | 1,553E-10 |
| TcCLB.506697.80 | Present in the outer mitochondrial membrane proteome 2 | 0,8178 | 9,815E-07 |
| TcCLB.509029.40 | hypothetical protein, conserved | 0,8176 | 1,213E-12 |
| TcCLB.504137.80 | adenylyl cyclase-associated protein, putative | 0,8176 | 1,035E-10 |
| TcCLB.509879.40 | fumarate hydratase, putative | 0,8176 | 1,129E-08 |
| TcCLB.510309.40 | 60S acidic ribosomal protein P2, putative | 0,8172 | 0,0000613 |
| TcCLB.507245.10 | retrotransposon hot spot protein (RHS, pseudogene), putat | 0,8168 | 5,324E-07 |
| TcCLB.508539.160 | mucin-associated surface protein (MASP), putative | 0,8163 | 0,01775 |
| TcCLB.508493.50 | mucin-associated surface protein (MASP, pseudogene), pu | 0,8163 | 0,02226 |
| TcCLB.511647.70 | hypothetical protein, conserved | 0,8161 | 5,454E-09 |
| TcCLB.503957.10 | trans-sialidase, putative | 0,8158 | 0,0002354 |
| TcCLB.511285.40 | hypothetical protein, conserved | 0,8152 | 3,285E-08 |
| TcCLB.506343.30 | retrotransposon hot spot protein (RHS, pseudogene), putat | 0,8151 | 1,044E-15 |
| TcCLB.505365.60 | trans-sialidase, Group VIII, putative | 0,815 | 0,3279 |
| TcCLB.505843.10 | glutamate dehydrogenase, putative | 0,8149 | 2,014E-17 |
| TcCLB.509211.10 | hypothetical protein, conserved | 0,8148 | 1,31E-09 |
| TcCLB.510303.204 | hypothetical protein, conserved | 0,8147 | 7,53E-11 |
| TcCLB.506401.80 | hypothetical protein, conserved | 0,8144 | 1,109E-18 |
| TcCLB.510533.50 | hypothetical protein, conserved | 0,8143 | 0,0001812 |
| TcCLB.510729.230 | hypothetical protein, conserved | 0,8142 | 7,768E-13 |
| TcCLB.506485.60 | hypothetical protein, conserved | 0,814 | 6,416E-13 |
| TcCLB.510373.70 | mucin TcMUCII, putative | 0,814 | 0,3433 |
| TcCLB.506281.50 | mucin TcMUCII, putative | 0,8139 | 0,00171 |
| TcCLB.507251.30 | hypothetical protein, conserved | 0,8138 | 1,217E-12 |
| TcCLB.506529.260 | chaperone DNAJ protein, putative | 0,8137 | 4,262E-11 |
| TcCLB.509905.10 | trans-sialidase (pseudogene), putative | 0,8137 | 7,795E-09 |

| | | | |
|---|---|---|---|
| TcCLB.505165.10 | RNA-binding protein, putative | 0,8134 | 0,1469 |
| TcCLB.510039.60 | hypothetical protein, conserved | 0,8133 | 2,832E-18 |
| TcCLB.506649.4 | hypothetical protein, conserved | 0,8132 | 0,00006246 |
| TcCLB.507167.259 | trans-sialidase (pseudogene), putative | 0,8132 | 0,02034 |
| TcCLB.511817.190 | hypothetical protein, conserved | 0,8129 | 1,316E-10 |
| TcCLB.506963.70 | hypothetical protein, conserved | 0,8127 | 3,592E-07 |
| TcCLB.509601.70 | vacuolar proton translocating ATPase subunit A, putative (\ | 0,8126 | 4,026E-17 |
| TcCLB.510861.70 | hypothetical protein, conserved | 0,8123 | 9,844E-09 |
| TcCLB.478463.10 | hypothetical protein | 0,8123 | 0,4678 |
| TcCLB.507501.10 | retrotransposon hot spot (RHS) protein, putative | 0,8121 | 1,994E-51 |
| TcCLB.510945.60 | hypothetical protein, conserved | 0,8116 | 1,047E-14 |
| TcCLB.507755.10 | hypothetical protein | 0,8116 | 1,347E-07 |
| TcCLB.506773.70 | transcription modulator/accessory protein, putative | 0,8106 | 5,145E-19 |
| TcCLB.509997.20 | tryparedoxin 2, putative (TXN2) | 0,8103 | 9,608E-12 |
| TcCLB.511303.64 | hypothetical protein, conserved | 0,8102 | 2,262E-10 |
| TcCLB.506593.40 | hypothetical protein, conserved | 0,8102 | 7,337E-08 |
| TcCLB.511259.80 | mucin TcMUCII, putative | 0,8101 | 0,1133 |
| TcCLB.511727.250 | hypothetical protein, conserved | 0,81 | 4,633E-17 |
| TcCLB.510101.290 | hypothetical protein, conserved | 0,8096 | 3,308E-29 |
| TcCLB.503609.60 | mucin-associated surface protein (MASP), putative | 0,8093 | 0,1144 |
| TcCLB.504339.10 | hypothetical protein, conserved | 0,8092 | 0,008848 |
| TcCLB.509149.20 | zinc finger protein family member, putative (POMP35) | 0,809 | 3,301E-08 |
| TcCLB.510101.410 | hypothetical protein, conserved | 0,8087 | 3,465E-12 |
| TcCLB.507641.250 | hypothetical protein | 0,8087 | 0,0000083 |
| TcCLB.505333.30 | hypothetical protein, conserved (pseudogene) | 0,8086 | 0,03716 |
| TcCLB.506245.210 | mucin TcMUCII, putative | 0,8083 | 0,1138 |
| TcCLB.505763.10 | proton motive ATPase 1, putative | 0,8082 | 4,482E-34 |
| TcCLB.504149.190 | hypothetical protein, conserved (pseudogene) | 0,8081 | 1,452E-16 |
| TcCLB.506779.44 | hypothetical protein, conserved | 0,8081 | 0,0004679 |
| TcCLB.509525.220 | mucin TcMUCII, putative | 0,8081 | 0,4003 |
| TcCLB.508999.120 | trans-sialidase (pseudogene), putative | 0,8079 | 0,03661 |
| TcCLB.504057.150 | hypothetical protein, conserved | 0,8076 | 1,814E-10 |
| TcCLB.507275.50 | hypothetical protein, conserved | 0,8076 | 3,866E-10 |
| TcCLB.507085.60 | mucin TcMUCII, putative | 0,8072 | 0,02007 |
| TcCLB.506863.20 | hypothetical protein | 0,8071 | 0,0000013 |
| TcCLB.510533.70 | protein kinase, putative | 0,807 | 6,016E-07 |
| TcCLB.510643.140 | 60S acidic ribosomal protein P2, putative | 0,8066 | 5,849E-16 |
| TcCLB.511393.79 | hypothetical protein, conserved | 0,8066 | 2,62E-07 |
| TcCLB.508641.240 | hypothetical protein, conserved | 0,8065 | 1,972E-08 |
| TcCLB.506499.220 | 90 kDa surface protein, putative | 0,8064 | 0,06201 |
| TcCLB.510303.210 | 2,4-dienoyl-coa reductase-like protein, putative | 0,8063 | 2,728E-23 |
| TcCLB.506331.50 | trans-sialidase (pseudogene), putative | 0,8062 | 0,03394 |
| TcCLB.506529.508 | glucose-6-phosphate isomerase, glycosomal, putative | 0,8061 | 3,354E-18 |
| TcCLB.509905.170 | trans-sialidase, Group VIII, putative | 0,8058 | 0,0001189 |
| TcCLB.509943.11 | cAMP specific phosphodiesterase, putative | 0,8057 | 0,0612 |

| | | | |
|---|---|---|---|
| TcCLB.503665.40 | surface protease GP63 (pseudogene), putative | 0,8056 | 0,002532 |
| TcCLB.506243.94 | retrotransposon hot spot protein (RHS, pseudogene), putat | 0,8055 | 0,00004179 |
| TcCLB.506999.100 | hypothetical protein, conserved | 0,8046 | 0,00000736 |
| TcCLB.505955.30 | surface protease GP63 (pseudogene), putative | 0,8041 | 0,008589 |
| TcCLB.510621.60 | mucin-associated surface protein (MASP), putative | 0,8039 | 0,1838 |
| TcCLB.508017.20 | UDP-Gal or UDP-GlcNAc-dependent glycosyltransferase (p | 0,8037 | 0,02741 |
| TcCLB.508389.70 | trans-sialidase (pseudogene), putative | 0,8036 | 0,03481 |
| TcCLB.504081.141 | syntaxin binding protein (pseudogene), putative | 0,8034 | 0,1484 |
| TcCLB.504109.160 | electron transfer flavoprotein, putative | 0,803 | 1,018E-18 |
| TcCLB.511773.10 | retrotransposon hot spot protein (RHS, pseudogene), putat | 0,803 | 7,47E-08 |
| TcCLB.504425.10 | trans-sialidase, Group III, putative | 0,8029 | 8,078E-08 |
| TcCLB.510889.4 | hypothetical protein, conserved | 0,8029 | 0,07455 |
| TcCLB.503613.30 | DNA repair protein, putative | 0,8025 | 4,198E-08 |
| TcCLB.510347.60 | protein kinase, putative | 0,8022 | 1,838E-08 |
| TcCLB.510939.25 | mucin TcMUCI, putative | 0,8021 | 0,02787 |
| TcCLB.507085.80 | trans-sialidase (pseudogene), putative | 0,8019 | 9,074E-06 |
| TcCLB.508043.40 | kinesin, putative | 0,8014 | 1,617E-14 |
| TcCLB.506813.130 | beta galactofuranosyl glycosyltransferase (pseudogene), p | 0,8014 | 0,1536 |
| TcCLB.507669.230 | hypothetical protein, conserved | 0,8013 | 1,336E-15 |
| TcCLB.508231.190 | ABC transporter, putative | 0,8013 | 2,109E-12 |
| TcCLB.508799.130 | hypothetical protein, conserved | 0,8009 | 6,108E-16 |
| TcCLB.507009.10 | Gim5A protein, putative | 0,8009 | 8,043E-08 |
| TcCLB.506015.20 | surface protease GP63, putative | 0,8009 | 0,00003877 |
| TcCLB.506813.5 | hypothetical protein | 0,8009 | 0,02491 |
| TcCLB.506741.150 | surface protease GP63 (pseudogene), putative | 0,8009 | 0,3388 |
| TcCLB.507625.99 | gamma-glutamylcysteine synthetase, putative | 0,8003 | 1,656E-12 |
| TcCLB.508163.280 | mucin TcMUCII, putative | 0,8003 | 0,1584 |
| TcCLB.506753.15 | dispersed gene family protein 1 (DGF-1, pseudogene), puta | 0,8002 | 0,4988 |
| TcCLB.509207.140 | hypothetical protein, conserved | 0,8 | 1,139E-13 |
| TcCLB.506885.413 | glyceraldehyde-3-phosphate dehydrogenase, putative | 0,7999 | 3,851E-08 |
| TcCLB.506289.94 | mucin-like glycoprotein, putative | 0,7999 | 0,03908 |
| TcCLB.505025.30 | mucin TcMUCII, putative | 0,7996 | 0,1622 |
| TcCLB.507093.160 | hypothetical protein, conserved | 0,7995 | 1,892E-09 |
| TcCLB.507079.20 | dynein heavy chain, cytosolic, putative | 0,7995 | 5,331E-08 |
| TcCLB.506769.90 | surface protease GP63, putative | 0,7992 | 0,09597 |
| TcCLB.511567.20 | retrotransposon hot spot protein (RHS, pseudogene), putat | 0,799 | 0,001815 |
| TcCLB.506825.120 | hypothetical protein, conserved | 0,7986 | 7,093E-24 |
| TcCLB.506123.60 | hypothetical protein, conserved | 0,7986 | 4,464E-08 |
| TcCLB.507857.60 | Dynein intermediate chain 1, axonemal (DNAI1) | 0,7985 | 1,229E-08 |
| TcCLB.507307.30 | peptidylprolyl isomerase (pseudogene), putative | 0,7982 | 0,2361 |
| TcCLB.506853.70 | hypothetical protein, conserved | 0,7981 | 5,321E-09 |
| TcCLB.510927.20 | trans-sialidase (pseudogene), putative | 0,7979 | 0,0001541 |
| TcCLB.505007.10 | RNA-binding protein, putative | 0,7978 | 0,00001903 |
| TcCLB.506885.40 | hypothetical protein, conserved | 0,7968 | 1,222E-11 |
| TcCLB.511259.130 | mucin TcMUCII, putative | 0,7965 | 0,06831 |

| | | | |
|---|---|---|---|
| TcCLB.509875.140 | retrotransposon hot spot protein (RHS, pseudogene), putat | 0,7965 | 0,08158 |
| TcCLB.421619.10 | hypothetical protein | 0,7964 | 9,241E-11 |
| TcCLB.506351.20 | hypothetical protein, conserved | 0,7961 | 5,501E-06 |
| TcCLB.511531.50 | glucosamine-6-phosphate isomerase, putative | 0,7959 | 4,143E-18 |
| TcCLB.505945.69 | hypothetical protein, conserved | 0,7953 | 0,002619 |
| TcCLB.509733.40 | hypothetical protein, conserved | 0,7948 | 3,603E-09 |
| TcCLB.506363.70 | amidohydrolase, putative | 0,7942 | 0,00001527 |
| TcCLB.504579.20 | trans-sialidase (pseudogene), putative | 0,7941 | 1,058E-07 |
| TcCLB.509549.140 | trans-sialidase, putative | 0,794 | 0,0003032 |
| TcCLB.504255.20 | hypothetical protein | 0,7934 | 1,55E-12 |
| TcCLB.506763.50 | mucin-associated surface protein (MASP), putative | 0,7931 | 0,1384 |
| TcCLB.503447.50 | trans-sialidase, Group IV, putative | 0,7929 | 0,003961 |
| TcCLB.503417.50 | hypothetical protein | 0,7929 | 0,01418 |
| TcCLB.507221.50 | hypothetical protein, conserved | 0,7928 | 8,372E-06 |
| TcCLB.510197.210 | mucin TcMUCII, putative | 0,7927 | 0,1512 |
| TcCLB.506869.54 | hypothetical protein, conserved | 0,7926 | 4,079E-06 |
| TcCLB.509461.30 | hypothetical protein, conserved | 0,7925 | 1,32E-09 |
| TcCLB.509725.126 | N-acetyltransferase complex ARD1 subunit (pseudogene), | 0,7922 | 0,04111 |
| TcCLB.511101.19 | retrotransposon hot spot (RHS) protein, putative | 0,792 | 0,00008483 |
| TcCLB.510693.176 | mucin TcMUC (pseudogene), putative | 0,7918 | 0,02846 |
| TcCLB.507473.10 | hypothetical protein | 0,7915 | 1,346E-13 |
| TcCLB.506337.100 | surface antigen TASV, putative | 0,7915 | 1,419E-10 |
| TcCLB.510089.100 | hypothetical protein | 0,7909 | 4,167E-08 |
| TcCLB.510275.405 | hypothetical protein | 0,7908 | 0,05632 |
| TcCLB.509153.60 | lipin, putative | 0,7907 | 1,971E-15 |
| TcCLB.510207.10 | mucin TcMUCII, putative | 0,7907 | 0,1912 |
| TcCLB.504153.120 | hypothetical protein, conserved | 0,7905 | 1,718E-20 |
| TcCLB.511819.10 | hypothetical protein, conserved | 0,7904 | 3,901E-10 |
| TcCLB.506623.20 | trans-sialidase (pseudogene), putative | 0,7904 | 0,05831 |
| TcCLB.509225.11 | trans-sialidase, putative | 0,7903 | 0,09994 |
| TcCLB.510187.520 | hypothetical protein, conserved | 0,7901 | 3,878E-08 |
| TcCLB.507777.30 | retrotransposon hot spot protein (RHS, pseudogene), putat | 0,7899 | 5,016E-08 |
| TcCLB.452699.10 | retrotransposon hot spot protein (RHS, pseudogene), putat | 0,7899 | 1,608E-06 |
| TcCLB.507811.40 | hypothetical protein, conserved | 0,7898 | 9,631E-13 |
| TcCLB.511257.110 | mucin-like glycoprotein, putative | 0,7898 | 0,04764 |
| TcCLB.508859.40 | leucine-rich repeat protein (LRRP), putative | 0,7895 | 7,225E-07 |
| TcCLB.506223.20 | hypothetical protein, conserved | 0,7894 | 0,00002259 |
| TcCLB.413293.20 | hypothetical protein | 0,7894 | 0,1336 |
| TcCLB.510719.210 | hypothetical protein, conserved | 0,7891 | 1,371E-08 |
| TcCLB.504213.90 | inositol polyphosphate kinase-like protein, putative | 0,789 | 0,06403 |
| TcCLB.509531.10 | trans-sialidase, putative (fragment) | 0,7889 | 0,08377 |
| TcCLB.511809.99 | 40S ribosomal protein S15, putative | 0,7888 | 3,341E-08 |
| TcCLB.506227.205 | hypothetical protein, conserved | 0,7888 | 0,00003147 |
| TcCLB.509223.60 | beta galactofuranosyl glycosyltransferase, putative | 0,7888 | 0,005011 |
| TcCLB.511277.110 | citrate synthase, putative | 0,7887 | 0,08069 |

| | | | |
|---|---|---|---|
| TcCLB.508277.310 | hypothetical protein | 0,7882 | 7,235E-06 |
| TcCLB.508561.10 | dispersed gene family protein 1 (DGF-1), putative | 0,7882 | 0,02405 |
| TcCLB.508789.100 | trans-sialidase (pseudogene), putative | 0,7882 | 0,05544 |
| TcCLB.504105.100 | hypothetical protein | 0,788 | 6,219E-10 |
| TcCLB.509331.110 | hypothetical protein, conserved | 0,7878 | 9,955E-13 |
| TcCLB.510187.120 | hypothetical protein, conserved | 0,7878 | 5,712E-09 |
| TcCLB.503635.10 | protein kinase, putative | 0,7872 | 8,316E-13 |
| TcCLB.505655.20 | hypothetical protein | 0,7872 | 0,07703 |
| TcCLB.506825.30 | hypothetical protein, conserved | 0,7869 | 5,994E-11 |
| TcCLB.504185.30 | hypothetical protein, conserved | 0,7865 | 2,144E-06 |
| TcCLB.503685.5 | RNA-binding protein, putative (fragment) | 0,7859 | 0,1605 |
| TcCLB.506519.60 | Protein kinase domain, putative | 0,7858 | 1,207E-07 |
| TcCLB.504797.40 | thioredoxin, putative | 0,7855 | 4,313E-07 |
| TcCLB.511537.69 | hypothetical protein, conserved | 0,7854 | 1,383E-13 |
| TcCLB.510337.20 | calpain-like cysteine peptidase, putative | 0,7847 | 3,856E-06 |
| TcCLB.503837.30 | hypothetical protein, conserved | 0,7844 | 0,00000661 |
| TcCLB.503831.20 | oxidoreductase, putative | 0,7842 | 0,00001823 |
| TcCLB.507957.229 | mucin TcMUCII, putative | 0,7842 | 0,08436 |
| TcCLB.508929.10 | UDP-Gal or UDP-GlcNAc-dependent glycosyltransferase, pu | 0,7841 | 0,003552 |
| TcCLB.504247.50 | hypothetical protein, conserved | 0,784 | 0,00000227 |
| TcCLB.511641.10 | hypothetical protein, conserved | 0,7832 | 0,00001084 |
| TcCLB.508045.10 | hypothetical protein, conserved | 0,7831 | 3,488E-11 |
| TcCLB.504643.30 | ubiquitin-conjugating enzyme E2, putative | 0,7825 | 5,12E-14 |
| TcCLB.506057.20 | retrotransposon hot spot protein (RHS, pseudogene), putat | 0,7824 | 0,0001104 |
| TcCLB.511649.94 | hypothetical protein, conserved | 0,7823 | 2,774E-07 |
| TcCLB.506321.10 | mucin TcMUCII, putative | 0,7823 | 0,1065 |
| TcCLB.510503.50 | trans-sialidase, Group V, putative | 0,7821 | 0,001265 |
| TcCLB.503665.30 | mucin-associated surface protein (MASP), putative | 0,7821 | 0,04083 |
| TcCLB.511517.130 | hypothetical protein, conserved | 0,782 | 7,427E-12 |
| TcCLB.511649.50 | hypothetical protein, conserved | 0,782 | 1,707E-07 |
| TcCLB.510431.30 | amino acid permease, putative | 0,7819 | 8,392E-09 |
| TcCLB.510719.74 | hypothetical protein, conserved | 0,7819 | 6,887E-06 |
| TcCLB.504231.5 | hypothetical protein, conserved (pseudogene) | 0,7818 | 0,01396 |
| TcCLB.510421.180 | casein kinase I, putative | 0,7817 | 2,412E-06 |
| TcCLB.510203.19 | surface protease GP63 (pseudogene), putative | 0,7817 | 0,1902 |
| TcCLB.506829.30 | hypothetical protein, conserved | 0,7814 | 0,00007205 |
| TcCLB.503747.30 | hypothetical protein | 0,781 | 0,0004637 |
| TcCLB.511277.100 | hypothetical protein, conserved | 0,7809 | 2,852E-06 |
| TcCLB.508175.220 | hypothetical protein, conserved | 0,7804 | 4,833E-11 |
| TcCLB.508543.90 | plasma membrane Ca2 ATPase | 0,7803 | 7,436E-15 |
| TcCLB.506925.64 | hypothetical protein, conserved | 0,7801 | 1,16E-13 |
| TcCLB.510497.114 | mucin-associated surface protein (MASP, pseudogene), pu | 0,7801 | 0,2685 |
| TcCLB.509233.160 | alpha-tubulin N-acetyltransferase, putative | 0,78 | 7,377E-07 |
| TcCLB.510135.10 | retrotransposon hot spot protein (RHS, pseudogene), putat | 0,7798 | 0,00003352 |
| TcCLB.504217.19 | mitochondrial carrier protein, putative | 0,7795 | 1,358E-06 |

| | | | |
|---|---|---|---|
| TcCLB.511427.19 | conserved protein, unknown function (fragment) | 0,7795 | 0,07063 |
| TcCLB.508221.490 | mucin-associated surface protein (MASP), putative | 0,7794 | 0,3869 |
| TcCLB.507237.60 | surface protease GP63 (pseudogene), putative | 0,7787 | 0,01182 |
| TcCLB.511315.56 | hypothetical protein, conserved | 0,7787 | 0,05332 |
| TcCLB.507881.20 | retrotransposon hot spot (RHS) protein, putative | 0,7783 | 1,22E-10 |
| TcCLB.510879.190 | hypothetical protein, conserved | 0,7781 | 8,496E-08 |
| TcCLB.506357.120 | ethanolamine phosphotransferase, putative | 0,7781 | 4,005E-07 |
| TcCLB.511771.10 | retrotransposon hot spot protein (RHS, pseudogene), putat | 0,7779 | 1,481E-10 |
| TcCLB.507849.70 | hypothetical protein | 0,7778 | 1,235E-16 |
| TcCLB.510679.30 | metal-ion transporter, putative | 0,7778 | 6,219E-12 |
| TcCLB.427247.10 | serine/threonine protein kinase, putative | 0,7778 | 0,02329 |
| TcCLB.506671.10 | mucin-associated surface protein (MASP), putative | 0,7778 | 0,0597 |
| TcCLB.506421.10 | retrotransposon hot spot (RHS) protein, putative | 0,7776 | 2,867E-18 |
| TcCLB.510375.10 | mucin-associated surface protein (MASP), putative | 0,7776 | 0,207 |
| TcCLB.506695.40 | hypothetical protein, conserved | 0,7775 | 5,542E-06 |
| TcCLB.511877.10 | surface antigen TASV-B2 (TASV-B2) | 0,7773 | 0,0004472 |
| TcCLB.506035.4 | trans-sialidase, putative (fragment) | 0,7773 | 0,002561 |
| TcCLB.509233.60 | hypothetical protein, conserved | 0,7771 | 1,603E-12 |
| TcCLB.508041.10 | SKP1-like protein | 0,7769 | 0,00554 |
| TcCLB.509551.20 | Putrescine-cadaverine transporter 1, putative (POT1.2) | 0,7766 | 8,422E-12 |
| TcCLB.504423.20 | electon transport protein SCO1/SCO2, putative | 0,7765 | 1,15E-11 |
| TcCLB.503925.3 | hypothetical protein, conserved | 0,7764 | 0,00007053 |
| TcCLB.510571.39 | actin 2, putative (fragment) | 0,7762 | 2,475E-07 |
| TcCLB.413133.10 | mucin TcMUCII (pseudogene), putative | 0,7762 | 0,05679 |
| TcCLB.509157.80 | DnaJ homolog, putative | 0,7759 | 9,873E-07 |
| TcCLB.511321.40 | hypothetical protein, conserved | 0,7757 | 3,447E-10 |
| TcCLB.511393.88 | hypothetical protein, conserved (fragment) | 0,7755 | 3,297E-10 |
| TcCLB.507547.9 | mucin TcMUCI, putative | 0,7753 | 0,0001531 |
| TcCLB.509933.20 | hypothetical protein, conserved | 0,7746 | 6,053E-08 |
| TcCLB.509231.10 | cytochrome P450, putative | 0,7745 | 4,449E-13 |
| TcCLB.461515.10 | mucin TcMUCI, putative (fragment) | 0,7742 | 1,528E-09 |
| TcCLB.511655.60 | hypothetical protein, conserved | 0,7735 | 0,0007979 |
| TcCLB.511395.90 | serine/threonine protein kinase, putative | 0,7732 | 1,594E-09 |
| TcCLB.508157.50 | trans-sialidase, putative | 0,7732 | 0,0005791 |
| TcCLB.510275.50 | mucin TcMUCII, putative | 0,7729 | 0,128 |
| TcCLB.510349.90 | Lys-63-specific deubiquitinase BRCC36, putative | 0,7724 | 8,286E-08 |
| TcCLB.506559.30 | hypothetical protein, conserved | 0,7724 | 3,624E-07 |
| TcCLB.509029.140 | hypothetical protein, conserved | 0,7723 | 5,077E-09 |
| TcCLB.507221.30 | hypothetical protein, conserved | 0,7723 | 0,00003168 |
| TcCLB.422921.10 | hypothetical protein, conserved | 0,7722 | 0,0003959 |
| TcCLB.510047.100 | mucin-associated surface protein (MASP, pseudogene), pu | 0,7722 | 0,02949 |
| TcCLB.510303.140 | surface protease GP63, putative | 0,772 | 3,164E-12 |
| TcCLB.422867.10 | trans-sialidase, Group II, putative | 0,772 | 0,007265 |
| TcCLB.510101.200 | zinc finger ccch and cchc domain-containing protein | 0,7719 | 1,753E-12 |
| TcCLB.507095.10 | hypothetical protein (fragment) | 0,7719 | 0,005398 |

| | | | |
|---|---|---|---|
| TcCLB.508665.14 | heat shock protein, putative | 0,7716 | 1,518E-17 |
| TcCLB.510963.50 | 6-phosphofructo-2-kinase/fructose-2,6-bisphosphatas, puta | 0,7716 | 3,555E-09 |
| TcCLB.508895.40 | hypothetical protein, conserved | 0,7713 | 4,904E-08 |
| TcCLB.509227.10 | mucin TcMUCI, putative | 0,7712 | 0,02109 |
| TcCLB.507957.50 | mucin TcMUCII, putative | 0,7708 | 0,1035 |
| TcCLB.510791.60 | mucin-associated surface protein (MASP, pseudogene), pu | 0,7707 | 0,001864 |
| TcCLB.503423.5 | retrotransposon hot spot (RHS) protein, putative (fragmen | 0,7707 | 0,1487 |
| TcCLB.509753.110 | mucin TcMUCII, putative | 0,7707 | 0,2974 |
| TcCLB.506713.70 | retrotransposon hot spot protein (RHS, pseudogene), putat | 0,77 | 2,299E-08 |
| TcCLB.510661.129 | hypothetical protein, conserved | 0,7692 | 2,167E-06 |
| TcCLB.510345.30 | hypothetical protein | 0,7692 | 0,001523 |
| TcCLB.508023.120 | hypothetical protein, conserved (pseudogene) | 0,7691 | 1,263E-13 |
| TcCLB.507229.29 | dispersed gene family protein 1 (DGF-1), putative | 0,7691 | 0,09847 |
| TcCLB.504249.111 | mucin-associated surface protein (MASP), putative | 0,769 | 0,07296 |
| TcCLB.508503.70 | hypothetical protein, conserved | 0,7689 | 1,708E-08 |
| TcCLB.506191.20 | retrotransposon hot spot protein (RHS, pseudogene), putat | 0,7684 | 0,0000296 |
| TcCLB.510155.190 | hypothetical protein, conserved | 0,7682 | 5,85E-10 |
| TcCLB.511693.90 | electron-transfer-flavoprotein, alpha polypeptide, putative | 0,7681 | 4,018E-14 |
| TcCLB.506535.10 | retrotransposon hot spot (RHS) protein, putative (fragmen | 0,7679 | 5,599E-06 |
| TcCLB.485335.10 | retrotransposon hot spot protein (RHS, pseudogene), putat | 0,7676 | 0,0002157 |
| TcCLB.505163.50 | hypothetical protein | 0,7676 | 0,09389 |
| TcCLB.511767.30 | retrotransposon hot spot protein (RHS, pseudogene), putat | 0,7671 | 1,287E-09 |
| TcCLB.509897.250 | retrotransposon hot spot protein (RHS, pseudogene), putat | 0,766 | 5,917E-06 |
| TcCLB.506743.110 | protein tyrosine phosphatase, putative | 0,7658 | 0,00004093 |
| TcCLB.508047.80 | mucin TcMUCII (pseudogene), putative | 0,7658 | 0,08104 |
| TcCLB.507253.70 | N-acetyltransferase complex ARD1 subunit (pseudogene), | 0,7658 | 0,2271 |
| TcCLB.508547.201 | ubiquitin-protein ligase, putative (fragment) | 0,7653 | 0,009148 |
| TcCLB.511401.40 | mucin-associated surface protein (MASP, pseudogene), pu | 0,7653 | 0,0198 |
| TcCLB.508869.59 | mucin-associated surface protein (MASP), putative | 0,7649 | 0,2705 |
| TcCLB.511577.120 | hypothetical protein, conserved | 0,7645 | 2,335E-06 |
| TcCLB.508383.10 | Present in the outer mitochondrial membrane proteome 3 | 0,7643 | 0,0001649 |
| TcCLB.506525.162 | hypothetical protein, conserved | 0,7641 | 0,002312 |
| TcCLB.508563.20 | trans-sialidase, Group II, putative | 0,764 | 0,003241 |
| TcCLB.509099.140 | hypothetical protein, conserved | 0,7638 | 0,00001439 |
| TcCLB.509337.15 | hypothetical protein, conserved | 0,7633 | 3,313E-12 |
| TcCLB.503681.20 | ethanolamine phosphotransferase, putative | 0,7632 | 5,14E-08 |
| TcCLB.511823.79 | hypothetical protein, conserved | 0,7632 | 2,514E-06 |
| TcCLB.508707.310 | leucine rich repeat protein, putative | 0,7631 | 8,85E-20 |
| TcCLB.508277.220 | hypothetical protein | 0,7626 | 4,848E-22 |
| TcCLB.511153.60 | palmitoyl acyltransferase 9, putative | 0,7626 | 1,458E-06 |
| TcCLB.510155.170 | hypothetical protein, conserved | 0,7626 | 0,00008586 |
| TcCLB.506683.140 | beta galactofuranosyl glycosyltransferase (pseudogene), p | 0,7621 | 1,338E-06 |
| TcCLB.508193.45 | trans-sialidase (pseudogene), putative | 0,762 | 0,0003607 |
| TcCLB.506265.130 | hypothetical protein, conserved | 0,7619 | 1,217E-10 |
| TcCLB.509149.70 | hypothetical protein, conserved | 0,7618 | 4,529E-10 |

| | | | |
|---|---|---|---|
| TcCLB.510755.65 | hypothetical protein | 0,7618 | 0,0003807 |
| TcCLB.511523.70 | Present in the outer mitochondrial membrane proteome 3: | 0,7617 | 0,0002506 |
| TcCLB.509769.63 | Ran-binding protein, putative | 0,7616 | 2,876E-07 |
| TcCLB.511389.120 | hypothetical protein, conserved | 0,7613 | 3,134E-07 |
| TcCLB.505193.20 | hypothetical protein, conserved | 0,7613 | 0,00000232 |
| TcCLB.508503.100 | hypothetical protein, conserved | 0,7613 | 0,000314 |
| TcCLB.507907.10 | trans-sialidase, putative | 0,7613 | 0,2747 |
| TcCLB.508315.5 | mucin TcMUCII, putative (fragment) | 0,7612 | 0,2039 |
| TcCLB.507317.40 | hypothetical protein, conserved | 0,7611 | 8,188E-09 |
| TcCLB.504183.20 | trans-sialidase, Group II, putative | 0,7608 | 0,0002671 |
| TcCLB.511155.10 | hypothetical protein, conserved (pseudogene) | 0,7601 | 0,00003476 |
| TcCLB.507667.20 | hypothetical protein, conserved | 0,7599 | 1,999E-19 |
| TcCLB.503633.11 | mucin-associated surface protein (MASP), putative (fragm | 0,7591 | 0,1636 |
| TcCLB.508913.10 | trans-sialidase (pseudogene), putative | 0,7589 | 0,00006618 |
| TcCLB.508239.20 | hypothetical protein, conserved | 0,7588 | 1,028E-16 |
| TcCLB.511361.30 | hypothetical protein, conserved | 0,7586 | 0,0004498 |
| TcCLB.510713.90 | trans-sialidase, Group V, putative | 0,7585 | 0,001162 |
| TcCLB.506443.100 | trans-sialidase, Group IV, putative | 0,7583 | 0,00007806 |
| TcCLB.504233.70 | mucin TcMUCI, putative | 0,7583 | 0,2746 |
| TcCLB.506633.80 | hypothetical protein, conserved | 0,758 | 4,681E-12 |
| TcCLB.510529.30 | hypothetical protein, conserved | 0,7574 | 8,051E-16 |
| TcCLB.511283.140 | hypothetical protein, conserved | 0,7569 | 1,867E-14 |
| TcCLB.510323.40 | heat shock protein 20, putative | 0,7568 | 7,977E-10 |
| TcCLB.506777.100 | hypothetical protein, conserved | 0,7567 | 0,00009773 |
| TcCLB.508741.20 | hypothetical protein, conserved | 0,7565 | 4,919E-07 |
| TcCLB.507691.50 | retrotransposon hot spot (RHS) protein, putative | 0,7565 | 6,56E-07 |
| TcCLB.506199.10 | hypothetical protein, conserved | 0,7565 | 0,0001064 |
| TcCLB.508677.120 | hypothetical protein, conserved | 0,7564 | 0,0005863 |
| TcCLB.511077.130 | mucin TcMUCII, putative | 0,7563 | 0,2002 |
| TcCLB.508881.50 | hypothetical protein, conserved | 0,756 | 0,00000111 |
| TcCLB.509115.60 | mucin-like glycoprotein, putative | 0,7559 | 0,03813 |
| TcCLB.509179.130 | hypothetical protein, conserved | 0,7558 | 3,913E-07 |
| TcCLB.506821.20 | ATP-dependent Clp protease subunit, heat shock protein 1( | 0,7557 | 8,493E-18 |
| TcCLB.511807.140 | hypothetical protein, conserved | 0,7555 | 3,926E-12 |
| TcCLB.507993.60 | hypothetical protein, conserved | 0,7555 | 1,873E-06 |
| TcCLB.509087.10 | retrotransposon hot spot protein (RHS, pseudogene), putat | 0,7555 | 0,008832 |
| TcCLB.506767.280 | retrotransposon hot spot protein (RHS, pseudogene), putat | 0,7555 | 0,2274 |
| TcCLB.508163.20 | mucin TcMUCII, putative | 0,7555 | 0,2917 |
| TcCLB.508165.100 | mucin TcMUCII, putative | 0,7554 | 0,1509 |
| TcCLB.511067.30 | transketolase, putative | 0,7546 | 2,22E-17 |
| TcCLB.507893.9 | dispersed gene family protein 1 (DGF-1), putative (fragme | 0,7542 | 0,008161 |
| TcCLB.511739.10 | hypothetical protein, conserved | 0,7541 | 3,813E-07 |
| TcCLB.510361.200 | mucin TcMUCII, putative | 0,7539 | 0,2907 |
| TcCLB.507035.80 | trans-sialidase (pseudogene), putative | 0,7538 | 1,201E-07 |
| TcCLB.507853.20 | dynein beta chain, flagellar outer arm (pseudogene), putat | 0,7538 | 0,00002709 |

| | | | |
|---|---|---|---|
| TcCLB.511807.120 | hypothetical protein, conserved | 0,7537 | 5,112E-11 |
| TcCLB.508461.10 | Present in the outer mitochondrial membrane proteome 6 | 0,7537 | 2,652E-08 |
| TcCLB.511367.70 | dihydrolipoamide acetyltransferase, putative | 0,7536 | 4,153E-10 |
| TcCLB.509141.20 | hypothetical protein | 0,7533 | 0,00003979 |
| TcCLB.503665.10 | trans-sialidase (pseudogene), putative | 0,7533 | 0,1088 |
| TcCLB.510417.40 | mucin-associated surface protein (MASP), putative (fragm | 0,753 | 0,4055 |
| TcCLB.507587.24 | dispersed gene family protein 1 (DGF-1), putative (fragme | 0,7528 | 0,00001221 |
| TcCLB.508389.114 | mucin TcMUCII, putative | 0,7526 | 0,284 |
| TcCLB.509005.10 | axoneme central apparatus protein (fragment) (PF16) | 0,7521 | 0,0001465 |
| TcCLB.511855.10 | trans-sialidase, putative | 0,752 | 0,002364 |
| TcCLB.509791.180 | hypothetical protein, conserved | 0,7518 | 1,81E-07 |
| TcCLB.508119.160 | hypothetical protein, conserved | 0,7515 | 2,78E-08 |
| TcCLB.510985.20 | retrotransposon hot spot protein (RHS, pseudogene), putat | 0,7512 | 6,326E-12 |
| TcCLB.506401.360 | hypothetical protein, conserved | 0,7509 | 0,0007878 |
| TcCLB.506481.30 | hypothetical protein, conserved | 0,7503 | 2,41E-08 |
| TcCLB.504025.10 | hypothetical protein, conserved | 0,7496 | 0,0006968 |
| TcCLB.506845.40 | hypothetical protein, conserved (pseudogene) | 0,7496 | 0,000834 |
| TcCLB.509725.60 | glycine dehydrogenase [decarboxylating] (pseudogene), pu | 0,7495 | 8,976E-07 |
| TcCLB.508153.560 | hypothetical protein, conserved | 0,7495 | 0,00001711 |
| TcCLB.508641.200 | hypothetical protein, conserved | 0,7494 | 9,164E-09 |
| TcCLB.509871.80 | dispersed gene family protein 1 (DGF-1), putative | 0,7494 | 0,00001244 |
| TcCLB.508293.160 | trans-sialidase (pseudogene), putative | 0,749 | 0,3473 |
| TcCLB.510565.130 | hypothetical protein | 0,7486 | 2,134E-08 |
| TcCLB.509633.60 | serine/threonine-protein phosphatase PP1, putative | 0,7486 | 0,001401 |
| TcCLB.508173.47 | autophagy-related protein 8 (ATG8) | 0,7485 | 8,787E-09 |
| TcCLB.420293.31 | hypothetical protein | 0,7485 | 0,05332 |
| TcCLB.506001.40 | trans-sialidase, Group V, putative | 0,748 | 0,01345 |
| TcCLB.507065.108 | mucin-associated surface protein (MASP, pseudogene), pu | 0,7479 | 0,1014 |
| TcCLB.508237.9 | protein kinase, putative | 0,7477 | 0,0005104 |
| TcCLB.506285.40 | mucin-associated surface protein (MASP, pseudogene), pu | 0,7477 | 0,01734 |
| TcCLB.511231.34 | hypothetical protein, conserved | 0,7476 | 5,815E-07 |
| TcCLB.506133.68 | mucin TcMUCII (pseudogene), putative | 0,7476 | 0,1729 |
| TcCLB.510155.20 | KREL1 | 0,7474 | 8,442E-12 |
| TcCLB.507559.100 | hypothetical protein, conserved | 0,7473 | 1,735E-18 |
| TcCLB.504017.54 | hypothetical protein, conserved | 0,7473 | 3,105E-08 |
| TcCLB.508331.30 | Checkpoint protein HUS1, putative | 0,7473 | 0,00001771 |
| TcCLB.503447.20 | trans-sialidase, Group II, putative | 0,7472 | 0,008435 |
| TcCLB.508873.420 | Present in the outer mitochondrial membrane proteome 1( | 0,747 | 1,169E-11 |
| TcCLB.509109.30 | hypothetical protein, conserved | 0,7457 | 5,163E-06 |
| TcCLB.504039.240 | mucin TcMUCII, putative | 0,7457 | 0,2055 |
| TcCLB.506657.40 | ADP/ATP translocase, putative | 0,7456 | 4,841E-11 |
| TcCLB.508501.270 | hypothetical protein, conserved | 0,7456 | 4,443E-09 |
| TcCLB.410589.10 | AMP deaminase, putative | 0,7454 | 0,0000948 |
| TcCLB.508959.4 | cytochrome c, putative | 0,7453 | 1,596E-09 |
| TcCLB.511105.29 | dispersed gene family protein 1 (DGF-1), putative | 0,7452 | 0,004116 |

| | | | |
|---|---|---|---|
| TcCLB.510497.50 | mucin-associated surface protein (MASP), putative | 0,7452 | 0,04428 |
| TcCLB.507699.37 | mucin-associated surface protein (MASP, pseudogene), pu | 0,745 | 0,174 |
| TcCLB.510061.20 | hypothetical protein | 0,745 | 0,4744 |
| TcCLB.506053.5 | amino acid transporter, putative (fragment) | 0,7443 | 1,132E-09 |
| TcCLB.506169.10 | glycine dehydrogenase [decarboxylating] (pseudogene), pu | 0,7443 | 0,009441 |
| TcCLB.508775.40 | trans-sialidase (pseudogene), putative | 0,7443 | 0,02046 |
| TcCLB.510791.11 | hypothetical protein | 0,7441 | 0,007788 |
| TcCLB.503715.10 | splicing factor TSR1, putative | 0,7438 | 6,167E-14 |
| TcCLB.507847.10 | mucin TcMUCII (pseudogene), putative | 0,7438 | 0,00514 |
| TcCLB.508983.20 | trans-sialidase (pseudogene), putative | 0,7437 | 0,00425 |
| TcCLB.507049.190 | hypothetical protein, conserved | 0,7435 | 1,789E-12 |
| TcCLB.510349.70 | hypothetical protein, conserved | 0,7434 | 0,0001002 |
| TcCLB.507849.60 | hypothetical protein, conserved | 0,7432 | 1,188E-11 |
| TcCLB.506301.50 | succinyl-coA:3-ketoacid-coenzyme A transferase, mitochor | 0,7429 | 1,476E-11 |
| TcCLB.507835.60 | retrotransposon hot spot protein (RHS, pseudogene), putat | 0,7429 | 0,00009351 |
| TcCLB.507953.200 | 90 kDa surface protein, putative | 0,7429 | 0,06601 |
| TcCLB.506735.80 | hypothetical protein, conserved | 0,7422 | 1,442E-10 |
| TcCLB.511257.70 | surface protease GP63, putative | 0,7422 | 0,00001088 |
| TcCLB.508569.70 | hypothetical protein, conserved | 0,7421 | 1,457E-13 |
| TcCLB.510039.100 | hypothetical protein, conserved | 0,7419 | 0,0002144 |
| TcCLB.509067.70 | pyrroline-5-carboxylate synthetase-like protein, putative | 0,7418 | 3,459E-13 |
| TcCLB.508241.40 | hypothetical protein | 0,7417 | 1,709E-09 |
| TcCLB.508325.60 | retrotransposon hot spot (RHS) protein, putative | 0,7417 | 1,539E-06 |
| TcCLB.510625.109 | mucin-associated surface protein (MASP), putative | 0,7417 | 0,4717 |
| TcCLB.510577.30 | hypothetical protein | 0,7415 | 1,445E-08 |
| TcCLB.511349.110 | retrotransposon hot spot protein (RHS, pseudogene), putat | 0,7415 | 0,0316 |
| TcCLB.510181.100 | acid phosphatase, putative (AP) | 0,7414 | 0,00005399 |
| TcCLB.510175.125 | hypothetical protein | 0,7414 | 0,00393 |
| TcCLB.507015.50 | retrotransposon hot spot protein (RHS, pseudogene), putat | 0,7408 | 2,484E-06 |
| TcCLB.506259.30 | retrotransposon hot spot protein (RHS, pseudogene), putat | 0,7407 | 2,586E-08 |
| TcCLB.508015.30 | hypothetical protein | 0,7407 | 0,004669 |
| TcCLB.509445.39 | glutamate dehydrogenase, putative | 0,7405 | 4,098E-14 |
| TcCLB.506773.120 | mRNA cap guanine-N7 methyltransferase, putative | 0,7405 | 9,264E-10 |
| TcCLB.508027.120 | hypothetical protein, conserved | 0,7403 | 1,371E-09 |
| TcCLB.421555.9 | mucin-associated surface protein (MASP), putative | 0,7403 | 0,0004173 |
| TcCLB.504427.40 | hypothetical protein, conserved | 0,74 | 0,0001091 |
| TcCLB.508041.30 | 40S ribosomal protein S15A, putative | 0,7398 | 3,241E-25 |
| TcCLB.503649.20 | mucin TcMUC, putative | 0,7397 | 0,4132 |
| TcCLB.509533.30 | retrotransposon hot spot protein (RHS, pseudogene), putat | 0,7396 | 2,959E-06 |
| TcCLB.511589.180 | hypothetical protein, conserved | 0,7395 | 5,366E-08 |
| TcCLB.510861.60 | hypothetical protein, conserved | 0,7394 | 7,596E-15 |
| TcCLB.505763.19 | P-type H -ATPase, putative | 0,7393 | 4,299E-25 |
| TcCLB.510717.20 | surface antigen TASV, putative | 0,7392 | 4,321E-10 |
| TcCLB.503565.29 | hypothetical protein, conserved | 0,7392 | 8,883E-09 |
| TcCLB.504425.100 | hypothetical protein, conserved | 0,7386 | 0,00001896 |

| | | | |
|---|---|---|---|
| TcCLB.510613.20 | kinesin, putative | 0,7385 | 8,011E-09 |
| TcCLB.504239.410 | mucin TcMUCII, putative | 0,7385 | 0,1393 |
| TcCLB.509105.4 | Generative cell specific 1 protein, putative (GCS1) | 0,7384 | 0,00001733 |
| TcCLB.506777.20 | cytoplasmic translation machinery associated protein, puta | 0,7381 | 4,705E-08 |
| TcCLB.509753.250 | mucin-associated surface protein (MASP), putative | 0,7381 | 0,07536 |
| TcCLB.511613.60 | mucin-associated surface protein (MASP), putative | 0,738 | 0,0163 |
| TcCLB.503821.19 | hypothetical protein, conserved | 0,7379 | 1,364E-06 |
| TcCLB.511529.210 | hypothetical protein, conserved | 0,7375 | 0,00000149 |
| TcCLB.510285.74 | hypothetical protein, conserved | 0,7375 | 0,0001048 |
| TcCLB.509549.90 | alpha-adaptin-like (pseudogene), putative | 0,7371 | 2,072E-17 |
| TcCLB.507047.130 | hypothetical protein, conserved | 0,7371 | 6,536E-07 |
| TcCLB.505919.20 | trans-sialidase, Group II, putative | 0,7371 | 0,0002524 |
| TcCLB.510275.60 | mucin-associated surface protein (MASP), putative | 0,7371 | 0,117 |
| TcCLB.505193.60 | hypothetical protein, conserved | 0,737 | 0,00000926 |
| TcCLB.507317.20 | hypothetical protein | 0,737 | 0,00001401 |
| TcCLB.511871.170 | trans-sialidase (pseudogene), putative | 0,737 | 0,001214 |
| TcCLB.511121.11 | retrotransposon hot spot (RHS) protein, putative (fragmen | 0,7367 | 0,0004474 |
| TcCLB.508165.30 | mucin-associated surface protein (MASP), putative | 0,7367 | 0,124 |
| TcCLB.508587.50 | hypothetical protein | 0,7366 | 0,1024 |
| TcCLB.507669.104 | transcription elongation factor, putative | 0,7363 | 9,097E-07 |
| TcCLB.507253.30 | hypothetical protein, conserved (pseudogene) | 0,7361 | 0,0001519 |
| TcCLB.509775.9 | protein kinase, putative (fragment) | 0,7359 | 0,001069 |
| TcCLB.506331.80 | beta galactofuranosyl glycosyltransferase (pseudogene), p | 0,7357 | 0,06138 |
| TcCLB.509549.100 | hypothetical protein, conserved | 0,7352 | 4,212E-14 |
| TcCLB.510791.30 | mucin-associated surface protein (MASP, pseudogene), pu | 0,7352 | 1,345E-07 |
| TcCLB.503559.70 | cystinosin, putative | 0,7351 | 0,000321 |
| TcCLB.510155.80 | mitochondrial-processing peptidase subunit beta, putative | 0,7348 | 1,744E-14 |
| TcCLB.507981.61 | hypothetical protein | 0,7347 | 0,1069 |
| TcCLB.506855.20 | hypothetical protein, conserved | 0,7346 | 9,607E-07 |
| TcCLB.510947.44 | hypothetical protein, conserved | 0,7346 | 0,00005987 |
| TcCLB.510887.70 | hypothetical protein, conserved | 0,7344 | 2,492E-07 |
| TcCLB.460625.9 | trans-sialidase (pseudogene), putative | 0,734 | 0,2377 |
| TcCLB.506529.80 | hypothetical protein, conserved | 0,7339 | 1,224E-15 |
| TcCLB.506671.50 | trans-sialidase (pseudogene), putative | 0,7338 | 0,02537 |
| TcCLB.506759.20 | mucin-associated surface protein (MASP), putative | 0,7338 | 0,119 |
| TcCLB.507615.60 | hypothetical protein, conserved | 0,7337 | 1,086E-06 |
| TcCLB.510241.100 | hypothetical protein, conserved | 0,7335 | 9,061E-06 |
| TcCLB.508543.60 | hypothetical protein, conserved | 0,7329 | 7,492E-08 |
| TcCLB.509331.40 | hypothetical protein, conserved | 0,7326 | 0,00002729 |
| TcCLB.510461.20 | trans-sialidase (pseudogene), putative | 0,7325 | 0,2837 |
| TcCLB.506401.230 | hypothetical protein, conserved | 0,7324 | 2,313E-14 |
| TcCLB.508285.40 | retrotransposon hot spot protein (RHS, pseudogene), putat | 0,7319 | 0,07978 |
| TcCLB.511817.230 | hypothetical protein, conserved | 0,7318 | 8,091E-14 |
| TcCLB.507213.60 | retrotransposon hot spot protein (RHS, pseudogene), putat | 0,7318 | 5,498E-06 |
| TcCLB.503839.39 | hypothetical protein, conserved | 0,7317 | 0,0002883 |

| | | | |
|---|---|---|---|
| TcCLB.431889.10 | hypothetical protein, conserved (fragment) | 0,7315 | 0,1545 |
| TcCLB.506697.20 | cyclic nucleotide phosphodiesterase | 0,7311 | 0,00002305 |
| TcCLB.508173.100 | Monooxygenase, putative | 0,7308 | 1,502E-11 |
| TcCLB.510947.10 | GDP-mannose pyrophosphorylase | 0,7307 | 1,63E-11 |
| TcCLB.509765.30 | retrotransposon hot spot protein (RHS, pseudogene), putat | 0,7307 | 0,0002706 |
| TcCLB.509207.50 | hypothetical protein, conserved | 0,7304 | 2,652E-15 |
| TcCLB.504797.34 | N-acetyltransferase, putative | 0,7301 | 1,104E-14 |
| TcCLB.510205.50 | mucin-associated surface protein (MASP), putative | 0,73 | 0,273 |
| TcCLB.505193.40 | SUMO1/Ulp2, putative | 0,7297 | 4,52E-10 |
| TcCLB.508621.30 | choline/carnitine O-acyltransferase (pseudogene), putative | 0,7295 | 3,837E-12 |
| TcCLB.507499.56 | N-acetyltransferase complex ARD1 subunit (pseudogene), | 0,7293 | 0,2115 |
| TcCLB.506349.53 | retrotransposon hot spot protein (RHS, pseudogene), putat | 0,7291 | 2,647E-13 |
| TcCLB.506529.314 | hypothetical protein, conserved | 0,7286 | 0,001702 |
| TcCLB.506269.70 | mucin TcMUCII, putative | 0,7285 | 0,1507 |
| TcCLB.508509.30 | surface protease GP63 (pseudogene), putative | 0,7283 | 0,09405 |
| TcCLB.508539.110 | mucin-associated surface protein (MASP), putative | 0,7281 | 0,07766 |
| TcCLB.506399.10 | sigma-adaptin 3, putative | 0,7281 | 0,1485 |
| TcCLB.503505.14 | PTP1-interacting protein, 39 kDa, putative | 0,7279 | 0,00003561 |
| TcCLB.509601.30 | hypothetical protein | 0,7277 | 1,083E-07 |
| TcCLB.509943.20 | cAMP specific phosphodiesterase, putative | 0,7277 | 9,072E-07 |
| TcCLB.510359.610 | trans-sialidase, Group II, putative | 0,7277 | 0,03096 |
| TcCLB.506743.69 | hypothetical protein, conserved | 0,7275 | 0,0004615 |
| TcCLB.434931.10 | paraxonemal rod protein PAR2, putative | 0,7273 | 5,494E-11 |
| TcCLB.510243.10 | hypothetical protein, conserved | 0,7273 | 0,000058 |
| TcCLB.508205.39 | dynein light chain, putative | 0,7271 | 9,112E-06 |
| TcCLB.510889.130 | hypothetical protein | 0,7266 | 0,008811 |
| TcCLB.510503.160 | mucin TcMUCII, putative | 0,7265 | 0,02403 |
| TcCLB.506485.80 | oxidoreductase, putative | 0,7264 | 1,726E-12 |
| TcCLB.507957.140 | trans-sialidase (pseudogene), putative | 0,7263 | 0,002225 |
| TcCLB.510715.90 | trans-sialidase, Group V, putative | 0,7259 | 0,0575 |
| TcCLB.510853.96 | trans-sialidase (pseudogene), putative | 0,7258 | 1,249E-07 |
| TcCLB.509611.150 | hypothetical protein, conserved | 0,7254 | 1,411E-08 |
| TcCLB.506847.30 | retrotransposon hot spot protein (RHS, pseudogene), putat | 0,7254 | 3,639E-06 |
| TcCLB.509979.100 | hypothetical protein, conserved | 0,7254 | 0,00001111 |
| TcCLB.510403.30 | trans-sialidase, Group II, putative | 0,7251 | 0,08349 |
| TcCLB.508601.20 | hypothetical protein, conserved | 0,7249 | 6,457E-08 |
| TcCLB.510153.5 | hypothetical protein, conserved | 0,7244 | 0,05593 |
| TcCLB.507875.159 | 60S acidic ribosomal protein P2 (pseudogene), putative | 0,7241 | 3,309E-13 |
| TcCLB.507663.80 | hypothetical protein, conserved | 0,724 | 8,031E-08 |
| TcCLB.511571.10 | ATP-dependent DEAD/H RNA helicase, putative | 0,7239 | 0,01308 |
| TcCLB.511491.9 | hypothetical protein, conserved | 0,7237 | 0,001179 |
| TcCLB.510381.40 | hypothetical protein, conserved | 0,7236 | 6,739E-10 |
| TcCLB.509571.40 | hypothetical protein, conserved | 0,7236 | 5,766E-09 |
| TcCLB.509153.139 | hypothetical protein, conserved | 0,7235 | 8,41E-10 |
| TcCLB.511385.100 | hypothetical protein, conserved | 0,7235 | 2,492E-08 |

| | | | |
|---|---|---|---|
| TcCLB.511255.370 | mucin TcMUCII, putative | 0,7233 | 0,2592 |
| TcCLB.511825.80 | I/6 autoantigen, putative | 0,7232 | 2,958E-10 |
| TcCLB.511509.40 | hypothetical protein, conserved | 0,7232 | 0,0007347 |
| TcCLB.503697.140 | hypothetical protein | 0,7231 | 0,0002096 |
| TcCLB.508207.160 | hypothetical protein, conserved | 0,7229 | 0,0001027 |
| TcCLB.506065.10 | retrotransposon hot spot protein (RHS, pseudogene), putat | 0,7229 | 0,03568 |
| TcCLB.510197.10 | trans-sialidase (pseudogene), putative | 0,7226 | 0,00004005 |
| TcCLB.506457.4 | hypothetical protein, conserved | 0,7226 | 0,07659 |
| TcCLB.509897.80 | mucin-associated surface protein (MASP, pseudogene), pu | 0,7221 | 0,1879 |
| TcCLB.503925.60 | hypothetical protein | 0,722 | 2,43E-08 |
| TcCLB.506499.105 | mucin-associated surface protein (MASP, pseudogene), pu | 0,7219 | 0,1619 |
| TcCLB.508325.160 | retrotransposon hot spot (RHS) protein, putative | 0,7218 | 4,836E-14 |
| TcCLB.503889.20 | mucin TcMUC (pseudogene), putative | 0,7218 | 0,0001424 |
| TcCLB.503979.20 | hypothetical protein, conserved | 0,7217 | 3,751E-06 |
| TcCLB.506601.51 | mucin TcMUCII (pseudogene), putative | 0,7216 | 0,2003 |
| TcCLB.508575.20 | hypothetical protein | 0,7214 | 0,1376 |
| TcCLB.511417.80 | phosphate-repressible phosphate permease, putative | 0,7211 | 1,924E-11 |
| TcCLB.510989.69 | hypothetical protein, conserved | 0,7211 | 0,00728 |
| TcCLB.503587.47 | hypothetical protein | 0,7211 | 0,1165 |
| TcCLB.507471.31 | elongation factor 1-gamma (EF-1-gamma, pseudogene), p | 0,7204 | 0,429 |
| TcCLB.504115.10 | trans-sialidase, Group VIII, putative | 0,7203 | 1,161E-06 |
| TcCLB.504035.41 | mucin-associated surface protein (MASP, pseudogene), pu | 0,7202 | 0,6909 |
| TcCLB.510235.6 | hypothetical protein | 0,72 | 0,05927 |
| TcCLB.506345.100 | beta galactofuranosyl glycosyltransferase (pseudogene), p | 0,7198 | 0,00007082 |
| TcCLB.506493.90 | cysteine peptidase, Clan CA, family C2, putative | 0,7197 | 1,598E-13 |
| TcCLB.510565.160 | hypothetical protein, conserved | 0,7196 | 0,00007051 |
| TcCLB.511225.20 | retrotransposon hot spot protein (RHS, pseudogene), putat | 0,7195 | 4,824E-12 |
| TcCLB.503905.10 | myosin heavy chain, putative | 0,7195 | 8,345E-09 |
| TcCLB.508207.120 | hypothetical protein, conserved | 0,7195 | 0,006869 |
| TcCLB.509855.10 | hypothetical protein, conserved | 0,7194 | 3,039E-13 |
| TcCLB.510713.170 | mucin-associated surface protein (MASP), putative | 0,7193 | 0,1584 |
| TcCLB.504427.270 | hypothetical protein, conserved | 0,7192 | 1,723E-08 |
| TcCLB.509033.20 | beta galactofuranosyl glycosyltransferase, putative | 0,7186 | 0,0001018 |
| TcCLB.503955.50 | hypothetical protein, conserved | 0,7185 | 3,586E-07 |
| TcCLB.511349.80 | retrotransposon hot spot protein (RHS, pseudogene), putat | 0,7185 | 0,2688 |
| TcCLB.511923.80 | mucin-associated surface protein (MASP, pseudogene), pu | 0,7182 | 0,4261 |
| TcCLB.511785.5 | hypothetical protein | 0,7182 | 0,5343 |
| TcCLB.506721.30 | calpain-like cysteine peptidase, putative | 0,7181 | 4,47E-11 |
| TcCLB.507641.236 | hypothetical protein, conserved | 0,7181 | 5,897E-09 |
| TcCLB.508965.30 | hypothetical protein | 0,718 | 4,605E-07 |
| TcCLB.511585.50 | hypothetical protein, conserved | 0,7178 | 1,587E-07 |
| TcCLB.507997.14 | trans-sialidase, putative (fragment) | 0,7178 | 0,001615 |
| TcCLB.510891.30 | retrotransposon hot spot protein (RHS, pseudogene), putat | 0,7177 | 8,846E-07 |
| TcCLB.504155.10 | mucin-associated surface protein (MASP), putative | 0,7177 | 0,4079 |
| TcCLB.509213.160 | protein kinase, putative | 0,7173 | 5,93E-11 |

| | | | |
|---|---|---|---|
| TcCLB.505343.10 | trans-sialidase (pseudogene), putative | 0,7173 | 0,1255 |
| TcCLB.505333.10 | N-acetyltransferase complex ARD1 subunit (pseudogene), | 0,7171 | 0,05283 |
| TcCLB.503955.40 | trans-sialidase, Group II, putative | 0,7169 | 2,455E-09 |
| TcCLB.511363.4 | hypothetical protein, conserved | 0,7168 | 2,905E-07 |
| TcCLB.510773.50 | hypothetical protein | 0,7168 | 0,005218 |
| TcCLB.503613.50 | PA domain containing protein, putative (ESP7) | 0,7167 | 1,885E-06 |
| TcCLB.506401.100 | DNA polymerase sigma, putative | 0,7165 | 7,297E-15 |
| TcCLB.508061.30 | retrotransposon hot spot protein (RHS, pseudogene), putat | 0,7165 | 0,0002699 |
| TcCLB.511861.50 | trans-sialidase, Group VIII, putative | 0,7159 | 0,03876 |
| TcCLB.511469.70 | prohibitin, putative | 0,7155 | 4,566E-11 |
| TcCLB.506989.70 | flagellar radial spoke protein-like, putative | 0,7155 | 1,44E-07 |
| TcCLB.506939.70 | retrotransposon hot spot protein (RHS, pseudogene), putat | 0,7153 | 0,02401 |
| TcCLB.511603.330 | mucin TcMUCII, putative | 0,7153 | 0,2193 |
| TcCLB.504069.60 | GPI transamidase component GAA1, putative | 0,7152 | 1,374E-07 |
| TcCLB.506779.10 | dynein regulatory complex subunit, putative | 0,7151 | 0,00009831 |
| TcCLB.506625.80 | cAMP-specific phosphodiesterase, putative | 0,715 | 1,691E-16 |
| TcCLB.509035.20 | surface protease GP63, putative | 0,7149 | 0,1162 |
| TcCLB.509525.380 | mucin TcMUCII, putative | 0,7148 | 0,3637 |
| TcCLB.510477.10 | surface protease GP63, putative | 0,7147 | 0,0005686 |
| TcCLB.507031.110 | hypothetical protein, conserved | 0,7146 | 8,785E-11 |
| TcCLB.506891.10 | Metal-binding domain-containing protein | 0,7144 | 0,00000333 |
| TcCLB.508577.150 | hypothetical protein, conserved | 0,7144 | 0,004001 |
| TcCLB.507093.250 | RNA-binding protein, putative | 0,7138 | 8,186E-14 |
| TcCLB.508567.100 | hypothetical protein, conserved | 0,7137 | 1,132E-14 |
| TcCLB.506327.60 | hypothetical protein, conserved | 0,7137 | 0,00001371 |
| TcCLB.508153.140 | protein kinase, putative | 0,7135 | 0,000663 |
| TcCLB.511649.160 | hypothetical protein, conserved | 0,7134 | 1,728E-08 |
| TcCLB.511535.30 | hypothetical protein, conserved | 0,7133 | 6,52E-12 |
| TcCLB.511663.40 | trans-sialidase (pseudogene), putative | 0,7133 | 0,0006908 |
| TcCLB.506265.45 | chaperone DNAJ protein, putative | 0,7124 | 0,0004518 |
| TcCLB.508119.40 | hypothetical protein | 0,7123 | 0,1517 |
| TcCLB.508951.90 | hypothetical protein, conserved | 0,7117 | 0,00005016 |
| TcCLB.508241.30 | hypothetical protein, conserved | 0,7115 | 1,374E-11 |
| TcCLB.508273.10 | AMP deaminase, putative | 0,7113 | 1,458E-11 |
| TcCLB.503837.5 | mucin TcMUCI, putative (fragment) | 0,7113 | 0,01137 |
| TcCLB.441729.9 | hypothetical protein (fragment) | 0,711 | 0,01995 |
| TcCLB.503909.84 | calpain-like cysteine peptidase, putative | 0,7109 | 6,693E-08 |
| TcCLB.510311.180 | hypothetical protein, conserved | 0,7104 | 3,116E-07 |
| TcCLB.511603.70 | mucin TcMUCII, putative | 0,7104 | 0,06718 |
| TcCLB.511773.35 | hypothetical protein, conserved (pseudogene) | 0,7102 | 0,001975 |
| TcCLB.507747.160 | mucin TcMUCII, putative | 0,7102 | 0,07154 |
| TcCLB.506321.300 | hypothetical protein, conserved | 0,7101 | 6,043E-10 |
| TcCLB.507071.160 | mucin TcMUCII, putative | 0,7101 | 0,2931 |
| TcCLB.507219.9 | trans-sialidase, putative | 0,71 | 0,02784 |
| TcCLB.508221.30 | mucin-associated surface protein (MASP), putative | 0,7097 | 0,2815 |

| | | | |
|---|---|---|---|
| TcCLB.506401.200 | hypothetical protein, conserved | 0,7093 | 0,00002854 |
| TcCLB.506281.90 | mucin TcMUCII, putative | 0,7092 | 0,3491 |
| TcCLB.509601.50 | hypothetical protein, conserved | 0,7088 | 7,381E-12 |
| TcCLB.511889.20 | retrotransposon hot spot protein (RHS, pseudogene), putat | 0,7087 | 6,984E-08 |
| TcCLB.506375.10 | tryptophanyl-tRNA synthetase, putative | 0,7086 | 0,1189 |
| TcCLB.510435.50 | AMP deaminase (pseudogene), putative | 0,7084 | 1,772E-06 |
| TcCLB.511807.80 | RNA capping enzyme, nuclear (Cgm1) | 0,7083 | 8,703E-17 |
| TcCLB.510411.40 | trans-sialidase, putative | 0,7083 | 0,06494 |
| TcCLB.508777.190 | hypothetical protein, conserved | 0,7081 | 0,0004399 |
| TcCLB.511849.20 | mucin TcMUCI, putative | 0,708 | 0,01008 |
| TcCLB.511801.50 | hypothetical protein, conserved | 0,7079 | 5,539E-08 |
| TcCLB.459793.10 | mucin TcMUCI, putative | 0,7078 | 0,2123 |
| TcCLB.504827.150 | hypothetical protein, conserved | 0,7076 | 2,958E-06 |
| TcCLB.506759.210 | mucin TcMUCII, putative | 0,7073 | 0,345 |
| TcCLB.503953.30 | protein kinase, putative | 0,707 | 1,959E-07 |
| TcCLB.508731.60 | adenylosuccinate synthetase, putative | 0,7068 | 1,798E-07 |
| TcCLB.508975.4 | hypothetical protein, conserved (fragment) | 0,7066 | 0,0002807 |
| TcCLB.511603.130 | hypothetical protein, conserved | 0,7063 | 0,3382 |
| TcCLB.508241.90 | hypothetical protein, conserved | 0,7059 | 1,059E-11 |
| TcCLB.503865.50 | ankyrin, putative | 0,7059 | 1,627E-10 |
| TcCLB.511127.429 | hypothetical protein, conserved | 0,7059 | 0,0007014 |
| TcCLB.506265.90 | hypothetical protein, conserved | 0,7058 | 0,0001548 |
| TcCLB.508137.40 | hypothetical protein, conserved | 0,7058 | 0,1063 |
| TcCLB.510089.5 | mucin TcMUCII (pseudogene), putative | 0,7056 | 0,4476 |
| TcCLB.511903.120 | transketolase, putative | 0,7055 | 1,689E-18 |
| TcCLB.508771.34 | hypothetical protein, conserved | 0,7055 | 0,01522 |
| TcCLB.510359.480 | mucin-associated surface protein (MASP, pseudogene), pu | 0,7055 | 0,4482 |
| TcCLB.509399.20 | metacaspase, putative | 0,7052 | 0,0003435 |
| TcCLB.508153.130 | enoyl-CoA hydratase, mitochondrial precursor, putative | 0,7051 | 9,828E-10 |
| TcCLB.504081.40 | mucin-associated surface protein (MASP), putative | 0,7051 | 0,09437 |
| TcCLB.510349.20 | hypothetical protein, conserved | 0,7048 | 4,233E-06 |
| TcCLB.506401.160 | hypothetical protein, conserved | 0,7046 | 0,00005637 |
| TcCLB.503541.5 | cAMP-specific phosphodiesterase, putative | 0,7045 | 0,1195 |
| TcCLB.509753.202 | hypothetical protein | 0,7044 | 0,3039 |
| TcCLB.506249.70 | ABC transporter, putative | 0,7043 | 6,568E-10 |
| TcCLB.511415.30 | retrotransposon hot spot protein (RHS, pseudogene), putat | 0,7043 | 1,537E-07 |
| TcCLB.507237.210 | mucin TcMUCII, putative | 0,7043 | 0,02598 |
| TcCLB.506763.35 | hypothetical protein, conserved (pseudogene) | 0,7043 | 0,235 |
| TcCLB.508015.50 | hypothetical protein, conserved | 0,7042 | 0,00001733 |
| TcCLB.509699.200 | mucin TcMUCII, putative | 0,7042 | 0,3709 |
| TcCLB.506767.60 | hypothetical protein, conserved (pseudogene) | 0,704 | 0,2624 |
| TcCLB.511213.70 | trans-sialidase (pseudogene), putative | 0,7038 | 0,00005899 |
| TcCLB.511643.70 | trans-sialidase, Group VI, putative | 0,7038 | 0,03824 |
| TcCLB.506297.350 | SKP1-like protein | 0,7036 | 0,09986 |
| TcCLB.510275.219 | mucin TcMUCII, putative | 0,7036 | 0,348 |

| | | | |
|---|---|---|---|
| TcCLB.511887.50 | hypothetical protein, conserved | 0,7033 | 0,04583 |
| TcCLB.506691.60 | hypothetical protein, conserved | 0,7032 | 0,0001771 |
| TcCLB.508779.9 | hypothetical protein, conserved (fragment) | 0,7031 | 1,008E-06 |
| TcCLB.506737.90 | trans-sialidase, Group V, putative | 0,703 | 5,349E-08 |
| TcCLB.508551.20 | 40S ribosomal protein S12, putative | 0,7028 | 2,642E-20 |
| TcCLB.508705.10 | retrotransposon hot spot protein (RHS, pseudogene), putat | 0,7026 | 6,304E-07 |
| TcCLB.506265.80 | hypothetical protein, conserved | 0,7025 | 0,001195 |
| TcCLB.509275.10 | elongation factor 1-gamma (EF-1-gamma, pseudogene), p | 0,7025 | 0,07088 |
| TcCLB.504149.70 | hypothetical protein, conserved | 0,7018 | 5,479E-08 |
| TcCLB.506035.20 | dispersed gene family protein 1 (DGF-1, pseudogene), puta | 0,7018 | 0,00002012 |
| TcCLB.511825.63 | peptidyl-prolyl cis-trans isomerase, putative | 0,7017 | 5,553E-07 |
| TcCLB.504125.100 | sugar transporter (pseudogene), putative | 0,7016 | 0,0001745 |
| TcCLB.506977.70 | ATPase subunit 9, putative | 0,7015 | 2,471E-11 |
| TcCLB.506277.3 | mucin TcMUCII, putative (fragment) | 0,7014 | 0,2933 |
| TcCLB.508257.160 | hypothetical protein, conserved | 0,7009 | 2,611E-09 |
| TcCLB.506009.40 | hypothetical protein | 0,7009 | 0,01373 |
| TcCLB.507555.50 | trans-sialidase, Group VIII, putative | 0,7007 | 0,00012 |
| TcCLB.511815.40 | guide RNA associated protein, GAP2, putative | 0,7004 | 2,056E-21 |
| TcCLB.511301.170 | retrotransposon hot spot (RHS) protein, putative | 0,6998 | 9,502E-07 |
| TcCLB.506779.30 | hypothetical protein, conserved (pseudogene) | 0,6996 | 7,609E-11 |
| TcCLB.510529.39 | hypothetical protein, conserved | 0,6994 | 0,003914 |
| TcCLB.507497.11 | trans-sialidase, putative (fragment) | 0,6993 | 0,05363 |
| TcCLB.508153.1074 | hypothetical protein, conserved | 0,6992 | 6,768E-08 |
| TcCLB.506513.130 | kinesin (pseudogene), putative | 0,6991 | 1,312E-14 |
| TcCLB.506545.5 | mucin TcMUCI, putative | 0,699 | 0,03101 |
| TcCLB.506173.50 | mucin TcMUCII (pseudogene), putative | 0,6989 | 0,1472 |
| TcCLB.511255.410 | surface protease GP63 (pseudogene), putative | 0,6988 | 0,1785 |
| TcCLB.509849.20 | hypothetical protein, conserved | 0,6987 | 1,068E-07 |
| TcCLB.508501.210 | hypothetical protein, conserved | 0,6986 | 0,00001478 |
| TcCLB.504149.90 | hypothetical protein, conserved | 0,6985 | 6,339E-16 |
| TcCLB.508293.130 | hypothetical protein | 0,6983 | 0,4493 |
| TcCLB.506773.180 | regulator of chromatin condensation, putative | 0,6982 | 2,421E-07 |
| TcCLB.507041.71 | mucin TcMUCI, putative | 0,6981 | 0,1513 |
| TcCLB.509151.50 | hypothetical protein, conserved | 0,6979 | 1,565E-14 |
| TcCLB.509111.28 | fucose kinase, putative | 0,6978 | 0,04131 |
| TcCLB.509157.60 | 1-acyl-sn-glycerol-3-phosphate acyltransferase, putative | 0,6977 | 1,054E-06 |
| TcCLB.509641.10 | hypothetical protein, conserved | 0,6976 | 0,000996 |
| TcCLB.504035.70 | cyclophilin, putative | 0,6971 | 0,000011 |
| TcCLB.511807.50 | hypothetical protein, conserved | 0,697 | 1,681E-15 |
| TcCLB.509437.170 | hypothetical protein | 0,697 | 0,008503 |
| TcCLB.511181.150 | hypothetical protein, conserved | 0,6969 | 6,445E-06 |
| TcCLB.511069.4 | transketolase, putative | 0,6969 | 0,0001608 |
| TcCLB.506133.80 | mucin-associated surface protein (MASP, pseudogene), pu | 0,6967 | 0,08079 |
| TcCLB.507681.200 | serine carboxypeptidase S28, putative | 0,6966 | 6,567E-10 |
| TcCLB.508903.30 | retrotransposon hot spot protein (RHS, pseudogene), putat | 0,6966 | 0,0004389 |

| | | | |
|---|---|---|---|
| TcCLB.504199.10 | hypothetical protein, conserved | 0,6964 | 1,038E-14 |
| TcCLB.511603.190 | mucin-associated surface protein (MASP), putative | 0,6964 | 0,1804 |
| TcCLB.510111.20 | trans-sialidase, Group II, putative | 0,696 | 0,01995 |
| TcCLB.510225.20 | hypothetical protein, conserved | 0,6959 | 0,00004801 |
| TcCLB.504801.10 | beta galactofuranosyl glycosyltransferase (pseudogene), pu | 0,6959 | 0,02976 |
| TcCLB.506129.50 | trans-sialidase, Group IV, putative | 0,6956 | 1,245E-07 |
| TcCLB.508669.21 | hypothetical protein | 0,6955 | 0,1918 |
| TcCLB.510005.11 | endosomal integral membrane protein (pseudogene), puta | 0,6954 | 0,00006537 |
| TcCLB.510865.10 | retrotransposon hot spot protein (RHS, pseudogene), putat | 0,6954 | 0,000676 |
| TcCLB.504055.30 | hypothetical protein | 0,6951 | 9,281E-06 |
| TcCLB.508767.20 | surface protein TolT (TolT) | 0,6951 | 0,001155 |
| TcCLB.504081.30 | trans-sialidase (pseudogene), putative | 0,6949 | 0,01471 |
| TcCLB.456097.30 | glycerophosphoryl diester phosphodiesterase, putative | 0,6948 | 0,00004589 |
| TcCLB.504249.30 | mucin-associated surface protein (MASP), putative | 0,6947 | 0,09107 |
| TcCLB.507573.10 | hypothetical protein, conserved (pseudogene) | 0,6937 | 1,091E-06 |
| TcCLB.508461.260 | Bem46-like serine peptidase, putative | 0,6933 | 4,261E-09 |
| TcCLB.506675.20 | hypothetical protein, conserved (pseudogene) | 0,6931 | 1,182E-08 |
| TcCLB.510933.90 | mucin-associated surface protein (MASP), putative | 0,6931 | 0,09239 |
| TcCLB.508891.60 | serine/threonine protein phosphatase, putative | 0,693 | 0,0006705 |
| TcCLB.508341.10 | mucin-associated surface protein (MASP), putative | 0,693 | 0,367 |
| TcCLB.510403.50 | retrotransposon hot spot protein (RHS, pseudogene), putat | 0,6927 | 3,753E-12 |
| TcCLB.509615.40 | alpha-adaptin-like (pseudogene), putative | 0,6925 | 6,909E-07 |
| TcCLB.508569.130 | 6-phosphofructo-2-kinase/fructose-2,6-biphosphatase, put | 0,6923 | 2,381E-13 |
| TcCLB.510939.30 | mucin TcMUCI, putative | 0,6921 | 0,06163 |
| TcCLB.508799.104 | hypothetical protein, conserved | 0,692 | 0,0002225 |
| TcCLB.506981.10 | dispersed gene family protein 1 (DGF-1), putative (fragme | 0,6918 | 0,1016 |
| TcCLB.482425.10 | retrotransposon hot spot (RHS) protein, putative | 0,6918 | 0,1052 |
| TcCLB.510037.20 | surface protease GP63 (pseudogene), putative | 0,6918 | 0,1105 |
| TcCLB.508555.60 | Calpain-like protein 1 (CAP5.5) | 0,6916 | 2,775E-09 |
| TcCLB.510661.140 | hypothetical protein, conserved | 0,6913 | 1,427E-09 |
| TcCLB.508347.50 | hypothetical protein, conserved | 0,6913 | 0,001749 |
| TcCLB.509429.30 | retrotransposon hot spot protein (RHS, pseudogene), putat | 0,6912 | 1,313E-08 |
| TcCLB.507681.210 | Flagellar Member 2 (FLAM2) | 0,6911 | 8,583E-24 |
| TcCLB.509479.20 | mucin TcMUCII, putative | 0,691 | 0,148 |
| TcCLB.510561.40 | mucin TcMUCII, putative | 0,6909 | 0,005895 |
| TcCLB.506775.50 | hypothetical protein, conserved | 0,6908 | 2,607E-14 |
| TcCLB.511809.20 | methylenetetrahydrofolate dehydrogenase-like protein, pu | 0,6904 | 5,269E-07 |
| TcCLB.506211.20 | hypothetical protein, conserved | 0,6901 | 0,00001114 |
| TcCLB.503905.40 | hypothetical protein, conserved | 0,69 | 0,00004467 |
| TcCLB.503717.20 | mucin-associated surface protein (MASP), putative | 0,69 | 0,04691 |
| TcCLB.510507.20 | 3-ketoacyl-CoA thiolase, putative | 0,6899 | 9,026E-16 |
| TcCLB.511809.140 | histone H2A, putative | 0,6899 | 5,736E-15 |
| TcCLB.503563.29 | hypothetical protein | 0,6899 | 0,04219 |
| TcCLB.511693.120 | hypothetical protein | 0,6898 | 4,247E-09 |
| TcCLB.510361.30 | mucin-associated surface protein (MASP), putative | 0,6894 | 0,2582 |

| | | | |
|---|---|---|---|
| TcCLB.509761.20 | 3,2-trans-enoyl-CoA isomerase, mitochondrial precursor (p | 0,689 | 1,29E-11 |
| TcCLB.509993.40 | mucin TcMUCI, putative | 0,689 | 0,0584 |
| TcCLB.508409.210 | Hsc70-interacting protein, putative (HIP) | 0,6887 | 6,634E-20 |
| TcCLB.507835.10 | trans-sialidase (pseudogene), putative | 0,6886 | 0,00495 |
| TcCLB.506899.40 | mucin TcMUCII, putative | 0,6886 | 0,1775 |
| TcCLB.509275.20 | mucin-associated surface protein (MASP), putative (fragm | 0,6886 | 0,1865 |
| TcCLB.508367.70 | trans-sialidase (pseudogene), putative | 0,6885 | 0,09232 |
| TcCLB.503989.30 | hypothetical protein, conserved (pseudogene) | 0,6882 | 9,48E-16 |
| TcCLB.506501.40 | mucin TcMUCII, putative | 0,6882 | 0,1511 |
| TcCLB.511389.150 | thiolase protein-like protein, putative | 0,6881 | 2,783E-17 |
| TcCLB.511903.170 | hypothetical protein, conserved | 0,6881 | 1,424E-10 |
| TcCLB.503581.20 | PPIase, putative | 0,688 | 0,0005254 |
| TcCLB.506811.120 | hypothetical protein, conserved | 0,6879 | 3,348E-11 |
| TcCLB.506247.370 | hypothetical protein, conserved | 0,6879 | 7,89E-11 |
| TcCLB.507523.80 | mucin-associated surface protein (MASP), putative | 0,6878 | 0,1093 |
| TcCLB.510741.20 | hypothetical protein, conserved | 0,6877 | 0,00002954 |
| TcCLB.503689.30 | flagellar protofilament ribbon protein, putative | 0,6876 | 1,191E-09 |
| TcCLB.507641.240 | hypothetical protein | 0,6875 | 0,09359 |
| TcCLB.509959.40 | beta galactofuranosyl glycosyltransferase, putative | 0,6875 | 0,2847 |
| TcCLB.503861.50 | beta galactofuranosyl glycosyltransferase, putative | 0,6873 | 1,2E-09 |
| TcCLB.508365.4 | mucin-associated surface protein (MASP), putative | 0,6873 | 0,3897 |
| TcCLB.507053.110 | hypothetical protein, conserved | 0,6872 | 1,413E-11 |
| TcCLB.430061.9 | hypothetical protein, conserved | 0,6868 | 0,06998 |
| TcCLB.510849.30 | hypothetical protein, conserved (pseudogene) | 0,6863 | 0,00004376 |
| TcCLB.506765.10 | retrotransposon hot spot protein (RHS, pseudogene), putat | 0,6863 | 0,04424 |
| TcCLB.511367.129 | hypothetical protein, conserved | 0,6862 | 8,569E-06 |
| TcCLB.510039.109 | sec1 family transport protein, putative (fragment) | 0,6862 | 0,001844 |
| TcCLB.509207.11 | histone H2A, putative | 0,6861 | 2,669E-19 |
| TcCLB.509219.50 | hypothetical protein, conserved (pseudogene) | 0,686 | 0,0001463 |
| TcCLB.510583.80 | surface protease GP63 (pseudogene), putative | 0,686 | 0,1807 |
| TcCLB.511007.10 | dispersed gene family protein 1 (DGF-1), putative | 0,6859 | 0,001592 |
| TcCLB.506067.198 | mucin-associated surface protein (MASP, pseudogene), pu | 0,6858 | 0,3312 |
| TcCLB.510053.70 | hypothetical protein, conserved | 0,6856 | 3,422E-12 |
| TcCLB.510755.80 | hypothetical protein, conserved | 0,6855 | 0,00002119 |
| TcCLB.506219.30 | exosome component CSL4, putative (CSL4) | 0,6852 | 0,00001289 |
| TcCLB.506599.430 | mucin TcMUCII, putative | 0,6852 | 0,2176 |
| TcCLB.508503.80 | hypothetical protein | 0,6851 | 0,0003289 |
| TcCLB.508293.80 | mucin-associated surface protein (MASP), putative | 0,685 | 0,3107 |
| TcCLB.504003.14 | hypothetical protein, conserved | 0,6849 | 0,00006613 |
| TcCLB.508221.680 | serine-alanine-and proline-rich protein (pseudogene), puta | 0,6848 | 0,3408 |
| TcCLB.508173.54 | hypothetical protein, conserved | 0,6846 | 1,836E-12 |
| TcCLB.506361.50 | mucin-associated surface protein (MASP), putative | 0,6845 | 0,1216 |
| TcCLB.506993.63 | mucin TcMUCI, putative | 0,6844 | 0,309 |
| TcCLB.511725.230 | protein kinase, putative | 0,6842 | 1,129E-12 |
| TcCLB.511809.135 | histone H2A, putative | 0,6841 | 2,397E-14 |

| | | | |
|---|---|---|---|
| TcCLB.503415.30 | hypothetical protein, conserved | 0,6838 | 3,266E-11 |
| TcCLB.511523.100 | aldehyde dehydrogenase, putative | 0,6837 | 6,899E-10 |
| TcCLB.509897.279 | mucin-associated surface protein (MASP), putative | 0,6837 | 0,2018 |
| TcCLB.511791.10 | mucin-associated surface protein (MASP), putative | 0,6837 | 0,4923 |
| TcCLB.510857.30 | hypothetical protein, conserved | 0,6835 | 8,349E-10 |
| TcCLB.506801.40 | mucin-associated surface protein (MASP), putative | 0,6834 | 0,008531 |
| TcCLB.508165.60 | mucin TcMUCII, putative | 0,6834 | 0,4253 |
| TcCLB.506987.10 | retrotransposon hot spot protein (RHS, pseudogene), putat | 0,6831 | 0,05897 |
| TcCLB.509373.21 | elongation factor 1-gamma (EF-1-gamma, pseudogene), p | 0,6831 | 0,08177 |
| TcCLB.510575.20 | hypothetical protein, conserved | 0,6829 | 0,0002912 |
| TcCLB.509429.210 | leucine-rich repeat protein (LRRP), putative | 0,6828 | 3,568E-11 |
| TcCLB.506063.59 | dispersed gene family protein 1 (DGF-1), putative | 0,6827 | 0,007651 |
| TcCLB.505975.41 | retrotransposon hot spot protein (RHS, pseudogene), putat | 0,6825 | 3,247E-06 |
| TcCLB.511239.49 | adenosine 5'-monophosphoramidase, putative | 0,6825 | 0,0001353 |
| TcCLB.509897.150 | mucin-associated surface protein (MASP), putative | 0,6825 | 0,0001398 |
| TcCLB.509993.10 | mucin TcMUCI, putative | 0,6824 | 0,1814 |
| TcCLB.510749.50 | Flagellar Member 5 (FLAM5) | 0,6822 | 9,661E-09 |
| TcCLB.507185.26 | mucin-associated surface protein (MASP, pseudogene), pu | 0,6822 | 0,002533 |
| TcCLB.509759.19 | pumilio/PUF RNA binding protein 1, putative (PUF1) | 0,6821 | 8,04E-11 |
| TcCLB.504213.100 | hypothetical protein, conserved | 0,6819 | 0,003245 |
| TcCLB.504149.55 | acyl-CoA thioesterase, putative | 0,6817 | 1,849E-10 |
| TcCLB.508593.20 | trans-sialidase, Group IV, putative | 0,6816 | 0,003922 |
| TcCLB.506773.150 | hypothetical protein | 0,6814 | 0,00000068 |
| TcCLB.507293.20 | dispersed gene family protein 1 (DGF-1, pseudogene), puta | 0,6814 | 0,00003696 |
| TcCLB.509049.30 | hypothetical protein, conserved | 0,6813 | 5,471E-13 |
| TcCLB.511815.10 | hypothetical protein | 0,6813 | 1,141E-12 |
| TcCLB.510439.30 | hypothetical protein, conserved | 0,6813 | 2,304E-10 |
| TcCLB.510103.10 | hypothetical protein | 0,6812 | 0,0006353 |
| TcCLB.507625.150 | hypothetical protein, conserved | 0,6807 | 0,0001316 |
| TcCLB.504639.60 | trans-sialidase, putative | 0,6802 | 0,01501 |
| TcCLB.506297.340 | hypothetical protein, conserved | 0,6801 | 2,745E-06 |
| TcCLB.506295.190 | glycine dehydrogenase [decarboxylating] (pseudogene), pu | 0,6799 | 0,02363 |
| TcCLB.510283.80 | hypothetical protein, conserved | 0,6795 | 0,00007304 |
| TcCLB.511801.34 | hypothetical protein, conserved | 0,6793 | 7,616E-06 |
| TcCLB.508647.210 | dynein arm light chain, putative | 0,6792 | 1,931E-13 |
| TcCLB.506009.50 | hypothetical protein, conserved | 0,679 | 9,477E-09 |
| TcCLB.510465.110 | mucin-associated surface protein (MASP, pseudogene), pu | 0,6787 | 0,09285 |
| TcCLB.510543.140 | retrotransposon hot spot protein (RHS, pseudogene), putat | 0,6784 | 0,0002616 |
| TcCLB.506759.160 | mucin TcMUCII, putative | 0,6781 | 0,5575 |
| TcCLB.504797.60 | hypothetical protein, conserved | 0,6777 | 2,622E-08 |
| TcCLB.510061.40 | retrotransposon hot spot protein (RHS, pseudogene), putat | 0,6775 | 0,0001161 |
| TcCLB.510495.20 | mucin-associated surface protein (MASP), putative | 0,6775 | 0,4915 |
| TcCLB.506211.40 | intraflagellar transport protein 52, putative (IFT52) | 0,6774 | 7,73E-13 |
| TcCLB.507811.107 | hypothetical protein, conserved | 0,6774 | 0,0001145 |
| TcCLB.511589.140 | phosphoenolpyruvate mutase, putative | 0,6771 | 4,106E-10 |

| | | | |
|---|---|---|---|
| TcCLB.511469.60 | Trk system potassium uptake protein (HKT1) | 0,677 | 8,242E-06 |
| TcCLB.506131.50 | hypothetical protein | 0,6769 | 0,438 |
| TcCLB.509211.20 | hypothetical protein, conserved | 0,6768 | 1,894E-13 |
| TcCLB.511659.9 | hypothetical protein, conserved (fragment) | 0,6768 | 0,00000116 |
| TcCLB.508237.18 | SNF2 DNA repair protein, putative | 0,6767 | 0,0000017 |
| TcCLB.511621.120 | hypothetical protein, conserved | 0,6764 | 0,00008881 |
| TcCLB.506781.30 | mucin TcMUCII, putative | 0,6764 | 0,1318 |
| TcCLB.510165.30 | hypothetical protein, conserved | 0,6761 | 4,253E-09 |
| TcCLB.505201.20 | dispersed gene family protein 1 (DGF-1, pseudogene), puta | 0,6761 | 0,0007883 |
| TcCLB.508837.120 | prohibitin, putative | 0,676 | 1,119E-11 |
| TcCLB.508173.210 | hypothetical protein, conserved | 0,676 | 5,85E-10 |
| TcCLB.511521.20 | hypothetical protein, conserved | 0,676 | 0,00005872 |
| TcCLB.506289.200 | hypothetical protein | 0,676 | 0,1575 |
| TcCLB.506729.80 | hypothetical protein, conserved | 0,6759 | 0,00001854 |
| TcCLB.506223.70 | hypothetical protein | 0,6759 | 0,003609 |
| TcCLB.510377.420 | surface protease GP63, putative | 0,6758 | 0,5133 |
| TcCLB.506201.60 | hypothetical protein | 0,6755 | 0,001649 |
| TcCLB.509295.5 | retrotransposon hot spot protein (RHS, pseudogene), putat | 0,6754 | 2,899E-08 |
| TcCLB.509699.85 | surface protease GP63 (pseudogene), putative | 0,6754 | 0,09389 |
| TcCLB.506789.240 | hypothetical protein, conserved | 0,6752 | 0,00002071 |
| TcCLB.503939.84 | hypothetical protein, conserved | 0,6752 | 0,001281 |
| TcCLB.511215.119 | 69 kDa paraflagellar rod protein, putative | 0,6751 | 1,473E-14 |
| TcCLB.506595.65 | hypothetical protein | 0,6751 | 0,002168 |
| TcCLB.511843.35 | hypothetical protein | 0,6749 | 0,07494 |
| TcCLB.508777.30 | hypothetical protein, conserved | 0,6746 | 4,626E-13 |
| TcCLB.508113.20 | trans-sialidase (pseudogene), putative | 0,6746 | 0,001157 |
| TcCLB.507777.41 | trans-sialidase (pseudogene), putative | 0,6744 | 0,03474 |
| TcCLB.503717.80 | mucin-associated surface protein (MASP, pseudogene), pu | 0,6744 | 0,1949 |
| TcCLB.506147.190 | hypothetical protein, conserved | 0,6743 | 2,08E-09 |
| TcCLB.510535.64 | hypothetical protein, conserved | 0,6742 | 5,861E-08 |
| TcCLB.507787.40 | hypothetical protein, conserved | 0,6742 | 0,0002655 |
| TcCLB.507953.110 | retrotransposon hot spot protein (RHS, pseudogene), putat | 0,6741 | 0,2177 |
| TcCLB.511217.170 | hypothetical protein, conserved | 0,674 | 1,813E-06 |
| TcCLB.504099.50 | trans-sialidase, Group II, putative | 0,674 | 0,0004407 |
| TcCLB.510915.10 | mucin-associated surface protein (MASP), putative | 0,6738 | 0,0001111 |
| TcCLB.506753.200 | hypothetical protein, conserved | 0,6735 | 9,096E-07 |
| TcCLB.509755.70 | mucin TcMUCII, putative | 0,6735 | 0,4606 |
| TcCLB.506405.140 | hypothetical protein, conserved | 0,6733 | 3,856E-09 |
| TcCLB.507063.150 | hypothetical protein, conserved | 0,6729 | 8,318E-07 |
| TcCLB.508177.64 | hypothetical protein, conserved | 0,6729 | 0,00001465 |
| TcCLB.509725.139 | damage-specific DNA binding protein, putative | 0,6728 | 0,00005016 |
| TcCLB.507669.70 | Dynein intermediate chain 1, axonemal (DNAI1) | 0,6727 | 8,39E-08 |
| TcCLB.507085.130 | hypothetical protein | 0,6726 | 0,2126 |
| TcCLB.507615.70 | deoxyhypusine hydroxylase, putative | 0,6722 | 0,0001133 |
| TcCLB.509023.150 | integral membrane transport protein, putative | 0,6718 | 0,0002623 |

| | | | |
|---|---|---|---|
| TcCLB.508265.10 | glutathione-S-transferase/glutaredoxin, putative | 0,6714 | 3,158E-06 |
| TcCLB.508153.540 | hypothetical protein, conserved | 0,6713 | 1,86E-10 |
| TcCLB.506459.230 | trans-sialidase, Group V, putative | 0,6709 | 0,005404 |
| TcCLB.506139.100 | mucin TcMUCII, putative | 0,6697 | 0,268 |
| TcCLB.508647.110 | Present in the outer mitochondrial membrane proteome 1 | 0,6696 | 6,493E-10 |
| TcCLB.509859.20 | hypothetical protein, conserved | 0,6696 | 5,368E-06 |
| TcCLB.506405.80 | hypothetical protein, conserved | 0,6695 | 0,0001324 |
| TcCLB.473111.10 | lathosterol oxidase, putative | 0,6692 | 5,201E-06 |
| TcCLB.506625.35 | trans-sialidase (pseudogene), putative | 0,6689 | 0,1039 |
| TcCLB.507383.10 | hypothetical protein, conserved | 0,6682 | 0,1336 |
| TcCLB.507809.60 | hypothetical protein, conserved | 0,6681 | 2,049E-09 |
| TcCLB.503729.10 | hypothetical protein, conserved | 0,668 | 0,0002937 |
| TcCLB.509229.110 | hypothetical protein, conserved | 0,6678 | 2,905E-11 |
| TcCLB.503575.50 | fatty acyl CoA synthetase, putative | 0,6676 | 4,708E-08 |
| TcCLB.509237.130 | Calpain-like protein 1 (CAP5.5) | 0,6675 | 2,252E-18 |
| TcCLB.511609.10 | retrotransposon hot spot protein (RHS, pseudogene), putat | 0,6674 | 0,07819 |
| TcCLB.511409.50 | hypothetical protein, conserved | 0,6671 | 8,748E-08 |
| TcCLB.506401.50 | malonyl-CoA decarboxylase, mitochondrial precursor, puta | 0,6669 | 4,401E-09 |
| TcCLB.508003.30 | hypothetical protein, conserved (pseudogene) | 0,6669 | 0,0009689 |
| TcCLB.510065.30 | translationally controlled tumor protein (TCTP), putative | 0,6665 | 2,28E-22 |
| TcCLB.506575.70 | Present in the outer mitochondrial membrane proteome 4 | 0,6664 | 6,857E-08 |
| TcCLB.510055.110 | fructosamine kinase, putative | 0,6662 | 0,0000439 |
| TcCLB.508045.60 | hypothetical protein, conserved | 0,6661 | 1,321E-09 |
| TcCLB.510307.140 | retrotransposon hot spot protein (RHS, pseudogene), putat | 0,666 | 2,019E-07 |
| TcCLB.506629.50 | hypothetical protein, conserved | 0,666 | 1,538E-06 |
| TcCLB.508547.180 | hypothetical protein, conserved | 0,6656 | 2,73E-10 |
| TcCLB.506907.30 | hypothetical protein, conserved | 0,6655 | 0,0001156 |
| TcCLB.506605.90 | hypothetical protein, conserved | 0,6652 | 0,0001011 |
| TcCLB.506943.4 | aldehyde dehydrogenase, putative | 0,6648 | 1,821E-12 |
| TcCLB.504427.110 | acetyl-CoA synthetase, putative | 0,6647 | 5,143E-08 |
| TcCLB.503433.10 | hypothetical protein, conserved | 0,6647 | 0,0001682 |
| TcCLB.508365.130 | mucin TcMUCII, putative | 0,6647 | 0,2713 |
| TcCLB.508221.40 | mucin TcMUCII, putative | 0,6647 | 0,2924 |
| TcCLB.506697.10 | hypothetical protein, conserved | 0,6646 | 4,476E-08 |
| TcCLB.397937.10 | pumilio/PUF RNA binding protein 1, putative (PUF1) | 0,6645 | 9,374E-06 |
| TcCLB.511579.9 | fatty acyl CoA syntetase 1, putative | 0,6645 | 0,00544 |
| TcCLB.509905.60 | trans-sialidase (pseudogene), putative | 0,6645 | 0,01202 |
| TcCLB.511681.20 | histone H4, putative | 0,6642 | 0,0018 |
| TcCLB.509507.10 | hypothetical protein, conserved | 0,6638 | 2,977E-10 |
| TcCLB.506677.30 | hypothetical protein, conserved | 0,6627 | 0,00000386 |
| TcCLB.510983.19 | retrotransposon hot spot (RHS) protein, putative | 0,6626 | 0,0003463 |
| TcCLB.507671.10 | hypothetical protein | 0,6623 | 0,0008047 |
| TcCLB.511013.10 | retrotransposon hot spot protein (RHS, pseudogene), putat | 0,6621 | 0,005626 |
| TcCLB.510421.310 | hypothetical protein, conserved | 0,662 | 5,793E-06 |
| TcCLB.510583.140 | trans-sialidase (pseudogene), putative | 0,662 | 0,09588 |

| | | | |
|---|---|---|---|
| TcCLB.509195.30 | mucin-associated surface protein (MASP), putative | 0,6615 | 0,3494 |
| TcCLB.510859.50 | hypothetical protein, conserved | 0,6614 | 0,00002202 |
| TcCLB.511635.60 | hypothetical protein, conserved | 0,6612 | 1,019E-06 |
| TcCLB.507509.50 | polypeptide deformylase-like protein, putative | 0,6611 | 3,982E-06 |
| TcCLB.509715.150 | hypothetical protein | 0,661 | 0,005498 |
| TcCLB.504149.150 | hypothetical protein, conserved | 0,6608 | 5,548E-12 |
| TcCLB.510853.30 | trans-sialidase (pseudogene), putative | 0,6607 | 0,0005062 |
| TcCLB.511671.130 | hypothetical protein, conserved | 0,6597 | 5,443E-08 |
| TcCLB.506761.40 | mucin-associated surface protein (MASP, pseudogene), pu | 0,6597 | 0,1123 |
| TcCLB.510377.330 | trans-sialidase, Group V, putative | 0,6591 | 0,0008778 |
| TcCLB.508175.280 | hypothetical protein, conserved | 0,6589 | 1,403E-09 |
| TcCLB.506295.100 | hypothetical protein, conserved | 0,6587 | 3,063E-13 |
| TcCLB.508323.84 | peptidyl-prolyl cis-trans isomerase, putative | 0,6587 | 0,002124 |
| TcCLB.508541.130 | mucin-associated surface protein (MASP), putative | 0,6586 | 0,004802 |
| TcCLB.504109.150 | elongation factor, putative | 0,6585 | 4,569E-12 |
| TcCLB.507083.30 | apurinic/apyrimidinic endonuclease, putative | 0,6583 | 0,0000052 |
| TcCLB.509177.59 | hypothetical protein, conserved | 0,6583 | 0,00001463 |
| TcCLB.507957.150 | mucin-associated surface protein (MASP), putative | 0,6582 | 0,04388 |
| TcCLB.504239.420 | mucin-associated surface protein (MASP), putative | 0,6578 | 0,1574 |
| TcCLB.503399.10 | hypothetical protein, conserved | 0,6577 | 0,0004143 |
| TcCLB.509787.30 | retrotransposon hot spot protein (RHS, pseudogene), putat | 0,6576 | 0,0001891 |
| TcCLB.507731.10 | mucin-associated surface protein (MASP), putative | 0,6575 | 0,08775 |
| TcCLB.508647.264 | hypothetical protein, conserved | 0,6568 | 6,887E-10 |
| TcCLB.506779.130 | hypothetical protein | 0,6567 | 2,766E-07 |
| TcCLB.506693.40 | hypothetical protein, conserved | 0,6563 | 0,009207 |
| TcCLB.506683.180 | dispersed gene family protein 1 (DGF-1), putative | 0,6562 | 0,00001985 |
| TcCLB.507975.10 | retrotransposon hot spot protein (RHS, pseudogene), putat | 0,6559 | 5,795E-11 |
| TcCLB.510241.90 | hypothetical protein, conserved | 0,6556 | 3,478E-10 |
| TcCLB.510885.130 | hypothetical protein, conserved | 0,6556 | 0,00004759 |
| TcCLB.511003.150 | hypothetical protein | 0,6555 | 2,581E-09 |
| TcCLB.506691.90 | serine/threonine protein kinase, putative | 0,6555 | 0,215 |
| TcCLB.510769.190 | retrotransposon hot spot protein (RHS, pseudogene), putat | 0,6554 | 0,00003064 |
| TcCLB.511255.299 | mucin TcMUCII, putative | 0,6554 | 0,1988 |
| TcCLB.506025.40 | hypothetical protein, conserved | 0,6552 | 2,67E-10 |
| TcCLB.511173.340 | mucin TcMUCII (pseudogene), putative | 0,6547 | 0,0545 |
| TcCLB.510155.200 | hypothetical protein, conserved | 0,6545 | 5,897E-06 |
| TcCLB.507045.5 | trans-sialidase (pseudogene), putative | 0,6545 | 0,2359 |
| TcCLB.511313.30 | formin, putative | 0,6543 | 0,0119 |
| TcCLB.506263.30 | D-isomer specific 2-hydroxyacid dehydrogenase-protein, pu | 0,6542 | 0,02436 |
| TcCLB.510431.274 | 60S ribosomal protein L37, putative | 0,654 | 7,38E-15 |
| TcCLB.506753.240 | hypothetical protein, conserved | 0,654 | 0,001722 |
| TcCLB.508045.70 | hypothetical protein, conserved | 0,6537 | 6,84E-17 |
| TcCLB.510279.330 | mucin-associated surface protein (MASP), putative | 0,6537 | 0,1126 |
| TcCLB.503733.70 | phosphoacetylglucosamine mutase, putative | 0,6535 | 7,327E-06 |
| TcCLB.402857.10 | hypothetical protein, conserved (fragment) | 0,6531 | 4,899E-06 |

| | | | |
|---|---|---|---|
| TcCLB.504625.80 | hypothetical protein, conserved | 0,6531 | 0,0000068 |
| TcCLB.507641.180 | endochitinase, putative | 0,6531 | 0,0003115 |
| TcCLB.450489.10 | histone acetyltransferase, putative | 0,653 | 0,0001646 |
| TcCLB.506931.20 | hypothetical protein, conserved | 0,6529 | 2,568E-07 |
| TcCLB.510641.10 | retrotransposon hot spot protein (RHS, pseudogene), putat | 0,6529 | 0,02013 |
| TcCLB.508119.150 | hypothetical protein, conserved | 0,6528 | 0,00006865 |
| TcCLB.507443.53 | hypothetical protein | 0,6528 | 0,1086 |
| TcCLB.507221.80 | 5'-3' exoribonuclease C, putative (fragment) (XRNC) | 0,6527 | 6,413E-06 |
| TcCLB.505977.26 | 60S acidic ribosomal protein P2, putative | 0,6524 | 6,958E-13 |
| TcCLB.504797.10 | acidocalcisomal exopolyphosphatase, putative | 0,6524 | 2,834E-06 |
| TcCLB.503957.120 | retrotransposon hot spot (RHS) protein, putative | 0,6515 | 0,00000301 |
| TcCLB.510659.130 | hypothetical protein, conserved | 0,6513 | 3,027E-16 |
| TcCLB.504219.50 | hypothetical protein, conserved (pseudogene) | 0,6512 | 0,4601 |
| TcCLB.506825.80 | hypothetical protein, conserved | 0,6509 | 6,778E-11 |
| TcCLB.506109.29 | dispersed gene family protein 1 (DGF-1), putative | 0,6509 | 0,001014 |
| TcCLB.509053.140 | succinate dehydrogenase subunit | 0,6508 | 7,784E-11 |
| TcCLB.509209.10 | endoplasmatic reticulum retrieval protein, putative | 0,6507 | 4,34E-12 |
| TcCLB.504045.100 | hypothetical protein, conserved | 0,6505 | 1,827E-07 |
| TcCLB.510577.10 | hypothetical protein, conserved | 0,6502 | 0,134 |
| TcCLB.510983.5 | trans-sialidase (pseudogene), putative | 0,6501 | 0,08619 |
| TcCLB.508501.230 | monothiol glutaredoxin, putative | 0,65 | 0,0003173 |
| TcCLB.503437.21 | N-acetyltransferase complex ARD1 subunit (pseudogene), | 0,65 | 0,222 |
| TcCLB.506741.30 | mucin-associated surface protein (MASP, pseudogene), pu | 0,65 | 0,46 |
| TcCLB.511183.590 | mucin TcMUCII, putative | 0,6497 | 0,07181 |
| TcCLB.508525.30 | hypothetical protein, conserved | 0,6495 | 6,434E-09 |
| TcCLB.506171.50 | retrotransposon hot spot protein (RHS, pseudogene), putat | 0,6495 | 0,0001645 |
| TcCLB.508257.190 | hypothetical protein, conserved | 0,6494 | 0,0005678 |
| TcCLB.511739.30 | hypothetical protein, conserved | 0,649 | 0,0008379 |
| TcCLB.510503.130 | mucin TcMUCII, putative | 0,649 | 0,2717 |
| TcCLB.504429.70 | retrotransposon hot spot protein (RHS, pseudogene), putat | 0,6487 | 3,83E-11 |
| TcCLB.503681.29 | 30S ribosomal protein S8, putative | 0,6486 | 0,2935 |
| TcCLB.511737.90 | hypothetical protein, conserved | 0,6484 | 9,217E-07 |
| TcCLB.511807.270 | hypothetical protein, conserved | 0,6482 | 6,461E-11 |
| TcCLB.506297.190 | pyruvate phosphate dikinase, putative | 0,6479 | 1,282E-11 |
| TcCLB.506873.10 | hypothetical protein, conserved | 0,6479 | 2,112E-08 |
| TcCLB.506251.20 | hypothetical protein, conserved | 0,6479 | 0,002603 |
| TcCLB.511009.9 | dispersed gene family protein 1 (DGF-1), putative (fragme | 0,6475 | 0,03799 |
| TcCLB.508163.90 | trans-sialidase (pseudogene), putative | 0,6474 | 0,1724 |
| TcCLB.510377.410 | mucin TcMUCII, putative | 0,6473 | 0,2978 |
| TcCLB.510303.190 | cell cycle associated protein MOB1, putative | 0,6471 | 1,595E-07 |
| TcCLB.504351.30 | hypothetical protein, conserved (pseudogene) | 0,6471 | 0,001481 |
| TcCLB.508053.5 | hypothetical protein, conserved (fragment) | 0,6469 | 0,1844 |
| TcCLB.508503.30 | hypothetical protein | 0,6468 | 0,00000505 |
| TcCLB.506021.20 | trans-sialidase, Group II, putative | 0,6465 | 0,02952 |
| TcCLB.509437.70 | hypothetical protein, conserved | 0,6463 | 2,511E-14 |

| | | | |
|---|---|---|---|
| TcCLB.510643.150 | ribosomal P protein AGP2beta-1 (pseudogene), putative | 0,6463 | 0,000654 |
| TcCLB.503903.70 | hypothetical protein, conserved | 0,6461 | 0,00003675 |
| TcCLB.506393.30 | beta galactofuranosyl glycosyltransferase, putative | 0,646 | 0,002853 |
| TcCLB.506887.100 | protein kinase, putative | 0,6459 | 0,0003173 |
| TcCLB.510565.40 | hypothetical protein | 0,6458 | 0,00002833 |
| TcCLB.507615.110 | Present in the outer mitochondrial membrane proteome 3 | 0,6456 | 4,001E-06 |
| TcCLB.511815.70 | hypothetical protein, conserved | 0,6455 | 0,01075 |
| TcCLB.506063.31 | ATP-dependent DEAD/H RNA helicase (pseudogene), putat | 0,6454 | 0,3009 |
| TcCLB.507959.80 | mucin-associated surface protein (MASP), putative | 0,6453 | 0,04465 |
| TcCLB.505297.70 | mucin TcMUCII, putative | 0,6451 | 0,01684 |
| TcCLB.506375.120 | hypothetical protein, conserved | 0,6449 | 2,412E-06 |
| TcCLB.511175.50 | mucin TcMUC (pseudogene), putative | 0,6446 | 0,00002312 |
| TcCLB.509817.50 | trans-sialidase, Group I, putative | 0,6446 | 0,02757 |
| TcCLB.506855.200 | hypothetical protein | 0,6446 | 0,03983 |
| TcCLB.510349.64 | hypothetical protein, conserved | 0,6444 | 8,288E-06 |
| TcCLB.507517.80 | hypothetical protein, conserved | 0,6443 | 0,00008458 |
| TcCLB.510377.50 | trans-sialidase, Group V, putative | 0,6442 | 0,2854 |
| TcCLB.510643.190 | retrotransposon hot spot (RHS) protein, putative | 0,6441 | 1,759E-08 |
| TcCLB.503483.9 | retrotransposon hot spot (RHS) protein, putative | 0,644 | 1,986E-21 |
| TcCLB.507615.19 | microtubule-associated protein, putative, (fragment) (GB4 | 0,644 | 4,324E-13 |
| TcCLB.507153.40 | tryptophanyl-tRNA synthetase (pseudogene), putative | 0,644 | 0,02752 |
| TcCLB.442427.10 | trans-sialidase (pseudogene), putative | 0,6439 | 0,005976 |
| TcCLB.508441.94 | mucin TcMUCII, putative | 0,6438 | 0,005774 |
| TcCLB.511817.280 | hypothetical protein, conserved | 0,6437 | 4,355E-12 |
| TcCLB.507063.130 | protein kinase, putative | 0,6436 | 4,329E-09 |
| TcCLB.503609.24 | mucin-associated surface protein (MASP, pseudogene), pu | 0,6434 | 0,1597 |
| TcCLB.510803.19 | trans-sialidase, putative | 0,6431 | 0,4943 |
| TcCLB.504077.50 | hypothetical protein, conserved | 0,643 | 7,419E-06 |
| TcCLB.511163.24 | hypothetical protein, conserved | 0,6426 | 0,001478 |
| TcCLB.510021.10 | mucin TcMUCII, putative | 0,6426 | 0,2498 |
| TcCLB.511817.319 | hypothetical protein, conserved | 0,6424 | 0,0008133 |
| TcCLB.510655.60 | hypothetical protein, conserved | 0,6423 | 0,0001129 |
| TcCLB.510115.50 | retrotransposon hot spot protein (RHS, pseudogene), putat | 0,6419 | 0,00003807 |
| TcCLB.508919.70 | protein kinase, putative | 0,6413 | 3,295E-09 |
| TcCLB.504277.20 | hypothetical protein | 0,6411 | 0,0004399 |
| TcCLB.509029.90 | hypothetical protein, conserved | 0,6411 | 0,0005339 |
| TcCLB.510381.5 | SNF2 DNA repair protein (pseudogene), putative | 0,6411 | 0,001418 |
| TcCLB.507079.30 | multidrug resistance protein E, putative | 0,6408 | 0,01428 |
| TcCLB.510857.10 | hypothetical protein, conserved | 0,6404 | 3,201E-08 |
| TcCLB.510483.224 | mucin TcMUCII (pseudogene), putative | 0,6403 | 0,2987 |
| TcCLB.507063.40 | hypothetical protein, conserved | 0,6398 | 0,0001012 |
| TcCLB.509745.40 | hypothetical protein, conserved | 0,6395 | 7,909E-07 |
| TcCLB.510021.30 | mucin TcMUCII, putative | 0,6394 | 0,3469 |
| TcCLB.510435.10 | hypothetical protein, conserved | 0,6393 | 0,001701 |
| TcCLB.511613.40 | mucin-associated surface protein (MASP), putative | 0,6393 | 0,06539 |

| | | | |
|---|---|---|---|
| TcCLB.506401.240 | hypothetical protein, conserved | 0,6391 | 1,842E-12 |
| TcCLB.504115.110 | trans-sialidase (pseudogene), putative | 0,6391 | 0,006605 |
| TcCLB.509995.20 | hypothetical protein, conserved | 0,6386 | 2,227E-08 |
| TcCLB.508415.30 | hypothetical protein | 0,6385 | 8,548E-08 |
| TcCLB.507087.14 | hypothetical protein, conserved | 0,6385 | 0,02489 |
| TcCLB.506067.169 | mucin-associated surface protein (MASP), putative | 0,6385 | 0,1666 |
| TcCLB.511089.10 | mucin-associated surface protein (MASP), putative | 0,6377 | 0,442 |
| TcCLB.505115.20 | hypothetical protein | 0,6374 | 0,1535 |
| TcCLB.509735.70 | retrotransposon hot spot protein (RHS, pseudogene), putat | 0,6373 | 3,876E-07 |
| TcCLB.508719.20 | hypothetical protein, conserved | 0,6373 | 0,0004819 |
| TcCLB.506129.89 | hypothetical protein, conserved | 0,6372 | 8,91E-11 |
| TcCLB.511317.60 | hypothetical protein, conserved | 0,6371 | 0,004851 |
| TcCLB.508125.140 | mucin-associated surface protein (MASP), putative | 0,6371 | 0,1587 |
| TcCLB.506357.199 | hypothetical protein, conserved | 0,6369 | 2,501E-09 |
| TcCLB.507653.110 | trans-sialidase (pseudogene), putative | 0,6368 | 0,0000613 |
| TcCLB.511817.130 | hypothetical protein, conserved | 0,6366 | 0,0005261 |
| TcCLB.508515.40 | protein kinase, putative | 0,6364 | 0,00169 |
| TcCLB.503613.20 | proteasome alpha 7 subunit, putative | 0,6362 | 3,699E-07 |
| TcCLB.511733.90 | hypothetical protein, conserved | 0,6361 | 0,00003359 |
| TcCLB.510773.110 | hypothetical protein, conserved | 0,6361 | 0,001317 |
| TcCLB.506595.70 | trans-sialidase (pseudogene), putative | 0,6361 | 0,002336 |
| TcCLB.506321.120 | mucin-associated surface protein (MASP), putative | 0,6361 | 0,003751 |
| TcCLB.508499.39 | trans-sialidase (pseudogene), putative | 0,636 | 0,0769 |
| TcCLB.506357.10 | hypothetical protein, conserved | 0,6359 | 0,0004431 |
| TcCLB.509443.39 | hypothetical protein, conserved | 0,6359 | 0,002222 |
| TcCLB.509161.60 | hypothetical protein, conserved | 0,6358 | 2,312E-08 |
| TcCLB.507173.30 | mucin-associated surface protein (MASP, pseudogene), pu | 0,6356 | 0,1202 |
| TcCLB.506401.280 | hypothetical protein, conserved | 0,6353 | 6,675E-07 |
| TcCLB.504081.342 | hypothetical protein (pseudogene) | 0,6353 | 0,3137 |
| TcCLB.506627.130 | dispersed gene family protein 1 (DGF-1, pseudogene), puta | 0,635 | 0,1076 |
| TcCLB.511001.60 | protein kinase, putative | 0,6349 | 7,163E-06 |
| TcCLB.506147.110 | hypothetical protein, conserved | 0,6348 | 8,528E-08 |
| TcCLB.509203.14 | hypothetical protein | 0,6348 | 0,0178 |
| TcCLB.506335.55 | elongation factor 1-gamma (EF-1-gamma, pseudogene), p | 0,6346 | 0,05217 |
| TcCLB.399373.9 | retrotransposon hot spot (RHS) protein, putative (fragmen | 0,6342 | 4,336E-08 |
| TcCLB.511581.10 | fatty acyl CoA synthetase 2, putative | 0,6342 | 0,0001391 |
| TcCLB.507071.20 | mucin-associated surface protein (MASP), putative | 0,6342 | 0,06866 |
| TcCLB.511391.10 | retrotransposon hot spot protein (RHS, pseudogene), putat | 0,6338 | 1,063E-10 |
| TcCLB.507905.39 | mucin-associated surface protein (MASP), putative | 0,6337 | 1,211E-08 |
| TcCLB.506001.50 | mucin-associated surface protein (MASP), putative | 0,6337 | 0,009844 |
| TcCLB.503973.57 | mucin TcMUCII (pseudogene), putative | 0,6336 | 0,1871 |
| TcCLB.503789.60 | hypothetical protein, conserved | 0,6335 | 4,495E-06 |
| TcCLB.507627.20 | proton motive ATPase 1, putative | 0,6333 | 1,159E-09 |
| TcCLB.510105.200 | hypothetical protein, conserved | 0,6331 | 1,069E-14 |
| TcCLB.508619.10 | retrotransposon hot spot protein (RHS, pseudogene), putat | 0,633 | 5,194E-06 |

| | | | |
|---|---|---|---|
| TcCLB.506843.50 | glycine dehydrogenase [decarboxylating] (pseudogene), pu | 0,6327 | 0,1199 |
| TcCLB.511281.20 | transporter, putative | 0,6325 | 1,596E-07 |
| TcCLB.504431.50 | hypothetical protein, conserved | 0,6324 | 5,69E-08 |
| TcCLB.509141.30 | hypothetical protein | 0,6323 | 0,0002022 |
| TcCLB.507885.10 | RNA-binding protein, putative | 0,6322 | 0,001236 |
| TcCLB.509717.9 | hypothetical protein (pseudogene) | 0,6322 | 0,03908 |
| TcCLB.503901.9 | hypothetical protein, conserved (pseudogene) | 0,6321 | 0,0006682 |
| TcCLB.504003.50 | hypothetical protein, conserved | 0,632 | 0,0006914 |
| TcCLB.507505.10 | trans-sialidase, putative | 0,632 | 0,3792 |
| TcCLB.506801.50 | surface protease GP63 (pseudogene), putative | 0,6318 | 0,00314 |
| TcCLB.510149.120 | flagellar radial spoke protein-like, putative | 0,6316 | 1,185E-06 |
| TcCLB.504427.120 | hypothetical protein, conserved | 0,6312 | 0,00004191 |
| TcCLB.511863.4 | retrotransposon hot spot (RHS) protein, putative (fragmen | 0,6312 | 0,00008551 |
| TcCLB.509029.60 | hypothetical protein, conserved | 0,6311 | 2,298E-12 |
| TcCLB.508351.40 | cytochrome c oxidase assembly factor, putative | 0,6311 | 0,00003175 |
| TcCLB.511173.360 | mucin-associated surface protein (MASP), putative | 0,631 | 0,1253 |
| TcCLB.508087.10 | hypothetical protein | 0,6308 | 0,00003535 |
| TcCLB.504797.120 | hypothetical protein, conserved | 0,6307 | 0,0005469 |
| TcCLB.503909.90 | hypothetical protein | 0,6305 | 0,00191 |
| TcCLB.509667.31 | hypothetical protein, conserved (pseudogene) | 0,6303 | 0,03854 |
| TcCLB.509611.80 | cation-transporting ATPase, putative | 0,6299 | 3,731E-14 |
| TcCLB.507911.15 | retrotransposon hot spot protein (RHS, pseudogene), putat | 0,6291 | 0,02719 |
| TcCLB.510555.41 | hypothetical protein, conserved (pseudogene) | 0,6291 | 0,2251 |
| TcCLB.510651.20 | hypothetical protein, conserved | 0,629 | 0,0007595 |
| TcCLB.509735.150 | retrotransposon hot spot protein (RHS, pseudogene), putat | 0,6289 | 1,703E-07 |
| TcCLB.508161.10 | mucin-associated surface protein (MASP), putative | 0,6289 | 0,374 |
| TcCLB.510699.110 | retrotransposon hot spot protein (RHS, pseudogene), putat | 0,6288 | 0,003159 |
| TcCLB.506717.30 | dispersed gene family protein 1 (DGF-1, pseudogene), puta | 0,6287 | 0,005963 |
| TcCLB.506885.180 | hypothetical protein, conserved | 0,6286 | 1,493E-06 |
| TcCLB.506357.170 | serine/threonine protein phosphatase, putative | 0,6286 | 0,00009842 |
| TcCLB.503893.30 | hypothetical protein, conserved | 0,6284 | 4,325E-06 |
| TcCLB.506435.410 | uracil phosphoribosyltransferase, putative | 0,6283 | 1,239E-07 |
| TcCLB.508815.130 | hypothetical protein, conserved | 0,6281 | 1,492E-09 |
| TcCLB.504119.29 | deoxyhypusine synthase, putative | 0,6281 | 1,772E-07 |
| TcCLB.506267.60 | mucin TcMUCII, putative | 0,628 | 0,2265 |
| TcCLB.506763.40 | hypothetical protein | 0,628 | 0,2616 |
| TcCLB.509581.35 | hypothetical protein, conserved (pseudogene) | 0,628 | 0,4064 |
| TcCLB.504155.150 | surface protease GP63, putative | 0,6278 | 0,01849 |
| TcCLB.510611.60 | nitroreductase | 0,6276 | 3,802E-06 |
| TcCLB.506859.10 | ubiquitin-conjugating enzyme E2, putative | 0,6276 | 8,549E-06 |
| TcCLB.508515.50 | glycogenin glucosyltransferase, putative | 0,6272 | 0,001446 |
| TcCLB.510155.60 | hypothetical protein, conserved | 0,6271 | 0,00005826 |
| TcCLB.430737.20 | hypothetical protein, conserved (fragment) | 0,6268 | 0,00225 |
| TcCLB.511875.29 | mucin-associated surface protein (MASP, pseudogene), pu | 0,6268 | 0,01784 |
| TcCLB.503823.84 | DNA-directed RNA polymerase I subunit, putative | 0,6267 | 0,0001424 |

| | | | |
|---|---|---|---|
| TcCLB.508539.80 | mucin-associated surface protein (MASP), putative | 0,6267 | 0,03996 |
| TcCLB.511469.40 | hypothetical protein, conserved | 0,6263 | 2,447E-09 |
| TcCLB.510851.40 | trans-sialidase (pseudogene), putative | 0,6263 | 0,002138 |
| TcCLB.507167.169 | dispersed gene family protein 1 (DGF-1), putative | 0,6263 | 0,002821 |
| TcCLB.511809.130 | 40S ribosomal protein S15, putative | 0,6262 | 7,662E-06 |
| TcCLB.508261.50 | mucin-associated surface protein (MASP), putative | 0,6258 | 0,2399 |
| TcCLB.504149.130 | hypothetical protein, conserved | 0,6254 | 1,157E-13 |
| TcCLB.504099.10 | retrotransposon hot spot (RHS) protein, putative | 0,6254 | 0,0001507 |
| TcCLB.507959.180 | surface protease GP63 (pseudogene), putative | 0,6254 | 0,27 |
| TcCLB.509905.100 | retrotransposon hot spot protein (RHS, pseudogene), putat | 0,625 | 0,00002559 |
| TcCLB.510089.170 | casein kinase I, putative | 0,6248 | 0,0004468 |
| TcCLB.506559.434 | hypothetical protein, conserved | 0,6247 | 0,0000139 |
| TcCLB.504001.3 | flagellar radial spoke component, putative (fragment) | 0,6246 | 0,000641 |
| TcCLB.511625.149 | mucin TcMUCII (pseudogene), putative | 0,6246 | 0,2332 |
| TcCLB.504035.120 | hypothetical protein, conserved | 0,6245 | 3,89E-10 |
| TcCLB.506991.10 | hypothetical protein, conserved | 0,6245 | 7,576E-06 |
| TcCLB.506975.63 | hypothetical protein, conserved | 0,6244 | 0,0001861 |
| TcCLB.510153.10 | hypothetical protein, conserved | 0,6241 | 4,036E-07 |
| TcCLB.508567.89 | hypothetical protein, conserved | 0,6239 | 0,00000312 |
| TcCLB.506501.50 | mucin-associated surface protein (MASP, pseudogene), pu | 0,6239 | 0,313 |
| TcCLB.508377.9 | fucose kinase, putative (fragment) | 0,6238 | 2,419E-07 |
| TcCLB.509965.310 | hypothetical protein, conserved (pseudogene) | 0,6237 | 1,207E-07 |
| TcCLB.507507.40 | ubiquitin hydrolase, putative | 0,6233 | 0,001381 |
| TcCLB.510275.370 | mucin-associated surface protein (MASP), putative | 0,6231 | 0,04055 |
| TcCLB.511131.30 | hypothetical protein, conserved | 0,623 | 4,173E-09 |
| TcCLB.506769.65 | trans-sialidase (pseudogene), putative | 0,6229 | 0,5472 |
| TcCLB.506959.110 | hypothetical protein, conserved | 0,6226 | 4,566E-07 |
| TcCLB.510357.100 | hypothetical protein, conserved | 0,6226 | 0,00002434 |
| TcCLB.506487.50 | trans-sialidase (pseudogene), putative | 0,6226 | 0,001317 |
| TcCLB.507649.90 | hypothetical protein, conserved | 0,6222 | 5,066E-08 |
| TcCLB.509127.90 | hypothetical protein, conserved | 0,6222 | 0,00000115 |
| TcCLB.505171.30 | hypothetical protein, conserved | 0,622 | 7,168E-08 |
| TcCLB.397923.10 | mucin-associated surface protein (MASP), putative | 0,622 | 0,1237 |
| TcCLB.508521.20 | trans-sialidase, Group VIII, putative | 0,6219 | 0,01974 |
| TcCLB.504147.160 | ADP-ribosylation factor-like protein, putative | 0,6217 | 8,738E-08 |
| TcCLB.506207.50 | translationally controlled tumor protein (TCTP), putative | 0,6216 | 6,32E-19 |
| TcCLB.509059.30 | calcium-binding protein, putative | 0,6213 | 0,0003869 |
| TcCLB.508741.360 | hypothetical protein, conserved | 0,6212 | 3,065E-07 |
| TcCLB.503717.50 | mucin TcMUCII (pseudogene), putative | 0,6212 | 0,1554 |
| TcCLB.508739.70 | protein kinase, putative | 0,6211 | 8,036E-19 |
| TcCLB.509231.23 | hypothetical protein, conserved | 0,621 | 4,444E-07 |
| TcCLB.508221.920 | mucin-associated surface protein (MASP, pseudogene), pu | 0,6209 | 0,0533 |
| TcCLB.509805.230 | hypothetical protein, conserved | 0,6204 | 0,001005 |
| TcCLB.511245.230 | hypothetical protein, conserved | 0,6201 | 2,402E-06 |
| TcCLB.503947.11 | surface protease GP63, putative (fragment) | 0,6201 | 0,2494 |

| | | | |
|---|---|---|---|
| TcCLB.468041.30 | retrotransposon hot spot protein (RHS, pseudogene), putat | 0,6196 | 3,264E-10 |
| TcCLB.504077.30 | hypothetical protein, conserved | 0,6194 | 2,375E-06 |
| TcCLB.510643.90 | mucin-associated surface protein (MASP, pseudogene), pu | 0,6193 | 0,007608 |
| TcCLB.509581.5 | retrotransposon hot spot (RHS) protein, putative (fragmen | 0,6189 | 0,008961 |
| TcCLB.510377.150 | mucin TcMUCII, putative | 0,6188 | 0,5623 |
| TcCLB.508375.30 | aspartate carbamoyltransferase, putative | 0,6187 | 1,403E-06 |
| TcCLB.509295.90 | hypothetical protein | 0,6187 | 0,004884 |
| TcCLB.506471.30 | trans-sialidase, Group II, putative | 0,6185 | 0,0006034 |
| TcCLB.510299.40 | hypothetical protein, conserved | 0,6183 | 1,344E-06 |
| TcCLB.510753.20 | hypothetical protein, conserved | 0,6182 | 0,00006146 |
| TcCLB.507091.120 | mucin-associated surface protein (MASP), putative | 0,6178 | 0,3881 |
| TcCLB.503467.20 | hypothetical protein, conserved | 0,6177 | 9,718E-08 |
| TcCLB.509769.66 | hypothetical protein, conserved | 0,6177 | 0,00008586 |
| TcCLB.508781.80 | hypothetical protein, conserved | 0,6174 | 1,512E-06 |
| TcCLB.422955.10 | hypothetical protein, conserved (pseudogene) | 0,6174 | 4,086E-06 |
| TcCLB.504741.4 | hypothetical protein, conserved | 0,6173 | 1,993E-06 |
| TcCLB.507487.10 | mucin TcMUCII, putative | 0,6173 | 0,2345 |
| TcCLB.503977.40 | mucin TcMUCII, putative | 0,6173 | 0,267 |
| TcCLB.510525.130 | Present in the outer mitochondrial membrane proteome 3 | 0,6172 | 6,482E-07 |
| TcCLB.503689.10 | small GTP-binding protein Rab7, putative | 0,6169 | 0,0000518 |
| TcCLB.508207.240 | C-1-tetrahydrofolate synthase, cytoplasmic, putative | 0,6166 | 3,348E-06 |
| TcCLB.503613.79 | hypothetical protein | 0,6166 | 0,0006008 |
| TcCLB.511733.10 | aldehyde dehydrogenase, putative | 0,6165 | 4,432E-07 |
| TcCLB.510731.90 | protein kinase, putative | 0,6164 | 0,0007961 |
| TcCLB.509545.40 | trans-sialidase (pseudogene), putative | 0,6163 | 0,04675 |
| TcCLB.507053.160 | hypothetical protein, conserved | 0,6161 | 1,198E-12 |
| TcCLB.504081.420 | mucin-associated surface protein (MASP), putative | 0,616 | 0,327 |
| TcCLB.458153.10 | serine/threonine protein phosphatase, putative (fragment) | 0,6156 | 0,00894 |
| TcCLB.510101.70 | protein transport protein SEC61 subunit alpha, putative | 0,6155 | 1,633E-12 |
| TcCLB.508173.70 | hypothetical protein, conserved | 0,6155 | 5,717E-07 |
| TcCLB.511517.164 | hypothetical protein, conserved | 0,6155 | 0,00004522 |
| TcCLB.507639.20 | universal minicircle sequence binding protein, putative (UN | 0,6154 | 0,0004408 |
| TcCLB.508317.80 | mannosyl-oligosaccharide 1,2-alpha-mannosidase IB, puta | 0,6151 | 5,282E-08 |
| TcCLB.506599.395 | elongation factor 1-gamma (EF-1-gamma, pseudogene), p | 0,6149 | 0,2708 |
| TcCLB.507809.30 | ribose-phosphate pyrophosphokinase, putative | 0,6148 | 2,409E-11 |
| TcCLB.509569.140 | hypothetical protein, conserved | 0,6146 | 1,041E-07 |
| TcCLB.507739.50 | hypothetical protein, conserved | 0,6146 | 0,004166 |
| TcCLB.508909.10 | OSM3-like kinesin, putative | 0,6145 | 5,185E-08 |
| TcCLB.507787.80 | hypothetical protein, conserved | 0,6145 | 0,0009311 |
| TcCLB.503635.20 | hypothetical protein, conserved | 0,614 | 0,00001235 |
| TcCLB.510797.39 | hypothetical protein, conserved | 0,6135 | 0,0002134 |
| TcCLB.420293.40 | beta galactofuranosyl glycosyltransferase (pseudogene), p | 0,6135 | 0,006641 |
| TcCLB.511303.40 | carnitine O-acetyltransferase, putative | 0,6134 | 6,236E-06 |
| TcCLB.509769.90 | mitochondrial carrier protein, putative | 0,6133 | 5,011E-07 |
| TcCLB.511603.110 | mucin TcMUCII, putative | 0,6133 | 0,3695 |

| | | | |
|---|---|---|---|
| TcCLB.506417.60 | terbinafine resistance locus protein (yip1), putative | 0,6129 | 1,107E-06 |
| TcCLB.509599.157 | serine/threonine protein kinase, putative | 0,6128 | 8,852E-07 |
| TcCLB.506375.50 | dispersed gene family protein 1 (DGF-1), putative | 0,6128 | 0,0009623 |
| TcCLB.506951.60 | mucin-associated surface protein (MASP), putative | 0,6126 | 0,009825 |
| TcCLB.511899.40 | 2-amino-3-ketobutyrate coenzyme A ligase, putative | 0,6123 | 2,118E-07 |
| TcCLB.506595.104 | ribosomal P protein AGP2beta-1, putative | 0,6123 | 0,009768 |
| TcCLB.507723.30 | retrotransposon hot spot protein (RHS, pseudogene), putat | 0,6122 | 0,00005466 |
| TcCLB.511771.75 | hypothetical protein | 0,6122 | 0,02998 |
| TcCLB.511807.160 | hypothetical protein, conserved | 0,6121 | 0,0008438 |
| TcCLB.505971.20 | hypothetical protein | 0,6118 | 0,0117 |
| TcCLB.503995.40 | hypothetical protein, conserved | 0,6117 | 0,0004151 |
| TcCLB.511755.19 | hypothetical protein, conserved | 0,6117 | 0,08112 |
| TcCLB.510271.30 | trans-sialidase (pseudogene), putative | 0,6113 | 0,05156 |
| TcCLB.511217.120 | peptidyl-prolyl cis-trans isomerase, putative | 0,6108 | 0,0001202 |
| TcCLB.507899.10 | retrotransposon hot spot protein (RHS, pseudogene), putat | 0,6108 | 0,0004162 |
| TcCLB.510371.50 | mucin-associated surface protein (MASP), putative | 0,6107 | 0,1233 |
| TcCLB.509967.40 | hypothetical protein, conserved | 0,6106 | 9,689E-06 |
| TcCLB.510409.10 | protein kinase, putative | 0,6106 | 0,05804 |
| TcCLB.509525.234 | mucin TcMUCII, putative | 0,6105 | 0,2997 |
| TcCLB.506705.20 | adenosine kinase, putative | 0,6104 | 0,002212 |
| TcCLB.506025.60 | 2-oxoglutarate dehydrogenase, E2 component, dihydrolipo | 0,6101 | 2,656E-11 |
| TcCLB.510021.80 | 90 kDa surface protein, putative | 0,6096 | 0,2119 |
| TcCLB.511863.20 | hypothetical protein, conserved | 0,6093 | 0,0000012 |
| TcCLB.506265.140 | hypothetical protein, conserved | 0,6093 | 0,00005174 |
| TcCLB.508379.10 | fucose kinase, putative | 0,6093 | 0,003054 |
| TcCLB.506759.31 | surface protease GP63 (pseudogene), putative | 0,6093 | 0,1394 |
| TcCLB.506999.160 | hypothetical protein | 0,6086 | 0,02137 |
| TcCLB.511727.310 | Leucine-rich repeat-containing protein ODA7 (ODA7) | 0,6083 | 1,978E-07 |
| TcCLB.510997.40 | enoyl-CoA hydratase/isomerase family protein, putative | 0,6083 | 3,307E-07 |
| TcCLB.510533.180 | microtubule-associated protein 1A/1B, light chain 3, putati | 0,6082 | 0,03129 |
| TcCLB.504975.10 | mucin-associated surface protein (MASP, pseudogene), pu | 0,6081 | 0,1558 |
| TcCLB.511575.110 | hypothetical protein, conserved | 0,6076 | 4,873E-08 |
| TcCLB.508731.50 | dynein light chain Tctex-type, putative | 0,6074 | 0,0005364 |
| TcCLB.509259.123 | hypothetical protein (pseudogene) | 0,6073 | 0,007395 |
| TcCLB.509217.170 | hypothetical protein | 0,6073 | 0,1885 |
| TcCLB.510933.20 | mucin-associated surface protein (MASP), putative | 0,6073 | 0,1934 |
| TcCLB.503903.60 | hypothetical protein, conserved | 0,6072 | 2,734E-08 |
| TcCLB.506821.80 | hypothetical protein, conserved | 0,6069 | 0,007273 |
| TcCLB.506331.104 | retrotransposon hot spot protein (RHS, pseudogene), putat | 0,6069 | 0,009033 |
| TcCLB.506911.30 | trans-sialidase, Group V, putative | 0,6069 | 0,03298 |
| TcCLB.508641.100 | hypothetical protein, conserved | 0,6067 | 1,392E-06 |
| TcCLB.503507.50 | mucin TcMUCII, putative | 0,6064 | 0,2508 |
| TcCLB.510395.30 | GTPase, putative | 0,6061 | 0,000122 |
| TcCLB.510361.90 | surface protease GP63, putative | 0,606 | 0,1631 |
| TcCLB.506159.5 | trans-sialidase, putative (fragment) | 0,6059 | 0,4524 |

| | | | |
|---|---|---|---|
| TcCLB.503619.10 | meiotic recombination protein SPO11, putative | 0,6056 | 0,01041 |
| TcCLB.507995.20 | trans-sialidase, putative | 0,6056 | 0,2455 |
| TcCLB.508879.150 | hypothetical protein, conserved | 0,6054 | 0,000298 |
| TcCLB.424383.9 | trans-sialidase (pseudogene), putative | 0,6053 | 0,06162 |
| TcCLB.509519.9 | dispersed gene family protein 1 (DGF-1), putative (fragme | 0,6052 | 0,3375 |
| TcCLB.511089.140 | mucin-associated surface protein (MASP), putative | 0,605 | 0,1933 |
| TcCLB.509889.9 | dispersed gene family protein 1 (DGF-1), putative | 0,6049 | 0,04943 |
| TcCLB.508879.110 | hypothetical protein, conserved | 0,6048 | 2,664E-06 |
| TcCLB.508293.120 | hypothetical protein | 0,6048 | 0,2652 |
| TcCLB.508893.30 | hypothetical protein, conserved | 0,6047 | 0,002264 |
| TcCLB.510323.50 | hypothetical protein, conserved | 0,6046 | 0,0006426 |
| TcCLB.507167.50 | hypothetical protein, conserved (pseudogene) | 0,6046 | 0,001195 |
| TcCLB.503429.39 | trans-sialidase, putative | 0,604 | 0,2285 |
| TcCLB.506801.120 | protein kinase, putative | 0,6039 | 3,927E-06 |
| TcCLB.503599.70 | hypothetical protein, conserved | 0,6039 | 0,0002041 |
| TcCLB.457547.9 | dispersed gene family protein 1 (DGF-1), putative | 0,6037 | 0,04108 |
| TcCLB.506777.120 | protein kinase, putative | 0,6035 | 0,000107 |
| TcCLB.508859.20 | hypothetical protein, conserved | 0,6035 | 0,0004996 |
| TcCLB.506579.119 | hypothetical protein, conserved | 0,6034 | 0,02778 |
| TcCLB.510345.20 | hypothetical protein | 0,6032 | 0,005239 |
| TcCLB.507497.20 | retrotransposon hot spot protein (RHS, pseudogene), putat | 0,6032 | 0,07597 |
| TcCLB.507233.10 | trans-sialidase, Group III, putative | 0,6032 | 0,2037 |
| TcCLB.510259.40 | hypothetical protein, conserved | 0,6031 | 0,02822 |
| TcCLB.506213.90 | hypothetical protein, conserved | 0,603 | 1,229E-06 |
| TcCLB.506563.200 | calpain cysteine peptidase, putative | 0,6029 | 3,172E-09 |
| TcCLB.508305.50 | mucin-associated surface protein (MASP), putative | 0,6025 | 0,1259 |
| TcCLB.506761.11 | mucin-associated surface protein (MASP), putative (fragm | 0,6024 | 0,4611 |
| TcCLB.510863.20 | hypothetical protein | 0,602 | 3,174E-06 |
| TcCLB.506741.80 | mucin TcMUCII, putative | 0,602 | 0,4637 |
| TcCLB.504029.101 | hypothetical protein, conserved | 0,6019 | 0,0001832 |
| TcCLB.507093.44 | dynein light chain, putative | 0,6018 | 0,001087 |
| TcCLB.509629.10 | trans-sialidase, Group III, putative | 0,6014 | 0,1415 |
| TcCLB.508971.10 | UDP-Gal or UDP-GlcNAc-dependent glycosyltransferase, pu | 0,6012 | 0,002009 |
| TcCLB.510155.180 | eukaryotic initiation factor 4a, putative | 0,6011 | 4,44E-11 |
| TcCLB.507653.40 | dispersed gene family protein 1 (DGF-1), putative | 0,6008 | 0,009234 |
| TcCLB.511817.240 | hypothetical protein, conserved | 0,6006 | 0,00002935 |
| TcCLB.510307.284 | trans-sialidase, Group IV, putative | 0,6005 | 0,00000617 |
| TcCLB.506605.210 | leucine-rich repeat protein (LRRP), putative | 0,6004 | 4,972E-08 |
| TcCLB.511711.80 | Ras-related protein RAB2B, putative (RAB2B) | 0,6 | 4,084E-06 |
| TcCLB.511873.30 | trans-sialidase (pseudogene), putative | 0,6 | 0,004525 |
| TcCLB.508461.210 | hypothetical protein, conserved | 0,5999 | 3,598E-06 |
| TcCLB.510861.80 | hypothetical protein | 0,5999 | 0,00009578 |
| TcCLB.511023.19 | Tetratricopeptide-like helical domain containing protein (fr | 0,5999 | 0,0004901 |
| TcCLB.408547.10 | mucin-associated surface protein (MASP), putative | 0,5998 | 0,1288 |
| TcCLB.510187.370 | hypothetical protein, conserved | 0,5997 | 0,0008817 |

| | | | |
|---|---|---|---|
| TcCLB.511821.30 | Mu-adaptin 1, putative | 0,5996 | 1,364E-06 |
| TcCLB.506193.70 | glucose-6-phosphate dehydrogenase (pseudogene), putativ | 0,5996 | 0,00003261 |
| TcCLB.506763.234 | mucin-associated surface protein (MASP, pseudogene), pu | 0,5996 | 0,1918 |
| TcCLB.503899.110 | glutathione peroxidase-like protein, putative | 0,5994 | 0,00000542 |
| TcCLB.510101.480 | hypothetical protein, conserved | 0,5991 | 0,00002179 |
| TcCLB.504031.30 | mucin-associated surface protein (MASP), putative | 0,5991 | 0,06304 |
| TcCLB.506851.20 | choline dehydrogenase, putative | 0,599 | 1,542E-09 |
| TcCLB.407477.10 | thiolase protein-like protein, putative | 0,599 | 0,00001597 |
| TcCLB.511191.19 | ras-related protein Rab21, putative | 0,5989 | 3,087E-06 |
| TcCLB.511693.100 | long-chain-fatty-acid-CoA ligase, putative | 0,5988 | 1,171E-09 |
| TcCLB.511817.310 | hypothetical protein, conserved | 0,5986 | 9,693E-10 |
| TcCLB.508785.10 | Present in the outer mitochondrial membrane proteome 1! | 0,5986 | 0,00002898 |
| TcCLB.509857.20 | hypothetical protein, conserved | 0,5986 | 0,00004631 |
| TcCLB.510349.40 | hypothetical protein, conserved | 0,5985 | 2,251E-11 |
| TcCLB.510853.40 | trans-sialidase, Group IV, putative | 0,5982 | 0,00004983 |
| TcCLB.508899.100 | hypothetical protein, conserved | 0,5982 | 0,001483 |
| TcCLB.508441.30 | protein kinase, putative | 0,5978 | 0,01149 |
| TcCLB.509993.60 | mucin TcMUCI, putative | 0,5978 | 0,2801 |
| TcCLB.506035.10 | retrotransposon hot spot protein (RHS, pseudogene), putat | 0,5977 | 0,0001414 |
| TcCLB.507275.10 | protein kinase, putative | 0,5977 | 0,001573 |
| TcCLB.507121.11 | trans-sialidase, putative (fragment) | 0,5976 | 0,3623 |
| TcCLB.506895.40 | Kinesin-13 1, putative (KIN13-1) | 0,5975 | 2,091E-08 |
| TcCLB.506763.280 | mucin-associated surface protein (MASP), putative | 0,5975 | 0,07502 |
| TcCLB.506773.170 | hypothetical protein, conserved | 0,5974 | 3,901E-06 |
| TcCLB.504201.40 | mucin-associated surface protein (MASP), putative | 0,5974 | 0,1738 |
| TcCLB.511843.90 | hypothetical protein, conserved (pseudogene) | 0,5973 | 0,006224 |
| TcCLB.504239.434 | trans-sialidase, putative (fragment) | 0,5973 | 0,3563 |
| TcCLB.507609.150 | hypothetical protein (pseudogene) | 0,5972 | 0,07814 |
| TcCLB.509917.19 | retrotransposon hot spot (RHS) protein, putative (fragmen | 0,5971 | 1,158E-10 |
| TcCLB.506041.50 | cytochrome c oxidase assembly protein, putative | 0,5969 | 0,00008529 |
| TcCLB.508993.10 | cytidine triphosphate synthase, putative | 0,5968 | 2,302E-11 |
| TcCLB.510993.19 | hypothetical protein | 0,5968 | 0,00744 |
| TcCLB.506269.45 | trans-sialidase (pseudogene), putative | 0,5967 | 0,4309 |
| TcCLB.508501.310 | hypothetical protein, conserved | 0,5966 | 3,254E-06 |
| TcCLB.506599.100 | mucin-associated surface protein (MASP), putative | 0,5965 | 0,2707 |
| TcCLB.506499.10 | hypothetical protein, conserved (pseudogene) | 0,5964 | 0,4616 |
| TcCLB.506323.10 | dispersed gene family protein 1 (DGF-1, pseudogene), puta | 0,5963 | 0,002914 |
| TcCLB.503677.4 | hypothetical protein, conserved | 0,5961 | 0,2784 |
| TcCLB.511167.40 | hypothetical protein, conserved | 0,5959 | 0,003768 |
| TcCLB.506803.30 | retrotransposon hot spot protein (RHS, pseudogene), putat | 0,5959 | 0,1061 |
| TcCLB.507993.290 | hypothetical protein, conserved | 0,5958 | 0,0003025 |
| TcCLB.508539.150 | surface protease GP63 (pseudogene), putative | 0,5958 | 0,1171 |
| TcCLB.507747.110 | mucin TcMUCII, putative | 0,5957 | 0,1374 |
| TcCLB.510579.40 | hypothetical protein, conserved | 0,5955 | 0,000246 |
| TcCLB.508613.10 | hypothetical protein, conserved | 0,5951 | 3,95E-07 |

| | | | |
|---|---|---|---|
| TcCLB.511757.70 | trans-sialidase, Group II, putative | 0,5951 | 0,00522 |
| TcCLB.506147.80 | Present in the outer mitochondrial membrane proteome 6 | 0,5948 | 0,00001658 |
| TcCLB.507969.10 | vesicular transport-associated repeat protein, putative | 0,5947 | 1,708E-09 |
| TcCLB.506527.4 | hypothetical protein, conserved (fragment) | 0,5945 | 1,055E-06 |
| TcCLB.483183.10 | trans-sialidase, putative (fragment) | 0,5945 | 0,4009 |
| TcCLB.507317.50 | hypothetical protein, conserved | 0,5944 | 1,093E-08 |
| TcCLB.457101.10 | hypothetical protein, conserved | 0,5943 | 0,00000015 |
| TcCLB.510205.40 | trans-sialidase, Group V, putative | 0,5943 | 0,02884 |
| TcCLB.511539.9 | hypothetical protein, conserved | 0,5939 | 1,832E-07 |
| TcCLB.506679.80 | SNF1-related protein kinases, putative | 0,5936 | 3,077E-07 |
| TcCLB.506513.150 | hypothetical protein, conserved | 0,5936 | 0,0003816 |
| TcCLB.509537.70 | hypothetical protein, conserved | 0,5934 | 0,00004709 |
| TcCLB.506661.140 | hypothetical protein, conserved | 0,5931 | 1,236E-07 |
| TcCLB.507615.90 | hypothetical protein, conserved | 0,5928 | 5,701E-13 |
| TcCLB.507645.59 | hypothetical protein, conserved | 0,5928 | 0,006303 |
| TcCLB.510691.10 | 1,2-Dihydroxy-3-keto-5-methylthiopentene dioxygenase, pu | 0,5927 | 7,803E-06 |
| TcCLB.506649.110 | hypothetical protein, conserved | 0,5927 | 0,004835 |
| TcCLB.510027.20 | transporter, putative | 0,5926 | 0,0006066 |
| TcCLB.511077.160 | mucin TcMUCII, putative | 0,5926 | 0,3361 |
| TcCLB.510575.220 | hypothetical protein, conserved | 0,5923 | 0,00006222 |
| TcCLB.509601.130 | hypothetical protein, conserved | 0,5922 | 0,00672 |
| TcCLB.511383.50 | prohibitin, putative | 0,5918 | 3,756E-08 |
| TcCLB.505171.40 | hypothetical protein, conserved | 0,5917 | 0,00005633 |
| TcCLB.506757.129 | mucin-associated surface protein (MASP, pseudogene), pu | 0,5917 | 0,01376 |
| TcCLB.511815.24 | hypothetical protein, conserved | 0,5915 | 2,232E-08 |
| TcCLB.433033.10 | retrotransposon hot spot protein (RHS, pseudogene), putat | 0,5914 | 0,0004043 |
| TcCLB.507071.250 | mucin TcMUCII, putative | 0,5913 | 0,2949 |
| TcCLB.510079.49 | dispersed gene family protein 1 (DGF-1), putative (fragme | 0,591 | 0,01471 |
| TcCLB.503619.25 | hypothetical protein, conserved | 0,5908 | 0,003281 |
| TcCLB.504153.330 | hypothetical protein, conserved | 0,5907 | 2,075E-11 |
| TcCLB.510901.20 | hypothetical protein | 0,5907 | 0,007484 |
| TcCLB.504039.130 | mucin-associated surface protein (MASP), putative | 0,5905 | 0,2491 |
| TcCLB.510105.230 | glyceraldehyde 3-phosphate dehydrogenase, cytosolic, puta | 0,5904 | 0,00009824 |
| TcCLB.510421.60 | hypothetical protein, conserved | 0,5903 | 6,388E-06 |
| TcCLB.511181.50 | metallo-peptidase, Clan ME, Family M16 | 0,5902 | 5,066E-11 |
| TcCLB.506595.120 | trans-sialidase (pseudogene), putative | 0,5901 | 0,0001283 |
| TcCLB.509797.60 | hypothetical protein | 0,59 | 0,00002566 |
| TcCLB.506375.60 | tryptophanyl-tRNA synthetase, putative | 0,5899 | 0,0009714 |
| TcCLB.505267.20 | trans-sialidase (pseudogene), putative | 0,5899 | 0,133 |
| TcCLB.507681.50 | hypothetical protein, conserved | 0,5898 | 1,047E-09 |
| TcCLB.509217.140 | hypothetical protein, conserved (pseudogene) | 0,5898 | 0,005427 |
| TcCLB.506675.90 | retrotransposon hot spot protein (RHS, pseudogene), putat | 0,5897 | 0,002804 |
| TcCLB.510085.50 | trans-sialidase (pseudogene), putative | 0,5897 | 0,1025 |
| TcCLB.508999.60 | mucin TcMUCII, putative | 0,5897 | 0,1202 |
| TcCLB.509947.10 | mucin TcMUCII, putative | 0,5895 | 0,05416 |

| | | | |
|---|---|---|---|
| TcCLB.507773.34 | 60S ribosomal protein L37, putative | 0,5894 | 3,289E-09 |
| TcCLB.506961.50 | dispersed gene family protein 1 (DGF-1), putative | 0,5891 | 0,00005574 |
| TcCLB.506773.40 | 3-methylcrotonoyl-CoA carboxylase beta subunit, putative | 0,5887 | 5,441E-13 |
| TcCLB.503763.30 | retrotransposon hot spot protein (RHS, pseudogene), putat | 0,5886 | 2,199E-08 |
| TcCLB.438059.10 | serine/threonine-protein phosphatase PP1 alpha | 0,5886 | 0,00735 |
| TcCLB.511689.10 | hypothetical protein | 0,5883 | 0,007293 |
| TcCLB.503899.130 | glutathione peroxidase-like protein, putative | 0,5881 | 0,0004422 |
| TcCLB.510497.130 | retrotransposon hot spot protein (RHS, pseudogene), putat | 0,5878 | 0,009665 |
| TcCLB.508289.20 | mucin TcMUCII, putative | 0,5877 | 0,5239 |
| TcCLB.422867.20 | retrotransposon hot spot protein (RHS, pseudogene), putat | 0,5876 | 0,000192 |
| TcCLB.504231.10 | hypothetical protein, conserved | 0,5872 | 7,316E-06 |
| TcCLB.511557.10 | serine peptidase, clan SC, family S9A-like protein (OPB) | 0,5871 | 6,509E-12 |
| TcCLB.509419.49 | mucin-associated surface protein (MASP, pseudogene), pu | 0,5871 | 0,4965 |
| TcCLB.511585.320 | retrotransposon hot spot (RHS) protein, putative | 0,5869 | 5,281E-06 |
| TcCLB.510697.105 | hypothetical protein, conserved (pseudogene) | 0,5866 | 0,1918 |
| TcCLB.511293.80 | hypothetical protein, conserved | 0,5862 | 0,0000435 |
| TcCLB.503829.10 | hypothetical protein, conserved | 0,5861 | 0,0007241 |
| TcCLB.510693.314 | mucin TcMUCII (pseudogene), putative | 0,5859 | 0,0539 |
| TcCLB.509151.120 | hypothetical protein, conserved | 0,5858 | 8,846E-07 |
| TcCLB.506765.74 | mucin-associated surface protein (MASP), putative | 0,5857 | 0,2321 |
| TcCLB.509053.170 | DREV methyltransferase, putative | 0,5856 | 0,0000656 |
| TcCLB.511769.10 | hypothetical protein | 0,5856 | 0,02413 |
| TcCLB.511211.200 | hypothetical protein, conserved | 0,5855 | 5,971E-06 |
| TcCLB.507491.90 | protein kinase, putative | 0,5853 | 0,0004576 |
| TcCLB.508221.780 | hypothetical protein | 0,5852 | 0,1098 |
| TcCLB.511871.60 | hypothetical protein, conserved (pseudogene) | 0,5846 | 0,00005941 |
| TcCLB.506885.124 | Enriched in surface-labeled proteome protein 19 (SLS3) | 0,5842 | 5,148E-06 |
| TcCLB.506825.219 | hypothetical protein, conserved | 0,584 | 2,23E-08 |
| TcCLB.506839.10 | serine/threonine kinase, putative | 0,5837 | 0,7031 |
| TcCLB.508461.142 | hypothetical protein | 0,5835 | 0,0000195 |
| TcCLB.509643.90 | serine peptidase, putative | 0,5834 | 8,927E-06 |
| TcCLB.510635.10 | trans-sialidase, Group VII, putative | 0,5832 | 0,1262 |
| TcCLB.510151.10 | fatty acyl CoA syntetase 1, putative | 0,5831 | 0,00003903 |
| TcCLB.508389.154 | mucin-associated surface protein (MASP), putative | 0,583 | 0,2252 |
| TcCLB.504575.20 | hypothetical protein, conserved | 0,5828 | 0,0001969 |
| TcCLB.511261.30 | NADH-ubiquinone oxidoreductase complex I subunit, putati | 0,5827 | 3,868E-07 |
| TcCLB.507049.50 | ABC transporter, putative | 0,5826 | 0,00003762 |
| TcCLB.506955.212 | mucin-associated surface protein (MASP), putative | 0,5826 | 0,1256 |
| TcCLB.508169.110 | hypothetical protein, conserved | 0,5825 | 0,001741 |
| TcCLB.507623.80 | hypothetical protein, conserved | 0,5824 | 0,00006198 |
| TcCLB.511183.360 | mucin-associated surface protein (MASP), putative | 0,5821 | 0,08668 |
| TcCLB.509819.20 | hypothetical protein, conserved | 0,582 | 1,347E-11 |
| TcCLB.508263.30 | kinesin, putative | 0,582 | 2,172E-06 |
| TcCLB.511357.9 | hypothetical protein, conserved | 0,582 | 5,316E-06 |
| TcCLB.418221.20 | hypothetical protein, conserved | 0,5818 | 1,827E-08 |

| | | | |
|---|---|---|---|
| TcCLB.511821.90 | hypothetical protein, conserved | 0,5815 | 0,001257 |
| TcCLB.506443.110 | retrotransposon hot spot protein (RHS, pseudogene), putat | 0,5814 | 0,0006718 |
| TcCLB.506455.30 | trans-sialidase, Group II, putative | 0,5814 | 0,06213 |
| TcCLB.506529.535 | hypothetical protein, conserved | 0,5814 | 0,3688 |
| TcCLB.509657.40 | mucin TcMUCII, putative | 0,5811 | 0,1367 |
| TcCLB.507069.50 | mucin-associated surface protein (MASP, pseudogene), pu | 0,581 | 0,2444 |
| TcCLB.509207.30 | hypothetical protein, conserved | 0,5809 | 0,0007565 |
| TcCLB.506965.10 | mucin TcMUCII, putative | 0,5809 | 0,283 |
| TcCLB.506463.10 | Lys-63-specific deubiquitinase BRCC36, putative | 0,5808 | 0,00008889 |
| TcCLB.507017.20 | Present in the outer mitochondrial membrane proteome 3( | 0,5803 | 0,00005395 |
| TcCLB.508173.260 | hypothetical protein, conserved | 0,5802 | 0,00000122 |
| TcCLB.511649.150 | phosphoglycerate mutase protein, putative | 0,58 | 0,0001179 |
| TcCLB.506351.9 | 5'-3' exoribonuclease A, putative (fragment) (XRNA) | 0,5797 | 0,0003777 |
| TcCLB.509217.146 | N-acetyltransferase complex ARD1 subunit (pseudogene), | 0,5797 | 0,3715 |
| TcCLB.506659.10 | immunodominant antigen, putative | 0,5795 | 1,307E-10 |
| TcCLB.507601.130 | hypothetical protein, conserved | 0,5795 | 7,002E-07 |
| TcCLB.510307.10 | surface protease GP63, putative | 0,5794 | 0,1666 |
| TcCLB.510519.50 | hypothetical protein, conserved | 0,5793 | 0,0009643 |
| TcCLB.503429.30 | mucin-associated surface protein (MASP), putative | 0,5791 | 0,2991 |
| TcCLB.511727.270 | RNA-binding protein, putative | 0,5785 | 1,538E-15 |
| TcCLB.506153.10 | amino acid transporter, putative | 0,5785 | 3,426E-12 |
| TcCLB.507765.110 | hypothetical protein, conserved | 0,5785 | 1,215E-08 |
| TcCLB.509151.40 | inorganic polyphosphate/ATP-NAD kinase, putative | 0,5783 | 1,956E-12 |
| TcCLB.510371.35 | hypothetical protein, conserved (pseudogene) | 0,5782 | 0,4104 |
| TcCLB.511919.20 | trans-sialidase, Group V, putative | 0,578 | 0,002951 |
| TcCLB.506767.127 | hypothetical protein, conserved (pseudogene) | 0,5778 | 0,4531 |
| TcCLB.506495.9 | hypothetical protein, conserved | 0,5777 | 5,092E-06 |
| TcCLB.506999.50 | hypothetical protein, conserved | 0,5776 | 2,658E-06 |
| TcCLB.510373.60 | mucin-associated surface protein (MASP), putative | 0,5774 | 0,09308 |
| TcCLB.510019.10 | mucin TcMUCII, putative | 0,5772 | 0,3733 |
| TcCLB.509825.20 | hypothetical protein, conserved | 0,577 | 0,00007428 |
| TcCLB.508869.20 | mucin-associated surface protein (MASP), putative | 0,577 | 0,2381 |
| TcCLB.506227.150 | Flagellum matrix protein 458 (PKA-R) | 0,5768 | 6,162E-06 |
| TcCLB.509167.110 | S-adenosylmethionine decarboxylase proenzyme, putative | 0,5768 | 8,289E-06 |
| TcCLB.508799.120 | amino acid permease/transporter, putative | 0,5763 | 9,19E-08 |
| TcCLB.508629.26 | retrotransposon hot spot protein (RHS, pseudogene), putat | 0,576 | 0,004912 |
| TcCLB.508173.230 | hypothetical protein, conserved | 0,5759 | 6,397E-08 |
| TcCLB.508469.30 | hypothetical protein, conserved | 0,5759 | 0,0001485 |
| TcCLB.511411.30 | arginine permease (AAAP411) | 0,5758 | 0,2708 |
| TcCLB.511017.50 | glycosomal membrane protein, putative | 0,5757 | 2,844E-08 |
| TcCLB.510155.120 | ubiquitin activating enzyme, putative | 0,5757 | 2,031E-06 |
| TcCLB.503403.20 | trans-sialidase, Group II, putative | 0,5757 | 0,1066 |
| TcCLB.510583.70 | mucin TcMUCII, putative | 0,5757 | 0,2794 |
| TcCLB.509871.10 | retrotransposon hot spot protein (RHS, pseudogene), putat | 0,5755 | 2,146E-07 |
| TcCLB.508893.20 | hypothetical protein, conserved | 0,5753 | 3,047E-08 |

| | | | |
|---|---|---|---|
| TcCLB.506743.200 | 3-Beta-hydroxysteroid-delta(8), delta(7)-isomerase, putati | 0,5752 | 0,004483 |
| TcCLB.506843.80 | trans-sialidase (pseudogene), putative | 0,5752 | 0,02559 |
| TcCLB.508559.90 | retrotransposon hot spot (RHS) protein, putative | 0,5749 | 0,0002716 |
| TcCLB.506635.90 | ubiquitin-protein ligase, putative | 0,5747 | 0,001355 |
| TcCLB.507559.90 | hypothetical protein | 0,5743 | 7,616E-10 |
| TcCLB.506335.50 | mucin-associated surface protein (MASP), putative | 0,5741 | 0,066 |
| TcCLB.506537.240 | hypothetical protein, conserved | 0,5739 | 0,005151 |
| TcCLB.510749.40 | ubiquitin hydrolase, putative | 0,5738 | 0,1025 |
| TcCLB.508909.210 | hypothetical protein, conserved | 0,5732 | 1,746E-08 |
| TcCLB.509979.14 | peroxisomal membrane protein 4, putative | 0,5732 | 0,00002259 |
| TcCLB.506247.24 | hypothetical protein, conserved | 0,5732 | 0,03259 |
| TcCLB.503855.50 | hypothetical protein, conserved | 0,573 | 3,348E-13 |
| TcCLB.507875.210 | retrotransposon hot spot protein (RHS, pseudogene), putat | 0,573 | 1,255E-08 |
| TcCLB.507049.65 | hypothetical protein, conserved (pseudogene) | 0,5729 | 0,06723 |
| TcCLB.510857.40 | RNA-editing complex protein, putative | 0,5724 | 2,089E-06 |
| TcCLB.511277.570 | 60S ribosomal protein L7, putative | 0,5721 | 7,456E-07 |
| TcCLB.453445.20 | ubiquitin-conjugating enzyme E2, putative | 0,5721 | 5,651E-06 |
| TcCLB.504081.350 | mucin-associated surface protein (MASP), putative | 0,5716 | 0,1814 |
| TcCLB.504023.10 | hypothetical protein, conserved | 0,5714 | 0,00224 |
| TcCLB.509925.10 | mucin-associated surface protein (MASP), putative | 0,5713 | 0,08219 |
| TcCLB.509223.10 | dispersed gene family protein 1 (DGF-1, pseudogene), puta | 0,5712 | 0,000551 |
| TcCLB.503487.30 | hypothetical protein, conserved | 0,5711 | 0,0002606 |
| TcCLB.510303.270 | hypothetical protein, conserved | 0,5706 | 3,357E-13 |
| TcCLB.508375.89 | fucose kinase, putative (fragment) | 0,5705 | 0,1215 |
| TcCLB.506435.400 | hypothetical protein, conserved | 0,5704 | 2,288E-08 |
| TcCLB.508463.29 | hypothetical protein, conserved | 0,5704 | 0,0001689 |
| TcCLB.511371.10 | retrotransposon hot spot (RHS) protein, putative | 0,5703 | 0,00003324 |
| TcCLB.510731.20 | hypothetical protein, conserved | 0,5701 | 0,00001527 |
| TcCLB.507121.20 | trans-sialidase, Group VII, putative | 0,5701 | 0,003797 |
| TcCLB.503715.20 | hypothetical protein | 0,5694 | 0,000218 |
| TcCLB.508401.10 | trans-sialidase, putative | 0,5694 | 0,2595 |
| TcCLB.511697.30 | hypothetical protein | 0,5692 | 0,1303 |
| TcCLB.508793.14 | dynein regulatory complex subunit, putative (fragment) | 0,5691 | 1,257E-06 |
| TcCLB.509029.70 | leucine-rich repeat protein, putative | 0,5691 | 0,0001272 |
| TcCLB.508971.50 | ubiquitin-protein ligase-like, putative | 0,569 | 4,718E-11 |
| TcCLB.507559.30 | hypothetical protein, conserved | 0,569 | 0,00001243 |
| TcCLB.508153.270 | heat shock protein 20, putative | 0,569 | 0,00001582 |
| TcCLB.510317.7 | Present in the outer mitochondrial membrane proteome 2 | 0,5689 | 2,973E-08 |
| TcCLB.509569.160 | hypothetical protein, conserved | 0,5688 | 0,00001467 |
| TcCLB.506525.150 | hypothetical protein, conserved | 0,5685 | 2,924E-08 |
| TcCLB.507669.90 | hypothetical protein, conserved | 0,5685 | 5,825E-06 |
| TcCLB.506759.170 | mucin-associated surface protein (MASP), putative | 0,5685 | 0,1978 |
| TcCLB.510683.20 | hypothetical protein, conserved | 0,5683 | 0,00009154 |
| TcCLB.506289.150 | mucin-associated surface protein (MASP), putative | 0,5678 | 0,07042 |
| TcCLB.508415.20 | hypothetical protein, conserved | 0,5677 | 0,00005321 |

| | | | |
|---|---|---|---|
| TcCLB.506401.300 | hypothetical protein, conserved | 0,5675 | 0,00001859 |
| TcCLB.509911.74 | hypothetical protein, conserved | 0,5674 | 0,00001529 |
| TcCLB.508247.100 | hypothetical protein (pseudogene) | 0,5674 | 0,1415 |
| TcCLB.511171.50 | mucin-associated surface protein (MASP, pseudogene), pu | 0,5674 | 0,1826 |
| TcCLB.509635.9 | hypothetical protein, conserved | 0,5673 | 0,00003395 |
| TcCLB.504131.40 | hypothetical protein, conserved | 0,5672 | 0,00001783 |
| TcCLB.508493.60 | trans-sialidase (pseudogene), putative | 0,5669 | 0,0007478 |
| TcCLB.509809.39 | P-type H -ATPase, putative | 0,5668 | 1,401E-11 |
| TcCLB.509971.60 | surface protease GP63, putative | 0,5666 | 0,2742 |
| TcCLB.509177.40 | hypothetical protein, conserved | 0,5664 | 0,0001955 |
| TcCLB.509683.20 | hypothetical protein, conserved | 0,5663 | 1,749E-07 |
| TcCLB.503881.20 | GDP-mannose 4,6 dehydratase, putative | 0,5661 | 0,00000397 |
| TcCLB.509899.100 | trans-sialidase, Group V, putative | 0,5661 | 0,00689 |
| TcCLB.509229.70 | hypothetical protein, conserved | 0,566 | 0,0002729 |
| TcCLB.508879.60 | hypothetical protein, conserved | 0,5658 | 4,859E-10 |
| TcCLB.507099.50 | programmed cell death 6 protein-like, putative | 0,5658 | 0,00001526 |
| TcCLB.453917.9 | hypothetical protein, conserved | 0,5657 | 0,0008031 |
| TcCLB.508977.50 | mucin-associated surface protein (MASP), putative | 0,5653 | 0,07978 |
| TcCLB.510121.170 | cysteine peptidase, Clan CA, family C2, putative | 0,5652 | 1,738E-11 |
| TcCLB.411337.10 | retrotransposon hot spot protein (RHS, pseudogene), putat | 0,5649 | 0,01247 |
| TcCLB.511657.79 | hypothetical protein, conserved | 0,5646 | 0,06478 |
| TcCLB.511621.90 | esterase, putative | 0,5645 | 0,0013 |
| TcCLB.506401.150 | hypothetical protein, conserved | 0,5637 | 0,0007591 |
| TcCLB.509633.50 | serine/threonine-protein phosphatase PP1, putative | 0,5636 | 0,0716 |
| TcCLB.504137.160 | hypothetical protein, conserved | 0,5635 | 0,005313 |
| TcCLB.506145.10 | hypothetical protein, conserved | 0,5634 | 6,359E-12 |
| TcCLB.511323.60 | hypothetical protein, conserved | 0,5634 | 0,0000101 |
| TcCLB.506479.130 | cytidine triphosphate synthase, putative | 0,5631 | 0,0001254 |
| TcCLB.506993.40 | mucin TcMUCII, putative | 0,5631 | 0,3803 |
| TcCLB.504557.40 | hypothetical protein | 0,563 | 0,2018 |
| TcCLB.508865.30 | immunodominant antigen, putative | 0,5625 | 1,849E-15 |
| TcCLB.510533.210 | hypothetical protein, conserved (pseudogene) | 0,5623 | 0,00003729 |
| TcCLB.400945.10 | trans-sialidase (pseudogene), putative | 0,5623 | 0,2299 |
| TcCLB.511725.30 | hypothetical protein, conserved | 0,5622 | 0,005466 |
| TcCLB.511175.40 | mucin TcMUCII, putative | 0,562 | 0,1284 |
| TcCLB.508761.280 | mucin TcMUCII, putative | 0,562 | 0,254 |
| TcCLB.421951.10 | trans-sialidase, putative (fragment) | 0,562 | 0,4853 |
| TcCLB.506465.10 | hypothetical protein, conserved | 0,5618 | 0,00002465 |
| TcCLB.503823.120 | hypothetical protein, conserved | 0,5617 | 0,0006065 |
| TcCLB.506605.150 | 60S ribosomal protein, putative | 0,5616 | 0,0007576 |
| TcCLB.508265.100 | hypothetical protein, conserved | 0,5615 | 7,615E-10 |
| TcCLB.509799.110 | hypothetical protein, conserved | 0,5613 | 0,007517 |
| TcCLB.506311.30 | hypothetical protein, conserved | 0,5612 | 0,000176 |
| TcCLB.508029.20 | ubiquitin-protein ligase, putative | 0,5612 | 0,002718 |
| TcCLB.503939.40 | hypothetical protein, conserved | 0,5611 | 8,24E-08 |

| | | | |
|---|---|---|---|
| TcCLB.507087.80 | 5'-3' exoribonuclease B, putative (XRNB) | 0,561 | 1,234E-06 |
| TcCLB.505945.80 | hypothetical protein, conserved | 0,561 | 0,00001365 |
| TcCLB.510783.30 | dispersed gene family protein 1 (DGF-1, pseudogene), puta | 0,561 | 0,1829 |
| TcCLB.511553.5 | trans-sialidase (pseudogene), putative | 0,561 | 0,4004 |
| TcCLB.503627.20 | protein kinase, putative | 0,5607 | 2,981E-09 |
| TcCLB.504425.70 | hypothetical protein, conserved | 0,5607 | 4,107E-06 |
| TcCLB.503973.80 | mucin-associated surface protein (MASP), putative | 0,5607 | 0,1883 |
| TcCLB.504057.90 | hypothetical protein, conserved | 0,5605 | 1,595E-07 |
| TcCLB.508325.230 | trans-sialidase, Group IV, putative | 0,5604 | 3,078E-08 |
| TcCLB.507669.80 | protein phosphotase, putative | 0,5604 | 4,047E-07 |
| TcCLB.505989.90 | hypothetical protein, conserved | 0,5602 | 1,634E-06 |
| TcCLB.506825.50 | hypothetical protein, conserved | 0,5601 | 5,446E-08 |
| TcCLB.511321.20 | hypothetical protein, conserved | 0,5599 | 3,23E-08 |
| TcCLB.504155.280 | trans-sialidase (pseudogene), putative | 0,5599 | 0,03744 |
| TcCLB.506337.50 | hypothetical protein, conserved | 0,5598 | 9,867E-08 |
| TcCLB.507907.30 | retrotransposon hot spot (RHS) protein, putative | 0,5598 | 0,07874 |
| TcCLB.508601.90 | Small Surface Antigen (LDK) | 0,5597 | 0,00002042 |
| TcCLB.503825.19 | protein kinase, putative | 0,5597 | 0,06129 |
| TcCLB.511261.4 | P-type H -ATPase, putative (fragment) | 0,5596 | 2,016E-10 |
| TcCLB.504149.210 | N-acetyltransferase complex ARD1 subunit (pseudogene), | 0,5595 | 0,06473 |
| TcCLB.508417.20 | hypothetical protein | 0,5594 | 0,2515 |
| TcCLB.511813.10 | hypothetical protein, conserved | 0,5588 | 0,0001612 |
| TcCLB.510381.20 | ABC transporter, putative | 0,5588 | 0,0001855 |
| TcCLB.510367.10 | dispersed gene family protein 1 (DGF-1), putative | 0,5588 | 0,02046 |
| TcCLB.460747.30 | hypothetical protein, conserved | 0,5586 | 0,03135 |
| TcCLB.508153.750 | hypothetical protein, conserved | 0,5581 | 0,00009131 |
| TcCLB.503749.5 | pyrroline-5-carboxylate synthetase-like protein, putative (f | 0,5581 | 0,0006032 |
| TcCLB.506923.10 | trans-sialidase (pseudogene), putative | 0,5581 | 0,09639 |
| TcCLB.511277.400 | UDP-galactose transporter | 0,5579 | 0,0001359 |
| TcCLB.510377.400 | mucin TcMUCII, putative | 0,5578 | 0,3407 |
| TcCLB.511809.90 | hypothetical protein, conserved | 0,5577 | 1,111E-06 |
| TcCLB.509753.290 | hypothetical protein, conserved (pseudogene) | 0,5577 | 0,1528 |
| TcCLB.508449.20 | retrotransposon hot spot protein (RHS, pseudogene), putat | 0,5575 | 0,001714 |
| TcCLB.508831.80 | hypothetical protein, conserved | 0,5574 | 0,01223 |
| TcCLB.511887.10 | retrotransposon hot spot protein (RHS, pseudogene), putat | 0,5573 | 0,002269 |
| TcCLB.510359.290 | hypothetical protein, conserved | 0,557 | 0,032 |
| TcCLB.510575.239 | hypothetical protein, conserved | 0,557 | 0,2123 |
| TcCLB.508465.30 | hypothetical protein, conserved | 0,5568 | 2,457E-06 |
| TcCLB.505959.10 | mucin TcMUCII, putative | 0,5568 | 0,2536 |
| TcCLB.509767.90 | alanine racemase, putative | 0,5567 | 1,711E-06 |
| TcCLB.511925.30 | fucose kinase (pseudogene), putative | 0,5565 | 1,037E-06 |
| TcCLB.506573.81 | ribosomal protein L39, putative | 0,5565 | 0,0004925 |
| TcCLB.509065.140 | hypothetical protein, conserved | 0,5564 | 0,004536 |
| TcCLB.509733.150 | hypothetical protein, conserved | 0,5562 | 4,943E-07 |
| TcCLB.504777.50 | mucin-associated surface protein (MASP, pseudogene), pu | 0,5562 | 0,05659 |

| | | | |
|---|---|---|---|
| TcCLB.507981.30 | hypothetical protein | 0,5562 | 0,5266 |
| TcCLB.503801.30 | DNA repair protein RAD51, putative (RAD51) | 0,5561 | 1,119E-09 |
| TcCLB.507641.200 | hypothetical protein, conserved | 0,5561 | 0,005148 |
| TcCLB.508741.220 | hypothetical protein | 0,556 | 2,121E-07 |
| TcCLB.507757.26 | hypothetical protein | 0,556 | 0,3562 |
| TcCLB.503893.120 | calcium uniporter protein, mitochondrial, putative | 0,5559 | 0,001733 |
| TcCLB.504147.50 | amino acid transporter (pseudogene), putative | 0,5558 | 0,000124 |
| TcCLB.508981.30 | hypothetical protein, conserved | 0,5557 | 0,000882 |
| TcCLB.509393.30 | hypothetical protein, conserved | 0,5554 | 5,844E-06 |
| TcCLB.511577.100 | hypothetical protein, conserved | 0,5554 | 0,0006234 |
| TcCLB.506773.30 | hypothetical protein, conserved | 0,5552 | 2,182E-08 |
| TcCLB.507875.220 | trans-sialidase, Group II, putative | 0,5552 | 0,008818 |
| TcCLB.506757.150 | mucin-associated surface protein (MASP), putative | 0,5551 | 0,08129 |
| TcCLB.503661.11 | trans-sialidase, putative (fragment) | 0,5549 | 0,005692 |
| TcCLB.511259.18 | mucin-associated surface protein (MASP, pseudogene), pu | 0,5549 | 0,1805 |
| TcCLB.505063.30 | mucin-associated surface protein (MASP, pseudogene), pu | 0,5548 | 0,0004645 |
| TcCLB.507519.260 | retrotransposon hot spot protein (RHS, pseudogene), putat | 0,5547 | 3,63E-07 |
| TcCLB.507851.40 | hypothetical protein, conserved | 0,5546 | 0,002892 |
| TcCLB.510025.90 | mucin TcMUCII, putative | 0,5546 | 0,2833 |
| TcCLB.507159.60 | hypothetical protein, conserved | 0,5545 | 3,827E-11 |
| TcCLB.412419.10 | mucin-associated surface protein (MASP, pseudogene), pu | 0,5545 | 0,1649 |
| TcCLB.511045.29 | hypothetical protein, conserved | 0,5544 | 0,00003071 |
| TcCLB.506051.20 | hypothetical protein | 0,5543 | 0,2773 |
| TcCLB.508409.329 | hypothetical protein, conserved | 0,5541 | 0,0001945 |
| TcCLB.511071.50 | 3-hydroxy-3-methylglutaryl-CoA synthase, putative | 0,5539 | 3,058E-13 |
| TcCLB.504593.10 | trans-sialidase, putative (CRP10) | 0,5539 | 0,08371 |
| TcCLB.511431.40 | glutaredoxin, putative | 0,5538 | 0,001379 |
| TcCLB.510105.210 | metallo-peptidase, Clan MA(E) Family M41 | 0,5536 | 4,316E-08 |
| TcCLB.508777.60 | hypothetical protein, conserved | 0,5536 | 0,005312 |
| TcCLB.508859.30 | hypothetical protein, conserved | 0,5535 | 0,00006778 |
| TcCLB.504201.44 | mucin TcMUCII, putative | 0,5534 | 0,3678 |
| TcCLB.508761.200 | trans-sialidase (pseudogene), putative | 0,5533 | 0,1507 |
| TcCLB.509459.70 | hypothetical protein, conserved | 0,5532 | 4,851E-06 |
| TcCLB.506757.10 | phospholipid:diacylglycerol acyltransferase, putative | 0,5531 | 4,088E-08 |
| TcCLB.511535.20 | hypothetical protein, conserved | 0,5531 | 0,0001107 |
| TcCLB.509023.160 | hypothetical protein, conserved | 0,553 | 0,00004454 |
| TcCLB.506799.90 | surface protease GP63 (pseudogene), putative | 0,5525 | 0,003769 |
| TcCLB.508461.440 | hypothetical protein, conserved | 0,5523 | 3,983E-06 |
| TcCLB.509801.30 | citrate synthase, putative | 0,5523 | 0,1596 |
| TcCLB.505007.30 | hypothetical protein, conserved | 0,5522 | 0,000938 |
| TcCLB.508153.460 | Present in the outer mitochondrial membrane proteome 2 | 0,5521 | 3,442E-10 |
| TcCLB.398751.10 | hypothetical protein, conserved | 0,5519 | 0,2899 |
| TcCLB.508153.670 | hypothetical protein, conserved | 0,5518 | 0,04069 |
| TcCLB.506401.350 | hypothetical protein, conserved | 0,5517 | 1,477E-06 |
| TcCLB.508251.31 | mucin-associated surface protein (MASP, pseudogene), pu | 0,5512 | 0,1057 |

| | | | |
|---|---|---|---|
| TcCLB.508961.79 | 69 kDa paraflagellar rod protein, putative | 0,5511 | 3,227E-08 |
| TcCLB.504165.10 | hypothetical protein, conserved | 0,551 | 0,00009853 |
| TcCLB.509537.20 | hypothetical protein, conserved | 0,5501 | 8,977E-06 |
| TcCLB.509353.20 | Present in the outer mitochondrial membrane proteome 4 | 0,55 | 0,003761 |
| TcCLB.506559.490 | hypothetical protein, conserved | 0,5499 | 0,0313 |
| TcCLB.511255.490 | trans-sialidase (pseudogene), putative | 0,5499 | 0,2454 |
| TcCLB.508555.70 | calpain-like cysteine peptidase, putative | 0,5497 | 0,0007783 |
| TcCLB.506203.20 | dynein light chain, putative | 0,5496 | 0,00003655 |
| TcCLB.508727.50 | hypothetical protein, conserved | 0,5495 | 0,00047 |
| TcCLB.511259.149 | mucin-associated surface protein (MASP, pseudogene), pu | 0,5495 | 0,27 |
| TcCLB.509053.160 | hypothetical protein, conserved | 0,5494 | 0,0003699 |
| TcCLB.511517.99 | hypothetical protein, conserved | 0,5493 | 0,004229 |
| TcCLB.506487.24 | dispersed gene family protein 1 (DGF-1), putative | 0,5493 | 0,01606 |
| TcCLB.508515.100 | hypothetical protein, conserved | 0,5489 | 0,002556 |
| TcCLB.504427.240 | metallo-beta-lactamase-like protein, putative | 0,5488 | 0,002118 |
| TcCLB.510655.20 | hypothetical protein, conserved | 0,5487 | 0,00007205 |
| TcCLB.511833.10 | trans-sialidase, Group VIII, putative | 0,5487 | 0,009461 |
| TcCLB.508973.50 | ATP-dependent DEAD/H RNA helicase, putative | 0,5486 | 1,265E-08 |
| TcCLB.511101.37 | retrotransposon hot spot protein (RHS, pseudogene), putat | 0,5486 | 0,0002679 |
| TcCLB.509195.61 | mucin TcMUCII, putative | 0,5483 | 0,3422 |
| TcCLB.510099.110 | hypothetical protein, conserved | 0,5482 | 3,356E-09 |
| TcCLB.504769.89 | hypothetical protein, conserved | 0,5481 | 0,001573 |
| TcCLB.506767.360 | mucin TcMUCII, putative | 0,5481 | 0,2737 |
| TcCLB.509287.170 | trans-sialidase (pseudogene), putative | 0,548 | 0,00142 |
| TcCLB.510633.9 | dispersed gene family protein 1 (DGF-1), putative (fragme | 0,548 | 0,08189 |
| TcCLB.503791.20 | hypothetical protein, conserved | 0,5479 | 0,04568 |
| TcCLB.509253.30 | 3-Beta-hydroxysteroid-delta(8), delta(7)-isomerase, putati | 0,5478 | 0,0008471 |
| TcCLB.510851.50 | hypothetical protein, conserved (pseudogene) | 0,5474 | 0,0001271 |
| TcCLB.506253.39 | trans-sialidase, putative (CRP10) | 0,5474 | 0,09268 |
| TcCLB.508613.74 | mucin-associated surface protein (MASP), putative | 0,5471 | 0,0236 |
| TcCLB.507093.69 | hypothetical protein, conserved | 0,5469 | 0,0121 |
| TcCLB.510661.120 | hypothetical protein, conserved | 0,5467 | 0,0000959 |
| TcCLB.511469.100 | 2-oxoisovalerate dehydrogenase beta subunit, mitochondri | 0,5466 | 0,00004183 |
| TcCLB.506843.75 | hypothetical protein, conserved (pseudogene) | 0,5466 | 0,1797 |
| TcCLB.511705.20 | putative prolyl carboxypeptidase, putative | 0,5465 | 0,007274 |
| TcCLB.509611.20 | peptidyl-prolyl cis-trans isomerase (PIN1) | 0,5464 | 0,01232 |
| TcCLB.503623.10 | dispersed gene family protein 1 (DGF-1), putative | 0,5464 | 0,3803 |
| TcCLB.508299.70 | peroxisome targeting signal 1 receptor, putative (PEX5) | 0,5463 | 1,932E-07 |
| TcCLB.509885.10 | V-type proton ATPase subunit D, putative (VAd) | 0,5463 | 3,608E-06 |
| TcCLB.510955.40 | axoneme central apparatus protein (PF16) | 0,5459 | 4,539E-06 |
| TcCLB.508325.80 | trans-sialidase (pseudogene), putative | 0,5459 | 0,00005923 |
| TcCLB.507529.30 | hypothetical protein, conserved | 0,5458 | 2,903E-07 |
| TcCLB.511173.210 | mucin-associated surface protein (MASP), putative | 0,5457 | 0,308 |
| TcCLB.510697.10 | mucin-associated surface protein (MASP), putative | 0,5455 | 0,2842 |
| TcCLB.503809.120 | hypothetical protein, conserved | 0,5454 | 0,009619 |

| | | | |
|---|---|---|---|
| TcCLB.506667.130 | surface protease GP63 (pseudogene), putative | 0,5454 | 0,3777 |
| TcCLB.509113.10 | mucin-associated surface protein (MASP, pseudogene), pu | 0,5453 | 0,1404 |
| TcCLB.510431.250 | phosphoribosylpyrophosphate synthetase, putative | 0,5452 | 2,872E-10 |
| TcCLB.506537.160 | glycine dehydrogenase [decarboxylating] (pseudogene), pu | 0,5452 | 0,001243 |
| TcCLB.509529.40 | trans-sialidase, Group VII, putative | 0,5451 | 0,04889 |
| TcCLB.507959.210 | mucin-associated surface protein (MASP), putative | 0,5449 | 0,231 |
| TcCLB.509053.120 | hypothetical protein, conserved | 0,5448 | 0,0001699 |
| TcCLB.508991.30 | trans-sialidase (pseudogene), putative | 0,5448 | 0,00811 |
| TcCLB.506401.70 | vacuolar protein sorting protein 18, putative | 0,5444 | 6,103E-09 |
| TcCLB.506301.40 | hypothetical protein, conserved | 0,5444 | 0,00001704 |
| TcCLB.506359.40 | hypothetical protein, conserved | 0,5443 | 0,00004021 |
| TcCLB.506047.16 | N-acetyltransferase complex ARD1 subunit (pseudogene), | 0,5442 | 0,1293 |
| TcCLB.511873.20 | hypothetical protein | 0,544 | 0,1677 |
| TcCLB.510025.20 | mucin TcMUCII, putative | 0,544 | 0,271 |
| TcCLB.503667.10 | trans-sialidase, putative | 0,5439 | 0,0648 |
| TcCLB.506611.40 | mucin-associated surface protein (MASP), putative | 0,5435 | 0,0778 |
| TcCLB.510105.20 | Ribosomal RNA processing protein 41A (RRP41A) | 0,5434 | 8,363E-08 |
| TcCLB.510235.10 | dispersed gene family protein 1 (DGF-1), putative | 0,5434 | 0,0201 |
| TcCLB.508711.10 | Tetratricopeptide-like helical domain containing protein (fı | 0,5433 | 2,483E-14 |
| TcCLB.507615.80 | hypothetical protein, conserved | 0,5429 | 2,994E-07 |
| TcCLB.506195.50 | hypothetical protein, conserved | 0,5429 | 0,003349 |
| TcCLB.510361.80 | trans-sialidase, Group V, putative | 0,5425 | 0,1058 |
| TcCLB.509733.60 | hypothetical protein | 0,5424 | 0,00007782 |
| TcCLB.509573.30 | flagellar protofilament ribbon protein, putative | 0,5422 | 3,211E-08 |
| TcCLB.507087.10 | hydrolase-like protein (pseudogene), putative | 0,5421 | 0,01539 |
| TcCLB.511313.10 | hypothetical protein, conserved | 0,5419 | 0,01231 |
| TcCLB.507043.10 | mucin TcMUCI, putative | 0,5418 | 0,1648 |
| TcCLB.510819.30 | hypothetical protein | 0,5416 | 0,0003273 |
| TcCLB.509571.79 | Ras-related protein RabX1, putative (RABX1) | 0,5414 | 0,000146 |
| TcCLB.510237.70 | mucin-associated surface protein (MASP), putative | 0,5414 | 0,3185 |
| TcCLB.506661.20 | fatty acid elongase, putative | 0,5412 | 3,972E-06 |
| TcCLB.511867.130 | hypothetical protein, conserved | 0,5411 | 3,856E-08 |
| TcCLB.511807.180 | hypothetical protein, conserved | 0,5409 | 3,037E-13 |
| TcCLB.506333.10 | plasma-membrane proton-efflux P-type ATPase, putative | 0,5406 | 4,809E-16 |
| TcCLB.510535.30 | hypothetical protein | 0,5406 | 0,001043 |
| TcCLB.508607.40 | retrotransposon hot spot protein (RHS, pseudogene), putat | 0,5403 | 0,00003308 |
| TcCLB.505183.50 | hypothetical protein, conserved | 0,5402 | 0,00002519 |
| TcCLB.510483.190 | mucin TcMUCII, putative | 0,5402 | 0,1294 |
| TcCLB.504881.50 | ABC transporter, putative | 0,54 | 0,005535 |
| TcCLB.506755.30 | hypothetical protein, conserved | 0,5398 | 0,0001687 |
| TcCLB.506995.70 | mucin-associated surface protein (MASP), putative | 0,5397 | 0,1532 |
| TcCLB.508881.20 | hypothetical protein, conserved | 0,5395 | 0,00006909 |
| TcCLB.508351.10 | fructose-1,6-bisphosphatase, cytosolic, putative | 0,5392 | 0,00003145 |
| TcCLB.506139.60 | trans-sialidase (pseudogene), putative | 0,5392 | 0,03138 |
| TcCLB.510571.30 | actin 1, putative | 0,5391 | 7,216E-12 |

| | | | |
|---|---|---|---|
| TcCLB.511127.410 | hypothetical protein, conserved | 0,5388 | 5,403E-07 |
| TcCLB.454391.10 | trans-sialidase (pseudogene), putative | 0,5388 | 0,0001328 |
| TcCLB.507069.130 | mucin-associated surface protein (MASP, pseudogene), pu | 0,5387 | 0,1951 |
| TcCLB.506449.10 | trans-sialidase, Group III, putative | 0,5385 | 0,6446 |
| TcCLB.503713.20 | hypothetical protein, conserved | 0,5379 | 0,0137 |
| TcCLB.506715.20 | Lys-63-specific deubiquitinase BRCC36, putative | 0,5378 | 0,01799 |
| TcCLB.508163.400 | surface antigen TASV-B1, putative | 0,5378 | 0,1666 |
| TcCLB.507869.20 | trans-sialidase (pseudogene), putative | 0,5376 | 7,684E-06 |
| TcCLB.503689.4 | Ras-related protein RabX1, putative (RABX1) | 0,5371 | 0,0008699 |
| TcCLB.506423.10 | mucin-associated surface protein (MASP), putative | 0,5371 | 0,4164 |
| TcCLB.503955.100 | protein kinase, putative | 0,537 | 1,102E-06 |
| TcCLB.509443.60 | hypothetical protein, conserved | 0,537 | 0,02742 |
| TcCLB.511141.20 | choline dehydrogenase, putative | 0,5369 | 4,064E-10 |
| TcCLB.506485.70 | hypothetical protein, conserved | 0,5369 | 8,772E-06 |
| TcCLB.506529.220 | hypothetical protein, conserved | 0,5369 | 0,0002594 |
| TcCLB.508147.159 | dispersed gene family protein 1 (DGF-1), putative | 0,5366 | 0,1908 |
| TcCLB.510877.180 | GPI transamidase subunit 16, putative (GPI16) | 0,5365 | 1,132E-08 |
| TcCLB.509029.20 | hypothetical protein, conserved | 0,5365 | 1,008E-06 |
| TcCLB.508589.60 | hypothetical protein, conserved (pseudogene) | 0,5362 | 0,01172 |
| TcCLB.511473.10 | retrotransposon hot spot protein (RHS, pseudogene), putat | 0,536 | 0,001568 |
| TcCLB.510155.40 | DNA-damage inducible protein DDI1-like protein, putative | 0,5358 | 2,255E-07 |
| TcCLB.506703.74 | mucin-associated surface protein (MASP, pseudogene), pu | 0,5358 | 0,5329 |
| TcCLB.506973.90 | mucin-associated surface protein (MASP, pseudogene), pu | 0,5352 | 0,1627 |
| TcCLB.510303.320 | dihydrofolate reductase-thymidylate synthase, putative | 0,5351 | 0,0008355 |
| TcCLB.506835.60 | calmodulin, putative | 0,535 | 0,00587 |
| TcCLB.511089.90 | mucin-associated surface protein (MASP), putative | 0,535 | 0,5922 |
| TcCLB.478283.10 | hypothetical protein, conserved (pseudogene) | 0,5348 | 0,004014 |
| TcCLB.506955.90 | trans-sialidase (pseudogene), putative | 0,5348 | 0,1752 |
| TcCLB.503761.40 | mucin-associated surface protein (MASP), putative | 0,5346 | 0,05606 |
| TcCLB.506409.200 | mucin TcMUCII, putative | 0,5346 | 0,2321 |
| TcCLB.506939.220 | mucin-associated surface protein (MASP), putative | 0,5346 | 0,2691 |
| TcCLB.505051.10 | hypothetical protein, conserved | 0,5345 | 1,07E-20 |
| TcCLB.511287.20 | hypothetical protein, conserved | 0,5345 | 9,659E-06 |
| TcCLB.511099.210 | mucin TcMUCII, putative | 0,5345 | 0,4097 |
| TcCLB.506337.210 | signal recognition particle receptor like protein, putative | 0,5344 | 3,033E-07 |
| TcCLB.505987.60 | hypothetical protein | 0,5343 | 5,56E-10 |
| TcCLB.511087.19 | trans-sialidase, putative (fragment) | 0,5343 | 0,2842 |
| TcCLB.506539.10 | trans-sialidase (pseudogene), putative | 0,534 | 0,115 |
| TcCLB.511851.5 | trans-sialidase, putative | 0,5333 | 0,2831 |
| TcCLB.510025.100 | mucin-associated surface protein (MASP), putative | 0,533 | 0,05873 |
| TcCLB.510623.80 | trans-sialidase (pseudogene), putative | 0,533 | 0,1497 |
| TcCLB.505931.30 | trans-sialidase, Group I, putative | 0,5329 | 0,009646 |
| TcCLB.509529.60 | dispersed gene family protein 1 (DGF-1), putative | 0,5327 | 0,0006738 |
| TcCLB.509109.140 | hypothetical protein, conserved | 0,5324 | 0,00003841 |
| TcCLB.510359.30 | hypothetical protein, conserved | 0,5324 | 0,002771 |

| | | | |
|---|---|---|---|
| TcCLB.511711.40 | hypothetical protein, conserved | 0,5324 | 0,004498 |
| TcCLB.507851.20 | hypothetical protein, conserved | 0,5321 | 0,001217 |
| TcCLB.508799.200 | hypothetical protein, conserved | 0,532 | 0,05853 |
| TcCLB.509979.40 | hypothetical protein, conserved | 0,5319 | 0,00006167 |
| TcCLB.508895.80 | intraflagellar transport protein 52, putative (IFT52) | 0,5318 | 3,724E-07 |
| TcCLB.511165.54 | pterin-4-alpha-carbinolamine dehydratase, putative | 0,5318 | 0,0009013 |
| TcCLB.509307.10 | retrotransposon hot spot protein (RHS, pseudogene), putat | 0,5317 | 0,001329 |
| TcCLB.509867.60 | mucin-associated surface protein (MASP), putative | 0,5316 | 0,1713 |
| TcCLB.508961.40 | hypothetical protein, conserved | 0,5315 | 0,00006097 |
| TcCLB.511185.114 | mucin TcMUCII (pseudogene), putative | 0,5315 | 0,5186 |
| TcCLB.510741.150 | Present in the outer mitochondrial membrane proteome 7 | 0,5313 | 0,0001432 |
| TcCLB.508435.10 | mucin-associated surface protein (MASP), putative | 0,5312 | 0,3044 |
| TcCLB.510323.60 | hypothetical protein | 0,5311 | 0,001707 |
| TcCLB.511127.250 | hypothetical protein, conserved | 0,531 | 0,003685 |
| TcCLB.504253.40 | hypothetical protein, conserved | 0,5309 | 0,0001804 |
| TcCLB.508223.30 | retrotransposon hot spot protein (RHS, pseudogene), putat | 0,5309 | 0,1375 |
| TcCLB.504163.20 | hypothetical protein, conserved | 0,5308 | 0,00002245 |
| TcCLB.511667.80 | trans-sialidase (pseudogene), putative | 0,5308 | 0,01459 |
| TcCLB.504221.9 | membrane associated protein, putative | 0,5307 | 0,00005389 |
| TcCLB.511063.120 | mucin TcMUCII, putative | 0,5307 | 0,5692 |
| TcCLB.508881.150 | hypothetical protein, conserved | 0,5304 | 0,0003002 |
| TcCLB.509963.60 | hypothetical protein, conserved | 0,5302 | 0,01611 |
| TcCLB.503823.104 | UDP-N-acetylglucosamine-dolichyl-phosphate N-acetylgluc | 0,5301 | 0,0003426 |
| TcCLB.504035.130 | hypothetical protein, conserved | 0,5301 | 0,00122 |
| TcCLB.506057.61 | elongation factor 1-gamma (EF-1-gamma, pseudogene), p | 0,5301 | 0,1803 |
| TcCLB.510307.320 | retrotransposon hot spot protein (RHS, pseudogene), putat | 0,5298 | 0,002694 |
| TcCLB.510417.30 | mucin TcMUCII, putative | 0,5298 | 0,2575 |
| TcCLB.504229.70 | retrotransposon hot spot (RHS) protein, putative | 0,5297 | 0,002104 |
| TcCLB.510713.70 | retrotransposon hot spot protein (RHS, pseudogene), putat | 0,5296 | 0,005588 |
| TcCLB.511555.30 | hypothetical protein, conserved | 0,5294 | 0,003197 |
| TcCLB.506195.230 | hypothetical protein, conserved | 0,5289 | 0,0001127 |
| TcCLB.506223.60 | hypothetical protein, conserved | 0,5289 | 0,00158 |
| TcCLB.507711.330 | serine peptidase, putative | 0,5288 | 0,0001758 |
| TcCLB.506871.20 | hypothetical protein, conserved | 0,5287 | 8,177E-07 |
| TcCLB.507825.40 | hypothetical protein, conserved | 0,5287 | 0,0001172 |
| TcCLB.504137.30 | hypothetical protein | 0,5284 | 0,00000913 |
| TcCLB.507163.70 | trans-sialidase, Group V, putative | 0,5284 | 0,008484 |
| TcCLB.508231.230 | protein transport protein Sec23A, putative | 0,5283 | 0,002139 |
| TcCLB.510025.14 | hypothetical protein, conserved (pseudogene) | 0,5282 | 0,5585 |
| TcCLB.506999.140 | hypothetical protein, conserved | 0,5277 | 3,712E-07 |
| TcCLB.410337.10 | trans-sialidase (pseudogene), putative | 0,5275 | 0,342 |
| TcCLB.508521.30 | beta galactofuranosyl glycosyltransferase (pseudogene), p | 0,5274 | 0,022 |
| TcCLB.510275.10 | hypothetical protein | 0,5273 | 0,008964 |
| TcCLB.508879.210 | hypothetical protein, conserved | 0,5272 | 0,0001307 |
| TcCLB.511361.40 | actin-related protein 2, putative | 0,5272 | 0,002161 |

| | | | |
|---|---|---|---|
| TcCLB.506823.100 | dispersed gene family protein 1 (DGF-1), putative | 0,527 | 0,168 |
| TcCLB.510377.260 | mucin-associated surface protein (MASP), putative | 0,527 | 0,5668 |
| TcCLB.506009.20 | hypothetical protein, conserved | 0,5268 | 1,132E-08 |
| TcCLB.511807.200 | hypothetical protein, conserved | 0,5267 | 8,321E-13 |
| TcCLB.510101.340 | hypothetical protein, conserved | 0,5267 | 2,305E-12 |
| TcCLB.505317.20 | hypothetical protein | 0,5267 | 0,002199 |
| TcCLB.505075.30 | hypothetical protein, conserved | 0,5266 | 0,002654 |
| TcCLB.503893.130 | hypothetical protein, conserved | 0,5265 | 0,0005944 |
| TcCLB.504797.130 | hypothetical protein, conserved | 0,5265 | 0,01096 |
| TcCLB.507949.210 | trans-sialidase, Group II, putative | 0,5265 | 0,01965 |
| TcCLB.508285.20 | hypothetical protein, conserved (pseudogene) | 0,5259 | 0,008732 |
| TcCLB.509099.30 | paraflagellar rod protein 5, putative (PFR5) | 0,5258 | 0,00004346 |
| TcCLB.445557.10 | mucin TcMUCII, putative | 0,5258 | 0,482 |
| TcCLB.511395.70 | hypothetical protein, conserved | 0,5256 | 0,0004678 |
| TcCLB.507677.80 | hypothetical protein, conserved | 0,5256 | 0,05656 |
| TcCLB.511089.74 | mucin TcMUCII (pseudogene), putative | 0,5254 | 0,436 |
| TcCLB.506279.19 | mucin-associated surface protein (MASP, pseudogene), pu | 0,5253 | 0,06492 |
| TcCLB.506301.20 | Golgi reassembly stacking protein (GRASP homologue), pu | 0,5251 | 0,00005507 |
| TcCLB.510287.10 | dynein light chain, putative | 0,5249 | 0,001818 |
| TcCLB.505183.124 | hypothetical protein, conserved | 0,5249 | 0,002729 |
| TcCLB.506825.90 | hypothetical protein, conserved | 0,5248 | 6,88E-09 |
| TcCLB.510819.4 | NADH-cytochrome b5 reductase, putative | 0,5248 | 0,0006653 |
| TcCLB.510155.30 | hypothetical protein, conserved | 0,5247 | 8,971E-08 |
| TcCLB.506529.425 | hypothetical protein | 0,5246 | 0,295 |
| TcCLB.508227.30 | mucin TcMUCII, putative | 0,5243 | 0,3513 |
| TcCLB.508153.650 | hypothetical protein, conserved (pseudogene) | 0,5241 | 4,556E-06 |
| TcCLB.508909.190 | hypothetical protein, conserved | 0,524 | 0,00007748 |
| TcCLB.506629.180 | hypothetical protein, conserved | 0,5238 | 0,003751 |
| TcCLB.511317.79 | OSM3-like kinesin, putative | 0,5238 | 0,01105 |
| TcCLB.510197.190 | mucin-associated surface protein (MASP), putative | 0,5238 | 0,223 |
| TcCLB.508081.10 | mucin-associated surface protein (MASP), putative | 0,5238 | 0,7259 |
| TcCLB.506977.49 | ribosomal protein L36, putative | 0,5237 | 2,985E-10 |
| TcCLB.509527.200 | hypothetical protein | 0,5236 | 0,1897 |
| TcCLB.511843.40 | dispersed gene family protein 1 (DGF-1), putative | 0,5234 | 0,04605 |
| TcCLB.507547.59 | hypothetical protein, conserved | 0,5233 | 0,02938 |
| TcCLB.507835.35 | trans-sialidase (pseudogene), putative | 0,5232 | 0,1727 |
| TcCLB.510741.80 | riboflavin kinase, putative | 0,5231 | 0,003681 |
| TcCLB.510945.30 | actin, putative | 0,5229 | 5,584E-06 |
| TcCLB.508973.110 | hypothetical protein, conserved | 0,5229 | 0,0004156 |
| TcCLB.506735.20 | hypothetical protein, conserved | 0,5228 | 0,0009093 |
| TcCLB.507445.9 | dispersed gene family protein 1 (DGF-1), putative (fragme | 0,5228 | 0,03039 |
| TcCLB.505949.100 | trans-sialidase (pseudogene), putative | 0,5226 | 0,008982 |
| TcCLB.503897.170 | citrate transporter, putative | 0,5225 | 0,001084 |
| TcCLB.508761.9 | mucin-associated surface protein (MASP, pseudogene), pu | 0,5225 | 0,2522 |
| TcCLB.508241.100 | hypothetical protein | 0,5224 | 1,293E-09 |

| | | | |
|---|---|---|---|
| TcCLB.506375.110 | (H )-ATPase G subunit, putative | 0,5224 | 2,471E-06 |
| TcCLB.416605.10 | mucin-associated surface protein (MASP), putative | 0,5222 | 0,4046 |
| TcCLB.509815.50 | beta galactofuranosyl glycosyltransferase, putative | 0,522 | 0,02098 |
| TcCLB.510807.10 | mucin TcMUCII, putative | 0,5218 | 0,4112 |
| TcCLB.509237.60 | UDP-Gal or UDP-GlcNAc-dependent glycosyltransferase, pu | 0,5217 | 0,002304 |
| TcCLB.508075.50 | hypothetical protein, conserved | 0,5216 | 0,0001262 |
| TcCLB.510227.20 | hypothetical protein | 0,5216 | 0,0009275 |
| TcCLB.510533.190 | hypothetical protein, conserved | 0,5215 | 0,005854 |
| TcCLB.507991.20 | protein kinase, putative | 0,5213 | 0,001227 |
| TcCLB.508221.52 | hypothetical protein | 0,5213 | 0,3948 |
| TcCLB.437349.10 | hypothetical protein, conserved | 0,5212 | 0,01006 |
| TcCLB.510371.120 | mucin-associated surface protein (MASP), putative | 0,5212 | 0,3567 |
| TcCLB.506789.200 | ubiquitin-conjugating enzyme E2, putative | 0,5211 | 4,381E-06 |
| TcCLB.510381.70 | hypothetical protein, conserved | 0,5211 | 0,005359 |
| TcCLB.506955.99 | mucin-associated surface protein (MASP), putative | 0,5211 | 0,2283 |
| TcCLB.506833.60 | UV excision repair RAD23-like protein, putative | 0,521 | 3,918E-07 |
| TcCLB.508479.110 | glutathionylspermidine synthase, putative | 0,5208 | 2,224E-08 |
| TcCLB.510283.140 | UDP-N-acetylglucosamine-dolichyl-phosphate N-acetylgluc | 0,5206 | 0,00008383 |
| TcCLB.506775.160 | hypothetical protein, conserved | 0,5206 | 0,0001379 |
| TcCLB.506459.240 | elongation factor 1-gamma (EF-1-gamma), putative | 0,5206 | 0,008636 |
| TcCLB.510053.60 | cytidine deaminase-like protein, putative | 0,5202 | 0,002855 |
| TcCLB.506673.20 | hypothetical protein, conserved | 0,5202 | 0,01592 |
| TcCLB.506645.9 | dispersed gene family protein 1 (DGF-1), putative | 0,5202 | 0,01601 |
| TcCLB.506529.516 | ATP synthase subunit, putative | 0,5202 | 0,02126 |
| TcCLB.508081.90 | mucin-associated surface protein (MASP), putative | 0,52 | 0,1469 |
| TcCLB.511693.40 | hypothetical protein, conserved | 0,5195 | 1,456E-08 |
| TcCLB.503591.10 | retrotransposon hot spot protein (RHS, pseudogene), putat | 0,5195 | 0,01779 |
| TcCLB.509875.80 | trans-sialidase, Group II, putative | 0,5192 | 0,1457 |
| TcCLB.505997.190 | retrotransposon hot spot protein (RHS, pseudogene), putat | 0,5189 | 0,001031 |
| TcCLB.506965.20 | mucin-associated surface protein (MASP), putative | 0,5189 | 0,3835 |
| TcCLB.506269.90 | mucin TcMUCII, putative | 0,5187 | 0,0475 |
| TcCLB.510055.60 | hypothetical protein, conserved | 0,5186 | 9,355E-09 |
| TcCLB.511267.24 | hypothetical protein, conserved | 0,5185 | 0,000258 |
| TcCLB.510879.80 | serine acetyltransferase, putative (SAT) | 0,5184 | 0,0002632 |
| TcCLB.503907.10 | trans-sialidase, Group VIII, putative | 0,5182 | 0,1372 |
| TcCLB.508991.10 | retrotransposon hot spot protein (RHS, pseudogene), putat | 0,5181 | 0,0004391 |
| TcCLB.509259.80 | hypothetical protein, conserved (pseudogene) | 0,518 | 0,001935 |
| TcCLB.504005.20 | N-Acetyl-D-glucosaminylphosphatidylinositol de-N-acetyla: | 0,518 | 0,003325 |
| TcCLB.507881.90 | trans-sialidase (pseudogene), putative | 0,518 | 0,00477 |
| TcCLB.506195.300 | deoxyhypusine synthase, putative | 0,5177 | 9,773E-06 |
| TcCLB.503537.18 | retrotransposon hot spot protein (RHS, pseudogene), putat | 0,5173 | 0,1482 |
| TcCLB.511605.90 | mucin TcMUCII, putative | 0,517 | 0,5102 |
| TcCLB.504899.20 | retrotransposon hot spot protein (RHS, pseudogene), putat | 0,5169 | 0,001669 |
| TcCLB.508479.220 | hypothetical protein, conserved | 0,5169 | 0,004459 |
| TcCLB.510625.88 | mucin-associated surface protein (MASP), putative | 0,5169 | 0,5319 |

| | | | |
|---|---|---|---|
| TcCLB.508973.90 | Present in the outer mitochondrial membrane proteome 3 | 0,5168 | 9,673E-06 |
| TcCLB.510025.170 | mucin TcMUCII, putative | 0,5163 | 0,3354 |
| TcCLB.511367.159 | hypothetical protein, conserved (pseudogene) | 0,5162 | 0,007514 |
| TcCLB.506699.6 | hypothetical protein, conserved | 0,5162 | 0,01292 |
| TcCLB.508789.80 | mucin-associated surface protein (MASP), putative | 0,5161 | 0,3581 |
| TcCLB.507091.4 | fucose kinase, putative | 0,516 | 0,03547 |
| TcCLB.510835.30 | hypothetical protein, conserved | 0,5158 | 0,001935 |
| TcCLB.509991.80 | hypothetical protein, conserved | 0,5153 | 0,000853 |
| TcCLB.510557.30 | hypothetical protein | 0,5153 | 0,3883 |
| TcCLB.508879.120 | hypothetical protein, conserved | 0,5152 | 0,0000188 |
| TcCLB.510761.76 | UDP-glucoronosyl and UDP-glucosyl transferase, putative | 0,5152 | 0,0001388 |
| TcCLB.510635.30 | dispersed gene family protein 1 (DGF-1, pseudogene), puta | 0,5152 | 0,2186 |
| TcCLB.508977.99 | mucin-associated surface protein (MASP, pseudogene), pu | 0,5152 | 0,3785 |
| TcCLB.503663.20 | hypothetical protein, conserved | 0,5151 | 6,889E-07 |
| TcCLB.511827.110 | trans-sialidase, Group II, putative | 0,5151 | 0,00005903 |
| TcCLB.510729.290 | Enriched in surface-labeled proteome protein 19 (SLS3) | 0,5149 | 0,00001053 |
| TcCLB.506615.75 | surface protease GP63 (pseudogene), putative | 0,5149 | 0,0001256 |
| TcCLB.510663.40 | hypothetical protein, conserved | 0,5148 | 0,0001386 |
| TcCLB.508889.19 | hypothetical protein | 0,5146 | 0,001495 |
| TcCLB.506543.30 | hypothetical protein, conserved | 0,5146 | 0,1036 |
| TcCLB.508421.30 | tryptophanyl-tRNA synthetase (pseudogene), putative | 0,5145 | 3,868E-10 |
| TcCLB.508083.10 | Present in the outer mitochondrial membrane proteome 1 | 0,5145 | 5,858E-07 |
| TcCLB.508261.130 | mucin-associated surface protein (MASP), putative | 0,5145 | 0,2857 |
| TcCLB.463901.10 | trans-sialidase, putative (fragment) | 0,5144 | 0,06375 |
| TcCLB.510729.115 | hypothetical protein | 0,5143 | 0,0004664 |
| TcCLB.506127.150 | hypothetical protein, conserved | 0,5142 | 2,97E-07 |
| TcCLB.511733.50 | hypothetical protein, conserved | 0,5141 | 0,00000239 |
| TcCLB.504149.117 | hypothetical protein, conserved | 0,5139 | 0,0001157 |
| TcCLB.511731.70 | hypothetical protein, conserved | 0,5135 | 0,006183 |
| TcCLB.507611.50 | hypothetical protein, conserved | 0,5134 | 0,366 |
| TcCLB.507159.70 | hypothetical protein, conserved | 0,5133 | 0,00006359 |
| TcCLB.511151.54 | surface protease GP63, putative | 0,5133 | 0,0003807 |
| TcCLB.511395.50 | hypothetical protein, conserved | 0,5133 | 0,0006028 |
| TcCLB.481099.9 | dispersed gene family protein 1 (DGF-1), putative (fragme | 0,5133 | 0,1294 |
| TcCLB.506913.10 | hypothetical protein, conserved | 0,513 | 0,02579 |
| TcCLB.506775.80 | 14-3-3 protein 2, putative (14-3-3-II) | 0,5127 | 2,635E-12 |
| TcCLB.511029.10 | enolase, putative | 0,5124 | 0,0004664 |
| TcCLB.507725.9 | dispersed gene family protein 1 (DGF-1, pseudogene), puta | 0,5124 | 0,003514 |
| TcCLB.511593.70 | mucin TcMUCII, putative | 0,5121 | 0,3964 |
| TcCLB.506851.10 | NIMA-related kinase, putative | 0,512 | 6,895E-09 |
| TcCLB.508813.80 | hypothetical protein, conserved | 0,512 | 6,052E-06 |
| TcCLB.509877.10 | hypothetical protein, conserved | 0,512 | 0,000699 |
| TcCLB.506475.110 | hypothetical protein | 0,5119 | 0,04923 |
| TcCLB.511509.50 | hypothetical protein, conserved | 0,5116 | 0,00001164 |
| TcCLB.507071.310 | mucin-associated surface protein (MASP), putative | 0,5116 | 0,3425 |

| | | | |
|---|---|---|---|
| TcCLB.506999.30 | hypothetical protein, conserved | 0,5113 | 3,674E-06 |
| TcCLB.511827.10 | hypothetical protein, conserved | 0,5112 | 0,08645 |
| TcCLB.506135.70 | hypothetical protein, conserved | 0,511 | 0,0004996 |
| TcCLB.507511.50 | hypothetical protein | 0,511 | 0,02513 |
| TcCLB.503587.80 | mucin-associated surface protein (MASP, pseudogene), pu | 0,5109 | 0,302 |
| TcCLB.506801.20 | mucin-like glycoprotein, putative | 0,5107 | 0,09821 |
| TcCLB.509601.44 | hypothetical protein, conserved | 0,5105 | 0,00000224 |
| TcCLB.504125.64 | hypothetical protein | 0,5105 | 0,00007001 |
| TcCLB.510105.70 | hypothetical protein, conserved | 0,5105 | 0,005995 |
| TcCLB.507039.20 | GDP-mannose 4,6 dehydratase (pseudogene), putative | 0,5104 | 0,00453 |
| TcCLB.509615.10 | hypothetical protein, conserved | 0,5101 | 0,006784 |
| TcCLB.510515.20 | hypothetical protein, conserved (pseudogene) | 0,51 | 8,601E-06 |
| TcCLB.510411.10 | trans-sialidase, Group II, putative | 0,5099 | 0,05175 |
| TcCLB.503645.10 | mucin-associated surface protein (MASP), putative | 0,5098 | 0,2594 |
| TcCLB.511451.59 | mucin-associated surface protein (MASP), putative | 0,5098 | 0,4064 |
| TcCLB.509207.110 | hypothetical protein, conserved | 0,5097 | 0,001666 |
| TcCLB.410943.10 | hypothetical protein, conserved | 0,5096 | 0,1457 |
| TcCLB.511731.5 | cationic amino acid transporter (pseudogene), putative | 0,5095 | 0,1622 |
| TcCLB.508171.10 | trans-sialidase, Group VIII, putative | 0,5094 | 0,1222 |
| TcCLB.506829.90 | conserved protein | 0,509 | 3,643E-10 |
| TcCLB.510055.160 | protein kinase, putative | 0,509 | 0,09678 |
| TcCLB.511255.630 | mucin-associated surface protein (MASP), putative | 0,509 | 0,3637 |
| TcCLB.504149.110 | GTP-binding protein, putative | 0,5089 | 0,00001375 |
| TcCLB.511625.30 | hypothetical protein, conserved | 0,5088 | 0,002221 |
| TcCLB.508515.10 | hypothetical protein, conserved | 0,5086 | 0,0109 |
| TcCLB.506449.30 | retrotransposon hot spot protein (RHS, pseudogene), putat | 0,5086 | 0,1432 |
| TcCLB.506267.70 | mucin-associated surface protein (MASP), putative | 0,5085 | 0,3091 |
| TcCLB.506925.450 | hypothetical protein, conserved | 0,5083 | 4,873E-06 |
| TcCLB.506653.50 | elongation factor 1-gamma (EF-1-gamma, pseudogene), p | 0,5083 | 0,2823 |
| TcCLB.504149.40 | tatD related deoxyribonuclease, putative | 0,5081 | 1,098E-07 |
| TcCLB.507059.80 | glutamine-dependent carbamoyl-phosphate synthetase, pu | 0,508 | 0,00001414 |
| TcCLB.508723.50 | hypothetical protein, conserved | 0,5078 | 0,00003325 |
| TcCLB.507851.10 | hypothetical protein, conserved | 0,5076 | 0,00003632 |
| TcCLB.509631.100 | hypothetical protein, conserved (pseudogene) | 0,5074 | 0,3935 |
| TcCLB.510839.11 | hypothetical protein | 0,507 | 0,02466 |
| TcCLB.508231.20 | 40S ribosomal protein S12, putative | 0,5069 | 4,95E-11 |
| TcCLB.504469.20 | hypothetical protein (fragment) | 0,5069 | 0,09279 |
| TcCLB.506779.50 | proteasome beta 3 subunit, putative | 0,5068 | 0,00001894 |
| TcCLB.506321.50 | mucin-associated surface protein (MASP), putative | 0,5068 | 0,005981 |
| TcCLB.403153.10 | receptor-type adenylate cyclase (pseudogene), putative | 0,5067 | 0,07519 |
| TcCLB.511221.20 | hypothetical protein | 0,5067 | 0,1375 |
| TcCLB.511807.10 | hypothetical protein, conserved (pseudogene) | 0,506 | 5,544E-08 |
| TcCLB.506501.184 | mucin-associated surface protein (MASP, pseudogene), pu | 0,5059 | 0,3986 |
| TcCLB.510303.70 | hypothetical protein, conserved | 0,5057 | 0,00001502 |
| TcCLB.506757.60 | trans-sialidase, Group V, putative | 0,5057 | 0,0002947 |

| | | | |
|---|---|---|---|
| TcCLB.511717.110 | hypothetical protein, conserved | 0,5055 | 0,001464 |
| TcCLB.448661.20 | mucin TcMUCII, putative | 0,5055 | 0,363 |
| TcCLB.507067.50 | mucin TcMUCII (pseudogene), putative | 0,5055 | 0,4213 |
| TcCLB.509437.140 | retrotransposon hot spot protein (RHS, pseudogene), putat | 0,5054 | 0,05357 |
| TcCLB.509467.20 | hypothetical protein, conserved (pseudogene) | 0,5054 | 0,2354 |
| TcCLB.508153.100 | acyltransferase, putative | 0,5049 | 0,0003938 |
| TcCLB.511593.60 | mucin-associated surface protein (MASP), putative | 0,5049 | 0,1302 |
| TcCLB.418613.20 | mucin-associated surface protein (MASP), putative (fragm | 0,5049 | 0,4785 |
| TcCLB.509011.60 | hypothetical protein, conserved | 0,5048 | 0,00001384 |
| TcCLB.511245.30 | ion transport protein, putative | 0,5047 | 0,0004615 |
| TcCLB.511817.70 | hypothetical protein, conserved | 0,5047 | 0,008136 |
| TcCLB.504239.10 | hypothetical protein, conserved (pseudogene) | 0,5047 | 0,4493 |
| TcCLB.511185.100 | mucin-associated surface protein (MASP), putative | 0,5047 | 0,6013 |
| TcCLB.511001.50 | hypothetical protein, conserved | 0,5046 | 0,0003816 |
| TcCLB.504343.30 | retrotransposon hot spot (RHS) protein, putative | 0,5046 | 0,01153 |
| TcCLB.510123.70 | vesicle-associated membrane protein, putative | 0,5042 | 0,0002166 |
| TcCLB.506599.240 | mucin-associated surface protein (MASP), putative | 0,5042 | 0,07693 |
| TcCLB.510861.150 | hypothetical protein | 0,5041 | 1,388E-08 |
| TcCLB.506173.150 | hypothetical protein | 0,5041 | 0,1981 |
| TcCLB.506559.300 | twinfilin, putative | 0,5037 | 1,925E-08 |
| TcCLB.507559.50 | kinesin, putative | 0,5034 | 0,0004174 |
| TcCLB.510157.10 | mucin-associated surface protein (MASP), putative | 0,5029 | 0,2063 |
| TcCLB.508051.30 | protein kinase, putative | 0,5028 | 0,00009538 |
| TcCLB.510311.30 | hypothetical protein, conserved | 0,5027 | 8,566E-06 |
| TcCLB.511221.35 | retrotransposon hot spot protein (RHS, pseudogene), putat | 0,5027 | 0,004953 |
| TcCLB.509763.20 | hypothetical protein | 0,5023 | 0,01416 |
| TcCLB.511557.20 | hypothetical protein, conserved | 0,5022 | 0,003097 |
| TcCLB.506989.309 | hypothetical protein, conserved | 0,502 | 5,012E-06 |
| TcCLB.510353.20 | hypothetical protein, conserved | 0,5018 | 0,00003115 |
| TcCLB.510421.200 | hypothetical protein, conserved | 0,5017 | 0,0001093 |
| TcCLB.507253.10 | retrotransposon hot spot (RHS) protein, putative | 0,5017 | 0,007548 |
| TcCLB.508221.199 | elongation factor 1-gamma (EF-1-gamma), putative | 0,5017 | 0,324 |
| TcCLB.484097.10 | trans-sialidase, putative (fragment) | 0,5016 | 0,03882 |
| TcCLB.509001.40 | calpain-like cysteine peptidase, putative | 0,5015 | 9,647E-08 |
| TcCLB.506583.50 | hypothetical protein, conserved | 0,5015 | 0,0009256 |
| TcCLB.504031.20 | surface protease GP63 (pseudogene), putative | 0,5013 | 0,2808 |
| TcCLB.508403.20 | hypothetical protein, conserved (pseudogene) | 0,5012 | 0,00002559 |
| TcCLB.508699.80 | hypothetical protein, conserved | 0,501 | 0,0002855 |
| TcCLB.503903.79 | hypothetical protein | 0,501 | 0,001381 |
| TcCLB.508413.20 | hypothetical protein, conserved | 0,5006 | 0,0001114 |
| TcCLB.508827.40 | acyl-CoA dehydrogenase, putative | 0,5003 | 0,0008433 |
| TcCLB.508593.10 | trans-sialidase, Group IV, putative | 0,5002 | 0,01119 |
| TcCLB.508331.20 | hypothetical protein, conserved | 0,4999 | 8,665E-10 |
| TcCLB.506743.20 | hypothetical protein, conserved | 0,4999 | 0,04279 |
| TcCLB.506791.70 | nitroreductase | 0,4998 | 0,0003286 |

| | | | |
|---|---|---|---|
| TcCLB.503923.20 | hypothetical protein, conserved | 0,4998 | 0,008194 |
| TcCLB.508625.150 | L-ribulokinase, putative | 0,4997 | 0,008643 |
| TcCLB.508479.250 | membrane-bound acid phosphatase 2, putative | 0,4996 | 2,399E-06 |
| TcCLB.506403.9 | hypothetical protein, conserved (fragment) | 0,4995 | 2,342E-06 |
| TcCLB.509957.20 | expression site-associated gene (ESAG-like) protein, putati | 0,4995 | 0,4284 |
| TcCLB.507643.30 | retrotransposon hot spot protein (RHS, pseudogene), putat | 0,4993 | 0,0006005 |
| TcCLB.504717.30 | hypothetical protein, conserved | 0,4993 | 0,001834 |
| TcCLB.510091.50 | hypothetical protein, conserved | 0,4991 | 0,0003555 |
| TcCLB.503465.49 | mitochondrial processing peptidase, beta subunit, putative | 0,4991 | 0,002924 |
| TcCLB.506811.180 | protein kinase, putative | 0,4991 | 0,04447 |
| TcCLB.508121.56 | mucin-associated surface protein (MASP, pseudogene), pu | 0,499 | 0,3365 |
| TcCLB.504445.20 | retrotransposon hot spot protein (RHS, pseudogene), putat | 0,4986 | 0,001916 |
| TcCLB.510359.60 | hypothetical protein, conserved | 0,4986 | 0,004221 |
| TcCLB.507957.270 | mucin TcMUCI, putative | 0,4985 | 0,2761 |
| TcCLB.510901.90 | ATPase protein, putative | 0,4983 | 0,00402 |
| TcCLB.506609.80 | mucin TcMUCII, putative | 0,4981 | 0,4172 |
| TcCLB.507295.61 | elongation factor 1-gamma (EF-1-gamma, pseudogene), p | 0,498 | 0,2924 |
| TcCLB.511045.50 | hypothetical protein, conserved | 0,4977 | 0,006725 |
| TcCLB.508917.30 | kinesin, putative | 0,4975 | 2,602E-06 |
| TcCLB.511603.20 | mucin TcMUCII, putative | 0,4974 | 0,4005 |
| TcCLB.506171.20 | hypothetical protein, conserved (pseudogene) | 0,4973 | 0,007768 |
| TcCLB.511915.20 | trans-sialidase (pseudogene), putative | 0,4971 | 0,02544 |
| TcCLB.504033.170 | tyrosine phosphatase, putative | 0,497 | 0,001239 |
| TcCLB.508221.610 | elongation factor 1-gamma (EF-1-gamma), putative | 0,497 | 0,4194 |
| TcCLB.507019.10 | protein kinase, putative | 0,4968 | 0,0002057 |
| TcCLB.503479.60 | surface protease GP63, putative | 0,4967 | 0,01142 |
| TcCLB.510715.30 | mucin-associated surface protein (MASP), putative | 0,4967 | 0,1067 |
| TcCLB.506833.70 | cationic amino acid transporter, putative | 0,4966 | 0,0003537 |
| TcCLB.505941.40 | hypothetical protein, conserved | 0,4965 | 0,004478 |
| TcCLB.508421.20 | dispersed gene family protein 1 (DGF-1, pseudogene), puta | 0,4965 | 0,07249 |
| TcCLB.507517.10 | hypothetical protein, conserved | 0,4962 | 0,0001751 |
| TcCLB.511231.40 | hypothetical protein, conserved | 0,4961 | 0,03836 |
| TcCLB.507669.190 | hypothetical protein, conserved | 0,496 | 0,0008875 |
| TcCLB.503439.40 | trans-sialidase (pseudogene), putative | 0,4959 | 0,003227 |
| TcCLB.506517.149 | hypothetical protein, conserved | 0,4958 | 0,000101 |
| TcCLB.511385.110 | hypothetical protein | 0,4958 | 0,006258 |
| TcCLB.507071.220 | mucin-associated surface protein (MASP, pseudogene), pu | 0,4957 | 0,3281 |
| TcCLB.506559.210 | hypothetical protein, conserved | 0,4956 | 0,0003816 |
| TcCLB.503977.79 | elongation factor 1-gamma (EF-1-gamma), putative | 0,4954 | 0,4858 |
| TcCLB.510857.50 | lipoate-protein ligase, putative | 0,4953 | 0,0007537 |
| TcCLB.510687.30 | hypothetical protein, conserved | 0,4952 | 2,144E-08 |
| TcCLB.508515.170 | retrotransposon hot spot protein (RHS, pseudogene), putat | 0,4952 | 0,001464 |
| TcCLB.510535.50 | hypothetical protein | 0,4952 | 0,01421 |
| TcCLB.430895.10 | hypothetical protein, conserved | 0,4952 | 0,02835 |
| TcCLB.508011.90 | trans-sialidase (pseudogene), putative | 0,4952 | 0,1574 |

| | | | |
|---|---|---|---|
| TcCLB.510477.100 | mucin-associated surface protein (MASP), putative | 0,495 | 0,2774 |
| TcCLB.506787.19 | mucin-associated surface protein (MASP, pseudogene), pu | 0,4948 | 0,5332 |
| TcCLB.507795.10 | membrane associated protein, putative | 0,4946 | 1,887E-06 |
| TcCLB.507503.70 | retrotransposon hot spot protein (RHS, pseudogene), putat | 0,4946 | 0,005133 |
| TcCLB.509875.230 | hypothetical protein, conserved | 0,4944 | 0,000521 |
| TcCLB.511661.120 | protein kinase, putative | 0,494 | 2,192E-07 |
| TcCLB.510947.20 | hypothetical protein, conserved | 0,494 | 0,00004093 |
| TcCLB.511071.140 | 2-amino-3-ketobutyrate coenzyme A ligase, putative | 0,4935 | 0,0009115 |
| TcCLB.506925.480 | ribokinase, putative | 0,4934 | 0,0002573 |
| TcCLB.508919.130 | hypothetical protein, conserved | 0,4933 | 0,0006314 |
| TcCLB.510459.20 | mucin-associated surface protein (MASP), putative | 0,4932 | 0,2479 |
| TcCLB.511825.90 | hypothetical protein, conserved | 0,4931 | 0,009293 |
| TcCLB.508001.10 | dispersed gene family protein 1 (DGF-1), putative | 0,4931 | 0,05071 |
| TcCLB.506855.160 | hypothetical protein, conserved | 0,4928 | 3,888E-08 |
| TcCLB.506753.210 | hypothetical protein, conserved | 0,4927 | 0,00902 |
| TcCLB.506559.550 | C2 domain containing protein, putative | 0,4924 | 3,182E-06 |
| TcCLB.511221.50 | mucin-associated surface protein (MASP), putative | 0,4924 | 0,08797 |
| TcCLB.510561.50 | mucin-associated surface protein (MASP), putative | 0,4924 | 0,6407 |
| TcCLB.509891.60 | hypothetical protein, conserved | 0,4923 | 0,0009013 |
| TcCLB.506939.10 | mucin-associated surface protein (MASP, pseudogene), pu | 0,4923 | 0,5134 |
| TcCLB.503769.20 | trans-sialidase (pseudogene), putative | 0,4921 | 0,3235 |
| TcCLB.506829.39 | ribosomal protein S7, putative | 0,4911 | 6,315E-12 |
| TcCLB.506869.10 | hypothetical protein | 0,491 | 0,004185 |
| TcCLB.504411.5 | hypothetical protein, conserved (fragment) | 0,491 | 0,5166 |
| TcCLB.509349.20 | retrotransposon hot spot (RHS) protein, putative | 0,4909 | 0,0001104 |
| TcCLB.509645.30 | hypothetical protein, conserved | 0,4908 | 0,00147 |
| TcCLB.510275.80 | trans-sialidase (pseudogene), putative | 0,4907 | 0,1765 |
| TcCLB.509161.40 | centrin, putative | 0,4906 | 0,002683 |
| TcCLB.481211.30 | mucin TcMUCII, putative | 0,4905 | 0,07238 |
| TcCLB.506289.130 | hypothetical protein | 0,4904 | 0,1762 |
| TcCLB.508109.14 | mucin TcMUCII, putative | 0,4904 | 0,302 |
| TcCLB.506891.30 | leucine-rich repeat-containing protein | 0,4902 | 0,01581 |
| TcCLB.506501.177 | hypothetical protein, conserved (pseudogene) | 0,4902 | 0,4985 |
| TcCLB.506915.20 | hypothetical protein | 0,4901 | 0,00003967 |
| TcCLB.504741.40 | hypothetical protein, conserved | 0,49 | 0,0014 |
| TcCLB.507691.31 | hypothetical protein | 0,4899 | 0,04016 |
| TcCLB.507237.120 | mucin-associated surface protein (MASP), putative | 0,4899 | 0,2922 |
| TcCLB.510095.40 | retrotransposon hot spot protein (RHS, pseudogene), putat | 0,4898 | 0,009855 |
| TcCLB.505365.80 | retrotransposon hot spot protein (RHS, pseudogene), putat | 0,4897 | 0,04643 |
| TcCLB.511337.10 | hypothetical protein, conserved | 0,4894 | 0,000268 |
| TcCLB.506529.370 | hypothetical protein | 0,4894 | 0,02125 |
| TcCLB.504189.10 | beta galactofuranosyl glycosyltransferase (pseudogene), p | 0,4894 | 0,3094 |
| TcCLB.505183.60 | hypothetical protein, conserved | 0,4889 | 0,00003127 |
| TcCLB.507085.110 | hypothetical protein | 0,4889 | 0,1516 |
| TcCLB.503879.50 | hypothetical protein, conserved | 0,4888 | 0,001205 |

| | | | |
|---|---|---|---|
| TcCLB.506213.60 | 40S ribosomal protein S2, putative | 0,4884 | 0,2681 |
| TcCLB.510475.100 | mucin-associated surface protein (MASP), putative | 0,4882 | 0,1263 |
| TcCLB.510483.30 | trans-sialidase (pseudogene), putative | 0,488 | 0,1608 |
| TcCLB.507991.70 | hypothetical protein, conserved | 0,4878 | 0,00233 |
| TcCLB.504229.4 | retrotransposon hot spot (RHS) protein, putative | 0,4877 | 0,009599 |
| TcCLB.510877.160 | hypothetical protein, conserved | 0,4876 | 0,0001126 |
| TcCLB.509967.50 | hypothetical protein, conserved | 0,4875 | 0,01517 |
| TcCLB.510487.79 | mucin-associated surface protein (MASP), putative | 0,4873 | 0,2934 |
| TcCLB.510643.110 | beta galactofuranosyl glycosyltransferase, putative | 0,4871 | 0,01918 |
| TcCLB.506393.80 | trans-sialidase (pseudogene), putative | 0,4871 | 0,4495 |
| TcCLB.506279.160 | mucin-associated surface protein (MASP, pseudogene), pu | 0,4869 | 0,3482 |
| TcCLB.508649.20 | hypothetical protein, conserved | 0,4868 | 1,289E-08 |
| TcCLB.509163.110 | retrotransposon hot spot (RHS) protein, putative | 0,4868 | 0,06482 |
| TcCLB.506885.204 | hypothetical protein, conserved | 0,4865 | 2,715E-07 |
| TcCLB.510155.130 | hypothetical protein, conserved | 0,4863 | 1,707E-07 |
| TcCLB.511649.90 | hypothetical protein, conserved | 0,4861 | 0,0002291 |
| TcCLB.509265.110 | trans-sialidase, Group IV, putative | 0,486 | 0,04355 |
| TcCLB.506455.10 | retrotransposon hot spot protein (RHS, pseudogene), putat | 0,4853 | 0,06728 |
| TcCLB.508779.30 | synaptobrevin, putative | 0,485 | 0,0001967 |
| TcCLB.511167.20 | hypothetical protein, conserved | 0,485 | 0,0005125 |
| TcCLB.507045.10 | retrotransposon hot spot protein (RHS, pseudogene), putat | 0,485 | 0,01036 |
| TcCLB.505997.40 | hypothetical protein, conserved (pseudogene) | 0,4847 | 0,004907 |
| TcCLB.506419.30 | hypothetical protein, conserved | 0,4847 | 0,02256 |
| TcCLB.506685.5 | trans-sialidase (pseudogene), putative | 0,4841 | 0,01354 |
| TcCLB.507059.20 | trans-sialidase, Group V, putative | 0,4841 | 0,05746 |
| TcCLB.506295.120 | Trk system potassium uptake protein (HKT1) | 0,484 | 0,002683 |
| TcCLB.510357.160 | hypothetical protein, conserved (fragment) | 0,484 | 0,1418 |
| TcCLB.511181.160 | choline/ethanolamine kinase (pseudogene), putative | 0,4838 | 0,001079 |
| TcCLB.504003.80 | hypothetical protein, conserved | 0,4837 | 0,133 |
| TcCLB.506437.40 | tuzin | 0,4835 | 1,087E-07 |
| TcCLB.505363.19 | trans-sialidase, putative | 0,4834 | 0,00811 |
| TcCLB.506579.60 | hypothetical protein, conserved | 0,483 | 0,002728 |
| TcCLB.508271.20 | hypothetical protein, conserved | 0,4829 | 0,002519 |
| TcCLB.510557.73 | mucin-associated surface protein (MASP, pseudogene), pu | 0,4829 | 0,2581 |
| TcCLB.506795.44 | hypothetical protein, conserved | 0,4827 | 3,107E-06 |
| TcCLB.506865.70 | hypothetical protein, conserved | 0,4827 | 7,463E-06 |
| TcCLB.511107.5 | mucin TcMUCII, putative | 0,4826 | 0,0671 |
| TcCLB.507075.40 | mucin-associated surface protein (MASP), putative (fragm | 0,4825 | 0,1838 |
| TcCLB.511249.34 | hypothetical protein, conserved | 0,4824 | 0,002529 |
| TcCLB.509093.10 | mucin-associated surface protein (MASP), putative | 0,4823 | 0,399 |
| TcCLB.506057.81 | mucin-associated surface protein (MASP, pseudogene), pu | 0,4822 | 0,00735 |
| TcCLB.503779.30 | hypothetical protein, conserved | 0,482 | 0,007078 |
| TcCLB.511907.30 | hypothetical protein, conserved | 0,4818 | 8,801E-06 |
| TcCLB.510025.250 | trans-sialidase (pseudogene), putative | 0,4817 | 0,1059 |
| TcCLB.510359.630 | retrotransposon hot spot protein (RHS, pseudogene), putat | 0,4817 | 0,2103 |

| | | | |
|---|---|---|---|
| TcCLB.504899.30 | hypothetical protein | 0,4815 | 0,0253 |
| TcCLB.508969.30 | retrotransposon hot spot protein (RHS, pseudogene), putat | 0,4809 | 0,05178 |
| TcCLB.506789.120 | hypothetical protein, conserved | 0,4808 | 4,094E-06 |
| TcCLB.507891.20 | flagellar calcium-binding-like protein, putative | 0,4808 | 0,001033 |
| TcCLB.504165.20 | hypothetical protein, conserved | 0,4808 | 0,002763 |
| TcCLB.508649.10 | beta propeller protein, putative | 0,4806 | 0,00003642 |
| TcCLB.504167.20 | mucin TcMUCI, putative | 0,4803 | 0,02456 |
| TcCLB.511807.110 | GMP synthase [glutamine-hydrolysing] (Gmps) | 0,48 | 4,936E-09 |
| TcCLB.508039.90 | phosphoinositide-specific phospholipase C (pseudogene), p | 0,48 | 0,008417 |
| TcCLB.506139.80 | mucin TcMUCII, putative | 0,4798 | 0,5052 |
| TcCLB.506965.100 | mucin-associated surface protein (MASP), putative | 0,4797 | 0,322 |
| TcCLB.510661.90 | ATP-dependent RNA helicase, putative | 0,4796 | 4,766E-07 |
| TcCLB.509411.10 | trans-sialidase, Group II, putative | 0,4796 | 0,371 |
| TcCLB.458015.10 | hypothetical protein, conserved | 0,4795 | 0,00003492 |
| TcCLB.511401.130 | mucin-associated surface protein (MASP, pseudogene), pu | 0,4795 | 0,2254 |
| TcCLB.510657.50 | hypothetical protein, conserved | 0,4794 | 0,005082 |
| TcCLB.506389.20 | trans-sialidase (pseudogene), putative | 0,4794 | 0,03894 |
| TcCLB.504149.20 | ATP-binding cassette transporter ABCA1, putative (ABCA1) | 0,4793 | 0,0003983 |
| TcCLB.509979.140 | hypothetical protein, conserved | 0,4793 | 0,0005663 |
| TcCLB.511127.300 | hypothetical protein, conserved | 0,479 | 0,194 |
| TcCLB.511393.20 | rhomboid-like protein, putative | 0,4789 | 0,0001868 |
| TcCLB.506661.10 | fatty acid elongase, putative | 0,4788 | 0,005445 |
| TcCLB.511385.90 | hypothetical protein, conserved | 0,4785 | 4,491E-07 |
| TcCLB.510861.14 | hypothetical protein, conserved (fragment) | 0,4785 | 2,918E-06 |
| TcCLB.506289.70 | hypothetical protein, conserved | 0,4785 | 0,0000158 |
| TcCLB.508501.165 | U6 snRNA-associated Sm-like protein LSm6p | 0,4781 | 0,08951 |
| TcCLB.506513.50 | protein kinase, putative | 0,478 | 0,0001916 |
| TcCLB.509027.50 | hypothetical protein, conserved | 0,4779 | 0,01305 |
| TcCLB.511311.20 | trans-sialidase, Group II, putative | 0,4777 | 0,08136 |
| TcCLB.507731.60 | trans-sialidase, putative | 0,4777 | 0,2965 |
| TcCLB.508119.90 | surface protease GP63 (pseudogene), putative | 0,4774 | 0,00003953 |
| TcCLB.509337.28 | hypothetical protein, conserved (pseudogene) | 0,4773 | 0,0004723 |
| TcCLB.509461.20 | hypothetical protein, conserved | 0,4773 | 0,009561 |
| TcCLB.503815.10 | alkyl-dihydroxyacetone phosphate synthase, putative | 0,4772 | 1,027E-09 |
| TcCLB.509153.40 | hypothetical protein, conserved | 0,4772 | 3,252E-07 |
| TcCLB.508649.14 | hypothetical protein, conserved | 0,4772 | 0,000502 |
| TcCLB.508161.23 | hypothetical protein | 0,4771 | 0,5264 |
| TcCLB.510751.10 | poly(A)-specific ribonuclease PARN, putative | 0,477 | 0,00009734 |
| TcCLB.503401.11 | retrotransposon hot spot (RHS) protein, putative (fragmen | 0,477 | 0,3992 |
| TcCLB.506613.40 | trans-sialidase, Group V, putative | 0,4767 | 0,1218 |
| TcCLB.511683.30 | hypothetical protein, conserved | 0,4766 | 0,0001678 |
| TcCLB.507647.39 | hypothetical protein, conserved | 0,4766 | 0,0274 |
| TcCLB.508097.81 | mucin TcMUCII, putative | 0,4765 | 0,3179 |
| TcCLB.507895.160 | hypothetical protein, conserved | 0,4764 | 0,0105 |
| TcCLB.506159.10 | retrotransposon hot spot protein (RHS, pseudogene), putat | 0,4764 | 0,01275 |

| | | | |
|---|---|---|---|
| TcCLB.503715.40 | protein kinase, putative | 0,4763 | 0,1056 |
| TcCLB.511401.100 | mucin-associated surface protein (MASP), putative | 0,4754 | 0,1022 |
| TcCLB.510809.10 | mucin TcMUCII, putative | 0,4749 | 0,3165 |
| TcCLB.508713.20 | hypothetical protein, conserved | 0,4748 | 0,001513 |
| TcCLB.510713.110 | mucin-associated surface protein (MASP), putative | 0,4745 | 0,351 |
| TcCLB.510887.64 | hypothetical protein, conserved | 0,4744 | 0,02749 |
| TcCLB.506425.50 | hypothetical protein, conserved | 0,4743 | 0,0001243 |
| TcCLB.510241.5 | hypothetical protein | 0,4743 | 0,0003562 |
| TcCLB.508307.134 | hypothetical protein, conserved | 0,4743 | 0,003733 |
| TcCLB.510351.50 | hypothetical protein, conserved | 0,4741 | 0,0002675 |
| TcCLB.504259.10 | hypothetical protein, conserved | 0,4741 | 0,1489 |
| TcCLB.511071.10 | hypothetical protein, conserved | 0,474 | 0,00006868 |
| TcCLB.510719.20 | short-chain dehydrogenase, putative | 0,4738 | 0,0009712 |
| TcCLB.503889.5 | hypothetical protein | 0,4737 | 0,07613 |
| TcCLB.511111.30 | hypothetical protein, conserved | 0,4736 | 0,0001823 |
| TcCLB.506303.100 | hypothetical protein, conserved | 0,4736 | 0,003155 |
| TcCLB.511077.30 | retrotransposon hot spot protein (RHS, pseudogene), putat | 0,4735 | 0,09018 |
| TcCLB.506789.180 | glutaminyl cyclase, putative | 0,4734 | 0,00001345 |
| TcCLB.503595.19 | ATP-dependent DEAD/H RNA helicase, putative (fragment) | 0,4734 | 0,00609 |
| TcCLB.511283.160 | hypothetical protein, conserved | 0,4733 | 2,129E-07 |
| TcCLB.506885.14 | ribosomal protein L37, putative | 0,4731 | 0,00006603 |
| TcCLB.511283.60 | hypothetical protein, conserved | 0,4731 | 0,0003157 |
| TcCLB.509213.150 | hypothetical protein, conserved | 0,473 | 0,001909 |
| TcCLB.508567.80 | DNA repair and recombination helicase protein PIF7, putat | 0,4729 | 4,875E-06 |
| TcCLB.507959.270 | mucin TcMUCII, putative | 0,4728 | 0,3577 |
| TcCLB.506845.10 | retrotransposon hot spot protein (RHS, pseudogene), putat | 0,4727 | 0,09134 |
| TcCLB.504099.70 | retrotransposon hot spot protein (RHS, pseudogene), putat | 0,4726 | 0,1965 |
| TcCLB.507679.20 | retrotransposon hot spot protein (RHS, pseudogene), putat | 0,4724 | 0,2557 |
| TcCLB.506959.94 | hypothetical protein, conserved | 0,4723 | 0,004109 |
| TcCLB.503441.10 | retrotransposon hot spot protein (RHS, pseudogene), putat | 0,472 | 0,0001249 |
| TcCLB.419469.30 | hypothetical protein, conserved | 0,472 | 0,001027 |
| TcCLB.508245.70 | mucin-associated surface protein (MASP), putative | 0,472 | 0,2943 |
| TcCLB.510857.20 | hypothetical protein, conserved | 0,4719 | 0,02529 |
| TcCLB.506173.100 | mucin-associated surface protein (MASP, pseudogene), pu | 0,4719 | 0,26 |
| TcCLB.507519.60 | Silent information regulator 2 related protein 1, putative (S | 0,4718 | 0,002046 |
| TcCLB.507949.219 | mucin-associated surface protein (MASP, pseudogene), pu | 0,4718 | 0,02401 |
| TcCLB.511183.430 | trans-sialidase, Group V, putative | 0,4718 | 0,1902 |
| TcCLB.510361.50 | mucin-associated surface protein (MASP, pseudogene), pu | 0,4717 | 0,1367 |
| TcCLB.508509.20 | mucin TcMUCII, putative | 0,4717 | 0,3943 |
| TcCLB.506751.116 | elongation factor 1-gamma (EF-1-gamma, pseudogene), p | 0,4716 | 0,3965 |
| TcCLB.509237.30 | glycosomal transporter (GAT1), putative (GAT1) | 0,4712 | 0,00005107 |
| TcCLB.504213.80 | threonine synthase, putative | 0,4711 | 0,00006149 |
| TcCLB.509427.60 | trans-sialidase (pseudogene), putative | 0,4711 | 0,1351 |
| TcCLB.509237.10 | mevalonate kinase, putative | 0,4708 | 0,00001367 |
| TcCLB.510283.160 | hypothetical protein, conserved | 0,4708 | 0,006451 |

| | | | |
|---|---|---|---|
| TcCLB.510629.420 | protein phosphatase 1, regulatory subunit, putative | 0,4706 | 0,0009003 |
| TcCLB.507471.20 | mucin-associated surface protein (MASP, pseudogene), pu | 0,4706 | 0,1452 |
| TcCLB.506939.110 | mucin-associated surface protein (MASP), putative | 0,4705 | 0,4111 |
| TcCLB.511575.20 | hypothetical protein, conserved | 0,4704 | 0,01394 |
| TcCLB.511231.69 | hypothetical protein, conserved | 0,47 | 0,0002018 |
| TcCLB.507089.270 | dihydrolipoyl dehydrogenase, putative (GCVL-2) | 0,4699 | 3,014E-08 |
| TcCLB.511871.130 | retrotransposon hot spot (RHS) protein, putative | 0,4699 | 0,006402 |
| TcCLB.511635.80 | hypothetical protein, conserved | 0,4696 | 0,002515 |
| TcCLB.509237.100 | hypothetical protein, conserved | 0,4696 | 0,005475 |
| TcCLB.508001.60 | trans-sialidase (pseudogene), putative | 0,4696 | 0,1639 |
| TcCLB.511533.19 | dynein light intermediate chain, putative | 0,4695 | 0,02846 |
| TcCLB.507547.80 | protein kinase, putative | 0,4695 | 0,03119 |
| TcCLB.508669.30 | hypothetical protein | 0,4693 | 0,0007348 |
| TcCLB.510569.10 | trans-sialidase, putative | 0,4693 | 0,2067 |
| TcCLB.508831.66 | hypothetical protein, conserved | 0,4692 | 0,0001247 |
| TcCLB.506173.170 | retrotransposon hot spot protein (RHS, pseudogene), putat | 0,4692 | 0,002825 |
| TcCLB.503869.9 | axoneme central apparatus protein, putative | 0,4691 | 0,001921 |
| TcCLB.509127.20 | hypothetical protein, conserved | 0,469 | 0,003656 |
| TcCLB.506895.30 | hypothetical protein, conserved | 0,469 | 0,01005 |
| TcCLB.511575.40 | hypothetical protein, conserved | 0,4688 | 4,044E-06 |
| TcCLB.509051.50 | hypothetical protein, conserved | 0,4687 | 0,0006299 |
| TcCLB.511401.95 | hypothetical protein (pseudogene) | 0,4687 | 0,08586 |
| TcCLB.508125.65 | elongation factor 1-gamma (EF-1-gamma, pseudogene), p | 0,4686 | 0,2348 |
| TcCLB.508741.30 | hypothetical protein, conserved | 0,4685 | 0,0001351 |
| TcCLB.511247.18 | 50S ribosomal protein L16, putative | 0,4684 | 0,003556 |
| TcCLB.509265.60 | retrotransposon hot spot protein (RHS, pseudogene), putat | 0,4684 | 0,02222 |
| TcCLB.506757.70 | retrotransposon hot spot protein (RHS, pseudogene), putat | 0,4684 | 0,3054 |
| TcCLB.508999.230 | calpain-like cysteine peptidase (pseudogene), putative | 0,4683 | 0,00003786 |
| TcCLB.510879.220 | hypothetical protein, conserved | 0,4682 | 0,0005118 |
| TcCLB.507641.50 | hypothetical protein, conserved | 0,4681 | 1,119E-06 |
| TcCLB.507837.30 | hypothetical protein, conserved | 0,4681 | 0,00005768 |
| TcCLB.508629.42 | retrotransposon hot spot protein (RHS, pseudogene), putat | 0,4681 | 0,0001038 |
| TcCLB.507691.10 | hypothetical protein, conserved | 0,4681 | 0,3107 |
| TcCLB.508975.20 | pre-rrna-processing protein tsr1 homolog | 0,4679 | 1,904E-06 |
| TcCLB.505401.40 | dispersed gene family protein 1 (DGF-1), putative | 0,4678 | 0,5208 |
| TcCLB.503441.19 | retrotransposon hot spot protein (RHS, pseudogene), putat | 0,4675 | 0,0054 |
| TcCLB.510665.70 | acyl transferase-like protein, putative | 0,4674 | 0,09389 |
| TcCLB.507701.4 | mucin-associated surface protein (MASP), putative (fragm | 0,4673 | 0,3569 |
| TcCLB.503605.10 | retrotransposon hot spot protein (RHS, pseudogene), putat | 0,4672 | 0,00001214 |
| TcCLB.511761.40 | hypothetical protein, conserved | 0,4672 | 0,1404 |
| TcCLB.506755.70 | hypothetical protein, conserved | 0,4671 | 0,0052 |
| TcCLB.504867.129 | hypothetical protein, conserved | 0,467 | 0,002036 |
| TcCLB.511727.290 | RNA-binding protein, putative | 0,4669 | 3,743E-13 |
| TcCLB.511903.230 | hypothetical protein, conserved | 0,4669 | 0,00347 |
| TcCLB.503947.20 | mucin-associated surface protein (MASP), putative | 0,4669 | 0,07721 |

| | | | |
|---|---|---|---|
| TcCLB.503823.50 | hypothetical protein, conserved | 0,4668 | 0,0005073 |
| TcCLB.510857.54 | hypothetical protein, conserved | 0,4668 | 0,004961 |
| TcCLB.506757.100 | retrotransposon hot spot protein (RHS, pseudogene), putat | 0,4668 | 0,1138 |
| TcCLB.429185.20 | trans-sialidase (pseudogene), putative | 0,4668 | 0,3521 |
| TcCLB.506855.250 | hypothetical protein, conserved | 0,4666 | 0,00001167 |
| TcCLB.511019.80 | retrotransposon hot spot (RHS) protein, putative | 0,4666 | 0,001636 |
| TcCLB.507857.50 | protein phosphotase, putative | 0,4664 | 0,0001353 |
| TcCLB.510309.30 | PTP1-interacting protein, 39 kDa, putative | 0,4662 | 0,009057 |
| TcCLB.509769.148 | dynein heavy chain, putative | 0,4661 | 0,004707 |
| TcCLB.506563.110 | calpain-like cysteine peptidase (pseudogene), putative | 0,4657 | 5,184E-07 |
| TcCLB.509723.20 | glycine dehydrogenase (pseudogene), putative | 0,4657 | 0,2901 |
| TcCLB.510797.30 | hypothetical protein, conserved | 0,4654 | 0,0002091 |
| TcCLB.506285.90 | mucin TcMUCII, putative | 0,4654 | 0,2921 |
| TcCLB.506909.150 | mucin TcMUCII (pseudogene), putative | 0,4654 | 0,5524 |
| TcCLB.506591.130 | hypothetical protein, conserved | 0,465 | 0,003034 |
| TcCLB.507959.170 | mucin-associated surface protein (MASP), putative | 0,4649 | 0,232 |
| TcCLB.510889.270 | hypothetical protein, conserved | 0,4648 | 0,0002192 |
| TcCLB.504239.120 | trans-sialidase (pseudogene), putative | 0,4646 | 0,4507 |
| TcCLB.506767.50 | mucin TcMUCII, putative | 0,4646 | 0,4599 |
| TcCLB.510101.110 | hypothetical protein, conserved | 0,4644 | 5,542E-06 |
| TcCLB.511127.180 | hypothetical protein, conserved | 0,4642 | 0,000217 |
| TcCLB.511001.90 | hypothetical protein, conserved | 0,4641 | 5,311E-09 |
| TcCLB.506577.60 | hypothetical protein, conserved | 0,4641 | 0,00003885 |
| TcCLB.509153.80 | Present in the outer mitochondrial membrane proteome 3( | 0,464 | 0,008369 |
| TcCLB.511715.90 | hypothetical protein, conserved | 0,4637 | 0,0006457 |
| TcCLB.510671.20 | tryptophanyl-tRNA synthetase, putative | 0,4635 | 8,985E-09 |
| TcCLB.504159.70 | hypothetical protein, conserved | 0,4632 | 0,002284 |
| TcCLB.509825.7 | hypothetical protein, conserved | 0,4632 | 0,4486 |
| TcCLB.506401.370 | hypothetical protein, conserved | 0,4629 | 0,003136 |
| TcCLB.509695.30 | hypothetical protein, conserved | 0,4628 | 0,01379 |
| TcCLB.504277.11 | surface protein TolT, putative | 0,4626 | 0,072 |
| TcCLB.511237.70 | hypothetical protein, conserved | 0,4625 | 0,008198 |
| TcCLB.511219.20 | retrotransposon hot spot protein (RHS, pseudogene), putat | 0,4625 | 0,009842 |
| TcCLB.511871.80 | frataxin-like, mitochondrial precursor, putative | 0,4622 | 0,0236 |
| TcCLB.508761.90 | mucin-associated surface protein (MASP), putative | 0,4619 | 0,632 |
| TcCLB.511291.50 | hypothetical protein, conserved | 0,4617 | 0,007485 |
| TcCLB.508977.130 | mucin-associated surface protein (MASP), putative | 0,4617 | 0,2051 |
| TcCLB.510515.100 | coronin, putative (CRN12) | 0,4616 | 0,00001853 |
| TcCLB.511277.486 | hypothetical protein, conserved | 0,4616 | 0,01202 |
| TcCLB.504183.10 | retrotransposon hot spot protein (RHS, pseudogene), putat | 0,4614 | 0,07816 |
| TcCLB.511237.50 | 40S ribosomal protein S33, putative | 0,4612 | 2,203E-06 |
| TcCLB.508277.320 | hypothetical protein, conserved | 0,4609 | 0,001031 |
| TcCLB.511607.110 | mucin-associated surface protein (MASP), putative | 0,4608 | 0,1846 |
| TcCLB.508629.58 | hypothetical protein, conserved (pseudogene) | 0,4607 | 0,0004695 |
| TcCLB.507445.50 | myosin heavy chain, putative | 0,4606 | 1,549E-06 |

| | | | |
|---|---|---|---|
| TcCLB.511409.40 | hypothetical protein, conserved | 0,4606 | 0,0005956 |
| TcCLB.511727.145 | hypothetical protein | 0,4605 | 6,913E-06 |
| TcCLB.507941.50 | hypothetical protein, conserved | 0,4603 | 0,0002018 |
| TcCLB.507993.36 | hypothetical protein, conserved | 0,4603 | 0,02308 |
| TcCLB.510729.140 | NIMA-related kinase, putative | 0,4603 | 0,06359 |
| TcCLB.508153.1130 | hypothetical protein | 0,4603 | 0,07579 |
| TcCLB.505919.40 | protein kinase, putative | 0,4602 | 0,1908 |
| TcCLB.510759.150 | hypothetical protein, conserved | 0,46 | 0,00002994 |
| TcCLB.511283.290 | hypothetical protein, conserved | 0,4599 | 5,117E-08 |
| TcCLB.511149.80 | hypothetical protein | 0,4599 | 0,3342 |
| TcCLB.507985.30 | trans-sialidase (pseudogene), putative | 0,4598 | 0,6802 |
| TcCLB.510657.20 | ubiquitin hydrolase, putative | 0,4597 | 0,01514 |
| TcCLB.509539.20 | receptor-type adenylate cyclase, putative | 0,4596 | 0,00404 |
| TcCLB.511771.143 | retrotransposon hot spot protein (RHS, pseudogene), putat | 0,4595 | 0,2426 |
| TcCLB.510425.19 | 40S ribosomal protein SA, putative | 0,4591 | 6,702E-08 |
| TcCLB.418185.70 | trans-sialidase, Group II, putative | 0,459 | 0,1629 |
| TcCLB.507783.20 | AGP2beta-2, putative | 0,4589 | 0,0001009 |
| TcCLB.506147.200 | hypothetical protein, conserved | 0,4588 | 0,0009239 |
| TcCLB.506517.14 | protein kinase, putative | 0,4588 | 0,1424 |
| TcCLB.510173.100 | hypothetical protein, conserved | 0,4587 | 0,0109 |
| TcCLB.511771.120 | hypothetical protein, conserved (pseudogene) | 0,4586 | 0,01247 |
| TcCLB.504161.59 | transporter, putative | 0,4585 | 0,01012 |
| TcCLB.510879.60 | hypothetical protein, conserved | 0,4585 | 0,01111 |
| TcCLB.507057.4 | myosin heavy chain, putative (fragment) | 0,4585 | 0,01329 |
| TcCLB.506573.10 | hypothetical protein, conserved | 0,4584 | 0,0003108 |
| TcCLB.510889.30 | kinase, putative | 0,4584 | 0,00157 |
| TcCLB.506145.50 | hypothetical protein, conserved | 0,4584 | 0,1047 |
| TcCLB.507021.20 | ESAG8-associated protein, putative (PIE8) | 0,4583 | 0,2302 |
| TcCLB.509525.244 | hypothetical protein | 0,4583 | 0,536 |
| TcCLB.506693.50 | ribonuclease inhibitor-like protein, putative | 0,4582 | 0,002015 |
| TcCLB.506563.60 | hypothetical protein, conserved | 0,4581 | 0,009866 |
| TcCLB.503891.50 | hypothetical protein, conserved | 0,458 | 0,0009435 |
| TcCLB.507501.30 | hypothetical protein, conserved | 0,4579 | 0,188 |
| TcCLB.506367.180 | vesicular transport protein (CDC48 homologue), putative | 0,4578 | 0,0001577 |
| TcCLB.511755.28 | hypothetical protein, conserved | 0,4578 | 0,08822 |
| TcCLB.506845.60 | retrotransposon hot spot (RHS) protein, putative | 0,4578 | 0,2453 |
| TcCLB.510423.70 | hypothetical protein, conserved | 0,4576 | 0,02832 |
| TcCLB.509209.20 | hypothetical protein, conserved | 0,4576 | 0,06917 |
| TcCLB.506195.25 | hypothetical protein, conserved | 0,4575 | 0,2178 |
| TcCLB.509745.10 | hypothetical protein, conserved | 0,4574 | 0,00003905 |
| TcCLB.510489.44 | mucin TcMUCII (pseudogene), putative | 0,4574 | 0,361 |
| TcCLB.511301.80 | choline/carnitine O-acyltransferase (pseudogene), putative | 0,4572 | 0,00001253 |
| TcCLB.510181.150 | caltractin, putative | 0,4571 | 0,02437 |
| TcCLB.508011.110 | mucin-associated surface protein (MASP), putative | 0,4571 | 0,225 |
| TcCLB.503537.9 | retrotransposon hot spot protein (RHS, pseudogene), putat | 0,457 | 0,0001672 |

| | | | |
|---|---|---|---|
| TcCLB.506459.299 | mucin-associated surface protein (MASP, pseudogene), pu | 0,4569 | 0,00003554 |
| TcCLB.507499.60 | retrotransposon hot spot protein (RHS, pseudogene), putat | 0,4568 | 1,224E-07 |
| TcCLB.511025.110 | dihydrolipoyl dehydrogenase, putative (GCVL-2) | 0,4568 | 6,228E-06 |
| TcCLB.510543.10 | retrotransposon hot spot protein (RHS, pseudogene), putat | 0,4567 | 0,01298 |
| TcCLB.506683.250 | retrotransposon hot spot protein (RHS, pseudogene), putat | 0,4567 | 0,1219 |
| TcCLB.511553.200 | mucin-associated surface protein (MASP), putative | 0,4567 | 0,3224 |
| TcCLB.506651.9 | dispersed gene family protein 1 (DGF-1), putative | 0,4566 | 0,1199 |
| TcCLB.510379.20 | hypothetical protein (fragment) | 0,4566 | 0,3596 |
| TcCLB.509429.124 | hypothetical protein, conserved | 0,4565 | 0,07144 |
| TcCLB.510017.59 | mucin-associated surface protein (MASP), putative | 0,4564 | 0,06549 |
| TcCLB.506975.90 | trans-sialidase, Group VIII, putative | 0,4564 | 0,2077 |
| TcCLB.504147.210 | hypothetical protein, conserved | 0,4563 | 0,009672 |
| TcCLB.511469.110 | dispersed gene family protein 1 (DGF-1), putative | 0,4559 | 0,06774 |
| TcCLB.511877.20 | mucin-associated surface protein (MASP), putative | 0,4559 | 0,07761 |
| TcCLB.509699.210 | mucin-associated surface protein (MASP), putative | 0,4558 | 0,3055 |
| TcCLB.511649.140 | hypothetical protein, conserved | 0,4555 | 0,005207 |
| TcCLB.503479.10 | hypothetical protein, conserved | 0,4553 | 0,001749 |
| TcCLB.510667.20 | hypothetical protein, conserved | 0,4552 | 0,01415 |
| TcCLB.508817.50 | DNA repair protein RAD51, putative (RAD51) | 0,4551 | 2,627E-07 |
| TcCLB.507583.60 | hypothetical protein, conserved | 0,4551 | 0,4351 |
| TcCLB.509109.80 | hypothetical protein, conserved | 0,4548 | 5,187E-09 |
| TcCLB.509105.20 | palmitoyl acyltransferase 2, putative | 0,4548 | 0,001332 |
| TcCLB.511821.120 | hypothetical protein, conserved | 0,4547 | 0,0009392 |
| TcCLB.510165.60 | expression site-associated gene (ESAG-like) protein, putati | 0,4547 | 0,1367 |
| TcCLB.508973.80 | hypothetical protein, conserved | 0,4546 | 8,878E-09 |
| TcCLB.510583.74 | mucin-associated surface protein (MASP, pseudogene), pu | 0,4543 | 0,3049 |
| TcCLB.506591.120 | receptor-type adenylate cyclase (pseudogene), putative | 0,4542 | 0,00008981 |
| TcCLB.506201.84 | hypothetical protein, conserved | 0,4541 | 0,02821 |
| TcCLB.506757.90 | trans-sialidase, Group II, putative | 0,4537 | 0,108 |
| TcCLB.507231.20 | trans-sialidase (pseudogene), putative | 0,4535 | 0,1818 |
| TcCLB.511607.100 | mucin TcMUCII, putative | 0,4534 | 0,1792 |
| TcCLB.506765.40 | trans-sialidase, Group II, putative | 0,4533 | 0,3421 |
| TcCLB.511367.50 | hypothetical protein, conserved | 0,4532 | 0,00005029 |
| TcCLB.509571.50 | hypothetical protein, conserved | 0,4531 | 0,00006438 |
| TcCLB.509857.10 | hypothetical protein, conserved | 0,453 | 0,0009573 |
| TcCLB.506725.40 | hypothetical protein, conserved | 0,453 | 0,001537 |
| TcCLB.508381.40 | retrotransposon hot spot protein (RHS, pseudogene), putat | 0,4526 | 0,09589 |
| TcCLB.509979.200 | mucin-associated surface protein (MASP), putative | 0,4524 | 0,1453 |
| TcCLB.508523.40 | dispersed gene family protein 1 (DGF-1), putative | 0,4522 | 0,0007565 |
| TcCLB.511621.30 | hypothetical protein, conserved | 0,4522 | 0,01042 |
| TcCLB.509331.9 | 5'-3' exoribonuclease C, putative (fragment) (XRNC) | 0,4521 | 0,06971 |
| TcCLB.504863.10 | retrotransposon hot spot (RHS) protein, putative | 0,4519 | 0,00006645 |
| TcCLB.504003.90 | hypothetical protein, conserved | 0,4516 | 0,003431 |
| TcCLB.511507.24 | hypothetical protein, conserved | 0,4514 | 0,04597 |
| TcCLB.507623.60 | hypothetical protein, conserved | 0,4513 | 5,862E-06 |

| | | | |
|---|---|---|---|
| TcCLB.508887.30 | hypothetical protein | 0,4513 | 0,2646 |
| TcCLB.508639.10 | trans-sialidase, putative | 0,4511 | 0,1185 |
| TcCLB.507429.6 | hypothetical protein | 0,4511 | 0,5211 |
| TcCLB.511183.200 | mucin-associated surface protein (MASP, pseudogene), pu | 0,4509 | 0,07039 |
| TcCLB.509897.210 | mucin-associated surface protein (MASP, pseudogene), pu | 0,4508 | 0,02847 |
| TcCLB.506419.69 | hypothetical protein, conserved | 0,4507 | 0,03494 |
| TcCLB.508209.100 | 10 kDa heat shock protein, putative | 0,4506 | 0,006287 |
| TcCLB.506683.19 | GMP synthase (pseudogene), putative | 0,4505 | 0,0005911 |
| TcCLB.509525.20 | trans-sialidase (pseudogene), putative | 0,4505 | 0,02967 |
| TcCLB.506173.70 | trans-sialidase, Group V, putative | 0,4504 | 0,09783 |
| TcCLB.509207.130 | peptidyl-prolyl cis-trans isomerase (PAR45) | 0,4503 | 0,0001423 |
| TcCLB.508731.30 | calmodulin, putative | 0,4503 | 0,003173 |
| TcCLB.511449.40 | mucin TcMUCII (pseudogene), putative | 0,4503 | 0,3186 |
| TcCLB.509291.10 | hypothetical protein | 0,4503 | 0,3686 |
| TcCLB.410243.10 | hypothetical protein, conserved (fragment) | 0,4502 | 0,04395 |
| TcCLB.509749.39 | short-chain dehydrogenase, putative | 0,4502 | 0,3442 |
| TcCLB.509099.10 | GTP-binding protein, putative | 0,45 | 0,0004522 |
| TcCLB.508285.60 | trans-sialidase, Group II, putative | 0,45 | 0,1087 |
| TcCLB.506559.360 | hypothetical protein, conserved | 0,4497 | 0,00007371 |
| TcCLB.506537.60 | dispersed gene family protein 1 (DGF-1, pseudogene), puta | 0,4496 | 0,003376 |
| TcCLB.508175.370 | glutamine synthetase, putative | 0,4496 | 0,003463 |
| TcCLB.508873.10 | mucin-associated surface protein (MASP), putative | 0,4496 | 0,1036 |
| TcCLB.505915.40 | beta galactofuranosyl glycosyltransferase (pseudogene), p | 0,4495 | 0,1647 |
| TcCLB.510493.20 | hypothetical protein, conserved | 0,4491 | 0,0467 |
| TcCLB.504207.20 | hypothetical protein, conserved | 0,4488 | 2,716E-06 |
| TcCLB.507939.70 | hypothetical protein | 0,4487 | 0,2537 |
| TcCLB.510347.10 | hypothetical protein | 0,4486 | 0,00002988 |
| TcCLB.509899.20 | retrotransposon hot spot protein (RHS, pseudogene), putat | 0,4486 | 0,01765 |
| TcCLB.504313.10 | trans-sialidase, Group IV, putative | 0,4486 | 0,3543 |
| TcCLB.508325.220 | beta galactofuranosyl glycosyltransferase (pseudogene), p | 0,4485 | 0,001522 |
| TcCLB.510483.120 | mucin-associated surface protein (MASP), putative | 0,4482 | 0,4954 |
| TcCLB.506531.60 | hypothetical protein, conserved | 0,448 | 0,0001775 |
| TcCLB.507933.29 | hypothetical protein, conserved (fragment) | 0,4476 | 0,1143 |
| TcCLB.503633.40 | trans-sialidase (pseudogene), putative | 0,4476 | 0,141 |
| TcCLB.511815.60 | prefoldin, putative | 0,4475 | 0,0001438 |
| TcCLB.510101.280 | kinesin, putative | 0,4474 | 7,842E-09 |
| TcCLB.511557.64 | hypothetical protein | 0,4474 | 0,006364 |
| TcCLB.507159.50 | hypothetical protein, conserved | 0,4467 | 0,0005743 |
| TcCLB.509059.10 | hypothetical protein | 0,4465 | 0,05659 |
| TcCLB.511183.550 | mucin TcMUCII, putative | 0,4463 | 0,5578 |
| TcCLB.510463.120 | mucin-associated surface protein (MASP), putative | 0,4463 | 0,7709 |
| TcCLB.510579.10 | tyrosine phospatase-like protein, putative | 0,4462 | 0,002924 |
| TcCLB.509511.10 | dispersed gene family protein 1 (DGF-1), putative | 0,446 | 0,0289 |
| TcCLB.507765.60 | 40S ribosomal protein S21, putative | 0,4457 | 8,157E-10 |
| TcCLB.432443.10 | mucin-associated surface protein (MASP), putative | 0,4456 | 0,3673 |

| | | | |
|---|---|---|---|
| TcCLB.511621.130 | hypothetical protein, conserved | 0,4453 | 0,000155 |
| TcCLB.506443.70 | glutathione-S-transferase/glutaredoxin, putative | 0,4453 | 0,001976 |
| TcCLB.509937.150 | serine hydroxymethyltransferase, putative | 0,4452 | 0,01802 |
| TcCLB.511903.45 | tuzin-like (pseudogene), putative | 0,4451 | 0,003167 |
| TcCLB.509437.183 | hypothetical protein, conserved (pseudogene) | 0,4451 | 0,004175 |
| TcCLB.503563.14 | ADG1, putative | 0,4451 | 0,008142 |
| TcCLB.506871.120 | hypothetical protein, conserved | 0,445 | 0,0003854 |
| TcCLB.404843.10 | hypothetical protein, conserved | 0,4449 | 0,004454 |
| TcCLB.508121.160 | mucin-associated surface protein (MASP, pseudogene), pu | 0,4449 | 0,3971 |
| TcCLB.506435.110 | hypothetical protein, conserved | 0,4445 | 0,02649 |
| TcCLB.507953.70 | 90 kDa surface protein, putative | 0,4445 | 0,395 |
| TcCLB.510759.170 | hypothetical protein, conserved | 0,4443 | 0,00006097 |
| TcCLB.511907.120 | mucin-like glycoprotein, putative | 0,4443 | 0,1562 |
| TcCLB.510687.120 | snoRNP protein GAR1, putative | 0,4439 | 0,004129 |
| TcCLB.508919.94 | hypothetical protein, conserved | 0,4438 | 0,01573 |
| TcCLB.507957.280 | mucin-associated surface protein (MASP), putative | 0,4438 | 0,2782 |
| TcCLB.509247.50 | methionyl-tRNA synthetase, putative | 0,4437 | 4,726E-08 |
| TcCLB.507087.60 | hypothetical protein, conserved | 0,4437 | 0,003309 |
| TcCLB.508173.90 | hypothetical protein, conserved | 0,4436 | 0,03267 |
| TcCLB.511797.120 | mucin-associated surface protein (MASP), putative | 0,4432 | 0,2791 |
| TcCLB.506833.50 | hypothetical protein, conserved | 0,4431 | 0,00005069 |
| TcCLB.510699.40 | mucin TcMUCII, putative | 0,4431 | 0,3851 |
| TcCLB.506895.5 | hypothetical protein | 0,4429 | 0,5052 |
| TcCLB.507053.40 | hypothetical protein, conserved | 0,4425 | 6,884E-07 |
| TcCLB.509529.19 | dispersed gene family protein 1 (DGF-1), putative (fragme | 0,4425 | 0,01542 |
| TcCLB.508289.70 | mucin-associated surface protein (MASP, pseudogene), pu | 0,4424 | 0,4523 |
| TcCLB.506139.10 | mucin-associated surface protein (MASP), putative | 0,4424 | 0,4557 |
| TcCLB.506175.50 | hypothetical protein, conserved | 0,4423 | 0,0001669 |
| TcCLB.506857.10 | glycogen synthase kinase-3 alpha, putative | 0,4422 | 0,002503 |
| TcCLB.508045.79 | spliceosomal U5 snRNP-specific protein, putative | 0,4421 | 0,001919 |
| TcCLB.504157.130 | surface protein TolT, putative (TolT) | 0,442 | 0,06304 |
| TcCLB.509717.140 | hypothetical protein, conserved (pseudogene) | 0,442 | 0,1519 |
| TcCLB.398915.5 | trans-sialidase, putative (fragment) | 0,4419 | 0,000452 |
| TcCLB.508585.10 | hypothetical protein, conserved | 0,4419 | 0,008538 |
| TcCLB.504125.90 | protein kinase, putative | 0,4417 | 0,00003786 |
| TcCLB.508061.60 | retrotransposon hot spot protein (RHS, pseudogene), putat | 0,4417 | 0,003407 |
| TcCLB.509907.60 | trans-sialidase (pseudogene), putative | 0,4416 | 0,1949 |
| TcCLB.506605.40 | hypothetical protein, conserved | 0,4411 | 0,002175 |
| TcCLB.506139.130 | hypothetical protein | 0,4411 | 0,5307 |
| TcCLB.511815.4 | proteasome regulatory ATPase subunit 5, putative (fragme | 0,441 | 0,001576 |
| TcCLB.508277.269 | hypothetical protein, conserved (pseudogene) | 0,441 | 0,005766 |
| TcCLB.506401.180 | hypothetical protein, conserved | 0,4407 | 8,65E-07 |
| TcCLB.510737.30 | mitochondrial carrier protein, putative | 0,4406 | 0,004625 |
| TcCLB.511399.24 | trans-sialidase, putative | 0,4406 | 0,2003 |
| TcCLB.506147.150 | hypothetical protein, conserved | 0,4403 | 0,0003198 |

| | | | |
|---|---|---|---|
| TcCLB.510041.10 | sec1 family transport protein, putative | 0,4403 | 0,07078 |
| TcCLB.511523.14 | succinate dehydrogenase cytochrome B subunit | 0,4402 | 0,00487 |
| TcCLB.511583.10 | hypothetical protein, conserved | 0,44 | 0,006998 |
| TcCLB.511609.30 | mucin-associated surface protein (MASP), putative | 0,44 | 0,5765 |
| TcCLB.509233.50 | hypothetical protein, conserved | 0,4399 | 9,94E-07 |
| TcCLB.509337.10 | hypothetical protein, conserved | 0,4399 | 0,0001895 |
| TcCLB.508897.70 | ADP-ribosylation factor family, putative | 0,4399 | 0,01244 |
| TcCLB.509561.20 | flagellum-adhesion glycoprotein, putative | 0,4397 | 0,00007367 |
| TcCLB.507047.50 | hypothetical protein, conserved | 0,4397 | 0,00169 |
| TcCLB.508577.110 | hypothetical protein, conserved | 0,4397 | 0,01418 |
| TcCLB.509753.210 | mucin-associated surface protein (MASP), putative | 0,4397 | 0,1818 |
| TcCLB.510773.100 | Flagellar Member 4 (FLAM4) | 0,4395 | 0,0002282 |
| TcCLB.509455.114 | Protein of unknown function (DUF1295), putative (POMP23 | 0,4395 | 0,0009987 |
| TcCLB.511851.37 | mucin TcMUCI, putative | 0,4395 | 0,2401 |
| TcCLB.509151.110 | hypothetical protein, conserved | 0,4392 | 0,00001715 |
| TcCLB.508365.10 | mucin TcMUCII, putative | 0,4391 | 0,4606 |
| TcCLB.506605.130 | hypothetical protein, conserved | 0,439 | 0,008226 |
| TcCLB.510515.90 | hypothetical protein, conserved | 0,4389 | 0,01582 |
| TcCLB.508525.5 | cAMP response protein 1, putative (pseudogene) | 0,4387 | 0,1646 |
| TcCLB.511245.140 | fatty acid elongase, putative | 0,4386 | 0,00003475 |
| TcCLB.503783.50 | surface protease GP63 (pseudogene), putative | 0,4386 | 0,3444 |
| TcCLB.507159.80 | hypothetical protein, conserved | 0,4385 | 0,0000378 |
| TcCLB.506727.10 | hypothetical protein | 0,4382 | 0,0007514 |
| TcCLB.508441.110 | mucin TcMUCI, putative | 0,4382 | 0,009775 |
| TcCLB.511807.70 | heat shock protein DNAJ, putative | 0,4381 | 0,000147 |
| TcCLB.509831.5 | vesicular transport protein (CDC48 homologue), putative (f | 0,4381 | 0,0253 |
| TcCLB.508231.180 | hypothetical protein, conserved | 0,438 | 0,0006113 |
| TcCLB.506269.59 | hypothetical protein (pseudogene) | 0,4379 | 0,1225 |
| TcCLB.509785.120 | hypothetical protein, conserved (pseudogene) | 0,4374 | 0,387 |
| TcCLB.508555.10 | hypothetical protein, conserved | 0,4372 | 0,05277 |
| TcCLB.511801.44 | DNA-dependent RNA polymerases, putative | 0,4368 | 0,01128 |
| TcCLB.511635.69 | hypothetical protein, conserved | 0,4368 | 0,1058 |
| TcCLB.505949.50 | mucin-associated surface protein (MASP, pseudogene), pu | 0,4366 | 0,5494 |
| TcCLB.510113.20 | dispersed gene family protein 1 (DGF-1, pseudogene), puta | 0,4365 | 0,02124 |
| TcCLB.510729.210 | hypothetical protein, conserved | 0,4364 | 0,00000985 |
| TcCLB.509885.20 | hypothetical protein, conserved | 0,4364 | 0,02194 |
| TcCLB.509399.180 | hypothetical protein, conserved | 0,4363 | 0,08413 |
| TcCLB.508831.120 | glucosamine 6-phosphate n-acetyltransferase, putative | 0,4362 | 0,02538 |
| TcCLB.506955.170 | trans-sialidase (pseudogene), putative | 0,4361 | 0,1131 |
| TcCLB.504151.20 | hypothetical protein, conserved | 0,4358 | 0,01364 |
| TcCLB.511899.10 | 2-amino-3-ketobutyrate coenzyme A ligase, putative | 0,4358 | 0,07469 |
| TcCLB.511255.540 | mucin-associated surface protein (MASP), putative | 0,4357 | 0,4312 |
| TcCLB.508173.264 | hypothetical protein, conserved | 0,4353 | 2,714E-06 |
| TcCLB.506581.20 | hypothetical protein, conserved | 0,4353 | 0,00898 |
| TcCLB.504131.100 | DnaJ chaperone protein, putative | 0,4352 | 3,006E-06 |

| | | | |
|---|---|---|---|
| TcCLB.507491.59 | ubiquitin fold modifier protein, putative (UFM1) | 0,4352 | 0,05694 |
| TcCLB.509023.90 | hypothetical protein, conserved | 0,4351 | 0,02456 |
| TcCLB.507211.10 | choline/carnitine O-acyltransferase, putative | 0,4348 | 0,0003146 |
| TcCLB.510243.50 | adenylate kinase, putative | 0,4348 | 0,00145 |
| TcCLB.506629.70 | cysteine desulfurase, putative | 0,4348 | 0,002888 |
| TcCLB.506529.250 | hypothetical protein, conserved | 0,4347 | 4,757E-07 |
| TcCLB.510081.30 | retrotransposon hot spot protein (RHS, pseudogene), putat | 0,4347 | 0,002459 |
| TcCLB.508355.210 | hypothetical protein, conserved | 0,4346 | 0,01436 |
| TcCLB.508013.70 | mucin TcMUCII, putative | 0,4346 | 0,5765 |
| TcCLB.508719.70 | flagellum targeting protein kharon1, putative (KH1) | 0,4343 | 0,00002486 |
| TcCLB.509067.40 | NADH dehydrogenase subunit NI8M, putative | 0,4343 | 0,005178 |
| TcCLB.510741.70 | protein kinase, putative | 0,4342 | 0,06817 |
| TcCLB.511277.115 | hypothetical protein, conserved | 0,434 | 0,00001704 |
| TcCLB.509207.160 | proteasome regulatory ATPase subunit 1, putative | 0,4337 | 4,788E-06 |
| TcCLB.506173.120 | mucin TcMUCI, putative | 0,4336 | 0,419 |
| TcCLB.508273.30 | hypothetical protein, conserved | 0,4335 | 0,0007576 |
| TcCLB.507521.100 | hypothetical protein, conserved | 0,4334 | 0,02515 |
| TcCLB.504371.20 | dispersed gene family protein 1 (DGF-1, pseudogene), puta | 0,4333 | 0,05801 |
| TcCLB.509391.10 | Flagellar calcium-binding protein 24, putative | 0,4333 | 0,5201 |
| TcCLB.506811.190 | UDP-galactose 4-epimerase (GalE) | 0,4331 | 0,00001279 |
| TcCLB.508317.10 | cysteine proteinase, putative | 0,433 | 0,0007722 |
| TcCLB.509319.109 | paraflagellar rod protein, putative (fragment) | 0,433 | 0,09613 |
| TcCLB.510733.50 | hypothetical protein, conserved | 0,4329 | 6,899E-08 |
| TcCLB.510187.410 | hypothetical protein, conserved | 0,4329 | 0,001219 |
| TcCLB.505999.160 | leucine-rich repeat-containing protein | 0,4329 | 0,006994 |
| TcCLB.506961.10 | retrotransposon hot spot (RHS) protein, putative | 0,4327 | 0,003908 |
| TcCLB.510237.80 | trans-sialidase, Group V, putative | 0,4327 | 0,1364 |
| TcCLB.506603.35 | trans-sialidase (pseudogene), putative | 0,4327 | 0,1408 |
| TcCLB.504717.10 | divalent cation tolerance protein, putative | 0,4325 | 0,4826 |
| TcCLB.506525.50 | hypothetical protein, conserved | 0,4324 | 0,002557 |
| TcCLB.510155.210 | calcium/potassium channel (CAKC), putative | 0,432 | 0,0005057 |
| TcCLB.507009.130 | hypothetical protein, conserved | 0,4319 | 0,0001172 |
| TcCLB.511167.30 | hypothetical protein, conserved | 0,4318 | 0,002853 |
| TcCLB.503521.89 | mannosyltransferase-II, putative | 0,4317 | 0,00115 |
| TcCLB.508979.40 | hypothetical protein | 0,4316 | 0,568 |
| TcCLB.511735.30 | hypothetical protein, conserved | 0,4315 | 0,03277 |
| TcCLB.507017.159 | hypothetical protein, conserved | 0,4311 | 0,02778 |
| TcCLB.508775.20 | retrotransposon hot spot protein (RHS, pseudogene), putat | 0,4311 | 0,04767 |
| TcCLB.507521.60 | hypothetical protein, conserved | 0,4311 | 0,06472 |
| TcCLB.509843.10 | retrotransposon hot spot (RHS) protein, putative | 0,431 | 0,0004121 |
| TcCLB.504111.10 | hypothetical protein, conserved | 0,4307 | 0,0001104 |
| TcCLB.504213.40 | hypothetical protein, conserved | 0,4305 | 0,0009772 |
| TcCLB.508239.30 | neurobeachin/beige-like protein, putative | 0,4302 | 7,099E-09 |
| TcCLB.509879.20 | hypothetical protein, conserved | 0,4302 | 0,005972 |
| TcCLB.510719.270 | hypothetical protein, conserved | 0,4302 | 0,0387 |

| | | | |
|---|---|---|---|
| TcCLB.508939.40 | hypothetical protein | 0,4302 | 0,4489 |
| TcCLB.510719.35 | ribosomal protein L29, putative | 0,4301 | 0,007242 |
| TcCLB.511209.30 | hypothetical protein, conserved (pseudogene) | 0,43 | 0,0001088 |
| TcCLB.506345.30 | retrotransposon hot spot protein (RHS, pseudogene), putat | 0,43 | 0,0001255 |
| TcCLB.504505.10 | retrotransposon hot spot protein (RHS, pseudogene), putat | 0,4298 | 0,001932 |
| TcCLB.511151.30 | hypothetical protein, conserved | 0,4297 | 0,04284 |
| TcCLB.511003.190 | hypothetical protein, conserved | 0,4296 | 0,01105 |
| TcCLB.505097.20 | hypothetical protein, conserved (pseudogene) | 0,4296 | 0,04109 |
| TcCLB.509265.120 | trans-sialidase, putative | 0,4296 | 0,3301 |
| TcCLB.508087.20 | retrotransposon hot spot protein (RHS, pseudogene), putat | 0,4295 | 0,0001541 |
| TcCLB.508507.24 | hypothetical protein, conserved | 0,4295 | 0,007581 |
| TcCLB.509791.189 | flagellum targeting protein kharon1, putative (fragment) ( | 0,429 | 0,000858 |
| TcCLB.504153.250 | Present in the outer mitochondrial membrane proteome 1 | 0,429 | 0,006804 |
| TcCLB.508823.140 | ribosomal protein S20, putative | 0,4287 | 6,742E-06 |
| TcCLB.409625.10 | trans-sialidase, putative (fragment) | 0,4287 | 0,3263 |
| TcCLB.506927.30 | hypothetical protein, conserved (fragment) | 0,4283 | 0,0144 |
| TcCLB.509993.20 | mucin TcMUCI, putative | 0,4283 | 0,2433 |
| TcCLB.466823.10 | Checkpoint protein HUS1, putative | 0,4281 | 0,05497 |
| TcCLB.509081.20 | mucin-associated surface protein (MASP), putative | 0,4281 | 0,1687 |
| TcCLB.506717.60 | retrotransposon hot spot protein (RHS, pseudogene), putat | 0,428 | 0,0009531 |
| TcCLB.509715.60 | hypothetical protein, conserved | 0,428 | 0,02091 |
| TcCLB.506435.250 | hypothetical protein, conserved | 0,428 | 0,03488 |
| TcCLB.511585.280 | retrotransposon hot spot protein (RHS, pseudogene), putat | 0,4278 | 0,001392 |
| TcCLB.506611.20 | mucin TcMUCII, putative | 0,4278 | 0,5782 |
| TcCLB.508759.40 | mucin-associated surface protein (MASP), putative | 0,4277 | 0,2143 |
| TcCLB.511811.30 | hypothetical protein, conserved | 0,4272 | 0,000934 |
| TcCLB.507831.10 | hypothetical protein | 0,4272 | 0,1451 |
| TcCLB.510101.440 | hypothetical protein, conserved | 0,427 | 0,007051 |
| TcCLB.504147.180 | hypothetical protein, conserved | 0,427 | 0,02276 |
| TcCLB.509205.110 | elongation factor G2-like protein, putative | 0,4269 | 0,3814 |
| TcCLB.503797.20 | centrin (pseudogene), putative | 0,4268 | 0,05245 |
| TcCLB.511875.110 | trans-sialidase (pseudogene), putative | 0,4268 | 0,06374 |
| TcCLB.511173.100 | mucin-associated surface protein (MASP), putative | 0,4268 | 0,2252 |
| TcCLB.508233.60 | heat shock protein 100 (clp protein), putative | 0,4267 | 0,1335 |
| TcCLB.509167.60 | hypothetical protein, conserved | 0,4266 | 0,00001931 |
| TcCLB.508355.140 | hypothetical protein, conserved | 0,4266 | 0,000629 |
| TcCLB.507573.6 | ATP-dependent DEAD/H RNA helicase (pseudogene), putat | 0,4266 | 0,1546 |
| TcCLB.506775.90 | PIF1 helicase-like protein, putative | 0,4262 | 4,111E-06 |
| TcCLB.507491.10 | hypothetical protein, conserved | 0,4262 | 0,002934 |
| TcCLB.507765.120 | hypothetical protein, conserved | 0,4261 | 0,00001371 |
| TcCLB.511421.150 | leucine-rich repeat-containing protein | 0,426 | 0,00005787 |
| TcCLB.506487.60 | endosomal integral membrane protein, putative (fragmen | 0,4259 | 0,1429 |
| TcCLB.503439.80 | glycine dehydrogenase [decarboxylating] (pseudogene), pu | 0,4257 | 0,05338 |
| TcCLB.509381.20 | 40S ribosomal protein S15A, putative | 0,4256 | 1,392E-06 |
| TcCLB.506529.590 | hypothetical protein, conserved | 0,4255 | 0,001409 |

| | | | |
|---|---|---|---|
| TcCLB.503431.80 | hypothetical protein, conserved | 0,4255 | 0,003621 |
| TcCLB.508071.80 | trans-sialidase (pseudogene), putative | 0,4255 | 0,004343 |
| TcCLB.511081.60 | mucin-associated surface protein (MASP), putative | 0,4254 | 0,4733 |
| TcCLB.454403.10 | trans-sialidase (pseudogene), putative | 0,4253 | 0,000878 |
| TcCLB.507241.39 | ABC transporter, putative | 0,4251 | 0,0000616 |
| TcCLB.509753.270 | trans-sialidase (pseudogene), putative | 0,4251 | 0,1719 |
| TcCLB.506605.139 | hypothetical protein, conserved | 0,4248 | 0,0143 |
| TcCLB.509683.70 | dynein light chain Tctex-type, putative | 0,4247 | 0,007169 |
| TcCLB.504173.30 | hypothetical protein, conserved | 0,4247 | 0,1448 |
| TcCLB.506127.9 | phosphoglycerate kinase, putative | 0,4247 | 0,1981 |
| TcCLB.510165.54 | hypothetical protein, conserved | 0,4243 | 0,0192 |
| TcCLB.503885.30 | hypothetical protein, conserved (pseudogene) | 0,4242 | 0,0179 |
| TcCLB.506249.20 | protein transport protein Sec23A, putative | 0,4241 | 0,00001022 |
| TcCLB.508719.60 | kinetoplast DNA-associated protein, putative | 0,4241 | 0,02691 |
| TcCLB.506803.14 | hypothetical protein | 0,4239 | 0,05073 |
| TcCLB.506991.19 | hypothetical protein, conserved | 0,4238 | 0,05022 |
| TcCLB.511393.30 | formin homology 2 (FH2) domain protein, putative | 0,4237 | 1,431E-06 |
| TcCLB.510759.70 | hypothetical protein, conserved | 0,4237 | 0,00001553 |
| TcCLB.510091.60 | hypothetical protein, conserved | 0,4237 | 0,001047 |
| TcCLB.506355.150 | hypothetical protein, conserved | 0,4237 | 0,01406 |
| TcCLB.510397.5 | GTP-binding protein, putative (fragment) | 0,4235 | 0,003569 |
| TcCLB.399685.10 | trans-sialidase, putative (fragment) | 0,4235 | 0,1703 |
| TcCLB.503735.49 | hypothetical protein, conserved | 0,4234 | 0,06075 |
| TcCLB.507163.11 | mucin TcMUCII, putative (fragment) | 0,4234 | 0,5485 |
| TcCLB.488767.20 | retrotransposon hot spot protein (RHS, pseudogene), putat | 0,4233 | 0,02376 |
| TcCLB.510631.11 | retrotransposon hot spot protein (RHS, pseudogene), putat | 0,4231 | 0,001808 |
| TcCLB.506213.20 | hypothetical protein, conserved | 0,4231 | 0,03621 |
| TcCLB.507641.100 | hypothetical protein, conserved | 0,423 | 0,0002038 |
| TcCLB.504009.70 | succinate dehydrogenase cytochrome B subunit | 0,4229 | 0,001158 |
| TcCLB.510289.10 | hypothetical protein, conserved | 0,4229 | 0,03479 |
| TcCLB.510519.60 | calcineurin B subunit, putative (CNB) | 0,4228 | 0,01578 |
| TcCLB.503667.39 | dispersed gene family protein 1 (DGF-1), putative (fragme | 0,4228 | 0,1157 |
| TcCLB.504073.10 | endonuclease/exonuclease/phosphatase, putative | 0,4227 | 0,0008056 |
| TcCLB.508839.20 | hypothetical protein, conserved | 0,4227 | 0,002813 |
| TcCLB.509885.30 | SPFH domain / Band 7 family protein, putative | 0,4227 | 0,00317 |
| TcCLB.506599.70 | mucin TcMUCII, putative | 0,4227 | 0,5407 |
| TcCLB.510733.20 | hypothetical protein, conserved | 0,4224 | 0,0008924 |
| TcCLB.510583.180 | mucin-associated surface protein (MASP, pseudogene), pu | 0,4224 | 0,4351 |
| TcCLB.504069.50 | peroxin 14, putative | 0,4223 | 0,00005249 |
| TcCLB.503745.30 | ascorbate peroxidase (APX) | 0,4222 | 0,00912 |
| TcCLB.508071.50 | surface protease GP63, putative | 0,4221 | 0,5829 |
| TcCLB.508501.220 | hypothetical protein, conserved | 0,422 | 0,001419 |
| TcCLB.506451.35 | hypothetical protein | 0,422 | 0,2445 |
| TcCLB.508689.30 | hypothetical protein, conserved | 0,4218 | 0,001607 |
| TcCLB.506613.60 | mucin TcMUCII (pseudogene), putative | 0,4218 | 0,3736 |

| | | | |
|---|---|---|---|
| TcCLB.507093.100 | hypothetical protein, conserved | 0,4216 | 0,0007489 |
| TcCLB.508453.10 | mucin-associated surface protein (MASP), putative | 0,4216 | 0,1386 |
| TcCLB.511747.14 | hypothetical protein, conserved | 0,4215 | 0,03784 |
| TcCLB.509685.40 | trans-sialidase, Group V, putative | 0,4215 | 0,06295 |
| TcCLB.506773.130 | pumilio-repeat, RNA-binding protein, putative | 0,4213 | 2,521E-06 |
| TcCLB.509905.150 | trans-sialidase, Group II, putative | 0,4211 | 0,00001806 |
| TcCLB.510283.120 | hypothetical protein, conserved | 0,4211 | 0,001833 |
| TcCLB.507049.170 | Axonemal inner arm dynein light chain, putative | 0,4211 | 0,002641 |
| TcCLB.508633.120 | trans-sialidase (pseudogene), putative | 0,4209 | 0,1147 |
| TcCLB.508853.30 | hypothetical protein | 0,4209 | 0,2707 |
| TcCLB.508209.130 | hypothetical protein, conserved | 0,4208 | 0,000546 |
| TcCLB.458759.14 | hypothetical protein, conserved | 0,4207 | 0,000634 |
| TcCLB.503999.60 | phosphoprotein phosphatase, putative | 0,4205 | 0,0003933 |
| TcCLB.511655.10 | hypothetical protein, conserved | 0,4205 | 0,0545 |
| TcCLB.510033.30 | trans-sialidase, Group V, putative | 0,4201 | 0,1393 |
| TcCLB.506279.30 | mucin TcMUCII, putative | 0,4201 | 0,3159 |
| TcCLB.511239.30 | hypothetical protein, conserved | 0,4197 | 0,01424 |
| TcCLB.509055.54 | hypothetical protein, conserved | 0,4197 | 0,1057 |
| TcCLB.507091.10 | hypothetical protein, conserved | 0,4195 | 0,3568 |
| TcCLB.504057.50 | hypothetical protein, conserved | 0,4194 | 0,0005376 |
| TcCLB.510361.260 | mucin-associated surface protein (MASP), putative | 0,4194 | 0,5194 |
| TcCLB.509453.40 | RNA polymerase subunit RPB10, putative (RPB10) | 0,4193 | 0,05546 |
| TcCLB.509103.30 | hypothetical protein, conserved | 0,4192 | 0,1189 |
| TcCLB.508919.150 | retrotransposon hot spot protein (RHS, pseudogene), putat | 0,4191 | 0,08196 |
| TcCLB.510009.19 | hypothetical protein, conserved (fragment) | 0,4191 | 0,1645 |
| TcCLB.508365.140 | hypothetical protein | 0,419 | 0,3274 |
| TcCLB.507023.150 | hypothetical protein | 0,4187 | 0,002942 |
| TcCLB.506615.150 | trans-sialidase, Group V, putative | 0,4186 | 0,009331 |
| TcCLB.506321.270 | hypothetical protein, conserved | 0,4185 | 0,018 |
| TcCLB.509419.20 | mucin-associated surface protein (MASP), putative | 0,4185 | 0,6622 |
| TcCLB.507539.10 | hypothetical protein, conserved | 0,4184 | 0,09842 |
| TcCLB.506965.140 | mucin-associated surface protein (MASP), putative | 0,4184 | 0,2088 |
| TcCLB.504239.365 | surface protease GP63 (pseudogene), putative | 0,4183 | 0,6145 |
| TcCLB.510981.20 | trans-sialidase, putative | 0,4181 | 0,02895 |
| TcCLB.508173.84 | hypothetical protein, conserved | 0,418 | 0,0004194 |
| TcCLB.506797.60 | hypothetical protein, conserved | 0,4177 | 0,004133 |
| TcCLB.511151.100 | Unconventional myosin | 0,4175 | 0,00003526 |
| TcCLB.511259.170 | trans-sialidase (pseudogene), putative | 0,4175 | 0,2285 |
| TcCLB.511077.150 | hypothetical protein | 0,4174 | 0,5874 |
| TcCLB.509525.360 | mucin-associated surface protein (MASP, pseudogene), pu | 0,4173 | 0,4182 |
| TcCLB.507053.90 | conserved protein, unknown function | 0,417 | 0,002012 |
| TcCLB.509079.20 | mucin-associated surface protein (MASP), putative | 0,417 | 0,4691 |
| TcCLB.506697.70 | hypothetical protein, conserved | 0,4169 | 0,003462 |
| TcCLB.510121.50 | calmodulin, putative | 0,4169 | 0,07184 |
| TcCLB.511553.120 | surface protease GP63, putative | 0,4169 | 0,298 |

| | | | |
|---|---|---|---|
| TcCLB.509509.20 | C-1-tetrahydrofolate synthase, cytoplasmic, putative | 0,4168 | 0,01052 |
| TcCLB.510565.70 | protein kinase, putative | 0,4165 | 8,179E-06 |
| TcCLB.510877.55 | acyl-CoA binding protein, putative | 0,4164 | 0,002401 |
| TcCLB.504029.70 | hypothetical protein, conserved | 0,4163 | 0,005034 |
| TcCLB.507937.20 | mucin-associated surface protein (MASP, pseudogene), pu | 0,4159 | 0,4057 |
| TcCLB.510041.20 | zinc carboxypeptidase, putative | 0,4157 | 0,1292 |
| TcCLB.506939.40 | mucin TcMUCII, putative | 0,4157 | 0,4595 |
| TcCLB.509743.40 | trans-sialidase, putative | 0,4157 | 0,4913 |
| TcCLB.509899.10 | trans-sialidase (pseudogene), putative | 0,4156 | 0,0203 |
| TcCLB.508475.10 | ribosomal protein S20, putative | 0,4155 | 0,0002927 |
| TcCLB.506653.40 | trans-sialidase (pseudogene), putative | 0,4155 | 0,02423 |
| TcCLB.509127.30 | hypothetical protein, conserved | 0,4155 | 0,1047 |
| TcCLB.510901.200 | differentiation inhibitory kinase, putative (ZFK) | 0,4154 | 0,00005749 |
| TcCLB.503587.10 | mucin-associated surface protein (MASP), putative | 0,4154 | 0,6507 |
| TcCLB.511389.10 | hypothetical protein, conserved | 0,4153 | 0,001861 |
| TcCLB.507927.4 | rac serine-threonine kinase, putative | 0,4152 | 0,03178 |
| TcCLB.505533.10 | dispersed gene family protein 1 (DGF-1, pseudogene), puta | 0,4152 | 0,307 |
| TcCLB.508387.120 | glutaminyl cyclase, putative | 0,4151 | 0,00002496 |
| TcCLB.508679.7 | hypothetical protein, conserved (pseudogene) | 0,4151 | 0,3331 |
| TcCLB.453767.50 | mucin-associated surface protein (MASP), putative | 0,4151 | 0,4777 |
| TcCLB.508737.110 | hypothetical protein, conserved | 0,415 | 0,0006011 |
| TcCLB.509237.140 | calpain-like cysteine peptidase, putative | 0,415 | 0,008329 |
| TcCLB.508347.24 | hypothetical protein, conserved | 0,415 | 0,03224 |
| TcCLB.509303.10 | retrotransposon hot spot protein (RHS, pseudogene), putat | 0,4148 | 0,07469 |
| TcCLB.511257.80 | hypothetical protein | 0,4147 | 0,129 |
| TcCLB.511257.60 | surface protease GP63, putative | 0,4144 | 0,02482 |
| TcCLB.510989.10 | fatty acid elongase, putative | 0,4143 | 0,01103 |
| TcCLB.507295.30 | mucin TcMUCII, putative | 0,4141 | 0,1429 |
| TcCLB.509723.10 | protein kinase, putative | 0,4141 | 0,1962 |
| TcCLB.508633.30 | mucin-associated surface protein (MASP, pseudogene), pu | 0,414 | 0,4204 |
| TcCLB.506885.10 | alpha-1,2-mannosyltransferase, putative | 0,4139 | 0,0008251 |
| TcCLB.511903.270 | hypothetical protein, conserved | 0,4138 | 0,09744 |
| TcCLB.510275.250 | mucin-associated surface protein (MASP), putative | 0,4138 | 0,1458 |
| TcCLB.506303.80 | hypothetical protein, conserved | 0,4137 | 1,142E-06 |
| TcCLB.503959.10 | hypothetical protein, conserved | 0,4135 | 0,0003576 |
| TcCLB.510587.10 | hypothetical protein | 0,4135 | 0,6715 |
| TcCLB.510355.150 | retrotransposon hot spot protein (RHS, pseudogene), putat | 0,4134 | 0,002919 |
| TcCLB.511807.210 | hypothetical protein, conserved (pseudogene) | 0,4133 | 0,0000402 |
| TcCLB.509837.10 | hypothetical protein, conserved | 0,4133 | 0,0002357 |
| TcCLB.511445.40 | hypothetical protein, conserved | 0,4132 | 0,002143 |
| TcCLB.504285.10 | retrotransposon hot spot protein (RHS, pseudogene), putat | 0,4131 | 0,05069 |
| TcCLB.508727.18 | centrin, putative | 0,4129 | 0,01784 |
| TcCLB.509099.170 | hypothetical protein, conserved | 0,4128 | 0,08313 |
| TcCLB.508233.4 | protein transport protein Sec23A, putative (fragment) | 0,4127 | 0,001766 |
| TcCLB.509967.70 | katanin, putative | 0,4127 | 0,2007 |

| | | | |
|---|---|---|---|
| TcCLB.510907.10 | trans-sialidase, Group VIII, putative | 0,4127 | 0,3699 |
| TcCLB.508355.290 | hypothetical protein, conserved | 0,4125 | 0,003285 |
| TcCLB.506553.10 | hypothetical protein, conserved | 0,4123 | 0,0655 |
| TcCLB.511657.10 | protein kinase, putative | 0,4123 | 0,1149 |
| TcCLB.503431.60 | acid phosphatase, putative (AP) | 0,4122 | 0,002907 |
| TcCLB.504239.140 | mucin-associated surface protein (MASP, pseudogene), pu | 0,4122 | 0,278 |
| TcCLB.508805.30 | choline/ethanolamine kinase (pseudogene), putative | 0,412 | 0,006566 |
| TcCLB.507803.10 | mucin-associated surface protein (MASP), putative | 0,412 | 0,3064 |
| TcCLB.508301.20 | metallo-peptidase, Clan ME, Family M16 | 0,4118 | 0,0004882 |
| TcCLB.510693.60 | retrotransposon hot spot protein (RHS, pseudogene), putat | 0,4118 | 0,4326 |
| TcCLB.508741.200 | hypothetical protein, conserved | 0,4117 | 6,348E-06 |
| TcCLB.508787.20 | hypothetical protein | 0,4117 | 0,0003514 |
| TcCLB.508535.20 | nuclear transmembrane protein, putative (NP) | 0,4117 | 0,0008922 |
| TcCLB.503507.35 | mucin-associated surface protein (MASP, pseudogene), pu | 0,4112 | 0,2 |
| TcCLB.510873.10 | NAD(P)-dependent steroid dehydrogenase protein, putative | 0,4111 | 1,031E-06 |
| TcCLB.506799.100 | mucin-like glycoprotein, putative | 0,4107 | 0,2679 |
| TcCLB.506831.30 | hypothetical protein, conserved | 0,4105 | 0,02547 |
| TcCLB.506755.80 | hypothetical protein | 0,4102 | 0,009765 |
| TcCLB.511641.5 | fucose kinase, putative | 0,4099 | 0,1882 |
| TcCLB.506499.80 | mucin-associated surface protein (MASP), putative | 0,4099 | 0,5866 |
| TcCLB.510121.70 | hypothetical protein, conserved | 0,4098 | 0,0001345 |
| TcCLB.511249.110 | kinesin K39 (pseudogene), putative | 0,4096 | 0,0002408 |
| TcCLB.510307.220 | retrotransposon hot spot (RHS) protein, putative | 0,4095 | 0,007433 |
| TcCLB.506627.80 | hypothetical protein, conserved | 0,4094 | 0,0004566 |
| TcCLB.507641.70 | hypothetical protein, conserved | 0,4094 | 0,002117 |
| TcCLB.511255.210 | mucin TcMUCII, putative | 0,4094 | 0,6445 |
| TcCLB.504231.20 | hypothetical protein, conserved | 0,4093 | 0,0000278 |
| TcCLB.509539.10 | trans-sialidase (pseudogene), putative | 0,4093 | 0,05689 |
| TcCLB.507035.110 | cAMP response protein 1, putative (CARP1) | 0,4092 | 0,004329 |
| TcCLB.506337.4 | short-chain dehydrogenase, putative (fragment) | 0,4092 | 0,03983 |
| TcCLB.506925.10 | hypothetical protein | 0,4092 | 0,1227 |
| TcCLB.505071.70 | hypothetical protein, conserved | 0,409 | 0,0312 |
| TcCLB.508139.80 | trans-sialidase (pseudogene), putative | 0,4088 | 0,02894 |
| TcCLB.508541.40 | surface protease GP63 (pseudogene), putative | 0,4088 | 0,3771 |
| TcCLB.510275.272 | hypothetical protein | 0,4088 | 0,4277 |
| TcCLB.509897.30 | mucin-associated surface protein (MASP, pseudogene), pu | 0,4087 | 0,3761 |
| TcCLB.506025.50 | Vacuolar proton pump subunit B, putative | 0,4086 | 0,00001054 |
| TcCLB.510661.80 | hypothetical protein | 0,4086 | 0,00008433 |
| TcCLB.506131.20 | mucin TcMUCII, putative | 0,4084 | 0,5454 |
| TcCLB.504479.20 | retrotransposon hot spot protein (RHS, pseudogene), putat | 0,4083 | 0,07262 |
| TcCLB.506329.20 | mucin TcMUCII, putative | 0,4082 | 0,5119 |
| TcCLB.508175.309 | 60S ribosomal protein L19, putative | 0,4079 | 2,586E-06 |
| TcCLB.507851.30 | hypothetical protein, conserved | 0,4079 | 0,00003643 |
| TcCLB.511369.10 | elongation factor 1-alpha, putative | 0,4079 | 0,1369 |
| TcCLB.506955.14 | mucin-associated surface protein (MASP, pseudogene), pu | 0,4079 | 0,3106 |

| | | | |
|---|---|---|---|
| TcCLB.508143.80 | mucin-associated surface protein (MASP), putative | 0,4079 | 0,3801 |
| TcCLB.509921.9 | dispersed gene family protein 1 (DGF-1), putative | 0,4078 | 0,3405 |
| TcCLB.506209.60 | retrotransposon hot spot protein (RHS, pseudogene), putat | 0,4075 | 0,007019 |
| TcCLB.509551.90 | hypothetical protein, conserved | 0,4074 | 0,00713 |
| TcCLB.506885.360 | hypothetical protein, conserved | 0,4072 | 0,001984 |
| TcCLB.505071.140 | protein kinase, putative | 0,4071 | 0,001373 |
| TcCLB.507485.140 | tuzin | 0,4071 | 0,001446 |
| TcCLB.503809.100 | GUP1, putative | 0,407 | 0,0003849 |
| TcCLB.503647.9 | hypothetical protein, conserved | 0,407 | 0,0006531 |
| TcCLB.503981.70 | hypothetical protein, conserved | 0,4069 | 0,002656 |
| TcCLB.508839.50 | hypothetical protein, conserved | 0,4068 | 0,0001939 |
| TcCLB.508273.70 | hypothetical protein, conserved | 0,4067 | 0,02741 |
| TcCLB.509785.40 | retrotransposon hot spot protein (RHS, pseudogene), putat | 0,4066 | 0,06089 |
| TcCLB.510101.20 | phosphatidylinositol-4-phosphate 5-kinase, putative | 0,4063 | 0,004793 |
| TcCLB.507071.70 | mucin-associated surface protein (MASP, pseudogene), pu | 0,4063 | 0,3252 |
| TcCLB.506599.189 | mucin TcMUCII, putative | 0,4063 | 0,3323 |
| TcCLB.508389.10 | hypothetical protein, conserved | 0,4058 | 0,0001797 |
| TcCLB.508659.30 | hypothetical protein, conserved | 0,4053 | 0,001112 |
| TcCLB.503525.30 | hypothetical protein, conserved | 0,4052 | 0,0008972 |
| TcCLB.511849.30 | mucin-associated surface protein (MASP), putative | 0,4052 | 0,2834 |
| TcCLB.511661.30 | hypothetical protein, conserved | 0,4051 | 7,117E-08 |
| TcCLB.506435.260 | hypothetical protein | 0,4051 | 0,005071 |
| TcCLB.510275.400 | mucin-associated surface protein (MASP), putative | 0,405 | 0,3503 |
| TcCLB.509599.50 | hypothetical protein, conserved | 0,4049 | 0,001431 |
| TcCLB.510003.9 | hypothetical protein, conserved | 0,4049 | 0,003726 |
| TcCLB.507067.70 | hypothetical protein | 0,4048 | 0,5556 |
| TcCLB.510509.20 | hypothetical protein, conserved | 0,4047 | 0,003128 |
| TcCLB.511281.30 | RNA-binding protein 27, putative (RBP27) | 0,4046 | 0,005398 |
| TcCLB.507649.10 | hypothetical protein, conserved | 0,4046 | 0,01299 |
| TcCLB.504015.11 | mucin-associated surface protein (MASP), putative (fragm | 0,4046 | 0,4615 |
| TcCLB.508647.90 | hypothetical protein, conserved | 0,4045 | 0,007944 |
| TcCLB.511103.9 | dispersed gene family protein 1 (DGF-1), putative (fragme | 0,4044 | 0,5305 |
| TcCLB.507837.100 | hypothetical protein, conserved | 0,4043 | 0,0001278 |
| TcCLB.509265.90 | trans-sialidase, Group II, putative | 0,4041 | 0,03124 |
| TcCLB.510303.10 | surface protease GP63, putative | 0,4041 | 0,1374 |
| TcCLB.508099.30 | hypothetical protein | 0,4037 | 0,4827 |
| TcCLB.511585.10 | hypothetical protein, conserved | 0,4035 | 0,0005757 |
| TcCLB.508927.20 | glycosomal transporter (GAT1), putative (GAT1) | 0,4035 | 0,003571 |
| TcCLB.506605.59 | adenosine 5'-monophosphoramidase, putative | 0,4035 | 0,01966 |
| TcCLB.503897.150 | hypothetical protein, conserved | 0,4034 | 0,007289 |
| TcCLB.511469.80 | hypothetical protein, conserved | 0,4031 | 0,004943 |
| TcCLB.508229.60 | trans-sialidase, Group VII, putative | 0,403 | 0,6261 |
| TcCLB.511369.20 | elongation factor 1-alpha, putative | 0,4029 | 0,076 |
| TcCLB.510007.50 | hypothetical protein, conserved | 0,4027 | 0,0007397 |
| TcCLB.508989.70 | hypothetical protein, conserved | 0,4025 | 0,001517 |

| | | | |
|---|---|---|---|
| TcCLB.511773.60 | glycine dehydrogenase [decarboxylating] (pseudogene), pu | 0,4023 | 0,0001115 |
| TcCLB.508645.30 | dynein light chain, putative | 0,4019 | 0,00302 |
| TcCLB.511217.70 | hypothetical protein, conserved | 0,4018 | 0,0001414 |
| TcCLB.506559.59 | ubiquitin, putative | 0,4017 | 0,01081 |
| TcCLB.509959.160 | hypothetical protein, conserved (pseudogene) | 0,4017 | 0,04896 |
| TcCLB.503593.30 | hypothetical protein, conserved | 0,4016 | 0,0005257 |
| TcCLB.510431.10 | isopentenyl-diphosphate delta-isomerase (fragment) (idi1) | 0,4016 | 0,004617 |
| TcCLB.507795.40 | hypothetical protein, conserved | 0,4016 | 0,1917 |
| TcCLB.511003.80 | hypothetical protein, conserved | 0,4015 | 0,0008025 |
| TcCLB.506733.80 | hypothetical protein, conserved | 0,4015 | 0,008589 |
| TcCLB.507801.80 | hypothetical protein, conserved | 0,4014 | 0,0552 |
| TcCLB.504797.70 | DNA excision/repair protein SNF2, putative | 0,4013 | 6,315E-06 |
| TcCLB.507027.70 | hypothetical protein, conserved | 0,4013 | 0,002534 |
| TcCLB.508851.59 | ATP synthase, putative | 0,4011 | 0,00438 |
| TcCLB.510759.120 | rieske iron-sulfur protein, mitochondrial precursor, putative | 0,4007 | 7,489E-06 |
| TcCLB.511865.40 | hypothetical protein, conserved | 0,4006 | 0,001332 |
| TcCLB.508173.80 | hypothetical protein, conserved | 0,4003 | 0,0001149 |
| TcCLB.509957.30 | retrotransposon hot spot protein (RHS, pseudogene), putat | 0,4001 | 0,1258 |
| TcCLB.511825.69 | cyclophilin, putative | 0,3998 | 0,0314 |
| TcCLB.506567.60 | ABC transporter, putative | 0,3996 | 0,0004757 |
| TcCLB.509237.80 | hypothetical protein, conserved | 0,3995 | 0,0001721 |
| TcCLB.463155.10 | retrotransposon hot spot protein (RHS, pseudogene), putat | 0,3993 | 0,01955 |
| TcCLB.511821.50 | hypothetical protein, conserved | 0,3989 | 0,02231 |
| TcCLB.504639.40 | trans-sialidase, putative | 0,3984 | 0,2554 |
| TcCLB.510209.9 | trans-sialidase, putative | 0,3983 | 0,5447 |
| TcCLB.448329.10 | trans-sialidase, putative | 0,3982 | 0,5621 |
| TcCLB.506205.40 | protein kinase, putative | 0,3981 | 0,007472 |
| TcCLB.506919.40 | retrotransposon hot spot protein (RHS, pseudogene), putat | 0,3981 | 0,3347 |
| TcCLB.506865.50 | Present in the outer mitochondrial membrane proteome 4 | 0,398 | 0,02894 |
| TcCLB.508573.90 | retrotransposon hot spot protein (RHS, pseudogene), putat | 0,398 | 0,09336 |
| TcCLB.508165.310 | surface protease GP63, putative | 0,3975 | 0,2161 |
| TcCLB.506553.30 | GTP-binding protein, putative | 0,3974 | 0,001686 |
| TcCLB.508695.30 | hypothetical protein (fragment) | 0,3974 | 0,02831 |
| TcCLB.504741.230 | hypothetical protein, conserved | 0,3967 | 0,0009313 |
| TcCLB.511691.30 | ubiquitin activating enzyme, putative | 0,3966 | 0,01766 |
| TcCLB.511643.50 | mucin-associated surface protein (MASP, pseudogene), pu | 0,3965 | 0,3424 |
| TcCLB.510713.50 | beta galactofuranosyl glycosyltransferase (pseudogene), p | 0,3964 | 0,3042 |
| TcCLB.507855.9 | dispersed gene family protein 1 (DGF-1), putative (fragme | 0,3962 | 0,1769 |
| TcCLB.511471.150 | hypothetical protein, conserved (pseudogene) | 0,3962 | 0,1838 |
| TcCLB.511755.100 | hypothetical protein, conserved | 0,3962 | 0,1965 |
| TcCLB.506237.10 | mismatch repair protein MSH5, putative | 0,3961 | 0,01448 |
| TcCLB.509979.174 | mucin-associated surface protein (MASP), putative | 0,3961 | 0,1639 |
| TcCLB.509013.19 | calpain-like cysteine peptidase, putative (fragment) | 0,3961 | 0,1786 |
| TcCLB.508437.5 | mucin-like glycoprotein (pseudogene), putative | 0,396 | 0,4763 |
| TcCLB.506727.4 | hypothetical protein, conserved | 0,3959 | 0,1867 |

| | | | |
|---|---|---|---|
| TcCLB.455171.9 | trans-sialidase, putative | 0,3959 | 0,2672 |
| TcCLB.507227.9 | hypothetical protein, conserved (fragment) | 0,3956 | 0,002107 |
| TcCLB.505807.150 | Leucine-rich repeat, ribonuclease inhibitor subtype domain | 0,3954 | 0,006445 |
| TcCLB.507529.50 | hypothetical protein, conserved | 0,3953 | 4,809E-07 |
| TcCLB.510489.20 | mucin-associated surface protein (MASP), putative | 0,3953 | 0,3281 |
| TcCLB.506163.40 | vacuolar type h ATPase subunit, putative | 0,3953 | 0,3842 |
| TcCLB.503665.15 | trans-sialidase (pseudogene), putative | 0,3952 | 0,04264 |
| TcCLB.510431.290 | hypothetical protein | 0,3952 | 0,1304 |
| TcCLB.506799.20 | mucin-like glycoprotein, putative | 0,3952 | 0,2961 |
| TcCLB.506925.380 | hypothetical protein | 0,395 | 0,007155 |
| TcCLB.503757.20 | hypothetical protein, conserved | 0,395 | 0,01872 |
| TcCLB.506425.80 | phosphatidylinositol kinase, putative | 0,3949 | 0,002332 |
| TcCLB.509999.60 | lipase domain protein, putative | 0,3947 | 9,476E-06 |
| TcCLB.504769.80 | hypothetical protein, conserved | 0,3946 | 0,0001169 |
| TcCLB.509665.40 | hypothetical protein | 0,3945 | 0,514 |
| TcCLB.509671.130 | trypanin-like protein (pseudogene), putative | 0,3944 | 0,005425 |
| TcCLB.510951.19 | hypothetical protein, conserved | 0,3944 | 0,06281 |
| TcCLB.506435.150 | hypothetical protein, conserved | 0,3942 | 0,0009414 |
| TcCLB.508931.30 | UDP-Gal or UDP-GlcNAc-dependent glycosyltransferase, pu | 0,3939 | 0,09155 |
| TcCLB.511809.80 | hypothetical protein, conserved | 0,3937 | 0,00001218 |
| TcCLB.510425.10 | hypothetical protein, conserved | 0,3937 | 0,04544 |
| TcCLB.508001.90 | hypothetical protein, conserved (pseudogene) | 0,3934 | 0,2247 |
| TcCLB.503911.10 | hypothetical protein | 0,3934 | 0,3498 |
| TcCLB.412723.9 | dispersed gene family protein 1 (DGF-1), putative (fragme | 0,3933 | 0,2356 |
| TcCLB.504105.160 | splicing factor ptsr1 interacting protein, putative | 0,3931 | 0,0001745 |
| TcCLB.507603.200 | proteasome regulatory ATPase subunit 3, putative | 0,3929 | 0,0003976 |
| TcCLB.504143.20 | hypothetical protein, conserved | 0,3922 | 0,01047 |
| TcCLB.511887.40 | hypothetical protein, conserved | 0,3922 | 0,2105 |
| TcCLB.510243.80 | hypothetical protein, conserved | 0,3921 | 0,006875 |
| TcCLB.504137.90 | hypothetical protein, conserved | 0,3916 | 0,002297 |
| TcCLB.506155.60 | aldehyde dehydrogenase, putative | 0,3915 | 0,0005419 |
| TcCLB.509229.10 | hypothetical protein, conserved | 0,3915 | 0,003479 |
| TcCLB.509977.15 | surface protease GP63 (pseudogene), putative | 0,3915 | 0,2877 |
| TcCLB.509349.30 | hypothetical protein | 0,3914 | 0,2297 |
| TcCLB.507071.340 | hypothetical protein | 0,3914 | 0,2961 |
| TcCLB.508081.80 | hypothetical protein | 0,3914 | 0,5717 |
| TcCLB.511277.180 | hypothetical protein, conserved | 0,3913 | 0,0001319 |
| TcCLB.506329.59 | mucin-associated surface protein (MASP, pseudogene), pu | 0,3913 | 0,2674 |
| TcCLB.509599.40 | hypothetical protein, conserved | 0,3908 | 0,03521 |
| TcCLB.511561.11 | hypothetical protein, conserved (pseudogene) | 0,3908 | 0,04268 |
| TcCLB.508269.20 | hypothetical protein, conserved | 0,3908 | 0,05164 |
| TcCLB.511817.110 | hypothetical protein, conserved | 0,3907 | 0,002489 |
| TcCLB.509591.60 | phospholipid-transporting ATPase 1-like protein, putative | 0,3905 | 0,5357 |
| TcCLB.503487.70 | hypothetical protein, conserved | 0,3904 | 0,0002725 |
| TcCLB.507641.239 | hypothetical protein, conserved | 0,3903 | 0,001726 |

| | | | |
|---|---|---|---|
| TcCLB.510303.20 | surface protease GP63, putative | 0,39 | 0,2278 |
| TcCLB.510741.190 | hypothetical protein, conserved (pseudogene) | 0,3899 | 0,006525 |
| TcCLB.508001.106 | serine/threonine protein kinase (pseudogene), putative | 0,3898 | 0,3395 |
| TcCLB.510303.230 | hypothetical protein, conserved | 0,3897 | 0,0007284 |
| TcCLB.504787.47 | elongation factor 1-gamma (EF-1-gamma, pseudogene), p | 0,3896 | 0,546 |
| TcCLB.510759.134 | chaperone DNAJ protein, putative | 0,3895 | 0,003458 |
| TcCLB.511867.20 | hypothetical protein, conserved | 0,3895 | 0,03428 |
| TcCLB.508883.48 | trans-sialidase, putative | 0,3894 | 0,1878 |
| TcCLB.510335.30 | hypothetical protein, conserved (pseudogene) | 0,3893 | 0,00001914 |
| TcCLB.504089.50 | flagellar radial spoke component, putative | 0,3893 | 0,0004198 |
| TcCLB.507669.14 | hypothetical protein, conserved | 0,3893 | 0,004927 |
| TcCLB.510429.30 | hypothetical protein, conserved | 0,3893 | 0,01092 |
| TcCLB.509967.180 | hypothetical protein, conserved | 0,3892 | 0,008852 |
| TcCLB.510373.119 | mucin-associated surface protein (MASP), putative | 0,3891 | 0,4736 |
| TcCLB.506617.20 | surface protein TolT (TolT) | 0,3889 | 0,0149 |
| TcCLB.509699.60 | mucin-associated surface protein (MASP, pseudogene), pu | 0,3887 | 0,1207 |
| TcCLB.425597.10 | trans-sialidase (pseudogene), putative | 0,3887 | 0,3781 |
| TcCLB.508247.149 | dispersed gene family protein 1 (DGF-1), putative | 0,3886 | 0,5326 |
| TcCLB.506823.65 | trans-sialidase (pseudogene), putative | 0,3884 | 0,2031 |
| TcCLB.510241.130 | tyrosine phospatase-like protein (pseudogene), putative | 0,3881 | 0,007931 |
| TcCLB.506147.220 | hypothetical protein, conserved | 0,3879 | 0,00145 |
| TcCLB.510583.34 | mucin-associated surface protein (MASP, pseudogene), pu | 0,3879 | 0,3574 |
| TcCLB.510267.10 | hypothetical protein, conserved | 0,3878 | 0,002108 |
| TcCLB.503987.4 | Ras-related protein RAB2B, putative (fragment) (RAB2B) | 0,3877 | 0,04474 |
| TcCLB.503687.58 | AMP deaminase, putative | 0,3876 | 0,09171 |
| TcCLB.507055.10 | hypothetical protein, conserved | 0,3875 | 0,1525 |
| TcCLB.506789.310 | hypothetical protein, conserved | 0,387 | 0,005613 |
| TcCLB.509561.9 | hypothetical protein, conserved | 0,3869 | 0,02397 |
| TcCLB.511707.10 | hypothetical protein | 0,3869 | 0,05744 |
| TcCLB.401041.10 | retrotransposon hot spot protein (RHS, pseudogene), putat | 0,3868 | 0,001761 |
| TcCLB.507411.10 | trans-sialidase, Group V, putative | 0,3868 | 0,08844 |
| TcCLB.511065.36 | hypothetical protein, conserved | 0,3867 | 0,0005762 |
| TcCLB.508939.10 | trans-sialidase (pseudogene), putative | 0,3867 | 0,128 |
| TcCLB.506529.190 | hypothetical protein, conserved | 0,3863 | 0,01295 |
| TcCLB.509747.70 | actin-like protein, putative | 0,3863 | 0,01437 |
| TcCLB.508325.150 | glycine dehydrogenase (pseudogene), putative | 0,3861 | 0,0001431 |
| TcCLB.508179.30 | hypothetical protein, conserved | 0,3861 | 0,004285 |
| TcCLB.511127.70 | hypothetical protein, conserved | 0,3859 | 0,003654 |
| TcCLB.509127.97 | hypothetical protein, conserved | 0,3859 | 0,02066 |
| TcCLB.506459.40 | dispersed gene family protein 1 (DGF-1, pseudogene), puta | 0,3858 | 0,4714 |
| TcCLB.506559.330 | clathrin coat assembly protein AP17, putative | 0,3857 | 0,02976 |
| TcCLB.506351.50 | hypothetical protein | 0,3855 | 0,01851 |
| TcCLB.510901.40 | hypothetical protein, conserved | 0,3855 | 0,03046 |
| TcCLB.487739.20 | choline/ethanolamine kinase (pseudogene), putative | 0,3849 | 0,07063 |
| TcCLB.510997.20 | hypothetical protein, conserved | 0,3848 | 0,02898 |

| | | | |
|---|---|---|---|
| TcCLB.503511.10 | hypothetical protein, conserved | 0,3846 | 0,006571 |
| TcCLB.511765.40 | retrotransposon hot spot protein (RHS, pseudogene), putat | 0,3846 | 0,02199 |
| TcCLB.510421.90 | protein kinase, putative | 0,3845 | 0,005576 |
| TcCLB.509081.10 | elongation factor 1-gamma (EF-1-gamma), putative | 0,3845 | 0,1015 |
| TcCLB.510191.70 | mucin-associated surface protein (MASP), putative | 0,3841 | 0,4644 |
| TcCLB.509221.10 | retrotransposon hot spot protein (RHS, pseudogene), putat | 0,3839 | 0,002489 |
| TcCLB.510079.40 | dispersed gene family protein 1 (DGF-1, pseudogene), puta | 0,3837 | 0,3056 |
| TcCLB.503755.20 | hypothetical protein | 0,3836 | 0,002777 |
| TcCLB.510657.80 | hypothetical protein, conserved | 0,3835 | 0,00171 |
| TcCLB.511185.10 | hypothetical protein | 0,3835 | 0,03718 |
| TcCLB.481211.20 | hypothetical protein | 0,3835 | 0,143 |
| TcCLB.510197.180 | trans-sialidase, Group VI, putative | 0,3831 | 0,2474 |
| TcCLB.510759.160 | metacaspase 5, putative | 0,3829 | 0,0009335 |
| TcCLB.506323.40 | trans-sialidase (pseudogene), putative | 0,3829 | 0,03666 |
| TcCLB.506067.140 | mucin-associated surface protein (MASP), putative | 0,3827 | 0,4851 |
| TcCLB.509525.150 | mucin TcMUCII, putative | 0,3827 | 0,4995 |
| TcCLB.511245.160 | fatty acid elongase, putative | 0,3825 | 0,2019 |
| TcCLB.507071.60 | mucin TcMUCII, putative | 0,3824 | 0,605 |
| TcCLB.508915.9 | hypothetical protein, conserved | 0,3823 | 0,02456 |
| TcCLB.508735.30 | mucin-associated surface protein (MASP), putative | 0,3823 | 0,04896 |
| TcCLB.506999.40 | metacaspase 5, putative | 0,3822 | 0,000696 |
| TcCLB.506201.170 | flagellar attachment zone protein, putative (FAZ1) | 0,3821 | 0,00002085 |
| TcCLB.415331.10 | trans-sialidase (pseudogene), putative | 0,382 | 0,01042 |
| TcCLB.505267.100 | retrotransposon hot spot protein (RHS, pseudogene), putat | 0,3819 | 0,006411 |
| TcCLB.503787.20 | mucin TcMUCII, putative | 0,3819 | 0,4342 |
| TcCLB.511653.30 | hypothetical protein, conserved | 0,3813 | 0,002368 |
| TcCLB.504933.20 | hypothetical protein | 0,3812 | 0,001601 |
| TcCLB.507849.40 | hypothetical protein, conserved | 0,3811 | 0,0000844 |
| TcCLB.508153.310 | protein kinase, putative | 0,3809 | 0,006281 |
| TcCLB.402863.9 | dynein heavy chain, putative (fragment) | 0,3808 | 0,0001031 |
| TcCLB.508909.214 | hypothetical protein, conserved | 0,3808 | 0,00141 |
| TcCLB.506139.160 | mucin-associated surface protein (MASP), putative | 0,3808 | 0,3935 |
| TcCLB.511287.124 | peroxin 13, putative (PEX13) | 0,3807 | 0,007602 |
| TcCLB.511083.30 | mucin-associated surface protein (MASP), putative | 0,3806 | 0,5587 |
| TcCLB.511391.200 | pitrilysin-like metalloprotease, putative | 0,3805 | 1,376E-07 |
| TcCLB.509217.143 | ATP-dependent DEAD/H RNA helicase (pseudogene), putat | 0,3805 | 0,4026 |
| TcCLB.511061.20 | mucin-associated surface protein (MASP, pseudogene), pu | 0,3804 | 0,5995 |
| TcCLB.507313.20 | trans-sialidase, Group VIII, putative | 0,3803 | 0,2725 |
| TcCLB.511277.150 | hypothetical protein | 0,3799 | 0,00009823 |
| TcCLB.506227.110 | hypothetical protein, conserved | 0,3798 | 0,0201 |
| TcCLB.508871.51 | serine-alanine-and proline-rich protein (pseudogene), puta | 0,3797 | 0,1819 |
| TcCLB.511851.20 | retrotransposon hot spot protein (RHS, pseudogene), putat | 0,3794 | 0,1462 |
| TcCLB.511239.80 | hypothetical protein, conserved | 0,3793 | 0,0141 |
| TcCLB.503929.30 | hypothetical protein, conserved | 0,3792 | 0,0004252 |
| TcCLB.506343.51 | retrotransposon hot spot protein (RHS, pseudogene), putat | 0,3791 | 0,2219 |

| | | | |
|---|---|---|---|
| TcCLB.504741.240 | hypothetical protein, conserved | 0,379 | 0,02913 |
| TcCLB.507625.120 | hypothetical protein, conserved | 0,379 | 0,05181 |
| TcCLB.506735.50 | hypothetical protein, conserved | 0,3788 | 0,0003707 |
| TcCLB.508589.30 | retrotransposon hot spot (RHS) protein, putative | 0,3788 | 0,002101 |
| TcCLB.511351.41 | choline/carnitine O-acyltransferase, putative | 0,3788 | 0,1849 |
| TcCLB.509643.120 | hypothetical protein, conserved | 0,3786 | 0,005467 |
| TcCLB.506737.114 | mucin TcMUCI, putative | 0,3786 | 0,2851 |
| TcCLB.510623.20 | mucin-associated surface protein (MASP), putative | 0,3785 | 0,4415 |
| TcCLB.508021.10 | trans-sialidase, putative | 0,3785 | 0,7912 |
| TcCLB.511079.40 | surface protease GP63 (pseudogene), putative | 0,3784 | 0,6456 |
| TcCLB.510985.10 | trans-sialidase, putative (fragment) | 0,3784 | 0,6909 |
| TcCLB.511727.150 | hypothetical protein, conserved | 0,3783 | 0,0001513 |
| TcCLB.511807.290 | hypothetical protein, conserved | 0,3783 | 0,0003139 |
| TcCLB.508541.140 | surface protease GP63 (pseudogene), putative | 0,3779 | 0,2499 |
| TcCLB.504239.195 | mucin-associated surface protein (MASP, pseudogene), pu | 0,3779 | 0,5119 |
| TcCLB.506985.50 | hypothetical protein, conserved | 0,3778 | 0,07159 |
| TcCLB.506933.10 | hypothetical protein, conserved | 0,3777 | 0,01541 |
| TcCLB.511023.10 | pyrroline-5-carboxylate synthetase-like protein, putative | 0,3776 | 0,001074 |
| TcCLB.506605.229 | SNF2 DNA repair protein, putative | 0,3775 | 0,0006754 |
| TcCLB.510007.10 | hypothetical protein, conserved | 0,3774 | 0,09618 |
| TcCLB.506537.120 | hypothetical protein, conserved | 0,3774 | 0,2551 |
| TcCLB.510483.270 | hypothetical protein | 0,3774 | 0,519 |
| TcCLB.505945.50 | ATP-dependent chaperone, putative | 0,3773 | 0,002562 |
| TcCLB.506945.280 | hypothetical protein, conserved | 0,3769 | 8,691E-06 |
| TcCLB.510091.7 | diacylglycerol acyltransferase, putative (fragment) | 0,3769 | 0,03523 |
| TcCLB.507671.30 | 25 kDa translation elongation factor 1-beta (EF-1-beta) | 0,3767 | 0,00002867 |
| TcCLB.510359.300 | hypothetical protein, conserved | 0,3766 | 0,0005775 |
| TcCLB.510425.28 | hypothetical protein, conserved | 0,3762 | 0,2597 |
| TcCLB.508999.110 | hypothetical protein | 0,3762 | 0,697 |
| TcCLB.507601.110 | prolyl oligopeptidase (pseudogene), putative | 0,3761 | 0,002089 |
| TcCLB.508723.24 | hypothetical protein, conserved | 0,3761 | 0,004246 |
| TcCLB.510835.24 | mitochondrial carrier protein, putative | 0,376 | 0,02122 |
| TcCLB.510741.124 | hypothetical protein, conserved | 0,376 | 0,02422 |
| TcCLB.507819.20 | retrotransposon hot spot protein (RHS, pseudogene), putat | 0,3757 | 0,02796 |
| TcCLB.508723.100 | rab-like GTPase activating protein, putative | 0,3756 | 0,005243 |
| TcCLB.509527.130 | hypothetical protein | 0,3756 | 0,4343 |
| TcCLB.503939.114 | hypothetical protein, conserved | 0,3755 | 0,02293 |
| TcCLB.506779.170 | haloacid dehalogenase hydrolase, putative | 0,3752 | 0,008844 |
| TcCLB.504189.5 | trans-sialidase, putative (fragment) | 0,3751 | 0,6213 |
| TcCLB.510887.30 | hypothetical protein, conserved | 0,3748 | 0,02473 |
| TcCLB.508419.30 | tryptophanyl-tRNA synthetase (pseudogene), putative | 0,3747 | 0,04556 |
| TcCLB.510657.10 | Flagellar Member 5 (fragment) (FLAM5) | 0,3746 | 0,01169 |
| TcCLB.508313.90 | mucin TcMUCII, putative | 0,3744 | 0,199 |
| TcCLB.506459.290 | elongation factor 1-gamma (EF-1-gamma, pseudogene), p | 0,3742 | 0,00002357 |
| TcCLB.503861.110 | retrotransposon hot spot protein (RHS, pseudogene), putat | 0,3741 | 0,01811 |

| | | | |
|---|---|---|---|
| TcCLB.509249.20 | trans-sialidase (pseudogene), putative | 0,3741 | 0,1285 |
| TcCLB.506629.230 | retrotransposon hot spot protein (RHS, pseudogene), putat | 0,374 | 0,001685 |
| TcCLB.511277.175 | ribosomal protein S27, putative | 0,3738 | 0,0002302 |
| TcCLB.506925.14 | hypothetical protein, conserved | 0,3737 | 0,00008052 |
| TcCLB.510053.30 | hypothetical protein, conserved | 0,3737 | 0,005319 |
| TcCLB.511813.30 | proteasome regulatory ATPase subunit 5, putative | 0,3737 | 0,01419 |
| TcCLB.511301.50 | hypothetical protein, conserved | 0,3735 | 0,0004327 |
| TcCLB.510125.20 | trans-sialidase, Group VIII, putative | 0,3735 | 0,5449 |
| TcCLB.511165.80 | protein kinase, putative | 0,3734 | 0,7351 |
| TcCLB.509997.60 | hypothetical protein, conserved | 0,373 | 0,00005289 |
| TcCLB.511439.50 | zinc transporter, putative | 0,373 | 0,004434 |
| TcCLB.508831.250 | dispersed gene family protein 1 (DGF-1), putative | 0,373 | 0,07069 |
| TcCLB.509777.40 | beta galactofuranosyl glycosyltransferase, putative | 0,373 | 0,1139 |
| TcCLB.409407.10 | trans-sialidase (pseudogene), putative | 0,3729 | 0,3507 |
| TcCLB.510199.10 | GP85-like protein, putative | 0,3729 | 0,5431 |
| TcCLB.511255.313 | syntaxin binding protein (pseudogene), putative | 0,3728 | 0,4398 |
| TcCLB.510735.40 | hypothetical protein, conserved | 0,3727 | 0,001756 |
| TcCLB.511625.100 | mucin TcMUCII, putative | 0,3726 | 0,4202 |
| TcCLB.508163.210 | trans-sialidase (pseudogene), putative | 0,3724 | 0,2856 |
| TcCLB.507467.110 | hypothetical protein, conserved | 0,3722 | 0,0001839 |
| TcCLB.433733.10 | trans-sialidase, putative | 0,3721 | 0,1694 |
| TcCLB.508903.110 | trans-sialidase, putative | 0,3716 | 0,3079 |
| TcCLB.508023.20 | trans-sialidase (pseudogene), putative | 0,3715 | 0,1708 |
| TcCLB.510483.180 | mucin-associated surface protein (MASP), putative | 0,3715 | 0,485 |
| TcCLB.508917.20 | hypothetical protein, conserved | 0,3714 | 0,01569 |
| TcCLB.511841.10 | trans-sialidase (pseudogene), putative | 0,3712 | 0,496 |
| TcCLB.510293.54 | hypothetical protein, conserved | 0,3711 | 0,002731 |
| TcCLB.506695.30 | protein kinase, putative | 0,371 | 0,03071 |
| TcCLB.503839.10 | hypothetical protein, conserved | 0,3705 | 0,005402 |
| TcCLB.467287.10 | alpha-tubulin N-acetyltransferase, putative | 0,3705 | 0,02994 |
| TcCLB.508767.10 | surface protein TolT | 0,3704 | 0,05114 |
| TcCLB.506469.10 | hypothetical protein, conserved | 0,3702 | 0,02702 |
| TcCLB.506793.30 | kinesin, putative | 0,37 | 0,0007181 |
| TcCLB.510609.40 | ATP synthase F1, alpha subunit, putative (pseudogene) | 0,3699 | 5,867E-07 |
| TcCLB.503527.29 | hypothetical protein, conserved | 0,3698 | 0,01259 |
| TcCLB.463323.10 | trans-sialidase, Group II, putative | 0,3698 | 0,1128 |
| TcCLB.508043.30 | ubiquitin-activating enzyme E1, putative | 0,3696 | 8,737E-07 |
| TcCLB.507817.80 | 5'-3' exoribonuclease A, putative (fragment) (XRNA) | 0,3696 | 4,826E-06 |
| TcCLB.509549.20 | mucin-associated surface protein (MASP), putative | 0,3696 | 0,2366 |
| TcCLB.404073.10 | hypothetical protein, conserved | 0,3694 | 0,2742 |
| TcCLB.506465.50 | serine/threonine protein phosphatase-like protein, putative | 0,3693 | 0,1729 |
| TcCLB.507821.40 | trans-sialidase (pseudogene), putative | 0,3692 | 0,1219 |
| TcCLB.507381.41 | trans-sialidase (pseudogene), putative | 0,3692 | 0,1378 |
| TcCLB.508257.210 | hypothetical protein, conserved | 0,3691 | 0,003911 |
| TcCLB.508799.60 | DNA ligase, putative | 0,3691 | 0,01642 |

| | | | |
|---|---|---|---|
| TcCLB.511731.10 | UV excision repair RAD23-like protein, putative | 0,369 | 0,0006113 |
| TcCLB.503783.70 | mucin TcMUCII, putative | 0,369 | 0,4137 |
| TcCLB.510089.180 | hypothetical protein, conserved | 0,3688 | 0,006976 |
| TcCLB.509109.100 | hypothetical protein, conserved | 0,3687 | 0,145 |
| TcCLB.511373.30 | mucin-associated surface protein (MASP), putative | 0,3685 | 0,672 |
| TcCLB.506859.170 | hypothetical protein, conserved | 0,3684 | 0,0009815 |
| TcCLB.511587.100 | retrotransposon hot spot (RHS) protein, putative | 0,3683 | 0,002571 |
| TcCLB.508165.110 | mucin-associated surface protein (MASP), putative | 0,3681 | 0,5487 |
| TcCLB.506401.24 | hypothetical protein, conserved | 0,3678 | 0,005771 |
| TcCLB.509965.134 | hypothetical protein, conserved | 0,3677 | 0,1813 |
| TcCLB.508461.234 | hypothetical protein, conserved | 0,3674 | 0,05365 |
| TcCLB.504085.70 | Zinc finger protein, C3H1 type-like (ZC3H46) | 0,3674 | 0,5838 |
| TcCLB.508165.430 | trans-sialidase, putative | 0,3669 | 0,3183 |
| TcCLB.508815.50 | hypothetical protein, conserved | 0,3668 | 0,005308 |
| TcCLB.508175.9 | hypothetical protein, conserved (fragment) | 0,3666 | 0,002874 |
| TcCLB.506187.40 | trans-sialidase (pseudogene), putative | 0,3666 | 0,2624 |
| TcCLB.511003.140 | vacuolar transport protein 4A, putative | 0,3665 | 0,005299 |
| TcCLB.506635.30 | hypothetical protein, conserved | 0,3664 | 0,002466 |
| TcCLB.508461.550 | hypothetical protein, conserved | 0,3664 | 0,003421 |
| TcCLB.506855.130 | hypothetical protein, conserved | 0,3664 | 0,03776 |
| TcCLB.511087.4 | mucin TcMUCII (pseudogene), putative | 0,3664 | 0,4236 |
| TcCLB.488591.10 | trans-sialidase, putative (fragment) | 0,3663 | 0,1487 |
| TcCLB.503897.140 | hypothetical protein, conserved | 0,3662 | 0,01333 |
| TcCLB.511257.10 | DNAK protein, putative | 0,3659 | 0,0004041 |
| TcCLB.509561.130 | hypothetical protein, conserved | 0,3657 | 0,005344 |
| TcCLB.506127.40 | hypothetical protein, conserved | 0,3657 | 0,036 |
| TcCLB.504163.100 | hypothetical protein, conserved | 0,3654 | 0,001699 |
| TcCLB.509259.129 | hypothetical protein, conserved (pseudogene) | 0,3654 | 0,2102 |
| TcCLB.503871.10 | trans-sialidase (pseudogene), putative | 0,3654 | 0,2792 |
| TcCLB.509105.10 | hypothetical protein, conserved | 0,3653 | 0,00728 |
| TcCLB.510933.80 | trans-sialidase, Group V, putative | 0,3653 | 0,3486 |
| TcCLB.510253.10 | hypothetical protein, conserved | 0,3652 | 0,01665 |
| TcCLB.509809.10 | NADH-ubiquinone oxidoreductase complex I subunit, putati | 0,3651 | 0,004076 |
| TcCLB.510719.220 | hypothetical protein | 0,3651 | 0,03971 |
| TcCLB.509063.2 | hypothetical protein, conserved (fragment) | 0,3651 | 0,4625 |
| TcCLB.504161.10 | hypothetical protein | 0,365 | 0,01501 |
| TcCLB.507959.260 | mucin-associated surface protein (MASP), putative | 0,365 | 0,446 |
| TcCLB.507949.4 | cytidine triphosphate synthase, putative | 0,3649 | 0,0284 |
| TcCLB.508911.50 | fructosamine kinase, putative | 0,3649 | 0,05444 |
| TcCLB.506409.100 | mucin TcMUCII, putative | 0,3647 | 0,3341 |
| TcCLB.503587.64 | retrotransposon hot spot protein (RHS, pseudogene), putat | 0,3644 | 0,3651 |
| TcCLB.504153.230 | hypothetical protein, conserved | 0,3643 | 0,0128 |
| TcCLB.510355.90 | dispersed gene family protein 1 (DGF-1), putative | 0,3642 | 0,05953 |
| TcCLB.506767.340 | mucin-associated surface protein (MASP), putative | 0,3642 | 0,2633 |
| TcCLB.510239.40 | mucin-associated surface protein (MASP, pseudogene), pu | 0,3642 | 0,3145 |

| | | | |
|---|---|---|---|
| TcCLB.509805.84 | hypothetical protein, conserved | 0,3641 | 0,07147 |
| TcCLB.508369.10 | hypothetical protein, conserved | 0,3641 | 0,08008 |
| TcCLB.508877.30 | retrotransposon hot spot protein (RHS, pseudogene), putat | 0,3639 | 0,2582 |
| TcCLB.506763.290 | mucin TcMUCII, putative | 0,3639 | 0,5033 |
| TcCLB.506685.20 | hypothetical protein, conserved | 0,3637 | 0,004683 |
| TcCLB.509669.70 | hypothetical protein, conserved | 0,3635 | 0,0000299 |
| TcCLB.511209.54 | ribosomal protein S29, putative | 0,3634 | 0,0009798 |
| TcCLB.509533.5 | retrotransposon hot spot protein (RHS, pseudogene), putat | 0,3634 | 0,002095 |
| TcCLB.506617.10 | surface protein TolT (TolT) | 0,3634 | 0,009627 |
| TcCLB.511391.170 | hypothetical protein, conserved | 0,3632 | 0,007251 |
| TcCLB.506629.240 | retrotransposon hot spot (RHS) protein, putative (fragmen | 0,3632 | 0,5349 |
| TcCLB.410923.40 | trans-sialidase (pseudogene), putative | 0,363 | 0,03586 |
| TcCLB.511449.30 | mucin-associated surface protein (MASP), putative | 0,363 | 0,3148 |
| TcCLB.507163.40 | mucin-associated surface protein (MASP), putative | 0,3629 | 0,462 |
| TcCLB.510431.150 | glycosomal transporter (GAT2), putative (GAT2) | 0,3628 | 6,843E-06 |
| TcCLB.509611.70 | hypothetical protein, conserved | 0,3628 | 0,06254 |
| TcCLB.506969.10 | mucin-associated surface protein (MASP, pseudogene), pu | 0,3628 | 0,1144 |
| TcCLB.511145.50 | vacuolar ATP synthase, putative | 0,3627 | 0,00008231 |
| TcCLB.506195.40 | hypothetical protein, conserved | 0,3626 | 0,05919 |
| TcCLB.504191.19 | sterol 24-c-methyltransferase, putative | 0,3626 | 0,07346 |
| TcCLB.506779.60 | DNA-direcetd RNA polymerase II, subunit 9, putative | 0,3625 | 0,03488 |
| TcCLB.511921.150 | trans-sialidase, Group V, putative | 0,3625 | 0,1788 |
| TcCLB.504081.490 | mucin-associated surface protein (MASP), putative | 0,3625 | 0,4434 |
| TcCLB.506517.158 | hypothetical protein, conserved | 0,3622 | 0,03267 |
| TcCLB.507015.10 | retrotransposon hot spot (RHS) protein, putative | 0,3621 | 0,00001451 |
| TcCLB.506155.40 | hypothetical protein, conserved | 0,3618 | 0,0004429 |
| TcCLB.506513.170 | hypothetical protein, conserved (pseudogene) | 0,3617 | 0,003212 |
| TcCLB.504239.260 | trans-sialidase, Group V, putative | 0,3617 | 0,4165 |
| TcCLB.510603.40 | hypothetical protein, conserved | 0,3616 | 0,001669 |
| TcCLB.506529.470 | hypothetical protein, conserved | 0,3615 | 0,00006826 |
| TcCLB.508153.240 | hypothetical protein, conserved | 0,3613 | 0,004577 |
| TcCLB.511575.50 | hypothetical protein, conserved | 0,3611 | 0,00003311 |
| TcCLB.510997.10 | 3-keto-dihydrosphingosine reductase | 0,3611 | 0,004742 |
| TcCLB.510361.130 | serine-alanine-and proline-rich protein, putative | 0,3611 | 0,6787 |
| TcCLB.511847.40 | hypothetical protein, conserved | 0,3608 | 0,1322 |
| TcCLB.506613.100 | mucin TcMUCII (pseudogene), putative | 0,3607 | 0,4494 |
| TcCLB.510191.80 | mucin TcMUCII, putative | 0,3606 | 0,5935 |
| TcCLB.506883.9 | isopentenyl-diphosphate delta-isomerase (type II), putative | 0,3605 | 0,03729 |
| TcCLB.510565.170 | hypothetical protein, conserved | 0,3605 | 0,4532 |
| TcCLB.510483.240 | mucin-associated surface protein (MASP, pseudogene), pu | 0,3601 | 0,324 |
| TcCLB.508145.10 | RNA-binding protein, putative | 0,3598 | 0,5655 |
| TcCLB.511825.40 | centrin-4, putative (Centrin4) | 0,3597 | 0,0008161 |
| TcCLB.511001.30 | hypothetical protein, conserved | 0,3596 | 0,004408 |
| TcCLB.509615.4 | hypothetical protein, conserved (pseudogene) | 0,3596 | 0,2611 |
| TcCLB.506309.80 | mucin TcMUCII, putative | 0,3596 | 0,6843 |

| | | | |
|---|---|---|---|
| TcCLB.511717.150 | hypothetical protein, conserved | 0,3593 | 0,01118 |
| TcCLB.510077.40 | small GTP-binding protein RAB6, putative | 0,3593 | 0,03764 |
| TcCLB.509979.350 | mucin-associated surface protein (MASP), putative | 0,3593 | 0,1505 |
| TcCLB.506955.70 | mucin-associated surface protein (MASP), putative | 0,3593 | 0,2368 |
| TcCLB.511159.30 | Present in the outer mitochondrial membrane proteome 1! | 0,3592 | 0,01317 |
| TcCLB.509767.80 | hypothetical protein, conserved | 0,3589 | 0,07332 |
| TcCLB.508141.20 | hypothetical protein, conserved | 0,3586 | 0,002187 |
| TcCLB.510359.580 | mucin TcMUCII, putative | 0,3585 | 0,5985 |
| TcCLB.506317.10 | hypothetical protein, conserved | 0,3584 | 0,002267 |
| TcCLB.508979.90 | hypothetical protein | 0,3584 | 0,059 |
| TcCLB.511825.260 | ribosomal RNA processing protein 45, putative (RRP45) | 0,3583 | 0,02396 |
| TcCLB.506173.60 | mucin-associated surface protein (MASP), putative | 0,3583 | 0,3116 |
| TcCLB.506797.30 | hypothetical protein | 0,3582 | 0,0001967 |
| TcCLB.508433.120 | mucin-associated surface protein (MASP, pseudogene), pu | 0,3582 | 0,5116 |
| TcCLB.511393.40 | hypothetical protein, conserved | 0,3581 | 0,01627 |
| TcCLB.506503.4 | transporter, putative | 0,358 | 0,01076 |
| TcCLB.506329.65 | trans-sialidase (pseudogene), putative | 0,3579 | 0,6104 |
| TcCLB.503521.80 | hypothetical protein, conserved | 0,3576 | 0,0116 |
| TcCLB.503579.80 | hypothetical protein, conserved | 0,3575 | 0,005446 |
| TcCLB.508547.30 | acyl-CoA dehydrogenase, mitochondrial precursor, putative | 0,3574 | 0,001091 |
| TcCLB.509831.30 | hypothetical protein, conserved | 0,3574 | 0,002512 |
| TcCLB.508981.39 | trifunctional enzyme alpha subunit, mitochondrial precurso | 0,3574 | 0,003141 |
| TcCLB.511167.50 | hypothetical protein, conserved | 0,3573 | 0,002487 |
| TcCLB.510943.150 | glycerol-3-phosphate acyltransferase, putative | 0,3572 | 0,007617 |
| TcCLB.508221.941 | surface protease GP63 (pseudogene), putative | 0,3572 | 0,4685 |
| TcCLB.506755.250 | mitochondrial import inner membrane translocase subunit | 0,3571 | 0,008517 |
| TcCLB.506529.380 | ser/thr protein phosphatase, putative | 0,3567 | 0,00001889 |
| TcCLB.511919.10 | mucin-associated surface protein (MASP), putative | 0,3565 | 0,1778 |
| TcCLB.508221.830 | hypothetical protein, conserved (pseudogene) | 0,3562 | 0,4654 |
| TcCLB.509769.134 | hypothetical protein, conserved | 0,3561 | 0,002114 |
| TcCLB.510099.60 | hypothetical protein | 0,356 | 0,003365 |
| TcCLB.506805.10 | mucin-associated surface protein (MASP), putative | 0,3559 | 0,4925 |
| TcCLB.511063.110 | mucin-associated surface protein (MASP, pseudogene), pu | 0,3558 | 0,5189 |
| TcCLB.448653.10 | trans-sialidase, putative (fragment) | 0,3557 | 0,728 |
| TcCLB.504149.140 | hypothetical protein, conserved | 0,3556 | 0,0006075 |
| TcCLB.511725.280 | hypothetical protein, conserved | 0,3555 | 0,001313 |
| TcCLB.504113.41 | hypothetical protein, conserved (fragment) | 0,3555 | 0,3082 |
| TcCLB.510163.20 | elongation factor 1-gamma (EF-1-gamma), putative | 0,3554 | 0,001786 |
| TcCLB.511921.90 | mucin-associated surface protein (MASP, pseudogene), pu | 0,355 | 0,3372 |
| TcCLB.506425.150 | hypothetical protein | 0,3548 | 0,1036 |
| TcCLB.508051.10 | hypothetical protein, conserved | 0,3547 | 0,003821 |
| TcCLB.508165.380 | hypothetical protein | 0,3547 | 0,5665 |
| TcCLB.510885.160 | hypothetical protein, conserved | 0,3543 | 0,00003969 |
| TcCLB.510649.24 | membrane associated protein, putative (fragment) | 0,3543 | 0,0002357 |
| TcCLB.506163.60 | hypothetical protein | 0,3542 | 0,002327 |

| | | | |
|---|---|---|---|
| TcCLB.509459.40 | hypothetical protein, conserved | 0,3542 | 0,003456 |
| TcCLB.503439.61 | retrotransposon hot spot protein (RHS, pseudogene), putat | 0,3542 | 0,03493 |
| TcCLB.510013.310 | mucin-associated surface protein (MASP), putative | 0,3542 | 0,2983 |
| TcCLB.509745.20 | glycosyl transferase-like, putative | 0,3539 | 0,01517 |
| TcCLB.510359.490 | trans-sialidase (pseudogene), putative | 0,3538 | 0,1715 |
| TcCLB.504111.30 | hypothetical protein, conserved | 0,3537 | 0,007352 |
| TcCLB.503603.10 | mitochondrial RNA binding complex 1 subunit, putative | 0,3536 | 0,02081 |
| TcCLB.506469.100 | hypothetical protein, conserved | 0,3535 | 0,0003424 |
| TcCLB.508119.11 | surface protease GP63 (pseudogene), putative | 0,3535 | 0,1352 |
| TcCLB.504019.6 | hypothetical protein | 0,3534 | 0,0005203 |
| TcCLB.504223.20 | amino acid permease, putative | 0,3533 | 0,2031 |
| TcCLB.505183.110 | hypothetical protein | 0,3533 | 0,2244 |
| TcCLB.510287.4 | Mutt-Nudix-related hydrolase, putative (fragment) | 0,3532 | 0,4612 |
| TcCLB.505071.90 | protein phosphatase 2A, regulatory subunit B, putative | 0,3531 | 0,003309 |
| TcCLB.511283.320 | hypothetical protein, conserved | 0,353 | 0,02491 |
| TcCLB.508443.10 | hypothetical protein, conserved | 0,3529 | 0,03143 |
| TcCLB.510283.110 | hypothetical protein, conserved | 0,3529 | 0,08809 |
| TcCLB.410269.10 | trans-sialidase, putative (fragment) | 0,3529 | 0,3573 |
| TcCLB.511237.60 | 40S ribosomal protein S33, putative | 0,3528 | 0,0003439 |
| TcCLB.510565.50 | hypothetical protein, conserved | 0,3528 | 0,01689 |
| TcCLB.506279.200 | mucin TcMUCII, putative | 0,3527 | 0,5516 |
| TcCLB.506861.10 | ATP-dependent DEAD/H RNA helicase, putative | 0,3526 | 0,004112 |
| TcCLB.508165.150 | mucin-associated surface protein (MASP), putative | 0,3526 | 0,3191 |
| TcCLB.510013.240 | mucin TcMUCII, putative | 0,3526 | 0,6586 |
| TcCLB.509289.20 | tuzin, putative | 0,3525 | 2,322E-07 |
| TcCLB.510719.130 | hypothetical protein, conserved | 0,3523 | 0,01053 |
| TcCLB.506853.10 | hypothetical protein, conserved | 0,3521 | 0,00004778 |
| TcCLB.507521.150 | dispersed gene family protein 1 (DGF-1), putative | 0,3521 | 0,05022 |
| TcCLB.507641.210 | T-lymphocyte triggering factor, putative | 0,352 | 0,0006702 |
| TcCLB.506745.10 | hypothetical protein, conserved | 0,352 | 0,0596 |
| TcCLB.503945.30 | hypothetical protein, conserved | 0,352 | 0,07913 |
| TcCLB.506133.10 | syntaxin binding protein (pseudogene), putative | 0,3518 | 0,5426 |
| TcCLB.508511.40 | mucin-associated surface protein (MASP, pseudogene), pu | 0,3518 | 0,5601 |
| TcCLB.511417.30 | hypothetical protein, conserved | 0,3517 | 0,01972 |
| TcCLB.510643.20 | retrotransposon hot spot protein (RHS, pseudogene), putat | 0,3517 | 0,05017 |
| TcCLB.507973.10 | mucin TcMUCI, putative | 0,3517 | 0,4925 |
| TcCLB.504131.70 | hypothetical protein, conserved | 0,3516 | 0,00005757 |
| TcCLB.505807.90 | hypothetical protein, conserved | 0,3516 | 0,0002379 |
| TcCLB.511217.100 | hypothetical protein, conserved | 0,3516 | 0,00167 |
| TcCLB.508779.40 | hypothetical protein, conserved | 0,3516 | 0,04305 |
| TcCLB.508515.60 | hypothetical protein, conserved | 0,3516 | 0,04953 |
| TcCLB.507047.134 | hypothetical protein, conserved | 0,3516 | 0,2741 |
| TcCLB.511647.30 | meiotic recombination protein SPO11, putative | 0,3515 | 0,1949 |
| TcCLB.511665.21 | trans-sialidase (pseudogene), putative | 0,3513 | 0,05529 |
| TcCLB.510943.20 | hypothetical protein | 0,3512 | 0,172 |

| | | | |
|---|---|---|---|
| TcCLB.508815.150 | hypothetical protein, conserved | 0,3511 | 0,08799 |
| TcCLB.507559.40 | hypothetical protein, conserved | 0,351 | 0,01107 |
| TcCLB.511135.10 | hypothetical protein, conserved (pseudogene) | 0,351 | 0,02742 |
| TcCLB.510503.89 | mucin-associated surface protein (MASP, pseudogene), pu | 0,3507 | 0,6595 |
| TcCLB.511907.180 | surface protease GP63 (pseudogene), putative | 0,3503 | 0,2382 |
| TcCLB.508277.130 | hypothetical protein, conserved | 0,3502 | 0,08537 |
| TcCLB.511825.10 | hypothetical protein, conserved | 0,3501 | 0,007577 |
| TcCLB.511907.340 | hypothetical protein, conserved | 0,35 | 0,1414 |
| TcCLB.511767.130 | expression site-associated gene (ESAG-like) protein, putati | 0,35 | 0,4174 |
| TcCLB.510879.110 | ribosomal protein L3, putative | 0,3499 | 0,000171 |
| TcCLB.507623.20 | hypothetical protein, conserved | 0,3498 | 0,004943 |
| TcCLB.511459.40 | hypothetical protein, conserved | 0,3498 | 0,007773 |
| TcCLB.506705.30 | hypothetical protein, conserved | 0,3498 | 0,07317 |
| TcCLB.511611.20 | mucin-associated surface protein (MASP), putative | 0,3498 | 0,6538 |
| TcCLB.510021.120 | trans-sialidase, Group V, putative | 0,3497 | 0,3237 |
| TcCLB.507689.25 | hypothetical protein, conserved | 0,3496 | 0,4267 |
| TcCLB.509233.190 | ribosomal protein S25, putative | 0,3494 | 8,624E-06 |
| TcCLB.505987.10 | hypothetical protein, conserved (pseudogene) | 0,3494 | 0,03782 |
| TcCLB.503561.10 | hypothetical protein, conserved | 0,3494 | 0,03996 |
| TcCLB.511603.40 | hypothetical protein | 0,3494 | 0,5373 |
| TcCLB.439803.9 | trans-sialidase (pseudogene), putative | 0,3493 | 0,1771 |
| TcCLB.509859.60 | hypothetical protein, conserved | 0,3491 | 0,001714 |
| TcCLB.511283.100 | hypothetical protein, conserved | 0,3491 | 0,008043 |
| TcCLB.505297.80 | trans-sialidase (pseudogene), putative | 0,3491 | 0,09488 |
| TcCLB.511367.230 | NADPH:adrenodoxin oxidoreductase, mitochondrial, putativ | 0,3489 | 0,003341 |
| TcCLB.510193.10 | mucin-associated surface protein (MASP), putative | 0,3488 | 0,3513 |
| TcCLB.503593.50 | hypothetical protein, conserved | 0,3484 | 0,0005594 |
| TcCLB.506223.30 | hypothetical protein, conserved | 0,3483 | 0,02294 |
| TcCLB.510713.130 | mucin-associated surface protein (MASP), putative | 0,3483 | 0,4502 |
| TcCLB.506667.20 | hypothetical protein | 0,3481 | 0,5667 |
| TcCLB.509733.180 | adenylate kinase, putative | 0,348 | 0,1844 |
| TcCLB.511835.10 | trans-sialidase, Group VIII, putative | 0,3476 | 0,3139 |
| TcCLB.508819.10 | hypothetical protein, conserved | 0,3475 | 0,0356 |
| TcCLB.503503.40 | mucin-associated surface protein (MASP), putative | 0,3475 | 0,4681 |
| TcCLB.506559.205 | hypothetical protein | 0,3474 | 0,08132 |
| TcCLB.508221.894 | mucin-associated surface protein (MASP), putative | 0,3473 | 0,4828 |
| TcCLB.510091.119 | calpain-like cysteine peptidase (pseudogene), putative | 0,3472 | 0,0427 |
| TcCLB.506819.20 | hypothetical protein | 0,3471 | 0,06963 |
| TcCLB.506835.20 | hypothetical protein | 0,347 | 0,003139 |
| TcCLB.503911.30 | hypothetical protein, conserved | 0,3469 | 0,001447 |
| TcCLB.508823.120 | ribosomal protein S20, putative | 0,3469 | 0,001675 |
| TcCLB.511765.20 | dispersed gene family protein 1 (DGF-1), putative | 0,3469 | 0,1098 |
| TcCLB.511843.70 | trans-sialidase, Group V, putative | 0,3469 | 0,3683 |
| TcCLB.507023.130 | hypothetical protein, conserved | 0,3468 | 0,02952 |
| TcCLB.509611.170 | hypothetical protein, conserved | 0,3467 | 0,009707 |

| | | | |
|---|---|---|---|
| TcCLB.511595.59 | trans-sialidase, putative (fragment) | 0,3466 | 0,7486 |
| TcCLB.508041.60 | eukaryotic release factor 3, putative | 0,3465 | 0,00006332 |
| TcCLB.509915.10 | trans-sialidase (pseudogene), putative | 0,3465 | 0,3189 |
| TcCLB.508153.760 | hypothetical protein, conserved | 0,3464 | 0,003751 |
| TcCLB.509967.80 | hypothetical protein, conserved | 0,3464 | 0,03229 |
| TcCLB.509767.70 | V-type proton ATPase subunit a, putative (VAa) | 0,3463 | 0,0007576 |
| TcCLB.505183.100 | hypothetical protein, conserved | 0,3463 | 0,01446 |
| TcCLB.510719.110 | actin-like protein, putative | 0,3462 | 0,03135 |
| TcCLB.509643.100 | hypothetical protein, conserved | 0,3461 | 0,04938 |
| TcCLB.503587.3 | mucin TcMUC (pseudogene), putative | 0,3461 | 0,6547 |
| TcCLB.511839.40 | trans-sialidase, Group V, putative | 0,346 | 0,3478 |
| TcCLB.508881.120 | prefoldin, putative | 0,3455 | 0,004122 |
| TcCLB.505807.250 | hypothetical protein, conserved | 0,3455 | 0,04059 |
| TcCLB.506289.180 | mucin-associated surface protein (MASP, pseudogene), pu | 0,3454 | 0,3971 |
| TcCLB.510101.130 | inositol-1,4,5-trisphosphate (IP3) 5-phosphatase, putative | 0,3453 | 0,0008986 |
| TcCLB.508817.110 | hypothetical protein, conserved | 0,3453 | 0,06328 |
| TcCLB.506615.21 | surface protease GP63 (pseudogene), putative | 0,3453 | 0,5124 |
| TcCLB.508319.60 | hypothetical protein, conserved | 0,3452 | 0,003253 |
| TcCLB.510363.30 | retrotransposon hot spot protein (RHS, pseudogene), putat | 0,3452 | 0,4366 |
| TcCLB.479253.9 | hypothetical protein, conserved (fragment) | 0,3451 | 0,1044 |
| TcCLB.507951.100 | hypothetical protein, conserved | 0,345 | 0,00762 |
| TcCLB.508153.260 | hypothetical protein, conserved | 0,345 | 0,07635 |
| TcCLB.506625.100 | hypothetical protein, conserved | 0,3448 | 0,001714 |
| TcCLB.504193.30 | retrotransposon hot spot protein (RHS, pseudogene), putat | 0,3448 | 0,004901 |
| TcCLB.506965.180 | mucin-associated surface protein (MASP), putative | 0,3447 | 0,6962 |
| TcCLB.506835.50 | hypothetical protein | 0,3445 | 0,002309 |
| TcCLB.510337.11 | hypothetical protein, conserved (fragment) | 0,3445 | 0,009665 |
| TcCLB.504411.40 | hypothetical protein, conserved | 0,3445 | 0,02234 |
| TcCLB.508805.120 | UDP-Gal or UDP-GlcNAc-dependent glycosyltransferase, pu | 0,3445 | 0,2601 |
| TcCLB.506825.60 | hypothetical protein, conserved | 0,3442 | 0,0005653 |
| TcCLB.507089.200 | chloride channel protein, putative | 0,3442 | 0,001057 |
| TcCLB.508355.200 | vacuolar protein sorting protein 16, putative | 0,3441 | 0,0002172 |
| TcCLB.506563.190 | calpain-like cysteine peptidase, putative | 0,3441 | 0,0008341 |
| TcCLB.508175.290 | hypothetical protein, conserved | 0,3441 | 0,003013 |
| TcCLB.509399.9 | helicase, putative | 0,3439 | 0,00331 |
| TcCLB.506327.40 | hypothetical protein, conserved | 0,3438 | 0,0001343 |
| TcCLB.510887.14 | chaperone DNAJ protein, putative | 0,3437 | 0,01616 |
| TcCLB.506963.90 | calmodulin, putative | 0,3435 | 0,02031 |
| TcCLB.511089.30 | mucin-associated surface protein (MASP), putative | 0,3434 | 0,5264 |
| TcCLB.507957.100 | mucin-associated surface protein (MASP, pseudogene), pu | 0,3433 | 0,3917 |
| TcCLB.511755.40 | hypothetical protein, conserved | 0,3429 | 0,06052 |
| TcCLB.510525.70 | hypothetical protein, conserved | 0,3428 | 0,02498 |
| TcCLB.510859.10 | nucleolar RNA-binding protein, putative | 0,3427 | 0,00003942 |
| TcCLB.508577.50 | hypothetical protein | 0,3427 | 0,1293 |
| TcCLB.511089.130 | mucin-associated surface protein (MASP), putative | 0,3426 | 0,5997 |

| | | | |
|---|---|---|---|
| TcCLB.509735.120 | retrotransposon hot spot (RHS) protein, putative | 0,3425 | 0,003596 |
| TcCLB.507295.50 | mucin-associated surface protein (MASP), putative | 0,3425 | 0,2301 |
| TcCLB.506345.90 | trans-sialidase, Group II, putative | 0,3424 | 0,2014 |
| TcCLB.506227.190 | hypothetical protein | 0,3424 | 0,2586 |
| TcCLB.506113.80 | hypothetical protein, conserved (pseudogene) | 0,3423 | 0,01767 |
| TcCLB.506179.20 | hypothetical protein, conserved | 0,3423 | 0,05443 |
| TcCLB.511853.21 | mucin-associated surface protein (MASP, pseudogene), pu | 0,3423 | 0,3444 |
| TcCLB.508963.10 | Flagellar Member 3 (pseudogene) | 0,3422 | 0,0009431 |
| TcCLB.508313.30 | mucin-associated surface protein (MASP), putative | 0,3422 | 0,5737 |
| TcCLB.506267.90 | mucin-associated surface protein (MASP), putative | 0,3421 | 0,2712 |
| TcCLB.511685.50 | hypothetical protein, conserved | 0,342 | 0,1044 |
| TcCLB.511665.100 | retrotransposon hot spot protein (RHS, pseudogene), putat | 0,342 | 0,1321 |
| TcCLB.510241.60 | nascent polypeptide associated complex subunit, putative | 0,3418 | 2,658E-06 |
| TcCLB.510275.120 | hypothetical protein | 0,3417 | 0,6191 |
| TcCLB.508529.30 | retrotransposon hot spot protein (RHS, pseudogene), putat | 0,3416 | 0,003754 |
| TcCLB.506513.60 | ADP-ribosylation factor, putative (ARL-1) | 0,3416 | 0,04168 |
| TcCLB.511105.60 | trans-sialidase, putative | 0,3415 | 0,1198 |
| TcCLB.506399.89 | Succinate dehydrogenase [ubiquinone] flavoprotein subuni | 0,3414 | 0,02427 |
| TcCLB.506537.100 | trans-sialidase (pseudogene), putative | 0,3412 | 0,0314 |
| TcCLB.508823.40 | hypothetical protein, conserved | 0,3412 | 0,07643 |
| TcCLB.506801.10 | surface protease GP63 (pseudogene), putative | 0,3412 | 0,4699 |
| TcCLB.511825.170 | serine/threonine protein phosphatase, putative | 0,3409 | 0,01632 |
| TcCLB.506789.210 | paraflagellar rod protein, putative | 0,3407 | 0,003995 |
| TcCLB.509925.50 | retrotransposon hot spot protein (RHS, pseudogene), putat | 0,3407 | 0,1714 |
| TcCLB.505655.10 | trans-sialidase (pseudogene), putative | 0,3403 | 0,3469 |
| TcCLB.509769.120 | hypothetical protein, conserved | 0,3402 | 0,00013 |
| TcCLB.506529.110 | hypothetical protein, conserved | 0,3401 | 0,02906 |
| TcCLB.507085.120 | hypothetical protein | 0,3401 | 0,6064 |
| TcCLB.506409.30 | mucin-associated surface protein (MASP), putative | 0,3399 | 0,4313 |
| TcCLB.504039.90 | mucin-associated surface protein (MASP, pseudogene), pu | 0,3395 | 0,5231 |
| TcCLB.507509.40 | hypothetical protein, conserved | 0,3394 | 0,001833 |
| TcCLB.511153.84 | hypothetical protein, conserved | 0,3394 | 0,03543 |
| TcCLB.504443.20 | hypothetical protein, conserved | 0,3394 | 0,05662 |
| TcCLB.506489.20 | monoglyceride lipase, putative | 0,3391 | 0,00005073 |
| TcCLB.504103.6 | hypothetical protein, conserved | 0,3391 | 0,1035 |
| TcCLB.508175.329 | 60S ribosomal protein L19, putative | 0,3389 | 0,001266 |
| TcCLB.506519.140 | hypothetical protein, conserved (pseudogene) | 0,3389 | 0,04828 |
| TcCLB.511717.200 | hypothetical protein, conserved | 0,3388 | 0,01153 |
| TcCLB.507875.90 | dispersed gene family protein 1 (DGF-1, pseudogene), puta | 0,3386 | 0,06787 |
| TcCLB.507993.100 | hypothetical protein, conserved | 0,3386 | 0,1416 |
| TcCLB.504049.20 | retrotransposon hot spot protein (RHS, pseudogene), putat | 0,3384 | 0,003323 |
| TcCLB.509023.40 | hypothetical protein, conserved | 0,3382 | 0,02332 |
| TcCLB.507257.90 | hypothetical protein, conserved | 0,3382 | 0,06683 |
| TcCLB.506947.110 | myosin heavy chain (pseudogene), putative | 0,3381 | 0,003639 |
| TcCLB.508521.100 | trans-sialidase, Group VIII, putative | 0,338 | 0,4801 |

| | | | |
|---|---|---|---|
| TcCLB.506295.70 | hypothetical protein, conserved | 0,3379 | 0,002153 |
| TcCLB.510303.170 | hypothetical protein, conserved | 0,3378 | 0,00008018 |
| TcCLB.503639.20 | hypothetical protein, conserved | 0,3374 | 0,1271 |
| TcCLB.511559.40 | hypothetical protein, conserved | 0,3373 | 0,01529 |
| TcCLB.503887.60 | glycerolphosphate mutase, putative | 0,3373 | 0,01654 |
| TcCLB.510339.60 | hypothetical protein, conserved | 0,3373 | 0,0248 |
| TcCLB.511717.170 | Metalloprotease M41 FtsH, putative | 0,3372 | 0,00501 |
| TcCLB.506529.320 | hypothetical protein, conserved | 0,3372 | 0,01317 |
| TcCLB.506557.10 | hypothetical protein, conserved | 0,3371 | 0,02572 |
| TcCLB.509893.20 | hypothetical protein, conserved | 0,3369 | 0,001516 |
| TcCLB.508661.34 | hypothetical protein, conserved | 0,3368 | 0,01256 |
| TcCLB.510163.30 | mucin-associated surface protein (MASP), putative | 0,3366 | 0,01044 |
| TcCLB.511929.20 | dispersed gene family protein 1 (DGF-1, pseudogene), puta | 0,3366 | 0,1288 |
| TcCLB.510715.40 | mucin-associated surface protein (MASP), putative | 0,3366 | 0,5727 |
| TcCLB.510311.80 | hypothetical protein, conserved | 0,3365 | 0,007741 |
| TcCLB.506907.14 | hypothetical protein, conserved | 0,3365 | 0,01753 |
| TcCLB.509373.10 | trans-sialidase (pseudogene), putative | 0,3365 | 0,2829 |
| TcCLB.508479.274 | hypothetical protein, conserved | 0,3364 | 0,002586 |
| TcCLB.503579.140 | hypothetical protein, conserved | 0,3364 | 0,009847 |
| TcCLB.507437.30 | trans-sialidase (pseudogene), putative | 0,3364 | 0,2271 |
| TcCLB.510261.41 | mucin TcMUC (pseudogene), putative | 0,3364 | 0,7031 |
| TcCLB.511577.44 | hypothetical protein, conserved | 0,3363 | 0,1656 |
| TcCLB.509057.10 | hypothetical protein, conserved | 0,3361 | 0,1931 |
| TcCLB.511153.110 | hypothetical protein, conserved | 0,3359 | 0,0007017 |
| TcCLB.503453.70 | hypothetical protein, conserved | 0,3359 | 0,0009662 |
| TcCLB.511163.10 | seryl-tRNA synthetase, putative (fragment) | 0,3358 | 0,0009898 |
| TcCLB.511875.64 | hypothetical protein, conserved (pseudogene) | 0,3358 | 0,169 |
| TcCLB.511127.340 | tyrosine phosphatase, putative | 0,3357 | 0,002449 |
| TcCLB.509791.20 | hypothetical protein, conserved | 0,3357 | 0,01152 |
| TcCLB.508089.50 | hypothetical protein, conserved | 0,3356 | 0,0147 |
| TcCLB.507699.200 | mucin TcMUCII, putative | 0,3356 | 0,4935 |
| TcCLB.511323.10 | trans-sialidase, Group I, putative | 0,3353 | 0,04953 |
| TcCLB.506147.210 | hypothetical protein, conserved | 0,3353 | 0,06619 |
| TcCLB.503833.40 | 40S ribosomal protein S2, putative | 0,3352 | 0,0001719 |
| TcCLB.506743.180 | recombination initiation protein NBS1, putative | 0,335 | 0,001419 |
| TcCLB.510979.10 | mucin-associated surface protein (MASP), putative | 0,335 | 0,3315 |
| TcCLB.506331.200 | trans-sialidase (pseudogene), putative | 0,335 | 0,4223 |
| TcCLB.509541.10 | hypothetical protein, conserved | 0,3349 | 0,00161 |
| TcCLB.506637.10 | hypothetical protein, conserved | 0,3348 | 0,004454 |
| TcCLB.511401.30 | trans-sialidase, Group V, putative | 0,3347 | 0,1713 |
| TcCLB.511067.20 | 60S ribosomal protein L26, putative | 0,3345 | 0,00004759 |
| TcCLB.506795.70 | hypothetical protein, conserved | 0,3344 | 0,118 |
| TcCLB.506153.6 | hypothetical protein, conserved | 0,3342 | 0,06028 |
| TcCLB.459061.10 | trans-sialidase, putative | 0,3342 | 0,3282 |
| TcCLB.503909.76 | ferric reductase transmembrane protein, putative | 0,3342 | 0,3484 |

| | | | |
|---|---|---|---|
| TcCLB.510375.40 | mucin-associated surface protein (MASP), putative | 0,3342 | 0,6619 |
| TcCLB.510543.110 | hypothetical protein, conserved (pseudogene) | 0,3341 | 0,1962 |
| TcCLB.509695.50 | katanin, putative | 0,3339 | 0,1986 |
| TcCLB.511521.24 | hypothetical protein, conserved | 0,3339 | 0,3261 |
| TcCLB.509717.90 | short chain 3-hydroxyacyl-CoA dehydrogenase, putative | 0,3337 | 0,0003536 |
| TcCLB.510901.70 | mitochondrial RNA binding complex 1 subunit, putative | 0,3337 | 0,001759 |
| TcCLB.511647.4 | glutamine-dependent carbamoyl-phosphate synthetase, pu | 0,3337 | 0,005619 |
| TcCLB.511315.30 | hypothetical protein, conserved | 0,3337 | 0,1043 |
| TcCLB.506811.140 | hypothetical protein, conserved | 0,3336 | 0,01406 |
| TcCLB.503757.10 | 40S ribosomal protein SA, putative | 0,3332 | 0,0009996 |
| TcCLB.506821.60 | mucin TcMUCI, putative | 0,3332 | 0,3678 |
| TcCLB.504081.400 | mucin-associated surface protein (MASP), putative | 0,3332 | 0,4647 |
| TcCLB.510531.40 | hypothetical protein, conserved | 0,3329 | 0,002961 |
| TcCLB.511529.90 | enolase, putative | 0,3328 | 0,04498 |
| TcCLB.505297.60 | mucin-associated surface protein (MASP), putative | 0,3327 | 0,2351 |
| TcCLB.507765.70 | 40S ribosomal protein S21, putative | 0,3325 | 4,408E-07 |
| TcCLB.510897.70 | glycerophosphoryl diester phosphodiesterase (pseudogene | 0,3325 | 0,04108 |
| TcCLB.511673.9 | hypothetical protein, conserved | 0,3325 | 0,2434 |
| TcCLB.510013.110 | mucin TcMUCII, putative | 0,3325 | 0,7023 |
| TcCLB.511529.150 | hypothetical protein, conserved | 0,3324 | 0,006167 |
| TcCLB.506705.10 | hypothetical protein | 0,3324 | 0,02957 |
| TcCLB.506761.47 | mucin TcMUCII, putative | 0,3323 | 0,6967 |
| TcCLB.506201.39 | 25 kDa translation elongation factor 1-beta, putative (EF-1 | 0,3322 | 6,865E-06 |
| TcCLB.508641.330 | pre-mRNA splicing factor, putative | 0,3322 | 0,2127 |
| TcCLB.510031.20 | NADH-ubiquinone oxidoreductase complex I subunit, putati | 0,3321 | 0,07958 |
| TcCLB.506695.7 | hypothetical protein, conserved | 0,3321 | 0,08472 |
| TcCLB.511183.49 | mucin TcMUCII (pseudogene), putative | 0,3319 | 0,6405 |
| TcCLB.510805.5 | trans-sialidase, putative | 0,3318 | 0,6484 |
| TcCLB.509545.10 | mucin-associated surface protein (MASP), putative | 0,3317 | 0,2708 |
| TcCLB.506953.30 | protein kinase, putative | 0,3314 | 0,05787 |
| TcCLB.416809.10 | dispersed gene family protein 1 (DGF-1), putative | 0,3313 | 0,08619 |
| TcCLB.511751.80 | hypothetical protein | 0,3312 | 0,01901 |
| TcCLB.506941.50 | hypothetical protein, conserved | 0,331 | 0,009319 |
| TcCLB.507009.90 | iron-sulfur cluster assembly protein, putative | 0,331 | 0,02103 |
| TcCLB.511499.30 | RNA decapping enzyme, putative (Dcp2) | 0,331 | 0,1087 |
| TcCLB.504147.190 | hypothetical protein, conserved | 0,3309 | 0,002367 |
| TcCLB.511367.290 | hypothetical protein, conserved | 0,3309 | 0,002731 |
| TcCLB.505941.20 | surface protease GP63 (pseudogene), putative | 0,3309 | 0,2586 |
| TcCLB.506839.80 | phosphoribosylpyrophosphate synthetase, putative | 0,3308 | 0,0001774 |
| TcCLB.508433.180 | mucin-associated surface protein (MASP), putative | 0,3308 | 0,3172 |
| TcCLB.510465.150 | dispersed gene family protein 1 (DGF-1), putative | 0,3307 | 0,443 |
| TcCLB.511809.70 | DNA primase small subunit, putative | 0,3306 | 0,04004 |
| TcCLB.511709.19 | phosphatidylinositol 3-kinase catalytic subunit, putative | 0,3306 | 0,09648 |
| TcCLB.508055.25 | hypothetical protein | 0,3304 | 0,5263 |
| TcCLB.508425.20 | tryptophanyl-tRNA synthetase (pseudogene), putative | 0,3303 | 0,1326 |

| | | | |
|---|---|---|---|
| TcCLB.511649.60 | COP9 signalosome complex subunit 5, putative | 0,3302 | 0,004972 |
| TcCLB.510361.180 | trans-sialidase (pseudogene), putative | 0,3302 | 0,571 |
| TcCLB.511745.10 | heat shock 70 kDa protein, mitochondrial precursor, putativ | 0,33 | 0,001145 |
| TcCLB.506245.302 | mucin-associated surface protein (MASP), putative | 0,3299 | 0,3797 |
| TcCLB.503835.39 | hypothetical protein, conserved | 0,3298 | 0,01714 |
| TcCLB.511291.20 | hypothetical protein, conserved | 0,3298 | 0,0478 |
| TcCLB.511751.114 | hypothetical protein, conserved | 0,3298 | 0,1832 |
| TcCLB.507121.40 | beta galactofuranosyl glycosyltransferase, putative | 0,3298 | 0,2565 |
| TcCLB.506767.196 | surface protease GP63 (pseudogene), putative | 0,3298 | 0,6867 |
| TcCLB.503719.20 | 40S ribosomal protein SA, putative | 0,3297 | 0,0002357 |
| TcCLB.510191.31 | trans-sialidase (pseudogene), putative | 0,3297 | 0,5878 |
| TcCLB.509693.50 | metallo-peptidase, Clan MG, Family M24 (MetAP2) | 0,3293 | 0,001462 |
| TcCLB.508299.60 | 60S ribosomal protein L2, putative | 0,3291 | 7,671E-06 |
| TcCLB.508387.130 | ATPase, putative | 0,3291 | 0,02741 |
| TcCLB.510101.460 | hypothetical protein, conserved | 0,329 | 0,009571 |
| TcCLB.507699.210 | hypothetical protein | 0,329 | 0,4855 |
| TcCLB.511239.120 | hypothetical protein, conserved | 0,3289 | 0,1554 |
| TcCLB.510761.10 | mismatch repair protein PMS1, putative | 0,3288 | 0,0004863 |
| TcCLB.504151.30 | hypothetical protein, conserved | 0,3288 | 0,05796 |
| TcCLB.507203.9 | dispersed gene family protein 1 (DGF-1), putative | 0,3288 | 0,3539 |
| TcCLB.508027.30 | hypothetical protein, conserved | 0,3287 | 0,06671 |
| TcCLB.507081.130 | 2,4-dihydroxyhept-2-ene-1,7-dioic acid aldolase, putative | 0,3286 | 0,007406 |
| TcCLB.511859.70 | hypothetical protein, conserved | 0,3285 | 0,04318 |
| TcCLB.510101.310 | hypothetical protein | 0,3284 | 0,001993 |
| TcCLB.508153.790 | hypothetical protein, conserved | 0,3284 | 0,08984 |
| TcCLB.508027.90 | hypothetical protein, conserved | 0,3283 | 0,008551 |
| TcCLB.503545.20 | retrotransposon hot spot protein (RHS, pseudogene), putat | 0,3282 | 0,1816 |
| TcCLB.511319.20 | hypothetical protein, conserved | 0,3281 | 0,001189 |
| TcCLB.507491.119 | hypothetical protein, conserved | 0,3281 | 0,004375 |
| TcCLB.511827.60 | CCR4 associated factor, putative | 0,3279 | 0,001645 |
| TcCLB.508989.50 | hypothetical protein, conserved | 0,3277 | 0,004017 |
| TcCLB.509073.110 | hypothetical protein, conserved | 0,3275 | 0,001599 |
| TcCLB.506605.180 | hypothetical protein, conserved | 0,3275 | 0,02541 |
| TcCLB.507485.120 | tuzin | 0,3275 | 0,1554 |
| TcCLB.508175.50 | hypothetical protein, conserved | 0,3274 | 0,001697 |
| TcCLB.511903.90 | hypothetical protein, conserved | 0,3273 | 0,01302 |
| TcCLB.506951.70 | surface protease GP63 (pseudogene), putative | 0,3273 | 0,4106 |
| TcCLB.453995.10 | retrotransposon hot spot protein (RHS, pseudogene), putat | 0,3272 | 0,03608 |
| TcCLB.505997.60 | retrotransposon hot spot (RHS) protein, putative | 0,3272 | 0,1303 |
| TcCLB.510943.40 | protein kinase, putative | 0,3271 | 0,004082 |
| TcCLB.510125.10 | pumilio/PUF RNA binding protein 6, putative (PUF6) | 0,3265 | 0,01607 |
| TcCLB.510437.30 | hypothetical protein, conserved | 0,3264 | 0,2494 |
| TcCLB.505155.10 | retrotransposon hot spot protein (RHS, pseudogene), putat | 0,3263 | 0,0922 |
| TcCLB.511169.10 | hypothetical protein, conserved | 0,3263 | 0,1112 |
| TcCLB.511277.554 | hypothetical protein, conserved | 0,3261 | 0,05942 |

| | | | |
|---|---|---|---|
| TcCLB.503721.39 | heat shock 70 kDa protein, putative | 0,326 | 0,003264 |
| TcCLB.506885.354 | hypothetical protein, conserved | 0,3258 | 0,1837 |
| TcCLB.508883.30 | mucin TcMUCII, putative | 0,3256 | 0,5952 |
| TcCLB.508737.170 | hypothetical protein, conserved | 0,3255 | 0,04193 |
| TcCLB.503761.20 | surface protease GP63 (pseudogene), putative | 0,3255 | 0,3942 |
| TcCLB.506957.120 | hypothetical protein, conserved | 0,3253 | 0,0005901 |
| TcCLB.506277.30 | mucin-associated surface protein (MASP, pseudogene), pu | 0,3253 | 0,3122 |
| TcCLB.509429.190 | hypothetical protein, conserved | 0,3252 | 0,002599 |
| TcCLB.510525.50 | hypothetical protein, conserved | 0,3252 | 0,006763 |
| TcCLB.507993.80 | glycogen synthase kinase 3, putative | 0,3252 | 0,1313 |
| TcCLB.508387.60 | hypothetical protein, conserved | 0,3251 | 0,06608 |
| TcCLB.510889.80 | protein phosphatase with EF-Hand domains (PPEF) | 0,325 | 0,0001932 |
| TcCLB.503787.30 | hypothetical protein | 0,325 | 0,4138 |
| TcCLB.509805.120 | hypothetical protein, conserved | 0,3249 | 0,119 |
| TcCLB.503509.10 | hypothetical protein, conserved | 0,3245 | 0,0000801 |
| TcCLB.506617.5 | surface protein TolT, putative (fragment) | 0,3243 | 0,06175 |
| TcCLB.508387.100 | ubiquitin-conjugating enzyme E2, putative | 0,3242 | 0,02364 |
| TcCLB.503849.51 | mucin TcMUCII (pseudogene), putative | 0,3242 | 0,5064 |
| TcCLB.511871.100 | hypothetical protein, conserved | 0,324 | 0,01679 |
| TcCLB.507949.260 | Cell cycle checkpoint protein RAD1-like, putative | 0,3239 | 0,1717 |
| TcCLB.510579.110 | hypothetical protein, conserved | 0,3238 | 0,0006744 |
| TcCLB.507723.160 | hypothetical protein, conserved | 0,3238 | 0,003775 |
| TcCLB.511195.10 | retrotransposon hot spot protein (RHS, pseudogene), putat | 0,3238 | 0,4916 |
| TcCLB.510083.30 | retrotransposon hot spot protein (RHS, pseudogene), putat | 0,3237 | 0,002822 |
| TcCLB.510835.40 | 60S ribosomal protein L7a, putative | 0,3236 | 0,00002027 |
| TcCLB.511465.20 | hypothetical protein, conserved | 0,3236 | 0,004847 |
| TcCLB.510087.30 | dihydroceramide synthase, putative (DHCS) | 0,3234 | 0,008702 |
| TcCLB.506777.70 | cdc2-related kinase 8, putative (CRK8) | 0,3232 | 0,0104 |
| TcCLB.507013.20 | hypothetical protein, conserved | 0,323 | 0,0172 |
| TcCLB.509003.20 | calpain-like cysteine peptidase, putative | 0,323 | 0,1298 |
| TcCLB.507957.10 | mucin-associated surface protein (MASP), putative | 0,3227 | 0,2701 |
| TcCLB.503571.19 | hypothetical protein, conserved | 0,3226 | 0,00623 |
| TcCLB.509913.24 | dynein light chain 2B, cytoplasmic, putative | 0,3226 | 0,02952 |
| TcCLB.506657.79 | 50S ribosomal protein L16, putative | 0,3226 | 0,04661 |
| TcCLB.511643.110 | trans-sialidase, Group V, putative | 0,3225 | 0,1503 |
| TcCLB.510187.300 | hypothetical protein, conserved | 0,3223 | 0,04064 |
| TcCLB.511705.5 | trans-sialidase (pseudogene), putative | 0,3223 | 0,5489 |
| TcCLB.511469.20 | hypothetical protein, conserved | 0,3222 | 0,003645 |
| TcCLB.511413.19 | trans-sialidase, putative | 0,3218 | 0,2652 |
| TcCLB.506859.20 | proteasome regulatory ATPase subunit 5, putative | 0,3217 | 0,002242 |
| TcCLB.509047.60 | hypothetical protein, conserved | 0,3217 | 0,008191 |
| TcCLB.506529.224 | hypothetical protein, conserved | 0,3215 | 0,0867 |
| TcCLB.506559.500 | hypothetical protein, conserved | 0,3214 | 0,003473 |
| TcCLB.507921.9 | dispersed gene family protein 1 (DGF-1), putative | 0,3213 | 0,2049 |
| TcCLB.510621.49 | mucin-associated surface protein (MASP), putative | 0,3212 | 0,6225 |

| | | | |
|---|---|---|---|
| TcCLB.510899.50 | palmitoyl acyltransferase 4, putative | 0,3211 | 0,02397 |
| TcCLB.508707.120 | hypothetical protein, conserved | 0,321 | 0,005378 |
| TcCLB.511199.20 | hypothetical protein, conserved | 0,321 | 0,0197 |
| TcCLB.504149.50 | hypothetical protein, conserved | 0,3209 | 0,003943 |
| TcCLB.507669.120 | ATPase domain protein, putative | 0,3208 | 0,03613 |
| TcCLB.507165.59 | hypothetical protein, conserved | 0,3207 | 0,02227 |
| TcCLB.510519.140 | hypothetical protein, conserved | 0,3207 | 0,1684 |
| TcCLB.508579.4 | hypothetical protein, conserved | 0,3206 | 0,1689 |
| TcCLB.507883.10 | glucose-6-phosphate 1-dehydrogenase (pseudogene), puta | 0,3205 | 0,2157 |
| TcCLB.506975.34 | aminoacylase, putative | 0,3204 | 0,267 |
| TcCLB.510737.10 | 6-phosphogluconate dehydrogenase, decarboxylating, puta | 0,3203 | 0,09842 |
| TcCLB.510429.59 | hypothetical protein, conserved | 0,3203 | 0,2545 |
| TcCLB.509151.80 | hypothetical protein, conserved | 0,32 | 0,07517 |
| TcCLB.508411.30 | hypothetical protein, conserved | 0,3199 | 0,002514 |
| TcCLB.506279.120 | mucin TcMUCII, putative | 0,3199 | 0,6474 |
| TcCLB.506821.150 | hypothetical protein, conserved | 0,3194 | 0,0165 |
| TcCLB.506485.20 | hypothetical protein, conserved | 0,3194 | 0,03872 |
| TcCLB.507857.30 | hypothetical protein, conserved | 0,3193 | 0,007738 |
| TcCLB.509803.10 | hypothetical protein | 0,3188 | 0,1916 |
| TcCLB.510565.125 | hypothetical protein, conserved | 0,3188 | 0,236 |
| TcCLB.506637.70 | hypothetical protein, conserved | 0,3187 | 0,009702 |
| TcCLB.506931.70 | hypothetical protein, conserved | 0,3184 | 0,008387 |
| TcCLB.506977.60 | hypothetical protein, conserved | 0,3181 | 0,05749 |
| TcCLB.510143.24 | Hsc70-interacting protein (Hip), putative | 0,318 | 0,0003169 |
| TcCLB.506599.120 | surface protease GP63 (pseudogene), putative | 0,3175 | 0,6929 |
| TcCLB.511173.150 | mucin-associated surface protein (MASP), putative | 0,3173 | 0,4138 |
| TcCLB.509455.30 | hypothetical protein, conserved | 0,3172 | 0,02989 |
| TcCLB.510661.100 | hypothetical protein | 0,3172 | 0,04662 |
| TcCLB.510525.20 | hypothetical protein, conserved | 0,3172 | 0,06873 |
| TcCLB.506989.270 | hypothetical protein, conserved | 0,3171 | 0,6379 |
| TcCLB.508221.350 | hypothetical protein | 0,317 | 0,4314 |
| TcCLB.507517.120 | hypothetical protein, conserved | 0,3168 | 0,1004 |
| TcCLB.506297.94 | 60S ribosomal protein L24, putative | 0,3167 | 0,01519 |
| TcCLB.506789.140 | MORN repeat-containing protein | 0,3166 | 0,02637 |
| TcCLB.511857.20 | hypothetical protein, conserved | 0,3165 | 0,009679 |
| TcCLB.503893.70 | reductase, putative | 0,3164 | 0,01912 |
| TcCLB.510121.20 | hexokinase, putative | 0,3163 | 0,0004197 |
| TcCLB.504003.60 | chaperonin/T-complex protein 1 gamma subunit, putative | 0,316 | 0,00001362 |
| TcCLB.504817.9 | hypothetical protein, conserved (fragment) | 0,316 | 0,3538 |
| TcCLB.510667.70 | iron-sulfur cluster assembly protein, putative | 0,3158 | 0,03349 |
| TcCLB.509259.45 | hypothetical protein | 0,3158 | 0,2982 |
| TcCLB.509971.44 | mucin-associated surface protein (MASP, pseudogene), pu | 0,3158 | 0,5953 |
| TcCLB.511657.30 | hypothetical protein, conserved | 0,3155 | 0,01173 |
| TcCLB.506929.30 | hypothetical protein, conserved | 0,3155 | 0,01666 |
| TcCLB.504103.60 | hypothetical protein, conserved | 0,3155 | 0,1332 |

| | | | |
|---|---|---|---|
| TcCLB.506943.27 | hypothetical protein, conserved | 0,3155 | 0,2186 |
| TcCLB.511875.80 | mucin-associated surface protein (MASP), putative | 0,3155 | 0,254 |
| TcCLB.505949.70 | mucin TcMUCII, putative | 0,3154 | 0,4177 |
| TcCLB.505949.120 | mucin TcMUCII, putative | 0,3153 | 0,538 |
| TcCLB.511169.29 | tubulin-tyrsoine ligase-like protein, putative | 0,3149 | 0,002521 |
| TcCLB.503741.20 | mucin-associated surface protein (MASP, pseudogene), pu | 0,3149 | 0,3033 |
| TcCLB.505807.110 | alkyl-dihydroxyacetone phosphate synthase, putative | 0,3145 | 0,00006902 |
| TcCLB.511577.86 | N-acetyltransferase, putative | 0,3145 | 0,007392 |
| TcCLB.510221.39 | adenosine monophosphate deaminase, putative (fragment | 0,3145 | 0,0255 |
| TcCLB.508125.149 | trans-sialidase, putative (fragment) | 0,3144 | 0,342 |
| TcCLB.506955.261 | hypothetical protein, conserved (pseudogene) | 0,3144 | 0,4511 |
| TcCLB.507701.10 | mucin TcMUCII, putative | 0,3143 | 0,1978 |
| TcCLB.511875.9 | mucin-associated surface protein (MASP), putative | 0,3143 | 0,2379 |
| TcCLB.508645.59 | dynein light chain, putative | 0,3139 | 0,01714 |
| TcCLB.504827.80 | hypothetical protein, conserved | 0,3139 | 0,1019 |
| TcCLB.506799.110 | hypothetical protein | 0,3139 | 0,5777 |
| TcCLB.508153.1060 | hypothetical protein, conserved | 0,3138 | 0,2754 |
| TcCLB.511181.80 | threonyl-tRNA synthetase, putative | 0,3137 | 0,00007088 |
| TcCLB.505183.140 | protein kinase, putative | 0,3135 | 0,01014 |
| TcCLB.509735.30 | retrotransposon hot spot protein (RHS, pseudogene), putat | 0,3133 | 0,01005 |
| TcCLB.506925.460 | hypothetical protein, conserved | 0,3133 | 0,08782 |
| TcCLB.506391.50 | polyubiquitin (pseudogene), putative | 0,3132 | 0,0313 |
| TcCLB.509169.20 | hypothetical protein, conserved | 0,3131 | 0,01213 |
| TcCLB.510431.70 | hypothetical protein, conserved | 0,3128 | 0,07151 |
| TcCLB.506845.31 | ATP-dependent DEAD/H RNA helicase (pseudogene), putat | 0,3128 | 0,575 |
| TcCLB.506755.270 | hypothetical protein, conserved | 0,3125 | 0,4234 |
| TcCLB.511553.160 | 90 kDa surface protein, putative | 0,3125 | 0,5954 |
| TcCLB.503905.30 | hypothetical protein, conserved | 0,3124 | 0,01922 |
| TcCLB.508233.30 | mucin TcSMUGS, putative | 0,3123 | 0,1619 |
| TcCLB.509805.240 | hypothetical protein, conserved (pseudogene) | 0,3121 | 0,08256 |
| TcCLB.509471.30 | hypothetical protein, conserved | 0,3117 | 0,06335 |
| TcCLB.507907.60 | retrotransposon hot spot (RHS) protein, putative | 0,3115 | 0,1931 |
| TcCLB.510737.60 | hypothetical protein, conserved | 0,3114 | 0,03863 |
| TcCLB.508349.10 | hypothetical protein, conserved | 0,3114 | 0,07746 |
| TcCLB.511133.20 | thymidine kinase, putative | 0,3114 | 0,09513 |
| TcCLB.506211.80 | hypothetical protein, conserved | 0,3113 | 0,05012 |
| TcCLB.504021.74 | hypothetical protein, conserved | 0,311 | 0,1088 |
| TcCLB.511733.100 | hypothetical protein, conserved | 0,3109 | 0,02334 |
| TcCLB.506457.10 | metallo-peptidase, Clan MG, Family M24 (MetAP2) | 0,3108 | 0,002521 |
| TcCLB.505807.50 | hypothetical protein, conserved | 0,3105 | 0,05283 |
| TcCLB.506407.80 | intraflagellar transport protein 81, putative (IFT81) | 0,3103 | 0,0001552 |
| TcCLB.507477.20 | hypothetical protein, conserved (pseudogene) | 0,3102 | 0,001756 |
| TcCLB.508277.330 | actin-like protein, putative | 0,3095 | 0,01611 |
| TcCLB.506267.30 | mucin TcMUCII, putative | 0,3095 | 0,4714 |
| TcCLB.506667.80 | mucin-associated surface protein (MASP), putative | 0,3095 | 0,605 |

| | | | |
|---|---|---|---|
| TcCLB.507651.10 | hypothetical protein, conserved (pseudogene) | 0,3094 | 0,002232 |
| TcCLB.507023.210 | hypothetical protein, conserved | 0,3094 | 0,2049 |
| TcCLB.510515.130 | S-adenosyl-methyltransferase mraW-like protein, putative | 0,3093 | 0,02123 |
| TcCLB.508823.100 | hypothetical protein, conserved | 0,3093 | 0,0314 |
| TcCLB.511233.130 | mucin-associated surface protein (MASP), putative | 0,3093 | 0,7281 |
| TcCLB.508589.10 | hypothetical protein | 0,3086 | 0,3761 |
| TcCLB.511867.140 | Acyl carrier protein, mitochondrial, putative (ACP) | 0,3085 | 0,004081 |
| TcCLB.510945.20 | hypothetical protein, conserved | 0,3085 | 0,01584 |
| TcCLB.506999.90 | rieske iron-sulfur protein, mitochondrial precursor, putative | 0,3082 | 0,000694 |
| TcCLB.509805.60 | Ras-related protein Rab5A, putative (RAB5A) | 0,3082 | 0,004285 |
| TcCLB.510659.90 | hypothetical protein, conserved | 0,3082 | 0,05278 |
| TcCLB.509607.10 | ubiquitin-conjugating enzyme E2, putative | 0,3081 | 0,01784 |
| TcCLB.422507.10 | glycine dehydrogenase [decarboxylating], putative | 0,3081 | 0,1877 |
| TcCLB.508461.350 | hypothetical protein, conserved | 0,308 | 0,008349 |
| TcCLB.506977.30 | hypothetical protein, conserved | 0,308 | 0,02765 |
| TcCLB.509417.30 | retrotransposon hot spot protein (RHS, pseudogene), putat | 0,308 | 0,3433 |
| TcCLB.507835.80 | trans-sialidase, putative | 0,3079 | 0,4916 |
| TcCLB.511543.30 | amino acid transporter, putative | 0,3078 | 0,2515 |
| TcCLB.510671.31 | hypothetical protein, conserved | 0,3077 | 0,02294 |
| TcCLB.509027.70 | kinetoplastid kinetochore protein 19, putative (kkt19) | 0,3077 | 0,02882 |
| TcCLB.507993.40 | hypothetical protein, conserved | 0,3076 | 0,01193 |
| TcCLB.506147.100 | hypothetical protein, conserved | 0,3075 | 0,007044 |
| TcCLB.506503.53 | hypothetical protein | 0,3075 | 0,06327 |
| TcCLB.508899.140 | hypothetical protein | 0,3073 | 0,0418 |
| TcCLB.510227.10 | hypothetical protein, conserved | 0,3071 | 0,1555 |
| TcCLB.510373.10 | mucin-associated surface protein (MASP), putative | 0,3071 | 0,3883 |
| TcCLB.508063.9 | dispersed gene family protein 1 (DGF-1), putative (fragme | 0,307 | 0,3411 |
| TcCLB.506195.210 | hypothetical protein, conserved | 0,3069 | 0,00458 |
| TcCLB.510355.70 | retrotransposon hot spot protein (RHS, pseudogene), putat | 0,3067 | 0,04649 |
| TcCLB.503733.10 | hypothetical protein, conserved | 0,3067 | 0,1167 |
| TcCLB.504427.80 | hypothetical protein, conserved | 0,3066 | 0,07068 |
| TcCLB.506517.40 | hypothetical protein | 0,3066 | 0,3757 |
| TcCLB.405165.19 | hypothetical protein, conserved (fragment) | 0,3065 | 0,05238 |
| TcCLB.508647.50 | hypothetical protein, conserved (pseudogene) | 0,3064 | 0,001028 |
| TcCLB.508411.50 | hypothetical protein, conserved | 0,3063 | 0,01864 |
| TcCLB.415187.19 | hypothetical protein, conserved (fragment) | 0,3063 | 0,3771 |
| TcCLB.508665.40 | hypothetical protein, conserved | 0,3061 | 0,2048 |
| TcCLB.506559.40 | cardiolipin synthetase, putative (cls) | 0,306 | 0,001742 |
| TcCLB.508601.30 | DNA-directed RNA polymerase I/III subunit, putative | 0,3059 | 0,09174 |
| TcCLB.506945.150 | hypothetical protein | 0,3058 | 0,08412 |
| TcCLB.509669.14 | thioredoxin, putative | 0,3058 | 0,1395 |
| TcCLB.510385.10 | dispersed gene family protein 1 (DGF-1), putative | 0,3057 | 0,3051 |
| TcCLB.508265.30 | hypothetical protein | 0,3056 | 0,01889 |
| TcCLB.509081.150 | mucin-associated surface protein (MASP), putative | 0,3055 | 0,4303 |
| TcCLB.503599.10 | protein kinase, putative | 0,3053 | 0,1097 |

| | | | |
|---|---|---|---|
| TcCLB.509897.260 | hypothetical protein | 0,3053 | 0,3587 |
| TcCLB.503575.60 | protein kinase, putative | 0,3052 | 0,01313 |
| TcCLB.507209.50 | hypothetical protein, conserved | 0,3052 | 0,04049 |
| TcCLB.506537.230 | hypothetical protein, conserved | 0,3052 | 0,3365 |
| TcCLB.509081.5 | mucin-associated surface protein (MASP), putative | 0,3052 | 0,5429 |
| TcCLB.507023.60 | hypothetical protein, conserved | 0,3049 | 0,05203 |
| TcCLB.506437.20 | tuzin, putative | 0,3048 | 0,00005819 |
| TcCLB.511211.104 | hypothetical protein, conserved | 0,3048 | 0,1471 |
| TcCLB.508431.5 | hypothetical protein | 0,3047 | 0,5476 |
| TcCLB.511715.70 | hypothetical protein, conserved | 0,3045 | 0,04379 |
| TcCLB.505193.84 | hypothetical protein, conserved | 0,3045 | 0,07457 |
| TcCLB.504153.320 | Golgi reassembly stacking protein (GRASP homologue), pu | 0,3043 | 0,04707 |
| TcCLB.510329.370 | hypothetical protein, conserved | 0,3043 | 0,1162 |
| TcCLB.503839.5 | chloride channel protein, putative | 0,3043 | 0,1971 |
| TcCLB.506763.150 | mucin-associated surface protein (MASP), putative | 0,3043 | 0,7065 |
| TcCLB.504097.30 | hypothetical protein, conserved | 0,3042 | 0,00137 |
| TcCLB.509525.420 | mucin TcMUCII, putative | 0,3042 | 0,7219 |
| TcCLB.511865.19 | frataxin-like, mitochondrial precursor, putative | 0,304 | 0,09567 |
| TcCLB.506967.27 | syntaxin binding protein (pseudogene), putative | 0,3037 | 0,6657 |
| TcCLB.509945.30 | hypothetical protein | 0,3036 | 0,001997 |
| TcCLB.510853.120 | hypothetical protein, conserved | 0,3034 | 0,002255 |
| TcCLB.503927.9 | dynein light intermediate chain, putative | 0,3034 | 0,02529 |
| TcCLB.505999.60 | hypothetical protein | 0,3034 | 0,2136 |
| TcCLB.423205.10 | trans-sialidase, putative (fragment) | 0,3034 | 0,8343 |
| TcCLB.506113.40 | hypothetical protein, conserved (pseudogene) | 0,3033 | 0,08367 |
| TcCLB.511695.10 | hypothetical protein | 0,3033 | 0,3861 |
| TcCLB.504153.20 | phosphoglycerate kinase, putative | 0,3032 | 0,001427 |
| TcCLB.508859.70 | hypothetical protein, conserved | 0,3032 | 0,06328 |
| TcCLB.511303.80 | hypothetical protein, conserved | 0,3031 | 0,07279 |
| TcCLB.511049.40 | leucyl-tRNA synthetase, putative | 0,3029 | 0,00006688 |
| TcCLB.413893.10 | trans-sialidase (pseudogene), putative | 0,3029 | 0,5356 |
| TcCLB.506479.30 | hypothetical protein, conserved | 0,3028 | 0,06633 |
| TcCLB.506737.10 | hypothetical protein, conserved | 0,3026 | 0,008982 |
| TcCLB.506247.400 | hypothetical protein, conserved | 0,3025 | 0,1087 |
| TcCLB.509419.230 | trans-sialidase (pseudogene), putative | 0,3024 | 0,2091 |
| TcCLB.508593.100 | hypothetical protein, conserved | 0,3024 | 0,2446 |
| TcCLB.507071.190 | trans-sialidase, Group V, putative | 0,3024 | 0,4672 |
| TcCLB.509525.120 | mucin TcMUCII, putative | 0,3024 | 0,6164 |
| TcCLB.498829.10 | mucin TcMUC, putative | 0,3023 | 0,2808 |
| TcCLB.508227.10 | hypothetical protein | 0,3023 | 0,7462 |
| TcCLB.510307.240 | trans-sialidase, Group VIII, putative | 0,3022 | 0,2429 |
| TcCLB.509895.40 | hypothetical protein, conserved | 0,3021 | 0,004776 |
| TcCLB.511849.10 | mucin TcMUCI, putative | 0,3021 | 0,3044 |
| TcCLB.506209.19 | dispersed gene family protein 1 (DGF-1), putative (fragme | 0,3021 | 0,5298 |
| TcCLB.509769.70 | DNA polymerase theta (helicase domain only), putative | 0,302 | 0,0006546 |

| | | | |
|---|---|---|---|
| TcCLB.511301.120 | hypothetical protein, conserved | 0,302 | 0,003456 |
| TcCLB.503655.30 | mucin-like glycoprotein, putative | 0,3019 | 0,4476 |
| TcCLB.509339.4 | poly(A) export protein, putative | 0,3017 | 0,05497 |
| TcCLB.511425.19 | hypothetical protein, conserved (pseudogene) | 0,3016 | 0,08299 |
| TcCLB.510297.50 | hypothetical protein, conserved | 0,3015 | 0,03057 |
| TcCLB.506145.40 | hypothetical protein, conserved | 0,3014 | 0,0003576 |
| TcCLB.511173.24 | mucin-associated surface protein (MASP), putative | 0,3014 | 0,482 |
| TcCLB.503819.30 | eukaryotic translation initiation factor 2 subunit, putative | 0,3013 | 0,0008148 |
| TcCLB.506985.60 | inositol polyphosphate kinase-like protein, putative | 0,3013 | 0,0872 |
| TcCLB.510689.70 | hypothetical protein, conserved | 0,301 | 0,2649 |
| TcCLB.506777.80 | seryl-tRNA synthetase, putative | 0,3009 | 9,057E-06 |
| TcCLB.509245.29 | hypothetical protein, conserved | 0,3009 | 0,1419 |
| TcCLB.511821.110 | translation initiation factor IF-2, putative | 0,3008 | 0,004542 |
| TcCLB.504491.20 | trans-sialidase, putative | 0,3008 | 0,5233 |
| TcCLB.504069.70 | vacuolar ATP synthase, putative | 0,3007 | 0,001028 |
| TcCLB.505073.10 | hypothetical protein | 0,3007 | 0,002033 |
| TcCLB.506591.30 | hypothetical protein, conserved | 0,3007 | 0,01493 |
| TcCLB.511751.50 | protein kinase, putative | 0,3007 | 0,02031 |
| TcCLB.507723.41 | trans-sialidase (pseudogene), putative | 0,3005 | 0,3359 |
| TcCLB.508247.30 | hypothetical protein, conserved | 0,3005 | 0,6475 |
| TcCLB.508805.90 | hypothetical protein, conserved | 0,3003 | 0,3184 |
| TcCLB.511467.4 | hypothetical protein, conserved | 0,3002 | 0,04737 |
| TcCLB.503693.29 | hypothetical protein | 0,3 | 0,03071 |
| TcCLB.503611.50 | hypothetical protein, conserved | 0,3 | 0,1271 |
| TcCLB.511399.50 | mucin-associated surface protein (MASP), putative | 0,2999 | 0,4142 |
| TcCLB.506559.220 | hypothetical protein, conserved | 0,2997 | 0,02609 |
| TcCLB.510315.4 | hypothetical protein, conserved (fragment) | 0,2997 | 0,1179 |
| TcCLB.509777.80 | hypothetical protein, conserved | 0,2995 | 0,04114 |
| TcCLB.508247.110 | serine-alanine-and proline-rich protein, putative | 0,2995 | 0,4522 |
| TcCLB.509393.10 | hypothetical protein, conserved | 0,2993 | 0,02208 |
| TcCLB.507663.30 | hypothetical protein, conserved | 0,2993 | 0,03421 |
| TcCLB.508881.90 | hypothetical protein, conserved | 0,2993 | 0,0484 |
| TcCLB.511189.30 | 40S ribosomal protein S13, putative | 0,2991 | 0,0006339 |
| TcCLB.511585.125 | hypothetical protein, conserved | 0,2991 | 0,0485 |
| TcCLB.509025.10 | hypothetical protein, conserved | 0,299 | 0,2404 |
| TcCLB.508577.70 | hypothetical protein | 0,2989 | 0,00611 |
| TcCLB.511217.145 | ribosomal protein L39, putative | 0,2988 | 0,005118 |
| TcCLB.508807.10 | ATP-dependent Clp protease subunit, heat shock protein 1( | 0,2988 | 0,01338 |
| TcCLB.511907.210 | hypothetical protein, conserved | 0,2988 | 0,07624 |
| TcCLB.507825.49 | hypothetical protein, conserved (fragment) | 0,2988 | 0,2168 |
| TcCLB.503531.30 | COP9 signalosome complex subunit 5, putative (pseudoger | 0,2987 | 0,04277 |
| TcCLB.508595.35 | hypothetical protein (pseudogene) | 0,2987 | 0,52 |
| TcCLB.507255.10 | surface protease GP63 (pseudogene), putative | 0,2987 | 0,5838 |
| TcCLB.503413.19 | hypothetical protein | 0,2986 | 0,06328 |
| TcCLB.510355.230 | hypothetical protein, conserved | 0,2985 | 0,09097 |

| | | | |
|---|---|---|---|
| TcCLB.503499.20 | hypothetical protein, conserved | 0,2981 | 0,04401 |
| TcCLB.503607.7 | glycine dehydrogenase (pseudogene), putative | 0,2981 | 0,09672 |
| TcCLB.511431.20 | hypothetical protein, conserved | 0,2977 | 0,0001512 |
| TcCLB.503935.20 | 2-oxoglutarate dehydrogenase, E2 component, dihydrolipo | 0,2975 | 0,00116 |
| TcCLB.511069.10 | 40S ribosomal protein S8, putative | 0,2975 | 0,008945 |
| TcCLB.508207.150 | Silent information regulator 2 related protein 1, putative ( | 0,2975 | 0,04216 |
| TcCLB.506401.220 | electron transfer flavoprotein-ubiquinone oxidoreductase, | 0,2974 | 0,03957 |
| TcCLB.509573.10 | small GTP-binding protein Rab7, putative | 0,2973 | 0,06544 |
| TcCLB.509979.270 | mucin-associated surface protein (MASP), putative | 0,2971 | 0,1631 |
| TcCLB.507647.30 | hypothetical protein, conserved | 0,2971 | 0,2639 |
| TcCLB.506475.130 | hypothetical protein, conserved | 0,2969 | 0,002695 |
| TcCLB.510477.80 | mucin-associated surface protein (MASP), putative | 0,2969 | 0,4424 |
| TcCLB.506629.80 | vesicular transport protein (CDC48 homologue), putative | 0,2967 | 0,0002102 |
| TcCLB.508977.40 | trans-sialidase, Group V, putative | 0,2967 | 0,1364 |
| TcCLB.511921.140 | mucin-associated surface protein (MASP), putative | 0,2966 | 0,6816 |
| TcCLB.510323.70 | hypothetical protein, conserved | 0,2964 | 0,008876 |
| TcCLB.504105.190 | hypothetical protein, conserved | 0,2963 | 0,04024 |
| TcCLB.509693.60 | hypothetical protein, conserved | 0,2963 | 0,1147 |
| TcCLB.506765.43 | hypothetical protein | 0,2962 | 0,2473 |
| TcCLB.507005.4 | acyl transferase-like protein | 0,296 | 0,04105 |
| TcCLB.507503.20 | glycine dehydrogenase [decarboxylating] (pseudogene), pu | 0,2959 | 0,03411 |
| TcCLB.507603.110 | hypothetical protein, conserved | 0,2958 | 0,04452 |
| TcCLB.504505.20 | retrotransposon hot spot (RHS) protein, putative | 0,2957 | 0,1553 |
| TcCLB.507611.380 | hypothetical protein | 0,2955 | 0,04118 |
| TcCLB.511741.50 | C-1-tetrahydrofolate synthase, cytoplasmic, putative | 0,2955 | 0,06152 |
| TcCLB.420495.10 | hypothetical protein, conserved (pseudogene) | 0,2954 | 0,1253 |
| TcCLB.508351.30 | hypothetical protein, conserved | 0,295 | 0,03375 |
| TcCLB.510743.60 | phospholipase A1, putative | 0,295 | 0,06861 |
| TcCLB.506581.69 | Mutt-Nudix-related hydrolase 5, putative | 0,2949 | 0,1629 |
| TcCLB.506993.130 | mucin-associated surface protein (MASP, pseudogene), pu | 0,2949 | 0,2565 |
| TcCLB.504105.80 | hypothetical protein, conserved | 0,2942 | 0,003801 |
| TcCLB.506265.170 | hypothetical protein, conserved | 0,2941 | 0,08271 |
| TcCLB.504013.11 | ribosomal protein L13, putative (fragment) | 0,2939 | 0,00001404 |
| TcCLB.507499.90 | trans-sialidase (pseudogene), putative | 0,2938 | 0,04883 |
| TcCLB.509055.69 | hypothetical protein, conserved (fragment) | 0,2937 | 0,3063 |
| TcCLB.511839.25 | elongation factor 1-gamma (EF-1-gamma, pseudogene), p | 0,2934 | 0,6111 |
| TcCLB.507625.130 | hypothetical protein, conserved | 0,2932 | 0,04554 |
| TcCLB.508629.74 | retrotransposon hot spot protein (RHS, pseudogene), putat | 0,2932 | 0,0658 |
| TcCLB.510395.10 | ATP synthase F1, alpha subunit, putative (ATPF1A) | 0,2931 | 0,006858 |
| TcCLB.506297.10 | hypothetical protein | 0,2931 | 0,007827 |
| TcCLB.507203.5 | trans-sialidase (pseudogene), putative | 0,2931 | 0,09219 |
| TcCLB.506117.9 | dispersed gene family protein 1 (DGF-1), putative | 0,2931 | 0,1029 |
| TcCLB.508857.130 | hypothetical protein, conserved | 0,293 | 0,1155 |
| TcCLB.508543.20 | protein kinase, putative | 0,2929 | 0,004621 |
| TcCLB.506855.150 | hypothetical protein, conserved | 0,2928 | 0,01858 |

| | | | |
|---|---|---|---|
| TcCLB.510661.279 | hypothetical protein, conserved | 0,2928 | 0,133 |
| TcCLB.504017.50 | hypothetical protein, conserved | 0,2927 | 0,2024 |
| TcCLB.508153.90 | hypothetical protein | 0,2923 | 0,07636 |
| TcCLB.509229.80 | mitochondrial processing peptidase alpha subunit, putative | 0,2922 | 0,001436 |
| TcCLB.506247.170 | ATPase, putative | 0,2922 | 0,0144 |
| TcCLB.510421.320 | pyruvate dehydrogenase E1 beta subunit, putative | 0,2922 | 0,01472 |
| TcCLB.510359.220 | hypothetical protein, conserved | 0,2922 | 0,01748 |
| TcCLB.503957.30 | retrotransposon hot spot (RHS) protein, putative | 0,2922 | 0,1077 |
| TcCLB.507753.10 | trans-sialidase, Group III, putative | 0,2922 | 0,3655 |
| TcCLB.506629.20 | hypothetical protein, conserved | 0,292 | 0,05852 |
| TcCLB.511621.259 | hypothetical protein, conserved | 0,2918 | 0,1926 |
| TcCLB.510785.20 | pteridine reductase (pseudogene), putative | 0,2917 | 0,001451 |
| TcCLB.509717.50 | hypothetical protein, conserved | 0,2915 | 0,0006969 |
| TcCLB.508177.30 | hypothetical protein, conserved | 0,2915 | 0,01605 |
| TcCLB.508603.10 | trans-sialidase, putative | 0,2915 | 0,3735 |
| TcCLB.511407.30 | hypothetical protein | 0,2914 | 0,005364 |
| TcCLB.511817.151 | histone H2A, putative | 0,2913 | 0,5533 |
| TcCLB.507801.70 | protein kinase, putative | 0,2912 | 0,2205 |
| TcCLB.508279.10 | trans-sialidase (pseudogene), putative | 0,2911 | 0,3135 |
| TcCLB.503995.50 | serine peptidase, clan SC, family S9A-like protein (OPB) | 0,291 | 0,007524 |
| TcCLB.509171.54 | hypothetical protein, conserved | 0,2906 | 0,13 |
| TcCLB.503973.290 | mucin TcMUCII, putative | 0,2906 | 0,6445 |
| TcCLB.504009.60 | hypothetical protein, conserved | 0,2904 | 0,007501 |
| TcCLB.509471.10 | hypothetical protein, conserved | 0,2904 | 0,07348 |
| TcCLB.506211.30 | intraflagellar transport protein 80, putative (IFT80) | 0,2903 | 0,02075 |
| TcCLB.504147.240 | hypothetical protein, conserved | 0,2903 | 0,04012 |
| TcCLB.511585.269 | retrotransposon hot spot protein (RHS, pseudogene), putat | 0,2903 | 0,1949 |
| TcCLB.507001.80 | acyl-CoA dehydrogenase, putative | 0,29 | 0,02947 |
| TcCLB.510311.120 | hypothetical protein | 0,2899 | 0,1058 |
| TcCLB.506171.35 | ATP-dependent DEAD/H RNA helicase (pseudogene), putat | 0,2899 | 0,5278 |
| TcCLB.506183.20 | dispersed gene family protein 1 (DGF-1), putative | 0,2898 | 0,2144 |
| TcCLB.510495.30 | trans-sialidase (pseudogene), putative | 0,2898 | 0,4649 |
| TcCLB.506511.60 | flavoprotein monooxygenase, putative | 0,2895 | 0,003776 |
| TcCLB.398009.10 | retrotransposon hot spot protein (RHS, pseudogene), putat | 0,2894 | 0,05757 |
| TcCLB.506417.50 | hypothetical protein, conserved | 0,2894 | 0,1119 |
| TcCLB.510661.244 | hypothetical protein, conserved | 0,2893 | 0,1096 |
| TcCLB.507777.20 | hypothetical protein, conserved | 0,2893 | 0,1204 |
| TcCLB.504033.110 | hypothetical protein, conserved | 0,2892 | 0,04471 |
| TcCLB.503973.300 | surface protease GP63 (pseudogene), putative | 0,2892 | 0,6196 |
| TcCLB.510281.5 | mucin TcMUCII (pseudogene), putative | 0,2892 | 0,6586 |
| TcCLB.511035.10 | surface protease GP63, putative | 0,2891 | 0,004651 |
| TcCLB.510659.28 | hypothetical protein, conserved | 0,2891 | 0,05365 |
| TcCLB.505949.160 | mucin TcMUCII, putative | 0,2891 | 0,6043 |
| TcCLB.509127.59 | hypothetical protein, conserved | 0,2889 | 0,3811 |
| TcCLB.507069.220 | mucin-associated surface protein (MASP), putative | 0,2889 | 0,5481 |

| | | | |
|---|---|---|---|
| TcCLB.511209.10 | V-type proton ATPase subunit B, putative | 0,2888 | 0,003939 |
| TcCLB.508139.40 | retrotransposon hot spot protein (RHS, pseudogene), putat | 0,2886 | 0,04058 |
| TcCLB.504149.30 | hypothetical protein, conserved | 0,2885 | 0,04335 |
| TcCLB.506409.140 | mucin TcMUCII, putative | 0,2885 | 0,6912 |
| TcCLB.510281.20 | surface protease GP63, putative | 0,2884 | 0,4926 |
| TcCLB.510737.50 | hypothetical protein, conserved | 0,2882 | 0,004264 |
| TcCLB.506669.10 | hypothetical protein, conserved | 0,2882 | 0,1656 |
| TcCLB.508415.10 | hypothetical protein, conserved (pseudogene) | 0,2878 | 0,2288 |
| TcCLB.508201.20 | surface protease GP63 (pseudogene), putative | 0,2878 | 0,4626 |
| TcCLB.508873.580 | mucin-associated surface protein (MASP), putative | 0,2876 | 0,5662 |
| TcCLB.503841.60 | hypothetical protein, conserved | 0,2873 | 0,04692 |
| TcCLB.508539.40 | hypothetical protein, conserved | 0,2873 | 0,08279 |
| TcCLB.506151.4 | hypothetical protein, conserved (fragment) | 0,2872 | 0,0912 |
| TcCLB.506247.190 | nuclear lim interactor-interacting factor, putative | 0,287 | 0,3336 |
| TcCLB.511801.14 | protein kinase, putative (LDK) | 0,2869 | 0,04232 |
| TcCLB.511603.304 | mucin TcMUCII, putative | 0,2867 | 0,6214 |
| TcCLB.506431.20 | retrotransposon hot spot protein (RHS, pseudogene), putat | 0,2866 | 0,02832 |
| TcCLB.507035.75 | retrotransposon hot spot protein (RHS, pseudogene), putat | 0,2863 | 0,07101 |
| TcCLB.511471.20 | trans-sialidase (pseudogene), putative | 0,2863 | 0,2295 |
| TcCLB.459619.10 | hypothetical protein, conserved (pseudogene) | 0,2863 | 0,5004 |
| TcCLB.506307.10 | mucin-associated surface protein (MASP), putative | 0,2863 | 0,6097 |
| TcCLB.511311.7 | trans-sialidase, putative (fragment) | 0,2862 | 0,2722 |
| TcCLB.509795.40 | hypothetical protein, conserved | 0,2859 | 0,06888 |
| TcCLB.506151.20 | hypothetical protein, conserved | 0,2855 | 0,001046 |
| TcCLB.503787.50 | mucin TcMUCII, putative | 0,2855 | 0,5612 |
| TcCLB.504009.20 | hypothetical protein, conserved | 0,2852 | 0,04923 |
| TcCLB.509029.120 | ATP-dependent DNA helicase, putative | 0,285 | 0,02585 |
| TcCLB.485889.10 | hypothetical protein, conserved | 0,2848 | 0,007583 |
| TcCLB.508041.80 | protein transport protein Sec31, putative (pseudogene) | 0,2846 | 0,002029 |
| TcCLB.507623.101 | surface protease GP63 (pseudogene), putative | 0,2845 | 0,1246 |
| TcCLB.506233.20 | retrotransposon hot spot protein (RHS, pseudogene), putat | 0,2845 | 0,4213 |
| TcCLB.430605.29 | 40S ribosomal protein S3, putative | 0,2844 | 0,00288 |
| TcCLB.508221.502 | hypothetical protein, conserved (pseudogene) | 0,2843 | 0,4163 |
| TcCLB.510463.190 | retrotransposon hot spot (RHS) protein, putative | 0,2842 | 0,2106 |
| TcCLB.506637.20 | hypothetical protein, conserved | 0,284 | 0,09539 |
| TcCLB.508309.5 | trans-sialidase, putative (fragment) | 0,284 | 0,6324 |
| TcCLB.504069.30 | 60S ribosomal protein L32, putative | 0,2839 | 0,01069 |
| TcCLB.510015.20 | mucin TcMUCII, putative | 0,2839 | 0,5846 |
| TcCLB.507765.80 | hypothetical protein, conserved | 0,2838 | 0,03208 |
| TcCLB.509799.5 | hypothetical protein, conserved | 0,2838 | 0,06643 |
| TcCLB.506409.220 | hypothetical protein | 0,2837 | 0,11 |
| TcCLB.508169.129 | Mutt-Nudix-related hydrolase 5, putative (Nudix5) | 0,2837 | 0,1855 |
| TcCLB.507515.30 | hypothetical protein, conserved | 0,2836 | 0,2144 |
| TcCLB.504639.19 | dispersed gene family protein 1 (DGF-1), putative (fragme | 0,2836 | 0,292 |
| TcCLB.509289.5 | tuzin (fragment) | 0,2835 | 0,00004416 |

| | | | |
|---|---|---|---|
| TcCLB.509699.35 | hypothetical protein, conserved (pseudogene) | 0,2834 | 0,4119 |
| TcCLB.507093.320 | hypothetical protein, conserved | 0,2831 | 0,0378 |
| TcCLB.511859.50 | hypothetical protein, conserved | 0,2829 | 0,0597 |
| TcCLB.511667.30 | trans-sialidase, Group I, putative | 0,2829 | 0,6756 |
| TcCLB.506025.14 | ribosomal protein S29, putative | 0,2828 | 0,03787 |
| TcCLB.506345.70 | retrotransposon hot spot protein (RHS, pseudogene), putat | 0,2828 | 0,1808 |
| TcCLB.511277.210 | peroxisomal enoyl-coa hydratase, putative | 0,2826 | 0,002208 |
| TcCLB.507547.40 | enoyl-CoA hydratase/Enoyl-CoA isomerase/3-hydroxyacyl-( | 0,2826 | 0,01641 |
| TcCLB.507671.20 | hypothetical protein, conserved | 0,2826 | 0,1861 |
| TcCLB.508539.155 | mucin TcMUCII, putative | 0,2825 | 0,5593 |
| TcCLB.510197.220 | mucin-associated surface protein (MASP, pseudogene), pu | 0,2824 | 0,4484 |
| TcCLB.508953.19 | hypothetical protein, conserved | 0,2822 | 0,04004 |
| TcCLB.511559.50 | hypothetical protein, conserved | 0,2821 | 0,009528 |
| TcCLB.510275.21 | surface protease GP63 (pseudogene), putative | 0,2821 | 0,665 |
| TcCLB.508117.30 | elongation factor 1-gamma (EF-1-gamma, pseudogene), p | 0,282 | 0,3289 |
| TcCLB.509525.240 | mucin-associated surface protein (MASP), putative | 0,2819 | 0,6876 |
| TcCLB.510971.30 | trans-sialidase, Group VIII, putative | 0,2818 | 0,4533 |
| TcCLB.508173.219 | hypothetical protein, conserved | 0,2816 | 0,004476 |
| TcCLB.503977.50 | surface protease GP63 (pseudogene), putative | 0,2816 | 0,6078 |
| TcCLB.509167.30 | hypothetical protein, conserved | 0,2812 | 0,1499 |
| TcCLB.510679.90 | expression site-associated gene (ESAG-like, pseudogene), | 0,2812 | 0,5065 |
| TcCLB.508479.330 | hypothetical protein, conserved | 0,2811 | 0,004081 |
| TcCLB.506737.151 | surface protease GP63 (pseudogene), putative | 0,281 | 0,1038 |
| TcCLB.511921.100 | trans-sialidase (pseudogene), putative | 0,281 | 0,2742 |
| TcCLB.505999.140 | Metal-binding domain-containing protein | 0,2809 | 0,06211 |
| TcCLB.506225.20 | trans-sialidase (pseudogene), putative | 0,2807 | 0,3572 |
| TcCLB.506529.90 | puromycin-sensitive aminopeptidase-like protein, putative | 0,2806 | 0,06903 |
| TcCLB.508027.80 | hypothetical protein, conserved | 0,2805 | 0,008739 |
| TcCLB.510885.20 | hypothetical protein, conserved | 0,2804 | 0,04644 |
| TcCLB.506207.20 | hypothetical protein, conserved | 0,2802 | 0,00356 |
| TcCLB.503751.30 | hypothetical protein, conserved (pseudogene) | 0,2801 | 0,04568 |
| TcCLB.510329.120 | Present in the outer mitochondrial membrane proteome 2( | 0,2799 | 0,03399 |
| TcCLB.506391.10 | calmodulin (CALA2) | 0,2798 | 0,05446 |
| TcCLB.508737.100 | ATP-dependent Clp protease subunit, heat shock protein 78 | 0,2795 | 0,004844 |
| TcCLB.507467.90 | diacylglycerol acyltransferase, putative | 0,2795 | 0,02609 |
| TcCLB.508577.30 | retrotransposon hot spot protein (RHS, pseudogene), putat | 0,2795 | 0,04196 |
| TcCLB.511821.10 | Cytoplasmic dynein 1 heavy chain (DYNC1H1), putative | 0,2794 | 0,003227 |
| TcCLB.511079.60 | mucin-associated surface protein (MASP), putative | 0,2794 | 0,633 |
| TcCLB.506611.55 | elongation factor 1-gamma (EF-1-gamma, pseudogene), p | 0,2794 | 0,6872 |
| TcCLB.506135.9 | heat shock 70 kDa protein, putative (fragment) | 0,2793 | 0,002677 |
| TcCLB.509733.140 | hypothetical protein, conserved | 0,2793 | 0,008017 |
| TcCLB.510693.200 | mucin TcMUCII, putative | 0,2793 | 0,6244 |
| TcCLB.506309.40 | mucin-associated surface protein (MASP), putative | 0,2793 | 0,7025 |
| TcCLB.511421.110 | developmentally regulated phosphoprotein, putative | 0,2791 | 0,01735 |
| TcCLB.510305.40 | hypothetical protein, conserved | 0,2791 | 0,05705 |

| | | | |
|---|---|---|---|
| TcCLB.507675.30 | hypothetical protein, conserved | 0,2791 | 0,254 |
| TcCLB.508139.150 | hypothetical protein, conserved | 0,279 | 0,3152 |
| TcCLB.504081.409 | mucin TcMUCII (pseudogene), putative | 0,279 | 0,7123 |
| TcCLB.511401.120 | trans-sialidase, Group V, putative | 0,2788 | 0,2476 |
| TcCLB.506331.30 | hypothetical protein | 0,2787 | 0,595 |
| TcCLB.511213.60 | mucin-associated surface protein (MASP), putative | 0,2786 | 0,2945 |
| TcCLB.508081.110 | trans-sialidase (pseudogene), putative | 0,2785 | 0,3688 |
| TcCLB.507991.123 | hypothetical protein, conserved | 0,2784 | 0,08895 |
| TcCLB.504071.40 | hypothetical protein | 0,2779 | 0,1023 |
| TcCLB.511665.30 | retrotransposon hot spot protein (RHS, pseudogene), putat | 0,2779 | 0,2176 |
| TcCLB.503783.80 | mucin-associated surface protein (MASP), putative | 0,2779 | 0,345 |
| TcCLB.507053.20 | hypothetical protein, conserved | 0,2778 | 0,04038 |
| TcCLB.511907.230 | hypothetical protein, conserved | 0,2778 | 0,05815 |
| TcCLB.511713.20 | hypothetical protein, conserved | 0,2777 | 0,03407 |
| TcCLB.507715.34 | hypothetical protein, conserved | 0,2776 | 0,009711 |
| TcCLB.504867.90 | glycerol-3-phosphate acyltransferase, putative | 0,2776 | 0,03223 |
| TcCLB.508547.50 | maoC-like dehydratase, putative | 0,2774 | 0,03913 |
| TcCLB.511865.80 | hypothetical protein, conserved | 0,2774 | 0,2429 |
| TcCLB.506667.120 | mucin-associated surface protein (MASP), putative | 0,2773 | 0,5022 |
| TcCLB.509937.50 | hypothetical protein, conserved | 0,2772 | 0,007745 |
| TcCLB.508051.20 | hypothetical protein, conserved | 0,2772 | 0,01294 |
| TcCLB.511217.60 | hypothetical protein | 0,2772 | 0,05287 |
| TcCLB.504161.20 | hypothetical protein, conserved | 0,277 | 0,01344 |
| TcCLB.503465.20 | hypothetical protein, conserved (pseudogene) | 0,277 | 0,1137 |
| TcCLB.509747.50 | hypothetical protein, conserved | 0,2769 | 0,03086 |
| TcCLB.504175.40 | dual specificity protein phosphatase, putative | 0,2766 | 0,008806 |
| TcCLB.511269.10 | hypothetical protein, conserved | 0,2766 | 0,03078 |
| TcCLB.508405.60 | hypothetical protein, conserved | 0,2765 | 0,03428 |
| TcCLB.509717.20 | dihydrolipoamide acetyltransferase precursor, putative | 0,2763 | 0,002386 |
| TcCLB.509099.160 | hypothetical protein, conserved | 0,2761 | 0,008169 |
| TcCLB.510973.9 | hypothetical protein, conserved | 0,2761 | 0,102 |
| TcCLB.511511.130 | hypothetical protein, conserved | 0,2759 | 0,01804 |
| TcCLB.510377.100 | mucin-associated surface protein (MASP), putative | 0,2759 | 0,753 |
| TcCLB.508011.30 | mucin TcMUCII, putative | 0,2756 | 0,7038 |
| TcCLB.509733.80 | hypothetical protein, conserved | 0,2754 | 0,107 |
| TcCLB.511143.6 | protein kinase, putative (fragment) | 0,2754 | 0,3948 |
| TcCLB.511681.10 | hypothetical protein | 0,2754 | 0,5518 |
| TcCLB.511395.60 | serine/threonine protein kinase, putative | 0,2753 | 0,01106 |
| TcCLB.508445.10 | rab11B GTPase, putative | 0,2753 | 0,0466 |
| TcCLB.509073.80 | hypothetical protein, conserved | 0,2752 | 0,0648 |
| TcCLB.503841.70 | aspartate aminotransferase, putative | 0,2751 | 0,3103 |
| TcCLB.511173.430 | mucin-associated surface protein (MASP), putative | 0,2751 | 0,4967 |
| TcCLB.504151.69 | hypothetical protein, conserved | 0,2748 | 0,04178 |
| TcCLB.511517.60 | C-1-tetrahydrofolate synthase, cytoplasmic, putative | 0,2746 | 0,04334 |
| TcCLB.507733.10 | trans-sialidase (pseudogene), putative | 0,2746 | 0,3027 |

| | | | |
|---|---|---|---|
| TcCLB.511821.100 | hypothetical protein, conserved | 0,2745 | 0,02831 |
| TcCLB.508061.154 | trans-sialidase, putative (fragment) | 0,2744 | 0,2143 |
| TcCLB.510119.40 | hypothetical protein, conserved | 0,2743 | 0,2851 |
| TcCLB.509317.10 | hypothetical protein, conserved | 0,2742 | 0,03502 |
| TcCLB.508815.90 | hypothetical protein, conserved | 0,274 | 0,04766 |
| TcCLB.511807.150 | hypothetical protein, conserved | 0,2737 | 0,02867 |
| TcCLB.503687.20 | glyceraldehyde-3-phosphate dehydrogenase, putative | 0,2737 | 0,06093 |
| TcCLB.504035.110 | hypothetical protein | 0,2737 | 0,07003 |
| TcCLB.508071.60 | mucin-associated surface protein (MASP), putative | 0,2736 | 0,2081 |
| TcCLB.506769.60 | mucin-associated surface protein (MASP, pseudogene), pu | 0,2736 | 0,5741 |
| TcCLB.510013.300 | mucin TcMUCII, putative | 0,2736 | 0,7352 |
| TcCLB.432997.10 | trans-sialidase, putative | 0,2733 | 0,4518 |
| TcCLB.508941.10 | retrotransposon hot spot protein (RHS, pseudogene), putat | 0,2733 | 0,609 |
| TcCLB.510643.160 | AGP2beta-2, putative | 0,2732 | 0,01975 |
| TcCLB.507095.30 | trans-sialidase, putative (fragment) | 0,2732 | 0,4947 |
| TcCLB.503571.10 | flagellum-adhesion glycoprotein, putative | 0,2729 | 0,01613 |
| TcCLB.504153.280 | hypothetical protein, conserved | 0,2729 | 0,04425 |
| TcCLB.511675.10 | trans-sialidase (pseudogene), putative | 0,2729 | 0,0948 |
| TcCLB.506855.170 | hypothetical protein, conserved | 0,2728 | 0,1647 |
| TcCLB.511001.70 | phosphatidylinositol-4-phosphate 5-kinase-like, putative | 0,2727 | 0,01208 |
| TcCLB.401661.10 | trans-sialidase (pseudogene), putative | 0,2727 | 0,06876 |
| TcCLB.510741.110 | hypothetical protein, conserved | 0,2727 | 0,08539 |
| TcCLB.509643.60 | hypothetical protein, conserved | 0,2727 | 0,1096 |
| TcCLB.508059.4 | hypothetical protein, conserved | 0,2727 | 0,1642 |
| TcCLB.511281.10 | hypothetical protein, conserved | 0,2726 | 0,1397 |
| TcCLB.507091.30 | succinate dehydrogenase subunit | 0,2726 | 0,3321 |
| TcCLB.439307.19 | hypothetical protein, conserved | 0,2726 | 0,5578 |
| TcCLB.508799.250 | hypothetical protein, conserved | 0,2724 | 0,009309 |
| TcCLB.511801.60 | DEAD-box ATP-dependent RNA helicase, mitochondrial (M | 0,2723 | 0,00276 |
| TcCLB.510509.30 | hypothetical protein, conserved | 0,2721 | 0,1917 |
| TcCLB.505193.50 | hypothetical protein | 0,2721 | 0,4501 |
| TcCLB.506767.324 | mucin TcMUCII, putative | 0,2719 | 0,7177 |
| TcCLB.509937.170 | hypothetical protein | 0,2716 | 0,1327 |
| TcCLB.504209.10 | conserved protein, unknown function | 0,2716 | 0,1559 |
| TcCLB.510073.80 | hypothetical protein, conserved | 0,2714 | 0,01408 |
| TcCLB.506195.90 | adenylate kinase 2 | 0,2713 | 0,03926 |
| TcCLB.504159.10 | hypothetical protein, conserved | 0,2713 | 0,045 |
| TcCLB.504427.150 | oxidoreductase-like protein, putative | 0,2713 | 0,09766 |
| TcCLB.507959.350 | mucin-associated surface protein (MASP), putative | 0,2713 | 0,3058 |
| TcCLB.511211.30 | hypothetical protein | 0,2712 | 0,1293 |
| TcCLB.506883.130 | hypothetical protein, conserved (pseudogene) | 0,2709 | 0,5556 |
| TcCLB.506679.70 | eukaryotic translation initiation factor 6 (eIF-6), putative | 0,2706 | 0,006446 |
| TcCLB.508137.30 | ubiquitin-conjugating enzyme E2, putative | 0,2706 | 0,06795 |
| TcCLB.506409.50 | mucin TcMUCII, putative | 0,2706 | 0,5409 |
| TcCLB.510275.360 | mucin TcMUCII (pseudogene), putative | 0,2705 | 0,6908 |

| | | | |
|---|---|---|---|
| TcCLB.510409.30 | hypothetical protein, conserved | 0,2704 | 0,07911 |
| TcCLB.503479.49 | U6 snRNA-associated Sm-like protein LSm7p | 0,2704 | 0,1175 |
| TcCLB.507519.100 | hypothetical protein, conserved | 0,2704 | 0,3533 |
| TcCLB.506525.90 | hypothetical protein, conserved | 0,2703 | 0,06901 |
| TcCLB.508965.90 | hypothetical protein, conserved | 0,2702 | 0,01257 |
| TcCLB.509215.19 | hypothetical protein, conserved | 0,2702 | 0,1194 |
| TcCLB.511717.80 | DNA repair and recombination helicase protein PIF4, putat | 0,2702 | 0,1657 |
| TcCLB.510749.20 | hypothetical protein, conserved | 0,27 | 0,06988 |
| TcCLB.510661.70 | hypothetical protein, conserved | 0,27 | 0,08159 |
| TcCLB.506573.30 | tyrosyl or methionyl-tRNA synthetase, putative | 0,27 | 0,1464 |
| TcCLB.506859.110 | 3'a2rel-related protein, putative | 0,2698 | 0,2713 |
| TcCLB.506909.46 | hypothetical protein | 0,2697 | 0,5016 |
| TcCLB.507537.40 | metacaspase, putative | 0,2696 | 0,4267 |
| TcCLB.511625.160 | hypothetical protein | 0,2695 | 0,4583 |
| TcCLB.508083.5 | elongation factor 1-gamma (EF-1-gamma, pseudogene), p | 0,2695 | 0,5662 |
| TcCLB.506989.210 | U-box domain protein, putative | 0,2693 | 0,006209 |
| TcCLB.509753.60 | mucin-associated surface protein (MASP), putative | 0,2693 | 0,6601 |
| TcCLB.508175.189 | hypothetical protein, conserved | 0,2692 | 0,1234 |
| TcCLB.509293.10 | mucin TcMUCII, putative | 0,2692 | 0,3616 |
| TcCLB.408485.20 | retrotransposon hot spot protein (RHS, pseudogene), putat | 0,2687 | 0,316 |
| TcCLB.508789.160 | mucin TcMUCII, putative | 0,2686 | 0,7137 |
| TcCLB.508909.310 | hypothetical protein | 0,2685 | 0,2572 |
| TcCLB.511603.160 | mucin-associated surface protein (MASP), putative | 0,2684 | 0,5665 |
| TcCLB.506857.80 | kinesin, putative | 0,2683 | 0,02195 |
| TcCLB.507675.39 | hypothetical protein, conserved | 0,2683 | 0,09376 |
| TcCLB.503427.9 | trans-sialidase, putative (fragment) | 0,2683 | 0,4339 |
| TcCLB.504881.14 | hypothetical protein, conserved | 0,2681 | 0,349 |
| TcCLB.509437.100 | retrotransposon hot spot protein (RHS, pseudogene), putat | 0,268 | 0,2134 |
| TcCLB.510005.50 | dispersed gene family protein 1 (DGF-1), putative | 0,268 | 0,2554 |
| TcCLB.510359.80 | hypothetical protein | 0,268 | 0,2637 |
| TcCLB.506055.40 | dispersed gene family protein 1 (DGF-1), putative | 0,2678 | 0,2453 |
| TcCLB.506753.170 | ubiquinone biosynthesis protein-like protein, putative | 0,2677 | 0,06014 |
| TcCLB.503399.29 | hydrolase-like protein | 0,2677 | 0,2782 |
| TcCLB.511683.20 | hypothetical protein, conserved | 0,2676 | 0,08978 |
| TcCLB.508075.14 | hypothetical protein, conserved | 0,2675 | 0,07228 |
| TcCLB.503973.181 | surface protease GP63 (pseudogene), putative | 0,2675 | 0,6815 |
| TcCLB.425311.9 | hypothetical protein, conserved (fragment) | 0,2674 | 0,04271 |
| TcCLB.510567.10 | trans-sialidase, putative (fragment) | 0,2674 | 0,7654 |
| TcCLB.509799.100 | hypothetical protein, conserved | 0,2673 | 0,01872 |
| TcCLB.510759.130 | hypothetical protein, conserved | 0,2671 | 0,09702 |
| TcCLB.510833.20 | mucin-associated surface protein (MASP), putative | 0,2671 | 0,461 |
| TcCLB.510807.40 | mucin TcMUCII, putative | 0,2671 | 0,6851 |
| TcCLB.509693.140 | hypothetical protein, conserved | 0,267 | 0,016 |
| TcCLB.509721.20 | hypothetical protein, conserved | 0,2669 | 0,1085 |
| TcCLB.506525.70 | hypothetical protein, conserved | 0,2668 | 0,008508 |

| | | | |
|---|---|---|---|
| TcCLB.506753.70 | hypothetical protein, conserved | 0,2668 | 0,1614 |
| TcCLB.509179.140 | myo-inositol-1(or 4)-monophosphatase 1, putative | 0,2667 | 0,242 |
| TcCLB.506135.30 | hypothetical protein, conserved | 0,2666 | 0,08906 |
| TcCLB.507283.39 | trans-sialidase (pseudogene), putative | 0,2666 | 0,1076 |
| TcCLB.507873.34 | hypothetical protein, conserved | 0,2666 | 0,2965 |
| TcCLB.506559.110 | hypothetical protein | 0,2665 | 0,04456 |
| TcCLB.507649.70 | nucleolar RNA-binding protein, putative | 0,2663 | 0,0006132 |
| TcCLB.509769.110 | Elongator-like Protein 3b, putative | 0,2663 | 0,0117 |
| TcCLB.503809.40 | protein kinase, putative | 0,2662 | 0,1676 |
| TcCLB.508161.20 | trans-sialidase, Group V, putative | 0,2662 | 0,4999 |
| TcCLB.511819.14 | small nuclear ribonucleoprotein, putative | 0,2661 | 0,09912 |
| TcCLB.510307.110 | hypothetical protein, conserved (pseudogene) | 0,266 | 0,01799 |
| TcCLB.511217.20 | Present in the outer mitochondrial membrane proteome 4 | 0,266 | 0,05709 |
| TcCLB.506599.40 | hypothetical protein | 0,266 | 0,7007 |
| TcCLB.509419.40 | trans-sialidase, Group V, putative | 0,266 | 0,8672 |
| TcCLB.506885.390 | AMP deaminase, putative | 0,2659 | 0,02749 |
| TcCLB.508793.30 | palmitoyl acyltransferase 8, putative | 0,2658 | 0,272 |
| TcCLB.460111.10 | trans-sialidase, putative (fragment) | 0,2658 | 0,7353 |
| TcCLB.505071.130 | hypothetical protein, conserved | 0,2657 | 0,0545 |
| TcCLB.509965.270 | hypothetical protein, conserved | 0,2655 | 0,03499 |
| TcCLB.505939.50 | hypothetical protein | 0,2655 | 0,1019 |
| TcCLB.509833.10 | hypothetical protein, conserved | 0,2655 | 0,1893 |
| TcCLB.511665.110 | glycine dehydrogenase [decarboxylating] (pseudogene), pu | 0,2655 | 0,3152 |
| TcCLB.511903.110 | 40S ribosomal protein S8, putative | 0,2654 | 0,0006868 |
| TcCLB.509455.100 | hypothetical protein, conserved | 0,2654 | 0,02273 |
| TcCLB.510513.80 | bilobe protein, conserved | 0,2653 | 0,02903 |
| TcCLB.503531.40 | hypothetical protein, conserved | 0,2652 | 0,2234 |
| TcCLB.504737.10 | trans-sialidase (pseudogene), putative | 0,2651 | 0,5525 |
| TcCLB.510715.100 | mucin-associated surface protein (MASP, pseudogene), pu | 0,265 | 0,5683 |
| TcCLB.508265.110 | hypothetical protein, conserved | 0,2648 | 0,01361 |
| TcCLB.508081.70 | mucin-associated surface protein (MASP), putative | 0,2648 | 0,6269 |
| TcCLB.509085.20 | retrotransposon hot spot protein (RHS, pseudogene), putat | 0,2646 | 0,687 |
| TcCLB.506943.80 | hypothetical protein, conserved | 0,2644 | 0,006354 |
| TcCLB.506303.130 | hypothetical protein, conserved | 0,2643 | 0,02097 |
| TcCLB.510407.70 | hypothetical protein | 0,2642 | 0,07916 |
| TcCLB.509631.140 | flagellar attachment zone protein, putative (FAZ1) | 0,2641 | 0,005517 |
| TcCLB.472777.20 | hypothetical protein, conserved | 0,2641 | 0,0462 |
| TcCLB.509937.210 | hypothetical protein, conserved | 0,2639 | 0,03582 |
| TcCLB.507519.170 | hypothetical protein, conserved | 0,2638 | 0,08196 |
| TcCLB.509991.9 | protein kinase, putative | 0,2638 | 0,2348 |
| TcCLB.471403.9 | dispersed gene family protein 1 (DGF-1), putative (fragme | 0,2635 | 0,1364 |
| TcCLB.509683.50 | calmodulin, putative | 0,2632 | 0,1613 |
| TcCLB.510001.59 | hypothetical protein, conserved (fragment) | 0,263 | 0,01286 |
| TcCLB.503675.40 | hypothetical protein, conserved | 0,2629 | 0,07153 |
| TcCLB.509599.60 | hypothetical protein | 0,2628 | 0,07506 |

| | | | |
|---|---|---|---|
| TcCLB.510973.30 | 40S ribosomal protein S13, putative | 0,2626 | 0,0004636 |
| TcCLB.510879.50 | regulatory subunit of protein kinase a-like protein, putative | 0,2626 | 0,002694 |
| TcCLB.509767.220 | hypothetical protein, conserved | 0,2625 | 0,01514 |
| TcCLB.504635.20 | hypothetical protein, conserved | 0,2624 | 0,1364 |
| TcCLB.507897.2 | hypothetical protein, conserved | 0,2624 | 0,1468 |
| TcCLB.510697.130 | mucin-associated surface protein (MASP), putative | 0,2624 | 0,6223 |
| TcCLB.510359.260 | mitochondrial import inner membrane translocase subunit | 0,2622 | 0,08182 |
| TcCLB.508325.340 | glycine dehydrogenase [decarboxylating] (pseudogene), pu | 0,2622 | 0,08343 |
| TcCLB.506733.130 | hypothetical protein, conserved | 0,262 | 0,02379 |
| TcCLB.509669.75 | hypothetical protein | 0,262 | 0,1656 |
| TcCLB.511027.20 | hypothetical protein, conserved | 0,2618 | 0,09514 |
| TcCLB.503973.320 | mucin TcMUCII, putative | 0,2616 | 0,7457 |
| TcCLB.511735.40 | hypothetical protein, conserved | 0,2614 | 0,2565 |
| TcCLB.511173.64 | mucin-associated surface protein (MASP), putative | 0,2613 | 0,7972 |
| TcCLB.509461.60 | mitochondrial RNA binding protein, putative | 0,2612 | 0,005054 |
| TcCLB.509505.20 | hypothetical protein, conserved | 0,2612 | 0,04554 |
| TcCLB.504213.50 | hypothetical protein, conserved | 0,2611 | 0,2218 |
| TcCLB.509937.20 | hypothetical protein, conserved | 0,2606 | 0,002881 |
| TcCLB.508723.59 | hypothetical protein, conserved | 0,2604 | 0,1029 |
| TcCLB.510921.30 | mucin-associated surface protein (MASP), putative | 0,2602 | 0,5383 |
| TcCLB.507959.140 | mucin TcMUCII, putative | 0,2602 | 0,8251 |
| TcCLB.507053.60 | hypothetical protein, conserved | 0,2599 | 0,02414 |
| TcCLB.506559.290 | hypothetical protein, conserved | 0,2597 | 0,02659 |
| TcCLB.507711.250 | hypothetical protein, conserved | 0,2597 | 0,02879 |
| TcCLB.511115.30 | hypothetical protein, conserved | 0,2596 | 0,05187 |
| TcCLB.507041.100 | ATP-dependent protease binding subunit (pseudogene), pu | 0,2596 | 0,08462 |
| TcCLB.510659.110 | hypothetical protein, conserved | 0,2595 | 0,04308 |
| TcCLB.507035.20 | retrotransposon hot spot protein (RHS, pseudogene), putat | 0,2595 | 0,06361 |
| TcCLB.506667.110 | trans-sialidase (pseudogene), putative | 0,2594 | 0,4177 |
| TcCLB.506339.40 | mucin-associated surface protein (MASP), putative | 0,259 | 0,5998 |
| TcCLB.508465.130 | hypothetical protein, conserved | 0,2587 | 0,03056 |
| TcCLB.479721.10 | hypothetical protein, conserved (pseudogene) | 0,2584 | 0,6654 |
| TcCLB.509419.90 | trans-sialidase, Group V, putative | 0,2583 | 0,8011 |
| TcCLB.508637.66 | hypothetical protein, conserved | 0,2582 | 0,09921 |
| TcCLB.509157.170 | trans-sialidase, Group VIII, putative | 0,2581 | 0,7477 |
| TcCLB.510531.10 | hypothetical protein, conserved | 0,2578 | 0,01136 |
| TcCLB.506435.190 | hypothetical protein, conserved | 0,2578 | 0,1977 |
| TcCLB.509211.39 | hypothetical protein, conserved (fragment) | 0,2577 | 0,09824 |
| TcCLB.509965.20 | hypothetical protein | 0,2576 | 0,1457 |
| TcCLB.508175.264 | hypothetical protein, conserved (pseudogene) | 0,2576 | 0,2438 |
| TcCLB.506227.200 | hypothetical protein | 0,2574 | 0,06842 |
| TcCLB.506245.270 | mucin-associated surface protein (MASP), putative | 0,2574 | 0,5213 |
| TcCLB.510025.180 | mucin-associated surface protein (MASP), putative | 0,2574 | 0,5284 |
| TcCLB.510647.10 | hypothetical protein, conserved | 0,2572 | 0,1933 |
| TcCLB.423707.10 | trans-sialidase, putative (fragment) | 0,2572 | 0,652 |

| | | | |
|---|---|---|---|
| TcCLB.509733.95 | hypothetical protein | 0,2571 | 0,05694 |
| TcCLB.511649.30 | hypothetical protein, conserved | 0,2566 | 0,1012 |
| TcCLB.508519.20 | retrotransposon hot spot protein (RHS, pseudogene), putat | 0,2565 | 0,08039 |
| TcCLB.506599.330 | mucin-associated surface protein (MASP), putative | 0,2565 | 0,4696 |
| TcCLB.510241.70 | heat shock protein, putative | 0,2564 | 0,1184 |
| TcCLB.508389.160 | trans-sialidase (pseudogene), putative | 0,2564 | 0,4626 |
| TcCLB.509971.90 | trans-sialidase, putative (fragment) | 0,2563 | 0,5982 |
| TcCLB.508999.127 | mucin-associated surface protein (MASP, pseudogene), pu | 0,2563 | 0,6015 |
| TcCLB.510661.110 | Present in the outer mitochondrial membrane proteome 7 | 0,2562 | 0,02634 |
| TcCLB.510729.60 | hypothetical protein, conserved | 0,256 | 0,2808 |
| TcCLB.508515.30 | RNA-binding protein 38, putative (RBP38) | 0,2559 | 0,001136 |
| TcCLB.503685.20 | hypothetical protein, conserved | 0,2559 | 0,17 |
| TcCLB.511345.30 | trans-sialidase (pseudogene), putative | 0,2559 | 0,2515 |
| TcCLB.506247.180 | hypothetical protein, conserved | 0,2558 | 0,01646 |
| TcCLB.444777.20 | hypothetical protein, conserved | 0,2558 | 0,07484 |
| TcCLB.510999.19 | hypothetical protein, conserved | 0,2558 | 0,1877 |
| TcCLB.508527.20 | retrotransposon hot spot protein (RHS, pseudogene), putat | 0,2557 | 0,01187 |
| TcCLB.508637.10 | trans-sialidase, Group III, putative | 0,2557 | 0,5744 |
| TcCLB.508029.50 | kinesin-like protein, putative | 0,2556 | 0,176 |
| TcCLB.506961.40 | retrotransposon hot spot protein (RHS, pseudogene), putat | 0,2555 | 0,0344 |
| TcCLB.506559.50 | hypothetical protein, conserved | 0,2553 | 0,04449 |
| TcCLB.506493.100 | hypothetical protein, conserved | 0,2553 | 0,1227 |
| TcCLB.509999.10 | hypothetical protein, conserved | 0,2551 | 0,1138 |
| TcCLB.508193.15 | trans-sialidase (pseudogene), putative | 0,255 | 0,02815 |
| TcCLB.506579.40 | hypothetical protein, conserved | 0,2548 | 0,001576 |
| TcCLB.511849.50 | surface protease GP63 (pseudogene), putative | 0,2548 | 0,2158 |
| TcCLB.442383.9 | 40S ribosomal protein SA, putative | 0,2547 | 0,01082 |
| TcCLB.510719.230 | hypothetical protein, conserved | 0,2547 | 0,02894 |
| TcCLB.503823.90 | hypothetical protein, conserved | 0,2547 | 0,04917 |
| TcCLB.508245.40 | mucin-associated surface protein (MASP), putative | 0,2547 | 0,5272 |
| TcCLB.506941.139 | zinc finger protein, putative | 0,2546 | 0,03654 |
| TcCLB.506393.90 | dispersed gene family protein 1 (DGF-1), putative | 0,2546 | 0,113 |
| TcCLB.510885.100 | hypothetical protein, conserved | 0,2545 | 0,1067 |
| TcCLB.508521.70 | mucin-associated surface protein (MASP, pseudogene), pu | 0,2545 | 0,3671 |
| TcCLB.511871.20 | hypothetical protein, conserved | 0,2543 | 0,09672 |
| TcCLB.509429.10 | trans-sialidase (pseudogene), putative | 0,2543 | 0,2225 |
| TcCLB.511089.19 | mucin-associated surface protein (MASP), putative | 0,2542 | 0,7086 |
| TcCLB.506477.14 | hypothetical protein, conserved | 0,254 | 0,1368 |
| TcCLB.507681.40 | hypothetical protein | 0,2539 | 0,06199 |
| TcCLB.508175.30 | hypothetical protein, conserved | 0,2539 | 0,1592 |
| TcCLB.507915.5 | glutaminyl-tRNA synthetase, putative | 0,2539 | 0,626 |
| TcCLB.504587.10 | surface protease GP63 (pseudogene), putative | 0,2538 | 0,5562 |
| TcCLB.509999.20 | hypothetical protein, conserved | 0,2537 | 0,09408 |
| TcCLB.507551.9 | dispersed gene family protein 1 (DGF-1), putative | 0,2537 | 0,1032 |
| TcCLB.511733.70 | hypothetical protein, conserved | 0,2536 | 0,1916 |

| | | | |
|---|---|---|---|
| TcCLB.508397.79 | hypothetical protein, conserved | 0,2536 | 0,4563 |
| TcCLB.511577.170 | hypothetical protein, conserved (pseudogene) | 0,2535 | 0,09297 |
| TcCLB.507899.5 | trans-sialidase (pseudogene), putative | 0,2533 | 0,09291 |
| TcCLB.507467.104 | hypothetical protein, conserved | 0,2533 | 0,2851 |
| TcCLB.511267.20 | hypothetical protein, conserved | 0,2533 | 0,338 |
| TcCLB.506597.30 | mucin TcMUCII, putative | 0,2533 | 0,7161 |
| TcCLB.503409.10 | hypothetical protein, conserved | 0,2531 | 0,1739 |
| TcCLB.504069.4 | hypothetical protein, conserved (fragment) | 0,2529 | 0,1509 |
| TcCLB.510247.20 | casein kinase I, putative | 0,2529 | 0,4795 |
| TcCLB.509791.30 | hypothetical protein, conserved | 0,2527 | 0,02743 |
| TcCLB.506227.170 | hypothetical protein, conserved | 0,2526 | 0,1912 |
| TcCLB.511255.430 | trans-sialidase (pseudogene), putative | 0,2524 | 0,5393 |
| TcCLB.509777.64 | ADG1-like protein, putative | 0,2523 | 0,1887 |
| TcCLB.510761.44 | cyclophilin-like protein, putative | 0,2522 | 0,008382 |
| TcCLB.479883.10 | hypothetical protein, conserved | 0,2521 | 0,1858 |
| TcCLB.506267.10 | mucin-associated surface protein (MASP), putative | 0,2519 | 0,765 |
| TcCLB.510797.20 | hypothetical protein, conserved | 0,2517 | 0,2144 |
| TcCLB.503525.10 | hypothetical protein, conserved | 0,2516 | 0,07529 |
| TcCLB.511755.24 | hypothetical protein, conserved (pseudogene) | 0,2515 | 0,2594 |
| TcCLB.510465.24 | mucin-associated surface protein (MASP, pseudogene), pu | 0,2515 | 0,7651 |
| TcCLB.510359.330 | hypothetical protein, conserved | 0,2512 | 0,04599 |
| TcCLB.509919.9 | dispersed gene family protein 1 (DGF-1), putative (fragme | 0,2512 | 0,1725 |
| TcCLB.509047.14 | hypothetical protein, conserved | 0,2511 | 0,1418 |
| TcCLB.506129.80 | retrotransposon hot spot protein (RHS, pseudogene), putat | 0,251 | 0,1418 |
| TcCLB.503913.29 | hypothetical protein, conserved | 0,251 | 0,3865 |
| TcCLB.508433.210 | mucin TcMUCII (pseudogene), putative | 0,2509 | 0,6334 |
| TcCLB.507517.40 | hypothetical protein, conserved | 0,2507 | 0,0196 |
| TcCLB.504817.20 | hypothetical protein, conserved | 0,2507 | 0,09529 |
| TcCLB.508837.170 | hypothetical protein, conserved | 0,2506 | 0,03239 |
| TcCLB.508277.100 | cAMP specific phosphodiesterase, putative | 0,2506 | 0,1319 |
| TcCLB.508197.39 | 60S ribosomal protein L11, putative | 0,2503 | 0,06323 |
| TcCLB.510377.370 | mucin TcMUCII, putative | 0,2503 | 0,6106 |
| TcCLB.510265.80 | hypothetical protein, conserved | 0,2502 | 0,04108 |
| TcCLB.509455.40 | hypothetical protein, conserved | 0,2502 | 0,08247 |
| TcCLB.504131.60 | hypothetical protein, conserved | 0,2501 | 0,06003 |
| TcCLB.506685.30 | thymidylate kinase, putative | 0,2501 | 0,139 |
| TcCLB.507637.170 | surface protease GP63, putative | 0,2501 | 0,4106 |
| TcCLB.511289.70 | Mitochondrial ADP/ATP carrier protein 5, putative (MCP5) | 0,25 | 0,004926 |
| TcCLB.508213.49 | hypothetical protein, conserved | 0,25 | 0,277 |
| TcCLB.509717.60 | metallo-peptidase, Clan MA(E) Family M41 | 0,2497 | 0,02427 |
| TcCLB.504085.50 | ATP-dependent DEAD/H RNA helicase, putative | 0,2497 | 0,05511 |
| TcCLB.506045.11 | retrotransposon hot spot protein (RHS, pseudogene), putat | 0,2497 | 0,2219 |
| TcCLB.503397.10 | retrotransposon hot spot protein (RHS, pseudogene), putat | 0,2496 | 0,04848 |
| TcCLB.510391.30 | mucin TcMUCII, putative | 0,2496 | 0,6098 |
| TcCLB.511697.40 | hypothetical protein, conserved (pseudogene) | 0,2494 | 0,2154 |

| | | | |
|---|---|---|---|
| TcCLB.511367.170 | vacuolar protein sorting-associated protein 35, putative | 0,2493 | 0,004168 |
| TcCLB.511307.3 | membrane transporter protein, putative (fragment) | 0,2492 | 0,2004 |
| TcCLB.510611.30 | kinesin, putative | 0,2491 | 0,02491 |
| TcCLB.511797.110 | trans-sialidase, Group V, putative | 0,249 | 0,3843 |
| TcCLB.510355.200 | hypothetical protein, conserved | 0,2489 | 0,1486 |
| TcCLB.510689.80 | hypothetical protein, conserved | 0,2488 | 0,2115 |
| TcCLB.506955.244 | mucin-associated surface protein (MASP), putative | 0,2488 | 0,5471 |
| TcCLB.506529.400 | hypothetical protein, conserved | 0,2487 | 0,1516 |
| TcCLB.511211.240 | hypothetical protein, conserved | 0,2487 | 0,1608 |
| TcCLB.508429.10 | hypothetical protein | 0,2487 | 0,5892 |
| TcCLB.509977.61 | surface protease GP63 (pseudogene), putative | 0,2486 | 0,3202 |
| TcCLB.504153.340 | hypothetical protein, conserved | 0,2482 | 0,04479 |
| TcCLB.509979.281 | elongation factor 1-gamma (EF-1-gamma, pseudogene), p | 0,2481 | 0,3819 |
| TcCLB.508999.134 | mucin TcMUCII, putative | 0,248 | 0,6511 |
| TcCLB.504017.10 | Nascent polypeptide-associated complex subunit beta, put | 0,2478 | 0,03016 |
| TcCLB.506649.60 | hypothetical protein, conserved | 0,2477 | 0,04007 |
| TcCLB.503995.10 | Present in the outer mitochondrial membrane proteome 1 | 0,2476 | 0,1611 |
| TcCLB.510693.260 | trans-sialidase (pseudogene), putative | 0,2476 | 0,596 |
| TcCLB.510099.50 | hypothetical protein, conserved | 0,2475 | 0,08465 |
| TcCLB.504087.10 | hypothetical protein, conserved | 0,2475 | 0,5078 |
| TcCLB.510363.80 | mucin-associated surface protein (MASP), putative | 0,2475 | 0,7462 |
| TcCLB.510285.60 | ubiquinone biosynthesis methyltransferase, putative | 0,2474 | 0,03744 |
| TcCLB.510431.140 | dual-specificity protein kinase, putative | 0,2473 | 0,02455 |
| TcCLB.504137.50 | hypothetical protein, conserved | 0,2472 | 0,1558 |
| TcCLB.506559.420 | dolichyl-P-Man:GDP-Man1GlcNAc2-PP-dolichyl alpha-1,3-n | 0,2471 | 0,001976 |
| TcCLB.511253.31 | kinesin-like protein, putative (fragment) | 0,2471 | 0,6453 |
| TcCLB.510877.190 | hypothetical protein, conserved | 0,247 | 0,3746 |
| TcCLB.509231.26 | hypothetical protein, conserved | 0,2468 | 0,07584 |
| TcCLB.509749.10 | Biotin--acetyl-CoA-carboxylase ligase, putative | 0,2468 | 0,08195 |
| TcCLB.509777.10 | hypothetical protein | 0,2468 | 0,4767 |
| TcCLB.509897.124 | mucin-associated surface protein (MASP, pseudogene), pu | 0,2467 | 0,4222 |
| TcCLB.507993.280 | hypothetical protein, conserved | 0,2466 | 0,0253 |
| TcCLB.504175.20 | hypothetical protein, conserved | 0,2466 | 0,1836 |
| TcCLB.510129.20 | surface glycoprotein, putative | 0,2465 | 0,04258 |
| TcCLB.507007.76 | receptor protein kinase, putative | 0,2465 | 0,2118 |
| TcCLB.506525.120 | hypothetical protein, conserved | 0,2465 | 0,3742 |
| TcCLB.508523.80 | cAMP response protein 1 (CARP1) | 0,2461 | 0,1455 |
| TcCLB.510239.70 | mucin-associated surface protein (MASP), putative | 0,246 | 0,5489 |
| TcCLB.506309.140 | mucin-associated surface protein (MASP), putative | 0,2458 | 0,8266 |
| TcCLB.510291.20 | hypothetical protein, conserved | 0,2457 | 0,1058 |
| TcCLB.508153.490 | peptidyl-prolyl cis-trans isomerase NIMA-interacting 4, put | 0,2457 | 0,3091 |
| TcCLB.503789.20 | hypothetical protein, conserved | 0,2456 | 0,03384 |
| TcCLB.506203.25 | hypothetical protein | 0,2456 | 0,06537 |
| TcCLB.506939.180 | mucin-associated surface protein (MASP), putative | 0,2456 | 0,4563 |
| TcCLB.508247.50 | mucin-associated surface protein (MASP, pseudogene), pu | 0,2456 | 0,6318 |

| | | | |
|---|---|---|---|
| TcCLB.506727.110 | hypothetical protein, conserved | 0,2455 | 0,06178 |
| TcCLB.508781.70 | palmitoyl acyltransferase 3, putative | 0,2454 | 0,07549 |
| TcCLB.509595.44 | hypothetical protein, conserved | 0,2454 | 0,1751 |
| TcCLB.511771.171 | hypothetical protein | 0,2454 | 0,5365 |
| TcCLB.510889.190 | hypothetical protein, conserved | 0,2453 | 0,07782 |
| TcCLB.503999.10 | squalene monooxygenase, putative | 0,245 | 0,01399 |
| TcCLB.421057.10 | adrenodoxin precursor, putative | 0,2449 | 0,2927 |
| TcCLB.510165.110 | retrotransposon hot spot (RHS) protein, putative | 0,2448 | 0,1856 |
| TcCLB.504105.87 | hypothetical protein, conserved | 0,2445 | 0,2051 |
| TcCLB.506581.4 | hypothetical protein, conserved | 0,2445 | 0,5329 |
| TcCLB.507803.30 | hypothetical protein | 0,2444 | 0,07218 |
| TcCLB.508385.40 | dual specificity protein phosphatase, putative | 0,2444 | 0,137 |
| TcCLB.508673.20 | UDP-Gal or UDP-GlcNAc-dependent glycosyltransferase, pu | 0,2444 | 0,2441 |
| TcCLB.506223.80 | Metalloprotease M41 FtsH, putative | 0,2443 | 0,02247 |
| TcCLB.509429.270 | proteasome regulatory ATPase subunit 3, putative | 0,2442 | 0,01527 |
| TcCLB.508529.40 | trans-sialidase (pseudogene), putative | 0,2442 | 0,1078 |
| TcCLB.508187.20 | hypothetical protein (pseudogene) | 0,2442 | 0,3957 |
| TcCLB.503827.5 | mucin-associated surface protein (MASP), putative | 0,2442 | 0,781 |
| TcCLB.506705.50 | hypothetical protein | 0,2441 | 0,02614 |
| TcCLB.510101.160 | hypothetical protein, conserved | 0,2441 | 0,1765 |
| TcCLB.511587.75 | hypothetical protein | 0,2438 | 0,5879 |
| TcCLB.511569.14 | dispersed gene family protein 1 (DGF-1), putative | 0,2437 | 0,2782 |
| TcCLB.503669.10 | hypothetical protein, conserved | 0,2436 | 0,02949 |
| TcCLB.509221.50 | trans-sialidase (pseudogene), putative | 0,2436 | 0,2831 |
| TcCLB.506871.40 | hypothetical protein, conserved | 0,2434 | 0,06523 |
| TcCLB.510943.206 | trans-sialidase, putative | 0,2434 | 0,8227 |
| TcCLB.508881.140 | hypothetical protein | 0,2433 | 0,03547 |
| TcCLB.508573.39 | dispersed gene family protein 1 (DGF-1), putative | 0,2433 | 0,2756 |
| TcCLB.506157.40 | trans-sialidase, putative | 0,2432 | 0,5283 |
| TcCLB.507709.50 | 60S ribosomal protein L6, putative | 0,2428 | 0,004541 |
| TcCLB.507483.50 | calmodulin (CALA2) | 0,2427 | 0,3425 |
| TcCLB.476725.9 | dispersed gene family protein 1 (DGF-1), putative (fragme | 0,2427 | 0,4398 |
| TcCLB.508047.70 | mucin-associated surface protein (MASP), putative | 0,2427 | 0,469 |
| TcCLB.508441.50 | hypothetical protein, conserved | 0,2426 | 0,327 |
| TcCLB.507049.140 | hypothetical protein, conserved | 0,2425 | 0,03021 |
| TcCLB.505807.220 | ribosomal RNA adenine dimethylase family protein, putati | 0,2424 | 0,04767 |
| TcCLB.506635.40 | hypothetical protein, conserved | 0,2424 | 0,09126 |
| TcCLB.506417.20 | hypothetical protein, conserved | 0,2424 | 0,3951 |
| TcCLB.506777.60 | NGG1 interacting factor 3, putative | 0,2422 | 0,2183 |
| TcCLB.510289.44 | hypothetical protein, conserved | 0,2422 | 0,2754 |
| TcCLB.506599.400 | trans-sialidase (pseudogene), putative | 0,2422 | 0,7362 |
| TcCLB.511285.100 | hypothetical protein | 0,2421 | 0,05645 |
| TcCLB.509805.140 | hypothetical protein | 0,2421 | 0,09147 |
| TcCLB.510165.20 | DNA repair protein, putative | 0,2421 | 0,09413 |
| TcCLB.511071.171 | 60S ribosomal protein L12, putative | 0,2419 | 0,001256 |

| | | | |
|---|---|---|---|
| TcCLB.509365.50 | hypothetical protein, conserved | 0,2419 | 0,6253 |
| TcCLB.507081.80 | hypothetical protein, conserved | 0,2418 | 0,05437 |
| TcCLB.506619.49 | ras-related protein Rab21, putative | 0,2418 | 0,075 |
| TcCLB.511181.20 | hypothetical protein, conserved | 0,2417 | 0,07569 |
| TcCLB.507521.44 | hypothetical protein, conserved | 0,2416 | 0,3326 |
| TcCLB.510879.210 | hypothetical protein, conserved | 0,2415 | 0,1281 |
| TcCLB.507071.180 | mucin-associated surface protein (MASP), putative | 0,2414 | 0,4916 |
| TcCLB.507791.20 | N-acetyltransferase complex ARD1 subunit (pseudogene), | 0,2413 | 0,5838 |
| TcCLB.506945.10 | hypothetical protein, conserved | 0,241 | 0,2166 |
| TcCLB.506949.30 | hypothetical protein, conserved | 0,2409 | 0,5008 |
| TcCLB.509835.10 | hypothetical protein, conserved | 0,2408 | 0,5685 |
| TcCLB.505949.169 | mucin-associated surface protein (MASP), putative (fragm | 0,2408 | 0,7373 |
| TcCLB.511391.110 | histone-lysine N-methyltransferase, putative (DOT1A) | 0,2406 | 0,03114 |
| TcCLB.506413.40 | hypothetical protein, conserved | 0,2405 | 0,224 |
| TcCLB.509819.30 | septum formation inhibitor, Maf-like protein, putative | 0,2402 | 0,2123 |
| TcCLB.505965.50 | phosphonopyruvate decarboxylase, putative | 0,2401 | 0,0506 |
| TcCLB.510621.30 | mucin TcMUCII, putative | 0,2401 | 0,7334 |
| TcCLB.506727.50 | hypothetical protein, conserved | 0,2399 | 0,04873 |
| TcCLB.508487.20 | dispersed gene family protein 1 (DGF-1, pseudogene), puta | 0,2399 | 0,3491 |
| TcCLB.504061.10 | retrotransposon hot spot protein (RHS, pseudogene), putat | 0,2398 | 0,0887 |
| TcCLB.510089.160 | hypothetical protein, conserved (pseudogene) | 0,2395 | 0,0147 |
| TcCLB.508461.60 | synaptojanin (N-terminal domain), putative | 0,2393 | 0,06929 |
| TcCLB.508461.100 | Bardet-Biedl syndrome 1 protein homolog (BBS1-like prote | 0,2393 | 0,06973 |
| TcCLB.507809.110 | hypothetical protein, conserved | 0,239 | 0,04097 |
| TcCLB.511305.40 | hypothetical protein, conserved | 0,239 | 0,2086 |
| TcCLB.511595.30 | mucin-associated surface protein (MASP), putative | 0,2389 | 0,6017 |
| TcCLB.508707.100 | hypothetical protein, conserved | 0,2387 | 0,0259 |
| TcCLB.503757.30 | hypothetical protein, conserved | 0,2386 | 0,1499 |
| TcCLB.506635.10 | hypothetical protein (pseudogene) | 0,2386 | 0,2814 |
| TcCLB.511173.309 | mucin-associated surface protein (MASP, pseudogene), pu | 0,2385 | 0,6935 |
| TcCLB.508175.230 | hypothetical protein, conserved | 0,2383 | 0,06995 |
| TcCLB.510543.80 | hypothetical protein | 0,2382 | 0,5133 |
| TcCLB.508445.90 | hypothetical protein, conserved | 0,2381 | 0,08312 |
| TcCLB.508181.124 | hypothetical protein, conserved | 0,2379 | 0,08668 |
| TcCLB.511135.20 | hypothetical protein, conserved | 0,2378 | 0,005842 |
| TcCLB.511907.320 | hypothetical protein, conserved | 0,2378 | 0,07059 |
| TcCLB.509903.10 | dispersed gene family protein 1 (DGF-1, pseudogene), puta | 0,2378 | 0,1413 |
| TcCLB.511245.240 | serine/threonine protein kinase, putative | 0,2378 | 0,2706 |
| TcCLB.508479.500 | retrotransposon hot spot (RHS) protein, putative | 0,2377 | 0,0181 |
| TcCLB.509251.30 | trans-sialidase (pseudogene), putative | 0,2377 | 0,3885 |
| TcCLB.506155.20 | hypothetical protein, conserved | 0,2376 | 0,0502 |
| TcCLB.511729.9 | calmodulin, putative | 0,2375 | 0,2756 |
| TcCLB.506767.10 | hypothetical protein, conserved (pseudogene) | 0,2374 | 0,7212 |
| TcCLB.511661.60 | hypothetical protein, conserved | 0,2373 | 0,1094 |
| TcCLB.509769.130 | DNA polymerase zeta catalytic subunit, putative | 0,2372 | 0,004931 |

| | | | |
|---|---|---|---|
| TcCLB.503899.50 | RNA-editing complex protein MP100, putative | 0,2372 | 0,02714 |
| TcCLB.433061.10 | hypothetical protein | 0,2372 | 0,7219 |
| TcCLB.506721.20 | radial spoke protein 3, putative (RSP3) | 0,2371 | 0,04915 |
| TcCLB.506047.10 | hypothetical protein, conserved (pseudogene) | 0,237 | 0,4399 |
| TcCLB.509993.31 | mucin-associated surface protein (MASP, pseudogene), pu | 0,2369 | 0,5482 |
| TcCLB.506799.130 | mucin-associated surface protein (MASP), putative | 0,2368 | 0,3505 |
| TcCLB.508789.120 | trans-sialidase, Group VI, putative | 0,2368 | 0,4554 |
| TcCLB.508165.185 | mucin TcMUCII (pseudogene), putative | 0,2368 | 0,8348 |
| TcCLB.509913.10 | 3-methylcrotonyl-CoA carboxylase, putative | 0,2367 | 0,05903 |
| TcCLB.405179.10 | dynein heavy chain, putative (fragment) | 0,2367 | 0,1918 |
| TcCLB.503491.29 | surface protease GP63 (pseudogene), putative | 0,2366 | 0,7236 |
| TcCLB.508325.90 | hypothetical protein | 0,2365 | 0,4709 |
| TcCLB.510213.80 | mucin TcMUCII, putative | 0,2365 | 0,7163 |
| TcCLB.510279.240 | hypothetical protein | 0,2364 | 0,4653 |
| TcCLB.508411.40 | hypothetical protein, conserved | 0,2363 | 0,0993 |
| TcCLB.506247.380 | hypothetical protein, conserved | 0,2362 | 0,09435 |
| TcCLB.508523.10 | retrotransposon hot spot protein (RHS, pseudogene), putat | 0,2362 | 0,209 |
| TcCLB.509047.10 | hypothetical protein | 0,2361 | 0,07059 |
| TcCLB.509025.29 | DNA photolyase, putative | 0,236 | 0,2288 |
| TcCLB.509701.10 | trifunctional enzyme alpha subunit, mitochondrial precurs | 0,2359 | 0,02497 |
| TcCLB.507927.90 | hypothetical protein, conserved | 0,2359 | 0,2462 |
| TcCLB.507623.110 | surface protease GP63, putative | 0,2358 | 0,09513 |
| TcCLB.511021.70 | ABC transporter of the mitochondrion, putative | 0,2357 | 0,07915 |
| TcCLB.510065.60 | hypothetical protein, conserved | 0,2357 | 0,223 |
| TcCLB.506133.130 | trans-sialidase, Group V, putative | 0,2356 | 0,5414 |
| TcCLB.503717.130 | trans-sialidase (pseudogene), putative | 0,2356 | 0,578 |
| TcCLB.509229.40 | hypothetical protein, conserved | 0,2355 | 0,04032 |
| TcCLB.507179.20 | dispersed gene family protein 1 (DGF-1), putative | 0,2355 | 0,1867 |
| TcCLB.508739.60 | histone H4, putative | 0,2355 | 0,2454 |
| TcCLB.504099.40 | retrotransposon hot spot protein (RHS, pseudogene), putat | 0,2355 | 0,4766 |
| TcCLB.508461.146 | hypothetical protein, conserved | 0,2354 | 0,08531 |
| TcCLB.506513.120 | hypothetical protein, conserved | 0,2353 | 0,08002 |
| TcCLB.508737.130 | hypothetical protein, conserved | 0,2353 | 0,247 |
| TcCLB.508457.30 | lipoic acid containing carrier protein, putative (GCVH) | 0,2352 | 0,07619 |
| TcCLB.511571.49 | nucleoside phosphatase, putative | 0,2351 | 0,0109 |
| TcCLB.507515.80 | hypothetical protein, conserved | 0,2351 | 0,135 |
| TcCLB.503981.40 | DnaJ chaperone protein, putative | 0,235 | 0,007047 |
| TcCLB.506591.20 | U-box domain protein, putative | 0,235 | 0,02268 |
| TcCLB.508777.70 | hypothetical protein, conserved | 0,235 | 0,1507 |
| TcCLB.506839.30 | hypothetical protein, conserved | 0,2349 | 0,08761 |
| TcCLB.506067.300 | trans-sialidase, Group V, putative | 0,2349 | 0,3743 |
| TcCLB.510433.20 | surface protein TolT, putative | 0,2348 | 0,1468 |
| TcCLB.511545.110 | hypothetical protein | 0,2347 | 0,2103 |
| TcCLB.473313.10 | mucin-associated surface protein (MASP), putative (fragm | 0,2345 | 0,3626 |
| TcCLB.510015.10 | hypothetical protein | 0,2345 | 0,6863 |

| | | | |
|---|---|---|---|
| TcCLB.506223.110 | hypothetical protein, conserved | 0,2344 | 0,04178 |
| TcCLB.506275.40 | hypothetical protein, conserved | 0,2343 | 0,07161 |
| TcCLB.508235.30 | mucin TcMUCI, putative | 0,2343 | 0,3528 |
| TcCLB.510101.10 | protein transport protein Sec31, putative (pseudogene) | 0,2342 | 0,1531 |
| TcCLB.511019.60 | hypothetical protein, conserved (pseudogene) | 0,2342 | 0,1574 |
| TcCLB.511213.15 | mucin-associated surface protein (MASP), putative | 0,2342 | 0,5384 |
| TcCLB.503891.54 | nucleolar RNA-binding protein, putative | 0,2341 | 0,1945 |
| TcCLB.508207.170 | hypothetical protein | 0,234 | 0,07897 |
| TcCLB.510101.230 | ATPase (pseudogene), putative | 0,2339 | 0,01247 |
| TcCLB.510187.310 | hypothetical protein, conserved | 0,2338 | 0,04917 |
| TcCLB.508069.5 | trans-sialidase, putative (fragment) | 0,2338 | 0,5673 |
| TcCLB.503855.20 | spermidine synthase, putative | 0,2337 | 0,0304 |
| TcCLB.509721.70 | hypothetical protein, conserved | 0,2337 | 0,1531 |
| TcCLB.506445.10 | actin-like protein, putative | 0,2337 | 0,2225 |
| TcCLB.511015.10 | mannosyl-oligosaccharide glucosidase, putative | 0,2336 | 0,226 |
| TcCLB.508211.40 | hypothetical protein, conserved | 0,2335 | 0,07213 |
| TcCLB.511887.45 | hypothetical protein, conserved (pseudogene) | 0,2335 | 0,6873 |
| TcCLB.511509.30 | protein kinase, putative | 0,2334 | 0,09087 |
| TcCLB.509753.310 | mucin-associated surface protein (MASP), putative | 0,2334 | 0,5641 |
| TcCLB.503823.160 | ubiquinone biosynthesis methyltransferase, putative | 0,2333 | 0,07978 |
| TcCLB.509979.110 | hypothetical protein, conserved | 0,2333 | 0,2051 |
| TcCLB.508479.5 | glycine dehydrogenase (pseudogene), putative | 0,233 | 0,5378 |
| TcCLB.511771.199 | hypothetical protein | 0,233 | 0,6969 |
| TcCLB.510663.30 | 60S ribosomal subunit protein L31, putative | 0,2328 | 0,007546 |
| TcCLB.511903.40 | 3-hydroxy-3-methylglutaryl-CoA synthase, putative (HMGS | 0,2328 | 0,008888 |
| TcCLB.510407.40 | hypothetical protein, conserved | 0,2328 | 0,09448 |
| TcCLB.510749.60 | hypothetical protein, conserved | 0,2328 | 0,1545 |
| TcCLB.510609.90 | hypothetical protein, conserved | 0,2325 | 0,0486 |
| TcCLB.507513.40 | hypothetical protein, conserved | 0,2325 | 0,2575 |
| TcCLB.509167.160 | hypothetical protein | 0,2322 | 0,04773 |
| TcCLB.507213.10 | hypothetical protein, conserved | 0,2321 | 0,04503 |
| TcCLB.508851.50 | hypothetical protein, conserved | 0,2321 | 0,09905 |
| TcCLB.508081.60 | hypothetical protein | 0,2321 | 0,645 |
| TcCLB.507993.230 | hypothetical protein, conserved (pseudogene) | 0,232 | 0,8206 |
| TcCLB.507035.70 | hypothetical protein | 0,2319 | 0,3763 |
| TcCLB.506605.194 | hypothetical protein, conserved | 0,2318 | 0,1766 |
| TcCLB.508411.80 | hypothetical protein, conserved | 0,2316 | 0,1458 |
| TcCLB.503805.20 | hypothetical protein, conserved | 0,2316 | 0,1853 |
| TcCLB.508433.200 | trans-sialidase (pseudogene), putative | 0,2315 | 0,545 |
| TcCLB.506563.90 | calpain-like cysteine peptidase (pseudogene), putative | 0,2313 | 0,2724 |
| TcCLB.482971.9 | dispersed gene family protein 1 (DGF-1), putative (fragme | 0,2313 | 0,6349 |
| TcCLB.511879.9 | hypothetical protein, conserved | 0,2311 | 0,05115 |
| TcCLB.510823.70 | 60S acidic ribosomal protein, putative | 0,2309 | 0,05512 |
| TcCLB.511451.80 | trans-sialidase, Group V, putative | 0,2307 | 0,2575 |
| TcCLB.507061.20 | ribosomal protein L2, putative | 0,2306 | 0,05626 |

| | | | |
|---|---|---|---|
| TcCLB.510303.290 | acyl-CoA dehydrogenase, putative | 0,2306 | 0,09538 |
| TcCLB.510595.40 | hypothetical protein, conserved | 0,2305 | 0,1589 |
| TcCLB.503643.3 | 60S ribosomal protein L4, putative (fragment) | 0,2302 | 0,008832 |
| TcCLB.506885.420 | hypothetical protein | 0,2301 | 0,2615 |
| TcCLB.510655.159 | hypothetical protein, conserved | 0,2301 | 0,4107 |
| TcCLB.508129.9 | adenosine monophosphate deaminase, putative (fragment | 0,2299 | 0,0953 |
| TcCLB.508917.40 | hypothetical protein, conserved | 0,2297 | 0,08887 |
| TcCLB.506435.100 | hypothetical protein, conserved | 0,2297 | 0,3066 |
| TcCLB.503829.40 | hypothetical protein, conserved | 0,2294 | 0,04146 |
| TcCLB.508711.20 | hypothetical protein, conserved | 0,2294 | 0,05869 |
| TcCLB.511389.70 | hypothetical protein, conserved | 0,2294 | 0,06891 |
| TcCLB.506471.50 | trans-sialidase (pseudogene), putative | 0,2293 | 0,1251 |
| TcCLB.511507.50 | mannosyltransferase, putative | 0,2293 | 0,1352 |
| TcCLB.511417.60 | hypothetical protein, conserved | 0,2293 | 0,1696 |
| TcCLB.509695.60 | hypothetical protein, conserved | 0,2293 | 0,1702 |
| TcCLB.510089.219 | diacylglycerol acyltransferase, putative | 0,2293 | 0,2329 |
| TcCLB.511663.19 | dispersed gene family protein 1 (DGF-1), putative (fragme | 0,2292 | 0,3818 |
| TcCLB.508347.20 | O-6 methyl-guanine alkyl transferase, putative | 0,2291 | 0,2337 |
| TcCLB.503973.255 | hypothetical protein, conserved (pseudogene) | 0,2291 | 0,679 |
| TcCLB.509867.70 | trans-sialidase, Group VI, putative | 0,229 | 0,6343 |
| TcCLB.506389.39 | elongation factor 1-gamma (EF-1-gamma, pseudogene), p | 0,2289 | 0,0722 |
| TcCLB.506389.79 | calmodulin, putative | 0,2289 | 0,7875 |
| TcCLB.506167.9 | hypothetical protein, conserved | 0,2288 | 0,02015 |
| TcCLB.507017.70 | ribosomal RNA adenine dimethylase family protein, putati | 0,2288 | 0,06127 |
| TcCLB.511071.80 | hypothetical protein, conserved | 0,2288 | 0,2799 |
| TcCLB.510119.20 | elongation factor 1-alpha, putative | 0,2287 | 0,006895 |
| TcCLB.508525.10 | hypothetical protein, conserved | 0,2287 | 0,1916 |
| TcCLB.505989.100 | hypothetical protein, conserved | 0,2286 | 0,2048 |
| TcCLB.509161.100 | hypothetical protein, conserved | 0,2285 | 0,05822 |
| TcCLB.509901.170 | apurinic/apyrimidinic endonuclease (TCAP) | 0,2285 | 0,1034 |
| TcCLB.507957.90 | mucin TcMUCII, putative | 0,2285 | 0,6584 |
| TcCLB.508909.30 | hypothetical protein, conserved | 0,2283 | 0,09166 |
| TcCLB.507831.70 | pyruvate dehydrogenase E1 component alpha subunit, puta | 0,2282 | 0,09024 |
| TcCLB.508957.40 | hypothetical protein, conserved (pseudogene) | 0,2281 | 0,02714 |
| TcCLB.503747.20 | hypothetical protein | 0,2281 | 0,3122 |
| TcCLB.507523.70 | trans-sialidase, Group V, putative | 0,2281 | 0,5653 |
| TcCLB.510799.70 | hypothetical protein, conserved | 0,228 | 0,08711 |
| TcCLB.509877.50 | glycosomal membrane protein, putative | 0,228 | 0,2425 |
| TcCLB.506679.130 | hypothetical protein, conserved | 0,2279 | 0,1119 |
| TcCLB.506529.530 | Present in the outer mitochondrial membrane proteome 1: | 0,2279 | 0,2899 |
| TcCLB.509905.190 | elongation factor 1-gamma (EF-1-gamma, pseudogene), p | 0,2279 | 0,5184 |
| TcCLB.506125.30 | phosphoglycerate kinase, putative | 0,2278 | 0,1692 |
| TcCLB.511499.69 | hypothetical protein (fragment) | 0,2277 | 0,3183 |
| TcCLB.506211.190 | hypothetical protein, conserved | 0,2276 | 0,03235 |
| TcCLB.509717.100 | membrane-bound acid phosphatase 2, putative | 0,2276 | 0,05259 |

| | | | |
|---|---|---|---|
| TcCLB.511179.130 | hypothetical protein, conserved | 0,2276 | 0,1109 |
| TcCLB.509543.50 | DNAK protein, putative | 0,2275 | 0,02498 |
| TcCLB.508397.20 | hypothetical protein, conserved | 0,2275 | 0,2734 |
| TcCLB.508221.650 | trans-sialidase, Group V, putative | 0,2274 | 0,5244 |
| TcCLB.507927.100 | hypothetical protein | 0,2273 | 0,02442 |
| TcCLB.507049.179 | hypothetical protein, conserved | 0,2273 | 0,06231 |
| TcCLB.506301.30 | hypothetical protein, conserved | 0,2272 | 0,01999 |
| TcCLB.506337.10 | vacuolar protein sorting-associated protein-like, putative | 0,2272 | 0,08586 |
| TcCLB.508813.70 | hypothetical protein, conserved | 0,2272 | 0,1016 |
| TcCLB.508165.14 | trans-sialidase, putative (fragment) | 0,2272 | 0,7374 |
| TcCLB.504033.160 | hypothetical protein | 0,227 | 0,107 |
| TcCLB.506821.10 | hypothetical protein | 0,2268 | 0,1591 |
| TcCLB.510759.100 | hypothetical protein, conserved | 0,2267 | 0,03779 |
| TcCLB.508557.20 | amino acid transporter, putative | 0,2267 | 0,3075 |
| TcCLB.506883.69 | hypothetical protein, conserved | 0,2266 | 0,3785 |
| TcCLB.504159.50 | aquaporin-like protein, putative | 0,2264 | 0,06297 |
| TcCLB.510303.310 | arginine N-methyltransferase, type III, putative (PRMT7) | 0,2262 | 0,03136 |
| TcCLB.507941.30 | PrimPol-like protein 1, putative (ppl1) | 0,2262 | 0,1245 |
| TcCLB.508235.20 | mucin TcMUCIII, putative | 0,2262 | 0,4902 |
| TcCLB.511505.10 | RNA pseudouridylate synthase, putative | 0,2261 | 0,1065 |
| TcCLB.510761.14 | 60S ribosomal protein L26, putative | 0,226 | 0,01108 |
| TcCLB.511661.100 | protein kinase, putative | 0,226 | 0,0147 |
| TcCLB.506297.160 | 40S ribosomal protein S5, putative | 0,226 | 0,02768 |
| TcCLB.510307.200 | hypothetical protein, conserved | 0,2259 | 0,1952 |
| TcCLB.507911.60 | retrotransposon hot spot protein (RHS, pseudogene), putat | 0,2259 | 0,6306 |
| TcCLB.511407.40 | hypothetical protein, conserved | 0,2258 | 0,1294 |
| TcCLB.509551.120 | hypothetical protein | 0,2256 | 0,2861 |
| TcCLB.503951.10 | retrotransposon hot spot protein (RHS, pseudogene), putat | 0,2256 | 0,5617 |
| TcCLB.508745.30 | trans-sialidase, Group V, putative | 0,2255 | 0,6197 |
| TcCLB.510157.20 | hypothetical protein, conserved | 0,2254 | 0,6842 |
| TcCLB.510575.130 | chloride channel protein, putative | 0,2253 | 0,02877 |
| TcCLB.508173.30 | trans-sialidase (pseudogene), putative | 0,2253 | 0,2132 |
| TcCLB.511751.22 | hypothetical protein, conserved | 0,2253 | 0,3735 |
| TcCLB.511417.20 | retrotransposon hot spot protein (RHS, pseudogene), putat | 0,2252 | 0,4946 |
| TcCLB.509317.29 | hypothetical protein | 0,2251 | 0,4297 |
| TcCLB.509259.21 | elongation factor 1-gamma (EF-1-gamma, pseudogene), p | 0,2251 | 0,7178 |
| TcCLB.506797.90 | short chain dehydrogenase/reductase, putative | 0,225 | 0,1058 |
| TcCLB.491081.10 | retrotransposon hot spot protein (RHS, pseudogene), putat | 0,225 | 0,3397 |
| TcCLB.508165.330 | trans-sialidase (pseudogene), putative | 0,2246 | 0,5142 |
| TcCLB.509207.20 | histone H2A, putative | 0,2243 | 0,7278 |
| TcCLB.506405.10 | phosphomannomutase-like protein, putative | 0,2242 | 0,01968 |
| TcCLB.510645.30 | 3,2-trans-enoyl-CoA isomerase, mitochondrial precursor, p | 0,2242 | 0,04575 |
| TcCLB.506179.30 | hypothetical protein, conserved | 0,2242 | 0,1684 |
| TcCLB.510579.119 | hypothetical protein, conserved | 0,2242 | 0,1884 |
| TcCLB.506407.100 | hypothetical protein, conserved | 0,2241 | 0,03549 |

| | | | |
|---|---|---|---|
| TcCLB.504147.70 | hypothetical protein, conserved | 0,2241 | 0,1207 |
| TcCLB.507669.130 | hypothetical protein, conserved | 0,2241 | 0,1294 |
| TcCLB.505997.100 | retrotransposon hot spot protein (RHS, pseudogene), putat | 0,224 | 0,2378 |
| TcCLB.509165.30 | AGP2beta-2, putative | 0,2239 | 0,03908 |
| TcCLB.511837.29 | iron-sulfur cluster assembly protein, putative | 0,2239 | 0,1552 |
| TcCLB.510299.70 | hypothetical protein, conserved | 0,2239 | 0,2304 |
| TcCLB.506131.30 | mucin-associated surface protein (MASP), putative | 0,2239 | 0,7366 |
| TcCLB.507711.300 | Archaic translocase of outer membrane 11 kDa subunit, pu | 0,2237 | 0,2292 |
| TcCLB.506335.60 | mucin TcMUCII (pseudogene), putative | 0,2237 | 0,418 |
| TcCLB.508165.300 | mucin-associated surface protein (MASP), putative | 0,2237 | 0,5309 |
| TcCLB.503665.21 | elongation factor 1-gamma (EF-1-gamma, pseudogene), p | 0,2237 | 0,5314 |
| TcCLB.508479.370 | hypothetical protein, conserved | 0,2236 | 0,1224 |
| TcCLB.507073.20 | hypothetical protein, conserved | 0,2236 | 0,3588 |
| TcCLB.508717.30 | phosphoribosylpyrophosphate synthetase, putative | 0,2235 | 0,02822 |
| TcCLB.508461.19 | hypothetical protein, conserved | 0,2235 | 0,1696 |
| TcCLB.507777.10 | hypothetical protein, conserved | 0,2235 | 0,254 |
| TcCLB.511031.10 | hypothetical protein, conserved | 0,2235 | 0,3881 |
| TcCLB.508165.34 | mucin TcMUC (pseudogene), putative | 0,2235 | 0,6589 |
| TcCLB.508241.110 | hypothetical protein, conserved | 0,2234 | 0,1051 |
| TcCLB.507677.70 | hypothetical protein, conserved | 0,223 | 0,1684 |
| TcCLB.511885.10 | hypothetical protein, conserved (pseudogene) | 0,2228 | 0,02405 |
| TcCLB.511253.20 | kinesin-like protein, putative | 0,2228 | 0,303 |
| TcCLB.507063.120 | hypothetical protein, conserved | 0,2226 | 0,2932 |
| TcCLB.510025.220 | mucin-associated surface protein (MASP), putative | 0,2226 | 0,6525 |
| TcCLB.511889.50 | retrotransposon hot spot protein (RHS, pseudogene), putat | 0,2225 | 0,1224 |
| TcCLB.510307.250 | trans-sialidase (pseudogene), putative | 0,2225 | 0,2303 |
| TcCLB.511577.40 | 35 kDa cyclophilin, putative (CYP35) | 0,2225 | 0,2903 |
| TcCLB.503703.50 | copper-transporting ATPase-like protein, putative | 0,2223 | 0,03815 |
| TcCLB.508777.40 | leucine-rich repeat protein, putative | 0,2222 | 0,0869 |
| TcCLB.508071.30 | beta galactofuranosyl glycosyltransferase, putative | 0,2221 | 0,5175 |
| TcCLB.511537.60 | hypothetical protein, conserved | 0,2218 | 0,07577 |
| TcCLB.509599.70 | hypothetical protein, conserved | 0,2218 | 0,119 |
| TcCLB.509163.100 | hypothetical protein, conserved (pseudogene) | 0,2218 | 0,2838 |
| TcCLB.508977.20 | mucin-associated surface protein (MASP), putative | 0,2218 | 0,6821 |
| TcCLB.504203.50 | hypothetical protein, conserved | 0,2217 | 0,245 |
| TcCLB.506727.150 | hypothetical protein, conserved | 0,2217 | 0,2722 |
| TcCLB.509055.60 | hypothetical protein, conserved | 0,2216 | 0,3727 |
| TcCLB.506973.14 | mucin TcMUCII (pseudogene), putative | 0,2215 | 0,4775 |
| TcCLB.508539.95 | surface protease GP63 (pseudogene), putative | 0,2215 | 0,7044 |
| TcCLB.506009.90 | hypothetical protein, conserved | 0,2213 | 0,07147 |
| TcCLB.506727.30 | hypothetical protein, conserved | 0,2212 | 0,1916 |
| TcCLB.457101.20 | dynein heavy chain, cytosolic, putative | 0,2209 | 0,04165 |
| TcCLB.506855.360 | hypothetical protein, conserved | 0,2207 | 0,02427 |
| TcCLB.509967.30 | aspartyl-tRNA synthetase, putative | 0,2207 | 0,1217 |
| TcCLB.510373.100 | hypothetical protein, conserved (pseudogene) | 0,2204 | 0,6976 |

| | | | |
|---|---|---|---|
| TcCLB.503611.20 | ribosomal protein L24, putative | 0,2202 | 0,03079 |
| TcCLB.510517.90 | hypothetical protein, conserved | 0,2202 | 0,181 |
| TcCLB.510091.80 | pyruvate dehydrogenase E1 beta subunit, putative | 0,2201 | 0,03077 |
| TcCLB.510885.120 | hypothetical protein, conserved | 0,2201 | 0,1138 |
| TcCLB.511689.19 | hypothetical protein, conserved (fragment) | 0,22 | 0,1744 |
| TcCLB.504035.90 | translation initiation factor IF-2, putative | 0,2199 | 0,04702 |
| TcCLB.508535.30 | hypothetical protein, conserved | 0,2199 | 0,1288 |
| TcCLB.509877.89 | hypothetical protein, conserved | 0,2199 | 0,3089 |
| TcCLB.510519.90 | hypothetical protein, conserved | 0,2199 | 0,4821 |
| TcCLB.510309.50 | 60S acidic ribosomal protein P2, putative | 0,2197 | 0,04621 |
| TcCLB.510879.160 | tyrosine phosphatase, putative | 0,2195 | 0,01711 |
| TcCLB.509715.90 | UDP-glucose/GDP-mannose dehydrogenase family, NAD bi | 0,2195 | 0,2003 |
| TcCLB.510659.74 | hypothetical protein, conserved | 0,2195 | 0,4765 |
| TcCLB.509937.100 | ubiquitin/ribosomal protein S27a, putative | 0,2194 | 0,002199 |
| TcCLB.510719.120 | hypothetical protein, conserved | 0,2194 | 0,06867 |
| TcCLB.506727.20 | hypothetical protein, conserved | 0,2193 | 0,05626 |
| TcCLB.509567.60 | hypothetical protein | 0,2192 | 0,1072 |
| TcCLB.509509.30 | 6-phosphofructo-2-kinase/fructose-2,6-biphosphatase, put | 0,2192 | 0,1134 |
| TcCLB.505025.40 | mucin-associated surface protein (MASP), putative | 0,2192 | 0,6561 |
| TcCLB.507485.34 | DNA-directed RNA polymerase III subunit, putative | 0,2192 | 0,7343 |
| TcCLB.506575.40 | hypothetical protein | 0,2191 | 0,1037 |
| TcCLB.507357.35 | hypothetical protein | 0,219 | 0,6708 |
| TcCLB.511001.190 | vacuolar ATP synthase subunit d, putative | 0,2188 | 0,09618 |
| TcCLB.463323.20 | retrotransposon hot spot protein (RHS, pseudogene), putat | 0,2188 | 0,3399 |
| TcCLB.508111.10 | trans-sialidase (pseudogene), putative | 0,2188 | 0,5134 |
| TcCLB.506599.170 | mucin-associated surface protein (MASP), putative | 0,2187 | 0,5047 |
| TcCLB.510535.20 | hypothetical protein, conserved | 0,2186 | 0,04134 |
| TcCLB.506265.114 | Enriched in surface-labeled proteome protein 11, putative | 0,2186 | 0,1134 |
| TcCLB.509317.90 | hypothetical protein, conserved | 0,2186 | 0,1791 |
| TcCLB.510519.10 | hypothetical protein | 0,2183 | 0,05081 |
| TcCLB.511127.160 | integral membrane transport protein, putative | 0,2183 | 0,6039 |
| TcCLB.410199.7 | hypothetical protein | 0,2182 | 0,6418 |
| TcCLB.507021.30 | hypothetical protein, conserved | 0,218 | 0,2844 |
| TcCLB.506705.40 | hypothetical protein, conserved | 0,2179 | 0,1325 |
| TcCLB.503827.50 | trans-sialidase (pseudogene), putative | 0,2178 | 0,6892 |
| TcCLB.504133.45 | hypothetical protein, conserved | 0,2177 | 0,5308 |
| TcCLB.511385.120 | hypothetical protein, conserved | 0,2177 | 0,5331 |
| TcCLB.506933.60 | mitochondrial RNA binding protein, putative | 0,2176 | 0,01299 |
| TcCLB.508781.40 | hypothetical protein, conserved | 0,2175 | 0,1369 |
| TcCLB.510513.40 | hypothetical protein, conserved | 0,2175 | 0,1794 |
| TcCLB.507759.10 | trans-sialidase, Group VIII, putative | 0,2175 | 0,6586 |
| TcCLB.511629.30 | hypothetical protein, conserved | 0,2174 | 0,2101 |
| TcCLB.511295.40 | tRNA pseudouridine synthase A-like protein, putative | 0,2173 | 0,08628 |
| TcCLB.510769.10 | hypothetical protein, conserved | 0,2173 | 0,1386 |
| TcCLB.511859.140 | hypothetical protein | 0,2172 | 0,2984 |

| | | | |
|---|---|---|---|
| TcCLB.510191.10 | mucin-associated surface protein (MASP), putative | 0,2172 | 0,6557 |
| TcCLB.508451.10 | trans-sialidase (pseudogene), putative | 0,217 | 0,3702 |
| TcCLB.503817.10 | trans-sialidase (pseudogene), putative | 0,2166 | 0,6242 |
| TcCLB.511761.89 | hypothetical protein, conserved (fragment) | 0,2164 | 0,3587 |
| TcCLB.504249.100 | trans-sialidase (pseudogene), putative | 0,2163 | 0,3984 |
| TcCLB.506959.64 | 3-keto-dihydrosphingosine reductase | 0,2162 | 0,13 |
| TcCLB.510047.120 | mucin-associated surface protein (MASP), putative | 0,2162 | 0,7029 |
| TcCLB.511301.160 | trans-sialidase, Group II, putative | 0,2161 | 0,2446 |
| TcCLB.504575.10 | hypothetical protein, conserved | 0,2161 | 0,5134 |
| TcCLB.410589.19 | hypothetical protein, conserved | 0,2158 | 0,4026 |
| TcCLB.508163.250 | trans-sialidase, Group VI, putative | 0,2158 | 0,5386 |
| TcCLB.506959.140 | hypothetical protein, conserved | 0,2157 | 0,2857 |
| TcCLB.509507.40 | hypothetical protein, conserved | 0,2155 | 0,1922 |
| TcCLB.509569.94 | hypothetical protein, conserved | 0,2154 | 0,2801 |
| TcCLB.506857.30 | hypothetical protein, conserved | 0,2153 | 0,1661 |
| TcCLB.507817.70 | hypothetical protein, conserved | 0,2152 | 0,1951 |
| TcCLB.507513.90 | RNA-editing complex protein MP100, putative | 0,215 | 0,03604 |
| TcCLB.508859.50 | hypothetical protein, conserved | 0,2148 | 0,3055 |
| TcCLB.508125.130 | mucin TcMUCII, putative | 0,2148 | 0,6851 |
| TcCLB.507809.80 | hypothetical protein, conserved | 0,2147 | 0,003774 |
| TcCLB.504229.20 | glycine dehydrogenase [decarboxylating] (pseudogene), pu | 0,2142 | 0,05269 |
| TcCLB.509733.190 | hypothetical protein, conserved | 0,2142 | 0,1223 |
| TcCLB.511421.190 | ESCO acetyltransferase domain-containing protein | 0,214 | 0,1786 |
| TcCLB.503773.20 | hypothetical protein, conserved | 0,214 | 0,4529 |
| TcCLB.506857.50 | hypothetical protein, conserved | 0,2139 | 0,06323 |
| TcCLB.510943.180 | hypothetical protein, conserved | 0,2138 | 0,1916 |
| TcCLB.508873.520 | mucin-associated surface protein (MASP, pseudogene), pu | 0,2138 | 0,6692 |
| TcCLB.511283.260 | hypothetical protein, conserved | 0,2137 | 0,01504 |
| TcCLB.507787.70 | hypothetical protein, conserved | 0,2136 | 0,2595 |
| TcCLB.510421.280 | hypothetical protein, conserved | 0,2135 | 0,1259 |
| TcCLB.508647.30 | hypothetical protein, conserved | 0,2135 | 0,1422 |
| TcCLB.504787.80 | mucin TcMUCII, putative | 0,2135 | 0,5622 |
| TcCLB.503683.20 | hypothetical protein, conserved | 0,2134 | 0,0552 |
| TcCLB.509671.84 | hypothetical protein, conserved | 0,2134 | 0,1692 |
| TcCLB.406115.10 | trans-sialidase, putative (fragment) | 0,2134 | 0,5665 |
| TcCLB.509537.90 | hypothetical protein, conserved | 0,2129 | 0,03461 |
| TcCLB.510659.36 | hypothetical protein, conserved | 0,2129 | 0,1672 |
| TcCLB.509133.10 | trans-sialidase (pseudogene), putative | 0,2128 | 0,4899 |
| TcCLB.504129.40 | hypothetical protein, conserved | 0,2126 | 0,2041 |
| TcCLB.506587.70 | hypothetical protein, conserved | 0,2125 | 0,088 |
| TcCLB.506635.130 | hypothetical protein, conserved | 0,2124 | 0,08085 |
| TcCLB.506529.480 | hypothetical protein, conserved | 0,2124 | 0,1143 |
| TcCLB.511269.4 | Ras-related protein Rab5A, putative (RAB5A) | 0,2124 | 0,1254 |
| TcCLB.509911.110 | hypothetical protein, conserved | 0,2123 | 0,2596 |
| TcCLB.504153.129 | hypothetical protein, conserved | 0,2122 | 0,1073 |

| | | | |
|---|---|---|---|
| TcCLB.506559.270 | sarcoplasmic reticulum glycoprotein, putative | 0,2121 | 0,03283 |
| TcCLB.506021.10 | retrotransposon hot spot protein (RHS, pseudogene), putat | 0,212 | 0,1594 |
| TcCLB.509733.20 | hypothetical protein | 0,2119 | 0,4583 |
| TcCLB.504929.10 | casein kinase, delta isoform, putative | 0,2119 | 0,7343 |
| TcCLB.510423.60 | hypothetical protein, conserved | 0,2118 | 0,09365 |
| TcCLB.508541.230 | casein kinase, putative | 0,2118 | 0,5166 |
| TcCLB.506425.130 | hypothetical protein, conserved | 0,2117 | 0,1874 |
| TcCLB.510279.70 | hypothetical protein | 0,2117 | 0,6988 |
| TcCLB.503885.20 | hypothetical protein, conserved | 0,2116 | 0,05027 |
| TcCLB.506971.70 | mucin TcMUCII, putative | 0,2114 | 0,7943 |
| TcCLB.503939.10 | acetyltransferase, putative | 0,2112 | 0,1765 |
| TcCLB.510207.40 | mucin-associated surface protein (MASP), putative | 0,2112 | 0,561 |
| TcCLB.507061.10 | hypothetical protein | 0,2111 | 0,3677 |
| TcCLB.508741.250 | hypothetical protein | 0,211 | 0,07511 |
| TcCLB.511507.20 | hypothetical protein, conserved | 0,211 | 0,1883 |
| TcCLB.511401.90 | trans-sialidase, Group V, putative | 0,211 | 0,3562 |
| TcCLB.507013.30 | hypothetical protein, conserved | 0,2109 | 0,325 |
| TcCLB.506445.30 | cation transporter, putative | 0,2105 | 0,0742 |
| TcCLB.506543.90 | cytochrome c oxidase assembly protein, putative | 0,2105 | 0,1715 |
| TcCLB.507509.70 | hypothetical protein, conserved | 0,2104 | 0,127 |
| TcCLB.509965.280 | hypothetical protein, conserved | 0,2102 | 0,2297 |
| TcCLB.506275.30 | hypothetical protein, conserved | 0,2102 | 0,3526 |
| TcCLB.511411.8 | hypothetical protein, conserved (fragment) | 0,2101 | 0,2679 |
| TcCLB.510275.90 | hypothetical protein | 0,2101 | 0,4962 |
| TcCLB.508647.290 | DNA repair and recombination protein RAD54, putative | 0,21 | 0,01646 |
| TcCLB.506783.20 | trans-sialidase (pseudogene), putative | 0,2099 | 0,4421 |
| TcCLB.503651.10 | hypothetical protein, conserved | 0,2097 | 0,09277 |
| TcCLB.510067.9 | hypothetical protein, conserved | 0,2096 | 0,1433 |
| TcCLB.507357.71 | hypothetical protein | 0,2096 | 0,7118 |
| TcCLB.506531.50 | metacaspase, putative | 0,2092 | 0,219 |
| TcCLB.509859.30 | hypothetical protein, conserved | 0,2091 | 0,114 |
| TcCLB.506977.110 | hypothetical protein, conserved (pseudogene) | 0,2091 | 0,1572 |
| TcCLB.503957.90 | dispersed gene family protein 1 (DGF-1, pseudogene), puta | 0,2089 | 0,3821 |
| TcCLB.506625.120 | hypothetical protein, conserved | 0,2088 | 0,08054 |
| TcCLB.509749.36 | ribosomal protein L29, putative | 0,2088 | 0,08994 |
| TcCLB.511603.80 | mucin-associated surface protein (MASP), putative | 0,2088 | 0,6453 |
| TcCLB.506887.20 | hypothetical protein, conserved | 0,2087 | 0,1392 |
| TcCLB.508185.10 | enoyl-CoA hydratase, mitochondrial precursor, putative | 0,2087 | 0,1394 |
| TcCLB.509777.70 | Calcium ATPase SERCA-like | 0,2085 | 0,006446 |
| TcCLB.503703.100 | hypothetical protein, conserved | 0,2084 | 0,1233 |
| TcCLB.506303.150 | hypothetical protein, conserved | 0,2084 | 0,1368 |
| TcCLB.509663.25 | hypothetical protein, conserved (pseudogene) | 0,2084 | 0,4853 |
| TcCLB.508153.420 | hypothetical protein, conserved | 0,2083 | 0,07081 |
| TcCLB.507513.100 | hypothetical protein, conserved | 0,2082 | 0,1005 |
| TcCLB.483009.9 | exonuclease, putative | 0,2081 | 0,1787 |

| | | | |
|---|---|---|---|
| TcCLB.507953.40 | mucin-associated surface protein (MASP), putative | 0,2081 | 0,7339 |
| TcCLB.508771.60 | retrotransposon hot spot protein (RHS, pseudogene), putat | 0,208 | 0,2084 |
| TcCLB.510889.251 | Present in the outer mitochondrial membrane proteome 1: | 0,208 | 0,3591 |
| TcCLB.511027.10 | chloride channel protein, putative | 0,2079 | 0,127 |
| TcCLB.510173.110 | hypothetical protein, conserved | 0,2078 | 0,1031 |
| TcCLB.506529.280 | hypothetical protein, conserved | 0,2077 | 0,1734 |
| TcCLB.506573.50 | hypothetical protein, conserved | 0,2077 | 0,2666 |
| TcCLB.506147.60 | 3-demethylubiquinone-9 3-methyltransferase, putative | 0,2076 | 0,1842 |
| TcCLB.507429.60 | hypothetical protein | 0,2075 | 0,5003 |
| TcCLB.507989.10 | hypothetical protein, conserved | 0,2072 | 0,1793 |
| TcCLB.511303.50 | hypothetical protein, conserved | 0,2072 | 0,3056 |
| TcCLB.505807.29 | hypothetical protein, conserved | 0,2072 | 0,5487 |
| TcCLB.508365.100 | surface protease GP63, putative | 0,2072 | 0,6642 |
| TcCLB.509727.20 | retrotransposon hot spot protein (RHS, pseudogene), putat | 0,2072 | 0,7184 |
| TcCLB.508233.70 | ATP-dependent protease binding subunit (pseudogene), pu | 0,2071 | 0,4266 |
| TcCLB.509429.70 | hypothetical protein | 0,207 | 0,08705 |
| TcCLB.447255.4 | ubiquitin, putative | 0,207 | 0,2496 |
| TcCLB.511353.60 | hypothetical protein, conserved | 0,207 | 0,2512 |
| TcCLB.506759.150 | hypothetical protein, conserved (pseudogene) | 0,207 | 0,6777 |
| TcCLB.503487.20 | hypothetical protein | 0,2069 | 0,09709 |
| TcCLB.509671.99 | hypothetical protein, conserved | 0,2069 | 0,3963 |
| TcCLB.511729.30 | hypothetical protein, conserved | 0,2068 | 0,196 |
| TcCLB.503437.31 | ATP-dependent DEAD/H RNA helicase (pseudogene), putat | 0,2066 | 0,6746 |
| TcCLB.504111.20 | metallo- peptidase, Clan MG, Family M24 | 0,2063 | 0,09396 |
| TcCLB.509399.170 | elongation factor 1-gamma (EF-1-gamma), putative | 0,2062 | 0,2128 |
| TcCLB.509777.4 | ribosomal protein L2, putative | 0,2062 | 0,2277 |
| TcCLB.509253.50 | recombination initiation protein NBS1, putative | 0,2061 | 0,04697 |
| TcCLB.506925.120 | eukaryotic initiation factor 5a, putative | 0,2061 | 0,08049 |
| TcCLB.506335.69 | mucin-associated surface protein (MASP, pseudogene), pu | 0,206 | 0,595 |
| TcCLB.510147.9 | hypothetical protein, conserved | 0,2057 | 0,4452 |
| TcCLB.509251.70 | mucin-associated surface protein (MASP, pseudogene), pu | 0,2056 | 0,3973 |
| TcCLB.510885.110 | hypothetical protein, conserved | 0,2055 | 0,1109 |
| TcCLB.504157.70 | hypothetical protein, conserved | 0,2054 | 0,06638 |
| TcCLB.510087.40 | hypothetical protein | 0,2053 | 0,6982 |
| TcCLB.511907.200 | cysteine peptidase inhibitor, putative | 0,2052 | 0,06282 |
| TcCLB.511593.51 | elongation factor 1-gamma (EF-1-gamma, pseudogene), p | 0,2051 | 0,4877 |
| TcCLB.511599.110 | mucin TcMUCII, putative | 0,2049 | 0,7251 |
| TcCLB.408547.5 | mucin TcMUCII, putative | 0,2049 | 0,7385 |
| TcCLB.509719.23 | hypothetical protein, conserved | 0,2048 | 0,3184 |
| TcCLB.510201.70 | hypothetical protein | 0,2048 | 0,652 |
| TcCLB.465323.10 | hypothetical protein (fragment) | 0,2048 | 0,7373 |
| TcCLB.508895.30 | Sarcalumenin, putative | 0,2047 | 0,03749 |
| TcCLB.506133.170 | mucin-associated surface protein (MASP, pseudogene), pu | 0,2047 | 0,573 |
| TcCLB.510013.50 | trans-sialidase, Group IV, putative | 0,2047 | 0,668 |
| TcCLB.511921.167 | hypothetical protein, conserved (pseudogene) | 0,2047 | 0,6839 |

| | | | |
|---|---|---|---|
| TcCLB.511553.100 | mucin-associated surface protein (MASP), putative | 0,2045 | 0,6251 |
| TcCLB.510949.10 | dual specificity protein phosphatase, putative | 0,2044 | 0,1444 |
| TcCLB.509769.100 | hypothetical protein, conserved | 0,2044 | 0,1589 |
| TcCLB.506751.70 | retrotransposon hot spot protein (RHS, pseudogene), putat | 0,2043 | 0,6104 |
| TcCLB.510879.120 | ribosomal protein L3, putative | 0,2041 | 0,003494 |
| TcCLB.506867.40 | surface protease GP63, putative | 0,2041 | 0,1374 |
| TcCLB.511861.14 | trans-sialidase (pseudogene), putative | 0,2039 | 0,3718 |
| TcCLB.505071.110 | hypothetical protein, conserved | 0,2037 | 0,08659 |
| TcCLB.509015.10 | copper-transporting ATPase, putative | 0,2037 | 0,5634 |
| TcCLB.510729.10 | AMP deaminase, putative | 0,2036 | 0,1082 |
| TcCLB.508831.63 | hypothetical protein | 0,2036 | 0,1511 |
| TcCLB.507711.340 | hypothetical protein, conserved | 0,2036 | 0,3006 |
| TcCLB.511491.120 | hypothetical protein, conserved | 0,2035 | 0,08592 |
| TcCLB.506955.240 | trans-sialidase, Group V, putative | 0,2035 | 0,4213 |
| TcCLB.503583.80 | dual specificity protein phosphatase or MAP kinase phosph | 0,2033 | 0,1683 |
| TcCLB.510309.60 | hypothetical protein, conserved | 0,2032 | 0,09837 |
| TcCLB.509769.140 | hypothetical protein, conserved | 0,2031 | 0,2456 |
| TcCLB.504069.56 | 60S ribosomal protein L37a, putative | 0,203 | 0,07305 |
| TcCLB.506573.40 | hypothetical protein, conserved | 0,203 | 0,1222 |
| TcCLB.507193.10 | dispersed gene family protein 1 (DGF-1), putative | 0,2029 | 0,481 |
| TcCLB.509647.70 | guanylate kinase, putative | 0,2027 | 0,1277 |
| TcCLB.507093.20 | hypothetical protein, conserved | 0,2027 | 0,2566 |
| TcCLB.511181.90 | peroxisome targeting signal 1 receptor, putative (PEX5) | 0,2026 | 0,07646 |
| TcCLB.508413.44 | 60S ribosomal protein L12, putative | 0,2025 | 0,01496 |
| TcCLB.506821.200 | hypothetical protein, conserved | 0,2025 | 0,1394 |
| TcCLB.507623.40 | hypothetical protein, conserved | 0,2024 | 0,2262 |
| TcCLB.506201.150 | hypothetical protein, conserved | 0,2023 | 0,4579 |
| TcCLB.510487.44 | elongation factor 1-gamma (EF-1-gamma, pseudogene), p | 0,2023 | 0,769 |
| TcCLB.509971.70 | hypothetical protein | 0,2022 | 0,6197 |
| TcCLB.509767.210 | vesicle transport v-SNARE 11, putative | 0,2021 | 0,2473 |
| TcCLB.510877.20 | hypothetical protein, conserved | 0,202 | 0,0495 |
| TcCLB.507867.100 | hypothetical protein | 0,2019 | 0,07575 |
| TcCLB.508853.10 | mucin-associated surface protein (MASP), putative | 0,2019 | 0,6101 |
| TcCLB.506637.80 | hypothetical protein | 0,2017 | 0,1278 |
| TcCLB.509205.70 | hypothetical protein, conserved | 0,2017 | 0,1962 |
| TcCLB.507089.70 | RNA-binding protein, putative | 0,2016 | 0,2809 |
| TcCLB.506567.77 | hypothetical protein, conserved (pseudogene) | 0,2016 | 0,5326 |
| TcCLB.507693.5 | tryptophanyl-tRNA synthetase, putative (fragment) | 0,2015 | 0,2889 |
| TcCLB.508895.20 | hypothetical protein, conserved | 0,2013 | 0,1354 |
| TcCLB.506829.20 | hypothetical protein, conserved | 0,2012 | 0,1 |
| TcCLB.510357.150 | hypothetical protein, conserved | 0,2012 | 0,2082 |
| TcCLB.506291.20 | hypothetical protein | 0,2012 | 0,7945 |
| TcCLB.509087.20 | dispersed gene family protein 1 (DGF-1), putative | 0,2011 | 0,3148 |
| TcCLB.506367.70 | hypothetical protein, conserved | 0,2009 | 0,2426 |
| TcCLB.416041.24 | retrotransposon hot spot protein (RHS, pseudogene), putat | 0,2009 | 0,3231 |

| | | | |
|---|---|---|---|
| TcCLB.506425.169 | Ras-related protein RAB2B, putative (fragment) (RAB2B) | 0,2007 | 0,3486 |
| TcCLB.508547.130 | NUDIX family hydrolase, putative | 0,2006 | 0,2063 |
| TcCLB.511649.120 | hypothetical protein, conserved | 0,2004 | 0,1952 |
| TcCLB.506291.31 | surface protease GP63 (pseudogene), putative | 0,2003 | 0,7242 |
| TcCLB.507491.130 | hypothetical protein, conserved | 0,2 | 0,211 |
| TcCLB.505807.20 | hypothetical protein, conserved | 0,1999 | 0,1525 |
| TcCLB.509725.50 | protein kinase, putative | 0,1998 | 0,6448 |
| TcCLB.509643.20 | hypothetical protein, conserved | 0,1995 | 0,1336 |
| TcCLB.511661.70 | hypothetical protein, conserved | 0,1995 | 0,2794 |
| TcCLB.507765.90 | hypothetical protein, conserved | 0,1992 | 0,1434 |
| TcCLB.508909.100 | hypothetical protein, conserved | 0,1992 | 0,1492 |
| TcCLB.504737.20 | mucin-associated surface protein (MASP, pseudogene), pu | 0,1992 | 0,6865 |
| TcCLB.507673.60 | hypothetical protein, conserved | 0,1991 | 0,5566 |
| TcCLB.510189.60 | trans-sialidase (pseudogene), putative | 0,199 | 0,5857 |
| TcCLB.510657.150 | hypothetical protein | 0,1989 | 0,1827 |
| TcCLB.509831.10 | cysteine desulfurase, putative | 0,1988 | 0,1893 |
| TcCLB.506727.60 | hypothetical protein, conserved | 0,1987 | 0,1124 |
| TcCLB.508851.150 | hypothetical protein, conserved | 0,1987 | 0,1807 |
| TcCLB.509985.10 | hypothetical protein, conserved (pseudogene) | 0,1987 | 0,2127 |
| TcCLB.511255.50 | mucin-associated surface protein (MASP), putative | 0,1987 | 0,6178 |
| TcCLB.510583.130 | mucin-associated surface protein (MASP), putative | 0,1986 | 0,7133 |
| TcCLB.511745.30 | E3 ubiquitin-protein ligase, putative | 0,1985 | 0,08697 |
| TcCLB.510445.40 | hypothetical protein | 0,1984 | 0,2476 |
| TcCLB.509747.30 | hypothetical protein, conserved | 0,1983 | 0,04122 |
| TcCLB.504255.30 | hypothetical protein, conserved | 0,1983 | 0,3948 |
| TcCLB.511693.60 | hypothetical protein, conserved | 0,1983 | 0,4302 |
| TcCLB.511825.66 | hypothetical protein, conserved | 0,1982 | 0,3851 |
| TcCLB.507093.310 | hypothetical protein, conserved | 0,1981 | 0,1435 |
| TcCLB.507603.100 | hypothetical protein, conserved | 0,198 | 0,2366 |
| TcCLB.506635.70 | hypothetical protein, conserved | 0,1979 | 0,09795 |
| TcCLB.508495.9 | mucin-associated surface protein (MASP, pseudogene), pu | 0,1979 | 0,7486 |
| TcCLB.510257.60 | 3-ketoacyl-CoA reductase, putative | 0,1978 | 0,1113 |
| TcCLB.506181.30 | hypothetical protein, conserved | 0,1978 | 0,4099 |
| TcCLB.511025.10 | dynein light intermediate chain, putative | 0,1977 | 0,1135 |
| TcCLB.509543.20 | hypothetical protein, conserved | 0,1977 | 0,1567 |
| TcCLB.504163.40 | fructose-bisphosphate aldolase, glycosomal, putative | 0,1976 | 0,1337 |
| TcCLB.508759.20 | trans-sialidase, Group V, putative | 0,1976 | 0,6608 |
| TcCLB.509297.30 | ADP-ribosylation factor, putative | 0,1976 | 0,7362 |
| TcCLB.509959.236 | N-acetyltransferase complex ARD1 subunit (pseudogene), | 0,1976 | 0,7869 |
| TcCLB.506999.170 | hypothetical protein | 0,1975 | 0,1492 |
| TcCLB.510477.30 | mucin TcMUCI, putative | 0,1974 | 0,7072 |
| TcCLB.507927.40 | hypothetical protein, conserved | 0,1973 | 0,05822 |
| TcCLB.430567.10 | trans-sialidase, putative (fragment) | 0,1973 | 0,4892 |
| TcCLB.506127.170 | eukaryotic translation initiation factor eIF-4E, putative | 0,1972 | 0,07597 |
| TcCLB.510187.210 | hypothetical protein, conserved | 0,1971 | 0,01759 |

| | | | |
|---|---|---|---|
| TcCLB.511557.29 | hypothetical protein | 0,1971 | 0,275 |
| TcCLB.506469.50 | hypothetical protein, conserved | 0,1971 | 0,2928 |
| TcCLB.507105.60 | hypothetical protein, conserved | 0,197 | 0,05788 |
| TcCLB.511553.30 | mucin-associated surface protein (MASP), putative | 0,197 | 0,523 |
| TcCLB.509885.64 | hypothetical protein, conserved | 0,1969 | 0,4747 |
| TcCLB.504081.450 | trans-sialidase (pseudogene), putative | 0,1969 | 0,4952 |
| TcCLB.511515.40 | heat shock 70 kDa protein, mitochondrial precursor, putativ | 0,1968 | 0,04441 |
| TcCLB.508691.10 | hypothetical protein, conserved | 0,1968 | 0,3956 |
| TcCLB.507669.160 | hypothetical protein, conserved | 0,1967 | 0,02583 |
| TcCLB.504021.80 | hypothetical protein, conserved | 0,1967 | 0,2344 |
| TcCLB.508647.174 | hypothetical protein, conserved | 0,1967 | 0,3163 |
| TcCLB.508765.30 | glycosylphosphatidylinositol-specific phospholipase C, puta | 0,1966 | 0,3051 |
| TcCLB.506037.10 | dispersed gene family protein 1 (DGF-1, pseudogene), puta | 0,1966 | 0,7055 |
| TcCLB.506733.120 | hypothetical protein, conserved | 0,1965 | 0,2133 |
| TcCLB.507653.10 | hypothetical protein, conserved | 0,1965 | 0,3754 |
| TcCLB.509979.290 | trans-sialidase (pseudogene), putative | 0,1963 | 0,405 |
| TcCLB.506245.200 | mucin-associated surface protein (MASP), putative | 0,1963 | 0,5703 |
| TcCLB.511603.200 | mucin TcMUCII, putative | 0,1961 | 0,7302 |
| TcCLB.509267.3 | receptor-type adenylate cyclase, putative | 0,1961 | 0,7454 |
| TcCLB.508379.20 | hypothetical protein, conserved | 0,1958 | 0,5119 |
| TcCLB.508811.10 | hypothetical protein | 0,1957 | 0,1737 |
| TcCLB.508837.74 | small GTP-binding protein Rab18, putative | 0,1957 | 0,1843 |
| TcCLB.506257.10 | DNA/RNA non-specific endonuclease protein-like, putative | 0,1955 | 0,32 |
| TcCLB.510693.280 | mucin-associated surface protein (MASP), putative | 0,1955 | 0,5078 |
| TcCLB.508125.22 | mucin TcMUCII (pseudogene), putative | 0,1955 | 0,6788 |
| TcCLB.510799.30 | hypothetical protein, conserved | 0,1954 | 0,1584 |
| TcCLB.511717.140 | hypothetical protein, conserved | 0,1954 | 0,2524 |
| TcCLB.508591.10 | dispersed gene family protein 1 (DGF-1), putative | 0,1954 | 0,4342 |
| TcCLB.507603.120 | hypothetical protein, conserved | 0,1953 | 0,08978 |
| TcCLB.504153.210 | hypothetical protein | 0,1953 | 0,3845 |
| TcCLB.508759.30 | mucin TcMUC (pseudogene), putative | 0,1952 | 0,7904 |
| TcCLB.505987.30 | hypothetical protein, conserved | 0,1951 | 0,1921 |
| TcCLB.506903.50 | dispersed gene family protein 1 (DGF-1, pseudogene), puta | 0,1951 | 0,3273 |
| TcCLB.508229.20 | hypothetical protein | 0,1951 | 0,7526 |
| TcCLB.511255.130 | mucin-associated surface protein (MASP), putative | 0,195 | 0,7141 |
| TcCLB.510911.40 | hypothetical protein, conserved | 0,1947 | 0,4087 |
| TcCLB.508355.230 | hypothetical protein | 0,1947 | 0,436 |
| TcCLB.507801.40 | hypothetical protein, conserved | 0,1947 | 0,4916 |
| TcCLB.509599.100 | hypothetical protein | 0,1946 | 0,4414 |
| TcCLB.503809.75 | hypothetical protein, conserved | 0,1945 | 0,3989 |
| TcCLB.506465.40 | protein kinase-like protein, putative | 0,1944 | 0,3502 |
| TcCLB.503905.50 | hypothetical protein, conserved | 0,1942 | 0,367 |
| TcCLB.510441.41 | trans-sialidase (pseudogene), putative | 0,1941 | 0,1887 |
| TcCLB.506755.120 | hypothetical protein, conserved | 0,194 | 0,0754 |
| TcCLB.506999.70 | chaperone DNAJ protein, putative | 0,194 | 0,1487 |

| | | | |
|---|---|---|---|
| TcCLB.503583.100 | hypothetical protein, conserved | 0,194 | 0,3227 |
| TcCLB.511863.30 | hypothetical protein, conserved | 0,1939 | 0,2178 |
| TcCLB.510409.39 | ubiquitin/ribosomal protein S27a, putative | 0,1938 | 0,0104 |
| TcCLB.504765.15 | succinyl-coA:3-ketoacid-coenzyme A transferase (pseudoge | 0,1934 | 0,4766 |
| TcCLB.506957.90 | 60S ribosomal protein L18a, putative | 0,1932 | 0,009229 |
| TcCLB.505009.40 | hypothetical protein, conserved | 0,1932 | 0,2005 |
| TcCLB.510303.30 | hypothetical protein, conserved | 0,193 | 0,07939 |
| TcCLB.507093.184 | hypothetical protein, conserved | 0,193 | 0,142 |
| TcCLB.511261.50 | hypothetical protein, conserved | 0,1928 | 0,2354 |
| TcCLB.510769.170 | trans-sialidase (pseudogene), putative | 0,1928 | 0,3678 |
| TcCLB.511305.20 | hypothetical protein, conserved | 0,1926 | 0,07094 |
| TcCLB.509103.20 | tubulin-tyrosine ligase, putative | 0,1925 | 0,2335 |
| TcCLB.511255.690 | mucin-associated surface protein (MASP), putative | 0,1925 | 0,6151 |
| TcCLB.508543.110 | hypothetical protein, conserved | 0,1924 | 0,1531 |
| TcCLB.511705.10 | receptor-type adenylate cyclase, putative | 0,1924 | 0,2256 |
| TcCLB.507765.100 | hypothetical protein, conserved | 0,1924 | 0,2798 |
| TcCLB.508799.110 | protein kinase, putative | 0,1924 | 0,2808 |
| TcCLB.507499.50 | hypothetical protein, conserved (pseudogene) | 0,1924 | 0,3493 |
| TcCLB.511383.89 | Flagellar Member 3 (FLAM3) | 0,1922 | 0,2267 |
| TcCLB.511807.3 | N-acetyltransferase complex ARD1 subunit (pseudogene), | 0,1922 | 0,4599 |
| TcCLB.509993.51 | mucin-associated surface protein (MASP, pseudogene), pu | 0,1922 | 0,6944 |
| TcCLB.507519.40 | hypothetical protein | 0,1921 | 0,1531 |
| TcCLB.510339.7 | hypothetical protein, conserved (fragment) | 0,1921 | 0,2253 |
| TcCLB.506767.123 | syntaxin binding protein (pseudogene), putative | 0,192 | 0,783 |
| TcCLB.507485.30 | hypothetical protein, conserved | 0,1919 | 0,4057 |
| TcCLB.508409.50 | hypothetical protein, conserved | 0,1917 | 0,1765 |
| TcCLB.508607.60 | beta galactofuranosyl glycosyltransferase, putative | 0,1916 | 0,5035 |
| TcCLB.505163.40 | hypothetical protein, conserved | 0,1913 | 0,3383 |
| TcCLB.508033.60 | mucin TcMUCII, putative | 0,1912 | 0,6891 |
| TcCLB.506961.90 | retrotransposon hot spot (RHS) protein, putative | 0,1911 | 0,1363 |
| TcCLB.511577.80 | hypothetical protein, conserved | 0,1911 | 0,1456 |
| TcCLB.511497.10 | hypothetical protein, conserved | 0,1911 | 0,4644 |
| TcCLB.509625.30 | hypothetical protein, conserved | 0,191 | 0,2848 |
| TcCLB.504229.44 | hypothetical protein, conserved (pseudogene) | 0,1909 | 0,3183 |
| TcCLB.511283.80 | hypothetical protein, conserved | 0,1908 | 0,04512 |
| TcCLB.510719.40 | hypothetical protein, conserved | 0,1907 | 0,3002 |
| TcCLB.510719.300 | metallo-peptidase, Clan MC, Family M14, putative | 0,1905 | 0,09842 |
| TcCLB.508955.10 | syntaxin, putative | 0,1904 | 0,3625 |
| TcCLB.506251.10 | hypothetical protein, conserved | 0,1903 | 0,2072 |
| TcCLB.508225.20 | retrotransposon hot spot protein (RHS, pseudogene), putat | 0,1903 | 0,6303 |
| TcCLB.511303.120 | hypothetical protein, conserved | 0,1901 | 0,04112 |
| TcCLB.508175.80 | protein kinase, putative | 0,19 | 0,0383 |
| TcCLB.509897.110 | trans-sialidase, Group V, putative | 0,19 | 0,4392 |
| TcCLB.511553.130 | mucin TcMUCII, putative | 0,1899 | 0,6192 |
| TcCLB.508149.10 | dispersed gene family protein 1 (DGF-1), putative | 0,1899 | 0,6204 |

| | | | |
|---|---|---|---|
| TcCLB.511001.120 | 60S ribosomal protein L18a, putative | 0,1898 | 0,006914 |
| TcCLB.511069.20 | 40S ribosomal protein S8, putative | 0,1897 | 0,07351 |
| TcCLB.511821.80 | hypothetical protein, conserved | 0,1897 | 0,2094 |
| TcCLB.504867.110 | hypothetical protein, conserved | 0,1896 | 0,2758 |
| TcCLB.510241.124 | hypothetical protein, conserved | 0,1895 | 0,09462 |
| TcCLB.507949.40 | hypothetical protein, conserved | 0,1895 | 0,1429 |
| TcCLB.506275.60 | zinc carboxypeptidase, putative | 0,1895 | 0,1468 |
| TcCLB.510553.20 | trans-sialidase (pseudogene), putative | 0,1895 | 0,6372 |
| TcCLB.509719.53 | hypothetical protein, conserved | 0,1893 | 0,1286 |
| TcCLB.509057.20 | importin alpha, putative | 0,1892 | 0,03071 |
| TcCLB.507079.10 | hypothetical protein, conserved | 0,1892 | 0,2262 |
| TcCLB.506573.70 | hypothetical protein, conserved | 0,1892 | 0,3408 |
| TcCLB.507711.190 | hypothetical protein, conserved | 0,189 | 0,02992 |
| TcCLB.510007.60 | hypothetical protein, conserved | 0,189 | 0,08997 |
| TcCLB.509157.279 | Chaperonin HSP60, mitochondrial (HSP60) | 0,189 | 0,4008 |
| TcCLB.508057.20 | mucin-associated surface protein (MASP, pseudogene), pu | 0,1889 | 0,437 |
| TcCLB.510107.30 | mucin-associated surface protein (MASP), putative | 0,1889 | 0,6351 |
| TcCLB.507081.20 | hypothetical protein, conserved | 0,1888 | 0,1375 |
| TcCLB.511431.10 | protein kinase, putative | 0,1888 | 0,4452 |
| TcCLB.404975.30 | trans-sialidase (pseudogene), putative | 0,1888 | 0,5755 |
| TcCLB.508461.30 | 3-demethylubiquinone-9 3-methyltransferase, putative | 0,1887 | 0,1832 |
| TcCLB.511183.60 | trans-sialidase, Group V, putative | 0,1887 | 0,5953 |
| TcCLB.503617.20 | kinetoplast DNA-associated protein, putative | 0,1885 | 0,046 |
| TcCLB.511353.4 | choline/carnitine O-acyltransferase, putative | 0,1885 | 0,2421 |
| TcCLB.506885.340 | Mitochondrial import receptor subunit ATOM40, putative ( | 0,1884 | 0,05932 |
| TcCLB.508323.20 | hypothetical protein, conserved | 0,188 | 0,06834 |
| TcCLB.506247.430 | hypothetical protein, conserved | 0,188 | 0,1902 |
| TcCLB.511051.80 | DNA/RNA non-specific endonuclease protein-like, putative | 0,1879 | 0,3305 |
| TcCLB.510657.30 | Sorting nexin-4 (ATG24) | 0,1877 | 0,08039 |
| TcCLB.507993.320 | RNA-binding protein 27, putative (RBP27) | 0,1877 | 0,209 |
| TcCLB.504257.9 | RNA-binding protein 42 (RNA-binding motif protein 42), pu | 0,1875 | 0,251 |
| TcCLB.506391.30 | ef-hand protein 5, putative | 0,1873 | 0,2394 |
| TcCLB.503909.19 | dispersed gene family protein 1 (DGF-1), putative (fragme | 0,1873 | 0,5274 |
| TcCLB.510767.20 | 60S ribosomal protein L36, putative | 0,1872 | 0,06861 |
| TcCLB.510359.604 | mucin TcMUCII (pseudogene), putative | 0,187 | 0,6268 |
| TcCLB.506297.150 | 40S ribosomal protein S5, putative | 0,1869 | 0,03815 |
| TcCLB.509979.51 | zinc carboxypeptidase, putative | 0,1869 | 0,6895 |
| TcCLB.509985.50 | PrimPol-like protein 1, putative (ppl1) | 0,1868 | 0,1688 |
| TcCLB.506195.270 | hypothetical protein, conserved | 0,1867 | 0,1555 |
| TcCLB.507521.70 | cyclophilin, putative | 0,1867 | 0,2112 |
| TcCLB.510221.30 | hypothetical protein, conserved | 0,1866 | 0,05803 |
| TcCLB.510121.180 | hypothetical protein, conserved | 0,1865 | 0,2314 |
| TcCLB.504777.30 | surface protease GP63 (pseudogene), putative | 0,1865 | 0,5635 |
| TcCLB.510377.390 | mucin-associated surface protein (MASP), putative | 0,1865 | 0,7318 |
| TcCLB.506559.400 | hypothetical protein, conserved | 0,1864 | 0,2353 |

| | | | |
|---|---|---|---|
| TcCLB.420495.19 | hypothetical protein, conserved (pseudogene) | 0,1864 | 0,5879 |
| TcCLB.507717.9 | hypothetical protein, conserved | 0,1863 | 0,361 |
| TcCLB.508661.40 | hypothetical protein, conserved | 0,1862 | 0,2798 |
| TcCLB.511593.80 | mucin-associated surface protein (MASP), putative | 0,186 | 0,5227 |
| TcCLB.508461.490 | 60S ribosomal protein L23, putative | 0,1859 | 0,01836 |
| TcCLB.508879.160 | hypothetical protein, conserved | 0,1859 | 0,09247 |
| TcCLB.508965.94 | hypothetical protein, conserved | 0,1859 | 0,2014 |
| TcCLB.508389.130 | mucin-associated surface protein (MASP), putative | 0,1859 | 0,8556 |
| TcCLB.507091.100 | trans-sialidase (pseudogene), putative | 0,1858 | 0,5396 |
| TcCLB.504125.10 | SWI/SNF-related matrix-associated actin-dependent regul | 0,1856 | 0,158 |
| TcCLB.504425.110 | permease-like protein, putative | 0,1856 | 0,319 |
| TcCLB.510667.90 | hypothetical protein, conserved | 0,1855 | 0,1755 |
| TcCLB.504069.40 | hypothetical protein, conserved | 0,1855 | 0,193 |
| TcCLB.510575.170 | hypothetical protein, conserved | 0,1855 | 0,2174 |
| TcCLB.511439.20 | lipase, putative | 0,1855 | 0,4696 |
| TcCLB.511757.100 | trans-sialidase, putative (fragment) | 0,1854 | 0,3116 |
| TcCLB.508153.184 | 60S ribosomal protein L39, putative | 0,1853 | 0,1088 |
| TcCLB.504137.60 | beta adaptin, putative | 0,1853 | 0,1457 |
| TcCLB.411235.9 | alpha tubulin, putative | 0,185 | 0,03028 |
| TcCLB.504187.21 | trans-sialidase (pseudogene), putative | 0,185 | 0,5457 |
| TcCLB.415187.10 | hypothetical protein, conserved | 0,1849 | 0,6178 |
| TcCLB.504037.30 | 60S ribosomal protein L12, putative | 0,1848 | 0,0384 |
| TcCLB.508207.10 | glycine dehydrogenase (pseudogene), putative | 0,1848 | 0,2787 |
| TcCLB.509611.30 | hypothetical protein | 0,1847 | 0,1727 |
| TcCLB.509147.21 | hypothetical protein, conserved (pseudogene) | 0,1847 | 0,569 |
| TcCLB.506963.20 | 60S ribosomal protein L34, putative | 0,1843 | 0,02424 |
| TcCLB.503923.40 | hypothetical protein, conserved | 0,1843 | 0,5417 |
| TcCLB.511903.130 | 60S ribosomal protein L26, putative | 0,1842 | 0,01547 |
| TcCLB.509067.50 | hypothetical protein, conserved | 0,1842 | 0,1041 |
| TcCLB.503925.10 | hypothetical protein, conserved | 0,1841 | 0,2557 |
| TcCLB.510901.250 | hypothetical protein, conserved | 0,184 | 0,204 |
| TcCLB.506901.60 | mucin TcMUCII, putative | 0,184 | 0,7936 |
| TcCLB.510119.70 | protein kinase, putative | 0,1837 | 0,3878 |
| TcCLB.506769.120 | mucin-associated surface protein (MASP), putative | 0,1837 | 0,6972 |
| TcCLB.511099.70 | mucin TcMUCII, putative | 0,1837 | 0,793 |
| TcCLB.510371.141 | hypothetical protein, conserved (pseudogene) | 0,1835 | 0,8508 |
| TcCLB.506925.60 | hypothetical protein, conserved | 0,1833 | 0,3467 |
| TcCLB.509437.125 | trans-sialidase (pseudogene), putative | 0,1833 | 0,6344 |
| TcCLB.510483.90 | trans-sialidase (pseudogene), putative | 0,1833 | 0,6466 |
| TcCLB.503727.10 | centrin, putative | 0,1831 | 0,2321 |
| TcCLB.411427.40 | hypothetical protein, conserved | 0,1831 | 0,4801 |
| TcCLB.511613.10 | mucin-associated surface protein (MASP), putative | 0,1831 | 0,6126 |
| TcCLB.506329.9 | mucin-associated surface protein (MASP, pseudogene), pu | 0,1831 | 0,7974 |
| TcCLB.508707.330 | chaperone DNAJ protein, putative | 0,183 | 0,2346 |
| TcCLB.507049.199 | pumilio protein, putative | 0,1829 | 0,6217 |

| | | | |
|---|---|---|---|
| TcCLB.506727.80 | hypothetical protein, conserved | 0,1827 | 0,2135 |
| TcCLB.505071.100 | phosphatidylglycerolphosphate synthase, mitochondrial, pu | 0,1826 | 0,1867 |
| TcCLB.504431.109 | cytochrome b5, putative | 0,1826 | 0,3365 |
| TcCLB.506247.310 | hypothetical protein, conserved | 0,1825 | 0,08163 |
| TcCLB.510895.9 | hypothetical protein, conserved (fragment) | 0,1824 | 0,07438 |
| TcCLB.509213.110 | hypothetical protein, conserved | 0,1824 | 0,1047 |
| TcCLB.511259.70 | mucin-associated surface protein (MASP, pseudogene), pu | 0,1824 | 0,6438 |
| TcCLB.504423.30 | hypothetical protein, conserved | 0,1823 | 0,2152 |
| TcCLB.506201.30 | serine/threonine-protein phosphatase PP1 beta | 0,1822 | 0,1844 |
| TcCLB.507687.70 | mucin-associated surface protein (MASP), putative | 0,182 | 0,7177 |
| TcCLB.510015.30 | mucin-associated surface protein (MASP), putative | 0,1819 | 0,3898 |
| TcCLB.509287.50 | glucose-6-phosphate 1-dehydrogenase, putative | 0,1816 | 0,1314 |
| TcCLB.508567.49 | hypothetical protein, conserved | 0,1816 | 0,4625 |
| TcCLB.507729.30 | trans-sialidase (pseudogene), putative | 0,1816 | 0,5027 |
| TcCLB.506945.240 | ATP synthase, epsilon chain, putative | 0,1815 | 0,1232 |
| TcCLB.504769.190 | trans-sialidase (pseudogene), putative | 0,1815 | 0,5434 |
| TcCLB.509461.90 | inositol 1,4,5-trisphosphate receptor, putative (IP3R) | 0,1814 | 0,01193 |
| TcCLB.511133.30 | hypothetical protein, conserved | 0,1814 | 0,08063 |
| TcCLB.507603.170 | hypothetical protein, conserved | 0,1814 | 0,1908 |
| TcCLB.509753.140 | mucin-associated surface protein (MASP), putative | 0,1814 | 0,6946 |
| TcCLB.511173.470 | trans-sialidase, Group V, putative | 0,181 | 0,6432 |
| TcCLB.511277.490 | hypothetical protein, conserved | 0,1809 | 0,0899 |
| TcCLB.511763.10 | hypothetical protein, conserved | 0,1809 | 0,3843 |
| TcCLB.506211.107 | hypothetical protein, conserved | 0,1809 | 0,4588 |
| TcCLB.507093.180 | transcription factor IIIb, putative | 0,1808 | 0,1635 |
| TcCLB.507023.180 | protein kinase, putative | 0,1808 | 0,1715 |
| TcCLB.503663.10 | septum formation inhibitor, Maf-like protein, putative | 0,1806 | 0,4189 |
| TcCLB.509941.85 | trans-sialidase (pseudogene), putative | 0,1804 | 0,3429 |
| TcCLB.506155.110 | hypothetical protein, conserved | 0,1803 | 0,2112 |
| TcCLB.511921.50 | trans-sialidase, Group VI, putative | 0,1803 | 0,5207 |
| TcCLB.506941.238 | hypothetical protein, conserved | 0,1802 | 0,293 |
| TcCLB.506067.290 | mucin-associated surface protein (MASP), putative | 0,1802 | 0,573 |
| TcCLB.506471.110 | retrotransposon hot spot protein (RHS, pseudogene), putat | 0,1798 | 0,2047 |
| TcCLB.509809.20 | chaperone DNAJ protein, putative | 0,1798 | 0,2706 |
| TcCLB.507961.10 | mucin-associated surface protein (MASP, pseudogene), pu | 0,1798 | 0,7118 |
| TcCLB.506865.60 | hypothetical protein, conserved | 0,1796 | 0,08342 |
| TcCLB.508465.20 | hypothetical protein, conserved | 0,1796 | 0,2417 |
| TcCLB.509569.40 | hypothetical protein, conserved | 0,1795 | 0,08263 |
| TcCLB.506559.517 | hypothetical protein, conserved | 0,1795 | 0,1497 |
| TcCLB.507907.40 | hypothetical protein, conserved | 0,1793 | 0,2245 |
| TcCLB.506187.50 | mucin-associated surface protein (MASP), putative | 0,1793 | 0,7267 |
| TcCLB.503979.10 | ethanolamine phosphotransferase, putative (fragment) | 0,1791 | 0,154 |
| TcCLB.504069.80 | ATP synthase F1 subunit gamma protein, putative (ATPF1G | 0,1789 | 0,1591 |
| TcCLB.509561.39 | hypothetical protein, conserved | 0,1789 | 0,282 |
| TcCLB.506173.219 | trans-sialidase (pseudogene), putative | 0,1788 | 0,6966 |

| | | | |
|---|---|---|---|
| TcCLB.503455.20 | hypothetical protein, conserved | 0,1786 | 0,264 |
| TcCLB.506637.120 | hypothetical protein, conserved | 0,1784 | 0,07424 |
| TcCLB.511277.200 | RNA-binding protein, putative | 0,1783 | 0,7792 |
| TcCLB.508919.60 | ADP-ribosylation factor, putative (ARL-1) | 0,1782 | 0,3611 |
| TcCLB.511647.44 | hypothetical protein, conserved | 0,1781 | 0,4599 |
| TcCLB.507513.70 | 40S ribosomal protein S16, putative | 0,178 | 0,04545 |
| TcCLB.504175.9 | hypothetical protein, conserved | 0,178 | 0,081 |
| TcCLB.506635.139 | kasal-type serpin (pseudogene), putative | 0,178 | 0,2183 |
| TcCLB.506777.30 | hypothetical protein, conserved | 0,1779 | 0,1051 |
| TcCLB.508515.150 | trans-sialidase, Group VIII, putative | 0,1779 | 0,5643 |
| TcCLB.507689.20 | hypothetical protein | 0,1776 | 0,3909 |
| TcCLB.509993.15 | mucin-associated surface protein (MASP, pseudogene), pu | 0,1776 | 0,4816 |
| TcCLB.508547.190 | hypothetical protein, conserved | 0,1775 | 0,2356 |
| TcCLB.511629.20 | hypothetical protein, conserved | 0,1775 | 0,2593 |
| TcCLB.509921.51 | hypothetical protein, conserved (pseudogene) | 0,1775 | 0,5639 |
| TcCLB.506735.10 | mitochondrial processing peptidase alpha subunit, putative | 0,1774 | 0,07659 |
| TcCLB.508897.100 | dynein light chain, putative | 0,1773 | 0,3442 |
| TcCLB.507047.120 | myo-inositol-1(or 4)-monophosphatase 1, putative | 0,1772 | 0,3539 |
| TcCLB.456021.10 | hypothetical protein, conserved (fragment) | 0,177 | 0,5713 |
| TcCLB.510889.100 | hypothetical protein, conserved | 0,1769 | 0,2724 |
| TcCLB.503479.20 | hypothetical protein, conserved (pseudogene) | 0,1768 | 0,4825 |
| TcCLB.509195.80 | trans-sialidase (pseudogene), putative | 0,1768 | 0,483 |
| TcCLB.511825.50 | centrin-4, putative (Centrin4) | 0,1766 | 0,117 |
| TcCLB.509693.70 | hypoxanthine-guanine phosphoribosyltransferase, putative | 0,1766 | 0,3225 |
| TcCLB.511529.30 | hypothetical protein, conserved | 0,1766 | 0,3438 |
| TcCLB.510579.70 | nascent polypeptide associated complex subunit, putative | 0,1763 | 0,04212 |
| TcCLB.511071.100 | Glycerophosphoryl diester phosphodiesterase family, putat | 0,1763 | 0,1696 |
| TcCLB.511491.100 | phosphatase 2C, putative | 0,1761 | 0,1699 |
| TcCLB.508827.79 | 40S ribosomal protein S17, putative | 0,176 | 0,04922 |
| TcCLB.510659.180 | hypothetical protein, conserved | 0,176 | 0,1957 |
| TcCLB.509549.15 | mucin TcMUCII (pseudogene), putative | 0,176 | 0,7696 |
| TcCLB.510749.10 | hypothetical protein, conserved | 0,1759 | 0,2134 |
| TcCLB.510463.90 | mucin-associated surface protein (MASP), putative | 0,1759 | 0,9141 |
| TcCLB.506609.15 | hypothetical protein (pseudogene) | 0,1758 | 0,7116 |
| TcCLB.511261.80 | hypothetical protein, conserved | 0,1756 | 0,1182 |
| TcCLB.511903.10 | hypothetical protein | 0,1756 | 0,5688 |
| TcCLB.506723.20 | trans-sialidase, Group VII, putative | 0,1755 | 0,4156 |
| TcCLB.506399.20 | hypothetical protein, conserved | 0,1754 | 0,2582 |
| TcCLB.506659.39 | protein phosphatase 2C, putative | 0,1754 | 0,5809 |
| TcCLB.506287.90 | hypothetical protein, conserved | 0,1753 | 0,2413 |
| TcCLB.510779.10 | tryptophanyl-tRNA synthetase (pseudogene), putative | 0,1753 | 0,4486 |
| TcCLB.503849.40 | mucin-associated surface protein (MASP), putative | 0,1752 | 0,6232 |
| TcCLB.506871.160 | RNA guanylyltransferase, putative | 0,175 | 0,1233 |
| TcCLB.505807.54 | hypothetical protein, conserved | 0,175 | 0,3612 |
| TcCLB.508175.320 | hypothetical protein, conserved | 0,1749 | 0,1422 |

| | | | |
|---|---|---|---|
| TcCLB.509303.20 | glycine dehydrogenase [decarboxylating] (pseudogene), pu | 0,1749 | 0,5038 |
| TcCLB.506175.20 | hypothetical protein, conserved | 0,1748 | 0,09268 |
| TcCLB.506993.190 | hypothetical protein | 0,1748 | 0,7922 |
| TcCLB.510339.14 | hypothetical protein, conserved | 0,1747 | 0,2379 |
| TcCLB.508761.270 | mucin-associated surface protein (MASP), putative | 0,1745 | 0,596 |
| TcCLB.509151.90 | hypothetical protein, conserved | 0,1744 | 0,19 |
| TcCLB.509587.20 | squalene monooxygenase, putative (fragment) | 0,1743 | 0,1922 |
| TcCLB.510733.40 | hypothetical protein, conserved | 0,1743 | 0,2477 |
| TcCLB.508641.40 | retrotransposon hot spot protein (RHS, pseudogene), putat | 0,1743 | 0,5997 |
| TcCLB.509443.30 | AUT2/APG4/ATG4 cysteine peptidase, putative | 0,1741 | 0,3906 |
| TcCLB.506321.140 | mucin-associated surface protein (MASP), putative | 0,174 | 0,5985 |
| TcCLB.486465.9 | cysteine peptidase, Clan CA, family C51, putative | 0,174 | 0,6045 |
| TcCLB.510577.50 | rhomboid-like protein, putative | 0,1739 | 0,05165 |
| TcCLB.511657.20 | protein kinase, putative | 0,1739 | 0,2259 |
| TcCLB.509419.150 | mucin-associated surface protein (MASP), putative | 0,1739 | 0,6543 |
| TcCLB.509825.30 | hypothetical protein, conserved | 0,1738 | 0,2567 |
| TcCLB.508399.49 | dispersed gene family protein 1 (DGF-1), putative | 0,1737 | 0,395 |
| TcCLB.424889.10 | UDP-Gal or UDP-GlcNAc-dependent glycosyltransferase, pu | 0,1736 | 0,8154 |
| TcCLB.511165.40 | acidocalcisomal pyrophosphatase, putative | 0,1735 | 0,1352 |
| TcCLB.503501.70 | mucin-associated surface protein (MASP), putative (fragm | 0,1735 | 0,6845 |
| TcCLB.508355.120 | hypothetical protein, conserved | 0,1734 | 0,1639 |
| TcCLB.511711.100 | hypothetical protein, conserved | 0,1734 | 0,2545 |
| TcCLB.509831.50 | intraflagellar transport protein 88, putative (fragment) (IF | 0,1733 | 0,3172 |
| TcCLB.509793.10 | kinetoplast-associated protein 4 (KAP4) | 0,1733 | 0,3618 |
| TcCLB.511725.40 | hypothetical protein, conserved | 0,1732 | 0,3317 |
| TcCLB.508151.5 | mucin-associated surface protein (MASP, pseudogene), pu | 0,1732 | 0,8483 |
| TcCLB.506937.30 | 60S ribosomal protein L14, putative | 0,173 | 0,08984 |
| TcCLB.508423.20 | hypothetical protein, conserved (pseudogene) | 0,173 | 0,1061 |
| TcCLB.473633.9 | dispersed gene family protein 1 (DGF-1), putative (fragme | 0,1729 | 0,6031 |
| TcCLB.511667.40 | retrotransposon hot spot (RHS) protein, putative | 0,1728 | 0,527 |
| TcCLB.511511.170 | ADP-ribosylation factor-like protein, putative | 0,1727 | 0,2816 |
| TcCLB.510731.40 | katanin, putative | 0,1727 | 0,3342 |
| TcCLB.507523.10 | trans-sialidase, putative | 0,1727 | 0,5634 |
| TcCLB.509033.100 | hypothetical protein, conserved | 0,1726 | 0,1874 |
| TcCLB.506309.60 | trans-sialidase, Group V, putative | 0,1726 | 0,866 |
| TcCLB.510355.110 | hypothetical protein, conserved (pseudogene) | 0,1725 | 0,5022 |
| TcCLB.511283.300 | hypothetical protein, conserved | 0,1724 | 0,2029 |
| TcCLB.506861.20 | hypothetical protein, conserved | 0,1721 | 0,2049 |
| TcCLB.503791.49 | vacuolar protein sorting-associated protein, putative (fragr | 0,1721 | 0,3719 |
| TcCLB.504787.90 | mucin-associated surface protein (MASP, pseudogene), pu | 0,1721 | 0,5591 |
| TcCLB.508157.20 | mucin-associated surface protein (MASP), putative | 0,1721 | 0,6227 |
| TcCLB.506799.50 | mucin-associated surface protein (MASP), putative | 0,172 | 0,4302 |
| TcCLB.508211.70 | hypothetical protein, conserved | 0,1719 | 0,2307 |
| TcCLB.478671.9 | hypothetical protein, conserved | 0,1718 | 0,176 |
| TcCLB.506297.320 | hypothetical protein | 0,1718 | 0,2401 |

| | | | |
|---|---|---|---|
| TcCLB.507993.200 | hypothetical protein, conserved | 0,1717 | 0,1236 |
| TcCLB.511589.250 | hypothetical protein, conserved | 0,1716 | 0,2615 |
| TcCLB.508873.250 | mucin-associated surface protein (MASP), putative | 0,1715 | 0,5489 |
| TcCLB.506131.170 | trans-sialidase, Group V, putative | 0,1715 | 0,6955 |
| TcCLB.506975.30 | amidohydrolase, putative | 0,1714 | 0,2213 |
| TcCLB.508605.20 | UDP-Gal or UDP-GlcNAc-dependent glycosyltransferase, pu | 0,1713 | 0,6237 |
| TcCLB.504261.30 | trans-sialidase (pseudogene), putative | 0,1713 | 0,7233 |
| TcCLB.509617.30 | mismatch repair protein MSH5, putative | 0,1711 | 0,324 |
| TcCLB.507515.100 | hypothetical protein, conserved | 0,1711 | 0,4289 |
| TcCLB.506245.160 | retrotransposon hot spot (RHS) protein, putative | 0,1711 | 0,6241 |
| TcCLB.504741.210 | hypothetical protein, conserved | 0,171 | 0,1566 |
| TcCLB.506123.40 | nucleolar RNA helicase II, putative | 0,1708 | 0,09123 |
| TcCLB.509899.40 | trans-sialidase (pseudogene), putative | 0,1708 | 0,4703 |
| TcCLB.506789.260 | hypothetical protein, conserved | 0,1708 | 0,4956 |
| TcCLB.506821.170 | aquaporin-like protein, putative | 0,1707 | 0,187 |
| TcCLB.503581.39 | coatomer delta subunit, putative | 0,1707 | 0,2462 |
| TcCLB.507991.40 | hypothetical protein | 0,1707 | 0,2907 |
| TcCLB.507017.150 | AUT2/APG4/ATG4 cysteine peptidase, putative | 0,1707 | 0,3382 |
| TcCLB.506101.20 | trans-sialidase (pseudogene), putative | 0,1707 | 0,5409 |
| TcCLB.509381.29 | hypothetical protein, conserved | 0,1706 | 0,1987 |
| TcCLB.510629.450 | hypothetical protein, conserved | 0,1706 | 0,3208 |
| TcCLB.476733.29 | serine-threonine dehydratase, putative | 0,1706 | 0,647 |
| TcCLB.511167.70 | DNA repair and recombination helicase protein PIF2, putat | 0,1705 | 0,3465 |
| TcCLB.509753.194 | mucin-associated surface protein (MASP), putative | 0,1704 | 0,8194 |
| TcCLB.510123.10 | hypothetical protein, conserved | 0,1703 | 0,2262 |
| TcCLB.507825.30 | Present in the outer mitochondrial membrane proteome 2( | 0,1701 | 0,1352 |
| TcCLB.506599.315 | hypothetical protein, conserved (pseudogene) | 0,1701 | 0,6847 |
| TcCLB.509295.110 | dispersed gene family protein 1 (DGF-1, pseudogene), puta | 0,1699 | 0,7371 |
| TcCLB.511149.199 | mucin TcMUCII (pseudogene), putative | 0,1698 | 0,8092 |
| TcCLB.503815.29 | hypothetical protein, conserved | 0,1697 | 0,5473 |
| TcCLB.504827.130 | eukaryotic translation initation factor 4 gamma, putative | 0,1696 | 0,1858 |
| TcCLB.506489.50 | serine/threonine protein kinase, putative | 0,1695 | 0,3713 |
| TcCLB.506413.20 | 40S ribosomal protein S33, putative | 0,1694 | 0,1114 |
| TcCLB.505943.50 | hypothetical protein, conserved | 0,1694 | 0,2358 |
| TcCLB.509267.60 | NADH-ubiquinone oxidoreductase complex I subunit, putati | 0,1694 | 0,3787 |
| TcCLB.507913.39 | glutaminyl-tRNA synthetase, putative | 0,1692 | 0,4386 |
| TcCLB.507797.18 | serine/threonine protein kinase, putative | 0,1691 | 0,376 |
| TcCLB.510911.14 | hypothetical protein, conserved | 0,169 | 0,2128 |
| TcCLB.509259.140 | glycine dehydrogenase [decarboxylating] (pseudogene), pu | 0,169 | 0,3108 |
| TcCLB.509161.80 | hypothetical protein, conserved | 0,1689 | 0,3159 |
| TcCLB.507007.40 | hypothetical protein, conserved | 0,1689 | 0,3377 |
| TcCLB.511521.9 | hypothetical protein, conserved | 0,1689 | 0,4959 |
| TcCLB.509979.365 | surface protease GP63 (pseudogene), putative | 0,1689 | 0,7462 |
| TcCLB.508461.510 | 60S ribosomal protein L27A/L29, putative | 0,1688 | 0,2134 |
| TcCLB.507711.290 | hypothetical protein, conserved | 0,1687 | 0,342 |

| | | | |
|---|---|---|---|
| TcCLB.511367.180 | hypothetical protein | 0,1682 | 0,154 |
| TcCLB.508355.100 | diacylglycerol acyltransferase, putative | 0,168 | 0,1073 |
| TcCLB.509927.10 | trans-sialidase, putative (fragment) | 0,168 | 0,4176 |
| TcCLB.503697.130 | hypothetical protein, conserved | 0,1677 | 0,1114 |
| TcCLB.506681.24 | hypothetical protein, conserved | 0,1677 | 0,2522 |
| TcCLB.511367.360 | elongation factor 1-alpha, putative | 0,1676 | 0,008648 |
| TcCLB.506569.50 | expression site-associated gene (ESAG-like) protein, putati | 0,1675 | 0,8456 |
| TcCLB.510351.60 | hypothetical protein, conserved | 0,1674 | 0,4371 |
| TcCLB.511287.120 | 40S ribosomal protein S2, putative | 0,1673 | 0,03823 |
| TcCLB.508911.60 | hypothetical protein, conserved | 0,1673 | 0,1274 |
| TcCLB.506821.120 | hypothetical protein, conserved | 0,1673 | 0,3797 |
| TcCLB.510897.6 | Dynein light chain LC6, flagellar outer arm, putative | 0,1672 | 0,3421 |
| TcCLB.505025.60 | 90 kDa surface protein, putative | 0,1672 | 0,7747 |
| TcCLB.507011.60 | structural maintenance of chromosome (SMC) family prote | 0,1671 | 0,07401 |
| TcCLB.511625.20 | hypothetical protein, conserved | 0,1671 | 0,3894 |
| TcCLB.511071.90 | hypothetical protein | 0,167 | 0,5957 |
| TcCLB.511871.120 | mucin-associated surface protein (MASP), putative | 0,167 | 0,6443 |
| TcCLB.503647.20 | hypothetical protein, conserved | 0,1669 | 0,5026 |
| TcCLB.508593.140 | hypothetical protein, conserved | 0,1668 | 0,1801 |
| TcCLB.506097.9 | dynein heavy chain, putative (fragment) | 0,1668 | 0,2358 |
| TcCLB.506555.9 | hypothetical protein, conserved (fragment) | 0,1668 | 0,3787 |
| TcCLB.508547.110 | ATP-dependent DEAD/H RNA helicase, putative | 0,1667 | 0,0977 |
| TcCLB.504427.140 | hypothetical protein, conserved | 0,1667 | 0,2275 |
| TcCLB.437575.18 | transporter, putative | 0,1666 | 0,5321 |
| TcCLB.510483.230 | mucin-associated surface protein (MASP), putative | 0,1664 | 0,7438 |
| TcCLB.510903.80 | retrotransposon hot spot protein (RHS, pseudogene), putat | 0,1663 | 0,05412 |
| TcCLB.511903.250 | hypothetical protein, conserved | 0,1663 | 0,3231 |
| TcCLB.507029.30 | heat shock 70 kDa protein, mitochondrial precursor, putativ | 0,1662 | 0,08374 |
| TcCLB.510329.160 | hypothetical protein, conserved | 0,1659 | 0,0749 |
| TcCLB.506217.10 | hypothetical protein | 0,1659 | 0,6547 |
| TcCLB.508469.59 | helicase, putative | 0,1657 | 0,2703 |
| TcCLB.506365.20 | retrotransposon hot spot protein (RHS, pseudogene), putat | 0,1656 | 0,08894 |
| TcCLB.504149.124 | conserved protein, unknown function | 0,1655 | 0,1882 |
| TcCLB.510359.10 | hypothetical protein, conserved | 0,1655 | 0,4097 |
| TcCLB.503679.10 | aspartate aminotransferase, putative | 0,1655 | 0,4641 |
| TcCLB.507769.40 | GTP-binding protein, putative | 0,1654 | 0,2096 |
| TcCLB.505989.70 | surface protease GP63, putative | 0,1653 | 0,3413 |
| TcCLB.510681.30 | lipase, putative | 0,1652 | 0,2899 |
| TcCLB.510731.10 | trans-sialidase, putative | 0,1652 | 0,8895 |
| TcCLB.507637.150 | mucin-associated surface protein (MASP, pseudogene), pu | 0,1651 | 0,5616 |
| TcCLB.511415.11 | retrotransposon hot spot (RHS) protein, putative (fragmen | 0,165 | 0,4275 |
| TcCLB.507071.230 | trans-sialidase, Group V, putative | 0,165 | 0,65 |
| TcCLB.508365.200 | mucin-associated surface protein (MASP), putative | 0,1648 | 0,8373 |
| TcCLB.506961.110 | hypothetical protein, conserved (pseudogene) | 0,1647 | 0,1524 |
| TcCLB.510047.10 | hypothetical protein, conserved | 0,1647 | 0,1914 |

| | | | |
|---|---|---|---|
| TcCLB.506743.190 | hypothetical protein, conserved | 0,1647 | 0,3746 |
| TcCLB.503583.40 | trans-splicing factor, putative | 0,1645 | 0,2921 |
| TcCLB.510205.80 | mucin-associated surface protein (MASP), putative | 0,1645 | 0,7284 |
| TcCLB.508363.10 | hypothetical protein | 0,1645 | 0,7911 |
| TcCLB.507513.50 | heat shock 70 kDa protein, putative | 0,1644 | 0,1231 |
| TcCLB.509171.90 | splicing factor 3a, putative | 0,1644 | 0,2064 |
| TcCLB.510899.19 | hypothetical protein, conserved | 0,1644 | 0,4797 |
| TcCLB.425839.10 | kinesin, putative | 0,1644 | 0,4937 |
| TcCLB.506767.240 | mucin-associated surface protein (MASP), putative | 0,1644 | 0,687 |
| TcCLB.511909.40 | Succinate dehydrogenase [ubiquinone] flavoprotein subuni | 0,1642 | 0,08094 |
| TcCLB.508299.80 | threonyl-tRNA synthetase, putative | 0,1642 | 0,1311 |
| TcCLB.508461.50 | hypothetical protein | 0,1641 | 0,4203 |
| TcCLB.506469.40 | hypothetical protein, conserved | 0,1641 | 0,4765 |
| TcCLB.506337.70 | 2-oxoglutarate dehydrogenase E1 component, putative | 0,164 | 0,02866 |
| TcCLB.509841.10 | trans-sialidase, putative | 0,164 | 0,799 |
| TcCLB.506529.60 | hypothetical protein, conserved | 0,1637 | 0,4438 |
| TcCLB.506905.59 | SNF2 DNA repair protein, putative (fragment) | 0,1635 | 0,1684 |
| TcCLB.507027.40 | hypothetical protein, conserved | 0,1635 | 0,3859 |
| TcCLB.510045.20 | ABC transporter, putative | 0,1635 | 0,4247 |
| TcCLB.509875.90 | retrotransposon hot spot protein (RHS, pseudogene), putat | 0,1635 | 0,5659 |
| TcCLB.504131.20 | hypothetical protein, conserved | 0,1634 | 0,09499 |
| TcCLB.478071.10 | trans-sialidase (pseudogene), putative | 0,1633 | 0,3575 |
| TcCLB.507099.120 | hypothetical protein, conserved | 0,1631 | 0,112 |
| TcCLB.510077.20 | trans-sialidase (pseudogene), putative | 0,1631 | 0,2034 |
| TcCLB.503501.30 | surface protease GP63 (pseudogene), putative | 0,1629 | 0,8181 |
| TcCLB.505023.10 | trans-sialidase (pseudogene), putative | 0,1628 | 0,4887 |
| TcCLB.511259.220 | mucin-associated surface protein (MASP), putative | 0,1628 | 0,7138 |
| TcCLB.506573.91 | coatomer delta subunit, putative | 0,1628 | 0,7222 |
| TcCLB.510629.460 | hypothetical protein | 0,1627 | 0,1584 |
| TcCLB.510575.150 | hypothetical protein, conserved | 0,1626 | 0,18 |
| TcCLB.509965.220 | hypothetical protein, conserved | 0,1624 | 0,1723 |
| TcCLB.506633.30 | hypothetical protein, conserved | 0,1624 | 0,2428 |
| TcCLB.509959.198 | ATP-dependent DEAD/H RNA helicase (pseudogene), putat | 0,1624 | 0,7455 |
| TcCLB.509671.10 | hypothetical protein, conserved | 0,1623 | 0,09317 |
| TcCLB.511277.340 | paralyzed flagella protein 20 (PF20) | 0,1623 | 0,1559 |
| TcCLB.509445.30 | solanesyl-diphosphate synthase (SPPS) | 0,1622 | 0,07351 |
| TcCLB.507059.30 | retrotransposon hot spot protein (RHS, pseudogene), putat | 0,1622 | 0,4174 |
| TcCLB.510573.10 | actin 2, putative (fragment) | 0,1621 | 0,1244 |
| TcCLB.508441.70 | enoyl-CoA hydratase/Enoyl-CoA isomerase/3-hydroxyacyl-( | 0,1621 | 0,1457 |
| TcCLB.507611.398 | RNA editing complex protein MP63, putative (fragment) | 0,1621 | 0,1499 |
| TcCLB.507831.40 | hypothetical protein, conserved | 0,1621 | 0,3089 |
| TcCLB.507001.120 | hypothetical protein, conserved | 0,1621 | 0,4174 |
| TcCLB.508631.40 | hypothetical protein, conserved (pseudogene) | 0,162 | 0,2713 |
| TcCLB.503945.20 | hypothetical protein, conserved | 0,162 | 0,5407 |
| TcCLB.506763.270 | hypothetical protein, conserved (pseudogene) | 0,162 | 0,6577 |

| | | | |
|---|---|---|---|
| TcCLB.508999.269 | hypothetical protein, conserved (fragment) | 0,1619 | 0,3929 |
| TcCLB.508273.50 | hypothetical protein, conserved | 0,1618 | 0,3461 |
| TcCLB.506501.141 | syntaxin binding protein (pseudogene), putative | 0,1618 | 0,7932 |
| TcCLB.507099.30 | Ran-binding protein 1, putative | 0,1617 | 0,1552 |
| TcCLB.508641.180 | RNA-binding protein 38, putative (RBP38) | 0,1616 | 0,1419 |
| TcCLB.508375.20 | trans-sialidase, Group V, putative | 0,1616 | 0,4476 |
| TcCLB.508161.50 | trans-sialidase, putative | 0,1615 | 0,7374 |
| TcCLB.510361.240 | mucin-associated surface protein (MASP, pseudogene), pu | 0,1615 | 0,7616 |
| TcCLB.508181.60 | hypothetical protein, conserved | 0,1612 | 0,1256 |
| TcCLB.506227.50 | hypothetical protein, conserved | 0,1612 | 0,375 |
| TcCLB.511685.40 | hypothetical protein | 0,1611 | 0,2884 |
| TcCLB.510729.100 | hypothetical protein, conserved | 0,1609 | 0,1902 |
| TcCLB.506113.90 | retrotransposon hot spot protein (RHS, pseudogene), putat | 0,1609 | 0,2356 |
| TcCLB.511367.370 | elongation factor 1-alpha, putative | 0,1608 | 0,01189 |
| TcCLB.508799.190 | hypothetical protein, conserved | 0,1608 | 0,1643 |
| TcCLB.509779.60 | dual-specificity protein phosphatase, putative | 0,1608 | 0,7837 |
| TcCLB.507641.233 | ribosomal protein L37, putative | 0,1605 | 0,1057 |
| TcCLB.507993.300 | hypothetical protein, conserved | 0,1605 | 0,376 |
| TcCLB.506223.40 | hypothetical protein, conserved | 0,1604 | 0,3155 |
| TcCLB.504131.30 | ubiquitin-activating enzyme, putative | 0,1603 | 0,27 |
| TcCLB.511129.80 | UDP-Gal or UDP-GlcNAc-dependent glycosyltransferase, pu | 0,1603 | 0,5601 |
| TcCLB.506679.250 | ribosomal protein L24, putative | 0,1602 | 0,08945 |
| TcCLB.507537.30 | cysteinyl-tRNA synthetase, putative | 0,1602 | 0,1035 |
| TcCLB.511301.10 | oxidoreductase-like protein, putative | 0,1602 | 0,5485 |
| TcCLB.510693.240 | trans-sialidase, Group VI, putative | 0,1602 | 0,6884 |
| TcCLB.508387.110 | hypothetical protein, conserved | 0,1601 | 0,3032 |
| TcCLB.506963.30 | hypothetical protein, conserved | 0,1601 | 0,396 |
| TcCLB.510729.280 | trans-sialidase, Group III, putative | 0,1601 | 0,4398 |
| TcCLB.511391.140 | alpha-adaptin-like, putative | 0,16 | 0,1942 |
| TcCLB.510535.80 | hypothetical protein, conserved | 0,1599 | 0,5184 |
| TcCLB.510275.340 | mucin-associated surface protein (MASP, pseudogene), pu | 0,1599 | 0,612 |
| TcCLB.506279.100 | hypothetical protein | 0,1598 | 0,5948 |
| TcCLB.503919.40 | hypothetical protein, conserved | 0,1597 | 0,158 |
| TcCLB.510855.4 | hypothetical protein, conserved | 0,1597 | 0,3088 |
| TcCLB.508457.40 | hypothetical protein | 0,1597 | 0,4164 |
| TcCLB.511311.30 | trans-sialidase, Group II, putative | 0,1596 | 0,5576 |
| TcCLB.503633.21 | mucin TcMUCII (pseudogene), putative | 0,1596 | 0,834 |
| TcCLB.509791.80 | hypothetical protein | 0,1595 | 0,108 |
| TcCLB.511589.10 | ATP synthase, putative | 0,1595 | 0,2315 |
| TcCLB.509805.150 | hypothetical protein, conserved | 0,1594 | 0,227 |
| TcCLB.508407.10 | membrane associated protein, putative | 0,1594 | 0,3518 |
| TcCLB.510025.260 | mucin-associated surface protein (MASP), putative | 0,1594 | 0,6483 |
| TcCLB.511607.90 | mucin-associated surface protein (MASP, pseudogene), pu | 0,1592 | 0,6423 |
| TcCLB.506625.180 | hypothetical protein, conserved | 0,1591 | 0,2672 |
| TcCLB.509017.20 | hypothetical protein, conserved | 0,1591 | 0,2987 |

| | | | |
|---|---|---|---|
| TcCLB.506053.60 | hypothetical protein (fragment) | 0,159 | 0,4057 |
| TcCLB.503771.10 | mucin-associated surface protein (MASP), putative | 0,159 | 0,8905 |
| TcCLB.511071.160 | hypothetical protein, conserved | 0,1589 | 0,3741 |
| TcCLB.510597.20 | hypothetical protein, conserved | 0,1589 | 0,4242 |
| TcCLB.507799.10 | mannosyl-oligosaccharide 1,2-alpha-mannosidase IB, puta | 0,1589 | 0,6706 |
| TcCLB.508859.90 | phosphatidylinositol 3-kinase 2, putative | 0,1588 | 0,2798 |
| TcCLB.503735.30 | hypothetical protein, conserved | 0,1588 | 0,284 |
| TcCLB.506349.40 | glycine dehydrogenase [decarboxylating] (pseudogene), pu | 0,1587 | 0,7453 |
| TcCLB.508547.140 | hypothetical protein, conserved | 0,1586 | 0,3454 |
| TcCLB.503453.40 | 60S ribosomal protein L30, putative | 0,1586 | 0,4902 |
| TcCLB.510955.20 | hypothetical protein, conserved | 0,1585 | 0,1982 |
| TcCLB.506419.60 | nucleoporin (NUP54/57), putative | 0,1585 | 0,2087 |
| TcCLB.510571.20 | leucine-rich repeat protein 1 (LRRP1), putative | 0,1585 | 0,45 |
| TcCLB.507057.20 | hypothetical protein, conserved | 0,1584 | 0,3486 |
| TcCLB.510257.70 | hypothetical protein, conserved | 0,1583 | 0,1655 |
| TcCLB.509399.90 | hypothetical protein | 0,1583 | 0,4246 |
| TcCLB.509575.4 | trans-sialidase, putative (fragment) | 0,1583 | 0,7066 |
| TcCLB.511261.150 | hypothetical protein, conserved | 0,1582 | 0,4297 |
| TcCLB.506605.220 | hypothetical protein, conserved | 0,1581 | 0,3048 |
| TcCLB.510021.50 | mucin-associated surface protein (MASP), putative | 0,1577 | 0,7838 |
| TcCLB.509797.9 | hypothetical protein, conserved (pseudogene) | 0,1576 | 0,3068 |
| TcCLB.505757.10 | hypothetical protein, conserved | 0,1576 | 0,4908 |
| TcCLB.506337.150 | hypothetical protein, conserved | 0,1575 | 0,3729 |
| TcCLB.511131.40 | hypothetical protein | 0,1575 | 0,4122 |
| TcCLB.511807.280 | hypothetical protein, conserved | 0,1574 | 0,03092 |
| TcCLB.509765.95 | mucin-associated surface protein (MASP, pseudogene), pu | 0,1572 | 0,5601 |
| TcCLB.507637.180 | mucin-associated surface protein (MASP, pseudogene), pu | 0,1572 | 0,5838 |
| TcCLB.511677.20 | trans-sialidase, putative | 0,1572 | 0,8004 |
| TcCLB.506741.140 | mucin-associated surface protein (MASP), putative | 0,1571 | 0,7532 |
| TcCLB.506267.19 | hypothetical protein | 0,157 | 0,7058 |
| TcCLB.509999.130 | hypothetical protein, conserved | 0,1568 | 0,2186 |
| TcCLB.506211.160 | Mitochondrial ADP/ATP carrier protein 5, putative (MCP5) | 0,1566 | 0,2356 |
| TcCLB.509569.60 | hypothetical protein, conserved | 0,1566 | 0,3209 |
| TcCLB.509243.20 | hypothetical protein, conserved | 0,1566 | 0,6551 |
| TcCLB.506525.130 | mitochondrial carrier protein, putative | 0,1565 | 0,368 |
| TcCLB.507009.20 | hypothetical protein, conserved | 0,1565 | 0,5015 |
| TcCLB.510105.100 | UDP-glucose dehydrogenase, putative | 0,1564 | 0,2672 |
| TcCLB.506513.24 | hypothetical protein, conserved | 0,1563 | 0,3687 |
| TcCLB.508875.10 | hypothetical protein (fragment) | 0,1563 | 0,7502 |
| TcCLB.418221.10 | hypothetical protein, conserved | 0,1562 | 0,1857 |
| TcCLB.508985.10 | mucin-associated surface protein (MASP, pseudogene), pu | 0,156 | 0,6002 |
| TcCLB.511257.40 | hypothetical protein, conserved | 0,1559 | 0,2565 |
| TcCLB.510091.30 | hypothetical protein, conserved | 0,1557 | 0,298 |
| TcCLB.509979.130 | hypothetical protein, conserved | 0,1556 | 0,1097 |
| TcCLB.506915.10 | hypothetical protein, conserved | 0,1556 | 0,1786 |

| | | | |
|---|---|---|---|
| TcCLB.511529.200 | hypothetical protein, conserved | 0,1555 | 0,265 |
| TcCLB.509167.190 | protein kinase-like protein, putative | 0,1554 | 0,1243 |
| TcCLB.511715.80 | hypothetical protein, conserved | 0,1554 | 0,2739 |
| TcCLB.508111.20 | hypothetical protein, conserved | 0,1552 | 0,2945 |
| TcCLB.511805.15 | ribosomal protein S29, putative | 0,1552 | 0,7333 |
| TcCLB.506679.230 | ribosomal protein L24, putative | 0,1551 | 0,1118 |
| TcCLB.503663.4 | hypothetical protein, conserved (fragment) | 0,1551 | 0,5197 |
| TcCLB.503981.9 | hypothetical protein, conserved (pseudogene) | 0,155 | 0,4665 |
| TcCLB.507771.114 | hypothetical protein, conserved | 0,155 | 0,6237 |
| TcCLB.506965.130 | mucin-associated surface protein (MASP), putative | 0,1549 | 0,6167 |
| TcCLB.503521.39 | mitochondrial carrier protein, putative | 0,1548 | 0,4437 |
| TcCLB.506443.20 | hypothetical protein, conserved | 0,1547 | 0,3971 |
| TcCLB.511627.159 | hypothetical protein, conserved (fragment) | 0,1547 | 0,5319 |
| TcCLB.503973.150 | mucin-associated surface protein (MASP), putative | 0,1547 | 0,7919 |
| TcCLB.511419.30 | hypothetical protein, conserved | 0,1545 | 0,4476 |
| TcCLB.435051.10 | trans-sialidase, putative (fragment) | 0,1545 | 0,8037 |
| TcCLB.504429.50 | trans-sialidase (pseudogene), putative | 0,1544 | 0,4995 |
| TcCLB.506523.9 | dispersed gene family protein 1 (DGF-1), putative | 0,1544 | 0,6762 |
| TcCLB.508013.50 | mucin TcMUCII, putative | 0,1544 | 0,8342 |
| TcCLB.511001.200 | hypothetical protein, conserved | 0,1543 | 0,3587 |
| TcCLB.509751.28 | mucin-associated surface protein (MASP, pseudogene), pu | 0,1541 | 0,2678 |
| TcCLB.506887.110 | hypothetical protein, conserved (pseudogene) | 0,1541 | 0,4269 |
| TcCLB.508197.10 | 60S ribosomal protein L11, putative | 0,1539 | 0,0578 |
| TcCLB.508227.40 | mucin-associated surface protein (MASP), putative | 0,1539 | 0,5563 |
| TcCLB.509177.20 | hypothetical protein | 0,1537 | 0,3743 |
| TcCLB.511505.30 | hypothetical protein, conserved | 0,1537 | 0,498 |
| TcCLB.506367.160 | hypothetical protein, conserved | 0,1536 | 0,3174 |
| TcCLB.506633.20 | hypothetical protein, conserved | 0,1535 | 0,1951 |
| TcCLB.509047.119 | hypothetical protein, conserved (fragment) | 0,1535 | 0,6199 |
| TcCLB.509885.70 | transporter, putative | 0,1534 | 0,227 |
| TcCLB.503891.110 | dolicholphosphate-mannose synthase, putative | 0,153 | 0,3943 |
| TcCLB.509911.130 | hypothetical protein, conserved | 0,1524 | 0,1711 |
| TcCLB.504141.20 | hypothetical protein, conserved | 0,1523 | 0,2843 |
| TcCLB.509767.120 | hypothetical protein, conserved | 0,1523 | 0,2872 |
| TcCLB.507049.5 | kynurenine aminotransferase (pseudogene), putative | 0,152 | 0,3621 |
| TcCLB.506919.60 | hypothetical protein | 0,152 | 0,7458 |
| TcCLB.511133.10 | hypothetical protein, conserved | 0,1518 | 0,5333 |
| TcCLB.510101.210 | hypothetical protein, conserved | 0,1516 | 0,2431 |
| TcCLB.504179.90 | hypothetical protein, conserved | 0,1516 | 0,2471 |
| TcCLB.506829.50 | hypothetical protein, conserved | 0,1516 | 0,4106 |
| TcCLB.506955.50 | trans-sialidase (pseudogene), putative | 0,1516 | 0,6178 |
| TcCLB.509399.80 | hypothetical protein, conserved | 0,1515 | 0,2691 |
| TcCLB.511277.170 | ATP-dependent zinc metallopeptidase, putative | 0,1514 | 0,1195 |
| TcCLB.511109.70 | hypothetical protein, conserved | 0,1514 | 0,2117 |
| TcCLB.506825.160 | hypothetical protein, conserved | 0,1513 | 0,3505 |

| | | | |
|---|---|---|---|
| TcCLB.510751.30 | hypothetical protein | 0,1512 | 0,3312 |
| TcCLB.441923.10 | UDP-Gal or UDP-GlcNAc-dependent glycosyltransferase, pu | 0,151 | 0,5217 |
| TcCLB.506599.320 | hypothetical protein | 0,151 | 0,8479 |
| TcCLB.508981.20 | hypothetical protein, conserved | 0,1509 | 0,4688 |
| TcCLB.511471.10 | hypothetical protein, conserved (pseudogene) | 0,1509 | 0,5248 |
| TcCLB.511445.160 | copper-transporting ATPase-like protein, putative | 0,1508 | 0,08115 |
| TcCLB.511421.200 | ARM-like helical domain-containing protein | 0,1508 | 0,1843 |
| TcCLB.507911.33 | retrotransposon hot spot protein (RHS, pseudogene), putat | 0,1506 | 0,7498 |
| TcCLB.511541.20 | hypothetical protein | 0,1505 | 0,2584 |
| TcCLB.511149.50 | mucin TcMUCII, putative | 0,1504 | 0,7932 |
| TcCLB.506435.460 | hypothetical protein | 0,1503 | 0,3563 |
| TcCLB.508471.10 | hypothetical protein, conserved | 0,1503 | 0,4487 |
| TcCLB.503701.9 | dispersed gene family protein 1 (DGF-1, pseudogene), puta | 0,1502 | 0,5908 |
| TcCLB.510303.180 | hypothetical protein, conserved | 0,1501 | 0,06841 |
| TcCLB.510297.90 | hypothetical protein, conserved | 0,15 | 0,159 |
| TcCLB.511671.30 | beta galactofuranosyl glycosyltransferase, putative | 0,1499 | 0,7087 |
| TcCLB.508977.111 | elongation factor 1-gamma (EF-1-gamma, pseudogene), p | 0,1496 | 0,6756 |
| TcCLB.503417.33 | hypothetical protein | 0,1495 | 0,8278 |
| TcCLB.509665.13 | trans-sialidase (pseudogene), putative | 0,1494 | 0,6572 |
| TcCLB.510065.20 | DNA repair protein, putative | 0,1493 | 0,1821 |
| TcCLB.509717.80 | glyceraldehyde 3-phosphate dehydrogenase, cytosolic, puta | 0,1493 | 0,3432 |
| TcCLB.511871.10 | hypothetical protein, conserved | 0,1493 | 0,5292 |
| TcCLB.511555.110 | hypothetical protein, conserved | 0,1492 | 0,1536 |
| TcCLB.506885.60 | hypothetical protein, conserved | 0,1492 | 0,262 |
| TcCLB.506205.20 | hypothetical protein | 0,1491 | 0,2433 |
| TcCLB.510687.140 | alkylated DNA repair protein, putative | 0,1491 | 0,2944 |
| TcCLB.510187.280 | hypothetical protein, conserved | 0,1491 | 0,3986 |
| TcCLB.503557.10 | hypothetical protein, conserved | 0,149 | 0,4849 |
| TcCLB.508977.10 | trans-sialidase, Group V, putative | 0,149 | 0,5472 |
| TcCLB.508325.310 | hypothetical protein, conserved | 0,149 | 0,5909 |
| TcCLB.509967.150 | hypothetical protein, conserved | 0,1489 | 0,2658 |
| TcCLB.511527.60 | hypothetical protein, conserved | 0,1489 | 0,3766 |
| TcCLB.510989.60 | hypothetical protein, conserved | 0,1488 | 0,2281 |
| TcCLB.506815.60 | 60S ribosomal protein L44 (RPL44) | 0,1487 | 0,09673 |
| TcCLB.503815.20 | lysyl-tRNA synthetase, putative | 0,1487 | 0,3507 |
| TcCLB.508163.120 | mucin-associated surface protein (MASP, pseudogene), pu | 0,1487 | 0,6213 |
| TcCLB.511183.420 | mucin-associated surface protein (MASP, pseudogene), pu | 0,1487 | 0,6747 |
| TcCLB.507001.14 | Present in the outer mitochondrial membrane proteome 8 | 0,1485 | 0,5116 |
| TcCLB.509023.10 | calcium binding protein EF hand, putative | 0,1482 | 0,1963 |
| TcCLB.507811.30 | ATP-dependent DEAD/H RNA helicase, putative | 0,1482 | 0,2375 |
| TcCLB.510115.10 | trans-sialidase (pseudogene), putative | 0,1482 | 0,5335 |
| TcCLB.504121.39 | 60S ribosomal protein L14, putative (fragment) | 0,1481 | 0,1368 |
| TcCLB.503703.30 | hypothetical protein, conserved | 0,148 | 0,3096 |
| TcCLB.511141.30 | protein kinase, putative | 0,148 | 0,5866 |
| TcCLB.507069.180 | mucin-associated surface protein (MASP), putative | 0,148 | 0,7834 |

| | | | |
|---|---|---|---|
| TcCLB.508641.230 | hypothetical protein, conserved | 0,1479 | 0,2357 |
| TcCLB.507165.20 | hypothetical protein, conserved | 0,1479 | 0,5907 |
| TcCLB.504051.30 | hypothetical protein, conserved | 0,1476 | 0,2956 |
| TcCLB.509561.50 | hypothetical protein, conserved | 0,1476 | 0,3809 |
| TcCLB.508011.120 | trans-sialidase (pseudogene), putative | 0,1475 | 0,7177 |
| TcCLB.506963.10 | 60S ribosomal protein L10a, putative | 0,1474 | 0,05363 |
| TcCLB.509167.40 | amino acid permease, putative | 0,1474 | 0,2869 |
| TcCLB.507047.160 | hypothetical protein, conserved | 0,1474 | 0,598 |
| TcCLB.506599.360 | mucin-associated surface protein (MASP, pseudogene), pu | 0,1474 | 0,7505 |
| TcCLB.507041.30 | hypothetical protein, conserved | 0,1472 | 0,3847 |
| TcCLB.510431.170 | hypothetical protein, conserved | 0,1471 | 0,2968 |
| TcCLB.511501.50 | GTPase activating protein, putative | 0,1471 | 0,3069 |
| TcCLB.511267.30 | cholinephosphate cytidylyltransferase A, putative | 0,147 | 0,1949 |
| TcCLB.508891.50 | hypothetical protein | 0,147 | 0,239 |
| TcCLB.507673.10 | serine/threonine-protein phosphatase PP1, putative | 0,147 | 0,5219 |
| TcCLB.506605.174 | hypothetical protein, conserved | 0,147 | 0,5276 |
| TcCLB.506609.70 | mucin-associated surface protein (MASP), putative | 0,147 | 0,7823 |
| TcCLB.506797.80 | hypothetical protein, conserved | 0,1469 | 0,4132 |
| TcCLB.508879.70 | hypothetical protein, conserved | 0,1468 | 0,4552 |
| TcCLB.507005.80 | hypothetical protein, conserved | 0,1467 | 0,536 |
| TcCLB.504493.10 | mucin-associated surface protein (MASP), putative (fragm | 0,1467 | 0,8021 |
| TcCLB.506321.320 | suppressive immunomodulating factor, putative (TSIF) | 0,1466 | 0,2232 |
| TcCLB.505997.200 | hypothetical protein, conserved | 0,1464 | 0,4123 |
| TcCLB.508815.100 | hypothetical protein, conserved | 0,1463 | 0,2326 |
| TcCLB.510119.9 | elongation factor 1-alpha, putative | 0,1462 | 0,03253 |
| TcCLB.510341.10 | hypothetical protein | 0,1462 | 0,612 |
| TcCLB.511653.60 | hypothetical protein, conserved | 0,1461 | 0,4772 |
| TcCLB.511289.50 | mitogen-activated protein kinase 9, putative (MPK9) | 0,1459 | 0,2949 |
| TcCLB.506131.190 | mucin-associated surface protein (MASP), putative | 0,1459 | 0,7901 |
| TcCLB.503909.68 | hypothetical protein, conserved | 0,1458 | 0,4335 |
| TcCLB.505097.49 | expression site-associated gene (ESAG-like) protein, putati | 0,1457 | 0,7606 |
| TcCLB.510055.10 | trans-sialidase, putative | 0,1456 | 0,5335 |
| TcCLB.509925.5 | trans-sialidase (pseudogene), putative | 0,1456 | 0,6915 |
| TcCLB.506375.70 | hypothetical protein, conserved | 0,1455 | 0,514 |
| TcCLB.506457.20 | hypothetical protein, conserved | 0,1454 | 0,4373 |
| TcCLB.506831.5 | hypothetical protein, conserved (fragment) | 0,1454 | 0,6574 |
| TcCLB.508405.40 | 40S ribosomal protein S11, putative | 0,1452 | 0,1335 |
| TcCLB.511167.90 | 14-3-3 protein 2, putative (14-3-3-II) | 0,1452 | 0,1796 |
| TcCLB.510487.50 | mucin-associated surface protein (MASP), putative | 0,1451 | 0,7701 |
| TcCLB.508349.20 | tRNA nucleotidyltransferase, putative | 0,1448 | 0,2757 |
| TcCLB.510615.10 | hypothetical protein, conserved | 0,1447 | 0,2936 |
| TcCLB.510603.110 | hypothetical protein, conserved | 0,1447 | 0,5369 |
| TcCLB.503575.34 | ribosomal protein L38, putative | 0,1446 | 0,1898 |
| TcCLB.505957.80 | retrotransposon hot spot protein (RHS, pseudogene), putat | 0,1446 | 0,4096 |
| TcCLB.506967.50 | mucin-associated surface protein (MASP), putative | 0,1443 | 0,8092 |

| | | | |
|---|---|---|---|
| TcCLB.511269.20 | Multisite-specific tRNA:(cytosine-C(5))-methyltransferase, | 0,1441 | 0,2086 |
| TcCLB.506503.60 | hypothetical protein, conserved | 0,1441 | 0,2965 |
| TcCLB.506313.10 | retrotransposon hot spot (RHS) protein, putative (fragmen | 0,144 | 0,2723 |
| TcCLB.511181.130 | hypothetical protein | 0,1438 | 0,385 |
| TcCLB.511745.60 | hypothetical protein | 0,1438 | 0,71 |
| TcCLB.511907.240 | protein kinase, putative | 0,1437 | 0,2274 |
| TcCLB.509171.29 | RNA pseudouridylate synthase, putative | 0,1436 | 0,3211 |
| TcCLB.510529.20 | hypothetical protein, conserved | 0,1436 | 0,333 |
| TcCLB.503953.10 | hypothetical protein, conserved | 0,1435 | 0,2294 |
| TcCLB.508231.204 | hypothetical protein, conserved | 0,1434 | 0,4974 |
| TcCLB.510643.130 | damage-specific DNA binding protein, putative | 0,1433 | 0,1559 |
| TcCLB.511277.410 | hypothetical protein, conserved | 0,1433 | 0,2742 |
| TcCLB.504741.160 | ribosomal protein L11, putative | 0,1433 | 0,4455 |
| TcCLB.510967.30 | hypothetical protein | 0,1432 | 0,6101 |
| TcCLB.505051.30 | tryptophanyl-tRNA synthetase, putative (fragment) | 0,1429 | 0,8313 |
| TcCLB.511859.40 | hypothetical protein, conserved | 0,1427 | 0,5785 |
| TcCLB.506871.90 | hypothetical protein, conserved | 0,1426 | 0,1707 |
| TcCLB.510405.9 | hypothetical protein, conserved (fragment) | 0,1426 | 0,253 |
| TcCLB.511577.188 | hypothetical protein, conserved | 0,1426 | 0,4569 |
| TcCLB.508127.9 | hypothetical protein, conserved | 0,1424 | 0,2782 |
| TcCLB.506479.60 | kinesin heavy chain, putative | 0,1424 | 0,3124 |
| TcCLB.506789.330 | hypothetical protein, conserved | 0,1424 | 0,3622 |
| TcCLB.504121.30 | 60S ribosomal protein L4, putative | 0,1423 | 0,1062 |
| TcCLB.509011.54 | ubiquinol-cytochrome c reductase, putative | 0,1421 | 0,552 |
| TcCLB.506925.150 | protein phosphatase, putative | 0,142 | 0,2545 |
| TcCLB.508153.1080 | hypothetical protein, conserved | 0,142 | 0,4008 |
| TcCLB.504081.540 | trans-sialidase, Group V, putative | 0,142 | 0,7288 |
| TcCLB.406933.10 | retrotransposon hot spot (RHS) protein, putative | 0,1419 | 0,5377 |
| TcCLB.504125.60 | Kinesin-13 3, putative (KIN13-3) | 0,1418 | 0,1212 |
| TcCLB.511797.167 | mucin-associated surface protein (MASP), putative | 0,1417 | 0,8329 |
| TcCLB.506473.4 | hypothetical protein, conserved (pseudogene) | 0,1417 | 0,8346 |
| TcCLB.506211.10 | lipoic acid synthetase, mitochondrial precursor, putative | 0,1416 | 0,4226 |
| TcCLB.506869.40 | mitochondrial RNA binding complex 1 subunit, putative | 0,1415 | 0,3205 |
| TcCLB.511927.30 | hypothetical protein, conserved | 0,1414 | 0,627 |
| TcCLB.511211.230 | hypothetical protein, conserved | 0,1412 | 0,2637 |
| TcCLB.506605.160 | histone acetyltransferase, putative | 0,1412 | 0,3221 |
| TcCLB.508889.10 | hypothetical protein, conserved | 0,1411 | 0,318 |
| TcCLB.509127.80 | hypothetical protein, conserved | 0,1411 | 0,5517 |
| TcCLB.507895.170 | hypothetical protein, conserved | 0,1409 | 0,3592 |
| TcCLB.511599.90 | mucin-associated surface protein (MASP), putative | 0,1409 | 0,7844 |
| TcCLB.510373.90 | mucin-associated surface protein (MASP), putative | 0,1408 | 0,7691 |
| TcCLB.508851.70 | hypothetical protein | 0,1407 | 0,404 |
| TcCLB.511277.550 | hypothetical protein, conserved | 0,1407 | 0,5639 |
| TcCLB.510279.160 | trans-sialidase (pseudogene), putative | 0,1405 | 0,765 |
| TcCLB.508827.70 | 40S ribosomal protein S17, putative | 0,1404 | 0,1692 |

| | | | |
|---|---|---|---|
| TcCLB.506529.200 | hypothetical protein, conserved | 0,1404 | 0,3337 |
| TcCLB.503563.10 | calcium-translocating P-type ATPase, putative | 0,1404 | 0,4098 |
| TcCLB.510089.210 | hypothetical protein, conserved | 0,1402 | 0,391 |
| TcCLB.507817.40 | hypothetical protein, conserved | 0,1402 | 0,494 |
| TcCLB.508819.50 | katanin, putative (fragment) | 0,14 | 0,5238 |
| TcCLB.511465.30 | hypothetical protein, conserved | 0,14 | 0,554 |
| TcCLB.447255.10 | hypothetical protein, conserved | 0,1399 | 0,333 |
| TcCLB.506537.280 | glycine dehydrogenase [decarboxylating] (pseudogene), pu | 0,1399 | 0,3486 |
| TcCLB.507929.10 | kinesin, putative (fragment) | 0,1398 | 0,3307 |
| TcCLB.511277.450 | GPI-anchor transamidase subunit 8, putative (GPI8) | 0,1396 | 0,3313 |
| TcCLB.504147.90 | hypothetical protein, conserved | 0,1396 | 0,3479 |
| TcCLB.510659.219 | hypothetical protein, conserved | 0,1396 | 0,4452 |
| TcCLB.506625.130 | hypothetical protein, conserved | 0,1396 | 0,6551 |
| TcCLB.506513.180 | hypothetical protein, conserved | 0,1395 | 0,3239 |
| TcCLB.508777.90 | ATP-dependent DNA helicase, putative | 0,1395 | 0,3255 |
| TcCLB.503827.94 | chitin binding-like protein (pseudogene), putative | 0,1395 | 0,8281 |
| TcCLB.507519.110 | 60S ribosomal protein L7, putative | 0,1394 | 0,08721 |
| TcCLB.511263.50 | hypothetical protein, conserved | 0,1394 | 0,6498 |
| TcCLB.510941.20 | hypothetical protein, conserved | 0,1393 | 0,3973 |
| TcCLB.503599.50 | hypothetical protein, conserved | 0,1393 | 0,5015 |
| TcCLB.506537.110 | hypothetical protein, conserved | 0,1393 | 0,5685 |
| TcCLB.506321.190 | mucin TcMUCII, putative | 0,1393 | 0,7821 |
| TcCLB.511901.30 | Glycerophosphoryl diester phosphodiesterase family, putat | 0,1392 | 0,3042 |
| TcCLB.508873.425 | elongation factor 1-gamma (EF-1-gamma, pseudogene), p | 0,1392 | 0,5045 |
| TcCLB.506351.70 | hypothetical protein, conserved | 0,1391 | 0,2712 |
| TcCLB.511277.614 | hypothetical protein | 0,1391 | 0,6895 |
| TcCLB.507911.50 | hypothetical protein, conserved | 0,139 | 0,4202 |
| TcCLB.508919.189 | dispersed gene family protein 1 (DGF-1), putative | 0,139 | 0,5193 |
| TcCLB.410797.10 | surface protease GP63, putative | 0,1389 | 0,4333 |
| TcCLB.506691.80 | hypothetical protein, conserved | 0,1388 | 0,2907 |
| TcCLB.510903.10 | tRNA pseudouridine synthase, putative | 0,1388 | 0,4755 |
| TcCLB.507629.30 | hypothetical protein, conserved | 0,1387 | 0,3198 |
| TcCLB.503697.100 | hypothetical protein, conserved | 0,1386 | 0,172 |
| TcCLB.511861.90 | retrotransposon hot spot (RHS) protein, putative | 0,1386 | 0,4447 |
| TcCLB.507031.120 | metallo- peptidase, Clan MG, Family M24 | 0,1385 | 0,2809 |
| TcCLB.507027.80 | hypothetical protein, conserved | 0,1385 | 0,2943 |
| TcCLB.509471.40 | hypothetical protein, conserved | 0,1384 | 0,4418 |
| TcCLB.511671.149 | phosphate transporter, putative (PHO91) | 0,1383 | 0,3675 |
| TcCLB.510719.10 | vacuolar protein sorting-associated protein-like, putative | 0,1382 | 0,2695 |
| TcCLB.448567.9 | dynein heavy chain, putative | 0,1382 | 0,4428 |
| TcCLB.509039.49 | hypothetical protein, conserved | 0,138 | 0,6255 |
| TcCLB.508471.19 | hypothetical protein, conserved (fragment) | 0,1379 | 0,2201 |
| TcCLB.506341.40 | hypothetical protein | 0,1379 | 0,7976 |
| TcCLB.504033.180 | hypothetical protein, conserved | 0,1376 | 0,1188 |
| TcCLB.510963.10 | DNA damage repair protein, putative | 0,1375 | 0,2715 |

| | | | |
|---|---|---|---|
| TcCLB.509259.130 | protein kinase, putative | 0,1375 | 0,8307 |
| TcCLB.506127.30 | phenylalanyl-tRNA synthetase, putative | 0,1372 | 0,1199 |
| TcCLB.503733.30 | hypothetical protein, conserved | 0,1372 | 0,3829 |
| TcCLB.506821.50 | hypothetical protein | 0,1371 | 0,5659 |
| TcCLB.506045.19 | dispersed gene family protein 1 (DGF-1), putative (fragme | 0,1371 | 0,595 |
| TcCLB.505035.10 | hypothetical protein | 0,137 | 0,4661 |
| TcCLB.510929.10 | trans-sialidase, putative | 0,137 | 0,8385 |
| TcCLB.504115.90 | retrotransposon hot spot protein (RHS, pseudogene), putat | 0,1369 | 0,5009 |
| TcCLB.509679.9 | leucyl-tRNA synthetase, putative | 0,1368 | 0,495 |
| TcCLB.506793.20 | hypothetical protein, conserved | 0,1367 | 0,2882 |
| TcCLB.511063.100 | trans-sialidase (pseudogene), putative | 0,1367 | 0,7825 |
| TcCLB.508699.20 | Translation initiation factor eIF-2B subunit epsilon, putativ | 0,1366 | 0,2063 |
| TcCLB.505999.50 | hypothetical protein, conserved | 0,1366 | 0,5294 |
| TcCLB.504035.149 | kinesin, putative | 0,1365 | 0,4034 |
| TcCLB.504105.110 | hypothetical protein, conserved | 0,1363 | 0,1495 |
| TcCLB.509831.36 | guide RNA-binding protein of 21 kDa (gBP21) | 0,1362 | 0,2317 |
| TcCLB.510089.190 | hypothetical protein, conserved | 0,1362 | 0,3067 |
| TcCLB.507205.10 | mucin-associated surface protein (MASP, pseudogene), pu | 0,1362 | 0,7861 |
| TcCLB.506355.80 | flagellum transition zone component, putative | 0,136 | 0,1042 |
| TcCLB.507083.10 | hypothetical protein, conserved | 0,1359 | 0,2067 |
| TcCLB.504059.30 | hypothetical protein, conserved | 0,1358 | 0,1784 |
| TcCLB.509053.60 | hypothetical protein, conserved | 0,1358 | 0,605 |
| TcCLB.509233.10 | hypothetical protein | 0,1357 | 0,8784 |
| TcCLB.460757.10 | receptor-type adenylate cyclase, putative | 0,1356 | 0,6263 |
| TcCLB.510595.58 | hypothetical protein, conserved | 0,1356 | 0,7063 |
| TcCLB.509165.40 | 60S acidic ribosomal protein P2 beta (H6.4), putative (P2-b | 0,1355 | 0,148 |
| TcCLB.507513.60 | 40S ribosomal protein S16, putative | 0,1355 | 0,1803 |
| TcCLB.506435.160 | hypothetical protein, conserved | 0,1355 | 0,2583 |
| TcCLB.510665.60 | hypothetical protein, conserved (pseudogene) | 0,1355 | 0,3585 |
| TcCLB.509287.100 | mucin TcMUCII, putative | 0,1353 | 0,6394 |
| TcCLB.508241.130 | hypothetical protein, conserved | 0,1352 | 0,161 |
| TcCLB.511391.80 | hypothetical protein | 0,1352 | 0,7226 |
| TcCLB.510489.36 | mucin-associated surface protein (MASP, pseudogene), pu | 0,1351 | 0,7622 |
| TcCLB.419637.9 | dispersed gene family protein 1 (DGF-1), putative | 0,1351 | 0,849 |
| TcCLB.438147.9 | hypothetical protein, conserved (fragment) | 0,1349 | 0,5163 |
| TcCLB.511523.60 | hypothetical protein, conserved | 0,1348 | 0,2857 |
| TcCLB.506337.30 | class-E vacuolar protein-sorting protein 24 (Vps24p), putat | 0,1348 | 0,4008 |
| TcCLB.507481.20 | retrotransposon hot spot protein (RHS, pseudogene), putat | 0,1348 | 0,5783 |
| TcCLB.506409.60 | hypothetical protein | 0,1346 | 0,8729 |
| TcCLB.508963.30 | trans-sialidase, Group VIII, putative | 0,1344 | 0,6432 |
| TcCLB.509967.120 | intraflagellar transport protein 172, putative | 0,1343 | 0,1628 |
| TcCLB.506961.70 | trans-sialidase (pseudogene), putative | 0,1342 | 0,6077 |
| TcCLB.504787.60 | mucin TcMUCII (pseudogene), putative | 0,1342 | 0,7493 |
| TcCLB.508461.270 | rab7 GTP binding protein, putative | 0,134 | 0,1672 |
| TcCLB.511727.240 | hypothetical protein | 0,134 | 0,3438 |

| | | | |
|---|---|---|---|
| TcCLB.509897.200 | trans-sialidase, Group V, putative | 0,134 | 0,5702 |
| TcCLB.510543.130 | ATP-dependent DEAD/H RNA helicase (pseudogene), putat | 0,134 | 0,7707 |
| TcCLB.506919.130 | retrotransposon hot spot protein (RHS, pseudogene), putat | 0,1338 | 0,7873 |
| TcCLB.508817.60 | ribosomal protein S26, putative | 0,1337 | 0,1473 |
| TcCLB.504741.50 | hypothetical protein, conserved | 0,1334 | 0,4738 |
| TcCLB.511911.98 | hypothetical protein, conserved | 0,1333 | 0,388 |
| TcCLB.508355.260 | 60S acidic ribosomal protein P0, putative (P0) | 0,1332 | 0,04637 |
| TcCLB.506973.10 | trans-sialidase, Group V, putative | 0,1332 | 0,7291 |
| TcCLB.503827.20 | mucin-associated surface protein (MASP), putative | 0,1332 | 0,7709 |
| TcCLB.507519.80 | eukaryotic peptide chain release factor subunit 1, putative | 0,133 | 0,4234 |
| TcCLB.506339.60 | trans-sialidase (pseudogene), putative | 0,133 | 0,71 |
| TcCLB.508693.180 | ankyrin, putative | 0,1329 | 0,2367 |
| TcCLB.507867.50 | hypothetical protein, conserved | 0,1329 | 0,5307 |
| TcCLB.509549.110 | hypothetical protein, conserved (pseudogene) | 0,1326 | 0,2884 |
| TcCLB.510087.50 | hypothetical protein, conserved | 0,1323 | 0,2919 |
| TcCLB.511107.41 | ATP-dependent Clp protease subunit heat shock protein 10 | 0,1323 | 0,5032 |
| TcCLB.509215.13 | hypothetical protein, conserved | 0,1322 | 0,71 |
| TcCLB.510483.130 | mucin TcMUCII, putative | 0,1321 | 0,848 |
| TcCLB.504113.20 | MYH7B protein, putative | 0,132 | 0,3941 |
| TcCLB.467795.19 | hypothetical protein, conserved | 0,1318 | 0,5209 |
| TcCLB.506221.40 | mitochondrial carrier protein, putative | 0,1316 | 0,415 |
| TcCLB.510181.140 | hypothetical protein, conserved | 0,1316 | 0,5276 |
| TcCLB.511391.220 | hypothetical protein, conserved | 0,1315 | 0,0648 |
| TcCLB.510721.10 | hypothetical protein, conserved | 0,1313 | 0,2135 |
| TcCLB.509525.270 | mucin TcMUCII, putative | 0,1313 | 0,8841 |
| TcCLB.509123.30 | mitochondrial RNA binding protein 1 | 0,1312 | 0,4551 |
| TcCLB.509719.56 | hypothetical protein, conserved | 0,131 | 0,2753 |
| TcCLB.506485.50 | hypothetical protein | 0,131 | 0,4165 |
| TcCLB.509233.180 | ATP synthase F1, beta subunit, putative (ATPB) | 0,1309 | 0,1547 |
| TcCLB.506529.150 | hypothetical protein, conserved | 0,1307 | 0,2453 |
| TcCLB.506919.10 | hypothetical protein | 0,1307 | 0,7114 |
| TcCLB.511391.210 | mitochondrial calcium uptake 1, putative | 0,1306 | 0,2587 |
| TcCLB.506195.110 | malate dehydrogenase, putative | 0,1305 | 0,1012 |
| TcCLB.509171.10 | hypothetical protein, conserved | 0,1305 | 0,6125 |
| TcCLB.511075.24 | zinc-finger multi-pass transmembrane protein, putative | 0,1304 | 0,4212 |
| TcCLB.508817.40 | hypothetical protein, conserved | 0,1301 | 0,3254 |
| TcCLB.510431.300 | trans-sialidase, Group II, putative | 0,1301 | 0,6447 |
| TcCLB.511683.10 | hypothetical protein | 0,1298 | 0,6023 |
| TcCLB.506207.40 | hypothetical protein, conserved | 0,1297 | 0,2413 |
| TcCLB.503925.80 | CDP-diacylglycerol--inositol 3-phosphatidyltransferase | 0,1296 | 0,5029 |
| TcCLB.510105.220 | transportin2-like protein, putative | 0,1295 | 0,2804 |
| TcCLB.509023.180 | kinesin, putative | 0,1294 | 0,2226 |
| TcCLB.506967.65 | mucin TcMUCII (pseudogene), putative | 0,1294 | 0,8137 |
| TcCLB.508839.60 | ADP-ribosylation factor, putative | 0,1293 | 0,5199 |
| TcCLB.510181.80 | hypothetical protein, conserved | 0,1292 | 0,4097 |

| | | | |
|---|---|---|---|
| TcCLB.508153.840 | hypothetical protein, conserved | 0,129 | 0,3869 |
| TcCLB.504137.140 | hypothetical protein, conserved | 0,129 | 0,3986 |
| TcCLB.414243.10 | hypothetical protein, conserved | 0,129 | 0,5643 |
| TcCLB.509003.50 | hypothetical protein, conserved | 0,129 | 0,6165 |
| TcCLB.511487.170 | mucin-associated surface protein (MASP), putative | 0,129 | 0,7105 |
| TcCLB.508221.840 | hypothetical protein | 0,1289 | 0,8341 |
| TcCLB.503497.10 | mucin-associated surface protein (MASP, pseudogene), pu | 0,1288 | 0,5974 |
| TcCLB.511279.10 | hypothetical protein, conserved | 0,1287 | 0,534 |
| TcCLB.510127.70 | leucine-rich repeat protein 1 (LRRP1), putative | 0,1287 | 0,5713 |
| TcCLB.506811.100 | hypothetical protein, conserved | 0,1284 | 0,3878 |
| TcCLB.511837.80 | hypothetical protein, conserved | 0,1284 | 0,4086 |
| TcCLB.506617.50 | hypothetical protein | 0,1284 | 0,4615 |
| TcCLB.506745.30 | hypothetical protein, conserved | 0,1283 | 0,808 |
| TcCLB.511655.50 | hypothetical protein, conserved (pseudogene) | 0,1282 | 0,367 |
| TcCLB.506177.70 | hypothetical protein | 0,1279 | 0,5794 |
| TcCLB.510099.131 | hypothetical protein, conserved (fragment) | 0,1278 | 0,3236 |
| TcCLB.509657.30 | mucin-associated surface protein (MASP), putative | 0,1278 | 0,7515 |
| TcCLB.506925.180 | protein kinase, putative | 0,1275 | 0,2799 |
| TcCLB.506155.99 | hypothetical protein, conserved | 0,1275 | 0,4935 |
| TcCLB.506181.130 | hypothetical protein | 0,1275 | 0,5135 |
| TcCLB.507801.50 | tubulin-tyrosine ligase, putative | 0,1273 | 0,4001 |
| TcCLB.511283.294 | b9 domain-containing protein 1 (b9d1) | 0,1272 | 0,283 |
| TcCLB.509127.110 | hypothetical protein, conserved | 0,1272 | 0,4171 |
| TcCLB.511301.40 | eukaryotic translation initiation factor 4E (eIF4E) interactir | 0,1272 | 0,4455 |
| TcCLB.508765.50 | mucin TcMUCI (pseudogene), putative | 0,1272 | 0,668 |
| TcCLB.508905.20 | hypothetical protein, conserved (pseudogene) | 0,1272 | 0,6681 |
| TcCLB.509995.10 | 60S ribosomal protein L4, putative | 0,1271 | 0,1697 |
| TcCLB.506905.40 | ABC transporter, putative | 0,1271 | 0,5607 |
| TcCLB.507711.60 | hypothetical protein, conserved | 0,127 | 0,3388 |
| TcCLB.506195.190 | intraflagellar transport protein 81, putative (IFT81) | 0,1269 | 0,1902 |
| TcCLB.506321.310 | phosphoribosylpyrophosphate synthetase, putative | 0,1269 | 0,1984 |
| TcCLB.506227.210 | hypothetical protein, conserved | 0,1268 | 0,4793 |
| TcCLB.425785.9 | hypothetical protein, conserved | 0,1266 | 0,5064 |
| TcCLB.511423.70 | glycerol-3-phosphate dehydrogenase (FAD-dependent), pu | 0,1265 | 0,4571 |
| TcCLB.506859.130 | hypothetical protein, conserved | 0,1263 | 0,6408 |
| TcCLB.504755.10 | surface protease GP63, putative | 0,1263 | 0,9106 |
| TcCLB.508277.350 | cation transporter, putative | 0,1262 | 0,1893 |
| TcCLB.503453.50 | hypothetical protein, conserved | 0,1261 | 0,4255 |
| TcCLB.509507.49 | mitotic cyclin, putative | 0,126 | 0,5709 |
| TcCLB.507505.30 | hypothetical protein | 0,1259 | 0,3923 |
| TcCLB.511131.60 | hypothetical protein, conserved | 0,1259 | 0,4396 |
| TcCLB.507275.30 | hypothetical protein, conserved | 0,1259 | 0,4592 |
| TcCLB.506363.80 | aminoacylase, putative | 0,1259 | 0,8243 |
| TcCLB.507625.170 | hypothetical protein, conserved | 0,1258 | 0,3118 |
| TcCLB.508857.120 | hypothetical protein, conserved | 0,1258 | 0,4608 |

| | | | |
|---|---|---|---|
| TcCLB.508593.80 | hypothetical protein, conserved | 0,1257 | 0,3282 |
| TcCLB.508799.10 | hypothetical protein, conserved | 0,1257 | 0,4684 |
| TcCLB.508443.40 | hypothetical protein | 0,1257 | 0,5255 |
| TcCLB.509999.120 | RNA-binding protein, putative | 0,1257 | 0,7226 |
| TcCLB.508333.9 | dispersed gene family protein 1 (DGF-1), putative (fragme | 0,1257 | 0,7711 |
| TcCLB.511145.40 | peroxin 14, putative | 0,1256 | 0,3497 |
| TcCLB.507997.30 | trans-sialidase, Group V, putative | 0,1256 | 0,6458 |
| TcCLB.511803.40 | Present in the outer mitochondrial membrane proteome 9 | 0,1255 | 0,1152 |
| TcCLB.509233.239 | hypothetical protein, conserved | 0,1255 | 0,6331 |
| TcCLB.506979.40 | hypothetical protein, conserved | 0,1254 | 0,4396 |
| TcCLB.507841.14 | retrotransposon hot spot (RHS) protein, putative (fragmen | 0,1254 | 0,4496 |
| TcCLB.508081.40 | hypothetical protein | 0,1254 | 0,8754 |
| TcCLB.510025.60 | hypothetical protein | 0,1254 | 0,9068 |
| TcCLB.506999.120 | hypothetical protein, conserved | 0,1253 | 0,3754 |
| TcCLB.505993.10 | trans-sialidase (pseudogene), putative | 0,1252 | 0,7029 |
| TcCLB.509599.80 | hypothetical protein, conserved | 0,1251 | 0,3987 |
| TcCLB.507991.130 | hypothetical protein, conserved | 0,1249 | 0,5109 |
| TcCLB.506613.24 | mucin TcMUCII, putative | 0,1249 | 0,8344 |
| TcCLB.511107.10 | hypothetical protein, conserved (pseudogene) | 0,1248 | 0,4047 |
| TcCLB.506989.120 | hypothetical protein, conserved | 0,1248 | 0,4563 |
| TcCLB.507935.10 | mucin-like glycoprotein (pseudogene), putative | 0,1248 | 0,8125 |
| TcCLB.506625.190 | hypothetical protein, conserved | 0,1247 | 0,5175 |
| TcCLB.503453.20 | hypothetical protein, conserved | 0,1246 | 0,2782 |
| TcCLB.504153.370 | hypothetical protein | 0,1244 | 0,3251 |
| TcCLB.509243.40 | hypothetical protein, conserved (pseudogene) | 0,1244 | 0,4705 |
| TcCLB.503721.10 | hypothetical protein, conserved | 0,124 | 0,4131 |
| TcCLB.507763.20 | hypothetical protein | 0,124 | 0,619 |
| TcCLB.506849.20 | dynein heavy chain (pseudogene), putative | 0,1239 | 0,431 |
| TcCLB.503909.120 | cyclosome subunit, putative | 0,1239 | 0,5976 |
| TcCLB.510013.190 | hypothetical protein | 0,1239 | 0,8438 |
| TcCLB.506309.30 | mucin-associated surface protein (MASP), putative | 0,1239 | 0,8669 |
| TcCLB.503887.70 | phosphoglycerate mutase, putative (iPGAM) | 0,1235 | 0,5786 |
| TcCLB.507649.5 | hypothetical protein, conserved | 0,1235 | 0,7594 |
| TcCLB.503431.90 | hypothetical protein, conserved | 0,1233 | 0,4212 |
| TcCLB.510303.260 | hypothetical protein, conserved | 0,123 | 0,4685 |
| TcCLB.506847.50 | trans-sialidase (pseudogene), putative | 0,123 | 0,918 |
| TcCLB.510531.60 | hypothetical protein, conserved | 0,1229 | 0,299 |
| TcCLB.503539.20 | Putative intraflagellar transport protein A1 (PIFTA1) | 0,1229 | 0,3534 |
| TcCLB.506757.50 | mucin-associated surface protein (MASP), putative | 0,1229 | 0,7228 |
| TcCLB.506501.170 | mucin TcMUCII, putative | 0,1229 | 0,8514 |
| TcCLB.508165.220 | mucin TcMUCII (pseudogene), putative | 0,1228 | 0,8252 |
| TcCLB.506309.100 | mucin-associated surface protein (MASP, pseudogene), pu | 0,1228 | 0,8904 |
| TcCLB.509429.170 | hypothetical protein, conserved | 0,1227 | 0,4989 |
| TcCLB.507011.230 | hypothetical protein, conserved (pseudogene) | 0,1226 | 0,3452 |
| TcCLB.510517.30 | hypothetical protein, conserved | 0,1226 | 0,3533 |

| | | | |
|---|---|---|---|
| TcCLB.504123.20 | hypothetical protein, conserved | 0,1226 | 0,4342 |
| TcCLB.503557.40 | Present in the outer mitochondrial membrane proteome 8 | 0,1226 | 0,5963 |
| TcCLB.506247.60 | hypothetical protein, conserved | 0,1225 | 0,3554 |
| TcCLB.507949.50 | SSU ribosomal protein, mitochondrial, putative (MRPS15) | 0,1222 | 0,3983 |
| TcCLB.510667.4 | hypothetical protein, conserved (fragment) | 0,1222 | 0,49 |
| TcCLB.506763.121 | elongation factor 1-gamma (EF-1-gamma, pseudogene), p | 0,1221 | 0,8603 |
| TcCLB.503415.10 | hypothetical protein (pseudogene) | 0,122 | 0,4458 |
| TcCLB.509647.180 | nicotinate phosphoribosyltransferase, putative | 0,1219 | 0,2479 |
| TcCLB.511233.20 | 60S ribosomal protein L34, putative | 0,1218 | 0,1958 |
| TcCLB.509023.60 | hypothetical protein, conserved | 0,1218 | 0,3853 |
| TcCLB.511057.40 | trans-sialidase, Group V, putative | 0,1218 | 0,6543 |
| TcCLB.511585.210 | hypothetical protein, conserved | 0,1216 | 0,496 |
| TcCLB.510647.50 | hypothetical protein, conserved | 0,1216 | 0,688 |
| TcCLB.506453.20 | mucin TcMUC (pseudogene), putative | 0,1215 | 0,6884 |
| TcCLB.509659.20 | dispersed gene family protein 1 (DGF-1, pseudogene), puta | 0,1215 | 0,7088 |
| TcCLB.506203.50 | DnaJ chaperone protein, putative | 0,1214 | 0,4451 |
| TcCLB.506425.120 | eRF1 methyltransferase catalytic subunit, putative | 0,1214 | 0,5925 |
| TcCLB.508541.190 | cell cycle sequence binding phosphoprotein (RBP33), putat | 0,1213 | 0,2816 |
| TcCLB.510089.80 | protein kinase, putative | 0,1213 | 0,406 |
| TcCLB.503617.40 | ATP synthase F1, alpha subunit, putative (fragment) (ATPF | 0,1212 | 0,2026 |
| TcCLB.509213.140 | mitochondrial RNA binding complex 1 subunit, putative | 0,1212 | 0,2592 |
| TcCLB.508211.30 | sphingosine kinase A, B, putative | 0,1212 | 0,3736 |
| TcCLB.508269.10 | hypothetical protein, conserved | 0,1212 | 0,4127 |
| TcCLB.506931.64 | RNA processing endonuclease 1, putative | 0,1212 | 0,4853 |
| TcCLB.504039.50 | retrotransposon hot spot protein (RHS, pseudogene), putat | 0,1211 | 0,6953 |
| TcCLB.504149.201 | ATP-dependent DEAD/H RNA helicase (pseudogene), putat | 0,1211 | 0,7038 |
| TcCLB.505009.4 | hypothetical protein, conserved | 0,1211 | 0,7653 |
| TcCLB.445869.10 | hypothetical protein, conserved | 0,121 | 0,4277 |
| TcCLB.507881.110 | trans-sialidase (pseudogene), putative | 0,121 | 0,4976 |
| TcCLB.509671.20 | hypothetical protein, conserved | 0,1208 | 0,3788 |
| TcCLB.509693.100 | 2-aminoethylphosphonate:pyruvateaminotransferase- like | 0,1208 | 0,5135 |
| TcCLB.511895.40 | hypothetical protein | 0,1208 | 0,5444 |
| TcCLB.510717.30 | 2-oxoglutarate dehydrogenase E1 component, putative | 0,1207 | 0,191 |
| TcCLB.509699.110 | mucin-associated surface protein (MASP), putative | 0,1206 | 0,7665 |
| TcCLB.510173.30 | hypothetical protein, conserved | 0,1205 | 0,5848 |
| TcCLB.507237.160 | mucin TcMUCII, putative | 0,1205 | 0,882 |
| TcCLB.509027.10 | DNA photolyase, putative | 0,1204 | 0,5209 |
| TcCLB.509647.50 | hypothetical protein, conserved | 0,1203 | 0,2003 |
| TcCLB.510743.120 | hypothetical protein, conserved | 0,1203 | 0,3665 |
| TcCLB.507923.40 | hypothetical protein, conserved | 0,1201 | 0,4974 |
| TcCLB.507711.260 | hypothetical protein, conserved | 0,12 | 0,3409 |
| TcCLB.506705.60 | hypothetical protein, conserved | 0,12 | 0,3499 |
| TcCLB.508523.21 | trans-sialidase (pseudogene), putative | 0,12 | 0,634 |
| TcCLB.504625.99 | protein tyrosine phosphatase, putative (fragment) | 0,12 | 0,6445 |
| TcCLB.506893.40 | hypothetical protein, conserved | 0,12 | 0,6847 |

| | | | |
|---|---|---|---|
| TcCLB.511229.30 | hypothetical protein, conserved | 0,1199 | 0,4218 |
| TcCLB.510575.210 | hypothetical protein, conserved | 0,1199 | 0,4349 |
| TcCLB.510063.20 | ATP-dependent DEAD/H DNA helicase recQ, putative | 0,1199 | 0,7074 |
| TcCLB.507821.20 | hypothetical protein | 0,1199 | 0,7972 |
| TcCLB.508741.240 | hypothetical protein, conserved | 0,1195 | 0,3451 |
| TcCLB.505071.30 | serine/threonine protein kinase, putative | 0,1195 | 0,4269 |
| TcCLB.509195.70 | mucin-associated surface protein (MASP), putative | 0,1195 | 0,6944 |
| TcCLB.506679.40 | mitogen-activated protein kinase kinase 5, putative (MKK5 | 0,1193 | 0,3478 |
| TcCLB.511239.70 | hypothetical protein, conserved | 0,1192 | 0,3956 |
| TcCLB.506879.100 | hypothetical protein, conserved | 0,1191 | 0,6589 |
| TcCLB.511245.60 | hypothetical protein, conserved | 0,1186 | 0,7454 |
| TcCLB.511185.120 | mucin-associated surface protein (MASP), putative | 0,1186 | 0,791 |
| TcCLB.511881.14 | RNA polymerase subunit RPB10, putative (RPB10) | 0,1185 | 0,569 |
| TcCLB.510355.41 | mucin-associated surface protein (MASP, pseudogene), pu | 0,1185 | 0,8065 |
| TcCLB.508479.90 | hypothetical protein, conserved (pseudogene) | 0,1184 | 0,4571 |
| TcCLB.504147.110 | asparaginase, putative | 0,1184 | 0,6156 |
| TcCLB.511469.105 | hypothetical protein | 0,1184 | 0,6322 |
| TcCLB.468217.14 | mucin-associated surface protein (MASP), putative | 0,1184 | 0,7173 |
| TcCLB.510465.100 | trans-sialidase, Group V, putative | 0,1184 | 0,8107 |
| TcCLB.507957.310 | hypothetical protein | 0,1184 | 0,8182 |
| TcCLB.430605.20 | PTP1-interacting protein, 39 kDa, putative | 0,1183 | 0,2759 |
| TcCLB.510723.10 | hypothetical protein, conserved | 0,1182 | 0,7054 |
| TcCLB.506201.90 | hypothetical protein, conserved | 0,1181 | 0,4551 |
| TcCLB.510371.90 | trans-sialidase, Group V, putative | 0,118 | 0,6184 |
| TcCLB.509695.10 | aspartyl-tRNA synthetase, putative | 0,1178 | 0,4257 |
| TcCLB.509063.10 | hypothetical protein, conserved | 0,1178 | 0,5515 |
| TcCLB.508347.80 | Present in the outer mitochondrial membrane proteome 2 | 0,1177 | 0,2695 |
| TcCLB.503909.40 | hypothetical protein | 0,1176 | 0,6749 |
| TcCLB.510311.140 | arginine N-methyltransferase, putative | 0,1175 | 0,3519 |
| TcCLB.504009.40 | Axonemal inner arm dynein light chain, putative | 0,1175 | 0,4818 |
| TcCLB.510275.110 | trans-sialidase (pseudogene), putative | 0,1175 | 0,7306 |
| TcCLB.506223.50 | hypothetical protein, conserved | 0,1174 | 0,4665 |
| TcCLB.503917.21 | hypothetical protein, conserved | 0,1174 | 0,5088 |
| TcCLB.504085.40 | hypothetical protein, conserved | 0,1173 | 0,4688 |
| TcCLB.509695.180 | 40S ribosomal protein S23, putative | 0,1172 | 0,1876 |
| TcCLB.507639.90 | hypothetical protein, conserved | 0,1171 | 0,658 |
| TcCLB.504081.460 | mucin-associated surface protein (MASP), putative | 0,1171 | 0,7527 |
| TcCLB.510359.50 | hypothetical protein, conserved | 0,117 | 0,6913 |
| TcCLB.432027.9 | hypothetical protein, conserved | 0,1169 | 0,596 |
| TcCLB.507979.30 | trans-sialidase, Group I, putative | 0,1168 | 0,6578 |
| TcCLB.511727.199 | CYC2-like cyclin 6, putative (CYC6) | 0,1167 | 0,4815 |
| TcCLB.510375.20 | trans-sialidase (pseudogene), putative | 0,1167 | 0,854 |
| TcCLB.429251.10 | mucin-associated surface protein (MASP), putative | 0,1167 | 0,9242 |
| TcCLB.509543.30 | hypothetical protein | 0,1166 | 0,5142 |
| TcCLB.508735.65 | trans-sialidase (pseudogene), putative | 0,1165 | 0,6014 |

| | | | |
|---|---|---|---|
| TcCLB.508207.60 | hypothetical protein, conserved | 0,1164 | 0,521 |
| TcCLB.507065.70 | mucin TcMUCII, putative | 0,1162 | 0,8761 |
| TcCLB.507001.34 | hypothetical protein, conserved | 0,1161 | 0,4803 |
| TcCLB.506281.100 | mucin-associated surface protein (MASP, pseudogene), pu | 0,116 | 0,7886 |
| TcCLB.503687.49 | hypothetical protein, conserved | 0,1159 | 0,4956 |
| TcCLB.511621.50 | RNA-binding protein, putative | 0,1158 | 0,2352 |
| TcCLB.508849.20 | retrotransposon hot spot protein (RHS, pseudogene), putat | 0,1158 | 0,4455 |
| TcCLB.506855.10 | palmitoyl acyltransferase 3, putative | 0,1158 | 0,5089 |
| TcCLB.507093.40 | hypothetical protein, conserved | 0,1157 | 0,2368 |
| TcCLB.507517.70 | hypothetical protein, conserved | 0,1157 | 0,4811 |
| TcCLB.510283.40 | phosphoglycerate mutase family member 5, putative | 0,1156 | 0,6251 |
| TcCLB.511211.10 | hypothetical protein, conserved | 0,1155 | 0,5052 |
| TcCLB.510525.60 | hypothetical protein, conserved | 0,1155 | 0,5278 |
| TcCLB.508689.50 | GPI transamidase subunit 16, putative (fragment) (GPI16) | 0,1155 | 0,6321 |
| TcCLB.504033.190 | EF hand, putative | 0,1153 | 0,4505 |
| TcCLB.509599.180 | hypothetical protein, conserved | 0,1153 | 0,5097 |
| TcCLB.510697.70 | elongation factor 1-gamma (EF-1-gamma, pseudogene), p | 0,1153 | 0,5378 |
| TcCLB.510359.380 | mucin-associated surface protein (MASP), putative | 0,1152 | 0,8206 |
| TcCLB.509693.200 | hypothetical protein, conserved | 0,1151 | 0,5323 |
| TcCLB.511127.360 | hypothetical protein, conserved | 0,1149 | 0,5189 |
| TcCLB.506003.10 | retrotransposon hot spot protein (RHS, pseudogene), putat | 0,1146 | 0,7616 |
| TcCLB.509055.10 | RNA-binding protein, putative | 0,1145 | 0,5365 |
| TcCLB.507093.300 | hypothetical protein SCD6.10 | 0,1144 | 0,3203 |
| TcCLB.509717.70 | transportin2-like protein, putative | 0,1143 | 0,3214 |
| TcCLB.511149.4 | mucin-associated surface protein (MASP, pseudogene), pu | 0,1143 | 0,8483 |
| TcCLB.510055.140 | hypothetical protein, conserved | 0,1142 | 0,5228 |
| TcCLB.510515.110 | hypothetical protein, conserved | 0,114 | 0,3942 |
| TcCLB.506303.160 | hypothetical protein | 0,114 | 0,4266 |
| TcCLB.507951.70 | hypothetical protein | 0,114 | 0,5957 |
| TcCLB.511383.10 | hypothetical protein, conserved | 0,114 | 0,6809 |
| TcCLB.506727.120 | hypothetical protein, conserved | 0,1139 | 0,285 |
| TcCLB.507957.179 | mucin-associated surface protein (MASP), putative | 0,1138 | 0,8431 |
| TcCLB.504029.50 | hypothetical protein, conserved | 0,1135 | 0,482 |
| TcCLB.508005.20 | trans-sialidase (pseudogene), putative | 0,1134 | 0,8601 |
| TcCLB.511389.80 | protein kinase, putative | 0,1131 | 0,3971 |
| TcCLB.509695.140 | hypothetical protein, conserved | 0,113 | 0,4438 |
| TcCLB.510437.20 | 60S ribosomal protein L14, putative | 0,1129 | 0,2219 |
| TcCLB.510403.89 | synaptojanin (N-terminal domain), putative | 0,1129 | 0,5071 |
| TcCLB.508479.270 | hypothetical protein, conserved | 0,1128 | 0,2925 |
| TcCLB.507925.10 | hypothetical protein, conserved (pseudogene) | 0,1128 | 0,7787 |
| TcCLB.511745.20 | AAA ATPase, putative | 0,1127 | 0,3382 |
| TcCLB.510149.9 | hypothetical protein, conserved (fragment) | 0,1126 | 0,5452 |
| TcCLB.508641.190 | hypothetical protein, conserved | 0,1126 | 0,596 |
| TcCLB.508707.300 | hypothetical protein, conserved | 0,1125 | 0,2711 |
| TcCLB.511145.30 | hypothetical protein, conserved | 0,1125 | 0,4106 |

| | | | |
|---|---|---|---|
| TcCLB.511109.30 | dynein intermediate chain, putative | 0,1125 | 0,4506 |
| TcCLB.510749.30 | Sorting nexin-4 (ATG24) | 0,1124 | 0,3165 |
| TcCLB.508355.104 | hypothetical protein, conserved | 0,1124 | 0,6756 |
| TcCLB.510767.10 | ribosomal protein L15, putative | 0,1123 | 0,203 |
| TcCLB.509669.80 | hypothetical protein, conserved | 0,1122 | 0,6838 |
| TcCLB.508165.120 | mucin-associated surface protein (MASP), putative | 0,1122 | 0,7659 |
| TcCLB.503823.70 | hypothetical protein, conserved | 0,1121 | 0,6919 |
| TcCLB.506435.280 | hypothetical protein, conserved | 0,112 | 0,4194 |
| TcCLB.508411.20 | hypothetical protein, conserved | 0,112 | 0,5267 |
| TcCLB.508257.170 | dynein heavy chain, putative | 0,1119 | 0,3776 |
| TcCLB.507957.220 | mucin-associated surface protein (MASP), putative | 0,1119 | 0,7599 |
| TcCLB.510901.260 | tRNA pseudouridine synthase (pseudogene), putative | 0,1118 | 0,5339 |
| TcCLB.506861.30 | 60S ribosomal protein L14, putative | 0,1117 | 0,2281 |
| TcCLB.506717.25 | trans-sialidase (pseudogene), putative | 0,1117 | 0,6575 |
| TcCLB.503439.14 | dispersed gene family protein 1 (DGF-1, pseudogene), puta | 0,1116 | 0,6573 |
| TcCLB.508873.370 | hypothetical protein | 0,1116 | 0,7932 |
| TcCLB.508261.120 | hypothetical protein | 0,1115 | 0,8073 |
| TcCLB.506227.40 | hypothetical protein, conserved | 0,1114 | 0,4725 |
| TcCLB.511515.9 | E3 ubiquitin-protein ligase, putative (fragment) | 0,1113 | 0,5453 |
| TcCLB.506559.470 | ribosomal protein l35a, putative | 0,1112 | 0,1469 |
| TcCLB.510631.20 | trans-sialidase (pseudogene), putative | 0,1111 | 0,5185 |
| TcCLB.511039.6 | hypothetical protein, conserved | 0,1111 | 0,5435 |
| TcCLB.505965.60 | hypothetical protein, conserved | 0,1111 | 0,5791 |
| TcCLB.452699.30 | dispersed gene family protein 1 (DGF-1), putative | 0,1111 | 0,837 |
| TcCLB.506679.94 | 40S ribosomal protein S18, putative | 0,111 | 0,2082 |
| TcCLB.509979.90 | 60S ribosomal protein L35, putative | 0,111 | 0,2757 |
| TcCLB.508977.120 | trans-sialidase, Group V, putative | 0,111 | 0,6067 |
| TcCLB.511455.10 | surface protease GP63 (pseudogene), putative | 0,111 | 0,6946 |
| TcCLB.507573.3 | N-acetyltransferase complex ARD1 subunit (pseudogene), | 0,1109 | 0,732 |
| TcCLB.511293.50 | kinesin, putative | 0,1107 | 0,4283 |
| TcCLB.509029.100 | hypothetical protein, conserved | 0,1107 | 0,5095 |
| TcCLB.508469.10 | hypothetical protein, conserved | 0,1107 | 0,7079 |
| TcCLB.506689.10 | trans-sialidase (pseudogene), putative | 0,1107 | 0,7738 |
| TcCLB.508205.20 | nucleolar RNA helicase II, putative | 0,1106 | 0,3665 |
| TcCLB.510823.100 | hypothetical protein, conserved | 0,1106 | 0,4283 |
| TcCLB.506241.10 | retrotransposon hot spot protein (RHS, pseudogene), putat | 0,1105 | 0,7114 |
| TcCLB.511545.140 | hypothetical protein, conserved | 0,1104 | 0,415 |
| TcCLB.510579.14 | hypothetical protein, conserved | 0,1103 | 0,4055 |
| TcCLB.508461.70 | hypothetical protein | 0,1103 | 0,6769 |
| TcCLB.508267.10 | hypothetical protein, conserved | 0,1102 | 0,3508 |
| TcCLB.506729.70 | hypothetical protein, conserved | 0,1102 | 0,4575 |
| TcCLB.511171.60 | surface protease GP63 (pseudogene), putative | 0,1101 | 0,8705 |
| TcCLB.503677.10 | phosphomannose isomerase, putative | 0,11 | 0,4109 |
| TcCLB.511737.9 | hypothetical protein, conserved | 0,11 | 0,7282 |
| TcCLB.506635.80 | Hydroxymethylglutaryl-CoA lyase, mitochondrial, putative | 0,1099 | 0,4065 |

| | | | |
|---|---|---|---|
| TcCLB.506961.160 | retrotransposon hot spot (RHS) protein, putative | 0,1098 | 0,4379 |
| TcCLB.509997.73 | hypothetical protein, conserved | 0,1098 | 0,5137 |
| TcCLB.509065.180 | protein kinase-like protein, putative | 0,1097 | 0,3528 |
| TcCLB.511197.40 | trans-sialidase (pseudogene), putative | 0,1097 | 0,7112 |
| TcCLB.503449.14 | hypothetical protein, conserved | 0,1096 | 0,5081 |
| TcCLB.510257.100 | mitochondrial RNA binding complex 1 subunit, putative | 0,1095 | 0,3089 |
| TcCLB.510359.360 | hypothetical protein, conserved | 0,1095 | 0,4832 |
| TcCLB.507027.50 | Present in the outer mitochondrial membrane proteome 3 | 0,1095 | 0,5189 |
| TcCLB.506559.314 | hypothetical protein, conserved | 0,1095 | 0,6125 |
| TcCLB.506289.230 | mucin-associated surface protein (MASP, pseudogene), pu | 0,1095 | 0,7281 |
| TcCLB.508265.4 | synaptojanin (N-terminal domain), putative (fragment) | 0,1094 | 0,6443 |
| TcCLB.511527.34 | 60S ribosomal protein L8, putative | 0,1093 | 0,1813 |
| TcCLB.505037.140 | retrotransposon hot spot protein (RHS, pseudogene), putat | 0,1093 | 0,5782 |
| TcCLB.511589.220 | hypothetical protein, conserved | 0,1093 | 0,6005 |
| TcCLB.511421.120 | E3 ubiquitin-protein ligase KCMF1, putative | 0,1091 | 0,3759 |
| TcCLB.511529.220 | hypothetical protein, conserved | 0,1091 | 0,6715 |
| TcCLB.511621.220 | esterase, putative | 0,109 | 0,703 |
| TcCLB.508795.19 | protein kinase, putative (fragment) | 0,1089 | 0,4382 |
| TcCLB.503455.10 | chaperone DNAJ protein, putative | 0,1089 | 0,5387 |
| TcCLB.504557.50 | dispersed gene family protein 1 (DGF-1), putative | 0,1089 | 0,7888 |
| TcCLB.508313.130 | mucin TcMUCII, putative | 0,1089 | 0,8469 |
| TcCLB.503925.119 | protein kinase, putative | 0,1087 | 0,5892 |
| TcCLB.508283.40 | trans-sialidase (pseudogene), putative | 0,1087 | 0,6465 |
| TcCLB.441061.9 | dynein heavy chain, putative (fragment) | 0,1084 | 0,5417 |
| TcCLB.506123.30 | protein kinase, putative | 0,1083 | 0,3422 |
| TcCLB.509763.10 | trans-sialidase, putative | 0,1083 | 0,8383 |
| TcCLB.509649.20 | hypothetical protein | 0,1082 | 0,4435 |
| TcCLB.509825.14 | 40S ribosomal protein SA, putative | 0,1082 | 0,5896 |
| TcCLB.506543.50 | hypothetical protein, conserved | 0,1082 | 0,8086 |
| TcCLB.509979.95 | 60S ribosomal protein L35, putative | 0,1081 | 0,2751 |
| TcCLB.507081.50 | hypothetical protein, conserved | 0,1078 | 0,4306 |
| TcCLB.503885.70 | hypothetical protein, conserved | 0,1078 | 0,6427 |
| TcCLB.511871.150 | trans-sialidase (pseudogene), putative | 0,1078 | 0,7036 |
| TcCLB.511145.20 | 60S ribosomal protein L32, putative | 0,1077 | 0,2878 |
| TcCLB.510275.283 | mucin TcMUCII (pseudogene), putative | 0,1077 | 0,8628 |
| TcCLB.509167.20 | 3-hydroxy-3-methylglutaryl-CoA reductase, putative | 0,1076 | 0,3231 |
| TcCLB.511621.160 | hypothetical protein, conserved | 0,1076 | 0,5114 |
| TcCLB.511239.20 | hypothetical protein, conserved | 0,1076 | 0,5202 |
| TcCLB.511449.10 | mucin-associated surface protein (MASP), putative | 0,1075 | 0,859 |
| TcCLB.511289.40 | hypothetical protein, conserved | 0,1074 | 0,3001 |
| TcCLB.508479.360 | hypothetical protein, conserved | 0,1074 | 0,4189 |
| TcCLB.506359.99 | hypothetical protein | 0,1074 | 0,6485 |
| TcCLB.508251.40 | mucin TcMUCII, putative | 0,1072 | 0,8782 |
| TcCLB.503679.30 | hypothetical protein, conserved | 0,1071 | 0,5593 |
| TcCLB.506393.120 | hypothetical protein, conserved (pseudogene) | 0,107 | 0,4954 |

| | | | |
|---|---|---|---|
| TcCLB.507711.170 | hypothetical protein, conserved | 0,107 | 0,5242 |
| TcCLB.509027.24 | hypothetical protein, conserved | 0,107 | 0,5766 |
| TcCLB.506759.220 | mucin-associated surface protein (MASP), putative | 0,107 | 0,8564 |
| TcCLB.510237.40 | mucin-associated surface protein (MASP, pseudogene), pu | 0,1069 | 0,7653 |
| TcCLB.508409.60 | hypothetical protein, conserved | 0,1068 | 0,4254 |
| TcCLB.510805.30 | mucin-associated surface protein (MASP, pseudogene), pu | 0,1067 | 0,7562 |
| TcCLB.511185.80 | trans-sialidase, Group V, putative | 0,1067 | 0,7888 |
| TcCLB.508209.159 | hypothetical protein, conserved | 0,1066 | 0,552 |
| TcCLB.511253.10 | kinesin-like protein, putative | 0,1065 | 0,6453 |
| TcCLB.511861.80 | UDP-Gal or UDP-GlcNAc-dependent glycosyltransferase, pu | 0,1064 | 0,5766 |
| TcCLB.507949.10 | chaperone DNAJ protein, putative | 0,1064 | 0,627 |
| TcCLB.511123.20 | retrotransposon hot spot protein (RHS, pseudogene), putat | 0,1064 | 0,6365 |
| TcCLB.507729.50 | mucin-associated surface protein (MASP), putative (fragm | 0,1064 | 0,7815 |
| TcCLB.510759.110 | hypothetical protein, conserved | 0,1063 | 0,4118 |
| TcCLB.511255.341 | trans-sialidase (pseudogene), putative | 0,1063 | 0,8075 |
| TcCLB.511421.180 | hypothetical protein, conserved | 0,1061 | 0,5639 |
| TcCLB.509047.110 | rac serine-threonine kinase, putative | 0,106 | 0,4001 |
| TcCLB.508389.100 | mucin-associated surface protein (MASP), putative | 0,106 | 0,7486 |
| TcCLB.506499.140 | mucin-associated surface protein (MASP), putative | 0,1059 | 0,7901 |
| TcCLB.506885.270 | tubulin tyrosine ligase, putative | 0,1057 | 0,4142 |
| TcCLB.511309.70 | hypothetical protein, conserved | 0,1056 | 0,4475 |
| TcCLB.508153.570 | protein kinase, putative | 0,1056 | 0,5599 |
| TcCLB.506755.74 | mitochondrial carrier protein, putative | 0,1055 | 0,5276 |
| TcCLB.511865.70 | hypothetical protein, conserved | 0,1055 | 0,5515 |
| TcCLB.506575.9 | hypothetical protein, conserved | 0,1053 | 0,3671 |
| TcCLB.511277.290 | aconitase, putative | 0,1053 | 0,4 |
| TcCLB.510575.140 | hypothetical protein, conserved | 0,1053 | 0,5364 |
| TcCLB.509115.20 | trans-sialidase, Group V, putative | 0,1053 | 0,7118 |
| TcCLB.503629.10 | hypothetical protein, conserved | 0,1051 | 0,4382 |
| TcCLB.509991.100 | Transcription elongation factor 1 domain-containing protei | 0,1051 | 0,5621 |
| TcCLB.509053.179 | RNA-binding protein, putative (fragment) | 0,105 | 0,5505 |
| TcCLB.510885.50 | hypothetical protein, conserved | 0,105 | 0,6481 |
| TcCLB.509051.40 | adenosine kinase, putative | 0,1048 | 0,3339 |
| TcCLB.508543.140 | electron transfer flavoprotein-ubiquinone oxidoreductase, | 0,1048 | 0,7253 |
| TcCLB.503527.10 | hypothetical protein, conserved | 0,1046 | 0,2374 |
| TcCLB.510311.40 | hypothetical protein, conserved | 0,1046 | 0,2882 |
| TcCLB.509429.180 | hypothetical protein, conserved | 0,1046 | 0,4404 |
| TcCLB.508911.70 | hypothetical protein | 0,1045 | 0,4946 |
| TcCLB.507821.140 | trans-sialidase (pseudogene), putative | 0,1045 | 0,7639 |
| TcCLB.508209.30 | hypothetical protein, conserved | 0,1044 | 0,3516 |
| TcCLB.503855.80 | hypothetical protein, conserved | 0,1044 | 0,4785 |
| TcCLB.510753.9 | helicase, putative | 0,1043 | 0,3466 |
| TcCLB.507597.10 | hypothetical protein | 0,1041 | 0,802 |
| TcCLB.505071.80 | hypothetical protein | 0,1039 | 0,4266 |
| TcCLB.506331.174 | dispersed gene family protein 1 (DGF-1, pseudogene), puta | 0,1039 | 0,6908 |

| | | | |
|---|---|---|---|
| TcCLB.507041.90 | hypothetical protein | 0,1038 | 0,6734 |
| TcCLB.508153.900 | hypothetical protein, conserved | 0,1037 | 0,3954 |
| TcCLB.509471.50 | hypothetical protein, conserved | 0,1037 | 0,4652 |
| TcCLB.508461.4 | hypothetical protein, conserved | 0,1035 | 0,4658 |
| TcCLB.510187.250 | hypothetical protein, conserved | 0,1035 | 0,5877 |
| TcCLB.511587.10 | hypothetical protein | 0,1035 | 0,8343 |
| TcCLB.510497.100 | mucin-associated surface protein (MASP), putative | 0,1033 | 0,8182 |
| TcCLB.510311.170 | hypothetical protein, conserved | 0,1032 | 0,5186 |
| TcCLB.506297.250 | palmitoyl acyltransferase 7, putative | 0,1031 | 0,415 |
| TcCLB.511633.50 | hypothetical protein, conserved | 0,1031 | 0,4298 |
| TcCLB.505843.20 | 60S ribosomal protein L6, putative | 0,1028 | 0,2737 |
| TcCLB.507447.10 | hypothetical protein, conserved | 0,1027 | 0,5917 |
| TcCLB.511071.45 | tuzin-like (pseudogene), putative | 0,1026 | 0,4451 |
| TcCLB.509671.60 | hypothetical protein | 0,1024 | 0,6445 |
| TcCLB.403875.19 | hypothetical protein, conserved (fragment) | 0,1022 | 0,7961 |
| TcCLB.511831.10 | trans-sialidase, putative (fragment) | 0,1021 | 0,691 |
| TcCLB.509115.30 | mucin-associated surface protein (MASP), putative | 0,1019 | 0,7425 |
| TcCLB.508919.180 | trans-sialidase (pseudogene), putative | 0,1018 | 0,3768 |
| TcCLB.509433.10 | hypothetical protein | 0,1018 | 0,8238 |
| TcCLB.509433.30 | hypothetical protein | 0,1017 | 0,7274 |
| TcCLB.511491.110 | rab6 GTPase activating protein, putative | 0,1014 | 0,5997 |
| TcCLB.511325.10 | amino acid permease (pseudogene), putative | 0,1014 | 0,8373 |
| TcCLB.506405.120 | V-type ATPase, A subunit, putative | 0,1011 | 0,2232 |
| TcCLB.510025.7 | syntaxin binding protein (pseudogene), putative | 0,1011 | 0,8619 |
| TcCLB.485683.10 | RNA-binding protein 42 (RNA-binding motif protein 42), pu | 0,101 | 0,3609 |
| TcCLB.508387.80 | hypothetical protein, conserved | 0,101 | 0,4746 |
| TcCLB.506635.104 | hypothetical protein | 0,101 | 0,6885 |
| TcCLB.506247.270 | ATPase, putative | 0,1007 | 0,3841 |
| TcCLB.506679.190 | hypothetical protein, conserved | 0,1007 | 0,4048 |
| TcCLB.508347.30 | hypothetical protein, conserved | 0,1007 | 0,4267 |
| TcCLB.510719.150 | hypothetical protein, conserved | 0,1005 | 0,3043 |
| TcCLB.510609.30 | GTPase, putative | 0,1003 | 0,487 |
| TcCLB.508207.54 | hypothetical protein, conserved | 0,1003 | 0,6453 |
| TcCLB.509805.220 | cholinephosphate cytidylyltransferase A, putative | 0,1002 | 0,3535 |
| TcCLB.504115.30 | beta galactofuranosyl glycosyltransferase, putative | 0,1002 | 0,5704 |
| TcCLB.508837.140 | U1 small nuclear ribonucleoprotein, putative | 0,1002 | 0,6185 |
| TcCLB.510007.20 | hypothetical protein, conserved | 0,1001 | 0,5396 |
| TcCLB.509665.20 | mucin-associated surface protein (MASP), putative | 0,1 | 0,788 |
| TcCLB.508823.90 | hypothetical protein | 0,09982 | 0,5407 |
| TcCLB.508465.149 | hypothetical protein, conserved (fragment) | 0,0998 | 0,6336 |
| TcCLB.506559.559 | antigenic protein, putative | 0,09949 | 0,342 |
| TcCLB.507993.5 | GPI-anchor transamidase subunit 8 (GPI8, pseudogene), pu | 0,09949 | 0,7654 |
| TcCLB.503775.41 | hypothetical protein, conserved (pseudogene) | 0,09946 | 0,8406 |
| TcCLB.509353.30 | 40S ribosomal protein S3, putative | 0,09944 | 0,2594 |
| TcCLB.506321.280 | hypothetical protein, conserved | 0,09942 | 0,2175 |

| | | | |
|---|---|---|---|
| TcCLB.510149.30 | hypothetical protein, conserved | 0,09932 | 0,4081 |
| TcCLB.511467.16 | hypothetical protein, conserved | 0,0993 | 0,5189 |
| TcCLB.507053.50 | hypothetical protein, conserved | 0,09925 | 0,4488 |
| TcCLB.511807.256 | hypothetical protein, conserved | 0,09919 | 0,5626 |
| TcCLB.510527.9 | kinesin, putative | 0,09914 | 0,5924 |
| TcCLB.508597.9 | retrotransposon hot spot (RHS) protein, putative | 0,09908 | 0,6504 |
| TcCLB.509033.60 | hypothetical protein, conserved | 0,09907 | 0,5232 |
| TcCLB.503449.10 | 60S ribosomal protein L17, putative | 0,09906 | 0,2453 |
| TcCLB.403869.10 | trans-sialidase, putative (fragment) | 0,09905 | 0,7835 |
| TcCLB.511423.130 | hypothetical protein, conserved (pseudogene) | 0,09901 | 0,5888 |
| TcCLB.510421.150 | protein kinase, putative | 0,09892 | 0,3251 |
| TcCLB.508059.10 | hypothetical protein, conserved | 0,09889 | 0,6764 |
| TcCLB.510933.140 | trans-sialidase (pseudogene), putative | 0,09889 | 0,8539 |
| TcCLB.507991.50 | kinesin, putative | 0,09887 | 0,3382 |
| TcCLB.506955.226 | elongation factor 1-gamma (EF-1-gamma, pseudogene), p | 0,09884 | 0,8973 |
| TcCLB.506755.150 | hypothetical protein, conserved | 0,09877 | 0,5985 |
| TcCLB.509695.230 | serine carboxypeptidase (CBP1), putative (fragment) | 0,09874 | 0,5637 |
| TcCLB.506247.90 | hypothetical protein | 0,09859 | 0,5721 |
| TcCLB.507715.100 | hypothetical protein, conserved | 0,09851 | 0,4932 |
| TcCLB.511627.150 | beta-ketoacyl-ACP reductase | 0,09848 | 0,5683 |
| TcCLB.510545.60 | hypothetical protein, conserved (pseudogene) | 0,09843 | 0,8729 |
| TcCLB.506409.90 | mucin-associated surface protein (MASP), putative | 0,0984 | 0,8984 |
| TcCLB.508207.230 | 6-phosphofructo-2-kinase/fructose-2,6-biphosphatase, put | 0,09839 | 0,4706 |
| TcCLB.510579.90 | hypothetical protein, conserved | 0,09838 | 0,5033 |
| TcCLB.506529.100 | hypothetical protein, conserved | 0,09836 | 0,4688 |
| TcCLB.506925.360 | hypothetical protein | 0,09818 | 0,5355 |
| TcCLB.510145.50 | hypothetical protein, conserved | 0,09817 | 0,3812 |
| TcCLB.509289.30 | hypothetical protein, conserved | 0,09808 | 0,3846 |
| TcCLB.508165.115 | mucin TcMUCII (pseudogene), putative | 0,09807 | 0,8618 |
| TcCLB.503479.70 | bilobe protein, conserved | 0,098 | 0,2931 |
| TcCLB.507519.20 | hypothetical protein, conserved | 0,09798 | 0,4364 |
| TcCLB.509799.70 | hypothetical protein, conserved | 0,09788 | 0,4837 |
| TcCLB.511837.129 | RNA-binding protein, putative | 0,09787 | 0,532 |
| TcCLB.507059.70 | dihydroorotase, putative | 0,09768 | 0,5462 |
| TcCLB.503583.30 | hypothetical protein, conserved | 0,09757 | 0,5846 |
| TcCLB.509451.70 | hypothetical protein, conserved | 0,09755 | 0,4177 |
| TcCLB.511215.14 | hypothetical protein, conserved | 0,09755 | 0,5957 |
| TcCLB.511167.10 | NADH-cytochrome b5 reductase, putative | 0,09755 | 0,6307 |
| TcCLB.509637.9 | hypothetical protein, conserved | 0,09753 | 0,635 |
| TcCLB.511277.610 | surface protease GP63, putative | 0,09748 | 0,6581 |
| TcCLB.503717.120 | trans-sialidase, Group V, putative | 0,09723 | 0,8111 |
| TcCLB.511459.20 | hypothetical protein, conserved | 0,09704 | 0,5053 |
| TcCLB.511847.80 | hypothetical protein, conserved | 0,09699 | 0,5183 |
| TcCLB.510889.20 | hypothetical protein, conserved | 0,09693 | 0,5836 |
| TcCLB.506575.54 | hypothetical protein, conserved | 0,09688 | 0,7173 |

| | | | |
|---|---|---|---|
| TcCLB.509733.30 | hypothetical protein, conserved | 0,09667 | 0,5102 |
| TcCLB.506303.90 | hypothetical protein, conserved | 0,09665 | 0,5137 |
| TcCLB.510737.100 | hypothetical protein, conserved | 0,09647 | 0,4212 |
| TcCLB.508857.70 | hypothetical protein, conserved | 0,0964 | 0,5219 |
| TcCLB.508933.40 | hypothetical protein | 0,09629 | 0,765 |
| TcCLB.510659.100 | hypothetical protein, conserved | 0,09621 | 0,6659 |
| TcCLB.510497.60 | mucin TcMUCII, putative | 0,09617 | 0,8626 |
| TcCLB.507987.10 | trans-sialidase, putative | 0,09609 | 0,8603 |
| TcCLB.507875.50 | hypothetical protein | 0,09608 | 0,7469 |
| TcCLB.508541.240 | casein kinase, putative | 0,09602 | 0,841 |
| TcCLB.506221.100 | hypothetical protein, conserved | 0,096 | 0,5198 |
| TcCLB.511001.230 | hypothetical protein, conserved | 0,09596 | 0,5156 |
| TcCLB.511585.310 | trans-sialidase (pseudogene), putative | 0,09593 | 0,6781 |
| TcCLB.507081.60 | helicase-like protein, putative | 0,0959 | 0,3747 |
| TcCLB.503499.10 | hypothetical protein, conserved | 0,09581 | 0,4315 |
| TcCLB.503433.20 | hypothetical protein, conserved | 0,09581 | 0,5562 |
| TcCLB.504431.20 | hypothetical protein | 0,09567 | 0,6359 |
| TcCLB.510437.70 | hypothetical protein, conserved | 0,0956 | 0,5804 |
| TcCLB.506855.140 | synaptobrevin, putative | 0,09551 | 0,5023 |
| TcCLB.506871.50 | ubiquitin hydrolase, putative | 0,09549 | 0,6448 |
| TcCLB.511895.10 | receptor-type adenylate cyclase, putative | 0,09547 | 0,6584 |
| TcCLB.507609.20 | hypothetical protein, conserved | 0,09536 | 0,7294 |
| TcCLB.511183.560 | mucin-associated surface protein (MASP), putative | 0,09496 | 0,832 |
| TcCLB.510763.100 | trypanin-like protein, putative | 0,09494 | 0,4828 |
| TcCLB.506985.40 | Polyamine transporter 1, putative (POT1.1) | 0,09478 | 0,4928 |
| TcCLB.509923.9 | dispersed gene family protein 1 (DGF-1), putative (fragme | 0,09478 | 0,7662 |
| TcCLB.507515.60 | hypothetical protein, conserved | 0,09471 | 0,6184 |
| TcCLB.508355.150 | proteasome regulatory non-ATPase subunit 3, putative | 0,09468 | 0,2973 |
| TcCLB.504255.50 | hypothetical protein, conserved | 0,09456 | 0,5623 |
| TcCLB.508409.230 | UBX domain-containing protein | 0,09416 | 0,4647 |
| TcCLB.509631.20 | mucin-associated surface protein (MASP), putative | 0,09408 | 0,8114 |
| TcCLB.510877.140 | hypothetical protein, conserved | 0,09404 | 0,5319 |
| TcCLB.508777.20 | hypothetical protein, conserved | 0,09399 | 0,6891 |
| TcCLB.507881.10 | retrotransposon hot spot protein (RHS, pseudogene), putat | 0,09392 | 0,6261 |
| TcCLB.511255.230 | mucin-associated surface protein (MASP, pseudogene), pu | 0,09388 | 0,8548 |
| TcCLB.506831.40 | 3-hydroxy-3-methylglutaryl-CoA reductase (HMGR) | 0,09382 | 0,3611 |
| TcCLB.509003.40 | calpain-like cysteine peptidase, putative | 0,09381 | 0,7221 |
| TcCLB.510889.300 | hypothetical protein | 0,09365 | 0,6311 |
| TcCLB.510939.20 | hypothetical protein, conserved | 0,09365 | 0,7188 |
| TcCLB.511397.10 | mucin-associated surface protein (MASP, pseudogene), pu | 0,09335 | 0,8561 |
| TcCLB.508737.187 | hypothetical protein, conserved | 0,09311 | 0,534 |
| TcCLB.510285.70 | kinetoplastid kinetochore protein 2, putative (kkt2) | 0,0931 | 0,5021 |
| TcCLB.508871.40 | mucin-associated surface protein (MASP), putative | 0,0931 | 0,8468 |
| TcCLB.506737.30 | hypothetical protein, conserved | 0,09304 | 0,5617 |
| TcCLB.508919.40 | hypothetical protein, conserved | 0,09302 | 0,489 |

| | | | |
|---|---|---|---|
| TcCLB.509203.50 | hypothetical protein | 0,09298 | 0,4482 |
| TcCLB.504867.20 | ABC transporter, putative | 0,09287 | 0,3761 |
| TcCLB.506147.50 | hypothetical protein, conserved | 0,09283 | 0,5618 |
| TcCLB.510201.15 | mucin TcMUCII (pseudogene), putative | 0,09275 | 0,805 |
| TcCLB.508697.10 | trans-sialidase, Group V, putative | 0,09272 | 0,7886 |
| TcCLB.506481.10 | tRNA pseudouridine synthase A-like protein (pseudogene), | 0,09259 | 0,6293 |
| TcCLB.506375.80 | hypothetical protein, conserved | 0,09254 | 0,701 |
| TcCLB.510603.130 | hypothetical protein, conserved | 0,09248 | 0,3469 |
| TcCLB.508881.40 | hypothetical protein, conserved | 0,09247 | 0,5227 |
| TcCLB.506241.170 | 40S ribosomal protein S6, putative | 0,09242 | 0,3085 |
| TcCLB.508375.60 | succinate dehydrogenase subunit | 0,09235 | 0,5638 |
| TcCLB.511001.220 | hypothetical protein, conserved | 0,0923 | 0,4986 |
| TcCLB.503453.64 | hypothetical protein, conserved | 0,09217 | 0,6018 |
| TcCLB.510187.380 | hypothetical protein, conserved | 0,09208 | 0,5414 |
| TcCLB.504161.40 | hypothetical protein, conserved | 0,09201 | 0,514 |
| TcCLB.506067.240 | mucin-associated surface protein (MASP), putative | 0,092 | 0,9128 |
| TcCLB.510021.90 | hypothetical protein, conserved (pseudogene) | 0,09195 | 0,8759 |
| TcCLB.509799.30 | adenylyl cyclase-associated protein, putative | 0,09182 | 0,5449 |
| TcCLB.504625.50 | dual specificity protein phosphatase or MAP kinase phosph | 0,09176 | 0,5598 |
| TcCLB.504131.130 | hypothetical protein, conserved | 0,0916 | 0,5446 |
| TcCLB.504427.60 | hypothetical protein, conserved | 0,09159 | 0,6323 |
| TcCLB.505363.10 | trans-sialidase, Group II, putative | 0,09127 | 0,809 |
| TcCLB.506753.80 | hypothetical protein, conserved | 0,09122 | 0,5623 |
| TcCLB.507667.11 | hypothetical protein, conserved (fragment) | 0,09115 | 0,62 |
| TcCLB.511109.40 | hypothetical protein, conserved | 0,09112 | 0,5417 |
| TcCLB.506959.60 | hypothetical protein, conserved | 0,09101 | 0,6438 |
| TcCLB.503635.89 | hypothetical protein, conserved | 0,09097 | 0,6355 |
| TcCLB.503801.20 | ribosomal protein S26, putative | 0,09089 | 0,3281 |
| TcCLB.506469.148 | hypothetical protein, conserved | 0,09078 | 0,5234 |
| TcCLB.509859.40 | cytosolic leucyl aminopeptidase, putative | 0,09059 | 0,4106 |
| TcCLB.510127.79 | actin 3, putative (fragment) | 0,09044 | 0,4893 |
| TcCLB.509799.40 | hypothetical protein, conserved | 0,09042 | 0,5427 |
| TcCLB.505267.30 | beta galactofuranosyl glycosyltransferase, putative | 0,09042 | 0,7946 |
| TcCLB.511529.140 | hypothetical protein, conserved | 0,09028 | 0,4414 |
| TcCLB.509017.60 | hypothetical protein, conserved | 0,09006 | 0,4418 |
| TcCLB.506899.30 | mucin-associated surface protein (MASP, pseudogene), pu | 0,09005 | 0,8934 |
| TcCLB.508851.19 | retrotransposon hot spot (RHS) protein, putative | 0,08994 | 0,5163 |
| TcCLB.506217.20 | beta galactofuranosyl glycosyltransferase (pseudogene), p | 0,08981 | 0,7486 |
| TcCLB.504053.10 | hypothetical protein, conserved (pseudogene) | 0,08972 | 0,4324 |
| TcCLB.511907.270 | ubiquitin-conjugating enzyme E2, putative | 0,08958 | 0,542 |
| TcCLB.506459.30 | trans-sialidase (pseudogene), putative | 0,08953 | 0,6954 |
| TcCLB.508723.110 | hypothetical protein | 0,08934 | 0,4681 |
| TcCLB.510941.6 | hypothetical protein, conserved | 0,08913 | 0,5417 |
| TcCLB.506577.30 | hypothetical protein, conserved | 0,0891 | 0,5414 |
| TcCLB.508153.1020 | hypothetical protein, conserved | 0,08904 | 0,6445 |

| | | | |
|---|---|---|---|
| TcCLB.503625.20 | hypothetical protein | 0,08904 | 0,7531 |
| TcCLB.506629.130 | hypothetical protein, conserved | 0,08901 | 0,6001 |
| TcCLB.414145.9 | hypothetical protein, conserved (fragment) | 0,08899 | 0,6703 |
| TcCLB.508233.19 | hypothetical protein, conserved | 0,08898 | 0,7281 |
| TcCLB.508239.10 | origin recognition complex subunit 1 (ORC1), putative | 0,08889 | 0,5771 |
| TcCLB.506973.50 | trans-sialidase (pseudogene), putative | 0,0888 | 0,816 |
| TcCLB.507975.50 | trans-sialidase (pseudogene), putative | 0,08879 | 0,3741 |
| TcCLB.509157.110 | hypothetical protein, conserved | 0,08878 | 0,7181 |
| TcCLB.506763.210 | hypothetical protein | 0,08875 | 0,8544 |
| TcCLB.506627.10 | AMP deaminase 2, putative | 0,08872 | 0,5156 |
| TcCLB.511011.30 | hypothetical protein, conserved (pseudogene) | 0,08849 | 0,7251 |
| TcCLB.510699.89 | mucin-associated surface protein (MASP), putative | 0,08836 | 0,8854 |
| TcCLB.510417.20 | mucin-associated surface protein (MASP, pseudogene), pu | 0,08825 | 0,8192 |
| TcCLB.506941.130 | hypothetical protein, conserved | 0,08818 | 0,5429 |
| TcCLB.506051.30 | dispersed gene family protein 1 (DGF-1), putative | 0,08818 | 0,829 |
| TcCLB.511239.140 | 60S ribosomal protein, putative | 0,08815 | 0,6692 |
| TcCLB.503973.240 | mucin-associated surface protein (MASP), putative | 0,08801 | 0,8536 |
| TcCLB.464807.20 | hypothetical protein, conserved (fragment) | 0,08799 | 0,5524 |
| TcCLB.506361.18 | mucin TcMUC (pseudogene), putative | 0,08778 | 0,8941 |
| TcCLB.509297.10 | hypothetical protein, conserved | 0,08776 | 0,6842 |
| TcCLB.508221.960 | mucin-associated surface protein (MASP), putative | 0,08749 | 0,8239 |
| TcCLB.506785.30 | mucin TcMUCII, putative | 0,08735 | 0,905 |
| TcCLB.511081.71 | hypothetical protein, conserved (pseudogene) | 0,08729 | 0,853 |
| TcCLB.506369.9 | hypothetical protein, conserved | 0,08723 | 0,765 |
| TcCLB.511887.30 | trans-sialidase (pseudogene), putative | 0,08721 | 0,7051 |
| TcCLB.511767.100 | hypothetical protein, conserved | 0,08717 | 0,8084 |
| TcCLB.509157.40 | hypothetical protein, conserved | 0,08707 | 0,6886 |
| TcCLB.511051.39 | 40S ribosomal protein S4, putative | 0,08704 | 0,3162 |
| TcCLB.506303.60 | protein kinase, putative | 0,08701 | 0,6336 |
| TcCLB.508615.10 | mucin-associated surface protein (MASP), putative | 0,087 | 0,8902 |
| TcCLB.510657.210 | hypothetical protein | 0,08679 | 0,7141 |
| TcCLB.507019.83 | 40S ribosomal protein S30, putative | 0,08673 | 0,417 |
| TcCLB.508397.30 | SPFH domain / Band 7 family protein, putative | 0,08652 | 0,5973 |
| TcCLB.504173.60 | hypothetical protein, conserved | 0,08645 | 0,5022 |
| TcCLB.508883.39 | mucin-associated surface protein (MASP, pseudogene), pu | 0,08645 | 0,7956 |
| TcCLB.510531.20 | hypothetical protein, conserved | 0,0864 | 0,6552 |
| TcCLB.511277.84 | dynein light chain Tctex-type, putative | 0,0863 | 0,71 |
| TcCLB.507867.31 | mucin TcMUCII (pseudogene), putative | 0,08628 | 0,7003 |
| TcCLB.510877.150 | hypothetical protein, conserved | 0,08623 | 0,4647 |
| TcCLB.506177.90 | hypothetical protein, conserved | 0,08597 | 0,7773 |
| TcCLB.510021.210 | mucin-associated surface protein (MASP, pseudogene), pu | 0,08591 | 0,9068 |
| TcCLB.509633.30 | hypothetical protein, conserved | 0,08584 | 0,5892 |
| TcCLB.506599.210 | mucin-associated surface protein (MASP), putative | 0,08575 | 0,8492 |
| TcCLB.510323.20 | hypothetical protein, conserved | 0,0857 | 0,3689 |
| TcCLB.506703.140 | mucin-associated surface protein (MASP), putative | 0,08559 | 0,9418 |

| | | | |
|---|---|---|---|
| TcCLB.509537.50 | Flagellar Member 1 (FLAM1) | 0,08557 | 0,4203 |
| TcCLB.510121.110 | hypothetical protein, conserved | 0,08555 | 0,433 |
| TcCLB.510663.19 | 60S ribosomal subunit protein L31, putative | 0,08538 | 0,5303 |
| TcCLB.509769.36 | palmitoyl acyltransferase 12, putative | 0,08538 | 0,5899 |
| TcCLB.510889.340 | hypothetical protein, conserved | 0,08537 | 0,7761 |
| TcCLB.511753.80 | GTPase activating protein, putative | 0,08528 | 0,614 |
| TcCLB.507019.86 | 40S ribosomal protein S30, putative | 0,08523 | 0,5268 |
| TcCLB.509611.189 | hypothetical protein, conserved | 0,08516 | 0,4224 |
| TcCLB.509551.40 | hypothetical protein, conserved | 0,08512 | 0,6784 |
| TcCLB.504575.40 | adrenodoxin precursor, putative | 0,08509 | 0,7971 |
| TcCLB.510175.200 | retrotransposon hot spot protein (RHS, pseudogene), putat | 0,08501 | 0,7759 |
| TcCLB.511239.90 | hypothetical protein | 0,08499 | 0,5659 |
| TcCLB.511435.40 | hypothetical protein, conserved | 0,08497 | 0,5417 |
| TcCLB.506413.30 | 40S ribosomal protein S33, putative | 0,08476 | 0,6867 |
| TcCLB.509177.68 | translation initiation factor, putative | 0,08475 | 0,6364 |
| TcCLB.504213.29 | hypothetical protein, conserved | 0,08467 | 0,6715 |
| TcCLB.508443.50 | hypothetical protein, conserved | 0,08448 | 0,6652 |
| TcCLB.506993.140 | hypothetical protein | 0,08431 | 0,8876 |
| TcCLB.511771.132 | ATP-dependent DEAD/H RNA helicase (pseudogene), putat | 0,08422 | 0,883 |
| TcCLB.508317.20 | hypothetical protein, conserved | 0,08414 | 0,6421 |
| TcCLB.509655.10 | trans-sialidase (pseudogene), putative | 0,08408 | 0,8125 |
| TcCLB.506413.60 | hypothetical protein, conserved | 0,08374 | 0,7998 |
| TcCLB.507875.60 | beta galactofuranosyl glycosyltransferase, putative | 0,08369 | 0,7282 |
| TcCLB.511189.60 | hypothetical protein, conserved | 0,08361 | 0,5927 |
| TcCLB.510409.20 | hypothetical protein, conserved (pseudogene) | 0,08349 | 0,7757 |
| TcCLB.507669.180 | hypothetical protein, conserved | 0,08345 | 0,5147 |
| TcCLB.510357.50 | hypothetical protein, conserved | 0,08336 | 0,674 |
| TcCLB.511291.120 | DNA-directed RNA polymerase III subunit C11, putative (RI | 0,0833 | 0,698 |
| TcCLB.506357.29 | dynein light chain Tctex-type, putative | 0,08328 | 0,7186 |
| TcCLB.504055.20 | hypothetical protein, conserved | 0,08314 | 0,6867 |
| TcCLB.507979.40 | hypothetical protein, conserved | 0,08313 | 0,7779 |
| TcCLB.508805.100 | hypothetical protein, conserved | 0,08294 | 0,5495 |
| TcCLB.509979.60 | sec1 family transport protein, putative | 0,08279 | 0,62 |
| TcCLB.506209.70 | dispersed gene family protein 1 (DGF-1), putative | 0,08265 | 0,8233 |
| TcCLB.508241.120 | rrp44p homologue, putative | 0,08264 | 0,3262 |
| TcCLB.511753.60 | ESAG8-associated protein, putative (PIE8) | 0,08255 | 0,6832 |
| TcCLB.509497.30 | agmatinase, putative | 0,08241 | 0,6628 |
| TcCLB.506227.30 | hypothetical protein, conserved | 0,08228 | 0,534 |
| TcCLB.460125.10 | hypothetical protein, conserved (fragment) | 0,08228 | 0,7606 |
| TcCLB.511259.200 | mucin-associated surface protein (MASP, pseudogene), pu | 0,08226 | 0,8761 |
| TcCLB.511025.50 | glucosamine-6-phosphate isomerase, putative | 0,08223 | 0,4087 |
| TcCLB.511145.46 | 60S ribosomal protein L37a, putative | 0,08222 | 0,4905 |
| TcCLB.509899.50 | mucin-associated surface protein (MASP, pseudogene), pu | 0,08197 | 0,7924 |
| TcCLB.509109.114 | mitochondrial carrier protein, putative | 0,08192 | 0,6521 |
| TcCLB.508879.40 | hypothetical protein, conserved | 0,08187 | 0,5472 |

| | | | |
|---|---|---|---|
| TcCLB.511121.20 | hypothetical protein | 0,08186 | 0,8504 |
| TcCLB.508593.69 | hypothetical protein, conserved | 0,08184 | 0,7899 |
| TcCLB.508109.60 | trans-sialidase, Group V, putative | 0,08167 | 0,8385 |
| TcCLB.511511.50 | hypothetical protein, conserved | 0,08165 | 0,5567 |
| TcCLB.506683.30 | hypothetical protein, conserved | 0,08162 | 0,7666 |
| TcCLB.508089.20 | UDP-glucoronosyl and UDP-glucosyl transferase, putative | 0,08155 | 0,6344 |
| TcCLB.509961.30 | hypothetical protein, conserved | 0,0815 | 0,5755 |
| TcCLB.509393.20 | hypothetical protein, conserved | 0,08143 | 0,6315 |
| TcCLB.511483.40 | prefoldin subunit 2, putative | 0,08136 | 0,6475 |
| TcCLB.461927.9 | adenosine monophosphate deaminase, putative | 0,08133 | 0,6568 |
| TcCLB.506633.109 | hypothetical protein (fragment) | 0,08124 | 0,6721 |
| TcCLB.506201.120 | hypothetical protein, conserved | 0,08119 | 0,5713 |
| TcCLB.504795.9 | dispersed gene family protein 1 (DGF-1), putative | 0,08119 | 0,7481 |
| TcCLB.511421.90 | hypothetical protein, conserved | 0,08094 | 0,615 |
| TcCLB.507723.140 | hypothetical protein, conserved | 0,08086 | 0,6823 |
| TcCLB.506357.100 | DNA-directed RNA polymerase I subunit RPB5z, putative (F | 0,08083 | 0,6614 |
| TcCLB.506965.70 | mucin-associated surface protein (MASP), putative | 0,08082 | 0,8188 |
| TcCLB.437575.9 | GDP-mannose 4,6 dehydratase, putative | 0,08081 | 0,8543 |
| TcCLB.509507.20 | hypothetical protein, conserved | 0,08072 | 0,5099 |
| TcCLB.511337.39 | hypothetical protein, conserved | 0,08071 | 0,7184 |
| TcCLB.503703.90 | hypothetical protein, conserved | 0,08065 | 0,6727 |
| TcCLB.503589.19 | hypothetical protein, conserved | 0,08062 | 0,7735 |
| TcCLB.506459.180 | trans-sialidase, Group V, putative | 0,0806 | 0,8087 |
| TcCLB.509805.50 | hypothetical protein, conserved | 0,08052 | 0,6886 |
| TcCLB.507223.20 | hypothetical protein, conserved | 0,08051 | 0,8346 |
| TcCLB.508761.30 | mucin-associated surface protein (MASP), putative | 0,0805 | 0,805 |
| TcCLB.511423.140 | hypothetical protein, conserved | 0,08033 | 0,5373 |
| TcCLB.510889.10 | hypothetical protein, conserved | 0,08029 | 0,7116 |
| TcCLB.503775.10 | surface protease GP63 (pseudogene), putative | 0,08029 | 0,8353 |
| TcCLB.507875.130 | trans-sialidase (pseudogene), putative | 0,08022 | 0,8085 |
| TcCLB.503925.90 | hypothetical protein, conserved | 0,08021 | 0,8155 |
| TcCLB.505071.104 | hypothetical protein, conserved | 0,08018 | 0,784 |
| TcCLB.507029.20 | DNA-J protein, putative | 0,08006 | 0,6512 |
| TcCLB.509747.60 | hypothetical protein, conserved | 0,07997 | 0,5472 |
| TcCLB.510055.30 | hypothetical protein, conserved | 0,07996 | 0,8182 |
| TcCLB.509897.160 | trans-sialidase (pseudogene), putative | 0,07989 | 0,7557 |
| TcCLB.503955.89 | hypothetical protein, conserved | 0,07973 | 0,6342 |
| TcCLB.506563.100 | hypothetical protein, conserved | 0,0797 | 0,4992 |
| TcCLB.511111.20 | hypothetical protein | 0,0797 | 0,6104 |
| TcCLB.466529.9 | hypothetical protein, conserved | 0,07961 | 0,7006 |
| TcCLB.510719.30 | ribosomal protein L29, putative | 0,07955 | 0,5393 |
| TcCLB.507001.20 | hypothetical protein, conserved | 0,07953 | 0,6976 |
| TcCLB.506567.20 | hypothetical protein | 0,07948 | 0,6495 |
| TcCLB.507233.60 | retrotransposon hot spot protein (RHS, pseudogene), putat | 0,07934 | 0,6598 |
| TcCLB.509897.170 | mucin-associated surface protein (MASP, pseudogene), pu | 0,07932 | 0,7919 |

| | | | |
|---|---|---|---|
| TcCLB.508295.40 | mucin-associated surface protein (MASP), putative | 0,07931 | 0,8985 |
| TcCLB.507513.110 | hypothetical protein | 0,07927 | 0,6195 |
| TcCLB.511217.140 | coatomer delta subunit, putative | 0,07926 | 0,4511 |
| TcCLB.508373.10 | glutamine dependent carbamoyl-phosphate synthase, puta | 0,07926 | 0,5065 |
| TcCLB.506503.140 | extracellular receptor, putative | 0,0792 | 0,5022 |
| TcCLB.510279.250 | mucin-associated surface protein (MASP), putative | 0,0792 | 0,9162 |
| TcCLB.507083.50 | hypothetical protein, conserved | 0,07919 | 0,6926 |
| TcCLB.509805.260 | dispersed gene family protein 1 (DGF-1, pseudogene), puta | 0,07911 | 0,6756 |
| TcCLB.508831.40 | hypothetical protein, conserved | 0,07909 | 0,7844 |
| TcCLB.509065.33 | hypothetical protein, conserved | 0,07898 | 0,7279 |
| TcCLB.509611.89 | hypothetical protein, conserved | 0,07898 | 0,7616 |
| TcCLB.506919.70 | hypothetical protein, conserved (pseudogene) | 0,07898 | 0,8499 |
| TcCLB.510359.680 | mucin-associated surface protein (MASP, pseudogene), pu | 0,07888 | 0,8854 |
| TcCLB.510293.60 | hypothetical protein, conserved | 0,07886 | 0,6867 |
| TcCLB.506581.60 | hypothetical protein, conserved | 0,07873 | 0,636 |
| TcCLB.511809.50 | hypothetical protein, conserved | 0,07868 | 0,6682 |
| TcCLB.511717.20 | hypothetical protein, conserved | 0,07863 | 0,5621 |
| TcCLB.511445.20 | radial spoke protein 3, putative (RSP3) | 0,07855 | 0,5827 |
| TcCLB.506855.120 | hypothetical protein, conserved | 0,07846 | 0,5563 |
| TcCLB.511557.50 | Present in the outer mitochondrial membrane proteome 1 | 0,0784 | 0,6867 |
| TcCLB.505999.189 | hypothetical protein, conserved (pseudogene) | 0,07837 | 0,7721 |
| TcCLB.506859.70 | guide RNA associated protein, GAP2, putative | 0,07832 | 0,4045 |
| TcCLB.510285.100 | eukaryotic translation initation factor 4 gamma, putative | 0,07811 | 0,4999 |
| TcCLB.508409.130 | protein transport protein Sec24A, putative | 0,07803 | 0,3744 |
| TcCLB.510359.371 | elongation factor 1-gamma (EF-1-gamma, pseudogene), p | 0,07803 | 0,8061 |
| TcCLB.506755.279 | hypothetical protein, conserved | 0,078 | 0,6293 |
| TcCLB.506147.30 | synaptojanin (N-terminal domain), putative | 0,07796 | 0,5382 |
| TcCLB.508365.30 | mucin TcMUCII, putative | 0,07792 | 0,8828 |
| TcCLB.509631.150 | hypothetical protein, conserved | 0,07783 | 0,8223 |
| TcCLB.504123.50 | hypothetical protein, conserved (pseudogene) | 0,07781 | 0,5012 |
| TcCLB.511539.18 | hypothetical protein, conserved | 0,07779 | 0,6549 |
| TcCLB.506025.70 | dolichyl-lipid chaperone, putative (rft1) | 0,07749 | 0,7281 |
| TcCLB.507001.130 | hypothetical protein, conserved | 0,07747 | 0,6301 |
| TcCLB.508139.120 | hypothetical protein | 0,07745 | 0,8752 |
| TcCLB.504215.50 | hypothetical protein, conserved | 0,07734 | 0,5537 |
| TcCLB.503899.10 | heat shock 70 kDa protein, putative | 0,07728 | 0,6092 |
| TcCLB.509799.10 | adaptin complex 4 subunit, putative | 0,07719 | 0,6411 |
| TcCLB.511291.10 | intraflagellar transport protein 80, putative (fragment) (IF | 0,07714 | 0,6872 |
| TcCLB.507017.100 | hypothetical protein, conserved | 0,07688 | 0,7533 |
| TcCLB.510123.50 | iron-sulfur cluster assembly protein, putative | 0,07683 | 0,6099 |
| TcCLB.509429.220 | dolichyl-P-Man:GDP-Man5GlcNAc2-PP-dolichyl alpha-1,2-n | 0,0768 | 0,5756 |
| TcCLB.511487.100 | retrotransposon hot spot protein (RHS, pseudogene), putat | 0,07677 | 0,7304 |
| TcCLB.510691.95 | hypothetical protein, conserved (fragment) | 0,07675 | 0,713 |
| TcCLB.503697.9 | proteasome regulatory non-ATPase subunit 11, putative (fr | 0,07672 | 0,6165 |
| TcCLB.511173.440 | trans-sialidase, Group V, putative | 0,07663 | 0,8252 |

| | | | |
|---|---|---|---|
| TcCLB.506493.30 | hypothetical protein, conserved | 0,0766 | 0,5926 |
| TcCLB.507089.260 | CYC2-like cyclin, putative | 0,07654 | 0,6271 |
| TcCLB.508927.10 | hypothetical protein, conserved | 0,07653 | 0,7394 |
| TcCLB.506459.100 | surface protease GP63 (pseudogene), putative | 0,07647 | 0,8885 |
| TcCLB.506443.60 | hypothetical protein | 0,07638 | 0,6011 |
| TcCLB.508633.100 | retrotransposon hot spot (RHS) protein, putative | 0,07636 | 0,7147 |
| TcCLB.508207.110 | 60S ribosomal protein L7, putative | 0,07633 | 0,3373 |
| TcCLB.510729.220 | hypothetical protein, conserved | 0,07632 | 0,3604 |
| TcCLB.511089.5 | trans-sialidase (pseudogene), putative | 0,07618 | 0,9228 |
| TcCLB.506925.520 | hypothetical protein, conserved | 0,07615 | 0,5928 |
| TcCLB.509569.50 | guanine nucleotide exchange factor subunit GCD7 | 0,07614 | 0,5695 |
| TcCLB.508257.204 | hypothetical protein, conserved | 0,07609 | 0,6442 |
| TcCLB.506265.70 | hypothetical protein, conserved | 0,07606 | 0,6067 |
| TcCLB.508541.225 | casein kinase, putative | 0,07601 | 0,6424 |
| TcCLB.510715.11 | hypothetical protein | 0,07598 | 0,7834 |
| TcCLB.510575.180 | adenylate kinase, putative | 0,07591 | 0,696 |
| TcCLB.506295.140 | hypothetical protein, conserved | 0,07571 | 0,6121 |
| TcCLB.506289.120 | mucin-associated surface protein (MASP), putative | 0,07562 | 0,7959 |
| TcCLB.510307.180 | glycine dehydrogenase [decarboxylating] (pseudogene), pu | 0,07557 | 0,5685 |
| TcCLB.503971.30 | hypothetical protein, conserved | 0,07552 | 0,7327 |
| TcCLB.506563.20 | exosome-associated protein 1, putative (EAP1) | 0,07537 | 0,6805 |
| TcCLB.505939.30 | hypothetical protein, conserved | 0,07535 | 0,4988 |
| TcCLB.507483.20 | ef-hand protein 5, putative | 0,07522 | 0,6909 |
| TcCLB.507911.40 | hypothetical protein, conserved (pseudogene) | 0,07507 | 0,7134 |
| TcCLB.506289.250 | surface protease GP63 (pseudogene), putative | 0,07507 | 0,8949 |
| TcCLB.506737.121 | mucin-associated surface protein (MASP, pseudogene), pu | 0,07481 | 0,6503 |
| TcCLB.510219.20 | hypothetical protein, conserved | 0,07465 | 0,7195 |
| TcCLB.511087.10 | mucin-associated surface protein (MASP, pseudogene), pu | 0,07462 | 0,8622 |
| TcCLB.511421.170 | hypothetical protein, conserved | 0,07461 | 0,6195 |
| TcCLB.507231.30 | mucin-associated surface protein (MASP), putative | 0,07456 | 0,8632 |
| TcCLB.511633.70 | protein kinase, putative | 0,07429 | 0,5284 |
| TcCLB.507811.114 | hypothetical protein, conserved | 0,07428 | 0,6306 |
| TcCLB.509105.110 | hypothetical protein, conserved | 0,0742 | 0,6116 |
| TcCLB.510105.170 | dihydrolipoamide acetyltransferase precursor, putative | 0,07416 | 0,4872 |
| TcCLB.511807.6 | ATP-dependent DEAD/H RNA helicase (pseudogene), putat | 0,07395 | 0,6652 |
| TcCLB.509203.79 | guanine-nucleotide-exchange-factor, putative (fragment) | 0,07384 | 0,7495 |
| TcCLB.508789.150 | mucin-associated surface protein (MASP), putative | 0,0737 | 0,8444 |
| TcCLB.511735.88 | hypothetical protein, conserved (fragment) | 0,07367 | 0,8385 |
| TcCLB.510259.12 | hypothetical protein, conserved | 0,07366 | 0,8238 |
| TcCLB.507847.20 | mucin-associated surface protein (MASP, pseudogene), pu | 0,07326 | 0,698 |
| TcCLB.490507.10 | hypothetical protein, conserved | 0,07323 | 0,6505 |
| TcCLB.503703.40 | hypothetical protein, conserved | 0,07319 | 0,654 |
| TcCLB.507527.4 | dolichyl-P-Man:GDP-Man5GlcNAc2-PP-dolichyl alpha-1,2-n | 0,07317 | 0,7908 |
| TcCLB.503647.40 | hypothetical protein | 0,07312 | 0,7007 |
| TcCLB.508527.30 | trans-sialidase (pseudogene), putative | 0,07311 | 0,7132 |

| | | | |
|---|---|---|---|
| TcCLB.503925.20 | hypothetical protein, conserved | 0,07295 | 0,5997 |
| TcCLB.510361.190 | mucin-associated surface protein (MASP), putative | 0,07289 | 0,8865 |
| TcCLB.508407.4 | hypothetical protein, conserved | 0,07273 | 0,8739 |
| TcCLB.509767.40 | hypothetical protein, conserved | 0,07272 | 0,6246 |
| TcCLB.448661.10 | mucin-associated surface protein (MASP), putative (fragm | 0,07266 | 0,8828 |
| TcCLB.509645.10 | hypothetical protein, conserved | 0,07264 | 0,5604 |
| TcCLB.510081.11 | hypothetical protein, conserved (pseudogene) | 0,0726 | 0,6483 |
| TcCLB.510061.30 | trans-sialidase, Group V, putative | 0,07254 | 0,8104 |
| TcCLB.506999.80 | hypothetical protein, conserved | 0,07248 | 0,6986 |
| TcCLB.503425.10 | hypothetical protein, conserved | 0,07245 | 0,632 |
| TcCLB.511907.160 | mucin-associated surface protein (MASP, pseudogene), pu | 0,07245 | 0,7597 |
| TcCLB.509011.70 | adrenodoxin precursor, putative | 0,0723 | 0,7715 |
| TcCLB.506629.110 | hypothetical protein, conserved | 0,07221 | 0,659 |
| TcCLB.505037.50 | small GTP-binding protein RAB6, putative | 0,0722 | 0,5515 |
| TcCLB.509695.184 | 40S ribosomal protein S23, putative | 0,0721 | 0,4524 |
| TcCLB.508277.340 | Eukaryotic translation initiation factor 4 gamma type 2, pu | 0,07202 | 0,4965 |
| TcCLB.506583.20 | hypothetical protein, conserved | 0,072 | 0,79 |
| TcCLB.510805.70 | mucin TcMUCII, putative | 0,07188 | 0,9075 |
| TcCLB.507985.20 | retrotransposon hot spot protein (RHS, pseudogene), putat | 0,07183 | 0,7599 |
| TcCLB.507011.130 | hypothetical protein, conserved | 0,07178 | 0,6354 |
| TcCLB.509195.10 | hypothetical protein | 0,07175 | 0,8922 |
| TcCLB.510667.100 | hypothetical protein, conserved | 0,07163 | 0,5097 |
| TcCLB.510613.30 | hypothetical protein | 0,07148 | 0,6279 |
| TcCLB.506767.220 | mucin TcMUCII, putative | 0,07142 | 0,9048 |
| TcCLB.506657.88 | hypothetical protein, conserved | 0,07141 | 0,912 |
| TcCLB.508277.110 | cAMP specific phosphodiesterase, putative | 0,07137 | 0,6579 |
| TcCLB.511217.200 | tyrosyl or methionyl-tRNA synthetase, putative | 0,07122 | 0,6504 |
| TcCLB.506041.40 | hypothetical protein, conserved | 0,07108 | 0,6521 |
| TcCLB.509161.50 | hypothetical protein, conserved | 0,07106 | 0,6707 |
| TcCLB.503999.110 | hypothetical protein, conserved (fragment) | 0,07098 | 0,8297 |
| TcCLB.510611.40 | hypothetical protein, conserved | 0,07096 | 0,614 |
| TcCLB.506797.10 | hypothetical protein, conserved | 0,07093 | 0,593 |
| TcCLB.511269.40 | cAMP phosphodiesterase A, putative (PDEA) | 0,07088 | 0,7486 |
| TcCLB.508287.10 | dispersed gene family protein 1 (DGF-1), putative | 0,07084 | 0,9026 |
| TcCLB.511647.42 | hypothetical protein, conserved | 0,07081 | 0,7837 |
| TcCLB.507797.9 | nuclear transcription factor, putative | 0,0708 | 0,7932 |
| TcCLB.510657.110 | hypothetical protein, conserved | 0,07075 | 0,6183 |
| TcCLB.506499.215 | elongation factor 1-gamma (EF-1-gamma, pseudogene), p | 0,07068 | 0,9069 |
| TcCLB.508881.110 | GTP-binding protein, putative | 0,07067 | 0,6739 |
| TcCLB.510295.10 | Enriched in surface-labeled proteome protein 18 (ESP18) | 0,07063 | 0,6123 |
| TcCLB.511423.100 | kinetoplast ribosomal PPR-repeat containing protein 3, put | 0,07063 | 0,6244 |
| TcCLB.412093.10 | antigenic protein (pseudogene), putative | 0,07063 | 0,8209 |
| TcCLB.503929.50 | hypothetical protein, conserved | 0,07061 | 0,7214 |
| TcCLB.504105.60 | hypothetical protein, conserved | 0,07027 | 0,6446 |
| TcCLB.511301.20 | hypothetical protein, conserved | 0,0702 | 0,6719 |

| | | | |
|---|---|---|---|
| TcCLB.511287.40 | 40S ribosomal protein S2, putative | 0,07018 | 0,3688 |
| TcCLB.504181.10 | 60S ribosomal protein L9, putative | 0,07018 | 0,4086 |
| TcCLB.416503.10 | trans-sialidase (pseudogene), putative | 0,07007 | 0,6507 |
| TcCLB.506585.90 | RNA processing endonuclease 1, putative | 0,07003 | 0,6908 |
| TcCLB.509085.120 | retrotransposon hot spot (RHS) protein, putative | 0,07002 | 0,7662 |
| TcCLB.511017.40 | hd phosphohydrolase family protein | 0,06984 | 0,8125 |
| TcCLB.506795.40 | phosphatidylserine synthase, putative | 0,06983 | 0,6708 |
| TcCLB.511107.20 | hypothetical protein, conserved (pseudogene) | 0,06969 | 0,6877 |
| TcCLB.503803.20 | hypothetical protein, conserved | 0,06969 | 0,7702 |
| TcCLB.507519.10 | hypothetical protein, conserved (pseudogene) | 0,0696 | 0,6775 |
| TcCLB.509167.50 | hypothetical protein | 0,06954 | 0,7284 |
| TcCLB.511339.10 | UDP-Gal or UDP-GlcNAc-dependent glycosyltransferase, pu | 0,06939 | 0,8741 |
| TcCLB.506941.170 | ADP-ribosylation factor-like protein, putative | 0,06935 | 0,7224 |
| TcCLB.508543.30 | 40S ribosomal protein S9, putative | 0,06929 | 0,4079 |
| TcCLB.508479.210 | hypothetical protein, conserved | 0,06916 | 0,6779 |
| TcCLB.509669.140 | hypothetical protein, conserved | 0,06911 | 0,7477 |
| TcCLB.504057.40 | NAD dependent epimerase/dehydratase family, putative (I | 0,06903 | 0,5462 |
| TcCLB.511585.90 | hypothetical protein, conserved | 0,06902 | 0,5665 |
| TcCLB.506567.130 | hypothetical protein, conserved | 0,06899 | 0,7844 |
| TcCLB.503847.80 | hypothetical protein, conserved | 0,06895 | 0,6581 |
| TcCLB.509247.20 | ras-related protein Rab21, putative (RAB21) | 0,06892 | 0,6909 |
| TcCLB.510579.20 | Peptide deformylase 2, putative (PDF2) | 0,0688 | 0,6521 |
| TcCLB.510903.70 | hypothetical protein, conserved | 0,06873 | 0,6547 |
| TcCLB.509965.120 | hypothetical protein, conserved | 0,06869 | 0,6665 |
| TcCLB.511423.90 | zeta tubulin, putative | 0,06865 | 0,6721 |
| TcCLB.508445.20 | iron superoxide dismutase, putative | 0,06863 | 0,6029 |
| TcCLB.507305.30 | casein kinase, putative | 0,0686 | 0,8274 |
| TcCLB.511537.50 | chaperone DNAJ protein, putative | 0,06859 | 0,6438 |
| TcCLB.509543.40 | hypothetical protein, conserved | 0,06856 | 0,6746 |
| TcCLB.507069.94 | mucin-associated surface protein (MASP), putative | 0,06845 | 0,8829 |
| TcCLB.508401.50 | hypothetical protein, conserved (pseudogene) | 0,06837 | 0,6483 |
| TcCLB.510943.90 | hypothetical protein, conserved | 0,06834 | 0,7982 |
| TcCLB.507709.40 | hypothetical protein, conserved | 0,06826 | 0,7112 |
| TcCLB.511033.20 | hypothetical protein | 0,06825 | 0,6542 |
| TcCLB.420369.40 | hypothetical protein, conserved | 0,06825 | 0,7009 |
| TcCLB.505051.20 | Checkpoint protein HUS1, putative (hus1) | 0,06823 | 0,665 |
| TcCLB.504159.40 | aquaporin-like protein, putative | 0,06817 | 0,6969 |
| TcCLB.510305.60 | hypothetical protein, conserved | 0,06806 | 0,5438 |
| TcCLB.506999.150 | vesicle-associated membrane protein, putative | 0,06793 | 0,6178 |
| TcCLB.509213.170 | transcription activator, putative | 0,06787 | 0,507 |
| TcCLB.507053.80 | hypothetical protein, conserved | 0,06786 | 0,633 |
| TcCLB.508547.40 | hypothetical protein, conserved | 0,06772 | 0,7561 |
| TcCLB.504035.100 | hypothetical protein, conserved | 0,06757 | 0,605 |
| TcCLB.508637.40 | hypothetical protein, conserved | 0,06743 | 0,7725 |
| TcCLB.507601.120 | hypothetical protein, conserved | 0,06742 | 0,7556 |

| | | | |
|---|---|---|---|
| TcCLB.506587.60 | target SNARE, putative | 0,06738 | 0,6925 |
| TcCLB.507519.154 | hypothetical protein, conserved | 0,06738 | 0,7126 |
| TcCLB.507041.40 | hypothetical protein, conserved | 0,06731 | 0,5602 |
| TcCLB.506733.90 | hypothetical protein, conserved | 0,0673 | 0,627 |
| TcCLB.508245.60 | mucin TcMUCII, putative | 0,06717 | 0,9185 |
| TcCLB.507993.70 | hypothetical protein, conserved | 0,06713 | 0,5816 |
| TcCLB.505267.60 | trans-sialidase (pseudogene), putative | 0,06712 | 0,5936 |
| TcCLB.503529.30 | hypothetical protein, conserved | 0,06706 | 0,7241 |
| TcCLB.506753.90 | hypothetical protein, conserved | 0,06704 | 0,5254 |
| TcCLB.508879.100 | hypothetical protein, conserved | 0,06704 | 0,5278 |
| TcCLB.503395.50 | 40S ribosomal protein S12, putative | 0,06697 | 0,5052 |
| TcCLB.507771.120 | trans-sialidase (pseudogene), putative | 0,06679 | 0,7881 |
| TcCLB.508465.60 | hypothetical protein, conserved | 0,06674 | 0,6219 |
| TcCLB.503939.70 | hypothetical protein | 0,0667 | 0,5081 |
| TcCLB.505025.20 | surface protease GP63 (pseudogene), putative | 0,06658 | 0,9129 |
| TcCLB.506163.80 | katanin, putative | 0,06654 | 0,6842 |
| TcCLB.510515.40 | hypothetical protein, conserved | 0,06653 | 0,8382 |
| TcCLB.510969.4 | hypothetical protein | 0,06636 | 0,8619 |
| TcCLB.503699.11 | mucin-associated surface protein (MASP, pseudogene), pu | 0,0663 | 0,9393 |
| TcCLB.504169.10 | dispersed gene family protein 1 (DGF-1), putative | 0,06625 | 0,7834 |
| TcCLB.511837.50 | DNA repair protein, putative | 0,06613 | 0,7232 |
| TcCLB.503971.10 | hypothetical protein, conserved | 0,06609 | 0,5081 |
| TcCLB.506661.130 | ion transport protein, putative | 0,06609 | 0,6807 |
| TcCLB.503843.30 | hypothetical protein, conserved | 0,06592 | 0,6398 |
| TcCLB.507103.29 | hypothetical protein, conserved (fragment) | 0,0659 | 0,7392 |
| TcCLB.506247.280 | hypothetical protein, conserved | 0,06588 | 0,6693 |
| TcCLB.509089.9 | dispersed gene family protein 1 (DGF-1), putative | 0,06585 | 0,7844 |
| TcCLB.506579.90 | hypothetical protein | 0,0658 | 0,582 |
| TcCLB.511349.100 | trans-sialidase, Group II, putative | 0,06559 | 0,8137 |
| TcCLB.508139.160 | expression site-associated gene (ESAG-like) protein, putati | 0,06558 | 0,8357 |
| TcCLB.506925.350 | hypothetical protein, conserved | 0,06556 | 0,5857 |
| TcCLB.508781.20 | ATP synthase, putative | 0,06554 | 0,5985 |
| TcCLB.507747.170 | mucin-associated surface protein (MASP), putative | 0,06551 | 0,8251 |
| TcCLB.436497.10 | hypothetical protein | 0,0654 | 0,6642 |
| TcCLB.510143.60 | hypothetical protein, conserved | 0,06534 | 0,7386 |
| TcCLB.511603.356 | syntaxin binding protein (pseudogene), putative | 0,06534 | 0,9203 |
| TcCLB.503903.10 | hypothetical protein | 0,065 | 0,7697 |
| TcCLB.508693.70 | hypothetical protein, conserved | 0,06491 | 0,6998 |
| TcCLB.510407.9 | protein kinase, putative | 0,06483 | 0,8507 |
| TcCLB.510475.90 | mucin TcMUCII, putative | 0,06483 | 0,9071 |
| TcCLB.422189.10 | nucleoside phosphatase, putative | 0,0647 | 0,7134 |
| TcCLB.507677.39 | 40S ribosomal protein S3, putative | 0,06457 | 0,4214 |
| TcCLB.510145.59 | exonuclease, putative | 0,06456 | 0,7173 |
| TcCLB.508243.10 | mucin-associated surface protein (MASP), putative | 0,06444 | 0,8674 |
| TcCLB.504431.80 | hypothetical protein | 0,06416 | 0,6807 |

| | | | |
|---|---|---|---|
| TcCLB.510121.100 | hypothetical protein | 0,06409 | 0,8341 |
| TcCLB.510063.39 | DNA repair protein, putative | 0,06403 | 0,6438 |
| TcCLB.506577.54 | 60S ribosomal protein L6, putative | 0,06392 | 0,5703 |
| TcCLB.506459.250 | hypothetical protein, conserved | 0,06392 | 0,7356 |
| TcCLB.506211.180 | mitogen-activated protein kinase 9, putative (MPK9) | 0,06391 | 0,6443 |
| TcCLB.508837.70 | trans-sialidase, Group II, putative | 0,0639 | 0,7405 |
| TcCLB.507873.50 | hypothetical protein, conserved | 0,06385 | 0,7842 |
| TcCLB.511021.60 | hypothetical protein, conserved | 0,06383 | 0,7006 |
| TcCLB.510665.10 | hypothetical protein, conserved | 0,06364 | 0,6521 |
| TcCLB.509505.30 | 10 kDa heat shock protein, putative | 0,06362 | 0,796 |
| TcCLB.506945.290 | ribosomal protein L15, putative | 0,06354 | 0,5474 |
| TcCLB.510761.80 | surface protease GP63, putative | 0,06352 | 0,8514 |
| TcCLB.509797.40 | isoleucyl-tRNA synthetase, putative | 0,06349 | 0,5211 |
| TcCLB.507827.10 | dispersed gene family protein 1 (DGF-1), putative | 0,06346 | 0,799 |
| TcCLB.506425.140 | hypothetical protein, conserved | 0,0634 | 0,703 |
| TcCLB.507015.59 | expression site-associated gene (ESAG-like, pseudogene), | 0,06338 | 0,7936 |
| TcCLB.506525.80 | N-myristoyl transferase, putative | 0,0633 | 0,691 |
| TcCLB.509073.40 | outer dynein arm docking complex protein (DC2) | 0,06329 | 0,6398 |
| TcCLB.507521.50 | proteasome beta 6 subunit, putative | 0,06319 | 0,6474 |
| TcCLB.510149.40 | hypothetical protein, conserved | 0,06312 | 0,7224 |
| TcCLB.507467.100 | hypothetical protein | 0,06308 | 0,6622 |
| TcCLB.510613.10 | hypothetical protein, conserved | 0,06308 | 0,7268 |
| TcCLB.510667.80 | hypothetical protein, conserved | 0,06304 | 0,7612 |
| TcCLB.504115.50 | retrotransposon hot spot protein (RHS, pseudogene), putat | 0,06292 | 0,7974 |
| TcCLB.509151.20 | hypothetical protein | 0,06289 | 0,7173 |
| TcCLB.507993.260 | hypothetical protein, conserved | 0,06288 | 0,6296 |
| TcCLB.503601.20 | hypothetical protein | 0,0628 | 0,9081 |
| TcCLB.509017.9 | hypothetical protein, conserved (fragment) | 0,06262 | 0,7919 |
| TcCLB.511495.11 | hypothetical protein, conserved (fragment) | 0,06255 | 0,8385 |
| TcCLB.411427.30 | hypothetical protein, conserved | 0,06242 | 0,6256 |
| TcCLB.504105.94 | ribosomal protein S25, putative | 0,06233 | 0,4925 |
| TcCLB.503877.19 | hypothetical protein, conserved | 0,06225 | 0,8114 |
| TcCLB.511903.220 | hypothetical protein, conserved | 0,0621 | 0,6157 |
| TcCLB.505997.160 | retrotransposon hot spot protein (RHS, pseudogene), putat | 0,06208 | 0,7839 |
| TcCLB.503981.60 | hypothetical protein, conserved | 0,06165 | 0,7023 |
| TcCLB.503465.30 | Iron-sulfur assembly protein 2, putative (Isa2) | 0,06165 | 0,848 |
| TcCLB.510275.330 | mucin TcMUCII, putative | 0,06162 | 0,9159 |
| TcCLB.508173.240 | hypothetical protein, conserved | 0,06143 | 0,744 |
| TcCLB.507825.20 | RNA helicase, putative | 0,06127 | 0,5408 |
| TcCLB.511603.30 | mucin-associated surface protein (MASP), putative | 0,06126 | 0,9139 |
| TcCLB.511885.50 | retrotransposon hot spot protein (RHS, pseudogene), putat | 0,06125 | 0,6962 |
| TcCLB.503609.32 | elongation factor 1-gamma (EF-1-gamma, pseudogene), p | 0,06121 | 0,9324 |
| TcCLB.508277.390 | hypothetical protein, conserved | 0,06116 | 0,608 |
| TcCLB.509205.50 | hypothetical protein, conserved | 0,06116 | 0,7252 |
| TcCLB.507033.29 | extracellular receptor, putative (fragment) | 0,06112 | 0,7333 |

| | | | |
|---|---|---|---|
| TcCLB.431849.20 | hypothetical protein, conserved (fragment) | 0,0611 | 0,84 |
| TcCLB.511299.79 | oxidoreductase, putative (fragment) | 0,06082 | 0,7629 |
| TcCLB.510085.60 | mucin-associated surface protein (MASP), putative | 0,06067 | 0,8529 |
| TcCLB.510295.50 | protein kinase, putative | 0,06065 | 0,7268 |
| TcCLB.507517.50 | hypothetical protein | 0,06065 | 0,7493 |
| TcCLB.507739.100 | hypothetical protein | 0,0606 | 0,7464 |
| TcCLB.506729.20 | hypothetical protein, conserved | 0,06057 | 0,7998 |
| TcCLB.506289.170 | surface protease GP63, putative | 0,0605 | 0,8614 |
| TcCLB.503819.10 | hypothetical protein, conserved | 0,06049 | 0,7029 |
| TcCLB.509353.60 | calmodulin, putative | 0,0604 | 0,5861 |
| TcCLB.510469.30 | trans-sialidase (pseudogene), putative | 0,06031 | 0,9122 |
| TcCLB.506755.260 | mitochondrial RNA-binding protein 2, putative | 0,06025 | 0,6679 |
| TcCLB.505007.39 | hypothetical protein, conserved | 0,06023 | 0,7922 |
| TcCLB.508507.30 | calcium uniporter protein, mitochondrial, putative | 0,06005 | 0,7747 |
| TcCLB.506857.110 | hypothetical protein, conserved | 0,05999 | 0,7455 |
| TcCLB.504057.130 | hypothetical protein, conserved | 0,05994 | 0,5659 |
| TcCLB.511367.340 | mitochondrial processing peptidase alpha subunit, putative | 0,05987 | 0,5478 |
| TcCLB.510165.50 | hypothetical protein, conserved | 0,05984 | 0,7932 |
| TcCLB.506755.40 | hypothetical protein, conserved | 0,05983 | 0,7869 |
| TcCLB.509203.30 | hd phosphohydrolase family protein | 0,05983 | 0,831 |
| TcCLB.507809.74 | signal peptidase type I, putative | 0,05971 | 0,7277 |
| TcCLB.507007.20 | hypothetical protein, conserved | 0,05962 | 0,644 |
| TcCLB.508347.110 | hypothetical protein, conserved | 0,05934 | 0,794 |
| TcCLB.506679.100 | 40S ribosomal protein S18, putative | 0,05926 | 0,5162 |
| TcCLB.511295.20 | hypothetical protein, conserved | 0,05906 | 0,8386 |
| TcCLB.506133.125 | hypothetical protein, conserved (pseudogene) | 0,05905 | 0,9473 |
| TcCLB.508641.170 | protein kinase, putative | 0,05885 | 0,782 |
| TcCLB.508569.90 | hypothetical protein, conserved | 0,05872 | 0,7112 |
| TcCLB.508661.20 | ubiquitin/ribosomal protein S27a, putative | 0,05869 | 0,5581 |
| TcCLB.511043.45 | trans-sialidase (pseudogene), putative | 0,05867 | 0,7397 |
| TcCLB.506697.50 | peptidyl-prolyl cis-trans isomerase NIMA-interacting 4, put | 0,05864 | 0,8048 |
| TcCLB.506947.90 | adenylate kinase, putative | 0,05862 | 0,7602 |
| TcCLB.510199.30 | mucin-associated surface protein (MASP), putative | 0,05858 | 0,8892 |
| TcCLB.509767.180 | exosome component CSL4, putative (CSL4) | 0,05854 | 0,812 |
| TcCLB.507255.50 | UDP-Gal or UDP-GlcNAc-dependent glycosyltransferase, pu | 0,05821 | 0,8666 |
| TcCLB.509613.10 | hypothetical protein, conserved | 0,0582 | 0,7687 |
| TcCLB.503489.10 | metallo-peptidase, Clan MC, Family M14, putative | 0,05813 | 0,6606 |
| TcCLB.510427.5 | hypothetical protein, conserved | 0,05806 | 0,8445 |
| TcCLB.509611.160 | proteasome regulatory non-ATPase subunit, putative | 0,05784 | 0,6716 |
| TcCLB.511883.49 | hypothetical protein, conserved (fragment) | 0,0578 | 0,745 |
| TcCLB.506499.170 | trans-sialidase, Group V, putative | 0,0578 | 0,8896 |
| TcCLB.508851.110 | hypothetical protein, conserved | 0,05776 | 0,8345 |
| TcCLB.508625.50 | 6-phosphofructo-2-kinase/fructose-2,6-bisphosphatas, puta | 0,05753 | 0,7697 |
| TcCLB.511383.70 | hypothetical protein, conserved | 0,05753 | 0,7922 |
| TcCLB.508469.50 | hypothetical protein, conserved (pseudogene) | 0,05744 | 0,7268 |

| | | | |
|---|---|---|---|
| TcCLB.506017.20 | hypothetical protein, conserved (pseudogene) | 0,05743 | 0,7055 |
| TcCLB.511507.120 | hypothetical protein, conserved | 0,05732 | 0,7561 |
| TcCLB.506471.90 | dispersed gene family protein 1 (DGF-1, pseudogene), puta | 0,0573 | 0,8065 |
| TcCLB.511237.120 | hypothetical protein | 0,05728 | 0,8729 |
| TcCLB.511711.90 | cop-coated vesicle membrane protein p24 precursor, putat | 0,05715 | 0,6717 |
| TcCLB.509229.120 | hypothetical protein, conserved | 0,057 | 0,7708 |
| TcCLB.507711.40 | RNA-binding protein, putative | 0,05694 | 0,7309 |
| TcCLB.510747.80 | Double RNA binding domain protein 9 (DRBD9) | 0,05692 | 0,7123 |
| TcCLB.508227.51 | surface protease GP63 (pseudogene), putative | 0,05687 | 0,9383 |
| TcCLB.509965.300 | hypothetical protein, conserved | 0,05673 | 0,835 |
| TcCLB.509897.70 | trans-sialidase, Group V, putative | 0,05667 | 0,8376 |
| TcCLB.511799.9 | hypothetical protein, conserved (fragment) | 0,05655 | 0,6637 |
| TcCLB.503573.9 | ATP-binding cassette transporter ABCA1, putative (ABCA1) | 0,05654 | 0,8057 |
| TcCLB.506123.10 | major vault protein, putative | 0,05652 | 0,7408 |
| TcCLB.510911.50 | glycine dehydrogenase [decarboxylating], putative | 0,05651 | 0,8677 |
| TcCLB.509169.39 | protein kinase, putative | 0,05649 | 0,8419 |
| TcCLB.509505.50 | 10 kDa heat shock protein, putative | 0,05639 | 0,7166 |
| TcCLB.506279.220 | mucin-associated surface protein (MASP, pseudogene), pu | 0,05622 | 0,8544 |
| TcCLB.508957.30 | hypothetical protein, conserved | 0,05612 | 0,6622 |
| TcCLB.510531.90 | hypothetical protein, conserved (pseudogene) | 0,0561 | 0,7454 |
| TcCLB.507021.50 | kinesin, putative | 0,05604 | 0,6318 |
| TcCLB.510579.60 | heat shock protein, putative | 0,05603 | 0,7282 |
| TcCLB.503823.20 | hypothetical protein, conserved | 0,05601 | 0,8076 |
| TcCLB.509607.40 | hypothetical protein | 0,05593 | 0,7313 |
| TcCLB.511383.30 | U1 small nuclear ribonucleoprotein, putative | 0,05592 | 0,7844 |
| TcCLB.511179.110 | hypothetical protein, conserved | 0,05579 | 0,7062 |
| TcCLB.510483.320 | retrotransposon hot spot protein (RHS, pseudogene), putat | 0,05578 | 0,8266 |
| TcCLB.508153.1070 | hypothetical protein, conserved | 0,05569 | 0,6959 |
| TcCLB.508717.53 | hypothetical protein, conserved | 0,0555 | 0,6685 |
| TcCLB.508387.30 | hypothetical protein, conserved | 0,05548 | 0,7156 |
| TcCLB.508553.10 | dispersed gene family protein 1 (DGF-1, pseudogene), puta | 0,05541 | 0,859 |
| TcCLB.503425.5 | mitochondrial carrier protein, putative | 0,0554 | 0,883 |
| TcCLB.510429.24 | hypothetical protein, conserved | 0,05527 | 0,8296 |
| TcCLB.508897.50 | hypothetical protein, conserved | 0,05525 | 0,6671 |
| TcCLB.507993.250 | hypothetical protein, conserved | 0,05516 | 0,7166 |
| TcCLB.506629.30 | guide RNA-binding protein of 21 kDa (gBP21) | 0,05504 | 0,6637 |
| TcCLB.404001.20 | hypothetical protein, conserved | 0,05482 | 0,6122 |
| TcCLB.510387.15 | syntaxin binding protein (pseudogene), putative | 0,05482 | 0,9139 |
| TcCLB.504033.53 | UDP-Gal or UDP-GlcNAc-dependent glycosyltransferase (p | 0,05474 | 0,9417 |
| TcCLB.507491.50 | hypothetical protein, conserved | 0,05458 | 0,7604 |
| TcCLB.507601.50 | hypothetical protein | 0,05457 | 0,7561 |
| TcCLB.506695.4 | Putative intraflagellar transport protein C3 (fragment) (PIF | 0,05456 | 0,7747 |
| TcCLB.511217.209 | hypothetical protein, conserved (fragment) | 0,05445 | 0,8239 |
| TcCLB.510565.120 | hypothetical protein, conserved | 0,05441 | 0,773 |
| TcCLB.503779.10 | hypothetical protein, conserved | 0,05439 | 0,8253 |

| | | | |
|---|---|---|---|
| TcCLB.510371.10 | mucin-associated surface protein (MASP), putative | 0,05426 | 0,8994 |
| TcCLB.510317.40 | hypothetical protein, conserved | 0,05425 | 0,7186 |
| TcCLB.506967.100 | mucin-associated surface protein (MASP), putative | 0,05423 | 0,9171 |
| TcCLB.510073.10 | cysteine peptidase (N-terminal), putative | 0,05418 | 0,8613 |
| TcCLB.504047.10 | hypothetical protein, conserved | 0,05407 | 0,8467 |
| TcCLB.511309.20 | hydrolase-like protein, putative | 0,054 | 0,7985 |
| TcCLB.508909.330 | protein kinase, putative | 0,05399 | 0,7015 |
| TcCLB.511277.600 | UDP-galactopyranose mutase (GLF) | 0,05378 | 0,6635 |
| TcCLB.506435.270 | dihydroxyacetone phosphate acyltransferase, putative (DA | 0,05375 | 0,6574 |
| TcCLB.507059.50 | trans-sialidase, Group V, putative | 0,05367 | 0,8475 |
| TcCLB.509245.20 | hypothetical protein, conserved | 0,05361 | 0,8617 |
| TcCLB.503917.40 | hypothetical protein, conserved | 0,05359 | 0,7312 |
| TcCLB.509157.260 | hypothetical protein | 0,05354 | 0,7725 |
| TcCLB.506163.35 | hypothetical protein, conserved | 0,05346 | 0,8252 |
| TcCLB.508257.180 | hypothetical protein, conserved | 0,05328 | 0,7227 |
| TcCLB.506943.130 | hypothetical protein, conserved | 0,05325 | 0,7764 |
| TcCLB.511353.30 | hypothetical protein, conserved | 0,05325 | 0,7855 |
| TcCLB.511471.90 | glycine dehydrogenase [decarboxylating] (pseudogene), pu | 0,0532 | 0,8605 |
| TcCLB.508817.20 | hypothetical protein | 0,05317 | 0,703 |
| TcCLB.508409.338 | DNA repair protein, putative | 0,05307 | 0,802 |
| TcCLB.508921.20 | hypothetical protein, conserved | 0,05302 | 0,8151 |
| TcCLB.505939.70 | hypothetical protein, conserved | 0,05271 | 0,7251 |
| TcCLB.509733.100 | translation elongation factor 1-beta, putative | 0,05257 | 0,5098 |
| TcCLB.506783.30 | elongation factor 1-gamma (EF-1-gamma), putative | 0,05252 | 0,9119 |
| TcCLB.504131.190 | mitochondrial carrier protein, putative | 0,05248 | 0,7421 |
| TcCLB.507491.110 | hypothetical protein | 0,05246 | 0,6884 |
| TcCLB.416883.9 | hypothetical protein, conserved (fragment) | 0,05242 | 0,7972 |
| TcCLB.511163.40 | cell cycle sequence binding phosphoprotein (RBP45), putat | 0,05241 | 0,7121 |
| TcCLB.507519.50 | hypothetical protein, conserved | 0,05241 | 0,8032 |
| TcCLB.507737.10 | hypothetical protein, conserved | 0,05241 | 0,8849 |
| TcCLB.510301.50 | hypothetical protein, conserved | 0,0524 | 0,7928 |
| TcCLB.507167.31 | N-acetyltransferase complex ARD1 subunit (pseudogene), | 0,05231 | 0,9348 |
| TcCLB.509643.110 | hypothetical protein, conserved | 0,05228 | 0,7355 |
| TcCLB.511603.380 | mucin-associated surface protein (MASP), putative | 0,05223 | 0,9179 |
| TcCLB.505037.40 | hypothetical protein, conserved | 0,0522 | 0,7506 |
| TcCLB.506457.60 | 2-aminoethylphosphonate:pyruvateaminotransferase- like | 0,0522 | 0,7661 |
| TcCLB.511911.140 | hypothetical protein, conserved | 0,05198 | 0,8 |
| TcCLB.509719.28 | hypothetical protein, conserved | 0,05195 | 0,8707 |
| TcCLB.508221.1050 | trans-sialidase, putative | 0,05193 | 0,916 |
| TcCLB.509789.30 | rab-like GTPase activating protein, putative | 0,05183 | 0,7845 |
| TcCLB.509669.100 | hypothetical protein, conserved | 0,05182 | 0,7522 |
| TcCLB.510289.6 | nucleolar RNA-binding protein, putative | 0,05173 | 0,7844 |
| TcCLB.507907.55 | ATP-dependent DEAD/H RNA helicase (pseudogene), putat | 0,05173 | 0,8416 |
| TcCLB.508271.10 | phenylalanyl-tRNA synthetase, putative | 0,05165 | 0,6234 |
| TcCLB.510143.50 | hypothetical protein, conserved | 0,05163 | 0,7661 |

| | | | |
|---|---|---|---|
| TcCLB.509671.80 | 60S ribosomal protein L5, putative | 0,05162 | 0,5404 |
| TcCLB.508759.10 | mucin-associated surface protein (MASP), putative | 0,0516 | 0,945 |
| TcCLB.507105.120 | hypothetical protein, conserved (pseudogene) | 0,0515 | 0,7641 |
| TcCLB.506435.430 | hypothetical protein, conserved | 0,05141 | 0,6979 |
| TcCLB.510283.50 | hypothetical protein, conserved | 0,05138 | 0,824 |
| TcCLB.507519.210 | glycine dehydrogenase [decarboxylating] (pseudogene), pu | 0,05127 | 0,7834 |
| TcCLB.508385.30 | CYC2-like cyclin, putative | 0,05094 | 0,6725 |
| TcCLB.510175.30 | trans-sialidase (pseudogene), putative | 0,0506 | 0,8434 |
| TcCLB.510735.100 | hypothetical protein, conserved | 0,05057 | 0,793 |
| TcCLB.506575.30 | hypothetical protein, conserved (pseudogene) | 0,05033 | 0,6945 |
| TcCLB.510363.60 | trans-sialidase (pseudogene), putative | 0,05031 | 0,9061 |
| TcCLB.508417.10 | hypothetical protein, conserved | 0,0502 | 0,6884 |
| TcCLB.460747.9 | hypothetical protein, conserved (fragment) | 0,05019 | 0,7229 |
| TcCLB.508205.10 | hypothetical protein, conserved | 0,05019 | 0,8111 |
| TcCLB.507623.129 | vacuolar type h ATPase subunit, putative | 0,05018 | 0,7823 |
| TcCLB.507515.50 | hypothetical protein | 0,05018 | 0,787 |
| TcCLB.507071.140 | mucin-associated surface protein (MASP), putative | 0,05017 | 0,9312 |
| TcCLB.510497.190 | hypothetical protein | 0,05 | 0,9423 |
| TcCLB.509579.40 | hypothetical protein, conserved (pseudogene) | 0,04998 | 0,8513 |
| TcCLB.506945.350 | serine/threonine protein kinase, putative | 0,0498 | 0,5835 |
| TcCLB.509871.39 | hypothetical protein, conserved (pseudogene) | 0,04968 | 0,7168 |
| TcCLB.507669.200 | hypothetical protein, conserved | 0,04967 | 0,7797 |
| TcCLB.504427.260 | kinesin, putative | 0,04954 | 0,6787 |
| TcCLB.510719.170 | hypothetical protein, conserved | 0,04947 | 0,8285 |
| TcCLB.511139.20 | 40S ribosomal protein S11, putative | 0,04944 | 0,5611 |
| TcCLB.504015.21 | elongation factor 1-gamma (EF-1-gamma, pseudogene), p | 0,04943 | 0,9297 |
| TcCLB.507467.30 | hypothetical protein, conserved | 0,04941 | 0,8458 |
| TcCLB.511181.100 | 60S ribosomal protein L8, putative | 0,0494 | 0,5738 |
| TcCLB.505419.20 | retrotransposon hot spot protein (RHS, pseudogene), putat | 0,04939 | 0,7771 |
| TcCLB.506625.60 | inositol 5-phosphatase 1, putative | 0,04936 | 0,6121 |
| TcCLB.509683.40 | hypothetical protein, conserved | 0,04928 | 0,7629 |
| TcCLB.511655.20 | hypothetical protein, conserved | 0,04928 | 0,7927 |
| TcCLB.506501.390 | mucin TcMUCII, putative | 0,04928 | 0,9495 |
| TcCLB.506297.270 | 60S ribosomal protein L28, putative | 0,04915 | 0,5901 |
| TcCLB.504147.20 | 60S ribosomal protein L18, putative | 0,04904 | 0,5846 |
| TcCLB.507713.20 | hypothetical protein, conserved | 0,04904 | 0,7829 |
| TcCLB.509205.20 | hypothetical protein, conserved | 0,04903 | 0,81 |
| TcCLB.508797.29 | hypothetical protein, conserved | 0,04883 | 0,6508 |
| TcCLB.485845.9 | dispersed gene family protein 1 (DGF-1), putative (fragme | 0,04874 | 0,9471 |
| TcCLB.508879.50 | hypothetical protein | 0,0487 | 0,7618 |
| TcCLB.509337.19 | hypothetical protein, conserved | 0,04867 | 0,7949 |
| TcCLB.508747.5 | mucin-associated surface protein (MASP), putative | 0,0486 | 0,9335 |
| TcCLB.506905.20 | hypothetical protein, conserved | 0,04855 | 0,7588 |
| TcCLB.505333.20 | ATP-dependent DEAD/H RNA helicase (pseudogene), putat | 0,04828 | 0,9294 |
| TcCLB.503421.20 | hypothetical protein, conserved (pseudogene) | 0,0482 | 0,8441 |

| | | | |
|---|---|---|---|
| TcCLB.508837.154 | hypothetical protein, conserved | 0,04815 | 0,8389 |
| TcCLB.507963.70 | hypothetical protein, conserved | 0,04811 | 0,7347 |
| TcCLB.509497.20 | hypothetical protein, conserved | 0,04791 | 0,7583 |
| TcCLB.511045.10 | hypothetical protein, conserved | 0,04788 | 0,7661 |
| TcCLB.503579.60 | hypothetical protein | 0,04773 | 0,7796 |
| TcCLB.508303.50 | hypothetical protein, conserved | 0,04773 | 0,7977 |
| TcCLB.509471.86 | histone H3, putative | 0,04754 | 0,594 |
| TcCLB.508303.20 | hypothetical protein | 0,04751 | 0,7919 |
| TcCLB.509287.220 | hypothetical protein, conserved | 0,0475 | 0,7496 |
| TcCLB.510329.330 | hypothetical protein, conserved | 0,04749 | 0,7923 |
| TcCLB.510315.14 | hypothetical protein, conserved | 0,04749 | 0,8346 |
| TcCLB.506865.30 | hypothetical protein, conserved | 0,04744 | 0,7521 |
| TcCLB.509115.41 | elongation factor 1-gamma (EF-1-gamma, pseudogene), p | 0,04738 | 0,8822 |
| TcCLB.509045.29 | kinesin, putative | 0,04736 | 0,8142 |
| TcCLB.508207.100 | 60S ribosomal protein L7, putative | 0,04734 | 0,5766 |
| TcCLB.508727.54 | hypothetical protein, conserved | 0,04728 | 0,7863 |
| TcCLB.509671.70 | ribosomal protein L15, putative | 0,04725 | 0,6016 |
| TcCLB.503903.20 | hypothetical protein, conserved | 0,0471 | 0,8233 |
| TcCLB.503987.70 | ribosome biogenesis protein TSR3, putative (tsr3) | 0,04708 | 0,8453 |
| TcCLB.504157.100 | hypothetical protein, conserved | 0,04697 | 0,7704 |
| TcCLB.508887.10 | hypothetical protein, conserved (pseudogene) | 0,04685 | 0,9468 |
| TcCLB.506337.200 | hypothetical protein, conserved | 0,04663 | 0,6919 |
| TcCLB.506603.40 | trans-sialidase, Group V, putative | 0,04661 | 0,89 |
| TcCLB.508779.70 | hypothetical protein, conserved | 0,04648 | 0,732 |
| TcCLB.506679.240 | hypothetical protein, conserved | 0,0464 | 0,7945 |
| TcCLB.478711.10 | trans-sialidase (pseudogene), putative | 0,04637 | 0,7623 |
| TcCLB.504787.70 | mucin-associated surface protein (MASP), putative | 0,04617 | 0,9022 |
| TcCLB.507699.170 | surface protease GP63 (pseudogene), putative | 0,04594 | 0,8667 |
| TcCLB.507427.10 | trans-sialidase, Group I, putative | 0,04589 | 0,8137 |
| TcCLB.507093.90 | hypothetical protein, conserved | 0,04585 | 0,7567 |
| TcCLB.506329.35 | hypothetical protein, conserved (pseudogene) | 0,04575 | 0,9505 |
| TcCLB.506835.40 | hypothetical protein, conserved | 0,04573 | 0,7905 |
| TcCLB.509885.50 | hypothetical protein, conserved | 0,04571 | 0,7946 |
| TcCLB.509733.160 | amino acid tansporter, putative | 0,04566 | 0,7881 |
| TcCLB.508323.100 | I/6 autoantigen, putative | 0,04563 | 0,793 |
| TcCLB.511491.130 | dolichyl pyrophosphate phosphatase, putative | 0,04557 | 0,8482 |
| TcCLB.503919.50 | hypothetical protein, conserved | 0,04553 | 0,7562 |
| TcCLB.510489.50 | trans-sialidase, putative | 0,0455 | 0,9441 |
| TcCLB.508813.60 | Present in the outer mitochondrial membrane proteome 1 | 0,04549 | 0,6536 |
| TcCLB.509539.50 | small GTP-binding protein, putative | 0,04538 | 0,7173 |
| TcCLB.511657.50 | hypothetical protein, conserved | 0,04536 | 0,9136 |
| TcCLB.509693.130 | hypothetical protein | 0,04534 | 0,805 |
| TcCLB.510187.400 | serine/threonine protein phosphatase, putative | 0,04518 | 0,7271 |
| TcCLB.433273.10 | dynein heavy chain, putative (fragment) | 0,045 | 0,7671 |
| TcCLB.511653.10 | delta-adaptin, putative | 0,04498 | 0,7388 |

| | | | |
|---|---|---|---|
| TcCLB.510187.130 | Bardet-Biedl syndrome 4 protein, putative (BBS4) | 0,04492 | 0,7587 |
| TcCLB.506967.60 | trans-sialidase, Group VI, putative | 0,04489 | 0,9114 |
| TcCLB.506195.280 | hypothetical protein, conserved | 0,04485 | 0,6966 |
| TcCLB.506357.180 | hypothetical protein | 0,04484 | 0,7529 |
| TcCLB.508737.40 | hypothetical protein | 0,04479 | 0,7797 |
| TcCLB.510665.40 | DNA-repair protein, putative | 0,0446 | 0,7173 |
| TcCLB.511001.110 | hypothetical protein, conserved | 0,04456 | 0,8214 |
| TcCLB.511287.10 | hypothetical protein, conserved | 0,04454 | 0,79 |
| TcCLB.506667.40 | mucin-associated surface protein (MASP), putative | 0,04453 | 0,9398 |
| TcCLB.507875.70 | trans-sialidase, Group VII, putative | 0,04452 | 0,9168 |
| TcCLB.506163.70 | hypothetical protein, conserved | 0,04437 | 0,7614 |
| TcCLB.503487.60 | hypothetical protein, conserved | 0,04431 | 0,832 |
| TcCLB.511407.50 | hypothetical protein, conserved | 0,04427 | 0,7243 |
| TcCLB.506363.90 | amidohydrolase, putative | 0,0442 | 0,7661 |
| TcCLB.508967.20 | dispersed gene family protein 1 (DGF-1), putative | 0,04414 | 0,9133 |
| TcCLB.510661.200 | hypothetical protein, conserved | 0,04409 | 0,7921 |
| TcCLB.506885.120 | protein kinase, putative | 0,04394 | 0,8137 |
| TcCLB.506613.70 | hypothetical protein | 0,04394 | 0,9249 |
| TcCLB.510219.10 | hypothetical protein, conserved | 0,04385 | 0,8238 |
| TcCLB.511311.14 | retrotransposon hot spot protein (RHS, pseudogene), putat | 0,04383 | 0,8056 |
| TcCLB.503847.50 | hypothetical protein, conserved | 0,04351 | 0,8433 |
| TcCLB.506457.99 | hypothetical protein, conserved | 0,04347 | 0,7602 |
| TcCLB.503709.20 | 60S ribosomal protein L44, putative (fragment) | 0,04343 | 0,6966 |
| TcCLB.504741.250 | protein kinase, putative | 0,04341 | 0,7454 |
| TcCLB.510877.130 | hypothetical protein, conserved | 0,04336 | 0,7224 |
| TcCLB.507837.50 | 40S ribosomal protein S11, putative | 0,04324 | 0,9229 |
| TcCLB.510599.50 | hypothetical protein | 0,04323 | 0,7606 |
| TcCLB.508845.5 | retrotransposon hot spot protein (RHS, pseudogene), putat | 0,04321 | 0,941 |
| TcCLB.504741.200 | hypothetical protein, conserved | 0,0432 | 0,787 |
| TcCLB.506855.330 | hypothetical protein, conserved | 0,04317 | 0,7622 |
| TcCLB.510551.20 | trans-sialidase (pseudogene), putative | 0,04298 | 0,9103 |
| TcCLB.507631.15 | hypothetical protein, conserved (pseudogene) | 0,04281 | 0,8948 |
| TcCLB.504025.19 | hypothetical protein, conserved (fragment) | 0,04277 | 0,8677 |
| TcCLB.503493.40 | hypothetical protein, conserved | 0,04272 | 0,9456 |
| TcCLB.506499.210 | mucin-associated surface protein (MASP), putative | 0,04267 | 0,8948 |
| TcCLB.511131.89 | hypothetical protein, conserved | 0,04266 | 0,8472 |
| TcCLB.504881.40 | hypothetical protein, conserved | 0,04263 | 0,7504 |
| TcCLB.506425.30 | hypothetical protein, conserved | 0,04255 | 0,7467 |
| TcCLB.506067.34 | mucin-associated surface protein (MASP, pseudogene), pu | 0,04255 | 0,91 |
| TcCLB.508207.260 | hypothetical protein, conserved | 0,04249 | 0,8291 |
| TcCLB.508461.450 | hypothetical protein, conserved | 0,04245 | 0,793 |
| TcCLB.506767.400 | mucin-associated surface protein (MASP), putative | 0,04242 | 0,9499 |
| TcCLB.510383.9 | dispersed gene family protein 1 (DGF-1), putative | 0,04238 | 0,8906 |
| TcCLB.507159.65 | DNA-directed RNA polymerase subunit, putative | 0,04227 | 0,8041 |
| TcCLB.509243.60 | ATP-dependent DEAD/H DNA helicase recQ, putative | 0,04215 | 0,9071 |

| | | | |
|---|---|---|---|
| TcCLB.511555.20 | hypothetical protein, conserved | 0,04209 | 0,7855 |
| TcCLB.511515.20 | AAA ATPase (pseudogene), putative | 0,04204 | 0,7662 |
| TcCLB.504047.60 | protein kinase, putative | 0,04201 | 0,7424 |
| TcCLB.504239.290 | mucin-associated surface protein (MASP), putative | 0,04197 | 0,9249 |
| TcCLB.508355.310 | hypothetical protein, conserved | 0,04196 | 0,7576 |
| TcCLB.509921.40 | retrotransposon hot spot protein (RHS, pseudogene), putat | 0,04193 | 0,8321 |
| TcCLB.511761.70 | hypothetical protein, conserved | 0,04188 | 0,8178 |
| TcCLB.508541.110 | mucin-associated surface protein (MASP), putative | 0,04167 | 0,9344 |
| TcCLB.508541.170 | hypothetical protein, conserved | 0,04159 | 0,876 |
| TcCLB.509039.40 | hypothetical protein | 0,04156 | 0,837 |
| TcCLB.506367.60 | vacuolar protein sorting protein 16, putative | 0,04155 | 0,7209 |
| TcCLB.511621.100 | hypothetical protein, conserved | 0,04139 | 0,802 |
| TcCLB.510293.20 | cytosolic factor SEC14, putative | 0,04138 | 0,8307 |
| TcCLB.509105.94 | hypothetical protein, conserved | 0,0412 | 0,8722 |
| TcCLB.511725.180 | dynein, putative | 0,04118 | 0,7619 |
| TcCLB.506843.70 | protein kinase, putative | 0,04118 | 0,9516 |
| TcCLB.508797.10 | tubulin-tyrosine ligase-like protein, putative | 0,04112 | 0,7685 |
| TcCLB.511017.30 | cyclin dependent kinase-binding protein, putative | 0,0411 | 0,7418 |
| TcCLB.509767.160 | hypothetical protein, conserved | 0,04109 | 0,885 |
| TcCLB.511291.90 | 40S ribosomal protein S13, putative | 0,04105 | 0,6776 |
| TcCLB.506501.10 | mucin-associated surface protein (MASP), putative | 0,04101 | 0,9441 |
| TcCLB.511127.260 | DNA photolyase, putative | 0,04095 | 0,7464 |
| TcCLB.510597.40 | nucleoporin (NUP54/57), putative | 0,04087 | 0,8729 |
| TcCLB.506977.130 | ras-related protein Rab21, putative (RAB21) | 0,0407 | 0,796 |
| TcCLB.511407.60 | small GTP-binding protein Rab11, putative (Rab11) | 0,04068 | 0,8075 |
| TcCLB.507401.9 | dispersed gene family protein 1 (DGF-1), putative (fragme | 0,04063 | 0,8826 |
| TcCLB.503395.20 | hypothetical protein, conserved | 0,04051 | 0,8583 |
| TcCLB.503953.59 | mitochondrial DNA polymerase beta-PAK, putative | 0,0405 | 0,8373 |
| TcCLB.505995.10 | dispersed gene family protein 1 (DGF-1), putative | 0,04044 | 0,92 |
| TcCLB.506605.200 | hypothetical protein, conserved | 0,04041 | 0,8334 |
| TcCLB.511293.69 | glycosomal malate dehydrogenase, putative | 0,04034 | 0,6276 |
| TcCLB.510627.60 | mucin-associated surface protein (MASP), putative | 0,04029 | 0,9542 |
| TcCLB.508135.10 | hypothetical protein, conserved | 0,04023 | 0,8341 |
| TcCLB.509453.30 | GCN5-like protein, putative | 0,04015 | 0,8841 |
| TcCLB.508361.30 | mucin TcMUCII, putative | 0,04009 | 0,9377 |
| TcCLB.503763.10 | retrotransposon hot spot (RHS) protein, putative | 0,04002 | 0,8934 |
| TcCLB.511303.110 | hypothetical protein, conserved | 0,03998 | 0,7984 |
| TcCLB.509053.100 | protein kinase, putative | 0,03988 | 0,802 |
| TcCLB.503643.39 | hypothetical protein, conserved | 0,03971 | 0,864 |
| TcCLB.504103.90 | hypothetical protein, conserved | 0,03967 | 0,8362 |
| TcCLB.506613.30 | hypothetical protein | 0,03955 | 0,9471 |
| TcCLB.506475.116 | glutaredoxin, putative | 0,03949 | 0,8617 |
| TcCLB.419417.4 | ADP-ribosylation factor, putative (fragment) | 0,03943 | 0,7998 |
| TcCLB.421321.9 | exportin-T, putative (fragment) | 0,03941 | 0,8161 |
| TcCLB.507023.124 | hypothetical protein, conserved | 0,03939 | 0,9055 |

| | | | |
|---|---|---|---|
| TcCLB.438147.18 | hypothetical protein, conserved | 0,03924 | 0,8837 |
| TcCLB.508657.20 | sec1-like protein (pseudogene), putative | 0,0392 | 0,8094 |
| TcCLB.506201.140 | hypothetical protein, conserved | 0,03903 | 0,7515 |
| TcCLB.506945.310 | hypothetical protein, conserved | 0,039 | 0,8249 |
| TcCLB.509649.50 | hypothetical protein, conserved | 0,03896 | 0,7511 |
| TcCLB.510197.20 | mucin-associated surface protein (MASP), putative | 0,03877 | 0,8924 |
| TcCLB.507091.60 | trans-sialidase, Group V, putative | 0,03874 | 0,8942 |
| TcCLB.507669.110 | hypothetical protein, conserved | 0,03871 | 0,7542 |
| TcCLB.506817.10 | hypothetical protein, conserved | 0,03868 | 0,6981 |
| TcCLB.510901.170 | hypothetical protein, conserved | 0,03866 | 0,7415 |
| TcCLB.511751.40 | DNA-J protein, putative | 0,03861 | 0,8578 |
| TcCLB.508661.30 | DnaJ chaperone protein, putative | 0,03821 | 0,7592 |
| TcCLB.510407.80 | hypothetical protein, conserved | 0,03817 | 0,7666 |
| TcCLB.506841.8 | hypothetical protein, conserved | 0,03813 | 0,7767 |
| TcCLB.507003.60 | hypothetical protein, conserved | 0,03787 | 0,7919 |
| TcCLB.506301.60 | hypothetical protein | 0,03781 | 0,7565 |
| TcCLB.509897.155 | trans-sialidase (pseudogene), putative | 0,03778 | 0,8734 |
| TcCLB.506809.10 | trans-sialidase (pseudogene), putative | 0,03775 | 0,8941 |
| TcCLB.506485.109 | cyclic nucleotide-binding domain containing protein, putati | 0,03774 | 0,8012 |
| TcCLB.510207.30 | trans-sialidase, Group V, putative | 0,03768 | 0,9339 |
| TcCLB.507011.120 | acyl transferase-like protein, putative | 0,03767 | 0,8105 |
| TcCLB.506577.5 | hypothetical protein | 0,03765 | 0,8374 |
| TcCLB.504021.149 | hypothetical protein, conserved (fragment) | 0,03765 | 0,8552 |
| TcCLB.508577.10 | trans-sialidase (pseudogene), putative | 0,0376 | 0,9168 |
| TcCLB.506241.50 | hypothetical protein, conserved | 0,03758 | 0,8226 |
| TcCLB.507065.60 | mucin-associated surface protein (MASP), putative | 0,0375 | 0,9128 |
| TcCLB.402449.10 | trans-sialidase (pseudogene), putative | 0,03745 | 0,8961 |
| TcCLB.511895.50 | hypothetical protein, conserved | 0,03744 | 0,7887 |
| TcCLB.511661.40 | signal peptidase type I, putative | 0,03735 | 0,8654 |
| TcCLB.509895.110 | hypothetical protein, conserved | 0,03733 | 0,9279 |
| TcCLB.508821.50 | vacuolar type h ATPase subunit, putative | 0,03728 | 0,8372 |
| TcCLB.507029.10 | hypothetical protein, conserved | 0,03714 | 0,8221 |
| TcCLB.508815.20 | hypothetical protein, conserved | 0,03714 | 0,8305 |
| TcCLB.503999.100 | phospholipid-transporting ATPase 1-like protein, putative | 0,03711 | 0,8777 |
| TcCLB.506865.4 | hypothetical protein, conserved | 0,03706 | 0,8805 |
| TcCLB.511239.159 | inosine-guanine nucleoside hydrolase, putative | 0,03695 | 0,9409 |
| TcCLB.507681.10 | hypothetical protein, conserved | 0,03693 | 0,9168 |
| TcCLB.510279.340 | mucin TcMUCII, putative | 0,03688 | 0,9582 |
| TcCLB.504741.149 | hypothetical protein | 0,03678 | 0,92 |
| TcCLB.507105.80 | dispersed gene family protein 1 (DGF-1), putative | 0,03663 | 0,9376 |
| TcCLB.509627.10 | bilobe protein, conserved | 0,03662 | 0,7614 |
| TcCLB.507811.70 | hypothetical protein, conserved | 0,0366 | 0,8266 |
| TcCLB.511259.161 | elongation factor 1-gamma (EF-1-gamma, pseudogene), p | 0,03634 | 0,9483 |
| TcCLB.509599.90 | hypothetical protein, conserved | 0,03598 | 0,8129 |
| TcCLB.506359.60 | UTP-glucose-1-phosphate uridylyltransferase 2, putative | 0,03595 | 0,8535 |

| | | | |
|---|---|---|---|
| TcCLB.508165.130 | surface protease GP63, putative | 0,03592 | 0,9363 |
| TcCLB.507875.20 | glutamate dehydrogenase, putative | 0,03579 | 0,8457 |
| TcCLB.511433.20 | hypothetical protein, conserved | 0,03577 | 0,9197 |
| TcCLB.511249.90 | hypothetical protein, conserved | 0,03571 | 0,7948 |
| TcCLB.509525.230 | mucin-associated surface protein (MASP), putative | 0,03566 | 0,9296 |
| TcCLB.511751.90 | hypothetical protein | 0,0355 | 0,8657 |
| TcCLB.509757.20 | leucine carboxyl methyltransferase, putative | 0,03541 | 0,8452 |
| TcCLB.503495.5 | hypothetical protein | 0,03535 | 0,9229 |
| TcCLB.506135.40 | hypothetical protein, conserved | 0,03531 | 0,7891 |
| TcCLB.506811.160 | R27-2 protein, putative | 0,03525 | 0,7197 |
| TcCLB.509031.10 | surface protease GP63 (pseudogene), putative | 0,03515 | 0,9025 |
| TcCLB.508879.229 | chaperone DNAJ protein, putative | 0,03506 | 0,9199 |
| TcCLB.510255.10 | hypothetical protein, conserved | 0,03484 | 0,8631 |
| TcCLB.507959.310 | mucin TcMUCII, putative | 0,03477 | 0,9505 |
| TcCLB.511667.140 | trans-sialidase (pseudogene), putative | 0,03472 | 0,8656 |
| TcCLB.503881.4 | hypothetical protein, conserved | 0,03471 | 0,9402 |
| TcCLB.509641.30 | Ubiquitin-like modifier-activating enzyme ATG7, putative ( | 0,03457 | 0,8111 |
| TcCLB.511391.90 | endosomal integral membrane protein, putative | 0,03451 | 0,8241 |
| TcCLB.508831.100 | hypothetical protein, conserved | 0,03447 | 0,8393 |
| TcCLB.506925.230 | hypothetical protein, conserved | 0,03426 | 0,7503 |
| TcCLB.511577.30 | hypothetical protein, conserved | 0,03402 | 0,8285 |
| TcCLB.510817.30 | hypothetical protein, conserved (pseudogene) | 0,034 | 0,8752 |
| TcCLB.511165.4 | cytoplasmic translation machinery associated protein, puta | 0,03395 | 0,9136 |
| TcCLB.511589.20 | hypothetical protein | 0,0339 | 0,8636 |
| TcCLB.506685.10 | hypothetical protein, conserved | 0,03382 | 0,7847 |
| TcCLB.509515.20 | UDP-Gal or UDP-GlcNAc-dependent glycosyltransferase, pu | 0,03371 | 0,9483 |
| TcCLB.412509.10 | retrotransposon hot spot (RHS) protein, putative | 0,03368 | 0,919 |
| TcCLB.508741.340 | hypothetical protein, conserved | 0,03352 | 0,8329 |
| TcCLB.507883.25 | hypothetical protein, conserved | 0,03352 | 0,9381 |
| TcCLB.509847.10 | retrotransposon hot spot protein (RHS, pseudogene), putat | 0,03349 | 0,9249 |
| TcCLB.509551.130 | hypothetical protein, conserved | 0,03331 | 0,8107 |
| TcCLB.508525.60 | hypothetical protein, conserved | 0,03317 | 0,7956 |
| TcCLB.504229.100 | trans-sialidase (pseudogene), putative | 0,03307 | 0,9051 |
| TcCLB.506817.30 | ribosomal protein L27, putative | 0,03291 | 0,7901 |
| TcCLB.510437.10 | dynein heavy chain, putative | 0,03287 | 0,856 |
| TcCLB.511143.19 | hypothetical protein, conserved | 0,03285 | 0,9048 |
| TcCLB.506821.140 | histone deacetylase, putative | 0,03284 | 0,8438 |
| TcCLB.509777.30 | trans-sialidase, Group II, putative | 0,0328 | 0,8419 |
| TcCLB.506327.50 | hypothetical protein, conserved | 0,03274 | 0,8413 |
| TcCLB.504081.280 | mucin-associated surface protein (MASP, pseudogene), pu | 0,0326 | 0,9591 |
| TcCLB.511725.240 | hypothetical protein, conserved | 0,03251 | 0,7858 |
| TcCLB.509617.70 | subtilisin-like serine peptidase, putative | 0,03249 | 0,8081 |
| TcCLB.511665.60 | trans-sialidase (pseudogene), putative | 0,03241 | 0,8179 |
| TcCLB.506531.9 | hypothetical protein (fragment) | 0,03239 | 0,8849 |
| TcCLB.506945.210 | Zinc finger CCCH domain-containing protein 47 (zc3h47) | 0,03229 | 0,7462 |

| | | | |
|---|---|---|---|
| TcCLB.503917.7 | RNA-binding protein, putative | 0,03227 | 0,8388 |
| TcCLB.510731.120 | hypothetical protein, conserved | 0,03217 | 0,8619 |
| TcCLB.506501.160 | mucin-associated surface protein (MASP, pseudogene), pu | 0,03197 | 0,9499 |
| TcCLB.504143.30 | kinesin protein (pseudogene), putative | 0,03196 | 0,8734 |
| TcCLB.506315.40 | hypothetical protein, conserved | 0,03184 | 0,8215 |
| TcCLB.511755.90 | hypothetical protein, conserved | 0,03175 | 0,8285 |
| TcCLB.505987.20 | hypothetical protein | 0,03163 | 0,8103 |
| TcCLB.406725.10 | trans-sialidase (pseudogene), putative | 0,03151 | 0,9568 |
| TcCLB.509489.29 | proteasome regulatory non-ATP-ase subunit, putative (frag | 0,03145 | 0,878 |
| TcCLB.506855.80 | ATP synthase, putative | 0,03144 | 0,8139 |
| TcCLB.506649.100 | tRNA nucleotidyltransferase, putative | 0,03142 | 0,8259 |
| TcCLB.511785.10 | mucin-associated surface protein (MASP), putative | 0,03129 | 0,9569 |
| TcCLB.505025.137 | mucin TcMUCII, putative | 0,03118 | 0,9663 |
| TcCLB.509877.80 | ATPase domain protein, putative | 0,03116 | 0,8916 |
| TcCLB.509571.10 | hypothetical protein, conserved | 0,03092 | 0,8955 |
| TcCLB.511651.10 | ATP-dependent DEAD/H RNA helicase, putative | 0,03091 | 0,8879 |
| TcCLB.504029.60 | hypothetical protein, conserved | 0,03079 | 0,8447 |
| TcCLB.510397.39 | dispersed gene family protein 1 (DGF-1), putative | 0,03073 | 0,9293 |
| TcCLB.506587.90 | surface protease GP63, putative | 0,0307 | 0,8729 |
| TcCLB.511409.60 | hypothetical protein, conserved | 0,03067 | 0,8835 |
| TcCLB.506181.97 | hypothetical protein, conserved | 0,03063 | 0,8918 |
| TcCLB.511019.74 | retrotransposon hot spot (RHS) protein, putative | 0,03056 | 0,8466 |
| TcCLB.507491.100 | hypothetical protein, conserved | 0,03055 | 0,8348 |
| TcCLB.508647.260 | hypothetical protein, conserved | 0,03054 | 0,8751 |
| TcCLB.508479.140 | hypothetical protein, conserved | 0,03054 | 0,888 |
| TcCLB.508461.284 | mitochondrial carrier protein, putative | 0,03045 | 0,8756 |
| TcCLB.511353.34 | hypothetical protein, conserved | 0,0304 | 0,849 |
| TcCLB.510157.5 | trans-sialidase, putative (fragment) | 0,03025 | 0,9876 |
| TcCLB.511131.80 | hypothetical protein, conserved | 0,03022 | 0,7864 |
| TcCLB.503823.130 | hypothetical protein, conserved | 0,03018 | 0,7819 |
| TcCLB.510189.30 | surface protease GP63 (pseudogene), putative | 0,03018 | 0,9527 |
| TcCLB.503809.110 | phosphatidic acid phosphatase protein, putative | 0,03005 | 0,8376 |
| TcCLB.510533.20 | hypothetical protein, conserved | 0,02997 | 0,8847 |
| TcCLB.504137.150 | hypothetical protein, conserved | 0,02987 | 0,8209 |
| TcCLB.509251.80 | trans-sialidase, Group V, putative | 0,02976 | 0,9243 |
| TcCLB.507069.40 | trans-sialidase, Group VI, putative | 0,02976 | 0,9452 |
| TcCLB.507953.60 | hypothetical protein | 0,02972 | 0,9618 |
| TcCLB.508601.10 | mitochondrial DEAD box protein, putative | 0,02968 | 0,8849 |
| TcCLB.510687.110 | hypothetical protein, conserved | 0,02966 | 0,8583 |
| TcCLB.504179.100 | hypothetical protein, conserved | 0,02963 | 0,898 |
| TcCLB.511249.44 | vacuolar transporter chaperone, putative | 0,02962 | 0,8149 |
| TcCLB.507519.30 | Eukaryotic translation initiation factor 4E type 6, putative ( | 0,0296 | 0,8544 |
| TcCLB.507941.14 | hypothetical protein, conserved | 0,02948 | 0,9362 |
| TcCLB.509157.240 | hypothetical protein, conserved | 0,02927 | 0,9103 |
| TcCLB.505957.140 | trans-sialidase (pseudogene), putative | 0,02914 | 0,9313 |

| | | | |
|---|---|---|---|
| TcCLB.503899.20 | 40S ribosomal protein S16, putative | 0,02894 | 0,7736 |
| TcCLB.506869.34 | hypothetical protein, conserved | 0,02883 | 0,8786 |
| TcCLB.508865.4 | protein phosphatase 2C, putative | 0,02875 | 0,8643 |
| TcCLB.506855.240 | bilobe protein, conserved (pseudogene) | 0,02868 | 0,7841 |
| TcCLB.509693.110 | hypothetical protein, conserved | 0,02859 | 0,9033 |
| TcCLB.510943.70 | hypothetical protein, conserved | 0,02852 | 0,8859 |
| TcCLB.511155.4 | hypothetical protein, conserved | 0,02839 | 0,8934 |
| TcCLB.504397.9 | surface protease GP63, putative (fragment) | 0,02805 | 0,9429 |
| TcCLB.507957.260 | surface protease GP63 (pseudogene), putative | 0,02787 | 0,9645 |
| TcCLB.509599.120 | hypothetical protein, conserved | 0,02768 | 0,8937 |
| TcCLB.507793.40 | hypothetical protein, conserved | 0,02765 | 0,8849 |
| TcCLB.508173.200 | hypothetical protein, conserved | 0,02758 | 0,7735 |
| TcCLB.506247.30 | RNA editing complex protein MP63, putative | 0,02739 | 0,7869 |
| TcCLB.509429.310 | hypothetical protein, conserved | 0,02739 | 0,8507 |
| TcCLB.506297.70 | hypothetical protein, conserved | 0,02735 | 0,8425 |
| TcCLB.509081.110 | mucin-associated surface protein (MASP), putative | 0,02733 | 0,9513 |
| TcCLB.503391.10 | ATP-dependent DEAD/H DNA helicase recQ, putative | 0,02725 | 0,9565 |
| TcCLB.510431.110 | hypothetical protein | 0,02724 | 0,8109 |
| TcCLB.506873.20 | hypothetical protein, conserved | 0,02715 | 0,8952 |
| TcCLB.506289.210 | surface protease GP63, putative | 0,02697 | 0,9471 |
| TcCLB.408799.19 | isopentenyl-diphosphate delta-isomerase (fragment) (idi1 | 0,02696 | 0,9087 |
| TcCLB.511261.100 | chaperone DNAJ protein, putative | 0,02694 | 0,8709 |
| TcCLB.509453.50 | serine/threonine-protein phosphatase | 0,02689 | 0,8879 |
| TcCLB.511049.30 | hypothetical protein, conserved | 0,02687 | 0,9122 |
| TcCLB.508475.30 | surface protease GP63, putative | 0,02671 | 0,9163 |
| TcCLB.509643.40 | hypothetical protein, conserved | 0,02661 | 0,8963 |
| TcCLB.511217.39 | hypothetical protein, conserved | 0,02641 | 0,9146 |
| TcCLB.508707.190 | hypothetical protein, conserved | 0,02639 | 0,8248 |
| TcCLB.503501.50 | trans-sialidase (pseudogene), putative | 0,02637 | 0,9499 |
| TcCLB.508133.10 | hypothetical protein | 0,02626 | 0,8939 |
| TcCLB.509527.90 | trans-sialidase (pseudogene), putative | 0,02622 | 0,8685 |
| TcCLB.508951.50 | calmodulin, putative | 0,02605 | 0,928 |
| TcCLB.506529.310 | hypothetical protein, conserved | 0,02601 | 0,8752 |
| TcCLB.507237.170 | mucin-associated surface protein (MASP), putative | 0,02597 | 0,9853 |
| TcCLB.503973.140 | hypothetical protein, conserved (pseudogene) | 0,02581 | 0,9614 |
| TcCLB.508153.470 | hypothetical protein, conserved | 0,02569 | 0,882 |
| TcCLB.510785.15 | hypothetical protein, conserved | 0,02566 | 0,9398 |
| TcCLB.508895.89 | intraflagellar transport protein 80, putative (fragment) (IF | 0,02565 | 0,8869 |
| TcCLB.511747.7 | cyclosome subunit, putative (fragment) | 0,02561 | 0,8835 |
| TcCLB.504741.130 | hypothetical protein, conserved | 0,0254 | 0,8194 |
| TcCLB.503521.60 | hypothetical protein, conserved | 0,02528 | 0,9291 |
| TcCLB.503973.10 | hypothetical protein | 0,02525 | 0,9423 |
| TcCLB.508173.180 | nuclear transport factor (MTR2) | 0,02518 | 0,8389 |
| TcCLB.510085.54 | mucin TcMUCII (pseudogene), putative | 0,0251 | 0,9612 |
| TcCLB.503935.10 | dolichyl-lipid chaperone, putative (rft1) | 0,02509 | 0,896 |

| | | | |
|---|---|---|---|
| TcCLB.506297.130 | zinc finger ccch and cchc domain-containing protein | 0,02502 | 0,8457 |
| TcCLB.509657.61 | elongation factor 1-gamma (EF-1-gamma, pseudogene), p | 0,02493 | 0,9617 |
| TcCLB.504027.20 | hypothetical protein | 0,02487 | 0,875 |
| TcCLB.458241.10 | hypothetical protein, conserved | 0,02487 | 0,9329 |
| TcCLB.511179.120 | hypothetical protein, conserved | 0,02484 | 0,9016 |
| TcCLB.509335.20 | retrotransposon hot spot protein (RHS, pseudogene), putat | 0,02462 | 0,9225 |
| TcCLB.504241.80 | trans-sialidase (pseudogene), putative | 0,02449 | 0,8784 |
| TcCLB.507093.80 | hypothetical protein, conserved | 0,02449 | 0,8819 |
| TcCLB.510181.90 | hypothetical protein, conserved | 0,02449 | 0,8924 |
| TcCLB.508445.70 | hypothetical protein, conserved | 0,02448 | 0,8808 |
| TcCLB.510691.4 | hypothetical protein, conserved | 0,02444 | 0,9489 |
| TcCLB.508153.300 | hypothetical protein, conserved | 0,0242 | 0,8161 |
| TcCLB.510519.80 | hypothetical protein, conserved | 0,02419 | 0,906 |
| TcCLB.508467.19 | hypothetical protein, conserved (fragment) | 0,0241 | 0,943 |
| TcCLB.503679.20 | hypothetical protein, conserved | 0,02403 | 0,8872 |
| TcCLB.507857.4 | transcription elongation factor, putative | 0,02399 | 0,9005 |
| TcCLB.506685.50 | myosin heavy chain, putative (fragment) | 0,02379 | 0,9191 |
| TcCLB.511365.94 | anaphase promoting complex, subunit 10-like protein | 0,02377 | 0,9137 |
| TcCLB.504171.40 | hypothetical protein, conserved | 0,02367 | 0,9103 |
| TcCLB.510229.10 | hypothetical protein, conserved | 0,02361 | 0,8597 |
| TcCLB.511517.37 | reticulon domain protein, putative | 0,0236 | 0,8817 |
| TcCLB.506933.30 | hypothetical protein, conserved | 0,02354 | 0,8628 |
| TcCLB.510621.20 | mucin-associated surface protein (MASP), putative | 0,02352 | 0,9636 |
| TcCLB.510803.15 | mucin-associated surface protein (MASP, pseudogene), pu | 0,02349 | 0,9818 |
| TcCLB.511261.160 | pyruvate dehydrogenase E1 component alpha subunit, puta | 0,02344 | 0,8754 |
| TcCLB.506801.129 | hypothetical protein (fragment) | 0,02343 | 0,8775 |
| TcCLB.506245.230 | mucin-associated surface protein (MASP, pseudogene), pu | 0,02334 | 0,9714 |
| TcCLB.511541.9 | hypothetical protein, conserved (fragment) | 0,02326 | 0,885 |
| TcCLB.507023.270 | hypothetical protein, conserved | 0,0232 | 0,9126 |
| TcCLB.503829.80 | hypothetical potein, conserved | 0,02306 | 0,8412 |
| TcCLB.510647.60 | intraflagellar transport protein 27 (IFT27) | 0,023 | 0,8974 |
| TcCLB.511303.140 | hypothetical protein, conserved | 0,02296 | 0,8854 |
| TcCLB.507803.40 | hypothetical protein | 0,02274 | 0,8578 |
| TcCLB.503587.30 | trans-sialidase, Group V, putative | 0,02273 | 0,9652 |
| TcCLB.509791.170 | small GTPase, putative | 0,02263 | 0,8951 |
| TcCLB.503721.20 | hypothetical protein, conserved | 0,02262 | 0,9051 |
| TcCLB.506829.5 | hypothetical protein, conserved | 0,02257 | 0,9335 |
| TcCLB.506945.230 | 40S ribosomal protein S14, putative | 0,02251 | 0,842 |
| TcCLB.507747.80 | mucin TcMUCII, putative | 0,0225 | 0,9683 |
| TcCLB.504397.20 | surface protease GP63, putative | 0,02233 | 0,951 |
| TcCLB.509669.90 | hypothetical protein, conserved | 0,02216 | 0,8885 |
| TcCLB.508873.90 | trans-sialidase (pseudogene), putative | 0,02213 | 0,9614 |
| TcCLB.507963.60 | hypothetical protein, conserved | 0,02198 | 0,9416 |
| TcCLB.509055.40 | hypothetical protein, conserved | 0,02192 | 0,8872 |
| TcCLB.511261.90 | hypothetical protein, conserved | 0,02183 | 0,9015 |

| | | | |
|---|---|---|---|
| TcCLB.506435.30 | mitochondrial DNA polymerase I protein A, putative | 0,02167 | 0,8728 |
| TcCLB.506579.129 | hypothetical protein, conserved (fragment) | 0,02152 | 0,9275 |
| TcCLB.511751.260 | hypothetical protein, conserved | 0,02146 | 0,9383 |
| TcCLB.506637.110 | hypothetical protein, conserved | 0,02126 | 0,8448 |
| TcCLB.510743.130 | denn domain-containing protein (pseudogene) | 0,02109 | 0,9438 |
| TcCLB.508029.59 | kinesin, putative | 0,0208 | 0,9471 |
| TcCLB.510769.90 | 60S ribosomal protein L32, putative | 0,02074 | 0,8752 |
| TcCLB.508689.10 | hypothetical protein, conserved | 0,02052 | 0,9034 |
| TcCLB.507843.20 | trans-sialidase, putative (pseudogene) | 0,02036 | 0,9724 |
| TcCLB.504039.80 | surface protease GP63 (pseudogene), putative | 0,02035 | 0,9753 |
| TcCLB.508577.130 | hypothetical protein, conserved | 0,02022 | 0,8993 |
| TcCLB.511071.30 | hypothetical protein, conserved | 0,02019 | 0,93 |
| TcCLB.511277.420 | hypothetical protein, conserved | 0,0201 | 0,8663 |
| TcCLB.508929.20 | UDP-Gal or UDP-GlcNAc-dependent glycosyltransferase, pu | 0,01998 | 0,9495 |
| TcCLB.503651.40 | hypothetical protein | 0,01991 | 0,9097 |
| TcCLB.507063.280 | hypothetical protein | 0,01983 | 0,8717 |
| TcCLB.511643.61 | elongation factor 1-gamma (EF-1-gamma, pseudogene), p | 0,01981 | 0,9293 |
| TcCLB.509399.60 | hypothetical protein, conserved | 0,01979 | 0,8704 |
| TcCLB.505035.5 | hypothetical protein, conserved | 0,01979 | 0,9499 |
| TcCLB.509615.2 | endosomal integral membrane protein (pseudogene), puta | 0,01977 | 0,9233 |
| TcCLB.511389.20 | hypothetical protein, conserved | 0,01976 | 0,9033 |
| TcCLB.508487.11 | retrotransposon hot spot protein (RHS, pseudogene), putat | 0,01975 | 0,9351 |
| TcCLB.508831.60 | Low-affinity phosphate transporter, putative (PHO91) | 0,01973 | 0,8614 |
| TcCLB.509915.55 | trans-sialidase (pseudogene), putative | 0,01971 | 0,9627 |
| TcCLB.507001.60 | hypothetical protein, conserved | 0,01968 | 0,8822 |
| TcCLB.504157.20 | beta-ketoacyl synthase family protein, putative | 0,0196 | 0,9229 |
| TcCLB.506617.70 | variant-surface-glycoprotein phospholipase C (GPI-PLC) | 0,01958 | 0,9301 |
| TcCLB.504073.20 | hypothetical protein, conserved | 0,01947 | 0,9065 |
| TcCLB.509157.250 | hypothetical protein, conserved | 0,01937 | 0,8966 |
| TcCLB.511523.40 | hypothetical protein, conserved | 0,01931 | 0,9255 |
| TcCLB.508307.184 | hypothetical protein, conserved | 0,01921 | 0,9386 |
| TcCLB.508647.20 | hypothetical protein | 0,01916 | 0,8763 |
| TcCLB.511511.159 | zinc finger protein, putative | 0,01916 | 0,886 |
| TcCLB.503987.20 | hypothetical protein, conserved | 0,01916 | 0,9197 |
| TcCLB.506203.80 | formin-like protein | 0,01892 | 0,9068 |
| TcCLB.508489.30 | hypothetical protein, conserved | 0,01885 | 0,9171 |
| TcCLB.511127.200 | kinesin, putative | 0,01874 | 0,9075 |
| TcCLB.507265.15 | transcription activator, putative (fragment) | 0,01871 | 0,8642 |
| TcCLB.507277.20 | hypothetical protein, conserved | 0,01863 | 0,9513 |
| TcCLB.505987.41 | hypothetical protein, conserved | 0,01846 | 0,9402 |
| TcCLB.509935.20 | hypothetical protein, conserved | 0,01841 | 0,8869 |
| TcCLB.508153.364 | hypothetical protein, conserved | 0,01838 | 0,9346 |
| TcCLB.509005.40 | hypothetical protein, conserved | 0,01837 | 0,9059 |
| TcCLB.511291.30 | lipoic acid synthetase, mitochondrial precursor, putative | 0,0183 | 0,9217 |
| TcCLB.503813.30 | hypothetical protein, conserved | 0,0183 | 0,925 |

| | | | |
|---|---|---|---|
| TcCLB.511461.50 | nuclear transmembrane protein, putative (NP) | 0,01821 | 0,9165 |
| TcCLB.507949.20 | RWD domain-containing protein | 0,01817 | 0,9326 |
| TcCLB.504125.20 | hypothetical protein | 0,01811 | 0,9228 |
| TcCLB.503407.10 | hypothetical protein, conserved | 0,0181 | 0,9343 |
| TcCLB.506829.60 | hypothetical protein, conserved | 0,01805 | 0,929 |
| TcCLB.511727.40 | ubiquitin-conjugating enzyme E2, putative | 0,01789 | 0,885 |
| TcCLB.510165.40 | hypothetical protein, conserved | 0,01777 | 0,9027 |
| TcCLB.508799.140 | transcription modulator/accessory protein, putative | 0,01775 | 0,8858 |
| TcCLB.507395.10 | dihydroceramide synthase, putative (DHCS) | 0,01775 | 0,9103 |
| TcCLB.510531.70 | hypothetical protein, conserved | 0,01739 | 0,9471 |
| TcCLB.511555.10 | mucin-associated surface protein (MASP), putative | 0,01739 | 0,9565 |
| TcCLB.508753.40 | mucin-associated surface protein (MASP, pseudogene), pu | 0,01738 | 0,9536 |
| TcCLB.509607.70 | serine/threonine protein kinase, putative | 0,01718 | 0,9452 |
| TcCLB.504533.20 | retrotransposon hot spot protein (RHS, pseudogene), putat | 0,01715 | 0,9347 |
| TcCLB.506567.10 | hypothetical protein, conserved | 0,01707 | 0,9298 |
| TcCLB.511487.150 | serine-alanine-and proline-rich protein, putative | 0,01704 | 0,977 |
| TcCLB.504949.39 | DNA repair protein Rad9, putative (fragment) (Rad9) | 0,01702 | 0,9293 |
| TcCLB.507257.160 | hypothetical protein, conserved | 0,01683 | 0,9381 |
| TcCLB.511541.30 | hypothetical protein, conserved | 0,01683 | 0,9417 |
| TcCLB.506959.90 | hypothetical protein, conserved | 0,01673 | 0,8774 |
| TcCLB.508873.530 | hypothetical protein, conserved (pseudogene) | 0,01671 | 0,9579 |
| TcCLB.506789.220 | hypothetical protein, conserved | 0,01658 | 0,9137 |
| TcCLB.504021.109 | 40S ribosomal protein S3, putative | 0,01649 | 0,8608 |
| TcCLB.507671.39 | serine/threonine-protein phosphatase PP1 | 0,01649 | 0,9544 |
| TcCLB.510391.20 | mucin-associated surface protein (MASP), putative | 0,01637 | 0,9686 |
| TcCLB.509753.50 | retrotransposon hot spot protein (RHS, pseudogene), putat | 0,01616 | 0,9752 |
| TcCLB.510329.300 | hypothetical protein, conserved | 0,01608 | 0,9463 |
| TcCLB.510835.60 | hypothetical protein, conserved | 0,01606 | 0,9007 |
| TcCLB.503635.68 | 40S ribosomal protein S3A, putative | 0,01599 | 0,8827 |
| TcCLB.511585.220 | calcium/potassium channel (CAKC), putative | 0,01597 | 0,9008 |
| TcCLB.503931.30 | hypothetical protein | 0,01596 | 0,9299 |
| TcCLB.506327.70 | engulfment and cell motility domain 2, putative | 0,01588 | 0,9201 |
| TcCLB.504081.260 | surface protease GP63, putative | 0,01586 | 0,9811 |
| TcCLB.504109.24 | dynein light chain, putative | 0,01584 | 0,945 |
| TcCLB.506297.80 | hypothetical protein, conserved | 0,01577 | 0,9065 |
| TcCLB.503955.110 | protein kinase, putative | 0,01575 | 0,9314 |
| TcCLB.504041.10 | hypothetical protein, conserved | 0,01557 | 0,9266 |
| TcCLB.511745.40 | hypothetical protein, conserved | 0,01556 | 0,9293 |
| TcCLB.510757.30 | hypothetical protein, conserved | 0,01551 | 0,9505 |
| TcCLB.510421.220 | hypothetical protein, conserved | 0,01545 | 0,917 |
| TcCLB.511217.90 | hypothetical protein, conserved | 0,01543 | 0,8897 |
| TcCLB.511507.80 | ferric reductase transmembrane protein, putative | 0,01543 | 0,9665 |
| TcCLB.507063.144 | hypothetical protein, conserved | 0,01533 | 0,943 |
| TcCLB.510645.40 | hypothetical protein, conserved (pseudogene) | 0,01527 | 0,8644 |
| TcCLB.506401.290 | hypothetical protein, conserved | 0,0152 | 0,9597 |

| | | | |
|---|---|---|---|
| TcCLB.511165.20 | hypothetical protein | 0,01503 | 0,9273 |
| TcCLB.507911.30 | dispersed gene family protein 1 (DGF-1), putative | 0,01484 | 0,9563 |
| TcCLB.506529.546 | hypothetical protein | 0,01469 | 0,9788 |
| TcCLB.506151.30 | small GTP-binding protein Rab11, putative (TcRab11) | 0,01455 | 0,9246 |
| TcCLB.507033.20 | histone H3 variant, putative | 0,01452 | 0,9313 |
| TcCLB.511753.90 | hypothetical protein, conserved | 0,01452 | 0,9411 |
| TcCLB.507515.110 | hypothetical protein, conserved | 0,01444 | 0,9332 |
| TcCLB.506933.70 | hypothetical protein, conserved | 0,01443 | 0,9248 |
| TcCLB.508647.40 | replication factor C large subunit, putative | 0,01441 | 0,8906 |
| TcCLB.509719.80 | hypothetical protein, conserved | 0,01438 | 0,9445 |
| TcCLB.506831.66 | hypothetical protein, conserved | 0,0143 | 0,9597 |
| TcCLB.510421.190 | hypothetical protein, conserved | 0,01424 | 0,9389 |
| TcCLB.507715.40 | GTP-binding elongation factor Tu family, putative | 0,014 | 0,9064 |
| TcCLB.508505.10 | hypothetical protein, conserved | 0,014 | 0,9329 |
| TcCLB.503675.20 | Charged multivesicular body protein 3, putative | 0,01399 | 0,9479 |
| TcCLB.507071.100 | mucin-associated surface protein (MASP), putative | 0,01387 | 0,9777 |
| TcCLB.511667.120 | dispersed gene family protein 1 (DGF-1, pseudogene), puta | 0,01381 | 0,9567 |
| TcCLB.511421.210 | hypothetical protein, conserved | 0,0138 | 0,9154 |
| TcCLB.511229.100 | hypothetical protein, conserved | 0,01376 | 0,924 |
| TcCLB.510187.340 | hypothetical protein, conserved | 0,01352 | 0,9306 |
| TcCLB.510325.20 | hypothetical protein | 0,01351 | 0,9688 |
| TcCLB.507001.100 | transporter, putative | 0,01345 | 0,9535 |
| TcCLB.506295.150 | hypothetical protein | 0,01344 | 0,9525 |
| TcCLB.510251.19 | hypothetical protein, conserved | 0,01325 | 0,9413 |
| TcCLB.504881.20 | hypothetical protein, conserved | 0,01318 | 0,9051 |
| TcCLB.503593.60 | hypothetical protein, conserved | 0,01317 | 0,9452 |
| TcCLB.507677.4 | hypothetical protein, conserved | 0,01314 | 0,9612 |
| TcCLB.511071.70 | ubiquitin-conjugating enzyme E2, putative | 0,01309 | 0,9409 |
| TcCLB.507159.90 | DNA-directed RNA polymerase III subunit, putative | 0,01284 | 0,9413 |
| TcCLB.506637.40 | hypothetical protein, conserved | 0,01261 | 0,9568 |
| TcCLB.503929.10 | V-type ATPase, A subunit, putative | 0,01253 | 0,9125 |
| TcCLB.510949.20 | thioredoxin, putative | 0,0125 | 0,9509 |
| TcCLB.511065.50 | phosphatidylinositol 3-kinase, putative | 0,01243 | 0,9581 |
| TcCLB.511245.210 | hypothetical protein, conserved | 0,01241 | 0,9379 |
| TcCLB.510197.64 | mucin-associated surface protein (MASP, pseudogene), pu | 0,0123 | 0,9724 |
| TcCLB.507025.10 | hypothetical protein, conserved | 0,01227 | 0,9296 |
| TcCLB.508723.30 | bilobe protein, conserved | 0,01223 | 0,9426 |
| TcCLB.510243.119 | hypothetical protein, conserved | 0,01217 | 0,9529 |
| TcCLB.511837.10 | vesicle-associated membrane protein, putative | 0,01214 | 0,9458 |
| TcCLB.506943.160 | eukaryotic translation initiation factor 3 subunit 7-like prot | 0,01205 | 0,9032 |
| TcCLB.509779.20 | hypothetical protein, conserved | 0,0119 | 0,9627 |
| TcCLB.508465.80 | hypothetical protein, conserved | 0,01184 | 0,9607 |
| TcCLB.506211.50 | membrane protein, putative | 0,01181 | 0,9553 |
| TcCLB.511367.110 | hypothetical protein, conserved | 0,01167 | 0,9562 |
| TcCLB.506135.110 | mitochondrial RNA binding complex 1 subunit, putative | 0,01147 | 0,9524 |

| | | | |
|---|---|---|---|
| TcCLB.509683.117 | 40S ribosomal protein S4, putative | 0,01143 | 0,888 |
| TcCLB.508323.199 | ATP-dependent DEAD/H RNA helicase, putative (fragment) | 0,01139 | 0,9614 |
| TcCLB.504081.9 | trans-sialidase (pseudogene), putative | 0,01135 | 0,9856 |
| TcCLB.419469.14 | kinesin, putative | 0,01127 | 0,9784 |
| TcCLB.511281.60 | pyruvate kinase 2, putative | 0,01126 | 0,943 |
| TcCLB.510259.30 | hypothetical protein, conserved | 0,01121 | 0,9633 |
| TcCLB.511529.259 | hypothetical protein, conserved | 0,01116 | 0,977 |
| TcCLB.511367.138 | chaperone DNAJ protein, putative | 0,01115 | 0,9417 |
| TcCLB.509205.40 | hypothetical protein, conserved | 0,01113 | 0,9526 |
| TcCLB.511017.60 | hypothetical protein | 0,01112 | 0,9295 |
| TcCLB.506745.20 | hypothetical protein, conserved | 0,01105 | 0,9533 |
| TcCLB.511719.30 | hypothetical protein, conserved | 0,01103 | 0,9614 |
| TcCLB.506443.50 | hypothetical protein, conserved | 0,01101 | 0,9453 |
| TcCLB.506341.20 | hypothetical protein, conserved | 0,01082 | 0,9614 |
| TcCLB.508153.120 | ubiquitin hydrolase, putative | 0,01081 | 0,9389 |
| TcCLB.504113.10 | protein kinase, putative | 0,01081 | 0,9614 |
| TcCLB.507949.80 | kinetoplast ribosomal PPR-repeat containing protein 3, put | 0,01077 | 0,949 |
| TcCLB.511083.35 | mucin TcMUCII (pseudogene), putative | 0,0106 | 0,9833 |
| TcCLB.511419.10 | hypothetical protein | 0,01039 | 0,967 |
| TcCLB.511809.60 | hypothetical protein, conserved | 0,01035 | 0,9065 |
| TcCLB.490165.19 | hypothetical protein, conserved | 0,01035 | 0,9748 |
| TcCLB.503431.70 | hypothetical protein, conserved | 0,01033 | 0,9533 |
| TcCLB.506789.130 | double-strand-break repair protein rad21 homolog, putativ | 0,01029 | 0,9231 |
| TcCLB.506819.49 | hypothetical protein, conserved | 0,01021 | 0,9667 |
| TcCLB.443397.9 | NADH-cytochrome b5 reductase, putative (fragment) | 0,01011 | 0,9617 |
| TcCLB.506435.440 | hypothetical protein, conserved | 0,01001 | 0,9489 |
| TcCLB.508153.480 | hypothetical protein, conserved | 0,00996 | 0,9299 |
| TcCLB.503635.59 | 40S ribosomal protein S3A, putative | 0,009873 | 0,9287 |
| TcCLB.504029.21 | lipase, putative (fragment) | 0,009796 | 0,9453 |
| TcCLB.508461.520 | hypothetical protein, conserved | 0,009633 | 0,9255 |
| TcCLB.503815.4 | hypothetical protein, conserved | 0,009265 | 0,9772 |
| TcCLB.507017.4 | hypothetical protein, conserved | 0,009142 | 0,9802 |
| TcCLB.504255.4 | rhomboid-like protein | 0,009106 | 0,9809 |
| TcCLB.508399.40 | trans-sialidase (pseudogene), putative | 0,0091 | 0,9642 |
| TcCLB.509829.9 | retrotransposon hot spot (RHS) protein, putative (fragmen | 0,009001 | 0,9767 |
| TcCLB.510885.90 | hypothetical protein, conserved | 0,009 | 0,9744 |
| TcCLB.506801.90 | hypothetical protein, conserved | 0,008931 | 0,9627 |
| TcCLB.507765.130 | hypothetical protein, conserved | 0,00892 | 0,9257 |
| TcCLB.511927.10 | hypothetical protein, conserved | 0,008859 | 0,9771 |
| TcCLB.510289.60 | hypothetical protein | 0,00883 | 0,9736 |
| TcCLB.508151.20 | mucin-associated surface protein (MASP), putative | 0,008737 | 0,9855 |
| TcCLB.503917.50 | hypothetical protein, conserved | 0,008628 | 0,9679 |
| TcCLB.507819.40 | retrotransposon hot spot protein (RHS, pseudogene), putat | 0,008621 | 0,975 |
| TcCLB.492827.10 | trans-sialidase, putative (fragment) | 0,00862 | 0,9951 |
| TcCLB.511671.140 | hypothetical protein | 0,008609 | 0,9621 |

| | | | |
|---|---|---|---|
| TcCLB.510257.20 | hypothetical protein, conserved | 0,008607 | 0,9591 |
| TcCLB.503659.30 | trans-sialidase, putative | 0,008606 | 0,9836 |
| TcCLB.509191.9 | dispersed gene family protein 1 (DGF-1, pseudogene), puta | 0,008491 | 0,9841 |
| TcCLB.508827.130 | hypothetical protein, conserved | 0,008345 | 0,9571 |
| TcCLB.412435.10 | hypothetical protein, conserved | 0,008319 | 0,9642 |
| TcCLB.507049.130 | hypothetical protein, conserved (pseudogene) | 0,008227 | 0,9789 |
| TcCLB.399033.10 | hypothetical protein | 0,008222 | 0,9759 |
| TcCLB.508351.50 | rho-like GTP binding protein, putative | 0,008123 | 0,963 |
| TcCLB.508613.60 | hypothetical protein, conserved | 0,008078 | 0,9544 |
| TcCLB.508257.130 | hypothetical protein, conserved | 0,007873 | 0,9699 |
| TcCLB.511183.400 | trans-sialidase, Group VI, putative | 0,007697 | 0,9833 |
| TcCLB.507959.41 | mucin TcMUCII, putative | 0,007679 | 0,9924 |
| TcCLB.508933.10 | dihydrolipoamide dehydrogenase, putative | 0,007648 | 0,9652 |
| TcCLB.509897.95 | hypothetical protein, conserved (pseudogene) | 0,007613 | 0,9704 |
| TcCLB.511585.70 | heat shock protein 70 (hsp70), putative | 0,007543 | 0,9568 |
| TcCLB.506587.100 | surface protease GP63, putative | 0,007464 | 0,9652 |
| TcCLB.509471.20 | Present in the outer mitochondrial membrane proteome 3 | 0,007324 | 0,9513 |
| TcCLB.509741.30 | hypothetical protein, conserved | 0,007205 | 0,9698 |
| TcCLB.508221.697 | trans-sialidase (pseudogene), putative | 0,007187 | 0,9845 |
| TcCLB.506559.480 | hypothetical protein, conserved | 0,007123 | 0,9756 |
| TcCLB.509767.200 | hypothetical protein, conserved | 0,007116 | 0,97 |
| TcCLB.442287.10 | serine/threonine protein phosphatase (pseudogene), putat | 0,00711 | 0,9622 |
| TcCLB.511145.60 | ATP synthase F1 subunit gamma protein, putative (ATPF1G | 0,007008 | 0,9622 |
| TcCLB.510667.50 | hypothetical protein, conserved | 0,006992 | 0,9719 |
| TcCLB.510167.10 | phosphatidylinositol 3-kinase 2, putative | 0,006698 | 0,9614 |
| TcCLB.504197.20 | mucin-associated surface protein (MASP), putative | 0,006625 | 0,9871 |
| TcCLB.511729.20 | hypothetical protein | 0,006477 | 0,9663 |
| TcCLB.509999.50 | lipase domain protein, putative | 0,006474 | 0,9595 |
| TcCLB.511821.140 | hypothetical protein, conserved | 0,006467 | 0,9683 |
| TcCLB.507991.10 | hypothetical protein, conserved | 0,006446 | 0,9779 |
| TcCLB.506287.180 | hypothetical protein, conserved | 0,006432 | 0,9724 |
| TcCLB.511873.5 | hypothetical protein | 0,006281 | 0,9818 |
| TcCLB.508161.40 | mucin TcMUCII (pseudogene), putative | 0,006248 | 0,9913 |
| TcCLB.511827.130 | dispersed gene family protein 1 (DGF-1), putative | 0,006245 | 0,9674 |
| TcCLB.507957.36 | syntaxin binding protein (pseudogene), putative | 0,006135 | 0,9909 |
| TcCLB.503987.10 | cop-coated vesicle membrane protein p24 precursor, putat | 0,006098 | 0,9704 |
| TcCLB.506529.230 | hypothetical protein, conserved | 0,006087 | 0,9767 |
| TcCLB.506649.30 | hypothetical protein, conserved | 0,005969 | 0,9636 |
| TcCLB.507053.130 | hypothetical protein, conserved | 0,00578 | 0,9724 |
| TcCLB.432677.20 | heat shock 70 kDa protein, mitochondrial precursor, putativ | 0,005756 | 0,9569 |
| TcCLB.507991.90 | hypothetical protein | 0,005571 | 0,9732 |
| TcCLB.506667.140 | hypothetical protein | 0,00557 | 0,9929 |
| TcCLB.509803.60 | retrotransposon hot spot protein (RHS, pseudogene), putat | 0,005525 | 0,9895 |
| TcCLB.506747.20 | dihydroorotase, putative | 0,005499 | 0,9821 |
| TcCLB.507105.40 | 60S ribosomal protein L44 (RPL44) | 0,005408 | 0,9756 |

| | | | |
|---|---|---|---|
| TcCLB.506835.110 | protein mkt1, putative | 0,005352 | 0,9627 |
| TcCLB.509965.60 | RNA decapping enzyme, putative (Dcp2) | 0,005065 | 0,983 |
| TcCLB.506791.30 | hypothetical protein, conserved | 0,00503 | 0,9672 |
| TcCLB.510329.180 | hypothetical protein, conserved | 0,004983 | 0,9929 |
| TcCLB.510073.75 | hypothetical protein, conserved | 0,004957 | 0,9908 |
| TcCLB.507209.20 | hypothetical protein, conserved | 0,004855 | 0,9801 |
| TcCLB.507773.20 | hypothetical protein | 0,004846 | 0,9838 |
| TcCLB.511857.70 | hypothetical protein, conserved | 0,004823 | 0,9752 |
| TcCLB.506811.90 | hypothetical protein, conserved | 0,004719 | 0,9761 |
| TcCLB.506859.249 | hypothetical protein, conserved | 0,00441 | 0,9851 |
| TcCLB.508347.120 | hypothetical protein, conserved | 0,004374 | 0,9756 |
| TcCLB.504163.30 | 40S ribosomal protein S9, putative | 0,004272 | 0,9667 |
| TcCLB.507711.10 | intraflagellar transport protein 57/55, putative (IFT57/55) | 0,004112 | 0,9789 |
| TcCLB.509569.150 | hypothetical protein, conserved | 0,004025 | 0,9898 |
| TcCLB.508819.30 | hypothetical protein, conserved | 0,003998 | 0,9826 |
| TcCLB.510483.154 | mucin-associated surface protein (MASP), putative | 0,003852 | 0,9954 |
| TcCLB.509373.39 | surface protease GP63, putative (fragment) | 0,003841 | 0,9965 |
| TcCLB.507793.20 | cysteine synthase, putative | 0,003821 | 0,981 |
| TcCLB.508541.120 | trans-sialidase (pseudogene), putative | 0,003769 | 0,9959 |
| TcCLB.511753.100 | ABC transporter, putative | 0,003706 | 0,9832 |
| TcCLB.511189.84 | hypothetical protein, conserved | 0,003683 | 0,9883 |
| TcCLB.506883.20 | hypothetical protein, conserved | 0,003625 | 0,9824 |
| TcCLB.511537.8 | ABC transporter of the mitochondrion, putative (fragment) | 0,003597 | 0,9838 |
| TcCLB.508015.20 | hypothetical protein | 0,003585 | 0,9908 |
| TcCLB.510575.230 | hypothetical protein, conserved | 0,003539 | 0,9837 |
| TcCLB.506605.170 | inosine-guanine nucleoside hydrolase, putative (IG-NH) | 0,003463 | 0,9844 |
| TcCLB.507959.224 | surface protease GP63 (pseudogene), putative | 0,003438 | 0,9954 |
| TcCLB.508445.80 | hypothetical protein, conserved | 0,003352 | 0,9833 |
| TcCLB.511753.30 | kinesin, putative | 0,003286 | 0,9809 |
| TcCLB.504341.10 | trans-sialidase, Group IV, putative | 0,003196 | 0,9908 |
| TcCLB.509959.10 | retrotransposon hot spot protein (RHS, pseudogene), putat | 0,003173 | 0,9931 |
| TcCLB.507949.130 | glycerol-3-phosphate dehydrogenase (FAD-dependent), pu | 0,003111 | 0,9892 |
| TcCLB.509979.330 | surface protease GP63 (pseudogene), putative | 0,002945 | 0,9921 |
| TcCLB.504911.20 | mucin-associated surface protein (MASP, pseudogene), pu | 0,002814 | 0,9983 |
| TcCLB.507951.154 | cytochrome b5, putative | 0,002761 | 0,9892 |
| TcCLB.506289.140 | surface protease GP63, putative | 0,002757 | 0,9926 |
| TcCLB.508943.20 | protein kinase, putative | 0,002692 | 0,9905 |
| TcCLB.506597.40 | trans-sialidase, Group V, putative | 0,002626 | 0,9974 |
| TcCLB.508979.60 | mucin-associated surface protein (MASP), putative | 0,00253 | 0,9937 |
| TcCLB.503879.30 | hypothetical protein, conserved | 0,002316 | 0,9886 |
| TcCLB.508153.610 | Putative intraflagellar transport protein C3 (PIFTC3) | 0,002308 | 0,988 |
| TcCLB.511381.60 | hypothetical protein, conserved | 0,002207 | 0,9922 |
| TcCLB.503651.30 | hypothetical protein | 0,002206 | 0,9888 |
| TcCLB.510693.274 | mucin TcMUCII (pseudogene), putative | 0,002196 | 0,9954 |
| TcCLB.506215.19 | intraflagellar transport protein 140, putative (IFT140) | 0,002193 | 0,9831 |

| | | | |
|---|---|---|---|
| TcCLB.509561.60 | hypothetical protein, conserved | 0,00215 | 0,9927 |
| TcCLB.504249.15 | surface protease GP63 (pseudogene), putative | 0,002133 | 0,9961 |
| TcCLB.508629.120 | mucin-associated surface protein (MASP), putative | 0,002108 | 0,9908 |
| TcCLB.507771.50 | glycerolphosphate mutase, putative | 0,001903 | 0,9909 |
| TcCLB.507041.10 | hypothetical protein, conserved | 0,001851 | 0,9898 |
| TcCLB.510665.20 | hypothetical protein, conserved | 0,001799 | 0,9915 |
| TcCLB.508381.20 | hypothetical protein, conserved (pseudogene) | 0,001708 | 0,996 |
| TcCLB.510483.340 | beta galactofuranosyl glycosyltransferase (pseudogene), p | 0,001682 | 0,9965 |
| TcCLB.508177.50 | hypothetical protein | 0,001674 | 0,9941 |
| TcCLB.509879.30 | hypothetical protein, conserved | 0,001585 | 0,993 |
| TcCLB.507773.50 | hypothetical protein, conserved | 0,001556 | 0,992 |
| TcCLB.507993.130 | 60S ribosomal protein L7, putative | 0,001535 | 0,991 |
| TcCLB.503677.29 | hypothetical protein, conserved | 0,001467 | 0,994 |
| TcCLB.507513.20 | hypothetical protein, conserved | 0,001453 | 0,9926 |
| TcCLB.504153.160 | metallo-peptidase, Clan MA(E) Family M32 | 0,001297 | 0,989 |
| TcCLB.509591.40 | hypothetical protein, conserved | 0,001159 | 0,9926 |
| TcCLB.503811.45 | hypothetical protein, conserved | 0,001144 | 0,9969 |
| TcCLB.507601.80 | hypothetical protein, conserved | 0,001041 | 0,9964 |
| TcCLB.436521.9 | mevalonate kinase, putative (fragment) | 0,0009913 | 0,9956 |
| TcCLB.504239.20 | mucin TcMUCII, putative | 0,0009251 | 0,9987 |
| TcCLB.506559.10 | hypothetical protein | 0,0008556 | 0,996 |
| TcCLB.506559.320 | hypothetical protein, conserved | 0,0007599 | 0,9968 |
| TcCLB.506289.240 | hypothetical protein | 0,0007431 | 0,9989 |
| TcCLB.506405.100 | hypothetical protein, conserved | 0,0005796 | 0,9962 |
| TcCLB.509791.60 | hypothetical protein, conserved | 0,0005406 | 0,9969 |
| TcCLB.504425.90 | aldose 1-epimerase-like protein, putative | 0,0005398 | 0,9982 |
| TcCLB.506337.20 | hypothetical protein, conserved | 0,0005059 | 0,9975 |
| TcCLB.508153.280 | 60S ribosomal protein L13, putative | 0,0004814 | 0,9964 |
| TcCLB.510149.50 | tfiih basal transcription factor complex helicase xpb subuni | 0,0004579 | 0,9968 |
| TcCLB.508707.170 | outer dynein arm docking complex, putative (DC2) | 0,0004463 | 0,9975 |
| TcCLB.511423.80 | Transmembrane protein 231, putative | 0,0004257 | 0,9978 |
| TcCLB.510609.100 | hypothetical protein, conserved | 0,00007583 | 0,9997 |
| TcCLB.508261.150 | trans-sialidase (pseudogene), putative | 0,00006967 | 0,9998 |
| TcCLB.508193.30 | dispersed gene family protein 1 (DGF-1, pseudogene), puta | 0,00006735 | 0,9998 |
| TcCLB.508269.80 | retrotransposon hot spot protein (RHS, pseudogene), putat | -0,0001254 | 0,9997 |
| TcCLB.510099.20 | hypothetical protein, conserved | -0,0001308 | 0,9994 |
| TcCLB.506425.10 | serine carboxypeptidase S28, putative | -0,0002832 | 0,9986 |
| TcCLB.510359.460 | mucin-associated surface protein (MASP), putative | -0,0002844 | 0,9997 |
| TcCLB.511867.40 | hypothetical protein, conserved | -0,0005147 | 0,9982 |
| TcCLB.509179.70 | hypothetical protein, conserved | -0,0005766 | 0,9986 |
| TcCLB.507049.110 | hypothetical protein, conserved | -0,0006474 | 0,9982 |
| TcCLB.510533.150 | hypothetical protein | -0,000673 | 0,9964 |
| TcCLB.503761.50 | hypothetical protein | -0,0006794 | 0,9987 |
| TcCLB.509031.20 | trans-sialidase, putative (fragment) | -0,0007076 | 0,999 |
| TcCLB.508175.146 | 60S ribosomal protein L23a, putative | -0,0007455 | 0,9945 |

| | | | |
|---|---|---|---|
| TcCLB.429185.31 | trans-sialidase (pseudogene), putative | -0,0008439 | 0,9987 |
| TcCLB.506691.70 | hypothetical protein, conserved | -0,0008452 | 0,9962 |
| TcCLB.511081.41 | mucin-associated surface protein (MASP, pseudogene), pu | -0,0008479 | 0,9987 |
| TcCLB.511897.19 | zinc-finger multi-pass transmembrane protein, putative | -0,0009042 | 0,9964 |
| TcCLB.503885.60 | hypothetical protein, conserved | -0,0009236 | 0,9959 |
| TcCLB.510609.65 | U6 snRNA-associated Sm-like protein LSm3p | -0,0009995 | 0,998 |
| TcCLB.503781.30 | hypothetical protein, conserved | -0,001061 | 0,9975 |
| TcCLB.511751.170 | protein kinase, putative | -0,001102 | 0,9926 |
| TcCLB.503597.5 | hypothetical protein (fragment) | -0,001302 | 0,9954 |
| TcCLB.503999.50 | ubiquitin-conjugating enzyme, putative | -0,001395 | 0,9932 |
| TcCLB.506479.100 | hypothetical protein, conserved | -0,001475 | 0,995 |
| TcCLB.508747.59 | mucin-associated surface protein (MASP), putative | -0,001507 | 0,9974 |
| TcCLB.507485.10 | hypothetical protein, conserved | -0,001526 | 0,9964 |
| TcCLB.506405.149 | ribosomal protein L21E (60S), putative | -0,001629 | 0,9843 |
| TcCLB.508693.60 | hypothetical protein, conserved | -0,001674 | 0,9941 |
| TcCLB.412393.10 | trans-sialidase, putative (pseudogene) | -0,001693 | 0,9986 |
| TcCLB.506007.30 | Protein C21orf2 homolog, putative | -0,001843 | 0,9923 |
| TcCLB.510769.49 | 40S ribosomal protein S6, putative | -0,00195 | 0,9841 |
| TcCLB.508547.90 | hypothetical protein, conserved | -0,002039 | 0,9952 |
| TcCLB.504203.10 | hypothetical protein, conserved | -0,002232 | 0,9898 |
| TcCLB.504639.50 | retrotransposon hot spot protein (RHS, pseudogene), putat | -0,002331 | 0,9908 |
| TcCLB.508457.10 | Kinesin-13 6, putative (KIN13-6) | -0,002333 | 0,9895 |
| TcCLB.507229.25 | trans-sialidase (pseudogene), putative | -0,002369 | 0,9915 |
| TcCLB.508625.120 | leucine carboxyl methyltransferase, putative | -0,002433 | 0,9894 |
| TcCLB.511173.241 | mucin TcMUCII (pseudogene), putative | -0,002488 | 0,9964 |
| TcCLB.507053.120 | hypothetical protein, conserved | -0,002555 | 0,9892 |
| TcCLB.506925.210 | hypothetical protein, conserved | -0,002584 | 0,9917 |
| TcCLB.509965.429 | hypothetical protein, conserved | -0,002668 | 0,9957 |
| TcCLB.510333.10 | hypothetical protein, conserved | -0,002737 | 0,984 |
| TcCLB.509905.130 | dispersed gene family protein 1 (DGF-1, pseudogene), puta | -0,002841 | 0,9841 |
| TcCLB.510293.40 | ubiquitin/ribosomal protein S27a, putative | -0,002876 | 0,9931 |
| TcCLB.508737.80 | mannosyltransferase-like protein, putative | -0,002931 | 0,9832 |
| TcCLB.506993.31 | mucin-associated surface protein (MASP, pseudogene), pu | -0,002976 | 0,9965 |
| TcCLB.504239.40 | surface protease GP63 (pseudogene), putative | -0,002992 | 0,9973 |
| TcCLB.509039.20 | bystin, putative | -0,003005 | 0,981 |
| TcCLB.510463.180 | trans-sialidase (pseudogene), putative | -0,003074 | 0,9957 |
| TcCLB.511729.50 | hypothetical protein, conserved | -0,003089 | 0,9809 |
| TcCLB.504055.50 | hypothetical protein, conserved | -0,003196 | 0,9831 |
| TcCLB.509067.60 | Tetratricopeptide-like helical domain containing protein (F | -0,003422 | 0,9676 |
| TcCLB.471901.10 | hypothetical protein, conserved | -0,003453 | 0,9803 |
| TcCLB.506577.160 | hypothetical protein, conserved | -0,003472 | 0,9862 |
| TcCLB.506885.430 | lipoic acid containing carrier protein, putative (GCVH) | -0,003503 | 0,9892 |
| TcCLB.511127.90 | hypothetical protein, conserved | -0,003515 | 0,987 |
| TcCLB.507801.90 | beta-ketoacyl-ACP reductase | -0,003545 | 0,9865 |
| TcCLB.506563.70 | calpain-like cysteine peptidase (pseudogene), putative | -0,003614 | 0,9895 |

| | | | |
|---|---|---|---|
| TcCLB.508641.280 | hypothetical protein, conserved | -0,003638 | 0,9724 |
| TcCLB.506507.30 | beta galactofuranosyl glycosyltransferase, putative | -0,003739 | 0,9871 |
| TcCLB.510547.14 | mucin-associated surface protein (MASP, pseudogene), pu | -0,00374 | 0,995 |
| TcCLB.506869.30 | hypothetical protein, conserved | -0,003766 | 0,9898 |
| TcCLB.506559.454 | GTP-binding protein, putative | -0,00377 | 0,9803 |
| TcCLB.506909.155 | mucin-associated surface protein (MASP, pseudogene), pu | -0,003806 | 0,996 |
| TcCLB.508881.100 | hypothetical protein, conserved | -0,00382 | 0,9788 |
| TcCLB.503745.20 | hypothetical protein, conserved | -0,003861 | 0,9779 |
| TcCLB.503593.70 | eukaryotic peptide chain release factor subunit 1, putative | -0,003931 | 0,9813 |
| TcCLB.507515.20 | hypothetical protein, conserved | -0,003982 | 0,9731 |
| TcCLB.506833.30 | serine/threonine protein phosphatase, putative | -0,004219 | 0,9801 |
| TcCLB.511153.90 | phopshatase, putative | -0,00428 | 0,9779 |
| TcCLB.506155.10 | hypothetical protein, conserved | -0,00431 | 0,9808 |
| TcCLB.506707.20 | ATP-dependent DEAD/H RNA helicase (pseudogene), putat | -0,004431 | 0,9917 |
| TcCLB.503801.70 | hypothetical protein | -0,00447 | 0,9785 |
| TcCLB.509111.19 | hypothetical protein, conserved | -0,004549 | 0,9892 |
| TcCLB.508235.40 | protein kinase, putative (fragment) | -0,004665 | 0,987 |
| TcCLB.503903.50 | hypothetical protein, conserved | -0,00468 | 0,9752 |
| TcCLB.508413.40 | serine/threonine protein phosphatase, putative | -0,0047 | 0,9762 |
| TcCLB.509789.20 | hypothetical protein | -0,004757 | 0,9727 |
| TcCLB.510523.29 | hypothetical protein, conserved (fragment) | -0,004767 | 0,9862 |
| TcCLB.511555.90 | acyl-CoA binding protein, putative | -0,004784 | 0,9729 |
| TcCLB.509259.111 | N-acetyltransferase complex ARD1 subunit (pseudogene), | -0,004807 | 0,9921 |
| TcCLB.508153.290 | 60S ribosomal protein L13, putative | -0,00484 | 0,9618 |
| TcCLB.507009.80 | hypothetical protein, conserved | -0,004982 | 0,9831 |
| TcCLB.507831.90 | 60S ribosomal protein L34, putative | -0,005018 | 0,9789 |
| TcCLB.509693.30 | hypothetical protein, conserved | -0,005046 | 0,9818 |
| TcCLB.504055.60 | hypothetical protein, conserved | -0,005056 | 0,983 |
| TcCLB.510303.250 | hypothetical protein, conserved | -0,005103 | 0,9676 |
| TcCLB.506905.4 | proliferator-activated receptor-interacting protein (PRIP) ir | -0,005132 | 0,9868 |
| TcCLB.505025.70 | mucin-associated surface protein (MASP), putative | -0,005163 | 0,9909 |
| TcCLB.504427.250 | ubiquitin-activating enzyme E1, putative | -0,005184 | 0,9567 |
| TcCLB.506767.110 | mucin-associated surface protein (MASP), putative | -0,005305 | 0,9908 |
| TcCLB.509805.200 | acetyltransferase, putative | -0,005377 | 0,9729 |
| TcCLB.511293.10 | extracellular receptor, putative | -0,00557 | 0,9809 |
| TcCLB.510287.40 | hypothetical protein | -0,005781 | 0,9773 |
| TcCLB.511867.70 | hypothetical protein | -0,005823 | 0,975 |
| TcCLB.509867.20 | mucin-associated surface protein (MASP), putative | -0,005975 | 0,9929 |
| TcCLB.509317.60 | RNA-binding protein, putative | -0,006144 | 0,9627 |
| TcCLB.509999.100 | hypothetical protein, conserved | -0,006172 | 0,9781 |
| TcCLB.510027.10 | GDP-mannose 4,6 dehydratase, putative | -0,006177 | 0,9726 |
| TcCLB.508277.160 | glutaminyl-tRNA synthetase, putative | -0,006191 | 0,9495 |
| TcCLB.509901.130 | Iron-sulfur assembly protein 1, putative (Isa1) | -0,006253 | 0,9718 |
| TcCLB.504873.10 | UDP-Gal or UDP-GlcNAc-dependent glycosyltransferase, pu | -0,006268 | 0,9855 |
| TcCLB.511625.90 | mucin-associated surface protein (MASP), putative | -0,006337 | 0,9877 |

| | | | |
|---|---|---|---|
| TcCLB.507925.50 | hypothetical protein, conserved | -0,006341 | 0,9586 |
| TcCLB.506811.70 | hypothetical protein | -0,006353 | 0,9654 |
| TcCLB.506755.220 | hypothetical protein, conserved | -0,006361 | 0,9675 |
| TcCLB.510183.40 | hypothetical protein, conserved | -0,00637 | 0,9808 |
| TcCLB.503579.90 | hypothetical protein, conserved | -0,006402 | 0,9719 |
| TcCLB.511511.139 | zinc finger protein, putative | -0,006508 | 0,9639 |
| TcCLB.504507.10 | hypothetical protein, conserved | -0,006517 | 0,975 |
| TcCLB.507603.54 | hypothetical protein, conserved | -0,006534 | 0,9812 |
| TcCLB.511233.30 | hypothetical protein, conserved | -0,006622 | 0,9724 |
| TcCLB.507993.190 | serine/threonine protein phosphatase type 5, putative | -0,006663 | 0,9603 |
| TcCLB.503779.60 | hypothetical protein, conserved | -0,006744 | 0,9677 |
| TcCLB.511127.220 | cdc2-related kinase 12 (CRK12) | -0,006861 | 0,9627 |
| TcCLB.475705.9 | hypothetical protein, conserved | -0,006951 | 0,9731 |
| TcCLB.507107.30 | UDP-Gal or UDP-GlcNAc-dependent glycosyltransferase, pu | -0,006968 | 0,9898 |
| TcCLB.510257.24 | hypothetical protein, conserved | -0,00715 | 0,976 |
| TcCLB.507951.130 | hypothetical protein | -0,007239 | 0,9673 |
| TcCLB.508075.60 | hypothetical protein, conserved | -0,007275 | 0,9665 |
| TcCLB.508387.40 | hypothetical protein, conserved | -0,007317 | 0,9782 |
| TcCLB.508479.216 | hypothetical protein, conserved | -0,007329 | 0,9659 |
| TcCLB.508355.124 | hypothetical protein, conserved | -0,007357 | 0,977 |
| TcCLB.507603.250 | hypothetical protein, conserved | -0,00736 | 0,9584 |
| TcCLB.506661.60 | hypothetical protein, conserved | -0,007403 | 0,9678 |
| TcCLB.511345.10 | retrotransposon hot spot (RHS) protein, putative | -0,007561 | 0,977 |
| TcCLB.509471.77 | histone H3, putative | -0,007609 | 0,9491 |
| TcCLB.511529.68 | cyclin-dependent kinases regulatory subunit, putative (P12 | -0,007681 | 0,9639 |
| TcCLB.506629.120 | hypothetical protein, conserved | -0,00772 | 0,9622 |
| TcCLB.509231.5 | hypothetical protein, conserved (fragment) | -0,007741 | 0,9917 |
| TcCLB.510431.200 | LisH domain-containing protein FOPNL, putative (fopnl) | -0,007798 | 0,9678 |
| TcCLB.507603.150 | dolichyl-P-Man:GDP-Man5GlcNAc2-PP-dolichyl alpha-1,2-m | -0,007813 | 0,9615 |
| TcCLB.508027.100 | hypothetical protein | -0,007821 | 0,9612 |
| TcCLB.509901.140 | COP9 signalosome complex subunit 5, putative | -0,007833 | 0,9661 |
| TcCLB.508109.50 | retrotransposon hot spot protein (RHS, pseudogene), putat | -0,007931 | 0,9772 |
| TcCLB.510955.10 | hypothetical protein, conserved | -0,008078 | 0,9538 |
| TcCLB.510519.149 | hypothetical protein, conserved | -0,008138 | 0,9568 |
| TcCLB.510329.380 | hypothetical protein, conserved | -0,008247 | 0,9677 |
| TcCLB.508545.30 | hypothetical protein, conserved | -0,008424 | 0,9765 |
| TcCLB.508359.80 | mucin-associated surface protein (MASP), putative | -0,008444 | 0,9917 |
| TcCLB.506177.130 | hypothetical protein, conserved | -0,008529 | 0,9765 |
| TcCLB.508657.10 | hypothetical protein, conserved | -0,008533 | 0,9647 |
| TcCLB.506929.40 | target SNARE, putative | -0,008573 | 0,9617 |
| TcCLB.508577.160 | proteasome beta 6 subunit, putative | -0,008843 | 0,9487 |
| TcCLB.511907.300 | DNA repair and transcription factor protein, putative | -0,008846 | 0,9594 |
| TcCLB.511771.60 | hypothetical protein | -0,008849 | 0,9895 |
| TcCLB.508229.10 | trans-sialidase, putative | -0,008863 | 0,9893 |
| TcCLB.510285.110 | hypothetical protein, conserved | -0,008883 | 0,9617 |

| | | | |
|---|---|---|---|
| TcCLB.506513.100 | hypothetical protein, conserved | -0,008889 | 0,9485 |
| TcCLB.511285.80 | hypothetical protein | -0,00897 | 0,9443 |
| TcCLB.503421.7 | N-acetyltransferase complex ARD1 subunit (pseudogene), | -0,008979 | 0,9841 |
| TcCLB.507641.110 | hypothetical protein, conserved | -0,008988 | 0,9497 |
| TcCLB.508755.11 | trans-sialidase (pseudogene), putative | -0,008998 | 0,9761 |
| TcCLB.506605.30 | hypothetical protein, conserved | -0,009026 | 0,9638 |
| TcCLB.507611.199 | dispersed gene family protein 1 (DGF-1), putative | -0,009037 | 0,9766 |
| TcCLB.509253.40 | hypothetical protein, conserved | -0,009088 | 0,9639 |
| TcCLB.508011.20 | mucin-associated surface protein (MASP, pseudogene), pu | -0,009214 | 0,9922 |
| TcCLB.511071.20 | hypothetical protein, conserved | -0,00929 | 0,9692 |
| TcCLB.503649.10 | mucin-associated surface protein (MASP, pseudogene), pu | -0,009321 | 0,9862 |
| TcCLB.510105.36 | hypothetical protein, conserved | -0,009338 | 0,9619 |
| TcCLB.510299.10 | hypothetical protein, conserved | -0,009345 | 0,9596 |
| TcCLB.508899.110 | actin-related protein 2, putative | -0,009346 | 0,9614 |
| TcCLB.511475.24 | dispersed gene family protein 1 (DGF-1), putative (fragme | -0,009361 | 0,975 |
| TcCLB.506295.200 | retrotransposon hot spot protein (RHS, pseudogene), putat | -0,009396 | 0,9576 |
| TcCLB.511173.120 | mucin-associated surface protein (MASP, pseudogene), pu | -0,009466 | 0,979 |
| TcCLB.506933.46 | hypothetical protein, conserved | -0,009474 | 0,9675 |
| TcCLB.509875.30 | trans-sialidase (pseudogene), putative | -0,009474 | 0,9781 |
| TcCLB.506697.60 | hypothetical protein, conserved | -0,009491 | 0,9335 |
| TcCLB.510099.30 | hypothetical protein, conserved | -0,009503 | 0,9563 |
| TcCLB.503955.10 | mitochondrial DNA polymerase beta-PAK, putative | -0,009801 | 0,963 |
| TcCLB.511657.60 | hypothetical protein, conserved | -0,00984 | 0,9371 |
| TcCLB.510431.260 | hypothetical protein, conserved | -0,009866 | 0,9568 |
| TcCLB.511277.160 | 60S ribosomal protein L10a, putative | -0,009904 | 0,8904 |
| TcCLB.506151.10 | hypothetical protein, conserved | -0,009913 | 0,9498 |
| TcCLB.505943.30 | hypothetical protein, conserved | -0,009956 | 0,9542 |
| TcCLB.508163.270 | mucin-associated surface protein (MASP), putative | -0,009989 | 0,9833 |
| TcCLB.506187.80 | trans-sialidase (pseudogene), putative | -0,01007 | 0,9896 |
| TcCLB.504109.90 | cysteine peptidase, Clan CA, family C12, putative | -0,01008 | 0,9568 |
| TcCLB.508175.340 | hypothetical protein, conserved | -0,0102 | 0,9337 |
| TcCLB.506559.484 | hypothetical protein, conserved | -0,01021 | 0,9674 |
| TcCLB.507811.50 | hypothetical protein, conserved | -0,01025 | 0,9538 |
| TcCLB.508707.320 | hypothetical protein, conserved | -0,01029 | 0,9452 |
| TcCLB.503971.20 | hypothetical protein, conserved | -0,01031 | 0,959 |
| TcCLB.506025.30 | hypothetical protein, conserved | -0,01034 | 0,9612 |
| TcCLB.508385.20 | hypothetical protein, conserved | -0,01035 | 0,9496 |
| TcCLB.509007.20 | hypothetical protein, conserved | -0,01037 | 0,9483 |
| TcCLB.510031.30 | hypothetical protein | -0,01045 | 0,9461 |
| TcCLB.506213.100 | hypothetical protein, conserved | -0,01059 | 0,9495 |
| TcCLB.509267.6 | trans-sialidase (pseudogene), putative | -0,01063 | 0,9719 |
| TcCLB.506929.20 | hypothetical protein, conserved | -0,01071 | 0,9471 |
| TcCLB.503593.10 | eukaryotic translation initiation factor eIF-4E, putative | -0,01073 | 0,9416 |
| TcCLB.509201.15 | ribosomal protein S29, putative | -0,0109 | 0,951 |
| TcCLB.506287.150 | GTP-binding protein, putative | -0,01118 | 0,9432 |

| | | | |
|---|---|---|---|
| TcCLB.509125.20 | ATP-dependent Clp protease subunit, heat shock protein 78 | -0,01129 | 0,9325 |
| TcCLB.506211.220 | Associated kinase of Tb14-3-3, putative (AKB1) | -0,01134 | 0,9315 |
| TcCLB.506389.60 | hypothetical protein, conserved | -0,01138 | 0,9491 |
| TcCLB.509551.100 | hypothetical protein, conserved | -0,01141 | 0,9495 |
| TcCLB.506961.20 | dispersed gene family protein 1 (DGF-1), putative | -0,01141 | 0,9788 |
| TcCLB.505985.9 | calpain cysteine peptidase, putative (fragment) | -0,01146 | 0,9495 |
| TcCLB.504575.15 | ubiquinol-cytochrome-C-reductase, putative | -0,01149 | 0,9777 |
| TcCLB.508355.130 | hypothetical protein, conserved | -0,0115 | 0,9406 |
| TcCLB.506457.50 | serine/threonine protein kinase, putative | -0,01151 | 0,9439 |
| TcCLB.511753.130 | U3 small nuclear ribonucleoprotein (snRNP), putative | -0,01166 | 0,914 |
| TcCLB.506399.60 | hypothetical protein, conserved | -0,01191 | 0,9486 |
| TcCLB.506703.30 | mucin TcMUCII (pseudogene), putative | -0,01194 | 0,9844 |
| TcCLB.504099.30 | hypothetical protein, conserved (pseudogene) | -0,0121 | 0,9645 |
| TcCLB.507601.10 | phosphoprotein phosphatase, putative | -0,01213 | 0,9139 |
| TcCLB.510279.100 | hypothetical protein | -0,01217 | 0,983 |
| TcCLB.509965.190 | DREV methyltransferase, putative | -0,01224 | 0,9421 |
| TcCLB.508461.90 | hypothetical protein, conserved | -0,01229 | 0,9416 |
| TcCLB.508667.30 | hypothetical protein, conserved | -0,01238 | 0,9452 |
| TcCLB.507041.50 | hypothetical protein, conserved | -0,01243 | 0,9625 |
| TcCLB.506203.120 | hypothetical protein, conserved | -0,01244 | 0,9064 |
| TcCLB.506979.20 | hypothetical protein, conserved | -0,01246 | 0,9603 |
| TcCLB.504029.90 | hypothetical protein, conserved | -0,01256 | 0,9493 |
| TcCLB.509551.30 | mitochondrial phosphate transporter, putative | -0,01259 | 0,9325 |
| TcCLB.509717.40 | hypothetical protein, conserved | -0,01263 | 0,9368 |
| TcCLB.507529.10 | hypothetical Zinc-finger containing protein | -0,01283 | 0,9471 |
| TcCLB.506503.40 | hypothetical protein, conserved | -0,01286 | 0,9536 |
| TcCLB.479517.30 | hypothetical protein | -0,0129 | 0,9886 |
| TcCLB.507957.200 | mucin-associated surface protein (MASP), putative | -0,01291 | 0,9851 |
| TcCLB.506469.120 | hypothetical protein, conserved | -0,01295 | 0,9452 |
| TcCLB.508265.40 | hypothetical protein, conserved | -0,01298 | 0,9335 |
| TcCLB.506511.20 | hypothetical protein, conserved | -0,01301 | 0,9399 |
| TcCLB.508265.70 | hypothetical protein, conserved | -0,01301 | 0,9538 |
| TcCLB.509897.132 | hypothetical protein, conserved (pseudogene) | -0,01317 | 0,9287 |
| TcCLB.506243.110 | expression site-associated gene (ESAG-like) protein, putati | -0,01325 | 0,973 |
| TcCLB.509733.70 | hypothetical protein, conserved | -0,01327 | 0,9312 |
| TcCLB.506297.310 | hypothetical protein, conserved | -0,01335 | 0,9291 |
| TcCLB.506351.35 | hypothetical protein, conserved | -0,01335 | 0,9663 |
| TcCLB.511837.110 | hypothetical protein, conserved | -0,01343 | 0,9266 |
| TcCLB.510257.130 | protein kinase, putative | -0,0135 | 0,919 |
| TcCLB.507949.69 | DNA-directed RNA polymerase 2, RPB12 subunit, putative | -0,01359 | 0,9463 |
| TcCLB.503811.80 | hypothetical protein, conserved | -0,01366 | 0,9383 |
| TcCLB.504109.30 | hypothetical protein, conserved | -0,01371 | 0,9171 |
| TcCLB.510729.80 | Mitochondrial import receptor subunit ATOM40, putative ( | -0,01386 | 0,9069 |
| TcCLB.508335.9 | dispersed gene family protein 1 (DGF-1), putative | -0,01389 | 0,9642 |
| TcCLB.506955.198 | hypothetical protein | -0,01392 | 0,969 |

| | | | |
|---|---|---|---|
| TcCLB.506495.24 | hypothetical protein, conserved | -0,01394 | 0,9564 |
| TcCLB.505989.78 | hypothetical protein, conserved | -0,01397 | 0,9493 |
| TcCLB.510513.30 | Probable methyltransferase BTM2 homolog | -0,01413 | 0,9441 |
| TcCLB.504091.5 | hypothetical protein, conserved (fragment) | -0,01418 | 0,9817 |
| TcCLB.508331.41 | tryptophanyl-tRNA synthetase (pseudogene), putative | -0,01423 | 0,9752 |
| TcCLB.503617.15 | U6 snRNA-associated Sm-like protein LSm3p | -0,01428 | 0,9678 |
| TcCLB.509755.10 | trans-sialidase, Group V, putative | -0,0144 | 0,9721 |
| TcCLB.506621.30 | ribonucleoside-diphosphate reductase small chain, putativ | -0,01442 | 0,9121 |
| TcCLB.507511.71 | hypothetical protein (pseudogene) | -0,01443 | 0,9711 |
| TcCLB.508461.160 | hypothetical protein, conserved | -0,01444 | 0,9157 |
| TcCLB.511773.50 | protein kinase, putative | -0,01462 | 0,9389 |
| TcCLB.507631.10 | hypothetical protein, conserved | -0,01469 | 0,9659 |
| TcCLB.506505.10 | hypothetical protein, conserved | -0,01473 | 0,9568 |
| TcCLB.507953.140 | trans-sialidase, Group V, putative | -0,01473 | 0,978 |
| TcCLB.504137.10 | isoleucyl-tRNA synthetase, putative | -0,01478 | 0,9301 |
| TcCLB.506479.110 | hypothetical protein | -0,01499 | 0,9171 |
| TcCLB.508385.60 | hypothetical protein, conserved | -0,01514 | 0,939 |
| TcCLB.509231.20 | protein kinase, putative | -0,01515 | 0,9326 |
| TcCLB.510351.130 | hypothetical protein, conserved | -0,01519 | 0,9419 |
| TcCLB.508273.40 | hypothetical protein, conserved | -0,01524 | 0,9317 |
| TcCLB.503565.10 | hypothetical protein | -0,01533 | 0,9458 |
| TcCLB.510377.134 | mucin-associated surface protein (MASP), putative | -0,01534 | 0,9809 |
| TcCLB.510747.150 | hypothetical protein, conserved | -0,01543 | 0,9261 |
| TcCLB.509967.90 | hypothetical protein, conserved | -0,01543 | 0,9361 |
| TcCLB.506931.40 | hypothetical protein, conserved | -0,01556 | 0,915 |
| TcCLB.503939.50 | hypothetical protein, conserved | -0,01559 | 0,903 |
| TcCLB.506221.30 | protein phosphatase 2C, putative | -0,01565 | 0,9211 |
| TcCLB.511303.20 | hypothetical protein | -0,0157 | 0,9317 |
| TcCLB.508355.250 | 60S acidic ribosomal protein P0, putative (P0) | -0,01582 | 0,8532 |
| TcCLB.510747.34 | hypothetical protein, conserved | -0,01584 | 0,9423 |
| TcCLB.509767.100 | hypothetical protein, conserved | -0,01596 | 0,9421 |
| TcCLB.511277.320 | hypothetical protein, conserved | -0,01599 | 0,8995 |
| TcCLB.508879.180 | hypothetical protein, conserved | -0,01602 | 0,9465 |
| TcCLB.508059.40 | syntaxin, putative | -0,01604 | 0,943 |
| TcCLB.508199.90 | hypothetical protein, conserved | -0,01605 | 0,9257 |
| TcCLB.507669.100 | transcription elongation factor, putative | -0,01606 | 0,9243 |
| TcCLB.506949.50 | cytochrome c, putative | -0,0161 | 0,9068 |
| TcCLB.507807.20 | hypothetical protein, conserved | -0,0161 | 0,9495 |
| TcCLB.509027.40 | hypothetical protein, conserved | -0,01615 | 0,8926 |
| TcCLB.426071.10 | dispersed gene family protein 1 (DGF-1), putative (fragme | -0,01623 | 0,967 |
| TcCLB.507069.160 | trans-sialidase, Group V, putative | -0,01627 | 0,9656 |
| TcCLB.511923.40 | hypothetical protein, conserved (pseudogene) | -0,01632 | 0,9619 |
| TcCLB.508179.40 | hypothetical protein, conserved | -0,01635 | 0,9156 |
| TcCLB.511505.20 | hypothetical protein, conserved | -0,01641 | 0,9253 |
| TcCLB.511209.40 | hypothetical protein, conserved | -0,01646 | 0,9392 |

| | | | |
|---|---|---|---|
| TcCLB.507729.41 | elongation factor 1-gamma (EF-1-gamma, pseudogene), p | -0,01646 | 0,9784 |
| TcCLB.504047.40 | kinesin, putative | -0,01651 | 0,8744 |
| TcCLB.508221.518 | mucin TcMUCII, putative | -0,01651 | 0,9805 |
| TcCLB.505163.20 | hypothetical protein, conserved | -0,01657 | 0,9375 |
| TcCLB.510297.100 | ADP-ribosylation factor-like protein, putative | -0,01664 | 0,9251 |
| TcCLB.506469.60 | hypothetical protein | -0,01676 | 0,9591 |
| TcCLB.503985.10 | trans-sialidase (pseudogene), putative | -0,01685 | 0,9617 |
| TcCLB.511077.7 | trans-sialidase, putative (fragment) | -0,01694 | 0,9939 |
| TcCLB.503805.30 | hypothetical protein, conserved | -0,01695 | 0,9335 |
| TcCLB.506501.5 | trans-sialidase (pseudogene), putative | -0,01695 | 0,9801 |
| TcCLB.510515.120 | hypothetical protein, conserved | -0,01703 | 0,9656 |
| TcCLB.506529.650 | hypothetical protein, conserved | -0,01706 | 0,9161 |
| TcCLB.510011.9 | mucin-associated surface protein (MASP), putative | -0,01727 | 0,9719 |
| TcCLB.511153.100 | RNA-binding protein, putative | -0,01738 | 0,9146 |
| TcCLB.511257.30 | ras-related protein rab, putative | -0,0174 | 0,9406 |
| TcCLB.510105.250 | membrane-bound acid phosphatase 2, putative | -0,01748 | 0,9253 |
| TcCLB.510127.60 | hypothetical protein, conserved | -0,01748 | 0,9324 |
| TcCLB.509117.10 | ubiquitin-activating enzyme, putative | -0,01754 | 0,9287 |
| TcCLB.504883.40 | hypothetical protein, conserved | -0,01758 | 0,9006 |
| TcCLB.508371.10 | glutamine-dependent carbamoyl-phosphate synthetase (ps | -0,0176 | 0,9251 |
| TcCLB.509161.90 | hypothetical protein, conserved | -0,01761 | 0,9273 |
| TcCLB.511185.60 | trans-sialidase (pseudogene), putative | -0,01765 | 0,9614 |
| TcCLB.506863.40 | hypothetical protein, conserved | -0,01767 | 0,9453 |
| TcCLB.506245.76 | trans-sialidase (pseudogene), putative | -0,01768 | 0,9627 |
| TcCLB.506767.250 | trans-sialidase (pseudogene), putative | -0,01773 | 0,9702 |
| TcCLB.504313.30 | dispersed gene family protein 1 (DGF-1), putative | -0,01783 | 0,9308 |
| TcCLB.508593.30 | hypothetical protein, conserved | -0,01792 | 0,9172 |
| TcCLB.508587.40 | dispersed gene family protein 1 (DGF-1, pseudogene), puta | -0,01802 | 0,9472 |
| TcCLB.507159.44 | DNA-directed RNA polymerase III subunit, putative | -0,01817 | 0,9145 |
| TcCLB.511921.10 | hypothetical protein, conserved (pseudogene) | -0,0182 | 0,9581 |
| TcCLB.508209.150 | hypothetical protein, conserved | -0,01824 | 0,9049 |
| TcCLB.508779.50 | hypothetical protein, conserved | -0,01829 | 0,8995 |
| TcCLB.509589.50 | hypothetical protein, conserved | -0,01832 | 0,898 |
| TcCLB.509617.60 | thioredoxin, putative | -0,01834 | 0,9385 |
| TcCLB.507793.60 | hypothetical protein, conserved | -0,01837 | 0,9567 |
| TcCLB.503899.70 | hypothetical protein | -0,01845 | 0,9299 |
| TcCLB.509937.130 | protein kinase, putative | -0,01849 | 0,9812 |
| TcCLB.504161.50 | hypothetical protein, conserved | -0,01859 | 0,9233 |
| TcCLB.509217.160 | complement regulatory protein, putative | -0,01869 | 0,9634 |
| TcCLB.508321.40 | hypothetical protein, conserved | -0,01874 | 0,8496 |
| TcCLB.511059.10 | trans-sialidase, putative | -0,01874 | 0,9806 |
| TcCLB.509887.10 | hypothetical protein | -0,01876 | 0,943 |
| TcCLB.503687.10 | regulator of chromosome condensation (RCC1), putative | -0,01888 | 0,9176 |
| TcCLB.506331.139 | mucin-associated surface protein (MASP, pseudogene), pu | -0,01891 | 0,9469 |
| TcCLB.506247.290 | hypothetical protein, conserved | -0,01895 | 0,8661 |

| | | | |
|---|---|---|---|
| TcCLB.506507.60 | retrotransposon hot spot protein (RHS, pseudogene), putat | -0,019 | 0,9203 |
| TcCLB.509781.10 | hypothetical protein, conserved | -0,01906 | 0,9341 |
| TcCLB.507987.30 | mucin-associated surface protein (MASP), putative (fragm | -0,01907 | 0,9719 |
| TcCLB.503609.50 | trans-sialidase, Group VI, putative | -0,01913 | 0,9602 |
| TcCLB.508081.20 | hypothetical protein | -0,01924 | 0,9762 |
| TcCLB.504017.40 | aminomethyltransferase, mitochondrial, putative (GCV) | -0,01949 | 0,9229 |
| TcCLB.507811.44 | phosphoglycerate mutase protein, putative | -0,01964 | 0,9368 |
| TcCLB.510655.80 | hypothetical protein, conserved | -0,01972 | 0,9491 |
| TcCLB.510371.70 | mucin-associated surface protein (MASP), putative | -0,01972 | 0,9665 |
| TcCLB.511383.20 | kinesin, putative | -0,01979 | 0,8963 |
| TcCLB.507757.60 | dihydrolipoamide dehydrogenase, putative | -0,02002 | 0,9203 |
| TcCLB.511137.10 | hypothetical protein, conserved | -0,02003 | 0,9314 |
| TcCLB.509803.30 | hypothetical protein, conserved | -0,02006 | 0,8927 |
| TcCLB.503897.50 | hypothetical protein, conserved | -0,02006 | 0,9006 |
| TcCLB.503851.10 | Elongator-like Protein 3a, putative | -0,02007 | 0,9249 |
| TcCLB.509911.100 | chaperone DNAJ protein, putative | -0,02014 | 0,9173 |
| TcCLB.511867.180 | hypothetical protein, conserved | -0,02018 | 0,9102 |
| TcCLB.511747.20 | hypothetical protein, conserved | -0,02018 | 0,9114 |
| TcCLB.510205.20 | mucin-associated surface protein (MASP), putative | -0,02023 | 0,9614 |
| TcCLB.510089.110 | mitochondrial RNA binding complex 1 subunit, putative | -0,02036 | 0,8958 |
| TcCLB.508963.20 | small GTP-binding protein Rab18, putative | -0,02047 | 0,9125 |
| TcCLB.511907.10 | hypothetical protein, conserved | -0,02049 | 0,9071 |
| TcCLB.504625.10 | trans-splicing factor, putative | -0,02064 | 0,9075 |
| TcCLB.506941.250 | hypothetical protein, conserved | -0,02065 | 0,9012 |
| TcCLB.510741.180 | ATP-dependent RNA helicase, putative | -0,02074 | 0,8411 |
| TcCLB.511925.40 | hypothetical protein, conserved | -0,02087 | 0,9452 |
| TcCLB.510295.15 | hypothetical protein, conserved | -0,02088 | 0,9103 |
| TcCLB.507215.9 | dispersed gene family protein 1 (DGF-1), putative (fragme | -0,02095 | 0,9702 |
| TcCLB.508825.20 | hypothetical protein | -0,02099 | 0,9293 |
| TcCLB.506303.50 | UDP-galactose 4-epimerase, putative | -0,02101 | 0,8937 |
| TcCLB.508065.10 | UDP-Gal or UDP-GlcNAc-dependent glycosyltransferase, pu | -0,02103 | 0,9499 |
| TcCLB.511287.80 | hypothetical protein, conserved | -0,02109 | 0,9197 |
| TcCLB.508165.410 | mucin-associated surface protein (MASP), putative | -0,02128 | 0,9639 |
| TcCLB.511929.40 | trans-sialidase (pseudogene), putative | -0,02133 | 0,9617 |
| TcCLB.509065.120 | hypothetical protein | -0,02135 | 0,9296 |
| TcCLB.508815.70 | hypothetical protein, conserved | -0,02141 | 0,8345 |
| TcCLB.503657.20 | hypothetical protein, conserved | -0,02144 | 0,9141 |
| TcCLB.509059.80 | hypothetical protein, conserved | -0,02148 | 0,8725 |
| TcCLB.509767.10 | neutral sphingomyelinase activation associated factor-like | -0,02159 | 0,8769 |
| TcCLB.503563.20 | ADG2, putative | -0,02182 | 0,9042 |
| TcCLB.506869.4 | hypothetical protein, conserved | -0,02195 | 0,9005 |
| TcCLB.503879.100 | hypothetical protein, conserved | -0,02196 | 0,819 |
| TcCLB.507609.70 | hypothetical protein, conserved | -0,02216 | 0,9363 |
| TcCLB.511747.30 | hypothetical protein, conserved | -0,02218 | 0,9251 |
| TcCLB.506871.130 | hypothetical protein, conserved | -0,02222 | 0,8924 |

| | | | |
|---|---|---|---|
| TcCLB.506173.20 | hypothetical protein | -0,02225 | 0,9189 |
| TcCLB.506765.58 | trans-sialidase (pseudogene), putative | -0,02226 | 0,9609 |
| TcCLB.509999.30 | hypothetical protein, conserved | -0,02232 | 0,9125 |
| TcCLB.508085.10 | GMP synthase, putative | -0,02235 | 0,8906 |
| TcCLB.511797.240 | trans-sialidase (pseudogene), putative | -0,02238 | 0,9522 |
| TcCLB.506859.80 | hypothetical protein, conserved | -0,0225 | 0,8629 |
| TcCLB.505025.110 | surface protease GP63 (pseudogene), putative | -0,02252 | 0,9617 |
| TcCLB.504109.70 | mitochondrial carrier protein, putative | -0,02261 | 0,8608 |
| TcCLB.509647.190 | hypothetical protein, conserved | -0,02266 | 0,8667 |
| TcCLB.508735.10 | trans-sialidase (pseudogene), putative | -0,02268 | 0,91 |
| TcCLB.505965.80 | hypothetical protein, conserved | -0,02269 | 0,9148 |
| TcCLB.509561.120 | deaminase, putative | -0,02275 | 0,928 |
| TcCLB.510745.10 | hypothetical protein, conserved | -0,02279 | 0,8948 |
| TcCLB.507583.30 | hypothetical protein | -0,0228 | 0,9091 |
| TcCLB.510355.250 | hypothetical protein, conserved | -0,02288 | 0,8597 |
| TcCLB.506195.150 | hypothetical protein, conserved | -0,02302 | 0,8619 |
| TcCLB.506527.10 | hypothetical protein, conserved | -0,02327 | 0,8389 |
| TcCLB.509037.50 | aspartyl-tRNA synthetase, putative | -0,02333 | 0,8706 |
| TcCLB.507527.10 | hypothetical protein, conserved | -0,02334 | 0,8887 |
| TcCLB.507591.20 | beta galactofuranosyl glycosyltransferase (pseudogene), p | -0,02337 | 0,939 |
| TcCLB.511625.74 | hypothetical protein, conserved | -0,02343 | 0,8662 |
| TcCLB.506205.30 | hypothetical protein, conserved | -0,02349 | 0,9128 |
| TcCLB.510105.184 | hypothetical protein, conserved | -0,02351 | 0,917 |
| TcCLB.507793.50 | hypothetical protein, conserved | -0,02365 | 0,9394 |
| TcCLB.473755.9 | hypothetical protein, conserved | -0,02366 | 0,8912 |
| TcCLB.511825.110 | hypothetical protein | -0,0237 | 0,9683 |
| TcCLB.504051.40 | hypothetical protein, conserved | -0,02372 | 0,8631 |
| TcCLB.511923.130 | hypothetical protein, conserved | -0,02383 | 0,8895 |
| TcCLB.507069.139 | mucin TcMUCII (pseudogene), putative | -0,02391 | 0,9673 |
| TcCLB.508921.10 | hypothetical protein, conserved | -0,02399 | 0,9134 |
| TcCLB.503835.20 | hypothetical protein, conserved | -0,02402 | 0,9068 |
| TcCLB.507001.40 | hypothetical protein, conserved | -0,02406 | 0,8717 |
| TcCLB.503833.30 | peroxin 13, putative (PEX13) | -0,02415 | 0,883 |
| TcCLB.506321.350 | hypothetical protein, conserved | -0,02415 | 0,9075 |
| TcCLB.508741.190 | hypothetical protein, conserved | -0,02423 | 0,8759 |
| TcCLB.507011.260 | hypothetical protein, conserved | -0,02425 | 0,8831 |
| TcCLB.511719.20 | hypothetical protein, conserved | -0,02425 | 0,9093 |
| TcCLB.504881.30 | hypothetical protein, conserved | -0,02433 | 0,7653 |
| TcCLB.504427.30 | hypothetical protein, conserved | -0,02437 | 0,8587 |
| TcCLB.506789.110 | hypothetical protein | -0,02442 | 0,854 |
| TcCLB.511025.30 | mitogen-activated protein kinase, putative | -0,02442 | 0,873 |
| TcCLB.511923.71 | hypothetical protein, conserved (pseudogene) | -0,02442 | 0,9522 |
| TcCLB.509539.44 | hypothetical protein, conserved | -0,02444 | 0,9503 |
| TcCLB.506411.39 | CDP-DAG synthase, putative (fragment) (CDS) | -0,02448 | 0,9483 |
| TcCLB.509633.40 | hypothetical protein, conserved | -0,02455 | 0,913 |

| | | | |
|---|---|---|---|
| TcCLB.504021.10 | G1 cyclin CycE4, putative (CYC5) | -0,02459 | 0,8372 |
| TcCLB.511851.44 | mucin-associated surface protein (MASP, pseudogene), pu | -0,02459 | 0,8402 |
| TcCLB.503985.20 | retrotransposon hot spot protein (RHS, pseudogene), putat | -0,0246 | 0,9409 |
| TcCLB.506277.20 | mucin TcMUCII (pseudogene), putative | -0,0246 | 0,9708 |
| TcCLB.509915.80 | hypothetical protein, conserved (pseudogene) | -0,02464 | 0,9335 |
| TcCLB.508355.280 | hypothetical protein, conserved | -0,02465 | 0,8564 |
| TcCLB.510437.45 | hypothetical protein | -0,02474 | 0,9137 |
| TcCLB.504741.110 | hypothetical protein, conserved | -0,02478 | 0,9091 |
| TcCLB.506529.460 | hypothetical protein, conserved | -0,02481 | 0,8669 |
| TcCLB.511183.391 | surface protease GP63 (pseudogene), putative | -0,02482 | 0,9495 |
| TcCLB.511477.40 | dispersed gene family protein 1 (DGF-1, pseudogene), puta | -0,02483 | 0,9242 |
| TcCLB.510921.40 | trans-sialidase, putative (fragment) | -0,02492 | 0,9615 |
| TcCLB.506989.155 | hypothetical protein, conserved | -0,02493 | 0,9203 |
| TcCLB.508917.4 | hypothetical protein, conserved | -0,02505 | 0,8922 |
| TcCLB.508827.110 | leucine-rich repeat protein (LRRP), putative | -0,02507 | 0,8934 |
| TcCLB.508781.10 | structural maintenance of chromosome protein 4 (pseudog | -0,02515 | 0,9346 |
| TcCLB.506343.10 | serine/threonine protein kinase, putative | -0,0252 | 0,9139 |
| TcCLB.509455.90 | Putative intraflagellar transport protein A1 (PIFTA1) | -0,02522 | 0,8491 |
| TcCLB.509543.10 | hypothetical protein | -0,02525 | 0,951 |
| TcCLB.511653.40 | hypothetical protein, conserved | -0,02551 | 0,902 |
| TcCLB.511003.200 | hypothetical protein, conserved | -0,02562 | 0,9163 |
| TcCLB.507991.30 | hypothetical protein, conserved | -0,0258 | 0,8946 |
| TcCLB.511737.50 | hypothetical protein, conserved | -0,0258 | 0,9016 |
| TcCLB.505941.30 | hypothetical protein, conserved | -0,02582 | 0,8507 |
| TcCLB.507073.30 | hypothetical protein, conserved | -0,02584 | 0,8253 |
| TcCLB.509505.80 | hypothetical protein, conserved | -0,02595 | 0,8761 |
| TcCLB.506703.20 | mucin-associated surface protein (MASP), putative | -0,02616 | 0,965 |
| TcCLB.506297.220 | hypothetical protein, conserved | -0,02619 | 0,8111 |
| TcCLB.509157.100 | hypothetical protein | -0,02632 | 0,9128 |
| TcCLB.506475.120 | conserved protein, unknown function | -0,02633 | 0,8729 |
| TcCLB.506919.110 | glycine dehydrogenase [decarboxylating] (pseudogene), pu | -0,02633 | 0,9606 |
| TcCLB.510755.129 | 60S ribosomal protein L12, putative | -0,02636 | 0,9114 |
| TcCLB.509671.64 | ribosomal protein L36, putative | -0,02641 | 0,8605 |
| TcCLB.507071.390 | trans-sialidase, Group V, putative | -0,02654 | 0,9581 |
| TcCLB.508965.39 | hypothetical protein, conserved | -0,02655 | 0,8552 |
| TcCLB.508265.44 | mitochondrial pyruvate carrier protein 2, putative | -0,02668 | 0,8766 |
| TcCLB.510967.16 | hypothetical protein, conserved | -0,02673 | 0,8796 |
| TcCLB.506379.10 | UDP-Gal or UDP-GlcNAc-dependent glycosyltransferase, pu | -0,02678 | 0,9427 |
| TcCLB.511173.460 | mucin-associated surface protein (MASP), putative | -0,0268 | 0,9618 |
| TcCLB.508871.160 | hypothetical protein | -0,0268 | 0,9675 |
| TcCLB.504835.29 | hypothetical protein, conserved | -0,02681 | 0,8455 |
| TcCLB.507681.70 | hypothetical protein, conserved | -0,02689 | 0,8538 |
| TcCLB.507747.220 | protein farnesyltransferase, putative | -0,02691 | 0,8636 |
| TcCLB.511215.50 | hypothetical protein, conserved | -0,02697 | 0,8348 |
| TcCLB.510603.119 | hypothetical protein, conserved | -0,02704 | 0,9278 |

| | | | |
|---|---|---|---|
| TcCLB.506357.190 | hypothetical protein, conserved | -0,02708 | 0,8685 |
| TcCLB.508707.130 | hypothetical protein, conserved | -0,02708 | 0,8711 |
| TcCLB.505999.150 | wd40 repeat domain-containing protein | -0,0271 | 0,8766 |
| TcCLB.511803.30 | hypothetical protein, conserved | -0,02719 | 0,8858 |
| TcCLB.506967.10 | trans-sialidase, Group V, putative | -0,02722 | 0,9508 |
| TcCLB.503717.40 | mucin-associated surface protein (MASP, pseudogene), pu | -0,02731 | 0,9461 |
| TcCLB.506201.4 | hypothetical protein, conserved | -0,0274 | 0,9093 |
| TcCLB.506477.4 | hypothetical protein, conserved | -0,02747 | 0,9275 |
| TcCLB.507993.20 | hypothetical protein, conserved | -0,0276 | 0,9325 |
| TcCLB.504449.30 | hypothetical protein, conserved | -0,02761 | 0,79 |
| TcCLB.507093.339 | hypothetical protein, conserved | -0,02763 | 0,8575 |
| TcCLB.507089.140 | hypothetical protein, conserved | -0,02765 | 0,8492 |
| TcCLB.506767.381 | syntaxin binding protein (pseudogene), putative | -0,02767 | 0,9663 |
| TcCLB.509595.30 | hypothetical protein, conserved | -0,02773 | 0,8688 |
| TcCLB.506469.110 | hypothetical protein | -0,0282 | 0,8994 |
| TcCLB.506713.14 | hypothetical protein, conserved | -0,02837 | 0,9055 |
| TcCLB.504011.70 | hypothetical protein, conserved | -0,02841 | 0,9329 |
| TcCLB.509805.160 | hypothetical protein, conserved | -0,02844 | 0,8336 |
| TcCLB.510437.50 | hypothetical protein, conserved | -0,02845 | 0,8722 |
| TcCLB.511239.130 | hypothetical protein, conserved | -0,0285 | 0,8908 |
| TcCLB.506203.100 | hypothetical protein | -0,02861 | 0,8499 |
| TcCLB.503925.70 | hypothetical protein, conserved | -0,02863 | 0,7998 |
| TcCLB.509541.19 | major vault protein, putative | -0,02879 | 0,8252 |
| TcCLB.508405.110 | hypothetical protein, conserved | -0,02886 | 0,8739 |
| TcCLB.507991.80 | hypothetical protein, conserved | -0,029 | 0,8275 |
| TcCLB.507521.140 | hypothetical protein, conserved | -0,02904 | 0,8085 |
| TcCLB.511621.244 | hypothetical protein, conserved | -0,02904 | 0,904 |
| TcCLB.507521.130 | hypothetical protein | -0,02912 | 0,8011 |
| TcCLB.511127.50 | hypothetical protein, conserved | -0,02912 | 0,8649 |
| TcCLB.511815.150 | hypothetical protein | -0,02913 | 0,7761 |
| TcCLB.508559.80 | trans-sialidase (pseudogene), putative | -0,02913 | 0,9213 |
| TcCLB.511239.10 | hypothetical protein, conserved | -0,02914 | 0,8653 |
| TcCLB.509867.50 | hypothetical protein, conserved (pseudogene) | -0,02916 | 0,9471 |
| TcCLB.511221.60 | trans-sialidase, Group V, putative | -0,02919 | 0,9193 |
| TcCLB.504105.170 | hypothetical protein, conserved | -0,02926 | 0,7939 |
| TcCLB.504017.20 | hypothetical protein, conserved | -0,02938 | 0,8434 |
| TcCLB.507747.240 | hypothetical protein, conserved | -0,02956 | 0,8419 |
| TcCLB.511139.30 | hypothetical protein, conserved | -0,0296 | 0,8643 |
| TcCLB.506437.10 | amastin, putative | -0,0296 | 0,8826 |
| TcCLB.506367.130 | hypothetical protein, conserved | -0,02975 | 0,845 |
| TcCLB.506853.20 | hypothetical protein, conserved | -0,02976 | 0,8259 |
| TcCLB.511587.40 | hypothetical protein, conserved (pseudogene) | -0,02997 | 0,8061 |
| TcCLB.509485.70 | retrotransposon hot spot protein (RHS, pseudogene), putat | -0,03001 | 0,8749 |
| TcCLB.506811.110 | hypothetical protein, conserved | -0,03008 | 0,8238 |
| TcCLB.503651.20 | hypothetical protein | -0,03014 | 0,87 |

| | | | |
|---|---|---|---|
| TcCLB.511797.184 | mucin TcMUC (pseudogene), putative | -0,03022 | 0,9656 |
| TcCLB.506227.250 | hypothetical protein, conserved | -0,03038 | 0,8439 |
| TcCLB.508153.350 | hypothetical protein, conserved | -0,03046 | 0,8934 |
| TcCLB.510655.40 | hypothetical protein, conserved | -0,03047 | 0,8422 |
| TcCLB.504175.30 | hypothetical protein, conserved | -0,03049 | 0,8671 |
| TcCLB.506767.140 | mucin-associated surface protein (MASP), putative | -0,03052 | 0,9599 |
| TcCLB.511283.280 | Serine/threonine-protein kinase Nek1-related, putative | -0,03059 | 0,8337 |
| TcCLB.508771.30 | hypothetical protein, conserved | -0,03059 | 0,8773 |
| TcCLB.506635.110 | kinesin-like protein, putative | -0,03061 | 0,8816 |
| TcCLB.510663.10 | hypothetical protein, conserved | -0,03073 | 0,9122 |
| TcCLB.506855.209 | U6 snRNA-associated Sm-like protein LSm7p | -0,03075 | 0,879 |
| TcCLB.508741.10 | hypothetical protein | -0,03077 | 0,8086 |
| TcCLB.511421.230 | Cell cycle checkpoint protein RAD1-like, putative | -0,03088 | 0,8504 |
| TcCLB.510965.29 | hypothetical protein, conserved | -0,03089 | 0,8779 |
| TcCLB.508507.20 | hypothetical protein, conserved | -0,03095 | 0,8561 |
| TcCLB.510625.140 | elongation factor 1-gamma (EF-1-gamma), putative | -0,03101 | 0,8657 |
| TcCLB.511237.80 | hypothetical protein, conserved | -0,0311 | 0,8346 |
| TcCLB.506211.100 | Sarcalumenin, putative | -0,03113 | 0,8209 |
| TcCLB.511727.140 | phenylalanyl-tRNA synthetase alpha chain, putative | -0,03115 | 0,7267 |
| TcCLB.509747.20 | 60S ribosomal protein L22, putative | -0,03122 | 0,7571 |
| TcCLB.503673.9 | eukaryotic translation initiation factor 2-alpha kinase, puta | -0,03125 | 0,8854 |
| TcCLB.511451.90 | mucin-associated surface protein (MASP), putative | -0,03129 | 0,9542 |
| TcCLB.506959.120 | 60S ribosomal protein L30, putative | -0,03144 | 0,7612 |
| TcCLB.509227.20 | mucin-associated surface protein (MASP, pseudogene), pu | -0,03144 | 0,7966 |
| TcCLB.504013.55 | hypothetical protein, conserved | -0,03156 | 0,8784 |
| TcCLB.506513.80 | hypothetical protein, conserved | -0,03159 | 0,8552 |
| TcCLB.506241.100 | 60S ribosomal protein L32, putative | -0,0316 | 0,7521 |
| TcCLB.508541.30 | hypothetical protein | -0,03165 | 0,9647 |
| TcCLB.510761.18 | hypothetical protein, conserved | -0,0317 | 0,834 |
| TcCLB.510149.70 | hypothetical protein, conserved | -0,03173 | 0,7844 |
| TcCLB.511545.60 | hypothetical protein | -0,03173 | 0,919 |
| TcCLB.509693.170 | hypothetical protein, conserved | -0,03182 | 0,8496 |
| TcCLB.509081.140 | mucin TcMUCII, putative | -0,03184 | 0,9536 |
| TcCLB.506241.30 | trans-sialidase, Group II, putative | -0,03192 | 0,9171 |
| TcCLB.509917.12 | N-acetyltransferase complex ARD1 subunit (pseudogene), | -0,03192 | 0,9485 |
| TcCLB.504081.310 | syntaxin binding protein (pseudogene), putative | -0,03208 | 0,9536 |
| TcCLB.510971.69 | hypothetical protein, conserved (fragment) | -0,03217 | 0,8132 |
| TcCLB.507687.20 | dispersed gene family protein 1 (DGF-1), putative | -0,03238 | 0,919 |
| TcCLB.511303.160 | sodium/hydrogen exchanger, putative (FTP) | -0,03249 | 0,8465 |
| TcCLB.503995.30 | hypothetical protein | -0,03264 | 0,8895 |
| TcCLB.506155.30 | kinesin-like protein, putative | -0,03269 | 0,8074 |
| TcCLB.507787.20 | hypothetical protein, conserved | -0,03279 | 0,7876 |
| TcCLB.511445.100 | hypothetical protein, conserved | -0,03285 | 0,8061 |
| TcCLB.508277.30 | hypothetical protein, conserved | -0,03286 | 0,815 |
| TcCLB.508641.80 | hypothetical protein, conserved | -0,03292 | 0,8345 |

| | | | |
|---|---|---|---|
| TcCLB.506239.50 | palmitoyl acyltransferase 6, putative | -0,03294 | 0,8769 |
| TcCLB.511529.160 | enoyl-CoA hydratase/isomerase family protein, putative | -0,03313 | 0,8086 |
| TcCLB.509817.40 | trans-sialidase (pseudogene), putative | -0,0332 | 0,884 |
| TcCLB.503813.10 | hypothetical protein, conserved | -0,03328 | 0,82 |
| TcCLB.511517.20 | hypothetical protein, conserved | -0,0333 | 0,8256 |
| TcCLB.506717.100 | mucin-associated surface protein (MASP, pseudogene), pu | -0,03339 | 0,8974 |
| TcCLB.510213.20 | hypothetical protein | -0,03355 | 0,9639 |
| TcCLB.505171.10 | phospholipid transporting ATPase-like protein, putative | -0,03361 | 0,832 |
| TcCLB.510819.90 | hypothetical protein, conserved | -0,03363 | 0,8609 |
| TcCLB.510797.10 | 2-oxoglutarate dehydrogenase subunit, putative | -0,03383 | 0,8285 |
| TcCLB.509109.145 | hypothetical protein, conserved (pseudogene) | -0,03407 | 0,9 |
| TcCLB.507605.26 | mucin TcMUCII, putative | -0,03418 | 0,9747 |
| TcCLB.508645.10 | SNF2 family protein | -0,03419 | 0,7934 |
| TcCLB.508507.50 | hypothetical protein, conserved | -0,0342 | 0,8799 |
| TcCLB.452039.9 | hypothetical protein, conserved | -0,03427 | 0,79 |
| TcCLB.509807.29 | hypothetical protein, conserved | -0,03427 | 0,8752 |
| TcCLB.507031.50 | hypothetical protein, conserved | -0,03427 | 0,9091 |
| TcCLB.511127.100 | catalytic subunit of the vacuolar transporter chaperone 4, p | -0,03435 | 0,777 |
| TcCLB.506835.30 | 60S ribosomal protein L9, putative | -0,03436 | 0,6953 |
| TcCLB.506979.5 | hypothetical protein, conserved (pseudogene) | -0,03439 | 0,9306 |
| TcCLB.447135.10 | surface protease GP63, putative | -0,03453 | 0,9636 |
| TcCLB.506193.20 | hypothetical protein, conserved | -0,03456 | 0,8297 |
| TcCLB.507997.80 | hypothetical protein, conserved | -0,03456 | 0,9139 |
| TcCLB.508461.589 | hypothetical protein, conserved (fragment) | -0,03462 | 0,9093 |
| TcCLB.508317.40 | hypothetical protein, conserved | -0,03464 | 0,876 |
| TcCLB.505071.119 | hypothetical protein, conserved | -0,0347 | 0,854 |
| TcCLB.506413.70 | hypothetical protein, conserved | -0,03482 | 0,8268 |
| TcCLB.503829.89 | spliceosomal U5 snRNP-specific protein, putative | -0,03488 | 0,8928 |
| TcCLB.510603.150 | hypothetical protein, conserved | -0,03492 | 0,8589 |
| TcCLB.507087.70 | hypothetical protein, conserved | -0,03501 | 0,7993 |
| TcCLB.508075.20 | DNA repair protein, putative | -0,0351 | 0,8358 |
| TcCLB.504051.9 | hypothetical protein, conserved | -0,03516 | 0,8622 |
| TcCLB.506887.60 | small nuclear RNA gene activation protein (SNAP) 50, puta | -0,03519 | 0,7708 |
| TcCLB.506685.40 | hypothetical protein, conserved | -0,0354 | 0,807 |
| TcCLB.507585.10 | amino acid transporter, putative | -0,03544 | 0,6349 |
| TcCLB.506807.30 | hypothetical protein, conserved | -0,03546 | 0,8711 |
| TcCLB.506675.70 | dispersed gene family protein 1 (DGF-1), putative | -0,03552 | 0,8943 |
| TcCLB.506689.20 | hypothetical protein, conserved (pseudogene) | -0,03554 | 0,8535 |
| TcCLB.508325.180 | retrotransposon hot spot protein (RHS, pseudogene), putat | -0,03566 | 0,8078 |
| TcCLB.509201.10 | hypothetical protein, conserved | -0,03569 | 0,7984 |
| TcCLB.511737.40 | hypothetical protein, conserved | -0,03586 | 0,8175 |
| TcCLB.507519.180 | hypothetical protein, conserved | -0,03595 | 0,8473 |
| TcCLB.418185.46 | N-acetyltransferase complex ARD1 subunit (pseudogene), | -0,03606 | 0,9301 |
| TcCLB.509715.120 | phosphatidylinositol-4-phosphate 5-kinase, putative | -0,0362 | 0,8135 |
| TcCLB.511649.110 | hypothetical protein, conserved | -0,03631 | 0,7876 |

| | | | |
|---|---|---|---|
| TcCLB.511907.250 | hypothetical protein | -0,03644 | 0,7721 |
| TcCLB.506251.120 | hypothetical protein, conserved | -0,03646 | 0,88 |
| TcCLB.506951.10 | retrotransposon hot spot protein (RHS, pseudogene), putat | -0,03647 | 0,9296 |
| TcCLB.510735.20 | hypothetical protein, conserved | -0,03657 | 0,8355 |
| TcCLB.504153.260 | hypothetical protein, conserved | -0,0366 | 0,7462 |
| TcCLB.507625.80 | hypothetical protein, conserved | -0,03665 | 0,8092 |
| TcCLB.504159.109 | hypothetical protein, conserved | -0,03665 | 0,8859 |
| TcCLB.510091.110 | mitochondrial RNA binding complex 1 subunit, putative | -0,03679 | 0,82 |
| TcCLB.510231.10 | eukaryotic translation initiation factor 2-alpha kinase, puta | -0,03686 | 0,8438 |
| TcCLB.510543.200 | trans-sialidase (pseudogene), putative | -0,03697 | 0,9347 |
| TcCLB.507739.70 | hypothetical protein, conserved | -0,037 | 0,7985 |
| TcCLB.506247.210 | hypothetical protein, conserved | -0,03708 | 0,671 |
| TcCLB.510421.130 | mitochondrial RNA binding complex 1 subunit, putative | -0,03726 | 0,789 |
| TcCLB.511255.470 | retrotransposon hot spot protein (RHS, pseudogene), putat | -0,03738 | 0,922 |
| TcCLB.507687.30 | retrotransposon hot spot protein (RHS, pseudogene), putat | -0,03748 | 0,8668 |
| TcCLB.506559.364 | hypothetical protein, conserved | -0,0375 | 0,7006 |
| TcCLB.506763.130 | mucin-associated surface protein (MASP), putative | -0,03751 | 0,9346 |
| TcCLB.510295.40 | Protein C21orf2 homolog, putative | -0,03756 | 0,8181 |
| TcCLB.509235.10 | UDP-Gal or UDP-GlcNAc-dependent glycosyltransferase, pu | -0,03793 | 0,9051 |
| TcCLB.416143.10 | Lys-63-specific deubiquitinase BRCC36, putative | -0,03794 | 0,7858 |
| TcCLB.503635.40 | hypothetical protein | -0,03803 | 0,8171 |
| TcCLB.507681.180 | hypothetical protein, conserved | -0,03803 | 0,8176 |
| TcCLB.503987.39 | hypothetical protein, conserved | -0,03822 | 0,8784 |
| TcCLB.508899.80 | hypothetical protein, conserved | -0,0383 | 0,7861 |
| TcCLB.504125.80 | hypothetical protein, conserved | -0,03836 | 0,7765 |
| TcCLB.511881.20 | serine/threonine-protein phosphatase | -0,03842 | 0,8254 |
| TcCLB.511671.70 | glucosamine 6-phosphate n-acetyltransferase, putative | -0,03846 | 0,8661 |
| TcCLB.503967.19 | hypothetical protein | -0,03857 | 0,8897 |
| TcCLB.507011.40 | hypothetical protein, conserved | -0,03861 | 0,8052 |
| TcCLB.509795.60 | tRNA pseudouridine synthase TruD, putative | -0,03863 | 0,7915 |
| TcCLB.511837.60 | hypothetical protein, conserved | -0,03863 | 0,8118 |
| TcCLB.509213.20 | hypothetical protein, conserved | -0,03863 | 0,8465 |
| TcCLB.511215.100 | hypothetical protein | -0,03869 | 0,8783 |
| TcCLB.508871.170 | trans-sialidase, Group V, putative | -0,03883 | 0,9091 |
| TcCLB.510197.47 | mucin TcMUCII (pseudogene), putative | -0,03886 | 0,9188 |
| TcCLB.508141.30 | hypothetical protein, conserved | -0,03898 | 0,7788 |
| TcCLB.508385.10 | hypothetical protein, conserved | -0,03905 | 0,7486 |
| TcCLB.503395.40 | 60S ribosomal protein L18, putative | -0,03915 | 0,6987 |
| TcCLB.510329.80 | hypothetical protein, conserved | -0,03921 | 0,7537 |
| TcCLB.511421.40 | transmembrane protein, putative | -0,03921 | 0,8302 |
| TcCLB.506983.60 | DNA excision repair protein, putative | -0,03937 | 0,7241 |
| TcCLB.504081.50 | mucin-associated surface protein (MASP), putative | -0,03938 | 0,9171 |
| TcCLB.511557.70 | hypothetical protein, conserved | -0,0395 | 0,7545 |
| TcCLB.511277.4 | 30S ribosomal protein S8, putative (fragment) | -0,03958 | 0,8566 |
| TcCLB.506667.5 | surface protease GP63 (pseudogene), putative | -0,03968 | 0,9512 |

| | | | |
|---|---|---|---|
| TcCLB.507681.230 | hypothetical protein, conserved | -0,03974 | 0,7956 |
| TcCLB.509755.50 | trans-sialidase, Group V, putative | -0,0398 | 0,9163 |
| TcCLB.507795.90 | hypothetical protein, conserved | -0,03981 | 0,8575 |
| TcCLB.506777.24 | hypothetical protein, conserved | -0,03986 | 0,8504 |
| TcCLB.508645.20 | hypothetical protein | -0,03998 | 0,8016 |
| TcCLB.508277.210 | hypothetical protein, conserved | -0,04001 | 0,8074 |
| TcCLB.511707.30 | hypothetical protein, conserved | -0,04016 | 0,7477 |
| TcCLB.507669.210 | hypothetical protein, conserved | -0,04017 | 0,831 |
| TcCLB.506175.80 | hypothetical protein, conserved | -0,0402 | 0,8753 |
| TcCLB.508373.20 | dihydroorotase, putative | -0,04024 | 0,8041 |
| TcCLB.506883.34 | hypothetical protein, conserved | -0,04032 | 0,8246 |
| TcCLB.408437.29 | hypothetical protein (fragment) | -0,04041 | 0,7898 |
| TcCLB.511041.50 | flagellum transition zone component, putative | -0,04046 | 0,6661 |
| TcCLB.509453.4 | hypothetical protein, conserved | -0,04055 | 0,8469 |
| TcCLB.511167.60 | hypothetical protein, conserved | -0,04058 | 0,7489 |
| TcCLB.507519.140 | Adenine phosphoribosyltransferase, putative | -0,0406 | 0,7949 |
| TcCLB.504625.74 | hypothetical protein, conserved | -0,04061 | 0,8979 |
| TcCLB.508687.25 | hypothetical protein | -0,04061 | 0,9092 |
| TcCLB.511635.40 | protein tyrosine phosphatase, putative | -0,04077 | 0,6976 |
| TcCLB.482319.19 | hypothetical protein, conserved | -0,04077 | 0,8612 |
| TcCLB.510943.26 | ribosomal protein L38, putative | -0,04098 | 0,7844 |
| TcCLB.511649.130 | vesicle-associated membrane protein, putative | -0,041 | 0,7709 |
| TcCLB.507017.60 | hypothetical protein, conserved | -0,04104 | 0,7971 |
| TcCLB.461879.10 | retrotransposon hot spot protein (RHS, pseudogene), putat | -0,04107 | 0,8629 |
| TcCLB.504131.120 | hypothetical protein, conserved | -0,04108 | 0,809 |
| TcCLB.504827.120 | hypothetical protein, conserved | -0,0411 | 0,8151 |
| TcCLB.506709.10 | hypothetical protein, conserved | -0,04111 | 0,781 |
| TcCLB.511589.100 | hypothetical protein, conserved | -0,04112 | 0,8921 |
| TcCLB.510303.119 | hypothetical protein, conserved | -0,04114 | 0,8247 |
| TcCLB.510933.160 | mucin-associated surface protein (MASP, pseudogene), pu | -0,04123 | 0,918 |
| TcCLB.511529.50 | hypothetical protein, conserved | -0,04128 | 0,8812 |
| TcCLB.506603.20 | mucin-associated surface protein (MASP, pseudogene), pu | -0,04136 | 0,9314 |
| TcCLB.507811.60 | vesicle-associated membrane protein, putative | -0,04157 | 0,7872 |
| TcCLB.508717.36 | selenophosphate synthetase, putative | -0,04159 | 0,7791 |
| TcCLB.511545.20 | ribosomal protein L27, putative | -0,04174 | 0,6975 |
| TcCLB.509997.86 | serine/threonine protein phosphatase, putative (fragment) | -0,04185 | 0,7661 |
| TcCLB.510747.90 | hypothetical protein | -0,04203 | 0,7927 |
| TcCLB.503879.119 | protein transport protein Sec24C, putative | -0,04215 | 0,7826 |
| TcCLB.506979.30 | hypothetical protein | -0,0422 | 0,9298 |
| TcCLB.507071.89 | mucin TcMUCII (pseudogene), putative | -0,04224 | 0,9393 |
| TcCLB.509669.20 | subtilisin-like serine peptidase, putative | -0,04234 | 0,7186 |
| TcCLB.508647.220 | mitochondrial RNA binding protein 1 | -0,04236 | 0,6656 |
| TcCLB.506529.620 | hypothetical protein | -0,04236 | 0,8346 |
| TcCLB.509721.10 | MRB1 complex subunit MRB3010 (MRB3010) | -0,04238 | 0,8527 |
| TcCLB.506857.100 | endoplasmatic reticulum retrieval protein, putative | -0,04243 | 0,732 |

| | | | |
|---|---|---|---|
| TcCLB.509985.20 | hypothetical protein, conserved | -0,04244 | 0,8037 |
| TcCLB.508777.120 | hypothetical protein, conserved | -0,04246 | 0,8389 |
| TcCLB.509179.220 | hypothetical protein, conserved | -0,04248 | 0,8292 |
| TcCLB.508357.23 | hypothetical protein, conserved | -0,04252 | 0,8297 |
| TcCLB.506435.170 | hypothetical protein, conserved | -0,04252 | 0,8784 |
| TcCLB.508637.140 | hypothetical protein, conserved | -0,04254 | 0,7708 |
| TcCLB.507071.240 | mucin-associated surface protein (MASP), putative | -0,04269 | 0,9319 |
| TcCLB.506335.20 | trans-sialidase (pseudogene), putative | -0,04272 | 0,9141 |
| TcCLB.506661.100 | hypothetical protein, conserved | -0,0428 | 0,8984 |
| TcCLB.432995.9 | trans-sialidase, putative | -0,04298 | 0,9612 |
| TcCLB.506441.10 | hypothetical protein, conserved | -0,04307 | 0,7046 |
| TcCLB.510377.30 | mucin-associated surface protein (MASP, pseudogene), pu | -0,04317 | 0,9499 |
| TcCLB.510603.30 | hypothetical protein, conserved | -0,04318 | 0,9152 |
| TcCLB.503805.3 | kinesin heavy chain, putative | -0,0432 | 0,8705 |
| TcCLB.511829.10 | dispersed gene family protein 1 (DGF-1), putative | -0,0433 | 0,7595 |
| TcCLB.511211.180 | hypothetical protein, conserved | -0,04335 | 0,7681 |
| TcCLB.511647.60 | hypothetical protein, conserved | -0,04342 | 0,7462 |
| TcCLB.504867.30 | hypothetical protein, conserved | -0,04347 | 0,8957 |
| TcCLB.511261.40 | hypothetical protein | -0,04362 | 0,7932 |
| TcCLB.508461.480 | 60S ribosomal protein L23, putative | -0,04364 | 0,6016 |
| TcCLB.511717.30 | chromosomal passenger protein, putative (CPC1) | -0,0437 | 0,7943 |
| TcCLB.511279.60 | hypothetical protein, conserved | -0,04376 | 0,7243 |
| TcCLB.508307.10 | hypothetical protein, conserved | -0,04381 | 0,9042 |
| TcCLB.406999.10 | dispersed gene family protein 1 (DGF-1), putative (fragme | -0,04393 | 0,9542 |
| TcCLB.511581.20 | fatty acyl CoA syntetase 1 | -0,04394 | 0,7835 |
| TcCLB.508815.120 | hypothetical protein, conserved | -0,044 | 0,79 |
| TcCLB.506227.180 | ras-related protein RAB5B, putative (RAB5B) | -0,04409 | 0,7422 |
| TcCLB.511753.140 | vacuolar assembly protein vps41, putative | -0,0441 | 0,8544 |
| TcCLB.507633.110 | mucin-associated surface protein (MASP, pseudogene), pu | -0,04412 | 0,9172 |
| TcCLB.503973.260 | syntaxin binding protein (pseudogene), putative | -0,04412 | 0,9291 |
| TcCLB.508221.180 | trans-sialidase, Group V, putative | -0,04426 | 0,9273 |
| TcCLB.506629.10 | intraflagellar transport protein 88, putative (fragment) (IF | -0,04435 | 0,8406 |
| TcCLB.506831.10 | hypothetical protein | -0,04436 | 0,7805 |
| TcCLB.504057.178 | hypothetical protein, conserved | -0,04439 | 0,8045 |
| TcCLB.473709.10 | trans-sialidase, putative (fragment) | -0,04444 | 0,9553 |
| TcCLB.510325.50 | DNA replication complex GINS protein SLD5, putative | -0,04446 | 0,8491 |
| TcCLB.504137.120 | hypothetical protein, conserved | -0,04451 | 0,79 |
| TcCLB.503957.20 | retrotransposon hot spot (RHS) protein, putative | -0,04454 | 0,7869 |
| TcCLB.511649.70 | structural maintenance of chromosome 3 protein, putative | -0,04462 | 0,8008 |
| TcCLB.511733.40 | FUN14 family, putative (POMP32) | -0,04481 | 0,8489 |
| TcCLB.507615.120 | hypothetical protein, conserved | -0,04493 | 0,842 |
| TcCLB.507941.60 | hypothetical protein, conserved | -0,04498 | 0,8337 |
| TcCLB.508375.50 | dihydroorotate dehydrogenase (fumarate), putative (DHOD | -0,04503 | 0,8107 |
| TcCLB.506503.160 | hypothetical protein, conserved | -0,04506 | 0,6639 |
| TcCLB.506581.50 | hypothetical protein, conserved | -0,04523 | 0,8248 |

| | | | |
|---|---|---|---|
| TcCLB.504147.99 | hypothetical protein | -0,04528 | 0,6841 |
| TcCLB.510759.4 | hypothetical protein, conserved | -0,04538 | 0,8211 |
| TcCLB.506855.350 | hypothetical protein | -0,04538 | 0,8296 |
| TcCLB.511651.19 | delta-adaptin, putative | -0,04539 | 0,8752 |
| TcCLB.503953.14 | hypothetical protein, conserved | -0,04547 | 0,8229 |
| TcCLB.506485.40 | kinesin (pseudogene), putative | -0,04588 | 0,645 |
| TcCLB.508347.100 | hypothetical protein, conserved | -0,0459 | 0,7438 |
| TcCLB.511717.50 | hypothetical protein, conserved | -0,04595 | 0,7752 |
| TcCLB.508857.80 | hypothetical protein, conserved | -0,04595 | 0,8337 |
| TcCLB.511671.50 | hypothetical protein, conserved | -0,04611 | 0,6385 |
| TcCLB.504147.80 | metallo-peptidase, Clan MA(E) Family M3 | -0,04617 | 0,6023 |
| TcCLB.509753.100 | mucin-associated surface protein (MASP), putative | -0,04631 | 0,9325 |
| TcCLB.511907.60 | hypothetical protein, conserved | -0,04633 | 0,8076 |
| TcCLB.506753.50 | hypothetical protein, conserved | -0,04647 | 0,7454 |
| TcCLB.504019.3 | hypothetical protein (fragment) | -0,04659 | 0,8194 |
| TcCLB.511149.190 | mucin-associated surface protein (MASP, pseudogene), pu | -0,04663 | 0,8952 |
| TcCLB.507433.9 | ATP-dependent DEAD/H DNA helicase recQ, putative (fragı | -0,04666 | 0,7008 |
| TcCLB.509895.100 | hypothetical protein, conserved | -0,04671 | 0,798 |
| TcCLB.506869.70 | mitochondrial structure specific endonuclease I (SSE-1), pu | -0,04676 | 0,7881 |
| TcCLB.508153.770 | hypothetical protein, conserved | -0,04681 | 0,7268 |
| TcCLB.508163.170 | trans-sialidase (pseudogene), putative | -0,04689 | 0,9297 |
| TcCLB.507207.19 | FtsJ cell division protein, putative (fragment) | -0,04693 | 0,5886 |
| TcCLB.509691.10 | hypothetical protein, conserved | -0,04695 | 0,863 |
| TcCLB.508681.20 | hypothetical protein | -0,04712 | 0,9407 |
| TcCLB.510877.120 | NADPH-dependent FMN/FAD containing oxidoreductase, p | -0,04716 | 0,7304 |
| TcCLB.506599.80 | mucin-associated surface protein (MASP), putative | -0,04724 | 0,9417 |
| TcCLB.508291.10 | mucin-associated surface protein (MASP, pseudogene), pu | -0,04727 | 0,9103 |
| TcCLB.508379.29 | hypothetical protein, conserved | -0,04733 | 0,8762 |
| TcCLB.408573.9 | hypothetical protein, conserved | -0,04735 | 0,8214 |
| TcCLB.510799.9 | hypothetical protein, conserved | -0,04742 | 0,8253 |
| TcCLB.511019.90 | iron superoxide dismutase, putative | -0,04748 | 0,6825 |
| TcCLB.509455.50 | hypothetical protein, conserved | -0,04749 | 0,7747 |
| TcCLB.503583.10 | metallo-peptidase, Clan MC, Family M14, putative | -0,04749 | 0,7871 |
| TcCLB.507251.20 | ribosomal protein L21E (60S), putative | -0,04751 | 0,5278 |
| TcCLB.507711.180 | hypothetical protein, conserved | -0,04754 | 0,7517 |
| TcCLB.511657.70 | hypothetical protein, conserved | -0,04755 | 0,7433 |
| TcCLB.508207.20 | protein kinase, putative | -0,04763 | 0,8693 |
| TcCLB.511733.130 | chaperonin HSP60, putative (fragment) | -0,04764 | 0,8268 |
| TcCLB.506957.23 | hypothetical protein, conserved | -0,04769 | 0,6689 |
| TcCLB.510687.40 | serine/threonine protein phosphatase, putative | -0,04778 | 0,7209 |
| TcCLB.511621.150 | hypothetical protein, conserved | -0,04786 | 0,7128 |
| TcCLB.510885.30 | hypothetical protein | -0,04788 | 0,7387 |
| TcCLB.510149.80 | ABC transporter, putative | -0,0479 | 0,594 |
| TcCLB.508209.80 | hypothetical protein, conserved | -0,04802 | 0,7756 |
| TcCLB.504039.250 | surface protease GP63 (pseudogene), putative | -0,04802 | 0,9027 |

| | | | |
|---|---|---|---|
| TcCLB.468005.9 | pumilio/PUF RNA binding protein 3, putative (PUF3) | -0,04809 | 0,7166 |
| TcCLB.511615.10 | trans-sialidase, Group V, putative | -0,04826 | 0,925 |
| TcCLB.504077.60 | hypothetical protein, conserved (pseudogene) | -0,0483 | 0,8974 |
| TcCLB.503477.30 | hypothetical protein, conserved | -0,04833 | 0,8268 |
| TcCLB.510725.99 | Chaperonin HSP60, mitochondrial (HSP60) | -0,04841 | 0,873 |
| TcCLB.507669.170 | WD40/YVTN repeat-like-containing protein | -0,04845 | 0,6736 |
| TcCLB.511715.30 | hypothetical protein, conserved | -0,04849 | 0,8407 |
| TcCLB.509953.9 | ubiquitin carboxyl-terminal hydrolase, putative (fragment) | -0,0485 | 0,8364 |
| TcCLB.510321.20 | hypothetical protein, conserved | -0,04853 | 0,7617 |
| TcCLB.506559.370 | hypothetical protein, conserved | -0,0486 | 0,7535 |
| TcCLB.509985.30 | hypothetical protein, conserved | -0,04872 | 0,8007 |
| TcCLB.511099.220 | mucin-associated surface protein (MASP), putative | -0,04878 | 0,9376 |
| TcCLB.507843.10 | retrotransposon hot spot (RHS) protein, putative | -0,04883 | 0,79 |
| TcCLB.507907.20 | trans-sialidase, Group II, putative | -0,04886 | 0,8656 |
| TcCLB.508175.20 | hypothetical protein, conserved | -0,04888 | 0,6178 |
| TcCLB.504181.30 | 40S ribosomal protein S23, putative | -0,04888 | 0,8583 |
| TcCLB.511711.50 | hypothetical protein, conserved | -0,04895 | 0,7823 |
| TcCLB.506725.30 | hypothetical protein, conserved | -0,04914 | 0,6251 |
| TcCLB.511819.59 | dynein heavy chain, cytosolic, putative | -0,04917 | 0,7197 |
| TcCLB.510431.310 | hypothetical protein, conserved | -0,04924 | 0,8869 |
| TcCLB.503811.40 | PTS2 receptor, putative (PEX7) | -0,04958 | 0,8162 |
| TcCLB.509911.80 | hypothetical protein, conserved | -0,04963 | 0,7966 |
| TcCLB.509055.20 | hypothetical protein, conserved | -0,04964 | 0,7176 |
| TcCLB.509875.200 | copper homeostasis protein, putative | -0,04966 | 0,7482 |
| TcCLB.506279.10 | trans-sialidase, Group VI, putative | -0,04972 | 0,8735 |
| TcCLB.507915.10 | hypothetical protein, conserved | -0,04982 | 0,8895 |
| TcCLB.504425.80 | hypothetical protein, conserved | -0,05001 | 0,7536 |
| TcCLB.510257.120 | hypothetical protein, conserved | -0,05006 | 0,7639 |
| TcCLB.511797.210 | trans-sialidase (pseudogene), putative | -0,05006 | 0,8878 |
| TcCLB.509733.50 | hypothetical protein, conserved | -0,05007 | 0,7502 |
| TcCLB.509245.10 | hypothetical protein, conserved | -0,0501 | 0,7899 |
| TcCLB.506767.21 | syntaxin binding protein (pseudogene), putative | -0,05018 | 0,9236 |
| TcCLB.508207.194 | hypothetical protein, conserved | -0,05039 | 0,7037 |
| TcCLB.508177.60 | hypothetical protein, conserved | -0,05041 | 0,7757 |
| TcCLB.510761.73 | dynein-associated protein, putative | -0,05042 | 0,7902 |
| TcCLB.460757.20 | hypothetical protein (fragment) | -0,05043 | 0,926 |
| TcCLB.510279.90 | mucin-associated surface protein (MASP, pseudogene), pu | -0,05049 | 0,9215 |
| TcCLB.511015.20 | ribosomal protein S29, putative | -0,05055 | 0,8025 |
| TcCLB.511715.100 | pumilio/PUF RNA binding protein 7, putative (PUF7) | -0,05064 | 0,6955 |
| TcCLB.509069.14 | hypothetical protein, conserved | -0,05069 | 0,7769 |
| TcCLB.511445.90 | hypothetical protein, conserved | -0,05071 | 0,6706 |
| TcCLB.510229.20 | 8-oxoguanine DNA glycosylase, putative | -0,05072 | 0,7821 |
| TcCLB.509645.19 | hypothetical protein, conserved | -0,05078 | 0,7676 |
| TcCLB.503485.20 | hypothetical protein, conserved (pseudogene) | -0,05079 | 0,841 |
| TcCLB.503973.350 | hypothetical protein | -0,05081 | 0,9306 |

| | | | |
|---|---|---|---|
| TcCLB.506733.50 | actin-like protein, putative | -0,05085 | 0,7425 |
| TcCLB.507595.10 | dispersed gene family protein 1 (DGF-1, pseudogene), puta | -0,05096 | 0,9139 |
| TcCLB.510765.50 | hypothetical protein, conserved | -0,05102 | 0,79 |
| TcCLB.509429.140 | kinetoplastid kinetochore protein 6, putative (kkt6) | -0,05112 | 0,8061 |
| TcCLB.510431.240 | hypothetical protein, conserved | -0,05115 | 0,7405 |
| TcCLB.510955.50 | hypothetical protein, conserved | -0,05115 | 0,8313 |
| TcCLB.510655.110 | poly(A)-specific ribonuclease PARN, putative | -0,05129 | 0,7249 |
| TcCLB.509073.60 | hypothetical protein, conserved | -0,05144 | 0,7766 |
| TcCLB.504147.120 | 60S ribosomal protein L22, putative | -0,05146 | 0,551 |
| TcCLB.509037.40 | eukaryotic translation initiation factor 4E, putative | -0,05154 | 0,6661 |
| TcCLB.506281.14 | hypothetical protein | -0,05154 | 0,8348 |
| TcCLB.503493.10 | hypothetical protein, conserved | -0,05157 | 0,7462 |
| TcCLB.508589.41 | N-acetyltransferase complex ARD1 subunit (pseudogene), | -0,05174 | 0,9251 |
| TcCLB.507209.60 | hypothetical protein, conserved | -0,05182 | 0,6891 |
| TcCLB.509875.210 | 60S ribosomal protein L17, putative | -0,05183 | 0,5753 |
| TcCLB.474339.9 | hypothetical protein, conserved | -0,05185 | 0,7648 |
| TcCLB.511283.310 | hypothetical protein, conserved | -0,05191 | 0,6591 |
| TcCLB.508589.51 | ATP-dependent DEAD/H RNA helicase (pseudogene), putat | -0,05192 | 0,8933 |
| TcCLB.509551.50 | hypothetical protein | -0,05197 | 0,5999 |
| TcCLB.509707.20 | hypothetical protein, conserved | -0,05197 | 0,6385 |
| TcCLB.508539.169 | trans-sialidase (pseudogene), putative | -0,05203 | 0,8947 |
| TcCLB.506503.50 | hypothetical protein, conserved | -0,05211 | 0,6961 |
| TcCLB.507521.115 | hypothetical protein, conserved | -0,05214 | 0,8375 |
| TcCLB.511643.10 | hypothetical protein, conserved | -0,05238 | 0,7651 |
| TcCLB.509083.9 | ATP-dependent DEAD/H RNA helicase, putative | -0,0526 | 0,832 |
| TcCLB.507093.229 | U-rich RNA-binding protein UBP-2 | -0,05283 | 0,6555 |
| TcCLB.509641.50 | hypothetical protein, conserved | -0,05285 | 0,7333 |
| TcCLB.504055.40 | dihydroxyacetone phosphate acyltransferase, putative (DAT | -0,05291 | 0,6273 |
| TcCLB.507937.50 | mucin-associated surface protein (MASP), putative | -0,05307 | 0,8991 |
| TcCLB.510485.9 | dispersed gene family protein 1 (DGF-1), putative | -0,05314 | 0,8897 |
| TcCLB.509551.149 | phosphatidylserine decarboxylase, putative | -0,05318 | 0,8544 |
| TcCLB.506189.10 | nucleolar protein, putative | -0,05327 | 0,7305 |
| TcCLB.509395.80 | hypothetical protein, conserved | -0,05328 | 0,697 |
| TcCLB.509099.130 | hypothetical protein, conserved | -0,05333 | 0,671 |
| TcCLB.509445.10 | tryparedoxin peroxidase, putative | -0,05333 | 0,7945 |
| TcCLB.506335.150 | mucin-associated surface protein (MASP, pseudogene), pu | -0,05336 | 0,8952 |
| TcCLB.511495.20 | hypothetical protein, conserved | -0,05337 | 0,7509 |
| TcCLB.507711.270 | hypothetical protein, conserved | -0,05342 | 0,7703 |
| TcCLB.504203.20 | sec1-like protein, putative | -0,05345 | 0,653 |
| TcCLB.511249.4 | RNA editing 3' terminal uridylyl transferase 1, putative | -0,05347 | 0,8075 |
| TcCLB.504161.30 | hypothetical protein, conserved | -0,05354 | 0,797 |
| TcCLB.509267.20 | hypothetical protein | -0,05355 | 0,757 |
| TcCLB.503965.39 | hypothetical protein, conserved | -0,05361 | 0,8181 |
| TcCLB.508389.90 | surface protease GP63, putative | -0,05363 | 0,8985 |
| TcCLB.508873.103 | mucin-associated surface protein (MASP, pseudogene), pu | -0,05366 | 0,9312 |

| | | | |
|---|---|---|---|
| TcCLB.510743.40 | prenyl protein specific carboxyl methyltransferase, putative | -0,05368 | 0,7659 |
| TcCLB.507541.40 | casein kinase, putative | -0,05373 | 0,4937 |
| TcCLB.510719.160 | 60S ribosomal protein L22, putative | -0,05373 | 0,5368 |
| TcCLB.506703.50 | mucin-associated surface protein (MASP), putative | -0,0539 | 0,8737 |
| TcCLB.506769.40 | hypothetical protein | -0,05415 | 0,9471 |
| TcCLB.507035.120 | hypothetical protein, conserved | -0,05437 | 0,7577 |
| TcCLB.508175.39 | hypothetical protein, conserved | -0,05458 | 0,7687 |
| TcCLB.509253.20 | tubulin tyrosine ligase, putative | -0,05469 | 0,7652 |
| TcCLB.506989.250 | hypothetical protein, conserved | -0,05474 | 0,6908 |
| TcCLB.508027.50 | hypothetical protein, conserved | -0,05479 | 0,7904 |
| TcCLB.507007.30 | hypothetical protein | -0,05484 | 0,7576 |
| TcCLB.506321.101 | hypothetical protein, conserved (pseudogene) | -0,05491 | 0,9133 |
| TcCLB.508303.30 | hypothetical protein, conserved | -0,05503 | 0,7888 |
| TcCLB.506311.20 | palmitoyl acyltransferase 2, putative | -0,05504 | 0,739 |
| TcCLB.509799.120 | hypothetical protein, conserved | -0,05504 | 0,776 |
| TcCLB.506575.60 | diacylglycerol kinase, putative | -0,05512 | 0,7357 |
| TcCLB.506327.20 | hypothetical protein, conserved | -0,05513 | 0,7844 |
| TcCLB.511911.94 | hypothetical protein, conserved | -0,05516 | 0,6602 |
| TcCLB.479517.50 | surface protease GP63, putative | -0,05524 | 0,8928 |
| TcCLB.507499.53 | ATP-dependent DEAD/H RNA helicase (pseudogene), putat | -0,05528 | 0,8406 |
| TcCLB.509715.9 | Ribosomal RNA processing protein 41A (RRP41A) | -0,05533 | 0,7717 |
| TcCLB.510901.10 | hypothetical protein, conserved | -0,05539 | 0,7463 |
| TcCLB.509999.40 | COPII coat assembly protein sec16, putative (sec16) | -0,05542 | 0,5541 |
| TcCLB.506213.80 | 60S ribosomal protein L17, putative | -0,05544 | 0,7621 |
| TcCLB.509931.20 | trans-sialidase, Group VIII, putative | -0,05545 | 0,875 |
| TcCLB.511291.110 | hypothetical protein, conserved | -0,05546 | 0,7405 |
| TcCLB.507673.20 | 60S ribosomal protein L13a, putative | -0,05547 | 0,6318 |
| TcCLB.507761.10 | dispersed gene family protein 1 (DGF-1, pseudogene), puta | -0,05552 | 0,8683 |
| TcCLB.511633.79 | microtubule-associated protein, putative | -0,05573 | 0,7144 |
| TcCLB.506989.20 | hypothetical protein, conserved | -0,05578 | 0,6457 |
| TcCLB.506667.170 | hypothetical protein | -0,05581 | 0,9203 |
| TcCLB.506789.170 | ATPase, putative | -0,05583 | 0,7318 |
| TcCLB.508559.29 | dispersed gene family protein 1 (DGF-1), putative | -0,05583 | 0,7747 |
| TcCLB.508521.11 | trans-sialidase (pseudogene), putative | -0,05602 | 0,8393 |
| TcCLB.474887.9 | pseudouridine synthase TruD, putative (fragment) | -0,05604 | 0,7719 |
| TcCLB.508165.290 | mucin TcMUCII, putative | -0,05622 | 0,9474 |
| TcCLB.510029.70 | 40S ribosomal protein S13, putative | -0,0563 | 0,6164 |
| TcCLB.507093.140 | hypothetical protein, conserved | -0,05631 | 0,7191 |
| TcCLB.511585.190 | eukaryotic initiation factor 4a, putative | -0,05636 | 0,5503 |
| TcCLB.506631.10 | hypothetical protein, conserved | -0,05643 | 0,7855 |
| TcCLB.507041.65 | hypothetical protein, conserved (pseudogene) | -0,05645 | 0,907 |
| TcCLB.511735.10 | chaperonin HSP60/CNP60, putative | -0,05649 | 0,806 |
| TcCLB.506057.101 | surface protease GP63, putative (fragment) | -0,05651 | 0,963 |
| TcCLB.511291.100 | Vesicle-trafficking protein SEC22, putative (SEC22) | -0,05656 | 0,7639 |
| TcCLB.506591.81 | hypothetical protein, conserved (pseudogene) | -0,05661 | 0,7881 |

| | | | |
|---|---|---|---|
| TcCLB.506757.120 | trans-sialidase, Group V, putative | -0,05663 | 0,7249 |
| TcCLB.510219.30 | CPSF 30 kDa subunit, putative (CPSF30) | -0,05663 | 0,7272 |
| TcCLB.507011.160 | hypothetical protein, conserved | -0,05672 | 0,7112 |
| TcCLB.457109.9 | dispersed gene family protein 1 (DGF-1), putative (fragme | -0,05681 | 0,8029 |
| TcCLB.511557.40 | hypothetical protein, conserved | -0,05683 | 0,7457 |
| TcCLB.509199.40 | hypothetical protein, conserved | -0,05686 | 0,7726 |
| TcCLB.506245.70 | mucin-associated surface protein (MASP, pseudogene), pu | -0,05689 | 0,8918 |
| TcCLB.507099.60 | hypothetical protein, conserved | -0,05694 | 0,7009 |
| TcCLB.509911.30 | hypothetical protein | -0,05699 | 0,7438 |
| TcCLB.503789.30 | hypothetical protein, conserved | -0,05732 | 0,8186 |
| TcCLB.508319.70 | 40S ribosomal protein S11, putative | -0,05735 | 0,5455 |
| TcCLB.511483.20 | hypothetical protein, conserved | -0,05748 | 0,7132 |
| TcCLB.506959.30 | ATP-dependent DEAD/H RNA helicase, putative | -0,05756 | 0,6271 |
| TcCLB.508459.9 | hypothetical protein, conserved | -0,05757 | 0,7844 |
| TcCLB.505229.10 | Isoprenylcysteine alpha-carbonyl methylesterase, putative | -0,05771 | 0,6503 |
| TcCLB.510431.245 | hypothetical protein, conserved (pseudogene) | -0,05776 | 0,7956 |
| TcCLB.503723.70 | protein farnesyltransferase, putative | -0,05778 | 0,6989 |
| TcCLB.508647.80 | hypothetical protein, conserved | -0,05782 | 0,7123 |
| TcCLB.509631.160 | hypothetical protein, conserved | -0,05802 | 0,7545 |
| TcCLB.506559.524 | protein phosphatase 2C, putative | -0,05807 | 0,7509 |
| TcCLB.507591.9 | retrotransposon hot spot protein (RHS, pseudogene), putat | -0,05812 | 0,7313 |
| TcCLB.503759.10 | trans-sialidase, Group II, putative | -0,05827 | 0,8008 |
| TcCLB.505207.40 | dispersed gene family protein 1 (DGF-1), putative | -0,05827 | 0,8486 |
| TcCLB.510579.100 | hypothetical protein, conserved | -0,05829 | 0,7282 |
| TcCLB.506559.510 | hypothetical protein, conserved | -0,05831 | 0,7932 |
| TcCLB.509163.80 | glycine dehydrogenase [decarboxylating], putative | -0,05838 | 0,8389 |
| TcCLB.504239.330 | surface protease GP63 (pseudogene), putative | -0,05847 | 0,9101 |
| TcCLB.508815.80 | hypothetical protein, conserved | -0,05852 | 0,5353 |
| TcCLB.507053.170 | hypothetical protein, conserved | -0,05864 | 0,6591 |
| TcCLB.508307.110 | hypothetical protein, conserved | -0,05865 | 0,5407 |
| TcCLB.511401.111 | elongation factor 1-gamma (EF-1-gamma, pseudogene), p | -0,05866 | 0,8285 |
| TcCLB.506367.100 | hypothetical protein, conserved | -0,05872 | 0,7556 |
| TcCLB.509647.110 | hypothetical protein, conserved | -0,0588 | 0,6597 |
| TcCLB.508153.800 | hypothetical protein | -0,05885 | 0,7595 |
| TcCLB.506223.10 | hypothetical protein, conserved | -0,05891 | 0,767 |
| TcCLB.510575.190 | 60S ribosomal protein L10, putative | -0,05894 | 0,5664 |
| TcCLB.511751.110 | hypothetical protein, conserved | -0,05907 | 0,7135 |
| TcCLB.508413.84 | kinetoplastid membrane protein KMP-11 (fragment) | -0,05909 | 0,7119 |
| TcCLB.507811.120 | myosin heavy chain MYA2-related, putative | -0,05911 | 0,6586 |
| TcCLB.510077.30 | Mitogen-activated kinase kinase kinase, putative | -0,05914 | 0,8252 |
| TcCLB.506713.10 | hypothetical protein, conserved | -0,05928 | 0,7671 |
| TcCLB.509099.80 | hypothetical protein, conserved | -0,05933 | 0,5282 |
| TcCLB.509877.40 | hypothetical protein, conserved | -0,05951 | 0,5641 |
| TcCLB.510401.50 | trans-sialidase (pseudogene), putative | -0,05955 | 0,7988 |
| TcCLB.508625.90 | DNA damage repair protein, putative | -0,05958 | 0,6607 |

| | | | |
|---|---|---|---|
| TcCLB.506351.30 | hypothetical protein, conserved | -0,05962 | 0,7186 |
| TcCLB.506503.30 | hypothetical protein, conserved | -0,05964 | 0,6783 |
| TcCLB.508047.20 | mucin-associated surface protein (MASP), putative | -0,05967 | 0,8759 |
| TcCLB.510899.40 | hypothetical protein, conserved | -0,05973 | 0,8239 |
| TcCLB.506751.100 | mucin-associated surface protein (MASP), putative | -0,05981 | 0,9059 |
| TcCLB.504827.90 | hypothetical protein, conserved | -0,05984 | 0,7454 |
| TcCLB.507933.20 | hypothetical protein, conserved | -0,05985 | 0,7631 |
| TcCLB.511311.80 | hypothetical protein, conserved | -0,05989 | 0,5157 |
| TcCLB.503791.10 | TPR-repeat-containing chaperone protein DNAJ, putative (f | -0,05997 | 0,7462 |
| TcCLB.511603.90 | trans-sialidase, Group V, putative | -0,06015 | 0,8591 |
| TcCLB.503423.10 | glycine dehydrogenase [decarboxylating] (pseudogene), pu | -0,06027 | 0,802 |
| TcCLB.503897.90 | hypothetical protein, conserved | -0,06034 | 0,6592 |
| TcCLB.503893.90 | hypothetical protein, conserved | -0,06048 | 0,7482 |
| TcCLB.504141.31 | hypothetical protein, conserved (pseudogene) | -0,06054 | 0,834 |
| TcCLB.507275.20 | hypothetical protein, conserved | -0,06059 | 0,5836 |
| TcCLB.508799.170 | 3-methylcrotonoyl-CoA carboxylase beta subunit, putative | -0,0606 | 0,5673 |
| TcCLB.511923.30 | trans-sialidase, Group V, putative | -0,06067 | 0,8281 |
| TcCLB.506885.320 | hypothetical protein, conserved | -0,0607 | 0,6441 |
| TcCLB.509749.33 | ribosomal protein L29, putative | -0,06071 | 0,6407 |
| TcCLB.506175.60 | GTPase activator protein, putative | -0,06076 | 0,6839 |
| TcCLB.503785.30 | hypothetical protein, conserved | -0,06077 | 0,802 |
| TcCLB.508243.23 | mucin TcMUCII, putative | -0,06089 | 0,9134 |
| TcCLB.506247.340 | hypothetical protein, conserved | -0,06096 | 0,611 |
| TcCLB.506205.10 | hypothetical protein, conserved | -0,06101 | 0,6634 |
| TcCLB.511621.250 | hypothetical protein, conserved | -0,06114 | 0,7183 |
| TcCLB.510175.110 | trans-sialidase (pseudogene), putative | -0,06121 | 0,8179 |
| TcCLB.508667.20 | hypothetical protein | -0,06122 | 0,7712 |
| TcCLB.507063.100 | mitochondrial DNA polymerase beta-PAK, putative | -0,06132 | 0,65 |
| TcCLB.507959.250 | mucin TcMUCII, putative | -0,0614 | 0,9229 |
| TcCLB.504131.80 | hypothetical protein, conserved | -0,06145 | 0,7565 |
| TcCLB.508707.280 | hypothetical protein, conserved | -0,06151 | 0,7776 |
| TcCLB.504057.60 | hypothetical protein, conserved | -0,06163 | 0,6311 |
| TcCLB.511283.180 | hypothetical protein, conserved | -0,06189 | 0,5663 |
| TcCLB.506961.35 | trans-sialidase (pseudogene), putative | -0,06194 | 0,8252 |
| TcCLB.508209.60 | spliced leader RNA PSE-promoter transcription factor, puta | -0,06203 | 0,661 |
| TcCLB.508355.370 | hypothetical protein, conserved | -0,06203 | 0,7263 |
| TcCLB.506599.280 | mucin-associated surface protein (MASP, pseudogene), pu | -0,06213 | 0,9174 |
| TcCLB.511751.130 | small GTP-binding protein Rab28, putative | -0,06224 | 0,7393 |
| TcCLB.503735.10 | hypothetical protein, conserved | -0,06225 | 0,7853 |
| TcCLB.507681.80 | hypothetical protein | -0,06238 | 0,6341 |
| TcCLB.506181.140 | hypothetical protein | -0,06242 | 0,6792 |
| TcCLB.511309.14 | hypothetical protein, conserved | -0,06251 | 0,8391 |
| TcCLB.509437.109 | trans-sialidase (pseudogene), putative | -0,0626 | 0,8266 |
| TcCLB.511321.70 | hypothetical protein, conserved | -0,06261 | 0,6563 |
| TcCLB.507063.70 | trans-sialidase, Group II, putative | -0,06267 | 0,6884 |

| | | | |
|---|---|---|---|
| TcCLB.511811.20 | hypothetical protein, conserved | -0,06267 | 0,6916 |
| TcCLB.511649.169 | ATP-dependent DEAD/H RNA helicase, putative | -0,06269 | 0,6758 |
| TcCLB.509791.40 | hypothetical protein, conserved | -0,06288 | 0,5832 |
| TcCLB.508305.20 | mucin-associated surface protein (MASP), putative | -0,06289 | 0,8854 |
| TcCLB.509169.10 | hypothetical protein, conserved | -0,0629 | 0,6707 |
| TcCLB.511019.26 | trans-sialidase (pseudogene), putative | -0,06293 | 0,752 |
| TcCLB.506753.180 | hypothetical protein, conserved | -0,06294 | 0,7587 |
| TcCLB.510149.140 | Triple RNA binding domain protein 3 (TRRM3) | -0,06307 | 0,6314 |
| TcCLB.508221.730 | 90 kDa surface protein, putative | -0,06318 | 0,9288 |
| TcCLB.507805.9 | histone deacetylase 4, putative | -0,0632 | 0,8362 |
| TcCLB.511261.140 | 60S ribosomal protein L34, putative | -0,06327 | 0,5211 |
| TcCLB.506217.30 | hypothetical protein | -0,06328 | 0,8597 |
| TcCLB.509419.160 | mucin TcMUCII, putative | -0,06329 | 0,9441 |
| TcCLB.508139.230 | retrotransposon hot spot protein (RHS, pseudogene), putat | -0,06351 | 0,6835 |
| TcCLB.509399.190 | pumilio/PUF RNA binding protein 4, putative (PUF4) | -0,06361 | 0,783 |
| TcCLB.507769.30 | hypothetical protein, conserved | -0,06362 | 0,5692 |
| TcCLB.511255.80 | mucin-associated surface protein (MASP), putative | -0,06367 | 0,8577 |
| TcCLB.507927.10 | hypothetical protein, conserved | -0,06372 | 0,7666 |
| TcCLB.506955.161 | elongation factor 1-gamma (EF-1-gamma, pseudogene), p | -0,06373 | 0,8945 |
| TcCLB.508409.240 | hypothetical protein, conserved | -0,06394 | 0,6343 |
| TcCLB.511441.10 | calpain cysteine peptidase, putative | -0,06398 | 0,717 |
| TcCLB.510415.9 | dispersed gene family protein 1 (DGF-1), putative (fragme | -0,06416 | 0,8455 |
| TcCLB.507673.40 | hypothetical protein, conserved | -0,06417 | 0,5824 |
| TcCLB.422207.21 | retrotransposon hot spot protein (RHS, pseudogene), putat | -0,06432 | 0,8805 |
| TcCLB.508543.10 | DNA polymerase sigma, putative | -0,06433 | 0,5845 |
| TcCLB.503939.130 | hypothetical protein, conserved | -0,06435 | 0,6104 |
| TcCLB.504045.10 | hypothetical protein, conserved | -0,06436 | 0,6271 |
| TcCLB.506723.30 | hypothetical protein | -0,06446 | 0,8382 |
| TcCLB.509795.30 | hypothetical protein, conserved | -0,06464 | 0,6431 |
| TcCLB.507089.50 | ATP-dependent DEAD/H RNA helicase, putative | -0,06471 | 0,6512 |
| TcCLB.511585.80 | mitochondrial-processing peptidase subunit beta, putative | -0,06472 | 0,6152 |
| TcCLB.511861.130 | trans-sialidase (pseudogene), putative | -0,06515 | 0,7639 |
| TcCLB.510359.210 | hypothetical protein, conserved | -0,0652 | 0,6663 |
| TcCLB.510525.30 | hypothetical protein, conserved | -0,06524 | 0,6178 |
| TcCLB.511621.80 | hypothetical protein | -0,06538 | 0,8104 |
| TcCLB.503685.30 | carnitine/choline acetyltransferase, putative | -0,06546 | 0,7413 |
| TcCLB.508231.50 | legume-like lectin, putative | -0,06547 | 0,6838 |
| TcCLB.506945.70 | protein mkt1, putative | -0,06552 | 0,4468 |
| TcCLB.508297.20 | UDP-Gal or UDP-GlcNAc-dependent glycosyltransferase, pu | -0,06561 | 0,8462 |
| TcCLB.508723.10 | hypothetical protein, conserved | -0,06564 | 0,6734 |
| TcCLB.509047.100 | hypothetical protein, conserved | -0,06565 | 0,7343 |
| TcCLB.507357.110 | hypothetical protein, conserved (pseudogene) | -0,06573 | 0,9182 |
| TcCLB.511849.15 | mucin-associated surface protein (MASP, pseudogene), pu | -0,06579 | 0,6131 |
| TcCLB.511737.30 | hypothetical protein, conserved | -0,06598 | 0,6457 |
| TcCLB.506673.10 | hypothetical protein, conserved | -0,06601 | 0,7479 |

| | | | |
|---|---|---|---|
| TcCLB.506131.140 | mucin-associated surface protein (MASP), putative | -0,06604 | 0,8961 |
| TcCLB.511277.630 | serine/threonine protein phosphatase type 5, putative | -0,06605 | 0,6249 |
| TcCLB.503805.40 | hypothetical protein | -0,06612 | 0,6349 |
| TcCLB.506957.80 | hypothetical protein, conserved | -0,06614 | 0,5268 |
| TcCLB.510729.20 | hypothetical protein, conserved | -0,0663 | 0,6904 |
| TcCLB.503777.19 | phosphoribosylpyrophosphate synthetase, putative | -0,0663 | 0,7348 |
| TcCLB.506885.30 | hypothetical protein, conserved | -0,06635 | 0,7265 |
| TcCLB.507641.120 | ATP-dependent DEAD/H RNA helicase, putative | -0,06643 | 0,5746 |
| TcCLB.511309.7 | hypothetical protein | -0,06643 | 0,7921 |
| TcCLB.506471.40 | beta galactofuranosyl glycosyltransferase, putative | -0,06655 | 0,8658 |
| TcCLB.507689.5 | hypothetical protein, conserved (fragment) | -0,06663 | 0,8427 |
| TcCLB.511391.60 | trans-sialidase (pseudogene), putative | -0,06671 | 0,8728 |
| TcCLB.506933.89 | calmodulin, putative | -0,06678 | 0,738 |
| TcCLB.508445.40 | hypothetical protein, conserved | -0,06686 | 0,7455 |
| TcCLB.506517.50 | hypothetical protein, conserved | -0,0669 | 0,6939 |
| TcCLB.509033.109 | hypothetical protein, conserved | -0,06693 | 0,8125 |
| TcCLB.511167.119 | hypothetical protein, conserved (fragment) | -0,06698 | 0,61 |
| TcCLB.511215.60 | hypothetical protein, conserved | -0,06699 | 0,7341 |
| TcCLB.511773.30 | hypothetical protein | -0,06708 | 0,8115 |
| TcCLB.503657.50 | hypothetical protein, conserved | -0,06712 | 0,7126 |
| TcCLB.508675.50 | hypothetical protein, conserved | -0,06721 | 0,6037 |
| TcCLB.507841.34 | exosome-associated protein 4, putative | -0,06721 | 0,7537 |
| TcCLB.509525.160 | mucin-associated surface protein (MASP, pseudogene), pu | -0,06726 | 0,862 |
| TcCLB.506763.350 | hypothetical protein, conserved (pseudogene) | -0,0673 | 0,9312 |
| TcCLB.511439.10 | expression site-associated gene (ESAG-like) protein, putati | -0,06737 | 0,9249 |
| TcCLB.509791.90 | hypothetical protein, conserved | -0,06744 | 0,7173 |
| TcCLB.506613.50 | mucin-associated surface protein (MASP), putative | -0,06751 | 0,8018 |
| TcCLB.504769.41 | surface protease GP63 (pseudogene), putative | -0,06751 | 0,8237 |
| TcCLB.503723.40 | hypothetical protein, conserved | -0,06753 | 0,6622 |
| TcCLB.511145.49 | GPI transamidase component GAA1, putative | -0,06756 | 0,6266 |
| TcCLB.511753.50 | hypothetical protein, conserved | -0,06756 | 0,775 |
| TcCLB.511577.90 | pseudouridylate synthase-like protein, putative | -0,06759 | 0,6649 |
| TcCLB.507739.30 | hypothetical protein, conserved | -0,06765 | 0,5995 |
| TcCLB.507939.14 | hypothetical protein, conserved | -0,06778 | 0,7582 |
| TcCLB.503971.5 | hypothetical protein, conserved | -0,06784 | 0,8848 |
| TcCLB.503453.30 | 60S ribosomal protein L30, putative | -0,06822 | 0,5318 |
| TcCLB.506597.36 | hypothetical protein, conserved (pseudogene) | -0,06829 | 0,9342 |
| TcCLB.506181.50 | 60S ribosomal protein L18, putative | -0,0686 | 0,4572 |
| TcCLB.505949.80 | mucin-associated surface protein (MASP), putative | -0,06865 | 0,822 |
| TcCLB.506413.10 | CDP-DAG synthase, putative (fragment) (CDS) | -0,06871 | 0,7678 |
| TcCLB.511923.120 | dihydroorotate dehydrogenase, putative (DHODH) | -0,06882 | 0,7455 |
| TcCLB.510805.60 | surface protease GP63 (pseudogene), putative | -0,06884 | 0,9217 |
| TcCLB.506425.40 | hypothetical protein, conserved | -0,06885 | 0,3539 |
| TcCLB.504431.30 | Present in the outer mitochondrial membrane proteome 2 | -0,06887 | 0,6037 |
| TcCLB.508919.50 | hypothetical protein | -0,0689 | 0,6891 |

| | | | |
|---|---|---|---|
| TcCLB.507071.294 | surface protease GP63 (pseudogene), putative | -0,069 | 0,933 |
| TcCLB.508837.34 | dispersed gene family protein 1 (DGF-1), putative (fragme | -0,06902 | 0,7056 |
| TcCLB.503555.50 | hypothetical protein, conserved | -0,06903 | 0,7599 |
| TcCLB.510021.20 | mucin-associated surface protein (MASP), putative | -0,06912 | 0,8819 |
| TcCLB.509551.64 | small nuclear ribonucleoprotein Sm-F, putative | -0,06914 | 0,8132 |
| TcCLB.508205.30 | protein kinase, putative | -0,06928 | 0,598 |
| TcCLB.507801.100 | hypothetical protein, conserved | -0,06942 | 0,6172 |
| TcCLB.509837.20 | hypothetical protein, conserved | -0,06954 | 0,5309 |
| TcCLB.505037.100 | dispersed gene family protein 1 (DGF-1), putative | -0,06954 | 0,6102 |
| TcCLB.511321.10 | FG-GAP repeat protein, putative | -0,0696 | 0,5017 |
| TcCLB.508601.40 | hypothetical protein, conserved | -0,06962 | 0,7544 |
| TcCLB.508179.50 | hypothetical protein, conserved | -0,06964 | 0,6721 |
| TcCLB.508607.50 | trans-sialidase, Group II, putative | -0,06968 | 0,771 |
| TcCLB.506971.30 | trans-sialidase (pseudogene), putative | -0,06971 | 0,8656 |
| TcCLB.503833.60 | hypothetical protein, conserved | -0,06988 | 0,6629 |
| TcCLB.507663.40 | hypothetical protein, conserved | -0,07005 | 0,812 |
| TcCLB.507787.130 | hypothetical protein, conserved | -0,07011 | 0,605 |
| TcCLB.510407.20 | hypothetical protein, conserved | -0,07017 | 0,6095 |
| TcCLB.506739.140 | hypothetical protein, conserved | -0,07026 | 0,8092 |
| TcCLB.445849.10 | hypothetical protein, conserved | -0,07048 | 0,7194 |
| TcCLB.509109.90 | hypothetical protein, conserved | -0,07049 | 0,7671 |
| TcCLB.509007.14 | hypothetical protein, conserved | -0,0705 | 0,674 |
| TcCLB.506797.120 | hypothetical protein, conserved | -0,07053 | 0,5914 |
| TcCLB.511245.190 | hypothetical protein, conserved | -0,07053 | 0,7121 |
| TcCLB.508207.45 | hypothetical protein, conserved | -0,07054 | 0,6772 |
| TcCLB.511283.194 | hypothetical protein, conserved | -0,07069 | 0,6752 |
| TcCLB.506887.90 | hypothetical protein, conserved | -0,07074 | 0,5584 |
| TcCLB.504021.40 | hypothetical protein, conserved | -0,07082 | 0,59 |
| TcCLB.511377.30 | mucin-associated surface protein (MASP), putative | -0,07082 | 0,9081 |
| TcCLB.508999.70 | mucin-associated surface protein (MASP), putative | -0,07086 | 0,8346 |
| TcCLB.399033.19 | mitochondrial RNA binding protein 1 | -0,07097 | 0,5891 |
| TcCLB.504159.60 | hypothetical protein, conserved | -0,07099 | 0,6715 |
| TcCLB.507723.20 | hypothetical protein | -0,07103 | 0,8597 |
| TcCLB.503525.40 | FAD dependent oxidoreductase, putative (POMP20) | -0,07106 | 0,5612 |
| TcCLB.509805.70 | hypothetical protein, conserved | -0,07109 | 0,5563 |
| TcCLB.504867.50 | hypothetical protein, conserved | -0,0711 | 0,5974 |
| TcCLB.504109.189 | hypothetical protein | -0,07112 | 0,8967 |
| TcCLB.508539.50 | hypothetical protein, conserved | -0,07113 | 0,6406 |
| TcCLB.509897.51 | hypothetical protein, conserved (pseudogene) | -0,07119 | 0,7391 |
| TcCLB.509053.20 | hypothetical protein, conserved | -0,07137 | 0,6158 |
| TcCLB.506755.100 | hypothetical protein, conserved | -0,07138 | 0,7319 |
| TcCLB.511263.30 | hypothetical protein, conserved | -0,07147 | 0,5511 |
| TcCLB.505073.30 | mu-adaptin 4, putative | -0,07152 | 0,71 |
| TcCLB.505297.50 | trans-sialidase (pseudogene), putative | -0,07155 | 0,5685 |
| TcCLB.510669.30 | hypothetical protein, conserved | -0,07158 | 0,6126 |

| | | | |
|---|---|---|---|
| TcCLB.503823.10 | phosphoglycerate mutase family member 5, putative | -0,07159 | 0,7846 |
| TcCLB.508293.10 | trans-sialidase (pseudogene), putative | -0,07168 | 0,8604 |
| TcCLB.505171.109 | Present in the outer mitochondrial membrane proteome 1 | -0,07172 | 0,8296 |
| TcCLB.509099.110 | hypothetical protein, conserved | -0,07175 | 0,7972 |
| TcCLB.511855.50 | hypothetical protein, conserved | -0,07185 | 0,5314 |
| TcCLB.511621.180 | hypothetical protein, conserved | -0,07211 | 0,6403 |
| TcCLB.510021.130 | mucin-associated surface protein (MASP), putative | -0,07214 | 0,8049 |
| TcCLB.509965.160 | hypothetical protein, conserved | -0,07216 | 0,7235 |
| TcCLB.508821.30 | hypothetical protein | -0,07223 | 0,5192 |
| TcCLB.506295.60 | hypothetical protein, conserved | -0,07224 | 0,7678 |
| TcCLB.506863.60 | hypothetical protein, conserved | -0,07231 | 0,6604 |
| TcCLB.510483.70 | mucin-associated surface protein (MASP), putative | -0,07234 | 0,87 |
| TcCLB.507083.80 | hypothetical protein, conserved | -0,07238 | 0,7252 |
| TcCLB.508257.120 | hypothetical protein, conserved | -0,07247 | 0,6864 |
| TcCLB.509521.70 | hypothetical protein | -0,07251 | 0,9146 |
| TcCLB.508693.50 | amidinotransferase, putative | -0,07255 | 0,4309 |
| TcCLB.510575.160 | hypothetical protein | -0,07255 | 0,669 |
| TcCLB.507229.20 | dispersed gene family protein 1 (DGF-1, pseudogene), puta | -0,07257 | 0,7179 |
| TcCLB.470687.9 | hypothetical protein, conserved | -0,07276 | 0,7173 |
| TcCLB.509315.20 | trans-sialidase (pseudogene), putative | -0,07279 | 0,8858 |
| TcCLB.508303.40 | hypothetical protein, conserved | -0,07296 | 0,6873 |
| TcCLB.510975.10 | hypothetical protein, conserved (pseudogene) | -0,07297 | 0,8048 |
| TcCLB.508723.90 | hypothetical protein, conserved | -0,07298 | 0,6517 |
| TcCLB.510821.40 | hypothetical protein, conserved | -0,07301 | 0,596 |
| TcCLB.504071.50 | spliceosome-associated protein, putative | -0,07312 | 0,5996 |
| TcCLB.506289.80 | small GTP-binding protein Rab32, putative | -0,07325 | 0,6229 |
| TcCLB.508893.40 | hypothetical protein, conserved (fragment) | -0,07331 | 0,8474 |
| TcCLB.511309.30 | hydrolase, alpha/beta fold family, putative | -0,07332 | 0,6594 |
| TcCLB.511589.80 | hypothetical protein, conserved | -0,07336 | 0,6948 |
| TcCLB.508175.260 | inorganic polyphosphate/ATP-NAD kinase, putative | -0,07342 | 0,506 |
| TcCLB.509099.89 | hypothetical protein, conserved | -0,07344 | 0,6411 |
| TcCLB.508213.30 | hypothetical protein | -0,07355 | 0,6962 |
| TcCLB.505999.80 | hypothetical protein, conserved | -0,07359 | 0,7003 |
| TcCLB.507011.180 | hypothetical protein, conserved | -0,07363 | 0,589 |
| TcCLB.508707.234 | hypothetical protein, conserved | -0,07372 | 0,7118 |
| TcCLB.506739.190 | serine/threonine protein kinase, putative | -0,0738 | 0,571 |
| TcCLB.507979.10 | hypothetical protein, conserved | -0,07392 | 0,7187 |
| TcCLB.509319.70 | hypothetical protein, conserved | -0,07396 | 0,6835 |
| TcCLB.511277.130 | hypothetical protein, conserved | -0,07397 | 0,4931 |
| TcCLB.510899.59 | hypothetical protein, conserved | -0,07398 | 0,5454 |
| TcCLB.506529.330 | hypothetical protein, conserved | -0,07407 | 0,6902 |
| TcCLB.508867.9 | kinesin, putative (fragment) | -0,0742 | 0,8056 |
| TcCLB.506765.50 | mucin-associated surface protein (MASP), putative | -0,0742 | 0,8706 |
| TcCLB.511527.9 | hypothetical protein, conserved | -0,07421 | 0,7132 |
| TcCLB.503673.27 | 8-oxoguanine DNA glycosylase (pseudogene), putative | -0,0743 | 0,8147 |

| | | | |
|---|---|---|---|
| TcCLB.506321.230 | hypothetical protein | -0,0743 | 0,8464 |
| TcCLB.508909.250 | hypothetical protein, conserved | -0,07432 | 0,6534 |
| TcCLB.506859.30 | hypothetical protein | -0,07434 | 0,5436 |
| TcCLB.511287.100 | hypothetical protein, conserved | -0,07436 | 0,6344 |
| TcCLB.506723.41 | beta galactofuranosyl glycosyltransferase (pseudogene), pu | -0,0744 | 0,7154 |
| TcCLB.506355.170 | hypothetical protein, conserved | -0,07455 | 0,7088 |
| TcCLB.511727.120 | ethanolamine-phosphate cytidylyltransferase (ECT) | -0,07456 | 0,4595 |
| TcCLB.506789.190 | hypothetical protein, conserved | -0,07465 | 0,6299 |
| TcCLB.507887.30 | hypothetical protein, conserved | -0,07466 | 0,7355 |
| TcCLB.507957.304 | surface protease GP63 (pseudogene), putative | -0,07478 | 0,9214 |
| TcCLB.504021.120 | hypothetical protein, conserved | -0,07495 | 0,6042 |
| TcCLB.508283.10 | dispersed gene family protein 1 (DGF-1), putative | -0,075 | 0,7881 |
| TcCLB.503997.19 | hypothetical protein, conserved | -0,07504 | 0,6752 |
| TcCLB.510729.150 | tubulin tyrosine ligase, putative | -0,07514 | 0,5571 |
| TcCLB.506529.640 | hypothetical protein, conserved | -0,07521 | 0,4867 |
| TcCLB.507637.45 | mucin-associated surface protein (MASP, pseudogene), pu | -0,07521 | 0,8963 |
| TcCLB.510441.30 | trans-sialidase (pseudogene), putative | -0,07533 | 0,6976 |
| TcCLB.509141.10 | hypothetical protein | -0,07539 | 0,4954 |
| TcCLB.511289.14 | mitogen-activated protein kinase kinase kinase, putative ( | -0,07548 | 0,5335 |
| TcCLB.506363.110 | RNA editing ligase, putative | -0,0755 | 0,6064 |
| TcCLB.506587.30 | hypothetical protein, conserved | -0,07554 | 0,5957 |
| TcCLB.511013.3 | ATP-dependent DEAD/H RNA helicase (pseudogene), putat | -0,07554 | 0,8933 |
| TcCLB.507801.114 | aldo-keto reductase (pseudogene), putative | -0,07562 | 0,7147 |
| TcCLB.511735.50 | hypothetical protein, conserved | -0,07572 | 0,6189 |
| TcCLB.511179.140 | hypothetical protein, conserved | -0,07594 | 0,5262 |
| TcCLB.511925.60 | hypothetical protein, conserved | -0,07602 | 0,6771 |
| TcCLB.510583.30 | mucin TcMUCII, putative | -0,07603 | 0,9371 |
| TcCLB.507023.280 | hypothetical protein, conserved | -0,07604 | 0,6696 |
| TcCLB.503703.60 | hypothetical protein | -0,07606 | 0,6453 |
| TcCLB.511643.4 | hypothetical protein, conserved | -0,07611 | 0,8452 |
| TcCLB.510601.10 | hypothetical protein, conserved | -0,07627 | 0,7873 |
| TcCLB.508397.50 | hypothetical protein, conserved | -0,07634 | 0,6222 |
| TcCLB.509733.134 | hypothetical protein, conserved | -0,07643 | 0,7562 |
| TcCLB.510297.110 | cAMP binding protein, putative | -0,0766 | 0,4042 |
| TcCLB.508965.60 | rac serine-threonine kinase, putative | -0,07667 | 0,5536 |
| TcCLB.510723.20 | hypothetical protein | -0,07667 | 0,5684 |
| TcCLB.506389.50 | hypothetical protein, conserved | -0,07673 | 0,5599 |
| TcCLB.506975.49 | f-actin capping protein beta subunit, putative | -0,07677 | 0,7032 |
| TcCLB.508479.170 | hypothetical protein, conserved | -0,07693 | 0,6121 |
| TcCLB.511903.60 | hypothetical protein, conserved | -0,07693 | 0,7465 |
| TcCLB.511355.40 | GUP1, putative | -0,07702 | 0,6086 |
| TcCLB.511715.10 | iron superoxide dismutase, putative | -0,07707 | 0,5993 |
| TcCLB.503723.10 | hypothetical protein, conserved | -0,07711 | 0,7112 |
| TcCLB.506211.70 | RNA-binding protein, putative | -0,07724 | 0,5806 |
| TcCLB.510657.130 | hypothetical protein, conserved | -0,07727 | 0,7066 |

| | | | |
|---|---|---|---|
| TcCLB.431849.10 | hypothetical protein, conserved | -0,07732 | 0,8107 |
| TcCLB.510725.70 | hypothetical protein, conserved | -0,07744 | 0,6006 |
| TcCLB.507503.11 | protein kinase (pseudogene), putative | -0,07745 | 0,5112 |
| TcCLB.507611.30 | glycine dehydrogenase (pseudogene), putative | -0,07745 | 0,845 |
| TcCLB.511037.40 | hypothetical protein, conserved | -0,07756 | 0,8307 |
| TcCLB.507041.20 | hypothetical protein, conserved | -0,07758 | 0,6086 |
| TcCLB.506513.70 | hypothetical protein | -0,0776 | 0,6299 |
| TcCLB.510303.280 | hypothetical protein | -0,07762 | 0,6281 |
| TcCLB.510629.410 | hypothetical protein, conserved | -0,07766 | 0,7617 |
| TcCLB.506657.70 | RNA editing 3' terminal uridylyl transferase 1, putative | -0,07768 | 0,388 |
| TcCLB.507603.160 | hypothetical protein, conserved | -0,07769 | 0,7173 |
| TcCLB.506635.50 | hypothetical protein | -0,07775 | 0,5327 |
| TcCLB.422031.10 | solanesyl-diphosphate synthase, putative | -0,07775 | 0,6584 |
| TcCLB.506967.150 | trans-sialidase (pseudogene), putative | -0,07778 | 0,8359 |
| TcCLB.508067.10 | hypothetical protein, conserved | -0,07784 | 0,3092 |
| TcCLB.506885.260 | hypothetical protein, conserved | -0,07786 | 0,4779 |
| TcCLB.425435.10 | trans-sialidase, putative (fragment) | -0,07795 | 0,8594 |
| TcCLB.508897.10 | GDP-mannose 4,6 dehydratase, putative | -0,07809 | 0,6873 |
| TcCLB.509163.50 | hypothetical protein | -0,07823 | 0,8466 |
| TcCLB.508871.60 | mucin-associated surface protein (MASP), putative | -0,07827 | 0,8837 |
| TcCLB.511189.54 | hypothetical protein | -0,0783 | 0,6796 |
| TcCLB.511231.80 | Nascent polypeptide-associated complex subunit beta, put | -0,07833 | 0,6671 |
| TcCLB.511109.120 | beta-ketoacyl synthase family protein, putative | -0,07856 | 0,687 |
| TcCLB.506767.209 | syntaxin binding protein (pseudogene), putative | -0,0787 | 0,8892 |
| TcCLB.504153.360 | succinyl-coA:3-ketoacid-coenzyme A transferase, mitochor | -0,07877 | 0,3433 |
| TcCLB.509965.370 | adenosine kinase, putative | -0,07887 | 0,4909 |
| TcCLB.511415.50 | retrotransposon hot spot protein (RHS, pseudogene), putat | -0,07894 | 0,7597 |
| TcCLB.509379.10 | dihydrolipoamide dehydrogenase, putative (GCVL-1) | -0,07896 | 0,6529 |
| TcCLB.511231.50 | aminomethyltransferase, mitochondrial, putative (GCV) | -0,079 | 0,5985 |
| TcCLB.507077.9 | hypothetical protein, conserved | -0,07904 | 0,633 |
| TcCLB.510311.190 | hypothetical protein, conserved | -0,07905 | 0,6 |
| TcCLB.508277.260 | actin-related protein 3, putative | -0,07911 | 0,5498 |
| TcCLB.506267.80 | mucin-associated surface protein (MASP), putative | -0,07915 | 0,8763 |
| TcCLB.511173.270 | mucin-associated surface protein (MASP), putative | -0,07922 | 0,7932 |
| TcCLB.506871.140 | cleavage and polyadenylation specificity factor, putative | -0,07928 | 0,3883 |
| TcCLB.507941.130 | hypothetical protein, conserved | -0,07937 | 0,7018 |
| TcCLB.507491.140 | hypothetical protein, conserved | -0,07938 | 0,6437 |
| TcCLB.507723.150 | hypothetical protein, conserved | -0,07941 | 0,641 |
| TcCLB.508479.224 | hypothetical protein, conserved | -0,07943 | 0,7284 |
| TcCLB.510297.70 | hypothetical protein | -0,07951 | 0,6516 |
| TcCLB.440083.10 | trans-sialidase, putative (fragment) | -0,07956 | 0,8191 |
| TcCLB.510105.310 | mucin-associated surface protein (MASP), putative | -0,07957 | 0,8642 |
| TcCLB.511621.140 | hypothetical protein, conserved | -0,07969 | 0,6443 |
| TcCLB.510719.280 | hypothetical protein, conserved | -0,07974 | 0,6092 |
| TcCLB.509653.10 | mucin-associated surface protein (MASP), putative | -0,07984 | 0,919 |

| | | | |
|---|---|---|---|
| TcCLB.510821.60 | ribonucleoprotein p18, mitochondrial precursor, putative | -0,08001 | 0,4258 |
| TcCLB.506529.290 | hypothetical protein, conserved | -0,08001 | 0,6553 |
| TcCLB.470169.9 | hypothetical protein, conserved | -0,08004 | 0,697 |
| TcCLB.509805.170 | hypothetical protein, conserved | -0,08006 | 0,4657 |
| TcCLB.511521.40 | hypothetical protein, conserved | -0,08007 | 0,6483 |
| TcCLB.510531.50 | hypothetical protein, conserved | -0,0801 | 0,5544 |
| TcCLB.509429.290 | hydroxyacylglutathione hydrolase, putative | -0,08012 | 0,6878 |
| TcCLB.504181.20 | 40S ribosomal protein S23, putative | -0,08021 | 0,3294 |
| TcCLB.506565.12 | RNA-binding protein, putative | -0,08031 | 0,6346 |
| TcCLB.511469.10 | hypothetical protein, conserved | -0,08031 | 0,7307 |
| TcCLB.509269.4 | Tb-291 membrane associated protein, putative | -0,0804 | 0,5827 |
| TcCLB.507669.150 | glycosomal membrane protein, putative | -0,08041 | 0,7052 |
| TcCLB.510447.10 | dispersed gene family protein 1 (DGF-1), putative | -0,08041 | 0,922 |
| TcCLB.506177.60 | hypothetical protein | -0,08054 | 0,6116 |
| TcCLB.506789.20 | trans-sialidase, Group V, putative | -0,08056 | 0,873 |
| TcCLB.506147.184 | hypothetical protein, conserved | -0,08059 | 0,7219 |
| TcCLB.507053.150 | hypothetical protein, conserved | -0,08065 | 0,4801 |
| TcCLB.508065.40 | hypothetical protein, conserved | -0,08067 | 0,5827 |
| TcCLB.506885.364 | hypothetical protein, conserved | -0,08071 | 0,7032 |
| TcCLB.504079.6 | selenophosphate synthetase, putative | -0,08075 | 0,5307 |
| TcCLB.511421.50 | transmembrane protein, putative | -0,08079 | 0,4954 |
| TcCLB.510697.80 | mucin-associated surface protein (MASP), putative | -0,0808 | 0,7815 |
| TcCLB.507611.320 | hypothetical protein, conserved | -0,08084 | 0,6775 |
| TcCLB.509145.19 | hypothetical protein, conserved | -0,08095 | 0,701 |
| TcCLB.505183.90 | hypothetical protein, conserved | -0,08095 | 0,7511 |
| TcCLB.506227.160 | MOZ/SAS family acetyltransferase, putative | -0,08098 | 0,6104 |
| TcCLB.508277.70 | hypothetical protein, conserved | -0,08109 | 0,6017 |
| TcCLB.507023.40 | protein kinase, putative | -0,08118 | 0,4616 |
| TcCLB.508141.70 | Kinesin-13 4, putative (KIN13-4) | -0,0812 | 0,663 |
| TcCLB.506605.80 | hypothetical protein, conserved | -0,08134 | 0,5997 |
| TcCLB.504933.30 | hypothetical protein | -0,08145 | 0,3611 |
| TcCLB.508641.140 | hypothetical protein, conserved | -0,0815 | 0,7051 |
| TcCLB.508805.136 | UDP-Gal or UDP-GlcNAc-dependent glycosyltransferase, pu | -0,08152 | 0,7888 |
| TcCLB.503885.40 | hypothetical protein, conserved | -0,08153 | 0,6304 |
| TcCLB.510625.19 | mucin-associated surface protein (MASP), putative | -0,0816 | 0,9126 |
| TcCLB.506727.130 | hypothetical protein, conserved | -0,08164 | 0,6736 |
| TcCLB.511317.30 | regulator of nonsense transcripts UPF1, putative | -0,0818 | 0,3546 |
| TcCLB.509127.10 | ATP-dependent Clp protease subunit, heat shock protein 78 | -0,08181 | 0,5652 |
| TcCLB.506439.9 | myosin heavy chain, putative | -0,08191 | 0,5976 |
| TcCLB.504185.40 | hypothetical protein, conserved | -0,08194 | 0,6041 |
| TcCLB.507895.10 | trans-sialidase (pseudogene), putative | -0,08199 | 0,9101 |
| TcCLB.506537.30 | hypothetical protein, conserved (pseudogene) | -0,08211 | 0,5989 |
| TcCLB.511165.30 | hypothetical protein, conserved | -0,08211 | 0,7689 |
| TcCLB.508247.60 | hypothetical protein, conserved (pseudogene) | -0,08211 | 0,8431 |
| TcCLB.506235.10 | translation initiation factor IF-2, putative | -0,0822 | 0,3469 |

| | | | |
|---|---|---|---|
| TcCLB.507775.20 | hypothetical protein, conserved | -0,0822 | 0,632 |
| TcCLB.508111.30 | glutamate dehydrogenase, putative | -0,08221 | 0,4254 |
| TcCLB.503859.45 | mucin TcMUCII, putative | -0,08225 | 0,9007 |
| TcCLB.508125.80 | trans-sialidase (pseudogene), putative | -0,08256 | 0,8822 |
| TcCLB.509925.40 | trans-sialidase (pseudogene), putative | -0,0826 | 0,8174 |
| TcCLB.506457.130 | hypothetical protein, conserved | -0,08268 | 0,5601 |
| TcCLB.414243.20 | hypothetical protein, conserved | -0,08272 | 0,6546 |
| TcCLB.509479.10 | mucin TcMUCII, putative | -0,08272 | 0,8008 |
| TcCLB.506173.190 | dispersed gene family protein 1 (DGF-1), putative | -0,0828 | 0,802 |
| TcCLB.508737.50 | hypothetical protein, conserved | -0,0829 | 0,6823 |
| TcCLB.509937.110 | hypothetical protein, conserved | -0,08302 | 0,6087 |
| TcCLB.507049.10 | hypothetical protein, conserved | -0,08307 | 0,6383 |
| TcCLB.510001.20 | hypothetical protein, conserved | -0,08309 | 0,5005 |
| TcCLB.503755.50 | hypothetical protein, conserved | -0,08309 | 0,5562 |
| TcCLB.507991.120 | translation initiation factor IF-2, putative | -0,0831 | 0,5839 |
| TcCLB.506585.80 | hypothetical protein, conserved | -0,0831 | 0,634 |
| TcCLB.510607.70 | trans-sialidase (pseudogene), putative | -0,08311 | 0,7984 |
| TcCLB.509263.10 | UDP-Gal or UDP-GlcNAc-dependent glycosyltransferase, pu | -0,08337 | 0,832 |
| TcCLB.420533.9 | dihydrofolate reductase-thymidylate synthase, putative (fra | -0,08338 | 0,7244 |
| TcCLB.509989.10 | phosphatidylinositol 3-kinase, putative | -0,08383 | 0,7184 |
| TcCLB.509805.40 | Multisite-specific tRNA:(cytosine-C(5))-methyltransferase, | -0,08389 | 0,5351 |
| TcCLB.510635.60 | hypothetical protein, conserved | -0,0839 | 0,6486 |
| TcCLB.511001.150 | hypothetical protein, conserved | -0,08396 | 0,6199 |
| TcCLB.506607.19 | phosphoribosylpyrophosphate synthetase, putative | -0,08402 | 0,6443 |
| TcCLB.511577.70 | hypothetical protein, conserved | -0,08405 | 0,5846 |
| TcCLB.510743.100 | chaperone DNAJ protein, putative | -0,08412 | 0,4607 |
| TcCLB.504949.14 | 60S ribosomal protein L6, putative | -0,08414 | 0,4783 |
| TcCLB.504883.10 | protein kinase, putative | -0,08428 | 0,4455 |
| TcCLB.507011.90 | hypothetical protein, conserved | -0,08432 | 0,5067 |
| TcCLB.510729.50 | hypothetical protein, conserved | -0,08434 | 0,6734 |
| TcCLB.509699.40 | trans-sialidase, Group V, putative | -0,08441 | 0,8186 |
| TcCLB.510143.40 | UBX domain-containing protein | -0,08448 | 0,4883 |
| TcCLB.510073.50 | elongation factor 1-gamma (EF-1-gamma), putative | -0,08458 | 0,606 |
| TcCLB.507093.330 | hypothetical protein, conserved | -0,0846 | 0,7166 |
| TcCLB.506305.10 | hypothetical protein, conserved | -0,0847 | 0,4557 |
| TcCLB.506811.220 | hypothetical protein, conserved | -0,08476 | 0,4549 |
| TcCLB.510879.20 | ribosomal protein S19, putative | -0,08492 | 0,3394 |
| TcCLB.507953.185 | hypothetical protein, conserved (pseudogene) | -0,08499 | 0,8763 |
| TcCLB.508153.970 | AGC/RSK family serine/threonine kinase, putative | -0,08503 | 0,6624 |
| TcCLB.511215.10 | hypothetical protein, conserved | -0,08511 | 0,5612 |
| TcCLB.508883.15 | trans-sialidase (pseudogene), putative | -0,08511 | 0,9006 |
| TcCLB.511001.130 | hypothetical protein, conserved | -0,08513 | 0,3339 |
| TcCLB.504427.90 | ribosomal RNA methyltransferase, putative | -0,08521 | 0,6202 |
| TcCLB.504101.40 | trans-sialidase (pseudogene), putative | -0,08528 | 0,5901 |
| TcCLB.507811.90 | hypothetical protein, conserved | -0,08531 | 0,5318 |

| | | | |
|---|---|---|---|
| TcCLB.507003.50 | hypothetical protein, conserved | -0,08536 | 0,705 |
| TcCLB.508461.290 | 4E-interacting protein, putative | -0,08537 | 0,534 |
| TcCLB.507669.239 | hypothetical protein, conserved | -0,08545 | 0,5921 |
| TcCLB.506203.110 | hypothetical protein, conserved | -0,0855 | 0,5622 |
| TcCLB.511865.90 | dynein heavy chain, putative | -0,08551 | 0,5612 |
| TcCLB.506737.107 | mucin-associated surface protein (MASP, pseudogene), pu | -0,08568 | 0,7161 |
| TcCLB.504213.10 | hypothetical protein, conserved | -0,08574 | 0,6943 |
| TcCLB.509965.50 | hypothetical protein, conserved | -0,08579 | 0,7386 |
| TcCLB.511907.70 | short chain dehydrogenase/reductase, putative | -0,0858 | 0,5524 |
| TcCLB.510767.69 | hypothetical protein, conserved | -0,08584 | 0,7077 |
| TcCLB.506405.30 | actin-like protein, putative | -0,08591 | 0,6522 |
| TcCLB.506213.10 | hypothetical protein, conserved | -0,08602 | 0,6289 |
| TcCLB.511589.190 | hypothetical protein, conserved | -0,08604 | 0,6793 |
| TcCLB.506565.29 | hypothetical protein, conserved | -0,08604 | 0,7051 |
| TcCLB.509575.10 | retrotransposon hot spot (RHS) protein, putative | -0,08609 | 0,5621 |
| TcCLB.510533.60 | hypothetical protein, conserved | -0,0861 | 0,6721 |
| TcCLB.506507.10 | N-acetylglucosamine-6-phosphate deacetylase-like proteir | -0,08613 | 0,4791 |
| TcCLB.510257.139 | transcription activator, putative | -0,08614 | 0,5839 |
| TcCLB.508459.18 | phosphatidylserine decarboxylase, putative | -0,08617 | 0,8273 |
| TcCLB.510173.50 | hypothetical protein, conserved | -0,08618 | 0,7168 |
| TcCLB.507037.40 | hypothetical protein, conserved (pseudogene) | -0,08625 | 0,4159 |
| TcCLB.510903.40 | hypothetical protein, conserved | -0,08625 | 0,6095 |
| TcCLB.511303.24 | hypothetical protein, conserved | -0,08625 | 0,6185 |
| TcCLB.509891.20 | hypothetical protein, conserved (pseudogene) | -0,0863 | 0,6276 |
| TcCLB.504935.10 | eukaryotic translation initiation factor 4E, putative | -0,08644 | 0,658 |
| TcCLB.506159.30 | dispersed gene family protein 1 (DGF-1), putative | -0,08644 | 0,7933 |
| TcCLB.506529.440 | hypothetical protein, conserved | -0,08646 | 0,5454 |
| TcCLB.505999.120 | hypothetical protein | -0,08655 | 0,6693 |
| TcCLB.509965.140 | hypothetical protein, conserved | -0,08687 | 0,5863 |
| TcCLB.506503.90 | kinesin, putative | -0,0869 | 0,6734 |
| TcCLB.511349.20 | trans-sialidase (pseudogene), putative | -0,08692 | 0,7037 |
| TcCLB.507501.25 | hypothetical protein, conserved (pseudogene) | -0,08706 | 0,8608 |
| TcCLB.510755.138 | serine/threonine protein phosphatase, putative | -0,08715 | 0,5905 |
| TcCLB.509527.50 | mucin TcMUCII (pseudogene), putative | -0,08718 | 0,8922 |
| TcCLB.507603.80 | hypothetical protein, conserved | -0,0872 | 0,5441 |
| TcCLB.510297.20 | hypothetical protein, conserved | -0,08731 | 0,4736 |
| TcCLB.506835.120 | DNA ligase I, putative | -0,08734 | 0,4561 |
| TcCLB.506243.50 | dispersed gene family protein 1 (DGF-1, pseudogene), puta | -0,08735 | 0,7425 |
| TcCLB.508263.10 | hypothetical protein | -0,08739 | 0,5549 |
| TcCLB.436535.10 | hypothetical protein, conserved | -0,08742 | 0,7093 |
| TcCLB.508977.80 | trans-sialidase, Group V, putative | -0,08756 | 0,7467 |
| TcCLB.503899.139 | hypothetical protein, conserved | -0,08762 | 0,3962 |
| TcCLB.508207.180 | Eukaryotic translation initiation factor 4E type 6, putative ( | -0,08766 | 0,5302 |
| TcCLB.509875.260 | hypothetical protein, conserved | -0,08776 | 0,6729 |
| TcCLB.506977.120 | hypothetical protein, conserved | -0,08778 | 0,5621 |

| | | | |
|---|---|---|---|
| TcCLB.507511.10 | cleavage and polyadenylation specificity factor, putative | -0,08781 | 0,5119 |
| TcCLB.511529.20 | hypothetical protein, conserved | -0,08782 | 0,6375 |
| TcCLB.507093.220 | RNA-binding protein, putative | -0,08783 | 0,4693 |
| TcCLB.511423.60 | ribosomal RNA-processing protein 8, putative (RRP8) | -0,08795 | 0,5637 |
| TcCLB.506957.140 | phosphatidylinositol-4-phosphate 5-kinase-like, putative | -0,08796 | 0,4938 |
| TcCLB.507625.110 | chaperone DNAJ protein, putative | -0,08797 | 0,5156 |
| TcCLB.510517.110 | hypothetical protein | -0,08797 | 0,7555 |
| TcCLB.509791.50 | bilobe protein, conserved | -0,08801 | 0,5312 |
| TcCLB.506279.250 | trans-sialidase (pseudogene), putative | -0,08803 | 0,8285 |
| TcCLB.510181.40 | hypothetical protein, conserved | -0,08813 | 0,7542 |
| TcCLB.510519.30 | mitochondrial structure specific endonuclease I (SSE-1), pu | -0,08824 | 0,5449 |
| TcCLB.508951.70 | hypothetical protein, conserved | -0,0883 | 0,52 |
| TcCLB.511851.61 | mucin-associated surface protein (MASP, pseudogene), pu | -0,08844 | 0,8643 |
| TcCLB.509161.70 | Mg transporter, putative | -0,08849 | 0,5746 |
| TcCLB.506435.450 | hypothetical protein, conserved | -0,0885 | 0,5269 |
| TcCLB.510145.30 | hypothetical protein, conserved | -0,08863 | 0,5867 |
| TcCLB.505807.120 | lysyl-tRNA synthetase, putative | -0,08875 | 0,6111 |
| TcCLB.506337.190 | hypothetical protein, conserved | -0,08882 | 0,5443 |
| TcCLB.506925.170 | hypothetical protein, conserved | -0,08897 | 0,8762 |
| TcCLB.508741.260 | hypothetical protein, conserved | -0,08903 | 0,4657 |
| TcCLB.503899.60 | hypothetical protein, conserved | -0,08912 | 0,5772 |
| TcCLB.503893.10 | copper-transporting ATPase-like protein, putative | -0,08916 | 0,7747 |
| TcCLB.506473.10 | Pre-mRNA-splicing factor 38, putative | -0,08917 | 0,7112 |
| TcCLB.508147.79 | mucin TcMUCII, putative | -0,08922 | 0,895 |
| TcCLB.503851.54 | proteasome 26S non-ATPase subunit 9, putative | -0,08943 | 0,8247 |
| TcCLB.503939.60 | hypothetical protein, conserved | -0,08947 | 0,4976 |
| TcCLB.507681.60 | hypothetical protein, conserved | -0,08947 | 0,713 |
| TcCLB.508693.84 | hypothetical protein, conserved | -0,08949 | 0,6525 |
| TcCLB.507389.70 | peptidase, putative | -0,08968 | 0,3419 |
| TcCLB.510301.10 | fructose-bisphosphate aldolase, glycosomal, putative | -0,0899 | 0,4799 |
| TcCLB.503487.10 | DNA-directed RNA polymerase, mitochondrial (fragment) | -0,09001 | 0,6847 |
| TcCLB.504129.20 | hypothetical protein, conserved | -0,09017 | 0,3929 |
| TcCLB.506817.50 | ribosomal protein L27, putative | -0,0902 | 0,3862 |
| TcCLB.503829.50 | hypothetical protein, conserved | -0,09047 | 0,6292 |
| TcCLB.507903.19 | dispersed gene family protein 1 (DGF-1, pseudogene), puta | -0,09048 | 0,7791 |
| TcCLB.504155.80 | mucin-associated surface protein (MASP), putative | -0,09049 | 0,8729 |
| TcCLB.506513.90 | hypothetical protein, conserved | -0,09051 | 0,602 |
| TcCLB.510755.60 | ADP-ribosylation factor 1, putative (ARF1) | -0,09054 | 0,7106 |
| TcCLB.504109.50 | hypothetical protein, conserved | -0,09056 | 0,5655 |
| TcCLB.509569.110 | carrier protein, putative | -0,09058 | 0,5841 |
| TcCLB.511911.21 | hypothetical protein, conserved | -0,0906 | 0,8348 |
| TcCLB.503899.100 | hypothetical protein | -0,09069 | 0,5677 |
| TcCLB.509585.10 | dynein heavy chain, putative | -0,09082 | 0,3971 |
| TcCLB.507649.50 | hypothetical protein, conserved | -0,09085 | 0,3962 |
| TcCLB.508089.71 | hypothetical protein | -0,09086 | 0,7008 |

| | | | |
|---|---|---|---|
| TcCLB.506465.20 | hypothetical protein, conserved | -0,09092 | 0,6095 |
| TcCLB.506869.50 | calcineurin B subunit, putative (CNB) | -0,09095 | 0,674 |
| TcCLB.511467.13 | hypothetical protein, conserved | -0,09101 | 0,7212 |
| TcCLB.506163.10 | surface protease GP63, putative | -0,09123 | 0,5618 |
| TcCLB.506757.30 | hypothetical protein, conserved | -0,09124 | 0,4937 |
| TcCLB.503719.10 | hypothetical protein | -0,09137 | 0,4853 |
| TcCLB.509767.240 | hypothetical protein, conserved | -0,09137 | 0,6972 |
| TcCLB.508351.60 | hypothetical protein, conserved | -0,09146 | 0,4642 |
| TcCLB.510325.45 | hypothetical protein | -0,09154 | 0,5555 |
| TcCLB.511711.70 | hypothetical protein, conserved | -0,09155 | 0,6049 |
| TcCLB.510737.70 | 60S ribosomal subunit protein L31, putative | -0,09164 | 0,6244 |
| TcCLB.508293.111 | elongation factor 1-gamma (EF-1-gamma, pseudogene), p | -0,09166 | 0,8849 |
| TcCLB.511629.10 | hypothetical protein, conserved | -0,09183 | 0,5124 |
| TcCLB.508817.160 | Trichohyalin, putative | -0,09185 | 0,5183 |
| TcCLB.506569.10 | cysteine peptidase, Clan CA, family C19, putative | -0,09187 | 0,4971 |
| TcCLB.509767.215 | vesicle transport v-SNARE 11, putative | -0,09207 | 0,6589 |
| TcCLB.510275.355 | hypothetical protein | -0,09217 | 0,8744 |
| TcCLB.509409.10 | protein kinase, putative | -0,09222 | 0,5635 |
| TcCLB.509629.20 | DnaJ chaperone protein, putative | -0,0923 | 0,7041 |
| TcCLB.510725.30 | hypothetical protein, conserved | -0,09231 | 0,6394 |
| TcCLB.510765.60 | 60S ribosomal protein L5, putative | -0,09232 | 0,2421 |
| TcCLB.507939.40 | hypothetical protein, conserved | -0,09232 | 0,6954 |
| TcCLB.506367.170 | hypothetical protein | -0,09235 | 0,4762 |
| TcCLB.506107.10 | hypothetical protein, conserved | -0,09236 | 0,4976 |
| TcCLB.506795.60 | hypothetical protein, conserved | -0,09241 | 0,4451 |
| TcCLB.507883.109 | mitochondrial malate dehydrogenase, putative | -0,09242 | 0,3004 |
| TcCLB.418185.43 | ATP-dependent DEAD/H RNA helicase (pseudogene), putat | -0,09243 | 0,7635 |
| TcCLB.507801.144 | hypothetical protein, conserved | -0,09247 | 0,6859 |
| TcCLB.506335.80 | hypothetical protein | -0,09258 | 0,8879 |
| TcCLB.508209.20 | hypothetical protein, conserved | -0,09261 | 0,6 |
| TcCLB.507053.100 | ubiquitin hydrolase, putative | -0,09264 | 0,3726 |
| TcCLB.508415.40 | ADP-ribosylation factor 1, putative (ARF1) | -0,09264 | 0,4217 |
| TcCLB.509509.69 | hypothetical protein, conserved | -0,09265 | 0,6282 |
| TcCLB.504105.210 | Translation factor GUF1, mitochondrial, putative (GUF1) | -0,09266 | 0,4343 |
| TcCLB.505965.86 | tRNA pseudouridine synthase (pseudogene), putative | -0,09268 | 0,4804 |
| TcCLB.506289.50 | hypothetical protein, conserved | -0,09273 | 0,4644 |
| TcCLB.503539.10 | hypothetical protein, conserved | -0,09274 | 0,6966 |
| TcCLB.504119.10 | eukaryotic translation initiation factor 5, putative | -0,09281 | 0,3719 |
| TcCLB.509171.47 | hypothetical protein, conserved | -0,09301 | 0,5509 |
| TcCLB.506947.30 | hypothetical protein, conserved | -0,09302 | 0,6818 |
| TcCLB.507077.20 | palmitoyl acyltransferase 4, putative | -0,09304 | 0,469 |
| TcCLB.505297.10 | dispersed gene family protein 1 (DGF-1, pseudogene), puta | -0,09304 | 0,8499 |
| TcCLB.509717.34 | hypothetical protein, conserved | -0,09307 | 0,6301 |
| TcCLB.506351.80 | oligopeptidase B-like protein, putative | -0,09308 | 0,4935 |
| TcCLB.506587.80 | hypothetical protein, conserved | -0,09311 | 0,5444 |

| | | | |
|---|---|---|---|
| TcCLB.510359.600 | mucin-associated surface protein (MASP), putative | -0,09316 | 0,8166 |
| TcCLB.503901.18 | hypothetical protein, conserved (fragment) | -0,09318 | 0,5331 |
| TcCLB.511899.30 | hypothetical protein, conserved | -0,09324 | 0,4877 |
| TcCLB.509671.150 | hypothetical protein, conserved | -0,09333 | 0,6325 |
| TcCLB.509939.19 | hypothetical protein, conserved | -0,09377 | 0,3997 |
| TcCLB.506965.160 | hypothetical protein | -0,09392 | 0,8614 |
| TcCLB.506965.40 | trans-sialidase (pseudogene), putative | -0,09409 | 0,808 |
| TcCLB.504163.10 | hypothetical protein, conserved | -0,09412 | 0,4253 |
| TcCLB.508013.120 | elongation factor 1-alpha (ef-1-alpha), putative | -0,09422 | 0,848 |
| TcCLB.509563.40 | hypothetical protein, conserved | -0,09444 | 0,5461 |
| TcCLB.508723.20 | hypothetical protein, conserved | -0,09448 | 0,5551 |
| TcCLB.506269.40 | mucin-associated surface protein (MASP), putative | -0,0945 | 0,8061 |
| TcCLB.427091.10 | solanesyl-diphosphate synthase, putative | -0,09453 | 0,6343 |
| TcCLB.510887.40 | hypothetical protein, conserved | -0,09454 | 0,6192 |
| TcCLB.504427.100 | hypothetical protein, conserved | -0,09457 | 0,6534 |
| TcCLB.504741.124 | hypothetical protein, conserved | -0,09457 | 0,7503 |
| TcCLB.411427.10 | hypothetical protein, conserved | -0,09458 | 0,781 |
| TcCLB.510023.20 | mucin TcMUCII, putative | -0,09459 | 0,8485 |
| TcCLB.463955.20 | mucin-associated surface protein (MASP), putative (fragm | -0,09481 | 0,9152 |
| TcCLB.508413.68 | kinetoplastid membrane protein KMP-11 | -0,09487 | 0,5197 |
| TcCLB.506925.319 | isocitrate dehydrogenase, putative | -0,0949 | 0,6091 |
| TcCLB.507381.20 | trans-sialidase (pseudogene), putative | -0,0949 | 0,6663 |
| TcCLB.440101.20 | activated protein kinase C receptor (pseudogene), putative | -0,09491 | 0,7653 |
| TcCLB.511445.140 | hypothetical protein | -0,09496 | 0,5377 |
| TcCLB.506925.400 | hypothetical protein, conserved | -0,09499 | 0,4267 |
| TcCLB.506713.20 | hypothetical protein, conserved | -0,09508 | 0,5266 |
| TcCLB.509111.10 | hypothetical protein, conserved | -0,09509 | 0,5464 |
| TcCLB.508817.30 | hypothetical protein | -0,09511 | 0,5736 |
| TcCLB.503529.20 | hypothetical protein, conserved | -0,09533 | 0,4759 |
| TcCLB.511421.60 | deoxyhypusine synthase, putative | -0,09536 | 0,5702 |
| TcCLB.416891.9 | protein kinase, putative | -0,09539 | 0,6549 |
| TcCLB.511907.40 | hypothetical protein, conserved | -0,09548 | 0,6178 |
| TcCLB.506355.50 | chaperone DNAJ protein, putative | -0,09549 | 0,3516 |
| TcCLB.508827.90 | hypothetical protein, conserved | -0,09553 | 0,5974 |
| TcCLB.511517.156 | hypothetical protein, conserved | -0,09554 | 0,6125 |
| TcCLB.507883.50 | hypothetical protein, conserved | -0,09555 | 0,5952 |
| TcCLB.509967.10 | hypothetical protein, conserved | -0,09563 | 0,5741 |
| TcCLB.509797.50 | hypothetical protein, conserved | -0,0957 | 0,5909 |
| TcCLB.511269.50 | protein kinase A catalytic subunit, putative (PKAC3) | -0,09578 | 0,7118 |
| TcCLB.509455.9 | hypothetical protein, conserved | -0,09585 | 0,4832 |
| TcCLB.506553.20 | bromodomain factor 2 protein | -0,0959 | 0,4486 |
| TcCLB.508277.50 | presenilin-like aspartic peptidase, putative | -0,09601 | 0,4415 |
| TcCLB.510241.10 | hypothetical protein, conserved | -0,09605 | 0,4117 |
| TcCLB.510969.30 | hypothetical protein, conserved | -0,09613 | 0,727 |
| TcCLB.509797.69 | hypothetical protein, conserved (fragment) | -0,09621 | 0,5196 |

| | | | |
|---|---|---|---|
| TcCLB.506297.180 | beta-adaptin 1, putative | -0,09638 | 0,4269 |
| TcCLB.508897.40 | hypothetical protein, conserved | -0,09654 | 0,3358 |
| TcCLB.510187.500 | serine/threonine protein phosphatase 2B catalytic subunit | -0,09654 | 0,5287 |
| TcCLB.507809.90 | hypothetical protein, conserved | -0,0966 | 0,4486 |
| TcCLB.506941.210 | hypothetical protein, conserved | -0,09668 | 0,565 |
| TcCLB.506359.30 | hypothetical protein, conserved (pseudogene) | -0,0967 | 0,523 |
| TcCLB.508413.76 | kinetoplastid membrane protein KMP-11 | -0,0967 | 0,5276 |
| TcCLB.506333.19 | kinesin, putative (fragment) | -0,09673 | 0,3908 |
| TcCLB.509965.260 | protein kinase, putative | -0,09677 | 0,5098 |
| TcCLB.508899.30 | hypothetical protein, conserved | -0,09679 | 0,5838 |
| TcCLB.507945.30 | pseudouridine synthase TruD, putative (fragment) | -0,0968 | 0,4824 |
| TcCLB.506359.10 | hypothetical protein, conserved | -0,09688 | 0,6555 |
| TcCLB.507717.20 | hypothetical protein | -0,09693 | 0,4134 |
| TcCLB.509213.80 | hypothetical protein, conserved | -0,09703 | 0,6165 |
| TcCLB.507925.69 | hypothetical protein, conserved | -0,09706 | 0,6734 |
| TcCLB.509921.60 | dispersed gene family protein 1 (DGF-1), putative | -0,09706 | 0,7545 |
| TcCLB.505183.80 | signal recognition particle, putative | -0,09724 | 0,3687 |
| TcCLB.507071.277 | syntaxin binding protein (pseudogene), putative | -0,09725 | 0,859 |
| TcCLB.508147.90 | mucin-associated surface protein (MASP, pseudogene), pu | -0,09731 | 0,782 |
| TcCLB.511111.10 | translation initiation factor IF-2, putative | -0,09734 | 0,2855 |
| TcCLB.508879.80 | hypothetical protein, conserved | -0,09734 | 0,5504 |
| TcCLB.508337.10 | hypothetical protein, conserved (fragment) | -0,09738 | 0,4469 |
| TcCLB.506813.70 | mucin-associated surface protein (MASP, pseudogene), pu | -0,09739 | 0,85 |
| TcCLB.505169.10 | polyubiquitin (pseudogene), putative | -0,09758 | 0,5254 |
| TcCLB.509429.4 | retrotransposon hot spot (RHS) protein, putative (fragmen | -0,09768 | 0,3774 |
| TcCLB.510943.184 | tyrosine phosphatase, putative | -0,09786 | 0,5323 |
| TcCLB.507879.10 | trans-sialidase, Group II, putative | -0,09786 | 0,5576 |
| TcCLB.505025.100 | mucin TcMUCII (pseudogene), putative | -0,09792 | 0,8507 |
| TcCLB.510731.80 | hypothetical protein, conserved | -0,09797 | 0,3916 |
| TcCLB.510243.60 | hypothetical protein | -0,09797 | 0,4937 |
| TcCLB.506479.50 | rRNA biogenesis protein, putative | -0,09798 | 0,4426 |
| TcCLB.507831.110 | pumilio/PUF RNA binding protein 2, putative (PUF2) | -0,09799 | 0,9021 |
| TcCLB.509913.30 | hypothetical protein, conserved | -0,09804 | 0,342 |
| TcCLB.510517.20 | hypothetical protein, conserved | -0,09807 | 0,6651 |
| TcCLB.506559.260 | ATP-dependent RNA helicase, putative | -0,09816 | 0,3478 |
| TcCLB.503697.30 | bilobe protein, conserved | -0,09817 | 0,4004 |
| TcCLB.507681.240 | hypothetical protein, conserved | -0,09817 | 0,4882 |
| TcCLB.509769.4 | Meckel syndrome type 1 protein, putative (fragment) (mks | -0,09826 | 0,5064 |
| TcCLB.503431.30 | hypothetical protein | -0,09828 | 0,6534 |
| TcCLB.506329.70 | mucin TcMUCII, putative | -0,09833 | 0,8932 |
| TcCLB.510351.110 | hypothetical protein | -0,09842 | 0,5108 |
| TcCLB.509757.40 | hypothetical protein, conserved | -0,0985 | 0,6842 |
| TcCLB.503891.90 | hypothetical protein | -0,09851 | 0,6136 |
| TcCLB.508325.110 | hypothetical protein, conserved | -0,09862 | 0,7294 |
| TcCLB.411915.10 | glycine dehydrogenase [decarboxylating] (pseudogene), pu | -0,09862 | 0,8195 |

| | | | |
|---|---|---|---|
| TcCLB.509891.10 | hypothetical protein, conserved | -0,09877 | 0,6167 |
| TcCLB.504105.40 | hypothetical protein, conserved | -0,09882 | 0,3734 |
| TcCLB.508537.10 | glyceraldehyde 3-phosphate dehydrogenase, cytosolic, puta | -0,09893 | 0,7858 |
| TcCLB.507735.69 | hypothetical protein, conserved | -0,0991 | 0,634 |
| TcCLB.511165.60 | DNA repair protein, putative | -0,09911 | 0,5956 |
| TcCLB.509081.40 | surface protease GP63 (pseudogene), putative | -0,0994 | 0,8517 |
| TcCLB.508739.40 | hypothetical protein, conserved | -0,09941 | 0,3442 |
| TcCLB.511451.100 | trans-sialidase (pseudogene), putative | -0,09944 | 0,777 |
| TcCLB.510243.54 | hypothetical protein, conserved | -0,09953 | 0,5524 |
| TcCLB.506789.300 | hypothetical protein, conserved | -0,09956 | 0,6657 |
| TcCLB.474341.9 | dynein heavy chain, putative | -0,09962 | 0,4596 |
| TcCLB.504129.30 | hypothetical protein, conserved | -0,09967 | 0,4978 |
| TcCLB.511421.155 | hypothetical protein, conserved | -0,09967 | 0,5417 |
| TcCLB.504045.5 | hypothetical protein, conserved (fragment) | -0,09996 | 0,791 |
| TcCLB.510919.9 | trans-sialidase, putative (fragment) | -0,09999 | 0,7872 |
| TcCLB.508837.80 | Flagellar Member 3 (FLAM3) | -0,1 | 0,4476 |
| TcCLB.509453.70 | protein transport protein Sec24C, putative | -0,1001 | 0,316 |
| TcCLB.510533.40 | Mu-adaptin 1, putative | -0,1001 | 0,4277 |
| TcCLB.507083.60 | COP9 signalosome complex subunit 5, putative | -0,1001 | 0,6828 |
| TcCLB.504109.195 | N-acetyltransferase complex ARD1 subunit (pseudogene), | -0,1001 | 0,8343 |
| TcCLB.509317.109 | hypothetical protein, conserved (fragment) | -0,1002 | 0,5243 |
| TcCLB.503977.70 | mucin-associated surface protein (MASP, pseudogene), pu | -0,1002 | 0,8104 |
| TcCLB.509059.40 | hypothetical protein, conserved | -0,1003 | 0,3797 |
| TcCLB.509029.170 | palmitoyl acyltransferase 5, putative | -0,1003 | 0,5017 |
| TcCLB.507563.9 | dispersed gene family protein 1 (DGF-1), putative (fragme | -0,1003 | 0,5844 |
| TcCLB.509791.10 | hypothetical protein, conserved | -0,1003 | 0,7267 |
| TcCLB.508175.350 | zinc finger protein family member, putative (POMP35) | -0,1004 | 0,4961 |
| TcCLB.504741.70 | hypothetical protein, conserved | -0,1004 | 0,4966 |
| TcCLB.508355.270 | hypothetical protein | -0,1004 | 0,4974 |
| TcCLB.510697.50 | trans-sialidase, Group VI, putative | -0,1004 | 0,8024 |
| TcCLB.511751.60 | hypothetical protein, conserved | -0,1006 | 0,3857 |
| TcCLB.509695.70 | hypothetical protein, conserved | -0,1006 | 0,4502 |
| TcCLB.503847.70 | hypothetical protein, conserved | -0,1006 | 0,4597 |
| TcCLB.506475.113 | hypothetical protein, conserved | -0,1007 | 0,5011 |
| TcCLB.504313.40 | hypothetical protein, conserved (pseudogene) | -0,1007 | 0,5528 |
| TcCLB.506221.20 | hypothetical protein, conserved | -0,1007 | 0,6202 |
| TcCLB.506795.100 | hypothetical protein | -0,1007 | 0,8146 |
| TcCLB.511233.7 | 60S ribosomal protein L10a, putative | -0,1008 | 0,3628 |
| TcCLB.506821.130 | hypothetical protein, conserved | -0,1008 | 0,4009 |
| TcCLB.504017.30 | hypothetical protein, conserved | -0,1008 | 0,4462 |
| TcCLB.509717.30 | hypothetical protein, conserved | -0,1008 | 0,4704 |
| TcCLB.510741.60 | hypothetical protein, conserved | -0,1008 | 0,519 |
| TcCLB.509247.60 | hypothetical protein, conserved (pseudogene) | -0,1008 | 0,5403 |
| TcCLB.508827.30 | eukaryotic translation initiation factor, putative | -0,1009 | 0,3173 |
| TcCLB.506155.50 | vacuolar protein sorting-associated protein-like, putative | -0,1009 | 0,5356 |

| | | | |
|---|---|---|---|
| TcCLB.506289.20 | hypothetical protein, conserved | -0,1009 | 0,5616 |
| TcCLB.510659.4 | hypothetical protein, conserved | -0,101 | 0,6541 |
| TcCLB.506963.14 | 40S ribosomal protein S27, putative | -0,1012 | 0,363 |
| TcCLB.510747.40 | surface protease GP63, putative | -0,1012 | 0,6616 |
| TcCLB.510023.10 | hypothetical protein, conserved (pseudogene) | -0,1012 | 0,8705 |
| TcCLB.511463.10 | rRNA dimethyltransferase, putative | -0,1015 | 0,5489 |
| TcCLB.511755.119 | hypothetical protein, conserved (fragment) | -0,1016 | 0,3122 |
| TcCLB.505999.10 | phosphate-repressible phosphate permease, putative | -0,1017 | 0,4639 |
| TcCLB.508917.10 | protein kinase, putative | -0,1017 | 0,4699 |
| TcCLB.510445.20 | chromosomal passenger protein, putative (CPC1) | -0,1017 | 0,4976 |
| TcCLB.508979.80 | hypothetical protein, conserved | -0,1017 | 0,5804 |
| TcCLB.506529.340 | Protein kinase domain, putative | -0,1017 | 0,6127 |
| TcCLB.506177.110 | hypothetical protein, conserved | -0,1018 | 0,6785 |
| TcCLB.508971.30 | lysyl-tRNA synthetase, putative | -0,102 | 0,2088 |
| TcCLB.507073.40 | hypothetical protein, conserved | -0,102 | 0,2994 |
| TcCLB.506637.30 | hypothetical protein, conserved | -0,102 | 0,3807 |
| TcCLB.506839.34 | oxidoreductase, putative | -0,102 | 0,7121 |
| TcCLB.510629.440 | hypothetical protein, conserved | -0,1021 | 0,2923 |
| TcCLB.511811.10 | 40S ribosomal protein S15, putative | -0,1021 | 0,7411 |
| TcCLB.504031.35 | mucin TcMUCII (pseudogene), putative | -0,1022 | 0,8707 |
| TcCLB.508949.20 | hypothetical protein, conserved | -0,1023 | 0,7263 |
| TcCLB.508779.20 | hypothetical protein, conserved | -0,1024 | 0,4661 |
| TcCLB.510329.320 | hypothetical protein, conserved | -0,1026 | 0,6031 |
| TcCLB.511237.100 | hypothetical protein, conserved | -0,1027 | 0,4742 |
| TcCLB.511297.20 | anaphase promoting complex subunit 10, putative | -0,1027 | 0,6486 |
| TcCLB.504239.110 | mucin-associated surface protein (MASP), putative | -0,1027 | 0,8604 |
| TcCLB.506767.371 | hypothetical protein, conserved (pseudogene) | -0,1027 | 0,8991 |
| TcCLB.503897.70 | hypothetical protein, conserved | -0,1028 | 0,5354 |
| TcCLB.507963.50 | nuclear cap binding complex subunit CBP30, putative (CBP3 | -0,1028 | 0,5476 |
| TcCLB.504141.9 | calcium-translocating P-type ATPase, putative | -0,1028 | 0,6006 |
| TcCLB.506885.330 | hypothetical protein, conserved | -0,103 | 0,3729 |
| TcCLB.506479.90 | hypothetical protein, conserved | -0,103 | 0,5214 |
| TcCLB.511737.70 | hypothetical protein, conserved | -0,103 | 0,5892 |
| TcCLB.506585.100 | hypothetical protein | -0,103 | 0,7486 |
| TcCLB.510281.10 | hypothetical protein | -0,1031 | 0,8705 |
| TcCLB.444331.10 | mucin-associated surface protein (MASP), putative | -0,1031 | 0,8869 |
| TcCLB.506009.70 | hypothetical protein | -0,1032 | 0,5744 |
| TcCLB.510187.470 | hypothetical protein | -0,1032 | 0,6721 |
| TcCLB.509777.21 | hypothetical protein | -0,1032 | 0,7186 |
| TcCLB.510183.20 | dipeptidyl-peptidase 8-like serine peptidase | -0,1033 | 0,3315 |
| TcCLB.506675.99 | trans-sialidase (pseudogene), putative | -0,1033 | 0,8252 |
| TcCLB.503585.10 | mucin-associated surface protein (MASP), putative | -0,1033 | 0,883 |
| TcCLB.511063.140 | trans-sialidase (pseudogene), putative | -0,1034 | 0,8656 |
| TcCLB.511025.20 | hypothetical protein, conserved | -0,1035 | 0,4801 |
| TcCLB.503953.40 | hypothetical protein, conserved | -0,1036 | 0,6573 |

| | | | |
|---|---|---|---|
| TcCLB.506883.60 | hypothetical protein, conserved | -0,1038 | 0,5198 |
| TcCLB.504229.37 | hypothetical protein | -0,1038 | 0,8123 |
| TcCLB.510997.30 | hypothetical protein, conserved | -0,1039 | 0,4048 |
| TcCLB.511277.240 | vacuolar ATP synthase subunit c, putative | -0,1039 | 0,4239 |
| TcCLB.511039.39 | hypothetical protein, conserved | -0,1039 | 0,5292 |
| TcCLB.509497.9 | hypothetical protein, conserved | -0,1039 | 0,5958 |
| TcCLB.507047.110 | otubain cysteine peptidase, Clan CA, family C65, putative | -0,104 | 0,4343 |
| TcCLB.511507.39 | intraflagellar transport (IFT) protein, putative (IFT20) | -0,104 | 0,6072 |
| TcCLB.510451.21 | trans-sialidase (pseudogene), putative | -0,104 | 0,8606 |
| TcCLB.505169.21 | phospholipid transporting ATPase-like protein (pseudogene | -0,1041 | 0,809 |
| TcCLB.511871.40 | hypothetical protein, conserved | -0,1043 | 0,5671 |
| TcCLB.506813.10 | hypothetical protein | -0,1043 | 0,6903 |
| TcCLB.507817.30 | hypothetical protein, conserved | -0,1044 | 0,4781 |
| TcCLB.507041.55 | mucin-associated surface protein (MASP, pseudogene), pu | -0,1044 | 0,7972 |
| TcCLB.509831.33 | hypothetical protein, conserved | -0,1046 | 0,2722 |
| TcCLB.503913.20 | actin-related protein 3, putative | -0,1046 | 0,5053 |
| TcCLB.506961.150 | trans-sialidase, Group VIII, putative | -0,1046 | 0,5748 |
| TcCLB.509611.60 | hypothetical protein | -0,1047 | 0,6763 |
| TcCLB.506297.330 | 40S ribosomal protein S15A, putative | -0,1048 | 0,2384 |
| TcCLB.508089.60 | hypothetical protein, conserved | -0,1048 | 0,5604 |
| TcCLB.511281.53 | hypothetical protein | -0,105 | 0,8353 |
| TcCLB.509163.105 | ATP-dependent DEAD/H RNA helicase (pseudogene), putat | -0,105 | 0,8747 |
| TcCLB.510659.70 | protein kinase-like protein, putative | -0,1051 | 0,4665 |
| TcCLB.508307.170 | 60S acidic ribosomal protein, putative | -0,1051 | 0,5395 |
| TcCLB.510351.140 | hypothetical protein, conserved | -0,1051 | 0,6371 |
| TcCLB.511077.95 | mucin TcMUCII (pseudogene), putative | -0,1051 | 0,9033 |
| TcCLB.506999.110 | hypothetical protein, conserved | -0,1052 | 0,3797 |
| TcCLB.508891.10 | dolichyl-P-Man:GDP-ManGlcNAc2-PP-dolichyl beta-1,4-ma | -0,1052 | 0,6187 |
| TcCLB.504019.9 | hypothetical protein, conserved | -0,1052 | 0,653 |
| TcCLB.506717.4 | hypothetical protein, conserved (fragment) | -0,1052 | 0,7116 |
| TcCLB.508277.65 | hypothetical protein, conserved | -0,1053 | 0,4795 |
| TcCLB.509955.10 | ubiquitin carboxyl-terminal hydrolase, putative (fragment) | -0,1053 | 0,6867 |
| TcCLB.510167.37 | trans-sialidase (pseudogene), putative | -0,1054 | 0,842 |
| TcCLB.508181.20 | 6-phosphofructo-2-kinase 1 | -0,1057 | 0,3336 |
| TcCLB.509967.60 | hypothetical protein, conserved | -0,1057 | 0,5372 |
| TcCLB.507623.90 | hypothetical protein, conserved | -0,1058 | 0,4034 |
| TcCLB.507009.60 | hypothetical protein, conserved | -0,1059 | 0,3338 |
| TcCLB.503789.10 | hypothetical protein | -0,106 | 0,6529 |
| TcCLB.510315.40 | Putative intraflagellar transport protein C3 (fragment) (PIF | -0,1061 | 0,4502 |
| TcCLB.435519.10 | DNA replication licensing factor MCM6, putative | -0,1062 | 0,6873 |
| TcCLB.506763.340 | mucin TcMUCII, putative | -0,1062 | 0,8592 |
| TcCLB.510609.60 | hypothetical protein, conserved | -0,1064 | 0,3342 |
| TcCLB.507507.30 | hypothetical protein, conserved | -0,1064 | 0,4637 |
| TcCLB.508461.130 | elongation factor 1-alpha (ef-1-alpha), putative | -0,1065 | 0,2939 |
| TcCLB.510691.20 | hypothetical protein, conserved | -0,1065 | 0,3947 |

| | | | |
|---|---|---|---|
| TcCLB.510105.180 | hypothetical protein, conserved | -0,1065 | 0,4437 |
| TcCLB.508919.20 | hypothetical protein, conserved | -0,1066 | 0,3518 |
| TcCLB.506933.24 | hypothetical protein, conserved | -0,1066 | 0,5116 |
| TcCLB.508837.100 | hypothetical protein, conserved | -0,1066 | 0,6158 |
| TcCLB.424771.10 | retrotransposon hot spot (RHS) protein, putative | -0,1068 | 0,7588 |
| TcCLB.507017.110 | hypothetical protein, conserved | -0,1069 | 0,4439 |
| TcCLB.509827.4 | retrotransposon hot spot (RHS) protein, putative | -0,1069 | 0,6657 |
| TcCLB.450061.10 | hypothetical protein, conserved | -0,107 | 0,4198 |
| TcCLB.511247.9 | hypothetical protein, conserved | -0,1072 | 0,8428 |
| TcCLB.506925.330 | hypothetical protein, conserved | -0,1073 | 0,2581 |
| TcCLB.509455.60 | hypothetical protein, conserved | -0,1074 | 0,4972 |
| TcCLB.503783.34 | mucin-associated surface protein (MASP, pseudogene), pu | -0,1074 | 0,7887 |
| TcCLB.506303.171 | hypothetical protein, conserved | -0,1074 | 0,8191 |
| TcCLB.506167.30 | hypothetical protein, conserved | -0,1075 | 0,5976 |
| TcCLB.511517.153 | SH3 domain protein, putative | -0,1076 | 0,3971 |
| TcCLB.511421.100 | Enkuring domain-containig protein | -0,1078 | 0,3503 |
| TcCLB.510055.100 | hypothetical protein, conserved | -0,1079 | 0,3536 |
| TcCLB.504125.70 | hypothetical protein, conserved | -0,1079 | 0,5487 |
| TcCLB.510359.34 | hypothetical protein, conserved | -0,1079 | 0,6022 |
| TcCLB.507295.20 | mucin-associated surface protein (MASP), putative | -0,1079 | 0,7609 |
| TcCLB.510807.30 | hypothetical protein | -0,1079 | 0,7773 |
| TcCLB.506213.40 | hypothetical protein, conserved | -0,108 | 0,2584 |
| TcCLB.511859.20 | hypothetical protein, conserved | -0,108 | 0,5255 |
| TcCLB.503729.19 | hypothetical protein (fragment) | -0,108 | 0,5558 |
| TcCLB.506755.130 | hypothetical protein, conserved | -0,1081 | 0,6806 |
| TcCLB.509055.30 | hypothetical protein, conserved | -0,1082 | 0,4689 |
| TcCLB.507923.20 | U3 small nuclear ribonucleoprotein (snRNP), putative | -0,1083 | 0,2595 |
| TcCLB.510259.24 | hypothetical protein, conserved | -0,1083 | 0,5817 |
| TcCLB.507775.10 | hypothetical protein, conserved | -0,1084 | 0,523 |
| TcCLB.506839.89 | mitochondrial chaperone BCS1, putative (fragment) | -0,1084 | 0,6443 |
| TcCLB.506271.10 | ATP-dependent DEAD/H RNA helicase (pseudogene), putat | -0,1084 | 0,7171 |
| TcCLB.509813.10 | hypothetical protein (pseudogene) | -0,1085 | 0,821 |
| TcCLB.510517.70 | hypothetical protein, conserved | -0,1086 | 0,3055 |
| TcCLB.510241.20 | hypothetical protein, conserved | -0,1086 | 0,3758 |
| TcCLB.510121.9 | hypothetical protein, conserved | -0,1086 | 0,6875 |
| TcCLB.504179.40 | hypothetical protein, conserved | -0,1088 | 0,3305 |
| TcCLB.506989.100 | Triple RNA binding domain protein 3 | -0,1088 | 0,4577 |
| TcCLB.511923.49 | mucin-associated surface protein (MASP, pseudogene), pu | -0,1088 | 0,9196 |
| TcCLB.511545.170 | hypothetical protein, conserved | -0,1089 | 0,5478 |
| TcCLB.507023.190 | DNA-J protein, putative | -0,1089 | 0,561 |
| TcCLB.510329.350 | hypothetical protein, conserved | -0,1092 | 0,4688 |
| TcCLB.509251.20 | trans-sialidase, Group V, putative | -0,1092 | 0,6965 |
| TcCLB.506201.20 | 60S ribosomal protein L13a, putative | -0,1093 | 0,2991 |
| TcCLB.510231.29 | multidrug resistance-associated protein, putative | -0,1094 | 0,4297 |
| TcCLB.506265.10 | hypothetical protein, conserved | -0,1094 | 0,5727 |

| | | | |
|---|---|---|---|
| TcCLB.503777.10 | suppressive immunomodulating factor, putative (TSIF) | -0,1095 | 0,3768 |
| TcCLB.510339.50 | spermidine synthase, putative | -0,1095 | 0,4309 |
| TcCLB.511075.60 | protein kinase, putative | -0,1097 | 0,5507 |
| TcCLB.510623.50 | mucin-associated surface protein (MASP, pseudogene), pu | -0,1099 | 0,7724 |
| TcCLB.506769.4 | mucin-associated surface protein (MASP, pseudogene), pu | -0,1099 | 0,8329 |
| TcCLB.506007.20 | RNA editing complex protein MP18, putative | -0,11 | 0,5015 |
| TcCLB.507089.120 | hypothetical protein, conserved | -0,11 | 0,6573 |
| TcCLB.507159.30 | hypothetical protein, conserved | -0,1101 | 0,342 |
| TcCLB.510687.70 | hypothetical protein, conserved | -0,1101 | 0,6097 |
| TcCLB.508771.10 | cysteine peptidase, Clan CA, family C51, putative | -0,1102 | 0,7847 |
| TcCLB.510303.100 | hypothetical protein, conserved | -0,1103 | 0,4563 |
| TcCLB.510903.50 | phosphonopyruvate decarboxylase, putative | -0,1103 | 0,4586 |
| TcCLB.508777.150 | hypothetical protein | -0,1103 | 0,4779 |
| TcCLB.508207.135 | hypothetical protein, conserved | -0,1104 | 0,4955 |
| TcCLB.506813.220 | trans-sialidase, Group V, putative | -0,1104 | 0,7922 |
| TcCLB.503855.30 | adenylosuccinate lyase, putative (ADSL) | -0,1105 | 0,3237 |
| TcCLB.508689.20 | hypothetical protein, conserved | -0,1105 | 0,3466 |
| TcCLB.507001.140 | hypothetical protein, conserved | -0,1105 | 0,483 |
| TcCLB.509747.10 | hypothetical protein, conserved | -0,1105 | 0,6414 |
| TcCLB.508221.870 | trans-sialidase, Group V, putative | -0,1105 | 0,7636 |
| TcCLB.511649.80 | myosin heavy chain, putative | -0,1106 | 0,3442 |
| TcCLB.506127.110 | eukaryotic peptide chain release factor subunit 1, putative | -0,1107 | 0,2874 |
| TcCLB.511245.170 | hypothetical protein, conserved | -0,1107 | 0,3284 |
| TcCLB.511469.90 | hypothetical protein | -0,1107 | 0,5925 |
| TcCLB.504127.10 | surface protease GP63 (pseudogene), putative | -0,1107 | 0,7579 |
| TcCLB.510363.320 | mucin-associated surface protein (MASP), putative | -0,1108 | 0,6266 |
| TcCLB.507709.130 | hypothetical protein, conserved | -0,1109 | 0,5557 |
| TcCLB.506793.60 | hypothetical protein, conserved | -0,111 | 0,4765 |
| TcCLB.511171.90 | mucin-associated surface protein (MASP), putative | -0,111 | 0,8812 |
| TcCLB.504157.50 | DnaJ chaperone protein, putative | -0,1111 | 0,4841 |
| TcCLB.508227.20 | mucin-associated surface protein (MASP), putative | -0,1111 | 0,7638 |
| TcCLB.506203.129 | DNA-directed RNA polymerase, mitochondrial (fragment) | -0,1112 | 0,2965 |
| TcCLB.504103.109 | hypothetical protein, conserved | -0,1112 | 0,5364 |
| TcCLB.509051.60 | ubiquitin-protein ligase, putative (fragment) | -0,1112 | 0,7383 |
| TcCLB.503459.20 | hypothetical protein, conserved | -0,1113 | 0,5029 |
| TcCLB.509779.40 | hypothetical protein, conserved | -0,1114 | 0,573 |
| TcCLB.506855.310 | hypothetical protein, conserved | -0,1115 | 0,2609 |
| TcCLB.511869.9 | dynein heavy chain, putative (fragment) | -0,1115 | 0,4339 |
| TcCLB.504045.50 | hypothetical protein | -0,1117 | 0,4478 |
| TcCLB.509117.19 | hypothetical protein, conserved | -0,1118 | 0,5008 |
| TcCLB.508355.50 | hypothetical protein, conserved | -0,1118 | 0,5649 |
| TcCLB.508409.180 | hypothetical protein, conserved | -0,1119 | 0,3001 |
| TcCLB.509831.40 | hypothetical protein, conserved | -0,1119 | 0,5386 |
| TcCLB.510661.160 | hypothetical protein, conserved | -0,1119 | 0,5444 |
| TcCLB.507991.60 | mismatch repair protein MLH1, putative | -0,112 | 0,4518 |

| | | | |
|---|---|---|---|
| TcCLB.506885.300 | hypothetical protein, conserved | -0,112 | 0,5962 |
| TcCLB.510065.40 | hypothetical protein, conserved | -0,1122 | 0,2479 |
| TcCLB.511787.30 | hypothetical protein, conserved (pseudogene) | -0,1122 | 0,721 |
| TcCLB.508919.30 | hypothetical protein, conserved | -0,1123 | 0,5352 |
| TcCLB.511391.100 | hypothetical protein, conserved | -0,1123 | 0,5735 |
| TcCLB.510323.30 | 60S ribosomal protein L13, putative | -0,1124 | 0,2148 |
| TcCLB.506885.110 | hypothetical protein, conserved | -0,1125 | 0,2264 |
| TcCLB.509937.140 | hypothetical protein, conserved | -0,1125 | 0,5014 |
| TcCLB.506741.60 | mucin-associated surface protein (MASP), putative | -0,1125 | 0,8175 |
| TcCLB.508027.60 | hypothetical protein | -0,1126 | 0,3464 |
| TcCLB.509437.50 | AAA ATPase, putative | -0,1127 | 0,3906 |
| TcCLB.509901.110 | hypothetical protein, conserved | -0,1127 | 0,5759 |
| TcCLB.506489.30 | hypothetical protein, conserved | -0,1129 | 0,3582 |
| TcCLB.510325.10 | ubiquitin hydrolase, putative | -0,1129 | 0,4562 |
| TcCLB.511277.270 | tubulin cofactor C domain-containing protein RP2, putative | -0,113 | 0,3332 |
| TcCLB.509233.40 | hypothetical protein, conserved | -0,113 | 0,5369 |
| TcCLB.505071.50 | hypothetical protein, conserved | -0,113 | 0,5966 |
| TcCLB.508641.210 | hypothetical protein, conserved | -0,1132 | 0,5926 |
| TcCLB.510175.10 | retrotransposon hot spot protein (RHS, pseudogene), putat | -0,1132 | 0,703 |
| TcCLB.506997.9 | dispersed gene family protein 1 (DGF-1), putative | -0,1132 | 0,7788 |
| TcCLB.506397.10 | trans-sialidase, Group II, putative | -0,1132 | 0,9522 |
| TcCLB.508319.30 | hypothetical protein, conserved | -0,1133 | 0,1908 |
| TcCLB.507213.20 | protein kinase, putative | -0,1134 | 0,4986 |
| TcCLB.507505.20 | DnaJ chaperone protein (pseudogene), putative | -0,1135 | 0,4604 |
| TcCLB.511229.39 | hypothetical protein, conserved | -0,1136 | 0,6525 |
| TcCLB.508605.40 | retrotransposon hot spot protein (RHS, pseudogene), putat | -0,1137 | 0,6276 |
| TcCLB.506457.110 | hypothetical protein, conserved | -0,1138 | 0,4649 |
| TcCLB.509379.29 | hypothetical protein, conserved | -0,1138 | 0,6499 |
| TcCLB.508355.330 | hypothetical protein, conserved | -0,1139 | 0,2308 |
| TcCLB.506203.30 | SWI/SNF-related matrix-associated actin-dependent regul | -0,114 | 0,3165 |
| TcCLB.511585.60 | hypothetical protein, conserved | -0,114 | 0,5189 |
| TcCLB.511031.30 | hypothetical protein, conserved | -0,1141 | 0,5119 |
| TcCLB.503759.59 | hypothetical protein, conserved | -0,1141 | 0,5211 |
| TcCLB.506477.30 | tubulin-tyrosine ligase, putative | -0,1142 | 0,4345 |
| TcCLB.510601.20 | hypothetical protein, conserved | -0,1142 | 0,4763 |
| TcCLB.506265.190 | hypothetical protein, conserved | -0,1143 | 0,4765 |
| TcCLB.504153.10 | hypothetical protein, conserved | -0,1143 | 0,6198 |
| TcCLB.509937.120 | hypothetical protein, conserved | -0,1143 | 0,664 |
| TcCLB.509001.10 | hypothetical protein, conserved | -0,1144 | 0,3718 |
| TcCLB.506467.20 | hypothetical protein, conserved | -0,1144 | 0,3942 |
| TcCLB.511143.12 | hypothetical protein, conserved | -0,1146 | 0,3656 |
| TcCLB.507305.11 | casein kinase (pseudogene), putative | -0,1147 | 0,1448 |
| TcCLB.511725.160 | hypothetical protein, conserved | -0,1147 | 0,1926 |
| TcCLB.510885.60 | hypothetical protein, conserved | -0,1147 | 0,302 |
| TcCLB.511909.70 | hypothetical protein, conserved | -0,1147 | 0,5637 |

| | | | |
|---|---|---|---|
| TcCLB.511761.50 | sodium/hydrogen exchanger, putative (FTP) | -0,1148 | 0,4366 |
| TcCLB.440099.10 | hypothetical protein | -0,1148 | 0,5276 |
| TcCLB.511865.50 | hypothetical protein, conserved | -0,1148 | 0,5692 |
| TcCLB.506767.70 | mucin-associated surface protein (MASP), putative | -0,115 | 0,7884 |
| TcCLB.510431.210 | vesicular protein trafficking mediator, putative | -0,1151 | 0,4549 |
| TcCLB.503899.30 | 40S ribosomal protein S16, putative | -0,1151 | 0,5901 |
| TcCLB.511109.60 | hypothetical protein, conserved | -0,1152 | 0,2984 |
| TcCLB.506833.40 | hypothetical protein | -0,1154 | 0,4569 |
| TcCLB.509695.170 | 60S ribosomal protein L9, putative | -0,1155 | 0,09336 |
| TcCLB.503449.20 | copper homeostasis protein, putative | -0,1155 | 0,4642 |
| TcCLB.507095.20 | hypothetical protein | -0,1155 | 0,8085 |
| TcCLB.510647.40 | syntaxin, putative | -0,1156 | 0,4493 |
| TcCLB.503993.10 | trans-sialidase, Group I, putative | -0,1157 | 0,5312 |
| TcCLB.508539.71 | elongation factor 1-gamma (EF-1-gamma, pseudogene), p | -0,1157 | 0,7556 |
| TcCLB.507023.140 | hypothetical protein | -0,1158 | 0,5562 |
| TcCLB.509965.110 | importin alpha, putative | -0,1159 | 0,2221 |
| TcCLB.510327.10 | hypothetical protein | -0,1159 | 0,7934 |
| TcCLB.511847.60 | mitochondrial DNA polymerase I protein B, putative (POLIE | -0,116 | 0,2934 |
| TcCLB.506561.14 | hypothetical protein, conserved | -0,116 | 0,5985 |
| TcCLB.510021.160 | 90 kDa surface protein, putative | -0,116 | 0,7894 |
| TcCLB.506563.160 | hypothetical protein, conserved | -0,1162 | 0,3034 |
| TcCLB.509081.166 | trans-sialidase, putative (fragment) | -0,1163 | 0,8322 |
| TcCLB.506211.230 | hypothetical protein, conserved | -0,1164 | 0,4428 |
| TcCLB.509583.4 | 60S ribosomal protein L44, putative | -0,1165 | 0,2184 |
| TcCLB.508277.190 | ERAD-associated E3 ubiquitin-protein ligase HRD1, putative | -0,1165 | 0,2583 |
| TcCLB.507839.40 | trans-sialidase, Group II, putative | -0,1166 | 0,5528 |
| TcCLB.510967.10 | GPI inositol deacylase precursor, putative | -0,1166 | 0,5957 |
| TcCLB.506775.70 | hypothetical protein, conserved | -0,1167 | 0,4234 |
| TcCLB.442801.10 | mucin TcMUCII, putative | -0,1167 | 0,8439 |
| TcCLB.506835.99 | hypothetical protein, conserved | -0,1168 | 0,1856 |
| TcCLB.507049.20 | hypothetical protein | -0,1168 | 0,5223 |
| TcCLB.508307.50 | hypothetical protein, conserved | -0,1169 | 0,385 |
| TcCLB.508139.10 | retrotransposon hot spot (RHS) protein, putative | -0,1171 | 0,6187 |
| TcCLB.509671.168 | hypothetical protein, conserved | -0,1171 | 0,6311 |
| TcCLB.510197.80 | hypothetical protein, conserved | -0,1172 | 0,687 |
| TcCLB.506741.135 | mucin TcMUCII (pseudogene), putative | -0,1173 | 0,809 |
| TcCLB.428403.5 | hypothetical protein, conserved | -0,1174 | 0,5155 |
| TcCLB.511533.10 | hypothetical protein, conserved | -0,1175 | 0,4382 |
| TcCLB.508579.10 | hypothetical protein, conserved | -0,1176 | 0,5556 |
| TcCLB.506531.30 | hypothetical protein, conserved | -0,1177 | 0,4806 |
| TcCLB.506911.20 | mucin-associated surface protein (MASP), putative | -0,1177 | 0,6322 |
| TcCLB.507611.40 | protein kinase, putative | -0,1178 | 0,845 |
| TcCLB.508029.30 | ATP dependent DEAD-box helicase, putative (RH) | -0,118 | 0,2902 |
| TcCLB.506715.50 | SNARE domain-containing protein, putative (SNARE2.2) | -0,118 | 0,4482 |
| TcCLB.507709.80 | hypothetical protein, conserved | -0,118 | 0,4809 |

| | | | |
|---|---|---|---|
| TcCLB.511923.100 | trans-sialidase (pseudogene), putative | -0,118 | 0,6438 |
| TcCLB.507519.90 | hypothetical protein, conserved | -0,1181 | 0,5216 |
| TcCLB.510823.40 | hypothetical protein, conserved | -0,1182 | 0,2772 |
| TcCLB.511001.100 | SNF-7-like protein, putative | -0,1182 | 0,4016 |
| TcCLB.508323.80 | aquaporin, putative | -0,1182 | 0,5343 |
| TcCLB.507539.49 | hypothetical protein, conserved | -0,1183 | 0,6992 |
| TcCLB.508629.10 | dispersed gene family protein 1 (DGF-1), putative | -0,1184 | 0,4976 |
| TcCLB.506989.130 | hypothetical protein, conserved | -0,1185 | 0,2882 |
| TcCLB.503839.19 | hypothetical protein, conserved | -0,1185 | 0,4486 |
| TcCLB.508937.10 | dispersed gene family protein 1 (DGF-1, pseudogene), puta | -0,1185 | 0,7058 |
| TcCLB.508221.750 | trans-sialidase, Group V, putative | -0,1185 | 0,7582 |
| TcCLB.510579.30 | hypothetical protein, conserved | -0,1186 | 0,2443 |
| TcCLB.507949.29 | conserved protein, unknown function | -0,1188 | 0,5353 |
| TcCLB.503785.20 | hypothetical protein, conserved | -0,1188 | 0,5562 |
| TcCLB.511871.4 | dynein heavy chain, putative | -0,1188 | 0,6064 |
| TcCLB.508809.50 | hypothetical protein | -0,1188 | 0,7058 |
| TcCLB.506503.150 | histone H3 variant, putative | -0,1189 | 0,3586 |
| TcCLB.508833.10 | hypothetical protein | -0,1189 | 0,6443 |
| TcCLB.511115.20 | GTP-binding elongation factor Tu family, putative | -0,119 | 0,2768 |
| TcCLB.509967.160 | epsilon tubulin, putative | -0,119 | 0,3892 |
| TcCLB.511025.120 | CYC2-like cyclin, putative | -0,1192 | 0,3661 |
| TcCLB.511001.210 | hypothetical protein, conserved | -0,1192 | 0,4804 |
| TcCLB.509875.180 | hypothetical protein | -0,1192 | 0,545 |
| TcCLB.508777.170 | hypothetical protein, conserved | -0,1193 | 0,4817 |
| TcCLB.506525.10 | nuclear segregation protein, putative | -0,1194 | 0,124 |
| TcCLB.508289.80 | trans-sialidase (pseudogene), putative | -0,1194 | 0,8048 |
| TcCLB.511173.200 | hypothetical protein | -0,1194 | 0,8235 |
| TcCLB.510315.30 | hypothetical protein, conserved | -0,1196 | 0,5462 |
| TcCLB.511751.250 | hypothetical protein, conserved | -0,1197 | 0,5387 |
| TcCLB.507077.50 | hypothetical protein, conserved | -0,1197 | 0,5416 |
| TcCLB.503893.140 | hypothetical protein, conserved | -0,1198 | 0,4937 |
| TcCLB.510421.240 | diacylglycerol acyltransferase, putative | -0,1198 | 0,5243 |
| TcCLB.442495.10 | cyclic nucleotide-binding domain containing protein, putati | -0,1199 | 0,5277 |
| TcCLB.510073.40 | hypothetical protein, conserved | -0,1199 | 0,6851 |
| TcCLB.510265.70 | hypothetical protein, conserved | -0,12 | 0,2055 |
| TcCLB.508909.80 | hypothetical protein | -0,1201 | 0,5397 |
| TcCLB.507523.60 | mucin-associated surface protein (MASP), putative | -0,1201 | 0,833 |
| TcCLB.511627.140 | hypothetical protein, conserved | -0,1202 | 0,4662 |
| TcCLB.511255.529 | surface protease GP63 (pseudogene), putative | -0,1202 | 0,7545 |
| TcCLB.508307.20 | hypothetical protein, conserved | -0,1205 | 0,4903 |
| TcCLB.511647.50 | hypothetical protein, conserved | -0,1206 | 0,4618 |
| TcCLB.511727.70 | hypothetical protein, conserved | -0,1207 | 0,3342 |
| TcCLB.510617.10 | hypothetical protein, conserved | -0,1207 | 0,5621 |
| TcCLB.506943.10 | mitogen-activated protein kinase 5, putative | -0,1207 | 0,578 |
| TcCLB.506599.35 | hypothetical protein, conserved (pseudogene) | -0,1208 | 0,7949 |

| | | | |
|---|---|---|---|
| TcCLB.510339.20 | hypothetical protein | -0,1209 | 0,3784 |
| TcCLB.508777.180 | prefoldin subunit 2, putative | -0,1209 | 0,4343 |
| TcCLB.510747.140 | hypothetical protein, conserved | -0,1209 | 0,5169 |
| TcCLB.507019.80 | hypothetical protein, conserved | -0,121 | 0,1665 |
| TcCLB.510885.140 | hypothetical protein | -0,121 | 0,4862 |
| TcCLB.511115.49 | hypothetical protein, conserved | -0,121 | 0,7119 |
| TcCLB.506503.110 | DNA-directed RNA polymerases I and III subunit RPAC1, pu | -0,1211 | 0,4159 |
| TcCLB.506577.40 | hypothetical protein, conserved | -0,1211 | 0,4762 |
| TcCLB.507017.80 | hypothetical protein | -0,1211 | 0,5772 |
| TcCLB.509527.70 | mucin TcMUCII, putative | -0,1211 | 0,8239 |
| TcCLB.510889.170 | hypothetical protein, conserved | -0,1212 | 0,2132 |
| TcCLB.509039.30 | hypothetical protein, conserved | -0,1212 | 0,4065 |
| TcCLB.510731.110 | hypothetical protein, conserved | -0,1212 | 0,5282 |
| TcCLB.511355.24 | hypothetical protein, conserved | -0,1212 | 0,5558 |
| TcCLB.511167.110 | hypothetical protein, conserved | -0,1213 | 0,3881 |
| TcCLB.506925.220 | hypothetical protein, conserved | -0,1213 | 0,3882 |
| TcCLB.507257.170 | hypothetical protein, conserved | -0,1213 | 0,486 |
| TcCLB.508193.20 | retrotransposon hot spot protein (RHS, pseudogene), putat | -0,1214 | 0,239 |
| TcCLB.506363.40 | hypothetical protein, conserved | -0,1214 | 0,5156 |
| TcCLB.506599.420 | mucin-associated surface protein (MASP), putative | -0,1214 | 0,8341 |
| TcCLB.507541.30 | casein kinase, putative | -0,1215 | 0,1721 |
| TcCLB.509877.20 | hypothetical protein, conserved | -0,1217 | 0,3706 |
| TcCLB.506263.40 | hypothetical protein, conserved | -0,1217 | 0,4845 |
| TcCLB.507775.40 | hypothetical protein, conserved | -0,1217 | 0,5358 |
| TcCLB.509237.40 | hypothetical protein, conserved | -0,1217 | 0,5454 |
| TcCLB.508977.31 | elongation factor 1-gamma (EF-1-gamma, pseudogene), p | -0,1218 | 0,767 |
| TcCLB.505807.260 | hypothetical protein, conserved | -0,1219 | 0,4263 |
| TcCLB.508139.140 | dispersed gene family protein 1 (DGF-1), putative | -0,1219 | 0,4896 |
| TcCLB.511049.20 | hypothetical protein, conserved | -0,122 | 0,4976 |
| TcCLB.508283.60 | retrotransposon hot spot protein (RHS, pseudogene), putat | -0,122 | 0,7397 |
| TcCLB.510453.9 | dispersed gene family protein 1 (DGF-1), putative | -0,122 | 0,7839 |
| TcCLB.506761.19 | mucin TcMUCII, putative | -0,1221 | 0,8432 |
| TcCLB.503923.50 | GPI transamidase subunit 16, putative (fragment) (GPI16) | -0,1222 | 0,382 |
| TcCLB.507655.9 | dispersed gene family protein 1 (DGF-1), putative | -0,1222 | 0,7349 |
| TcCLB.508601.60 | hypothetical protein, conserved | -0,1223 | 0,4685 |
| TcCLB.503999.40 | hypothetical protein, conserved | -0,1224 | 0,3197 |
| TcCLB.504157.120 | hypothetical protein, conserved | -0,1224 | 0,5793 |
| TcCLB.510289.30 | phosphatidylinositol-4-phosphate 5-kinase, putative | -0,1225 | 0,5001 |
| TcCLB.510701.10 | mucin-associated surface protein (MASP), putative | -0,1225 | 0,7625 |
| TcCLB.503575.27 | hypothetical protein, conserved | -0,1226 | 0,347 |
| TcCLB.508897.90 | hypothetical protein, conserved | -0,1226 | 0,3548 |
| TcCLB.506763.110 | trans-sialidase, Group VII, putative | -0,1226 | 0,8407 |
| TcCLB.511731.20 | hypothetical protein, conserved | -0,1227 | 0,3131 |
| TcCLB.510529.10 | hypothetical protein, conserved | -0,1227 | 0,5157 |
| TcCLB.503655.20 | dispersed gene family protein 1 (DGF-1), putative | -0,1228 | 0,4469 |

| | | | |
|---|---|---|---|
| TcCLB.510355.97 | retrotransposon hot spot protein (RHS, pseudogene), putat | -0,123 | 0,8427 |
| TcCLB.506943.100 | hypothetical protein, conserved | -0,1231 | 0,25 |
| TcCLB.507677.60 | hypothetical protein | -0,1231 | 0,3084 |
| TcCLB.511423.40 | TFIIH basal transcription factor subunit, putative (TSP1) | -0,1231 | 0,4542 |
| TcCLB.504179.9 | pre-mRNA splicing factor ATP-dependent RNA helicase, pu | -0,1231 | 0,578 |
| TcCLB.507951.160 | succinate dehydrogenase flavoprotein subunit | -0,1232 | 0,1385 |
| TcCLB.506181.59 | 40S ribosomal protein S12, putative | -0,1232 | 0,1924 |
| TcCLB.508647.140 | DNA replication licensing factor MCM5, putative | -0,1232 | 0,2859 |
| TcCLB.509965.400 | hypothetical protein, conserved | -0,1232 | 0,3185 |
| TcCLB.408799.10 | hypothetical protein, conserved | -0,1233 | 0,5844 |
| TcCLB.511735.60 | superoxide dismutase, putative | -0,1234 | 0,4201 |
| TcCLB.508831.70 | U5 snRNP-specific 40 kDa protein, putative | -0,1235 | 0,4502 |
| TcCLB.510835.50 | hypothetical protein, conserved | -0,1235 | 0,5067 |
| TcCLB.507209.40 | cdc2-related kinase 9 (CRK9) | -0,1236 | 0,27 |
| TcCLB.508851.140 | hypothetical protein, conserved | -0,1236 | 0,4976 |
| TcCLB.506537.250 | hypothetical protein, conserved | -0,1236 | 0,629 |
| TcCLB.511731.80 | hypothetical protein, conserved | -0,1237 | 0,3878 |
| TcCLB.510667.30 | hypothetical protein, conserved | -0,1238 | 0,3118 |
| TcCLB.508817.5 | dynein heavy chain, putative | -0,1238 | 0,5453 |
| TcCLB.510105.150 | hypothetical protein, conserved | -0,1238 | 0,5918 |
| TcCLB.510903.20 | hypothetical protein, conserved | -0,1239 | 0,6181 |
| TcCLB.510513.60 | hypothetical protein, conserved | -0,124 | 0,5186 |
| TcCLB.504017.60 | hypothetical protein, conserved | -0,1241 | 0,4878 |
| TcCLB.511529.110 | hypothetical protein, conserved | -0,1241 | 0,5763 |
| TcCLB.503441.5 | trans-sialidase, putative (fragment) | -0,1242 | 0,7486 |
| TcCLB.509937.220 | protein kinase, putative | -0,1242 | 0,8305 |
| TcCLB.511487.19 | trans-sialidase (pseudogene), putative | -0,1244 | 0,264 |
| TcCLB.510187.448 | hypothetical protein, conserved | -0,1245 | 0,1784 |
| TcCLB.510355.190 | hypothetical protein | -0,1245 | 0,5256 |
| TcCLB.504411.30 | serine/threonine protein kinase, putative | -0,1245 | 0,6151 |
| TcCLB.506127.160 | hypothetical protein, conserved | -0,1246 | 0,3725 |
| TcCLB.508153.164 | hypothetical protein, conserved | -0,1247 | 0,4196 |
| TcCLB.506223.130 | hypothetical protein, conserved | -0,1247 | 0,4701 |
| TcCLB.509233.70 | Spastin, putative | -0,1248 | 0,302 |
| TcCLB.506303.111 | hypothetical protein, conserved | -0,1248 | 0,3098 |
| TcCLB.504031.60 | mucin-associated surface protein (MASP, pseudogene), pu | -0,1248 | 0,7773 |
| TcCLB.506367.20 | hypothetical protein, conserved | -0,1249 | 0,3169 |
| TcCLB.507951.90 | hypothetical protein, conserved | -0,1249 | 0,4324 |
| TcCLB.506303.140 | hypothetical protein, conserved | -0,125 | 0,3634 |
| TcCLB.511261.130 | hypothetical protein, conserved | -0,125 | 0,5148 |
| TcCLB.506691.40 | hypothetical protein, conserved | -0,1251 | 0,5087 |
| TcCLB.510885.40 | WD repeat and HMG-box DNA-binding protein, putative | -0,1252 | 0,2046 |
| TcCLB.503447.60 | beta galactofuranosyl glycosyltransferase, putative | -0,1252 | 0,8212 |
| TcCLB.511585.100 | hypothetical protein, conserved | -0,1253 | 0,5939 |
| TcCLB.510377.500 | trans-sialidase (pseudogene), putative | -0,1253 | 0,8343 |

| | | | |
|---|---|---|---|
| TcCLB.505843.30 | DNA repair protein Rad9, putative (Rad9) | -0,1254 | 0,2759 |
| TcCLB.509429.150 | hypothetical protein, conserved | -0,1254 | 0,3767 |
| TcCLB.504351.41 | ATP-dependent DEAD/H RNA helicase (pseudogene), putat | -0,1255 | 0,7773 |
| TcCLB.509251.98 | hypothetical protein, conserved (pseudogene) | -0,1257 | 0,7773 |
| TcCLB.509809.30 | hypothetical protein, conserved | -0,1258 | 0,2143 |
| TcCLB.503697.60 | nuclear cap binding complex subunit CBP110, putative (CBI | -0,1258 | 0,2592 |
| TcCLB.511217.160 | hypothetical protein, conserved | -0,1258 | 0,3726 |
| TcCLB.506755.10 | hypothetical protein, conserved | -0,1258 | 0,451 |
| TcCLB.508141.60 | hypothetical protein, conserved | -0,1258 | 0,4802 |
| TcCLB.511577.19 | hypothetical protein, conserved | -0,1258 | 0,6047 |
| TcCLB.506211.200 | metallo-peptidase, Clan MG, Family M24 | -0,1259 | 0,1771 |
| TcCLB.509877.70 | hypothetical protein, conserved | -0,1259 | 0,3611 |
| TcCLB.504041.20 | hypothetical protein, conserved | -0,1259 | 0,4545 |
| TcCLB.509453.90 | hypothetical protein, conserved | -0,1261 | 0,2698 |
| TcCLB.510421.260 | diacylglycerol acyltransferase, putative | -0,1262 | 0,4502 |
| TcCLB.506199.30 | hypothetical protein, conserved | -0,1262 | 0,5189 |
| TcCLB.508815.179 | dynein heavy chain, putative (fragment) | -0,1263 | 0,298 |
| TcCLB.510989.20 | hypothetical protein, conserved | -0,1263 | 0,3339 |
| TcCLB.508153.150 | hypothetical protein, conserved | -0,1264 | 0,382 |
| TcCLB.509171.80 | hypothetical protein, conserved | -0,1264 | 0,3986 |
| TcCLB.506767.74 | mucin TcMUCII (pseudogene), putative | -0,1264 | 0,8318 |
| TcCLB.510293.30 | Eukaryotic translation initiation factor 4E type 5, putative ( | -0,1265 | 0,2983 |
| TcCLB.504153.70 | hypothetical protein, conserved | -0,1265 | 0,3079 |
| TcCLB.503779.80 | hypothetical protein, conserved (pseudogene) | -0,1265 | 0,3453 |
| TcCLB.506529.324 | hypothetical protein, conserved | -0,1265 | 0,5447 |
| TcCLB.507007.34 | hypothetical protein, conserved | -0,1266 | 0,3027 |
| TcCLB.503987.50 | hypothetical protein, conserved | -0,1267 | 0,4171 |
| TcCLB.511587.90 | trans-sialidase, Group VIII, putative | -0,1268 | 0,4442 |
| TcCLB.509573.20 | hypothetical protein, conserved | -0,1268 | 0,655 |
| TcCLB.506941.260 | hypothetical protein, conserved | -0,127 | 0,3913 |
| TcCLB.506435.60 | hypothetical protein, conserved | -0,127 | 0,4156 |
| TcCLB.511721.10 | hypothetical protein, conserved | -0,1271 | 0,3793 |
| TcCLB.427789.30 | NAD synthase, putative | -0,1271 | 0,49 |
| TcCLB.510655.10 | hypothetical protein, conserved | -0,1271 | 0,5148 |
| TcCLB.511857.50 | actin-like protein, putative | -0,1272 | 0,3615 |
| TcCLB.507089.154 | hypothetical protein, conserved | -0,1273 | 0,166 |
| TcCLB.509827.20 | hypothetical protein, conserved | -0,1273 | 0,5772 |
| TcCLB.506175.90 | hypothetical protein, conserved | -0,1274 | 0,361 |
| TcCLB.469735.9 | hypothetical protein, conserved | -0,1274 | 0,471 |
| TcCLB.503671.20 | hypothetical protein, conserved | -0,1275 | 0,5122 |
| TcCLB.508479.450 | hypothetical protein, conserved (pseudogene) | -0,1275 | 0,5555 |
| TcCLB.398235.20 | ubiquitin fusion degardation protein, putative | -0,1275 | 0,7922 |
| TcCLB.511127.400 | serine/threonine protein phosphatase, putative | -0,1276 | 0,3293 |
| TcCLB.401469.20 | pumilio/PUF RNA binding protein 7, putative (PUF7) | -0,1278 | 0,4826 |
| TcCLB.511861.60 | beta galactofuranosyl glycosyltransferase (pseudogene), pı | -0,1278 | 0,596 |

| | | | |
|---|---|---|---|
| TcCLB.511585.30 | hypothetical protein, conserved | -0,1279 | 0,2854 |
| TcCLB.511621.110 | hypothetical protein | -0,1279 | 0,366 |
| TcCLB.508207.210 | hypothetical protein, conserved | -0,1279 | 0,5167 |
| TcCLB.511055.10 | retrotransposon hot spot protein (RHS, pseudogene), putat | -0,128 | 0,2894 |
| TcCLB.511417.90 | phosphate-repressible phosphate permease, putative | -0,128 | 0,564 |
| TcCLB.510431.80 | hypothetical protein, conserved | -0,1282 | 0,1966 |
| TcCLB.506925.470 | heat shock protein DNAJ, putative | -0,1282 | 0,2986 |
| TcCLB.511633.30 | hypothetical protein, conserved | -0,1282 | 0,4595 |
| TcCLB.510695.10 | surface protease GP63 (pseudogene), putative | -0,1282 | 0,6446 |
| TcCLB.506739.70 | hypothetical protein, conserved | -0,1283 | 0,3269 |
| TcCLB.509177.10 | hypothetical protein, conserved | -0,1283 | 0,4753 |
| TcCLB.508489.40 | hypothetical protein, conserved | -0,1285 | 0,4183 |
| TcCLB.507011.70 | hypothetical protein | -0,1286 | 0,4384 |
| TcCLB.506601.40 | mucin-associated surface protein (MASP), putative | -0,1286 | 0,818 |
| TcCLB.508179.10 | DnaJ chaperone protein, putative | -0,1288 | 0,5517 |
| TcCLB.505807.230 | hypothetical protein | -0,1289 | 0,5963 |
| TcCLB.506831.60 | hypothetical protein, conserved | -0,129 | 0,3762 |
| TcCLB.508033.20 | hypothetical protein | -0,129 | 0,8458 |
| TcCLB.468217.30 | trans-sialidase, putative | -0,1291 | 0,661 |
| TcCLB.508221.970 | mucin-associated surface protein (MASP), putative | -0,1291 | 0,735 |
| TcCLB.507809.14 | protein kinase, putative | -0,1293 | 0,4034 |
| TcCLB.511001.170 | hypothetical protein, conserved | -0,1294 | 0,2101 |
| TcCLB.509563.30 | hypothetical protein, conserved | -0,1294 | 0,3199 |
| TcCLB.510355.220 | GDP-mannose transporter, putative | -0,1294 | 0,5814 |
| TcCLB.505123.10 | hypothetical protein, conserved | -0,1294 | 0,6346 |
| TcCLB.508465.140 | hypothetical protein, conserved | -0,1296 | 0,2187 |
| TcCLB.506525.140 | MEKK-related kinase 1, putative (MRK1) | -0,1296 | 0,303 |
| TcCLB.511367.320 | hypothetical protein, conserved | -0,1296 | 0,3506 |
| TcCLB.506619.40 | tyrosyl-DNA Phosphodiesterase (Tdp1), putative | -0,1296 | 0,3762 |
| TcCLB.509005.30 | hypothetical protein, conserved | -0,1297 | 0,3754 |
| TcCLB.507089.220 | hypothetical protein, conserved | -0,1298 | 0,4169 |
| TcCLB.510239.60 | mucin TcMUCII, putative | -0,1299 | 0,8397 |
| TcCLB.506195.250 | hypothetical protein, conserved | -0,1301 | 0,2577 |
| TcCLB.508347.70 | Engulfment and cell motility domain-containing protein (P | -0,1301 | 0,3812 |
| TcCLB.510917.9 | mucin-associated surface protein (MASP), putative | -0,1301 | 0,7443 |
| TcCLB.503891.100 | 20S proteasome subunit | -0,1302 | 0,3805 |
| TcCLB.511183.280 | mucin-associated surface protein (MASP), putative | -0,1302 | 0,7616 |
| TcCLB.504007.20 | trans-sialidase, Group V, putative | -0,1302 | 0,7708 |
| TcCLB.510565.100 | hypothetical protein, conserved | -0,1303 | 0,4901 |
| TcCLB.508257.100 | hypothetical protein, conserved | -0,1303 | 0,5438 |
| TcCLB.509901.120 | hypothetical protein, conserved | -0,1304 | 0,4888 |
| TcCLB.509815.120 | dispersed gene family protein 1 (DGF-1, pseudogene), puta | -0,1304 | 0,5421 |
| TcCLB.510657.120 | hypothetical protein, conserved | -0,1304 | 0,6509 |
| TcCLB.507005.60 | structural maintenance of chromosome (SMC) family prote | -0,1305 | 0,1537 |
| TcCLB.511741.40 | RNA-binding protein, putative | -0,1305 | 0,3725 |

| | | | |
|---|---|---|---|
| TcCLB.511671.110 | hypothetical protein, conserved | -0,1305 | 0,5576 |
| TcCLB.509439.20 | UDP-Gal or UDP-GlcNAc-dependent glycosyltransferase, pu | -0,1306 | 0,7831 |
| TcCLB.509525.90 | mucin TcMUCII, putative | -0,1306 | 0,842 |
| TcCLB.510355.210 | hypothetical protein, conserved | -0,1307 | 0,2381 |
| TcCLB.506321.210 | mucin TcMUCII, putative | -0,1307 | 0,8207 |
| TcCLB.508811.55 | hypothetical protein | -0,1309 | 0,1517 |
| TcCLB.508737.200 | hypothetical protein, conserved | -0,1309 | 0,2294 |
| TcCLB.511907.110 | fatty acid transporter protein-like, putative | -0,1309 | 0,2403 |
| TcCLB.503893.170 | hypothetical protein, conserved | -0,1312 | 0,3116 |
| TcCLB.507821.10 | trans-sialidase, Group VI, putative | -0,1312 | 0,5617 |
| TcCLB.507941.70 | hypothetical protein, conserved | -0,1313 | 0,3077 |
| TcCLB.506743.120 | Elongator-like Protein 3a, putative | -0,1313 | 0,3723 |
| TcCLB.510047.109 | mucin TcMUCII, putative | -0,1313 | 0,7339 |
| TcCLB.507047.180 | hypothetical protein, conserved | -0,1314 | 0,5733 |
| TcCLB.508451.5 | mucin-associated surface protein (MASP, pseudogene), pu | -0,1314 | 0,5829 |
| TcCLB.510085.10 | mucin-associated surface protein (MASP, pseudogene), pu | -0,1315 | 0,7754 |
| TcCLB.508739.80 | hypothetical protein, conserved | -0,1316 | 0,1785 |
| TcCLB.504257.20 | hypothetical protein, conserved | -0,1317 | 0,2831 |
| TcCLB.507087.20 | hypothetical protein | -0,1317 | 0,3962 |
| TcCLB.510329.360 | hypothetical protein, conserved | -0,1317 | 0,4804 |
| TcCLB.508939.30 | mucin-associated surface protein (MASP), putative | -0,1317 | 0,7894 |
| TcCLB.510517.80 | hypothetical protein, conserved | -0,1318 | 0,6101 |
| TcCLB.507583.4 | hypothetical protein, conserved (fragment) | -0,1319 | 0,6276 |
| TcCLB.509761.10 | hypothetical protein, conserved | -0,132 | 0,1112 |
| TcCLB.506701.29 | hypothetical protein, conserved | -0,132 | 0,4896 |
| TcCLB.427743.9 | translation initiation factor eIF2B delta subunit, putative (f | -0,1323 | 0,6023 |
| TcCLB.506893.90 | hypothetical protein, conserved | -0,1324 | 0,605 |
| TcCLB.511183.40 | mucin-associated surface protein (MASP, pseudogene), pu | -0,1324 | 0,7952 |
| TcCLB.505999.130 | phosphatidylinositol-4-phosphate 5-kinase related | -0,1325 | 0,4929 |
| TcCLB.431357.10 | 3,2-trans-enoyl-CoA isomerase, mitochondrial precursor, p | -0,1325 | 0,5918 |
| TcCLB.506941.80 | tubulin-tyrosine ligase-like protein, putative | -0,1326 | 0,356 |
| TcCLB.509671.40 | hypothetical protein, conserved | -0,1327 | 0,4633 |
| TcCLB.510359.140 | hypothetical protein, conserved | -0,1327 | 0,6094 |
| TcCLB.510759.60 | vesicle-associated membrane protein, putative | -0,1328 | 0,4459 |
| TcCLB.503925.110 | hypothetical protein, conserved | -0,1328 | 0,5452 |
| TcCLB.508207.40 | hypothetical protein, conserved | -0,1328 | 0,5879 |
| TcCLB.504147.200 | Regulatory particle triple-A ATPase subunit 6 (RPT6) | -0,1329 | 0,2801 |
| TcCLB.507601.90 | hypothetical protein, conserved | -0,1331 | 0,3092 |
| TcCLB.508741.330 | hypothetical protein, conserved | -0,1331 | 0,3157 |
| TcCLB.510435.20 | hypothetical protein, conserved | -0,1331 | 0,4365 |
| TcCLB.506943.110 | hypothetical protein | -0,1331 | 0,5546 |
| TcCLB.508837.147 | kinesin, putative | -0,1332 | 0,298 |
| TcCLB.503981.50 | hypothetical protein, conserved | -0,1332 | 0,4343 |
| TcCLB.506567.100 | hypothetical protein, conserved | -0,1334 | 0,3442 |
| TcCLB.510087.20 | hypothetical protein, conserved | -0,1334 | 0,4225 |

| | | | |
|---|---|---|---|
| TcCLB.511903.134 | hypothetical protein, conserved | -0,1334 | 0,5306 |
| TcCLB.506943.30 | hypothetical protein, conserved | -0,1335 | 0,2497 |
| TcCLB.506475.14 | phosphatidylinositol 3-kinase, putative | -0,1335 | 0,311 |
| TcCLB.506755.210 | hypothetical protein, conserved | -0,1335 | 0,4523 |
| TcCLB.506729.30 | hypothetical protein, conserved | -0,1336 | 0,5578 |
| TcCLB.511787.20 | trans-sialidase (pseudogene), putative | -0,1336 | 0,6933 |
| TcCLB.508461.149 | hypothetical protein, conserved | -0,1337 | 0,2706 |
| TcCLB.511301.180 | trans-sialidase (pseudogene), putative | -0,1337 | 0,392 |
| TcCLB.436137.9 | dispersed gene family protein 1 (DGF-1), putative (fragme | -0,1337 | 0,5064 |
| TcCLB.506391.20 | calmodulin, putative | -0,1339 | 0,4274 |
| TcCLB.506247.70 | nuclear pore complex protein (NUP155), putative | -0,134 | 0,09755 |
| TcCLB.504227.10 | hypothetical protein, conserved | -0,134 | 0,405 |
| TcCLB.510943.140 | DNA primase large subunit, putative (pril) | -0,1341 | 0,285 |
| TcCLB.466965.10 | trans-sialidase (pseudogene), putative | -0,1343 | 0,5121 |
| TcCLB.510131.20 | hypothetical protein, conserved | -0,1343 | 0,5905 |
| TcCLB.507067.30 | mucin-associated surface protein (MASP, pseudogene), pu | -0,1343 | 0,6399 |
| TcCLB.507283.10 | trans-sialidase, Group V, putative | -0,1344 | 0,3842 |
| TcCLB.507601.30 | hypothetical protein, conserved | -0,1344 | 0,5326 |
| TcCLB.509695.90 | intraflagellar transport protein 172, putative (IFT172) | -0,1345 | 0,2342 |
| TcCLB.510099.100 | hypothetical protein, conserved | -0,1345 | 0,24 |
| TcCLB.506865.10 | ARP2/3 complex subunit, putative | -0,1345 | 0,4502 |
| TcCLB.506041.60 | hypothetical protein, conserved | -0,1346 | 0,2491 |
| TcCLB.511655.69 | ubiquitin-activating enzyme-like protein | -0,1346 | 0,4033 |
| TcCLB.477471.10 | trans-sialidase (pseudogene), putative | -0,1346 | 0,4995 |
| TcCLB.506929.60 | ATP-dependent DEAD/H RNA helicase (pseudogene), putat | -0,1346 | 0,5219 |
| TcCLB.507003.70 | hypothetical protein, conserved | -0,1346 | 0,5364 |
| TcCLB.507105.70 | ABC transporter, putative | -0,1347 | 0,2888 |
| TcCLB.503809.149 | TATA binding protein (TBP) | -0,1348 | 0,3802 |
| TcCLB.422921.20 | 50S ribosomal protein L17, putative | -0,1349 | 0,6156 |
| TcCLB.506759.70 | mucin TcMUCII, putative | -0,1349 | 0,8849 |
| TcCLB.510143.80 | RNA-binding protein, putative | -0,1351 | 0,2557 |
| TcCLB.504159.100 | hypothetical protein, conserved | -0,1351 | 0,5188 |
| TcCLB.506181.110 | hypothetical protein, conserved | -0,1351 | 0,632 |
| TcCLB.506925.140 | DNA repair protein, putative | -0,1352 | 0,2071 |
| TcCLB.508445.99 | pumilio/PUF RNA binding protein 7, putative (PUF7) | -0,1353 | 0,3715 |
| TcCLB.508741.180 | hypothetical protein, conserved | -0,1353 | 0,4182 |
| TcCLB.508175.200 | hypothetical protein, conserved | -0,1353 | 0,4323 |
| TcCLB.510279.310 | mucin-associated surface protein (MASP), putative | -0,1354 | 0,7186 |
| TcCLB.511491.20 | hypothetical protein, conserved | -0,1356 | 0,1276 |
| TcCLB.510091.130 | hypothetical protein, conserved | -0,1356 | 0,2594 |
| TcCLB.503611.10 | nucleoplasmin-like protein (NLP), putative | -0,1357 | 0,3076 |
| TcCLB.508219.170 | trans-sialidase (pseudogene), putative | -0,1357 | 0,7536 |
| TcCLB.509429.50 | retrotransposon hot spot protein (RHS, pseudogene), putat | -0,1358 | 0,2425 |
| TcCLB.510025.230 | hypothetical protein | -0,1358 | 0,8575 |
| TcCLB.503893.40 | hypothetical protein, conserved | -0,1359 | 0,4263 |

| | | | |
|---|---|---|---|
| TcCLB.509671.172 | hypothetical protein, conserved | -0,1359 | 0,5089 |
| TcCLB.507953.190 | hypothetical protein | -0,1359 | 0,8366 |
| TcCLB.511807.260 | hypothetical protein, conserved | -0,1361 | 0,1221 |
| TcCLB.507875.30 | hypothetical protein, conserved | -0,1361 | 0,4273 |
| TcCLB.510307.210 | trans-sialidase, Group II, putative | -0,1362 | 0,6522 |
| TcCLB.506133.190 | trans-sialidase, Group VI, putative | -0,1363 | 0,7902 |
| TcCLB.506627.40 | hypothetical protein, conserved | -0,1364 | 0,4205 |
| TcCLB.503763.50 | trans-sialidase (pseudogene), putative | -0,1364 | 0,4243 |
| TcCLB.511249.10 | ADP/ATP mitochondrial carrier protein, putative | -0,1365 | 0,2605 |
| TcCLB.511317.70 | RNA-binding protein, putative (RBP30) | -0,1366 | 0,3308 |
| TcCLB.506789.50 | mucin-associated surface protein (MASP), putative | -0,1366 | 0,7629 |
| TcCLB.506425.160 | hypothetical protein, conserved | -0,1367 | 0,42 |
| TcCLB.508153.910 | hypothetical protein, conserved | -0,1367 | 0,451 |
| TcCLB.508153.80 | DNA replication complex GINS protein SLD5, putative | -0,1369 | 0,4738 |
| TcCLB.509267.70 | hypothetical protein | -0,137 | 0,2201 |
| TcCLB.506577.140 | hypothetical protein, conserved | -0,137 | 0,5605 |
| TcCLB.506679.220 | nucleoplasmin-like protein (NLP), putative | -0,1371 | 0,1711 |
| TcCLB.510889.205 | hypothetical protein, conserved | -0,1371 | 0,3935 |
| TcCLB.506355.20 | receptor-type adenylate cyclase, putative | -0,1371 | 0,4502 |
| TcCLB.511923.60 | trans-sialidase, Group V, putative | -0,1371 | 0,598 |
| TcCLB.506813.29 | mucin-associated surface protein (MASP), putative | -0,1371 | 0,8214 |
| TcCLB.506649.40 | rho-like GTP binding protein, putative | -0,1372 | 0,4238 |
| TcCLB.506503.170 | hypothetical protein, conserved | -0,1372 | 0,6222 |
| TcCLB.507491.70 | hypothetical protein, conserved (pseudogene) | -0,1375 | 0,2346 |
| TcCLB.508199.70 | hypothetical protein | -0,1376 | 0,3267 |
| TcCLB.506297.280 | hypothetical protein, conserved | -0,1376 | 0,3739 |
| TcCLB.511577.130 | poly(A) export protein, putative | -0,1376 | 0,3858 |
| TcCLB.508461.390 | hypothetical protein, conserved | -0,1377 | 0,2149 |
| TcCLB.506929.10 | surface protease GP63, putative | -0,1378 | 0,3516 |
| TcCLB.509779.50 | hypothetical protein, conserved | -0,1378 | 0,3954 |
| TcCLB.507057.50 | hypothetical protein, conserved | -0,1379 | 0,3853 |
| TcCLB.472777.9 | hypothetical protein, conserved | -0,1379 | 0,71 |
| TcCLB.508813.50 | oxidoreductase-like protein, putative | -0,138 | 0,2292 |
| TcCLB.507467.94 | hypothetical protein, conserved | -0,138 | 0,5787 |
| TcCLB.510025.191 | surface protease GP63 (pseudogene), putative | -0,138 | 0,8312 |
| TcCLB.507927.20 | mitochondrial oligo_U binding protein TBRGG1, putative | -0,1381 | 0,3417 |
| TcCLB.507483.60 | translation-associated element 2, putative | -0,1382 | 0,2396 |
| TcCLB.511725.130 | hypothetical protein, conserved | -0,1383 | 0,1076 |
| TcCLB.507991.110 | hypothetical protein, conserved | -0,1384 | 0,2915 |
| TcCLB.511603.250 | mucin-associated surface protein (MASP), putative | -0,1385 | 0,7237 |
| TcCLB.503823.60 | hypothetical protein, conserved | -0,1386 | 0,3433 |
| TcCLB.504251.20 | hypothetical protein, conserved | -0,1386 | 0,4995 |
| TcCLB.503801.79 | hypothetical protein, conserved | -0,1387 | 0,5081 |
| TcCLB.506269.25 | hypothetical protein | -0,1387 | 0,7379 |
| TcCLB.506515.29 | trans-sialidase, putative | -0,1388 | 0,6872 |

| | | | |
|---|---|---|---|
| TcCLB.508277.360 | ubiquitin hydrolase, putative | -0,1389 | 0,1614 |
| TcCLB.508675.40 | hypothetical protein, conserved | -0,1389 | 0,3025 |
| TcCLB.509735.84 | dispersed gene family protein 1 (DGF-1), putative | -0,1389 | 0,3945 |
| TcCLB.508307.60 | hypothetical protein, conserved | -0,139 | 0,2371 |
| TcCLB.508461.400 | nucleoside diphosphate kinase (NDPK2) | -0,139 | 0,3015 |
| TcCLB.506209.50 | trans-sialidase (pseudogene), putative | -0,1392 | 0,6266 |
| TcCLB.504149.80 | hypothetical protein, conserved | -0,1393 | 0,2427 |
| TcCLB.510945.10 | ubiquitin carboxyl-terminal hydrolase, putative | -0,1393 | 0,3719 |
| TcCLB.511711.60 | hypothetical protein | -0,1393 | 0,5917 |
| TcCLB.447847.19 | retrotransposon hot spot protein (RHS, pseudogene), putat | -0,1395 | 0,5558 |
| TcCLB.507063.110 | hypothetical protein, conserved | -0,1396 | 0,1697 |
| TcCLB.507005.30 | hypothetical protein, conserved | -0,1398 | 0,3027 |
| TcCLB.503779.100 | hypothetical protein, conserved | -0,1398 | 0,3727 |
| TcCLB.506289.110 | surface protease GP63 (pseudogene), putative | -0,1398 | 0,5429 |
| TcCLB.506203.70 | pre-mRNA-splicing factor ATP-dependent RNA helicase, pu | -0,1399 | 0,1989 |
| TcCLB.508927.30 | hypothetical protein, conserved | -0,1399 | 0,4042 |
| TcCLB.467287.30 | ATPase beta subunit, putative | -0,1401 | 0,19 |
| TcCLB.503473.4 | proteasome 26S non-ATPase subunit 9, putative | -0,1401 | 0,6447 |
| TcCLB.508873.590 | retrotransposon hot spot protein (RHS, pseudogene), putat | -0,1403 | 0,6247 |
| TcCLB.508909.20 | guanosine monophosphate reductase, putative | -0,1404 | 0,1202 |
| TcCLB.511837.138 | RNA-binding protein, putative | -0,1404 | 0,2424 |
| TcCLB.506467.4 | hypothetical protein, conserved | -0,1404 | 0,4734 |
| TcCLB.506629.40 | hypothetical protein, conserved | -0,1405 | 0,1275 |
| TcCLB.511545.180 | hypothetical protein, conserved | -0,1405 | 0,2 |
| TcCLB.507611.70 | trans-sialidase (pseudogene), putative | -0,1405 | 0,6032 |
| TcCLB.507681.259 | Flagellar Member 1 (fragment) (FLAM1) | -0,1406 | 0,3678 |
| TcCLB.511875.25 | elongation factor 1-gamma (EF-1-gamma, pseudogene), p | -0,1407 | 0,7842 |
| TcCLB.511051.50 | 40S ribosomal protein S4, putative | -0,1408 | 0,07153 |
| TcCLB.482471.10 | hypothetical protein, conserved | -0,1408 | 0,3713 |
| TcCLB.506883.90 | hypothetical protein, conserved | -0,1408 | 0,3821 |
| TcCLB.507811.20 | delta-adaptin, putative | -0,141 | 0,2533 |
| TcCLB.510963.20 | hypothetical protein, conserved | -0,141 | 0,303 |
| TcCLB.506577.170 | hypothetical protein, conserved | -0,141 | 0,3545 |
| TcCLB.506267.75 | hypothetical protein, conserved (pseudogene) | -0,141 | 0,6764 |
| TcCLB.408825.5 | mucin-associated surface protein (MASP), putative (fragm | -0,141 | 0,816 |
| TcCLB.511727.210 | Repressor of differentiation kinase 1, putative (RDK1.1) | -0,1411 | 0,154 |
| TcCLB.504243.30 | hypothetical protein, conserved | -0,1411 | 0,1549 |
| TcCLB.509205.10 | guanine-nucleotide-exchange-factor, putative | -0,1412 | 0,454 |
| TcCLB.503457.4 | hypothetical protein, conserved | -0,1412 | 0,6351 |
| TcCLB.510963.80 | hypothetical protein, conserved | -0,1414 | 0,5842 |
| TcCLB.508541.90 | mucin-associated surface protein (MASP), putative | -0,1414 | 0,7029 |
| TcCLB.510243.30 | chaperone DNAJ protein, putative | -0,1415 | 0,243 |
| TcCLB.506893.60 | hypothetical protein, conserved | -0,1415 | 0,5457 |
| TcCLB.503919.10 | spliced leader RNA PSE-promoter transcription factor, puta | -0,1416 | 0,3069 |
| TcCLB.506007.40 | protein kinase, putative | -0,1417 | 0,2816 |

| | | | |
|---|---|---|---|
| TcCLB.504045.70 | hypothetical protein, conserved | -0,1417 | 0,5048 |
| TcCLB.506417.70 | hypothetical protein | -0,1418 | 0,4593 |
| TcCLB.509579.30 | retrotransposon hot spot protein (RHS, pseudogene), putat | -0,1419 | 0,4286 |
| TcCLB.413605.10 | kinesin, putative | -0,1421 | 0,2697 |
| TcCLB.508213.20 | hypothetical protein, conserved | -0,1421 | 0,5576 |
| TcCLB.506943.23 | hypothetical protein, conserved | -0,1422 | 0,4524 |
| TcCLB.458369.10 | trans-sialidase (pseudogene), putative | -0,1425 | 0,7639 |
| TcCLB.510323.10 | protein kinase, putative | -0,1426 | 0,3761 |
| TcCLB.506203.90 | hypothetical protein, conserved | -0,1428 | 0,25 |
| TcCLB.506467.10 | hypothetical protein, conserved | -0,1428 | 0,3634 |
| TcCLB.510075.60 | hypothetical protein | -0,1429 | 0,5574 |
| TcCLB.505163.10 | hypothetical protein, conserved | -0,143 | 0,6251 |
| TcCLB.511075.30 | TFIIH basal transcription factor complex helicase subunit, p | -0,1431 | 0,1058 |
| TcCLB.506627.90 | hypothetical protein | -0,1431 | 0,1494 |
| TcCLB.508647.100 | hypothetical protein, conserved | -0,1431 | 0,2229 |
| TcCLB.507237.220 | mucin-associated surface protein (MASP), putative | -0,1431 | 0,8344 |
| TcCLB.507897.40 | acetyl-CoA carboxylase, putative | -0,1432 | 0,2799 |
| TcCLB.503855.60 | hypothetical protein, conserved | -0,1432 | 0,3572 |
| TcCLB.509611.50 | hypothetical protein, conserved | -0,1432 | 0,423 |
| TcCLB.399173.10 | trans-sialidase, putative (fragment) | -0,1432 | 0,7195 |
| TcCLB.509937.190 | hypothetical protein, conserved | -0,1433 | 0,2692 |
| TcCLB.507031.60 | nuclear cap binding complex subunit CBP30, putative (CBP3 | -0,1433 | 0,4027 |
| TcCLB.507887.10 | dihydroorotase (pseudogene), putative | -0,1433 | 0,6412 |
| TcCLB.511769.20 | dispersed gene family protein 1 (DGF-1), putative | -0,1434 | 0,4892 |
| TcCLB.510143.129 | hypothetical protein, conserved | -0,1434 | 0,5268 |
| TcCLB.508647.120 | hypothetical protein, conserved | -0,1435 | 0,1902 |
| TcCLB.503643.10 | KU70 protein, putative | -0,1435 | 0,303 |
| TcCLB.504101.50 | trans-sialidase, Group V, putative | -0,1435 | 0,4717 |
| TcCLB.508679.20 | ribonuclease inhibitor, putative | -0,1435 | 0,4765 |
| TcCLB.506127.130 | hypothetical protein, conserved | -0,1436 | 0,3187 |
| TcCLB.506895.50 | hypothetical protein, conserved | -0,1436 | 0,3541 |
| TcCLB.504205.10 | hypothetical protein | -0,1437 | 0,5567 |
| TcCLB.505941.10 | surface protease GP63, putative | -0,1438 | 0,4644 |
| TcCLB.503953.20 | hypothetical protein, conserved | -0,1438 | 0,5368 |
| TcCLB.506297.30 | hypothetical protein, conserved | -0,1439 | 0,08537 |
| TcCLB.509721.79 | hypothetical protein, conserved (fragment) | -0,1439 | 0,2496 |
| TcCLB.510377.450 | mucin-associated surface protein (MASP), putative | -0,1439 | 0,7243 |
| TcCLB.510197.130 | trans-sialidase, Group VI, putative | -0,144 | 0,7816 |
| TcCLB.506989.80 | hypothetical protein, conserved | -0,1441 | 0,4901 |
| TcCLB.508409.320 | hypothetical protein, conserved | -0,1442 | 0,1734 |
| TcCLB.506303.120 | hypothetical protein, conserved | -0,1442 | 0,2428 |
| TcCLB.509641.70 | hypothetical protein, conserved | -0,1443 | 0,1301 |
| TcCLB.506131.114 | mucin TcMUC (pseudogene), putative | -0,1443 | 0,7824 |
| TcCLB.509753.220 | mucin TcMUCII, putative | -0,1444 | 0,7986 |
| TcCLB.507501.5 | trans-sialidase, putative (fragment) | -0,1444 | 0,8377 |

| | | | |
|---|---|---|---|
| TcCLB.503897.110 | plectin-like protein, putative | -0,1445 | 0,3497 |
| TcCLB.504625.90 | hypothetical protein, conserved | -0,1445 | 0,4217 |
| TcCLB.508699.100 | surface protease GP63, putative | -0,1446 | 0,5119 |
| TcCLB.509223.80 | trans-sialidase (pseudogene), putative | -0,1446 | 0,5136 |
| TcCLB.511213.81 | retrotransposon hot spot protein (RHS, pseudogene), putat | -0,1446 | 0,578 |
| TcCLB.506221.110 | chromosomal passenger protein, putative (CPC2) | -0,1447 | 0,3425 |
| TcCLB.511237.40 | CDP-DAG synthase, putative (CDS) | -0,1448 | 0,301 |
| TcCLB.511855.30 | hypothetical protein, conserved | -0,1448 | 0,6247 |
| TcCLB.506939.50 | mucin-associated surface protein (MASP), putative | -0,1448 | 0,8595 |
| TcCLB.503847.60 | hypothetical protein, conserved | -0,1449 | 0,2114 |
| TcCLB.511071.150 | hypothetical protein, conserved | -0,1449 | 0,406 |
| TcCLB.509693.194 | hypothetical protein, conserved | -0,1449 | 0,6958 |
| TcCLB.510507.60 | inositol 1,4,5-trisphosphate receptor, putative (fragment) ( | -0,145 | 0,1671 |
| TcCLB.511071.120 | hypothetical protein, conserved | -0,145 | 0,2115 |
| TcCLB.511517.70 | RNA-binding protein, putative | -0,1453 | 0,2719 |
| TcCLB.511171.25 | hypothetical protein, conserved (pseudogene) | -0,1453 | 0,5609 |
| TcCLB.506971.10 | mucin-associated surface protein (MASP), putative | -0,1453 | 0,7711 |
| TcCLB.507875.200 | trans-sialidase (pseudogene), putative | -0,1454 | 0,5457 |
| TcCLB.507099.110 | hypothetical protein, conserved | -0,1455 | 0,2746 |
| TcCLB.506959.130 | 60S ribosomal protein L30, putative | -0,1456 | 0,07432 |
| TcCLB.511637.9 | zinc carboxypeptidase, putative | -0,1456 | 0,1991 |
| TcCLB.509537.30 | hypothetical protein, conserved | -0,1456 | 0,3514 |
| TcCLB.508479.390 | RNA triphosphatase, putative | -0,1457 | 0,4218 |
| TcCLB.508707.60 | hypothetical protein, conserved | -0,1458 | 0,1528 |
| TcCLB.507083.20 | dispersed gene family protein 1 (DGF-1), putative | -0,1458 | 0,286 |
| TcCLB.509767.130 | hypothetical protein, conserved | -0,1458 | 0,601 |
| TcCLB.503667.20 | hypothetical protein | -0,1458 | 0,7636 |
| TcCLB.511597.50 | trans-sialidase, putative (fragment) | -0,1458 | 0,8604 |
| TcCLB.432677.10 | heat shock 70 kDa protein, mitochondrial precursor (pseud | -0,1459 | 0,1758 |
| TcCLB.504215.20 | dolichyl-P-Man:GDP-ManGlcNAc2-PP-dolichyl beta-1,4-ma | -0,1459 | 0,3512 |
| TcCLB.409117.20 | 40S ribosomal protein S14, putative | -0,146 | 0,1177 |
| TcCLB.509911.40 | hypothetical protein, conserved | -0,146 | 0,4899 |
| TcCLB.511803.80 | hypothetical protein, conserved (fragment) | -0,1461 | 0,7692 |
| TcCLB.508347.159 | microtubule-associated protein, putative, (fragment) (GB4 | -0,1462 | 0,3702 |
| TcCLB.510289.50 | hypothetical protein, conserved | -0,1463 | 0,3374 |
| TcCLB.509067.14 | hypothetical protein, conserved | -0,1464 | 0,5339 |
| TcCLB.508145.49 | hypothetical protein, conserved | -0,1466 | 0,1828 |
| TcCLB.510357.70 | ubiquinone biosynthesis protein-like protein, putative | -0,1466 | 0,366 |
| TcCLB.511281.56 | sjogren s syndrome nuclear autoantigen 1, putative | -0,1466 | 0,4684 |
| TcCLB.511725.10 | proteasome regulatory non-ATPase subunit, putative | -0,1467 | 0,3291 |
| TcCLB.506925.550 | calpain-like cysteine peptidase (pseudogene), putative | -0,1467 | 0,5044 |
| TcCLB.508415.50 | ADP-ribosylation factor 1, putative (ARF1) | -0,1468 | 0,2484 |
| TcCLB.510357.120 | hypothetical protein, conserved | -0,1468 | 0,7046 |
| TcCLB.508257.260 | AP-1 adapter complex gamma subunit, putative | -0,1469 | 0,1336 |
| TcCLB.508779.90 | structural maintenance of chromosome protein 4, putative | -0,1469 | 0,1761 |

| | | | |
|---|---|---|---|
| TcCLB.508321.30 | hypothetical protein, conserved | -0,1469 | 0,5815 |
| TcCLB.509171.100 | hypothetical protein, conserved | -0,1472 | 0,2717 |
| TcCLB.506435.10 | hypothetical protein, conserved | -0,1472 | 0,4987 |
| TcCLB.507677.30 | hypothetical protein, conserved | -0,1474 | 0,267 |
| TcCLB.511367.74 | hypothetical protein, conserved | -0,1474 | 0,4229 |
| TcCLB.506265.30 | mitochondrial DNA polymerase I protein A, putative | -0,1475 | 0,2167 |
| TcCLB.504085.30 | RNA-binding protein, putative | -0,1476 | 0,3179 |
| TcCLB.507703.10 | retrotransposon hot spot protein (RHS, pseudogene), putat | -0,1476 | 0,5216 |
| TcCLB.506925.260 | protein kinase, putative | -0,1477 | 0,3678 |
| TcCLB.504079.3 | mitochondrial chaperone BCS1, putative (fragment) | -0,1479 | 0,3464 |
| TcCLB.507951.30 | receptor-type adenylate cyclase (pseudogene), putative | -0,1481 | 0,2292 |
| TcCLB.506367.140 | hypothetical protein, conserved | -0,1482 | 0,2446 |
| TcCLB.505071.40 | hypothetical protein | -0,1482 | 0,3841 |
| TcCLB.509777.60 | ADG2, putative | -0,1483 | 0,3337 |
| TcCLB.508415.59 | hypothetical protein, conserved | -0,1483 | 0,6192 |
| TcCLB.506101.10 | mucin-associated surface protein (MASP), putative | -0,1483 | 0,6481 |
| TcCLB.508387.150 | hypothetical protein | -0,1485 | 0,3143 |
| TcCLB.508153.400 | serine/threonine-protein kinase, putative | -0,1486 | 0,1424 |
| TcCLB.510173.120 | hypothetical protein, conserved | -0,1486 | 0,4827 |
| TcCLB.511599.10 | trans-sialidase, putative | -0,1486 | 0,8362 |
| TcCLB.503861.80 | dispersed gene family protein 1 (DGF-1), putative | -0,1487 | 0,3127 |
| TcCLB.509007.80 | hypothetical protein, conserved | -0,1487 | 0,3393 |
| TcCLB.511611.11 | hypothetical protein | -0,1487 | 0,6693 |
| TcCLB.462617.10 | dynein heavy chain (pseudogene), putative | -0,1488 | 0,1798 |
| TcCLB.509965.180 | RNA-binding protein, putative | -0,1488 | 0,1947 |
| TcCLB.510163.60 | trans-sialidase, Group V, putative | -0,1488 | 0,2768 |
| TcCLB.509797.20 | hypothetical protein, conserved | -0,1488 | 0,3395 |
| TcCLB.509743.19 | dispersed gene family protein 1 (DGF-1), putative (fragme | -0,149 | 0,5525 |
| TcCLB.506513.160 | protein kinase, putative | -0,1491 | 0,2254 |
| TcCLB.510747.26 | hypothetical protein, conserved | -0,1491 | 0,6023 |
| TcCLB.510291.7 | TPR-repeat-containing chaperone protein DNAJ, putative (f | -0,1492 | 0,359 |
| TcCLB.506911.10 | mucin TcMUCII, putative | -0,1492 | 0,8186 |
| TcCLB.511585.160 | hypothetical protein, conserved | -0,1493 | 0,2761 |
| TcCLB.507623.50 | oxidoreductase, putative | -0,1493 | 0,3787 |
| TcCLB.511735.34 | hypothetical protein, conserved | -0,1493 | 0,4819 |
| TcCLB.503643.20 | cyclic nucleotide-binding domain containing protein, putati | -0,1494 | 0,1874 |
| TcCLB.511127.280 | hypothetical protein, conserved | -0,1494 | 0,4298 |
| TcCLB.503879.60 | hypothetical protein, conserved | -0,1495 | 0,3302 |
| TcCLB.510329.310 | hypothetical protein, conserved | -0,1496 | 0,3139 |
| TcCLB.511481.40 | N-Acetyl-D-glucosaminylphosphatidylinositol de-N-acetylas | -0,1496 | 0,4365 |
| TcCLB.506727.40 | hypothetical protein, conserved | -0,1496 | 0,5224 |
| TcCLB.504827.40 | AP-3 complex subunit beta, putative (AP3B) | -0,15 | 0,2239 |
| TcCLB.506821.180 | aquaporin-like protein, putative | -0,1501 | 0,411 |
| TcCLB.507927.80 | hypothetical protein | -0,1503 | 0,3649 |
| TcCLB.507867.90 | hypothetical protein, conserved | -0,1503 | 0,378 |

| | | | |
|---|---|---|---|
| TcCLB.510283.114 | DNA-directed RNA polymerase I subunit, putative | -0,1503 | 0,4226 |
| TcCLB.506625.150 | hypothetical protein, conserved | -0,1503 | 0,4335 |
| TcCLB.509769.10 | hypothetical protein, conserved | -0,1503 | 0,5838 |
| TcCLB.508703.25 | hypothetical protein | -0,1504 | 0,2915 |
| TcCLB.510351.40 | ATP-dependent RNA helicase, putative | -0,1507 | 0,1949 |
| TcCLB.421959.10 | eukaryotic translation initiation factor 4E, putative | -0,1507 | 0,2764 |
| TcCLB.511889.40 | dispersed gene family protein 1 (DGF-1), putative | -0,1507 | 0,4461 |
| TcCLB.511001.180 | hypothetical protein, conserved | -0,1508 | 0,2261 |
| TcCLB.506679.170 | hypothetical protein, conserved | -0,151 | 0,1185 |
| TcCLB.507031.179 | hypothetical protein, conserved | -0,151 | 0,356 |
| TcCLB.506195.180 | hypothetical protein, conserved | -0,151 | 0,4227 |
| TcCLB.511291.80 | hypothetical protein, conserved | -0,151 | 0,5552 |
| TcCLB.510307.19 | mucin-associated surface protein (MASP, pseudogene), pu | -0,1511 | 0,6212 |
| TcCLB.506631.50 | hypothetical protein, conserved (fragment) | -0,1511 | 0,7976 |
| TcCLB.509073.20 | hypothetical protein, conserved | -0,1512 | 0,1233 |
| TcCLB.510889.310 | hypothetical protein, conserved | -0,1512 | 0,1928 |
| TcCLB.509715.26 | hypothetical protein, conserved | -0,1512 | 0,3835 |
| TcCLB.509985.60 | hypothetical protein, conserved | -0,1513 | 0,1561 |
| TcCLB.507711.30 | hypothetical protein, conserved | -0,1513 | 0,3838 |
| TcCLB.425969.9 | cyclin-dependent kinases regulatory subunit, putative (P12 | -0,1514 | 0,4847 |
| TcCLB.510663.50 | hypothetical protein, conserved | -0,1515 | 0,2059 |
| TcCLB.508175.300 | hypothetical protein, conserved | -0,1515 | 0,3338 |
| TcCLB.510963.70 | ARP2/3 complex subunit, putative | -0,1517 | 0,4377 |
| TcCLB.511525.29 | hypothetical protein, conserved | -0,1518 | 0,2119 |
| TcCLB.508165.50 | mucin-associated surface protein (MASP), putative | -0,1519 | 0,8259 |
| TcCLB.510187.390 | Protein kinase domain, putative | -0,152 | 0,1695 |
| TcCLB.506793.90 | hypothetical protein, conserved | -0,152 | 0,3099 |
| TcCLB.511407.20 | hypothetical protein, conserved | -0,152 | 0,3546 |
| TcCLB.511431.30 | conserved protein, unknown function | -0,1521 | 0,2917 |
| TcCLB.506795.29 | hypothetical protein, conserved | -0,1521 | 0,3192 |
| TcCLB.510363.19 | trans-sialidase (pseudogene), putative | -0,1523 | 0,6663 |
| TcCLB.468405.9 | dispersed gene family protein 1 (DGF-1), putative | -0,1523 | 0,7764 |
| TcCLB.506221.60 | hypothetical protein, conserved (pseudogene) | -0,1524 | 0,1912 |
| TcCLB.509163.30 | trans-sialidase (pseudogene), putative | -0,1524 | 0,6062 |
| TcCLB.508141.50 | hypothetical protein, conserved | -0,1525 | 0,4647 |
| TcCLB.507895.60 | UDP-Gal or UDP-GlcNAc-dependent glycosyltransferase, pu | -0,1525 | 0,7019 |
| TcCLB.504089.60 | hypothetical protein, conserved | -0,1526 | 0,1946 |
| TcCLB.504257.30 | S-adenosylmethionine decarboxylase proenzyme, putative | -0,1526 | 0,2196 |
| TcCLB.510657.160 | Double RNA binding domain protein 9 (DRBD9) | -0,1526 | 0,3398 |
| TcCLB.505983.30 | hypothetical protein, conserved | -0,1528 | 0,1218 |
| TcCLB.510817.40 | hypothetical protein, conserved | -0,1528 | 0,6496 |
| TcCLB.509545.50 | mucin-associated surface protein (MASP), putative | -0,1529 | 0,7099 |
| TcCLB.504625.30 | hypothetical protein | -0,153 | 0,1835 |
| TcCLB.506543.100 | hypothetical protein, conserved | -0,153 | 0,3534 |
| TcCLB.511595.10 | trans-sialidase (pseudogene), putative | -0,153 | 0,7449 |

| | | | |
|---|---|---|---|
| TcCLB.508831.20 | oligomeric golgi complex component 8, putative | -0,1531 | 0,2831 |
| TcCLB.509399.70 | hypothetical protein, conserved | -0,1532 | 0,3059 |
| TcCLB.508189.20 | dispersed gene family protein 1 (DGF-1, pseudogene), puta | -0,1532 | 0,6151 |
| TcCLB.511903.260 | hypothetical protein, conserved | -0,1533 | 0,316 |
| TcCLB.507083.90 | hypothetical protein, conserved | -0,1533 | 0,405 |
| TcCLB.510017.50 | surface protease GP63 (pseudogene), putative | -0,1535 | 0,7888 |
| TcCLB.504163.90 | hypothetical protein, conserved | -0,1537 | 0,2756 |
| TcCLB.507041.80 | hypothetical protein | -0,1537 | 0,5565 |
| TcCLB.510519.110 | hypothetical protein | -0,1538 | 0,4936 |
| TcCLB.506931.30 | hypothetical protein, conserved | -0,1539 | 0,537 |
| TcCLB.420369.20 | trans-sialidase (pseudogene), putative | -0,1539 | 0,5766 |
| TcCLB.504163.50 | fructose-bisphosphate aldolase, glycosomal, putative | -0,154 | 0,3208 |
| TcCLB.507787.170 | retrotransposon hot spot protein (RHS, pseudogene), putat | -0,1541 | 0,1685 |
| TcCLB.511517.110 | hypothetical protein, conserved | -0,1541 | 0,2385 |
| TcCLB.509999.90 | carnitine/choline acetyltransferase, putative | -0,1543 | 0,09248 |
| TcCLB.507831.50 | hypothetical protein, conserved | -0,1545 | 0,1162 |
| TcCLB.508817.150 | hypothetical protein, conserved | -0,1546 | 0,239 |
| TcCLB.508397.64 | hypothetical protein, conserved | -0,1546 | 0,5034 |
| TcCLB.508355.80 | trans-sialidase, Group VIII, putative | -0,1546 | 0,5892 |
| TcCLB.506885.370 | hypothetical protein, conserved | -0,1547 | 0,4505 |
| TcCLB.510207.35 | mucin TcMUCII (pseudogene), putative | -0,1547 | 0,7015 |
| TcCLB.507099.10 | hypothetical protein, conserved | -0,1548 | 0,1281 |
| TcCLB.511189.10 | hypothetical protein, conserved (pseudogene) | -0,1548 | 0,4738 |
| TcCLB.509777.130 | inositol phosphosphingolipid phospholipase C-Like, putative | -0,1549 | 0,1707 |
| TcCLB.509551.110 | hypothetical protein, conserved | -0,1549 | 0,231 |
| TcCLB.511523.80 | hypothetical protein, conserved | -0,1549 | 0,2567 |
| TcCLB.506435.290 | hypothetical protein, conserved | -0,1549 | 0,4356 |
| TcCLB.506321.359 | MRB1-associated protein, putative | -0,1549 | 0,5034 |
| TcCLB.510733.30 | hypothetical protein, conserved | -0,155 | 0,5303 |
| TcCLB.510483.220 | mucin-associated surface protein (MASP), putative | -0,155 | 0,6446 |
| TcCLB.510519.40 | protein kinase, putative | -0,1551 | 0,2653 |
| TcCLB.507943.99 | hypothetical protein, conserved (fragment) | -0,1551 | 0,5846 |
| TcCLB.506625.70 | RNA-binding protein, putative | -0,1552 | 0,1223 |
| TcCLB.506821.160 | hypothetical protein, conserved | -0,1554 | 0,2993 |
| TcCLB.509429.200 | hypothetical protein, conserved | -0,1554 | 0,3506 |
| TcCLB.506425.110 | hypothetical protein, conserved | -0,1554 | 0,3973 |
| TcCLB.508913.25 | trans-sialidase, Group I, putative | -0,1554 | 0,7742 |
| TcCLB.458759.10 | profilin, putative | -0,1555 | 0,3502 |
| TcCLB.507521.40 | hypothetical protein, conserved | -0,1555 | 0,3854 |
| TcCLB.506629.210 | hypothetical protein, conserved | -0,1556 | 0,2901 |
| TcCLB.510165.10 | heat shock protein DNAJ, putative | -0,1558 | 0,2729 |
| TcCLB.510763.50 | hypothetical protein, conserved | -0,1558 | 0,4104 |
| TcCLB.508721.20 | hypothetical protein, conserved | -0,1559 | 0,3434 |
| TcCLB.506435.70 | hypothetical protein, conserved (pseudogene) | -0,156 | 0,1342 |
| TcCLB.510769.110 | hypothetical protein, conserved | -0,156 | 0,3339 |

| | | | |
|---|---|---|---|
| TcCLB.508765.40 | hypothetical protein | -0,156 | 0,3368 |
| TcCLB.506753.40 | hypothetical protein, conserved | -0,156 | 0,3961 |
| TcCLB.511043.20 | small G-protein, putative | -0,1561 | 0,132 |
| TcCLB.506793.100 | hypothetical protein, conserved | -0,1561 | 0,2344 |
| TcCLB.503635.80 | hypothetical protein, conserved | -0,1561 | 0,2879 |
| TcCLB.511725.120 | kinetoplast polyadenylation/uridylation factor 2, putative | -0,1562 | 0,1946 |
| TcCLB.504137.20 | hypothetical protein, conserved | -0,1562 | 0,2464 |
| TcCLB.504045.20 | hypothetical protein, conserved | -0,1563 | 0,1575 |
| TcCLB.511661.50 | small ubiquitin protein, putative | -0,1563 | 0,4691 |
| TcCLB.508175.70 | lanosterol synthase, putative | -0,1564 | 0,1449 |
| TcCLB.511517.5 | 70 kDa heat shock protein, putative (fragment) | -0,1564 | 0,5093 |
| TcCLB.510299.50 | GPI alpha-mannosyltransferase III (GPI10) | -0,1565 | 0,1242 |
| TcCLB.510173.40 | MRB1 complex subunit MRB3010 (MRB3010) | -0,1565 | 0,2004 |
| TcCLB.508775.70 | dispersed gene family protein 1 (DGF-1), putative | -0,1565 | 0,7562 |
| TcCLB.507089.90 | hypothetical protein, conserved | -0,1566 | 0,1938 |
| TcCLB.509573.60 | hypothetical protein, conserved | -0,1566 | 0,4947 |
| TcCLB.510397.30 | retrotransposon hot spot protein (RHS, pseudogene), putat | -0,1566 | 0,7132 |
| TcCLB.417797.9 | hypothetical protein, conserved | -0,1567 | 0,381 |
| TcCLB.508823.110 | hypothetical protein, conserved | -0,1568 | 0,07874 |
| TcCLB.504147.224 | RNA-editing-associated protein 1, putative (REAP-1) | -0,1568 | 0,2921 |
| TcCLB.506291.10 | mucin-associated surface protein (MASP), putative | -0,1568 | 0,6823 |
| TcCLB.510659.140 | hypothetical protein, conserved (pseudogene) | -0,157 | 0,1305 |
| TcCLB.503869.30 | hypothetical protein, conserved | -0,157 | 0,151 |
| TcCLB.508551.10 | target of rapamycin (TOR) kinase 1, putative | -0,1571 | 0,3078 |
| TcCLB.507053.70 | geranylgeranyl transferase type II beta subunit, putative | -0,1572 | 0,2782 |
| TcCLB.503789.34 | hypothetical protein, conserved | -0,1573 | 0,3116 |
| TcCLB.503909.44 | hypothetical protein, conserved | -0,1573 | 0,3316 |
| TcCLB.509579.34 | ATP-dependent DEAD/H RNA helicase (pseudogene), putat | -0,1573 | 0,7253 |
| TcCLB.508295.80 | mucin TcMUCII, putative | -0,1573 | 0,7512 |
| TcCLB.507105.10 | beta prime cop protein, putative | -0,1574 | 0,08617 |
| TcCLB.508299.40 | hypothetical protein | -0,1574 | 0,2793 |
| TcCLB.506247.250 | hypothetical protein, conserved | -0,1575 | 0,05904 |
| TcCLB.508659.9 | TPR-repeat-containing chaperone protein DNAJ, putative (f | -0,1575 | 0,3446 |
| TcCLB.507881.30 | glycine dehydrogenase [decarboxylating] (pseudogene), pu | -0,1575 | 0,6585 |
| TcCLB.510435.30 | hypothetical protein, conserved | -0,1577 | 0,1667 |
| TcCLB.507023.250 | hypothetical protein, conserved | -0,1577 | 0,3092 |
| TcCLB.506679.140 | 40S ribosomal protein S10, putative | -0,1578 | 0,2454 |
| TcCLB.506287.80 | hypothetical protein, conserved | -0,1579 | 0,3039 |
| TcCLB.509171.20 | hypothetical protein, conserved | -0,158 | 0,2594 |
| TcCLB.506955.130 | trans-sialidase (pseudogene), putative | -0,1581 | 0,617 |
| TcCLB.508987.20 | hypothetical protein, conserved | -0,1582 | 0,1889 |
| TcCLB.505183.130 | sugar transporter, putative | -0,1582 | 0,4289 |
| TcCLB.506679.200 | hypothetical protein, conserved | -0,1583 | 0,2945 |
| TcCLB.506247.355 | hypothetical protein, conserved | -0,1583 | 0,5436 |
| TcCLB.509767.50 | hypothetical protein, conserved | -0,1584 | 0,2238 |

| | | | |
|---|---|---|---|
| TcCLB.510943.30 | hypothetical protein, conserved | -0,1584 | 0,4371 |
| TcCLB.511361.70 | hypothetical protein, conserved | -0,1586 | 0,2315 |
| TcCLB.508643.9 | pre-mRNA-splicing factor ATP-dependent RNA helicase, pu | -0,1586 | 0,2771 |
| TcCLB.508515.80 | hypothetical protein, conserved | -0,1587 | 0,3042 |
| TcCLB.510753.30 | hypothetical protein, conserved | -0,1591 | 0,3271 |
| TcCLB.508181.50 | hypothetical protein, conserved | -0,1591 | 0,3776 |
| TcCLB.506405.70 | hypothetical protein, conserved | -0,1591 | 0,3826 |
| TcCLB.509203.20 | cyclin dependent kinase-binding protein, putative | -0,1593 | 0,1568 |
| TcCLB.506529.120 | hypothetical protein | -0,1593 | 0,2488 |
| TcCLB.508059.20 | hypothetical protein, conserved | -0,1594 | 0,3033 |
| TcCLB.510877.90 | hypothetical protein, conserved | -0,1595 | 0,1794 |
| TcCLB.506221.129 | hypothetical protein, conserved (fragment) | -0,1595 | 0,389 |
| TcCLB.507711.210 | hypothetical protein, conserved | -0,1596 | 0,1417 |
| TcCLB.504035.84 | succinate dehydrogenase subunit | -0,1596 | 0,3768 |
| TcCLB.508761.70 | mucin-associated surface protein (MASP), putative | -0,1596 | 0,5243 |
| TcCLB.506989.240 | hypothetical protein, conserved (pseudogene) | -0,1598 | 0,09078 |
| TcCLB.509999.110 | serine/threonine-protein kinase, putative | -0,1598 | 0,2896 |
| TcCLB.503811.70 | hypothetical protein, conserved | -0,1599 | 0,1743 |
| TcCLB.506925.200 | hypothetical protein, conserved | -0,16 | 0,1477 |
| TcCLB.507677.129 | hypothetical protein, conserved | -0,16 | 0,1565 |
| TcCLB.504689.30 | trans-sialidase (pseudogene), putative | -0,16 | 0,596 |
| TcCLB.503545.10 | trans-sialidase (pseudogene), putative | -0,16 | 0,6163 |
| TcCLB.510477.90 | mucin TcMUCII, putative | -0,16 | 0,7881 |
| TcCLB.511603.500 | mucin-associated surface protein (MASP), putative | -0,16 | 0,7984 |
| TcCLB.503717.31 | mucin TcMUCII, putative | -0,1602 | 0,7661 |
| TcCLB.510181.130 | hypothetical protein | -0,1603 | 0,4119 |
| TcCLB.510513.20 | hypothetical protein, conserved | -0,1605 | 0,2596 |
| TcCLB.511127.40 | hypothetical protein | -0,1606 | 0,4262 |
| TcCLB.511173.50 | mucin-associated surface protein (MASP), putative | -0,1606 | 0,5845 |
| TcCLB.505843.24 | succinate dehydrogenase, putative | -0,1607 | 0,1617 |
| TcCLB.509715.20 | clathrin coat assembly protein, putative | -0,1607 | 0,3165 |
| TcCLB.507883.80 | adenylate kinase, putative | -0,1607 | 0,4693 |
| TcCLB.510021.60 | hypothetical protein | -0,1607 | 0,8285 |
| TcCLB.511729.40 | 60S ribosomal protein L9, putative | -0,1608 | 0,3666 |
| TcCLB.508777.160 | hypothetical protein, conserved | -0,1609 | 0,19 |
| TcCLB.506297.210 | hypothetical protein, conserved | -0,161 | 0,1462 |
| TcCLB.503435.14 | hypothetical protein | -0,161 | 0,2145 |
| TcCLB.507029.60 | hypothetical protein, conserved | -0,161 | 0,2884 |
| TcCLB.414243.29 | inosine-guanine nucleoside hydrolase, putative (fragment) | -0,161 | 0,665 |
| TcCLB.511467.50 | hypothetical protein, conserved | -0,1611 | 0,3318 |
| TcCLB.509049.51 | hypothetical protein, conserved | -0,1611 | 0,4079 |
| TcCLB.509599.110 | hypothetical protein, conserved | -0,1612 | 0,3023 |
| TcCLB.506537.210 | trans-sialidase (pseudogene), putative | -0,1612 | 0,4411 |
| TcCLB.505999.20 | phosphate-repressible phosphate permease, putative | -0,1613 | 0,1258 |
| TcCLB.505231.10 | dynein heavy chain, putative | -0,1613 | 0,1804 |

| | | | |
|---|---|---|---|
| TcCLB.509669.60 | hypothetical protein, conserved | -0,1613 | 0,3824 |
| TcCLB.503853.10 | hypothetical protein, conserved | -0,1615 | 0,2192 |
| TcCLB.509047.30 | hypothetical protein, conserved | -0,1616 | 0,4612 |
| TcCLB.507949.140 | ribosomal RNA-processing protein 8, putative (RRP8) | -0,1617 | 0,2657 |
| TcCLB.510121.130 | serine/threonine protein kinase, putative | -0,1617 | 0,5351 |
| TcCLB.506047.13 | ATP-dependent DEAD/H RNA helicase (pseudogene), putat | -0,1617 | 0,7268 |
| TcCLB.504219.10 | hypothetical protein, conserved (pseudogene) | -0,1618 | 0,8025 |
| TcCLB.506943.150 | hypothetical protein, conserved (pseudogene) | -0,1619 | 0,2185 |
| TcCLB.510231.20 | hypothetical protein | -0,1619 | 0,2226 |
| TcCLB.510431.220 | ribosome assembly protein RRB1, putative | -0,162 | 0,2905 |
| TcCLB.407335.9 | phosphatidylserine decarboxylase, putative | -0,1621 | 0,5253 |
| TcCLB.507063.30 | hypothetical protein, conserved | -0,1622 | 0,3873 |
| TcCLB.510131.70 | DNA polymerase kappa, putative | -0,1622 | 0,4827 |
| TcCLB.506509.50 | hypothetical protein, conserved | -0,1623 | 0,1959 |
| TcCLB.511279.20 | hypothetical protein, conserved | -0,1623 | 0,2244 |
| TcCLB.508899.40 | hypothetical protein, conserved | -0,1624 | 0,08231 |
| TcCLB.503593.20 | hypothetical protein, conserved | -0,1624 | 0,1993 |
| TcCLB.509963.10 | hypothetical protein, conserved (fragment) | -0,1624 | 0,4314 |
| TcCLB.510213.9 | mucin-associated surface protein (MASP), putative | -0,1624 | 0,7387 |
| TcCLB.507659.10 | hypothetical protein, conserved | -0,1627 | 0,2262 |
| TcCLB.509179.160 | hypothetical protein, conserved | -0,1627 | 0,2469 |
| TcCLB.507053.190 | hypothetical protein, conserved | -0,1627 | 0,253 |
| TcCLB.511347.20 | mucin-associated surface protein (MASP, pseudogene), pu | -0,1627 | 0,4877 |
| TcCLB.507049.63 | hypothetical protein, conserved | -0,1627 | 0,525 |
| TcCLB.506499.100 | hypothetical protein | -0,1628 | 0,8656 |
| TcCLB.506743.210 | tubulin tyrosine ligase, putative | -0,1629 | 0,4482 |
| TcCLB.511749.9 | hypothetical protein, conserved (fragment) | -0,163 | 0,1003 |
| TcCLB.510161.10 | hypothetical protein, conserved | -0,163 | 0,2231 |
| TcCLB.506939.120 | trans-sialidase, Group V, putative | -0,163 | 0,5751 |
| TcCLB.507877.5 | elongation factor 1-gamma (EF-1-gamma, pseudogene), p | -0,1631 | 0,7712 |
| TcCLB.508647.160 | hypothetical protein, conserved | -0,1632 | 0,1218 |
| TcCLB.506681.30 | hypothetical protein, conserved | -0,1632 | 0,2833 |
| TcCLB.511711.20 | hypothetical protein, conserved | -0,1632 | 0,3182 |
| TcCLB.503727.20 | hypothetical protein, conserved | -0,1633 | 0,157 |
| TcCLB.511231.60 | hypothetical protein, conserved | -0,1633 | 0,2747 |
| TcCLB.508441.10 | NUP-1 protein, putative | -0,1634 | 0,009455 |
| TcCLB.508153.990 | hypothetical protein, conserved | -0,1635 | 0,3083 |
| TcCLB.511293.30 | DNA-directed RNA polymerases I and III subunit RPAC1, pu | -0,1636 | 0,2585 |
| TcCLB.507625.200 | hypothetical protein, conserved | -0,1637 | 0,2582 |
| TcCLB.506799.150 | surface protease GP63 (pseudogene), putative | -0,1637 | 0,6031 |
| TcCLB.510579.80 | 60S ribosomal protein L10, putative | -0,1638 | 0,06958 |
| TcCLB.511021.10 | protein phosphatase 2A catalytic subunit, putative | -0,1638 | 0,3147 |
| TcCLB.510279.60 | mucin-associated surface protein (MASP), putative | -0,1638 | 0,6525 |
| TcCLB.510303.160 | hypothetical protein, conserved | -0,164 | 0,1786 |
| TcCLB.506239.20 | hypothetical protein, conserved | -0,164 | 0,2218 |

| | | | |
|---|---|---|---|
| TcCLB.504009.10 | hypothetical protein, conserved | -0,1641 | 0,2142 |
| TcCLB.510867.19 | trans-sialidase (pseudogene), putative | -0,1641 | 0,3972 |
| TcCLB.503981.90 | SWI/SNF-related matrix-associated actin-dependent regul | -0,1641 | 0,669 |
| TcCLB.510625.30 | mucin-associated surface protein (MASP, pseudogene), pu | -0,1641 | 0,6902 |
| TcCLB.503547.20 | dispersed gene family protein 1 (DGF-1, pseudogene), puta | -0,1641 | 0,7001 |
| TcCLB.506933.50 | hypothetical protein, conserved | -0,1643 | 0,2018 |
| TcCLB.508899.90 | hypothetical protein | -0,1643 | 0,2542 |
| TcCLB.511075.50 | hypothetical protein | -0,1644 | 0,3382 |
| TcCLB.511803.20 | A/G-specific adenine glycosylase, putative | -0,1645 | 0,2587 |
| TcCLB.506123.20 | hypothetical protein, conserved | -0,1645 | 0,3514 |
| TcCLB.511727.100 | hypothetical protein, conserved | -0,1646 | 0,1577 |
| TcCLB.510645.20 | nuclear transcription factor, putative | -0,1646 | 0,1882 |
| TcCLB.509513.5 | hypothetical protein, conserved (pseudogene) | -0,1647 | 0,3856 |
| TcCLB.485889.4 | hypothetical protein, conserved | -0,1647 | 0,5122 |
| TcCLB.508153.500 | helicase, putative | -0,1648 | 0,08157 |
| TcCLB.506577.47 | hypothetical protein, conserved | -0,1648 | 0,2386 |
| TcCLB.510497.199 | dispersed gene family protein 1 (DGF-1), putative (fragme | -0,1648 | 0,7209 |
| TcCLB.504147.320 | hypothetical protein, conserved | -0,1649 | 0,09194 |
| TcCLB.511535.39 | hypothetical protein, conserved | -0,1649 | 0,5558 |
| TcCLB.511109.10 | surface protein TolT (pseudogene) (TolT) | -0,1649 | 0,6851 |
| TcCLB.504213.110 | Putrescine-cadaverine transporter (POT1.1) | -0,165 | 0,08811 |
| TcCLB.508647.190 | hypothetical protein, conserved | -0,165 | 0,2379 |
| TcCLB.511375.20 | hypothetical protein | -0,165 | 0,7353 |
| TcCLB.507071.349 | mucin-associated surface protein (MASP), putative | -0,165 | 0,8412 |
| TcCLB.509583.19 | dynein heavy chain, putative (fragment) | -0,1651 | 0,306 |
| TcCLB.503983.10 | hypothetical protein, conserved | -0,1651 | 0,4941 |
| TcCLB.511287.30 | 60S ribosomal protein L17, putative | -0,1652 | 0,04267 |
| TcCLB.503939.90 | hypothetical protein, conserved | -0,1652 | 0,1121 |
| TcCLB.508257.60 | hypothetical protein, conserved | -0,1655 | 0,4239 |
| TcCLB.510691.70 | hypothetical protein, conserved | -0,1656 | 0,4641 |
| TcCLB.508097.40 | trans-sialidase (pseudogene), putative | -0,1656 | 0,6649 |
| TcCLB.508577.60 | hypothetical protein, conserved | -0,1657 | 0,1312 |
| TcCLB.511277.350 | tubulin-specific chaperone, putative | -0,1657 | 0,1463 |
| TcCLB.507937.10 | mucin TcMUCII, putative | -0,1657 | 0,8181 |
| TcCLB.506493.70 | serine/threonine protein kinase, putative | -0,1658 | 0,1432 |
| TcCLB.511761.10 | UDP-sugar pyrophosphorylase (Sloppy) | -0,1658 | 0,2021 |
| TcCLB.511589.240 | hypothetical protein, conserved | -0,1658 | 0,254 |
| TcCLB.507049.80 | hypothetical protein, conserved | -0,1658 | 0,3453 |
| TcCLB.442629.9 | hypothetical protein, conserved | -0,1658 | 0,3878 |
| TcCLB.507927.50 | metallo-beta-lactamase superfamily, putative | -0,1659 | 0,3743 |
| TcCLB.506319.10 | hypothetical protein | -0,166 | 0,07218 |
| TcCLB.503461.30 | hypothetical protein, conserved | -0,166 | 0,1472 |
| TcCLB.509627.30 | proteasome regulatory non-ATPase subunit 11, putative (R | -0,166 | 0,1564 |
| TcCLB.511211.235 | hypothetical protein, conserved | -0,166 | 0,4106 |
| TcCLB.506569.60 | retrotransposon hot spot protein (RHS, pseudogene), putat | -0,166 | 0,5418 |

| | | | |
|---|---|---|---|
| TcCLB.507611.114 | retrotransposon hot spot protein (RHS, pseudogene), putat | -0,166 | 0,5692 |
| TcCLB.511589.160 | replication factor C, subunit 4, putative | -0,1662 | 0,26 |
| TcCLB.505183.120 | aldo/keto reductase, putative | -0,1662 | 0,298 |
| TcCLB.511507.90 | hypothetical protein, conserved | -0,1662 | 0,3246 |
| TcCLB.510837.30 | hypothetical protein, conserved | -0,1663 | 0,3034 |
| TcCLB.509489.10 | hypothetical protein, conserved | -0,1664 | 0,2047 |
| TcCLB.509605.10 | malonyl-CoA decarboxylase, mitochondrial precursor, puta | -0,1664 | 0,3971 |
| TcCLB.506567.90 | hypothetical protein, conserved | -0,1665 | 0,2916 |
| TcCLB.509567.30 | serine/threonine-protein kinase, putative | -0,1666 | 0,1365 |
| TcCLB.510889.110 | primase 2, putative | -0,1668 | 0,1918 |
| TcCLB.503955.60 | hypothetical protein, conserved | -0,1668 | 0,3135 |
| TcCLB.510263.30 | surface protease GP63, putative | -0,1668 | 0,4703 |
| TcCLB.510663.57 | transport protein particle (TRAPP) subunit, putative | -0,167 | 0,3503 |
| TcCLB.503973.120 | mucin-associated surface protein (MASP), putative | -0,167 | 0,5941 |
| TcCLB.510967.13 | hypothetical protein, conserved | -0,1671 | 0,5467 |
| TcCLB.436535.19 | Glucose-6-phosphate dehydrogenase, putative | -0,1671 | 0,6533 |
| TcCLB.503751.10 | hypothetical protein | -0,1672 | 0,2621 |
| TcCLB.508491.9 | hypothetical protein, conserved | -0,1672 | 0,5446 |
| TcCLB.511553.60 | hypothetical protein | -0,1672 | 0,7615 |
| TcCLB.508533.10 | hypothetical protein, conserved | -0,1673 | 0,1839 |
| TcCLB.506869.60 | protein kinase, putative | -0,1673 | 0,2099 |
| TcCLB.503847.30 | hypothetical protein, conserved | -0,1673 | 0,3089 |
| TcCLB.425851.9 | hypothetical protein, conserved | -0,1673 | 0,4544 |
| TcCLB.510015.50 | mucin TcMUC (pseudogene), putative | -0,1673 | 0,6212 |
| TcCLB.511867.110 | flap endonuclease-1 (FEN-1), putative | -0,1675 | 0,1656 |
| TcCLB.507837.80 | TBC1 domain family member 20/GTPase, putative | -0,1675 | 0,3869 |
| TcCLB.507989.40 | ATP-dependent DEAD/H DNA helicase recQ, putative | -0,1677 | 0,1034 |
| TcCLB.509647.80 | hypothetical protein, conserved | -0,1677 | 0,2398 |
| TcCLB.506459.280 | hypothetical protein, conserved | -0,1677 | 0,2851 |
| TcCLB.509179.170 | hypothetical protein | -0,1677 | 0,3052 |
| TcCLB.508521.112 | beta galactofuranosyl glycosyltransferase (pseudogene), pu | -0,1677 | 0,6324 |
| TcCLB.508273.130 | hypothetical protein | -0,1677 | 0,6966 |
| TcCLB.506297.240 | pretranslocation protein, alpha subunit, putative | -0,1678 | 0,05681 |
| TcCLB.510187.360 | hypothetical protein, conserved | -0,1678 | 0,1395 |
| TcCLB.425597.5 | hypothetical protein, conserved (pseudogene) | -0,1679 | 0,5487 |
| TcCLB.503487.40 | hypothetical protein, conserved | -0,168 | 0,1392 |
| TcCLB.505939.20 | oligopeptidase B-like protein, putative | -0,1683 | 0,07924 |
| TcCLB.510665.4 | hypothetical protein, conserved (fragment) | -0,1683 | 0,2686 |
| TcCLB.506315.50 | 60S ribosomal protein L13a, putative | -0,1684 | 0,0445 |
| TcCLB.510533.120 | translation initiation factor IF-2, putative | -0,1685 | 0,1609 |
| TcCLB.508479.20 | hypothetical protein, conserved (pseudogene) | -0,1685 | 0,486 |
| TcCLB.506557.30 | hypothetical protein, conserved | -0,1686 | 0,1749 |
| TcCLB.508165.280 | mucin-associated surface protein (MASP), putative | -0,1687 | 0,7155 |
| TcCLB.506573.60 | hypothetical protein, conserved | -0,1689 | 0,2655 |
| TcCLB.509801.10 | hypothetical protein, conserved | -0,1691 | 0,2102 |

| | | | |
|---|---|---|---|
| TcCLB.506893.70 | hypothetical protein, conserved | -0,1691 | 0,2897 |
| TcCLB.511753.110 | hypothetical protein, conserved | -0,1692 | 0,05164 |
| TcCLB.510431.270 | hypothetical protein, conserved | -0,1692 | 0,1771 |
| TcCLB.510483.234 | mucin TcMUC (pseudogene), putative | -0,1692 | 0,7116 |
| TcCLB.440099.28 | protein kinase, putative | -0,1694 | 0,4321 |
| TcCLB.507809.170 | protein kinase, putative | -0,1694 | 0,636 |
| TcCLB.511403.40 | mucin-associated surface protein (MASP, pseudogene), pu | -0,1694 | 0,7267 |
| TcCLB.511625.40 | protein phosphatase 2C, putative | -0,1695 | 0,3545 |
| TcCLB.510289.80 | hypothetical protein, conserved | -0,1695 | 0,3841 |
| TcCLB.509753.20 | trans-sialidase (pseudogene), putative | -0,1695 | 0,5356 |
| TcCLB.508689.5 | hypothetical protein, conserved (pseudogene) | -0,1696 | 0,4398 |
| TcCLB.507063.240 | hypothetical protein, conserved (pseudogene) | -0,1697 | 0,04562 |
| TcCLB.506147.140 | hypothetical protein, conserved | -0,1697 | 0,1656 |
| TcCLB.509179.200 | hypothetical protein, conserved | -0,1697 | 0,5417 |
| TcCLB.511249.30 | hypothetical protein, conserved | -0,1698 | 0,09529 |
| TcCLB.511067.4 | phosphatidylinositol 3-kinase, putative | -0,1699 | 0,3602 |
| TcCLB.507305.20 | casein kinase, putative | -0,17 | 0,09091 |
| TcCLB.503939.30 | ttaggg binding factor, putative | -0,17 | 0,218 |
| TcCLB.506863.50 | ribosomal RNA processing protein 45, putative (RRP45) | -0,17 | 0,4005 |
| TcCLB.412943.9 | hypothetical protein, conserved | -0,1701 | 0,4197 |
| TcCLB.508219.70 | mucin-associated surface protein (MASP), putative | -0,1702 | 0,7192 |
| TcCLB.506679.110 | dynein heavy chain, putative | -0,1703 | 0,09389 |
| TcCLB.510213.70 | mucin-associated surface protein (MASP), putative | -0,1703 | 0,6584 |
| TcCLB.504241.57 | retrotransposon hot spot protein (RHS, pseudogene), putat | -0,1703 | 0,7206 |
| TcCLB.511907.290 | hypothetical protein, conserved | -0,1705 | 0,2159 |
| TcCLB.506953.49 | glucose-6-phosphate 1-dehydrogenase, putative | -0,1706 | 0,6437 |
| TcCLB.509393.50 | hypothetical protein, conserved | -0,1707 | 0,1506 |
| TcCLB.509567.50 | hypothetical protein, conserved | -0,1707 | 0,2669 |
| TcCLB.508397.60 | methyltransferase, putative | -0,1707 | 0,3906 |
| TcCLB.507711.220 | hypothetical protein, conserved | -0,1708 | 0,5008 |
| TcCLB.509693.20 | plectin, putative | -0,1709 | 0,06042 |
| TcCLB.509965.240 | hypothetical protein, conserved | -0,1709 | 0,3116 |
| TcCLB.506635.100 | ATP dependent DEAD-box helicase, putative (RH) | -0,1711 | 0,2018 |
| TcCLB.509793.30 | kinetoplast DNA-associated protein, putative | -0,1712 | 0,249 |
| TcCLB.511301.44 | hypothetical protein, conserved | -0,1712 | 0,2661 |
| TcCLB.507167.190 | trans-sialidase (pseudogene), putative | -0,1712 | 0,5455 |
| TcCLB.508987.30 | serine/threonine-protein kinase, putative | -0,1714 | 0,1152 |
| TcCLB.508897.80 | hypothetical protein, conserved | -0,1714 | 0,1285 |
| TcCLB.503939.120 | hypothetical protein, conserved | -0,1715 | 0,1516 |
| TcCLB.510653.19 | hypothetical protein, conserved (fragment) | -0,1715 | 0,2881 |
| TcCLB.506195.130 | hypothetical protein, conserved | -0,1716 | 0,3478 |
| TcCLB.508277.60 | hypothetical protein, conserved | -0,1717 | 0,06608 |
| TcCLB.511233.14 | 40S ribosomal protein S27, putative | -0,1717 | 0,07009 |
| TcCLB.507087.40 | kinesin, putative | -0,1717 | 0,102 |
| TcCLB.503799.10 | DNA repair protein RAD50, putative | -0,1718 | 0,08086 |

| | | | |
|---|---|---|---|
| TcCLB.460127.20 | metallo- peptidase, Clan M- Family M48 (FACE1) | -0,1718 | 0,2368 |
| TcCLB.503579.100 | nucleotide-binding protein, putative | -0,1719 | 0,2624 |
| TcCLB.510305.20 | hypothetical protein, conserved | -0,1719 | 0,2949 |
| TcCLB.511851.30 | trans-sialidase (pseudogene), putative | -0,1719 | 0,3009 |
| TcCLB.508799.100 | hypothetical protein, conserved | -0,172 | 0,2051 |
| TcCLB.504239.350 | mucin-associated surface protein (MASP), putative | -0,1721 | 0,5197 |
| TcCLB.505997.70 | ATPase, putative | -0,1721 | 0,8084 |
| TcCLB.509875.250 | hypothetical protein, conserved | -0,1722 | 0,3145 |
| TcCLB.506435.200 | hypothetical protein, conserved | -0,1722 | 0,3934 |
| TcCLB.509967.189 | kinesin, putative (fragment) | -0,1722 | 0,4834 |
| TcCLB.506603.43 | hypothetical protein | -0,1722 | 0,5283 |
| TcCLB.510307.77 | N-acetyltransferase complex ARD1 subunit (pseudogene), | -0,1722 | 0,5605 |
| TcCLB.506811.80 | hypothetical protein, conserved | -0,1723 | 0,26 |
| TcCLB.507949.110 | zeta tubulin, putative | -0,1725 | 0,227 |
| TcCLB.507833.10 | mucin-associated surface protein (MASP), putative | -0,1727 | 0,7782 |
| TcCLB.511717.100 | hypothetical protein, conserved | -0,1728 | 0,2041 |
| TcCLB.511899.20 | hypothetical protein, conserved | -0,1728 | 0,2546 |
| TcCLB.507083.70 | Iron-sulfur assembly protein 1, putative (Isa1) | -0,1729 | 0,2552 |
| TcCLB.504147.30 | hypothetical protein, conserved | -0,1731 | 0,1763 |
| TcCLB.509695.40 | hypothetical protein, conserved | -0,1731 | 0,1813 |
| TcCLB.507685.20 | trans-sialidase (pseudogene), putative | -0,1731 | 0,5963 |
| TcCLB.508257.140 | Aquaporin 2 | -0,1732 | 0,1636 |
| TcCLB.506811.150 | hypothetical protein, conserved | -0,1732 | 0,2163 |
| TcCLB.511895.60 | hypothetical protein, conserved | -0,1732 | 0,312 |
| TcCLB.511283.170 | hypothetical protein, conserved | -0,1733 | 0,03309 |
| TcCLB.506435.120 | ATP-dependent RNA helicase, putative | -0,1733 | 0,07469 |
| TcCLB.508169.100 | hypothetical protein, conserved | -0,1733 | 0,2351 |
| TcCLB.510123.20 | mitogen activated protein kinase, putative | -0,1734 | 0,1099 |
| TcCLB.508387.170 | double-strand-break repair protein rad21 homolog, putativ | -0,1734 | 0,1335 |
| TcCLB.507649.40 | hypothetical protein, conserved | -0,1734 | 0,2724 |
| TcCLB.510431.190 | hypothetical protein | -0,1736 | 0,03877 |
| TcCLB.506445.40 | ubiquitin hydrolase (pseudogene), putative | -0,1736 | 0,1206 |
| TcCLB.508873.330 | trans-sialidase, Group V, putative | -0,1736 | 0,5365 |
| TcCLB.508727.40 | hypothetical protein, conserved | -0,1738 | 0,2764 |
| TcCLB.510823.50 | hypothetical protein, conserved | -0,1739 | 0,1715 |
| TcCLB.503419.20 | adenosine deaminase-like protein, putative (ADAT1h) | -0,1739 | 0,2322 |
| TcCLB.510911.58 | retrotransposon hot spot protein (RHS, pseudogene), putat | -0,1739 | 0,7272 |
| TcCLB.506925.100 | hypothetical protein, conserved | -0,174 | 0,2498 |
| TcCLB.506747.30 | ubiquitin hydrolase, putative | -0,1741 | 0,09918 |
| TcCLB.509695.120 | epsilon tubulin, putative | -0,1741 | 0,2376 |
| TcCLB.506287.209 | DNA ligase, putative | -0,1742 | 0,1885 |
| TcCLB.504243.40 | hypothetical protein, conserved | -0,1743 | 0,1094 |
| TcCLB.506817.60 | hypothetical protein, conserved | -0,1743 | 0,2709 |
| TcCLB.507967.10 | hypothetical protein | -0,1744 | 0,236 |
| TcCLB.504689.20 | mucin-associated surface protein (MASP, pseudogene), pu | -0,1744 | 0,4888 |

| | | | |
|---|---|---|---|
| TcCLB.503551.20 | hypothetical protein, conserved | -0,1744 | 0,8151 |
| TcCLB.506733.30 | hypothetical protein, conserved | -0,1745 | 0,09549 |
| TcCLB.506273.10 | trans-sialidase (pseudogene), putative | -0,1745 | 0,4081 |
| TcCLB.507951.80 | Present in the outer mitochondrial membrane proteome 2! | -0,1746 | 0,1612 |
| TcCLB.503793.10 | 2-oxoglutarate dehydrogenase subunit, putative | -0,1746 | 0,2752 |
| TcCLB.508355.350 | hypothetical protein, conserved | -0,1747 | 0,2281 |
| TcCLB.507257.150 | hypothetical protein, conserved | -0,1748 | 0,2356 |
| TcCLB.509109.190 | hypothetical protein, conserved | -0,1748 | 0,3637 |
| TcCLB.508153.950 | 6-phosphofructo-2-kinase 1 | -0,1749 | 0,04674 |
| TcCLB.506753.120 | DnaJ chaperone protein, putative | -0,1749 | 0,1591 |
| TcCLB.505807.200 | hypothetical protein, conserved | -0,1749 | 0,1727 |
| TcCLB.506417.30 | hypothetical protein, conserved | -0,175 | 0,3755 |
| TcCLB.509597.10 | hypothetical protein, conserved | -0,1751 | 0,1185 |
| TcCLB.504159.90 | hypothetical protein, conserved | -0,1751 | 0,1202 |
| TcCLB.510815.9 | dispersed gene family protein 1 (DGF-1), putative | -0,1751 | 0,2109 |
| TcCLB.506743.30 | 50S ribosomal protein L17, putative | -0,1751 | 0,3326 |
| TcCLB.511893.10 | ribonuclease inhibitor, putative | -0,1752 | 0,09636 |
| TcCLB.506459.260 | hypothetical protein, conserved | -0,1752 | 0,1057 |
| TcCLB.504741.220 | hypothetical protein, conserved | -0,1752 | 0,2769 |
| TcCLB.507677.140 | hypothetical protein, conserved | -0,1752 | 0,7992 |
| TcCLB.511239.150 | histone acetyltransferase, putative | -0,1754 | 0,2443 |
| TcCLB.506613.110 | mucin-associated surface protein (MASP, pseudogene), pu | -0,1754 | 0,6316 |
| TcCLB.509965.319 | hypothetical protein, conserved | -0,1755 | 0,1525 |
| TcCLB.508199.80 | EF hand, putative | -0,1756 | 0,1903 |
| TcCLB.509429.120 | Mitochondrial Ribosomal Protein of Small Subunit 29, puta | -0,1756 | 0,2253 |
| TcCLB.508707.290 | hypothetical protein | -0,1756 | 0,3116 |
| TcCLB.509775.34 | hypothetical protein, conserved | -0,1756 | 0,3419 |
| TcCLB.511323.20 | hypothetical protein, conserved | -0,1756 | 0,3897 |
| TcCLB.436761.20 | hypothetical protein, conserved (fragment) | -0,1756 | 0,7188 |
| TcCLB.504039.175 | hypothetical protein, conserved (pseudogene) | -0,1757 | 0,6932 |
| TcCLB.511803.15 | dynein light chain lc6, flagellar outer arm (pseudogene), pu | -0,176 | 0,4289 |
| TcCLB.511577.10 | hypothetical protein, conserved | -0,176 | 0,5301 |
| TcCLB.511707.39 | hypothetical protein, conserved | -0,1761 | 0,1029 |
| TcCLB.511529.100 | hypothetical protein, conserved | -0,1761 | 0,2903 |
| TcCLB.510175.170 | glycine dehydrogenase [decarboxylating] (pseudogene), pu | -0,1761 | 0,4438 |
| TcCLB.511127.110 | ATP-dependent protease ATPase subunit HslU2 (HslU2) | -0,1762 | 0,08104 |
| TcCLB.504103.80 | hypothetical protein, conserved | -0,1762 | 0,1906 |
| TcCLB.506955.150 | mucin-associated surface protein (MASP), putative | -0,1762 | 0,5733 |
| TcCLB.511421.24 | kelch domain-containing protein | -0,1764 | 0,2474 |
| TcCLB.510043.50 | hypothetical protein, conserved | -0,1765 | 0,05114 |
| TcCLB.511445.70 | hypothetical protein, conserved | -0,1765 | 0,1622 |
| TcCLB.508083.20 | hypothetical protein, conserved | -0,1767 | 0,3851 |
| TcCLB.511867.190 | tubulin gamma chain (TUBG) | -0,1768 | 0,125 |
| TcCLB.504021.130 | hypothetical protein, conserved | -0,1768 | 0,3503 |
| TcCLB.507735.50 | hypothetical protein, conserved (pseudogene) | -0,1769 | 0,04971 |

| | | | |
|---|---|---|---|
| TcCLB.506887.50 | katanin, putative | -0,177 | 0,255 |
| TcCLB.509985.40 | hypothetical protein, conserved | -0,177 | 0,2582 |
| TcCLB.508121.80 | mucin TcMUCII, putative | -0,177 | 0,7861 |
| TcCLB.510099.70 | hypothetical protein, conserved | -0,1771 | 0,111 |
| TcCLB.511739.20 | hypothetical protein, conserved | -0,1772 | 0,1662 |
| TcCLB.508805.20 | hypothetical protein, conserved | -0,1773 | 0,08214 |
| TcCLB.509399.160 | CCR4-NOT transcription complex subunit 2, putative | -0,1773 | 0,1604 |
| TcCLB.508525.40 | hypothetical protein, conserved | -0,1775 | 0,1902 |
| TcCLB.506357.140 | Periodic tryptophan protein 2 homolog, putative | -0,1776 | 0,1272 |
| TcCLB.511389.140 | hypothetical protein, conserved | -0,1776 | 0,2932 |
| TcCLB.510351.100 | hypothetical protein, conserved | -0,1776 | 0,3208 |
| TcCLB.503803.10 | rRNA biogenesis protein, putative | -0,1777 | 0,1708 |
| TcCLB.507093.260 | multidrug resistance protein, mitochondrial, putative (ABC | -0,1778 | 0,01904 |
| TcCLB.510999.39 | 40S ribosomal protein S3A, putative (fragment) | -0,1778 | 0,1163 |
| TcCLB.510759.50 | hypothetical protein | -0,1778 | 0,2145 |
| TcCLB.510517.60 | hypothetical protein, conserved | -0,1778 | 0,3159 |
| TcCLB.507515.70 | hypothetical protein, conserved | -0,1779 | 0,09933 |
| TcCLB.511285.70 | hypothetical protein, conserved | -0,1781 | 0,07749 |
| TcCLB.507089.80 | exosome-associated protein 2, putative (EAP2) | -0,1782 | 0,3051 |
| TcCLB.508257.104 | ribosomal protein L14, putative | -0,1782 | 0,4106 |
| TcCLB.509979.379 | hypothetical protein, conserved | -0,1783 | 0,2352 |
| TcCLB.506017.31 | ATP-dependent DEAD/H RNA helicase (pseudogene), putat | -0,1785 | 0,6112 |
| TcCLB.507467.50 | hypothetical protein, conserved | -0,1786 | 0,2642 |
| TcCLB.510395.20 | hypothetical protein, conserved | -0,1786 | 0,301 |
| TcCLB.504197.15 | surface protease GP63 (pseudogene), putative | -0,1786 | 0,705 |
| TcCLB.503847.10 | hypothetical protein, conserved (pseudogene) | -0,1787 | 0,1386 |
| TcCLB.507611.210 | hypothetical protein | -0,1787 | 0,7462 |
| TcCLB.511297.10 | hypothetical protein, conserved | -0,1788 | 0,04833 |
| TcCLB.506935.19 | hypothetical protein, conserved (fragment) | -0,1788 | 0,5195 |
| TcCLB.503973.271 | surface protease GP63 (pseudogene), putative | -0,1789 | 0,6053 |
| TcCLB.509549.120 | endosomal integral membrane protein (pseudogene), puta | -0,179 | 0,142 |
| TcCLB.508741.350 | hypothetical protein, conserved | -0,179 | 0,1709 |
| TcCLB.509837.30 | hypothetical protein | -0,179 | 0,3239 |
| TcCLB.511901.10 | Trypanosome basal body component protein (TBBC) | -0,179 | 0,357 |
| TcCLB.511001.40 | hypothetical protein | -0,1791 | 0,187 |
| TcCLB.511847.89 | hypothetical protein, conserved | -0,1791 | 0,4932 |
| TcCLB.508389.20 | hypothetical protein | -0,1792 | 0,1466 |
| TcCLB.510603.20 | hypothetical protein, conserved | -0,1792 | 0,3635 |
| TcCLB.506795.90 | hypothetical protein, conserved | -0,1793 | 0,07346 |
| TcCLB.506063.34 | hypothetical protein, conserved (pseudogene) | -0,1793 | 0,08819 |
| TcCLB.508409.150 | hypothetical protein, conserved | -0,1794 | 0,09087 |
| TcCLB.507993.89 | hypothetical protein, conserved | -0,1794 | 0,3783 |
| TcCLB.506625.19 | dispersed gene family protein 1 (DGF-1), putative (fragme | -0,1794 | 0,6141 |
| TcCLB.507007.74 | small nuclear ribonucleoprotein Sm-F, putative | -0,1795 | 0,5227 |
| TcCLB.509291.20 | mucin-associated surface protein (MASP), putative | -0,1796 | 0,7835 |

| | | | |
|---|---|---|---|
| TcCLB.511881.30 | hypothetical protein, conserved | -0,1797 | 0,1882 |
| TcCLB.511855.60 | hypothetical protein, conserved | -0,1797 | 0,2394 |
| TcCLB.511143.39 | hypothetical protein, conserved | -0,1797 | 0,4381 |
| TcCLB.506893.50 | hypothetical protein, conserved | -0,1797 | 0,5145 |
| TcCLB.508307.160 | hypothetical protein, conserved | -0,1799 | 0,1166 |
| TcCLB.504109.10 | hypothetical protein, conserved | -0,1799 | 0,1176 |
| TcCLB.506369.50 | hypothetical protein, conserved | -0,1799 | 0,2803 |
| TcCLB.506753.190 | hypothetical protein, conserved | -0,18 | 0,2914 |
| TcCLB.508775.35 | retrotransposon hot spot protein (RHS, pseudogene), putat | -0,1801 | 0,7226 |
| TcCLB.504741.80 | hypothetical protein, conserved | -0,1802 | 0,2534 |
| TcCLB.509715.80 | hypothetical protein, conserved | -0,1803 | 0,1531 |
| TcCLB.510525.120 | hypothetical protein, conserved | -0,1803 | 0,2253 |
| TcCLB.506839.20 | hypothetical protein | -0,1804 | 0,1059 |
| TcCLB.506499.40 | mucin-associated surface protein (MASP), putative | -0,1804 | 0,6988 |
| TcCLB.507071.34 | mucin TcMUCII, putative | -0,1804 | 0,8086 |
| TcCLB.509571.4 | PAB1-binding protein , putative | -0,1805 | 0,06659 |
| TcCLB.506647.10 | hypothetical protein, conserved | -0,1805 | 0,1462 |
| TcCLB.503455.30 | hypothetical protein, conserved | -0,1805 | 0,1548 |
| TcCLB.503891.30 | phosphatidylinositol-4-phosphate 5-kinase, putative | -0,1805 | 0,2872 |
| TcCLB.510359.69 | mitochondrial carrier protein, putative | -0,1806 | 0,2595 |
| TcCLB.508243.50 | mucin-associated surface protein (MASP), putative | -0,1806 | 0,6601 |
| TcCLB.509979.187 | surface protease GP63 (pseudogene), putative | -0,1808 | 0,6685 |
| TcCLB.508637.150 | ribonuclease L inhibitor, putative | -0,1809 | 0,0613 |
| TcCLB.511159.14 | hypothetical protein, conserved | -0,1809 | 0,09378 |
| TcCLB.507653.30 | hypothetical protein | -0,1809 | 0,5431 |
| TcCLB.509537.80 | hypothetical protein, conserved | -0,181 | 0,03334 |
| TcCLB.510031.40 | hypothetical protein | -0,181 | 0,3383 |
| TcCLB.509735.140 | trans-sialidase (pseudogene), putative | -0,181 | 0,3608 |
| TcCLB.507539.40 | cell cycle sequence binding phosphoprotein (RBP33), putat | -0,1811 | 0,09456 |
| TcCLB.509287.150 | retrotransposon hot spot protein (RHS, pseudogene), putat | -0,1811 | 0,2307 |
| TcCLB.510105.90 | hypothetical protein, conserved | -0,1812 | 0,179 |
| TcCLB.508699.60 | hypothetical protein, conserved | -0,1812 | 0,2419 |
| TcCLB.506859.120 | hypothetical protein | -0,1814 | 0,2021 |
| TcCLB.508533.40 | ecotin, putative | -0,1814 | 0,2816 |
| TcCLB.509395.50 | hypothetical protein, conserved | -0,1815 | 0,06785 |
| TcCLB.511825.130 | hypothetical protein | -0,1815 | 0,4438 |
| TcCLB.506181.20 | hypothetical protein, conserved | -0,1816 | 0,1446 |
| TcCLB.511127.20 | calcium binding protein EF hand, putative | -0,1817 | 0,07932 |
| TcCLB.463673.9 | hypothetical protein, conserved | -0,1817 | 0,3029 |
| TcCLB.510759.80 | hypothetical protein, conserved | -0,182 | 0,06656 |
| TcCLB.504131.170 | ubiquitin carboxyl-terminal hydrolase, putative | -0,182 | 0,1004 |
| TcCLB.508355.190 | choline kinase | -0,182 | 0,1149 |
| TcCLB.507257.110 | hypothetical protein, conserved | -0,182 | 0,3208 |
| TcCLB.507611.270 | hypothetical protein, conserved | -0,1822 | 0,3295 |
| TcCLB.506725.60 | hypothetical protein, conserved | -0,1822 | 0,4228 |

| | | | |
|---|---|---|---|
| TcCLB.511217.30 | diacylglycerol kinase, putative | -0,1823 | 0,1769 |
| TcCLB.508209.39 | hypothetical protein, conserved | -0,1823 | 0,2046 |
| TcCLB.509911.70 | mu-adaptin 4, putative | -0,1823 | 0,2474 |
| TcCLB.511163.59 | hypothetical protein, conserved (fragment) | -0,1824 | 0,4953 |
| TcCLB.509203.10 | 2,3-bisphosphoglycerate-independent phosphoglycerate m | -0,1825 | 0,5239 |
| TcCLB.508199.30 | hypothetical protein, conserved | -0,1826 | 0,2298 |
| TcCLB.510655.30 | helicase, putative | -0,1829 | 0,0253 |
| TcCLB.503959.60 | hypothetical protein, conserved | -0,183 | 0,1175 |
| TcCLB.507049.100 | mitochondrial RNA binding protein, putative | -0,183 | 0,1964 |
| TcCLB.510609.130 | hypothetical protein, conserved | -0,183 | 0,2025 |
| TcCLB.510743.90 | hypothetical protein, conserved | -0,183 | 0,2901 |
| TcCLB.511211.110 | hypothetical protein, conserved | -0,1831 | 0,07996 |
| TcCLB.506559.240 | hypothetical protein, conserved | -0,1831 | 0,09102 |
| TcCLB.510289.20 | hypothetical protein, conserved | -0,1831 | 0,1274 |
| TcCLB.504171.20 | hypothetical protein, conserved | -0,1831 | 0,3211 |
| TcCLB.508479.480 | N-acetyltransferase complex ARD1 subunit (pseudogene), | -0,1832 | 0,5195 |
| TcCLB.510261.10 | mucin-associated surface protein (MASP), putative | -0,1832 | 0,6093 |
| TcCLB.508181.110 | hypothetical protein, conserved | -0,1833 | 0,1155 |
| TcCLB.506275.70 | hypothetical protein, conserved | -0,1833 | 0,1672 |
| TcCLB.509747.40 | hypothetical protein, conserved | -0,1833 | 0,176 |
| TcCLB.510149.130 | hypothetical protein, conserved | -0,1833 | 0,2824 |
| TcCLB.507677.100 | hypothetical protein, conserved | -0,1834 | 0,2799 |
| TcCLB.506853.60 | hypothetical protein, conserved | -0,1834 | 0,3417 |
| TcCLB.504097.4 | nucleoplasmin-like protein (NLP), putative | -0,1835 | 0,1968 |
| TcCLB.504131.110 | hypothetical protein, conserved | -0,1835 | 0,2807 |
| TcCLB.510265.110 | hypothetical protein, conserved | -0,1835 | 0,7163 |
| TcCLB.510105.40 | cullin-like protein, putative | -0,1836 | 0,06367 |
| TcCLB.506743.140 | hypothetical protein, conserved | -0,1836 | 0,2134 |
| TcCLB.510241.30 | hypothetical protein, conserved | -0,1836 | 0,2365 |
| TcCLB.510241.50 | 60S ribosomal protein L10, putative | -0,1838 | 0,02756 |
| TcCLB.508153.980 | hypothetical protein, conserved | -0,1839 | 0,07401 |
| TcCLB.504883.50 | hypothetical protein, conserved | -0,1839 | 0,07615 |
| TcCLB.507913.10 | hypothetical protein, conserved | -0,184 | 0,5193 |
| TcCLB.507963.79 | nucleosome assembly protein, putative | -0,1841 | 0,05781 |
| TcCLB.503959.30 | hypothetical protein, conserved | -0,1841 | 0,06851 |
| TcCLB.507949.150 | conserved protein, unknown function | -0,1841 | 0,1258 |
| TcCLB.510381.60 | proliferator-activated receptor-interacting protein (PRIP) ir | -0,1841 | 0,3157 |
| TcCLB.507823.30 | hypothetical protein, conserved | -0,1842 | 0,1582 |
| TcCLB.506247.320 | methyltransferase, putative | -0,1843 | 0,1061 |
| TcCLB.508909.200 | hypothetical protein | -0,1843 | 0,1077 |
| TcCLB.511153.130 | DNA-direcetd RNA polymerase II, subunit 9, putative | -0,1843 | 0,3921 |
| TcCLB.509109.110 | hypothetical protein, conserved | -0,1843 | 0,4176 |
| TcCLB.510001.50 | hypothetical protein, conserved | -0,1845 | 0,4 |
| TcCLB.511183.170 | trans-sialidase (pseudogene), putative | -0,1845 | 0,6342 |
| TcCLB.509175.18 | hypothetical protein, conserved | -0,1846 | 0,3634 |

| | | | |
|---|---|---|---|
| TcCLB.510373.40 | hypothetical protein | -0,1846 | 0,7844 |
| TcCLB.507801.179 | vesicle-associated membrane protein | -0,1847 | 0,1994 |
| TcCLB.510021.180 | trans-sialidase, Group V, putative | -0,1848 | 0,5862 |
| TcCLB.506453.30 | mucin-associated surface protein (MASP), putative | -0,1848 | 0,7152 |
| TcCLB.508285.5 | trans-sialidase (pseudogene), putative | -0,185 | 0,5584 |
| TcCLB.398265.9 | trans-sialidase, putative | -0,185 | 0,7876 |
| TcCLB.506965.190 | mucin TcMUCII, putative | -0,185 | 0,8016 |
| TcCLB.511215.20 | hypothetical protein | -0,1851 | 0,03577 |
| TcCLB.507081.90 | hypothetical protein, conserved | -0,1851 | 0,1648 |
| TcCLB.510961.10 | mucin-associated surface protein (MASP), putative | -0,1851 | 0,6549 |
| TcCLB.504131.150 | SNF2 family protein | -0,1852 | 0,2356 |
| TcCLB.506247.300 | hypothetical protein, conserved | -0,1852 | 0,2824 |
| TcCLB.510655.70 | hypothetical protein, conserved | -0,1853 | 0,3314 |
| TcCLB.507913.30 | proliferative cell nuclear antigen (PCNA), putative | -0,1855 | 0,0743 |
| TcCLB.429257.10 | hypothetical protein, conserved | -0,1855 | 0,4974 |
| TcCLB.506737.40 | hypothetical protein, conserved | -0,1856 | 0,08544 |
| TcCLB.506979.50 | hypothetical protein, conserved | -0,1856 | 0,1094 |
| TcCLB.505171.44 | hypothetical protein, conserved | -0,1857 | 0,4772 |
| TcCLB.511529.180 | hypothetical protein, conserved | -0,1858 | 0,1744 |
| TcCLB.505231.20 | trans-sialidase, Group II, putative | -0,1858 | 0,3457 |
| TcCLB.508629.63 | ATP-dependent DEAD/H RNA helicase (pseudogene), putat | -0,1859 | 0,614 |
| TcCLB.505945.10 | DNA repair and recombination helicase protein PIF5, putat | -0,186 | 0,07909 |
| TcCLB.510769.150 | hypothetical protein, conserved | -0,1862 | 0,1209 |
| TcCLB.509121.20 | dispersed gene family protein 1 (DGF-1, pseudogene), puta | -0,1863 | 0,3213 |
| TcCLB.511275.20 | hypothetical protein, conserved | -0,1864 | 0,142 |
| TcCLB.510241.40 | hypothetical protein, conserved | -0,1865 | 0,2352 |
| TcCLB.510731.130 | hypothetical protein, conserved | -0,1865 | 0,5343 |
| TcCLB.511417.129 | hypothetical protein, conserved | -0,1865 | 0,6663 |
| TcCLB.509185.10 | retrotransposon hot spot protein (RHS, pseudogene), putat | -0,1865 | 0,737 |
| TcCLB.510421.170 | hypothetical protein, conserved | -0,1866 | 0,02816 |
| TcCLB.506963.110 | NUDIX hydrolase 3, putative (Nudix3) | -0,1866 | 0,3273 |
| TcCLB.508607.10 | retrotransposon hot spot (RHS) protein, putative | -0,1866 | 0,6943 |
| TcCLB.511621.40 | hypothetical protein, conserved | -0,1867 | 0,1964 |
| TcCLB.507047.90 | hypothetical protein | -0,1867 | 0,253 |
| TcCLB.506175.40 | hypothetical protein, conserved | -0,1868 | 0,1269 |
| TcCLB.504253.30 | hypothetical protein, conserved | -0,1871 | 0,1434 |
| TcCLB.510901.110 | hypothetical protein, conserved | -0,1871 | 0,2964 |
| TcCLB.508153.170 | ubiquinone biosynthesis protein COQ7 homolog, putative | -0,1871 | 0,4253 |
| TcCLB.507527.20 | hypothetical protein, conserved | -0,1872 | 0,109 |
| TcCLB.506989.50 | hypothetical protein, conserved | -0,1872 | 0,249 |
| TcCLB.511621.190 | dual specificity protein phosphatase, putative | -0,1874 | 0,3009 |
| TcCLB.427789.10 | hypothetical protein, conserved | -0,1874 | 0,4677 |
| TcCLB.509103.39 | dynein heavy chain, putative | -0,1875 | 0,1008 |
| TcCLB.506855.60 | hypothetical protein, conserved | -0,1875 | 0,1798 |
| TcCLB.509749.30 | hypothetical protein, conserved | -0,1876 | 0,2983 |

| | | | |
|---|---|---|---|
| TcCLB.509029.150 | hypothetical protein, conserved | -0,1877 | 0,3763 |
| TcCLB.510357.60 | hypothetical protein, conserved | -0,1877 | 0,4872 |
| TcCLB.511409.70 | hypothetical protein, conserved | -0,1878 | 0,2133 |
| TcCLB.511619.5 | FtsJ cell division protein, putative (fragment) | -0,188 | 0,1867 |
| TcCLB.509941.30 | retrotransposon hot spot protein (RHS, pseudogene), putat | -0,188 | 0,2761 |
| TcCLB.507023.230 | splicing factor 3a, putative | -0,1881 | 0,1464 |
| TcCLB.510959.30 | mucin-associated surface protein (MASP), putative | -0,1881 | 0,6026 |
| TcCLB.509483.10 | surface protease GP63, putative | -0,1881 | 0,7757 |
| TcCLB.510437.60 | hypothetical protein, conserved | -0,1882 | 0,3171 |
| TcCLB.508089.10 | trans-sialidase, Group I, putative | -0,1882 | 0,3605 |
| TcCLB.507771.20 | hypothetical protein, conserved | -0,1883 | 0,1803 |
| TcCLB.506223.100 | hypothetical protein, conserved | -0,1883 | 0,2037 |
| TcCLB.506933.40 | DNA replication licensing factor MCM2, putative | -0,1885 | 0,08803 |
| TcCLB.509791.110 | hypothetical protein, conserved | -0,1885 | 0,1557 |
| TcCLB.506247.80 | DNA-binding protein, putative | -0,1885 | 0,1852 |
| TcCLB.506123.50 | hypothetical protein, conserved | -0,1885 | 0,301 |
| TcCLB.506767.230 | mucin-associated surface protein (MASP), putative | -0,1885 | 0,5766 |
| TcCLB.507817.60 | Present in the outer mitochondrial membrane proteome 3 | -0,1886 | 0,06504 |
| TcCLB.509345.11 | retrotransposon hot spot protein (RHS, pseudogene), putat | -0,1886 | 0,1666 |
| TcCLB.506451.80 | mucin-associated surface protein (MASP, pseudogene), pu | -0,1886 | 0,5006 |
| TcCLB.504179.50 | hypothetical protein | -0,1887 | 0,1049 |
| TcCLB.510335.10 | hypothetical protein, conserved | -0,1887 | 0,1423 |
| TcCLB.511765.10 | trans-sialidase (pseudogene), putative | -0,1887 | 0,6102 |
| TcCLB.503465.10 | hypothetical protein, conserved | -0,1888 | 0,141 |
| TcCLB.507257.120 | hypothetical protein, conserved | -0,1888 | 0,3027 |
| TcCLB.508385.50 | hypothetical protein, conserved | -0,1889 | 0,2535 |
| TcCLB.507071.80 | trans-sialidase, Group VI, putative | -0,1889 | 0,6453 |
| TcCLB.510715.20 | trans-sialidase, Group V, putative | -0,189 | 0,5029 |
| TcCLB.506637.90 | hypothetical protein, conserved | -0,1891 | 0,07706 |
| TcCLB.508153.710 | hypothetical protein | -0,1893 | 0,2036 |
| TcCLB.508153.780 | hypothetical protein | -0,1894 | 0,06861 |
| TcCLB.510517.50 | hypothetical protein, conserved | -0,1894 | 0,1065 |
| TcCLB.510371.100 | mucin-associated surface protein (MASP), putative | -0,1894 | 0,6745 |
| TcCLB.511129.47 | hypothetical protein, conserved (pseudogene) | -0,1895 | 0,4139 |
| TcCLB.507611.389 | hypothetical protein, conserved | -0,1895 | 0,4883 |
| TcCLB.504089.14 | hypothetical protein, conserved | -0,1896 | 0,07706 |
| TcCLB.508257.244 | hypothetical protein, conserved | -0,1896 | 0,4213 |
| TcCLB.511001.9 | 40S ribosomal protein S3A, putative | -0,1897 | 0,2143 |
| TcCLB.510551.50 | trans-sialidase, putative | -0,1897 | 0,7735 |
| TcCLB.510659.210 | chaperone DNAJ protein, putative | -0,1898 | 0,06965 |
| TcCLB.510265.100 | hypothetical protein, conserved | -0,1898 | 0,109 |
| TcCLB.510213.39 | mucin-associated surface protein (MASP), putative | -0,1898 | 0,5357 |
| TcCLB.508739.11 | trans-sialidase (pseudogene), putative | -0,1898 | 0,5438 |
| TcCLB.507867.80 | hypothetical protein, conserved | -0,19 | 0,04653 |
| TcCLB.511165.100 | serine/threonine protein kinase, putative | -0,1901 | 0,2244 |

| | | | |
|---|---|---|---|
| TcCLB.508879.90 | hypothetical protein, conserved | -0,1901 | 0,3113 |
| TcCLB.508277.250 | hypothetical protein, conserved | -0,1902 | 0,09868 |
| TcCLB.510769.40 | hypothetical protein, conserved | -0,1902 | 0,2306 |
| TcCLB.509941.70 | dispersed gene family protein 1 (DGF-1, pseudogene), puta | -0,1902 | 0,4195 |
| TcCLB.507951.179 | hypothetical protein, conserved | -0,1904 | 0,1795 |
| TcCLB.507555.70 | trans-sialidase (pseudogene), putative | -0,1904 | 0,3842 |
| TcCLB.510827.40 | hypothetical protein | -0,1906 | 0,4476 |
| TcCLB.510283.60 | hypothetical protein, conserved | -0,1907 | 0,3469 |
| TcCLB.511585.130 | ubiquitin activating enzyme, putative | -0,1908 | 0,1474 |
| TcCLB.509213.50 | hypothetical protein, conserved | -0,1909 | 0,2674 |
| TcCLB.480975.10 | hypothetical protein, conserved | -0,191 | 0,2366 |
| TcCLB.506765.46 | mucin-associated surface protein (MASP, pseudogene), pu | -0,191 | 0,4029 |
| TcCLB.511283.130 | MEKK-related kinase 1, putative (MRK1) | -0,1911 | 0,1099 |
| TcCLB.506155.70 | SWI/SNF-related helicase, putative | -0,1912 | 0,0301 |
| TcCLB.510777.29 | eukaryotic translation initiation factor 4E, putative | -0,1912 | 0,4343 |
| TcCLB.503723.90 | glyceraldehyde 3-phosphate dehydrogenase, C-terminal do | -0,1912 | 0,579 |
| TcCLB.506167.20 | hypothetical protein, conserved | -0,1913 | 0,03579 |
| TcCLB.506355.60 | small G-protein, putative | -0,1913 | 0,06485 |
| TcCLB.510823.80 | cytoplasmic l-asparaginase i-like protein, putative | -0,1913 | 0,3298 |
| TcCLB.509665.16 | elongation factor 1-gamma (EF-1-gamma, pseudogene), p | -0,1913 | 0,7272 |
| TcCLB.503589.10 | GDP-mannose pyrophosphorylase | -0,1914 | 0,1605 |
| TcCLB.508165.140 | mucin TcMUCII (pseudogene), putative | -0,1914 | 0,7596 |
| TcCLB.511229.60 | hypothetical protein, conserved | -0,1915 | 0,5535 |
| TcCLB.508737.30 | mitochondrial RNA binding protein 1 | -0,1916 | 0,01482 |
| TcCLB.510445.30 | hypothetical protein, conserved | -0,1916 | 0,1665 |
| TcCLB.503691.9 | hypothetical protein, conserved (fragment) | -0,1916 | 0,2802 |
| TcCLB.510421.300 | hypothetical protein, conserved | -0,1916 | 0,298 |
| TcCLB.511543.10 | hypothetical protein, conserved | -0,1916 | 0,316 |
| TcCLB.506885.280 | NIMA-related kinase, putative | -0,1916 | 0,5552 |
| TcCLB.508047.30 | trans-sialidase (pseudogene), putative | -0,1916 | 0,6359 |
| TcCLB.511917.10 | trans-sialidase, putative | -0,1917 | 0,4602 |
| TcCLB.508175.250 | hypothetical protein, conserved | -0,1919 | 0,05794 |
| TcCLB.509023.110 | ATP-dependent protease ATPase subunit HslU2 (HslU2) | -0,1919 | 0,06733 |
| TcCLB.503579.150 | RNA editing 3' terminal uridylyl transferase 2, putative | -0,1919 | 0,2078 |
| TcCLB.509049.10 | hypothetical protein, conserved | -0,1919 | 0,3498 |
| TcCLB.507677.90 | hypothetical protein, conserved | -0,192 | 0,1474 |
| TcCLB.504109.120 | hypothetical protein, conserved | -0,192 | 0,2343 |
| TcCLB.507787.30 | hypothetical protein, conserved | -0,192 | 0,4091 |
| TcCLB.506201.110 | hypothetical protein, conserved | -0,1921 | 0,2074 |
| TcCLB.507057.10 | hypothetical protein, conserved | -0,1921 | 0,2119 |
| TcCLB.507771.80 | hypothetical protein, conserved | -0,1922 | 0,1126 |
| TcCLB.508153.634 | hypothetical protein, conserved | -0,1922 | 0,349 |
| TcCLB.508325.130 | protein kinase, putative | -0,1923 | 0,1878 |
| TcCLB.511671.90 | hypothetical protein, conserved | -0,1923 | 0,2285 |
| TcCLB.509487.20 | hypothetical protein, conserved | -0,1923 | 0,236 |

| | | | |
|---|---|---|---|
| TcCLB.511725.190 | vacuolar sorting protein, putative | -0,1923 | 0,2463 |
| TcCLB.509097.28 | hypothetical protein, conserved | -0,1923 | 0,4598 |
| TcCLB.506475.30 | hypothetical protein, conserved | -0,1924 | 0,1143 |
| TcCLB.506287.120 | hypothetical protein | -0,1924 | 0,1442 |
| TcCLB.508539.60 | mucin-associated surface protein (MASP), putative | -0,1924 | 0,6613 |
| TcCLB.506211.210 | mitogen-activated protein kinase kinase kinase, putative (( | -0,1925 | 0,07706 |
| TcCLB.509029.50 | hypothetical protein, conserved | -0,1925 | 0,3058 |
| TcCLB.504427.50 | AMPK subunit beta, putative (AMPKB) | -0,1926 | 0,1993 |
| TcCLB.508313.122 | trans-sialidase (pseudogene), putative | -0,1926 | 0,806 |
| TcCLB.510737.40 | transport protein particle (TRAPP) subunit, putative | -0,1927 | 0,2715 |
| TcCLB.506295.80 | ADP-ribosylation factor-like 2, arl2, putative (ARL2) | -0,1927 | 0,5479 |
| TcCLB.506067.179 | elongation factor 1-gamma (EF-1-gamma, pseudogene), p | -0,1927 | 0,7669 |
| TcCLB.424123.10 | hypothetical protein, conserved (pseudogene) | -0,1928 | 0,2027 |
| TcCLB.503619.29 | hypothetical protein, conserved | -0,1928 | 0,4205 |
| TcCLB.506367.40 | hypothetical protein, conserved | -0,1929 | 0,1175 |
| TcCLB.506661.90 | hypothetical protein, conserved | -0,193 | 0,1056 |
| TcCLB.509715.140 | hypothetical protein, conserved | -0,193 | 0,3324 |
| TcCLB.504057.145 | cytochrome b5-like Heme/Steroid binding domain containi | -0,1931 | 0,1968 |
| TcCLB.506627.70 | hypothetical protein | -0,1931 | 0,2136 |
| TcCLB.503809.24 | hypothetical protein, conserved | -0,1931 | 0,3811 |
| TcCLB.508219.30 | surface protease GP63 (pseudogene), putative | -0,1931 | 0,7562 |
| TcCLB.510121.150 | serine/threonine protein kinase, putative | -0,1932 | 0,1269 |
| TcCLB.507213.30 | potassium voltage-gated channel, putative | -0,1932 | 0,1397 |
| TcCLB.503637.20 | hypothetical protein, conserved | -0,1933 | 0,08666 |
| TcCLB.507677.150 | G1 cyclin CycE4, putative (CYC5) | -0,1933 | 0,09578 |
| TcCLB.503789.70 | hypothetical protein, conserved | -0,1933 | 0,106 |
| TcCLB.509549.10 | trans-sialidase, Group V, putative | -0,1933 | 0,5382 |
| TcCLB.511545.50 | hypothetical protein, conserved | -0,1934 | 0,1195 |
| TcCLB.510105.130 | phosphatidylinositol-4-phosphate 5-kinase type II beta, put | -0,1934 | 0,1587 |
| TcCLB.511013.6 | N-acetyltransferase complex ARD1 subunit (pseudogene), | -0,1934 | 0,7408 |
| TcCLB.506257.80 | hypothetical protein, conserved | -0,1935 | 0,2319 |
| TcCLB.509065.160 | eukaryotic translation initiation factor 3 subunit 7-like prot | -0,1936 | 0,01386 |
| TcCLB.508355.160 | hypothetical protein, conserved | -0,1937 | 0,06718 |
| TcCLB.509455.130 | cAMP response protein, putative (CARP4) | -0,1937 | 0,08118 |
| TcCLB.503717.71 | trans-sialidase (pseudogene), putative | -0,1937 | 0,7979 |
| TcCLB.503895.10 | hypothetical protein, conserved | -0,1939 | 0,143 |
| TcCLB.506357.160 | hypothetical protein, conserved | -0,1939 | 0,3534 |
| TcCLB.507565.10 | dispersed gene family protein 1 (DGF-1, pseudogene), puta | -0,194 | 0,5478 |
| TcCLB.505363.15 | elongation factor 1-gamma (EF-1-gamma, pseudogene), p | -0,194 | 0,6125 |
| TcCLB.507047.140 | hypothetical protein, conserved | -0,1941 | 0,1361 |
| TcCLB.509033.70 | hypothetical protein, conserved | -0,1941 | 0,2876 |
| TcCLB.506267.101 | surface protease GP63 (pseudogene), putative | -0,1941 | 0,6178 |
| TcCLB.508647.230 | synaptojanin (pseudogene), putative | -0,1942 | 0,02616 |
| TcCLB.506559.70 | hypothetical protein, conserved | -0,1942 | 0,1463 |
| TcCLB.506519.80 | hypothetical protein, conserved | -0,1942 | 0,2737 |

| | | | |
|---|---|---|---|
| TcCLB.511527.50 | AUT2/APG4/ATG4 cysteine peptidase, putative | -0,1943 | 0,1267 |
| TcCLB.506673.40 | hypothetical protein, conserved | -0,1943 | 0,1308 |
| TcCLB.506821.30 | glycosylphosphatidylinositol-specific phospholipase C | -0,1943 | 0,1749 |
| TcCLB.508873.439 | mucin TcMUCII (pseudogene), putative | -0,1943 | 0,7521 |
| TcCLB.506509.30 | hypothetical protein, conserved | -0,1945 | 0,1152 |
| TcCLB.468941.9 | ATP-dependent DEAD/H RNA helicase, putative (fragment) | -0,1946 | 0,1004 |
| TcCLB.509561.110 | hypothetical protein, conserved | -0,1947 | 0,1258 |
| TcCLB.510877.110 | hypothetical protein, conserved | -0,1948 | 0,01747 |
| TcCLB.507795.20 | hypothetical protein, conserved | -0,1948 | 0,21 |
| TcCLB.506493.20 | hypothetical protein | -0,1949 | 0,5558 |
| TcCLB.510941.10 | hypothetical protein, conserved | -0,195 | 0,138 |
| TcCLB.510685.20 | hypothetical protein | -0,195 | 0,2794 |
| TcCLB.506577.80 | trans-sialidase, Group VIII, putative | -0,1952 | 0,3119 |
| TcCLB.507057.40 | hypothetical protein, conserved | -0,1952 | 0,3567 |
| TcCLB.510659.279 | nucleic acid binding protein, putative | -0,1953 | 0,3199 |
| TcCLB.510279.50 | mucin TcMUCII (pseudogene), putative | -0,1954 | 0,7524 |
| TcCLB.510943.190 | myosin heavy chain MYA2-related, putative | -0,1956 | 0,06361 |
| TcCLB.506581.40 | hypothetical protein, conserved | -0,1957 | 0,1526 |
| TcCLB.509021.9 | dispersed gene family protein 1 (DGF-1), putative | -0,1957 | 0,7654 |
| TcCLB.511603.300 | mucin-associated surface protein (MASP), putative | -0,1958 | 0,5564 |
| TcCLB.511277.120 | hypothetical protein, conserved | -0,1959 | 0,09006 |
| TcCLB.507001.70 | syntaxin, putative | -0,196 | 0,2133 |
| TcCLB.505807.160 | hypothetical protein, conserved | -0,196 | 0,3212 |
| TcCLB.508331.50 | hypothetical protein, conserved | -0,1961 | 0,1983 |
| TcCLB.509385.19 | retrotransposon hot spot protein (RHS, pseudogene), putat | -0,1961 | 0,603 |
| TcCLB.508851.180 | 14-3-3 protein 1, putative (14-3-3-I) | -0,1962 | 0,02235 |
| TcCLB.511491.70 | endosomal integral membrane protein, putative | -0,1962 | 0,08346 |
| TcCLB.508301.10 | hypothetical protein, conserved | -0,1962 | 0,2494 |
| TcCLB.506587.44 | hypothetical protein | -0,1962 | 0,3782 |
| TcCLB.504013.100 | ribosomal protein S19, putative | -0,1963 | 0,0161 |
| TcCLB.507089.190 | hypothetical protein, conserved | -0,1963 | 0,1364 |
| TcCLB.509951.40 | hypothetical protein | -0,1963 | 0,3851 |
| TcCLB.511385.70 | hypothetical protein | -0,1963 | 0,5418 |
| TcCLB.505183.40 | DNA repair and recombination protein RAD54, putative | -0,1965 | 0,03645 |
| TcCLB.506241.190 | hypothetical protein, conserved | -0,1966 | 0,2677 |
| TcCLB.506519.130 | guanosine monophosphate reductase, putative | -0,1967 | 0,05266 |
| TcCLB.508153.70 | hypothetical protein, conserved | -0,1967 | 0,2111 |
| TcCLB.507209.30 | hypothetical protein, conserved | -0,1968 | 0,05261 |
| TcCLB.506135.100 | hypothetical protein, conserved | -0,1968 | 0,3343 |
| TcCLB.506893.25 | U6 snRNA-associated Sm-like protein LSm6p | -0,1968 | 0,6088 |
| TcCLB.504105.200 | hypothetical protein, conserved | -0,1969 | 0,02656 |
| TcCLB.506241.40 | hypothetical protein, conserved | -0,1969 | 0,1467 |
| TcCLB.511537.16 | hypothetical protein, conserved | -0,1969 | 0,1954 |
| TcCLB.510283.90 | hypothetical protein, conserved | -0,197 | 0,1413 |
| TcCLB.506815.50 | hypothetical protein, conserved | -0,197 | 0,1544 |

| | | | |
|---|---|---|---|
| TcCLB.506679.150 | 40S ribosomal protein S10, putative | -0,197 | 0,1784 |
| TcCLB.510943.80 | ABC transporter, putative | -0,1971 | 0,0304 |
| TcCLB.510353.10 | major vault protein, putative | -0,1972 | 0,0197 |
| TcCLB.504427.280 | hypothetical protein, conserved | -0,1974 | 0,1483 |
| TcCLB.509161.149 | hypothetical protein, conserved | -0,1974 | 0,4266 |
| TcCLB.506883.100 | GTP-binding protein, putative | -0,1975 | 0,1047 |
| TcCLB.506801.70 | hypothetical protein | -0,1975 | 0,1113 |
| TcCLB.510965.20 | mitochondrial inner membrane signal peptidase, putative | -0,1975 | 0,3948 |
| TcCLB.506443.80 | synaptojanin (N-terminal domain), putative | -0,1976 | 0,07108 |
| TcCLB.511807.250 | hypothetical protein, conserved | -0,1976 | 0,0874 |
| TcCLB.504199.30 | kinesin-like protein, putative | -0,1976 | 0,1451 |
| TcCLB.509719.69 | hypothetical protein, conserved | -0,1976 | 0,1829 |
| TcCLB.507801.230 | trans-sialidase (pseudogene), putative | -0,1976 | 0,4365 |
| TcCLB.509623.10 | hypothetical protein, conserved | -0,1977 | 0,1613 |
| TcCLB.510759.200 | hypothetical protein, conserved | -0,1978 | 0,1531 |
| TcCLB.503823.169 | kinetoplastid kinetochore protein 2, putative (fragment) (k | -0,1979 | 0,1796 |
| TcCLB.511323.30 | hypothetical protein, conserved | -0,1979 | 0,2234 |
| TcCLB.511237.110 | cyclophilin, putative | -0,198 | 0,1724 |
| TcCLB.510737.110 | replication factor C, subunit 3, putative | -0,198 | 0,1944 |
| TcCLB.503637.10 | hypothetical protein, conserved | -0,198 | 0,2705 |
| TcCLB.509509.50 | hypothetical protein, conserved | -0,1981 | 0,2117 |
| TcCLB.511577.4 | CLC-type chloride channel, putative (fragment) | -0,1981 | 0,5302 |
| TcCLB.507989.20 | sigma-adaptin 3, putative | -0,1982 | 0,2949 |
| TcCLB.510255.4 | hypothetical protein, conserved | -0,1982 | 0,6154 |
| TcCLB.510697.55 | mucin TcMUCII (pseudogene), putative | -0,1985 | 0,6615 |
| TcCLB.508125.14 | mucin-associated surface protein (MASP), putative (fragm | -0,1985 | 0,6909 |
| TcCLB.511021.20 | hypothetical protein | -0,1986 | 0,3853 |
| TcCLB.511181.10 | hypothetical protein, conserved | -0,1987 | 0,1328 |
| TcCLB.508999.40 | hypothetical protein, conserved | -0,1989 | 0,1786 |
| TcCLB.506591.69 | hypothetical protein, conserved | -0,199 | 0,1094 |
| TcCLB.508031.10 | dispersed gene family protein 1 (DGF-1, pseudogene), puta | -0,1991 | 0,5216 |
| TcCLB.508771.40 | hypothetical protein, conserved | -0,1992 | 0,3048 |
| TcCLB.509161.140 | hypothetical protein, conserved | -0,1993 | 0,1317 |
| TcCLB.509885.40 | hypothetical protein, conserved | -0,1993 | 0,1339 |
| TcCLB.507727.19 | hypothetical protein | -0,1993 | 0,5343 |
| TcCLB.508707.340 | serine/threonine protein phosphatase, putative (fragment) | -0,1994 | 0,415 |
| TcCLB.503965.30 | hypothetical protein, conserved | -0,1995 | 0,2452 |
| TcCLB.511531.59 | mitogen activated protein kinase, putative | -0,1995 | 0,2893 |
| TcCLB.508911.10 | hypothetical protein, conserved | -0,1996 | 0,02996 |
| TcCLB.506941.270 | DNA-J protein, putative | -0,1996 | 0,1893 |
| TcCLB.506767.389 | surface protease GP63 (pseudogene), putative | -0,1996 | 0,7218 |
| TcCLB.509067.30 | mitochondrial RNA binding protein, putative | -0,1997 | 0,06654 |
| TcCLB.509911.90 | peptidase, putative | -0,1998 | 0,01795 |
| TcCLB.503703.10 | hypothetical protein, conserved | -0,1998 | 0,1938 |
| TcCLB.510661.220 | hypothetical protein, conserved | -0,1998 | 0,2089 |

| | | | |
|---|---|---|---|
| TcCLB.511127.290 | hypothetical protein, conserved | -0,2 | 0,08007 |
| TcCLB.506885.394 | hypothetical protein, conserved | -0,2 | 0,2392 |
| TcCLB.510329.210 | SNF1-related protein kinase catalytic subunit alpha, putativ | -0,2001 | 0,03439 |
| TcCLB.503789.40 | signal recognition particle (pseudogene), putative | -0,2001 | 0,05801 |
| TcCLB.507813.10 | structural maintenance of chromosome 3 protein, putative | -0,2001 | 0,2719 |
| TcCLB.506279.60 | trans-sialidase, Group VI, putative | -0,2001 | 0,3546 |
| TcCLB.506591.50 | hypothetical protein, conserved (pseudogene) | -0,2002 | 0,02403 |
| TcCLB.506247.20 | hypothetical protein | -0,2002 | 0,1179 |
| TcCLB.511713.9 | hypothetical protein, conserved | -0,2003 | 0,4607 |
| TcCLB.507005.20 | hypothetical protein, conserved | -0,2004 | 0,1893 |
| TcCLB.503861.30 | trans-sialidase (pseudogene), putative | -0,2004 | 0,2824 |
| TcCLB.511671.100 | hypothetical protein, conserved | -0,2004 | 0,3607 |
| TcCLB.511255.200 | mucin-associated surface protein (MASP), putative | -0,2004 | 0,665 |
| TcCLB.508707.149 | hypothetical protein, conserved | -0,2005 | 0,2739 |
| TcCLB.510997.50 | ATP-dependent DEAD/H RNA helicase (pseudogene), putat | -0,2007 | 0,1528 |
| TcCLB.510075.40 | hypothetical protein, conserved | -0,2007 | 0,2453 |
| TcCLB.507093.130 | hypothetical protein, conserved | -0,2008 | 0,0668 |
| TcCLB.419703.10 | hypothetical protein | -0,2008 | 0,495 |
| TcCLB.508355.300 | hypothetical protein, conserved | -0,2009 | 0,1903 |
| TcCLB.509047.50 | metallo-beta-lactamase superfamily, putative | -0,2009 | 0,3113 |
| TcCLB.511153.80 | hypothetical protein | -0,201 | 0,238 |
| TcCLB.507673.30 | hypothetical protein, conserved | -0,2011 | 0,1868 |
| TcCLB.503579.130 | hypothetical protein, conserved | -0,2011 | 0,2362 |
| TcCLB.510187.199 | hypothetical protein, conserved | -0,2011 | 0,2713 |
| TcCLB.508955.20 | hypothetical protein, conserved | -0,2012 | 0,03636 |
| TcCLB.507969.20 | hypothetical protein, conserved | -0,2012 | 0,2394 |
| TcCLB.507603.180 | ubiquitin hydrolase, putative | -0,2013 | 0,06549 |
| TcCLB.504045.90 | hypothetical protein, conserved (pseudogene) | -0,2013 | 0,26 |
| TcCLB.503823.80 | hypothetical protein, conserved | -0,2013 | 0,3289 |
| TcCLB.506537.130 | protein kinase, putative | -0,2014 | 0,5373 |
| TcCLB.511727.129 | ribosomal protein L18, putative | -0,2015 | 0,02165 |
| TcCLB.510311.10 | hypothetical protein, conserved | -0,2015 | 0,08048 |
| TcCLB.506375.20 | hypothetical protein, conserved (pseudogene) | -0,2016 | 0,1125 |
| TcCLB.505807.100 | hypothetical protein, conserved | -0,2016 | 0,2162 |
| TcCLB.506343.20 | dispersed gene family protein 1 (DGF-1, pseudogene), puta | -0,2016 | 0,5289 |
| TcCLB.511897.10 | TFIIH basal transcription factor complex helicase subunit, p | -0,2017 | 0,1139 |
| TcCLB.508355.20 | hypothetical protein, conserved | -0,2017 | 0,3758 |
| TcCLB.507093.280 | hypothetical protein | -0,2018 | 0,02623 |
| TcCLB.509007.90 | hypothetical protein | -0,2018 | 0,2548 |
| TcCLB.507663.70 | hypothetical protein, conserved | -0,2019 | 0,228 |
| TcCLB.510897.80 | hypothetical protein, conserved | -0,2019 | 0,3359 |
| TcCLB.508461.470 | hypothetical protein, conserved | -0,202 | 0,1228 |
| TcCLB.504217.10 | coatomer subunit epsilon (COPE) | -0,202 | 0,1893 |
| TcCLB.511287.180 | Associated kinase of Tb14-3-3, putative (AKB1) | -0,202 | 0,3881 |
| TcCLB.507713.10 | hypothetical protein, conserved | -0,202 | 0,4672 |

| | | | |
|---|---|---|---|
| TcCLB.509597.4 | hypothetical protein, conserved | -0,2021 | 0,1202 |
| TcCLB.511277.220 | hypothetical protein, conserved | -0,2021 | 0,1574 |
| TcCLB.506289.30 | hypothetical protein, conserved | -0,2021 | 0,2533 |
| TcCLB.508121.90 | mucin-associated surface protein (MASP, pseudogene), pu | -0,2021 | 0,6474 |
| TcCLB.509237.20 | hypothetical protein, conserved | -0,2022 | 0,04333 |
| TcCLB.508233.40 | hypothetical protein | -0,2022 | 0,1463 |
| TcCLB.511277.380 | hypothetical protein, conserved | -0,2022 | 0,4925 |
| TcCLB.509551.95 | anaphase promoting complex subunit protein, putative | -0,2022 | 0,5382 |
| TcCLB.511183.210 | mucin TcMUCII, putative | -0,2022 | 0,5827 |
| TcCLB.511173.380 | mucin TcMUCII (pseudogene), putative | -0,2023 | 0,7562 |
| TcCLB.507521.80 | hypothetical protein, conserved | -0,2024 | 0,1017 |
| TcCLB.508909.230 | calcium-transporting ATPase (pseudogene), putative | -0,2024 | 0,1691 |
| TcCLB.509805.80 | hypothetical protein, conserved | -0,2025 | 0,1209 |
| TcCLB.506457.70 | hypothetical protein, conserved | -0,2025 | 0,3187 |
| TcCLB.509951.30 | hypothetical protein, conserved | -0,2026 | 0,08812 |
| TcCLB.506127.120 | hypothetical protein, conserved | -0,2027 | 0,1763 |
| TcCLB.508543.170 | hypothetical protein, conserved | -0,2028 | 0,06088 |
| TcCLB.510187.80 | endonuclease V, putative | -0,2029 | 0,2383 |
| TcCLB.506937.20 | hypothetical protein, conserved | -0,2029 | 0,4774 |
| TcCLB.510013.230 | mucin-associated surface protein (MASP), putative | -0,2029 | 0,6609 |
| TcCLB.507437.21 | elongation factor 1-gamma (EF-1-gamma, pseudogene), p | -0,203 | 0,718 |
| TcCLB.508375.5 | elongation factor 1-gamma (EF-1-gamma, pseudogene), p | -0,2031 | 0,4426 |
| TcCLB.510717.40 | hypothetical protein, conserved | -0,2033 | 0,09177 |
| TcCLB.506227.230 | N-acetyltransferase complex ARD1 subunit, putative | -0,2033 | 0,1361 |
| TcCLB.508521.18 | trans-sialidase, putative (pseudogene) | -0,2033 | 0,6536 |
| TcCLB.508169.90 | eukaryotic translation initiation factor 3 subunit L, putative | -0,2034 | 0,01509 |
| TcCLB.510311.110 | hypothetical protein, conserved | -0,2034 | 0,1554 |
| TcCLB.504057.80 | Present in the outer mitochondrial membrane proteome 2 | -0,2035 | 0,06772 |
| TcCLB.507063.270 | histone deacetylase, putative | -0,2035 | 0,1074 |
| TcCLB.511211.170 | heat shock protein 70 (hsp70), putative | -0,2037 | 0,0106 |
| TcCLB.508169.50 | hypothetical protein, conserved | -0,2037 | 0,1532 |
| TcCLB.505943.10 | hypothetical protein, conserved | -0,2037 | 0,362 |
| TcCLB.510747.100 | hypothetical protein, conserved | -0,2038 | 0,1604 |
| TcCLB.506445.80 | hypothetical protein, conserved | -0,2038 | 0,1842 |
| TcCLB.508539.30 | hypothetical protein, conserved | -0,2038 | 0,2317 |
| TcCLB.509539.30 | fatty acid elongase, putative | -0,2038 | 0,3851 |
| TcCLB.504001.10 | hypothetical protein, conserved | -0,2039 | 0,03102 |
| TcCLB.508207.250 | mitotic cyclin, putative | -0,2039 | 0,2566 |
| TcCLB.506393.40 | trans-sialidase (pseudogene), putative | -0,2039 | 0,7294 |
| TcCLB.510877.50 | AP-3 complex subunit mu, putative (AP3M) | -0,204 | 0,06792 |
| TcCLB.511435.30 | MORN repeat-containing protein | -0,204 | 0,06913 |
| TcCLB.509607.60 | hypothetical protein | -0,2041 | 0,0753 |
| TcCLB.509755.60 | hypothetical protein, conserved (pseudogene) | -0,2041 | 0,6092 |
| TcCLB.511429.30 | conserved protein, unknown function | -0,2042 | 0,04619 |
| TcCLB.507639.40 | proteasome subunit beta type-5, putative | -0,2042 | 0,05746 |

| | | | |
|---|---|---|---|
| TcCLB.510063.10 | sigma-adaptin 3, putative | -0,2043 | 0,2852 |
| TcCLB.504717.20 | hypothetical protein, conserved | -0,2044 | 0,03744 |
| TcCLB.504827.170 | kinetoplastid kinetochore protein 2, putative (fragment) (k | -0,2044 | 0,2289 |
| TcCLB.508593.90 | hypothetical protein, conserved | -0,2044 | 0,2563 |
| TcCLB.511507.100 | hypothetical protein, conserved | -0,2045 | 0,2825 |
| TcCLB.511365.4 | hypothetical protein, conserved (fragment) | -0,2045 | 0,3862 |
| TcCLB.508479.280 | hypothetical protein, conserved | -0,2046 | 0,1525 |
| TcCLB.507093.63 | RNA-binding protein, putative | -0,2046 | 0,1632 |
| TcCLB.508015.40 | hypothetical protein | -0,2046 | 0,4611 |
| TcCLB.507787.150 | hypothetical protein, conserved | -0,2048 | 0,04573 |
| TcCLB.511229.9 | replication factor C, subunit 4, putative | -0,2048 | 0,3943 |
| TcCLB.506711.10 | cAMP response protein, putative (CARP4) | -0,2049 | 0,04077 |
| TcCLB.507031.29 | nucleosome assembly protein, putative | -0,2049 | 0,04739 |
| TcCLB.506479.74 | Quinonprotein alcohol dehydrogenase-like protein, putative | -0,2049 | 0,1328 |
| TcCLB.510307.330 | dispersed gene family protein 1 (DGF-1), putative | -0,2049 | 0,2018 |
| TcCLB.507021.100 | hypothetical protein, conserved | -0,205 | 0,1058 |
| TcCLB.508073.20 | hypothetical protein | -0,205 | 0,2218 |
| TcCLB.510271.20 | dispersed gene family protein 1 (DGF-1, pseudogene), puta | -0,205 | 0,3947 |
| TcCLB.503865.80 | phosphatidylinositol 4-kinase, putative | -0,2051 | 0,2193 |
| TcCLB.508273.91 | hypothetical protein, conserved | -0,2051 | 0,5462 |
| TcCLB.507071.210 | mucin TcMUCII, putative | -0,2051 | 0,759 |
| TcCLB.503599.30 | hypothetical protein, conserved | -0,2052 | 0,2446 |
| TcCLB.506661.160 | hypothetical protein, conserved | -0,2052 | 0,5368 |
| TcCLB.504427.190 | hypothetical protein, conserved | -0,2053 | 0,1914 |
| TcCLB.506835.150 | zinc-binding protein (Yippee), putative | -0,2053 | 0,3209 |
| TcCLB.419417.10 | hypothetical protein, conserved | -0,2053 | 0,5585 |
| TcCLB.507197.20 | oxidoreductase-like protein, putative | -0,2054 | 0,0944 |
| TcCLB.509643.70 | Archaic translocase of outer membrane 11 kDa subunit, pu | -0,2055 | 0,1335 |
| TcCLB.506567.140 | hypothetical protein, conserved | -0,2055 | 0,4007 |
| TcCLB.511473.3 | hypothetical protein, conserved (pseudogene) | -0,2056 | 0,3742 |
| TcCLB.511521.30 | superoxide dismutase, putative | -0,2057 | 0,1138 |
| TcCLB.509231.30 | hypothetical protein | -0,2057 | 0,3835 |
| TcCLB.506315.90 | serine/threonine protein kinase, putative | -0,2058 | 0,06524 |
| TcCLB.507723.120 | U1A small nuclear ribonucleoprotein, putative | -0,2058 | 0,4108 |
| TcCLB.503981.20 | hypothetical protein, conserved | -0,2059 | 0,2759 |
| TcCLB.507951.290 | hypothetical protein, conserved | -0,2059 | 0,3486 |
| TcCLB.509053.10 | ubiquitin-protein ligase, putative | -0,206 | 0,1914 |
| TcCLB.510289.90 | hypothetical protein, conserved | -0,206 | 0,5193 |
| TcCLB.504029.80 | serine/threonine protein phosphatase, putative | -0,2061 | 0,1286 |
| TcCLB.506879.109 | protein kinase, putative | -0,2061 | 0,3611 |
| TcCLB.510279.220 | mucin TcMUC (pseudogene), putative | -0,2061 | 0,7219 |
| TcCLB.510611.20 | UDP-glucuronic acid/UDP-N-acetylgalactosamine transpor | -0,2062 | 0,2595 |
| TcCLB.509229.140 | hypothetical protein, conserved | -0,2065 | 0,06153 |
| TcCLB.508881.80 | DNA ligase, putative | -0,2065 | 0,2871 |
| TcCLB.511115.40 | transcription elongation factor s-II, putative | -0,2066 | 0,08627 |

| | | | |
|---|---|---|---|
| TcCLB.504765.9 | hypothetical protein, conserved (fragment) | -0,2066 | 0,09774 |
| TcCLB.509179.60 | hypothetical protein, conserved | -0,2066 | 0,1908 |
| TcCLB.509925.70 | dispersed gene family protein 1 (DGF-1), putative | -0,2066 | 0,3231 |
| TcCLB.506611.10 | mucin-associated surface protein (MASP), putative | -0,2066 | 0,6892 |
| TcCLB.506319.5 | hypothetical protein, conserved (fragment) | -0,2069 | 0,5716 |
| TcCLB.507029.41 | hypothetical protein | -0,207 | 0,527 |
| TcCLB.508357.30 | hypothetical protein, conserved | -0,2071 | 0,1916 |
| TcCLB.510393.10 | hypothetical protein, conserved | -0,2071 | 0,2585 |
| TcCLB.508831.4 | dynein heavy chain, putative | -0,2072 | 0,09252 |
| TcCLB.511859.100 | trans-sialidase (pseudogene), putative | -0,2073 | 0,1929 |
| TcCLB.506363.100 | hypothetical protein, conserved | -0,2075 | 0,1033 |
| TcCLB.507519.70 | hypothetical protein, conserved | -0,2075 | 0,1289 |
| TcCLB.510451.10 | trans-sialidase (pseudogene), putative | -0,2075 | 0,7557 |
| TcCLB.507997.89 | hypothetical protein, conserved | -0,2076 | 0,4821 |
| TcCLB.510359.240 | hypothetical protein, conserved | -0,2078 | 0,1843 |
| TcCLB.503493.30 | hypothetical protein, conserved | -0,2079 | 0,2278 |
| TcCLB.506945.100 | hypothetical protein, conserved | -0,2081 | 0,06246 |
| TcCLB.506465.30 | hypothetical protein, conserved | -0,2082 | 0,07802 |
| TcCLB.506661.50 | hypothetical protein, conserved | -0,2084 | 0,06475 |
| TcCLB.510901.190 | hypothetical protein, conserved | -0,2086 | 0,1087 |
| TcCLB.511875.50 | syntaxin binding protein (pseudogene), putative | -0,2086 | 0,1747 |
| TcCLB.506957.110 | SNF-7-like protein, putative | -0,2087 | 0,1264 |
| TcCLB.506755.240 | hypothetical protein, conserved | -0,2087 | 0,4988 |
| TcCLB.511859.170 | NADH-cytochrome b5 reductase, putative | -0,2088 | 0,09897 |
| TcCLB.510807.20 | hypothetical protein | -0,2088 | 0,7254 |
| TcCLB.511409.10 | PAB1-binding protein , putative | -0,209 | 0,005413 |
| TcCLB.509027.60 | kinetoplastid kinetochore protein 10, putative (kkt10) | -0,209 | 0,2432 |
| TcCLB.506407.110 | hypothetical protein, conserved | -0,209 | 0,3524 |
| TcCLB.506999.20 | hypothetical protein, conserved | -0,2091 | 0,04713 |
| TcCLB.460747.20 | hypothetical protein, conserved | -0,2091 | 0,08624 |
| TcCLB.509455.20 | hypothetical protein, conserved | -0,2091 | 0,1101 |
| TcCLB.507265.30 | DNA polymerase delta catalytic subunit, putative | -0,2091 | 0,258 |
| TcCLB.506207.10 | hypothetical protein, conserved | -0,2091 | 0,2979 |
| TcCLB.510435.40 | GPI transamidase component Tta1, putative (TTA1) | -0,2092 | 0,2134 |
| TcCLB.504157.80 | hypothetical protein, conserved | -0,2093 | 0,06759 |
| TcCLB.510359.180 | hypothetical protein, conserved | -0,2093 | 0,1973 |
| TcCLB.507937.40 | elongation factor 1-gamma (EF-1-gamma), putative | -0,2093 | 0,7303 |
| TcCLB.509453.10 | hypothetical protein, conserved | -0,2094 | 0,1104 |
| TcCLB.507583.10 | hypothetical protein, conserved | -0,2094 | 0,5417 |
| TcCLB.509171.40 | hypothetical protein, conserved | -0,2095 | 0,05346 |
| TcCLB.508731.20 | hypothetical protein, conserved | -0,2096 | 0,1756 |
| TcCLB.509023.200 | cdc2-related kinase 12 (CRK12) | -0,2097 | 0,06472 |
| TcCLB.510911.30 | Ras-related protein Rab4, putative (RAB4) | -0,2097 | 0,09568 |
| TcCLB.507641.90 | hypothetical protein, conserved | -0,2097 | 0,2118 |
| TcCLB.508293.140 | mucin-associated surface protein (MASP), putative | -0,2098 | 0,7762 |

| | | | |
|---|---|---|---|
| TcCLB.511367.280 | hypothetical protein, conserved | -0,21 | 0,1155 |
| TcCLB.506147.20 | hypothetical protein | -0,21 | 0,3776 |
| TcCLB.510553.11 | hypothetical protein, conserved (pseudogene) | -0,21 | 0,6039 |
| TcCLB.509799.20 | hypothetical protein, conserved | -0,2101 | 0,1206 |
| TcCLB.420719.9 | Associated kinase of Tb14-3-3, putative (AKB1) | -0,2101 | 0,2246 |
| TcCLB.510995.9 | dispersed gene family protein 1 (DGF-1), putative | -0,2101 | 0,5113 |
| TcCLB.510121.80 | hypothetical protein, conserved | -0,2102 | 0,06303 |
| TcCLB.506629.150 | hypothetical protein, conserved | -0,2102 | 0,155 |
| TcCLB.509965.10 | hypothetical protein, conserved (pseudogene) | -0,2103 | 0,1985 |
| TcCLB.506495.3 | hypothetical protein, conserved | -0,2103 | 0,427 |
| TcCLB.506543.20 | hypothetical protein, conserved | -0,2104 | 0,03855 |
| TcCLB.508153.620 | hypothetical protein, conserved | -0,2104 | 0,203 |
| TcCLB.510599.14 | acetyl-CoA carboxylase (fragment) | -0,2105 | 0,02268 |
| TcCLB.508185.19 | ubiquitin hydrolase, putative (fragment) | -0,2105 | 0,6035 |
| TcCLB.508873.450 | mucin-associated surface protein (MASP), putative | -0,2105 | 0,665 |
| TcCLB.507257.180 | hypothetical protein | -0,2106 | 0,07307 |
| TcCLB.503407.20 | SUMO1/Ulp2, putative | -0,2106 | 0,1246 |
| TcCLB.510763.80 | DNA polymerase epsilon subunit B, putative | -0,2106 | 0,1748 |
| TcCLB.511483.30 | hypothetical protein, conserved | -0,2106 | 0,176 |
| TcCLB.511585.150 | Meckel syndrome type 6 protein, putative (mks6) | -0,2107 | 0,03459 |
| TcCLB.508325.300 | hypothetical protein, conserved | -0,2107 | 0,406 |
| TcCLB.510275.260 | trans-sialidase (pseudogene), putative | -0,2107 | 0,5753 |
| TcCLB.508015.10 | hypothetical protein | -0,2108 | 0,4051 |
| TcCLB.504055.10 | hypothetical protein, conserved | -0,2109 | 0,2596 |
| TcCLB.506529.130 | WD repeat and HMG-box DNA-binding protein, putative | -0,2111 | 0,03585 |
| TcCLB.507013.10 | dolichol kinase, putative | -0,2111 | 0,119 |
| TcCLB.505183.5 | hypothetical protein, conserved (fragment) | -0,2111 | 0,2277 |
| TcCLB.507001.50 | hypothetical protein, conserved | -0,2112 | 0,05711 |
| TcCLB.510287.20 | dolicholphosphate-mannose synthase, putative | -0,2113 | 0,2037 |
| TcCLB.510363.310 | trans-sialidase (pseudogene), putative | -0,2113 | 0,6075 |
| TcCLB.510555.21 | hypothetical protein | -0,2114 | 0,6486 |
| TcCLB.506867.20 | vacuolar type h ATPase subunit, putative | -0,2115 | 0,102 |
| TcCLB.481729.20 | hypothetical protein, conserved | -0,2115 | 0,5097 |
| TcCLB.506857.70 | hypothetical protein, conserved | -0,2117 | 0,09621 |
| TcCLB.509053.150 | hypothetical protein, conserved | -0,2117 | 0,3179 |
| TcCLB.507023.289 | hypothetical protein, conserved (fragment) | -0,2117 | 0,3634 |
| TcCLB.507681.120 | hypothetical protein, conserved (pseudogene) | -0,2118 | 0,2192 |
| TcCLB.506133.179 | hypothetical protein | -0,2118 | 0,5512 |
| TcCLB.510351.150 | phosphoprotein phosphatase, putative | -0,2119 | 0,2852 |
| TcCLB.469113.10 | trans-sialidase (pseudogene), putative | -0,2119 | 0,4091 |
| TcCLB.510269.50 | mucin-associated surface protein (MASP), putative | -0,2119 | 0,4596 |
| TcCLB.511911.110 | RNA helicase, putative | -0,2122 | 0,07895 |
| TcCLB.511517.150 | hypothetical protein, conserved | -0,2122 | 0,2447 |
| TcCLB.508817.170 | hypothetical protein, conserved | -0,2122 | 0,2527 |
| TcCLB.511315.44 | hypothetical protein, conserved (pseudogene) | -0,2123 | 0,4071 |

| | | | |
|---|---|---|---|
| TcCLB.506227.100 | hypothetical protein, conserved | -0,2124 | 0,4008 |
| TcCLB.511303.60 | translation initiation factor, putative | -0,2126 | 0,004815 |
| TcCLB.511277.280 | hypothetical protein, conserved | -0,2126 | 0,04575 |
| TcCLB.510305.30 | lipin, putative | -0,2126 | 0,04808 |
| TcCLB.504013.110 | serine/threonine-protein phosphatase 2A, catalytic subunit | -0,2127 | 0,2014 |
| TcCLB.511019.108 | hypothetical protein, conserved | -0,2128 | 0,2193 |
| TcCLB.506681.70 | serine-threonine dehydratase, putative | -0,2128 | 0,2897 |
| TcCLB.508543.120 | hypothetical protein, conserved | -0,2128 | 0,3198 |
| TcCLB.510725.60 | hypothetical protein, conserved | -0,2128 | 0,3926 |
| TcCLB.511529.240 | hypothetical protein, conserved | -0,2129 | 0,04609 |
| TcCLB.509429.250 | ubiquitin hydrolase, putative | -0,2129 | 0,04706 |
| TcCLB.508461.190 | hypothetical protein, conserved | -0,2129 | 0,04834 |
| TcCLB.510531.30 | hypothetical protein, conserved | -0,213 | 0,05844 |
| TcCLB.506407.40 | hypothetical protein, conserved | -0,213 | 0,07059 |
| TcCLB.508409.270 | RNA-binding protein, putative | -0,213 | 0,08369 |
| TcCLB.503697.50 | hypothetical protein, conserved | -0,2131 | 0,0329 |
| TcCLB.503823.30 | hypothetical protein, conserved | -0,2132 | 0,1671 |
| TcCLB.508827.120 | hypothetical protein, conserved | -0,2132 | 0,1742 |
| TcCLB.504443.10 | cysteine peptidase, Clan CA, family C19, putative | -0,2134 | 0,08254 |
| TcCLB.509641.60 | hypothetical protein, conserved | -0,2135 | 0,2043 |
| TcCLB.511589.150 | hypothetical protein, conserved | -0,2135 | 0,27 |
| TcCLB.506559.430 | TPR-repeat protein, putative | -0,2137 | 0,01764 |
| TcCLB.511277.460 | hypothetical protein, conserved | -0,2137 | 0,1324 |
| TcCLB.510777.20 | aspartyl-tRNA synthetase, putative | -0,2138 | 0,02451 |
| TcCLB.511365.70 | proteasome regulatory non-ATP-ase subunit 9, putative (R | -0,2138 | 0,05951 |
| TcCLB.507773.30 | RNA editing complex protein MP46, putative | -0,2138 | 0,171 |
| TcCLB.510655.150 | hypothetical protein, conserved | -0,2138 | 0,1938 |
| TcCLB.503757.40 | Casein kinase II subunit beta, putative | -0,2139 | 0,1535 |
| TcCLB.509859.90 | NAD-dependent deacetylase, putative (fragment) | -0,2139 | 0,5766 |
| TcCLB.503909.50 | hypothetical protein, conserved (YCF45) | -0,214 | 0,1056 |
| TcCLB.503823.110 | hypothetical protein, conserved | -0,214 | 0,2292 |
| TcCLB.507259.10 | tryparedoxin peroxidase, putative | -0,2141 | 0,01435 |
| TcCLB.508781.30 | hypothetical protein, conserved | -0,2141 | 0,09389 |
| TcCLB.511467.60 | hypothetical protein, conserved | -0,2141 | 0,1339 |
| TcCLB.507105.30 | hypothetical protein, conserved | -0,2142 | 0,08891 |
| TcCLB.509123.5 | trans-sialidase (pseudogene), putative | -0,2142 | 0,8175 |
| TcCLB.511511.10 | endosomal trafficking protein RME-8, putative | -0,2143 | 0,03183 |
| TcCLB.509353.40 | PTP1-interacting protein, 39 kDa, putative | -0,2143 | 0,07776 |
| TcCLB.503923.30 | hypothetical protein, conserved | -0,2143 | 0,1673 |
| TcCLB.511287.60 | hypothetical protein, conserved | -0,2144 | 0,01547 |
| TcCLB.507895.124 | hypothetical protein, conserved | -0,2146 | 0,008073 |
| TcCLB.509261.30 | 3,2-trans-enoyl-CoA isomerase, mitochondrial precursor, p | -0,2146 | 0,02573 |
| TcCLB.507047.100 | hypothetical protein, conserved | -0,2146 | 0,08133 |
| TcCLB.511625.120 | hypothetical protein | -0,2147 | 0,6839 |
| TcCLB.511191.10 | tyrosyl-DNA Phosphodiesterase (Tdp1), putative | -0,2148 | 0,2134 |

| | | | |
|---|---|---|---|
| TcCLB.506265.209 | hypothetical protein, conserved (fragment) | -0,2148 | 0,6138 |
| TcCLB.503781.50 | hypothetical protein, conserved | -0,2149 | 0,1164 |
| TcCLB.508699.110 | elongation factor G2-like protein, putative | -0,2149 | 0,3044 |
| TcCLB.508181.4 | AGC/RSK family serine/threonine kinase, putative (fragme | -0,2149 | 0,5656 |
| TcCLB.463297.10 | hypothetical protein (fragment) | -0,2149 | 0,571 |
| TcCLB.510439.61 | heat shock 70 kDa protein, putative (fragment) | -0,2152 | 0,01894 |
| TcCLB.506937.40 | dynein heavy chain, putative | -0,2153 | 0,04906 |
| TcCLB.511925.50 | hypothetical protein, conserved | -0,2154 | 0,141 |
| TcCLB.509063.20 | hypothetical protein, conserved | -0,2154 | 0,1818 |
| TcCLB.506855.260 | thymidine kinase, putative | -0,2154 | 0,2271 |
| TcCLB.510021.185 | hypothetical protein | -0,2155 | 0,6962 |
| TcCLB.511727.230 | Repressor of differentiation kinase 1, putative (RDK1.2) | -0,2156 | 0,03304 |
| TcCLB.508521.90 | trans-sialidase (pseudogene), putative | -0,2156 | 0,3305 |
| TcCLB.509547.40 | mucin-associated surface protein (MASP), putative | -0,2156 | 0,6494 |
| TcCLB.508641.260 | DNA-directed RNA polymerase, mitochondrial (MTRNAP) | -0,2157 | 0,01125 |
| TcCLB.511801.30 | hypothetical protein, conserved | -0,2157 | 0,1412 |
| TcCLB.510767.40 | hypothetical protein, conserved | -0,2157 | 0,1877 |
| TcCLB.507699.230 | trans-sialidase, putative | -0,2157 | 0,4668 |
| TcCLB.506147.70 | hypothetical protein, conserved | -0,2158 | 0,1335 |
| TcCLB.508625.70 | mRNA capping methyltransferase, putative | -0,2158 | 0,2743 |
| TcCLB.510285.20 | hypothetical protein, conserved | -0,2161 | 0,03971 |
| TcCLB.509647.90 | chloride channel protein, putative | -0,2161 | 0,06301 |
| TcCLB.506533.70 | hypothetical protein, conserved | -0,2161 | 0,1431 |
| TcCLB.504179.20 | hypothetical protein, conserved | -0,2161 | 0,2403 |
| TcCLB.508827.100 | splicing factor 3B subunit 1, putative | -0,2163 | 0,0109 |
| TcCLB.510491.30 | hypothetical protein, conserved (pseudogene) | -0,2164 | 0,2035 |
| TcCLB.508859.80 | hypothetical protein, conserved | -0,2164 | 0,2065 |
| TcCLB.503525.50 | RNA helicase, putative | -0,2164 | 0,2381 |
| TcCLB.511643.40 | trans-sialidase, Group V, putative | -0,2164 | 0,3597 |
| TcCLB.464879.9 | ribonuclease L inhibitor, putative (fragment) | -0,2166 | 0,222 |
| TcCLB.510131.90 | hypothetical protein, conserved | -0,2167 | 0,07877 |
| TcCLB.503877.10 | hypothetical protein, conserved | -0,2167 | 0,232 |
| TcCLB.504625.60 | hydrogenase, putative | -0,2168 | 0,129 |
| TcCLB.508613.50 | hypothetical protein, conserved | -0,2169 | 0,02495 |
| TcCLB.510663.80 | hypothetical protein, conserved | -0,2169 | 0,1601 |
| TcCLB.508817.70 | DNA repair protein RAD50, putative | -0,217 | 0,003042 |
| TcCLB.509617.50 | hypothetical protein, conserved | -0,2171 | 0,1192 |
| TcCLB.511499.40 | hypothetical protein, conserved | -0,2171 | 0,2945 |
| TcCLB.508899.70 | hypothetical protein, conserved | -0,2172 | 0,1709 |
| TcCLB.507771.40 | phosphoglycerate mutase, putative (iPGAM) | -0,2173 | 0,2026 |
| TcCLB.508153.200 | kinesin, putative | -0,2174 | 0,09389 |
| TcCLB.506505.50 | hypothetical protein, conserved | -0,2175 | 0,04847 |
| TcCLB.509033.80 | hypothetical protein, conserved | -0,2175 | 0,06011 |
| TcCLB.506661.120 | hypothetical protein | -0,2175 | 0,1813 |
| TcCLB.505155.4 | trans-sialidase, putative (fragment) | -0,2175 | 0,6908 |

| | | | |
|---|---|---|---|
| TcCLB.510901.220 | hypothetical protein, conserved | -0,2176 | 0,08882 |
| TcCLB.508637.79 | DNA polymerase eta, putative | -0,2176 | 0,1287 |
| TcCLB.506459.10 | mucin-associated surface protein (MASP), putative | -0,2176 | 0,478 |
| TcCLB.508547.100 | hypothetical protein, conserved | -0,2177 | 0,1234 |
| TcCLB.511771.156 | retrotransposon hot spot protein (RHS, pseudogene), putat | -0,2177 | 0,1989 |
| TcCLB.511365.100 | hypothetical protein, conserved | -0,2177 | 0,2585 |
| TcCLB.506603.46 | hypothetical protein, conserved (pseudogene) | -0,2177 | 0,6936 |
| TcCLB.506399.40 | hypothetical protein, conserved | -0,2178 | 0,4112 |
| TcCLB.503667.31 | retrotransposon hot spot protein (RHS, pseudogene), putat | -0,2179 | 0,54 |
| TcCLB.511499.10 | hypothetical protein, conserved | -0,218 | 0,06209 |
| TcCLB.507641.160 | hypothetical protein, conserved | -0,2181 | 0,008112 |
| TcCLB.510901.210 | hypothetical protein, conserved | -0,2182 | 0,05187 |
| TcCLB.506289.74 | chaperone DNAJ protein, putative | -0,2182 | 0,1489 |
| TcCLB.506315.100 | protein phosphatase 2C, putative | -0,2183 | 0,06832 |
| TcCLB.506469.20 | hypothetical protein | -0,2183 | 0,09638 |
| TcCLB.504131.160 | hypothetical protein | -0,2184 | 0,152 |
| TcCLB.508257.50 | actin-like protein, putative | -0,2184 | 0,245 |
| TcCLB.504037.20 | hypothetical protein, conserved | -0,2186 | 0,07063 |
| TcCLB.506891.20 | wd40 repeat domain-containing protein | -0,2186 | 0,1643 |
| TcCLB.510877.10 | antigenic protein, putative | -0,2187 | 0,1405 |
| TcCLB.511713.40 | ribosome biogenesis protein TSR3, putative (tsr3) | -0,2187 | 0,2039 |
| TcCLB.507061.30 | iron superoxide dismutase, putative | -0,2187 | 0,2072 |
| TcCLB.507835.70 | dispersed gene family protein 1 (DGF-1), putative | -0,2187 | 0,2989 |
| TcCLB.507093.240 | hypothetical protein, conserved | -0,2188 | 0,02579 |
| TcCLB.445553.10 | hypothetical protein, conserved | -0,2188 | 0,1322 |
| TcCLB.506701.20 | Tob55, putative | -0,219 | 0,139 |
| TcCLB.510737.90 | hypothetical protein, conserved | -0,2191 | 0,2738 |
| TcCLB.511153.50 | haloacid dehalogenase hydrolase, putative | -0,2192 | 0,1999 |
| TcCLB.503983.20 | dihydroxyacetone kinase 1-like, putative | -0,2193 | 0,1004 |
| TcCLB.510291.30 | protein phosphatase 2C-like, putative | -0,2194 | 0,1869 |
| TcCLB.506943.114 | U6 snRNA-associated Sm-like protein LSm5p | -0,2194 | 0,3174 |
| TcCLB.511089.80 | trans-sialidase, Group V, putative | -0,2194 | 0,5938 |
| TcCLB.511127.369 | hypothetical protein, conserved | -0,2195 | 0,1355 |
| TcCLB.503925.50 | hypothetical protein, conserved | -0,2195 | 0,1951 |
| TcCLB.506821.110 | hypothetical protein, conserved | -0,2195 | 0,2285 |
| TcCLB.439301.9 | hypothetical protein, conserved | -0,2197 | 0,06707 |
| TcCLB.509105.60 | adenosine deaminase-like protein, putative (ADAT1h) | -0,2198 | 0,1502 |
| TcCLB.511279.30 | ARF-like 2-binding protein, putative | -0,2198 | 0,2816 |
| TcCLB.511469.24 | ADP-ribosylation factor-like 2, arl2, putative (ARL2) | -0,2198 | 0,4441 |
| TcCLB.507677.120 | hypothetical protein, conserved | -0,22 | 0,07214 |
| TcCLB.511855.70 | protein kinase, putative | -0,22 | 0,07922 |
| TcCLB.506801.100 | hypothetical protein, conserved | -0,2201 | 0,143 |
| TcCLB.511071.181 | hypothetical protein, conserved | -0,2202 | 0,08363 |
| TcCLB.510743.80 | hypothetical protein, conserved | -0,2202 | 0,1315 |
| TcCLB.510729.130 | DNA replication licensing factor MCM9, putative | -0,2203 | 0,06747 |

| | | | |
|---|---|---|---|
| TcCLB.510167.29 | exosome complex exonuclease, putative | -0,2203 | 0,1933 |
| TcCLB.506699.3 | hypothetical protein, conserved | -0,2203 | 0,2434 |
| TcCLB.510691.30 | hypothetical protein, conserved | -0,2206 | 0,09617 |
| TcCLB.508409.140 | hypothetical protein, conserved | -0,2207 | 0,1524 |
| TcCLB.511281.40 | mevalonate-diphosphate decarboxylase, putative | -0,2208 | 0,07008 |
| TcCLB.508209.70 | hypothetical protein, conserved | -0,2208 | 0,1443 |
| TcCLB.507167.41 | ATP-dependent DEAD/H RNA helicase (pseudogene), putat | -0,2209 | 0,6974 |
| TcCLB.504827.60 | hypothetical protein, conserved | -0,221 | 0,07438 |
| TcCLB.506631.20 | cardiolipin synthetase, putative (cls) | -0,2211 | 0,07935 |
| TcCLB.508169.80 | hypothetical protein, conserved | -0,2212 | 0,2325 |
| TcCLB.506963.100 | hypothetical protein, conserved | -0,2213 | 0,2818 |
| TcCLB.508261.25 | trans-sialidase (pseudogene), putative | -0,2214 | 0,06079 |
| TcCLB.506517.130 | hypothetical protein, conserved | -0,2214 | 0,08282 |
| TcCLB.506885.200 | hypothetical protein, conserved | -0,2215 | 0,007361 |
| TcCLB.508899.20 | hypothetical protein, conserved | -0,2215 | 0,02582 |
| TcCLB.507711.349 | hypothetical protein, conserved | -0,2215 | 0,3844 |
| TcCLB.503697.90 | Translation factor GUF1, mitochondrial, putative (GUF1) | -0,2216 | 0,03812 |
| TcCLB.506255.10 | hypothetical protein, conserved (pseudogene) | -0,2216 | 0,3844 |
| TcCLB.503809.20 | peroxin 12, putative (PEX12) | -0,2217 | 0,1299 |
| TcCLB.511717.120 | hypothetical protein, conserved | -0,2218 | 0,1113 |
| TcCLB.510181.120 | hypothetical protein, conserved | -0,2218 | 0,1276 |
| TcCLB.507769.10 | hypothetical protein, conserved | -0,2218 | 0,5243 |
| TcCLB.507073.10 | hypothetical protein, conserved | -0,2219 | 0,07259 |
| TcCLB.511319.10 | hypothetical protein, conserved | -0,222 | 0,03129 |
| TcCLB.510525.10 | protein kinase, putative | -0,2221 | 0,1628 |
| TcCLB.508027.40 | folate/pteridine transporter, putative | -0,2222 | 0,02239 |
| TcCLB.511589.90 | hypothetical protein, conserved | -0,2222 | 0,09869 |
| TcCLB.510407.100 | hypothetical protein, conserved | -0,2222 | 0,1187 |
| TcCLB.511903.290 | hypothetical protein, conserved | -0,2224 | 0,01696 |
| TcCLB.510463.259 | mucin-associated surface protein (MASP), putative (fragm | -0,2224 | 0,6112 |
| TcCLB.509965.340 | hypothetical protein, conserved | -0,2225 | 0,06745 |
| TcCLB.504115.5 | trans-sialidase (pseudogene), putative | -0,2226 | 0,2344 |
| TcCLB.507023.220 | hypothetical protein, conserved | -0,2227 | 0,01749 |
| TcCLB.504179.30 | hypothetical protein, conserved | -0,2228 | 0,1937 |
| TcCLB.510639.10 | dispersed gene family protein 1 (DGF-1, pseudogene), puta | -0,2228 | 0,3441 |
| TcCLB.511627.80 | hypothetical protein, conserved | -0,2229 | 0,3306 |
| TcCLB.504093.4 | hypothetical protein, conserved | -0,2229 | 0,3348 |
| TcCLB.503431.20 | hypothetical protein, conserved | -0,223 | 0,216 |
| TcCLB.506789.270 | hypothetical protein, conserved | -0,2231 | 0,04368 |
| TcCLB.509395.70 | hypothetical protein, conserved | -0,2232 | 0,06981 |
| TcCLB.510731.50 | small nuclear RNA gene activation protein (SNAP) 50, puta | -0,2233 | 0,05494 |
| TcCLB.504057.45 | hypothetical protein, conserved | -0,2233 | 0,0849 |
| TcCLB.504147.130 | hypothetical protein, conserved | -0,2234 | 0,09521 |
| TcCLB.508027.94 | hypothetical protein, conserved | -0,2234 | 0,156 |
| TcCLB.509317.80 | succinate dehydrogenase subunit | -0,2236 | 0,09393 |

| | | | |
|---|---|---|---|
| TcCLB.508461.143 | hypothetical protein, conserved | -0,2238 | 0,3176 |
| TcCLB.511873.10 | trans-sialidase, Group V, putative | -0,2238 | 0,3586 |
| TcCLB.508787.10 | hypothetical protein, conserved | -0,2239 | 0,1692 |
| TcCLB.508511.10 | mucin-associated surface protein (MASP, pseudogene), pu | -0,2239 | 0,6938 |
| TcCLB.503559.80 | hypothetical protein | -0,224 | 0,09162 |
| TcCLB.507519.160 | hypothetical protein, conserved | -0,2241 | 0,1892 |
| TcCLB.511731.24 | hypothetical protein | -0,2242 | 0,08998 |
| TcCLB.508369.50 | acetyl-CoA carboxylase (fragment) | -0,2243 | 0,01403 |
| TcCLB.511283.30 | hypothetical protein, conserved | -0,2243 | 0,05115 |
| TcCLB.504077.40 | translation elongation factor EF-2, putative | -0,2243 | 0,06918 |
| TcCLB.510301.40 | hypothetical protein, conserved | -0,2243 | 0,1183 |
| TcCLB.509099.150 | cdc2-related kinase 9 (CRK9) | -0,2245 | 0,05727 |
| TcCLB.506945.170 | thiosulfate sulfurtransferase, mitochondrial, putative | -0,2246 | 0,176 |
| TcCLB.506467.14 | hypothetical protein, conserved | -0,2246 | 0,2351 |
| TcCLB.510387.60 | trans-sialidase (pseudogene), putative | -0,2247 | 0,3814 |
| TcCLB.511283.109 | AP-4 complex subunit sigma-1, putative | -0,2249 | 0,2197 |
| TcCLB.507959.300 | mucin-associated surface protein (MASP), putative | -0,2249 | 0,4992 |
| TcCLB.511883.30 | dispersed gene family protein 1 (DGF-1, pseudogene), puta | -0,225 | 0,2746 |
| TcCLB.504057.170 | protein kinase, putative | -0,2251 | 0,0779 |
| TcCLB.508827.20 | hypothetical protein, conserved | -0,2251 | 0,1565 |
| TcCLB.511257.90 | mucin-associated surface protein (MASP), putative | -0,2251 | 0,4781 |
| TcCLB.511475.50 | dispersed gene family protein 1 (DGF-1), putative | -0,2254 | 0,3128 |
| TcCLB.509525.60 | mucin TcMUCII, putative | -0,2254 | 0,76 |
| TcCLB.507793.10 | hypothetical protein, conserved | -0,2256 | 0,01598 |
| TcCLB.509247.10 | hypothetical protein, conserved | -0,2256 | 0,1444 |
| TcCLB.511183.260 | mucin-associated surface protein (MASP), putative | -0,2256 | 0,5816 |
| TcCLB.509017.40 | hypothetical protein, conserved | -0,2257 | 0,08746 |
| TcCLB.506909.100 | trans-sialidase, Group V, putative | -0,2257 | 0,5064 |
| TcCLB.510243.70 | hypothetical protein, conserved | -0,2259 | 0,03019 |
| TcCLB.504129.10 | hypothetical protein, conserved | -0,2259 | 0,4364 |
| TcCLB.511901.20 | hypothetical protein, conserved | -0,226 | 0,03212 |
| TcCLB.503869.50 | exosome-associated protein 1, putative (EAP1) | -0,226 | 0,07309 |
| TcCLB.503575.54 | hypothetical protein, conserved | -0,226 | 0,1405 |
| TcCLB.511725.100 | phosphatidylinositol-4-phosphate 5-kinase, putative | -0,2261 | 0,04178 |
| TcCLB.506531.20 | hypothetical protein, conserved | -0,2261 | 0,06244 |
| TcCLB.509471.68 | histone H3, putative | -0,2263 | 0,02327 |
| TcCLB.509287.250 | retrotransposon hot spot protein (RHS, pseudogene), putat | -0,2263 | 0,1277 |
| TcCLB.508881.60 | hypothetical protein, conserved | -0,2263 | 0,1402 |
| TcCLB.509399.140 | hypothetical protein, conserved | -0,2263 | 0,1475 |
| TcCLB.507599.10 | Ribosome biogenesis protein Tsr3, putative (fragment) | -0,2263 | 0,4735 |
| TcCLB.504081.170 | mucin-associated surface protein (MASP), putative | -0,2263 | 0,6202 |
| TcCLB.509693.40 | hypothetical protein, conserved | -0,2264 | 0,2373 |
| TcCLB.510487.20 | dispersed gene family protein 1 (DGF-1, pseudogene), puta | -0,2265 | 0,4138 |
| TcCLB.506787.9 | dispersed gene family protein 1 (DGF-1), putative (fragme | -0,2265 | 0,6707 |
| TcCLB.510091.14 | hypothetical protein, conserved | -0,2266 | 0,3424 |

| | | | |
|---|---|---|---|
| TcCLB.506811.170 | hypothetical protein, conserved | -0,2269 | 0,03114 |
| TcCLB.510719.310 | hypothetical protein, conserved | -0,2269 | 0,0396 |
| TcCLB.510007.70 | ATP-dependent zinc metallopeptidase, putative | -0,2269 | 0,08456 |
| TcCLB.510265.14 | hypothetical protein, conserved | -0,2269 | 0,1452 |
| TcCLB.508273.60 | hypothetical protein, conserved | -0,227 | 0,04172 |
| TcCLB.509965.360 | hypothetical protein, conserved | -0,227 | 0,2205 |
| TcCLB.506147.170 | hypothetical protein, conserved | -0,2271 | 0,02345 |
| TcCLB.511421.80 | Midasin, putative (pseudogene) | -0,2272 | 0,02104 |
| TcCLB.507625.190 | hypothetical protein, conserved | -0,2273 | 0,1017 |
| TcCLB.510945.40 | hypothetical protein, conserved | -0,2274 | 0,04784 |
| TcCLB.504741.30 | protein phosphatase 2C, putative | -0,2274 | 0,1552 |
| TcCLB.504071.100 | hypothetical protein, conserved | -0,2275 | 0,02396 |
| TcCLB.504035.60 | hypothetical protein, conserved | -0,2275 | 0,2183 |
| TcCLB.507939.30 | hypothetical protein, conserved | -0,2276 | 0,1316 |
| TcCLB.511423.50 | conserved protein, unknown function | -0,2277 | 0,05821 |
| TcCLB.511277.520 | protein kinase (pseudogene), putative | -0,2277 | 0,2168 |
| TcCLB.506635.44 | hypothetical protein, conserved | -0,2277 | 0,2299 |
| TcCLB.511283.10 | nuclear segregation protein, putative | -0,2278 | 0,01005 |
| TcCLB.506857.60 | peptidyl-prolyl cis-trans isomerase, putative (PAR45) | -0,2278 | 0,05329 |
| TcCLB.506953.40 | hypothetical protein, conserved | -0,2278 | 0,08292 |
| TcCLB.504085.10 | hypothetical protein, conserved | -0,2278 | 0,09911 |
| TcCLB.507039.41 | hypothetical protein, conserved | -0,2278 | 0,297 |
| TcCLB.508871.150 | mucin-associated surface protein (MASP), putative | -0,2278 | 0,4397 |
| TcCLB.508965.50 | hypothetical protein, conserved | -0,2279 | 0,04826 |
| TcCLB.509643.30 | hypothetical protein, conserved | -0,2279 | 0,0578 |
| TcCLB.510029.50 | hypothetical protein, conserved | -0,2279 | 0,1035 |
| TcCLB.507829.10 | cysteine peptidase, Clan CA, family C2, putative | -0,2279 | 0,1598 |
| TcCLB.508707.20 | As/Sb Reductase, putative (ACR2) | -0,2279 | 0,2504 |
| TcCLB.511529.250 | hypothetical protein | -0,228 | 0,09059 |
| TcCLB.510459.30 | dispersed gene family protein 1 (DGF-1), putative | -0,2282 | 0,5738 |
| TcCLB.503831.40 | mitochondrial DNA primase, putative | -0,2283 | 0,0596 |
| TcCLB.511715.50 | hypothetical protein, conserved (pseudogene) | -0,2285 | 0,1645 |
| TcCLB.511311.60 | hypothetical protein, conserved | -0,2285 | 0,2358 |
| TcCLB.506265.60 | hypothetical protein, conserved | -0,2286 | 0,01612 |
| TcCLB.508153.40 | hypothetical protein, conserved | -0,2286 | 0,2112 |
| TcCLB.511919.30 | hypothetical protein, conserved (pseudogene) | -0,2286 | 0,4863 |
| TcCLB.503755.54 | hypothetical protein, conserved | -0,2287 | 0,08581 |
| TcCLB.511247.27 | RNA editing 3' terminal uridylyl transferase 1, putative | -0,2287 | 0,1646 |
| TcCLB.508461.420 | 30S Ribosomal protein S17, putative | -0,2288 | 0,07457 |
| TcCLB.506559.170 | 8-oxoguanine DNA glycosylase, putative | -0,2289 | 0,033 |
| TcCLB.509023.100 | catalytic subunit of the vacuolar transporter chaperone 4, p | -0,2292 | 0,06264 |
| TcCLB.509167.180 | kinesin, putative | -0,2294 | 0,03967 |
| TcCLB.509871.31 | ATP-dependent DEAD/H RNA helicase (pseudogene), putat | -0,2294 | 0,5637 |
| TcCLB.507801.150 | hypothetical protein, conserved | -0,2295 | 0,2781 |
| TcCLB.506941.9 | endosomal trafficking protein RME-8, putative | -0,2297 | 0,06748 |

| | | | |
|---|---|---|---|
| TcCLB.506193.10 | hypothetical protein, conserved | -0,2298 | 0,01833 |
| TcCLB.506755.200 | hypothetical protein, conserved | -0,2298 | 0,2085 |
| TcCLB.508409.30 | DNA-directed RNA polymerase II subunit 3, putative (RPB3 | -0,2299 | 0,03631 |
| TcCLB.508899.120 | hypothetical protein, conserved | -0,2299 | 0,3948 |
| TcCLB.506857.90 | proteasome regulatory ATPase subunit 1, putative | -0,23 | 0,0222 |
| TcCLB.507897.4 | inositol polyphosphate 1-phosphatase, putative | -0,23 | 0,08761 |
| TcCLB.508439.10 | hypothetical protein, conserved | -0,23 | 0,3567 |
| TcCLB.474937.9 | acetyl-CoA carboxylase, putative | -0,2304 | 0,04212 |
| TcCLB.510087.10 | hypothetical protein, conserved | -0,2304 | 0,1676 |
| TcCLB.511767.60 | trans-sialidase (pseudogene), putative | -0,2305 | 0,09286 |
| TcCLB.508799.20 | chromatin binding protein, putative | -0,2306 | 0,114 |
| TcCLB.507023.90 | hypothetical protein, conserved | -0,2307 | 0,06641 |
| TcCLB.510357.130 | hypothetical protein, conserved | -0,2307 | 0,1727 |
| TcCLB.509267.100 | methionyl-tRNA formyltransferase, putative | -0,2308 | 0,1275 |
| TcCLB.511503.9 | hypothetical protein, conserved (pseudogene) | -0,2308 | 0,1766 |
| TcCLB.511575.90 | hypothetical protein | -0,2308 | 0,2164 |
| TcCLB.511151.70 | hypothetical protein, conserved | -0,2309 | 0,03291 |
| TcCLB.503525.20 | hypothetical protein, conserved | -0,2309 | 0,03358 |
| TcCLB.509063.30 | chaperonin HSP60/CNP60, putative | -0,2309 | 0,07686 |
| TcCLB.510241.120 | Peptide deformylase 2, putative (PDF2) | -0,2309 | 0,0916 |
| TcCLB.504027.4 | kinesin, putative (fragment) | -0,2309 | 0,2095 |
| TcCLB.507031.40 | hypothetical protein, conserved | -0,231 | 0,07487 |
| TcCLB.506135.80 | hypothetical protein, conserved | -0,231 | 0,2498 |
| TcCLB.507603.30 | PITH domain-containing protein | -0,231 | 0,2551 |
| TcCLB.505983.20 | nucleosome assembly protein-like protein, putative | -0,2311 | 0,004237 |
| TcCLB.506503.10 | hypothetical protein, conserved | -0,2311 | 0,1398 |
| TcCLB.509875.25 | trans-sialidase (pseudogene), putative | -0,2311 | 0,6412 |
| TcCLB.510253.20 | hypothetical protein, conserved | -0,2312 | 0,01725 |
| TcCLB.509999.80 | hypothetical protein, conserved | -0,2312 | 0,03951 |
| TcCLB.508323.30 | hypothetical protein, conserved | -0,2312 | 0,1136 |
| TcCLB.508411.90 | tfiih basal transcription factor complex helicase xpb subuni | -0,2313 | 0,0134 |
| TcCLB.506709.30 | hypothetical protein, conserved | -0,2313 | 0,08342 |
| TcCLB.504427.180 | hypothetical protein | -0,2313 | 0,1797 |
| TcCLB.508815.170 | protein kinase, putative | -0,2314 | 0,003004 |
| TcCLB.511277.70 | Elongation factor Tu, mitochondrial, putative (EF-Tu) | -0,2314 | 0,01175 |
| TcCLB.507857.10 | transcription elongation factor, putative | -0,2314 | 0,1091 |
| TcCLB.504253.60 | hypothetical protein, conserved (fragment) | -0,2314 | 0,511 |
| TcCLB.503995.20 | hypothetical protein, conserved | -0,2315 | 0,1344 |
| TcCLB.504147.40 | Mitochondrial N(5)-glutamine methyltransferase MTQ1, pu | -0,2316 | 0,02913 |
| TcCLB.504075.6 | dual specificity protein phosphatase, putative | -0,2316 | 0,1087 |
| TcCLB.507721.10 | hypothetical protein, conserved | -0,2316 | 0,1766 |
| TcCLB.507823.20 | hypothetical protein | -0,2317 | 0,08521 |
| TcCLB.507297.20 | cysteinyl-tRNA synthetase, putative | -0,2318 | 0,007053 |
| TcCLB.507011.200 | dihydroxyacetone kinase 1-like, putative | -0,2318 | 0,09148 |
| TcCLB.510377.70 | mucin TcMUCII, putative | -0,2318 | 0,8399 |

| | | | |
|---|---|---|---|
| TcCLB.511301.140 | potassium voltage-gated channel | -0,2319 | 0,1116 |
| TcCLB.506173.140 | dispersed gene family protein 1 (DGF-1), putative | -0,2319 | 0,2893 |
| TcCLB.503829.70 | hypothetical protein, conserved | -0,232 | 0,06088 |
| TcCLB.503809.130 | hypothetical protein, conserved | -0,232 | 0,1158 |
| TcCLB.507035.30 | dispersed gene family protein 1 (DGF-1), putative | -0,2321 | 0,155 |
| TcCLB.506435.40 | hypothetical protein, conserved | -0,2321 | 0,272 |
| TcCLB.504137.70 | hypothetical protein, conserved | -0,2322 | 0,1123 |
| TcCLB.506999.130 | hypothetical protein, conserved | -0,2323 | 0,02151 |
| TcCLB.508779.80 | hypothetical protein, conserved | -0,2323 | 0,07078 |
| TcCLB.509917.6 | ATP-dependent DEAD/H RNA helicase (pseudogene), putat | -0,2323 | 0,3503 |
| TcCLB.507167.140 | trans-sialidase (pseudogene), putative | -0,2323 | 0,527 |
| TcCLB.503805.10 | hypothetical protein, conserved | -0,2324 | 0,1535 |
| TcCLB.510491.20 | dynein light chain, putative | -0,2324 | 0,2398 |
| TcCLB.508709.10 | hypothetical protein, conserved | -0,2325 | 0,2458 |
| TcCLB.504129.50 | hypothetical protein, conserved (pseudogene) | -0,2325 | 0,3617 |
| TcCLB.507485.60 | hypothetical protein, conserved | -0,2326 | 0,06042 |
| TcCLB.510609.80 | hypothetical protein, conserved | -0,2328 | 0,2544 |
| TcCLB.506895.10 | hypothetical protein, conserved | -0,233 | 0,1695 |
| TcCLB.507711.230 | hypothetical protein, conserved | -0,2332 | 0,05114 |
| TcCLB.504827.140 | hypothetical protein, conserved | -0,2332 | 0,09247 |
| TcCLB.508641.290 | hypothetical protein, conserved | -0,2332 | 0,105 |
| TcCLB.511725.174 | small nuclear ribonucleoprotein Sm-G, putative | -0,2332 | 0,111 |
| TcCLB.510119.60 | hypothetical protein, conserved | -0,2332 | 0,211 |
| TcCLB.506797.70 | lysophospholipase, putative | -0,2333 | 0,04216 |
| TcCLB.506207.30 | hypothetical protein, conserved | -0,2333 | 0,1775 |
| TcCLB.508533.50 | hypothetical protein, conserved | -0,2334 | 0,003115 |
| TcCLB.511217.150 | hypothetical protein, conserved | -0,2334 | 0,1857 |
| TcCLB.506363.60 | f-actin capping protein beta subunit, putative | -0,2335 | 0,1575 |
| TcCLB.506265.110 | hypothetical protein, conserved | -0,2336 | 0,1516 |
| TcCLB.511559.34 | hypothetical protein, conserved | -0,2337 | 0,2931 |
| TcCLB.511183.565 | surface protease GP63 (pseudogene), putative | -0,2337 | 0,7081 |
| TcCLB.508579.40 | hypothetical protein, conserved | -0,2339 | 0,19 |
| TcCLB.506479.67 | hypothetical protein, conserved | -0,234 | 0,06291 |
| TcCLB.506625.240 | hypothetical protein, conserved | -0,2342 | 0,01503 |
| TcCLB.503805.9 | Quinonprotein alcohol dehydrogenase-like protein, putative | -0,2342 | 0,09597 |
| TcCLB.506847.4 | hypothetical protein, conserved | -0,2342 | 0,3477 |
| TcCLB.508741.300 | proteasome regulatory non-ATPase subunit 6, putative | -0,2343 | 0,01573 |
| TcCLB.485889.16 | hypothetical protein (fragment) | -0,2343 | 0,3138 |
| TcCLB.506559.460 | hypothetical protein | -0,2344 | 0,09049 |
| TcCLB.506835.10 | hypothetical protein, conserved | -0,2345 | 0,01166 |
| TcCLB.505183.10 | hypothetical protein, conserved | -0,2345 | 0,0944 |
| TcCLB.511757.20 | hypothetical protein, conserved | -0,2345 | 0,2416 |
| TcCLB.509661.10 | retrotransposon hot spot protein (RHS, pseudogene), putat | -0,2347 | 0,1361 |
| TcCLB.509363.31 | hypothetical protein | -0,2347 | 0,6196 |
| TcCLB.511517.160 | hypothetical protein, conserved | -0,2348 | 0,06254 |

| | | | |
|---|---|---|---|
| TcCLB.508323.94 | cyclophilin, putative | -0,2348 | 0,2904 |
| TcCLB.509049.20 | hypothetical protein, conserved | -0,2349 | 0,1028 |
| TcCLB.508831.90 | hypothetical protein, conserved | -0,2349 | 0,2672 |
| TcCLB.510601.40 | tubulin tyrosine ligase (pseudogene), putative | -0,2351 | 0,1814 |
| TcCLB.507873.30 | Double RNA binding domain protein 7 (DRBD7) | -0,2352 | 0,02507 |
| TcCLB.511127.150 | hypothetical protein, conserved | -0,2352 | 0,117 |
| TcCLB.510747.120 | hypothetical protein, conserved | -0,2353 | 0,07172 |
| TcCLB.504151.50 | hypothetical protein, conserved | -0,2353 | 0,2741 |
| TcCLB.506489.40 | hypothetical protein, conserved | -0,2355 | 0,08185 |
| TcCLB.511559.30 | ras-related GTP-binding protein, putative | -0,2355 | 0,1475 |
| TcCLB.511419.80 | phosphatidylinositol-4-phosphate 5-kinase related | -0,2356 | 0,1863 |
| TcCLB.509965.200 | hypothetical protein, conserved | -0,2358 | 0,175 |
| TcCLB.511283.90 | N-myristoyl transferase, putative | -0,2359 | 0,07138 |
| TcCLB.509875.220 | hypothetical protein, conserved | -0,236 | 0,06226 |
| TcCLB.511737.60 | hypothetical protein, conserved | -0,236 | 0,1001 |
| TcCLB.508211.50 | hypothetical protein, conserved | -0,236 | 0,1518 |
| TcCLB.506835.131 | hypothetical protein, conserved | -0,236 | 0,3534 |
| TcCLB.507709.140 | hypothetical protein, conserved | -0,2361 | 0,1135 |
| TcCLB.503457.10 | hypothetical protein | -0,2361 | 0,2032 |
| TcCLB.508829.9 | dynein heavy chain, putative (fragment) | -0,2362 | 0,07033 |
| TcCLB.506945.300 | V-type ATPase, C subunit, putative | -0,2363 | 0,04718 |
| TcCLB.503909.60 | hypothetical protein, conserved | -0,2364 | 0,1168 |
| TcCLB.506649.80 | RNA-binding protein, putative (DRBD3) | -0,2367 | 0,002622 |
| TcCLB.504203.60 | TPR-repeat-containing chaperone protein DNAJ, putative (1 | -0,2367 | 0,06974 |
| TcCLB.509429.230 | hypothetical protein, conserved | -0,2367 | 0,2072 |
| TcCLB.507541.50 | casein kinase, putative (fragment) | -0,237 | 0,02422 |
| TcCLB.507099.100 | hypothetical protein, conserved | -0,237 | 0,1427 |
| TcCLB.504163.80 | hypothetical protein, conserved | -0,2371 | 0,07704 |
| TcCLB.511421.217 | hypothetical protein, conserved | -0,2371 | 0,1695 |
| TcCLB.509319.50 | hypothetical protein, conserved | -0,2372 | 0,002771 |
| TcCLB.511677.10 | dispersed gene family protein 1 (DGF-1), putative | -0,2372 | 0,1454 |
| TcCLB.508355.114 | hypothetical protein, conserved | -0,2373 | 0,251 |
| TcCLB.511419.20 | hypothetical protein, conserved | -0,2374 | 0,04997 |
| TcCLB.509767.30 | hypothetical protein | -0,2374 | 0,109 |
| TcCLB.509895.50 | hypothetical protein, conserved | -0,2375 | 0,02768 |
| TcCLB.504239.80 | syntaxin binding protein (pseudogene), putative | -0,2375 | 0,6437 |
| TcCLB.509507.30 | hypothetical protein, conserved | -0,2376 | 0,03678 |
| TcCLB.510323.90 | hypothetical protein, conserved | -0,2376 | 0,06396 |
| TcCLB.506007.10 | MP24, putative | -0,2376 | 0,2928 |
| TcCLB.417993.10 | dispersed gene family protein 1 (DGF-1, pseudogene), puta | -0,2376 | 0,7765 |
| TcCLB.511733.120 | hypothetical protein, conserved | -0,2377 | 0,02711 |
| TcCLB.506945.140 | hypothetical protein, conserved (pseudogene) | -0,2377 | 0,07728 |
| TcCLB.508169.40 | mitochondrial inner membrane signal peptidase, putative | -0,2377 | 0,2867 |
| TcCLB.510201.40 | trans-sialidase (pseudogene), putative | -0,2377 | 0,3337 |
| TcCLB.508437.20 | trans-sialidase (pseudogene), putative | -0,2377 | 0,5509 |

| | | | |
|---|---|---|---|
| TcCLB.508723.70 | hypothetical protein, conserved | -0,2378 | 0,004214 |
| TcCLB.511289.110 | hypothetical protein, conserved | -0,2378 | 0,1034 |
| TcCLB.404073.20 | actin, putative (fragment) | -0,2378 | 0,3732 |
| TcCLB.511449.80 | mucin-associated surface protein (MASP, pseudogene), pu | -0,2378 | 0,7383 |
| TcCLB.511661.20 | hypothetical protein, conserved | -0,238 | 0,04922 |
| TcCLB.504085.20 | exosome-associated protein 2, putative (EAP2) | -0,238 | 0,1094 |
| TcCLB.510187.220 | hypothetical protein, conserved | -0,238 | 0,1523 |
| TcCLB.508761.110 | trans-sialidase, Group VI, putative | -0,238 | 0,4297 |
| TcCLB.507681.150 | 40S ribosomal protein S24E, putative | -0,2382 | 0,04096 |
| TcCLB.504035.140 | mismatch repair protein MLH1, putative | -0,2383 | 0,06015 |
| TcCLB.510099.40 | hypothetical protein, conserved | -0,2383 | 0,1024 |
| TcCLB.511161.10 | hypothetical protein, conserved (pseudogene) | -0,2383 | 0,3375 |
| TcCLB.508367.20 | UDP-Gal or UDP-GlcNAc-dependent glycosyltransferase, pu | -0,2383 | 0,6022 |
| TcCLB.506925.500 | hypothetical protein, conserved | -0,2384 | 0,037 |
| TcCLB.506697.40 | helicase, putative | -0,2385 | 0,02134 |
| TcCLB.510689.20 | quinonoid dihydropteridine reductase, putative | -0,2385 | 0,07706 |
| TcCLB.511727.234 | hypothetical protein, conserved | -0,2385 | 0,08265 |
| TcCLB.505807.190 | polynucleotide kinase 3'-phosphatase, putative | -0,2385 | 0,1402 |
| TcCLB.503703.80 | hypothetical protein, conserved | -0,2385 | 0,1734 |
| TcCLB.511903.70 | hypothetical protein, conserved | -0,2385 | 0,243 |
| TcCLB.508647.250 | phosphotransferase, putative | -0,2386 | 0,02065 |
| TcCLB.505965.20 | hypothetical protein, conserved | -0,2386 | 0,1106 |
| TcCLB.504239.30 | mucin-associated surface protein (MASP), putative | -0,2386 | 0,6215 |
| TcCLB.504427.130 | 5'-3' exoribonuclease C, putative (XRNC) | -0,2387 | 0,06763 |
| TcCLB.507927.60 | hypothetical protein, conserved | -0,2389 | 0,008648 |
| TcCLB.509109.180 | hypothetical protein, conserved | -0,2389 | 0,1033 |
| TcCLB.508325.250 | trans-sialidase (pseudogene), putative | -0,239 | 0,2315 |
| TcCLB.510887.60 | hypothetical protein, conserved | -0,2391 | 0,17 |
| TcCLB.510657.90 | hypothetical protein, conserved | -0,2392 | 0,2125 |
| TcCLB.504253.10 | hypothetical protein, conserved | -0,2393 | 0,01947 |
| TcCLB.508163.140 | mucin-associated surface protein (MASP), putative | -0,2393 | 0,3231 |
| TcCLB.510443.40 | hypothetical protein, conserved | -0,2394 | 0,07165 |
| TcCLB.510731.30 | hypothetical protein, conserved | -0,2394 | 0,07343 |
| TcCLB.511753.70 | hypothetical protein, conserved | -0,2395 | 0,02018 |
| TcCLB.510423.40 | hypothetical protein, conserved | -0,2396 | 0,01319 |
| TcCLB.510301.60 | hypothetical protein, conserved | -0,2396 | 0,07137 |
| TcCLB.510317.30 | polynucleotide adenylyltransferase, putative (PAP) | -0,2396 | 0,2167 |
| TcCLB.510009.10 | hypothetical protein, conserved | -0,2396 | 0,4611 |
| TcCLB.506717.10 | fatty acid elongase, putative | -0,2398 | 0,1828 |
| TcCLB.463451.4 | hypothetical protein, conserved | -0,2399 | 0,3237 |
| TcCLB.510651.10 | hypothetical protein, conserved | -0,2399 | 0,347 |
| TcCLB.506941.90 | ATP-dependent DEAD/H DNA helicase recQ family, putativ | -0,24 | 0,02431 |
| TcCLB.508625.80 | hypothetical protein, conserved | -0,24 | 0,07336 |
| TcCLB.506559.129 | eukaryotic translation initiation factor 2-alpha kinase, puta | -0,2402 | 0,02279 |
| TcCLB.511643.80 | hypothetical protein, conserved (pseudogene) | -0,2403 | 0,03809 |

| | | | |
|---|---|---|---|
| TcCLB.508949.50 | protein kinase, putative | -0,2404 | 0,1527 |
| TcCLB.503879.90 | hypothetical protein, conserved | -0,2406 | 0,05237 |
| TcCLB.504867.80 | DNA primase large subunit, putative (pril) | -0,2406 | 0,07506 |
| TcCLB.506025.20 | hypothetical protein, conserved | -0,2406 | 0,1023 |
| TcCLB.503823.140 | hypothetical protein, conserved | -0,2406 | 0,12 |
| TcCLB.508247.40 | mucin TcMUCII, putative | -0,2406 | 0,691 |
| TcCLB.507019.30 | ribosome biogenesis regulatory protein (RRS1), putative | -0,2407 | 0,06499 |
| TcCLB.504109.40 | hypothetical protein, conserved | -0,2407 | 0,12 |
| TcCLB.508121.110 | mucin-associated surface protein (MASP), putative | -0,2407 | 0,4661 |
| TcCLB.506287.130 | hypothetical protein, conserved | -0,2408 | 0,108 |
| TcCLB.511507.10 | hypothetical protein, conserved | -0,2408 | 0,3207 |
| TcCLB.511353.10 | FK506-binding protein (FKBP)-type peptidyl-prolyl isomera: | -0,2409 | 0,06785 |
| TcCLB.504013.30 | hypothetical protein, conserved | -0,241 | 0,2313 |
| TcCLB.506559.280 | hypothetical protein, conserved | -0,2413 | 0,02561 |
| TcCLB.510761.30 | RNA re-capping enzyme, cytoplasmic, putative (Ce1) | -0,2413 | 0,03062 |
| TcCLB.510655.130 | hypothetical protein, conserved | -0,2413 | 0,3487 |
| TcCLB.507993.180 | hypothetical protein, conserved | -0,2414 | 0,06109 |
| TcCLB.511127.320 | kinetoplastid kinetochore protein 10, putative (kkt10) | -0,2414 | 0,1278 |
| TcCLB.506373.30 | retrotransposon hot spot protein (RHS, pseudogene), putat | -0,2414 | 0,1558 |
| TcCLB.511391.180 | Golgi/lysosome glycoprotein 1, putative (GLP1) | -0,2415 | 0,00507 |
| TcCLB.509173.9 | hypothetical protein, conserved | -0,2415 | 0,1083 |
| TcCLB.507977.9 | hypothetical protein, conserved (pseudogene) | -0,2415 | 0,2428 |
| TcCLB.508313.14 | trans-sialidase, putative | -0,2415 | 0,595 |
| TcCLB.510489.5 | mucin-associated surface protein (MASP), putative | -0,2415 | 0,9006 |
| TcCLB.507897.10 | hypothetical protein, conserved | -0,2416 | 0,2048 |
| TcCLB.509799.60 | hypothetical protein, conserved | -0,2417 | 0,01016 |
| TcCLB.510301.80 | N-acetyltransferase subunit Nat1, putative | -0,2417 | 0,03168 |
| TcCLB.510841.10 | Kinesin-13 4b, putative (fragment) (KIN13-4b) | -0,2417 | 0,3882 |
| TcCLB.508717.33 | mitochondrial chaperone BCS1, putative | -0,2418 | 0,03952 |
| TcCLB.506241.150 | hypothetical protein | -0,2418 | 0,08566 |
| TcCLB.506885.380 | hypothetical protein, conserved | -0,2418 | 0,09389 |
| TcCLB.511907.80 | beta-ketoacyl-ACP reductase 2, putative | -0,2418 | 0,1168 |
| TcCLB.508231.64 | hypothetical protein, conserved | -0,2418 | 0,1696 |
| TcCLB.506163.20 | surface protease GP63, putative | -0,2418 | 0,3864 |
| TcCLB.506637.50 | hypothetical protein, conserved | -0,2419 | 0,1358 |
| TcCLB.510283.130 | hypothetical protein, conserved | -0,2419 | 0,1944 |
| TcCLB.479481.10 | mucin-associated surface protein (MASP), putative (fragm | -0,2419 | 0,6431 |
| TcCLB.503719.30 | hypothetical protein, conserved | -0,242 | 0,1952 |
| TcCLB.506409.230 | hypothetical protein, conserved | -0,2421 | 0,09842 |
| TcCLB.511283.50 | hypothetical protein, conserved | -0,2421 | 0,2991 |
| TcCLB.506577.20 | hypothetical protein | -0,2423 | 0,03636 |
| TcCLB.506975.9 | REL2, putative | -0,2423 | 0,3515 |
| TcCLB.503977.30 | mucin-associated surface protein (MASP), putative | -0,2423 | 0,5323 |
| TcCLB.506985.30 | proteasome alpha 3 subunit, putative | -0,2424 | 0,04259 |
| TcCLB.511523.4 | hypothetical protein, conserved | -0,2424 | 0,08898 |

| | | | |
|---|---|---|---|
| TcCLB.506717.219 | GMP synthase (pseudogene), putative | -0,2424 | 0,1019 |
| TcCLB.505965.92 | hypothetical protein, conserved (pseudogene) | -0,2424 | 0,3094 |
| TcCLB.510961.30 | mucin-associated surface protein (MASP), putative | -0,2425 | 0,5148 |
| TcCLB.507723.90 | hypothetical protein, conserved | -0,2427 | 0,1654 |
| TcCLB.511719.41 | hypothetical protein, conserved (fragment) | -0,2428 | 0,1004 |
| TcCLB.510989.40 | hypothetical protein, conserved | -0,2428 | 0,1234 |
| TcCLB.507483.80 | hypothetical protein, conserved | -0,2429 | 0,1006 |
| TcCLB.511603.100 | mucin-associated surface protein (MASP), putative | -0,2429 | 0,5359 |
| TcCLB.506567.120 | mitochondrial RNA binding protein, putative | -0,2431 | 0,1267 |
| TcCLB.507735.30 | hypothetical protein, conserved | -0,2431 | 0,1333 |
| TcCLB.510303.60 | hypothetical protein, conserved | -0,2431 | 0,1413 |
| TcCLB.508153.890 | hypothetical protein, conserved | -0,2433 | 0,002534 |
| TcCLB.509243.50 | sigma-adaptin 3, putative | -0,2434 | 0,196 |
| TcCLB.511867.200 | hypothetical protein, conserved | -0,2435 | 0,05059 |
| TcCLB.509669.10 | hypothetical protein, conserved | -0,2435 | 0,2647 |
| TcCLB.508183.20 | hypothetical protein, conserved | -0,2437 | 0,08125 |
| TcCLB.506733.70 | glycosyl transferase-like protein, putative | -0,2437 | 0,1798 |
| TcCLB.506413.89 | trans-sialidase, putative | -0,2439 | 0,0376 |
| TcCLB.508153.600 | actin-like protein, putative | -0,2439 | 0,1376 |
| TcCLB.441967.9 | hypothetical protein, conserved | -0,2439 | 0,1415 |
| TcCLB.510889.352 | retrotransposon hot spot (RHS) protein, putative (fragmen | -0,244 | 0,571 |
| TcCLB.504171.59 | pseudouridine synthase TruD, putative | -0,2443 | 0,1516 |
| TcCLB.509823.10 | hypothetical protein, conserved | -0,2443 | 0,2154 |
| TcCLB.510747.114 | hypothetical protein, conserved | -0,2443 | 0,4206 |
| TcCLB.511717.130 | hypothetical protein, conserved | -0,2444 | 0,1199 |
| TcCLB.506895.60 | hypothetical protein, conserved (pseudogene) | -0,2446 | 0,07713 |
| TcCLB.510431.160 | nucleolar GTP-binding protein 1, putative | -0,2447 | 0,06831 |
| TcCLB.511911.120 | DNA polymerase eta, putative | -0,2447 | 0,07415 |
| TcCLB.506247.350 | hypothetical protein, conserved | -0,2447 | 0,08298 |
| TcCLB.510911.20 | hypothetical protein, conserved | -0,2448 | 0,1019 |
| TcCLB.507519.164 | hypothetical protein, conserved | -0,2449 | 0,2525 |
| TcCLB.506901.10 | mucin TcMUCI, putative | -0,2449 | 0,8052 |
| TcCLB.504105.130 | Flagellum surface protein 179 (FS179) | -0,245 | 0,01618 |
| TcCLB.509941.100 | 2,4-dienoyl-coa reductase fadh1, putative | -0,245 | 0,01648 |
| TcCLB.508413.50 | RNA-binding protein, putative | -0,245 | 0,01784 |
| TcCLB.511559.70 | phospholipid transporting ATPase-like protein (pseudogene | -0,245 | 0,03459 |
| TcCLB.511353.40 | eukaryotic translation initiation factor 4E (eIF4E) interactir | -0,245 | 0,0877 |
| TcCLB.505807.240 | hypothetical protein, conserved | -0,2451 | 0,1419 |
| TcCLB.479379.9 | dynein heavy chain, putative | -0,2452 | 0,02371 |
| TcCLB.503459.29 | hypothetical protein, conserved | -0,2452 | 0,1427 |
| TcCLB.511753.150 | hypothetical protein, conserved | -0,2452 | 0,1771 |
| TcCLB.510625.10 | dispersed gene family protein 1 (DGF-1), putative | -0,2452 | 0,4462 |
| TcCLB.510943.36 | hypothetical protein, conserved | -0,2453 | 0,06409 |
| TcCLB.508707.194 | hypothetical protein, conserved | -0,2454 | 0,01867 |
| TcCLB.503419.60 | Enriched in surface-labeled proteome protein 15 (ESP15) | -0,2454 | 0,03677 |

| | | | |
|---|---|---|---|
| TcCLB.510265.90 | hypothetical protein, conserved | -0,2454 | 0,06976 |
| TcCLB.508675.20 | DNA excision repair protein, putative | -0,2455 | 0,01731 |
| TcCLB.507061.34 | hypothetical protein, conserved | -0,2455 | 0,1348 |
| TcCLB.506743.220 | hypothetical protein, conserved | -0,2455 | 0,3472 |
| TcCLB.507681.130 | hypothetical protein, conserved | -0,2456 | 0,007902 |
| TcCLB.507039.10 | iron superoxide dismutase, putative | -0,2456 | 0,01131 |
| TcCLB.510187.540 | amino acid permease, putative | -0,2456 | 0,04938 |
| TcCLB.511439.90 | hypothetical protein, conserved | -0,2456 | 0,5156 |
| TcCLB.506581.10 | dolicholphosphate-mannose synthase, putative (DPMS) | -0,2457 | 0,1669 |
| TcCLB.508721.10 | small GTPase, putative | -0,2458 | 0,08849 |
| TcCLB.509073.119 | proteasome regulatory non-ATP-ase subunit 7, putative | -0,2458 | 0,1155 |
| TcCLB.510329.240 | hypothetical protein, conserved | -0,2458 | 0,1687 |
| TcCLB.511287.140 | hypothetical protein, conserved | -0,2459 | 0,0143 |
| TcCLB.507037.60 | hypothetical protein, conserved | -0,246 | 0,01274 |
| TcCLB.510897.3 | A/G-specific adenine glycosylase (pseudogene), putative | -0,246 | 0,5081 |
| TcCLB.506953.10 | hypothetical protein, conserved (pseudogene) | -0,246 | 0,5447 |
| TcCLB.510123.24 | hypothetical protein, conserved | -0,2461 | 0,07485 |
| TcCLB.509669.120 | hypothetical protein, conserved (pseudogene) | -0,2461 | 0,0765 |
| TcCLB.508475.40 | hypothetical protein | -0,2461 | 0,3099 |
| TcCLB.506459.160 | mucin TcMUCII (pseudogene), putative | -0,2461 | 0,605 |
| TcCLB.504039.70 | hypothetical protein | -0,2461 | 0,7195 |
| TcCLB.509059.20 | hypothetical protein, conserved | -0,2462 | 0,03576 |
| TcCLB.506177.150 | hypothetical protein, conserved | -0,2462 | 0,0484 |
| TcCLB.511909.10 | hypothetical protein, conserved | -0,2463 | 0,006563 |
| TcCLB.509525.410 | mucin-associated surface protein (MASP, pseudogene), pu | -0,2463 | 0,5612 |
| TcCLB.508349.39 | RNA-binding protein, putative (fragment) (DRBD3) | -0,2464 | 0,04515 |
| TcCLB.510455.10 | dispersed gene family protein 1 (DGF-1), putative | -0,2466 | 0,6702 |
| TcCLB.506297.40 | hypothetical protein, conserved | -0,2467 | 0,001251 |
| TcCLB.510657.70 | hypothetical protein, conserved (pseudogene) | -0,2468 | 0,2379 |
| TcCLB.506529.410 | primase 2, putative | -0,2469 | 0,0157 |
| TcCLB.510065.50 | hypothetical protein, conserved | -0,2469 | 0,1564 |
| TcCLB.510657.220 | hypothetical protein, conserved | -0,2469 | 0,1842 |
| TcCLB.511181.60 | hypothetical protein, conserved | -0,247 | 0,07974 |
| TcCLB.508479.310 | hypothetical protein, conserved | -0,247 | 0,08347 |
| TcCLB.509261.40 | UDP-Gal or UDP-GlcNAc-dependent glycosyltransferase, pu | -0,247 | 0,4181 |
| TcCLB.508541.80 | mucin TcMUCII, putative | -0,247 | 0,538 |
| TcCLB.508827.139 | Coatomer subunit gamma (fragment) (COPG) | -0,2471 | 0,01313 |
| TcCLB.507993.369 | sjogren s syndrome nuclear autoantigen 1, putative | -0,2471 | 0,1527 |
| TcCLB.506203.40 | hypothetical protein, conserved | -0,2472 | 0,00866 |
| TcCLB.511285.20 | hypothetical protein, conserved | -0,2472 | 0,01181 |
| TcCLB.508909.300 | hypothetical protein, conserved | -0,2472 | 0,07315 |
| TcCLB.511421.160 | hypothetical protein, conserved | -0,2473 | 0,1418 |
| TcCLB.510187.330 | hypothetical protein, conserved | -0,2474 | 0,02226 |
| TcCLB.503909.100 | mannosyltransferase, putative | -0,2474 | 0,09091 |
| TcCLB.509103.4 | protein kinase, putative | -0,2475 | 0,5086 |

| | | | |
|---|---|---|---|
| TcCLB.511263.20 | hypothetical protein, conserved | -0,2476 | 0,008438 |
| TcCLB.508297.30 | hypothetical protein, conserved | -0,2476 | 0,07176 |
| TcCLB.507949.120 | Transmembrane protein 231, putative | -0,2477 | 0,06641 |
| TcCLB.503555.20 | hypothetical protein, conserved | -0,2477 | 0,1209 |
| TcCLB.407477.40 | hypothetical protein, conserved | -0,2477 | 0,1595 |
| TcCLB.508321.60 | hypothetical protein, conserved | -0,2478 | 0,02583 |
| TcCLB.508347.150 | hypothetical protein, conserved | -0,2481 | 0,01681 |
| TcCLB.508857.90 | hypothetical protein, conserved | -0,2481 | 0,04995 |
| TcCLB.510519.20 | hypothetical protein, conserved | -0,2481 | 0,1226 |
| TcCLB.509753.70 | mucin-associated surface protein (MASP), putative | -0,2481 | 0,6534 |
| TcCLB.511603.450 | trans-sialidase, Group V, putative | -0,2481 | 0,6593 |
| TcCLB.509399.110 | hypothetical protein, conserved | -0,2482 | 0,06036 |
| TcCLB.509023.140 | hypothetical protein, conserved | -0,2482 | 0,1557 |
| TcCLB.503801.60 | hypothetical protein | -0,2482 | 0,1558 |
| TcCLB.509795.20 | hypothetical protein, conserved | -0,2482 | 0,1929 |
| TcCLB.507611.230 | trans-sialidase (pseudogene), putative | -0,2482 | 0,2226 |
| TcCLB.508199.60 | hypothetical protein, conserved | -0,2482 | 0,27 |
| TcCLB.507277.10 | hypothetical protein, conserved | -0,2483 | 0,1083 |
| TcCLB.509991.20 | conserved protein, unknown function | -0,2484 | 0,01886 |
| TcCLB.510121.60 | hypothetical protein, conserved | -0,2484 | 0,1067 |
| TcCLB.508029.40 | hypothetical protein | -0,2484 | 0,2494 |
| TcCLB.511277.510 | hypothetical protein, conserved | -0,2485 | 0,01347 |
| TcCLB.506621.70 | hypothetical protein, conserved | -0,2485 | 0,04248 |
| TcCLB.509559.40 | trans-sialidase (pseudogene), putative | -0,2485 | 0,2538 |
| TcCLB.509601.110 | hypothetical protein, conserved | -0,2486 | 0,1016 |
| TcCLB.511661.10 | hypothetical protein, conserved | -0,2487 | 0,00953 |
| TcCLB.510741.220 | hypothetical protein, conserved | -0,2487 | 0,03576 |
| TcCLB.511733.30 | hypothetical protein, conserved | -0,2488 | 0,1554 |
| TcCLB.503929.40 | hypothetical protein, conserved | -0,2488 | 0,2176 |
| TcCLB.509937.160 | hypothetical protein, conserved | -0,2489 | 0,01875 |
| TcCLB.509065.110 | hypothetical protein, conserved | -0,2489 | 0,02282 |
| TcCLB.508879.190 | hypothetical protein, conserved | -0,2489 | 0,07578 |
| TcCLB.508909.339 | hypothetical protein, conserved | -0,2489 | 0,08699 |
| TcCLB.510609.110 | ATP-dependent RNA helicase, putative | -0,2489 | 0,09291 |
| TcCLB.508307.40 | hypothetical protein, conserved | -0,2489 | 0,1853 |
| TcCLB.506405.20 | hypothetical protein, conserved | -0,2489 | 0,2729 |
| TcCLB.511173.280 | trans-sialidase, Group V, putative | -0,2489 | 0,4267 |
| TcCLB.504089.30 | hypothetical protein, conserved | -0,2491 | 0,02154 |
| TcCLB.507529.45 | hypothetical protein, conserved | -0,2491 | 0,2096 |
| TcCLB.416891.19 | hypothetical protein | -0,2491 | 0,3801 |
| TcCLB.509143.10 | hypothetical protein, conserved | -0,2492 | 0,01652 |
| TcCLB.507957.60 | mucin-associated surface protein (MASP, pseudogene), pu | -0,2492 | 0,5852 |
| TcCLB.504625.70 | kinetoplast DNA-associated protein, putative | -0,2493 | 0,01913 |
| TcCLB.503967.10 | hypothetical protein, conserved | -0,2493 | 0,08885 |
| TcCLB.509179.40 | hypothetical protein, conserved | -0,2493 | 0,1259 |

| | | | |
|---|---|---|---|
| TcCLB.509687.19 | dispersed gene family protein 1 (DGF-1), putative (fragme | -0,2494 | 0,4107 |
| TcCLB.510553.25 | elongation factor 1-gamma (EF-1-gamma, pseudogene), p | -0,2494 | 0,6871 |
| TcCLB.506481.39 | hypothetical protein, conserved | -0,2496 | 0,009574 |
| TcCLB.511743.20 | hypothetical protein, conserved | -0,2496 | 0,03513 |
| TcCLB.503745.10 | hypothetical protein, conserved | -0,2496 | 0,06202 |
| TcCLB.503881.30 | retrotransposon hot spot (RHS) protein, putative | -0,2496 | 0,3179 |
| TcCLB.417939.9 | dispersed gene family protein 1 (DGF-1), putative | -0,2496 | 0,5169 |
| TcCLB.511293.20 | hypothetical protein, conserved | -0,2497 | 0,004003 |
| TcCLB.511131.70 | hypothetical protein, conserved | -0,2497 | 0,1128 |
| TcCLB.511031.40 | hypothetical protein, conserved | -0,2497 | 0,134 |
| TcCLB.511861.7 | elongation factor 1-gamma (EF-1-gamma), putative | -0,2498 | 0,5867 |
| TcCLB.506245.60 | trans-sialidase, Group V, putative | -0,2499 | 0,6071 |
| TcCLB.506243.135 | hypothetical protein, conserved (pseudogene) | -0,2501 | 0,4421 |
| TcCLB.508465.120 | syntaxin, putative | -0,2502 | 0,08046 |
| TcCLB.509267.90 | ATP-dependent RNA helicase, putative | -0,2502 | 0,1187 |
| TcCLB.507641.150 | hypothetical protein, conserved | -0,2503 | 0,009753 |
| TcCLB.506871.150 | hypothetical protein, conserved | -0,2505 | 0,03965 |
| TcCLB.506009.100 | ADP-ribosylation factor-like protein, putative | -0,2505 | 0,1346 |
| TcCLB.506739.170 | hypothetical protein, conserved | -0,2506 | 0,03302 |
| TcCLB.509897.270 | trans-sialidase, Group V, putative | -0,2506 | 0,2815 |
| TcCLB.511291.129 | hypothetical protein, conserved | -0,2507 | 0,2822 |
| TcCLB.509789.59 | hypothetical protein, conserved | -0,2508 | 0,007328 |
| TcCLB.511421.10 | hypothetical protein, conserved | -0,2508 | 0,01179 |
| TcCLB.507023.10 | epsilon-adaptin, putative | -0,2509 | 0,006301 |
| TcCLB.506963.40 | hypothetical protein, conserved | -0,2509 | 0,03971 |
| TcCLB.507063.140 | hypothetical protein, conserved | -0,2509 | 0,2433 |
| TcCLB.507883.60 | hypothetical protein, conserved | -0,251 | 0,06849 |
| TcCLB.508231.220 | hypothetical protein, conserved | -0,251 | 0,3219 |
| TcCLB.509029.30 | hypothetical protein, conserved | -0,2511 | 0,1844 |
| TcCLB.511859.30 | hypothetical protein, conserved | -0,2512 | 0,1093 |
| TcCLB.510587.20 | mucin-associated surface protein (MASP, pseudogene), pu | -0,2513 | 0,6993 |
| TcCLB.507795.30 | intraflagellar transport protein 27 (IFT27) | -0,2514 | 0,07558 |
| TcCLB.508881.54 | hypothetical protein, conserved | -0,2514 | 0,1056 |
| TcCLB.510645.9 | serine/threonine protein kinase, putative | -0,2515 | 0,03507 |
| TcCLB.510755.50 | ADP-ribosylation factor 1, putative (ARF1) | -0,2516 | 0,007556 |
| TcCLB.505997.210 | DNA-directed RNA polymerase III subunit, putative | -0,2516 | 0,2161 |
| TcCLB.506139.110 | mucin-associated surface protein (MASP), putative | -0,2516 | 0,6814 |
| TcCLB.511283.40 | syntaxin 5, putative | -0,2517 | 0,06814 |
| TcCLB.510143.17 | hypothetical protein, conserved | -0,2517 | 0,1243 |
| TcCLB.506661.70 | hypothetical protein, conserved | -0,2517 | 0,145 |
| TcCLB.503717.5 | mucin TcMUCII (pseudogene), putative | -0,2517 | 0,7333 |
| TcCLB.511389.40 | hypothetical protein, conserved | -0,2518 | 0,005988 |
| TcCLB.506127.20 | Nuclear RNA export factor (MEX67) | -0,2518 | 0,04225 |
| TcCLB.506755.230 | hypothetical protein, conserved | -0,2518 | 0,1687 |
| TcCLB.510741.30 | hypothetical protein, conserved | -0,2519 | 0,02013 |

| | | | |
|---|---|---|---|
| TcCLB.508593.40 | hypothetical protein, conserved | -0,252 | 0,1284 |
| TcCLB.503957.25 | trans-sialidase (pseudogene), putative | -0,252 | 0,4329 |
| TcCLB.508785.4 | hypothetical protein, conserved (fragment) | -0,2521 | 0,02559 |
| TcCLB.503635.50 | hypothetical protein, conserved | -0,2521 | 0,064 |
| TcCLB.509893.29 | hypothetical protein, conserved | -0,2521 | 0,1887 |
| TcCLB.509525.320 | mucin TcMUCII, putative | -0,2522 | 0,6535 |
| TcCLB.504153.100 | Tubulin cofactor C domain-containing protein 1, putative (t | -0,2523 | 0,04178 |
| TcCLB.509023.50 | kinesin, putative | -0,2524 | 0,02136 |
| TcCLB.507591.15 | trans-sialidase (pseudogene), putative | -0,2526 | 0,1397 |
| TcCLB.509385.11 | retrotransposon hot spot protein (RHS, pseudogene), putat | -0,2526 | 0,3926 |
| TcCLB.506811.40 | hypothetical protein, conserved | -0,2529 | 0,02771 |
| TcCLB.508153.510 | serine/threonine-protein kinase Nek, putative | -0,2529 | 0,05573 |
| TcCLB.509757.30 | stress-inducible protein STI1-like, putative | -0,2529 | 0,1623 |
| TcCLB.509127.70 | hypothetical protein, conserved | -0,253 | 0,05089 |
| TcCLB.506855.270 | hypothetical protein, conserved | -0,253 | 0,253 |
| TcCLB.503757.49 | hypothetical protein, conserved (fragment) | -0,253 | 0,5409 |
| TcCLB.509569.10 | hypothetical protein, conserved | -0,2533 | 0,1683 |
| TcCLB.508409.250 | hypothetical protein, conserved | -0,2534 | 0,1494 |
| TcCLB.503659.20 | mucin TcMUCII (pseudogene), putative | -0,2534 | 0,4629 |
| TcCLB.506833.9 | hypothetical protein, conserved | -0,2536 | 0,07173 |
| TcCLB.509017.24 | hypothetical protein, conserved | -0,2537 | 0,05675 |
| TcCLB.509065.150 | hypothetical protein, conserved | -0,2537 | 0,0869 |
| TcCLB.507539.20 | hypothetical protein, conserved | -0,2538 | 0,3959 |
| TcCLB.508869.10 | hypothetical protein | -0,2538 | 0,6205 |
| TcCLB.506815.30 | beta prime cop protein, putative | -0,2539 | 0,01085 |
| TcCLB.504111.49 | hypothetical protein, conserved | -0,2539 | 0,02735 |
| TcCLB.506443.40 | mitochondrial pyruvate carrier protein 2, putative | -0,2539 | 0,04824 |
| TcCLB.503713.40 | hypothetical protein | -0,2539 | 0,4082 |
| TcCLB.507509.89 | cleavage and polyadenylation specificity factor-like protein | -0,254 | 0,03584 |
| TcCLB.506753.164 | hypothetical protein, conserved | -0,254 | 0,292 |
| TcCLB.511459.60 | hypothetical protein, conserved | -0,2541 | 0,2284 |
| TcCLB.507809.130 | hypothetical protein, conserved | -0,2542 | 0,008178 |
| TcCLB.506559.100 | multidrug resistance-associated protein, putative | -0,2542 | 0,01589 |
| TcCLB.511751.220 | hypothetical protein, conserved | -0,2542 | 0,04319 |
| TcCLB.508275.9 | dynein heavy chain, putative (fragment) | -0,2543 | 0,05277 |
| TcCLB.504153.150 | Unc104-like kinesin, putative | -0,2544 | 0,005387 |
| TcCLB.506803.20 | trans-sialidase (pseudogene), putative | -0,2544 | 0,1369 |
| TcCLB.504239.401 | hypothetical protein, conserved (pseudogene) | -0,2544 | 0,7007 |
| TcCLB.504103.20 | hypothetical protein, conserved | -0,2545 | 0,0035 |
| TcCLB.506671.70 | dispersed gene family protein 1 (DGF-1), putative | -0,2547 | 0,3783 |
| TcCLB.509643.10 | hypothetical protein, conserved | -0,2547 | 0,4288 |
| TcCLB.511255.360 | mucin-associated surface protein (MASP), putative | -0,2547 | 0,5538 |
| TcCLB.508207.190 | hypothetical protein, conserved | -0,2548 | 0,03889 |
| TcCLB.511383.40 | hypothetical protein, conserved | -0,2548 | 0,06021 |
| TcCLB.505183.160 | hypothetical protein, conserved | -0,2549 | 0,07713 |

| | | | |
|---|---|---|---|
| TcCLB.510195.9 | dispersed gene family protein 1 (DGF-1), putative | -0,2549 | 0,2241 |
| TcCLB.506025.10 | hypothetical protein, conserved | -0,255 | 0,08454 |
| TcCLB.509201.20 | hypothetical protein, conserved | -0,2551 | 0,518 |
| TcCLB.506863.80 | serine/threonine protein phosphatase, putative | -0,2552 | 0,06699 |
| TcCLB.506139.40 | mucin-associated surface protein (MASP), putative | -0,2552 | 0,545 |
| TcCLB.503823.150 | hypothetical protein, conserved | -0,2553 | 0,08345 |
| TcCLB.507389.50 | hypothetical protein, conserved | -0,2553 | 0,08349 |
| TcCLB.507641.140 | ATP-dependent DEAD/H RNA helicase, putative | -0,2554 | 0,003708 |
| TcCLB.507005.50 | hypothetical protein | -0,2554 | 0,03955 |
| TcCLB.511209.60 | hypothetical protein, conserved | -0,2554 | 0,06761 |
| TcCLB.506829.10 | hypothetical protein, conserved | -0,2555 | 0,01536 |
| TcCLB.511507.30 | cyclosome subunit, putative | -0,2555 | 0,01602 |
| TcCLB.507825.10 | hypothetical protein, conserved | -0,2555 | 0,1858 |
| TcCLB.509639.20 | hypothetical protein, conserved | -0,2556 | 0,1132 |
| TcCLB.509835.20 | guanine nucleotide exchange factor subunit GCD2 | -0,2557 | 0,06396 |
| TcCLB.509625.20 | hypothetical protein, conserved | -0,2557 | 0,3563 |
| TcCLB.508221.580 | mucin-associated surface protein (MASP), putative | -0,2557 | 0,639 |
| TcCLB.508297.14 | trans-sialidase (pseudogene), putative | -0,256 | 0,1218 |
| TcCLB.505171.80 | hypothetical protein, conserved | -0,2561 | 0,02504 |
| TcCLB.507723.60 | hypothetical protein | -0,2562 | 0,1858 |
| TcCLB.510329.280 | protein farnesyltransferase alpha subunit, putative | -0,2564 | 0,02329 |
| TcCLB.506711.30 | cyclin, putative | -0,2564 | 0,1227 |
| TcCLB.508649.5 | tryparedoxin peroxidase, putative (fragment) | -0,2565 | 0,0234 |
| TcCLB.506363.50 | hypothetical protein, conserved | -0,2566 | 0,009765 |
| TcCLB.508515.110 | hypothetical protein, conserved | -0,2569 | 0,01594 |
| TcCLB.506649.10 | hypothetical protein | -0,2569 | 0,1206 |
| TcCLB.509047.20 | hypothetical protein | -0,257 | 0,07405 |
| TcCLB.509259.30 | mucin-associated surface protein (MASP, pseudogene), pu | -0,257 | 0,3803 |
| TcCLB.510565.150 | surface protease GP63, putative | -0,2572 | 0,1294 |
| TcCLB.509777.100 | ubiquitin-activating enzyme, putative | -0,2574 | 0,03377 |
| TcCLB.507049.70 | hypothetical protein, conserved | -0,2574 | 0,1205 |
| TcCLB.484949.9 | AGC/RSK family serine/threonine kinase, putative (fragme | -0,2574 | 0,3509 |
| TcCLB.509215.10 | DNA polymerase delta catalytic subunit, putative | -0,2576 | 0,003086 |
| TcCLB.506565.4 | RNA-binding protein, putative | -0,2576 | 0,4789 |
| TcCLB.508229.40 | trans-sialidase, Group V, putative | -0,2577 | 0,1335 |
| TcCLB.506475.40 | MORN repeat-containing protein | -0,2578 | 0,01332 |
| TcCLB.506755.180 | hypothetical protein, conserved | -0,2578 | 0,1932 |
| TcCLB.507021.80 | hypothetical protein, conserved | -0,2579 | 0,09842 |
| TcCLB.510765.70 | 60S ribosomal protein L5, putative | -0,258 | 0,02503 |
| TcCLB.511257.20 | hypothetical protein, conserved | -0,258 | 0,05444 |
| TcCLB.508211.10 | oxysterol-binding protein, putative | -0,258 | 0,133 |
| TcCLB.508325.25 | trans-sialidase (pseudogene), putative | -0,258 | 0,2699 |
| TcCLB.413977.10 | guide RNA associated protein, GAP1, putative | -0,2581 | 0,04198 |
| TcCLB.504057.70 | hypothetical protein, conserved | -0,2582 | 0,08546 |
| TcCLB.508163.340 | trans-sialidase (pseudogene), putative | -0,2582 | 0,6256 |

| | | | |
|---|---|---|---|
| TcCLB.506885.140 | trans-sialidase (pseudogene), putative | -0,2583 | 0,231 |
| TcCLB.507603.40 | proteasome beta-1 subunit, putative | -0,2584 | 0,02349 |
| TcCLB.510141.4 | hypothetical protein, conserved (fragment) | -0,2584 | 0,1298 |
| TcCLB.508707.110 | hypothetical protein, conserved | -0,2584 | 0,2498 |
| TcCLB.506321.180 | hypothetical protein | -0,2585 | 0,733 |
| TcCLB.508461.360 | hypothetical protein, conserved | -0,2586 | 0,01075 |
| TcCLB.508891.20 | hypothetical protein, conserved | -0,2586 | 0,03196 |
| TcCLB.510283.150 | hypothetical protein, conserved | -0,2587 | 0,2318 |
| TcCLB.506473.14 | Protein transport protein yif1, putative | -0,2588 | 0,09171 |
| TcCLB.511245.40 | hypothetical protein | -0,259 | 0,1022 |
| TcCLB.511751.191 | hypothetical protein (pseudogene) | -0,259 | 0,1323 |
| TcCLB.507021.70 | hypothetical protein, conserved | -0,2591 | 0,03617 |
| TcCLB.506697.30 | serine/threonine-protein kinase Nek, putative | -0,2591 | 0,03855 |
| TcCLB.511127.190 | hypothetical protein, conserved | -0,2591 | 0,05147 |
| TcCLB.507081.120 | hypothetical protein, conserved | -0,2592 | 0,07347 |
| TcCLB.506945.180 | hypothetical protein, conserved | -0,2593 | 0,007445 |
| TcCLB.508641.90 | hypothetical protein, conserved | -0,2593 | 0,02067 |
| TcCLB.510575.40 | hypothetical protein, conserved | -0,2593 | 0,06759 |
| TcCLB.506855.70 | hypothetical protein, conserved | -0,2594 | 0,01395 |
| TcCLB.506407.60 | hypothetical protein, conserved | -0,2594 | 0,1015 |
| TcCLB.508633.10 | trans-sialidase (pseudogene), putative | -0,2594 | 0,5454 |
| TcCLB.508995.69 | pseudouridine synthase TruD, putative (fragment) | -0,2596 | 0,04069 |
| TcCLB.503527.20 | hypothetical protein, conserved | -0,2596 | 0,06365 |
| TcCLB.509099.59 | hypothetical protein, conserved | -0,2596 | 0,1008 |
| TcCLB.511109.20 | hypothetical protein, conserved | -0,2596 | 0,1878 |
| TcCLB.510565.141 | hypothetical protein, conserved | -0,2596 | 0,2124 |
| TcCLB.508355.110 | hypothetical protein | -0,2597 | 0,05821 |
| TcCLB.510747.110 | hypothetical protein, conserved | -0,2597 | 0,1185 |
| TcCLB.509033.44 | hypothetical protein, conserved | -0,2597 | 0,5315 |
| TcCLB.506971.80 | mucin-associated surface protein (MASP), putative (fragm | -0,2598 | 0,551 |
| TcCLB.508299.20 | hypothetical protein, conserved | -0,26 | 0,1264 |
| TcCLB.510257.40 | hypothetical protein, conserved | -0,2602 | 0,1177 |
| TcCLB.506511.80 | hypothetical protein, conserved | -0,2603 | 0,02186 |
| TcCLB.505025.130 | mucin-associated surface protein (MASP), putative | -0,2603 | 0,6547 |
| TcCLB.511281.59 | hypothetical protein, conserved | -0,2604 | 0,0492 |
| TcCLB.506359.90 | hypothetical protein, conserved | -0,2604 | 0,05641 |
| TcCLB.508231.210 | hypothetical protein, conserved | -0,2607 | 0,02448 |
| TcCLB.511135.40 | hypothetical protein, conserved | -0,2608 | 0,1294 |
| TcCLB.506267.39 | mucin-associated surface protein (MASP, pseudogene), pu | -0,2608 | 0,2764 |
| TcCLB.504157.110 | dynein intermediate chain, putative | -0,2609 | 0,08454 |
| TcCLB.509715.30 | cullin-like protein, putative | -0,2611 | 0,03474 |
| TcCLB.507677.20 | hypothetical protein, conserved | -0,2612 | 0,1458 |
| TcCLB.510689.89 | hypothetical protein, conserved | -0,2612 | 0,4563 |
| TcCLB.505123.20 | hypothetical protein, conserved | -0,2612 | 0,477 |
| TcCLB.509053.50 | hypothetical protein, conserved | -0,2613 | 0,3502 |

| | | | |
|---|---|---|---|
| TcCLB.504147.140 | hypothetical protein | -0,2614 | 0,02306 |
| TcCLB.507993.10 | hypothetical protein, conserved | -0,2614 | 0,04917 |
| TcCLB.504255.10 | hypothetical protein, conserved | -0,2614 | 0,1247 |
| TcCLB.510241.110 | hypothetical protein, conserved | -0,2615 | 0,00373 |
| TcCLB.508207.270 | hypothetical protein, conserved | -0,2616 | 0,01648 |
| TcCLB.510517.100 | hypothetical protein, conserved | -0,2617 | 0,1146 |
| TcCLB.507669.50 | hypothetical protein, conserved | -0,2618 | 0,00149 |
| TcCLB.509429.100 | PITH domain-containing protein | -0,2618 | 0,07623 |
| TcCLB.503521.10 | hypothetical protein | -0,2619 | 0,03592 |
| TcCLB.506531.40 | hypothetical protein, conserved | -0,2619 | 0,04305 |
| TcCLB.506585.50 | hypothetical protein, conserved | -0,2619 | 0,05854 |
| TcCLB.506979.10 | hypothetical protein, conserved | -0,2619 | 0,06373 |
| TcCLB.506655.7 | phospholipid-transporting ATPase, putative (fragment) | -0,2619 | 0,1524 |
| TcCLB.506317.30 | hypothetical protein, conserved (pseudogene) | -0,262 | 0,05873 |
| TcCLB.511079.25 | mucin-associated surface protein (MASP, pseudogene), pu | -0,262 | 0,5655 |
| TcCLB.510743.110 | hypothetical protein, conserved | -0,2621 | 0,09114 |
| TcCLB.482097.10 | hypothetical protein, conserved (fragment) | -0,2621 | 0,3885 |
| TcCLB.506671.41 | elongation factor 1-gamma (EF-1-gamma, pseudogene), p | -0,2621 | 0,7159 |
| TcCLB.506529.669 | hypothetical protein, conserved | -0,2622 | 0,2117 |
| TcCLB.506513.200 | histone H2A, putative | -0,2622 | 0,7155 |
| TcCLB.511151.50 | phosphoadenosine phosphosulfate reductase-like protein, | -0,2623 | 0,02931 |
| TcCLB.510687.20 | hypothetical protein, conserved | -0,2623 | 0,08919 |
| TcCLB.510595.4 | hypothetical protein, conserved | -0,2623 | 0,2773 |
| TcCLB.506599.310 | mucin-associated surface protein (MASP), putative | -0,2623 | 0,7026 |
| TcCLB.509719.30 | hypothetical protein, conserved | -0,2625 | 0,1041 |
| TcCLB.503765.10 | trans-sialidase (pseudogene), putative | -0,2625 | 0,2562 |
| TcCLB.510535.60 | CCR4 associated factor, putative | -0,2626 | 0,02091 |
| TcCLB.504253.20 | hypothetical protein, conserved | -0,2626 | 0,04061 |
| TcCLB.509695.80 | hypothetical protein, conserved | -0,2626 | 0,04418 |
| TcCLB.507603.130 | hypothetical protein, conserved | -0,2626 | 0,09018 |
| TcCLB.506435.180 | hypothetical protein, conserved | -0,2627 | 0,1874 |
| TcCLB.511145.89 | hypothetical protein, conserved | -0,2627 | 0,2169 |
| TcCLB.511099.190 | mucin-associated surface protein (MASP), putative | -0,2627 | 0,5247 |
| TcCLB.508633.20 | trans-sialidase, Group V, putative | -0,2627 | 0,5804 |
| TcCLB.507887.20 | ubiquitin hydrolase, putative | -0,2629 | 0,004155 |
| TcCLB.507801.140 | TPR-repeat protein, putative | -0,2629 | 0,03319 |
| TcCLB.511239.40 | hypothetical protein, conserved | -0,2629 | 0,0332 |
| TcCLB.506179.50 | hypothetical protein, conserved | -0,2631 | 0,08607 |
| TcCLB.511349.61 | retrotransposon hot spot protein (RHS, pseudogene), putat | -0,2631 | 0,6593 |
| TcCLB.509897.190 | hypothetical protein, conserved (pseudogene) | -0,2632 | 0,02606 |
| TcCLB.511355.7 | TFIID-like protein, putative (fragment) | -0,2632 | 0,1044 |
| TcCLB.507979.20 | hypothetical protein, conserved | -0,2632 | 0,1216 |
| TcCLB.507011.269 | 6-phosphogluconate dehydrogenase, decarboxylating (6PG | -0,2632 | 0,2388 |
| TcCLB.506739.150 | 60S ribosomal protein L13a, putative | -0,2633 | 0,000252 |
| TcCLB.506825.190 | hypothetical protein | -0,2633 | 0,01952 |

| | | | |
|---|---|---|---|
| TcCLB.506473.20 | DnaJ chaperone protein, putative | -0,2633 | 0,02828 |
| TcCLB.511417.120 | DNA repair protein BRCA2 (pseudogene), putative | -0,2633 | 0,04379 |
| TcCLB.511423.160 | RWD domain-containing protein | -0,2634 | 0,1565 |
| TcCLB.510299.60 | hypothetical protein, conserved | -0,2635 | 0,1833 |
| TcCLB.509637.27 | hypothetical protein, conserved | -0,2637 | 0,09945 |
| TcCLB.506529.14 | hypothetical protein, conserved | -0,2637 | 0,1624 |
| TcCLB.506445.121 | heat shock protein DNAJ, putative (fragment) | -0,2638 | 0,3042 |
| TcCLB.509671.164 | hypothetical protein, conserved | -0,2639 | 0,1037 |
| TcCLB.509029.110 | hypothetical protein, conserved | -0,264 | 0,04601 |
| TcCLB.504203.29 | tRNA-dihydrouridine synthase 2, putative | -0,2641 | 0,2083 |
| TcCLB.511491.80 | mitogen-activated protein kinase, putative | -0,2642 | 0,04465 |
| TcCLB.507257.60 | Fumarate hydratase class I, cytosolic (FHc) | -0,2642 | 0,1037 |
| TcCLB.503823.100 | hypothetical protein, conserved | -0,2643 | 0,08625 |
| TcCLB.509053.90 | hypothetical protein, conserved | -0,2644 | 0,08487 |
| TcCLB.509965.130 | hypothetical protein, conserved | -0,2644 | 0,2433 |
| TcCLB.508553.20 | hypothetical protein | -0,2645 | 0,5753 |
| TcCLB.504147.150 | ATP-dependent DEAD/H RNA helicase, putative | -0,2646 | 0,002263 |
| TcCLB.507101.20 | hypothetical protein (pseudogene) | -0,2646 | 0,3743 |
| TcCLB.511903.240 | hypothetical protein, conserved | -0,2647 | 0,03143 |
| TcCLB.511761.30 | hypothetical protein, conserved | -0,2647 | 0,1171 |
| TcCLB.507603.20 | translation initiation factor EIF-2b alpha subunit, putative | -0,2648 | 0,1405 |
| TcCLB.508355.170 | hypothetical protein, conserved | -0,2649 | 0,02705 |
| TcCLB.505931.10 | surface protease GP63, putative | -0,2649 | 0,132 |
| TcCLB.503917.35 | B9 domain-containing protein 2 (b9d2) | -0,2649 | 0,2345 |
| TcCLB.510685.30 | RNA polymerase IIA largest subunit, putative | -0,265 | 0,01509 |
| TcCLB.510329.270 | Fumarate hydratase class I, cytosolic (FHc) | -0,265 | 0,1887 |
| TcCLB.508269.50 | hypothetical protein, conserved | -0,2651 | 0,01546 |
| TcCLB.510625.40 | trans-sialidase, Group V, putative | -0,2651 | 0,5135 |
| TcCLB.506195.70 | PrimPol-like protein 2, putative (ppl2) | -0,2653 | 0,02621 |
| TcCLB.405737.14 | hypothetical protein | -0,2655 | 0,1227 |
| TcCLB.511419.60 | NADH-ubiquinone oxidoreductase complex I subunit, putati | -0,2656 | 0,06001 |
| TcCLB.504171.10 | hypothetical protein, conserved | -0,2656 | 0,1371 |
| TcCLB.506741.90 | mucin-associated surface protein (MASP), putative | -0,2656 | 0,4061 |
| TcCLB.506885.480 | trans-sialidase, Group V, putative | -0,2657 | 0,4047 |
| TcCLB.506473.6 | hypothetical protein, conserved | -0,2659 | 0,05741 |
| TcCLB.511003.210 | hypothetical protein, conserved | -0,2659 | 0,08677 |
| TcCLB.508895.70 | membrane protein, putative | -0,266 | 0,1332 |
| TcCLB.508257.200 | hypothetical protein, conserved | -0,2662 | 0,007328 |
| TcCLB.503885.90 | hypothetical protein, conserved | -0,2662 | 0,01573 |
| TcCLB.509639.10 | procyclic form surface phosphoprotein, putative | -0,2662 | 0,08377 |
| TcCLB.510227.29 | thioredoxin, putative (fragment) | -0,2663 | 0,003378 |
| TcCLB.511727.280 | hypothetical protein, conserved | -0,2663 | 0,004394 |
| TcCLB.510265.40 | hypothetical protein, conserved | -0,2663 | 0,005731 |
| TcCLB.506859.160 | hypothetical protein, conserved | -0,2663 | 0,01426 |
| TcCLB.508745.11 | trans-sialidase, putative (fragment) | -0,2663 | 0,5447 |

| | | | |
|---|---|---|---|
| TcCLB.509805.100 | hypothetical protein, conserved | -0,2664 | 0,01326 |
| TcCLB.510659.170 | denn domain-containing protein | -0,2664 | 0,08479 |
| TcCLB.511255.160 | trans-sialidase (pseudogene), putative | -0,2664 | 0,5846 |
| TcCLB.503841.20 | AMPK subunit gamma (AMPKG) | -0,2665 | 0,03285 |
| TcCLB.506503.100 | U3 snoRNA-associated protein UTP11, putative | -0,2665 | 0,09206 |
| TcCLB.511717.70 | hypothetical protein, conserved | -0,2666 | 0,08104 |
| TcCLB.510575.100 | hypothetical protein, conserved | -0,2668 | 0,09148 |
| TcCLB.508577.120 | hypothetical protein, conserved | -0,2668 | 0,1264 |
| TcCLB.506587.10 | hypothetical protein, conserved | -0,2668 | 0,1732 |
| TcCLB.503771.40 | syntaxin binding protein (pseudogene), putative | -0,2668 | 0,5899 |
| TcCLB.508627.21 | retrotransposon hot spot protein (RHS, pseudogene), putat | -0,2669 | 0,3818 |
| TcCLB.506781.50 | dispersed gene family protein 1 (DGF-1), putative | -0,2669 | 0,4128 |
| TcCLB.511667.10 | hypothetical protein, conserved | -0,267 | 0,5367 |
| TcCLB.506855.110 | hypothetical protein, conserved | -0,2671 | 0,03333 |
| TcCLB.503989.60 | protein phosphatase 2C, putative (fragment) | -0,2671 | 0,1565 |
| TcCLB.508173.120 | trans-sialidase, Group I, putative | -0,2672 | 0,002853 |
| TcCLB.503897.40 | hypothetical protein, conserved | -0,2672 | 0,01546 |
| TcCLB.507743.9 | tyrosine protein kinase, putative | -0,2672 | 0,04191 |
| TcCLB.510969.20 | hypothetical protein, conserved | -0,2673 | 0,08396 |
| TcCLB.506351.60 | hypothetical protein, conserved | -0,2675 | 0,06276 |
| TcCLB.507771.99 | hypothetical protein, conserved | -0,2675 | 0,2455 |
| TcCLB.447255.29 | multidrug resistance-associated protein, putative | -0,2675 | 0,2722 |
| TcCLB.510943.110 | hypothetical protein, conserved | -0,2676 | 0,02829 |
| TcCLB.511317.20 | hypothetical protein, conserved | -0,2678 | 0,04662 |
| TcCLB.505999.40 | DNA repair protein BRCA2, putative | -0,2678 | 0,06745 |
| TcCLB.510025.160 | mucin-associated surface protein (MASP), putative | -0,2678 | 0,6195 |
| TcCLB.503461.40 | succinyl-CoA:3-ketoacid-coenzyme-A transferase (pseudog | -0,268 | 0,4202 |
| TcCLB.504057.11 | phospholipid-translocating ATPase, putative | -0,2681 | 0,006043 |
| TcCLB.504221.20 | 26S proteasome regulatory non-ATPase subunit, putative | -0,2681 | 0,02449 |
| TcCLB.511231.30 | hypothetical protein, conserved | -0,2682 | 0,0461 |
| TcCLB.508431.80 | trans-sialidase (pseudogene), putative | -0,2682 | 0,3906 |
| TcCLB.508761.40 | trans-sialidase, Group V, putative | -0,2682 | 0,4279 |
| TcCLB.511361.60 | hypothetical protein | -0,2683 | 0,0253 |
| TcCLB.507957.160 | mucin-associated surface protein (MASP), putative | -0,2683 | 0,5096 |
| TcCLB.503671.10 | Aminopeptidase M1, putative (fragment) | -0,2684 | 0,04172 |
| TcCLB.506559.80 | Sir2-related protein 3, putative (SIR2rp3) | -0,2684 | 0,1332 |
| TcCLB.506415.10 | dispersed gene family protein 1 (DGF-1), putative | -0,2685 | 0,006563 |
| TcCLB.505807.60 | hypothetical protein, conserved | -0,2685 | 0,01596 |
| TcCLB.507541.10 | hypothetical protein, conserved | -0,2685 | 0,2063 |
| TcCLB.504881.10 | hypothetical protein, conserved | -0,2686 | 0,04913 |
| TcCLB.503923.10 | hypothetical protein, conserved | -0,2686 | 0,05561 |
| TcCLB.509157.210 | hypothetical protein, conserved | -0,2686 | 0,2698 |
| TcCLB.503809.68 | glycerol uptake protein (pseudogene), putative | -0,2687 | 0,407 |
| TcCLB.506625.220 | hypothetical protein, conserved | -0,2688 | 0,03666 |
| TcCLB.511109.110 | hypothetical protein, conserved | -0,2688 | 0,2513 |

| | | | |
|---|---|---|---|
| TcCLB.510769.140 | hypothetical protein, conserved | -0,2689 | 0,07392 |
| TcCLB.503935.4 | hypothetical protein, conserved (pseudogene) | -0,2689 | 0,2039 |
| TcCLB.507357.20 | hypothetical protein | -0,2689 | 0,5018 |
| TcCLB.508989.90 | Eukaryotic translation initiation factor 4 gamma type 5, pu | -0,269 | 0,001728 |
| TcCLB.503835.10 | hypothetical protein, conserved | -0,2691 | 0,1949 |
| TcCLB.506677.4 | hypothetical protein, conserved | -0,2691 | 0,5695 |
| TcCLB.504243.49 | hypothetical protein, conserved (fragment) | -0,2692 | 0,07249 |
| TcCLB.510359.310 | eukaryotic translation initiation factor 2 subunit, putative | -0,2693 | 0,006415 |
| TcCLB.509807.20 | hypothetical protein, conserved | -0,2694 | 0,02719 |
| TcCLB.508695.20 | phosphatidylinositol 4-kinase, putative | -0,2696 | 0,02299 |
| TcCLB.503863.9 | hypothetical protein, conserved | -0,2696 | 0,1125 |
| TcCLB.503881.39 | ribosomal protein L38, putative | -0,2698 | 0,02749 |
| TcCLB.506247.460 | hypothetical protein, conserved | -0,27 | 0,05933 |
| TcCLB.508153.110 | hypothetical protein | -0,2701 | 0,1502 |
| TcCLB.509081.60 | syntaxin binding protein (pseudogene), putative | -0,2701 | 0,4048 |
| TcCLB.508891.40 | hypothetical protein, conserved | -0,2702 | 0,04454 |
| TcCLB.506625.90 | hypothetical protein, conserved | -0,2702 | 0,1206 |
| TcCLB.511727.30 | hypothetical protein, conserved | -0,2703 | 0,005359 |
| TcCLB.506249.40 | hypothetical protein, conserved | -0,2703 | 0,03874 |
| TcCLB.508429.40 | mucin-associated surface protein (MASP, pseudogene), pu | -0,2703 | 0,6039 |
| TcCLB.504069.53 | proteasome subunit alpha type-5, putative | -0,2705 | 0,005385 |
| TcCLB.503449.40 | hypothetical protein, conserved | -0,2706 | 0,08196 |
| TcCLB.509551.80 | hypothetical protein, conserved | -0,2706 | 0,1064 |
| TcCLB.508479.150 | serine/threonine protein kinase, putative | -0,2707 | 0,00652 |
| TcCLB.503579.50 | hypothetical protein, conserved | -0,2707 | 0,007331 |
| TcCLB.507257.100 | hypothetical protein, conserved | -0,2707 | 0,05809 |
| TcCLB.508899.10 | proteasome regulatory non-ATP-ase subunit 9, putative (fr | -0,2707 | 0,0784 |
| TcCLB.507709.30 | hypothetical protein, conserved | -0,2707 | 0,1041 |
| TcCLB.479961.20 | retrotransposon hot spot protein (RHS, pseudogene), putat | -0,2707 | 0,1828 |
| TcCLB.510431.180 | Present in the outer mitochondrial membrane proteome 1 | -0,2708 | 0,03725 |
| TcCLB.508133.19 | hypothetical protein, conserved | -0,2708 | 0,1051 |
| TcCLB.511381.40 | DNA-directed rna polymerase I largest subunit, putative | -0,2709 | 0,001949 |
| TcCLB.510655.140 | hypothetical protein, conserved | -0,2709 | 0,0608 |
| TcCLB.510659.230 | hypothetical protein, conserved | -0,2709 | 0,1135 |
| TcCLB.505989.110 | amidinotransferase, putative | -0,271 | 0,0003939 |
| TcCLB.510363.160 | mucin-associated surface protein (MASP, pseudogene), pu | -0,271 | 0,4491 |
| TcCLB.508693.90 | peroxisome biogenesis factor 1, putative | -0,2711 | 0,004224 |
| TcCLB.508219.110 | mucin-associated surface protein (MASP), putative | -0,2712 | 0,4941 |
| TcCLB.503419.54 | hypothetical protein, conserved | -0,2713 | 0,1503 |
| TcCLB.511727.300 | GDP-L-fucose synthetase, putative | -0,2715 | 0,01736 |
| TcCLB.506367.80 | hypothetical protein | -0,2715 | 0,2692 |
| TcCLB.508307.130 | hypothetical protein, conserved | -0,2716 | 0,02708 |
| TcCLB.504827.70 | hypothetical protein, conserved | -0,2716 | 0,09897 |
| TcCLB.510357.90 | hypothetical protein, conserved | -0,2716 | 0,1392 |
| TcCLB.506557.20 | hypothetical protein, conserved | -0,2717 | 0,0252 |

| | | | |
|---|---|---|---|
| TcCLB.509073.90 | hypothetical protein, conserved | -0,2718 | 0,02456 |
| TcCLB.508317.30 | hypothetical protein | -0,2719 | 0,01033 |
| TcCLB.506813.160 | dispersed gene family protein 1 (DGF-1, pseudogene), puta | -0,272 | 0,3337 |
| TcCLB.508209.110 | hypothetical protein, conserved | -0,2721 | 0,005774 |
| TcCLB.410923.20 | retrotransposon hot spot (RHS) protein, putative | -0,2721 | 0,1081 |
| TcCLB.510463.240 | mucin-associated surface protein (MASP), putative | -0,2721 | 0,6092 |
| TcCLB.511249.80 | hypothetical protein, conserved | -0,2722 | 0,01226 |
| TcCLB.507521.110 | pumilio/PUF RNA binding protein 5, putative (PUF5) | -0,2722 | 0,02312 |
| TcCLB.508795.10 | SNF2/RAD54 related DNA helicase (pseudogene), putative | -0,2723 | 0,1563 |
| TcCLB.510699.100 | hypothetical protein | -0,2723 | 0,5147 |
| TcCLB.511389.130 | diphthine synthase, putative | -0,2726 | 0,08867 |
| TcCLB.511803.50 | hypothetical protein, conserved | -0,2726 | 0,1005 |
| TcCLB.509047.80 | mitochondrial oligo_U binding protein TBRGG1 (pseudoge | -0,2727 | 0,01509 |
| TcCLB.507639.100 | phosphatidylinositol N-acetylglucosaminyltransferase subu | -0,2727 | 0,1761 |
| TcCLB.506673.69 | hypothetical protein, conserved | -0,2727 | 0,1859 |
| TcCLB.506817.79 | kinteoplast poly(A) polymerase complex 1 subunit, putative | -0,2727 | 0,2084 |
| TcCLB.505773.10 | retrotransposon hot spot protein (RHS, pseudogene), putat | -0,2727 | 0,2132 |
| TcCLB.506559.230 | hypothetical protein, conserved | -0,2727 | 0,2248 |
| TcCLB.506955.115 | surface protease GP63 (pseudogene), putative | -0,2727 | 0,599 |
| TcCLB.509399.120 | hypothetical protein, conserved | -0,2728 | 0,01247 |
| TcCLB.510533.100 | hypothetical protein, conserved | -0,2728 | 0,1365 |
| TcCLB.504247.30 | transcription elongation factor 1 homolog (ELOF1) | -0,2728 | 0,1563 |
| TcCLB.507663.9 | hypothetical protein, conserved | -0,2728 | 0,2483 |
| TcCLB.510697.60 | mucin-associated surface protein (MASP), putative | -0,2728 | 0,3901 |
| TcCLB.504147.310 | hypothetical protein, conserved | -0,2729 | 0,01113 |
| TcCLB.508577.100 | pumilio/PUF RNA binding protein 5, putative (PUF5) | -0,2729 | 0,05568 |
| TcCLB.507511.81 | mucin TcMUCII, putative | -0,2731 | 0,08341 |
| TcCLB.508999.80 | mucin-associated surface protein (MASP), putative | -0,2731 | 0,6551 |
| TcCLB.511767.20 | dispersed gene family protein 1 (DGF-1), putative | -0,2732 | 0,1535 |
| TcCLB.507521.30 | hypothetical protein, conserved | -0,2732 | 0,1909 |
| TcCLB.420989.10 | protein phosphatase 2C, putative | -0,2732 | 0,3657 |
| TcCLB.507771.60 | hypothetical protein, conserved | -0,2733 | 0,01105 |
| TcCLB.508305.30 | hypothetical protein | -0,2733 | 0,7037 |
| TcCLB.507941.10 | hypothetical protein, conserved | -0,2734 | 0,008033 |
| TcCLB.506545.10 | hypothetical protein | -0,2735 | 0,02402 |
| TcCLB.510175.60 | dispersed gene family protein 1 (DGF-1), putative | -0,2735 | 0,1 |
| TcCLB.508027.70 | ATP-dependent RNA helicase, putative | -0,2736 | 0,01181 |
| TcCLB.509167.140 | RNA-binding protein 42 (RNA-binding motif protein 42), pu | -0,2736 | 0,02017 |
| TcCLB.506139.140 | trans-sialidase (pseudogene), putative | -0,2736 | 0,4983 |
| TcCLB.510893.9 | dispersed gene family protein 1 (DGF-1, pseudogene), puta | -0,2737 | 0,3781 |
| TcCLB.511183.190 | trans-sialidase, Group V, putative | -0,2738 | 0,4749 |
| TcCLB.511075.20 | notchless homolog, putative | -0,274 | 0,01009 |
| TcCLB.504643.10 | kinesin (pseudogene), putative | -0,2741 | 0,01276 |
| TcCLB.508295.30 | mucin TcMUCII, putative | -0,2741 | 0,5717 |
| TcCLB.506425.60 | hypothetical protein, conserved | -0,2742 | 0,05363 |

| | | | |
|---|---|---|---|
| TcCLB.507841.30 | hypothetical protein, conserved | -0,2742 | 0,2613 |
| TcCLB.506573.34 | hypothetical protein, conserved | -0,2742 | 0,4079 |
| TcCLB.508737.154 | hypothetical protein, conserved | -0,2745 | 0,2197 |
| TcCLB.506925.440 | Nuclear pore complex protein 158 (pseudogene) | -0,2746 | 0,0002553 |
| TcCLB.507257.50 | hypothetical protein, conserved | -0,2746 | 0,0004807 |
| TcCLB.509179.210 | hypothetical protein, conserved | -0,2746 | 0,06108 |
| TcCLB.511717.190 | hypothetical protein, conserved | -0,2746 | 0,08182 |
| TcCLB.503929.60 | hypothetical protein, conserved | -0,2746 | 0,1125 |
| TcCLB.509985.69 | hypothetical protein, conserved | -0,2748 | 0,2808 |
| TcCLB.510661.260 | hypothetical protein, conserved | -0,2749 | 0,007487 |
| TcCLB.508397.10 | V-type proton ATPase subunit D, putative (VAd) | -0,2749 | 0,03137 |
| TcCLB.509053.130 | DnaJ chaperone protein, putative | -0,2749 | 0,1171 |
| TcCLB.511171.10 | dispersed gene family protein 1 (DGF-1), putative | -0,2749 | 0,6012 |
| TcCLB.506795.50 | poly(A) polymerase, putative | -0,2752 | 0,008296 |
| TcCLB.509429.90 | translation initiation factor EIF-2b alpha subunit, putative | -0,2752 | 0,0905 |
| TcCLB.506513.30 | hypothetical protein, conserved | -0,2752 | 0,1646 |
| TcCLB.506177.10 | hypothetical protein, conserved | -0,2753 | 0,03205 |
| TcCLB.510719.140 | hypothetical protein, conserved | -0,2753 | 0,04219 |
| TcCLB.509581.10 | trans-sialidase, Group VII, putative | -0,2754 | 0,7084 |
| TcCLB.508741.370 | hypothetical protein, conserved | -0,2755 | 0,0004805 |
| TcCLB.507013.24 | poly(ADP-ribose) glycohydrolase, putative | -0,2755 | 0,07487 |
| TcCLB.507009.70 | hypothetical protein, conserved | -0,2757 | 0,02935 |
| TcCLB.508197.30 | hypothetical protein, conserved | -0,2758 | 0,007394 |
| TcCLB.508277.80 | hypothetical protein, conserved | -0,276 | 0,05903 |
| TcCLB.506265.160 | hypothetical protein, conserved | -0,276 | 0,1613 |
| TcCLB.503679.40 | actin-related protein 2, putative | -0,276 | 0,417 |
| TcCLB.503669.39 | hypothetical protein, conserved (fragment) | -0,2761 | 0,3509 |
| TcCLB.510287.30 | proteasome subunit beta type-2, putative | -0,2762 | 0,03648 |
| TcCLB.445011.10 | hypothetical protein, conserved | -0,2762 | 0,08542 |
| TcCLB.504013.90 | hypothetical protein, conserved | -0,2763 | 0,004927 |
| TcCLB.508989.20 | carrier protein, putative | -0,2764 | 0,08592 |
| TcCLB.504261.20 | trans-sialidase, Group VI, putative | -0,2765 | 0,4502 |
| TcCLB.507709.44 | hypothetical protein, conserved | -0,2766 | 0,04267 |
| TcCLB.506337.171 | ubiquitin (pseudogene), putative | -0,2766 | 0,2628 |
| TcCLB.503781.20 | phosphatidylinositol N-acetylglucosaminyltransferase subu | -0,2767 | 0,09975 |
| TcCLB.506925.290 | hypothetical protein | -0,2768 | 0,01507 |
| TcCLB.511165.70 | proteasome alpha 7 subunit, putative | -0,2768 | 0,02657 |
| TcCLB.509003.10 | hypothetical protein, conserved | -0,2769 | 0,003542 |
| TcCLB.508213.9 | regulator of chromosome condensation, putative | -0,2771 | 0,009382 |
| TcCLB.506419.20 | tousled-like kinase II, putative | -0,2771 | 0,03795 |
| TcCLB.510259.6 | DNA polymerase delta catalytic subunit, putative (fragmer | -0,2774 | 0,01159 |
| TcCLB.510745.40 | protein kinase, putative | -0,2774 | 0,02972 |
| TcCLB.504949.30 | succinate dehydrogenase iron-sulfur subunit (SDH2C) | -0,2774 | 0,06599 |
| TcCLB.511277.310 | hypothetical protein, conserved | -0,2775 | 0,01917 |
| TcCLB.506619.10 | sedoheptulose-1,7-bisphosphatase, putative | -0,2776 | 0,0706 |

| | | | |
|---|---|---|---|
| TcCLB.511737.80 | hypothetical protein | -0,2777 | 0,08124 |
| TcCLB.507513.10 | hypothetical protein, conserved | -0,2778 | 0,03569 |
| TcCLB.511751.163 | hypothetical protein, conserved | -0,2778 | 0,05455 |
| TcCLB.506529.170 | hypothetical protein, conserved | -0,2778 | 0,09383 |
| TcCLB.509937.30 | phosphatidylserine synthase, putative | -0,278 | 0,06502 |
| TcCLB.509527.80 | mucin-associated surface protein (MASP), putative | -0,2781 | 0,4202 |
| TcCLB.509161.30 | OSM3-like kinesin, putative | -0,2782 | 0,001076 |
| TcCLB.508895.60 | RNA-binding protein, putative | -0,2784 | 0,01346 |
| TcCLB.506287.140 | prefoldin, putative | -0,2784 | 0,05127 |
| TcCLB.507257.140 | hypothetical protein, conserved | -0,2784 | 0,07482 |
| TcCLB.505207.10 | trans-sialidase, Group VII, putative | -0,2784 | 0,2876 |
| TcCLB.503887.30 | hypothetical protein, conserved | -0,2785 | 0,02189 |
| TcCLB.508461.560 | hypothetical protein, conserved | -0,2787 | 0,03972 |
| TcCLB.506829.100 | fatty acyl CoA syntetase 1, putative | -0,2788 | 0,003911 |
| TcCLB.506857.40 | hypothetical protein, conserved | -0,2788 | 0,1162 |
| TcCLB.508707.70 | hypothetical protein, conserved | -0,2789 | 0,0007393 |
| TcCLB.509165.49 | damage-specific DNA binding protein, putative | -0,2789 | 0,04489 |
| TcCLB.510699.11 | surface protease GP63 (pseudogene), putative | -0,2789 | 0,6179 |
| TcCLB.506147.160 | hypothetical protein, conserved | -0,279 | 0,08689 |
| TcCLB.445011.19 | hypothetical protein, conserved | -0,279 | 0,09199 |
| TcCLB.510209.27 | mucin-associated surface protein (MASP), putative | -0,279 | 0,5993 |
| TcCLB.510749.74 | hypothetical protein, conserved | -0,2791 | 0,03191 |
| TcCLB.507063.80 | hypothetical protein, conserved | -0,2791 | 0,06433 |
| TcCLB.510879.30 | hypothetical protein, conserved | -0,2793 | 0,006762 |
| TcCLB.511245.50 | hypothetical protein, conserved | -0,2794 | 0,003004 |
| TcCLB.506985.10 | hypothetical protein, conserved | -0,2795 | 0,1124 |
| TcCLB.503395.10 | hypothetical protein, conserved | -0,2795 | 0,254 |
| TcCLB.506493.60 | hypothetical protein, conserved | -0,2796 | 0,01984 |
| TcCLB.506341.50 | trans-sialidase, Group VIII, putative | -0,2796 | 0,2609 |
| TcCLB.510201.10 | trans-sialidase, Group V, putative | -0,2796 | 0,3647 |
| TcCLB.506121.9 | dispersed gene family protein 1 (DGF-1), putative | -0,2796 | 0,4827 |
| TcCLB.506871.180 | G5-interacting protein, putative (G5-IP) | -0,2797 | 0,007198 |
| TcCLB.509109.40 | hypothetical protein, conserved | -0,2797 | 0,07867 |
| TcCLB.506393.166 | ATP-dependent DEAD/H RNA helicase (pseudogene), putat | -0,2798 | 0,4831 |
| TcCLB.511071.130 | Trypanosome basal body component protein (TBBC) | -0,2799 | 0,001376 |
| TcCLB.509195.50 | mucin-associated surface protein (MASP), putative | -0,2799 | 0,4619 |
| TcCLB.506983.10 | hypothetical protein, conserved | -0,28 | 0,08983 |
| TcCLB.511003.70 | hypothetical protein, conserved | -0,28 | 0,1377 |
| TcCLB.403875.10 | cysteine peptidase, Clan CA, family C51, putative | -0,2801 | 0,01259 |
| TcCLB.506305.5 | hypothetical protein, conserved | -0,2802 | 0,1367 |
| TcCLB.504121.10 | KU70 protein, putative | -0,2803 | 0,08659 |
| TcCLB.506703.110 | mucin-associated surface protein (MASP), putative | -0,2803 | 0,2967 |
| TcCLB.506131.10 | mucin TcMUCII, putative | -0,2803 | 0,6405 |
| TcCLB.511671.120 | U5 snRNP-specific 40 kDa protein, putative | -0,2805 | 0,07414 |
| TcCLB.507389.79 | hypothetical protein, conserved | -0,2805 | 0,128 |

| | | | |
|---|---|---|---|
| TcCLB.507913.20 | phosphomevalonate kinase protein, putative | -0,2807 | 0,04383 |
| TcCLB.511543.20 | As/Sb Reductase, putative (ACR2) | -0,2808 | 0,2267 |
| TcCLB.509437.80 | hypothetical protein, conserved | -0,2809 | 0,07062 |
| TcCLB.510757.20 | hypothetical protein, conserved | -0,2811 | 0,05653 |
| TcCLB.509823.4 | hypothetical protein, conserved | -0,2812 | 0,05882 |
| TcCLB.508821.10 | katanin, putative | -0,2812 | 0,14 |
| TcCLB.511725.270 | hypothetical protein, conserved | -0,2813 | 0,001687 |
| TcCLB.508919.90 | hypothetical protein, conserved | -0,2813 | 0,1138 |
| TcCLB.506583.4 | serine/threonine protein kinase, putative | -0,2813 | 0,1346 |
| TcCLB.508545.20 | hypothetical protein, conserved | -0,2814 | 0,2413 |
| TcCLB.508329.9 | potassium voltage-gated channel, putative | -0,2814 | 0,536 |
| TcCLB.506489.59 | divalent cation tolerance protein, putative | -0,2815 | 0,3234 |
| TcCLB.510187.190 | vacuolar protein sorting-associated protein, putative | -0,2816 | 0,002581 |
| TcCLB.507037.80 | hypothetical protein, conserved | -0,2816 | 0,0347 |
| TcCLB.510741.100 | hypothetical protein, conserved | -0,2816 | 0,06649 |
| TcCLB.509199.10 | Elongation factor G 2, mitochondrial, putative (EF-G2) | -0,2817 | 0,01219 |
| TcCLB.506245.73 | elongation factor 1-gamma (EF-1-gamma, pseudogene), p | -0,2817 | 0,5958 |
| TcCLB.503983.29 | hypothetical protein, conserved | -0,2818 | 0,1798 |
| TcCLB.511061.51 | mucin-associated surface protein (MASP), putative | -0,282 | 0,4173 |
| TcCLB.507787.120 | hypothetical protein, conserved | -0,2821 | 0,1755 |
| TcCLB.511449.70 | trans-sialidase, Group V, putative | -0,2821 | 0,459 |
| TcCLB.506175.10 | hypothetical protein, conserved | -0,2822 | 0,044 |
| TcCLB.510729.299 | protein kinase, putative | -0,2822 | 0,1878 |
| TcCLB.508637.120 | hypothetical protein, conserved | -0,2823 | 0,04466 |
| TcCLB.511599.120 | mucin-associated surface protein (MASP), putative | -0,2823 | 0,5312 |
| TcCLB.508623.10 | trans-sialidase (pseudogene), putative | -0,2824 | 0,00393 |
| TcCLB.508247.19 | syntaxin binding protein 1 (pseudogene), putative | -0,2824 | 0,5299 |
| TcCLB.503851.40 | hypothetical protein, conserved | -0,2826 | 0,04149 |
| TcCLB.503669.20 | hypothetical protein, conserved | -0,2826 | 0,08155 |
| TcCLB.432867.10 | hypothetical protein, conserved (fragment) | -0,2827 | 0,07744 |
| TcCLB.507517.130 | RNA editing 3' terminal uridylyl transferase 2, putative (RE | -0,2828 | 0,08272 |
| TcCLB.434945.4 | ADP-ribosylation factor, putative | -0,283 | 0,04102 |
| TcCLB.511109.90 | DnaJ chaperone protein, putative | -0,283 | 0,07617 |
| TcCLB.506959.100 | GTP-ase activating protein, putative | -0,2831 | 0,0632 |
| TcCLB.510879.229 | protein kinase, putative | -0,2832 | 0,2658 |
| TcCLB.506973.70 | dispersed gene family protein 1 (DGF-1), putative | -0,2832 | 0,6091 |
| TcCLB.507779.30 | dispersed gene family protein 1 (DGF-1), putative | -0,2833 | 0,1025 |
| TcCLB.507159.10 | Wee1-like protein kinase, putative (pseudogene, point mut | -0,2834 | 0,07413 |
| TcCLB.510729.160 | hypothetical protein, conserved | -0,2835 | 0,01248 |
| TcCLB.511511.80 | tubulin-tyrosine ligase-like protein, putative | -0,2835 | 0,03356 |
| TcCLB.511927.20 | hypothetical protein, conserved | -0,2835 | 0,1488 |
| TcCLB.506755.5 | hypothetical protein, conserved | -0,2835 | 0,3333 |
| TcCLB.507047.70 | hypothetical protein, conserved | -0,2836 | 0,2064 |
| TcCLB.509747.100 | hypothetical protein, conserved | -0,2837 | 0,001732 |
| TcCLB.506175.100 | hypothetical protein, conserved | -0,2837 | 0,01396 |

| | | | |
|---|---|---|---|
| TcCLB.507549.20 | trans-sialidase (pseudogene), putative | -0,2837 | 0,1524 |
| TcCLB.510293.50 | DnaJ chaperone protein, putative | -0,2838 | 0,01812 |
| TcCLB.509647.150 | calcium-transporting ATPase, putative | -0,2839 | 0,000473 |
| TcCLB.506155.120 | glycosyl hydrolase, putative | -0,2839 | 0,01228 |
| TcCLB.504171.30 | hypothetical protein, conserved | -0,284 | 0,01127 |
| TcCLB.509937.180 | tyrosyl-tRNA synthetase, putative | -0,2841 | 0,006489 |
| TcCLB.506331.120 | trans-sialidase (pseudogene), putative | -0,2841 | 0,2596 |
| TcCLB.503837.40 | hypothetical protein, conserved | -0,2843 | 0,2774 |
| TcCLB.506503.120 | hypothetical protein, conserved | -0,2847 | 0,0003285 |
| TcCLB.509119.39 | dynein heavy chain, putative | -0,2848 | 0,198 |
| TcCLB.510391.10 | dispersed gene family protein 1 (DGF-1), putative | -0,2848 | 0,3524 |
| TcCLB.506565.8 | RNA-binding protein, putative | -0,285 | 0,006773 |
| TcCLB.511109.80 | hypothetical protein, conserved | -0,285 | 0,1479 |
| TcCLB.503999.70 | ras-family member, GTP-binding protein, putative | -0,2851 | 0,01352 |
| TcCLB.503689.20 | hypothetical protein, conserved | -0,2851 | 0,3413 |
| TcCLB.511183.116 | trans-sialidase (pseudogene), putative | -0,2851 | 0,4857 |
| TcCLB.506591.90 | hypothetical protein, conserved | -0,2851 | 0,6618 |
| TcCLB.504155.293 | mucin-associated surface protein (MASP), putative | -0,2851 | 0,7431 |
| TcCLB.509777.90 | hypothetical protein, conserved | -0,2854 | 0,0207 |
| TcCLB.506241.80 | hypothetical protein, conserved | -0,2854 | 0,0795 |
| TcCLB.509965.420 | hypothetical protein, conserved | -0,2855 | 0,04259 |
| TcCLB.508163.330 | surface protease GP63, putative | -0,2855 | 0,5711 |
| TcCLB.511617.9 | NUP-1 protein (pseudogene), putative | -0,2857 | 0,00001108 |
| TcCLB.503991.30 | hypothetical protein, conserved | -0,2857 | 0,0166 |
| TcCLB.508789.130 | surface protease GP63 (pseudogene), putative | -0,2857 | 0,5078 |
| TcCLB.506781.19 | mucin-associated surface protein (MASP), putative | -0,2857 | 0,6027 |
| TcCLB.482369.10 | protein phosphatase 2A regulatory subunit, putative | -0,2858 | 0,01195 |
| TcCLB.506729.9 | heat shock 70 kDa protein, putative (fragment) | -0,2858 | 0,1677 |
| TcCLB.511001.18 | 40S ribosomal protein S3A, putative | -0,286 | 0,1355 |
| TcCLB.509947.20 | mucin-associated surface protein (MASP, pseudogene), pu | -0,286 | 0,7037 |
| TcCLB.503515.20 | Kinetoplastid RNA editing protein 1, putative (KREPA1) | -0,2863 | 0,006017 |
| TcCLB.506475.50 | Replication Factor C Subunit 1-related protein | -0,2863 | 0,02638 |
| TcCLB.506227.130 | calpain-like cysteine peptidase, putative | -0,2864 | 0,007694 |
| TcCLB.506127.70 | hypothetical protein | -0,2864 | 0,01485 |
| TcCLB.508789.220 | mucin TcMUCII, putative | -0,2864 | 0,5689 |
| TcCLB.506885.20 | hypothetical protein, conserved | -0,2865 | 0,008001 |
| TcCLB.508573.80 | trans-sialidase (pseudogene), putative | -0,2865 | 0,1763 |
| TcCLB.509099.70 | endo/exonuclease Mre11, putative | -0,2866 | 0,004139 |
| TcCLB.503685.10 | serine/threonine-protein kinase, putative | -0,2866 | 0,06228 |
| TcCLB.511423.110 | hypothetical protein, conserved | -0,2867 | 0,04831 |
| TcCLB.509799.90 | hypothetical protein, conserved | -0,2867 | 0,08542 |
| TcCLB.508873.280 | mucin-associated surface protein (MASP, pseudogene), pu | -0,2867 | 0,3068 |
| TcCLB.511019.99 | glutathione peroxidase, putative | -0,2868 | 0,16 |
| TcCLB.510523.10 | hypothetical protein, conserved (pseudogene) | -0,2869 | 0,02961 |
| TcCLB.509269.10 | hypothetical protein, conserved | -0,287 | 0,06365 |

| | | | |
|---|---|---|---|
| TcCLB.506695.20 | hypothetical protein, conserved | -0,287 | 0,07577 |
| TcCLB.506017.41 | N-acetyltransferase complex ARD1 subunit (pseudogene), | -0,2871 | 0,3369 |
| TcCLB.509437.106 | trans-sialidase (pseudogene), putative | -0,2871 | 0,5124 |
| TcCLB.506835.140 | hypothetical protein, conserved | -0,2872 | 0,006046 |
| TcCLB.506577.70 | hypothetical protein, conserved | -0,2872 | 0,03977 |
| TcCLB.510547.30 | trans-sialidase (pseudogene), putative | -0,2872 | 0,4085 |
| TcCLB.506563.10 | pumilio/PUF RNA binding protein 9, putative (PUF9) | -0,2873 | 0,02205 |
| TcCLB.507023.240 | hypothetical protein, conserved | -0,2873 | 0,05679 |
| TcCLB.510761.22 | hypothetical protein, conserved | -0,2874 | 0,01347 |
| TcCLB.511151.39 | hypothetical protein, conserved | -0,2874 | 0,07911 |
| TcCLB.508625.140 | hypothetical protein, conserved | -0,2874 | 0,1218 |
| TcCLB.508539.20 | hypothetical protein, conserved | -0,2874 | 0,2066 |
| TcCLB.505949.20 | mucin-associated surface protein (MASP), putative | -0,2874 | 0,3786 |
| TcCLB.510223.9 | hypothetical protein, conserved | -0,2875 | 0,5294 |
| TcCLB.504643.49 | hypothetical protein, conserved (fragment) | -0,2876 | 0,005125 |
| TcCLB.503637.30 | serine peptidase, Clan SC, Family S9B | -0,2876 | 0,006032 |
| TcCLB.507513.80 | hypothetical protein, conserved | -0,2876 | 0,01225 |
| TcCLB.506793.50 | hypothetical protein, conserved | -0,2877 | 0,0575 |
| TcCLB.509121.10 | retrotransposon hot spot protein (RHS, pseudogene), putat | -0,2877 | 0,3233 |
| TcCLB.506113.86 | N-acetyltransferase complex ARD1 subunit (pseudogene), | -0,2878 | 0,459 |
| TcCLB.508175.133 | hypothetical protein, conserved (pseudogene) | -0,2879 | 0,003477 |
| TcCLB.506127.140 | hypothetical protein | -0,2879 | 0,006071 |
| TcCLB.504213.70 | hypothetical protein, conserved | -0,2879 | 0,02544 |
| TcCLB.507737.30 | hypothetical protein, conserved | -0,2879 | 0,05533 |
| TcCLB.511111.50 | ubiquitin-conjugating enzyme, putative | -0,2879 | 0,5313 |
| TcCLB.507195.60 | hypothetical protein | -0,2879 | 0,6648 |
| TcCLB.511277.539 | hypothetical protein, conserved | -0,288 | 0,1359 |
| TcCLB.503989.50 | hypothetical protein, conserved | -0,2883 | 0,001752 |
| TcCLB.503653.30 | ribosomal RNA processing protein 41B, putative (RRP41B) | -0,2883 | 0,06348 |
| TcCLB.507549.40 | trans-sialidase (pseudogene), putative | -0,2883 | 0,1876 |
| TcCLB.511017.20 | 2,3-bisphosphoglycerate-independent phosphoglycerate m | -0,2884 | 0,01416 |
| TcCLB.507747.130 | trans-sialidase (pseudogene), putative | -0,2885 | 0,2917 |
| TcCLB.509537.40 | hypothetical protein, conserved | -0,2886 | 0,01274 |
| TcCLB.507757.70 | dihydrolipoamide dehydrogenase, putative | -0,2886 | 0,1324 |
| TcCLB.506525.20 | protein transport protein SEC13, putative | -0,2887 | 0,03059 |
| TcCLB.511627.100 | TPR-repeat protein, putative | -0,2888 | 0,02002 |
| TcCLB.511337.20 | UDP-Gal or UDP-GlcNAc-dependent glycosyltransferase, pu | -0,2888 | 0,5104 |
| TcCLB.507633.59 | mucin-associated surface protein (MASP, pseudogene), pu | -0,2888 | 0,5425 |
| TcCLB.503557.30 | hypothetical protein, conserved | -0,2889 | 0,05157 |
| TcCLB.510767.50 | hypothetical protein, conserved | -0,2889 | 0,09288 |
| TcCLB.507007.50 | hypothetical protein, conserved | -0,289 | 0,0358 |
| TcCLB.503583.70 | hypothetical protein, conserved | -0,289 | 0,0533 |
| TcCLB.508131.19 | hypothetical protein, conserved (fragment) | -0,289 | 0,06279 |
| TcCLB.510889.330 | hypothetical protein, conserved | -0,2891 | 0,1008 |
| TcCLB.426397.9 | kinesin, putative (fragment) | -0,2892 | 0,0341 |

| | | | |
|---|---|---|---|
| TcCLB.510313.20 | hypothetical protein, conserved | -0,2893 | 0,007929 |
| TcCLB.508293.60 | trans-sialidase, Group VI, putative | -0,2893 | 0,3995 |
| TcCLB.503955.70 | eukaryotic translation initiation factor 2 beta subunit, puta | -0,2894 | 0,02135 |
| TcCLB.508189.11 | retrotransposon hot spot protein (RHS, pseudogene), putat | -0,2894 | 0,5238 |
| TcCLB.506679.180 | hypothetical protein, conserved | -0,2895 | 0,007208 |
| TcCLB.509671.30 | hypothetical protein, conserved | -0,2895 | 0,06927 |
| TcCLB.511761.20 | hypothetical protein, conserved | -0,2896 | 0,1385 |
| TcCLB.510399.39 | dispersed gene family protein 1 (DGF-1), putative | -0,2896 | 0,3776 |
| TcCLB.510625.150 | mucin-associated surface protein (MASP), putative | -0,2898 | 0,7117 |
| TcCLB.511303.9 | hypothetical protein, conserved | -0,2899 | 0,08644 |
| TcCLB.507711.140 | hypothetical protein, conserved | -0,2899 | 0,103 |
| TcCLB.507959.240 | syntaxin binding protein (pseudogene), putative | -0,2899 | 0,5007 |
| TcCLB.506529.140 | hypothetical protein, conserved | -0,29 | 0,07867 |
| TcCLB.511183.103 | mucin-associated surface protein (MASP, pseudogene), pu | -0,2902 | 0,4593 |
| TcCLB.505949.139 | mucin-associated surface protein (MASP, pseudogene), pu | -0,2903 | 0,5357 |
| TcCLB.503959.20 | hypothetical protein, conserved | -0,2904 | 0,01263 |
| TcCLB.507521.120 | hypothetical protein, conserved | -0,2904 | 0,1451 |
| TcCLB.506133.54 | mucin-associated surface protein (MASP), putative | -0,2906 | 0,4686 |
| TcCLB.511439.120 | aminopeptidase, putative | -0,2907 | 0,004017 |
| TcCLB.507075.30 | trans-sialidase (pseudogene), putative | -0,2907 | 0,2254 |
| TcCLB.506961.25 | trans-sialidase, Group I, putative | -0,2908 | 0,108 |
| TcCLB.510133.9 | dispersed gene family protein 1 (DGF-1), putative (fragme | -0,2908 | 0,4234 |
| TcCLB.510259.18 | GlcNAc-PI synthesis protein (GPI3) | -0,2909 | 0,09539 |
| TcCLB.508409.80 | hypothetical protein, conserved | -0,291 | 0,0003756 |
| TcCLB.507031.150 | hypothetical protein, conserved | -0,291 | 0,001062 |
| TcCLB.507941.40 | hypothetical protein, conserved | -0,291 | 0,04025 |
| TcCLB.508547.60 | hypothetical protein, conserved | -0,2911 | 0,06685 |
| TcCLB.506737.140 | syntaxin binding protein 1, putative | -0,2912 | 0,00052 |
| TcCLB.504071.90 | glycosyltransferase family 28 protein, putative | -0,2912 | 0,02206 |
| TcCLB.504835.10 | hypothetical protein, conserved | -0,2912 | 0,04291 |
| TcCLB.510515.70 | hypothetical protein, conserved | -0,2912 | 0,2474 |
| TcCLB.509317.40 | ribosomal protein L11, putative | -0,2914 | 0,1031 |
| TcCLB.507021.90 | hypothetical protein, conserved | -0,2915 | 0,03876 |
| TcCLB.506537.50 | retrotransposon hot spot (RHS) protein, putative | -0,2915 | 0,4188 |
| TcCLB.511231.14 | hypothetical protein (fragment) | -0,2916 | 0,03386 |
| TcCLB.509253.10 | hypothetical protein, conserved | -0,2916 | 0,04446 |
| TcCLB.509595.70 | Trichohyalin, putative | -0,2919 | 0,01494 |
| TcCLB.504243.10 | RNA-binding protein, putative | -0,2919 | 0,02097 |
| TcCLB.507959.336 | surface protease GP63 (pseudogene), putative | -0,2919 | 0,3604 |
| TcCLB.469435.4 | ribosomal protein L14, putative | -0,292 | 0,1693 |
| TcCLB.510303.80 | kinesin, putative | -0,2921 | 0,02788 |
| TcCLB.503651.49 | hypothetical protein, conserved (pseudogene) | -0,2921 | 0,07978 |
| TcCLB.504923.10 | dispersed gene family protein 1 (DGF-1), putative | -0,2921 | 0,6139 |
| TcCLB.506347.10 | dynein heavy chain (pseudogene), putative | -0,2922 | 0,00007664 |
| TcCLB.504225.10 | hypothetical protein, conserved | -0,2922 | 0,02302 |

| | | | |
|---|---|---|---|
| TcCLB.508601.110 | hypothetical protein, conserved | -0,2923 | 0,002523 |
| TcCLB.510943.120 | hypothetical protein, conserved | -0,2923 | 0,00664 |
| TcCLB.503513.40 | proteasome regulatory non-ATPase subunit 6, putative | -0,2923 | 0,02865 |
| TcCLB.431357.20 | UDP-Gal or UDP-GlcNAc-dependent glycosyltransferase, pu | -0,2923 | 0,6017 |
| TcCLB.507007.80 | hypothetical protein | -0,2924 | 0,001524 |
| TcCLB.509905.180 | mucin-associated surface protein (MASP), putative | -0,2925 | 0,1983 |
| TcCLB.511367.90 | hypothetical protein, conserved | -0,2926 | 0,004985 |
| TcCLB.504153.300 | GTPase protein, putative | -0,2927 | 0,01006 |
| TcCLB.511419.50 | 3-phosphoglycerate kinase, glycosomal (PGKA) | -0,2927 | 0,1006 |
| TcCLB.508139.210 | dispersed gene family protein 1 (DGF-1), putative | -0,2927 | 0,1117 |
| TcCLB.469785.5 | hypothetical protein, conserved (fragment) | -0,2928 | 0,2245 |
| TcCLB.510763.40 | hypothetical protein, conserved | -0,293 | 0,08721 |
| TcCLB.506933.80 | RNA polymerase B subunit RPB8, putative | -0,2931 | 0,01397 |
| TcCLB.510475.120 | mucin-associated surface protein (MASP, pseudogene), pu | -0,2931 | 0,7404 |
| TcCLB.504431.118 | hypothetical protein, conserved | -0,2934 | 0,03432 |
| TcCLB.508271.50 | hypothetical protein | -0,2935 | 0,009674 |
| TcCLB.506895.20 | hypothetical protein, conserved | -0,2935 | 0,01032 |
| TcCLB.511071.190 | hypothetical protein, conserved | -0,2936 | 0,008463 |
| TcCLB.506933.20 | ribonucleoside-diphosphate reductase large chain, putative | -0,2937 | 0,007006 |
| TcCLB.510143.90 | hypothetical protein, conserved | -0,2938 | 0,08078 |
| TcCLB.507089.130 | hypothetical protein, conserved | -0,2939 | 0,0322 |
| TcCLB.511277.30 | hypothetical protein, conserved (pseudogene) | -0,294 | 0,00293 |
| TcCLB.508839.80 | hypothetical protein, conserved | -0,294 | 0,01008 |
| TcCLB.508349.30 | hypothetical protein, conserved | -0,2943 | 0,0002238 |
| TcCLB.510659.10 | hypothetical protein, conserved | -0,2943 | 0,01335 |
| TcCLB.509647.120 | hypothetical protein | -0,2944 | 0,04127 |
| TcCLB.511709.10 | hypothetical protein, conserved | -0,2945 | 0,04496 |
| TcCLB.511857.59 | katanin, putative | -0,2945 | 0,05631 |
| TcCLB.510183.30 | hypothetical protein, conserved | -0,2946 | 0,004542 |
| TcCLB.511461.10 | hypothetical protein, conserved | -0,2946 | 0,07964 |
| TcCLB.509965.350 | ubiquitin-protein ligase, putative | -0,2948 | 0,001562 |
| TcCLB.510187.110 | Mitochondrial ribosomal protein S18, putative (MRPS18) | -0,2949 | 0,003949 |
| TcCLB.509505.40 | hypothetical protein, conserved | -0,2949 | 0,03074 |
| TcCLB.509575.50 | Casein kinase II subunit beta, putative | -0,2949 | 0,1089 |
| TcCLB.507877.10 | mucin-associated surface protein (MASP, pseudogene), pu | -0,2949 | 0,5356 |
| TcCLB.506583.30 | ubiquitin-like protein, putative | -0,2952 | 0,0006914 |
| TcCLB.453767.20 | trans-sialidase (pseudogene), putative | -0,2952 | 0,3534 |
| TcCLB.506265.150 | hypothetical protein, conserved | -0,2953 | 0,1696 |
| TcCLB.507063.20 | eukaryotic translation initiation factor 2 beta subunit, puta | -0,2954 | 0,003876 |
| TcCLB.511277.480 | hypothetical protein, conserved | -0,2954 | 0,01779 |
| TcCLB.506407.20 | hypothetical protein, conserved | -0,2954 | 0,09443 |
| TcCLB.506975.69 | hypothetical protein, conserved | -0,2954 | 0,1277 |
| TcCLB.504167.7 | hypothetical protein, conserved | -0,2954 | 0,2138 |
| TcCLB.422843.10 | trans-sialidase (pseudogene), putative | -0,2955 | 0,09374 |
| TcCLB.506451.20 | trans-sialidase, Group V, putative | -0,2955 | 0,2955 |

| | | | |
|---|---|---|---|
| TcCLB.503893.181 | hypothetical protein, conserved (fragment) | -0,2955 | 0,4026 |
| TcCLB.504031.40 | hypothetical protein | -0,2955 | 0,5427 |
| TcCLB.508873.160 | mucin-associated surface protein (MASP), putative | -0,2955 | 0,6593 |
| TcCLB.506863.70 | hypothetical protein, conserved | -0,2957 | 0,008217 |
| TcCLB.509875.190 | clathrin coat assembly protein, putative | -0,2957 | 0,008269 |
| TcCLB.509945.24 | hypothetical protein, conserved | -0,2957 | 0,1066 |
| TcCLB.397937.5 | N-acetylglucosamine-6-phosphate deacetylase-like proteir | -0,2957 | 0,1119 |
| TcCLB.504125.50 | mitochondrial carrier protein, putative | -0,2959 | 0,02483 |
| TcCLB.510121.40 | hypothetical protein | -0,2959 | 0,07987 |
| TcCLB.509033.10 | trans-sialidase (pseudogene), putative | -0,2959 | 0,3519 |
| TcCLB.506859.90 | prefoldin, putative | -0,296 | 0,0213 |
| TcCLB.505939.10 | hypothetical protein | -0,296 | 0,5266 |
| TcCLB.510421.210 | hypothetical protein, conserved | -0,2961 | 0,004429 |
| TcCLB.506831.50 | hypothetical protein, conserved | -0,2961 | 0,004863 |
| TcCLB.510885.70 | hypothetical protein, conserved | -0,2961 | 0,04271 |
| TcCLB.511751.144 | hypothetical protein, conserved | -0,2962 | 0,01034 |
| TcCLB.507609.30 | ATP-dependent DEAD/H RNA helicase, putative | -0,2962 | 0,04004 |
| TcCLB.508153.1120 | DNA repair helicase, putative | -0,2964 | 0,01501 |
| TcCLB.511885.35 | trans-sialidase (pseudogene), putative | -0,2964 | 0,2729 |
| TcCLB.510377.439 | mucin TcMUCII, putative | -0,2964 | 0,5267 |
| TcCLB.509827.10 | hypothetical protein, conserved | -0,2965 | 0,02419 |
| TcCLB.507511.30 | RNA guanylyltransferase, putative | -0,2966 | 0,01756 |
| TcCLB.508817.140 | hypothetical protein, conserved | -0,2969 | 0,002791 |
| TcCLB.506627.50 | mitochondrial exoribonuclease DSS-1, putative | -0,2969 | 0,004683 |
| TcCLB.510661.40 | hypothetical protein, conserved | -0,2969 | 0,005119 |
| TcCLB.504071.60 | hypothetical protein, conserved | -0,2969 | 0,01603 |
| TcCLB.508207.279 | hypothetical protein, conserved | -0,2969 | 0,1083 |
| TcCLB.507041.85 | hypothetical protein, conserved (pseudogene) | -0,2969 | 0,1432 |
| TcCLB.509315.10 | hypothetical protein, conserved (pseudogene) | -0,2969 | 0,4441 |
| TcCLB.506865.40 | guanine nucleotide exchange factor subunit GCD7 | -0,2972 | 0,01086 |
| TcCLB.508409.20 | hypothetical protein, conserved | -0,2972 | 0,01296 |
| TcCLB.507747.260 | hypothetical protein, conserved | -0,2972 | 0,0184 |
| TcCLB.511625.50 | GTP-binding protein, putative | -0,2974 | 0,003407 |
| TcCLB.507019.40 | histidyl-tRNA synthetase | -0,2975 | 0,00116 |
| TcCLB.508153.960 | hypothetical protein, conserved | -0,2975 | 0,02282 |
| TcCLB.503751.20 | hypothetical protein | -0,2975 | 0,3586 |
| TcCLB.504103.40 | hypothetical protein, conserved | -0,2976 | 0,05004 |
| TcCLB.508463.10 | hypothetical protein, conserved | -0,2976 | 0,2236 |
| TcCLB.506909.120 | hypothetical protein | -0,2976 | 0,6313 |
| TcCLB.511157.9 | neurobeachin/beige protein, putative (fragment) | -0,2977 | 0,005895 |
| TcCLB.508879.220 | hypothetical protein, conserved | -0,2978 | 0,01624 |
| TcCLB.509979.119 | hypothetical protein, conserved | -0,2978 | 0,1006 |
| TcCLB.509933.10 | hypothetical protein, conserved | -0,2978 | 0,1944 |
| TcCLB.511303.30 | hypothetical protein, conserved | -0,298 | 0,03457 |
| TcCLB.506979.59 | hypothetical protein, conserved | -0,298 | 0,1012 |

| | | | |
|---|---|---|---|
| TcCLB.506361.44 | mucin TcMUC (pseudogene), putative | -0,298 | 0,342 |
| TcCLB.511491.40 | hypothetical protein, conserved | -0,2981 | 0,001683 |
| TcCLB.505123.4 | ATP-dependent RNA helicase, putative (fragment) | -0,2981 | 0,0125 |
| TcCLB.506493.110 | hypothetical protein, conserved | -0,2981 | 0,1684 |
| TcCLB.508107.30 | trans-sialidase, Group VI, putative | -0,2982 | 0,5576 |
| TcCLB.505163.30 | hypothetical protein, conserved | -0,2983 | 0,01557 |
| TcCLB.507823.10 | hypothetical protein, conserved | -0,2983 | 0,03775 |
| TcCLB.503489.30 | hypothetical protein, conserved | -0,2983 | 0,05567 |
| TcCLB.509715.130 | hypothetical protein, conserved | -0,2984 | 0,04414 |
| TcCLB.510205.15 | mucin TcMUCII (pseudogene), putative | -0,2984 | 0,5076 |
| TcCLB.508647.200 | triosephosphate isomerase, putative (TIM) | -0,2985 | 0,00002167 |
| TcCLB.507165.10 | hypothetical protein, conserved | -0,2985 | 0,07159 |
| TcCLB.511491.90 | hypothetical protein | -0,2986 | 0,07804 |
| TcCLB.506717.200 | hypothetical protein, conserved | -0,2986 | 0,1823 |
| TcCLB.506605.10 | hypothetical protein, conserved | -0,2987 | 0,1314 |
| TcCLB.507803.49 | histone deacetylase, putative | -0,2987 | 0,2477 |
| TcCLB.509395.10 | hypothetical protein, conserved | -0,2988 | 0,02424 |
| TcCLB.511229.90 | hypothetical protein, conserved | -0,2988 | 0,02526 |
| TcCLB.507735.60 | hypothetical protein, conserved | -0,2989 | 0,01107 |
| TcCLB.509455.110 | hypothetical protein, conserved | -0,2989 | 0,09774 |
| TcCLB.508593.120 | hypothetical protein, conserved | -0,2989 | 0,09947 |
| TcCLB.508857.140 | hypothetical protein, conserved | -0,2991 | 0,01953 |
| TcCLB.509161.20 | hypothetical protein, conserved | -0,2992 | 0,08824 |
| TcCLB.511235.10 | hypothetical protein, conserved (pseudogene) | -0,2992 | 0,4063 |
| TcCLB.506357.40 | Elongation factor Tu, mitochondrial, putative (EF-Tu) | -0,2993 | 0,001341 |
| TcCLB.510187.260 | hypothetical protein, conserved | -0,2993 | 0,03068 |
| TcCLB.507969.50 | actin 2, putative | -0,2993 | 0,1051 |
| TcCLB.508455.20 | trans-sialidase, Group V, putative | -0,2993 | 0,3425 |
| TcCLB.507019.100 | nuclear movement protein, putative | -0,2994 | 0,02607 |
| TcCLB.511859.161 | hypothetical protein | -0,2996 | 0,1836 |
| TcCLB.507001.4 | hypothetical protein, conserved | -0,2996 | 0,3263 |
| TcCLB.506959.50 | hypothetical protein, conserved | -0,2997 | 0,01715 |
| TcCLB.506887.40 | hypothetical protein, conserved | -0,2997 | 0,03112 |
| TcCLB.511289.59 | Multiprotein-bridging factor 1, putative (mbf1) | -0,2998 | 0,00884 |
| TcCLB.506753.220 | hypothetical protein, conserved | -0,2998 | 0,2638 |
| TcCLB.506525.30 | hypothetical protein, conserved | -0,3 | 0,01267 |
| TcCLB.509875.204 | hypothetical protein, conserved | -0,3001 | 0,05499 |
| TcCLB.480653.10 | hypothetical protein, conserved | -0,3002 | 0,09336 |
| TcCLB.508637.70 | RNA helicase, putative | -0,3003 | 0,006067 |
| TcCLB.509611.10 | DNA repair and recombination helicase protein PIF7, putat | -0,3004 | 0,06044 |
| TcCLB.509741.34 | hypothetical protein, conserved | -0,3005 | 0,1645 |
| TcCLB.506635.120 | kinesin, putative | -0,3006 | 0,01366 |
| TcCLB.508479.83 | N-acetyltransferase complex ARD1 subunit (pseudogene), | -0,3006 | 0,4383 |
| TcCLB.510329.260 | hypothetical protein, conserved | -0,3007 | 0,000169 |
| TcCLB.503885.50 | hypothetical protein, conserved | -0,3007 | 0,03612 |

| | | | |
|---|---|---|---|
| TcCLB.507839.35 | trans-sialidase, putative | -0,3007 | 0,3178 |
| TcCLB.506945.270 | cyclin, putative | -0,3008 | 0,01594 |
| TcCLB.506791.20 | hypothetical protein, conserved | -0,3009 | 0,01423 |
| TcCLB.510029.10 | hypothetical protein, conserved (pseudogene) | -0,3009 | 0,1153 |
| TcCLB.504051.49 | hypothetical protein, conserved | -0,3009 | 0,1238 |
| TcCLB.506529.160 | hypothetical protein, conserved | -0,301 | 0,02261 |
| TcCLB.507709.110 | hypothetical protein, conserved | -0,301 | 0,2 |
| TcCLB.503893.50 | hypothetical protein, conserved | -0,3011 | 0,04628 |
| TcCLB.503495.10 | dispersed gene family protein 1 (DGF-1), putative | -0,3011 | 0,2014 |
| TcCLB.510143.70 | hypothetical protein, conserved | -0,3013 | 0,0009447 |
| TcCLB.508995.30 | tubulin-tyrosine ligase, putative | -0,3013 | 0,03075 |
| TcCLB.507031.10 | hypothetical protein, conserved | -0,3013 | 0,03631 |
| TcCLB.434259.19 | hypothetical protein, conserved | -0,3013 | 0,0931 |
| TcCLB.506743.90 | hypothetical protein, conserved | -0,3013 | 0,236 |
| TcCLB.510301.20 | fructose-bisphosphate aldolase, glycosomal, putative | -0,3014 | 0,01162 |
| TcCLB.511283.190 | U2 small nuclear ribonucleoprotein 40K, putative | -0,3014 | 0,01284 |
| TcCLB.508637.160 | dual specificity protein phosphatase, putative | -0,3014 | 0,04437 |
| TcCLB.509213.60 | hypothetical protein, conserved | -0,3014 | 0,1334 |
| TcCLB.506525.45 | hypothetical protein, conserved (pseudogene) | -0,3014 | 0,2754 |
| TcCLB.510577.40 | hypothetical protein, conserved | -0,3016 | 0,04678 |
| TcCLB.507987.20 | hypothetical protein | -0,3016 | 0,6635 |
| TcCLB.510533.220 | hypothetical protein, conserved | -0,3017 | 0,01541 |
| TcCLB.510359.450 | hypothetical protein | -0,3017 | 0,58 |
| TcCLB.507517.110 | hypothetical protein, conserved | -0,3018 | 0,0636 |
| TcCLB.506495.10 | hypothetical protein, conserved | -0,3018 | 0,08759 |
| TcCLB.511211.120 | receptor for activated C kinase 1, putative | -0,3019 | 0,00002057 |
| TcCLB.508899.130 | hypothetical protein, conserved | -0,3019 | 0,1543 |
| TcCLB.505989.74 | peroxisome biogenesis factor 1, putative | -0,3021 | 0,004066 |
| TcCLB.510029.40 | DNA-directed RNA polymerase III subunit C11, putative (RI | -0,3021 | 0,09598 |
| TcCLB.509135.9 | hypothetical protein (fragment) | -0,3021 | 0,1594 |
| TcCLB.509569.90 | ARP2/3 complex subunit, putative | -0,3023 | 0,06824 |
| TcCLB.504185.9 | hypothetical protein, conserved | -0,3023 | 0,3233 |
| TcCLB.510149.110 | phosphate transporter, putative | -0,3024 | 0,01766 |
| TcCLB.506201.154 | hypothetical protein, conserved | -0,3024 | 0,113 |
| TcCLB.510013.150 | mucin-associated surface protein (MASP, pseudogene), pu | -0,3024 | 0,4058 |
| TcCLB.509627.40 | succinate dehydrogenase subunit (SDH9) | -0,3025 | 0,02409 |
| TcCLB.511589.29 | hypothetical protein, conserved | -0,3025 | 0,06173 |
| TcCLB.511179.114 | hypothetical protein, conserved | -0,3025 | 0,1229 |
| TcCLB.506131.161 | trans-sialidase (pseudogene), putative | -0,3025 | 0,4513 |
| TcCLB.511145.24 | hypothetical protein, conserved | -0,3026 | 0,2568 |
| TcCLB.508543.150 | hypothetical protein, conserved | -0,3027 | 0,001554 |
| TcCLB.507099.20 | hypothetical protein, conserved | -0,3028 | 0,006031 |
| TcCLB.510105.160 | hypothetical protein | -0,3029 | 0,07268 |
| TcCLB.503855.10 | hypothetical protein, conserved | -0,303 | 0,1072 |
| TcCLB.511417.94 | hypothetical protein, conserved | -0,3031 | 0,09171 |

| | | | |
|---|---|---|---|
| TcCLB.511643.20 | dihydroorotate dehydrogenase, putative (DHODH) | -0,3034 | 0,02806 |
| TcCLB.508165.40 | trans-sialidase (pseudogene), putative | -0,3034 | 0,3733 |
| TcCLB.510499.10 | dispersed gene family protein 1 (DGF-1), putative | -0,3034 | 0,4797 |
| TcCLB.510595.49 | hypothetical protein, conserved | -0,3035 | 0,02697 |
| TcCLB.511249.40 | hypothetical protein, conserved | -0,3035 | 0,0293 |
| TcCLB.508277.180 | mismatch repair protein MSH3, putative | -0,3036 | 0,0004932 |
| TcCLB.506709.40 | hypothetical protein, conserved | -0,3037 | 0,05889 |
| TcCLB.506363.20 | hypothetical protein, conserved | -0,3037 | 0,08962 |
| TcCLB.509319.100 | hypothetical protein, conserved | -0,304 | 0,03557 |
| TcCLB.511055.20 | retrotransposon hot spot (RHS) protein, putative | -0,3041 | 0,000459 |
| TcCLB.504047.70 | hypothetical protein, conserved | -0,3041 | 0,008206 |
| TcCLB.506479.80 | hypothetical protein, conserved | -0,3041 | 0,0358 |
| TcCLB.452895.10 | hypothetical protein, conserved | -0,3041 | 0,1086 |
| TcCLB.511507.60 | hypothetical protein | -0,3041 | 0,1103 |
| TcCLB.508719.40 | kinetoplast DNA-associated protein, putative | -0,3042 | 0,01873 |
| TcCLB.511577.144 | hypothetical protein, conserved | -0,3042 | 0,09758 |
| TcCLB.504239.380 | hypothetical protein | -0,3042 | 0,6291 |
| TcCLB.506625.200 | ribosomal RNA processing protein 40, putative (RRP40) | -0,3043 | 0,07103 |
| TcCLB.507011.110 | hypothetical protein | -0,3044 | 0,06262 |
| TcCLB.509395.59 | hypothetical protein, conserved | -0,3044 | 0,1577 |
| TcCLB.506627.100 | hypothetical protein, conserved | -0,3045 | 0,07779 |
| TcCLB.507011.80 | zinc finger protein, putative | -0,3046 | 0,03498 |
| TcCLB.507443.14 | RNA polymerase IIA largest subunit, putative (fragment) | -0,3046 | 0,04305 |
| TcCLB.508329.19 | trans-sialidase (pseudogene), putative | -0,3047 | 0,4495 |
| TcCLB.507093.60 | hypothetical protein, conserved | -0,3048 | 0,004683 |
| TcCLB.511203.20 | surface protease GP63, putative | -0,3048 | 0,1439 |
| TcCLB.506841.4 | hypothetical protein, conserved | -0,305 | 0,2158 |
| TcCLB.507837.10 | Peptidase C50 separase, putative | -0,3051 | 0,04305 |
| TcCLB.508815.40 | hypothetical protein, conserved | -0,3051 | 0,05593 |
| TcCLB.510663.64 | mitochondrial carrier protein, putative | -0,3051 | 0,08285 |
| TcCLB.481729.10 | hypothetical protein, conserved | -0,3052 | 0,06 |
| TcCLB.511369.5 | elongation factor 1-alpha, putative (fragment) | -0,3052 | 0,5197 |
| TcCLB.510955.60 | hypothetical protein, conserved | -0,3053 | 0,01824 |
| TcCLB.509895.60 | hypothetical protein, conserved | -0,3055 | 0,2158 |
| TcCLB.508257.110 | hypothetical protein, conserved | -0,3057 | 0,004724 |
| TcCLB.507875.180 | AGP2beta-2, putative | -0,3057 | 0,006216 |
| TcCLB.511277.620 | hypothetical protein, conserved | -0,3057 | 0,009633 |
| TcCLB.507677.10 | hypothetical protein, conserved | -0,3057 | 0,01296 |
| TcCLB.510289.99 | hypothetical protein, conserved | -0,3058 | 0,1091 |
| TcCLB.510659.80 | hypothetical protein, conserved | -0,3059 | 0,02238 |
| TcCLB.508153.220 | hypothetical protein, conserved | -0,306 | 0,02308 |
| TcCLB.506223.84 | hypothetical protein, conserved | -0,306 | 0,0427 |
| TcCLB.504027.10 | hypothetical protein, conserved | -0,306 | 0,07212 |
| TcCLB.508153.810 | hypothetical protein, conserved | -0,3061 | 0,09376 |
| TcCLB.457979.10 | mucin-associated surface protein (MASP), putative | -0,3061 | 0,6407 |

| | | | |
|---|---|---|---|
| TcCLB.507669.60 | hypothetical protein, conserved | -0,3062 | 0,02522 |
| TcCLB.511771.185 | hypothetical protein | -0,3062 | 0,3103 |
| TcCLB.507979.50 | hypothetical protein, conserved | -0,3063 | 0,005868 |
| TcCLB.511401.24 | hypothetical protein, conserved (pseudogene) | -0,3063 | 0,1215 |
| TcCLB.508181.70 | hypothetical protein, conserved | -0,3064 | 0,001086 |
| TcCLB.504149.60 | hypothetical protein | -0,3065 | 0,001849 |
| TcCLB.511367.190 | hypothetical protein, conserved | -0,3065 | 0,005524 |
| TcCLB.507023.170 | hypothetical protein, conserved | -0,3065 | 0,01264 |
| TcCLB.505037.157 | trans-sialidase (pseudogene), putative | -0,3065 | 0,03754 |
| TcCLB.508221.880 | hypothetical protein | -0,3065 | 0,6703 |
| TcCLB.511445.110 | hypothetical protein, conserved | -0,3066 | 0,07424 |
| TcCLB.510747.170 | hypothetical protein, conserved | -0,3066 | 0,1629 |
| TcCLB.508277.140 | phosphomevalonate kinase protein, putative | -0,3067 | 0,0007878 |
| TcCLB.511751.14 | hypothetical protein, conserved (fragment) | -0,3067 | 0,05397 |
| TcCLB.506175.30 | hypothetical protein, conserved | -0,3067 | 0,2183 |
| TcCLB.510509.50 | mitochondrial RNA binding protein, putative | -0,3068 | 0,01729 |
| TcCLB.508797.20 | hypothetical protein, conserved | -0,3068 | 0,1748 |
| TcCLB.510355.104 | ATP-dependent DEAD/H RNA helicase (pseudogene), putat | -0,3068 | 0,4119 |
| TcCLB.506975.60 | hypothetical protein, conserved | -0,3069 | 0,003306 |
| TcCLB.506337.140 | hypothetical protein, conserved | -0,3069 | 0,0112 |
| TcCLB.508531.10 | glycine dehydrogenase (pseudogene), putative | -0,3069 | 0,2965 |
| TcCLB.508543.80 | hypothetical protein, conserved | -0,307 | 0,05593 |
| TcCLB.508823.130 | hypothetical protein, conserved | -0,3071 | 0,0008967 |
| TcCLB.509171.70 | hypothetical protein, conserved | -0,3071 | 0,03724 |
| TcCLB.509395.40 | hypothetical protein, conserved | -0,3071 | 0,05417 |
| TcCLB.511461.40 | hypothetical protein, conserved | -0,3072 | 0,01784 |
| TcCLB.507649.60 | hypothetical protein, conserved | -0,3073 | 0,0291 |
| TcCLB.504867.114 | tyrosine phosphatase, putative | -0,3073 | 0,04399 |
| TcCLB.511439.80 | hypothetical protein, conserved | -0,3074 | 0,0009007 |
| TcCLB.505999.70 | hypothetical protein, conserved | -0,3076 | 0,01857 |
| TcCLB.510243.110 | hypothetical protein, conserved | -0,3078 | 0,05187 |
| TcCLB.510329.200 | UDP-GlcNAc:PI a1-6 GlcNAc-transferase, putative | -0,308 | 0,001341 |
| TcCLB.506679.60 | hypothetical protein, conserved | -0,308 | 0,02323 |
| TcCLB.511089.70 | mucin-associated surface protein (MASP, pseudogene), pu | -0,308 | 0,3205 |
| TcCLB.506575.50 | hypothetical protein, conserved | -0,3082 | 0,0008592 |
| TcCLB.506859.49 | hypothetical protein, conserved | -0,3082 | 0,004502 |
| TcCLB.510123.40 | hypothetical protein, conserved | -0,3082 | 0,2156 |
| TcCLB.510225.40 | hypothetical protein, conserved | -0,3084 | 0,006806 |
| TcCLB.504001.6 | protein kinase, putative | -0,3084 | 0,009481 |
| TcCLB.509033.40 | hypothetical protein, conserved | -0,3085 | 0,01834 |
| TcCLB.508251.50 | mucin-associated surface protein (MASP), putative (fragm | -0,3085 | 0,6663 |
| TcCLB.511389.50 | hypothetical protein, conserved | -0,3086 | 0,001181 |
| TcCLB.505365.50 | beta galactofuranosyl glycosyltransferase (pseudogene), p | -0,3086 | 0,2078 |
| TcCLB.510767.30 | hypothetical protein | -0,3087 | 0,1302 |
| TcCLB.511127.350 | hypothetical protein, conserved | -0,3088 | 0,01527 |

| | | | |
|---|---|---|---|
| TcCLB.511647.40 | hypothetical protein, conserved | -0,3088 | 0,02324 |
| TcCLB.510163.10 | hypothetical protein, conserved | -0,3088 | 0,02583 |
| TcCLB.506295.90 | coatomer subunit zeta, putative | -0,3089 | 0,09897 |
| TcCLB.508515.20 | translation factor sui1, putative | -0,3091 | 0,01731 |
| TcCLB.507963.30 | hypothetical protein, conserved | -0,3092 | 0,0004682 |
| TcCLB.511603.320 | mucin-associated surface protein (MASP), putative | -0,3092 | 0,5581 |
| TcCLB.508409.9 | 26S proteasome regulatory non-ATPase subunit, putative | -0,3093 | 0,00837 |
| TcCLB.506989.150 | hypothetical protein, conserved | -0,3095 | 0,01286 |
| TcCLB.509231.39 | hypothetical protein, conserved | -0,3095 | 0,132 |
| TcCLB.506357.110 | hypothetical protein, conserved | -0,3096 | 0,01895 |
| TcCLB.511425.40 | hypothetical protein, conserved | -0,3097 | 0,01203 |
| TcCLB.503431.40 | hypothetical protein, conserved | -0,3098 | 0,07447 |
| TcCLB.417091.10 | RNA pseudouridylate synthase, putative | -0,3098 | 0,1387 |
| TcCLB.504985.20 | UDP-Gal or UDP-GlcNAc-dependent glycosyltransferase, pu | -0,3098 | 0,4407 |
| TcCLB.511907.220 | hypothetical protein, conserved | -0,3099 | 0,02828 |
| TcCLB.510509.9 | inositol 1,4,5-trisphosphate receptor, putative (fragment) ( | -0,31 | 0,01658 |
| TcCLB.503925.100 | hypothetical protein, conserved | -0,3101 | 0,05114 |
| TcCLB.507683.34 | hypothetical protein, conserved | -0,3101 | 0,09024 |
| TcCLB.510359.190 | hypothetical protein, conserved | -0,3101 | 0,1158 |
| TcCLB.510189.40 | mucin-associated surface protein (MASP), putative | -0,3101 | 0,378 |
| TcCLB.416761.10 | surface protease GP63, putative | -0,3101 | 0,4447 |
| TcCLB.511109.50 | hypothetical protein, conserved (pseudogene) | -0,3103 | 0,07379 |
| TcCLB.504241.85 | hypothetical protein, conserved (pseudogene) | -0,3103 | 0,1827 |
| TcCLB.509571.70 | hypothetical protein, conserved | -0,3104 | 0,01831 |
| TcCLB.509569.180 | hypothetical protein, conserved | -0,3104 | 0,03211 |
| TcCLB.503697.110 | hypothetical protein, conserved | -0,3105 | 0,03178 |
| TcCLB.507795.50 | syntaxin, putative | -0,3105 | 0,03491 |
| TcCLB.508547.9 | UDP-Gal or UDP-GlcNAc-dependent glycosyltransferase, pu | -0,3105 | 0,06752 |
| TcCLB.506547.10 | dispersed gene family protein 1 (DGF-1, pseudogene), puta | -0,3107 | 0,3743 |
| TcCLB.507257.20 | hypothetical protein, conserved | -0,3108 | 0,01698 |
| TcCLB.503881.50 | hypothetical protein, conserved | -0,3109 | 0,2541 |
| TcCLB.511049.10 | hypothetical protein, conserved (pseudogene) | -0,3111 | 0,1065 |
| TcCLB.506925.160 | hypothetical protein, conserved | -0,3112 | 0,0512 |
| TcCLB.507093.120 | hypothetical protein, conserved | -0,3113 | 0,05216 |
| TcCLB.508605.10 | retrotransposon hot spot protein (RHS, pseudogene), putat | -0,3113 | 0,2259 |
| TcCLB.504411.20 | divalent cation tolerance protein, putative | -0,3115 | 0,3475 |
| TcCLB.504137.130 | hypothetical protein, conserved | -0,3116 | 0,01257 |
| TcCLB.507057.30 | arginine N-methyltransferase, type I (PRMT6) | -0,3116 | 0,09529 |
| TcCLB.510149.20 | hypothetical protein, conserved | -0,3119 | 0,02577 |
| TcCLB.506821.100 | mitochondrial RNA binding complex 1 subunit, putative (M | -0,312 | 0,007659 |
| TcCLB.506507.20 | hypothetical protein, conserved | -0,312 | 0,09729 |
| TcCLB.511021.99 | hypothetical protein, conserved | -0,312 | 0,1364 |
| TcCLB.508753.20 | trans-sialidase, Group V, putative | -0,312 | 0,3589 |
| TcCLB.511307.10 | kinesin, putative | -0,3121 | 0,01352 |
| TcCLB.510359.230 | hypothetical protein, conserved | -0,3124 | 0,006699 |

| | | | |
|---|---|---|---|
| TcCLB.506471.19 | hypothetical protein, conserved | -0,3124 | 0,1682 |
| TcCLB.508165.425 | mucin TcMUC (pseudogene), putative | -0,3124 | 0,5352 |
| TcCLB.509991.40 | transmembrane protein, conserved (pseudogene) | -0,3125 | 0,0000997 |
| TcCLB.504153.220 | DNA recombination and repair protein RAD51, putative | -0,3126 | 0,008838 |
| TcCLB.507641.230 | hypothetical protein, conserved | -0,3126 | 0,009454 |
| TcCLB.503499.24 | hypothetical protein | -0,3126 | 0,3364 |
| TcCLB.508559.40 | retrotransposon hot spot protein (RHS, pseudogene), putat | -0,3127 | 0,08357 |
| TcCLB.508731.40 | hypothetical protein, conserved | -0,3128 | 0,01852 |
| TcCLB.511137.20 | hypothetical protein, conserved | -0,3128 | 0,06967 |
| TcCLB.509179.90 | hypothetical protein, conserved | -0,3128 | 0,2908 |
| TcCLB.506741.100 | trans-sialidase, Group V, putative | -0,3128 | 0,411 |
| TcCLB.504213.60 | hypothetical protein, conserved | -0,3129 | 0,007443 |
| TcCLB.510173.10 | hypothetical protein, conserved | -0,3129 | 0,05546 |
| TcCLB.510279.314 | mucin TcMUCII (pseudogene), putative | -0,313 | 0,5684 |
| TcCLB.509561.70 | hypothetical protein, conserved | -0,3131 | 0,003056 |
| TcCLB.504247.10 | hypothetical protein, conserved | -0,3131 | 0,05198 |
| TcCLB.506201.10 | hypothetical protein, conserved | -0,3132 | 0,05033 |
| TcCLB.506653.10 | trans-sialidase, putative | -0,3132 | 0,6601 |
| TcCLB.503869.40 | pumilio/PUF RNA binding protein 9, putative (PUF9) | -0,3134 | 0,0381 |
| TcCLB.508325.320 | protein kinase, putative | -0,3134 | 0,165 |
| TcCLB.511621.240 | hypothetical protein, conserved | -0,3138 | 0,00121 |
| TcCLB.508549.10 | legume-like lectin, putative | -0,3138 | 0,009473 |
| TcCLB.507997.70 | hypothetical protein, conserved | -0,3138 | 0,01501 |
| TcCLB.509721.50 | hypothetical protein | -0,3138 | 0,0668 |
| TcCLB.505267.110 | dispersed gene family protein 1 (DGF-1), putative | -0,3138 | 0,178 |
| TcCLB.510983.10 | hypothetical protein | -0,3139 | 0,2271 |
| TcCLB.511051.70 | aminopeptidase, putative | -0,314 | 0,003149 |
| TcCLB.507515.140 | oxysterol-binding protein, putative | -0,314 | 0,01345 |
| TcCLB.507775.50 | proteasome alpha 7 subunit, putative | -0,314 | 0,02411 |
| TcCLB.508323.40 | hypothetical protein, conserved | -0,314 | 0,08757 |
| TcCLB.506945.260 | hypothetical protein, conserved | -0,3141 | 0,0006113 |
| TcCLB.509647.26 | hypothetical protein, conserved | -0,3141 | 0,01802 |
| TcCLB.507997.50 | hypothetical protein, conserved | -0,3141 | 0,03396 |
| TcCLB.506641.9 | dispersed gene family protein 1 (DGF-1), putative | -0,3141 | 0,3541 |
| TcCLB.510583.209 | mucin-associated surface protein (MASP, pseudogene), pu | -0,3141 | 0,7282 |
| TcCLB.506011.20 | hypothetical protein, conserved (pseudogene) | -0,3142 | 0,1789 |
| TcCLB.511211.160 | heat shock protein 70 (hsp70), putative | -0,3144 | 0,00004775 |
| TcCLB.509235.20 | C-5 sterol desaturase, putative | -0,3144 | 0,03584 |
| TcCLB.511057.60 | mucin-associated surface protein (MASP), putative | -0,3144 | 0,3769 |
| TcCLB.507611.300 | U1A small nuclear ribonucleoprotein, putative | -0,3145 | 0,1208 |
| TcCLB.506945.340 | hypothetical protein, conserved | -0,3146 | 0,001673 |
| TcCLB.511491.150 | hypothetical protein, conserved | -0,3147 | 0,02756 |
| TcCLB.509569.170 | hypothetical protein | -0,3148 | 0,001336 |
| TcCLB.506925.540 | hypothetical protein, conserved | -0,3148 | 0,004146 |
| TcCLB.510075.30 | hypothetical protein, conserved | -0,3149 | 0,002614 |

| | | | |
|---|---|---|---|
| TcCLB.490165.9 | DNA repair protein, putative | -0,3149 | 0,3165 |
| TcCLB.503733.40 | hypothetical protein, conserved | -0,3151 | 0,1303 |
| TcCLB.511481.50 | endonuclease III, putative | -0,3154 | 0,05586 |
| TcCLB.507805.20 | hypothetical protein | -0,3155 | 0,05071 |
| TcCLB.504137.40 | hypothetical protein, conserved | -0,3156 | 0,01792 |
| TcCLB.506941.194 | hypothetical protein, conserved | -0,3158 | 0,00007659 |
| TcCLB.510329.60 | RAB GDP dissociation inhibitor alpha, putative | -0,3158 | 0,0004586 |
| TcCLB.511077.40 | trans-sialidase (pseudogene), putative | -0,3158 | 0,4903 |
| TcCLB.511545.40 | ribosomal protein L27, putative | -0,3159 | 0,2084 |
| TcCLB.506743.160 | protein kinase, putative | -0,316 | 0,007082 |
| TcCLB.511299.60 | hypothetical protein, conserved | -0,3161 | 0,07033 |
| TcCLB.504123.40 | hypothetical protein, conserved | -0,3162 | 0,07417 |
| TcCLB.510311.20 | hypothetical protein, conserved | -0,3163 | 0,008158 |
| TcCLB.511867.80 | Vacuolar iron transporter 1, putative (VIT1) | -0,3163 | 0,02713 |
| TcCLB.511153.64 | histone H2B variant, putative | -0,3164 | 0,003126 |
| TcCLB.508871.100 | hypothetical protein | -0,3165 | 0,6877 |
| TcCLB.510661.240 | hypothetical protein, conserved | -0,3166 | 0,001791 |
| TcCLB.511181.140 | hypothetical protein | -0,3168 | 0,004589 |
| TcCLB.507159.41 | hypothetical protein, conserved | -0,3168 | 0,2384 |
| TcCLB.510737.119 | hypothetical protein, conserved | -0,3169 | 0,03729 |
| TcCLB.507467.119 | hypothetical protein, conserved | -0,3171 | 0,08309 |
| TcCLB.511249.100 | hypothetical protein, conserved | -0,3172 | 0,0002048 |
| TcCLB.504057.104 | phosphatidylinositol phosphate kinase alpha (PIPKA) | -0,3172 | 0,02223 |
| TcCLB.506351.90 | surface protease GP63, putative | -0,3172 | 0,09254 |
| TcCLB.510047.70 | hypothetical protein (pseudogene) | -0,3172 | 0,5301 |
| TcCLB.509061.20 | hypothetical protein, conserved | -0,3173 | 0,03093 |
| TcCLB.504035.80 | hypothetical protein, conserved | -0,3173 | 0,1631 |
| TcCLB.506945.200 | mago nashi-like protein, putative | -0,3175 | 0,001341 |
| TcCLB.508325.20 | retrotransposon hot spot protein (RHS, pseudogene), putat | -0,3175 | 0,1812 |
| TcCLB.506435.90 | hypothetical protein, conserved | -0,3176 | 0,01403 |
| TcCLB.507017.30 | hypothetical protein, conserved | -0,3176 | 0,08632 |
| TcCLB.506297.110 | ATPase, putative | -0,3177 | 0,0001514 |
| TcCLB.507611.330 | hypothetical protein, conserved | -0,3177 | 0,05185 |
| TcCLB.509065.40 | hypothetical protein, conserved | -0,3178 | 0,01933 |
| TcCLB.506459.210 | mucin TcMUCII, putative | -0,3178 | 0,5721 |
| TcCLB.507641.30 | hypothetical protein, conserved | -0,3179 | 0,002033 |
| TcCLB.508461.250 | ATP-dependent RNA helicase, putative | -0,3181 | 0,0007869 |
| TcCLB.507905.30 | trans-sialidase (pseudogene), putative | -0,3181 | 0,001515 |
| TcCLB.503583.60 | hypothetical protein | -0,3181 | 0,00478 |
| TcCLB.509695.190 | cdc2-related kinase 1, putative (CRK1) | -0,3181 | 0,01346 |
| TcCLB.508269.76 | N-acetyltransferase complex ARD1 subunit (pseudogene), | -0,3181 | 0,4036 |
| TcCLB.509419.260 | dispersed gene family protein 1 (DGF-1), putative | -0,3181 | 0,4227 |
| TcCLB.511881.40 | protein transport protein Sec24C, putative | -0,3182 | 0,01453 |
| TcCLB.509563.20 | hypothetical protein, conserved | -0,3183 | 0,03215 |
| TcCLB.503801.50 | hypothetical protein, conserved | -0,3184 | 0,0108 |

| | | | |
|---|---|---|---|
| TcCLB.505987.50 | adenosine kinase, putative | -0,3184 | 0,01606 |
| TcCLB.508479.160 | hypothetical protein, conserved | -0,3185 | 0,0008797 |
| TcCLB.509647.60 | hypothetical protein, conserved | -0,3185 | 0,00199 |
| TcCLB.506265.50 | hypothetical protein, conserved | -0,3187 | 0,03582 |
| TcCLB.509591.10 | ubiquitin-conjugating enzyme, putative | -0,3187 | 0,05045 |
| TcCLB.509793.40 | kinetoplast-associated protein (TcKAP4) | -0,3188 | 0,01619 |
| TcCLB.508659.40 | protein phosphatase 2C-like (pseudogene), putative | -0,3188 | 0,08864 |
| TcCLB.509749.20 | hypothetical protein, conserved | -0,3188 | 0,2037 |
| TcCLB.503453.4 | hypothetical protein, conserved | -0,3188 | 0,2607 |
| TcCLB.505999.30 | anti-silencing protein ASF 1, putative | -0,3189 | 0,1334 |
| TcCLB.506113.83 | ATP-dependent DEAD/H RNA helicase (pseudogene), putat | -0,319 | 0,2383 |
| TcCLB.508463.20 | hypothetical protein, conserved | -0,3191 | 0,0001172 |
| TcCLB.506195.60 | hypothetical protein, conserved | -0,3191 | 0,02755 |
| TcCLB.507033.4 | hypothetical protein, conserved | -0,3191 | 0,1143 |
| TcCLB.507011.190 | hypothetical protein, conserved | -0,3191 | 0,1387 |
| TcCLB.507787.140 | hypothetical protein, conserved | -0,3193 | 0,01422 |
| TcCLB.509693.160 | hypothetical protein, conserved | -0,3193 | 0,01618 |
| TcCLB.507801.240 | mucin-associated surface protein (MASP, pseudogene), pu | -0,3193 | 0,3154 |
| TcCLB.509699.140 | mucin-associated surface protein (MASP, pseudogene), pu | -0,3193 | 0,3371 |
| TcCLB.503899.40 | hypothetical protein, conserved | -0,3194 | 0,00577 |
| TcCLB.505009.30 | serine/threonine protein kinase, putative | -0,3194 | 0,1516 |
| TcCLB.506609.50 | hypothetical protein | -0,3195 | 0,5196 |
| TcCLB.511363.10 | hypothetical protein, conserved | -0,3196 | 0,001447 |
| TcCLB.506925.50 | hypothetical protein, conserved | -0,3196 | 0,003923 |
| TcCLB.511559.20 | hypothetical protein, conserved | -0,3197 | 0,001493 |
| TcCLB.509901.150 | hypothetical protein, conserved | -0,3197 | 0,1096 |
| TcCLB.510105.33 | RNA-binding protein, putative | -0,3197 | 0,2943 |
| TcCLB.506725.10 | hypothetical protein, conserved | -0,3198 | 0,03315 |
| TcCLB.509775.40 | iron superoxide dismutase, putative | -0,3198 | 0,05056 |
| TcCLB.503669.30 | hypothetical protein, conserved | -0,3199 | 0,04814 |
| TcCLB.504047.30 | kinesin, putative | -0,32 | 0,01975 |
| TcCLB.508777.80 | hypothetical protein, conserved | -0,3201 | 0,01187 |
| TcCLB.510755.40 | hypothetical protein, conserved | -0,3201 | 0,08864 |
| TcCLB.511245.70 | hypothetical protein, conserved | -0,3202 | 0,004455 |
| TcCLB.507817.50 | hypothetical protein, conserved | -0,3204 | 0,03826 |
| TcCLB.511377.25 | elongation factor 1-gamma (EF-1-gamma, pseudogene), p | -0,3204 | 0,5462 |
| TcCLB.510687.10 | dynein heavy chain (pseudogene), putative | -0,3206 | 0,00002326 |
| TcCLB.508629.100 | trans-sialidase (pseudogene), putative | -0,3206 | 0,02226 |
| TcCLB.503431.50 | hypothetical protein, conserved | -0,3206 | 0,07623 |
| TcCLB.507641.170 | hypothetical protein, conserved | -0,3207 | 0,00432 |
| TcCLB.508543.129 | hypothetical protein, conserved | -0,3207 | 0,01471 |
| TcCLB.508181.140 | inorganic pyrophosphatase, putative | -0,3207 | 0,03392 |
| TcCLB.504153.170 | hypothetical protein, conserved | -0,3209 | 4,652E-06 |
| TcCLB.511501.10 | hypothetical protein, conserved | -0,321 | 0,00003216 |
| TcCLB.511587.39 | ATP-dependent DEAD/H RNA helicase (pseudogene), putat | -0,321 | 0,2127 |

| | | | |
|---|---|---|---|
| TcCLB.506691.14 | ATP-dependent DEAD/H RNA helicase, putative (fragment) | -0,3211 | 0,005226 |
| TcCLB.510187.140 | hypothetical protein, conserved | -0,3214 | 0,0007199 |
| TcCLB.506755.60 | hypothetical protein, conserved | -0,3214 | 0,0972 |
| TcCLB.509163.10 | hypothetical protein | -0,3215 | 0,1329 |
| TcCLB.508307.200 | hypothetical protein, conserved | -0,3216 | 0,02808 |
| TcCLB.510517.10 | chaperonin/T-complex protein 1 gamma subunit, putative | -0,3217 | 0,002233 |
| TcCLB.511283.20 | protein transport protein SEC13, putative | -0,3217 | 0,01469 |
| TcCLB.503913.10 | hypothetical protein, conserved | -0,3217 | 0,02737 |
| TcCLB.507623.30 | hypothetical protein, conserved | -0,3218 | 0,001914 |
| TcCLB.508221.150 | mucin-associated surface protein (MASP), putative | -0,3218 | 0,5122 |
| TcCLB.511141.10 | hypothetical protein, conserved | -0,3219 | 0,1077 |
| TcCLB.506589.130 | trans-sialidase (pseudogene), putative | -0,322 | 0,1885 |
| TcCLB.511913.9 | ribonuclease L inhibitor (pseudogene), putative | -0,3221 | 0,004921 |
| TcCLB.506945.80 | DNA ligase I, putative | -0,3222 | 0,001032 |
| TcCLB.506315.10 | hypothetical protein | -0,3222 | 0,0368 |
| TcCLB.505949.30 | trans-sialidase, Group V, putative | -0,3222 | 0,4185 |
| TcCLB.508699.30 | hypothetical protein, conserved | -0,3223 | 0,07717 |
| TcCLB.504097.40 | hypothetical protein, conserved | -0,3224 | 0,005981 |
| TcCLB.506195.100 | malate dehydrogenase (pseudogene), putative | -0,3224 | 0,04378 |
| TcCLB.509751.10 | hypothetical protein, conserved | -0,3226 | 0,07079 |
| TcCLB.511003.180 | hypothetical protein, conserved | -0,3226 | 0,1121 |
| TcCLB.509831.20 | vesicular-fusion ATPase-like protein, putative | -0,3227 | 0,001286 |
| TcCLB.507711.150 | Ubiquitin-like modifier-activating enzyme ATG7, putative ( | -0,3227 | 0,008789 |
| TcCLB.509429.41 | trans-sialidase (pseudogene), putative | -0,3229 | 0,2291 |
| TcCLB.508153.1140 | hypothetical protein, conserved | -0,323 | 0,02567 |
| TcCLB.510729.90 | hypothetical protein, conserved | -0,3231 | 0,00825 |
| TcCLB.506287.170 | hypothetical protein, conserved | -0,3232 | 0,002271 |
| TcCLB.508799.40 | hypothetical protein, conserved | -0,3232 | 0,003679 |
| TcCLB.510943.170 | hypothetical protein, conserved | -0,3232 | 0,006021 |
| TcCLB.503841.40 | actin, putative | -0,3232 | 0,02839 |
| TcCLB.506817.70 | hypothetical protein | -0,3232 | 0,2574 |
| TcCLB.509779.30 | hypothetical protein, conserved | -0,3233 | 0,01571 |
| TcCLB.507809.50 | hypothetical protein, conserved | -0,3233 | 0,07613 |
| TcCLB.507831.100 | hypothetical protein, conserved | -0,3234 | 0,01777 |
| TcCLB.505615.10 | surface protease GP63, putative | -0,3234 | 0,06222 |
| TcCLB.503489.24 | hypothetical protein, conserved | -0,3235 | 0,0004344 |
| TcCLB.508061.50 | dispersed gene family protein 1 (DGF-1), putative | -0,3236 | 0,02893 |
| TcCLB.509027.30 | hypothetical protein, conserved | -0,3237 | 0,04739 |
| TcCLB.506413.50 | hypothetical protein, conserved | -0,3238 | 0,02632 |
| TcCLB.504167.40 | ubiquitin fusion degradation protein, putative | -0,3238 | 0,02896 |
| TcCLB.507087.50 | hypothetical protein, conserved | -0,3238 | 0,03749 |
| TcCLB.509037.30 | hypothetical protein, conserved | -0,3239 | 0,0325 |
| TcCLB.510407.60 | tyrosyl-tRNA synthetase (pseudogene), putative | -0,324 | 0,0001608 |
| TcCLB.507021.10 | hypothetical protein, conserved | -0,324 | 0,0136 |
| TcCLB.440363.10 | hypothetical protein, conserved | -0,324 | 0,1176 |

| | | | |
|---|---|---|---|
| TcCLB.508641.250 | hypothetical protein | -0,3241 | 0,002264 |
| TcCLB.508489.20 | hypothetical protein, conserved | -0,3242 | 0,02199 |
| TcCLB.506763.361 | syntaxin binding protein (pseudogene), putative | -0,3243 | 0,5323 |
| TcCLB.508153.20 | vacuolar protein sorting-associated protein, putative | -0,3244 | 0,0001968 |
| TcCLB.511067.10 | tubulin folding cofactor D, putative | -0,3244 | 0,0002033 |
| TcCLB.508357.9 | hypothetical protein, conserved (fragment) | -0,3244 | 0,003506 |
| TcCLB.506629.220 | isovaleryl-coA dehydrogenase, putative | -0,3244 | 0,005329 |
| TcCLB.503453.80 | hypothetical protein, conserved | -0,3244 | 0,0377 |
| TcCLB.511361.80 | hypothetical protein, conserved | -0,3244 | 0,03813 |
| TcCLB.508805.110 | hypothetical protein | -0,3244 | 0,3571 |
| TcCLB.510311.160 | hypothetical protein, conserved | -0,3245 | 0,02161 |
| TcCLB.511665.50 | dispersed gene family protein 1 (DGF-1), putative | -0,3245 | 0,07743 |
| TcCLB.509269.19 | sphingosine-1-phosphate phosphatase, putative | -0,3246 | 0,006508 |
| TcCLB.511517.170 | hypothetical protein, conserved | -0,3246 | 0,007346 |
| TcCLB.506945.120 | chromosomal passenger protein, putative (CPC1) | -0,3246 | 0,01649 |
| TcCLB.508577.80 | hypothetical protein, conserved | -0,3246 | 0,09851 |
| TcCLB.510377.310 | mucin-associated surface protein (MASP, pseudogene), pu | -0,3246 | 0,569 |
| TcCLB.511877.30 | surface protease GP63 (pseudogene), putative | -0,3247 | 0,2284 |
| TcCLB.506247.100 | hypothetical protein, conserved | -0,3248 | 0,08202 |
| TcCLB.510565.110 | hypothetical protein | -0,3248 | 0,1404 |
| TcCLB.507611.290 | acetyltransferase, putative | -0,3249 | 0,04733 |
| TcCLB.508469.20 | hypothetical protein, conserved | -0,3249 | 0,05198 |
| TcCLB.511423.170 | chaperone DNAJ protein, putative | -0,3249 | 0,08124 |
| TcCLB.506913.20 | protein kinase, putative (fragment) | -0,3249 | 0,2315 |
| TcCLB.511767.10 | trans-sialidase, putative | -0,325 | 0,3784 |
| TcCLB.511585.20 | KREL1 | -0,3252 | 0,001998 |
| TcCLB.503779.40 | protein-l-isoaspartate o-methyltransferase, putative | -0,3252 | 0,01177 |
| TcCLB.508479.350 | hypothetical protein, conserved | -0,3253 | 0,002142 |
| TcCLB.507467.20 | hypothetical protein, conserved | -0,3253 | 0,0151 |
| TcCLB.509617.89 | Zn-finger protein, putative (fragment) | -0,3254 | 0,02831 |
| TcCLB.457339.19 | hypothetical protein, conserved | -0,3254 | 0,1196 |
| TcCLB.510325.69 | Aminopeptidase M1, putative (fragment) | -0,3255 | 0,00147 |
| TcCLB.506681.20 | hypothetical protein, conserved | -0,3256 | 0,0007858 |
| TcCLB.510769.30 | hypothetical protein, conserved | -0,3256 | 0,003504 |
| TcCLB.509573.50 | hypothetical protein, conserved | -0,3257 | 0,000447 |
| TcCLB.510745.49 | hypothetical protein, conserved (fragment) | -0,3257 | 0,1799 |
| TcCLB.510243.100 | chloride channel protein, putative | -0,3258 | 0,002717 |
| TcCLB.503559.100 | long-chain-fatty-acid-CoA ligase, putative | -0,3258 | 0,004166 |
| TcCLB.503897.130 | hypothetical protein, conserved | -0,3258 | 0,04096 |
| TcCLB.507257.130 | hypothetical protein, conserved | -0,3258 | 0,05155 |
| TcCLB.506629.100 | hypothetical protein | -0,3259 | 0,004393 |
| TcCLB.510729.30 | hypothetical protein, conserved | -0,3259 | 0,06109 |
| TcCLB.509633.10 | hypothetical protein, conserved | -0,3259 | 0,08614 |
| TcCLB.511473.9 | N-acetyltransferase complex ARD1 subunit (pseudogene), | -0,326 | 0,2638 |
| TcCLB.511245.110 | hypothetical protein, conserved | -0,3261 | 0,003456 |

| | | | |
|---|---|---|---|
| TcCLB.506885.290 | DNA replication licensing factor MCM9, putative | -0,3261 | 0,005612 |
| TcCLB.511173.190 | trans-sialidase (pseudogene), putative | -0,3261 | 0,4036 |
| TcCLB.508409.100 | hypothetical protein, conserved | -0,3262 | 0,00007568 |
| TcCLB.506559.180 | hypothetical protein, conserved | -0,3262 | 0,003808 |
| TcCLB.506289.60 | hypothetical protein, conserved | -0,3262 | 0,04739 |
| TcCLB.511665.15 | dispersed gene family protein 1 (DGF-1), putative (fragme | -0,3262 | 0,3473 |
| TcCLB.506557.24 | hypothetical protein, conserved | -0,3263 | 0,1313 |
| TcCLB.511183.570 | syntaxin binding protein (pseudogene), putative | -0,3263 | 0,4819 |
| TcCLB.506165.10 | hypothetical protein, conserved | -0,3264 | 0,01319 |
| TcCLB.508405.20 | TBC1 domain family member 20/GTPase, putative | -0,3264 | 0,08857 |
| TcCLB.506897.10 | mucin TcMUCII, putative | -0,3264 | 0,5496 |
| TcCLB.504163.110 | N-acetyltransferase subunit Nat1, putative | -0,3265 | 0,0009643 |
| TcCLB.508949.40 | hypothetical protein, conserved | -0,3265 | 0,05239 |
| TcCLB.508277.300 | hypothetical protein, conserved | -0,3265 | 0,05392 |
| TcCLB.508707.64 | exonuclease, putative | -0,3265 | 0,09268 |
| TcCLB.508871.20 | mucin TcMUCII, putative | -0,3265 | 0,5846 |
| TcCLB.508257.40 | hypothetical protein, conserved | -0,3269 | 0,01185 |
| TcCLB.510659.194 | hypothetical protein, conserved | -0,3269 | 0,0207 |
| TcCLB.509321.19 | GTP-binding protein, putative | -0,3269 | 0,02391 |
| TcCLB.508405.50 | hypothetical protein, conserved | -0,3269 | 0,05144 |
| TcCLB.508647.130 | hypothetical protein, conserved | -0,327 | 0,01577 |
| TcCLB.505171.50 | ras-related GTP-binding protein, putative | -0,327 | 0,09943 |
| TcCLB.508479.240 | hypothetical protein, conserved | -0,3271 | 0,003829 |
| TcCLB.506227.140 | hypothetical protein, conserved | -0,3272 | 0,05741 |
| TcCLB.508661.10 | Eukaryotic translation initiation factor 4E type 5, putative ( | -0,3273 | 0,005922 |
| TcCLB.510729.180 | hypothetical protein | -0,3273 | 0,006441 |
| TcCLB.510299.79 | protein phosphatase 2C, putative | -0,3273 | 0,01893 |
| TcCLB.509793.60 | hypothetical protein, conserved | -0,3274 | 0,01668 |
| TcCLB.508409.220 | hypothetical protein, conserved | -0,3274 | 0,01733 |
| TcCLB.505939.40 | hypothetical protein, conserved | -0,3274 | 0,02501 |
| TcCLB.508303.65 | thioredoxin-like protein, putative | -0,3274 | 0,04187 |
| TcCLB.506589.140 | mucin-associated surface protein (MASP), putative | -0,3274 | 0,2858 |
| TcCLB.503897.80 | deaminase, putative | -0,3275 | 0,1026 |
| TcCLB.511523.30 | hypothetical protein | -0,3276 | 0,004019 |
| TcCLB.506599.270 | mucin TcMUCII, putative | -0,3276 | 0,5796 |
| TcCLB.508139.170 | retrotransposon hot spot protein (RHS, pseudogene), putat | -0,3279 | 0,01996 |
| TcCLB.511731.40 | hypothetical protein, conserved | -0,3281 | 0,004597 |
| TcCLB.511575.100 | protein transport protein SEC23, putative | -0,3282 | 0,0001548 |
| TcCLB.508059.30 | hypothetical protein, conserved | -0,3282 | 0,0105 |
| TcCLB.449247.4 | serine/threonine protein phosphatase, putative (fragment) | -0,3283 | 0,0312 |
| TcCLB.503401.20 | hypothetical protein, conserved | -0,3283 | 0,09845 |
| TcCLB.508827.44 | hypothetical protein, conserved | -0,3284 | 0,02408 |
| TcCLB.509593.59 | dispersed gene family protein 1 (DGF-1), putative | -0,3284 | 0,6426 |
| TcCLB.509607.30 | splicing factor TSR1, putative | -0,3285 | 0,002364 |
| TcCLB.510519.120 | hypothetical protein, conserved | -0,3285 | 0,007208 |

| | | | |
|---|---|---|---|
| TcCLB.506943.120 | hypothetical protein, conserved | -0,3286 | 0,0253 |
| TcCLB.503599.20 | hypothetical protein, conserved | -0,3286 | 0,09617 |
| TcCLB.428999.20 | receptor-type adenylate cyclase, putative | -0,3286 | 0,1401 |
| TcCLB.506253.21 | hypothetical protein, conserved | -0,3286 | 0,5513 |
| TcCLB.511255.65 | surface protease GP63 (pseudogene), putative | -0,3287 | 0,545 |
| TcCLB.511729.60 | MORN repeat-containing protein 1 (MORN1) | -0,3288 | 0,0007397 |
| TcCLB.509429.240 | hypothetical protein, conserved | -0,3288 | 0,01094 |
| TcCLB.510091.40 | Mad2-like HORMA domain containing protein (mad2) | -0,3288 | 0,01166 |
| TcCLB.511555.40 | CPSF 30 kDa subunit, putative (CPSF30) | -0,3288 | 0,01852 |
| TcCLB.508807.5 | glycosylphosphatidylinositol-specific phospholipase C, puta | -0,3288 | 0,09558 |
| TcCLB.507993.340 | hypothetical protein | -0,3288 | 0,2225 |
| TcCLB.507805.30 | hypothetical protein, conserved (pseudogene) | -0,3288 | 0,2308 |
| TcCLB.510659.120 | dolichol kinase, putative | -0,3289 | 0,001388 |
| TcCLB.509979.230 | hypothetical protein, conserved | -0,3289 | 0,04395 |
| TcCLB.510625.130 | hypothetical protein | -0,3289 | 0,6099 |
| TcCLB.402647.9 | hypothetical protein, conserved | -0,329 | 0,05318 |
| TcCLB.506989.199 | hypothetical protein, conserved | -0,329 | 0,1602 |
| TcCLB.507681.160 | 40S ribosomal protein S24E, putative | -0,3291 | 0,005486 |
| TcCLB.509683.100 | hypothetical protein, conserved | -0,3291 | 0,01805 |
| TcCLB.511553.140 | mucin-associated surface protein (MASP), putative | -0,3291 | 0,5202 |
| TcCLB.509261.10 | p-nitrophenylphosphatase, putative | -0,3292 | 0,02624 |
| TcCLB.510697.25 | surface protease GP63 (pseudogene), putative | -0,3294 | 0,3481 |
| TcCLB.510889.160 | cell division control protein (pseudogene), putative | -0,3295 | 0,01166 |
| TcCLB.509589.10 | mitochondrial DNA polymerase I protein D, putative | -0,3295 | 0,0171 |
| TcCLB.511043.60 | receptor-type adenylate cyclase, putative | -0,3295 | 0,03664 |
| TcCLB.506791.10 | CYC2-like cyclin 4, putative (CYC4) | -0,3296 | 0,006236 |
| TcCLB.506977.80 | hypothetical protein, conserved | -0,3297 | 0,00009489 |
| TcCLB.508647.270 | malic enzyme, putative | -0,3297 | 0,00009886 |
| TcCLB.507257.40 | hypothetical protein, conserved | -0,3297 | 0,1672 |
| TcCLB.508121.130 | mucin-associated surface protein (MASP, pseudogene), pu | -0,3297 | 0,2566 |
| TcCLB.511145.43 | proteasome subunit alpha type-5, putative | -0,3299 | 0,01265 |
| TcCLB.506297.140 | hypothetical protein, conserved | -0,3299 | 0,02832 |
| TcCLB.509049.40 | hypothetical protein, conserved | -0,3299 | 0,03536 |
| TcCLB.503395.30 | hypothetical protein, conserved | -0,33 | 0,006076 |
| TcCLB.506733.10 | hypothetical protein, conserved | -0,33 | 0,04228 |
| TcCLB.507997.54 | Alpha/beta hydrolase domain-containing protein | -0,3301 | 0,01075 |
| TcCLB.509213.10 | hypothetical protein, conserved | -0,3302 | 0,05818 |
| TcCLB.510421.80 | hypothetical protein | -0,3302 | 0,1175 |
| TcCLB.507629.20 | hypothetical protein, conserved | -0,3303 | 0,00005501 |
| TcCLB.511627.60 | vesicle-associated membrane protein | -0,3303 | 0,01319 |
| TcCLB.503931.46 | hypothetical protein, conserved (pseudogene) | -0,3303 | 0,294 |
| TcCLB.508951.4 | protein kinase, putative (fragment) | -0,3303 | 0,3124 |
| TcCLB.506739.50 | ATP-dependent protease ATPase subunit HslU1, putative (I | -0,3304 | 0,0001534 |
| TcCLB.510285.90 | hypothetical protein, conserved | -0,3304 | 0,009765 |
| TcCLB.511407.70 | hypothetical protein, conserved | -0,3306 | 0,002011 |

| | | | |
|---|---|---|---|
| TcCLB.510545.30 | mucin-associated surface protein (MASP, pseudogene), pu | -0,3306 | 0,4243 |
| TcCLB.507857.70 | hypothetical protein, conserved | -0,3307 | 0,00833 |
| TcCLB.503913.49 | hypothetical protein, conserved (fragment) | -0,3307 | 0,4242 |
| TcCLB.507793.30 | hypothetical protein, conserved | -0,3308 | 0,02993 |
| TcCLB.507673.65 | DNA-directed RNA polymerase subunit, putative | -0,3308 | 0,5515 |
| TcCLB.506973.20 | mucin-associated surface protein (MASP), putative | -0,3309 | 0,1931 |
| TcCLB.510693.270 | trans-sialidase, Group VI, putative | -0,3309 | 0,4197 |
| TcCLB.511877.15 | elongation factor 1-gamma (EF-1-gamma, pseudogene), p | -0,331 | 0,1269 |
| TcCLB.509453.20 | hypothetical protein, conserved | -0,3312 | 0,1783 |
| TcCLB.504133.30 | tyrosine protein kinase, putative | -0,3313 | 0,001383 |
| TcCLB.503843.10 | hypothetical protein, conserved | -0,3313 | 0,007048 |
| TcCLB.508425.31 | hypothetical protein, conserved (pseudogene) | -0,3313 | 0,04297 |
| TcCLB.507641.130 | hypothetical protein, conserved | -0,3313 | 0,05822 |
| TcCLB.504213.130 | hypothetical protein, conserved | -0,3314 | 0,06472 |
| TcCLB.506009.109 | hypothetical protein, conserved | -0,3314 | 0,08847 |
| TcCLB.510371.5 | elongation factor 1-gamma (EF-1-gamma, pseudogene), p | -0,3314 | 0,5423 |
| TcCLB.434259.10 | RNA polymerase-associated protein CTR9, putative | -0,3315 | 0,033 |
| TcCLB.511517.10 | hypothetical protein, conserved | -0,3315 | 0,03595 |
| TcCLB.508859.109 | exosome complex exonuclease, putative | -0,3315 | 0,08775 |
| TcCLB.509897.140 | trans-sialidase, Group V, putative | -0,3315 | 0,124 |
| TcCLB.503827.60 | mucin-associated surface protein (MASP), putative | -0,3316 | 0,3916 |
| TcCLB.508693.10 | cleavage and polyadenylation specificity factor, putative | -0,3318 | 0,0001544 |
| TcCLB.509733.130 | hypothetical protein, conserved | -0,3318 | 0,002273 |
| TcCLB.510821.20 | hypothetical protein, conserved | -0,3319 | 0,00004011 |
| TcCLB.510123.30 | DNA repair protein, putative | -0,3319 | 0,02845 |
| TcCLB.507757.21 | hypothetical protein, conserved | -0,332 | 0,1656 |
| TcCLB.506601.21 | syntaxin binding protein (pseudogene), putative | -0,332 | 0,5327 |
| TcCLB.511075.40 | cullin, putative | -0,3321 | 0,001101 |
| TcCLB.510115.30 | trans-sialidase, Group VI, putative | -0,3321 | 0,1136 |
| TcCLB.509381.10 | mitochondrial import receptor subunit, putative | -0,3322 | 0,004674 |
| TcCLB.506195.260 | hypothetical protein | -0,3322 | 0,01053 |
| TcCLB.510533.130 | hypothetical protein, conserved | -0,3323 | 0,009664 |
| TcCLB.506833.79 | hypothetical protein, conserved (fragment) | -0,3324 | 0,005085 |
| TcCLB.510519.70 | mitochondrial RNA binding complex 1 subunit, putative | -0,3324 | 0,01108 |
| TcCLB.508479.230 | peroxin-2, putative | -0,3324 | 0,02555 |
| TcCLB.507681.90 | hypothetical protein, conserved | -0,3325 | 0,006326 |
| TcCLB.511055.40 | glycine dehydrogenase [decarboxylating] (pseudogene), pu | -0,3325 | 0,04957 |
| TcCLB.506661.110 | hypothetical protein, conserved | -0,3326 | 0,0001908 |
| TcCLB.510359.130 | hypothetical protein, conserved | -0,3326 | 0,01949 |
| TcCLB.509023.190 | hypothetical protein, conserved | -0,3326 | 0,1685 |
| TcCLB.506269.50 | mucin-associated surface protein (MASP, pseudogene), pu | -0,3326 | 0,406 |
| TcCLB.507017.90 | hypothetical protein, conserved | -0,3327 | 0,04498 |
| TcCLB.509099.5 | NUP-1 protein, putative (fragment) | -0,3328 | 0,02953 |
| TcCLB.506111.10 | retrotransposon hot spot protein (RHS, pseudogene), putat | -0,3328 | 0,03341 |
| TcCLB.506733.40 | katanin-like protein, putative | -0,3328 | 0,03485 |

| | | | |
|---|---|---|---|
| TcCLB.510765.20 | hypothetical protein, conserved | -0,3328 | 0,1543 |
| TcCLB.508289.94 | hypothetical protein, conserved (pseudogene) | -0,3328 | 0,6033 |
| TcCLB.507099.140 | endomembrane protein, putative | -0,3329 | 0,0001866 |
| TcCLB.504057.21 | hypothetical protein, conserved | -0,3331 | 0,01665 |
| TcCLB.506315.80 | hypothetical protein, conserved | -0,3332 | 0,01216 |
| TcCLB.505963.9 | dispersed gene family protein 1 (DGF-1), putative (fragme | -0,3332 | 0,2104 |
| TcCLB.510181.69 | hypothetical protein, conserved | -0,3333 | 0,0309 |
| TcCLB.506257.70 | hypothetical protein, conserved | -0,3335 | 0,01868 |
| TcCLB.507807.10 | hypothetical protein, conserved | -0,3337 | 0,0001358 |
| TcCLB.508303.4 | ubiquitin-protein ligase, putative | -0,3337 | 0,01041 |
| TcCLB.507037.20 | RNA-binding protein, putative | -0,3337 | 0,01569 |
| TcCLB.503463.30 | protein kinase, putative | -0,3337 | 0,0287 |
| TcCLB.508911.20 | hypothetical protein, conserved | -0,3339 | 0,0548 |
| TcCLB.507047.170 | hypothetical protein, conserved | -0,3339 | 0,224 |
| TcCLB.511169.20 | hypothetical protein, conserved | -0,334 | 0,03046 |
| TcCLB.506309.110 | mucin TcMUCII, putative | -0,3341 | 0,5156 |
| TcCLB.511751.200 | epsilon-adaptin, putative | -0,3342 | 0,001463 |
| TcCLB.506839.14 | hypothetical protein, conserved | -0,3342 | 0,007353 |
| TcCLB.503759.30 | trans-sialidase, putative | -0,3342 | 0,447 |
| TcCLB.508269.60 | exosome-associated protein 4, putative (EAP4) | -0,3343 | 0,1255 |
| TcCLB.511585.140 | hypothetical protein, conserved | -0,3344 | 0,000739 |
| TcCLB.509965.90 | hypothetical protein, conserved | -0,3344 | 0,001072 |
| TcCLB.432787.10 | mitochondrial RNA binding complex 1 subunit, putative (fra | -0,3344 | 0,05299 |
| TcCLB.506741.120 | mucin-associated surface protein (MASP), putative | -0,3344 | 0,4537 |
| TcCLB.509499.14 | peroxidoxin | -0,3345 | 0,0001801 |
| TcCLB.510533.230 | palmitoyl acyltransferase 11, putative | -0,3346 | 0,04166 |
| TcCLB.504069.110 | hypothetical protein, conserved | -0,3346 | 0,1092 |
| TcCLB.506067.230 | trans-sialidase (pseudogene), putative | -0,3346 | 0,3388 |
| TcCLB.510965.5 | elongation factor 2, putative (fragment) | -0,3347 | 0,00006688 |
| TcCLB.509567.10 | hypothetical protein, conserved | -0,3348 | 0,02025 |
| TcCLB.510735.9 | hypothetical protein, conserved | -0,3349 | 0,001418 |
| TcCLB.509103.10 | hypothetical protein, conserved | -0,3349 | 0,01174 |
| TcCLB.507943.60 | hypothetical protein, conserved | -0,3349 | 0,01299 |
| TcCLB.504235.9 | hypothetical protein, conserved (fragment) | -0,335 | 0,1597 |
| TcCLB.511201.19 | hypothetical protein (fragment) | -0,3352 | 0,02031 |
| TcCLB.506925.104 | dynein light chain, putative | -0,3354 | 0,000135 |
| TcCLB.506855.340 | U4/U6 small nuclear ribonuclear protein, putative | -0,3354 | 0,01673 |
| TcCLB.510519.130 | hypothetical protein, conserved | -0,3354 | 0,02795 |
| TcCLB.510645.49 | ribosomal protein L3 mitochondrial, putative | -0,3355 | 0,1029 |
| TcCLB.507025.30 | hypothetical protein, conserved | -0,3357 | 0,05199 |
| TcCLB.510733.54 | hypothetical protein, conserved | -0,3357 | 0,07488 |
| TcCLB.511629.40 | hypothetical protein, conserved | -0,3357 | 0,08209 |
| TcCLB.432867.20 | hypothetical protein, conserved | -0,3357 | 0,2547 |
| TcCLB.508393.9 | dispersed gene family protein 1 (DGF-1), putative | -0,3358 | 0,6352 |
| TcCLB.509805.20 | cAMP phosphodiesterase A, putative (PDEA) | -0,3359 | 0,04373 |

| | | | |
|---|---|---|---|
| TcCLB.511367.354 | U1 small nuclear ribonucleoprotein C, putative | -0,336 | 0,05602 |
| TcCLB.508541.210 | hypothetical protein, conserved | -0,3361 | 0,006192 |
| TcCLB.506637.60 | hypothetical protein, conserved | -0,3361 | 0,03428 |
| TcCLB.510755.147 | hypothetical protein, conserved | -0,3361 | 0,07033 |
| TcCLB.509613.20 | trans-sialidase (pseudogene), putative | -0,3362 | 0,306 |
| TcCLB.508577.140 | cyclophilin, putative | -0,3363 | 0,001389 |
| TcCLB.467795.9 | diphthamide synthesis protein, putative | -0,3363 | 0,01121 |
| TcCLB.504081.295 | surface protease GP63 (pseudogene), putative | -0,3363 | 0,532 |
| TcCLB.508505.30 | hypothetical protein, conserved | -0,3365 | 0,006896 |
| TcCLB.509913.20 | hypothetical protein, conserved | -0,3365 | 0,007435 |
| TcCLB.508261.29 | mucin TcMUCII (pseudogene), putative | -0,3365 | 0,6033 |
| TcCLB.508897.20 | signal recognition particle protein, putative | -0,3366 | 0,00006265 |
| TcCLB.506679.160 | intraflagellar transport protein 122, putative (IFT122) | -0,3366 | 0,0002421 |
| TcCLB.511365.50 | hypothetical protein, conserved (pseudogene) | -0,3366 | 0,002599 |
| TcCLB.506419.10 | ribosomal protein L7/L12, putative | -0,3367 | 0,04873 |
| TcCLB.511867.209 | dynein heavy chain, putative (fragment) | -0,3368 | 0,004175 |
| TcCLB.508769.10 | hypothetical protein, conserved (pseudogene) | -0,3368 | 0,008174 |
| TcCLB.510637.9 | dispersed gene family protein 1 (DGF-1), putative | -0,3369 | 0,1238 |
| TcCLB.509683.60 | hypothetical protein, conserved | -0,337 | 0,01775 |
| TcCLB.507807.40 | hypothetical protein, conserved | -0,3371 | 0,004849 |
| TcCLB.510355.269 | cytochrome b5, putative | -0,3371 | 0,1563 |
| TcCLB.506649.64 | hypothetical protein, conserved | -0,3371 | 0,1935 |
| TcCLB.505807.140 | hypothetical protein, conserved | -0,3372 | 0,002273 |
| TcCLB.506265.200 | hypothetical protein, conserved | -0,3372 | 0,08251 |
| TcCLB.507639.60 | hypothetical protein, conserved | -0,3373 | 0,005964 |
| TcCLB.509991.90 | RNA-editing nuclease 1 (KREN1) | -0,3373 | 0,01529 |
| TcCLB.410199.10 | hypothetical protein | -0,3373 | 0,07438 |
| TcCLB.506341.10 | N-acetylglucosamine-6-phosphate deacetylase-like proteir | -0,3376 | 0,0199 |
| TcCLB.508857.110 | hypothetical protein, conserved | -0,3377 | 0,08833 |
| TcCLB.508707.220 | hypothetical protein | -0,3378 | 0,00989 |
| TcCLB.508461.200 | hypothetical protein, conserved | -0,3378 | 0,01264 |
| TcCLB.508277.401 | magnesium and cobalt transport protein, putative (fragme | -0,3378 | 0,01676 |
| TcCLB.508997.20 | phosphatidylinositol 3-kinase, putative | -0,338 | 0,0005001 |
| TcCLB.511733.80 | hypothetical protein | -0,338 | 0,00236 |
| TcCLB.510963.90 | elongation factor 2, putative | -0,3381 | 1,902E-06 |
| TcCLB.510131.10 | hypothetical protein, conserved | -0,3382 | 0,003894 |
| TcCLB.508933.20 | dihydrolipoamide dehydrogenase, putative | -0,3382 | 0,0316 |
| TcCLB.509685.20 | syntaxin binding protein (pseudogene), putative | -0,3383 | 0,4571 |
| TcCLB.503781.60 | tRNA pseudouridine synthase A, putative | -0,3385 | 0,008923 |
| TcCLB.507669.84 | ATP-dependent DEAD/H RNA helicase, putative | -0,3387 | 0,0009042 |
| TcCLB.509505.89 | hypothetical protein, conserved | -0,3387 | 0,02533 |
| TcCLB.510283.100 | hypothetical protein, conserved | -0,3387 | 0,1684 |
| TcCLB.508153.440 | polynucleotide adenylyltransferase, putative (PAP) | -0,3388 | 0,01047 |
| TcCLB.511893.20 | hypothetical protein, conserved | -0,3388 | 0,5921 |
| TcCLB.510877.30 | hypothetical protein, conserved | -0,339 | 0,00001351 |

| | | | |
|---|---|---|---|
| TcCLB.508479.213 | hypothetical protein, conserved | -0,339 | 0,1418 |
| TcCLB.508479.465 | ATP-dependent DEAD/H RNA helicase (pseudogene), putat | -0,339 | 0,3463 |
| TcCLB.508277.290 | hypothetical protein, conserved | -0,3391 | 8,853E-06 |
| TcCLB.510357.80 | hypothetical protein, conserved | -0,3391 | 0,01055 |
| TcCLB.508851.100 | cyclophilin-like protein, putative | -0,3391 | 0,0926 |
| TcCLB.511283.200 | hypothetical protein, conserved | -0,3392 | 0,0002142 |
| TcCLB.510719.180 | hypothetical protein, conserved | -0,3392 | 0,002745 |
| TcCLB.509065.30 | hypothetical protein, conserved | -0,3392 | 0,006098 |
| TcCLB.507445.20 | trans-sialidase (pseudogene), putative | -0,3392 | 0,02228 |
| TcCLB.506963.60 | hypothetical protein, conserved (pseudogene) | -0,3392 | 0,02998 |
| TcCLB.506855.100 | hypothetical protein, conserved | -0,3392 | 0,04067 |
| TcCLB.511217.180 | hypothetical protein, conserved | -0,3393 | 0,004331 |
| TcCLB.503395.59 | hypothetical protein, conserved (fragment) | -0,3393 | 0,2658 |
| TcCLB.506737.135 | hypothetical protein, conserved (pseudogene) | -0,3395 | 0,03417 |
| TcCLB.508777.200 | hypothetical protein, conserved | -0,3395 | 0,246 |
| TcCLB.509179.30 | hypothetical protein, conserved | -0,3396 | 0,01948 |
| TcCLB.505987.70 | phosphatidylinositol 4-kinase, putative | -0,3396 | 0,03037 |
| TcCLB.510431.230 | hypothetical protein, conserved | -0,3397 | 0,006516 |
| TcCLB.508303.44 | hypothetical protein, conserved | -0,3398 | 0,07177 |
| TcCLB.506649.90 | hypothetical protein, conserved | -0,3399 | 5,423E-07 |
| TcCLB.510719.80 | hypothetical protein, conserved | -0,3399 | 1,827E-06 |
| TcCLB.504867.120 | hypothetical protein | -0,3399 | 0,0002395 |
| TcCLB.504021.60 | hypothetical protein, conserved | -0,34 | 0,01929 |
| TcCLB.507731.50 | trans-sialidase (pseudogene), putative | -0,34 | 0,4986 |
| TcCLB.510377.405 | hypothetical protein, conserved (pseudogene) | -0,34 | 0,5892 |
| TcCLB.511255.280 | mucin-associated surface protein (MASP), putative | -0,3402 | 0,3338 |
| TcCLB.510573.19 | hypothetical protein, conserved | -0,3403 | 0,05522 |
| TcCLB.508445.50 | hypothetical protein, conserved | -0,3403 | 0,1029 |
| TcCLB.511737.3 | superoxide dismutase, putative | -0,3405 | 0,156 |
| TcCLB.509017.50 | hypothetical protein, conserved | -0,3407 | 0,0001488 |
| TcCLB.508919.100 | hypothetical protein, conserved | -0,3408 | 0,005951 |
| TcCLB.507129.40 | hypothetical protein, conserved | -0,3408 | 0,1074 |
| TcCLB.507053.209 | Cytoplasmic dynein 2 heavy chain (DYNC2H1), putative | -0,3409 | 0,00007252 |
| TcCLB.506181.40 | hypothetical protein, conserved | -0,3409 | 0,005281 |
| TcCLB.506865.79 | hypothetical protein, conserved (fragment) | -0,3411 | 0,4609 |
| TcCLB.504105.120 | hypothetical protein, conserved | -0,3414 | 0,00001069 |
| TcCLB.509937.10 | poly(A) polymerase, putative | -0,3414 | 0,01635 |
| TcCLB.404845.10 | trans-sialidase (pseudogene), putative | -0,3415 | 0,02904 |
| TcCLB.506583.10 | U2 small nuclear ribonucleoprotein 16.5K, putative | -0,3415 | 0,02999 |
| TcCLB.509735.100 | trans-sialidase (pseudogene), putative | -0,3415 | 0,0749 |
| TcCLB.506177.120 | hypothetical protein | -0,3416 | 0,03014 |
| TcCLB.511543.40 | hypothetical protein, conserved | -0,3418 | 0,002558 |
| TcCLB.508851.79 | hypothetical protein, conserved | -0,3418 | 0,03014 |
| TcCLB.509485.20 | retrotransposon hot spot protein (RHS, pseudogene), putat | -0,3419 | 0,124 |
| TcCLB.507611.340 | hypothetical protein, conserved | -0,342 | 0,003567 |

| | | | |
|---|---|---|---|
| TcCLB.510535.40 | hypothetical protein, conserved | -0,342 | 0,03149 |
| TcCLB.510187.430 | DNA mismatch repair protein MSH6, putative (MSH6) | -0,3422 | 0,00002253 |
| TcCLB.504103.100 | ABC1 protein, putative | -0,3423 | 0,006032 |
| TcCLB.510289.70 | hypothetical protein, conserved | -0,3424 | 0,03956 |
| TcCLB.511047.50 | proteasome regulatory ATPase subunit 2, putative | -0,3425 | 0,0006646 |
| TcCLB.511421.224 | hypothetical protein, conserved (pseudogene) | -0,3425 | 0,001929 |
| TcCLB.504215.10 | hypothetical protein, conserved | -0,3425 | 0,006017 |
| TcCLB.506941.60 | phospholipid-translocating P-type ATPase (flippase), putati | -0,3425 | 0,007053 |
| TcCLB.508409.90 | hypothetical protein, conserved | -0,3428 | 0,00001361 |
| TcCLB.507031.20 | hypothetical protein, conserved | -0,3429 | 0,002269 |
| TcCLB.509053.40 | hypothetical protein, conserved | -0,3429 | 0,005228 |
| TcCLB.503903.30 | hypothetical protein | -0,343 | 0,0103 |
| TcCLB.507883.100 | malate dehydrogenase, putative | -0,3432 | 0,01032 |
| TcCLB.508445.60 | hypothetical protein, conserved | -0,3433 | 0,01437 |
| TcCLB.410049.10 | hypothetical protein, conserved | -0,3434 | 0,5558 |
| TcCLB.508989.80 | hypothetical protein | -0,3435 | 0,0005113 |
| TcCLB.511365.10 | hypothetical protein, conserved | -0,3436 | 0,0109 |
| TcCLB.508295.70 | hypothetical protein | -0,3436 | 0,5028 |
| TcCLB.504153.190 | mitochondrial edited mRNA stability factor 1 subunit | -0,3438 | 0,0005067 |
| TcCLB.506743.150 | proteasome 26S non-ATPase subunit 9, putative | -0,3438 | 0,04675 |
| TcCLB.506537.260 | protein kinase, putative | -0,3438 | 0,3648 |
| TcCLB.506067.80 | mucin TcMUCII, putative | -0,3439 | 0,5567 |
| TcCLB.506137.40 | mucin TcMUCII, putative | -0,3439 | 0,7056 |
| TcCLB.504193.51 | ATP-dependent DEAD/H RNA helicase (pseudogene), putat | -0,344 | 0,0403 |
| TcCLB.503693.20 | hypothetical protein, conserved | -0,344 | 0,04248 |
| TcCLB.507159.37 | hypothetical protein, conserved | -0,3441 | 0,01499 |
| TcCLB.506925.490 | hypothetical protein, conserved | -0,3443 | 0,01164 |
| TcCLB.507993.120 | synaptotagmin, putative | -0,3443 | 0,05884 |
| TcCLB.503541.20 | hypothetical protein, conserved | -0,3444 | 0,03276 |
| TcCLB.503531.49 | structural maintenance of chromosome 3, putative | -0,3444 | 0,1299 |
| TcCLB.511215.30 | hypothetical protein | -0,3445 | 0,003386 |
| TcCLB.506725.50 | hypothetical protein, conserved | -0,3445 | 0,01116 |
| TcCLB.505555.40 | hypothetical protein, conserved | -0,3446 | 0,002033 |
| TcCLB.506629.60 | vesicular-fusion ATPase-like protein, putative | -0,3446 | 0,004291 |
| TcCLB.508141.40 | ATP-dependent RNA helicase, putative | -0,3447 | 3,377E-06 |
| TcCLB.511881.10 | GCN5-like protein, putative | -0,3448 | 0,1854 |
| TcCLB.509065.20 | mitogen-activated protein kinase 5, putative | -0,3449 | 0,04711 |
| TcCLB.509395.30 | tRNA exportin, putative | -0,345 | 0,0001716 |
| TcCLB.504089.40 | long-chain-fatty-acid-CoA ligase, putative | -0,345 | 0,0008972 |
| TcCLB.511479.30 | hypothetical protein, conserved | -0,345 | 0,02138 |
| TcCLB.509099.40 | hypothetical protein, conserved | -0,345 | 0,05322 |
| TcCLB.508613.100 | surface protease GP63 (pseudogene), putative | -0,345 | 0,2407 |
| TcCLB.508153.994 | cytochrome c oxidase copper chaperone, putative | -0,3452 | 0,1118 |
| TcCLB.509767.20 | hypothetical protein, conserved | -0,3452 | 0,1238 |
| TcCLB.511537.40 | protein phosphatase 2A catalytic subunit, putative | -0,3453 | 0,00442 |

| | | | |
|---|---|---|---|
| TcCLB.510533.90 | hypothetical protein, conserved | -0,3454 | 0,01374 |
| TcCLB.511423.149 | conserved protein, unknown function | -0,3454 | 0,04064 |
| TcCLB.509617.40 | hypothetical protein, conserved | -0,3455 | 0,001438 |
| TcCLB.511303.100 | hypothetical protein | -0,3455 | 0,02235 |
| TcCLB.511165.10 | ATP-dependent DEAD/H RNA helicase, putative | -0,3455 | 0,02741 |
| TcCLB.505025.80 | hypothetical protein, conserved | -0,3455 | 0,04103 |
| TcCLB.508637.130 | hypothetical protein, conserved | -0,3456 | 0,2908 |
| TcCLB.510013.120 | surface protease GP63, putative | -0,3458 | 0,6779 |
| TcCLB.511275.10 | hypothetical protein, conserved (pseudogene) | -0,3459 | 0,00002546 |
| TcCLB.508965.100 | hypothetical protein, conserved | -0,3459 | 0,0002848 |
| TcCLB.461165.10 | hypothetical protein, conserved | -0,3461 | 0,03907 |
| TcCLB.507739.120 | amastin, putative | -0,3461 | 0,3572 |
| TcCLB.509297.20 | hypothetical protein, conserved | -0,3463 | 0,004466 |
| TcCLB.426799.19 | serine/threonine protein kinase, putative | -0,3463 | 0,09488 |
| TcCLB.506627.110 | hypothetical protein, conserved | -0,3465 | 0,0005777 |
| TcCLB.506247.450 | hypothetical protein, conserved | -0,3465 | 0,002802 |
| TcCLB.503959.50 | hypothetical protein, conserved | -0,3465 | 0,01757 |
| TcCLB.508959.20 | hypothetical protein, conserved | -0,3465 | 0,01821 |
| TcCLB.506613.20 | mucin-associated surface protein (MASP, pseudogene), pu | -0,3465 | 0,3431 |
| TcCLB.506431.5 | trans-sialidase, putative (pseudogene) | -0,3466 | 0,5276 |
| TcCLB.508435.60 | mucin TcMUCII, putative | -0,3466 | 0,5661 |
| TcCLB.508989.30 | RNA-binding protein, putative | -0,3467 | 0,002827 |
| TcCLB.508269.30 | translation initiation factor 2 subunit, putative | -0,3467 | 0,01849 |
| TcCLB.506227.80 | mitochondrial DNA polymerase I protein B, putative (POLIE | -0,3468 | 0,0002409 |
| TcCLB.504741.10 | hypothetical protein, conserved | -0,3469 | 0,04424 |
| TcCLB.505073.14 | hypothetical protein, conserved | -0,3471 | 0,06833 |
| TcCLB.504173.50 | dihydrouridine synthase (Dus), putative | -0,3473 | 0,04122 |
| TcCLB.505009.49 | hypothetical protein, conserved | -0,3474 | 0,01283 |
| TcCLB.504073.30 | ribonucleoside-diphosphate reductase large chain, putative | -0,3475 | 0,1056 |
| TcCLB.511283.120 | hypothetical protein, conserved | -0,3475 | 0,5623 |
| TcCLB.508879.170 | mitochondrial RNA binding complex 1 subunit, putative | -0,3476 | 0,001306 |
| TcCLB.503651.4 | hypothetical protein, conserved (fragment) | -0,3476 | 0,008101 |
| TcCLB.507993.210 | hypothetical protein, conserved | -0,3477 | 0,002827 |
| TcCLB.511431.50 | hypothetical protein, conserved | -0,3477 | 0,005528 |
| TcCLB.510725.80 | hypothetical protein | -0,3477 | 0,02209 |
| TcCLB.508309.10 | mucin-associated surface protein (MASP), putative | -0,3477 | 0,483 |
| TcCLB.504081.80 | trans-sialidase, Group V, putative | -0,3478 | 0,3992 |
| TcCLB.508465.70 | hypothetical protein, conserved | -0,3479 | 0,009058 |
| TcCLB.506501.380 | mucin-associated surface protein (MASP), putative | -0,3479 | 0,3847 |
| TcCLB.508951.80 | hypothetical protein, conserved | -0,348 | 0,003087 |
| TcCLB.510603.100 | hypothetical protein, conserved | -0,348 | 0,07468 |
| TcCLB.503657.9 | hypothetical protein, conserved | -0,3481 | 0,006261 |
| TcCLB.508153.1030 | hypothetical protein, conserved | -0,3482 | 0,00013 |
| TcCLB.508911.30 | hypothetical protein, conserved (pseudogene) | -0,3482 | 0,007638 |
| TcCLB.506753.110 | cytochrome b5-like protein, putative | -0,3482 | 0,01414 |

| | | | |
|---|---|---|---|
| TcCLB.509671.180 | hypothetical protein, conserved | -0,3482 | 0,02475 |
| TcCLB.509563.50 | hypothetical protein, conserved | -0,3483 | 0,006026 |
| TcCLB.511003.10 | phospholipid-translocating ATPase, putative | -0,3484 | 0,002992 |
| TcCLB.511621.70 | ribosomal protein L11, putative | -0,3484 | 0,01342 |
| TcCLB.507641.80 | 5'-3' exoribonuclease D, putative (XRND) | -0,3485 | 0,00005633 |
| TcCLB.510089.64 | hypothetical protein, conserved | -0,3485 | 0,02281 |
| TcCLB.510175.140 | hypothetical protein, conserved (pseudogene) | -0,3485 | 0,05578 |
| TcCLB.510359.550 | mucin-associated surface protein (MASP), putative | -0,3485 | 0,4916 |
| TcCLB.506977.90 | proteasome regulatory non-ATP-ase subunit 5, putative (R | -0,3487 | 7,631E-07 |
| TcCLB.511211.150 | spindle pole body component alp6, putative | -0,3487 | 0,003172 |
| TcCLB.508741.310 | hypothetical protein | -0,3487 | 0,005754 |
| TcCLB.509033.50 | hypothetical protein, conserved | -0,3489 | 0,0004743 |
| TcCLB.507485.70 | hypothetical protein, conserved | -0,3489 | 0,03988 |
| TcCLB.507031.80 | hypothetical protein, conserved | -0,349 | 0,00004186 |
| TcCLB.511417.50 | hypothetical protein | -0,349 | 0,07338 |
| TcCLB.506871.110 | hypothetical protein, conserved | -0,3491 | 0,003785 |
| TcCLB.511307.6 | hypothetical protein, conserved | -0,3491 | 0,0407 |
| TcCLB.507963.90 | hypothetical protein, conserved | -0,3491 | 0,04824 |
| TcCLB.507485.90 | Wee1-like protein kinase, putative | -0,3492 | 0,001315 |
| TcCLB.503893.60 | hypothetical protein, conserved | -0,3492 | 0,008362 |
| TcCLB.507709.100 | protein phosphatase 2A regulatory subunit, putative | -0,3494 | 0,01258 |
| TcCLB.509877.30 | guanine nucleotide-binding beta subunit-like protein | -0,3495 | 0,0006969 |
| TcCLB.510953.49 | lipase-like protein, putative | -0,3495 | 0,06209 |
| TcCLB.508461.500 | 60S ribosomal protein L27A/L29, putative | -0,3496 | 0,03801 |
| TcCLB.506589.89 | mucin-associated surface protein (MASP, pseudogene), pu | -0,3496 | 0,3421 |
| TcCLB.510901.120 | protein kinase, putative | -0,35 | 0,0002162 |
| TcCLB.506941.70 | hypothetical protein, conserved | -0,35 | 0,0005296 |
| TcCLB.503925.30 | protein kinase, putative | -0,35 | 0,01589 |
| TcCLB.503599.40 | hypothetical protein, conserved | -0,3501 | 0,003929 |
| TcCLB.505939.89 | 5'-3' exoribonuclease A, putative (fragment) (XRNA) | -0,3501 | 0,006875 |
| TcCLB.510175.160 | protein kinase, putative | -0,3501 | 0,1164 |
| TcCLB.511249.60 | protein kinase, putative | -0,3506 | 0,000435 |
| TcCLB.509641.40 | hypothetical protein, conserved | -0,3507 | 0,00006512 |
| TcCLB.510575.200 | chaperone DNAJ protein, putative | -0,3507 | 0,0001788 |
| TcCLB.511127.380 | hypothetical protein, conserved | -0,3508 | 0,00001371 |
| TcCLB.506635.20 | ATP-dependent RNA helicase, putative | -0,3508 | 0,0005301 |
| TcCLB.510457.10 | mucin-associated surface protein (MASP), putative | -0,351 | 0,5996 |
| TcCLB.510647.69 | hypothetical protein, conserved | -0,3511 | 0,2731 |
| TcCLB.509647.140 | hypothetical protein, conserved | -0,3512 | 0,002036 |
| TcCLB.506587.20 | hypothetical protein | -0,3513 | 0,006345 |
| TcCLB.506513.40 | hypothetical protein, conserved | -0,3513 | 0,0312 |
| TcCLB.509959.60 | trans-sialidase (pseudogene), putative | -0,3513 | 0,534 |
| TcCLB.506519.120 | hypothetical protein, conserved | -0,3514 | 0,008754 |
| TcCLB.511301.130 | protein kinase, putative | -0,3514 | 0,03119 |
| TcCLB.504117.50 | hypothetical protein, conserved (pseudogene) | -0,3514 | 0,3522 |

| | | | |
|---|---|---|---|
| TcCLB.506505.40 | protein kinase, putative | -0,3515 | 0,0002277 |
| TcCLB.509455.70 | DNA polymerase delta subunit 2, putative | -0,3515 | 0,001655 |
| TcCLB.508723.40 | hypothetical protein, conserved | -0,3516 | 0,0005385 |
| TcCLB.508479.174 | hypothetical protein, conserved | -0,3517 | 0,009129 |
| TcCLB.506315.70 | hypothetical protein, conserved | -0,3518 | 0,02417 |
| TcCLB.511371.5 | trans-sialidase (pseudogene), putative | -0,3518 | 0,1497 |
| TcCLB.471901.5 | GDP-mannose 4,6 dehydratase, putative (fragment) | -0,3518 | 0,5278 |
| TcCLB.507483.30 | calmodulin, putative | -0,3519 | 0,003513 |
| TcCLB.509267.80 | hypothetical protein | -0,352 | 0,03875 |
| TcCLB.506755.140 | ATP-dependent helicase, putative | -0,3521 | 0,0007434 |
| TcCLB.510535.70 | hypothetical protein, conserved | -0,3521 | 0,007863 |
| TcCLB.509821.20 | trans-sialidase (pseudogene), putative | -0,3522 | 0,03608 |
| TcCLB.511925.10 | hypothetical protein, conserved | -0,3523 | 0,0033 |
| TcCLB.509647.160 | SKP1-like protein | -0,3525 | 0,01528 |
| TcCLB.507471.41 | trans-sialidase (pseudogene), putative | -0,3525 | 0,09898 |
| TcCLB.511245.79 | hypothetical protein, conserved | -0,3526 | 0,01738 |
| TcCLB.510203.49 | hypothetical protein, conserved (pseudogene) | -0,3527 | 0,06814 |
| TcCLB.508891.120 | periodic tryptophan protein 2 (pseudogene), putative | -0,3528 | 0,001001 |
| TcCLB.509595.40 | hypothetical protein, conserved | -0,3528 | 0,002791 |
| TcCLB.510237.129 | dispersed gene family protein 1 (DGF-1), putative (fragme | -0,3528 | 0,1567 |
| TcCLB.511233.100 | NUDIX hydrolase 3, putative (Nudix3) | -0,3529 | 0,0404 |
| TcCLB.506475.80 | WD40 repeat-containing protein | -0,353 | 0,0001595 |
| TcCLB.503845.60 | trans-sialidase (pseudogene), putative | -0,353 | 0,01013 |
| TcCLB.508987.9 | hypothetical protein, conserved | -0,3532 | 0,0001534 |
| TcCLB.510759.180 | hypothetical protein, conserved | -0,3532 | 0,000952 |
| TcCLB.511277.10 | DNA-directed RNA polymerase I subunit RPB5z, putative (F | -0,3533 | 0,02336 |
| TcCLB.507715.30 | transcription elongation factor-like protein, putative | -0,3534 | 0,002333 |
| TcCLB.506681.50 | hypothetical protein, conserved | -0,3536 | 0,0009524 |
| TcCLB.510303.40 | hypothetical protein | -0,3536 | 0,007444 |
| TcCLB.506989.60 | phosphate transporter, putative | -0,3536 | 0,01192 |
| TcCLB.508509.70 | mucin TcMUCII, putative | -0,3536 | 0,6008 |
| TcCLB.507031.130 | hypothetical protein, conserved | -0,3538 | 0,00168 |
| TcCLB.506425.20 | hypothetical protein | -0,3538 | 0,0284 |
| TcCLB.510661.150 | hypothetical protein, conserved | -0,3539 | 0,02115 |
| TcCLB.503909.110 | intraflagellar transport (IFT) protein, putative (IFT20) | -0,3539 | 0,03639 |
| TcCLB.508703.20 | trans-sialidase (pseudogene), putative | -0,354 | 0,3772 |
| TcCLB.511033.10 | hypothetical protein, conserved | -0,3541 | 0,0102 |
| TcCLB.506735.60 | hypothetical protein, conserved | -0,3542 | 0,09375 |
| TcCLB.506175.70 | hypothetical protein, conserved | -0,3543 | 0,004823 |
| TcCLB.508857.100 | hypothetical protein, conserved | -0,3544 | 0,001834 |
| TcCLB.511751.70 | hypothetical protein, conserved | -0,3545 | 7,114E-06 |
| TcCLB.507089.180 | hypothetical protein, conserved | -0,3545 | 0,001197 |
| TcCLB.506605.100 | hypothetical protein | -0,3546 | 0,006286 |
| TcCLB.506855.370 | hypothetical protein, conserved | -0,3546 | 0,007546 |
| TcCLB.511419.70 | hypothetical protein | -0,3546 | 0,03832 |

| | | | |
|---|---|---|---|
| TcCLB.508427.20 | mucin-associated surface protein (MASP), putative | -0,3546 | 0,2934 |
| TcCLB.504081.390 | trans-sialidase (pseudogene), putative | -0,3547 | 0,2183 |
| TcCLB.511211.20 | Coatomer subunit gamma (COPG) | -0,3552 | 0,00006661 |
| TcCLB.507809.70 | small ubiquitin protein, putative | -0,3554 | 0,0442 |
| TcCLB.507077.30 | hypothetical protein, conserved | -0,3555 | 0,1116 |
| TcCLB.444777.29 | intraflagellar transport protein 88, putative (fragment) (IF | -0,3556 | 0,01086 |
| TcCLB.503513.20 | hypothetical protein | -0,3556 | 0,01158 |
| TcCLB.509785.90 | trans-sialidase (pseudogene), putative | -0,3557 | 0,2252 |
| TcCLB.509011.50 | hypothetical protein, conserved | -0,3558 | 0,00005118 |
| TcCLB.510073.90 | hypothetical protein, conserved | -0,3558 | 0,008705 |
| TcCLB.506405.40 | hypothetical protein, conserved | -0,3558 | 0,01965 |
| TcCLB.509395.90 | hypothetical protein, conserved | -0,3558 | 0,04802 |
| TcCLB.510429.10 | hypothetical protein, conserved | -0,3558 | 0,07777 |
| TcCLB.508707.180 | ubiquitin hydrolase, putative | -0,3562 | 0,0001463 |
| TcCLB.511285.120 | ATP-dependent RNA helicase, putative | -0,3563 | 0,009261 |
| TcCLB.506265.40 | hypothetical protein, conserved | -0,3563 | 0,1326 |
| TcCLB.507389.10 | hypothetical protein, conserved | -0,3564 | 0,02472 |
| TcCLB.509019.30 | dispersed gene family protein 1 (DGF-1), putative | -0,3564 | 0,2397 |
| TcCLB.509733.10 | hypothetical protein, conserved | -0,3567 | 0,0095 |
| TcCLB.507603.230 | hydroxyacylglutathione hydrolase, putative | -0,3568 | 0,0503 |
| TcCLB.511237.90 | hypothetical protein, conserved | -0,3568 | 0,2043 |
| TcCLB.511277.330 | uracil-DNA glycosylase | -0,3569 | 0,00137 |
| TcCLB.506435.130 | hypothetical protein, conserved | -0,3569 | 0,00825 |
| TcCLB.506947.100 | hypothetical protein, conserved | -0,3569 | 0,0116 |
| TcCLB.508533.20 | hypothetical protein, conserved | -0,3569 | 0,0129 |
| TcCLB.508175.180 | hypothetical protein, conserved | -0,3571 | 0,0009164 |
| TcCLB.509007.60 | rac serine-threonine kinase, putative | -0,3571 | 0,003612 |
| TcCLB.509157.270 | hypothetical protein, conserved | -0,3572 | 0,0201 |
| TcCLB.503841.50 | hypothetical protein, conserved | -0,3574 | 0,035 |
| TcCLB.506947.60 | hypothetical protein, conserved | -0,3575 | 0,02372 |
| TcCLB.507507.49 | hypothetical protein, conserved | -0,3575 | 0,27 |
| TcCLB.504045.120 | hypothetical protein, conserved | -0,3576 | 0,0044 |
| TcCLB.510001.31 | hypothetical protein, conserved | -0,3576 | 0,03853 |
| TcCLB.510879.150 | hypothetical protein, conserved | -0,3577 | 0,004268 |
| TcCLB.506289.9 | DNA ligase, putative (fragment) | -0,3577 | 0,01676 |
| TcCLB.506853.30 | arginine N-methyltransferase, type II, putative (PRMT5) | -0,3578 | 0,003134 |
| TcCLB.508153.590 | hypothetical protein, conserved | -0,3579 | 0,01456 |
| TcCLB.511189.70 | hypothetical protein, conserved | -0,3579 | 0,4034 |
| TcCLB.503697.120 | hypothetical protein, conserved | -0,358 | 0,0009932 |
| TcCLB.509649.30 | 19S proteasome non-atpase subunit 8 (Rpn8) | -0,358 | 0,001178 |
| TcCLB.506177.40 | palmitoyl acyltransferase 6, putative | -0,358 | 0,08318 |
| TcCLB.506247.440 | hypothetical protein, conserved | -0,358 | 0,1143 |
| TcCLB.506699.9 | hypothetical protein, conserved | -0,3582 | 0,05751 |
| TcCLB.506241.19 | dispersed gene family protein 1 (DGF-1), putative | -0,3582 | 0,06738 |
| TcCLB.448387.9 | hypothetical protein, conserved | -0,3583 | 0,01919 |

| | | | |
|---|---|---|---|
| TcCLB.507395.20 | hypothetical protein, conserved | -0,3583 | 0,04125 |
| TcCLB.506355.35 | trans-sialidase (pseudogene), putative | -0,3584 | 0,03871 |
| TcCLB.504059.24 | hypothetical protein, conserved | -0,3585 | 0,07171 |
| TcCLB.511725.150 | exportin 1, putative (CRM1) | -0,3586 | 0,00003794 |
| TcCLB.511507.110 | hypothetical protein, conserved (YCF45) | -0,3586 | 0,006021 |
| TcCLB.510387.70 | hypothetical protein, conserved (pseudogene) | -0,3586 | 0,1555 |
| TcCLB.510747.130 | hypothetical protein, conserved | -0,3588 | 0,0006133 |
| TcCLB.506513.20 | hypothetical protein, conserved | -0,3588 | 0,04737 |
| TcCLB.506751.20 | mucin-associated surface protein (MASP, pseudogene), pu | -0,3589 | 0,4373 |
| TcCLB.506211.60 | RNA-binding protein, putative | -0,359 | 0,001544 |
| TcCLB.511173.80 | trans-sialidase, Group V, putative | -0,3591 | 0,3083 |
| TcCLB.504575.60 | surface protease GP63, putative (fragment) | -0,3591 | 0,6818 |
| TcCLB.511903.160 | phosphatidylinositol 3-kinase vps34-like | -0,3592 | 0,0001011 |
| TcCLB.509455.140 | cyclin, putative | -0,3592 | 0,01541 |
| TcCLB.508271.4 | hypothetical protein, conserved | -0,3592 | 0,07196 |
| TcCLB.510953.40 | hypothetical protein, conserved | -0,3593 | 0,01224 |
| TcCLB.507969.60 | C-14 sterol reductase, putative | -0,3594 | 0,004296 |
| TcCLB.508593.44 | serine peptidase, Clan SC, Family S9D | -0,3594 | 0,004563 |
| TcCLB.506583.40 | cdc2-related kinase 3, putative (CRK3) | -0,3594 | 0,02644 |
| TcCLB.511529.230 | hypothetical protein, conserved | -0,3595 | 0,0316 |
| TcCLB.507011.170 | hypothetical protein, conserved | -0,3596 | 0,003306 |
| TcCLB.507049.150 | hypothetical protein, conserved | -0,3596 | 0,01667 |
| TcCLB.507773.40 | hypothetical protein, conserved | -0,3597 | 0,004275 |
| TcCLB.503513.10 | casein kinase II, alpha chain, putative | -0,3597 | 0,007524 |
| TcCLB.510105.30 | clathrin coat assembly protein, putative | -0,3597 | 0,01465 |
| TcCLB.506959.80 | hypothetical protein, conserved | -0,3597 | 0,01996 |
| TcCLB.507547.30 | Mitochondrial-processing peptidase subunit alpha (MPPA) | -0,3598 | 0,000171 |
| TcCLB.506855.90 | structural maintenance of chromosome protein 4, putative | -0,36 | 0,00001433 |
| TcCLB.510769.120 | calcium motive p-type ATPase, putative | -0,36 | 0,00007686 |
| TcCLB.505057.10 | hypothetical protein, conserved | -0,36 | 0,004441 |
| TcCLB.509215.20 | hypothetical protein, conserved | -0,36 | 0,07147 |
| TcCLB.511063.80 | dispersed gene family protein 1 (DGF-1, pseudogene), puta | -0,3601 | 0,277 |
| TcCLB.503893.160 | hypothetical protein, conserved | -0,3602 | 0,004924 |
| TcCLB.510445.61 | hypothetical protein, conserved (fragment) | -0,3603 | 0,07413 |
| TcCLB.509069.40 | hypothetical protein | -0,3604 | 0,002459 |
| TcCLB.507763.10 | hypothetical protein, conserved | -0,3604 | 0,01492 |
| TcCLB.511733.60 | hypothetical protein, conserved | -0,3605 | 0,01997 |
| TcCLB.510265.30 | hypothetical protein, conserved | -0,3606 | 0,00000134 |
| TcCLB.508461.230 | kinetoplastid kinetochore protein 3, putative (kkt3) | -0,3606 | 0,002235 |
| TcCLB.511003.120 | hypothetical protein, conserved | -0,3606 | 0,04124 |
| TcCLB.509591.30 | ras-family member, GTP-binding protein, putative | -0,3607 | 0,001436 |
| TcCLB.506389.10 | trans-sialidase, putative | -0,3607 | 0,05998 |
| TcCLB.508883.20 | hypothetical protein | -0,3607 | 0,5256 |
| TcCLB.508231.150 | hypothetical protein | -0,3608 | 0,0007352 |
| TcCLB.508153.550 | hypothetical protein, conserved | -0,3609 | 0,000878 |

| | | | |
|---|---|---|---|
| TcCLB.511653.50 | hypothetical protein, conserved | -0,3609 | 0,009689 |
| TcCLB.508605.31 | trans-sialidase (pseudogene), putative | -0,3609 | 0,02097 |
| TcCLB.511771.70 | dispersed gene family protein 1 (DGF-1), putative | -0,3609 | 0,03815 |
| TcCLB.510749.90 | hypothetical protein, conserved | -0,3609 | 0,07461 |
| TcCLB.506797.100 | beta-ketoacyl-ACP reductase 2, putative | -0,361 | 0,004704 |
| TcCLB.506181.90 | F-actin capping protein, putative | -0,361 | 0,006765 |
| TcCLB.511867.120 | hypothetical protein, conserved | -0,3611 | 0,0105 |
| TcCLB.506639.50 | dispersed gene family protein 1 (DGF-1, pseudogene), puta | -0,3611 | 0,09791 |
| TcCLB.506605.214 | hypothetical protein, conserved | -0,3612 | 0,09336 |
| TcCLB.508277.20 | protein kinase, putative | -0,3613 | 0,001081 |
| TcCLB.509965.150 | hypothetical protein, conserved | -0,3614 | 0,0159 |
| TcCLB.507031.64 | hypothetical protein, conserved | -0,3615 | 0,02697 |
| TcCLB.511801.40 | hypothetical protein, conserved | -0,3615 | 0,02847 |
| TcCLB.418221.4 | hypothetical protein, conserved (fragment) | -0,3616 | 0,02128 |
| TcCLB.508207.31 | hypothetical protein | -0,3617 | 0,04692 |
| TcCLB.510307.94 | ATP-dependent DEAD/H RNA helicase (pseudogene), putat | -0,3617 | 0,139 |
| TcCLB.507883.70 | PrimPol-like protein 2, putative (ppl2) | -0,3619 | 0,001008 |
| TcCLB.508461.300 | hypothetical protein, conserved | -0,3619 | 0,001446 |
| TcCLB.509805.180 | ttaggg binding factor, putative | -0,3619 | 0,004668 |
| TcCLB.508641.320 | formin-like protein | -0,362 | 0,0005911 |
| TcCLB.511603.369 | surface protease GP63 (pseudogene), putative | -0,362 | 0,5412 |
| TcCLB.508803.5 | retrotransposon hot spot protein (RHS, pseudogene), putat | -0,362 | 0,558 |
| TcCLB.510741.40 | hypothetical protein, conserved | -0,3621 | 0,0001342 |
| TcCLB.508859.100 | hypothetical protein, conserved | -0,3622 | 0,03974 |
| TcCLB.506131.120 | trans-sialidase (pseudogene), putative | -0,3622 | 0,2614 |
| TcCLB.508221.990 | surface protease GP63 (pseudogene), putative | -0,3622 | 0,3883 |
| TcCLB.507053.10 | 50S ribosomal protein L7Ae, putative | -0,3623 | 0,0004174 |
| TcCLB.511367.310 | hypothetical protein, conserved | -0,3623 | 0,01284 |
| TcCLB.509985.78 | hypothetical protein, conserved | -0,3623 | 0,04424 |
| TcCLB.506953.20 | hypothetical protein, conserved | -0,3623 | 0,1334 |
| TcCLB.506297.50 | kinesin, putative | -0,3624 | 4,931E-06 |
| TcCLB.503577.20 | U2 splicing auxiliary factor, putative (u2af26) | -0,3624 | 0,00002577 |
| TcCLB.506559.530 | protein phosphatase 2C, putative | -0,3626 | 0,00724 |
| TcCLB.506579.110 | hypothetical protein, conserved | -0,3627 | 0,00000128 |
| TcCLB.511183.330 | dispersed gene family protein 1 (DGF-1), putative | -0,3628 | 0,2481 |
| TcCLB.503809.140 | cation transporter, putative | -0,3629 | 0,002077 |
| TcCLB.507243.9 | dispersed gene family protein 1 (DGF-1), putative | -0,363 | 0,08823 |
| TcCLB.510933.110 | mucin-associated surface protein (MASP, pseudogene), pu | -0,363 | 0,4048 |
| TcCLB.507941.20 | hypothetical protein, conserved | -0,3631 | 0,00004742 |
| TcCLB.503627.9 | hypothetical protein, conserved (fragment) | -0,3631 | 0,00386 |
| TcCLB.504103.70 | hypothetical protein, conserved | -0,3631 | 0,02214 |
| TcCLB.506701.10 | dihydrolipoamide dehydrogenase, putative (GCVL-1) | -0,3631 | 0,05997 |
| TcCLB.506529.40 | hypothetical protein, conserved | -0,3632 | 0,04909 |
| TcCLB.506335.100 | mucin-associated surface protein (MASP), putative | -0,3633 | 0,1034 |
| TcCLB.505939.65 | hypothetical protein, conserved | -0,3633 | 0,1995 |

| | | | |
|---|---|---|---|
| TcCLB.511545.70 | kinteoplast poly(A) polymerase complex 1 subunit, putative | -0,3634 | 0,00006927 |
| TcCLB.511847.10 | calpain-like cysteine peptidase, putative | -0,3634 | 0,002328 |
| TcCLB.510879.90 | hypothetical protein, conserved | -0,3634 | 0,04147 |
| TcCLB.508999.20 | Protein transport protein yif1, putative | -0,3636 | 0,03066 |
| TcCLB.511523.50 | hypothetical protein, conserved | -0,3637 | 0,01462 |
| TcCLB.469785.30 | hypothetical protein | -0,3637 | 0,01729 |
| TcCLB.508323.120 | hypothetical protein, conserved | -0,3638 | 0,03536 |
| TcCLB.511727.60 | hypothetical protein, conserved | -0,3639 | 0,00005239 |
| TcCLB.503841.30 | rRNA dimethyltransferase, putative | -0,3639 | 0,04943 |
| TcCLB.508413.60 | hypothetical protein, conserved | -0,364 | 0,01561 |
| TcCLB.508739.30 | hypothetical protein | -0,364 | 0,03416 |
| TcCLB.503887.10 | hypothetical protein, conserved | -0,364 | 0,04474 |
| TcCLB.511131.50 | U4/U6 small nuclear ribonuclear protein, putative | -0,3641 | 0,002613 |
| TcCLB.508641.310 | hypothetical protein, conserved | -0,3642 | 0,0006594 |
| TcCLB.506367.50 | hypothetical protein, conserved | -0,3642 | 0,05456 |
| TcCLB.506925.510 | ribosome production factor 1, putative | -0,3643 | 0,0004059 |
| TcCLB.508325.30 | dispersed gene family protein 1 (DGF-1), putative | -0,3643 | 0,01567 |
| TcCLB.510659.250 | phospholipase A1, putative | -0,3644 | 0,002244 |
| TcCLB.510297.40 | hypothetical protein | -0,3644 | 0,07548 |
| TcCLB.511509.5 | hypothetical protein, conserved | -0,3645 | 0,004185 |
| TcCLB.503851.24 | protein tyrosine phosphatase PRL | -0,3646 | 0,1039 |
| TcCLB.506569.45 | protein kinase (pseudogene), putative | -0,3646 | 0,1727 |
| TcCLB.507093.200 | hypothetical protein, conserved | -0,3647 | 0,0001092 |
| TcCLB.509261.20 | aminopeptidase, putative | -0,3647 | 0,0003303 |
| TcCLB.509287.60 | hypothetical protein, conserved | -0,3647 | 0,0004114 |
| TcCLB.511287.170 | hypothetical protein, conserved | -0,3647 | 0,004182 |
| TcCLB.509991.70 | hypothetical protein, conserved (pseudogene) | -0,3647 | 0,007701 |
| TcCLB.511295.50 | hypothetical protein, conserved | -0,3648 | 1,783E-06 |
| TcCLB.510167.4 | hypothetical protein, conserved | -0,3649 | 0,1172 |
| TcCLB.508817.90 | hypothetical protein, conserved (pseudogene) | -0,3651 | 0,008943 |
| TcCLB.508891.80 | hypothetical protein, conserved (pseudogene) | -0,3651 | 0,01163 |
| TcCLB.511527.82 | hypothetical protein, conserved | -0,3651 | 0,1324 |
| TcCLB.510607.80 | dispersed gene family protein 1 (DGF-1), putative | -0,3652 | 0,1392 |
| TcCLB.506885.4 | hypothetical protein, conserved (fragment) | -0,3654 | 0,006611 |
| TcCLB.508799.50 | hypothetical protein | -0,3655 | 0,01341 |
| TcCLB.504105.240 | hypothetical protein, conserved | -0,3656 | 0,0001577 |
| TcCLB.507081.30 | translation elongation factor EF-2, putative | -0,3656 | 0,0007304 |
| TcCLB.507795.100 | mitochondrial ribosomal protein L3, putative | -0,3657 | 0,0003849 |
| TcCLB.506195.309 | hypothetical protein, conserved | -0,3659 | 0,006545 |
| TcCLB.506945.105 | zinc-binding protein (Yippee), putative | -0,3659 | 0,04259 |
| TcCLB.506621.60 | acyl-CoA binding protein, putative | -0,366 | 0,00004772 |
| TcCLB.503801.9 | RAD50 DNA repair protein, putative (fragment) | -0,366 | 0,0003155 |
| TcCLB.511163.50 | hypothetical protein, conserved | -0,366 | 0,01188 |
| TcCLB.506885.250 | hypothetical protein | -0,3662 | 0,001919 |
| TcCLB.509073.100 | hypothetical protein, conserved | -0,3662 | 0,08209 |

| | | | |
|---|---|---|---|
| TcCLB.506517.110 | hypothetical protein, conserved | -0,3662 | 0,1163 |
| TcCLB.504181.50 | T-complex protein 1, delta subunit, putative | -0,3664 | 0,003201 |
| TcCLB.507039.30 | retrotransposon hot spot protein (RHS, pseudogene), putat | -0,3664 | 0,003653 |
| TcCLB.506733.110 | hypothetical protein, conserved | -0,3665 | 0,03025 |
| TcCLB.510889.210 | hypothetical protein, conserved | -0,3666 | 0,02533 |
| TcCLB.511125.9 | dispersed gene family protein 1 (DGF-1), putative | -0,3666 | 0,1729 |
| TcCLB.507883.90 | adenylate kinase, putative | -0,3667 | 0,04097 |
| TcCLB.511501.40 | hypothetical protein, conserved | -0,3667 | 0,05114 |
| TcCLB.506677.39 | Elongation factor G1, mitochondrial, putative (EF-G1) | -0,3669 | 0,00001828 |
| TcCLB.508977.65 | elongation factor 1-gamma (EF-1-gamma, pseudogene), p | -0,367 | 0,2915 |
| TcCLB.511837.70 | mitogen activated protein kinase, putative | -0,3671 | 0,0001242 |
| TcCLB.506195.140 | hypothetical protein, conserved | -0,3671 | 0,0009749 |
| TcCLB.511907.280 | hypothetical protein, conserved | -0,3671 | 0,002395 |
| TcCLB.506131.110 | mucin-associated surface protein (MASP), putative | -0,3671 | 0,448 |
| TcCLB.510659.50 | RNA polymerase-associated protein LEO1, putative | -0,3672 | 0,00002046 |
| TcCLB.511583.30 | hypothetical protein, conserved | -0,3672 | 0,01681 |
| TcCLB.506511.30 | hypothetical protein, conserved | -0,3673 | 0,000341 |
| TcCLB.510325.60 | hypothetical protein, conserved | -0,3673 | 0,003911 |
| TcCLB.504741.120 | hypothetical protein, conserved | -0,3673 | 0,01063 |
| TcCLB.511149.60 | mucin-associated surface protein (MASP), putative | -0,3674 | 0,5558 |
| TcCLB.511261.60 | hypothetical protein, conserved | -0,3675 | 0,000192 |
| TcCLB.495369.10 | hypothetical protein, conserved (fragment) | -0,3675 | 0,03868 |
| TcCLB.510741.210 | glycerol kinase, glycosomal (pseudogene), putative | -0,3676 | 0,002855 |
| TcCLB.510105.190 | hypothetical protein, conserved | -0,3676 | 0,01516 |
| TcCLB.503601.10 | trans-sialidase, Group V, putative | -0,3676 | 0,2405 |
| TcCLB.509805.110 | hypothetical protein, conserved | -0,3677 | 0,0004455 |
| TcCLB.511625.10 | transcription initiation protein, putative | -0,3677 | 0,0007296 |
| TcCLB.507711.160 | hypothetical protein, conserved | -0,3678 | 6,926E-06 |
| TcCLB.506739.180 | hypothetical protein, conserved | -0,3678 | 0,00007556 |
| TcCLB.504033.100 | hypothetical protein, conserved | -0,3679 | 0,001023 |
| TcCLB.506139.70 | mucin-associated surface protein (MASP), putative | -0,3679 | 0,3762 |
| TcCLB.427303.10 | 5'-3' exoribonuclease A, putative (fragment) (XRNA) | -0,368 | 0,000107 |
| TcCLB.508717.56 | hypothetical protein, conserved | -0,368 | 0,01002 |
| TcCLB.506459.140 | hypothetical protein | -0,368 | 0,4292 |
| TcCLB.508183.29 | hypothetical protein, conserved (fragment) | -0,3682 | 0,03302 |
| TcCLB.508995.20 | Tubulin/FtsZ family, putative | -0,3683 | 0,0006845 |
| TcCLB.510911.10 | profilin, putative | -0,3683 | 0,005051 |
| TcCLB.508851.210 | replication factor C, subunit 4, putative | -0,3683 | 0,04603 |
| TcCLB.511493.5 | hypothetical protein, conserved | -0,3683 | 0,2344 |
| TcCLB.507011.210 | hypothetical protein, conserved | -0,3684 | 0,00293 |
| TcCLB.511153.150 | hypothetical protein | -0,3684 | 0,03711 |
| TcCLB.506629.190 | hypothetical protein, conserved | -0,3687 | 0,001702 |
| TcCLB.509065.130 | hypothetical protein, conserved | -0,3687 | 0,009171 |
| TcCLB.508961.30 | trafficking protein particle complex subunit 3, putative | -0,3687 | 0,1143 |
| TcCLB.511019.120 | hypothetical protein, conserved | -0,3688 | 0,0003967 |

| | | | |
|---|---|---|---|
| TcCLB.509059.70 | hypothetical protein, conserved | -0,3689 | 0,000618 |
| TcCLB.508153.454 | hypothetical protein, conserved | -0,369 | 0,004769 |
| TcCLB.507925.20 | hypothetical protein, conserved | -0,3693 | 0,0006174 |
| TcCLB.503811.50 | hypothetical protein, conserved | -0,3693 | 0,003227 |
| TcCLB.507839.9 | Myotubularin-related protein, putative (fragment) | -0,3694 | 0,06713 |
| TcCLB.510835.20 | hypothetical protein, conserved | -0,3695 | 0,005634 |
| TcCLB.508131.10 | hypothetical protein, conserved (pseudogene) | -0,3696 | 0,00006279 |
| TcCLB.508465.50 | hypothetical protein, conserved | -0,3696 | 0,006739 |
| TcCLB.505183.20 | malic enzyme | -0,3697 | 0,00004812 |
| TcCLB.507993.160 | UDP-galactopyranose mutase (GLF) | -0,3698 | 0,002053 |
| TcCLB.506181.70 | hypothetical protein, conserved | -0,3698 | 0,002061 |
| TcCLB.508803.19 | dispersed gene family protein 1 (DGF-1), putative | -0,3698 | 0,1507 |
| TcCLB.504081.344 | elongation factor 1-gamma (EF-1-gamma, pseudogene), p | -0,3698 | 0,5139 |
| TcCLB.506477.20 | vesicular-fusion protein nsf, putative | -0,37 | 0,001619 |
| TcCLB.504039.25 | syntaxin binding protein (pseudogene), putative | -0,37 | 0,4722 |
| TcCLB.506587.40 | ATP-dependent RNA helicase FAL1, putative | -0,3701 | 0,0008456 |
| TcCLB.510755.19 | hypothetical protein, conserved | -0,3703 | 0,006801 |
| TcCLB.510091.90 | hypothetical protein, conserved | -0,3704 | 0,0001067 |
| TcCLB.508199.20 | hypothetical protein, conserved | -0,3704 | 0,008857 |
| TcCLB.507999.10 | dispersed gene family protein 1 (DGF-1), putative | -0,3705 | 0,002067 |
| TcCLB.508959.10 | hypothetical protein, conserved | -0,3705 | 0,1097 |
| TcCLB.508089.40 | hypothetical protein, conserved | -0,3706 | 0,01938 |
| TcCLB.405453.9 | dispersed gene family protein 1 (DGF-1), putative | -0,3706 | 0,1885 |
| TcCLB.506435.210 | hypothetical protein, conserved | -0,3707 | 0,005961 |
| TcCLB.503767.5 | potassium voltage-gated channel, putative | -0,3707 | 0,2285 |
| TcCLB.511857.10 | hypothetical protein, conserved | -0,3708 | 0,01062 |
| TcCLB.507023.110 | small GTP-binding protein Rab28, putative | -0,3708 | 0,01211 |
| TcCLB.507857.40 | ATP-dependent DEAD/H RNA helicase, putative | -0,371 | 0,0001737 |
| TcCLB.419703.20 | ubiquitin carboxyl-terminal hydrolase, putative | -0,371 | 0,01202 |
| TcCLB.511283.240 | hypothetical protein, conserved | -0,3712 | 5,664E-07 |
| TcCLB.508153.1010 | hypothetical protein, conserved | -0,3712 | 0,001481 |
| TcCLB.507517.20 | DNA polymerase sigma-like protein, putative | -0,3712 | 0,02042 |
| TcCLB.504005.10 | endonuclease III, putative | -0,3713 | 0,02206 |
| TcCLB.511403.30 | mucin TcMUCII, putative | -0,3714 | 0,4935 |
| TcCLB.510729.119 | hypothetical protein, conserved | -0,3715 | 0,03587 |
| TcCLB.508351.20 | hypothetical protein, conserved | -0,3715 | 0,09815 |
| TcCLB.510761.4 | hypothetical protein, conserved | -0,3716 | 0,003015 |
| TcCLB.507531.10 | hypothetical protein, conserved (pseudogene) | -0,3718 | 0,0000124 |
| TcCLB.504109.130 | hypothetical protein, conserved | -0,3718 | 0,00002772 |
| TcCLB.510359.200 | hypothetical protein, conserved | -0,3719 | 0,0003742 |
| TcCLB.503653.70 | hypothetical protein, conserved | -0,3719 | 0,02781 |
| TcCLB.508895.10 | C-terminal motor kinesin, putative | -0,3719 | 0,04679 |
| TcCLB.508799.70 | pumilio-repeat, RNA-binding protein, putative | -0,372 | 0,0003761 |
| TcCLB.503419.40 | hypothetical protein, conserved | -0,3722 | 0,01376 |
| TcCLB.510187.290 | ATP-dependent DEAD/H RNA helicase, putative (HEL64) | -0,3723 | 5,635E-06 |

| | | | |
|---|---|---|---|
| TcCLB.506943.20 | hypothetical protein, conserved | -0,3723 | 0,006956 |
| TcCLB.506623.30 | hypothetical protein | -0,3723 | 0,3528 |
| TcCLB.506767.119 | surface protease GP63 (pseudogene), putative | -0,3724 | 0,6296 |
| TcCLB.509319.20 | hypothetical protein, conserved | -0,3725 | 0,09115 |
| TcCLB.506435.140 | hypothetical protein, conserved | -0,3726 | 0,004377 |
| TcCLB.511837.20 | hypothetical protein, conserved | -0,373 | 0,0171 |
| TcCLB.507723.130 | eukaryotic translation initiation factor 3 subunit 8, putative | -0,3732 | 2,465E-07 |
| TcCLB.511211.220 | heat shock protein 70, putative | -0,3732 | 0,0005625 |
| TcCLB.507129.19 | hypothetical protein, conserved | -0,3733 | 0,00468 |
| TcCLB.507049.41 | hypothetical protein, conserved | -0,3735 | 0,1035 |
| TcCLB.510745.20 | hypothetical protein, conserved | -0,3737 | 0,008878 |
| TcCLB.506265.100 | ATP-dependent RNA helicase, putative | -0,3738 | 0,00002542 |
| TcCLB.510175.150 | hypothetical protein, conserved | -0,3738 | 0,1413 |
| TcCLB.509229.90 | hypothetical protein, conserved | -0,3739 | 0,0001929 |
| TcCLB.508461.140 | poly(A)-binding protein, putative | -0,374 | 7,531E-08 |
| TcCLB.508515.120 | hypothetical protein, conserved | -0,374 | 0,002332 |
| TcCLB.506925.410 | ATPase, putative | -0,374 | 0,004355 |
| TcCLB.506267.48 | hypothetical protein | -0,374 | 0,1889 |
| TcCLB.508667.49 | small nuclear ribonucleoprotein SmD2, putative | -0,3743 | 0,008155 |
| TcCLB.503421.14 | ATP-dependent DEAD/H RNA helicase (pseudogene), putat | -0,3743 | 0,2977 |
| TcCLB.510659.104 | poly(ADP-ribose) glycohydrolase, putative | -0,3744 | 0,009764 |
| TcCLB.507053.180 | hypothetical protein, conserved | -0,3746 | 2,091E-08 |
| TcCLB.508613.30 | peroxisomal membrane protein 4, putative | -0,3746 | 0,02623 |
| TcCLB.511735.20 | hypothetical protein, conserved | -0,3747 | 0,01777 |
| TcCLB.505071.60 | protein kinase, putative | -0,375 | 0,0002434 |
| TcCLB.507179.14 | hypothetical protein | -0,375 | 0,2286 |
| TcCLB.511509.21 | hypothetical protein, conserved | -0,3751 | 0,001164 |
| TcCLB.508909.240 | hypothetical protein, conserved | -0,3752 | 0,001714 |
| TcCLB.511091.9 | mucin-associated surface protein (MASP), putative | -0,3752 | 0,5765 |
| TcCLB.506241.70 | calcium motive p-type ATPase, putative | -0,3754 | 0,00002401 |
| TcCLB.508879.30 | dynein heavy chain, putative | -0,3755 | 9,379E-07 |
| TcCLB.511529.80 | kinetoplast-associated protein 3 (KAP3) | -0,3756 | 0,001752 |
| TcCLB.509693.80 | hypoxanthine-guanine phosphoribosyltransferase, putative | -0,3756 | 0,002016 |
| TcCLB.506885.50 | hypothetical protein, conserved | -0,3756 | 0,002069 |
| TcCLB.508711.49 | hypothetical protein, conserved | -0,3757 | 0,0001194 |
| TcCLB.509789.10 | hypothetical protein, conserved | -0,3757 | 0,0007038 |
| TcCLB.506883.80 | tRNA (guanine-N(7)-)-methyltransferase, putative | -0,3758 | 0,01519 |
| TcCLB.510731.104 | hypothetical protein, conserved | -0,3758 | 0,03461 |
| TcCLB.506735.70 | hypothetical protein, conserved | -0,3759 | 0,00005339 |
| TcCLB.506807.10 | tubulin delta chain (TUBD1) | -0,376 | 0,02091 |
| TcCLB.509979.320 | trans-sialidase, Group V, putative | -0,376 | 0,04235 |
| TcCLB.510687.130 | hypothetical protein, conserved | -0,3761 | 0,02641 |
| TcCLB.508409.160 | thymine-7-hydroxylase, putative | -0,3762 | 0,03352 |
| TcCLB.506709.50 | hypothetical protein, conserved | -0,3765 | 0,1234 |
| TcCLB.510311.50 | hypothetical protein, conserved | -0,3767 | 9,029E-06 |

| | | | |
|---|---|---|---|
| TcCLB.511491.60 | kinesin, putative | -0,3767 | 0,00004794 |
| TcCLB.508949.30 | hypothetical protein, conserved | -0,3767 | 0,0006877 |
| TcCLB.510719.260 | ATP-dependent protease subunit HslV, putative (HslV) | -0,3767 | 0,002193 |
| TcCLB.504045.30 | hypothetical protein | -0,3767 | 0,02364 |
| TcCLB.508873.410 | hypothetical protein, conserved | -0,3768 | 0,09529 |
| TcCLB.509895.20 | mitochondrial RNA binding protein, putative | -0,3769 | 5,584E-06 |
| TcCLB.507765.140 | hypothetical protein, conserved | -0,377 | 0,00002689 |
| TcCLB.509167.130 | hypothetical protein, conserved | -0,3771 | 0,0001572 |
| TcCLB.511855.80 | hypothetical protein, conserved | -0,3771 | 0,001427 |
| TcCLB.511077.110 | mucin-associated surface protein (MASP), putative | -0,3773 | 0,4711 |
| TcCLB.506925.270 | hypothetical protein, conserved | -0,3774 | 0,00005693 |
| TcCLB.509023.170 | hypothetical protein, conserved | -0,3774 | 0,004792 |
| TcCLB.507009.110 | hypothetical protein, conserved | -0,3775 | 0,02224 |
| TcCLB.509697.15 | mucin-associated surface protein (MASP, pseudogene), pu | -0,3775 | 0,5147 |
| TcCLB.510767.60 | hypothetical protein, conserved | -0,3776 | 0,02449 |
| TcCLB.506971.20 | surface protease GP63, putative | -0,3776 | 0,1495 |
| TcCLB.511511.90 | ATP-dependent DEAD/H DNA helicase recQ family, putativ | -0,3777 | 0,001296 |
| TcCLB.507941.80 | hypothetical protein, conserved | -0,3779 | 0,0253 |
| TcCLB.506269.79 | mucin-associated surface protein (MASP, pseudogene), pu | -0,3779 | 0,2405 |
| TcCLB.507107.40 | 3,2-trans-enoyl-CoA isomerase, mitochondrial precursor, p | -0,378 | 1,247E-06 |
| TcCLB.504119.20 | hypothetical protein, conserved | -0,3783 | 0,0009904 |
| TcCLB.507583.70 | ribonuclease inhibitor, putative | -0,3783 | 0,00391 |
| TcCLB.506529.524 | hypothetical protein, conserved | -0,3783 | 0,08761 |
| TcCLB.507069.120 | trans-sialidase (pseudogene), putative | -0,3784 | 0,2127 |
| TcCLB.507647.20 | hypothetical protein, conserved (pseudogene) | -0,3785 | 0,00004847 |
| TcCLB.510491.10 | hypothetical protein, conserved | -0,3785 | 0,08948 |
| TcCLB.506139.1 | mucin-associated surface protein (MASP), putative (fragm | -0,3785 | 0,6296 |
| TcCLB.511277.300 | hypothetical protein | -0,3786 | 0,00003848 |
| TcCLB.509045.20 | co-chaperone GrpE, putative | -0,3787 | 0,00009396 |
| TcCLB.506843.120 | trans-sialidase (pseudogene), putative | -0,3787 | 0,2152 |
| TcCLB.508873.130 | hypothetical protein | -0,3787 | 0,5052 |
| TcCLB.511211.60 | protein phosphatase 4 catalytic subunit, putative | -0,3788 | 0,0009988 |
| TcCLB.511249.50 | hypothetical protein, conserved | -0,3788 | 0,001035 |
| TcCLB.508153.410 | hypothetical protein, conserved | -0,3789 | 0,002709 |
| TcCLB.503723.80 | hypothetical protein, conserved | -0,3789 | 0,0072 |
| TcCLB.510181.30 | ATP-dependent DEAD/H RNA helicase, putative | -0,379 | 0,004739 |
| TcCLB.509715.23 | RNA-binding protein, putative | -0,379 | 0,1872 |
| TcCLB.508121.150 | trans-sialidase, Group V, putative | -0,379 | 0,2878 |
| TcCLB.506275.80 | hypothetical protein, conserved | -0,3791 | 0,0208 |
| TcCLB.509795.10 | hypothetical protein, conserved | -0,3793 | 0,002093 |
| TcCLB.506973.5 | mucin-associated surface protein (MASP), putative | -0,3794 | 0,5621 |
| TcCLB.509611.110 | RNA-editing complex protein, putative | -0,3796 | 0,00502 |
| TcCLB.507991.126 | hypothetical protein, conserved | -0,3796 | 0,09964 |
| TcCLB.504159.30 | hypothetical protein, conserved | -0,3797 | 0,006664 |
| TcCLB.507683.30 | hypothetical protein, conserved | -0,3798 | 0,0002388 |

| | | | |
|---|---|---|---|
| TcCLB.511229.120 | hypothetical protein | -0,3799 | 0,03249 |
| TcCLB.509581.20 | beta galactofuranosyl glycosyltransferase, putative | -0,3802 | 0,02199 |
| TcCLB.506633.100 | hypothetical protein, conserved | -0,3803 | 0,02437 |
| TcCLB.506411.25 | hypothetical protein, conserved (pseudogene) | -0,3803 | 0,0589 |
| TcCLB.508277.150 | proliferative cell nuclear antigen (PCNA), putative | -0,3804 | 3,443E-06 |
| TcCLB.508413.10 | hypothetical protein, conserved | -0,3804 | 0,001543 |
| TcCLB.504043.10 | hypothetical protein, conserved | -0,3805 | 0,0001712 |
| TcCLB.508699.50 | hypothetical protein, conserved | -0,3805 | 0,003119 |
| TcCLB.511279.50 | hypothetical protein, conserved | -0,3807 | 0,018 |
| TcCLB.510013.170 | mucin-associated surface protein (MASP), putative | -0,3807 | 0,5199 |
| TcCLB.510225.10 | histone H1, putative | -0,3808 | 1,932E-07 |
| TcCLB.508061.130 | trans-sialidase (pseudogene), putative | -0,3809 | 0,0004529 |
| TcCLB.504147.170 | gamma-tubulin complex subunit, putative | -0,381 | 0,00002531 |
| TcCLB.507025.40 | hypothetical protein | -0,381 | 0,01213 |
| TcCLB.508277.170 | hypothetical protein, conserved | -0,381 | 0,07308 |
| TcCLB.511153.144 | hypothetical protein, conserved | -0,381 | 0,2027 |
| TcCLB.507649.100 | hypothetical protein, conserved | -0,3811 | 0,0000893 |
| TcCLB.511215.40 | hypothetical protein, conserved | -0,3811 | 0,0002771 |
| TcCLB.503843.40 | chaperone DNAJ protein, putative (fragment) | -0,3811 | 0,0006568 |
| TcCLB.506817.20 | ABC transporter, putative | -0,3811 | 0,0009245 |
| TcCLB.511599.85 | elongation factor 1-gamma (EF-1-gamma, pseudogene), p | -0,3811 | 0,6202 |
| TcCLB.511603.170 | mucin TcMUCII, putative | -0,3812 | 0,504 |
| TcCLB.509161.110 | hypothetical protein, conserved | -0,3813 | 0,00002983 |
| TcCLB.507969.54 | hypothetical protein, conserved | -0,3813 | 0,006958 |
| TcCLB.508585.20 | hypothetical protein, conserved | -0,3814 | 0,01534 |
| TcCLB.511717.60 | hypothetical protein, conserved | -0,3814 | 0,0182 |
| TcCLB.511173.160 | trans-sialidase (pseudogene), putative | -0,3815 | 0,27 |
| TcCLB.510669.9 | hypothetical protein, conserved | -0,3816 | 0,02069 |
| TcCLB.506155.90 | hypothetical protein, conserved (pseudogene) | -0,3817 | 0,00001735 |
| TcCLB.503697.20 | hypothetical protein, conserved | -0,3817 | 0,00195 |
| TcCLB.511277.92 | hypothetical protein, conserved | -0,3817 | 0,001989 |
| TcCLB.510943.60 | U2 splicing auxiliary factor, putative (u2af26) | -0,3818 | 0,00004365 |
| TcCLB.508823.152 | hypothetical protein (fragment) | -0,3818 | 0,006528 |
| TcCLB.504887.10 | dispersed gene family protein 1 (DGF-1), putative | -0,3819 | 0,01556 |
| TcCLB.457977.10 | trans-sialidase, putative (fragment) | -0,3819 | 0,3569 |
| TcCLB.506637.100 | tyrosine phosphatase isoform, putative | -0,382 | 0,003902 |
| TcCLB.510487.37 | trans-sialidase (pseudogene), putative | -0,3821 | 0,2348 |
| TcCLB.510043.29 | hypothetical protein, conserved | -0,3822 | 0,09918 |
| TcCLB.510879.40 | Carboxypeptidase Nna1-like protein, putative | -0,3823 | 0,0001296 |
| TcCLB.508375.10 | mucin-associated surface protein (MASP), putative | -0,3823 | 0,1968 |
| TcCLB.507711.240 | hypothetical protein, conserved | -0,3825 | 0,001867 |
| TcCLB.510187.460 | hypothetical protein | -0,3825 | 0,003179 |
| TcCLB.508919.80 | hypothetical protein, conserved | -0,3825 | 0,01583 |
| TcCLB.511167.80 | PIF1 helicase-like protein, putative | -0,3827 | 0,0001342 |
| TcCLB.509039.4 | aspartyl-tRNA synthetase, putative | -0,3828 | 0,02741 |

| | | | |
|---|---|---|---|
| TcCLB.509069.49 | DNA replication licensing factor MCM7, putative (fragmen | -0,3828 | 0,0533 |
| TcCLB.506983.30 | hypothetical protein, conserved | -0,383 | 0,0005294 |
| TcCLB.457251.10 | 3-oxo-5-alpha-steroid 4-dehydrogenase, putative (fragmer | -0,383 | 0,0008904 |
| TcCLB.508461.540 | hypothetical protein, conserved | -0,383 | 0,004667 |
| TcCLB.509179.231 | trans-sialidase (pseudogene), putative | -0,383 | 0,2525 |
| TcCLB.511211.100 | hypothetical protein, conserved | -0,3832 | 0,002834 |
| TcCLB.508839.30 | trafficking protein particle complex subunit 3, putative | -0,3833 | 0,1344 |
| TcCLB.506729.50 | hypothetical protein, conserved | -0,3834 | 0,00005431 |
| TcCLB.506737.70 | ubiquitin-like protein, putative | -0,3835 | 1,168E-06 |
| TcCLB.506743.60 | hypothetical protein, conserved | -0,3836 | 0,0002907 |
| TcCLB.507709.20 | hypothetical protein | -0,3837 | 0,0006621 |
| TcCLB.509055.50 | hypothetical protein, conserved | -0,3837 | 0,006694 |
| TcCLB.510043.40 | hypothetical protein, conserved | -0,3838 | 0,002534 |
| TcCLB.509999.70 | hypothetical protein, conserved | -0,3839 | 0,0007927 |
| TcCLB.508547.70 | Present in the outer mitochondrial membrane proteome 3 | -0,3839 | 0,004971 |
| TcCLB.507993.380 | hypothetical protein, conserved | -0,3839 | 0,008635 |
| TcCLB.509567.4 | hypothetical protein, conserved | -0,3839 | 0,01282 |
| TcCLB.510445.50 | S-adenosylmethionine synthetase, putative (METK1) | -0,384 | 0,00001524 |
| TcCLB.407477.69 | hypothetical protein, conserved (fragment) | -0,384 | 0,01349 |
| TcCLB.508323.110 | hypothetical protein, conserved | -0,3841 | 0,0141 |
| TcCLB.509937.90 | hypothetical protein, conserved | -0,3842 | 0,003582 |
| TcCLB.416883.18 | proteasome alpha 2 subunit, putative | -0,3843 | 0,00432 |
| TcCLB.507771.110 | hypothetical protein, conserved | -0,3844 | 0,003912 |
| TcCLB.503869.20 | hypothetical protein, conserved | -0,3844 | 0,03968 |
| TcCLB.506989.300 | receptor-type adenylate cyclase (pseudogene), putative | -0,3845 | 0,0004681 |
| TcCLB.510055.130 | hypothetical protein, conserved (pseudogene) | -0,3847 | 0,004915 |
| TcCLB.503449.30 | clathrin coat assembly protein, putative | -0,3848 | 0,000203 |
| TcCLB.509875.160 | dispersed gene family protein 1 (DGF-1), putative | -0,385 | 0,03255 |
| TcCLB.510741.90 | hypothetical protein, conserved | -0,3851 | 0,0239 |
| TcCLB.506885.210 | trans-sialidase, Group III, putative | -0,3851 | 0,04526 |
| TcCLB.509371.10 | dispersed gene family protein 1 (DGF-1), putative | -0,3851 | 0,4119 |
| TcCLB.504105.230 | nuclear cap binding complex subunit CBP110, putative (CBI | -0,3852 | 0,0005057 |
| TcCLB.503915.9 | hypothetical protein, conserved (fragment) | -0,3852 | 0,002732 |
| TcCLB.506297.170 | hypothetical protein, conserved | -0,3852 | 0,04502 |
| TcCLB.510731.100 | hypothetical protein, conserved | -0,3853 | 0,01247 |
| TcCLB.506941.159 | zinc finger protein, putative | -0,3854 | 0,0007496 |
| TcCLB.511707.20 | hypothetical protein | -0,3854 | 0,01938 |
| TcCLB.511217.190 | hypothetical protein, conserved | -0,3854 | 0,2326 |
| TcCLB.508625.130 | stress-inducible protein STI1-like, putative | -0,3857 | 0,04913 |
| TcCLB.506559.148 | hypothetical protein, conserved | -0,3858 | 0,01861 |
| TcCLB.506671.20 | mucin TcMUC (pseudogene), putative | -0,3858 | 0,3675 |
| TcCLB.509491.10 | hypothetical protein, conserved | -0,386 | 0,009298 |
| TcCLB.508183.10 | ubiquinone biosynthesis protein COQ7 homolog, putative | -0,386 | 0,0116 |
| TcCLB.508825.10 | surface protease GP63, putative | -0,386 | 0,0147 |
| TcCLB.509737.9 | trans-sialidase, putative (fragment) | -0,386 | 0,2178 |

| | | | |
|---|---|---|---|
| TcCLB.507143.50 | hypothetical protein, conserved | -0,3861 | 0,0003901 |
| TcCLB.503467.30 | oligosaccharyl transferase subunit, putative | -0,3862 | 0,001574 |
| TcCLB.510719.290 | Golgi SNAP receptor complex member 1, putative | -0,3862 | 0,01718 |
| TcCLB.511127.120 | cysteine peptidase, Clan CA, family C19, putative | -0,3863 | 0,009168 |
| TcCLB.507237.10 | trans-sialidase, Group V, putative | -0,3864 | 0,2263 |
| TcCLB.506195.200 | hypothetical protein, conserved | -0,3865 | 0,00002636 |
| TcCLB.503565.20 | hypothetical protein, conserved | -0,3865 | 0,005842 |
| TcCLB.509569.44 | Present in the outer mitochondrial membrane proteome 4( | -0,3865 | 0,01259 |
| TcCLB.471901.20 | SNARE protein, putative | -0,3866 | 0,005621 |
| TcCLB.508625.30 | ARP2/3 complex subunit, putative | -0,3868 | 0,08047 |
| TcCLB.503521.20 | hypothetical protein, conserved | -0,3869 | 0,001986 |
| TcCLB.506485.90 | hypothetical protein, conserved | -0,387 | 0,0006898 |
| TcCLB.510143.30 | hypothetical protein, conserved | -0,387 | 0,002492 |
| TcCLB.510715.150 | trans-sialidase, Group VI, putative | -0,387 | 0,2429 |
| TcCLB.407477.20 | hypothetical protein, conserved | -0,3871 | 0,04032 |
| TcCLB.511041.10 | hypothetical protein, conserved | -0,3873 | 0,00316 |
| TcCLB.509109.170 | tubulin tyrosine ligase (pseudogene), putative | -0,3873 | 0,01102 |
| TcCLB.506457.149 | hypothetical protein, conserved (fragment) | -0,3873 | 0,04403 |
| TcCLB.506863.4 | dephospho-CoA kinase, putative | -0,3873 | 0,2156 |
| TcCLB.511895.30 | receptor-type adenylate cyclase, putative | -0,3873 | 0,2352 |
| TcCLB.510657.40 | hypothetical protein, conserved | -0,3874 | 0,002462 |
| TcCLB.511419.40 | phosphoglycerate kinase, putative | -0,3875 | 0,00009019 |
| TcCLB.509693.190 | ABC transporter, putative | -0,3875 | 0,007152 |
| TcCLB.510889.150 | hypothetical protein, conserved | -0,3875 | 0,01839 |
| TcCLB.506493.50 | serine/threonine protein kinase, putative | -0,3875 | 0,08638 |
| TcCLB.506813.200 | mucin-like glycoprotein (pseudogene), putative | -0,3876 | 0,5193 |
| TcCLB.510989.50 | hypothetical protein, conserved | -0,3877 | 1,121E-06 |
| TcCLB.508411.10 | cofilin/actin depolymerizing factor, putative | -0,3877 | 0,0004958 |
| TcCLB.506295.50 | hypothetical protein, conserved | -0,3877 | 0,05401 |
| TcCLB.511473.6 | ATP-dependent DEAD/H RNA helicase (pseudogene), putat | -0,3878 | 0,2713 |
| TcCLB.508717.10 | tyrosine specific protein phosphatase, putative | -0,3879 | 0,0002909 |
| TcCLB.511391.130 | hypothetical protein, conserved | -0,388 | 0,02396 |
| TcCLB.505807.209 | hypothetical protein, conserved | -0,388 | 0,0523 |
| TcCLB.511553.150 | mucin-associated surface protein (MASP), putative | -0,388 | 0,2959 |
| TcCLB.508153.860 | hypothetical protein, conserved | -0,3882 | 0,00008171 |
| TcCLB.509429.110 | proteasome beta-1 subunit, putative | -0,3883 | 0,0001623 |
| TcCLB.482319.10 | hypothetical protein, conserved | -0,3883 | 0,001268 |
| TcCLB.506871.100 | polypeptide deformylase-like protein, putative | -0,3883 | 0,003006 |
| TcCLB.508139.135 | retrotransposon hot spot protein (RHS, pseudogene), putat | -0,3883 | 0,01379 |
| TcCLB.511265.10 | dispersed gene family protein 1 (DGF-1, pseudogene), puta | -0,3883 | 0,02143 |
| TcCLB.511283.210 | hypothetical protein, conserved | -0,3884 | 0,00007507 |
| TcCLB.503473.10 | protein kinase, putative | -0,3884 | 0,002624 |
| TcCLB.505121.9 | dispersed gene family protein 1 (DGF-1), putative (fragme | -0,3884 | 0,0482 |
| TcCLB.508221.64 | syntaxin binding protein (pseudogene), putative | -0,3884 | 0,3231 |
| TcCLB.508347.60 | hypothetical protein, conserved | -0,3886 | 0,04355 |

| | | | |
|---|---|---|---|
| TcCLB.503885.80 | hypothetical protein, conserved | -0,3887 | 0,005383 |
| TcCLB.507129.10 | actin 2, putative | -0,3887 | 0,04943 |
| TcCLB.510221.10 | RNA pseudouridylate synthase, putative | -0,3888 | 0,002208 |
| TcCLB.511553.80 | hypothetical protein, conserved | -0,3888 | 0,5343 |
| TcCLB.504883.30 | hypothetical protein, conserved | -0,3889 | 0,0000563 |
| TcCLB.508747.50 | mucin TcMUCII, putative | -0,3889 | 0,2692 |
| TcCLB.510141.20 | hypothetical protein, conserved | -0,3891 | 0,00801 |
| TcCLB.510201.20 | mucin-associated surface protein (MASP), putative | -0,3891 | 0,4086 |
| TcCLB.509453.60 | hypothetical protein, conserved | -0,3893 | 0,0002381 |
| TcCLB.511127.274 | hypothetical protein, conserved | -0,3893 | 0,02894 |
| TcCLB.508745.70 | mucin-associated surface protein (MASP), putative (fragm | -0,3894 | 0,6826 |
| TcCLB.507257.30 | hypothetical protein, conserved | -0,3895 | 0,02157 |
| TcCLB.506415.30 | hypothetical protein, conserved | -0,3896 | 0,01705 |
| TcCLB.510597.9 | tousled-like kinase II, putative | -0,3899 | 0,006469 |
| TcCLB.503919.30 | hypothetical protein, conserved | -0,3899 | 0,01265 |
| TcCLB.507611.310 | eukaryotic translation initiation factor 3 subunit 8, putative | -0,3901 | 4,558E-06 |
| TcCLB.503809.10 | hypothetical protein, conserved | -0,3901 | 0,000179 |
| TcCLB.507011.140 | DNA-repair protein, putative | -0,3902 | 0,0007453 |
| TcCLB.507037.50 | exonuclease, putative | -0,3904 | 0,02856 |
| TcCLB.506811.30 | hypothetical protein, conserved | -0,3905 | 0,001517 |
| TcCLB.506973.105 | mucin-associated surface protein (MASP, pseudogene), pu | -0,3905 | 0,3292 |
| TcCLB.510421.314 | U6 snRNA-associated Sm-like protein LSm8p | -0,3906 | 0,05504 |
| TcCLB.510885.10 | hypothetical protein, conserved | -0,3907 | 0,003167 |
| TcCLB.508403.10 | hypothetical protein, conserved | -0,3908 | 0,0007778 |
| TcCLB.506297.120 | hypothetical protein, conserved | -0,3908 | 0,001741 |
| TcCLB.507837.40 | hypothetical protein, conserved | -0,3908 | 0,02523 |
| TcCLB.507485.45 | hypothetical protein, conserved | -0,391 | 0,09545 |
| TcCLB.506201.130 | hypothetical protein, conserved | -0,3913 | 0,002466 |
| TcCLB.507949.154 | TFIIH basal transcription factor subunit, putative (TSP1) | -0,3913 | 0,00779 |
| TcCLB.510607.75 | hypothetical protein, conserved (pseudogene) | -0,3913 | 0,1363 |
| TcCLB.510397.20 | trans-sialidase (pseudogene), putative | -0,3913 | 0,3862 |
| TcCLB.508245.50 | hypothetical protein | -0,3914 | 0,1298 |
| TcCLB.507237.100 | mucin-associated surface protein (MASP), putative | -0,3914 | 0,5586 |
| TcCLB.504151.39 | hypothetical protein, conserved | -0,3915 | 0,002212 |
| TcCLB.432067.20 | hypothetical protein, conserved (pseudogene) | -0,3915 | 0,1589 |
| TcCLB.508169.20 | elongation factor 2, putative | -0,3916 | 3,236E-10 |
| TcCLB.510661.60 | glycerol kinase, glycosomal, putative | -0,3917 | 0,0001575 |
| TcCLB.508647.170 | DNA polymerase theta (polymerase domain only), putative | -0,3917 | 0,0002824 |
| TcCLB.510187.234 | tyrosine specific protein phosphatase, putative | -0,3917 | 0,001095 |
| TcCLB.508001.30 | hypothetical protein, conserved (pseudogene) | -0,3918 | 0,02661 |
| TcCLB.511291.70 | hypothetical protein, conserved | -0,3919 | 0,0002617 |
| TcCLB.510317.14 | hypothetical protein, conserved | -0,3919 | 0,005441 |
| TcCLB.511921.71 | hypothetical protein, conserved (pseudogene) | -0,3919 | 0,2225 |
| TcCLB.509719.50 | protein kinase, putative | -0,392 | 0,003146 |
| TcCLB.508815.160 | ER lumen retaining receptor protein, putative | -0,3921 | 0,0002905 |

| | | | |
|---|---|---|---|
| TcCLB.506989.160 | PTS2 receptor, putative (PEX7) | -0,3921 | 0,001024 |
| TcCLB.508545.9 | hypothetical protein, conserved | -0,3922 | 0,03042 |
| TcCLB.511277.560 | synaptotagmin, putative | -0,3923 | 0,007587 |
| TcCLB.504079.10 | branch point binding protein, putative | -0,3927 | 0,0063 |
| TcCLB.510055.70 | hypothetical protein, conserved | -0,3929 | 0,001736 |
| TcCLB.505807.40 | hypothetical protein, conserved | -0,393 | 0,00001777 |
| TcCLB.509847.50 | trans-sialidase (pseudogene), putative | -0,393 | 0,2756 |
| TcCLB.511211.40 | molybdenum cofactor biosynthesis protein, putative | -0,3931 | 0,001136 |
| TcCLB.505183.84 | hypothetical protein, conserved | -0,3931 | 0,006143 |
| TcCLB.508305.9 | trans-sialidase, putative | -0,3932 | 0,2525 |
| TcCLB.506893.80 | monothiol glutaredoxin, putative | -0,3933 | 0,03995 |
| TcCLB.506647.20 | hypothetical protein, conserved | -0,3934 | 0,000345 |
| TcCLB.509685.27 | surface protease GP63 (pseudogene), putative | -0,3934 | 0,4793 |
| TcCLB.508707.80 | RNA-binding protein, putative | -0,3935 | 0,003037 |
| TcCLB.508139.155 | protein kinase (pseudogene), putative | -0,3937 | 0,03678 |
| TcCLB.507663.60 | hypothetical protein, conserved | -0,3938 | 0,004485 |
| TcCLB.510089.130 | protein kinase, putative | -0,3939 | 0,0000538 |
| TcCLB.507049.119 | hypothetical protein, conserved | -0,3939 | 0,0002724 |
| TcCLB.506577.130 | protein phosphatase 2A regulatory subunit, putative | -0,3939 | 0,002812 |
| TcCLB.510901.50 | C-terminal motor kinesin, putative | -0,3941 | 0,0002304 |
| TcCLB.507601.140 | ATP-dependent RNA helicase, putative | -0,3941 | 0,0003964 |
| TcCLB.510101.490 | anion-transporting ATPase-like, putative | -0,3942 | 0,0001107 |
| TcCLB.511803.70 | hypothetical protein, conserved | -0,3942 | 0,005657 |
| TcCLB.506249.30 | hypothetical protein, conserved | -0,3943 | 0,09194 |
| TcCLB.506179.59 | ATP-dependent RNA helicase, putative (fragment) | -0,3944 | 0,07878 |
| TcCLB.510877.60 | ruvB-like DNA helicase, putative (RUVBL) | -0,3946 | 5,766E-07 |
| TcCLB.504125.30 | ubiquitin carboxyl-terminal hydrolase, putative | -0,3946 | 0,00001034 |
| TcCLB.503611.30 | hypothetical protein, conserved | -0,3946 | 0,002777 |
| TcCLB.508197.20 | ADP-ribosylation factor, putative | -0,3946 | 0,02567 |
| TcCLB.508413.30 | hypothetical protein, conserved | -0,3947 | 0,00006481 |
| TcCLB.508065.60 | hypothetical protein, conserved | -0,3947 | 0,001584 |
| TcCLB.510979.40 | trans-sialidase, Group V, putative | -0,3947 | 0,05278 |
| TcCLB.507023.20 | hypothetical protein | -0,3948 | 0,01668 |
| TcCLB.508691.20 | hypothetical protein, conserved | -0,3949 | 0,007445 |
| TcCLB.510159.10 | hypothetical protein, conserved | -0,3949 | 0,009841 |
| TcCLB.509693.120 | hypothetical protein, conserved | -0,395 | 0,0004286 |
| TcCLB.506363.120 | hypothetical protein, conserved | -0,3951 | 0,01412 |
| TcCLB.506625.249 | RNA pseudouridylate synthase protein, putative | -0,3952 | 0,0004435 |
| TcCLB.508147.150 | mucin-associated surface protein (MASP), putative | -0,3953 | 0,3586 |
| TcCLB.509733.170 | Valosin-containing protein, putative (VCP) | -0,3955 | 2,172E-06 |
| TcCLB.504057.97 | hypothetical protein, conserved | -0,3955 | 0,003602 |
| TcCLB.510969.10 | hypothetical protein, conserved | -0,3955 | 0,007216 |
| TcCLB.508873.510 | mucin-associated surface protein (MASP), putative | -0,3955 | 0,3163 |
| TcCLB.507089.60 | hypothetical protein, conserved | -0,3956 | 0,002108 |
| TcCLB.506435.220 | hypothetical protein, conserved | -0,3956 | 0,03351 |

| | | | |
|---|---|---|---|
| TcCLB.506925.30 | hypothetical protein, conserved | -0,3957 | 6,337E-08 |
| TcCLB.506279.140 | trans-sialidase, Group V, putative | -0,3957 | 0,1387 |
| TcCLB.511003.170 | hypothetical protein, conserved | -0,3959 | 0,01564 |
| TcCLB.504703.10 | glycosyl hydrolase (pseudogene), putative | -0,396 | 0,00004885 |
| TcCLB.503955.30 | hypothetical protein, conserved | -0,3961 | 0,004701 |
| TcCLB.503417.40 | hypothetical protein | -0,3961 | 0,4781 |
| TcCLB.507901.30 | retrotransposon hot spot protein (RHS, pseudogene), putat | -0,3962 | 0,00979 |
| TcCLB.511127.240 | hypothetical protein, conserved | -0,3962 | 0,1897 |
| TcCLB.508421.40 | hypothetical protein, conserved (pseudogene) | -0,3963 | 0,3879 |
| TcCLB.511131.20 | hypothetical protein, conserved | -0,3965 | 0,001553 |
| TcCLB.510841.30 | hypothetical protein | -0,3965 | 0,1682 |
| TcCLB.511867.50 | proteasome alpha 7 subunit, putative | -0,3966 | 0,00124 |
| TcCLB.504021.90 | hypothetical protein | -0,3966 | 0,006589 |
| TcCLB.505807.254 | hypothetical protein | -0,3966 | 0,05601 |
| TcCLB.507063.160 | hypothetical protein, conserved | -0,3967 | 0,0002605 |
| TcCLB.503557.14 | hypothetical protein, conserved | -0,3967 | 0,002032 |
| TcCLB.508517.9 | dispersed gene family protein 1 (DGF-1), putative | -0,3967 | 0,1211 |
| TcCLB.509579.32 | N-acetyltransferase complex ARD1 subunit (pseudogene), | -0,3967 | 0,2863 |
| TcCLB.506247.150 | hypothetical protein, conserved | -0,3968 | 0,02086 |
| TcCLB.511017.69 | histone acetyltransferase, putative (fragment) | -0,3968 | 0,1192 |
| TcCLB.511145.80 | RNA polymerase II, putative | -0,3969 | 0,01239 |
| TcCLB.507625.183 | SNARE domain-containing protein, putative (SNARE2.1) | -0,3969 | 0,01851 |
| TcCLB.507953.100 | trans-sialidase, Group V, putative | -0,3969 | 0,209 |
| TcCLB.506363.159 | separase, putative | -0,397 | 0,3263 |
| TcCLB.504407.10 | dispersed gene family protein 1 (DGF-1, pseudogene), puta | -0,3971 | 0,2276 |
| TcCLB.507063.260 | hypothetical protein | -0,3972 | 0,2406 |
| TcCLB.511577.160 | hypothetical protein, conserved | -0,3974 | 0,002731 |
| TcCLB.508221.530 | trans-sialidase (pseudogene), putative | -0,3975 | 0,3339 |
| TcCLB.503653.104 | hypothetical protein, conserved | -0,3975 | 0,3755 |
| TcCLB.511233.40 | hypothetical protein, conserved | -0,3976 | 0,0002679 |
| TcCLB.511321.26 | vesicle transport v-SNARE protein, putative | -0,3976 | 0,01669 |
| TcCLB.507837.90 | cell division cycle protein, putative | -0,3977 | 0,008743 |
| TcCLB.510761.70 | ubiquitin hydrolase, putative | -0,3979 | 0,00003574 |
| TcCLB.504259.19 | hypothetical protein, conserved | -0,3979 | 0,006938 |
| TcCLB.506743.10 | hypothetical protein | -0,3979 | 0,04911 |
| TcCLB.509179.120 | hypothetical protein, conserved | -0,3979 | 0,06401 |
| TcCLB.504839.60 | hypothetical protein, conserved | -0,398 | 0,002201 |
| TcCLB.507049.30 | hypothetical protein, conserved | -0,398 | 0,01159 |
| TcCLB.505941.49 | hypothetical protein, conserved | -0,398 | 0,02129 |
| TcCLB.506287.200 | DNA ligase, putative | -0,398 | 0,02463 |
| TcCLB.427355.10 | oligosaccharyl transferase subunit, putative (fragment) | -0,3981 | 0,009094 |
| TcCLB.509611.120 | hypothetical protein, conserved | -0,3981 | 0,009165 |
| TcCLB.510763.90 | hypothetical protein, conserved | -0,3981 | 0,01939 |
| TcCLB.510269.60 | trans-sialidase (pseudogene), putative | -0,3981 | 0,1583 |
| TcCLB.408345.20 | 2-oxoglutarate dehydrogenase subunit, putative (fragment | -0,3983 | 0,01616 |

| | | | |
|---|---|---|---|
| TcCLB.510515.139 | T-complex protein 1, gamma subunit, putative | -0,3984 | 0,002443 |
| TcCLB.506221.80 | NAD synthase, putative | -0,3984 | 0,01707 |
| TcCLB.506769.30 | mucin-associated surface protein (MASP), putative | -0,3985 | 0,1914 |
| TcCLB.507019.130 | hypothetical protein | -0,3986 | 0,02318 |
| TcCLB.511529.190 | hypothetical protein, conserved | -0,3986 | 0,03447 |
| TcCLB.509991.110 | transcription elongation factor 1 homolog (ELOF1) | -0,3988 | 0,01002 |
| TcCLB.503733.20 | 6-phosphofructo-2-kinase/fructose-2,6-biphosphatase, put | -0,3989 | 0,001181 |
| TcCLB.508099.9 | mucin-associated surface protein (MASP, pseudogene), pu | -0,399 | 0,2967 |
| TcCLB.504011.40 | XPA-interacting protein, putative | -0,3992 | 0,001525 |
| TcCLB.506177.80 | hypothetical protein, conserved | -0,3992 | 0,09499 |
| TcCLB.510575.30 | hypothetical protein, conserved | -0,3994 | 0,00044 |
| TcCLB.507601.60 | hypothetical protein, conserved | -0,3994 | 0,01754 |
| TcCLB.506247.50 | CCT-theta, putative | -0,3995 | 1,273E-11 |
| TcCLB.510733.10 | p21-activated kinase 3, putative | -0,3995 | 6,481E-07 |
| TcCLB.511593.65 | hypothetical protein | -0,3995 | 0,3776 |
| TcCLB.508207.200 | small GTP-binding protein, putative | -0,3996 | 0,001084 |
| TcCLB.508153.360 | hypothetical protein, conserved | -0,3996 | 0,02497 |
| TcCLB.510659.60 | hypothetical protein, conserved | -0,3997 | 0,02396 |
| TcCLB.508467.10 | hypothetical protein, conserved | -0,3997 | 0,1066 |
| TcCLB.506973.61 | hypothetical protein, conserved (pseudogene) | -0,3997 | 0,1136 |
| TcCLB.506327.30 | hypothetical protein, conserved | -0,3998 | 0,00000241 |
| TcCLB.511627.120 | aldo/keto reductase, putative | -0,3998 | 0,004966 |
| TcCLB.469785.20 | hypothetical protein, conserved | -0,3998 | 0,02698 |
| TcCLB.510571.10 | hypothetical protein, conserved | -0,3998 | 0,0503 |
| TcCLB.504057.140 | ADP/ATP mitochondrial carrier protein, putative | -0,3999 | 0,0001111 |
| TcCLB.506693.7 | hypothetical protein, conserved | -0,3999 | 0,05332 |
| TcCLB.506275.50 | Golgi SNAP receptor complex member 1, putative | -0,4 | 0,01665 |
| TcCLB.508547.20 | pre-rrna-processing protein tsr1 homolog | -0,4001 | 0,0002576 |
| TcCLB.511755.50 | hypothetical protein, conserved | -0,4001 | 0,002236 |
| TcCLB.511201.58 | hypothetical protein | -0,4001 | 0,002876 |
| TcCLB.510963.60 | hypothetical protein, conserved | -0,4001 | 0,006104 |
| TcCLB.504021.70 | hypothetical protein, conserved | -0,4002 | 0,002534 |
| TcCLB.508979.30 | mucin TcMUCII, putative | -0,4002 | 0,192 |
| TcCLB.510279.110 | trans-sialidase, Group VI, putative | -0,4002 | 0,3318 |
| TcCLB.508347.130 | deoxyhypusine hydroxylase, putative | -0,4004 | 0,02645 |
| TcCLB.507517.90 | nucleotide-binding protein, putative | -0,4005 | 0,003511 |
| TcCLB.506657.50 | ADP/ATP mitochondrial carrier protein (pseudogene), puta | -0,4005 | 0,003679 |
| TcCLB.509875.120 | trans-sialidase, Group II, putative | -0,4006 | 0,01496 |
| TcCLB.506961.30 | hypothetical protein | -0,4006 | 0,1829 |
| TcCLB.506515.10 | FG-GAP repeat protein, putative | -0,4007 | 0,0003482 |
| TcCLB.511211.140 | hypothetical protein, conserved | -0,4008 | 0,0007712 |
| TcCLB.510355.10 | dispersed gene family protein 1 (DGF-1), putative | -0,4008 | 0,01192 |
| TcCLB.508123.10 | hypothetical protein, conserved | -0,4008 | 0,04647 |
| TcCLB.508787.30 | pumilio/PUF RNA binding protein 3, putative (PUF3) | -0,4009 | 0,0003469 |
| TcCLB.511907.100 | hypothetical protein, conserved | -0,401 | 0,001079 |

| | | | |
|---|---|---|---|
| TcCLB.503719.39 | hypothetical protein, conserved | -0,401 | 0,008731 |
| TcCLB.506839.60 | tyrosine specific protein phosphatase, putative | -0,4011 | 0,0002346 |
| TcCLB.510689.40 | phosphatidylinositol 3-kinase (tor2), putative (TOR2) | -0,4012 | 9,819E-07 |
| TcCLB.503541.40 | hypothetical protein, conserved | -0,4013 | 0,003162 |
| TcCLB.509627.20 | hypothetical protein, conserved | -0,4014 | 0,000651 |
| TcCLB.419833.10 | vacuolar protein sorting 18, putative | -0,4014 | 0,02046 |
| TcCLB.510431.340 | hypothetical protein, conserved | -0,4015 | 0,03956 |
| TcCLB.510611.70 | hypothetical protein, conserved | -0,4017 | 0,0005419 |
| TcCLB.510043.59 | hypothetical protein, conserved (fragment) | -0,4017 | 0,001129 |
| TcCLB.509595.60 | hypothetical protein, conserved | -0,4018 | 0,01239 |
| TcCLB.411427.49 | hypothetical protein, conserved (fragment) | -0,4018 | 0,1266 |
| TcCLB.504057.110 | ribosomal RNA processing protein 6, putative (RRP6) | -0,4019 | 0,00002068 |
| TcCLB.509029.130 | cyclin 1 (CYC1) | -0,4019 | 0,005707 |
| TcCLB.510939.9 | hypothetical protein, conserved | -0,402 | 0,1003 |
| TcCLB.508799.30 | hypothetical protein, conserved | -0,4021 | 0,002005 |
| TcCLB.503797.4 | hypothetical protein, conserved | -0,4022 | 0,0007717 |
| TcCLB.505607.9 | dispersed gene family protein 1 (DGF-1), putative | -0,4022 | 0,115 |
| TcCLB.508175.210 | hypothetical protein, conserved | -0,4023 | 0,04838 |
| TcCLB.506195.240 | hypothetical protein, conserved | -0,4025 | 0,0002426 |
| TcCLB.505075.20 | hypothetical protein | -0,4025 | 0,05452 |
| TcCLB.506133.120 | syntaxin binding protein (pseudogene), putative | -0,4025 | 0,4064 |
| TcCLB.511779.31 | hypothetical protein | -0,4025 | 0,424 |
| TcCLB.509719.26 | hypothetical protein, conserved | -0,4026 | 0,0003459 |
| TcCLB.508635.10 | hypothetical protein | -0,4026 | 0,1499 |
| TcCLB.510601.30 | hypothetical protein, conserved | -0,4027 | 0,001079 |
| TcCLB.508409.170 | hypothetical protein, conserved | -0,4027 | 0,00631 |
| TcCLB.511285.10 | hypothetical protein | -0,4028 | 0,00003534 |
| TcCLB.506983.20 | hypothetical protein, conserved | -0,4028 | 0,0002501 |
| TcCLB.510897.10 | cullin 2, putative | -0,4028 | 0,001565 |
| TcCLB.511529.130 | hypothetical protein, conserved | -0,4028 | 0,008701 |
| TcCLB.438897.10 | hypothetical protein, conserved | -0,4029 | 0,001607 |
| TcCLB.503579.30 | DNA polymerase sigma-like protein, putative | -0,4029 | 0,006313 |
| TcCLB.410199.4 | kinase, putative (fragment) | -0,4031 | 0,04554 |
| TcCLB.511317.50 | hypothetical protein, conserved | -0,4032 | 0,0009144 |
| TcCLB.509591.20 | phosphoprotein phosphatase, putative | -0,4032 | 0,5365 |
| TcCLB.507951.280 | ABC transporter, putative | -0,4033 | 0,0001367 |
| TcCLB.503987.60 | hypothetical protein, conserved | -0,4034 | 0,02141 |
| TcCLB.510285.30 | hypothetical protein, conserved | -0,4035 | 0,00148 |
| TcCLB.507017.140 | mitochondrial RNA binding protein 1, putative | -0,4037 | 0,00006271 |
| TcCLB.508831.140 | surface antigen 2 (CA-2), putative | -0,4037 | 0,000762 |
| TcCLB.507581.9 | dispersed gene family protein 1 (DGF-1), putative (fragme | -0,4037 | 0,08499 |
| TcCLB.508737.180 | hypothetical protein, conserved | -0,4038 | 2,235E-06 |
| TcCLB.507875.40 | trans-sialidase (pseudogene), putative | -0,4038 | 0,01668 |
| TcCLB.510603.90 | mitochondrial carrier protein, putative | -0,4039 | 0,03411 |
| TcCLB.508629.68 | N-acetyltransferase complex ARD1 subunit (pseudogene), | -0,4041 | 0,2023 |

| | | | |
|---|---|---|---|
| TcCLB.510421.290 | Mad2-like HORMA domain containing protein (mad2) | -0,4042 | 0,001376 |
| TcCLB.504033.200 | hypothetical protein | -0,4043 | 0,002179 |
| TcCLB.508881.10 | hypothetical protein, conserved | -0,4043 | 0,02127 |
| TcCLB.503535.9 | hypothetical protein, conserved | -0,4044 | 5,322E-06 |
| TcCLB.510181.110 | hypothetical protein, conserved | -0,4044 | 0,004011 |
| TcCLB.508839.70 | hypothetical protein, conserved | -0,4044 | 0,02613 |
| TcCLB.509539.40 | hypothetical protein, conserved | -0,4044 | 0,03889 |
| TcCLB.506265.120 | hypothetical protein, conserved | -0,4048 | 0,0002872 |
| TcCLB.476733.20 | hypothetical protein, conserved | -0,4049 | 0,00008721 |
| TcCLB.509499.10 | cyclophilin, putative (CYP) | -0,4049 | 0,0004819 |
| TcCLB.504239.100 | mucin TcMUCII, putative | -0,4049 | 0,369 |
| TcCLB.503939.4 | hypothetical protein, conserved | -0,4049 | 0,4502 |
| TcCLB.506425.100 | hypothetical protein, conserved | -0,405 | 0,0005424 |
| TcCLB.506789.150 | hypothetical protein | -0,405 | 0,001014 |
| TcCLB.506887.80 | hypothetical protein, conserved | -0,405 | 0,02917 |
| TcCLB.506789.249 | bilobe protein, conserved | -0,4051 | 0,0001203 |
| TcCLB.509819.40 | hypothetical protein, conserved | -0,4051 | 0,0004782 |
| TcCLB.408437.10 | hypothetical protein, conserved | -0,4051 | 0,000541 |
| TcCLB.404711.10 | trans-sialidase (pseudogene), putative | -0,4051 | 0,001765 |
| TcCLB.509979.70 | hypothetical protein, conserved | -0,4051 | 0,03928 |
| TcCLB.511327.20 | hypothetical protein (fragment) | -0,4051 | 0,04168 |
| TcCLB.510351.80 | hypothetical protein, conserved | -0,4053 | 0,02693 |
| TcCLB.508909.50 | hypothetical protein, conserved | -0,4053 | 0,04921 |
| TcCLB.510311.60 | hypothetical protein, conserved | -0,4054 | 1,333E-06 |
| TcCLB.506691.50 | hypothetical protein, conserved | -0,4054 | 0,002294 |
| TcCLB.507815.20 | trans-sialidase (pseudogene), putative | -0,4054 | 0,06524 |
| TcCLB.507239.10 | hypothetical protein, conserved | -0,4054 | 0,07917 |
| TcCLB.508641.184 | Protein translation factor SUI1 homolog, putative | -0,4056 | 0,01347 |
| TcCLB.506947.70 | hypothetical protein, conserved | -0,4059 | 0,06327 |
| TcCLB.504155.170 | mucin TcMUCII, putative | -0,4059 | 0,481 |
| TcCLB.506367.89 | hypothetical protein, conserved | -0,406 | 0,02649 |
| TcCLB.510243.20 | hypothetical protein, conserved | -0,4061 | 0,003179 |
| TcCLB.506679.210 | RNA editing complex protein MP61, putative | -0,4063 | 0,00152 |
| TcCLB.503399.20 | hydrolase, alpha/beta fold family, putative | -0,4064 | 0,007601 |
| TcCLB.507945.20 | hypothetical protein, conserved | -0,4064 | 0,01627 |
| TcCLB.510053.50 | vesicle transport v-SNARE protein, putative | -0,4064 | 0,03789 |
| TcCLB.504009.4 | hypothetical protein, conserved | -0,4065 | 0,04299 |
| TcCLB.507711.280 | hypothetical protein, conserved | -0,4066 | 0,007459 |
| TcCLB.504013.50 | selenocysteine-tRNA-specific elongation factor, putative | -0,4067 | 0,0005838 |
| TcCLB.511423.104 | DNA-directed RNA polymerase 2, RPB12 subunit, putative | -0,4068 | 0,02506 |
| TcCLB.509011.30 | hypothetical protein | -0,4069 | 0,01549 |
| TcCLB.510819.40 | hypothetical protein, conserved | -0,4071 | 0,0169 |
| TcCLB.508999.10 | DnaJ chaperone protein, putative | -0,4072 | 0,0001814 |
| TcCLB.510763.70 | hypothetical protein, conserved | -0,4072 | 0,002203 |
| TcCLB.506859.100 | hypothetical protein, conserved | -0,4072 | 0,1335 |

| | | | |
|---|---|---|---|
| TcCLB.509835.30 | hypothetical protein, conserved | -0,4073 | 0,008101 |
| TcCLB.504091.10 | hypothetical protein, conserved | -0,4074 | 0,0002299 |
| TcCLB.506759.230 | hypothetical protein | -0,4074 | 0,6421 |
| TcCLB.508851.170 | DNA topoisomerase III, putative | -0,4075 | 0,0004065 |
| TcCLB.507715.80 | ubiquitin-conjugating enzyme, putative | -0,4075 | 0,005655 |
| TcCLB.506529.180 | hypothetical protein, conserved | -0,4075 | 0,08965 |
| TcCLB.506297.20 | hypothetical protein | -0,4077 | 0,00002105 |
| TcCLB.504021.140 | hypothetical protein, conserved | -0,4078 | 0,0002524 |
| TcCLB.436083.9 | hypothetical protein, conserved | -0,4078 | 0,007245 |
| TcCLB.508879.200 | hypothetical protein, conserved | -0,4078 | 0,01237 |
| TcCLB.507625.84 | hypothetical protein, conserved | -0,4078 | 0,02845 |
| TcCLB.507197.10 | surface protease GP63, putative | -0,4079 | 0,00004221 |
| TcCLB.508199.50 | palmitoyl acyltransferase 1, putative | -0,4081 | 0,00003099 |
| TcCLB.510045.30 | protein kinase, putative | -0,4082 | 0,002834 |
| TcCLB.506495.20 | helicase-like protein, putative | -0,4083 | 0,0001064 |
| TcCLB.503879.10 | DNA polymerase delta subunit 2, putative | -0,4083 | 0,00114 |
| TcCLB.511367.30 | hypothetical protein | -0,4084 | 0,3324 |
| TcCLB.506925.70 | epsin, putative | -0,4085 | 0,0000013 |
| TcCLB.504243.20 | hypothetical protein, conserved | -0,4085 | 0,01164 |
| TcCLB.510533.160 | hypothetical protein, conserved | -0,4088 | 0,002618 |
| TcCLB.506993.160 | trans-sialidase (pseudogene), putative | -0,4088 | 0,1756 |
| TcCLB.506541.10 | hypothetical protein, conserved (pseudogene) | -0,4088 | 0,2987 |
| TcCLB.509669.50 | hypothetical protein, conserved | -0,4089 | 4,532E-06 |
| TcCLB.508921.30 | Kinesin-13 4, putative (KIN13-4) | -0,4089 | 0,03003 |
| TcCLB.504193.41 | N-acetyltransferase complex ARD1 subunit (pseudogene), | -0,4089 | 0,04716 |
| TcCLB.507963.44 | hypothetical protein, conserved | -0,409 | 0,005932 |
| TcCLB.506753.60 | GDP-mannose transporter, putative | -0,4093 | 0,02776 |
| TcCLB.511057.15 | hypothetical protein | -0,4094 | 0,3278 |
| TcCLB.511021.40 | hypothetical protein, conserved | -0,4095 | 0,00164 |
| TcCLB.506351.40 | ribonuclease H1, putative | -0,4095 | 0,02 |
| TcCLB.511403.15 | mucin TcMUC (pseudogene), putative | -0,4095 | 0,5796 |
| TcCLB.505115.10 | hypothetical protein, conserved | -0,4096 | 0,0001941 |
| TcCLB.507801.60 | hypothetical protein, conserved | -0,4096 | 0,0003298 |
| TcCLB.509893.10 | hypothetical protein | -0,4096 | 0,0009305 |
| TcCLB.504123.10 | hypothetical protein, conserved | -0,4096 | 0,003549 |
| TcCLB.503865.30 | hypothetical protein, conserved | -0,4097 | 0,0008977 |
| TcCLB.506657.30 | hypothetical protein, conserved | -0,4099 | 6,121E-06 |
| TcCLB.510719.240 | hypothetical protein, conserved | -0,4099 | 0,01387 |
| TcCLB.506491.10 | casein kinase I, putative | -0,4101 | 0,1788 |
| TcCLB.506813.140 | trans-sialidase, Group VII, putative | -0,4103 | 0,259 |
| TcCLB.504155.200 | syntaxin binding protein (pseudogene), putative | -0,4103 | 0,5166 |
| TcCLB.509805.130 | prolyl-tRNA synthetase, putative | -0,4104 | 3,585E-07 |
| TcCLB.506501.120 | mucin TcMUCII, putative | -0,4105 | 0,4762 |
| TcCLB.511861.111 | elongation factor 1-gamma (EF-1-gamma), putative | -0,4105 | 0,4804 |
| TcCLB.508693.160 | hypothetical protein, conserved | -0,4106 | 0,0008477 |

| | | | |
|---|---|---|---|
| TcCLB.508995.40 | vesicular-fusion protein nsf, putative | -0,4107 | 0,001811 |
| TcCLB.463359.9 | vacuolar assembly protein vps41, putative | -0,4107 | 0,0186 |
| TcCLB.511717.160 | hypothetical protein | -0,4109 | 0,01606 |
| TcCLB.506717.80 | trans-sialidase, Group VIII, putative | -0,4109 | 0,04772 |
| TcCLB.504047.34 | acylphosphatase, putative | -0,411 | 0,04053 |
| TcCLB.510055.150 | hypothetical protein, conserved | -0,4111 | 4,945E-07 |
| TcCLB.507257.70 | protein farnesyltransferase alpha subunit, putative | -0,4111 | 0,0002416 |
| TcCLB.504121.20 | hypothetical protein, conserved | -0,4111 | 0,005346 |
| TcCLB.506935.10 | hypothetical protein, conserved | -0,4112 | 0,0004274 |
| TcCLB.506147.40 | hypothetical protein | -0,4112 | 0,002082 |
| TcCLB.505999.170 | hypothetical protein, conserved | -0,4113 | 2,208E-07 |
| TcCLB.506989.260 | hypothetical protein, conserved (pseudogene) | -0,4113 | 0,01546 |
| TcCLB.511797.54 | dispersed gene family protein 1 (DGF-1), putative | -0,4113 | 0,158 |
| TcCLB.508465.10 | aminotransferase, putative | -0,4114 | 0,005389 |
| TcCLB.508523.50 | trans-sialidase, Group II, putative | -0,4115 | 0,05187 |
| TcCLB.508693.30 | hypothetical protein, conserved | -0,4117 | 0,00007215 |
| TcCLB.506561.10 | helicase-like protein, putative | -0,4118 | 0,0002011 |
| TcCLB.508909.70 | hypothetical protein, conserved | -0,4118 | 0,03219 |
| TcCLB.457251.19 | hypothetical protein, conserved (fragment) | -0,4118 | 0,05529 |
| TcCLB.508057.41 | trans-sialidase (pseudogene), putative | -0,4118 | 0,2692 |
| TcCLB.510973.20 | hypothetical protein, conserved | -0,4123 | 0,0002141 |
| TcCLB.509669.110 | GTPase activator protein, putative | -0,4123 | 0,0009102 |
| TcCLB.511293.40 | U3 snoRNA-associated protein UTP11, putative | -0,4124 | 0,002175 |
| TcCLB.507949.60 | hypothetical protein, conserved | -0,4124 | 0,002892 |
| TcCLB.508323.60 | centrin-4, putative (Centrin4) | -0,4125 | 0,00002346 |
| TcCLB.510431.40 | metallo- peptidase, Clan M- Family M48 (FACE1) | -0,4125 | 0,0004274 |
| TcCLB.505999.110 | NADH-ubiquinone oxidoreductase complex I subunit, putati | -0,4125 | 0,002623 |
| TcCLB.511771.210 | dispersed gene family protein 1 (DGF-1), putative | -0,4125 | 0,0146 |
| TcCLB.509695.110 | hypothetical protein, conserved | -0,4127 | 0,0007066 |
| TcCLB.511407.80 | hypothetical protein, conserved | -0,413 | 0,08953 |
| TcCLB.511543.50 | hypothetical protein, conserved | -0,4131 | 0,01463 |
| TcCLB.504115.60 | trans-sialidase, Group V, putative | -0,4131 | 0,09559 |
| TcCLB.511721.20 | hypothetical protein, conserved | -0,4131 | 0,3649 |
| TcCLB.507841.40 | hypothetical protein, conserved | -0,4132 | 0,0005768 |
| TcCLB.509965.229 | heat shock protein (pseudogene), putative | -0,4132 | 0,02046 |
| TcCLB.506247.160 | hypothetical protein, conserved | -0,4133 | 0,00003476 |
| TcCLB.511543.60 | glutathione peroxidase, putative | -0,4133 | 0,01023 |
| TcCLB.511537.21 | hypothetical protein, conserved | -0,4133 | 0,02408 |
| TcCLB.511803.10 | cullin 2, putative | -0,4134 | 0,003209 |
| TcCLB.508163.81 | hypothetical protein, conserved (pseudogene) | -0,4134 | 0,4183 |
| TcCLB.507957.300 | mucin-associated surface protein (MASP), putative | -0,4136 | 0,3702 |
| TcCLB.507519.130 | Mini-chromosome maintenance complex-binding protein ( | -0,4137 | 0,0007241 |
| TcCLB.507993.310 | transporter, putative | -0,4137 | 0,008113 |
| TcCLB.510377.475 | hypothetical protein, conserved (pseudogene) | -0,4137 | 0,577 |
| TcCLB.504867.99 | hypothetical protein, conserved | -0,4138 | 0,00009937 |

| | | | |
|---|---|---|---|
| TcCLB.508169.69 | peptidyl-prolyl cis-trans isomerase, putative | -0,4138 | 0,0007063 |
| TcCLB.508865.10 | glutathione synthetase, putative | -0,4138 | 0,001733 |
| TcCLB.507513.30 | hypothetical protein, conserved | -0,4139 | 0,09022 |
| TcCLB.507771.90 | hypothetical protein, conserved | -0,4141 | 1,754E-06 |
| TcCLB.509105.90 | MRB1-associated protein, putative (RGG2) | -0,4141 | 0,003054 |
| TcCLB.511727.260 | cyclin 9, putative (CYC9) | -0,4141 | 0,01569 |
| TcCLB.508257.80 | hypothetical protein, conserved | -0,4142 | 2,561E-06 |
| TcCLB.511671.60 | surface antigen 2 (CA-2), putative | -0,4142 | 0,00003216 |
| TcCLB.506407.70 | hypothetical protein, conserved | -0,4142 | 0,004978 |
| TcCLB.508505.20 | hypothetical protein, conserved | -0,4143 | 0,0003482 |
| TcCLB.509119.10 | oligomeric golgi complex component 8, putative | -0,4143 | 0,00242 |
| TcCLB.507065.29 | mucin-associated surface protein (MASP, pseudogene), pu | -0,4147 | 0,283 |
| TcCLB.508831.150 | hypothetical protein, conserved | -0,4149 | 0,00001163 |
| TcCLB.424123.30 | hypothetical protein, conserved | -0,4149 | 0,0053 |
| TcCLB.511003.30 | mitochondrial RNA binding complex 1 subunit, putative (RI | -0,4152 | 2,093E-07 |
| TcCLB.511909.20 | expression site-associated gene (ESAG-like) protein, putati | -0,4152 | 0,004077 |
| TcCLB.507547.70 | hypothetical protein, conserved | -0,4153 | 0,00005201 |
| TcCLB.503721.30 | hypothetical protein, conserved | -0,4154 | 0,00002764 |
| TcCLB.511339.30 | UDP-Gal or UDP-GlcNAc-dependent glycosyltransferase, pu | -0,4154 | 0,1451 |
| TcCLB.510361.10 | trans-sialidase, Group V, putative | -0,4154 | 0,2799 |
| TcCLB.503583.90 | hydrogenase, putative | -0,4155 | 0,001376 |
| TcCLB.506127.80 | molybdopterin synthase sulphurylase-like protein, putative | -0,4157 | 0,000352 |
| TcCLB.508989.60 | hypothetical protein, conserved | -0,4157 | 0,08481 |
| TcCLB.508821.20 | hypothetical protein, conserved | -0,4158 | 0,0001758 |
| TcCLB.506407.30 | hypothetical protein, conserved | -0,4158 | 0,0005658 |
| TcCLB.506543.70 | hypothetical protein, conserved | -0,4158 | 0,01585 |
| TcCLB.510187.74 | short-chain dehydrogenase, putative | -0,4159 | 0,005112 |
| TcCLB.506519.90 | hypothetical protein, conserved | -0,4159 | 0,009326 |
| TcCLB.511183.80 | mucin-associated surface protein (MASP), putative | -0,4159 | 0,309 |
| TcCLB.511583.20 | hypothetical protein, conserved | -0,416 | 0,002264 |
| TcCLB.508319.40 | ATP-dependent RNA helicase SUB2, putative (SUB2) | -0,4161 | 2,089E-07 |
| TcCLB.506337.130 | hypothetical protein, conserved | -0,4161 | 0,004504 |
| TcCLB.504047.50 | hypothetical protein, conserved | -0,4162 | 6,368E-06 |
| TcCLB.506247.130 | hypothetical protein, conserved | -0,4163 | 0,00003358 |
| TcCLB.509525.94 | mucin-associated surface protein (MASP, pseudogene), pu | -0,4163 | 0,1192 |
| TcCLB.511145.70 | centromere/microtubule binding protein cbf5, putative | -0,4164 | 0,00001442 |
| TcCLB.511269.30 | hypothetical protein, conserved | -0,4164 | 0,01323 |
| TcCLB.511797.200 | mucin-associated surface protein (MASP), putative | -0,4164 | 0,02752 |
| TcCLB.508307.90 | hypothetical protein, conserved | -0,4165 | 0,00003285 |
| TcCLB.506303.10 | hypothetical protein, conserved | -0,4166 | 0,00001221 |
| TcCLB.505077.9 | dispersed gene family protein 1 (DGF-1), putative (fragme | -0,4166 | 0,1846 |
| TcCLB.511183.140 | dispersed gene family protein 1 (DGF-1), putative | -0,4166 | 0,2133 |
| TcCLB.506411.20 | hypothetical protein, conserved | -0,4167 | 0,001812 |
| TcCLB.506847.70 | hypothetical protein, conserved | -0,4167 | 0,01759 |
| TcCLB.507729.10 | dispersed gene family protein 1 (DGF-1), putative | -0,4168 | 0,3525 |

| | | | |
|---|---|---|---|
| TcCLB.505009.20 | divalent cation tolerance protein, putative | -0,4169 | 0,2431 |
| TcCLB.506885.70 | polyadenylate-binding protein 1, putative | -0,417 | 0,00007218 |
| TcCLB.511537.24 | hypothetical protein, conserved | -0,4171 | 0,001045 |
| TcCLB.510001.40 | hypothetical protein, conserved | -0,4171 | 0,001487 |
| TcCLB.506419.40 | hypothetical protein, conserved | -0,4172 | 7,225E-06 |
| TcCLB.510329.230 | hypothetical protein, conserved | -0,4172 | 0,002454 |
| TcCLB.484299.4 | ATP-dependent DEAD/H RNA helicase, putative | -0,4174 | 0,1502 |
| TcCLB.506625.210 | hypothetical protein, conserved | -0,4177 | 0,001613 |
| TcCLB.510889.240 | hypothetical protein, conserved | -0,4177 | 0,0211 |
| TcCLB.506753.230 | hypothetical protein, conserved | -0,4178 | 0,0004038 |
| TcCLB.508461.110 | hypothetical protein, conserved | -0,4178 | 0,0008811 |
| TcCLB.506251.110 | hypothetical protein, conserved | -0,4178 | 0,002694 |
| TcCLB.508815.60 | hypothetical protein, conserved | -0,4178 | 0,008572 |
| TcCLB.472833.10 | hypothetical protein, conserved | -0,4183 | 0,3532 |
| TcCLB.508161.30 | mucin-associated surface protein (MASP), putative | -0,4184 | 0,1234 |
| TcCLB.508857.150 | hypothetical protein, conserved | -0,4185 | 0,00412 |
| TcCLB.511857.40 | N(2), N(2)-dimethylguanosine tRNA methyltransferase, pu | -0,4187 | 0,0003542 |
| TcCLB.503809.90 | glycerol uptake protein, putative | -0,4187 | 0,001134 |
| TcCLB.509023.120 | ubiquitin carboxyl-terminal hydrolase, putative | -0,4188 | 0,00005361 |
| TcCLB.509899.70 | hypothetical protein, conserved (pseudogene) | -0,4188 | 0,00008204 |
| TcCLB.510329.50 | hypothetical protein | -0,419 | 0,00001069 |
| TcCLB.511277.580 | RNA-binding protein 29, putative (RBP29) | -0,4191 | 4,227E-06 |
| TcCLB.508409.260 | hypothetical protein, conserved | -0,4191 | 0,0000192 |
| TcCLB.410923.30 | hypothetical protein | -0,4191 | 0,1649 |
| TcCLB.503453.90 | hypothetical protein, conserved | -0,4192 | 0,0001927 |
| TcCLB.511209.50 | hypothetical protein, conserved | -0,4192 | 0,00122 |
| TcCLB.509937.39 | hypothetical protein, conserved | -0,4194 | 0,007768 |
| TcCLB.510653.10 | hypothetical protein, conserved | -0,4194 | 0,05073 |
| TcCLB.504155.160 | mucin-associated surface protein (MASP), putative | -0,4194 | 0,2078 |
| TcCLB.506811.210 | cell division cycle protein 45 (CDC45), putative | -0,4195 | 0,0006763 |
| TcCLB.510421.70 | hypothetical protein, conserved | -0,4195 | 0,003575 |
| TcCLB.507001.10 | hypothetical protein, conserved | -0,4196 | 0,00005321 |
| TcCLB.413293.30 | retrotransposon hot spot protein (RHS, pseudogene), putat | -0,4196 | 0,001632 |
| TcCLB.504005.50 | hypothetical protein, conserved | -0,4197 | 0,001562 |
| TcCLB.511003.90 | hypothetical protein, conserved | -0,4198 | 0,0007326 |
| TcCLB.511151.20 | hypothetical protein, conserved | -0,4199 | 0,00007768 |
| TcCLB.506221.50 | coatomer subunit epsilon (COPE) | -0,4199 | 0,001781 |
| TcCLB.511737.20 | hypothetical protein, conserved | -0,42 | 0,00006957 |
| TcCLB.509603.10 | vacuolar protein sorting 18, putative | -0,42 | 0,01665 |
| TcCLB.507543.10 | dispersed gene family protein 1 (DGF-1), putative | -0,4201 | 0,3949 |
| TcCLB.508479.190 | peroxisome assembly protein, putative | -0,4202 | 0,0003329 |
| TcCLB.506067.70 | mucin-associated surface protein (MASP), putative | -0,4202 | 0,4508 |
| TcCLB.503887.50 | hypothetical protein, conserved | -0,4204 | 0,001177 |
| TcCLB.507943.20 | histone H4, putative | -0,4207 | 0,0000026 |
| TcCLB.508799.80 | mRNA cap guanine-N7 methyltransferase, putative | -0,4207 | 0,0003552 |

| | | | |
|---|---|---|---|
| TcCLB.509077.20 | trans-sialidase (pseudogene), putative | -0,4207 | 0,3238 |
| TcCLB.506679.30 | hypothetical protein, conserved | -0,4208 | 1,975E-06 |
| TcCLB.511491.50 | KU80 protein, putative | -0,421 | 0,0002942 |
| TcCLB.506727.70 | hypothetical protein, conserved | -0,421 | 0,001337 |
| TcCLB.506967.13 | hypothetical protein, conserved | -0,421 | 0,072 |
| TcCLB.507025.20 | hypothetical protein, conserved | -0,4214 | 0,002573 |
| TcCLB.511285.4 | intraflagellar transport protein 140, putative (fragment) (II | -0,4215 | 0,0007799 |
| TcCLB.509575.30 | hypothetical protein, conserved | -0,4215 | 0,0009292 |
| TcCLB.508299.89 | hypothetical protein, conserved | -0,4215 | 0,02952 |
| TcCLB.511143.30 | hypothetical protein, conserved | -0,4216 | 0,003567 |
| TcCLB.504079.20 | hypothetical protein, conserved | -0,4217 | 0,06246 |
| TcCLB.510731.68 | hypothetical protein, conserved | -0,4218 | 0,02825 |
| TcCLB.504097.10 | RNA editing complex protein MP61, putative | -0,422 | 0,0006422 |
| TcCLB.508871.180 | hypothetical protein | -0,422 | 0,2718 |
| TcCLB.509005.50 | hypothetical protein, conserved | -0,4223 | 0,00003249 |
| TcCLB.506257.60 | XPA-interacting protein, putative | -0,4223 | 0,002866 |
| TcCLB.509267.40 | hypothetical protein, conserved | -0,4226 | 0,0002482 |
| TcCLB.503753.10 | hypothetical protein, conserved | -0,4226 | 0,007669 |
| TcCLB.506389.70 | translation-associated element 2, putative | -0,4228 | 0,00004537 |
| TcCLB.511743.30 | hypothetical protein, conserved | -0,4229 | 0,0005221 |
| TcCLB.509561.80 | plectin-like protein, putative | -0,4229 | 0,001296 |
| TcCLB.510999.30 | hypothetical protein, conserved | -0,423 | 0,001096 |
| TcCLB.510777.10 | hypothetical protein, conserved | -0,423 | 0,01413 |
| TcCLB.503673.18 | hypothetical protein, conserved | -0,4234 | 0,009499 |
| TcCLB.507621.5 | trans-sialidase (pseudogene), putative | -0,4234 | 0,2254 |
| TcCLB.508457.20 | hypothetical protein, conserved | -0,4235 | 9,269E-06 |
| TcCLB.504071.20 | hypothetical protein, conserved | -0,4235 | 0,0001915 |
| TcCLB.506729.40 | hypothetical protein, conserved | -0,4238 | 0,0002782 |
| TcCLB.511861.20 | trans-sialidase, Group V, putative | -0,4238 | 0,02499 |
| TcCLB.507485.110 | hypothetical protein, conserved | -0,4239 | 0,00002206 |
| TcCLB.506855.300 | hypothetical protein, conserved | -0,4239 | 0,0002134 |
| TcCLB.511181.70 | hypothetical protein, conserved | -0,4239 | 0,01682 |
| TcCLB.510121.30 | actin-like protein, putative | -0,4239 | 0,01794 |
| TcCLB.506139.170 | trans-sialidase (pseudogene), putative | -0,4239 | 0,2104 |
| TcCLB.506975.20 | hypothetical protein, conserved | -0,4241 | 0,004011 |
| TcCLB.507007.45 | anaphase promoting complex subunit protein, putative | -0,4241 | 0,2139 |
| TcCLB.506247.240 | hypothetical protein, conserved | -0,4242 | 0,002175 |
| TcCLB.510253.29 | hypothetical protein, conserved (fragment) | -0,4243 | 0,07706 |
| TcCLB.505025.150 | trans-sialidase (pseudogene), putative | -0,4243 | 0,2194 |
| TcCLB.506211.114 | C-terminal motor kinesin, putative | -0,4247 | 5,688E-08 |
| TcCLB.509747.110 | hypothetical protein, conserved | -0,4247 | 0,0008258 |
| TcCLB.508717.50 | hypothetical protein, conserved | -0,4247 | 0,003227 |
| TcCLB.506577.150 | hypothetical protein, conserved | -0,4248 | 3,88E-08 |
| TcCLB.506629.90 | hypothetical protein, conserved | -0,4248 | 0,09741 |
| TcCLB.511083.39 | mucin-associated surface protein (MASP), putative | -0,4248 | 0,1974 |

| | | | |
|---|---|---|---|
| TcCLB.509053.80 | hypothetical protein, conserved | -0,4249 | 0,01051 |
| TcCLB.510285.40 | hypothetical protein, conserved | -0,425 | 0,01361 |
| TcCLB.505997.65 | trans-sialidase (pseudogene), putative | -0,4251 | 0,06033 |
| TcCLB.504109.110 | cleavage and polyadenylation specificity factor, putative | -0,4252 | 0,00009937 |
| TcCLB.506695.49 | hypothetical protein, conserved | -0,4254 | 0,00001316 |
| TcCLB.511003.50 | hypothetical protein, conserved | -0,4254 | 0,00003304 |
| TcCLB.509791.160 | hypothetical protein, conserved | -0,4256 | 5,111E-07 |
| TcCLB.506339.25 | hypothetical protein, conserved (pseudogene) | -0,4256 | 0,0582 |
| TcCLB.509757.10 | hypothetical protein, conserved | -0,4257 | 0,001952 |
| TcCLB.507993.33 | hypothetical protein, conserved | -0,4257 | 0,01523 |
| TcCLB.503499.30 | mitochondrial carrier protein (pseudogene), putative | -0,4258 | 0,02348 |
| TcCLB.506745.4 | hypothetical protein, conserved (fragment) | -0,4258 | 0,2941 |
| TcCLB.510329.40 | hypothetical protein, conserved | -0,4259 | 0,00004673 |
| TcCLB.511801.4 | hypothetical protein, conserved | -0,4259 | 0,001558 |
| TcCLB.510015.5 | hypothetical protein, conserved (pseudogene) | -0,4259 | 0,3468 |
| TcCLB.510391.40 | mucin TcMUCII, putative | -0,4261 | 0,3635 |
| TcCLB.510317.20 | hypothetical protein, conserved | -0,4262 | 0,00006602 |
| TcCLB.506741.130 | trans-sialidase, Group V, putative | -0,4262 | 0,2781 |
| TcCLB.503897.20 | hypothetical protein, conserved | -0,4264 | 0,0001258 |
| TcCLB.510781.30 | tryptophanyl-tRNA synthetase (pseudogene), putative | -0,4264 | 0,4275 |
| TcCLB.506407.90 | hypothetical protein, conserved | -0,4267 | 1,974E-08 |
| TcCLB.510889.320 | hypothetical protein, conserved | -0,4267 | 0,0001088 |
| TcCLB.508305.40 | trans-sialidase, Group V, putative | -0,4268 | 0,1813 |
| TcCLB.509105.100 | Enriched in surface-labeled proteome protein 15 (ESP15) | -0,427 | 0,0005516 |
| TcCLB.506737.60 | hypothetical protein, conserved | -0,427 | 0,06796 |
| TcCLB.508819.20 | hypothetical protein, conserved | -0,4272 | 0,000377 |
| TcCLB.421173.4 | trans-sialidase, putative (fragment) | -0,4272 | 0,07595 |
| TcCLB.506457.120 | hypothetical protein, conserved | -0,4274 | 0,000493 |
| TcCLB.510091.70 | U6 snRNA-associated Sm-like protein LSm8, putative | -0,4274 | 0,02916 |
| TcCLB.508231.200 | 40S ribosomal protein S8, putative | -0,4275 | 0,0006014 |
| TcCLB.507831.80 | hypothetical protein, conserved | -0,4276 | 0,001447 |
| TcCLB.504839.28 | tryparedoxin peroxidase, putative | -0,4277 | 3,066E-07 |
| TcCLB.508059.50 | hypothetical protein, conserved | -0,4277 | 5,335E-06 |
| TcCLB.504867.60 | hypothetical protein, conserved | -0,4277 | 0,0001273 |
| TcCLB.504157.60 | hypothetical protein, conserved | -0,4279 | 0,01926 |
| TcCLB.506945.250 | hypothetical protein, conserved | -0,428 | 8,804E-06 |
| TcCLB.510187.60 | glyceraldehyde 3-phosphate dehydrogenase, cytosolic, puta | -0,428 | 0,001489 |
| TcCLB.506771.18 | hypothetical protein, conserved | -0,428 | 0,002112 |
| TcCLB.508139.240 | trans-sialidase, Group II, putative | -0,428 | 0,004149 |
| TcCLB.509167.120 | S-adenosylmethionine decarboxylase proenzyme, putative | -0,4281 | 0,00007001 |
| TcCLB.511725.140 | mitochondrial carrier protein, putative | -0,4283 | 0,0009811 |
| TcCLB.398235.10 | protein kinase, putative (fragment) | -0,4283 | 0,09037 |
| TcCLB.440099.19 | hypothetical protein, conserved | -0,4284 | 0,004306 |
| TcCLB.506269.99 | mucin-associated surface protein (MASP), putative (fragm | -0,4284 | 0,01462 |
| TcCLB.508365.280 | mucin TcMUCII, putative | -0,4284 | 0,3469 |

| | | | |
|---|---|---|---|
| TcCLB.509569.30 | hypothetical protein, conserved | -0,4285 | 0,0002754 |
| TcCLB.503543.4 | hypothetical protein, conserved (fragment) | -0,4285 | 0,06062 |
| TcCLB.508409.300 | hypothetical protein, conserved | -0,4286 | 0,005996 |
| TcCLB.507711.120 | hypothetical protein, conserved | -0,4286 | 0,007397 |
| TcCLB.506405.110 | ribosomal protein L13, putative | -0,4286 | 0,03621 |
| TcCLB.511165.50 | Enriched in surface-labeled proteome protein 7 (ESP7) | -0,4288 | 0,008786 |
| TcCLB.506215.10 | hypothetical protein | -0,4288 | 0,1389 |
| TcCLB.508461.530 | hypothetical protein, conserved | -0,4289 | 0,0001594 |
| TcCLB.509181.9 | dispersed gene family protein 1 (DGF-1), putative | -0,4289 | 0,3107 |
| TcCLB.503423.30 | hypothetical protein, conserved | -0,429 | 0,1557 |
| TcCLB.508461.380 | calmodulin, putative | -0,4292 | 4,915E-07 |
| TcCLB.504827.110 | hypothetical protein, conserved | -0,4292 | 0,00002436 |
| TcCLB.508175.114 | hypothetical protein, conserved | -0,4292 | 0,0001617 |
| TcCLB.508461.120 | hypothetical protein, conserved | -0,4295 | 8,43E-07 |
| TcCLB.511313.20 | hypothetical protein, conserved | -0,4295 | 0,002564 |
| TcCLB.506559.200 | thioredoxin, putative | -0,4296 | 1,458E-07 |
| TcCLB.506249.10 | hypothetical protein, conserved | -0,4296 | 0,02156 |
| TcCLB.511255.580 | mucin-associated surface protein (MASP, pseudogene), pu | -0,4296 | 0,3908 |
| TcCLB.511589.130 | 14-3-3 protein 1, putative (14-3-3-I) | -0,4297 | 1,277E-06 |
| TcCLB.508299.30 | hypothetical protein | -0,4298 | 3,202E-06 |
| TcCLB.506211.120 | hypothetical protein, conserved | -0,4299 | 0,0009696 |
| TcCLB.511765.55 | trans-sialidase (pseudogene), putative | -0,43 | 0,3497 |
| TcCLB.510055.80 | hypothetical protein | -0,4301 | 0,002468 |
| TcCLB.506993.180 | mucin-associated surface protein (MASP), putative | -0,4301 | 0,2244 |
| TcCLB.506809.5 | retrotransposon hot spot (RHS) protein, putative | -0,4301 | 0,271 |
| TcCLB.503999.20 | Kinesin-13 5, putative (KIN13-5) | -0,4302 | 0,0000147 |
| TcCLB.509595.50 | DIS3-like exonuclease, putative | -0,4303 | 0,00001856 |
| TcCLB.506755.170 | hypothetical protein, conserved | -0,4303 | 0,00004936 |
| TcCLB.510167.20 | hypothetical protein, conserved | -0,4304 | 0,02082 |
| TcCLB.511897.28 | hypothetical protein, conserved | -0,4306 | 0,009934 |
| TcCLB.506195.220 | hypothetical protein, conserved | -0,4306 | 0,0108 |
| TcCLB.506721.10 | hypothetical protein, conserved (fragment) | -0,4306 | 0,1096 |
| TcCLB.508859.10 | hypothetical protein, conserved | -0,4307 | 0,003561 |
| TcCLB.510541.9 | dispersed gene family protein 1 (DGF-1), putative | -0,431 | 0,3946 |
| TcCLB.506331.20 | retrotransposon hot spot protein (RHS, pseudogene), putat | -0,4311 | 0,07786 |
| TcCLB.506357.80 | hypothetical protein, conserved | -0,4312 | 2,651E-06 |
| TcCLB.437545.10 | 3-oxo-5-alpha-steroid 4-dehydrogenase, putative | -0,4312 | 0,0004353 |
| TcCLB.508871.140 | mucin TcMUCII, putative | -0,4312 | 0,3642 |
| TcCLB.503809.30 | RNA-binding protein, putative | -0,4314 | 0,01193 |
| TcCLB.509721.30 | intraflagellar transport protein 122B, putative (IFT122B) | -0,4315 | 7,494E-06 |
| TcCLB.430603.10 | mucin-associated surface protein (MASP, pseudogene), pu | -0,4315 | 0,2467 |
| TcCLB.509695.200 | T-complex protein 1, delta subunit, putative | -0,4316 | 1,39E-08 |
| TcCLB.508153.380 | RNA polymerase-associated protein CTR9, putative | -0,4316 | 2,398E-06 |
| TcCLB.507511.91 | mucin TcMUCIII, putative | -0,4317 | 0,002383 |
| TcCLB.508441.60 | hypothetical protein, conserved | -0,4318 | 0,0006048 |

| | | | |
|---|---|---|---|
| TcCLB.510441.20 | dispersed gene family protein 1 (DGF-1), putative | -0,4318 | 0,04691 |
| TcCLB.509637.18 | hypothetical protein, conserved | -0,432 | 0,00604 |
| TcCLB.510359.250 | hypothetical protein, conserved | -0,432 | 0,1364 |
| TcCLB.509265.36 | ATP-dependent DEAD/H RNA helicase (pseudogene), putat | -0,432 | 0,2073 |
| TcCLB.511239.60 | hypothetical protein, conserved | -0,4321 | 5,541E-06 |
| TcCLB.511367.350 | hypothetical protein, conserved | -0,4322 | 0,0002831 |
| TcCLB.510359.150 | ATP-dependent helicase, putative | -0,4323 | 0,00001118 |
| TcCLB.509445.20 | hypothetical protein, conserved | -0,4323 | 0,0000266 |
| TcCLB.507715.20 | hypothetical protein, conserved | -0,4323 | 0,0004345 |
| TcCLB.506287.30 | hypothetical protein, conserved | -0,4324 | 0,01546 |
| TcCLB.508347.90 | hypothetical protein, conserved | -0,4325 | 2,014E-06 |
| TcCLB.508799.180 | hypothetical protein, conserved | -0,4325 | 0,0001115 |
| TcCLB.510029.30 | hypothetical protein, conserved | -0,4325 | 0,0002717 |
| TcCLB.506249.50 | hypothetical protein, conserved | -0,4325 | 0,03223 |
| TcCLB.504147.60 | chaperone DNAJ protein, putative | -0,4326 | 1,198E-06 |
| TcCLB.507787.90 | chaperone DNAJ protein, putative | -0,4327 | 1,164E-06 |
| TcCLB.508277.370 | mitochondrial DNA topoisomerase II, putative | -0,4329 | 8,666E-09 |
| TcCLB.509609.10 | hypothetical protein, conserved | -0,4329 | 0,005562 |
| TcCLB.504507.20 | DNA replication licensing factor MCM8, putative | -0,4329 | 0,06607 |
| TcCLB.511555.60 | ubiquitin ligase, putative | -0,433 | 1,63E-11 |
| TcCLB.509669.150 | hypothetical protein, conserved | -0,4331 | 0,001129 |
| TcCLB.510599.40 | hypothetical protein, conserved | -0,4335 | 0,00001329 |
| TcCLB.506319.40 | hypothetical protein | -0,4339 | 0,2119 |
| TcCLB.508295.50 | mucin TcMUCII, putative | -0,4339 | 0,2682 |
| TcCLB.506135.50 | hypothetical protein, conserved | -0,434 | 0,0006036 |
| TcCLB.508699.90 | surface protease GP63, putative | -0,4343 | 0,02741 |
| TcCLB.510575.120 | hypothetical protein, conserved | -0,4344 | 0,007856 |
| TcCLB.510825.20 | hypothetical protein, conserved | -0,4344 | 0,07161 |
| TcCLB.506239.30 | lectin, putative (AG38) | -0,4345 | 1,112E-06 |
| TcCLB.504137.110 | hypothetical protein, conserved | -0,4345 | 7,911E-06 |
| TcCLB.507007.37 | hypothetical protein, conserved | -0,4346 | 0,3932 |
| TcCLB.508641.120 | hypothetical protein, conserved | -0,4349 | 0,00275 |
| TcCLB.507007.90 | hypothetical protein, conserved | -0,4349 | 0,01017 |
| TcCLB.506621.14 | ubiquitin ligase, putative | -0,435 | 0,000473 |
| TcCLB.506597.51 | hypothetical protein | -0,435 | 0,547 |
| TcCLB.508707.90 | proteasome regulatory non-ATPase subunit 7, putative | -0,4351 | 0,00001168 |
| TcCLB.509215.40 | peptidyl-prolyl cis-trans isomerase, putative | -0,4352 | 2,114E-06 |
| TcCLB.507087.30 | cysteine peptidase, Clan CA, family C19, putative | -0,4352 | 0,00002007 |
| TcCLB.507031.170 | protein kinase (pseudogene), putative | -0,4353 | 0,00003582 |
| TcCLB.506559.310 | hypothetical protein, conserved | -0,4353 | 0,0000624 |
| TcCLB.506227.90 | hypothetical protein, conserved | -0,4353 | 0,001554 |
| TcCLB.510557.10 | trans-sialidase (pseudogene), putative | -0,4354 | 0,01731 |
| TcCLB.506589.100 | hypothetical protein | -0,4357 | 0,3833 |
| TcCLB.506743.170 | hypothetical protein, conserved | -0,4358 | 0,001494 |
| TcCLB.507649.80 | nucleolar RNA-binding protein, putative | -0,436 | 1,37E-07 |

| | | | |
|---|---|---|---|
| TcCLB.504247.40 | RNA-editing nuclease 1 (KREN1) | -0,436 | 0,001899 |
| TcCLB.508331.80 | dispersed gene family protein 1 (DGF-1, pseudogene), puta | -0,4361 | 0,0539 |
| TcCLB.509525.190 | hypothetical protein | -0,4361 | 0,5365 |
| TcCLB.506295.30 | hypothetical protein, conserved | -0,4362 | 0,001593 |
| TcCLB.507485.130 | amastin, putative | -0,4363 | 0,00007061 |
| TcCLB.507485.80 | hypothetical protein, conserved | -0,4363 | 0,02038 |
| TcCLB.510143.120 | hypothetical protein, conserved | -0,4363 | 0,02187 |
| TcCLB.508153.370 | hypothetical protein, conserved | -0,4364 | 0,0000389 |
| TcCLB.503749.20 | hypothetical protein | -0,4364 | 0,0005653 |
| TcCLB.503575.40 | hypothetical protein, conserved | -0,4364 | 0,02373 |
| TcCLB.509487.10 | hypothetical protein | -0,4365 | 0,01223 |
| TcCLB.507053.30 | hypothetical protein, conserved | -0,4366 | 0,00001841 |
| TcCLB.504037.40 | hypothetical protein, conserved | -0,4366 | 0,04395 |
| TcCLB.507061.40 | hypothetical protein, conserved (pseudogene) | -0,4367 | 0,00001048 |
| TcCLB.506011.30 | hypothetical protein, conserved | -0,4367 | 0,06213 |
| TcCLB.511055.60 | hypothetical protein, conserved | -0,4368 | 0,1328 |
| TcCLB.510073.30 | pumilio/PUF RNA binding protein 4, putative (PUF4) | -0,437 | 0,5323 |
| TcCLB.506929.50 | hypothetical protein, conserved | -0,4373 | 0,0003333 |
| TcCLB.503821.10 | DNA-directed RNA polymerase I subunit, putative | -0,4373 | 0,0228 |
| TcCLB.510363.170 | mucin TcMUCII, putative | -0,4373 | 0,6029 |
| TcCLB.506703.100 | mucin TcMUCII (pseudogene), putative | -0,4374 | 0,5275 |
| TcCLB.509969.20 | surface protease GP63 (pseudogene), putative | -0,4375 | 0,2049 |
| TcCLB.507627.9 | kinesin, putative | -0,4376 | 8,711E-06 |
| TcCLB.508837.160 | hypothetical protein, conserved | -0,4376 | 0,00009883 |
| TcCLB.511847.20 | hypothetical protein, conserved | -0,4376 | 0,000346 |
| TcCLB.509805.30 | hypothetical protein, conserved | -0,4376 | 0,009204 |
| TcCLB.505171.70 | ATP-dependent DEAD/H RNA helicase, putative | -0,4377 | 0,00001549 |
| TcCLB.509443.20 | mitochondrial RNA binding protein 1, putative | -0,4377 | 0,00003007 |
| TcCLB.508665.30 | hypothetical protein, conserved | -0,4377 | 0,00008603 |
| TcCLB.511511.6 | zinc finger protein ZFP1 | -0,4379 | 0,3224 |
| TcCLB.510225.30 | hypothetical protein, conserved | -0,4383 | 0,000431 |
| TcCLB.511301.60 | hypothetical protein, conserved | -0,4383 | 0,0006568 |
| TcCLB.510265.20 | 26S protease regulatory subunit, putative | -0,4384 | 0,00000585 |
| TcCLB.504167.14 | hypothetical protein | -0,4384 | 0,005895 |
| TcCLB.503579.40 | hypothetical protein, conserved | -0,4384 | 0,006309 |
| TcCLB.504741.60 | transcription initiation protein, putative | -0,4386 | 0,00003038 |
| TcCLB.506661.80 | hypothetical protein, conserved | -0,4387 | 0,001146 |
| TcCLB.505957.20 | dispersed gene family protein 1 (DGF-1, pseudogene), puta | -0,4388 | 0,02967 |
| TcCLB.509229.130 | hypothetical protein, conserved | -0,4391 | 0,00301 |
| TcCLB.511367.260 | Peptidase C19, ubiquitin carboxyl-terminal hydrolase, puta | -0,4392 | 9,089E-06 |
| TcCLB.510657.100 | hypothetical protein, conserved | -0,4393 | 0,00001748 |
| TcCLB.503847.90 | ABC1 protein, putative | -0,4395 | 0,0000767 |
| TcCLB.510197.110 | dispersed gene family protein 1 (DGF-1), putative | -0,4395 | 0,04013 |
| TcCLB.506583.60 | Elongation factor G1, mitochondrial, putative (EF-G1) | -0,4396 | 4,874E-08 |
| TcCLB.507527.30 | DNA replication licensing factor MCM6, putative | -0,4396 | 0,0002108 |

| | | | |
|---|---|---|---|
| TcCLB.509017.30 | vacuolar ATP synthase subunit d | -0,4396 | 0,0009218 |
| TcCLB.506113.50 | ATP-dependent DEAD/H RNA helicase (pseudogene), putat | -0,4396 | 0,1661 |
| TcCLB.509161.130 | ATP-dependent DEAD/H RNA helicase, putative | -0,4397 | 1,649E-06 |
| TcCLB.509965.40 | ubiquitin hydrolase, putative | -0,4399 | 0,0002112 |
| TcCLB.510431.320 | hypothetical protein, conserved | -0,4399 | 0,0002577 |
| TcCLB.508153.1090 | hypothetical protein, conserved | -0,4399 | 0,0003897 |
| TcCLB.511181.30 | Iron-sulfur assembly protein 2, putative (Isa2) | -0,44 | 0,05401 |
| TcCLB.504039.160 | mucin-associated surface protein (MASP), putative | -0,44 | 0,4701 |
| TcCLB.511445.120 | hypothetical protein, conserved | -0,4401 | 0,001107 |
| TcCLB.510329.220 | hypothetical protein, conserved | -0,4404 | 0,0002023 |
| TcCLB.506833.20 | hypothetical protein, conserved | -0,4404 | 0,01957 |
| TcCLB.511277.440 | tubulin tyrosine ligase protein, putative | -0,4405 | 0,003197 |
| TcCLB.504839.36 | hypothetical protein, conserved | -0,4407 | 0,01364 |
| TcCLB.509571.30 | hypothetical protein, conserved | -0,4408 | 4,086E-06 |
| TcCLB.506919.20 | trans-sialidase (pseudogene), putative | -0,4408 | 0,44 |
| TcCLB.507609.80 | hypothetical protein, conserved | -0,4409 | 0,02071 |
| TcCLB.508405.80 | cysteine peptidase, Clan CD, family C50, putative | -0,441 | 0,0001555 |
| TcCLB.508461.460 | hypothetical protein, conserved | -0,441 | 0,0009066 |
| TcCLB.510687.80 | hypothetical protein, conserved | -0,441 | 0,001479 |
| TcCLB.511577.140 | hypothetical protein, conserved | -0,441 | 0,004245 |
| TcCLB.504201.30 | trans-sialidase, Group V, putative | -0,4412 | 0,2072 |
| TcCLB.511019.47 | N-acetyltransferase complex ARD1 subunit (pseudogene), | -0,4413 | 0,1099 |
| TcCLB.504643.20 | hypothetical protein, conserved | -0,4415 | 0,006405 |
| TcCLB.504115.100 | dispersed gene family protein 1 (DGF-1), putative | -0,4415 | 0,01025 |
| TcCLB.511907.330 | retrotransposon hot spot (RHS) protein, putative | -0,4416 | 0,000239 |
| TcCLB.511355.20 | hypothetical protein, conserved | -0,4416 | 0,0009319 |
| TcCLB.504157.30 | hypothetical protein, conserved | -0,4416 | 0,01919 |
| TcCLB.510839.20 | hypothetical protein | -0,4418 | 0,0005376 |
| TcCLB.508153.250 | hypothetical protein | -0,4419 | 0,0138 |
| TcCLB.508715.10 | oxidoreductase, putative (fragment) | -0,442 | 0,009751 |
| TcCLB.510073.70 | CCR4-NOT transcription complex subunit 2, putative | -0,4422 | 0,0000716 |
| TcCLB.508411.60 | hypothetical protein, conserved | -0,4422 | 0,001092 |
| TcCLB.507801.19 | dynein heavy chain, putative | -0,4423 | 0,0004132 |
| TcCLB.511389.90 | hypothetical protein, conserved | -0,4423 | 0,001543 |
| TcCLB.510359.100 | hypothetical protein, conserved | -0,4425 | 0,02112 |
| TcCLB.508153.390 | hypothetical protein, conserved | -0,4426 | 0,0002243 |
| TcCLB.509683.24 | DNA-directed RNA polymerase I subunit, putative | -0,4427 | 0,008654 |
| TcCLB.506567.74 | hypothetical protein, conserved | -0,4427 | 0,02408 |
| TcCLB.506425.70 | hypothetical protein, conserved | -0,4428 | 0,00003187 |
| TcCLB.511513.30 | hypothetical protein, conserved | -0,4429 | 1,499E-07 |
| TcCLB.506627.60 | ADP-ribosylation factor, putative | -0,4429 | 0,00348 |
| TcCLB.510279.30 | hypothetical protein | -0,4429 | 0,4685 |
| TcCLB.510533.110 | hypothetical protein, conserved | -0,443 | 0,0003077 |
| TcCLB.506755.190 | hypothetical protein, conserved | -0,4432 | 0,0003637 |
| TcCLB.507077.40 | hypothetical protein, conserved | -0,4432 | 0,002331 |

| | | | |
|---|---|---|---|
| TcCLB.508217.9 | dynein heavy chain, putative | -0,4432 | 0,003189 |
| TcCLB.511229.70 | hypothetical protein, conserved | -0,4432 | 0,009665 |
| TcCLB.505943.20 | oxidoreductase, putative | -0,4433 | 0,006088 |
| TcCLB.507143.30 | hypothetical protein | -0,4434 | 7,321E-09 |
| TcCLB.511587.111 | N-acetyltransferase complex ARD1 subunit (pseudogene), | -0,4434 | 0,1608 |
| TcCLB.508873.390 | trans-sialidase, Group V, putative | -0,4435 | 0,1677 |
| TcCLB.507089.230 | hypothetical protein, conserved | -0,4436 | 0,00003696 |
| TcCLB.503653.90 | hypothetical protein, conserved | -0,4436 | 0,001619 |
| TcCLB.503969.10 | tubulin delta chain (TUBD1) | -0,4436 | 0,005739 |
| TcCLB.510819.60 | ATP-dependent protease ATPase subunit HslU1, putative (I | -0,4439 | 3,996E-07 |
| TcCLB.511037.30 | hypothetical protein, conserved | -0,4439 | 0,00000227 |
| TcCLB.508643.30 | DnaJ chaperone protein, putative | -0,4439 | 0,007106 |
| TcCLB.510543.30 | trans-sialidase (pseudogene), putative | -0,444 | 0,1955 |
| TcCLB.469785.10 | hypothetical protein, conserved | -0,4441 | 0,000108 |
| TcCLB.507541.20 | hypothetical protein, conserved | -0,4441 | 0,0001994 |
| TcCLB.507951.260 | aspartic peptidase, clan AD, family A22B, putative | -0,4442 | 3,743E-06 |
| TcCLB.504109.80 | Intraflagellar transport protein 144, putative (PIFTF6) | -0,4444 | 4,435E-06 |
| TcCLB.506605.190 | hypothetical protein, conserved | -0,4444 | 0,001128 |
| TcCLB.511429.10 | transmembrane protein, conserved (fragment) | -0,4445 | 3,117E-06 |
| TcCLB.506173.10 | hypothetical protein, conserved | -0,4446 | 0,01471 |
| TcCLB.511019.54 | ATP-dependent DEAD/H RNA helicase (pseudogene), putat | -0,4446 | 0,2565 |
| TcCLB.507015.30 | trans-sialidase (pseudogene), putative | -0,4447 | 0,007548 |
| TcCLB.511419.4 | hypothetical protein, conserved (fragment) | -0,4447 | 0,09834 |
| TcCLB.510819.20 | hypothetical protein, conserved | -0,4448 | 0,01141 |
| TcCLB.504217.4 | hypothetical protein, conserved (fragment) | -0,4448 | 0,02766 |
| TcCLB.511847.70 | hypothetical protein, conserved | -0,4449 | 0,001829 |
| TcCLB.506181.80 | hypothetical protein, conserved | -0,4449 | 0,01848 |
| TcCLB.511211.130 | receptor for activated C kinase 1, putative | -0,445 | 6,099E-08 |
| TcCLB.510421.330 | hypothetical protein, conserved (fragment) | -0,445 | 9,827E-07 |
| TcCLB.510895.20 | hypothetical protein, conserved | -0,445 | 0,0006705 |
| TcCLB.510173.80 | hypothetical protein | -0,4451 | 0,008705 |
| TcCLB.510659.44 | cysteine desulfurase, putative | -0,4452 | 0,0002701 |
| TcCLB.504185.20 | hypothetical protein, conserved | -0,4452 | 0,0008967 |
| TcCLB.506947.80 | arginine N-methyltransferase, type I (PRMT6) | -0,4454 | 0,01216 |
| TcCLB.510483.60 | trans-sialidase, Group V, putative | -0,4456 | 0,2025 |
| TcCLB.510735.50 | hypothetical protein, conserved | -0,4457 | 0,0001448 |
| TcCLB.510657.140 | hypothetical protein, conserved | -0,4457 | 0,001182 |
| TcCLB.503515.14 | ubiquitin-conjugating enzyme E2, putative | -0,4459 | 0,004146 |
| TcCLB.510013.180 | hypothetical protein | -0,446 | 0,3626 |
| TcCLB.476733.10 | hypothetical protein, conserved | -0,4461 | 0,00001606 |
| TcCLB.507017.10 | hypothetical protein, conserved | -0,4462 | 9,542E-06 |
| TcCLB.510339.74 | mitochondrial carrier protein, putative | -0,4462 | 0,0001266 |
| TcCLB.510737.79 | 60S ribosomal subunit protein L31, putative | -0,4462 | 0,2636 |
| TcCLB.506249.60 | 40S ribosomal protein S8, putative | -0,4463 | 0,00003012 |
| TcCLB.510943.100 | acetyltransferase, putative | -0,4463 | 0,005923 |

| | | | |
|---|---|---|---|
| TcCLB.508479.120 | pumilio/PUF RNA binding protein 8, putative (PUF8) | -0,4464 | 5,972E-08 |
| TcCLB.511421.20 | Non cytoplasmic domain containing protein, conserved | -0,4465 | 0,0002591 |
| TcCLB.509911.4 | hypothetical protein, conserved | -0,4465 | 0,01988 |
| TcCLB.508755.20 | dispersed gene family protein 1 (DGF-1, pseudogene), puta | -0,4466 | 0,1548 |
| TcCLB.509229.20 | hypothetical protein, conserved | -0,4468 | 2,256E-06 |
| TcCLB.509911.60 | hypothetical protein | -0,4468 | 0,00001386 |
| TcCLB.504041.4 | RNA polymerase I | -0,4468 | 0,03124 |
| TcCLB.506305.20 | hypothetical protein, conserved | -0,4471 | 0,0007924 |
| TcCLB.508207.64 | hypothetical protein, conserved | -0,4471 | 0,001805 |
| TcCLB.510275.306 | mucin-associated surface protein (MASP, pseudogene), pu | -0,4472 | 0,1067 |
| TcCLB.510089.70 | hypothetical protein | -0,4473 | 0,05109 |
| TcCLB.506303.30 | hypothetical protein, conserved | -0,4474 | 0,005382 |
| TcCLB.511471.60 | protein kinase, putative | -0,4476 | 0,2143 |
| TcCLB.463155.40 | glycine dehydrogenase [decarboxylating] (pseudogene), pu | -0,4477 | 0,002078 |
| TcCLB.511509.10 | hypothetical protein, conserved | -0,4477 | 0,007082 |
| TcCLB.506599.150 | mucin-associated surface protein (MASP), putative | -0,4478 | 0,1814 |
| TcCLB.511283.124 | mitochondrial carrier protein, putative | -0,448 | 0,0002774 |
| TcCLB.506519.110 | hypothetical protein, conserved | -0,4481 | 0,000298 |
| TcCLB.511003.221 | cleavage and polyadenylation specificity factor, putative (f | -0,4482 | 0,00006969 |
| TcCLB.510597.30 | hypothetical protein, conserved | -0,4483 | 3,913E-06 |
| TcCLB.447255.20 | Sir2-related protein 3, putative (SIR2rp3) | -0,4483 | 0,004834 |
| TcCLB.510275.160 | trans-sialidase (pseudogene), putative | -0,4483 | 0,2123 |
| TcCLB.508207.80 | Mini-chromosome maintenance complex-binding protein ( | -0,4484 | 0,000549 |
| TcCLB.507257.10 | hypothetical protein, conserved | -0,4484 | 0,007288 |
| TcCLB.504213.120 | proteasome alpha 3 subunit, putative | -0,4485 | 3,049E-06 |
| TcCLB.504117.20 | sedoheptulose-1,7-bisphosphatase, putative | -0,4486 | 0,01416 |
| TcCLB.511193.9 | dispersed gene family protein 1 (DGF-1, pseudogene), puta | -0,4486 | 0,03788 |
| TcCLB.506499.160 | mucin-associated surface protein (MASP), putative | -0,4487 | 0,1492 |
| TcCLB.507641.40 | hypothetical protein, conserved | -0,4488 | 2,109E-08 |
| TcCLB.507395.30 | hypothetical protein | -0,4488 | 0,03957 |
| TcCLB.503891.40 | hypothetical protein, conserved | -0,4489 | 0,00009888 |
| TcCLB.508641.300 | hypothetical protein | -0,4489 | 0,0002605 |
| TcCLB.510901.140 | hypothetical protein, conserved | -0,4489 | 0,002508 |
| TcCLB.509965.170 | hypothetical protein, conserved | -0,449 | 0,00005464 |
| TcCLB.506841.12 | hypothetical protein, conserved | -0,449 | 0,002612 |
| TcCLB.506957.46 | hypothetical protein, conserved | -0,4491 | 0,003671 |
| TcCLB.503897.30 | hypothetical protein, conserved | -0,4492 | 0,0005302 |
| TcCLB.508059.24 | hypothetical protein, conserved | -0,4494 | 0,0003547 |
| TcCLB.508153.880 | hypothetical protein, conserved | -0,4495 | 0,00003043 |
| TcCLB.511859.180 | proteasome regulatory ATPase subunit 2, putative | -0,4496 | 2,172E-06 |
| TcCLB.503643.6 | hypothetical protein, conserved | -0,4497 | 0,0005296 |
| TcCLB.506337.40 | glycosyl transferase-like protein, putative | -0,4497 | 0,007827 |
| TcCLB.442495.19 | KU70 protein, putative (fragment) | -0,4497 | 0,04626 |
| TcCLB.508725.4 | hypothetical protein, conserved | -0,4498 | 0,003912 |
| TcCLB.504033.210 | hypothetical protein, conserved | -0,4498 | 0,04561 |

| | | | |
|---|---|---|---|
| TcCLB.506945.50 | hypothetical protein, conserved | -0,4502 | 0,00004905 |
| TcCLB.510099.10 | hypothetical protein, conserved | -0,4502 | 0,0005934 |
| TcCLB.511317.40 | hypothetical protein, conserved | -0,4502 | 0,0006803 |
| TcCLB.510431.280 | RNA editing complex protein MP46, putative | -0,4502 | 0,001765 |
| TcCLB.506479.20 | hypothetical protein, conserved | -0,4505 | 0,001868 |
| TcCLB.506657.10 | hypothetical protein, conserved | -0,4506 | 0,04766 |
| TcCLB.509065.79 | hypothetical protein, conserved | -0,4507 | 0,0005693 |
| TcCLB.506195.80 | adenylate kinase, putative | -0,4507 | 0,005592 |
| TcCLB.511045.20 | hypothetical protein, conserved | -0,4507 | 0,01403 |
| TcCLB.510237.21 | hypothetical protein, conserved (pseudogene) | -0,4508 | 0,1392 |
| TcCLB.506321.260 | hypothetical protein, conserved | -0,4509 | 0,01519 |
| TcCLB.508817.10 | hypothetical protein, conserved | -0,451 | 0,02115 |
| TcCLB.510329.140 | hypothetical protein, conserved | -0,4512 | 1,858E-09 |
| TcCLB.505937.70 | trans-sialidase (pseudogene), putative | -0,4512 | 0,05605 |
| TcCLB.507943.40 | histone H4, putative | -0,4513 | 3,595E-08 |
| TcCLB.508169.60 | dolicholphosphate-mannose synthase, putative (DPMS) | -0,4513 | 0,001109 |
| TcCLB.511555.50 | hypothetical protein, conserved | -0,4514 | 0,002344 |
| TcCLB.506975.70 | hypothetical protein | -0,4514 | 0,04715 |
| TcCLB.506509.60 | hypothetical protein, conserved | -0,4515 | 0,02528 |
| TcCLB.503593.40 | hypothetical protein | -0,4517 | 0,00005928 |
| TcCLB.507997.40 | hypothetical protein, conserved | -0,4517 | 0,001145 |
| TcCLB.508321.50 | hypothetical protein, conserved | -0,4518 | 6,468E-07 |
| TcCLB.510121.160 | DNA topoisomerase IA, putative | -0,4518 | 1,034E-06 |
| TcCLB.508881.130 | hypothetical protein, conserved | -0,4518 | 0,003356 |
| TcCLB.405165.10 | hypothetical protein, conserved | -0,4519 | 0,01474 |
| TcCLB.508387.50 | bilobe protein, conserved | -0,452 | 0,00008627 |
| TcCLB.510377.350 | mucin-associated surface protein (MASP), putative | -0,452 | 0,1783 |
| TcCLB.508165.420 | mucin-associated surface protein (MASP), putative | -0,4524 | 0,03045 |
| TcCLB.510295.20 | MP24, putative | -0,4524 | 0,03289 |
| TcCLB.508777.140 | palmitoyl acyltransferase 5, putative | -0,4525 | 0,00008872 |
| TcCLB.511027.29 | hypothetical protein, conserved (pseudogene) | -0,4525 | 0,003006 |
| TcCLB.509179.50 | hypothetical protein | -0,4526 | 0,002833 |
| TcCLB.506495.6 | hypothetical protein, conserved | -0,4526 | 0,005498 |
| TcCLB.506529.541 | hypothetical protein | -0,4526 | 0,0487 |
| TcCLB.510131.60 | hypothetical protein | -0,4527 | 0,001956 |
| TcCLB.508939.60 | hypothetical protein | -0,4527 | 0,02473 |
| TcCLB.506975.66 | hypothetical protein, conserved | -0,453 | 0,00002195 |
| TcCLB.503787.65 | hypothetical protein (pseudogene) | -0,4531 | 0,3249 |
| TcCLB.504045.114 | hypothetical protein, conserved | -0,4532 | 0,03914 |
| TcCLB.507081.70 | hypothetical protein, conserved | -0,4533 | 0,0000472 |
| TcCLB.507017.104 | hypothetical protein | -0,4533 | 0,01006 |
| TcCLB.510173.60 | intraflagellar transport protein 122B, putative (IFT122B) | -0,4535 | 2,256E-06 |
| TcCLB.507511.20 | hypothetical protein, conserved | -0,4535 | 0,00004498 |
| TcCLB.507159.20 | hypothetical protein, conserved | -0,4535 | 0,002033 |
| TcCLB.510963.30 | mRNA capping methyltransferase, putative | -0,4535 | 0,005187 |

| | | | |
|---|---|---|---|
| TcCLB.511875.40 | surface protease GP63 (pseudogene), putative | -0,4536 | 0,005286 |
| TcCLB.511731.35 | hypothetical protein, conserved | -0,4537 | 0,004023 |
| TcCLB.508269.70 | hypothetical protein, conserved | -0,4537 | 0,06119 |
| TcCLB.510031.50 | ATP-dependent RNA helicase, putative | -0,4538 | 0,001469 |
| TcCLB.509029.10 | serine/threonine protein phosphatase, putative | -0,4539 | 0,001089 |
| TcCLB.506811.10 | kinesin, putative | -0,454 | 0,0001277 |
| TcCLB.507071.10 | trans-sialidase, Group V, putative | -0,454 | 0,1578 |
| TcCLB.510409.50 | hypothetical protein, conserved | -0,4542 | 0,00131 |
| TcCLB.510089.120 | hypothetical protein, conserved | -0,4542 | 0,002525 |
| TcCLB.504213.20 | hypothetical protein, conserved | -0,4543 | 1,476E-06 |
| TcCLB.467287.20 | hypothetical protein, conserved | -0,4543 | 0,1984 |
| TcCLB.509875.20 | dispersed gene family protein 1 (DGF-1), putative | -0,4544 | 0,03766 |
| TcCLB.506511.10 | hypothetical protein, conserved | -0,4545 | 0,03561 |
| TcCLB.408345.10 | hypothetical protein, conserved | -0,4546 | 0,08102 |
| TcCLB.511217.110 | hypothetical protein, conserved | -0,4547 | 5,847E-06 |
| TcCLB.504179.69 | phosphoprotein lepp12, putative | -0,4547 | 0,003338 |
| TcCLB.510025.200 | syntaxin binding protein (pseudogene), putative | -0,4547 | 0,249 |
| TcCLB.508231.30 | target of rapamycin (TOR) kinase 1, putative (TOR1) | -0,4548 | 1,444E-13 |
| TcCLB.506329.79 | mucin-associated surface protein (MASP, pseudogene), pu | -0,4549 | 0,4932 |
| TcCLB.508409.40 | Putative intraflagellar transport protein B2 (PIFTB2) | -0,4553 | 5,613E-06 |
| TcCLB.509161.34 | hypothetical protein, conserved | -0,4553 | 0,0005388 |
| TcCLB.511183.460 | dispersed gene family protein 1 (DGF-1), putative | -0,4554 | 0,06682 |
| TcCLB.506331.64 | hypothetical protein | -0,4554 | 0,1342 |
| TcCLB.506211.130 | hypothetical protein, conserved | -0,4555 | 0,00005699 |
| TcCLB.511611.40 | hypothetical protein | -0,4555 | 0,3298 |
| TcCLB.508723.120 | hypothetical protein, conserved | -0,4556 | 0,004449 |
| TcCLB.507009.100 | cyclophilin, putative | -0,4557 | 0,005323 |
| TcCLB.509167.170 | hypothetical protein, conserved | -0,4557 | 0,008068 |
| TcCLB.511127.5 | RNA helicase, putative | -0,4559 | 0,1663 |
| TcCLB.510299.30 | hypothetical protein, conserved | -0,4561 | 0,001921 |
| TcCLB.503453.60 | GTP-ase activating protein, putative | -0,4561 | 0,004652 |
| TcCLB.503939.100 | hypothetical protein, conserved | -0,4562 | 0,0000013 |
| TcCLB.508549.30 | target of rapamycin (TOR) kinase 1, putative | -0,4562 | 0,0006435 |
| TcCLB.511797.215 | hypothetical protein, conserved (pseudogene) | -0,4562 | 0,08747 |
| TcCLB.506945.30 | CAS/CSE/importin domain protein, putative | -0,4563 | 1,021E-06 |
| TcCLB.507521.90 | hypothetical protein, conserved | -0,4564 | 0,01682 |
| TcCLB.507485.150 | amastin, putative | -0,4566 | 0,00001956 |
| TcCLB.504867.10 | hypothetical protein, conserved | -0,4566 | 0,002342 |
| TcCLB.509771.10 | trans-sialidase, Group II, putative | -0,4566 | 0,3956 |
| TcCLB.509099.120 | hypothetical protein, conserved | -0,4567 | 0,004283 |
| TcCLB.507605.13 | mucin-associated surface protein (MASP), putative (fragm | -0,4567 | 0,5289 |
| TcCLB.508153.1050 | hypothetical protein, conserved | -0,4568 | 3,51E-09 |
| TcCLB.505229.5 | hypothetical protein, conserved (fragment) | -0,4569 | 0,001845 |
| TcCLB.510121.190 | hypothetical protein, conserved | -0,4569 | 0,009171 |
| TcCLB.504431.40 | hypothetical protein, conserved | -0,457 | 0,0005363 |

| | | | |
|---|---|---|---|
| TcCLB.508741.280 | ubiquitin-conjugating enzyme E2, putative | -0,457 | 0,001157 |
| TcCLB.506691.64 | cytochrome c oxidase copper chaperone, putative | -0,4571 | 0,03804 |
| TcCLB.510355.260 | hypothetical protein, conserved | -0,4572 | 0,05885 |
| TcCLB.507007.99 | mitochondrial carrier protein (pseudogene), putative | -0,4573 | 0,0002849 |
| TcCLB.509205.90 | surface protease GP63 (pseudogene), putative | -0,4576 | 0,02794 |
| TcCLB.504069.90 | centromere/microtubule binding protein cbf5, putative | -0,4577 | 8,846E-06 |
| TcCLB.508441.80 | Mitochondrial-processing peptidase subunit alpha (MPPA) | -0,4578 | 1,667E-07 |
| TcCLB.511587.33 | hypothetical protein | -0,4578 | 0,07757 |
| TcCLB.508909.270 | hypothetical protein, conserved | -0,4579 | 0,00002113 |
| TcCLB.509287.20 | protein kinase, putative | -0,4579 | 0,0001835 |
| TcCLB.506227.240 | hypothetical protein, conserved | -0,458 | 1,044E-06 |
| TcCLB.506241.90 | hypothetical protein, conserved | -0,458 | 0,0003723 |
| TcCLB.511725.90 | hypothetical protein, conserved | -0,458 | 0,00109 |
| TcCLB.506559.4 | hypothetical protein, conserved (fragment) | -0,458 | 0,001719 |
| TcCLB.509657.20 | dispersed gene family protein 1 (DGF-1), putative | -0,458 | 0,01996 |
| TcCLB.507185.40 | trans-sialidase (pseudogene), putative | -0,458 | 0,145 |
| TcCLB.506517.140 | hypothetical protein, conserved | -0,4581 | 0,000175 |
| TcCLB.508347.140 | hypothetical protein, conserved | -0,4581 | 0,0001764 |
| TcCLB.506515.20 | O-sialoglycoprotein endopeptidase, putative | -0,4581 | 0,0007192 |
| TcCLB.508235.10 | hypothetical protein, conserved (pseudogene) | -0,4582 | 0,0019 |
| TcCLB.511229.20 | oxidoreductase, putative | -0,4582 | 0,01365 |
| TcCLB.509163.6 | hypothetical protein | -0,4583 | 0,4566 |
| TcCLB.511731.30 | serine/threonine protein phosphatase, putative | -0,4589 | 0,0001404 |
| TcCLB.506977.10 | dual specificity protein phosphatase, putative | -0,459 | 0,004617 |
| TcCLB.508903.90 | dispersed gene family protein 1 (DGF-1, pseudogene), puta | -0,4592 | 0,0109 |
| TcCLB.506717.20 | receptor-type adenylate cyclase, putative | -0,4594 | 0,00003136 |
| TcCLB.511847.50 | hypothetical protein, conserved | -0,4597 | 0,00003738 |
| TcCLB.511245.20 | hypothetical protein, conserved | -0,4598 | 0,00836 |
| TcCLB.507023.200 | queuine tRNA-ribosyltransferase, putative | -0,4599 | 0,000163 |
| TcCLB.508783.10 | hypothetical protein, conserved (pseudogene) | -0,4599 | 0,0001992 |
| TcCLB.509967.7 | hypothetical protein, conserved | -0,4599 | 0,06356 |
| TcCLB.503827.100 | trans-sialidase, Group VI, putative | -0,4599 | 0,2824 |
| TcCLB.509789.50 | rab-like GTPase activating protein, putative | -0,46 | 0,0002084 |
| TcCLB.508307.44 | hypothetical protein, conserved | -0,4601 | 0,00005911 |
| TcCLB.509795.70 | hypothetical protein, conserved | -0,4602 | 0,00005237 |
| TcCLB.506739.10 | Eukaryotic translation initiation factor 4 gamma type 1, pu | -0,4603 | 0,00009477 |
| TcCLB.503919.20 | hypothetical protein, conserved | -0,4603 | 0,036 |
| TcCLB.503875.40 | hypothetical protein | -0,4603 | 0,2615 |
| TcCLB.508139.250 | trans-sialidase, Group II, putative | -0,4604 | 0,002363 |
| TcCLB.449759.9 | hypothetical protein, conserved | -0,4604 | 0,0109 |
| TcCLB.511459.70 | hypothetical protein, conserved | -0,4604 | 0,2285 |
| TcCLB.508243.60 | mucin TcMUCII, putative | -0,4605 | 0,3503 |
| TcCLB.508247.80 | mucin-associated surface protein (MASP), putative | -0,4606 | 0,3713 |
| TcCLB.510835.10 | hypothetical protein, conserved | -0,4608 | 0,1436 |
| TcCLB.510089.200 | hypothetical protein, conserved | -0,4609 | 0,00000319 |

| | | | |
|---|---|---|---|
| TcCLB.504109.60 | hypothetical protein, conserved | -0,4609 | 0,00001096 |
| TcCLB.509837.40 | histone H1, putative | -0,4609 | 0,02573 |
| TcCLB.510687.90 | hypothetical protein, conserved | -0,461 | 0,0002905 |
| TcCLB.511667.50 | mucin-associated surface protein (MASP), putative | -0,4611 | 0,08432 |
| TcCLB.508711.40 | mitochondrial RNA binding protein, putative | -0,4614 | 4,636E-06 |
| TcCLB.508871.110 | trans-sialidase, Group V, putative | -0,4614 | 0,1963 |
| TcCLB.511437.20 | expression site-associated gene (ESAG-like) protein, putati | -0,4614 | 0,2126 |
| TcCLB.511727.110 | hypothetical protein, conserved | -0,4615 | 0,0003203 |
| TcCLB.506735.40 | hypothetical protein | -0,4617 | 0,00117 |
| TcCLB.509793.69 | hypothetical protein, conserved | -0,4618 | 6,826E-06 |
| TcCLB.508895.50 | RNA-binding protein, putative | -0,462 | 0,00002458 |
| TcCLB.504105.10 | hypothetical protein, conserved | -0,4621 | 0,002367 |
| TcCLB.508219.10 | mucin-associated surface protein (MASP, pseudogene), pu | -0,4621 | 0,3067 |
| TcCLB.507027.59 | pre-mRNA cleavage complex II Clp1 protein, putative | -0,4622 | 0,001175 |
| TcCLB.511589.209 | hypothetical protein, conserved | -0,4622 | 0,0749 |
| TcCLB.503619.20 | hypothetical protein, conserved | -0,4623 | 0,0004353 |
| TcCLB.507629.39 | hypothetical protein, conserved (fragment) | -0,4624 | 9,993E-08 |
| TcCLB.507993.30 | hypothetical protein, conserved | -0,4624 | 0,000174 |
| TcCLB.504219.70 | mucin TcMUC, putative (fragment) | -0,4624 | 0,1364 |
| TcCLB.508347.10 | hypothetical protein, conserved | -0,4625 | 0,00004312 |
| TcCLB.510955.4 | lipase-like protein, putative | -0,4625 | 0,0003161 |
| TcCLB.511151.60 | hypothetical protein | -0,4626 | 0,0005003 |
| TcCLB.503997.10 | hypothetical protein, conserved | -0,4626 | 0,005004 |
| TcCLB.509695.150 | kinesin, putative | -0,4627 | 6,129E-13 |
| TcCLB.511173.250 | mucin-associated surface protein (MASP, pseudogene), pu | -0,4627 | 0,1129 |
| TcCLB.508525.20 | hypothetical protein, conserved | -0,4631 | 3,592E-07 |
| TcCLB.511907.260 | cell division cycle protein 20, putative (CDC20) | -0,4634 | 8,169E-08 |
| TcCLB.508637.110 | nucleotide-binding protein, putative | -0,4634 | 0,002071 |
| TcCLB.511293.60 | kinesin, putative | -0,4635 | 1,039E-06 |
| TcCLB.506355.160 | hypothetical protein, conserved | -0,4635 | 0,0003849 |
| TcCLB.510979.50 | mucin-associated surface protein (MASP), putative | -0,4635 | 0,03739 |
| TcCLB.506319.50 | hypothetical protein, conserved | -0,4637 | 0,0000852 |
| TcCLB.509795.89 | phosphoprotein lepp12, putative | -0,4637 | 0,001829 |
| TcCLB.508249.10 | dispersed gene family protein 1 (DGF-1), putative | -0,4637 | 0,04166 |
| TcCLB.506529.660 | hypothetical protein, conserved | -0,4638 | 0,01036 |
| TcCLB.509897.20 | trans-sialidase, Group V, putative | -0,4638 | 0,07538 |
| TcCLB.511119.10 | dispersed gene family protein 1 (DGF-1), putative (fragme | -0,4638 | 0,1173 |
| TcCLB.509819.10 | hypothetical protein, conserved | -0,4639 | 0,0001267 |
| TcCLB.509127.40 | hypothetical protein, conserved | -0,4639 | 0,0006142 |
| TcCLB.510891.40 | trans-sialidase (pseudogene), putative | -0,4639 | 0,0052 |
| TcCLB.511865.30 | hypothetical protein, conserved | -0,4639 | 0,005992 |
| TcCLB.510713.30 | trans-sialidase, Group VII, putative | -0,464 | 0,09151 |
| TcCLB.509775.49 | ribosomal protein L2, putative | -0,4641 | 0,0725 |
| TcCLB.508435.40 | mucin-associated surface protein (MASP, pseudogene), pu | -0,4641 | 0,204 |
| TcCLB.510105.140 | hypothetical protein, conserved (pseudogene) | -0,4642 | 0,0007723 |

| | | | |
|---|---|---|---|
| TcCLB.510879.200 | hypothetical protein, conserved | -0,4643 | 0,004174 |
| TcCLB.506337.120 | hypothetical protein, conserved | -0,4643 | 0,007935 |
| TcCLB.508163.70 | dispersed gene family protein 1 (DGF-1), putative | -0,4643 | 0,09944 |
| TcCLB.508355.320 | arginyl-tRNA synthetase, putative | -0,4645 | 1,047E-07 |
| TcCLB.511287.110 | hypothetical protein, conserved | -0,4645 | 1,137E-07 |
| TcCLB.506409.255 | surface protease GP63, putative | -0,4645 | 0,5861 |
| TcCLB.508219.20 | trans-sialidase, Group V, putative | -0,4646 | 0,1763 |
| TcCLB.511491.30 | hypothetical protein, conserved | -0,4648 | 0,0002042 |
| TcCLB.510769.70 | hypothetical protein | -0,4648 | 0,00037 |
| TcCLB.509265.50 | hypothetical protein, conserved (pseudogene) | -0,4648 | 0,002016 |
| TcCLB.509023.30 | hypothetical protein | -0,4649 | 0,01275 |
| TcCLB.511523.90 | hypothetical protein, conserved | -0,465 | 0,0000129 |
| TcCLB.468041.10 | hypothetical protein | -0,465 | 0,0269 |
| TcCLB.507093.75 | B9 domain-containing protein 2 (b9d2) | -0,4653 | 0,007627 |
| TcCLB.511277.20 | DNA-directed RNA polymerase II/III subunit, putative | -0,4654 | 0,001499 |
| TcCLB.510915.21 | surface protease GP63 (pseudogene), putative | -0,4655 | 0,06831 |
| TcCLB.510693.310 | syntaxin binding protein (pseudogene), putative | -0,4655 | 0,2898 |
| TcCLB.508269.73 | ATP-dependent DEAD/H RNA helicase (pseudogene), putat | -0,4656 | 0,03653 |
| TcCLB.507395.40 | hypothetical protein | -0,4656 | 0,509 |
| TcCLB.511713.30 | hypothetical protein, conserved | -0,4657 | 0,002615 |
| TcCLB.509589.30 | Kinesin-13 5, putative (KIN13-5) | -0,4661 | 0,00000082 |
| TcCLB.509105.80 | hypothetical protein, conserved | -0,4661 | 0,001959 |
| TcCLB.506245.140 | trans-sialidase (pseudogene), putative | -0,4661 | 0,5118 |
| TcCLB.508741.100 | hypothetical protein, conserved | -0,4662 | 0,00005502 |
| TcCLB.507023.160 | hypothetical protein, conserved | -0,4663 | 3,257E-09 |
| TcCLB.503809.50 | hypothetical protein, conserved | -0,4663 | 0,009562 |
| TcCLB.507099.40 | hypothetical protein, conserved | -0,4664 | 2,638E-12 |
| TcCLB.506925.130 | eukaryotic initiation factor 5a, putative | -0,4665 | 0,0004509 |
| TcCLB.506213.120 | ATP-dependent RNA helicase, putative | -0,4666 | 0,0005454 |
| TcCLB.511491.140 | hypothetical protein, conserved | -0,4667 | 0,00001766 |
| TcCLB.503527.50 | hypothetical protein, conserved | -0,4668 | 0,003929 |
| TcCLB.507715.90 | hypothetical protein, conserved | -0,4669 | 0,00000141 |
| TcCLB.506553.40 | ATP-dependent RNA helicase (pseudogene), putative | -0,4669 | 0,00003901 |
| TcCLB.508951.60 | hypothetical protein, conserved | -0,4669 | 0,0008403 |
| TcCLB.507445.30 | trans-sialidase (pseudogene), putative | -0,467 | 0,02188 |
| TcCLB.506369.40 | hypothetical protein, conserved | -0,4671 | 2,723E-06 |
| TcCLB.509395.120 | histone deacetylase, putative | -0,4671 | 7,196E-06 |
| TcCLB.509671.120 | rRNA processing protein, putative | -0,4672 | 0,0006942 |
| TcCLB.511367.100 | hypothetical protein, conserved | -0,4672 | 0,001366 |
| TcCLB.507035.130 | hypothetical protein, conserved | -0,4673 | 8,872E-06 |
| TcCLB.504179.80 | hypothetical protein, conserved | -0,4674 | 0,00004301 |
| TcCLB.509561.100 | hypothetical protein, conserved | -0,4675 | 0,00001006 |
| TcCLB.506613.150 | trans-sialidase (pseudogene), putative | -0,4675 | 0,1851 |
| TcCLB.508127.5 | hypothetical protein, conserved | -0,4676 | 0,04225 |
| TcCLB.509179.110 | hypothetical protein, conserved | -0,4677 | 0,00005875 |

| | | | |
|---|---|---|---|
| TcCLB.508177.40 | hypothetical protein, conserved | -0,4678 | 2,621E-07 |
| TcCLB.508693.20 | DNA topoisomerase IB, large subunit | -0,4681 | 4,435E-06 |
| TcCLB.510119.50 | hypothetical protein, conserved | -0,4683 | 0,0001747 |
| TcCLB.504867.40 | acetyltransferase, putative | -0,4684 | 0,001368 |
| TcCLB.509735.60 | trans-sialidase (pseudogene), putative | -0,4684 | 0,02789 |
| TcCLB.511153.140 | proteasome beta 3 subunit, putative | -0,4685 | 0,001524 |
| TcCLB.511289.100 | hypothetical protein, conserved | -0,4686 | 2,366E-07 |
| TcCLB.511429.20 | cyclophilin 15, putative | -0,4686 | 0,0007467 |
| TcCLB.507081.139 | hypothetical protein, conserved (fragment) | -0,4687 | 0,00001952 |
| TcCLB.504163.70 | hypothetical protein, conserved | -0,4687 | 0,000349 |
| TcCLB.509957.10 | hypothetical protein | -0,4688 | 0,1432 |
| TcCLB.507071.120 | mucin-associated surface protein (MASP), putative | -0,4688 | 0,1554 |
| TcCLB.509259.100 | ATP-dependent DEAD/H RNA helicase (pseudogene), putat | -0,469 | 0,1978 |
| TcCLB.510131.50 | DNA polymerase kappa, putative | -0,4691 | 0,001494 |
| TcCLB.508719.50 | hypothetical protein, conserved | -0,4691 | 0,006895 |
| TcCLB.510337.40 | spermidine synthase, putative | -0,4692 | 2,935E-07 |
| TcCLB.511669.20 | hypothetical protein | -0,4692 | 0,1535 |
| TcCLB.510841.20 | hypothetical protein, conserved | -0,4694 | 0,008305 |
| TcCLB.510123.60 | hypothetical protein, conserved | -0,4696 | 0,0003959 |
| TcCLB.506135.20 | hypothetical protein, conserved | -0,4698 | 3,478E-07 |
| TcCLB.511625.60 | hypothetical protein, conserved | -0,4698 | 0,01141 |
| TcCLB.511517.90 | hypothetical protein, conserved | -0,4699 | 0,00008357 |
| TcCLB.504075.3 | calmodulin, putative | -0,4699 | 0,2164 |
| TcCLB.511511.60 | phospholipid-translocating P-type ATPase (flippase), putati | -0,47 | 0,0003818 |
| TcCLB.509657.70 | trans-sialidase (pseudogene), putative | -0,4701 | 0,3227 |
| TcCLB.511511.20 | tRNA modification enzyme, putative | -0,4702 | 0,007696 |
| TcCLB.510431.330 | hypothetical protein, conserved | -0,4702 | 0,008668 |
| TcCLB.504425.50 | hypothetical protein, conserved | -0,4702 | 0,04831 |
| TcCLB.507923.30 | vacuolar assembly protein vps41, putative | -0,4703 | 4,112E-06 |
| TcCLB.510879.100 | hypothetical protein | -0,4703 | 0,001539 |
| TcCLB.506053.20 | hypothetical protein, conserved | -0,4703 | 0,002667 |
| TcCLB.509569.120 | RNA-binding protein, putative | -0,4705 | 9,779E-06 |
| TcCLB.509895.10 | mitochondrial RNA binding complex 1 subunit, putative | -0,4706 | 1,209E-13 |
| TcCLB.509003.79 | coatomer beta subunit, putative (fragment) | -0,4707 | 0,000165 |
| TcCLB.510119.30 | hypothetical protein, conserved | -0,4707 | 0,002982 |
| TcCLB.503973.64 | hypothetical protein, conserved (pseudogene) | -0,4707 | 0,3275 |
| TcCLB.511907.50 | lysophospholipase, putative | -0,4708 | 8,258E-06 |
| TcCLB.509767.44 | hypothetical protein, conserved | -0,4708 | 0,0002133 |
| TcCLB.506211.170 | Multiprotein-bridging factor 1, putative (mbf1) | -0,4711 | 0,00001221 |
| TcCLB.503945.10 | eukaryotic translation initiation factor 1A, putative | -0,4711 | 0,0007189 |
| TcCLB.509069.20 | ATP binding protein-like protein, putative | -0,4712 | 0,001515 |
| TcCLB.509647.100 | hypothetical protein, conserved | -0,4713 | 0,0006515 |
| TcCLB.507723.110 | acetyltransferase, putative | -0,4713 | 0,0008558 |
| TcCLB.506369.30 | hypothetical protein, conserved | -0,4713 | 0,005794 |
| TcCLB.506699.10 | hypothetical protein, conserved | -0,4713 | 0,005808 |

| | | | |
|---|---|---|---|
| TcCLB.506915.30 | hypothetical protein | -0,4714 | 1,997E-06 |
| TcCLB.510279.10 | trans-sialidase (pseudogene), putative | -0,4715 | 0,2199 |
| TcCLB.508547.80 | hypothetical protein, conserved | -0,4716 | 2,297E-06 |
| TcCLB.506679.20 | hypothetical protein, conserved | -0,4717 | 0,08663 |
| TcCLB.510199.20 | elongation factor 1-gamma (EF-1-gamma, pseudogene), p | -0,4717 | 0,3212 |
| TcCLB.509939.10 | hypothetical protein, conserved | -0,4718 | 0,0000031 |
| TcCLB.511761.80 | hypothetical protein | -0,4718 | 0,0001544 |
| TcCLB.504433.14 | ADP-ribosylation factor-like protein 3A, putative | -0,4718 | 0,0007429 |
| TcCLB.505939.60 | ribonuclease H1, putative | -0,4718 | 0,003345 |
| TcCLB.509707.25 | hypothetical protein, conserved | -0,4718 | 0,02719 |
| TcCLB.507547.50 | hypothetical protein, conserved | -0,4719 | 0,0001914 |
| TcCLB.508277.230 | nucleolar RNA binding protein, putative | -0,472 | 1,769E-07 |
| TcCLB.505931.50 | histone H3, putative | -0,472 | 0,09911 |
| TcCLB.511541.40 | hypothetical protein, conserved | -0,4721 | 0,00002036 |
| TcCLB.509997.30 | tryparedoxin 1, putative (TXN1) | -0,4721 | 0,0001765 |
| TcCLB.511275.30 | actin-related protein 2/3 complex subunit 1, putative (ARP | -0,4721 | 0,001223 |
| TcCLB.428999.10 | hypothetical protein | -0,4721 | 0,002661 |
| TcCLB.503829.30 | hypothetical protein, conserved | -0,4721 | 0,005721 |
| TcCLB.511577.83 | thioredoxin, putative | -0,4721 | 0,0201 |
| TcCLB.508479.490 | hypothetical protein | -0,4721 | 0,05284 |
| TcCLB.510275.350 | trans-sialidase, Group V, putative | -0,4721 | 0,1444 |
| TcCLB.511551.10 | trans-sialidase, Group II, putative | -0,4721 | 0,2396 |
| TcCLB.504935.20 | hypothetical protein, conserved | -0,4722 | 0,0008595 |
| TcCLB.503749.10 | hypothetical protein, conserved | -0,4723 | 0,0002836 |
| TcCLB.507823.39 | hypothetical protein, conserved | -0,4723 | 0,02067 |
| TcCLB.510363.260 | mucin-associated surface protein (MASP), putative | -0,4723 | 0,3837 |
| TcCLB.507019.50 | hypothetical protein, conserved | -0,4724 | 0,001415 |
| TcCLB.509693.150 | hypothetical protein, conserved | -0,4727 | 0,0001296 |
| TcCLB.506559.440 | hypothetical protein, conserved | -0,4729 | 1,985E-09 |
| TcCLB.511903.30 | ubiquitin-conjugating enzyme E2, putative | -0,4729 | 0,00004863 |
| TcCLB.508169.120 | hypothetical protein, conserved | -0,4729 | 0,001116 |
| TcCLB.507093.30 | hypothetical protein, conserved | -0,473 | 0,00003351 |
| TcCLB.511111.40 | hypothetical protein, conserved | -0,473 | 0,0005218 |
| TcCLB.511727.50 | hypothetical protein, conserved | -0,4731 | 5,782E-09 |
| TcCLB.508625.10 | elongation factor 2, putative | -0,4731 | 8,04E-07 |
| TcCLB.508979.15 | mucin-associated surface protein (MASP, pseudogene), pu | -0,4731 | 0,4642 |
| TcCLB.509587.10 | mitochondrial DNA polymerase I protein D, putative | -0,4732 | 2,094E-11 |
| TcCLB.510729.70 | proteasome alpha 1 subunit, putative | -0,4732 | 6,344E-07 |
| TcCLB.509525.330 | hypothetical protein | -0,4732 | 0,3351 |
| TcCLB.504033.220 | palmitoyl acyltransferase 1, putative | -0,4733 | 1,347E-06 |
| TcCLB.507103.10 | DnaJ chaperone protein, putative | -0,4733 | 0,0001839 |
| TcCLB.509229.50 | hypothetical protein | -0,4733 | 0,04077 |
| TcCLB.508837.130 | hypothetical protein, conserved | -0,4735 | 0,00007033 |
| TcCLB.435519.19 | hypothetical protein, conserved (fragment) | -0,4735 | 0,0612 |
| TcCLB.509321.10 | hypothetical protein, conserved | -0,4736 | 3,922E-06 |

| | | | |
|---|---|---|---|
| TcCLB.509965.30 | small nuclear ribonucleoprotein Sm-E, putative | -0,4736 | 0,03027 |
| TcCLB.510483.360 | trans-sialidase, Group VII, putative | -0,4736 | 0,07956 |
| TcCLB.506181.120 | hypothetical protein, conserved | -0,4737 | 0,03686 |
| TcCLB.511229.80 | eukaryotic translation initiation factor 3 subunit 2, putative | -0,4738 | 1,141E-06 |
| TcCLB.504427.200 | poly(A)-specific ribonuclease PARN, putative | -0,4739 | 8,387E-06 |
| TcCLB.510975.30 | hypothetical protein, conserved | -0,474 | 0,0000648 |
| TcCLB.511865.60 | hypothetical protein, conserved | -0,4741 | 4,272E-06 |
| TcCLB.507297.30 | metacaspase, putative | -0,4741 | 0,0003189 |
| TcCLB.503473.20 | hypothetical protein, conserved | -0,4741 | 0,001708 |
| TcCLB.511245.100 | hypothetical protein, conserved | -0,4742 | 0,0001261 |
| TcCLB.507005.70 | Pre-rRNA-processing protein PNO1, putative (PNO1) | -0,4742 | 0,0004599 |
| TcCLB.505999.100 | 3-phosphoglycerate kinase, glycosomal (PGKA) | -0,4743 | 0,00001975 |
| TcCLB.506973.30 | mucin-associated surface protein (MASP), putative | -0,4744 | 0,1903 |
| TcCLB.510629.430 | prefoldin subunit, putative | -0,4745 | 0,00975 |
| TcCLB.506227.120 | hypothetical protein, conserved | -0,4747 | 0,0002258 |
| TcCLB.504157.90 | hypothetical protein, conserved | -0,4749 | 0,0007905 |
| TcCLB.507993.330 | mevalonate-diphosphate decarboxylase, putative | -0,475 | 0,00006094 |
| TcCLB.503973.280 | mucin-associated surface protein (MASP), putative | -0,475 | 0,2007 |
| TcCLB.504071.70 | hypothetical protein, conserved | -0,4751 | 1,357E-08 |
| TcCLB.503425.29 | DNA replication licensing factor MCM8, putative | -0,4751 | 0,001708 |
| TcCLB.508777.130 | hypothetical protein, conserved | -0,4752 | 0,00000461 |
| TcCLB.509145.10 | hypothetical protein, conserved | -0,4752 | 0,00006957 |
| TcCLB.506727.100 | enoyl-CoA hydratase/isomerase family protein, putative | -0,4753 | 0,00004093 |
| TcCLB.511075.54 | hypothetical protein, conserved | -0,4753 | 0,002112 |
| TcCLB.510661.230 | hypothetical protein, conserved | -0,4754 | 7,048E-07 |
| TcCLB.507357.140 | hypothetical protein | -0,4754 | 0,3341 |
| TcCLB.510725.90 | hypothetical protein, conserved | -0,4755 | 0,007583 |
| TcCLB.509885.60 | methyltransferase, putative | -0,4756 | 0,004444 |
| TcCLB.427789.39 | hypothetical protein, conserved | -0,4756 | 0,0097 |
| TcCLB.508221.120 | hypothetical protein | -0,4757 | 0,2317 |
| TcCLB.506883.30 | ribosome-interacting GTPase 2, putative | -0,4759 | 0,00001225 |
| TcCLB.504147.330 | hypothetical protein, conserved | -0,476 | 1,097E-11 |
| TcCLB.510687.94 | Unc104-like kinesin, putative | -0,476 | 0,0001382 |
| TcCLB.508389.174 | hypothetical protein, conserved (pseudogene) | -0,4761 | 0,081 |
| TcCLB.507009.120 | DNA repair protein RAD2, putative | -0,4762 | 0,0002256 |
| TcCLB.510719.50 | hypothetical protein, conserved | -0,4762 | 0,04474 |
| TcCLB.511151.11 | hypothetical protein (fragment) | -0,4763 | 1,951E-07 |
| TcCLB.511001.140 | hypothetical protein, conserved | -0,4763 | 0,0001372 |
| TcCLB.510913.10 | dispersed gene family protein 1 (DGF-1), putative (fragme | -0,4763 | 0,008084 |
| TcCLB.510359.530 | trans-sialidase, Group V, putative | -0,4763 | 0,06633 |
| TcCLB.508989.110 | hypothetical protein, conserved | -0,4765 | 0,00001192 |
| TcCLB.508051.40 | hypothetical protein, conserved | -0,4767 | 0,003281 |
| TcCLB.506275.20 | ATP-dependent protease subunit HslV, putative (HslV) | -0,4772 | 0,0001865 |
| TcCLB.509671.160 | DNA polymerase epsilon subunit B, putative | -0,4774 | 0,000239 |
| TcCLB.509319.30 | hypothetical protein, conserved | -0,4775 | 0,00009038 |

| | | | |
|---|---|---|---|
| TcCLB.507047.60 | hypothetical protein, conserved | -0,4775 | 0,001031 |
| TcCLB.506799.10 | fatty acid transporter protein-like, putative | -0,4776 | 2,602E-06 |
| TcCLB.504839.44 | tryparedoxin peroxidase | -0,4777 | 1,306E-10 |
| TcCLB.506791.60 | hypothetical protein, conserved | -0,4777 | 3,544E-06 |
| TcCLB.506765.80 | mucin TcMUCII, putative | -0,4778 | 0,4905 |
| TcCLB.507003.24 | hypothetical protein, conserved (fragment) | -0,4779 | 2,935E-09 |
| TcCLB.506525.104 | AP-4 complex subunit sigma-1, putative | -0,4779 | 0,01216 |
| TcCLB.509715.70 | folylpolyglutamate synthase, putative (FPGS) | -0,478 | 0,0008739 |
| TcCLB.506579.70 | hypothetical protein, conserved | -0,4782 | 5,704E-10 |
| TcCLB.506221.90 | hypothetical protein, conserved | -0,4783 | 5,784E-06 |
| TcCLB.509551.10 | protein kinase, putative | -0,4783 | 0,0004822 |
| TcCLB.508781.90 | hypothetical protein, conserved | -0,4783 | 0,009355 |
| TcCLB.506819.10 | hypothetical protein, conserved | -0,4783 | 0,02925 |
| TcCLB.504105.150 | importin beta-1 subunit, putative | -0,4784 | 1,044E-11 |
| TcCLB.504247.34 | Transcription elongation factor 1 domain-containing protei | -0,4785 | 0,001724 |
| TcCLB.506943.69 | hypothetical protein, conserved | -0,4786 | 0,0003383 |
| TcCLB.511867.100 | hypothetical protein, conserved | -0,4788 | 0,0001562 |
| TcCLB.505171.60 | hypothetical protein, conserved | -0,4789 | 2,173E-06 |
| TcCLB.507019.60 | hypothetical protein, conserved | -0,4791 | 3,389E-06 |
| TcCLB.507019.45 | hypothetical protein | -0,4791 | 0,001048 |
| TcCLB.510143.110 | hypothetical protein, conserved | -0,4793 | 0,000363 |
| TcCLB.508365.114 | mucin-associated surface protein (MASP, pseudogene), pu | -0,4793 | 0,08761 |
| TcCLB.506445.100 | E2-like ubiquitin-conjugation enzyme, putative (UFC1) | -0,4794 | 1,225E-07 |
| TcCLB.506905.10 | hypothetical protein, conserved | -0,4794 | 0,005763 |
| TcCLB.511127.390 | hypothetical protein, conserved | -0,4795 | 1,188E-10 |
| TcCLB.506625.50 | hypothetical protein | -0,4796 | 0,187 |
| TcCLB.509967.100 | hypothetical protein, conserved (pseudogene) | -0,4797 | 0,006036 |
| TcCLB.508943.40 | hypothetical protein, conserved | -0,4797 | 0,01913 |
| TcCLB.504109.170 | hypothetical protein, conserved | -0,4801 | 3,371E-07 |
| TcCLB.510007.40 | hypothetical protein, conserved | -0,4801 | 0,0002778 |
| TcCLB.509525.390 | mucin-associated surface protein (MASP), putative | -0,4803 | 0,1075 |
| TcCLB.503583.24 | hypothetical protein, conserved | -0,4804 | 9,17E-09 |
| TcCLB.509693.90 | serine/threonine protein kinase, putative | -0,4804 | 0,00001388 |
| TcCLB.470399.10 | cyclophilin, putative (fragment) | -0,4804 | 0,001715 |
| TcCLB.511189.90 | hypothetical protein, conserved | -0,4805 | 3,624E-06 |
| TcCLB.506567.134 | hypothetical protein, conserved | -0,4805 | 0,0001608 |
| TcCLB.504131.140 | hypothetical protein, conserved | -0,4806 | 1,282E-09 |
| TcCLB.507207.10 | hypothetical protein, conserved | -0,4807 | 0,00002115 |
| TcCLB.510145.20 | cofilin/actin depolymerizing factor, putative | -0,4809 | 2,833E-07 |
| TcCLB.504069.10 | proteasome alpha 2 subunit, putative | -0,4809 | 0,00007602 |
| TcCLB.457339.10 | U6 snRNA-associated Sm-like protein LSm4, putative | -0,481 | 0,01284 |
| TcCLB.414517.9 | hypothetical protein, conserved | -0,481 | 0,01774 |
| TcCLB.511899.50 | Trypanosome basal body component protein (fragment) (T | -0,4811 | 2,244E-06 |
| TcCLB.508409.190 | hypothetical protein, conserved | -0,4812 | 0,01638 |
| TcCLB.509393.40 | hypothetical protein | -0,4813 | 1,009E-06 |

| | | | |
|---|---|---|---|
| TcCLB.503435.30 | hypothetical protein, conserved | -0,4814 | 1,068E-06 |
| TcCLB.509745.70 | hypothetical protein, conserved | -0,4818 | 0,00002803 |
| TcCLB.506127.50 | hypothetical protein, conserved | -0,4818 | 0,0001883 |
| TcCLB.508233.10 | hypothetical protein, conserved | -0,4818 | 0,003565 |
| TcCLB.510593.49 | kinetoplast DNA-associated protein, putative | -0,4819 | 1,039E-06 |
| TcCLB.503775.30 | syntaxin binding protein (pseudogene), putative | -0,4819 | 0,0116 |
| TcCLB.506297.300 | eukaryotic release factor 3, putative | -0,482 | 1,456E-09 |
| TcCLB.508475.20 | hypothetical protein | -0,482 | 0,00000129 |
| TcCLB.506863.10 | alanyl-tRNA synthetase, putative | -0,4821 | 4,535E-12 |
| TcCLB.507007.60 | hypothetical protein, conserved | -0,4821 | 0,003327 |
| TcCLB.510551.15 | hypothetical protein | -0,4821 | 0,1588 |
| TcCLB.508175.170 | hypothetical protein, conserved | -0,4823 | 1,991E-06 |
| TcCLB.510693.251 | mucin TcMUC (pseudogene), putative | -0,4823 | 0,2312 |
| TcCLB.511173.40 | mucin TcMUCII (pseudogene), putative | -0,4825 | 0,2805 |
| TcCLB.510181.60 | chromatin assembly factor 1 subunit B, putative | -0,4828 | 7,235E-06 |
| TcCLB.507093.270 | guanine nucleotide releasing protein, putative | -0,4828 | 0,0002636 |
| TcCLB.507091.20 | hypothetical protein, conserved | -0,4829 | 0,00009144 |
| TcCLB.507619.10 | hypothetical protein, conserved (fragment) | -0,4829 | 0,000113 |
| TcCLB.507093.150 | hypothetical protein, conserved (pseudogene) | -0,4831 | 2,383E-06 |
| TcCLB.509287.110 | mucin-associated surface protein (MASP), putative | -0,4831 | 0,03333 |
| TcCLB.508153.730 | eukaryotic initiation factor 2a, putative | -0,4833 | 4,205E-10 |
| TcCLB.507933.10 | plectin, putative | -0,4833 | 9,65E-10 |
| TcCLB.511159.20 | origin recognition complex subunit 1 (ORC1), putative | -0,4835 | 0,005951 |
| TcCLB.508777.100 | cyclin 1 (CYC1) | -0,4838 | 0,0003965 |
| TcCLB.508207.130 | hypothetical protein, conserved | -0,484 | 0,004726 |
| TcCLB.510877.100 | hypothetical protein, conserved | -0,4841 | 0,00003833 |
| TcCLB.508281.9 | dispersed gene family protein 1 (DGF-1), putative | -0,4841 | 0,2249 |
| TcCLB.508401.70 | ATP-dependent DEAD/H RNA helicase (pseudogene), putat | -0,4844 | 0,03222 |
| TcCLB.510275.255 | hypothetical protein | -0,4844 | 0,4812 |
| TcCLB.508741.380 | hypothetical protein, conserved | -0,4845 | 1,732E-10 |
| TcCLB.509007.70 | hypothetical protein, conserved | -0,4845 | 1,343E-06 |
| TcCLB.508881.70 | DNA ligase, putative | -0,4846 | 2,444E-07 |
| TcCLB.509719.60 | hypothetical protein | -0,4846 | 0,01933 |
| TcCLB.510029.80 | hypothetical protein, conserved | -0,4847 | 3,258E-06 |
| TcCLB.508909.320 | hypothetical protein | -0,4847 | 0,00001817 |
| TcCLB.508307.190 | hypothetical protein, conserved | -0,4848 | 0,003799 |
| TcCLB.511559.10 | Present in the outer mitochondrial membrane proteome 1 | -0,4848 | 0,3772 |
| TcCLB.506495.30 | retrotransposon hot spot protein (RHS, pseudogene), putat | -0,4849 | 0,01556 |
| TcCLB.509683.110 | nucleobase transporter, putative | -0,485 | 0,0004212 |
| TcCLB.506815.40 | mitochondrial RNA binding complex 1 subunit, putative | -0,485 | 0,004708 |
| TcCLB.508277.380 | hypothetical protein, conserved | -0,4851 | 5,795E-10 |
| TcCLB.509443.48 | hypothetical protein, conserved | -0,4851 | 0,02025 |
| TcCLB.506225.30 | hypothetical protein, conserved (pseudogene) | -0,4852 | 0,1052 |
| TcCLB.505025.90 | mucin-associated surface protein (MASP), putative | -0,4852 | 0,3075 |
| TcCLB.507093.210 | myosin heavy chain, putative | -0,4853 | 1,907E-09 |

| | | | |
|---|---|---|---|
| TcCLB.508075.7 | hypothetical protein, conserved | -0,4853 | 0,01193 |
| TcCLB.506367.10 | isovaleryl-coA dehydrogenase, putative (fragment) | -0,4855 | 0,04044 |
| TcCLB.511233.140 | trans-sialidase, Group V, putative | -0,4855 | 0,4774 |
| TcCLB.511741.20 | hypothetical protein, conserved | -0,4859 | 0,0001031 |
| TcCLB.510071.30 | UDP-Gal or UDP-GlcNAc-dependent glycosyltransferase, pu | -0,4859 | 0,07053 |
| TcCLB.506499.110 | mucin TcMUCII (pseudogene), putative | -0,4859 | 0,3015 |
| TcCLB.509039.10 | hypothetical protein, conserved | -0,4861 | 2,914E-08 |
| TcCLB.510943.23 | hypothetical protein, conserved | -0,4862 | 0,0002525 |
| TcCLB.511545.190 | GPI transamidase component Tta1, putative (TTA1) | -0,4862 | 0,002293 |
| TcCLB.503891.70 | ribonuclease HII, putative | -0,4863 | 0,0002707 |
| TcCLB.509287.5 | hypothetical protein, conserved (pseudogene) | -0,4863 | 0,0616 |
| TcCLB.506501.280 | mucin TcMUCII, putative | -0,4863 | 0,4969 |
| TcCLB.511483.10 | hypothetical protein | -0,4864 | 0,00009154 |
| TcCLB.510693.320 | mucin-associated surface protein (MASP), putative | -0,4864 | 0,1222 |
| TcCLB.508461.370 | hypothetical protein, conserved | -0,4865 | 0,00002463 |
| TcCLB.506887.10 | trans-sialidase, putative | -0,4865 | 0,2887 |
| TcCLB.505983.40 | hypothetical protein, conserved | -0,4866 | 7,294E-10 |
| TcCLB.506327.10 | proteasome subunit alpha type-1, putative (fragment) (PS/ | -0,4868 | 0,001469 |
| TcCLB.507739.80 | hypothetical protein, conserved | -0,4869 | 0,0000302 |
| TcCLB.507029.50 | hypothetical protein, conserved | -0,4869 | 0,04449 |
| TcCLB.511577.150 | hypothetical protein, conserved | -0,487 | 0,00009742 |
| TcCLB.507031.70 | hypothetical protein, conserved | -0,487 | 0,0009438 |
| TcCLB.511907.190 | hypothetical protein | -0,4871 | 2,921E-06 |
| TcCLB.508505.40 | hypothetical protein, conserved | -0,4871 | 0,0000932 |
| TcCLB.508707.140 | phosphatidate cytidylyltransferase-like protein, putative | -0,4872 | 0,0005376 |
| TcCLB.507357.10 | surface protease GP63, putative | -0,4872 | 0,21 |
| TcCLB.506679.50 | pre-mRNA splicing factor ATP-dependent RNA helicase, pu | -0,4873 | 2,43E-08 |
| TcCLB.504507.5 | trypanothione reductase, putative (fragment) | -0,4873 | 0,0001484 |
| TcCLB.507943.50 | hypothetical protein, conserved | -0,4873 | 0,009491 |
| TcCLB.507519.167 | hypothetical protein | -0,4873 | 0,037 |
| TcCLB.508461.340 | hypothetical protein, conserved | -0,4874 | 1,042E-08 |
| TcCLB.511773.40 | hypothetical protein, conserved | -0,4874 | 0,1282 |
| TcCLB.507951.15 | hypothetical protein, conserved (fragment) | -0,4876 | 1,41E-07 |
| TcCLB.510121.140 | hypothetical protein, conserved | -0,4876 | 0,00003878 |
| TcCLB.510053.79 | FG-GAP repeat protein, putative | -0,4876 | 0,02059 |
| TcCLB.506127.90 | hypothetical protein, conserved | -0,4877 | 3,938E-11 |
| TcCLB.506533.34 | phosphatidylinositol kinase related protein, putative | -0,4877 | 6,192E-10 |
| TcCLB.508479.340 | succinyl-CoA synthetase alpha subunit, putative | -0,4879 | 7,749E-12 |
| TcCLB.510255.20 | hypothetical protein, conserved | -0,4885 | 0,001826 |
| TcCLB.419417.19 | hypothetical protein, conserved | -0,4885 | 0,02082 |
| TcCLB.509399.130 | hypothetical protein, conserved | -0,4886 | 0,0001255 |
| TcCLB.508559.20 | trans-sialidase (pseudogene), putative | -0,4886 | 0,0002304 |
| TcCLB.507389.60 | chaperone DNAJ protein, putative | -0,4886 | 0,01469 |
| TcCLB.511911.102 | hypothetical protein, conserved | -0,4887 | 0,001679 |
| TcCLB.507929.30 | hypothetical protein, conserved | -0,4887 | 0,01148 |

| | | | |
|---|---|---|---|
| TcCLB.508307.80 | mitochondrial RNA binding complex 1 subunit, putative | -0,4888 | 3,309E-06 |
| TcCLB.506681.60 | hypothetical protein, conserved | -0,4888 | 3,465E-06 |
| TcCLB.508541.200 | hypothetical protein, conserved | -0,4888 | 0,00736 |
| TcCLB.503493.20 | hypothetical protein, conserved | -0,4889 | 0,007788 |
| TcCLB.511903.20 | hypothetical protein, conserved | -0,4891 | 0,01243 |
| TcCLB.511753.9 | hypothetical protein, conserved | -0,4893 | 0,01046 |
| TcCLB.510833.40 | trans-sialidase (pseudogene), putative | -0,4893 | 0,07231 |
| TcCLB.508993.20 | homocysteine S-methyltransferase, putative | -0,4894 | 0,0005704 |
| TcCLB.510303.90 | hypothetical protein, conserved | -0,4895 | 0,0007701 |
| TcCLB.511501.30 | ABC transporter, putative | -0,4897 | 0,001284 |
| TcCLB.507017.50 | polynucleotide kinase 3'-phosphatase, putative | -0,4898 | 0,0003626 |
| TcCLB.511589.50 | cyclophilin-like protein, putative | -0,4898 | 0,000462 |
| TcCLB.508325.101 | hypothetical protein, conserved | -0,49 | 0,2217 |
| TcCLB.504131.10 | hypothetical protein, conserved (pseudogene) | -0,4902 | 0,00005285 |
| TcCLB.503693.11 | hypothetical protein, conserved (fragment) | -0,4902 | 0,07837 |
| TcCLB.506811.50 | hypothetical protein, conserved | -0,4903 | 0,2118 |
| TcCLB.508601.130 | Elongation factor G 2, mitochondrial, putative (EF-G2) | -0,4905 | 1,588E-06 |
| TcCLB.511269.70 | cysteine peptidase, Clan CA, family C2, putative | -0,4905 | 0,003451 |
| TcCLB.508479.86 | ATP-dependent DEAD/H RNA helicase (pseudogene), putat | -0,4905 | 0,09211 |
| TcCLB.510257.50 | hypothetical protein, conserved | -0,4906 | 0,003681 |
| TcCLB.508815.4 | RNA polymerase I, putative | -0,4907 | 0,08445 |
| TcCLB.504153.290 | hypothetical protein, conserved | -0,4908 | 0,00003694 |
| TcCLB.506469.139 | hypothetical protein, conserved | -0,4908 | 0,03358 |
| TcCLB.508593.50 | hypothetical protein, conserved | -0,491 | 0,00001838 |
| TcCLB.511269.60 | Vacuolar protein sorting-associated protein 16 homolog, p | -0,491 | 0,0001699 |
| TcCLB.506367.120 | hypothetical protein, conserved | -0,4911 | 1,539E-07 |
| TcCLB.506459.215 | hypothetical protein, conserved (pseudogene) | -0,4911 | 0,2672 |
| TcCLB.511675.30 | retrotransposon hot spot protein (RHS, pseudogene), putat | -0,4912 | 0,3321 |
| TcCLB.510421.264 | hypothetical protein, conserved | -0,4913 | 0,02175 |
| TcCLB.508401.90 | N-acetyltransferase complex ARD1 subunit (pseudogene), | -0,4913 | 0,05114 |
| TcCLB.508153.940 | hypothetical protein, conserved (pseudogene) | -0,4914 | 1,731E-09 |
| TcCLB.506501.110 | mucin-associated surface protein (MASP), putative | -0,4916 | 0,2567 |
| TcCLB.508231.70 | hypothetical protein, conserved | -0,4918 | 0,0007659 |
| TcCLB.509215.16 | GlcNAc-PI synthesis protein (GPI3) | -0,4918 | 0,00253 |
| TcCLB.503835.5 | rac serine threonine protein kinase, putative | -0,4918 | 0,1372 |
| TcCLB.511289.119 | C-terminal motor kinesin, putative (fragment) | -0,4919 | 0,0008737 |
| TcCLB.503897.100 | hypothetical protein, conserved | -0,4921 | 0,00001031 |
| TcCLB.507959.110 | mucin-associated surface protein (MASP), putative | -0,4922 | 0,2761 |
| TcCLB.511237.20 | hypothetical protein, conserved | -0,4923 | 0,0001069 |
| TcCLB.511725.110 | hypothetical protein, conserved | -0,4923 | 0,006544 |
| TcCLB.511797.220 | dispersed gene family protein 1 (DGF-1), putative | -0,4924 | 0,09642 |
| TcCLB.507639.50 | tRNA pseudouridine synthase A, putative | -0,4928 | 0,0004174 |
| TcCLB.508405.30 | Suppressor of G2 allele of SKP1, putative | -0,4928 | 0,001334 |
| TcCLB.509557.10 | hypothetical protein | -0,4929 | 0,0007501 |
| TcCLB.510143.10 | hypothetical protein, conserved | -0,4929 | 0,003097 |

| | | | |
|---|---|---|---|
| TcCLB.511589.230 | eukaryotic translation initiation factor 3 subunit 2, putative | -0,493 | 7,192E-07 |
| TcCLB.503965.10 | hypothetical protein, conserved | -0,493 | 0,001467 |
| TcCLB.503557.20 | ankyrin repeat protein, putative | -0,493 | 0,0185 |
| TcCLB.510799.80 | hypothetical protein, conserved | -0,4932 | 0,00004953 |
| TcCLB.510187.230 | formin, putative | -0,4932 | 0,00005498 |
| TcCLB.508951.10 | hypothetical protein, conserved | -0,4932 | 0,00006359 |
| TcCLB.508357.20 | hypothetical protein, conserved | -0,4932 | 0,01894 |
| TcCLB.421173.14 | trans-sialidase, putative | -0,4932 | 0,409 |
| TcCLB.506457.30 | hypoxanthine-guanine phosphoribosyltransferase, putative | -0,4933 | 0,01571 |
| TcCLB.511353.69 | oxidoreductase-protein, putative | -0,4933 | 0,02815 |
| TcCLB.510719.70 | bromodomain-containing protein 3 (BDF3) | -0,4934 | 0,00006487 |
| TcCLB.511245.90 | hypothetical protein, conserved | -0,4934 | 0,0008462 |
| TcCLB.507895.140 | hypothetical protein | -0,4935 | 2,974E-06 |
| TcCLB.508215.10 | hypothetical protein, conserved | -0,4935 | 0,0001856 |
| TcCLB.507739.90 | hypothetical protein, conserved | -0,4937 | 0,0005774 |
| TcCLB.508693.170 | hypothetical protein, conserved | -0,4937 | 0,001368 |
| TcCLB.511857.30 | glycosyl transferase-like protein, putative | -0,4938 | 0,004394 |
| TcCLB.510317.34 | unspecified product | -0,4939 | 0,09491 |
| TcCLB.508299.50 | hypothetical protein, conserved | -0,4941 | 3,284E-10 |
| TcCLB.511293.90 | hypothetical protein, conserved | -0,4941 | 0,0002083 |
| TcCLB.510609.34 | hypothetical protein, conserved | -0,4941 | 0,0003967 |
| TcCLB.507775.30 | hypothetical protein, conserved | -0,4941 | 0,0006515 |
| TcCLB.511283.230 | hypothetical protein, conserved | -0,4942 | 9,789E-10 |
| TcCLB.503703.70 | Mitochondrial SSU ribosomal protein, putative | -0,4942 | 3,044E-07 |
| TcCLB.510667.60 | peptidyl-prolyl cis-trans isomerase (cyclophilin), putative | -0,4942 | 0,001403 |
| TcCLB.511183.230 | trans-sialidase, Group V, putative | -0,4942 | 0,2175 |
| TcCLB.504215.40 | actin-related protein 2/3 complex subunit 1, putative (ARP | -0,4943 | 0,0003404 |
| TcCLB.436761.10 | hypothetical protein, conserved | -0,4945 | 0,008627 |
| TcCLB.510187.10 | hypothetical protein, conserved | -0,4945 | 0,1057 |
| TcCLB.504985.10 | hypothetical protein | -0,4945 | 0,1844 |
| TcCLB.510339.70 | hypothetical protein, conserved | -0,4946 | 0,00002073 |
| TcCLB.508739.50 | hypothetical protein | -0,4947 | 0,001287 |
| TcCLB.511633.20 | hypothetical protein, conserved | -0,4948 | 0,0001814 |
| TcCLB.505171.100 | hypothetical protein, conserved | -0,4948 | 0,005591 |
| TcCLB.508583.10 | hypothetical protein, conserved | -0,4949 | 0,00379 |
| TcCLB.505977.40 | inositol/phosphatidylinositol phosphatase (pseudogene), p | -0,495 | 4,886E-10 |
| TcCLB.510609.20 | hypothetical protein, conserved | -0,4953 | 2,801E-06 |
| TcCLB.503425.20 | monothiol glutaredoxin, putative | -0,4954 | 0,00009671 |
| TcCLB.509429.59 | hypothetical protein, conserved | -0,4955 | 1,107E-06 |
| TcCLB.506435.230 | hypothetical protein, conserved (pseudogene) | -0,4955 | 0,001884 |
| TcCLB.511043.10 | hypothetical protein, conserved | -0,4956 | 2,119E-08 |
| TcCLB.509081.80 | hypothetical protein, conserved (pseudogene) | -0,4956 | 0,2543 |
| TcCLB.511277.483 | hypothetical protein, conserved | -0,4957 | 0,003388 |
| TcCLB.506177.20 | lectin, putative | -0,4959 | 0,00000729 |
| TcCLB.508805.10 | pre-mRNA splicing factor ATP-dependent RNA helicase (ps | -0,4959 | 0,000102 |

| | | | |
|---|---|---|---|
| TcCLB.509643.50 | hypothetical protein, conserved | -0,4959 | 0,0001044 |
| TcCLB.506167.40 | proteasome subunit alpha type-1, putative (PSA1) | -0,4959 | 0,0001436 |
| TcCLB.511173.230 | trans-sialidase (pseudogene), putative | -0,4959 | 0,1094 |
| TcCLB.507017.120 | ubiquitin hydrolase, putative | -0,496 | 3,347E-09 |
| TcCLB.511217.80 | hypothetical protein, conserved | -0,496 | 0,06326 |
| TcCLB.506925.190 | hypothetical protein, conserved | -0,4962 | 0,0002914 |
| TcCLB.506679.10 | hypothetical protein, conserved | -0,4962 | 0,01717 |
| TcCLB.506845.21 | N-acetyltransferase complex ARD1 subunit (pseudogene), | -0,4962 | 0,4052 |
| TcCLB.508441.40 | hypothetical protein, conserved | -0,4963 | 7,143E-06 |
| TcCLB.504021.30 | hypothetical protein, conserved | -0,4965 | 1,509E-07 |
| TcCLB.508613.20 | hypothetical protein, conserved | -0,4967 | 0,0006638 |
| TcCLB.506349.83 | retrotransposon hot spot protein (RHS, pseudogene), putat | -0,4968 | 0,007583 |
| TcCLB.511563.30 | retrotransposon hot spot protein (RHS, pseudogene), putat | -0,4969 | 0,3382 |
| TcCLB.510833.10 | trans-sialidase, putative | -0,4973 | 0,2901 |
| TcCLB.507993.140 | RNA-binding protein 29, putative (RBP29) | -0,4975 | 8,964E-07 |
| TcCLB.508039.50 | dispersed gene family protein 1 (DGF-1, pseudogene), puta | -0,4976 | 0,02563 |
| TcCLB.508277.90 | hypothetical protein, conserved | -0,4977 | 0,0006 |
| TcCLB.507011.100 | hypothetical protein, conserved | -0,4978 | 0,0001541 |
| TcCLB.507993.350 | surface protease GP63, putative | -0,4978 | 0,04268 |
| TcCLB.508247.140 | hypothetical protein | -0,4979 | 0,1202 |
| TcCLB.509671.50 | hypothetical protein, conserved | -0,4981 | 0,00334 |
| TcCLB.511287.160 | hypothetical protein, conserved | -0,4983 | 2,448E-06 |
| TcCLB.508181.80 | hypothetical protein, conserved | -0,4983 | 8,449E-06 |
| TcCLB.510025.205 | hypothetical protein, conserved (pseudogene) | -0,4984 | 0,2458 |
| TcCLB.505943.40 | hypothetical protein, conserved | -0,4985 | 6,013E-08 |
| TcCLB.511445.130 | Mitochondrial SSU ribosomal protein, putative | -0,4985 | 4,479E-07 |
| TcCLB.503797.7 | hypothetical protein, conserved (pseudogene) | -0,4985 | 0,0001435 |
| TcCLB.504071.80 | nuclear cap binding protein, putative | -0,4985 | 0,01162 |
| TcCLB.511295.70 | hypothetical protein, conserved (fragment) | -0,4985 | 0,2398 |
| TcCLB.508307.150 | hypothetical protein, conserved | -0,4986 | 0,00004408 |
| TcCLB.509615.60 | hypothetical protein | -0,4988 | 0,3732 |
| TcCLB.511727.90 | vacuolar proton-ATPase-like protein, putative | -0,499 | 5,352E-10 |
| TcCLB.507957.29 | surface protease GP63 (pseudogene), putative | -0,499 | 0,4583 |
| TcCLB.506375.100 | hypothetical protein | -0,4992 | 0,0007465 |
| TcCLB.510261.31 | mucin-associated surface protein (MASP, pseudogene), pu | -0,4992 | 0,1821 |
| TcCLB.507033.10 | hypothetical protein, conserved | -0,4997 | 4,164E-09 |
| TcCLB.506539.21 | hypothetical protein, conserved (pseudogene) | -0,4997 | 0,1003 |
| TcCLB.504033.90 | mitochondrial carrier protein, putative | -0,4998 | 2,91E-07 |
| TcCLB.510667.14 | membrane transporter protein, putative | -0,4998 | 0,0001273 |
| TcCLB.506613.130 | dispersed gene family protein 1 (DGF-1), putative | -0,4998 | 0,03511 |
| TcCLB.511511.70 | hypothetical protein, conserved | -0,4999 | 5,893E-08 |
| TcCLB.508951.40 | acyl-CoA dehydrogenase, putative | -0,4999 | 0,002297 |
| TcCLB.507757.50 | serine/threonine-protein phosphatase PP1, putative | -0,5002 | 2,946E-06 |
| TcCLB.511921.130 | hypothetical protein, conserved (pseudogene) | -0,5003 | 0,0897 |
| TcCLB.507951.270 | LETM1 and EF-hand domain-containing protein 1, putative | -0,5008 | 4,538E-07 |

| | | | |
|---|---|---|---|
| TcCLB.511355.14 | cation transporter, putative | -0,5009 | 0,00000821 |
| TcCLB.507093.190 | hypothetical protein, conserved | -0,5012 | 0,003345 |
| TcCLB.506737.100 | hypothetical protein, conserved | -0,5013 | 0,0002064 |
| TcCLB.506793.80 | ATP-dependent RNA helicase, putative | -0,5015 | 0,00002174 |
| TcCLB.508397.40 | succinate dehydrogenase subunit | -0,5016 | 0,0001227 |
| TcCLB.410961.10 | phosphatidylinositol kinase related protein, putative | -0,5017 | 0,1096 |
| TcCLB.510543.210 | retrotransposon hot spot protein (RHS, pseudogene), putat | -0,5018 | 0,07376 |
| TcCLB.509525.40 | hypothetical protein, conserved | -0,5018 | 0,5198 |
| TcCLB.505967.19 | dispersed gene family protein 1 (DGF-1), putative (fragme | -0,502 | 0,09389 |
| TcCLB.510575.10 | hypothetical protein | -0,5024 | 0,0008626 |
| TcCLB.510075.50 | hypothetical protein, conserved | -0,5025 | 3,732E-07 |
| TcCLB.509525.10 | mucin-associated surface protein (MASP, pseudogene), pu | -0,5026 | 0,2629 |
| TcCLB.504787.54 | mucin-associated surface protein (MASP, pseudogene), pu | -0,5027 | 0,03333 |
| TcCLB.511127.140 | DNA replication licensing factor MCM4, putative | -0,5029 | 0,00001699 |
| TcCLB.510361.110 | mucin-associated surface protein (MASP), putative | -0,5029 | 0,1525 |
| TcCLB.506733.100 | hypothetical protein, conserved | -0,503 | 2,347E-06 |
| TcCLB.507857.80 | hypothetical protein, conserved (pseudogene) | -0,503 | 0,00003398 |
| TcCLB.511751.166 | hypothetical protein, conserved | -0,503 | 0,0001214 |
| TcCLB.509011.90 | lipase, putative | -0,5031 | 0,004488 |
| TcCLB.506933.107 | hypothetical protein, conserved | -0,5035 | 0,0007293 |
| TcCLB.507509.60 | hypothetical protein, conserved | -0,5037 | 0,00001994 |
| TcCLB.504239.171 | mucin TcMUCII, putative | -0,5038 | 0,2911 |
| TcCLB.508573.120 | dispersed gene family protein 1 (DGF-1), putative | -0,504 | 0,09943 |
| TcCLB.510553.40 | trans-sialidase, Group V, putative | -0,5042 | 0,09393 |
| TcCLB.504827.21 | hypothetical protein, conserved (fragment) | -0,5043 | 3,968E-08 |
| TcCLB.509729.10 | hypothetical protein, conserved | -0,5043 | 5,707E-06 |
| TcCLB.508255.10 | trans-sialidase (pseudogene), putative | -0,5043 | 0,005556 |
| TcCLB.510013.59 | trans-sialidase (pseudogene), putative | -0,5043 | 0,3045 |
| TcCLB.443683.9 | DNA-directed rna polymerase I largest subunit, putative (fr | -0,5044 | 0,001908 |
| TcCLB.503583.20 | hypothetical protein, conserved | -0,5045 | 0,0007751 |
| TcCLB.507711.80 | hypothetical protein, conserved | -0,5047 | 0,0004412 |
| TcCLB.505317.9 | trans-sialidase, putative (fragment) | -0,5047 | 0,003351 |
| TcCLB.508153.190 | kinesin, putative | -0,5051 | 3,452E-06 |
| TcCLB.508637.69 | hypothetical protein, conserved | -0,5051 | 0,001377 |
| TcCLB.507975.80 | hypothetical protein, conserved | -0,5052 | 3,322E-09 |
| TcCLB.507895.164 | hypothetical protein, conserved | -0,5052 | 0,0001164 |
| TcCLB.510691.80 | hypothetical protein, conserved | -0,5054 | 0,003238 |
| TcCLB.509247.30 | hypothetical protein, conserved | -0,5055 | 1,956E-11 |
| TcCLB.504103.50 | hypothetical protein, conserved | -0,5056 | 0,00007054 |
| TcCLB.511435.49 | phosphatidylinositol 3-kinase, putative | -0,5057 | 0,07527 |
| TcCLB.511525.11 | hypothetical protein, conserved (fragment) | -0,5057 | 0,1117 |
| TcCLB.511751.180 | pseudouridine synthase A-like protein, putative | -0,5058 | 0,00007406 |
| TcCLB.508405.100 | hypothetical protein, conserved | -0,5058 | 0,001329 |
| TcCLB.506605.50 | hypothetical protein, conserved | -0,5059 | 3,639E-06 |
| TcCLB.419043.10 | hypothetical protein, conserved | -0,5059 | 0,005886 |

| | | | |
|---|---|---|---|
| TcCLB.509609.19 | DNA repair and recombination helicase protein PIF7, putat | -0,5059 | 0,007621 |
| TcCLB.508771.55 | hypothetical protein, conserved (pseudogene) | -0,506 | 0,002898 |
| TcCLB.507907.50 | hypothetical protein, conserved | -0,5063 | 1,196E-12 |
| TcCLB.508261.70 | mucin-associated surface protein (MASP, pseudogene), pu | -0,5063 | 0,1492 |
| TcCLB.508479.260 | hypothetical protein, conserved | -0,5064 | 0,00002149 |
| TcCLB.511589.120 | DNA topoisomerase III, putative | -0,5064 | 0,00008019 |
| TcCLB.511435.10 | WD40 repeat-containing protein | -0,5065 | 1,429E-06 |
| TcCLB.510901.150 | proteasome regulatory non-ATPase subunit 2, putative | -0,5066 | 5,262E-11 |
| TcCLB.510575.50 | hypothetical protein, conserved | -0,5066 | 0,04252 |
| TcCLB.506265.180 | hypothetical protein, conserved | -0,507 | 0,01082 |
| TcCLB.508909.60 | hypothetical protein, conserved | -0,5071 | 0,0006118 |
| TcCLB.503973.20 | trans-sialidase (pseudogene), putative | -0,5071 | 0,03913 |
| TcCLB.507625.70 | hypothetical protein, conserved | -0,5072 | 1,716E-06 |
| TcCLB.469435.10 | hypothetical protein, conserved | -0,5072 | 0,00252 |
| TcCLB.503777.4 | hypothetical protein, conserved | -0,5072 | 0,08401 |
| TcCLB.505809.10 | dispersed gene family protein 1 (DGF-1, pseudogene), puta | -0,5073 | 0,4243 |
| TcCLB.511439.100 | p-nitrophenylphosphatase, putative | -0,5075 | 0,00267 |
| TcCLB.506739.209 | hypothetical protein, conserved | -0,5076 | 0,001061 |
| TcCLB.504247.70 | AATF protein, putative (pseudogene) | -0,5077 | 0,0003792 |
| TcCLB.505999.180 | Non cytoplasmic domain containing protein, conserved | -0,5078 | 0,00006584 |
| TcCLB.511499.59 | small nuclear ribonucleoprotein Sm-E, putative | -0,5078 | 0,003167 |
| TcCLB.503981.80 | hypothetical protein, conserved | -0,508 | 5,481E-11 |
| TcCLB.507041.132 | hypothetical protein (fragment) | -0,508 | 0,04071 |
| TcCLB.511063.40 | trans-sialidase, Group V, putative | -0,5081 | 0,2276 |
| TcCLB.504055.81 | ubiquitin-protein ligase, putative | -0,5083 | 0,0002311 |
| TcCLB.508355.10 | hypothetical protein, conserved | -0,5085 | 0,0003065 |
| TcCLB.508231.120 | trans-sialidase (pseudogene), putative | -0,5085 | 0,3404 |
| TcCLB.508357.80 | cation transporter protein, putative | -0,5086 | 0,00005923 |
| TcCLB.506459.270 | hypothetical protein, conserved | -0,5088 | 2,297E-09 |
| TcCLB.511181.110 | hypothetical protein, conserved | -0,5089 | 2,498E-07 |
| TcCLB.511727.160 | hypothetical protein, conserved | -0,5091 | 1,493E-11 |
| TcCLB.407477.60 | hypothetical protein, conserved | -0,5091 | 0,0006833 |
| TcCLB.506479.10 | hypothetical protein, conserved | -0,5092 | 0,0001138 |
| TcCLB.511415.40 | dispersed gene family protein 1 (DGF-1), putative | -0,5092 | 0,0008318 |
| TcCLB.507023.120 | aldehyde dehydrogenase family, putative | -0,5096 | 0,0001688 |
| TcCLB.433677.19 | hypothetical protein, conserved | -0,5097 | 0,04183 |
| TcCLB.510029.90 | hypothetical protein, conserved | -0,5098 | 8,152E-06 |
| TcCLB.509167.90 | hypothetical protein, conserved | -0,5098 | 0,0001829 |
| TcCLB.507509.30 | RNA polymerase-like protein, putative | -0,5098 | 0,003079 |
| TcCLB.508873.40 | mucin TcMUCII (pseudogene), putative | -0,5098 | 0,1543 |
| TcCLB.510295.30 | RNA editing complex protein MP18, putative | -0,5103 | 0,00003248 |
| TcCLB.508181.130 | peter pan protein, putative | -0,5103 | 0,00004751 |
| TcCLB.511289.30 | metallo-peptidase, Clan MG, Family M24 | -0,5104 | 2,197E-10 |
| TcCLB.510329.110 | hypothetical protein | -0,5105 | 6,752E-07 |
| TcCLB.508741.210 | hypothetical protein | -0,5106 | 6,432E-12 |

| | | | |
|---|---|---|---|
| TcCLB.506739.200 | protein phosphatase 2C, putative | -0,5108 | 3,801E-11 |
| TcCLB.418113.9 | dispersed gene family protein 1 (DGF-1), putative | -0,5109 | 0,3693 |
| TcCLB.503833.10 | hypothetical protein, conserved | -0,511 | 9,159E-07 |
| TcCLB.507017.170 | hypothetical protein, conserved | -0,5111 | 0,01047 |
| TcCLB.506405.90 | hypothetical protein, conserved | -0,5114 | 0,001215 |
| TcCLB.507027.30 | hypothetical protein, conserved (pseudogene) | -0,5116 | 2,975E-09 |
| TcCLB.511021.51 | hypothetical protein, conserved | -0,5117 | 0,004545 |
| TcCLB.510087.100 | GINS complex subunit Psf3, putative | -0,5118 | 0,03439 |
| TcCLB.508479.200 | palmitoyl acyltransferase 10, putative | -0,512 | 4,742E-07 |
| TcCLB.510275.190 | mucin-associated surface protein (MASP), putative | -0,512 | 0,246 |
| TcCLB.508153.920 | choline/ethanolamine phosphotransferase, putative | -0,5121 | 4,831E-06 |
| TcCLB.508593.110 | arginine N-methyltransferase, putative | -0,5121 | 0,0000475 |
| TcCLB.510089.39 | CAAX amino terminal protease, putative | -0,5122 | 0,00482 |
| TcCLB.503655.40 | surface protease GP63 (pseudogene), putative | -0,5122 | 0,05763 |
| TcCLB.509553.30 | ribonuclease mar1, putative | -0,5123 | 0,03447 |
| TcCLB.506493.80 | DNA topoisomerase IA, putative | -0,5124 | 1,146E-09 |
| TcCLB.511867.170 | hypothetical protein, conserved | -0,5124 | 2,116E-06 |
| TcCLB.506733.60 | N(2), N(2)-dimethylguanosine tRNA methyltransferase, pu | -0,5124 | 0,00001215 |
| TcCLB.511303.130 | hypothetical protein | -0,5125 | 0,00008073 |
| TcCLB.504067.10 | dispersed gene family protein 1 (DGF-1), putative | -0,5125 | 0,1742 |
| TcCLB.510243.128 | hypothetical protein, conserved | -0,5127 | 0,03563 |
| TcCLB.509229.30 | hypothetical protein, conserved | -0,5129 | 0,02244 |
| TcCLB.511589.60 | hypothetical protein, conserved | -0,513 | 0,01684 |
| TcCLB.509745.60 | telomerase reverse transcriptase, putative (TERT) | -0,5133 | 7,852E-06 |
| TcCLB.506413.80 | cyclophilin, putative | -0,5134 | 0,0005608 |
| TcCLB.511911.71 | hypothetical protein, conserved | -0,5135 | 0,2388 |
| TcCLB.511153.124 | hypothetical protein, conserved | -0,5136 | 0,005442 |
| TcCLB.506671.30 | mucin-associated surface protein (MASP, pseudogene), pu | -0,5136 | 0,02107 |
| TcCLB.511801.20 | hypothetical protein, conserved | -0,5137 | 0,0003805 |
| TcCLB.506247.200 | adaptin complex 1 subunit, putative | -0,5138 | 5,769E-11 |
| TcCLB.507037.10 | proteasome regulatory non-ATPase subunit 7 (pseudogene | -0,5142 | 0,00003295 |
| TcCLB.509611.140 | hypothetical protein, conserved | -0,5143 | 0,00004436 |
| TcCLB.511127.30 | hypothetical protein, conserved | -0,5144 | 0,0006634 |
| TcCLB.507005.40 | zinc finger protein, putative | -0,5144 | 0,0006651 |
| TcCLB.506711.4 | peptide chain release factor 1, putative (fragment) | -0,5144 | 0,01664 |
| TcCLB.505555.61 | cytoplasmic l-asparaginase i-like protein, putative | -0,5144 | 0,02262 |
| TcCLB.510745.30 | hypothetical protein, conserved | -0,5149 | 0,03787 |
| TcCLB.511733.20 | hypothetical protein, conserved | -0,5152 | 6,579E-06 |
| TcCLB.506231.40 | retrotransposon hot spot protein (RHS, pseudogene), putat | -0,5152 | 0,002534 |
| TcCLB.508489.10 | hypothetical protein, conserved | -0,5155 | 0,001151 |
| TcCLB.510593.40 | kinetoplastid kinetochore protein 8, putative (kkt8) | -0,5156 | 0,00004495 |
| TcCLB.507155.9 | dispersed gene family protein 1 (DGF-1, pseudogene), puta | -0,5156 | 0,02677 |
| TcCLB.506247.260 | mitochondrial carrier protein, putative | -0,5157 | 0,008993 |
| TcCLB.504239.161 | mucin-associated surface protein (MASP, pseudogene), pu | -0,5159 | 0,2055 |
| TcCLB.509073.10 | hypothetical protein, conserved | -0,5161 | 9,061E-06 |

| | | | |
|---|---|---|---|
| TcCLB.509875.110 | hypothetical protein | -0,5161 | 0,2894 |
| TcCLB.509525.260 | hypothetical protein | -0,5163 | 0,03909 |
| TcCLB.506321.330 | hypothetical protein, conserved | -0,5164 | 0,00001266 |
| TcCLB.511565.15 | hypothetical protein | -0,5164 | 0,2287 |
| TcCLB.509895.80 | ATP-dependent zinc metallopeptidase (pseudogene), putat | -0,5165 | 3,546E-07 |
| TcCLB.510823.20 | conserved protein, unknown function | -0,5166 | 3,544E-06 |
| TcCLB.511517.80 | replication factor C, subunit 2, putative | -0,5168 | 0,0000419 |
| TcCLB.510817.10 | protein kinase, putative | -0,5168 | 0,09933 |
| TcCLB.507997.60 | hypothetical protein, conserved | -0,5169 | 4,225E-06 |
| TcCLB.506739.40 | hypothetical protein, conserved | -0,5169 | 0,00003535 |
| TcCLB.507895.110 | hypothetical protein, conserved | -0,5169 | 0,0001598 |
| TcCLB.509125.10 | mannosyltransferase-like protein, putative | -0,517 | 2,916E-07 |
| TcCLB.511039.3 | choline/ethanolamine phosphotransferase, putative (CEPT | -0,5171 | 0,07835 |
| TcCLB.506925.280 | hypothetical protein, conserved | -0,5172 | 0,0002341 |
| TcCLB.510761.60 | casein kinase II, putative | -0,5176 | 7,742E-07 |
| TcCLB.508387.70 | hypothetical protein, conserved | -0,5178 | 1,492E-10 |
| TcCLB.509979.249 | syntaxin binding protein 1 (pseudogene), putative | -0,5178 | 0,003244 |
| TcCLB.508461.174 | hypothetical protein, conserved | -0,5178 | 0,03746 |
| TcCLB.511577.110 | exopolyphosphatase | -0,5181 | 0,002647 |
| TcCLB.509023.20 | hypothetical protein, conserved | -0,5181 | 0,007693 |
| TcCLB.507603.50 | Mitochondrial Ribosomal Protein of Small Subunit 29, puta | -0,5183 | 0,00002242 |
| TcCLB.508933.5 | serine/threonine protein phosphatase PP1 (fragment) | -0,5184 | 0,01805 |
| TcCLB.511133.4 | DNA repair and recombination helicase protein PIF6, putat | -0,5185 | 0,0004012 |
| TcCLB.507959.323 | syntaxin binding protein (pseudogene), putative | -0,5187 | 0,1131 |
| TcCLB.510187.240 | hypothetical protein, conserved | -0,5188 | 1,528E-09 |
| TcCLB.510237.95 | hypothetical protein, conserved (pseudogene) | -0,5188 | 0,2308 |
| TcCLB.503835.30 | hypothetical protein, conserved | -0,5189 | 0,00003249 |
| TcCLB.510687.50 | hypothetical protein, conserved | -0,519 | 1,761E-06 |
| TcCLB.506135.90 | hypothetical protein, conserved | -0,519 | 0,0001211 |
| TcCLB.508707.227 | ATP binding protein-like protein, putative | -0,5192 | 0,001968 |
| TcCLB.511001.80 | hypothetical protein, conserved | -0,5193 | 2,95E-07 |
| TcCLB.508931.50 | hypothetical protein | -0,5193 | 0,00002485 |
| TcCLB.511001.160 | hypothetical protein, conserved | -0,5194 | 4,328E-09 |
| TcCLB.506529.500 | hypothetical protein, conserved | -0,5194 | 0,00003365 |
| TcCLB.504005.29 | hypothetical protein, conserved | -0,5194 | 0,001969 |
| TcCLB.511131.98 | hypothetical protein, conserved | -0,5194 | 0,003902 |
| TcCLB.509473.10 | hypothetical protein (pseudogene) | -0,5195 | 8,723E-06 |
| TcCLB.509995.29 | hypothetical protein, conserved | -0,5196 | 0,003926 |
| TcCLB.508355.240 | hypothetical protein, conserved | -0,5196 | 0,003941 |
| TcCLB.506363.150 | soluble N-ethylmaleimide sensitive factor (NSF) attachme | -0,5197 | 0,0009557 |
| TcCLB.508839.89 | DNA-directed rna polymerase I largest subunit, putative (fr | -0,5198 | 0,0001099 |
| TcCLB.507089.40 | CMP-sialic acid transporter, putative | -0,5199 | 0,0003982 |
| TcCLB.506479.120 | cytidine triphosphate synthase, putative | -0,5199 | 0,005091 |
| TcCLB.506595.21 | hypothetical protein | -0,5201 | 0,002865 |
| TcCLB.506961.104 | ATP-dependent DEAD/H RNA helicase (pseudogene), putat | -0,5201 | 0,02027 |

| | | | |
|---|---|---|---|
| TcCLB.511313.40 | ATP-dependent RNA helicase, putative | -0,5202 | 6,344E-08 |
| TcCLB.509791.150 | choline/ethanolamine phosphotransferase, putative (CEPT | -0,5202 | 2,493E-07 |
| TcCLB.510031.9 | U5 snrnp-specific protein, putative | -0,5202 | 0,0000438 |
| TcCLB.507967.30 | hypothetical protein, conserved (fragment) | -0,5202 | 0,0004347 |
| TcCLB.507093.72 | hypothetical protein, conserved | -0,5203 | 4,431E-08 |
| TcCLB.506959.74 | hypothetical protein, conserved | -0,5204 | 4,501E-06 |
| TcCLB.510337.50 | hypothetical protein, conserved (pseudogene) | -0,5204 | 0,005072 |
| TcCLB.506691.22 | hypothetical protein, conserved | -0,5206 | 1,936E-06 |
| TcCLB.424123.40 | RNA-binding protein, putative | -0,5208 | 0,001083 |
| TcCLB.506499.30 | trans-sialidase (pseudogene), putative | -0,5209 | 0,188 |
| TcCLB.504105.20 | eukaryotic translation initiation factor 5, putative | -0,521 | 2,172E-09 |
| TcCLB.509287.30 | hypothetical protein, conserved | -0,5212 | 5,658E-09 |
| TcCLB.511903.280 | hypothetical protein, conserved | -0,5212 | 0,0001773 |
| TcCLB.508355.420 | cation transporter protein, putative | -0,5213 | 0,00002177 |
| TcCLB.506241.200 | hypothetical protein, conserved | -0,5214 | 1,601E-07 |
| TcCLB.503813.20 | hypothetical protein, conserved | -0,5214 | 5,493E-06 |
| TcCLB.509005.20 | hypothetical protein, conserved | -0,5214 | 0,0000916 |
| TcCLB.503671.39 | dynamin-1-like protein (fragment) | -0,5215 | 0,0000243 |
| TcCLB.508827.60 | hypothetical protein, conserved | -0,5217 | 1,935E-06 |
| TcCLB.511511.120 | hypothetical protein, conserved | -0,5217 | 0,00003521 |
| TcCLB.504011.60 | hypothetical protein, conserved | -0,5219 | 0,0000417 |
| TcCLB.506871.30 | ADP-ribosylation factor GTPase activating protein 1, putati | -0,522 | 8,722E-06 |
| TcCLB.508153.720 | hypothetical protein, conserved | -0,5221 | 0,01194 |
| TcCLB.511179.160 | ubiquitin-protein ligase, putative | -0,5223 | 0,0005192 |
| TcCLB.508741.90 | hypothetical protein | -0,5227 | 7,747E-06 |
| TcCLB.508999.49 | hypothetical protein | -0,5227 | 0,2302 |
| TcCLB.506297.60 | hypothetical protein, conserved | -0,5228 | 0,00004893 |
| TcCLB.509589.40 | hypothetical protein, conserved | -0,523 | 1,313E-11 |
| TcCLB.508479.180 | hypothetical protein, conserved | -0,5232 | 8,119E-11 |
| TcCLB.504229.50 | trans-sialidase (pseudogene), putative | -0,5234 | 0,02025 |
| TcCLB.508707.50 | hypothetical protein, conserved | -0,5237 | 0,00001321 |
| TcCLB.510025.139 | hypothetical protein, conserved (pseudogene) | -0,5237 | 0,4126 |
| TcCLB.507939.50 | hypothetical protein, conserved | -0,5238 | 9,093E-06 |
| TcCLB.506563.30 | coatomer beta subunit, putative | -0,5239 | 2,053E-14 |
| TcCLB.508153.340 | ATP-dependent 6-phosphofructokinase, glycosomal (PFK) | -0,5241 | 8,087E-06 |
| TcCLB.508953.40 | hypothetical protein, conserved | -0,5241 | 0,000195 |
| TcCLB.510515.50 | hypothetical protein, conserved | -0,5241 | 0,0002246 |
| TcCLB.506871.80 | RNA polymerase-like protein, putative | -0,5243 | 0,001903 |
| TcCLB.510535.109 | trans-sialidase, putative | -0,5243 | 0,009547 |
| TcCLB.508165.170 | mucin-associated surface protein (MASP), putative | -0,5243 | 0,1584 |
| TcCLB.511479.20 | hypothetical protein, conserved | -0,5244 | 8,565E-06 |
| TcCLB.507709.10 | hypothetical protein, conserved | -0,5249 | 0,007245 |
| TcCLB.510535.10 | hypothetical protein, conserved | -0,5251 | 0,00002374 |
| TcCLB.509941.140 | hypothetical protein, conserved | -0,5254 | 0,001496 |
| TcCLB.509127.104 | ARP2/3 complex subunit, putative | -0,5254 | 0,01452 |

| | | | |
|---|---|---|---|
| TcCLB.508815.10 | hypothetical protein, conserved | -0,5256 | 0,0003736 |
| TcCLB.506925.420 | hypothetical protein, conserved | -0,5256 | 0,0008271 |
| TcCLB.508741.270 | Kinetoplastid RNA editing protein 1, putative (KREPA1) | -0,5257 | 3,494E-08 |
| TcCLB.511715.40 | hypothetical protein, conserved | -0,5257 | 0,01845 |
| TcCLB.511851.10 | dispersed gene family protein 1 (DGF-1), putative | -0,5258 | 0,0005057 |
| TcCLB.509287.240 | dispersed gene family protein 1 (DGF-1), putative | -0,5258 | 0,0007593 |
| TcCLB.506957.70 | hypothetical protein, conserved | -0,5262 | 7,917E-06 |
| TcCLB.506629.140 | hypothetical protein, conserved | -0,5264 | 0,00000287 |
| TcCLB.509005.70 | hypothetical protein, conserved | -0,5264 | 0,0001344 |
| TcCLB.511355.30 | phosphatidic acid phosphatase protein, putative | -0,5266 | 0,00001568 |
| TcCLB.507711.70 | hypothetical protein, conserved | -0,5268 | 0,0005974 |
| TcCLB.510429.40 | Present in the outer mitochondrial membrane proteome 5 | -0,5269 | 0,0001043 |
| TcCLB.508909.150 | hypothetical protein, conserved | -0,5269 | 0,006443 |
| TcCLB.507957.320 | mucin-associated surface protein (MASP), putative | -0,5271 | 0,1157 |
| TcCLB.507641.190 | kinetoplastid kinetochore protein 1, putative (kkt1) | -0,5272 | 9,599E-08 |
| TcCLB.509577.9 | hypothetical protein, conserved (fragment) | -0,5272 | 0,0008592 |
| TcCLB.506925.40 | RNA polymerase I second largest subunit, putative | -0,5273 | 3,81E-13 |
| TcCLB.460061.10 | hypothetical protein (fragment) | -0,5274 | 0,04123 |
| TcCLB.509621.10 | hypothetical protein, conserved | -0,5275 | 8,625E-06 |
| TcCLB.508897.30 | ABC transport system ATP-binding protein, putative | -0,5276 | 1,276E-12 |
| TcCLB.507625.180 | kinesin, putative | -0,5277 | 7,331E-07 |
| TcCLB.505037.20 | ribosomal RNA processing protein 41B, putative (RRP41B) | -0,5277 | 6,139E-06 |
| TcCLB.506619.30 | Flagellar Member 8 (FLAM8) | -0,5277 | 0,2462 |
| TcCLB.509429.80 | hypothetical protein, conserved | -0,5278 | 0,00002097 |
| TcCLB.510901.80 | hypothetical protein, conserved | -0,528 | 1,384E-08 |
| TcCLB.503713.10 | hypothetical protein, conserved | -0,528 | 0,002052 |
| TcCLB.506369.70 | histone H1, putative | -0,5282 | 1,078E-17 |
| TcCLB.510755.120 | RNA-binding protein, putative | -0,5282 | 1,162E-07 |
| TcCLB.506829.71 | hypothetical protein, conserved (pseudogene) | -0,5282 | 0,03621 |
| TcCLB.510901.160 | hypothetical protein, conserved | -0,5284 | 0,002884 |
| TcCLB.506405.60 | hypothetical protein, conserved | -0,5285 | 0,0009815 |
| TcCLB.504215.59 | hypothetical protein, conserved | -0,5286 | 1,771E-06 |
| TcCLB.507515.120 | sphingosine kinase A, B, putative | -0,5287 | 5,578E-06 |
| TcCLB.510827.10 | hypothetical protein, conserved | -0,5287 | 0,003373 |
| TcCLB.503867.10 | hypothetical protein, conserved | -0,5288 | 0,0001642 |
| TcCLB.507711.50 | hypothetical protein, conserved | -0,5288 | 0,001422 |
| TcCLB.509575.20 | hypothetical protein, conserved | -0,529 | 0,0005658 |
| TcCLB.503999.80 | hypothetical protein, conserved | -0,5291 | 9,104E-10 |
| TcCLB.506235.20 | guide RNA associated protein 1, putative (GAP1) | -0,5292 | 3,534E-11 |
| TcCLB.507763.30 | anion-transporting ATPase-like, putative | -0,5292 | 4,14E-07 |
| TcCLB.506587.50 | hypothetical protein, conserved | -0,5292 | 5,239E-06 |
| TcCLB.503785.10 | hypothetical protein, conserved (pseudogene) | -0,5292 | 0,00002923 |
| TcCLB.509203.60 | histone acetyltransferase, putative | -0,5292 | 0,0000551 |
| TcCLB.506367.174 | hypothetical protein, conserved | -0,5292 | 0,008116 |
| TcCLB.511321.4 | O-sialoglycoprotein endopeptidase, putative | -0,5293 | 0,07391 |

| | | | |
|---|---|---|---|
| TcCLB.511867.160 | hypothetical protein, conserved | -0,5294 | 0,004495 |
| TcCLB.509237.50 | hypothetical protein, conserved | -0,5295 | 1,826E-08 |
| TcCLB.503839.30 | hypothetical protein, conserved | -0,5296 | 3,984E-07 |
| TcCLB.508405.10 | cell division cycle protein, putative | -0,5296 | 9,545E-06 |
| TcCLB.503867.20 | trans-sialidase (pseudogene), putative | -0,5297 | 0,1255 |
| TcCLB.511385.14 | hypothetical protein, conserved (fragment) | -0,5298 | 3,016E-07 |
| TcCLB.511307.20 | Peptidase C19, ubiquitin carboxyl-terminal hydrolase, puta | -0,5299 | 7,505E-06 |
| TcCLB.506561.20 | retrotransposon hot spot protein (RHS, pseudogene), putat | -0,53 | 0,001273 |
| TcCLB.505965.70 | helicase (pseudogene), putative | -0,5302 | 1,431E-06 |
| TcCLB.508543.160 | hypothetical protein, conserved | -0,5303 | 3,152E-07 |
| TcCLB.508479.300 | hypothetical protein, conserved | -0,5303 | 0,00005199 |
| TcCLB.510121.120 | hypothetical protein, conserved | -0,5305 | 0,0001244 |
| TcCLB.508479.50 | trans-sialidase (pseudogene), putative | -0,5307 | 0,0004492 |
| TcCLB.504741.180 | hypothetical protein, conserved | -0,5309 | 0,0001116 |
| TcCLB.506337.220 | hypothetical protein | -0,5309 | 0,0008389 |
| TcCLB.510311.70 | 26S protease regulatory subunit, putative | -0,5312 | 8,772E-08 |
| TcCLB.509625.10 | ATP-dependent RNA helicase, putative (fragment) | -0,5314 | 3,913E-06 |
| TcCLB.506493.40 | hypothetical protein, conserved | -0,5314 | 0,00001626 |
| TcCLB.508137.20 | hypothetical protein, conserved | -0,5316 | 1,099E-08 |
| TcCLB.508949.10 | hypothetical protein, conserved | -0,5317 | 0,0004042 |
| TcCLB.506927.10 | hypothetical protein, conserved (pseudogene) | -0,5318 | 0,01416 |
| TcCLB.506067.189 | trans-sialidase (pseudogene), putative | -0,5318 | 0,1298 |
| TcCLB.506559.250 | hypothetical protein, conserved | -0,532 | 4,617E-06 |
| TcCLB.508873.300 | hypothetical protein | -0,532 | 0,3103 |
| TcCLB.503689.40 | hypothetical protein, conserved | -0,5321 | 2,306E-08 |
| TcCLB.509879.10 | hypothetical protein, conserved | -0,5322 | 0,003115 |
| TcCLB.508737.160 | hypothetical protein, conserved | -0,5323 | 2,905E-07 |
| TcCLB.508365.210 | mucin TcMUCI, putative | -0,5323 | 0,4144 |
| TcCLB.507037.70 | hypothetical protein, conserved | -0,5324 | 0,00000106 |
| TcCLB.508541.220 | casein kinase, putative | -0,5324 | 0,02061 |
| TcCLB.511675.6 | hypothetical protein, conserved (pseudogene) | -0,5324 | 0,3324 |
| TcCLB.510403.71 | trans-sialidase (pseudogene), putative | -0,5325 | 2,086E-06 |
| TcCLB.510201.50 | dispersed gene family protein 1 (DGF-1), putative | -0,5325 | 0,003761 |
| TcCLB.511379.9 | hypothetical protein, conserved (fragment) | -0,5326 | 0,0001934 |
| TcCLB.510739.10 | hypothetical protein, conserved | -0,5328 | 0,001739 |
| TcCLB.506529.30 | hypothetical protein, conserved | -0,533 | 0,0103 |
| TcCLB.509619.10 | Zn-finger protein, putative | -0,5331 | 0,00001004 |
| TcCLB.511409.20 | hypothetical protein, conserved | -0,5331 | 0,004938 |
| TcCLB.506247.420 | hypothetical protein, conserved | -0,5331 | 0,01415 |
| TcCLB.507237.310 | surface protease GP63 (pseudogene), putative | -0,5331 | 0,2051 |
| TcCLB.510279.276 | hypothetical protein, conserved (pseudogene) | -0,5336 | 0,1016 |
| TcCLB.511355.50 | glycerol uptake protein, putative | -0,5338 | 1,924E-06 |
| TcCLB.506213.70 | hypothetical protein, conserved | -0,5338 | 0,00005197 |
| TcCLB.508359.70 | hypothetical protein | -0,5338 | 0,2066 |
| TcCLB.503713.50 | trans-sialidase, putative (fragment) | -0,5338 | 0,2968 |

| | | | |
|---|---|---|---|
| TcCLB.509911.10 | hypothetical protein, conserved | -0,5339 | 1,389E-08 |
| TcCLB.508409.110 | hypothetical protein, conserved | -0,5339 | 1,529E-06 |
| TcCLB.510285.140 | hypothetical protein, conserved (fragment) | -0,5339 | 0,2483 |
| TcCLB.508289.41 | syntaxin binding protein (pseudogene), putative | -0,534 | 0,1078 |
| TcCLB.510687.100 | Unc104-like kinesin, putative | -0,5341 | 1,272E-08 |
| TcCLB.509109.50 | hypothetical protein | -0,5342 | 2,891E-08 |
| TcCLB.506739.30 | hypothetical protein | -0,5343 | 1,402E-08 |
| TcCLB.510181.50 | nucleoporin interacting component (NUP93), putative | -0,5343 | 5,461E-06 |
| TcCLB.508909.160 | hypothetical protein, conserved | -0,5343 | 0,005069 |
| TcCLB.504431.90 | U2 small nuclear ribonucleoprotein B, putative | -0,5343 | 0,006337 |
| TcCLB.511285.90 | 19S proteasome non-atpase subunit 8 (Rpn8) | -0,5344 | 9,258E-07 |
| TcCLB.508277.200 | hypothetical protein, conserved | -0,5344 | 0,00004228 |
| TcCLB.508257.90 | hypothetical protein, conserved | -0,5345 | 4,952E-06 |
| TcCLB.507041.76 | hypothetical protein, conserved (pseudogene) | -0,5345 | 0,002846 |
| TcCLB.506253.30 | hypothetical protein | -0,5345 | 0,1421 |
| TcCLB.509059.60 | hypothetical protein, conserved | -0,5347 | 0,04245 |
| TcCLB.507895.31 | retrotransposon hot spot protein (RHS, pseudogene), putat | -0,5347 | 0,05591 |
| TcCLB.506813.80 | trans-sialidase, Group V, putative | -0,5347 | 0,16 |
| TcCLB.509455.120 | Peptide chain release factor 1, mitochondrial, putative (RF | -0,5349 | 9,227E-06 |
| TcCLB.508821.60 | hypothetical protein, conserved | -0,5349 | 0,08659 |
| TcCLB.507519.190 | protein kinase (pseudogene), putative | -0,5349 | 0,114 |
| TcCLB.508365.284 | mucin-associated surface protein (MASP, pseudogene), pu | -0,5349 | 0,1239 |
| TcCLB.504741.170 | dual specificity protein phosphatase, putative | -0,535 | 0,00137 |
| TcCLB.504009.30 | hypothetical protein, conserved | -0,5351 | 0,0000159 |
| TcCLB.510029.60 | Vesicle-trafficking protein SEC22, putative (SEC22) | -0,5352 | 0,0001872 |
| TcCLB.510515.80 | hypothetical protein, conserved | -0,5353 | 4,323E-07 |
| TcCLB.511003.100 | hypothetical protein, conserved | -0,5353 | 2,325E-06 |
| TcCLB.510155.270 | dispersed gene family protein 1 (DGF-1, pseudogene), puta | -0,5354 | 9,675E-06 |
| TcCLB.509319.40 | hypothetical protein, conserved | -0,5357 | 0,00002569 |
| TcCLB.509669.40 | Zn-finger protein, putative | -0,5358 | 1,249E-11 |
| TcCLB.510329.190 | hypothetical protein, conserved | -0,5358 | 2,55E-08 |
| TcCLB.503675.10 | glycosyl transferase-like protein, putative | -0,536 | 0,004939 |
| TcCLB.509161.120 | hypothetical protein, conserved | -0,5361 | 4,473E-06 |
| TcCLB.511481.20 | hypothetical protein, conserved | -0,5361 | 0,003611 |
| TcCLB.510355.240 | hypothetical protein, conserved | -0,5362 | 0,0001936 |
| TcCLB.508221.1010 | mucin-associated surface protein (MASP), putative | -0,5363 | 0,01841 |
| TcCLB.509317.100 | hypothetical protein, conserved | -0,5367 | 0,0003755 |
| TcCLB.503541.30 | hypothetical protein, conserved | -0,5368 | 0,008942 |
| TcCLB.503973.340 | trans-sialidase (pseudogene), putative | -0,5368 | 0,2478 |
| TcCLB.507715.70 | hypothetical protein, conserved | -0,5369 | 4,45E-08 |
| TcCLB.510363.270 | mucin TcMUCII, putative | -0,5369 | 0,6796 |
| TcCLB.506147.180 | DNA polymerase epsilon catalytic subunit, putative | -0,537 | 1,965E-08 |
| TcCLB.508409.199 | hypothetical protein, conserved | -0,537 | 0,0007396 |
| TcCLB.510247.10 | R27-2 protein, putative | -0,5371 | 0,00001219 |
| TcCLB.509819.49 | Myotubularin-related protein, putative (fragment) | -0,5371 | 0,000664 |

| | | | |
|---|---|---|---|
| TcCLB.509755.80 | mucin-associated surface protein (MASP, pseudogene), pu | -0,5371 | 0,3706 |
| TcCLB.503653.50 | histone deacetylase, putative | -0,5372 | 3,628E-06 |
| TcCLB.509987.10 | hypothetical protein, conserved | -0,5372 | 0,0006052 |
| TcCLB.503781.70 | proteasome subunit beta type-5, putative | -0,5373 | 1,039E-06 |
| TcCLB.509167.10 | hypothetical protein, conserved | -0,5373 | 1,408E-06 |
| TcCLB.504039.140 | surface protease GP63 (pseudogene), putative | -0,5373 | 0,3078 |
| TcCLB.506869.20 | hypothetical protein, conserved | -0,5374 | 0,0001269 |
| TcCLB.508637.100 | hypothetical protein, conserved | -0,5374 | 0,001266 |
| TcCLB.508659.50 | serine/threonine protein kinase, putative | -0,5374 | 0,007995 |
| TcCLB.506949.10 | DNA-directed rna polymerase I largest subunit, putative | -0,5374 | 0,03739 |
| TcCLB.509059.89 | vacuolar assembly protein vps41, putative (fragment) | -0,5375 | 0,0507 |
| TcCLB.509177.30 | hypothetical protein, conserved | -0,5376 | 1,925E-07 |
| TcCLB.401469.10 | zinc finger protein 2, putative | -0,5376 | 5,163E-06 |
| TcCLB.511545.150 | hypothetical protein, conserved | -0,5376 | 0,00001512 |
| TcCLB.511127.420 | hypothetical protein, conserved | -0,5376 | 0,0001981 |
| TcCLB.509611.130 | hypothetical protein, conserved | -0,538 | 0,009403 |
| TcCLB.503833.69 | ribosome biogenesis protein, putative (fragment) | -0,5381 | 0,0005073 |
| TcCLB.504449.40 | hypothetical protein | -0,5383 | 0,00002979 |
| TcCLB.508405.90 | soluble N-ethylmaleimide sensitive factor (NSF) attachme | -0,5386 | 0,0006668 |
| TcCLB.504835.20 | 3-methylcrotonyl-CoA carboxylase, putative | -0,5387 | 5,788E-08 |
| TcCLB.509799.130 | hypothetical protein, conserved | -0,539 | 2,533E-08 |
| TcCLB.510873.20 | surface protease GP63, putative | -0,539 | 0,0001286 |
| TcCLB.510439.20 | hypothetical protein, conserved | -0,5394 | 0,01593 |
| TcCLB.508247.90 | trans-sialidase, Group V, putative | -0,5394 | 0,4818 |
| TcCLB.506813.190 | trans-sialidase, Group VI, putative | -0,5395 | 0,0411 |
| TcCLB.510105.9 | ATP-dependent DEAD/H RNA helicase, putative | -0,5396 | 0,00000643 |
| TcCLB.511511.100 | hypothetical protein, conserved | -0,5398 | 1,997E-10 |
| TcCLB.511385.80 | kinesin-like protein, putative | -0,5398 | 0,00009773 |
| TcCLB.508543.70 | hypothetical protein, conserved | -0,5399 | 0,0003084 |
| TcCLB.506405.4 | hypothetical protein, conserved (fragment) | -0,5402 | 8,508E-06 |
| TcCLB.506247.360 | hypothetical protein, conserved | -0,5403 | 0,00008186 |
| TcCLB.504041.29 | DNA repair protein, putative | -0,5403 | 0,0002299 |
| TcCLB.507259.4 | hypothetical protein, conserved | -0,5403 | 0,001418 |
| TcCLB.510081.21 | ATP-dependent DEAD/H RNA helicase (pseudogene), putat | -0,5404 | 0,03438 |
| TcCLB.508317.60 | hypothetical protein, conserved | -0,5407 | 0,001358 |
| TcCLB.511107.30 | hypothetical protein, conserved | -0,5409 | 3,294E-07 |
| TcCLB.507925.30 | hypothetical protein, conserved | -0,5409 | 0,01825 |
| TcCLB.503431.100 | chromatin assembly factor 1 subunit B, putative | -0,5411 | 6,813E-07 |
| TcCLB.507677.110 | hypothetical protein | -0,5411 | 8,911E-07 |
| TcCLB.510431.90 | hypothetical protein, conserved | -0,5412 | 0,001824 |
| TcCLB.506483.69 | mitogen activated protein kinase 4, putative (MPK4) | -0,5413 | 0,0004522 |
| TcCLB.507675.20 | hypothetical protein, conserved | -0,5414 | 3,263E-07 |
| TcCLB.506175.110 | hypothetical protein, conserved | -0,5415 | 7,634E-08 |
| TcCLB.504033.140 | hypothetical protein, conserved (pseudogene) | -0,5415 | 1,668E-07 |
| TcCLB.511577.60 | DNA excision/repair protein SNF2, putative | -0,5415 | 3,479E-07 |

| | | | |
|---|---|---|---|
| TcCLB.503703.20 | hypothetical protein, conserved | -0,5415 | 0,00006548 |
| TcCLB.507011.20 | hypothetical protein, conserved | -0,5417 | 0,00005682 |
| TcCLB.506577.110 | hypothetical protein, conserved | -0,5417 | 0,004847 |
| TcCLB.509213.90 | hypothetical protein, conserved | -0,5418 | 0,002537 |
| TcCLB.507625.186 | hypothetical protein, conserved | -0,542 | 0,002718 |
| TcCLB.506241.60 | kinesin, putative | -0,5421 | 2,685E-07 |
| TcCLB.511553.180 | hypothetical protein | -0,5422 | 0,4134 |
| TcCLB.508095.20 | methyltransferase, putative | -0,5423 | 1,217E-11 |
| TcCLB.510611.10 | hypothetical protein, conserved | -0,5423 | 0,00001122 |
| TcCLB.509289.10 | amastin, putative | -0,5423 | 0,00001327 |
| TcCLB.506945.90 | hypothetical protein, conserved | -0,5423 | 0,02866 |
| TcCLB.509965.70 | hypothetical protein, conserved | -0,5424 | 2,373E-06 |
| TcCLB.507585.20 | hypothetical protein, conserved | -0,5424 | 0,00001227 |
| TcCLB.506113.55 | N-acetyltransferase complex ARD1 subunit (pseudogene), | -0,5424 | 0,08492 |
| TcCLB.503885.10 | hypothetical protein, conserved | -0,5425 | 0,0002481 |
| TcCLB.508231.170 | hypothetical protein, conserved | -0,5427 | 1,468E-08 |
| TcCLB.506303.20 | cell division cycle protein 45 (CDC45), putative | -0,5427 | 0,0000315 |
| TcCLB.507995.10 | trans-sialidase (pseudogene), putative | -0,5427 | 0,0004425 |
| TcCLB.508461.344 | hypothetical protein, conserved | -0,543 | 0,0006022 |
| TcCLB.507047.20 | dispersed gene family protein 1 (DGF-1, pseudogene), puta | -0,543 | 0,0018 |
| TcCLB.506817.40 | hypothetical protein, conserved | -0,5431 | 4,142E-07 |
| TcCLB.511857.80 | AMP deaminase, putative | -0,5432 | 0,0000025 |
| TcCLB.508543.40 | hypothetical protein, conserved | -0,5432 | 0,008049 |
| TcCLB.510131.99 | hypothetical protein, conserved | -0,5434 | 0,0002836 |
| TcCLB.508409.310 | hypothetical protein, conserved | -0,5436 | 0,0001316 |
| TcCLB.510729.240 | hypothetical protein, conserved | -0,5438 | 0,0002376 |
| TcCLB.508177.100 | ubiquitin activating enzyme, putative | -0,5439 | 0,001536 |
| TcCLB.506363.30 | hypothetical protein, conserved | -0,544 | 6,272E-08 |
| TcCLB.509073.50 | hypothetical protein, conserved | -0,544 | 0,00002439 |
| TcCLB.508593.60 | hypothetical protein, conserved | -0,544 | 0,00007732 |
| TcCLB.511199.9 | ATP-dependent RNA helicase FAL1, putative (fragment) | -0,544 | 0,003071 |
| TcCLB.506917.27 | hypothetical protein, conserved (pseudogene) | -0,544 | 0,02578 |
| TcCLB.470827.20 | trans-sialidase, putative | -0,5442 | 0,0001564 |
| TcCLB.506567.50 | hypothetical protein, conserved | -0,5442 | 0,0002497 |
| TcCLB.506629.160 | hypothetical protein, conserved | -0,5443 | 0,01072 |
| TcCLB.506247.110 | hypothetical protein, conserved | -0,5444 | 0,0002457 |
| TcCLB.508255.20 | hypothetical protein, conserved (pseudogene) | -0,5445 | 0,0006718 |
| TcCLB.503847.20 | myosin heavy chain, putative | -0,5446 | 7,2E-08 |
| TcCLB.510689.60 | Double RNA binding domain protein 7 (DRBD7) | -0,5446 | 9,816E-07 |
| TcCLB.509031.6 | hypothetical protein, conserved | -0,5446 | 0,005729 |
| TcCLB.506941.100 | hypothetical protein, conserved | -0,5447 | 1,283E-09 |
| TcCLB.508181.90 | hypothetical protein, conserved (pseudogene) | -0,5447 | 2,333E-06 |
| TcCLB.503993.30 | surface protease GP63, putative | -0,5449 | 0,00379 |
| TcCLB.504119.4 | DNA-directed RNA polymerase III subunit C1, putative (frag | -0,5449 | 0,016 |
| TcCLB.509733.120 | hypothetical protein, conserved | -0,5451 | 1,165E-09 |

| | | | |
|---|---|---|---|
| TcCLB.511867.60 | hypothetical protein, conserved | -0,5453 | 0,0002933 |
| TcCLB.511589.170 | oxidoreductase, putative | -0,5453 | 0,001317 |
| TcCLB.509525.350 | mucin TcMUCII, putative | -0,5454 | 0,2943 |
| TcCLB.506677.20 | cdc2-related kinase 3, putative (CRK3) | -0,5456 | 0,004229 |
| TcCLB.509471.59 | histone H3, putative | -0,5457 | 0,02375 |
| TcCLB.508817.180 | DIS3-like exonuclease, putative (pseudogene) | -0,546 | 1,036E-07 |
| TcCLB.507047.80 | zinc-finger protein ZPR1, putative | -0,5462 | 0,00003413 |
| TcCLB.508199.10 | ADP-ribosylation factor, putative | -0,5466 | 0,0001544 |
| TcCLB.507943.70 | hypothetical protein, conserved | -0,5467 | 0,0001478 |
| TcCLB.510047.129 | mucin TcMUCII, putative | -0,5467 | 0,4882 |
| TcCLB.504251.9 | peroxisome assembly protein, putative (fragment) | -0,5468 | 3,666E-06 |
| TcCLB.506247.40 | hypothetical protein, conserved | -0,5468 | 0,00001393 |
| TcCLB.504089.54 | protein kinase, putative | -0,547 | 2,314E-07 |
| TcCLB.511751.240 | hypothetical protein, conserved (pseudogene) | -0,547 | 0,00001115 |
| TcCLB.506445.60 | mitochondrial DNA topoisomerase II, putative | -0,5473 | 4,6E-12 |
| TcCLB.507093.10 | hypothetical protein, conserved | -0,5473 | 8,203E-08 |
| TcCLB.510187.180 | hypothetical protein, conserved | -0,5475 | 3,261E-13 |
| TcCLB.506483.60 | hypothetical protein, conserved | -0,5477 | 0,0003448 |
| TcCLB.510661.19 | ubiquitin-activating enzyme E1, putative (fragment) | -0,5478 | 0,0000155 |
| TcCLB.511531.44 | hypothetical protein | -0,5478 | 0,0004606 |
| TcCLB.504207.5 | hypothetical protein, conserved | -0,5478 | 0,137 |
| TcCLB.506297.290 | protein transport protein SEC31, putative | -0,5479 | 4,384E-15 |
| TcCLB.504105.260 | bilobe protein, conserved | -0,5479 | 0,1282 |
| TcCLB.507975.40 | dispersed gene family protein 1 (DGF-1), putative | -0,5483 | 0,0001506 |
| TcCLB.509167.100 | valyl-tRNA synthetase, putative | -0,5484 | 4,251E-16 |
| TcCLB.508013.10 | mucin-associated surface protein (MASP, pseudogene), pu | -0,5486 | 0,3639 |
| TcCLB.511233.50 | hypothetical protein, conserved | -0,5487 | 0,00001431 |
| TcCLB.508717.40 | branch point binding protein, putative | -0,5488 | 0,00006846 |
| TcCLB.511599.61 | hypothetical protein, conserved (pseudogene) | -0,5489 | 0,006256 |
| TcCLB.510361.100 | mucin TcMUCII, putative | -0,5489 | 0,2085 |
| TcCLB.506989.180 | U3 small nucleolar ribonucleoprotein protein MPP10, putat | -0,549 | 2,899E-08 |
| TcCLB.510435.5 | hypothetical protein, conserved (pseudogene) | -0,549 | 0,002251 |
| TcCLB.509753.180 | hypothetical protein, conserved (pseudogene) | -0,549 | 0,0666 |
| TcCLB.510219.40 | hypothetical protein, conserved | -0,5491 | 0,0001185 |
| TcCLB.511163.30 | NGG1 interacting factor 3, putative | -0,5491 | 0,004325 |
| TcCLB.507765.10 | hypothetical protein, conserved | -0,5491 | 0,01703 |
| TcCLB.510357.10 | DnaJ chaperone protein, putative | -0,5493 | 2,046E-09 |
| TcCLB.511287.130 | hypothetical protein, conserved | -0,5494 | 1,346E-07 |
| TcCLB.507809.119 | hypothetical protein, conserved | -0,5496 | 1,857E-07 |
| TcCLB.507969.70 | hypothetical protein, conserved | -0,5496 | 0,001766 |
| TcCLB.507105.20 | mitochondrial RNA binding complex 1 subunit, putative | -0,5497 | 0,0001711 |
| TcCLB.509525.50 | hypothetical protein | -0,55 | 0,3725 |
| TcCLB.508707.270 | hypothetical protein, conserved | -0,5501 | 8,953E-08 |
| TcCLB.506479.39 | serine/threonine protein phosphatase 2A regulatory subun | -0,5503 | 0,00001883 |
| TcCLB.510429.20 | tRNA (guanine-N(7)-)-methyltransferase, putative | -0,5503 | 0,002711 |

| | | | |
|---|---|---|---|
| TcCLB.510547.7 | surface protease GP63 (pseudogene), putative | -0,5504 | 0,175 |
| TcCLB.507757.10 | hypothetical protein | -0,5505 | 0,0003879 |
| TcCLB.504109.174 | hypothetical protein, conserved | -0,5506 | 0,00002562 |
| TcCLB.510693.230 | mucin-associated surface protein (MASP), putative | -0,5506 | 0,2583 |
| TcCLB.507969.30 | sphingosine-1-phosphate phosphatase, putative | -0,5507 | 6,876E-07 |
| TcCLB.506127.100 | hypothetical protein, conserved | -0,5507 | 0,0001832 |
| TcCLB.504239.390 | syntaxin binding protein (pseudogene), putative | -0,5508 | 0,2863 |
| TcCLB.508707.30 | hypothetical protein, conserved | -0,551 | 4,075E-10 |
| TcCLB.510895.29 | hypothetical protein, conserved | -0,5511 | 0,006289 |
| TcCLB.507143.10 | hypothetical protein, conserved | -0,5512 | 4,209E-10 |
| TcCLB.509951.20 | hypothetical protein, conserved | -0,5512 | 0,04776 |
| TcCLB.503869.60 | coatomer beta subunit (pseudogene), putative | -0,5514 | 4,749E-10 |
| TcCLB.507603.60 | hypothetical protein, conserved | -0,5514 | 0,00004853 |
| TcCLB.511277.260 | hypothetical protein, conserved | -0,5515 | 0,00002727 |
| TcCLB.510001.10 | hypothetical Zinc-finger containing protein | -0,5515 | 0,000377 |
| TcCLB.510743.50 | phospholipase A2-like protein, putative | -0,5516 | 0,0001202 |
| TcCLB.506335.130 | trans-sialidase, Group V, putative | -0,5518 | 0,1747 |
| TcCLB.508851.200 | hypothetical protein, conserved | -0,5519 | 0,0002607 |
| TcCLB.461165.4 | hypothetical protein, conserved (fragment) | -0,5519 | 0,04599 |
| TcCLB.508609.20 | hypothetical protein | -0,5521 | 0,001543 |
| TcCLB.507163.90 | hypothetical protein, conserved | -0,5522 | 0,009949 |
| TcCLB.509539.59 | hypothetical protein, conserved | -0,5523 | 0,0134 |
| TcCLB.507037.89 | hypothetical protein, conserved | -0,5523 | 0,01552 |
| TcCLB.509349.10 | trans-sialidase (pseudogene), putative | -0,5523 | 0,2532 |
| TcCLB.507083.109 | hypothetical protein, conserved (fragment) | -0,5525 | 9,688E-11 |
| TcCLB.506563.4 | hypothetical protein, conserved | -0,5526 | 0,009842 |
| TcCLB.510359.270 | mitochondrial RNA-binding protein 2, putative | -0,5527 | 1,671E-06 |
| TcCLB.511389.100 | hypothetical protein, conserved | -0,5528 | 0,0001015 |
| TcCLB.506949.20 | hypothetical protein, conserved | -0,5529 | 1,069E-12 |
| TcCLB.503419.50 | MRB1-associated protein, putative (RGG2) | -0,553 | 0,00007162 |
| TcCLB.504427.170 | hypothetical protein, conserved | -0,553 | 0,0006102 |
| TcCLB.507001.30 | ankyrin repeat protein, putative | -0,553 | 0,007131 |
| TcCLB.507389.20 | hypothetical protein, conserved | -0,5532 | 0,002401 |
| TcCLB.504441.10 | trans-sialidase, putative (fragment) | -0,5533 | 0,2452 |
| TcCLB.509775.30 | hypothetical protein, conserved | -0,5534 | 1,443E-11 |
| TcCLB.508815.140 | protein transport protein SEC13, putative | -0,5534 | 7,229E-06 |
| TcCLB.506945.220 | serine/arginine-rich protein specific kinase SRPK, putative | -0,5535 | 7,656E-15 |
| TcCLB.511275.40 | hypothetical protein, conserved | -0,5535 | 0,001508 |
| TcCLB.506223.90 | hypothetical protein, conserved | -0,5536 | 0,00001314 |
| TcCLB.507739.20 | metallo-peptidase, Clan MA(E) Family M41 | -0,5539 | 7,384E-08 |
| TcCLB.508771.50 | hypothetical protein, conserved | -0,554 | 1,097E-06 |
| TcCLB.511711.10 | hypothetical protein, conserved | -0,5541 | 8,427E-06 |
| TcCLB.509949.9 | dispersed gene family protein 1 (DGF-1), putative | -0,5547 | 0,02636 |
| TcCLB.503919.4 | hypothetical protein, conserved | -0,5547 | 0,0634 |
| TcCLB.508727.44 | hypothetical protein, conserved | -0,555 | 1,942E-07 |

| | | | |
|---|---|---|---|
| TcCLB.510659.264 | prenyl protein specific carboxyl methyltransferase, putative | -0,5551 | 0,000333 |
| TcCLB.506201.160 | hypothetical protein, conserved | -0,5553 | 0,02481 |
| TcCLB.410199.30 | trans-sialidase, Group VI, putative | -0,5554 | 0,004648 |
| TcCLB.506629.170 | hypothetical protein | -0,5556 | 0,02452 |
| TcCLB.509287.10 | hypothetical protein, conserved | -0,5558 | 0,001392 |
| TcCLB.511859.90 | hypothetical protein | -0,5559 | 0,002694 |
| TcCLB.509179.80 | hypothetical protein, conserved | -0,5559 | 0,009119 |
| TcCLB.511867.10 | hypothetical protein, conserved | -0,556 | 2,512E-06 |
| TcCLB.510091.20 | hypothetical protein, conserved | -0,556 | 0,004139 |
| TcCLB.506939.209 | mucin TcMUCII (pseudogene), putative | -0,5561 | 0,09371 |
| TcCLB.505997.174 | dispersed gene family protein 1 (DGF-1), putative | -0,5562 | 0,003081 |
| TcCLB.507875.150 | damage-specific DNA binding protein (pseudogene), putati | -0,5564 | 0,0007878 |
| TcCLB.506579.100 | hypothetical protein, conserved | -0,5568 | 0,00003195 |
| TcCLB.510421.140 | hypothetical protein, conserved | -0,5569 | 0,0003607 |
| TcCLB.503639.30 | hypothetical protein, conserved | -0,557 | 0,00005771 |
| TcCLB.506525.40 | syntaxin 5, putative | -0,5571 | 5,476E-06 |
| TcCLB.511039.20 | hypothetical protein, conserved | -0,5571 | 8,403E-06 |
| TcCLB.508355.180 | hypothetical protein, conserved | -0,5572 | 0,00003837 |
| TcCLB.437999.9 | hypothetical protein, conserved (fragment) | -0,5572 | 0,0003074 |
| TcCLB.509157.230 | hypothetical protein, conserved | -0,5573 | 8,548E-08 |
| TcCLB.508911.80 | hypothetical protein, conserved | -0,5576 | 0,0002117 |
| TcCLB.506357.69 | hypothetical protein, conserved | -0,5576 | 0,05069 |
| TcCLB.511041.20 | hypothetical protein, conserved | -0,5577 | 5,015E-10 |
| TcCLB.503733.80 | hypothetical protein, conserved | -0,5577 | 0,00000022 |
| TcCLB.508455.30 | hypothetical protein | -0,5578 | 0,3014 |
| TcCLB.507103.20 | hypothetical protein | -0,5581 | 9,917E-12 |
| TcCLB.509603.19 | malonyl-CoA decarboxylase, mitochondrial precursor, puta | -0,5584 | 0,02102 |
| TcCLB.510595.10 | hypothetical protein | -0,5587 | 2,469E-06 |
| TcCLB.507603.10 | hypothetical protein, conserved (pseudogene) | -0,5587 | 0,0003369 |
| TcCLB.509047.40 | hypothetical protein, conserved | -0,5589 | 2,937E-11 |
| TcCLB.510599.30 | hypothetical protein, conserved | -0,5589 | 2,06E-09 |
| TcCLB.508579.20 | hypothetical protein, conserved | -0,5589 | 0,01593 |
| TcCLB.430281.10 | retrotransposon hot spot protein (RHS, pseudogene), putat | -0,5589 | 0,07033 |
| TcCLB.506269.10 | mucin-associated surface protein (MASP), putative | -0,5591 | 0,05431 |
| TcCLB.507437.10 | mucin-associated surface protein (MASP), putative | -0,5592 | 0,07083 |
| TcCLB.508741.390 | hypothetical protein, conserved | -0,5594 | 9,533E-10 |
| TcCLB.507159.14 | hypothetical protein, conserved | -0,5594 | 0,003481 |
| TcCLB.509187.10 | trans-sialidase, Group II, putative | -0,5594 | 0,551 |
| TcCLB.511051.30 | nucleobase transporter, putative | -0,5595 | 3,232E-07 |
| TcCLB.506753.160 | hypothetical protein, conserved | -0,5596 | 0,002084 |
| TcCLB.507819.11 | N-acetyltransferase complex ARD1 subunit (pseudogene), | -0,5598 | 0,02148 |
| TcCLB.506337.180 | hypothetical protein, conserved | -0,5599 | 2,877E-07 |
| TcCLB.510013.290 | mucin-associated surface protein (MASP), putative | -0,5599 | 0,3516 |
| TcCLB.506355.130 | hypothetical protein, conserved | -0,56 | 2,866E-06 |
| TcCLB.511031.49 | hypothetical protein, conserved | -0,56 | 0,00009096 |

| | | | |
|---|---|---|---|
| TcCLB.511029.20 | kinetoplast-associated protein 3 (KAP3) | -0,56 | 0,0001332 |
| TcCLB.511635.50 | hypothetical protein, conserved | -0,5601 | 0,0008649 |
| TcCLB.508875.20 | mucin-associated surface protein (MASP), putative (fragm | -0,5601 | 0,004367 |
| TcCLB.511585.40 | DNA-damage inducible protein DDI1-like protein, putative | -0,5602 | 9,278E-07 |
| TcCLB.507951.114 | high mobility group protein, putative | -0,5604 | 1,034E-07 |
| TcCLB.511391.24 | dispersed gene family protein 1 (DGF-1, pseudogene), puta | -0,5604 | 0,0002318 |
| TcCLB.506519.100 | hypothetical protein, conserved | -0,5604 | 0,002506 |
| TcCLB.503809.60 | glycerol uptake protein (pseudogene), putative | -0,5605 | 0,00647 |
| TcCLB.503833.20 | hypothetical protein, conserved | -0,5606 | 3,338E-08 |
| TcCLB.510867.9 | dispersed gene family protein 1 (DGF-1), putative | -0,5606 | 0,007547 |
| TcCLB.510367.20 | retrotransposon hot spot protein (RHS, pseudogene), putat | -0,5606 | 0,009025 |
| TcCLB.511321.50 | hypothetical protein, conserved | -0,5607 | 4,469E-08 |
| TcCLB.504207.29 | hypothetical protein, conserved | -0,5607 | 0,00002397 |
| TcCLB.506445.20 | Eukaryotic translation initiation factor 4 gamma type 2, pu | -0,5611 | 5,45E-09 |
| TcCLB.506221.120 | hypothetical protein, conserved | -0,5611 | 0,0008623 |
| TcCLB.511911.81 | hypothetical protein, conserved (pseudogene) | -0,5611 | 0,01448 |
| TcCLB.506871.10 | hypothetical protein, conserved | -0,5612 | 0,00000984 |
| TcCLB.504105.30 | DNA-directed RNA polymerase III subunit C1, putative (RP( | -0,5613 | 9,325E-18 |
| TcCLB.511277.50 | hypothetical protein, conserved | -0,5615 | 3,21E-09 |
| TcCLB.504047.20 | hypothetical protein, conserved | -0,5615 | 6,878E-06 |
| TcCLB.508891.4 | hypothetical protein, conserved | -0,5617 | 0,0004401 |
| TcCLB.507611.120 | expression site-associated gene (ESAG-like) protein, putati | -0,5617 | 0,2943 |
| TcCLB.511603.260 | mucin TcMUCII, putative | -0,5618 | 0,3432 |
| TcCLB.511751.30 | queuine tRNA-ribosyltransferase, putative | -0,5621 | 8,416E-08 |
| TcCLB.503561.20 | hypothetical protein, conserved | -0,5622 | 0,0002423 |
| TcCLB.507633.51 | trans-sialidase (pseudogene), putative | -0,5622 | 0,2722 |
| TcCLB.508153.450 | hypothetical protein, conserved | -0,5623 | 2,981E-07 |
| TcCLB.507629.10 | hypothetical protein, conserved | -0,5624 | 3,865E-11 |
| TcCLB.509751.4 | hypothetical protein, conserved (fragment) | -0,5624 | 0,03844 |
| TcCLB.507235.10 | dispersed gene family protein 1 (DGF-1), putative | -0,5625 | 0,177 |
| TcCLB.507011.150 | hypothetical protein, conserved | -0,5626 | 0,0002569 |
| TcCLB.509669.160 | hypothetical protein, conserved | -0,5628 | 3,187E-08 |
| TcCLB.508313.60 | hypothetical protein | -0,5628 | 0,4138 |
| TcCLB.511301.110 | inosine-5'-monophosphate dehydrogenase, putative | -0,5629 | 2,146E-16 |
| TcCLB.507663.50 | ATP-dependent RNA helicase, putative | -0,563 | 4,626E-13 |
| TcCLB.510719.200 | telomerase reverse transcriptase, putative (TERT) | -0,563 | 0,00000312 |
| TcCLB.511625.4 | hypothetical protein, conserved | -0,5631 | 0,007748 |
| TcCLB.510003.20 | hypothetical protein, conserved | -0,5632 | 4,631E-13 |
| TcCLB.508533.30 | metalloprotease, putative | -0,5636 | 5,486E-11 |
| TcCLB.511021.90 | hypothetical protein | -0,5636 | 0,00002336 |
| TcCLB.511255.618 | syntaxin binding protein (pseudogene), putative | -0,5636 | 0,3053 |
| TcCLB.507875.140 | retrotransposon hot spot protein (RHS, pseudogene), putat | -0,5637 | 0,02727 |
| TcCLB.508815.30 | DNA repair protein, putative | -0,5639 | 6,095E-10 |
| TcCLB.511511.30 | hypothetical protein, conserved | -0,5641 | 4,72E-08 |
| TcCLB.506793.70 | hypothetical protein, conserved | -0,5642 | 4,052E-09 |

| | | | |
|---|---|---|---|
| TcCLB.510531.54 | small nuclear ribonucleoprotein, putative | -0,5642 | 5,234E-06 |
| TcCLB.509753.126 | hypothetical protein, conserved (pseudogene) | -0,5642 | 0,1096 |
| TcCLB.509379.20 | Tob55, putative | -0,5643 | 8,949E-06 |
| TcCLB.511461.4 | hypothetical protein, conserved (fragment) | -0,5643 | 0,005729 |
| TcCLB.511623.10 | hypothetical protein, conserved | -0,565 | 2,562E-08 |
| TcCLB.510351.70 | hypothetical protein, conserved | -0,565 | 0,00003739 |
| TcCLB.511865.54 | protein transport protein Sec61 gamma subunit, putative | -0,5651 | 0,00001169 |
| TcCLB.508439.30 | hypothetical protein, conserved | -0,5654 | 3,555E-06 |
| TcCLB.510713.20 | dispersed gene family protein 1 (DGF-1), putative | -0,5654 | 0,001203 |
| TcCLB.511255.10 | trans-sialidase, Group V, putative | -0,5654 | 0,1176 |
| TcCLB.508207.50 | hypothetical protein | -0,5655 | 0,002665 |
| TcCLB.508661.70 | Enriched in surface-labeled proteome protein 18 (ESP18) | -0,5656 | 0,00002533 |
| TcCLB.510593.30 | hypothetical protein, conserved | -0,5656 | 0,0000398 |
| TcCLB.508595.30 | hypothetical protein (pseudogene) | -0,5658 | 0,001746 |
| TcCLB.507951.170 | ADP-ribosylation factor-like protein 3, putative (arl3) | -0,5659 | 7,572E-06 |
| TcCLB.504109.180 | hypothetical protein, conserved | -0,5659 | 0,002525 |
| TcCLB.447705.9 | hypothetical protein, conserved | -0,566 | 0,02829 |
| TcCLB.507513.130 | hypothetical protein, conserved (pseudogene) | -0,5662 | 2,252E-11 |
| TcCLB.509729.20 | hypothetical protein, conserved | -0,5662 | 6,754E-07 |
| TcCLB.511367.60 | la RNA binding protein, putative | -0,5663 | 8,342E-11 |
| TcCLB.506425.5 | receptor-type adenylate cyclase, putative (fragment) | -0,5666 | 0,04583 |
| TcCLB.508865.20 | mitochondrial DNA polymerase I protein C, putative | -0,5669 | 1,547E-14 |
| TcCLB.508963.40 | trans-sialidase (pseudogene), putative | -0,5673 | 0,0001877 |
| TcCLB.503557.49 | hypothetical protein, conserved (fragment) | -0,5674 | 0,02974 |
| TcCLB.506581.30 | eukaryotic translation initiation factor 3 subunit L, putative | -0,5675 | 2,836E-12 |
| TcCLB.410199.40 | hypothetical protein | -0,5676 | 0,07679 |
| TcCLB.506517.100 | hypothetical protein, conserved | -0,5677 | 7,454E-08 |
| TcCLB.511233.150 | hypothetical protein | -0,5677 | 0,2888 |
| TcCLB.508307.120 | hypothetical protein, conserved | -0,5678 | 1,849E-10 |
| TcCLB.511381.50 | hypothetical protein, conserved | -0,5678 | 1,241E-06 |
| TcCLB.504213.140 | hypothetical protein, conserved | -0,5679 | 0,008551 |
| TcCLB.510741.50 | hypothetical protein, conserved | -0,568 | 9,321E-07 |
| TcCLB.510755.110 | hypothetical protein, conserved | -0,568 | 0,000535 |
| TcCLB.509537.10 | pre-mRNA splicing factor ATP-dependent RNA helicase, pu | -0,5681 | 0,00002764 |
| TcCLB.506985.20 | hypothetical protein, conserved | -0,5681 | 0,0002566 |
| TcCLB.507765.30 | hypothetical protein, conserved | -0,5682 | 0,00000314 |
| TcCLB.430605.10 | hypothetical protein, conserved | -0,5686 | 4,303E-11 |
| TcCLB.510765.10 | hypothetical protein, conserved | -0,5687 | 9,686E-06 |
| TcCLB.509353.10 | E1-like ubiquitin-activating enzyme, putative (UBA5) | -0,5687 | 0,009712 |
| TcCLB.506673.60 | AP-3 complex subunit beta, putative (AP3B) | -0,5689 | 8,286E-08 |
| TcCLB.509167.70 | hypothetical protein, conserved | -0,5689 | 0,00003599 |
| TcCLB.510421.29 | CAAX prenyl protease 2, putative | -0,5691 | 0,0004753 |
| TcCLB.507967.4 | hypothetical protein, conserved | -0,5694 | 0,004826 |
| TcCLB.503555.10 | DNA replication licensing factor MCM8, putative | -0,5695 | 0,00009158 |
| TcCLB.511893.40 | receptor-type adenlyate cyclase, putative (fragment) | -0,5696 | 0,04893 |

| | | | |
|---|---|---|---|
| TcCLB.508163.105 | elongation factor 1-gamma (EF-1-gamma, pseudogene), p | -0,5697 | 0,241 |
| TcCLB.506885.440 | hypothetical protein, conserved | -0,5699 | 9,797E-09 |
| TcCLB.508347.40 | hypothetical protein, conserved | -0,5704 | 4,517E-13 |
| TcCLB.509073.30 | ubiquitin hydrolase, putative | -0,5704 | 2,859E-10 |
| TcCLB.508283.69 | dispersed gene family protein 1 (DGF-1), putative (fragme | -0,5704 | 0,006621 |
| TcCLB.511463.20 | AMPK subunit gamma (AMPKG) | -0,5708 | 0,00007392 |
| TcCLB.511519.18 | hypothetical protein, conserved | -0,5708 | 0,003082 |
| TcCLB.510889.200 | hypothetical protein, conserved | -0,5711 | 1,045E-07 |
| TcCLB.508257.240 | hypothetical protein, conserved | -0,5711 | 0,00004728 |
| TcCLB.511527.20 | TFIIH basal transcription factor complex helicase XPB subu | -0,5714 | 4,264E-08 |
| TcCLB.503555.40 | ATP-dependent RNA helicase, putative | -0,5715 | 6,232E-09 |
| TcCLB.511287.90 | ribosome biogenesis protein, putative | -0,5716 | 1,285E-08 |
| TcCLB.509631.60 | dispersed gene family protein 1 (DGF-1), putative | -0,5716 | 0,01397 |
| TcCLB.510953.30 | hypothetical protein, conserved | -0,5717 | 3,283E-06 |
| TcCLB.504155.230 | mucin-associated surface protein (MASP), putative | -0,5717 | 0,1485 |
| TcCLB.511529.40 | hypothetical protein, conserved | -0,5718 | 0,00005987 |
| TcCLB.511109.100 | DNA replication licensing factor MCM3, putative | -0,5719 | 0,0003648 |
| TcCLB.506477.40 | Tubulin/FtsZ family, putative | -0,572 | 8,185E-08 |
| TcCLB.511587.36 | N-acetyltransferase complex ARD1 subunit (pseudogene), | -0,572 | 0,07609 |
| TcCLB.509105.150 | heat shock protein 85, putative | -0,5722 | 2,793E-08 |
| TcCLB.503957.60 | trans-sialidase (pseudogene), putative | -0,5723 | 0,00005955 |
| TcCLB.505009.10 | hypothetical protein, conserved | -0,5725 | 0,00004119 |
| TcCLB.506287.20 | hypothetical protein, conserved | -0,5725 | 0,001596 |
| TcCLB.507709.70 | hypothetical protein, conserved | -0,5726 | 1,063E-06 |
| TcCLB.507641.220 | hypothetical protein | -0,5726 | 2,172E-06 |
| TcCLB.507281.9 | dispersed gene family protein 1 (DGF-1, pseudogene), puta | -0,5726 | 0,1037 |
| TcCLB.509047.70 | hypothetical protein, conserved | -0,5727 | 0,0005355 |
| TcCLB.511589.260 | glycyl-tRNA synthetase, putative | -0,5727 | 0,001951 |
| TcCLB.511417.40 | DNA-directed RNA polymerase III, putative | -0,5728 | 1,103E-07 |
| TcCLB.508035.10 | dispersed gene family protein 1 (DGF-1, pseudogene), puta | -0,5728 | 0,1463 |
| TcCLB.511855.20 | ubiquitin-like protein, putative | -0,5729 | 3,483E-12 |
| TcCLB.506925.80 | hypothetical protein, conserved | -0,5732 | 1,091E-08 |
| TcCLB.503689.34 | hypothetical protein, conserved | -0,5733 | 0,02659 |
| TcCLB.506579.30 | myo-inositol-1 phosphatase, putative | -0,5734 | 1,13E-10 |
| TcCLB.510727.9 | hypothetical protein, conserved (fragment) | -0,5734 | 0,001409 |
| TcCLB.508857.60 | hypothetical protein, conserved | -0,5735 | 0,00006246 |
| TcCLB.507993.220 | ARF-like 2-binding protein, putative | -0,5735 | 0,007945 |
| TcCLB.508277.10 | condensin subunit 1, putative | -0,5736 | 3,237E-10 |
| TcCLB.462761.4 | hypothetical protein, conserved | -0,5736 | 0,004457 |
| TcCLB.510551.10 | dispersed gene family protein 1 (DGF-1, pseudogene), puta | -0,5737 | 0,05121 |
| TcCLB.504239.270 | hypothetical protein | -0,5738 | 0,3458 |
| TcCLB.506783.10 | hypothetical protein, conserved (pseudogene) | -0,574 | 0,04077 |
| TcCLB.508153.1040 | hypothetical protein, conserved | -0,5741 | 4,71E-07 |
| TcCLB.510901.60 | replication factor A, 51kDa subunit, putative | -0,5742 | 3,39E-08 |
| TcCLB.507099.130 | hypothetical protein, conserved | -0,5742 | 0,0001013 |

| | | | |
|---|---|---|---|
| TcCLB.508153.830 | peter pan protein, putative | -0,5743 | 0,00004709 |
| TcCLB.511183.371 | hypothetical protein, conserved (pseudogene) | -0,5745 | 0,2287 |
| TcCLB.504195.9 | dispersed gene family protein 1 (DGF-1, pseudogene), puta | -0,5746 | 0,06121 |
| TcCLB.511421.30 | hypothetical protein, conserved | -0,5747 | 0,00001995 |
| TcCLB.507795.80 | hypothetical protein, conserved | -0,5748 | 7,171E-07 |
| TcCLB.508261.166 | hypothetical protein, conserved (pseudogene) | -0,5748 | 0,04504 |
| TcCLB.439991.10 | trans-sialidase (pseudogene), putative | -0,5749 | 0,05705 |
| TcCLB.510307.191 | protein kinase (pseudogene), putative | -0,5751 | 3,908E-08 |
| TcCLB.507623.125 | hypothetical protein, conserved | -0,5751 | 0,06831 |
| TcCLB.506471.10 | hypothetical protein, conserved | -0,5752 | 5,52E-07 |
| TcCLB.506439.18 | hypothetical protein, conserved | -0,5753 | 7,972E-06 |
| TcCLB.503609.10 | trans-sialidase (pseudogene), putative | -0,5753 | 0,347 |
| TcCLB.506297.200 | inositol-1,4,5-trisphosphate (IP3) 5-phosphatase, putative | -0,5755 | 5,647E-08 |
| TcCLB.507165.40 | hypothetical protein, conserved | -0,5755 | 0,0006369 |
| TcCLB.511315.20 | hypothetical protein, conserved | -0,5756 | 0,00000059 |
| TcCLB.507099.90 | hypothetical protein, conserved | -0,5757 | 1,573E-09 |
| TcCLB.510105.80 | folylpolyglutamate synthetase (FPGS) | -0,5757 | 0,0000134 |
| TcCLB.509255.10 | hypothetical protein, conserved | -0,5758 | 9,874E-09 |
| TcCLB.511233.90 | hypothetical protein, conserved | -0,5759 | 0,003455 |
| TcCLB.508257.250 | hypothetical protein, conserved | -0,576 | 1,148E-19 |
| TcCLB.506339.100 | dispersed gene family protein 1 (DGF-1), putative | -0,5762 | 0,004046 |
| TcCLB.511585.200 | hypothetical protein, conserved | -0,5763 | 2,854E-06 |
| TcCLB.404001.10 | hypothetical protein, conserved | -0,5764 | 1,395E-07 |
| TcCLB.508221.310 | trans-sialidase (pseudogene), putative | -0,5764 | 0,1419 |
| TcCLB.509267.10 | hypothetical protein, conserved | -0,5765 | 3,268E-09 |
| TcCLB.506945.330 | hypothetical protein, conserved | -0,5767 | 1,932E-14 |
| TcCLB.504157.40 | DNA replication licensing factor MCM3, putative | -0,5769 | 0,00003046 |
| TcCLB.506821.190 | hypothetical protein, conserved | -0,5771 | 1,058E-06 |
| TcCLB.439125.10 | Peptide chain release factor 1, mitochondrial, putative (RF | -0,5771 | 0,0002136 |
| TcCLB.510879.140 | hypothetical protein, conserved | -0,5772 | 3,363E-06 |
| TcCLB.510699.27 | elongation factor 1-gamma (EF-1-gamma, pseudogene), p | -0,5772 | 0,3005 |
| TcCLB.508141.80 | hypothetical protein, conserved | -0,5774 | 4,105E-10 |
| TcCLB.509965.290 | p22 protein precursor, putative | -0,5776 | 7,334E-09 |
| TcCLB.507089.100 | hypothetical protein, conserved | -0,5778 | 0,00001221 |
| TcCLB.503827.80 | syntaxin binding protein (pseudogene), putative | -0,5778 | 0,0562 |
| TcCLB.503717.10 | trans-sialidase, Group V, putative | -0,5779 | 0,05568 |
| TcCLB.508857.50 | hypothetical protein, conserved | -0,578 | 8,387E-15 |
| TcCLB.508355.340 | hypothetical protein, conserved | -0,578 | 0,0003807 |
| TcCLB.506679.120 | hypothetical protein | -0,5781 | 0,0001004 |
| TcCLB.508951.29 | actin-like protein, putative | -0,5782 | 0,00005087 |
| TcCLB.404001.4 | metalloprotease, putative (fragment) | -0,5783 | 1,625E-10 |
| TcCLB.508999.30 | Pre-mRNA-splicing factor 38, putative | -0,5784 | 0,01059 |
| TcCLB.508153.820 | inorganic pyrophosphatase, putative | -0,5788 | 0,00007249 |
| TcCLB.507091.150 | hypothetical protein, conserved (pseudogene) | -0,5788 | 0,0005135 |
| TcCLB.506945.160 | S-adenosylmethionine synthetase, putative (METK1) | -0,5792 | 1,574E-13 |

| | | | |
|---|---|---|---|
| TcCLB.508909.260 | hypothetical protein, conserved | -0,5793 | 0,009746 |
| TcCLB.508165.350 | mucin-associated surface protein (MASP), putative | -0,5793 | 0,1772 |
| TcCLB.511425.10 | serine/threonine protein phosphatase 2A regulatory subun | -0,5794 | 0,002173 |
| TcCLB.509319.10 | hypothetical protein, conserved | -0,5798 | 0,00005699 |
| TcCLB.510963.5 | hypothetical protein, conserved (fragment) | -0,5798 | 0,007171 |
| TcCLB.511051.20 | hypothetical protein, conserved | -0,5803 | 0,00008924 |
| TcCLB.510303.299 | hypothetical protein | -0,5805 | 0,00002384 |
| TcCLB.507005.10 | hypothetical protein | -0,5805 | 0,0198 |
| TcCLB.506175.5 | hypothetical protein, conserved | -0,5808 | 0,002508 |
| TcCLB.505983.9 | tryparedoxin peroxidase, putative | -0,5812 | 6,272E-08 |
| TcCLB.508909.40 | hypothetical protein, conserved | -0,5812 | 1,138E-06 |
| TcCLB.504251.30 | hypothetical protein, conserved | -0,5813 | 6,416E-07 |
| TcCLB.510475.110 | mucin TcMUCII, putative | -0,5813 | 0,2596 |
| TcCLB.506855.280 | DNA repair and recombination helicase protein PIF6, putat | -0,5814 | 3,757E-08 |
| TcCLB.473619.9 | dispersed gene family protein 1 (DGF-1), putative (fragme | -0,5815 | 0,3286 |
| TcCLB.506767.215 | hypothetical protein, conserved (pseudogene) | -0,5815 | 0,4261 |
| TcCLB.510081.40 | trans-sialidase (pseudogene), putative | -0,5816 | 0,6063 |
| TcCLB.510607.29 | dispersed gene family protein 1 (DGF-1), putative (fragme | -0,5817 | 0,0107 |
| TcCLB.506469.70 | hypothetical protein | -0,5818 | 0,01549 |
| TcCLB.510189.51 | elongation factor 1-gamma (EF-1-gamma, pseudogene), p | -0,5818 | 0,2768 |
| TcCLB.506643.10 | mucin-associated surface protein (MASP), putative | -0,5819 | 0,5051 |
| TcCLB.511315.9 | ATP-dependent RNA helicase, putative | -0,5824 | 0,001191 |
| TcCLB.506625.140 | hypothetical protein, conserved | -0,5826 | 0,00001827 |
| TcCLB.506739.20 | hypothetical protein, conserved | -0,5828 | 9,797E-09 |
| TcCLB.504167.30 | protein kinase, putative | -0,5829 | 0,00006969 |
| TcCLB.507897.20 | squalene synthase, putative | -0,5831 | 1,83E-20 |
| TcCLB.510689.50 | cell differentiation protein, putative | -0,5831 | 7,347E-07 |
| TcCLB.511529.9 | hypothetical protein, conserved | -0,5832 | 2,191E-08 |
| TcCLB.503797.10 | hypothetical protein, conserved | -0,5835 | 0,002791 |
| TcCLB.508409.280 | hypothetical protein, conserved | -0,5836 | 0,0001165 |
| TcCLB.507689.10 | dihydrouridine synthase (Dus, pseudogene), putative | -0,5836 | 0,01397 |
| TcCLB.504039.170 | mucin-associated surface protein (MASP, pseudogene), pu | -0,5837 | 0,258 |
| TcCLB.509205.30 | eukaryotic translation initiation factor, putative | -0,5838 | 1,871E-07 |
| TcCLB.510259.50 | 21 kDa cyclophilin, putative (CYP21) | -0,5839 | 7,149E-08 |
| TcCLB.503527.4 | hypothetical protein, conserved (pseudogene) | -0,5839 | 1,201E-07 |
| TcCLB.507507.20 | ADP-ribosylation factor GTPase activating protein 1, putati | -0,5839 | 1,727E-06 |
| TcCLB.511481.10 | hypothetical protein, conserved | -0,5839 | 0,0003854 |
| TcCLB.503859.25 | surface protease GP63 (pseudogene), putative | -0,5839 | 0,3338 |
| TcCLB.506821.90 | hypothetical protein, conserved | -0,5841 | 0,00000211 |
| TcCLB.510421.50 | hypothetical protein, conserved | -0,5844 | 1,066E-06 |
| TcCLB.503865.40 | hypothetical protein, conserved | -0,5844 | 0,00008309 |
| TcCLB.509681.10 | hypothetical protein, conserved | -0,5844 | 0,2386 |
| TcCLB.504179.60 | hypothetical protein, conserved | -0,5847 | 2,898E-07 |
| TcCLB.506655.20 | polyubiquitin (pseudogene), putative | -0,5847 | 0,04424 |
| TcCLB.506883.50 | Present in the outer mitochondrial membrane proteome 5 | -0,5848 | 3,682E-07 |

| | | | |
|---|---|---|---|
| TcCLB.440363.19 | Vacuolar iron transporter 1, putative (fragment) (VIT1) | -0,5848 | 0,000256 |
| TcCLB.509657.50 | mucin-associated surface protein (MASP), putative | -0,5848 | 0,281 |
| TcCLB.509105.140 | heat shock protein 85, putative | -0,5849 | 1,043E-08 |
| TcCLB.508479.380 | hypothetical protein, conserved | -0,5849 | 4,334E-07 |
| TcCLB.511353.20 | hypothetical protein, conserved | -0,585 | 1,569E-07 |
| TcCLB.508999.170 | surface protease GP63, putative | -0,5851 | 0,7842 |
| TcCLB.511493.10 | quiescin sulfhydryl oxidase, putative (QSOX) | -0,5855 | 0,0000312 |
| TcCLB.509395.20 | phosphatidylinositol kinase related protein, putative | -0,5857 | 7,324E-13 |
| TcCLB.510329.30 | hypothetical protein, conserved | -0,5857 | 1,479E-07 |
| TcCLB.505037.60 | NIMA/Nek Serine/threonine-protein kinase family, putative | -0,5859 | 9,278E-07 |
| TcCLB.508693.80 | ruvB-like DNA helicase, putative | -0,586 | 7,953E-13 |
| TcCLB.509023.130 | DNA replication licensing factor MCM4, putative (MCM4) | -0,5862 | 3,244E-07 |
| TcCLB.510667.40 | DNA repair protein RAD2, putative | -0,5867 | 3,649E-07 |
| TcCLB.506251.40 | hypothetical protein | -0,5871 | 1,75E-09 |
| TcCLB.508431.10 | dispersed gene family protein 1 (DGF-1, pseudogene), puta | -0,5877 | 0,001235 |
| TcCLB.507789.9 | dispersed gene family protein 1 (DGF-1), putative (fragme | -0,5877 | 0,03041 |
| TcCLB.510013.70 | mucin-associated surface protein (MASP), putative | -0,5877 | 0,3121 |
| TcCLB.508153.30 | Protein tilB homolog, putative (LRTP) | -0,5881 | 0,00005876 |
| TcCLB.506595.10 | dispersed gene family protein 1 (DGF-1), putative | -0,5881 | 0,02103 |
| TcCLB.504431.10 | hypothetical protein, conserved | -0,5885 | 2,525E-07 |
| TcCLB.508431.60 | hypothetical protein, conserved (pseudogene) | -0,5886 | 0,04009 |
| TcCLB.509345.20 | trans-sialidase (pseudogene), putative | -0,5887 | 0,04097 |
| TcCLB.508153.850 | hypothetical protein, conserved | -0,5889 | 3,763E-10 |
| TcCLB.509619.20 | hypothetical protein, conserved (pseudogene) | -0,589 | 0,003347 |
| TcCLB.511745.50 | hypothetical protein, conserved | -0,589 | 0,003599 |
| TcCLB.511183.409 | mucin TcMUC (pseudogene), putative | -0,5892 | 0,1656 |
| TcCLB.507483.4 | polyubiquitin (pseudogene), putative | -0,5894 | 0,0164 |
| TcCLB.509503.9 | dispersed gene family protein 1 (DGF-1), putative (fragme | -0,5894 | 0,09837 |
| TcCLB.508953.28 | hypothetical protein, conserved | -0,5896 | 0,0002709 |
| TcCLB.508163.181 | hypothetical protein, conserved (pseudogene) | -0,5896 | 0,1339 |
| TcCLB.504449.14 | hypothetical protein, conserved (fragment) | -0,5898 | 3,359E-14 |
| TcCLB.508823.50 | 40S ribosomal protein S14, putative | -0,59 | 2,241E-15 |
| TcCLB.507211.40 | inosine-5'-monophosphate dehydrogenase, putative | -0,59 | 1,402E-14 |
| TcCLB.511041.30 | hypothetical protein, conserved | -0,5903 | 1,698E-11 |
| TcCLB.510257.30 | ribulose-5-phosphate 3-epimerase, putative | -0,5903 | 0,0001048 |
| TcCLB.508727.24 | hypothetical protein | -0,5904 | 2,89E-10 |
| TcCLB.507011.50 | Pre-rRNA-processing protein PNO1, putative (PNO1) | -0,5906 | 3,398E-06 |
| TcCLB.508625.100 | hypothetical protein, conserved (pseudogene) | -0,5909 | 5,17E-07 |
| TcCLB.507053.140 | hypothetical protein, conserved | -0,5909 | 0,00008807 |
| TcCLB.510661.270 | replication factor C, subunit 3, putative | -0,5911 | 0,00001578 |
| TcCLB.504261.100 | mucin-associated surface protein (MASP, pseudogene), pu | -0,5913 | 0,1599 |
| TcCLB.503819.20 | DNA-directed RNA polymerase III subunit, putative | -0,5914 | 0,004018 |
| TcCLB.505163.80 | oligosaccharyl transferase subunit, putative | -0,5916 | 8,978E-22 |
| TcCLB.508851.154 | hypothetical protein, conserved | -0,5917 | 0,02275 |
| TcCLB.504207.10 | hypothetical protein, conserved | -0,5918 | 0,00001179 |

| | | | |
|---|---|---|---|
| TcCLB.508209.10 | hypothetical protein, conserved (pseudogene) | -0,5918 | 0,02069 |
| TcCLB.507785.20 | proteasome subunit alpha type-1, putative (fragment) (PS/ | -0,5918 | 0,2017 |
| TcCLB.506227.220 | hypothetical protein, conserved | -0,5921 | 2,596E-08 |
| TcCLB.507621.10 | retrotransposon hot spot protein (RHS, pseudogene), putat | -0,5921 | 0,04263 |
| TcCLB.506275.90 | hypothetical protein, conserved | -0,5922 | 0,00001007 |
| TcCLB.506305.40 | kinesin, putative | -0,5922 | 0,01342 |
| TcCLB.507757.40 | hypothetical protein | -0,5926 | 2,705E-08 |
| TcCLB.511459.50 | hypothetical protein, conserved | -0,5932 | 0,0002209 |
| TcCLB.506245.110 | dispersed gene family protein 1 (DGF-1, pseudogene), puta | -0,5933 | 0,1032 |
| TcCLB.511755.60 | ATP-dependent RNA helicase, putative | -0,5934 | 5,365E-14 |
| TcCLB.503897.120 | hypothetical protein, conserved | -0,5934 | 4,795E-08 |
| TcCLB.510303.50 | deoxyribose-phosphate aldolase, putative | -0,5936 | 5,995E-07 |
| TcCLB.510163.40 | surface protease GP63 (pseudogene), putative | -0,5936 | 0,00002739 |
| TcCLB.510465.140 | mucin-associated surface protein (MASP), putative | -0,5936 | 0,1275 |
| TcCLB.510311.130 | DNA repair helicase, putative | -0,5939 | 5,189E-09 |
| TcCLB.507681.250 | hypothetical protein, conserved | -0,5941 | 4,831E-06 |
| TcCLB.506445.70 | hypothetical protein, conserved | -0,5942 | 8,005E-06 |
| TcCLB.508175.270 | hypothetical protein | -0,5945 | 0,0002175 |
| TcCLB.510129.30 | procyclic form surface phosphoprotein, putative | -0,5946 | 1,916E-08 |
| TcCLB.511619.10 | hypothetical protein, conserved | -0,5946 | 0,0002091 |
| TcCLB.506501.210 | surface protease GP63 (pseudogene), putative | -0,5946 | 0,3612 |
| TcCLB.510971.60 | protein kinase, putative | -0,5948 | 1,508E-06 |
| TcCLB.510879.130 | hypothetical protein, conserved | -0,5948 | 7,382E-06 |
| TcCLB.506885.489 | dispersed gene family protein 1 (DGF-1), putative | -0,595 | 0,006769 |
| TcCLB.506585.40 | glucose-regulated protein 78, putative | -0,5953 | 1,084E-16 |
| TcCLB.511295.30 | hypothetical protein, conserved | -0,5953 | 6,484E-12 |
| TcCLB.509161.135 | hypothetical protein, conserved | -0,5954 | 0,00005502 |
| TcCLB.509915.20 | dispersed gene family protein 1 (DGF-1), putative | -0,5955 | 0,002621 |
| TcCLB.511805.29 | 2,3-bisphosphoglycerate-independent phosphoglycerate m | -0,5957 | 0,009087 |
| TcCLB.508601.70 | hypothetical protein, conserved | -0,5959 | 0,00005908 |
| TcCLB.509265.23 | N-acetyltransferase complex ARD1 subunit (pseudogene), | -0,5959 | 0,06064 |
| TcCLB.507515.130 | hypothetical protein, conserved | -0,596 | 0,000124 |
| TcCLB.511197.10 | trans-sialidase, putative | -0,5961 | 0,07755 |
| TcCLB.506805.20 | hypothetical protein | -0,5961 | 0,1202 |
| TcCLB.509399.100 | hypothetical protein, conserved | -0,5963 | 5,244E-10 |
| TcCLB.504883.70 | hypothetical protein, conserved (pseudogene) | -0,5964 | 6,65E-17 |
| TcCLB.504261.40 | mucin-associated surface protein (MASP), putative | -0,5965 | 0,1187 |
| TcCLB.507839.30 | dispersed gene family protein 1 (DGF-1), putative | -0,5965 | 0,2367 |
| TcCLB.508157.10 | serine-alanine-and proline-rich protein (pseudogene), puta | -0,5968 | 0,04548 |
| TcCLB.506961.97 | N-acetyltransferase complex ARD1 subunit (pseudogene), | -0,5968 | 0,09944 |
| TcCLB.506789.290 | hypothetical protein, conserved | -0,5973 | 8,234E-10 |
| TcCLB.506357.150 | hypothetical protein, conserved | -0,5974 | 3,218E-08 |
| TcCLB.509109.150 | hypothetical protein, conserved | -0,5975 | 0,0003088 |
| TcCLB.508873.460 | mucin-associated surface protein (MASP), putative | -0,5975 | 0,002771 |
| TcCLB.507929.20 | co-chaperone GrpE, putative | -0,5976 | 1,217E-08 |

| | | | |
|---|---|---|---|
| TcCLB.511499.50 | ubiquitin hydrolase, putative | -0,5977 | 1,787E-08 |
| TcCLB.506179.40 | hypothetical protein, conserved | -0,5977 | 0,00001081 |
| TcCLB.507063.90 | mitochondrial DNA polymerase beta, putative | -0,5979 | 3,856E-11 |
| TcCLB.504229.30 | protein kinase, putative | -0,598 | 9,231E-08 |
| TcCLB.511211.80 | hypothetical protein, conserved | -0,598 | 9,552E-06 |
| TcCLB.508707.210 | DNA replication licensing factor MCM7, putative | -0,5981 | 1,132E-08 |
| TcCLB.511281.50 | surface protease GP63, putative | -0,5982 | 0,001505 |
| TcCLB.506509.20 | hypothetical protein, conserved | -0,5984 | 0,0004754 |
| TcCLB.511277.370 | phosphatidic acid phosphatase, putative | -0,5984 | 0,0006814 |
| TcCLB.509179.180 | zinc-finger protein ZPR1, putative | -0,5987 | 1,092E-07 |
| TcCLB.511189.80 | small nuclear ribonucleoprotein SmD2, putative | -0,5987 | 2,037E-06 |
| TcCLB.504045.130 | kinesin, putative (fragment) | -0,5987 | 0,03924 |
| TcCLB.510443.20 | hypothetical protein, conserved | -0,5989 | 0,00001197 |
| TcCLB.508719.9 | hypothetical protein, conserved | -0,599 | 7,401E-13 |
| TcCLB.510053.20 | hypothetical protein, conserved | -0,599 | 5,454E-08 |
| TcCLB.509671.110 | hypothetical protein, conserved | -0,5991 | 2,862E-06 |
| TcCLB.508815.110 | serine/threonine protein phosphatase catalytic subunit, pu | -0,5992 | 1,789E-11 |
| TcCLB.511267.40 | hypothetical protein, conserved | -0,5992 | 0,04291 |
| TcCLB.506941.229 | pre-mRNA cleavage complex II Clp1 protein, putative | -0,5994 | 5,136E-07 |
| TcCLB.507023.30 | pseudouridine synthase A-like protein, putative | -0,5996 | 1,713E-07 |
| TcCLB.508903.10 | trans-sialidase (pseudogene), putative | -0,5997 | 0,04698 |
| TcCLB.503541.10 | hypothetical protein, conserved | -0,5998 | 0,00005816 |
| TcCLB.508825.30 | surface protease GP63 (pseudogene), putative | -0,6001 | 0,00003429 |
| TcCLB.506945.359 | damage-specific DNA binding protein, putative | -0,6006 | 2,713E-08 |
| TcCLB.509233.170 | hypothetical protein, conserved | -0,6007 | 0,0005483 |
| TcCLB.510449.20 | hypothetical protein, conserved (pseudogene) | -0,6009 | 0,1949 |
| TcCLB.509259.10 | trans-sialidase (pseudogene), putative | -0,6012 | 0,04538 |
| TcCLB.510059.10 | dispersed gene family protein 1 (DGF-1), putative | -0,6012 | 0,1535 |
| TcCLB.407477.44 | hypothetical protein, conserved | -0,6013 | 0,01223 |
| TcCLB.510735.70 | hypothetical protein, conserved | -0,6014 | 0,00001338 |
| TcCLB.472777.30 | protein disulfide isomerase, putative | -0,6014 | 0,00277 |
| TcCLB.506481.20 | hypothetical protein, conserved | -0,6022 | 7,251E-19 |
| TcCLB.510089.90 | hypothetical protein, conserved | -0,6026 | 4,146E-06 |
| TcCLB.509509.40 | hypothetical protein, conserved | -0,6029 | 0,00745 |
| TcCLB.506691.30 | hypothetical protein, conserved | -0,603 | 3,759E-15 |
| TcCLB.511733.110 | GTPase activating protein of Rab-like GTPase, putative | -0,6033 | 4,357E-06 |
| TcCLB.509007.10 | hypothetical protein, conserved | -0,6034 | 1,047E-08 |
| TcCLB.511499.20 | hypothetical protein, conserved | -0,6034 | 2,119E-06 |
| TcCLB.510679.14 | hypothetical protein, conserved | -0,6034 | 0,008993 |
| TcCLB.511299.70 | mitogen-activated protein kinase, putative | -0,6035 | 8,955E-08 |
| TcCLB.503755.10 | DNA polymerase kappa, putative | -0,6038 | 2,972E-07 |
| TcCLB.506363.130 | hypothetical protein, conserved | -0,6039 | 0,00001312 |
| TcCLB.507927.30 | hypothetical protein, conserved | -0,6042 | 0,0002781 |
| TcCLB.508181.30 | hypothetical protein, conserved | -0,6043 | 3,553E-14 |
| TcCLB.511365.80 | mitochondrial carrier protein, putative | -0,6046 | 2,911E-06 |

| | | | |
|---|---|---|---|
| TcCLB.507735.10 | hypothetical protein, conserved | -0,6047 | 0,004708 |
| TcCLB.510729.40 | hypothetical protein, conserved | -0,6051 | 0,0001351 |
| TcCLB.511283.220 | hypothetical protein, conserved | -0,6052 | 1,2E-11 |
| TcCLB.507395.14 | hypothetical protein, conserved | -0,6052 | 7,625E-06 |
| TcCLB.508121.155 | mucin TcMUCII (pseudogene), putative | -0,6053 | 0,1792 |
| TcCLB.511903.200 | hypothetical protein, conserved | -0,6055 | 6,58E-07 |
| TcCLB.506737.20 | protein kinase, putative | -0,6056 | 5,783E-08 |
| TcCLB.503861.100 | trans-sialidase (pseudogene), putative | -0,6057 | 0,0009145 |
| TcCLB.510329.100 | leucine-rich repeat protein, putative | -0,6058 | 1,113E-07 |
| TcCLB.508187.10 | vacuolar protein sorting-associated protein, putative (fragr | -0,606 | 1,785E-06 |
| TcCLB.507873.20 | cell differentiation protein, putative | -0,606 | 4,718E-06 |
| TcCLB.507017.64 | hypothetical protein, conserved | -0,606 | 0,0006376 |
| TcCLB.503779.70 | hypothetical protein, conserved | -0,6061 | 0,007113 |
| TcCLB.510819.10 | Eukaryotic translation initiation factor 4 gamma type 1, pu | -0,6062 | 4,087E-09 |
| TcCLB.511903.140 | tubulin folding cofactor D (pseudogene), putative | -0,6065 | 1,965E-12 |
| TcCLB.509751.20 | hypothetical protein, conserved | -0,6068 | 9,272E-08 |
| TcCLB.503489.20 | hypothetical protein, conserved | -0,6068 | 0,00002559 |
| TcCLB.509573.40 | hypothetical protein, conserved | -0,6068 | 0,01234 |
| TcCLB.507381.31 | retrotransposon hot spot protein (RHS, pseudogene), putat | -0,6071 | 1,659E-06 |
| TcCLB.410199.20 | hypothetical protein | -0,6071 | 0,02276 |
| TcCLB.509527.10 | mucin TcMUCII, putative | -0,6072 | 0,186 |
| TcCLB.511189.100 | hypothetical protein, conserved | -0,6075 | 1,856E-06 |
| TcCLB.504113.30 | hypothetical protein, conserved | -0,6075 | 0,001046 |
| TcCLB.506753.100 | hypothetical protein, conserved | -0,6076 | 0,003368 |
| TcCLB.504021.20 | hypothetical protein, conserved | -0,6076 | 0,08306 |
| TcCLB.504203.4 | hypothetical protein, conserved | -0,6077 | 6,478E-06 |
| TcCLB.507211.30 | hypothetical protein, conserved | -0,608 | 0,00005948 |
| TcCLB.508831.30 | hypothetical protein, conserved | -0,608 | 0,00008414 |
| TcCLB.506955.79 | surface protease GP63 (pseudogene), putative | -0,6082 | 0,2127 |
| TcCLB.506577.10 | hypothetical protein, conserved | -0,6083 | 0,0007624 |
| TcCLB.506173.179 | trans-sialidase (pseudogene), putative | -0,6084 | 0,02104 |
| TcCLB.402857.20 | hypothetical protein | -0,6085 | 4,459E-13 |
| TcCLB.506885.170 | hypothetical protein, conserved | -0,6085 | 0,00003576 |
| TcCLB.510743.10 | ubiquitin-activating enzyme E1, putative | -0,6089 | 1,043E-10 |
| TcCLB.511043.30 | DnaJ chaperone protein, putative (TcJ6) | -0,6089 | 5,167E-10 |
| TcCLB.507643.50 | dispersed gene family protein 1 (DGF-1), putative | -0,6089 | 0,0000474 |
| TcCLB.503971.59 | hypothetical protein, conserved (fragment) | -0,6089 | 0,00007025 |
| TcCLB.510769.100 | hypothetical protein, conserved | -0,6091 | 1,127E-07 |
| TcCLB.508307.140 | hypothetical protein, conserved | -0,6092 | 1,832E-07 |
| TcCLB.507711.320 | DNA mismatch repair protein MSH2, putative (MSH2) | -0,6093 | 2,195E-12 |
| TcCLB.506713.40 | N-acetyltransferase complex ARD1 subunit, putative | -0,6093 | 0,002637 |
| TcCLB.511603.141 | syntaxin binding protein (pseudogene), putative | -0,6093 | 0,2004 |
| TcCLB.510089.10 | mucin-associated surface protein (MASP), putative | -0,6094 | 0,06973 |
| TcCLB.508879.10 | hypothetical protein, conserved | -0,6095 | 3,945E-07 |
| TcCLB.505073.20 | hypothetical protein | -0,6097 | 1,145E-10 |

| | | | |
|---|---|---|---|
| TcCLB.510647.20 | hypothetical protein, conserved | -0,6097 | 2,195E-08 |
| TcCLB.507837.60 | Suppressor of G2 allele of SKP1, putative | -0,6098 | 6,992E-06 |
| TcCLB.508233.25 | hypothetical protein | -0,6098 | 0,00009996 |
| TcCLB.510635.50 | zinc-binding phosphatase, putative | -0,6099 | 1,865E-07 |
| TcCLB.507787.10 | kinesin-like protein (pseudogene), putative | -0,6099 | 0,00002764 |
| TcCLB.408573.5 | hypothetical protein | -0,6099 | 0,2039 |
| TcCLB.506131.151 | elongation factor 1-gamma (EF-1-gamma, pseudogene), p | -0,61 | 0,312 |
| TcCLB.510291.40 | serine/threonine protein kinase, putative | -0,6102 | 0,003949 |
| TcCLB.507633.90 | hypothetical protein | -0,6103 | 0,1274 |
| TcCLB.508761.130 | hypothetical protein | -0,6103 | 0,3473 |
| TcCLB.509945.10 | ERAD-associated E3 ubiquitin-protein ligase HRD1, putativ | -0,6104 | 2,199E-10 |
| TcCLB.511715.20 | protein kinase, putative | -0,6106 | 9,512E-07 |
| TcCLB.511867.30 | hypothetical protein, conserved | -0,6106 | 0,00006569 |
| TcCLB.503579.10 | hypothetical protein, conserved | -0,611 | 2,833E-07 |
| TcCLB.504081.530 | mucin-associated surface protein (MASP), putative | -0,611 | 0,2247 |
| TcCLB.506725.80 | hypothetical protein, conserved | -0,6111 | 8,176E-07 |
| TcCLB.508221.420 | mucin-associated surface protein (MASP), putative | -0,6113 | 0,2035 |
| TcCLB.511185.130 | dispersed gene family protein 1 (DGF-1), putative | -0,6115 | 0,07157 |
| TcCLB.506665.10 | hypothetical protein | -0,6116 | 0,2133 |
| TcCLB.508707.40 | hypothetical protein, conserved | -0,6118 | 2,374E-09 |
| TcCLB.510437.79 | heat shock protein 70, putative | -0,6119 | 0,0003619 |
| TcCLB.507895.50 | dispersed gene family protein 1 (DGF-1, pseudogene), puta | -0,6119 | 0,1535 |
| TcCLB.506679.260 | hypothetical protein, conserved | -0,6122 | 9,326E-09 |
| TcCLB.508817.80 | protein kinase, putative | -0,6124 | 7,566E-06 |
| TcCLB.509065.124 | U6 snRNA-associated Sm-like protein LSm5p | -0,6124 | 0,002983 |
| TcCLB.511229.110 | glycyl-tRNA synthetase, putative | -0,6126 | 6,681E-13 |
| TcCLB.507941.110 | hypothetical protein, conserved | -0,6127 | 2,974E-06 |
| TcCLB.508579.30 | kinesin, putative | -0,6128 | 0,00001429 |
| TcCLB.421621.35 | surface protease GP63 (pseudogene), putative | -0,6128 | 0,2736 |
| TcCLB.509965.250 | ubiquitin fusion degradation protein 2, putative | -0,6129 | 4,714E-13 |
| TcCLB.508699.10 | DNA topoisomerase II, putative | -0,613 | 9,008E-15 |
| TcCLB.510311.150 | mitochondrial protein 99, putative (MP99) | -0,613 | 1,779E-08 |
| TcCLB.506947.20 | hypothetical protein, conserved | -0,6132 | 2,668E-08 |
| TcCLB.507237.89 | mucin TcMUCII, putative | -0,6133 | 0,3913 |
| TcCLB.511425.30 | hypothetical protein, conserved | -0,6137 | 4,825E-06 |
| TcCLB.510821.50 | replication Factor A 28 kDa subunit, putative | -0,6138 | 9,01E-11 |
| TcCLB.509719.20 | ATP-dependent DEAD/H RNA helicase, putative | -0,6142 | 4,163E-09 |
| TcCLB.442297.10 | ARP2/3 complex 16kDa subunit, putative | -0,6145 | 0,0001402 |
| TcCLB.508221.20 | trans-sialidase (pseudogene), putative | -0,6146 | 0,01154 |
| TcCLB.510275.180 | elongation factor 1-gamma (EF-1-gamma), putative | -0,6146 | 0,2987 |
| TcCLB.510351.120 | Nudix hydrolase 2, putative (Nudix2) | -0,6147 | 3,608E-06 |
| TcCLB.507517.100 | pseudouridylate synthase I, putative | -0,6147 | 9,155E-06 |
| TcCLB.509965.380 | dTDP-glucose 4,6-dehydratase, putative | -0,6148 | 5,833E-09 |
| TcCLB.510733.60 | DNA polymerase epsilon catalytic subunit, putative | -0,6149 | 1,708E-12 |
| TcCLB.509991.29 | cyclophilin 15, putative | -0,615 | 1,338E-06 |

| | | | |
|---|---|---|---|
| TcCLB.407477.30 | diphthine synthase, putative | -0,6151 | 0,00003046 |
| TcCLB.510693.50 | trans-sialidase (pseudogene), putative | -0,6151 | 0,07106 |
| TcCLB.508693.40 | hypothetical protein, conserved | -0,6152 | 2,351E-10 |
| TcCLB.510275.390 | mucin TcMUCII (pseudogene), putative | -0,6154 | 0,2603 |
| TcCLB.506255.20 | hypothetical protein, conserved | -0,6155 | 0,0006117 |
| TcCLB.504147.10 | hypothetical protein, conserved | -0,6157 | 1,759E-11 |
| TcCLB.508293.100 | chitin-binding like protein, putative | -0,6157 | 0,2423 |
| TcCLB.508661.50 | ribonuclease H, putative | -0,6158 | 0,00006618 |
| TcCLB.510021.220 | trans-sialidase, Group V, putative | -0,6158 | 0,01243 |
| TcCLB.506247.140 | hypothetical protein, conserved | -0,6159 | 5,196E-09 |
| TcCLB.511115.10 | hypothetical protein, conserved | -0,6161 | 1,101E-09 |
| TcCLB.511255.330 | mucin-associated surface protein (MASP), putative | -0,6161 | 0,2708 |
| TcCLB.507687.10 | trans-sialidase, putative | -0,6162 | 0,01519 |
| TcCLB.507019.140 | CDC16, putative | -0,6164 | 5,962E-07 |
| TcCLB.506753.30 | hypothetical protein | -0,6164 | 0,0001149 |
| TcCLB.504741.100 | hypothetical protein, conserved | -0,6165 | 3,465E-11 |
| TcCLB.508845.9 | dispersed gene family protein 1 (DGF-1), putative | -0,6165 | 0,05927 |
| TcCLB.509525.280 | hypothetical protein | -0,6165 | 0,2602 |
| TcCLB.508891.130 | hypothetical protein, conserved | -0,6166 | 1,208E-10 |
| TcCLB.505935.30 | dispersed gene family protein 1 (DGF-1), putative | -0,6166 | 0,1218 |
| TcCLB.511633.60 | structural maintenance of chromosome (SMC), putative | -0,617 | 1,191E-16 |
| TcCLB.507063.250 | hypothetical protein | -0,6171 | 0,003957 |
| TcCLB.508741.110 | hypothetical protein, conserved | -0,6172 | 3,216E-06 |
| TcCLB.507099.70 | hypothetical protein, conserved | -0,6173 | 1,593E-11 |
| TcCLB.510689.30 | proteasome beta 7 subunit, putative | -0,6173 | 9,849E-11 |
| TcCLB.506363.140 | hypothetical protein, conserved | -0,6176 | 0,00009197 |
| TcCLB.510769.130 | kinesin, putative | -0,6181 | 9,471E-09 |
| TcCLB.504147.260 | hypothetical protein, conserved | -0,6185 | 3,125E-08 |
| TcCLB.507681.189 | hypothetical protein, conserved | -0,6186 | 0,00005388 |
| TcCLB.508147.70 | trans-sialidase (pseudogene), putative | -0,6186 | 0,1131 |
| TcCLB.508365.40 | mucin-associated surface protein (MASP), putative | -0,6186 | 0,2285 |
| TcCLB.511435.20 | Replication Factor C Subunit 1-related protein | -0,6187 | 7,47E-08 |
| TcCLB.504059.20 | endomembrane protein, putative | -0,6188 | 1,691E-10 |
| TcCLB.506357.20 | hypothetical protein, conserved | -0,6189 | 2,576E-08 |
| TcCLB.509643.139 | Heat shock protein 83, putative | -0,619 | 5,995E-09 |
| TcCLB.509099.20 | hypothetical protein, conserved | -0,6193 | 1,143E-10 |
| TcCLB.506393.143 | ATP-dependent DEAD/H RNA helicase (pseudogene), putat | -0,6194 | 0,02168 |
| TcCLB.506355.120 | hypothetical protein, conserved | -0,6196 | 1,543E-11 |
| TcCLB.511197.60 | mucin-associated surface protein (MASP), putative | -0,62 | 0,02428 |
| TcCLB.508607.30 | dispersed gene family protein 1 (DGF-1), putative | -0,6201 | 0,00478 |
| TcCLB.509065.100 | hypothetical protein, conserved | -0,6203 | 0,0002391 |
| TcCLB.510373.110 | hypothetical protein | -0,6203 | 0,328 |
| TcCLB.511481.30 | hypothetical protein, conserved | -0,6204 | 0,00009136 |
| TcCLB.508737.140 | hypothetical protein, conserved | -0,6206 | 1,166E-08 |
| TcCLB.509005.60 | hypothetical protein, conserved | -0,6207 | 3,439E-07 |

| | | | |
|---|---|---|---|
| TcCLB.506743.100 | hypothetical protein, conserved | -0,6209 | 0,01473 |
| TcCLB.511389.114 | hypothetical protein, conserved | -0,621 | 0,01443 |
| TcCLB.511353.50 | hypothetical protein, conserved | -0,6212 | 0,0001581 |
| TcCLB.508317.70 | hypothetical protein | -0,6217 | 2,692E-12 |
| TcCLB.510901.100 | DNA topoisomerase III, putative | -0,6217 | 1,686E-11 |
| TcCLB.508285.30 | ATP-dependent DEAD/H RNA helicase (pseudogene), putat | -0,6218 | 0,2287 |
| TcCLB.510493.10 | hypothetical protein, conserved | -0,6221 | 0,001695 |
| TcCLB.508741.290 | ribosomal RNA methyltransferase, putative | -0,6223 | 8,023E-10 |
| TcCLB.510161.20 | hypothetical protein, conserved | -0,6224 | 9,466E-15 |
| TcCLB.507951.250 | tRNA isopentenyltransferase, putative | -0,6224 | 4,336E-07 |
| TcCLB.511285.50 | hypothetical protein, conserved | -0,6225 | 5,378E-14 |
| TcCLB.506677.10 | ubiquitin-like protein, putative | -0,6226 | 3,927E-13 |
| TcCLB.509509.60 | small GTP-binding protein, putative | -0,6226 | 5,576E-08 |
| TcCLB.506303.70 | hypothetical protein, conserved | -0,6229 | 5,409E-09 |
| TcCLB.511031.20 | hypothetical protein, conserved | -0,6229 | 0,00000742 |
| TcCLB.506659.20 | mitochondrial DNA polymerase I protein C, putative | -0,6231 | 1,886E-17 |
| TcCLB.508643.40 | hypothetical protein, conserved | -0,6232 | 5,709E-14 |
| TcCLB.472253.10 | vacuolar protein sorting complex subunit (pseudogene), pu | -0,6232 | 0,0001011 |
| TcCLB.506165.4 | hypothetical protein, conserved | -0,6235 | 7,494E-06 |
| TcCLB.507253.50 | ATP-dependent DEAD/H RNA helicase (pseudogene), putat | -0,6235 | 0,2541 |
| TcCLB.506435.470 | hypothetical protein, conserved | -0,6238 | 0,003137 |
| TcCLB.506355.110 | hypothetical protein, conserved | -0,6239 | 6,753E-12 |
| TcCLB.511179.100 | monothiol glutaredoxin, putative | -0,6241 | 4,476E-07 |
| TcCLB.507611.260 | trans-sialidase (pseudogene), putative | -0,6241 | 0,105 |
| TcCLB.506789.230 | hypothetical protein, conserved | -0,6244 | 8,079E-18 |
| TcCLB.511417.100 | anti-silencing protein ASF 1, putative | -0,6244 | 0,004059 |
| TcCLB.510089.50 | hypothetical protein, conserved | -0,6246 | 5,505E-08 |
| TcCLB.508445.30 | protein kinase, putative | -0,625 | 1,031E-06 |
| TcCLB.511725.134 | mitochondrial carrier protein, putative | -0,6252 | 6,702E-08 |
| TcCLB.507937.6 | trans-sialidase (pseudogene), putative | -0,6254 | 0,2529 |
| TcCLB.510973.50 | hypothetical protein, conserved | -0,6255 | 6,819E-06 |
| TcCLB.508461.430 | proteasome beta 2 subunit, putative | -0,6256 | 5,731E-13 |
| TcCLB.503917.28 | hypothetical protein, conserved | -0,6258 | 3,609E-10 |
| TcCLB.506941.280 | heat shock 70 kDa protein, mitochondrial precursor, putativ | -0,6258 | 0,001409 |
| TcCLB.506195.170 | hypothetical protein, conserved | -0,6263 | 7,277E-10 |
| TcCLB.509649.59 | hypothetical protein, conserved | -0,6263 | 3,46E-07 |
| TcCLB.508885.30 | hypothetical protein | -0,6263 | 0,2998 |
| TcCLB.511531.40 | chaperonin alpha subunit, putative | -0,6266 | 8,148E-19 |
| TcCLB.507765.50 | hypothetical protein, conserved | -0,6267 | 2,513E-06 |
| TcCLB.398345.10 | hypothetical protein, conserved | -0,6268 | 3,591E-07 |
| TcCLB.506955.270 | trans-sialidase, Group V, putative | -0,627 | 0,01398 |
| TcCLB.510285.130 | hypothetical protein, conserved | -0,6273 | 0,00001429 |
| TcCLB.507709.120 | hypothetical protein, conserved | -0,6274 | 5,091E-18 |
| TcCLB.506183.109 | dispersed gene family protein 1 (DGF-1, pseudogene), puta | -0,6274 | 0,3904 |
| TcCLB.511523.10 | GTPase activating protein of Rab-like GTPase, putative | -0,6275 | 0,00001244 |

| | | | |
|---|---|---|---|
| TcCLB.510975.40 | hypothetical protein, conserved | -0,6276 | 4,311E-13 |
| TcCLB.508177.20 | carbohydrate kinase, thermoresistant glucokinase, putative | -0,6279 | 0,02187 |
| TcCLB.509643.130 | heat shock protein 85, putative | -0,6282 | 1,172E-09 |
| TcCLB.510693.130 | mucin-associated surface protein (MASP), putative | -0,6282 | 0,1531 |
| TcCLB.509895.70 | hypothetical protein, conserved | -0,6283 | 0,0309 |
| TcCLB.505977.13 | protein kinase, putative (fragment) | -0,6286 | 3,847E-09 |
| TcCLB.507517.30 | hypothetical protein, conserved | -0,6286 | 0,0001053 |
| TcCLB.504013.70 | metallo-peptidase, Clan MC, Family M14 | -0,6289 | 1,972E-09 |
| TcCLB.508681.31 | hypothetical protein | -0,6289 | 0,08279 |
| TcCLB.506459.190 | hypothetical protein | -0,6291 | 0,04392 |
| TcCLB.504021.50 | hypothetical protein | -0,6296 | 4,063E-09 |
| TcCLB.507943.80 | hypothetical protein, conserved (pseudogene) | -0,6296 | 6,245E-06 |
| TcCLB.506503.56 | hypothetical protein, conserved | -0,6298 | 6,142E-07 |
| TcCLB.506491.20 | myosin heavy chain, putative | -0,6299 | 7,839E-16 |
| TcCLB.511159.7 | neurobeachin/beige protein, putative | -0,6299 | 7,117E-09 |
| TcCLB.508675.60 | hypothetical protein, conserved | -0,63 | 8,846E-09 |
| TcCLB.503911.20 | pre-mRNA splicing factor ATP-dependent RNA helicase, pu | -0,6302 | 3,59E-08 |
| TcCLB.509971.80 | hypothetical protein, conserved | -0,6304 | 0,3168 |
| TcCLB.418405.30 | trans-sialidase, putative | -0,6304 | 0,427 |
| TcCLB.508857.15 | trans-sialidase, putative (fragment) | -0,6306 | 0,00221 |
| TcCLB.503653.9 | hypothetical protein, conserved | -0,6307 | 0,005525 |
| TcCLB.507521.10 | hypothetical protein, conserved | -0,6308 | 2,098E-07 |
| TcCLB.504159.80 | histone deacetylase, putative | -0,6311 | 1,882E-07 |
| TcCLB.506529.600 | metacyclin II, putative | -0,6312 | 0,1535 |
| TcCLB.506139.4 | mucin TcMUCII, putative | -0,6313 | 0,2958 |
| TcCLB.511287.4 | hypothetical protein | -0,6314 | 5,097E-09 |
| TcCLB.504101.10 | hypothetical protein | -0,6315 | 0,001977 |
| TcCLB.508175.110 | hypothetical protein, conserved | -0,6318 | 5,949E-09 |
| TcCLB.507683.10 | hypothetical protein, conserved | -0,6319 | 0,0006047 |
| TcCLB.508221.320 | mucin-associated surface protein (MASP, pseudogene), pu | -0,6322 | 0,1647 |
| TcCLB.506355.70 | hypothetical protein, conserved | -0,6323 | 2,141E-12 |
| TcCLB.508307.100 | hypothetical protein, conserved | -0,6323 | 0,0007838 |
| TcCLB.504069.100 | RNA polymerase II, putative | -0,6324 | 1,216E-06 |
| TcCLB.510225.50 | hypothetical protein, conserved | -0,6324 | 0,00004109 |
| TcCLB.510371.61 | surface protease GP63 (pseudogene), putative | -0,6326 | 0,1377 |
| TcCLB.511621.20 | hypothetical protein, conserved | -0,6327 | 0,00000384 |
| TcCLB.510699.34 | trans-sialidase (pseudogene), putative | -0,6328 | 0,02789 |
| TcCLB.510359.160 | hypothetical protein, conserved | -0,6329 | 0,0004212 |
| TcCLB.507911.36 | ATP-dependent DEAD/H RNA helicase (pseudogene), putat | -0,633 | 0,09981 |
| TcCLB.503795.10 | hypothetical protein, conserved | -0,6331 | 0,1368 |
| TcCLB.506925.90 | hypothetical protein, conserved | -0,6334 | 6,811E-10 |
| TcCLB.503555.30 | trypanothione reductase, putative | -0,6334 | 3,694E-09 |
| TcCLB.509789.40 | hypothetical protein, conserved | -0,6334 | 1,961E-06 |
| TcCLB.505931.20 | surface protease GP63, putative | -0,6335 | 0,006071 |
| TcCLB.508319.79 | hypothetical protein, conserved | -0,6337 | 0,1166 |

| | | | |
|---|---|---|---|
| TcCLB.504029.40 | COPII coat assembly protein sec16, putative (sec16) | -0,6338 | 2,485E-08 |
| TcCLB.504069.34 | hypothetical protein, conserved | -0,6338 | 0,004447 |
| TcCLB.409101.9 | hypothetical protein, conserved (fragment) | -0,634 | 0,002747 |
| TcCLB.510765.30 | COP-coated vesicle membrane protein gp25L precursor, pu | -0,6341 | 2,205E-07 |
| TcCLB.475169.10 | DNA-directed RNA polymerase II subunit 2 (pseudogene), p | -0,6342 | 0,0008741 |
| TcCLB.504741.20 | GTP-binding protein, putative | -0,6344 | 2,433E-10 |
| TcCLB.504017.70 | hypothetical protein | -0,6344 | 2,933E-07 |
| TcCLB.508539.130 | trans-sialidase (pseudogene), putative | -0,6345 | 0,00466 |
| TcCLB.508153.870 | hypothetical protein, conserved | -0,6346 | 2,001E-09 |
| TcCLB.504105.180 | hypothetical protein, conserved | -0,6347 | 1,263E-08 |
| TcCLB.508207.140 | hypothetical protein, conserved | -0,6347 | 9,852E-07 |
| TcCLB.503847.99 | hypothetical protein, conserved (pseudogene) | -0,6347 | 0,0009447 |
| TcCLB.503929.70 | hypothetical protein, conserved | -0,6348 | 0,00002672 |
| TcCLB.511173.130 | trans-sialidase, Group V, putative | -0,635 | 0,009544 |
| TcCLB.508181.120 | hypothetical protein, conserved | -0,6351 | 2,32E-11 |
| TcCLB.511129.54 | hypothetical protein, conserved (pseudogene) | -0,6351 | 0,001797 |
| TcCLB.511183.381 | syntaxin binding protein 1, (pseudogene), putative | -0,6351 | 0,05655 |
| TcCLB.511725.200 | hypothetical protein, conserved | -0,6354 | 4,334E-09 |
| TcCLB.510431.100 | hypothetical protein, conserved | -0,6355 | 2,332E-07 |
| TcCLB.510057.20 | hypothetical protein, conserved | -0,6355 | 0,0002456 |
| TcCLB.509773.9 | protein kinase, putative | -0,6356 | 0,00005398 |
| TcCLB.511895.69 | C-5 sterol desaturase, putative | -0,6358 | 0,001164 |
| TcCLB.509233.230 | hypothetical protein, conserved | -0,6359 | 2,146E-09 |
| TcCLB.510159.19 | hypothetical protein, conserved (fragment) | -0,636 | 0,0005468 |
| TcCLB.509487.4 | DNA-directed RNA polymerase III, putative | -0,6362 | 0,00633 |
| TcCLB.510823.10 | hypothetical protein, conserved | -0,6368 | 2,687E-11 |
| TcCLB.503929.20 | ribosomal protein L13, putative | -0,6368 | 0,0006599 |
| TcCLB.504057.120 | lipophosphoglycan biosynthetic protein 2, putative | -0,6371 | 3,105E-06 |
| TcCLB.508625.60 | hypothetical protein, conserved | -0,6371 | 0,00002119 |
| TcCLB.511517.140 | hypothetical protein | -0,6373 | 2,693E-06 |
| TcCLB.504827.100 | hypothetical protein, conserved | -0,6373 | 4,642E-06 |
| TcCLB.507023.50 | hypothetical protein, conserved | -0,6374 | 3,846E-07 |
| TcCLB.507681.20 | succinyl-CoA ligase [GDP-forming] beta-chain, putative | -0,6377 | 2,023E-19 |
| TcCLB.508645.50 | hypothetical protein, conserved | -0,6377 | 7,745E-10 |
| TcCLB.508479.290 | vacuolar sorting protein, putative | -0,6378 | 0,00004874 |
| TcCLB.510475.40 | mucin TcMUCII, putative | -0,6381 | 0,2862 |
| TcCLB.506567.34 | hypothetical protein, conserved | -0,6385 | 7,493E-09 |
| TcCLB.506513.14 | hypothetical protein, conserved | -0,6385 | 0,001943 |
| TcCLB.510483.250 | trans-sialidase (pseudogene), putative | -0,6385 | 0,02124 |
| TcCLB.507983.10 | dispersed gene family protein 1 (DGF-1), putative (fragme | -0,6388 | 0,07049 |
| TcCLB.507049.90 | hypothetical protein, conserved | -0,6391 | 0,00000107 |
| TcCLB.504411.10 | hypothetical protein, conserved | -0,6392 | 2,994E-07 |
| TcCLB.506057.40 | trans-sialidase (pseudogene), putative | -0,6392 | 0,00005057 |
| TcCLB.506255.30 | hypothetical protein, conserved | -0,6392 | 0,009406 |
| TcCLB.508257.150 | small nuclear ribonucleoprotein SmD3, putative | -0,6394 | 1,203E-06 |

| | | | |
|---|---|---|---|
| TcCLB.504075.10 | RNA polymerase I, putative | -0,6395 | 1,289E-06 |
| TcCLB.509203.70 | DNA topoisomerase II, putative | -0,6402 | 3,063E-13 |
| TcCLB.510565.179 | hypothetical protein, conserved | -0,6404 | 3,169E-15 |
| TcCLB.510773.9 | hypothetical protein, conserved (fragment) | -0,6407 | 0,0002446 |
| TcCLB.509955.50 | hypothetical protein, conserved | -0,6407 | 0,2446 |
| TcCLB.510217.70 | trans-sialidase (pseudogene), putative | -0,6408 | 0,1203 |
| TcCLB.507981.10 | trans-sialidase (pseudogene), putative | -0,6409 | 0,07721 |
| TcCLB.508163.320 | mucin-associated surface protein (MASP), putative | -0,6413 | 0,09413 |
| TcCLB.509451.50 | hypothetical protein, conserved | -0,6415 | 2,515E-10 |
| TcCLB.509085.60 | dispersed gene family protein 1 (DGF-1), putative | -0,6415 | 0,001092 |
| TcCLB.510339.40 | hypothetical protein, conserved | -0,6416 | 1,151E-07 |
| TcCLB.506529.490 | hypothetical protein, conserved | -0,6416 | 1,292E-07 |
| TcCLB.505943.95 | trans-sialidase (pseudogene), putative | -0,6416 | 0,00007883 |
| TcCLB.504089.79 | retrotransposon hot spot protein (RHS, pseudogene), putat | -0,6417 | 0,1241 |
| TcCLB.511383.60 | hypothetical protein, conserved | -0,6418 | 0,0001401 |
| TcCLB.508683.9 | dispersed gene family protein 1 (DGF-1), putative | -0,642 | 0,09538 |
| TcCLB.504443.30 | DNA-directed RNA polymerase II subunit 2, putative (pseu | -0,6421 | 7,424E-14 |
| TcCLB.511189.20 | hypothetical protein, conserved | -0,6421 | 1,126E-07 |
| TcCLB.510087.110 | hypothetical protein, conserved | -0,6422 | 8,91E-11 |
| TcCLB.415789.9 | dispersed gene family protein 1 (DGF-1), putative (fragme | -0,6422 | 0,00002161 |
| TcCLB.422041.9 | pre-mRNA splicing factor ATP-dependent RNA helicase, pu | -0,6424 | 0,0003544 |
| TcCLB.508365.220 | hypothetical protein, conserved (pseudogene) | -0,6424 | 0,09372 |
| TcCLB.504557.60 | hypothetical protein, conserved | -0,6425 | 0,00006737 |
| TcCLB.507529.40 | hypothetical protein, conserved | -0,6426 | 4,253E-08 |
| TcCLB.508153.60 | metallo-peptidase, Clan MA(E) Family M1 | -0,6427 | 1,418E-13 |
| TcCLB.509233.200 | hypothetical protein, conserved | -0,6431 | 2,452E-12 |
| TcCLB.507639.70 | hypothetical protein, conserved | -0,6431 | 0,00016 |
| TcCLB.511463.4 | actin, putative | -0,6433 | 0,01309 |
| TcCLB.507049.160 | hypothetical protein, conserved | -0,6434 | 2,052E-06 |
| TcCLB.510019.30 | mucin-associated surface protein (MASP), putative | -0,6434 | 0,076 |
| TcCLB.506957.130 | hypothetical protein, conserved | -0,644 | 1,385E-08 |
| TcCLB.510519.100 | hypothetical protein, conserved | -0,644 | 0,00005489 |
| TcCLB.506815.20 | mucin-associated surface protein (MASP, pseudogene), pu | -0,6443 | 0,00005322 |
| TcCLB.510823.90 | hypothetical protein | -0,6447 | 5,843E-10 |
| TcCLB.503783.17 | mucin TcMUCII, putative | -0,6447 | 0,2216 |
| TcCLB.510289.3 | hypothetical protein, conserved | -0,6449 | 0,004237 |
| TcCLB.508405.119 | hypothetical protein, conserved | -0,6449 | 0,03495 |
| TcCLB.511217.50 | hypothetical protein, conserved | -0,645 | 1,947E-15 |
| TcCLB.508357.26 | hypothetical protein, conserved | -0,6452 | 9,487E-06 |
| TcCLB.503999.90 | RNA-binding protein 34, putative (RBP34) | -0,6453 | 9,069E-06 |
| TcCLB.503845.20 | dispersed gene family protein 1 (DGF-1, pseudogene), puta | -0,6456 | 0,009808 |
| TcCLB.510099.90 | tyrosine phosphatase isoform, putative | -0,6458 | 2,267E-08 |
| TcCLB.511867.150 | hypothetical protein, conserved | -0,6459 | 5,864E-06 |
| TcCLB.506319.20 | hypothetical protein, conserved | -0,646 | 4,8E-08 |
| TcCLB.510363.70 | mucin-associated surface protein (MASP), putative | -0,6461 | 0,2546 |

| | | | |
|---|---|---|---|
| TcCLB.510323.120 | kinesin, putative | -0,6462 | 2,091E-07 |
| TcCLB.509525.210 | hypothetical protein | -0,6462 | 0,06465 |
| TcCLB.440343.9 | dispersed gene family protein 1 (DGF-1), putative (fragme | -0,6463 | 0,2118 |
| TcCLB.506559.540 | hypothetical protein, conserved | -0,6464 | 2,913E-07 |
| TcCLB.504147.250 | hypothetical protein, conserved | -0,6465 | 2,874E-15 |
| TcCLB.508541.10 | elongation factor 1-gamma (EF-1-gamma, pseudogene), p | -0,6468 | 0,05098 |
| TcCLB.508209.50 | hypothetical protein, conserved | -0,6472 | 0,001322 |
| TcCLB.508213.40 | hypothetical protein, conserved | -0,6473 | 0,0002141 |
| TcCLB.503781.10 | hypothetical protein, conserved | -0,6474 | 3,073E-16 |
| TcCLB.506855.320 | hypothetical protein, conserved | -0,6474 | 0,00002331 |
| TcCLB.511627.90 | hypothetical protein, conserved | -0,6474 | 0,0003604 |
| TcCLB.509419.140 | mucin-associated surface protein (MASP), putative | -0,6475 | 0,02809 |
| TcCLB.506409.110 | hypothetical protein | -0,6476 | 0,01833 |
| TcCLB.508211.19 | hypothetical protein, conserved | -0,648 | 0,0002577 |
| TcCLB.508761.60 | hypothetical protein (pseudogene) | -0,648 | 0,09827 |
| TcCLB.508775.60 | hypothetical protein, conserved (pseudogene) | -0,6482 | 0,02446 |
| TcCLB.506335.15 | hypothetical protein | -0,6485 | 0,03938 |
| TcCLB.503973.50 | mucin-associated surface protein (MASP), putative | -0,6488 | 0,08081 |
| TcCLB.507601.20 | hypothetical protein, conserved | -0,649 | 0,00319 |
| TcCLB.510239.10 | dispersed gene family protein 1 (DGF-1), putative | -0,649 | 0,1004 |
| TcCLB.509641.20 | hypothetical protein, conserved | -0,6491 | 0,00003475 |
| TcCLB.504031.50 | trans-sialidase, Group V, putative | -0,6492 | 0,03358 |
| TcCLB.510747.18 | hypothetical protein, conserved | -0,6495 | 0,00001692 |
| TcCLB.507357.50 | hypothetical protein | -0,6496 | 0,08158 |
| TcCLB.507681.220 | pre-mRNA splicing factor ATP-dependent RNA helicase, pu | -0,6497 | 7,847E-13 |
| TcCLB.506453.10 | trans-sialidase, putative | -0,6497 | 0,05147 |
| TcCLB.511741.30 | replication factor C, subunit 2, putative | -0,6498 | 5,386E-09 |
| TcCLB.510901.30 | hypothetical protein, conserved | -0,6498 | 5,634E-07 |
| TcCLB.506319.30 | hypothetical protein, conserved | -0,6498 | 1,248E-06 |
| TcCLB.505075.10 | hypothetical protein, conserved | -0,6501 | 0,00001295 |
| TcCLB.507017.130 | hypothetical protein | -0,6501 | 0,001664 |
| TcCLB.503833.50 | hypothetical protein, conserved | -0,6503 | 7,965E-16 |
| TcCLB.511479.10 | trans-sialidase (pseudogene), putative | -0,6508 | 0,3611 |
| TcCLB.511401.80 | hypothetical protein, conserved (pseudogene) | -0,6509 | 0,00007233 |
| TcCLB.509803.40 | 5-oxoprolinase, putative | -0,6511 | 2,698E-08 |
| TcCLB.506667.10 | mucin-associated surface protein (MASP), putative | -0,6511 | 0,1906 |
| TcCLB.504117.9 | hypothetical protein, conserved | -0,6512 | 1,208E-09 |
| TcCLB.507093.50 | hypothetical protein, conserved | -0,6513 | 3,238E-14 |
| TcCLB.509457.10 | hypothetical protein | -0,6514 | 1,693E-12 |
| TcCLB.508987.39 | hypothetical protein, conserved | -0,6514 | 3,033E-08 |
| TcCLB.505807.270 | ubiquitin hydrolase, putative | -0,6515 | 0,00001368 |
| TcCLB.506591.10 | hypothetical protein, conserved | -0,6519 | 0,005152 |
| TcCLB.416511.9 | mismatch repair protein MSH3, putative | -0,652 | 4,821E-06 |
| TcCLB.504155.140 | mucin TcMUCII, putative | -0,652 | 0,6097 |
| TcCLB.505999.24 | hypothetical protein, conserved | -0,6523 | 0,0001148 |

| | | | |
|---|---|---|---|
| TcCLB.507237.24 | mucin TcMUCII (pseudogene), putative | -0,6523 | 0,4504 |
| TcCLB.509845.9 | dispersed gene family protein 1 (DGF-1), putative | -0,6524 | 0,01472 |
| TcCLB.510279.290 | dispersed gene family protein 1 (DGF-1, pseudogene), puta | -0,6526 | 0,01009 |
| TcCLB.511303.90 | hypothetical protein, conserved | -0,6529 | 1,353E-09 |
| TcCLB.509525.70 | hypothetical protein | -0,653 | 0,0744 |
| TcCLB.510305.9 | Present in the outer mitochondrial membrane proteome 3 | -0,6531 | 0,00002086 |
| TcCLB.503671.30 | Protein tilB homolog, putative (LRTP) | -0,6533 | 0,00001078 |
| TcCLB.508145.40 | hypothetical protein, conserved | -0,6533 | 0,00009001 |
| TcCLB.510713.120 | surface protease GP63, putative | -0,6533 | 0,2172 |
| TcCLB.506317.20 | hypothetical protein, conserved | -0,6536 | 1,358E-13 |
| TcCLB.510149.90 | Golgi vesicular membrane trafficking protein, putative | -0,6536 | 0,0009017 |
| TcCLB.510091.100 | hypothetical protein, conserved | -0,6541 | 0,00001557 |
| TcCLB.509499.4 | hypothetical protein, conserved | -0,6541 | 0,00007307 |
| TcCLB.510513.50 | hypothetical protein, conserved | -0,6544 | 8,766E-06 |
| TcCLB.505023.40 | mucin-associated surface protein (MASP), putative | -0,6544 | 0,01417 |
| TcCLB.508259.20 | dispersed gene family protein 1 (DGF-1, pseudogene), puta | -0,6546 | 0,02223 |
| TcCLB.509911.120 | hypothetical protein | -0,6547 | 2,193E-15 |
| TcCLB.504643.40 | U6 snRNA-associated Sm-like protein LSm4, putative | -0,6552 | 0,00006793 |
| TcCLB.507575.10 | dispersed gene family protein 1 (DGF-1), putative | -0,6552 | 0,02978 |
| TcCLB.510897.99 | hypothetical protein, conserved (fragment) | -0,6556 | 0,002277 |
| TcCLB.504057.30 | hypothetical protein, conserved | -0,6558 | 9,602E-17 |
| TcCLB.506519.70 | hypothetical protein | -0,6559 | 0,00002739 |
| TcCLB.509231.50 | hypothetical protein, conserved | -0,6565 | 0,00002388 |
| TcCLB.511593.40 | trans-sialidase (pseudogene), putative | -0,657 | 1,234E-07 |
| TcCLB.510085.4 | mucin TcMUCII (pseudogene), putative | -0,6572 | 0,2135 |
| TcCLB.509105.130 | ATP-binding cassette sub-family F member 1, putative (AB | -0,6573 | 3,526E-16 |
| TcCLB.507019.110 | hypothetical protein | -0,6575 | 0,00005654 |
| TcCLB.506357.60 | hypothetical protein, conserved | -0,6577 | 8,724E-16 |
| TcCLB.511211.210 | hypothetical protein, conserved | -0,6581 | 0,004199 |
| TcCLB.503543.10 | presenilin-like aspartic peptidase, putative | -0,6582 | 9,048E-09 |
| TcCLB.509353.50 | hypothetical protein, conserved | -0,6583 | 3,805E-09 |
| TcCLB.509319.60 | endo/exonuclease Mre11, putative | -0,6589 | 2,367E-13 |
| TcCLB.509791.70 | hypothetical protein, conserved | -0,6589 | 0,0000735 |
| TcCLB.510285.120 | hypothetical protein, conserved | -0,659 | 3,412E-11 |
| TcCLB.507711.235 | deoxyribodipyrimidine photo-lyase, putative | -0,659 | 4,366E-06 |
| TcCLB.509941.110 | condensin subunit 1, putative | -0,6591 | 1,951E-15 |
| TcCLB.509213.30 | hypothetical protein, conserved | -0,6591 | 0,00001286 |
| TcCLB.510719.250 | RNA polymerase-associated protein CDC73, putative | -0,6592 | 1,398E-08 |
| TcCLB.507611.131 | protein kinase (pseudogene), putative | -0,6592 | 0,0001929 |
| TcCLB.510321.10 | RNA polymerase-associated protein CTR9, putative | -0,6594 | 0,00002547 |
| TcCLB.403789.9 | DNA-directed rna polymerase I largest subunit, putative | -0,6595 | 0,0008006 |
| TcCLB.510763.109 | rRNA processing protein, putative | -0,6597 | 6,794E-06 |
| TcCLB.503791.40 | dihydrouridine synthase, putative | -0,6598 | 0,0006752 |
| TcCLB.506695.10 | actin-like protein, putative | -0,6599 | 0,00004709 |
| TcCLB.507685.10 | hypothetical protein, conserved | -0,6601 | 1,874E-06 |

| | | | |
|---|---|---|---|
| TcCLB.503749.30 | hypothetical protein, conserved | -0,6603 | 4,247E-09 |
| TcCLB.508461.40 | hypothetical protein, conserved | -0,6603 | 0,00001072 |
| TcCLB.508741.320 | casein kinase II, alpha chain, putative | -0,6605 | 5,617E-10 |
| TcCLB.506511.50 | hypothetical protein, conserved | -0,6607 | 1,509E-17 |
| TcCLB.506435.300 | ubiquitin-protein ligase, putative | -0,6607 | 0,00001658 |
| TcCLB.484311.10 | calpain-like cysteine peptidase, putative (fragment) | -0,6609 | 0,0392 |
| TcCLB.511113.9 | hypothetical protein, conserved (fragment) | -0,661 | 4,56E-11 |
| TcCLB.509167.200 | hypothetical protein, conserved | -0,6613 | 1,663E-09 |
| TcCLB.506835.80 | CAS/CSE/importin domain protein, putative | -0,6618 | 3,136E-13 |
| TcCLB.508325.290 | trans-sialidase (pseudogene), putative | -0,6618 | 3,471E-11 |
| TcCLB.510879.70 | selenocysteine-tRNA-specific elongation factor, putative | -0,6621 | 4,314E-09 |
| TcCLB.511725.210 | hypothetical protein, conserved | -0,6621 | 4,568E-07 |
| TcCLB.506213.30 | hypothetical protein, conserved | -0,6621 | 0,0001721 |
| TcCLB.504625.20 | hypothetical protein, conserved | -0,6624 | 6,228E-06 |
| TcCLB.509665.30 | trans-sialidase, Group V, putative | -0,6624 | 0,06641 |
| TcCLB.510257.90 | autophagocytosis associated protein, putative | -0,6628 | 0,00005302 |
| TcCLB.504431.64 | high mobility group protein, putative | -0,6629 | 1,732E-10 |
| TcCLB.506315.60 | hypothetical protein, conserved | -0,6632 | 0,0000013 |
| TcCLB.508207.5 | retrotransposon hot spot protein (RHS, pseudogene), putat | -0,6633 | 6,952E-10 |
| TcCLB.504103.3 | hypothetical protein, conserved (fragment) | -0,6634 | 0,00000764 |
| TcCLB.508837.180 | DNA polymerase I alpha catalytic subunit, putative | -0,6635 | 1,975E-17 |
| TcCLB.508355.70 | diacylglycerol acyltransferase, putative | -0,6635 | 1,51E-11 |
| TcCLB.508707.160 | hypothetical protein, conserved | -0,6636 | 1,161E-06 |
| TcCLB.510173.20 | hypothetical protein, conserved | -0,6636 | 0,00001164 |
| TcCLB.509967.3 | hypothetical protein, conserved (pseudogene) | -0,6636 | 0,004924 |
| TcCLB.509499.38 | hypothetical protein, conserved | -0,6637 | 7,175E-11 |
| TcCLB.507687.60 | trans-sialidase (pseudogene), putative | -0,6638 | 0,03481 |
| TcCLB.506667.160 | trans-sialidase, Group V, putative | -0,6643 | 0,0544 |
| TcCLB.510755.98 | kinetoplastid membrane protein KMP-11, putative | -0,6645 | 0,0009323 |
| TcCLB.510315.20 | mitochondrial carrier protein, putative | -0,6646 | 4,991E-07 |
| TcCLB.503879.20 | hypothetical protein, conserved | -0,6648 | 3,841E-08 |
| TcCLB.510643.180 | trans-sialidase (pseudogene), putative | -0,6652 | 0,00001435 |
| TcCLB.506155.80 | hypothetical protein, conserved | -0,6654 | 9,238E-11 |
| TcCLB.504103.10 | hypothetical protein, conserved | -0,6654 | 0,0001496 |
| TcCLB.511599.40 | dispersed gene family protein 1 (DGF-1), putative | -0,6654 | 0,0002053 |
| TcCLB.506709.20 | hypothetical protein, conserved | -0,6657 | 3,729E-12 |
| TcCLB.509165.10 | trans-sialidase (pseudogene), putative | -0,6661 | 0,00001559 |
| TcCLB.510475.70 | mucin TcMUCII, putative | -0,6661 | 0,1156 |
| TcCLB.506671.90 | mucin-associated surface protein (MASP), putative | -0,6663 | 0,1693 |
| TcCLB.506989.40 | Golgi vesicular membrane trafficking protein, putative | -0,6665 | 0,001044 |
| TcCLB.511003.130 | hypothetical protein, conserved | -0,6666 | 0,0000277 |
| TcCLB.506767.183 | mucin-associated surface protein (MASP, pseudogene), pu | -0,6667 | 0,005 |
| TcCLB.511301.100 | hypothetical protein, conserved | -0,6668 | 2,069E-08 |
| TcCLB.510357.110 | hypothetical protein, conserved | -0,6668 | 0,00001957 |
| TcCLB.504109.100 | hypothetical protein, conserved | -0,6672 | 8,516E-14 |

| | | | |
|---|---|---|---|
| TcCLB.508221.539 | hypothetical protein, conserved | -0,6673 | 0,22 |
| TcCLB.508479.320 | hypothetical protein, conserved | -0,6675 | 0,0009237 |
| TcCLB.510359.320 | DNA-directed RNA polymerase III subunit, putative | -0,6675 | 0,004164 |
| TcCLB.511139.10 | hypothetical protein, conserved (pseudogene) | -0,6678 | 5,395E-06 |
| TcCLB.506489.10 | hypothetical protein, conserved | -0,6682 | 1,502E-17 |
| TcCLB.511585.180 | hypothetical protein, conserved | -0,6682 | 0,0006014 |
| TcCLB.506253.11 | hypothetical protein, conserved (pseudogene) | -0,6682 | 0,009264 |
| TcCLB.506567.110 | hypothetical protein, conserved | -0,6686 | 1,861E-08 |
| TcCLB.434931.20 | hypothetical protein, conserved | -0,6686 | 9,865E-07 |
| TcCLB.507625.160 | hypothetical protein, conserved | -0,6686 | 0,0004344 |
| TcCLB.510943.198 | hypothetical protein, conserved | -0,6687 | 0,00002825 |
| TcCLB.510999.10 | hypothetical protein, conserved | -0,6688 | 3,012E-09 |
| TcCLB.504827.50 | apurinic/apyrimidinic endonuclease, putative | -0,6688 | 0,00001329 |
| TcCLB.506807.39 | hypothetical protein, conserved (fragment) | -0,6691 | 0,00001523 |
| TcCLB.510609.120 | hypothetical protein, conserved | -0,6692 | 1,275E-10 |
| TcCLB.506239.10 | hypothetical protein, conserved | -0,6695 | 7,788E-06 |
| TcCLB.458759.20 | hypothetical protein, conserved | -0,6695 | 0,0001151 |
| TcCLB.508699.40 | eukaryotic translation initiation factor, putative | -0,6696 | 5,479E-10 |
| TcCLB.507767.10 | succinyl-CoA ligase [GDP-forming] beta-chain, putative | -0,6697 | 1,77E-19 |
| TcCLB.506941.30 | hypothetical protein, conserved | -0,6699 | 7,667E-14 |
| TcCLB.509157.50 | hypothetical protein | -0,6699 | 1,463E-11 |
| TcCLB.509741.20 | hypothetical protein, conserved | -0,6699 | 2,509E-06 |
| TcCLB.504779.10 | UDP-Gal or UDP-GlcNAc-dependent glycosyltransferase, pu | -0,6699 | 0,05122 |
| TcCLB.503811.20 | U3 small nucleolar ribonucleoprotein protein MPP10, putat | -0,6703 | 3,463E-12 |
| TcCLB.510327.5 | surface protease GP63 (pseudogene), putative | -0,6703 | 0,07528 |
| TcCLB.510689.10 | coatomer alpha subunit (pseudogene), putative | -0,6707 | 1,929E-16 |
| TcCLB.506435.20 | hypothetical protein, conserved | -0,6713 | 6,337E-16 |
| TcCLB.510713.101 | surface protease GP63 (pseudogene), putative | -0,6714 | 0,2295 |
| TcCLB.506727.90 | enoyl-CoA hydratase/isomerase family protein, putative | -0,6718 | 4,786E-08 |
| TcCLB.508625.20 | hypothetical protein, conserved | -0,6719 | 0,0003407 |
| TcCLB.510821.10 | hypothetical protein, conserved | -0,6722 | 7,581E-09 |
| TcCLB.418069.10 | heat shock protein 70 (hsp70), putative (fragment) | -0,6723 | 2,734E-10 |
| TcCLB.511871.30 | hypothetical protein, conserved | -0,6728 | 6,289E-11 |
| TcCLB.511003.40 | hypothetical protein, conserved | -0,6729 | 1,27E-11 |
| TcCLB.507713.30 | heat shock protein 85, putative | -0,6729 | 0,05948 |
| TcCLB.463955.10 | mucin TcMUCII, putative | -0,6731 | 0,1116 |
| TcCLB.506579.20 | hypothetical protein, conserved | -0,6732 | 1,743E-09 |
| TcCLB.511367.80 | hypothetical protein, conserved | -0,6732 | 1,838E-06 |
| TcCLB.505555.20 | hypothetical protein, conserved | -0,6733 | 0,00003675 |
| TcCLB.506703.90 | trans-sialidase, Group VI, putative | -0,6733 | 0,06996 |
| TcCLB.506659.30 | glutathione synthetase, putative | -0,6735 | 1,899E-07 |
| TcCLB.509443.10 | hypothetical protein | -0,6739 | 0,0007643 |
| TcCLB.506247.10 | bloodstream- specific protein 2 precursor (fragment) (PDI2 | -0,674 | 1,278E-12 |
| TcCLB.506211.259 | hypothetical protein, conserved | -0,674 | 0,0000959 |
| TcCLB.453445.10 | hypothetical protein, conserved | -0,6744 | 1,997E-13 |

| | | | |
|---|---|---|---|
| TcCLB.506247.74 | hypothetical protein, conserved | -0,6744 | 0,002491 |
| TcCLB.504001.20 | hypothetical protein, conserved | -0,6748 | 9,749E-09 |
| TcCLB.511625.140 | mucin-associated surface protein (MASP, pseudogene), pu | -0,675 | 0,0001668 |
| TcCLB.506435.390 | metallo-peptidase, Clan MA(E) Family M41 | -0,6755 | 1,625E-10 |
| TcCLB.511663.50 | dispersed gene family protein 1 (DGF-1), putative | -0,6756 | 0,1205 |
| TcCLB.508599.10 | hypothetical protein, conserved (pseudogene) | -0,6758 | 0,1496 |
| TcCLB.508909.170 | hypothetical protein, conserved | -0,676 | 2,254E-11 |
| TcCLB.511351.20 | hypothetical protein, conserved | -0,6762 | 2,151E-07 |
| TcCLB.509935.10 | hypothetical protein | -0,6763 | 0,01148 |
| TcCLB.511469.30 | coatomer subunit zeta, putative | -0,6769 | 0,00001934 |
| TcCLB.510327.20 | surface protease GP63 (pseudogene), putative | -0,6778 | 0,01352 |
| TcCLB.505989.82 | ruvB-like DNA helicase, putative | -0,6781 | 3,543E-11 |
| TcCLB.509429.300 | hypothetical protein, conserved | -0,6782 | 1,118E-07 |
| TcCLB.509643.80 | DNA mismatch repair protein MSH2, putative (MSH2) | -0,6785 | 6,905E-15 |
| TcCLB.511731.89 | hypothetical protein, conserved | -0,6786 | 0,00001701 |
| TcCLB.508479.10 | protein kinase, putative | -0,6788 | 0,0333 |
| TcCLB.511545.30 | hypothetical protein, conserved | -0,6789 | 6,795E-09 |
| TcCLB.507521.20 | kinesin, putative | -0,6789 | 0,00001393 |
| TcCLB.507583.40 | hypothetical protein, conserved | -0,6792 | 6,349E-12 |
| TcCLB.511245.200 | hypothetical protein, conserved | -0,6793 | 3,888E-17 |
| TcCLB.507163.80 | hypothetical protein, conserved | -0,6796 | 8,403E-06 |
| TcCLB.509071.10 | hypothetical protein, conserved (pseudogene) | -0,6797 | 0,003524 |
| TcCLB.506457.40 | hypoxanthine-guanine phosphoribosyltransferase, putative | -0,6798 | 9,355E-09 |
| TcCLB.511127.210 | hypothetical protein, conserved | -0,6798 | 0,0004553 |
| TcCLB.510665.30 | hypothetical protein, conserved | -0,6799 | 3,133E-06 |
| TcCLB.508461.320 | PRP8 protein homologue, putative | -0,6803 | 7,8E-27 |
| TcCLB.510629.10 | mucin-associated surface protein (MASP), putative | -0,6805 | 0,268 |
| TcCLB.503939.80 | prolyl-tRNA synthetase, putative | -0,6806 | 1,518E-19 |
| TcCLB.508283.25 | hypothetical protein, conserved (pseudogene) | -0,6809 | 0,01694 |
| TcCLB.510057.10 | hypothetical protein, conserved | -0,681 | 0,009693 |
| TcCLB.507747.210 | hypothetical protein, conserved | -0,6811 | 3,949E-07 |
| TcCLB.510463.211 | trans-sialidase (pseudogene), putative | -0,6811 | 0,0795 |
| TcCLB.510659.257 | phospholipase A2-like protein, putative | -0,6812 | 5,038E-08 |
| TcCLB.506599.130 | mucin TcMUCII (pseudogene), putative | -0,6812 | 0,01072 |
| TcCLB.505843.40 | E1-like ubiquitin-activating enzyme, putative (UBA5) | -0,6813 | 4,989E-13 |
| TcCLB.511481.60 | RNA helicase, putative | -0,6818 | 6,853E-19 |
| TcCLB.510899.30 | hypothetical protein, conserved | -0,6819 | 1,875E-06 |
| TcCLB.511789.10 | dispersed gene family protein 1 (DGF-1, pseudogene), puta | -0,6819 | 0,08482 |
| TcCLB.509437.40 | heat shock protein DNAJ, putative | -0,682 | 1,92E-15 |
| TcCLB.510025.126 | syntaxin binding protein (pseudogene), putative | -0,6827 | 0,0689 |
| TcCLB.503781.40 | hypothetical protein, conserved | -0,6828 | 0,0000844 |
| TcCLB.509829.20 | isovaleryl-coA dehydrogenase, putative | -0,6829 | 0,00001929 |
| TcCLB.510421.100 | hypothetical protein, conserved | -0,6831 | 8,549E-09 |
| TcCLB.506989.30 | ABC transporter, putative | -0,6833 | 2,568E-21 |
| TcCLB.510933.180 | syntaxin binding protein (pseudogene), putative | -0,6835 | 0,1106 |

| | | | |
|---|---|---|---|
| TcCLB.508939.50 | hypothetical protein, conserved | -0,6836 | 0,0008538 |
| TcCLB.508921.40 | hypothetical protein, conserved | -0,6836 | 0,001765 |
| TcCLB.510643.121 | trans-sialidase (pseudogene), putative | -0,6838 | 0,0008421 |
| TcCLB.509429.130 | hypothetical protein, conserved | -0,684 | 9,161E-07 |
| TcCLB.511255.516 | syntaxin binding protein (pseudogene), putative | -0,6841 | 0,09915 |
| TcCLB.507753.3 | hypothetical protein, conserved (pseudogene) | -0,6843 | 0,01221 |
| TcCLB.508365.50 | mucin TcMUCII, putative | -0,6846 | 0,2968 |
| TcCLB.503779.50 | hypothetical protein, conserved | -0,685 | 0,001069 |
| TcCLB.506295.40 | hypothetical protein, conserved | -0,6851 | 1,627E-06 |
| TcCLB.510889.280 | metacyclin II, putative | -0,6852 | 0,163 |
| TcCLB.506517.30 | hypothetical protein | -0,6854 | 1,173E-11 |
| TcCLB.508153.1110 | arginine N-methyltransferase, putative | -0,6856 | 5,042E-09 |
| TcCLB.506883.40 | Lsm12 protein, putative | -0,6858 | 8,602E-09 |
| TcCLB.504239.220 | mucin-associated surface protein (MASP52) | -0,6858 | 0,09228 |
| TcCLB.511801.70 | hypothetical protein, conserved | -0,6863 | 1,3E-08 |
| TcCLB.509669.170 | hypothetical protein, conserved | -0,6864 | 5,427E-12 |
| TcCLB.510685.10 | calreticulin, putative | -0,6867 | 7,869E-21 |
| TcCLB.507945.10 | ATP-dependent RNA helicase, putative | -0,6867 | 1,644E-12 |
| TcCLB.509063.4 | hypothetical protein, conserved | -0,6867 | 0,0005209 |
| TcCLB.506693.4 | elongation initiation factor 2 alpha subunit, putative | -0,687 | 4,133E-08 |
| TcCLB.510769.202 | trans-sialidase (pseudogene), putative | -0,6872 | 0,06882 |
| TcCLB.506631.40 | hypothetical protein, conserved | -0,6875 | 2,911E-11 |
| TcCLB.511495.30 | hypothetical protein, conserved | -0,6877 | 0,00004644 |
| TcCLB.507069.150 | mucin-associated surface protein (MASP), putative | -0,6877 | 0,03706 |
| TcCLB.511409.30 | hypothetical protein, conserved | -0,6878 | 1,203E-11 |
| TcCLB.509287.80 | trans-sialidase, Group VII, putative | -0,6882 | 1,592E-08 |
| TcCLB.511605.30 | mucin TcMUCII, putative | -0,6882 | 0,3348 |
| TcCLB.506447.19 | hypothetical protein, conserved | -0,6886 | 1,818E-08 |
| TcCLB.511215.110 | hypothetical protein, conserved | -0,689 | 4,792E-12 |
| TcCLB.510965.10 | hypothetical protein, conserved | -0,6892 | 2,037E-07 |
| TcCLB.506811.200 | hypothetical protein, conserved | -0,6893 | 8,566E-06 |
| TcCLB.511563.9 | dispersed gene family protein 1 (DGF-1), putative | -0,6893 | 0,0545 |
| TcCLB.509945.20 | hypothetical protein, conserved | -0,6894 | 2,584E-06 |
| TcCLB.504103.30 | myosin heavy chain, putative | -0,6902 | 6,184E-09 |
| TcCLB.508375.70 | hypothetical protein, conserved | -0,6902 | 8,37E-07 |
| TcCLB.510315.10 | IAG-nucleoside hydrolase, putative (IAGNH) | -0,6903 | 3,389E-12 |
| TcCLB.481435.9 | clathrin heavy chain, putative | -0,6904 | 5,441E-11 |
| TcCLB.507611.370 | protein disulfide isomerase 2 (PDI2) | -0,6907 | 1,852E-28 |
| TcCLB.510713.160 | trans-sialidase, Group V, putative | -0,6907 | 0,04013 |
| TcCLB.503805.6 | hypothetical protein, conserved | -0,6908 | 8,327E-06 |
| TcCLB.507785.10 | clathrin heavy chain, putative | -0,6914 | 5,831E-16 |
| TcCLB.506941.20 | tRNA modification enzyme, putative | -0,6915 | 0,00000306 |
| TcCLB.506581.14 | peptidyl-prolyl cis-trans isomerase, putative | -0,6916 | 2,894E-06 |
| TcCLB.506977.40 | hypothetical protein, conserved | -0,6918 | 4,456E-07 |
| TcCLB.509029.160 | hypothetical protein, conserved | -0,692 | 4,337E-10 |

| | | | |
|---|---|---|---|
| TcCLB.503859.10 | syntaxin binding protein (pseudogene), putative | -0,6923 | 0,1882 |
| TcCLB.510491.40 | hypothetical protein, conserved | -0,6925 | 6,409E-06 |
| TcCLB.511099.180 | trans-sialidase (pseudogene), putative | -0,693 | 0,09446 |
| TcCLB.509805.10 | protein kinase A catalytic subunit, putative | -0,6932 | 0,01055 |
| TcCLB.503583.50 | hypothetical protein, conserved | -0,6936 | 1,042E-06 |
| TcCLB.510305.50 | hypothetical protein, conserved | -0,6941 | 1,001E-14 |
| TcCLB.430603.20 | mucin TcMUCII, putative | -0,6942 | 0,078 |
| TcCLB.508143.30 | hypothetical protein, conserved (pseudogene) | -0,6945 | 2,422E-06 |
| TcCLB.511255.270 | trans-sialidase (pseudogene), putative | -0,6945 | 0,05789 |
| TcCLB.507941.120 | hypothetical protein, conserved | -0,6949 | 0,00001542 |
| TcCLB.506975.97 | hypothetical protein, conserved (fragment) | -0,6949 | 0,0372 |
| TcCLB.510885.80 | hypothetical protein, conserved | -0,695 | 8,571E-07 |
| TcCLB.504133.15 | hypothetical protein, conserved (fragment) | -0,695 | 0,001584 |
| TcCLB.509841.20 | hypothetical protein | -0,6952 | 0,01202 |
| TcCLB.510475.60 | hypothetical protein | -0,696 | 0,4182 |
| TcCLB.507943.30 | histone H4, putative | -0,6961 | 2,092E-10 |
| TcCLB.511407.10 | nuclear transport factor 2, putative | -0,6963 | 3,765E-10 |
| TcCLB.506507.40 | trans-sialidase (pseudogene), putative | -0,6965 | 7,388E-08 |
| TcCLB.509733.110 | hypothetical protein, conserved | -0,6967 | 8,111E-16 |
| TcCLB.506977.139 | hypothetical protein, conserved | -0,6968 | 6,507E-09 |
| TcCLB.508293.27 | mucin-associated surface protein (MASP, pseudogene), pu | -0,697 | 0,1484 |
| TcCLB.510681.41 | surface protease GP63, putative | -0,6971 | 0,06888 |
| TcCLB.508175.160 | deoxyuridine triphosphatase, putative | -0,698 | 1,025E-08 |
| TcCLB.510047.60 | mucin-associated surface protein (MASP, pseudogene), pu | -0,698 | 0,002173 |
| TcCLB.506965.110 | hypothetical protein | -0,6981 | 0,1629 |
| TcCLB.506201.100 | hypothetical protein, conserved | -0,6992 | 4,559E-09 |
| TcCLB.511203.10 | surface protease GP63, putative | -0,7 | 0,008643 |
| TcCLB.510963.39 | hypothetical protein, conserved | -0,7002 | 0,00001332 |
| TcCLB.511893.30 | hypothetical protein, conserved (pseudogene) | -0,7004 | 0,00005287 |
| TcCLB.507485.50 | hypothetical protein, conserved | -0,7005 | 5,739E-11 |
| TcCLB.507771.9 | kinesin, putative (fragment) | -0,7008 | 0,004297 |
| TcCLB.509127.50 | mitochondrial carrier protein, putative | -0,701 | 5,486E-11 |
| TcCLB.503891.60 | hypothetical protein, conserved | -0,701 | 1,422E-07 |
| TcCLB.508641.10 | dispersed gene family protein 1 (DGF-1), putative | -0,701 | 0,04172 |
| TcCLB.511871.34 | protein transport protein Sec61 gamma subunit, putative | -0,7011 | 8,437E-07 |
| TcCLB.510483.20 | mucin-associated surface protein (MASP), putative | -0,7013 | 0,01462 |
| TcCLB.507241.10 | protein phosphatase 2A regulatory subunit, putative | -0,7014 | 1,293E-11 |
| TcCLB.511189.50 | hypothetical protein, conserved (pseudogene) | -0,7017 | 2,259E-09 |
| TcCLB.511487.71 | hypothetical protein | -0,7017 | 3,454E-07 |
| TcCLB.511717.174 | hypothetical protein, conserved | -0,702 | 1,614E-08 |
| TcCLB.511321.30 | hypothetical protein, conserved | -0,7023 | 0,00005693 |
| TcCLB.506357.130 | hypothetical protein, conserved | -0,7026 | 5,071E-26 |
| TcCLB.508221.600 | mucin-associated surface protein (MASP), putative | -0,7027 | 0,1794 |
| TcCLB.503953.50 | hypothetical protein, conserved | -0,7028 | 2,187E-12 |
| TcCLB.507809.100 | hypothetical protein, conserved | -0,7029 | 7,051E-20 |

| | | | |
|---|---|---|---|
| TcCLB.506759.90 | trans-sialidase (pseudogene), putative | -0,7032 | 0,0549 |
| TcCLB.510141.10 | thymine-7-hydroxylase, putative | -0,7033 | 0,02031 |
| TcCLB.510409.60 | hypothetical protein, conserved | -0,7034 | 8,88E-10 |
| TcCLB.510819.70 | hypothetical protein, conserved | -0,7035 | 2,703E-07 |
| TcCLB.510307.352 | retrotransposon hot spot protein (RHS, pseudogene), putat | -0,7036 | 0,00004825 |
| TcCLB.506665.20 | hypothetical protein | -0,7043 | 0,01041 |
| TcCLB.508511.15 | trans-sialidase (pseudogene), putative | -0,7048 | 0,3417 |
| TcCLB.509715.40 | fibrillarin, putative | -0,7049 | 6,452E-10 |
| TcCLB.506443.130 | trans-sialidase (pseudogene), putative | -0,7049 | 9,22E-10 |
| TcCLB.508387.10 | hypothetical protein, conserved | -0,7049 | 0,0003849 |
| TcCLB.504033.120 | hypothetical protein, conserved | -0,705 | 1,365E-12 |
| TcCLB.511025.70 | chaperonin alpha subunit, putative | -0,7052 | 2,459E-24 |
| TcCLB.508357.90 | T-complex protein 1, beta subunit, putative | -0,7053 | 4,621E-24 |
| TcCLB.504059.11 | hypothetical protein, conserved (fragment) | -0,7055 | 0,00003387 |
| TcCLB.504005.40 | hypothetical protein, conserved | -0,7061 | 0,00000617 |
| TcCLB.507237.150 | hypothetical protein | -0,7061 | 0,1983 |
| TcCLB.508167.10 | mucin-associated surface protein (MASP, pseudogene), pu | -0,7064 | 0,007936 |
| TcCLB.508879.130 | telomerase component (pseudogene), putative | -0,7067 | 4,821E-27 |
| TcCLB.510439.10 | heat shock protein 70, putative | -0,7074 | 3,429E-07 |
| TcCLB.504049.30 | trans-sialidase (pseudogene), putative | -0,7076 | 0,02916 |
| TcCLB.510553.50 | UDP-Gal or UDP-GlcNAc-dependent glycosyltransferase, pu | -0,7078 | 0,02819 |
| TcCLB.507739.110 | myosin IB heavy chain, putative | -0,7079 | 4,34E-19 |
| TcCLB.511737.10 | small glutamine-rich tetratricopeptide repeat protein, puta | -0,708 | 3,477E-13 |
| TcCLB.506205.49 | hypothetical protein, conserved | -0,7084 | 0,07201 |
| TcCLB.510389.10 | hypothetical protein, conserved (pseudogene) | -0,7085 | 0,0007044 |
| TcCLB.507011.250 | hypothetical protein, conserved | -0,7093 | 8,108E-21 |
| TcCLB.504081.90 | mucin-associated surface protein (MASP), putative | -0,7096 | 0,02702 |
| TcCLB.509233.210 | hypothetical protein, conserved | -0,7097 | 6,945E-11 |
| TcCLB.511445.30 | hypothetical protein, conserved | -0,7105 | 2,193E-10 |
| TcCLB.420091.10 | dispersed gene family protein 1 (DGF-1), putative | -0,7106 | 0,003227 |
| TcCLB.511611.5 | mucin TcMUCII (pseudogene), putative | -0,7107 | 0,2576 |
| TcCLB.506925.250 | hypothetical protein, conserved (pseudogene) | -0,7109 | 1,045E-07 |
| TcCLB.506147.130 | hypothetical protein, conserved | -0,711 | 0,001476 |
| TcCLB.503989.10 | zinc finger protein 2, putative | -0,7111 | 3,447E-10 |
| TcCLB.511021.80 | hypothetical protein, conserved | -0,7116 | 3,23E-11 |
| TcCLB.510421.10 | surface protease GP63 (pseudogene), putative | -0,712 | 0,01817 |
| TcCLB.506859.180 | hypothetical protein, conserved | -0,7121 | 0,0006865 |
| TcCLB.509911.124 | hypothetical protein, conserved | -0,7122 | 3,427E-06 |
| TcCLB.511553.71 | syntaxin binding protein (pseudogene), putative | -0,7122 | 0,05101 |
| TcCLB.510101.470 | PUF nine target 1 | -0,7123 | 1,057E-19 |
| TcCLB.508153.1100 | mitochondrial protein 99, putative (MP99) | -0,7123 | 1,36E-11 |
| TcCLB.507993.390 | pyruvate kinase 2, putative | -0,7125 | 1,699E-09 |
| TcCLB.401473.9 | DNA-directed rna polymerase I largest subunit, putative | -0,7127 | 0,00000985 |
| TcCLB.510397.10 | hypothetical protein, conserved | -0,7131 | 3,403E-12 |
| TcCLB.482471.19 | ATP-dependent RNA helicase SUB2, putative (fragment) (S | -0,7131 | 4,698E-09 |

| | | | |
|---|---|---|---|
| TcCLB.511255.625 | surface protease GP63 (pseudogene), putative | -0,7132 | 0,2401 |
| TcCLB.507041.60 | syntaxin binding protein (pseudogene), putative | -0,7133 | 0,0007382 |
| TcCLB.510269.55 | elongation factor 1-gamma (EF-1-gamma, pseudogene), p | -0,7133 | 0,1365 |
| TcCLB.508637.30 | hypothetical protein, conserved (pseudogene) | -0,7143 | 0,01033 |
| TcCLB.507881.60 | protein kinase, putative | -0,7144 | 0,001339 |
| TcCLB.507603.240 | hypothetical protein, conserved | -0,7149 | 3,249E-09 |
| TcCLB.511287.34 | hypothetical protein, conserved | -0,715 | 1,613E-08 |
| TcCLB.511139.40 | ATP-dependent RNA helicase SUB2, putative (fragment) (S | -0,7152 | 0,00000601 |
| TcCLB.507639.110 | hypothetical protein, conserved | -0,7154 | 1,376E-09 |
| TcCLB.506559.20 | hypothetical protein, conserved | -0,7156 | 5,125E-14 |
| TcCLB.503775.60 | hypothetical protein | -0,7159 | 0,004408 |
| TcCLB.503707.9 | dispersed gene family protein 1 (DGF-1, pseudogene), puta | -0,7162 | 0,0001602 |
| TcCLB.511717.10 | hypothetical protein, conserved (pseudogene) | -0,7162 | 0,005804 |
| TcCLB.508995.10 | protein kinase, putative | -0,7168 | 0,00000783 |
| TcCLB.511453.10 | dispersed gene family protein 1 (DGF-1), putative | -0,7169 | 0,006513 |
| TcCLB.510149.100 | hypothetical protein, conserved | -0,717 | 5,298E-07 |
| TcCLB.508507.9 | hypothetical protein, conserved | -0,7172 | 0,00004054 |
| TcCLB.510817.50 | trans-sialidase, Group II, putative | -0,7172 | 0,00005868 |
| TcCLB.511731.50 | ATP-dependent RNA helicase, putative | -0,7173 | 7,622E-10 |
| TcCLB.506181.150 | hypothetical protein (fragment) | -0,7175 | 7,023E-16 |
| TcCLB.508717.20 | NADH dehydrogenase, putative | -0,7176 | 2,769E-13 |
| TcCLB.510017.30 | mucin-associated surface protein (MASP, pseudogene), pu | -0,7176 | 0,09905 |
| TcCLB.503581.10 | hypothetical protein, conserved | -0,7179 | 1,481E-13 |
| TcCLB.508167.29 | mucin-associated surface protein (MASP), putative (fragm | -0,7183 | 0,08375 |
| TcCLB.508169.9 | elongation factor 2, putative (fragment) | -0,7184 | 8,156E-17 |
| TcCLB.508153.630 | mitochondrial carrier protein, putative | -0,7184 | 7,225E-06 |
| TcCLB.509419.245 | hypothetical protein, conserved (pseudogene) | -0,7186 | 0,004491 |
| TcCLB.506623.60 | dispersed gene family protein 1 (DGF-1), putative | -0,7188 | 0,01221 |
| TcCLB.507603.70 | kinetoplastid kinetochore protein 6, putative (kkt6) | -0,7189 | 0,0001964 |
| TcCLB.505229.20 | conserved protein, unknown function | -0,7191 | 2,545E-16 |
| TcCLB.509937.80 | hypothetical protein, conserved | -0,7191 | 1,021E-08 |
| TcCLB.510401.31 | retrotransposon hot spot protein (RHS, pseudogene), putat | -0,7191 | 0,1864 |
| TcCLB.511531.9 | chaperone DNAJ protein, putative | -0,7193 | 1,246E-06 |
| TcCLB.511089.15 | hypothetical protein | -0,7193 | 0,09275 |
| TcCLB.510877.40 | hypothetical protein, conserved | -0,7195 | 1,574E-18 |
| TcCLB.409139.9 | trans-sialidase, putative | -0,7195 | 0,1565 |
| TcCLB.507093.170 | hypothetical protein, conserved | -0,7196 | 5,371E-22 |
| TcCLB.506437.30 | amastin, putative | -0,7198 | 9,675E-10 |
| TcCLB.511751.120 | aldehyde dehydrogenase family, putative | -0,7201 | 9,214E-08 |
| TcCLB.510725.50 | hypothetical protein, conserved | -0,7205 | 2,991E-13 |
| TcCLB.506203.10 | nucleoside transporter 1, putative | -0,7212 | 6,463E-09 |
| TcCLB.506357.90 | DNA-directed RNA polymerase II/III subunit, putative | -0,7212 | 6,532E-08 |
| TcCLB.509941.120 | protein kinase, putative | -0,7213 | 8,448E-10 |
| TcCLB.506601.31 | surface protease GP63 (pseudogene), putative | -0,7213 | 0,2179 |
| TcCLB.506673.50 | apurinic/apyrimidinic endonuclease, putative | -0,7214 | 0,00006118 |

| | | | |
|---|---|---|---|
| TcCLB.509171.60 | hypothetical protein, conserved | -0,722 | 1,733E-09 |
| TcCLB.506917.60 | mucin TcMUCII, putative | -0,722 | 0,1616 |
| TcCLB.508781.129 | arginine N-methyltransferase, type II, putative | -0,7223 | 0,00001274 |
| TcCLB.509611.40 | hypothetical protein, conserved | -0,7223 | 0,01334 |
| TcCLB.506247.120 | hypothetical protein, conserved | -0,7225 | 2,061E-14 |
| TcCLB.509611.100 | lipoate-protein ligase, putative | -0,7225 | 1,797E-06 |
| TcCLB.510287.60 | ribonuclease HII, putative | -0,7226 | 4,431E-08 |
| TcCLB.407335.18 | hypothetical protein, conserved | -0,7227 | 4,134E-06 |
| TcCLB.504257.70 | valyl-tRNA synthetase, putative | -0,7229 | 3,961E-18 |
| TcCLB.503727.29 | hypothetical protein | -0,7229 | 1,501E-09 |
| TcCLB.507971.29 | hypothetical protein, conserved | -0,7229 | 0,00006705 |
| TcCLB.503451.9 | hypothetical protein, conserved (fragment) | -0,723 | 2,668E-19 |
| TcCLB.506147.120 | hypothetical protein, conserved | -0,7232 | 6,099E-08 |
| TcCLB.506411.10 | thimet oligopeptidase, putative | -0,7235 | 1,012E-16 |
| TcCLB.506315.20 | hypothetical protein, conserved | -0,7236 | 1,074E-08 |
| TcCLB.433717.10 | dispersed gene family protein 1 (DGF-1, pseudogene), puta | -0,7236 | 0,01265 |
| TcCLB.511255.190 | trans-sialidase, Group V, putative | -0,7238 | 0,02401 |
| TcCLB.504201.50 | mucin-associated surface protein (MASP), putative | -0,7241 | 0,05794 |
| TcCLB.506729.90 | hypothetical protein, conserved | -0,7242 | 5,471E-11 |
| TcCLB.508153.680 | RNA-binding protein 6, putative | -0,7243 | 0,01075 |
| TcCLB.506925.390 | MP44, putative | -0,7244 | 1,095E-08 |
| TcCLB.508637.114 | histone deacetylase 1, putative | -0,725 | 0,00004285 |
| TcCLB.505765.10 | hypothetical protein, conserved (pseudogene) | -0,7259 | 0,00003835 |
| TcCLB.508215.6 | hypothetical protein, conserved | -0,7264 | 5,95E-10 |
| TcCLB.508577.40 | dispersed gene family protein 1 (DGF-1), putative | -0,7264 | 6,339E-06 |
| TcCLB.510497.40 | trans-sialidase, Group V, putative | -0,7265 | 0,033 |
| TcCLB.511467.70 | hypothetical protein, conserved | -0,7266 | 0,0003964 |
| TcCLB.509525.100 | trans-sialidase (pseudogene), putative | -0,7271 | 0,006051 |
| TcCLB.508647.10 | hypothetical protein, conserved | -0,7273 | 7,114E-12 |
| TcCLB.510421.270 | hypothetical protein, conserved | -0,7273 | 0,0002036 |
| TcCLB.508165.200 | trans-sialidase (pseudogene), putative | -0,7273 | 0,06516 |
| TcCLB.510187.270 | transmembrane emp24 domain-containing protein | -0,7274 | 2,981E-09 |
| TcCLB.508681.40 | dispersed gene family protein 1 (DGF-1), putative | -0,7275 | 0,1287 |
| TcCLB.507625.90 | hypothetical protein, conserved | -0,7278 | 6,166E-06 |
| TcCLB.503641.10 | hypothetical protein, conserved | -0,7281 | 3,093E-11 |
| TcCLB.508813.40 | surface protease GP63, putative | -0,7281 | 0,00002209 |
| TcCLB.506443.30 | hypothetical protein, conserved | -0,7283 | 2,461E-10 |
| TcCLB.507927.70 | hypothetical protein, conserved | -0,7283 | 0,0000186 |
| TcCLB.506865.20 | hypothetical protein | -0,7284 | 2,134E-08 |
| TcCLB.509233.80 | heat shock protein DNAJ, putative | -0,7287 | 1,799E-12 |
| TcCLB.510545.73 | syntaxin binding protein (pseudogene), putative | -0,729 | 0,01446 |
| TcCLB.503931.14 | hypothetical protein, conserved | -0,7299 | 1,424E-08 |
| TcCLB.510043.20 | hypothetical protein, conserved (pseudogene) | -0,73 | 4,732E-18 |
| TcCLB.506579.80 | hypothetical protein, conserved | -0,7305 | 1,215E-09 |
| TcCLB.507951.140 | U2 small nuclear ribonucleoprotein B, putative | -0,7311 | 0,001467 |

| | | | |
|---|---|---|---|
| TcCLB.505937.29 | dispersed gene family protein 1 (DGF-1), putative | -0,7312 | 0,01787 |
| TcCLB.510013.200 | 90 kDa surface protein, putative | -0,7315 | 0,03267 |
| TcCLB.510359.90 | hypothetical protein, conserved | -0,7318 | 3,959E-06 |
| TcCLB.503603.20 | hypothetical protein, conserved | -0,7323 | 2,857E-09 |
| TcCLB.506831.20 | amino acid permease, putative | -0,7324 | 9,873E-07 |
| TcCLB.511439.60 | hypothetical protein, conserved | -0,7324 | 0,00000252 |
| TcCLB.510297.60 | zinc transporter, putative | -0,7325 | 2,36E-09 |
| TcCLB.511911.150 | nucleotide-binding protein, putative | -0,7326 | 4,225E-06 |
| TcCLB.508625.40 | hypothetical protein, conserved | -0,7327 | 0,00001591 |
| TcCLB.511529.120 | hypothetical protein, conserved | -0,7329 | 3,934E-06 |
| TcCLB.511725.250 | chaperonin containing t-complex protein, putative | -0,733 | 1,34E-36 |
| TcCLB.503687.40 | 40 kDa cyclophilin, putative (CYP40) | -0,7339 | 7,603E-18 |
| TcCLB.508277.120 | ribosomal protein S6, putative | -0,734 | 8,832E-09 |
| TcCLB.450155.9 | hypothetical protein, conserved | -0,7341 | 0,00001314 |
| TcCLB.509213.70 | ribulose-5-phosphate 3-epimerase, putative | -0,7342 | 5,066E-08 |
| TcCLB.506321.340 | hypothetical protein, conserved | -0,7342 | 1,514E-07 |
| TcCLB.508961.20 | hypothetical protein, conserved | -0,7349 | 1,057E-06 |
| TcCLB.508231.60 | ubiquitin-protein ligase, putative | -0,7351 | 3,446E-29 |
| TcCLB.506501.341 | surface protease GP63 (pseudogene), putative | -0,7351 | 0,1733 |
| TcCLB.505555.50 | hypothetical protein | -0,7352 | 1,236E-20 |
| TcCLB.510515.60 | cytosolic leucyl aminopeptidase, putative | -0,7352 | 1,888E-11 |
| TcCLB.508187.30 | surface protease GP63, putative (fragment) | -0,7352 | 0,04075 |
| TcCLB.510969.40 | hypothetical protein, conserved | -0,7353 | 3,997E-10 |
| TcCLB.506999.10 | hypothetical protein, conserved | -0,7359 | 5,802E-07 |
| TcCLB.473703.10 | surface protease GP63, putative | -0,7359 | 0,1882 |
| TcCLB.508461.150 | hypothetical protein, conserved | -0,736 | 1,904E-07 |
| TcCLB.508047.25 | elongation factor 1-gamma (EF-1-gamma, pseudogene), p | -0,7362 | 0,1725 |
| TcCLB.510421.250 | hypothetical protein, conserved | -0,7368 | 2,754E-14 |
| TcCLB.511533.5 | mitogen activated protein kinase (pseudogene), putative | -0,7368 | 0,07378 |
| TcCLB.510249.14 | hypothetical protein, conserved | -0,7368 | 0,2383 |
| TcCLB.507091.80 | mucin-associated surface protein (MASP), putative | -0,7369 | 0,2043 |
| TcCLB.504155.221 | surface protease GP63 (pseudogene), putative | -0,7369 | 0,3469 |
| TcCLB.510757.9 | hypothetical protein, conserved (fragment) | -0,7375 | 2,51E-10 |
| TcCLB.508725.10 | CCR4-NOT transcription complex subunit 10, putative | -0,7377 | 4,69E-13 |
| TcCLB.510323.4 | katanin, putative | -0,7379 | 1,571E-11 |
| TcCLB.508919.140 | hypothetical protein, conserved | -0,738 | 1,117E-25 |
| TcCLB.510279.349 | mucin-associated surface protein (MASP), putative | -0,7381 | 0,0736 |
| TcCLB.510817.21 | hypothetical protein | -0,7385 | 0,0004402 |
| TcCLB.504233.50 | hypothetical protein, conserved | -0,7385 | 0,2501 |
| TcCLB.511587.120 | Octanoyltransferase, putative | -0,7391 | 0,007346 |
| TcCLB.506321.200 | mucin-associated surface protein (MASP), putative | -0,7393 | 0,07751 |
| TcCLB.506933.98 | dual specificity protein phosphatase, putative | -0,7394 | 8,011E-10 |
| TcCLB.510053.40 | hypothetical protein, conserved | -0,7394 | 0,00001523 |
| TcCLB.506925.300 | cyclophilin a, putative (CYPA) | -0,7399 | 3,412E-11 |
| TcCLB.507515.40 | hypothetical protein, conserved | -0,7401 | 2,218E-06 |

| | | | |
|---|---|---|---|
| TcCLB.506749.10 | flagellar calcium-binding-like protein, putative | -0,7402 | 1,146E-15 |
| TcCLB.508607.70 | trans-sialidase (pseudogene), putative | -0,7403 | 1,383E-06 |
| TcCLB.510223.5 | guanine nucleotide exchange factor subunit GCD2 | -0,7406 | 0,05473 |
| TcCLB.510681.20 | myosin heavy chain kinase A, putative | -0,7407 | 6,394E-08 |
| TcCLB.504109.140 | DNA-directed RNA polymerase I subunit, putative | -0,7409 | 1,379E-07 |
| TcCLB.508175.60 | hypothetical protein, conserved | -0,7412 | 4,855E-07 |
| TcCLB.503917.14 | hypothetical protein, conserved | -0,7415 | 4,892E-09 |
| TcCLB.510629.200 | retrotransposon hot spot protein (RHS, pseudogene), putat | -0,7415 | 0,005903 |
| TcCLB.504171.50 | hypothetical protein, conserved | -0,7416 | 5,107E-22 |
| TcCLB.506341.30 | beta galactofuranosyl glycosyltransferase, putative | -0,7418 | 2,072E-10 |
| TcCLB.507611.180 | hypothetical protein | -0,742 | 0,02819 |
| TcCLB.504181.40 | cdc2-related kinase 1, putative (TzCRK1) | -0,7426 | 1,516E-09 |
| TcCLB.507583.20 | nucleolar protein, putative | -0,7431 | 4,063E-09 |
| TcCLB.506855.290 | hypothetical protein, conserved | -0,7435 | 5,067E-08 |
| TcCLB.506469.90 | hypothetical protein, conserved | -0,7437 | 0,000432 |
| TcCLB.507829.19 | Vacuolar protein sorting-associated protein 16 homolog, p | -0,7439 | 0,00007882 |
| TcCLB.509803.50 | hypothetical protein, conserved | -0,744 | 2,305E-12 |
| TcCLB.507459.10 | hypothetical protein, conserved (pseudogene) | -0,744 | 0,02756 |
| TcCLB.506321.290 | stress-induced protein sti1, putative | -0,7441 | 2,743E-25 |
| TcCLB.432067.10 | ATP-dependent RNA helicase, putative | -0,7441 | 0,00002023 |
| TcCLB.484299.10 | trypanothione reductase, putative | -0,7442 | 2,774E-09 |
| TcCLB.510279.230 | trans-sialidase, Group VI, putative | -0,7444 | 0,05734 |
| TcCLB.508813.15 | heat shock protein 70, putative (fragment) | -0,7445 | 0,00001392 |
| TcCLB.511287.150 | fibrillarin, putative | -0,7449 | 4,466E-10 |
| TcCLB.510359.170 | AAA ATPase, putative | -0,7453 | 1,805E-24 |
| TcCLB.508897.110 | macrophage infectivity potentiator, precursor, putative (M | -0,7456 | 1,233E-12 |
| TcCLB.508163.190 | dispersed gene family protein 1 (DGF-1), putative | -0,7456 | 0,00314 |
| TcCLB.509647.40 | hypothetical protein, conserved | -0,7459 | 1,462E-08 |
| TcCLB.506625.160 | hypothetical protein, conserved | -0,746 | 3,296E-23 |
| TcCLB.508013.40 | trans-sialidase, Group VI, putative | -0,746 | 0,006766 |
| TcCLB.510533.10 | Cytoplasmic dynein 1 heavy chain (DYNC1H1), putative | -0,7462 | 1,657E-16 |
| TcCLB.430727.9 | inositol phosphatidylinositol phosphatase, putative | -0,7472 | 0,00000318 |
| TcCLB.510595.20 | hypothetical protein, conserved | -0,7474 | 5,774E-08 |
| TcCLB.511635.20 | histone H2B, putative | -0,7477 | 4,277E-19 |
| TcCLB.509795.80 | hypothetical protein, conserved | -0,7477 | 1,234E-13 |
| TcCLB.503735.40 | hypothetical protein | -0,7478 | 4,775E-08 |
| TcCLB.509525.180 | mucin TcMUCII, putative | -0,7478 | 0,1547 |
| TcCLB.503717.90 | surface protease GP63, putative | -0,7479 | 0,1101 |
| TcCLB.509045.10 | hypothetical protein, conserved | -0,7484 | 6,434E-06 |
| TcCLB.505035.20 | DNA-directed RNA polymerase II subunit 2 (pseudogene), ı | -0,7489 | 1,146E-07 |
| TcCLB.487507.10 | tryparedoxin peroxidase, putative | -0,7492 | 3,868E-07 |
| TcCLB.504703.20 | hypothetical protein, conserved | -0,7499 | 2,58E-15 |
| TcCLB.506393.189 | retrotransposon hot spot (RHS) protein, putative (fragmen | -0,7499 | 0,0005215 |
| TcCLB.511381.19 | hypothetical protein, conserved (fragment) | -0,7502 | 1,1E-11 |
| TcCLB.506609.30 | trans-sialidase, Group V, putative | -0,7502 | 0,2552 |

| | | | |
|---|---|---|---|
| TcCLB.504109.20 | hypothetical protein, conserved | -0,7503 | 1,303E-15 |
| TcCLB.507007.70 | hypothetical protein, conserved | -0,7503 | 2,107E-11 |
| TcCLB.511067.14 | hypothetical protein, conserved | -0,7511 | 0,00001331 |
| TcCLB.505997.31 | ATP-dependent DEAD/H RNA helicase (pseudogene), putat | -0,7513 | 0,01477 |
| TcCLB.511025.100 | chaperone DNAJ protein, putative | -0,7515 | 5,431E-14 |
| TcCLB.510323.80 | hypothetical protein, conserved | -0,7519 | 2,07E-14 |
| TcCLB.506165.16 | protein kinase, putative | -0,7519 | 0,00571 |
| TcCLB.510763.60 | hypothetical protein, conserved | -0,7522 | 1,499E-06 |
| TcCLB.507649.20 | hypothetical protein, conserved | -0,7526 | 1,262E-22 |
| TcCLB.510443.9 | valyl-tRNA synthetase, putative | -0,7528 | 1,339E-18 |
| TcCLB.507801.160 | hypothetical protein, conserved | -0,7528 | 2,492E-08 |
| TcCLB.510187.510 | hypothetical protein, conserved | -0,753 | 7,63E-10 |
| TcCLB.482369.20 | hypothetical protein, conserved | -0,7531 | 1,637E-20 |
| TcCLB.509471.95 | histone H3, putative | -0,7531 | 5,986E-08 |
| TcCLB.508535.40 | hypothetical protein, conserved | -0,7532 | 0,0001494 |
| TcCLB.510933.10 | trans-sialidase, Group V, putative | -0,7532 | 0,01083 |
| TcCLB.509951.10 | nucleolar protein, putative | -0,7534 | 0,00001379 |
| TcCLB.508221.460 | mucin-associated surface protein (MASP, pseudogene), pu | -0,7537 | 0,03898 |
| TcCLB.506529.50 | arginine N-methyltransferase, putative | -0,7539 | 7,688E-12 |
| TcCLB.507941.100 | hypothetical protein, conserved | -0,7541 | 3,541E-10 |
| TcCLB.507093.66 | hypothetical protein, conserved | -0,7543 | 7,685E-09 |
| TcCLB.507735.20 | hypothetical protein, conserved | -0,7546 | 1,144E-21 |
| TcCLB.510319.9 | hypothetical protein, conserved | -0,7546 | 3,719E-08 |
| TcCLB.504039.57 | mucin-associated surface protein (MASP, pseudogene), pu | -0,7549 | 0,3509 |
| TcCLB.510539.10 | mucin-associated surface protein (MASP), putative | -0,7551 | 0,3586 |
| TcCLB.503515.7 | ribosomal RNA methyltransferase, putative (fragment) | -0,7553 | 0,00006569 |
| TcCLB.506575.20 | hypothetical protein, conserved | -0,7554 | 0,00358 |
| TcCLB.506947.50 | hypothetical protein, conserved | -0,7555 | 1,544E-08 |
| TcCLB.508257.230 | target of rapamycin kinase (TOR) kinase 3, putative | -0,7556 | 2,441E-29 |
| TcCLB.510819.110 | ATP-dependent RNA helicase, putative | -0,7557 | 1,318E-16 |
| TcCLB.507201.9 | dispersed gene family protein 1 (DGF-1), putative | -0,7566 | 1,877E-06 |
| TcCLB.506885.350 | proteasome alpha 1 subunit, putative | -0,7568 | 1,988E-13 |
| TcCLB.508693.150 | hypothetical protein, conserved | -0,7569 | 6,577E-10 |
| TcCLB.509993.5 | hypothetical protein, conserved (pseudogene) | -0,7569 | 3,222E-06 |
| TcCLB.508153.160 | hypothetical protein, conserved | -0,7572 | 1,997E-06 |
| TcCLB.508661.60 | GPI transamidase component Tta2, putative (TTA2) | -0,7576 | 3,309E-12 |
| TcCLB.508365.171 | elongation factor 1-gamma (EF-1-gamma, pseudogene), p | -0,7576 | 0,07033 |
| TcCLB.503999.30 | hypothetical protein, conserved | -0,7579 | 5,459E-22 |
| TcCLB.504013.20 | hypothetical protein | -0,758 | 1,167E-06 |
| TcCLB.505567.31 | mucin-associated surface protein (MASP, pseudogene), pu | -0,7582 | 0,3597 |
| TcCLB.511149.180 | trans-sialidase, Group VI, putative | -0,7586 | 0,008479 |
| TcCLB.504433.10 | ADP-ribosylation factor-like protein 3, putative (arl3) | -0,7593 | 8,188E-09 |
| TcCLB.507867.70 | prefoldin subunit, putative | -0,7593 | 0,00001804 |
| TcCLB.508221.360 | mucin-associated surface protein (MASP, pseudogene), pu | -0,7595 | 0,144 |
| TcCLB.504839.50 | nucleosome assembly protein-like protein, putative | -0,7597 | 2,799E-17 |

| | | | |
|---|---|---|---|
| TcCLB.510293.79 | GPI transamidase component Tta2, putative (fragment) (T | -0,7597 | 5,638E-08 |
| TcCLB.464415.10 | hypothetical protein | -0,7598 | 0,05633 |
| TcCLB.510901.130 | hypothetical protein, conserved | -0,76 | 1,161E-08 |
| TcCLB.506963.80 | hypothetical protein, conserved | -0,7601 | 4,997E-18 |
| TcCLB.511557.60 | hypothetical protein, conserved | -0,7604 | 0,00002245 |
| TcCLB.509583.10 | chaperonin TCP20, putative | -0,7606 | 3,447E-21 |
| TcCLB.510617.19 | hypothetical protein, conserved | -0,7607 | 4,47E-08 |
| TcCLB.506943.90 | hypothetical protein, conserved | -0,7607 | 0,0000232 |
| TcCLB.506769.10 | hypothetical protein | -0,7607 | 0,1427 |
| TcCLB.509233.140 | aldehyde dehydrogenase (pseudogene), putative | -0,761 | 0,004773 |
| TcCLB.511633.10 | hypothetical protein, conserved | -0,7614 | 0,1471 |
| TcCLB.508637.63 | hypothetical protein, conserved | -0,7615 | 3,06E-14 |
| TcCLB.508737.194 | ARP2/3 complex subunit, putative | -0,7616 | 3,519E-07 |
| TcCLB.510491.50 | hypothetical protein, conserved | -0,7617 | 0,001037 |
| TcCLB.511233.70 | hypothetical protein, conserved | -0,7625 | 3,941E-20 |
| TcCLB.506335.110 | trans-sialidase (pseudogene), putative | -0,7625 | 0,0009801 |
| TcCLB.506309.130 | trans-sialidase (pseudogene), putative | -0,7629 | 0,2234 |
| TcCLB.508181.149 | hypothetical protein, conserved | -0,7632 | 0,002158 |
| TcCLB.510655.90 | Importin 1 (IMP1) | -0,7633 | 9,218E-30 |
| TcCLB.508839.40 | hypothetical protein, conserved | -0,7641 | 8,958E-08 |
| TcCLB.510187.320 | hypothetical protein | -0,7643 | 1,125E-11 |
| TcCLB.504247.20 | ATP-dependent RNA helicase, putative | -0,7645 | 1,668E-17 |
| TcCLB.506107.20 | dihydrouridine synthase (Dus), putative | -0,7645 | 0,0002022 |
| TcCLB.508253.20 | trans-sialidase (pseudogene), putative | -0,7648 | 0,05477 |
| TcCLB.510759.190 | hypothetical protein, conserved | -0,7649 | 7,491E-15 |
| TcCLB.509591.50 | RNA-binding protein 34, putative (RBP34) | -0,7656 | 1,815E-10 |
| TcCLB.507609.10 | nucleoporin interacting component (NUP93), putative | -0,7659 | 5,139E-09 |
| TcCLB.506275.9 | RNA polymerase-associated protein CDC73, putative | -0,7659 | 1,328E-06 |
| TcCLB.511759.20 | trans-sialidase (pseudogene), putative | -0,7665 | 1,493E-09 |
| TcCLB.507023.260 | hypothetical protein, conserved | -0,767 | 1,035E-10 |
| TcCLB.509891.50 | hypothetical protein, conserved | -0,7672 | 7,853E-06 |
| TcCLB.510221.20 | hypothetical protein, conserved | -0,7677 | 1,913E-09 |
| TcCLB.507259.16 | protein kinase, putative (fragment) | -0,7679 | 0,08998 |
| TcCLB.510269.73 | hypothetical protein, conserved (pseudogene) | -0,7681 | 0,0004412 |
| TcCLB.511437.10 | surface protease GP63, putative | -0,7686 | 0,00088 |
| TcCLB.508847.9 | dispersed gene family protein 1 (DGF-1), putative | -0,7686 | 0,1017 |
| TcCLB.511339.20 | C-5 sterol desaturase, putative | -0,7687 | 1,044E-08 |
| TcCLB.511283.184 | hypothetical protein, conserved | -0,7687 | 1,053E-06 |
| TcCLB.503803.4 | serine/threonine protein phosphatase 2A regulatory subun | -0,7687 | 0,00009309 |
| TcCLB.505999.4 | hypothetical protein, conserved | -0,7688 | 1,794E-06 |
| TcCLB.510771.9 | hypothetical protein, conserved (fragment) | -0,769 | 2,649E-21 |
| TcCLB.508159.40 | dispersed gene family protein 1 (DGF-1), putative | -0,7695 | 0,002272 |
| TcCLB.446067.9 | hypothetical protein, conserved (fragment) | -0,7696 | 7,628E-10 |
| TcCLB.466593.19 | hypothetical protein, conserved | -0,7706 | 1,62E-09 |
| TcCLB.509555.10 | hypothetical protein (fragment) | -0,7708 | 5,254E-10 |

| | | | |
|---|---|---|---|
| TcCLB.510499.20 | hypothetical protein | -0,7708 | 0,0302 |
| TcCLB.508075.30 | hypothetical protein, conserved (pseudogene) | -0,7709 | 5,876E-08 |
| TcCLB.504215.30 | hypothetical protein, conserved | -0,7712 | 2,605E-09 |
| TcCLB.510387.30 | surface protease GP63 (pseudogene), putative | -0,7713 | 0,05435 |
| TcCLB.511875.5 | elongation factor 1-gamma (EF-1-gamma, pseudogene), p | -0,7714 | 0,03142 |
| TcCLB.508139.20 | dispersed gene family protein 1 (DGF-1), putative | -0,7715 | 7,22E-09 |
| TcCLB.508007.10 | trans-sialidase, putative | -0,7723 | 0,4989 |
| TcCLB.508813.30 | surface protease GP63 (pseudogene), putative | -0,7726 | 0,004007 |
| TcCLB.503491.19 | mucin-associated surface protein (MASP), putative | -0,7728 | 0,02507 |
| TcCLB.508143.65 | elongation factor 1-gamma (EF-1-gamma, pseudogene), p | -0,7735 | 0,06365 |
| TcCLB.506939.200 | trans-sialidase (pseudogene), putative | -0,774 | 0,00674 |
| TcCLB.504741.90 | hypothetical protein, conserved | -0,7741 | 5,808E-07 |
| TcCLB.510487.70 | trans-sialidase (pseudogene), putative | -0,7742 | 0,003389 |
| TcCLB.507165.50 | cysteine synthase, putative | -0,7744 | 3,803E-07 |
| TcCLB.508775.30 | dispersed gene family protein 1 (DGF-1), putative | -0,7744 | 0,004027 |
| TcCLB.509551.140 | hypothetical protein, conserved | -0,7747 | 2,961E-17 |
| TcCLB.509733.90 | hypothetical protein, conserved | -0,7751 | 1,214E-19 |
| TcCLB.508873.344 | mucin-associated surface protein (MASP, pseudogene), pu | -0,7751 | 0,01665 |
| TcCLB.509799.80 | hypothetical protein, conserved | -0,7752 | 9,651E-12 |
| TcCLB.508727.30 | hypothetical protein, conserved | -0,7753 | 0,00001812 |
| TcCLB.508957.14 | hypothetical protein, conserved (fragment) | -0,7757 | 4,704E-08 |
| TcCLB.506753.249 | hypothetical protein, conserved | -0,7761 | 0,1277 |
| TcCLB.509741.50 | kinesin-like protein (pseudogene), putative | -0,7763 | 0,000212 |
| TcCLB.510955.30 | hypothetical protein, conserved | -0,7766 | 6,322E-12 |
| TcCLB.507129.30 | C-14 sterol reductase, putative | -0,777 | 8,534E-14 |
| TcCLB.508461.410 | T-complex protein 1, eta subunit, putative | -0,7771 | 4,568E-38 |
| TcCLB.511273.59 | 5-oxoprolinase, putative | -0,7771 | 2,742E-12 |
| TcCLB.503899.90 | asparagine synthetase a, putative | -0,7772 | 5,395E-06 |
| TcCLB.510741.229 | ubiquitin-activating enzyme E1, putative | -0,7775 | 1,804E-13 |
| TcCLB.509551.70 | hypothetical protein, conserved | -0,7775 | 3,52E-13 |
| TcCLB.509683.130 | aminopeptidase, putative | -0,7776 | 2,296E-12 |
| TcCLB.506359.50 | hypothetical protein, conserved | -0,7777 | 2,161E-06 |
| TcCLB.506465.5 | hypothetical protein | -0,7777 | 0,00002903 |
| TcCLB.510773.70 | dynein heavy chain, putative | -0,7781 | 8,182E-10 |
| TcCLB.504625.40 | hypothetical protein, conserved | -0,7782 | 2,421E-12 |
| TcCLB.505931.40 | histone H3, putative | -0,7784 | 1,945E-06 |
| TcCLB.510275.130 | dispersed gene family protein 1 (DGF-1), putative | -0,7785 | 0,0005614 |
| TcCLB.506763.370 | hypothetical protein | -0,7785 | 0,2004 |
| TcCLB.510463.230 | dispersed gene family protein 1 (DGF-1), putative | -0,7787 | 0,02398 |
| TcCLB.509567.34 | hypothetical protein, conserved | -0,7788 | 2,116E-10 |
| TcCLB.510061.11 | mucin-associated surface protein (MASP), putative | -0,7789 | 0,008383 |
| TcCLB.508881.30 | hypothetical protein, conserved | -0,779 | 1,89E-09 |
| TcCLB.509871.21 | N-acetyltransferase complex ARD1 subunit (pseudogene), | -0,7796 | 0,1927 |
| TcCLB.508595.20 | UDP-Gal or UDP-GlcNAc-dependent glycosyltransferase, pu | -0,7798 | 0,002479 |
| TcCLB.506459.235 | hypothetical protein, conserved | -0,7806 | 0,0002129 |

| | | | |
|---|---|---|---|
| TcCLB.509065.170 | hypothetical protein, conserved | -0,7811 | 9,913E-14 |
| TcCLB.417203.9 | dispersed gene family protein 1 (DGF-1), putative | -0,7811 | 0,005474 |
| TcCLB.510431.120 | hypothetical protein, conserved | -0,7818 | 2,862E-10 |
| TcCLB.508823.60 | hypothetical protein, conserved | -0,782 | 8,746E-08 |
| TcCLB.506679.90 | RNA editing complex protein MP67, putative | -0,7825 | 1,231E-15 |
| TcCLB.510699.20 | mucin-associated surface protein (MASP, pseudogene), pu | -0,7825 | 0,0004349 |
| TcCLB.506321.60 | surface protease GP63 (pseudogene), putative | -0,7826 | 0,02071 |
| TcCLB.511801.10 | hypothetical protein, conserved | -0,7828 | 1,138E-19 |
| TcCLB.508219.90 | 90 kDa surface protein, putative | -0,7831 | 0,01769 |
| TcCLB.511545.100 | hypothetical protein, conserved | -0,7835 | 3,035E-19 |
| TcCLB.511741.10 | hypothetical protein, conserved | -0,7836 | 1,257E-06 |
| TcCLB.503955.20 | mitochondrial DNA polymerase beta, putative | -0,7846 | 6,519E-23 |
| TcCLB.511337.30 | UDP-Gal or UDP-GlcNAc-dependent glycosyltransferase, pu | -0,7846 | 0,01169 |
| TcCLB.506965.120 | hypothetical protein | -0,7848 | 0,1081 |
| TcCLB.511519.9 | small glutamine-rich tetratricopeptide repeat protein, puta | -0,7868 | 3,486E-10 |
| TcCLB.503681.9 | hypothetical protein, conserved | -0,7869 | 2,4E-15 |
| TcCLB.506743.4 | eukaryotic translation initiation factor 1A, putative (fragme | -0,7872 | 6,16E-10 |
| TcCLB.503967.4 | hypothetical protein, conserved | -0,7873 | 0,002753 |
| TcCLB.411383.10 | surface protease GP63 (pseudogene), putative | -0,7874 | 0,1825 |
| TcCLB.507511.40 | G5-interacting protein, putative (G5-IP) | -0,7885 | 7,315E-16 |
| TcCLB.511467.20 | hypothetical protein, conserved | -0,7889 | 1,317E-37 |
| TcCLB.506009.60 | hypothetical protein | -0,7892 | 7,255E-09 |
| TcCLB.508645.40 | nucleoside transporter 1, putative | -0,7897 | 4,396E-38 |
| TcCLB.507915.19 | mismatch repair protein MSH3, putative | -0,7906 | 0,0001637 |
| TcCLB.507951.60 | hypothetical protein, conserved | -0,7911 | 8,905E-15 |
| TcCLB.505025.10 | mucin-associated surface protein (MASP), putative | -0,7918 | 0,08221 |
| TcCLB.511485.20 | retrotransposon hot spot protein (RHS, pseudogene), putat | -0,7919 | 2,636E-06 |
| TcCLB.508323.9 | farnesyl pyrophosphate synthase, putative | -0,7921 | 3,961E-13 |
| TcCLB.508427.30 | surface protease GP63 (pseudogene), putative | -0,7924 | 0,02713 |
| TcCLB.504017.79 | glycyl-tRNA synthetase, putative | -0,7925 | 8,97E-12 |
| TcCLB.511099.20 | trans-sialidase (pseudogene), putative | -0,7925 | 0,09837 |
| TcCLB.511871.70 | hypothetical protein, conserved | -0,7926 | 2,556E-07 |
| TcCLB.511477.10 | trans-sialidase (pseudogene), putative | -0,7929 | 0,1406 |
| TcCLB.508271.30 | hypothetical protein, conserved | -0,793 | 6,355E-12 |
| TcCLB.508601.100 | hypothetical protein, conserved | -0,7933 | 1,098E-18 |
| TcCLB.503893.150 | hypothetical protein, conserved | -0,7938 | 2,578E-10 |
| TcCLB.509053.110 | ubiquitin fusion degradation protein 2, putative | -0,794 | 5,98E-27 |
| TcCLB.510105.50 | fibrillarin, putative | -0,7943 | 9,097E-15 |
| TcCLB.505997.21 | N-acetyltransferase complex ARD1 subunit (pseudogene), | -0,7944 | 0,07627 |
| TcCLB.509267.50 | U5 small nuclear ribonucleoprotein component, putative | -0,7945 | 2,663E-19 |
| TcCLB.504005.60 | hypothetical protein, conserved | -0,7951 | 0,000051 |
| TcCLB.510337.30 | adenylosuccinate lyase, putative (ADSL) | -0,7953 | 2,077E-16 |
| TcCLB.511445.80 | hypothetical protein, conserved | -0,7957 | 1,035E-10 |
| TcCLB.511795.10 | hypothetical protein, conserved (pseudogene) | -0,7958 | 0,0109 |
| TcCLB.507143.70 | hypothetical protein, conserved | -0,7961 | 9,259E-17 |

| | | | |
|---|---|---|---|
| TcCLB.486901.10 | endonuclease/exonuclease/phosphatase, putative | -0,7963 | 0,00001128 |
| TcCLB.511261.120 | pumilio/PUF RNA binding protein 2, putative (PUF2) | -0,7963 | 0,000213 |
| TcCLB.506885.410 | hypothetical protein, conserved | -0,7975 | 1,269E-08 |
| TcCLB.511837.40 | hypothetical protein, conserved | -0,7975 | 8,186E-07 |
| TcCLB.509157.70 | hypothetical protein, conserved | -0,7976 | 0,000493 |
| TcCLB.510209.18 | mucin TcMUCII (pseudogene), putative | -0,7983 | 0,09462 |
| TcCLB.508943.30 | RNA-binding protein, putative | -0,7984 | 8,228E-09 |
| TcCLB.506947.40 | peroxisome assembly protein, putative | -0,7989 | 5,596E-15 |
| TcCLB.508073.10 | surface protease GP63, putative | -0,7989 | 0,0001814 |
| TcCLB.508163.390 | hypothetical protein, conserved | -0,7989 | 0,001551 |
| TcCLB.508391.10 | dispersed gene family protein 1 (DGF-1), putative | -0,7997 | 0,004979 |
| TcCLB.508365.160 | mucin-associated surface protein (MASP), putative | -0,7997 | 0,02719 |
| TcCLB.507089.280 | chaperone DnaJ protein (pseudogene), putative | -0,8005 | 0,000053 |
| TcCLB.507277.30 | hypothetical protein, conserved | -0,8006 | 1,919E-09 |
| TcCLB.511649.39 | hypothetical protein, conserved | -0,8007 | 6,822E-06 |
| TcCLB.503973.40 | dispersed gene family protein 1 (DGF-1), putative | -0,801 | 0,00161 |
| TcCLB.506459.120 | mucin TcMUCII, putative | -0,8017 | 0,06757 |
| TcCLB.506503.69 | glycosomal malate dehydrogenase, putative | -0,8023 | 1,016E-23 |
| TcCLB.511357.5 | glycerol uptake protein, putative (fragment) | -0,8024 | 6,527E-06 |
| TcCLB.507209.10 | small nuclear ribonucleoprotein protein, putative | -0,8026 | 1,864E-11 |
| TcCLB.511045.40 | subtilisin peptidase | -0,8027 | 1,337E-16 |
| TcCLB.510693.120 | trans-sialidase (pseudogene), putative | -0,803 | 0,01076 |
| TcCLB.508207.220 | hypothetical protein, conserved | -0,8031 | 0,0001949 |
| TcCLB.510385.5 | hypothetical protein, conserved (pseudogene) | -0,8031 | 0,001092 |
| TcCLB.506661.150 | hypothetical protein, conserved | -0,8032 | 2,763E-06 |
| TcCLB.508853.20 | trans-sialidase, Group V, putative | -0,8044 | 0,0006502 |
| TcCLB.411059.18 | clathrin heavy chain, putative | -0,8047 | 1,295E-07 |
| TcCLB.507839.18 | hypothetical protein, conserved | -0,8048 | 5,709E-10 |
| TcCLB.506625.84 | hypothetical protein, conserved | -0,8049 | 3,468E-07 |
| TcCLB.450019.9 | dispersed gene family protein 1 (DGF-1), putative (fragme | -0,8054 | 7,136E-07 |
| TcCLB.508181.10 | hypothetical protein, conserved | -0,8055 | 5E-10 |
| TcCLB.503899.80 | hypothetical protein, conserved | -0,8057 | 5,597E-14 |
| TcCLB.506363.10 | hypothetical protein | -0,8059 | 0,00001086 |
| TcCLB.508355.380 | cysteine peptidase, Clan CA, family C19, putative | -0,8062 | 9,973E-06 |
| TcCLB.511173.10 | trans-sialidase, putative | -0,8064 | 0,09817 |
| TcCLB.503423.20 | protein kinase, putative | -0,8072 | 0,1068 |
| TcCLB.509527.154 | mucin-associated surface protein (MASP, pseudogene), pu | -0,8077 | 0,01279 |
| TcCLB.506885.450 | Kinesin-13 6, putative (KIN13-6) | -0,8081 | 5,464E-10 |
| TcCLB.510187.420 | chaperonin HSP60, mitochondrial precursor (pseudogene) ( | -0,8082 | 5,452E-11 |
| TcCLB.446785.10 | trans-sialidase, putative (fragment) | -0,8083 | 0,00001808 |
| TcCLB.509805.210 | hypothetical protein, conserved | -0,8085 | 2,373E-06 |
| TcCLB.506315.30 | protein phosphatase 1 catalitic subunit, putative | -0,8088 | 4,522E-13 |
| TcCLB.471467.10 | trans-sialidase, putative (fragment) | -0,8093 | 0,002087 |
| TcCLB.511575.60 | isocitrate dehydrogenase [NADP], mitochondrial precursor, | -0,8096 | 4,581E-10 |
| TcCLB.511255.350 | trans-sialidase, Group V, putative | -0,8097 | 0,02083 |

| | | | |
|---|---|---|---|
| TcCLB.508153.330 | vacuolar protein sorting-associated protein 4, putative (VP! | -0,81 | 5,244E-26 |
| TcCLB.509505.70 | 10 kDa heat shock protein, putative | -0,81 | 0,2195 |
| TcCLB.506137.60 | mucin-associated surface protein (MASP), putative | -0,8106 | 0,2 |
| TcCLB.511367.20 | hypothetical protein | -0,8111 | 0,09627 |
| TcCLB.507859.120 | dispersed gene family protein 1 (DGF-1), putative | -0,8111 | 0,1321 |
| TcCLB.511163.20 | cdc2-related kinase 8, putative (CRK8) | -0,8119 | 2,188E-10 |
| TcCLB.506925.370 | MP44 (pseudogene), putative | -0,8122 | 0,002956 |
| TcCLB.434945.10 | hypothetical protein, conserved | -0,8123 | 0,005699 |
| TcCLB.504057.65 | hypothetical protein, conserved | -0,8127 | 6,523E-08 |
| TcCLB.508251.10 | hypothetical protein | -0,8127 | 0,0612 |
| TcCLB.508891.90 | hypothetical protein, conserved | -0,8128 | 6,694E-15 |
| TcCLB.507851.70 | hypothetical protein | -0,813 | 1,899E-12 |
| TcCLB.510825.10 | hypothetical protein, conserved | -0,8138 | 1,133E-09 |
| TcCLB.511911.159 | histone deacetylase 1, putative | -0,8141 | 1,699E-06 |
| TcCLB.503687.30 | hypothetical protein, conserved | -0,8144 | 0,00000229 |
| TcCLB.506193.5 | glucose-6-phosphate 1-dehydrogenase (pseudogene), puta | -0,8146 | 0,0004034 |
| TcCLB.510823.60 | hypothetical protein, conserved | -0,815 | 1,296E-17 |
| TcCLB.511351.9 | inosine-5'-monophosphate dehydrogenase, putative | -0,8151 | 2,03E-13 |
| TcCLB.509551.60 | hypothetical protein, conserved | -0,8153 | 6,225E-07 |
| TcCLB.508873.260 | hypothetical protein | -0,8154 | 0,1129 |
| TcCLB.506755.160 | AAA ATPase, putative | -0,8156 | 1,179E-31 |
| TcCLB.503945.40 | 6-phosphogluconolactonase, putative | -0,8156 | 1,037E-06 |
| TcCLB.506605.70 | hypothetical protein, conserved | -0,8157 | 1,913E-16 |
| TcCLB.507069.10 | mucin-associated surface protein (MASP), putative | -0,816 | 0,009497 |
| TcCLB.511183.130 | hypothetical protein, conserved (pseudogene) | -0,816 | 0,05817 |
| TcCLB.511547.10 | mucin-associated surface protein (MASP), putative | -0,816 | 0,08202 |
| TcCLB.506831.70 | glutamyl-tRNA synthetase, putative | -0,8162 | 9,941E-16 |
| TcCLB.425785.18 | hypothetical protein, conserved | -0,8166 | 0,002297 |
| TcCLB.510709.41 | hypothetical protein, conserved | -0,8183 | 0,07424 |
| TcCLB.506831.79 | pre-mRNA splicing factor ATP-dependent RNA helicase, pu | -0,8192 | 0,0001019 |
| TcCLB.511573.58 | Nucleolar protein 56, putative (fragment) (NOP56) | -0,8197 | 8,452E-19 |
| TcCLB.509499.29 | hypothetical protein, conserved | -0,8202 | 7,637E-07 |
| TcCLB.506265.20 | hypothetical protein, conserved | -0,8205 | 4,257E-21 |
| TcCLB.508259.11 | hypothetical protein, conserved (pseudogene) | -0,821 | 0,001908 |
| TcCLB.511229.50 | S-adenosylhomocysteine hydrolase, putative | -0,8211 | 5,427E-16 |
| TcCLB.510525.40 | hypothetical protein, conserved | -0,8211 | 1,142E-07 |
| TcCLB.426897.4 | hypothetical protein, conserved (fragment) | -0,8211 | 0,0002178 |
| TcCLB.443971.10 | hypothetical protein, conserved (fragment) | -0,8212 | 0,06131 |
| TcCLB.508365.290 | mucin TcMUCII, putative | -0,8218 | 0,05734 |
| TcCLB.511635.10 | histone H2B, putative | -0,822 | 4,498E-26 |
| TcCLB.506699.34 | ATP-dependent 6-phosphofructokinase, glycosomal (pseud | -0,822 | 1,519E-15 |
| TcCLB.511867.90 | hypothetical protein, conserved | -0,822 | 5,266E-09 |
| TcCLB.508965.70 | hypothetical protein, conserved | -0,8227 | 0,1095 |
| TcCLB.509729.29 | neurobeachin/beige protein, putative (fragment) | -0,8229 | 1,206E-13 |
| TcCLB.510799.18 | hypothetical protein, conserved | -0,823 | 1,293E-09 |

| | | | |
|---|---|---|---|
| TcCLB.504155.30 | retrotransposon hot spot protein (RHS, pseudogene), putat | -0,8234 | 0,02178 |
| TcCLB.507389.30 | hypothetical protein | -0,8235 | 4,079E-16 |
| TcCLB.507063.180 | hypothetical protein, conserved | -0,8238 | 7,556E-07 |
| TcCLB.511255.420 | mucin-associated surface protein (MASP, pseudogene), pu | -0,8239 | 0,006001 |
| TcCLB.504099.15 | hypothetical protein | -0,824 | 0,0147 |
| TcCLB.466007.9 | dispersed gene family protein 1 (DGF-1), putative (fragme | -0,8242 | 0,05565 |
| TcCLB.508831.180 | beta galactofuranosyl glycosyltransferase (pseudogene), p | -0,8244 | 0,05681 |
| TcCLB.507963.40 | hypothetical protein, conserved | -0,8245 | 1,241E-08 |
| TcCLB.470521.10 | protein kinase, putative (fragment) | -0,8246 | 1,559E-06 |
| TcCLB.506451.13 | hypothetical protein (pseudogene) | -0,825 | 0,00002039 |
| TcCLB.509213.130 | autophagocytosis associated protein, putative | -0,8252 | 2,199E-08 |
| TcCLB.511367.220 | hypothetical protein, conserved | -0,8253 | 3,665E-16 |
| TcCLB.505773.20 | dispersed gene family protein 1 (DGF-1), putative | -0,8259 | 0,007709 |
| TcCLB.508969.40 | dispersed gene family protein 1 (DGF-1), putative | -0,8263 | 0,03032 |
| TcCLB.511717.90 | phosphomannose isomerase, putative | -0,8266 | 2,981E-12 |
| TcCLB.511485.11 | retrotransposon hot spot protein (RHS, pseudogene), putat | -0,827 | 0,02515 |
| TcCLB.507809.140 | hypothetical protein, conserved | -0,8277 | 4,354E-07 |
| TcCLB.509011.20 | RNA polymerase IIA largest subunit, putative | -0,8278 | 1,145E-26 |
| TcCLB.508163.360 | dispersed gene family protein 1 (DGF-1), putative | -0,8289 | 0,0002585 |
| TcCLB.503897.10 | hypothetical protein | -0,829 | 8,784E-18 |
| TcCLB.508871.30 | surface protease GP63 (pseudogene), putative | -0,8294 | 0,003721 |
| TcCLB.507769.50 | ATP-dependent DEAD/H RNA helicase (pseudogene), putat | -0,8297 | 5,257E-06 |
| TcCLB.510877.70 | ATP-dependent RNA and DNA helicase, mitochondrial, puta | -0,83 | 2,893E-17 |
| TcCLB.508265.60 | hypothetical protein, conserved (pseudogene) | -0,83 | 3,098E-15 |
| TcCLB.510187.490 | hypothetical protein, conserved | -0,83 | 2,825E-08 |
| TcCLB.508539.10 | hypothetical protein, conserved | -0,8302 | 0,003216 |
| TcCLB.504741.190 | hypothetical protein, conserved | -0,8313 | 9,683E-07 |
| TcCLB.509793.20 | hypothetical protein, conserved | -0,8316 | 1,201E-10 |
| TcCLB.510687.149 | coatomer alpha subunit, putative | -0,8319 | 6,231E-16 |
| TcCLB.511731.60 | glutamyl-tRNA synthetase, putative | -0,8328 | 6,937E-18 |
| TcCLB.509767.110 | hypothetical protein, conserved | -0,8328 | 5,259E-10 |
| TcCLB.510187.160 | developmentally regulated GTP-binding protein, putative | -0,8338 | 9,923E-25 |
| TcCLB.506321.160 | syntaxin binding protein (pseudogene), putative | -0,8339 | 0,00007399 |
| TcCLB.510297.30 | TPR-repeat-containing chaperone protein DNAJ, putative | -0,8341 | 9,453E-18 |
| TcCLB.506137.27 | mucin-associated surface protein (MASP, pseudogene), pu | -0,8342 | 0,01342 |
| TcCLB.511803.60 | hypothetical protein, conserved | -0,8344 | 6,449E-07 |
| TcCLB.511717.180 | hypothetical protein, conserved | -0,8347 | 1,505E-11 |
| TcCLB.509345.30 | trans-sialidase (pseudogene), putative | -0,8348 | 3,042E-10 |
| TcCLB.511301.30 | hypothetical protein, conserved | -0,8348 | 1,417E-06 |
| TcCLB.508943.10 | hypothetical protein, conserved | -0,8353 | 1,811E-08 |
| TcCLB.507007.77 | hypothetical protein, conserved | -0,8356 | 0,0003127 |
| TcCLB.506211.250 | fibrillarin, putative | -0,8362 | 5,419E-12 |
| TcCLB.509331.134 | trypanothione synthetase, putative | -0,8365 | 8,955E-08 |
| TcCLB.510371.110 | mucin TcMUCII, putative | -0,8369 | 0,1039 |
| TcCLB.510215.10 | NADH-dependent fumarate reductase, putative | -0,8372 | 7,737E-11 |

| | | | |
|---|---|---|---|
| TcCLB.511255.110 | trans-sialidase (pseudogene), putative | -0,8381 | 0,0165 |
| TcCLB.510603.140 | hypothetical protein | -0,8382 | 2,549E-21 |
| TcCLB.508873.360 | retrotransposon hot spot protein (RHS, pseudogene), putat | -0,8382 | 0,001317 |
| TcCLB.506859.150 | hypothetical protein | -0,8385 | 4,319E-22 |
| TcCLB.510329.250 | hypothetical protein, conserved | -0,8385 | 3,161E-07 |
| TcCLB.510361.209 | mucin-associated surface protein (MASP, pseudogene), pu | -0,8386 | 0,02879 |
| TcCLB.506009.80 | DNA repair and transcription factor protein, putative | -0,8391 | 1,136E-10 |
| TcCLB.511611.16 | mucin TcMUCII (pseudogene), putative | -0,8395 | 0,1144 |
| TcCLB.509633.20 | hypothetical protein, conserved | -0,84 | 8,078E-15 |
| TcCLB.509901.90 | hypothetical protein, conserved (pseudogene) | -0,8402 | 7,35E-25 |
| TcCLB.511589.200 | S-adenosylhomocysteine hydrolase, putative | -0,8406 | 1,047E-14 |
| TcCLB.503959.40 | hypothetical protein | -0,8407 | 6,038E-12 |
| TcCLB.505025.147 | hypothetical protein | -0,8407 | 0,003697 |
| TcCLB.509123.20 | hypothetical protein | -0,841 | 0,1177 |
| TcCLB.506537.71 | hypothetical protein, conserved (pseudogene) | -0,8414 | 1,979E-11 |
| TcCLB.421621.20 | syntaxin binding protein (pseudogene), putative | -0,8416 | 0,007738 |
| TcCLB.504241.90 | dispersed gene family protein 1 (DGF-1), putative | -0,8424 | 0,08943 |
| TcCLB.511367.10 | hypothetical protein | -0,8425 | 0,09517 |
| TcCLB.511061.40 | trans-sialidase, Group V, putative | -0,8432 | 0,006693 |
| TcCLB.511717.209 | phosphatidylinositol 3-related kinase, putative (fragment) | -0,8435 | 0,008857 |
| TcCLB.511277.250 | hypothetical protein, conserved | -0,8437 | 4,123E-24 |
| TcCLB.510463.221 | hypothetical protein, conserved (pseudogene) | -0,8439 | 0,1606 |
| TcCLB.506967.140 | mucin-associated surface protein (MASP, pseudogene), pu | -0,8441 | 0,09035 |
| TcCLB.509031.3 | trans-sialidase, putative (fragment) | -0,8442 | 0,01872 |
| TcCLB.510477.70 | mucin TcMUCII, putative | -0,8446 | 0,04796 |
| TcCLB.510429.50 | Lsm12 protein, putative | -0,845 | 1,125E-11 |
| TcCLB.505555.10 | lysosomal/endosomal membrane protein p67 (pseudogene | -0,8453 | 8,249E-22 |
| TcCLB.462761.10 | surface protease GP63, putative | -0,8459 | 9,008E-10 |
| TcCLB.510491.80 | hypothetical protein, conserved | -0,8461 | 2,22E-13 |
| TcCLB.506941.120 | hypothetical protein, conserved | -0,8462 | 1,508E-11 |
| TcCLB.510449.10 | dispersed gene family protein 1 (DGF-1), putative | -0,8468 | 9,377E-06 |
| TcCLB.509205.100 | surface protease GP63, putative | -0,8473 | 9,268E-07 |
| TcCLB.506229.60 | hypothetical protein, conserved | -0,848 | 7,756E-13 |
| TcCLB.511277.190 | ras-related GTP-binding protein, putative | -0,8491 | 1,381E-12 |
| TcCLB.511477.25 | hypothetical protein, conserved (pseudogene) | -0,8495 | 0,000364 |
| TcCLB.508771.20 | cysteine peptidase, Clan CA, family C51, putative | -0,8497 | 1,533E-17 |
| TcCLB.508909.140 | hypothetical protein, conserved | -0,8503 | 2,065E-11 |
| TcCLB.511633.40 | dynein light chain, putative | -0,8504 | 6,6E-08 |
| TcCLB.510623.45 | surface protease GP63 (pseudogene), putative | -0,8506 | 0,006064 |
| TcCLB.508231.74 | poly(A)-specific ribonuclease PARN, putative | -0,8509 | 7,323E-09 |
| TcCLB.505965.10 | surface protease GP63, putative | -0,8509 | 0,02104 |
| TcCLB.509951.49 | DNA-directed RNA polymerase II subunit 2, putative | -0,8515 | 9,3E-14 |
| TcCLB.509065.90 | hypothetical protein, conserved | -0,8517 | 2,024E-19 |
| TcCLB.505943.79 | retrotransposon hot spot protein (RHS, pseudogene), putat | -0,8517 | 0,07721 |
| TcCLB.511173.370 | trans-sialidase, Group VI, putative | -0,8519 | 0,009286 |

| | | | |
|---|---|---|---|
| TcCLB.507953.10 | trans-sialidase, Group V, putative | -0,8519 | 0,1215 |
| TcCLB.509715.104 | protein kinase, putative | -0,8523 | 1,235E-14 |
| TcCLB.509163.60 | hypothetical protein, conserved | -0,8528 | 2,834E-11 |
| TcCLB.509167.80 | hypothetical protein, conserved | -0,8529 | 1,57E-17 |
| TcCLB.508773.9 | dispersed gene family protein 1 (DGF-1), putative | -0,853 | 0,1144 |
| TcCLB.508543.100 | hypothetical protein, conserved | -0,8531 | 1,506E-24 |
| TcCLB.509105.120 | protein kinase, putative | -0,8534 | 6,435E-11 |
| TcCLB.510329.170 | hypothetical protein, conserved | -0,8535 | 8,95E-22 |
| TcCLB.507019.120 | hypothetical protein, conserved | -0,8541 | 1,23E-09 |
| TcCLB.504839.70 | hypothetical protein, conserved | -0,8542 | 2,414E-24 |
| TcCLB.508667.40 | hypothetical protein, conserved | -0,8547 | 9,983E-18 |
| TcCLB.503865.20 | hypothetical protein, conserved | -0,8557 | 2,59E-19 |
| TcCLB.504247.79 | transmembrane protein, conserved (fragment) | -0,8562 | 1,104E-08 |
| TcCLB.507255.63 | UDP-Gal or UDP-GlcNAc-dependent glycosyltransferase, pu | -0,8563 | 0,005672 |
| TcCLB.506447.10 | hypothetical protein | -0,8566 | 7,067E-08 |
| TcCLB.506009.30 | TPR-repeat-containing chaperone protein DNAJ, putative | -0,8579 | 2,802E-15 |
| TcCLB.509795.50 | hypothetical protein, conserved | -0,8581 | 1,079E-22 |
| TcCLB.506321.150 | surface protease GP63 (pseudogene), putative | -0,8585 | 0,02149 |
| TcCLB.509051.30 | dTDP-glucose 4,6-dehydratase, putative | -0,859 | 1,141E-13 |
| TcCLB.508667.9 | hypothetical protein, conserved | -0,859 | 0,001155 |
| TcCLB.508851.40 | Octanoyltransferase, putative | -0,8592 | 0,0002063 |
| TcCLB.511597.25 | hypothetical protein | -0,8592 | 0,105 |
| TcCLB.506743.130 | protein tyrosine phosphatase-like protein, putative | -0,8593 | 0,0003342 |
| TcCLB.511855.40 | hypothetical protein, conserved | -0,86 | 3,455E-11 |
| TcCLB.509741.40 | hypothetical protein, conserved | -0,86 | 8,567E-08 |
| TcCLB.504153.270 | hypothetical protein, conserved | -0,8601 | 1,867E-23 |
| TcCLB.508103.30 | trans-sialidase, Group V, putative | -0,8602 | 0,06688 |
| TcCLB.509767.170 | hypothetical protein, conserved | -0,8604 | 1,062E-26 |
| TcCLB.507723.70 | hypothetical protein, conserved (pseudogene) | -0,8607 | 4,294E-25 |
| TcCLB.510031.60 | methionyl-tRNA formyltransferase, putative | -0,8607 | 7,986E-14 |
| TcCLB.508707.224 | tubulin binding cofactor A-like protein, putative | -0,861 | 1,787E-07 |
| TcCLB.509569.80 | hypothetical protein | -0,8616 | 1,547E-10 |
| TcCLB.506925.340 | hypothetical protein, conserved | -0,8618 | 2,377E-10 |
| TcCLB.508727.9 | CCR4-NOT transcription complex subunit 10, putative | -0,8629 | 5,697E-19 |
| TcCLB.507723.189 | bloodstream- specific protein 2 precursor (fragment) | -0,8632 | 6,345E-12 |
| TcCLB.507641.14 | neurobeachin/beige protein, putative | -0,8635 | 7,059E-21 |
| TcCLB.508261.140 | elongation factor 1-gamma (EF-1-gamma), putative | -0,8638 | 0,04659 |
| TcCLB.506245.80 | hypothetical protein, conserved (pseudogene) | -0,8641 | 0,0008625 |
| TcCLB.506875.10 | mucin TcMUCII (pseudogene), putative | -0,8646 | 0,0718 |
| TcCLB.510657.200 | surface protease GP63, putative | -0,8648 | 3,771E-07 |
| TcCLB.511273.50 | hypothetical protein, conserved | -0,8651 | 2,463E-13 |
| TcCLB.427237.9 | dispersed gene family protein 1 (DGF-1), putative | -0,8654 | 0,04564 |
| TcCLB.510627.110 | mucin-associated surface protein (MASP), putative | -0,8658 | 0,3469 |
| TcCLB.503971.40 | translation initiation factor 2 subunit, putative | -0,8661 | 2,283E-09 |
| TcCLB.511757.10 | hypothetical protein, conserved | -0,8664 | 7,822E-23 |

| | | | |
|---|---|---|---|
| TcCLB.510431.130 | hypothetical protein, conserved | -0,8668 | 1,142E-10 |
| TcCLB.507821.101 | hypothetical protein, conserved (pseudogene) | -0,8668 | 0,00001503 |
| TcCLB.506321.90 | syntaxin binding protein (pseudogene), putative | -0,8668 | 0,00045 |
| TcCLB.508427.40 | mucin-associated surface protein (MASP), putative | -0,8677 | 0,0003235 |
| TcCLB.509545.25 | elongation factor 1-gamma (EF-1-gamma, pseudogene), p | -0,8677 | 0,1985 |
| TcCLB.507817.9 | histone H3, putative | -0,8682 | 4,138E-06 |
| TcCLB.509965.210 | hypothetical protein, conserved | -0,8687 | 0,003688 |
| TcCLB.510465.10 | dispersed gene family protein 1 (DGF-1), putative | -0,8687 | 0,04526 |
| TcCLB.507007.9 | hypothetical protein, conserved (pseudogene) | -0,8691 | 2,229E-18 |
| TcCLB.506727.140 | hypothetical protein, conserved | -0,8702 | 1,376E-09 |
| TcCLB.511205.10 | surface protease GP63 (pseudogene), putative | -0,8704 | 1,46E-08 |
| TcCLB.509979.261 | surface protease GP63 (pseudogene), putative | -0,8704 | 0,0004424 |
| TcCLB.503973.31 | hypothetical protein, conserved (pseudogene) | -0,8714 | 0,00008498 |
| TcCLB.504057.160 | hypothetical protein, conserved | -0,8722 | 3,763E-10 |
| TcCLB.507195.40 | trans-sialidase (pseudogene), putative | -0,8722 | 0,04013 |
| TcCLB.511323.40 | histone H2A, putative | -0,8734 | 4,704E-08 |
| TcCLB.505555.30 | hypothetical protein, conserved | -0,8738 | 1,193E-26 |
| TcCLB.506501.329 | mucin-associated surface protein (MASP, pseudogene), pu | -0,8741 | 0,07007 |
| TcCLB.509719.9 | N-acetyltransferase complex ARD1 subunit, putative (fragr | -0,8746 | 0,0007207 |
| TcCLB.506739.160 | hypothetical protein, conserved | -0,8748 | 8,554E-16 |
| TcCLB.510625.210 | trans-sialidase (pseudogene), putative | -0,8749 | 0,02593 |
| TcCLB.506193.60 | ascorbate peroxidase, putative (APX) | -0,8751 | 1,723E-07 |
| TcCLB.507753.6 | hypothetical protein, conserved (pseudogene) | -0,8751 | 0,0001673 |
| TcCLB.511875.20 | trans-sialidase, Group VII, putative | -0,8754 | 2,831E-08 |
| TcCLB.511183.315 | hypothetical protein, conserved (pseudogene) | -0,8762 | 0,04026 |
| TcCLB.506909.131 | elongation factor 1-gamma (EF-1-gamma, pseudogene), p | -0,8774 | 0,0659 |
| TcCLB.506167.50 | clathrin heavy chain, putative | -0,8776 | 3,801E-38 |
| TcCLB.507875.10 | hypothetical protein, conserved | -0,8776 | 3,542E-14 |
| TcCLB.503529.10 | hypothetical protein, conserved (fragment) | -0,8777 | 0,06114 |
| TcCLB.508273.80 | syntaxin binding protein, putative | -0,878 | 0,0008286 |
| TcCLB.506739.60 | hypothetical protein, conserved | -0,8783 | 3,221E-15 |
| TcCLB.509163.54 | zinc finger protein, putative | -0,8784 | 0,1668 |
| TcCLB.508153.210 | hypothetical protein, conserved | -0,8797 | 2,572E-07 |
| TcCLB.508365.180 | hypothetical protein | -0,8798 | 0,1499 |
| TcCLB.506443.10 | hypothetical protein | -0,8802 | 5,367E-13 |
| TcCLB.508645.4 | hypothetical protein, conserved | -0,8803 | 0,0001184 |
| TcCLB.511215.70 | kinesin, putative | -0,8826 | 9,394E-13 |
| TcCLB.510003.30 | phosphatidylinositol 4-kinase alpha, putative | -0,883 | 7,4E-25 |
| TcCLB.510057.4 | hypothetical protein, conserved | -0,8831 | 3,085E-09 |
| TcCLB.506509.40 | hypothetical protein, conserved | -0,8836 | 3,788E-09 |
| TcCLB.508469.4 | hypothetical protein, conserved (fragment) | -0,8843 | 1,106E-12 |
| TcCLB.509069.30 | tubulin binding cofactor A-like protein, putative | -0,8844 | 1,364E-07 |
| TcCLB.510239.30 | hypothetical protein, conserved (pseudogene) | -0,8844 | 0,001326 |
| TcCLB.506251.70 | trans-sialidase (pseudogene), putative | -0,8846 | 0,2772 |
| TcCLB.506693.30 | RNA-binding protein 6, putative | -0,8854 | 0,05537 |

| | | | |
|---|---|---|---|
| TcCLB.507509.10 | hypothetical protein, conserved (pseudogene) | -0,8858 | 0,00000012 |
| TcCLB.507817.18 | histone H3, putative | -0,8859 | 1,017E-08 |
| TcCLB.508809.30 | ABC transporter family-like protein | -0,8865 | 2,992E-17 |
| TcCLB.511873.41 | hypothetical protein, conserved (pseudogene) | -0,8866 | 0,01362 |
| TcCLB.511859.60 | subtilisin peptidase | -0,8868 | 2,13E-13 |
| TcCLB.506693.10 | hypothetical protein | -0,8873 | 1,355E-10 |
| TcCLB.506739.110 | hypothetical protein | -0,8877 | 2,755E-12 |
| TcCLB.510625.225 | hypothetical protein, conserved (pseudogene) | -0,8878 | 0,00524 |
| TcCLB.421621.11 | hypothetical protein, conserved (pseudogene) | -0,888 | 0,102 |
| TcCLB.510823.109 | hypothetical protein, conserved | -0,8881 | 4,647E-08 |
| TcCLB.508243.81 | syntaxin binding protein (pseudogene), putative | -0,8881 | 0,004218 |
| TcCLB.510807.50 | mucin-associated surface protein (MASP), putative | -0,8883 | 0,03325 |
| TcCLB.510293.70 | ribonuclease H, putative | -0,8886 | 4,929E-08 |
| TcCLB.510751.40 | Importin 1 (fragment) (IMP1) | -0,8891 | 1,351E-23 |
| TcCLB.507379.9 | dispersed gene family protein 1 (DGF-1), putative (fragme | -0,8909 | 0,000239 |
| TcCLB.508851.31 | N-acetyltransferase complex ARD1 subunit (pseudogene), | -0,8913 | 0,0008645 |
| TcCLB.511055.50 | protein kinase, putative | -0,8914 | 0,0006311 |
| TcCLB.507241.20 | hypothetical protein, conserved | -0,8919 | 2,558E-19 |
| TcCLB.509085.31 | trans-sialidase (pseudogene), putative | -0,892 | 0,0006963 |
| TcCLB.507245.40 | dispersed gene family protein 1 (DGF-1), putative | -0,8926 | 2,418E-08 |
| TcCLB.508147.45 | hypothetical protein, conserved (pseudogene) | -0,8935 | 0,003378 |
| TcCLB.510021.190 | hypothetical protein | -0,8939 | 0,2387 |
| TcCLB.510257.10 | hypothetical protein, conserved | -0,8947 | 2,008E-34 |
| TcCLB.511711.30 | eRF1 methyltransferase catalytic subunit, putative | -0,8948 | 6,823E-11 |
| TcCLB.510749.4 | ubiquitin-protein ligase, putative | -0,8948 | 0,1088 |
| TcCLB.507771.85 | hypothetical protein | -0,8949 | 4,499E-07 |
| TcCLB.511079.80 | mucin TcMUCII, putative | -0,8957 | 0,1874 |
| TcCLB.504153.90 | Cytoplasmic dynein 2 heavy chain (DYNC2H1), putative | -0,8961 | 1,235E-50 |
| TcCLB.506733.140 | hypothetical protein, conserved | -0,8964 | 2,271E-17 |
| TcCLB.509085.40 | retrotransposon hot spot protein (RHS, pseudogene), putat | -0,8964 | 0,003677 |
| TcCLB.503887.20 | hypothetical protein | -0,8974 | 2,044E-08 |
| TcCLB.503847.40 | hypothetical protein, conserved | -0,8975 | 1,034E-14 |
| TcCLB.510443.30 | hypothetical protein, conserved | -0,8989 | 2,058E-17 |
| TcCLB.505037.70 | trans-sialidase (pseudogene), putative | -0,8991 | 2,79E-08 |
| TcCLB.510773.60 | hypothetical protein | -0,8992 | 0,001465 |
| TcCLB.511261.10 | hypothetical protein, conserved | -0,8999 | 1,908E-27 |
| TcCLB.506737.50 | hypothetical protein, conserved | -0,9009 | 4,718E-11 |
| TcCLB.511367.40 | hypothetical protein | -0,9014 | 0,009038 |
| TcCLB.505025.144 | hypothetical protein, conserved | -0,9017 | 0,08522 |
| TcCLB.508153.640 | IAG-nucleoside hydrolase, putative (IAGNH) | -0,9024 | 8,626E-29 |
| TcCLB.508215.9 | protein kinase C substrate protein, heavy chain, putative | -0,9025 | 7,518E-23 |
| TcCLB.506899.10 | trans-sialidase, putative | -0,9028 | 0,08272 |
| TcCLB.510903.30 | helicase-like protein, putative | -0,9029 | 1,611E-15 |
| TcCLB.510899.10 | surface protease GP63, putative | -0,9032 | 2,573E-16 |
| TcCLB.507787.110 | hypothetical protein, conserved | -0,9038 | 2,572E-08 |

| | | | |
|---|---|---|---|
| TcCLB.510419.11 | mucin-associated surface protein (MASP, pseudogene), pu | -0,9045 | 0,2019 |
| TcCLB.511549.20 | trans-sialidase (pseudogene), putative | -0,905 | 0,0002679 |
| TcCLB.399389.10 | hypothetical protein, conserved | -0,9054 | 1,085E-29 |
| TcCLB.510057.30 | hypothetical protein, conserved | -0,9061 | 6,893E-13 |
| TcCLB.509915.35 | hypothetical protein, conserved (pseudogene) | -0,9064 | 5,594E-06 |
| TcCLB.511299.10 | Triglyceride lipase, putative (fragment) | -0,9067 | 0,0005755 |
| TcCLB.511861.100 | mucin-associated surface protein (MASP, pseudogene), pu | -0,9072 | 1,833E-06 |
| TcCLB.506251.100 | poly(A)-specific ribonuclease PARN, putative | -0,9077 | 3,308E-14 |
| TcCLB.508837.110 | hypothetical protein, conserved | -0,908 | 8,85E-13 |
| TcCLB.511287.64 | hypothetical protein, conserved | -0,908 | 5,828E-08 |
| TcCLB.511821.70 | hypothetical protein, conserved | -0,9091 | 1,627E-12 |
| TcCLB.506797.40 | hypothetical protein, conserved | -0,9091 | 0,0002147 |
| TcCLB.503913.4 | nucleolar RNA binding protein (pseudogene), putative | -0,9096 | 6,712E-11 |
| TcCLB.504153.310 | heat shock protein, putative | -0,9099 | 1,729E-49 |
| TcCLB.506619.90 | ABC transporter family-like protein | -0,9117 | 3,954E-12 |
| TcCLB.510933.190 | mucin TcMUCII, putative | -0,9139 | 0,02097 |
| TcCLB.510635.40 | hypothetical protein, conserved | -0,914 | 2,149E-12 |
| TcCLB.507145.84 | mucin-associated surface protein (MASP, pseudogene), pu | -0,914 | 0,00003725 |
| TcCLB.508873.170 | retrotransposon hot spot protein (RHS, pseudogene), putat | -0,914 | 0,005816 |
| TcCLB.511311.70 | hypothetical protein, conserved | -0,9144 | 4,405E-22 |
| TcCLB.506213.110 | hypothetical protein | -0,9147 | 1,018E-23 |
| TcCLB.511635.30 | ARP2/3 complex 16kDa subunit, putative | -0,9154 | 3,83E-09 |
| TcCLB.508727.60 | hypothetical protein, conserved | -0,9157 | 5,662E-15 |
| TcCLB.459389.10 | hypothetical protein, conserved (fragment) | -0,9162 | 7,443E-11 |
| TcCLB.504005.3 | RNA helicase, putative | -0,9163 | 2,499E-15 |
| TcCLB.510307.245 | elongation factor 1-gamma (EF-1-gamma, pseudogene), p | -0,9169 | 0,02213 |
| TcCLB.511255.660 | mucin TcMUCII, putative | -0,9176 | 0,01523 |
| TcCLB.506229.40 | leucine-rich repeat protein, putative | -0,9179 | 0,0001065 |
| TcCLB.508835.30 | trans-sialidase (pseudogene), putative | -0,9182 | 2,549E-07 |
| TcCLB.506499.200 | hypothetical protein | -0,9182 | 0,01189 |
| TcCLB.506223.120 | phosphatidylinositol 3-related kinase, putative | -0,92 | 5,311E-31 |
| TcCLB.508873.210 | mucin-associated surface protein (MASP), putative | -0,9204 | 0,005909 |
| TcCLB.509099.100 | hypothetical protein, conserved | -0,9207 | 2,132E-11 |
| TcCLB.510329.130 | UDP-glucose:glycoprotein glucosyltransferase (tcgt1) | -0,9217 | 2,383E-20 |
| TcCLB.509011.80 | surface protease GP63, putative | -0,9225 | 8,351E-07 |
| TcCLB.508855.10 | trans-sialidase, putative | -0,9228 | 1,227E-08 |
| TcCLB.503893.100 | hypothetical protein, conserved | -0,9229 | 0,00001947 |
| TcCLB.506457.80 | hypothetical protein, conserved | -0,9241 | 2,596E-15 |
| TcCLB.511451.110 | hypothetical protein, conserved (pseudogene) | -0,9252 | 0,01769 |
| TcCLB.506599.390 | mucin-associated surface protein (MASP), putative | -0,9265 | 0,01831 |
| TcCLB.504787.30 | mucin-associated surface protein (MASP), putative | -0,9269 | 0,06465 |
| TcCLB.506925.310 | hypothetical protein, conserved | -0,9271 | 2,489E-53 |
| TcCLB.507077.60 | surface protease GP63, putative | -0,9275 | 2,018E-10 |
| TcCLB.421621.50 | mucin-associated surface protein (MASP), putative | -0,9276 | 0,07688 |
| TcCLB.508707.200 | nucleoside diphosphate kinase (NDPK1) | -0,9289 | 2,762E-25 |

| | | | |
|---|---|---|---|
| TcCLB.504081.26 | hypothetical protein | -0,9296 | 0,0001582 |
| TcCLB.510297.80 | DNA repair and transcription factor protein, putative | -0,9298 | 7,393E-13 |
| TcCLB.511717.40 | hypothetical protein, conserved | -0,9301 | 2,314E-08 |
| TcCLB.507359.30 | hypothetical protein, conserved (pseudogene) | -0,9319 | 0,101 |
| TcCLB.505387.9 | dispersed gene family protein 1 (DGF-1), putative (fragme | -0,933 | 2,036E-09 |
| TcCLB.511239.110 | hypothetical protein, conserved | -0,9335 | 2,434E-22 |
| TcCLB.511127.230 | hypothetical protein, conserved | -0,9338 | 4,887E-11 |
| TcCLB.510021.109 | mucin-associated surface protein (MASP, pseudogene), pu | -0,9341 | 0,0001773 |
| TcCLB.506901.85 | mucin-associated surface protein (MASP, pseudogene), pu | -0,9341 | 0,07144 |
| TcCLB.508355.360 | hypothetical protein, conserved | -0,9342 | 4,622E-08 |
| TcCLB.506095.9 | dispersed gene family protein 1 (DGF-1), putative (fragme | -0,9346 | 2,614E-10 |
| TcCLB.506625.110 | DNA topoisomerase type IB small subunit, putative | -0,9347 | 2,324E-13 |
| TcCLB.511277.39 | hypothetical protein, conserved | -0,9348 | 0,00004272 |
| TcCLB.505949.110 | hypothetical protein, conserved (pseudogene) | -0,9352 | 0,03861 |
| TcCLB.504245.9 | hypothetical protein, conserved | -0,9354 | 6,581E-14 |
| TcCLB.506349.10 | protein kinase, putative | -0,9357 | 0,02168 |
| TcCLB.510953.10 | surface protease GP63 (pseudogene), putative | -0,936 | 0,0006846 |
| TcCLB.507677.50 | hypothetical protein, conserved | -0,9362 | 5,696E-13 |
| TcCLB.508723.80 | disgorgin, putative | -0,9362 | 9,848E-12 |
| TcCLB.507143.60 | hypothetical protein | -0,9367 | 0,1025 |
| TcCLB.480997.10 | hypothetical protein, conserved | -0,937 | 0,104 |
| TcCLB.511479.48 | hypothetical protein, conserved (fragment) | -0,9372 | 0,0004292 |
| TcCLB.511417.70 | histone-lysine N-methyltransferase, putative (DOT1B) | -0,9378 | 4,696E-22 |
| TcCLB.505101.50 | GMP synthetase (pseudogene), putative | -0,9379 | 6,249E-06 |
| TcCLB.510779.20 | hypothetical protein (fragment) | -0,9382 | 5,692E-06 |
| TcCLB.506733.20 | AMP deaminase, putative | -0,9389 | 4,466E-28 |
| TcCLB.504011.30 | surface protease GP63 (pseudogene), putative | -0,9397 | 0,01009 |
| TcCLB.511859.10 | AMP deaminase, putative | -0,9398 | 0,00002133 |
| TcCLB.506343.41 | trans-sialidase (pseudogene), putative | -0,9403 | 0,0007081 |
| TcCLB.510329.340 | hypothetical protein, conserved | -0,9408 | 2,332E-14 |
| TcCLB.508877.10 | trans-sialidase (pseudogene), putative | -0,9408 | 0,0005693 |
| TcCLB.511277.470 | hypothetical protein, conserved | -0,9413 | 3,222E-07 |
| TcCLB.506739.130 | protein phosphatase 1 catalitic subunit, putative | -0,9423 | 1,204E-18 |
| TcCLB.508369.20 | squalene synthase, putative | -0,9429 | 5,589E-47 |
| TcCLB.426761.10 | retrotransposon hot spot protein (RHS, pseudogene), putat | -0,9433 | 0,0507 |
| TcCLB.506445.90 | magnesium and cobalt transport protein, putative | -0,9442 | 9,28E-16 |
| TcCLB.509671.90 | COP-coated vesicle membrane protein gp25L precursor, pu | -0,9449 | 6,033E-16 |
| TcCLB.508879.4 | protein kinase C substrate protein, heavy chain, putative | -0,9451 | 2,05E-12 |
| TcCLB.510657.170 | cytochrome P450 reductase, putative | -0,9455 | 5,013E-11 |
| TcCLB.504797.80 | endo-beta-N-acetylglucosaminidase, putative | -0,9458 | 9,002E-27 |
| TcCLB.506229.70 | ATP-dependent DEAD/H RNA helicase, putative | -0,9458 | 2,809E-11 |
| TcCLB.509565.9 | hypothetical protein (fragment) | -0,9465 | 5,878E-34 |
| TcCLB.506739.120 | hypothetical protein, conserved | -0,9466 | 4,989E-17 |
| TcCLB.510903.60 | prenyltransferase, putative | -0,9466 | 3,7E-13 |
| TcCLB.508523.35 | hypothetical protein, conserved (pseudogene) | -0,9472 | 1,116E-13 |

| | | | |
|---|---|---|---|
| TcCLB.510255.29 | hypothetical protein, conserved (fragment) | -0,9473 | 1,334E-09 |
| TcCLB.510047.80 | trans-sialidase, Group V, putative | -0,9475 | 0,0005483 |
| TcCLB.503467.9 | NADH-dependent fumarate reductase, putative (fragment) | -0,948 | 2,225E-08 |
| TcCLB.511719.10 | phosphatidylinositol 3-related kinase, putative | -0,9487 | 6,422E-30 |
| TcCLB.506319.70 | hypothetical protein | -0,9495 | 0,02713 |
| TcCLB.507053.200 | hypothetical protein, conserved | -0,9498 | 2,551E-16 |
| TcCLB.510825.30 | phospholipase B-like, putative (p67) | -0,9512 | 2,028E-41 |
| TcCLB.511271.10 | dispersed gene family protein 1 (DGF-1, pseudogene), puta | -0,9516 | 6,451E-06 |
| TcCLB.509599.20 | hypothetical protein, conserved (fragment) | -0,952 | 3,336E-15 |
| TcCLB.510681.9 | surface protease GP63, putative | -0,9526 | 0,007438 |
| TcCLB.506155.130 | hypothetical protein, conserved | -0,9533 | 0,0004146 |
| TcCLB.511183.505 | trans-sialidase (pseudogene), putative | -0,9534 | 0,005813 |
| TcCLB.503579.110 | pseudouridylate synthase I, putative | -0,9538 | 3,312E-12 |
| TcCLB.509451.24 | hypothetical protein, conserved (fragment) | -0,954 | 4,303E-18 |
| TcCLB.511183.300 | trans-sialidase (pseudogene), putative | -0,954 | 0,03514 |
| TcCLB.506959.20 | hypothetical protein, conserved | -0,9553 | 2,421E-32 |
| TcCLB.510441.10 | hypothetical protein | -0,9556 | 0,0001683 |
| TcCLB.510439.40 | hypothetical protein | -0,956 | 1,866E-10 |
| TcCLB.507257.80 | diacylglycerol kinase-like protein, putative | -0,9569 | 2,085E-08 |
| TcCLB.510747.60 | cytochrome P450 reductase, putative | -0,9574 | 9,951E-15 |
| TcCLB.508121.15 | hypothetical protein, conserved (pseudogene) | -0,9586 | 4,452E-07 |
| TcCLB.503755.30 | DNA polymerase kappa, putative | -0,9609 | 0,00001591 |
| TcCLB.509875.60 | hypothetical protein | -0,963 | 0,01353 |
| TcCLB.509011.100 | myosin heavy chain kinase A, putative | -0,9632 | 6,509E-17 |
| TcCLB.511727.190 | hypothetical protein, conserved | -0,9639 | 9,939E-29 |
| TcCLB.507601.40 | Nudix hydrolase 2, putative (Nudix2) | -0,9641 | 4,163E-13 |
| TcCLB.508889.4 | 5-oxoprolinase, putative | -0,9657 | 0,0001565 |
| TcCLB.509485.30 | hypothetical protein, conserved (pseudogene) | -0,9672 | 0,02148 |
| TcCLB.504277.30 | hypothetical protein | -0,9693 | 0,00006138 |
| TcCLB.508365.240 | mucin TcMUCII, putative | -0,9697 | 0,09306 |
| TcCLB.508153.690 | hypothetical protein | -0,9705 | 0,08462 |
| TcCLB.510017.15 | surface protease GP63 (pseudogene), putative | -0,9731 | 0,02573 |
| TcCLB.507531.19 | phosphatidylinositol 4-kinase alpha, putative | -0,9732 | 3,29E-21 |
| TcCLB.507681.30 | hypothetical protein, conserved | -0,9736 | 2,2E-28 |
| TcCLB.508357.100 | hypothetical protein, conserved | -0,974 | 2,861E-34 |
| TcCLB.503431.109 | nucleoporin interacting component (NUP93), putative | -0,9745 | 0,00004765 |
| TcCLB.511605.20 | mucin-associated surface protein (MASP), putative | -0,9748 | 0,0019 |
| TcCLB.511511.5 | hypothetical protein | -0,9748 | 0,001984 |
| TcCLB.511631.9 | hypothetical protein, conserved (fragment) | -0,9753 | 8,899E-08 |
| TcCLB.447847.10 | trans-sialidase (pseudogene), putative | -0,976 | 0,03237 |
| TcCLB.506247.230 | prolyl endopeptidase (POP) | -0,9762 | 2,06E-30 |
| TcCLB.503641.19 | peroxin 12, putative (PEX12) | -0,9765 | 1,049E-07 |
| TcCLB.510201.45 | hypothetical protein, conserved (pseudogene) | -0,9784 | 6,337E-07 |
| TcCLB.511127.60 | kinesin, putative | -0,9794 | 3,265E-25 |
| TcCLB.510329.290 | diacylglycerol kinase-like protein, putative | -0,9797 | 7,751E-08 |

| | | | |
|---|---|---|---|
| TcCLB.506071.9 | dispersed gene family protein 1 (DGF-1), putative (fragme | -0,9799 | 0,0003338 |
| TcCLB.503895.19 | hypothetical protein | -0,9801 | 1,461E-32 |
| TcCLB.506591.4 | heat shock protein 90, putative (LPG3) | -0,9812 | 4,848E-22 |
| TcCLB.508221.590 | trans-sialidase, Group V, putative | -0,9813 | 0,007563 |
| TcCLB.510399.10 | dispersed gene family protein 1 (DGF-1), putative | -0,9815 | 0,05114 |
| TcCLB.511211.49 | hypothetical protein, conserved | -0,9817 | 2,169E-16 |
| TcCLB.510359.520 | mucin-associated surface protein (MASP), putative | -0,9819 | 0,4451 |
| TcCLB.508149.60 | trans-sialidase (pseudogene), putative | -0,9834 | 0,001041 |
| TcCLB.511735.70 | hypothetical protein, conserved | -0,9838 | 5,591E-16 |
| TcCLB.510191.40 | trans-sialidase (pseudogene), putative | -0,9845 | 0,0253 |
| TcCLB.506585.60 | hypothetical protein, conserved | -0,9853 | 8,921E-15 |
| TcCLB.511623.20 | hypothetical protein, conserved | -0,9858 | 7,385E-11 |
| TcCLB.508355.431 | chaperonin containing t-complex protein, putative (fragme | -0,9865 | 0,008682 |
| TcCLB.509053.70 | p22 protein precursor, putative | -0,9878 | 5,706E-18 |
| TcCLB.508357.40 | cysteine peptidase, Clan CA, family C19, putative | -0,988 | 2,886E-11 |
| TcCLB.511837.100 | amino acid permease-like protein, putative | -0,9898 | 8,444E-19 |
| TcCLB.506251.130 | ubiquitin-protein ligase (pseudogene), putative | -0,9902 | 1,587E-22 |
| TcCLB.506725.20 | hypothetical protein, conserved | -0,9906 | 2,278E-46 |
| TcCLB.507533.10 | phosphatidylinositol 4-kinase alpha, putative | -0,9909 | 2,107E-21 |
| TcCLB.504021.100 | hypothetical protein, conserved | -0,9909 | 1,123E-17 |
| TcCLB.479481.5 | elongation factor 1-gamma (EF-1-gamma, pseudogene), p | -0,9911 | 0,02106 |
| TcCLB.506511.70 | hypothetical protein, conserved | -0,992 | 2,3E-18 |
| TcCLB.506871.60 | hypothetical protein, conserved (pseudogene) | -0,9926 | 9,485E-16 |
| TcCLB.507143.40 | hypothetical protein, conserved | -0,994 | 1,772E-34 |
| TcCLB.506301.11 | heat shock protein, putative (fragment) | -0,9944 | 6,05E-30 |
| TcCLB.508323.190 | serine/threonine protein phosphatase, putative | -0,9944 | 1,078E-12 |
| TcCLB.506559.380 | centrin, putative | -0,9955 | 2,454E-18 |
| TcCLB.506177.50 | hypothetical protein, conserved | -0,9956 | 8,8E-38 |
| TcCLB.511483.50 | hypothetical protein, conserved | -0,9981 | 2,099E-08 |
| TcCLB.507895.150 | hypothetical protein, conserved | -0,9982 | 8,184E-12 |
| TcCLB.509073.70 | phosphatidate cytidylyltransferase-like protein, putative | -0,9988 | 1,73E-16 |
| TcCLB.510187.394 | hypothetical protein, conserved | -0,9993 | 2,097E-16 |
| TcCLB.508209.120 | 10 kDa heat shock protein, putative | -0,9999 | 0,001003 |
| TcCLB.510743.20 | hypothetical protein, conserved | -1 | 9,289E-11 |
| TcCLB.506965.170 | trans-sialidase, Group V, putative | -1 | 0,0003021 |
| TcCLB.509563.60 | hypothetical protein | -1,001 | 1,169E-07 |
| TcCLB.511621.170 | hypothetical protein, conserved | -1,001 | 1,589E-07 |
| TcCLB.510475.51 | mucin-associated surface protein (MASP, pseudogene), pu | -1,001 | 0,04518 |
| TcCLB.508873.50 | hypothetical protein | -1,003 | 0,06031 |
| TcCLB.506229.64 | hypothetical protein, conserved | -1,004 | 6,297E-09 |
| TcCLB.508481.30 | trans-sialidase (pseudogene), putative | -1,004 | 0,013 |
| TcCLB.503875.30 | mucin-associated surface protein (MASP), putative | -1,005 | 9,801E-07 |
| TcCLB.508737.150 | mitochondrial carrier protein, putative | -1,007 | 1,332E-33 |
| TcCLB.506755.90 | hypothetical protein, conserved | -1,007 | 1,396E-11 |
| TcCLB.510275.150 | hypothetical protein, conserved (pseudogene) | -1,007 | 0,0002391 |

| | | | |
|---|---|---|---|
| TcCLB.509943.30 | ribosomal protein S6, putative | -1,008 | 2,444E-14 |
| TcCLB.509567.40 | nuclear transport factor 2, putative | -1,009 | 1,26E-16 |
| TcCLB.511079.100 | mucin-associated surface protein (MASP), putative (fragm | -1,009 | 0,2089 |
| TcCLB.510187.480 | phosphomannomutase, putative | -1,01 | 4,954E-25 |
| TcCLB.507711.110 | procyclic form surface phosphoprotein, putative | -1,01 | 2,556E-13 |
| TcCLB.506499.60 | hypothetical protein | -1,01 | 0,08526 |
| TcCLB.506839.70 | NADH dehydrogenase, putative | -1,013 | 5,927E-36 |
| TcCLB.507747.10 | surface protease GP63, putative | -1,013 | 0,00002254 |
| TcCLB.507447.19 | microtubule-associated protein, putative (fragment) | -1,014 | 0,0211 |
| TcCLB.511627.110 | heat shock protein DNAJ, putative | -1,015 | 2,526E-62 |
| TcCLB.510013.13 | syntaxin binding protein (pseudogene), putative | -1,016 | 0,01355 |
| TcCLB.510667.10 | hypothetical protein, conserved | -1,017 | 7,307E-13 |
| TcCLB.503811.10 | heat shock protein 90, putative (LPG3) | -1,018 | 1,854E-28 |
| TcCLB.507895.100 | hypothetical protein, conserved (pseudogene) | -1,018 | 6,111E-15 |
| TcCLB.508181.40 | choline/ethanolamine phosphotransferase, putative | -1,019 | 2,072E-20 |
| TcCLB.447801.10 | hypothetical protein, conserved (fragment) | -1,019 | 2,262E-09 |
| TcCLB.508291.71 | hypothetical protein | -1,02 | 4,327E-06 |
| TcCLB.507607.10 | surface protease GP63 (pseudogene), putative | -1,021 | 0,07639 |
| TcCLB.508111.40 | hypothetical protein, conserved | -1,023 | 2,705E-31 |
| TcCLB.506477.60 | protein kinase, putative | -1,023 | 8,2E-24 |
| TcCLB.506763.330 | mucin-associated surface protein (MASP), putative | -1,024 | 0,004454 |
| TcCLB.510461.25 | elongation factor 1-gamma (EF-1-gamma, pseudogene), p | -1,024 | 0,04106 |
| TcCLB.506053.35 | hypothetical protein, conserved (pseudogene) | -1,025 | 1,473E-13 |
| TcCLB.506457.90 | hypothetical protein | -1,025 | 2,383E-12 |
| TcCLB.509369.20 | retrotransposon hot spot protein (RHS, pseudogene), putat | -1,025 | 0,009141 |
| TcCLB.509097.36 | surface protease GP63 (pseudogene), putative | -1,025 | 0,1236 |
| TcCLB.511415.21 | trans-sialidase (pseudogene), putative | -1,026 | 0,00002515 |
| TcCLB.463269.10 | eukaryotic translation initiation factor 1A, putative (fragme | -1,026 | 0,0005031 |
| TcCLB.510013.40 | mucin-associated surface protein (MASP), putative | -1,026 | 0,007258 |
| TcCLB.509437.30 | hypothetical protein, conserved | -1,027 | 5,96E-42 |
| TcCLB.506671.61 | hypothetical protein, conserved (pseudogene) | -1,027 | 2,464E-06 |
| TcCLB.508145.30 | RNA-binding protein, putative | -1,028 | 0,001886 |
| TcCLB.508169.30 | hypothetical protein, conserved | -1,029 | 5,469E-14 |
| TcCLB.510877.80 | phospholipid-transporting ATPase-like protein, putative | -1,03 | 8,348E-37 |
| TcCLB.508327.10 | mucin-associated surface protein (MASP), putative | -1,03 | 0,07476 |
| TcCLB.503463.20 | hypothetical protein, conserved | -1,031 | 1,962E-19 |
| TcCLB.509233.220 | hypothetical protein, conserved | -1,033 | 4,911E-28 |
| TcCLB.505481.10 | hypothetical protein | -1,033 | 0,0002648 |
| TcCLB.510359.40 | hypothetical protein, conserved | -1,034 | 7,505E-06 |
| TcCLB.511179.60 | hypothetical protein, conserved | -1,034 | 0,004289 |
| TcCLB.510547.46 | hypothetical protein, conserved (pseudogene) | -1,035 | 0,0538 |
| TcCLB.510769.80 | ankyrin repeat protein, putative | -1,036 | 6,86E-31 |
| TcCLB.511491.170 | quiescin sulfhydryl oxidase, putative (QSOX) | -1,036 | 3,265E-25 |
| TcCLB.510105.120 | protein kinase, putative | -1,036 | 1,502E-16 |
| TcCLB.509505.10 | protein disulfide isomerase, putative | -1,037 | 2,238E-18 |

| | | | |
|---|---|---|---|
| TcCLB.507237.260 | mucin TcMUCII, putative | -1,038 | 0,04958 |
| TcCLB.506501.150 | surface protease GP63 (pseudogene), putative | -1,039 | 0,06015 |
| TcCLB.511529.170 | enoyl-CoA hydratase/isomerase family protein, putative | -1,04 | 2,009E-08 |
| TcCLB.506739.90 | ATP-dependent RNA helicase, putative | -1,041 | 6,856E-37 |
| TcCLB.508173.129 | hypothetical protein, conserved | -1,042 | 3,427E-25 |
| TcCLB.506713.30 | ATP-dependent DEAD/H RNA helicase, putative | -1,042 | 9,89E-25 |
| TcCLB.506747.10 | orotidine-5-phosphate decarboxylase/orotate phosphoribo: | -1,044 | 1,663E-07 |
| TcCLB.510607.51 | retrotransposon hot spot protein (RHS, pseudogene), putat | -1,044 | 0,004317 |
| TcCLB.508655.10 | DNA-directed RNA polymerase II subunit 2 (pseudogene), | | -1,044 | 0,01333 |
| TcCLB.509605.20 | hypothetical protein, conserved | -1,045 | 1,733E-29 |
| TcCLB.507021.60 | p1/s1 nuclease, putative | -1,046 | 7,101E-23 |
| TcCLB.410275.18 | trans-sialidase (pseudogene), putative | -1,046 | 0,05437 |
| TcCLB.508757.10 | trans-sialidase (pseudogene), putative | -1,046 | 0,1765 |
| TcCLB.506459.130 | mucin-associated surface protein (MASP), putative | -1,047 | 0,005114 |
| TcCLB.510187.404 | dolichyl-P-Man:GDP-Man5GlcNAc2-PP-dolichyl alpha-1,3-n | -1,048 | 3,863E-19 |
| TcCLB.398009.20 | trans-sialidase (pseudogene), putative | -1,048 | 3,023E-14 |
| TcCLB.506857.120 | hypothetical protein, conserved | -1,048 | 0,03084 |
| TcCLB.511219.10 | dispersed gene family protein 1 (DGF-1), putative | -1,049 | 0,02203 |
| TcCLB.507801.130 | heat shock protein DnaJ (pseudogene), putative | -1,05 | 4,756E-49 |
| TcCLB.510197.5 | hypothetical protein, conserved (pseudogene) | -1,05 | 0,00001392 |
| TcCLB.506133.30 | surface protease GP63 (pseudogene), putative | -1,05 | 0,03314 |
| TcCLB.510061.60 | dispersed gene family protein 1 (DGF-1), putative | -1,05 | 0,0688 |
| TcCLB.507467.70 | diacylglycerol acyltransferase, putative | -1,051 | 1,073E-26 |
| TcCLB.511413.11 | hypothetical protein, conserved (pseudogene) | -1,051 | 1,897E-14 |
| TcCLB.509097.10 | hypothetical protein | -1,053 | 0,06702 |
| TcCLB.506503.80 | kinesin, putative | -1,054 | 3,76E-28 |
| TcCLB.510339.54 | hypothetical protein, conserved | -1,054 | 9,065E-27 |
| TcCLB.508221.381 | mucin-associated surface protein (MASP, pseudogene), pu | -1,055 | 0,07379 |
| TcCLB.443899.10 | trans-sialidase, putative (fragment) | -1,055 | 0,1902 |
| TcCLB.508961.70 | hypothetical protein, conserved | -1,056 | 2,236E-19 |
| TcCLB.511753.20 | class I nuclease-like protein, putative | -1,056 | 9,015E-15 |
| TcCLB.509085.136 | trans-sialidase (pseudogene), putative | -1,056 | 0,005621 |
| TcCLB.511577.104 | hypothetical protein, conserved | -1,057 | 3,834E-09 |
| TcCLB.509007.40 | hypothetical protein, conserved | -1,057 | 0,125 |
| TcCLB.511439.70 | hypothetical protein, conserved | -1,058 | 3,336E-22 |
| TcCLB.510721.21 | hypothetical protein | -1,058 | 0,004886 |
| TcCLB.511063.61 | hypothetical protein, conserved (pseudogene) | -1,058 | 0,007544 |
| TcCLB.510701.31 | hypothetical protein | -1,058 | 0,05567 |
| TcCLB.506989.190 | heat shock protein 90, putative (LPG3) | -1,059 | 4,578E-44 |
| TcCLB.506241.124 | ankyrin repeat protein, putative | -1,059 | 3,071E-37 |
| TcCLB.510251.10 | amino acid permease-like protein, putative | -1,06 | 3,379E-26 |
| TcCLB.509647.170 | hypothetical protein, conserved | -1,061 | 7,956E-32 |
| TcCLB.510701.50 | mucin TcMUC (pseudogene), putative | -1,061 | 0,005677 |
| TcCLB.510497.170 | trans-sialidase (pseudogene), putative | -1,061 | 0,03804 |
| TcCLB.511375.10 | hypothetical protein (pseudogene) | -1,061 | 0,1031 |

| | | | |
|---|---|---|---|
| TcCLB.511301.70 | FK506-binding protein (FKBP)-type peptidyl-prolyl isomera: | -1,062 | 1,532E-33 |
| TcCLB.510329.20 | hypothetical protein, conserved | -1,062 | 1,091E-16 |
| TcCLB.504197.10 | hypothetical protein | -1,063 | 0,03289 |
| TcCLB.510487.30 | hypothetical protein, conserved (pseudogene) | -1,065 | 2,605E-09 |
| TcCLB.510971.10 | dispersed gene family protein 1 (DGF-1), putative | -1,065 | 1,334E-07 |
| TcCLB.509527.120 | trans-sialidase, Group V, putative | -1,065 | 0,00004373 |
| TcCLB.506509.10 | hypothetical protein, conserved | -1,067 | 4,553E-16 |
| TcCLB.510773.30 | hypothetical protein | -1,068 | 0,001233 |
| TcCLB.510211.10 | trans-sialidase (pseudogene), putative | -1,068 | 0,1004 |
| TcCLB.508719.30 | hypothetical protein, conserved | -1,07 | 2,068E-42 |
| TcCLB.504089.70 | hypothetical protein, conserved | -1,071 | 3,713E-15 |
| TcCLB.507831.60 | heat shock protein 110, putative | -1,072 | 3,529E-50 |
| TcCLB.511875.123 | hypothetical protein, conserved (pseudogene) | -1,072 | 0,02909 |
| TcCLB.508209.140 | protein disulfide isomerase, putative | -1,073 | 2,359E-50 |
| TcCLB.506469.130 | SNARE domain-containing protein, putative | -1,073 | 2,373E-14 |
| TcCLB.511025.60 | hypothetical protein | -1,075 | 1,237E-12 |
| TcCLB.506211.240 | hypothetical protein, conserved | -1,078 | 2,111E-33 |
| TcCLB.507979.4 | histone H2A, putative | -1,079 | 3,091E-07 |
| TcCLB.509657.11 | hypothetical protein, conserved (pseudogene) | -1,079 | 7,684E-06 |
| TcCLB.503799.4 | protein kinase, putative (fragment) | -1,082 | 8,57E-11 |
| TcCLB.509753.10 | hypothetical protein, conserved (pseudogene) | -1,082 | 0,003703 |
| TcCLB.506567.80 | hypothetical protein | -1,084 | 1,218E-16 |
| TcCLB.508913.39 | O-sialoglycoprotein endopeptidase, putative | -1,084 | 6,772E-09 |
| TcCLB.508221.400 | trans-sialidase, Group V, putative | -1,084 | 0,002948 |
| TcCLB.506605.120 | hypothetical protein, conserved | -1,085 | 4,798E-30 |
| TcCLB.506679.280 | endonuclease/exonuclease/phosphatase, putative | -1,087 | 3,171E-15 |
| TcCLB.509525.340 | hypothetical protein | -1,087 | 0,01712 |
| TcCLB.510501.40 | trans-sialidase, putative | -1,088 | 0,02268 |
| TcCLB.506885.400 | 40 kDa cyclophilin, putative (CYP40) | -1,089 | 2,299E-25 |
| TcCLB.508221.130 | mucin TcMUCII (pseudogene), putative | -1,089 | 0,005966 |
| TcCLB.509075.50 | trans-sialidase, Group II, putative | -1,09 | 2,825E-07 |
| TcCLB.508153.180 | aspartyl aminopeptidase, putative | -1,091 | 4,57E-41 |
| TcCLB.511751.140 | hypothetical protein | -1,092 | 2,619E-13 |
| TcCLB.503479.40 | hypothetical protein, conserved | -1,093 | 9,878E-37 |
| TcCLB.506191.39 | dispersed gene family protein 1 (DGF-1), putative | -1,094 | 1,282E-11 |
| TcCLB.430281.20 | hypothetical protein (fragment) | -1,095 | 0,00914 |
| TcCLB.508209.144 | hypothetical protein, conserved | -1,096 | 1,382E-14 |
| TcCLB.511183.600 | mucin-associated surface protein (MASP, pseudogene), pu | -1,096 | 5,907E-07 |
| TcCLB.509663.10 | myosin heavy chain, putative | -1,097 | 4,892E-31 |
| TcCLB.510031.70 | heat shock protein Hsp20, putative | -1,097 | 5,285E-14 |
| TcCLB.503417.20 | hypothetical protein | -1,097 | 0,00468 |
| TcCLB.511185.150 | hypothetical protein, conserved (pseudogene) | -1,099 | 1,037E-14 |
| TcCLB.504155.250 | trans-sialidase (pseudogene), putative | -1,099 | 0,007279 |
| TcCLB.506855.190 | hypothetical protein, conserved | -1,103 | 1,561E-29 |
| TcCLB.506257.5 | aminopeptidase, putative (fragment) | -1,103 | 2,17E-10 |

| | | | |
|---|---|---|---|
| TcCLB.508865.2 | hypothetical protein, conserved (fragment) | -1,105 | 0,004991 |
| TcCLB.508099.50 | trans-sialidase, putative | -1,105 | 0,1659 |
| TcCLB.506793.40 | UDP-glucuronic acid/UDP-N-acetylgalactosamine transpor | -1,106 | 3,166E-18 |
| TcCLB.506111.20 | dispersed gene family protein 1 (DGF-1), putative | -1,106 | 5,822E-13 |
| TcCLB.511367.330 | hypothetical protein, conserved | -1,106 | 2,801E-12 |
| TcCLB.507163.75 | hypothetical protein | -1,107 | 0,0244 |
| TcCLB.511233.170 | elongation factor 1-gamma (EF-1-gamma, pseudogene), p | -1,108 | 0,02213 |
| TcCLB.509213.40 | hypothetical protein, conserved | -1,109 | 3,728E-46 |
| TcCLB.507765.20 | PUF nine target 1 | -1,109 | 2,121E-40 |
| TcCLB.510021.100 | mucin TcMUCII, putative | -1,109 | 0,001516 |
| TcCLB.503531.10 | hypothetical protein, conserved | -1,11 | 0,0003254 |
| TcCLB.506229.10 | Mitogen-activated protein kinase 10, putative (MPK10) | -1,111 | 1,471E-12 |
| TcCLB.508933.50 | hypothetical protein, conserved | -1,112 | 1,761E-18 |
| TcCLB.507633.80 | trans-sialidase, Group V, putative | -1,112 | 0,0001895 |
| TcCLB.506719.10 | phosphatidylinositol (3,5) kinase, putative | -1,114 | 4,766E-19 |
| TcCLB.508189.31 | hypothetical protein, conserved (pseudogene) | -1,114 | 0,0003047 |
| TcCLB.510419.20 | trans-sialidase, Group VI, putative | -1,115 | 0,0005092 |
| TcCLB.510363.250 | trans-sialidase, Group V, putative | -1,115 | 0,002372 |
| TcCLB.510479.40 | trans-sialidase, putative (fragment) | -1,117 | 0,006046 |
| TcCLB.508101.9 | trans-sialidase, putative (fragment) | -1,119 | 0,003491 |
| TcCLB.503989.20 | hypothetical protein, conserved | -1,12 | 5,356E-19 |
| TcCLB.504241.34 | dispersed gene family protein 1 (DGF-1), putative (fragme | -1,121 | 1,688E-07 |
| TcCLB.509233.120 | hypothetical protein, conserved | -1,122 | 6,567E-50 |
| TcCLB.510735.60 | hypothetical protein, conserved | -1,122 | 4,164E-13 |
| TcCLB.507641.260 | chaperonin HSP60, mitochondrial precursor (pseudogene) | -1,123 | 5,885E-19 |
| TcCLB.511203.29 | surface protease GP63, putative (fragment) | -1,123 | 2,51E-08 |
| TcCLB.506859.4 | N-acetyltransferase, putative (fragment) | -1,123 | 0,00001337 |
| TcCLB.509469.10 | dispersed gene family protein 1 (DGF-1, pseudogene), puta | -1,123 | 0,00002269 |
| TcCLB.457113.10 | trans-sialidase, putative (fragment) | -1,123 | 0,244 |
| TcCLB.503867.5 | hypothetical protein, conserved | -1,125 | 0,00001508 |
| TcCLB.510365.10 | trans-sialidase (pseudogene), putative | -1,125 | 0,01211 |
| TcCLB.508245.21 | surface protease GP63 (pseudogene), putative | -1,125 | 0,05581 |
| TcCLB.506953.45 | ascorbate-dependent peroxidase (pseudogene), putative | -1,126 | 4,018E-09 |
| TcCLB.503733.50 | hypothetical protein, conserved (pseudogene) | -1,127 | 6,875E-15 |
| TcCLB.510817.60 | hypothetical protein, conserved | -1,128 | 2,94E-47 |
| TcCLB.511237.130 | trans-sialidase, putative (fragment) | -1,128 | 6,482E-07 |
| TcCLB.508251.21 | hypothetical protein, conserved (pseudogene) | -1,13 | 9,161E-06 |
| TcCLB.503533.40 | mucin-associated surface protein (MASP), putative | -1,13 | 0,01287 |
| TcCLB.511385.40 | hypothetical protein | -1,131 | 1,074E-07 |
| TcCLB.506483.10 | Triglyceride lipase, putative | -1,132 | 3,958E-15 |
| TcCLB.507641.60 | aldehyde dehydrogenase, putative | -1,133 | 1,404E-36 |
| TcCLB.503849.80 | NADH-dependent fumarate reductase, putative | -1,137 | 1,544E-10 |
| TcCLB.504533.10 | dispersed gene family protein 1 (DGF-1), putative | -1,138 | 3,705E-09 |
| TcCLB.506459.200 | hypothetical protein | -1,139 | 0,01011 |
| TcCLB.504147.280 | trypanothione synthetase-like protein, putative | -1,14 | 1,448E-41 |

| | | | |
|---|---|---|---|
| TcCLB.507057.60 | hypothetical protein, conserved | -1,14 | 5,548E-13 |
| TcCLB.439653.10 | NADH-dependent fumarate reductase, putative | -1,14 | 0,00001422 |
| TcCLB.510531.109 | dynein heavy chain, cytosolic, putative (fragment) | -1,143 | 3,053E-13 |
| TcCLB.510279.263 | trans-sialidase (pseudogene), putative | -1,143 | 0,0003173 |
| TcCLB.507957.334 | mucin-associated surface protein (MASP), putative | -1,145 | 0,2392 |
| TcCLB.509085.90 | retrotransposon hot spot protein (RHS, pseudogene), putat | -1,148 | 2,96E-07 |
| TcCLB.510819.100 | mitochondrial carrier protein, putative | -1,149 | 3,489E-21 |
| TcCLB.506683.10 | hypothetical protein, conserved (pseudogene) | -1,15 | 0,0138 |
| TcCLB.510713.11 | hypothetical protein, conserved (pseudogene) | -1,153 | 0,00000418 |
| TcCLB.504069.140 | trans-sialidase (pseudogene), putative | -1,154 | 9,604E-12 |
| TcCLB.508731.9 | hypothetical protein, conserved (fragment) | -1,155 | 1,408E-20 |
| TcCLB.508373.29 | orotidine-5-phosphate decarboxylase/orotate phosphoribo: | -1,155 | 6,772E-20 |
| TcCLB.507993.150 | hypothetical protein, conserved | -1,156 | 3,858E-36 |
| TcCLB.506229.30 | hypothetical protein | -1,159 | 8,286E-23 |
| TcCLB.506501.350 | mucin-associated surface protein (MASP), putative | -1,159 | 0,008178 |
| TcCLB.506251.30 | hypothetical protein, conserved | -1,16 | 5,976E-39 |
| TcCLB.506699.40 | katanin-like protein, putative | -1,161 | 1,639E-15 |
| TcCLB.507611.190 | retrotransposon hot spot protein (RHS, pseudogene), putat | -1,161 | 0,00000103 |
| TcCLB.510363.220 | mucin-associated surface protein (MASP), putative | -1,161 | 0,004734 |
| TcCLB.503875.10 | mucin-associated surface protein (MASP), putative | -1,161 | 0,02257 |
| TcCLB.511667.90 | hypothetical protein, conserved (pseudogene) | -1,164 | 3,426E-12 |
| TcCLB.498829.21 | hypothetical protein | -1,166 | 0,01451 |
| TcCLB.510629.120 | dispersed gene family protein 1 (DGF-1, pseudogene), puta | -1,168 | 0,009522 |
| TcCLB.510187.551 | chaperonin HSP60, mitochondrial precursor, putative (fragı | -1,171 | 4,715E-24 |
| TcCLB.511627.70 | hypothetical protein, conserved | -1,172 | 1,508E-18 |
| TcCLB.506263.49 | hypothetical protein, conserved (fragment) | -1,173 | 9,529E-16 |
| TcCLB.503809.158 | Triglyceride lipase, putative (fragment) | -1,173 | 1,429E-06 |
| TcCLB.506171.60 | trans-sialidase, Group II, putative | -1,179 | 0,00006106 |
| TcCLB.503919.60 | hypothetical protein, conserved | -1,181 | 4,933E-21 |
| TcCLB.507065.127 | mucin-associated surface protein (MASP, pseudogene), pu | -1,181 | 9,163E-08 |
| TcCLB.508221.660 | mucin-associated surface protein (MASP), putative | -1,181 | 0,005736 |
| TcCLB.506579.50 | hypothetical protein, conserved | -1,183 | 1,165E-29 |
| TcCLB.511365.90 | hypothetical protein, conserved | -1,183 | 3,619E-24 |
| TcCLB.399373.20 | dispersed gene family protein 1 (DGF-1), putative | -1,184 | 2,374E-06 |
| TcCLB.507729.20 | hypothetical protein, conserved (pseudogene) | -1,185 | 4,442E-08 |
| TcCLB.508265.80 | hypothetical protein | -1,186 | 5,451E-25 |
| TcCLB.511127.10 | RNA-binding protein 5, putative | -1,187 | 0,1076 |
| TcCLB.503463.10 | protein kinase C substrate protein, heavy chain, putative | -1,188 | 3,53E-21 |
| TcCLB.503539.30 | GTP-binding nuclear protein rtb2, putative | -1,189 | 8,81E-62 |
| TcCLB.506501.290 | mucin-associated surface protein (MASP), putative | -1,194 | 0,00003675 |
| TcCLB.505965.30 | prenyltransferase, putative | -1,195 | 7,105E-20 |
| TcCLB.509663.60 | retrotransposon hot spot protein (RHS, pseudogene), putat | -1,195 | 4,165E-17 |
| TcCLB.511487.130 | trans-sialidase (pseudogene), putative | -1,196 | 9,283E-07 |
| TcCLB.506239.60 | hypothetical protein, conserved | -1,197 | 2,136E-15 |
| TcCLB.503695.10 | retrotransposon hot spot protein (RHS, pseudogene), putat | -1,198 | 7,494E-06 |

| | | | |
|---|---|---|---|
| TcCLB.504929.5 | zinc finger CCCH domain containing protein 11 (ZC3H11) | -1,198 | 0,02118 |
| TcCLB.511575.70 | kinetoplastid kinetochore protein 4, putative | -1,199 | 2,825E-25 |
| TcCLB.511467.30 | hypothetical protein, conserved | -1,2 | 1,996E-19 |
| TcCLB.511263.9 | heat shock protein 110, putative (fragment) | -1,202 | 3,393E-57 |
| TcCLB.511215.90 | hypothetical protein, conserved | -1,205 | 0,0000402 |
| TcCLB.511723.10 | surface protease GP63, putative | -1,206 | 7,049E-16 |
| TcCLB.508313.140 | mucin-associated surface protein (MASP), putative | -1,206 | 0,00995 |
| TcCLB.508645.70 | nucleoside transporter 1, putative | -1,207 | 5,478E-17 |
| TcCLB.508159.26 | hypothetical protein, conserved (pseudogene) | -1,209 | 1,595E-07 |
| TcCLB.510629.80 | trans-sialidase (pseudogene), putative | -1,211 | 0,01192 |
| TcCLB.507061.49 | hypothetical protein | -1,212 | 1,882E-21 |
| TcCLB.422207.10 | UDP-Gal or UDP-GlcNAc-dependent glycosyltransferase, pu | -1,212 | 0,121 |
| TcCLB.507859.80 | trans-sialidase (pseudogene), putative | -1,213 | 0,01883 |
| TcCLB.508323.70 | centrin-4, putative (Centrin4) | -1,219 | 6,092E-10 |
| TcCLB.506501.220 | mucin-associated surface protein (MASP), putative | -1,219 | 0,001746 |
| TcCLB.508221.440 | trans-sialidase (pseudogene), putative | -1,22 | 0,0000302 |
| TcCLB.511277.590 | hypothetical protein, conserved | -1,221 | 8,466E-43 |
| TcCLB.509023.209 | hypothetical protein, conserved | -1,221 | 6,068E-34 |
| TcCLB.511751.160 | hypothetical protein, conserved | -1,221 | 1,597E-15 |
| TcCLB.508365.190 | trans-sialidase, Group VI, putative | -1,223 | 1,741E-06 |
| TcCLB.506835.90 | hypothetical protein, conserved | -1,224 | 1,315E-34 |
| TcCLB.511039.10 | kinetoplast DNA-associated protein, putative | -1,224 | 4,706E-31 |
| TcCLB.509505.60 | hypothetical protein | -1,225 | 3,733E-22 |
| TcCLB.508367.50 | hypothetical protein, conserved (pseudogene) | -1,226 | 4,605E-07 |
| TcCLB.510217.10 | hypothetical protein | -1,229 | 0,06443 |
| TcCLB.460061.20 | trans-sialidase, putative (fragment) | -1,229 | 0,09326 |
| TcCLB.503821.5 | hypothetical protein, conserved | -1,232 | 9,069E-09 |
| TcCLB.507641.280 | chaperonin HSP60, mitochondrial precursor (HSP60) | -1,234 | 6,096E-26 |
| TcCLB.510197.30 | trans-sialidase, Group V, putative | -1,235 | 1,803E-07 |
| TcCLB.510627.200 | trans-sialidase (pseudogene), putative | -1,235 | 0,02944 |
| TcCLB.511237.10 | thimet oligopeptidase, putative | -1,236 | 4,975E-47 |
| TcCLB.506739.80 | mitochondrial carrier protein, putative | -1,236 | 2,356E-33 |
| TcCLB.503793.20 | phosphatidylinositol (3,5) kinase, putative (fragment) | -1,236 | 2,037E-08 |
| TcCLB.504233.10 | surface protease GP63, putative | -1,237 | 7,186E-10 |
| TcCLB.506451.16 | hypothetical protein, conserved (pseudogene) | -1,237 | 0,0002098 |
| TcCLB.503753.20 | hypothetical protein, conserved | -1,239 | 4,858E-20 |
| TcCLB.509521.84 | retrotransposon hot spot protein (RHS, pseudogene), putat | -1,24 | 7,225E-06 |
| TcCLB.505101.30 | hypothetical protein, conserved | -1,24 | 0,001761 |
| TcCLB.508943.4 | glycerol uptake protein, putative | -1,24 | 0,009053 |
| TcCLB.509553.40 | ribonuclease mar1, putative | -1,242 | 3,112E-16 |
| TcCLB.506613.141 | hypothetical protein, conserved (pseudogene) | -1,244 | 7,953E-13 |
| TcCLB.504445.10 | dispersed gene family protein 1 (DGF-1), putative | -1,244 | 4,535E-07 |
| TcCLB.510413.10 | hypothetical protein, conserved (pseudogene) | -1,245 | 1,049E-10 |
| TcCLB.506667.90 | trans-sialidase, Group V, putative | -1,246 | 0,0001996 |
| TcCLB.510289.40 | GPI inositol deacylase precursor, putative | -1,247 | 3,662E-27 |

| | | | |
|---|---|---|---|
| TcCLB.506501.401 | hypothetical protein, conserved (pseudogene) | -1,247 | 0,06456 |
| TcCLB.506559.450 | hypothetical protein | -1,249 | 4,307E-21 |
| TcCLB.503755.60 | surface protease GP63 (pseudogene), putative | -1,25 | 0,0003604 |
| TcCLB.508737.20 | hypothetical protein | -1,251 | 1,552E-16 |
| TcCLB.507733.40 | hypothetical protein, conserved (pseudogene) | -1,251 | 0,000104 |
| TcCLB.509683.30 | hypothetical protein, conserved | -1,252 | 5,514E-34 |
| TcCLB.476911.10 | trans-sialidase, putative (fragment) | -1,252 | 0,02696 |
| TcCLB.508595.11 | hypothetical protein | -1,253 | 0,03028 |
| TcCLB.508483.13 | trans-sialidase (pseudogene), putative | -1,254 | 0,0000664 |
| TcCLB.510797.4 | phosphatidylinositol (3,5) kinase, putative (fragment) | -1,256 | 1,259E-16 |
| TcCLB.503825.10 | protein kinase (pseudogene), putative | -1,256 | 1,504E-14 |
| TcCLB.506965.60 | elongation factor 1-gamma (EF-1-gamma, pseudogene), p | -1,256 | 0,01865 |
| TcCLB.511183.480 | hypothetical protein, conserved (pseudogene) | -1,257 | 0,0007178 |
| TcCLB.425969.5 | hypothetical protein | -1,259 | 1,891E-21 |
| TcCLB.508297.7 | trans-sialidase, putative (fragment) | -1,261 | 5,831E-08 |
| TcCLB.506945.40 | hypothetical protein, conserved | -1,264 | 2,669E-33 |
| TcCLB.506867.10 | endonuclease G, putative | -1,264 | 7,334E-17 |
| TcCLB.506755.50 | hypothetical protein, conserved | -1,267 | 7,038E-24 |
| TcCLB.506667.50 | trans-sialidase (pseudogene), putative | -1,268 | 0,000136 |
| TcCLB.511517.95 | hypothetical protein, conserved | -1,269 | 8,415E-09 |
| TcCLB.509891.70 | hypothetical protein, conserved (pseudogene) | -1,27 | 5,215E-29 |
| TcCLB.509791.120 | kinetoplast-associated protein (KAP6) | -1,271 | 1,391E-34 |
| TcCLB.510481.90 | hypothetical protein, conserved | -1,275 | 3,701E-06 |
| TcCLB.507237.250 | mucin-associated surface protein (MASP, pseudogene), pu | -1,282 | 9,527E-09 |
| TcCLB.509499.20 | beta propeller protein, putative | -1,285 | 1,283E-35 |
| TcCLB.508277.280 | hypothetical protein | -1,287 | 1,391E-19 |
| TcCLB.506211.150 | polyprenyl synthase, putative | -1,293 | 7,265E-28 |
| TcCLB.507011.30 | hypothetical protein, conserved | -1,294 | 3,181E-34 |
| TcCLB.510693.150 | mucin-associated surface protein (MASP), putative | -1,296 | 0,1037 |
| TcCLB.405737.4 | RNA polymerase-associated protein CDC73, putative (fragr | -1,297 | 0,001052 |
| TcCLB.506251.50 | Cell division control protein 48, putative | -1,3 | 1,914E-18 |
| TcCLB.509085.50 | trans-sialidase (pseudogene), putative | -1,302 | 1,634E-09 |
| TcCLB.503709.10 | hypothetical protein | -1,305 | 0,00204 |
| TcCLB.507105.50 | hypothetical protein | -1,306 | 1,883E-81 |
| TcCLB.510503.40 | surface protease GP63 (pseudogene), putative | -1,306 | 0,001025 |
| TcCLB.510831.10 | mucin-associated surface protein (MASP, pseudogene), pu | -1,308 | 0,005824 |
| TcCLB.507059.60 | orotidine-5-phosphate decarboxylase/orotate phosphoribo: | -1,309 | 1,448E-36 |
| TcCLB.505935.10 | hypothetical protein, conserved (pseudogene) | -1,309 | 1,586E-07 |
| TcCLB.508163.371 | hypothetical protein, conserved (pseudogene) | -1,309 | 3,212E-07 |
| TcCLB.510111.30 | hypothetical protein, conserved (pseudogene) | -1,311 | 6,618E-21 |
| TcCLB.510627.179 | syntaxin binding protein 1 (pseudogene), putative | -1,313 | 0,001851 |
| TcCLB.504427.10 | trypanothione synthetase, putative | -1,315 | 3,241E-15 |
| TcCLB.508461.330 | hypothetical protein | -1,317 | 1,599E-28 |
| TcCLB.510197.95 | hypothetical protein, conserved (pseudogene) | -1,317 | 4,241E-11 |
| TcCLB.509085.110 | trans-sialidase (pseudogene), putative | -1,317 | 4,262E-09 |

| | | | |
|---|---|---|---|
| TcCLB.509553.20 | ATP-dependent chaperone, putative | -1,318 | 1,995E-22 |
| TcCLB.508835.10 | hypothetical protein, conserved (pseudogene) | -1,318 | 3,882E-15 |
| TcCLB.508153.230 | hypothetical protein, conserved | -1,321 | 1,266E-42 |
| TcCLB.511561.20 | dispersed gene family protein 1 (DGF-1), putative | -1,321 | 7,56E-17 |
| TcCLB.506445.110 | nucleobase transporter, putative | -1,325 | 1,964E-64 |
| TcCLB.507081.100 | hypothetical protein, conserved | -1,325 | 2,637E-20 |
| TcCLB.508183.4 | aspartyl aminopeptidase, putative (fragment) | -1,327 | 1,924E-20 |
| TcCLB.508323.90 | hypothetical protein, conserved | -1,327 | 2,843E-12 |
| TcCLB.508219.50 | mucin TcMUCII, putative | -1,329 | 0,004648 |
| TcCLB.510647.30 | tryptophanyl-tRNA synthetase, putative | -1,33 | 7,621E-44 |
| TcCLB.507641.104 | small GTPase, putative | -1,33 | 5,351E-22 |
| TcCLB.508873.60 | trans-sialidase, Group V, putative | -1,331 | 0,0001149 |
| TcCLB.510279.320 | trans-sialidase, Group V, putative | -1,332 | 2,361E-07 |
| TcCLB.506501.270 | syntaxin binding protein (pseudogene), putative | -1,334 | 0,01459 |
| TcCLB.511297.30 | hypothetical protein, conserved (fragment) | -1,338 | 1,559E-30 |
| TcCLB.506813.170 | hypothetical protein, conserved (pseudogene) | -1,338 | 3,746E-10 |
| TcCLB.506929.54 | hypothetical protein | -1,339 | 1,214E-11 |
| TcCLB.504035.21 | hypothetical protein | -1,339 | 0,001254 |
| TcCLB.506459.195 | hypothetical protein, conserved | -1,34 | 0,0000921 |
| TcCLB.511553.45 | surface protease GP63 (pseudogene), putative | -1,34 | 0,002845 |
| TcCLB.506881.10 | hypothetical protein, conserved (pseudogene) | -1,343 | 2,602E-06 |
| TcCLB.509655.20 | hypothetical protein, conserved (pseudogene) | -1,346 | 8,402E-09 |
| TcCLB.508209.90 | 10 kDa heat shock protein, putative | -1,347 | 0,001687 |
| TcCLB.509455.80 | GTP-binding nuclear protein rtb2, putative | -1,35 | 1,824E-47 |
| TcCLB.509899.110 | orotidine-5-phosphate decarboxylase/orotate phosphoribo: | -1,351 | 4,021E-41 |
| TcCLB.507089.170 | hypothetical protein, conserved | -1,353 | 2,664E-39 |
| TcCLB.511481.55 | RNA-binding protein, putative | -1,364 | 2,133E-19 |
| TcCLB.510323.110 | hypothetical protein, conserved | -1,367 | 4,681E-12 |
| TcCLB.509011.40 | calreticulin, putative | -1,373 | 5,86E-117 |
| TcCLB.510401.10 | dispersed gene family protein 1 (DGF-1), putative | -1,375 | 0,0204 |
| TcCLB.505999.90 | phosphoglycerate kinase, putative | -1,38 | 2,013E-59 |
| TcCLB.510533.80 | hypothetical protein, conserved | -1,38 | 1,559E-22 |
| TcCLB.503991.10 | hypothetical protein, conserved | -1,38 | 9,186E-19 |
| TcCLB.510493.4 | hypothetical protein, conserved (fragment) | -1,382 | 4,006E-09 |
| TcCLB.506133.207 | mucin-associated surface protein (MASP, pseudogene), pu | -1,385 | 0,0001541 |
| TcCLB.510363.196 | syntaxin binding protein (pseudogene), putative | -1,387 | 0,005162 |
| TcCLB.507795.60 | tryptophanyl-tRNA synthetase (pseudogene), putative | -1,39 | 1,073E-39 |
| TcCLB.506973.25 | hypothetical protein | -1,39 | 1,241E-07 |
| TcCLB.509287.140 | dispersed gene family protein 1 (DGF-1), putative | -1,393 | 1,654E-16 |
| TcCLB.508221.414 | elongation factor 1-gamma (EF-1-gamma), putative | -1,394 | 0,0002341 |
| TcCLB.508175.90 | hypothetical protein | -1,395 | 5,451E-77 |
| TcCLB.510629.101 | hypothetical protein, conserved (pseudogene) | -1,395 | 0,01109 |
| TcCLB.503471.19 | expression site-associated gene (ESAG-like) protein, putati | -1,397 | 2,17E-18 |
| TcCLB.511167.100 | hypothetical protein, conserved | -1,404 | 1,793E-33 |
| TcCLB.510489.10 | hypothetical protein | -1,417 | 1,769E-07 |

| | | | |
|---|---|---|---|
| TcCLB.507049.66 | hypothetical protein | -1,421 | 6,894E-17 |
| TcCLB.504239.150 | hypothetical protein | -1,421 | 0,04291 |
| TcCLB.507771.30 | hypothetical protein, conserved | -1,422 | 3,925E-30 |
| TcCLB.510291.14 | mitochondrial carrier protein, putative | -1,422 | 6,099E-27 |
| TcCLB.508149.40 | hypothetical protein, conserved (pseudogene) | -1,423 | 1,16E-08 |
| TcCLB.508369.40 | serine/threonine-protein kinase A (pseudogene), putative | -1,432 | 1,804E-42 |
| TcCLB.433673.10 | trans-sialidase, putative (fragment) | -1,432 | 6,353E-33 |
| TcCLB.509369.40 | dispersed gene family protein 1 (DGF-1, pseudogene), puta | -1,433 | 1,598E-08 |
| TcCLB.511529.59 | hypothetical protein | -1,436 | 3,573E-26 |
| TcCLB.509793.50 | hypothetical protein, conserved | -1,437 | 6,26E-86 |
| TcCLB.507859.101 | hypothetical protein, conserved (pseudogene) | -1,45 | 0,0004901 |
| TcCLB.510711.10 | trans-sialidase (pseudogene), putative | -1,452 | 0,0002048 |
| TcCLB.503631.9 | hypothetical protein, conserved | -1,454 | 1,025E-30 |
| TcCLB.511385.50 | hypothetical protein | -1,457 | 1,214E-18 |
| TcCLB.510329.70 | cdc2-related kinase 10, putative (CRK10) | -1,458 | 2,147E-22 |
| TcCLB.505479.10 | mucin-associated surface protein (MASP, pseudogene), pu | -1,458 | 0,0007304 |
| TcCLB.507055.20 | hypothetical protein, conserved (fragment) | -1,459 | 3,77E-23 |
| TcCLB.507641.290 | chaperonin HSP60, mitochondrial precursor (HSP60) | -1,46 | 0,00002417 |
| TcCLB.508395.9 | dispersed gene family protein 1 (DGF-1), putative (fragme | -1,467 | 9,627E-06 |
| TcCLB.508637.19 | hypothetical protein, conserved | -1,468 | 0,008348 |
| TcCLB.506501.230 | mucin TcMUCII, putative | -1,468 | 0,009001 |
| TcCLB.504081.23 | hypothetical protein, conserved | -1,47 | 0,008005 |
| TcCLB.511575.120 | hypothetical protein, conserved | -1,473 | 2,359E-50 |
| TcCLB.511071.40 | amastin, putative | -1,475 | 1,922E-31 |
| TcCLB.508433.80 | hypothetical protein | -1,479 | 0,002097 |
| TcCLB.506765.49 | hypothetical protein | -1,487 | 0,01375 |
| TcCLB.510537.40 | trans-sialidase (pseudogene), putative | -1,488 | 0,003349 |
| TcCLB.506599.300 | hypothetical protein | -1,489 | 0,01629 |
| TcCLB.507041.120 | variant-surface-glycoprotein phospholipase C, putative | -1,49 | 3,482E-30 |
| TcCLB.508919.110 | hypothetical protein, conserved | -1,492 | 1,908E-20 |
| TcCLB.507863.6 | hypothetical protein, conserved (pseudogene) | -1,492 | 0,006691 |
| TcCLB.467465.10 | surface protease GP63, putative | -1,495 | 0,0004864 |
| TcCLB.503887.79 | hypothetical protein, conserved | -1,496 | 7,442E-30 |
| TcCLB.510625.240 | cell-surface protein (cysteine-rich repeat motif) (pseudoge | -1,502 | 0,001601 |
| TcCLB.507579.30 | surface protease GP63 (pseudogene), putative | -1,511 | 8,461E-08 |
| TcCLB.506851.30 | hypothetical protein, conserved | -1,512 | 8,878E-09 |
| TcCLB.507973.20 | hypothetical protein | -1,513 | 1,556E-08 |
| TcCLB.511173.140 | hypothetical protein | -1,513 | 0,008488 |
| TcCLB.503533.30 | trans-sialidase, Group V, putative | -1,515 | 0,002945 |
| TcCLB.509663.5 | dehydrogenase (pseudogene), putative | -1,517 | 3,327E-09 |
| TcCLB.508319.10 | hypothetical protein | -1,517 | 0,000526 |
| TcCLB.511273.10 | dispersed gene family protein 1 (DGF-1, pseudogene), puta | -1,523 | 0,00001222 |
| TcCLB.511449.95 | hypothetical protein, conserved (pseudogene) | -1,525 | 0,003563 |
| TcCLB.509663.47 | trans-sialidase (pseudogene), putative | -1,526 | 5,754E-09 |
| TcCLB.509631.45 | hypothetical protein, conserved (pseudogene) | -1,531 | 4,694E-11 |

| | | | |
|---|---|---|---|
| TcCLB.508741.415 | hypothetical protein, conserved (pseudogene) | -1,532 | 4,434E-25 |
| TcCLB.511547.14 | mucin-associated surface protein (MASP), putative | -1,537 | 0,00003807 |
| TcCLB.504039.210 | hypothetical protein | -1,543 | 0,02347 |
| TcCLB.509065.70 | glyceraldehyde 3-phosphate dehydrogenase, putative | -1,545 | 7,179E-67 |
| TcCLB.506461.10 | hypothetical protein, conserved | -1,547 | 0,003172 |
| TcCLB.507949.190 | trans-sialidase (pseudogene), putative | -1,549 | 7,268E-29 |
| TcCLB.503973.190 | mucin-associated surface protein (MASP), putative | -1,552 | 0,00001446 |
| TcCLB.508307.180 | hypothetical protein, conserved | -1,574 | 3,903E-54 |
| TcCLB.511483.60 | hypothetical protein, conserved | -1,574 | 3,492E-10 |
| TcCLB.506435.370 | surface protease GP63, putative | -1,577 | 3,458E-31 |
| TcCLB.509607.50 | protein kinase, putative | -1,582 | 6,763E-11 |
| TcCLB.504037.10 | hypothetical protein, conserved | -1,587 | 1,892E-38 |
| TcCLB.509065.60 | glyceraldehyde 3-phosphate dehydrogenase, putative | -1,588 | 3,824E-101 |
| TcCLB.510071.20 | hypothetical protein | -1,595 | 0,00043 |
| TcCLB.503625.10 | asparagine synthetase a, putative | -1,596 | 9,66E-44 |
| TcCLB.503773.30 | hypothetical protein, conserved | -1,596 | 1,054E-11 |
| TcCLB.504689.60 | dispersed gene family protein 1 (DGF-1), putative | -1,607 | 7,52E-14 |
| TcCLB.507251.10 | hypothetical protein, conserved | -1,609 | 1,703E-42 |
| TcCLB.510627.215 | hypothetical protein, conserved (pseudogene) | -1,614 | 0,006355 |
| TcCLB.511537.30 | hypothetical protein, conserved | -1,621 | 1,374E-74 |
| TcCLB.507953.170 | hypothetical protein | -1,622 | 0,001599 |
| TcCLB.506859.230 | hypothetical protein, conserved | -1,622 | 0,002011 |
| TcCLB.507947.10 | conserved protein, unknown function | -1,637 | 2,37E-14 |
| TcCLB.507661.10 | retrotransposon hot spot protein (RHS, pseudogene), putat | -1,638 | 0,0003399 |
| TcCLB.424171.10 | trans-sialidase (pseudogene), putative | -1,638 | 0,0004008 |
| TcCLB.506229.20 | hypothetical protein, conserved | -1,64 | 3,054E-13 |
| TcCLB.509775.20 | protein kinase, putative | -1,641 | 9,349E-76 |
| TcCLB.510377.340 | hypothetical protein | -1,643 | 0,003344 |
| TcCLB.507237.280 | mucin TcMUCII, putative | -1,643 | 0,05157 |
| TcCLB.511211.190 | hypothetical protein | -1,647 | 4,537E-25 |
| TcCLB.507681.100 | ATP-binding cassette protein, putative | -1,651 | 6,949E-42 |
| TcCLB.508049.40 | hypothetical protein, conserved | -1,655 | 0,0001295 |
| TcCLB.508659.18 | mitochondrial carrier protein, putative | -1,656 | 3,371E-37 |
| TcCLB.511605.5 | mucin-associated surface protein (MASP, pseudogene), pu | -1,67 | 8,683E-10 |
| TcCLB.507895.69 | hypothetical protein, conserved | -1,683 | 2,807E-18 |
| TcCLB.506039.10 | retrotransposon hot spot protein (RHS, pseudogene), putat | -1,683 | 4,4E-18 |
| TcCLB.425855.9 | hypothetical protein, conserved (fragment) | -1,683 | 0,00003756 |
| TcCLB.507897.30 | serine/threonine-protein kinase a, putative | -1,686 | 1,309E-47 |
| TcCLB.507023.100 | hypothetical protein | -1,687 | 4,169E-32 |
| TcCLB.506247.330 | 2,3-bisphosphoglycerate-independent phosphoglycerate m | -1,689 | 8,85E-54 |
| TcCLB.511461.30 | hypothetical protein, conserved | -1,69 | 2,576E-16 |
| TcCLB.511577.50 | endo-beta-N-acetylglucosaminidase, putative | -1,697 | 2,77E-53 |
| TcCLB.508747.10 | hypothetical protein | -1,7 | 0,00308 |
| TcCLB.507025.60 | RNA-binding protein, putative | -1,704 | 1,132E-08 |
| TcCLB.506943.60 | glyceraldehyde 3-phosphate dehydrogenase, putative | -1,712 | 9,958E-25 |

| | | | |
|---|---|---|---|
| TcCLB.508717.60 | trans-sialidase, Group I, putative | -1,722 | 1,252E-22 |
| TcCLB.506705.70 | beta-fructofuranosidase-like protein (fragment) | -1,723 | 2,465E-44 |
| TcCLB.511487.30 | dispersed gene family protein 1 (DGF-1), putative | -1,723 | 4,46E-17 |
| TcCLB.508355.40 | choline/carnitine O-acetyltransferase, putative | -1,726 | 1E-31 |
| TcCLB.508433.90 | mucin-associated surface protein (MASP), putative | -1,726 | 0,002995 |
| TcCLB.511565.30 | mucin TcMUCII, putative | -1,729 | 3,001E-09 |
| TcCLB.507859.31 | hypothetical protein | -1,732 | 1,709E-09 |
| TcCLB.508221.410 | hypothetical protein | -1,736 | 2,513E-06 |
| TcCLB.510593.10 | hypothetical protein, conserved | -1,738 | 2,748E-18 |
| TcCLB.510419.30 | hypothetical protein | -1,74 | 0,00134 |
| TcCLB.511725.80 | hypothetical protein | -1,745 | 1,451E-78 |
| TcCLB.506501.100 | hypothetical protein | -1,748 | 0,002743 |
| TcCLB.510537.68 | syntaxin binding protein (pseudogene), putative | -1,749 | 0,00003358 |
| TcCLB.503913.40 | hypothetical protein | -1,751 | 2,952E-21 |
| TcCLB.509553.10 | hypothetical protein, conserved | -1,752 | 6,825E-21 |
| TcCLB.509663.30 | thymidylate kinase, putative | -1,754 | 2,719E-21 |
| TcCLB.511863.10 | mucin-associated surface protein (MASP), putative | -1,755 | 3,753E-12 |
| TcCLB.508221.329 | mucin-associated surface protein (MASP, pseudogene), pu | -1,758 | 0,003529 |
| TcCLB.456627.10 | trans-sialidase (pseudogene), putative | -1,762 | 0,02921 |
| TcCLB.511183.610 | hypothetical protein | -1,763 | 0,00001488 |
| TcCLB.511255.320 | surface protease GP63, putative | -1,766 | 0,004127 |
| TcCLB.511289.80 | polyprenyl synthase, putative | -1,783 | 2,127E-72 |
| TcCLB.508693.100 | surface protease GP63, putative | -1,783 | 4,52E-49 |
| TcCLB.508231.130 | Cell division control protein 48, putative | -1,788 | 3,996E-24 |
| TcCLB.511099.140 | mucin-associated surface protein (MASP), putative | -1,789 | 0,001233 |
| TcCLB.511427.10 | conserved protein, unknown function (fragment) | -1,789 | 0,007259 |
| TcCLB.506943.50 | glyceraldehyde 3-phosphate dehydrogenase, putative | -1,794 | 1,419E-68 |
| TcCLB.510329.90 | C-8 sterol isomerase, putative | -1,795 | 5,612E-59 |
| TcCLB.505567.20 | surface protease GP63, putative | -1,8 | 4,512E-06 |
| TcCLB.511309.80 | trans-sialidase (pseudogene), putative | -1,803 | 7,148E-39 |
| TcCLB.507143.90 | hypothetical protein | -1,81 | 0,0172 |
| TcCLB.508089.30 | hypothetical protein, conserved | -1,818 | 1,055E-83 |
| TcCLB.511277.359 | phosphatidic acid phosphatase, putative | -1,826 | 9,649E-61 |
| TcCLB.511487.51 | hypothetical protein | -1,829 | 6,218E-10 |
| TcCLB.506067.110 | mucin-associated surface protein (MASP, pseudogene), pu | -1,834 | 0,00001981 |
| TcCLB.508365.80 | hypothetical protein | -1,846 | 0,008377 |
| TcCLB.507281.19 | retrotransposon hot spot protein (RHS, pseudogene), putat | -1,848 | 1,481E-13 |
| TcCLB.509697.29 | hypothetical protein | -1,85 | 0,0001608 |
| TcCLB.510629.210 | dispersed gene family protein 1 (DGF-1), putative | -1,856 | 2,333E-09 |
| TcCLB.506501.300 | hypothetical protein | -1,856 | 0,0003789 |
| TcCLB.510615.4 | hypothetical protein, conserved | -1,857 | 1,953E-29 |
| TcCLB.508743.5 | hypothetical protein, conserved (pseudogene) | -1,861 | 1,786E-12 |
| TcCLB.508233.50 | variant-surface-glycoprotein phospholipase C, putative | -1,866 | 1,057E-45 |
| TcCLB.506257.50 | surface protease GP63, putative | -1,874 | 7,039E-21 |
| TcCLB.508907.10 | hypothetical protein, conserved | -1,883 | 7,93E-41 |

| | | | |
|---|---|---|---|
| TcCLB.510629.70 | mucin-associated surface protein (MASP), putative | -1,884 | 0,00001471 |
| TcCLB.509907.50 | hypothetical protein, conserved (pseudogene) | -1,891 | 0,00003311 |
| TcCLB.508433.50 | retrotransposon hot spot protein (RHS, pseudogene), putat | -1,894 | 1,45E-08 |
| TcCLB.509663.50 | trans-sialidase, Group VIII, putative | -1,896 | 2,477E-18 |
| TcCLB.437805.9 | trans-sialidase (pseudogene), putative | -1,896 | 0,0004935 |
| TcCLB.507305.40 | zinc finger CCCH domain containing protein 11 (ZC3H11) | -1,904 | 0,0005112 |
| TcCLB.504005.6 | RNA-binding protein, putative | -1,905 | 9,9E-39 |
| TcCLB.507623.140 | endonuclease G, putative | -1,911 | 7,088E-44 |
| TcCLB.507467.40 | choline/carnitine O-acetyltransferase (pseudogene), putati | -1,912 | 7,605E-32 |
| TcCLB.503773.40 | hypothetical protein, conserved | -1,913 | 8,736E-28 |
| TcCLB.508209.94 | hypothetical protein | -1,919 | 4,207E-37 |
| TcCLB.507099.80 | ABC transporter, putative | -1,939 | 1,064E-136 |
| TcCLB.508857.40 | UDP-glucoronosyl and UDP-glucosyl transferase, putative | -1,944 | 2,284E-51 |
| TcCLB.507237.356 | syntaxin binding protein (pseudogene), putative | -1,946 | 0,0000414 |
| TcCLB.511547.20 | trans-sialidase, Group IV, putative | -1,95 | 0,00003091 |
| TcCLB.510709.60 | mucin-associated surface protein (MASP), putative | -1,95 | 0,000981 |
| TcCLB.507859.15 | hypothetical protein | -1,971 | 1,516E-20 |
| TcCLB.510709.50 | mucin TcMUCII (pseudogene), putative | -1,973 | 0,001379 |
| TcCLB.511911.10 | trans-sialidase, Group II, putative | -1,995 | 3,256E-15 |
| TcCLB.510481.30 | hypothetical protein, conserved | -1,997 | 0,00002662 |
| TcCLB.510261.21 | hypothetical protein | -2,005 | 0,0001307 |
| TcCLB.507641.300 | chaperonin HSP60, mitochondrial precursor (HSP60) | -2,014 | 0,0001136 |
| TcCLB.510629.270 | syntaxin binding protein (pseudogene), putative | -2,051 | 0,00001498 |
| TcCLB.506229.50 | nucleoside phosphatase, putative | -2,053 | 5,249E-83 |
| TcCLB.511605.10 | hypothetical protein | -2,054 | 0,0002872 |
| TcCLB.510701.40 | mucin-associated surface protein (MASP, pseudogene), pu | -2,058 | 7,994E-06 |
| TcCLB.428403.10 | hypothetical protein (fragment) | -2,069 | 1,074E-08 |
| TcCLB.509553.60 | mitogen-activated protein kinase 3, putative | -2,083 | 1,451E-50 |
| TcCLB.509137.20 | hypothetical protein | -2,087 | 5,044E-86 |
| TcCLB.510773.40 | hypothetical protein | -2,096 | 2,012E-29 |
| TcCLB.507069.20 | hypothetical protein | -2,097 | 0,001826 |
| TcCLB.508777.210 | hypothetical protein | -2,098 | 2,283E-08 |
| TcCLB.510705.10 | mucin-associated surface protein (MASP), putative | -2,099 | 0,00004131 |
| TcCLB.506495.40 | hypothetical protein | -2,102 | 4,412E-102 |
| TcCLB.507705.20 | hypothetical protein, conserved | -2,112 | 3,91E-21 |
| TcCLB.507397.9 | dispersed gene family protein 1 (DGF-1), putative | -2,114 | 0,000196 |
| TcCLB.510089.20 | surface protease GP63, putative | -2,12 | 0,00315 |
| TcCLB.510629.290 | mucin-associated surface protein (MASP, pseudogene), pu | -2,121 | 1,484E-15 |
| TcCLB.507705.10 | hypothetical protein, conserved | -2,126 | 4,864E-30 |
| TcCLB.503639.10 | inositol-3-phosphate synthase, putative | -2,128 | 2,726E-65 |
| TcCLB.509313.9 | trans-sialidase (pseudogene), putative | -2,142 | 0,0005228 |
| TcCLB.510197.160 | mucin-associated surface protein (MASP, pseudogene), pu | -2,147 | 9,681E-47 |
| TcCLB.507959.290 | hypothetical protein | -2,147 | 0,001236 |
| TcCLB.509075.20 | dispersed gene family protein 1 (DGF-1, pseudogene), puta | -2,165 | 1,514E-07 |
| TcCLB.506039.30 | dispersed gene family protein 1 (DGF-1), putative | -2,175 | 4,516E-16 |

| | | | |
|---|---|---|---|
| TcCLB.510181.9 | inositol-3-phosphate synthase, putative (fragment) | -2,179 | 3,421E-79 |
| TcCLB.509475.10 | mitogen-activated protein kinase 3, putative | -2,222 | 2,201E-30 |
| TcCLB.511547.40 | retrotransposon hot spot protein (RHS, pseudogene), putat | -2,247 | 3,658E-31 |
| TcCLB.511521.27 | hypothetical protein, conserved | -2,252 | 7,885E-72 |
| TcCLB.508481.10 | trans-sialidase, Group IV, putative | -2,257 | 0,003125 |
| TcCLB.511021.30 | hypothetical protein, conserved | -2,274 | 7,34E-111 |
| TcCLB.506683.240 | trans-sialidase, Group II, putative | -2,277 | 1,025E-138 |
| TcCLB.509731.10 | poly-zinc finger protein 2, putative | -2,314 | 4,763E-13 |
| TcCLB.507609.60 | inositol-3-phosphate synthase, putative | -2,323 | 3,061E-83 |
| TcCLB.509197.39 | cation transporter, putative | -2,333 | 2,09E-162 |
| TcCLB.503773.24 | hypothetical protein, conserved | -2,351 | 1,569E-31 |
| TcCLB.508243.36 | hypothetical protein | -2,36 | 0,00006498 |
| TcCLB.503773.9 | hypothetical protein, conserved (fragment) | -2,382 | 9,282E-41 |
| TcCLB.506789.334 | hypothetical protein, conserved | -2,385 | 1,714E-53 |
| TcCLB.509663.20 | hypothetical protein, conserved | -2,389 | 8,349E-50 |
| TcCLB.505945.5 | mitogen-activated protein kinase 3, putative (fragment) | -2,396 | 2,386E-15 |
| TcCLB.508741.130 | hypothetical protein | -2,397 | 5,919E-150 |
| TcCLB.504039.200 | mucin-associated surface protein (MASP), putative | -2,4 | 0,0001342 |
| TcCLB.509075.31 | hypothetical protein, conserved (pseudogene) | -2,408 | 0,002532 |
| TcCLB.506115.9 | dispersed gene family protein 1 (DGF-1), putative | -2,425 | 2,46E-20 |
| TcCLB.511903.50 | amastin, putative | -2,431 | 8,584E-83 |
| TcCLB.506841.20 | trans-sialidase, Group I, putative | -2,445 | 3,002E-47 |
| TcCLB.510363.280 | mucin-associated surface protein (MASP), putative | -2,448 | 1,514E-12 |
| TcCLB.510171.10 | trans-sialidase, putative (fragment) | -2,453 | 2,523E-83 |
| TcCLB.469785.40 | RNA-binding protein, putative | -2,49 | 2,783E-65 |
| TcCLB.509325.10 | mucin-associated surface protein (MASP), putative | -2,503 | 0,00007217 |
| TcCLB.506149.20 | hypothetical protein, conserved | -2,52 | 5,52E-34 |
| TcCLB.511565.10 | mucin TcMUCII, putative (fragment) | -2,52 | 6,115E-09 |
| TcCLB.509075.70 | retrotransposon hot spot protein (RHS, pseudogene), putat | -2,538 | 0,00003049 |
| TcCLB.510709.31 | syntaxin binding protein (pseudogene), putative | -2,541 | 2,562E-09 |
| TcCLB.507025.50 | RNA-binding protein, putative | -2,553 | 1,585E-48 |
| TcCLB.507859.60 | mucin-associated surface protein (MASP), putative | -2,561 | 6,455E-06 |
| TcCLB.510709.70 | trans-sialidase, Group V, putative | -2,565 | 3,025E-09 |
| TcCLB.508433.30 | mucin-associated surface protein (MASP), putative | -2,567 | 4,062E-06 |
| TcCLB.507237.359 | surface protease GP63 (pseudogene), putative | -2,575 | 0,00001597 |
| TcCLB.509663.37 | hypothetical protein, conserved | -2,586 | 5,633E-93 |
| TcCLB.509881.9 | dispersed gene family protein 1 (DGF-1), putative (fragme | -2,592 | 2,645E-07 |
| TcCLB.506149.29 | hypothetical protein, conserved | -2,603 | 1,026E-17 |
| TcCLB.503705.11 | UDP-Gal or UDP-GlcNAc-dependent glycosyltransferase (p: | -2,621 | 1,126E-07 |
| TcCLB.438923.10 | hypothetical protein (fragment) | -2,633 | 1,63E-108 |
| TcCLB.508647.280 | malic enzyme | -2,654 | 3,893E-250 |
| TcCLB.416041.14 | trans-sialidase, putative (fragment) | -2,67 | 8,07E-167 |
| TcCLB.402255.10 | retrotransposon hot spot protein (RHS, pseudogene), putat | -2,732 | 2,828E-26 |
| TcCLB.510831.21 | trans-sialidase, putative (fragment) | -2,739 | 9,239E-10 |
| TcCLB.508433.10 | mucin-associated surface protein (MASP), putative | -2,743 | 7,009E-06 |

| | | | |
|---|---|---|---|
| TcCLB.504405.10 | dispersed gene family protein 1 (DGF-1), putative | -2,747 | 0,00004793 |
| TcCLB.503475.10 | dispersed gene family protein 1 (DGF-1, pseudogene), puta | -2,747 | 0,00005469 |
| TcCLB.504105.140 | enolase | -2,777 | 2,551E-198 |
| TcCLB.510875.10 | dispersed gene family protein 1 (DGF-1, pseudogene), puta | -2,778 | 4,466E-13 |
| TcCLB.504733.10 | beta galactofuranosyl glycosyltransferase (pseudogene), p | -2,786 | 7,602E-15 |
| TcCLB.507613.10 | dispersed gene family protein 1 (DGF-1), putative | -2,804 | 1,481E-11 |
| TcCLB.511563.20 | trans-sialidase (pseudogene), putative | -2,844 | 1,046E-12 |
| TcCLB.506067.100 | mucin TcMUCII, putative | -2,86 | 5,606E-08 |
| TcCLB.450161.10 | cation transporter, putative | -2,861 | 0 |
| TcCLB.503773.50 | Nucleolar protein 56, putative (fragment) (NOP56) | -2,871 | 5,659E-89 |
| TcCLB.508701.10 | ZIP Zn transporter, putative | -2,891 | 9,552E-92 |
| TcCLB.508699.130 | cation transporter, putative (fragment) | -2,908 | 3,123E-219 |
| TcCLB.509241.10 | surface protease GP63, putative (fragment) | -2,933 | 1,492E-64 |
| TcCLB.511907.350 | surface protease GP63 (pseudogene), putative | -2,944 | 4,977E-53 |
| TcCLB.509197.30 | cation transporter, putative | -2,945 | 1,812E-105 |
| TcCLB.511755.10 | hypothetical protein, conserved | -2,947 | 6,075E-103 |
| TcCLB.511551.40 | retrotransposon hot spot protein (RHS, pseudogene), putat | -2,964 | 4,498E-16 |
| TcCLB.507489.9 | dispersed gene family protein 1 (DGF-1), putative (fragme | -2,971 | 1,189E-12 |
| TcCLB.509663.44 | hypothetical protein, conserved | -2,979 | 3,934E-121 |
| TcCLB.508905.10 | hypothetical protein, conserved (pseudogene) | -2,983 | 4,567E-113 |
| TcCLB.509267.110 | heat shock protein Hsp20, putative | -2,999 | 2,051E-109 |
| TcCLB.413489.10 | retrotransposon hot spot protein (RHS, pseudogene), putat | -3,04 | 1,063E-12 |
| TcCLB.505183.30 | malic enzyme, putative | -3,069 | 0 |
| TcCLB.507867.10 | mucin-associated surface protein (MASP), putative | -3,07 | 1,677E-10 |
| TcCLB.505425.9 | dispersed gene family protein 1 (DGF-1, pseudogene), puta | -3,152 | 1,122E-14 |
| TcCLB.464391.10 | retrotransposon hot spot protein (RHS, pseudogene), putat | -3,162 | 3,124E-12 |
| TcCLB.509553.50 | DNA repair and recombination helicase protein PIF5, putat | -3,201 | 3,06E-166 |
| TcCLB.504145.10 | trans-sialidase, Group II, putative | -3,207 | 1,099E-07 |
| TcCLB.506495.50 | hypothetical protein | -3,246 | 8,202E-164 |
| TcCLB.506087.10 | retrotransposon hot spot protein (RHS, pseudogene), putat | -3,251 | 2,639E-12 |
| TcCLB.508061.10 | hypothetical protein | -3,305 | 9,713E-193 |
| TcCLB.507711.100 | procyclic form surface phosphoprotein, putative | -3,308 | 1,192E-192 |
| TcCLB.510363.290 | hypothetical protein | -3,354 | 2,012E-06 |
| TcCLB.510481.20 | dispersed gene family protein 1 (DGF-1, pseudogene), puta | -3,358 | 1,341E-30 |
| TcCLB.509325.16 | elongation factor 1-gamma (EF-1-gamma, pseudogene), p | -3,385 | 3,708E-14 |
| TcCLB.509325.4 | surface protease GP63, putative (fragment) | -3,393 | 0,08195 |
| TcCLB.507951.150 | guanine deaminase, putative | -3,394 | 0 |
| TcCLB.511549.10 | retrotransposon hot spot protein (RHS, pseudogene), putat | -3,426 | 2,167E-24 |
| TcCLB.508481.20 | mucin-associated surface protein (MASP), putative | -3,443 | 3,606E-06 |
| TcCLB.507705.30 | hypothetical protein, conserved | -3,473 | 2,242E-100 |
| TcCLB.504431.100 | guanine deaminase, putative | -3,533 | 4,749E-239 |
| TcCLB.508485.30 | retrotransposon hot spot protein (RHS, pseudogene), putat | -3,539 | 8,201E-32 |
| TcCLB.508581.30 | beta galactofuranosyl glycosyltransferase (pseudogene), p | -3,546 | 2,894E-09 |
| TcCLB.507639.10 | poly-zinc finger protein 2, putative | -3,551 | 1,046E-82 |
| TcCLB.466325.10 | trans-sialidase (pseudogene), putative | -3,872 | 1,168E-41 |

| | | | |
|---|---|---|---|
| TcCLB.507145.70 | mucin-associated surface protein (MASP, pseudogene), pu | -3,877 | 5,456E-25 |
| TcCLB.508581.10 | trans-sialidase, Group VII, putative | -3,882 | 2,571E-33 |
| TcCLB.511551.50 | trans-sialidase (pseudogene), putative | -4,191 | 5,831E-10 |
| TcCLB.508485.10 | trans-sialidase, Group II, putative | -4,218 | 2,434E-14 |
| TcCLB.509075.40 | mucin-associated surface protein (MASP), putative | -4,342 | 0,0009103 |
| TcCLB.507145.79 | mucin TcMUCII, putative | -4,452 | 2,839E-07 |
| TcCLB.507867.5 | trans-sialidase, putative (fragment) | -4,584 | 0,00002407 |
| TcCLB.508483.40 | retrotransposon hot spot protein (RHS, pseudogene), putat | -4,673 | 2,268E-28 |
| TcCLB.506231.10 | trans-sialidase, putative | -4,865 | 2,241E-213 |
| TcCLB.509663.69 | trans-sialidase, putative (fragment) | -4,895 | 2,523E-11 |
| TcCLB.508483.26 | trans-sialidase (pseudogene), putative | -5,217 | 3,514E-16 |
| TcCLB.508581.40 | trans-sialidase, Group II, putative | -5,48 | 5,666E-09 |

epi_over_tryp

| geneid | genedescription | deseq_logfc | deseq_adjp |
|---|---|---|---|
| TcCLB.506357.50 | alcohol dehydrogenase, putative | 6,298 | 0 |
| TcCLB.511391.120 | glutamamyl carboxypeptidase, putative | 5,936 | 0 |
| TcCLB.511277.60 | alcohol dehydrogenase, putative | 5,81 | 0 |
| TcCLB.467287.20 | hypothetical protein, conserved | 5,528 | 2,967E-43 |
| TcCLB.510103.24 | hypothetical protein, conserved | 5,437 | 0 |
| TcCLB.509713.30 | hypothetical protein, conserved | 5,412 | 0 |
| TcCLB.509213.120 | cytosolic nonspecific dipeptidase, putative | 5,393 | 0 |
| TcCLB.510535.100 | cysteine peptidase C (CPC), putative (TCCB) | 5,333 | 0 |
| TcCLB.509233.170 | hypothetical protein, conserved | 5,207 | 6,839E-100 |
| TcCLB.509105.70 | thiol-dependent reductase 1, putative (TcAc2) | 5,134 | 0 |
| TcCLB.508999.260 | calpain-like cysteine peptidase (pseudogene), putative | 5,089 | 0 |
| TcCLB.510257.80 | cytosolic nonspecific dipeptidase, putative | 5,072 | 0 |
| TcCLB.506563.170 | calpain-like cysteine peptidase (pseudogene), putative | 4,92 | 0 |
| TcCLB.510357.20 | hypothetical protein, conserved | 4,874 | 6,743E-103 |
| TcCLB.511497.19 | amastin, putative | 4,753 | 1,345E-193 |
| TcCLB.511073.10 | fatty acid desaturase, putative | 4,739 | 3,974E-292 |
| TcCLB.503419.30 | thiol-dependent reductase 1, putative (TcAc2) | 4,71 | 0 |
| TcCLB.511003.160 | hypothetical protein, conserved | 4,704 | 2,003E-179 |
| TcCLB.511075.9 | fatty acid desaturase, putative (fragment) | 4,686 | 8,306E-207 |
| TcCLB.510855.10 | peptide methionine sulfoxide reductase, putative | 4,641 | 0 |
| TcCLB.510837.9 | acetylornithine deacetylase, putative (fragment) | 4,442 | 1,141E-104 |
| TcCLB.507853.10 | lathosterol oxidase, putative | 4,42 | 5,482E-163 |
| TcCLB.510357.30 | hypothetical protein, conserved | 4,372 | 3,562E-141 |
| TcCLB.504425.60 | NADP-dependent alcohol hydrogenase, putative | 4,301 | 2,153E-244 |
| TcCLB.507891.29 | flagellar calcium-binding protein, putative | 4,295 | 1,466E-12 |
| TcCLB.511201.49 | hypothetical protein, conserved | 4,243 | 0 |
| TcCLB.508017.30 | amino acid transporter, putative (fragment) | 4,202 | 2,027E-295 |
| TcCLB.510679.100 | lipase, putative | 4,201 | 6,705E-234 |
| TcCLB.511859.200 | hypothetical protein, conserved | 4,182 | 0 |
| TcCLB.506773.50 | nucleoside transporter-like, putative | 4,174 | 0 |
| TcCLB.506559.410 | hypothetical protein, conserved | 4,163 | 3,535E-234 |
| TcCLB.509003.30 | calpain-like cysteine peptidase, putative | 4,135 | 5,324E-257 |
| TcCLB.506661.30 | fatty acid elongase, putative | 4,125 | 0 |
| TcCLB.509965.394 | amastin, putative | 4,112 | 9,014E-137 |
| TcCLB.503837.10 | hypothetical protein, conserved | 4,046 | 1,655E-247 |
| TcCLB.507949.250 | hypothetical protein, conserved | 4,01 | 0 |
| TcCLB.508111.30 | glutamate dehydrogenase, putative | 3,992 | 0 |
| TcCLB.506789.352 | hypothetical protein, conserved | 3,987 | 3,613E-66 |
| TcCLB.507689.40 | glutamamyl carboxypeptidase, putative | 3,985 | 0 |
| TcCLB.509239.10 | fatty acid desaturase, putative | 3,974 | 9,73E-298 |
| TcCLB.508065.70 | protein associated with differentiation 8, putative | 3,971 | 4,557E-143 |
| TcCLB.506563.79 | calpain-like cysteine peptidase (pseudogene), putative | 3,95 | 7,728E-247 |
| TcCLB.511245.130 | fatty acid elongase, putative | 3,91 | 3,403E-287 |

| | | | |
|---|---|---|---|
| TcCLB.504045.60 | thermostable carboxypeptidase 2, putative | 3,907 | 0 |
| TcCLB.507047.150 | D-arabinono-1,4-lactone oxidase, putative (ALO) | 3,903 | 5,233E-227 |
| TcCLB.426435.9 | fatty acid desaturase, putative | 3,868 | 9,065E-288 |
| TcCLB.511439.40 | hypothetical protein | 3,867 | 3,599E-293 |
| TcCLB.511685.20 | mucin TcSMUGS, putative | 3,848 | 1,403E-284 |
| TcCLB.511687.19 | voltage-dependent anion-selective channel, putative (frag | 3,848 | 1,516E-28 |
| TcCLB.509023.80 | hypothetical protein, conserved | 3,84 | 0 |
| TcCLB.509157.220 | hypothetical protein, conserved | 3,837 | 6,147E-200 |
| TcCLB.506825.200 | hypothetical protein | 3,833 | 1,35E-302 |
| TcCLB.508545.40 | surface protease GP63, putative | 3,809 | 0 |
| TcCLB.506367.30 | 1,2-Dihydroxy-3-keto-5-methylthiopentene dioxygenase, p | 3,804 | 0 |
| TcCLB.508209.100 | 10 kDa heat shock protein, putative | 3,803 | 4,509E-101 |
| TcCLB.506789.340 | hypothetical protein, conserved | 3,796 | 4,061E-206 |
| TcCLB.509197.50 | hypothetical protein, conserved | 3,784 | 1,558E-154 |
| TcCLB.504533.50 | hypothetical protein, conserved | 3,751 | 1,221E-181 |
| TcCLB.506551.10 | protein associated with differentiation 4, putative | 3,72 | 3,592E-279 |
| TcCLB.507689.50 | metallo-peptidase, clan MH, family M18, putative (fragm | 3,716 | 0 |
| TcCLB.510613.40 | hypothetical protein, conserved (fragment) | 3,704 | 2,244E-63 |
| TcCLB.508923.10 | amino acid permease, putative | 3,7 | 9,624E-184 |
| TcCLB.509965.390 | amastin, putative | 3,693 | 5,035E-234 |
| TcCLB.506753.130 | hypothetical protein, conserved | 3,691 | 5,237E-138 |
| TcCLB.509205.120 | hypothetical protein, conserved | 3,684 | 0 |
| TcCLB.506247.220 | histidine ammonia-lyase, putative | 3,683 | 0 |
| TcCLB.506401.380 | surface protease GP63, putative | 3,671 | 0 |
| TcCLB.503413.4 | Present in the outer mitochondrial membrane proteome 2 | 3,633 | 5,652E-112 |
| TcCLB.508735.100 | hypothetical protein, conserved | 3,627 | 9,555E-208 |
| TcCLB.508741.80 | mucin TcSMUGS, putative | 3,614 | 4,53E-48 |
| TcCLB.503747.10 | glutamine aminotransferase, putative (fragment) (GlnAT) | 3,604 | 5,541E-193 |
| TcCLB.508263.5 | hypothetical protein, conserved (fragment) | 3,594 | 2,513E-135 |
| TcCLB.511303.34 | Macro domain containing protein, putative | 3,588 | 6,843E-163 |
| TcCLB.510667.120 | galactokinase, putative | 3,583 | 1,245E-285 |
| TcCLB.511685.10 | mucin TcSMUGS, putative | 3,58 | 4,941E-172 |
| TcCLB.510725.40 | hypothetical protein, conserved | 3,574 | 6,996E-166 |
| TcCLB.510811.20 | protein associated with differentiation 4, putative | 3,56 | 2,174E-202 |
| TcCLB.509051.20 | amastin, putative | 3,548 | 5,717E-189 |
| TcCLB.508741.70 | mucin TcSMUGS, putative | 3,547 | 3,928E-81 |
| TcCLB.506227.10 | amino acid transporter, putative | 3,539 | 2,152E-178 |
| TcCLB.506629.200 | 1,2-Dihydroxy-3-keto-5-methylthiopentene dioxygenase, p | 3,532 | 7,992E-273 |
| TcCLB.509569.100 | nucleoside phosphorylase, putative | 3,529 | 8,766E-259 |
| TcCLB.510187.30 | tyrosine aminotransferase | 3,524 | 6,788E-58 |
| TcCLB.508173.160 | hypothetical protein, conserved | 3,499 | 3,013E-254 |
| TcCLB.511679.10 | mucin TcSMUGS, putative | 3,49 | 3,729E-102 |
| TcCLB.511809.120 | hypothetical protein, conserved | 3,488 | 1,663E-231 |
| TcCLB.506563.40 | beta tubulin, putative | 3,483 | 2,526E-137 |
| TcCLB.508693.50 | amidinotransferase, putative | 3,478 | 0 |

| | | | |
|---|---|---|---|
| TcCLB.511391.110 | histone-lysine N-methyltransferase, putative (DOT1A) | 3,478 | 7,732E-208 |
| TcCLB.507359.10 | hypothetical protein, conserved | 3,472 | 2,435E-64 |
| TcCLB.511385.30 | Pyrophosphate-energized vacuolar membrane proton pum | 3,461 | 0 |
| TcCLB.511693.10 | hypothetical protein | 3,461 | 4,385E-247 |
| TcCLB.510311.90 | quinone oxidoreductase, putative | 3,45 | 1,662E-265 |
| TcCLB.509107.9 | hypothetical protein, conserved (fragment) | 3,446 | 2,432E-69 |
| TcCLB.511685.30 | mucin TcSMUGS, putative | 3,443 | 2,946E-69 |
| TcCLB.504047.5 | urocanate hydratase, putative (fragment) | 3,439 | 2,229E-16 |
| TcCLB.508503.20 | cytochrome c oxidase subunit V, putative | 3,436 | 0 |
| TcCLB.509683.10 | hypothetical protein | 3,432 | 0 |
| TcCLB.511817.40 | NADH-cytochrome b5 reductase, putative | 3,416 | 8,664E-151 |
| TcCLB.440101.10 | fatty acid desaturase, putative (fragment) | 3,409 | 2,114E-64 |
| TcCLB.507187.9 | dispersed gene family protein 1 (DGF-1), putative | 3,406 | 2,607E-133 |
| TcCLB.506937.10 | cytosolic malate dehydrogenase, putative | 3,401 | 2,41E-126 |
| TcCLB.510661.250 | cystathione gamma lyase, putative | 3,397 | 0 |
| TcCLB.510679.40 | hypothetical protein | 3,392 | 3,387E-183 |
| TcCLB.506739.99 | zinc finger (CCCH type) protein, putative | 3,388 | 0 |
| TcCLB.511051.10 | hypothetical protein | 3,383 | 3,074E-64 |
| TcCLB.437545.10 | 3-oxo-5-alpha-steroid 4-dehydrogenase, putative | 3,375 | 4,69E-146 |
| TcCLB.510187.40 | tyrosine aminotransferase | 3,372 | 4,413E-235 |
| TcCLB.510837.20 | glutamamyl carboxypeptidase, putative | 3,371 | 7,58E-62 |
| TcCLB.509895.30 | hypothetical protein, conserved | 3,364 | 0 |
| TcCLB.503411.30 | hypothetical protein, conserved | 3,363 | 2,299E-53 |
| TcCLB.509041.10 | tyrosine aminotransferase, putative | 3,36 | 4,963E-11 |
| TcCLB.509237.151 | calpain-like cysteine peptidase, putative (fragment) | 3,358 | 9,494E-117 |
| TcCLB.511717.30 | chromosomal passenger protein, putative (CPC1) | 3,344 | 1,465E-112 |
| TcCLB.511909.50 | hypothetical protein, conserved | 3,341 | 2,849E-126 |
| TcCLB.507657.20 | glutamamyl carboxypeptidase, putative | 3,336 | 5,569E-305 |
| TcCLB.505989.110 | amidinotransferase, putative | 3,333 | 0 |
| TcCLB.508741.60 | mucin TcSMUGS, putative | 3,331 | 2,014E-82 |
| TcCLB.507689.30 | glutamamyl carboxypeptidase, putative | 3,33 | 0 |
| TcCLB.509707.10 | protein associated with differentiation 8, putative | 3,33 | 6,349E-74 |
| TcCLB.507657.10 | glutamamyl carboxypeptidase, putative | 3,31 | 0 |
| TcCLB.510187.70 | tyrosine aminotransferase, putative | 3,306 | 6,908E-229 |
| TcCLB.503925.6 | hypothetical protein, conserved | 3,293 | 8,463E-26 |
| TcCLB.503939.20 | hypothetical protein, conserved | 3,289 | 8,038E-169 |
| TcCLB.509805.190 | hypothetical protein, conserved | 3,289 | 2,841E-122 |
| TcCLB.508741.229 | Mitochondrial outer membrane protein porin, putative (VI | 3,284 | 1,085E-255 |
| TcCLB.510249.30 | hypothetical protein, conserved | 3,272 | 5,406E-36 |
| TcCLB.506885.310 | meiotic recombination protein DMC1, putative (dmc1) | 3,266 | 1,28E-140 |
| TcCLB.508737.210 | Mitochondrial outer membrane protein porin, putative (VI | 3,262 | 1,197E-241 |
| TcCLB.411235.9 | alpha tubulin, putative | 3,257 | 0 |
| TcCLB.506513.110 | peptidase t, putative | 3,248 | 8,573E-199 |
| TcCLB.510187.50 | tyrosine aminotransferase | 3,245 | 0 |
| TcCLB.508917.49 | peptidase t, putative | 3,239 | 2,65E-87 |

| | | | |
|---|---|---|---|
| TcCLB.462761.16 | hypothetical protein, conserved (fragment) | 3,228 | 4,828E-69 |
| TcCLB.508799.270 | protein associated with differentiation 4, putative | 3,223 | 7,674E-288 |
| TcCLB.508919.10 | peptidase T (pseudogene), putative | 3,22 | 2,12E-99 |
| TcCLB.504765.20 | protein associated with differentiation 8, putative | 3,2 | 3,306E-28 |
| TcCLB.509003.70 | beta tubulin (pseudogene), putative | 3,187 | 2,64E-235 |
| TcCLB.511287.49 | aldo-keto reductase (AKR) | 3,187 | 3,153E-130 |
| TcCLB.510565.30 | cytochrome c oxidase subunit V, putative | 3,184 | 2,332E-290 |
| TcCLB.504045.110 | urocanate hydratase, putative | 3,171 | 0 |
| TcCLB.508741.40 | mucin TcSMUGS, putative | 3,171 | 1,744E-227 |
| TcCLB.511129.60 | hypothetical protein, conserved | 3,151 | 1,198E-134 |
| TcCLB.510773.20 | Pyrophosphate-energized vacuolar membrane proton pum | 3,15 | 8,25E-247 |
| TcCLB.508001.70 | retrotransposon hot spot (RHS) protein, putative | 3,148 | 6,716E-145 |
| TcCLB.506823.50 | retrotransposon hot spot protein (RHS, pseudogene), puta | 3,147 | 9,389E-179 |
| TcCLB.510795.10 | tyrosine aminotransferase (2.6.1.5) | 3,144 | 8,507E-256 |
| TcCLB.511687.10 | Mitochondrial outer membrane protein porin, putative (VI | 3,141 | 2,901E-158 |
| TcCLB.506257.98 | hypothetical protein, conserved (pseudogene) | 3,133 | 4,54E-134 |
| TcCLB.507009.40 | galactokinase, putative | 3,128 | 2,381E-195 |
| TcCLB.510039.40 | nitrilase, putative | 3,123 | 2,166E-241 |
| TcCLB.508535.50 | tyrosine aminotransferase | 3,118 | 2,628E-132 |
| TcCLB.506789.280 | methylthioadenosine phosphorylase, putative | 3,112 | 2,344E-146 |
| TcCLB.511367.240 | argonaute-like protein, putative | 3,107 | 3,26E-211 |
| TcCLB.509179.100 | L-galactonolactone oxidase (ALO) | 3,1 | 1,066E-176 |
| TcCLB.508823.10 | hypothetical protein, conserved | 3,099 | 1,846E-153 |
| TcCLB.511479.39 | hypothetical protein, conserved | 3,098 | 5,096E-81 |
| TcCLB.506931.4 | hypothetical protein, conserved | 3,096 | 5,433E-196 |
| TcCLB.503837.20 | amino acid transporter, putative | 3,088 | 9,659E-155 |
| TcCLB.503713.30 | 6-phosphogluconolactonase, putative | 3,081 | 7,013E-127 |
| TcCLB.503415.40 | hypothetical protein | 3,08 | 2,543E-74 |
| TcCLB.504933.10 | calpain-like cysteine peptidase, putative | 3,079 | 3,328E-66 |
| TcCLB.511807.220 | hypothetical protein | 3,075 | 1,38E-101 |
| TcCLB.511825.240 | hypothetical protein, conserved | 3,058 | 3,803E-71 |
| TcCLB.503617.10 | protein kinase, putative | 3,057 | 4,258E-187 |
| TcCLB.506243.10 | retrotransposon hot spot protein (RHS, pseudogene), puta | 3,056 | 3,433E-127 |
| TcCLB.507031.90 | arginase, putative | 3,051 | 1,866E-167 |
| TcCLB.510437.40 | cytosolic malate dehydrogenase, putative | 3,047 | 5,952E-165 |
| TcCLB.504137.100 | hypothetical protein, conserved | 3,045 | 9,297E-161 |
| TcCLB.509207.90 | pyrroline-5-carboxylate reductase, putative (P5CR) | 3,04 | 7,075E-208 |
| TcCLB.511325.40 | amino acid permease, putative | 3,04 | 2,262E-170 |
| TcCLB.503975.100 | Present in the outer mitochondrial membrane proteome 2 | 3,04 | 2,313E-122 |
| TcCLB.509997.10 | hypothetical protein, conserved | 3,04 | 2,218E-46 |
| TcCLB.509767.170 | hypothetical protein, conserved | 3,038 | 1,208E-225 |
| TcCLB.506773.44 | cytochrome b5, putative | 3,037 | 1,04E-154 |
| TcCLB.506863.30 | hypothetical protein, conserved | 3,037 | 1,286E-67 |
| TcCLB.508823.20 | hypothetical protein, conserved | 3,032 | 4,67E-243 |
| TcCLB.509141.40 | Mitochondrial outer membrane protein porin, putative (VI | 3,029 | 2,21E-124 |

| | | | |
|---|---|---|---|
| TcCLB.508805.80 | short chain dehydrogenase, putative | 3,026 | 7,205E-98 |
| TcCLB.507603.220 | hypothetical protein | 3,023 | 1,477E-86 |
| TcCLB.508811.40 | hypothetical protein | 3,018 | 5,626E-125 |
| TcCLB.510187.20 | tyrosine aminotransferase, putative | 3,011 | 3,08E-227 |
| TcCLB.510187.100 | glucokinase | 3,004 | 8,632E-154 |
| TcCLB.473111.10 | lathosterol oxidase, putative | 3,004 | 2,948E-88 |
| TcCLB.503975.40 | hypothetical protein, conserved | 3,003 | 7,502E-151 |
| TcCLB.509043.10 | tyrosine aminotransferase (pseudogene), putative | 3,002 | 2,6E-155 |
| TcCLB.506893.100 | UMP-CMP kinase, mitochondrial, putative | 3,002 | 8,299E-86 |
| TcCLB.503445.9 | retrotransposon hot spot (RHS) protein, putative | 2,997 | 1,953E-94 |
| TcCLB.510669.49 | Gim5A protein, putative | 2,992 | 5,843E-151 |
| TcCLB.509463.30 | 3-ketoacyl-CoA thiolase, putative | 2,989 | 4,092E-276 |
| TcCLB.510339.80 | hypothetical protein, conserved | 2,983 | 3,748E-276 |
| TcCLB.511329.10 | Calpain-like protein 2 (CALP2) | 2,979 | 7,27E-136 |
| TcCLB.503653.60 | cytochrome b-domain protein, putative | 2,979 | 1,015E-88 |
| TcCLB.510565.11 | tyrosine aminotransferase, putative (fragment) | 2,978 | 2,973E-210 |
| TcCLB.506399.80 | hypothetical protein, conserved | 2,968 | 3,078E-97 |
| TcCLB.510609.70 | protein kinase, putative | 2,967 | 1,42E-185 |
| TcCLB.511047.70 | hypothetical protein, conserved | 2,959 | 1,03E-207 |
| TcCLB.506011.10 | hypothetical protein, conserved | 2,958 | 2,203E-131 |
| TcCLB.509129.10 | Mitochondrial outer membrane protein porin, putative (VI | 2,955 | 1,325E-213 |
| TcCLB.507101.10 | amino acid permease, putative | 2,953 | 1,022E-183 |
| TcCLB.509961.9 | cytochrome c oxidase subunit 10, putative | 2,95 | 7,879E-153 |
| TcCLB.507247.14 | dispersed gene family protein 1 (DGF-1), putative (fragme | 2,945 | 4,999E-77 |
| TcCLB.507963.20 | arginase, putative | 2,934 | 3,941E-140 |
| TcCLB.511245.140 | fatty acid elongase, putative | 2,924 | 5,145E-156 |
| TcCLB.508741.50 | mucin TcSMUGS, putative | 2,919 | 2,423E-264 |
| TcCLB.509331.74 | hypothetical protein, conserved | 2,919 | 2,735E-77 |
| TcCLB.504087.20 | hypothetical protein | 2,904 | 6,306E-225 |
| TcCLB.506563.180 | calpain-like cysteine peptidase (pseudogene), putative | 2,896 | 7,938E-140 |
| TcCLB.511911.90 | hypothetical protein, conserved | 2,887 | 9,267E-157 |
| TcCLB.510977.9 | hypothetical protein, conserved (fragment) | 2,887 | 3,846E-127 |
| TcCLB.505789.20 | hypothetical protein, conserved | 2,886 | 9,267E-157 |
| TcCLB.509197.20 | cation transporter, putative | 2,872 | 9,133E-09 |
| TcCLB.508643.20 | hypothetical protein, conserved | 2,869 | 1,544E-122 |
| TcCLB.506325.10 | retrotransposon hot spot (RHS) protein, putative | 2,869 | 3,002E-72 |
| TcCLB.504045.55 | hypothetical protein, conserved | 2,868 | 4,62E-202 |
| TcCLB.508651.20 | cytochrome c oxidase subunit 10, putative | 2,86 | 3,762E-142 |
| TcCLB.508675.29 | calpain-like cysteine peptidase, putative | 2,86 | 1,71E-131 |
| TcCLB.505945.20 | ribonuclease mar1, putative | 2,855 | 1,078E-124 |
| TcCLB.506221.110 | chromosomal passenger protein, putative (CPC2) | 2,85 | 5,778E-86 |
| TcCLB.508799.290 | hypothetical protein, conserved (fragment) | 2,847 | 6,142E-155 |
| TcCLB.506459.60 | retrotransposon hot spot protein (RHS, pseudogene), puta | 2,846 | 4,436E-92 |
| TcCLB.506593.60 | nicotinamidase, putative (PNC1) | 2,845 | 2,8E-179 |
| TcCLB.507527.50 | cation transporter, putative | 2,844 | 2,134E-09 |

| | | | |
|---|---|---|---|
| TcCLB.509641.79 | hypothetical protein, conserved (fragment) | 2,843 | 1,446E-83 |
| TcCLB.506213.50 | prostaglandin F synthase (pseudogene), putative (PGFS) | 2,842 | 1,783E-115 |
| TcCLB.504033.80 | hypothetical protein, conserved | 2,841 | 2,452E-243 |
| TcCLB.509505.50 | 10 kDa heat shock protein, putative | 2,84 | 8,381E-100 |
| TcCLB.508999.250 | calpain-like cysteine peptidase (pseudogene), putative | 2,837 | 4,967E-202 |
| TcCLB.506621.30 | ribonucleoside-diphosphate reductase small chain, putativ | 2,829 | 8,698E-157 |
| TcCLB.509197.10 | cation transporter, putative | 2,826 | 5,166E-09 |
| TcCLB.509965.330 | hypothetical protein, conserved | 2,821 | 1,281E-252 |
| TcCLB.504203.40 | hypothetical protein, conserved | 2,818 | 5,069E-180 |
| TcCLB.506661.20 | fatty acid elongase, putative | 2,818 | 2,841E-122 |
| TcCLB.506983.48 | calpain-like cysteine peptidase, putative | 2,816 | 6,989E-72 |
| TcCLB.509611.180 | peptide methionine sulfoxide reductase, putative | 2,813 | 5,532E-82 |
| TcCLB.509059.50 | pyridoxal kinase, putative | 2,809 | 1,499E-174 |
| TcCLB.508863.10 | hypothetical protein, conserved | 2,801 | 4,791E-140 |
| TcCLB.504223.10 | amino acid permease, putative | 2,797 | 1,324E-163 |
| TcCLB.506199.20 | hypothetical protein, conserved | 2,795 | 3,424E-57 |
| TcCLB.509203.40 | glycosomal membrane protein, putative | 2,793 | 1,168E-236 |
| TcCLB.508799.150 | nucleoside transporter-like, putative | 2,792 | 5,552E-133 |
| TcCLB.507515.4 | trypanothione/tryparedoxin dependent peroxidase 2, puta | 2,786 | 1,6E-188 |
| TcCLB.506529.360 | cytochrome oxidase subunit IV, putative (COXIV) | 2,783 | 8,685E-289 |
| TcCLB.506931.10 | haloacid dehalogenase-like hydrolase, putative | 2,783 | 5,101E-92 |
| TcCLB.506177.100 | actyltransferase-like, putative | 2,782 | 6,666E-76 |
| TcCLB.509237.120 | calpain-like cysteine peptidase, putative | 2,778 | 1,63E-209 |
| TcCLB.508719.30 | hypothetical protein, conserved | 2,771 | 2,774E-205 |
| TcCLB.504013.60 | regulatory subunit of protein kinase a-like protein, putativ | 2,767 | 2,894E-113 |
| TcCLB.508501.250 | UMP-CMP kinase, mitochondrial, putative | 2,767 | 7,197E-85 |
| TcCLB.508827.50 | DNA repair and recombination helicase protein PIF3, puta | 2,764 | 9,058E-139 |
| TcCLB.508999.200 | cysteine peptidase, Clan CA, family C2, putative | 2,763 | 3,053E-120 |
| TcCLB.510521.10 | NADH-cytochrome b5 reductase (pseudogene), putative | 2,758 | 1,116E-114 |
| TcCLB.470399.10 | cyclophilin, putative (fragment) | 2,75 | 6,493E-58 |
| TcCLB.511437.30 | lipase, putative | 2,749 | 4,602E-75 |
| TcCLB.508823.70 | P27 protein, putative (P27) | 2,746 | 0 |
| TcCLB.506853.50 | 2-oxoisovalerate dehydrogenase alpha subunit, putative | 2,742 | 2,278E-148 |
| TcCLB.504223.30 | hypothetical protein, conserved (fragment) | 2,739 | 2,649E-21 |
| TcCLB.511575.30 | hypothetical protein, conserved | 2,735 | 1,096E-108 |
| TcCLB.506201.39 | 25 kDa translation elongation factor 1-beta, putative (EF-: | 2,734 | 1,003E-286 |
| TcCLB.510121.20 | hexokinase, putative | 2,727 | 2,384E-197 |
| TcCLB.503861.10 | retrotransposon hot spot (RHS) protein, putative | 2,724 | 9,773E-191 |
| TcCLB.508387.20 | methylthioadenosine phosphorylase, putative | 2,717 | 7,478E-107 |
| TcCLB.510629.460 | hypothetical protein | 2,713 | 6,238E-136 |
| TcCLB.510879.50 | regulatory subunit of protein kinase a-like protein, putativ | 2,71 | 2,538E-207 |
| TcCLB.511697.40 | hypothetical protein, conserved (pseudogene) | 2,709 | 6,313E-48 |
| TcCLB.509295.80 | retrotransposon hot spot (RHS) protein, putative | 2,706 | 3,036E-126 |
| TcCLB.507707.20 | hypothetical protein, conserved | 2,704 | 4,362E-97 |
| TcCLB.510905.10 | dispersed gene family protein 1 (DGF-1), putative | 2,703 | 1,296E-13 |

| | | | |
|---|---|---|---|
| TcCLB.510101.140 | pyruvate phosphate dikinase, putative | 2,701 | 4,81E-116 |
| TcCLB.506773.90 | amino acid permease/transporter, putative | 2,7 | 3,06E-161 |
| TcCLB.508677.80 | NADP-dependent alcohol hydrogenase, putative | 2,696 | 3,13E-88 |
| TcCLB.511245.180 | guanin nucleotide-binding protein, putative | 2,695 | 2,012E-128 |
| TcCLB.506319.60 | hypothetical protein, conserved | 2,689 | 1,197E-76 |
| TcCLB.508601.141 | dipeptidyl-peptidase, putative (fragment) | 2,688 | 9,123E-204 |
| TcCLB.510783.49 | cytochrome c oxidase subunit 10, putative | 2,685 | 3,048E-30 |
| TcCLB.510259.50 | 21 kDa cyclophilin, putative (CYP21) | 2,684 | 7,251E-108 |
| TcCLB.511135.20 | hypothetical protein, conserved | 2,679 | 1,797E-211 |
| TcCLB.511389.110 | cytochrome c oxidase VIII (COX VIII), putative | 2,675 | 2,213E-150 |
| TcCLB.510741.10 | cystathione gamma lyase, putative | 2,673 | 4,737E-52 |
| TcCLB.508677.50 | beta galactofuranosyl glycosyltransferase, putative | 2,669 | 1,167E-92 |
| TcCLB.508699.120 | dipeptidyl-peptidase, putative | 2,667 | 3,325E-254 |
| TcCLB.510565.20 | fatty acid desaturase, putative | 2,665 | 8,004E-124 |
| TcCLB.503791.30 | hypothetical protein, conserved | 2,66 | 6,426E-200 |
| TcCLB.506635.60 | pyroglutamyl-peptidase I, putative (PPI) | 2,658 | 5,814E-116 |
| TcCLB.511325.50 | amino acid permease, putative | 2,657 | 9,851E-47 |
| TcCLB.503393.10 | amino acid permease, putative (fragment) | 2,657 | 1,945E-39 |
| TcCLB.506219.40 | hypothetical protein, conserved | 2,654 | 2,392E-207 |
| TcCLB.509053.30 | hypothetical protein, conserved | 2,652 | 7,425E-174 |
| TcCLB.509719.40 | cytochrome P450, putative | 2,652 | 8,804E-89 |
| TcCLB.506857.20 | pyrroline-5-carboxylate reductase, putative (P5CR) | 2,65 | 2,046E-97 |
| TcCLB.509601.140 | hypothetical protein | 2,648 | 4,422E-194 |
| TcCLB.507677.160 | hypothetical protein, conserved | 2,639 | 1,711E-154 |
| TcCLB.507167.70 | retrotransposon hot spot (RHS) protein, putative | 2,639 | 5,574E-101 |
| TcCLB.508303.10 | alanine racemase 2, putative | 2,636 | 1,126E-140 |
| TcCLB.506779.200 | glycerol-3-phosphate dehydrogenase (pseudogene), putat | 2,628 | 6,654E-188 |
| TcCLB.506947.10 | hypothetical protein, conserved | 2,625 | 6,124E-127 |
| TcCLB.508821.40 | hypothetical protein, conserved | 2,622 | 2,471E-222 |
| TcCLB.510943.50 | delta-1-pyrroline-5-carboxylate dehydrogenase, putative | 2,619 | 5,032E-135 |
| TcCLB.506753.140 | hypothetical protein, conserved | 2,619 | 2,482E-88 |
| TcCLB.508955.20 | hypothetical protein, conserved | 2,618 | 3,544E-163 |
| TcCLB.503769.40 | cytochrome oxidase subunit VII (COXVII) | 2,618 | 1,35E-160 |
| TcCLB.509199.24 | ribose 5-phosphate isomerase type B | 2,617 | 1,02E-100 |
| TcCLB.510357.40 | hypothetical protein, conserved | 2,609 | 1,622E-57 |
| TcCLB.509683.90 | hypothetical protein | 2,606 | 1,968E-86 |
| TcCLB.505373.10 | retrotransposon hot spot (RHS) protein, putative | 2,603 | 2,095E-100 |
| TcCLB.509331.210 | NADP-dependent alcohol hydrogenase, putative | 2,602 | 2,758E-59 |
| TcCLB.511545.80 | L-Lysine transport protein | 2,598 | 6,486E-119 |
| TcCLB.506529.508 | glucose-6-phosphate isomerase, glycosomal, putative | 2,596 | 1,231E-153 |
| TcCLB.506925.300 | cyclophilin a, putative (CYPA) | 2,591 | 6,11E-102 |
| TcCLB.506563.210 | cysteine peptidase, Clan CA, family C2, putative | 2,59 | 2,306E-272 |
| TcCLB.508951.20 | hexokinase, putative | 2,589 | 9,055E-171 |
| TcCLB.506203.60 | hypothetical protein, conserved | 2,586 | 2,962E-73 |
| TcCLB.509793.50 | hypothetical protein, conserved | 2,584 | 1,922E-183 |

| | | | |
|---|---|---|---|
| TcCLB.511207.10 | hypothetical protein, conserved | 2,583 | 2,811E-128 |
| TcCLB.507105.50 | hypothetical protein | 2,581 | 3,651E-216 |
| TcCLB.511811.30 | hypothetical protein, conserved | 2,579 | 4,554E-92 |
| TcCLB.510739.19 | cystathione gamma lyase, putative (fragment) | 2,577 | 3,062E-37 |
| TcCLB.504081.220 | retrotransposon hot spot protein (RHS, pseudogene), puta | 2,571 | 4,637E-85 |
| TcCLB.508535.10 | NADH-dependent fumarate reductase, putative | 2,569 | 1,214E-285 |
| TcCLB.507129.5 | hypothetical protein, conserved (fragment) | 2,569 | 4,383E-25 |
| TcCLB.508029.10 | Hydroxymethylglutaryl-CoA lyase, mitochondrial, putative | 2,568 | 1,926E-74 |
| TcCLB.511691.39 | hypothetical protein, conserved | 2,562 | 6,659E-190 |
| TcCLB.510605.10 | hypothetical protein, conserved | 2,562 | 4,557E-87 |
| TcCLB.507709.90 | sterol C-24 reductase, putative | 2,559 | 1,99E-90 |
| TcCLB.509197.39 | cation transporter, putative | 2,556 | 2,651E-140 |
| TcCLB.504867.70 | cytochrome c oxidase assembly protein, putative | 2,552 | 2,073E-115 |
| TcCLB.509589.20 | squalene monooxygenase, putative | 2,551 | 8,386E-132 |
| TcCLB.504339.19 | NADP-dependent alcohol hydrogenase, putative | 2,55 | 5,495E-12 |
| TcCLB.508701.20 | cation transporter, putative | 2,537 | 4,421E-07 |
| TcCLB.511333.10 | calpain-like cysteine peptidase, putative | 2,529 | 3,909E-23 |
| TcCLB.510265.10 | quinone oxidoreductase, putative | 2,528 | 1,303E-81 |
| TcCLB.508241.10 | glycerol-3-phosphate dehydrogenase, putative | 2,525 | 1,448E-119 |
| TcCLB.506843.100 | retrotransposon hot spot (RHS) protein, putative | 2,523 | 1,137E-169 |
| TcCLB.506529.80 | hypothetical protein, conserved | 2,52 | 9,334E-163 |
| TcCLB.508027.129 | Hydroxymethylglutaryl-CoA lyase, mitochondrial, putative | 2,502 | 1,854E-44 |
| TcCLB.506147.9 | prostaglandin F2alpha synthase (TcPGFS) | 2,501 | 9,529E-238 |
| TcCLB.507867.100 | hypothetical protein | 2,498 | 4,304E-123 |
| TcCLB.508033.39 | mucin-associated surface protein (MASP, pseudogene), pu | 2,497 | 6,138E-54 |
| TcCLB.507559.80 | deoxyribose-phosphate aldolase, putative | 2,494 | 1,522E-187 |
| TcCLB.510245.10 | amino acid permease, putative | 2,492 | 4,773E-166 |
| TcCLB.509351.10 | delta-1-pyrroline-5-carboxylate dehydrogenase, putative | 2,492 | 1,445E-54 |
| TcCLB.508799.160 | cytochrome b5-like protein, putative | 2,49 | 5,544E-160 |
| TcCLB.506181.104 | actyltransferase-like protein | 2,49 | 2,316E-86 |
| TcCLB.507049.60 | NAD(P)-dependent oxidoreductase, putative | 2,489 | 7,022E-129 |
| TcCLB.505057.20 | hypothetical protein | 2,482 | 5,358E-61 |
| TcCLB.505807.180 | 2-hydroxy-3-oxopropionate reductase, putative | 2,474 | 5,846E-159 |
| TcCLB.509331.70 | 3-oxo-5-alpha-steroid 4-dehydrogenase, putative | 2,471 | 8,571E-90 |
| TcCLB.511613.100 | elongation factor 1-gamma (EF-1-gamma), putative | 2,469 | 2,701E-06 |
| TcCLB.509705.10 | cellulosomal scaffoldin anchoring protein, putative | 2,464 | 8,133E-91 |
| TcCLB.506983.39 | calpain-like cysteine peptidase, putative | 2,458 | 2,85E-76 |
| TcCLB.507081.130 | 2,4-dihydroxyhept-2-ene-1,7-dioic acid aldolase, putative | 2,457 | 1,887E-96 |
| TcCLB.505171.30 | hypothetical protein, conserved | 2,456 | 7,47E-95 |
| TcCLB.450161.10 | cation transporter, putative | 2,455 | 5,829E-116 |
| TcCLB.508781.110 | 2-oxoisovalerate dehydrogenase alpha subunit, putative | 2,453 | 4,741E-93 |
| TcCLB.506975.40 | amidohydrolase, putative | 2,449 | 8,12E-154 |
| TcCLB.511335.9 | calpain-like cysteine peptidase, putative (fragment) | 2,445 | 4,191E-26 |
| TcCLB.504105.220 | hypothetical protein, conserved | 2,44 | 8,007E-51 |
| TcCLB.482097.20 | hypothetical protein, conserved | 2,438 | 2,16E-09 |

| | | | |
|---|---|---|---|
| TcCLB.509045.20 | co-chaperone GrpE, putative | 2,437 | 6,755E-127 |
| TcCLB.511867.140 | Acyl carrier protein, mitochondrial, putative (ACP) | 2,434 | 3,259E-121 |
| TcCLB.507517.60 | ATPase subunit 9, putative | 2,43 | 2,061E-98 |
| TcCLB.509233.150 | cytochrome oxidase subunit VII (COXVII) | 2,428 | 9,581E-147 |
| TcCLB.509395.100 | cytochrome b-domain protein, putative | 2,423 | 3,009E-56 |
| TcCLB.509179.20 | Macro domain containing protein, putative | 2,422 | 4,471E-102 |
| TcCLB.507317.34 | hypothetical protein, conserved | 2,422 | 7,637E-79 |
| TcCLB.508173.150 | protein disulfide isomerase, putative | 2,415 | 2,098E-98 |
| TcCLB.509265.10 | retrotransposon hot spot (RHS) protein, putative | 2,415 | 3,111E-76 |
| TcCLB.508873.420 | Present in the outer mitochondrial membrane proteome 1 | 2,414 | 1,105E-110 |
| TcCLB.484311.10 | calpain-like cysteine peptidase, putative (fragment) | 2,412 | 9,136E-13 |
| TcCLB.441401.10 | imidazolonepropienase, putative | 2,407 | 1,564E-81 |
| TcCLB.503899.119 | trypanothione/tryparedoxin dependent peroxidase 2, puta | 2,406 | 9,94E-126 |
| TcCLB.508461.80 | prostaglandin F2alpha synthase (TcPGFS) | 2,402 | 7,837E-234 |
| TcCLB.508257.220 | expression site-associated gene (ESAG-like) protein, puta | 2,4 | 2,037E-103 |
| TcCLB.503753.29 | NADPH--cytochrome P450 reductase, putative | 2,399 | 5,094E-97 |
| TcCLB.509769.40 | hypothetical protein, conserved | 2,398 | 1,993E-87 |
| TcCLB.505193.100 | Present in the outer mitochondrial membrane proteome 1 | 2,396 | 1,754E-40 |
| TcCLB.506169.21 | retrotransposon hot spot (RHS) protein, putative (fragmer | 2,39 | 3,46E-30 |
| TcCLB.503697.70 | hypothetical protein, conserved | 2,388 | 9,672E-78 |
| TcCLB.510507.20 | 3-ketoacyl-CoA thiolase, putative | 2,387 | 7,284E-166 |
| TcCLB.503461.20 | per-hexamer repeat protein 5, putative | 2,386 | 1,001E-73 |
| TcCLB.438313.9 | dispersed gene family protein 1 (DGF-1), putative | 2,383 | 4,2E-71 |
| TcCLB.508897.119 | FKBP-type peptidylprolyl cis-trans isomerase, putative | 2,382 | 3,612E-98 |
| TcCLB.508045.100 | mitochondrial DNA primase, putative (PRI1) | 2,379 | 2,326E-116 |
| TcCLB.506789.130 | double-strand-break repair protein rad21 homolog, putativ | 2,377 | 5,913E-181 |
| TcCLB.509137.30 | imidazolonepropionase, putative | 2,375 | 7,925E-147 |
| TcCLB.504983.10 | hypothetical protein | 2,375 | 3,269E-81 |
| TcCLB.506417.5 | endoribonuclease L-PSP (pb5), putative | 2,371 | 6,94E-110 |
| TcCLB.510063.30 | hypothetical protein, conserved | 2,371 | 1,145E-98 |
| TcCLB.509799.50 | hypothetical protein, conserved | 2,369 | 3,808E-103 |
| TcCLB.511573.20 | hypothetical protein | 2,367 | 5,378E-173 |
| TcCLB.508741.140 | imidazolonepropionase, putative | 2,361 | 1,13E-147 |
| TcCLB.510517.40 | hypothetical protein | 2,361 | 4,968E-98 |
| TcCLB.503991.39 | 2,4-dihydroxyhept-2-ene-1,7-dioic acid aldolase, putative | 2,36 | 5,614E-68 |
| TcCLB.511145.10 | cytochrome c oxidase subunit VI, putative | 2,357 | 1,93E-65 |
| TcCLB.511827.100 | cysteine peptidase C (CPC), putative (TCCB) | 2,35 | 1,074E-87 |
| TcCLB.507929.20 | co-chaperone GrpE, putative | 2,344 | 4,743E-95 |
| TcCLB.509331.30 | acetyl-CoA synthetase, putative | 2,344 | 1,647E-75 |
| TcCLB.509979.150 | nitrilase, putative | 2,343 | 5,782E-88 |
| TcCLB.506325.40 | retrotransposon hot spot protein (RHS, pseudogene), puta | 2,332 | 2,283E-120 |
| TcCLB.508647.180 | hypothetical protein, conserved | 2,331 | 5,552E-103 |
| TcCLB.504257.60 | S-adenosylmethionine decarboxylase proenzyme, putative | 2,33 | 8,524E-34 |
| TcCLB.506297.190 | pyruvate phosphate dikinase, putative | 2,328 | 1,596E-116 |
| TcCLB.507617.9 | prostaglandin F2alpha synthase (TcPGFS) | 2,326 | 8,586E-209 |

| | | | |
|---|---|---|---|
| TcCLB.507611.280 | cytochrome oxidase subunit IX, putative (COXIX) | 2,325 | 3,816E-102 |
| TcCLB.509587.20 | squalene monooxygenase, putative (fragment) | 2,316 | 7,218E-84 |
| TcCLB.511017.50 | glycosomal membrane protein, putative | 2,313 | 5,235E-105 |
| TcCLB.504069.20 | cytochrome c oxidase subunit VI, putative | 2,313 | 4,009E-81 |
| TcCLB.509215.40 | peptidyl-prolyl cis-trans isomerase, putative | 2,305 | 2,736E-124 |
| TcCLB.428403.10 | hypothetical protein (fragment) | 2,3 | 7,877E-06 |
| TcCLB.503579.70 | ATPase subunit 9, putative | 2,298 | 6,957E-87 |
| TcCLB.506853.10 | hypothetical protein, conserved | 2,297 | 9,191E-154 |
| TcCLB.507875.20 | glutamate dehydrogenase, putative | 2,297 | 1,576E-52 |
| TcCLB.507837.20 | protein kinase, putative | 2,296 | 2,639E-84 |
| TcCLB.511639.20 | hypothetical protein, conserved | 2,292 | 4,994E-149 |
| TcCLB.508699.130 | cation transporter, putative (fragment) | 2,287 | 2,982E-81 |
| TcCLB.508555.50 | calpain-like cysteine peptidase, putative | 2,286 | 1,589E-63 |
| TcCLB.510729.110 | meiotic recombination protein DMC1, putative (dmc1) | 2,285 | 1,39E-119 |
| TcCLB.509267.30 | hypothetical protein, conserved | 2,282 | 1,558E-32 |
| TcCLB.507219.20 | retrotransposon hot spot protein (RHS, pseudogene), puta | 2,28 | 4,865E-87 |
| TcCLB.506467.29 | methyltransferase domain containing protein, putative (P | 2,276 | 1,888E-106 |
| TcCLB.509787.10 | retrotransposon hot spot (RHS) protein, putative | 2,276 | 9,327E-104 |
| TcCLB.511393.49 | hypothetical protein, conserved | 2,276 | 2,91E-63 |
| TcCLB.510131.44 | hypothetical protein, conserved | 2,27 | 3,158E-104 |
| TcCLB.509703.10 | cellulosomal scaffoldin anchoring protein, putative | 2,264 | 1,811E-74 |
| TcCLB.511545.90 | hypothetical protein | 2,259 | 4,937E-79 |
| TcCLB.510101.80 | hypothetical protein, conserved | 2,258 | 1,327E-68 |
| TcCLB.507711.200 | hypothetical protein, conserved | 2,256 | 3,465E-281 |
| TcCLB.508555.20 | hypothetical protein, conserved | 2,255 | 1,588E-48 |
| TcCLB.455721.9 | cytochrome c oxidase subunit 10, putative | 2,253 | 1,924E-55 |
| TcCLB.503855.40 | calpain-like cysteine peptidase, putative | 2,251 | 7,301E-71 |
| TcCLB.506183.70 | beta galactofuranosyl glycosyltransferase, putative | 2,25 | 1,329E-67 |
| TcCLB.511309.60 | hypothetical protein, conserved | 2,25 | 1,655E-52 |
| TcCLB.510769.10 | hypothetical protein, conserved | 2,245 | 3,497E-64 |
| TcCLB.507723.100 | cytochrome oxidase subunit IX, putative (COXIX) | 2,243 | 2,107E-87 |
| TcCLB.510977.20 | hypothetical protein, conserved (pseudogene) | 2,239 | 1,078E-60 |
| TcCLB.507739.40 | uracil phosphoribosyltransferase, putative | 2,237 | 3,666E-104 |
| TcCLB.503577.9 | delta-1-pyrroline-5-carboxylate dehydrogenase (pseudoge | 2,236 | 1,405E-45 |
| TcCLB.510069.20 | protein associated with differentiation 4, putative | 2,235 | 4,018E-126 |
| TcCLB.510309.20 | MFS transporter, putative | 2,233 | 5,071E-96 |
| TcCLB.508153.740 | hypothetical protein | 2,229 | 1,144E-93 |
| TcCLB.508549.20 | hypothetical protein, conserved | 2,227 | 2,161E-132 |
| TcCLB.509223.20 | retrotransposon hot spot (RHS) protein, putative | 2,227 | 6,007E-98 |
| TcCLB.510795.30 | fatty acid desaturase (pseudogene), putative | 2,226 | 4,634E-79 |
| TcCLB.503999.10 | squalene monooxygenase, putative | 2,225 | 4,669E-120 |
| TcCLB.504005.54 | hypothetical protein, conserved | 2,225 | 8,256E-51 |
| TcCLB.511639.9 | hypothetical protein, conserved | 2,224 | 1,562E-66 |
| TcCLB.503989.30 | hypothetical protein, conserved (pseudogene) | 2,223 | 3,664E-150 |
| TcCLB.508471.19 | hypothetical protein, conserved (fragment) | 2,223 | 1,324E-104 |

| | | | |
|---|---|---|---|
| TcCLB.509329.20 | retrotransposon hot spot protein (RHS, pseudogene), puta | 2,223 | 1,174E-82 |
| TcCLB.506625.230 | hypothetical protein, conserved | 2,216 | 2,139E-43 |
| TcCLB.509023.70 | hypothetical protein, conserved | 2,211 | 2,948E-22 |
| TcCLB.510943.130 | cytochrome c oxidase assembly protein, putative | 2,21 | 7,03E-87 |
| TcCLB.510749.99 | aminopeptidase P, putative | 2,208 | 1,821E-26 |
| TcCLB.511151.90 | glycerol-3-phosphate dehydrogenase, putative | 2,204 | 6,41E-149 |
| TcCLB.506505.30 | hypothetical protein, conserved | 2,201 | 2,827E-92 |
| TcCLB.511903.40 | 3-hydroxy-3-methylglutaryl-CoA synthase, putative (HMGS | 2,196 | 2,367E-139 |
| TcCLB.504069.120 | amino acid permease 24, putative (AAP24) | 2,194 | 4,418E-58 |
| TcCLB.507771.70 | hypothetical protein, conserved | 2,192 | 1,172E-104 |
| TcCLB.505647.40 | endoribonuclease L-PSP (pb5), putative | 2,19 | 7,124E-164 |
| TcCLB.509937.200 | hypothetical protein, conserved | 2,184 | 1,097E-57 |
| TcCLB.508221.414 | elongation factor 1-gamma (EF-1-gamma), putative | 2,181 | 2,726E-06 |
| TcCLB.506945.190 | cytochrome p450-like protein, putative | 2,171 | 7,176E-127 |
| TcCLB.509427.50 | retrotransposon hot spot protein (RHS, pseudogene), puta | 2,171 | 2,574E-77 |
| TcCLB.510747.10 | hypothetical protein, conserved | 2,171 | 1,273E-55 |
| TcCLB.508799.120 | amino acid permease/transporter, putative | 2,166 | 2,232E-86 |
| TcCLB.508567.60 | hypothetical protein | 2,165 | 1,454E-80 |
| TcCLB.505647.60 | hypothetical protein, conserved | 2,164 | 2,056E-83 |
| TcCLB.506375.110 | (H )-ATPase G subunit, putative | 2,159 | 1,021E-90 |
| TcCLB.509367.30 | dispersed gene family protein 1 (DGF-1, pseudogene), put | 2,159 | 7,096E-23 |
| TcCLB.506731.10 | hypothetical protein | 2,158 | 4,346E-25 |
| TcCLB.510759.40 | Protein kinase domain, putative | 2,156 | 1,675E-99 |
| TcCLB.503745.25 | hypothetical protein, conserved | 2,155 | 5,954E-32 |
| TcCLB.510853.10 | dispersed gene family protein 1 (DGF-1, pseudogene), put | 2,146 | 4,766E-46 |
| TcCLB.503841.80 | NADH-dependent fumarate reductase, putative | 2,145 | 2,531E-31 |
| TcCLB.507115.20 | dispersed gene family protein 1 (DGF-1), putative | 2,141 | 5,574E-42 |
| TcCLB.508147.159 | dispersed gene family protein 1 (DGF-1), putative | 2,141 | 1,16E-09 |
| TcCLB.511575.130 | folate/pteridine transporter, putative | 2,14 | 3,965E-90 |
| TcCLB.509287.230 | trans-sialidase, Group II, putative | 2,139 | 1,446E-87 |
| TcCLB.511277.500 | hypothetical protein, conserved | 2,136 | 1,099E-69 |
| TcCLB.510143.24 | Hsc70-interacting protein (Hip), putative | 2,126 | 1,753E-134 |
| TcCLB.508023.80 | retrotransposon hot spot protein (RHS, pseudogene), puta | 2,126 | 1,355E-42 |
| TcCLB.503975.70 | hypothetical protein, conserved | 2,124 | 8,023E-90 |
| TcCLB.508971.40 | acetylornithine deacetylase-like, putative | 2,123 | 6,654E-180 |
| TcCLB.510301.30 | hypothetical protein, conserved | 2,121 | 2,745E-61 |
| TcCLB.507671.30 | 25 kDa translation elongation factor 1-beta (EF-1-beta) | 2,12 | 1,012E-125 |
| TcCLB.507515.90 | hypothetical protein, conserved | 2,119 | 1,056E-61 |
| TcCLB.503453.100 | serine palmitoyltransferase, putative | 2,115 | 4,632E-96 |
| TcCLB.506999.180 | hypothetical protein | 2,114 | 1,201E-106 |
| TcCLB.508465.40 | hypothetical protein, conserved | 2,111 | 5,428E-86 |
| TcCLB.511903.120 | transketolase, putative | 2,11 | 3,094E-142 |
| TcCLB.511667.20 | retrotransposon hot spot (RHS) protein, putative | 2,105 | 3,575E-47 |
| TcCLB.506129.89 | hypothetical protein, conserved | 2,104 | 1,627E-106 |
| TcCLB.507641.64 | hypothetical protein, conserved | 2,104 | 1,354E-83 |

| | | | |
|---|---|---|---|
| TcCLB.510213.110 | NADH-dependent fumarate reductase, putative | 2,103 | 2,729E-40 |
| TcCLB.508503.10 | fatty acid desaturase, putative | 2,1 | 1,046E-48 |
| TcCLB.511337.4 | hypothetical protein, conserved | 2,095 | 1,452E-21 |
| TcCLB.508059.50 | hypothetical protein, conserved | 2,094 | 8,163E-102 |
| TcCLB.510889.50 | cytochrome oxidase subunit IV, putative (COXIV) | 2,093 | 3,093E-99 |
| TcCLB.506931.80 | p450 reductase, putative | 2,093 | 8,484E-89 |
| TcCLB.506789.160 | hypothetical protein, conserved | 2,092 | 1,273E-71 |
| TcCLB.511773.10 | retrotransposon hot spot protein (RHS, pseudogene), puta | 2,092 | 6,651E-57 |
| TcCLB.507153.30 | dispersed gene family protein 1 (DGF-1), putative | 2,089 | 1,444E-41 |
| TcCLB.507233.110 | mismatch repair protein MSH4, putative | 2,088 | 6,863E-127 |
| TcCLB.508139.130 | trans-sialidase, Group II, putative | 2,086 | 1,581E-104 |
| TcCLB.510407.30 | hypothetical protein, conserved | 2,083 | 7,763E-47 |
| TcCLB.508809.40 | hypothetical protein, conserved | 2,083 | 2,341E-33 |
| TcCLB.506171.10 | retrotransposon hot spot (RHS) protein, putative | 2,083 | 6,435E-27 |
| TcCLB.508231.40 | hypothetical protein, conserved | 2,081 | 4,147E-117 |
| TcCLB.509671.10 | hypothetical protein, conserved | 2,075 | 3,486E-112 |
| TcCLB.407477.50 | cytochrome c oxidase VIII (COX VIII), putative | 2,074 | 1,145E-93 |
| TcCLB.511909.30 | hypothetical protein, conserved | 2,07 | 1,594E-44 |
| TcCLB.507019.20 | nitrate reductase, putative | 2,068 | 1,822E-108 |
| TcCLB.510105.100 | UDP-glucose dehydrogenase, putative | 2,066 | 5,804E-60 |
| TcCLB.510993.10 | (H )-ATPase G subunit | 2,061 | 2,642E-76 |
| TcCLB.507897.20 | squalene synthase, putative | 2,06 | 7,56E-201 |
| TcCLB.509331.160 | hypothetical protein, conserved | 2,06 | 6,143E-53 |
| TcCLB.506865.2 | nucleoside phosphorylase, putative | 2,06 | 3,048E-45 |
| TcCLB.509003.60 | hypothetical protein, conserved | 2,058 | 1,02E-87 |
| TcCLB.511895.20 | hypothetical protein | 2,057 | 8,64E-72 |
| TcCLB.511295.10 | FKBP-type peptidylprolyl cis-trans isomerase, putative | 2,056 | 3,761E-74 |
| TcCLB.506391.30 | ef-hand protein 5, putative | 2,052 | 3,105E-47 |
| TcCLB.506773.120 | mRNA cap guanine-N7 methyltransferase, putative | 2,051 | 7,772E-69 |
| TcCLB.511485.11 | retrotransposon hot spot protein (RHS, pseudogene), puta | 2,051 | 1,409E-07 |
| TcCLB.510743.70 | lactoylglutathione lyase-like protein, putative | 2,048 | 9,934E-100 |
| TcCLB.511303.64 | hypothetical protein, conserved | 2,045 | 4,114E-71 |
| TcCLB.511815.170 | hypothetical protein, conserved | 2,045 | 6,434E-60 |
| TcCLB.506811.160 | R27-2 protein, putative | 2,044 | 3,252E-140 |
| TcCLB.426897.10 | hypothetical protein | 2,043 | 5,291E-75 |
| TcCLB.506989.140 | Elongation factor Ts, mitochondrial, putative (EF-Ts) | 2,042 | 3,758E-71 |
| TcCLB.511589.70 | ADP-ribosylation factor GTPase activating protein, putativ | 2,039 | 1,54E-69 |
| TcCLB.511131.80 | hypothetical protein, conserved | 2,035 | 1,728E-109 |
| TcCLB.509693.209 | mismatch repair protein, putative | 2,033 | 1,382E-58 |
| TcCLB.510861.140 | serine/threonine protein kinase, putative | 2,031 | 1,862E-123 |
| TcCLB.506125.20 | hypothetical protein, conserved | 2,029 | 1,51E-84 |
| TcCLB.509767.59 | protoheme IX farnesyltransferase, putative | 2,027 | 8,681E-85 |
| TcCLB.506855.310 | hypothetical protein, conserved | 2,026 | 1,599E-108 |
| TcCLB.511037.20 | L-2-hydroxyglutarate dehydrogenase, mitochondrial, putat | 2,025 | 1,65E-87 |
| TcCLB.508881.70 | DNA ligase, putative | 2,023 | 3,895E-93 |

| | | | |
|---|---|---|---|
| TcCLB.507719.9 | dispersed gene family protein 1 (DGF-1), putative | 2,021 | 8,944E-47 |
| TcCLB.506323.30 | retrotransposon hot spot (RHS) protein, putative | 2,019 | 1,814E-75 |
| TcCLB.510677.9 | dispersed gene family protein 1 (DGF-1), putative (fragme | 2,018 | 1,31E-35 |
| TcCLB.503411.10 | hypothetical protein | 2,015 | 5,46E-58 |
| TcCLB.511807.120 | hypothetical protein, conserved | 2,007 | 5,623E-73 |
| TcCLB.506355.5 | hexose transporter, putative | 2,007 | 1,154E-37 |
| TcCLB.504003.30 | hypothetical protein | 2,006 | 1,336E-67 |
| TcCLB.504033.130 | spermidine synthase, putative | 2,006 | 1,236E-19 |
| TcCLB.511759.40 | hypothetical protein, conserved | 2,004 | 7,179E-23 |
| TcCLB.508405.70 | protein kinase, putative | 2,001 | 1,147E-76 |
| TcCLB.508141.90 | hypothetical protein, conserved | 1,999 | 1,784E-62 |
| TcCLB.510603.169 | hypothetical protein, conserved | 1,997 | 1,207E-22 |
| TcCLB.507089.210 | hypothetical protein, conserved | 1,995 | 7,576E-71 |
| TcCLB.510971.5 | retrotransposon hot spot protein (RHS, pseudogene), puta | 1,994 | 3,173E-46 |
| TcCLB.506885.204 | hypothetical protein, conserved | 1,993 | 6,928E-99 |
| TcCLB.511469.70 | prohibitin, putative | 1,988 | 6,652E-91 |
| TcCLB.511485.20 | retrotransposon hot spot protein (RHS, pseudogene), puta | 1,988 | 4,011E-28 |
| TcCLB.505207.20 | beta galactofuranosyl glycosyltransferase, putative | 1,986 | 2,256E-20 |
| TcCLB.509029.140 | hypothetical protein, conserved | 1,985 | 1,086E-55 |
| TcCLB.507117.9 | dispersed gene family protein 1 (DGF-1), putative | 1,985 | 1,188E-32 |
| TcCLB.503739.10 | beta galactofuranosyl glycosyltransferase (pseudogene), p | 1,985 | 3,159E-09 |
| TcCLB.510669.20 | Gim5A protein, putative | 1,982 | 1,417E-83 |
| TcCLB.509601.59 | protoheme IX farnesyltransferase, putative | 1,981 | 1,945E-93 |
| TcCLB.510005.41 | retrotransposon hot spot (RHS) protein, putative | 1,98 | 5,428E-86 |
| TcCLB.511661.80 | hypothetical protein, conserved | 1,98 | 3,71E-48 |
| TcCLB.506263.10 | D-isomer specific 2-hydroxyacid dehydrogenase-protein, p | 1,979 | 3,095E-138 |
| TcCLB.505971.10 | UDP-Gal or UDP-GlcNAc-dependent glycosyltransferase, p | 1,979 | 1,541E-17 |
| TcCLB.509295.10 | dispersed gene family protein 1 (DGF-1), putative | 1,979 | 2,192E-17 |
| TcCLB.504741.4 | hypothetical protein, conserved | 1,978 | 2,147E-63 |
| TcCLB.509327.10 | retrotransposon hot spot protein (RHS, pseudogene), puta | 1,978 | 1,419E-40 |
| TcCLB.509551.30 | mitochondrial phosphate transporter, putative | 1,975 | 1,391E-64 |
| TcCLB.507895.124 | hypothetical protein, conserved | 1,971 | 1,312E-136 |
| TcCLB.507967.20 | hypothetical protein, conserved | 1,97 | 1,857E-49 |
| TcCLB.508211.60 | hypothetical protein, conserved | 1,965 | 5,351E-52 |
| TcCLB.510655.120 | aminopeptidase P, putative | 1,962 | 4,194E-78 |
| TcCLB.503975.60 | hypothetical protein | 1,956 | 3,269E-75 |
| TcCLB.510065.5 | hypothetical protein, conserved (fragment) | 1,952 | 4,007E-38 |
| TcCLB.511707.10 | hypothetical protein | 1,951 | 1,966E-28 |
| TcCLB.504427.75 | hypothetical protein, conserved (pseudogene) | 1,95 | 1,681E-35 |
| TcCLB.506001.90 | retrotransposon hot spot (RHS) protein, putative | 1,946 | 1,251E-71 |
| TcCLB.511237.30 | proline oxidase (pseudogene), putative | 1,946 | 1,908E-59 |
| TcCLB.506811.190 | UDP-galactose 4-epimerase (GalE) | 1,945 | 1,075E-99 |
| TcCLB.508003.10 | retrotransposon hot spot (RHS) protein, putative | 1,945 | 1,677E-51 |
| TcCLB.508837.90 | hypothetical protein, conserved | 1,944 | 1,061E-63 |
| TcCLB.511691.10 | cystathionine beta-synthase (CBS) | 1,941 | 9,033E-27 |

| | | | |
|---|---|---|---|
| TcCLB.503823.40 | glycosyl hydrolase-like protein, putative | 1,94 | 6,186E-98 |
| TcCLB.503959.78 | hypothetical protein, conserved | 1,939 | 8,939E-81 |
| TcCLB.509429.280 | hypothetical protein | 1,939 | 2,074E-77 |
| TcCLB.507625.10 | retrotransposon hot spot (RHS) protein, putative | 1,935 | 1,468E-60 |
| TcCLB.506999.60 | hypothetical protein, conserved | 1,933 | 6,541E-55 |
| TcCLB.510149.80 | ABC transporter, putative | 1,929 | 2,633E-136 |
| TcCLB.504933.30 | hypothetical protein | 1,929 | 1,208E-127 |
| TcCLB.510605.19 | hypothetical protein, conserved | 1,929 | 8,504E-111 |
| TcCLB.510573.10 | actin 2, putative (fragment) | 1,928 | 3,296E-89 |
| TcCLB.508851.120 | ADP-ribosylation factor GTPase activating protein, putativ | 1,928 | 2,004E-63 |
| TcCLB.509363.20 | retrotransposon hot spot protein (RHS, pseudogene), puta | 1,928 | 2,013E-06 |
| TcCLB.507167.20 | retrotransposon hot spot (RHS) protein, putative | 1,927 | 1,884E-121 |
| TcCLB.509875.50 | beta galactofuranosyl glycosyltransferase, putative | 1,926 | 1,708E-16 |
| TcCLB.508401.20 | retrotransposon hot spot (RHS) protein, putative | 1,925 | 1,379E-77 |
| TcCLB.458015.4 | hypothetical protein, conserved | 1,925 | 3,618E-45 |
| TcCLB.508567.40 | hypothetical protein, conserved | 1,924 | 6,433E-66 |
| TcCLB.508541.160 | hypothetical protein, conserved | 1,924 | 4,598E-25 |
| TcCLB.503613.40 | pterin-4-alpha-carbinolamine dehydratase, putative | 1,922 | 1,039E-60 |
| TcCLB.506585.70 | p450 reductase, putative | 1,92 | 1,257E-126 |
| TcCLB.510359.340 | phospholipid:diacylglycerol acyltransferase, putative | 1,919 | 5,037E-87 |
| TcCLB.511461.60 | NADH-dependent fumarate reductase (pseudogene), puta | 1,917 | 9,854E-107 |
| TcCLB.503845.30 | retrotransposon hot spot protein (RHS, pseudogene), puta | 1,917 | 2,201E-62 |
| TcCLB.510889.221 | glucose-6-phosphate isomerase, glycosomal, putative | 1,917 | 1,149E-61 |
| TcCLB.507081.110 | aminopeptidase P1, putative | 1,917 | 1,442E-58 |
| TcCLB.506549.9 | dispersed gene family protein 1 (DGF-1), putative (fragme | 1,916 | 1,658E-39 |
| TcCLB.419637.9 | dispersed gene family protein 1 (DGF-1), putative | 1,913 | 0,0004567 |
| TcCLB.510347.20 | hypothetical protein, conserved | 1,912 | 1,602E-65 |
| TcCLB.506563.120 | hypothetical protein, conserved | 1,911 | 3,701E-57 |
| TcCLB.510571.39 | actin 2, putative (fragment) | 1,91 | 2,859E-47 |
| TcCLB.511011.10 | retrotransposon hot spot (RHS) protein, putative | 1,908 | 1,465E-59 |
| TcCLB.504421.19 | hypothetical protein, conserved | 1,906 | 6,793E-76 |
| TcCLB.510299.20 | hypothetical protein, conserved | 1,906 | 2,464E-34 |
| TcCLB.430539.10 | hypothetical protein, conserved (fragment) | 1,905 | 4,771E-14 |
| TcCLB.510669.40 | hypothetical protein | 1,903 | 4,956E-74 |
| TcCLB.507519.150 | Adenine phosphoribosyltransferase, putative | 1,903 | 3,554E-59 |
| TcCLB.506567.70 | NAD(P)-dependent oxidoreductase, putative | 1,901 | 7,715E-83 |
| TcCLB.510751.4 | aminopeptidase P, putative | 1,901 | 2,084E-71 |
| TcCLB.508387.140 | hypothetical protein, conserved | 1,899 | 3,106E-56 |
| TcCLB.508241.140 | cystathionine beta-synthase, putative (CBS) | 1,899 | 1,331E-36 |
| TcCLB.503835.39 | hypothetical protein, conserved | 1,896 | 1,004E-52 |
| TcCLB.511009.9 | dispersed gene family protein 1 (DGF-1), putative (fragme | 1,896 | 4,138E-13 |
| TcCLB.508507.40 | hypothetical protein, conserved | 1,892 | 5,518E-26 |
| TcCLB.508297.50 | short chain dehydrogenase, putative | 1,891 | 6,353E-58 |
| TcCLB.503975.50 | hypothetical protein, conserved | 1,889 | 1,443E-95 |
| TcCLB.507167.200 | beta galactofuranosyl glycosyltransferase, putative | 1,888 | 3,205E-12 |

| | | | |
|---|---|---|---|
| TcCLB.506303.80 | hypothetical protein, conserved | 1,884 | 3,183E-116 |
| TcCLB.506989.220 | hypothetical protein, conserved | 1,884 | 5,121E-52 |
| TcCLB.508569.70 | hypothetical protein, conserved | 1,88 | 8,536E-80 |
| TcCLB.509259.50 | retrotransposon hot spot protein (RHS, pseudogene), puta | 1,88 | 8,112E-46 |
| TcCLB.442383.19 | hypothetical protein | 1,877 | 1,818E-39 |
| TcCLB.510819.119 | zinc finger (CCCH type) protein, putative | 1,876 | 8,11E-70 |
| TcCLB.510127.79 | actin 3, putative (fragment) | 1,876 | 4,581E-62 |
| TcCLB.511911.94 | hypothetical protein, conserved | 1,873 | 2,045E-76 |
| TcCLB.509245.29 | hypothetical protein, conserved | 1,872 | 1,098E-27 |
| TcCLB.506773.140 | DNA ligase, putative | 1,87 | 6,068E-57 |
| TcCLB.511039.30 | hypothetical protein, conserved | 1,87 | 6,633E-41 |
| TcCLB.510667.140 | ribulose-phosphate 3-epimerase, putative | 1,87 | 1,294E-39 |
| TcCLB.506793.40 | UDP-glucuronic acid/UDP-N-acetylgalactosamine transpor | 1,869 | 2,669E-39 |
| TcCLB.505615.20 | lipase (pseudogene), putative | 1,869 | 1,643E-08 |
| TcCLB.509109.14 | hypothetical protein, conserved (fragment) | 1,868 | 1,189E-41 |
| TcCLB.508149.10 | dispersed gene family protein 1 (DGF-1), putative | 1,868 | 1,174E-08 |
| TcCLB.511071.50 | 3-hydroxy-3-methylglutaryl-CoA synthase, putative | 1,867 | 5,74E-135 |
| TcCLB.508601.119 | ribose 5-phosphate isomerase, putative | 1,864 | 3,062E-48 |
| TcCLB.511229.50 | S-adenosylhomocysteine hydrolase, putative | 1,863 | 5,488E-64 |
| TcCLB.511421.185 | hypothetical protein, conserved | 1,863 | 1,445E-34 |
| TcCLB.503903.60 | hypothetical protein, conserved | 1,862 | 8,097E-79 |
| TcCLB.509597.20 | carbonic anhydrase-like protein, putative | 1,861 | 3,123E-73 |
| TcCLB.506381.10 | retrotransposon hot spot protein (RHS, pseudogene), puta | 1,86 | 5,144E-32 |
| TcCLB.511515.40 | heat shock 70 kDa protein, mitochondrial precursor, putati | 1,859 | 1,337E-89 |
| TcCLB.402375.10 | retrotransposon hot spot protein (RHS, pseudogene), puta | 1,858 | 4,721E-66 |
| TcCLB.508945.10 | hypothetical protein, conserved | 1,857 | 8,102E-47 |
| TcCLB.509331.120 | hypothetical protein, conserved | 1,856 | 2,365E-60 |
| TcCLB.507221.20 | poly(A)-specific ribonuclease PARN, putative | 1,854 | 7,5E-46 |
| TcCLB.506421.10 | retrotransposon hot spot (RHS) protein, putative | 1,852 | 4,782E-92 |
| TcCLB.506789.240 | hypothetical protein, conserved | 1,85 | 1,304E-43 |
| TcCLB.508177.90 | hypothetical protein, conserved | 1,845 | 9,758E-86 |
| TcCLB.510911.14 | hypothetical protein, conserved | 1,845 | 1,319E-55 |
| TcCLB.503657.70 | tyrosine phosphatase, putative | 1,845 | 2,177E-34 |
| TcCLB.508409.210 | Hsc70-interacting protein, putative (HIP) | 1,844 | 3,907E-139 |
| TcCLB.510131.40 | haloacid dehalogenase-like hydrolase, putative | 1,842 | 1,105E-47 |
| TcCLB.443403.9 | retrotransposon hot spot (RHS) protein, putative | 1,84 | 4,422E-39 |
| TcCLB.508461.570 | Gim5A protein, putative | 1,838 | 2,239E-122 |
| TcCLB.509721.60 | poly(ADP-ribose) polymerase, putative | 1,837 | 7,402E-74 |
| TcCLB.503703.40 | hypothetical protein, conserved | 1,837 | 2,568E-42 |
| TcCLB.503411.4 | hypothetical protein, conserved | 1,837 | 4,177E-13 |
| TcCLB.509733.100 | translation elongation factor 1-beta, putative | 1,834 | 2,279E-164 |
| TcCLB.508801.10 | protein associated with differentiation 4, putative | 1,834 | 1,753E-69 |
| TcCLB.458759.14 | hypothetical protein, conserved | 1,834 | 1,711E-58 |
| TcCLB.510541.9 | dispersed gene family protein 1 (DGF-1), putative | 1,833 | 0,00004621 |
| TcCLB.509463.10 | transmembrane amino acid transporter, putative | 1,832 | 7,075E-75 |

| | | | |
|---|---|---|---|
| TcCLB.508319.20 | Present in the outer mitochondrial membrane proteome 1 | 1,832 | 2,325E-65 |
| TcCLB.509903.5 | retrotransposon hot spot (RHS) protein, putative (fragmer | 1,83 | 6,305E-72 |
| TcCLB.425969.9 | cyclin-dependent kinases regulatory subunit, putative (P12 | 1,83 | 2,772E-24 |
| TcCLB.507889.20 | hypothetical protein, conserved | 1,828 | 8,128E-33 |
| TcCLB.510879.160 | tyrosine phosphatase, putative | 1,827 | 5,598E-106 |
| TcCLB.511627.130 | hypothetical protein, conserved | 1,827 | 1,505E-56 |
| TcCLB.510857.4 | hypothetical protein, conserved | 1,824 | 2,48E-63 |
| TcCLB.510611.20 | UDP-glucuronic acid/UDP-N-acetylgalactosamine transpo | 1,822 | 2,434E-28 |
| TcCLB.508221.87 | retrotransposon hot spot protein (RHS, pseudogene), puta | 1,82 | 9,643E-12 |
| TcCLB.511007.10 | dispersed gene family protein 1 (DGF-1), putative | 1,819 | 8,423E-22 |
| TcCLB.509965.429 | hypothetical protein, conserved | 1,818 | 1,032E-07 |
| TcCLB.508425.10 | dispersed gene family protein 1 (DGF-1), putative | 1,817 | 3,152E-11 |
| TcCLB.509167.110 | S-adenosylmethionine decarboxylase proenzyme, putative | 1,816 | 7,502E-54 |
| TcCLB.507641.250 | hypothetical protein | 1,816 | 3,898E-35 |
| TcCLB.509791.100 | hypothetical protein, conserved | 1,815 | 7,271E-44 |
| TcCLB.509963.40 | dispersed gene family protein 1 (DGF-1), putative | 1,808 | 9,167E-34 |
| TcCLB.503925.3 | hypothetical protein, conserved | 1,807 | 2,048E-26 |
| TcCLB.399997.10 | retrotransposon hot spot (RHS) protein, putative | 1,805 | 4,213E-44 |
| TcCLB.503477.10 | hypothetical protein, conserved | 1,804 | 1,181E-41 |
| TcCLB.506831.63 | hypothetical protein, conserved | 1,804 | 1,771E-41 |
| TcCLB.511245.150 | fatty acid elongase, putative | 1,803 | 3,394E-42 |
| TcCLB.511513.14 | hypothetical protein, conserved | 1,803 | 5,788E-15 |
| TcCLB.508931.10 | hypothetical protein | 1,801 | 2,291E-47 |
| TcCLB.508265.10 | glutathione-S-transferase/glutaredoxin, putative | 1,801 | 1,452E-36 |
| TcCLB.511053.10 | hypothetical protein, conserved | 1,801 | 5,47E-34 |
| TcCLB.509683.124 | hypothetical protein, conserved | 1,8 | 4,269E-85 |
| TcCLB.509463.41 | retrotransposon hot spot (RHS) protein, putative (fragmer | 1,797 | 3,067E-59 |
| TcCLB.510153.19 | tricarboxylate carrier, putative | 1,797 | 2,093E-29 |
| TcCLB.509721.40 | MIX protein, putative | 1,796 | 1,972E-43 |
| TcCLB.510847.20 | beta galactofuranosyl glycosyltransferase, putative | 1,796 | 5,688E-30 |
| TcCLB.506047.20 | retrotransposon hot spot (RHS) protein, putative | 1,795 | 5,189E-45 |
| TcCLB.511573.49 | hypothetical protein | 1,793 | 1,571E-60 |
| TcCLB.509457.50 | retrotransposon hot spot (RHS) protein, putative | 1,793 | 1,017E-30 |
| TcCLB.509877.40 | hypothetical protein, conserved | 1,792 | 7,348E-96 |
| TcCLB.508931.40 | hypothetical protein, conserved | 1,791 | 4,879E-29 |
| TcCLB.504207.5 | hypothetical protein, conserved | 1,791 | 2,229E-07 |
| TcCLB.506109.29 | dispersed gene family protein 1 (DGF-1), putative | 1,79 | 1,223E-23 |
| TcCLB.506751.116 | elongation factor 1-gamma (EF-1-gamma, pseudogene), | 1,789 | 0,00001895 |
| TcCLB.507675.10 | hypothetical protein, conserved | 1,788 | 7,262E-110 |
| TcCLB.511531.30 | hypothetical protein, conserved | 1,787 | 7,044E-17 |
| TcCLB.506183.80 | hypothetical protein | 1,786 | 8,298E-06 |
| TcCLB.506375.90 | hypothetical protein, conserved | 1,784 | 8,12E-43 |
| TcCLB.508557.29 | hypothetical protein, conserved | 1,783 | 1,498E-74 |
| TcCLB.506261.10 | fatty acyl CoA synthetase 2, putative | 1,783 | 3,471E-59 |
| TcCLB.507687.30 | retrotransposon hot spot protein (RHS, pseudogene), puta | 1,783 | 1,796E-26 |

| | | | |
|---|---|---|---|
| TcCLB.510571.30 | actin 1, putative | 1,781 | 5,808E-119 |
| TcCLB.506633.90 | folate/pteridine transporter, putative | 1,781 | 2,718E-89 |
| TcCLB.507053.40 | hypothetical protein, conserved | 1,78 | 8,279E-104 |
| TcCLB.509237.10 | mevalonate kinase, putative | 1,779 | 2,757E-69 |
| TcCLB.418185.21 | retrotransposon hot spot protein (RHS, pseudogene), puta | 1,777 | 6,138E-16 |
| TcCLB.506905.50 | cystathionine beta-synthase (CBS) | 1,776 | 4,813E-113 |
| TcCLB.506017.51 | retrotransposon hot spot (RHS) protein, putative (fragmer | 1,775 | 5,224E-94 |
| TcCLB.506811.120 | hypothetical protein, conserved | 1,772 | 8,524E-80 |
| TcCLB.507395.40 | hypothetical protein | 1,771 | 0,004052 |
| TcCLB.506933.60 | mitochondrial RNA binding protein, putative | 1,768 | 4,909E-102 |
| TcCLB.446325.9 | retrotransposon hot spot (RHS) protein, putative (fragmer | 1,766 | 8,016E-33 |
| TcCLB.509397.9 | dispersed gene family protein 1 (DGF-1), putative (fragme | 1,766 | 2,512E-17 |
| TcCLB.503887.40 | hypothetical protein, conserved | 1,765 | 2,959E-74 |
| TcCLB.506605.59 | adenosine 5'-monophosphoramidase, putative | 1,762 | 3,75E-32 |
| TcCLB.506521.9 | dispersed gene family protein 1 (DGF-1), putative | 1,762 | 3,567E-31 |
| TcCLB.503605.20 | retrotransposon hot spot (RHS) protein, putative | 1,762 | 1,027E-19 |
| TcCLB.508241.20 | hypothetical protein, conserved | 1,761 | 1,008E-79 |
| TcCLB.504427.64 | hypothetical protein, conserved | 1,761 | 1,071E-35 |
| TcCLB.506195.120 | lysosomal alpha-mannosidase precursor, putative | 1,761 | 1,217E-30 |
| TcCLB.511837.90 | protein kinase, putative | 1,76 | 4,067E-54 |
| TcCLB.508355.127 | hypothetical protein, conserved | 1,759 | 2,14E-61 |
| TcCLB.507655.9 | dispersed gene family protein 1 (DGF-1), putative | 1,759 | 2,735E-10 |
| TcCLB.508869.20 | mucin-associated surface protein (MASP), putative | 1,759 | 1,734E-06 |
| TcCLB.509539.30 | fatty acid elongase, putative | 1,758 | 2,346E-17 |
| TcCLB.507669.160 | hypothetical protein, conserved | 1,757 | 4,305E-108 |
| TcCLB.506959.70 | serine palmitoyltransferase, putative | 1,756 | 3,379E-87 |
| TcCLB.509859.10 | hypothetical protein, conserved | 1,754 | 3,883E-111 |
| TcCLB.508177.110 | cystathionine beta-synthase | 1,754 | 7,092E-91 |
| TcCLB.505201.9 | retrotransposon hot spot (RHS) protein, putative | 1,754 | 3,741E-29 |
| TcCLB.504125.100 | sugar transporter (pseudogene), putative | 1,754 | 1,805E-20 |
| TcCLB.510945.70 | aspartate aminotransferase, mitochondrial, putative | 1,753 | 5,996E-58 |
| TcCLB.508641.240 | hypothetical protein, conserved | 1,753 | 3,414E-50 |
| TcCLB.508285.50 | retrotransposon hot spot protein (RHS, pseudogene), puta | 1,753 | 1,492E-49 |
| TcCLB.511277.490 | hypothetical protein, conserved | 1,752 | 1,111E-78 |
| TcCLB.506175.100 | hypothetical protein, conserved | 1,749 | 5,481E-55 |
| TcCLB.504153.120 | hypothetical protein, conserved | 1,748 | 1,628E-90 |
| TcCLB.509959.90 | retrotransposon hot spot protein (RHS, pseudogene), puta | 1,748 | 2,539E-37 |
| TcCLB.509715.30 | cullin-like protein, putative | 1,747 | 3,987E-48 |
| TcCLB.507849.70 | hypothetical protein | 1,746 | 1,997E-77 |
| TcCLB.507099.30 | Ran-binding protein 1, putative | 1,746 | 2,052E-65 |
| TcCLB.511531.20 | hypothetical protein, conserved | 1,746 | 2,191E-32 |
| TcCLB.511211.70 | cytochrome c oxidase assembly protein (COX15) | 1,745 | 1,218E-74 |
| TcCLB.508387.120 | glutaminyl cyclase, putative | 1,744 | 6,937E-80 |
| TcCLB.511693.40 | hypothetical protein, conserved | 1,744 | 6,025E-79 |
| TcCLB.509319.90 | trypanothione synthetase (TRYS) | 1,743 | 2,056E-45 |

| | | | |
|---|---|---|---|
| TcCLB.509431.10 | hypothetical protein, conserved (pseudogene) | 1,743 | 1,304E-39 |
| TcCLB.509937.20 | hypothetical protein, conserved | 1,74 | 1,028E-100 |
| TcCLB.486465.9 | cysteine peptidase, Clan CA, family C51, putative | 1,739 | 7,04E-11 |
| TcCLB.507923.10 | L-threonine 3-dehydrogenase, putative | 1,737 | 3,366E-105 |
| TcCLB.504427.60 | hypothetical protein, conserved | 1,737 | 3,742E-28 |
| TcCLB.509109.130 | protein tyrosine phosphatase-like protein, putative | 1,736 | 3,442E-36 |
| TcCLB.432677.20 | heat shock 70 kDa protein, mitochondrial precursor, putati | 1,733 | 6,856E-108 |
| TcCLB.506425.50 | hypothetical protein, conserved | 1,733 | 1,577E-54 |
| TcCLB.509437.70 | hypothetical protein, conserved | 1,731 | 5,676E-95 |
| TcCLB.504427.70 | 3-oxo-5-alpha-steroid 4-dehydrogenase, putative | 1,728 | 1,466E-40 |
| TcCLB.511277.175 | ribosomal protein S27, putative | 1,727 | 3,254E-72 |
| TcCLB.504427.110 | acetyl-CoA synthetase, putative | 1,727 | 9,984E-51 |
| TcCLB.510361.160 | retrotransposon hot spot protein (RHS, pseudogene), puta | 1,727 | 1,208E-18 |
| TcCLB.476421.9 | protein kinase, putative | 1,726 | 1,424E-38 |
| TcCLB.506753.150 | hypothetical protein, conserved | 1,724 | 9,114E-31 |
| TcCLB.511907.260 | cell division cycle protein 20, putative (CDC20) | 1,723 | 1,323E-79 |
| TcCLB.511753.120 | L-threonine 3-dehydrogenase, putative | 1,721 | 3,053E-128 |
| TcCLB.504313.30 | dispersed gene family protein 1 (DGF-1), putative | 1,721 | 1,009E-32 |
| TcCLB.509223.10 | dispersed gene family protein 1 (DGF-1, pseudogene), put | 1,721 | 4,832E-31 |
| TcCLB.511773.90 | retrotransposon hot spot protein (RHS, pseudogene), puta | 1,72 | 1,54E-53 |
| TcCLB.511655.30 | hypothetical protein | 1,719 | 2,424E-86 |
| TcCLB.507017.40 | 2-hydroxy-3-oxopropionate reductase, putative | 1,718 | 2,534E-76 |
| TcCLB.510445.50 | S-adenosylmethionine synthetase, putative (METK1) | 1,717 | 6,831E-80 |
| TcCLB.507031.140 | hypothetical protein, conserved (pseudogene) | 1,716 | 1,38E-36 |
| TcCLB.509231.10 | cytochrome P450, putative | 1,715 | 3,568E-67 |
| TcCLB.506721.50 | hypothetical protein | 1,714 | 4,456E-14 |
| TcCLB.508177.70 | hypothetical protein | 1,712 | 3,783E-74 |
| TcCLB.510595.4 | hypothetical protein, conserved | 1,712 | 3,58E-16 |
| TcCLB.509329.9 | retrotransposon hot spot (RHS) protein, putative | 1,711 | 2,983E-23 |
| TcCLB.506465.14 | hypothetical protein, conserved | 1,706 | 2,48E-48 |
| TcCLB.508451.30 | retrotransposon hot spot protein (RHS, pseudogene), puta | 1,706 | 4,434E-17 |
| TcCLB.510801.10 | calpain-like cysteine peptidase, putative (fragment) | 1,704 | 3,049E-21 |
| TcCLB.511555.80 | ribonucleoside-diphosphate reductase small chain, putativ | 1,703 | 4,741E-33 |
| TcCLB.507603.260 | cysteine peptidase, putative | 1,703 | 0,00003611 |
| TcCLB.510879.180 | hypothetical protein | 1,702 | 3,019E-33 |
| TcCLB.503811.60 | Elongation factor Ts, mitochondrial, putative (EF-Ts) | 1,701 | 1,229E-64 |
| TcCLB.507059.80 | glutamine-dependent carbamoyl-phosphate synthetase, p | 1,701 | 2,631E-56 |
| TcCLB.509505.30 | 10 kDa heat shock protein, putative | 1,701 | 1,133E-16 |
| TcCLB.507569.10 | hypothetical protein, conserved | 1,699 | 2,815E-42 |
| TcCLB.510633.9 | dispersed gene family protein 1 (DGF-1), putative (fragme | 1,699 | 6,408E-12 |
| TcCLB.505047.9 | dispersed gene family protein 1 (DGF-1), putative | 1,699 | 1,16E-09 |
| TcCLB.511647.4 | glutamine-dependent carbamoyl-phosphate synthetase, p | 1,698 | 1,297E-51 |
| TcCLB.510543.60 | retrotransposon hot spot protein (RHS, pseudogene), puta | 1,697 | 1,061E-41 |
| TcCLB.511475.24 | dispersed gene family protein 1 (DGF-1), putative (fragme | 1,697 | 2,52E-16 |
| TcCLB.511391.160 | cytochrome c1, heme protein, mitochondrial, putative (CY | 1,696 | 3,047E-105 |

| | | | |
|---|---|---|---|
| TcCLB.510861.125 | hypothetical protein, conserved | 1,693 | 1,04E-47 |
| TcCLB.507009.10 | Gim5A protein, putative | 1,693 | 2,799E-41 |
| TcCLB.506925.14 | hypothetical protein, conserved | 1,692 | 5,702E-84 |
| TcCLB.504131.180 | ATPase subunit 9, putative | 1,691 | 1,09E-72 |
| TcCLB.507473.40 | retrotransposon hot spot protein (RHS, pseudogene), puta | 1,691 | 1,352E-50 |
| TcCLB.506835.159 | casein kinase 1 isoform 2, putative (fragment) | 1,688 | 1,519E-16 |
| TcCLB.508461.84 | hypothetical protein | 1,686 | 1,28E-42 |
| TcCLB.507777.20 | hypothetical protein, conserved | 1,685 | 3,11E-26 |
| TcCLB.506811.20 | hypothetical protein, conserved | 1,684 | 8,529E-39 |
| TcCLB.506675.110 | retrotransposon hot spot (RHS) protein, putative | 1,684 | 2,896E-32 |
| TcCLB.509457.60 | retrotransposon hot spot (RHS) protein, putative | 1,682 | 3,932E-19 |
| TcCLB.508177.120 | cystathionine beta-synthase (CBS) | 1,681 | 1,134E-88 |
| TcCLB.510079.49 | dispersed gene family protein 1 (DGF-1), putative (fragme | 1,679 | 3,415E-16 |
| TcCLB.511277.210 | peroxisomal enoyl-coa hydratase, putative | 1,677 | 1,739E-88 |
| TcCLB.510301.80 | N-acetyltransferase subunit Nat1, putative | 1,677 | 1,732E-53 |
| TcCLB.509331.80 | hypothetical protein, conserved | 1,677 | 1,046E-29 |
| TcCLB.507091.50 | aspartate carbamoyltransferase, putative | 1,676 | 3,572E-46 |
| TcCLB.509725.80 | retrotransposon hot spot protein (RHS, pseudogene), puta | 1,675 | 8,408E-100 |
| TcCLB.511817.20 | hypothetical protein, conserved | 1,674 | 7,608E-54 |
| TcCLB.507517.80 | hypothetical protein, conserved | 1,674 | 1,069E-33 |
| TcCLB.506469.10 | hypothetical protein, conserved | 1,674 | 1,552E-32 |
| TcCLB.505807.110 | alkyl-dihydroxyacetone phosphate synthase, putative | 1,672 | 4,197E-116 |
| TcCLB.510989.30 | guanin nucleotide-binding protein, putative | 1,672 | 1,616E-37 |
| TcCLB.510445.10 | casein kinase 1 isoform 2, putative | 1,672 | 3,014E-23 |
| TcCLB.511217.145 | ribosomal protein L39, putative | 1,671 | 1,526E-69 |
| TcCLB.507225.19 | hypothetical protein | 1,671 | 1,214E-50 |
| TcCLB.503991.20 | Xaa-Pro aminopeptidase, putative | 1,67 | 1,136E-55 |
| TcCLB.509885.80 | hypothetical protein, conserved | 1,67 | 7,371E-15 |
| TcCLB.507167.169 | dispersed gene family protein 1 (DGF-1), putative | 1,669 | 1,312E-17 |
| TcCLB.507555.80 | retrotransposon hot spot (RHS) protein, putative | 1,666 | 9,462E-75 |
| TcCLB.504797.70 | DNA excision/repair protein SNF2, putative | 1,665 | 5,713E-87 |
| TcCLB.509917.19 | retrotransposon hot spot (RHS) protein, putative (fragmer | 1,662 | 3,472E-70 |
| TcCLB.509167.150 | homoserine kinase, putative | 1,662 | 5,853E-50 |
| TcCLB.507485.20 | hypothetical protein, conserved | 1,661 | 4,951E-34 |
| TcCLB.507611.50 | hypothetical protein, conserved | 1,66 | 0,00007973 |
| TcCLB.506949.50 | cytochrome c, putative | 1,659 | 8,625E-55 |
| TcCLB.457251.10 | 3-oxo-5-alpha-steroid 4-dehydrogenase, putative (fragme | 1,658 | 7,796E-46 |
| TcCLB.509831.36 | guide RNA-binding protein of 21 kDa (gBP21) | 1,657 | 5,765E-66 |
| TcCLB.510283.70 | glycosyl hydrolase-like protein, putative | 1,657 | 4,824E-53 |
| TcCLB.511745.10 | heat shock 70 kDa protein, mitochondrial precursor, putati | 1,655 | 3,01E-68 |
| TcCLB.507691.20 | dispersed gene family protein 1 (DGF-1), putative | 1,655 | 7,193E-36 |
| TcCLB.511077.20 | beta galactofuranosyl glycosyltransferase (pseudogene), p | 1,655 | 1,468E-06 |
| TcCLB.507631.20 | hypothetical protein, conserved | 1,654 | 4,716E-39 |
| TcCLB.503923.40 | hypothetical protein, conserved | 1,654 | 7,945E-13 |
| TcCLB.507105.100 | retrotransposon hot spot protein (RHS, pseudogene), puta | 1,652 | 3,394E-77 |

| | | | |
|---|---|---|---|
| TcCLB.508831.110 | serine/threonine kinase, putative | 1,652 | 7,102E-35 |
| TcCLB.511589.200 | S-adenosylhomocysteine hydrolase, putative | 1,651 | 2,144E-44 |
| TcCLB.511025.90 | hypothetical protein, conserved | 1,65 | 1,075E-32 |
| TcCLB.505919.70 | glycine dehydrogenase [decarboxylating] (pseudogene), pı | 1,65 | 5,234E-20 |
| TcCLB.504425.100 | hypothetical protein, conserved | 1,649 | 2,032E-28 |
| TcCLB.483389.10 | retrotransposon hot spot protein (RHS, pseudogene), puta | 1,648 | 1,568E-54 |
| TcCLB.503965.20 | hypothetical protein, conserved | 1,648 | 3,691E-53 |
| TcCLB.510181.20 | delta-4 fatty acid desaturase, putative | 1,648 | 8,343E-25 |
| TcCLB.503893.30 | hypothetical protein, conserved | 1,644 | 2,907E-39 |
| TcCLB.507875.110 | retrotransposon hot spot protein (RHS, pseudogene), puta | 1,644 | 5,069E-32 |
| TcCLB.509717.90 | short chain 3-hydroxyacyl-CoA dehydrogenase, putative | 1,642 | 1,263E-79 |
| TcCLB.459199.10 | retrotransposon hot spot (RHS) protein, putative | 1,641 | 7,656E-94 |
| TcCLB.511179.100 | monothiol glutaredoxin, putative | 1,641 | 2,487E-39 |
| TcCLB.506635.80 | Hydroxymethylglutaryl-CoA lyase, mitochondrial, putative | 1,639 | 1,599E-46 |
| TcCLB.511635.40 | protein tyrosine phosphatase, putative | 1,638 | 3,973E-81 |
| TcCLB.509185.10 | retrotransposon hot spot protein (RHS, pseudogene), puta | 1,637 | 0,0001164 |
| TcCLB.506945.160 | S-adenosylmethionine synthetase, putative (METK1) | 1,636 | 1,402E-87 |
| TcCLB.507687.20 | dispersed gene family protein 1 (DGF-1), putative | 1,636 | 3,037E-13 |
| TcCLB.509601.90 | alanine racemase, putative | 1,635 | 3,926E-53 |
| TcCLB.509733.60 | hypothetical protein | 1,633 | 2,738E-36 |
| TcCLB.506577.120 | sterol C-24 reductase, putative | 1,632 | 7,875E-33 |
| TcCLB.507167.110 | retrotransposon hot spot protein (RHS, pseudogene), puta | 1,63 | 6,781E-42 |
| TcCLB.509615.2 | endosomal integral membrane protein (pseudogene), put | 1,63 | 2,199E-29 |
| TcCLB.507653.50 | retrotransposon hot spot protein (RHS, pseudogene), puta | 1,63 | 1,458E-22 |
| TcCLB.508461.360 | hypothetical protein, conserved | 1,629 | 2,038E-63 |
| TcCLB.508669.10 | hypothetical protein, conserved | 1,627 | 3,123E-30 |
| TcCLB.506661.10 | fatty acid elongase, putative | 1,626 | 4,331E-27 |
| TcCLB.507041.10 | hypothetical protein, conserved | 1,625 | 4,168E-56 |
| TcCLB.507757.30 | hypothetical protein, conserved | 1,625 | 1,661E-33 |
| TcCLB.509163.80 | glycine dehydrogenase [decarboxylating], putative | 1,625 | 2,356E-14 |
| TcCLB.507007.108 | amino acid transporter, putative | 1,624 | 2,798E-39 |
| TcCLB.504177.10 | fatty acyl CoA synthetase 2, putative | 1,624 | 2,92E-30 |
| TcCLB.509229.80 | mitochondrial processing peptidase alpha subunit, putativ | 1,623 | 2,424E-78 |
| TcCLB.507485.40 | hypothetical protein, conserved | 1,622 | 1,012E-58 |
| TcCLB.505975.30 | beta galactofuranosyl glycosyltransferase (pseudogene), ı | 1,622 | 9,104E-17 |
| TcCLB.506835.70 | PAS-domain containing phosphoglycerate kinase, putative | 1,62 | 4,38E-59 |
| TcCLB.507609.40 | delta-4 fatty acid desaturase, putative | 1,62 | 4,768E-22 |
| TcCLB.506925.380 | hypothetical protein | 1,619 | 5,738E-36 |
| TcCLB.511529.80 | kinetoplast-associated protein 3 (KAP3) | 1,618 | 2,869E-45 |
| TcCLB.508827.10 | membrane-bound acid phosphatase, putative | 1,618 | 5,234E-34 |
| TcCLB.510515.30 | hypothetical protein, conserved | 1,617 | 8,339E-63 |
| TcCLB.507715.10 | cysteine peptidase, Clan CA, family C51, putative | 1,617 | 1,506E-54 |
| TcCLB.511613.189 | mucin TcMUCII, putative | 1,617 | 0,00008789 |
| TcCLB.510065.10 | hypothetical protein, conserved | 1,616 | 3,286E-70 |
| TcCLB.511233.80 | calmodulin, putative | 1,616 | 3,591E-43 |

| | | | |
|---|---|---|---|
| TcCLB.508937.10 | dispersed gene family protein 1 (DGF-1, pseudogene), put | 1,616 | 2,861E-11 |
| TcCLB.510039.64 | hypothetical protein, conserved | 1,615 | 1,816E-39 |
| TcCLB.507293.20 | dispersed gene family protein 1 (DGF-1, pseudogene), put | 1,614 | 9,247E-25 |
| TcCLB.506325.36 | N-acetyltransferase complex ARD1 subunit (pseudogene), | 1,613 | 7,312E-07 |
| TcCLB.510355.150 | retrotransposon hot spot protein (RHS, pseudogene), puta | 1,612 | 1,02E-40 |
| TcCLB.506183.109 | dispersed gene family protein 1 (DGF-1, pseudogene), put | 1,612 | 0,01559 |
| TcCLB.504165.10 | hypothetical protein, conserved | 1,611 | 1,481E-37 |
| TcCLB.510593.49 | kinetoplast DNA-associated protein, putative | 1,608 | 7,841E-58 |
| TcCLB.510359.350 | hypothetical protein, conserved | 1,607 | 6,969E-74 |
| TcCLB.509205.60 | hypothetical protein, conserved | 1,607 | 3,264E-61 |
| TcCLB.503849.60 | NADH-dependent fumarate reductase, putative | 1,606 | 6,089E-49 |
| TcCLB.507465.10 | receptor-type adenylate cyclase, putative (fragment) | 1,606 | 2,378E-12 |
| TcCLB.508277.220 | hypothetical protein | 1,605 | 6,61E-107 |
| TcCLB.503597.10 | homoserine kinase, putative | 1,605 | 1,211E-48 |
| TcCLB.509259.120 | retrotransposon hot spot (RHS) protein, putative | 1,605 | 2,754E-29 |
| TcCLB.507159.80 | hypothetical protein, conserved | 1,603 | 3,614E-61 |
| TcCLB.507499.30 | retrotransposon hot spot protein (RHS, pseudogene), puta | 1,601 | 2,032E-30 |
| TcCLB.507855.9 | dispersed gene family protein 1 (DGF-1), putative (fragme | 1,601 | 7,31E-11 |
| TcCLB.510355.10 | dispersed gene family protein 1 (DGF-1), putative | 1,6 | 4,596E-27 |
| TcCLB.509289.39 | hypothetical protein, conserved (pseudogene) | 1,598 | 3,814E-26 |
| TcCLB.504137.10 | isoleucyl-tRNA synthetase, putative | 1,597 | 4,165E-41 |
| TcCLB.505945.69 | hypothetical protein, conserved | 1,595 | 5,303E-15 |
| TcCLB.504041.10 | hypothetical protein, conserved | 1,594 | 1,783E-39 |
| TcCLB.503607.4 | retrotransposon hot spot (RHS) protein, putative | 1,594 | 6,144E-38 |
| TcCLB.506169.10 | glycine dehydrogenase [decarboxylating] (pseudogene), p | 1,594 | 1,214E-13 |
| TcCLB.508461.580 | hypothetical protein | 1,593 | 7,487E-70 |
| TcCLB.509169.30 | hypothetical protein, conserved | 1,592 | 2,966E-32 |
| TcCLB.507443.30 | UDP-Gal or UDP-GlcNAc-dependent glycosyltransferase, p | 1,592 | 3,107E-13 |
| TcCLB.503815.10 | alkyl-dihydroxyacetone phosphate synthase, putative | 1,591 | 5,515E-101 |
| TcCLB.508177.129 | cystathionine beta-synthase, putative | 1,591 | 5,302E-91 |
| TcCLB.511025.80 | hypothetical protein, conserved | 1,591 | 3,055E-18 |
| TcCLB.506625.60 | inositol 5-phosphatase 1, putative | 1,59 | 9,28E-80 |
| TcCLB.506771.9 | hypothetical protein, conserved | 1,59 | 1,428E-43 |
| TcCLB.503661.11 | trans-sialidase, putative (fragment) | 1,59 | 2,591E-23 |
| TcCLB.506795.44 | hypothetical protein, conserved | 1,589 | 1,286E-55 |
| TcCLB.504109.24 | dynein light chain, putative | 1,589 | 4,252E-24 |
| TcCLB.509177.59 | hypothetical protein, conserved | 1,588 | 2,24E-31 |
| TcCLB.508273.30 | hypothetical protein, conserved | 1,585 | 9,303E-44 |
| TcCLB.511529.200 | hypothetical protein, conserved | 1,585 | 3,543E-42 |
| TcCLB.509471.30 | hypothetical protein, conserved | 1,584 | 1,678E-31 |
| TcCLB.507393.9 | dispersed gene family protein 1 (DGF-1), putative | 1,583 | 8,997E-27 |
| TcCLB.511383.50 | prohibitin, putative | 1,582 | 3,893E-60 |
| TcCLB.506779.90 | hypothetical protein, conserved | 1,581 | 8,813E-82 |
| TcCLB.506537.180 | retrotransposon hot spot protein (RHS, pseudogene), puta | 1,581 | 1,944E-57 |
| TcCLB.511051.59 | hypothetical protein, conserved | 1,58 | 5,222E-83 |

| | | | |
|---|---|---|---|
| TcCLB.508621.10 | protein kinase, putative | 1,579 | 8,911E-57 |
| TcCLB.510105.240 | short chain 3-hydroxyacyl-CoA dehydrogenase, putative | 1,579 | 1,447E-52 |
| TcCLB.510859.10 | nucleolar RNA-binding protein, putative | 1,577 | 2,005E-84 |
| TcCLB.436521.9 | mevalonate kinase, putative (fragment) | 1,577 | 1,038E-36 |
| TcCLB.511671.80 | serine/threonine kinase, putative | 1,577 | 7,022E-26 |
| TcCLB.511277.390 | hypothetical protein, conserved | 1,576 | 5,651E-66 |
| TcCLB.509713.10 | protein associated with differentiation 4, putative | 1,575 | 1,485E-141 |
| TcCLB.505945.90 | folate/pteridine transporter, putative | 1,575 | 2,017E-46 |
| TcCLB.508153.700 | hypothetical protein, conserved | 1,575 | 4,067E-33 |
| TcCLB.463155.20 | retrotransposon hot spot (RHS) protein, putative | 1,574 | 2,206E-85 |
| TcCLB.508677.150 | glycosyl transferase, putative | 1,574 | 9,589E-84 |
| TcCLB.507809.39 | hypothetical protein, conserved | 1,574 | 2,642E-29 |
| TcCLB.507477.10 | dispersed gene family protein 1 (DGF-1), putative | 1,574 | 1,428E-24 |
| TcCLB.506559.300 | twinfilin, putative | 1,573 | 2,949E-81 |
| TcCLB.506163.50 | hypothetical protein, conserved | 1,572 | 3,92E-44 |
| TcCLB.510083.10 | retrotransposon hot spot protein (RHS, pseudogene), puta | 1,572 | 2,519E-37 |
| TcCLB.507093.44 | dynein light chain, putative | 1,571 | 1,446E-25 |
| TcCLB.506823.20 | retrotransposon hot spot protein (RHS, pseudogene), puta | 1,571 | 4,476E-22 |
| TcCLB.505807.220 | ribosomal RNA adenine dimethylase family protein, putat | 1,57 | 5,376E-48 |
| TcCLB.508221.381 | mucin-associated surface protein (MASP, pseudogene), pu | 1,569 | 0,009171 |
| TcCLB.509295.50 | beta galactofuranosyl glycosyltransferase, putative | 1,568 | 1,218E-14 |
| TcCLB.503929.10 | V-type ATPase, A subunit, putative | 1,567 | 6,473E-75 |
| TcCLB.511139.40 | ATP-dependent RNA helicase SUB2, putative (fragment) ( | 1,567 | 1,497E-22 |
| TcCLB.511727.190 | hypothetical protein, conserved | 1,566 | 2,334E-62 |
| TcCLB.511573.30 | hypothetical protein, conserved | 1,566 | 2,457E-30 |
| TcCLB.504025.10 | hypothetical protein, conserved | 1,565 | 1,329E-19 |
| TcCLB.509237.130 | Calpain-like protein 1 (CAP5.5) | 1,564 | 8,836E-91 |
| TcCLB.511047.40 | NADH-cytochrome b5 reductase, putative | 1,564 | 4,341E-69 |
| TcCLB.506305.29 | hypothetical protein, conserved | 1,564 | 8,512E-48 |
| TcCLB.504153.240 | hypothetical protein, conserved | 1,563 | 1,087E-38 |
| TcCLB.507969.40 | hypothetical protein, conserved | 1,563 | 4,135E-33 |
| TcCLB.511487.100 | retrotransposon hot spot protein (RHS, pseudogene), puta | 1,563 | 7,617E-22 |
| TcCLB.507063.90 | mitochondrial DNA polymerase beta, putative | 1,562 | 3,103E-62 |
| TcCLB.506513.7 | hypothetical protein, conserved | 1,561 | 6,849E-40 |
| TcCLB.506459.40 | dispersed gene family protein 1 (DGF-1, pseudogene), put | 1,561 | 0,00006351 |
| TcCLB.509177.50 | hypothetical protein, conserved | 1,56 | 2,526E-41 |
| TcCLB.428999.30 | hypothetical protein | 1,56 | 3,195E-36 |
| TcCLB.510091.110 | mitochondrial RNA binding complex 1 subunit, putative | 1,56 | 6,277E-36 |
| TcCLB.507411.20 | retrotransposon hot spot (RHS) protein, putative | 1,56 | 2,063E-11 |
| TcCLB.508661.10 | Eukaryotic translation initiation factor 4E type 5, putative | 1,557 | 1,1E-41 |
| TcCLB.508045.131 | retrotransposon hot spot protein (RHS, pseudogene), puta | 1,556 | 4,497E-10 |
| TcCLB.503831.40 | mitochondrial DNA primase, putative | 1,555 | 1,986E-45 |
| TcCLB.510669.5 | ribulose-phosphate 3-epimerase, putative (fragment) | 1,555 | 2,164E-07 |
| TcCLB.506175.25 | hypothetical protein | 1,553 | 3,648E-43 |
| TcCLB.509809.10 | NADH-ubiquinone oxidoreductase complex I subunit, puta | 1,553 | 1,465E-39 |

| | | | |
|---|---|---|---|
| TcCLB.508601.50 | NADH dehydrogenase subunit NB6M, putative | 1,553 | 3,786E-27 |
| TcCLB.506865.50 | Present in the outer mitochondrial membrane proteome 4 | 1,553 | 4,647E-25 |
| TcCLB.507569.4 | electon transport protein SCO1/SCO2, putative | 1,552 | 1,94E-30 |
| TcCLB.504109.200 | retrotransposon hot spot (RHS) protein, putative | 1,551 | 9,462E-54 |
| TcCLB.510359.270 | mitochondrial RNA-binding protein 2, putative | 1,55 | 6,469E-41 |
| TcCLB.509717.20 | dihydrolipoamide acetyltransferase precursor, putative | 1,549 | 4,45E-77 |
| TcCLB.510729.210 | hypothetical protein, conserved | 1,549 | 7,896E-58 |
| TcCLB.507739.100 | hypothetical protein | 1,549 | 1,976E-27 |
| TcCLB.504557.10 | UDP-Gal or UDP-GlcNAc-dependent glycosyltransferase, p | 1,549 | 3,383E-25 |
| TcCLB.506129.60 | beta galactofuranosyl glycosyltransferase, putative | 1,549 | 2,015E-10 |
| TcCLB.506779.140 | hypothetical protein, conserved | 1,548 | 1,565E-62 |
| TcCLB.509601.70 | vacuolar proton translocating ATPase subunit A, putative ( | 1,548 | 1,592E-58 |
| TcCLB.508257.30 | hypothetical protein, conserved | 1,547 | 4,028E-32 |
| TcCLB.506795.80 | proline racemase | 1,547 | 2,246E-31 |
| TcCLB.511621.9 | hypothetical protein, conserved | 1,547 | 1,061E-29 |
| TcCLB.507029.30 | heat shock 70 kDa protein, mitochondrial precursor, putati | 1,546 | 2,098E-61 |
| TcCLB.507929.10 | kinesin, putative (fragment) | 1,546 | 9,062E-34 |
| TcCLB.508071.90 | retrotransposon hot spot (RHS) protein, putative | 1,545 | 1,514E-42 |
| TcCLB.510479.20 | dispersed gene family protein 1 (DGF-1, pseudogene), put | 1,545 | 5,763E-23 |
| TcCLB.511301.44 | hypothetical protein, conserved | 1,543 | 5,744E-32 |
| TcCLB.510347.50 | hypothetical protein, conserved | 1,542 | 6,44E-43 |
| TcCLB.510421.110 | hypothetical protein | 1,542 | 1,817E-36 |
| TcCLB.511395.40 | hypothetical protein | 1,54 | 6,894E-76 |
| TcCLB.506623.40 | retrotransposon hot spot protein (RHS, pseudogene), puta | 1,539 | 9,344E-39 |
| TcCLB.506925.360 | hypothetical protein | 1,539 | 3,567E-33 |
| TcCLB.511155.10 | hypothetical protein, conserved (pseudogene) | 1,538 | 5,59E-24 |
| TcCLB.508387.170 | double-strand-break repair protein rad21 homolog, putativ | 1,537 | 3,307E-50 |
| TcCLB.509153.20 | hypothetical protein, conserved | 1,537 | 8,786E-25 |
| TcCLB.510359.300 | hypothetical protein, conserved | 1,536 | 5,358E-61 |
| TcCLB.511395.70 | hypothetical protein, conserved | 1,535 | 9,725E-34 |
| TcCLB.506025.4 | membrane-bound acid phosphatase, putative (fragment) | 1,535 | 2,551E-29 |
| TcCLB.504111.40 | hypothetical protein, conserved | 1,533 | 3,367E-31 |
| TcCLB.506943.4 | aldehyde dehydrogenase, putative | 1,532 | 9,061E-73 |
| TcCLB.510643.140 | 60S acidic ribosomal protein P2, putative | 1,532 | 1,215E-63 |
| TcCLB.510687.60 | hypothetical protein, conserved | 1,532 | 2,305E-59 |
| TcCLB.503995.50 | serine peptidase, clan SC, family S9A-like protein (OPB) | 1,531 | 3,624E-49 |
| TcCLB.506295.160 | 2-oxoisovalerate dehydrogenase beta subunit, mitochondr | 1,531 | 1,039E-41 |
| TcCLB.509229.60 | hypothetical protein, conserved | 1,531 | 1,128E-19 |
| TcCLB.509427.30 | dispersed gene family protein 1 (DGF-1, pseudogene), put | 1,53 | 6,758E-08 |
| TcCLB.511817.260 | hypothetical protein, conserved | 1,529 | 7,497E-60 |
| TcCLB.509725.139 | damage-specific DNA binding protein, putative | 1,528 | 1,235E-29 |
| TcCLB.511815.40 | guide RNA associated protein, GAP2, putative | 1,526 | 1,298E-103 |
| TcCLB.506147.60 | 3-demethylubiquinone-9 3-methyltransferase, putative | 1,525 | 1,272E-30 |
| TcCLB.504071.30 | hypothetical protein, conserved | 1,524 | 2,157E-97 |
| TcCLB.511475.50 | dispersed gene family protein 1 (DGF-1), putative | 1,524 | 7,258E-16 |

| | | | |
|---|---|---|---|
| TcCLB.506649.50 | cytochrome c oxidase assembly factor, putative | 1,523 | 1,7E-45 |
| TcCLB.509717.176 | retrotransposon hot spot (RHS) protein, putative (fragmer | 1,523 | 8,019E-10 |
| TcCLB.506003.10 | retrotransposon hot spot protein (RHS, pseudogene), puta | 1,523 | 2,688E-08 |
| TcCLB.432677.10 | heat shock 70 kDa protein, mitochondrial precursor (pseuc | 1,521 | 2,364E-54 |
| TcCLB.510507.40 | transmembrane amino acid transporter, putative | 1,521 | 1,442E-46 |
| TcCLB.509029.100 | hypothetical protein, conserved | 1,521 | 3,916E-33 |
| TcCLB.507991.129 | cyclophilin, putative | 1,521 | 4,471E-25 |
| TcCLB.506401.110 | protein kinase, putative | 1,52 | 6,121E-82 |
| TcCLB.505163.70 | Selenoprotein T, putative (SelT) | 1,52 | 5,349E-56 |
| TcCLB.511589.180 | hypothetical protein, conserved | 1,52 | 4,011E-39 |
| TcCLB.503903.79 | hypothetical protein | 1,52 | 2,245E-24 |
| TcCLB.504069.70 | vacuolar ATP synthase, putative | 1,519 | 1,104E-74 |
| TcCLB.506775.90 | PIF1 helicase-like protein, putative | 1,519 | 1,196E-70 |
| TcCLB.509153.90 | dihydrofolate reductase-thymidylate synthase | 1,519 | 2,538E-68 |
| TcCLB.509045.29 | kinesin, putative | 1,518 | 8,281E-22 |
| TcCLB.426799.9 | hypothetical protein, conserved (fragment) | 1,518 | 1,825E-11 |
| TcCLB.503841.10 | proteasome activator protein pa26, putative | 1,517 | 5,343E-32 |
| TcCLB.507625.60 | dispersed gene family protein 1 (DGF-1, pseudogene), put | 1,517 | 9,965E-25 |
| TcCLB.506147.190 | hypothetical protein, conserved | 1,516 | 1,177E-41 |
| TcCLB.510759.140 | hypothetical protein, conserved | 1,515 | 1,383E-41 |
| TcCLB.503683.11 | RNA-binding protein, putative | 1,515 | 2,996E-12 |
| TcCLB.509611.30 | hypothetical protein | 1,514 | 1,059E-37 |
| TcCLB.505999.4 | hypothetical protein, conserved | 1,513 | 2,099E-20 |
| TcCLB.505945.10 | DNA repair and recombination helicase protein PIF5, puta | 1,511 | 3,166E-56 |
| TcCLB.509337.5 | hypothetical protein, conserved (pseudogene) | 1,511 | 2,038E-36 |
| TcCLB.511909.60 | hypothetical protein, conserved | 1,51 | 3,207E-29 |
| TcCLB.509237.90 | hypothetical protein, conserved | 1,51 | 3,414E-21 |
| TcCLB.508799.240 | aminopeptidase, putative | 1,509 | 1,352E-59 |
| TcCLB.503829.80 | hypothetical potein, conserved | 1,508 | 9,812E-66 |
| TcCLB.508317.30 | hypothetical protein | 1,508 | 1,739E-49 |
| TcCLB.506777.50 | cell cycle sequence binding phosphoprotein (RBP45), puta | 1,508 | 6,814E-45 |
| TcCLB.508333.9 | dispersed gene family protein 1 (DGF-1), putative (fragme | 1,508 | 3,404E-06 |
| TcCLB.503939.114 | hypothetical protein, conserved | 1,506 | 3,945E-28 |
| TcCLB.426071.10 | dispersed gene family protein 1 (DGF-1), putative (fragme | 1,506 | 2,88E-08 |
| TcCLB.510133.9 | dispersed gene family protein 1 (DGF-1), putative (fragme | 1,506 | 1,325E-06 |
| TcCLB.511625.74 | hypothetical protein, conserved | 1,505 | 1,144E-44 |
| TcCLB.508641.200 | hypothetical protein, conserved | 1,505 | 1,023E-34 |
| TcCLB.510453.9 | dispersed gene family protein 1 (DGF-1), putative | 1,505 | 5,358E-06 |
| TcCLB.507611.10 | retrotransposon hot spot (RHS) protein, putative | 1,504 | 1,165E-60 |
| TcCLB.504161.20 | hypothetical protein, conserved | 1,504 | 8,344E-54 |
| TcCLB.503599.60 | hypothetical protein, conserved | 1,504 | 7,514E-24 |
| TcCLB.506675.70 | dispersed gene family protein 1 (DGF-1), putative | 1,504 | 1,027E-15 |
| TcCLB.510101.90 | hypothetical protein, conserved | 1,503 | 5,65E-79 |
| TcCLB.510723.29 | 1-acyl-sn-glycerol-3-phosphate acyltransferase, putative ( | 1,503 | 1,418E-36 |
| TcCLB.506405.43 | hypothetical protein, conserved | 1,503 | 2,176E-29 |

| | | | |
|---|---|---|---|
| TcCLB.508023.140 | dispersed gene family protein 1 (DGF-1), putative | 1,503 | 5,723E-21 |
| TcCLB.508369.20 | squalene synthase, putative | 1,501 | 4,493E-98 |
| TcCLB.509047.10 | hypothetical protein | 1,501 | 4,621E-38 |
| TcCLB.511827.64 | hypothetical protein, conserved | 1,5 | 3,027E-45 |
| TcCLB.444777.10 | drug resistance protein, putative | 1,5 | 8,332E-12 |
| TcCLB.511261.20 | chaperone DNAJ protein, putative | 1,499 | 1,419E-44 |
| TcCLB.508257.70 | phosphomannomutase-like protein, putative | 1,498 | 5,864E-40 |
| TcCLB.505757.20 | transcriptional regulator, putative | 1,498 | 6,421E-26 |
| TcCLB.508043.20 | metallo-beta-lactamase-like protein, putative | 1,495 | 5,471E-44 |
| TcCLB.508799.230 | hypothetical protein, conserved | 1,495 | 3,454E-40 |
| TcCLB.506405.120 | V-type ATPase, A subunit, putative | 1,494 | 1,998E-98 |
| TcCLB.507167.240 | retrotransposon hot spot (RHS) protein, putative | 1,493 | 3,301E-25 |
| TcCLB.510347.29 | hypothetical protein, conserved | 1,493 | 2,042E-14 |
| TcCLB.507659.20 | amino acid transporter, putative | 1,491 | 1,834E-103 |
| TcCLB.508989.9 | nucleoside phosphorylase, putative | 1,491 | 1,164E-63 |
| TcCLB.509649.20 | hypothetical protein | 1,49 | 8,171E-37 |
| TcCLB.510307.60 | retrotransposon hot spot protein (RHS, pseudogene), puta | 1,489 | 7,849E-68 |
| TcCLB.511733.10 | aldehyde dehydrogenase, putative | 1,488 | 2,591E-40 |
| TcCLB.506337.50 | hypothetical protein, conserved | 1,487 | 6,925E-54 |
| TcCLB.507221.60 | oxidoreductase-like protein (pseudogene), putative | 1,487 | 1,857E-29 |
| TcCLB.508195.10 | UDP-Gal or UDP-GlcNAc-dependent glycosyltransferase, p | 1,487 | 9,649E-25 |
| TcCLB.506693.20 | hypothetical protein, conserved | 1,485 | 2,679E-35 |
| TcCLB.504625.99 | protein tyrosine phosphatase, putative (fragment) | 1,485 | 1,376E-14 |
| TcCLB.505207.30 | retrotransposon hot spot (RHS) protein, putative | 1,484 | 7,43E-46 |
| TcCLB.506631.30 | hypothetical protein, conserved | 1,484 | 1,151E-43 |
| TcCLB.507927.100 | hypothetical protein | 1,483 | 5,633E-55 |
| TcCLB.508153.130 | enoyl-CoA hydratase, mitochondrial precursor, putative | 1,482 | 1,164E-48 |
| TcCLB.506687.10 | retrotransposon hot spot protein (RHS, pseudogene), puta | 1,482 | 1,179E-35 |
| TcCLB.507063.10 | hypothetical protein | 1,482 | 1,454E-30 |
| TcCLB.503971.10 | hypothetical protein, conserved | 1,481 | 2,795E-68 |
| TcCLB.506821.210 | cytochrome-b5 reductase, putative | 1,481 | 1,594E-51 |
| TcCLB.504557.20 | UDP-Gal or UDP-GlcNAc-dependent glycosyltransferase, p | 1,479 | 1,269E-24 |
| TcCLB.509797.40 | isoleucyl-tRNA synthetase, putative | 1,478 | 1,071E-65 |
| TcCLB.508975.10 | acyl-CoA dehydrogenase, mitochondrial precursor, putativ | 1,478 | 6,923E-54 |
| TcCLB.510357.140 | hypothetical protein, conserved | 1,477 | 6,483E-21 |
| TcCLB.507935.10 | mucin-like glycoprotein (pseudogene), putative | 1,477 | 0,0001097 |
| TcCLB.506287.209 | DNA ligase, putative | 1,476 | 1,043E-39 |
| TcCLB.510659.240 | lactoylglutathione lyase-like protein, putative | 1,476 | 2,907E-32 |
| TcCLB.509715.90 | UDP-glucose/GDP-mannose dehydrogenase family, NAD b | 1,476 | 8,63E-22 |
| TcCLB.508219.140 | retrotransposon hot spot protein (RHS, pseudogene), puta | 1,476 | 3,042E-09 |
| TcCLB.511417.70 | histone-lysine N-methyltransferase, putative (DOT1B) | 1,475 | 2,147E-45 |
| TcCLB.504077.30 | hypothetical protein, conserved | 1,472 | 2,966E-32 |
| TcCLB.510581.20 | receptor-type adenylate cyclase, putative | 1,47 | 5,27E-23 |
| TcCLB.510509.50 | mitochondrial RNA binding protein, putative | 1,469 | 1,011E-34 |
| TcCLB.511843.80 | retrotransposon hot spot (RHS) protein, putative | 1,469 | 2,968E-27 |

| | | | |
|---|---|---|---|
| TcCLB.504741.140 | esterase, putative | 1,467 | 1,32E-26 |
| TcCLB.508837.189 | endoplasmic reticulum oxidoreductin, putative | 1,466 | 1,695E-33 |
| TcCLB.507001.110 | galactokinase-like protein, putative | 1,466 | 3,143E-32 |
| TcCLB.504153.370 | hypothetical protein | 1,465 | 7,324E-44 |
| TcCLB.506777.64 | hypothetical protein, conserved | 1,465 | 6,287E-25 |
| TcCLB.504065.10 | retrotransposon hot spot protein (RHS, pseudogene), puta | 1,464 | 6,328E-36 |
| TcCLB.482471.19 | ATP-dependent RNA helicase SUB2, putative (fragment) ( | 1,464 | 5,329E-32 |
| TcCLB.506195.210 | hypothetical protein, conserved | 1,463 | 5,385E-52 |
| TcCLB.507681.210 | Flagellar Member 2 (FLAM2) | 1,461 | 1,008E-97 |
| TcCLB.506223.40 | hypothetical protein, conserved | 1,461 | 1,674E-30 |
| TcCLB.511005.10 | retrotransposon hot spot (RHS) protein, putative (fragmer | 1,461 | 1,648E-27 |
| TcCLB.503843.20 | hypothetical protein, conserved | 1,46 | 1,182E-55 |
| TcCLB.468405.9 | dispersed gene family protein 1 (DGF-1), putative | 1,46 | 0,000296 |
| TcCLB.506469.148 | hypothetical protein, conserved | 1,459 | 1,239E-40 |
| TcCLB.503499.20 | hypothetical protein, conserved | 1,459 | 1,099E-28 |
| TcCLB.510091.80 | pyruvate dehydrogenase E1 beta subunit, putative | 1,458 | 6,413E-62 |
| TcCLB.503721.39 | heat shock 70 kDa protein, putative | 1,458 | 5,562E-47 |
| TcCLB.509911.74 | hypothetical protein, conserved | 1,458 | 5,377E-38 |
| TcCLB.508609.10 | surface protease GP63, putative | 1,458 | 9,546E-34 |
| TcCLB.506873.4 | ATP-dependent DNA DEAD/DEAH box helicase, putative (f | 1,457 | 6,412E-18 |
| TcCLB.506945.20 | PAS-domain containing phosphoglycerate kinase, putative | 1,455 | 1,107E-49 |
| TcCLB.511179.150 | alanine racemase, putative | 1,455 | 2,135E-27 |
| TcCLB.506529.516 | ATP synthase subunit, putative | 1,455 | 1,692E-16 |
| TcCLB.507695.14 | dispersed gene family protein 1 (DGF-1), putative | 1,454 | 0,00009188 |
| TcCLB.508079.10 | conserved protein | 1,453 | 6,734E-65 |
| TcCLB.507645.59 | hypothetical protein, conserved | 1,453 | 1,713E-16 |
| TcCLB.511025.110 | dihydrolipoyl dehydrogenase, putative (GCVL-2) | 1,452 | 1,898E-46 |
| TcCLB.504207.20 | hypothetical protein, conserved | 1,45 | 3,027E-65 |
| TcCLB.503847.80 | hypothetical protein, conserved | 1,45 | 5,098E-34 |
| TcCLB.504867.129 | hypothetical protein, conserved | 1,45 | 9,238E-30 |
| TcCLB.508817.130 | carbonic anhydrase-like protein, putative | 1,449 | 3,677E-64 |
| TcCLB.507821.70 | dispersed gene family protein 1 (DGF-1, pseudogene), put | 1,448 | 1,854E-13 |
| TcCLB.508479.500 | retrotransposon hot spot (RHS) protein, putative | 1,447 | 9,347E-54 |
| TcCLB.506779.20 | hypothetical protein, conserved | 1,447 | 2,689E-49 |
| TcCLB.511641.10 | hypothetical protein, conserved | 1,447 | 1,997E-23 |
| TcCLB.505401.19 | dispersed gene family protein 1 (DGF-1), putative (fragme | 1,447 | 5,515E-14 |
| TcCLB.511755.119 | hypothetical protein, conserved (fragment) | 1,446 | 4,732E-66 |
| TcCLB.509875.170 | pyruvate dehydrogenase (lipoamide) kinase, putative | 1,446 | 6,471E-42 |
| TcCLB.511151.80 | hypothetical protein, conserved | 1,446 | 1,485E-40 |
| TcCLB.508569.80 | phosphoacetylglucosamine mutase, putative | 1,446 | 5,245E-35 |
| TcCLB.505373.30 | retrotransposon hot spot protein (RHS, pseudogene), puta | 1,446 | 6,746E-18 |
| TcCLB.511887.10 | retrotransposon hot spot protein (RHS, pseudogene), puta | 1,442 | 8,071E-20 |
| TcCLB.510449.10 | dispersed gene family protein 1 (DGF-1), putative | 1,441 | 2,612E-13 |
| TcCLB.511825.200 | 60S ribosomal protein L14, putative | 1,44 | 5,255E-68 |
| TcCLB.510121.160 | DNA topoisomerase IA, putative | 1,44 | 9,524E-53 |

| | | | |
|---|---|---|---|
| TcCLB.508677.160 | hypothetical protein, conserved | 1,44 | 3,31E-38 |
| TcCLB.510099.120 | D-isomer specific 2-hydroxyacid dehydrogenase-protein, p | 1,439 | 1,156E-117 |
| TcCLB.509335.10 | retrotransposon hot spot (RHS) protein, putative | 1,439 | 1,127E-61 |
| TcCLB.508027.110 | pyroglutamyl-peptidase I, putative (PPI) | 1,439 | 2,014E-37 |
| TcCLB.510581.9 | receptor-type adenylate cyclase, putative (fragment) | 1,439 | 3,763E-21 |
| TcCLB.507221.40 | hypothetical protein | 1,439 | 1,899E-20 |
| TcCLB.508677.134 | hypothetical protein, conserved | 1,438 | 4,758E-21 |
| TcCLB.506537.299 | retrotransposon hot spot (RHS) protein, putative | 1,436 | 5,38E-57 |
| TcCLB.507709.60 | hypothetical protein, conserved | 1,435 | 7,243E-55 |
| TcCLB.508965.14 | multidrug resistance protein E, putative (MRPE) | 1,435 | 1,59E-48 |
| TcCLB.509799.5 | hypothetical protein, conserved | 1,435 | 4,672E-31 |
| TcCLB.506583.70 | cytosolic factor SEC14, putative | 1,435 | 5,288E-23 |
| TcCLB.511431.54 | hypothetical protein | 1,435 | 1,027E-09 |
| TcCLB.507089.240 | hypothetical protein, conserved | 1,434 | 2,496E-26 |
| TcCLB.507143.80 | hypothetical protein, conserved | 1,433 | 2,706E-16 |
| TcCLB.507093.69 | hypothetical protein, conserved | 1,432 | 9,746E-17 |
| TcCLB.507565.10 | dispersed gene family protein 1 (DGF-1, pseudogene), put | 1,432 | 8,633E-09 |
| TcCLB.509647.13 | hypothetical protein, conserved (fragment) | 1,431 | 1,446E-62 |
| TcCLB.510759.120 | rieske iron-sulfur protein, mitochondrial precursor, putativ | 1,43 | 4,049E-63 |
| TcCLB.503599.70 | hypothetical protein, conserved | 1,43 | 6,958E-25 |
| TcCLB.503527.4 | hypothetical protein, conserved (pseudogene) | 1,429 | 9,501E-39 |
| TcCLB.473633.9 | dispersed gene family protein 1 (DGF-1), putative (fragme | 1,429 | 4,199E-08 |
| TcCLB.511827.30 | hypothetical protein | 1,428 | 2,412E-39 |
| TcCLB.503505.20 | MFS transporter, putative | 1,428 | 2,615E-24 |
| TcCLB.507023.70 | hypothetical protein, conserved | 1,428 | 9,026E-19 |
| TcCLB.506925.310 | hypothetical protein, conserved | 1,427 | 1,97E-103 |
| TcCLB.506247.310 | hypothetical protein, conserved | 1,427 | 3,492E-57 |
| TcCLB.509067.70 | pyrroline-5-carboxylate synthetase-like protein, putative | 1,426 | 2,211E-45 |
| TcCLB.510345.9 | protein kinase, putative (fragment) | 1,426 | 2,83E-12 |
| TcCLB.506977.49 | ribosomal protein L36, putative | 1,425 | 2,404E-74 |
| TcCLB.508375.89 | fucose kinase, putative (fragment) | 1,425 | 2,577E-07 |
| TcCLB.511067.30 | transketolase, putative | 1,424 | 1,153E-56 |
| TcCLB.506517.149 | hypothetical protein, conserved | 1,424 | 1,342E-34 |
| TcCLB.507517.50 | hypothetical protein | 1,424 | 2,748E-19 |
| TcCLB.506417.83 | multidrug resistance-associated protein (pseudogene), pu | 1,423 | 3,226E-36 |
| TcCLB.507711.90 | procyclic form surface phosphoprotein, putative | 1,423 | 2,63E-31 |
| TcCLB.510349.10 | hypothetical protein, conserved | 1,423 | 4,688E-25 |
| TcCLB.506713.24 | hypothetical protein, conserved | 1,423 | 4,42E-13 |
| TcCLB.508175.100 | hypothetical protein | 1,422 | 2,398E-41 |
| TcCLB.511153.70 | hypothetical protein, conserved | 1,422 | 2,283E-33 |
| TcCLB.509153.10 | hypothetical protein, conserved | 1,421 | 5,213E-60 |
| TcCLB.508637.63 | hypothetical protein, conserved | 1,421 | 1,509E-42 |
| TcCLB.506823.80 | retrotransposon hot spot protein (RHS, pseudogene), puta | 1,421 | 2,86E-21 |
| TcCLB.511217.200 | tyrosyl or methionyl-tRNA synthetase, putative | 1,42 | 6,674E-34 |
| TcCLB.508335.9 | dispersed gene family protein 1 (DGF-1), putative | 1,42 | 3,973E-11 |

| | | | |
|---|---|---|---|
| TcCLB.507195.60 | hypothetical protein | 1,42 | 0,006706 |
| TcCLB.509461.60 | mitochondrial RNA binding protein, putative | 1,419 | 2,995E-65 |
| TcCLB.508637.140 | hypothetical protein, conserved | 1,419 | 1,646E-39 |
| TcCLB.511809.10 | hypothetical protein | 1,419 | 3,679E-32 |
| TcCLB.508417.29 | dispersed gene family protein 1 (DGF-1), putative (fragm | 1,419 | 2,528E-19 |
| TcCLB.507677.80 | hypothetical protein, conserved | 1,417 | 1,202E-10 |
| TcCLB.511761.60 | hypothetical protein, conserved | 1,415 | 4,464E-24 |
| TcCLB.508927.20 | glycosomal transporter (GAT1), putative (GAT1) | 1,414 | 3,97E-28 |
| TcCLB.506683.180 | dispersed gene family protein 1 (DGF-1), putative | 1,414 | 1,518E-20 |
| TcCLB.504137.50 | hypothetical protein, conserved | 1,413 | 1,137E-21 |
| TcCLB.511523.100 | aldehyde dehydrogenase, putative | 1,412 | 3,595E-49 |
| TcCLB.506147.100 | hypothetical protein, conserved | 1,412 | 1,159E-43 |
| TcCLB.509159.10 | ATP-dependent DNA DEAD/DEAH box helicase, putative (f | 1,412 | 7,261E-43 |
| TcCLB.511471.120 | retrotransposon hot spot protein (RHS, pseudogene), puta | 1,412 | 1,02E-41 |
| TcCLB.507009.70 | hypothetical protein, conserved | 1,412 | 4,665E-31 |
| TcCLB.511573.40 | Mitogen-activated protein kinase 10, putative (MPK10) | 1,41 | 6,89E-40 |
| TcCLB.509463.20 | hypothetical protein, conserved | 1,409 | 6,395E-33 |
| TcCLB.506363.70 | amidohydrolase, putative | 1,409 | 1,011E-18 |
| TcCLB.508373.10 | glutamine dependent carbamoyl-phosphate synthase, put | 1,408 | 4,303E-43 |
| TcCLB.508567.70 | peptidyl-prolyl cis-trans isomerase (PIN1) | 1,408 | 4,02E-36 |
| TcCLB.510577.50 | rhomboid-like protein, putative | 1,407 | 2,096E-70 |
| TcCLB.503939.110 | hypothetical protein, conserved | 1,407 | 4,523E-41 |
| TcCLB.508461.10 | Present in the outer mitochondrial membrane proteome 6 | 1,407 | 1,534E-33 |
| TcCLB.508319.40 | ATP-dependent RNA helicase SUB2, putative (SUB2) | 1,406 | 9,648E-68 |
| TcCLB.507659.30 | amino acid transporter, putative (fragment) | 1,406 | 1,235E-64 |
| TcCLB.510901.230 | hypothetical protein, conserved | 1,405 | 5,498E-35 |
| TcCLB.507993.290 | hypothetical protein, conserved | 1,405 | 1,794E-26 |
| TcCLB.506843.40 | retrotransposon hot spot (RHS) protein, putative | 1,403 | 7,85E-77 |
| TcCLB.506053.20 | hypothetical protein, conserved | 1,403 | 3,82E-22 |
| TcCLB.506571.10 | dispersed gene family protein 1 (DGF-1), putative | 1,403 | 1,146E-13 |
| TcCLB.510039.90 | 60S ribosomal protein L35, putative | 1,402 | 3,825E-85 |
| TcCLB.508399.49 | dispersed gene family protein 1 (DGF-1), putative | 1,402 | 1,394E-15 |
| TcCLB.506529.430 | alanine aminotransferase, putative | 1,402 | 6,025E-10 |
| TcCLB.510065.30 | translationally controlled tumor protein (TCTP), putative | 1,4 | 4,523E-100 |
| TcCLB.511809.30 | hypothetical protein, conserved | 1,4 | 4,469E-46 |
| TcCLB.510811.10 | protein associated with differentiation 4, putative | 1,4 | 4,788E-21 |
| TcCLB.507669.220 | hypothetical protein, conserved | 1,399 | 1,832E-36 |
| TcCLB.511803.15 | dynein light chain lc6, flagellar outer arm (pseudogene), p | 1,399 | 2,261E-15 |
| TcCLB.509365.20 | dispersed gene family protein 1 (DGF-1, pseudogene), put | 1,398 | 5,494E-16 |
| TcCLB.506529.300 | hypothetical protein, conserved | 1,397 | 2,444E-59 |
| TcCLB.510345.50 | hypothetical protein, conserved | 1,397 | 4,453E-44 |
| TcCLB.510843.10 | beta galactofuranosyl glycosyltransferase (pseudogene), p | 1,397 | 1,719E-37 |
| TcCLB.508815.20 | hypothetical protein, conserved | 1,397 | 2,139E-28 |
| TcCLB.511213.89 | dispersed gene family protein 1 (DGF-1), putative (fragm | 1,397 | 5,923E-09 |
| TcCLB.509769.60 | succinate dehydrogenase iron-sulfur subunit (SDH2N) | 1,396 | 2,603E-57 |

| | | | |
|---|---|---|---|
| TcCLB.506401.120 | 40S ribosomal protein S9, putative | 1,395 | 5,949E-71 |
| TcCLB.509999.140 | RNA-binding protein, putative | 1,395 | 9,643E-11 |
| TcCLB.507159.60 | hypothetical protein, conserved | 1,394 | 8,237E-71 |
| TcCLB.511649.70 | structural maintenance of chromosome 3 protein, putativ | 1,394 | 8,228E-21 |
| TcCLB.508507.24 | hypothetical protein, conserved | 1,393 | 9,13E-20 |
| TcCLB.510105.20 | Ribosomal RNA processing protein 41A (RRP41A) | 1,392 | 3,599E-56 |
| TcCLB.507875.159 | 60S acidic ribosomal protein P2 (pseudogene), putative | 1,391 | 2,078E-48 |
| TcCLB.506853.70 | hypothetical protein, conserved | 1,391 | 3,354E-32 |
| TcCLB.508909.214 | hypothetical protein, conserved | 1,39 | 1,131E-42 |
| TcCLB.511531.50 | glucosamine-6-phosphate isomerase, putative | 1,388 | 5,048E-50 |
| TcCLB.503873.10 | cytochrome-b5 reductase, putative | 1,388 | 2,821E-46 |
| TcCLB.506777.80 | seryl-tRNA synthetase, putative | 1,387 | 6,699E-97 |
| TcCLB.509099.10 | GTP-binding protein, putative | 1,387 | 6,996E-38 |
| TcCLB.511643.30 | aspartate carbamoyltransferase, putative | 1,387 | 5,749E-37 |
| TcCLB.503749.5 | pyrroline-5-carboxylate synthetase-like protein, putative ( | 1,386 | 6,012E-22 |
| TcCLB.509667.10 | dispersed gene family protein 1 (DGF-1), putative | 1,386 | 4,361E-15 |
| TcCLB.511825.230 | hypothetical protein | 1,385 | 2,414E-28 |
| TcCLB.507831.70 | pyruvate dehydrogenase E1 component alpha subunit, put | 1,385 | 6,093E-27 |
| TcCLB.504153.330 | hypothetical protein, conserved | 1,384 | 3,247E-62 |
| TcCLB.506247.370 | hypothetical protein, conserved | 1,384 | 3,479E-48 |
| TcCLB.503543.20 | divalent cation transporter, putative (fragment) | 1,383 | 1,799E-45 |
| TcCLB.504423.20 | electon transport protein SCO1/SCO2, putative | 1,383 | 4,231E-43 |
| TcCLB.505919.31 | hypothetical protein, conserved (pseudogene) | 1,383 | 3,554E-21 |
| TcCLB.510131.30 | hypothetical protein, conserved | 1,382 | 7,584E-45 |
| TcCLB.511649.90 | hypothetical protein, conserved | 1,382 | 7,353E-36 |
| TcCLB.511389.120 | hypothetical protein, conserved | 1,381 | 7,714E-31 |
| TcCLB.511383.80 | hypothetical protein, conserved | 1,381 | 4,527E-23 |
| TcCLB.430605.40 | ERV/ALR sulfhydryl oxidase domain-containing protein (P | 1,38 | 1,091E-34 |
| TcCLB.508027.60 | hypothetical protein | 1,379 | 3,722E-42 |
| TcCLB.507003.60 | hypothetical protein, conserved | 1,378 | 4,72E-32 |
| TcCLB.507167.50 | hypothetical protein, conserved (pseudogene) | 1,378 | 2,083E-19 |
| TcCLB.507857.70 | hypothetical protein, conserved | 1,377 | 3,315E-34 |
| TcCLB.509455.114 | Protein of unknown function (DUF1295), putative (POMP2 | 1,377 | 3,225E-30 |
| TcCLB.504147.160 | ADP-ribosylation factor-like protein, putative | 1,376 | 1,53E-41 |
| TcCLB.510421.320 | pyruvate dehydrogenase E1 beta subunit, putative | 1,376 | 2,053E-37 |
| TcCLB.511529.68 | cyclin-dependent kinases regulatory subunit, putative (P1 | 1,376 | 1,912E-32 |
| TcCLB.510309.10 | hypothetical protein, conserved | 1,376 | 9,022E-30 |
| TcCLB.503479.30 | adenylate kinase 1 | 1,375 | 2,652E-28 |
| TcCLB.510265.110 | hypothetical protein, conserved | 1,375 | 0,0002624 |
| TcCLB.506925.104 | dynein light chain, putative | 1,374 | 1,583E-57 |
| TcCLB.509441.10 | hypothetical protein, conserved | 1,374 | 9,792E-41 |
| TcCLB.510657.40 | hypothetical protein, conserved | 1,374 | 2,664E-29 |
| TcCLB.511105.29 | dispersed gene family protein 1 (DGF-1), putative | 1,374 | 1,117E-09 |
| TcCLB.503615.10 | UDP-Gal or UDP-GlcNAc-dependent glycosyltransferase, p | 1,371 | 5,289E-16 |
| TcCLB.511585.120 | hypothetical protein, conserved | 1,37 | 7,63E-40 |

| | | | |
|---|---|---|---|
| TcCLB.510173.90 | poly(ADP-ribose) polymerase, putative | 1,37 | 5,434E-35 |
| TcCLB.508399.10 | NADH-ubiquinone oxidoreductase, mitochondrial, putative | 1,369 | 8,816E-38 |
| TcCLB.503697.40 | hypothetical protein | 1,369 | 4,377E-17 |
| TcCLB.504255.4 | rhomboid-like protein | 1,369 | 3,992E-08 |
| TcCLB.507723.90 | hypothetical protein, conserved | 1,367 | 1,458E-17 |
| TcCLB.508433.50 | retrotransposon hot spot protein (RHS, pseudogene), puta | 1,367 | 0,0005453 |
| TcCLB.510103.30 | protein associated with differentiation 4, putative | 1,366 | 1,264E-53 |
| TcCLB.506945.240 | ATP synthase, epsilon chain, putative | 1,366 | 1,459E-38 |
| TcCLB.506979.40 | hypothetical protein, conserved | 1,366 | 3,267E-26 |
| TcCLB.511443.10 | calpain-like cysteine peptidase (pseudogene), putative | 1,366 | 2,106E-13 |
| TcCLB.506299.9 | dispersed gene family protein 1 (DGF-1), putative (fragme | 1,366 | 2,123E-13 |
| TcCLB.511867.130 | hypothetical protein, conserved | 1,365 | 3,779E-58 |
| TcCLB.507479.29 | retrotransposon hot spot protein (RHS, pseudogene), puta | 1,365 | 2,371E-16 |
| TcCLB.509805.90 | hypothetical protein, conserved | 1,364 | 1,292E-44 |
| TcCLB.508909.320 | hypothetical protein | 1,364 | 2,054E-34 |
| TcCLB.506859.170 | hypothetical protein, conserved | 1,363 | 1,159E-35 |
| TcCLB.509551.20 | Putrescine-cadaverine transporter 1, putative (POT1.2) | 1,362 | 1,901E-35 |
| TcCLB.510883.10 | retrotransposon hot spot (RHS) protein, putative | 1,362 | 1,67E-27 |
| TcCLB.510095.50 | dispersed gene family protein 1 (DGF-1), putative | 1,362 | 1,897E-13 |
| TcCLB.508541.190 | cell cycle sequence binding phosphoprotein (RBP33), puta | 1,361 | 2,538E-49 |
| TcCLB.506147.80 | Present in the outer mitochondrial membrane proteome 6 | 1,361 | 2,224E-28 |
| TcCLB.509037.50 | aspartyl-tRNA synthetase, putative | 1,36 | 2,341E-40 |
| TcCLB.507445.9 | dispersed gene family protein 1 (DGF-1), putative (fragme | 1,36 | 7,497E-12 |
| TcCLB.507467.10 | receptor-type adenylate cyclase, putative (fragment) | 1,36 | 3,766E-11 |
| TcCLB.506487.60 | endosomal integral membrane protein, putative (fragmer | 1,36 | 1,905E-09 |
| TcCLB.508591.10 | dispersed gene family protein 1 (DGF-1), putative | 1,359 | 1,287E-11 |
| TcCLB.509747.20 | 60S ribosomal protein L22, putative | 1,358 | 5,292E-64 |
| TcCLB.510307.140 | retrotransposon hot spot protein (RHS, pseudogene), puta | 1,358 | 1,371E-33 |
| TcCLB.511335.19 | hypothetical protein, conserved (pseudogene) | 1,358 | 1,82E-18 |
| TcCLB.508257.10 | serine-palmitoyl-CoA transferase, putative | 1,358 | 1,567E-15 |
| TcCLB.508077.9 | conserved protein (fragment) | 1,357 | 3,921E-50 |
| TcCLB.508739.21 | mucin TcSMUGS (pseudogene), putative | 1,357 | 6,824E-34 |
| TcCLB.506535.10 | retrotransposon hot spot (RHS) protein, putative (fragmer | 1,357 | 1,518E-23 |
| TcCLB.506295.130 | prohibitin, putative | 1,356 | 7,45E-33 |
| TcCLB.506063.31 | ATP-dependent DEAD/H RNA helicase (pseudogene), puta | 1,355 | 0,001201 |
| TcCLB.503431.60 | acid phosphatase, putative (AP) | 1,354 | 1,537E-24 |
| TcCLB.509065.9 | aldehyde dehydrogenase, putative (fragment) | 1,353 | 1,149E-48 |
| TcCLB.509437.60 | hypothetical protein, conserved | 1,353 | 6,35E-47 |
| TcCLB.511469.20 | hypothetical protein, conserved | 1,353 | 4,71E-46 |
| TcCLB.509191.9 | dispersed gene family protein 1 (DGF-1, pseudogene), put | 1,353 | 9,626E-06 |
| TcCLB.507669.10 | fumarate hydratase, putative | 1,351 | 1,974E-32 |
| TcCLB.511249.44 | vacuolar transporter chaperone, putative | 1,349 | 1,142E-42 |
| TcCLB.508647.110 | Present in the outer mitochondrial membrane proteome 1 | 1,349 | 2,861E-40 |
| TcCLB.506405.50 | serine-palmitoyl-CoA transferase, putative | 1,348 | 5,667E-58 |
| TcCLB.511771.80 | retrotransposon hot spot protein (RHS, pseudogene), puta | 1,348 | 1,782E-27 |

| | | | |
|---|---|---|---|
| TcCLB.503469.9 | dispersed gene family protein 1 (DGF-1), putative (fragme | 1,348 | 4,102E-16 |
| TcCLB.506417.10 | multidrug resistance protein A (pseudogene), putative | 1,347 | 3,363E-86 |
| TcCLB.506517.30 | hypothetical protein | 1,347 | 2,97E-39 |
| TcCLB.509791.120 | kinetoplast-associated protein (KAP6) | 1,347 | 1,179E-32 |
| TcCLB.509445.10 | tryparedoxin peroxidase, putative | 1,347 | 1,397E-19 |
| TcCLB.507611.270 | hypothetical protein, conserved | 1,347 | 4,576E-16 |
| TcCLB.507649.80 | nucleolar RNA-binding protein, putative | 1,346 | 3,169E-54 |
| TcCLB.507801.110 | hypothetical protein, conserved | 1,345 | 8,055E-38 |
| TcCLB.418185.50 | retrotransposon hot spot protein (RHS, pseudogene), puta | 1,345 | 1,093E-29 |
| TcCLB.511579.9 | fatty acyl CoA syntetase 1, putative | 1,345 | 5,236E-12 |
| TcCLB.505843.49 | ERV/ALR sulfhydryl oxidase domain-containing protein (P( | 1,345 | 1,247E-09 |
| TcCLB.507993.330 | mevalonate-diphosphate decarboxylase, putative | 1,344 | 2,836E-30 |
| TcCLB.506571.20 | retrotransposon hot spot protein (RHS, pseudogene), puta | 1,344 | 3,742E-20 |
| TcCLB.506041.50 | cytochrome c oxidase assembly protein, putative | 1,343 | 6,829E-25 |
| TcCLB.510355.200 | hypothetical protein, conserved | 1,343 | 9,183E-23 |
| TcCLB.506573.81 | ribosomal protein L39, putative | 1,343 | 1,291E-19 |
| TcCLB.510673.10 | dispersed gene family protein 1 (DGF-1, pseudogene), put | 1,343 | 9,026E-17 |
| TcCLB.508277.40 | divalent cation transporter, putative | 1,342 | 1,026E-70 |
| TcCLB.503955.55 | hypothetical protein, conserved | 1,342 | 1,055E-21 |
| TcCLB.506825.150 | hypothetical protein, conserved | 1,34 | 7,129E-63 |
| TcCLB.508303.65 | thioredoxin-like protein, putative | 1,339 | 1,348E-20 |
| TcCLB.506977.70 | ATPase subunit 9, putative | 1,337 | 5,589E-46 |
| TcCLB.510235.10 | dispersed gene family protein 1 (DGF-1), putative | 1,337 | 9,275E-10 |
| TcCLB.506295.190 | glycine dehydrogenase [decarboxylating] (pseudogene), pu | 1,337 | 2,789E-09 |
| TcCLB.508473.10 | retrotransposon hot spot (RHS) protein, putative | 1,336 | 4,497E-37 |
| TcCLB.510355.190 | hypothetical protein | 1,336 | 3,58E-18 |
| TcCLB.506401.320 | 60S ribosomal protein L7a, putative | 1,335 | 2,819E-58 |
| TcCLB.509167.140 | RNA-binding protein 42 (RNA-binding motif protein 42), p | 1,335 | 4,634E-30 |
| TcCLB.509901.160 | hypothetical protein, conserved | 1,335 | 2,49E-20 |
| TcCLB.506691.64 | cytochrome c oxidase copper chaperone, putative | 1,335 | 1,184E-11 |
| TcCLB.506411.10 | thimet oligopeptidase, putative | 1,334 | 1,224E-46 |
| TcCLB.504427.200 | poly(A)-specific ribonuclease PARN, putative | 1,334 | 7,051E-36 |
| TcCLB.510355.90 | dispersed gene family protein 1 (DGF-1), putative | 1,334 | 7,502E-19 |
| TcCLB.506651.9 | dispersed gene family protein 1 (DGF-1), putative | 1,334 | 1,543E-08 |
| TcCLB.506945.110 | casein kinase 1 isoform 2, putative | 1,333 | 1,018E-42 |
| TcCLB.509279.10 | hypothetical protein, conserved | 1,333 | 2,681E-13 |
| TcCLB.506629.30 | guide RNA-binding protein of 21 kDa (gBP21) | 1,331 | 2,648E-44 |
| TcCLB.508119.120 | GDP-mannose 4,6 dehydratase, putative | 1,331 | 3,645E-41 |
| TcCLB.506635.139 | kasal-type serpin (pseudogene), putative | 1,331 | 1,057E-26 |
| TcCLB.510101.330 | hypothetical protein, conserved | 1,33 | 1,087E-41 |
| TcCLB.506999.50 | hypothetical protein, conserved | 1,33 | 2,441E-34 |
| TcCLB.511657.40 | hypothetical protein, conserved | 1,33 | 1,824E-25 |
| TcCLB.506675.30 | retrotransposon hot spot protein (RHS, pseudogene), puta | 1,329 | 9,393E-37 |
| TcCLB.504163.110 | N-acetyltransferase subunit Nat1, putative | 1,327 | 1,591E-42 |
| TcCLB.511353.10 | FK506-binding protein (FKBP)-type peptidyl-prolyl isomera | 1,327 | 8,014E-27 |

| | | | |
|---|---|---|---|
| TcCLB.508153.994 | cytochrome c oxidase copper chaperone, putative | 1,327 | 2,352E-12 |
| TcCLB.508247.149 | dispersed gene family protein 1 (DGF-1), putative | 1,327 | 0,003828 |
| TcCLB.509207.40 | hypothetical protein, conserved | 1,326 | 3,781E-63 |
| TcCLB.507867.60 | protein phosphatase 1, regulatory subunit, putative | 1,326 | 3,451E-33 |
| TcCLB.508411.10 | cofilin/actin depolymerizing factor, putative | 1,325 | 9,008E-34 |
| TcCLB.511847.30 | hypothetical protein, conserved | 1,325 | 7,911E-29 |
| TcCLB.506537.60 | dispersed gene family protein 1 (DGF-1, pseudogene), put | 1,325 | 1,127E-19 |
| TcCLB.511625.50 | GTP-binding protein, putative | 1,324 | 3,091E-44 |
| TcCLB.508503.30 | hypothetical protein | 1,324 | 1,65E-23 |
| TcCLB.481099.9 | dispersed gene family protein 1 (DGF-1), putative (fragme | 1,322 | 5,356E-07 |
| TcCLB.417993.10 | dispersed gene family protein 1 (DGF-1, pseudogene), put | 1,322 | 0,05335 |
| TcCLB.506195.290 | hypothetical protein, conserved | 1,321 | 3,422E-30 |
| TcCLB.506779.80 | pantothenate kinase subunit, putative | 1,321 | 4,784E-29 |
| TcCLB.503823.84 | DNA-directed RNA polymerase I subunit, putative | 1,321 | 1,594E-20 |
| TcCLB.510381.10 | cystathionine beta-synthase, putative | 1,321 | 9,964E-08 |
| TcCLB.511145.50 | vacuolar ATP synthase, putative | 1,32 | 4,461E-57 |
| TcCLB.506357.40 | Elongation factor Tu, mitochondrial, putative (EF-Tu) | 1,32 | 3,693E-48 |
| TcCLB.508241.50 | surface protease GP63, putative | 1,319 | 1,44E-33 |
| TcCLB.506189.20 | hypothetical protein, conserved | 1,319 | 4,917E-20 |
| TcCLB.504055.40 | dihydroxyacetone phosphate acyltransferase, putative (DA | 1,318 | 1,252E-48 |
| TcCLB.511825.180 | ATP-dependent DEAD/H RNA helicase, putative | 1,318 | 7,835E-46 |
| TcCLB.510301.10 | fructose-bisphosphate aldolase, glycosomal, putative | 1,318 | 2,795E-38 |
| TcCLB.508823.100 | hypothetical protein, conserved | 1,318 | 2,184E-24 |
| TcCLB.506195.110 | malate dehydrogenase, putative | 1,317 | 5,836E-76 |
| TcCLB.510101.50 | lanosterol 14-alpha-demethylase, putative | 1,316 | 1,509E-78 |
| TcCLB.509331.110 | hypothetical protein, conserved | 1,316 | 2,158E-32 |
| TcCLB.506327.99 | serine/threonine protein kinase, putative (fragment) | 1,316 | 6,499E-32 |
| TcCLB.510225.20 | hypothetical protein, conserved | 1,316 | 5,004E-20 |
| TcCLB.510105.170 | dihydrolipoamide acetyltransferase precursor, putative | 1,315 | 2,916E-51 |
| TcCLB.510821.60 | ribonucleoprotein p18, mitochondrial precursor, putative | 1,314 | 7,069E-51 |
| TcCLB.503715.10 | splicing factor TSR1, putative | 1,314 | 2,092E-50 |
| TcCLB.504269.20 | GPR1/FUN34/yaaH family, putative | 1,314 | 3,05E-36 |
| TcCLB.506863.20 | hypothetical protein | 1,314 | 4,149E-23 |
| TcCLB.506443.70 | glutathione-S-transferase/glutaredoxin, putative | 1,314 | 5,103E-22 |
| TcCLB.510785.20 | pteridine reductase (pseudogene), putative | 1,313 | 8,978E-58 |
| TcCLB.510853.73 | retrotransposon hot spot protein (pseudogene), putative | 1,313 | 1,136E-55 |
| TcCLB.506485.80 | oxidoreductase, putative | 1,313 | 1,006E-42 |
| TcCLB.511217.130 | hypothetical protein, conserved | 1,313 | 2,709E-41 |
| TcCLB.508371.10 | glutamine-dependent carbamoyl-phosphate synthetase (p | 1,313 | 4,858E-21 |
| TcCLB.510603.60 | protein tyrosine phosphatase-like protein, putative | 1,313 | 2,174E-15 |
| TcCLB.506147.50 | hypothetical protein, conserved | 1,312 | 3,659E-26 |
| TcCLB.505035.10 | hypothetical protein | 1,312 | 1,296E-17 |
| TcCLB.510105.40 | cullin-like protein, putative | 1,311 | 1,23E-46 |
| TcCLB.503809.140 | cation transporter, putative | 1,311 | 7,609E-32 |
| TcCLB.509713.20 | protein associated with differentiation 4, putative | 1,309 | 1,432E-102 |

| | | | |
|---|---|---|---|
| TcCLB.503411.20 | hypothetical protein, conserved | 1,309 | 4,66E-25 |
| TcCLB.509769.66 | hypothetical protein, conserved | 1,309 | 5,396E-25 |
| TcCLB.507479.51 | beta galactofuranosyl glycosyltransferase, putative | 1,309 | 7,706E-06 |
| TcCLB.508023.120 | hypothetical protein, conserved (pseudogene) | 1,308 | 1,058E-46 |
| TcCLB.508089.50 | hypothetical protein, conserved | 1,308 | 1,1E-30 |
| TcCLB.506399.70 | hypothetical protein, conserved | 1,308 | 3,388E-18 |
| TcCLB.457101.30 | multidrug resistance protein E, putative | 1,307 | 2,461E-24 |
| TcCLB.511439.30 | lipase, putative | 1,307 | 5,325E-15 |
| TcCLB.506669.4 | MFS transporter, putative | 1,307 | 9,275E-08 |
| TcCLB.508351.40 | cytochrome c oxidase assembly factor, putative | 1,306 | 5,755E-26 |
| TcCLB.506717.10 | fatty acid elongase, putative | 1,306 | 2,408E-17 |
| TcCLB.511817.270 | hypothetical protein, conserved | 1,305 | 6,014E-41 |
| TcCLB.508153.270 | heat shock protein 20, putative | 1,305 | 3,313E-27 |
| TcCLB.511441.20 | calpain-like cysteine peptidase, putative (fragment) | 1,305 | 0,006618 |
| TcCLB.508535.30 | hypothetical protein, conserved | 1,304 | 4,724E-29 |
| TcCLB.511815.100 | hypothetical protein, conserved | 1,304 | 5,273E-21 |
| TcCLB.509549.100 | hypothetical protein, conserved | 1,303 | 4,153E-49 |
| TcCLB.509747.40 | hypothetical protein, conserved | 1,303 | 1,297E-30 |
| TcCLB.511211.30 | hypothetical protein | 1,303 | 1,862E-20 |
| TcCLB.504163.60 | hypothetical protein, conserved | 1,302 | 4,58E-33 |
| TcCLB.436137.9 | dispersed gene family protein 1 (DGF-1), putative (fragme | 1,302 | 1,018E-15 |
| TcCLB.504033.21 | UDP-Gal or UDP-GlcNAc-dependent glycosyltransferase, p | 1,302 | 0,00000263 |
| TcCLB.432867.20 | hypothetical protein, conserved | 1,301 | 1,312E-07 |
| TcCLB.506211.140 | hypothetical protein, conserved | 1,3 | 1,626E-49 |
| TcCLB.508231.9 | hexose transporter, putative | 1,3 | 4,727E-17 |
| TcCLB.507121.40 | beta galactofuranosyl glycosyltransferase, putative | 1,3 | 1,001E-08 |
| TcCLB.507831.20 | hypothetical protein, conserved | 1,3 | 0,00266 |
| TcCLB.509237.30 | glycosomal transporter (GAT1), putative (GAT1) | 1,299 | 8,054E-34 |
| TcCLB.503453.64 | hypothetical protein, conserved | 1,299 | 7,361E-23 |
| TcCLB.506925.64 | hypothetical protein, conserved | 1,297 | 2,247E-43 |
| TcCLB.511417.80 | phosphate-repressible phosphate permease, putative | 1,297 | 4,767E-38 |
| TcCLB.506629.50 | hypothetical protein, conserved | 1,297 | 2,932E-29 |
| TcCLB.511771.132 | ATP-dependent DEAD/H RNA helicase (pseudogene), puta | 1,297 | 0,0006505 |
| TcCLB.511311.80 | hypothetical protein, conserved | 1,296 | 4,641E-68 |
| TcCLB.508119.200 | hypothetical protein | 1,295 | 3,405E-36 |
| TcCLB.511367.70 | dihydrolipoamide acetyltransferase, putative | 1,295 | 1,774E-30 |
| TcCLB.511181.150 | hypothetical protein, conserved | 1,295 | 1,01E-25 |
| TcCLB.506005.10 | dispersed gene family protein 1 (DGF-1, pseudogene), put | 1,295 | 1,416E-19 |
| TcCLB.509449.5 | receptor-type adenylate cyclase, putative (fragment) | 1,295 | 0,003889 |
| TcCLB.406999.10 | dispersed gene family protein 1 (DGF-1), putative (fragme | 1,295 | 0,0234 |
| TcCLB.511825.40 | centrin-4, putative (Centrin4) | 1,294 | 8,865E-42 |
| TcCLB.507383.10 | hypothetical protein, conserved | 1,294 | 0,0000806 |
| TcCLB.509601.100 | hypothetical protein, conserved | 1,293 | 6,082E-43 |
| TcCLB.510329.10 | hypothetical protein, conserved | 1,293 | 2,65E-41 |
| TcCLB.510323.40 | heat shock protein 20, putative | 1,293 | 3,141E-33 |

| | | | |
|---|---|---|---|
| TcCLB.508001.10 | dispersed gene family protein 1 (DGF-1), putative | 1,293 | 3,55E-09 |
| TcCLB.510101.170 | 40S ribosomal protein S5, putative | 1,292 | 2,519E-47 |
| TcCLB.506735.10 | mitochondrial processing peptidase alpha subunit, putativ | 1,292 | 1,644E-46 |
| TcCLB.508637.90 | phosphoglucomutase | 1,292 | 1,581E-30 |
| TcCLB.506605.150 | 60S ribosomal protein, putative | 1,292 | 1,017E-21 |
| TcCLB.505201.20 | dispersed gene family protein 1 (DGF-1, pseudogene), put | 1,292 | 2,354E-15 |
| TcCLB.510629.200 | retrotransposon hot spot protein (RHS, pseudogene), puta | 1,292 | 6,239E-07 |
| TcCLB.510577.30 | hypothetical protein | 1,291 | 1,132E-30 |
| TcCLB.511469.100 | 2-oxoisovalerate dehydrogenase beta subunit, mitochondr | 1,291 | 1,174E-24 |
| TcCLB.503929.79 | serine-palmitoyl-CoA transferase, putative | 1,291 | 1,471E-19 |
| TcCLB.506925.530 | glycosomal transporter (GAT3), putative | 1,29 | 2,374E-63 |
| TcCLB.511283.290 | hypothetical protein, conserved | 1,29 | 1,826E-51 |
| TcCLB.511733.80 | hypothetical protein | 1,29 | 2,741E-32 |
| TcCLB.503891.80 | hypothetical protein, conserved | 1,29 | 2,53E-27 |
| TcCLB.507587.24 | dispersed gene family protein 1 (DGF-1), putative (fragme | 1,29 | 5,919E-15 |
| TcCLB.507259.10 | tryparedoxin peroxidase, putative | 1,289 | 3,001E-49 |
| TcCLB.509153.100 | arginine N-methyltransferase, type III, putative (PRMT7) | 1,289 | 3,479E-48 |
| TcCLB.508153.760 | hypothetical protein, conserved | 1,289 | 6,231E-36 |
| TcCLB.510847.40 | retrotransposon hot spot protein (RHS, pseudogene), puta | 1,289 | 1,78E-26 |
| TcCLB.508799.280 | protein associated with differentiation 4, putative | 1,288 | 1,29E-37 |
| TcCLB.511819.30 | hypothetical protein, conserved | 1,288 | 5,56E-25 |
| TcCLB.509687.19 | dispersed gene family protein 1 (DGF-1), putative (fragme | 1,288 | 2,391E-07 |
| TcCLB.508461.30 | 3-demethylubiquinone-9 3-methyltransferase, putative | 1,287 | 2,377E-29 |
| TcCLB.505171.60 | hypothetical protein, conserved | 1,286 | 1,454E-38 |
| TcCLB.509295.110 | dispersed gene family protein 1 (DGF-1, pseudogene), put | 1,285 | 0,0004958 |
| TcCLB.510529.30 | hypothetical protein, conserved | 1,283 | 1,455E-44 |
| TcCLB.509825.14 | 40S ribosomal protein SA, putative | 1,283 | 8,505E-18 |
| TcCLB.504639.19 | dispersed gene family protein 1 (DGF-1), putative (fragme | 1,283 | 1,28E-09 |
| TcCLB.510643.190 | retrotransposon hot spot (RHS) protein, putative | 1,281 | 1,061E-33 |
| TcCLB.510397.39 | dispersed gene family protein 1 (DGF-1), putative | 1,281 | 6,783E-07 |
| TcCLB.507159.30 | hypothetical protein, conserved | 1,28 | 1,198E-40 |
| TcCLB.508961.10 | hypothetical protein, conserved | 1,28 | 1,334E-32 |
| TcCLB.511209.69 | membrane-bound acid phosphatase, putative (fragment) | 1,28 | 2,377E-21 |
| TcCLB.511575.20 | hypothetical protein, conserved | 1,28 | 4,117E-18 |
| TcCLB.506401.170 | plasma membrane Ca2 ATPase | 1,279 | 1,797E-66 |
| TcCLB.506301.11 | heat shock protein, putative (fragment) | 1,279 | 5,926E-43 |
| TcCLB.507673.70 | hypothetical protein, conserved | 1,279 | 5,342E-31 |
| TcCLB.511217.10 | hypothetical protein | 1,279 | 2,78E-21 |
| TcCLB.508193.60 | UDP-Gal or UDP-GlcNAc-dependent glycosyltransferase, p | 1,279 | 2,834E-21 |
| TcCLB.506755.260 | mitochondrial RNA-binding protein 2, putative | 1,278 | 3,116E-32 |
| TcCLB.511285.100 | hypothetical protein | 1,278 | 2,068E-30 |
| TcCLB.510089.100 | hypothetical protein | 1,278 | 2,169E-25 |
| TcCLB.510357.70 | ubiquinone biosynthesis protein-like protein, putative | 1,278 | 7,349E-22 |
| TcCLB.511469.110 | dispersed gene family protein 1 (DGF-1), putative | 1,278 | 1,795E-08 |
| TcCLB.506773.160 | hypothetical protein, conserved | 1,277 | 3,224E-34 |

| | | | |
|---|---|---|---|
| TcCLB.506931.20 | hypothetical protein, conserved | 1,277 | 2,034E-29 |
| TcCLB.507781.10 | hypothetical protein, conserved (pseudogene) | 1,277 | 1,483E-13 |
| TcCLB.511817.280 | hypothetical protein, conserved | 1,276 | 2,733E-51 |
| TcCLB.506369.20 | transporter, putative | 1,276 | 4,393E-13 |
| TcCLB.505533.10 | dispersed gene family protein 1 (DGF-1, pseudogene), put | 1,276 | 0,00004343 |
| TcCLB.506649.20 | P-type H -ATPase, putative | 1,275 | 4,2E-35 |
| TcCLB.511529.210 | hypothetical protein, conserved | 1,275 | 4,22E-20 |
| TcCLB.510089.210 | hypothetical protein, conserved | 1,275 | 9,405E-20 |
| TcCLB.424937.10 | glucose transporter, putative | 1,275 | 8,027E-14 |
| TcCLB.511803.40 | Present in the outer mitochondrial membrane proteome ! | 1,274 | 2,994E-69 |
| TcCLB.503955.20 | mitochondrial DNA polymerase beta, putative | 1,274 | 1,152E-53 |
| TcCLB.506241.160 | hypothetical protein, conserved | 1,274 | 2,278E-45 |
| TcCLB.510329.150 | hypothetical protein, conserved | 1,274 | 3,436E-41 |
| TcCLB.506407.100 | hypothetical protein, conserved | 1,273 | 2,971E-45 |
| TcCLB.506829.80 | nicotinamidase, putative (PNC1) | 1,272 | 2,072E-26 |
| TcCLB.507653.40 | dispersed gene family protein 1 (DGF-1), putative | 1,272 | 6,268E-11 |
| TcCLB.510879.60 | hypothetical protein, conserved | 1,271 | 3,409E-19 |
| TcCLB.508395.9 | dispersed gene family protein 1 (DGF-1), putative (fragme | 1,271 | 0,000455 |
| TcCLB.509023.10 | calcium binding protein EF hand, putative | 1,27 | 2,068E-39 |
| TcCLB.505807.60 | hypothetical protein, conserved | 1,27 | 1,478E-35 |
| TcCLB.511301.90 | protein kinase, putative | 1,27 | 2,432E-30 |
| TcCLB.511257.10 | DNAK protein, putative | 1,269 | 5,061E-43 |
| TcCLB.511511.40 | hypothetical protein, conserved | 1,269 | 2,288E-35 |
| TcCLB.510661.174 | riboflavin kinase, putative | 1,269 | 4,636E-14 |
| TcCLB.511167.80 | PIF1 helicase-like protein, putative | 1,268 | 1,367E-38 |
| TcCLB.506337.160 | NADH-ubiquinone oxidoreductase complex I subunit, puta | 1,267 | 5,236E-31 |
| TcCLB.507543.30 | retrotransposon hot spot protein (RHS, pseudogene), puta | 1,265 | 9,919E-49 |
| TcCLB.511729.60 | MORN repeat-containing protein 1 (MORN1) | 1,264 | 1,011E-39 |
| TcCLB.506053.5 | amino acid transporter, putative (fragment) | 1,264 | 2,089E-35 |
| TcCLB.510755.89 | kinetoplastid membrane protein KMP-11, putative | 1,264 | 2,421E-31 |
| TcCLB.509725.30 | hypothetical protein, conserved (pseudogene) | 1,263 | 7,466E-15 |
| TcCLB.506829.110 | fatty acyl CoA synthetase 2, putative | 1,262 | 3,786E-37 |
| TcCLB.509331.40 | hypothetical protein, conserved | 1,262 | 3,629E-17 |
| TcCLB.506559.110 | hypothetical protein | 1,261 | 2,926E-30 |
| TcCLB.506825.140 | hypothetical protein, conserved | 1,261 | 4,197E-23 |
| TcCLB.511355.24 | hypothetical protein, conserved | 1,261 | 3,653E-15 |
| TcCLB.507919.10 | surface protease GP63, putative | 1,26 | 2,245E-26 |
| TcCLB.511277.70 | Elongation factor Tu, mitochondrial, putative (EF-Tu) | 1,259 | 1,47E-48 |
| TcCLB.503603.10 | mitochondrial RNA binding complex 1 subunit, putative | 1,259 | 4,03E-24 |
| TcCLB.504153.30 | hypothetical protein, conserved | 1,258 | 2,158E-33 |
| TcCLB.506823.100 | dispersed gene family protein 1 (DGF-1), putative | 1,258 | 0,00000708 |
| TcCLB.510357.10 | DnaJ chaperone protein, putative | 1,257 | 4,559E-44 |
| TcCLB.509459.60 | Na/H antiporter-like protein, putative | 1,257 | 1,676E-39 |
| TcCLB.509937.210 | hypothetical protein, conserved | 1,257 | 4,843E-31 |
| TcCLB.511655.40 | hypothetical protein, conserved | 1,257 | 1,336E-27 |

| | | | |
|---|---|---|---|
| TcCLB.503521.30 | Dynein light chain roadblock-type protein, putative | 1,257 | 1,734E-25 |
| TcCLB.508831.80 | hypothetical protein, conserved | 1,257 | 6,417E-13 |
| TcCLB.506153.20 | amino acid transporter, putative | 1,256 | 6,913E-47 |
| TcCLB.511391.10 | retrotransposon hot spot protein (RHS, pseudogene), puta | 1,256 | 4,765E-37 |
| TcCLB.506475.105 | hypothetical protein, conserved | 1,256 | 5,442E-19 |
| TcCLB.509509.20 | C-1-tetrahydrofolate synthase, cytoplasmic, putative | 1,256 | 1,516E-17 |
| TcCLB.506625.80 | cAMP-specific phosphodiesterase, putative | 1,255 | 8,605E-53 |
| TcCLB.509317.20 | esterase, putative | 1,255 | 5,383E-20 |
| TcCLB.506925.319 | isocitrate dehydrogenase, putative | 1,255 | 2,312E-14 |
| TcCLB.506529.420 | alanine aminotransferase, putative | 1,254 | 4,303E-42 |
| TcCLB.510861.110 | hypothetical protein | 1,254 | 2,785E-37 |
| TcCLB.506843.20 | retrotransposon hot spot protein (RHS, pseudogene), puta | 1,254 | 3,415E-20 |
| TcCLB.509775.34 | hypothetical protein, conserved | 1,254 | 2,586E-17 |
| TcCLB.508323.70 | centrin-4, putative (Centrin4) | 1,254 | 1,596E-09 |
| TcCLB.506593.30 | ribosomal protein S7, putative | 1,253 | 2,972E-67 |
| TcCLB.508153.605 | hypothetical protein, conserved | 1,253 | 1,244E-25 |
| TcCLB.504427.150 | oxidoreductase-like protein, putative | 1,253 | 5,386E-19 |
| TcCLB.511621.60 | Asparagine-linked glycosylation protein 12 (ALG12) | 1,253 | 1,213E-07 |
| TcCLB.509233.50 | hypothetical protein, conserved | 1,252 | 5,05E-48 |
| TcCLB.509233.180 | ATP synthase F1, beta subunit, putative (ATPB) | 1,252 | 1,443E-47 |
| TcCLB.507869.10 | NADH-ubiquinone oxidoreductase, mitochondrial, putative | 1,251 | 1,316E-39 |
| TcCLB.507993.40 | hypothetical protein, conserved | 1,251 | 9,737E-31 |
| TcCLB.504137.80 | adenylyl cyclase-associated protein, putative | 1,25 | 4,771E-30 |
| TcCLB.506529.550 | cysteine peptidase, putative | 1,249 | 1,565E-39 |
| TcCLB.507427.20 | dispersed gene family protein 1 (DGF-1, pseudogene), put | 1,249 | 7,487E-10 |
| TcCLB.507943.40 | histone H4, putative | 1,248 | 7,025E-50 |
| TcCLB.510877.190 | hypothetical protein, conserved | 1,248 | 4,669E-08 |
| TcCLB.510421.220 | hypothetical protein, conserved | 1,247 | 6,031E-33 |
| TcCLB.503401.40 | glycosyl transferase, putative | 1,246 | 6,061E-51 |
| TcCLB.510479.30 | hypothetical protein | 1,246 | 7,038E-08 |
| TcCLB.504153.310 | heat shock protein, putative | 1,245 | 5,937E-78 |
| TcCLB.506417.60 | terbinafine resistance locus protein (yip1), putative | 1,245 | 4,512E-31 |
| TcCLB.507717.20 | hypothetical protein | 1,244 | 4,985E-36 |
| TcCLB.510667.90 | hypothetical protein, conserved | 1,243 | 1,776E-24 |
| TcCLB.510827.40 | hypothetical protein | 1,243 | 8,845E-11 |
| TcCLB.503617.31 | nuclear receptor binding factor, putative | 1,241 | 9,083E-36 |
| TcCLB.506435.410 | uracil phosphoribosyltransferase, putative | 1,241 | 5,732E-29 |
| TcCLB.509611.20 | peptidyl-prolyl cis-trans isomerase (PIN1) | 1,241 | 6,849E-14 |
| TcCLB.510839.11 | hypothetical protein | 1,241 | 7,695E-13 |
| TcCLB.424795.10 | hypothetical protein, conserved | 1,24 | 2,345E-22 |
| TcCLB.510343.15 | protein kinase, putative | 1,24 | 8,951E-09 |
| TcCLB.503715.20 | hypothetical protein | 1,238 | 2,87E-21 |
| TcCLB.509013.10 | calpain-like cysteine peptidase, putative | 1,238 | 2,854E-16 |
| TcCLB.510925.9 | dispersed gene family protein 1 (DGF-1), putative (fragme | 1,238 | 0,001382 |
| TcCLB.445553.10 | hypothetical protein, conserved | 1,237 | 5,898E-23 |

| | | | |
|---|---|---|---|
| TcCLB.508349.30 | hypothetical protein, conserved | 1,236 | 6,442E-54 |
| TcCLB.507539.40 | cell cycle sequence binding phosphoprotein (RBP33), puta | 1,236 | 1,284E-40 |
| TcCLB.511517.60 | C-1-tetrahydrofolate synthase, cytoplasmic, putative | 1,236 | 9,223E-25 |
| TcCLB.508507.59 | NADH dehydrogenase [ubiquinone] iron-sulfur protein 7, r | 1,235 | 1,346E-27 |
| TcCLB.509535.9 | electon transport protein SCO1/SCO2, putative | 1,235 | 1,264E-17 |
| TcCLB.508419.20 | dispersed gene family protein 1 (DGF-1, pseudogene), put | 1,235 | 6,278E-14 |
| TcCLB.508119.144 | ribosomal protein L38, putative | 1,234 | 4,985E-34 |
| TcCLB.504839.44 | tryparedoxin peroxidase | 1,233 | 1,834E-54 |
| TcCLB.508741.340 | hypothetical protein, conserved | 1,233 | 1,266E-21 |
| TcCLB.508221.840 | hypothetical protein | 1,233 | 0,004223 |
| TcCLB.510575.5 | proton motive ATPase, putative | 1,232 | 4,444E-26 |
| TcCLB.507817.50 | hypothetical protein, conserved | 1,232 | 1,921E-19 |
| TcCLB.507221.50 | hypothetical protein, conserved | 1,232 | 6,179E-18 |
| TcCLB.505071.50 | hypothetical protein, conserved | 1,232 | 1,921E-14 |
| TcCLB.507907.55 | ATP-dependent DEAD/H RNA helicase (pseudogene), puta | 1,232 | 2,899E-11 |
| TcCLB.509151.130 | deoxyuridine triphosphatase, putative | 1,231 | 2,241E-45 |
| TcCLB.508567.100 | hypothetical protein, conserved | 1,231 | 7,524E-41 |
| TcCLB.511261.160 | pyruvate dehydrogenase E1 component alpha subunit, put | 1,231 | 4,055E-28 |
| TcCLB.447255.4 | ubiquitin, putative | 1,231 | 2,257E-17 |
| TcCLB.510511.9 | Na/H antiporter-like protein, putative | 1,231 | 2,608E-16 |
| TcCLB.510041.20 | zinc carboxypeptidase, putative | 1,23 | 1,129E-09 |
| TcCLB.510431.274 | 60S ribosomal protein L37, putative | 1,229 | 1,189E-57 |
| TcCLB.506295.70 | hypothetical protein, conserved | 1,229 | 2,402E-39 |
| TcCLB.510339.50 | spermidine synthase, putative | 1,229 | 4,511E-26 |
| TcCLB.503531.40 | hypothetical protein, conserved | 1,229 | 4,107E-12 |
| TcCLB.504907.9 | dispersed gene family protein 1 (DGF-1), putative (fragme | 1,229 | 8,791E-12 |
| TcCLB.506685.40 | hypothetical protein, conserved | 1,228 | 1,611E-31 |
| TcCLB.509963.30 | tryptophanyl-tRNA synthetase (pseudogene), putative | 1,228 | 0,002128 |
| TcCLB.504149.160 | phosphoinositide-specific phospholipase C, putative | 1,227 | 4,604E-41 |
| TcCLB.507473.20 | hypothetical protein | 1,227 | 3,566E-20 |
| TcCLB.511293.69 | glycosomal malate dehydrogenase, putative | 1,226 | 1,513E-75 |
| TcCLB.503483.9 | retrotransposon hot spot (RHS) protein, putative | 1,226 | 1,398E-70 |
| TcCLB.509875.240 | hypothetical protein, conserved | 1,226 | 7,057E-41 |
| TcCLB.509213.160 | protein kinase, putative | 1,226 | 4,202E-37 |
| TcCLB.511759.30 | hypothetical protein, conserved | 1,226 | 1,071E-31 |
| TcCLB.510095.70 | retrotransposon hot spot protein (RHS, pseudogene), puta | 1,226 | 2,204E-23 |
| TcCLB.447255.10 | hypothetical protein, conserved | 1,226 | 2,326E-22 |
| TcCLB.506743.110 | protein tyrosine phosphatase, putative | 1,226 | 3,122E-17 |
| TcCLB.509039.4 | aspartyl-tRNA synthetase, putative | 1,225 | 4,9E-16 |
| TcCLB.510309.40 | 60S acidic ribosomal protein P2, putative | 1,225 | 1,424E-14 |
| TcCLB.507603.250 | hypothetical protein, conserved | 1,224 | 5,226E-36 |
| TcCLB.463451.10 | hypothetical protein, conserved | 1,223 | 6,301E-33 |
| TcCLB.506999.90 | rieske iron-sulfur protein, mitochondrial precursor, putativ | 1,222 | 1,661E-48 |
| TcCLB.504149.175 | hypothetical protein, conserved | 1,222 | 1,158E-39 |
| TcCLB.504041.20 | hypothetical protein, conserved | 1,222 | 6,72E-21 |

| | | | |
|---|---|---|---|
| TcCLB.510385.10 | dispersed gene family protein 1 (DGF-1), putative | 1,222 | 2,264E-06 |
| TcCLB.509373.10 | trans-sialidase (pseudogene), putative | 1,221 | 1,411E-07 |
| TcCLB.509761.10 | hypothetical protein, conserved | 1,22 | 2,042E-59 |
| TcCLB.509945.30 | hypothetical protein | 1,22 | 2,427E-48 |
| TcCLB.507667.20 | hypothetical protein, conserved | 1,22 | 2,175E-46 |
| TcCLB.508799.80 | mRNA cap guanine-N7 methyltransferase, putative | 1,22 | 5,483E-25 |
| TcCLB.458759.10 | profilin, putative | 1,22 | 1,176E-16 |
| TcCLB.506053.10 | amino acid transporter, putative | 1,218 | 4,559E-68 |
| TcCLB.511825.100 | hypothetical protein, conserved | 1,218 | 1,746E-25 |
| TcCLB.509237.70 | UDP-Gal or UDP-GlcNAc-dependent glycosyltransferase, p | 1,217 | 3,193E-11 |
| TcCLB.507445.50 | myosin heavy chain, putative | 1,216 | 1,165E-42 |
| TcCLB.511621.90 | esterase, putative | 1,215 | 1,028E-17 |
| TcCLB.506487.24 | dispersed gene family protein 1 (DGF-1), putative | 1,215 | 1,068E-10 |
| TcCLB.508837.120 | prohibitin, putative | 1,214 | 3,451E-45 |
| TcCLB.509033.80 | hypothetical protein, conserved | 1,214 | 4,312E-33 |
| TcCLB.507603.270 | cruzipain precursor, putative | 1,214 | 1,974E-28 |
| TcCLB.507769.30 | hypothetical protein, conserved | 1,213 | 1,033E-41 |
| TcCLB.509237.100 | hypothetical protein, conserved | 1,213 | 2,298E-19 |
| TcCLB.508069.10 | retrotransposon hot spot protein (RHS, pseudogene), puta | 1,213 | 1,842E-14 |
| TcCLB.503617.40 | ATP synthase F1, alpha subunit, putative (fragment) (ATP | 1,212 | 9,683E-46 |
| TcCLB.506941.180 | hypothetical protein, conserved | 1,212 | 3,708E-28 |
| TcCLB.510305.70 | hypothetical protein, conserved | 1,212 | 4,607E-23 |
| TcCLB.511069.4 | transketolase, putative | 1,212 | 1,135E-14 |
| TcCLB.507501.30 | hypothetical protein, conserved | 1,212 | 2,888E-06 |
| TcCLB.506925.550 | calpain-like cysteine peptidase (pseudogene), putative | 1,211 | 4,842E-11 |
| TcCLB.507943.20 | histone H4, putative | 1,21 | 5,878E-40 |
| TcCLB.511647.60 | hypothetical protein, conserved | 1,21 | 3,11E-32 |
| TcCLB.508265.44 | mitochondrial pyruvate carrier protein 2, putative | 1,21 | 8,215E-23 |
| TcCLB.509583.4 | 60S ribosomal protein L44, putative | 1,209 | 9,41E-50 |
| TcCLB.506775.170 | NADH-cytochrome b5 reductase, putative | 1,209 | 9,35E-31 |
| TcCLB.504427.240 | metallo-beta-lactamase-like protein, putative | 1,209 | 1,555E-18 |
| TcCLB.503527.40 | GPI alpha-mannosyltransferase III (GPI10) | 1,209 | 5,351E-16 |
| TcCLB.511353.34 | hypothetical protein, conserved | 1,208 | 1,918E-25 |
| TcCLB.510359.260 | mitochondrial import inner membrane translocase subuni | 1,208 | 2,263E-24 |
| TcCLB.504795.9 | dispersed gene family protein 1 (DGF-1), putative | 1,208 | 3,984E-10 |
| TcCLB.506219.10 | hypothetical protein, conserved | 1,207 | 3,875E-32 |
| TcCLB.510139.50 | retrotransposon hot spot protein (RHS, pseudogene), puta | 1,207 | 6,037E-16 |
| TcCLB.511001.240 | proteasome activator protein pa26, putative | 1,207 | 4,841E-12 |
| TcCLB.504717.10 | divalent cation tolerance protein, putative | 1,207 | 0,005899 |
| TcCLB.510627.179 | syntaxin binding protein 1 (pseudogene), putative | 1,207 | 0,006425 |
| TcCLB.506321.280 | hypothetical protein, conserved | 1,206 | 4,431E-63 |
| TcCLB.511581.10 | fatty acyl CoA synthetase 2, putative | 1,206 | 5,885E-20 |
| TcCLB.510877.160 | hypothetical protein, conserved | 1,204 | 2,556E-28 |
| TcCLB.508169.69 | peptidyl-prolyl cis-trans isomerase, putative | 1,204 | 3,459E-27 |
| TcCLB.505115.20 | hypothetical protein | 1,204 | 0,0001661 |

| | | | |
|---|---|---|---|
| TcCLB.508175.360 | cystathionine beta-synthase (CBS) | 1,204 | 0,001557 |
| TcCLB.509669.160 | hypothetical protein, conserved | 1,203 | 2,749E-34 |
| TcCLB.509179.150 | otubain cysteine peptidase, Clan CA, family C65, putative | 1,203 | 2,882E-31 |
| TcCLB.511817.80 | protein kinase, putative | 1,203 | 1,673E-25 |
| TcCLB.507167.150 | retrotransposon hot spot protein (RHS, pseudogene), puta | 1,203 | 2,158E-18 |
| TcCLB.505983.9 | tryparedoxin peroxidase, putative | 1,202 | 3,746E-27 |
| TcCLB.509141.30 | hypothetical protein | 1,202 | 4,203E-19 |
| TcCLB.508283.69 | dispersed gene family protein 1 (DGF-1), putative (fragme | 1,202 | 1,162E-09 |
| TcCLB.511571.20 | hypothetical protein, conserved | 1,201 | 2,745E-35 |
| TcCLB.511925.20 | hypothetical protein, conserved | 1,201 | 8,404E-26 |
| TcCLB.509961.70 | dispersed gene family protein 1 (DGF-1, pseudogene), put | 1,201 | 9,971E-17 |
| TcCLB.507089.270 | dihydrolipoyl dehydrogenase, putative (GCVL-2) | 1,2 | 2,375E-48 |
| TcCLB.509767.70 | V-type proton ATPase subunit a, putative (VAa) | 1,2 | 7,175E-35 |
| TcCLB.511127.80 | hypothetical protein, conserved | 1,2 | 1,262E-31 |
| TcCLB.508647.174 | hypothetical protein, conserved | 1,2 | 5,3E-15 |
| TcCLB.509543.50 | DNAK protein, putative | 1,199 | 1,341E-42 |
| TcCLB.508785.10 | Present in the outer mitochondrial membrane proteome 1 | 1,199 | 3,388E-23 |
| TcCLB.506025.50 | Vacuolar proton pump subunit B, putative | 1,198 | 1,547E-44 |
| TcCLB.508839.50 | hypothetical protein, conserved | 1,197 | 7,216E-38 |
| TcCLB.509549.120 | endosomal integral membrane protein (pseudogene), put | 1,197 | 2,675E-27 |
| TcCLB.511277.160 | 60S ribosomal protein L10a, putative | 1,196 | 3,131E-101 |
| TcCLB.509099.50 | trypanothione synthetase (TRYS) | 1,196 | 6,735E-50 |
| TcCLB.505163.20 | hypothetical protein, conserved | 1,196 | 1,369E-17 |
| TcCLB.506247.340 | hypothetical protein, conserved | 1,195 | 4,407E-39 |
| TcCLB.506355.140 | hypothetical protein, conserved | 1,195 | 1,799E-35 |
| TcCLB.510659.20 | hypothetical protein, conserved | 1,195 | 9,747E-19 |
| TcCLB.510101.470 | PUF nine target 1 | 1,194 | 1,093E-46 |
| TcCLB.509965.290 | p22 protein precursor, putative | 1,194 | 8,722E-33 |
| TcCLB.511277.486 | hypothetical protein, conserved | 1,194 | 1,219E-16 |
| TcCLB.506845.40 | hypothetical protein, conserved (pseudogene) | 1,194 | 3,129E-11 |
| TcCLB.508023.50 | hypothetical protein | 1,194 | 6,91E-10 |
| TcCLB.508465.90 | galactokinase, putative | 1,193 | 2,287E-27 |
| TcCLB.504579.10 | retrotransposon hot spot protein (RHS, pseudogene), puta | 1,193 | 9,276E-16 |
| TcCLB.507873.50 | hypothetical protein, conserved | 1,193 | 1,484E-12 |
| TcCLB.507401.9 | dispersed gene family protein 1 (DGF-1), putative (fragme | 1,193 | 8,371E-09 |
| TcCLB.506595.90 | damage-specific DNA binding protein (pseudogene), puta | 1,192 | 7,781E-26 |
| TcCLB.509331.50 | ribosomal RNA methyltransferase, putative | 1,191 | 7,458E-21 |
| TcCLB.506593.19 | ribosomal protein S7, putative | 1,19 | 8,479E-58 |
| TcCLB.510769.60 | hypothetical protein, conserved (pseudogene) | 1,19 | 4,558E-32 |
| TcCLB.505985.9 | calpain cysteine peptidase, putative (fragment) | 1,189 | 1,411E-16 |
| TcCLB.506129.10 | dispersed gene family protein 1 (DGF-1, pseudogene), put | 1,189 | 2,365E-09 |
| TcCLB.510079.40 | dispersed gene family protein 1 (DGF-1, pseudogene), put | 1,189 | 0,0007892 |
| TcCLB.511233.7 | 60S ribosomal protein L10a, putative | 1,188 | 6,012E-37 |
| TcCLB.506351.70 | hypothetical protein, conserved | 1,188 | 3,117E-27 |
| TcCLB.507195.40 | trans-sialidase (pseudogene), putative | 1,188 | 0,003076 |

| | | | |
|---|---|---|---|
| TcCLB.511295.60 | hypothetical protein, conserved (pseudogene) | 1,187 | 5,032E-34 |
| TcCLB.503505.10 | 60S acidic ribosomal protein P2, putative | 1,187 | 1,666E-26 |
| TcCLB.508973.69 | hypothetical protein, conserved | 1,187 | 5,042E-20 |
| TcCLB.503751.20 | hypothetical protein | 1,187 | 4,863E-06 |
| TcCLB.510293.30 | Eukaryotic translation initiation factor 4E type 5, putative | 1,186 | 1,747E-32 |
| TcCLB.506721.30 | calpain-like cysteine peptidase, putative | 1,186 | 9,427E-27 |
| TcCLB.508815.150 | hypothetical protein, conserved | 1,186 | 9,106E-14 |
| TcCLB.507047.180 | hypothetical protein, conserved | 1,186 | 4,106E-12 |
| TcCLB.510325.20 | hypothetical protein | 1,186 | 1,112E-06 |
| TcCLB.506207.50 | translationally controlled tumor protein (TCTP), putative | 1,184 | 3,313E-66 |
| TcCLB.509151.140 | 60S ribosomal protein L23a, putative | 1,183 | 6,926E-44 |
| TcCLB.503801.30 | DNA repair protein RAD51, putative (RAD51) | 1,183 | 1,221E-42 |
| TcCLB.509967.20 | mismatch repair protein MSH4, putative | 1,183 | 1,536E-22 |
| TcCLB.506641.9 | dispersed gene family protein 1 (DGF-1), putative | 1,183 | 0,00002928 |
| TcCLB.507537.20 | cysteine peptidase, putative | 1,182 | 6,702E-38 |
| TcCLB.511461.40 | hypothetical protein, conserved | 1,182 | 1,303E-24 |
| TcCLB.503839.39 | hypothetical protein, conserved | 1,182 | 1,655E-12 |
| TcCLB.510397.5 | GTP-binding protein, putative (fragment) | 1,181 | 5,822E-20 |
| TcCLB.508859.74 | SNF2 DNA repair protein, putative | 1,18 | 6,623E-48 |
| TcCLB.503521.50 | UTP-glucose-1-phosphate uridylyltransferase 2, putative | 1,18 | 6,553E-18 |
| TcCLB.511751.205 | hypothetical protein | 1,179 | 7,086E-32 |
| TcCLB.507537.10 | cruzipain precursor (pseudogene), putative | 1,178 | 1,85E-42 |
| TcCLB.510325.40 | acyltransferase, putative | 1,178 | 2,408E-25 |
| TcCLB.430895.16 | hypothetical protein, conserved (fragment) | 1,178 | 3,139E-10 |
| TcCLB.504001.10 | hypothetical protein, conserved | 1,176 | 2,337E-44 |
| TcCLB.508479.370 | hypothetical protein, conserved | 1,176 | 2,857E-21 |
| TcCLB.510755.65 | hypothetical protein | 1,176 | 3,11E-12 |
| TcCLB.510847.70 | dispersed gene family protein 1 (DGF-1), putative | 1,176 | 8,327E-08 |
| TcCLB.505943.110 | retrotransposon hot spot protein (RHS, pseudogene), puta | 1,175 | 2,881E-18 |
| TcCLB.506777.20 | cytoplasmic translation machinery associated protein, put | 1,174 | 5,572E-25 |
| TcCLB.510079.10 | retrotransposon hot spot protein (RHS, pseudogene), puta | 1,174 | 8,779E-23 |
| TcCLB.508387.60 | hypothetical protein, conserved | 1,174 | 5,874E-16 |
| TcCLB.504099.10 | retrotransposon hot spot (RHS) protein, putative | 1,174 | 1,4E-14 |
| TcCLB.505995.10 | dispersed gene family protein 1 (DGF-1), putative | 1,174 | 0,00007033 |
| TcCLB.508779.90 | structural maintenance of chromosome protein 4, putativ | 1,173 | 2,494E-35 |
| TcCLB.511261.30 | NADH-ubiquinone oxidoreductase complex I subunit, puta | 1,173 | 6,367E-30 |
| TcCLB.508523.40 | dispersed gene family protein 1 (DGF-1), putative | 1,173 | 2,103E-21 |
| TcCLB.510135.10 | retrotransposon hot spot protein (RHS, pseudogene), puta | 1,173 | 1,542E-14 |
| TcCLB.506997.9 | dispersed gene family protein 1 (DGF-1), putative | 1,173 | 0,0001527 |
| TcCLB.507015.10 | retrotransposon hot spot (RHS) protein, putative | 1,172 | 1,945E-46 |
| TcCLB.511393.70 | hypothetical protein, conserved | 1,172 | 2,624E-33 |
| TcCLB.510643.10 | dispersed gene family protein 1 (DGF-1), putative | 1,172 | 1,277E-10 |
| TcCLB.509029.60 | hypothetical protein, conserved | 1,171 | 1,345E-46 |
| TcCLB.511289.90 | hypothetical protein, conserved | 1,171 | 9,047E-32 |
| TcCLB.506591.30 | hypothetical protein, conserved | 1,17 | 1,374E-31 |

| | | | |
|---|---|---|---|
| TcCLB.503893.130 | hypothetical protein, conserved | 1,17 | 1,647E-15 |
| TcCLB.504199.10 | hypothetical protein, conserved | 1,169 | 1,01E-42 |
| TcCLB.505939.30 | hypothetical protein, conserved | 1,169 | 4,587E-37 |
| TcCLB.510609.40 | ATP synthase F1, alpha subunit, putative (pseudogene) | 1,167 | 1,99E-52 |
| TcCLB.511865.60 | hypothetical protein, conserved | 1,167 | 3,242E-33 |
| TcCLB.508017.10 | hypothetical protein, conserved | 1,167 | 0,00006874 |
| TcCLB.506945.230 | 40S ribosomal protein S14, putative | 1,166 | 1,051E-39 |
| TcCLB.510777.20 | aspartyl-tRNA synthetase, putative | 1,166 | 5,94E-39 |
| TcCLB.511527.70 | Unconventional myosin | 1,166 | 7,901E-26 |
| TcCLB.506941.280 | heat shock 70 kDa protein, mitochondrial precursor, putati | 1,166 | 1,637E-10 |
| TcCLB.509857.60 | hypothetical protein, conserved | 1,165 | 4,558E-43 |
| TcCLB.509749.10 | Biotin--acetyl-CoA-carboxylase ligase, putative | 1,165 | 1,276E-24 |
| TcCLB.510395.20 | hypothetical protein, conserved | 1,165 | 9,527E-15 |
| TcCLB.509437.103 | dispersed gene family protein 1 (DGF-1, pseudogene), put | 1,165 | 0,0004461 |
| TcCLB.507711.190 | hypothetical protein, conserved | 1,164 | 1,591E-52 |
| TcCLB.506403.9 | hypothetical protein, conserved (fragment) | 1,164 | 1,839E-33 |
| TcCLB.507061.34 | hypothetical protein, conserved | 1,164 | 3,206E-17 |
| TcCLB.510073.75 | hypothetical protein, conserved | 1,164 | 0,00003389 |
| TcCLB.509693.80 | hypoxanthine-guanine phosphoribosyltransferase, putative | 1,163 | 6,488E-26 |
| TcCLB.503721.10 | hypothetical protein, conserved | 1,163 | 4,102E-24 |
| TcCLB.505763.10 | proton motive ATPase 1, putative | 1,162 | 4,174E-77 |
| TcCLB.509023.180 | kinesin, putative | 1,162 | 2,973E-41 |
| TcCLB.507669.60 | hypothetical protein, conserved | 1,162 | 8,887E-22 |
| TcCLB.508375.80 | hypothetical protein, conserved | 1,162 | 1,281E-17 |
| TcCLB.507223.10 | dynein heavy chain, cytosolic, putative | 1,162 | 4,027E-08 |
| TcCLB.509695.100 | hypothetical protein, conserved | 1,161 | 2,001E-24 |
| TcCLB.508487.20 | dispersed gene family protein 1 (DGF-1, pseudogene), put | 1,161 | 2,814E-09 |
| TcCLB.509921.9 | dispersed gene family protein 1 (DGF-1), putative | 1,161 | 0,0002949 |
| TcCLB.504153.160 | metallo-peptidase, Clan MA(E) Family M32 | 1,16 | 1,915E-69 |
| TcCLB.507381.41 | trans-sialidase (pseudogene), putative | 1,16 | 2,676E-10 |
| TcCLB.509011.10 | UDP-Gal or UDP-GlcNAc-dependent glycosyltransferase (p | 1,16 | 4,086E-09 |
| TcCLB.507693.10 | dispersed gene family protein 1 (DGF-1), putative | 1,16 | 4,607E-08 |
| TcCLB.509859.40 | cytosolic leucyl aminopeptidase, putative | 1,159 | 9,947E-39 |
| TcCLB.503881.20 | GDP-mannose 4,6 dehydratase, putative | 1,159 | 1,554E-28 |
| TcCLB.509231.23 | hypothetical protein, conserved | 1,159 | 4,98E-25 |
| TcCLB.432027.9 | hypothetical protein, conserved | 1,159 | 2,481E-12 |
| TcCLB.509895.20 | mitochondrial RNA binding protein, putative | 1,158 | 5,988E-45 |
| TcCLB.509151.70 | hypothetical protein, conserved | 1,158 | 1,614E-40 |
| TcCLB.507773.34 | 60S ribosomal protein L37, putative | 1,158 | 1,12E-34 |
| TcCLB.510851.21 | retrotransposon hot spot protein (RHS, pseudogene), puta | 1,158 | 1,023E-33 |
| TcCLB.509599.130 | serine/threonine protein kinase, putative | 1,158 | 1,158E-28 |
| TcCLB.507521.110 | pumilio/PUF RNA binding protein 5, putative (PUF5) | 1,158 | 2,657E-25 |
| TcCLB.508981.20 | hypothetical protein, conserved | 1,158 | 1,537E-14 |
| TcCLB.508389.50 | mucin-associated surface protein (MASP), putative | 1,158 | 0,00002892 |
| TcCLB.509473.10 | hypothetical protein (pseudogene) | 1,157 | 3,408E-21 |

| | | | |
|---|---|---|---|
| TcCLB.510173.130 | hypothetical protein, conserved | 1,157 | 6,815E-19 |
| TcCLB.503413.19 | hypothetical protein | 1,157 | 1,119E-17 |
| TcCLB.510325.69 | Aminopeptidase M1, putative (fragment) | 1,156 | 1,272E-33 |
| TcCLB.508699.70 | hypothetical protein, conserved | 1,156 | 3,508E-31 |
| TcCLB.507927.90 | hypothetical protein, conserved | 1,156 | 2,197E-14 |
| TcCLB.503433.30 | hypothetical protein, conserved | 1,155 | 1,703E-28 |
| TcCLB.510643.150 | ribosomal P protein AGP2beta-1 (pseudogene), putative | 1,155 | 1,378E-14 |
| TcCLB.510451.21 | trans-sialidase (pseudogene), putative | 1,155 | 0,004707 |
| TcCLB.509429.59 | hypothetical protein, conserved | 1,154 | 3,724E-32 |
| TcCLB.507603.140 | leucine-rich repeat protein (LRRP), putative | 1,154 | 3,439E-26 |
| TcCLB.510719.74 | hypothetical protein, conserved | 1,154 | 9,152E-16 |
| TcCLB.506493.80 | DNA topoisomerase IA, putative | 1,153 | 5,537E-41 |
| TcCLB.506351.9 | 5'-3' exoribonuclease A, putative (fragment) (XRNA) | 1,153 | 5,618E-18 |
| TcCLB.415789.9 | dispersed gene family protein 1 (DGF-1), putative (fragme | 1,153 | 6,053E-15 |
| TcCLB.506435.270 | dihydroxyacetone phosphate acyltransferase, putative (DA | 1,152 | 3,399E-30 |
| TcCLB.509633.50 | serine/threonine-protein phosphatase PP1, putative | 1,152 | 7,675E-07 |
| TcCLB.506987.10 | retrotransposon hot spot protein (RHS, pseudogene), puta | 1,152 | 0,0000138 |
| TcCLB.511465.10 | proteasome activator protein pa26, putative | 1,151 | 1,553E-19 |
| TcCLB.506559.480 | hypothetical protein, conserved | 1,151 | 5,291E-13 |
| TcCLB.506559.350 | asparaginyl-tRNA synthetase, putative | 1,15 | 5,125E-59 |
| TcCLB.509381.20 | 40S ribosomal protein S15A, putative | 1,15 | 1,513E-44 |
| TcCLB.511587.30 | retrotransposon hot spot protein (RHS, pseudogene), puta | 1,15 | 1,471E-39 |
| TcCLB.510101.240 | 60S ribosomal protein L24, putative | 1,15 | 4,569E-20 |
| TcCLB.507925.40 | pyridoxal kinase, putative | 1,15 | 1,312E-13 |
| TcCLB.506961.130 | retrotransposon hot spot protein (RHS, pseudogene), puta | 1,149 | 4,634E-29 |
| TcCLB.507663.30 | hypothetical protein, conserved | 1,149 | 2,194E-23 |
| TcCLB.504769.170 | retrotransposon hot spot protein (RHS, pseudogene), puta | 1,149 | 1,852E-21 |
| TcCLB.508799.220 | hypothetical protein, conserved (pseudogene) | 1,149 | 0,0004239 |
| TcCLB.509649.50 | hypothetical protein, conserved | 1,148 | 1,464E-33 |
| TcCLB.504089.60 | hypothetical protein, conserved | 1,148 | 4,181E-29 |
| TcCLB.510237.129 | dispersed gene family protein 1 (DGF-1), putative (fragme | 1,148 | 5,76E-07 |
| TcCLB.504163.30 | 40S ribosomal protein S9, putative | 1,147 | 3,651E-52 |
| TcCLB.508547.30 | acyl-CoA dehydrogenase, mitochondrial precursor, putativ | 1,147 | 1,044E-30 |
| TcCLB.510603.80 | protein tyrosine phosphatase-like protein, putative | 1,146 | 5,666E-19 |
| TcCLB.504033.90 | mitochondrial carrier protein, putative | 1,145 | 1,87E-33 |
| TcCLB.506503.80 | kinesin, putative | 1,145 | 1,739E-27 |
| TcCLB.511237.50 | 40S ribosomal protein S33, putative | 1,144 | 6,464E-40 |
| TcCLB.510381.30 | hypothetical protein, conserved | 1,144 | 1,069E-30 |
| TcCLB.504051.20 | ATP-dependent DNA DEAD/DEAH box helicase, putative | 1,144 | 3,509E-25 |
| TcCLB.509449.10 | receptor-type adenylate cyclase, putative | 1,144 | 4,807E-17 |
| TcCLB.504037.30 | 60S ribosomal protein L12, putative | 1,143 | 3,806E-44 |
| TcCLB.509219.20 | retrotransposon hot spot (RHS) protein, putative | 1,143 | 2,037E-37 |
| TcCLB.510101.30 | 60S ribosomal protein L28, putative | 1,143 | 8,826E-36 |
| TcCLB.510759.100 | hypothetical protein, conserved | 1,143 | 2,452E-33 |
| TcCLB.507093.220 | RNA-binding protein, putative | 1,143 | 2,271E-30 |

| | | | |
|---|---|---|---|
| TcCLB.504125.40 | ATPase subunit 9, putative | 1,143 | 5,817E-27 |
| TcCLB.506025.14 | ribosomal protein S29, putative | 1,143 | 1,359E-24 |
| TcCLB.510359.240 | hypothetical protein, conserved | 1,143 | 3,464E-19 |
| TcCLB.503461.30 | hypothetical protein, conserved | 1,142 | 7,999E-33 |
| TcCLB.508461.60 | synaptojanin (N-terminal domain), putative | 1,142 | 2,688E-25 |
| TcCLB.508995.60 | electron transport protein SCO1/2, putative | 1,142 | 4,328E-17 |
| TcCLB.506147.40 | hypothetical protein | 1,141 | 3,737E-20 |
| TcCLB.510225.59 | transporter, putative | 1,141 | 2,613E-10 |
| TcCLB.510431.110 | hypothetical protein | 1,14 | 1,972E-38 |
| TcCLB.508811.10 | hypothetical protein | 1,14 | 2,352E-19 |
| TcCLB.509207.30 | hypothetical protein, conserved | 1,14 | 1,67E-16 |
| TcCLB.504139.10 | retrotransposon hot spot protein (RHS, pseudogene), puta | 1,14 | 5,28E-13 |
| TcCLB.508543.30 | 40S ribosomal protein S9, putative | 1,139 | 2,114E-58 |
| TcCLB.506595.100 | 60S acidic ribosomal protein P2, putative | 1,139 | 6,051E-39 |
| TcCLB.509725.100 | hypothetical protein, conserved (pseudogene) | 1,139 | 2,158E-18 |
| TcCLB.509767.240 | hypothetical protein, conserved | 1,139 | 1,197E-11 |
| TcCLB.510497.199 | dispersed gene family protein 1 (DGF-1), putative (fragm | 1,139 | 0,001868 |
| TcCLB.507649.70 | nucleolar RNA-binding protein, putative | 1,138 | 1,766E-54 |
| TcCLB.508817.50 | DNA repair protein RAD51, putative (RAD51) | 1,138 | 5,741E-44 |
| TcCLB.504741.20 | GTP-binding protein, putative | 1,138 | 2,285E-29 |
| TcCLB.507669.20 | hypothetical protein, conserved | 1,138 | 1,441E-26 |
| TcCLB.509743.19 | dispersed gene family protein 1 (DGF-1), putative (fragm | 1,138 | 1,867E-08 |
| TcCLB.508283.10 | dispersed gene family protein 1 (DGF-1), putative | 1,138 | 2,006E-07 |
| TcCLB.508441.20 | Phosphoenolpyruvate carboxykinase [ATP], glycosomal | 1,137 | 6,689E-48 |
| TcCLB.503781.90 | universal minicircle sequence binding protein 2, putative ( | 1,137 | 3,316E-34 |
| TcCLB.507547.90 | Phosphoenolpyruvate carboxykinase [ATP], glycosomal | 1,136 | 7,279E-41 |
| TcCLB.437121.9 | hypothetical protein, conserved (fragment) | 1,136 | 7,182E-22 |
| TcCLB.506715.44 | hypothetical protein, conserved | 1,135 | 7,045E-14 |
| TcCLB.510155.20 | KREL1 | 1,134 | 2,216E-31 |
| TcCLB.506633.109 | hypothetical protein (fragment) | 1,134 | 4,63E-17 |
| TcCLB.418221.30 | hypothetical protein, conserved | 1,133 | 4,828E-45 |
| TcCLB.506829.39 | ribosomal protein S7, putative | 1,132 | 8,137E-59 |
| TcCLB.511145.46 | 60S ribosomal protein L37a, putative | 1,132 | 1,192E-34 |
| TcCLB.508173.100 | Monooxygenase, putative | 1,132 | 1,445E-24 |
| TcCLB.508909.190 | hypothetical protein, conserved | 1,132 | 2,278E-21 |
| TcCLB.506209.19 | dispersed gene family protein 1 (DGF-1), putative (fragm | 1,132 | 0,000879 |
| TcCLB.511559.50 | hypothetical protein, conserved | 1,131 | 5,403E-30 |
| TcCLB.509749.36 | ribosomal protein L29, putative | 1,131 | 5,41E-28 |
| TcCLB.507179.20 | dispersed gene family protein 1 (DGF-1), putative | 1,131 | 3,686E-14 |
| TcCLB.509023.30 | hypothetical protein | 1,131 | 1,1E-11 |
| TcCLB.503823.130 | hypothetical protein, conserved | 1,13 | 5,037E-43 |
| TcCLB.508777.110 | hypothetical protein, conserved | 1,13 | 1,165E-37 |
| TcCLB.506727.60 | hypothetical protein, conserved | 1,13 | 5,23E-28 |
| TcCLB.511655.20 | hypothetical protein, conserved | 1,13 | 1,341E-18 |
| TcCLB.509903.10 | dispersed gene family protein 1 (DGF-1, pseudogene), put | 1,13 | 1,51E-17 |

| | | | |
|---|---|---|---|
| TcCLB.509887.19 | dispersed gene family protein 1 (DGF-1), putative | 1,13 | 1,569E-08 |
| TcCLB.510487.20 | dispersed gene family protein 1 (DGF-1, pseudogene), put | 1,13 | 2,227E-06 |
| TcCLB.510877.55 | acyl-CoA binding protein, putative | 1,129 | 4,375E-22 |
| TcCLB.510299.10 | hypothetical protein, conserved | 1,129 | 3,81E-18 |
| TcCLB.509749.30 | hypothetical protein, conserved | 1,129 | 1,537E-14 |
| TcCLB.507645.11 | hypothetical protein, conserved (fragment) | 1,129 | 5,873E-13 |
| TcCLB.509979.90 | 60S ribosomal protein L35, putative | 1,128 | 1,011E-43 |
| TcCLB.506683.140 | beta galactofuranosyl glycosyltransferase (pseudogene), p | 1,128 | 4,969E-19 |
| TcCLB.507539.10 | hypothetical protein, conserved | 1,128 | 8,581E-09 |
| TcCLB.504339.10 | hypothetical protein, conserved | 1,128 | 9,948E-07 |
| TcCLB.503903.50 | hypothetical protein, conserved | 1,127 | 7,165E-27 |
| TcCLB.506325.33 | ATP-dependent DEAD/H RNA helicase (pseudogene), puta | 1,127 | 0,0004555 |
| TcCLB.508577.70 | hypothetical protein | 1,126 | 9,448E-31 |
| TcCLB.507681.170 | hypothetical protein, conserved | 1,126 | 2,4E-21 |
| TcCLB.509551.60 | hypothetical protein, conserved | 1,126 | 2,117E-12 |
| TcCLB.509331.180 | aldose 1-epimerase-like protein, putative | 1,126 | 2,292E-12 |
| TcCLB.509805.84 | hypothetical protein, conserved | 1,126 | 1,436E-11 |
| TcCLB.506639.50 | dispersed gene family protein 1 (DGF-1, pseudogene), put | 1,126 | 5,267E-09 |
| TcCLB.509149.40 | 60S ribosomal protein L19, putative | 1,125 | 4,218E-76 |
| TcCLB.506989.30 | ABC transporter, putative | 1,125 | 4,255E-48 |
| TcCLB.507969.10 | vesicular transport-associated repeat protein, putative | 1,125 | 8,813E-33 |
| TcCLB.510095.10 | hypothetical protein | 1,125 | 0,002229 |
| TcCLB.457101.20 | dynein heavy chain, cytosolic, putative | 1,124 | 7,66E-32 |
| TcCLB.509163.90 | retrotransposon hot spot protein (RHS, pseudogene), puta | 1,124 | 9,33E-26 |
| TcCLB.503733.80 | hypothetical protein, conserved | 1,124 | 2,486E-24 |
| TcCLB.510431.90 | hypothetical protein, conserved | 1,124 | 9,302E-13 |
| TcCLB.508813.15 | heat shock protein 70, putative (fragment) | 1,124 | 8,781E-12 |
| TcCLB.508803.19 | dispersed gene family protein 1 (DGF-1), putative | 1,124 | 1,822E-06 |
| TcCLB.506247.210 | hypothetical protein, conserved | 1,123 | 1,202E-61 |
| TcCLB.508173.10 | retrotransposon hot spot protein (RHS, pseudogene), puta | 1,123 | 1,263E-38 |
| TcCLB.507093.320 | hypothetical protein, conserved | 1,123 | 4,707E-24 |
| TcCLB.506795.29 | hypothetical protein, conserved | 1,123 | 6,313E-19 |
| TcCLB.510151.19 | fatty acyl CoA syntetase 1, putative | 1,123 | 1,808E-18 |
| TcCLB.506325.30 | hypothetical protein, conserved (pseudogene) | 1,123 | 4,236E-18 |
| TcCLB.508177.40 | hypothetical protein, conserved | 1,122 | 1,658E-33 |
| TcCLB.503471.10 | protein tyrosine phosphatase, putative | 1,122 | 3,45E-21 |
| TcCLB.509047.14 | hypothetical protein, conserved | 1,122 | 1,016E-17 |
| TcCLB.510113.20 | dispersed gene family protein 1 (DGF-1, pseudogene), put | 1,122 | 5,877E-12 |
| TcCLB.510357.100 | hypothetical protein, conserved | 1,121 | 1,035E-21 |
| TcCLB.509895.10 | mitochondrial RNA binding complex 1 subunit, putative | 1,12 | 4,126E-67 |
| TcCLB.506931.70 | hypothetical protein, conserved | 1,12 | 2,676E-28 |
| TcCLB.503439.14 | dispersed gene family protein 1 (DGF-1, pseudogene), put | 1,12 | 2,705E-08 |
| TcCLB.508153.100 | acyltransferase, putative | 1,119 | 6,212E-20 |
| TcCLB.506327.90 | serine/threonine protein kinase, putative | 1,118 | 6,862E-39 |
| TcCLB.507787.50 | membrane transporter protein (pseudogene), putative | 1,118 | 4,696E-13 |

| | | | |
|---|---|---|---|
| TcCLB.471403.9 | dispersed gene family protein 1 (DGF-1), putative (fragme | 1,118 | 3,981E-12 |
| TcCLB.511477.40 | dispersed gene family protein 1 (DGF-1, pseudogene), put | 1,118 | 2,837E-10 |
| TcCLB.507091.4 | fucose kinase, putative | 1,118 | 8,785E-10 |
| TcCLB.509769.63 | Ran-binding protein, putative | 1,117 | 1,108E-20 |
| TcCLB.511649.50 | hypothetical protein, conserved | 1,117 | 4,56E-19 |
| TcCLB.506443.150 | retrotransposon hot spot (RHS) protein, putative | 1,116 | 1,883E-53 |
| TcCLB.511647.70 | hypothetical protein, conserved | 1,115 | 3,354E-23 |
| TcCLB.506553.20 | bromodomain factor 2 protein | 1,114 | 8,005E-28 |
| TcCLB.506219.20 | hypothetical protein | 1,114 | 1,921E-12 |
| TcCLB.506153.10 | amino acid transporter, putative | 1,113 | 6,481E-39 |
| TcCLB.507019.86 | 40S ribosomal protein S30, putative | 1,113 | 1,078E-22 |
| TcCLB.507253.30 | hypothetical protein, conserved (pseudogene) | 1,113 | 5,383E-14 |
| TcCLB.504035.84 | succinate dehydrogenase subunit | 1,113 | 2,034E-13 |
| TcCLB.507563.9 | dispersed gene family protein 1 (DGF-1), putative (fragme | 1,113 | 2,667E-13 |
| TcCLB.506323.10 | dispersed gene family protein 1 (DGF-1, pseudogene), put | 1,113 | 2,092E-08 |
| TcCLB.507979.4 | histone H2A, putative | 1,113 | 2,332E-07 |
| TcCLB.509457.10 | hypothetical protein | 1,112 | 1,932E-31 |
| TcCLB.508811.30 | hypothetical protein, conserved | 1,112 | 5,27E-29 |
| TcCLB.508223.30 | retrotransposon hot spot protein (RHS, pseudogene), puta | 1,112 | 0,00001926 |
| TcCLB.509499.14 | peroxidoxin | 1,111 | 1,25E-35 |
| TcCLB.503893.80 | NADH dehydrogenase [ubiquinone] iron-sulfur protein 7, r | 1,111 | 1,929E-22 |
| TcCLB.506885.410 | hypothetical protein, conserved | 1,111 | 5,022E-16 |
| TcCLB.506859.70 | guide RNA associated protein, GAP2, putative | 1,11 | 5,422E-47 |
| TcCLB.506149.10 | folate/pteridine transporter, putative | 1,11 | 3,805E-17 |
| TcCLB.503735.20 | hypothetical protein | 1,11 | 1,795E-15 |
| TcCLB.410243.10 | hypothetical protein, conserved (fragment) | 1,11 | 9,232E-11 |
| TcCLB.505945.30 | ribonuclease mar1, putative | 1,109 | 1,074E-22 |
| TcCLB.511773.40 | hypothetical protein, conserved | 1,109 | 0,00005443 |
| TcCLB.510243.10 | hypothetical protein, conserved | 1,108 | 1,034E-14 |
| TcCLB.511291.90 | 40S ribosomal protein S13, putative | 1,107 | 4,292E-47 |
| TcCLB.506135.40 | hypothetical protein, conserved | 1,107 | 6,492E-30 |
| TcCLB.510173.70 | MIX protein, putative | 1,107 | 2,202E-15 |
| TcCLB.487507.10 | tryparedoxin peroxidase, putative | 1,107 | 3,491E-13 |
| TcCLB.510439.30 | hypothetical protein, conserved | 1,106 | 3,224E-34 |
| TcCLB.509001.40 | calpain-like cysteine peptidase, putative | 1,105 | 4,914E-40 |
| TcCLB.506459.270 | hypothetical protein, conserved | 1,105 | 5,127E-38 |
| TcCLB.506519.130 | guanosine monophosphate reductase, putative | 1,105 | 1,599E-33 |
| TcCLB.510339.20 | hypothetical protein | 1,105 | 1,423E-22 |
| TcCLB.504229.4 | retrotransposon hot spot (RHS) protein, putative | 1,105 | 2,045E-12 |
| TcCLB.509175.9 | hypothetical protein, conserved | 1,105 | 0,00001856 |
| TcCLB.511653.69 | DNA repair protein, putative (fragment) | 1,104 | 5,11E-40 |
| TcCLB.397937.10 | pumilio/PUF RNA binding protein 1, putative (PUF1) | 1,104 | 5,264E-18 |
| TcCLB.510671.10 | dispersed gene family protein 1 (DGF-1), putative | 1,104 | 1,128E-10 |
| TcCLB.503779.20 | hypothetical protein, conserved | 1,103 | 2,15E-21 |
| TcCLB.504313.40 | hypothetical protein, conserved (pseudogene) | 1,103 | 1,057E-15 |

| | | | |
|---|---|---|---|
| TcCLB.509871.70 | retrotransposon hot spot protein (RHS, pseudogene), puta | 1,103 | 1,52E-13 |
| TcCLB.508893.30 | hypothetical protein, conserved | 1,103 | 7,827E-13 |
| TcCLB.511807.60 | hypothetical protein, conserved | 1,102 | 2,114E-28 |
| TcCLB.509455.80 | GTP-binding nuclear protein rtb2, putative | 1,102 | 4,686E-27 |
| TcCLB.511357.9 | hypothetical protein, conserved | 1,102 | 1,094E-24 |
| TcCLB.506401.360 | hypothetical protein, conserved | 1,102 | 1,008E-10 |
| TcCLB.509275.10 | elongation factor 1-gamma (EF-1-gamma, pseudogene), | 1,102 | 0,0002399 |
| TcCLB.510661.80 | hypothetical protein | 1,101 | 6,355E-32 |
| TcCLB.510363.30 | retrotransposon hot spot protein (RHS, pseudogene), puta | 1,101 | 0,0003164 |
| TcCLB.506959.40 | enoyl-CoA hydratase/isomerase family protein, putative | 1,1 | 6,112E-30 |
| TcCLB.508881.54 | hypothetical protein, conserved | 1,1 | 1,443E-17 |
| TcCLB.511517.5 | 70 kDa heat shock protein, putative (fragment) | 1,1 | 6,142E-09 |
| TcCLB.508587.40 | dispersed gene family protein 1 (DGF-1, pseudogene), put | 1,1 | 3,152E-08 |
| TcCLB.506263.20 | D-isomer specific 2-hydroxyacid dehydrogenase-protein, p | 1,099 | 3,988E-33 |
| TcCLB.504057.97 | hypothetical protein, conserved | 1,099 | 6,429E-17 |
| TcCLB.511261.50 | hypothetical protein, conserved | 1,099 | 9,515E-16 |
| TcCLB.509557.10 | hypothetical protein | 1,099 | 3,619E-15 |
| TcCLB.507517.104 | hypothetical protein, conserved | 1,099 | 1,715E-12 |
| TcCLB.506679.230 | ribosomal protein L24, putative | 1,098 | 1,624E-41 |
| TcCLB.509791.80 | hypothetical protein | 1,098 | 5,639E-38 |
| TcCLB.510155.110 | hypothetical protein, conserved | 1,098 | 6,258E-21 |
| TcCLB.511007.20 | retrotransposon hot spot (RHS) protein, putative | 1,098 | 8,71E-12 |
| TcCLB.506831.40 | 3-hydroxy-3-methylglutaryl-CoA reductase (HMGR) | 1,097 | 2,26E-38 |
| TcCLB.511285.80 | hypothetical protein | 1,097 | 3,713E-35 |
| TcCLB.506495.9 | hypothetical protein, conserved | 1,097 | 5,339E-22 |
| TcCLB.511391.90 | endosomal integral membrane protein, putative | 1,097 | 9,912E-21 |
| TcCLB.511821.90 | hypothetical protein, conserved | 1,097 | 2,04E-14 |
| TcCLB.511809.130 | 40S ribosomal protein S15, putative | 1,097 | 3,461E-14 |
| TcCLB.506239.10 | hypothetical protein, conserved | 1,097 | 1,824E-12 |
| TcCLB.511825.50 | centrin-4, putative (Centrin4) | 1,096 | 5,259E-31 |
| TcCLB.510743.100 | chaperone DNAJ protein, putative | 1,095 | 1,352E-31 |
| TcCLB.510073.17 | cysteine peptidase (C-terminal), putative | 1,095 | 1,987E-31 |
| TcCLB.509815.90 | retrotransposon hot spot protein (RHS, pseudogene), puta | 1,094 | 3,249E-17 |
| TcCLB.511323.50 | NADH-ubiquinone oxidoreductase, mitochondrial, putative | 1,094 | 6,668E-13 |
| TcCLB.511393.30 | formin homology 2 (FH2) domain protein, putative | 1,093 | 2,005E-41 |
| TcCLB.511303.120 | hypothetical protein, conserved | 1,093 | 5,991E-41 |
| TcCLB.508649.5 | tryparedoxin peroxidase, putative (fragment) | 1,093 | 1,443E-26 |
| TcCLB.509559.20 | retrotransposon hot spot protein (RHS, pseudogene), puta | 1,093 | 4,025E-21 |
| TcCLB.507221.80 | 5'-3' exoribonuclease C, putative (fragment) (XRNC) | 1,093 | 7,409E-18 |
| TcCLB.508515.20 | translation factor sui1, putative | 1,092 | 1,24E-21 |
| TcCLB.511575.10 | hypothetical protein, conserved | 1,092 | 3,859E-18 |
| TcCLB.509157.279 | Chaperonin HSP60, mitochondrial (HSP60) | 1,092 | 4,778E-11 |
| TcCLB.506563.110 | calpain-like cysteine peptidase (pseudogene), putative | 1,091 | 1,316E-39 |
| TcCLB.511303.70 | hypothetical protein, conserved | 1,091 | 1,802E-19 |
| TcCLB.508771.10 | cysteine peptidase, Clan CA, family C51, putative | 1,091 | 0,0001686 |

| | | | |
|---|---|---|---|
| TcCLB.503757.10 | 40S ribosomal protein SA, putative | 1,09 | 2,386E-31 |
| TcCLB.506833.79 | hypothetical protein, conserved (fragment) | 1,09 | 6,818E-25 |
| TcCLB.511899.10 | 2-amino-3-ketobutyrate coenzyme A ligase, putative | 1,09 | 2,879E-08 |
| TcCLB.510823.70 | 60S acidic ribosomal protein, putative | 1,089 | 2,04E-23 |
| TcCLB.509979.14 | peroxisomal membrane protein 4, putative | 1,089 | 3,357E-22 |
| TcCLB.509919.9 | dispersed gene family protein 1 (DGF-1), putative (fragme | 1,089 | 2,984E-13 |
| TcCLB.506649.90 | hypothetical protein, conserved | 1,088 | 5,244E-57 |
| TcCLB.507515.10 | glutathione peroxidase-like protein, putative | 1,088 | 1,355E-19 |
| TcCLB.503415.10 | hypothetical protein (pseudogene) | 1,088 | 2,454E-16 |
| TcCLB.506129.20 | dispersed gene family protein 1 (DGF-1, pseudogene), put | 1,088 | 0,0003144 |
| TcCLB.494675.10 | fatty acyl CoA synthetase 2, putative | 1,087 | 6,496E-27 |
| TcCLB.507603.90 | hypothetical protein | 1,087 | 1,591E-24 |
| TcCLB.506375.120 | hypothetical protein, conserved | 1,087 | 1,108E-20 |
| TcCLB.482971.9 | dispersed gene family protein 1 (DGF-1), putative (fragme | 1,087 | 0,002544 |
| TcCLB.510645.40 | hypothetical protein, conserved (pseudogene) | 1,086 | 1,623E-57 |
| TcCLB.511421.170 | hypothetical protein, conserved | 1,086 | 1,316E-24 |
| TcCLB.510785.10 | hypothetical protein | 1,086 | 2,591E-23 |
| TcCLB.503401.29 | hypothetical protein, conserved | 1,086 | 5,795E-14 |
| TcCLB.508815.130 | hypothetical protein, conserved | 1,085 | 1,166E-32 |
| TcCLB.509289.30 | hypothetical protein, conserved | 1,085 | 1,311E-31 |
| TcCLB.507969.60 | C-14 sterol reductase, putative | 1,085 | 1,582E-18 |
| TcCLB.510535.30 | hypothetical protein | 1,085 | 2,743E-16 |
| TcCLB.510347.40 | hypothetical protein, conserved | 1,084 | 1,908E-27 |
| TcCLB.511127.74 | hypothetical protein, conserved | 1,084 | 1,099E-12 |
| TcCLB.509633.60 | serine/threonine-protein phosphatase PP1, putative | 1,084 | 1,608E-09 |
| TcCLB.506537.250 | hypothetical protein, conserved | 1,084 | 1,698E-08 |
| TcCLB.507651.10 | hypothetical protein, conserved (pseudogene) | 1,082 | 6,006E-31 |
| TcCLB.511809.90 | hypothetical protein, conserved | 1,082 | 3,96E-29 |
| TcCLB.508547.50 | maoC-like dehydratase, putative | 1,082 | 2,321E-21 |
| TcCLB.510243.80 | hypothetical protein, conserved | 1,082 | 7,095E-19 |
| TcCLB.508409.140 | hypothetical protein, conserved | 1,082 | 8,709E-18 |
| TcCLB.509747.104 | hypothetical protein, conserved | 1,082 | 2,766E-17 |
| TcCLB.511289.70 | Mitochondrial ADP/ATP carrier protein 5, putative (MCP5) | 1,081 | 5,392E-37 |
| TcCLB.508909.200 | hypothetical protein | 1,081 | 1,298E-27 |
| TcCLB.503955.80 | hypothetical protein | 1,081 | 1,071E-16 |
| TcCLB.511767.30 | retrotransposon hot spot protein (RHS, pseudogene), puta | 1,079 | 2,354E-23 |
| TcCLB.507023.150 | hypothetical protein | 1,079 | 3,714E-21 |
| TcCLB.506443.60 | hypothetical protein | 1,078 | 4,927E-22 |
| TcCLB.508879.229 | chaperone DNAJ protein, putative | 1,078 | 1,713E-06 |
| TcCLB.508221.714 | retrotransposon hot spot protein (RHS, pseudogene), puta | 1,078 | 0,01045 |
| TcCLB.510355.70 | retrotransposon hot spot protein (RHS, pseudogene), puta | 1,077 | 6,583E-18 |
| TcCLB.511665.15 | dispersed gene family protein 1 (DGF-1), putative (fragme | 1,077 | 0,0001632 |
| TcCLB.510835.40 | 60S ribosomal protein L7a, putative | 1,076 | 8,247E-45 |
| TcCLB.508277.65 | hypothetical protein, conserved | 1,076 | 1,479E-16 |
| TcCLB.511417.90 | phosphate-repressible phosphate permease, putative | 1,076 | 3,623E-10 |

| | | | |
|---|---|---|---|
| TcCLB.508543.90 | plasma membrane Ca2 ATPase | 1,075 | 4,442E-30 |
| TcCLB.503671.10 | Aminopeptidase M1, putative (fragment) | 1,075 | 4,68E-19 |
| TcCLB.510267.20 | 60S acidic ribosomal protein P2, putative | 1,075 | 3,233E-18 |
| TcCLB.508063.9 | dispersed gene family protein 1 (DGF-1), putative (fragme | 1,075 | 0,00002135 |
| TcCLB.504839.28 | tryparedoxin peroxidase, putative | 1,074 | 4,038E-34 |
| TcCLB.511583.40 | tricarboxylate carrier, putative | 1,074 | 2,423E-24 |
| TcCLB.509009.10 | dynein heavy chain, cytosolic, putative | 1,074 | 1,906E-18 |
| TcCLB.507009.30 | ribulose-phosphate 3-epimerase, putative | 1,074 | 3,936E-14 |
| TcCLB.509649.40 | hypothetical protein | 1,073 | 3,139E-26 |
| TcCLB.509967.170 | dynein arm light chain, putative | 1,073 | 2,542E-25 |
| TcCLB.507883.109 | mitochondrial malate dehydrogenase, putative | 1,072 | 4,847E-44 |
| TcCLB.506937.30 | 60S ribosomal protein L14, putative | 1,072 | 2,144E-36 |
| TcCLB.510337.40 | spermidine synthase, putative | 1,072 | 6,565E-35 |
| TcCLB.511431.40 | glutaredoxin, putative | 1,072 | 9,399E-16 |
| TcCLB.511819.14 | small nuclear ribonucleoprotein, putative | 1,072 | 2,014E-15 |
| TcCLB.508023.40 | retrotransposon hot spot (RHS) protein, putative | 1,072 | 1,497E-11 |
| TcCLB.510629.120 | dispersed gene family protein 1 (DGF-1, pseudogene), put | 1,072 | 0,0212 |
| TcCLB.506195.300 | deoxyhypusine synthase, putative | 1,071 | 9,078E-26 |
| TcCLB.510733.50 | hypothetical protein, conserved | 1,07 | 2,524E-41 |
| TcCLB.509795.40 | hypothetical protein, conserved | 1,07 | 3,092E-18 |
| TcCLB.503599.10 | protein kinase, putative | 1,07 | 8,703E-11 |
| TcCLB.510359.630 | retrotransposon hot spot protein (RHS, pseudogene), puta | 1,07 | 0,00008731 |
| TcCLB.508221.440 | trans-sialidase (pseudogene), putative | 1,07 | 0,0003773 |
| TcCLB.504119.29 | deoxyhypusine synthase, putative | 1,069 | 6,852E-26 |
| TcCLB.506183.9 | pyruvate dehydrogenase (lipoamide) kinase, putative (frag | 1,069 | 3,183E-12 |
| TcCLB.507537.30 | cysteinyl-tRNA synthetase, putative | 1,068 | 4,235E-37 |
| TcCLB.503997.19 | hypothetical protein, conserved | 1,068 | 2,345E-16 |
| TcCLB.506487.37 | hypothetical protein | 1,068 | 0,0001265 |
| TcCLB.509141.20 | hypothetical protein | 1,067 | 1,055E-10 |
| TcCLB.508137.40 | hypothetical protein, conserved | 1,067 | 0,001329 |
| TcCLB.508041.30 | 40S ribosomal protein S15A, putative | 1,066 | 8,222E-58 |
| TcCLB.506595.149 | retrotransposon hot spot (RHS) protein, putative | 1,066 | 2,937E-43 |
| TcCLB.511023.10 | pyrroline-5-carboxylate synthetase-like protein, putative | 1,066 | 4,59E-24 |
| TcCLB.508569.20 | ubiquitin-conjugating enzyme E2, putative | 1,066 | 1,379E-23 |
| TcCLB.509569.44 | Present in the outer mitochondrial membrane proteome 4 | 1,066 | 2,935E-14 |
| TcCLB.509295.90 | hypothetical protein | 1,066 | 2,818E-10 |
| TcCLB.511287.120 | 40S ribosomal protein S2, putative | 1,065 | 4,331E-44 |
| TcCLB.511667.120 | dispersed gene family protein 1 (DGF-1, pseudogene), put | 1,065 | 2,823E-08 |
| TcCLB.508221.199 | elongation factor 1-gamma (EF-1-gamma), putative | 1,065 | 0,002215 |
| TcCLB.511911.30 | mismatch repair protein MSH4 (pseudogene), putative | 1,064 | 4,237E-21 |
| TcCLB.509725.126 | N-acetyltransferase complex ARD1 subunit (pseudogene), | 1,064 | 0,0001224 |
| TcCLB.506369.60 | hypothetical protein, conserved | 1,063 | 4,372E-12 |
| TcCLB.509891.60 | hypothetical protein, conserved | 1,062 | 4,759E-15 |
| TcCLB.507485.60 | hypothetical protein, conserved | 1,06 | 2,638E-22 |
| TcCLB.510359.120 | hypothetical protein, conserved (pseudogene) | 1,06 | 1,455E-20 |

| | | | |
|---|---|---|---|
| TcCLB.511577.100 | hypothetical protein, conserved | 1,06 | 3,163E-16 |
| TcCLB.510101.180 | 40S ribosomal protein S5, putative | 1,06 | 6,571E-06 |
| TcCLB.506843.137 | retrotransposon hot spot protein (RHS, pseudogene), puta | 1,06 | 0,001092 |
| TcCLB.506443.40 | mitochondrial pyruvate carrier protein 2, putative | 1,059 | 2,41E-21 |
| TcCLB.508953.10 | hypothetical protein, conserved | 1,059 | 5,083E-10 |
| TcCLB.510101.420 | 40S ribosomal protein S21, putative | 1,058 | 2,525E-45 |
| TcCLB.506989.190 | heat shock protein 90, putative (LPG3) | 1,058 | 7,874E-37 |
| TcCLB.510187.530 | hypothetical protein, conserved | 1,058 | 6,102E-26 |
| TcCLB.511929.30 | retrotransposon hot spot (RHS) protein, putative | 1,058 | 8,522E-16 |
| TcCLB.508153.300 | hypothetical protein, conserved | 1,057 | 5,731E-38 |
| TcCLB.511237.60 | 40S ribosomal protein S33, putative | 1,057 | 1,633E-36 |
| TcCLB.503575.60 | protein kinase, putative | 1,057 | 6,941E-26 |
| TcCLB.511827.50 | hypothetical protein | 1,057 | 9,366E-13 |
| TcCLB.510279.290 | dispersed gene family protein 1 (DGF-1, pseudogene), put | 1,057 | 0,0000173 |
| TcCLB.506135.9 | heat shock 70 kDa protein, putative (fragment) | 1,056 | 3,36E-37 |
| TcCLB.509053.140 | succinate dehydrogenase subunit | 1,056 | 1,377E-30 |
| TcCLB.508799.60 | DNA ligase, putative | 1,056 | 7,132E-16 |
| TcCLB.509891.40 | hypothetical protein, conserved | 1,055 | 1,495E-16 |
| TcCLB.507711.20 | paraflagellar rod component, putative (PFC16) | 1,055 | 4,218E-11 |
| TcCLB.511817.151 | histone H2A, putative | 1,055 | 0,001904 |
| TcCLB.510711.10 | trans-sialidase (pseudogene), putative | 1,055 | 0,01144 |
| TcCLB.511041.40 | hexose transporter, putative | 1,054 | 0,0001417 |
| TcCLB.506843.75 | hypothetical protein, conserved (pseudogene) | 1,054 | 0,0003609 |
| TcCLB.510293.54 | hypothetical protein, conserved | 1,053 | 3,418E-25 |
| TcCLB.507659.10 | hypothetical protein, conserved | 1,053 | 9,125E-22 |
| TcCLB.511239.90 | hypothetical protein | 1,053 | 1,271E-19 |
| TcCLB.511229.30 | hypothetical protein, conserved | 1,052 | 2,015E-20 |
| TcCLB.507619.10 | hypothetical protein, conserved (fragment) | 1,052 | 4,467E-17 |
| TcCLB.510849.11 | retrotransposon hot spot protein (RHS, pseudogene), puta | 1,052 | 5,865E-07 |
| TcCLB.507105.40 | 60S ribosomal protein L44 (RPL44) | 1,051 | 1,511E-19 |
| TcCLB.509019.21 | retrotransposon hot spot protein (RHS, pseudogene), puta | 1,051 | 1,209E-16 |
| TcCLB.508677.60 | hypothetical protein | 1,051 | 9,621E-08 |
| TcCLB.511805.15 | ribosomal protein S29, putative | 1,051 | 0,0008715 |
| TcCLB.506503.50 | hypothetical protein, conserved | 1,05 | 2,155E-27 |
| TcCLB.509849.30 | hypothetical protein | 1,049 | 1,443E-39 |
| TcCLB.510323.20 | hypothetical protein, conserved | 1,049 | 1,673E-36 |
| TcCLB.507927.60 | hypothetical protein, conserved | 1,049 | 8,8E-35 |
| TcCLB.511909.40 | Succinate dehydrogenase [ubiquinone] flavoprotein subun | 1,049 | 2,79E-33 |
| TcCLB.504069.80 | ATP synthase F1 subunit gamma protein, putative (ATPF1 | 1,049 | 7,914E-21 |
| TcCLB.511431.60 | hypothetical protein, conserved | 1,049 | 7,147E-11 |
| TcCLB.425785.18 | hypothetical protein, conserved | 1,049 | 0,00004267 |
| TcCLB.510507.29 | hypothetical protein, conserved | 1,048 | 9,451E-14 |
| TcCLB.509013.19 | calpain-like cysteine peptidase, putative (fragment) | 1,048 | 4,781E-06 |
| TcCLB.508737.100 | ATP-dependent Clp protease subunit, heat shock protein 7 | 1,047 | 9,417E-27 |
| TcCLB.510941.3 | protein kinase, putative (fragment) | 1,047 | 3,641E-20 |

| | | | |
|---|---|---|---|
| TcCLB.504049.40 | retrotransposon hot spot protein (RHS, pseudogene), puta | 1,047 | 1,828E-16 |
| TcCLB.511825.220 | alanyl-tRNA synthetase, putative | 1,046 | 7,945E-50 |
| TcCLB.506829.90 | conserved protein | 1,046 | 2,182E-37 |
| TcCLB.511239.110 | hypothetical protein, conserved | 1,046 | 9,489E-25 |
| TcCLB.470169.9 | hypothetical protein, conserved | 1,046 | 8,922E-13 |
| TcCLB.508637.40 | hypothetical protein, conserved | 1,046 | 8,517E-10 |
| TcCLB.510667.100 | hypothetical protein, conserved | 1,045 | 1,444E-32 |
| TcCLB.508083.10 | Present in the outer mitochondrial membrane proteome 1 | 1,045 | 6,66E-31 |
| TcCLB.511001.40 | hypothetical protein | 1,045 | 1,449E-21 |
| TcCLB.511233.59 | hypothetical protein, conserved | 1,045 | 2,455E-18 |
| TcCLB.510855.20 | hypothetical protein, conserved | 1,044 | 8,412E-15 |
| TcCLB.506563.200 | calpain cysteine peptidase, putative | 1,043 | 3,836E-27 |
| TcCLB.508787.20 | hypothetical protein | 1,043 | 1,95E-25 |
| TcCLB.509913.10 | 3-methylcrotonyl-CoA carboxylase, putative | 1,043 | 3,207E-24 |
| TcCLB.510257.100 | mitochondrial RNA binding complex 1 subunit, putative | 1,042 | 1,528E-33 |
| TcCLB.506331.174 | dispersed gene family protein 1 (DGF-1, pseudogene), put | 1,042 | 7,082E-07 |
| TcCLB.506627.20 | hypothetical protein, conserved | 1,041 | 1,085E-37 |
| TcCLB.509047.40 | hypothetical protein, conserved | 1,041 | 1,096E-34 |
| TcCLB.506675.20 | hypothetical protein, conserved (pseudogene) | 1,041 | 1,47E-22 |
| TcCLB.507943.30 | histone H4, putative | 1,041 | 3,296E-21 |
| TcCLB.510101.430 | 40S ribosomal protein S21, putative | 1,041 | 2,812E-18 |
| TcCLB.507551.9 | dispersed gene family protein 1 (DGF-1), putative | 1,041 | 4,749E-15 |
| TcCLB.438059.10 | serine/threonine-protein phosphatase PP1 alpha | 1,041 | 2,937E-09 |
| TcCLB.509505.70 | 10 kDa heat shock protein, putative | 1,041 | 0,1265 |
| TcCLB.510395.10 | ATP synthase F1, alpha subunit, putative (ATPF1A) | 1,04 | 3,788E-25 |
| TcCLB.507711.300 | Archaic translocase of outer membrane 11 kDa subunit, pu | 1,04 | 4,006E-13 |
| TcCLB.506629.40 | hypothetical protein, conserved | 1,039 | 7,755E-37 |
| TcCLB.508501.290 | Kinesin-13 1, putative (KIN13-1) | 1,039 | 7,012E-36 |
| TcCLB.509109.50 | hypothetical protein | 1,039 | 4,112E-28 |
| TcCLB.510861.30 | hypothetical protein, conserved | 1,039 | 4,252E-24 |
| TcCLB.510661.70 | hypothetical protein, conserved | 1,039 | 1,879E-16 |
| TcCLB.506925.400 | hypothetical protein, conserved | 1,038 | 9,354E-29 |
| TcCLB.511209.54 | ribosomal protein S29, putative | 1,038 | 7,353E-26 |
| TcCLB.505945.80 | hypothetical protein, conserved | 1,038 | 9,106E-21 |
| TcCLB.508257.180 | hypothetical protein, conserved | 1,038 | 1,076E-19 |
| TcCLB.506695.7 | hypothetical protein, conserved | 1,038 | 2,086E-11 |
| TcCLB.510447.10 | dispersed gene family protein 1 (DGF-1), putative | 1,038 | 0,08473 |
| TcCLB.508909.20 | guanosine monophosphate reductase, putative | 1,037 | 1,458E-40 |
| TcCLB.506363.90 | amidohydrolase, putative | 1,037 | 3,686E-21 |
| TcCLB.506831.10 | hypothetical protein | 1,037 | 3,589E-20 |
| TcCLB.509549.110 | hypothetical protein, conserved (pseudogene) | 1,036 | 8,517E-23 |
| TcCLB.506859.240 | hypothetical protein, conserved | 1,036 | 0,0001209 |
| TcCLB.509007.99 | P-glycoprotein (pseudogene) (tcpgp1) | 1,035 | 6,711E-32 |
| TcCLB.510101.360 | hypothetical protein, conserved | 1,035 | 8,113E-29 |
| TcCLB.506855.250 | hypothetical protein, conserved | 1,035 | 2,072E-25 |

| | | | |
|---|---|---|---|
| TcCLB.511001.150 | hypothetical protein, conserved | 1,035 | 6,25E-15 |
| TcCLB.503579.60 | hypothetical protein | 1,035 | 8,93E-14 |
| TcCLB.508475.10 | ribosomal protein S20, putative | 1,034 | 6,084E-20 |
| TcCLB.508153.164 | hypothetical protein, conserved | 1,034 | 3,006E-16 |
| TcCLB.507529.10 | hypothetical Zinc-finger containing protein | 1,034 | 6,707E-13 |
| TcCLB.505957.20 | dispersed gene family protein 1 (DGF-1, pseudogene), put | 1,034 | 4,481E-08 |
| TcCLB.510483.280 | 90 kDa surface protein, putative | 1,034 | 1,349E-06 |
| TcCLB.511049.40 | leucyl-tRNA synthetase, putative | 1,033 | 5,359E-44 |
| TcCLB.511727.270 | RNA-binding protein, putative | 1,033 | 2,381E-42 |
| TcCLB.506485.10 | hypothetical protein, conserved | 1,033 | 1,054E-11 |
| TcCLB.506839.89 | mitochondrial chaperone BCS1, putative (fragment) | 1,033 | 1,587E-09 |
| TcCLB.504011.70 | hypothetical protein, conserved | 1,033 | 4,097E-06 |
| TcCLB.503539.30 | GTP-binding nuclear protein rtb2, putative | 1,032 | 9,928E-41 |
| TcCLB.509695.184 | 40S ribosomal protein S23, putative | 1,032 | 2,827E-38 |
| TcCLB.511903.170 | hypothetical protein, conserved | 1,032 | 5,606E-27 |
| TcCLB.509149.60 | 60S ribosomal protein L19, putative | 1,032 | 2,066E-19 |
| TcCLB.511165.54 | pterin-4-alpha-carbinolamine dehydratase, putative | 1,032 | 9,931E-17 |
| TcCLB.511801.34 | hypothetical protein, conserved | 1,032 | 4,22E-16 |
| TcCLB.511051.80 | DNA/RNA non-specific endonuclease protein-like, putative | 1,031 | 1,736E-12 |
| TcCLB.507783.10 | ribosomal P protein AGP2beta-1, putative | 1,031 | 4,611E-08 |
| TcCLB.506271.30 | retrotransposon hot spot (RHS) protein, putative (fragmer | 1,031 | 3,227E-07 |
| TcCLB.508189.20 | dispersed gene family protein 1 (DGF-1, pseudogene), put | 1,031 | 0,00003451 |
| TcCLB.504001.20 | hypothetical protein, conserved | 1,03 | 7,718E-18 |
| TcCLB.508677.90 | hypothetical protein, conserved | 1,03 | 2,07E-08 |
| TcCLB.508125.65 | elongation factor 1-gamma (EF-1-gamma, pseudogene), | 1,03 | 0,0003845 |
| TcCLB.504181.20 | 40S ribosomal protein S23, putative | 1,029 | 6,807E-50 |
| TcCLB.503395.50 | 40S ribosomal protein S12, putative | 1,029 | 2,099E-36 |
| TcCLB.510815.9 | dispersed gene family protein 1 (DGF-1), putative | 1,029 | 1,933E-16 |
| TcCLB.511109.130 | heterogeneous nuclear ribonucleoprotein H/F, putative | 1,028 | 6,899E-21 |
| TcCLB.509437.183 | hypothetical protein, conserved (pseudogene) | 1,028 | 1,011E-16 |
| TcCLB.507501.10 | retrotransposon hot spot (RHS) protein, putative | 1,027 | 1,847E-75 |
| TcCLB.504069.56 | 60S ribosomal protein L37a, putative | 1,027 | 9,113E-27 |
| TcCLB.507721.20 | clathrin assembly protein (pseudogene), putative | 1,027 | 1,301E-24 |
| TcCLB.503467.20 | hypothetical protein, conserved | 1,027 | 1,598E-24 |
| TcCLB.503905.50 | hypothetical protein, conserved | 1,027 | 7,208E-09 |
| TcCLB.506999.191 | hypothetical protein, conserved (fragment) | 1,027 | 0,00003585 |
| TcCLB.511817.180 | histone H2A, putative | 1,027 | 0,001973 |
| TcCLB.507019.40 | histidyl-tRNA synthetase | 1,026 | 1,219E-32 |
| TcCLB.506775.180 | hypothetical protein, conserved | 1,026 | 2,318E-31 |
| TcCLB.506779.190 | Unconventional myosin | 1,026 | 4,8E-21 |
| TcCLB.509875.120 | trans-sialidase, Group II, putative | 1,026 | 1,167E-12 |
| TcCLB.509779.4 | hypothetical protein, conserved | 1,025 | 3,52E-13 |
| TcCLB.507681.150 | 40S ribosomal protein S24E, putative | 1,024 | 1,474E-19 |
| TcCLB.510359.371 | elongation factor 1-gamma (EF-1-gamma, pseudogene), | 1,024 | 2,271E-06 |
| TcCLB.507039.10 | iron superoxide dismutase, putative | 1,023 | 1,238E-29 |

| | | | |
|---|---|---|---|
| TcCLB.506945.320 | hypothetical protein, conserved | 1,023 | 7,978E-23 |
| TcCLB.507521.150 | dispersed gene family protein 1 (DGF-1), putative | 1,023 | 7,777E-10 |
| TcCLB.510145.20 | cofilin/actin depolymerizing factor, putative | 1,022 | 3,154E-29 |
| TcCLB.506775.14 | hypothetical protein, conserved | 1,022 | 1,329E-20 |
| TcCLB.510839.20 | hypothetical protein | 1,022 | 1,445E-15 |
| TcCLB.506581.14 | peptidyl-prolyl cis-trans isomerase, putative | 1,022 | 3,809E-12 |
| TcCLB.508781.10 | structural maintenance of chromosome protein 4 (pseudo | 1,022 | 2,088E-06 |
| TcCLB.507007.70 | hypothetical protein, conserved | 1,021 | 2,251E-17 |
| TcCLB.507993.36 | hypothetical protein, conserved | 1,021 | 4,062E-10 |
| TcCLB.506679.250 | ribosomal protein L24, putative | 1,02 | 1,514E-37 |
| TcCLB.505807.20 | hypothetical protein, conserved | 1,02 | 2,71E-21 |
| TcCLB.503555.30 | trypanothione reductase, putative | 1,02 | 1,89E-19 |
| TcCLB.510665.60 | hypothetical protein, conserved (pseudogene) | 1,02 | 2,961E-19 |
| TcCLB.507523.75 | mucin TcMUCII (pseudogene), putative | 1,02 | 0,001608 |
| TcCLB.508355.420 | cation transporter protein, putative | 1,019 | 1,658E-16 |
| TcCLB.504769.89 | hypothetical protein, conserved | 1,019 | 5,303E-14 |
| TcCLB.511487.130 | trans-sialidase (pseudogene), putative | 1,019 | 0,00004793 |
| TcCLB.508561.10 | dispersed gene family protein 1 (DGF-1), putative | 1,019 | 0,0001121 |
| TcCLB.509857.50 | bilobe protein, conserved | 1,018 | 3,495E-29 |
| TcCLB.510431.30 | amino acid permease, putative | 1,018 | 9,497E-18 |
| TcCLB.508975.4 | hypothetical protein, conserved (fragment) | 1,018 | 7,927E-12 |
| TcCLB.506659.35 | hypothetical protein, conserved | 1,018 | 5,841E-09 |
| TcCLB.507871.9 | dispersed gene family protein 1 (DGF-1), putative | 1,018 | 0,003351 |
| TcCLB.508319.10 | hypothetical protein | 1,018 | 0,02988 |
| TcCLB.511067.20 | 60S ribosomal protein L26, putative | 1,017 | 2,874E-38 |
| TcCLB.503395.40 | 60S ribosomal protein L18, putative | 1,017 | 2,209E-34 |
| TcCLB.508457.20 | hypothetical protein, conserved | 1,017 | 1,032E-25 |
| TcCLB.510719.140 | hypothetical protein, conserved | 1,017 | 1,927E-18 |
| TcCLB.508547.160 | hypothetical protein, conserved | 1,017 | 6,295E-18 |
| TcCLB.507519.154 | hypothetical protein, conserved | 1,017 | 2,189E-14 |
| TcCLB.506063.20 | retrotransposon hot spot protein (RHS, pseudogene), puta | 1,017 | 1,604E-13 |
| TcCLB.509087.20 | dispersed gene family protein 1 (DGF-1), putative | 1,017 | 2,079E-09 |
| TcCLB.505647.10 | hypothetical protein, conserved | 1,016 | 1,124E-34 |
| TcCLB.506825.190 | hypothetical protein | 1,016 | 5,788E-20 |
| TcCLB.508879.120 | hypothetical protein, conserved | 1,016 | 1,166E-19 |
| TcCLB.507573.10 | hypothetical protein, conserved (pseudogene) | 1,016 | 1,084E-17 |
| TcCLB.511111.20 | hypothetical protein | 1,016 | 4,778E-17 |
| TcCLB.507725.9 | dispersed gene family protein 1 (DGF-1, pseudogene), put | 1,016 | 1,613E-11 |
| TcCLB.503965.39 | hypothetical protein, conserved | 1,016 | 1,285E-09 |
| TcCLB.510769.49 | 40S ribosomal protein S6, putative | 1,015 | 1,713E-44 |
| TcCLB.507873.30 | Double RNA binding domain protein 7 (DRBD7) | 1,015 | 5,02E-28 |
| TcCLB.508827.70 | 40S ribosomal protein S17, putative | 1,015 | 2,372E-27 |
| TcCLB.510943.40 | protein kinase, putative | 1,015 | 4,894E-26 |
| TcCLB.506637.10 | hypothetical protein, conserved | 1,015 | 2,105E-23 |
| TcCLB.506367.170 | hypothetical protein | 1,015 | 2,861E-22 |

| | | | |
|---|---|---|---|
| TcCLB.504797.20 | hypothetical protein, conserved | 1,015 | 6,719E-21 |
| TcCLB.503891.50 | hypothetical protein, conserved | 1,015 | 4,008E-19 |
| TcCLB.506157.30 | retrotransposon hot spot protein (RHS, pseudogene), puta | 1,015 | 3,449E-14 |
| TcCLB.509065.36 | hypothetical protein, conserved | 1,015 | 1,935E-12 |
| TcCLB.506375.80 | hypothetical protein, conserved | 1,015 | 4,769E-09 |
| TcCLB.503967.19 | hypothetical protein | 1,015 | 5,942E-08 |
| TcCLB.509845.9 | dispersed gene family protein 1 (DGF-1), putative | 1,015 | 0,00003913 |
| TcCLB.506963.10 | 60S ribosomal protein L10a, putative | 1,014 | 1,17E-44 |
| TcCLB.507991.123 | hypothetical protein, conserved | 1,014 | 1,117E-14 |
| TcCLB.508183.29 | hypothetical protein, conserved (fragment) | 1,014 | 1,1E-11 |
| TcCLB.511769.20 | dispersed gene family protein 1 (DGF-1), putative | 1,014 | 7,322E-09 |
| TcCLB.509437.140 | retrotransposon hot spot protein (RHS, pseudogene), puta | 1,014 | 2,299E-07 |
| TcCLB.511487.30 | dispersed gene family protein 1 (DGF-1), putative | 1,014 | 9,785E-06 |
| TcCLB.510625.240 | cell-surface protein (cysteine-rich repeat motif) (pseudog | 1,014 | 0,0461 |
| TcCLB.510311.10 | hypothetical protein, conserved | 1,013 | 3,141E-25 |
| TcCLB.506223.60 | hypothetical protein, conserved | 1,013 | 5,13E-14 |
| TcCLB.511809.99 | 40S ribosomal protein S15, putative | 1,013 | 2,376E-11 |
| TcCLB.508001.106 | serine/threonine protein kinase (pseudogene), putative | 1,013 | 0,0004238 |
| TcCLB.503905.10 | myosin heavy chain, putative | 1,012 | 3,496E-23 |
| TcCLB.511589.10 | ATP synthase, putative | 1,012 | 8,962E-21 |
| TcCLB.506845.21 | N-acetyltransferase complex ARD1 subunit (pseudogene), | 1,012 | 0,03862 |
| TcCLB.509607.40 | hypothetical protein | 1,011 | 3,019E-18 |
| TcCLB.506559.70 | hypothetical protein, conserved | 1,011 | 1,111E-16 |
| TcCLB.503607.7 | glycine dehydrogenase (pseudogene), putative | 1,011 | 5,551E-13 |
| TcCLB.511753.9 | hypothetical protein, conserved | 1,011 | 2,44E-09 |
| TcCLB.504195.9 | dispersed gene family protein 1 (DGF-1, pseudogene), put | 1,011 | 0,0003213 |
| TcCLB.510425.19 | 40S ribosomal protein SA, putative | 1,01 | 5,594E-32 |
| TcCLB.510407.20 | hypothetical protein, conserved | 1,01 | 1,095E-22 |
| TcCLB.509099.89 | hypothetical protein, conserved | 1,01 | 1,124E-17 |
| TcCLB.507831.90 | 60S ribosomal protein L34, putative | 1,01 | 1,08E-14 |
| TcCLB.507801.200 | dispersed gene family protein 1 (DGF-1), putative | 1,01 | 0,01449 |
| TcCLB.510357.80 | hypothetical protein, conserved | 1,009 | 5,751E-18 |
| TcCLB.511621.120 | hypothetical protein, conserved | 1,009 | 7,33E-10 |
| TcCLB.508293.100 | chitin-binding like protein, putative | 1,009 | 0,02973 |
| TcCLB.511051.39 | 40S ribosomal protein S4, putative | 1,008 | 6,12E-40 |
| TcCLB.511237.10 | thimet oligopeptidase, putative | 1,008 | 9,637E-27 |
| TcCLB.508461.480 | 60S ribosomal protein L23, putative | 1,007 | 2,357E-46 |
| TcCLB.511671.149 | phosphate transporter, putative (PHO91) | 1,007 | 4,637E-19 |
| TcCLB.505999.10 | phosphate-repressible phosphate permease, putative | 1,007 | 6,062E-19 |
| TcCLB.506275.20 | ATP-dependent protease subunit HslV, putative (HslV) | 1,007 | 3,856E-17 |
| TcCLB.507021.40 | hypothetical protein, conserved | 1,007 | 9,286E-11 |
| TcCLB.510101.40 | 60S ribosomal protein L28, putative | 1,006 | 1,525E-40 |
| TcCLB.508827.79 | 40S ribosomal protein S17, putative | 1,006 | 2,455E-34 |
| TcCLB.511751.260 | hypothetical protein, conserved | 1,006 | 2,916E-08 |
| TcCLB.506963.20 | 60S ribosomal protein L34, putative | 1,005 | 1,215E-40 |

| | | | |
|---|---|---|---|
| TcCLB.505843.10 | glutamate dehydrogenase, putative | 1,005 | 6,735E-26 |
| TcCLB.509799.30 | adenylyl cyclase-associated protein, putative | 1,005 | 3,489E-17 |
| TcCLB.511001.120 | 60S ribosomal protein L18a, putative | 1,004 | 2,714E-52 |
| TcCLB.509695.150 | kinesin, putative | 1,004 | 1,523E-51 |
| TcCLB.506181.59 | 40S ribosomal protein S12, putative | 1,004 | 9,728E-34 |
| TcCLB.511807.253 | hypothetical protein, conserved | 1,004 | 1,724E-24 |
| TcCLB.510609.50 | nuclear receptor binding factor, putative | 1,004 | 5,404E-17 |
| TcCLB.463155.10 | retrotransposon hot spot protein (RHS, pseudogene), puta | 1,004 | 1,883E-13 |
| TcCLB.508197.10 | 60S ribosomal protein L11, putative | 1,003 | 2,145E-37 |
| TcCLB.506727.50 | hypothetical protein, conserved | 1,003 | 1,131E-21 |
| TcCLB.505037.40 | hypothetical protein, conserved | 1,003 | 1,156E-17 |
| TcCLB.507009.50 | hypothetical protein, conserved | 1,003 | 6,968E-15 |
| TcCLB.503599.30 | hypothetical protein, conserved | 1,003 | 1,089E-12 |
| TcCLB.511133.20 | thymidine kinase, putative | 1,003 | 2,204E-10 |
| TcCLB.457109.9 | dispersed gene family protein 1 (DGF-1), putative (fragm | 1,003 | 5,684E-10 |
| TcCLB.511569.14 | dispersed gene family protein 1 (DGF-1), putative | 1,003 | 1,322E-08 |
| TcCLB.510219.30 | CPSF 30 kDa subunit, putative (CPSF30) | 1,002 | 2,717E-18 |
| TcCLB.507895.110 | hypothetical protein, conserved | 1,002 | 4,123E-15 |
| TcCLB.510195.9 | dispersed gene family protein 1 (DGF-1), putative | 1,002 | 1,251E-08 |
| TcCLB.511173.483 | mucin TcMUCII (pseudogene), putative | 1,002 | 0,004105 |
| TcCLB.504181.10 | 60S ribosomal protein L9, putative | 1,001 | 3,983E-42 |
| TcCLB.506739.170 | hypothetical protein, conserved | 1,001 | 1,584E-22 |
| TcCLB.510999.39 | 40S ribosomal protein S3A, putative (fragment) | 1,001 | 6,378E-20 |
| TcCLB.506903.50 | dispersed gene family protein 1 (DGF-1, pseudogene), pu | 1 | 3,47E-10 |
| TcCLB.508707.310 | leucine rich repeat protein, putative | 0,9999 | 1,379E-29 |
| TcCLB.508479.216 | hypothetical protein, conserved | 0,9999 | 1,951E-18 |
| TcCLB.510285.20 | hypothetical protein, conserved | 0,9996 | 8,884E-25 |
| TcCLB.510117.10 | retrotransposon hot spot protein (RHS, pseudogene), puta | 0,9996 | 4,949E-16 |
| TcCLB.508981.30 | hypothetical protein, conserved | 0,9995 | 7,577E-13 |
| TcCLB.509725.113 | ATP-dependent DEAD/H RNA helicase (pseudogene), puta | 0,9992 | 0,00007478 |
| TcCLB.506585.40 | glucose-regulated protein 78, putative | 0,999 | 9,005E-38 |
| TcCLB.511663.19 | dispersed gene family protein 1 (DGF-1), putative (fragm | 0,999 | 1,148E-06 |
| TcCLB.506817.10 | hypothetical protein, conserved | 0,9987 | 1,559E-38 |
| TcCLB.510565.40 | hypothetical protein | 0,9982 | 4,25E-14 |
| TcCLB.509157.90 | hypothetical protein, conserved | 0,9981 | 7,249E-41 |
| TcCLB.508041.50 | hypothetical protein, conserved | 0,9979 | 2,068E-27 |
| TcCLB.506529.410 | primase 2, putative | 0,9978 | 6,768E-27 |
| TcCLB.511717.130 | hypothetical protein, conserved | 0,9975 | 1,659E-13 |
| TcCLB.509569.94 | hypothetical protein, conserved | 0,9975 | 2,235E-11 |
| TcCLB.508501.230 | monothiol glutaredoxin, putative | 0,997 | 1,776E-13 |
| TcCLB.506543.90 | cytochrome c oxidase assembly protein, putative | 0,9969 | 2,448E-16 |
| TcCLB.510535.64 | hypothetical protein, conserved | 0,9966 | 1,176E-19 |
| TcCLB.503847.50 | hypothetical protein, conserved | 0,9959 | 2,2E-11 |
| TcCLB.508045.70 | hypothetical protein, conserved | 0,9958 | 3,245E-45 |
| TcCLB.509105.90 | MRB1-associated protein, putative (RGG2) | 0,9957 | 2,805E-13 |

| | | | |
|---|---|---|---|
| TcCLB.506405.10 | phosphomannomutase-like protein, putative | 0,9955 | 8,313E-29 |
| TcCLB.506181.50 | 60S ribosomal protein L18, putative | 0,9954 | 1,641E-38 |
| TcCLB.508479.110 | glutathionylspermidine synthase, putative | 0,9954 | 1,523E-33 |
| TcCLB.504257.9 | RNA-binding protein 42 (RNA-binding motif protein 42), p | 0,9954 | 9,628E-14 |
| TcCLB.511395.80 | hypothetical protein | 0,9952 | 3,851E-19 |
| TcCLB.503719.20 | 40S ribosomal protein SA, putative | 0,9951 | 7,229E-29 |
| TcCLB.509241.10 | surface protease GP63, putative (fragment) | 0,9949 | 5,571E-06 |
| TcCLB.511727.145 | hypothetical protein | 0,9944 | 4,497E-24 |
| TcCLB.506919.40 | retrotransposon hot spot protein (RHS, pseudogene), puta | 0,9943 | 0,0008883 |
| TcCLB.508331.30 | Checkpoint protein HUS1, putative | 0,9942 | 1,537E-12 |
| TcCLB.509659.20 | dispersed gene family protein 1 (DGF-1, pseudogene), put | 0,9942 | 0,00003196 |
| TcCLB.510759.150 | hypothetical protein, conserved | 0,9934 | 9,722E-28 |
| TcCLB.503751.10 | hypothetical protein | 0,9932 | 5,43E-16 |
| TcCLB.510339.74 | mitochondrial carrier protein, putative | 0,9926 | 2,192E-20 |
| TcCLB.506755.110 | hypothetical protein, conserved | 0,9922 | 7,654E-29 |
| TcCLB.510857.30 | hypothetical protein, conserved | 0,992 | 1,126E-24 |
| TcCLB.504427.20 | hypothetical protein, conserved | 0,992 | 9,428E-12 |
| TcCLB.511183.140 | dispersed gene family protein 1 (DGF-1), putative | 0,9917 | 0,000396 |
| TcCLB.511751.80 | hypothetical protein | 0,9913 | 4,018E-16 |
| TcCLB.505193.110 | serine/threonine protein kinase, putative | 0,9912 | 1,573E-31 |
| TcCLB.511545.40 | ribosomal protein L27, putative | 0,9912 | 8,423E-07 |
| TcCLB.507001.80 | acyl-CoA dehydrogenase, putative | 0,9907 | 1,904E-17 |
| TcCLB.510761.44 | cyclophilin-like protein, putative | 0,9906 | 1,225E-31 |
| TcCLB.510943.198 | hypothetical protein, conserved | 0,9902 | 2,885E-10 |
| TcCLB.510901.60 | replication factor A, 51kDa subunit, putative | 0,9901 | 2,587E-20 |
| TcCLB.507255.30 | retrotransposon hot spot protein (RHS, pseudogene), puta | 0,9899 | 2,931E-14 |
| TcCLB.508249.10 | dispersed gene family protein 1 (DGF-1), putative | 0,9899 | 2,776E-06 |
| TcCLB.509757.49 | L-ribulokinase, putative | 0,9898 | 0,0002574 |
| TcCLB.511883.30 | dispersed gene family protein 1 (DGF-1, pseudogene), put | 0,9897 | 1,641E-08 |
| TcCLB.510187.439 | hypothetical protein | 0,989 | 8,108E-34 |
| TcCLB.506357.120 | ethanolamine phosphotransferase, putative | 0,989 | 5,103E-16 |
| TcCLB.508391.10 | dispersed gene family protein 1 (DGF-1), putative | 0,9889 | 0,0004384 |
| TcCLB.506843.50 | glycine dehydrogenase [decarboxylating] (pseudogene), pu | 0,9887 | 0,0006415 |
| TcCLB.503487.30 | hypothetical protein, conserved | 0,9886 | 3,165E-15 |
| TcCLB.509759.19 | pumilio/PUF RNA binding protein 1, putative (PUF1) | 0,9879 | 2,814E-29 |
| TcCLB.510091.50 | hypothetical protein, conserved | 0,9879 | 5,32E-19 |
| TcCLB.507623.129 | vacuolar type h ATPase subunit, putative | 0,9878 | 1,428E-14 |
| TcCLB.509779.10 | DNA repair protein, putative | 0,9876 | 3,251E-24 |
| TcCLB.511577.144 | hypothetical protein, conserved | 0,9875 | 5,092E-10 |
| TcCLB.507711.40 | RNA-binding protein, putative | 0,9873 | 3,887E-16 |
| TcCLB.510911.50 | glycine dehydrogenase [decarboxylating], putative | 0,9873 | 0,0000104 |
| TcCLB.511303.150 | hypothetical protein, conserved | 0,9872 | 3,23E-14 |
| TcCLB.510087.80 | hypothetical protein, conserved | 0,9871 | 6,112E-26 |
| TcCLB.508619.10 | retrotransposon hot spot protein (RHS, pseudogene), puta | 0,987 | 3,547E-18 |
| TcCLB.511727.290 | RNA-binding protein, putative | 0,9869 | 4,803E-49 |

| | | | |
|---|---|---|---|
| TcCLB.504147.120 | 60S ribosomal protein L22, putative | 0,9869 | 8,232E-46 |
| TcCLB.507125.30 | retrotransposon hot spot protein (RHS, pseudogene), puta | 0,9868 | 0,0001027 |
| TcCLB.507245.20 | trans-sialidase (pseudogene), putative | 0,986 | 7,405E-10 |
| TcCLB.510437.20 | 60S ribosomal protein L14, putative | 0,9858 | 8,241E-34 |
| TcCLB.507229.29 | dispersed gene family protein 1 (DGF-1), putative | 0,9855 | 0,004995 |
| TcCLB.503575.34 | ribosomal protein L38, putative | 0,9854 | 3,173E-24 |
| TcCLB.507079.20 | dynein heavy chain, cytosolic, putative | 0,9853 | 2,75E-17 |
| TcCLB.508821.50 | vacuolar type h ATPase subunit, putative | 0,984 | 5,038E-15 |
| TcCLB.506393.30 | beta galactofuranosyl glycosyltransferase, putative | 0,9837 | 6,361E-09 |
| TcCLB.509331.189 | hypothetical protein, conserved | 0,983 | 1,116E-10 |
| TcCLB.508567.30 | hypothetical protein | 0,983 | 1,122E-10 |
| TcCLB.445281.9 | calpain-like cysteine peptidase, putative | 0,9829 | 0,0001238 |
| TcCLB.509979.95 | 60S ribosomal protein L35, putative | 0,9828 | 1,807E-32 |
| TcCLB.511245.24 | hypothetical protein, conserved | 0,9827 | 4,46E-32 |
| TcCLB.504351.30 | hypothetical protein, conserved (pseudogene) | 0,9827 | 2,002E-10 |
| TcCLB.509733.40 | hypothetical protein, conserved | 0,9822 | 4,757E-20 |
| TcCLB.507253.70 | N-acetyltransferase complex ARD1 subunit (pseudogene), | 0,9817 | 0,02272 |
| TcCLB.507017.70 | ribosomal RNA adenine dimethylase family protein, putat | 0,9815 | 3,284E-24 |
| TcCLB.508207.240 | C-1-tetrahydrofolate synthase, cytoplasmic, putative | 0,9814 | 1,224E-16 |
| TcCLB.508851.59 | ATP synthase, putative | 0,9814 | 7,024E-16 |
| TcCLB.507533.21 | clathrin assembly protein (pseudogene), putative | 0,9811 | 7,745E-27 |
| TcCLB.511903.130 | 60S ribosomal protein L26, putative | 0,9805 | 2,626E-44 |
| TcCLB.506825.24 | hypothetical protein, conserved | 0,9804 | 2,167E-22 |
| TcCLB.509033.10 | trans-sialidase (pseudogene), putative | 0,9801 | 0,0002825 |
| TcCLB.506985.30 | proteasome alpha 3 subunit, putative | 0,9799 | 1,362E-20 |
| TcCLB.503667.39 | dispersed gene family protein 1 (DGF-1), putative (fragme | 0,9798 | 4,005E-06 |
| TcCLB.508041.40 | hypothetical protein | 0,9792 | 1,345E-22 |
| TcCLB.507491.39 | hypothetical protein, conserved | 0,9788 | 2,72E-13 |
| TcCLB.511589.110 | hypothetical protein, conserved | 0,9787 | 1,541E-27 |
| TcCLB.511899.40 | 2-amino-3-ketobutyrate coenzyme A ligase, putative | 0,9784 | 1,122E-21 |
| TcCLB.511249.10 | ADP/ATP mitochondrial carrier protein, putative | 0,9782 | 1,349E-22 |
| TcCLB.511319.30 | hypothetical protein | 0,9782 | 1,836E-12 |
| TcCLB.506205.20 | hypothetical protein | 0,978 | 9,187E-23 |
| TcCLB.509109.30 | hypothetical protein, conserved | 0,9778 | 6,166E-10 |
| TcCLB.510095.80 | hypothetical protein | 0,9777 | 0,001413 |
| TcCLB.508321.40 | hypothetical protein, conserved | 0,9774 | 1,177E-34 |
| TcCLB.507519.110 | 60S ribosomal protein L7, putative | 0,9773 | 3,092E-37 |
| TcCLB.510479.11 | retrotransposon hot spot (RHS) protein, putative (fragmer | 0,9773 | 9,859E-19 |
| TcCLB.508465.110 | acyl-CoA dehydrogenase, putative | 0,9772 | 1,349E-19 |
| TcCLB.507529.20 | hypothetical protein, conserved | 0,9771 | 7,894E-30 |
| TcCLB.508125.40 | hypothetical protein | 0,9771 | 0,01129 |
| TcCLB.510355.230 | hypothetical protein, conserved | 0,9761 | 2,491E-13 |
| TcCLB.508539.30 | hypothetical protein, conserved | 0,9757 | 1,112E-11 |
| TcCLB.505965.50 | phosphonopyruvate decarboxylase, putative | 0,9755 | 3,129E-21 |
| TcCLB.510293.20 | cytosolic factor SEC14, putative | 0,9754 | 4,892E-13 |

| | | | |
|---|---|---|---|
| TcCLB.508159.13 | trans-sialidase (pseudogene), putative | 0,9752 | 0,00007741 |
| TcCLB.506301.60 | hypothetical protein | 0,975 | 6,041E-27 |
| TcCLB.509599.60 | hypothetical protein | 0,9749 | 1,268E-13 |
| TcCLB.511773.60 | glycine dehydrogenase [decarboxylating] (pseudogene), pu | 0,9744 | 4,406E-28 |
| TcCLB.509615.40 | alpha-adaptin-like (pseudogene), putative | 0,9744 | 1,028E-17 |
| TcCLB.504149.100 | hypothetical protein, conserved | 0,9742 | 5,778E-27 |
| TcCLB.510337.20 | calpain-like cysteine peptidase, putative | 0,9741 | 2,753E-09 |
| TcCLB.510659.140 | hypothetical protein, conserved (pseudogene) | 0,974 | 6,806E-27 |
| TcCLB.507081.50 | hypothetical protein, conserved | 0,9739 | 7,502E-20 |
| TcCLB.506583.30 | ubiquitin-like protein, putative | 0,9734 | 6,648E-32 |
| TcCLB.506577.54 | 60S ribosomal protein L6, putative | 0,9726 | 2,398E-24 |
| TcCLB.511541.30 | hypothetical protein, conserved | 0,9724 | 3,589E-11 |
| TcCLB.507645.50 | hypothetical protein, conserved | 0,9719 | 1,092E-15 |
| TcCLB.508799.260 | hypothetical protein, conserved | 0,9717 | 1,142E-28 |
| TcCLB.511469.40 | hypothetical protein, conserved | 0,9717 | 5,621E-26 |
| TcCLB.504431.118 | hypothetical protein, conserved | 0,9717 | 8,554E-16 |
| TcCLB.508197.39 | 60S ribosomal protein L11, putative | 0,9715 | 5,184E-16 |
| TcCLB.509849.10 | ammonium transporter, putative | 0,9713 | 1,137E-17 |
| TcCLB.506817.30 | ribosomal protein L27, putative | 0,971 | 3,102E-25 |
| TcCLB.508207.90 | hypothetical protein, conserved | 0,971 | 1,002E-16 |
| TcCLB.418185.40 | hypothetical protein, conserved (pseudogene) | 0,9706 | 1,874E-15 |
| TcCLB.504109.160 | electron transfer flavoprotein, putative | 0,9703 | 1,25E-28 |
| TcCLB.509247.50 | methionyl-tRNA synthetase, putative | 0,9702 | 2,739E-35 |
| TcCLB.507521.130 | hypothetical protein | 0,9702 | 8,018E-29 |
| TcCLB.511133.30 | hypothetical protein, conserved | 0,9699 | 9,935E-26 |
| TcCLB.510155.100 | hypothetical protein | 0,9697 | 7,568E-26 |
| TcCLB.503447.60 | beta galactofuranosyl glycosyltransferase, putative | 0,9692 | 0,01157 |
| TcCLB.510761.14 | 60S ribosomal protein L26, putative | 0,969 | 6,095E-31 |
| TcCLB.506289.9 | DNA ligase, putative (fragment) | 0,9687 | 8,031E-14 |
| TcCLB.510611.60 | nitroreductase | 0,968 | 3,034E-18 |
| TcCLB.503809.120 | hypothetical protein, conserved | 0,968 | 6,645E-09 |
| TcCLB.509683.117 | 40S ribosomal protein S4, putative | 0,9678 | 8,378E-53 |
| TcCLB.504269.10 | protein kinase (pseudogene), putative | 0,9676 | 4,451E-19 |
| TcCLB.506343.30 | retrotransposon hot spot protein (RHS, pseudogene), puta | 0,9673 | 1,379E-23 |
| TcCLB.506301.30 | hypothetical protein, conserved | 0,967 | 3,814E-29 |
| TcCLB.511647.10 | hypothetical protein, conserved | 0,967 | 8,14E-26 |
| TcCLB.470827.10 | retrotransposon hot spot protein (RHS, pseudogene), puta | 0,9668 | 1,13E-10 |
| TcCLB.509949.9 | dispersed gene family protein 1 (DGF-1), putative | 0,9668 | 0,00002661 |
| TcCLB.510719.160 | 60S ribosomal protein L22, putative | 0,9667 | 4,085E-39 |
| TcCLB.507019.83 | 40S ribosomal protein S30, putative | 0,9666 | 1,26E-26 |
| TcCLB.510945.50 | hypothetical protein, conserved | 0,9662 | 2,61E-30 |
| TcCLB.467287.30 | ATPase beta subunit, putative | 0,9662 | 1,517E-25 |
| TcCLB.510659.210 | chaperone DNAJ protein, putative | 0,9662 | 1,666E-25 |
| TcCLB.511385.60 | hypothetical protein | 0,9659 | 1,545E-11 |
| TcCLB.506435.160 | hypothetical protein, conserved | 0,9657 | 1,291E-25 |

| | | | |
|---|---|---|---|
| TcCLB.507585.20 | hypothetical protein, conserved | 0,9657 | 2,692E-17 |
| TcCLB.507509.80 | hypothetical protein, conserved | 0,9654 | 7,259E-24 |
| TcCLB.506883.9 | isopentenyl-diphosphate delta-isomerase (type II), putativ | 0,9654 | 1,908E-11 |
| TcCLB.506345.100 | beta galactofuranosyl glycosyltransferase (pseudogene), p | 0,9653 | 2,849E-12 |
| TcCLB.509007.30 | hypothetical protein, conserved | 0,9652 | 1,054E-12 |
| TcCLB.510005.30 | retrotransposon hot spot protein (RHS, pseudogene), puta | 0,9652 | 3,037E-09 |
| TcCLB.506503.53 | hypothetical protein | 0,9648 | 2,566E-13 |
| TcCLB.507711.50 | hypothetical protein, conserved | 0,9641 | 2,452E-10 |
| TcCLB.508163.360 | dispersed gene family protein 1 (DGF-1), putative | 0,9638 | 0,00001225 |
| TcCLB.511483.20 | hypothetical protein, conserved | 0,9637 | 1,435E-15 |
| TcCLB.508851.160 | hypothetical protein, conserved | 0,9632 | 2,074E-27 |
| TcCLB.506417.50 | hypothetical protein, conserved | 0,9632 | 1,457E-11 |
| TcCLB.506513.190 | NADH-ubiquinone oxidoreductase, mitochondrial, putative | 0,9624 | 2,048E-16 |
| TcCLB.509001.30 | hypothetical protein, conserved | 0,9624 | 7,845E-14 |
| TcCLB.507963.79 | nucleosome assembly protein, putative | 0,9623 | 7,936E-30 |
| TcCLB.507819.30 | trans-sialidase, Group II, putative | 0,962 | 5,074E-13 |
| TcCLB.504769.80 | hypothetical protein, conserved | 0,9616 | 1,511E-22 |
| TcCLB.511585.110 | hypothetical protein | 0,9615 | 1,197E-23 |
| TcCLB.511599.40 | dispersed gene family protein 1 (DGF-1), putative | 0,96 | 6,758E-08 |
| TcCLB.508897.50 | hypothetical protein, conserved | 0,9599 | 3,898E-21 |
| TcCLB.508463.20 | hypothetical protein, conserved | 0,9598 | 5,406E-36 |
| TcCLB.505607.9 | dispersed gene family protein 1 (DGF-1), putative | 0,9598 | 0,00005052 |
| TcCLB.511893.10 | ribonuclease inhibitor, putative | 0,9597 | 6,415E-27 |
| TcCLB.508631.20 | retrotransposon hot spot protein (RHS, pseudogene), puta | 0,9595 | 7,106E-29 |
| TcCLB.485683.10 | RNA-binding protein 42 (RNA-binding motif protein 42), p | 0,9595 | 8,135E-22 |
| TcCLB.503445.20 | trans-sialidase (pseudogene), putative | 0,9592 | 2,17E-15 |
| TcCLB.511613.151 | syntaxin binding protein (pseudogene), putative | 0,9592 | 0,03086 |
| TcCLB.506679.100 | 40S ribosomal protein S18, putative | 0,9583 | 1,293E-36 |
| TcCLB.511355.14 | cation transporter, putative | 0,9582 | 1,976E-19 |
| TcCLB.510359.330 | hypothetical protein, conserved | 0,9581 | 9,307E-19 |
| TcCLB.508153.540 | hypothetical protein, conserved | 0,9577 | 1,023E-26 |
| TcCLB.508815.160 | ER lumen retaining receptor protein, putative | 0,9577 | 2,052E-21 |
| TcCLB.506791.50 | Present in the outer mitochondrial membrane proteome 3 | 0,957 | 1,255E-14 |
| TcCLB.509967.140 | hypothetical protein, conserved | 0,957 | 1,706E-13 |
| TcCLB.509419.260 | dispersed gene family protein 1 (DGF-1), putative | 0,957 | 0,002869 |
| TcCLB.508911.50 | fructosamine kinase, putative | 0,9569 | 3,644E-10 |
| TcCLB.508357.90 | T-complex protein 1, beta subunit, putative | 0,9568 | 4,917E-38 |
| TcCLB.506733.130 | hypothetical protein, conserved | 0,9563 | 1,291E-19 |
| TcCLB.510943.26 | ribosomal protein L38, putative | 0,9563 | 2,684E-15 |
| TcCLB.506989.110 | NADH-ubiquinone oxidoreductase complex I subunit, puta | 0,955 | 3,148E-17 |
| TcCLB.503823.160 | ubiquinone biosynthesis methyltransferase, putative | 0,955 | 1,337E-16 |
| TcCLB.510383.9 | dispersed gene family protein 1 (DGF-1), putative | 0,955 | 0,00001311 |
| TcCLB.503557.14 | hypothetical protein, conserved | 0,9548 | 1,344E-16 |
| TcCLB.506203.120 | hypothetical protein, conserved | 0,9547 | 1,203E-34 |
| TcCLB.509259.180 | retrotransposon hot spot (RHS) protein, putative | 0,9546 | 1,115E-17 |

| | | | |
|---|---|---|---|
| TcCLB.503833.40 | 40S ribosomal protein S2, putative | 0,9545 | 8,69E-26 |
| TcCLB.506945.290 | ribosomal protein L15, putative | 0,9544 | 5,87E-28 |
| TcCLB.507007.99 | mitochondrial carrier protein (pseudogene), putative | 0,9544 | 1,744E-13 |
| TcCLB.506773.40 | 3-methylcrotonoyl-CoA carboxylase beta subunit, putative | 0,9543 | 1,102E-36 |
| TcCLB.504797.14 | hypothetical protein, conserved | 0,9542 | 1,372E-11 |
| TcCLB.511665.110 | glycine dehydrogenase [decarboxylating] (pseudogene), p | 0,9541 | 9,306E-07 |
| TcCLB.511867.20 | hypothetical protein, conserved | 0,954 | 2,083E-09 |
| TcCLB.511019.40 | retrotransposon hot spot protein (RHS, pseudogene), puta | 0,9538 | 6,623E-42 |
| TcCLB.510729.80 | Mitochondrial import receptor subunit ATOM40, putative | 0,9538 | 1,495E-31 |
| TcCLB.507887.10 | dihydroorotase (pseudogene), putative | 0,9538 | 0,00003922 |
| TcCLB.506399.89 | Succinate dehydrogenase [ubiquinone] flavoprotein subun | 0,9536 | 8,635E-15 |
| TcCLB.508169.90 | eukaryotic translation initiation factor 3 subunit L, putativ | 0,953 | 1,388E-36 |
| TcCLB.511523.30 | hypothetical protein | 0,9528 | 5,486E-18 |
| TcCLB.506715.30 | hypothetical protein | 0,9525 | 6,23E-17 |
| TcCLB.507089.260 | CYC2-like cyclin, putative | 0,9523 | 2,596E-16 |
| TcCLB.509695.180 | 40S ribosomal protein S23, putative | 0,9516 | 1,272E-34 |
| TcCLB.509027.70 | kinetoplastid kinetochore protein 19, putative (kkt19) | 0,9516 | 3,297E-17 |
| TcCLB.509701.10 | trifunctional enzyme alpha subunit, mitochondrial precurs | 0,9513 | 2,541E-23 |
| TcCLB.507503.70 | retrotransposon hot spot protein (RHS, pseudogene), puta | 0,9513 | 3,404E-11 |
| TcCLB.509123.30 | mitochondrial RNA binding protein 1 | 0,9512 | 1,32E-13 |
| TcCLB.509719.23 | hypothetical protein, conserved | 0,9509 | 5,622E-10 |
| TcCLB.442383.9 | 40S ribosomal protein SA, putative | 0,9504 | 8,14E-29 |
| TcCLB.505183.60 | hypothetical protein, conserved | 0,9504 | 1,594E-22 |
| TcCLB.506705.50 | hypothetical protein | 0,9503 | 2,282E-22 |
| TcCLB.506679.94 | 40S ribosomal protein S18, putative | 0,9502 | 1,049E-33 |
| TcCLB.510761.18 | hypothetical protein, conserved | 0,9499 | 7,693E-19 |
| TcCLB.510431.40 | metallo- peptidase, Clan M- Family M48 (FACE1) | 0,9497 | 1,066E-16 |
| TcCLB.509067.40 | NADH dehydrogenase subunit NI8M, putative | 0,9494 | 9,612E-13 |
| TcCLB.510535.50 | hypothetical protein | 0,9491 | 5,956E-09 |
| TcCLB.409117.20 | 40S ribosomal protein S14, putative | 0,949 | 3,552E-31 |
| TcCLB.508661.34 | hypothetical protein, conserved | 0,9489 | 9,304E-18 |
| TcCLB.506981.10 | dispersed gene family protein 1 (DGF-1), putative (fragme | 0,9487 | 0,001721 |
| TcCLB.503595.10 | hypothetical protein, conserved | 0,9485 | 2,268E-25 |
| TcCLB.506559.420 | dolichyl-P-Man:GDP-Man1GlcNAc2-PP-dolichyl alpha-1,3- | 0,9482 | 4,476E-40 |
| TcCLB.508413.44 | 60S ribosomal protein L12, putative | 0,9482 | 6,34E-36 |
| TcCLB.509233.190 | ribosomal protein S25, putative | 0,9481 | 3,018E-35 |
| TcCLB.509295.5 | retrotransposon hot spot protein (RHS, pseudogene), puta | 0,948 | 1,066E-20 |
| TcCLB.511813.20 | ubiquitin-conjugating enzyme E2, putative | 0,9477 | 1,055E-25 |
| TcCLB.504213.80 | threonine synthase, putative | 0,9476 | 5,218E-23 |
| TcCLB.503415.30 | hypothetical protein, conserved | 0,9475 | 7,849E-24 |
| TcCLB.507603.80 | hypothetical protein, conserved | 0,9461 | 5,936E-16 |
| TcCLB.509561.130 | hypothetical protein, conserved | 0,9458 | 2,91E-18 |
| TcCLB.508461.490 | 60S ribosomal protein L23, putative | 0,9453 | 6,594E-43 |
| TcCLB.511253.31 | kinesin-like protein, putative (fragment) | 0,9452 | 0,01156 |
| TcCLB.399033.19 | mitochondrial RNA binding protein 1 | 0,9451 | 2,895E-21 |

| | | | |
|---|---|---|---|
| TcCLB.506177.30 | hypothetical protein, conserved | 0,9451 | 8,371E-14 |
| TcCLB.508387.80 | hypothetical protein, conserved | 0,9446 | 3,301E-19 |
| TcCLB.510101.34 | hypothetical protein, conserved | 0,944 | 1,66E-17 |
| TcCLB.509747.80 | hypothetical protein, conserved | 0,9436 | 6,451E-11 |
| TcCLB.511883.49 | hypothetical protein, conserved (fragment) | 0,9426 | 1,936E-12 |
| TcCLB.508153.184 | 60S ribosomal protein L39, putative | 0,9425 | 2,874E-21 |
| TcCLB.511239.49 | adenosine 5'-monophosphoramidase, putative | 0,9425 | 1,082E-10 |
| TcCLB.507057.4 | myosin heavy chain, putative (fragment) | 0,9424 | 8,054E-10 |
| TcCLB.504029.60 | hypothetical protein, conserved | 0,9423 | 7,296E-18 |
| TcCLB.508179.70 | GPR1/FUN34/yaaH family, putative | 0,9411 | 2,4E-13 |
| TcCLB.503511.10 | hypothetical protein, conserved | 0,9411 | 8,675E-12 |
| TcCLB.511807.256 | hypothetical protein, conserved | 0,941 | 8,217E-14 |
| TcCLB.511305.40 | hypothetical protein, conserved | 0,9404 | 1,715E-09 |
| TcCLB.511309.7 | hypothetical protein | 0,9404 | 1,846E-07 |
| TcCLB.509599.150 | serine/threonine protein kinase, putative | 0,9399 | 3,147E-13 |
| TcCLB.509923.9 | dispersed gene family protein 1 (DGF-1), putative (fragm | 0,9392 | 0,0000123 |
| TcCLB.506885.14 | ribosomal protein L37, putative | 0,938 | 5,935E-20 |
| TcCLB.511653.40 | hypothetical protein, conserved | 0,9377 | 1,519E-11 |
| TcCLB.511771.120 | hypothetical protein, conserved (pseudogene) | 0,9377 | 1,642E-10 |
| TcCLB.509213.170 | transcription activator, putative | 0,9376 | 1,018E-28 |
| TcCLB.511209.10 | V-type proton ATPase subunit B, putative | 0,9376 | 1,069E-25 |
| TcCLB.510963.90 | elongation factor 2, putative | 0,9374 | 3,979E-36 |
| TcCLB.506683.30 | hypothetical protein, conserved | 0,9374 | 0,00001497 |
| TcCLB.508737.60 | hypothetical protein, conserved | 0,9372 | 2,306E-12 |
| TcCLB.506775.80 | 14-3-3 protein 2, putative (14-3-3-II) | 0,9371 | 5,22E-41 |
| TcCLB.509815.120 | dispersed gene family protein 1 (DGF-1, pseudogene), put | 0,9371 | 7,296E-08 |
| TcCLB.509027.60 | kinetoplastid kinetochore protein 10, putative (kkt10) | 0,9367 | 7,969E-11 |
| TcCLB.510227.29 | thioredoxin, putative (fragment) | 0,9366 | 4,148E-30 |
| TcCLB.418069.10 | heat shock protein 70 (hsp70), putative (fragment) | 0,9366 | 1,025E-17 |
| TcCLB.504079.3 | mitochondrial chaperone BCS1, putative (fragment) | 0,9366 | 2,779E-14 |
| TcCLB.504069.50 | peroxin 14, putative | 0,9363 | 3,573E-26 |
| TcCLB.509213.140 | mitochondrial RNA binding complex 1 subunit, putative | 0,9359 | 3,973E-28 |
| TcCLB.510395.30 | GTPase, putative | 0,9359 | 1,497E-10 |
| TcCLB.510893.9 | dispersed gene family protein 1 (DGF-1, pseudogene), put | 0,9359 | 0,0002686 |
| TcCLB.510755.129 | 60S ribosomal protein L12, putative | 0,9358 | 1,584E-09 |
| TcCLB.509509.50 | hypothetical protein, conserved | 0,9355 | 2,221E-13 |
| TcCLB.509317.10 | hypothetical protein, conserved | 0,9353 | 1,156E-18 |
| TcCLB.510719.260 | ATP-dependent protease subunit HslV, putative (HslV) | 0,9353 | 4,166E-17 |
| TcCLB.509109.150 | hypothetical protein, conserved | 0,9353 | 6,33E-10 |
| TcCLB.504009.70 | succinate dehydrogenase cytochrome B subunit | 0,9352 | 1,641E-18 |
| TcCLB.509437.170 | hypothetical protein | 0,9349 | 1,444E-06 |
| TcCLB.506825.170 | lanosterol synthase, putative | 0,9348 | 1,193E-20 |
| TcCLB.504769.70 | hypothetical protein, conserved | 0,9345 | 3,884E-30 |
| TcCLB.507671.10 | hypothetical protein | 0,9343 | 2,321E-09 |
| TcCLB.508031.10 | dispersed gene family protein 1 (DGF-1, pseudogene), put | 0,9343 | 0,0001723 |

| | | | |
|---|---|---|---|
| TcCLB.506773.130 | pumilio-repeat, RNA-binding protein, putative | 0,934 | 3,096E-32 |
| TcCLB.509889.9 | dispersed gene family protein 1 (DGF-1), putative | 0,934 | 0,0001215 |
| TcCLB.506817.50 | ribosomal protein L27, putative | 0,9337 | 8,8E-27 |
| TcCLB.506503.69 | glycosomal malate dehydrogenase, putative | 0,9336 | 3,675E-29 |
| TcCLB.511415.30 | retrotransposon hot spot protein (RHS, pseudogene), puta | 0,9336 | 7,484E-16 |
| TcCLB.508463.29 | hypothetical protein, conserved | 0,9336 | 3,288E-14 |
| TcCLB.508981.39 | trifunctional enzyme alpha subunit, mitochondrial precurs | 0,9335 | 1,794E-17 |
| TcCLB.511255.299 | mucin TcMUCII, putative | 0,9334 | 0,006819 |
| TcCLB.510985.20 | retrotransposon hot spot protein (RHS, pseudogene), puta | 0,9333 | 1,019E-21 |
| TcCLB.511261.140 | 60S ribosomal protein L34, putative | 0,9329 | 3,513E-31 |
| TcCLB.509109.120 | protein tyrosine phosphatase, putative | 0,9328 | 3,769E-15 |
| TcCLB.503709.20 | 60S ribosomal protein L44, putative (fragment) | 0,9327 | 5,071E-29 |
| TcCLB.506975.30 | amidohydrolase, putative | 0,9326 | 7,381E-16 |
| TcCLB.507093.80 | hypothetical protein, conserved | 0,9322 | 3,107E-13 |
| TcCLB.508823.120 | ribosomal protein S20, putative | 0,9321 | 5,053E-18 |
| TcCLB.507723.30 | retrotransposon hot spot protein (RHS, pseudogene), puta | 0,9319 | 2,645E-14 |
| TcCLB.505035.5 | hypothetical protein, conserved | 0,9319 | 4,058E-06 |
| TcCLB.511529.160 | enoyl-CoA hydratase/isomerase family protein, putative | 0,9316 | 3,295E-18 |
| TcCLB.503909.120 | cyclosome subunit, putative | 0,9315 | 1,121E-07 |
| TcCLB.510899.19 | hypothetical protein, conserved | 0,9311 | 8,264E-08 |
| TcCLB.511367.340 | mitochondrial processing peptidase alpha subunit, putativ | 0,931 | 4,194E-32 |
| TcCLB.506925.120 | eukaryotic initiation factor 5a, putative | 0,9308 | 1,98E-16 |
| TcCLB.511653.30 | hypothetical protein, conserved | 0,9304 | 3,673E-19 |
| TcCLB.507037.80 | hypothetical protein, conserved | 0,9302 | 3,355E-15 |
| TcCLB.511815.150 | hypothetical protein | 0,9299 | 4,316E-33 |
| TcCLB.510575.220 | hypothetical protein, conserved | 0,9298 | 6E-15 |
| TcCLB.510301.50 | hypothetical protein, conserved | 0,9297 | 3,209E-09 |
| TcCLB.511183.460 | dispersed gene family protein 1 (DGF-1), putative | 0,9296 | 0,00002025 |
| TcCLB.506919.120 | retrotransposon hot spot protein (RHS, pseudogene), puta | 0,9289 | 3,333E-06 |
| TcCLB.431357.10 | 3,2-trans-enoyl-CoA isomerase, mitochondrial precursor, p | 0,9285 | 4,046E-07 |
| TcCLB.508093.29 | hypothetical protein, conserved (pseudogene) | 0,9285 | 0,002019 |
| TcCLB.511823.70 | farnesyl pyrophosphate synthase | 0,9284 | 1,21E-19 |
| TcCLB.507773.60 | phosphoribosylpyrophosphate synthetase, putative | 0,9282 | 8,628E-22 |
| TcCLB.507653.10 | hypothetical protein, conserved | 0,9281 | 5,068E-09 |
| TcCLB.506241.170 | 40S ribosomal protein S6, putative | 0,9279 | 2,431E-31 |
| TcCLB.509865.9 | retrotransposon hot spot (RHS) protein, putative | 0,9279 | 5,413E-18 |
| TcCLB.507023.40 | protein kinase, putative | 0,9278 | 4,409E-27 |
| TcCLB.503943.10 | serine/threonine protein kinase, putative (fragment) | 0,9277 | 1,855E-26 |
| TcCLB.509641.70 | hypothetical protein, conserved | 0,9274 | 1,186E-27 |
| TcCLB.508241.100 | hypothetical protein | 0,9273 | 1,104E-29 |
| TcCLB.509021.9 | dispersed gene family protein 1 (DGF-1), putative | 0,9271 | 0,0545 |
| TcCLB.507589.30 | Checkpoint protein HUS1, putative | 0,9269 | 6,059E-23 |
| TcCLB.508823.140 | ribosomal protein S20, putative | 0,9263 | 7,093E-24 |
| TcCLB.506375.60 | tryptophanyl-tRNA synthetase, putative | 0,9263 | 1,128E-10 |
| TcCLB.509331.20 | hypothetical protein, conserved | 0,926 | 2,219E-15 |

| | | | |
|---|---|---|---|
| TcCLB.507625.140 | membrane transporter protein, putative | 0,9256 | 4,585E-20 |
| TcCLB.508587.10 | retrotransposon hot spot protein (RHS, pseudogene), puta | 0,9256 | 3,732E-13 |
| TcCLB.503529.20 | hypothetical protein, conserved | 0,9252 | 8,01E-17 |
| TcCLB.511903.110 | 40S ribosomal protein S8, putative | 0,925 | 1,428E-32 |
| TcCLB.508881.120 | prefoldin, putative | 0,925 | 1,203E-21 |
| TcCLB.509165.40 | 60S acidic ribosomal protein P2 beta (H6.4), putative (P2-I | 0,9247 | 3,743E-32 |
| TcCLB.510911.10 | profilin, putative | 0,9243 | 1,145E-13 |
| TcCLB.504097.30 | hypothetical protein, conserved | 0,9241 | 8,996E-29 |
| TcCLB.506297.230 | hypothetical protein, conserved | 0,9234 | 7,848E-14 |
| TcCLB.460747.9 | hypothetical protein, conserved (fragment) | 0,923 | 1,004E-19 |
| TcCLB.504229.110 | amino acid permease 24, putative (AAP24) | 0,923 | 4,337E-10 |
| TcCLB.510659.270 | hypothetical protein, conserved | 0,9226 | 5,745E-14 |
| TcCLB.506855.20 | hypothetical protein, conserved | 0,922 | 2,786E-15 |
| TcCLB.509905.100 | retrotransposon hot spot protein (RHS, pseudogene), puta | 0,9215 | 2,083E-15 |
| TcCLB.510005.50 | dispersed gene family protein 1 (DGF-1), putative | 0,9214 | 0,00000479 |
| TcCLB.510001.20 | hypothetical protein, conserved | 0,9212 | 1,923E-20 |
| TcCLB.507875.90 | dispersed gene family protein 1 (DGF-1, pseudogene), put | 0,9212 | 2,651E-09 |
| TcCLB.509429.210 | leucine-rich repeat protein (LRRP), putative | 0,921 | 1,403E-26 |
| TcCLB.508173.20 | retrotransposon hot spot protein (RHS, pseudogene), puta | 0,9209 | 2,771E-44 |
| TcCLB.508807.10 | ATP-dependent Clp protease subunit, heat shock protein 1 | 0,9205 | 5,89E-19 |
| TcCLB.506511.20 | hypothetical protein, conserved | 0,9205 | 3,801E-16 |
| TcCLB.507951.160 | succinate dehydrogenase flavoprotein subunit | 0,9203 | 2,916E-37 |
| TcCLB.509937.39 | hypothetical protein, conserved | 0,9201 | 4,625E-11 |
| TcCLB.511773.110 | retrotransposon hot spot (RHS) protein, putative | 0,9197 | 2,34E-25 |
| TcCLB.511621.50 | RNA-binding protein, putative | 0,9196 | 2,352E-26 |
| TcCLB.511135.40 | hypothetical protein, conserved | 0,9196 | 6,598E-11 |
| TcCLB.504003.60 | chaperonin/T-complex protein 1 gamma subunit, putative | 0,9195 | 6,777E-40 |
| TcCLB.506357.100 | DNA-directed RNA polymerase I subunit RPB5z, putative ( | 0,9189 | 1,057E-11 |
| TcCLB.504507.5 | trypanothione reductase, putative (fragment) | 0,9185 | 4,136E-12 |
| TcCLB.510551.10 | dispersed gene family protein 1 (DGF-1, pseudogene), put | 0,9184 | 0,0007267 |
| TcCLB.511621.220 | esterase, putative | 0,9183 | 3,412E-06 |
| TcCLB.507251.20 | ribosomal protein L21E (60S), putative | 0,9182 | 1,167E-47 |
| TcCLB.510101.10 | protein transport protein Sec31, putative (pseudogene) | 0,9182 | 3,3E-12 |
| TcCLB.511647.20 | hypothetical protein, conserved | 0,9181 | 6,849E-11 |
| TcCLB.511913.9 | ribonuclease L inhibitor (pseudogene), putative | 0,9178 | 4,498E-19 |
| TcCLB.506885.340 | Mitochondrial import receptor subunit ATOM40, putative | 0,9177 | 5,828E-28 |
| TcCLB.505989.90 | hypothetical protein, conserved | 0,9175 | 9,497E-20 |
| TcCLB.511763.10 | hypothetical protein, conserved | 0,9173 | 3,177E-08 |
| TcCLB.416041.24 | retrotransposon hot spot protein (RHS, pseudogene), puta | 0,9172 | 3,349E-09 |
| TcCLB.507595.10 | dispersed gene family protein 1 (DGF-1, pseudogene), put | 0,917 | 0,005596 |
| TcCLB.505647.50 | hypothetical protein, conserved | 0,9167 | 3,387E-07 |
| TcCLB.506359.30 | hypothetical protein, conserved (pseudogene) | 0,9164 | 1,556E-15 |
| TcCLB.508647.270 | malic enzyme, putative | 0,9163 | 3,546E-30 |
| TcCLB.506941.40 | hypothetical protein, conserved | 0,9163 | 4,079E-16 |
| TcCLB.509643.70 | Archaic translocase of outer membrane 11 kDa subunit, pu | 0,9162 | 3,414E-16 |

| | | | |
|---|---|---|---|
| TcCLB.511771.70 | dispersed gene family protein 1 (DGF-1), putative | 0,9161 | 8,451E-09 |
| TcCLB.511885.10 | hypothetical protein, conserved (pseudogene) | 0,9155 | 2,271E-23 |
| TcCLB.506581.30 | eukaryotic translation initiation factor 3 subunit L, putativ | 0,915 | 1,774E-29 |
| TcCLB.509725.60 | glycine dehydrogenase [decarboxylating] (pseudogene), p | 0,915 | 1,7E-14 |
| TcCLB.509049.30 | hypothetical protein, conserved | 0,9146 | 3,093E-25 |
| TcCLB.507925.60 | calcium-binding protein, putative | 0,9144 | 8,058E-11 |
| TcCLB.506413.30 | 40S ribosomal protein S33, putative | 0,9144 | 2,523E-08 |
| TcCLB.511445.10 | calpain-like cysteine peptidase, putative | 0,9141 | 1,693E-06 |
| TcCLB.511287.40 | 40S ribosomal protein S2, putative | 0,9138 | 6,009E-41 |
| TcCLB.509029.80 | hypothetical protein | 0,9138 | 1,468E-10 |
| TcCLB.504155.221 | surface protease GP63 (pseudogene), putative | 0,9137 | 0,1883 |
| TcCLB.506633.74 | hypothetical protein, conserved (pseudogene) | 0,9135 | 4,465E-12 |
| TcCLB.506657.40 | ADP/ATP translocase, putative | 0,9134 | 1,576E-22 |
| TcCLB.505229.5 | hypothetical protein, conserved (fragment) | 0,9134 | 3,881E-12 |
| TcCLB.511361.50 | hypothetical protein, conserved | 0,9134 | 2,404E-08 |
| TcCLB.511817.100 | hypothetical protein, conserved | 0,9133 | 4,698E-25 |
| TcCLB.506657.79 | 50S ribosomal protein L16, putative | 0,9131 | 1,819E-13 |
| TcCLB.510359.310 | eukaryotic translation initiation factor 2 subunit, putative | 0,913 | 6,581E-23 |
| TcCLB.508173.60 | cis-prenyltransferase, putative | 0,913 | 2,309E-13 |
| TcCLB.511131.40 | hypothetical protein | 0,913 | 2,907E-10 |
| TcCLB.507709.50 | 60S ribosomal protein L6, putative | 0,9128 | 1,46E-28 |
| TcCLB.511847.80 | hypothetical protein, conserved | 0,9127 | 2,765E-17 |
| TcCLB.511825.80 | I/6 autoantigen, putative | 0,9127 | 3,645E-14 |
| TcCLB.508163.70 | dispersed gene family protein 1 (DGF-1), putative | 0,9127 | 0,000508 |
| TcCLB.509201.15 | ribosomal protein S29, putative | 0,9124 | 1,033E-13 |
| TcCLB.510485.9 | dispersed gene family protein 1 (DGF-1), putative | 0,9123 | 0,0007276 |
| TcCLB.504255.20 | hypothetical protein | 0,9118 | 7,28E-23 |
| TcCLB.510349.20 | hypothetical protein, conserved | 0,9111 | 1,429E-13 |
| TcCLB.510095.5 | retrotransposon hot spot protein (RHS, pseudogene), puta | 0,911 | 0,01517 |
| TcCLB.509589.10 | mitochondrial DNA polymerase I protein D, putative | 0,9109 | 2,717E-14 |
| TcCLB.510499.10 | dispersed gene family protein 1 (DGF-1), putative | 0,9109 | 0,008987 |
| TcCLB.509453.79 | hypothetical protein, conserved | 0,9106 | 6,501E-27 |
| TcCLB.506275.121 | hypothetical protein | 0,9105 | 0,09897 |
| TcCLB.509601.120 | hypothetical protein, conserved | 0,9104 | 1,543E-26 |
| TcCLB.506475.100 | conserved protein, unknown function | 0,9104 | 2,739E-15 |
| TcCLB.505193.84 | hypothetical protein, conserved | 0,9104 | 4,061E-12 |
| TcCLB.510359.180 | hypothetical protein, conserved | 0,9104 | 9,896E-11 |
| TcCLB.464879.9 | ribonuclease L inhibitor, putative (fragment) | 0,9099 | 8,392E-10 |
| TcCLB.511751.210 | phosphopantothenoylcysteine decarboxylase, putative (PP | 0,9098 | 1,136E-18 |
| TcCLB.507715.34 | hypothetical protein, conserved | 0,9095 | 7,708E-22 |
| TcCLB.508377.9 | fucose kinase, putative (fragment) | 0,9094 | 1,034E-20 |
| TcCLB.510123.10 | hypothetical protein, conserved | 0,9092 | 2,679E-16 |
| TcCLB.509023.60 | hypothetical protein, conserved | 0,909 | 1,967E-17 |
| TcCLB.509695.20 | hypothetical protein, conserved | 0,9086 | 3,351E-12 |
| TcCLB.506393.90 | dispersed gene family protein 1 (DGF-1), putative | 0,9086 | 1,438E-09 |

| | | | |
|---|---|---|---|
| TcCLB.509127.50 | mitochondrial carrier protein, putative | 0,9084 | 7,611E-18 |
| TcCLB.506331.104 | retrotransposon hot spot protein (RHS, pseudogene), puta | 0,9084 | 5,181E-07 |
| TcCLB.503441.19 | retrotransposon hot spot protein (RHS, pseudogene), puta | 0,9083 | 1,532E-11 |
| TcCLB.503579.90 | hypothetical protein, conserved | 0,9076 | 3,523E-14 |
| TcCLB.505171.90 | hypothetical protein | 0,9073 | 1,764E-14 |
| TcCLB.511323.40 | histone H2A, putative | 0,9072 | 9,63E-09 |
| TcCLB.503563.14 | ADG1, putative | 0,9071 | 1,747E-12 |
| TcCLB.504153.340 | hypothetical protein, conserved | 0,9068 | 1,333E-19 |
| TcCLB.506925.110 | hypothetical protein, conserved | 0,9064 | 1,025E-29 |
| TcCLB.509695.170 | 60S ribosomal protein L9, putative | 0,9062 | 1,728E-46 |
| TcCLB.509693.200 | hypothetical protein, conserved | 0,9062 | 8,089E-12 |
| TcCLB.510863.20 | hypothetical protein | 0,9061 | 3,928E-16 |
| TcCLB.506715.40 | hypothetical protein, conserved | 0,9055 | 5,688E-24 |
| TcCLB.507873.60 | 1,2-Dihydroxy-3-keto-5-methylthiopentene dioxygenase, p | 0,9054 | 3,013E-07 |
| TcCLB.508625.160 | pumilio/PUF RNA binding protein 1, putative (PUF1) | 0,9051 | 4,76E-21 |
| TcCLB.510515.60 | cytosolic leucyl aminopeptidase, putative | 0,905 | 2,883E-16 |
| TcCLB.508859.30 | hypothetical protein, conserved | 0,9046 | 4,225E-16 |
| TcCLB.506469.80 | protein kinase, putative | 0,9046 | 9,426E-10 |
| TcCLB.511255.470 | retrotransposon hot spot protein (RHS, pseudogene), puta | 0,9042 | 0,0003447 |
| TcCLB.510661.140 | hypothetical protein, conserved | 0,904 | 4,032E-20 |
| TcCLB.511817.240 | hypothetical protein, conserved | 0,904 | 7,011E-12 |
| TcCLB.506295.100 | hypothetical protein, conserved | 0,9038 | 1,442E-29 |
| TcCLB.506925.230 | hypothetical protein, conserved | 0,9037 | 9,406E-30 |
| TcCLB.507581.9 | dispersed gene family protein 1 (DGF-1), putative (fragme | 0,9033 | 0,00003359 |
| TcCLB.511419.30 | hypothetical protein, conserved | 0,903 | 3,7E-10 |
| TcCLB.507529.50 | hypothetical protein, conserved | 0,9029 | 4,173E-33 |
| TcCLB.509331.60 | hypothetical protein, conserved | 0,9029 | 3,403E-10 |
| TcCLB.507681.160 | 40S ribosomal protein S24E, putative | 0,9021 | 4,331E-14 |
| TcCLB.511771.210 | dispersed gene family protein 1 (DGF-1), putative | 0,9021 | 1,8E-08 |
| TcCLB.506797.30 | hypothetical protein | 0,9015 | 3,82E-26 |
| TcCLB.511071.171 | 60S ribosomal protein L12, putative | 0,9012 | 1,436E-37 |
| TcCLB.508319.70 | 40S ribosomal protein S11, putative | 0,9003 | 6,156E-29 |
| TcCLB.509599.100 | hypothetical protein | 0,9003 | 3,383E-07 |
| TcCLB.506467.50 | serine/threonine protein kinase, putative | 0,9002 | 5,93E-25 |
| TcCLB.509153.120 | acyl-CoA dehydrogenase, putative | 0,9 | 2,189E-20 |
| TcCLB.508065.40 | hypothetical protein, conserved | 0,9 | 9,334E-15 |
| TcCLB.508975.20 | pre-rrna-processing protein tsr1 homolog | 0,8998 | 3,219E-25 |
| TcCLB.509767.230 | hypothetical protein, conserved | 0,8997 | 5,077E-12 |
| TcCLB.509647.180 | nicotinate phosphoribosyltransferase, putative | 0,8994 | 2,286E-24 |
| TcCLB.510359.604 | mucin TcMUCII (pseudogene), putative | 0,8991 | 0,0008967 |
| TcCLB.509461.10 | actin interacting protein-like protein, putative | 0,8985 | 2,665E-20 |
| TcCLB.485335.10 | retrotransposon hot spot protein (RHS, pseudogene), puta | 0,8985 | 5,126E-09 |
| TcCLB.510713.50 | beta galactofuranosyl glycosyltransferase (pseudogene), p | 0,8985 | 0,001581 |
| TcCLB.506469.30 | protein kinase, putative | 0,8983 | 8,177E-08 |
| TcCLB.505035.20 | DNA-directed RNA polymerase II subunit 2 (pseudogene), | 0,8982 | 1,66E-10 |

| | | | |
|---|---|---|---|
| TcCLB.503893.70 | reductase, putative | 0,898 | 1,778E-16 |
| TcCLB.511585.260 | dispersed gene family protein 1 (DGF-1, pseudogene), put | 0,8975 | 5,476E-10 |
| TcCLB.509611.150 | hypothetical protein, conserved | 0,8974 | 3,076E-15 |
| TcCLB.508269.20 | hypothetical protein, conserved | 0,8968 | 5,366E-09 |
| TcCLB.508873.370 | hypothetical protein | 0,8967 | 0,00138 |
| TcCLB.511753.60 | ESAG8-associated protein, putative (PIE8) | 0,8966 | 2,154E-10 |
| TcCLB.504021.74 | hypothetical protein, conserved | 0,8965 | 5,683E-09 |
| TcCLB.503603.20 | hypothetical protein, conserved | 0,8962 | 8,743E-14 |
| TcCLB.508911.70 | hypothetical protein | 0,8958 | 1,649E-15 |
| TcCLB.506435.400 | hypothetical protein, conserved | 0,8954 | 2,006E-21 |
| TcCLB.506495.3 | hypothetical protein, conserved | 0,8953 | 0,00001023 |
| TcCLB.504949.14 | 60S ribosomal protein L6, putative | 0,8951 | 5,24E-18 |
| TcCLB.508059.4 | hypothetical protein, conserved | 0,8949 | 1,42E-09 |
| TcCLB.511887.45 | hypothetical protein, conserved (pseudogene) | 0,8945 | 0,02419 |
| TcCLB.508687.20 | retrotransposon hot spot protein (RHS, pseudogene), puta | 0,894 | 2,207E-10 |
| TcCLB.506751.70 | retrotransposon hot spot protein (RHS, pseudogene), puta | 0,8939 | 0,001797 |
| TcCLB.508325.160 | retrotransposon hot spot (RHS) protein, putative | 0,8937 | 2,272E-21 |
| TcCLB.511245.30 | ion transport protein, putative | 0,8936 | 4E-15 |
| TcCLB.511725.130 | hypothetical protein, conserved | 0,8933 | 5,82E-32 |
| TcCLB.508831.120 | glucosamine 6-phosphate n-acetyltransferase, putative | 0,8926 | 3,167E-09 |
| TcCLB.510055.140 | hypothetical protein, conserved | 0,8923 | 4,232E-12 |
| TcCLB.508737.70 | cytochrome b5-dependent oleate desaturase | 0,8918 | 4,405E-17 |
| TcCLB.508379.10 | fucose kinase, putative | 0,8917 | 1,949E-08 |
| TcCLB.506401.240 | hypothetical protein, conserved | 0,8916 | 7,382E-28 |
| TcCLB.508647.200 | triosephosphate isomerase, putative (TIM) | 0,8915 | 1,186E-40 |
| TcCLB.508415.50 | ADP-ribosylation factor 1, putative (ARF1) | 0,8915 | 3,108E-16 |
| TcCLB.508729.18 | hypothetical protein, conserved | 0,8911 | 4,061E-15 |
| TcCLB.510187.150 | guanylate kinase, putative | 0,8908 | 1,137E-14 |
| TcCLB.510719.40 | hypothetical protein, conserved | 0,8908 | 4,419E-11 |
| TcCLB.511249.20 | ADP/ATP translocase, putative | 0,8907 | 4,208E-20 |
| TcCLB.509529.60 | dispersed gene family protein 1 (DGF-1), putative | 0,8907 | 9,605E-12 |
| TcCLB.508719.40 | kinetoplast DNA-associated protein, putative | 0,8904 | 1,614E-14 |
| TcCLB.511291.40 | protein kinase, putative | 0,8902 | 2,262E-20 |
| TcCLB.505945.60 | hypothetical protein, conserved | 0,8901 | 2,023E-15 |
| TcCLB.506895.40 | Kinesin-13 1, putative (KIN13-1) | 0,89 | 1,433E-18 |
| TcCLB.506183.40 | retrotransposon hot spot protein (RHS, pseudogene), puta | 0,8897 | 4,094E-08 |
| TcCLB.510947.40 | agmatinase, putative | 0,8895 | 3,935E-12 |
| TcCLB.511367.150 | hypothetical protein, conserved | 0,8892 | 1,005E-14 |
| TcCLB.508045.90 | oxidoreductase, putative | 0,889 | 3,565E-15 |
| TcCLB.504181.30 | 40S ribosomal protein S23, putative | 0,889 | 9,086E-07 |
| TcCLB.508325.270 | retrotransposon hot spot protein (RHS, pseudogene), puta | 0,8887 | 2,694E-12 |
| TcCLB.508351.60 | hypothetical protein, conserved | 0,8884 | 1,671E-17 |
| TcCLB.503935.20 | 2-oxoglutarate dehydrogenase, E2 component, dihydrolipo | 0,888 | 1,122E-27 |
| TcCLB.438147.18 | hypothetical protein, conserved | 0,8879 | 1,723E-06 |
| TcCLB.506821.150 | hypothetical protein, conserved | 0,8874 | 1,146E-16 |

| | | | |
|---|---|---|---|
| TcCLB.510101.300 | hypothetical protein, conserved | 0,8866 | 1,575E-28 |
| TcCLB.508355.260 | 60S acidic ribosomal protein P0, putative (P0) | 0,8864 | 2,886E-45 |
| TcCLB.504151.80 | Na/H antiporter-like protein, putative | 0,8859 | 5,785E-11 |
| TcCLB.508027.120 | hypothetical protein, conserved | 0,8858 | 1,53E-16 |
| TcCLB.506619.40 | tyrosyl-DNA Phosphodiesterase (Tdp1), putative | 0,8858 | 9,184E-13 |
| TcCLB.504081.30 | trans-sialidase (pseudogene), putative | 0,8856 | 0,00001111 |
| TcCLB.510123.50 | iron-sulfur cluster assembly protein, putative | 0,8854 | 4,161E-16 |
| TcCLB.506117.9 | dispersed gene family protein 1 (DGF-1), putative | 0,8852 | 1,162E-09 |
| TcCLB.504427.170 | hypothetical protein, conserved | 0,8846 | 1,132E-09 |
| TcCLB.511301.80 | choline/carnitine O-acyltransferase (pseudogene), putativ | 0,8844 | 1,994E-19 |
| TcCLB.511771.143 | retrotransposon hot spot protein (RHS, pseudogene), puta | 0,8841 | 0,001357 |
| TcCLB.506977.60 | hypothetical protein, conserved | 0,884 | 3,043E-10 |
| TcCLB.509029.90 | hypothetical protein, conserved | 0,884 | 8,674E-10 |
| TcCLB.507779.30 | dispersed gene family protein 1 (DGF-1), putative | 0,884 | 2,872E-08 |
| TcCLB.508175.309 | 60S ribosomal protein L19, putative | 0,8834 | 1,729E-23 |
| TcCLB.511159.30 | Present in the outer mitochondrial membrane proteome 1 | 0,8831 | 3,482E-13 |
| TcCLB.510149.150 | NADH-ubiquinone oxidoreductase complex I subunit, puta | 0,8831 | 2,455E-10 |
| TcCLB.506887.100 | protein kinase, putative | 0,8829 | 4,941E-10 |
| TcCLB.510089.60 | hypothetical protein, conserved | 0,8828 | 4,1E-09 |
| TcCLB.510105.220 | transportin2-like protein, putative | 0,8826 | 1,366E-17 |
| TcCLB.508693.80 | ruvB-like DNA helicase, putative | 0,8825 | 1,088E-27 |
| TcCLB.509671.64 | ribosomal protein L36, putative | 0,8824 | 1,907E-14 |
| TcCLB.510299.50 | GPI alpha-mannosyltransferase III (GPI10) | 0,8823 | 5,187E-24 |
| TcCLB.509331.100 | hypothetical protein, conserved | 0,8822 | 1,567E-09 |
| TcCLB.506645.9 | dispersed gene family protein 1 (DGF-1), putative | 0,8822 | 0,00000212 |
| TcCLB.510303.320 | dihydrofolate reductase-thymidylate synthase, putative | 0,8818 | 2,727E-12 |
| TcCLB.507063.170 | hypothetical protein | 0,8815 | 8,647E-06 |
| TcCLB.510287.50 | hypothetical protein, conserved | 0,8814 | 6,922E-11 |
| TcCLB.508551.20 | 40S ribosomal protein S12, putative | 0,8813 | 3,4E-30 |
| TcCLB.507809.160 | hypothetical protein, conserved | 0,8813 | 6,357E-16 |
| TcCLB.505419.20 | retrotransposon hot spot protein (RHS, pseudogene), puta | 0,8813 | 1,46E-13 |
| TcCLB.507275.40 | hypothetical protein, conserved | 0,881 | 2,224E-10 |
| TcCLB.507049.190 | hypothetical protein, conserved | 0,8806 | 3,2E-23 |
| TcCLB.508461.440 | hypothetical protein, conserved | 0,8806 | 9,356E-15 |
| TcCLB.510763.34 | hypothetical protein, conserved | 0,8803 | 2,724E-11 |
| TcCLB.510735.50 | hypothetical protein, conserved | 0,8802 | 2,985E-15 |
| TcCLB.510661.90 | ATP-dependent RNA helicase, putative | 0,8798 | 2,158E-20 |
| TcCLB.504103.60 | hypothetical protein, conserved | 0,8794 | 9,844E-09 |
| TcCLB.506675.90 | retrotransposon hot spot protein (RHS, pseudogene), puta | 0,8793 | 3,371E-08 |
| TcCLB.507501.25 | hypothetical protein, conserved (pseudogene) | 0,8793 | 0,008811 |
| TcCLB.510999.10 | hypothetical protein, conserved | 0,879 | 1,611E-15 |
| TcCLB.456021.10 | hypothetical protein, conserved (fragment) | 0,879 | 0,00009005 |
| TcCLB.506405.149 | ribosomal protein L21E (60S), putative | 0,8789 | 1,517E-45 |
| TcCLB.507101.20 | hypothetical protein (pseudogene) | 0,8787 | 0,0002607 |
| TcCLB.506657.20 | hypothetical protein, conserved | 0,8786 | 4,39E-15 |

| | | | |
|---|---|---|---|
| TcCLB.506735.30 | hypothetical protein, conserved | 0,8785 | 3,974E-14 |
| TcCLB.508299.60 | 60S ribosomal protein L2, putative | 0,8783 | 3,876E-36 |
| TcCLB.505999.20 | phosphate-repressible phosphate permease, putative | 0,8783 | 3,873E-20 |
| TcCLB.508641.280 | hypothetical protein, conserved | 0,8776 | 2,05E-31 |
| TcCLB.510349.30 | phosphatidylinositol-4-phosphate 5-kinase, putative | 0,8776 | 2,513E-12 |
| TcCLB.508479.330 | hypothetical protein, conserved | 0,8775 | 2,172E-24 |
| TcCLB.506773.104 | hypothetical protein, conserved | 0,8774 | 7,479E-15 |
| TcCLB.507735.50 | hypothetical protein, conserved (pseudogene) | 0,8772 | 4,34E-25 |
| TcCLB.509991.80 | hypothetical protein, conserved | 0,8772 | 1,785E-13 |
| TcCLB.506855.180 | adenylate kinase, putative | 0,877 | 1,888E-10 |
| TcCLB.511441.10 | calpain cysteine peptidase, putative | 0,877 | 2,313E-09 |
| TcCLB.511117.10 | retrotransposon hot spot protein (RHS, pseudogene), puta | 0,8767 | 5,17E-12 |
| TcCLB.506257.5 | aminopeptidase, putative (fragment) | 0,8766 | 1,291E-06 |
| TcCLB.509339.10 | hypothetical protein, conserved | 0,8765 | 1,69E-12 |
| TcCLB.508221.410 | hypothetical protein | 0,8765 | 0,03184 |
| TcCLB.507615.110 | Present in the outer mitochondrial membrane proteome 3 | 0,876 | 1,406E-13 |
| TcCLB.505171.100 | hypothetical protein, conserved | 0,8759 | 1,411E-08 |
| TcCLB.504035.70 | cyclophilin, putative | 0,8756 | 9,728E-12 |
| TcCLB.509331.9 | 5'-3' exoribonuclease C, putative (fragment) (XRNC) | 0,8756 | 7,246E-06 |
| TcCLB.506055.40 | dispersed gene family protein 1 (DGF-1), putative | 0,8755 | 3,338E-06 |
| TcCLB.506263.40 | hypothetical protein, conserved | 0,8753 | 1,477E-10 |
| TcCLB.510687.30 | hypothetical protein, conserved | 0,8752 | 2,015E-30 |
| TcCLB.504177.30 | hypothetical protein, conserved | 0,8752 | 7,92E-14 |
| TcCLB.509563.50 | hypothetical protein, conserved | 0,875 | 7,545E-12 |
| TcCLB.506475.90 | hypothetical protein, conserved | 0,8749 | 1,961E-14 |
| TcCLB.511545.120 | superoxide dismutase, putative | 0,8748 | 1,827E-14 |
| TcCLB.510301.20 | fructose-bisphosphate aldolase, glycosomal, putative | 0,8745 | 8,337E-15 |
| TcCLB.503801.20 | ribosomal protein S26, putative | 0,8744 | 2,222E-29 |
| TcCLB.503669.39 | hypothetical protein, conserved (fragment) | 0,8742 | 0,0001478 |
| TcCLB.505987.20 | hypothetical protein | 0,874 | 2,687E-18 |
| TcCLB.511307.3 | membrane transporter protein, putative (fragment) | 0,874 | 1,738E-09 |
| TcCLB.510519.10 | hypothetical protein | 0,8739 | 6,865E-20 |
| TcCLB.510903.40 | hypothetical protein, conserved | 0,8738 | 2,608E-12 |
| TcCLB.506707.20 | ATP-dependent DEAD/H RNA helicase (pseudogene), puta | 0,8738 | 0,001165 |
| TcCLB.510659.130 | hypothetical protein, conserved | 0,8734 | 1,246E-29 |
| TcCLB.506679.150 | 40S ribosomal protein S10, putative | 0,873 | 3,791E-10 |
| TcCLB.503611.40 | ribosomal protein L24, putative | 0,8729 | 0,002468 |
| TcCLB.508909.310 | hypothetical protein | 0,8728 | 3,545E-07 |
| TcCLB.509337.28 | hypothetical protein, conserved (pseudogene) | 0,8727 | 7,149E-16 |
| TcCLB.511411.24 | hypothetical protein, conserved | 0,8726 | 7,216E-11 |
| TcCLB.507857.50 | protein phosphotase, putative | 0,8725 | 2,132E-17 |
| TcCLB.509759.10 | L-ribulokinase, putative | 0,8721 | 6,242E-07 |
| TcCLB.509233.60 | hypothetical protein, conserved | 0,8718 | 5,943E-18 |
| TcCLB.506407.10 | lysosomal alpha-mannosidase precursor, putative | 0,8716 | 3,29E-16 |
| TcCLB.510101.410 | hypothetical protein, conserved | 0,8715 | 2,51E-19 |

| | | | |
|---|---|---|---|
| TcCLB.471901.10 | hypothetical protein, conserved | 0,871 | 1,354E-20 |
| TcCLB.510403.20 | retrotransposon hot spot protein (RHS, pseudogene), puta | 0,871 | 3,725E-13 |
| TcCLB.509029.20 | hypothetical protein, conserved | 0,8707 | 3,351E-19 |
| TcCLB.511819.40 | hypothetical protein, conserved | 0,8706 | 1,407E-16 |
| TcCLB.510055.110 | fructosamine kinase, putative | 0,8705 | 8,24E-12 |
| TcCLB.508153.290 | 60S ribosomal protein L13, putative | 0,8704 | 8,145E-31 |
| TcCLB.507975.10 | retrotransposon hot spot protein (RHS, pseudogene), puta | 0,8704 | 3,497E-23 |
| TcCLB.511809.135 | histone H2A, putative | 0,8697 | 1,461E-20 |
| TcCLB.506941.159 | zinc finger protein, putative | 0,8695 | 1,317E-16 |
| TcCLB.507573.6 | ATP-dependent DEAD/H RNA helicase (pseudogene), puta | 0,8695 | 0,00008202 |
| TcCLB.509911.20 | hypothetical protein | 0,8694 | 0,00004264 |
| TcCLB.506529.530 | Present in the outer mitochondrial membrane proteome 1 | 0,8693 | 1,823E-07 |
| TcCLB.507573.3 | N-acetyltransferase complex ARD1 subunit (pseudogene), | 0,8693 | 0,00007332 |
| TcCLB.509859.50 | hypothetical protein, conserved | 0,8692 | 9,002E-13 |
| TcCLB.509151.100 | hypothetical protein, conserved | 0,869 | 1,652E-15 |
| TcCLB.503899.130 | glutathione peroxidase-like protein, putative | 0,8688 | 5,16E-10 |
| TcCLB.508269.10 | hypothetical protein, conserved | 0,8686 | 7,739E-15 |
| TcCLB.511231.34 | hypothetical protein, conserved | 0,8686 | 3,306E-14 |
| TcCLB.509109.145 | hypothetical protein, conserved (pseudogene) | 0,8685 | 1,074E-06 |
| TcCLB.511715.10 | iron superoxide dismutase, putative | 0,8684 | 1,284E-12 |
| TcCLB.509555.10 | hypothetical protein (fragment) | 0,8683 | 6,767E-11 |
| TcCLB.509065.50 | hypothetical protein, conserved | 0,8682 | 2,293E-14 |
| TcCLB.507559.100 | hypothetical protein, conserved | 0,8678 | 2,791E-29 |
| TcCLB.509965.30 | small nuclear ribonucleoprotein Sm-E, putative | 0,8675 | 0,00000461 |
| TcCLB.506621.60 | acyl-CoA binding protein, putative | 0,867 | 3,791E-25 |
| TcCLB.508909.230 | calcium-transporting ATPase (pseudogene), putative | 0,8665 | 4,724E-13 |
| TcCLB.506679.140 | 40S ribosomal protein S10, putative | 0,8663 | 8,432E-12 |
| TcCLB.510857.60 | cation-transporting ATPase, putative | 0,8661 | 6,348E-14 |
| TcCLB.507649.90 | hypothetical protein, conserved | 0,866 | 6,013E-19 |
| TcCLB.509337.19 | hypothetical protein, conserved | 0,8657 | 4,05E-10 |
| TcCLB.510529.39 | hypothetical protein, conserved | 0,8657 | 1,391E-06 |
| TcCLB.508479.340 | succinyl-CoA synthetase alpha subunit, putative | 0,8656 | 2,719E-32 |
| TcCLB.509911.100 | chaperone DNAJ protein, putative | 0,8656 | 9,157E-12 |
| TcCLB.509819.20 | hypothetical protein, conserved | 0,8653 | 7,208E-28 |
| TcCLB.510733.5 | hypothetical protein, conserved (fragment) | 0,8651 | 3,181E-07 |
| TcCLB.510655.150 | hypothetical protein, conserved | 0,8649 | 2,43E-10 |
| TcCLB.417939.9 | dispersed gene family protein 1 (DGF-1), putative | 0,8648 | 0,003643 |
| TcCLB.511511.159 | zinc finger protein, putative | 0,8645 | 8,472E-21 |
| TcCLB.511211.200 | hypothetical protein, conserved | 0,8645 | 5,249E-16 |
| TcCLB.504137.30 | hypothetical protein | 0,8641 | 1,45E-18 |
| TcCLB.509647.130 | hypothetical protein, conserved | 0,864 | 6,15E-15 |
| TcCLB.509367.15 | tryptophanyl-tRNA synthetase (pseudogene), putative | 0,864 | 0,0002053 |
| TcCLB.503677.20 | DNA repair and recombination helicase protein PIF4, puta | 0,8636 | 2,516E-08 |
| TcCLB.466007.9 | dispersed gene family protein 1 (DGF-1), putative (fragme | 0,8631 | 0,03907 |
| TcCLB.511001.30 | hypothetical protein, conserved | 0,8629 | 2,687E-15 |

| | | | |
|---|---|---|---|
| TcCLB.509455.100 | hypothetical protein, conserved | 0,8624 | 2,099E-18 |
| TcCLB.504019.6 | hypothetical protein | 0,8622 | 1,45E-23 |
| TcCLB.510949.20 | thioredoxin, putative | 0,8621 | 5,522E-11 |
| TcCLB.506775.50 | hypothetical protein, conserved | 0,8619 | 6,617E-25 |
| TcCLB.508857.70 | hypothetical protein, conserved | 0,8618 | 1,315E-15 |
| TcCLB.511101.37 | retrotransposon hot spot protein (RHS, pseudogene), puta | 0,8618 | 7,731E-11 |
| TcCLB.509611.80 | cation-transporting ATPase, putative | 0,8615 | 4,414E-23 |
| TcCLB.503703.90 | hypothetical protein, conserved | 0,8613 | 3,094E-08 |
| TcCLB.507817.80 | 5'-3' exoribonuclease A, putative (fragment) (XRNA) | 0,8612 | 1,092E-30 |
| TcCLB.510119.40 | hypothetical protein, conserved | 0,861 | 3,618E-06 |
| TcCLB.506825.30 | hypothetical protein, conserved | 0,8609 | 1,137E-13 |
| TcCLB.508043.30 | ubiquitin-activating enzyme E1, putative | 0,8606 | 1,559E-31 |
| TcCLB.506975.69 | hypothetical protein, conserved | 0,8604 | 8,898E-08 |
| TcCLB.510997.40 | enoyl-CoA hydratase/isomerase family protein, putative | 0,8598 | 7,672E-19 |
| TcCLB.507735.10 | hypothetical protein, conserved | 0,8598 | 6,231E-06 |
| TcCLB.510391.10 | dispersed gene family protein 1 (DGF-1), putative | 0,8595 | 0,002081 |
| TcCLB.507775.20 | hypothetical protein, conserved | 0,8594 | 1,556E-11 |
| TcCLB.506629.130 | hypothetical protein, conserved | 0,8592 | 1,751E-11 |
| TcCLB.507063.20 | eukaryotic translation initiation factor 2 beta subunit, puta | 0,8589 | 1,31E-19 |
| TcCLB.509179.70 | hypothetical protein, conserved | 0,8585 | 0,00007588 |
| TcCLB.504165.20 | hypothetical protein, conserved | 0,8581 | 7,778E-12 |
| TcCLB.504575.20 | hypothetical protein, conserved | 0,8581 | 2,719E-10 |
| TcCLB.508153.1070 | hypothetical protein, conserved | 0,8579 | 8,848E-16 |
| TcCLB.509187.20 | retrotransposon hot spot protein (RHS, pseudogene), puta | 0,8579 | 0,01272 |
| TcCLB.511003.110 | hypothetical protein, conserved | 0,8578 | 1,577E-10 |
| TcCLB.511635.69 | hypothetical protein, conserved | 0,8577 | 7,391E-06 |
| TcCLB.504669.40 | mucin-associated surface protein (MASP, pseudogene), pu | 0,8576 | 7,853E-14 |
| TcCLB.511907.350 | surface protease GP63 (pseudogene), putative | 0,8576 | 0,0004303 |
| TcCLB.510879.20 | ribosomal protein S19, putative | 0,8574 | 6,032E-29 |
| TcCLB.510355.110 | hypothetical protein, conserved (pseudogene) | 0,8574 | 1,853E-06 |
| TcCLB.511471.90 | glycine dehydrogenase [decarboxylating] (pseudogene), pu | 0,8574 | 0,00002766 |
| TcCLB.510307.330 | dispersed gene family protein 1 (DGF-1), putative | 0,8572 | 1,644E-08 |
| TcCLB.507503.20 | glycine dehydrogenase [decarboxylating] (pseudogene), pu | 0,857 | 4,947E-14 |
| TcCLB.509459.79 | actin interacting protein-like protein, putative | 0,8568 | 4,987E-16 |
| TcCLB.505115.10 | hypothetical protein, conserved | 0,8566 | 1,3E-16 |
| TcCLB.509011.54 | ubiquinol-cytochrome c reductase, putative | 0,8566 | 0,00000929 |
| TcCLB.508355.240 | hypothetical protein, conserved | 0,8565 | 7,364E-08 |
| TcCLB.508965.80 | hypothetical protein, conserved | 0,8561 | 6,005E-15 |
| TcCLB.509693.140 | hypothetical protein, conserved | 0,856 | 8,711E-20 |
| TcCLB.511333.4 | Calpain-like protein 1 (CAP5.5) | 0,8559 | 1,228E-11 |
| TcCLB.509805.220 | cholinephosphate cytidylyltransferase A, putative | 0,8554 | 3,952E-24 |
| TcCLB.510755.50 | ADP-ribosylation factor 1, putative (ARF1) | 0,8552 | 2,858E-23 |
| TcCLB.484299.10 | trypanothione reductase, putative | 0,8551 | 1,722E-11 |
| TcCLB.507281.9 | dispersed gene family protein 1 (DGF-1, pseudogene), put | 0,855 | 0,008111 |
| TcCLB.509207.50 | hypothetical protein, conserved | 0,8546 | 4,633E-25 |

| | | | |
|---|---|---|---|
| TcCLB.504161.10 | hypothetical protein | 0,8544 | 4,33E-13 |
| TcCLB.509837.10 | hypothetical protein, conserved | 0,8542 | 2,109E-14 |
| TcCLB.511871.30 | hypothetical protein, conserved | 0,8537 | 1,204E-17 |
| TcCLB.509207.11 | histone H2A, putative | 0,8534 | 1,348E-27 |
| TcCLB.506635.10 | hypothetical protein (pseudogene) | 0,8534 | 7,795E-07 |
| TcCLB.510435.30 | hypothetical protein, conserved | 0,853 | 3,294E-17 |
| TcCLB.507641.50 | hypothetical protein, conserved | 0,8527 | 2,338E-23 |
| TcCLB.466593.10 | hypothetical protein, conserved | 0,8527 | 4,554E-10 |
| TcCLB.507875.170 | ribosomal P protein AGP2beta-1, putative | 0,8524 | 2,476E-06 |
| TcCLB.509393.10 | hypothetical protein, conserved | 0,8523 | 1,965E-16 |
| TcCLB.504123.30 | map kinase, putative | 0,8523 | 8,907E-14 |
| TcCLB.505401.31 | trans-sialidase (pseudogene), putative | 0,8519 | 1,001E-09 |
| TcCLB.507081.20 | hypothetical protein, conserved | 0,8517 | 6,181E-16 |
| TcCLB.508501.165 | U6 snRNA-associated Sm-like protein LSm6p | 0,8511 | 0,0000361 |
| TcCLB.503745.20 | hypothetical protein, conserved | 0,8508 | 2,351E-19 |
| TcCLB.507021.110 | phosphopantothenoylcysteine decarboxylase, putative (PP | 0,8505 | 2,176E-14 |
| TcCLB.510719.60 | Biotin--acetyl-CoA-carboxylase ligase, putative | 0,8504 | 9,044E-14 |
| TcCLB.508271.10 | phenylalanyl-tRNA synthetase, putative | 0,8503 | 6,937E-25 |
| TcCLB.510013.50 | trans-sialidase, Group IV, putative | 0,8502 | 0,01047 |
| TcCLB.506529.324 | hypothetical protein, conserved | 0,8501 | 2,09E-07 |
| TcCLB.508169.20 | elongation factor 2, putative | 0,8499 | 7,318E-38 |
| TcCLB.511295.50 | hypothetical protein, conserved | 0,8499 | 3,743E-30 |
| TcCLB.507829.10 | cysteine peptidase, Clan CA, family C2, putative | 0,8499 | 1,963E-08 |
| TcCLB.503763.10 | retrotransposon hot spot (RHS) protein, putative | 0,8497 | 0,00003796 |
| TcCLB.506727.10 | hypothetical protein | 0,8496 | 1,372E-14 |
| TcCLB.511807.230 | hypothetical protein | 0,8494 | 1,249E-19 |
| TcCLB.511391.200 | pitrilysin-like metalloprotease, putative | 0,8492 | 1,717E-34 |
| TcCLB.511665.50 | dispersed gene family protein 1 (DGF-1), putative | 0,8492 | 1,456E-07 |
| TcCLB.506959.120 | 60S ribosomal protein L30, putative | 0,8488 | 6,453E-28 |
| TcCLB.503781.80 | universal minicircle sequence binding protein 1 (UMSBP1) | 0,8488 | 2,089E-12 |
| TcCLB.506595.30 | retrotransposon hot spot protein (RHS, pseudogene), puta | 0,8487 | 5,537E-12 |
| TcCLB.504081.450 | trans-sialidase (pseudogene), putative | 0,8487 | 0,00003663 |
| TcCLB.508657.10 | hypothetical protein, conserved | 0,8486 | 8,622E-10 |
| TcCLB.511577.60 | DNA excision/repair protein SNF2, putative | 0,848 | 6,073E-16 |
| TcCLB.503899.110 | glutathione peroxidase-like protein, putative | 0,8479 | 2,343E-13 |
| TcCLB.509799.140 | L-2-hydroxyglutarate dehydrogenase, mitochondrial, putat | 0,8479 | 6,136E-13 |
| TcCLB.506063.50 | retrotransposon hot spot protein (RHS, pseudogene), puta | 0,8475 | 8,916E-09 |
| TcCLB.424123.40 | RNA-binding protein, putative | 0,8469 | 1,866E-08 |
| TcCLB.506055.30 | retrotransposon hot spot protein (RHS, pseudogene), puta | 0,8468 | 0,00004308 |
| TcCLB.506289.30 | hypothetical protein, conserved | 0,8466 | 2,906E-09 |
| TcCLB.508717.33 | mitochondrial chaperone BCS1, putative | 0,8465 | 2,387E-17 |
| TcCLB.506167.20 | hypothetical protein, conserved | 0,8464 | 8,776E-24 |
| TcCLB.510675.9 | dispersed gene family protein 1 (DGF-1), putative (fragme | 0,846 | 0,0003814 |
| TcCLB.509719.53 | hypothetical protein, conserved | 0,8458 | 3,384E-16 |
| TcCLB.507639.20 | universal minicircle sequence binding protein, putative (U | 0,8458 | 5,528E-08 |

| | | | |
|---|---|---|---|
| TcCLB.506375.50 | dispersed gene family protein 1 (DGF-1), putative | 0,8458 | 4,754E-07 |
| TcCLB.507683.40 | protein kinase, putative | 0,8455 | 5,951E-18 |
| TcCLB.506391.20 | calmodulin, putative | 0,8455 | 3,253E-08 |
| TcCLB.506247.190 | nuclear lim interactor-interacting factor, putative | 0,8453 | 0,00007651 |
| TcCLB.510731.90 | protein kinase, putative | 0,845 | 8,054E-09 |
| TcCLB.506547.10 | dispersed gene family protein 1 (DGF-1, pseudogene), put | 0,845 | 0,004068 |
| TcCLB.508993.10 | cytidine triphosphate synthase, putative | 0,8449 | 2,362E-26 |
| TcCLB.508823.90 | hypothetical protein | 0,8449 | 1,212E-09 |
| TcCLB.506529.541 | hypothetical protein | 0,8449 | 0,00001228 |
| TcCLB.510795.4 | CLC-type chloride channel protein (pseudogene), putative | 0,8449 | 0,009632 |
| TcCLB.507513.60 | 40S ribosomal protein S16, putative | 0,8446 | 3,189E-21 |
| TcCLB.511195.10 | retrotransposon hot spot protein (RHS, pseudogene), puta | 0,8446 | 0,01042 |
| TcCLB.504171.40 | hypothetical protein, conserved | 0,8444 | 9,155E-10 |
| TcCLB.507765.149 | hypothetical protein, conserved | 0,8441 | 5,014E-12 |
| TcCLB.506519.51 | hypothetical protein, conserved | 0,8439 | 2,804E-16 |
| TcCLB.508741.170 | glycerate kinase, putative | 0,8436 | 2,094E-35 |
| TcCLB.441241.10 | microtubule-associated protein Gb4, putative (fragment) | 0,8434 | 2,238E-12 |
| TcCLB.510741.150 | Present in the outer mitochondrial membrane proteome 7 | 0,8432 | 1,851E-11 |
| TcCLB.506671.70 | dispersed gene family protein 1 (DGF-1), putative | 0,8428 | 0,0007897 |
| TcCLB.510323.60 | hypothetical protein | 0,8426 | 2,588E-09 |
| TcCLB.509805.140 | hypothetical protein | 0,8421 | 4,354E-10 |
| TcCLB.508319.30 | hypothetical protein, conserved | 0,8419 | 1,127E-29 |
| TcCLB.509769.120 | hypothetical protein, conserved | 0,8419 | 2,61E-27 |
| TcCLB.508039.70 | hypothetical protein, conserved | 0,8419 | 5,966E-22 |
| TcCLB.452699.10 | retrotransposon hot spot protein (RHS, pseudogene), puta | 0,8418 | 8,023E-11 |
| TcCLB.506529.120 | hypothetical protein | 0,8417 | 3,738E-13 |
| TcCLB.510767.10 | ribosomal protein L15, putative | 0,8416 | 1,345E-26 |
| TcCLB.507723.189 | bloodstream- specific protein 2 precursor (fragment) | 0,8415 | 3,038E-11 |
| TcCLB.509317.60 | RNA-binding protein, putative | 0,8414 | 3,547E-17 |
| TcCLB.507615.19 | microtubule-associated protein, putative, (fragment) (GB4 | 0,8411 | 6,712E-23 |
| TcCLB.509511.10 | dispersed gene family protein 1 (DGF-1), putative | 0,8411 | 5,392E-07 |
| TcCLB.510265.100 | hypothetical protein, conserved | 0,8409 | 1,722E-17 |
| TcCLB.510431.10 | isopentenyl-diphosphate delta-isomerase (fragment) (idi1 | 0,8409 | 7,479E-13 |
| TcCLB.504147.20 | 60S ribosomal protein L18, putative | 0,8405 | 9,02E-33 |
| TcCLB.504153.20 | phosphoglycerate kinase, putative | 0,8404 | 5,761E-21 |
| TcCLB.506223.110 | hypothetical protein, conserved | 0,8401 | 2,22E-16 |
| TcCLB.509895.40 | hypothetical protein, conserved | 0,8399 | 6,973E-18 |
| TcCLB.508689.50 | GPI transamidase subunit 16, putative (fragment) (GPI16) | 0,8397 | 7,622E-07 |
| TcCLB.509331.221 | hypothetical protein, conserved | 0,8397 | 0,003718 |
| TcCLB.508461.510 | 60S ribosomal protein L27A/L29, putative | 0,8396 | 2,825E-11 |
| TcCLB.507603.70 | kinetoplastid kinetochore protein 6, putative (kkt6) | 0,8396 | 2,376E-06 |
| TcCLB.507153.40 | tryptophanyl-tRNA synthetase (pseudogene), putative | 0,8393 | 0,0001897 |
| TcCLB.506893.40 | hypothetical protein, conserved | 0,8392 | 0,00005899 |
| TcCLB.508891.30 | hypothetical protein, conserved | 0,839 | 6,444E-23 |
| TcCLB.511233.20 | 60S ribosomal protein L34, putative | 0,8388 | 4,316E-26 |

| | | | |
|---|---|---|---|
| TcCLB.505037.30 | hypothetical protein, conserved | 0,8388 | 0,003065 |
| TcCLB.506679.70 | eukaryotic translation initiation factor 6 (eIF-6), putative | 0,8387 | 1,537E-21 |
| TcCLB.511051.50 | 40S ribosomal protein S4, putative | 0,8386 | 1,109E-31 |
| TcCLB.506857.100 | endoplasmatic reticulum retrieval protein, putative | 0,8385 | 1,184E-19 |
| TcCLB.510515.20 | hypothetical protein, conserved (pseudogene) | 0,8384 | 1,243E-14 |
| TcCLB.503635.40 | hypothetical protein | 0,8383 | 2,885E-12 |
| TcCLB.508637.150 | ribonuclease L inhibitor, putative | 0,8382 | 1,189E-21 |
| TcCLB.505183.40 | DNA repair and recombination protein RAD54, putative | 0,838 | 2,337E-21 |
| TcCLB.506861.30 | 60S ribosomal protein L14, putative | 0,8379 | 6,175E-25 |
| TcCLB.503897.20 | hypothetical protein, conserved | 0,8373 | 1,987E-13 |
| TcCLB.506529.110 | hypothetical protein, conserved | 0,8372 | 4,464E-12 |
| TcCLB.509601.50 | hypothetical protein, conserved | 0,8371 | 1,799E-22 |
| TcCLB.511239.140 | 60S ribosomal protein, putative | 0,8369 | 1,557E-08 |
| TcCLB.506477.10 | cytochrome c oxidase assembly factor, putative | 0,8365 | 2,26E-13 |
| TcCLB.506457.20 | hypothetical protein, conserved | 0,8365 | 7,08E-10 |
| TcCLB.508405.40 | 40S ribosomal protein S11, putative | 0,8363 | 4,791E-22 |
| TcCLB.506959.64 | 3-keto-dihydrosphingosine reductase | 0,8363 | 1,19E-13 |
| TcCLB.508045.30 | hypothetical protein, conserved | 0,8361 | 1,732E-16 |
| TcCLB.509671.189 | hypothetical protein, conserved | 0,8359 | 1,055E-10 |
| TcCLB.504185.30 | hypothetical protein, conserved | 0,8357 | 5,814E-10 |
| TcCLB.504105.94 | ribosomal protein S25, putative | 0,8356 | 1,142E-28 |
| TcCLB.506489.59 | divalent cation tolerance protein, putative | 0,8356 | 0,0002189 |
| TcCLB.507851.60 | hypothetical protein, conserved | 0,8354 | 2,482E-16 |
| TcCLB.510767.20 | 60S ribosomal protein L36, putative | 0,8353 | 7,566E-21 |
| TcCLB.509793.40 | kinetoplast-associated protein (TcKAP4) | 0,8352 | 2,586E-12 |
| TcCLB.506559.30 | hypothetical protein, conserved | 0,8352 | 2,915E-12 |
| TcCLB.509499.10 | cyclophilin, putative (CYP) | 0,835 | 6,507E-15 |
| TcCLB.510643.110 | beta galactofuranosyl glycosyltransferase, putative | 0,835 | 1,423E-07 |
| TcCLB.508267.60 | hypothetical protein, conserved (fragment) | 0,8346 | 9,277E-09 |
| TcCLB.507851.50 | hypothetical protein, conserved | 0,8343 | 2,923E-20 |
| TcCLB.503701.9 | dispersed gene family protein 1 (DGF-1, pseudogene), put | 0,8343 | 0,00008908 |
| TcCLB.507059.70 | dihydroorotase, putative | 0,8336 | 2,768E-12 |
| TcCLB.506457.140 | hypothetical protein, conserved | 0,8336 | 1,595E-06 |
| TcCLB.507559.20 | hypothetical protein, conserved | 0,8333 | 1,428E-13 |
| TcCLB.506929.30 | hypothetical protein, conserved | 0,833 | 4,117E-13 |
| TcCLB.506701.20 | Tob55, putative | 0,8326 | 9,719E-12 |
| TcCLB.511291.120 | DNA-directed RNA polymerase III subunit C11, putative (R | 0,832 | 1,45E-08 |
| TcCLB.511697.30 | hypothetical protein | 0,832 | 0,002975 |
| TcCLB.511827.130 | dispersed gene family protein 1 (DGF-1), putative | 0,8319 | 1,785E-12 |
| TcCLB.505057.10 | hypothetical protein, conserved | 0,8318 | 3,485E-12 |
| TcCLB.510575.100 | hypothetical protein, conserved | 0,8316 | 1,015E-10 |
| TcCLB.507641.233 | ribosomal protein L37, putative | 0,8315 | 3,364E-20 |
| TcCLB.510663.30 | 60S ribosomal subunit protein L31, putative | 0,8314 | 1,84E-22 |
| TcCLB.507653.30 | hypothetical protein | 0,8312 | 0,0001506 |
| TcCLB.509593.59 | dispersed gene family protein 1 (DGF-1), putative | 0,8312 | 0,1382 |

| | | | |
|---|---|---|---|
| TcCLB.508817.60 | ribosomal protein S26, putative | 0,831 | 8,311E-26 |
| TcCLB.507243.9 | dispersed gene family protein 1 (DGF-1), putative | 0,831 | 0,00003755 |
| TcCLB.508211.30 | sphingosine kinase A, B, putative | 0,8307 | 1,745E-14 |
| TcCLB.505915.40 | beta galactofuranosyl glycosyltransferase (pseudogene), p | 0,8307 | 0,0003236 |
| TcCLB.506743.69 | hypothetical protein, conserved | 0,8304 | 4,324E-08 |
| TcCLB.508505.4 | hypothetical protein, conserved | 0,8304 | 3,268E-06 |
| TcCLB.443397.9 | NADH-cytochrome b5 reductase, putative (fragment) | 0,8298 | 3,945E-09 |
| TcCLB.511829.10 | dispersed gene family protein 1 (DGF-1), putative | 0,8297 | 3,477E-13 |
| TcCLB.511911.130 | phosphoglucomutase | 0,8297 | 2,372E-12 |
| TcCLB.508649.20 | hypothetical protein, conserved | 0,8294 | 1,806E-25 |
| TcCLB.509761.20 | 3,2-trans-enoyl-CoA isomerase, mitochondrial precursor ( | 0,8294 | 8,474E-20 |
| TcCLB.510965.5 | elongation factor 2, putative (fragment) | 0,8293 | 1,383E-24 |
| TcCLB.511771.30 | beta galactofuranosyl glycosyltransferase (pseudogene), p | 0,8293 | 0,00004222 |
| TcCLB.511619.5 | FtsJ cell division protein, putative (fragment) | 0,829 | 4,238E-13 |
| TcCLB.510857.54 | hypothetical protein, conserved | 0,8289 | 8,425E-10 |
| TcCLB.503611.20 | ribosomal protein L24, putative | 0,8281 | 1,286E-18 |
| TcCLB.511001.190 | vacuolar ATP synthase subunit d, putative | 0,8278 | 8,042E-16 |
| TcCLB.504251.9 | peroxisome assembly protein, putative (fragment) | 0,8277 | 5,803E-13 |
| TcCLB.507669.200 | hypothetical protein, conserved | 0,8277 | 1,306E-11 |
| TcCLB.504539.10 | mucin TcSMUGL, putative | 0,8277 | 0,002167 |
| TcCLB.505919.40 | protein kinase, putative | 0,8275 | 0,0005745 |
| TcCLB.511587.60 | retrotransposon hot spot protein (RHS, pseudogene), puta | 0,8273 | 4,782E-11 |
| TcCLB.508297.41 | hypothetical protein, conserved | 0,8273 | 2,073E-06 |
| TcCLB.509965.270 | hypothetical protein, conserved | 0,827 | 6,076E-17 |
| TcCLB.508231.20 | 40S ribosomal protein S12, putative | 0,8268 | 1,912E-26 |
| TcCLB.504827.60 | hypothetical protein, conserved | 0,8264 | 2,514E-15 |
| TcCLB.509027.50 | hypothetical protein, conserved | 0,8264 | 1,464E-08 |
| TcCLB.510859.17 | nucleolar RNA-binding protein, putative | 0,8262 | 1,223E-30 |
| TcCLB.422507.10 | glycine dehydrogenase [decarboxylating], putative | 0,8262 | 2,617E-06 |
| TcCLB.507787.20 | hypothetical protein, conserved | 0,8261 | 1,543E-20 |
| TcCLB.506333.10 | plasma-membrane proton-efflux P-type ATPase, putative | 0,8255 | 5,76E-36 |
| TcCLB.504863.10 | retrotransposon hot spot (RHS) protein, putative | 0,8255 | 4,705E-18 |
| TcCLB.511181.130 | hypothetical protein | 0,8253 | 1,087E-10 |
| TcCLB.503647.40 | hypothetical protein | 0,8253 | 1,9E-09 |
| TcCLB.504153.140 | metallo-peptidase, Clan MF, Family M17 | 0,8252 | 1,335E-22 |
| TcCLB.508221.539 | hypothetical protein, conserved | 0,8246 | 0,07826 |
| TcCLB.507105.60 | hypothetical protein, conserved | 0,8245 | 4,624E-18 |
| TcCLB.509209.10 | endoplasmatic reticulum retrieval protein, putative | 0,8239 | 4,829E-22 |
| TcCLB.507017.20 | Present in the outer mitochondrial membrane proteome 3 | 0,8238 | 9,635E-11 |
| TcCLB.503453.70 | hypothetical protein, conserved | 0,8237 | 9,806E-20 |
| TcCLB.511069.20 | 40S ribosomal protein S8, putative | 0,8237 | 1,427E-16 |
| TcCLB.511765.40 | retrotransposon hot spot protein (RHS, pseudogene), puta | 0,8229 | 3,682E-09 |
| TcCLB.511807.130 | hypothetical protein, conserved | 0,8224 | 0,0004962 |
| TcCLB.511529.50 | hypothetical protein, conserved | 0,8222 | 0,0000111 |
| TcCLB.419043.10 | hypothetical protein, conserved | 0,8219 | 9,115E-07 |

| | | | |
|---|---|---|---|
| TcCLB.509951.30 | hypothetical protein, conserved | 0,8214 | 3,583E-16 |
| TcCLB.508157.50 | trans-sialidase, putative | 0,8213 | 4,872E-06 |
| TcCLB.510997.10 | 3-keto-dihydrosphingosine reductase | 0,8211 | 5,858E-15 |
| TcCLB.503869.40 | pumilio/PUF RNA binding protein 9, putative (PUF9) | 0,8211 | 4,262E-09 |
| TcCLB.510343.5 | hypothetical protein, conserved (fragment) | 0,821 | 1,904E-09 |
| TcCLB.506327.30 | hypothetical protein, conserved | 0,8206 | 1,819E-22 |
| TcCLB.511809.140 | histone H2A, putative | 0,8203 | 4,587E-19 |
| TcCLB.508185.10 | enoyl-CoA hydratase, mitochondrial precursor, putative | 0,8197 | 2,551E-13 |
| TcCLB.506303.111 | hypothetical protein, conserved | 0,8196 | 6,867E-16 |
| TcCLB.457547.9 | dispersed gene family protein 1 (DGF-1), putative | 0,8195 | 0,0008609 |
| TcCLB.509875.210 | 60S ribosomal protein L17, putative | 0,8193 | 1,007E-26 |
| TcCLB.511655.10 | hypothetical protein, conserved | 0,8191 | 3,037E-06 |
| TcCLB.508537.10 | glyceraldehyde 3-phosphate dehydrogenase, cytosolic, put | 0,819 | 0,001272 |
| TcCLB.503531.20 | hypothetical protein, conserved | 0,8189 | 2,169E-14 |
| TcCLB.510241.50 | 60S ribosomal protein L10, putative | 0,8188 | 1,727E-24 |
| TcCLB.508299.20 | hypothetical protein, conserved | 0,8188 | 2,671E-09 |
| TcCLB.508441.70 | enoyl-CoA hydratase/Enoyl-CoA isomerase/3-hydroxyacyl- | 0,8182 | 6,065E-19 |
| TcCLB.510289.6 | nucleolar RNA-binding protein, putative | 0,8182 | 3,962E-09 |
| TcCLB.511063.80 | dispersed gene family protein 1 (DGF-1, pseudogene), put | 0,8182 | 0,005398 |
| TcCLB.510773.120 | hypothetical protein | 0,8181 | 0,05937 |
| TcCLB.510575.190 | 60S ribosomal protein L10, putative | 0,8176 | 1,236E-20 |
| TcCLB.510861.14 | hypothetical protein, conserved (fragment) | 0,8173 | 5,253E-15 |
| TcCLB.424195.5 | hypothetical protein, conserved | 0,8168 | 0,02189 |
| TcCLB.504159.70 | hypothetical protein, conserved | 0,8166 | 9,628E-11 |
| TcCLB.504073.30 | ribonucleoside-diphosphate reductase large chain, putativ | 0,8161 | 3,618E-06 |
| TcCLB.506529.50 | arginine N-methyltransferase, putative | 0,816 | 1,834E-13 |
| TcCLB.509587.10 | mitochondrial DNA polymerase I protein D, putative | 0,8159 | 4,681E-29 |
| TcCLB.510823.30 | Isoprenylcysteine alpha-carbonyl methylesterase, putative | 0,8152 | 6,062E-19 |
| TcCLB.508785.4 | hypothetical protein, conserved (fragment) | 0,8149 | 6,132E-17 |
| TcCLB.504929.10 | casein kinase, delta isoform, putative | 0,8146 | 0,04986 |
| TcCLB.505071.80 | hypothetical protein | 0,8145 | 7,5E-16 |
| TcCLB.509261.30 | 3,2-trans-enoyl-CoA isomerase, mitochondrial precursor, p | 0,8144 | 3,863E-22 |
| TcCLB.503425.20 | monothiol glutaredoxin, putative | 0,8144 | 2,63E-11 |
| TcCLB.507515.80 | hypothetical protein, conserved | 0,8143 | 2,082E-10 |
| TcCLB.503925.60 | hypothetical protein | 0,8142 | 3,976E-14 |
| TcCLB.508567.80 | DNA repair and recombination helicase protein PIF7, puta | 0,8139 | 7,264E-19 |
| TcCLB.509567.60 | hypothetical protein | 0,8139 | 3,312E-14 |
| TcCLB.410241.10 | retrotransposon hot spot protein (RHS, pseudogene), puta | 0,8138 | 0,03758 |
| TcCLB.511303.60 | translation initiation factor, putative | 0,8137 | 2,646E-30 |
| TcCLB.510187.320 | hypothetical protein | 0,8137 | 9,689E-14 |
| TcCLB.511811.10 | 40S ribosomal protein S15, putative | 0,8137 | 0,00321 |
| TcCLB.510243.119 | hypothetical protein, conserved | 0,8136 | 2,073E-09 |
| TcCLB.508415.40 | ADP-ribosylation factor 1, putative (ARF1) | 0,8131 | 1,18E-18 |
| TcCLB.506413.20 | 40S ribosomal protein S33, putative | 0,8131 | 5,454E-18 |
| TcCLB.510149.130 | hypothetical protein, conserved | 0,8131 | 1,72E-09 |

| | | | |
|---|---|---|---|
| TcCLB.511467.60 | hypothetical protein, conserved | 0,8128 | 2,418E-10 |
| TcCLB.505077.9 | dispersed gene family protein 1 (DGF-1), putative (fragm | 0,8125 | 0,00305 |
| TcCLB.511825.260 | ribosomal RNA processing protein 45, putative (RRP45) | 0,8124 | 3,823E-11 |
| TcCLB.505773.10 | retrotransposon hot spot protein (RHS, pseudogene), puta | 0,812 | 0,00001696 |
| TcCLB.508641.184 | Protein translation factor SUI1 homolog, putative | 0,8117 | 9,874E-09 |
| TcCLB.507739.50 | hypothetical protein, conserved | 0,8115 | 6,618E-06 |
| TcCLB.507245.40 | dispersed gene family protein 1 (DGF-1), putative | 0,811 | 1,588E-07 |
| TcCLB.509331.200 | ABC transporter, putative | 0,811 | 4,648E-07 |
| TcCLB.506855.80 | ATP synthase, putative | 0,8109 | 1,465E-16 |
| TcCLB.506583.40 | cdc2-related kinase 3, putative (CRK3) | 0,8108 | 1,285E-07 |
| TcCLB.507589.14 | tryptophanyl-tRNA synthetase, putative | 0,8104 | 4,155E-37 |
| TcCLB.506375.10 | tryptophanyl-tRNA synthetase, putative | 0,8104 | 0,00776 |
| TcCLB.506247.390 | hypothetical protein, conserved | 0,8101 | 2,471E-17 |
| TcCLB.510719.30 | ribosomal protein L29, putative | 0,8093 | 3,061E-17 |
| TcCLB.510531.10 | hypothetical protein, conserved | 0,8092 | 1,06E-20 |
| TcCLB.510547.30 | trans-sialidase (pseudogene), putative | 0,8089 | 0,00253 |
| TcCLB.511809.20 | methylenetetrahydrofolate dehydrogenase-like protein, pu | 0,8087 | 6,189E-13 |
| TcCLB.506435.134 | Enriched in surface-labeled proteome protein 11, putative | 0,8086 | 4,017E-10 |
| TcCLB.508461.40 | hypothetical protein, conserved | 0,8086 | 5,564E-08 |
| TcCLB.508041.80 | protein transport protein Sec31, putative (pseudogene) | 0,8083 | 1,827E-22 |
| TcCLB.506537.160 | glycine dehydrogenase [decarboxylating] (pseudogene), pu | 0,8079 | 3,958E-10 |
| TcCLB.510725.99 | Chaperonin HSP60, mitochondrial (HSP60) | 0,8079 | 0,0001819 |
| TcCLB.510687.120 | snoRNP protein GAR1, putative | 0,8078 | 7,069E-11 |
| TcCLB.506743.200 | 3-Beta-hydroxysteroid-delta(8), delta(7)-isomerase, putat | 0,8078 | 7,565E-08 |
| TcCLB.511425.30 | hypothetical protein, conserved | 0,8074 | 9,628E-11 |
| TcCLB.504055.30 | hypothetical protein | 0,8073 | 1,192E-09 |
| TcCLB.507625.170 | hypothetical protein, conserved | 0,8069 | 2,089E-14 |
| TcCLB.506213.60 | 40S ribosomal protein S2, putative | 0,8069 | 0,01042 |
| TcCLB.507927.20 | mitochondrial oligo_U binding protein TBRGG1, putative | 0,8068 | 7,447E-10 |
| TcCLB.507873.34 | hypothetical protein, conserved | 0,8068 | 0,00000603 |
| TcCLB.506963.14 | 40S ribosomal protein S27, putative | 0,8067 | 1,223E-17 |
| TcCLB.504157.10 | heterogeneous nuclear ribonucleoprotein H/F, putative | 0,8066 | 7,202E-12 |
| TcCLB.511807.64 | DNA repair protein, putative | 0,8064 | 5,726E-14 |
| TcCLB.508973.9 | hypothetical protein, conserved (fragment) | 0,8064 | 1,116E-08 |
| TcCLB.511901.30 | Glycerophosphoryl diester phosphodiesterase family, puta | 0,8061 | 1,139E-14 |
| TcCLB.511837.80 | hypothetical protein, conserved | 0,8059 | 6,308E-11 |
| TcCLB.505163.50 | hypothetical protein | 0,8059 | 0,01209 |
| TcCLB.460757.20 | hypothetical protein (fragment) | 0,8058 | 0,03043 |
| TcCLB.505317.9 | trans-sialidase, putative (fragment) | 0,8053 | 4,89E-07 |
| TcCLB.509025.10 | hypothetical protein, conserved | 0,8053 | 0,00001499 |
| TcCLB.509695.200 | T-complex protein 1, delta subunit, putative | 0,8052 | 5,837E-25 |
| TcCLB.510761.73 | dynein-associated protein, putative | 0,8052 | 6,667E-09 |
| TcCLB.509723.20 | glycine dehydrogenase (pseudogene), putative | 0,8048 | 0,01009 |
| TcCLB.510357.120 | hypothetical protein, conserved | 0,8044 | 0,009328 |
| TcCLB.508461.4 | hypothetical protein, conserved | 0,8042 | 2,511E-14 |

| | | | |
|---|---|---|---|
| TcCLB.506925.460 | hypothetical protein, conserved | 0,8039 | 7,543E-09 |
| TcCLB.504059.30 | hypothetical protein, conserved | 0,8033 | 1,6E-21 |
| TcCLB.506213.80 | 60S ribosomal protein L17, putative | 0,8032 | 1,407E-09 |
| TcCLB.506063.59 | dispersed gene family protein 1 (DGF-1), putative | 0,8031 | 0,000562 |
| TcCLB.511657.30 | hypothetical protein, conserved | 0,8027 | 1,626E-15 |
| TcCLB.511277.170 | ATP-dependent zinc metallopeptidase, putative | 0,8019 | 5,781E-22 |
| TcCLB.511829.20 | retrotransposon hot spot protein (RHS, pseudogene), puta | 0,8019 | 0,00000432 |
| TcCLB.508323.60 | centrin-4, putative (Centrin4) | 0,8018 | 3,778E-19 |
| TcCLB.503763.30 | retrotransposon hot spot protein (RHS, pseudogene), puta | 0,8017 | 9,655E-16 |
| TcCLB.511417.30 | hypothetical protein, conserved | 0,8016 | 4,135E-12 |
| TcCLB.505807.209 | hypothetical protein, conserved | 0,8016 | 1,177E-06 |
| TcCLB.507513.70 | 40S ribosomal protein S16, putative | 0,8015 | 7,442E-24 |
| TcCLB.506151.4 | hypothetical protein, conserved (fragment) | 0,801 | 1,006E-09 |
| TcCLB.511671.110 | hypothetical protein, conserved | 0,801 | 7,442E-07 |
| TcCLB.504443.30 | DNA-directed RNA polymerase II subunit 2, putative (pseu | 0,8009 | 2,179E-19 |
| TcCLB.506943.80 | hypothetical protein, conserved | 0,8006 | 3,728E-23 |
| TcCLB.511821.110 | translation initiation factor IF-2, putative | 0,8006 | 8,855E-19 |
| TcCLB.503583.24 | hypothetical protein, conserved | 0,8001 | 1,057E-22 |
| TcCLB.504797.90 | hypothetical protein, conserved | 0,7999 | 1,213E-08 |
| TcCLB.509215.30 | hypothetical protein, conserved | 0,7998 | 6,785E-07 |
| TcCLB.511019.13 | retrotransposon hot spot (RHS) protein, putative (fragmen | 0,7993 | 1,573E-14 |
| TcCLB.503453.40 | 60S ribosomal protein L30, putative | 0,7992 | 6,771E-07 |
| TcCLB.509089.9 | dispersed gene family protein 1 (DGF-1), putative | 0,7987 | 0,00001412 |
| TcCLB.509921.51 | hypothetical protein, conserved (pseudogene) | 0,7985 | 0,0003126 |
| TcCLB.504163.100 | hypothetical protein, conserved | 0,7979 | 2,774E-15 |
| TcCLB.511211.130 | receptor for activated C kinase 1, putative | 0,7978 | 1,432E-20 |
| TcCLB.510755.60 | ADP-ribosylation factor 1, putative (ARF1) | 0,7978 | 5,977E-06 |
| TcCLB.509365.11 | tryptophanyl-tRNA synthetase (pseudogene), putative | 0,7974 | 0,00002668 |
| TcCLB.510683.20 | hypothetical protein, conserved | 0,7972 | 1,137E-08 |
| TcCLB.504089.70 | hypothetical protein, conserved | 0,797 | 5,947E-08 |
| TcCLB.503909.19 | dispersed gene family protein 1 (DGF-1), putative (fragme | 0,797 | 0,0005047 |
| TcCLB.511421.10 | hypothetical protein, conserved | 0,7965 | 1,691E-19 |
| TcCLB.510517.10 | chaperonin/T-complex protein 1 gamma subunit, putative | 0,7965 | 2,755E-17 |
| TcCLB.509177.68 | translation initiation factor, putative | 0,7958 | 9,9E-10 |
| TcCLB.506989.10 | hypothetical protein, conserved | 0,7958 | 3,229E-08 |
| TcCLB.503579.119 | hypothetical protein, conserved | 0,7958 | 0,00003612 |
| TcCLB.503571.10 | flagellum-adhesion glycoprotein, putative | 0,7957 | 5,537E-17 |
| TcCLB.508421.20 | dispersed gene family protein 1 (DGF-1, pseudogene), put | 0,7956 | 0,0000781 |
| TcCLB.511389.140 | hypothetical protein, conserved | 0,7955 | 4,999E-09 |
| TcCLB.509165.34 | ribosomal P protein AGP2beta-1, putative | 0,7954 | 5,707E-11 |
| TcCLB.510533.100 | hypothetical protein, conserved | 0,7954 | 1,997E-07 |
| TcCLB.511145.20 | 60S ribosomal protein L32, putative | 0,7953 | 8,935E-20 |
| TcCLB.511145.60 | ATP synthase F1 subunit gamma protein, putative (ATPF1 | 0,7953 | 7,488E-14 |
| TcCLB.456097.30 | glycerophosphoryl diester phosphodiesterase, putative | 0,7953 | 7,007E-10 |
| TcCLB.510735.100 | hypothetical protein, conserved | 0,795 | 9,864E-09 |

| | | | |
|---|---|---|---|
| TcCLB.506503.160 | hypothetical protein, conserved | 0,7948 | 1,193E-23 |
| TcCLB.509805.150 | hypothetical protein, conserved | 0,7948 | 2,042E-12 |
| TcCLB.506861.10 | ATP-dependent DEAD/H RNA helicase, putative | 0,7946 | 5,08E-13 |
| TcCLB.507673.20 | 60S ribosomal protein L13a, putative | 0,7944 | 2,424E-19 |
| TcCLB.506885.40 | hypothetical protein, conserved | 0,7942 | 9,373E-16 |
| TcCLB.509233.160 | alpha-tubulin N-acetyltransferase, putative | 0,7942 | 1,959E-10 |
| TcCLB.507265.15 | transcription activator, putative (fragment) | 0,7935 | 8,111E-23 |
| TcCLB.509461.70 | hypothetical protein, conserved | 0,7933 | 6,895E-15 |
| TcCLB.508153.1040 | hypothetical protein, conserved | 0,7933 | 4,799E-12 |
| TcCLB.510731.120 | hypothetical protein, conserved | 0,7932 | 5,181E-11 |
| TcCLB.508003.30 | hypothetical protein, conserved (pseudogene) | 0,7932 | 1,539E-07 |
| TcCLB.508501.280 | hypothetical protein, conserved | 0,7932 | 1,89E-07 |
| TcCLB.511017.25 | hypothetical protein | 0,7931 | 0,00005098 |
| TcCLB.507231.20 | trans-sialidase (pseudogene), putative | 0,7931 | 0,001302 |
| TcCLB.506789.180 | glutaminyl cyclase, putative | 0,7929 | 1,186E-17 |
| TcCLB.506819.40 | hypothetical protein, conserved | 0,7922 | 1,331E-10 |
| TcCLB.510699.110 | retrotransposon hot spot protein (RHS, pseudogene), puta | 0,7922 | 2,653E-06 |
| TcCLB.503973.255 | hypothetical protein, conserved (pseudogene) | 0,792 | 0,03419 |
| TcCLB.511545.20 | ribosomal protein L27, putative | 0,7916 | 2,971E-19 |
| TcCLB.506773.80 | hypothetical protein, conserved | 0,7916 | 2,2E-15 |
| TcCLB.509209.20 | hypothetical protein, conserved | 0,7914 | 0,000021 |
| TcCLB.509395.10 | hypothetical protein, conserved | 0,7912 | 3,804E-11 |
| TcCLB.510901.70 | mitochondrial RNA binding complex 1 subunit, putative | 0,791 | 8,307E-17 |
| TcCLB.504423.30 | hypothetical protein, conserved | 0,791 | 2,002E-08 |
| TcCLB.508153.170 | ubiquinone biosynthesis protein COQ7 homolog, putative | 0,791 | 0,0000868 |
| TcCLB.506139.120 | trans-sialidase, Group V, putative | 0,7909 | 8,329E-06 |
| TcCLB.506435.420 | hypothetical protein, conserved | 0,7908 | 2,652E-07 |
| TcCLB.508787.14 | hypothetical protein, conserved | 0,7902 | 0,00002465 |
| TcCLB.511729.50 | hypothetical protein, conserved | 0,7901 | 1,228E-18 |
| TcCLB.509637.36 | procyclic form surface glycoprotein, putative (fragment) | 0,7901 | 2,564E-10 |
| TcCLB.510637.9 | dispersed gene family protein 1 (DGF-1), putative | 0,7901 | 0,00003805 |
| TcCLB.506265.130 | hypothetical protein, conserved | 0,79 | 5,349E-13 |
| TcCLB.506489.40 | hypothetical protein, conserved | 0,7899 | 1,168E-11 |
| TcCLB.511419.70 | hypothetical protein | 0,7899 | 6,767E-08 |
| TcCLB.507053.170 | hypothetical protein, conserved | 0,7897 | 1,93E-15 |
| TcCLB.511771.10 | retrotransposon hot spot protein (RHS, pseudogene), puta | 0,7897 | 2,704E-15 |
| TcCLB.505915.10 | retrotransposon hot spot protein (RHS, pseudogene), puta | 0,7896 | 7,596E-08 |
| TcCLB.504883.50 | hypothetical protein, conserved | 0,7891 | 2,43E-18 |
| TcCLB.506503.60 | hypothetical protein, conserved | 0,7889 | 6,548E-14 |
| TcCLB.506041.10 | hypothetical protein, conserved (fragment) | 0,7889 | 1,611E-08 |
| TcCLB.511003.30 | mitochondrial RNA binding complex 1 subunit, putative (R | 0,7886 | 4,953E-21 |
| TcCLB.507615.130 | hypothetical protein, conserved | 0,7883 | 5,535E-08 |
| TcCLB.506469.40 | hypothetical protein, conserved | 0,7883 | 1,868E-06 |
| TcCLB.506835.30 | 60S ribosomal protein L9, putative | 0,7882 | 1,09E-28 |
| TcCLB.506565.20 | hypothetical protein, conserved | 0,7882 | 3,008E-09 |

| | | | |
|---|---|---|---|
| TcCLB.510407.60 | tyrosyl-tRNA synthetase (pseudogene), putative | 0,788 | 5,05E-23 |
| TcCLB.510257.139 | transcription activator, putative | 0,7874 | 8,252E-12 |
| TcCLB.506739.150 | 60S ribosomal protein L13a, putative | 0,7873 | 2,721E-27 |
| TcCLB.507091.10 | hypothetical protein, conserved | 0,7871 | 0,01592 |
| TcCLB.506685.20 | hypothetical protein, conserved | 0,787 | 1,722E-11 |
| TcCLB.511211.120 | receptor for activated C kinase 1, putative | 0,7868 | 2,969E-28 |
| TcCLB.511523.70 | Present in the outer mitochondrial membrane proteome 3 | 0,7865 | 1,316E-06 |
| TcCLB.507087.80 | 5'-3' exoribonuclease B, putative (XRNB) | 0,7859 | 8,573E-15 |
| TcCLB.504071.40 | hypothetical protein | 0,7859 | 2,851E-08 |
| TcCLB.504109.70 | mitochondrial carrier protein, putative | 0,7855 | 1,475E-18 |
| TcCLB.503453.30 | 60S ribosomal protein L30, putative | 0,7855 | 3,245E-17 |
| TcCLB.510307.200 | hypothetical protein, conserved | 0,7855 | 1,702E-09 |
| TcCLB.506359.80 | Dynein light chain roadblock-type protein, putative | 0,7855 | 8,171E-09 |
| TcCLB.509735.70 | retrotransposon hot spot protein (RHS, pseudogene), puta | 0,7853 | 1,409E-13 |
| TcCLB.507467.110 | hypothetical protein, conserved | 0,785 | 4,585E-19 |
| TcCLB.505193.90 | hypothetical protein, conserved | 0,7848 | 1,424E-14 |
| TcCLB.509645.30 | hypothetical protein, conserved | 0,7846 | 6,648E-11 |
| TcCLB.510603.140 | hypothetical protein | 0,7845 | 2,294E-18 |
| TcCLB.507625.200 | hypothetical protein, conserved | 0,7843 | 1,932E-11 |
| TcCLB.508461.550 | hypothetical protein, conserved | 0,7843 | 6,558E-10 |
| TcCLB.508501.240 | hypothetical protein, conserved | 0,7842 | 8,111E-06 |
| TcCLB.511163.10 | seryl-tRNA synthetase, putative (fragment) | 0,7838 | 6,018E-21 |
| TcCLB.509733.140 | hypothetical protein, conserved | 0,7838 | 4,001E-17 |
| TcCLB.508965.30 | hypothetical protein | 0,7837 | 1,152E-12 |
| TcCLB.503613.30 | DNA repair protein, putative | 0,7833 | 2,058E-11 |
| TcCLB.511469.90 | hypothetical protein | 0,783 | 1,156E-07 |
| TcCLB.509163.10 | hypothetical protein | 0,7825 | 0,00000934 |
| TcCLB.509891.20 | hypothetical protein, conserved (pseudogene) | 0,7824 | 1,79E-08 |
| TcCLB.503687.30 | hypothetical protein, conserved | 0,7824 | 3,743E-06 |
| TcCLB.506789.310 | hypothetical protein, conserved | 0,7819 | 7,069E-11 |
| TcCLB.508027.40 | folate/pteridine transporter, putative | 0,7818 | 4,863E-18 |
| TcCLB.511717.150 | hypothetical protein, conserved | 0,7815 | 2,437E-12 |
| TcCLB.482471.10 | hypothetical protein, conserved | 0,7815 | 2,688E-10 |
| TcCLB.511035.10 | surface protease GP63, putative | 0,7813 | 6,265E-20 |
| TcCLB.503981.40 | DnaJ chaperone protein, putative | 0,781 | 8,52E-23 |
| TcCLB.511247.27 | RNA editing 3' terminal uridylyl transferase 1, putative | 0,7809 | 1,512E-08 |
| TcCLB.506657.50 | ADP/ATP mitochondrial carrier protein (pseudogene), puta | 0,7808 | 5,308E-10 |
| TcCLB.511529.130 | hypothetical protein, conserved | 0,7808 | 4,724E-09 |
| TcCLB.508355.250 | 60S acidic ribosomal protein P0, putative (P0) | 0,7805 | 6,646E-30 |
| TcCLB.508897.30 | ABC transport system ATP-binding protein, putative | 0,7803 | 1,734E-25 |
| TcCLB.511221.35 | retrotransposon hot spot protein (RHS, pseudogene), puta | 0,7798 | 1,415E-08 |
| TcCLB.509307.10 | retrotransposon hot spot protein (RHS, pseudogene), puta | 0,7798 | 1,979E-08 |
| TcCLB.509177.20 | hypothetical protein | 0,7794 | 1,461E-09 |
| TcCLB.509527.200 | hypothetical protein | 0,7792 | 0,02121 |
| TcCLB.506417.20 | hypothetical protein, conserved | 0,779 | 0,00007204 |

| | | | |
|---|---|---|---|
| TcCLB.503431.70 | hypothetical protein, conserved | 0,7781 | 8,581E-12 |
| TcCLB.504017.20 | hypothetical protein, conserved | 0,7779 | 5,707E-14 |
| TcCLB.506581.20 | hypothetical protein, conserved | 0,7779 | 1,824E-09 |
| TcCLB.509667.20 | tryptophanyl-tRNA synthetase (pseudogene), putative | 0,7773 | 2,209E-06 |
| TcCLB.507081.40 | hypothetical protein, conserved | 0,7771 | 6,625E-11 |
| TcCLB.508139.210 | dispersed gene family protein 1 (DGF-1), putative | 0,7771 | 5,036E-06 |
| TcCLB.506649.70 | fructose-1,6-bisphosphatase, cytosolic, putative | 0,7769 | 6,159E-15 |
| TcCLB.503431.30 | hypothetical protein | 0,7762 | 2,018E-06 |
| TcCLB.503811.10 | heat shock protein 90, putative (LPG3) | 0,7759 | 2,405E-15 |
| TcCLB.510053.60 | cytidine deaminase-like protein, putative | 0,7759 | 5,766E-09 |
| TcCLB.509749.33 | ribosomal protein L29, putative | 0,7758 | 2,217E-15 |
| TcCLB.507521.44 | hypothetical protein, conserved | 0,7755 | 0,00001953 |
| TcCLB.506685.10 | hypothetical protein, conserved | 0,7752 | 9,911E-17 |
| TcCLB.508147.45 | hypothetical protein, conserved (pseudogene) | 0,7748 | 0,008841 |
| TcCLB.504069.30 | 60S ribosomal protein L32, putative | 0,7747 | 6,098E-14 |
| TcCLB.511327.20 | hypothetical protein (fragment) | 0,7746 | 0,00001532 |
| TcCLB.509257.10 | trans-sialidase, Group II, putative | 0,7745 | 1,204E-13 |
| TcCLB.511825.69 | cyclophilin, putative | 0,7742 | 2,795E-08 |
| TcCLB.503861.50 | beta galactofuranosyl glycosyltransferase, putative | 0,7741 | 8,251E-17 |
| TcCLB.511871.10 | hypothetical protein, conserved | 0,7741 | 5,561E-06 |
| TcCLB.506351.50 | hypothetical protein | 0,774 | 1,922E-08 |
| TcCLB.509561.20 | flagellum-adhesion glycoprotein, putative | 0,7739 | 8,992E-15 |
| TcCLB.508017.20 | UDP-Gal or UDP-GlcNAc-dependent glycosyltransferase (p | 0,7739 | 0,003467 |
| TcCLB.510431.250 | phosphoribosylpyrophosphate synthetase, putative | 0,7737 | 4,71E-20 |
| TcCLB.508555.34 | hypothetical protein, conserved | 0,7737 | 4,355E-06 |
| TcCLB.507801.160 | hypothetical protein, conserved | 0,7734 | 5,847E-09 |
| TcCLB.510733.20 | hypothetical protein, conserved | 0,7731 | 7,86E-14 |
| TcCLB.426397.9 | kinesin, putative (fragment) | 0,7731 | 3,203E-11 |
| TcCLB.506815.60 | 60S ribosomal protein L44 (RPL44) | 0,773 | 6,091E-23 |
| TcCLB.506475.110 | hypothetical protein | 0,773 | 0,0001219 |
| TcCLB.507993.60 | hypothetical protein, conserved | 0,7727 | 2,808E-10 |
| TcCLB.511797.54 | dispersed gene family protein 1 (DGF-1), putative | 0,7727 | 0,004813 |
| TcCLB.510155.10 | hypothetical protein, conserved | 0,7726 | 4,805E-29 |
| TcCLB.503847.10 | hypothetical protein, conserved (pseudogene) | 0,7726 | 1,227E-14 |
| TcCLB.504443.10 | cysteine peptidase, Clan CA, family C19, putative | 0,7723 | 1,374E-12 |
| TcCLB.505169.21 | phospholipid transporting ATPase-like protein (pseudogen | 0,7722 | 0,01066 |
| TcCLB.506595.110 | AGP2beta-2, putative | 0,7721 | 5,771E-10 |
| TcCLB.508859.90 | phosphatidylinositol 3-kinase 2, putative | 0,7718 | 1,183E-09 |
| TcCLB.511277.150 | hypothetical protein | 0,7717 | 1,782E-19 |
| TcCLB.509613.10 | hypothetical protein, conserved | 0,771 | 1,852E-08 |
| TcCLB.510543.110 | hypothetical protein, conserved (pseudogene) | 0,7707 | 0,00002647 |
| TcCLB.507499.50 | hypothetical protein, conserved (pseudogene) | 0,7705 | 5,828E-07 |
| TcCLB.511755.19 | hypothetical protein, conserved | 0,7705 | 0,001864 |
| TcCLB.506413.80 | cyclophilin, putative | 0,7703 | 1,149E-08 |
| TcCLB.506869.10 | hypothetical protein | 0,77 | 2,228E-09 |

| | | | |
|---|---|---|---|
| TcCLB.506819.30 | superoxide dismutase, putative | 0,7696 | 1,375E-11 |
| TcCLB.508707.40 | hypothetical protein, conserved | 0,7695 | 4,623E-15 |
| TcCLB.506945.210 | Zinc finger CCCH domain-containing protein 47 (zc3h47) | 0,7694 | 4,821E-25 |
| TcCLB.511537.69 | hypothetical protein, conserved | 0,7694 | 5,381E-15 |
| TcCLB.506243.50 | dispersed gene family protein 1 (DGF-1, pseudogene), put | 0,7692 | 0,0001565 |
| TcCLB.511017.60 | hypothetical protein | 0,7691 | 1,774E-19 |
| TcCLB.503925.40 | hypothetical protein | 0,769 | 8,466E-08 |
| TcCLB.509229.110 | hypothetical protein, conserved | 0,7687 | 1,281E-16 |
| TcCLB.511825.30 | hypothetical protein, conserved | 0,7684 | 2,937E-09 |
| TcCLB.509787.30 | retrotransposon hot spot protein (RHS, pseudogene), puta | 0,7683 | 3,031E-08 |
| TcCLB.508515.90 | hypothetical protein, conserved | 0,7683 | 3,313E-06 |
| TcCLB.507777.41 | trans-sialidase (pseudogene), putative | 0,768 | 0,002732 |
| TcCLB.510891.30 | retrotransposon hot spot protein (RHS, pseudogene), puta | 0,7679 | 1,306E-11 |
| TcCLB.506567.110 | hypothetical protein, conserved | 0,7679 | 4,268E-11 |
| TcCLB.511843.40 | dispersed gene family protein 1 (DGF-1), putative | 0,7679 | 0,000279 |
| TcCLB.506247.410 | protein-l-isoaspartate o-methyltransferase, putative | 0,7677 | 6,269E-11 |
| TcCLB.509631.60 | dispersed gene family protein 1 (DGF-1), putative | 0,7676 | 0,0005044 |
| TcCLB.508267.40 | hypothetical protein, conserved | 0,7675 | 1,331E-14 |
| TcCLB.511747.7 | cyclosome subunit, putative (fragment) | 0,7675 | 9,632E-10 |
| TcCLB.508153.460 | Present in the outer mitochondrial membrane proteome 2 | 0,7671 | 1,651E-17 |
| TcCLB.511267.30 | cholinephosphate cytidylyltransferase A, putative | 0,7669 | 4,955E-16 |
| TcCLB.507157.10 | mucin TcSMUGL, putative | 0,7669 | 0,00003323 |
| TcCLB.506489.20 | monoglyceride lipase, putative | 0,7668 | 3,717E-20 |
| TcCLB.508087.10 | hypothetical protein | 0,7668 | 5,411E-10 |
| TcCLB.511713.9 | hypothetical protein, conserved | 0,7668 | 0,000135 |
| TcCLB.509615.4 | hypothetical protein, conserved (pseudogene) | 0,7668 | 0,0005236 |
| TcCLB.509815.50 | beta galactofuranosyl glycosyltransferase, putative | 0,7665 | 8,686E-06 |
| TcCLB.506631.20 | cardiolipin synthetase, putative (cls) | 0,7663 | 1,306E-12 |
| TcCLB.510773.110 | hypothetical protein, conserved | 0,7663 | 3,118E-07 |
| TcCLB.503681.20 | ethanolamine phosphotransferase, putative | 0,7661 | 1,098E-11 |
| TcCLB.511761.50 | sodium/hydrogen exchanger, putative (FTP) | 0,7661 | 1,706E-11 |
| TcCLB.510823.40 | hypothetical protein, conserved | 0,7658 | 1,989E-17 |
| TcCLB.508027.100 | hypothetical protein | 0,7658 | 7,029E-12 |
| TcCLB.508707.200 | nucleoside diphosphate kinase (NDPK1) | 0,7657 | 1,24E-15 |
| TcCLB.506617.50 | hypothetical protein | 0,7656 | 2,008E-09 |
| TcCLB.506975.63 | hypothetical protein, conserved | 0,7656 | 2,751E-09 |
| TcCLB.507849.50 | cysteine proteinase, putative | 0,7655 | 2,152E-20 |
| TcCLB.506559.470 | ribosomal protein l35a, putative | 0,7654 | 9,394E-29 |
| TcCLB.503917.40 | hypothetical protein, conserved | 0,7654 | 5,963E-10 |
| TcCLB.511815.50 | hypothetical protein, conserved | 0,7653 | 8,425E-21 |
| TcCLB.506529.669 | hypothetical protein, conserved | 0,7652 | 2,925E-06 |
| TcCLB.508317.50 | ammonium transporter, putative | 0,7651 | 3,442E-14 |
| TcCLB.504097.40 | hypothetical protein, conserved | 0,7649 | 1,452E-13 |
| TcCLB.509059.30 | calcium-binding protein, putative | 0,7649 | 8,273E-07 |
| TcCLB.510773.50 | hypothetical protein | 0,7647 | 0,00003268 |

| | | | |
|---|---|---|---|
| TcCLB.506407.80 | intraflagellar transport protein 81, putative (IFT81) | 0,7643 | 6,993E-26 |
| TcCLB.511003.150 | hypothetical protein | 0,7639 | 2,76E-15 |
| TcCLB.505977.26 | 60S acidic ribosomal protein P2, putative | 0,7638 | 1,241E-18 |
| TcCLB.510945.30 | actin, putative | 0,7638 | 1,125E-12 |
| TcCLB.508461.380 | calmodulin, putative | 0,7637 | 2,538E-19 |
| TcCLB.506773.70 | transcription modulator/accessory protein, putative | 0,7636 | 1,065E-17 |
| TcCLB.504213.120 | proteasome alpha 3 subunit, putative | 0,7635 | 5,694E-18 |
| TcCLB.506297.4 | hypothetical protein, conserved (pseudogene) | 0,7635 | 3,073E-08 |
| TcCLB.507991.40 | hypothetical protein | 0,7632 | 5,501E-10 |
| TcCLB.503449.10 | 60S ribosomal protein L17, putative | 0,7628 | 1,215E-23 |
| TcCLB.510007.70 | ATP-dependent zinc metallopeptidase, putative | 0,7628 | 1,147E-10 |
| TcCLB.511183.330 | dispersed gene family protein 1 (DGF-1), putative | 0,7627 | 0,002408 |
| TcCLB.506295.150 | hypothetical protein | 0,7626 | 1,111E-07 |
| TcCLB.511661.90 | ribose-phosphate pyrophosphokinase, putative | 0,7625 | 1,967E-21 |
| TcCLB.504137.140 | hypothetical protein, conserved | 0,7624 | 4,046E-11 |
| TcCLB.508153.1080 | hypothetical protein, conserved | 0,7624 | 5,709E-08 |
| TcCLB.503973.10 | hypothetical protein | 0,7624 | 0,0006894 |
| TcCLB.510889.231 | ATP synthase subunit, putative | 0,7623 | 6,551E-06 |
| TcCLB.506973.70 | dispersed gene family protein 1 (DGF-1), putative | 0,7623 | 0,1424 |
| TcCLB.506943.27 | hypothetical protein, conserved | 0,7619 | 0,00003349 |
| TcCLB.510543.90 | retrotransposon hot spot (RHS) protein, putative | 0,7619 | 0,0001745 |
| TcCLB.506885.416 | regulator of chromosome condensation (RCC1), putative | 0,7618 | 5,911E-13 |
| TcCLB.508651.10 | dispersed gene family protein 1 (DGF-1, pseudogene), put | 0,7616 | 0,02674 |
| TcCLB.509583.10 | chaperonin TCP20, putative | 0,7614 | 1,972E-18 |
| TcCLB.510187.290 | ATP-dependent DEAD/H RNA helicase, putative (HEL64) | 0,7608 | 2,509E-21 |
| TcCLB.511249.4 | RNA editing 3' terminal uridylyl transferase 1, putative | 0,7608 | 3,824E-07 |
| TcCLB.507895.170 | hypothetical protein, conserved | 0,7607 | 7,845E-11 |
| TcCLB.508397.79 | hypothetical protein, conserved | 0,7607 | 0,001097 |
| TcCLB.509445.39 | glutamate dehydrogenase, putative | 0,7606 | 7,18E-14 |
| TcCLB.506401.150 | hypothetical protein, conserved | 0,7606 | 9,89E-09 |
| TcCLB.510311.190 | hypothetical protein, conserved | 0,7603 | 4,443E-12 |
| TcCLB.509913.24 | dynein light chain 2B, cytoplasmic, putative | 0,7602 | 5,878E-11 |
| TcCLB.503437.31 | ATP-dependent DEAD/H RNA helicase (pseudogene), puta | 0,7601 | 0,02016 |
| TcCLB.509443.20 | mitochondrial RNA binding protein 1, putative | 0,76 | 7,706E-15 |
| TcCLB.507477.20 | hypothetical protein, conserved (pseudogene) | 0,7592 | 6,976E-16 |
| TcCLB.509551.70 | hypothetical protein, conserved | 0,7592 | 1,903E-11 |
| TcCLB.470827.20 | trans-sialidase, putative | 0,7591 | 1,217E-08 |
| TcCLB.412723.9 | dispersed gene family protein 1 (DGF-1), putative (fragme | 0,7591 | 0,01761 |
| TcCLB.511825.63 | peptidyl-prolyl cis-trans isomerase, putative | 0,7588 | 5,487E-11 |
| TcCLB.504105.110 | hypothetical protein, conserved | 0,7585 | 4,312E-22 |
| TcCLB.511807.200 | hypothetical protein, conserved | 0,7583 | 5,839E-24 |
| TcCLB.509541.19 | major vault protein, putative | 0,7582 | 3,494E-14 |
| TcCLB.504669.30 | surface protease GP63, putative | 0,7581 | 2,016E-10 |
| TcCLB.506511.30 | hypothetical protein, conserved | 0,7579 | 1,063E-15 |
| TcCLB.510315.14 | hypothetical protein, conserved | 0,7578 | 4,364E-07 |

| | | | |
|---|---|---|---|
| TcCLB.468041.30 | retrotransposon hot spot protein (RHS, pseudogene), puta | 0,7577 | 7,039E-18 |
| TcCLB.507897.10 | hypothetical protein, conserved | 0,7575 | 3,547E-07 |
| TcCLB.507913.20 | phosphomevalonate kinase protein, putative | 0,7574 | 7,999E-10 |
| TcCLB.506297.260 | lanosterol 14-alpha-demethylase, putative (CYP51) | 0,7572 | 4,53E-19 |
| TcCLB.504181.50 | T-complex protein 1, delta subunit, putative | 0,7571 | 7,729E-11 |
| TcCLB.507891.56 | flagellar calcium-binding protein, putative (fragment) | 0,7566 | 5,739E-09 |
| TcCLB.509965.20 | hypothetical protein | 0,7566 | 1,366E-06 |
| TcCLB.510341.4 | Monooxygenase, putative (fragment) | 0,7562 | 5,923E-07 |
| TcCLB.511233.14 | 40S ribosomal protein S27, putative | 0,7561 | 1,512E-19 |
| TcCLB.510087.50 | hypothetical protein, conserved | 0,7561 | 3,006E-15 |
| TcCLB.510155.160 | protein kinase, putative | 0,756 | 3,748E-18 |
| TcCLB.507875.140 | retrotransposon hot spot protein (RHS, pseudogene), puta | 0,7557 | 0,0006685 |
| TcCLB.511649.80 | myosin heavy chain, putative | 0,7556 | 1,168E-15 |
| TcCLB.506677.20 | cdc2-related kinase 3, putative (CRK3) | 0,7553 | 0,00007763 |
| TcCLB.511281.40 | mevalonate-diphosphate decarboxylase, putative | 0,7552 | 1,547E-11 |
| TcCLB.509047.80 | mitochondrial oligo_U binding protein TBRGG1 (pseudoge | 0,7549 | 5,797E-12 |
| TcCLB.508703.25 | hypothetical protein | 0,7543 | 3,014E-10 |
| TcCLB.509455.140 | cyclin, putative | 0,7541 | 5,662E-09 |
| TcCLB.460127.20 | metallo- peptidase, Clan M- Family M48 (FACE1) | 0,7541 | 8,704E-09 |
| TcCLB.504741.240 | hypothetical protein, conserved | 0,7541 | 2,088E-06 |
| TcCLB.507529.30 | hypothetical protein, conserved | 0,7537 | 2,302E-15 |
| TcCLB.506907.30 | hypothetical protein, conserved | 0,7535 | 7,118E-09 |
| TcCLB.506677.39 | Elongation factor G1, mitochondrial, putative (EF-G1) | 0,7533 | 4,216E-21 |
| TcCLB.506943.40 | hypothetical protein, conserved | 0,7532 | 2,672E-10 |
| TcCLB.511031.10 | hypothetical protein, conserved | 0,753 | 0,00007648 |
| TcCLB.507779.20 | trans-sialidase (pseudogene), putative | 0,7528 | 0,0002003 |
| TcCLB.507049.65 | hypothetical protein, conserved (pseudogene) | 0,7528 | 0,001494 |
| TcCLB.507677.39 | 40S ribosomal protein S3, putative | 0,7526 | 5,839E-28 |
| TcCLB.510731.20 | hypothetical protein, conserved | 0,7523 | 1,051E-12 |
| TcCLB.503855.20 | spermidine synthase, putative | 0,7515 | 2,937E-16 |
| TcCLB.507611.310 | eukaryotic translation initiation factor 3 subunit 8, putativ | 0,7511 | 3,612E-18 |
| TcCLB.510101.390 | hypothetical protein, conserved | 0,7511 | 4,654E-13 |
| TcCLB.510719.35 | ribosomal protein L29, putative | 0,7511 | 4,139E-08 |
| TcCLB.509965.50 | hypothetical protein, conserved | 0,751 | 0,00002549 |
| TcCLB.511277.240 | vacuolar ATP synthase subunit c, putative | 0,7501 | 5,337E-12 |
| TcCLB.506303.171 | hypothetical protein, conserved | 0,75 | 0,0219 |
| TcCLB.509789.20 | hypothetical protein | 0,7498 | 2,337E-15 |
| TcCLB.504013.55 | hypothetical protein, conserved | 0,7496 | 3,511E-08 |
| TcCLB.504013.100 | ribosomal protein S19, putative | 0,7494 | 2,165E-22 |
| TcCLB.511283.124 | mitochondrial carrier protein, putative | 0,7492 | 1,251E-11 |
| TcCLB.503939.70 | hypothetical protein | 0,749 | 2,658E-20 |
| TcCLB.510877.180 | GPI transamidase subunit 16, putative (GPI16) | 0,749 | 1,887E-17 |
| TcCLB.506391.10 | calmodulin (CALA2) | 0,749 | 6,581E-08 |
| TcCLB.506241.90 | hypothetical protein, conserved | 0,7489 | 8,86E-10 |
| TcCLB.504153.200 | hypothetical protein, conserved | 0,7488 | 9,721E-08 |

| | | | |
|---|---|---|---|
| TcCLB.508323.100 | I/6 autoantigen, putative | 0,7486 | 2,747E-07 |
| TcCLB.510415.9 | dispersed gene family protein 1 (DGF-1), putative (fragme | 0,7485 | 0,002635 |
| TcCLB.509033.20 | beta galactofuranosyl glycosyltransferase, putative | 0,7484 | 2,074E-07 |
| TcCLB.509797.60 | hypothetical protein | 0,7483 | 3,033E-09 |
| TcCLB.509611.170 | hypothetical protein, conserved | 0,7482 | 2,97E-11 |
| TcCLB.507513.10 | hypothetical protein, conserved | 0,7482 | 1,297E-10 |
| TcCLB.506357.170 | serine/threonine protein phosphatase, putative | 0,7481 | 2,262E-09 |
| TcCLB.503419.50 | MRB1-associated protein, putative (RGG2) | 0,7478 | 1,093E-07 |
| TcCLB.511351.30 | protein kinase, putative | 0,7475 | 1,218E-17 |
| TcCLB.510257.20 | hypothetical protein, conserved | 0,7473 | 5,326E-12 |
| TcCLB.509859.70 | hypothetical protein, conserved | 0,7473 | 1,841E-11 |
| TcCLB.509153.70 | exosome-associated protein 3, putative (EAP3) | 0,7472 | 5,236E-12 |
| TcCLB.506775.150 | hypothetical protein, conserved | 0,7472 | 1,3E-10 |
| TcCLB.510005.11 | endosomal integral membrane protein (pseudogene), put | 0,7472 | 1,545E-08 |
| TcCLB.414243.10 | hypothetical protein, conserved | 0,7472 | 1,381E-06 |
| TcCLB.506961.50 | dispersed gene family protein 1 (DGF-1), putative | 0,7471 | 5,443E-08 |
| TcCLB.511821.179 | hypothetical protein, conserved | 0,7469 | 7,032E-13 |
| TcCLB.510357.60 | hypothetical protein, conserved | 0,7469 | 0,0001434 |
| TcCLB.504797.100 | 35 kDa cyclophilin, putative (CYP35) | 0,7467 | 1,537E-08 |
| TcCLB.507693.5 | tryptophanyl-tRNA synthetase, putative (fragment) | 0,7467 | 2,948E-07 |
| TcCLB.510053.10 | hypothetical protein, conserved | 0,7466 | 0,00000865 |
| TcCLB.507077.50 | hypothetical protein, conserved | 0,7465 | 2,035E-07 |
| TcCLB.511807.280 | hypothetical protein, conserved | 0,7464 | 5,357E-31 |
| TcCLB.510285.74 | hypothetical protein, conserved | 0,7464 | 4,557E-07 |
| TcCLB.508267.30 | hypothetical protein, conserved | 0,746 | 1,089E-11 |
| TcCLB.508307.70 | hypothetical protein, conserved | 0,7459 | 3,956E-09 |
| TcCLB.504107.10 | calpain-like cysteine peptidase, putative (fragment) | 0,7459 | 0,06242 |
| TcCLB.506227.110 | hypothetical protein, conserved | 0,7458 | 8,397E-09 |
| TcCLB.508267.20 | hypothetical protein, conserved | 0,7453 | 5,461E-19 |
| TcCLB.507129.30 | C-14 sterol reductase, putative | 0,7452 | 6,674E-12 |
| TcCLB.508683.9 | dispersed gene family protein 1 (DGF-1), putative | 0,7451 | 0,02536 |
| TcCLB.504147.99 | hypothetical protein | 0,7443 | 1,973E-17 |
| TcCLB.511889.20 | retrotransposon hot spot protein (RHS, pseudogene), puta | 0,7443 | 4,284E-10 |
| TcCLB.508919.130 | hypothetical protein, conserved | 0,7442 | 1,037E-10 |
| TcCLB.452699.30 | dispersed gene family protein 1 (DGF-1), putative | 0,744 | 0,03943 |
| TcCLB.506825.10 | RNA-binding protein, putative | 0,744 | 0,09059 |
| TcCLB.509529.19 | dispersed gene family protein 1 (DGF-1), putative (fragme | 0,7438 | 0,00000127 |
| TcCLB.510575.130 | chloride channel protein, putative | 0,7432 | 1,269E-17 |
| TcCLB.509125.20 | ATP-dependent Clp protease subunit, heat shock protein 7 | 0,7431 | 3,318E-15 |
| TcCLB.508881.140 | hypothetical protein | 0,7429 | 5,898E-14 |
| TcCLB.511385.100 | hypothetical protein, conserved | 0,7429 | 1,429E-12 |
| TcCLB.504061.10 | retrotransposon hot spot protein (RHS, pseudogene), puta | 0,7429 | 1,765E-11 |
| TcCLB.505193.70 | hypothetical protein, conserved | 0,7428 | 1,047E-10 |
| TcCLB.506349.47 | retrotransposon hot spot protein (RHS, pseudogene), puta | 0,7427 | 0,001767 |
| TcCLB.504163.80 | hypothetical protein, conserved | 0,7424 | 9,43E-11 |

| | | | |
|---|---|---|---|
| TcCLB.506405.100 | hypothetical protein, conserved | 0,742 | 1,262E-19 |
| TcCLB.506561.10 | helicase-like protein, putative | 0,742 | 1,218E-13 |
| TcCLB.503827.94 | chitin binding-like protein (pseudogene), putative | 0,742 | 0,0753 |
| TcCLB.503757.20 | hypothetical protein, conserved | 0,7419 | 1,168E-06 |
| TcCLB.510307.180 | glycine dehydrogenase [decarboxylating] (pseudogene), pu | 0,7417 | 4,551E-12 |
| TcCLB.510303.210 | 2,4-dienoyl-coa reductase-like protein, putative | 0,7414 | 1,45E-23 |
| TcCLB.508287.10 | dispersed gene family protein 1 (DGF-1), putative | 0,7413 | 0,05787 |
| TcCLB.511765.20 | dispersed gene family protein 1 (DGF-1), putative | 0,7412 | 0,00002884 |
| TcCLB.509937.100 | ubiquitin/ribosomal protein S27a, putative | 0,7411 | 1,086E-29 |
| TcCLB.511661.60 | hypothetical protein, conserved | 0,7411 | 7,682E-09 |
| TcCLB.511077.30 | retrotransposon hot spot protein (RHS, pseudogene), puta | 0,7411 | 0,0005033 |
| TcCLB.485845.9 | dispersed gene family protein 1 (DGF-1), putative (fragme | 0,7411 | 0,134 |
| TcCLB.510309.50 | 60S acidic ribosomal protein P2, putative | 0,7408 | 4,259E-14 |
| TcCLB.508845.9 | dispersed gene family protein 1 (DGF-1), putative | 0,7408 | 0,01791 |
| TcCLB.510465.150 | dispersed gene family protein 1 (DGF-1), putative | 0,7408 | 0,01994 |
| TcCLB.503639.10 | inositol-3-phosphate synthase, putative | 0,7407 | 1,966E-07 |
| TcCLB.504189.10 | beta galactofuranosyl glycosyltransferase (pseudogene), p | 0,7405 | 0,02487 |
| TcCLB.510001.59 | hypothetical protein, conserved (fragment) | 0,7404 | 2,285E-16 |
| TcCLB.506171.50 | retrotransposon hot spot protein (RHS, pseudogene), puta | 0,7404 | 1,316E-07 |
| TcCLB.507835.70 | dispersed gene family protein 1 (DGF-1), putative | 0,7403 | 0,00003448 |
| TcCLB.509597.29 | hypothetical protein, conserved | 0,7402 | 1,053E-08 |
| TcCLB.507583.4 | hypothetical protein, conserved (fragment) | 0,7402 | 0,0001391 |
| TcCLB.506401.70 | vacuolar protein sorting protein 18, putative | 0,7401 | 2,465E-16 |
| TcCLB.509643.120 | hypothetical protein, conserved | 0,7401 | 1,17E-10 |
| TcCLB.507055.10 | hypothetical protein, conserved | 0,7401 | 0,00009294 |
| TcCLB.505999.50 | hypothetical protein, conserved | 0,7399 | 1,598E-06 |
| TcCLB.503493.10 | hypothetical protein, conserved | 0,7397 | 2,352E-09 |
| TcCLB.504797.10 | acidocalcisomal exopolyphosphatase, putative | 0,7391 | 4,121E-09 |
| TcCLB.508457.30 | lipoic acid containing carrier protein, putative (GCVH) | 0,7389 | 4,975E-12 |
| TcCLB.507499.56 | N-acetyltransferase complex ARD1 subunit (pseudogene), | 0,7387 | 0,05734 |
| TcCLB.509671.70 | ribosomal protein L15, putative | 0,7386 | 5,583E-25 |
| TcCLB.506825.210 | hypothetical protein, conserved | 0,7385 | 1,76E-17 |
| TcCLB.511733.70 | hypothetical protein, conserved | 0,7385 | 1,018E-06 |
| TcCLB.509017.60 | hypothetical protein, conserved | 0,7382 | 1,741E-13 |
| TcCLB.506553.30 | GTP-binding protein, putative | 0,7379 | 2,238E-12 |
| TcCLB.508163.190 | dispersed gene family protein 1 (DGF-1), putative | 0,7379 | 0,002736 |
| TcCLB.510533.120 | translation initiation factor IF-2, putative | 0,7378 | 2,316E-13 |
| TcCLB.508741.220 | hypothetical protein | 0,7377 | 5,302E-14 |
| TcCLB.456139.10 | hypothetical protein, conserved | 0,7376 | 0,0002305 |
| TcCLB.511871.40 | hypothetical protein, conserved | 0,7372 | 8,157E-08 |
| TcCLB.509615.10 | hypothetical protein, conserved | 0,7372 | 1,152E-07 |
| TcCLB.510629.410 | hypothetical protein, conserved | 0,737 | 0,00006869 |
| TcCLB.508153.280 | 60S ribosomal protein L13, putative | 0,7368 | 2,018E-21 |
| TcCLB.507963.30 | hypothetical protein, conserved | 0,7367 | 2,115E-19 |
| TcCLB.508299.40 | hypothetical protein | 0,7362 | 3,593E-10 |

| | | | |
|---|---|---|---|
| TcCLB.510685.30 | RNA polymerase IIA largest subunit, putative | 0,7358 | 2,677E-14 |
| TcCLB.506507.60 | retrotransposon hot spot protein (RHS, pseudogene), puta | 0,7358 | 2,422E-09 |
| TcCLB.507211.20 | protein kinase, putative | 0,7358 | 0,002617 |
| TcCLB.508949.20 | hypothetical protein, conserved | 0,7352 | 0,000485 |
| TcCLB.510629.420 | protein phosphatase 1, regulatory subunit, putative | 0,7347 | 5,96E-10 |
| TcCLB.507079.30 | multidrug resistance protein E, putative | 0,7347 | 0,0002692 |
| TcCLB.511693.30 | hypothetical protein, conserved | 0,7341 | 3,552E-19 |
| TcCLB.508087.20 | retrotransposon hot spot protein (RHS, pseudogene), puta | 0,734 | 3,964E-14 |
| TcCLB.510579.80 | 60S ribosomal protein L10, putative | 0,7337 | 7,234E-18 |
| TcCLB.507443.14 | RNA polymerase IIA largest subunit, putative (fragment) | 0,7337 | 3,045E-08 |
| TcCLB.505975.10 | hypothetical protein | 0,7337 | 0,1332 |
| TcCLB.510323.30 | 60S ribosomal protein L13, putative | 0,7335 | 2,037E-18 |
| TcCLB.504225.10 | hypothetical protein, conserved | 0,7334 | 6,704E-11 |
| TcCLB.506791.70 | nitroreductase | 0,7333 | 3,467E-11 |
| TcCLB.503449.14 | hypothetical protein, conserved | 0,7333 | 2,874E-09 |
| TcCLB.511139.20 | 40S ribosomal protein S11, putative | 0,7326 | 6,02E-24 |
| TcCLB.506957.90 | 60S ribosomal protein L18a, putative | 0,7323 | 2,012E-27 |
| TcCLB.509221.10 | retrotransposon hot spot protein (RHS, pseudogene), puta | 0,7323 | 9,594E-12 |
| TcCLB.511479.20 | hypothetical protein, conserved | 0,7323 | 1,584E-11 |
| TcCLB.506171.20 | hypothetical protein, conserved (pseudogene) | 0,7323 | 3,195E-07 |
| TcCLB.511293.80 | hypothetical protein, conserved | 0,7322 | 3,002E-09 |
| TcCLB.510459.30 | dispersed gene family protein 1 (DGF-1), putative | 0,7321 | 0,02929 |
| TcCLB.508393.9 | dispersed gene family protein 1 (DGF-1), putative | 0,7321 | 0,2099 |
| TcCLB.510689.14 | NADH-ubiquinone oxidoreductase complex I subunit, puta | 0,732 | 3,451E-10 |
| TcCLB.511821.160 | microtubial binding protein, putative | 0,732 | 0,00003542 |
| TcCLB.510055.70 | hypothetical protein, conserved | 0,7315 | 3,019E-11 |
| TcCLB.506865.4 | hypothetical protein, conserved | 0,7315 | 9,726E-06 |
| TcCLB.508153.490 | peptidyl-prolyl cis-trans isomerase NIMA-interacting 4, pu | 0,7314 | 0,00007382 |
| TcCLB.504797.40 | thioredoxin, putative | 0,7313 | 7,743E-10 |
| TcCLB.505763.19 | P-type H -ATPase, putative | 0,7312 | 1,149E-22 |
| TcCLB.509767.150 | hypothetical protein, conserved | 0,7312 | 1,958E-11 |
| TcCLB.506495.20 | helicase-like protein, putative | 0,7307 | 8,128E-14 |
| TcCLB.504769.160 | dispersed gene family protein 1 (DGF-1), putative | 0,7306 | 0,00009283 |
| TcCLB.511517.120 | hypothetical protein, conserved | 0,7305 | 2,859E-15 |
| TcCLB.475169.10 | DNA-directed RNA polymerase II subunit 2 (pseudogene), | 0,7305 | 0,00006943 |
| TcCLB.511277.270 | tubulin cofactor C domain-containing protein RP2, putativ | 0,7297 | 4,797E-13 |
| TcCLB.504797.140 | CLC-type chloride channel, putative | 0,7296 | 7,131E-20 |
| TcCLB.508285.40 | retrotransposon hot spot protein (RHS, pseudogene), puta | 0,7296 | 0,0139 |
| TcCLB.509533.30 | retrotransposon hot spot protein (RHS, pseudogene), puta | 0,7295 | 1,05E-08 |
| TcCLB.511181.100 | 60S ribosomal protein L8, putative | 0,7291 | 1,318E-24 |
| TcCLB.508541.170 | hypothetical protein, conserved | 0,7288 | 0,000023 |
| TcCLB.504021.109 | 40S ribosomal protein S3, putative | 0,7285 | 1,224E-23 |
| TcCLB.503865.50 | ankyrin, putative | 0,7284 | 1,816E-13 |
| TcCLB.509127.10 | ATP-dependent Clp protease subunit, heat shock protein 7 | 0,7281 | 4,015E-10 |
| TcCLB.510857.50 | lipoate-protein ligase, putative | 0,7281 | 4,651E-09 |

| | | | |
|---|---|---|---|
| TcCLB.503951.10 | retrotransposon hot spot protein (RHS, pseudogene), puta | 0,7281 | 0,006183 |
| TcCLB.510233.20 | hypothetical protein | 0,7279 | 0,02926 |
| TcCLB.509353.30 | 40S ribosomal protein S3, putative | 0,7277 | 5,253E-20 |
| TcCLB.504817.9 | hypothetical protein, conserved (fragment) | 0,7269 | 0,002796 |
| TcCLB.510293.40 | ubiquitin/ribosomal protein S27a, putative | 0,7268 | 0,000435 |
| TcCLB.510897.80 | hypothetical protein, conserved | 0,7267 | 0,00002121 |
| TcCLB.506691.40 | hypothetical protein, conserved | 0,7266 | 2,783E-07 |
| TcCLB.506779.44 | hypothetical protein, conserved | 0,7265 | 0,00001829 |
| TcCLB.506163.35 | hypothetical protein, conserved | 0,7262 | 0,00001238 |
| TcCLB.508141.50 | hypothetical protein, conserved | 0,7261 | 0,00001012 |
| TcCLB.509303.20 | glycine dehydrogenase [decarboxylating] (pseudogene), pu | 0,726 | 0,00006452 |
| TcCLB.506533.106 | mucin TcSMUGL, putative | 0,7258 | 2,55E-10 |
| TcCLB.503931.30 | hypothetical protein | 0,7257 | 3,048E-09 |
| TcCLB.511417.100 | anti-silencing protein ASF 1, putative | 0,7257 | 0,0001825 |
| TcCLB.509253.30 | 3-Beta-hydroxysteroid-delta(8), delta(7)-isomerase, putat | 0,7253 | 7,211E-08 |
| TcCLB.506201.80 | serine/threonine-protein phosphatase PP1, putative | 0,7253 | 0,0008207 |
| TcCLB.506933.20 | ribonucleoside-diphosphate reductase large chain, putativ | 0,725 | 3,463E-14 |
| TcCLB.506435.460 | hypothetical protein | 0,725 | 1,999E-09 |
| TcCLB.504949.30 | succinate dehydrogenase iron-sulfur subunit (SDH2C) | 0,7249 | 7,436E-08 |
| TcCLB.511421.190 | ESCO acetyltransferase domain-containing protein | 0,7247 | 2,813E-09 |
| TcCLB.508851.130 | hypothetical protein | 0,7241 | 5,115E-16 |
| TcCLB.507011.150 | hypothetical protein, conserved | 0,724 | 2,865E-07 |
| TcCLB.505207.40 | dispersed gene family protein 1 (DGF-1), putative | 0,724 | 0,0003386 |
| TcCLB.510091.119 | calpain-like cysteine peptidase (pseudogene), putative | 0,7239 | 4,108E-07 |
| TcCLB.508221.830 | hypothetical protein, conserved (pseudogene) | 0,7231 | 0,03418 |
| TcCLB.510359.220 | hypothetical protein, conserved | 0,7229 | 1,036E-12 |
| TcCLB.508307.200 | hypothetical protein, conserved | 0,7227 | 5,018E-09 |
| TcCLB.508153.60 | metallo-peptidase, Clan MA(E) Family M1 | 0,7226 | 3,404E-15 |
| TcCLB.507063.120 | hypothetical protein, conserved | 0,7224 | 2,235E-06 |
| TcCLB.507739.59 | hypothetical protein, conserved | 0,7218 | 0,00001381 |
| TcCLB.509127.97 | hypothetical protein, conserved | 0,7216 | 4,109E-08 |
| TcCLB.507519.260 | retrotransposon hot spot protein (RHS, pseudogene), puta | 0,7215 | 1,073E-13 |
| TcCLB.508153.454 | hypothetical protein, conserved | 0,7215 | 2,928E-09 |
| TcCLB.511727.140 | phenylalanyl-tRNA synthetase alpha chain, putative | 0,7213 | 8,754E-26 |
| TcCLB.506779.160 | palmitoyl acyltransferase 9, putative | 0,7213 | 4,985E-11 |
| TcCLB.506025.60 | 2-oxoglutarate dehydrogenase, E2 component, dihydrolipo | 0,721 | 3,233E-18 |
| TcCLB.511635.10 | histone H2B, putative | 0,7209 | 3,332E-17 |
| TcCLB.506975.70 | hypothetical protein | 0,7208 | 0,0002935 |
| TcCLB.511189.30 | 40S ribosomal protein S13, putative | 0,7206 | 4,042E-18 |
| TcCLB.503465.20 | hypothetical protein, conserved (pseudogene) | 0,7205 | 3,662E-06 |
| TcCLB.509871.80 | dispersed gene family protein 1 (DGF-1), putative | 0,7204 | 0,000015 |
| TcCLB.508879.40 | hypothetical protein, conserved | 0,7202 | 5,706E-12 |
| TcCLB.447483.10 | hypothetical protein, conserved | 0,7198 | 4,344E-11 |
| TcCLB.510873.10 | NAD(P)-dependent steroid dehydrogenase protein, putativ | 0,7194 | 4,118E-20 |
| TcCLB.510403.50 | retrotransposon hot spot protein (RHS, pseudogene), puta | 0,7194 | 1,93E-15 |

| | | | |
|---|---|---|---|
| TcCLB.506591.4 | heat shock protein 90, putative (LPG3) | 0,7189 | 1,241E-11 |
| TcCLB.509807.29 | hypothetical protein, conserved | 0,7188 | 1,134E-06 |
| TcCLB.506813.120 | retrotransposon hot spot protein (RHS, pseudogene), puta | 0,7188 | 0,0005332 |
| TcCLB.508647.264 | hypothetical protein, conserved | 0,7182 | 2,177E-15 |
| TcCLB.504003.70 | S-adenosyl-methyltransferase mraW-like protein, putative | 0,7181 | 1,017E-14 |
| TcCLB.507875.10 | hypothetical protein, conserved | 0,7181 | 9,352E-09 |
| TcCLB.504121.30 | 60S ribosomal protein L4, putative | 0,7179 | 9,968E-18 |
| TcCLB.510285.60 | ubiquinone biosynthesis methyltransferase, putative | 0,7177 | 2,353E-13 |
| TcCLB.506989.20 | hypothetical protein, conserved | 0,7177 | 2,92E-12 |
| TcCLB.510769.100 | hypothetical protein, conserved | 0,7177 | 3,342E-10 |
| TcCLB.508577.130 | hypothetical protein, conserved | 0,7176 | 3,688E-11 |
| TcCLB.509437.100 | retrotransposon hot spot protein (RHS, pseudogene), puta | 0,7175 | 4,973E-06 |
| TcCLB.506401.250 | hypothetical protein, conserved | 0,7173 | 3,064E-20 |
| TcCLB.506211.40 | intraflagellar transport protein 52, putative (IFT52) | 0,7173 | 9,732E-19 |
| TcCLB.504867.20 | ABC transporter, putative | 0,7173 | 1,354E-16 |
| TcCLB.510205.70 | surface protease GP63, putative | 0,7172 | 0,04457 |
| TcCLB.507083.50 | hypothetical protein, conserved | 0,7169 | 5,771E-07 |
| TcCLB.511263.40 | hypothetical protein, conserved | 0,7167 | 0,08686 |
| TcCLB.508153.770 | hypothetical protein, conserved | 0,7165 | 5,519E-13 |
| TcCLB.510741.190 | hypothetical protein, conserved (pseudogene) | 0,716 | 7,365E-10 |
| TcCLB.506683.100 | retrotransposon hot spot protein (RHS, pseudogene), puta | 0,7159 | 0,00006221 |
| TcCLB.506859.80 | hypothetical protein, conserved | 0,7158 | 5,816E-15 |
| TcCLB.510303.220 | hypothetical protein, conserved | 0,7156 | 1,02E-15 |
| TcCLB.503665.10 | trans-sialidase (pseudogene), putative | 0,7154 | 0,03229 |
| TcCLB.509105.10 | hypothetical protein, conserved | 0,7147 | 1,016E-10 |
| TcCLB.510749.20 | hypothetical protein, conserved | 0,7146 | 8,036E-10 |
| TcCLB.510663.19 | 60S ribosomal subunit protein L31, putative | 0,7144 | 3,961E-09 |
| TcCLB.509177.10 | hypothetical protein, conserved | 0,7139 | 1,622E-07 |
| TcCLB.511499.40 | hypothetical protein, conserved | 0,7139 | 6,245E-06 |
| TcCLB.510513.80 | bilobe protein, conserved | 0,7137 | 1,265E-12 |
| TcCLB.506635.50 | hypothetical protein | 0,7133 | 3,24E-14 |
| TcCLB.508919.189 | dispersed gene family protein 1 (DGF-1), putative | 0,7133 | 0,00000464 |
| TcCLB.507167.259 | trans-sialidase (pseudogene), putative | 0,7131 | 0,00325 |
| TcCLB.509979.51 | zinc carboxypeptidase, putative | 0,713 | 0,03456 |
| TcCLB.510609.34 | hypothetical protein, conserved | 0,7128 | 6,724E-08 |
| TcCLB.507167.31 | N-acetyltransferase complex ARD1 subunit (pseudogene), | 0,7126 | 0,07872 |
| TcCLB.508207.100 | 60S ribosomal protein L7, putative | 0,7124 | 7,026E-25 |
| TcCLB.507813.10 | structural maintenance of chromosome 3 protein, putative | 0,7117 | 9,017E-06 |
| TcCLB.508661.20 | ubiquitin/ribosomal protein S27a, putative | 0,7116 | 7,903E-18 |
| TcCLB.506295.200 | retrotransposon hot spot protein (RHS, pseudogene), puta | 0,7114 | 7,189E-09 |
| TcCLB.508179.60 | protein kinase, putative | 0,7113 | 1,037E-10 |
| TcCLB.506529.130 | WD repeat and HMG-box DNA-binding protein, putative | 0,7108 | 3,009E-15 |
| TcCLB.506041.20 | hypothetical protein, conserved | 0,7107 | 1,743E-10 |
| TcCLB.507603.110 | hypothetical protein, conserved | 0,7106 | 3,892E-09 |
| TcCLB.506351.80 | oligopeptidase B-like protein, putative | 0,7101 | 2,107E-10 |

| | | | |
|---|---|---|---|
| TcCLB.510187.460 | hypothetical protein | 0,7096 | 6,864E-08 |
| TcCLB.509139.20 | hypothetical protein, conserved | 0,7094 | 6,602E-11 |
| TcCLB.506355.50 | chaperone DNAJ protein, putative | 0,7093 | 7,726E-17 |
| TcCLB.504427.250 | ubiquitin-activating enzyme E1, putative | 0,7089 | 2,815E-24 |
| TcCLB.509803.60 | retrotransposon hot spot protein (RHS, pseudogene), puta | 0,7088 | 0,007665 |
| TcCLB.504049.20 | retrotransposon hot spot protein (RHS, pseudogene), puta | 0,7086 | 1,366E-13 |
| TcCLB.510903.50 | phosphonopyruvate decarboxylase, putative | 0,7086 | 1,298E-09 |
| TcCLB.507513.50 | heat shock 70 kDa protein, putative | 0,7083 | 1,022E-16 |
| TcCLB.510963.50 | 6-phosphofructo-2-kinase/fructose-2,6-bisphosphatas, pu | 0,7083 | 2,497E-11 |
| TcCLB.504605.10 | retrotransposon hot spot (RHS) protein, putative (fragmer | 0,7083 | 5,143E-07 |
| TcCLB.509167.50 | hypothetical protein | 0,7082 | 2,568E-06 |
| TcCLB.508881.50 | hypothetical protein, conserved | 0,7079 | 1,746E-08 |
| TcCLB.505809.10 | dispersed gene family protein 1 (DGF-1, pseudogene), pu | 0,7077 | 0,166 |
| TcCLB.508461.500 | 60S ribosomal protein L27A/L29, putative | 0,7076 | 0,00003258 |
| TcCLB.507681.10 | hypothetical protein, conserved | 0,7073 | 0,001837 |
| TcCLB.508221.610 | elongation factor 1-gamma (EF-1-gamma), putative | 0,7072 | 0,0876 |
| TcCLB.511019.90 | iron superoxide dismutase, putative | 0,7071 | 1,719E-13 |
| TcCLB.503665.15 | trans-sialidase (pseudogene), putative | 0,7071 | 7,725E-07 |
| TcCLB.511661.50 | small ubiquitin protein, putative | 0,7067 | 0,00000859 |
| TcCLB.506363.100 | hypothetical protein, conserved | 0,7065 | 2,793E-11 |
| TcCLB.509505.80 | hypothetical protein, conserved | 0,7065 | 2,084E-10 |
| TcCLB.508153.1140 | hypothetical protein, conserved | 0,7065 | 2,467E-08 |
| TcCLB.508781.20 | ATP synthase, putative | 0,7061 | 6,427E-14 |
| TcCLB.503879.110 | hypothetical protein, conserved | 0,706 | 1,136E-14 |
| TcCLB.506947.110 | myosin heavy chain (pseudogene), putative | 0,706 | 7,286E-12 |
| TcCLB.506635.70 | hypothetical protein, conserved | 0,7059 | 1,09E-13 |
| TcCLB.511499.59 | small nuclear ribonucleoprotein Sm-E, putative | 0,7058 | 2,452E-06 |
| TcCLB.509151.10 | hypothetical protein, conserved | 0,7057 | 4,023E-09 |
| TcCLB.506357.199 | hypothetical protein, conserved | 0,7055 | 1,577E-15 |
| TcCLB.503525.10 | hypothetical protein, conserved | 0,7055 | 7,295E-10 |
| TcCLB.504797.110 | hypothetical protein, conserved | 0,7054 | 2,182E-15 |
| TcCLB.506747.20 | dihydroorotase, putative | 0,7054 | 0,00000658 |
| TcCLB.509023.100 | catalytic subunit of the vacuolar transporter chaperone 4, | 0,7048 | 1,745E-10 |
| TcCLB.506559.59 | ubiquitin, putative | 0,7048 | 2,279E-08 |
| TcCLB.508737.187 | hypothetical protein, conserved | 0,7045 | 1,665E-09 |
| TcCLB.509875.260 | hypothetical protein, conserved | 0,7045 | 0,00001343 |
| TcCLB.506633.80 | hypothetical protein, conserved | 0,7044 | 4,971E-12 |
| TcCLB.504933.20 | hypothetical protein | 0,7042 | 5,488E-12 |
| TcCLB.460127.30 | amino acid permease, putative | 0,7042 | 0,0005601 |
| TcCLB.507991.130 | hypothetical protein, conserved | 0,704 | 8,186E-08 |
| TcCLB.509671.120 | rRNA processing protein, putative | 0,7037 | 1,152E-08 |
| TcCLB.506559.290 | hypothetical protein, conserved | 0,7036 | 1,127E-12 |
| TcCLB.506411.30 | proline oxidase, putative | 0,7035 | 6,137E-08 |
| TcCLB.510759.54 | hypothetical protein | 0,7035 | 0,00006147 |
| TcCLB.504105.130 | Flagellum surface protein 179 (FS179) | 0,7033 | 6,579E-12 |

| | | | |
|---|---|---|---|
| TcCLB.508265.30 | hypothetical protein | 0,7033 | 2,047E-10 |
| TcCLB.503487.70 | hypothetical protein, conserved | 0,7031 | 3,231E-15 |
| TcCLB.503967.10 | hypothetical protein, conserved | 0,7031 | 7,977E-08 |
| TcCLB.508859.80 | hypothetical protein, conserved | 0,7031 | 6,83E-07 |
| TcCLB.508409.199 | hypothetical protein, conserved | 0,7028 | 6,78E-07 |
| TcCLB.505843.20 | 60S ribosomal protein L6, putative | 0,702 | 1,379E-16 |
| TcCLB.510167.10 | phosphatidylinositol 3-kinase 2, putative | 0,7018 | 5,93E-14 |
| TcCLB.508439.40 | hypothetical protein | 0,7017 | 0,131 |
| TcCLB.509331.150 | permease-like protein, putative | 0,7016 | 0,0001585 |
| TcCLB.509693.180 | hypothetical protein, conserved | 0,7015 | 1,801E-06 |
| TcCLB.507627.20 | proton motive ATPase 1, putative | 0,7014 | 1,825E-16 |
| TcCLB.508421.30 | tryptophanyl-tRNA synthetase (pseudogene), putative | 0,7013 | 2,684E-21 |
| TcCLB.504141.9 | calcium-translocating P-type ATPase, putative | 0,7013 | 4,005E-06 |
| TcCLB.510855.40 | proteasome regulatory non-ATPase subunit, putative | 0,7012 | 8,139E-12 |
| TcCLB.510155.90 | hypothetical protein, conserved | 0,7011 | 2,185E-19 |
| TcCLB.508873.90 | trans-sialidase (pseudogene), putative | 0,7011 | 0,01462 |
| TcCLB.510721.10 | hypothetical protein, conserved | 0,701 | 8,006E-17 |
| TcCLB.510575.200 | chaperone DNAJ protein, putative | 0,7008 | 1,196E-15 |
| TcCLB.510859.50 | hypothetical protein, conserved | 0,7007 | 4,599E-08 |
| TcCLB.508325.340 | glycine dehydrogenase [decarboxylating] (pseudogene), p | 0,7006 | 8,486E-10 |
| TcCLB.508713.10 | hypothetical protein, conserved | 0,7004 | 1,186E-18 |
| TcCLB.510431.150 | glycosomal transporter (GAT2), putative (GAT2) | 0,7002 | 1,146E-22 |
| TcCLB.507143.20 | hypothetical protein, conserved | 0,6999 | 5,833E-09 |
| TcCLB.509161.140 | hypothetical protein, conserved | 0,6998 | 6,611E-10 |
| TcCLB.511361.4 | hypothetical protein, conserved | 0,6998 | 0,02706 |
| TcCLB.507709.120 | hypothetical protein, conserved | 0,6997 | 1,723E-20 |
| TcCLB.503683.20 | hypothetical protein, conserved | 0,6994 | 5,792E-13 |
| TcCLB.509611.189 | hypothetical protein, conserved | 0,6992 | 2,838E-14 |
| TcCLB.433677.19 | hypothetical protein, conserved | 0,6992 | 0,0009224 |
| TcCLB.506921.20 | hypothetical protein, conserved | 0,6991 | 4,868E-12 |
| TcCLB.506045.19 | dispersed gene family protein 1 (DGF-1), putative (fragm | 0,699 | 0,0001254 |
| TcCLB.507089.110 | hypothetical protein, conserved | 0,6989 | 2,427E-09 |
| TcCLB.511737.70 | hypothetical protein, conserved | 0,6985 | 2,771E-06 |
| TcCLB.506171.35 | ATP-dependent DEAD/H RNA helicase (pseudogene), puta | 0,6982 | 0,02497 |
| TcCLB.510105.200 | hypothetical protein, conserved | 0,6975 | 4,199E-21 |
| TcCLB.503733.70 | phosphoacetylglucosamine mutase, putative | 0,6971 | 4,568E-07 |
| TcCLB.504797.34 | N-acetyltransferase, putative | 0,697 | 4,892E-17 |
| TcCLB.503431.50 | hypothetical protein, conserved | 0,6959 | 0,00001449 |
| TcCLB.509879.20 | hypothetical protein, conserved | 0,6958 | 2,936E-07 |
| TcCLB.507275.24 | ubiquitin fold modifier protein, putative (UFM1) | 0,6958 | 6,189E-06 |
| TcCLB.506717.140 | retrotransposon hot spot protein (RHS, pseudogene), puta | 0,6958 | 0,008506 |
| TcCLB.510857.40 | RNA-editing complex protein, putative | 0,6957 | 1,995E-12 |
| TcCLB.507615.40 | hypothetical protein, conserved | 0,6954 | 4,872E-16 |
| TcCLB.507491.30 | hypothetical protein, conserved | 0,6954 | 2,905E-09 |
| TcCLB.510201.50 | dispersed gene family protein 1 (DGF-1), putative | 0,6952 | 0,000112 |

| | | | |
|---|---|---|---|
| TcCLB.504147.80 | metallo-peptidase, Clan MA(E) Family M3 | 0,6949 | 5,985E-22 |
| TcCLB.510101.190 | hypothetical protein, conserved | 0,6949 | 2,384E-09 |
| TcCLB.508707.280 | hypothetical protein, conserved | 0,6948 | 0,00000343 |
| TcCLB.510339.60 | hypothetical protein, conserved | 0,6947 | 2,156E-07 |
| TcCLB.508817.119 | hypothetical protein, conserved | 0,6947 | 2,511E-07 |
| TcCLB.511185.130 | dispersed gene family protein 1 (DGF-1), putative | 0,6947 | 0,03393 |
| TcCLB.509767.120 | hypothetical protein, conserved | 0,6946 | 2,232E-10 |
| TcCLB.510719.210 | hypothetical protein, conserved | 0,6946 | 4,349E-08 |
| TcCLB.510729.100 | hypothetical protein, conserved | 0,6945 | 2,749E-13 |
| TcCLB.453445.20 | ubiquitin-conjugating enzyme E2, putative | 0,6945 | 1,828E-09 |
| TcCLB.510719.20 | short-chain dehydrogenase, putative | 0,6945 | 5,236E-09 |
| TcCLB.510423.50 | hypothetical protein, conserved | 0,6944 | 4,366E-12 |
| TcCLB.508567.20 | hypothetical protein, conserved | 0,694 | 3,497E-06 |
| TcCLB.510241.60 | nascent polypeptide associated complex subunit, putative | 0,6935 | 5,888E-24 |
| TcCLB.507997.80 | hypothetical protein, conserved | 0,6935 | 0,0007682 |
| TcCLB.506441.20 | hypothetical protein, conserved | 0,6931 | 4,173E-11 |
| TcCLB.511825.210 | dephospho-CoA kinase, putative | 0,6929 | 5,821E-09 |
| TcCLB.503447.40 | retrotransposon hot spot (RHS) protein, putative | 0,6926 | 0,00008106 |
| TcCLB.507999.10 | dispersed gene family protein 1 (DGF-1), putative | 0,6925 | 1,564E-08 |
| TcCLB.511459.60 | hypothetical protein, conserved | 0,6925 | 0,00002735 |
| TcCLB.507011.10 | hypothetical protein, conserved | 0,6924 | 1,759E-10 |
| TcCLB.509741.20 | hypothetical protein, conserved | 0,6922 | 5,012E-07 |
| TcCLB.506195.170 | hypothetical protein, conserved | 0,6919 | 8,293E-13 |
| TcCLB.504421.10 | serine/threonine protein kinase, putative | 0,6917 | 4,891E-24 |
| TcCLB.507317.40 | hypothetical protein, conserved | 0,6917 | 1,104E-09 |
| TcCLB.504469.20 | hypothetical protein (fragment) | 0,6915 | 0,00154 |
| TcCLB.511293.60 | kinesin, putative | 0,6914 | 2,856E-12 |
| TcCLB.509235.20 | C-5 sterol desaturase, putative | 0,6914 | 1,733E-08 |
| TcCLB.508545.20 | hypothetical protein, conserved | 0,6914 | 0,0002896 |
| TcCLB.511573.10 | leucine-rich repeat protein, putative | 0,6913 | 0,00007209 |
| TcCLB.420533.9 | dihydrofolate reductase-thymidylate synthase, putative (fr | 0,6911 | 0,00003258 |
| TcCLB.508461.260 | Bem46-like serine peptidase, putative | 0,6908 | 5,64E-11 |
| TcCLB.506691.70 | hypothetical protein, conserved | 0,6908 | 6,173E-10 |
| TcCLB.511577.140 | hypothetical protein, conserved | 0,6906 | 4,839E-07 |
| TcCLB.511395.14 | hypothetical protein, conserved | 0,6906 | 0,00006839 |
| TcCLB.503491.29 | surface protease GP63 (pseudogene), putative | 0,6905 | 0,153 |
| TcCLB.507063.290 | hypothetical protein | 0,6904 | 2,78E-10 |
| TcCLB.508461.50 | hypothetical protein | 0,6903 | 2,133E-06 |
| TcCLB.509341.10 | hypothetical protein, conserved | 0,6901 | 5,768E-13 |
| TcCLB.508865.30 | immunodominant antigen, putative | 0,6898 | 3,635E-24 |
| TcCLB.507063.280 | hypothetical protein | 0,6898 | 4,123E-15 |
| TcCLB.508851.189 | phosphoenolpyruvate mutase, putative | 0,6896 | 4,683E-13 |
| TcCLB.510743.80 | hypothetical protein, conserved | 0,6895 | 2,243E-09 |
| TcCLB.506755.279 | hypothetical protein, conserved | 0,6895 | 3,632E-09 |
| TcCLB.507807.30 | hypothetical protein | 0,6891 | 0,0029 |

| | | | |
|---|---|---|---|
| TcCLB.509805.230 | hypothetical protein, conserved | 0,6886 | 0,00001074 |
| TcCLB.509537.40 | hypothetical protein, conserved | 0,6885 | 4,878E-12 |
| TcCLB.508461.410 | T-complex protein 1, eta subunit, putative | 0,6884 | 7,613E-27 |
| TcCLB.511815.10 | hypothetical protein | 0,6881 | 5,854E-16 |
| TcCLB.508053.5 | hypothetical protein, conserved (fragment) | 0,6881 | 0,0264 |
| TcCLB.511309.40 | hydrolase-like protein, putative | 0,6877 | 0,00004298 |
| TcCLB.508241.130 | hypothetical protein, conserved | 0,6875 | 3,299E-15 |
| TcCLB.509715.40 | fibrillarin, putative | 0,6875 | 1,492E-09 |
| TcCLB.508321.50 | hypothetical protein, conserved | 0,6871 | 2,837E-15 |
| TcCLB.506943.160 | eukaryotic translation initiation factor 3 subunit 7-like pro | 0,687 | 1,129E-21 |
| TcCLB.511287.30 | 60S ribosomal protein L17, putative | 0,687 | 1,909E-20 |
| TcCLB.506211.160 | Mitochondrial ADP/ATP carrier protein 5, putative (MCP5) | 0,6869 | 2,881E-08 |
| TcCLB.508267.50 | hypothetical protein, conserved | 0,6867 | 3,113E-10 |
| TcCLB.508851.180 | 14-3-3 protein 1, putative (14-3-3-I) | 0,6866 | 3,565E-18 |
| TcCLB.503655.20 | dispersed gene family protein 1 (DGF-1), putative | 0,6864 | 1,749E-06 |
| TcCLB.506577.150 | hypothetical protein, conserved | 0,6856 | 1,947E-18 |
| TcCLB.511907.310 | hypothetical protein, conserved | 0,6856 | 1,97E-15 |
| TcCLB.506257.80 | hypothetical protein, conserved | 0,6856 | 4,617E-08 |
| TcCLB.506315.10 | hypothetical protein | 0,6855 | 2,192E-06 |
| TcCLB.507951.170 | ADP-ribosylation factor-like protein 3, putative (arl3) | 0,6853 | 1,142E-08 |
| TcCLB.510065.40 | hypothetical protein, conserved | 0,6852 | 9,264E-18 |
| TcCLB.509777.70 | Calcium ATPase SERCA-like | 0,6851 | 4,167E-22 |
| TcCLB.506513.160 | protein kinase, putative | 0,6849 | 2,574E-11 |
| TcCLB.510349.60 | hypothetical protein, conserved | 0,6849 | 0,0001862 |
| TcCLB.506605.200 | hypothetical protein, conserved | 0,6847 | 8,996E-08 |
| TcCLB.511145.43 | proteasome subunit alpha type-5, putative | 0,6846 | 5,334E-08 |
| TcCLB.509727.20 | retrotransposon hot spot protein (RHS, pseudogene), puta | 0,6846 | 0,08207 |
| TcCLB.510643.130 | damage-specific DNA binding protein, putative | 0,6839 | 1,122E-16 |
| TcCLB.511071.190 | hypothetical protein, conserved | 0,6836 | 7,567E-11 |
| TcCLB.506839.80 | phosphoribosylpyrophosphate synthetase, putative | 0,6835 | 4,263E-18 |
| TcCLB.506941.220 | Present in the outer mitochondrial membrane proteome 3 | 0,6835 | 2,061E-06 |
| TcCLB.511771.75 | hypothetical protein | 0,6832 | 0,0007598 |
| TcCLB.508813.60 | Present in the outer mitochondrial membrane proteome 1 | 0,6831 | 5,816E-18 |
| TcCLB.510759.10 | hypothetical protein, conserved | 0,6831 | 3,404E-10 |
| TcCLB.508831.250 | dispersed gene family protein 1 (DGF-1), putative | 0,683 | 0,0003226 |
| TcCLB.508479.250 | membrane-bound acid phosphatase 2, putative | 0,6828 | 5,496E-14 |
| TcCLB.418185.43 | ATP-dependent DEAD/H RNA helicase (pseudogene), puta | 0,6827 | 0,001221 |
| TcCLB.508143.50 | trans-sialidase (pseudogene), putative | 0,6825 | 8,559E-07 |
| TcCLB.506959.94 | hypothetical protein, conserved | 0,682 | 9,543E-08 |
| TcCLB.511539.9 | hypothetical protein, conserved | 0,6819 | 2,456E-10 |
| TcCLB.511021.70 | ABC transporter of the mitochondrion, putative | 0,6819 | 1,373E-08 |
| TcCLB.511311.14 | retrotransposon hot spot protein (RHS, pseudogene), puta | 0,6819 | 2,769E-08 |
| TcCLB.508207.110 | 60S ribosomal protein L7, putative | 0,6818 | 5,06E-24 |
| TcCLB.506511.40 | hypothetical protein, conserved | 0,6818 | 2,76E-08 |
| TcCLB.510355.220 | GDP-mannose transporter, putative | 0,6818 | 0,0001291 |

| | | | |
|---|---|---|---|
| TcCLB.511421.50 | transmembrane protein, putative | 0,6817 | 1,267E-12 |
| TcCLB.508989.50 | hypothetical protein, conserved | 0,6816 | 1,49E-10 |
| TcCLB.509887.10 | hypothetical protein | 0,6815 | 0,00006738 |
| TcCLB.509795.60 | tRNA pseudouridine synthase TruD, putative | 0,6811 | 1,056E-10 |
| TcCLB.506407.60 | hypothetical protein, conserved | 0,6811 | 2,454E-06 |
| TcCLB.509831.40 | hypothetical protein, conserved | 0,681 | 5,537E-07 |
| TcCLB.510741.80 | riboflavin kinase, putative | 0,6809 | 0,00002019 |
| TcCLB.507485.110 | hypothetical protein, conserved | 0,6807 | 5,985E-13 |
| TcCLB.506247.290 | hypothetical protein, conserved | 0,6806 | 1,183E-16 |
| TcCLB.511825.170 | serine/threonine protein phosphatase, putative | 0,6805 | 4,24E-08 |
| TcCLB.505937.80 | retrotransposon hot spot protein (RHS, pseudogene), puta | 0,6803 | 0,000144 |
| TcCLB.504407.10 | dispersed gene family protein 1 (DGF-1, pseudogene), put | 0,6803 | 0,01839 |
| TcCLB.506679.270 | hypothetical protein, conserved | 0,6802 | 8,337E-13 |
| TcCLB.508625.130 | stress-inducible protein STI1-like, putative | 0,68 | 0,00008975 |
| TcCLB.504103.20 | hypothetical protein, conserved | 0,6799 | 2,024E-16 |
| TcCLB.511731.74 | hypothetical protein, conserved | 0,6798 | 2,836E-07 |
| TcCLB.509065.120 | hypothetical protein | 0,6798 | 6,909E-06 |
| TcCLB.511743.10 | hypothetical protein, conserved | 0,6797 | 1,969E-06 |
| TcCLB.511751.191 | hypothetical protein (pseudogene) | 0,6794 | 3,855E-06 |
| TcCLB.510579.70 | nascent polypeptide associated complex subunit, putative | 0,6788 | 1,497E-17 |
| TcCLB.510181.100 | acid phosphatase, putative (AP) | 0,6788 | 0,0002056 |
| TcCLB.510355.260 | hypothetical protein, conserved | 0,6785 | 0,0009949 |
| TcCLB.507049.50 | ABC transporter, putative | 0,6784 | 9,229E-08 |
| TcCLB.511545.160 | hypothetical protein, conserved | 0,6783 | 2,214E-07 |
| TcCLB.504003.50 | hypothetical protein, conserved | 0,6783 | 0,00001689 |
| TcCLB.509915.20 | dispersed gene family protein 1 (DGF-1), putative | 0,678 | 0,0003071 |
| TcCLB.511821.40 | hypothetical protein, conserved | 0,6774 | 2,875E-06 |
| TcCLB.506337.150 | hypothetical protein, conserved | 0,6773 | 1,694E-07 |
| TcCLB.506659.20 | mitochondrial DNA polymerase I protein C, putative | 0,677 | 2,343E-19 |
| TcCLB.508827.90 | hypothetical protein, conserved | 0,6769 | 2,727E-07 |
| TcCLB.509395.20 | phosphatidylinositol kinase related protein, putative | 0,6767 | 1,245E-14 |
| TcCLB.508559.29 | dispersed gene family protein 1 (DGF-1), putative | 0,6767 | 0,00001005 |
| TcCLB.507881.10 | retrotransposon hot spot protein (RHS, pseudogene), puta | 0,6766 | 6,123E-07 |
| TcCLB.506401.160 | hypothetical protein, conserved | 0,6766 | 0,00003502 |
| TcCLB.504131.100 | DnaJ chaperone protein, putative | 0,6763 | 5,126E-14 |
| TcCLB.510721.21 | hypothetical protein | 0,6756 | 0,06117 |
| TcCLB.510519.14 | hypothetical protein, conserved | 0,6754 | 0,00003395 |
| TcCLB.507663.60 | hypothetical protein, conserved | 0,6753 | 1,814E-08 |
| TcCLB.510099.60 | hypothetical protein | 0,6752 | 8,621E-12 |
| TcCLB.506635.90 | ubiquitin-protein ligase, putative | 0,675 | 2,125E-06 |
| TcCLB.503945.20 | hypothetical protein, conserved | 0,675 | 0,0006376 |
| TcCLB.504013.40 | serine acetyltransferase, putative (SAT) | 0,6748 | 1,898E-09 |
| TcCLB.510257.130 | protein kinase, putative | 0,6745 | 7,102E-14 |
| TcCLB.509915.60 | retrotransposon hot spot (RHS) protein, putative | 0,6742 | 4,084E-06 |
| TcCLB.507053.20 | hypothetical protein, conserved | 0,674 | 1,361E-10 |

| | | | |
|---|---|---|---|
| TcCLB.508851.150 | hypothetical protein, conserved | 0,674 | 1,114E-07 |
| TcCLB.506773.20 | hypothetical protein, conserved | 0,6739 | 1,053E-12 |
| TcCLB.507049.90 | hypothetical protein, conserved | 0,6737 | 1,14E-07 |
| TcCLB.506297.330 | 40S ribosomal protein S15A, putative | 0,6734 | 2,755E-19 |
| TcCLB.449247.10 | hypothetical protein, conserved | 0,6734 | 0,00000374 |
| TcCLB.507179.7 | trans-sialidase (pseudogene), putative | 0,6732 | 3,163E-06 |
| TcCLB.504111.20 | metallo- peptidase, Clan MG, Family M24 | 0,673 | 7,484E-11 |
| TcCLB.507233.70 | trans-sialidase (pseudogene), putative | 0,673 | 2,027E-10 |
| TcCLB.503851.54 | proteasome 26S non-ATPase subunit 9, putative | 0,673 | 0,01606 |
| TcCLB.511181.20 | hypothetical protein, conserved | 0,6729 | 1,282E-08 |
| TcCLB.508119.150 | hypothetical protein, conserved | 0,6729 | 1,921E-06 |
| TcCLB.503439.80 | glycine dehydrogenase [decarboxylating] (pseudogene), pu | 0,6729 | 0,00005517 |
| TcCLB.511125.9 | dispersed gene family protein 1 (DGF-1), putative | 0,6724 | 0,005712 |
| TcCLB.507009.60 | hypothetical protein, conserved | 0,6723 | 1,235E-13 |
| TcCLB.504103.90 | hypothetical protein, conserved | 0,6723 | 6,639E-07 |
| TcCLB.508987.20 | hypothetical protein, conserved | 0,6722 | 1,961E-12 |
| TcCLB.506147.120 | hypothetical protein, conserved | 0,672 | 1,166E-07 |
| TcCLB.506389.60 | hypothetical protein, conserved | 0,6718 | 1,366E-08 |
| TcCLB.510691.95 | hypothetical protein, conserved (fragment) | 0,6718 | 1,727E-06 |
| TcCLB.503635.68 | 40S ribosomal protein S3A, putative | 0,6717 | 4,293E-15 |
| TcCLB.506737.10 | hypothetical protein, conserved | 0,6717 | 1,563E-11 |
| TcCLB.504433.10 | ADP-ribosylation factor-like protein 3, putative (arl3) | 0,6716 | 9,741E-08 |
| TcCLB.503689.4 | Ras-related protein RabX1, putative (RABX1) | 0,6715 | 7,325E-07 |
| TcCLB.511071.140 | 2-amino-3-ketobutyrate coenzyme A ligase, putative | 0,6711 | 1,022E-06 |
| TcCLB.510121.70 | hypothetical protein, conserved | 0,671 | 5,101E-11 |
| TcCLB.505789.10 | hypothetical protein, conserved | 0,6706 | 3,286E-06 |
| TcCLB.506587.70 | hypothetical protein, conserved | 0,6704 | 2,951E-11 |
| TcCLB.503973.40 | dispersed gene family protein 1 (DGF-1), putative | 0,6703 | 0,008409 |
| TcCLB.508741.330 | hypothetical protein, conserved | 0,6702 | 4,604E-09 |
| TcCLB.503905.30 | hypothetical protein, conserved | 0,6699 | 1,799E-08 |
| TcCLB.510609.30 | GTPase, putative | 0,6696 | 1,167E-08 |
| TcCLB.511895.69 | C-5 sterol desaturase, putative | 0,6694 | 0,0002724 |
| TcCLB.505647.30 | hypothetical protein, conserved | 0,6693 | 1,044E-10 |
| TcCLB.511767.20 | dispersed gene family protein 1 (DGF-1), putative | 0,669 | 0,00005712 |
| TcCLB.509717.50 | hypothetical protein, conserved | 0,6686 | 9,618E-18 |
| TcCLB.506015.30 | mucin-associated surface protein (MASP, pseudogene), pu | 0,6686 | 0,00001634 |
| TcCLB.511577.188 | hypothetical protein, conserved | 0,6685 | 1,222E-06 |
| TcCLB.509961.51 | tryptophanyl-tRNA synthetase (pseudogene), putative | 0,6685 | 0,004818 |
| TcCLB.505123.20 | hypothetical protein, conserved | 0,6684 | 0,01704 |
| TcCLB.510029.70 | 40S ribosomal protein S13, putative | 0,6682 | 1,072E-14 |
| TcCLB.510671.20 | tryptophanyl-tRNA synthetase, putative | 0,6681 | 4,487E-20 |
| TcCLB.509867.50 | hypothetical protein, conserved (pseudogene) | 0,6681 | 0,01941 |
| TcCLB.510359.210 | hypothetical protein, conserved | 0,6679 | 1,661E-09 |
| TcCLB.507615.73 | hypothetical protein, conserved | 0,6677 | 4,589E-11 |
| TcCLB.511039.10 | kinetoplast DNA-associated protein, putative | 0,6677 | 7,497E-10 |

| | | | |
|---|---|---|---|
| TcCLB.508959.4 | cytochrome c, putative | 0,6677 | 1,477E-08 |
| TcCLB.509167.20 | 3-hydroxy-3-methylglutaryl-CoA reductase, putative | 0,6671 | 1,492E-13 |
| TcCLB.511731.35 | hypothetical protein, conserved | 0,6671 | 1,141E-06 |
| TcCLB.504425.90 | aldose 1-epimerase-like protein, putative | 0,667 | 0,0001144 |
| TcCLB.510073.40 | hypothetical protein, conserved | 0,667 | 0,001623 |
| TcCLB.508589.60 | hypothetical protein, conserved (pseudogene) | 0,6666 | 0,00002775 |
| TcCLB.511649.94 | hypothetical protein, conserved | 0,6664 | 3,743E-08 |
| TcCLB.510519.60 | calcineurin B subunit, putative (CNB) | 0,6663 | 7,128E-07 |
| TcCLB.506753.240 | hypothetical protein, conserved | 0,6663 | 0,00004369 |
| TcCLB.511585.90 | hypothetical protein, conserved | 0,6662 | 3,827E-12 |
| TcCLB.509171.90 | splicing factor 3a, putative | 0,6661 | 2,545E-10 |
| TcCLB.504149.110 | GTP-binding protein, putative | 0,6661 | 2,893E-10 |
| TcCLB.507017.64 | hypothetical protein, conserved | 0,6659 | 0,00002459 |
| TcCLB.508723.110 | hypothetical protein | 0,6658 | 4,652E-12 |
| TcCLB.509429.329 | cruzipain precursor, putative (fragment) | 0,6657 | 8,038E-25 |
| TcCLB.510973.30 | 40S ribosomal protein S13, putative | 0,6657 | 1,845E-22 |
| TcCLB.431889.10 | hypothetical protein, conserved (fragment) | 0,6655 | 0,05153 |
| TcCLB.508649.14 | hypothetical protein, conserved | 0,6653 | 1,97E-10 |
| TcCLB.506211.130 | hypothetical protein, conserved | 0,6653 | 6,037E-09 |
| TcCLB.511289.100 | hypothetical protein, conserved | 0,6652 | 1,576E-13 |
| TcCLB.505171.44 | hypothetical protein, conserved | 0,6644 | 0,0004354 |
| TcCLB.511649.14 | hypothetical protein, conserved (fragment) | 0,6641 | 5,243E-06 |
| TcCLB.507883.40 | hypothetical protein, conserved | 0,6632 | 1,175E-06 |
| TcCLB.509321.19 | GTP-binding protein, putative | 0,6631 | 1,001E-07 |
| TcCLB.506163.70 | hypothetical protein, conserved | 0,663 | 9,489E-09 |
| TcCLB.503965.30 | hypothetical protein, conserved | 0,663 | 1,263E-06 |
| TcCLB.508813.80 | hypothetical protein, conserved | 0,6625 | 9,635E-11 |
| TcCLB.506129.80 | retrotransposon hot spot protein (RHS, pseudogene), puta | 0,6622 | 6,851E-07 |
| TcCLB.509471.86 | histone H3, putative | 0,6621 | 2,14E-17 |
| TcCLB.509625.30 | hypothetical protein, conserved | 0,662 | 0,0000043 |
| TcCLB.509611.60 | hypothetical protein | 0,662 | 0,0002381 |
| TcCLB.511303.110 | hypothetical protein, conserved | 0,6619 | 6,667E-09 |
| TcCLB.507245.10 | retrotransposon hot spot protein (RHS, pseudogene), puta | 0,6617 | 8,651E-08 |
| TcCLB.511309.30 | hydrolase, alpha/beta fold family, putative | 0,6617 | 8,534E-07 |
| TcCLB.508145.40 | hypothetical protein, conserved | 0,6617 | 0,00003339 |
| TcCLB.508593.110 | arginine N-methyltransferase, putative | 0,6613 | 1,864E-08 |
| TcCLB.509589.40 | hypothetical protein, conserved | 0,6611 | 9,676E-18 |
| TcCLB.506621.14 | ubiquitin ligase, putative | 0,6608 | 2,657E-07 |
| TcCLB.511865.50 | hypothetical protein, conserved | 0,6605 | 0,00002013 |
| TcCLB.511613.90 | mucin-associated surface protein (MASP), putative | 0,6604 | 0,01086 |
| TcCLB.507021.70 | hypothetical protein, conserved | 0,6603 | 3,348E-09 |
| TcCLB.510357.50 | hypothetical protein, conserved | 0,6603 | 7,898E-07 |
| TcCLB.503729.19 | hypothetical protein (fragment) | 0,6602 | 4,215E-07 |
| TcCLB.503987.39 | hypothetical protein, conserved | 0,6601 | 0,00006878 |
| TcCLB.508779.80 | hypothetical protein, conserved | 0,66 | 3,574E-10 |

| | | | |
|---|---|---|---|
| TcCLB.507007.77 | hypothetical protein, conserved | 0,66 | 0,002647 |
| TcCLB.507531.10 | hypothetical protein, conserved (pseudogene) | 0,6598 | 8,939E-17 |
| TcCLB.506363.10 | hypothetical protein | 0,6598 | 0,0002151 |
| TcCLB.508967.20 | dispersed gene family protein 1 (DGF-1), putative | 0,6597 | 0,0182 |
| TcCLB.506529.100 | hypothetical protein, conserved | 0,6595 | 6,002E-11 |
| TcCLB.468941.9 | ATP-dependent DEAD/H RNA helicase, putative (fragment | 0,6595 | 8,357E-11 |
| TcCLB.506363.110 | RNA editing ligase, putative | 0,6594 | 1,322E-09 |
| TcCLB.508373.20 | dihydroorotase, putative | 0,6593 | 3,629E-09 |
| TcCLB.504827.120 | hypothetical protein, conserved | 0,6591 | 7,686E-08 |
| TcCLB.511011.30 | hypothetical protein, conserved (pseudogene) | 0,6591 | 0,0001071 |
| TcCLB.509785.120 | hypothetical protein, conserved (pseudogene) | 0,6582 | 0,04761 |
| TcCLB.509769.140 | hypothetical protein, conserved | 0,6581 | 3,254E-07 |
| TcCLB.511561.11 | hypothetical protein, conserved (pseudogene) | 0,6581 | 0,00001081 |
| TcCLB.506435.260 | hypothetical protein | 0,658 | 2,678E-08 |
| TcCLB.510719.100 | hypothetical protein, conserved | 0,6578 | 0,000204 |
| TcCLB.511003.60 | 3-ketoacyl-CoA thiolase, putative (ACAT) | 0,6576 | 1,123E-09 |
| TcCLB.508611.30 | surface protease GP63, putative | 0,6576 | 3,38E-07 |
| TcCLB.506417.70 | hypothetical protein | 0,6576 | 3,273E-06 |
| TcCLB.511819.20 | hypothetical protein, conserved | 0,6573 | 5,116E-13 |
| TcCLB.503843.10 | hypothetical protein, conserved | 0,6571 | 1,432E-08 |
| TcCLB.436083.9 | hypothetical protein, conserved | 0,6569 | 3,714E-07 |
| TcCLB.506401.200 | hypothetical protein, conserved | 0,6568 | 1,141E-06 |
| TcCLB.508173.264 | hypothetical protein, conserved | 0,6567 | 6,683E-16 |
| TcCLB.506779.50 | proteasome beta 3 subunit, putative | 0,6566 | 1,629E-11 |
| TcCLB.511395.20 | phosphatidylglycerolphosphate synthase, mitochondrial, p | 0,6565 | 2,72E-09 |
| TcCLB.508539.20 | hypothetical protein, conserved | 0,6561 | 0,0003124 |
| TcCLB.509693.60 | hypothetical protein, conserved | 0,656 | 0,00001824 |
| TcCLB.429257.20 | cytochrome b5-dependent oleate desaturase | 0,6559 | 3,408E-11 |
| TcCLB.508705.10 | retrotransposon hot spot protein (RHS, pseudogene), puta | 0,6558 | 1,853E-08 |
| TcCLB.506271.20 | N-acetyltransferase complex ARD1 subunit (pseudogene), | 0,6558 | 0,1091 |
| TcCLB.507817.18 | histone H3, putative | 0,6557 | 8,675E-06 |
| TcCLB.504233.4 | trifunctional enzyme alpha subunit, mitochondrial precurs | 0,6555 | 0,0505 |
| TcCLB.506249.70 | ABC transporter, putative | 0,6553 | 6,416E-11 |
| TcCLB.511421.110 | developmentally regulated phosphoprotein, putative | 0,6552 | 4,105E-11 |
| TcCLB.510659.180 | hypothetical protein, conserved | 0,6547 | 2,243E-09 |
| TcCLB.510543.10 | retrotransposon hot spot protein (RHS, pseudogene), puta | 0,6546 | 3,206E-06 |
| TcCLB.506359.60 | UTP-glucose-1-phosphate uridylyltransferase 2, putative | 0,6544 | 0,00001768 |
| TcCLB.506481.20 | hypothetical protein, conserved | 0,6543 | 1,52E-21 |
| TcCLB.510755.80 | hypothetical protein, conserved | 0,6541 | 1,241E-07 |
| TcCLB.507615.100 | Present in the outer mitochondrial membrane proteome 2 | 0,6539 | 3,974E-14 |
| TcCLB.510819.4 | NADH-cytochrome b5 reductase, putative | 0,6539 | 9,329E-07 |
| TcCLB.509207.140 | hypothetical protein, conserved | 0,6537 | 1,679E-11 |
| TcCLB.510685.10 | calreticulin, putative | 0,6536 | 3,367E-17 |
| TcCLB.511827.10 | hypothetical protein, conserved | 0,6536 | 0,005418 |
| TcCLB.509965.140 | hypothetical protein, conserved | 0,6535 | 5,901E-08 |

| | | | |
|---|---|---|---|
| TcCLB.510313.20 | hypothetical protein, conserved | 0,6534 | 6,273E-12 |
| TcCLB.509149.80 | hypothetical protein, conserved | 0,6533 | 2,123E-09 |
| TcCLB.511179.110 | hypothetical protein, conserved | 0,6532 | 1,426E-10 |
| TcCLB.510311.140 | arginine N-methyltransferase, putative | 0,6531 | 4,219E-11 |
| TcCLB.511277.260 | hypothetical protein, conserved | 0,6531 | 3,76E-07 |
| TcCLB.507053.50 | hypothetical protein, conserved | 0,6528 | 1,232E-09 |
| TcCLB.511001.9 | 40S ribosomal protein S3A, putative | 0,6528 | 1,015E-06 |
| TcCLB.510105.90 | hypothetical protein, conserved | 0,6524 | 2,188E-09 |
| TcCLB.508139.140 | dispersed gene family protein 1 (DGF-1), putative | 0,6523 | 0,00001016 |
| TcCLB.509831.30 | hypothetical protein, conserved | 0,6522 | 1,017E-10 |
| TcCLB.505317.20 | hypothetical protein | 0,6521 | 3,813E-07 |
| TcCLB.510359.80 | hypothetical protein | 0,6521 | 0,0007195 |
| TcCLB.508479.180 | hypothetical protein, conserved | 0,6515 | 5,063E-15 |
| TcCLB.504141.20 | hypothetical protein, conserved | 0,651 | 6,264E-09 |
| TcCLB.509053.160 | hypothetical protein, conserved | 0,651 | 1,491E-07 |
| TcCLB.504159.40 | aquaporin-like protein, putative | 0,6509 | 7,586E-08 |
| TcCLB.509719.28 | hypothetical protein, conserved | 0,6508 | 0,001778 |
| TcCLB.506595.60 | beta galactofuranosyl glycosyltransferase (pseudogene), p | 0,6505 | 4,031E-07 |
| TcCLB.509053.179 | RNA-binding protein, putative (fragment) | 0,6504 | 1,273E-06 |
| TcCLB.503431.10 | caltractin, putative | 0,6504 | 0,003436 |
| TcCLB.507835.60 | retrotransposon hot spot protein (RHS, pseudogene), puta | 0,6501 | 7,809E-06 |
| TcCLB.508865.20 | mitochondrial DNA polymerase I protein C, putative | 0,65 | 8,535E-18 |
| TcCLB.511729.40 | 60S ribosomal protein L9, putative | 0,65 | 1,857E-06 |
| TcCLB.510003.20 | hypothetical protein, conserved | 0,6499 | 7,421E-18 |
| TcCLB.510089.110 | mitochondrial RNA binding complex 1 subunit, putative | 0,6499 | 7,666E-08 |
| TcCLB.421057.10 | adrenodoxin precursor, putative | 0,6499 | 0,0001548 |
| TcCLB.507517.40 | hypothetical protein, conserved | 0,6498 | 2,34E-13 |
| TcCLB.508647.290 | DNA repair and recombination protein RAD54, putative | 0,6497 | 3,973E-15 |
| TcCLB.508355.150 | proteasome regulatory non-ATPase subunit 3, putative | 0,6495 | 4,647E-19 |
| TcCLB.510657.150 | hypothetical protein | 0,6495 | 1,722E-08 |
| TcCLB.511305.10 | 5'-3' exoribonuclease B, putative | 0,6494 | 5,301E-08 |
| TcCLB.509815.70 | retrotransposon hot spot protein (RHS, pseudogene), puta | 0,6493 | 4,33E-07 |
| TcCLB.508241.104 | hypothetical protein, conserved | 0,6492 | 9,355E-14 |
| TcCLB.510003.9 | hypothetical protein, conserved | 0,6491 | 1,622E-08 |
| TcCLB.508055.20 | retrotransposon hot spot protein (RHS, pseudogene), puta | 0,6489 | 0,0002458 |
| TcCLB.507611.199 | dispersed gene family protein 1 (DGF-1), putative | 0,6489 | 0,002239 |
| TcCLB.506657.70 | RNA editing 3' terminal uridylyl transferase 1, putative | 0,6486 | 2,764E-18 |
| TcCLB.511759.20 | trans-sialidase (pseudogene), putative | 0,6486 | 1,655E-07 |
| TcCLB.510323.4 | katanin, putative | 0,6485 | 9,107E-10 |
| TcCLB.507611.370 | protein disulfide isomerase 2 (PDI2) | 0,6483 | 3,414E-23 |
| TcCLB.509571.79 | Ras-related protein RabX1, putative (RABX1) | 0,6483 | 1,917E-07 |
| TcCLB.506975.80 | trans-sialidase, Group I, putative | 0,6483 | 0,0000331 |
| TcCLB.511367.60 | la RNA binding protein, putative | 0,6482 | 1,197E-13 |
| TcCLB.509109.140 | hypothetical protein, conserved | 0,6482 | 2,445E-08 |
| TcCLB.509715.150 | hypothetical protein | 0,6482 | 0,0001489 |

| | | | |
|---|---|---|---|
| TcCLB.508299.70 | peroxisome targeting signal 1 receptor, putative (PEX5) | 0,6481 | 1,347E-10 |
| TcCLB.504037.10 | hypothetical protein, conserved | 0,6481 | 1,688E-06 |
| TcCLB.506559.170 | 8-oxoguanine DNA glycosylase, putative | 0,6478 | 2,822E-12 |
| TcCLB.510737.30 | mitochondrial carrier protein, putative | 0,6478 | 3,289E-07 |
| TcCLB.508637.120 | hypothetical protein, conserved | 0,6477 | 8,06E-08 |
| TcCLB.510463.190 | retrotransposon hot spot (RHS) protein, putative | 0,6476 | 0,0001573 |
| TcCLB.507831.10 | hypothetical protein | 0,6473 | 0,002245 |
| TcCLB.427355.10 | oligosaccharyl transferase subunit, putative (fragment) | 0,6471 | 6,073E-07 |
| TcCLB.511275.10 | hypothetical protein, conserved (pseudogene) | 0,647 | 1,114E-15 |
| TcCLB.511643.10 | hypothetical protein, conserved | 0,647 | 3,62E-07 |
| TcCLB.509611.50 | hypothetical protein, conserved | 0,647 | 0,00000305 |
| TcCLB.506739.200 | protein phosphatase 2C, putative | 0,6469 | 3,144E-17 |
| TcCLB.508035.10 | dispersed gene family protein 1 (DGF-1, pseudogene), put | 0,6468 | 0,05914 |
| TcCLB.508577.100 | pumilio/PUF RNA binding protein 5, putative (PUF5) | 0,6466 | 1,341E-06 |
| TcCLB.408799.19 | isopentenyl-diphosphate delta-isomerase (fragment) (idi1 | 0,6465 | 0,00004702 |
| TcCLB.507063.70 | trans-sialidase, Group II, putative | 0,6464 | 1,541E-07 |
| TcCLB.503557.30 | hypothetical protein, conserved | 0,6462 | 6,469E-07 |
| TcCLB.510293.60 | hypothetical protein, conserved | 0,6459 | 9,549E-06 |
| TcCLB.504445.20 | retrotransposon hot spot protein (RHS, pseudogene), puta | 0,6458 | 3,671E-06 |
| TcCLB.507547.30 | Mitochondrial-processing peptidase subunit alpha (MPPA) | 0,6455 | 1,609E-13 |
| TcCLB.503605.10 | retrotransposon hot spot protein (RHS, pseudogene), puta | 0,6455 | 3,352E-12 |
| TcCLB.505121.9 | dispersed gene family protein 1 (DGF-1), putative (fragme | 0,6454 | 0,0004048 |
| TcCLB.511237.110 | cyclophilin, putative | 0,6453 | 2,839E-08 |
| TcCLB.507787.170 | retrotransposon hot spot protein (RHS, pseudogene), puta | 0,6452 | 1,331E-12 |
| TcCLB.511135.10 | hypothetical protein, conserved (pseudogene) | 0,645 | 1,83E-07 |
| TcCLB.510781.10 | dispersed gene family protein 1 (DGF-1, pseudogene), put | 0,645 | 0,0006914 |
| TcCLB.506489.30 | hypothetical protein, conserved | 0,6449 | 4,807E-11 |
| TcCLB.504133.15 | hypothetical protein, conserved (fragment) | 0,6449 | 0,001241 |
| TcCLB.508569.100 | hypothetical protein, conserved | 0,6448 | 1,307E-06 |
| TcCLB.505807.10 | cyclophilin, putative | 0,6447 | 8,424E-06 |
| TcCLB.506689.10 | trans-sialidase (pseudogene), putative | 0,6443 | 0,01165 |
| TcCLB.506853.40 | hypothetical protein, conserved | 0,6442 | 8,402E-07 |
| TcCLB.511711.40 | hypothetical protein, conserved | 0,6442 | 6,307E-06 |
| TcCLB.507625.183 | SNARE domain-containing protein, putative (SNARE2.1) | 0,6438 | 0,00001629 |
| TcCLB.506425.130 | hypothetical protein, conserved | 0,6437 | 1,031E-07 |
| TcCLB.509147.50 | mucin TcSMUGL, putative | 0,6436 | 4,82E-08 |
| TcCLB.503707.9 | dispersed gene family protein 1 (DGF-1, pseudogene), put | 0,6436 | 0,0003421 |
| TcCLB.510995.9 | dispersed gene family protein 1 (DGF-1), putative | 0,6436 | 0,007777 |
| TcCLB.508999.220 | calpain cysteine peptidase, putative | 0,6432 | 0,00000607 |
| TcCLB.511825.60 | aquaporin, putative | 0,6431 | 8,068E-10 |
| TcCLB.509213.50 | hypothetical protein, conserved | 0,6431 | 3,507E-06 |
| TcCLB.507921.9 | dispersed gene family protein 1 (DGF-1), putative | 0,643 | 0,0004992 |
| TcCLB.401469.20 | pumilio/PUF RNA binding protein 7, putative (PUF7) | 0,6429 | 2,955E-06 |
| TcCLB.508307.134 | hypothetical protein, conserved | 0,6429 | 0,00001222 |
| TcCLB.510345.40 | hypothetical protein, conserved | 0,6425 | 1,633E-07 |

| | | | |
|---|---|---|---|
| TcCLB.506193.10 | hypothetical protein, conserved | 0,6422 | 9,476E-13 |
| TcCLB.509109.160 | hypothetical protein, conserved | 0,6422 | 0,0005862 |
| TcCLB.507203.9 | dispersed gene family protein 1 (DGF-1), putative | 0,6421 | 0,02554 |
| TcCLB.508285.30 | ATP-dependent DEAD/H RNA helicase (pseudogene), puta | 0,6419 | 0,1428 |
| TcCLB.510181.9 | inositol-3-phosphate synthase, putative (fragment) | 0,6418 | 1,159E-06 |
| TcCLB.504431.50 | hypothetical protein, conserved | 0,6417 | 1,372E-10 |
| TcCLB.410923.40 | trans-sialidase (pseudogene), putative | 0,6415 | 7,666E-07 |
| TcCLB.509777.140 | hypothetical protein, conserved | 0,6415 | 1,377E-06 |
| TcCLB.509965.180 | RNA-binding protein, putative | 0,6412 | 1,314E-10 |
| TcCLB.510037.5 | elongation factor 1-gamma (EF-1-gamma), putative | 0,6412 | 0,00004813 |
| TcCLB.509319.40 | hypothetical protein, conserved | 0,6409 | 1,014E-07 |
| TcCLB.505939.70 | hypothetical protein, conserved | 0,6407 | 1,83E-09 |
| TcCLB.510749.10 | hypothetical protein, conserved | 0,6405 | 1,807E-09 |
| TcCLB.509995.10 | 60S ribosomal protein L4, putative | 0,6404 | 5,66E-16 |
| TcCLB.509805.160 | hypothetical protein, conserved | 0,6404 | 1,36E-10 |
| TcCLB.503809.110 | phosphatidic acid phosphatase protein, putative | 0,6403 | 6,189E-10 |
| TcCLB.507559.30 | hypothetical protein, conserved | 0,6403 | 1,611E-09 |
| TcCLB.506813.160 | dispersed gene family protein 1 (DGF-1, pseudogene), put | 0,6403 | 0,006484 |
| TcCLB.510519.40 | protein kinase, putative | 0,6402 | 4,025E-09 |
| TcCLB.510359.360 | hypothetical protein, conserved | 0,6398 | 2,077E-07 |
| TcCLB.506605.210 | leucine-rich repeat protein (LRRP), putative | 0,6397 | 4,201E-12 |
| TcCLB.505071.130 | hypothetical protein, conserved | 0,6396 | 6,881E-09 |
| TcCLB.506155.20 | hypothetical protein, conserved | 0,6395 | 1,943E-10 |
| TcCLB.508953.19 | hypothetical protein, conserved | 0,6395 | 1,626E-08 |
| TcCLB.508641.110 | hypothetical protein, conserved | 0,6392 | 8,944E-06 |
| TcCLB.504017.54 | hypothetical protein, conserved | 0,639 | 1,558E-09 |
| TcCLB.511887.50 | hypothetical protein, conserved | 0,639 | 0,01433 |
| TcCLB.508747.40 | mucin TcMUCII, putative | 0,6389 | 0,002884 |
| TcCLB.503635.59 | 40S ribosomal protein S3A, putative | 0,6382 | 6,673E-14 |
| TcCLB.510431.80 | hypothetical protein, conserved | 0,6381 | 5,878E-14 |
| TcCLB.506727.100 | enoyl-CoA hydratase/isomerase family protein, putative | 0,6377 | 2,634E-09 |
| TcCLB.508917.10 | protein kinase, putative | 0,6377 | 1,03E-08 |
| TcCLB.510421.60 | hypothetical protein, conserved | 0,6374 | 5,033E-09 |
| TcCLB.507585.10 | amino acid transporter, putative | 0,637 | 5,82E-24 |
| TcCLB.509099.80 | hypothetical protein, conserved | 0,637 | 1,248E-16 |
| TcCLB.510285.110 | hypothetical protein, conserved | 0,637 | 1,297E-07 |
| TcCLB.504099.15 | hypothetical protein | 0,637 | 0,03637 |
| TcCLB.511211.220 | heat shock protein 70, putative | 0,6366 | 5,905E-10 |
| TcCLB.508153.920 | choline/ethanolamine phosphotransferase, putative | 0,6366 | 4,769E-09 |
| TcCLB.506559.210 | hypothetical protein, conserved | 0,6363 | 2,688E-08 |
| TcCLB.507875.40 | trans-sialidase (pseudogene), putative | 0,6363 | 0,00001441 |
| TcCLB.506163.40 | vacuolar type h ATPase subunit, putative | 0,6363 | 0,05232 |
| TcCLB.510039.60 | hypothetical protein, conserved | 0,6362 | 5,726E-14 |
| TcCLB.508737.80 | mannosyltransferase-like protein, putative | 0,636 | 3,533E-09 |
| TcCLB.509055.10 | RNA-binding protein, putative | 0,636 | 0,00001154 |

| | | | |
|---|---|---|---|
| TcCLB.511301.50 | hypothetical protein, conserved | 0,6358 | 1,475E-12 |
| TcCLB.507583.30 | hypothetical protein | 0,6358 | 5,705E-06 |
| TcCLB.508153.730 | eukaryotic initiation factor 2a, putative | 0,6355 | 3,585E-16 |
| TcCLB.509087.10 | retrotransposon hot spot protein (RHS, pseudogene), puta | 0,6355 | 0,005789 |
| TcCLB.507803.30 | hypothetical protein | 0,6354 | 8,633E-10 |
| TcCLB.506859.10 | ubiquitin-conjugating enzyme E2, putative | 0,6353 | 1,966E-07 |
| TcCLB.511529.220 | hypothetical protein, conserved | 0,6351 | 0,0003671 |
| TcCLB.508071.40 | hypothetical protein | 0,6349 | 0,04164 |
| TcCLB.504241.80 | trans-sialidase (pseudogene), putative | 0,6348 | 1,756E-08 |
| TcCLB.507009.20 | hypothetical protein, conserved | 0,6347 | 0,0002784 |
| TcCLB.506893.80 | monothiol glutaredoxin, putative | 0,6344 | 0,00007498 |
| TcCLB.507295.61 | elongation factor 1-gamma (EF-1-gamma, pseudogene), | 0,6344 | 0,06257 |
| TcCLB.504231.20 | hypothetical protein, conserved | 0,6343 | 2,064E-12 |
| TcCLB.508547.150 | hypothetical protein, conserved | 0,634 | 3E-12 |
| TcCLB.510603.130 | hypothetical protein, conserved | 0,6339 | 1,801E-13 |
| TcCLB.507811.114 | hypothetical protein, conserved | 0,6339 | 1,216E-08 |
| TcCLB.510421.130 | mitochondrial RNA binding complex 1 subunit, putative | 0,6338 | 7,593E-09 |
| TcCLB.504105.150 | importin beta-1 subunit, putative | 0,6335 | 1,005E-18 |
| TcCLB.506737.70 | ubiquitin-like protein, putative | 0,6333 | 1,321E-16 |
| TcCLB.511201.40 | hypothetical protein, conserved | 0,6329 | 4,913E-06 |
| TcCLB.511733.130 | chaperonin HSP60, putative (fragment) | 0,6329 | 0,0000144 |
| TcCLB.511511.130 | hypothetical protein, conserved | 0,6328 | 1,564E-10 |
| TcCLB.511163.24 | hypothetical protein, conserved | 0,6326 | 0,00001549 |
| TcCLB.509967.189 | kinesin, putative (fragment) | 0,6325 | 0,0006145 |
| TcCLB.504009.20 | hypothetical protein, conserved | 0,6322 | 2,658E-08 |
| TcCLB.510105.230 | glyceraldehyde 3-phosphate dehydrogenase, cytosolic, pu | 0,6322 | 0,00000301 |
| TcCLB.506625.70 | RNA-binding protein, putative | 0,632 | 1,526E-13 |
| TcCLB.504229.20 | glycine dehydrogenase [decarboxylating] (pseudogene), p | 0,632 | 4,99E-12 |
| TcCLB.509453.60 | hypothetical protein, conserved | 0,6319 | 1,423E-10 |
| TcCLB.506583.60 | Elongation factor G1, mitochondrial, putative (EF-G1) | 0,6318 | 2,46E-16 |
| TcCLB.508595.40 | cysteine peptidase, putative | 0,6318 | 4,539E-07 |
| TcCLB.507083.10 | hypothetical protein, conserved | 0,6316 | 3,568E-12 |
| TcCLB.511823.79 | hypothetical protein, conserved | 0,6316 | 2,532E-06 |
| TcCLB.416809.10 | dispersed gene family protein 1 (DGF-1), putative | 0,6313 | 0,0003008 |
| TcCLB.509223.60 | beta galactofuranosyl glycosyltransferase, putative | 0,6312 | 0,002382 |
| TcCLB.504077.60 | hypothetical protein, conserved (pseudogene) | 0,6312 | 0,01245 |
| TcCLB.508283.60 | retrotransposon hot spot protein (RHS, pseudogene), puta | 0,6312 | 0,01344 |
| TcCLB.504765.9 | hypothetical protein, conserved (fragment) | 0,6308 | 1,49E-09 |
| TcCLB.510149.50 | tfiih basal transcription factor complex helicase xpb subur | 0,6307 | 2,944E-15 |
| TcCLB.506145.70 | hypothetical protein, conserved | 0,6306 | 9,375E-07 |
| TcCLB.510127.60 | hypothetical protein, conserved | 0,6304 | 0,00000153 |
| TcCLB.510615.10 | hypothetical protein, conserved | 0,6303 | 9,758E-10 |
| TcCLB.504047.40 | kinesin, putative | 0,6302 | 2,1E-16 |
| TcCLB.511635.20 | histone H2B, putative | 0,6299 | 3,059E-12 |
| TcCLB.510455.10 | dispersed gene family protein 1 (DGF-1), putative | 0,6299 | 0,1597 |

| | | | |
|---|---|---|---|
| TcCLB.507093.100 | hypothetical protein, conserved | 0,6296 | 2,557E-09 |
| TcCLB.509139.10 | glycerate kinase, putative | 0,6292 | 3,778E-16 |
| TcCLB.506007.20 | RNA editing complex protein MP18, putative | 0,6292 | 3,182E-07 |
| TcCLB.511261.4 | P-type H -ATPase, putative (fragment) | 0,629 | 8,719E-15 |
| TcCLB.509207.70 | hypothetical protein, conserved | 0,6289 | 2,156E-08 |
| TcCLB.506685.30 | thymidylate kinase, putative | 0,6289 | 5,772E-07 |
| TcCLB.511303.160 | sodium/hydrogen exchanger, putative (FTP) | 0,6288 | 5,006E-08 |
| TcCLB.506247.24 | hypothetical protein, conserved | 0,6287 | 0,001041 |
| TcCLB.508689.30 | hypothetical protein, conserved | 0,6286 | 3,1E-09 |
| TcCLB.511069.10 | 40S ribosomal protein S8, putative | 0,6284 | 2,522E-08 |
| TcCLB.508543.20 | protein kinase, putative | 0,6283 | 1,59E-10 |
| TcCLB.506781.50 | dispersed gene family protein 1 (DGF-1), putative | 0,6283 | 0,02615 |
| TcCLB.510119.9 | elongation factor 1-alpha, putative | 0,628 | 2,973E-20 |
| TcCLB.509271.10 | trans-sialidase, putative | 0,6279 | 4,552E-07 |
| TcCLB.503531.49 | structural maintenance of chromosome 3, putative | 0,6274 | 0,002105 |
| TcCLB.503939.60 | hypothetical protein, conserved | 0,6272 | 3,858E-10 |
| TcCLB.506989.130 | hypothetical protein, conserved | 0,6271 | 5,706E-12 |
| TcCLB.509683.100 | hypothetical protein, conserved | 0,6271 | 1,331E-07 |
| TcCLB.510087.30 | dihydroceramide synthase, putative (DHCS) | 0,627 | 4,762E-09 |
| TcCLB.511625.20 | hypothetical protein, conserved | 0,6267 | 7,559E-06 |
| TcCLB.505919.80 | retrotransposon hot spot (RHS) protein, putative | 0,6266 | 1,683E-08 |
| TcCLB.511545.180 | hypothetical protein, conserved | 0,6265 | 1,061E-10 |
| TcCLB.509245.10 | hypothetical protein, conserved | 0,6265 | 1,354E-06 |
| TcCLB.507951.299 | 5-histidylcysteine sulfoxide synthase | 0,6264 | 1,831E-12 |
| TcCLB.509965.420 | hypothetical protein, conserved | 0,6264 | 7,374E-08 |
| TcCLB.427237.9 | dispersed gene family protein 1 (DGF-1), putative | 0,6263 | 0,1156 |
| TcCLB.507007.76 | receptor protein kinase, putative | 0,6261 | 0,00001851 |
| TcCLB.503779.109 | hypothetical protein, conserved (fragment) | 0,6258 | 0,00000199 |
| TcCLB.504625.74 | hypothetical protein, conserved | 0,6258 | 0,002316 |
| TcCLB.506389.79 | calmodulin, putative | 0,6257 | 0,4371 |
| TcCLB.507809.30 | ribose-phosphate pyrophosphokinase, putative | 0,6256 | 1,238E-13 |
| TcCLB.510529.10 | hypothetical protein, conserved | 0,6256 | 0,0000644 |
| TcCLB.506173.150 | hypothetical protein | 0,6254 | 0,03855 |
| TcCLB.507017.140 | mitochondrial RNA binding protein 1, putative | 0,6253 | 8,984E-12 |
| TcCLB.511523.14 | succinate dehydrogenase cytochrome B subunit | 0,6247 | 5,823E-06 |
| TcCLB.504035.120 | hypothetical protein, conserved | 0,6246 | 1,024E-12 |
| TcCLB.506855.120 | hypothetical protein, conserved | 0,6242 | 3,117E-10 |
| TcCLB.510859.30 | hypothetical protein, conserved | 0,624 | 3,382E-17 |
| TcCLB.506789.120 | hypothetical protein, conserved | 0,624 | 2,759E-11 |
| TcCLB.506869.40 | mitochondrial RNA binding complex 1 subunit, putative | 0,624 | 7,556E-08 |
| TcCLB.508461.470 | hypothetical protein, conserved | 0,6239 | 1,436E-08 |
| TcCLB.506957.23 | hypothetical protein, conserved | 0,6237 | 4,227E-11 |
| TcCLB.511837.29 | iron-sulfur cluster assembly protein, putative | 0,6237 | 9,833E-08 |
| TcCLB.509803.30 | hypothetical protein, conserved | 0,6236 | 1,18E-09 |
| TcCLB.507379.9 | dispersed gene family protein 1 (DGF-1), putative (fragme | 0,6236 | 0,009597 |

| | | | |
|---|---|---|---|
| TcCLB.503687.40 | 40 kDa cyclophilin, putative (CYP40) | 0,6235 | 4,737E-13 |
| TcCLB.509005.60 | hypothetical protein, conserved | 0,6234 | 5,327E-08 |
| TcCLB.507769.20 | hypothetical protein, conserved | 0,6232 | 0,0001273 |
| TcCLB.504105.20 | eukaryotic translation initiation factor 5, putative | 0,6231 | 3,193E-14 |
| TcCLB.504029.90 | hypothetical protein, conserved | 0,6225 | 0,00000385 |
| TcCLB.510483.340 | beta galactofuranosyl glycosyltransferase (pseudogene), p | 0,6224 | 0,01149 |
| TcCLB.506523.9 | dispersed gene family protein 1 (DGF-1), putative | 0,6224 | 0,02012 |
| TcCLB.508847.9 | dispersed gene family protein 1 (DGF-1), putative | 0,6222 | 0,1614 |
| TcCLB.504393.10 | hypothetical protein | 0,622 | 0,1051 |
| TcCLB.503663.20 | hypothetical protein, conserved | 0,6217 | 6,131E-11 |
| TcCLB.508169.110 | hypothetical protein, conserved | 0,6215 | 0,00004004 |
| TcCLB.506047.10 | hypothetical protein, conserved (pseudogene) | 0,6214 | 0,004809 |
| TcCLB.511477.10 | trans-sialidase (pseudogene), putative | 0,6212 | 0,1969 |
| TcCLB.509267.40 | hypothetical protein, conserved | 0,6211 | 2,275E-09 |
| TcCLB.509607.70 | serine/threonine protein kinase, putative | 0,621 | 0,0003631 |
| TcCLB.506927.20 | hypothetical protein, conserved | 0,6209 | 0,00001793 |
| TcCLB.511563.9 | dispersed gene family protein 1 (DGF-1), putative | 0,6209 | 0,04509 |
| TcCLB.504109.150 | elongation factor, putative | 0,6208 | 9,248E-13 |
| TcCLB.506533.70 | hypothetical protein, conserved | 0,6208 | 1,461E-07 |
| TcCLB.505999.80 | hypothetical protein, conserved | 0,6208 | 2,476E-06 |
| TcCLB.510897.10 | cullin 2, putative | 0,6202 | 1,041E-06 |
| TcCLB.506835.120 | DNA ligase I, putative | 0,6201 | 7,798E-12 |
| TcCLB.508153.990 | hypothetical protein, conserved | 0,6201 | 1,689E-06 |
| TcCLB.506625.240 | hypothetical protein, conserved | 0,62 | 4,372E-12 |
| TcCLB.508041.60 | eukaryotic release factor 3, putative | 0,6199 | 5,776E-14 |
| TcCLB.510437.79 | heat shock protein 70, putative | 0,6199 | 0,00005922 |
| TcCLB.507017.10 | hypothetical protein, conserved | 0,6198 | 2,63E-11 |
| TcCLB.508331.90 | retrotransposon hot spot protein (RHS, pseudogene), puta | 0,6196 | 9,527E-07 |
| TcCLB.508175.329 | 60S ribosomal protein L19, putative | 0,6195 | 7,118E-09 |
| TcCLB.507809.150 | hypothetical protein, conserved | 0,6194 | 1,58E-07 |
| TcCLB.509011.50 | hypothetical protein, conserved | 0,6188 | 5,553E-14 |
| TcCLB.510087.110 | hypothetical protein, conserved | 0,6188 | 1,654E-10 |
| TcCLB.511807.160 | hypothetical protein, conserved | 0,6188 | 0,0001944 |
| TcCLB.506933.40 | DNA replication licensing factor MCM2, putative | 0,6185 | 4,071E-10 |
| TcCLB.506677.10 | ubiquitin-like protein, putative | 0,6183 | 1,227E-12 |
| TcCLB.508517.9 | dispersed gene family protein 1 (DGF-1), putative | 0,6183 | 0,007884 |
| TcCLB.507811.40 | hypothetical protein, conserved | 0,6182 | 1,55E-10 |
| TcCLB.407477.10 | thiolase protein-like protein, putative | 0,6182 | 1,313E-06 |
| TcCLB.510689.30 | proteasome beta 7 subunit, putative | 0,6181 | 1,358E-11 |
| TcCLB.509033.30 | cyclophilin, putative | 0,6181 | 6,595E-06 |
| TcCLB.508323.199 | ATP-dependent DEAD/H RNA helicase, putative (fragment | 0,6181 | 0,00009006 |
| TcCLB.509901.120 | hypothetical protein, conserved | 0,6179 | 6,669E-06 |
| TcCLB.511209.60 | hypothetical protein, conserved | 0,6176 | 6,645E-08 |
| TcCLB.506559.200 | thioredoxin, putative | 0,6174 | 1,237E-15 |
| TcCLB.511535.20 | hypothetical protein, conserved | 0,6174 | 1,095E-07 |

| | | | |
|---|---|---|---|
| TcCLB.510329.330 | hypothetical protein, conserved | 0,6173 | 3,643E-07 |
| TcCLB.511145.70 | centromere/microtubule binding protein cbf5, putative | 0,6172 | 9,494E-12 |
| TcCLB.509317.90 | hypothetical protein, conserved | 0,617 | 4,29E-07 |
| TcCLB.506125.30 | phosphoglycerate kinase, putative | 0,6169 | 8,493E-07 |
| TcCLB.511001.18 | 40S ribosomal protein S3A, putative | 0,6169 | 0,0005051 |
| TcCLB.509631.30 | mucin TcMUCII (pseudogene), putative | 0,6168 | 0,08395 |
| TcCLB.509429.160 | hypothetical protein | 0,6167 | 3,208E-11 |
| TcCLB.506241.100 | 60S ribosomal protein L32, putative | 0,6165 | 1,236E-15 |
| TcCLB.506629.100 | hypothetical protein | 0,6163 | 5,234E-09 |
| TcCLB.509153.110 | hypothetical protein | 0,6161 | 1,294E-10 |
| TcCLB.507031.29 | nucleosome assembly protein, putative | 0,6159 | 7,228E-11 |
| TcCLB.507643.50 | dispersed gene family protein 1 (DGF-1), putative | 0,6155 | 0,00007203 |
| TcCLB.507603.40 | proteasome beta-1 subunit, putative | 0,6153 | 7,389E-10 |
| TcCLB.510899.59 | hypothetical protein, conserved | 0,6152 | 7,133E-11 |
| TcCLB.508241.60 | phosphoadenosine phosphosulfate reductase-like protein, | 0,6151 | 1,012E-10 |
| TcCLB.508629.10 | dispersed gene family protein 1 (DGF-1), putative | 0,6151 | 0,00005433 |
| TcCLB.506977.30 | hypothetical protein, conserved | 0,615 | 9,654E-09 |
| TcCLB.509337.15 | hypothetical protein, conserved | 0,6149 | 1,267E-10 |
| TcCLB.503661.20 | retrotransposon hot spot (RHS) protein, putative | 0,6148 | 0,03556 |
| TcCLB.506129.30 | trans-sialidase, Group II, putative | 0,6147 | 1,122E-07 |
| TcCLB.508153.1120 | DNA repair helicase, putative | 0,6144 | 1,369E-07 |
| TcCLB.511889.50 | retrotransposon hot spot protein (RHS, pseudogene), puta | 0,6143 | 1,965E-07 |
| TcCLB.510155.170 | hypothetical protein, conserved | 0,6143 | 0,00001665 |
| TcCLB.506633.10 | protein kinase, putative | 0,6143 | 0,0001933 |
| TcCLB.511395.30 | protein phosphatase 2A, regulatory subunit B, putative | 0,6139 | 1,913E-15 |
| TcCLB.503723.20 | hypothetical protein, conserved | 0,6139 | 0,00001118 |
| TcCLB.508507.50 | hypothetical protein, conserved | 0,6136 | 0,00007846 |
| TcCLB.510283.114 | DNA-directed RNA polymerase I subunit, putative | 0,6135 | 0,00001299 |
| TcCLB.510733.10 | p21-activated kinase 3, putative | 0,6133 | 2,351E-15 |
| TcCLB.506813.130 | beta galactofuranosyl glycosyltransferase (pseudogene), p | 0,6133 | 0,1084 |
| TcCLB.506469.100 | hypothetical protein, conserved | 0,6132 | 2,306E-12 |
| TcCLB.510659.264 | prenyl protein specific carboxyl methyltransferase, putativ | 0,6132 | 0,00001388 |
| TcCLB.510409.39 | ubiquitin/ribosomal protein S27a, putative | 0,6131 | 5,255E-18 |
| TcCLB.506127.30 | phenylalanyl-tRNA synthetase, putative | 0,613 | 3,186E-16 |
| TcCLB.511577.44 | hypothetical protein, conserved | 0,613 | 0,0003389 |
| TcCLB.504109.189 | hypothetical protein | 0,613 | 0,09464 |
| TcCLB.511575.40 | hypothetical protein, conserved | 0,6129 | 5,489E-12 |
| TcCLB.507969.30 | sphingosine-1-phosphate phosphatase, putative | 0,6126 | 1,656E-09 |
| TcCLB.511683.10 | hypothetical protein | 0,6126 | 0,0004932 |
| TcCLB.508521.30 | beta galactofuranosyl glycosyltransferase (pseudogene), p | 0,6122 | 0,0002246 |
| TcCLB.503565.10 | hypothetical protein | 0,6121 | 0,00001466 |
| TcCLB.508741.160 | hypothetical protein, conserved | 0,612 | 1,134E-11 |
| TcCLB.504257.20 | hypothetical protein, conserved | 0,6116 | 3,598E-09 |
| TcCLB.504017.10 | Nascent polypeptide-associated complex subunit beta, pu | 0,6116 | 1,175E-08 |
| TcCLB.510197.110 | dispersed gene family protein 1 (DGF-1), putative | 0,6115 | 0,002902 |

| | | | |
|---|---|---|---|
| TcCLB.506947.40 | peroxisome assembly protein, putative | 0,6114 | 2,522E-09 |
| TcCLB.508281.9 | dispersed gene family protein 1 (DGF-1), putative | 0,6113 | 0,07891 |
| TcCLB.511247.18 | 50S ribosomal protein L16, putative | 0,6108 | 2,078E-06 |
| TcCLB.511667.30 | trans-sialidase, Group I, putative | 0,6107 | 0,1986 |
| TcCLB.507099.120 | hypothetical protein, conserved | 0,6105 | 3,651E-12 |
| TcCLB.511309.70 | hypothetical protein, conserved | 0,6105 | 1,319E-09 |
| TcCLB.511621.30 | hypothetical protein, conserved | 0,6105 | 0,00001512 |
| TcCLB.507207.19 | FtsJ cell division protein, putative (fragment) | 0,6104 | 8,97E-19 |
| TcCLB.511279.10 | hypothetical protein, conserved | 0,6103 | 0,00001759 |
| TcCLB.510595.30 | hypothetical protein | 0,61 | 0,0004172 |
| TcCLB.510897.70 | glycerophosphoryl diester phosphodiesterase (pseudogene | 0,6099 | 9,612E-06 |
| TcCLB.511575.90 | hypothetical protein | 0,6099 | 0,0000442 |
| TcCLB.506529.60 | hypothetical protein, conserved | 0,6099 | 0,00006284 |
| TcCLB.509997.20 | tryparedoxin 2, putative (TXN2) | 0,6098 | 3,079E-11 |
| TcCLB.509025.20 | hypothetical protein, conserved | 0,6097 | 0,0001761 |
| TcCLB.511389.40 | hypothetical protein, conserved | 0,6096 | 1,082E-12 |
| TcCLB.510377.270 | retrotransposon hot spot protein (RHS, pseudogene), puta | 0,6095 | 0,01258 |
| TcCLB.504075.6 | dual specificity protein phosphatase, putative | 0,6093 | 1,481E-06 |
| TcCLB.506787.9 | dispersed gene family protein 1 (DGF-1), putative (fragme | 0,6093 | 0,1546 |
| TcCLB.505183.130 | sugar transporter, putative | 0,6088 | 0,0008805 |
| TcCLB.511231.50 | aminomethyltransferase, mitochondrial, putative (GCV) | 0,6086 | 8,625E-08 |
| TcCLB.507937.40 | elongation factor 1-gamma (EF-1-gamma), putative | 0,6086 | 0,1619 |
| TcCLB.510741.229 | ubiquitin-activating enzyme E1, putative | 0,6084 | 7,981E-09 |
| TcCLB.510797.39 | hypothetical protein, conserved | 0,6083 | 0,0001506 |
| TcCLB.511929.20 | dispersed gene family protein 1 (DGF-1, pseudogene), put | 0,6081 | 0,001944 |
| TcCLB.507027.50 | Present in the outer mitochondrial membrane proteome 3 | 0,608 | 5,981E-07 |
| TcCLB.510295.30 | RNA editing complex protein MP18, putative | 0,6079 | 5,672E-08 |
| TcCLB.504011.80 | hypothetical protein, conserved | 0,6079 | 0,0001862 |
| TcCLB.508503.100 | hypothetical protein, conserved | 0,6078 | 0,00007881 |
| TcCLB.507473.10 | hypothetical protein | 0,6073 | 3,949E-12 |
| TcCLB.510119.20 | elongation factor 1-alpha, putative | 0,6072 | 1,96E-14 |
| TcCLB.510421.80 | hypothetical protein | 0,6072 | 0,0004168 |
| TcCLB.506811.170 | hypothetical protein, conserved | 0,6071 | 2,613E-11 |
| TcCLB.506467.40 | serine/threonine protein kinase, putative | 0,6068 | 1,632E-09 |
| TcCLB.509997.30 | tryparedoxin 1, putative (TXN1) | 0,6068 | 4,958E-06 |
| TcCLB.511089.5 | trans-sialidase (pseudogene), putative | 0,6062 | 0,2446 |
| TcCLB.511525.20 | hypothetical protein, conserved | 0,6061 | 7,151E-10 |
| TcCLB.509029.130 | cyclin 1 (CYC1) | 0,6061 | 1,421E-06 |
| TcCLB.510151.10 | fatty acyl CoA syntetase 1, putative | 0,6061 | 1,709E-06 |
| TcCLB.511003.180 | hypothetical protein, conserved | 0,606 | 0,0002506 |
| TcCLB.506531.50 | metacaspase, putative | 0,6055 | 2,254E-06 |
| TcCLB.506015.20 | surface protease GP63, putative | 0,6054 | 0,0000181 |
| TcCLB.507047.80 | zinc-finger protein ZPR1, putative | 0,6053 | 2,194E-06 |
| TcCLB.506173.140 | dispersed gene family protein 1 (DGF-1), putative | 0,6053 | 0,0008414 |
| TcCLB.511211.180 | hypothetical protein, conserved | 0,6051 | 6,086E-09 |

| | | | |
|---|---|---|---|
| TcCLB.509905.130 | dispersed gene family protein 1 (DGF-1, pseudogene), put | 0,605 | 1,704E-09 |
| TcCLB.511389.150 | thiolase protein-like protein, putative | 0,6047 | 2,271E-14 |
| TcCLB.506057.20 | retrotransposon hot spot protein (RHS, pseudogene), puta | 0,6046 | 0,00008635 |
| TcCLB.511521.40 | hypothetical protein, conserved | 0,6044 | 1,569E-06 |
| TcCLB.508813.70 | hypothetical protein, conserved | 0,6043 | 2,239E-08 |
| TcCLB.506559.190 | dual specificity protein phosphatase, putative | 0,6042 | 2,014E-08 |
| TcCLB.506557.10 | hypothetical protein, conserved | 0,604 | 4,185E-07 |
| TcCLB.508323.94 | cyclophilin, putative | 0,604 | 0,0003353 |
| TcCLB.506113.60 | retrotransposon hot spot (RHS) protein, putative | 0,6039 | 3,479E-16 |
| TcCLB.508601.10 | mitochondrial DEAD box protein, putative | 0,6039 | 0,00004751 |
| TcCLB.507033.4 | hypothetical protein, conserved | 0,6037 | 0,0006032 |
| TcCLB.511191.19 | ras-related protein Rab21, putative | 0,6036 | 2,379E-07 |
| TcCLB.506789.110 | hypothetical protein | 0,6035 | 3,719E-09 |
| TcCLB.508837.147 | kinesin, putative | 0,6034 | 1,542E-08 |
| TcCLB.511507.120 | hypothetical protein, conserved | 0,6034 | 2,671E-06 |
| TcCLB.508857.160 | threonine synthase, putative | 0,6032 | 5,754E-10 |
| TcCLB.507609.30 | ATP-dependent DEAD/H RNA helicase, putative | 0,6032 | 2,153E-06 |
| TcCLB.504143.30 | kinesin protein (pseudogene), putative | 0,6032 | 8,795E-06 |
| TcCLB.508307.184 | hypothetical protein, conserved | 0,6032 | 0,0001302 |
| TcCLB.505945.50 | ATP-dependent chaperone, putative | 0,6031 | 4,036E-09 |
| TcCLB.418113.9 | dispersed gene family protein 1 (DGF-1), putative | 0,6029 | 0,203 |
| TcCLB.506563.10 | pumilio/PUF RNA binding protein 9, putative (PUF9) | 0,6028 | 3,851E-08 |
| TcCLB.509875.160 | dispersed gene family protein 1 (DGF-1), putative | 0,6028 | 0,0001954 |
| TcCLB.508831.63 | hypothetical protein | 0,6027 | 1,486E-07 |
| TcCLB.510165.70 | retrotransposon hot spot protein (RHS, pseudogene), puta | 0,6027 | 0,004211 |
| TcCLB.510397.10 | hypothetical protein, conserved | 0,6025 | 1,891E-09 |
| TcCLB.511111.30 | hypothetical protein, conserved | 0,6025 | 9,608E-08 |
| TcCLB.506337.200 | hypothetical protein, conserved | 0,6023 | 2,38E-12 |
| TcCLB.509741.40 | hypothetical protein, conserved | 0,6022 | 0,00007196 |
| TcCLB.508221.284 | retrotransposon hot spot protein (RHS, pseudogene), puta | 0,6022 | 0,02914 |
| TcCLB.503981.30 | hypothetical protein, conserved | 0,6021 | 3,06E-08 |
| TcCLB.510359.130 | hypothetical protein, conserved | 0,6021 | 4,727E-06 |
| TcCLB.508325.220 | beta galactofuranosyl glycosyltransferase (pseudogene), p | 0,602 | 7,644E-08 |
| TcCLB.511391.170 | hypothetical protein, conserved | 0,6016 | 9,282E-08 |
| TcCLB.507809.70 | small ubiquitin protein, putative | 0,6016 | 0,00003537 |
| TcCLB.506417.30 | hypothetical protein, conserved | 0,6016 | 0,0000622 |
| TcCLB.510359.160 | hypothetical protein, conserved | 0,6016 | 0,0001863 |
| TcCLB.511731.60 | glutamyl-tRNA synthetase, putative | 0,6011 | 3,452E-09 |
| TcCLB.510667.110 | hypothetical protein, conserved | 0,6011 | 3,213E-06 |
| TcCLB.509677.10 | hypothetical protein, conserved | 0,601 | 2,194E-07 |
| TcCLB.503527.20 | hypothetical protein, conserved | 0,6008 | 2,896E-07 |
| TcCLB.507011.130 | hypothetical protein, conserved | 0,6007 | 1,853E-07 |
| TcCLB.511649.150 | phosphoglycerate mutase protein, putative | 0,6006 | 2,075E-07 |
| TcCLB.508607.40 | retrotransposon hot spot protein (RHS, pseudogene), puta | 0,6003 | 1,253E-08 |
| TcCLB.485889.4 | hypothetical protein, conserved | 0,6001 | 0,0009556 |

| | | | |
|---|---|---|---|
| TcCLB.509029.10 | serine/threonine protein phosphatase, putative | 0,5997 | 2,212E-06 |
| TcCLB.508593.140 | hypothetical protein, conserved | 0,5996 | 1,1E-08 |
| TcCLB.511825.4 | hypothetical protein, conserved (fragment) | 0,5991 | 1,199E-07 |
| TcCLB.503999.60 | phosphoprotein phosphatase, putative | 0,599 | 1,555E-08 |
| TcCLB.503925.50 | hypothetical protein, conserved | 0,5985 | 5,733E-06 |
| TcCLB.503823.104 | UDP-N-acetylglucosamine-dolichyl-phosphate N-acetylglu | 0,5984 | 2,619E-06 |
| TcCLB.507891.47 | flagellar calcium-binding protein, putative | 0,5983 | 1,201E-07 |
| TcCLB.508567.89 | hypothetical protein, conserved | 0,5981 | 3,685E-07 |
| TcCLB.510719.10 | vacuolar protein sorting-associated protein-like, putative | 0,598 | 5,557E-09 |
| TcCLB.511909.10 | hypothetical protein, conserved | 0,5978 | 8,121E-14 |
| TcCLB.509203.50 | hypothetical protein | 0,5976 | 8,486E-10 |
| TcCLB.503899.70 | hypothetical protein | 0,5973 | 0,0001963 |
| TcCLB.511665.30 | retrotransposon hot spot protein (RHS, pseudogene), puta | 0,5972 | 0,0003317 |
| TcCLB.508357.80 | cation transporter protein, putative | 0,597 | 2,283E-06 |
| TcCLB.503893.90 | hypothetical protein, conserved | 0,597 | 7,898E-06 |
| TcCLB.509843.10 | retrotransposon hot spot (RHS) protein, putative | 0,5964 | 3,838E-08 |
| TcCLB.508665.14 | heat shock protein, putative | 0,5963 | 3,83E-11 |
| TcCLB.507645.40 | hypothetical protein, conserved | 0,5963 | 1,179E-10 |
| TcCLB.510165.30 | hypothetical protein, conserved | 0,5963 | 8,954E-10 |
| TcCLB.508731.20 | hypothetical protein, conserved | 0,5963 | 4,197E-06 |
| TcCLB.410199.20 | hypothetical protein | 0,5963 | 0,01278 |
| TcCLB.507253.50 | ATP-dependent DEAD/H RNA helicase (pseudogene), puta | 0,5963 | 0,1983 |
| TcCLB.482319.10 | hypothetical protein, conserved | 0,5961 | 9,738E-08 |
| TcCLB.510259.24 | hypothetical protein, conserved | 0,5961 | 0,00009646 |
| TcCLB.506989.150 | hypothetical protein, conserved | 0,596 | 1,785E-08 |
| TcCLB.511353.30 | hypothetical protein, conserved | 0,596 | 0,0001337 |
| TcCLB.506303.50 | UDP-galactose 4-epimerase, putative | 0,5958 | 7,556E-08 |
| TcCLB.511289.59 | Multiprotein-bridging factor 1, putative (mbf1) | 0,5955 | 7,165E-09 |
| TcCLB.511737.9 | hypothetical protein, conserved | 0,5955 | 0,007093 |
| TcCLB.506331.80 | beta galactofuranosyl glycosyltransferase (pseudogene), p | 0,5955 | 0,0301 |
| TcCLB.508221.52 | hypothetical protein | 0,5954 | 0,1518 |
| TcCLB.510729.290 | Enriched in surface-labeled proteome protein 19 (SLS3) | 0,5953 | 1,902E-08 |
| TcCLB.509875.20 | dispersed gene family protein 1 (DGF-1), putative | 0,5952 | 0,002062 |
| TcCLB.511797.220 | dispersed gene family protein 1 (DGF-1), putative | 0,5952 | 0,03518 |
| TcCLB.511807.40 | hypothetical protein, conserved | 0,5951 | 5,403E-07 |
| TcCLB.509951.40 | hypothetical protein | 0,5951 | 0,002986 |
| TcCLB.508641.10 | dispersed gene family protein 1 (DGF-1), putative | 0,595 | 0,06897 |
| TcCLB.505975.20 | trans-sialidase, Group IV, putative | 0,5949 | 0,00001759 |
| TcCLB.509651.10 | hypothetical protein, conserved | 0,5949 | 0,00007956 |
| TcCLB.507971.20 | syntaxin binding protein (pseudogene), putative | 0,5948 | 5,584E-12 |
| TcCLB.506485.40 | kinesin (pseudogene), putative | 0,5947 | 1,676E-14 |
| TcCLB.506635.130 | hypothetical protein, conserved | 0,5945 | 4,082E-08 |
| TcCLB.506037.10 | dispersed gene family protein 1 (DGF-1, pseudogene), put | 0,5945 | 0,1109 |
| TcCLB.505989.82 | ruvB-like DNA helicase, putative | 0,5943 | 7,718E-09 |
| TcCLB.506939.70 | retrotransposon hot spot protein (RHS, pseudogene), puta | 0,5942 | 0,01328 |

| | | | |
|---|---|---|---|
| TcCLB.511255.313 | syntaxin binding protein (pseudogene), putative | 0,5942 | 0,06826 |
| TcCLB.510307.110 | hypothetical protein, conserved (pseudogene) | 0,5941 | 3,147E-10 |
| TcCLB.511015.20 | ribosomal protein S29, putative | 0,594 | 0,00001059 |
| TcCLB.511715.100 | pumilio/PUF RNA binding protein 7, putative (PUF7) | 0,5935 | 1,334E-09 |
| TcCLB.506697.60 | hypothetical protein, conserved | 0,5933 | 7,444E-14 |
| TcCLB.509857.30 | hypothetical protein, conserved | 0,5928 | 4,032E-07 |
| TcCLB.506605.100 | hypothetical protein | 0,5928 | 5,704E-07 |
| TcCLB.506559.50 | hypothetical protein, conserved | 0,5927 | 1,748E-09 |
| TcCLB.503495.20 | retrotransposon hot spot protein (RHS, pseudogene), puta | 0,5926 | 0,002333 |
| TcCLB.511419.50 | 3-phosphoglycerate kinase, glycosomal (PGKA) | 0,5925 | 0,00004592 |
| TcCLB.509671.80 | 60S ribosomal protein L5, putative | 0,5922 | 5,229E-16 |
| TcCLB.504003.90 | hypothetical protein, conserved | 0,5922 | 3,085E-07 |
| TcCLB.507611.30 | glycine dehydrogenase (pseudogene), putative | 0,5922 | 0,02335 |
| TcCLB.506559.550 | C2 domain containing protein, putative | 0,5921 | 3,376E-08 |
| TcCLB.507483.30 | calmodulin, putative | 0,592 | 3,795E-07 |
| TcCLB.503891.54 | nucleolar RNA-binding protein, putative | 0,592 | 0,00006181 |
| TcCLB.506173.190 | dispersed gene family protein 1 (DGF-1), putative | 0,5919 | 0,01412 |
| TcCLB.507641.110 | hypothetical protein, conserved | 0,5917 | 5,704E-10 |
| TcCLB.505999.100 | 3-phosphoglycerate kinase, glycosomal (PGKA) | 0,5916 | 1,792E-08 |
| TcCLB.511025.120 | CYC2-like cyclin, putative | 0,5915 | 4,203E-09 |
| TcCLB.511827.139 | retrotransposon hot spot protein (RHS, pseudogene), puta | 0,5914 | 0,01251 |
| TcCLB.507513.110 | hypothetical protein | 0,5913 | 3,296E-06 |
| TcCLB.402857.10 | hypothetical protein, conserved (fragment) | 0,5912 | 5,199E-07 |
| TcCLB.507851.40 | hypothetical protein, conserved | 0,5911 | 0,00005158 |
| TcCLB.511301.70 | FK506-binding protein (FKBP)-type peptidyl-prolyl isomera | 0,591 | 5,475E-11 |
| TcCLB.507907.40 | hypothetical protein, conserved | 0,591 | 3,311E-07 |
| TcCLB.506725.20 | hypothetical protein, conserved | 0,5909 | 1,563E-15 |
| TcCLB.510737.70 | 60S ribosomal subunit protein L31, putative | 0,5908 | 0,0004913 |
| TcCLB.509813.10 | hypothetical protein (pseudogene) | 0,5907 | 0,07764 |
| TcCLB.506357.140 | Periodic tryptophan protein 2 homolog, putative | 0,5906 | 3,577E-09 |
| TcCLB.508037.10 | retrotransposon hot spot (RHS) protein, putative | 0,5906 | 5,283E-07 |
| TcCLB.510143.17 | hypothetical protein, conserved | 0,5903 | 6,167E-06 |
| TcCLB.510515.90 | hypothetical protein, conserved | 0,5903 | 0,00003232 |
| TcCLB.506779.100 | RNA-binding protein, putative | 0,5902 | 6,986E-06 |
| TcCLB.507023.10 | epsilon-adaptin, putative | 0,59 | 9,121E-13 |
| TcCLB.508515.120 | hypothetical protein, conserved | 0,5899 | 5,863E-07 |
| TcCLB.507737.40 | hypothetical protein, conserved (fragment) | 0,5899 | 0,09238 |
| TcCLB.509471.68 | histone H3, putative | 0,5898 | 3,175E-09 |
| TcCLB.506153.6 | hypothetical protein, conserved | 0,5896 | 0,00001164 |
| TcCLB.509965.144 | hypothetical protein, conserved | 0,5895 | 1,175E-06 |
| TcCLB.510155.80 | mitochondrial-processing peptidase subunit beta, putative | 0,5893 | 1,171E-12 |
| TcCLB.508153.1074 | hypothetical protein, conserved | 0,5893 | 2,064E-08 |
| TcCLB.504431.20 | hypothetical protein | 0,5893 | 0,00001995 |
| TcCLB.507149.20 | retrotransposon hot spot protein (RHS, pseudogene), puta | 0,5892 | 0,03187 |
| TcCLB.508569.140 | hypothetical protein, conserved | 0,5891 | 3,385E-06 |

| | | | |
|---|---|---|---|
| TcCLB.507167.41 | ATP-dependent DEAD/H RNA helicase (pseudogene), puta | 0,5891 | 0,148 |
| TcCLB.511571.40 | hypothetical protein, conserved | 0,5889 | 4,682E-10 |
| TcCLB.506567.60 | ABC transporter, putative | 0,5889 | 1,496E-09 |
| TcCLB.506625.19 | dispersed gene family protein 1 (DGF-1), putative (fragm | 0,5886 | 0,03035 |
| TcCLB.503521.89 | mannosyltransferase-II, putative | 0,5885 | 3,203E-08 |
| TcCLB.503583.80 | dual specificity protein phosphatase or MAP kinase phosp | 0,5885 | 7,155E-07 |
| TcCLB.506335.60 | mucin TcMUCII (pseudogene), putative | 0,5884 | 0,002892 |
| TcCLB.506941.130 | hypothetical protein, conserved | 0,5883 | 3,822E-08 |
| TcCLB.511693.70 | hypothetical protein, conserved | 0,5883 | 0,00001768 |
| TcCLB.511865.80 | hypothetical protein, conserved | 0,5882 | 0,0007933 |
| TcCLB.511385.70 | hypothetical protein | 0,5881 | 0,01087 |
| TcCLB.508307.150 | hypothetical protein, conserved | 0,5877 | 1,381E-07 |
| TcCLB.510879.220 | hypothetical protein, conserved | 0,5874 | 1,162E-06 |
| TcCLB.506419.69 | hypothetical protein, conserved | 0,5874 | 0,0003879 |
| TcCLB.508139.40 | retrotransposon hot spot protein (RHS, pseudogene), puta | 0,5871 | 2,924E-07 |
| TcCLB.510531.54 | small nuclear ribonucleoprotein, putative | 0,5869 | 3,94E-07 |
| TcCLB.452283.10 | amino acid permease (pseudogene), putative | 0,5869 | 0,000522 |
| TcCLB.507011.240 | hypothetical protein, conserved | 0,5868 | 0,0001248 |
| TcCLB.507911.40 | hypothetical protein, conserved (pseudogene) | 0,5867 | 0,00004485 |
| TcCLB.511255.341 | trans-sialidase (pseudogene), putative | 0,5867 | 0,04174 |
| TcCLB.511847.50 | hypothetical protein, conserved | 0,5865 | 4,733E-09 |
| TcCLB.510641.10 | retrotransposon hot spot protein (RHS, pseudogene), puta | 0,5865 | 0,002999 |
| TcCLB.506857.110 | hypothetical protein, conserved | 0,586 | 0,00002521 |
| TcCLB.506739.50 | ATP-dependent protease ATPase subunit HslU1, putative ( | 0,5859 | 1,383E-12 |
| TcCLB.509055.44 | hypothetical protein, conserved | 0,5859 | 5,94E-08 |
| TcCLB.506227.190 | hypothetical protein | 0,5858 | 0,01064 |
| TcCLB.511213.70 | trans-sialidase (pseudogene), putative | 0,5857 | 0,000166 |
| TcCLB.510645.30 | 3,2-trans-enoyl-CoA isomerase, mitochondrial precursor, | 0,5856 | 1,533E-10 |
| TcCLB.503955.50 | hypothetical protein, conserved | 0,5854 | 0,00001667 |
| TcCLB.508817.20 | hypothetical protein | 0,5853 | 4,769E-09 |
| TcCLB.511217.60 | hypothetical protein | 0,5852 | 2,116E-07 |
| TcCLB.509467.20 | hypothetical protein, conserved (pseudogene) | 0,5852 | 0,04153 |
| TcCLB.509471.59 | histone H3, putative | 0,5851 | 0,004304 |
| TcCLB.510759.114 | hypothetical protein, conserved | 0,585 | 3,198E-06 |
| TcCLB.509855.10 | hypothetical protein, conserved | 0,5849 | 2,878E-12 |
| TcCLB.506127.150 | hypothetical protein, conserved | 0,5847 | 1,281E-09 |
| TcCLB.510105.130 | phosphatidylinositol-4-phosphate 5-kinase type II beta, pu | 0,5847 | 5,019E-07 |
| TcCLB.508777.30 | hypothetical protein, conserved | 0,5846 | 4,258E-13 |
| TcCLB.508045.79 | spliceosomal U5 snRNP-specific protein, putative | 0,5843 | 6,557E-07 |
| TcCLB.509679.9 | leucyl-tRNA synthetase, putative | 0,5843 | 0,0003495 |
| TcCLB.511717.90 | phosphomannose isomerase, putative | 0,5842 | 2,079E-06 |
| TcCLB.508957.40 | hypothetical protein, conserved (pseudogene) | 0,5841 | 7,32E-11 |
| TcCLB.510759.130 | hypothetical protein, conserved | 0,5841 | 0,00001067 |
| TcCLB.507827.10 | dispersed gene family protein 1 (DGF-1), putative | 0,5841 | 0,002166 |
| TcCLB.507159.10 | Wee1-like protein kinase, putative (pseudogene, point mu | 0,5839 | 7,866E-06 |

| | | | |
|---|---|---|---|
| TcCLB.507073.40 | hypothetical protein, conserved | 0,5838 | 6,027E-13 |
| TcCLB.508839.70 | hypothetical protein, conserved | 0,5833 | 0,0003547 |
| TcCLB.507769.10 | hypothetical protein, conserved | 0,5828 | 0,02782 |
| TcCLB.510887.50 | hypothetical protein, conserved | 0,5827 | 3,791E-10 |
| TcCLB.510885.20 | hypothetical protein, conserved | 0,5827 | 1,075E-07 |
| TcCLB.508679.20 | ribonuclease inhibitor, putative | 0,5827 | 0,0001054 |
| TcCLB.411427.30 | hypothetical protein, conserved | 0,5824 | 2,709E-09 |
| TcCLB.509735.84 | dispersed gene family protein 1 (DGF-1), putative | 0,5824 | 0,00002588 |
| TcCLB.507963.44 | hypothetical protein, conserved | 0,5823 | 6,815E-06 |
| TcCLB.505171.40 | hypothetical protein, conserved | 0,5822 | 5,281E-07 |
| TcCLB.511731.70 | hypothetical protein, conserved | 0,5822 | 0,00002899 |
| TcCLB.507609.60 | inositol-3-phosphate synthase, putative | 0,5821 | 0,00002648 |
| TcCLB.504371.20 | dispersed gene family protein 1 (DGF-1, pseudogene), put | 0,5821 | 0,00577 |
| TcCLB.504423.10 | hypothetical protein, conserved | 0,582 | 0,0003871 |
| TcCLB.511807.150 | hypothetical protein, conserved | 0,5819 | 4,252E-09 |
| TcCLB.511865.54 | protein transport protein Sec61 gamma subunit, putative | 0,5819 | 1,545E-06 |
| TcCLB.507639.30 | universal minicircle sequence binding protein 1 (pdz5) | 0,5818 | 0,0002086 |
| TcCLB.474937.9 | acetyl-CoA carboxylase, putative | 0,5817 | 6,162E-09 |
| TcCLB.511907.10 | hypothetical protein, conserved | 0,5817 | 3,171E-07 |
| TcCLB.508555.10 | hypothetical protein, conserved | 0,5817 | 0,0004813 |
| TcCLB.511855.50 | hypothetical protein, conserved | 0,5815 | 2,358E-10 |
| TcCLB.504163.50 | fructose-bisphosphate aldolase, glycosomal, putative | 0,5814 | 6,943E-06 |
| TcCLB.511289.40 | hypothetical protein, conserved | 0,5812 | 4,566E-12 |
| TcCLB.506207.40 | hypothetical protein, conserved | 0,5812 | 2,174E-11 |
| TcCLB.510971.10 | dispersed gene family protein 1 (DGF-1), putative | 0,5812 | 0,004629 |
| TcCLB.509885.10 | V-type proton ATPase subunit D, putative (VAd) | 0,581 | 2,261E-09 |
| TcCLB.509033.60 | hypothetical protein, conserved | 0,5809 | 4,519E-06 |
| TcCLB.458241.10 | hypothetical protein, conserved | 0,5809 | 0,002312 |
| TcCLB.506629.90 | hypothetical protein, conserved | 0,5809 | 0,009688 |
| TcCLB.511667.40 | retrotransposon hot spot (RHS) protein, putative | 0,5808 | 0,001779 |
| TcCLB.506797.20 | hypothetical protein | 0,5807 | 1,429E-13 |
| TcCLB.506529.650 | hypothetical protein, conserved | 0,5806 | 8,975E-08 |
| TcCLB.510639.10 | dispersed gene family protein 1 (DGF-1, pseudogene), put | 0,5806 | 0,003791 |
| TcCLB.508625.120 | leucine carboxyl methyltransferase, putative | 0,5805 | 2,648E-07 |
| TcCLB.504129.10 | hypothetical protein, conserved | 0,5805 | 0,008669 |
| TcCLB.505401.40 | dispersed gene family protein 1 (DGF-1), putative | 0,5805 | 0,3082 |
| TcCLB.503865.20 | hypothetical protein, conserved | 0,5804 | 1,546E-08 |
| TcCLB.506867.20 | vacuolar type h ATPase subunit, putative | 0,5803 | 3,593E-08 |
| TcCLB.504153.180 | UDP-N-acetylglucosamine pyrophosphorylase, putative | 0,5799 | 8,056E-09 |
| TcCLB.507949.4 | cytidine triphosphate synthase, putative | 0,5799 | 0,00002108 |
| TcCLB.511423.104 | DNA-directed RNA polymerase 2, RPB12 subunit, putative | 0,5799 | 0,0001741 |
| TcCLB.506575.9 | hypothetical protein, conserved | 0,5797 | 1,047E-09 |
| TcCLB.508079.20 | fatty acyl CoA syntetase 1 (pseudogene), putative | 0,5794 | 1,457E-08 |
| TcCLB.511107.41 | ATP-dependent Clp protease subunit heat shock protein 1( | 0,5789 | 0,0001868 |
| TcCLB.506825.219 | hypothetical protein, conserved | 0,5786 | 5,939E-11 |

| | | | |
|---|---|---|---|
| TcCLB.503899.30 | 40S ribosomal protein S16, putative | 0,5785 | 0,0002629 |
| TcCLB.506959.130 | 60S ribosomal protein L30, putative | 0,5784 | 2,192E-16 |
| TcCLB.506711.10 | cAMP response protein, putative (CARP4) | 0,5783 | 2,883E-10 |
| TcCLB.506543.30 | hypothetical protein, conserved | 0,5782 | 0,01086 |
| TcCLB.506123.40 | nucleolar RNA helicase II, putative | 0,5779 | 2,962E-10 |
| TcCLB.508413.84 | kinetoplastid membrane protein KMP-11 (fragment) | 0,5778 | 0,00002198 |
| TcCLB.511277.550 | hypothetical protein, conserved | 0,5778 | 0,0008022 |
| TcCLB.505297.10 | dispersed gene family protein 1 (DGF-1, pseudogene), put | 0,5778 | 0,0815 |
| TcCLB.504445.10 | dispersed gene family protein 1 (DGF-1), putative | 0,5777 | 0,0252 |
| TcCLB.511283.100 | hypothetical protein, conserved | 0,577 | 2,56E-08 |
| TcCLB.506593.50 | hypothetical protein, conserved | 0,577 | 5,108E-08 |
| TcCLB.510187.420 | chaperonin HSP60, mitochondrial precursor (pseudogene) | 0,577 | 0,00002432 |
| TcCLB.507275.50 | hypothetical protein, conserved | 0,5769 | 1,39E-07 |
| TcCLB.503861.80 | dispersed gene family protein 1 (DGF-1), putative | 0,5769 | 0,00002364 |
| TcCLB.509955.10 | ubiquitin carboxyl-terminal hydrolase, putative (fragment | 0,5769 | 0,001498 |
| TcCLB.506155.60 | aldehyde dehydrogenase, putative | 0,5768 | 2,208E-08 |
| TcCLB.506943.23 | hypothetical protein, conserved | 0,5767 | 0,00002487 |
| TcCLB.508815.70 | hypothetical protein, conserved | 0,5764 | 8,498E-14 |
| TcCLB.511517.153 | SH3 domain protein, putative | 0,5764 | 2,11E-08 |
| TcCLB.506147.140 | hypothetical protein, conserved | 0,5763 | 9,982E-09 |
| TcCLB.507691.10 | hypothetical protein, conserved | 0,5762 | 0,06503 |
| TcCLB.506743.180 | recombination initiation protein NBS1, putative | 0,5758 | 5,527E-10 |
| TcCLB.509799.60 | hypothetical protein, conserved | 0,5755 | 1,028E-11 |
| TcCLB.506683.210 | retrotransposon hot spot protein (RHS, pseudogene), puta | 0,5755 | 0,0002584 |
| TcCLB.510497.130 | retrotransposon hot spot protein (RHS, pseudogene), puta | 0,5755 | 0,001119 |
| TcCLB.504177.20 | fatty acyl CoA synthetase 2, putative | 0,5754 | 6,713E-09 |
| TcCLB.508047.25 | elongation factor 1-gamma (EF-1-gamma, pseudogene), | 0,5754 | 0,2225 |
| TcCLB.510525.130 | Present in the outer mitochondrial membrane proteome 3 | 0,5753 | 9,71E-09 |
| TcCLB.510241.130 | tyrosine phospatase-like protein (pseudogene), putative | 0,5752 | 3,635E-07 |
| TcCLB.507107.40 | 3,2-trans-enoyl-CoA isomerase, mitochondrial precursor, | 0,5751 | 1,225E-14 |
| TcCLB.508285.10 | retrotransposon hot spot (RHS) protein, putative | 0,575 | 0,0005241 |
| TcCLB.445777.10 | retrotransposon hot spot (RHS) protein, putative, (fragme | 0,5747 | 1,678E-07 |
| TcCLB.508159.40 | dispersed gene family protein 1 (DGF-1), putative | 0,5747 | 0,02325 |
| TcCLB.509029.170 | palmitoyl acyltransferase 5, putative | 0,5746 | 1,345E-06 |
| TcCLB.507053.160 | hypothetical protein, conserved | 0,5745 | 4,65E-14 |
| TcCLB.506605.120 | hypothetical protein, conserved | 0,574 | 7,47E-09 |
| TcCLB.507913.30 | proliferative cell nuclear antigen (PCNA), putative | 0,5737 | 4,339E-11 |
| TcCLB.506297.310 | hypothetical protein, conserved | 0,5737 | 2,827E-08 |
| TcCLB.508909.30 | hypothetical protein, conserved | 0,5737 | 2,903E-08 |
| TcCLB.507023.280 | hypothetical protein, conserved | 0,5732 | 3,887E-06 |
| TcCLB.509339.4 | poly(A) export protein, putative | 0,5727 | 0,00001977 |
| TcCLB.508607.30 | dispersed gene family protein 1 (DGF-1), putative | 0,5727 | 0,008003 |
| TcCLB.510853.120 | hypothetical protein, conserved | 0,5726 | 1,712E-11 |
| TcCLB.504103.6 | hypothetical protein, conserved | 0,5726 | 0,0002048 |
| TcCLB.506209.60 | retrotransposon hot spot protein (RHS, pseudogene), puta | 0,5723 | 2,738E-06 |

| | | | |
|---|---|---|---|
| TcCLB.510769.90 | 60S ribosomal protein L32, putative | 0,5722 | 2,395E-09 |
| TcCLB.511145.40 | peroxin 14, putative | 0,5721 | 1,989E-08 |
| TcCLB.507895.164 | hypothetical protein, conserved | 0,5721 | 1,414E-06 |
| TcCLB.504131.190 | mitochondrial carrier protein, putative | 0,5718 | 1,051E-06 |
| TcCLB.508895.80 | intraflagellar transport protein 52, putative (IFT52) | 0,5716 | 1,058E-10 |
| TcCLB.511751.90 | hypothetical protein | 0,5715 | 0,00009149 |
| TcCLB.508431.10 | dispersed gene family protein 1 (DGF-1, pseudogene), put | 0,5715 | 0,001082 |
| TcCLB.511181.50 | metallo-peptidase, Clan ME, Family M16 | 0,5714 | 2,857E-11 |
| TcCLB.506127.50 | hypothetical protein, conserved | 0,5714 | 0,0000016 |
| TcCLB.511821.10 | Cytoplasmic dynein 1 heavy chain (DYNC1H1), putative | 0,571 | 1,66E-11 |
| TcCLB.506425.40 | hypothetical protein, conserved | 0,5707 | 3,919E-18 |
| TcCLB.510139.10 | mucin-associated surface protein (MASP, pseudogene), pu | 0,5707 | 0,01126 |
| TcCLB.506389.50 | hypothetical protein, conserved | 0,5706 | 6,072E-09 |
| TcCLB.509767.90 | alanine racemase, putative | 0,5704 | 1,84E-08 |
| TcCLB.511115.30 | hypothetical protein, conserved | 0,5704 | 1,375E-06 |
| TcCLB.506283.10 | retrotransposon hot spot protein (RHS, pseudogene), puta | 0,5704 | 0,0001146 |
| TcCLB.511255.65 | surface protease GP63 (pseudogene), putative | 0,5703 | 0,1742 |
| TcCLB.508503.60 | protein kinase, putative | 0,5701 | 1,323E-13 |
| TcCLB.509151.60 | hypothetical protein, conserved | 0,5699 | 4,83E-09 |
| TcCLB.508277.370 | mitochondrial DNA topoisomerase II, putative | 0,5698 | 2,789E-13 |
| TcCLB.504231.10 | hypothetical protein, conserved | 0,5694 | 6,631E-08 |
| TcCLB.504121.39 | 60S ribosomal protein L14, putative (fragment) | 0,569 | 6,272E-11 |
| TcCLB.509977.100 | hypothetical protein, conserved | 0,5689 | 2,304E-10 |
| TcCLB.506599.395 | elongation factor 1-gamma (EF-1-gamma, pseudogene), | 0,5689 | 0,1515 |
| TcCLB.507723.130 | eukaryotic translation initiation factor 3 subunit 8, putativ | 0,5688 | 6,615E-16 |
| TcCLB.510731.80 | hypothetical protein, conserved | 0,5687 | 4,798E-10 |
| TcCLB.508641.160 | glycogenin glucosyltransferase, putative | 0,5687 | 5,466E-06 |
| TcCLB.506845.10 | retrotransposon hot spot protein (RHS, pseudogene), puta | 0,5687 | 0,003861 |
| TcCLB.505037.100 | dispersed gene family protein 1 (DGF-1), putative | 0,5686 | 1,433E-06 |
| TcCLB.511769.10 | hypothetical protein | 0,5681 | 0,001738 |
| TcCLB.506471.10 | hypothetical protein, conserved | 0,5679 | 8,79E-08 |
| TcCLB.509211.39 | hypothetical protein, conserved (fragment) | 0,5677 | 1,482E-06 |
| TcCLB.510127.70 | leucine-rich repeat protein 1 (LRRP1), putative | 0,5677 | 0,001672 |
| TcCLB.509023.160 | hypothetical protein, conserved | 0,5675 | 6,157E-07 |
| TcCLB.503557.40 | Present in the outer mitochondrial membrane proteome 8 | 0,5675 | 0,0005138 |
| TcCLB.506629.20 | hypothetical protein, conserved | 0,5674 | 3,593E-06 |
| TcCLB.510865.10 | retrotransposon hot spot protein (RHS, pseudogene), puta | 0,5674 | 0,0002264 |
| TcCLB.503643.3 | 60S ribosomal protein L4, putative (fragment) | 0,5673 | 5,933E-13 |
| TcCLB.508891.50 | hypothetical protein | 0,5673 | 1,18E-07 |
| TcCLB.511479.30 | hypothetical protein, conserved | 0,5673 | 0,00002217 |
| TcCLB.507023.110 | small GTP-binding protein Rab28, putative | 0,5672 | 7,365E-06 |
| TcCLB.509395.30 | tRNA exportin, putative | 0,567 | 6,091E-10 |
| TcCLB.511907.110 | fatty acid transporter protein-like, putative | 0,567 | 3,652E-09 |
| TcCLB.506459.260 | hypothetical protein, conserved | 0,567 | 2,233E-08 |
| TcCLB.504443.20 | hypothetical protein, conserved | 0,5669 | 0,00005773 |

| | | | |
|---|---|---|---|
| TcCLB.511725.170 | hypothetical protein, conserved | 0,5665 | 1,204E-08 |
| TcCLB.503905.40 | hypothetical protein, conserved | 0,5665 | 0,00003881 |
| TcCLB.503979.20 | hypothetical protein, conserved | 0,5664 | 0,00001145 |
| TcCLB.503653.70 | hypothetical protein, conserved | 0,5663 | 0,00009074 |
| TcCLB.509429.310 | hypothetical protein, conserved | 0,5661 | 3,491E-08 |
| TcCLB.509179.10 | choline/carnitine O-acyltransferase (pseudogene), putativ | 0,5661 | 0,004028 |
| TcCLB.503665.21 | elongation factor 1-gamma (EF-1-gamma, pseudogene), | 0,566 | 0,0188 |
| TcCLB.504125.20 | hypothetical protein | 0,5658 | 0,00001933 |
| TcCLB.510105.250 | membrane-bound acid phosphatase 2, putative | 0,5657 | 0,00004884 |
| TcCLB.509029.110 | hypothetical protein, conserved | 0,5656 | 8,104E-07 |
| TcCLB.510741.170 | hypothetical protein | 0,5655 | 1,07E-08 |
| TcCLB.511537.8 | ABC transporter of the mitochondrion, putative (fragment | 0,5654 | 6,276E-07 |
| TcCLB.511389.130 | diphthine synthase, putative | 0,5653 | 0,00001132 |
| TcCLB.508241.70 | hypothetical protein, conserved | 0,5651 | 1,097E-07 |
| TcCLB.507047.110 | otubain cysteine peptidase, Clan CA, family C65, putative | 0,565 | 2,795E-08 |
| TcCLB.504069.90 | centromere/microtubule binding protein cbf5, putative | 0,5649 | 1,391E-08 |
| TcCLB.510763.50 | hypothetical protein, conserved | 0,5649 | 0,0003154 |
| TcCLB.506661.140 | hypothetical protein, conserved | 0,5646 | 1,098E-09 |
| TcCLB.509725.50 | protein kinase, putative | 0,5645 | 0,04902 |
| TcCLB.503715.30 | ras-related protein RAB1A, putative (Rab1A) | 0,5643 | 1,032E-13 |
| TcCLB.511545.130 | hypothetical protein, conserved | 0,5638 | 3,853E-07 |
| TcCLB.507875.30 | hypothetical protein, conserved | 0,5637 | 0,00001624 |
| TcCLB.506835.10 | hypothetical protein, conserved | 0,5636 | 2,543E-11 |
| TcCLB.509875.204 | hypothetical protein, conserved | 0,5636 | 0,00001965 |
| TcCLB.509011.20 | RNA polymerase IIA largest subunit, putative | 0,5635 | 2,168E-11 |
| TcCLB.503971.30 | hypothetical protein, conserved | 0,5634 | 0,000251 |
| TcCLB.509769.80 | hypothetical protein, conserved | 0,5633 | 3,787E-17 |
| TcCLB.511807.180 | hypothetical protein, conserved | 0,5633 | 7,167E-16 |
| TcCLB.507061.20 | ribosomal protein L2, putative | 0,5633 | 1,104E-08 |
| TcCLB.509171.47 | hypothetical protein, conserved | 0,5631 | 1,162E-06 |
| TcCLB.508879.110 | hypothetical protein, conserved | 0,5631 | 0,0000016 |
| TcCLB.506179.50 | hypothetical protein, conserved | 0,5631 | 7,586E-06 |
| TcCLB.508089.20 | UDP-glucoronosyl and UDP-glucosyl transferase, putative | 0,5628 | 5,261E-06 |
| TcCLB.508831.10 | hypothetical protein, conserved | 0,5624 | 5,128E-08 |
| TcCLB.508873.590 | retrotransposon hot spot protein (RHS, pseudogene), puta | 0,5623 | 0,007248 |
| TcCLB.508301.20 | metallo-peptidase, Clan ME, Family M16 | 0,5622 | 1,409E-08 |
| TcCLB.509935.29 | proline racemase (fragment) | 0,562 | 0,00003356 |
| TcCLB.506459.240 | elongation factor 1-gamma (EF-1-gamma), putative | 0,562 | 0,0001263 |
| TcCLB.510647.50 | hypothetical protein, conserved | 0,5617 | 0,006063 |
| TcCLB.507795.30 | intraflagellar transport protein 27 (IFT27) | 0,5616 | 1,944E-06 |
| TcCLB.506941.170 | ADP-ribosylation factor-like protein, putative | 0,5615 | 0,0002542 |
| TcCLB.507521.10 | hypothetical protein, conserved | 0,5614 | 8,235E-07 |
| TcCLB.506195.230 | hypothetical protein, conserved | 0,561 | 1,374E-06 |
| TcCLB.510943.180 | hypothetical protein, conserved | 0,5609 | 0,00005083 |
| TcCLB.438897.10 | hypothetical protein, conserved | 0,5608 | 8,715E-07 |

| | | | |
|---|---|---|---|
| TcCLB.504057.90 | hypothetical protein, conserved | 0,5607 | 3,489E-10 |
| TcCLB.504151.60 | actin interacting protein-like protein, putative | 0,56 | 4,27E-09 |
| TcCLB.511801.50 | hypothetical protein, conserved | 0,56 | 4,81E-07 |
| TcCLB.506537.280 | glycine dehydrogenase [decarboxylating] (pseudogene), pu | 0,5597 | 3,778E-07 |
| TcCLB.508331.20 | hypothetical protein, conserved | 0,5596 | 2,684E-13 |
| TcCLB.504077.50 | hypothetical protein, conserved | 0,5596 | 5,052E-07 |
| TcCLB.506727.130 | hypothetical protein, conserved | 0,5596 | 0,00007148 |
| TcCLB.506935.19 | hypothetical protein, conserved (fragment) | 0,5595 | 0,006286 |
| TcCLB.510901.20 | hypothetical protein | 0,5591 | 0,0005977 |
| TcCLB.509287.60 | hypothetical protein, conserved | 0,5586 | 1,811E-09 |
| TcCLB.510355.269 | cytochrome b5, putative | 0,5586 | 0,004235 |
| TcCLB.511817.120 | hypothetical protein, conserved | 0,5578 | 9,63E-09 |
| TcCLB.510829.10 | hypothetical protein, conserved | 0,5577 | 1,357E-06 |
| TcCLB.504429.70 | retrotransposon hot spot protein (RHS, pseudogene), puta | 0,5576 | 6,702E-09 |
| TcCLB.508859.40 | leucine-rich repeat protein (LRRP), putative | 0,5575 | 4,062E-06 |
| TcCLB.504131.160 | hypothetical protein | 0,5575 | 0,00001637 |
| TcCLB.507703.10 | retrotransposon hot spot protein (RHS, pseudogene), puta | 0,5575 | 0,005327 |
| TcCLB.504105.250 | hypothetical protein | 0,5574 | 0,0002105 |
| TcCLB.503843.30 | hypothetical protein, conserved | 0,5573 | 4,08E-08 |
| TcCLB.508445.99 | pumilio/PUF RNA binding protein 7, putative (PUF7) | 0,5572 | 1,477E-06 |
| TcCLB.508173.140 | hypothetical protein | 0,5571 | 0,00001374 |
| TcCLB.511577.170 | hypothetical protein, conserved (pseudogene) | 0,5569 | 6,191E-06 |
| TcCLB.511317.60 | hypothetical protein, conserved | 0,5569 | 0,002193 |
| TcCLB.507133.10 | trans-sialidase, putative | 0,5569 | 0,004407 |
| TcCLB.509471.77 | histone H3, putative | 0,5568 | 3,662E-09 |
| TcCLB.511227.20 | retrotransposon hot spot protein (RHS, pseudogene), puta | 0,5567 | 3,819E-10 |
| TcCLB.507297.10 | cruzipain precursor, putative | 0,5565 | 2,045E-13 |
| TcCLB.510679.30 | metal-ion transporter, putative | 0,5565 | 3,4E-09 |
| TcCLB.509259.160 | retrotransposon hot spot (RHS) protein, putative | 0,5565 | 2,864E-06 |
| TcCLB.503745.10 | hypothetical protein, conserved | 0,5565 | 3,597E-06 |
| TcCLB.503925.10 | hypothetical protein, conserved | 0,5565 | 0,00004246 |
| TcCLB.509999.100 | hypothetical protein, conserved | 0,5565 | 0,00006406 |
| TcCLB.511473.3 | hypothetical protein, conserved (pseudogene) | 0,5564 | 0,001401 |
| TcCLB.506201.120 | hypothetical protein, conserved | 0,5563 | 1,423E-07 |
| TcCLB.511807.284 | carnitine O-palmitoyltransferase II, putative | 0,5562 | 1,123E-07 |
| TcCLB.507019.80 | hypothetical protein, conserved | 0,5561 | 6,565E-13 |
| TcCLB.508271.30 | hypothetical protein, conserved | 0,556 | 6,927E-07 |
| TcCLB.507975.70 | retrotransposon hot spot protein (RHS, pseudogene), puta | 0,5559 | 5,361E-09 |
| TcCLB.511471.150 | hypothetical protein, conserved (pseudogene) | 0,5559 | 0,01076 |
| TcCLB.510631.11 | retrotransposon hot spot protein (RHS, pseudogene), puta | 0,5558 | 1,837E-07 |
| TcCLB.510167.4 | hypothetical protein, conserved | 0,5557 | 0,003073 |
| TcCLB.507789.9 | dispersed gene family protein 1 (DGF-1), putative (fragme | 0,5555 | 0,02937 |
| TcCLB.503959.10 | hypothetical protein, conserved | 0,5553 | 5,805E-09 |
| TcCLB.509661.21 | ferric reductase (pseudogene), putative | 0,5553 | 0,0001165 |
| TcCLB.508821.20 | hypothetical protein, conserved | 0,555 | 1,987E-07 |

| | | | |
|---|---|---|---|
| TcCLB.508601.30 | DNA-directed RNA polymerase I/III subunit, putative | 0,5548 | 0,00004005 |
| TcCLB.510285.70 | kinetoplastid kinetochore protein 2, putative (kkt2) | 0,554 | 6,161E-07 |
| TcCLB.508661.40 | hypothetical protein, conserved | 0,5537 | 0,00005393 |
| TcCLB.507129.40 | hypothetical protein, conserved | 0,5537 | 0,002117 |
| TcCLB.511589.90 | hypothetical protein, conserved | 0,5534 | 3,488E-07 |
| TcCLB.508873.360 | retrotransposon hot spot protein (RHS, pseudogene), puta | 0,5533 | 0,02086 |
| TcCLB.509841.20 | hypothetical protein | 0,5532 | 0,02689 |
| TcCLB.508871.180 | hypothetical protein | 0,5532 | 0,07208 |
| TcCLB.506591.20 | U-box domain protein, putative | 0,5531 | 1,027E-09 |
| TcCLB.511289.14 | mitogen-activated protein kinase kinase kinase, putative ( | 0,553 | 1,262E-09 |
| TcCLB.508891.60 | serine/threonine protein phosphatase, putative | 0,553 | 0,000286 |
| TcCLB.507831.60 | heat shock protein 110, putative | 0,5527 | 2,969E-12 |
| TcCLB.510943.20 | hypothetical protein | 0,5527 | 0,00255 |
| TcCLB.504257.70 | valyl-tRNA synthetase, putative | 0,5526 | 4,403E-11 |
| TcCLB.509401.20 | cysteine peptidase (pseudogene), putative | 0,5525 | 9,627E-13 |
| TcCLB.508269.30 | translation initiation factor 2 subunit, putative | 0,5524 | 0,00001469 |
| TcCLB.506517.40 | hypothetical protein | 0,5524 | 0,02131 |
| TcCLB.509913.20 | hypothetical protein, conserved | 0,5523 | 9,533E-07 |
| TcCLB.509601.150 | hypothetical protein, conserved | 0,5523 | 3,497E-06 |
| TcCLB.508625.140 | hypothetical protein, conserved | 0,5522 | 0,0001884 |
| TcCLB.510659.80 | hypothetical protein, conserved | 0,5521 | 1,047E-06 |
| TcCLB.508711.30 | NADH dehydrogenase subunit NI8M, putative | 0,5521 | 6,665E-06 |
| TcCLB.509023.20 | hypothetical protein, conserved | 0,5521 | 0,0009279 |
| TcCLB.509549.90 | alpha-adaptin-like (pseudogene), putative | 0,5518 | 1,776E-12 |
| TcCLB.508857.50 | hypothetical protein, conserved | 0,5517 | 9,719E-14 |
| TcCLB.507519.20 | hypothetical protein, conserved | 0,5517 | 1,705E-08 |
| TcCLB.463323.20 | retrotransposon hot spot protein (RHS, pseudogene), puta | 0,5517 | 0,0008962 |
| TcCLB.506747.10 | orotidine-5-phosphate decarboxylase/orotate phosphoribo | 0,5517 | 0,004281 |
| TcCLB.506691.14 | ATP-dependent DEAD/H RNA helicase, putative (fragment | 0,5516 | 7,554E-07 |
| TcCLB.506605.229 | SNF2 DNA repair protein, putative | 0,5515 | 3,099E-08 |
| TcCLB.507603.200 | proteasome regulatory ATPase subunit 3, putative | 0,5514 | 2,282E-09 |
| TcCLB.503421.50 | retrotransposon hot spot protein (RHS, pseudogene), puta | 0,551 | 6,962E-09 |
| TcCLB.507519.210 | glycine dehydrogenase [decarboxylating] (pseudogene), p | 0,551 | 0,00002699 |
| TcCLB.510241.5 | hypothetical protein | 0,5509 | 1,395E-07 |
| TcCLB.508515.30 | RNA-binding protein 38, putative (RBP38) | 0,5508 | 1,315E-15 |
| TcCLB.507215.9 | dispersed gene family protein 1 (DGF-1), putative (fragme | 0,5508 | 0,1776 |
| TcCLB.511577.120 | hypothetical protein, conserved | 0,5507 | 0,00001945 |
| TcCLB.507091.30 | succinate dehydrogenase subunit | 0,5506 | 0,005955 |
| TcCLB.422189.20 | leucine-rich repeat protein, putative | 0,5503 | 0,000299 |
| TcCLB.508175.146 | 60S ribosomal protein L23a, putative | 0,5501 | 8,023E-13 |
| TcCLB.506247.230 | prolyl endopeptidase (POP) | 0,55 | 2,81E-09 |
| TcCLB.503499.30 | mitochondrial carrier protein (pseudogene), putative | 0,55 | 0,0005577 |
| TcCLB.511163.40 | cell cycle sequence binding phosphoprotein (RBP45), puta | 0,5499 | 1,142E-08 |
| TcCLB.503897.120 | hypothetical protein, conserved | 0,5498 | 1,82E-07 |
| TcCLB.506559.40 | cardiolipin synthetase, putative (cls) | 0,5495 | 5,636E-11 |

| | | | |
|---|---|---|---|
| TcCLB.509937.180 | tyrosyl-tRNA synthetase, putative | 0,5494 | 2,488E-08 |
| TcCLB.510101.400 | hypothetical protein, conserved | 0,5492 | 1,549E-10 |
| TcCLB.510861.60 | hypothetical protein, conserved | 0,5492 | 3,828E-10 |
| TcCLB.503499.24 | hypothetical protein | 0,5488 | 0,02787 |
| TcCLB.506739.110 | hypothetical protein | 0,5487 | 0,00002997 |
| TcCLB.509551.149 | phosphatidylserine decarboxylase, putative | 0,5486 | 0,005444 |
| TcCLB.505171.10 | phospholipid transporting ATPase-like protein, putative | 0,5484 | 1,157E-06 |
| TcCLB.503647.9 | hypothetical protein, conserved | 0,5483 | 1,975E-07 |
| TcCLB.508347.70 | Engulfment and cell motility domain-containing protein (F | 0,5482 | 1,857E-06 |
| TcCLB.504247.50 | hypothetical protein, conserved | 0,5481 | 0,00001214 |
| TcCLB.504179.100 | hypothetical protein, conserved | 0,5473 | 0,0001894 |
| TcCLB.509965.370 | adenosine kinase, putative | 0,5472 | 6,139E-10 |
| TcCLB.504161.59 | transporter, putative | 0,547 | 0,0001614 |
| TcCLB.503999.30 | hypothetical protein, conserved | 0,5468 | 9,857E-12 |
| TcCLB.505935.30 | dispersed gene family protein 1 (DGF-1), putative | 0,5466 | 0,126 |
| TcCLB.506173.200 | retrotransposon hot spot protein (RHS, pseudogene), puta | 0,5465 | 0,0005413 |
| TcCLB.504213.40 | hypothetical protein, conserved | 0,5464 | 2,133E-07 |
| TcCLB.506885.420 | hypothetical protein | 0,5464 | 0,0005274 |
| TcCLB.505807.250 | hypothetical protein, conserved | 0,5463 | 0,00002315 |
| TcCLB.508815.10 | hypothetical protein, conserved | 0,5462 | 0,00006738 |
| TcCLB.506445.60 | mitochondrial DNA topoisomerase II, putative | 0,5461 | 1,01E-11 |
| TcCLB.509117.10 | ubiquitin-activating enzyme, putative | 0,5461 | 0,00002009 |
| TcCLB.510665.30 | hypothetical protein, conserved | 0,5459 | 0,00005269 |
| TcCLB.510311.110 | hypothetical protein, conserved | 0,5456 | 1,235E-06 |
| TcCLB.510029.40 | DNA-directed RNA polymerase III subunit C11, putative (R | 0,5456 | 0,0002103 |
| TcCLB.511303.50 | hypothetical protein, conserved | 0,5454 | 0,0008894 |
| TcCLB.506507.10 | N-acetylglucosamine-6-phosphate deacetylase-like protei | 0,5452 | 2,353E-08 |
| TcCLB.506975.20 | hypothetical protein, conserved | 0,5445 | 0,00009021 |
| TcCLB.503669.10 | hypothetical protein, conserved | 0,5443 | 3,095E-09 |
| TcCLB.509721.70 | hypothetical protein, conserved | 0,5443 | 0,0001071 |
| TcCLB.506529.470 | hypothetical protein, conserved | 0,5442 | 3,523E-12 |
| TcCLB.510799.9 | hypothetical protein, conserved | 0,5442 | 0,0001272 |
| TcCLB.506401.90 | centrin, putative | 0,544 | 2,134E-06 |
| TcCLB.506529.524 | hypothetical protein, conserved | 0,5439 | 0,003796 |
| TcCLB.510629.210 | dispersed gene family protein 1 (DGF-1), putative | 0,5439 | 0,1157 |
| TcCLB.506963.70 | hypothetical protein, conserved | 0,5436 | 0,00003613 |
| TcCLB.480561.10 | trans-sialidase, putative | 0,5435 | 0,00115 |
| TcCLB.509109.114 | mitochondrial carrier protein, putative | 0,5434 | 0,00001214 |
| TcCLB.510187.250 | hypothetical protein, conserved | 0,5434 | 0,0002297 |
| TcCLB.508015.30 | hypothetical protein | 0,5434 | 0,008602 |
| TcCLB.506631.10 | hypothetical protein, conserved | 0,5431 | 0,0001616 |
| TcCLB.510531.70 | hypothetical protein, conserved | 0,543 | 0,002149 |
| TcCLB.510357.90 | hypothetical protein, conserved | 0,5429 | 0,0004156 |
| TcCLB.507037.40 | hypothetical protein, conserved (pseudogene) | 0,5428 | 7,392E-10 |
| TcCLB.506859.249 | hypothetical protein, conserved | 0,5426 | 0,0004929 |

| | | | |
|---|---|---|---|
| TcCLB.506315.50 | 60S ribosomal protein L13a, putative | 0,5425 | 5,58E-12 |
| TcCLB.510359.30 | hypothetical protein, conserved | 0,5424 | 0,00007014 |
| TcCLB.509999.130 | hypothetical protein, conserved | 0,5423 | 1,903E-07 |
| TcCLB.509011.60 | hypothetical protein, conserved | 0,5423 | 3,548E-07 |
| TcCLB.503931.46 | hypothetical protein, conserved (pseudogene) | 0,5421 | 0,02628 |
| TcCLB.506595.65 | hypothetical protein | 0,542 | 0,001179 |
| TcCLB.507907.60 | retrotransposon hot spot (RHS) protein, putative | 0,542 | 0,002744 |
| TcCLB.509429.320 | cysteine peptidase, putative | 0,5419 | 2,073E-14 |
| TcCLB.511283.140 | hypothetical protein, conserved | 0,5419 | 1,053E-10 |
| TcCLB.510609.20 | hypothetical protein, conserved | 0,5416 | 5,316E-08 |
| TcCLB.508425.20 | tryptophanyl-tRNA synthetase (pseudogene), putative | 0,5415 | 0,0009084 |
| TcCLB.507519.164 | hypothetical protein, conserved | 0,5414 | 0,001123 |
| TcCLB.503995.40 | hypothetical protein, conserved | 0,5413 | 0,0006351 |
| TcCLB.509395.90 | hypothetical protein, conserved | 0,5412 | 0,0006855 |
| TcCLB.506953.30 | protein kinase, putative | 0,541 | 0,0001282 |
| TcCLB.506529.595 | hypothetical protein | 0,5408 | 0,00000177 |
| TcCLB.509765.30 | retrotransposon hot spot protein (RHS, pseudogene), puta | 0,5408 | 0,0003182 |
| TcCLB.506513.130 | kinesin (pseudogene), putative | 0,5407 | 3,043E-10 |
| TcCLB.509561.70 | hypothetical protein, conserved | 0,5405 | 1,052E-08 |
| TcCLB.507591.15 | trans-sialidase (pseudogene), putative | 0,5404 | 0,0001425 |
| TcCLB.436535.19 | Glucose-6-phosphate dehydrogenase, putative | 0,5402 | 0,04207 |
| TcCLB.503465.49 | mitochondrial processing peptidase, beta subunit, putativ | 0,5401 | 0,00002366 |
| TcCLB.510747.90 | hypothetical protein | 0,5399 | 0,00000105 |
| TcCLB.511649.120 | hypothetical protein, conserved | 0,5399 | 3,916E-06 |
| TcCLB.510121.180 | hypothetical protein, conserved | 0,5398 | 6,352E-06 |
| TcCLB.511585.20 | KREL1 | 0,5397 | 5,207E-09 |
| TcCLB.509259.80 | hypothetical protein, conserved (pseudogene) | 0,5396 | 0,00003569 |
| TcCLB.506337.4 | short-chain dehydrogenase, putative (fragment) | 0,5395 | 0,0003209 |
| TcCLB.509181.9 | dispersed gene family protein 1 (DGF-1), putative | 0,5393 | 0,1169 |
| TcCLB.511589.130 | 14-3-3 protein 1, putative (14-3-3-I) | 0,5392 | 2,904E-09 |
| TcCLB.509719.69 | hypothetical protein, conserved | 0,5392 | 0,0000192 |
| TcCLB.504253.10 | hypothetical protein, conserved | 0,5385 | 1,701E-08 |
| TcCLB.509215.10 | DNA polymerase delta catalytic subunit, putative | 0,5383 | 3,349E-11 |
| TcCLB.504427.80 | hypothetical protein, conserved | 0,5383 | 0,00003954 |
| TcCLB.511727.220 | hypothetical protein, conserved | 0,5382 | 3,087E-06 |
| TcCLB.506439.9 | myosin heavy chain, putative | 0,5381 | 0,00005959 |
| TcCLB.511393.88 | hypothetical protein, conserved (fragment) | 0,5377 | 1,244E-07 |
| TcCLB.508409.60 | hypothetical protein, conserved | 0,5376 | 2,52E-07 |
| TcCLB.504033.160 | hypothetical protein | 0,5376 | 0,00000405 |
| TcCLB.504055.100 | retrotransposon hot spot protein (RHS, pseudogene), puta | 0,5374 | 0,0587 |
| TcCLB.507897.40 | acetyl-CoA carboxylase, putative | 0,5372 | 2,838E-07 |
| TcCLB.506617.90 | ATP-dependent Clp protease subunit, heat shock protein 1 | 0,5368 | 0,00002169 |
| TcCLB.506925.100 | hypothetical protein, conserved | 0,5367 | 0,00007925 |
| TcCLB.508065.50 | phosphatidylinositol-4-phosphate 5-kinase-like protein, pu | 0,5365 | 8,087E-09 |
| TcCLB.507997.50 | hypothetical protein, conserved | 0,5362 | 0,00002824 |

| | | | |
|---|---|---|---|
| TcCLB.507679.20 | retrotransposon hot spot protein (RHS, pseudogene), puta | 0,5362 | 0,06588 |
| TcCLB.508207.120 | hypothetical protein, conserved | 0,5361 | 0,009435 |
| TcCLB.509039.20 | bystin, putative | 0,5357 | 6,714E-10 |
| TcCLB.511799.9 | hypothetical protein, conserved (fragment) | 0,5355 | 4,572E-08 |
| TcCLB.510303.200 | glucosamine-fructose-6-phosphate aminotransferase, put | 0,5354 | 0,00002199 |
| TcCLB.505333.10 | N-acetyltransferase complex ARD1 subunit (pseudogene), | 0,5353 | 0,04683 |
| TcCLB.510105.184 | hypothetical protein, conserved | 0,5352 | 0,0006476 |
| TcCLB.510515.139 | T-complex protein 1, gamma subunit, putative | 0,5351 | 9,549E-06 |
| TcCLB.511819.50 | hypothetical protein, conserved | 0,535 | 1,726E-10 |
| TcCLB.508501.260 | hypothetical protein, conserved | 0,5348 | 6,021E-06 |
| TcCLB.511661.40 | signal peptidase type I, putative | 0,5348 | 0,001133 |
| TcCLB.505047.19 | trans-sialidase (pseudogene), putative | 0,5348 | 0,1679 |
| TcCLB.508999.40 | hypothetical protein, conserved | 0,5345 | 7,818E-06 |
| TcCLB.476125.9 | dispersed gene family protein 1 (DGF-1), putative | 0,5344 | 0,02276 |
| TcCLB.508411.30 | hypothetical protein, conserved | 0,5343 | 1,563E-08 |
| TcCLB.511367.360 | elongation factor 1-alpha, putative | 0,5341 | 1,461E-16 |
| TcCLB.503405.10 | glutamine synthetase, putative | 0,5339 | 1,463E-09 |
| TcCLB.508547.40 | hypothetical protein, conserved | 0,5339 | 0,0002204 |
| TcCLB.507093.250 | RNA-binding protein, putative | 0,5336 | 3,329E-11 |
| TcCLB.506211.200 | metallo-peptidase, Clan MG, Family M24 | 0,5336 | 3,882E-11 |
| TcCLB.510255.10 | hypothetical protein, conserved | 0,5336 | 0,00007607 |
| TcCLB.504149.40 | tatD related deoxyribonuclease, putative | 0,5334 | 4,071E-11 |
| TcCLB.511739.10 | hypothetical protein, conserved | 0,5334 | 2,572E-06 |
| TcCLB.509409.10 | protein kinase, putative | 0,5334 | 0,0000208 |
| TcCLB.509391.30 | flagellar calcium-binding protein, putative | 0,5334 | 0,2665 |
| TcCLB.511141.20 | choline dehydrogenase, putative | 0,5332 | 3,434E-12 |
| TcCLB.504055.60 | hypothetical protein, conserved | 0,5332 | 0,0003554 |
| TcCLB.503899.20 | 40S ribosomal protein S16, putative | 0,5331 | 5,166E-12 |
| TcCLB.503437.21 | N-acetyltransferase complex ARD1 subunit (pseudogene), | 0,5331 | 0,1398 |
| TcCLB.510301.70 | hypothetical protein, conserved | 0,533 | 4,962E-08 |
| TcCLB.511817.30 | hypothetical protein, conserved | 0,5329 | 0,0003012 |
| TcCLB.506605.30 | hypothetical protein, conserved | 0,5326 | 0,00006009 |
| TcCLB.506147.220 | hypothetical protein, conserved | 0,5325 | 8,172E-08 |
| TcCLB.503575.27 | hypothetical protein, conserved | 0,5325 | 2,396E-07 |
| TcCLB.511365.80 | mitochondrial carrier protein, putative | 0,5325 | 9,603E-06 |
| TcCLB.506779.70 | hypothetical protein, conserved | 0,5323 | 0,00004566 |
| TcCLB.508351.20 | hypothetical protein, conserved | 0,5321 | 0,003672 |
| TcCLB.506573.30 | tyrosyl or methionyl-tRNA synthetase, putative | 0,5319 | 0,0001093 |
| TcCLB.506147.30 | synaptojanin (N-terminal domain), putative | 0,5315 | 1,421E-08 |
| TcCLB.507093.229 | U-rich RNA-binding protein UBP-2 | 0,5315 | 2,596E-08 |
| TcCLB.508337.10 | hypothetical protein, conserved (fragment) | 0,5315 | 7,999E-07 |
| TcCLB.507061.40 | hypothetical protein, conserved (pseudogene) | 0,5314 | 2,924E-07 |
| TcCLB.511763.19 | hypothetical protein, conserved | 0,5313 | 0,00011 |
| TcCLB.506957.46 | hypothetical protein, conserved | 0,5312 | 0,0001443 |
| TcCLB.506537.260 | protein kinase, putative | 0,5311 | 0,08092 |

| | | | |
|---|---|---|---|
| TcCLB.508221.64 | syntaxin binding protein (pseudogene), putative | 0,5311 | 0,08786 |
| TcCLB.506127.170 | eukaryotic translation initiation factor eIF-4E, putative | 0,5308 | 2,457E-08 |
| TcCLB.505123.10 | hypothetical protein, conserved | 0,5308 | 0,007387 |
| TcCLB.503431.109 | nucleoporin interacting component (NUP93), putative | 0,5304 | 0,02568 |
| TcCLB.511419.40 | phosphoglycerate kinase, putative | 0,5303 | 7,1E-08 |
| TcCLB.508503.90 | hypothetical protein, conserved | 0,5303 | 0,00001218 |
| TcCLB.508153.330 | vacuolar protein sorting-associated protein 4, putative (VI | 0,5301 | 9,969E-12 |
| TcCLB.511731.80 | hypothetical protein, conserved | 0,5301 | 3,909E-06 |
| TcCLB.508501.190 | hypothetical protein, conserved | 0,5301 | 0,0004738 |
| TcCLB.510643.50 | retrotransposon hot spot protein (RHS, pseudogene), puta | 0,5299 | 1,904E-06 |
| TcCLB.509281.30 | hypothetical protein | 0,5299 | 0,0007822 |
| TcCLB.505071.40 | hypothetical protein | 0,5296 | 0,000205 |
| TcCLB.511063.180 | trans-sialidase (pseudogene), putative | 0,5296 | 0,03645 |
| TcCLB.506201.170 | flagellar attachment zone protein, putative (FAZ1) | 0,5294 | 2,673E-09 |
| TcCLB.510149.70 | hypothetical protein, conserved | 0,5293 | 1,638E-08 |
| TcCLB.508669.30 | hypothetical protein | 0,5292 | 4,459E-06 |
| TcCLB.508895.30 | Sarcalumenin, putative | 0,5291 | 1,618E-10 |
| TcCLB.507527.10 | hypothetical protein, conserved | 0,5291 | 1,709E-06 |
| TcCLB.509767.50 | hypothetical protein, conserved | 0,5287 | 7,51E-07 |
| TcCLB.506251.120 | hypothetical protein, conserved | 0,5283 | 0,001606 |
| TcCLB.425311.9 | hypothetical protein, conserved (fragment) | 0,5282 | 2,161E-06 |
| TcCLB.509023.120 | ubiquitin carboxyl-terminal hydrolase, putative | 0,5281 | 2,18E-07 |
| TcCLB.506435.150 | hypothetical protein, conserved | 0,5281 | 2,378E-07 |
| TcCLB.506155.50 | vacuolar protein sorting-associated protein-like, putative | 0,5279 | 0,00002367 |
| TcCLB.503971.20 | hypothetical protein, conserved | 0,5279 | 0,00005669 |
| TcCLB.509349.20 | retrotransposon hot spot (RHS) protein, putative | 0,5273 | 3,381E-07 |
| TcCLB.508319.60 | hypothetical protein, conserved | 0,5272 | 6,34E-08 |
| TcCLB.508693.150 | hypothetical protein, conserved | 0,5272 | 0,00007226 |
| TcCLB.510785.15 | hypothetical protein, conserved | 0,527 | 0,01327 |
| TcCLB.511815.160 | hypothetical protein, conserved | 0,5268 | 2,191E-09 |
| TcCLB.508911.10 | hypothetical protein, conserved | 0,5267 | 4,013E-10 |
| TcCLB.504231.5 | hypothetical protein, conserved (pseudogene) | 0,5261 | 0,01637 |
| TcCLB.509855.20 | Probable methyltransferase BTM2 homolog | 0,5258 | 5,214E-07 |
| TcCLB.508741.350 | hypothetical protein, conserved | 0,5258 | 5,24E-07 |
| TcCLB.508459.18 | phosphatidylserine decarboxylase, putative | 0,5258 | 0,04916 |
| TcCLB.511707.39 | hypothetical protein, conserved | 0,5257 | 9,895E-09 |
| TcCLB.508231.170 | hypothetical protein, conserved | 0,5256 | 5,796E-08 |
| TcCLB.508169.9 | elongation factor 2, putative (fragment) | 0,5255 | 1,407E-08 |
| TcCLB.510713.70 | retrotransposon hot spot protein (RHS, pseudogene), puta | 0,5255 | 0,0002278 |
| TcCLB.509551.64 | small nuclear ribonucleoprotein Sm-F, putative | 0,5255 | 0,009183 |
| TcCLB.508411.90 | tfiih basal transcription factor complex helicase xpb subur | 0,5254 | 4,857E-10 |
| TcCLB.508181.140 | inorganic pyrophosphatase, putative | 0,5253 | 0,00003951 |
| TcCLB.508781.60 | hypothetical protein, conserved | 0,5253 | 0,00005627 |
| TcCLB.511717.200 | hypothetical protein, conserved | 0,5252 | 0,00001325 |
| TcCLB.503823.120 | hypothetical protein, conserved | 0,5252 | 0,0002432 |

| | | | |
|---|---|---|---|
| TcCLB.506529.590 | hypothetical protein, conserved | 0,5251 | 0,00006754 |
| TcCLB.509921.60 | dispersed gene family protein 1 (DGF-1), putative | 0,525 | 0,02135 |
| TcCLB.507773.30 | RNA editing complex protein MP46, putative | 0,5249 | 0,00006239 |
| TcCLB.511491.110 | rab6 GTPase activating protein, putative | 0,5249 | 0,0001205 |
| TcCLB.503749.20 | hypothetical protein | 0,5248 | 3,154E-06 |
| TcCLB.509891.10 | hypothetical protein, conserved | 0,5248 | 0,0003273 |
| TcCLB.508573.39 | dispersed gene family protein 1 (DGF-1), putative | 0,5247 | 0,002434 |
| TcCLB.509753.290 | hypothetical protein, conserved (pseudogene) | 0,5246 | 0,05029 |
| TcCLB.508285.20 | hypothetical protein, conserved (pseudogene) | 0,5245 | 0,0006055 |
| TcCLB.506245.110 | dispersed gene family protein 1 (DGF-1, pseudogene), put | 0,5245 | 0,1321 |
| TcCLB.508461.370 | hypothetical protein, conserved | 0,5244 | 3,339E-06 |
| TcCLB.511567.20 | retrotransposon hot spot protein (RHS, pseudogene), puta | 0,5244 | 0,005309 |
| TcCLB.506051.30 | dispersed gene family protein 1 (DGF-1), putative | 0,5242 | 0,0624 |
| TcCLB.509167.100 | valyl-tRNA synthetase, putative | 0,5241 | 7,99E-14 |
| TcCLB.507803.40 | hypothetical protein | 0,524 | 1,465E-08 |
| TcCLB.506147.130 | hypothetical protein, conserved | 0,5238 | 0,0139 |
| TcCLB.509027.20 | ankyrin repeat protein, putative | 0,5237 | 0,0002568 |
| TcCLB.509303.10 | retrotransposon hot spot protein (RHS, pseudogene), puta | 0,5236 | 0,002192 |
| TcCLB.504431.64 | high mobility group protein, putative | 0,5235 | 2,082E-06 |
| TcCLB.506743.100 | hypothetical protein, conserved | 0,5231 | 0,024 |
| TcCLB.509393.30 | hypothetical protein, conserved | 0,523 | 4,436E-07 |
| TcCLB.503593.10 | eukaryotic translation initiation factor eIF-4E, putative | 0,523 | 1,326E-06 |
| TcCLB.510101.380 | hypothetical protein, conserved | 0,5229 | 7,939E-06 |
| TcCLB.447705.9 | hypothetical protein, conserved | 0,5229 | 0,0232 |
| TcCLB.506999.140 | hypothetical protein, conserved | 0,5227 | 1,29E-08 |
| TcCLB.507841.34 | exosome-associated protein 4, putative | 0,5227 | 0,0003426 |
| TcCLB.508831.100 | hypothetical protein, conserved | 0,5226 | 0,00000342 |
| TcCLB.510901.10 | hypothetical protein, conserved | 0,5225 | 0,00001301 |
| TcCLB.508347.159 | microtubule-associated protein, putative, (fragment) (GB4 | 0,5224 | 0,00004088 |
| TcCLB.504013.90 | hypothetical protein, conserved | 0,5223 | 2,306E-09 |
| TcCLB.510533.130 | hypothetical protein, conserved | 0,5223 | 1,939E-06 |
| TcCLB.507507.10 | hypothetical protein, conserved | 0,5223 | 6,039E-06 |
| TcCLB.511817.50 | hypothetical protein, conserved | 0,5223 | 0,00002013 |
| TcCLB.504081.344 | elongation factor 1-gamma (EF-1-gamma, pseudogene), | 0,5223 | 0,2393 |
| TcCLB.420293.40 | beta galactofuranosyl glycosyltransferase (pseudogene), | 0,5222 | 0,002853 |
| TcCLB.511827.120 | retrotransposon hot spot protein (RHS, pseudogene), puta | 0,5215 | 9,006E-09 |
| TcCLB.504119.10 | eukaryotic translation initiation factor 5, putative | 0,5214 | 1,432E-09 |
| TcCLB.510435.10 | hypothetical protein, conserved | 0,5214 | 0,0007177 |
| TcCLB.506367.20 | hypothetical protein, conserved | 0,5213 | 1,647E-07 |
| TcCLB.511671.70 | glucosamine 6-phosphate n-acetyltransferase, putative | 0,5212 | 0,000685 |
| TcCLB.511903.45 | tuzin-like (pseudogene), putative | 0,5211 | 0,00004511 |
| TcCLB.511277.600 | UDP-galactopyranose mutase (GLF) | 0,521 | 1,406E-08 |
| TcCLB.510659.230 | hypothetical protein, conserved | 0,521 | 0,0002053 |
| TcCLB.506795.90 | hypothetical protein, conserved | 0,5209 | 4,671E-09 |
| TcCLB.510897.6 | Dynein light chain LC6, flagellar outer arm, putative | 0,5208 | 0,00004695 |

| | | | |
|---|---|---|---|
| TcCLB.511219.60 | retrotransposon hot spot protein (RHS, pseudogene), puta | 0,5208 | 0,0002086 |
| TcCLB.508969.30 | retrotransposon hot spot protein (RHS, pseudogene), puta | 0,5208 | 0,003751 |
| TcCLB.511889.40 | dispersed gene family protein 1 (DGF-1), putative | 0,5207 | 0,001127 |
| TcCLB.503657.20 | hypothetical protein, conserved | 0,5205 | 0,000156 |
| TcCLB.507031.120 | metallo- peptidase, Clan MG, Family M24 | 0,5203 | 9,592E-07 |
| TcCLB.508921.10 | hypothetical protein, conserved | 0,5203 | 0,0002684 |
| TcCLB.511051.70 | aminopeptidase, putative | 0,5201 | 0,00000182 |
| TcCLB.505365.20 | retrotransposon hot spot protein (RHS, pseudogene), puta | 0,52 | 0,00003879 |
| TcCLB.508905.30 | hypothetical protein, conserved (pseudogene) | 0,5199 | 0,00001478 |
| TcCLB.507485.90 | Wee1-like protein kinase, putative | 0,5198 | 2,066E-07 |
| TcCLB.507033.29 | extracellular receptor, putative (fragment) | 0,5198 | 0,00001398 |
| TcCLB.508877.30 | retrotransposon hot spot protein (RHS, pseudogene), puta | 0,5197 | 0,02476 |
| TcCLB.479253.9 | hypothetical protein, conserved (fragment) | 0,5196 | 0,001925 |
| TcCLB.508973.20 | hypothetical protein, conserved | 0,5195 | 1,007E-06 |
| TcCLB.511827.115 | hypothetical protein | 0,5195 | 0,002336 |
| TcCLB.468005.9 | pumilio/PUF RNA binding protein 3, putative (PUF3) | 0,5193 | 6,298E-08 |
| TcCLB.503613.20 | proteasome alpha 7 subunit, putative | 0,5192 | 6,973E-06 |
| TcCLB.509051.50 | hypothetical protein, conserved | 0,5191 | 4,429E-06 |
| TcCLB.509885.70 | transporter, putative | 0,519 | 3,152E-07 |
| TcCLB.507063.110 | hypothetical protein, conserved | 0,5189 | 4,339E-10 |
| TcCLB.503653.9 | hypothetical protein, conserved | 0,5189 | 0,0126 |
| TcCLB.507769.40 | GTP-binding protein, putative | 0,5188 | 7,312E-07 |
| TcCLB.508331.80 | dispersed gene family protein 1 (DGF-1, pseudogene), put | 0,5188 | 0,01051 |
| TcCLB.506459.290 | elongation factor 1-gamma (EF-1-gamma, pseudogene), | 0,5187 | 1,944E-08 |
| TcCLB.510665.10 | hypothetical protein, conserved | 0,5186 | 4,185E-07 |
| TcCLB.509157.60 | 1-acyl-sn-glycerol-3-phosphate acyltransferase, putative | 0,5186 | 3,445E-06 |
| TcCLB.507391.20 | hypothetical protein, conserved (pseudogene) | 0,5185 | 6,565E-07 |
| TcCLB.505051.10 | hypothetical protein, conserved | 0,5184 | 1,72E-19 |
| TcCLB.507777.30 | retrotransposon hot spot protein (RHS, pseudogene), puta | 0,5184 | 2,477E-06 |
| TcCLB.509055.54 | hypothetical protein, conserved | 0,5184 | 0,004915 |
| TcCLB.510657.80 | hypothetical protein, conserved | 0,5183 | 4,196E-07 |
| TcCLB.510747.80 | Double RNA binding domain protein 9 (DRBD9) | 0,5182 | 2,787E-06 |
| TcCLB.511589.50 | cyclophilin-like protein, putative | 0,5182 | 0,00002713 |
| TcCLB.506351.30 | hypothetical protein, conserved | 0,5181 | 0,00003244 |
| TcCLB.508241.30 | hypothetical protein, conserved | 0,518 | 2,276E-07 |
| TcCLB.506431.20 | retrotransposon hot spot protein (RHS, pseudogene), puta | 0,5179 | 0,00002344 |
| TcCLB.510271.20 | dispersed gene family protein 1 (DGF-1, pseudogene), put | 0,5177 | 0,01052 |
| TcCLB.507087.60 | hypothetical protein, conserved | 0,5176 | 0,00001281 |
| TcCLB.511747.30 | hypothetical protein, conserved | 0,5176 | 0,001348 |
| TcCLB.506239.40 | hypothetical protein, conserved | 0,5174 | 0,0006724 |
| TcCLB.503625.20 | hypothetical protein | 0,5174 | 0,008299 |
| TcCLB.508153.620 | hypothetical protein, conserved | 0,5169 | 0,00004991 |
| TcCLB.430605.29 | 40S ribosomal protein S3, putative | 0,5168 | 1,247E-09 |
| TcCLB.511277.10 | DNA-directed RNA polymerase I subunit RPB5z, putative ( | 0,5167 | 0,0001166 |
| TcCLB.504175.9 | hypothetical protein, conserved | 0,5166 | 2,105E-08 |

| | | | |
|---|---|---|---|
| TcCLB.507677.90 | hypothetical protein, conserved | 0,5166 | 4,464E-06 |
| TcCLB.506401.50 | malonyl-CoA decarboxylase, mitochondrial precursor, puta | 0,5165 | 5,239E-08 |
| TcCLB.508175.50 | hypothetical protein, conserved | 0,5164 | 4,044E-08 |
| TcCLB.504109.10 | hypothetical protein, conserved | 0,5164 | 6,068E-08 |
| TcCLB.504427.90 | ribosomal RNA methyltransferase, putative | 0,5164 | 0,00002247 |
| TcCLB.509967.30 | aspartyl-tRNA synthetase, putative | 0,5163 | 9,961E-06 |
| TcCLB.504065.50 | trans-sialidase (pseudogene), putative | 0,5162 | 2,133E-08 |
| TcCLB.509719.26 | hypothetical protein, conserved | 0,5158 | 1,685E-07 |
| TcCLB.505939.20 | oligopeptidase B-like protein, putative | 0,5157 | 2,746E-10 |
| TcCLB.507093.140 | hypothetical protein, conserved | 0,5154 | 5,134E-06 |
| TcCLB.507049.140 | hypothetical protein, conserved | 0,5153 | 1,52E-08 |
| TcCLB.507081.60 | helicase-like protein, putative | 0,5152 | 1,207E-09 |
| TcCLB.503563.10 | calcium-translocating P-type ATPase, putative | 0,5151 | 0,00003529 |
| TcCLB.509109.80 | hypothetical protein, conserved | 0,515 | 2,884E-12 |
| TcCLB.467287.10 | alpha-tubulin N-acetyltransferase, putative | 0,5148 | 0,0001626 |
| TcCLB.506637.80 | hypothetical protein | 0,5147 | 1,541E-06 |
| TcCLB.503917.21 | hypothetical protein, conserved | 0,5146 | 0,00004506 |
| TcCLB.511337.39 | hypothetical protein, conserved | 0,5143 | 0,0009407 |
| TcCLB.510283.110 | hypothetical protein, conserved | 0,514 | 0,001212 |
| TcCLB.503619.25 | hypothetical protein, conserved | 0,514 | 0,003821 |
| TcCLB.508465.30 | hypothetical protein, conserved | 0,5139 | 4,293E-07 |
| TcCLB.503529.30 | hypothetical protein, conserved | 0,5138 | 0,0003118 |
| TcCLB.509671.20 | hypothetical protein, conserved | 0,5137 | 0,00001186 |
| TcCLB.509165.30 | AGP2beta-2, putative | 0,5135 | 3,469E-08 |
| TcCLB.509581.5 | retrotransposon hot spot (RHS) protein, putative (fragmer | 0,5135 | 0,006177 |
| TcCLB.508397.70 | transporter, putative | 0,5133 | 0,000478 |
| TcCLB.511103.9 | dispersed gene family protein 1 (DGF-1), putative (fragme | 0,5132 | 0,2584 |
| TcCLB.503453.10 | hypothetical protein, conserved | 0,5131 | 2,728E-06 |
| TcCLB.506127.20 | Nuclear RNA export factor (MEX67) | 0,513 | 9,285E-07 |
| TcCLB.507007.74 | small nuclear ribonucleoprotein Sm-F, putative | 0,5127 | 0,01969 |
| TcCLB.510761.10 | mismatch repair protein PMS1, putative | 0,5126 | 1,413E-09 |
| TcCLB.509099.140 | hypothetical protein, conserved | 0,5126 | 0,000382 |
| TcCLB.509179.120 | hypothetical protein, conserved | 0,5124 | 0,00326 |
| TcCLB.509733.190 | hypothetical protein, conserved | 0,5123 | 0,0000102 |
| TcCLB.510857.10 | hypothetical protein, conserved | 0,5122 | 3,411E-07 |
| TcCLB.511065.36 | hypothetical protein, conserved | 0,5121 | 1,936E-08 |
| TcCLB.506467.14 | hypothetical protein, conserved | 0,5121 | 0,0009187 |
| TcCLB.510943.80 | ABC transporter, putative | 0,5118 | 1,805E-10 |
| TcCLB.509791.20 | hypothetical protein, conserved | 0,5118 | 1,306E-06 |
| TcCLB.511907.100 | hypothetical protein, conserved | 0,5116 | 0,0000024 |
| TcCLB.504801.10 | beta galactofuranosyl glycosyltransferase (pseudogene), p | 0,5115 | 0,02145 |
| TcCLB.510889.261 | cruzipain precursor, putative | 0,5112 | 0,0005866 |
| TcCLB.508555.60 | Calpain-like protein 1 (CAP5.5) | 0,5109 | 1,351E-06 |
| TcCLB.508899.140 | hypothetical protein | 0,5108 | 0,00008276 |
| TcCLB.510173.30 | hypothetical protein, conserved | 0,5107 | 0,0009015 |

| | | | |
|---|---|---|---|
| TcCLB.507001.14 | Present in the outer mitochondrial membrane proteome 8 | 0,5107 | 0,00175 |
| TcCLB.506311.30 | hypothetical protein, conserved | 0,5105 | 8,116E-06 |
| TcCLB.511889.11 | trans-sialidase (pseudogene), putative | 0,51 | 3,896E-06 |
| TcCLB.503855.50 | hypothetical protein, conserved | 0,5097 | 2,331E-10 |
| TcCLB.506673.30 | hypothetical protein, conserved | 0,5096 | 0,0004402 |
| TcCLB.507603.30 | PITH domain-containing protein | 0,5095 | 0,002321 |
| TcCLB.509959.10 | retrotransposon hot spot protein (RHS, pseudogene), puta | 0,5094 | 0,03302 |
| TcCLB.511859.170 | NADH-cytochrome b5 reductase, putative | 0,5091 | 4,435E-06 |
| TcCLB.507801.144 | hypothetical protein, conserved | 0,5089 | 0,001863 |
| TcCLB.509915.80 | hypothetical protein, conserved (pseudogene) | 0,5089 | 0,005486 |
| TcCLB.510543.130 | ATP-dependent DEAD/H RNA helicase (pseudogene), puta | 0,5089 | 0,1206 |
| TcCLB.503687.49 | hypothetical protein, conserved | 0,5088 | 0,00004599 |
| TcCLB.511473.10 | retrotransposon hot spot protein (RHS, pseudogene), puta | 0,5088 | 0,00007008 |
| TcCLB.508899.100 | hypothetical protein, conserved | 0,5088 | 0,0005922 |
| TcCLB.505939.89 | 5'-3' exoribonuclease A, putative (fragment) (XRNA) | 0,5086 | 0,00002088 |
| TcCLB.508851.200 | hypothetical protein, conserved | 0,5086 | 0,0002629 |
| TcCLB.506885.360 | hypothetical protein, conserved | 0,5082 | 1,986E-06 |
| TcCLB.508413.5 | hypothetical protein, conserved | 0,508 | 0,01149 |
| TcCLB.508441.80 | Mitochondrial-processing peptidase subunit alpha (MPPA) | 0,5079 | 3,796E-10 |
| TcCLB.511003.170 | hypothetical protein, conserved | 0,5075 | 0,0002857 |
| TcCLB.511771.60 | hypothetical protein | 0,5075 | 0,2335 |
| TcCLB.506811.90 | hypothetical protein, conserved | 0,5073 | 4,45E-07 |
| TcCLB.506123.10 | major vault protein, putative | 0,5072 | 0,00001706 |
| TcCLB.506401.210 | hypothetical protein, conserved | 0,5072 | 0,00009232 |
| TcCLB.511665.21 | trans-sialidase (pseudogene), putative | 0,5071 | 0,0002627 |
| TcCLB.510889.120 | alanine aminotransferase, putative | 0,507 | 3,433E-09 |
| TcCLB.509033.40 | hypothetical protein, conserved | 0,507 | 4,374E-06 |
| TcCLB.430895.4 | hypothetical protein, conserved | 0,5068 | 0,2188 |
| TcCLB.509733.20 | hypothetical protein | 0,5067 | 0,009279 |
| TcCLB.507837.50 | 40S ribosomal protein S11, putative | 0,5067 | 0,08013 |
| TcCLB.506737.40 | hypothetical protein, conserved | 0,5066 | 1,616E-07 |
| TcCLB.508461.520 | hypothetical protein, conserved | 0,5065 | 5,313E-12 |
| TcCLB.508211.70 | hypothetical protein, conserved | 0,5065 | 3,444E-06 |
| TcCLB.511761.10 | UDP-sugar pyrophosphorylase (Sloppy) | 0,5065 | 0,00001067 |
| TcCLB.507227.9 | hypothetical protein, conserved (fragment) | 0,5064 | 7,956E-07 |
| TcCLB.508139.230 | retrotransposon hot spot protein (RHS, pseudogene), puta | 0,5062 | 8,102E-06 |
| TcCLB.509901.150 | hypothetical protein, conserved | 0,5062 | 0,002126 |
| TcCLB.508625.10 | elongation factor 2, putative | 0,506 | 8,449E-08 |
| TcCLB.510187.120 | hypothetical protein, conserved | 0,506 | 0,0001013 |
| TcCLB.510689.60 | Double RNA binding domain protein 7 (DRBD7) | 0,5059 | 5,689E-06 |
| TcCLB.509693.20 | plectin, putative | 0,5058 | 1,072E-08 |
| TcCLB.508695.10 | hypothetical protein, conserved | 0,5058 | 0,005551 |
| TcCLB.508805.20 | hypothetical protein, conserved | 0,5055 | 7,487E-09 |
| TcCLB.504343.30 | retrotransposon hot spot (RHS) protein, putative | 0,5054 | 0,0007919 |
| TcCLB.507063.100 | mitochondrial DNA polymerase beta-PAK, putative | 0,5053 | 9,327E-07 |

| | | | |
|---|---|---|---|
| TcCLB.511621.100 | hypothetical protein, conserved | 0,5052 | 0,00001038 |
| TcCLB.511367.370 | elongation factor 1-alpha, putative | 0,505 | 5,899E-15 |
| TcCLB.504169.10 | dispersed gene family protein 1 (DGF-1), putative | 0,5049 | 0,007297 |
| TcCLB.507485.140 | tuzin | 0,5048 | 8,402E-07 |
| TcCLB.510335.30 | hypothetical protein, conserved (pseudogene) | 0,5047 | 2,689E-08 |
| TcCLB.508355.290 | hypothetical protein, conserved | 0,5047 | 0,0001065 |
| TcCLB.511751.170 | protein kinase, putative | 0,5046 | 3,429E-10 |
| TcCLB.507087.20 | hypothetical protein | 0,5046 | 0,00007196 |
| TcCLB.506295.120 | Trk system potassium uptake protein (HKT1) | 0,5044 | 0,0001484 |
| TcCLB.510749.60 | hypothetical protein, conserved | 0,5037 | 0,0000379 |
| TcCLB.508413.20 | hypothetical protein, conserved | 0,5036 | 1,977E-06 |
| TcCLB.507033.20 | histone H3 variant, putative | 0,5036 | 0,00005388 |
| TcCLB.506391.50 | polyubiquitin (pseudogene), putative | 0,5035 | 0,00003698 |
| TcCLB.509715.70 | folylpolyglutamate synthase, putative (FPGS) | 0,5035 | 0,0001332 |
| TcCLB.509623.19 | clathrin coat assembly protein ap19, putative (ClaATc) | 0,5033 | 4,66E-17 |
| TcCLB.511019.84 | GDP-mannose 4,6 dehydratase, putative | 0,5033 | 0,00002775 |
| TcCLB.506831.70 | glutamyl-tRNA synthetase, putative | 0,5031 | 0,00000306 |
| TcCLB.508519.20 | retrotransposon hot spot protein (RHS, pseudogene), puta | 0,5031 | 0,0000211 |
| TcCLB.410589.10 | AMP deaminase, putative | 0,5029 | 0,002096 |
| TcCLB.504163.40 | fructose-bisphosphate aldolase, glycosomal, putative | 0,5028 | 0,00002929 |
| TcCLB.508397.40 | succinate dehydrogenase subunit | 0,5027 | 0,00006106 |
| TcCLB.503929.30 | hypothetical protein, conserved | 0,5026 | 1,832E-07 |
| TcCLB.506211.170 | Multiprotein-bridging factor 1, putative (mbf1) | 0,5026 | 8,424E-07 |
| TcCLB.503457.4 | hypothetical protein, conserved | 0,5024 | 0,01761 |
| TcCLB.510359.170 | AAA ATPase, putative | 0,5023 | 4,366E-11 |
| TcCLB.416503.10 | trans-sialidase (pseudogene), putative | 0,5022 | 0,00001179 |
| TcCLB.509601.160 | hypothetical protein, conserved | 0,5022 | 0,002229 |
| TcCLB.509353.60 | calmodulin, putative | 0,5021 | 7,987E-09 |
| TcCLB.506355.100 | hexose transporter (pseudogene), putative | 0,5019 | 0,0005053 |
| TcCLB.503957.120 | retrotransposon hot spot (RHS) protein, putative | 0,5018 | 3,999E-06 |
| TcCLB.504131.20 | hypothetical protein, conserved | 0,5016 | 1,594E-09 |
| TcCLB.506699.40 | katanin-like protein, putative | 0,5016 | 0,0007567 |
| TcCLB.506561.14 | hypothetical protein, conserved | 0,5015 | 0,001474 |
| TcCLB.507011.60 | structural maintenance of chromosome (SMC) family prot | 0,5014 | 8,916E-09 |
| TcCLB.504243.40 | hypothetical protein, conserved | 0,5012 | 5,905E-08 |
| TcCLB.503539.10 | hypothetical protein, conserved | 0,5012 | 0,003432 |
| TcCLB.508799.70 | pumilio-repeat, RNA-binding protein, putative | 0,5011 | 1,545E-07 |
| TcCLB.511393.10 | nonspecific lipid-transfer protein, putative | 0,5011 | 3,962E-06 |
| TcCLB.511277.290 | aconitase, putative | 0,5011 | 0,00001369 |
| TcCLB.509661.10 | retrotransposon hot spot protein (RHS, pseudogene), puta | 0,5011 | 0,0003779 |
| TcCLB.503909.44 | hypothetical protein, conserved | 0,501 | 0,00008051 |
| TcCLB.504797.130 | hypothetical protein, conserved | 0,501 | 0,001184 |
| TcCLB.417203.9 | dispersed gene family protein 1 (DGF-1), putative | 0,501 | 0,06147 |
| TcCLB.508325.310 | hypothetical protein, conserved | 0,5007 | 0,008425 |
| TcCLB.508183.10 | ubiquinone biosynthesis protein COQ7 homolog, putative | 0,5005 | 0,0003133 |

| | | | |
|---|---|---|---|
| TcCLB.506525.150 | hypothetical protein, conserved | 0,5004 | 1,185E-08 |
| TcCLB.510531.50 | hypothetical protein, conserved | 0,5002 | 1,698E-06 |
| TcCLB.506937.20 | hypothetical protein, conserved | 0,5002 | 0,01569 |
| TcCLB.508755.20 | dispersed gene family protein 1 (DGF-1, pseudogene), put | 0,5002 | 0,05031 |
| TcCLB.508173.110 | cysteine protease, putative | 0,5 | 0,00006437 |
| TcCLB.510003.41 | clathrin coat assembly protein ap19, putative | 0,4999 | 8,939E-17 |
| TcCLB.508321.60 | hypothetical protein, conserved | 0,4996 | 5,15E-07 |
| TcCLB.509965.220 | hypothetical protein, conserved | 0,4995 | 1,723E-07 |
| TcCLB.506991.10 | hypothetical protein, conserved | 0,4994 | 0,00001058 |
| TcCLB.506831.30 | hypothetical protein, conserved | 0,4993 | 0,0003021 |
| TcCLB.511925.30 | fucose kinase (pseudogene), putative | 0,4992 | 5,477E-07 |
| TcCLB.509767.100 | hypothetical protein, conserved | 0,4992 | 0,0005876 |
| TcCLB.509745.20 | glycosyl transferase-like, putative | 0,499 | 0,00001979 |
| TcCLB.506753.110 | cytochrome b5-like protein, putative | 0,4989 | 0,00007169 |
| TcCLB.503909.68 | hypothetical protein, conserved | 0,4989 | 0,0003193 |
| TcCLB.509317.80 | succinate dehydrogenase subunit | 0,4988 | 7,181E-06 |
| TcCLB.506247.50 | CCT-theta, putative | 0,4987 | 1,962E-16 |
| TcCLB.503583.60 | hypothetical protein | 0,4987 | 2,732E-07 |
| TcCLB.508781.120 | hypothetical protein, conserved | 0,4984 | 0,0002857 |
| TcCLB.503893.100 | hypothetical protein, conserved | 0,498 | 0,0135 |
| TcCLB.509805.260 | dispersed gene family protein 1 (DGF-1, pseudogene), put | 0,4979 | 0,001307 |
| TcCLB.511507.50 | mannosyltransferase, putative | 0,4978 | 0,00002658 |
| TcCLB.506905.30 | hypothetical protein, conserved | 0,4978 | 0,00006384 |
| TcCLB.508015.50 | hypothetical protein, conserved | 0,4975 | 0,00006469 |
| TcCLB.510877.10 | antigenic protein, putative | 0,4975 | 0,0002625 |
| TcCLB.508389.154 | mucin-associated surface protein (MASP), putative | 0,4974 | 0,06885 |
| TcCLB.509569.140 | hypothetical protein, conserved | 0,4971 | 3,743E-06 |
| TcCLB.506697.80 | Present in the outer mitochondrial membrane proteome 2 | 0,4969 | 0,0002224 |
| TcCLB.506235.10 | translation initiation factor IF-2, putative | 0,4967 | 7,475E-11 |
| TcCLB.426799.19 | serine/threonine protein kinase, putative | 0,4967 | 0,005218 |
| TcCLB.511167.90 | 14-3-3 protein 2, putative (14-3-3-II) | 0,4964 | 2,659E-07 |
| TcCLB.508741.100 | hypothetical protein, conserved | 0,4962 | 6,353E-06 |
| TcCLB.506183.20 | dispersed gene family protein 1 (DGF-1), putative | 0,496 | 0,009356 |
| TcCLB.510283.140 | UDP-N-acetylglucosamine-dolichyl-phosphate N-acetylglu | 0,4959 | 1,799E-06 |
| TcCLB.511627.70 | hypothetical protein, conserved | 0,4959 | 0,0004486 |
| TcCLB.509207.80 | glycogen synthase kinase-3 alpha, putative | 0,4958 | 0,0000311 |
| TcCLB.510849.30 | hypothetical protein, conserved (pseudogene) | 0,4957 | 0,0002085 |
| TcCLB.507031.64 | hypothetical protein, conserved | 0,4957 | 0,0003417 |
| TcCLB.508579.4 | hypothetical protein, conserved | 0,4957 | 0,00371 |
| TcCLB.509153.130 | hypothetical protein | 0,4955 | 4,568E-07 |
| TcCLB.508469.59 | helicase, putative | 0,4954 | 0,00001114 |
| TcCLB.508737.150 | mitochondrial carrier protein, putative | 0,4953 | 8,765E-09 |
| TcCLB.510889.140 | alanine aminotransferase, putative | 0,4953 | 5,459E-07 |
| TcCLB.508891.130 | hypothetical protein, conserved | 0,4952 | 8,956E-08 |
| TcCLB.479847.20 | retrotransposon hot spot (RHS) protein, putative | 0,4952 | 0,05132 |

| | | | |
|---|---|---|---|
| TcCLB.510149.140 | Triple RNA binding domain protein 3 (TRRM3) | 0,4951 | 4,328E-07 |
| TcCLB.511837.129 | RNA-binding protein, putative | 0,4951 | 0,00001878 |
| TcCLB.511819.59 | dynein heavy chain, cytosolic, putative | 0,4951 | 0,00002302 |
| TcCLB.506257.10 | DNA/RNA non-specific endonuclease protein-like, putative | 0,495 | 0,001067 |
| TcCLB.508577.50 | hypothetical protein | 0,4948 | 0,004043 |
| TcCLB.503973.271 | surface protease GP63 (pseudogene), putative | 0,4947 | 0,04585 |
| TcCLB.511303.100 | hypothetical protein | 0,4946 | 0,0001007 |
| TcCLB.509809.39 | P-type H -ATPase, putative | 0,4945 | 1,74E-10 |
| TcCLB.509331.90 | AMPK subunit beta, putative (AMPKB) | 0,4945 | 3,108E-06 |
| TcCLB.510515.130 | S-adenosyl-methyltransferase mraW-like protein, putative | 0,4945 | 4,498E-06 |
| TcCLB.508231.64 | hypothetical protein, conserved | 0,4945 | 0,0006399 |
| TcCLB.508971.30 | lysyl-tRNA synthetase, putative | 0,4944 | 3,831E-12 |
| TcCLB.506297.150 | 40S ribosomal protein S5, putative | 0,4944 | 4,859E-09 |
| TcCLB.510543.80 | hypothetical protein | 0,4943 | 0,04102 |
| TcCLB.509825.40 | hypothetical protein, conserved | 0,494 | 1,415E-07 |
| TcCLB.506795.40 | phosphatidylserine synthase, putative | 0,494 | 0,00003577 |
| TcCLB.509537.60 | meiosis recombination protein SPO11, putative | 0,4939 | 2,239E-06 |
| TcCLB.503973.260 | syntaxin binding protein (pseudogene), putative | 0,4939 | 0,1465 |
| TcCLB.511025.50 | glucosamine-6-phosphate isomerase, putative | 0,4938 | 1,463E-09 |
| TcCLB.506559.320 | hypothetical protein, conserved | 0,4937 | 0,0004178 |
| TcCLB.511317.30 | regulator of nonsense transcripts UPF1, putative | 0,4936 | 7,917E-11 |
| TcCLB.509023.50 | kinesin, putative | 0,4935 | 2,897E-07 |
| TcCLB.509905.20 | retrotransposon hot spot protein (RHS, pseudogene), puta | 0,4935 | 0,005554 |
| TcCLB.510997.20 | hypothetical protein, conserved | 0,4934 | 0,0001856 |
| TcCLB.508241.80 | hypothetical protein, conserved | 0,4933 | 0,00008693 |
| TcCLB.503509.10 | hypothetical protein, conserved | 0,4932 | 1,396E-11 |
| TcCLB.509245.20 | hypothetical protein, conserved | 0,493 | 0,01523 |
| TcCLB.509055.50 | hypothetical protein, conserved | 0,4927 | 0,00008245 |
| TcCLB.507053.180 | hypothetical protein, conserved | 0,4926 | 2,509E-13 |
| TcCLB.506595.140 | hypothetical protein | 0,4926 | 0,001593 |
| TcCLB.511179.60 | hypothetical protein, conserved | 0,4922 | 0,1498 |
| TcCLB.506387.9 | chaperone DNAJ protein, putative | 0,492 | 0,01039 |
| TcCLB.509179.180 | zinc-finger protein ZPR1, putative | 0,4918 | 5,516E-06 |
| TcCLB.509157.270 | hypothetical protein, conserved | 0,4918 | 0,0002524 |
| TcCLB.510533.20 | hypothetical protein, conserved | 0,4918 | 0,0004782 |
| TcCLB.491081.10 | retrotransposon hot spot protein (RHS, pseudogene), puta | 0,4916 | 0,005433 |
| TcCLB.510057.10 | hypothetical protein, conserved | 0,4913 | 0,03606 |
| TcCLB.510851.50 | hypothetical protein, conserved (pseudogene) | 0,4911 | 9,464E-06 |
| TcCLB.506733.120 | hypothetical protein, conserved | 0,4911 | 0,00003992 |
| TcCLB.509999.20 | hypothetical protein, conserved | 0,4909 | 0,0001367 |
| TcCLB.503957.90 | dispersed gene family protein 1 (DGF-1, pseudogene), put | 0,4909 | 0,01872 |
| TcCLB.510165.54 | hypothetical protein, conserved | 0,4908 | 0,0002423 |
| TcCLB.508375.30 | aspartate carbamoyltransferase, putative | 0,4907 | 5,962E-06 |
| TcCLB.510317.40 | hypothetical protein, conserved | 0,4906 | 0,00001274 |
| TcCLB.511751.160 | hypothetical protein, conserved | 0,4905 | 0,002374 |

| | | | |
|---|---|---|---|
| TcCLB.503829.40 | hypothetical protein, conserved | 0,4904 | 1,143E-07 |
| TcCLB.511907.230 | hypothetical protein, conserved | 0,4904 | 8,174E-06 |
| TcCLB.510225.30 | hypothetical protein, conserved | 0,4904 | 0,00001248 |
| TcCLB.504021.70 | hypothetical protein, conserved | 0,4904 | 0,00005388 |
| TcCLB.507765.60 | 40S ribosomal protein S21, putative | 0,4902 | 2,397E-11 |
| TcCLB.509023.110 | ATP-dependent protease ATPase subunit HslU2 (HslU2) | 0,49 | 3,139E-08 |
| TcCLB.510859.24 | hypothetical protein, conserved | 0,4898 | 1,592E-07 |
| TcCLB.510903.60 | prenyltransferase, putative | 0,4896 | 0,0001282 |
| TcCLB.506401.180 | hypothetical protein, conserved | 0,4893 | 3,141E-09 |
| TcCLB.509793.10 | kinetoplast-associated protein 4 (KAP4) | 0,4892 | 0,0005852 |
| TcCLB.510289.99 | hypothetical protein, conserved | 0,489 | 0,00126 |
| TcCLB.509149.30 | hypothetical protein, conserved | 0,4889 | 1,368E-08 |
| TcCLB.511259.161 | elongation factor 1-gamma (EF-1-gamma, pseudogene), | 0,4889 | 0,1799 |
| TcCLB.511727.250 | hypothetical protein, conserved | 0,4888 | 4,98E-09 |
| TcCLB.508589.30 | retrotransposon hot spot (RHS) protein, putative | 0,4887 | 1,203E-06 |
| TcCLB.505267.30 | beta galactofuranosyl glycosyltransferase, putative | 0,4887 | 0,03834 |
| TcCLB.510357.130 | hypothetical protein, conserved | 0,4885 | 0,002565 |
| TcCLB.509599.80 | hypothetical protein, conserved | 0,4884 | 0,00002631 |
| TcCLB.511471.10 | hypothetical protein, conserved (pseudogene) | 0,4884 | 0,009855 |
| TcCLB.509979.119 | hypothetical protein, conserved | 0,4881 | 0,001308 |
| TcCLB.408437.29 | hypothetical protein (fragment) | 0,488 | 0,00000378 |
| TcCLB.509995.20 | hypothetical protein, conserved | 0,4878 | 0,00000199 |
| TcCLB.508551.30 | hexose transporter, putative | 0,4878 | 0,001031 |
| TcCLB.503881.50 | hypothetical protein, conserved | 0,4878 | 0,02292 |
| TcCLB.509611.160 | proteasome regulatory non-ATPase subunit, putative | 0,4874 | 3,568E-06 |
| TcCLB.511439.50 | zinc transporter, putative | 0,4874 | 5,048E-06 |
| TcCLB.506321.350 | hypothetical protein, conserved | 0,4874 | 0,0002581 |
| TcCLB.507221.30 | hypothetical protein, conserved | 0,4874 | 0,003165 |
| TcCLB.511029.20 | kinetoplast-associated protein 3 (KAP3) | 0,4873 | 0,0001882 |
| TcCLB.509057.10 | hypothetical protein, conserved | 0,4871 | 0,009185 |
| TcCLB.507787.140 | hypothetical protein, conserved | 0,487 | 0,00006973 |
| TcCLB.506919.110 | glycine dehydrogenase [decarboxylating] (pseudogene), p | 0,4869 | 0,1586 |
| TcCLB.507711.30 | hypothetical protein, conserved | 0,4867 | 0,0007795 |
| TcCLB.506321.320 | suppressive immunomodulating factor, putative (TSIF) | 0,4866 | 1,449E-06 |
| TcCLB.511121.20 | hypothetical protein | 0,4866 | 0,07237 |
| TcCLB.511423.140 | hypothetical protein, conserved | 0,4865 | 5,823E-07 |
| TcCLB.509491.20 | hypothetical protein, conserved (pseudogene) | 0,4864 | 0,00001829 |
| TcCLB.507485.80 | hypothetical protein, conserved | 0,4864 | 0,003147 |
| TcCLB.510741.20 | hypothetical protein, conserved | 0,4862 | 0,0001238 |
| TcCLB.511381.60 | hypothetical protein, conserved | 0,4862 | 0,0004723 |
| TcCLB.510697.70 | elongation factor 1-gamma (EF-1-gamma, pseudogene), | 0,4861 | 0,0003458 |
| TcCLB.508831.60 | Low-affinity phosphate transporter, putative (PHO91) | 0,4858 | 1,197E-09 |
| TcCLB.508547.20 | pre-rrna-processing protein tsr1 homolog | 0,4858 | 1,834E-06 |
| TcCLB.508307.80 | mitochondrial RNA binding complex 1 subunit, putative | 0,4857 | 2,277E-06 |
| TcCLB.508771.55 | hypothetical protein, conserved (pseudogene) | 0,4855 | 0,001057 |

| | | | |
|---|---|---|---|
| TcCLB.506247.360 | hypothetical protein, conserved | 0,4852 | 0,0001794 |
| TcCLB.511469.60 | Trk system potassium uptake protein (HKT1) | 0,4851 | 0,00004059 |
| TcCLB.508769.10 | hypothetical protein, conserved (pseudogene) | 0,4849 | 9,763E-06 |
| TcCLB.506457.110 | hypothetical protein, conserved | 0,4849 | 0,00002867 |
| TcCLB.507559.70 | hypothetical protein, conserved | 0,4849 | 0,00004589 |
| TcCLB.503843.40 | chaperone DNAJ protein, putative (fragment) | 0,4846 | 3,642E-06 |
| TcCLB.506529.230 | hypothetical protein, conserved | 0,4846 | 0,0006886 |
| TcCLB.508629.42 | retrotransposon hot spot protein (RHS, pseudogene), puta | 0,4845 | 1,617E-06 |
| TcCLB.506845.31 | ATP-dependent DEAD/H RNA helicase (pseudogene), puta | 0,4844 | 0,1911 |
| TcCLB.511827.20 | hypothetical protein, conserved | 0,4843 | 1,682E-07 |
| TcCLB.503525.40 | FAD dependent oxidoreductase, putative (POMP20) | 0,4842 | 1,901E-07 |
| TcCLB.507491.162 | flagellar calcium-binding protein, putative (fragment) | 0,4842 | 0,00004524 |
| TcCLB.511181.140 | hypothetical protein | 0,4841 | 9,025E-07 |
| TcCLB.506679.40 | mitogen-activated protein kinase kinase 5, putative (MKK! | 0,484 | 0,00000317 |
| TcCLB.511483.30 | hypothetical protein, conserved | 0,4839 | 0,00008835 |
| TcCLB.506843.70 | protein kinase, putative | 0,4839 | 0,2621 |
| TcCLB.506989.100 | Triple RNA binding domain protein 3 | 0,4838 | 0,00001575 |
| TcCLB.511681.10 | hypothetical protein | 0,4838 | 0,128 |
| TcCLB.510519.30 | mitochondrial structure specific endonuclease I (SSE-1), p | 0,4837 | 0,00001983 |
| TcCLB.511527.34 | 60S ribosomal protein L8, putative | 0,4835 | 1,556E-11 |
| TcCLB.503955.70 | eukaryotic translation initiation factor 2 beta subunit, puta | 0,4834 | 0,00005047 |
| TcCLB.510655.20 | hypothetical protein, conserved | 0,4831 | 0,00004582 |
| TcCLB.503591.10 | retrotransposon hot spot protein (RHS, pseudogene), puta | 0,4831 | 0,005226 |
| TcCLB.511751.166 | hypothetical protein, conserved | 0,483 | 0,00002974 |
| TcCLB.507011.230 | hypothetical protein, conserved (pseudogene) | 0,4829 | 2,376E-06 |
| TcCLB.510819.60 | ATP-dependent protease ATPase subunit HslU1, putative ( | 0,4827 | 5,9E-09 |
| TcCLB.506925.440 | Nuclear pore complex protein 158 (pseudogene) | 0,4826 | 7,245E-11 |
| TcCLB.509459.70 | hypothetical protein, conserved | 0,4825 | 0,00000459 |
| TcCLB.508535.20 | nuclear transmembrane protein, putative (NP) | 0,4825 | 0,00001793 |
| TcCLB.504797.30 | pseudouridylate synthase-like protein, putative | 0,4824 | 0,00001529 |
| TcCLB.409117.10 | hypothetical protein, conserved | 0,4824 | 0,002534 |
| TcCLB.510861.94 | hypothetical protein, conserved | 0,4821 | 1,369E-09 |
| TcCLB.510603.20 | hypothetical protein, conserved | 0,4819 | 0,001878 |
| TcCLB.508823.50 | 40S ribosomal protein S14, putative | 0,4817 | 4,072E-11 |
| TcCLB.510753.9 | helicase, putative | 0,4817 | 5,888E-08 |
| TcCLB.506885.430 | lipoic acid containing carrier protein, putative (GCVH) | 0,4817 | 0,007589 |
| TcCLB.506871.40 | hypothetical protein, conserved | 0,4816 | 0,00001801 |
| TcCLB.509933.20 | hypothetical protein, conserved | 0,4813 | 0,00001183 |
| TcCLB.508479.360 | hypothetical protein, conserved | 0,4812 | 0,00001579 |
| TcCLB.507603.10 | hypothetical protein, conserved (pseudogene) | 0,4812 | 0,0009571 |
| TcCLB.509259.140 | glycine dehydrogenase [decarboxylating] (pseudogene), p | 0,4808 | 0,00008482 |
| TcCLB.508779.70 | hypothetical protein, conserved | 0,4806 | 8,13E-07 |
| TcCLB.509917.12 | N-acetyltransferase complex ARD1 subunit (pseudogene), | 0,4804 | 0,135 |
| TcCLB.510129.11 | surface glycoprotein, putative | 0,4803 | 0,00004843 |
| TcCLB.507903.19 | dispersed gene family protein 1 (DGF-1, pseudogene), put | 0,4803 | 0,04241 |

| | | | |
|---|---|---|---|
| TcCLB.505773.20 | dispersed gene family protein 1 (DGF-1), putative | 0,4802 | 0,1047 |
| TcCLB.511463.20 | AMPK subunit gamma (AMPKG) | 0,48 | 0,0002283 |
| TcCLB.505699.20 | retrotransposon hot spot protein (RHS, pseudogene), puta | 0,48 | 0,004038 |
| TcCLB.416761.10 | surface protease GP63, putative | 0,4798 | 0,136 |
| TcCLB.505807.140 | hypothetical protein, conserved | 0,4797 | 0,00000165 |
| TcCLB.445011.19 | hypothetical protein, conserved | 0,4796 | 0,0003214 |
| TcCLB.506235.20 | guide RNA associated protein 1, putative (GAP1) | 0,4795 | 2,03E-09 |
| TcCLB.509453.70 | protein transport protein Sec24C, putative | 0,4795 | 1,399E-08 |
| TcCLB.506145.20 | hypothetical protein, conserved | 0,4793 | 2,57E-08 |
| TcCLB.506581.50 | hypothetical protein, conserved | 0,4793 | 0,0007767 |
| TcCLB.506437.10 | amastin, putative | 0,4792 | 0,0003474 |
| TcCLB.511419.60 | NADH-ubiquinone oxidoreductase complex I subunit, puta | 0,4789 | 0,00005681 |
| TcCLB.511285.70 | hypothetical protein, conserved | 0,4788 | 2,828E-08 |
| TcCLB.511115.10 | hypothetical protein, conserved | 0,4787 | 1,512E-06 |
| TcCLB.511255.430 | trans-sialidase (pseudogene), putative | 0,4786 | 0,08786 |
| TcCLB.510625.10 | dispersed gene family protein 1 (DGF-1), putative | 0,4784 | 0,0973 |
| TcCLB.508449.20 | retrotransposon hot spot protein (RHS, pseudogene), puta | 0,4782 | 0,0003421 |
| TcCLB.506211.190 | hypothetical protein, conserved | 0,4781 | 1,617E-07 |
| TcCLB.511279.60 | hypothetical protein, conserved | 0,4781 | 5,015E-06 |
| TcCLB.508629.58 | hypothetical protein, conserved (pseudogene) | 0,4781 | 8,362E-06 |
| TcCLB.508153.480 | hypothetical protein, conserved | 0,478 | 1,569E-09 |
| TcCLB.511827.90 | hypothetical protein, conserved | 0,4779 | 0,01862 |
| TcCLB.510877.30 | hypothetical protein, conserved | 0,4778 | 8,05E-10 |
| TcCLB.510661.110 | Present in the outer mitochondrial membrane proteome 7 | 0,4778 | 1,173E-06 |
| TcCLB.510757.30 | hypothetical protein, conserved | 0,4776 | 0,002907 |
| TcCLB.506887.80 | hypothetical protein, conserved | 0,4775 | 0,002145 |
| TcCLB.506153.3 | hypothetical protein, conserved | 0,4774 | 0,0005168 |
| TcCLB.510363.19 | trans-sialidase (pseudogene), putative | 0,4774 | 0,05578 |
| TcCLB.506513.200 | histone H2A, putative | 0,4774 | 0,3818 |
| TcCLB.507993.200 | hypothetical protein, conserved | 0,4773 | 1,066E-07 |
| TcCLB.503623.10 | dispersed gene family protein 1 (DGF-1), putative | 0,4772 | 0,2847 |
| TcCLB.507467.104 | hypothetical protein, conserved | 0,4771 | 0,004094 |
| TcCLB.510187.280 | hypothetical protein, conserved | 0,4768 | 0,0003136 |
| TcCLB.509969.20 | surface protease GP63 (pseudogene), putative | 0,4768 | 0,09522 |
| TcCLB.510317.7 | Present in the outer mitochondrial membrane proteome 2 | 0,4767 | 3,462E-06 |
| TcCLB.508589.51 | ATP-dependent DEAD/H RNA helicase (pseudogene), puta | 0,4767 | 0,06381 |
| TcCLB.503881.39 | ribosomal protein L38, putative | 0,4765 | 0,00001939 |
| TcCLB.510239.10 | dispersed gene family protein 1 (DGF-1), putative | 0,4764 | 0,1995 |
| TcCLB.506789.210 | paraflagellar rod protein, putative | 0,4762 | 8,384E-06 |
| TcCLB.503909.100 | mannosyltransferase, putative | 0,4762 | 0,00005365 |
| TcCLB.508901.10 | hypothetical protein, conserved | 0,4762 | 0,0002634 |
| TcCLB.509751.20 | hypothetical protein, conserved | 0,4761 | 8,802E-06 |
| TcCLB.506885.20 | hypothetical protein, conserved | 0,476 | 5,779E-07 |
| TcCLB.511491.70 | endosomal integral membrane protein, putative | 0,476 | 0,00001282 |
| TcCLB.508909.339 | hypothetical protein, conserved | 0,4757 | 0,00005693 |

| | | | |
|---|---|---|---|
| TcCLB.509671.60 | hypothetical protein | 0,4756 | 0,002411 |
| TcCLB.511421.90 | hypothetical protein, conserved | 0,4755 | 0,0001697 |
| TcCLB.511133.4 | DNA repair and recombination helicase protein PIF6, puta | 0,4755 | 0,0002333 |
| TcCLB.506957.100 | hypothetical protein, conserved | 0,4755 | 0,0009976 |
| TcCLB.509427.80 | hypothetical protein | 0,4754 | 0,214 |
| TcCLB.507649.10 | hypothetical protein, conserved | 0,4753 | 0,0009857 |
| TcCLB.511725.174 | small nuclear ribonucleoprotein Sm-G, putative | 0,4752 | 0,0001442 |
| TcCLB.511895.40 | hypothetical protein | 0,4751 | 0,0007507 |
| TcCLB.507019.30 | ribosome biogenesis regulatory protein (RRS1), putative | 0,4749 | 0,00002381 |
| TcCLB.509459.50 | hypothetical protein, conserved | 0,4748 | 0,00003724 |
| TcCLB.506821.20 | ATP-dependent Clp protease subunit, heat shock protein 1 | 0,4747 | 1,367E-08 |
| TcCLB.507911.60 | retrotransposon hot spot protein (RHS, pseudogene), puta | 0,4747 | 0,1377 |
| TcCLB.503811.70 | hypothetical protein, conserved | 0,4745 | 0,00000176 |
| TcCLB.510517.90 | hypothetical protein, conserved | 0,4745 | 0,0007233 |
| TcCLB.506303.160 | hypothetical protein | 0,4744 | 0,0000278 |
| TcCLB.510259.6 | DNA polymerase delta catalytic subunit, putative (fragme | 0,4743 | 5,171E-06 |
| TcCLB.506529.380 | ser/thr protein phosphatase, putative | 0,4742 | 1,605E-10 |
| TcCLB.510421.250 | hypothetical protein, conserved | 0,4741 | 7,168E-07 |
| TcCLB.511353.60 | hypothetical protein, conserved | 0,474 | 0,0008165 |
| TcCLB.508737.30 | mitochondrial RNA binding protein 1 | 0,4739 | 1,706E-11 |
| TcCLB.503651.30 | hypothetical protein | 0,4739 | 1,714E-06 |
| TcCLB.509237.60 | UDP-Gal or UDP-GlcNAc-dependent glycosyltransferase, p | 0,4733 | 0,0009752 |
| TcCLB.506503.170 | hypothetical protein, conserved | 0,4731 | 0,0182 |
| TcCLB.509391.10 | Flagellar calcium-binding protein 24, putative | 0,4731 | 0,2793 |
| TcCLB.508839.40 | hypothetical protein, conserved | 0,473 | 0,0007291 |
| TcCLB.415187.19 | hypothetical protein, conserved (fragment) | 0,473 | 0,04911 |
| TcCLB.510077.40 | small GTP-binding protein RAB6, putative | 0,4729 | 0,0007781 |
| TcCLB.511013.10 | retrotransposon hot spot protein (RHS, pseudogene), puta | 0,4728 | 0,008631 |
| TcCLB.509617.80 | ABC transporter, putative | 0,4725 | 1,438E-06 |
| TcCLB.408437.20 | hypothetical protein, conserved | 0,4721 | 0,006827 |
| TcCLB.510759.70 | hypothetical protein, conserved | 0,472 | 6,831E-08 |
| TcCLB.507809.74 | signal peptidase type I, putative | 0,472 | 0,00009429 |
| TcCLB.503437.10 | retrotransposon hot spot protein (RHS, pseudogene), puta | 0,4719 | 0,002156 |
| TcCLB.510579.110 | hypothetical protein, conserved | 0,4716 | 3,258E-08 |
| TcCLB.509153.40 | hypothetical protein, conserved | 0,4716 | 2,404E-07 |
| TcCLB.506529.370 | hypothetical protein | 0,4716 | 0,002213 |
| TcCLB.506623.60 | dispersed gene family protein 1 (DGF-1), putative | 0,4715 | 0,09367 |
| TcCLB.506575.40 | hypothetical protein | 0,4714 | 4,446E-06 |
| TcCLB.507601.110 | prolyl oligopeptidase (pseudogene), putative | 0,4714 | 4,674E-06 |
| TcCLB.507063.144 | hypothetical protein, conserved | 0,4714 | 0,001206 |
| TcCLB.507791.20 | N-acetyltransferase complex ARD1 subunit (pseudogene), | 0,4714 | 0,1136 |
| TcCLB.506563.190 | calpain-like cysteine peptidase, putative | 0,4713 | 2,715E-07 |
| TcCLB.509029.40 | hypothetical protein, conserved | 0,4713 | 0,00000611 |
| TcCLB.507515.60 | hypothetical protein, conserved | 0,4713 | 0,0002915 |
| TcCLB.506953.10 | hypothetical protein, conserved (pseudogene) | 0,4713 | 0,1161 |

| | | | |
|---|---|---|---|
| TcCLB.506035.20 | dispersed gene family protein 1 (DGF-1, pseudogene), put | 0,4711 | 0,00009739 |
| TcCLB.511801.44 | DNA-dependent RNA polymerases, putative | 0,4711 | 0,0004204 |
| TcCLB.511817.300 | hypothetical protein, conserved | 0,471 | 7,003E-06 |
| TcCLB.506945.300 | V-type ATPase, C subunit, putative | 0,4709 | 6,442E-06 |
| TcCLB.507033.10 | hypothetical protein, conserved | 0,4707 | 7E-09 |
| TcCLB.503687.10 | regulator of chromosome condensation (RCC1), putative | 0,4704 | 0,0001475 |
| TcCLB.510741.180 | ATP-dependent RNA helicase, putative | 0,4702 | 1,216E-08 |
| TcCLB.510969.40 | hypothetical protein, conserved | 0,4702 | 0,00003104 |
| TcCLB.510063.39 | DNA repair protein, putative | 0,4701 | 8,685E-06 |
| TcCLB.506743.30 | 50S ribosomal protein L17, putative | 0,4701 | 0,001107 |
| TcCLB.505957.80 | retrotransposon hot spot protein (RHS, pseudogene), puta | 0,4695 | 0,0003123 |
| TcCLB.511499.69 | hypothetical protein (fragment) | 0,4695 | 0,007445 |
| TcCLB.505931.50 | histone H3, putative | 0,4695 | 0,05015 |
| TcCLB.507093.339 | hypothetical protein, conserved | 0,4694 | 8,763E-06 |
| TcCLB.503527.29 | hypothetical protein, conserved | 0,4693 | 0,00008035 |
| TcCLB.509965.310 | hypothetical protein, conserved (pseudogene) | 0,4691 | 0,00001348 |
| TcCLB.511663.40 | trans-sialidase (pseudogene), putative | 0,4691 | 0,001937 |
| TcCLB.508153.780 | hypothetical protein | 0,469 | 2,606E-07 |
| TcCLB.410337.10 | trans-sialidase (pseudogene), putative | 0,469 | 0,2378 |
| TcCLB.508607.60 | beta galactofuranosyl glycosyltransferase, putative | 0,4689 | 0,0256 |
| TcCLB.511367.290 | hypothetical protein, conserved | 0,4687 | 6,351E-06 |
| TcCLB.510525.10 | protein kinase, putative | 0,4687 | 0,0004041 |
| TcCLB.507057.10 | hypothetical protein, conserved | 0,4685 | 0,0002173 |
| TcCLB.503749.30 | hypothetical protein, conserved | 0,4683 | 0,00001623 |
| TcCLB.509065.160 | eukaryotic translation initiation factor 3 subunit 7-like pro | 0,4679 | 4,43E-11 |
| TcCLB.511267.24 | hypothetical protein, conserved | 0,4679 | 0,0001472 |
| TcCLB.504411.20 | divalent cation tolerance protein, putative | 0,4679 | 0,06909 |
| TcCLB.508851.90 | hypothetical protein, conserved | 0,4678 | 0,0006914 |
| TcCLB.511115.49 | hypothetical protein, conserved | 0,4678 | 0,04284 |
| TcCLB.507933.10 | plectin, putative | 0,4677 | 1,963E-09 |
| TcCLB.510301.60 | hypothetical protein, conserved | 0,4676 | 0,00003548 |
| TcCLB.506189.10 | nucleolar protein, putative | 0,4675 | 0,0002366 |
| TcCLB.506127.9 | phosphoglycerate kinase, putative | 0,4675 | 0,05532 |
| TcCLB.506479.130 | cytidine triphosphate synthase, putative | 0,4671 | 0,00007262 |
| TcCLB.509965.190 | DREV methyltransferase, putative | 0,467 | 0,0002254 |
| TcCLB.507513.90 | RNA-editing complex protein MP100, putative | 0,4669 | 4,446E-08 |
| TcCLB.510877.150 | hypothetical protein, conserved | 0,4667 | 8,565E-07 |
| TcCLB.506357.130 | hypothetical protein, conserved | 0,4665 | 3,72E-11 |
| TcCLB.509055.20 | hypothetical protein, conserved | 0,4664 | 5,948E-06 |
| TcCLB.507603.230 | hydroxyacylglutathione hydrolase, putative | 0,4663 | 0,004528 |
| TcCLB.508443.20 | heat shock protein-like protein, putative | 0,4659 | 0,0008236 |
| TcCLB.504045.130 | kinesin, putative (fragment) | 0,4657 | 0,06073 |
| TcCLB.507023.120 | aldehyde dehydrogenase family, putative | 0,4656 | 0,0002947 |
| TcCLB.511789.10 | dispersed gene family protein 1 (DGF-1, pseudogene), put | 0,4656 | 0,2112 |
| TcCLB.510577.40 | hypothetical protein, conserved | 0,4655 | 0,0002161 |

| | | | |
|---|---|---|---|
| TcCLB.507427.30 | retrotransposon hot spot protein (RHS, pseudogene), puta | 0,4654 | 0,001795 |
| TcCLB.508821.30 | hypothetical protein | 0,4653 | 3,719E-08 |
| TcCLB.506247.470 | hypothetical protein, conserved | 0,4651 | 3,961E-08 |
| TcCLB.505183.14 | hypothetical protein, conserved | 0,4651 | 0,0007238 |
| TcCLB.511249.34 | hypothetical protein, conserved | 0,4651 | 0,0009315 |
| TcCLB.510001.10 | hypothetical Zinc-finger containing protein | 0,4651 | 0,002563 |
| TcCLB.507883.100 | malate dehydrogenase, putative | 0,465 | 0,00003803 |
| TcCLB.507253.10 | retrotransposon hot spot (RHS) protein, putative | 0,465 | 0,001163 |
| TcCLB.506959.20 | hypothetical protein, conserved | 0,4649 | 1,517E-08 |
| TcCLB.509161.110 | hypothetical protein, conserved | 0,4649 | 2,961E-08 |
| TcCLB.507491.151 | flagellar calcium-binding protein, putative | 0,4648 | 0,0000543 |
| TcCLB.504143.10 | hypothetical protein, conserved | 0,4648 | 0,001015 |
| TcCLB.506559.80 | Sir2-related protein 3, putative (SIR2rp3) | 0,4646 | 0,001586 |
| TcCLB.509941.70 | dispersed gene family protein 1 (DGF-1, pseudogene), put | 0,4645 | 0,02409 |
| TcCLB.506389.39 | elongation factor 1-gamma (EF-1-gamma, pseudogene), | 0,464 | 6,677E-06 |
| TcCLB.508881.30 | hypothetical protein, conserved | 0,4639 | 0,0002482 |
| TcCLB.507475.30 | retrotransposon hot spot protein (RHS, pseudogene), puta | 0,4639 | 0,0005051 |
| TcCLB.511807.80 | RNA capping enzyme, nuclear (Cgm1) | 0,4638 | 1,623E-09 |
| TcCLB.508741.320 | casein kinase II, alpha chain, putative | 0,4638 | 0,0000177 |
| TcCLB.507019.50 | hypothetical protein, conserved | 0,4638 | 0,0004152 |
| TcCLB.510149.90 | Golgi vesicular membrane trafficking protein, putative | 0,4638 | 0,008665 |
| TcCLB.510241.10 | hypothetical protein, conserved | 0,4636 | 3,531E-07 |
| TcCLB.508761.260 | mucin TcMUCII, putative | 0,4636 | 0,009523 |
| TcCLB.506683.19 | GMP synthase (pseudogene), putative | 0,4635 | 0,00003599 |
| TcCLB.506435.424 | hypothetical protein, conserved | 0,4634 | 0,001986 |
| TcCLB.505073.10 | hypothetical protein | 0,4633 | 3,403E-07 |
| TcCLB.509371.10 | dispersed gene family protein 1 (DGF-1), putative | 0,4633 | 0,2318 |
| TcCLB.508539.40 | hypothetical protein, conserved | 0,4632 | 0,001217 |
| TcCLB.509337.10 | hypothetical protein, conserved | 0,463 | 0,0000337 |
| TcCLB.507001.20 | hypothetical protein, conserved | 0,463 | 0,003218 |
| TcCLB.506975.9 | REL2, putative | 0,4629 | 0,02119 |
| TcCLB.506945.150 | hypothetical protein | 0,4628 | 0,001663 |
| TcCLB.509153.50 | hypothetical protein, conserved | 0,4626 | 0,0001318 |
| TcCLB.510755.98 | kinetoplastid membrane protein KMP-11, putative | 0,4626 | 0,02317 |
| TcCLB.510181.90 | hypothetical protein, conserved | 0,4624 | 0,000132 |
| TcCLB.503909.90 | hypothetical protein | 0,462 | 0,002627 |
| TcCLB.506195.10 | glucose-6-phosphate dehydrogenase (pseudogene), putati | 0,462 | 0,004133 |
| TcCLB.505983.20 | nucleosome assembly protein-like protein, putative | 0,4619 | 5,001E-09 |
| TcCLB.509891.70 | hypothetical protein, conserved (pseudogene) | 0,4618 | 0,00028 |
| TcCLB.506933.30 | hypothetical protein, conserved | 0,4617 | 5,707E-06 |
| TcCLB.510105.80 | folylpolyglutamate synthetase (FPGS) | 0,4617 | 0,0000938 |
| TcCLB.510045.20 | ABC transporter, putative | 0,4615 | 0,004627 |
| TcCLB.510719.310 | hypothetical protein, conserved | 0,4614 | 1,633E-06 |
| TcCLB.507001.34 | hypothetical protein, conserved | 0,4614 | 0,0003829 |
| TcCLB.506195.190 | intraflagellar transport protein 81, putative (IFT81) | 0,4613 | 1,606E-08 |

| | | | |
|---|---|---|---|
| TcCLB.508589.41 | N-acetyltransferase complex ARD1 subunit (pseudogene), | 0,4612 | 0,203 |
| TcCLB.511725.280 | hypothetical protein, conserved | 0,461 | 1,447E-06 |
| TcCLB.511257.40 | hypothetical protein, conserved | 0,461 | 0,00004528 |
| TcCLB.503439.61 | retrotransposon hot spot protein (RHS, pseudogene), puta | 0,461 | 0,0003685 |
| TcCLB.508401.90 | N-acetyltransferase complex ARD1 subunit (pseudogene), | 0,461 | 0,0301 |
| TcCLB.511017.40 | hd phosphohydrolase family protein | 0,4609 | 0,01973 |
| TcCLB.506431.5 | trans-sialidase, putative (pseudogene) | 0,4606 | 0,2653 |
| TcCLB.509051.40 | adenosine kinase, putative | 0,4605 | 4,082E-08 |
| TcCLB.507093.60 | hypothetical protein, conserved | 0,4604 | 1,502E-06 |
| TcCLB.511395.10 | hypothetical protein, conserved | 0,4604 | 0,0003743 |
| TcCLB.511807.36 | hypothetical protein | 0,4603 | 0,00004367 |
| TcCLB.508659.30 | hypothetical protein, conserved | 0,4601 | 7,566E-06 |
| TcCLB.506155.120 | glycosyl hydrolase, putative | 0,4601 | 0,00001144 |
| TcCLB.507601.70 | dihydrolipoamide branched chain transacylase, putative | 0,46 | 0,00002829 |
| TcCLB.511149.210 | mucin TcMUCII, putative | 0,46 | 0,1495 |
| TcCLB.509027.24 | hypothetical protein, conserved | 0,4599 | 0,0005084 |
| TcCLB.508547.180 | hypothetical protein, conserved | 0,4598 | 1,422E-06 |
| TcCLB.507583.70 | ribonuclease inhibitor, putative | 0,4598 | 0,0001594 |
| TcCLB.506847.30 | retrotransposon hot spot protein (RHS, pseudogene), puta | 0,4598 | 0,0002915 |
| TcCLB.510863.30 | hypothetical protein, conserved | 0,4597 | 0,0002736 |
| TcCLB.511629.30 | hypothetical protein, conserved | 0,4596 | 0,0006713 |
| TcCLB.510279.276 | hypothetical protein, conserved (pseudogene) | 0,4596 | 0,09492 |
| TcCLB.506265.20 | hypothetical protein, conserved | 0,4595 | 7,904E-07 |
| TcCLB.511773.35 | hypothetical protein, conserved (pseudogene) | 0,4591 | 0,008095 |
| TcCLB.504131.90 | hypothetical protein, conserved | 0,4588 | 0,00001793 |
| TcCLB.510879.150 | hypothetical protein, conserved | 0,4587 | 0,00002507 |
| TcCLB.506533.142 | mucin TcSMUGL, putative | 0,4587 | 0,001075 |
| TcCLB.503991.30 | hypothetical protein, conserved | 0,4584 | 9,024E-06 |
| TcCLB.511713.40 | ribosome biogenesis protein TSR3, putative (tsr3) | 0,4584 | 0,001667 |
| TcCLB.506247.100 | hypothetical protein, conserved | 0,458 | 0,004494 |
| TcCLB.507603.170 | hypothetical protein, conserved | 0,4576 | 0,00001049 |
| TcCLB.508895.40 | hypothetical protein, conserved | 0,4576 | 0,00008015 |
| TcCLB.508971.50 | ubiquitin-protein ligase-like, putative | 0,4575 | 1,017E-06 |
| TcCLB.508831.66 | hypothetical protein, conserved | 0,4574 | 3,925E-06 |
| TcCLB.511727.129 | ribosomal protein L18, putative | 0,4573 | 3,435E-08 |
| TcCLB.511393.20 | rhomboid-like protein, putative | 0,4573 | 0,00001217 |
| TcCLB.506321.290 | stress-induced protein sti1, putative | 0,4572 | 3,274E-09 |
| TcCLB.508873.430 | trans-sialidase (pseudogene), putative | 0,4568 | 0,01137 |
| TcCLB.510311.170 | hypothetical protein, conserved | 0,4566 | 0,0002873 |
| TcCLB.506259.30 | retrotransposon hot spot protein (RHS, pseudogene), puta | 0,4564 | 0,00002775 |
| TcCLB.511863.4 | retrotransposon hot spot (RHS) protein, putative (fragmer | 0,4564 | 0,0003553 |
| TcCLB.508857.30 | trans-sialidase, Group I, putative | 0,4563 | 0,0001207 |
| TcCLB.511293.10 | extracellular receptor, putative | 0,4562 | 0,001782 |
| TcCLB.511249.60 | protein kinase, putative | 0,4561 | 5,169E-06 |
| TcCLB.511055.40 | glycine dehydrogenase [decarboxylating] (pseudogene), pu | 0,4561 | 0,002733 |

| | | | |
|---|---|---|---|
| TcCLB.509019.30 | dispersed gene family protein 1 (DGF-1), putative | 0,4561 | 0,08165 |
| TcCLB.504835.20 | 3-methylcrotonyl-CoA carboxylase, putative | 0,4559 | 1,202E-06 |
| TcCLB.508299.80 | threonyl-tRNA synthetase, putative | 0,4558 | 9,273E-06 |
| TcCLB.509053.170 | DREV methyltransferase, putative | 0,4558 | 0,0003685 |
| TcCLB.510053.70 | hypothetical protein, conserved | 0,4552 | 6,426E-08 |
| TcCLB.511211.104 | hypothetical protein, conserved | 0,4551 | 0,003313 |
| TcCLB.510309.30 | PTP1-interacting protein, 39 kDa, putative | 0,4551 | 0,003446 |
| TcCLB.511903.90 | hypothetical protein, conserved | 0,4549 | 0,00003681 |
| TcCLB.510121.50 | calmodulin, putative | 0,4547 | 0,009733 |
| TcCLB.506797.70 | lysophospholipase, putative | 0,4544 | 8,042E-06 |
| TcCLB.508823.40 | hypothetical protein, conserved | 0,4544 | 0,001967 |
| TcCLB.508269.76 | N-acetyltransferase complex ARD1 subunit (pseudogene), | 0,4542 | 0,1315 |
| TcCLB.506425.150 | hypothetical protein | 0,4541 | 0,003643 |
| TcCLB.506533.34 | phosphatidylinositol kinase related protein, putative | 0,454 | 1,193E-08 |
| TcCLB.508303.50 | hypothetical protein, conserved | 0,4539 | 0,00115 |
| TcCLB.507559.40 | hypothetical protein, conserved | 0,4538 | 0,00002713 |
| TcCLB.511837.60 | hypothetical protein, conserved | 0,4534 | 0,0001006 |
| TcCLB.507519.120 | hypothetical protein, conserved | 0,4534 | 0,001463 |
| TcCLB.510943.30 | hypothetical protein, conserved | 0,4534 | 0,002365 |
| TcCLB.511151.100 | Unconventional myosin | 0,4533 | 2,473E-07 |
| TcCLB.506245.160 | retrotransposon hot spot (RHS) protein, putative | 0,4533 | 0,06515 |
| TcCLB.507625.130 | hypothetical protein, conserved | 0,4532 | 0,0001528 |
| TcCLB.508133.10 | hypothetical protein | 0,4531 | 0,0006617 |
| TcCLB.510421.210 | hypothetical protein, conserved | 0,4529 | 1,392E-06 |
| TcCLB.509111.19 | hypothetical protein, conserved | 0,4529 | 0,02721 |
| TcCLB.509879.40 | fumarate hydratase, putative | 0,4527 | 0,001406 |
| TcCLB.508823.130 | hypothetical protein, conserved | 0,4525 | 6,594E-08 |
| TcCLB.509489.20 | hypothetical protein, conserved | 0,4523 | 0,00004047 |
| TcCLB.508817.10 | hypothetical protein, conserved | 0,4523 | 0,006985 |
| TcCLB.510007.50 | hypothetical protein, conserved | 0,4522 | 5,934E-06 |
| TcCLB.508641.290 | hypothetical protein, conserved | 0,4522 | 0,0002044 |
| TcCLB.511245.190 | hypothetical protein, conserved | 0,4522 | 0,0007536 |
| TcCLB.509497.20 | hypothetical protein, conserved | 0,4521 | 0,0006297 |
| TcCLB.506435.60 | hypothetical protein, conserved | 0,452 | 0,000171 |
| TcCLB.510287.4 | Mutt-Nudix-related hydrolase, putative (fragment) | 0,452 | 0,1761 |
| TcCLB.506771.18 | hypothetical protein, conserved | 0,4519 | 0,000333 |
| TcCLB.506017.41 | N-acetyltransferase complex ARD1 subunit (pseudogene), | 0,4518 | 0,05109 |
| TcCLB.506457.10 | metallo-peptidase, Clan MG, Family M24 (MetAP2) | 0,4517 | 3,914E-06 |
| TcCLB.503809.75 | hypothetical protein, conserved | 0,4517 | 0,008103 |
| TcCLB.508153.390 | hypothetical protein, conserved | 0,4516 | 0,00008277 |
| TcCLB.506193.20 | hypothetical protein, conserved | 0,4516 | 0,0001215 |
| TcCLB.511461.50 | nuclear transmembrane protein, putative (NP) | 0,4516 | 0,0004735 |
| TcCLB.510227.10 | hypothetical protein, conserved | 0,4516 | 0,00521 |
| TcCLB.510359.100 | hypothetical protein, conserved | 0,4514 | 0,006551 |
| TcCLB.511649.110 | hypothetical protein, conserved | 0,4512 | 2,751E-06 |

| | | | |
|---|---|---|---|
| TcCLB.509023.150 | integral membrane transport protein, putative | 0,4511 | 0,0008071 |
| TcCLB.507023.140 | hypothetical protein | 0,4511 | 0,001516 |
| TcCLB.510443.9 | valyl-tRNA synthetase, putative | 0,451 | 1,529E-07 |
| TcCLB.509551.10 | protein kinase, putative | 0,451 | 0,0002001 |
| TcCLB.504175.40 | dual specificity protein phosphatase, putative | 0,4509 | 8,601E-07 |
| TcCLB.503487.50 | UDP-Gal or UDP-GlcNAc-dependent glycosyltransferase, p | 0,4509 | 0,00008276 |
| TcCLB.508317.10 | cysteine proteinase, putative | 0,4507 | 0,00007315 |
| TcCLB.507485.150 | amastin, putative | 0,4503 | 0,00001307 |
| TcCLB.504047.20 | hypothetical protein, conserved | 0,4502 | 0,0001666 |
| TcCLB.511421.200 | ARM-like helical domain-containing protein | 0,4501 | 0,00001449 |
| TcCLB.507521.70 | cyclophilin, putative | 0,4501 | 0,0008363 |
| TcCLB.508051.30 | protein kinase, putative | 0,4499 | 0,0001898 |
| TcCLB.510173.100 | hypothetical protein, conserved | 0,4499 | 0,003141 |
| TcCLB.508531.10 | glycine dehydrogenase (pseudogene), putative | 0,4499 | 0,05092 |
| TcCLB.506201.50 | hypothetical protein, conserved | 0,4498 | 0,005545 |
| TcCLB.509769.70 | DNA polymerase theta (helicase domain only), putative | 0,4496 | 1,171E-08 |
| TcCLB.509797.30 | hypothetical protein, conserved | 0,4496 | 0,00005644 |
| TcCLB.510183.20 | dipeptidyl-peptidase 8-like serine peptidase | 0,4493 | 2,249E-07 |
| TcCLB.505097.20 | hypothetical protein, conserved (pseudogene) | 0,4493 | 0,005073 |
| TcCLB.504213.140 | hypothetical protein, conserved | 0,4493 | 0,02754 |
| TcCLB.511479.48 | hypothetical protein, conserved (fragment) | 0,4493 | 0,07178 |
| TcCLB.511229.9 | replication factor C, subunit 4, putative | 0,4491 | 0,01091 |
| TcCLB.506743.150 | proteasome 26S non-ATPase subunit 9, putative | 0,449 | 0,002032 |
| TcCLB.503809.100 | GUP1, putative | 0,4489 | 0,00000194 |
| TcCLB.511075.20 | notchless homolog, putative | 0,4488 | 1,365E-06 |
| TcCLB.508207.54 | hypothetical protein, conserved | 0,4488 | 0,004331 |
| TcCLB.504047.10 | hypothetical protein, conserved | 0,4488 | 0,0135 |
| TcCLB.507993.190 | serine/threonine protein phosphatase type 5, putative | 0,4485 | 7,489E-07 |
| TcCLB.506175.110 | hypothetical protein, conserved | 0,4485 | 1,744E-06 |
| TcCLB.504431.30 | Present in the outer mitochondrial membrane proteome 2 | 0,4484 | 0,00001233 |
| TcCLB.509769.90 | mitochondrial carrier protein, putative | 0,4484 | 0,00003474 |
| TcCLB.450489.10 | histone acetyltransferase, putative | 0,4484 | 0,0009233 |
| TcCLB.427303.10 | 5'-3' exoribonuclease A, putative (fragment) (XRNA) | 0,4482 | 4,84E-07 |
| TcCLB.509233.20 | retrotransposon hot spot protein (RHS, pseudogene), puta | 0,4482 | 0,004569 |
| TcCLB.506163.60 | hypothetical protein | 0,4481 | 6,661E-06 |
| TcCLB.503681.9 | hypothetical protein, conserved | 0,4481 | 0,00002293 |
| TcCLB.509715.9 | Ribosomal RNA processing protein 41A (RRP41A) | 0,4478 | 0,0005205 |
| TcCLB.510187.199 | hypothetical protein, conserved | 0,4478 | 0,001119 |
| TcCLB.510439.10 | heat shock protein 70, putative | 0,4477 | 0,0005081 |
| TcCLB.506649.30 | hypothetical protein, conserved | 0,4476 | 7,818E-07 |
| TcCLB.510735.70 | hypothetical protein, conserved | 0,4476 | 0,0003962 |
| TcCLB.511671.90 | hypothetical protein, conserved | 0,4475 | 0,0003823 |
| TcCLB.507063.160 | hypothetical protein, conserved | 0,4474 | 5,386E-06 |
| TcCLB.510087.20 | hypothetical protein, conserved | 0,4474 | 0,000769 |
| TcCLB.434259.19 | hypothetical protein, conserved | 0,4474 | 0,002865 |

| | | | |
|---|---|---|---|
| TcCLB.509893.10 | hypothetical protein | 0,4472 | 0,0001517 |
| TcCLB.506887.20 | hypothetical protein, conserved | 0,4471 | 0,0003591 |
| TcCLB.503767.5 | potassium voltage-gated channel, putative | 0,4471 | 0,07657 |
| TcCLB.503823.50 | hypothetical protein, conserved | 0,447 | 0,00005103 |
| TcCLB.511369.20 | elongation factor 1-alpha, putative | 0,447 | 0,008936 |
| TcCLB.510765.60 | 60S ribosomal protein L5, putative | 0,4469 | 7,199E-10 |
| TcCLB.508273.70 | hypothetical protein, conserved | 0,4469 | 0,002234 |
| TcCLB.506629.220 | isovaleryl-coA dehydrogenase, putative | 0,4468 | 0,00002415 |
| TcCLB.510853.50 | beta galactofuranosyl glycosyltransferase, putative | 0,4467 | 0,2246 |
| TcCLB.507951.260 | aspartic peptidase, clan AD, family A22B, putative | 0,4466 | 1,848E-06 |
| TcCLB.508899.90 | hypothetical protein | 0,4465 | 0,0002979 |
| TcCLB.511049.30 | hypothetical protein, conserved | 0,4461 | 0,003053 |
| TcCLB.506713.20 | hypothetical protein, conserved | 0,446 | 0,00003138 |
| TcCLB.507639.40 | proteasome subunit beta type-5, putative | 0,4459 | 2,942E-06 |
| TcCLB.503545.20 | retrotransposon hot spot protein (RHS, pseudogene), puta | 0,4452 | 0,01066 |
| TcCLB.507509.50 | polypeptide deformylase-like protein, putative | 0,445 | 0,00006579 |
| TcCLB.510039.70 | hypothetical protein, conserved | 0,4448 | 0,00153 |
| TcCLB.511071.100 | Glycerophosphoryl diester phosphodiesterase family, puta | 0,4446 | 9,726E-06 |
| TcCLB.506727.80 | hypothetical protein, conserved | 0,4446 | 0,0001257 |
| TcCLB.509965.229 | heat shock protein (pseudogene), putative | 0,4445 | 0,004052 |
| TcCLB.506689.20 | hypothetical protein, conserved (pseudogene) | 0,4444 | 0,001514 |
| TcCLB.506529.314 | hypothetical protein, conserved | 0,444 | 0,01133 |
| TcCLB.506297.220 | hypothetical protein, conserved | 0,4435 | 3,23E-08 |
| TcCLB.511825.130 | hypothetical protein | 0,4435 | 0,01441 |
| TcCLB.505939.50 | hypothetical protein | 0,4433 | 0,0004992 |
| TcCLB.510111.11 | trans-sialidase (pseudogene), putative | 0,4433 | 0,01885 |
| TcCLB.508727.50 | hypothetical protein, conserved | 0,443 | 0,0004885 |
| TcCLB.506315.70 | hypothetical protein, conserved | 0,443 | 0,0009199 |
| TcCLB.506661.130 | ion transport protein, putative | 0,4429 | 0,0001117 |
| TcCLB.511167.30 | hypothetical protein, conserved | 0,4428 | 0,00007882 |
| TcCLB.511685.40 | hypothetical protein | 0,4427 | 0,00009818 |
| TcCLB.506595.104 | ribosomal P protein AGP2beta-1, putative | 0,4426 | 0,009096 |
| TcCLB.504215.59 | hypothetical protein, conserved | 0,4425 | 0,00004108 |
| TcCLB.511897.19 | zinc-finger multi-pass transmembrane protein, putative | 0,4424 | 0,0004087 |
| TcCLB.508153.180 | aspartyl aminopeptidase, putative | 0,442 | 1,136E-07 |
| TcCLB.504159.80 | histone deacetylase, putative | 0,442 | 0,0001475 |
| TcCLB.508307.170 | 60S acidic ribosomal protein, putative | 0,4419 | 0,001342 |
| TcCLB.511707.30 | hypothetical protein, conserved | 0,4418 | 2,221E-06 |
| TcCLB.503657.50 | hypothetical protein, conserved | 0,4418 | 0,0008212 |
| TcCLB.509497.30 | agmatinase, putative | 0,4418 | 0,0009402 |
| TcCLB.509533.5 | retrotransposon hot spot protein (RHS, pseudogene), puta | 0,4416 | 3,028E-06 |
| TcCLB.511317.50 | hypothetical protein, conserved | 0,4416 | 0,0001241 |
| TcCLB.507083.40 | hypothetical protein, conserved | 0,4416 | 0,009155 |
| TcCLB.509237.80 | hypothetical protein, conserved | 0,4412 | 2,504E-06 |
| TcCLB.509063.10 | hypothetical protein, conserved | 0,4412 | 0,003858 |

| | | | |
|---|---|---|---|
| TcCLB.507485.100 | hypothetical protein, conserved | 0,441 | 0,0001149 |
| TcCLB.508837.110 | hypothetical protein, conserved | 0,441 | 0,0003628 |
| TcCLB.509695.130 | dynein arm light chain, putative | 0,441 | 0,0005533 |
| TcCLB.510101.320 | hypothetical protein | 0,4409 | 1,607E-06 |
| TcCLB.509069.30 | tubulin binding cofactor A-like protein, putative | 0,4408 | 0,00652 |
| TcCLB.510347.10 | hypothetical protein | 0,4406 | 8,898E-07 |
| TcCLB.509965.250 | ubiquitin fusion degradation protein 2, putative | 0,4403 | 1,698E-07 |
| TcCLB.506943.130 | hypothetical protein, conserved | 0,4402 | 0,000388 |
| TcCLB.509219.50 | hypothetical protein, conserved (pseudogene) | 0,4399 | 0,0007981 |
| TcCLB.511865.19 | frataxin-like, mitochondrial precursor, putative | 0,4395 | 0,00115 |
| TcCLB.506625.150 | hypothetical protein, conserved | 0,4395 | 0,002737 |
| TcCLB.506493.100 | hypothetical protein, conserved | 0,4393 | 0,0005013 |
| TcCLB.504285.10 | retrotransposon hot spot protein (RHS, pseudogene), puta | 0,4392 | 0,009223 |
| TcCLB.511655.69 | ubiquitin-activating enzyme-like protein | 0,4391 | 0,0003761 |
| TcCLB.510717.40 | hypothetical protein, conserved | 0,439 | 0,00001689 |
| TcCLB.506357.190 | hypothetical protein, conserved | 0,439 | 0,00007792 |
| TcCLB.506363.40 | hypothetical protein, conserved | 0,439 | 0,001148 |
| TcCLB.485889.16 | hypothetical protein (fragment) | 0,439 | 0,01225 |
| TcCLB.506925.20 | hypothetical protein, conserved | 0,4389 | 0,0000108 |
| TcCLB.511055.60 | hypothetical protein, conserved | 0,4389 | 0,07446 |
| TcCLB.510149.120 | flagellar radial spoke protein-like, putative | 0,4386 | 0,000629 |
| TcCLB.511191.10 | tyrosyl-DNA Phosphodiesterase (Tdp1), putative | 0,4386 | 0,00666 |
| TcCLB.508805.100 | hypothetical protein, conserved | 0,4384 | 0,00001132 |
| TcCLB.509733.170 | Valosin-containing protein, putative (VCP) | 0,4383 | 7,901E-07 |
| TcCLB.510061.40 | retrotransposon hot spot protein (RHS, pseudogene), puta | 0,4383 | 0,0009685 |
| TcCLB.506979.10 | hypothetical protein, conserved | 0,4382 | 0,0001671 |
| TcCLB.507591.9 | retrotransposon hot spot protein (RHS, pseudogene), puta | 0,4382 | 0,0002429 |
| TcCLB.433033.10 | retrotransposon hot spot protein (RHS, pseudogene), puta | 0,4382 | 0,0007656 |
| TcCLB.511171.40 | trans-sialidase (pseudogene), putative | 0,4381 | 0,0154 |
| TcCLB.506435.280 | hypothetical protein, conserved | 0,4379 | 0,00002697 |
| TcCLB.510243.30 | chaperone DNAJ protein, putative | 0,4379 | 0,00003433 |
| TcCLB.508045.4 | hypothetical protein, conserved | 0,4379 | 0,0005773 |
| TcCLB.510599.50 | hypothetical protein | 0,4378 | 0,00001503 |
| TcCLB.506577.60 | hypothetical protein, conserved | 0,4378 | 0,00008102 |
| TcCLB.506167.30 | hypothetical protein, conserved | 0,4378 | 0,007428 |
| TcCLB.510659.90 | hypothetical protein, conserved | 0,4377 | 0,0008243 |
| TcCLB.510989.40 | hypothetical protein, conserved | 0,4376 | 0,0005577 |
| TcCLB.506807.30 | hypothetical protein, conserved | 0,4375 | 0,002313 |
| TcCLB.511613.180 | mucin-associated surface protein (MASP), putative | 0,4375 | 0,1992 |
| TcCLB.511657.79 | hypothetical protein, conserved | 0,4374 | 0,05176 |
| TcCLB.510719.180 | hypothetical protein, conserved | 0,4371 | 0,00001216 |
| TcCLB.508445.20 | iron superoxide dismutase, putative | 0,437 | 7,388E-06 |
| TcCLB.506223.130 | hypothetical protein, conserved | 0,4369 | 0,0006648 |
| TcCLB.511181.80 | threonyl-tRNA synthetase, putative | 0,4368 | 2,207E-09 |
| TcCLB.503613.60 | acidocalcisomal pyrophosphatase, putative | 0,4368 | 6,643E-06 |

| | | | |
|---|---|---|---|
| TcCLB.511735.20 | hypothetical protein, conserved | 0,4368 | 0,001203 |
| TcCLB.505943.30 | hypothetical protein, conserved | 0,4367 | 0,0001037 |
| TcCLB.506789.300 | hypothetical protein, conserved | 0,4366 | 0,006918 |
| TcCLB.509499.20 | beta propeller protein, putative | 0,4365 | 0,00006942 |
| TcCLB.511817.110 | hypothetical protein, conserved | 0,4365 | 0,00008811 |
| TcCLB.510747.140 | hypothetical protein, conserved | 0,4365 | 0,001272 |
| TcCLB.508717.30 | phosphoribosylpyrophosphate synthetase, putative | 0,4364 | 2,857E-06 |
| TcCLB.508109.50 | retrotransposon hot spot protein (RHS, pseudogene), puta | 0,4364 | 0,0109 |
| TcCLB.506945.70 | protein mkt1, putative | 0,4363 | 2,706E-10 |
| TcCLB.511369.10 | elongation factor 1-alpha, putative | 0,4362 | 0,03041 |
| TcCLB.509875.60 | hypothetical protein | 0,4362 | 0,2269 |
| TcCLB.509039.30 | hypothetical protein, conserved | 0,4361 | 0,00008167 |
| TcCLB.503527.10 | hypothetical protein, conserved | 0,4355 | 1,414E-09 |
| TcCLB.509151.50 | hypothetical protein, conserved | 0,4355 | 1,751E-07 |
| TcCLB.508559.90 | retrotransposon hot spot (RHS) protein, putative | 0,4355 | 0,00137 |
| TcCLB.511389.114 | hypothetical protein, conserved | 0,4354 | 0,06084 |
| TcCLB.507053.80 | hypothetical protein, conserved | 0,4353 | 0,00001498 |
| TcCLB.509269.19 | sphingosine-1-phosphate phosphatase, putative | 0,4349 | 0,00009429 |
| TcCLB.510407.90 | serine hydroxymethyltransferase, putative | 0,4348 | 0,0001176 |
| TcCLB.508347.20 | O-6 methyl-guanine alkyl transferase, putative | 0,4348 | 0,001749 |
| TcCLB.509965.380 | dTDP-glucose 4,6-dehydratase, putative | 0,4346 | 0,00001155 |
| TcCLB.508233.4 | protein transport protein Sec23A, putative (fragment) | 0,4346 | 0,0002186 |
| TcCLB.508677.120 | hypothetical protein, conserved | 0,4345 | 0,02269 |
| TcCLB.511907.70 | short chain dehydrogenase/reductase, putative | 0,4344 | 0,00003789 |
| TcCLB.509023.170 | hypothetical protein, conserved | 0,4343 | 0,000227 |
| TcCLB.508723.10 | hypothetical protein, conserved | 0,4341 | 0,0005436 |
| TcCLB.506503.140 | extracellular receptor, putative | 0,434 | 2,858E-06 |
| TcCLB.507817.30 | hypothetical protein, conserved | 0,4337 | 0,00008955 |
| TcCLB.511659.9 | hypothetical protein, conserved (fragment) | 0,4337 | 0,0003435 |
| TcCLB.510105.50 | fibrillarin, putative | 0,4334 | 0,0000432 |
| TcCLB.506567.90 | hypothetical protein, conserved | 0,4332 | 0,0005267 |
| TcCLB.508169.80 | hypothetical protein, conserved | 0,4332 | 0,00337 |
| TcCLB.506953.49 | glucose-6-phosphate 1-dehydrogenase, putative | 0,4331 | 0,114 |
| TcCLB.503891.100 | 20S proteasome subunit | 0,433 | 0,0002365 |
| TcCLB.510733.30 | hypothetical protein, conserved | 0,433 | 0,02202 |
| TcCLB.511817.319 | hypothetical protein, conserved | 0,4328 | 0,003998 |
| TcCLB.503837.40 | hypothetical protein, conserved | 0,4327 | 0,03375 |
| TcCLB.511209.30 | hypothetical protein, conserved (pseudogene) | 0,4325 | 6,523E-06 |
| TcCLB.504147.60 | chaperone DNAJ protein, putative | 0,4324 | 1,939E-07 |
| TcCLB.511301.120 | hypothetical protein, conserved | 0,4324 | 3,65E-07 |
| TcCLB.511621.180 | hypothetical protein, conserved | 0,4324 | 0,0001715 |
| TcCLB.507521.30 | hypothetical protein, conserved | 0,4324 | 0,008666 |
| TcCLB.508095.20 | methyltransferase, putative | 0,4323 | 2,753E-08 |
| TcCLB.511585.10 | hypothetical protein, conserved | 0,4323 | 0,00001401 |
| TcCLB.509829.20 | isovaleryl-coA dehydrogenase, putative | 0,4321 | 0,003683 |

| | | | |
|---|---|---|---|
| TcCLB.507625.99 | gamma-glutamylcysteine synthetase, putative | 0,4318 | 0,0001416 |
| TcCLB.510041.30 | hypothetical protein, conserved (fragment) | 0,4318 | 0,02711 |
| TcCLB.509967.120 | intraflagellar transport protein 172, putative | 0,4315 | 2,899E-07 |
| TcCLB.510355.210 | hypothetical protein, conserved | 0,4313 | 4,094E-06 |
| TcCLB.507715.100 | hypothetical protein, conserved | 0,4312 | 0,00003854 |
| TcCLB.510663.70 | 6-phosphogluconate dehydrogenase, decarboxylating, puta | 0,4312 | 0,001825 |
| TcCLB.506121.9 | dispersed gene family protein 1 (DGF-1), putative | 0,4312 | 0,1505 |
| TcCLB.506585.80 | hypothetical protein, conserved | 0,4309 | 0,002492 |
| TcCLB.506871.140 | cleavage and polyadenylation specificity factor, putative | 0,4308 | 9,853E-08 |
| TcCLB.506963.40 | hypothetical protein, conserved | 0,4307 | 0,00004561 |
| TcCLB.511823.30 | hypothetical protein, conserved | 0,4306 | 0,00004569 |
| TcCLB.511801.60 | DEAD-box ATP-dependent RNA helicase, mitochondrial (M | 0,4305 | 1,516E-07 |
| TcCLB.509167.160 | hypothetical protein | 0,4305 | 6,293E-06 |
| TcCLB.503609.32 | elongation factor 1-gamma (EF-1-gamma, pseudogene), | 0,4305 | 0,3623 |
| TcCLB.510187.480 | phosphomannomutase, putative | 0,4304 | 0,0000192 |
| TcCLB.506673.40 | hypothetical protein, conserved | 0,4303 | 0,00005978 |
| TcCLB.510943.36 | hypothetical protein, conserved | 0,4303 | 0,00008975 |
| TcCLB.503431.20 | hypothetical protein, conserved | 0,4302 | 0,002536 |
| TcCLB.506605.130 | hypothetical protein, conserved | 0,43 | 0,001008 |
| TcCLB.511321.23 | cytidine deaminase-like protein | 0,4297 | 0,00128 |
| TcCLB.506247.10 | bloodstream- specific protein 2 precursor (fragment) (PDI | 0,4295 | 7,323E-06 |
| TcCLB.506265.120 | hypothetical protein, conserved | 0,4295 | 0,0000153 |
| TcCLB.508873.344 | mucin-associated surface protein (MASP, pseudogene), pu | 0,4295 | 0,1508 |
| TcCLB.504055.20 | hypothetical protein, conserved | 0,4294 | 0,002154 |
| TcCLB.504163.90 | hypothetical protein, conserved | 0,4293 | 0,0001207 |
| TcCLB.508331.41 | tryptophanyl-tRNA synthetase (pseudogene), putative | 0,4293 | 0,1378 |
| TcCLB.505999.90 | phosphoglycerate kinase, putative | 0,429 | 2,374E-06 |
| TcCLB.509353.20 | Present in the outer mitochondrial membrane proteome 4 | 0,4289 | 0,004214 |
| TcCLB.511815.60 | prefoldin, putative | 0,4288 | 0,00001314 |
| TcCLB.503703.100 | hypothetical protein, conserved | 0,4288 | 0,0001275 |
| TcCLB.511211.100 | hypothetical protein, conserved | 0,4288 | 0,0001325 |
| TcCLB.510579.10 | tyrosine phospatase-like protein, putative | 0,4286 | 0,0001531 |
| TcCLB.510879.30 | hypothetical protein, conserved | 0,4282 | 3,594E-06 |
| TcCLB.506359.70 | hypothetical protein, conserved | 0,4282 | 0,002445 |
| TcCLB.510105.210 | metallo-peptidase, Clan MA(E) Family M41 | 0,4279 | 1,387E-06 |
| TcCLB.506753.120 | DnaJ chaperone protein, putative | 0,4279 | 0,0001297 |
| TcCLB.414243.29 | inosine-guanine nucleoside hydrolase, putative (fragment | 0,4278 | 0,1178 |
| TcCLB.510123.20 | mitogen activated protein kinase, putative | 0,4273 | 1,927E-06 |
| TcCLB.505229.10 | Isoprenylcysteine alpha-carbonyl methylesterase, putative | 0,4273 | 0,00001105 |
| TcCLB.504213.29 | hypothetical protein, conserved | 0,4273 | 0,00298 |
| TcCLB.511857.70 | hypothetical protein, conserved | 0,427 | 0,0000763 |
| TcCLB.507541.20 | hypothetical protein, conserved | 0,427 | 0,0001026 |
| TcCLB.511045.10 | hypothetical protein, conserved | 0,4269 | 0,0001207 |
| TcCLB.504143.20 | hypothetical protein, conserved | 0,4266 | 0,0009446 |
| TcCLB.509695.70 | hypothetical protein, conserved | 0,4265 | 0,00001856 |

| | | | |
|---|---|---|---|
| TcCLB.509669.140 | hypothetical protein, conserved | 0,4265 | 0,003664 |
| TcCLB.510747.150 | hypothetical protein, conserved | 0,4262 | 0,0002368 |
| TcCLB.506799.40 | hypothetical protein | 0,4262 | 0,02484 |
| TcCLB.511807.22 | retrotransposon hot spot protein (RHS, pseudogene), puta | 0,426 | 0,0002333 |
| TcCLB.511545.199 | AMP deaminase, putative | 0,426 | 0,005958 |
| TcCLB.510877.130 | hypothetical protein, conserved | 0,4258 | 3,819E-06 |
| TcCLB.504115.30 | beta galactofuranosyl glycosyltransferase, putative | 0,4257 | 0,0008445 |
| TcCLB.510307.94 | ATP-dependent DEAD/H RNA helicase (pseudogene), puta | 0,4256 | 0,03595 |
| TcCLB.504835.10 | hypothetical protein, conserved | 0,4255 | 0,0004845 |
| TcCLB.506341.10 | N-acetylglucosamine-6-phosphate deacetylase-like protei | 0,4253 | 0,002167 |
| TcCLB.506825.130 | hypothetical protein, conserved | 0,4252 | 0,00002872 |
| TcCLB.510431.70 | hypothetical protein, conserved | 0,4252 | 0,001862 |
| TcCLB.510089.180 | hypothetical protein, conserved | 0,425 | 0,0004987 |
| TcCLB.506821.170 | aquaporin-like protein, putative | 0,4249 | 0,00002159 |
| TcCLB.504425.70 | hypothetical protein, conserved | 0,4249 | 0,000206 |
| TcCLB.510311.180 | hypothetical protein, conserved | 0,4249 | 0,0009709 |
| TcCLB.509757.30 | stress-inducible protein STI1-like, putative | 0,4249 | 0,003968 |
| TcCLB.511181.40 | hypothetical protein | 0,4248 | 0,002806 |
| TcCLB.507641.280 | chaperonin HSP60, mitochondrial precursor (HSP60) | 0,4246 | 0,00153 |
| TcCLB.511541.40 | hypothetical protein, conserved | 0,4245 | 0,00009578 |
| TcCLB.504881.40 | hypothetical protein, conserved | 0,4244 | 0,00004085 |
| TcCLB.503781.70 | proteasome subunit beta type-5, putative | 0,4244 | 0,000106 |
| TcCLB.508413.68 | kinetoplastid membrane protein KMP-11 | 0,4242 | 0,001544 |
| TcCLB.510849.20 | trans-sialidase (pseudogene), putative | 0,4241 | 0,005714 |
| TcCLB.508501.200 | hypothetical protein, conserved | 0,4241 | 0,04649 |
| TcCLB.511823.14 | hypothetical protein, conserved | 0,424 | 0,04524 |
| TcCLB.510509.30 | hypothetical protein, conserved | 0,4238 | 0,0135 |
| TcCLB.505937.29 | dispersed gene family protein 1 (DGF-1), putative | 0,4237 | 0,15 |
| TcCLB.511809.60 | hypothetical protein, conserved | 0,4236 | 7,947E-11 |
| TcCLB.508939.50 | hypothetical protein, conserved | 0,4236 | 0,02327 |
| TcCLB.507783.20 | AGP2beta-2, putative | 0,4234 | 6,964E-06 |
| TcCLB.410199.10 | hypothetical protein | 0,4234 | 0,006205 |
| TcCLB.418221.20 | hypothetical protein, conserved | 0,4231 | 0,00004401 |
| TcCLB.453917.9 | hypothetical protein, conserved | 0,4231 | 0,007001 |
| TcCLB.507047.50 | hypothetical protein, conserved | 0,423 | 0,0003002 |
| TcCLB.506717.30 | dispersed gene family protein 1 (DGF-1, pseudogene), put | 0,423 | 0,009644 |
| TcCLB.511725.150 | exportin 1, putative (CRM1) | 0,4228 | 1,224E-06 |
| TcCLB.511113.9 | hypothetical protein, conserved (fragment) | 0,4228 | 0,00006301 |
| TcCLB.506543.40 | hypothetical protein, conserved | 0,4228 | 0,002715 |
| TcCLB.509265.60 | retrotransposon hot spot protein (RHS, pseudogene), puta | 0,4228 | 0,009595 |
| TcCLB.511181.90 | peroxisome targeting signal 1 receptor, putative (PEX5) | 0,4224 | 0,00005674 |
| TcCLB.508479.320 | hypothetical protein, conserved | 0,4224 | 0,03189 |
| TcCLB.509147.21 | hypothetical protein, conserved (pseudogene) | 0,4223 | 0,06176 |
| TcCLB.511301.40 | eukaryotic translation initiation factor 4E (eIF4E) interacti | 0,4222 | 0,0007947 |
| TcCLB.507691.50 | retrotransposon hot spot (RHS) protein, putative | 0,4221 | 0,0007055 |

| | | | |
|---|---|---|---|
| TcCLB.510965.29 | hypothetical protein, conserved | 0,4218 | 0,002938 |
| TcCLB.510055.150 | hypothetical protein, conserved | 0,4217 | 2,554E-07 |
| TcCLB.506191.39 | dispersed gene family protein 1 (DGF-1), putative | 0,4216 | 0,01264 |
| TcCLB.509733.150 | hypothetical protein, conserved | 0,4213 | 9,666E-06 |
| TcCLB.510743.10 | ubiquitin-activating enzyme E1, putative | 0,4212 | 0,00001428 |
| TcCLB.507671.20 | hypothetical protein, conserved | 0,4212 | 0,009757 |
| TcCLB.509871.10 | retrotransposon hot spot protein (RHS, pseudogene), puta | 0,4211 | 8,094E-06 |
| TcCLB.510165.20 | DNA repair protein, putative | 0,4211 | 0,0001399 |
| TcCLB.479517.50 | surface protease GP63, putative | 0,4209 | 0,1231 |
| TcCLB.507717.9 | hypothetical protein, conserved | 0,4208 | 0,0058 |
| TcCLB.506419.30 | hypothetical protein, conserved | 0,4206 | 0,006858 |
| TcCLB.511621.40 | hypothetical protein, conserved | 0,4205 | 0,0003071 |
| TcCLB.507761.10 | dispersed gene family protein 1 (DGF-1, pseudogene), put | 0,4204 | 0,05524 |
| TcCLB.510643.160 | AGP2beta-2, putative | 0,4203 | 5,629E-06 |
| TcCLB.503489.24 | hypothetical protein, conserved | 0,42 | 9,567E-07 |
| TcCLB.506999.120 | hypothetical protein, conserved | 0,4198 | 0,0001024 |
| TcCLB.506481.30 | hypothetical protein, conserved | 0,4198 | 0,0002863 |
| TcCLB.511295.40 | tRNA pseudouridine synthase A-like protein, putative | 0,4193 | 0,00003301 |
| TcCLB.506537.120 | hypothetical protein, conserved | 0,4192 | 0,0719 |
| TcCLB.506977.80 | hypothetical protein, conserved | 0,4191 | 1,968E-07 |
| TcCLB.511481.50 | endonuclease III, putative | 0,4191 | 0,001863 |
| TcCLB.510731.104 | hypothetical protein, conserved | 0,4191 | 0,006663 |
| TcCLB.506447.10 | hypothetical protein | 0,4191 | 0,009235 |
| TcCLB.506025.30 | hypothetical protein, conserved | 0,419 | 0,001999 |
| TcCLB.508409.240 | hypothetical protein, conserved | 0,4189 | 0,0001605 |
| TcCLB.413977.10 | guide RNA associated protein, GAP1, putative | 0,4189 | 0,0002226 |
| TcCLB.504427.100 | hypothetical protein, conserved | 0,4187 | 0,01041 |
| TcCLB.507485.120 | tuzin | 0,4187 | 0,01852 |
| TcCLB.507715.80 | ubiquitin-conjugating enzyme, putative | 0,4185 | 0,001043 |
| TcCLB.506945.350 | serine/threonine protein kinase, putative | 0,4183 | 6,806E-09 |
| TcCLB.509207.150 | kinesin, putative | 0,4182 | 1,322E-08 |
| TcCLB.506775.60 | hypothetical protein, conserved | 0,4182 | 9,823E-06 |
| TcCLB.511165.4 | cytoplasmic translation machinery associated protein, put | 0,4181 | 0,04741 |
| TcCLB.503677.10 | phosphomannose isomerase, putative | 0,4179 | 0,0001133 |
| TcCLB.508827.40 | acyl-CoA dehydrogenase, putative | 0,4175 | 0,008129 |
| TcCLB.509785.40 | retrotransposon hot spot protein (RHS, pseudogene), puta | 0,4173 | 0,009231 |
| TcCLB.505939.80 | hypothetical protein, conserved | 0,4172 | 0,01222 |
| TcCLB.507765.70 | 40S ribosomal protein S21, putative | 0,417 | 2,052E-10 |
| TcCLB.511253.10 | kinesin-like protein, putative | 0,417 | 0,01051 |
| TcCLB.507485.130 | amastin, putative | 0,4169 | 0,0001032 |
| TcCLB.507677.100 | hypothetical protein, conserved | 0,4169 | 0,001892 |
| TcCLB.511821.170 | hypothetical protein, conserved | 0,4169 | 0,00228 |
| TcCLB.511277.420 | hypothetical protein, conserved | 0,4168 | 2,997E-06 |
| TcCLB.510877.20 | hypothetical protein, conserved | 0,4168 | 7,558E-06 |
| TcCLB.503657.9 | hypothetical protein, conserved | 0,4168 | 0,0001318 |

| | | | |
|---|---|---|---|
| TcCLB.511693.20 | hypothetical protein, conserved | 0,4168 | 0,008982 |
| TcCLB.506569.10 | cysteine peptidase, Clan CA, family C19, putative | 0,4166 | 0,00005368 |
| TcCLB.511383.40 | hypothetical protein, conserved | 0,4166 | 0,0007024 |
| TcCLB.507771.80 | hypothetical protein, conserved | 0,4164 | 0,00003524 |
| TcCLB.511039.6 | hypothetical protein, conserved | 0,4164 | 0,004158 |
| TcCLB.506297.270 | 60S ribosomal protein L28, putative | 0,4162 | 1,215E-08 |
| TcCLB.510575.230 | hypothetical protein, conserved | 0,4161 | 0,000257 |
| TcCLB.505071.90 | protein phosphatase 2A, regulatory subunit B, putative | 0,416 | 0,00003195 |
| TcCLB.510435.50 | AMP deaminase (pseudogene), putative | 0,416 | 0,0007013 |
| TcCLB.504887.10 | dispersed gene family protein 1 (DGF-1), putative | 0,4159 | 0,006231 |
| TcCLB.504069.53 | proteasome subunit alpha type-5, putative | 0,4158 | 1,058E-06 |
| TcCLB.507005.60 | structural maintenance of chromosome (SMC) family prot | 0,4157 | 7,283E-07 |
| TcCLB.509641.10 | hypothetical protein, conserved | 0,4155 | 0,01257 |
| TcCLB.445635.10 | receptor-type adenylate cyclase (pseudogene), putative | 0,415 | 0,0003772 |
| TcCLB.503473.4 | proteasome 26S non-ATPase subunit 9, putative | 0,415 | 0,05757 |
| TcCLB.511633.40 | dynein light chain, putative | 0,4149 | 0,005502 |
| TcCLB.507265.30 | DNA polymerase delta catalytic subunit, putative | 0,4147 | 0,007247 |
| TcCLB.510763.30 | hypothetical protein, conserved | 0,4146 | 0,0237 |
| TcCLB.507729.10 | dispersed gene family protein 1 (DGF-1), putative | 0,4144 | 0,2776 |
| TcCLB.506735.80 | hypothetical protein, conserved | 0,4143 | 0,00001133 |
| TcCLB.407477.40 | hypothetical protein, conserved | 0,4143 | 0,00335 |
| TcCLB.507041.30 | hypothetical protein, conserved | 0,4142 | 0,0006654 |
| TcCLB.506869.30 | hypothetical protein, conserved | 0,4142 | 0,02616 |
| TcCLB.506725.70 | hypothetical protein, conserved | 0,4139 | 0,0003312 |
| TcCLB.403985.10 | retrotransposon hot spot (RHS) protein, putative, (fragme | 0,4137 | 0,0006962 |
| TcCLB.511261.71 | hypothetical protein, conserved (pseudogene) | 0,4137 | 0,002916 |
| TcCLB.509171.54 | hypothetical protein, conserved | 0,4137 | 0,007961 |
| TcCLB.511661.30 | hypothetical protein, conserved | 0,4136 | 3,892E-08 |
| TcCLB.509127.90 | hypothetical protein, conserved | 0,4136 | 0,00003232 |
| TcCLB.508569.90 | hypothetical protein, conserved | 0,4136 | 0,0004093 |
| TcCLB.506945.310 | hypothetical protein, conserved | 0,4136 | 0,0004188 |
| TcCLB.509395.80 | hypothetical protein, conserved | 0,4135 | 0,0001442 |
| TcCLB.510761.30 | RNA re-capping enzyme, cytoplasmic, putative (Ce1) | 0,4134 | 0,00001031 |
| TcCLB.508001.90 | hypothetical protein, conserved (pseudogene) | 0,4134 | 0,07426 |
| TcCLB.509875.90 | retrotransposon hot spot protein (RHS, pseudogene), puta | 0,4133 | 0,04876 |
| TcCLB.508837.180 | DNA polymerase I alpha catalytic subunit, putative | 0,4132 | 4,006E-07 |
| TcCLB.508277.140 | phosphomevalonate kinase protein, putative | 0,413 | 9,547E-07 |
| TcCLB.507013.30 | hypothetical protein, conserved | 0,4128 | 0,01063 |
| TcCLB.511209.40 | hypothetical protein, conserved | 0,4126 | 0,005538 |
| TcCLB.510285.100 | eukaryotic translation initation factor 4 gamma, putative | 0,4124 | 0,00000759 |
| TcCLB.507099.60 | hypothetical protein, conserved | 0,4124 | 0,0001222 |
| TcCLB.504147.50 | amino acid transporter (pseudogene), putative | 0,4123 | 0,002881 |
| TcCLB.508845.5 | retrotransposon hot spot protein (RHS, pseudogene), puta | 0,4123 | 0,2649 |
| TcCLB.510155.190 | hypothetical protein, conserved | 0,4121 | 0,0001119 |
| TcCLB.509111.28 | fucose kinase, putative | 0,4121 | 0,08167 |

| | | | |
|---|---|---|---|
| TcCLB.506829.10 | hypothetical protein, conserved | 0,4118 | 0,00002105 |
| TcCLB.510307.250 | trans-sialidase (pseudogene), putative | 0,4118 | 0,003895 |
| TcCLB.508551.39 | hexose transporter, putative | 0,4118 | 0,0166 |
| TcCLB.508355.190 | choline kinase | 0,4117 | 0,00001542 |
| TcCLB.508881.90 | hypothetical protein, conserved | 0,4117 | 0,0002844 |
| TcCLB.504131.60 | hypothetical protein, conserved | 0,4114 | 0,0000782 |
| TcCLB.510279.240 | hypothetical protein | 0,4114 | 0,0726 |
| TcCLB.504883.60 | hypothetical protein, conserved | 0,411 | 0,00000109 |
| TcCLB.509429.4 | retrotransposon hot spot (RHS) protein, putative (fragmen | 0,4109 | 3,276E-06 |
| TcCLB.507491.70 | hypothetical protein, conserved (pseudogene) | 0,4109 | 9,189E-06 |
| TcCLB.507021.20 | ESAG8-associated protein, putative (PIE8) | 0,4108 | 0,1609 |
| TcCLB.504203.4 | hypothetical protein, conserved | 0,4106 | 0,0008535 |
| TcCLB.511627.150 | beta-ketoacyl-ACP reductase | 0,4104 | 0,0005494 |
| TcCLB.509437.160 | trans-sialidase, Group IV, putative | 0,4104 | 0,00124 |
| TcCLB.506195.160 | hypothetical protein, conserved | 0,4104 | 0,001339 |
| TcCLB.506525.90 | hypothetical protein, conserved | 0,4102 | 0,0004467 |
| TcCLB.509461.80 | hypothetical protein, conserved | 0,4098 | 0,0001977 |
| TcCLB.511661.120 | protein kinase, putative | 0,4097 | 1,268E-06 |
| TcCLB.508741.200 | hypothetical protein, conserved | 0,4096 | 2,659E-06 |
| TcCLB.506191.20 | retrotransposon hot spot protein (RHS, pseudogene), puta | 0,4096 | 0,004052 |
| TcCLB.507603.150 | dolichyl-P-Man:GDP-Man5GlcNAc2-PP-dolichyl alpha-1,2-r | 0,4094 | 0,0001262 |
| TcCLB.507083.80 | hypothetical protein, conserved | 0,4094 | 0,00458 |
| TcCLB.511415.50 | retrotransposon hot spot protein (RHS, pseudogene), puta | 0,4092 | 0,02372 |
| TcCLB.511827.110 | trans-sialidase, Group II, putative | 0,4091 | 0,0002438 |
| TcCLB.511821.120 | hypothetical protein, conserved | 0,409 | 0,0001558 |
| TcCLB.504137.60 | beta adaptin, putative | 0,409 | 0,0003721 |
| TcCLB.506797.60 | hypothetical protein, conserved | 0,409 | 0,0005461 |
| TcCLB.507211.30 | hypothetical protein, conserved | 0,4089 | 0,003172 |
| TcCLB.510155.120 | ubiquitin activating enzyme, putative | 0,4087 | 0,00005948 |
| TcCLB.508401.50 | hypothetical protein, conserved (pseudogene) | 0,4087 | 0,0001922 |
| TcCLB.511745.40 | hypothetical protein, conserved | 0,4087 | 0,0003259 |
| TcCLB.510747.100 | hypothetical protein, conserved | 0,4086 | 0,000737 |
| TcCLB.510737.50 | hypothetical protein, conserved | 0,4085 | 1,806E-06 |
| TcCLB.507089.50 | ATP-dependent DEAD/H RNA helicase, putative | 0,4083 | 0,00009524 |
| TcCLB.504213.10 | hypothetical protein, conserved | 0,4082 | 0,009247 |
| TcCLB.503643.10 | KU70 protein, putative | 0,4081 | 0,000281 |
| TcCLB.509215.19 | hypothetical protein, conserved | 0,408 | 0,003672 |
| TcCLB.508543.129 | hypothetical protein, conserved | 0,4079 | 0,0002191 |
| TcCLB.506265.40 | hypothetical protein, conserved | 0,4079 | 0,03193 |
| TcCLB.511587.40 | hypothetical protein, conserved (pseudogene) | 0,4078 | 3,924E-06 |
| TcCLB.508325.150 | glycine dehydrogenase (pseudogene), putative | 0,4078 | 9,159E-06 |
| TcCLB.508273.10 | AMP deaminase, putative | 0,4078 | 0,00006659 |
| TcCLB.511821.20 | hypothetical protein, conserved | 0,4078 | 0,001326 |
| TcCLB.506855.350 | hypothetical protein | 0,4078 | 0,003796 |
| TcCLB.508221.20 | trans-sialidase (pseudogene), putative | 0,4078 | 0,05887 |

| | | | |
|---|---|---|---|
| TcCLB.511391.24 | dispersed gene family protein 1 (DGF-1, pseudogene), put | 0,4076 | 0,006233 |
| TcCLB.507951.114 | high mobility group protein, putative | 0,4074 | 0,0002574 |
| TcCLB.424123.20 | paraflagellar rod component, putative (PFC16) | 0,4074 | 0,01486 |
| TcCLB.511867.110 | flap endonuclease-1 (FEN-1), putative | 0,4073 | 0,0001408 |
| TcCLB.504051.49 | hypothetical protein, conserved | 0,4073 | 0,01046 |
| TcCLB.510761.76 | UDP-glucoronosyl and UDP-glucosyl transferase, putative | 0,4072 | 0,001521 |
| TcCLB.503685.20 | hypothetical protein, conserved | 0,4072 | 0,002686 |
| TcCLB.482425.10 | retrotransposon hot spot (RHS) protein, putative | 0,4071 | 0,1796 |
| TcCLB.506297.94 | 60S ribosomal protein L24, putative | 0,4067 | 0,0003515 |
| TcCLB.508153.1020 | hypothetical protein, conserved | 0,4065 | 0,002463 |
| TcCLB.507711.150 | Ubiquitin-like modifier-activating enzyme ATG7, putative | 0,4064 | 0,0004542 |
| TcCLB.510819.30 | hypothetical protein | 0,4064 | 0,00418 |
| TcCLB.511807.70 | heat shock protein DNAJ, putative | 0,4063 | 0,00005779 |
| TcCLB.511867.150 | hypothetical protein, conserved | 0,4063 | 0,004026 |
| TcCLB.506797.90 | short chain dehydrogenase/reductase, putative | 0,4061 | 0,0002152 |
| TcCLB.507603.20 | translation initiation factor EIF-2b alpha subunit, putative | 0,4061 | 0,007629 |
| TcCLB.508711.10 | Tetratricopeptide-like helical domain containing protein (1 | 0,406 | 6,667E-09 |
| TcCLB.404001.20 | hypothetical protein, conserved | 0,406 | 0,00001343 |
| TcCLB.507517.10 | hypothetical protein, conserved | 0,4058 | 0,00008889 |
| TcCLB.508153.710 | hypothetical protein | 0,4057 | 0,0008229 |
| TcCLB.508211.40 | hypothetical protein, conserved | 0,4056 | 0,0000508 |
| TcCLB.507609.80 | hypothetical protein, conserved | 0,4055 | 0,02854 |
| TcCLB.506831.66 | hypothetical protein, conserved | 0,4054 | 0,02272 |
| TcCLB.507053.10 | 50S ribosomal protein L7Ae, putative | 0,4053 | 0,00002405 |
| TcCLB.506925.140 | DNA repair protein, putative | 0,4053 | 0,00003994 |
| TcCLB.506895.30 | hypothetical protein, conserved | 0,4053 | 0,004767 |
| TcCLB.510659.250 | phospholipase A1, putative | 0,4052 | 0,0001332 |
| TcCLB.503755.20 | hypothetical protein | 0,405 | 0,0001097 |
| TcCLB.506471.19 | hypothetical protein, conserved | 0,405 | 0,02632 |
| TcCLB.511827.80 | hypothetical protein, conserved | 0,4046 | 0,001499 |
| TcCLB.422955.10 | hypothetical protein, conserved (pseudogene) | 0,4044 | 0,0008873 |
| TcCLB.503903.30 | hypothetical protein | 0,4044 | 0,0009136 |
| TcCLB.510345.20 | hypothetical protein | 0,4043 | 0,01139 |
| TcCLB.511827.40 | hypothetical protein, conserved | 0,4042 | 0,0001893 |
| TcCLB.508693.199 | hypothetical protein, conserved | 0,4042 | 0,0003745 |
| TcCLB.511321.20 | hypothetical protein, conserved | 0,404 | 3,595E-06 |
| TcCLB.509695.90 | intraflagellar transport protein 172, putative (IFT172) | 0,4038 | 0,00008332 |
| TcCLB.418185.46 | N-acetyltransferase complex ARD1 subunit (pseudogene), | 0,4037 | 0,1301 |
| TcCLB.510465.10 | dispersed gene family protein 1 (DGF-1), putative | 0,4037 | 0,335 |
| TcCLB.509455.70 | DNA polymerase delta subunit 2, putative | 0,4035 | 0,00009315 |
| TcCLB.507397.9 | dispersed gene family protein 1 (DGF-1), putative | 0,4035 | 0,5342 |
| TcCLB.507951.70 | hypothetical protein | 0,4034 | 0,007369 |
| TcCLB.503955.40 | trans-sialidase, Group II, putative | 0,403 | 0,0000389 |
| TcCLB.507923.40 | hypothetical protein, conserved | 0,4028 | 0,00134 |
| TcCLB.506855.280 | DNA repair and recombination helicase protein PIF6, puta | 0,4027 | 0,00007117 |

| | | | |
|---|---|---|---|
| TcCLB.506471.110 | retrotransposon hot spot protein (RHS, pseudogene), puta | 0,4027 | 0,0003384 |
| TcCLB.509175.18 | hypothetical protein, conserved | 0,4027 | 0,0122 |
| TcCLB.504839.50 | nucleosome assembly protein-like protein, putative | 0,4026 | 0,00001842 |
| TcCLB.511045.20 | hypothetical protein, conserved | 0,4025 | 0,01123 |
| TcCLB.511923.130 | hypothetical protein, conserved | 0,4021 | 0,0003871 |
| TcCLB.510187.60 | glyceraldehyde 3-phosphate dehydrogenase, cytosolic, put | 0,402 | 0,002259 |
| TcCLB.509669.70 | hypothetical protein, conserved | 0,4019 | 5,881E-08 |
| TcCLB.509485.70 | retrotransposon hot spot protein (RHS, pseudogene), puta | 0,4019 | 0,003162 |
| TcCLB.410961.10 | phosphatidylinositol kinase related protein, putative | 0,4019 | 0,1205 |
| TcCLB.457101.10 | hypothetical protein, conserved | 0,4018 | 0,00002202 |
| TcCLB.506227.30 | hypothetical protein, conserved | 0,4017 | 0,0001147 |
| TcCLB.508445.10 | rab11B GTPase, putative | 0,4017 | 0,0003069 |
| TcCLB.508837.100 | hypothetical protein, conserved | 0,4016 | 0,009227 |
| TcCLB.503847.20 | myosin heavy chain, putative | 0,4012 | 0,00004317 |
| TcCLB.511283.320 | hypothetical protein, conserved | 0,4012 | 0,004043 |
| TcCLB.504067.10 | dispersed gene family protein 1 (DGF-1), putative | 0,4012 | 0,214 |
| TcCLB.509925.70 | dispersed gene family protein 1 (DGF-1), putative | 0,4011 | 0,02429 |
| TcCLB.510431.245 | hypothetical protein, conserved (pseudogene) | 0,401 | 0,009725 |
| TcCLB.510661.19 | ubiquitin-activating enzyme E1, putative (fragment) | 0,4008 | 0,001831 |
| TcCLB.508897.100 | dynein light chain, putative | 0,4006 | 0,003203 |
| TcCLB.509643.40 | hypothetical protein, conserved | 0,4006 | 0,005157 |
| TcCLB.511727.40 | ubiquitin-conjugating enzyme E2, putative | 0,4005 | 7,672E-06 |
| TcCLB.508921.20 | hypothetical protein, conserved | 0,4004 | 0,009536 |
| TcCLB.511001.220 | hypothetical protein, conserved | 0,4003 | 0,00007006 |
| TcCLB.510087.90 | hypothetical protein, conserved | 0,4003 | 0,02313 |
| TcCLB.511557.10 | serine peptidase, clan SC, family S9A-like protein (OPB) | 0,4 | 5,02E-07 |
| TcCLB.508173.210 | hypothetical protein, conserved | 0,4 | 0,00005273 |
| TcCLB.508089.71 | hypothetical protein | 0,3999 | 0,01557 |
| TcCLB.506529.440 | hypothetical protein, conserved | 0,3997 | 0,0005172 |
| TcCLB.507873.40 | hypothetical protein, conserved | 0,3997 | 0,01159 |
| TcCLB.511301.100 | hypothetical protein, conserved | 0,3996 | 0,0005651 |
| TcCLB.511367.190 | hypothetical protein, conserved | 0,3995 | 0,0000811 |
| TcCLB.511131.70 | hypothetical protein, conserved | 0,3995 | 0,002336 |
| TcCLB.507711.270 | hypothetical protein, conserved | 0,3994 | 0,004723 |
| TcCLB.509103.30 | hypothetical protein, conserved | 0,3994 | 0,03588 |
| TcCLB.504741.160 | ribosomal protein L11, putative | 0,3992 | 0,002786 |
| TcCLB.503527.50 | hypothetical protein, conserved | 0,3992 | 0,004532 |
| TcCLB.504149.210 | N-acetyltransferase complex ARD1 subunit (pseudogene), | 0,399 | 0,06526 |
| TcCLB.508837.34 | dispersed gene family protein 1 (DGF-1), putative (fragm | 0,3988 | 0,004861 |
| TcCLB.508523.10 | retrotransposon hot spot protein (RHS, pseudogene), puta | 0,3986 | 0,007058 |
| TcCLB.510305.79 | hypothetical protein, conserved | 0,3984 | 8,329E-07 |
| TcCLB.506843.80 | trans-sialidase (pseudogene), putative | 0,3983 | 0,03068 |
| TcCLB.408485.20 | retrotransposon hot spot protein (RHS, pseudogene), puta | 0,3983 | 0,04192 |
| TcCLB.509373.21 | elongation factor 1-gamma (EF-1-gamma, pseudogene), | 0,3983 | 0,1464 |
| TcCLB.510299.40 | hypothetical protein, conserved | 0,3982 | 0,00009878 |

| | | | |
|---|---|---|---|
| TcCLB.506789.330 | hypothetical protein, conserved | 0,3982 | 0,0008516 |
| TcCLB.506857.10 | glycogen synthase kinase-3 alpha, putative | 0,3982 | 0,0027 |
| TcCLB.506127.40 | hypothetical protein, conserved | 0,398 | 0,002858 |
| TcCLB.508897.20 | signal recognition particle protein, putative | 0,3979 | 0,00000092 |
| TcCLB.507011.170 | hypothetical protein, conserved | 0,3978 | 0,0004097 |
| TcCLB.509261.20 | aminopeptidase, putative | 0,3973 | 0,00001643 |
| TcCLB.507213.10 | hypothetical protein, conserved | 0,3973 | 0,00001672 |
| TcCLB.511751.60 | hypothetical protein, conserved | 0,397 | 0,00003602 |
| TcCLB.509053.130 | DnaJ chaperone protein, putative | 0,397 | 0,004563 |
| TcCLB.506419.60 | nucleoporin (NUP54/57), putative | 0,3969 | 0,0000794 |
| TcCLB.510153.10 | hypothetical protein, conserved | 0,3964 | 0,0002786 |
| TcCLB.508805.30 | choline/ethanolamine kinase (pseudogene), putative | 0,3964 | 0,0008609 |
| TcCLB.510187.500 | serine/threonine protein phosphatase 2B catalytic subunit | 0,3964 | 0,0008897 |
| TcCLB.506977.90 | proteasome regulatory non-ATP-ase subunit 5, putative (F | 0,3962 | 7,135E-09 |
| TcCLB.511807.290 | hypothetical protein, conserved | 0,3959 | 0,00003543 |
| TcCLB.509809.20 | chaperone DNAJ protein, putative | 0,3959 | 0,000945 |
| TcCLB.510253.10 | hypothetical protein, conserved | 0,3959 | 0,002763 |
| TcCLB.505997.10 | retrotransposon hot spot (RHS) protein, putative | 0,3957 | 0,005753 |
| TcCLB.506513.60 | ADP-ribosylation factor, putative (ARL-1) | 0,3956 | 0,001879 |
| TcCLB.510765.70 | 60S ribosomal protein L5, putative | 0,3955 | 0,0005731 |
| TcCLB.510579.20 | Peptide deformylase 2, putative (PDF2) | 0,3954 | 0,0003815 |
| TcCLB.506563.100 | hypothetical protein, conserved | 0,3951 | 0,00005558 |
| TcCLB.509453.50 | serine/threonine-protein phosphatase | 0,3951 | 0,003582 |
| TcCLB.508231.70 | hypothetical protein, conserved | 0,3951 | 0,00521 |
| TcCLB.508461.190 | hypothetical protein, conserved | 0,3948 | 0,0000276 |
| TcCLB.506529.546 | hypothetical protein | 0,3948 | 0,2751 |
| TcCLB.506885.440 | hypothetical protein, conserved | 0,3947 | 0,0001046 |
| TcCLB.409139.9 | trans-sialidase, putative | 0,3947 | 0,3718 |
| TcCLB.511127.330 | kinetoplastid kinetochore protein 19, putative (kkt19) | 0,3946 | 0,04179 |
| TcCLB.508239.40 | palmitoyl acyltransferase 8, putative | 0,3943 | 0,003657 |
| TcCLB.509065.33 | hypothetical protein, conserved | 0,3943 | 0,01207 |
| TcCLB.508355.431 | chaperonin containing t-complex protein, putative (fragm | 0,3941 | 0,2601 |
| TcCLB.511693.60 | hypothetical protein, conserved | 0,394 | 0,02371 |
| TcCLB.508547.201 | ubiquitin-protein ligase, putative (fragment) | 0,394 | 0,07016 |
| TcCLB.510661.60 | glycerol kinase, glycosomal, putative | 0,3938 | 0,0000706 |
| TcCLB.509105.94 | hypothetical protein, conserved | 0,3938 | 0,02551 |
| TcCLB.507213.60 | retrotransposon hot spot protein (RHS, pseudogene), puta | 0,3934 | 0,002402 |
| TcCLB.508181.60 | hypothetical protein, conserved | 0,3933 | 6,519E-06 |
| TcCLB.508221.502 | hypothetical protein, conserved (pseudogene) | 0,3933 | 0,1093 |
| TcCLB.506743.160 | protein kinase, putative | 0,3929 | 0,0001273 |
| TcCLB.506717.60 | retrotransposon hot spot protein (RHS, pseudogene), puta | 0,3928 | 0,0001412 |
| TcCLB.507907.30 | retrotransposon hot spot (RHS) protein, putative | 0,3928 | 0,1015 |
| TcCLB.413293.9 | trans-sialidase, putative (fragment) | 0,3928 | 0,2055 |
| TcCLB.507499.60 | retrotransposon hot spot protein (RHS, pseudogene), puta | 0,3926 | 2,814E-06 |
| TcCLB.509729.10 | hypothetical protein, conserved | 0,3925 | 0,000128 |

| | | | |
|---|---|---|---|
| TcCLB.510091.100 | hypothetical protein, conserved | 0,3925 | 0,007683 |
| TcCLB.508179.50 | hypothetical protein, conserved | 0,3924 | 0,001472 |
| TcCLB.508413.76 | kinetoplastid membrane protein KMP-11 | 0,3924 | 0,005112 |
| TcCLB.447255.20 | Sir2-related protein 3, putative (SIR2rp3) | 0,3923 | 0,004863 |
| TcCLB.508909.300 | hypothetical protein, conserved | 0,3922 | 0,000649 |
| TcCLB.506025.40 | hypothetical protein, conserved | 0,3919 | 0,00004528 |
| TcCLB.506851.20 | choline dehydrogenase, putative | 0,3918 | 0,0000176 |
| TcCLB.506605.180 | hypothetical protein, conserved | 0,3918 | 0,0006416 |
| TcCLB.509229.120 | hypothetical protein, conserved | 0,3917 | 0,003204 |
| TcCLB.508173.170 | protein kinase, putative | 0,3914 | 0,00002552 |
| TcCLB.506401.304 | mitochondrial carrier protein, putative | 0,3914 | 0,0001277 |
| TcCLB.506835.110 | protein mkt1, putative | 0,3912 | 3,365E-06 |
| TcCLB.506573.70 | hypothetical protein, conserved | 0,3912 | 0,01105 |
| TcCLB.506457.60 | 2-aminoethylphosphonate:pyruvateaminotransferase- like | 0,3911 | 0,001634 |
| TcCLB.511239.20 | hypothetical protein, conserved | 0,391 | 0,00132 |
| TcCLB.506795.70 | hypothetical protein, conserved | 0,391 | 0,02068 |
| TcCLB.509911.110 | hypothetical protein, conserved | 0,3909 | 0,004235 |
| TcCLB.507041.40 | hypothetical protein, conserved | 0,3907 | 0,00001449 |
| TcCLB.504175.20 | hypothetical protein, conserved | 0,3906 | 0,01265 |
| TcCLB.507023.210 | hypothetical protein, conserved | 0,3906 | 0,03461 |
| TcCLB.509643.90 | serine peptidase, putative | 0,3905 | 0,0004084 |
| TcCLB.507949.50 | SSU ribosomal protein, mitochondrial, putative (MRPS15) | 0,3904 | 0,0003181 |
| TcCLB.506999.20 | hypothetical protein, conserved | 0,3903 | 0,0000194 |
| TcCLB.506869.50 | calcineurin B subunit, putative (CNB) | 0,3902 | 0,01878 |
| TcCLB.506979.30 | hypothetical protein | 0,3902 | 0,2005 |
| TcCLB.511071.10 | hypothetical protein, conserved | 0,3901 | 0,0001509 |
| TcCLB.506883.30 | ribosome-interacting GTPase 2, putative | 0,3901 | 0,0003019 |
| TcCLB.508973.30 | NUDIX family hydrolase, putative | 0,3901 | 0,004158 |
| TcCLB.507875.150 | damage-specific DNA binding protein (pseudogene), puta | 0,3899 | 0,009616 |
| TcCLB.510571.20 | leucine-rich repeat protein 1 (LRRP1), putative | 0,3899 | 0,01842 |
| TcCLB.509047.20 | hypothetical protein | 0,3897 | 0,001245 |
| TcCLB.511821.30 | Mu-adaptin 1, putative | 0,3896 | 0,0006178 |
| TcCLB.509393.40 | hypothetical protein | 0,3895 | 0,00002727 |
| TcCLB.511695.10 | hypothetical protein | 0,3895 | 0,1169 |
| TcCLB.503877.19 | hypothetical protein, conserved | 0,3894 | 0,03005 |
| TcCLB.504071.90 | glycosyltransferase family 28 protein, putative | 0,3889 | 0,000249 |
| TcCLB.508501.210 | hypothetical protein, conserved | 0,3889 | 0,00228 |
| TcCLB.510903.80 | retrotransposon hot spot protein (RHS, pseudogene), puta | 0,3887 | 4,291E-06 |
| TcCLB.506473.6 | hypothetical protein, conserved | 0,3882 | 0,0005835 |
| TcCLB.510953.40 | hypothetical protein, conserved | 0,3882 | 0,00213 |
| TcCLB.506303.20 | cell division cycle protein 45 (CDC45), putative | 0,3881 | 0,001724 |
| TcCLB.506305.5 | hypothetical protein, conserved | 0,3881 | 0,009099 |
| TcCLB.506925.420 | hypothetical protein, conserved | 0,3879 | 0,007255 |
| TcCLB.508231.190 | ABC transporter, putative | 0,3878 | 0,0002859 |
| TcCLB.510599.14 | acetyl-CoA carboxylase (fragment) | 0,3872 | 9,493E-06 |

| | | | |
|---|---|---|---|
| TcCLB.506691.80 | hypothetical protein, conserved | 0,3872 | 0,0002193 |
| TcCLB.506515.10 | FG-GAP repeat protein, putative | 0,3872 | 0,0002859 |
| TcCLB.510947.30 | hypothetical protein, conserved | 0,3872 | 0,01108 |
| TcCLB.510737.110 | replication factor C, subunit 3, putative | 0,3871 | 0,00127 |
| TcCLB.508515.40 | protein kinase, putative | 0,3871 | 0,02242 |
| TcCLB.506337.210 | signal recognition particle receptor like protein, putative | 0,3869 | 0,00009675 |
| TcCLB.509391.20 | flagellar calcium-binding protein, putative | 0,3869 | 0,4988 |
| TcCLB.418221.10 | hypothetical protein, conserved | 0,3868 | 0,00001537 |
| TcCLB.508173.180 | nuclear transport factor (MTR2) | 0,3867 | 8,307E-06 |
| TcCLB.505193.10 | serine/threonine protein phosphatase-like protein, putativ | 0,3867 | 0,002106 |
| TcCLB.511487.90 | trans-sialidase, Group V, putative | 0,3867 | 0,1716 |
| TcCLB.507513.30 | hypothetical protein, conserved | 0,3866 | 0,05422 |
| TcCLB.506435.20 | hypothetical protein, conserved | 0,3863 | 9,828E-06 |
| TcCLB.510691.10 | 1,2-Dihydroxy-3-keto-5-methylthiopentene dioxygenase, p | 0,3863 | 0,001667 |
| TcCLB.509161.135 | hypothetical protein, conserved | 0,3861 | 0,005294 |
| TcCLB.510647.30 | tryptophanyl-tRNA synthetase, putative | 0,386 | 0,0001979 |
| TcCLB.508851.100 | cyclophilin-like protein, putative | 0,386 | 0,02251 |
| TcCLB.507201.9 | dispersed gene family protein 1 (DGF-1), putative | 0,3859 | 0,02166 |
| TcCLB.510381.40 | hypothetical protein, conserved | 0,3858 | 0,0002278 |
| TcCLB.510565.125 | hypothetical protein, conserved | 0,3856 | 0,05547 |
| TcCLB.510737.100 | hypothetical protein, conserved | 0,3855 | 0,00003783 |
| TcCLB.504147.90 | hypothetical protein, conserved | 0,3854 | 0,001251 |
| TcCLB.505963.9 | dispersed gene family protein 1 (DGF-1), putative (fragme | 0,3854 | 0,08955 |
| TcCLB.511807.3 | N-acetyltransferase complex ARD1 subunit (pseudogene), | 0,3848 | 0,0372 |
| TcCLB.510439.50 | hypothetical protein, conserved | 0,3843 | 0,0001727 |
| TcCLB.506321.300 | hypothetical protein, conserved | 0,3843 | 0,0004239 |
| TcCLB.503713.10 | hypothetical protein, conserved | 0,3843 | 0,009555 |
| TcCLB.511171.25 | hypothetical protein, conserved (pseudogene) | 0,384 | 0,03768 |
| TcCLB.510187.551 | chaperonin HSP60, mitochondrial precursor, putative (frag | 0,3837 | 0,003874 |
| TcCLB.503831.20 | oxidoreductase, putative | 0,3837 | 0,004329 |
| TcCLB.508173.120 | trans-sialidase, Group I, putative | 0,3836 | 3,103E-06 |
| TcCLB.510515.110 | hypothetical protein, conserved | 0,3833 | 0,0001929 |
| TcCLB.511249.50 | hypothetical protein, conserved | 0,3832 | 0,0003865 |
| TcCLB.510121.110 | hypothetical protein, conserved | 0,383 | 0,00003044 |
| TcCLB.511425.40 | hypothetical protein, conserved | 0,383 | 0,001141 |
| TcCLB.507709.80 | hypothetical protein, conserved | 0,383 | 0,002051 |
| TcCLB.508023.150 | trans-sialidase (pseudogene), putative | 0,3829 | 0,001063 |
| TcCLB.511589.220 | hypothetical protein, conserved | 0,3829 | 0,007848 |
| TcCLB.507485.34 | DNA-directed RNA polymerase III subunit, putative | 0,3829 | 0,3858 |
| TcCLB.506943.100 | hypothetical protein, conserved | 0,3828 | 6,624E-06 |
| TcCLB.506885.80 | hypothetical protein | 0,3827 | 0,0002815 |
| TcCLB.508839.20 | hypothetical protein, conserved | 0,3827 | 0,0007995 |
| TcCLB.510655.90 | Importin 1 (IMP1) | 0,3826 | 5,375E-08 |
| TcCLB.509747.110 | hypothetical protein, conserved | 0,3824 | 0,0008775 |
| TcCLB.508871.160 | hypothetical protein | 0,3824 | 0,3911 |

| | | | |
|---|---|---|---|
| TcCLB.506831.60 | hypothetical protein, conserved | 0,3823 | 0,001023 |
| TcCLB.407335.9 | phosphatidylserine decarboxylase, putative | 0,3823 | 0,05101 |
| TcCLB.509341.20 | hypothetical protein, conserved | 0,3822 | 3,076E-06 |
| TcCLB.507997.54 | Alpha/beta hydrolase domain-containing protein | 0,3822 | 0,0008811 |
| TcCLB.505965.60 | hypothetical protein, conserved | 0,382 | 0,01009 |
| TcCLB.507519.190 | protein kinase (pseudogene), putative | 0,3819 | 0,1779 |
| TcCLB.507089.200 | chloride channel protein, putative | 0,3818 | 0,00001907 |
| TcCLB.511127.320 | kinetoplastid kinetochore protein 10, putative (kkt10) | 0,3818 | 0,004507 |
| TcCLB.510315.30 | hypothetical protein, conserved | 0,3818 | 0,007651 |
| TcCLB.507967.30 | hypothetical protein, conserved (fragment) | 0,3815 | 0,006475 |
| TcCLB.504015.21 | elongation factor 1-gamma (EF-1-gamma, pseudogene), | 0,3807 | 0,304 |
| TcCLB.506661.60 | hypothetical protein, conserved | 0,3806 | 0,001451 |
| TcCLB.503925.80 | CDP-diacylglycerol--inositol 3-phosphatidyltransferase | 0,3806 | 0,008292 |
| TcCLB.510661.279 | hypothetical protein, conserved | 0,3806 | 0,009719 |
| TcCLB.508805.110 | hypothetical protein | 0,3806 | 0,1651 |
| TcCLB.506413.70 | hypothetical protein, conserved | 0,3804 | 0,001142 |
| TcCLB.506009.90 | hypothetical protein, conserved | 0,3799 | 0,0001887 |
| TcCLB.507027.70 | hypothetical protein, conserved | 0,3798 | 0,0003972 |
| TcCLB.504427.130 | 5'-3' exoribonuclease C, putative (XRNC) | 0,3798 | 0,000607 |
| TcCLB.508273.40 | hypothetical protein, conserved | 0,3798 | 0,001066 |
| TcCLB.429185.20 | trans-sialidase (pseudogene), putative | 0,3798 | 0,282 |
| TcCLB.511429.30 | conserved protein, unknown function | 0,3797 | 0,00002859 |
| TcCLB.509599.120 | hypothetical protein, conserved | 0,3797 | 0,004022 |
| TcCLB.510031.20 | NADH-ubiquinone oxidoreductase complex I subunit, puta | 0,3795 | 0,007115 |
| TcCLB.510493.20 | hypothetical protein, conserved | 0,3795 | 0,02651 |
| TcCLB.508569.120 | hypothetical protein, conserved | 0,3792 | 0,00005 |
| TcCLB.510947.50 | cyclophilin, putative (CYP) | 0,3792 | 0,005502 |
| TcCLB.506425.120 | eRF1 methyltransferase catalytic subunit, putative | 0,3789 | 0,03186 |
| TcCLB.506789.220 | hypothetical protein, conserved | 0,3788 | 0,0002295 |
| TcCLB.507609.20 | hypothetical protein, conserved | 0,3788 | 0,04623 |
| TcCLB.508529.10 | retrotransposon hot spot (RHS) protein, putative, (fragme | 0,3787 | 0,004159 |
| TcCLB.511211.80 | hypothetical protein, conserved | 0,3786 | 0,004585 |
| TcCLB.510667.80 | hypothetical protein, conserved | 0,3786 | 0,006667 |
| TcCLB.510533.180 | microtubule-associated protein 1A/1B, light chain 3, puta | 0,3785 | 0,05618 |
| TcCLB.503935.4 | hypothetical protein, conserved (pseudogene) | 0,3784 | 0,02574 |
| TcCLB.506583.10 | U2 small nuclear ribonucleoprotein 16.5K, putative | 0,3783 | 0,006768 |
| TcCLB.506247.320 | methyltransferase, putative | 0,3781 | 0,0001004 |
| TcCLB.510359.34 | hypothetical protein, conserved | 0,378 | 0,01276 |
| TcCLB.511807.10 | hypothetical protein, conserved (pseudogene) | 0,3777 | 7,288E-06 |
| TcCLB.507167.140 | trans-sialidase (pseudogene), putative | 0,3776 | 0,1806 |
| TcCLB.508461.400 | nucleoside diphosphate kinase (NDPK2) | 0,3775 | 0,000809 |
| TcCLB.509205.70 | hypothetical protein, conserved | 0,3774 | 0,002056 |
| TcCLB.508121.30 | trans-sialidase (pseudogene), putative | 0,3774 | 0,01394 |
| TcCLB.508857.110 | hypothetical protein, conserved | 0,3774 | 0,01833 |
| TcCLB.510691.30 | hypothetical protein, conserved | 0,3773 | 0,0004686 |

| | | | |
|---|---|---|---|
| TcCLB.511239.30 | hypothetical protein, conserved | 0,3771 | 0,01565 |
| TcCLB.510611.79 | Present in the outer mitochondrial membrane proteome 3 | 0,3769 | 0,01175 |
| TcCLB.511593.51 | elongation factor 1-gamma (EF-1-gamma, pseudogene), | 0,3769 | 0,0733 |
| TcCLB.510655.30 | helicase, putative | 0,3764 | 1,595E-06 |
| TcCLB.506945.270 | cyclin, putative | 0,3763 | 0,0006055 |
| TcCLB.510943.90 | hypothetical protein, conserved | 0,3763 | 0,03534 |
| TcCLB.510941.6 | hypothetical protein, conserved | 0,376 | 0,0005668 |
| TcCLB.509717.30 | hypothetical protein, conserved | 0,376 | 0,0007451 |
| TcCLB.510437.10 | dynein heavy chain, putative | 0,376 | 0,004548 |
| TcCLB.508259.20 | dispersed gene family protein 1 (DGF-1, pseudogene), put | 0,376 | 0,2001 |
| TcCLB.508717.36 | selenophosphate synthetase, putative | 0,3756 | 0,0002862 |
| TcCLB.507625.70 | hypothetical protein, conserved | 0,3753 | 0,0001348 |
| TcCLB.504867.120 | hypothetical protein | 0,3749 | 0,00004237 |
| TcCLB.504147.260 | hypothetical protein, conserved | 0,3749 | 0,0005096 |
| TcCLB.503395.20 | hypothetical protein, conserved | 0,3749 | 0,0179 |
| TcCLB.511545.60 | hypothetical protein | 0,3749 | 0,06004 |
| TcCLB.508163.105 | elongation factor 1-gamma (EF-1-gamma, pseudogene), | 0,3748 | 0,361 |
| TcCLB.503839.5 | chloride channel protein, putative | 0,3746 | 0,02785 |
| TcCLB.507949.69 | DNA-directed RNA polymerase 2, RPB12 subunit, putative | 0,3744 | 0,003799 |
| TcCLB.506959.90 | hypothetical protein, conserved | 0,374 | 1,509E-06 |
| TcCLB.508307.50 | hypothetical protein, conserved | 0,3738 | 0,0002853 |
| TcCLB.508479.120 | pumilio/PUF RNA binding protein 8, putative (PUF8) | 0,3737 | 5,282E-06 |
| TcCLB.509717.9 | hypothetical protein (pseudogene) | 0,3737 | 0,09823 |
| TcCLB.506775.30 | tubulin-tyrosine ligase-like protein, putative | 0,3736 | 0,00009692 |
| TcCLB.511829.5 | trans-sialidase (pseudogene), putative | 0,3736 | 0,009684 |
| TcCLB.509669.75 | hypothetical protein | 0,3736 | 0,01467 |
| TcCLB.511369.5 | elongation factor 1-alpha, putative (fragment) | 0,3734 | 0,297 |
| TcCLB.506559.220 | hypothetical protein, conserved | 0,3733 | 0,0009016 |
| TcCLB.511585.230 | trans-sialidase, Group II, putative | 0,3732 | 0,005151 |
| TcCLB.506559.314 | hypothetical protein, conserved | 0,3731 | 0,01855 |
| TcCLB.506513.170 | hypothetical protein, conserved (pseudogene) | 0,373 | 0,0002042 |
| TcCLB.507689.20 | hypothetical protein | 0,3729 | 0,01789 |
| TcCLB.507221.70 | hypothetical protein, conserved | 0,3724 | 0,03043 |
| TcCLB.507133.20 | trans-sialidase, putative (fragment) | 0,3724 | 0,04278 |
| TcCLB.508741.190 | hypothetical protein, conserved | 0,3722 | 0,0004905 |
| TcCLB.508567.49 | hypothetical protein, conserved | 0,3722 | 0,03227 |
| TcCLB.507793.50 | hypothetical protein, conserved | 0,3721 | 0,06659 |
| TcCLB.510879.140 | hypothetical protein, conserved | 0,372 | 0,001145 |
| TcCLB.511221.20 | hypothetical protein | 0,372 | 0,124 |
| TcCLB.506297.160 | 40S ribosomal protein S5, putative | 0,3719 | 0,0002028 |
| TcCLB.508649.10 | beta propeller protein, putative | 0,3718 | 0,0001904 |
| TcCLB.463451.4 | hypothetical protein, conserved | 0,3717 | 0,0457 |
| TcCLB.508381.20 | hypothetical protein, conserved (pseudogene) | 0,3717 | 0,07238 |
| TcCLB.505971.20 | hypothetical protein | 0,3715 | 0,04923 |
| TcCLB.509683.40 | hypothetical protein, conserved | 0,3712 | 0,001076 |

| | | | |
|---|---|---|---|
| TcCLB.509427.10 | trans-sialidase, Group II, putative | 0,3712 | 0,1867 |
| TcCLB.504009.10 | hypothetical protein, conserved | 0,3709 | 0,0006684 |
| TcCLB.511907.50 | lysophospholipase, putative | 0,3708 | 0,0001581 |
| TcCLB.509569.120 | RNA-binding protein, putative | 0,3708 | 0,0002467 |
| TcCLB.506959.60 | hypothetical protein, conserved | 0,3705 | 0,006259 |
| TcCLB.509905.190 | elongation factor 1-gamma (EF-1-gamma, pseudogene), | 0,3703 | 0,1249 |
| TcCLB.510745.20 | hypothetical protein, conserved | 0,3702 | 0,002739 |
| TcCLB.511459.40 | hypothetical protein, conserved | 0,3701 | 0,0006886 |
| TcCLB.503735.10 | hypothetical protein, conserved | 0,37 | 0,0201 |
| TcCLB.506945.80 | DNA ligase I, putative | 0,3699 | 0,00003548 |
| TcCLB.480653.10 | hypothetical protein, conserved | 0,3695 | 0,01864 |
| TcCLB.511735.10 | chaperonin HSP60/CNP60, putative | 0,3695 | 0,0314 |
| TcCLB.507993.220 | ARF-like 2-binding protein, putative | 0,3695 | 0,06179 |
| TcCLB.508647.40 | replication factor C large subunit, putative | 0,3693 | 7,087E-07 |
| TcCLB.509885.40 | hypothetical protein, conserved | 0,3693 | 0,0005518 |
| TcCLB.511903.190 | STOP axonemal protein, putative | 0,3692 | 0,003086 |
| TcCLB.511471.60 | protein kinase, putative | 0,3692 | 0,2245 |
| TcCLB.511731.24 | hypothetical protein | 0,3691 | 0,001044 |
| TcCLB.506961.110 | hypothetical protein, conserved (pseudogene) | 0,369 | 0,00008615 |
| TcCLB.508415.30 | hypothetical protein | 0,369 | 0,0004199 |
| TcCLB.507993.89 | hypothetical protein, conserved | 0,369 | 0,01927 |
| TcCLB.507063.240 | hypothetical protein, conserved (pseudogene) | 0,3689 | 2,999E-06 |
| TcCLB.508409.110 | hypothetical protein, conserved | 0,3689 | 0,0005465 |
| TcCLB.511817.90 | hypothetical protein, conserved | 0,3689 | 0,002689 |
| TcCLB.503995.20 | hypothetical protein, conserved | 0,3689 | 0,00323 |
| TcCLB.508153.1050 | hypothetical protein, conserved | 0,3687 | 8,098E-06 |
| TcCLB.511517.70 | RNA-binding protein, putative | 0,3687 | 0,000369 |
| TcCLB.506869.34 | hypothetical protein, conserved | 0,3686 | 0,003543 |
| TcCLB.506727.40 | hypothetical protein, conserved | 0,3685 | 0,02591 |
| TcCLB.506859.150 | hypothetical protein | 0,3684 | 0,00003069 |
| TcCLB.506567.20 | hypothetical protein | 0,3682 | 0,008078 |
| TcCLB.504867.110 | hypothetical protein, conserved | 0,3681 | 0,008241 |
| TcCLB.510143.50 | hypothetical protein, conserved | 0,3681 | 0,009568 |
| TcCLB.509445.30 | solanesyl-diphosphate synthase (SPPS) | 0,3676 | 5,298E-06 |
| TcCLB.510065.20 | DNA repair protein, putative | 0,3674 | 0,00005941 |
| TcCLB.506295.140 | hypothetical protein, conserved | 0,3673 | 0,0009298 |
| TcCLB.511445.140 | hypothetical protein | 0,3673 | 0,001715 |
| TcCLB.511825.10 | hypothetical protein, conserved | 0,3672 | 0,0005768 |
| TcCLB.506581.10 | dolicholphosphate-mannose synthase, putative (DPMS) | 0,3672 | 0,02481 |
| TcCLB.508621.30 | choline/carnitine O-acyltransferase (pseudogene), putativ | 0,3671 | 0,00001617 |
| TcCLB.507951.80 | Present in the outer mitochondrial membrane proteome | 0,3671 | 0,0005138 |
| TcCLB.506855.90 | structural maintenance of chromosome protein 4, putativ | 0,367 | 5,456E-06 |
| TcCLB.406933.10 | retrotransposon hot spot (RHS) protein, putative | 0,367 | 0,0244 |
| TcCLB.508153.420 | hypothetical protein, conserved | 0,3669 | 0,00006822 |
| TcCLB.503939.50 | hypothetical protein, conserved | 0,3668 | 0,00004885 |

| | | | |
|---|---|---|---|
| TcCLB.504741.250 | protein kinase, putative | 0,3666 | 0,0001042 |
| TcCLB.506679.130 | hypothetical protein, conserved | 0,3666 | 0,001079 |
| TcCLB.509823.10 | hypothetical protein, conserved | 0,3666 | 0,01613 |
| TcCLB.511467.10 | hypothetical protein, conserved | 0,3664 | 0,001654 |
| TcCLB.505333.30 | hypothetical protein, conserved (pseudogene) | 0,3664 | 0,17 |
| TcCLB.511367.320 | hypothetical protein, conserved | 0,3663 | 0,0005745 |
| TcCLB.509911.70 | mu-adaptin 4, putative | 0,3663 | 0,003399 |
| TcCLB.508199.20 | hypothetical protein, conserved | 0,3661 | 0,006896 |
| TcCLB.508973.50 | ATP-dependent DEAD/H RNA helicase, putative | 0,366 | 0,0001139 |
| TcCLB.503541.10 | hypothetical protein, conserved | 0,3659 | 0,007413 |
| TcCLB.510747.40 | surface protease GP63, putative | 0,3659 | 0,0382 |
| TcCLB.507275.20 | hypothetical protein, conserved | 0,3658 | 0,00001248 |
| TcCLB.511391.180 | Golgi/lysosome glycoprotein 1, putative (GLP1) | 0,3657 | 0,00002847 |
| TcCLB.508355.170 | hypothetical protein, conserved | 0,3657 | 0,0004649 |
| TcCLB.506173.170 | retrotransposon hot spot protein (RHS, pseudogene), puta | 0,3656 | 0,003307 |
| TcCLB.508239.30 | neurobeachin/beige-like protein, putative | 0,3654 | 7,127E-07 |
| TcCLB.511649.130 | vesicle-associated membrane protein, putative | 0,3654 | 0,0001954 |
| TcCLB.509769.148 | dynein heavy chain, putative | 0,3654 | 0,007811 |
| TcCLB.509717.34 | hypothetical protein, conserved | 0,3654 | 0,009884 |
| TcCLB.509775.30 | hypothetical protein, conserved | 0,3652 | 0,00002626 |
| TcCLB.507503.11 | protein kinase (pseudogene), putative | 0,3652 | 0,00008082 |
| TcCLB.508181.110 | hypothetical protein, conserved | 0,3652 | 0,0001732 |
| TcCLB.506619.49 | ras-related protein Rab21, putative | 0,3651 | 0,0007535 |
| TcCLB.509419.230 | trans-sialidase (pseudogene), putative | 0,3649 | 0,03362 |
| TcCLB.509671.99 | hypothetical protein, conserved | 0,3649 | 0,04858 |
| TcCLB.508577.140 | cyclophilin, putative | 0,3648 | 0,0002206 |
| TcCLB.504021.80 | hypothetical protein, conserved | 0,3647 | 0,006082 |
| TcCLB.504171.59 | pseudouridine synthase TruD, putative | 0,3647 | 0,01512 |
| TcCLB.503431.40 | hypothetical protein, conserved | 0,3647 | 0,0155 |
| TcCLB.508139.10 | retrotransposon hot spot (RHS) protein, putative | 0,3646 | 0,04387 |
| TcCLB.509631.160 | hypothetical protein, conserved | 0,3645 | 0,003887 |
| TcCLB.507437.30 | trans-sialidase (pseudogene), putative | 0,364 | 0,1303 |
| TcCLB.511281.59 | hypothetical protein, conserved | 0,3637 | 0,0008282 |
| TcCLB.509895.80 | ATP-dependent zinc metallopeptidase (pseudogene), puta | 0,3635 | 0,0001773 |
| TcCLB.507867.50 | hypothetical protein, conserved | 0,3635 | 0,01261 |
| TcCLB.506485.20 | hypothetical protein, conserved | 0,3634 | 0,002212 |
| TcCLB.510895.9 | hypothetical protein, conserved (fragment) | 0,3632 | 0,00001095 |
| TcCLB.505009.40 | hypothetical protein, conserved | 0,3632 | 0,002165 |
| TcCLB.506729.80 | hypothetical protein, conserved | 0,3632 | 0,007457 |
| TcCLB.510943.33 | fatty acyl CoA synthetase, putative | 0,3631 | 0,003224 |
| TcCLB.511761.89 | hypothetical protein, conserved (fragment) | 0,3631 | 0,04687 |
| TcCLB.511231.69 | hypothetical protein, conserved | 0,3629 | 0,0004115 |
| TcCLB.507985.20 | retrotransposon hot spot protein (RHS, pseudogene), puta | 0,3628 | 0,0389 |
| TcCLB.415331.10 | trans-sialidase (pseudogene), putative | 0,3627 | 0,003251 |
| TcCLB.509203.10 | 2,3-bisphosphoglycerate-independent phosphoglycerate m | 0,3625 | 0,09133 |

| | | | |
|---|---|---|---|
| TcCLB.506871.100 | polypeptide deformylase-like protein, putative | 0,3624 | 0,001678 |
| TcCLB.510577.10 | hypothetical protein, conserved | 0,3624 | 0,3094 |
| TcCLB.511535.30 | hypothetical protein, conserved | 0,3623 | 0,0001235 |
| TcCLB.510347.60 | protein kinase, putative | 0,3622 | 0,002882 |
| TcCLB.508689.5 | hypothetical protein, conserved (pseudogene) | 0,3621 | 0,04137 |
| TcCLB.506919.10 | hypothetical protein | 0,362 | 0,1516 |
| TcCLB.506915.10 | hypothetical protein, conserved | 0,3619 | 0,00008364 |
| TcCLB.510101.290 | hypothetical protein, conserved | 0,3616 | 1,516E-07 |
| TcCLB.510055.60 | hypothetical protein, conserved | 0,3616 | 0,00002695 |
| TcCLB.504505.10 | retrotransposon hot spot protein (RHS, pseudogene), puta | 0,3615 | 0,002412 |
| TcCLB.511277.539 | hypothetical protein, conserved | 0,3615 | 0,01953 |
| TcCLB.511231.80 | Nascent polypeptide-associated complex subunit beta, pu | 0,3614 | 0,02404 |
| TcCLB.511421.210 | hypothetical protein, conserved | 0,3612 | 0,00006592 |
| TcCLB.508181.80 | hypothetical protein, conserved | 0,3612 | 0,0005094 |
| TcCLB.507499.53 | ATP-dependent DEAD/H RNA helicase (pseudogene), puta | 0,3612 | 0,06768 |
| TcCLB.511641.5 | fucose kinase, putative | 0,3612 | 0,1183 |
| TcCLB.509627.40 | succinate dehydrogenase subunit (SDH9) | 0,3611 | 0,003073 |
| TcCLB.509287.240 | dispersed gene family protein 1 (DGF-1), putative | 0,3611 | 0,02058 |
| TcCLB.505807.50 | hypothetical protein, conserved | 0,361 | 0,01413 |
| TcCLB.509607.30 | splicing factor TSR1, putative | 0,3607 | 0,0001743 |
| TcCLB.445849.10 | hypothetical protein, conserved | 0,3607 | 0,009174 |
| TcCLB.507521.80 | hypothetical protein, conserved | 0,3603 | 0,0004402 |
| TcCLB.510187.80 | endonuclease V, putative | 0,3601 | 0,008948 |
| TcCLB.506739.30 | hypothetical protein | 0,36 | 0,0001735 |
| TcCLB.508141.60 | hypothetical protein, conserved | 0,3599 | 0,005749 |
| TcCLB.510955.10 | hypothetical protein, conserved | 0,3598 | 0,0001794 |
| TcCLB.506729.50 | hypothetical protein, conserved | 0,3595 | 0,00003335 |
| TcCLB.508479.350 | hypothetical protein, conserved | 0,3595 | 0,0002317 |
| TcCLB.508071.30 | beta galactofuranosyl glycosyltransferase, putative | 0,3595 | 0,155 |
| TcCLB.508613.60 | hypothetical protein, conserved | 0,3593 | 0,0001799 |
| TcCLB.506817.20 | ABC transporter, putative | 0,3593 | 0,001039 |
| TcCLB.507023.270 | hypothetical protein, conserved | 0,3592 | 0,0075 |
| TcCLB.504199.4 | hypothetical protein, conserved | 0,3591 | 0,1062 |
| TcCLB.511871.60 | hypothetical protein, conserved (pseudogene) | 0,3587 | 0,003533 |
| TcCLB.510647.10 | hypothetical protein, conserved | 0,3586 | 0,01399 |
| TcCLB.504153.230 | hypothetical protein, conserved | 0,3584 | 0,002346 |
| TcCLB.506359.40 | hypothetical protein, conserved | 0,3584 | 0,005169 |
| TcCLB.507615.76 | hypothetical protein, conserved | 0,3584 | 0,005278 |
| TcCLB.507791.10 | retrotransposon hot spot (RHS) protein, putative (fragmer | 0,3584 | 0,3091 |
| TcCLB.508713.20 | hypothetical protein, conserved | 0,3582 | 0,00591 |
| TcCLB.507501.20 | hypothetical protein, conserved | 0,3582 | 0,2935 |
| TcCLB.504161.40 | hypothetical protein, conserved | 0,3581 | 0,0003675 |
| TcCLB.510355.240 | hypothetical protein, conserved | 0,3581 | 0,006532 |
| TcCLB.511521.9 | hypothetical protein, conserved | 0,3578 | 0,04016 |
| TcCLB.507257.150 | hypothetical protein, conserved | 0,3577 | 0,004523 |

| | | | |
|---|---|---|---|
| TcCLB.506979.20 | hypothetical protein, conserved | 0,3576 | 0,03001 |
| TcCLB.506433.10 | retrotransposon hot spot protein (RHS, pseudogene), puta | 0,3575 | 0,005177 |
| TcCLB.508153.634 | hypothetical protein, conserved | 0,3575 | 0,02903 |
| TcCLB.508183.4 | aspartyl aminopeptidase, putative (fragment) | 0,3574 | 0,01624 |
| TcCLB.506775.140 | hypothetical protein, conserved | 0,3573 | 0,007792 |
| TcCLB.508355.114 | hypothetical protein, conserved | 0,3573 | 0,02776 |
| TcCLB.510181.130 | hypothetical protein | 0,3572 | 0,01575 |
| TcCLB.506071.9 | dispersed gene family protein 1 (DGF-1), putative (fragm | 0,3572 | 0,1779 |
| TcCLB.510971.30 | trans-sialidase, Group VIII, putative | 0,3572 | 0,1903 |
| TcCLB.507641.236 | hypothetical protein, conserved | 0,3571 | 0,0003646 |
| TcCLB.506529.220 | hypothetical protein, conserved | 0,3571 | 0,003122 |
| TcCLB.506985.10 | hypothetical protein, conserved | 0,3571 | 0,01106 |
| TcCLB.509693.30 | hypothetical protein, conserved | 0,3571 | 0,01348 |
| TcCLB.508323.50 | hypothetical protein, conserved | 0,3567 | 0,015 |
| TcCLB.504427.180 | hypothetical protein | 0,3567 | 0,01769 |
| TcCLB.509379.20 | Tob55, putative | 0,3566 | 0,003913 |
| TcCLB.508737.180 | hypothetical protein, conserved | 0,3564 | 0,00001253 |
| TcCLB.504027.20 | hypothetical protein | 0,3563 | 0,001163 |
| TcCLB.508181.40 | choline/ethanolamine phosphotransferase, putative | 0,3563 | 0,002101 |
| TcCLB.458015.10 | hypothetical protein, conserved | 0,3562 | 0,0003069 |
| TcCLB.509795.20 | hypothetical protein, conserved | 0,3562 | 0,02283 |
| TcCLB.510603.30 | hypothetical protein, conserved | 0,3561 | 0,1998 |
| TcCLB.511421.80 | Midasin, putative (pseudogene) | 0,356 | 0,0004009 |
| TcCLB.508207.64 | hypothetical protein, conserved | 0,356 | 0,005437 |
| TcCLB.508919.140 | hypothetical protein, conserved | 0,3559 | 9,152E-07 |
| TcCLB.511229.80 | eukaryotic translation initiation factor 3 subunit 2, putativ | 0,3559 | 0,0001144 |
| TcCLB.511303.80 | hypothetical protein, conserved | 0,3559 | 0,007604 |
| TcCLB.508231.210 | hypothetical protein, conserved | 0,3558 | 0,0002937 |
| TcCLB.503703.60 | hypothetical protein | 0,3556 | 0,004313 |
| TcCLB.481435.9 | clathrin heavy chain, putative | 0,3554 | 0,001102 |
| TcCLB.506745.10 | hypothetical protein, conserved | 0,3553 | 0,01256 |
| TcCLB.507939.70 | hypothetical protein | 0,3553 | 0,1961 |
| TcCLB.509053.120 | hypothetical protein, conserved | 0,3552 | 0,003779 |
| TcCLB.508647.120 | hypothetical protein, conserved | 0,3551 | 0,00006344 |
| TcCLB.506123.70 | hypothetical protein, conserved | 0,3551 | 0,08536 |
| TcCLB.506613.130 | dispersed gene family protein 1 (DGF-1), putative | 0,3551 | 0,1445 |
| TcCLB.510081.30 | retrotransposon hot spot protein (RHS, pseudogene), puta | 0,3548 | 0,005349 |
| TcCLB.506227.40 | hypothetical protein, conserved | 0,3547 | 0,002101 |
| TcCLB.505009.49 | hypothetical protein, conserved | 0,3547 | 0,003087 |
| TcCLB.507035.20 | retrotransposon hot spot protein (RHS, pseudogene), puta | 0,3546 | 0,002619 |
| TcCLB.508645.70 | nucleoside transporter 1, putative | 0,3545 | 0,02091 |
| TcCLB.503923.50 | GPI transamidase subunit 16, putative (fragment) (GPI16) | 0,3544 | 0,003194 |
| TcCLB.509799.100 | hypothetical protein, conserved | 0,3539 | 0,0005774 |
| TcCLB.506729.9 | heat shock 70 kDa protein, putative (fragment) | 0,3539 | 0,02952 |
| TcCLB.506629.230 | retrotransposon hot spot protein (RHS, pseudogene), puta | 0,3537 | 0,00166 |

| | | | |
|---|---|---|---|
| TcCLB.511855.20 | ubiquitin-like protein, putative | 0,3536 | 0,00001425 |
| TcCLB.507105.70 | ABC transporter, putative | 0,3536 | 0,001023 |
| TcCLB.506369.50 | hypothetical protein, conserved | 0,3536 | 0,01095 |
| TcCLB.505171.50 | ras-related GTP-binding protein, putative | 0,3536 | 0,02729 |
| TcCLB.506999.80 | hypothetical protein, conserved | 0,3535 | 0,01116 |
| TcCLB.506635.44 | hypothetical protein, conserved | 0,3535 | 0,02117 |
| TcCLB.508153.900 | hypothetical protein, conserved | 0,3534 | 0,0002593 |
| TcCLB.503809.149 | TATA binding protein (TBP) | 0,3533 | 0,002588 |
| TcCLB.508479.210 | hypothetical protein, conserved | 0,3531 | 0,002882 |
| TcCLB.503633.30 | surface protease GP63, putative | 0,3531 | 0,3077 |
| TcCLB.507939.14 | hypothetical protein, conserved | 0,353 | 0,01696 |
| TcCLB.511279.30 | ARF-like 2-binding protein, putative | 0,3529 | 0,04387 |
| TcCLB.510719.70 | bromodomain-containing protein 3 (BDF3) | 0,3527 | 0,001297 |
| TcCLB.506697.50 | peptidyl-prolyl cis-trans isomerase NIMA-interacting 4, pu | 0,3526 | 0,02602 |
| TcCLB.509799.10 | adaptin complex 4 subunit, putative | 0,3521 | 0,0106 |
| TcCLB.508355.120 | hypothetical protein, conserved | 0,3518 | 0,0007849 |
| TcCLB.508025.10 | retrotransposon hot spot (RHS) protein, putative (fragmer | 0,3518 | 0,09733 |
| TcCLB.511301.20 | hypothetical protein, conserved | 0,3517 | 0,002666 |
| TcCLB.503575.54 | hypothetical protein, conserved | 0,3517 | 0,004708 |
| TcCLB.510091.40 | Mad2-like HORMA domain containing protein (mad2) | 0,3515 | 0,002197 |
| TcCLB.509611.10 | DNA repair and recombination helicase protein PIF7, puta | 0,3514 | 0,009768 |
| TcCLB.508419.30 | tryptophanyl-tRNA synthetase (pseudogene), putative | 0,3514 | 0,01248 |
| TcCLB.508205.20 | nucleolar RNA helicase II, putative | 0,3512 | 0,001533 |
| TcCLB.510769.190 | retrotransposon hot spot protein (RHS, pseudogene), puta | 0,3512 | 0,007903 |
| TcCLB.408345.20 | 2-oxoglutarate dehydrogenase subunit, putative (fragmen | 0,3511 | 0,02862 |
| TcCLB.509971.80 | hypothetical protein, conserved | 0,351 | 0,5093 |
| TcCLB.508169.40 | mitochondrial inner membrane signal peptidase, putative | 0,3509 | 0,04453 |
| TcCLB.508175.220 | hypothetical protein, conserved | 0,3508 | 0,0006667 |
| TcCLB.503703.30 | hypothetical protein, conserved | 0,3508 | 0,002771 |
| TcCLB.511531.40 | chaperonin alpha subunit, putative | 0,3506 | 7,452E-07 |
| TcCLB.511907.340 | hypothetical protein, conserved | 0,3506 | 0,04107 |
| TcCLB.507165.59 | hypothetical protein, conserved | 0,3505 | 0,003983 |
| TcCLB.503575.50 | fatty acyl CoA synthetase, putative | 0,3504 | 0,002934 |
| TcCLB.505931.40 | histone H3, putative | 0,3501 | 0,02202 |
| TcCLB.507975.40 | dispersed gene family protein 1 (DGF-1), putative | 0,35 | 0,01683 |
| TcCLB.509693.110 | hypothetical protein, conserved | 0,3498 | 0,02094 |
| TcCLB.511131.30 | hypothetical protein, conserved | 0,3497 | 0,0008126 |
| TcCLB.398009.10 | retrotransposon hot spot protein (RHS, pseudogene), puta | 0,3497 | 0,003632 |
| TcCLB.507715.70 | hypothetical protein, conserved | 0,3495 | 0,0007007 |
| TcCLB.510431.230 | hypothetical protein, conserved | 0,3494 | 0,00184 |
| TcCLB.505807.150 | Leucine-rich repeat, ribonuclease inhibitor subtype domai | 0,3494 | 0,002883 |
| TcCLB.507027.40 | hypothetical protein, conserved | 0,3494 | 0,01991 |
| TcCLB.511871.90 | hypothetical protein, conserved | 0,3493 | 0,004244 |
| TcCLB.511277.350 | tubulin-specific chaperone, putative | 0,3492 | 0,0001573 |
| TcCLB.507645.20 | hypothetical protein, conserved | 0,3492 | 0,001709 |

| | | | |
|---|---|---|---|
| TcCLB.510241.110 | hypothetical protein, conserved | 0,3491 | 0,00004207 |
| TcCLB.508257.150 | small nuclear ribonucleoprotein SmD3, putative | 0,3489 | 0,004339 |
| TcCLB.509551.50 | hypothetical protein | 0,3487 | 4,964E-06 |
| TcCLB.510659.120 | dolichol kinase, putative | 0,3485 | 0,0002128 |
| TcCLB.511211.90 | surface protease GP63, putative | 0,3483 | 0,02189 |
| TcCLB.506051.20 | hypothetical protein | 0,3483 | 0,3207 |
| TcCLB.509551.130 | hypothetical protein, conserved | 0,3482 | 0,001165 |
| TcCLB.511735.50 | hypothetical protein, conserved | 0,3482 | 0,001431 |
| TcCLB.509997.73 | hypothetical protein, conserved | 0,3478 | 0,003883 |
| TcCLB.511689.19 | hypothetical protein, conserved (fragment) | 0,3475 | 0,007678 |
| TcCLB.509645.10 | hypothetical protein, conserved | 0,3473 | 0,0004788 |
| TcCLB.504057.50 | hypothetical protein, conserved | 0,3473 | 0,00106 |
| TcCLB.506401.330 | hypothetical protein, conserved | 0,3473 | 0,002215 |
| TcCLB.510115.50 | retrotransposon hot spot protein (RHS, pseudogene), puta | 0,3473 | 0,004449 |
| TcCLB.504159.10 | hypothetical protein, conserved | 0,3472 | 0,002714 |
| TcCLB.507801.150 | hypothetical protein, conserved | 0,3472 | 0,05727 |
| TcCLB.507179.14 | hypothetical protein | 0,3472 | 0,1607 |
| TcCLB.504741.200 | hypothetical protein, conserved | 0,347 | 0,001679 |
| TcCLB.507063.80 | hypothetical protein, conserved | 0,3469 | 0,005896 |
| TcCLB.445011.10 | hypothetical protein, conserved | 0,3469 | 0,008045 |
| TcCLB.511509.40 | hypothetical protein, conserved | 0,3469 | 0,04586 |
| TcCLB.507811.70 | hypothetical protein, conserved | 0,3468 | 0,002298 |
| TcCLB.511691.30 | ubiquitin activating enzyme, putative | 0,3467 | 0,006156 |
| TcCLB.510689.20 | quinonoid dihydropteridine reductase, putative | 0,3466 | 0,002924 |
| TcCLB.511023.19 | Tetratricopeptide-like helical domain containing protein (f | 0,3465 | 0,01068 |
| TcCLB.510911.20 | hypothetical protein, conserved | 0,3464 | 0,004792 |
| TcCLB.508681.40 | dispersed gene family protein 1 (DGF-1), putative | 0,3464 | 0,4147 |
| TcCLB.508323.20 | hypothetical protein, conserved | 0,3463 | 0,00004144 |
| TcCLB.506573.40 | hypothetical protein, conserved | 0,3463 | 0,0006902 |
| TcCLB.505193.80 | hypothetical protein, conserved | 0,3463 | 0,002656 |
| TcCLB.511151.54 | surface protease GP63, putative | 0,3463 | 0,006889 |
| TcCLB.506485.50 | hypothetical protein | 0,3462 | 0,004742 |
| TcCLB.509513.5 | hypothetical protein, conserved (pseudogene) | 0,3462 | 0,01587 |
| TcCLB.509207.120 | hypothetical protein, conserved | 0,3461 | 0,00002062 |
| TcCLB.503599.20 | hypothetical protein, conserved | 0,346 | 0,03216 |
| TcCLB.510165.50 | hypothetical protein, conserved | 0,3459 | 0,02489 |
| TcCLB.511019.80 | retrotransposon hot spot (RHS) protein, putative | 0,3458 | 0,002491 |
| TcCLB.509753.180 | hypothetical protein, conserved (pseudogene) | 0,3458 | 0,1923 |
| TcCLB.509027.10 | DNA photolyase, putative | 0,3457 | 0,01355 |
| TcCLB.507993.340 | hypothetical protein | 0,3456 | 0,1415 |
| TcCLB.509599.170 | hypothetical protein, conserved | 0,3455 | 0,007564 |
| TcCLB.510661.40 | hypothetical protein, conserved | 0,3454 | 0,0002174 |
| TcCLB.506145.24 | hypothetical protein, conserved | 0,3454 | 0,004484 |
| TcCLB.506777.40 | hypothetical protein, conserved | 0,3453 | 0,0001639 |
| TcCLB.506885.250 | hypothetical protein | 0,3453 | 0,001384 |

| | | | |
|---|---|---|---|
| TcCLB.507825.30 | Present in the outer mitochondrial membrane proteome 2 | 0,3452 | 0,0001414 |
| TcCLB.510657.110 | hypothetical protein, conserved | 0,3452 | 0,0006779 |
| TcCLB.503805.9 | Quinonprotein alcohol dehydrogenase-like protein, putativ | 0,345 | 0,003186 |
| TcCLB.503643.30 | hypothetical protein, conserved | 0,3447 | 0,005367 |
| TcCLB.506459.250 | hypothetical protein, conserved | 0,3447 | 0,007583 |
| TcCLB.510835.60 | hypothetical protein, conserved | 0,3446 | 0,0002361 |
| TcCLB.504411.40 | hypothetical protein, conserved | 0,3446 | 0,006167 |
| TcCLB.511719.30 | hypothetical protein, conserved | 0,3441 | 0,02054 |
| TcCLB.422207.21 | retrotransposon hot spot protein (RHS, pseudogene), puta | 0,3439 | 0,2612 |
| TcCLB.507029.60 | hypothetical protein, conserved | 0,3437 | 0,004488 |
| TcCLB.506251.110 | hypothetical protein, conserved | 0,3437 | 0,007704 |
| TcCLB.506887.90 | hypothetical protein, conserved | 0,3436 | 0,0003869 |
| TcCLB.509059.70 | hypothetical protein, conserved | 0,3436 | 0,0007773 |
| TcCLB.511041.10 | hypothetical protein, conserved | 0,3436 | 0,003366 |
| TcCLB.506017.20 | hypothetical protein, conserved (pseudogene) | 0,3432 | 0,001103 |
| TcCLB.503613.70 | hypothetical protein, conserved | 0,3432 | 0,03111 |
| TcCLB.504351.41 | ATP-dependent DEAD/H RNA helicase (pseudogene), puta | 0,3431 | 0,2682 |
| TcCLB.511445.130 | Mitochondrial SSU ribosomal protein, putative | 0,3429 | 0,0004839 |
| TcCLB.507005.4 | acyl transferase-like protein | 0,3428 | 0,002159 |
| TcCLB.509229.130 | hypothetical protein, conserved | 0,3428 | 0,01344 |
| TcCLB.503793.20 | phosphatidylinositol (3,5) kinase, putative (fragment) | 0,3428 | 0,1261 |
| TcCLB.507093.300 | hypothetical protein SCD6.10 | 0,3426 | 0,000906 |
| TcCLB.507505.30 | hypothetical protein | 0,3426 | 0,002273 |
| TcCLB.507785.10 | clathrin heavy chain, putative | 0,3425 | 0,00004109 |
| TcCLB.505807.90 | hypothetical protein, conserved | 0,3423 | 0,00009816 |
| TcCLB.508547.90 | hypothetical protein, conserved | 0,3423 | 0,08531 |
| TcCLB.506063.34 | hypothetical protein, conserved (pseudogene) | 0,3421 | 0,0002419 |
| TcCLB.509599.70 | hypothetical protein, conserved | 0,3421 | 0,006943 |
| TcCLB.506265.90 | hypothetical protein, conserved | 0,3421 | 0,01655 |
| TcCLB.508269.73 | ATP-dependent DEAD/H RNA helicase (pseudogene), puta | 0,3421 | 0,07661 |
| TcCLB.511283.300 | hypothetical protein, conserved | 0,342 | 0,0007811 |
| TcCLB.510121.150 | serine/threonine protein kinase, putative | 0,3418 | 0,002478 |
| TcCLB.504575.15 | ubiquinol-cytochrome-C-reductase, putative | 0,3418 | 0,1944 |
| TcCLB.511277.280 | hypothetical protein, conserved | 0,3417 | 0,000196 |
| TcCLB.511217.90 | hypothetical protein, conserved | 0,3416 | 0,00002688 |
| TcCLB.510751.40 | Importin 1 (fragment) (IMP1) | 0,3416 | 0,0003206 |
| TcCLB.509167.190 | protein kinase-like protein, putative | 0,3414 | 0,0002247 |
| TcCLB.506859.160 | hypothetical protein, conserved | 0,3414 | 0,0003825 |
| TcCLB.510989.60 | hypothetical protein, conserved | 0,3414 | 0,0005691 |
| TcCLB.466823.10 | Checkpoint protein HUS1, putative | 0,3414 | 0,03511 |
| TcCLB.506349.40 | glycine dehydrogenase [decarboxylating] (pseudogene), pu | 0,3413 | 0,2746 |
| TcCLB.507899.10 | retrotransposon hot spot protein (RHS, pseudogene), puta | 0,3407 | 0,009955 |
| TcCLB.511733.50 | hypothetical protein, conserved | 0,3406 | 0,0002723 |
| TcCLB.507603.50 | Mitochondrial Ribosomal Protein of Small Subunit 29, put | 0,3406 | 0,002332 |
| TcCLB.506193.70 | glucose-6-phosphate dehydrogenase (pseudogene), putati | 0,3405 | 0,0039 |

| | | | |
|---|---|---|---|
| TcCLB.508541.220 | casein kinase, putative | 0,3405 | 0,0869 |
| TcCLB.509167.30 | hypothetical protein, conserved | 0,3403 | 0,01796 |
| TcCLB.506933.80 | RNA polymerase B subunit RPB8, putative | 0,3402 | 0,001163 |
| TcCLB.510535.10 | hypothetical protein, conserved | 0,3402 | 0,003503 |
| TcCLB.509027.30 | hypothetical protein, conserved | 0,3402 | 0,01328 |
| TcCLB.508919.94 | hypothetical protein, conserved | 0,3401 | 0,01762 |
| TcCLB.508799.190 | hypothetical protein, conserved | 0,3399 | 0,0002352 |
| TcCLB.506999.170 | hypothetical protein | 0,3399 | 0,001265 |
| TcCLB.506933.107 | hypothetical protein, conserved | 0,3398 | 0,009642 |
| TcCLB.510837.30 | hypothetical protein, conserved | 0,3397 | 0,02624 |
| TcCLB.506977.130 | ras-related protein Rab21, putative (RAB21) | 0,3395 | 0,00304 |
| TcCLB.511065.23 | hypothetical protein, conserved | 0,3393 | 0,0166 |
| TcCLB.506355.80 | flagellum transition zone component, putative | 0,3392 | 2,064E-06 |
| TcCLB.505999.170 | hypothetical protein, conserved | 0,3389 | 6,385E-06 |
| TcCLB.506239.30 | lectin, putative (AG38) | 0,3388 | 0,0001287 |
| TcCLB.509921.20 | trans-sialidase (pseudogene), putative | 0,3387 | 0,04949 |
| TcCLB.510311.80 | hypothetical protein, conserved | 0,3386 | 0,002859 |
| TcCLB.510867.9 | dispersed gene family protein 1 (DGF-1), putative | 0,3384 | 0,09069 |
| TcCLB.510437.30 | hypothetical protein, conserved | 0,3384 | 0,1292 |
| TcCLB.510275.10 | hypothetical protein | 0,3383 | 0,03046 |
| TcCLB.511507.60 | hypothetical protein | 0,3383 | 0,03871 |
| TcCLB.507529.45 | hypothetical protein, conserved | 0,3381 | 0,03018 |
| TcCLB.510759.180 | hypothetical protein, conserved | 0,338 | 0,000484 |
| TcCLB.506679.190 | hypothetical protein, conserved | 0,338 | 0,0005575 |
| TcCLB.511421.60 | deoxyhypusine synthase, putative | 0,338 | 0,01025 |
| TcCLB.507795.40 | hypothetical protein, conserved | 0,338 | 0,1249 |
| TcCLB.506695.49 | hypothetical protein, conserved | 0,3379 | 0,0002953 |
| TcCLB.510219.40 | hypothetical protein, conserved | 0,3378 | 0,008836 |
| TcCLB.507519.30 | Eukaryotic translation initiation factor 4E type 6, putative | 0,3377 | 0,003234 |
| TcCLB.508375.60 | succinate dehydrogenase subunit | 0,3377 | 0,005357 |
| TcCLB.510299.30 | hypothetical protein, conserved | 0,3376 | 0,0125 |
| TcCLB.510119.60 | hypothetical protein, conserved | 0,3374 | 0,02427 |
| TcCLB.511843.90 | hypothetical protein, conserved (pseudogene) | 0,3369 | 0,03346 |
| TcCLB.504033.110 | hypothetical protein, conserved | 0,3368 | 0,004944 |
| TcCLB.458759.20 | hypothetical protein, conserved | 0,3368 | 0,03941 |
| TcCLB.506179.5 | ferric reductase transmembrane protein, putative (fragme | 0,3368 | 0,3437 |
| TcCLB.510947.10 | GDP-mannose pyrophosphorylase | 0,3363 | 0,0003289 |
| TcCLB.508547.170 | hypothetical protein, conserved | 0,3361 | 0,000029 |
| TcCLB.503431.80 | hypothetical protein, conserved | 0,336 | 0,005408 |
| TcCLB.507049.170 | Axonemal inner arm dynein light chain, putative | 0,336 | 0,00563 |
| TcCLB.509023.200 | cdc2-related kinase 12 (CRK12) | 0,3358 | 0,0004448 |
| TcCLB.510359.570 | mucin-associated surface protein (MASP), putative | 0,3358 | 0,06055 |
| TcCLB.504021.60 | hypothetical protein, conserved | 0,3357 | 0,00663 |
| TcCLB.510565.160 | hypothetical protein, conserved | 0,3357 | 0,02272 |
| TcCLB.511219.10 | dispersed gene family protein 1 (DGF-1), putative | 0,3357 | 0,4407 |

| | | | |
|---|---|---|---|
| TcCLB.507001.130 | hypothetical protein, conserved | 0,3356 | 0,00435 |
| TcCLB.506435.110 | hypothetical protein, conserved | 0,3355 | 0,03677 |
| TcCLB.503593.50 | hypothetical protein, conserved | 0,3352 | 0,0002149 |
| TcCLB.504125.50 | mitochondrial carrier protein, putative | 0,3351 | 0,004431 |
| TcCLB.506251.100 | poly(A)-specific ribonuclease PARN, putative | 0,3349 | 0,009047 |
| TcCLB.507165.20 | hypothetical protein, conserved | 0,3349 | 0,09257 |
| TcCLB.506493.30 | hypothetical protein, conserved | 0,3348 | 0,004924 |
| TcCLB.508879.210 | hypothetical protein, conserved | 0,3345 | 0,003637 |
| TcCLB.510155.60 | hypothetical protein, conserved | 0,3345 | 0,008903 |
| TcCLB.506791.30 | hypothetical protein, conserved | 0,3343 | 0,0001086 |
| TcCLB.505647.20 | hypothetical protein, conserved | 0,334 | 0,00005112 |
| TcCLB.507483.20 | ef-hand protein 5, putative | 0,334 | 0,01723 |
| TcCLB.510283.120 | hypothetical protein, conserved | 0,3339 | 0,005092 |
| TcCLB.508775.20 | retrotransposon hot spot protein (RHS, pseudogene), puta | 0,3339 | 0,03358 |
| TcCLB.506305.40 | kinesin, putative | 0,3339 | 0,107 |
| TcCLB.504191.10 | sterol 24-c-methyltransferase, putative | 0,3338 | 0,00005691 |
| TcCLB.510307.220 | retrotransposon hot spot (RHS) protein, putative | 0,3338 | 0,01635 |
| TcCLB.504923.10 | dispersed gene family protein 1 (DGF-1), putative | 0,3337 | 0,4922 |
| TcCLB.503925.90 | hypothetical protein, conserved | 0,3336 | 0,1579 |
| TcCLB.507257.120 | hypothetical protein, conserved | 0,3335 | 0,01689 |
| TcCLB.507159.37 | hypothetical protein, conserved | 0,3334 | 0,01035 |
| TcCLB.509151.29 | hypothetical protein, conserved (pseudogene) | 0,3334 | 0,09335 |
| TcCLB.511419.4 | hypothetical protein, conserved (fragment) | 0,3334 | 0,1311 |
| TcCLB.506559.10 | hypothetical protein | 0,3333 | 0,002029 |
| TcCLB.506999.100 | hypothetical protein, conserved | 0,3333 | 0,0154 |
| TcCLB.510593.20 | hypothetical protein, conserved | 0,3332 | 0,001153 |
| TcCLB.506303.70 | hypothetical protein, conserved | 0,3329 | 0,001195 |
| TcCLB.508461.140 | poly(A)-binding protein, putative | 0,3326 | 6,915E-06 |
| TcCLB.510121.190 | hypothetical protein, conserved | 0,3326 | 0,02895 |
| TcCLB.510647.60 | intraflagellar transport protein 27 (IFT27) | 0,3325 | 0,005136 |
| TcCLB.507641.40 | hypothetical protein, conserved | 0,3323 | 0,00002197 |
| TcCLB.509715.120 | phosphatidylinositol-4-phosphate 5-kinase, putative | 0,3323 | 0,002918 |
| TcCLB.509025.29 | DNA photolyase, putative | 0,3323 | 0,026 |
| TcCLB.506537.240 | hypothetical protein, conserved | 0,3323 | 0,03946 |
| TcCLB.508645.40 | nucleoside transporter 1, putative | 0,3321 | 5,942E-07 |
| TcCLB.510751.10 | poly(A)-specific ribonuclease PARN, putative | 0,3321 | 0,0009975 |
| TcCLB.506831.5 | hypothetical protein, conserved (fragment) | 0,3319 | 0,1386 |
| TcCLB.511477.25 | hypothetical protein, conserved (pseudogene) | 0,3318 | 0,1417 |
| TcCLB.506859.30 | hypothetical protein | 0,3317 | 0,0004255 |
| TcCLB.506245.73 | elongation factor 1-gamma (EF-1-gamma, pseudogene), | 0,3315 | 0,421 |
| TcCLB.508257.100 | hypothetical protein, conserved | 0,3312 | 0,03432 |
| TcCLB.511263.9 | heat shock protein 110, putative (fragment) | 0,3311 | 0,00006488 |
| TcCLB.511453.10 | dispersed gene family protein 1 (DGF-1), putative | 0,331 | 0,173 |
| TcCLB.509599.164 | engulfment and cell motility domain 2, putative | 0,3309 | 0,0009955 |
| TcCLB.508659.40 | protein phosphatase 2C-like (pseudogene), putative | 0,3309 | 0,04737 |

| | | | |
|---|---|---|---|
| TcCLB.509859.80 | coronin, putative (CRN12) | 0,3308 | 0,0000321 |
| TcCLB.511215.30 | hypothetical protein | 0,3308 | 0,001808 |
| TcCLB.506775.100 | DNA repair and recombination helicase protein PIF2, puta | 0,3308 | 0,02806 |
| TcCLB.503953.59 | mitochondrial DNA polymerase beta-PAK, putative | 0,3305 | 0,01312 |
| TcCLB.510565.70 | protein kinase, putative | 0,3304 | 0,0001514 |
| TcCLB.508475.30 | surface protease GP63, putative | 0,3304 | 0,04627 |
| TcCLB.508449.31 | trans-sialidase, putative | 0,3304 | 0,4282 |
| TcCLB.506297.290 | protein transport protein SEC31, putative | 0,3303 | 4,481E-06 |
| TcCLB.506581.60 | hypothetical protein, conserved | 0,3301 | 0,005167 |
| TcCLB.504575.10 | hypothetical protein, conserved | 0,3301 | 0,1478 |
| TcCLB.511907.40 | hypothetical protein, conserved | 0,33 | 0,02036 |
| TcCLB.506531.60 | hypothetical protein, conserved | 0,3299 | 0,0009664 |
| TcCLB.508269.50 | hypothetical protein, conserved | 0,3299 | 0,002053 |
| TcCLB.503987.60 | hypothetical protein, conserved | 0,3299 | 0,02608 |
| TcCLB.508471.10 | hypothetical protein, conserved | 0,3299 | 0,03462 |
| TcCLB.510743.40 | prenyl protein specific carboxyl methyltransferase, putativ | 0,3297 | 0,007945 |
| TcCLB.506821.110 | hypothetical protein, conserved | 0,3297 | 0,02175 |
| TcCLB.511277.4 | 30S ribosomal protein S8, putative (fragment) | 0,3295 | 0,02148 |
| TcCLB.510307.77 | N-acetyltransferase complex ARD1 subunit (pseudogene), | 0,3294 | 0,125 |
| TcCLB.509657.20 | dispersed gene family protein 1 (DGF-1), putative | 0,3291 | 0,09216 |
| TcCLB.510007.30 | RNA-binding protein, putative | 0,329 | 0,002503 |
| TcCLB.509695.60 | hypothetical protein, conserved | 0,329 | 0,01228 |
| TcCLB.507011.269 | 6-phosphogluconate dehydrogenase, decarboxylating (6PG | 0,3288 | 0,09768 |
| TcCLB.508265.100 | hypothetical protein, conserved | 0,3287 | 0,0007838 |
| TcCLB.510603.160 | hypothetical protein, conserved | 0,3287 | 0,01172 |
| TcCLB.508723.59 | hypothetical protein, conserved | 0,3284 | 0,008509 |
| TcCLB.507229.20 | dispersed gene family protein 1 (DGF-1, pseudogene), put | 0,3284 | 0,04693 |
| TcCLB.510729.70 | proteasome alpha 1 subunit, putative | 0,3283 | 0,0002598 |
| TcCLB.510779.10 | tryptophanyl-tRNA synthetase (pseudogene), putative | 0,328 | 0,04428 |
| TcCLB.506801.90 | hypothetical protein, conserved | 0,3278 | 0,01042 |
| TcCLB.397937.5 | N-acetylglucosamine-6-phosphate deacetylase-like protei | 0,3276 | 0,03996 |
| TcCLB.509023.190 | hypothetical protein, conserved | 0,3274 | 0,08882 |
| TcCLB.508595.50 | cysteine peptidase, putative | 0,3273 | 0,00008576 |
| TcCLB.507771.50 | glycerolphosphate mutase, putative | 0,3273 | 0,00249 |
| TcCLB.508401.70 | ATP-dependent DEAD/H RNA helicase (pseudogene), puta | 0,3273 | 0,09657 |
| TcCLB.503559.109 | electron-transfer-flavoprotein, alpha polypeptide, putative | 0,3272 | 0,001492 |
| TcCLB.507511.10 | cleavage and polyadenylation specificity factor, putative | 0,3272 | 0,001808 |
| TcCLB.510155.200 | hypothetical protein, conserved | 0,3271 | 0,004524 |
| TcCLB.410943.10 | hypothetical protein, conserved | 0,3271 | 0,1986 |
| TcCLB.509053.70 | p22 protein precursor, putative | 0,3269 | 0,007477 |
| TcCLB.506745.30 | hypothetical protein, conserved | 0,3269 | 0,3657 |
| TcCLB.507467.50 | hypothetical protein, conserved | 0,3268 | 0,01083 |
| TcCLB.503579.20 | hypothetical protein, conserved | 0,3264 | 0,002844 |
| TcCLB.506833.30 | serine/threonine protein phosphatase, putative | 0,3264 | 0,003616 |
| TcCLB.509631.150 | hypothetical protein, conserved | 0,3263 | 0,1743 |

| | | | |
|---|---|---|---|
| TcCLB.506925.470 | heat shock protein DNAJ, putative | 0,3262 | 0,0007922 |
| TcCLB.507609.10 | nucleoporin interacting component (NUP93), putative | 0,3262 | 0,01684 |
| TcCLB.508479.310 | hypothetical protein, conserved | 0,3261 | 0,009147 |
| TcCLB.479721.10 | hypothetical protein, conserved (pseudogene) | 0,3261 | 0,4378 |
| TcCLB.509097.28 | hypothetical protein, conserved | 0,3259 | 0,103 |
| TcCLB.416605.20 | mucin TcMUCII (pseudogene), putative | 0,3259 | 0,285 |
| TcCLB.507257.30 | hypothetical protein, conserved | 0,3257 | 0,02519 |
| TcCLB.504221.9 | membrane associated protein, putative | 0,3255 | 0,002865 |
| TcCLB.506209.70 | dispersed gene family protein 1 (DGF-1), putative | 0,3255 | 0,1918 |
| TcCLB.510853.96 | trans-sialidase (pseudogene), putative | 0,3253 | 0,00381 |
| TcCLB.509755.89 | hypothetical protein | 0,325 | 0,2798 |
| TcCLB.490507.10 | hypothetical protein, conserved | 0,3246 | 0,005245 |
| TcCLB.508873.530 | hypothetical protein, conserved (pseudogene) | 0,3244 | 0,106 |
| TcCLB.511737.10 | small glutamine-rich tetratricopeptide repeat protein, put | 0,3241 | 0,002223 |
| TcCLB.504131.140 | hypothetical protein, conserved | 0,3238 | 0,0000319 |
| TcCLB.507047.130 | hypothetical protein, conserved | 0,3238 | 0,008621 |
| TcCLB.510519.80 | hypothetical protein, conserved | 0,3234 | 0,01923 |
| TcCLB.509953.9 | ubiquitin carboxyl-terminal hydrolase, putative (fragment | 0,3233 | 0,04525 |
| TcCLB.506529.150 | hypothetical protein, conserved | 0,323 | 0,0002863 |
| TcCLB.511133.10 | hypothetical protein, conserved | 0,323 | 0,06131 |
| TcCLB.509765.60 | retrotransposon hot spot protein (RHS, pseudogene), puta | 0,3229 | 0,004152 |
| TcCLB.511589.260 | glycyl-tRNA synthetase, putative | 0,3228 | 0,04654 |
| TcCLB.507819.11 | N-acetyltransferase complex ARD1 subunit (pseudogene), | 0,3228 | 0,1409 |
| TcCLB.510969.20 | hypothetical protein, conserved | 0,3227 | 0,01081 |
| TcCLB.509505.89 | hypothetical protein, conserved | 0,3227 | 0,01141 |
| TcCLB.509763.20 | hypothetical protein | 0,3226 | 0,04617 |
| TcCLB.503925.20 | hypothetical protein, conserved | 0,3225 | 0,003068 |
| TcCLB.508699.80 | hypothetical protein, conserved | 0,3222 | 0,003051 |
| TcCLB.509059.80 | hypothetical protein, conserved | 0,3221 | 0,0004299 |
| TcCLB.505807.200 | hypothetical protein, conserved | 0,3221 | 0,001967 |
| TcCLB.508707.270 | hypothetical protein, conserved | 0,322 | 0,001266 |
| TcCLB.507211.40 | inosine-5'-monophosphate dehydrogenase, putative | 0,3219 | 0,00003965 |
| TcCLB.509567.10 | hypothetical protein, conserved | 0,3219 | 0,01236 |
| TcCLB.454787.20 | elongation factor 1-gamma (EF-1-gamma, pseudogene), | 0,3217 | 0,418 |
| TcCLB.511367.170 | vacuolar protein sorting-associated protein 35, putative | 0,3215 | 0,00006708 |
| TcCLB.506319.10 | hypothetical protein | 0,3213 | 0,00006069 |
| TcCLB.511745.20 | AAA ATPase, putative | 0,3213 | 0,001091 |
| TcCLB.509733.199 | Extradiol ring-cleavage dioxygenase, putative | 0,3213 | 0,00296 |
| TcCLB.508851.210 | replication factor C, subunit 4, putative | 0,3213 | 0,03919 |
| TcCLB.507395.14 | hypothetical protein, conserved | 0,3212 | 0,01367 |
| TcCLB.508919.40 | hypothetical protein, conserved | 0,3211 | 0,0012 |
| TcCLB.506485.100 | hypothetical protein, conserved | 0,3211 | 0,004702 |
| TcCLB.507691.31 | hypothetical protein | 0,3209 | 0,08058 |
| TcCLB.504881.20 | hypothetical protein, conserved | 0,3208 | 0,00004301 |
| TcCLB.508871.10 | trans-sialidase, Group V, putative | 0,3208 | 0,1934 |

| | | | |
|---|---|---|---|
| TcCLB.510743.30 | hypothetical protein, conserved | 0,3207 | 0,00188 |
| TcCLB.508383.10 | Present in the outer mitochondrial membrane proteome 3 | 0,3207 | 0,05659 |
| TcCLB.506495.24 | hypothetical protein, conserved | 0,3206 | 0,04404 |
| TcCLB.511803.10 | cullin 2, putative | 0,3203 | 0,01483 |
| TcCLB.503847.70 | hypothetical protein, conserved | 0,3202 | 0,002195 |
| TcCLB.511021.90 | hypothetical protein | 0,3202 | 0,01157 |
| TcCLB.507645.30 | hypothetical protein, conserved | 0,3202 | 0,01327 |
| TcCLB.509777.21 | hypothetical protein | 0,3202 | 0,1158 |
| TcCLB.511469.80 | hypothetical protein, conserved | 0,3201 | 0,005232 |
| TcCLB.509693.50 | metallo-peptidase, Clan MG, Family M24 (MetAP2) | 0,3199 | 0,0007281 |
| TcCLB.506791.79 | hypothetical protein, conserved | 0,3199 | 0,02127 |
| TcCLB.511269.70 | cysteine peptidase, Clan CA, family C2, putative | 0,3197 | 0,0601 |
| TcCLB.506167.9 | hypothetical protein, conserved | 0,3196 | 0,0002836 |
| TcCLB.511361.10 | kinesin-like protein (pseudogene), putative | 0,3196 | 0,004535 |
| TcCLB.504091.10 | hypothetical protein, conserved | 0,3195 | 0,002195 |
| TcCLB.511277.630 | serine/threonine protein phosphatase type 5, putative | 0,3195 | 0,003548 |
| TcCLB.508465.100 | hypothetical protein | 0,3194 | 0,05226 |
| TcCLB.504099.40 | retrotransposon hot spot protein (RHS, pseudogene), puta | 0,3194 | 0,155 |
| TcCLB.506289.40 | hypothetical protein, conserved | 0,3189 | 0,008542 |
| TcCLB.504205.10 | hypothetical protein | 0,3189 | 0,08321 |
| TcCLB.504147.140 | hypothetical protein | 0,3188 | 0,00165 |
| TcCLB.511435.20 | Replication Factor C Subunit 1-related protein | 0,3188 | 0,006871 |
| TcCLB.507011.260 | hypothetical protein, conserved | 0,3187 | 0,004117 |
| TcCLB.503811.50 | hypothetical protein, conserved | 0,3186 | 0,004886 |
| TcCLB.506123.24 | dynein light chain, putative | 0,3186 | 0,03862 |
| TcCLB.511585.80 | mitochondrial-processing peptidase subunit beta, putative | 0,3183 | 0,001348 |
| TcCLB.506479.20 | hypothetical protein, conserved | 0,3182 | 0,01488 |
| TcCLB.506201.60 | hypothetical protein | 0,3182 | 0,05865 |
| TcCLB.506221.40 | mitochondrial carrier protein, putative | 0,3181 | 0,01093 |
| TcCLB.507993.130 | 60S ribosomal protein L7, putative | 0,318 | 0,0003258 |
| TcCLB.510687.40 | serine/threonine protein phosphatase, putative | 0,3179 | 0,001087 |
| TcCLB.509645.5 | hypothetical protein, conserved (pseudogene) | 0,3177 | 0,01022 |
| TcCLB.511881.30 | hypothetical protein, conserved | 0,3175 | 0,008002 |
| TcCLB.508771.34 | hypothetical protein, conserved | 0,3175 | 0,1265 |
| TcCLB.504253.60 | hypothetical protein, conserved (fragment) | 0,3174 | 0,2356 |
| TcCLB.511033.29 | hypothetical protein | 0,3172 | 0,05018 |
| TcCLB.507077.9 | hypothetical protein, conserved | 0,3167 | 0,01325 |
| TcCLB.504255.10 | hypothetical protein, conserved | 0,3165 | 0,03341 |
| TcCLB.509911.130 | hypothetical protein, conserved | 0,3164 | 0,0005442 |
| TcCLB.510531.60 | hypothetical protein, conserved | 0,3164 | 0,0005846 |
| TcCLB.509539.67 | paraflagellar rod component par4, putative (fragment) | 0,3164 | 0,1113 |
| TcCLB.511819.10 | hypothetical protein, conserved | 0,3163 | 0,003636 |
| TcCLB.509997.86 | serine/threonine protein phosphatase, putative (fragment | 0,3162 | 0,001894 |
| TcCLB.506885.394 | hypothetical protein, conserved | 0,3162 | 0,0242 |
| TcCLB.508577.30 | retrotransposon hot spot protein (RHS, pseudogene), puta | 0,3161 | 0,004605 |

| | | | |
|---|---|---|---|
| TcCLB.509149.20 | zinc finger protein family member, putative (POMP35) | 0,3161 | 0,02189 |
| TcCLB.507817.40 | hypothetical protein, conserved | 0,316 | 0,04811 |
| TcCLB.511773.50 | protein kinase, putative | 0,3159 | 0,009725 |
| TcCLB.504881.50 | ABC transporter, putative | 0,3159 | 0,08245 |
| TcCLB.510743.120 | hypothetical protein, conserved | 0,3158 | 0,002324 |
| TcCLB.509179.130 | hypothetical protein, conserved | 0,3158 | 0,0121 |
| TcCLB.510729.220 | hypothetical protein, conserved | 0,3157 | 0,00001448 |
| TcCLB.504827.130 | eukaryotic translation initation factor 4 gamma, putative | 0,3157 | 0,003687 |
| TcCLB.511499.10 | hypothetical protein, conserved | 0,3155 | 0,001982 |
| TcCLB.509669.30 | ABC transporter, putative | 0,3155 | 0,002239 |
| TcCLB.511837.20 | hypothetical protein, conserved | 0,3155 | 0,02112 |
| TcCLB.509217.140 | hypothetical protein, conserved (pseudogene) | 0,3154 | 0,05026 |
| TcCLB.510435.20 | hypothetical protein, conserved | 0,3152 | 0,01929 |
| TcCLB.509625.20 | hypothetical protein, conserved | 0,3152 | 0,1485 |
| TcCLB.507077.40 | hypothetical protein, conserved | 0,3151 | 0,01732 |
| TcCLB.510105.70 | hypothetical protein, conserved | 0,3151 | 0,02421 |
| TcCLB.509795.80 | hypothetical protein, conserved | 0,3149 | 0,001282 |
| TcCLB.510845.10 | retrotransposon hot spot (RHS) protein, putative (fragmer | 0,3149 | 0,06813 |
| TcCLB.510187.470 | hypothetical protein | 0,3149 | 0,08338 |
| TcCLB.511807.190 | hypothetical protein, conserved | 0,3146 | 0,004132 |
| TcCLB.511219.20 | retrotransposon hot spot protein (RHS, pseudogene), puta | 0,3146 | 0,02451 |
| TcCLB.511635.60 | hypothetical protein, conserved | 0,3144 | 0,005625 |
| TcCLB.505183.20 | malic enzyme | 0,3142 | 0,0002579 |
| TcCLB.508815.80 | hypothetical protein, conserved | 0,314 | 0,00004521 |
| TcCLB.510441.20 | dispersed gene family protein 1 (DGF-1), putative | 0,3139 | 0,1587 |
| TcCLB.506545.10 | hypothetical protein | 0,3136 | 0,003318 |
| TcCLB.508911.80 | hypothetical protein, conserved | 0,3136 | 0,02442 |
| TcCLB.511809.70 | DNA primase small subunit, putative | 0,3136 | 0,04697 |
| TcCLB.511411.16 | hypothetical protein, conserved | 0,3135 | 0,002053 |
| TcCLB.510349.50 | CDP-diacylglycerol--inositol 3-phosphatidyltransferase, pu | 0,3133 | 0,05986 |
| TcCLB.511817.250 | hypothetical protein, conserved | 0,3131 | 0,001641 |
| TcCLB.507023.20 | hypothetical protein | 0,3131 | 0,03786 |
| TcCLB.508973.80 | hypothetical protein, conserved | 0,3129 | 9,637E-06 |
| TcCLB.506301.40 | hypothetical protein, conserved | 0,3129 | 0,006104 |
| TcCLB.504109.130 | hypothetical protein, conserved | 0,3128 | 0,0002593 |
| TcCLB.510289.80 | hypothetical protein, conserved | 0,3127 | 0,0313 |
| TcCLB.506217.20 | beta galactofuranosyl glycosyltransferase (pseudogene), p | 0,3127 | 0,09154 |
| TcCLB.511051.20 | hypothetical protein, conserved | 0,3126 | 0,02553 |
| TcCLB.508817.80 | protein kinase, putative | 0,3125 | 0,01845 |
| TcCLB.511755.28 | hypothetical protein, conserved | 0,3125 | 0,1216 |
| TcCLB.482097.10 | hypothetical protein, conserved (fragment) | 0,3125 | 0,1976 |
| TcCLB.507641.190 | kinetoplastid kinetochore protein 1, putative (kkt1) | 0,3124 | 0,002599 |
| TcCLB.510735.60 | hypothetical protein, conserved | 0,3124 | 0,04164 |
| TcCLB.506925.130 | eukaryotic initiation factor 5a, putative | 0,3122 | 0,02883 |
| TcCLB.506615.111 | elongation factor 1-gamma (EF-1-gamma, pseudogene), | | 0,3122 | 0,09726 |

| | | | |
|---|---|---|---|
| TcCLB.508153.650 | hypothetical protein, conserved (pseudogene) | 0,312 | 0,003044 |
| TcCLB.405453.9 | dispersed gene family protein 1 (DGF-1), putative | 0,312 | 0,1967 |
| TcCLB.506693.4 | elongation initiation factor 2 alpha subunit, putative | 0,3115 | 0,009055 |
| TcCLB.504575.40 | adrenodoxin precursor, putative | 0,3114 | 0,1665 |
| TcCLB.510411.5 | retrotransposon hot spot protein (RHS, pseudogene), puta | 0,3112 | 0,3781 |
| TcCLB.506691.22 | hypothetical protein, conserved | 0,3111 | 0,005004 |
| TcCLB.511465.30 | hypothetical protein, conserved | 0,311 | 0,06245 |
| TcCLB.509059.10 | hypothetical protein | 0,311 | 0,1291 |
| TcCLB.440101.20 | activated protein kinase C receptor (pseudogene), putative | 0,311 | 0,1583 |
| TcCLB.508479.20 | hypothetical protein, conserved (pseudogene) | 0,3107 | 0,1434 |
| TcCLB.506833.40 | hypothetical protein | 0,3106 | 0,009644 |
| TcCLB.509967.80 | hypothetical protein, conserved | 0,3106 | 0,03035 |
| TcCLB.507023.230 | splicing factor 3a, putative | 0,3105 | 0,003862 |
| TcCLB.509965.120 | hypothetical protein, conserved | 0,3104 | 0,005673 |
| TcCLB.507087.50 | hypothetical protein, conserved | 0,3104 | 0,02058 |
| TcCLB.511471.20 | trans-sialidase (pseudogene), putative | 0,3103 | 0,131 |
| TcCLB.510697.105 | hypothetical protein, conserved (pseudogene) | 0,3103 | 0,3271 |
| TcCLB.509027.40 | hypothetical protein, conserved | 0,3102 | 0,0001296 |
| TcCLB.507105.120 | hypothetical protein, conserved (pseudogene) | 0,3102 | 0,01083 |
| TcCLB.507047.70 | hypothetical protein, conserved | 0,3101 | 0,0835 |
| TcCLB.507003.24 | hypothetical protein, conserved (fragment) | 0,3099 | 0,0001412 |
| TcCLB.510547.7 | surface protease GP63 (pseudogene), putative | 0,3099 | 0,3756 |
| TcCLB.503969.20 | hypothetical protein, conserved | 0,3099 | 0,4926 |
| TcCLB.510269.55 | elongation factor 1-gamma (EF-1-gamma, pseudogene), | 0,3098 | 0,4635 |
| TcCLB.507615.90 | hypothetical protein, conserved | 0,3097 | 0,00007559 |
| TcCLB.504449.40 | hypothetical protein | 0,3097 | 0,009485 |
| TcCLB.506357.180 | hypothetical protein | 0,3096 | 0,004088 |
| TcCLB.511415.40 | dispersed gene family protein 1 (DGF-1), putative | 0,3095 | 0,03247 |
| TcCLB.503493.40 | hypothetical protein, conserved | 0,3095 | 0,453 |
| TcCLB.504159.50 | aquaporin-like protein, putative | 0,3094 | 0,00153 |
| TcCLB.506885.90 | hypothetical protein, conserved | 0,3094 | 0,001708 |
| TcCLB.505071.110 | hypothetical protein, conserved | 0,3092 | 0,002346 |
| TcCLB.507975.80 | hypothetical protein, conserved | 0,309 | 0,000235 |
| TcCLB.506205.30 | hypothetical protein, conserved | 0,309 | 0,0345 |
| TcCLB.510131.90 | hypothetical protein, conserved | 0,3087 | 0,004446 |
| TcCLB.508257.160 | hypothetical protein, conserved | 0,3087 | 0,005056 |
| TcCLB.511249.110 | kinesin K39 (pseudogene), putative | 0,3087 | 0,00831 |
| TcCLB.511171.10 | dispersed gene family protein 1 (DGF-1), putative | 0,3086 | 0,4575 |
| TcCLB.503671.30 | Protein tilB homolog, putative (LRTP) | 0,3085 | 0,02998 |
| TcCLB.511741.60 | hypothetical protein, conserved | 0,3084 | 0,006571 |
| TcCLB.506839.70 | NADH dehydrogenase, putative | 0,3083 | 0,0003519 |
| TcCLB.509745.70 | hypothetical protein, conserved | 0,3083 | 0,003523 |
| TcCLB.508257.120 | hypothetical protein, conserved | 0,3083 | 0,01969 |
| TcCLB.507979.20 | hypothetical protein, conserved | 0,3083 | 0,02679 |
| TcCLB.508027.70 | ATP-dependent RNA helicase, putative | 0,308 | 0,001216 |

| | | | |
|---|---|---|---|
| TcCLB.510817.30 | hypothetical protein, conserved (pseudogene) | 0,308 | 0,04687 |
| TcCLB.485889.10 | hypothetical protein, conserved | 0,3077 | 0,0006696 |
| TcCLB.511807.140 | hypothetical protein, conserved | 0,3077 | 0,002798 |
| TcCLB.507065.150 | mucin TcMUCII, putative | 0,3077 | 0,2333 |
| TcCLB.509599.157 | serine/threonine protein kinase, putative | 0,3076 | 0,009374 |
| TcCLB.511281.60 | pyruvate kinase 2, putative | 0,3075 | 0,01405 |
| TcCLB.509933.10 | hypothetical protein, conserved | 0,3074 | 0,09974 |
| TcCLB.511407.30 | hypothetical protein | 0,3073 | 0,0004579 |
| TcCLB.506493.110 | hypothetical protein, conserved | 0,3073 | 0,07433 |
| TcCLB.510431.300 | trans-sialidase, Group II, putative | 0,3073 | 0,1295 |
| TcCLB.509769.134 | hypothetical protein, conserved | 0,3068 | 0,001367 |
| TcCLB.510519.70 | mitochondrial RNA binding complex 1 subunit, putative | 0,3067 | 0,0091 |
| TcCLB.511391.130 | hypothetical protein, conserved | 0,3065 | 0,04824 |
| TcCLB.511235.31 | surface protease GP63, putative | 0,3064 | 0,5326 |
| TcCLB.508291.10 | mucin-associated surface protein (MASP, pseudogene), pu | 0,3061 | 0,2683 |
| TcCLB.511287.110 | hypothetical protein, conserved | 0,3059 | 0,0006168 |
| TcCLB.506503.10 | hypothetical protein, conserved | 0,3058 | 0,02126 |
| TcCLB.506779.30 | hypothetical protein, conserved (pseudogene) | 0,3057 | 0,0005354 |
| TcCLB.508551.10 | target of rapamycin (TOR) kinase 1, putative | 0,3057 | 0,009532 |
| TcCLB.511507.110 | hypothetical protein, conserved (YCF45) | 0,3055 | 0,007814 |
| TcCLB.506475.116 | glutaredoxin, putative | 0,3054 | 0,04779 |
| TcCLB.503637.30 | serine peptidase, Clan SC, Family S9B | 0,3053 | 0,001734 |
| TcCLB.509827.20 | hypothetical protein, conserved | 0,3053 | 0,07312 |
| TcCLB.509261.10 | p-nitrophenylphosphatase, putative | 0,3052 | 0,01337 |
| TcCLB.506465.20 | hypothetical protein, conserved | 0,3051 | 0,03039 |
| TcCLB.408573.9 | hypothetical protein, conserved | 0,3051 | 0,04134 |
| TcCLB.508369.10 | hypothetical protein, conserved | 0,3051 | 0,06275 |
| TcCLB.504147.30 | hypothetical protein, conserved | 0,305 | 0,005672 |
| TcCLB.508271.20 | hypothetical protein, conserved | 0,3049 | 0,01679 |
| TcCLB.508153.790 | hypothetical protein, conserved | 0,3049 | 0,04192 |
| TcCLB.509259.21 | elongation factor 1-gamma (EF-1-gamma, pseudogene), | | 0,3049 | 0,4594 |
| TcCLB.508919.70 | protein kinase, putative | 0,3048 | 0,001401 |
| TcCLB.506145.30 | O-6 methyl-guanine alkyl transferase, putative | 0,3047 | 0,075 |
| TcCLB.510355.97 | retrotransposon hot spot protein (RHS, pseudogene), puta | 0,3047 | 0,481 |
| TcCLB.507895.50 | dispersed gene family protein 1 (DGF-1, pseudogene), put | 0,3042 | 0,4186 |
| TcCLB.509067.30 | mitochondrial RNA binding protein, putative | 0,3039 | 0,0008509 |
| TcCLB.510349.64 | hypothetical protein, conserved | 0,3038 | 0,01928 |
| TcCLB.504179.69 | phosphoprotein lepp12, putative | 0,3038 | 0,02563 |
| TcCLB.510659.60 | hypothetical protein, conserved | 0,3038 | 0,04203 |
| TcCLB.506743.135 | trans-sialidase (pseudogene), putative | 0,3038 | 0,1639 |
| TcCLB.508655.10 | DNA-directed RNA polymerase II subunit 2 (pseudogene), | 0,3038 | 0,4584 |
| TcCLB.506435.430 | hypothetical protein, conserved | 0,3037 | 0,001814 |
| TcCLB.507623.80 | hypothetical protein, conserved | 0,3036 | 0,01635 |
| TcCLB.508443.30 | hypothetical protein, conserved | 0,3034 | 0,000373 |
| TcCLB.504433.14 | ADP-ribosylation factor-like protein 3A, putative | 0,3028 | 0,02117 |

| | | | |
|---|---|---|---|
| TcCLB.506685.5 | trans-sialidase (pseudogene), putative | 0,3028 | 0,04462 |
| TcCLB.511055.20 | retrotransposon hot spot (RHS) protein, putative | 0,3027 | 0,0003393 |
| TcCLB.506821.190 | hypothetical protein, conserved | 0,3027 | 0,005644 |
| TcCLB.508269.70 | hypothetical protein, conserved | 0,3027 | 0,1719 |
| TcCLB.510657.50 | hypothetical protein, conserved | 0,3025 | 0,03706 |
| TcCLB.511827.70 | hypothetical protein, conserved | 0,3024 | 0,01831 |
| TcCLB.506177.110 | hypothetical protein, conserved | 0,3024 | 0,08387 |
| TcCLB.508461.160 | hypothetical protein, conserved | 0,3023 | 0,00191 |
| TcCLB.510687.110 | hypothetical protein, conserved | 0,3021 | 0,009185 |
| TcCLB.506135.70 | hypothetical protein, conserved | 0,3021 | 0,02147 |
| TcCLB.504047.30 | kinesin, putative | 0,3019 | 0,0233 |
| TcCLB.509063.30 | chaperonin HSP60/CNP60, putative | 0,3018 | 0,01076 |
| TcCLB.507993.390 | pyruvate kinase 2, putative | 0,3018 | 0,01991 |
| TcCLB.505097.35 | retrotransposon hot spot protein (RHS, pseudogene), puta | 0,3017 | 0,02462 |
| TcCLB.511469.105 | hypothetical protein | 0,3017 | 0,07655 |
| TcCLB.508015.20 | hypothetical protein | 0,3017 | 0,1035 |
| TcCLB.508903.90 | dispersed gene family protein 1 (DGF-1, pseudogene), put | 0,3015 | 0,0889 |
| TcCLB.511671.60 | surface antigen 2 (CA-2), putative | 0,3014 | 0,001974 |
| TcCLB.506803.30 | retrotransposon hot spot protein (RHS, pseudogene), puta | 0,3014 | 0,2826 |
| TcCLB.511303.90 | hypothetical protein, conserved | 0,3013 | 0,003557 |
| TcCLB.504089.79 | retrotransposon hot spot protein (RHS, pseudogene), puta | 0,3013 | 0,4097 |
| TcCLB.510679.14 | hypothetical protein, conserved | 0,3011 | 0,1338 |
| TcCLB.509519.9 | dispersed gene family protein 1 (DGF-1), putative (fragme | 0,3011 | 0,5142 |
| TcCLB.508837.170 | hypothetical protein, conserved | 0,301 | 0,009035 |
| TcCLB.506817.70 | hypothetical protein | 0,301 | 0,1986 |
| TcCLB.507951.100 | hypothetical protein, conserved | 0,3009 | 0,004041 |
| TcCLB.509769.33 | palmitoyl acyltransferase 12, putative | 0,3009 | 0,01624 |
| TcCLB.510357.110 | hypothetical protein, conserved | 0,3009 | 0,0373 |
| TcCLB.507977.9 | hypothetical protein, conserved (pseudogene) | 0,3009 | 0,06592 |
| TcCLB.506711.4 | peptide chain release factor 1, putative (fragment) | 0,3008 | 0,1022 |
| TcCLB.504005.20 | N-Acetyl-D-glucosaminylphosphatidylinositol de-N-acetyla | 0,3006 | 0,04777 |
| TcCLB.507013.10 | dolichol kinase, putative | 0,3005 | 0,005912 |
| TcCLB.504149.20 | ATP-binding cassette transporter ABCA1, putative (ABCA1 | 0,3005 | 0,0113 |
| TcCLB.506885.489 | dispersed gene family protein 1 (DGF-1), putative | 0,3005 | 0,1535 |
| TcCLB.506801.110 | hypothetical protein, conserved | 0,3003 | 0,02761 |
| TcCLB.509167.130 | hypothetical protein, conserved | 0,3002 | 0,001004 |
| TcCLB.507049.70 | hypothetical protein, conserved | 0,3 | 0,02679 |
| TcCLB.509163.100 | hypothetical protein, conserved (pseudogene) | 0,2998 | 0,03968 |
| TcCLB.511431.30 | conserved protein, unknown function | 0,2997 | 0,009624 |
| TcCLB.503397.10 | retrotransposon hot spot protein (RHS, pseudogene), puta | 0,2994 | 0,00376 |
| TcCLB.507519.60 | Silent information regulator 2 related protein 1, putative ( | 0,2992 | 0,01676 |
| TcCLB.510463.230 | dispersed gene family protein 1 (DGF-1), putative | 0,2992 | 0,385 |
| TcCLB.511237.100 | hypothetical protein, conserved | 0,2991 | 0,006239 |
| TcCLB.509717.60 | metallo-peptidase, Clan MA(E) Family M41 | 0,299 | 0,001933 |
| TcCLB.506705.40 | hypothetical protein, conserved | 0,299 | 0,007705 |

| | | | |
|---|---|---|---|
| TcCLB.510329.120 | Present in the outer mitochondrial membrane proteome 2 | 0,299 | 0,01027 |
| TcCLB.509065.110 | hypothetical protein, conserved | 0,2989 | 0,002063 |
| TcCLB.417091.10 | RNA pseudouridylate synthase, putative | 0,2989 | 0,08168 |
| TcCLB.507063.30 | hypothetical protein, conserved | 0,2987 | 0,04127 |
| TcCLB.507745.10 | mucin-associated surface protein (MASP, pseudogene), pu | 0,2987 | 0,2475 |
| TcCLB.507951.179 | hypothetical protein, conserved | 0,2986 | 0,008212 |
| TcCLB.506559.400 | hypothetical protein, conserved | 0,2986 | 0,01853 |
| TcCLB.507575.10 | dispersed gene family protein 1 (DGF-1), putative | 0,2986 | 0,2893 |
| TcCLB.509317.109 | hypothetical protein, conserved (fragment) | 0,2984 | 0,0102 |
| TcCLB.510649.24 | membrane associated protein, putative (fragment) | 0,2982 | 0,002183 |
| TcCLB.507213.30 | potassium voltage-gated channel, putative | 0,2981 | 0,004203 |
| TcCLB.510101.70 | protein transport protein SEC61 subunit alpha, putative | 0,298 | 0,0008648 |
| TcCLB.511419.10 | hypothetical protein | 0,298 | 0,07829 |
| TcCLB.508797.29 | hypothetical protein, conserved | 0,2978 | 0,0004595 |
| TcCLB.508257.104 | ribosomal protein L14, putative | 0,2976 | 0,06856 |
| TcCLB.509161.149 | hypothetical protein, conserved | 0,2976 | 0,1289 |
| TcCLB.508297.61 | choline ethanolamine kinase, putative | 0,2976 | 0,138 |
| TcCLB.510877.170 | hypothetical protein, conserved | 0,2975 | 0,001211 |
| TcCLB.510353.20 | hypothetical protein, conserved | 0,2975 | 0,006255 |
| TcCLB.510039.109 | sec1 family transport protein, putative (fragment) | 0,2975 | 0,08241 |
| TcCLB.506629.70 | cysteine desulfurase, putative | 0,2974 | 0,01032 |
| TcCLB.507943.99 | hypothetical protein, conserved (fragment) | 0,2974 | 0,179 |
| TcCLB.507681.70 | hypothetical protein, conserved | 0,2973 | 0,00458 |
| TcCLB.509205.20 | hypothetical protein, conserved | 0,2973 | 0,04267 |
| TcCLB.504637.21 | trans-sialidase (pseudogene), putative | 0,2973 | 0,06415 |
| TcCLB.508543.80 | hypothetical protein, conserved | 0,2972 | 0,04433 |
| TcCLB.511361.60 | hypothetical protein | 0,2971 | 0,005444 |
| TcCLB.508871.20 | mucin TcMUCII, putative | 0,2971 | 0,5185 |
| TcCLB.510735.20 | hypothetical protein, conserved | 0,297 | 0,01135 |
| TcCLB.506393.143 | ATP-dependent DEAD/H RNA helicase (pseudogene), puta | 0,297 | 0,2203 |
| TcCLB.510627.200 | trans-sialidase (pseudogene), putative | 0,2967 | 0,603 |
| TcCLB.511907.80 | beta-ketoacyl-ACP reductase 2, putative | 0,2966 | 0,02735 |
| TcCLB.505163.40 | hypothetical protein, conserved | 0,2964 | 0,04361 |
| TcCLB.511025.70 | chaperonin alpha subunit, putative | 0,2962 | 0,00003436 |
| TcCLB.509213.40 | hypothetical protein, conserved | 0,2961 | 0,0003833 |
| TcCLB.508657.20 | sec1-like protein (pseudogene), putative | 0,296 | 0,008734 |
| TcCLB.511529.20 | hypothetical protein, conserved | 0,296 | 0,02655 |
| TcCLB.511717.70 | hypothetical protein, conserved | 0,2959 | 0,01784 |
| TcCLB.508207.10 | glycine dehydrogenase (pseudogene), putative | 0,2959 | 0,01935 |
| TcCLB.511595.10 | trans-sialidase (pseudogene), putative | 0,2959 | 0,381 |
| TcCLB.508369.50 | acetyl-CoA carboxylase (fragment) | 0,2957 | 0,0004004 |
| TcCLB.506227.90 | hypothetical protein, conserved | 0,2957 | 0,01631 |
| TcCLB.504057.45 | hypothetical protein, conserved | 0,2956 | 0,006461 |
| TcCLB.509895.100 | hypothetical protein, conserved | 0,2956 | 0,02523 |
| TcCLB.506327.20 | hypothetical protein, conserved | 0,2956 | 0,03271 |

| | | | |
|---|---|---|---|
| TcCLB.507969.70 | hypothetical protein, conserved | 0,2955 | 0,05786 |
| TcCLB.504155.30 | retrotransposon hot spot protein (RHS, pseudogene), puta | 0,2955 | 0,4045 |
| TcCLB.511385.90 | hypothetical protein, conserved | 0,2954 | 0,0009318 |
| TcCLB.508351.10 | fructose-1,6-bisphosphatase, cytosolic, putative | 0,2954 | 0,01506 |
| TcCLB.509753.126 | hypothetical protein, conserved (pseudogene) | 0,2951 | 0,3268 |
| TcCLB.509057.20 | importin alpha, putative | 0,2946 | 0,0002878 |
| TcCLB.506725.30 | hypothetical protein, conserved | 0,2946 | 0,0002978 |
| TcCLB.506679.180 | hypothetical protein, conserved | 0,2945 | 0,00284 |
| TcCLB.511015.10 | mannosyl-oligosaccharide glucosidase, putative | 0,2945 | 0,04301 |
| TcCLB.510137.40 | retrotransposon hot spot protein (RHS, pseudogene), puta | 0,2944 | 0,005458 |
| TcCLB.503441.10 | retrotransposon hot spot protein (RHS, pseudogene), puta | 0,2944 | 0,006482 |
| TcCLB.509297.20 | hypothetical protein, conserved | 0,2942 | 0,007406 |
| TcCLB.509017.9 | hypothetical protein, conserved (fragment) | 0,2941 | 0,06481 |
| TcCLB.507519.10 | hypothetical protein, conserved (pseudogene) | 0,294 | 0,01882 |
| TcCLB.508411.70 | hypothetical protein, conserved | 0,2939 | 0,02982 |
| TcCLB.510175.60 | dispersed gene family protein 1 (DGF-1), putative | 0,2939 | 0,05782 |
| TcCLB.507771.99 | hypothetical protein, conserved | 0,2939 | 0,1073 |
| TcCLB.506527.10 | hypothetical protein, conserved | 0,2937 | 0,0002575 |
| TcCLB.508115.10 | hypothetical protein | 0,2936 | 0,2265 |
| TcCLB.511153.110 | hypothetical protein, conserved | 0,2934 | 0,0007568 |
| TcCLB.511627.80 | hypothetical protein, conserved | 0,2934 | 0,1278 |
| TcCLB.508215.10 | hypothetical protein, conserved | 0,2932 | 0,01635 |
| TcCLB.503973.31 | hypothetical protein, conserved (pseudogene) | 0,293 | 0,1554 |
| TcCLB.510399.39 | dispersed gene family protein 1 (DGF-1), putative | 0,293 | 0,2944 |
| TcCLB.436761.10 | hypothetical protein, conserved | 0,2929 | 0,07682 |
| TcCLB.420091.10 | dispersed gene family protein 1 (DGF-1), putative | 0,2929 | 0,2034 |
| TcCLB.507711.320 | DNA mismatch repair protein MSH2, putative (MSH2) | 0,2928 | 0,00062 |
| TcCLB.509437.40 | heat shock protein DNAJ, putative | 0,2928 | 0,0008448 |
| TcCLB.415803.10 | hypothetical protein, conserved | 0,2928 | 0,09974 |
| TcCLB.506779.110 | phopshatase, putative | 0,2925 | 0,001151 |
| TcCLB.504179.40 | hypothetical protein, conserved | 0,2925 | 0,001345 |
| TcCLB.508541.225 | casein kinase, putative | 0,2925 | 0,01514 |
| TcCLB.510761.80 | surface protease GP63, putative | 0,2925 | 0,2444 |
| TcCLB.510289.54 | hypothetical protein, conserved | 0,2924 | 0,06146 |
| TcCLB.510731.68 | hypothetical protein, conserved | 0,2923 | 0,06891 |
| TcCLB.507793.10 | hypothetical protein, conserved | 0,2922 | 0,000411 |
| TcCLB.507765.80 | hypothetical protein, conserved | 0,2922 | 0,01096 |
| TcCLB.506699.3 | hypothetical protein, conserved | 0,2922 | 0,05183 |
| TcCLB.511739.30 | hypothetical protein, conserved | 0,292 | 0,05095 |
| TcCLB.509225.21 | hypothetical protein | 0,292 | 0,2224 |
| TcCLB.506503.120 | hypothetical protein, conserved | 0,2919 | 0,0001631 |
| TcCLB.508503.80 | hypothetical protein | 0,2918 | 0,04991 |
| TcCLB.509831.10 | cysteine desulfurase, putative | 0,2916 | 0,0114 |
| TcCLB.508163.240 | hypothetical protein | 0,2916 | 0,4795 |
| TcCLB.508085.10 | GMP synthase, putative | 0,2915 | 0,007397 |

| | | | |
|---|---|---|---|
| TcCLB.509967.10 | hypothetical protein, conserved | 0,2915 | 0,02849 |
| TcCLB.509003.10 | hypothetical protein, conserved | 0,2914 | 0,0009904 |
| TcCLB.503919.40 | hypothetical protein, conserved | 0,2913 | 0,001529 |
| TcCLB.510321.20 | hypothetical protein, conserved | 0,2913 | 0,01761 |
| TcCLB.508859.118 | trans-sialidase, putative | 0,2913 | 0,09213 |
| TcCLB.506367.130 | hypothetical protein, conserved | 0,2912 | 0,004852 |
| TcCLB.510855.4 | hypothetical protein, conserved | 0,2912 | 0,01932 |
| TcCLB.509733.185 | hypothetical protein, conserved (pseudogene) | 0,2909 | 0,01879 |
| TcCLB.506821.200 | hypothetical protein, conserved | 0,2907 | 0,01073 |
| TcCLB.509599.50 | hypothetical protein, conserved | 0,2906 | 0,004515 |
| TcCLB.511725.230 | protein kinase, putative | 0,2905 | 0,0003396 |
| TcCLB.506493.60 | hypothetical protein, conserved | 0,2905 | 0,004276 |
| TcCLB.507509.40 | hypothetical protein, conserved | 0,2905 | 0,006716 |
| TcCLB.506719.10 | phosphatidylinositol (3,5) kinase, putative | 0,2905 | 0,02382 |
| TcCLB.507009.110 | hypothetical protein, conserved | 0,2905 | 0,04218 |
| TcCLB.508989.120 | hypothetical protein, conserved | 0,2904 | 0,02546 |
| TcCLB.508231.74 | poly(A)-specific ribonuclease PARN, putative | 0,2904 | 0,06584 |
| TcCLB.508613.30 | peroxisomal membrane protein 4, putative | 0,2903 | 0,0404 |
| TcCLB.511231.30 | hypothetical protein, conserved | 0,2902 | 0,01126 |
| TcCLB.510029.50 | hypothetical protein, conserved | 0,2902 | 0,01205 |
| TcCLB.511589.250 | hypothetical protein, conserved | 0,2902 | 0,02745 |
| TcCLB.505171.80 | hypothetical protein, conserved | 0,2901 | 0,002962 |
| TcCLB.506339.100 | dispersed gene family protein 1 (DGF-1), putative | 0,2901 | 0,1588 |
| TcCLB.506435.250 | hypothetical protein, conserved | 0,2899 | 0,06153 |
| TcCLB.506287.9 | hypothetical protein, conserved | 0,2897 | 0,02258 |
| TcCLB.509003.79 | coatomer beta subunit, putative (fragment) | 0,2895 | 0,01585 |
| TcCLB.507609.150 | hypothetical protein (pseudogene) | 0,2895 | 0,2467 |
| TcCLB.510741.210 | glycerol kinase, glycosomal (pseudogene), putative | 0,2894 | 0,0101 |
| TcCLB.508659.9 | TPR-repeat-containing chaperone protein DNAJ, putative ( | 0,2893 | 0,02109 |
| TcCLB.509831.20 | vesicular-fusion ATPase-like protein, putative | 0,2892 | 0,002145 |
| TcCLB.504137.150 | hypothetical protein, conserved | 0,289 | 0,002298 |
| TcCLB.511577.90 | pseudouridylate synthase-like protein, putative | 0,2887 | 0,009183 |
| TcCLB.511571.10 | ATP-dependent DEAD/H RNA helicase, putative | 0,2885 | 0,1738 |
| TcCLB.508193.15 | trans-sialidase (pseudogene), putative | 0,2884 | 0,003218 |
| TcCLB.506211.100 | Sarcalumenin, putative | 0,2884 | 0,007016 |
| TcCLB.510047.10 | hypothetical protein, conserved | 0,2883 | 0,005423 |
| TcCLB.508577.40 | dispersed gene family protein 1 (DGF-1), putative | 0,2882 | 0,08414 |
| TcCLB.511671.130 | hypothetical protein, conserved | 0,2881 | 0,004865 |
| TcCLB.508451.10 | trans-sialidase (pseudogene), putative | 0,2881 | 0,1143 |
| TcCLB.509721.10 | MRB1 complex subunit MRB3010 (MRB3010) | 0,288 | 0,06052 |
| TcCLB.507681.200 | serine carboxypeptidase S28, putative | 0,2879 | 0,005156 |
| TcCLB.511459.50 | hypothetical protein, conserved | 0,2879 | 0,0646 |
| TcCLB.510431.210 | vesicular protein trafficking mediator, putative | 0,2878 | 0,01863 |
| TcCLB.508689.20 | hypothetical protein, conserved | 0,2876 | 0,004853 |
| TcCLB.508911.20 | hypothetical protein, conserved | 0,2876 | 0,0414 |

| | | | |
|---|---|---|---|
| TcCLB.504035.80 | hypothetical protein, conserved | 0,2873 | 0,1443 |
| TcCLB.510967.30 | hypothetical protein | 0,2872 | 0,1593 |
| TcCLB.507795.10 | membrane associated protein, putative | 0,2871 | 0,002786 |
| TcCLB.510947.20 | hypothetical protein, conserved | 0,2868 | 0,003723 |
| TcCLB.506683.240 | trans-sialidase, Group II, putative | 0,2867 | 0,005718 |
| TcCLB.508239.20 | hypothetical protein, conserved | 0,2866 | 0,0009807 |
| TcCLB.511055.10 | retrotransposon hot spot protein (RHS, pseudogene), puta | 0,2866 | 0,005433 |
| TcCLB.511529.180 | hypothetical protein, conserved | 0,2865 | 0,01194 |
| TcCLB.509779.40 | hypothetical protein, conserved | 0,2865 | 0,04891 |
| TcCLB.508175.39 | hypothetical protein, conserved | 0,2864 | 0,04038 |
| TcCLB.507007.34 | hypothetical protein, conserved | 0,2863 | 0,002868 |
| TcCLB.506221.100 | hypothetical protein, conserved | 0,2862 | 0,01229 |
| TcCLB.511303.30 | hypothetical protein, conserved | 0,2862 | 0,01522 |
| TcCLB.510449.20 | hypothetical protein, conserved (pseudogene) | 0,2861 | 0,4805 |
| TcCLB.509877.60 | hypothetical protein, conserved | 0,286 | 0,008573 |
| TcCLB.510311.150 | mitochondrial protein 99, putative (MP99) | 0,286 | 0,01079 |
| TcCLB.508153.880 | hypothetical protein, conserved | 0,2859 | 0,003828 |
| TcCLB.510329.380 | hypothetical protein, conserved | 0,2859 | 0,03712 |
| TcCLB.510359.320 | DNA-directed RNA polymerase III subunit, putative | 0,2858 | 0,1735 |
| TcCLB.509177.30 | hypothetical protein, conserved | 0,2857 | 0,00314 |
| TcCLB.511081.71 | hypothetical protein, conserved (pseudogene) | 0,2857 | 0,4179 |
| TcCLB.509561.30 | citrate transporter, putative | 0,2855 | 0,02322 |
| TcCLB.510949.10 | dual specificity protein phosphatase, putative | 0,2854 | 0,01594 |
| TcCLB.420719.9 | Associated kinase of Tb14-3-3, putative (AKB1) | 0,2853 | 0,03611 |
| TcCLB.507485.50 | hypothetical protein, conserved | 0,2852 | 0,005822 |
| TcCLB.506113.80 | hypothetical protein, conserved (pseudogene) | 0,2851 | 0,01237 |
| TcCLB.508815.140 | protein transport protein SEC13, putative | 0,285 | 0,01728 |
| TcCLB.506425.169 | Ras-related protein RAB2B, putative (fragment) (RAB2B) | 0,285 | 0,08897 |
| TcCLB.504033.40 | hypothetical protein | 0,2849 | 0,0018 |
| TcCLB.508349.39 | RNA-binding protein, putative (fragment) (DRBD3) | 0,2849 | 0,00775 |
| TcCLB.509715.60 | hypothetical protein, conserved | 0,2849 | 0,05807 |
| TcCLB.511321.10 | FG-GAP repeat protein, putative | 0,2848 | 0,0007678 |
| TcCLB.509769.110 | Elongator-like Protein 3b, putative | 0,2848 | 0,001044 |
| TcCLB.511483.40 | prefoldin subunit 2, putative | 0,2848 | 0,02297 |
| TcCLB.506733.110 | hypothetical protein, conserved | 0,2848 | 0,07824 |
| TcCLB.510053.79 | FG-GAP repeat protein, putative | 0,2847 | 0,129 |
| TcCLB.507809.60 | hypothetical protein, conserved | 0,2846 | 0,004265 |
| TcCLB.503579.140 | hypothetical protein, conserved | 0,2845 | 0,01813 |
| TcCLB.506315.20 | hypothetical protein, conserved | 0,2844 | 0,02915 |
| TcCLB.510289.10 | hypothetical protein, conserved | 0,2844 | 0,06903 |
| TcCLB.509237.50 | hypothetical protein, conserved | 0,2842 | 0,001285 |
| TcCLB.506863.4 | dephospho-CoA kinase, putative | 0,2842 | 0,2519 |
| TcCLB.510841.20 | hypothetical protein, conserved | 0,2841 | 0,06131 |
| TcCLB.511867.190 | tubulin gamma chain (TUBG) | 0,284 | 0,003626 |
| TcCLB.503917.7 | RNA-binding protein, putative | 0,284 | 0,01284 |

| | | | |
|---|---|---|---|
| TcCLB.510381.20 | ABC transporter, putative | 0,284 | 0,02409 |
| TcCLB.511315.20 | hypothetical protein, conserved | 0,2839 | 0,01053 |
| TcCLB.504149.130 | hypothetical protein, conserved | 0,2834 | 0,0002636 |
| TcCLB.504153.129 | hypothetical protein, conserved | 0,2834 | 0,007316 |
| TcCLB.508039.10 | retrotransposon hot spot protein (RHS, pseudogene), puta | 0,2833 | 0,007892 |
| TcCLB.510565.130 | hypothetical protein | 0,2832 | 0,01346 |
| TcCLB.504147.70 | hypothetical protein, conserved | 0,2832 | 0,02728 |
| TcCLB.506961.20 | dispersed gene family protein 1 (DGF-1), putative | 0,2831 | 0,3075 |
| TcCLB.510719.270 | hypothetical protein, conserved | 0,2829 | 0,07498 |
| TcCLB.510175.150 | hypothetical protein, conserved | 0,2828 | 0,1729 |
| TcCLB.508479.480 | N-acetyltransferase complex ARD1 subunit (pseudogene), | 0,2827 | 0,1698 |
| TcCLB.511585.280 | retrotransposon hot spot protein (RHS, pseudogene), puta | 0,2826 | 0,01959 |
| TcCLB.504427.120 | hypothetical protein, conserved | 0,2826 | 0,02324 |
| TcCLB.507155.9 | dispersed gene family protein 1 (DGF-1, pseudogene), put | 0,2826 | 0,1914 |
| TcCLB.509507.49 | mitotic cyclin, putative | 0,2825 | 0,06383 |
| TcCLB.510575.239 | hypothetical protein, conserved | 0,2824 | 0,3636 |
| TcCLB.511423.40 | TFIIH basal transcription factor subunit, putative (TSP1) | 0,2823 | 0,0274 |
| TcCLB.506863.10 | alanyl-tRNA synthetase, putative | 0,2822 | 0,00004744 |
| TcCLB.504149.30 | hypothetical protein, conserved | 0,2822 | 0,01855 |
| TcCLB.508139.20 | dispersed gene family protein 1 (DGF-1), putative | 0,2821 | 0,0487 |
| TcCLB.507993.100 | hypothetical protein, conserved | 0,282 | 0,1029 |
| TcCLB.508131.10 | hypothetical protein, conserved (pseudogene) | 0,2819 | 0,001554 |
| TcCLB.507105.20 | mitochondrial RNA binding complex 1 subunit, putative | 0,2819 | 0,03138 |
| TcCLB.511445.90 | hypothetical protein, conserved | 0,2818 | 0,001484 |
| TcCLB.511581.20 | fatty acyl CoA syntetase 1 | 0,2818 | 0,01465 |
| TcCLB.504021.40 | hypothetical protein, conserved | 0,2817 | 0,003881 |
| TcCLB.504167.40 | ubiquitin fusion degradation protein, putative | 0,2816 | 0,02501 |
| TcCLB.421619.10 | hypothetical protein | 0,2815 | 0,01314 |
| TcCLB.508903.30 | retrotransposon hot spot protein (RHS, pseudogene), puta | 0,2815 | 0,06638 |
| TcCLB.411337.10 | retrotransposon hot spot protein (RHS, pseudogene), puta | 0,2812 | 0,1037 |
| TcCLB.506587.60 | target SNARE, putative | 0,2811 | 0,02668 |
| TcCLB.504153.170 | hypothetical protein, conserved | 0,281 | 0,000039 |
| TcCLB.506179.10 | ferric reductase transmembrane protein, putative (FR1) | 0,2809 | 0,00007882 |
| TcCLB.506631.50 | hypothetical protein, conserved (fragment) | 0,2809 | 0,5047 |
| TcCLB.510287.30 | proteasome subunit beta type-2, putative | 0,2808 | 0,01214 |
| TcCLB.505965.80 | hypothetical protein, conserved | 0,2808 | 0,04708 |
| TcCLB.509769.100 | hypothetical protein, conserved | 0,2807 | 0,01789 |
| TcCLB.511713.30 | hypothetical protein, conserved | 0,2806 | 0,03561 |
| TcCLB.506893.25 | U6 snRNA-associated Sm-like protein LSm6p | 0,2806 | 0,3398 |
| TcCLB.508207.150 | Silent information regulator 2 related protein 1, putative ( | 0,2805 | 0,01817 |
| TcCLB.507517.120 | hypothetical protein, conserved | 0,2805 | 0,04749 |
| TcCLB.507025.30 | hypothetical protein, conserved | 0,2805 | 0,06816 |
| TcCLB.506155.40 | hypothetical protein, conserved | 0,2803 | 0,002451 |
| TcCLB.510307.320 | retrotransposon hot spot protein (RHS, pseudogene), puta | 0,2803 | 0,03543 |
| TcCLB.511837.120 | hypothetical protein, conserved | 0,2803 | 0,2308 |

| | | | |
|---|---|---|---|
| TcCLB.510411.30 | retrotransposon hot spot protein (RHS, pseudogene), puta | 0,2802 | 0,1509 |
| TcCLB.507643.15 | beta galactofuranosyl glycosyltransferase, putative (fragm | 0,2802 | 0,2075 |
| TcCLB.510699.100 | hypothetical protein | 0,28 | 0,3951 |
| TcCLB.507711.10 | intraflagellar transport protein 57/55, putative (IFT57/55) | 0,2798 | 0,01106 |
| TcCLB.506211.210 | mitogen-activated protein kinase kinase kinase, putative ( | 0,2796 | 0,003091 |
| TcCLB.511511.10 | endosomal trafficking protein RME-8, putative | 0,2795 | 0,005932 |
| TcCLB.506405.70 | hypothetical protein, conserved | 0,2795 | 0,04812 |
| TcCLB.511215.10 | hypothetical protein, conserved | 0,2791 | 0,03023 |
| TcCLB.511211.110 | hypothetical protein, conserved | 0,279 | 0,003291 |
| TcCLB.507811.120 | myosin heavy chain MYA2-related, putative | 0,279 | 0,008963 |
| TcCLB.508039.90 | phosphoinositide-specific phospholipase C (pseudogene), | 0,279 | 0,04578 |
| TcCLB.510121.140 | hypothetical protein, conserved | 0,2786 | 0,01197 |
| TcCLB.510349.40 | hypothetical protein, conserved | 0,2785 | 0,0009036 |
| TcCLB.507547.40 | enoyl-CoA hydratase/Enoyl-CoA isomerase/3-hydroxyacyl- | 0,2784 | 0,007136 |
| TcCLB.507993.250 | hypothetical protein, conserved | 0,2784 | 0,01013 |
| TcCLB.511649.140 | hypothetical protein, conserved | 0,2784 | 0,05057 |
| TcCLB.508717.10 | tyrosine specific protein phosphatase, putative | 0,2783 | 0,004697 |
| TcCLB.509733.95 | hypothetical protein | 0,2782 | 0,009293 |
| TcCLB.511577.80 | hypothetical protein, conserved | 0,2781 | 0,009536 |
| TcCLB.507389.60 | chaperone DNAJ protein, putative | 0,2781 | 0,1411 |
| TcCLB.509073.110 | hypothetical protein, conserved | 0,278 | 0,003323 |
| TcCLB.503581.30 | hypothetical protein, conserved | 0,278 | 0,1298 |
| TcCLB.509551.110 | hypothetical protein, conserved | 0,2779 | 0,007704 |
| TcCLB.503653.30 | ribosomal RNA processing protein 41B, putative (RRP41B) | 0,2779 | 0,03112 |
| TcCLB.511189.84 | hypothetical protein, conserved | 0,2777 | 0,06777 |
| TcCLB.506895.50 | hypothetical protein, conserved | 0,2776 | 0,0317 |
| TcCLB.452039.9 | hypothetical protein, conserved | 0,2775 | 0,002333 |
| TcCLB.508737.170 | hypothetical protein, conserved | 0,2775 | 0,02673 |
| TcCLB.506943.110 | hypothetical protein | 0,2773 | 0,09361 |
| TcCLB.511233.40 | hypothetical protein, conserved | 0,2772 | 0,004251 |
| TcCLB.504231.30 | hypothetical protein, conserved | 0,2771 | 0,007658 |
| TcCLB.511139.30 | hypothetical protein, conserved | 0,2771 | 0,02111 |
| TcCLB.506775.190 | serine/threonine protein kinase, putative | 0,277 | 0,11 |
| TcCLB.508381.40 | retrotransposon hot spot protein (RHS, pseudogene), puta | 0,2769 | 0,1709 |
| TcCLB.508067.10 | hypothetical protein, conserved | 0,2768 | 0,00002568 |
| TcCLB.508241.90 | hypothetical protein, conserved | 0,2768 | 0,00191 |
| TcCLB.511649.60 | COP9 signalosome complex subunit 5, putative | 0,2767 | 0,006968 |
| TcCLB.507507.30 | hypothetical protein, conserved | 0,2766 | 0,01855 |
| TcCLB.509591.40 | hypothetical protein, conserved | 0,2765 | 0,001056 |
| TcCLB.510537.68 | syntaxin binding protein (pseudogene), putative | 0,2765 | 0,5505 |
| TcCLB.509541.10 | hypothetical protein, conserved | 0,2764 | 0,002797 |
| TcCLB.508153.640 | IAG-nucleoside hydrolase, putative (IAGNH) | 0,2763 | 0,0009155 |
| TcCLB.508787.30 | pumilio/PUF RNA binding protein 3, putative (PUF3) | 0,2763 | 0,006316 |
| TcCLB.506303.140 | hypothetical protein, conserved | 0,2761 | 0,008788 |
| TcCLB.509429.140 | kinetoplastid kinetochore protein 6, putative (kkt6) | 0,2761 | 0,06164 |

| | | | |
|---|---|---|---|
| TcCLB.511751.22 | hypothetical protein, conserved | 0,2761 | 0,1369 |
| TcCLB.472777.9 | hypothetical protein, conserved | 0,2761 | 0,2937 |
| TcCLB.506503.150 | histone H3 variant, putative | 0,276 | 0,01196 |
| TcCLB.508163.40 | mucin TcMUCII (pseudogene), putative | 0,2754 | 0,4604 |
| TcCLB.511897.28 | hypothetical protein, conserved | 0,2752 | 0,06733 |
| TcCLB.511541.20 | hypothetical protein | 0,2751 | 0,01267 |
| TcCLB.510851.40 | trans-sialidase (pseudogene), putative | 0,275 | 0,08046 |
| TcCLB.510101.310 | hypothetical protein | 0,2749 | 0,002881 |
| TcCLB.504125.90 | protein kinase, putative | 0,2749 | 0,005444 |
| TcCLB.506577.30 | hypothetical protein, conserved | 0,2749 | 0,01319 |
| TcCLB.510469.30 | trans-sialidase (pseudogene), putative | 0,2748 | 0,4554 |
| TcCLB.510241.120 | Peptide deformylase 2, putative (PDF2) | 0,2747 | 0,01868 |
| TcCLB.509647.120 | hypothetical protein | 0,2747 | 0,02273 |
| TcCLB.508897.40 | hypothetical protein, conserved | 0,2746 | 0,002479 |
| TcCLB.511277.570 | 60S ribosomal protein L7, putative | 0,2742 | 0,01563 |
| TcCLB.503919.30 | hypothetical protein, conserved | 0,2742 | 0,05765 |
| TcCLB.508137.30 | ubiquitin-conjugating enzyme E2, putative | 0,2737 | 0,04408 |
| TcCLB.507817.9 | histone H3, putative | 0,2736 | 0,149 |
| TcCLB.508707.224 | tubulin binding cofactor A-like protein, putative | 0,2735 | 0,09474 |
| TcCLB.509065.140 | hypothetical protein, conserved | 0,2733 | 0,08538 |
| TcCLB.509755.60 | hypothetical protein, conserved (pseudogene) | 0,2732 | 0,3585 |
| TcCLB.506223.80 | Metalloprotease M41 FtsH, putative | 0,273 | 0,002023 |
| TcCLB.504045.20 | hypothetical protein, conserved | 0,273 | 0,003711 |
| TcCLB.506575.70 | Present in the outer mitochondrial membrane proteome 4 | 0,2728 | 0,006753 |
| TcCLB.507029.20 | DNA-J protein, putative | 0,2728 | 0,02941 |
| TcCLB.511741.40 | RNA-binding protein, putative | 0,2727 | 0,01501 |
| TcCLB.511245.100 | hypothetical protein, conserved | 0,2727 | 0,0203 |
| TcCLB.510593.40 | kinetoplastid kinetochore protein 8, putative (kkt8) | 0,2726 | 0,01722 |
| TcCLB.509777.4 | ribosomal protein L2, putative | 0,2726 | 0,03452 |
| TcCLB.509825.20 | hypothetical protein, conserved | 0,2726 | 0,06143 |
| TcCLB.510687.50 | hypothetical protein, conserved | 0,2725 | 0,00692 |
| TcCLB.507669.80 | protein phosphotase, putative | 0,2724 | 0,002487 |
| TcCLB.509543.20 | hypothetical protein, conserved | 0,2724 | 0,02927 |
| TcCLB.511523.40 | hypothetical protein, conserved | 0,2722 | 0,04521 |
| TcCLB.506893.90 | hypothetical protein, conserved | 0,2722 | 0,1341 |
| TcCLB.510651.10 | hypothetical protein, conserved | 0,2722 | 0,1735 |
| TcCLB.507089.40 | CMP-sialic acid transporter, putative | 0,2721 | 0,04943 |
| TcCLB.510687.140 | alkylated DNA repair protein, putative | 0,272 | 0,0229 |
| TcCLB.505267.100 | retrotransposon hot spot protein (RHS, pseudogene), puta | 0,2719 | 0,02209 |
| TcCLB.508461.420 | 30S Ribosomal protein S17, putative | 0,2718 | 0,01425 |
| TcCLB.508543.100 | hypothetical protein, conserved | 0,2717 | 0,001213 |
| TcCLB.511189.80 | small nuclear ribonucleoprotein SmD2, putative | 0,2717 | 0,01945 |
| TcCLB.511851.10 | dispersed gene family protein 1 (DGF-1), putative | 0,2717 | 0,09055 |
| TcCLB.506425.10 | serine carboxypeptidase S28, putative | 0,2714 | 0,03173 |
| TcCLB.511529.120 | hypothetical protein, conserved | 0,2714 | 0,06843 |

| | | | |
|---|---|---|---|
| TcCLB.509635.9 | hypothetical protein, conserved | 0,2713 | 0,04318 |
| TcCLB.504141.31 | hypothetical protein, conserved (pseudogene) | 0,2712 | 0,1889 |
| TcCLB.507023.50 | hypothetical protein, conserved | 0,2707 | 0,02061 |
| TcCLB.508781.80 | hypothetical protein, conserved | 0,2706 | 0,01099 |
| TcCLB.509461.20 | hypothetical protein, conserved | 0,2704 | 0,04594 |
| TcCLB.508469.50 | hypothetical protein, conserved (pseudogene) | 0,2702 | 0,02068 |
| TcCLB.511903.220 | hypothetical protein, conserved | 0,27 | 0,003243 |
| TcCLB.503727.10 | centrin, putative | 0,2699 | 0,02253 |
| TcCLB.509157.80 | DnaJ homolog, putative | 0,2699 | 0,0551 |
| TcCLB.507641.80 | 5'-3' exoribonuclease D, putative (XRND) | 0,2691 | 0,001204 |
| TcCLB.507077.20 | palmitoyl acyltransferase 4, putative | 0,2691 | 0,007325 |
| TcCLB.509319.30 | hypothetical protein, conserved | 0,269 | 0,01491 |
| TcCLB.508409.100 | hypothetical protein, conserved | 0,2688 | 0,00127 |
| TcCLB.506961.10 | retrotransposon hot spot (RHS) protein, putative | 0,2688 | 0,02582 |
| TcCLB.509693.170 | hypothetical protein, conserved | 0,2688 | 0,02874 |
| TcCLB.507057.20 | hypothetical protein, conserved | 0,2688 | 0,03186 |
| TcCLB.507543.10 | dispersed gene family protein 1 (DGF-1), putative | 0,2687 | 0,5182 |
| TcCLB.507513.80 | hypothetical protein, conserved | 0,2686 | 0,008537 |
| TcCLB.509585.10 | dynein heavy chain, putative | 0,2685 | 0,00898 |
| TcCLB.504153.250 | Present in the outer mitochondrial membrane proteome 1 | 0,2684 | 0,0425 |
| TcCLB.470521.10 | protein kinase, putative (fragment) | 0,2684 | 0,1005 |
| TcCLB.509959.198 | ATP-dependent DEAD/H RNA helicase (pseudogene), puta | 0,2684 | 0,4207 |
| TcCLB.478671.9 | hypothetical protein, conserved | 0,2683 | 0,01665 |
| TcCLB.510041.10 | sec1 family transport protein, putative | 0,2683 | 0,1421 |
| TcCLB.509029.150 | hypothetical protein, conserved | 0,2681 | 0,1121 |
| TcCLB.510183.30 | hypothetical protein, conserved | 0,268 | 0,005061 |
| TcCLB.503723.60 | tartrate-resistant acid phosphatase type 5 precursor, puta | 0,2678 | 0,03375 |
| TcCLB.508741.270 | Kinetoplastid RNA editing protein 1, putative (KREPA1) | 0,2673 | 0,005161 |
| TcCLB.511649.160 | hypothetical protein, conserved | 0,2673 | 0,01623 |
| TcCLB.506047.16 | N-acetyltransferase complex ARD1 subunit (pseudogene), | 0,2672 | 0,2753 |
| TcCLB.506733.80 | hypothetical protein, conserved | 0,2671 | 0,02855 |
| TcCLB.511233.30 | hypothetical protein, conserved | 0,267 | 0,03234 |
| TcCLB.506925.370 | MP44 (pseudogene), putative | 0,2669 | 0,3037 |
| TcCLB.509141.10 | hypothetical protein | 0,2668 | 0,001644 |
| TcCLB.510289.60 | hypothetical protein | 0,2668 | 0,1192 |
| TcCLB.507681.20 | succinyl-CoA ligase [GDP-forming] beta-chain, putative | 0,2667 | 0,0004001 |
| TcCLB.504203.10 | hypothetical protein, conserved | 0,2667 | 0,02147 |
| TcCLB.506727.120 | hypothetical protein, conserved | 0,2666 | 0,001287 |
| TcCLB.508505.20 | hypothetical protein, conserved | 0,2665 | 0,01852 |
| TcCLB.509601.130 | hypothetical protein, conserved | 0,2664 | 0,1133 |
| TcCLB.508817.70 | DNA repair protein RAD50, putative | 0,2663 | 0,0001461 |
| TcCLB.508479.274 | hypothetical protein, conserved | 0,2663 | 0,003481 |
| TcCLB.507911.50 | hypothetical protein, conserved | 0,2663 | 0,08201 |
| TcCLB.503879.50 | hypothetical protein, conserved | 0,2662 | 0,02638 |
| TcCLB.506683.250 | retrotransposon hot spot protein (RHS, pseudogene), puta | 0,2661 | 0,2147 |

| | | | |
|---|---|---|---|
| TcCLB.511337.10 | hypothetical protein, conserved | 0,2659 | 0,01698 |
| TcCLB.507673.65 | DNA-directed RNA polymerase subunit, putative | 0,2659 | 0,5386 |
| TcCLB.506365.20 | retrotransposon hot spot protein (RHS, pseudogene), puta | 0,2658 | 0,001216 |
| TcCLB.506529.320 | hypothetical protein, conserved | 0,2657 | 0,02257 |
| TcCLB.509393.20 | hypothetical protein, conserved | 0,2656 | 0,02986 |
| TcCLB.506909.131 | elongation factor 1-gamma (EF-1-gamma, pseudogene), | 0,2656 | 0,5498 |
| TcCLB.510965.20 | mitochondrial inner membrane signal peptidase, putative | 0,2653 | 0,138 |
| TcCLB.510525.30 | hypothetical protein, conserved | 0,265 | 0,005984 |
| TcCLB.511817.200 | Present in the outer mitochondrial membrane proteome 3 | 0,265 | 0,0134 |
| TcCLB.510181.150 | caltractin, putative | 0,2649 | 0,1004 |
| TcCLB.511767.100 | hypothetical protein, conserved | 0,2646 | 0,2771 |
| TcCLB.506779.130 | hypothetical protein | 0,2645 | 0,01393 |
| TcCLB.504241.34 | dispersed gene family protein 1 (DGF-1), putative (fragme | 0,2645 | 0,2198 |
| TcCLB.511237.80 | hypothetical protein, conserved | 0,2643 | 0,009681 |
| TcCLB.509979.230 | hypothetical protein, conserved | 0,2643 | 0,05784 |
| TcCLB.510187.160 | developmentally regulated GTP-binding protein, putative | 0,2642 | 0,001934 |
| TcCLB.506829.100 | fatty acyl CoA syntetase 1, putative | 0,2642 | 0,002732 |
| TcCLB.511181.160 | choline/ethanolamine kinase (pseudogene), putative | 0,2642 | 0,02827 |
| TcCLB.508347.150 | hypothetical protein, conserved | 0,2641 | 0,004102 |
| TcCLB.510091.7 | diacylglycerol acyltransferase, putative (fragment) | 0,2641 | 0,06177 |
| TcCLB.506459.280 | hypothetical protein, conserved | 0,264 | 0,03356 |
| TcCLB.503897.170 | citrate transporter, putative | 0,2639 | 0,07087 |
| TcCLB.507875.210 | retrotransposon hot spot protein (RHS, pseudogene), puta | 0,2638 | 0,00698 |
| TcCLB.508355.20 | hypothetical protein, conserved | 0,2637 | 0,1236 |
| TcCLB.511885.50 | retrotransposon hot spot protein (RHS, pseudogene), puta | 0,2636 | 0,03114 |
| TcCLB.509747.70 | actin-like protein, putative | 0,2636 | 0,05275 |
| TcCLB.510103.17 | hypothetical protein, conserved | 0,2634 | 0,02506 |
| TcCLB.510749.90 | hypothetical protein, conserved | 0,2634 | 0,12 |
| TcCLB.511445.40 | hypothetical protein, conserved | 0,2633 | 0,01693 |
| TcCLB.510155.50 | hypothetical protein, conserved | 0,2633 | 0,04478 |
| TcCLB.504079.6 | selenophosphate synthetase, putative | 0,2631 | 0,007187 |
| TcCLB.506265.114 | Enriched in surface-labeled proteome protein 11, putative | 0,2631 | 0,01302 |
| TcCLB.510101.130 | inositol-1,4,5-trisphosphate (IP3) 5-phosphatase, putative | 0,2629 | 0,003748 |
| TcCLB.506289.50 | hypothetical protein, conserved | 0,2628 | 0,005895 |
| TcCLB.505843.24 | succinate dehydrogenase, putative | 0,2627 | 0,008156 |
| TcCLB.511351.9 | inosine-5'-monophosphate dehydrogenase, putative | 0,2626 | 0,0171 |
| TcCLB.505387.9 | dispersed gene family protein 1 (DGF-1), putative (fragme | 0,2626 | 0,1286 |
| TcCLB.510121.120 | hypothetical protein, conserved | 0,2624 | 0,03924 |
| TcCLB.503453.4 | hypothetical protein, conserved | 0,2624 | 0,2765 |
| TcCLB.511225.20 | retrotransposon hot spot protein (RHS, pseudogene), puta | 0,2623 | 0,003783 |
| TcCLB.506745.20 | hypothetical protein, conserved | 0,2623 | 0,03439 |
| TcCLB.507883.80 | adenylate kinase, putative | 0,2623 | 0,1128 |
| TcCLB.508169.60 | dolicholphosphate-mannose synthase, putative (DPMS) | 0,2622 | 0,04361 |
| TcCLB.510445.40 | hypothetical protein | 0,262 | 0,04298 |
| TcCLB.509105.110 | hypothetical protein, conserved | 0,2618 | 0,03093 |

| | | | |
|---|---|---|---|
| TcCLB.504827.110 | hypothetical protein, conserved | 0,2616 | 0,006733 |
| TcCLB.508013.90 | hypothetical protein | 0,2616 | 0,5012 |
| TcCLB.506559.230 | hypothetical protein, conserved | 0,2615 | 0,1445 |
| TcCLB.511861.60 | beta galactofuranosyl glycosyltransferase (pseudogene), p | 0,2615 | 0,1456 |
| TcCLB.509951.10 | nucleolar protein, putative | 0,2614 | 0,1077 |
| TcCLB.506855.60 | hypothetical protein, conserved | 0,2613 | 0,0236 |
| TcCLB.504129.40 | hypothetical protein, conserved | 0,2613 | 0,05122 |
| TcCLB.413293.20 | hypothetical protein | 0,2613 | 0,4773 |
| TcCLB.508919.30 | hypothetical protein, conserved | 0,2612 | 0,07403 |
| TcCLB.506823.65 | trans-sialidase (pseudogene), putative | 0,2612 | 0,2478 |
| TcCLB.503401.11 | retrotransposon hot spot (RHS) protein, putative (fragmer | 0,2609 | 0,516 |
| TcCLB.511111.10 | translation initiation factor IF-2, putative | 0,2608 | 0,001199 |
| TcCLB.507583.20 | nucleolar protein, putative | 0,2608 | 0,04123 |
| TcCLB.506773.30 | hypothetical protein, conserved | 0,2606 | 0,001871 |
| TcCLB.507211.10 | choline/carnitine O-acyltransferase, putative | 0,2606 | 0,01177 |
| TcCLB.504427.270 | hypothetical protein, conserved | 0,2604 | 0,03084 |
| TcCLB.507317.20 | hypothetical protein | 0,2602 | 0,04708 |
| TcCLB.508033.20 | hypothetical protein | 0,2602 | 0,5806 |
| TcCLB.510543.140 | retrotransposon hot spot protein (RHS, pseudogene), puta | 0,2601 | 0,06816 |
| TcCLB.508061.30 | retrotransposon hot spot protein (RHS, pseudogene), puta | 0,2601 | 0,1057 |
| TcCLB.506337.10 | vacuolar protein sorting-associated protein-like, putative | 0,26 | 0,02309 |
| TcCLB.506271.10 | ATP-dependent DEAD/H RNA helicase (pseudogene), puta | 0,26 | 0,2302 |
| TcCLB.508221.780 | hypothetical protein | 0,26 | 0,3182 |
| TcCLB.461879.10 | retrotransposon hot spot protein (RHS, pseudogene), puta | 0,2598 | 0,1476 |
| TcCLB.507859.120 | dispersed gene family protein 1 (DGF-1), putative | 0,2598 | 0,6056 |
| TcCLB.509055.60 | hypothetical protein, conserved | 0,2595 | 0,1554 |
| TcCLB.506493.70 | serine/threonine protein kinase, putative | 0,2594 | 0,009634 |
| TcCLB.507143.50 | hypothetical protein, conserved | 0,2593 | 0,01623 |
| TcCLB.508209.144 | hypothetical protein, conserved | 0,2593 | 0,08273 |
| TcCLB.511809.80 | hypothetical protein, conserved | 0,2591 | 0,001692 |
| TcCLB.509037.40 | eukaryotic translation initiation factor 4E, putative | 0,2591 | 0,003773 |
| TcCLB.510143.80 | RNA-binding protein, putative | 0,259 | 0,008334 |
| TcCLB.510659.257 | phospholipase A2-like protein, putative | 0,259 | 0,02684 |
| TcCLB.503939.40 | hypothetical protein, conserved | 0,2589 | 0,01103 |
| TcCLB.503689.30 | flagellar protofilament ribbon protein, putative | 0,2589 | 0,01475 |
| TcCLB.510519.140 | hypothetical protein, conserved | 0,2588 | 0,13 |
| TcCLB.503541.40 | hypothetical protein, conserved | 0,2586 | 0,05085 |
| TcCLB.506833.20 | hypothetical protein, conserved | 0,2586 | 0,1257 |
| TcCLB.506855.70 | hypothetical protein, conserved | 0,2582 | 0,004465 |
| TcCLB.508461.390 | hypothetical protein, conserved | 0,2581 | 0,007037 |
| TcCLB.510879.80 | serine acetyltransferase, putative (SAT) | 0,258 | 0,02758 |
| TcCLB.504179.50 | hypothetical protein | 0,2579 | 0,01209 |
| TcCLB.503729.10 | hypothetical protein, conserved | 0,2577 | 0,1011 |
| TcCLB.510655.40 | hypothetical protein, conserved | 0,2576 | 0,0201 |
| TcCLB.508111.20 | hypothetical protein, conserved | 0,2576 | 0,02192 |

| | | | |
|---|---|---|---|
| TcCLB.506925.190 | hypothetical protein, conserved | 0,2576 | 0,06966 |
| TcCLB.508951.70 | hypothetical protein, conserved | 0,2574 | 0,02809 |
| TcCLB.503719.30 | hypothetical protein, conserved | 0,2574 | 0,1109 |
| TcCLB.508283.40 | trans-sialidase (pseudogene), putative | 0,2574 | 0,1329 |
| TcCLB.503755.54 | hypothetical protein, conserved | 0,2573 | 0,01821 |
| TcCLB.510879.110 | ribosomal protein L3, putative | 0,2572 | 0,009851 |
| TcCLB.506345.30 | retrotransposon hot spot protein (RHS, pseudogene), puta | 0,2571 | 0,007573 |
| TcCLB.508741.10 | hypothetical protein | 0,257 | 0,006554 |
| TcCLB.509859.90 | NAD-dependent deacetylase, putative (fragment) | 0,257 | 0,3645 |
| TcCLB.507153.10 | hypothetical protein, conserved | 0,257 | 0,3917 |
| TcCLB.508061.60 | retrotransposon hot spot protein (RHS, pseudogene), puta | 0,2569 | 0,03527 |
| TcCLB.506989.199 | hypothetical protein, conserved | 0,2567 | 0,1772 |
| TcCLB.504741.130 | hypothetical protein, conserved | 0,2566 | 0,001569 |
| TcCLB.506327.40 | hypothetical protein, conserved | 0,2566 | 0,002655 |
| TcCLB.509177.40 | hypothetical protein, conserved | 0,2566 | 0,03464 |
| TcCLB.508441.30 | protein kinase, putative | 0,2564 | 0,1829 |
| TcCLB.510421.290 | Mad2-like HORMA domain containing protein (mad2) | 0,2563 | 0,02245 |
| TcCLB.510691.20 | hypothetical protein, conserved | 0,2562 | 0,007874 |
| TcCLB.508693.180 | ankyrin, putative | 0,2562 | 0,008297 |
| TcCLB.510337.11 | hypothetical protein, conserved (fragment) | 0,2562 | 0,03587 |
| TcCLB.504161.50 | hypothetical protein, conserved | 0,2562 | 0,05189 |
| TcCLB.511865.10 | hypothetical protein, conserved | 0,256 | 0,01102 |
| TcCLB.506045.11 | retrotransposon hot spot protein (RHS, pseudogene), puta | 0,256 | 0,0948 |
| TcCLB.503801.9 | RAD50 DNA repair protein, putative (fragment) | 0,2559 | 0,00785 |
| TcCLB.509617.10 | paraflagellar rod protein 3, putative (pseudogene) (PAR3) | 0,2559 | 0,03585 |
| TcCLB.509965.170 | hypothetical protein, conserved | 0,2556 | 0,01597 |
| TcCLB.510975.10 | hypothetical protein, conserved (pseudogene) | 0,2556 | 0,1922 |
| TcCLB.508355.200 | vacuolar protein sorting protein 16, putative | 0,2555 | 0,002005 |
| TcCLB.509039.10 | hypothetical protein, conserved | 0,2555 | 0,002475 |
| TcCLB.508221.880 | hypothetical protein | 0,2555 | 0,636 |
| TcCLB.510247.10 | R27-2 protein, putative | 0,2554 | 0,05097 |
| TcCLB.511613.40 | mucin-associated surface protein (MASP), putative | 0,2553 | 0,3504 |
| TcCLB.510425.10 | hypothetical protein, conserved | 0,2552 | 0,1356 |
| TcCLB.503805.10 | hypothetical protein, conserved | 0,2551 | 0,06159 |
| TcCLB.508641.170 | protein kinase, putative | 0,2551 | 0,1029 |
| TcCLB.510629.430 | prefoldin subunit, putative | 0,2551 | 0,1182 |
| TcCLB.506437.40 | tuzin | 0,255 | 0,004438 |
| TcCLB.508897.80 | hypothetical protein, conserved | 0,255 | 0,00742 |
| TcCLB.506777.120 | protein kinase, putative | 0,2549 | 0,0474 |
| TcCLB.506559.364 | hypothetical protein, conserved | 0,2548 | 0,0005941 |
| TcCLB.506855.140 | synaptobrevin, putative | 0,2548 | 0,02322 |
| TcCLB.511261.150 | hypothetical protein, conserved | 0,2548 | 0,08834 |
| TcCLB.507891.20 | flagellar calcium-binding-like protein, putative | 0,2547 | 0,05858 |
| TcCLB.510767.30 | hypothetical protein | 0,2547 | 0,1285 |
| TcCLB.506389.20 | trans-sialidase (pseudogene), putative | 0,2547 | 0,1656 |

| | | | |
|---|---|---|---|
| TcCLB.507665.30 | ferric reductase transmembrane protein, putative (FR1) | 0,2546 | 0,004297 |
| TcCLB.510007.60 | hypothetical protein, conserved | 0,2546 | 0,004596 |
| TcCLB.503987.4 | Ras-related protein RAB2B, putative (fragment) (RAB2B) | 0,2546 | 0,09573 |
| TcCLB.510431.280 | RNA editing complex protein MP46, putative | 0,2545 | 0,0495 |
| TcCLB.504625.30 | hypothetical protein | 0,2544 | 0,007489 |
| TcCLB.511825.20 | hypothetical protein, conserved | 0,2544 | 0,0526 |
| TcCLB.511377.25 | elongation factor 1-gamma (EF-1-gamma, pseudogene), | 0,2543 | 0,5348 |
| TcCLB.507873.20 | cell differentiation protein, putative | 0,2542 | 0,0559 |
| TcCLB.509539.50 | small GTP-binding protein, putative | 0,2541 | 0,007246 |
| TcCLB.506635.40 | hypothetical protein, conserved | 0,2541 | 0,02575 |
| TcCLB.506933.24 | hypothetical protein, conserved | 0,2541 | 0,03855 |
| TcCLB.506457.70 | hypothetical protein, conserved | 0,2541 | 0,1112 |
| TcCLB.503799.10 | DNA repair protein RAD50, putative | 0,254 | 0,002351 |
| TcCLB.508241.110 | hypothetical protein, conserved | 0,254 | 0,01689 |
| TcCLB.505987.10 | hypothetical protein, conserved (pseudogene) | 0,254 | 0,09631 |
| TcCLB.510903.20 | hypothetical protein, conserved | 0,254 | 0,191 |
| TcCLB.507583.40 | hypothetical protein, conserved | 0,2539 | 0,008181 |
| TcCLB.503909.60 | hypothetical protein, conserved | 0,2539 | 0,04262 |
| TcCLB.506247.90 | hypothetical protein | 0,2539 | 0,0582 |
| TcCLB.510657.90 | hypothetical protein, conserved | 0,2534 | 0,1028 |
| TcCLB.503431.100 | chromatin assembly factor 1 subunit B, putative | 0,2527 | 0,0123 |
| TcCLB.506479.30 | hypothetical protein, conserved | 0,2527 | 0,04453 |
| TcCLB.509155.30 | hypothetical protein, conserved | 0,2525 | 0,01177 |
| TcCLB.506821.180 | aquaporin-like protein, putative | 0,2525 | 0,07613 |
| TcCLB.507623.40 | hypothetical protein, conserved | 0,2524 | 0,06397 |
| TcCLB.510167.20 | hypothetical protein, conserved | 0,2523 | 0,1386 |
| TcCLB.503815.20 | lysyl-tRNA synthetase, putative | 0,2521 | 0,03976 |
| TcCLB.507003.50 | hypothetical protein, conserved | 0,2521 | 0,1096 |
| TcCLB.508549.10 | legume-like lectin, putative | 0,2518 | 0,02658 |
| TcCLB.509063.20 | hypothetical protein, conserved | 0,2518 | 0,04725 |
| TcCLB.509029.50 | hypothetical protein, conserved | 0,2514 | 0,08941 |
| TcCLB.511339.20 | C-5 sterol desaturase, putative | 0,2513 | 0,04665 |
| TcCLB.507875.60 | beta galactofuranosyl glycosyltransferase, putative | 0,2512 | 0,1261 |
| TcCLB.504055.50 | hypothetical protein, conserved | 0,2511 | 0,01292 |
| TcCLB.510311.20 | hypothetical protein, conserved | 0,251 | 0,02792 |
| TcCLB.511185.10 | hypothetical protein | 0,251 | 0,08845 |
| TcCLB.508323.9 | farnesyl pyrophosphate synthase, putative | 0,2509 | 0,02282 |
| TcCLB.509105.4 | Generative cell specific 1 protein, putative (GCS1) | 0,2509 | 0,1233 |
| TcCLB.508207.180 | Eukaryotic translation initiation factor 4E type 6, putative | 0,2507 | 0,01799 |
| TcCLB.504075.10 | RNA polymerase I, putative | 0,2506 | 0,06229 |
| TcCLB.506989.40 | Golgi vesicular membrane trafficking protein, putative | 0,2506 | 0,1671 |
| TcCLB.503723.40 | hypothetical protein, conserved | 0,2503 | 0,02867 |
| TcCLB.511165.109 | hypothetical protein, conserved | 0,2503 | 0,08708 |
| TcCLB.507047.140 | hypothetical protein, conserved | 0,2502 | 0,03551 |
| TcCLB.506331.64 | hypothetical protein | 0,2501 | 0,3244 |

| | | | |
|---|---|---|---|
| TcCLB.503919.4 | hypothetical protein, conserved | 0,2499 | 0,3186 |
| TcCLB.511533.10 | hypothetical protein, conserved | 0,2496 | 0,03944 |
| TcCLB.504741.180 | hypothetical protein, conserved | 0,2495 | 0,06144 |
| TcCLB.503987.20 | hypothetical protein, conserved | 0,2495 | 0,08021 |
| TcCLB.510789.20 | mucin TcMUC (pseudogene), putative | 0,2495 | 0,2232 |
| TcCLB.510035.10 | trans-sialidase (pseudogene), putative | 0,2494 | 0,007431 |
| TcCLB.510763.109 | rRNA processing protein, putative | 0,2494 | 0,06564 |
| TcCLB.507895.160 | hypothetical protein, conserved | 0,2493 | 0,07825 |
| TcCLB.509719.60 | hypothetical protein | 0,2491 | 0,1746 |
| TcCLB.505183.84 | hypothetical protein, conserved | 0,2489 | 0,04339 |
| TcCLB.511655.50 | hypothetical protein, conserved (pseudogene) | 0,2489 | 0,05396 |
| TcCLB.508045.40 | hypothetical protein, conserved | 0,2488 | 0,01297 |
| TcCLB.506693.50 | ribonuclease inhibitor-like protein, putative | 0,2487 | 0,06577 |
| TcCLB.506943.114 | U6 snRNA-associated Sm-like protein LSm5p | 0,2486 | 0,1446 |
| TcCLB.445869.10 | hypothetical protein, conserved | 0,2485 | 0,02853 |
| TcCLB.504045.70 | hypothetical protein, conserved | 0,2485 | 0,1126 |
| TcCLB.506963.30 | hypothetical protein, conserved | 0,2485 | 0,1184 |
| TcCLB.440363.10 | hypothetical protein, conserved | 0,2485 | 0,1344 |
| TcCLB.511583.10 | hypothetical protein, conserved | 0,2483 | 0,07033 |
| TcCLB.506519.80 | hypothetical protein, conserved | 0,2483 | 0,07073 |
| TcCLB.503929.50 | hypothetical protein, conserved | 0,2483 | 0,08448 |
| TcCLB.507807.40 | hypothetical protein, conserved | 0,2482 | 0,02536 |
| TcCLB.506321.359 | MRB1-associated protein, putative | 0,2481 | 0,1516 |
| TcCLB.504333.20 | trans-sialidase (pseudogene), putative | 0,248 | 0,02524 |
| TcCLB.508799.10 | hypothetical protein, conserved | 0,248 | 0,06372 |
| TcCLB.511725.250 | chaperonin containing t-complex protein, putative | 0,2479 | 0,00008171 |
| TcCLB.510655.60 | hypothetical protein, conserved | 0,2479 | 0,08733 |
| TcCLB.507787.80 | hypothetical protein, conserved | 0,2479 | 0,1769 |
| TcCLB.508163.181 | hypothetical protein, conserved (pseudogene) | 0,2479 | 0,4655 |
| TcCLB.508459.9 | hypothetical protein, conserved | 0,2478 | 0,09098 |
| TcCLB.508893.10 | hypothetical protein, conserved | 0,2477 | 0,02981 |
| TcCLB.506495.6 | hypothetical protein, conserved | 0,2476 | 0,1037 |
| TcCLB.503899.10 | heat shock 70 kDa protein, putative | 0,2475 | 0,026 |
| TcCLB.511577.150 | hypothetical protein, conserved | 0,2475 | 0,03416 |
| TcCLB.510187.380 | hypothetical protein, conserved | 0,2474 | 0,03419 |
| TcCLB.504827.159 | hypothetical protein, conserved | 0,2471 | 0,1192 |
| TcCLB.510911.40 | hypothetical protein, conserved | 0,2471 | 0,1569 |
| TcCLB.509507.20 | hypothetical protein, conserved | 0,2467 | 0,00824 |
| TcCLB.511367.280 | hypothetical protein, conserved | 0,2467 | 0,02443 |
| TcCLB.507927.80 | hypothetical protein | 0,2466 | 0,06807 |
| TcCLB.506363.20 | hypothetical protein, conserved | 0,2466 | 0,1052 |
| TcCLB.506711.30 | cyclin, putative | 0,2466 | 0,1086 |
| TcCLB.508231.174 | hypothetical protein, conserved | 0,2466 | 0,1469 |
| TcCLB.511021.60 | hypothetical protein, conserved | 0,2465 | 0,03766 |
| TcCLB.510533.50 | hypothetical protein, conserved | 0,2465 | 0,1185 |

| | | | |
|---|---|---|---|
| TcCLB.510101.110 | hypothetical protein, conserved | 0,2464 | 0,006654 |
| TcCLB.506155.80 | hypothetical protein, conserved | 0,2464 | 0,01803 |
| TcCLB.511237.40 | CDP-DAG synthase, putative (CDS) | 0,2461 | 0,02551 |
| TcCLB.511725.120 | kinetoplast polyadenylation/uridylation factor 2, putative | 0,246 | 0,02087 |
| TcCLB.509641.50 | hypothetical protein, conserved | 0,2459 | 0,02658 |
| TcCLB.504125.120 | hypothetical protein, conserved | 0,2459 | 0,1208 |
| TcCLB.510073.50 | elongation factor 1-gamma (EF-1-gamma), putative | 0,2458 | 0,04025 |
| TcCLB.507669.180 | hypothetical protein, conserved | 0,2457 | 0,01372 |
| TcCLB.506793.10 | hypothetical protein, conserved (pseudogene) | 0,2457 | 0,04334 |
| TcCLB.507991.80 | hypothetical protein, conserved | 0,2455 | 0,01053 |
| TcCLB.510799.70 | hypothetical protein, conserved | 0,2452 | 0,03197 |
| TcCLB.506713.14 | hypothetical protein, conserved | 0,2452 | 0,1743 |
| TcCLB.510737.10 | 6-phosphogluconate dehydrogenase, decarboxylating, puta | 0,2452 | 0,183 |
| TcCLB.506915.20 | hypothetical protein | 0,2451 | 0,02161 |
| TcCLB.506435.440 | hypothetical protein, conserved | 0,245 | 0,01718 |
| TcCLB.511867.180 | hypothetical protein, conserved | 0,245 | 0,06771 |
| TcCLB.506529.250 | hypothetical protein, conserved | 0,2448 | 0,004118 |
| TcCLB.506445.30 | cation transporter, putative | 0,2448 | 0,01853 |
| TcCLB.511711.60 | hypothetical protein | 0,2448 | 0,2216 |
| TcCLB.510879.120 | ribosomal protein L3, putative | 0,2446 | 0,0005787 |
| TcCLB.507817.70 | hypothetical protein, conserved | 0,2445 | 0,0639 |
| TcCLB.510731.59 | dual specificity protein phosphatase, putative | 0,2444 | 0,06555 |
| TcCLB.508711.40 | mitochondrial RNA binding protein, putative | 0,2442 | 0,01149 |
| TcCLB.508325.180 | retrotransposon hot spot protein (RHS, pseudogene), puta | 0,2442 | 0,0182 |
| TcCLB.506773.150 | hypothetical protein | 0,2442 | 0,031 |
| TcCLB.503651.40 | hypothetical protein | 0,244 | 0,04594 |
| TcCLB.511481.10 | hypothetical protein, conserved | 0,2439 | 0,108 |
| TcCLB.508387.40 | hypothetical protein, conserved | 0,2439 | 0,1524 |
| TcCLB.503895.10 | hypothetical protein, conserved | 0,2438 | 0,02374 |
| TcCLB.511589.60 | hypothetical protein, conserved | 0,2437 | 0,1911 |
| TcCLB.506525.104 | AP-4 complex subunit sigma-1, putative | 0,2436 | 0,1657 |
| TcCLB.509767.220 | hypothetical protein, conserved | 0,2435 | 0,008415 |
| TcCLB.503955.10 | mitochondrial DNA polymerase beta-PAK, putative | 0,2435 | 0,07076 |
| TcCLB.507897.2 | hypothetical protein, conserved | 0,2432 | 0,09362 |
| TcCLB.507811.107 | hypothetical protein, conserved | 0,2432 | 0,1292 |
| TcCLB.503423.5 | retrotransposon hot spot (RHS) protein, putative (fragmer | 0,2429 | 0,5094 |
| TcCLB.511621.240 | hypothetical protein, conserved | 0,2427 | 0,006643 |
| TcCLB.511877.15 | elongation factor 1-gamma (EF-1-gamma, pseudogene), | 0,2427 | 0,1829 |
| TcCLB.510243.50 | adenylate kinase, putative | 0,2426 | 0,02399 |
| TcCLB.507143.10 | hypothetical protein, conserved | 0,2425 | 0,004167 |
| TcCLB.504159.109 | hypothetical protein, conserved | 0,2424 | 0,1526 |
| TcCLB.506637.110 | hypothetical protein, conserved | 0,2423 | 0,003563 |
| TcCLB.509791.140 | hypothetical protein, conserved | 0,2423 | 0,07286 |
| TcCLB.510963.80 | hypothetical protein, conserved | 0,2423 | 0,2234 |
| TcCLB.508873.50 | hypothetical protein | 0,2423 | 0,6316 |

| | | | |
|---|---|---|---|
| TcCLB.511817.230 | hypothetical protein, conserved | 0,2422 | 0,004965 |
| TcCLB.511807.6 | ATP-dependent DEAD/H RNA helicase (pseudogene), puta | 0,2422 | 0,04117 |
| TcCLB.503789.34 | hypothetical protein, conserved | 0,2421 | 0,0515 |
| TcCLB.506389.70 | translation-associated element 2, putative | 0,2419 | 0,01299 |
| TcCLB.508209.80 | hypothetical protein, conserved | 0,2419 | 0,04437 |
| TcCLB.503521.80 | hypothetical protein, conserved | 0,2419 | 0,07945 |
| TcCLB.508547.70 | Present in the outer mitochondrial membrane proteome 3 | 0,2418 | 0,04832 |
| TcCLB.508239.10 | origin recognition complex subunit 1 (ORC1), putative | 0,2418 | 0,05807 |
| TcCLB.507603.130 | hypothetical protein, conserved | 0,2417 | 0,06676 |
| TcCLB.506265.140 | hypothetical protein, conserved | 0,2416 | 0,07057 |
| TcCLB.402863.9 | dynein heavy chain, putative (fragment) | 0,2415 | 0,01879 |
| TcCLB.508255.20 | hypothetical protein, conserved (pseudogene) | 0,2415 | 0,1196 |
| TcCLB.511507.10 | hypothetical protein, conserved | 0,2415 | 0,2362 |
| TcCLB.506457.99 | hypothetical protein, conserved | 0,2414 | 0,01959 |
| TcCLB.506123.50 | hypothetical protein, conserved | 0,2414 | 0,0887 |
| TcCLB.504071.60 | hypothetical protein, conserved | 0,241 | 0,04638 |
| TcCLB.506885.170 | hypothetical protein, conserved | 0,2409 | 0,07388 |
| TcCLB.510431.190 | hypothetical protein | 0,2407 | 0,002201 |
| TcCLB.506221.120 | hypothetical protein, conserved | 0,2407 | 0,1107 |
| TcCLB.511651.19 | delta-adaptin, putative | 0,2407 | 0,2397 |
| TcCLB.510741.220 | hypothetical protein, conserved | 0,2404 | 0,01796 |
| TcCLB.509927.10 | trans-sialidase, putative (fragment) | 0,2403 | 0,1118 |
| TcCLB.503945.30 | hypothetical protein, conserved | 0,2403 | 0,1284 |
| TcCLB.506243.77 | retrotransposon hot spot protein (RHS, pseudogene), puta | 0,2403 | 0,3175 |
| TcCLB.510089.200 | hypothetical protein, conserved | 0,2402 | 0,01243 |
| TcCLB.506979.50 | hypothetical protein, conserved | 0,2402 | 0,0166 |
| TcCLB.506925.450 | hypothetical protein, conserved | 0,2402 | 0,01836 |
| TcCLB.509571.4 | PAB1-binding protein , putative | 0,2397 | 0,004278 |
| TcCLB.503835.20 | hypothetical protein, conserved | 0,2397 | 0,07267 |
| TcCLB.510149.60 | hypothetical protein, conserved | 0,2396 | 0,1043 |
| TcCLB.509155.10 | hypothetical protein, conserved | 0,2395 | 0,002322 |
| TcCLB.508879.60 | hypothetical protein, conserved | 0,2395 | 0,004633 |
| TcCLB.503957.20 | retrotransposon hot spot (RHS) protein, putative | 0,2393 | 0,04968 |
| TcCLB.511847.60 | mitochondrial DNA polymerase I protein B, putative (POLI | 0,2392 | 0,01021 |
| TcCLB.509213.20 | hypothetical protein, conserved | 0,2392 | 0,07748 |
| TcCLB.509509.69 | hypothetical protein, conserved | 0,2392 | 0,1086 |
| TcCLB.510745.49 | hypothetical protein, conserved (fragment) | 0,2392 | 0,2324 |
| TcCLB.505967.19 | dispersed gene family protein 1 (DGF-1), putative (fragme | 0,2389 | 0,3694 |
| TcCLB.506349.53 | retrotransposon hot spot protein (RHS, pseudogene), puta | 0,2388 | 0,007022 |
| TcCLB.508397.10 | V-type proton ATPase subunit D, putative (VAd) | 0,2382 | 0,03071 |
| TcCLB.509353.50 | hypothetical protein, conserved | 0,2382 | 0,03902 |
| TcCLB.511647.44 | hypothetical protein, conserved | 0,2381 | 0,1623 |
| TcCLB.511367.20 | hypothetical protein | 0,2381 | 0,5884 |
| TcCLB.511257.60 | surface protease GP63, putative | 0,2379 | 0,1023 |
| TcCLB.466965.10 | trans-sialidase (pseudogene), putative | 0,2378 | 0,1184 |

| | | | |
|---|---|---|---|
| TcCLB.508575.20 | hypothetical protein | 0,2378 | 0,5118 |
| TcCLB.509807.11 | hypothetical protein, conserved (pseudogene) | 0,2377 | 0,1135 |
| TcCLB.506227.170 | hypothetical protein, conserved | 0,2376 | 0,1809 |
| TcCLB.510431.270 | hypothetical protein, conserved | 0,2374 | 0,02173 |
| TcCLB.508487.11 | retrotransposon hot spot protein (RHS, pseudogene), puta | 0,2374 | 0,1279 |
| TcCLB.503421.14 | ATP-dependent DEAD/H RNA helicase (pseudogene), puta | 0,2373 | 0,42 |
| TcCLB.507519.160 | hypothetical protein, conserved | 0,2372 | 0,08451 |
| TcCLB.511577.40 | 35 kDa cyclophilin, putative (CYP35) | 0,2372 | 0,1537 |
| TcCLB.509965.110 | importin alpha, putative | 0,2371 | 0,00652 |
| TcCLB.508039.30 | retrotransposon hot spot (RHS) protein, putative | 0,2371 | 0,02031 |
| TcCLB.511811.20 | hypothetical protein, conserved | 0,2371 | 0,06597 |
| TcCLB.511577.86 | N-acetyltransferase, putative | 0,2369 | 0,02478 |
| TcCLB.506579.40 | hypothetical protein, conserved | 0,2368 | 0,002171 |
| TcCLB.510719.240 | hypothetical protein, conserved | 0,2368 | 0,1161 |
| TcCLB.507643.90 | beta galactofuranosyl glycosyltransferase (pseudogene), p | 0,2367 | 0,05442 |
| TcCLB.508739.70 | protein kinase, putative | 0,2365 | 0,0005153 |
| TcCLB.509647.70 | guanylate kinase, putative | 0,2365 | 0,02163 |
| TcCLB.506829.30 | hypothetical protein, conserved | 0,2365 | 0,1178 |
| TcCLB.506335.55 | elongation factor 1-gamma (EF-1-gamma, pseudogene), | 0,2365 | 0,3313 |
| TcCLB.509611.120 | hypothetical protein, conserved | 0,2362 | 0,1062 |
| TcCLB.508951.40 | acyl-CoA dehydrogenase, putative | 0,2362 | 0,1176 |
| TcCLB.510963.10 | DNA damage repair protein, putative | 0,2361 | 0,01637 |
| TcCLB.506871.130 | hypothetical protein, conserved | 0,236 | 0,03836 |
| TcCLB.511737.60 | hypothetical protein, conserved | 0,2359 | 0,07486 |
| TcCLB.505999.30 | anti-silencing protein ASF 1, putative | 0,2359 | 0,1676 |
| TcCLB.503953.40 | hypothetical protein, conserved | 0,2359 | 0,1706 |
| TcCLB.510821.50 | replication Factor A 28 kDa subunit, putative | 0,2358 | 0,01135 |
| TcCLB.506401.310 | hypothetical protein, conserved | 0,2354 | 0,09394 |
| TcCLB.510571.10 | hypothetical protein, conserved | 0,2351 | 0,1906 |
| TcCLB.509289.20 | tuzin, putative | 0,235 | 0,0004691 |
| TcCLB.504427.30 | hypothetical protein, conserved | 0,235 | 0,03051 |
| TcCLB.508909.100 | hypothetical protein, conserved | 0,2348 | 0,02721 |
| TcCLB.504741.40 | hypothetical protein, conserved | 0,2345 | 0,08857 |
| TcCLB.510665.70 | acyl transferase-like protein, putative | 0,2345 | 0,2433 |
| TcCLB.510735.9 | hypothetical protein, conserved | 0,2343 | 0,02124 |
| TcCLB.510895.20 | hypothetical protein, conserved | 0,2342 | 0,05615 |
| TcCLB.508501.300 | hypothetical protein, conserved | 0,2342 | 0,07369 |
| TcCLB.509753.202 | hypothetical protein | 0,2342 | 0,6298 |
| TcCLB.398235.20 | ubiquitin fusion degardation protein, putative | 0,234 | 0,4991 |
| TcCLB.504097.20 | hypothetical protein, conserved | 0,2339 | 0,0721 |
| TcCLB.511807.270 | hypothetical protein, conserved | 0,2338 | 0,0118 |
| TcCLB.511751.50 | protein kinase, putative | 0,2338 | 0,03583 |
| TcCLB.511725.210 | hypothetical protein, conserved | 0,2338 | 0,06894 |
| TcCLB.504057.70 | hypothetical protein, conserved | 0,2338 | 0,07612 |
| TcCLB.510099.131 | hypothetical protein, conserved (fragment) | 0,2337 | 0,02908 |

| | | | |
|---|---|---|---|
| TcCLB.507883.60 | hypothetical protein, conserved | 0,2337 | 0,04458 |
| TcCLB.508001.30 | hypothetical protein, conserved (pseudogene) | 0,2337 | 0,1403 |
| TcCLB.506683.10 | hypothetical protein, conserved (pseudogene) | 0,2337 | 0,6319 |
| TcCLB.510901.210 | hypothetical protein, conserved | 0,2336 | 0,01949 |
| TcCLB.507615.80 | hypothetical protein, conserved | 0,2335 | 0,008079 |
| TcCLB.511291.110 | hypothetical protein, conserved | 0,2335 | 0,04943 |
| TcCLB.507497.20 | retrotransposon hot spot protein (RHS, pseudogene), puta | 0,2335 | 0,3238 |
| TcCLB.508257.140 | Aquaporin 2 | 0,2334 | 0,0222 |
| TcCLB.508189.11 | retrotransposon hot spot protein (RHS, pseudogene), puta | 0,2333 | 0,4905 |
| TcCLB.509831.33 | hypothetical protein, conserved | 0,2331 | 0,003808 |
| TcCLB.509805.40 | Multisite-specific tRNA:(cytosine-C(5))-methyltransferase | 0,2331 | 0,03819 |
| TcCLB.503597.5 | hypothetical protein (fragment) | 0,2331 | 0,08789 |
| TcCLB.509671.172 | hypothetical protein, conserved | 0,2331 | 0,1717 |
| TcCLB.509875.110 | hypothetical protein | 0,2331 | 0,5925 |
| TcCLB.504151.69 | hypothetical protein, conserved | 0,2325 | 0,02848 |
| TcCLB.506025.10 | hypothetical protein, conserved | 0,2325 | 0,0609 |
| TcCLB.508271.4 | hypothetical protein, conserved | 0,2325 | 0,19 |
| TcCLB.511467.70 | hypothetical protein, conserved | 0,2325 | 0,2518 |
| TcCLB.509971.60 | surface protease GP63, putative | 0,2324 | 0,5265 |
| TcCLB.511821.100 | hypothetical protein, conserved | 0,2323 | 0,02045 |
| TcCLB.508539.71 | elongation factor 1-gamma (EF-1-gamma, pseudogene), | 0,2322 | 0,3665 |
| TcCLB.506067.179 | elongation factor 1-gamma (EF-1-gamma, pseudogene), | 0,2322 | 0,636 |
| TcCLB.511151.11 | hypothetical protein (fragment) | 0,2321 | 0,007557 |
| TcCLB.508645.10 | SNF2 family protein | 0,2321 | 0,01644 |
| TcCLB.507757.26 | hypothetical protein | 0,2321 | 0,5732 |
| TcCLB.510055.50 | hypothetical protein, conserved | 0,232 | 0,01095 |
| TcCLB.510005.20 | trans-sialidase, Group II, putative | 0,2318 | 0,1476 |
| TcCLB.504149.190 | hypothetical protein, conserved (pseudogene) | 0,2317 | 0,004851 |
| TcCLB.510039.80 | hypothetical protein, conserved | 0,2317 | 0,1136 |
| TcCLB.508153.560 | hypothetical protein, conserved | 0,2317 | 0,1449 |
| TcCLB.510089.190 | hypothetical protein, conserved | 0,2316 | 0,0431 |
| TcCLB.422921.20 | 50S ribosomal protein L17, putative | 0,2316 | 0,2596 |
| TcCLB.510303.190 | cell cycle associated protein MOB1, putative | 0,2315 | 0,04227 |
| TcCLB.507711.160 | hypothetical protein, conserved | 0,2314 | 0,003237 |
| TcCLB.511647.40 | hypothetical protein, conserved | 0,2314 | 0,04387 |
| TcCLB.506553.10 | hypothetical protein, conserved | 0,2314 | 0,19 |
| TcCLB.505071.104 | hypothetical protein, conserved | 0,2313 | 0,2462 |
| TcCLB.508479.5 | glycine dehydrogenase (pseudogene), putative | 0,2312 | 0,3817 |
| TcCLB.508647.210 | dynein arm light chain, putative | 0,2309 | 0,003472 |
| TcCLB.508479.170 | hypothetical protein, conserved | 0,2309 | 0,04241 |
| TcCLB.509319.50 | hypothetical protein, conserved | 0,2305 | 0,002237 |
| TcCLB.509979.10 | hypothetical protein, conserved | 0,2305 | 0,004139 |
| TcCLB.505193.60 | hypothetical protein, conserved | 0,2305 | 0,09005 |
| TcCLB.506195.140 | hypothetical protein, conserved | 0,2304 | 0,02721 |
| TcCLB.509671.40 | hypothetical protein, conserved | 0,2303 | 0,09249 |

| | | | |
|---|---|---|---|
| TcCLB.507547.59 | hypothetical protein, conserved | 0,2301 | 0,2256 |
| TcCLB.505163.80 | oligosaccharyl transferase subunit, putative | 0,2298 | 0,0003353 |
| TcCLB.511303.24 | hypothetical protein, conserved | 0,2298 | 0,07823 |
| TcCLB.503785.30 | hypothetical protein, conserved | 0,2298 | 0,1741 |
| TcCLB.443971.10 | hypothetical protein, conserved (fragment) | 0,2298 | 0,571 |
| TcCLB.509029.70 | leucine-rich repeat protein, putative | 0,2297 | 0,1294 |
| TcCLB.509715.140 | hypothetical protein, conserved | 0,2295 | 0,1402 |
| TcCLB.507993.260 | hypothetical protein, conserved | 0,2294 | 0,04939 |
| TcCLB.507711.330 | serine peptidase, putative | 0,2292 | 0,0453 |
| TcCLB.508355.160 | hypothetical protein, conserved | 0,2289 | 0,01635 |
| TcCLB.508027.90 | hypothetical protein, conserved | 0,2289 | 0,02426 |
| TcCLB.504625.50 | dual specificity protein phosphatase or MAP kinase phospl | 0,2289 | 0,05097 |
| TcCLB.510967.20 | hypothetical protein, conserved | 0,2285 | 0,1319 |
| TcCLB.509317.40 | ribosomal protein L11, putative | 0,2282 | 0,1401 |
| TcCLB.510959.15 | elongation factor 1-gamma (EF-1-gamma, pseudogene), | 0,2281 | 0,6175 |
| TcCLB.510149.30 | hypothetical protein, conserved | 0,228 | 0,0191 |
| TcCLB.511647.42 | hypothetical protein, conserved | 0,228 | 0,2199 |
| TcCLB.510421.300 | hypothetical protein, conserved | 0,2279 | 0,1349 |
| TcCLB.506201.70 | serine/threonine-protein phosphatase PP1 alpha | 0,2279 | 0,2207 |
| TcCLB.509875.165 | trans-sialidase (pseudogene), putative | 0,2279 | 0,2465 |
| TcCLB.508279.10 | trans-sialidase (pseudogene), putative | 0,2279 | 0,2874 |
| TcCLB.508741.240 | hypothetical protein, conserved | 0,2278 | 0,02647 |
| TcCLB.507925.20 | hypothetical protein, conserved | 0,2278 | 0,03179 |
| TcCLB.508461.240 | chaperone protein DNAJ, putatative | 0,2278 | 0,05687 |
| TcCLB.506579.129 | hypothetical protein, conserved (fragment) | 0,2277 | 0,1319 |
| TcCLB.505023.10 | trans-sialidase (pseudogene), putative | 0,2277 | 0,248 |
| TcCLB.503467.30 | oligosaccharyl transferase subunit, putative | 0,2276 | 0,03209 |
| TcCLB.511135.30 | hypothetical protein, conserved | 0,2276 | 0,05733 |
| TcCLB.508119.100 | hypothetical protein, conserved | 0,2275 | 0,02685 |
| TcCLB.509055.69 | hypothetical protein, conserved (fragment) | 0,2275 | 0,278 |
| TcCLB.508999.20 | Protein transport protein yif1, putative | 0,2271 | 0,1458 |
| TcCLB.507045.10 | retrotransposon hot spot protein (RHS, pseudogene), puta | 0,2268 | 0,1179 |
| TcCLB.508413.50 | RNA-binding protein, putative | 0,2267 | 0,01625 |
| TcCLB.508397.30 | SPFH domain / Band 7 family protein, putative | 0,2265 | 0,06899 |
| TcCLB.511283.160 | hypothetical protein, conserved | 0,2264 | 0,01317 |
| TcCLB.511523.60 | hypothetical protein, conserved | 0,2263 | 0,01773 |
| TcCLB.503823.169 | kinetoplastid kinetochore protein 2, putative (fragment) (l | 0,2263 | 0,07068 |
| TcCLB.507159.14 | hypothetical protein, conserved | 0,2262 | 0,2037 |
| TcCLB.506975.97 | hypothetical protein, conserved (fragment) | 0,2262 | 0,4472 |
| TcCLB.511417.94 | hypothetical protein, conserved | 0,2261 | 0,1542 |
| TcCLB.504221.20 | 26S proteasome regulatory non-ATPase subunit, putative | 0,2259 | 0,04854 |
| TcCLB.511199.20 | hypothetical protein, conserved | 0,2259 | 0,05765 |
| TcCLB.510121.130 | serine/threonine protein kinase, putative | 0,2259 | 0,2291 |
| TcCLB.503805.30 | hypothetical protein, conserved | 0,2257 | 0,1148 |
| TcCLB.508717.56 | hypothetical protein, conserved | 0,2255 | 0,07322 |

| | | | |
|---|---|---|---|
| TcCLB.509575.10 | retrotransposon hot spot (RHS) protein, putative | 0,2255 | 0,08561 |
| TcCLB.511825.190 | hypothetical protein, conserved | 0,2252 | 0,008988 |
| TcCLB.507925.50 | hypothetical protein, conserved | 0,2251 | 0,007635 |
| TcCLB.508169.120 | hypothetical protein, conserved | 0,2251 | 0,09032 |
| TcCLB.508827.30 | eukaryotic translation initiation factor, putative | 0,225 | 0,007455 |
| TcCLB.511589.40 | hypothetical protein, conserved | 0,225 | 0,1378 |
| TcCLB.511533.19 | dynein light intermediate chain, putative | 0,225 | 0,1857 |
| TcCLB.508409.9 | 26S proteasome regulatory non-ATPase subunit, putative | 0,2249 | 0,04221 |
| TcCLB.509231.30 | hypothetical protein | 0,2249 | 0,2068 |
| TcCLB.503999.100 | phospholipid-transporting ATPase 1-like protein, putative | 0,2249 | 0,2372 |
| TcCLB.510187.110 | Mitochondrial ribosomal protein S18, putative (MRPS18) | 0,2248 | 0,01794 |
| TcCLB.509715.80 | hypothetical protein, conserved | 0,2248 | 0,02864 |
| TcCLB.510667.30 | hypothetical protein, conserved | 0,2248 | 0,04614 |
| TcCLB.510431.330 | hypothetical protein, conserved | 0,2248 | 0,1514 |
| TcCLB.506529.640 | hypothetical protein, conserved | 0,2247 | 0,01155 |
| TcCLB.510749.30 | Sorting nexin-4 (ATG24) | 0,2247 | 0,01634 |
| TcCLB.511715.30 | hypothetical protein, conserved | 0,2247 | 0,2294 |
| TcCLB.473619.9 | dispersed gene family protein 1 (DGF-1), putative (fragme | 0,2247 | 0,6597 |
| TcCLB.510187.410 | hypothetical protein, conserved | 0,2246 | 0,04017 |
| TcCLB.509265.50 | hypothetical protein, conserved (pseudogene) | 0,2246 | 0,09816 |
| TcCLB.509979.281 | elongation factor 1-gamma (EF-1-gamma, pseudogene), | 0,2244 | 0,2896 |
| TcCLB.511109.60 | hypothetical protein, conserved | 0,2243 | 0,0128 |
| TcCLB.510317.14 | hypothetical protein, conserved | 0,2243 | 0,08055 |
| TcCLB.506941.270 | DNA-J protein, putative | 0,2241 | 0,06796 |
| TcCLB.506469.139 | hypothetical protein, conserved | 0,2241 | 0,2683 |
| TcCLB.511215.50 | hypothetical protein, conserved | 0,2239 | 0,0217 |
| TcCLB.506885.320 | hypothetical protein, conserved | 0,2239 | 0,02632 |
| TcCLB.510729.299 | protein kinase, putative | 0,2239 | 0,2089 |
| TcCLB.507949.40 | hypothetical protein, conserved | 0,2238 | 0,02968 |
| TcCLB.511517.110 | hypothetical protein, conserved | 0,2238 | 0,04249 |
| TcCLB.507603.160 | hypothetical protein, conserved | 0,2237 | 0,1674 |
| TcCLB.511653.10 | delta-adaptin, putative | 0,2235 | 0,01864 |
| TcCLB.454403.30 | retrotransposon hot spot protein (RHS, pseudogene), puta | 0,2234 | 0,1272 |
| TcCLB.506437.20 | tuzin, putative | 0,2231 | 0,002779 |
| TcCLB.511859.50 | hypothetical protein, conserved | 0,2231 | 0,08019 |
| TcCLB.503995.30 | hypothetical protein | 0,223 | 0,1612 |
| TcCLB.506629.60 | vesicular-fusion ATPase-like protein, putative | 0,2229 | 0,06298 |
| TcCLB.511693.110 | hypothetical protein, conserved | 0,2228 | 0,1706 |
| TcCLB.506789.140 | MORN repeat-containing protein | 0,2227 | 0,1208 |
| TcCLB.508727.18 | centrin, putative | 0,2227 | 0,1298 |
| TcCLB.506065.10 | retrotransposon hot spot protein (RHS, pseudogene), puta | 0,2227 | 0,3758 |
| TcCLB.510519.110 | hypothetical protein | 0,2225 | 0,1916 |
| TcCLB.509875.200 | copper homeostasis protein, putative | 0,2223 | 0,05456 |
| TcCLB.503617.20 | kinetoplast DNA-associated protein, putative | 0,2222 | 0,007431 |
| TcCLB.510661.129 | hypothetical protein, conserved | 0,2222 | 0,06504 |

| | | | |
|---|---|---|---|
| TcCLB.504041.4 | RNA polymerase I | 0,222 | 0,207 |
| TcCLB.508851.70 | hypothetical protein | 0,2219 | 0,07566 |
| TcCLB.506343.10 | serine/threonine protein kinase, putative | 0,2216 | 0,1601 |
| TcCLB.508089.40 | hypothetical protein, conserved | 0,2213 | 0,1286 |
| TcCLB.506577.90 | retrotransposon hot spot protein (RHS, pseudogene), puta | 0,2212 | 0,04961 |
| TcCLB.508741.210 | hypothetical protein | 0,2211 | 0,004063 |
| TcCLB.511585.130 | ubiquitin activating enzyme, putative | 0,221 | 0,04321 |
| TcCLB.509571.10 | hypothetical protein, conserved | 0,221 | 0,1564 |
| TcCLB.506303.90 | hypothetical protein, conserved | 0,2209 | 0,04589 |
| TcCLB.511817.70 | hypothetical protein, conserved | 0,2208 | 0,1202 |
| TcCLB.510767.50 | hypothetical protein, conserved | 0,2208 | 0,1286 |
| TcCLB.506777.100 | hypothetical protein, conserved | 0,2208 | 0,1313 |
| TcCLB.503893.170 | hypothetical protein, conserved | 0,2207 | 0,05153 |
| TcCLB.506177.130 | hypothetical protein, conserved | 0,2207 | 0,2479 |
| TcCLB.504033.180 | hypothetical protein, conserved | 0,2205 | 0,003311 |
| TcCLB.503885.50 | hypothetical protein, conserved | 0,2205 | 0,07074 |
| TcCLB.503395.30 | hypothetical protein, conserved | 0,2203 | 0,04871 |
| TcCLB.506933.98 | dual specificity protein phosphatase, putative | 0,2203 | 0,0578 |
| TcCLB.504123.50 | hypothetical protein, conserved (pseudogene) | 0,2202 | 0,01103 |
| TcCLB.510579.90 | hypothetical protein, conserved | 0,2202 | 0,04267 |
| TcCLB.506961.40 | retrotransposon hot spot protein (RHS, pseudogene), puta | 0,2201 | 0,02586 |
| TcCLB.506435.40 | hypothetical protein, conserved | 0,2201 | 0,1891 |
| TcCLB.509815.10 | retrotransposon hot spot (RHS) protein, putative | 0,22 | 0,178 |
| TcCLB.511021.80 | hypothetical protein, conserved | 0,2199 | 0,03583 |
| TcCLB.508645.20 | hypothetical protein | 0,2199 | 0,05069 |
| TcCLB.511211.60 | protein phosphatase 4 catalytic subunit, putative | 0,2198 | 0,03589 |
| TcCLB.509785.70 | retrotransposon hot spot (RHS) protein, putative | 0,2196 | 0,1265 |
| TcCLB.503893.40 | hypothetical protein, conserved | 0,2195 | 0,09438 |
| TcCLB.506321.270 | hypothetical protein, conserved | 0,2195 | 0,1554 |
| TcCLB.510001.31 | hypothetical protein, conserved | 0,2195 | 0,1591 |
| TcCLB.509023.140 | hypothetical protein, conserved | 0,2194 | 0,141 |
| TcCLB.508973.90 | Present in the outer mitochondrial membrane proteome 3 | 0,2193 | 0,0318 |
| TcCLB.421321.9 | exportin-T, putative (fragment) | 0,2193 | 0,06858 |
| TcCLB.509999.90 | carnitine/choline acetyltransferase, putative | 0,2192 | 0,009155 |
| TcCLB.510329.240 | hypothetical protein, conserved | 0,2192 | 0,1268 |
| TcCLB.509395.50 | hypothetical protein, conserved | 0,2191 | 0,01378 |
| TcCLB.509951.49 | DNA-directed RNA polymerase II subunit 2, putative | 0,2191 | 0,06783 |
| TcCLB.506679.200 | hypothetical protein, conserved | 0,2191 | 0,07044 |
| TcCLB.511351.20 | hypothetical protein, conserved | 0,2191 | 0,08479 |
| TcCLB.505999.120 | hypothetical protein | 0,2191 | 0,1185 |
| TcCLB.510603.10 | hypothetical protein, conserved | 0,2191 | 0,2378 |
| TcCLB.511809.110 | hypothetical protein, conserved | 0,2189 | 0,02535 |
| TcCLB.507257.20 | hypothetical protein, conserved | 0,2189 | 0,05373 |
| TcCLB.508533.10 | hypothetical protein, conserved | 0,2188 | 0,0832 |
| TcCLB.509539.59 | hypothetical protein, conserved | 0,2188 | 0,2871 |

| | | | |
|---|---|---|---|
| TcCLB.506367.10 | isovaleryl-coA dehydrogenase, putative (fragment) | 0,2184 | 0,2828 |
| TcCLB.506859.49 | hypothetical protein, conserved | 0,2183 | 0,03131 |
| TcCLB.509735.150 | retrotransposon hot spot protein (RHS, pseudogene), puta | 0,2182 | 0,02913 |
| TcCLB.510689.89 | hypothetical protein, conserved | 0,2182 | 0,461 |
| TcCLB.510359.20 | paraflagellar rod component, putative | 0,2182 | 0,4845 |
| TcCLB.509103.20 | tubulin-tyrosine ligase, putative | 0,2181 | 0,08678 |
| TcCLB.505941.30 | hypothetical protein, conserved | 0,218 | 0,0313 |
| TcCLB.509711.20 | retrotransposon hot spot protein (RHS, pseudogene), puta | 0,218 | 0,5029 |
| TcCLB.506885.180 | hypothetical protein, conserved | 0,2178 | 0,07921 |
| TcCLB.420495.10 | hypothetical protein, conserved (pseudogene) | 0,2178 | 0,1661 |
| TcCLB.506245.76 | trans-sialidase (pseudogene), putative | 0,2177 | 0,3925 |
| TcCLB.507951.280 | ABC transporter, putative | 0,2176 | 0,03204 |
| TcCLB.508409.280 | hypothetical protein, conserved | 0,2175 | 0,112 |
| TcCLB.506625.220 | hypothetical protein, conserved | 0,2174 | 0,04883 |
| TcCLB.507993.369 | sjogren s syndrome nuclear autoantigen 1, putative | 0,2172 | 0,1212 |
| TcCLB.511353.4 | choline/carnitine O-acyltransferase, putative | 0,2171 | 0,08551 |
| TcCLB.507089.140 | hypothetical protein, conserved | 0,217 | 0,04814 |
| TcCLB.508961.20 | hypothetical protein, conserved | 0,217 | 0,141 |
| TcCLB.511481.30 | hypothetical protein, conserved | 0,2168 | 0,1674 |
| TcCLB.511487.19 | trans-sialidase (pseudogene), putative | 0,2166 | 0,0441 |
| TcCLB.506577.40 | hypothetical protein, conserved | 0,2166 | 0,08612 |
| TcCLB.511727.90 | vacuolar proton-ATPase-like protein, putative | 0,2164 | 0,006229 |
| TcCLB.511507.90 | hypothetical protein, conserved | 0,2164 | 0,09515 |
| TcCLB.509267.60 | NADH-ubiquinone oxidoreductase complex I subunit, puta | 0,2164 | 0,1294 |
| TcCLB.509717.70 | transportin2-like protein, putative | 0,2163 | 0,0202 |
| TcCLB.511287.124 | peroxin 13, putative (PEX13) | 0,2163 | 0,09279 |
| TcCLB.506369.70 | histone H1, putative | 0,2162 | 0,001526 |
| TcCLB.507801.60 | hypothetical protein, conserved | 0,2162 | 0,03732 |
| TcCLB.508059.20 | hypothetical protein, conserved | 0,2162 | 0,1011 |
| TcCLB.506885.10 | alpha-1,2-mannosyltransferase, putative | 0,2161 | 0,04332 |
| TcCLB.507941.30 | PrimPol-like protein 1, putative (ppl1) | 0,216 | 0,08834 |
| TcCLB.503939.84 | hypothetical protein, conserved | 0,216 | 0,171 |
| TcCLB.511815.24 | hypothetical protein, conserved | 0,2159 | 0,03795 |
| TcCLB.508779.50 | hypothetical protein, conserved | 0,2158 | 0,02985 |
| TcCLB.504105.80 | hypothetical protein, conserved | 0,2157 | 0,01123 |
| TcCLB.510297.90 | hypothetical protein, conserved | 0,2156 | 0,01179 |
| TcCLB.511509.30 | protein kinase, putative | 0,2155 | 0,09429 |
| TcCLB.509153.139 | hypothetical protein, conserved | 0,2154 | 0,04567 |
| TcCLB.508527.39 | retrotransposon hot spot (RHS) protein, putative, (fragme | 0,2153 | 0,08 |
| TcCLB.503515.14 | ubiquitin-conjugating enzyme E2, putative | 0,2153 | 0,1329 |
| TcCLB.510341.10 | hypothetical protein | 0,2153 | 0,2885 |
| TcCLB.510155.70 | heat shock protein 70 (hsp70), putative | 0,2152 | 0,01213 |
| TcCLB.507209.10 | small nuclear ribonucleoprotein protein, putative | 0,2151 | 0,07849 |
| TcCLB.508273.50 | hypothetical protein, conserved | 0,215 | 0,09286 |
| TcCLB.508775.40 | trans-sialidase (pseudogene), putative | 0,2149 | 0,3278 |

| | | | |
|---|---|---|---|
| TcCLB.506503.30 | hypothetical protein, conserved | 0,2147 | 0,03745 |
| TcCLB.511143.19 | hypothetical protein, conserved | 0,2147 | 0,2336 |
| TcCLB.504173.30 | hypothetical protein, conserved | 0,2147 | 0,3117 |
| TcCLB.510055.80 | hypothetical protein | 0,2146 | 0,09106 |
| TcCLB.507823.20 | hypothetical protein | 0,2146 | 0,09112 |
| TcCLB.507787.100 | gamma-glutamylcysteine synthetase, putative | 0,2144 | 0,09772 |
| TcCLB.511277.400 | UDP-galactose transporter | 0,2143 | 0,07388 |
| TcCLB.509967.40 | hypothetical protein, conserved | 0,2142 | 0,06921 |
| TcCLB.504229.44 | hypothetical protein, conserved (pseudogene) | 0,2142 | 0,1541 |
| TcCLB.507093.75 | B9 domain-containing protein 2 (b9d2) | 0,214 | 0,1651 |
| TcCLB.510285.40 | hypothetical protein, conserved | 0,2139 | 0,1893 |
| TcCLB.506659.10 | immunodominant antigen, putative | 0,2138 | 0,01707 |
| TcCLB.511211.170 | heat shock protein 70 (hsp70), putative | 0,2137 | 0,01188 |
| TcCLB.506577.170 | hypothetical protein, conserved | 0,2137 | 0,0958 |
| TcCLB.508741.360 | hypothetical protein, conserved | 0,2136 | 0,09429 |
| TcCLB.507823.30 | hypothetical protein, conserved | 0,2135 | 0,06304 |
| TcCLB.509017.30 | vacuolar ATP synthase subunit d | 0,2135 | 0,07223 |
| TcCLB.506219.30 | exosome component CSL4, putative (CSL4) | 0,2135 | 0,1069 |
| TcCLB.503899.50 | RNA-editing complex protein MP100, putative | 0,2134 | 0,01698 |
| TcCLB.506941.238 | hypothetical protein, conserved | 0,2134 | 0,09969 |
| TcCLB.511383.60 | hypothetical protein, conserved | 0,2134 | 0,1808 |
| TcCLB.508013.60 | hypothetical protein | 0,2134 | 0,5855 |
| TcCLB.506247.350 | hypothetical protein, conserved | 0,2133 | 0,08046 |
| TcCLB.506247.280 | hypothetical protein, conserved | 0,2133 | 0,08342 |
| TcCLB.507531.19 | phosphatidylinositol 4-kinase alpha, putative | 0,2131 | 0,04403 |
| TcCLB.511839.25 | elongation factor 1-gamma (EF-1-gamma, pseudogene), | 0,2131 | 0,6064 |
| TcCLB.506175.20 | hypothetical protein, conserved | 0,213 | 0,02106 |
| TcCLB.509233.200 | hypothetical protein, conserved | 0,213 | 0,02724 |
| TcCLB.506919.70 | hypothetical protein, conserved (pseudogene) | 0,213 | 0,4339 |
| TcCLB.507011.90 | hypothetical protein, conserved | 0,2129 | 0,03156 |
| TcCLB.506605.160 | histone acetyltransferase, putative | 0,2129 | 0,06441 |
| TcCLB.504057.150 | hypothetical protein, conserved | 0,2127 | 0,09632 |
| TcCLB.507277.20 | hypothetical protein, conserved | 0,2127 | 0,2856 |
| TcCLB.503453.20 | hypothetical protein, conserved | 0,2125 | 0,02307 |
| TcCLB.507013.20 | hypothetical protein, conserved | 0,2125 | 0,1054 |
| TcCLB.510879.170 | hypothetical protein, conserved | 0,2125 | 0,1607 |
| TcCLB.504491.10 | hypothetical protein | 0,2125 | 0,3417 |
| TcCLB.508153.1000 | hypothetical protein, conserved | 0,2124 | 0,1349 |
| TcCLB.506989.80 | hypothetical protein, conserved | 0,2124 | 0,1692 |
| TcCLB.508851.110 | hypothetical protein, conserved | 0,2123 | 0,279 |
| TcCLB.510759.134 | chaperone DNAJ protein, putative | 0,2121 | 0,06552 |
| TcCLB.509777.64 | ADG1-like protein, putative | 0,2121 | 0,1554 |
| TcCLB.511589.160 | replication factor C, subunit 4, putative | 0,2119 | 0,07426 |
| TcCLB.504115.90 | retrotransposon hot spot protein (RHS, pseudogene), puta | 0,2119 | 0,1659 |
| TcCLB.510279.70 | hypothetical protein | 0,2119 | 0,5914 |

| | | | |
|---|---|---|---|
| TcCLB.506871.110 | hypothetical protein, conserved | 0,2117 | 0,04888 |
| TcCLB.506563.50 | calpain-like cysteine peptidase (pseudogene), putative | 0,2117 | 0,07228 |
| TcCLB.509911.10 | hypothetical protein, conserved | 0,2115 | 0,02209 |
| TcCLB.507795.100 | mitochondrial ribosomal protein L3, putative | 0,2115 | 0,03419 |
| TcCLB.511423.130 | hypothetical protein, conserved (pseudogene) | 0,2115 | 0,1154 |
| TcCLB.511599.61 | hypothetical protein, conserved (pseudogene) | 0,2115 | 0,2366 |
| TcCLB.511859.80 | hypothetical protein | 0,2112 | 0,1149 |
| TcCLB.510445.20 | chromosomal passenger protein, putative (CPC1) | 0,2111 | 0,08677 |
| TcCLB.504029.80 | serine/threonine protein phosphatase, putative | 0,211 | 0,06873 |
| TcCLB.509979.140 | hypothetical protein, conserved | 0,2109 | 0,08101 |
| TcCLB.506693.40 | hypothetical protein, conserved | 0,2108 | 0,2889 |
| TcCLB.510899.50 | palmitoyl acyltransferase 4, putative | 0,2107 | 0,07192 |
| TcCLB.510033.22 | surface protease GP63 (pseudogene), putative | 0,2102 | 0,1305 |
| TcCLB.507093.30 | hypothetical protein, conserved | 0,2101 | 0,04555 |
| TcCLB.511043.30 | DnaJ chaperone protein, putative (TcJ6) | 0,21 | 0,0427 |
| TcCLB.505037.20 | ribosomal RNA processing protein 41B, putative (RRP41B | 0,21 | 0,0573 |
| TcCLB.503643.39 | hypothetical protein, conserved | 0,2099 | 0,1856 |
| TcCLB.506469.20 | hypothetical protein | 0,2097 | 0,05132 |
| TcCLB.509097.36 | surface protease GP63 (pseudogene), putative | 0,2097 | 0,7519 |
| TcCLB.504935.20 | hypothetical protein, conserved | 0,2096 | 0,1064 |
| TcCLB.509997.40 | hypothetical protein, conserved | 0,2096 | 0,1182 |
| TcCLB.509499.38 | hypothetical protein, conserved | 0,2095 | 0,05245 |
| TcCLB.506201.20 | 60S ribosomal protein L13a, putative | 0,2094 | 0,01796 |
| TcCLB.508411.60 | hypothetical protein, conserved | 0,2094 | 0,09209 |
| TcCLB.509471.40 | hypothetical protein, conserved | 0,2094 | 0,1028 |
| TcCLB.509179.50 | hypothetical protein | 0,2094 | 0,1333 |
| TcCLB.504741.110 | hypothetical protein, conserved | 0,2094 | 0,219 |
| TcCLB.509109.180 | hypothetical protein, conserved | 0,2093 | 0,09404 |
| TcCLB.508479.90 | hypothetical protein, conserved (pseudogene) | 0,2091 | 0,07103 |
| TcCLB.410049.20 | hypothetical protein, conserved | 0,2091 | 0,2773 |
| TcCLB.508247.60 | hypothetical protein, conserved (pseudogene) | 0,2091 | 0,4859 |
| TcCLB.509259.129 | hypothetical protein, conserved (pseudogene) | 0,209 | 0,3406 |
| TcCLB.510689.10 | coatomer alpha subunit (pseudogene), putative | 0,2088 | 0,0157 |
| TcCLB.470687.9 | hypothetical protein, conserved | 0,2088 | 0,1861 |
| TcCLB.511675.6 | hypothetical protein, conserved (pseudogene) | 0,2087 | 0,6456 |
| TcCLB.508799.170 | 3-methylcrotonoyl-CoA carboxylase beta subunit, putative | 0,2086 | 0,01485 |
| TcCLB.509733.110 | hypothetical protein, conserved | 0,2086 | 0,02037 |
| TcCLB.509287.250 | retrotransposon hot spot protein (RHS, pseudogene), puta | 0,2086 | 0,1091 |
| TcCLB.510689.70 | hypothetical protein, conserved | 0,2086 | 0,2842 |
| TcCLB.507317.50 | hypothetical protein, conserved | 0,2084 | 0,05458 |
| TcCLB.506621.70 | hypothetical protein, conserved | 0,2084 | 0,09739 |
| TcCLB.506337.70 | 2-oxoglutarate dehydrogenase E1 component, putative | 0,2083 | 0,003312 |
| TcCLB.506517.50 | hypothetical protein, conserved | 0,2083 | 0,08674 |
| TcCLB.508547.140 | hypothetical protein, conserved | 0,2081 | 0,0954 |
| TcCLB.511303.20 | hypothetical protein | 0,208 | 0,08252 |

| | | | |
|---|---|---|---|
| TcCLB.508869.10 | hypothetical protein | 0,208 | 0,5919 |
| TcCLB.506625.100 | hypothetical protein, conserved | 0,2079 | 0,02534 |
| TcCLB.511391.100 | hypothetical protein, conserved | 0,2079 | 0,1542 |
| TcCLB.509211.20 | hypothetical protein, conserved | 0,2078 | 0,008841 |
| TcCLB.506289.60 | hypothetical protein, conserved | 0,2078 | 0,1565 |
| TcCLB.509871.39 | hypothetical protein, conserved (pseudogene) | 0,2076 | 0,04222 |
| TcCLB.503409.20 | hypothetical protein, conserved | 0,2075 | 0,1443 |
| TcCLB.510145.10 | hypothetical protein, conserved | 0,2075 | 0,1999 |
| TcCLB.511717.170 | Metalloprotease M41 FtsH, putative | 0,2074 | 0,05602 |
| TcCLB.511153.100 | RNA-binding protein, putative | 0,2074 | 0,06159 |
| TcCLB.508323.80 | aquaporin, putative | 0,2073 | 0,1749 |
| TcCLB.511727.100 | hypothetical protein, conserved | 0,2072 | 0,0433 |
| TcCLB.511211.160 | heat shock protein 70 (hsp70), putative | 0,2068 | 0,01562 |
| TcCLB.510943.184 | tyrosine phosphatase, putative | 0,2068 | 0,09686 |
| TcCLB.503955.60 | hypothetical protein, conserved | 0,2067 | 0,1087 |
| TcCLB.507765.20 | PUF nine target 1 | 0,2065 | 0,02798 |
| TcCLB.504081.250 | 90 kDa surface protein, putative | 0,2065 | 0,4635 |
| TcCLB.506181.97 | hypothetical protein, conserved | 0,2064 | 0,1735 |
| TcCLB.509735.120 | retrotransposon hot spot (RHS) protein, putative | 0,2063 | 0,03459 |
| TcCLB.508645.30 | dynein light chain, putative | 0,2059 | 0,08708 |
| TcCLB.511577.4 | CLC-type chloride channel, putative (fragment) | 0,2059 | 0,3832 |
| TcCLB.504217.19 | mitochondrial carrier protein, putative | 0,2057 | 0,1343 |
| TcCLB.503839.10 | hypothetical protein, conserved | 0,2056 | 0,08255 |
| TcCLB.504259.19 | hypothetical protein, conserved | 0,2056 | 0,1088 |
| TcCLB.507275.10 | protein kinase, putative | 0,2056 | 0,1864 |
| TcCLB.506351.20 | hypothetical protein, conserved | 0,2056 | 0,2527 |
| TcCLB.507875.130 | trans-sialidase (pseudogene), putative | 0,2056 | 0,3728 |
| TcCLB.506999.130 | hypothetical protein, conserved | 0,2053 | 0,02567 |
| TcCLB.399373.9 | retrotransposon hot spot (RHS) protein, putative (fragmer | 0,2053 | 0,05745 |
| TcCLB.508199.90 | hypothetical protein, conserved | 0,2051 | 0,1221 |
| TcCLB.510523.10 | hypothetical protein, conserved (pseudogene) | 0,205 | 0,1044 |
| TcCLB.509005.40 | hypothetical protein, conserved | 0,2048 | 0,05693 |
| TcCLB.511923.110 | aspartate carbamoyltransferase, putative | 0,2048 | 0,08532 |
| TcCLB.503779.30 | hypothetical protein, conserved | 0,2048 | 0,1835 |
| TcCLB.510401.10 | dispersed gene family protein 1 (DGF-1), putative | 0,2047 | 0,7407 |
| TcCLB.511621.190 | dual specificity protein phosphatase, putative | 0,2046 | 0,1885 |
| TcCLB.507831.50 | hypothetical protein, conserved | 0,2045 | 0,01487 |
| TcCLB.510659.28 | hypothetical protein, conserved | 0,2045 | 0,1343 |
| TcCLB.504115.100 | dispersed gene family protein 1 (DGF-1), putative | 0,2044 | 0,2193 |
| TcCLB.510729.240 | hypothetical protein, conserved | 0,2043 | 0,1309 |
| TcCLB.511277.330 | uracil-DNA glycosylase | 0,2042 | 0,0383 |
| TcCLB.511837.50 | DNA repair protein, putative | 0,2042 | 0,1102 |
| TcCLB.511275.20 | hypothetical protein, conserved | 0,2041 | 0,05764 |
| TcCLB.510283.130 | hypothetical protein, conserved | 0,2041 | 0,1657 |
| TcCLB.508113.7 | trans-sialidase (pseudogene), putative | 0,2041 | 0,2736 |

| | | | |
|---|---|---|---|
| TcCLB.508815.30 | DNA repair protein, putative | 0,204 | 0,02561 |
| TcCLB.506203.50 | DnaJ chaperone protein, putative | 0,2039 | 0,0788 |
| TcCLB.511519.9 | small glutamine-rich tetratricopeptide repeat protein, put | 0,2039 | 0,1552 |
| TcCLB.506181.30 | hypothetical protein, conserved | 0,2039 | 0,2771 |
| TcCLB.509199.40 | hypothetical protein, conserved | 0,2037 | 0,1715 |
| TcCLB.507049.63 | hypothetical protein, conserved | 0,2037 | 0,3138 |
| TcCLB.508045.10 | hypothetical protein, conserved | 0,2036 | 0,04221 |
| TcCLB.511903.250 | hypothetical protein, conserved | 0,2036 | 0,1263 |
| TcCLB.483009.9 | exonuclease, putative | 0,2035 | 0,08731 |
| TcCLB.511911.98 | hypothetical protein, conserved | 0,2035 | 0,09664 |
| TcCLB.506471.90 | dispersed gene family protein 1 (DGF-1, pseudogene), put | 0,2033 | 0,2195 |
| TcCLB.509121.20 | dispersed gene family protein 1 (DGF-1, pseudogene), put | 0,2032 | 0,167 |
| TcCLB.505365.80 | retrotransposon hot spot protein (RHS, pseudogene), puta | 0,2032 | 0,2973 |
| TcCLB.510155.140 | Meckel syndrome type 6 protein, putative (mks6) | 0,2031 | 0,01134 |
| TcCLB.511307.6 | hypothetical protein, conserved | 0,2031 | 0,1774 |
| TcCLB.503847.90 | ABC1 protein, putative | 0,203 | 0,06247 |
| TcCLB.509287.50 | glucose-6-phosphate 1-dehydrogenase, putative | 0,2029 | 0,0613 |
| TcCLB.509171.29 | RNA pseudouridylate synthase, putative | 0,2029 | 0,06342 |
| TcCLB.511515.9 | E3 ubiquitin-protein ligase, putative (fragment) | 0,2029 | 0,1687 |
| TcCLB.509967.160 | epsilon tubulin, putative | 0,2027 | 0,06459 |
| TcCLB.507933.20 | hypothetical protein, conserved | 0,2026 | 0,162 |
| TcCLB.511587.33 | hypothetical protein | 0,2026 | 0,3504 |
| TcCLB.511237.20 | hypothetical protein, conserved | 0,2025 | 0,08448 |
| TcCLB.511727.150 | hypothetical protein, conserved | 0,2023 | 0,02165 |
| TcCLB.503667.31 | retrotransposon hot spot protein (RHS, pseudogene), puta | 0,2023 | 0,4575 |
| TcCLB.510841.30 | hypothetical protein | 0,2022 | 0,4001 |
| TcCLB.508379.29 | hypothetical protein, conserved | 0,2021 | 0,3247 |
| TcCLB.506481.10 | tRNA pseudouridine synthase A-like protein (pseudogene) | 0,202 | 0,1401 |
| TcCLB.506513.90 | hypothetical protein, conserved | 0,202 | 0,1483 |
| TcCLB.508633.100 | retrotransposon hot spot (RHS) protein, putative | 0,202 | 0,1489 |
| TcCLB.506339.81 | elongation factor 1-gamma (EF-1-gamma, pseudogene), | 0,2019 | 0,5118 |
| TcCLB.508857.120 | hypothetical protein, conserved | 0,2017 | 0,1304 |
| TcCLB.506435.290 | hypothetical protein, conserved | 0,2017 | 0,1823 |
| TcCLB.508777.190 | hypothetical protein, conserved | 0,2017 | 0,2096 |
| TcCLB.506625.249 | RNA pseudouridylate synthase protein, putative | 0,2016 | 0,05456 |
| TcCLB.504185.20 | hypothetical protein, conserved | 0,2015 | 0,1004 |
| TcCLB.509419.245 | hypothetical protein, conserved (pseudogene) | 0,2015 | 0,3814 |
| TcCLB.507849.40 | hypothetical protein, conserved | 0,2014 | 0,0196 |
| TcCLB.509119.30 | hypothetical protein, conserved | 0,2014 | 0,08423 |
| TcCLB.511577.160 | hypothetical protein, conserved | 0,2013 | 0,1173 |
| TcCLB.504137.120 | hypothetical protein, conserved | 0,2012 | 0,1125 |
| TcCLB.511903.180 | hypothetical protein, conserved | 0,2012 | 0,131 |
| TcCLB.511217.20 | Present in the outer mitochondrial membrane proteome 4 | 0,2011 | 0,09159 |
| TcCLB.506401.350 | hypothetical protein, conserved | 0,2011 | 0,1106 |
| TcCLB.503829.10 | hypothetical protein, conserved | 0,2008 | 0,1343 |

| | | | |
|---|---|---|---|
| TcCLB.505009.20 | divalent cation tolerance protein, putative | 0,2008 | 0,5002 |
| TcCLB.510669.9 | hypothetical protein, conserved | 0,2007 | 0,1686 |
| TcCLB.508777.180 | prefoldin subunit 2, putative | 0,2006 | 0,09622 |
| TcCLB.511747.14 | hypothetical protein, conserved | 0,2006 | 0,1989 |
| TcCLB.479883.10 | hypothetical protein, conserved | 0,2004 | 0,165 |
| TcCLB.506355.10 | glucose transporter | 0,2003 | 0,2117 |
| TcCLB.507007.20 | hypothetical protein, conserved | 0,2002 | 0,05033 |
| TcCLB.508501.310 | hypothetical protein, conserved | 0,2002 | 0,1162 |
| TcCLB.508023.20 | trans-sialidase (pseudogene), putative | 0,2001 | 0,3397 |
| TcCLB.504057.145 | cytochrome b5-like Heme/Steroid binding domain contain | 0,2 | 0,1094 |
| TcCLB.509607.10 | ubiquitin-conjugating enzyme E2, putative | 0,1999 | 0,0729 |
| TcCLB.508457.40 | hypothetical protein | 0,1999 | 0,1786 |
| TcCLB.509105.130 | ATP-binding cassette sub-family F member 1, putative (A | 0,1997 | 0,01386 |
| TcCLB.509579.40 | hypothetical protein, conserved (pseudogene) | 0,1997 | 0,2628 |
| TcCLB.508879.170 | mitochondrial RNA binding complex 1 subunit, putative | 0,1996 | 0,05944 |
| TcCLB.506147.110 | hypothetical protein, conserved | 0,1996 | 0,1057 |
| TcCLB.508601.20 | hypothetical protein, conserved | 0,1996 | 0,1087 |
| TcCLB.507993.270 | hypothetical protein, conserved | 0,1996 | 0,1159 |
| TcCLB.506789.260 | hypothetical protein, conserved | 0,1996 | 0,2651 |
| TcCLB.442495.19 | KU70 protein, putative (fragment) | 0,1994 | 0,3038 |
| TcCLB.506181.80 | hypothetical protein, conserved | 0,1993 | 0,2294 |
| TcCLB.510661.200 | hypothetical protein, conserved | 0,1991 | 0,08834 |
| TcCLB.506859.110 | 3'a2rel-related protein, putative | 0,1991 | 0,2837 |
| TcCLB.511365.94 | anaphase promoting complex, subunit 10-like protein | 0,1989 | 0,1672 |
| TcCLB.506171.60 | trans-sialidase, Group II, putative | 0,1989 | 0,5035 |
| TcCLB.506531.30 | hypothetical protein, conserved | 0,1985 | 0,1233 |
| TcCLB.510517.80 | hypothetical protein, conserved | 0,1985 | 0,2973 |
| TcCLB.509551.95 | anaphase promoting complex subunit protein, putative | 0,1985 | 0,4312 |
| TcCLB.511545.170 | hypothetical protein, conserved | 0,1984 | 0,1582 |
| TcCLB.511797.130 | mucin TcMUCII (pseudogene), putative | 0,1982 | 0,3675 |
| TcCLB.508873.260 | hypothetical protein | 0,1982 | 0,6844 |
| TcCLB.503513.10 | casein kinase II, alpha chain, putative | 0,1981 | 0,1248 |
| TcCLB.508521.112 | beta galactofuranosyl glycosyltransferase (pseudogene), p | 0,1979 | 0,4449 |
| TcCLB.508385.10 | hypothetical protein, conserved | 0,1978 | 0,03772 |
| TcCLB.509153.60 | lipin, putative | 0,1977 | 0,02294 |
| TcCLB.510329.80 | hypothetical protein, conserved | 0,1975 | 0,03131 |
| TcCLB.511693.100 | long-chain-fatty-acid-CoA ligase, putative | 0,1974 | 0,02156 |
| TcCLB.506839.20 | hypothetical protein | 0,1974 | 0,04198 |
| TcCLB.508355.270 | hypothetical protein | 0,1974 | 0,08375 |
| TcCLB.506685.50 | myosin heavy chain, putative (fragment) | 0,1974 | 0,1912 |
| TcCLB.510579.40 | hypothetical protein, conserved | 0,1972 | 0,1614 |
| TcCLB.511771.156 | retrotransposon hot spot protein (RHS, pseudogene), puta | 0,1972 | 0,1655 |
| TcCLB.511281.56 | sjogren s syndrome nuclear autoantigen 1, putative | 0,1969 | 0,2035 |
| TcCLB.510719.170 | hypothetical protein, conserved | 0,1969 | 0,2088 |
| TcCLB.509641.30 | Ubiquitin-like modifier-activating enzyme ATG7, putative | 0,1967 | 0,06483 |

| | | | |
|---|---|---|---|
| TcCLB.509607.60 | hypothetical protein | 0,1966 | 0,04916 |
| TcCLB.507257.170 | hypothetical protein, conserved | 0,1966 | 0,1378 |
| TcCLB.511801.10 | hypothetical protein, conserved | 0,1965 | 0,02568 |
| TcCLB.509099.70 | endo/exonuclease Mre11, putative | 0,1964 | 0,03244 |
| TcCLB.507681.50 | hypothetical protein, conserved | 0,1963 | 0,03037 |
| TcCLB.508647.170 | DNA polymerase theta (polymerase domain only), putativ | 0,1962 | 0,06272 |
| TcCLB.509693.100 | 2-aminoethylphosphonate:pyruvateaminotransferase- like | 0,1961 | 0,1462 |
| TcCLB.506543.50 | hypothetical protein, conserved | 0,1959 | 0,5186 |
| TcCLB.506247.240 | hypothetical protein, conserved | 0,1958 | 0,1161 |
| TcCLB.507715.20 | hypothetical protein, conserved | 0,1956 | 0,08729 |
| TcCLB.506201.150 | hypothetical protein, conserved | 0,1952 | 0,3456 |
| TcCLB.504103.80 | hypothetical protein, conserved | 0,1951 | 0,09913 |
| TcCLB.507853.20 | dynein beta chain, flagellar outer arm (pseudogene), puta | 0,1951 | 0,2058 |
| TcCLB.506637.120 | hypothetical protein, conserved | 0,195 | 0,04317 |
| TcCLB.510575.120 | hypothetical protein, conserved | 0,195 | 0,1866 |
| TcCLB.506367.174 | hypothetical protein, conserved | 0,195 | 0,2803 |
| TcCLB.506713.40 | N-acetyltransferase complex ARD1 subunit, putative | 0,1949 | 0,295 |
| TcCLB.510315.40 | Putative intraflagellar transport protein C3 (fragment) (PI | 0,1948 | 0,07074 |
| TcCLB.509507.10 | hypothetical protein, conserved | 0,1943 | 0,04253 |
| TcCLB.506587.30 | hypothetical protein, conserved | 0,1943 | 0,06463 |
| TcCLB.508933.10 | dihydrolipoamide dehydrogenase, putative | 0,1942 | 0,1118 |
| TcCLB.505807.120 | lysyl-tRNA synthetase, putative | 0,1942 | 0,146 |
| TcCLB.503537.9 | retrotransposon hot spot protein (RHS, pseudogene), puta | 0,1939 | 0,05267 |
| TcCLB.508641.190 | hypothetical protein, conserved | 0,1938 | 0,2281 |
| TcCLB.504173.40 | hypothetical protein | 0,1937 | 0,2529 |
| TcCLB.504579.20 | trans-sialidase (pseudogene), putative | 0,1936 | 0,1075 |
| TcCLB.511903.260 | hypothetical protein, conserved | 0,1933 | 0,1432 |
| TcCLB.511585.60 | hypothetical protein, conserved | 0,1932 | 0,1525 |
| TcCLB.511217.140 | coatomer delta subunit, putative | 0,193 | 0,0191 |
| TcCLB.506201.84 | hypothetical protein, conserved | 0,193 | 0,217 |
| TcCLB.503459.20 | hypothetical protein, conserved | 0,1928 | 0,1116 |
| TcCLB.508169.129 | Mutt-Nudix-related hydrolase 5, putative (Nudix5) | 0,1928 | 0,2274 |
| TcCLB.507775.50 | proteasome alpha 7 subunit, putative | 0,1927 | 0,1249 |
| TcCLB.508645.59 | dynein light chain, putative | 0,1926 | 0,06485 |
| TcCLB.508909.70 | hypothetical protein, conserved | 0,1926 | 0,2467 |
| TcCLB.511543.30 | amino acid transporter, putative | 0,1925 | 0,3927 |
| TcCLB.508241.120 | rrp44p homologue, putative | 0,1922 | 0,009627 |
| TcCLB.508873.410 | hypothetical protein, conserved | 0,1921 | 0,3245 |
| TcCLB.509023.130 | DNA replication licensing factor MCM4, putative (MCM4) | 0,1918 | 0,08219 |
| TcCLB.509499.29 | hypothetical protein, conserved | 0,1918 | 0,2497 |
| TcCLB.510099.20 | hypothetical protein, conserved | 0,1917 | 0,1118 |
| TcCLB.510181.110 | hypothetical protein, conserved | 0,1917 | 0,1394 |
| TcCLB.511807.110 | GMP synthase [glutamine-hydrolysing] (Gmps) | 0,1916 | 0,0143 |
| TcCLB.511507.30 | cyclosome subunit, putative | 0,1916 | 0,05877 |
| TcCLB.508873.425 | elongation factor 1-gamma (EF-1-gamma, pseudogene), | 0,1915 | 0,1934 |

| | | | |
|---|---|---|---|
| TcCLB.511577.10 | hypothetical protein, conserved | 0,1913 | 0,3486 |
| TcCLB.506241.40 | hypothetical protein, conserved | 0,1912 | 0,107 |
| TcCLB.508461.110 | hypothetical protein, conserved | 0,1911 | 0,1033 |
| TcCLB.506323.40 | trans-sialidase (pseudogene), putative | 0,1911 | 0,1554 |
| TcCLB.511371.10 | retrotransposon hot spot (RHS) protein, putative | 0,1909 | 0,1227 |
| TcCLB.507881.20 | retrotransposon hot spot (RHS) protein, putative | 0,1908 | 0,05074 |
| TcCLB.508693.160 | hypothetical protein, conserved | 0,1908 | 0,08811 |
| TcCLB.509775.49 | ribosomal protein L2, putative | 0,1907 | 0,384 |
| TcCLB.510731.130 | hypothetical protein, conserved | 0,1906 | 0,4282 |
| TcCLB.508599.10 | hypothetical protein, conserved (pseudogene) | 0,1906 | 0,6517 |
| TcCLB.504137.20 | hypothetical protein, conserved | 0,1905 | 0,07642 |
| TcCLB.503833.69 | ribosome biogenesis protein, putative (fragment) | 0,1905 | 0,1935 |
| TcCLB.511585.150 | Meckel syndrome type 6 protein, putative (mks6) | 0,1904 | 0,04089 |
| TcCLB.510121.60 | hypothetical protein, conserved | 0,1904 | 0,1384 |
| TcCLB.511127.340 | tyrosine phosphatase, putative | 0,1903 | 0,06664 |
| TcCLB.503499.10 | hypothetical protein, conserved | 0,1902 | 0,05148 |
| TcCLB.509601.40 | hypothetical protein, conserved | 0,1902 | 0,06568 |
| TcCLB.503935.10 | dolichyl-lipid chaperone, putative (rft1) | 0,1902 | 0,1379 |
| TcCLB.449247.20 | hypothetical protein, conserved | 0,1902 | 0,3098 |
| TcCLB.511621.140 | hypothetical protein, conserved | 0,1901 | 0,1353 |
| TcCLB.506227.80 | mitochondrial DNA polymerase I protein B, putative (POLI | 0,19 | 0,03167 |
| TcCLB.509085.120 | retrotransposon hot spot (RHS) protein, putative | 0,19 | 0,2393 |
| TcCLB.509265.36 | ATP-dependent DEAD/H RNA helicase (pseudogene), puta | 0,1899 | 0,5082 |
| TcCLB.510933.180 | syntaxin binding protein (pseudogene), putative | 0,1899 | 0,6224 |
| TcCLB.507047.120 | myo-inositol-1(or 4)-monophosphatase 1, putative | 0,1898 | 0,1819 |
| TcCLB.507941.50 | hypothetical protein, conserved | 0,1897 | 0,07114 |
| TcCLB.509429.100 | PITH domain-containing protein | 0,1896 | 0,1244 |
| TcCLB.507875.50 | hypothetical protein | 0,1896 | 0,3683 |
| TcCLB.503431.90 | hypothetical protein, conserved | 0,1895 | 0,1269 |
| TcCLB.508951.50 | calmodulin, putative | 0,1895 | 0,3139 |
| TcCLB.424771.10 | retrotransposon hot spot (RHS) protein, putative | 0,1894 | 0,4826 |
| TcCLB.510729.230 | hypothetical protein, conserved | 0,1893 | 0,05828 |
| TcCLB.506989.160 | PTS2 receptor, putative (PEX7) | 0,1893 | 0,08808 |
| TcCLB.510657.160 | Double RNA binding domain protein 9 (DRBD9) | 0,1891 | 0,1642 |
| TcCLB.503703.70 | Mitochondrial SSU ribosomal protein, putative | 0,189 | 0,04757 |
| TcCLB.511353.50 | hypothetical protein, conserved | 0,189 | 0,2097 |
| TcCLB.506557.24 | hypothetical protein, conserved | 0,189 | 0,2938 |
| TcCLB.506825.90 | hypothetical protein, conserved | 0,1888 | 0,03444 |
| TcCLB.507851.70 | hypothetical protein | 0,1888 | 0,09223 |
| TcCLB.503903.10 | hypothetical protein | 0,1888 | 0,2234 |
| TcCLB.510317.30 | polynucleotide adenylyltransferase, putative (PAP) | 0,1888 | 0,2801 |
| TcCLB.511881.20 | serine/threonine-protein phosphatase | 0,1887 | 0,1306 |
| TcCLB.503703.50 | copper-transporting ATPase-like protein, putative | 0,1886 | 0,03653 |
| TcCLB.510359.10 | hypothetical protein, conserved | 0,1886 | 0,3111 |
| TcCLB.504045.114 | hypothetical protein, conserved | 0,1886 | 0,3169 |

| | | | |
|---|---|---|---|
| TcCLB.511677.10 | dispersed gene family protein 1 (DGF-1), putative | 0,1884 | 0,2192 |
| TcCLB.510575.20 | hypothetical protein, conserved | 0,1884 | 0,2464 |
| TcCLB.511153.120 | pantothenate kinase subunit, putative | 0,1883 | 0,09855 |
| TcCLB.509395.70 | hypothetical protein, conserved | 0,1883 | 0,1246 |
| TcCLB.504797.50 | hypothetical protein, conserved | 0,1882 | 0,08528 |
| TcCLB.507979.40 | hypothetical protein, conserved | 0,1882 | 0,3728 |
| TcCLB.511511.80 | tubulin-tyrosine ligase-like protein, putative | 0,1881 | 0,1023 |
| TcCLB.425597.10 | trans-sialidase (pseudogene), putative | 0,1881 | 0,5398 |
| TcCLB.503903.20 | hypothetical protein, conserved | 0,188 | 0,1772 |
| TcCLB.507257.90 | hypothetical protein, conserved | 0,1879 | 0,1875 |
| TcCLB.510153.5 | hypothetical protein, conserved | 0,1876 | 0,4766 |
| TcCLB.511495.20 | hypothetical protein, conserved | 0,1875 | 0,1302 |
| TcCLB.508273.130 | hypothetical protein | 0,1875 | 0,5484 |
| TcCLB.503515.20 | Kinetoplastid RNA editing protein 1, putative (KREPA1) | 0,1874 | 0,04022 |
| TcCLB.450019.9 | dispersed gene family protein 1 (DGF-1), putative (fragme | 0,1874 | 0,2759 |
| TcCLB.507519.180 | hypothetical protein, conserved | 0,1873 | 0,1399 |
| TcCLB.508675.20 | DNA excision repair protein, putative | 0,1871 | 0,04914 |
| TcCLB.507005.30 | hypothetical protein, conserved | 0,187 | 0,1158 |
| TcCLB.507509.89 | cleavage and polyadenylation specificity factor-like protei | 0,1869 | 0,07521 |
| TcCLB.508029.20 | ubiquitin-protein ligase, putative | 0,1869 | 0,2054 |
| TcCLB.508641.330 | pre-mRNA splicing factor, putative | 0,1868 | 0,33 |
| TcCLB.509211.30 | serine/threonine protein phosphatase, putative | 0,1866 | 0,1924 |
| TcCLB.506349.10 | protein kinase, putative | 0,1866 | 0,6153 |
| TcCLB.510353.10 | major vault protein, putative | 0,1865 | 0,01959 |
| TcCLB.510657.140 | hypothetical protein, conserved | 0,1865 | 0,1508 |
| TcCLB.511445.100 | hypothetical protein, conserved | 0,1863 | 0,07068 |
| TcCLB.510181.30 | ATP-dependent DEAD/H RNA helicase, putative | 0,1863 | 0,1404 |
| TcCLB.506693.10 | hypothetical protein | 0,1863 | 0,1696 |
| TcCLB.509965.300 | hypothetical protein, conserved | 0,1862 | 0,3138 |
| TcCLB.503487.20 | hypothetical protein | 0,1861 | 0,08807 |
| TcCLB.507467.30 | hypothetical protein, conserved | 0,1861 | 0,269 |
| TcCLB.506755.20 | paraflagellar rod component, putative | 0,1858 | 0,1076 |
| TcCLB.510719.200 | telomerase reverse transcriptase, putative (TERT) | 0,1857 | 0,1308 |
| TcCLB.503667.20 | hypothetical protein | 0,1857 | 0,5965 |
| TcCLB.510645.49 | ribosomal protein L3 mitochondrial, putative | 0,1856 | 0,2847 |
| TcCLB.503485.20 | hypothetical protein, conserved (pseudogene) | 0,1855 | 0,2955 |
| TcCLB.509207.100 | hypothetical protein, conserved | 0,1853 | 0,09908 |
| TcCLB.506887.110 | hypothetical protein, conserved (pseudogene) | 0,1853 | 0,2197 |
| TcCLB.511439.60 | hypothetical protein, conserved | 0,1853 | 0,2263 |
| TcCLB.510769.120 | calcium motive p-type ATPase, putative | 0,1852 | 0,04428 |
| TcCLB.510613.10 | hypothetical protein, conserved | 0,1852 | 0,157 |
| TcCLB.510735.40 | hypothetical protein, conserved | 0,1851 | 0,06486 |
| TcCLB.507047.134 | hypothetical protein, conserved | 0,185 | 0,4373 |
| TcCLB.511817.310 | hypothetical protein, conserved | 0,1849 | 0,02954 |
| TcCLB.408437.10 | hypothetical protein, conserved | 0,1849 | 0,09701 |

| | | | |
|---|---|---|---|
| TcCLB.511821.140 | hypothetical protein, conserved | 0,1849 | 0,1263 |
| TcCLB.507851.20 | hypothetical protein, conserved | 0,1849 | 0,151 |
| TcCLB.504105.210 | Translation factor GUF1, mitochondrial, putative (GUF1) | 0,1846 | 0,05063 |
| TcCLB.510359.420 | mucin-associated surface protein (MASP, pseudogene), pu | 0,1845 | 0,3223 |
| TcCLB.510461.25 | elongation factor 1-gamma (EF-1-gamma, pseudogene), | 0,1845 | 0,7002 |
| TcCLB.508153.980 | hypothetical protein, conserved | 0,1843 | 0,0385 |
| TcCLB.509331.134 | trypanothione synthetase, putative | 0,1843 | 0,2278 |
| TcCLB.507007.60 | hypothetical protein, conserved | 0,1843 | 0,2295 |
| TcCLB.509937.30 | phosphatidylserine synthase, putative | 0,1842 | 0,1758 |
| TcCLB.511245.70 | hypothetical protein, conserved | 0,184 | 0,09723 |
| TcCLB.506629.10 | intraflagellar transport protein 88, putative (fragment) (IF | 0,1839 | 0,2284 |
| TcCLB.505183.140 | protein kinase, putative | 0,1838 | 0,07153 |
| TcCLB.506529.260 | chaperone DNAJ protein, putative | 0,1838 | 0,08288 |
| TcCLB.508827.120 | hypothetical protein, conserved | 0,1838 | 0,1652 |
| TcCLB.510101.340 | hypothetical protein, conserved | 0,1837 | 0,01635 |
| TcCLB.511733.100 | hypothetical protein, conserved | 0,1836 | 0,09069 |
| TcCLB.508641.80 | hypothetical protein, conserved | 0,1836 | 0,1586 |
| TcCLB.510429.24 | hypothetical protein, conserved | 0,1836 | 0,3279 |
| TcCLB.509207.110 | hypothetical protein, conserved | 0,1835 | 0,1858 |
| TcCLB.508543.170 | hypothetical protein, conserved | 0,1834 | 0,04831 |
| TcCLB.509573.30 | flagellar protofilament ribbon protein, putative | 0,1833 | 0,05022 |
| TcCLB.508827.20 | hypothetical protein, conserved | 0,1833 | 0,1893 |
| TcCLB.510663.10 | hypothetical protein, conserved | 0,183 | 0,3238 |
| TcCLB.506337.120 | hypothetical protein, conserved | 0,1829 | 0,2418 |
| TcCLB.509151.120 | hypothetical protein, conserved | 0,1826 | 0,1311 |
| TcCLB.506931.40 | hypothetical protein, conserved | 0,1825 | 0,08019 |
| TcCLB.508541.210 | hypothetical protein, conserved | 0,1825 | 0,09567 |
| TcCLB.511575.100 | protein transport protein SEC23, putative | 0,1824 | 0,03392 |
| TcCLB.453445.10 | hypothetical protein, conserved | 0,1824 | 0,04718 |
| TcCLB.511505.10 | RNA pseudouridylate synthase, putative | 0,1823 | 0,09323 |
| TcCLB.509799.90 | hypothetical protein, conserved | 0,1822 | 0,2081 |
| TcCLB.507003.70 | hypothetical protein, conserved | 0,1822 | 0,2618 |
| TcCLB.504081.26 | hypothetical protein | 0,1822 | 0,4669 |
| TcCLB.508139.150 | hypothetical protein, conserved | 0,1821 | 0,4249 |
| TcCLB.510297.60 | zinc transporter, putative | 0,1819 | 0,1521 |
| TcCLB.510507.60 | inositol 1,4,5-trisphosphate receptor, putative (fragment) | 0,1818 | 0,05703 |
| TcCLB.510095.40 | retrotransposon hot spot protein (RHS, pseudogene), puta | 0,1817 | 0,2243 |
| TcCLB.506653.50 | elongation factor 1-gamma (EF-1-gamma, pseudogene), | 0,1817 | 0,5883 |
| TcCLB.507711.260 | hypothetical protein, conserved | 0,1816 | 0,0662 |
| TcCLB.511249.90 | hypothetical protein, conserved | 0,1816 | 0,09284 |
| TcCLB.511727.199 | CYC2-like cyclin 6, putative (CYC6) | 0,1815 | 0,1454 |
| TcCLB.511621.259 | hypothetical protein, conserved | 0,1815 | 0,3332 |
| TcCLB.506525.80 | N-myristoyl transferase, putative | 0,1813 | 0,1767 |
| TcCLB.507009.100 | cyclophilin, putative | 0,1813 | 0,2185 |
| TcCLB.506831.50 | hypothetical protein, conserved | 0,1812 | 0,08623 |

| | | | |
|---|---|---|---|
| TcCLB.507641.260 | chaperonin HSP60, mitochondrial precursor (pseudogene) | 0,1811 | 0,2311 |
| TcCLB.511711.90 | cop-coated vesicle membrane protein p24 precursor, puta | 0,181 | 0,05828 |
| TcCLB.508503.40 | hypothetical protein, conserved | 0,181 | 0,08924 |
| TcCLB.511925.40 | hypothetical protein, conserved | 0,1809 | 0,3787 |
| TcCLB.506563.130 | calpain-like cysteine peptidase (pseudogene), putative | 0,1807 | 0,1329 |
| TcCLB.507395.10 | dihydroceramide synthase, putative (DHCS) | 0,1806 | 0,113 |
| TcCLB.504185.9 | hypothetical protein, conserved | 0,1806 | 0,4598 |
| TcCLB.504689.60 | dispersed gene family protein 1 (DGF-1), putative | 0,1806 | 0,4695 |
| TcCLB.506513.80 | hypothetical protein, conserved | 0,1805 | 0,1575 |
| TcCLB.507857.60 | Dynein intermediate chain 1, axonemal (DNAI1) | 0,1805 | 0,1963 |
| TcCLB.511507.39 | intraflagellar transport (IFT) protein, putative (IFT20) | 0,1805 | 0,2236 |
| TcCLB.509127.40 | hypothetical protein, conserved | 0,1804 | 0,148 |
| TcCLB.506649.100 | tRNA nucleotidyltransferase, putative | 0,1803 | 0,08577 |
| TcCLB.510259.40 | hypothetical protein, conserved | 0,1802 | 0,4051 |
| TcCLB.507519.130 | Mini-chromosome maintenance complex-binding protein | 0,1801 | 0,1015 |
| TcCLB.506845.60 | retrotransposon hot spot (RHS) protein, putative | 0,1799 | 0,5191 |
| TcCLB.508389.10 | hypothetical protein, conserved | 0,1798 | 0,06524 |
| TcCLB.506895.10 | hypothetical protein, conserved | 0,1798 | 0,2073 |
| TcCLB.507767.10 | succinyl-CoA ligase [GDP-forming] beta-chain, putative | 0,1797 | 0,02152 |
| TcCLB.511425.19 | hypothetical protein, conserved (pseudogene) | 0,1797 | 0,2072 |
| TcCLB.508415.10 | hypothetical protein, conserved (pseudogene) | 0,1797 | 0,3176 |
| TcCLB.510073.10 | cysteine peptidase (N-terminal), putative | 0,1796 | 0,3884 |
| TcCLB.506821.90 | hypothetical protein, conserved | 0,1795 | 0,1238 |
| TcCLB.506195.100 | malate dehydrogenase (pseudogene), putative | 0,1794 | 0,1972 |
| TcCLB.511767.60 | trans-sialidase (pseudogene), putative | 0,1793 | 0,1155 |
| TcCLB.508413.60 | hypothetical protein, conserved | 0,1793 | 0,1816 |
| TcCLB.511693.50 | hypothetical protein, conserved | 0,1792 | 0,08133 |
| TcCLB.503495.5 | hypothetical protein | 0,1792 | 0,4398 |
| TcCLB.506401.80 | hypothetical protein, conserved | 0,1791 | 0,03634 |
| TcCLB.511727.70 | hypothetical protein, conserved | 0,179 | 0,06894 |
| TcCLB.509053.20 | hypothetical protein, conserved | 0,179 | 0,1036 |
| TcCLB.508357.23 | hypothetical protein, conserved | 0,1789 | 0,2346 |
| TcCLB.511737.6 | hypothetical protein, conserved | 0,1789 | 0,2828 |
| TcCLB.511041.50 | flagellum transition zone component, putative | 0,1788 | 0,01872 |
| TcCLB.506529.310 | hypothetical protein, conserved | 0,1788 | 0,1166 |
| TcCLB.509831.50 | intraflagellar transport protein 88, putative (fragment) (IF | 0,1788 | 0,1883 |
| TcCLB.506705.20 | adenosine kinase, putative | 0,1788 | 0,353 |
| TcCLB.508405.110 | hypothetical protein, conserved | 0,1787 | 0,1686 |
| TcCLB.509379.10 | dihydrolipoamide dehydrogenase, putative (GCVL-1) | 0,1787 | 0,1756 |
| TcCLB.510167.29 | exosome complex exonuclease, putative | 0,1787 | 0,2044 |
| TcCLB.508641.180 | RNA-binding protein 38, putative (RBP38) | 0,1786 | 0,0835 |
| TcCLB.506629.120 | hypothetical protein, conserved | 0,1786 | 0,1284 |
| TcCLB.509169.39 | protein kinase, putative | 0,1786 | 0,3759 |
| TcCLB.506435.50 | hypothetical protein, conserved | 0,1785 | 0,2518 |
| TcCLB.508995.20 | Tubulin/FtsZ family, putative | 0,1781 | 0,07659 |

| | | | |
|---|---|---|---|
| TcCLB.511249.40 | hypothetical protein, conserved | 0,178 | 0,169 |
| TcCLB.507735.60 | hypothetical protein, conserved | 0,1779 | 0,09862 |
| TcCLB.506713.70 | retrotransposon hot spot protein (RHS, pseudogene), puta | 0,1779 | 0,1826 |
| TcCLB.505555.20 | hypothetical protein, conserved | 0,1779 | 0,2473 |
| TcCLB.506367.80 | hypothetical protein | 0,1778 | 0,4059 |
| TcCLB.505037.140 | retrotransposon hot spot protein (RHS, pseudogene), puta | 0,1777 | 0,2456 |
| TcCLB.510299.79 | protein phosphatase 2C, putative | 0,1776 | 0,146 |
| TcCLB.508153.860 | hypothetical protein, conserved | 0,1775 | 0,05411 |
| TcCLB.506517.130 | hypothetical protein, conserved | 0,1773 | 0,09148 |
| TcCLB.508515.80 | hypothetical protein, conserved | 0,1773 | 0,1588 |
| TcCLB.511045.29 | hypothetical protein, conserved | 0,1772 | 0,1015 |
| TcCLB.508679.7 | hypothetical protein, conserved (pseudogene) | 0,1772 | 0,5635 |
| TcCLB.504883.40 | hypothetical protein, conserved | 0,1771 | 0,07628 |
| TcCLB.504867.114 | tyrosine phosphatase, putative | 0,1771 | 0,1915 |
| TcCLB.510143.90 | hypothetical protein, conserved | 0,1771 | 0,2025 |
| TcCLB.509569.160 | hypothetical protein, conserved | 0,177 | 0,1401 |
| TcCLB.511719.20 | hypothetical protein, conserved | 0,1769 | 0,2381 |
| TcCLB.511869.9 | dynein heavy chain, putative (fragment) | 0,1768 | 0,1425 |
| TcCLB.510145.50 | hypothetical protein, conserved | 0,1766 | 0,05292 |
| TcCLB.509891.50 | hypothetical protein, conserved | 0,1766 | 0,3006 |
| TcCLB.510359.250 | hypothetical protein, conserved | 0,1766 | 0,493 |
| TcCLB.506737.60 | hypothetical protein, conserved | 0,1765 | 0,3957 |
| TcCLB.506717.200 | hypothetical protein, conserved | 0,1765 | 0,4159 |
| TcCLB.508461.530 | hypothetical protein, conserved | 0,1764 | 0,09632 |
| TcCLB.507653.110 | trans-sialidase (pseudogene), putative | 0,1764 | 0,173 |
| TcCLB.506777.110 | hypothetical protein, conserved | 0,1763 | 0,1639 |
| TcCLB.505997.40 | hypothetical protein, conserved (pseudogene) | 0,1763 | 0,1695 |
| TcCLB.511705.20 | putative prolyl carboxypeptidase, putative | 0,1763 | 0,3298 |
| TcCLB.509247.10 | hypothetical protein, conserved | 0,1762 | 0,1991 |
| TcCLB.508543.70 | hypothetical protein, conserved | 0,1762 | 0,2236 |
| TcCLB.507047.100 | hypothetical protein, conserved | 0,1759 | 0,09095 |
| TcCLB.506449.30 | retrotransposon hot spot protein (RHS, pseudogene), puta | 0,1758 | 0,4831 |
| TcCLB.511899.30 | hypothetical protein, conserved | 0,1756 | 0,09424 |
| TcCLB.511859.30 | hypothetical protein, conserved | 0,1756 | 0,1979 |
| TcCLB.506491.20 | myosin heavy chain, putative | 0,1755 | 0,04489 |
| TcCLB.509747.10 | hypothetical protein, conserved | 0,1754 | 0,306 |
| TcCLB.507539.20 | hypothetical protein, conserved | 0,1754 | 0,4653 |
| TcCLB.507521.50 | proteasome beta 6 subunit, putative | 0,175 | 0,1178 |
| TcCLB.507099.50 | programmed cell death 6 protein-like, putative | 0,1749 | 0,127 |
| TcCLB.510289.40 | GPI inositol deacylase precursor, putative | 0,1749 | 0,1495 |
| TcCLB.509233.239 | hypothetical protein, conserved | 0,1747 | 0,3762 |
| TcCLB.509595.50 | DIS3-like exonuclease, putative | 0,1746 | 0,08115 |
| TcCLB.506705.10 | hypothetical protein | 0,1746 | 0,1511 |
| TcCLB.506989.50 | hypothetical protein, conserved | 0,1746 | 0,1759 |
| TcCLB.504057.140 | ADP/ATP mitochondrial carrier protein, putative | 0,1745 | 0,0726 |

| | | | |
|---|---|---|---|
| TcCLB.504147.40 | Mitochondrial N(5)-glutamine methyltransferase MTQ1, p | 0,1745 | 0,07692 |
| TcCLB.510009.10 | hypothetical protein, conserved | 0,1742 | 0,4857 |
| TcCLB.508139.135 | retrotransposon hot spot protein (RHS, pseudogene), puta | 0,174 | 0,2158 |
| TcCLB.506629.190 | hypothetical protein, conserved | 0,1739 | 0,1149 |
| TcCLB.510861.40 | hypothetical protein, conserved | 0,1738 | 0,1777 |
| TcCLB.506343.20 | dispersed gene family protein 1 (DGF-1, pseudogene), put | 0,1738 | 0,4922 |
| TcCLB.503879.30 | hypothetical protein, conserved | 0,1737 | 0,122 |
| TcCLB.509019.10 | trans-sialidase (pseudogene), putative | 0,1737 | 0,1355 |
| TcCLB.508169.50 | hypothetical protein, conserved | 0,1735 | 0,1373 |
| TcCLB.508059.30 | hypothetical protein, conserved | 0,1734 | 0,1411 |
| TcCLB.511317.40 | hypothetical protein, conserved | 0,1734 | 0,1483 |
| TcCLB.507837.60 | Suppressor of G2 allele of SKP1, putative | 0,1734 | 0,1819 |
| TcCLB.509053.110 | ubiquitin fusion degradation protein 2, putative | 0,1733 | 0,02374 |
| TcCLB.508667.49 | small nuclear ribonucleoprotein SmD2, putative | 0,1733 | 0,1871 |
| TcCLB.507907.50 | hypothetical protein, conserved | 0,1732 | 0,01959 |
| TcCLB.506537.130 | protein kinase, putative | 0,1732 | 0,4899 |
| TcCLB.509429.110 | proteasome beta-1 subunit, putative | 0,1731 | 0,07526 |
| TcCLB.504037.20 | hypothetical protein, conserved | 0,1731 | 0,09721 |
| TcCLB.414145.9 | hypothetical protein, conserved (fragment) | 0,1731 | 0,2476 |
| TcCLB.508609.20 | hypothetical protein | 0,1731 | 0,2856 |
| TcCLB.442287.10 | serine/threonine protein phosphatase (pseudogene), puta | 0,173 | 0,08872 |
| TcCLB.511861.90 | retrotransposon hot spot (RHS) protein, putative | 0,1729 | 0,2026 |
| TcCLB.504003.40 | hypothetical protein, conserved | 0,1728 | 0,08445 |
| TcCLB.504033.120 | hypothetical protein, conserved | 0,1728 | 0,09152 |
| TcCLB.503887.30 | hypothetical protein, conserved | 0,1727 | 0,1074 |
| TcCLB.503943.20 | hypothetical protein, conserved | 0,1726 | 0,04973 |
| TcCLB.506977.120 | hypothetical protein, conserved | 0,1726 | 0,1289 |
| TcCLB.507747.260 | hypothetical protein, conserved | 0,1726 | 0,143 |
| TcCLB.511751.200 | epsilon-adaptin, putative | 0,1725 | 0,07526 |
| TcCLB.509853.10 | mannosyl-oligosaccharide 1,2-alpha-mannosidase IB, puta | 0,1722 | 0,1225 |
| TcCLB.507991.30 | hypothetical protein, conserved | 0,1722 | 0,1939 |
| TcCLB.509213.110 | hypothetical protein, conserved | 0,1721 | 0,06155 |
| TcCLB.511911.120 | DNA polymerase eta, putative | 0,1719 | 0,1358 |
| TcCLB.507711.180 | hypothetical protein, conserved | 0,1719 | 0,1678 |
| TcCLB.503487.60 | hypothetical protein, conserved | 0,1719 | 0,2314 |
| TcCLB.508175.160 | deoxyuridine triphosphatase, putative | 0,1718 | 0,1792 |
| TcCLB.504635.20 | hypothetical protein, conserved | 0,1718 | 0,2236 |
| TcCLB.509663.25 | hypothetical protein, conserved (pseudogene) | 0,1717 | 0,4375 |
| TcCLB.507773.50 | hypothetical protein, conserved | 0,1716 | 0,09978 |
| TcCLB.511857.50 | actin-like protein, putative | 0,1714 | 0,1392 |
| TcCLB.510187.260 | hypothetical protein, conserved | 0,1714 | 0,161 |
| TcCLB.507587.50 | hypothetical protein | 0,1712 | 0,7084 |
| TcCLB.508879.10 | hypothetical protein, conserved | 0,1711 | 0,1544 |
| TcCLB.508837.80 | Flagellar Member 3 (FLAM3) | 0,171 | 0,09797 |
| TcCLB.507811.90 | hypothetical protein, conserved | 0,171 | 0,1237 |

| | | | |
|---|---|---|---|
| TcCLB.510359.190 | hypothetical protein, conserved | 0,171 | 0,3027 |
| TcCLB.511825.250 | hypothetical protein, conserved | 0,1709 | 0,2817 |
| TcCLB.504013.11 | ribosomal protein L13, putative (fragment) | 0,1708 | 0,0196 |
| TcCLB.511773.30 | hypothetical protein | 0,1708 | 0,3839 |
| TcCLB.507663.40 | hypothetical protein, conserved | 0,1708 | 0,4116 |
| TcCLB.508951.60 | hypothetical protein, conserved | 0,1707 | 0,175 |
| TcCLB.508541.30 | hypothetical protein | 0,1705 | 0,736 |
| TcCLB.509149.70 | hypothetical protein, conserved | 0,1703 | 0,1148 |
| TcCLB.506863.50 | ribosomal RNA processing protein 45, putative (RRP45) | 0,1703 | 0,2887 |
| TcCLB.506275.70 | hypothetical protein, conserved | 0,1701 | 0,1731 |
| TcCLB.506321.310 | phosphoribosylpyrophosphate synthetase, putative | 0,17 | 0,04344 |
| TcCLB.469435.4 | ribosomal protein L14, putative | 0,1699 | 0,3484 |
| TcCLB.507997.89 | hypothetical protein, conserved | 0,1699 | 0,439 |
| TcCLB.508857.90 | hypothetical protein, conserved | 0,1696 | 0,1207 |
| TcCLB.511517.140 | hypothetical protein | 0,1696 | 0,1989 |
| TcCLB.510797.30 | hypothetical protein, conserved | 0,1696 | 0,2077 |
| TcCLB.510777.10 | hypothetical protein, conserved | 0,1696 | 0,269 |
| TcCLB.420495.19 | hypothetical protein, conserved (pseudogene) | 0,1696 | 0,4894 |
| TcCLB.508891.40 | hypothetical protein, conserved | 0,1695 | 0,1366 |
| TcCLB.511847.89 | hypothetical protein, conserved | 0,1695 | 0,4007 |
| TcCLB.510089.219 | diacylglycerol acyltransferase, putative | 0,1693 | 0,2533 |
| TcCLB.510101.440 | hypothetical protein, conserved | 0,1692 | 0,1916 |
| TcCLB.509643.80 | DNA mismatch repair protein MSH2, putative (MSH2) | 0,1689 | 0,05852 |
| TcCLB.510325.45 | hypothetical protein | 0,1689 | 0,1487 |
| TcCLB.508515.50 | glycogenin glucosyltransferase, putative | 0,1689 | 0,3722 |
| TcCLB.508027.94 | hypothetical protein, conserved | 0,1688 | 0,1947 |
| TcCLB.509099.170 | hypothetical protein, conserved | 0,1687 | 0,3395 |
| TcCLB.506321.230 | hypothetical protein | 0,1687 | 0,5327 |
| TcCLB.506357.110 | hypothetical protein, conserved | 0,1683 | 0,1517 |
| TcCLB.506625.84 | hypothetical protein, conserved | 0,168 | 0,267 |
| TcCLB.508641.210 | hypothetical protein, conserved | 0,1679 | 0,288 |
| TcCLB.508879.20 | hypothetical protein | 0,1677 | 0,3573 |
| TcCLB.510311.130 | DNA repair helicase, putative | 0,1676 | 0,109 |
| TcCLB.506633.100 | hypothetical protein, conserved | 0,1676 | 0,2572 |
| TcCLB.510517.20 | hypothetical protein, conserved | 0,1675 | 0,3154 |
| TcCLB.508879.100 | hypothetical protein, conserved | 0,1674 | 0,05622 |
| TcCLB.504193.41 | N-acetyltransferase complex ARD1 subunit (pseudogene), | 0,1674 | 0,3436 |
| TcCLB.504163.20 | hypothetical protein, conserved | 0,1672 | 0,1403 |
| TcCLB.508355.310 | hypothetical protein, conserved | 0,1671 | 0,1033 |
| TcCLB.509745.60 | telomerase reverse transcriptase, putative (TERT) | 0,1666 | 0,1437 |
| TcCLB.511279.20 | hypothetical protein, conserved | 0,1665 | 0,1604 |
| TcCLB.509537.80 | hypothetical protein, conserved | 0,1664 | 0,0345 |
| TcCLB.511391.140 | alpha-adaptin-like, putative | 0,1664 | 0,1148 |
| TcCLB.508579.40 | hypothetical protein, conserved | 0,1662 | 0,266 |
| TcCLB.504505.20 | retrotransposon hot spot (RHS) protein, putative | 0,1661 | 0,3391 |

| | | | |
|---|---|---|---|
| TcCLB.506507.30 | beta galactofuranosyl glycosyltransferase, putative | 0,166 | 0,2683 |
| TcCLB.503815.29 | hypothetical protein, conserved | 0,1659 | 0,4593 |
| TcCLB.506829.5 | hypothetical protein, conserved | 0,1658 | 0,3348 |
| TcCLB.510217.10 | hypothetical protein | 0,1658 | 0,8085 |
| TcCLB.508231.230 | protein transport protein Sec23A, putative | 0,1657 | 0,2295 |
| TcCLB.511511.139 | zinc finger protein, putative | 0,1654 | 0,1236 |
| TcCLB.511229.100 | hypothetical protein, conserved | 0,1652 | 0,0922 |
| TcCLB.506147.180 | DNA polymerase epsilon catalytic subunit, putative | 0,1652 | 0,1069 |
| TcCLB.510039.50 | hypothetical protein, conserved | 0,1652 | 0,175 |
| TcCLB.510101.150 | adaptin-related protein-like, putative | 0,1651 | 0,09019 |
| TcCLB.506627.50 | mitochondrial exoribonuclease DSS-1, putative | 0,1651 | 0,1021 |
| TcCLB.504105.100 | hypothetical protein | 0,1651 | 0,1465 |
| TcCLB.511621.70 | ribosomal protein L11, putative | 0,1651 | 0,1727 |
| TcCLB.511585.50 | hypothetical protein, conserved | 0,1651 | 0,2012 |
| TcCLB.507053.110 | hypothetical protein, conserved | 0,165 | 0,1057 |
| TcCLB.505997.190 | retrotransposon hot spot protein (RHS, pseudogene), puta | 0,1649 | 0,206 |
| TcCLB.510219.10 | hypothetical protein, conserved | 0,1648 | 0,2597 |
| TcCLB.504157.120 | hypothetical protein, conserved | 0,1648 | 0,3818 |
| TcCLB.506925.390 | MP44, putative | 0,1647 | 0,1893 |
| TcCLB.511171.60 | surface protease GP63 (pseudogene), putative | 0,1647 | 0,7288 |
| TcCLB.504479.20 | retrotransposon hot spot protein (RHS, pseudogene), puta | 0,1646 | 0,3318 |
| TcCLB.507297.20 | cysteinyl-tRNA synthetase, putative | 0,1645 | 0,04198 |
| TcCLB.509029.120 | ATP-dependent DNA helicase, putative | 0,1644 | 0,1786 |
| TcCLB.511537.60 | hypothetical protein, conserved | 0,1643 | 0,1232 |
| TcCLB.511847.70 | hypothetical protein, conserved | 0,1643 | 0,1938 |
| TcCLB.504241.90 | dispersed gene family protein 1 (DGF-1), putative | 0,1643 | 0,7247 |
| TcCLB.503541.20 | hypothetical protein, conserved | 0,1642 | 0,2522 |
| TcCLB.506187.20 | trans-sialidase, Group V, putative | 0,1642 | 0,3117 |
| TcCLB.507801.50 | tubulin-tyrosine ligase, putative | 0,1641 | 0,1494 |
| TcCLB.511807.210 | hypothetical protein, conserved (pseudogene) | 0,164 | 0,06733 |
| TcCLB.504213.130 | hypothetical protein, conserved | 0,1639 | 0,2879 |
| TcCLB.510951.19 | hypothetical protein, conserved | 0,1637 | 0,3163 |
| TcCLB.510427.5 | hypothetical protein, conserved | 0,1637 | 0,4082 |
| TcCLB.507083.100 | hypothetical protein, conserved | 0,1636 | 0,04999 |
| TcCLB.507991.60 | mismatch repair protein MLH1, putative | 0,1636 | 0,2196 |
| TcCLB.510943.120 | hypothetical protein, conserved | 0,1634 | 0,09739 |
| TcCLB.511017.20 | 2,3-bisphosphoglycerate-independent phosphoglycerate m | 0,1634 | 0,1218 |
| TcCLB.510687.10 | dynein heavy chain (pseudogene), putative | 0,1633 | 0,0273 |
| TcCLB.510769.30 | hypothetical protein, conserved | 0,1633 | 0,1155 |
| TcCLB.506203.100 | hypothetical protein | 0,1632 | 0,1345 |
| TcCLB.508629.26 | retrotransposon hot spot protein (RHS, pseudogene), puta | 0,1632 | 0,3021 |
| TcCLB.506195.70 | PrimPol-like protein 2, putative (ppl2) | 0,1629 | 0,136 |
| TcCLB.511291.30 | lipoic acid synthetase, mitochondrial precursor, putative | 0,1629 | 0,1827 |
| TcCLB.503757.49 | hypothetical protein, conserved (fragment) | 0,1629 | 0,6316 |
| TcCLB.506743.90 | hypothetical protein, conserved | 0,1628 | 0,4615 |

| | | | |
|---|---|---|---|
| TcCLB.503599.40 | hypothetical protein, conserved | 0,1627 | 0,1478 |
| TcCLB.507049.20 | hypothetical protein | 0,1627 | 0,2856 |
| TcCLB.510043.59 | hypothetical protein, conserved (fragment) | 0,1626 | 0,1571 |
| TcCLB.506181.40 | hypothetical protein, conserved | 0,1622 | 0,1397 |
| TcCLB.508873.170 | retrotransposon hot spot protein (RHS, pseudogene), puta | 0,1622 | 0,6238 |
| TcCLB.507841.40 | hypothetical protein, conserved | 0,1621 | 0,1934 |
| TcCLB.510885.40 | WD repeat and HMG-box DNA-binding protein, putative | 0,162 | 0,05705 |
| TcCLB.509073.50 | hypothetical protein, conserved | 0,162 | 0,1963 |
| TcCLB.509959.160 | hypothetical protein, conserved (pseudogene) | 0,162 | 0,2898 |
| TcCLB.506249.60 | 40S ribosomal protein S8, putative | 0,1619 | 0,1095 |
| TcCLB.509179.40 | hypothetical protein, conserved | 0,1619 | 0,263 |
| TcCLB.511355.40 | GUP1, putative | 0,1618 | 0,1935 |
| TcCLB.510663.80 | hypothetical protein, conserved | 0,1618 | 0,2249 |
| TcCLB.507611.40 | protein kinase, putative | 0,1618 | 0,6996 |
| TcCLB.509609.19 | DNA repair and recombination helicase protein PIF7, puta | 0,1617 | 0,3457 |
| TcCLB.504427.190 | hypothetical protein, conserved | 0,1616 | 0,2376 |
| TcCLB.510759.110 | hypothetical protein, conserved | 0,1615 | 0,1258 |
| TcCLB.509033.100 | hypothetical protein, conserved | 0,1615 | 0,1891 |
| TcCLB.506485.60 | hypothetical protein, conserved | 0,1614 | 0,1943 |
| TcCLB.509243.30 | hypothetical protein, conserved | 0,1612 | 0,342 |
| TcCLB.509617.20 | paraflagellar rod protein 3, putative (PAR3) | 0,161 | 0,192 |
| TcCLB.507547.80 | protein kinase, putative | 0,161 | 0,3568 |
| TcCLB.503733.40 | hypothetical protein, conserved | 0,161 | 0,364 |
| TcCLB.507641.140 | ATP-dependent DEAD/H RNA helicase, putative | 0,1609 | 0,04585 |
| TcCLB.503637.20 | hypothetical protein, conserved | 0,1608 | 0,1259 |
| TcCLB.504103.100 | ABC1 protein, putative | 0,1607 | 0,1916 |
| TcCLB.508641.120 | hypothetical protein, conserved | 0,1607 | 0,2202 |
| TcCLB.506983.60 | DNA excision repair protein, putative | 0,1606 | 0,05606 |
| TcCLB.509857.40 | hypothetical protein, conserved | 0,1606 | 0,09032 |
| TcCLB.510533.160 | hypothetical protein, conserved | 0,1605 | 0,2078 |
| TcCLB.507515.50 | hypothetical protein | 0,1604 | 0,1995 |
| TcCLB.503847.30 | hypothetical protein, conserved | 0,1604 | 0,2128 |
| TcCLB.508143.65 | elongation factor 1-gamma (EF-1-gamma, pseudogene), | 0,1604 | 0,6715 |
| TcCLB.506989.250 | hypothetical protein, conserved | 0,16 | 0,1221 |
| TcCLB.503719.10 | hypothetical protein | 0,1599 | 0,1207 |
| TcCLB.508479.390 | RNA triphosphatase, putative | 0,1599 | 0,2651 |
| TcCLB.509123.20 | hypothetical protein | 0,1599 | 0,7525 |
| TcCLB.505183.50 | hypothetical protein, conserved | 0,1598 | 0,1155 |
| TcCLB.506301.50 | succinyl-coA:3-ketoacid-coenzyme A transferase, mitocho | 0,1598 | 0,1897 |
| TcCLB.503897.10 | hypothetical protein | 0,1597 | 0,1068 |
| TcCLB.507533.10 | phosphatidylinositol 4-kinase alpha, putative | 0,1597 | 0,1366 |
| TcCLB.503909.50 | hypothetical protein, conserved (YCF45) | 0,1596 | 0,1516 |
| TcCLB.507711.130 | hypothetical protein, conserved | 0,1593 | 0,3023 |
| TcCLB.510741.70 | protein kinase, putative | 0,1593 | 0,3808 |
| TcCLB.511065.10 | STOP axonemal protein, putative | 0,1591 | 0,1695 |

| | | | |
|---|---|---|---|
| TcCLB.506527.4 | hypothetical protein, conserved (fragment) | 0,1589 | 0,1192 |
| TcCLB.508209.39 | hypothetical protein, conserved | 0,1589 | 0,1681 |
| TcCLB.506941.210 | hypothetical protein, conserved | 0,1589 | 0,2044 |
| TcCLB.508153.1100 | mitochondrial protein 99, putative (MP99) | 0,1588 | 0,1569 |
| TcCLB.504111.30 | hypothetical protein, conserved | 0,1588 | 0,2125 |
| TcCLB.510355.250 | hypothetical protein, conserved | 0,1587 | 0,1203 |
| TcCLB.510423.70 | hypothetical protein, conserved | 0,1587 | 0,4149 |
| TcCLB.511515.20 | AAA ATPase (pseudogene), putative | 0,1586 | 0,1254 |
| TcCLB.511693.120 | hypothetical protein | 0,1586 | 0,1417 |
| TcCLB.507681.60 | hypothetical protein, conserved | 0,1586 | 0,3621 |
| TcCLB.506743.4 | eukaryotic translation initiation factor 1A, putative (fragm | 0,1585 | 0,2312 |
| TcCLB.508881.40 | hypothetical protein, conserved | 0,1584 | 0,1519 |
| TcCLB.506855.110 | hypothetical protein, conserved | 0,1584 | 0,1582 |
| TcCLB.511867.40 | hypothetical protein, conserved | 0,1583 | 0,2852 |
| TcCLB.507001.60 | hypothetical protein, conserved | 0,1582 | 0,08577 |
| TcCLB.508973.110 | hypothetical protein, conserved | 0,1581 | 0,1912 |
| TcCLB.511899.20 | hypothetical protein, conserved | 0,158 | 0,2015 |
| TcCLB.504147.210 | hypothetical protein, conserved | 0,1578 | 0,3092 |
| TcCLB.508979.80 | hypothetical protein, conserved | 0,1577 | 0,275 |
| TcCLB.510283.80 | hypothetical protein, conserved | 0,1576 | 0,2774 |
| TcCLB.504017.40 | aminomethyltransferase, mitochondrial, putative (GCV) | 0,1576 | 0,296 |
| TcCLB.508209.70 | hypothetical protein, conserved | 0,1575 | 0,2665 |
| TcCLB.506405.110 | ribosomal protein L13, putative | 0,1575 | 0,3756 |
| TcCLB.508799.130 | hypothetical protein, conserved | 0,1574 | 0,08804 |
| TcCLB.506537.10 | retrotransposon hot spot protein (RHS, pseudogene), puta | 0,1574 | 0,1758 |
| TcCLB.511183.116 | trans-sialidase (pseudogene), putative | 0,1573 | 0,6178 |
| TcCLB.511907.210 | hypothetical protein, conserved | 0,157 | 0,2365 |
| TcCLB.511585.269 | retrotransposon hot spot protein (RHS, pseudogene), puta | 0,157 | 0,3604 |
| TcCLB.509233.80 | heat shock protein DNAJ, putative | 0,1569 | 0,1579 |
| TcCLB.511621.160 | hypothetical protein, conserved | 0,1569 | 0,1903 |
| TcCLB.511301.140 | potassium voltage-gated channel | 0,1569 | 0,2291 |
| TcCLB.509591.30 | ras-family member, GTP-binding protein, putative | 0,1568 | 0,1378 |
| TcCLB.509965.200 | hypothetical protein, conserved | 0,1568 | 0,281 |
| TcCLB.511217.39 | hypothetical protein, conserved | 0,1568 | 0,3427 |
| TcCLB.511417.20 | retrotransposon hot spot protein (RHS, pseudogene), puta | 0,1568 | 0,548 |
| TcCLB.506147.184 | hypothetical protein, conserved | 0,1567 | 0,3301 |
| TcCLB.508687.25 | hypothetical protein | 0,1567 | 0,5084 |
| TcCLB.506239.20 | hypothetical protein, conserved | 0,1566 | 0,2226 |
| TcCLB.428403.5 | hypothetical protein, conserved | 0,1565 | 0,2759 |
| TcCLB.490165.19 | hypothetical protein, conserved | 0,1564 | 0,4766 |
| TcCLB.510399.10 | dispersed gene family protein 1 (DGF-1), putative | 0,1564 | 0,7486 |
| TcCLB.503847.60 | hypothetical protein, conserved | 0,1563 | 0,112 |
| TcCLB.511545.110 | hypothetical protein | 0,1563 | 0,3624 |
| TcCLB.503829.89 | spliceosomal U5 snRNP-specific protein, putative | 0,1561 | 0,3803 |
| TcCLB.504247.10 | hypothetical protein, conserved | 0,156 | 0,2606 |

| | | | |
|---|---|---|---|
| TcCLB.506825.20 | hypothetical protein, conserved | 0,1559 | 0,06296 |
| TcCLB.509991.110 | transcription elongation factor 1 homolog (ELOF1) | 0,1558 | 0,2604 |
| TcCLB.508443.50 | hypothetical protein, conserved | 0,1558 | 0,2653 |
| TcCLB.503521.39 | mitochondrial carrier protein, putative | 0,1558 | 0,2825 |
| TcCLB.507787.150 | hypothetical protein, conserved | 0,1557 | 0,1589 |
| TcCLB.507005.50 | hypothetical protein | 0,1556 | 0,1448 |
| TcCLB.511429.20 | cyclophilin 15, putative | 0,1556 | 0,2194 |
| TcCLB.510741.60 | hypothetical protein, conserved | 0,1556 | 0,23 |
| TcCLB.503841.20 | AMPK subunit gamma (AMPKG) | 0,1555 | 0,1973 |
| TcCLB.509259.45 | hypothetical protein | 0,1554 | 0,475 |
| TcCLB.504163.10 | hypothetical protein, conserved | 0,1553 | 0,09069 |
| TcCLB.510687.149 | coatomer alpha subunit, putative | 0,1553 | 0,1512 |
| TcCLB.508899.54 | hypothetical protein, conserved | 0,1552 | 0,2779 |
| TcCLB.510099.30 | hypothetical protein, conserved | 0,1551 | 0,1912 |
| TcCLB.510761.70 | ubiquitin hydrolase, putative | 0,1549 | 0,136 |
| TcCLB.508207.60 | hypothetical protein, conserved | 0,1549 | 0,2538 |
| TcCLB.504069.60 | GPI transamidase component GAA1, putative | 0,1549 | 0,2711 |
| TcCLB.508357.9 | hypothetical protein, conserved (fragment) | 0,1546 | 0,1235 |
| TcCLB.504741.210 | hypothetical protein, conserved | 0,1546 | 0,1834 |
| TcCLB.511671.50 | hypothetical protein, conserved | 0,1541 | 0,04711 |
| TcCLB.511001.180 | hypothetical protein, conserved | 0,1541 | 0,1303 |
| TcCLB.508045.14 | hypothetical protein, conserved | 0,1541 | 0,1787 |
| TcCLB.508849.20 | retrotransposon hot spot protein (RHS, pseudogene), puta | 0,1541 | 0,1853 |
| TcCLB.508153.120 | ubiquitin hydrolase, putative | 0,154 | 0,152 |
| TcCLB.507063.40 | hypothetical protein, conserved | 0,1539 | 0,2596 |
| TcCLB.509011.40 | calreticulin, putative | 0,1538 | 0,02273 |
| TcCLB.508355.50 | hypothetical protein, conserved | 0,1537 | 0,3241 |
| TcCLB.511185.150 | hypothetical protein, conserved (pseudogene) | 0,1536 | 0,316 |
| TcCLB.507049.5 | kynurenine aminotransferase (pseudogene), putative | 0,1533 | 0,2572 |
| TcCLB.507711.220 | hypothetical protein, conserved | 0,1532 | 0,4352 |
| TcCLB.508409.230 | UBX domain-containing protein | 0,153 | 0,1311 |
| TcCLB.503869.60 | coatomer beta subunit (pseudogene), putative | 0,1529 | 0,1043 |
| TcCLB.507713.20 | hypothetical protein, conserved | 0,1529 | 0,2159 |
| TcCLB.506925.290 | hypothetical protein | 0,1528 | 0,1386 |
| TcCLB.506835.140 | hypothetical protein, conserved | 0,1526 | 0,1064 |
| TcCLB.506925.50 | hypothetical protein, conserved | 0,1526 | 0,1493 |
| TcCLB.510381.5 | SNF2 DNA repair protein (pseudogene), putative | 0,1526 | 0,316 |
| TcCLB.503821.10 | DNA-directed RNA polymerase I subunit, putative | 0,1526 | 0,3823 |
| TcCLB.506287.120 | hypothetical protein | 0,1523 | 0,2175 |
| TcCLB.510659.110 | hypothetical protein, conserved | 0,1522 | 0,2118 |
| TcCLB.506017.31 | ATP-dependent DEAD/H RNA helicase (pseudogene), puta | 0,1522 | 0,5904 |
| TcCLB.511117.60 | mucin-associated surface protein (MASP, pseudogene), pu | 0,1521 | 0,6313 |
| TcCLB.511003.40 | hypothetical protein, conserved | 0,1519 | 0,135 |
| TcCLB.507073.49 | hypothetical protein, conserved | 0,1518 | 0,228 |
| TcCLB.506147.160 | hypothetical protein, conserved | 0,1518 | 0,2767 |

| | | | |
|---|---|---|---|
| TcCLB.503791.49 | vacuolar protein sorting-associated protein, putative (frag | 0,1516 | 0,298 |
| TcCLB.509733.50 | hypothetical protein, conserved | 0,1515 | 0,2214 |
| TcCLB.503975.20 | hypothetical protein | 0,1515 | 0,3076 |
| TcCLB.508409.220 | hypothetical protein, conserved | 0,1514 | 0,2166 |
| TcCLB.507491.50 | hypothetical protein, conserved | 0,1511 | 0,2342 |
| TcCLB.506529.620 | hypothetical protein | 0,1511 | 0,2898 |
| TcCLB.507079.10 | hypothetical protein, conserved | 0,1508 | 0,2294 |
| TcCLB.504257.30 | S-adenosylmethionine decarboxylase proenzyme, putative | 0,1506 | 0,151 |
| TcCLB.508461.90 | hypothetical protein, conserved | 0,1505 | 0,1889 |
| TcCLB.506529.190 | hypothetical protein, conserved | 0,1504 | 0,2157 |
| TcCLB.508965.94 | hypothetical protein, conserved | 0,1504 | 0,2258 |
| TcCLB.506435.450 | hypothetical protein, conserved | 0,1503 | 0,1553 |
| TcCLB.509455.130 | cAMP response protein, putative (CARP4) | 0,1501 | 0,1557 |
| TcCLB.504171.20 | hypothetical protein, conserved | 0,1501 | 0,3288 |
| TcCLB.511529.140 | hypothetical protein, conserved | 0,1499 | 0,1018 |
| TcCLB.503897.110 | plectin-like protein, putative | 0,1499 | 0,2568 |
| TcCLB.506095.9 | dispersed gene family protein 1 (DGF-1), putative (fragme | 0,1499 | 0,356 |
| TcCLB.510635.40 | hypothetical protein, conserved | 0,1498 | 0,2569 |
| TcCLB.509779.50 | hypothetical protein, conserved | 0,1498 | 0,2583 |
| TcCLB.508325.130 | protein kinase, putative | 0,1497 | 0,226 |
| TcCLB.506475.50 | Replication Factor C Subunit 1-related protein | 0,1497 | 0,228 |
| TcCLB.507591.20 | beta galactofuranosyl glycosyltransferase (pseudogene), p | 0,1497 | 0,4854 |
| TcCLB.508175.90 | hypothetical protein | 0,1496 | 0,07941 |
| TcCLB.420369.40 | hypothetical protein, conserved | 0,1496 | 0,3094 |
| TcCLB.510165.40 | hypothetical protein, conserved | 0,1495 | 0,1577 |
| TcCLB.507093.310 | hypothetical protein, conserved | 0,1495 | 0,2224 |
| TcCLB.505931.30 | trans-sialidase, Group I, putative | 0,1495 | 0,4006 |
| TcCLB.460125.10 | hypothetical protein, conserved (fragment) | 0,1494 | 0,4676 |
| TcCLB.511481.40 | N-Acetyl-D-glucosaminylphosphatidylinositol de-N-acetyla | 0,1492 | 0,3368 |
| TcCLB.511309.20 | hydrolase-like protein, putative | 0,1492 | 0,3475 |
| TcCLB.510039.100 | hypothetical protein, conserved | 0,1491 | 0,3633 |
| TcCLB.509085.60 | dispersed gene family protein 1 (DGF-1), putative | 0,1491 | 0,4843 |
| TcCLB.507541.30 | casein kinase, putative | 0,149 | 0,08093 |
| TcCLB.503781.20 | phosphatidylinositol N-acetylglucosaminyltransferase sub | 0,1489 | 0,2797 |
| TcCLB.510241.100 | hypothetical protein, conserved | 0,1489 | 0,2937 |
| TcCLB.508109.20 | mucin TcMUCII, putative | 0,1489 | 0,6181 |
| TcCLB.503629.10 | hypothetical protein, conserved | 0,1484 | 0,1879 |
| TcCLB.508479.150 | serine/threonine protein kinase, putative | 0,1483 | 0,102 |
| TcCLB.507041.20 | hypothetical protein, conserved | 0,1483 | 0,1947 |
| TcCLB.507521.40 | hypothetical protein, conserved | 0,1483 | 0,2934 |
| TcCLB.511269.20 | Multisite-specific tRNA:(cytosine-C(5))-methyltransferase | 0,1482 | 0,1289 |
| TcCLB.509791.180 | hypothetical protein, conserved | 0,1481 | 0,303 |
| TcCLB.504425.50 | hypothetical protein, conserved | 0,1481 | 0,4846 |
| TcCLB.508799.250 | hypothetical protein, conserved | 0,148 | 0,1179 |
| TcCLB.504017.79 | glycyl-tRNA synthetase, putative | 0,1477 | 0,2112 |

| | | | |
|---|---|---|---|
| TcCLB.509645.19 | hypothetical protein, conserved | 0,1477 | 0,2268 |
| TcCLB.506661.100 | hypothetical protein, conserved | 0,1477 | 0,5159 |
| TcCLB.503999.90 | RNA-binding protein 34, putative (RBP34) | 0,1475 | 0,2946 |
| TcCLB.511893.20 | hypothetical protein, conserved | 0,1475 | 0,7664 |
| TcCLB.509911.90 | peptidase, putative | 0,1474 | 0,0587 |
| TcCLB.506531.20 | hypothetical protein, conserved | 0,1474 | 0,1544 |
| TcCLB.506581.4 | hypothetical protein, conserved | 0,1473 | 0,5891 |
| TcCLB.508891.120 | periodic tryptophan protein 2 (pseudogene), putative | 0,1472 | 0,163 |
| TcCLB.506833.9 | hypothetical protein, conserved | 0,1471 | 0,2219 |
| TcCLB.506821.30 | glycosylphosphatidylinositol-specific phospholipase C | 0,1471 | 0,2289 |
| TcCLB.508479.260 | hypothetical protein, conserved | 0,147 | 0,2075 |
| TcCLB.503479.10 | hypothetical protein, conserved | 0,147 | 0,2683 |
| TcCLB.472777.20 | hypothetical protein, conserved | 0,1469 | 0,1922 |
| TcCLB.506627.80 | hypothetical protein, conserved | 0,1469 | 0,1938 |
| TcCLB.503953.20 | hypothetical protein, conserved | 0,1469 | 0,4196 |
| TcCLB.510323.70 | hypothetical protein, conserved | 0,1468 | 0,1544 |
| TcCLB.506529.14 | hypothetical protein, conserved | 0,1468 | 0,3391 |
| TcCLB.504229.70 | retrotransposon hot spot (RHS) protein, putative | 0,1467 | 0,2712 |
| TcCLB.506243.94 | retrotransposon hot spot protein (RHS, pseudogene), puta | 0,1467 | 0,3087 |
| TcCLB.506665.20 | hypothetical protein | 0,1466 | 0,5619 |
| TcCLB.508489.40 | hypothetical protein, conserved | 0,1465 | 0,2602 |
| TcCLB.511315.56 | hypothetical protein, conserved | 0,1465 | 0,6195 |
| TcCLB.506969.10 | mucin-associated surface protein (MASP, pseudogene), pu | 0,1464 | 0,4192 |
| TcCLB.510175.170 | glycine dehydrogenase [decarboxylating] (pseudogene), pu | 0,1463 | 0,4014 |
| TcCLB.506557.20 | hypothetical protein, conserved | 0,1462 | 0,1728 |
| TcCLB.511003.140 | vacuolar transport protein 4A, putative | 0,1462 | 0,2437 |
| TcCLB.508153.820 | inorganic pyrophosphatase, putative | 0,1462 | 0,3041 |
| TcCLB.509235.10 | UDP-Gal or UDP-GlcNAc-dependent glycosyltransferase, p | 0,1462 | 0,5053 |
| TcCLB.511779.31 | hypothetical protein | 0,1462 | 0,7205 |
| TcCLB.506459.190 | hypothetical protein | 0,1461 | 0,6146 |
| TcCLB.504033.170 | tyrosine phosphatase, putative | 0,146 | 0,2802 |
| TcCLB.510627.70 | hypothetical protein, conserved (pseudogene) | 0,146 | 0,7426 |
| TcCLB.507689.25 | hypothetical protein, conserved | 0,1458 | 0,6512 |
| TcCLB.511537.21 | hypothetical protein, conserved | 0,1457 | 0,3728 |
| TcCLB.506479.80 | hypothetical protein, conserved | 0,1456 | 0,242 |
| TcCLB.510333.10 | hypothetical protein, conserved | 0,1455 | 0,1331 |
| TcCLB.510237.95 | hypothetical protein, conserved (pseudogene) | 0,1455 | 0,6908 |
| TcCLB.503537.18 | retrotransposon hot spot protein (RHS, pseudogene), puta | 0,1454 | 0,5712 |
| TcCLB.506155.70 | SWI/SNF-related helicase, putative | 0,1452 | 0,07986 |
| TcCLB.509167.40 | amino acid permease, putative | 0,1452 | 0,2371 |
| TcCLB.511633.30 | hypothetical protein, conserved | 0,1452 | 0,2762 |
| TcCLB.510901.100 | DNA topoisomerase III, putative | 0,1451 | 0,1199 |
| TcCLB.508699.100 | surface protease GP63, putative | 0,1451 | 0,3903 |
| TcCLB.508541.120 | trans-sialidase (pseudogene), putative | 0,1449 | 0,7427 |
| TcCLB.509797.20 | hypothetical protein, conserved | 0,1447 | 0,2504 |

| | | | |
|---|---|---|---|
| TcCLB.507911.30 | dispersed gene family protein 1 (DGF-1), putative | 0,1446 | 0,4498 |
| TcCLB.508741.415 | hypothetical protein, conserved (pseudogene) | 0,1445 | 0,3971 |
| TcCLB.459619.10 | hypothetical protein, conserved (pseudogene) | 0,1445 | 0,6244 |
| TcCLB.506871.20 | hypothetical protein, conserved | 0,1444 | 0,1319 |
| TcCLB.510101.250 | hypothetical protein, conserved | 0,1444 | 0,2134 |
| TcCLB.509721.50 | hypothetical protein | 0,1442 | 0,3179 |
| TcCLB.509683.130 | aminopeptidase, putative | 0,1441 | 0,2547 |
| TcCLB.509179.200 | hypothetical protein, conserved | 0,1441 | 0,5135 |
| TcCLB.506737.30 | hypothetical protein, conserved | 0,1439 | 0,2331 |
| TcCLB.507681.80 | hypothetical protein | 0,1438 | 0,1537 |
| TcCLB.503929.20 | ribosomal protein L13, putative | 0,1438 | 0,4121 |
| TcCLB.511663.50 | dispersed gene family protein 1 (DGF-1), putative | 0,1438 | 0,7305 |
| TcCLB.510759.4 | hypothetical protein, conserved | 0,1437 | 0,296 |
| TcCLB.503465.40 | hypothetical protein | 0,1437 | 0,3083 |
| TcCLB.506053.60 | hypothetical protein (fragment) | 0,1435 | 0,3125 |
| TcCLB.504035.130 | hypothetical protein, conserved | 0,1432 | 0,2743 |
| TcCLB.508569.10 | hypothetical protein, conserved | 0,1432 | 0,6819 |
| TcCLB.457251.19 | hypothetical protein, conserved (fragment) | 0,143 | 0,4347 |
| TcCLB.509205.110 | elongation factor G2-like protein, putative | 0,143 | 0,6709 |
| TcCLB.508285.5 | trans-sialidase (pseudogene), putative | 0,1428 | 0,5612 |
| TcCLB.506757.10 | phospholipid:diacylglycerol acyltransferase, putative | 0,1427 | 0,1019 |
| TcCLB.508469.10 | hypothetical protein, conserved | 0,1427 | 0,4832 |
| TcCLB.504867.30 | hypothetical protein, conserved | 0,1427 | 0,5365 |
| TcCLB.511229.110 | glycyl-tRNA synthetase, putative | 0,1426 | 0,1041 |
| TcCLB.509943.11 | cAMP specific phosphodiesterase, putative | 0,1426 | 0,6433 |
| TcCLB.503733.10 | hypothetical protein, conserved | 0,1425 | 0,3471 |
| TcCLB.503399.10 | hypothetical protein, conserved | 0,1424 | 0,3308 |
| TcCLB.511575.50 | hypothetical protein, conserved | 0,1422 | 0,07862 |
| TcCLB.510769.150 | hypothetical protein, conserved | 0,1422 | 0,1673 |
| TcCLB.506885.60 | hypothetical protein, conserved | 0,1422 | 0,1764 |
| TcCLB.509225.30 | trans-sialidase (pseudogene), putative | 0,1421 | 0,2692 |
| TcCLB.511627.60 | vesicle-associated membrane protein | 0,1419 | 0,2232 |
| TcCLB.510791.30 | mucin-associated surface protein (MASP, pseudogene), pu | 0,1418 | 0,2159 |
| TcCLB.510439.61 | heat shock 70 kDa protein, putative (fragment) | 0,1415 | 0,1562 |
| TcCLB.508387.90 | paraflagellar rod protein, putative | 0,1414 | 0,1493 |
| TcCLB.511805.29 | 2,3-bisphosphoglycerate-independent phosphoglycerate m | 0,1414 | 0,517 |
| TcCLB.506457.130 | hypothetical protein, conserved | 0,1412 | 0,1948 |
| TcCLB.506561.20 | retrotransposon hot spot protein (RHS, pseudogene), puta | 0,1412 | 0,3532 |
| TcCLB.506529.460 | hypothetical protein, conserved | 0,1411 | 0,1867 |
| TcCLB.507225.10 | retrotransposon hot spot (RHS) protein, putative | 0,1408 | 0,2124 |
| TcCLB.508355.300 | hypothetical protein, conserved | 0,1408 | 0,2841 |
| TcCLB.510311.30 | hypothetical protein, conserved | 0,1407 | 0,1895 |
| TcCLB.510595.20 | hypothetical protein, conserved | 0,1407 | 0,3241 |
| TcCLB.508297.30 | hypothetical protein, conserved | 0,1406 | 0,2525 |
| TcCLB.506821.10 | hypothetical protein | 0,1405 | 0,2713 |

| | | | |
|---|---|---|---|
| TcCLB.506835.20 | hypothetical protein | 0,1403 | 0,1564 |
| TcCLB.509069.40 | hypothetical protein | 0,1402 | 0,2128 |
| TcCLB.506537.230 | hypothetical protein, conserved | 0,1402 | 0,5457 |
| TcCLB.510661.270 | replication factor C, subunit 3, putative | 0,14 | 0,2929 |
| TcCLB.511469.10 | hypothetical protein, conserved | 0,14 | 0,4513 |
| TcCLB.509611.110 | RNA-editing complex protein, putative | 0,1399 | 0,2656 |
| TcCLB.509493.120 | hypothetical protein | 0,1397 | 0,4893 |
| TcCLB.476725.9 | dispersed gene family protein 1 (DGF-1), putative (fragme | 0,1397 | 0,5825 |
| TcCLB.508799.104 | hypothetical protein, conserved | 0,1396 | 0,3852 |
| TcCLB.504033.220 | palmitoyl acyltransferase 1, putative | 0,1395 | 0,146 |
| TcCLB.506885.110 | hypothetical protein, conserved | 0,1393 | 0,09593 |
| TcCLB.511813.30 | proteasome regulatory ATPase subunit 5, putative | 0,1393 | 0,2572 |
| TcCLB.509289.5 | tuzin (fragment) | 0,1392 | 0,04354 |
| TcCLB.510431.170 | hypothetical protein, conserved | 0,1392 | 0,1991 |
| TcCLB.510163.20 | elongation factor 1-gamma (EF-1-gamma), putative | 0,1392 | 0,249 |
| TcCLB.507049.100 | mitochondrial RNA binding protein, putative | 0,139 | 0,2484 |
| TcCLB.508267.10 | hypothetical protein, conserved | 0,1389 | 0,1348 |
| TcCLB.506927.30 | hypothetical protein, conserved (fragment) | 0,1389 | 0,3889 |
| TcCLB.504425.110 | permease-like protein, putative | 0,1387 | 0,3931 |
| TcCLB.504071.100 | hypothetical protein, conserved | 0,1386 | 0,1416 |
| TcCLB.510029.10 | hypothetical protein, conserved (pseudogene) | 0,1386 | 0,4145 |
| TcCLB.503527.60 | hypothetical protein, conserved | 0,1384 | 0,3004 |
| TcCLB.507991.10 | hypothetical protein, conserved | 0,1384 | 0,3829 |
| TcCLB.511837.70 | mitogen activated protein kinase, putative | 0,1383 | 0,1293 |
| TcCLB.510797.4 | phosphatidylinositol (3,5) kinase, putative (fragment) | 0,1383 | 0,3957 |
| TcCLB.506801.30 | hypothetical protein | 0,1383 | 0,4031 |
| TcCLB.508153.340 | ATP-dependent 6-phosphofructokinase, glycosomal (PFK) | 0,1382 | 0,2849 |
| TcCLB.508899.10 | proteasome regulatory non-ATP-ase subunit 9, putative (f | 0,1382 | 0,3 |
| TcCLB.510283.160 | hypothetical protein, conserved | 0,138 | 0,3292 |
| TcCLB.510299.60 | hypothetical protein, conserved | 0,1379 | 0,4068 |
| TcCLB.444777.29 | intraflagellar transport protein 88, putative (fragment) (IF | 0,1378 | 0,2692 |
| TcCLB.509935.20 | hypothetical protein, conserved | 0,1377 | 0,1844 |
| TcCLB.507603.54 | hypothetical protein, conserved | 0,1374 | 0,4371 |
| TcCLB.506819.20 | hypothetical protein | 0,1371 | 0,4737 |
| TcCLB.503627.20 | protein kinase, putative | 0,1368 | 0,1246 |
| TcCLB.506999.30 | hypothetical protein, conserved | 0,1368 | 0,1462 |
| TcCLB.511283.220 | hypothetical protein, conserved | 0,1368 | 0,1645 |
| TcCLB.507019.60 | hypothetical protein, conserved | 0,1367 | 0,1741 |
| TcCLB.507663.80 | hypothetical protein, conserved | 0,1367 | 0,2249 |
| TcCLB.506525.162 | hypothetical protein, conserved | 0,1367 | 0,4513 |
| TcCLB.510345.30 | hypothetical protein | 0,1367 | 0,4734 |
| TcCLB.508317.70 | hypothetical protein | 0,1366 | 0,1271 |
| TcCLB.507691.61 | trans-sialidase (pseudogene), putative | 0,1366 | 0,4353 |
| TcCLB.509127.70 | hypothetical protein, conserved | 0,1365 | 0,2259 |
| TcCLB.506211.30 | intraflagellar transport protein 80, putative (IFT80) | 0,1365 | 0,2613 |

| | | | |
|---|---|---|---|
| TcCLB.508879.50 | hypothetical protein | 0,1365 | 0,2664 |
| TcCLB.508961.30 | trafficking protein particle complex subunit 3, putative | 0,1365 | 0,4833 |
| TcCLB.506797.120 | hypothetical protein, conserved | 0,1364 | 0,1862 |
| TcCLB.506241.50 | hypothetical protein, conserved | 0,1364 | 0,2459 |
| TcCLB.511351.41 | choline/carnitine O-acyltransferase, putative | 0,1364 | 0,5292 |
| TcCLB.504105.120 | hypothetical protein, conserved | 0,136 | 0,08653 |
| TcCLB.504741.50 | hypothetical protein, conserved | 0,1359 | 0,3038 |
| TcCLB.511739.20 | hypothetical protein, conserved | 0,1358 | 0,2162 |
| TcCLB.505333.20 | ATP-dependent DEAD/H RNA helicase (pseudogene), puta | 0,1358 | 0,7065 |
| TcCLB.511727.50 | hypothetical protein, conserved | 0,1357 | 0,1107 |
| TcCLB.511507.70 | calpain-like cysteine peptidase, putative | 0,1357 | 0,3519 |
| TcCLB.508479.213 | hypothetical protein, conserved | 0,1354 | 0,4836 |
| TcCLB.509875.140 | retrotransposon hot spot protein (RHS, pseudogene), puta | 0,1354 | 0,6653 |
| TcCLB.508913.25 | trans-sialidase, Group I, putative | 0,1354 | 0,7259 |
| TcCLB.508681.20 | hypothetical protein | 0,1354 | 0,7576 |
| TcCLB.509207.20 | histone H2A, putative | 0,1354 | 0,7697 |
| TcCLB.505231.10 | dynein heavy chain, putative | 0,1353 | 0,208 |
| TcCLB.510791.60 | mucin-associated surface protein (MASP, pseudogene), pu | 0,1352 | 0,4598 |
| TcCLB.511717.20 | hypothetical protein, conserved | 0,1351 | 0,1922 |
| TcCLB.509461.30 | hypothetical protein, conserved | 0,1351 | 0,2291 |
| TcCLB.508307.110 | hypothetical protein, conserved | 0,135 | 0,07506 |
| TcCLB.511633.60 | structural maintenance of chromosome (SMC), putative | 0,135 | 0,09422 |
| TcCLB.506887.69 | dual specificity protein phosphatase, putative | 0,135 | 0,2889 |
| TcCLB.507927.50 | metallo-beta-lactamase superfamily, putative | 0,135 | 0,3743 |
| TcCLB.507031.80 | hypothetical protein, conserved | 0,1349 | 0,09871 |
| TcCLB.508643.9 | pre-mRNA-splicing factor ATP-dependent RNA helicase, p | 0,1349 | 0,247 |
| TcCLB.506457.40 | hypoxanthine-guanine phosphoribosyltransferase, putative | 0,1349 | 0,2497 |
| TcCLB.504045.120 | hypothetical protein, conserved | 0,1348 | 0,24 |
| TcCLB.509767.210 | vesicle transport v-SNARE 11, putative | 0,1348 | 0,304 |
| TcCLB.507937.14 | mucin TcMUCII (pseudogene), putative | 0,1346 | 0,7261 |
| TcCLB.503525.4 | hypothetical protein, conserved | 0,1345 | 0,1963 |
| TcCLB.507857.30 | hypothetical protein, conserved | 0,1345 | 0,1994 |
| TcCLB.506201.30 | serine/threonine-protein phosphatase PP1 beta | 0,1345 | 0,2237 |
| TcCLB.510509.20 | hypothetical protein, conserved | 0,1345 | 0,2336 |
| TcCLB.507017.60 | hypothetical protein, conserved | 0,1344 | 0,2664 |
| TcCLB.506217.10 | hypothetical protein | 0,1342 | 0,6013 |
| TcCLB.511487.51 | hypothetical protein | 0,1342 | 0,6904 |
| TcCLB.504427.50 | AMPK subunit beta, putative (AMPKB) | 0,1341 | 0,2838 |
| TcCLB.505075.30 | hypothetical protein, conserved | 0,1341 | 0,3744 |
| TcCLB.507017.30 | hypothetical protein, conserved | 0,134 | 0,3919 |
| TcCLB.510889.110 | primase 2, putative | 0,1339 | 0,2481 |
| TcCLB.504081.342 | hypothetical protein (pseudogene) | 0,1339 | 0,7517 |
| TcCLB.508207.210 | hypothetical protein, conserved | 0,1338 | 0,3584 |
| TcCLB.511571.49 | nucleoside phosphatase, putative | 0,1337 | 0,09215 |
| TcCLB.507519.170 | hypothetical protein, conserved | 0,1337 | 0,2568 |

| | | | |
|---|---|---|---|
| TcCLB.506531.40 | hypothetical protein, conserved | 0,1337 | 0,3196 |
| TcCLB.510597.20 | hypothetical protein, conserved | 0,1336 | 0,3569 |
| TcCLB.508989.70 | hypothetical protein, conserved | 0,1334 | 0,2262 |
| TcCLB.511817.60 | hypothetical protein, conserved | 0,1334 | 0,3504 |
| TcCLB.503595.19 | ATP-dependent DEAD/H RNA helicase, putative (fragment | 0,1333 | 0,3226 |
| TcCLB.507257.140 | hypothetical protein, conserved | 0,1331 | 0,3458 |
| TcCLB.508881.10 | hypothetical protein, conserved | 0,1331 | 0,438 |
| TcCLB.503653.20 | hypothetical protein, conserved | 0,1331 | 0,5058 |
| TcCLB.503593.70 | eukaryotic peptide chain release factor subunit 1, putative | 0,1329 | 0,2527 |
| TcCLB.511737.80 | hypothetical protein | 0,1329 | 0,3552 |
| TcCLB.507059.60 | orotidine-5-phosphate decarboxylase/orotate phosphoribo | 0,1328 | 0,2543 |
| TcCLB.509695.120 | epsilon tubulin, putative | 0,1328 | 0,2993 |
| TcCLB.422921.10 | hypothetical protein, conserved | 0,1328 | 0,4341 |
| TcCLB.504261.30 | trans-sialidase (pseudogene), putative | 0,1328 | 0,7109 |
| TcCLB.506529.535 | hypothetical protein, conserved | 0,1326 | 0,7863 |
| TcCLB.506871.80 | RNA polymerase-like protein, putative | 0,1325 | 0,4011 |
| TcCLB.509723.10 | protein kinase, putative | 0,1324 | 0,5732 |
| TcCLB.511881.40 | protein transport protein Sec24C, putative | 0,1321 | 0,2464 |
| TcCLB.506485.70 | hypothetical protein, conserved | 0,1319 | 0,2557 |
| TcCLB.507511.30 | RNA guanylyltransferase, putative | 0,1319 | 0,2697 |
| TcCLB.503915.9 | hypothetical protein, conserved (fragment) | 0,1319 | 0,2753 |
| TcCLB.508641.230 | hypothetical protein, conserved | 0,1318 | 0,2061 |
| TcCLB.507505.20 | DnaJ chaperone protein (pseudogene), putative | 0,1318 | 0,2895 |
| TcCLB.507805.20 | hypothetical protein | 0,1318 | 0,35 |
| TcCLB.504147.180 | hypothetical protein, conserved | 0,1318 | 0,4267 |
| TcCLB.511577.130 | poly(A) export protein, putative | 0,1317 | 0,3367 |
| TcCLB.506297.10 | hypothetical protein | 0,1316 | 0,1827 |
| TcCLB.509895.110 | hypothetical protein, conserved | 0,1316 | 0,6294 |
| TcCLB.506159.30 | dispersed gene family protein 1 (DGF-1), putative | 0,1316 | 0,6299 |
| TcCLB.509237.140 | calpain-like cysteine peptidase, putative | 0,1315 | 0,3258 |
| TcCLB.509401.30 | cysteine peptidase, putative | 0,1314 | 0,0777 |
| TcCLB.508349.20 | tRNA nucleotidyltransferase, putative | 0,1314 | 0,2181 |
| TcCLB.508625.30 | ARP2/3 complex subunit, putative | 0,1314 | 0,5048 |
| TcCLB.511233.70 | hypothetical protein, conserved | 0,1312 | 0,1385 |
| TcCLB.510265.20 | 26S protease regulatory subunit, putative | 0,1312 | 0,1611 |
| TcCLB.508347.120 | hypothetical protein, conserved | 0,1312 | 0,2015 |
| TcCLB.510121.90 | hypothetical protein, conserved | 0,1312 | 0,2153 |
| TcCLB.503983.10 | hypothetical protein, conserved | 0,1311 | 0,4914 |
| TcCLB.506925.170 | hypothetical protein, conserved | 0,1311 | 0,7632 |
| TcCLB.511585.320 | retrotransposon hot spot (RHS) protein, putative | 0,1309 | 0,2483 |
| TcCLB.503793.10 | 2-oxoglutarate dehydrogenase subunit, putative | 0,1307 | 0,3836 |
| TcCLB.507093.160 | hypothetical protein, conserved | 0,1306 | 0,2713 |
| TcCLB.506857.30 | hypothetical protein, conserved | 0,1306 | 0,374 |
| TcCLB.507677.70 | hypothetical protein, conserved | 0,1303 | 0,3061 |
| TcCLB.506697.40 | helicase, putative | 0,1302 | 0,1777 |

| | | | |
|---|---|---|---|
| TcCLB.508141.40 | ATP-dependent RNA helicase, putative | 0,1301 | 0,07543 |
| TcCLB.509213.80 | hypothetical protein, conserved | 0,1301 | 0,3741 |
| TcCLB.508719.60 | kinetoplast DNA-associated protein, putative | 0,1301 | 0,4718 |
| TcCLB.503781.30 | hypothetical protein, conserved | 0,1301 | 0,5933 |
| TcCLB.511423.90 | zeta tubulin, putative | 0,13 | 0,2711 |
| TcCLB.508409.30 | DNA-directed RNA polymerase II subunit 3, putative (RPB: | 0,1298 | 0,188 |
| TcCLB.510877.140 | hypothetical protein, conserved | 0,1298 | 0,3298 |
| TcCLB.511003.120 | hypothetical protein, conserved | 0,1298 | 0,4168 |
| TcCLB.510377.210 | trans-sialidase (pseudogene), putative | 0,1298 | 0,7082 |
| TcCLB.506661.90 | hypothetical protein, conserved | 0,1297 | 0,24 |
| TcCLB.507603.190 | hypothetical protein, conserved | 0,1297 | 0,2687 |
| TcCLB.507841.14 | retrotransposon hot spot (RHS) protein, putative (fragmer | 0,1295 | 0,354 |
| TcCLB.503697.50 | hypothetical protein, conserved | 0,1294 | 0,1469 |
| TcCLB.441923.10 | UDP-Gal or UDP-GlcNAc-dependent glycosyltransferase, p | 0,1294 | 0,444 |
| TcCLB.511903.290 | hypothetical protein, conserved | 0,1292 | 0,1238 |
| TcCLB.479961.20 | retrotransposon hot spot protein (RHS, pseudogene), puta | 0,129 | 0,457 |
| TcCLB.503797.20 | centrin (pseudogene), putative | 0,1287 | 0,4943 |
| TcCLB.511229.70 | hypothetical protein, conserved | 0,1286 | 0,4037 |
| TcCLB.511635.80 | hypothetical protein, conserved | 0,1284 | 0,4008 |
| TcCLB.511643.4 | hypothetical protein, conserved | 0,1284 | 0,637 |
| TcCLB.509985.50 | PrimPol-like protein 1, putative (ppl1) | 0,1283 | 0,2311 |
| TcCLB.507623.90 | hypothetical protein, conserved | 0,1282 | 0,2173 |
| TcCLB.509979.60 | sec1 family transport protein, putative | 0,128 | 0,3081 |
| TcCLB.503495.10 | dispersed gene family protein 1 (DGF-1), putative | 0,128 | 0,5344 |
| TcCLB.509259.130 | protein kinase, putative | 0,128 | 0,7698 |
| TcCLB.508257.250 | hypothetical protein, conserved | 0,1279 | 0,06353 |
| TcCLB.511867.209 | dynein heavy chain, putative (fragment) | 0,1279 | 0,2598 |
| TcCLB.508741.250 | hypothetical protein | 0,1278 | 0,1914 |
| TcCLB.510579.14 | hypothetical protein, conserved | 0,1277 | 0,2684 |
| TcCLB.507723.110 | acetyltransferase, putative | 0,1277 | 0,3258 |
| TcCLB.506619.10 | sedoheptulose-1,7-bisphosphatase, putative | 0,1277 | 0,3495 |
| TcCLB.511001.110 | hypothetical protein, conserved | 0,1277 | 0,3944 |
| TcCLB.508209.90 | 10 kDa heat shock protein, putative | 0,1277 | 0,776 |
| TcCLB.510297.70 | hypothetical protein | 0,1276 | 0,3235 |
| TcCLB.503801.79 | hypothetical protein, conserved | 0,1276 | 0,4394 |
| TcCLB.510517.100 | hypothetical protein, conserved | 0,1275 | 0,363 |
| TcCLB.508199.80 | EF hand, putative | 0,1274 | 0,2527 |
| TcCLB.475705.9 | hypothetical protein, conserved | 0,1274 | 0,4013 |
| TcCLB.469435.10 | hypothetical protein, conserved | 0,1274 | 0,4114 |
| TcCLB.507519.40 | hypothetical protein | 0,1273 | 0,2291 |
| TcCLB.508205.10 | hypothetical protein, conserved | 0,1273 | 0,3826 |
| TcCLB.504797.120 | hypothetical protein, conserved | 0,1273 | 0,4238 |
| TcCLB.509917.6 | ATP-dependent DEAD/H RNA helicase (pseudogene), puta | 0,1273 | 0,53 |
| TcCLB.510595.58 | hypothetical protein, conserved | 0,1273 | 0,6192 |
| TcCLB.507723.160 | hypothetical protein, conserved | 0,1272 | 0,1975 |

| | | | |
|---|---|---|---|
| TcCLB.507795.90 | hypothetical protein, conserved | 0,1272 | 0,4462 |
| TcCLB.509109.190 | hypothetical protein, conserved | 0,1271 | 0,4301 |
| TcCLB.509503.9 | dispersed gene family protein 1 (DGF-1), putative (fragme | 0,1271 | 0,6893 |
| TcCLB.506247.70 | nuclear pore complex protein (NUP155), putative | 0,127 | 0,1037 |
| TcCLB.508965.100 | hypothetical protein, conserved | 0,127 | 0,1984 |
| TcCLB.506213.100 | hypothetical protein, conserved | 0,1268 | 0,3234 |
| TcCLB.506925.260 | protein kinase, putative | 0,1268 | 0,3248 |
| TcCLB.508989.30 | RNA-binding protein, putative | 0,1267 | 0,2397 |
| TcCLB.511421.20 | Non cytoplasmic domain containing protein, conserved | 0,1266 | 0,2759 |
| TcCLB.509233.40 | hypothetical protein, conserved | 0,1266 | 0,3891 |
| TcCLB.506227.180 | ras-related protein RAB5B, putative (RAB5B) | 0,1265 | 0,2117 |
| TcCLB.507951.270 | LETM1 and EF-hand domain-containing protein 1, putative | 0,1265 | 0,234 |
| TcCLB.508407.10 | membrane associated protein, putative | 0,1265 | 0,3593 |
| TcCLB.510329.340 | hypothetical protein, conserved | 0,1264 | 0,3127 |
| TcCLB.509037.30 | hypothetical protein, conserved | 0,1263 | 0,3471 |
| TcCLB.508421.40 | hypothetical protein, conserved (pseudogene) | 0,1263 | 0,7401 |
| TcCLB.506577.110 | hypothetical protein, conserved | 0,1262 | 0,4996 |
| TcCLB.507867.70 | prefoldin subunit, putative | 0,1261 | 0,4618 |
| TcCLB.508349.10 | hypothetical protein, conserved | 0,126 | 0,3683 |
| TcCLB.511871.4 | dynein heavy chain, putative | 0,1258 | 0,4673 |
| TcCLB.503999.70 | ras-family member, GTP-binding protein, putative | 0,1257 | 0,221 |
| TcCLB.503425.10 | hypothetical protein, conserved | 0,1254 | 0,3111 |
| TcCLB.503421.20 | hypothetical protein, conserved (pseudogene) | 0,1254 | 0,4565 |
| TcCLB.510221.30 | hypothetical protein, conserved | 0,1253 | 0,1377 |
| TcCLB.510101.200 | zinc finger ccch and cchc domain-containing protein | 0,1253 | 0,2309 |
| TcCLB.510969.4 | hypothetical protein | 0,1253 | 0,6309 |
| TcCLB.509769.130 | DNA polymerase zeta catalytic subunit, putative | 0,125 | 0,1109 |
| TcCLB.508257.80 | hypothetical protein, conserved | 0,125 | 0,1523 |
| TcCLB.506241.150 | hypothetical protein | 0,125 | 0,3 |
| TcCLB.511265.10 | dispersed gene family protein 1 (DGF-1, pseudogene), put | 0,125 | 0,4615 |
| TcCLB.509805.130 | prolyl-tRNA synthetase, putative | 0,1249 | 0,1363 |
| TcCLB.510303.230 | hypothetical protein, conserved | 0,1247 | 0,213 |
| TcCLB.508741.300 | proteasome regulatory non-ATPase subunit 6, putative | 0,1246 | 0,2157 |
| TcCLB.503885.60 | hypothetical protein, conserved | 0,1246 | 0,2993 |
| TcCLB.511389.90 | hypothetical protein, conserved | 0,1246 | 0,3839 |
| TcCLB.509151.80 | hypothetical protein, conserved | 0,1246 | 0,3939 |
| TcCLB.419417.19 | hypothetical protein, conserved | 0,1245 | 0,5881 |
| TcCLB.506175.80 | hypothetical protein, conserved | 0,1243 | 0,5007 |
| TcCLB.508547.110 | ATP-dependent DEAD/H RNA helicase, putative | 0,1242 | 0,19 |
| TcCLB.509369.40 | dispersed gene family protein 1 (DGF-1, pseudogene), put | 0,1242 | 0,6572 |
| TcCLB.510861.80 | hypothetical protein | 0,1241 | 0,3226 |
| TcCLB.509999.110 | serine/threonine-protein kinase, putative | 0,1241 | 0,3332 |
| TcCLB.506445.80 | hypothetical protein, conserved | 0,1241 | 0,3418 |
| TcCLB.509695.10 | aspartyl-tRNA synthetase, putative | 0,124 | 0,2876 |
| TcCLB.506925.500 | hypothetical protein, conserved | 0,1239 | 0,2335 |

| | | | |
|---|---|---|---|
| TcCLB.508625.100 | hypothetical protein, conserved (pseudogene) | 0,1238 | 0,2889 |
| TcCLB.511633.20 | hypothetical protein, conserved | 0,1238 | 0,3121 |
| TcCLB.504899.30 | hypothetical protein | 0,1238 | 0,4519 |
| TcCLB.506015.50 | hypothetical protein (pseudogene) | 0,1238 | 0,7129 |
| TcCLB.509795.89 | phosphoprotein lepp12, putative | 0,1236 | 0,361 |
| TcCLB.506829.20 | hypothetical protein, conserved | 0,1235 | 0,2349 |
| TcCLB.505807.190 | polynucleotide kinase 3'-phosphatase, putative | 0,1235 | 0,3753 |
| TcCLB.507515.20 | hypothetical protein, conserved | 0,1234 | 0,1517 |
| TcCLB.507087.40 | kinesin, putative | 0,1234 | 0,1936 |
| TcCLB.511727.280 | hypothetical protein, conserved | 0,1233 | 0,1642 |
| TcCLB.480975.10 | hypothetical protein, conserved | 0,1233 | 0,3529 |
| TcCLB.511145.80 | RNA polymerase II, putative | 0,1233 | 0,408 |
| TcCLB.510431.200 | LisH domain-containing protein FOPNL, putative (fopnl) | 0,1232 | 0,3596 |
| TcCLB.511049.20 | hypothetical protein, conserved | 0,1232 | 0,372 |
| TcCLB.508543.120 | hypothetical protein, conserved | 0,1232 | 0,499 |
| TcCLB.505193.40 | SUMO1/Ulp2, putative | 0,1231 | 0,2306 |
| TcCLB.504147.340 | hypothetical protein, conserved | 0,123 | 0,1597 |
| TcCLB.506559.270 | sarcoplasmic reticulum glycoprotein, putative | 0,1229 | 0,1492 |
| TcCLB.506945.120 | chromosomal passenger protein, putative (CPC1) | 0,1228 | 0,3074 |
| TcCLB.509253.10 | hypothetical protein, conserved | 0,1228 | 0,3404 |
| TcCLB.508831.90 | hypothetical protein, conserved | 0,1228 | 0,489 |
| TcCLB.509163.50 | hypothetical protein | 0,1228 | 0,6693 |
| TcCLB.503865.60 | hypothetical protein, conserved | 0,1227 | 0,2861 |
| TcCLB.510221.10 | RNA pseudouridylate synthase, putative | 0,1227 | 0,2997 |
| TcCLB.474341.9 | dynein heavy chain, putative | 0,1226 | 0,2845 |
| TcCLB.507681.250 | hypothetical protein, conserved | 0,1225 | 0,3504 |
| TcCLB.503433.20 | hypothetical protein, conserved | 0,1224 | 0,3522 |
| TcCLB.510717.30 | 2-oxoglutarate dehydrogenase E1 component, putative | 0,1223 | 0,1684 |
| TcCLB.511803.70 | hypothetical protein, conserved | 0,1223 | 0,3859 |
| TcCLB.511001.100 | SNF-7-like protein, putative | 0,1221 | 0,3176 |
| TcCLB.509047.70 | hypothetical protein, conserved | 0,122 | 0,432 |
| TcCLB.509011.70 | adrenodoxin precursor, putative | 0,122 | 0,4869 |
| TcCLB.511255.625 | surface protease GP63 (pseudogene), putative | 0,122 | 0,8261 |
| TcCLB.511863.20 | hypothetical protein, conserved | 0,1219 | 0,3145 |
| TcCLB.508257.200 | hypothetical protein, conserved | 0,1218 | 0,2017 |
| TcCLB.506811.70 | hypothetical protein | 0,1216 | 0,2323 |
| TcCLB.510879.200 | hypothetical protein, conserved | 0,1216 | 0,4204 |
| TcCLB.508689.40 | hypothetical protein, conserved | 0,1215 | 0,1953 |
| TcCLB.506155.110 | hypothetical protein, conserved | 0,1213 | 0,3119 |
| TcCLB.509877.20 | hypothetical protein, conserved | 0,1212 | 0,2928 |
| TcCLB.507857.40 | ATP-dependent DEAD/H RNA helicase, putative | 0,121 | 0,1993 |
| TcCLB.504121.10 | KU70 protein, putative | 0,1208 | 0,4145 |
| TcCLB.506919.130 | retrotransposon hot spot protein (RHS, pseudogene), puta | 0,1208 | 0,7208 |
| TcCLB.506493.40 | hypothetical protein, conserved | 0,1207 | 0,2963 |
| TcCLB.507843.10 | retrotransposon hot spot (RHS) protein, putative | 0,1207 | 0,4274 |

| | | | |
|---|---|---|---|
| TcCLB.511537.50 | chaperone DNAJ protein, putative | 0,1206 | 0,2734 |
| TcCLB.466529.9 | hypothetical protein, conserved | 0,1206 | 0,4771 |
| TcCLB.507119.20 | hypothetical protein, conserved (pseudogene) | 0,1205 | 0,2483 |
| TcCLB.505123.4 | ATP-dependent RNA helicase, putative (fragment) | 0,1205 | 0,2721 |
| TcCLB.511071.45 | tuzin-like (pseudogene), putative | 0,1205 | 0,2979 |
| TcCLB.507739.70 | hypothetical protein, conserved | 0,1204 | 0,2783 |
| TcCLB.511589.20 | hypothetical protein | 0,1204 | 0,3883 |
| TcCLB.510357.150 | hypothetical protein, conserved | 0,1203 | 0,3925 |
| TcCLB.506369.9 | hypothetical protein, conserved | 0,1202 | 0,5482 |
| TcCLB.504089.50 | flagellar radial spoke component, putative | 0,12 | 0,2881 |
| TcCLB.511367.354 | U1 small nuclear ribonucleoprotein C, putative | 0,1198 | 0,4179 |
| TcCLB.506885.124 | Enriched in surface-labeled proteome protein 19 (SLS3) | 0,1197 | 0,295 |
| TcCLB.508987.39 | hypothetical protein, conserved | 0,1197 | 0,3196 |
| TcCLB.507895.80 | hypothetical protein, conserved | 0,1195 | 0,3053 |
| TcCLB.506337.220 | hypothetical protein | 0,1195 | 0,4492 |
| TcCLB.508857.80 | hypothetical protein, conserved | 0,1194 | 0,4937 |
| TcCLB.509571.60 | hypothetical protein, conserved | 0,1192 | 0,3411 |
| TcCLB.507601.20 | hypothetical protein, conserved | 0,1192 | 0,5593 |
| TcCLB.507677.110 | hypothetical protein | 0,1191 | 0,2569 |
| TcCLB.508205.39 | dynein light chain, putative | 0,1191 | 0,3406 |
| TcCLB.507603.100 | hypothetical protein, conserved | 0,119 | 0,381 |
| TcCLB.509717.100 | membrane-bound acid phosphatase 2, putative | 0,1188 | 0,2467 |
| TcCLB.508479.240 | hypothetical protein, conserved | 0,1188 | 0,2854 |
| TcCLB.511353.40 | eukaryotic translation initiation factor 4E (eIF4E) interacti | 0,1188 | 0,3438 |
| TcCLB.510767.69 | hypothetical protein, conserved | 0,1186 | 0,53 |
| TcCLB.460061.10 | hypothetical protein (fragment) | 0,1186 | 0,5979 |
| TcCLB.437575.18 | transporter, putative | 0,1185 | 0,5475 |
| TcCLB.511727.310 | Leucine-rich repeat-containing protein ODA7 (ODA7) | 0,1184 | 0,2966 |
| TcCLB.508949.40 | hypothetical protein, conserved | 0,1184 | 0,4133 |
| TcCLB.504149.150 | hypothetical protein, conserved | 0,1182 | 0,1691 |
| TcCLB.508889.19 | hypothetical protein | 0,1181 | 0,3889 |
| TcCLB.505975.41 | retrotransposon hot spot protein (RHS, pseudogene), puta | 0,118 | 0,3219 |
| TcCLB.510181.40 | hypothetical protein, conserved | 0,118 | 0,5736 |
| TcCLB.509199.10 | Elongation factor G 2, mitochondrial, putative (EF-G2) | 0,1179 | 0,2645 |
| TcCLB.506113.10 | retrotransposon hot spot protein (RHS, pseudogene), puta | 0,1179 | 0,3597 |
| TcCLB.511419.80 | phosphatidylinositol-4-phosphate 5-kinase related | 0,1179 | 0,4302 |
| TcCLB.506743.190 | hypothetical protein, conserved | 0,1179 | 0,4524 |
| TcCLB.509667.31 | hypothetical protein, conserved (pseudogene) | 0,1179 | 0,5999 |
| TcCLB.509849.20 | hypothetical protein, conserved | 0,1177 | 0,2684 |
| TcCLB.506925.280 | hypothetical protein, conserved | 0,1177 | 0,3791 |
| TcCLB.506893.70 | hypothetical protein, conserved | 0,1176 | 0,3995 |
| TcCLB.511217.70 | hypothetical protein, conserved | 0,1175 | 0,2059 |
| TcCLB.508817.40 | hypothetical protein, conserved | 0,1175 | 0,2512 |
| TcCLB.506529.330 | hypothetical protein, conserved | 0,1175 | 0,4674 |
| TcCLB.505065.30 | carnitine O-palmitoyltransferase (pseudogene), putative | 0,1175 | 0,7366 |

| | | | |
|---|---|---|---|
| TcCLB.511741.50 | C-1-tetrahydrofolate synthase, cytoplasmic, putative | 0,1173 | 0,4283 |
| TcCLB.510797.10 | 2-oxoglutarate dehydrogenase subunit, putative | 0,1172 | 0,4082 |
| TcCLB.510091.130 | hypothetical protein, conserved | 0,117 | 0,2494 |
| TcCLB.508317.60 | hypothetical protein, conserved | 0,117 | 0,4657 |
| TcCLB.508479.450 | hypothetical protein, conserved (pseudogene) | 0,1169 | 0,4858 |
| TcCLB.504867.50 | hypothetical protein, conserved | 0,1168 | 0,2659 |
| TcCLB.507009.130 | hypothetical protein, conserved | 0,1167 | 0,2813 |
| TcCLB.507723.140 | hypothetical protein, conserved | 0,1167 | 0,4152 |
| TcCLB.511217.150 | hypothetical protein, conserved | 0,1166 | 0,4341 |
| TcCLB.510123.60 | hypothetical protein, conserved | 0,1164 | 0,3552 |
| TcCLB.508347.100 | hypothetical protein, conserved | 0,1163 | 0,2685 |
| TcCLB.509877.10 | hypothetical protein, conserved | 0,1163 | 0,4032 |
| TcCLB.503399.29 | hydrolase-like protein | 0,1163 | 0,5748 |
| TcCLB.504159.30 | hypothetical protein, conserved | 0,1162 | 0,3831 |
| TcCLB.509979.40 | hypothetical protein, conserved | 0,1161 | 0,4123 |
| TcCLB.507837.40 | hypothetical protein, conserved | 0,1161 | 0,4499 |
| TcCLB.509069.14 | hypothetical protein, conserved | 0,116 | 0,3716 |
| TcCLB.510659.100 | hypothetical protein, conserved | 0,1159 | 0,4796 |
| TcCLB.510311.60 | hypothetical protein, conserved | 0,1156 | 0,1905 |
| TcCLB.505989.78 | hypothetical protein, conserved | 0,1156 | 0,4338 |
| TcCLB.508173.230 | hypothetical protein, conserved | 0,1155 | 0,2954 |
| TcCLB.508667.30 | hypothetical protein, conserved | 0,1154 | 0,3701 |
| TcCLB.506825.180 | protein kinase, putative | 0,1153 | 0,1793 |
| TcCLB.507601.90 | hypothetical protein, conserved | 0,1153 | 0,3175 |
| TcCLB.510579.60 | heat shock protein, putative | 0,1153 | 0,3472 |
| TcCLB.507579.30 | surface protease GP63 (pseudogene), putative | 0,1153 | 0,7197 |
| TcCLB.506111.20 | dispersed gene family protein 1 (DGF-1), putative | 0,1152 | 0,5295 |
| TcCLB.507257.180 | hypothetical protein | 0,115 | 0,2625 |
| TcCLB.511409.50 | hypothetical protein, conserved | 0,1149 | 0,3543 |
| TcCLB.506265.45 | chaperone DNAJ protein, putative | 0,1148 | 0,4865 |
| TcCLB.508117.30 | elongation factor 1-gamma (EF-1-gamma, pseudogene), | 0,1146 | 0,632 |
| TcCLB.506241.70 | calcium motive p-type ATPase, putative | 0,1143 | 0,2143 |
| TcCLB.508113.14 | mucin TcMUC (pseudogene), putative | 0,1143 | 0,5903 |
| TcCLB.503455.30 | hypothetical protein, conserved | 0,1142 | 0,3157 |
| TcCLB.507045.5 | trans-sialidase (pseudogene), putative | 0,1141 | 0,7841 |
| TcCLB.506595.120 | trans-sialidase (pseudogene), putative | 0,1139 | 0,3994 |
| TcCLB.506067.189 | trans-sialidase (pseudogene), putative | 0,1139 | 0,7127 |
| TcCLB.508139.120 | hypothetical protein | 0,1139 | 0,7339 |
| TcCLB.510609.90 | hypothetical protein, conserved | 0,1138 | 0,2625 |
| TcCLB.507611.320 | hypothetical protein, conserved | 0,1138 | 0,4321 |
| TcCLB.506215.10 | hypothetical protein | 0,1137 | 0,6505 |
| TcCLB.509167.60 | hypothetical protein, conserved | 0,1135 | 0,2581 |
| TcCLB.511449.40 | mucin TcMUCII (pseudogene), putative | 0,1135 | 0,7225 |
| TcCLB.511277.480 | hypothetical protein, conserved | 0,1132 | 0,3146 |
| TcCLB.506203.20 | dynein light chain, putative | 0,1132 | 0,3607 |

| | | | |
|---|---|---|---|
| TcCLB.508357.30 | hypothetical protein, conserved | 0,1131 | 0,4241 |
| TcCLB.506175.60 | GTPase activator protein, putative | 0,1129 | 0,312 |
| TcCLB.506811.140 | hypothetical protein, conserved | 0,1129 | 0,3315 |
| TcCLB.507277.10 | hypothetical protein, conserved | 0,1129 | 0,3931 |
| TcCLB.507881.30 | glycine dehydrogenase [decarboxylating] (pseudogene), pu | 0,1126 | 0,6853 |
| TcCLB.508887.30 | hypothetical protein | 0,1126 | 0,699 |
| TcCLB.503617.15 | U6 snRNA-associated Sm-like protein LSm3p | 0,1119 | 0,6484 |
| TcCLB.509161.60 | hypothetical protein, conserved | 0,1118 | 0,2736 |
| TcCLB.510099.100 | hypothetical protein, conserved | 0,1117 | 0,2761 |
| TcCLB.508013.120 | elongation factor 1-alpha (ef-1-alpha), putative | 0,1116 | 0,756 |
| TcCLB.510877.60 | ruvB-like DNA helicase, putative (RUVBL) | 0,1114 | 0,1879 |
| TcCLB.510073.80 | hypothetical protein, conserved | 0,1114 | 0,255 |
| TcCLB.504507.10 | hypothetical protein, conserved | 0,1114 | 0,4267 |
| TcCLB.507519.90 | hypothetical protein, conserved | 0,1113 | 0,4363 |
| TcCLB.511313.20 | hypothetical protein, conserved | 0,1112 | 0,4029 |
| TcCLB.506211.50 | membrane protein, putative | 0,1112 | 0,449 |
| TcCLB.506999.40 | metacaspase 5, putative | 0,111 | 0,2557 |
| TcCLB.506627.110 | hypothetical protein, conserved | 0,111 | 0,27 |
| TcCLB.505989.70 | surface protease GP63, putative | 0,111 | 0,4091 |
| TcCLB.504171.30 | hypothetical protein, conserved | 0,1109 | 0,2693 |
| TcCLB.504251.20 | hypothetical protein, conserved | 0,1106 | 0,4817 |
| TcCLB.507513.100 | hypothetical protein, conserved | 0,1105 | 0,3194 |
| TcCLB.511321.40 | hypothetical protein, conserved | 0,1105 | 0,3389 |
| TcCLB.504109.40 | hypothetical protein, conserved | 0,1105 | 0,3961 |
| TcCLB.511295.70 | hypothetical protein, conserved (fragment) | 0,1105 | 0,7616 |
| TcCLB.511577.70 | hypothetical protein, conserved | 0,1104 | 0,3903 |
| TcCLB.509213.100 | 3-ketoacyl-CoA reductase, putative | 0,1102 | 0,2804 |
| TcCLB.511587.75 | hypothetical protein | 0,1102 | 0,7516 |
| TcCLB.508127.9 | hypothetical protein, conserved | 0,1101 | 0,3196 |
| TcCLB.511751.180 | pseudouridine synthase A-like protein, putative | 0,11 | 0,3799 |
| TcCLB.503789.10 | hypothetical protein | 0,11 | 0,5381 |
| TcCLB.503927.9 | dynein light intermediate chain, putative | 0,1098 | 0,3299 |
| TcCLB.506941.9 | endosomal trafficking protein RME-8, putative | 0,1098 | 0,3596 |
| TcCLB.506223.120 | phosphatidylinositol 3-related kinase, putative | 0,1097 | 0,2122 |
| TcCLB.504431.90 | U2 small nuclear ribonucleoprotein B, putative | 0,1097 | 0,5396 |
| TcCLB.508237.18 | SNF2 DNA repair protein, putative | 0,1096 | 0,3816 |
| TcCLB.506739.10 | Eukaryotic translation initiation factor 4 gamma type 1, pu | 0,1096 | 0,4223 |
| TcCLB.510491.20 | dynein light chain, putative | 0,1096 | 0,5185 |
| TcCLB.510877.120 | NADPH-dependent FMN/FAD containing oxidoreductase, p | 0,1095 | 0,3088 |
| TcCLB.511801.70 | hypothetical protein, conserved | 0,1094 | 0,3823 |
| TcCLB.510733.60 | DNA polymerase epsilon catalytic subunit, putative | 0,1093 | 0,2492 |
| TcCLB.509805.100 | hypothetical protein, conserved | 0,1093 | 0,3054 |
| TcCLB.506529.610 | hypothetical protein | 0,1092 | 0,6625 |
| TcCLB.511481.20 | hypothetical protein, conserved | 0,1091 | 0,5459 |
| TcCLB.510575.150 | hypothetical protein, conserved | 0,1089 | 0,2945 |

| | | | |
|---|---|---|---|
| TcCLB.507611.110 | retrotransposon hot spot (RHS) protein, putative | 0,1089 | 0,4114 |
| TcCLB.506401.190 | hypothetical protein, conserved | 0,1089 | 0,4536 |
| TcCLB.508355.40 | choline/carnitine O-acetyltransferase, putative | 0,1089 | 0,5371 |
| TcCLB.511439.80 | hypothetical protein, conserved | 0,1087 | 0,24 |
| TcCLB.506203.25 | hypothetical protein | 0,1087 | 0,3359 |
| TcCLB.509899.110 | orotidine-5-phosphate decarboxylase/orotate phosphoribc | 0,1086 | 0,322 |
| TcCLB.504149.201 | ATP-dependent DEAD/H RNA helicase (pseudogene), puta | 0,1086 | 0,6424 |
| TcCLB.506629.150 | hypothetical protein, conserved | 0,1083 | 0,3801 |
| TcCLB.507519.100 | hypothetical protein, conserved | 0,1083 | 0,6013 |
| TcCLB.506479.67 | hypothetical protein, conserved | 0,1082 | 0,3304 |
| TcCLB.506777.70 | cdc2-related kinase 8, putative (CRK8) | 0,1081 | 0,3221 |
| TcCLB.503887.10 | hypothetical protein, conserved | 0,1079 | 0,5118 |
| TcCLB.508465.130 | hypothetical protein, conserved | 0,1078 | 0,354 |
| TcCLB.511395.50 | hypothetical protein, conserved | 0,1076 | 0,3933 |
| TcCLB.506773.170 | hypothetical protein, conserved | 0,1074 | 0,3823 |
| TcCLB.511291.100 | Vesicle-trafficking protein SEC22, putative (SEC22) | 0,1074 | 0,4188 |
| TcCLB.508181.20 | 6-phosphofructo-2-kinase 1 | 0,1071 | 0,2827 |
| TcCLB.505965.30 | prenyltransferase, putative | 0,1071 | 0,4468 |
| TcCLB.507031.130 | hypothetical protein, conserved | 0,1069 | 0,3421 |
| TcCLB.508501.220 | hypothetical protein, conserved | 0,1069 | 0,3489 |
| TcCLB.506367.100 | hypothetical protein, conserved | 0,1069 | 0,4727 |
| TcCLB.506513.50 | protein kinase, putative | 0,1068 | 0,4082 |
| TcCLB.506503.4 | transporter, putative | 0,1068 | 0,4418 |
| TcCLB.506145.40 | hypothetical protein, conserved | 0,1066 | 0,2444 |
| TcCLB.507165.30 | hypothetical protein, conserved | 0,1065 | 0,4327 |
| TcCLB.504109.90 | cysteine peptidase, Clan CA, family C12, putative | 0,1064 | 0,4746 |
| TcCLB.509127.104 | ARP2/3 complex subunit, putative | 0,1064 | 0,6075 |
| TcCLB.506319.5 | hypothetical protein, conserved (fragment) | 0,1064 | 0,6959 |
| TcCLB.509995.29 | hypothetical protein, conserved | 0,1063 | 0,5298 |
| TcCLB.509911.80 | hypothetical protein, conserved | 0,1062 | 0,4387 |
| TcCLB.506507.40 | trans-sialidase (pseudogene), putative | 0,106 | 0,4179 |
| TcCLB.503669.20 | hypothetical protein, conserved | 0,1057 | 0,465 |
| TcCLB.511529.30 | hypothetical protein, conserved | 0,1057 | 0,4739 |
| TcCLB.510913.10 | dispersed gene family protein 1 (DGF-1), putative (fragme | 0,1057 | 0,5833 |
| TcCLB.510421.240 | diacylglycerol acyltransferase, putative | 0,1056 | 0,507 |
| TcCLB.510329.300 | hypothetical protein, conserved | 0,1056 | 0,5082 |
| TcCLB.507057.50 | hypothetical protein, conserved | 0,1055 | 0,3968 |
| TcCLB.509631.100 | hypothetical protein, conserved (pseudogene) | 0,1054 | 0,8155 |
| TcCLB.510535.70 | hypothetical protein, conserved | 0,1053 | 0,3876 |
| TcCLB.507023.170 | hypothetical protein, conserved | 0,1053 | 0,3909 |
| TcCLB.511127.410 | hypothetical protein, conserved | 0,1052 | 0,2664 |
| TcCLB.506223.90 | hypothetical protein, conserved | 0,1052 | 0,4037 |
| TcCLB.507675.30 | hypothetical protein, conserved | 0,1051 | 0,571 |
| TcCLB.508257.210 | hypothetical protein, conserved | 0,105 | 0,398 |
| TcCLB.506729.90 | hypothetical protein, conserved | 0,1049 | 0,3841 |

| | | | |
|---|---|---|---|
| TcCLB.508515.110 | hypothetical protein, conserved | 0,1048 | 0,3075 |
| TcCLB.506241.19 | dispersed gene family protein 1 (DGF-1), putative | 0,1048 | 0,5955 |
| TcCLB.439301.9 | hypothetical protein, conserved | 0,1047 | 0,3227 |
| TcCLB.507089.154 | hypothetical protein, conserved | 0,1046 | 0,2224 |
| TcCLB.510121.80 | hypothetical protein, conserved | 0,1045 | 0,2994 |
| TcCLB.509103.10 | hypothetical protein, conserved | 0,1044 | 0,4143 |
| TcCLB.508461.350 | hypothetical protein, conserved | 0,1043 | 0,3382 |
| TcCLB.507053.120 | hypothetical protein, conserved | 0,1043 | 0,426 |
| TcCLB.509335.20 | retrotransposon hot spot protein (RHS, pseudogene), puta | 0,1043 | 0,5777 |
| TcCLB.506401.100 | DNA polymerase sigma, putative | 0,1042 | 0,2214 |
| TcCLB.506959.74 | hypothetical protein, conserved | 0,1042 | 0,3443 |
| TcCLB.511349.61 | retrotransposon hot spot protein (RHS, pseudogene), puta | 0,1041 | 0,8268 |
| TcCLB.436535.10 | hypothetical protein, conserved | 0,104 | 0,5559 |
| TcCLB.511843.35 | hypothetical protein | 0,104 | 0,6974 |
| TcCLB.510315.4 | hypothetical protein, conserved (fragment) | 0,1035 | 0,53 |
| TcCLB.511323.20 | hypothetical protein, conserved | 0,1035 | 0,5563 |
| TcCLB.507305.20 | casein kinase, putative | 0,1034 | 0,2657 |
| TcCLB.508699.110 | elongation factor G2-like protein, putative | 0,1034 | 0,5456 |
| TcCLB.510265.80 | hypothetical protein, conserved | 0,1033 | 0,3608 |
| TcCLB.506393.166 | ATP-dependent DEAD/H RNA helicase (pseudogene), puta | 0,1033 | 0,749 |
| TcCLB.504255.50 | hypothetical protein, conserved | 0,1032 | 0,4168 |
| TcCLB.511871.130 | retrotransposon hot spot (RHS) protein, putative | 0,1032 | 0,4433 |
| TcCLB.507081.120 | hypothetical protein, conserved | 0,1031 | 0,4513 |
| TcCLB.507649.50 | hypothetical protein, conserved | 0,1029 | 0,2559 |
| TcCLB.503713.20 | hypothetical protein, conserved | 0,1028 | 0,539 |
| TcCLB.510783.30 | dispersed gene family protein 1 (DGF-1, pseudogene), put | 0,1028 | 0,7527 |
| TcCLB.509537.70 | hypothetical protein, conserved | 0,1027 | 0,4151 |
| TcCLB.509109.40 | hypothetical protein, conserved | 0,1027 | 0,4607 |
| TcCLB.511867.90 | hypothetical protein, conserved | 0,1027 | 0,472 |
| TcCLB.508779.9 | hypothetical protein, conserved (fragment) | 0,1023 | 0,4663 |
| TcCLB.437349.10 | hypothetical protein, conserved | 0,1021 | 0,5413 |
| TcCLB.511189.100 | hypothetical protein, conserved | 0,102 | 0,4373 |
| TcCLB.506679.80 | SNF1-related protein kinases, putative | 0,1019 | 0,3397 |
| TcCLB.511903.270 | hypothetical protein, conserved | 0,1019 | 0,6204 |
| TcCLB.508409.130 | protein transport protein Sec24A, putative | 0,1018 | 0,202 |
| TcCLB.503791.10 | TPR-repeat-containing chaperone protein DNAJ, putative ( | 0,1018 | 0,4446 |
| TcCLB.477935.9 | calpain-like cysteine peptidase (pseudogene), putative | 0,1018 | 0,5215 |
| TcCLB.508247.140 | hypothetical protein | 0,1018 | 0,7167 |
| TcCLB.507601.80 | hypothetical protein, conserved | 0,1017 | 0,5232 |
| TcCLB.507715.40 | GTP-binding elongation factor Tu family, putative | 0,1016 | 0,2518 |
| TcCLB.511817.220 | kinesin, putative | 0,1016 | 0,3354 |
| TcCLB.510595.49 | hypothetical protein, conserved | 0,1015 | 0,4083 |
| TcCLB.511735.30 | hypothetical protein, conserved | 0,1015 | 0,5945 |
| TcCLB.510747.34 | hypothetical protein, conserved | 0,1014 | 0,4995 |
| TcCLB.507019.120 | hypothetical protein, conserved | 0,1013 | 0,5166 |

| | | | |
|---|---|---|---|
| TcCLB.508707.70 | hypothetical protein, conserved | 0,1012 | 0,2127 |
| TcCLB.504343.10 | trans-sialidase, Group II, putative | 0,1012 | 0,5443 |
| TcCLB.510119.30 | hypothetical protein, conserved | 0,1011 | 0,5009 |
| TcCLB.510283.90 | hypothetical protein, conserved | 0,101 | 0,3874 |
| TcCLB.507991.70 | hypothetical protein, conserved | 0,1009 | 0,4547 |
| TcCLB.510181.140 | hypothetical protein, conserved | 0,1008 | 0,5051 |
| TcCLB.506883.69 | hypothetical protein, conserved | 0,1008 | 0,637 |
| TcCLB.511321.60 | hypothetical protein, conserved | 0,1008 | 0,6923 |
| TcCLB.510231.20 | hypothetical protein | 0,1007 | 0,3935 |
| TcCLB.508409.40 | Putative intraflagellar transport protein B2 (PIFTB2) | 0,1004 | 0,3522 |
| TcCLB.509951.20 | hypothetical protein, conserved | 0,1004 | 0,6882 |
| TcCLB.506627.90 | hypothetical protein | 0,1003 | 0,2528 |
| TcCLB.509229.140 | hypothetical protein, conserved | 0,1003 | 0,3201 |
| TcCLB.508623.10 | trans-sialidase (pseudogene), putative | 0,1003 | 0,3396 |
| TcCLB.506701.29 | hypothetical protein, conserved | 0,1003 | 0,557 |
| TcCLB.503717.130 | trans-sialidase (pseudogene), putative | 0,1003 | 0,7404 |
| TcCLB.503943.14 | hypothetical protein, conserved | 0,1002 | 0,4549 |
| TcCLB.511727.160 | hypothetical protein, conserved | 0,1001 | 0,232 |
| TcCLB.511909.70 | hypothetical protein, conserved | 0,1001 | 0,5594 |
| TcCLB.511733.40 | FUN14 family, putative (POMP32) | 0,1001 | 0,5876 |
| TcCLB.511673.9 | hypothetical protein, conserved | 0,09985 | 0,6422 |
| TcCLB.510291.30 | protein phosphatase 2C-like, putative | 0,09982 | 0,4898 |
| TcCLB.511109.30 | dynein intermediate chain, putative | 0,09969 | 0,4574 |
| TcCLB.510975.40 | hypothetical protein, conserved | 0,09954 | 0,2598 |
| TcCLB.511001.160 | hypothetical protein, conserved | 0,09953 | 0,3313 |
| TcCLB.511589.150 | hypothetical protein, conserved | 0,09953 | 0,5882 |
| TcCLB.509837.20 | hypothetical protein, conserved | 0,09948 | 0,2602 |
| TcCLB.509167.10 | hypothetical protein, conserved | 0,09932 | 0,3927 |
| TcCLB.511277.180 | hypothetical protein, conserved | 0,09931 | 0,3518 |
| TcCLB.510613.20 | kinesin, putative | 0,09924 | 0,4262 |
| TcCLB.510517.60 | hypothetical protein, conserved | 0,09921 | 0,4856 |
| TcCLB.509179.160 | hypothetical protein, conserved | 0,09913 | 0,4226 |
| TcCLB.511525.29 | hypothetical protein, conserved | 0,0991 | 0,321 |
| TcCLB.506277.10 | mucin-associated surface protein (MASP), putative | 0,09909 | 0,6935 |
| TcCLB.509243.50 | sigma-adaptin 3, putative | 0,09889 | 0,5364 |
| TcCLB.507491.90 | protein kinase, putative | 0,09885 | 0,4922 |
| TcCLB.509911.60 | hypothetical protein | 0,09879 | 0,3343 |
| TcCLB.511925.60 | hypothetical protein, conserved | 0,09872 | 0,4891 |
| TcCLB.504149.55 | acyl-CoA thioesterase, putative | 0,09869 | 0,2807 |
| TcCLB.511711.80 | Ras-related protein RAB2B, putative (RAB2B) | 0,09863 | 0,4328 |
| TcCLB.506677.30 | hypothetical protein, conserved | 0,09861 | 0,3948 |
| TcCLB.510661.160 | hypothetical protein, conserved | 0,09855 | 0,4883 |
| TcCLB.509641.60 | hypothetical protein, conserved | 0,09848 | 0,5536 |
| TcCLB.511321.26 | vesicle transport v-SNARE protein, putative | 0,09845 | 0,528 |
| TcCLB.508445.40 | hypothetical protein, conserved | 0,09837 | 0,5214 |

| | | | |
|---|---|---|---|
| TcCLB.508815.90 | hypothetical protein, conserved | 0,09829 | 0,4509 |
| TcCLB.506925.330 | hypothetical protein, conserved | 0,09816 | 0,2512 |
| TcCLB.507381.31 | retrotransposon hot spot protein (RHS, pseudogene), puta | 0,0981 | 0,4383 |
| TcCLB.511283.294 | b9 domain-containing protein 1 (b9d1) | 0,09806 | 0,3349 |
| TcCLB.504149.117 | hypothetical protein, conserved | 0,098 | 0,38 |
| TcCLB.510859.40 | hypothetical protein, conserved | 0,09797 | 0,5104 |
| TcCLB.510835.30 | hypothetical protein, conserved | 0,09785 | 0,5023 |
| TcCLB.506825.120 | hypothetical protein, conserved | 0,0977 | 0,2631 |
| TcCLB.510659.44 | cysteine desulfurase, putative | 0,09763 | 0,3921 |
| TcCLB.511725.110 | hypothetical protein, conserved | 0,09746 | 0,5747 |
| TcCLB.509165.49 | damage-specific DNA binding protein, putative | 0,09731 | 0,4682 |
| TcCLB.511587.100 | retrotransposon hot spot (RHS) protein, putative | 0,09713 | 0,373 |
| TcCLB.510155.115 | hypothetical protein, conserved | 0,0969 | 0,4017 |
| TcCLB.511661.100 | protein kinase, putative | 0,09676 | 0,2771 |
| TcCLB.506789.170 | ATPase, putative | 0,09674 | 0,4423 |
| TcCLB.508441.50 | hypothetical protein, conserved | 0,09668 | 0,6122 |
| TcCLB.510659.219 | hypothetical protein, conserved | 0,09663 | 0,4966 |
| TcCLB.508717.20 | NADH dehydrogenase, putative | 0,09661 | 0,3895 |
| TcCLB.503689.10 | small GTP-binding protein Rab7, putative | 0,09657 | 0,4321 |
| TcCLB.511685.50 | hypothetical protein, conserved | 0,09634 | 0,5518 |
| TcCLB.507559.90 | hypothetical protein | 0,09628 | 0,2664 |
| TcCLB.506649.80 | RNA-binding protein, putative (DRBD3) | 0,09622 | 0,2129 |
| TcCLB.440099.10 | hypothetical protein | 0,09622 | 0,5216 |
| TcCLB.481729.20 | hypothetical protein, conserved | 0,09615 | 0,7086 |
| TcCLB.510351.130 | hypothetical protein, conserved | 0,09603 | 0,5067 |
| TcCLB.510187.430 | DNA mismatch repair protein MSH6, putative (MSH6) | 0,09595 | 0,2356 |
| TcCLB.508963.10 | Flagellar Member 3 (pseudogene) | 0,09594 | 0,3204 |
| TcCLB.503685.30 | carnitine/choline acetyltransferase, putative | 0,09594 | 0,5177 |
| TcCLB.504013.30 | hypothetical protein, conserved | 0,09578 | 0,5823 |
| TcCLB.508851.50 | hypothetical protein, conserved | 0,09573 | 0,4914 |
| TcCLB.507951.250 | tRNA isopentenyltransferase, putative | 0,09571 | 0,4631 |
| TcCLB.508193.20 | retrotransposon hot spot protein (RHS, pseudogene), puta | 0,09565 | 0,3059 |
| TcCLB.506821.140 | histone deacetylase, putative | 0,0956 | 0,4432 |
| TcCLB.510681.20 | myosin heavy chain kinase A, putative | 0,09547 | 0,4843 |
| TcCLB.503401.20 | hypothetical protein, conserved | 0,09542 | 0,5961 |
| TcCLB.508515.170 | retrotransposon hot spot protein (RHS, pseudogene), puta | 0,09541 | 0,4681 |
| TcCLB.511025.10 | dynein light intermediate chain, putative | 0,0953 | 0,4 |
| TcCLB.504741.220 | hypothetical protein, conserved | 0,09516 | 0,4868 |
| TcCLB.508379.20 | hypothetical protein, conserved | 0,09511 | 0,671 |
| TcCLB.503811.80 | hypothetical protein, conserved | 0,0951 | 0,4626 |
| TcCLB.504717.20 | hypothetical protein, conserved | 0,09501 | 0,2856 |
| TcCLB.506517.100 | hypothetical protein, conserved | 0,09479 | 0,362 |
| TcCLB.509149.50 | hypothetical protein, conserved | 0,09472 | 0,3272 |
| TcCLB.506147.150 | hypothetical protein, conserved | 0,09466 | 0,3993 |
| TcCLB.504085.50 | ATP-dependent DEAD/H RNA helicase, putative | 0,09458 | 0,4029 |

| | | | |
|---|---|---|---|
| TcCLB.509145.10 | hypothetical protein, conserved | 0,09448 | 0,4398 |
| TcCLB.509563.30 | hypothetical protein, conserved | 0,09438 | 0,3902 |
| TcCLB.511511.150 | sphingosine 1-phosphate lyase, putative (SPL) | 0,09433 | 0,4966 |
| TcCLB.404843.20 | hypothetical protein, conserved | 0,09426 | 0,4516 |
| TcCLB.506339.25 | hypothetical protein, conserved (pseudogene) | 0,09424 | 0,6352 |
| TcCLB.508153.90 | hypothetical protein | 0,09421 | 0,5062 |
| TcCLB.508289.41 | syntaxin binding protein (pseudogene), putative | 0,09419 | 0,7478 |
| TcCLB.511269.10 | hypothetical protein, conserved | 0,09418 | 0,3889 |
| TcCLB.507467.100 | hypothetical protein | 0,09418 | 0,3935 |
| TcCLB.508355.350 | hypothetical protein, conserved | 0,09403 | 0,4495 |
| TcCLB.510173.40 | MRB1 complex subunit MRB3010 (MRB3010) | 0,09402 | 0,3722 |
| TcCLB.511255.618 | syntaxin binding protein (pseudogene), putative | 0,09401 | 0,846 |
| TcCLB.506539.10 | trans-sialidase (pseudogene), putative | 0,094 | 0,7255 |
| TcCLB.509827.4 | retrotransposon hot spot (RHS) protein, putative | 0,09388 | 0,6248 |
| TcCLB.507819.20 | retrotransposon hot spot protein (RHS, pseudogene), puta | 0,09377 | 0,5029 |
| TcCLB.434931.10 | paraxonemal rod protein PAR2, putative | 0,09372 | 0,4493 |
| TcCLB.510719.110 | actin-like protein, putative | 0,09367 | 0,5009 |
| TcCLB.508153.370 | hypothetical protein, conserved | 0,09364 | 0,3712 |
| TcCLB.506405.60 | hypothetical protein, conserved | 0,09353 | 0,5478 |
| TcCLB.506331.30 | hypothetical protein | 0,09352 | 0,7974 |
| TcCLB.507047.20 | dispersed gene family protein 1 (DGF-1, pseudogene), put | 0,0935 | 0,6026 |
| TcCLB.508541.240 | casein kinase, putative | 0,09341 | 0,7739 |
| TcCLB.504109.140 | DNA-directed RNA polymerase I subunit, putative | 0,09324 | 0,5069 |
| TcCLB.511217.160 | hypothetical protein, conserved | 0,0932 | 0,417 |
| TcCLB.433273.10 | dynein heavy chain, putative (fragment) | 0,09315 | 0,474 |
| TcCLB.508931.50 | hypothetical protein | 0,09308 | 0,4601 |
| TcCLB.510713.20 | dispersed gene family protein 1 (DGF-1), putative | 0,09308 | 0,6109 |
| TcCLB.510329.210 | SNF1-related protein kinase catalytic subunit alpha, putat | 0,09304 | 0,3364 |
| TcCLB.511585.220 | calcium/potassium channel (CAKC), putative | 0,09299 | 0,3353 |
| TcCLB.503451.9 | hypothetical protein, conserved (fragment) | 0,09291 | 0,2986 |
| TcCLB.508027.50 | hypothetical protein, conserved | 0,0929 | 0,5209 |
| TcCLB.509287.140 | dispersed gene family protein 1 (DGF-1), putative | 0,09262 | 0,6517 |
| TcCLB.503521.60 | hypothetical protein, conserved | 0,09257 | 0,6327 |
| TcCLB.510889.240 | hypothetical protein, conserved | 0,09252 | 0,5611 |
| TcCLB.511071.30 | hypothetical protein, conserved | 0,09239 | 0,5733 |
| TcCLB.511109.20 | hypothetical protein, conserved | 0,09231 | 0,6242 |
| TcCLB.509857.10 | hypothetical protein, conserved | 0,09225 | 0,4534 |
| TcCLB.510007.10 | hypothetical protein, conserved | 0,09221 | 0,5924 |
| TcCLB.509631.140 | flagellar attachment zone protein, putative (FAZ1) | 0,0922 | 0,3098 |
| TcCLB.506227.50 | hypothetical protein, conserved | 0,09217 | 0,5159 |
| TcCLB.508515.60 | hypothetical protein, conserved | 0,09217 | 0,5421 |
| TcCLB.503847.40 | hypothetical protein, conserved | 0,09216 | 0,4635 |
| TcCLB.416883.18 | proteasome alpha 2 subunit, putative | 0,0919 | 0,4666 |
| TcCLB.509967.7 | hypothetical protein, conserved | 0,09174 | 0,6852 |
| TcCLB.510533.210 | hypothetical protein, conserved (pseudogene) | 0,09173 | 0,5124 |

| | | | |
|---|---|---|---|
| TcCLB.424383.9 | trans-sialidase (pseudogene), putative | 0,09168 | 0,7107 |
| TcCLB.506885.350 | proteasome alpha 1 subunit, putative | 0,09157 | 0,4058 |
| TcCLB.419417.4 | ADP-ribosylation factor, putative (fragment) | 0,09157 | 0,4489 |
| TcCLB.511823.50 | palmitoyl acyltransferase 11, putative | 0,09147 | 0,4647 |
| TcCLB.506289.70 | hypothetical protein, conserved | 0,09127 | 0,3503 |
| TcCLB.510421.150 | protein kinase, putative | 0,09125 | 0,2967 |
| TcCLB.510329.350 | hypothetical protein, conserved | 0,09111 | 0,4431 |
| TcCLB.510145.59 | exonuclease, putative | 0,09095 | 0,4849 |
| TcCLB.508815.4 | RNA polymerase I, putative | 0,09095 | 0,7207 |
| TcCLB.511501.50 | GTPase activating protein, putative | 0,09061 | 0,4313 |
| TcCLB.506717.219 | GMP synthase (pseudogene), putative | 0,09059 | 0,516 |
| TcCLB.511277.580 | RNA-binding protein 29, putative (RBP29) | 0,09053 | 0,3358 |
| TcCLB.506181.130 | hypothetical protein | 0,09052 | 0,571 |
| TcCLB.503987.70 | ribosome biogenesis protein TSR3, putative (tsr3) | 0,09038 | 0,5955 |
| TcCLB.452699.5 | trans-sialidase (pseudogene), putative | 0,09027 | 0,8279 |
| TcCLB.506885.70 | polyadenylate-binding protein 1, putative | 0,0902 | 0,4522 |
| TcCLB.511277.410 | hypothetical protein, conserved | 0,09003 | 0,4228 |
| TcCLB.509049.51 | hypothetical protein, conserved | 0,08991 | 0,571 |
| TcCLB.506211.10 | lipoic acid synthetase, mitochondrial precursor, putative | 0,08982 | 0,5033 |
| TcCLB.508601.90 | Small Surface Antigen (LDK) | 0,08966 | 0,4438 |
| TcCLB.504241.57 | retrotransposon hot spot protein (RHS, pseudogene), puta | 0,08965 | 0,7989 |
| TcCLB.511735.70 | hypothetical protein, conserved | 0,08961 | 0,4891 |
| TcCLB.508153.1060 | hypothetical protein, conserved | 0,08961 | 0,669 |
| TcCLB.511535.39 | hypothetical protein, conserved | 0,08951 | 0,6952 |
| TcCLB.507599.10 | Ribosome biogenesis protein Tsr3, putative (fragment) | 0,08948 | 0,728 |
| TcCLB.511409.60 | hypothetical protein, conserved | 0,08946 | 0,5457 |
| TcCLB.511751.150 | hypothetical protein, conserved | 0,08938 | 0,5667 |
| TcCLB.510941.10 | hypothetical protein, conserved | 0,08937 | 0,4657 |
| TcCLB.504099.30 | hypothetical protein, conserved (pseudogene) | 0,08936 | 0,6229 |
| TcCLB.508179.30 | hypothetical protein, conserved | 0,08914 | 0,5265 |
| TcCLB.504131.40 | hypothetical protein, conserved | 0,08892 | 0,4266 |
| TcCLB.506963.90 | calmodulin, putative | 0,08892 | 0,4602 |
| TcCLB.506857.80 | kinesin, putative | 0,08882 | 0,3983 |
| TcCLB.506839.60 | tyrosine specific protein phosphatase, putative | 0,08875 | 0,4168 |
| TcCLB.503929.5 | hypothetical protein, conserved | 0,08859 | 0,6577 |
| TcCLB.503667.10 | trans-sialidase, putative | 0,08852 | 0,7031 |
| TcCLB.506443.50 | hypothetical protein, conserved | 0,08826 | 0,4534 |
| TcCLB.510719.220 | hypothetical protein | 0,08826 | 0,5494 |
| TcCLB.509099.110 | hypothetical protein, conserved | 0,08808 | 0,7059 |
| TcCLB.511235.10 | hypothetical protein, conserved (pseudogene) | 0,08795 | 0,7852 |
| TcCLB.507735.20 | hypothetical protein, conserved | 0,08791 | 0,3457 |
| TcCLB.506401.230 | hypothetical protein, conserved | 0,08782 | 0,3839 |
| TcCLB.415187.10 | hypothetical protein, conserved | 0,08781 | 0,7371 |
| TcCLB.506287.180 | hypothetical protein, conserved | 0,08766 | 0,488 |
| TcCLB.509803.40 | 5-oxoprolinase, putative | 0,08764 | 0,5082 |

| | | | |
|---|---|---|---|
| TcCLB.511725.270 | hypothetical protein, conserved | 0,0876 | 0,3345 |
| TcCLB.507953.110 | retrotransposon hot spot protein (RHS, pseudogene), puta | 0,0876 | 0,8383 |
| TcCLB.508781.90 | hypothetical protein, conserved | 0,08737 | 0,6048 |
| TcCLB.503635.80 | hypothetical protein, conserved | 0,0873 | 0,4837 |
| TcCLB.509595.44 | hypothetical protein, conserved | 0,08706 | 0,6165 |
| TcCLB.509647.50 | hypothetical protein, conserved | 0,087 | 0,2975 |
| TcCLB.506529.90 | puromycin-sensitive aminopeptidase-like protein, putative | 0,087 | 0,5294 |
| TcCLB.509067.60 | Tetratricopeptide-like helical domain containing protein (I | 0,08691 | 0,189 |
| TcCLB.510235.6 | hypothetical protein | 0,08685 | 0,753 |
| TcCLB.503889.20 | mucin TcMUC (pseudogene), putative | 0,0867 | 0,5636 |
| TcCLB.511587.10 | hypothetical protein | 0,08669 | 0,7989 |
| TcCLB.511901.10 | Trypanosome basal body component protein (TBBC) | 0,08658 | 0,5943 |
| TcCLB.509053.90 | hypothetical protein, conserved | 0,0865 | 0,5412 |
| TcCLB.506363.50 | hypothetical protein, conserved | 0,08645 | 0,3509 |
| TcCLB.510053.50 | vesicle transport v-SNARE protein, putative | 0,08642 | 0,6505 |
| TcCLB.505987.30 | hypothetical protein, conserved | 0,08621 | 0,4735 |
| TcCLB.509757.20 | leucine carboxyl methyltransferase, putative | 0,08609 | 0,5215 |
| TcCLB.510663.64 | mitochondrial carrier protein, putative | 0,08597 | 0,6119 |
| TcCLB.506829.60 | hypothetical protein, conserved | 0,08593 | 0,5228 |
| TcCLB.506989.240 | hypothetical protein, conserved (pseudogene) | 0,08591 | 0,3215 |
| TcCLB.511503.9 | hypothetical protein, conserved (pseudogene) | 0,08584 | 0,5691 |
| TcCLB.511907.270 | ubiquitin-conjugating enzyme E2, putative | 0,08574 | 0,4429 |
| TcCLB.511349.110 | retrotransposon hot spot protein (RHS, pseudogene), puta | 0,08571 | 0,7458 |
| TcCLB.508139.110 | hypothetical protein | 0,08566 | 0,7892 |
| TcCLB.509925.50 | retrotransposon hot spot protein (RHS, pseudogene), puta | 0,08564 | 0,6516 |
| TcCLB.503757.30 | hypothetical protein, conserved | 0,08546 | 0,6097 |
| TcCLB.510575.170 | hypothetical protein, conserved | 0,08536 | 0,5011 |
| TcCLB.510253.20 | hypothetical protein, conserved | 0,08535 | 0,3517 |
| TcCLB.507519.80 | eukaryotic peptide chain release factor subunit 1, putative | 0,08535 | 0,4952 |
| TcCLB.432787.10 | mitochondrial RNA binding complex 1 subunit, putative (fr | 0,08523 | 0,5906 |
| TcCLB.506709.20 | hypothetical protein, conserved | 0,08494 | 0,3975 |
| TcCLB.508207.194 | hypothetical protein, conserved | 0,08467 | 0,4384 |
| TcCLB.506817.40 | hypothetical protein, conserved | 0,08466 | 0,4236 |
| TcCLB.509857.20 | hypothetical protein, conserved | 0,08453 | 0,5129 |
| TcCLB.508265.80 | hypothetical protein | 0,08442 | 0,5066 |
| TcCLB.505937.70 | trans-sialidase (pseudogene), putative | 0,08416 | 0,701 |
| TcCLB.510359.150 | ATP-dependent helicase, putative | 0,08415 | 0,4025 |
| TcCLB.508593.80 | hypothetical protein, conserved | 0,08406 | 0,4423 |
| TcCLB.510755.70 | hypothetical protein, conserved | 0,08391 | 0,5295 |
| TcCLB.507821.130 | trans-sialidase, Group II, putative | 0,08379 | 0,7177 |
| TcCLB.511473.6 | ATP-dependent DEAD/H RNA helicase (pseudogene), puta | 0,08378 | 0,7873 |
| TcCLB.511859.70 | hypothetical protein, conserved | 0,08369 | 0,5684 |
| TcCLB.511517.150 | hypothetical protein, conserved | 0,08366 | 0,6125 |
| TcCLB.508693.20 | DNA topoisomerase IB, large subunit | 0,08365 | 0,4762 |
| TcCLB.504769.140 | trans-sialidase, Group VIII, putative | 0,08349 | 0,7809 |

| | | | |
|---|---|---|---|
| TcCLB.507611.389 | hypothetical protein, conserved | 0,08341 | 0,704 |
| TcCLB.506147.210 | hypothetical protein, conserved | 0,08339 | 0,5871 |
| TcCLB.509049.40 | hypothetical protein, conserved | 0,08333 | 0,5572 |
| TcCLB.504867.60 | hypothetical protein, conserved | 0,08329 | 0,4655 |
| TcCLB.509033.109 | hypothetical protein, conserved | 0,08328 | 0,6791 |
| TcCLB.504153.110 | hypothetical protein, conserved | 0,08324 | 0,4727 |
| TcCLB.507775.40 | hypothetical protein, conserved | 0,08323 | 0,6093 |
| TcCLB.510483.320 | retrotransposon hot spot protein (RHS, pseudogene), puta | 0,08322 | 0,642 |
| TcCLB.508269.60 | exosome-associated protein 4, putative (EAP4) | 0,08308 | 0,6554 |
| TcCLB.504103.30 | myosin heavy chain, putative | 0,08297 | 0,4916 |
| TcCLB.511217.120 | peptidyl-prolyl cis-trans isomerase, putative | 0,08297 | 0,5716 |
| TcCLB.506303.150 | hypothetical protein, conserved | 0,0825 | 0,4761 |
| TcCLB.510773.60 | hypothetical protein | 0,08246 | 0,7706 |
| TcCLB.503899.100 | hypothetical protein | 0,08238 | 0,5106 |
| TcCLB.509597.10 | hypothetical protein, conserved | 0,08218 | 0,4374 |
| TcCLB.509213.150 | hypothetical protein, conserved | 0,08207 | 0,5439 |
| TcCLB.503453.90 | hypothetical protein, conserved | 0,08192 | 0,4707 |
| TcCLB.506195.60 | hypothetical protein, conserved | 0,08192 | 0,5424 |
| TcCLB.508153.960 | hypothetical protein, conserved | 0,08186 | 0,5064 |
| TcCLB.506775.130 | hypothetical protein, conserved | 0,08185 | 0,4721 |
| TcCLB.503945.10 | eukaryotic translation initiation factor 1A, putative | 0,08182 | 0,5443 |
| TcCLB.510647.40 | syntaxin, putative | 0,08176 | 0,504 |
| TcCLB.510243.110 | hypothetical protein, conserved | 0,08175 | 0,5563 |
| TcCLB.508347.90 | hypothetical protein, conserved | 0,08172 | 0,4034 |
| TcCLB.511867.100 | hypothetical protein, conserved | 0,08162 | 0,5032 |
| TcCLB.511867.170 | hypothetical protein, conserved | 0,08152 | 0,4427 |
| TcCLB.508775.70 | dispersed gene family protein 1 (DGF-1), putative | 0,08142 | 0,8368 |
| TcCLB.510181.120 | hypothetical protein, conserved | 0,08133 | 0,5201 |
| TcCLB.508451.5 | mucin-associated surface protein (MASP, pseudogene), pu | 0,08133 | 0,6712 |
| TcCLB.503983.29 | hypothetical protein, conserved | 0,08125 | 0,6621 |
| TcCLB.511391.220 | hypothetical protein, conserved | 0,08124 | 0,2474 |
| TcCLB.508277.230 | nucleolar RNA binding protein, putative | 0,08122 | 0,4372 |
| TcCLB.508915.9 | hypothetical protein, conserved | 0,081 | 0,5622 |
| TcCLB.509913.30 | hypothetical protein, conserved | 0,08098 | 0,3988 |
| TcCLB.503521.20 | hypothetical protein, conserved | 0,08095 | 0,4835 |
| TcCLB.510311.160 | hypothetical protein, conserved | 0,08081 | 0,542 |
| TcCLB.509683.70 | dynein light chain Tctex-type, putative | 0,08069 | 0,5395 |
| TcCLB.508405.50 | hypothetical protein, conserved | 0,08056 | 0,5801 |
| TcCLB.511183.130 | hypothetical protein, conserved (pseudogene) | 0,08048 | 0,8401 |
| TcCLB.506415.30 | hypothetical protein, conserved | 0,08042 | 0,6156 |
| TcCLB.511293.20 | hypothetical protein, conserved | 0,08034 | 0,379 |
| TcCLB.510249.14 | hypothetical protein, conserved | 0,08031 | 0,8886 |
| TcCLB.507755.10 | hypothetical protein | 0,07976 | 0,5344 |
| TcCLB.509769.4 | Meckel syndrome type 1 protein, putative (fragment) (mk | 0,07963 | 0,5147 |
| TcCLB.510305.40 | hypothetical protein, conserved | 0,07957 | 0,5106 |

| | | | |
|---|---|---|---|
| TcCLB.510691.70 | hypothetical protein, conserved | 0,07938 | 0,6547 |
| TcCLB.509541.4 | paraflagellar rod component par4, putative | 0,07937 | 0,6611 |
| TcCLB.506871.150 | hypothetical protein, conserved | 0,07935 | 0,4762 |
| TcCLB.508153.610 | Putative intraflagellar transport protein C3 (PIFTC3) | 0,07922 | 0,5069 |
| TcCLB.506743.10 | hypothetical protein | 0,07918 | 0,6799 |
| TcCLB.510901.40 | hypothetical protein, conserved | 0,07912 | 0,66 |
| TcCLB.507621.10 | retrotransposon hot spot protein (RHS, pseudogene), puta | 0,07907 | 0,7878 |
| TcCLB.507093.120 | hypothetical protein, conserved | 0,07899 | 0,5806 |
| TcCLB.506313.10 | retrotransposon hot spot (RHS) protein, putative (fragmer | 0,07897 | 0,4889 |
| TcCLB.509471.20 | Present in the outer mitochondrial membrane proteome 3 | 0,07896 | 0,3665 |
| TcCLB.509597.4 | hypothetical protein, conserved | 0,07896 | 0,5058 |
| TcCLB.506885.400 | 40 kDa cyclophilin, putative (CYP40) | 0,07894 | 0,5123 |
| TcCLB.510291.7 | TPR-repeat-containing chaperone protein DNAJ, putative ( | 0,07892 | 0,5592 |
| TcCLB.511121.11 | retrotransposon hot spot (RHS) protein, putative (fragmer | 0,07891 | 0,619 |
| TcCLB.508153.350 | hypothetical protein, conserved | 0,07888 | 0,6153 |
| TcCLB.509573.40 | hypothetical protein, conserved | 0,07883 | 0,7429 |
| TcCLB.507035.75 | retrotransposon hot spot protein (RHS, pseudogene), puta | 0,07866 | 0,5649 |
| TcCLB.505965.86 | tRNA pseudouridine synthase (pseudogene), putative | 0,07865 | 0,496 |
| TcCLB.506913.20 | protein kinase, putative (fragment) | 0,07856 | 0,7351 |
| TcCLB.503435.14 | hypothetical protein | 0,07854 | 0,5148 |
| TcCLB.504001.3 | flagellar radial spoke component, putative (fragment) | 0,07849 | 0,6483 |
| TcCLB.504203.60 | TPR-repeat-containing chaperone protein DNAJ, putative ( | 0,07811 | 0,5032 |
| TcCLB.508197.20 | ADP-ribosylation factor, putative | 0,07801 | 0,654 |
| TcCLB.509829.9 | retrotransposon hot spot (RHS) protein, putative (fragmer | 0,07794 | 0,7128 |
| TcCLB.511245.50 | hypothetical protein, conserved | 0,07788 | 0,4035 |
| TcCLB.510599.40 | hypothetical protein, conserved | 0,07786 | 0,4486 |
| TcCLB.506975.34 | aminoacylase, putative | 0,07773 | 0,7237 |
| TcCLB.510963.5 | hypothetical protein, conserved (fragment) | 0,07771 | 0,699 |
| TcCLB.511807.50 | hypothetical protein, conserved | 0,07764 | 0,3476 |
| TcCLB.509769.36 | palmitoyl acyltransferase 12, putative | 0,07746 | 0,5369 |
| TcCLB.506739.180 | hypothetical protein, conserved | 0,07736 | 0,423 |
| TcCLB.504167.14 | hypothetical protein | 0,07716 | 0,5997 |
| TcCLB.511859.161 | hypothetical protein | 0,07712 | 0,6799 |
| TcCLB.508373.29 | orotidine-5-phosphate decarboxylase/orotate phosphoribo | 0,07691 | 0,5686 |
| TcCLB.506851.10 | NIMA-related kinase, putative | 0,07679 | 0,3788 |
| TcCLB.506959.30 | ATP-dependent DEAD/H RNA helicase, putative | 0,07677 | 0,4309 |
| TcCLB.510533.10 | Cytoplasmic dynein 1 heavy chain (DYNC1H1), putative | 0,07671 | 0,4652 |
| TcCLB.506577.20 | hypothetical protein | 0,07665 | 0,4741 |
| TcCLB.506287.100 | hypothetical protein, conserved | 0,07662 | 0,5485 |
| TcCLB.506133.75 | mucin TcMUCII (pseudogene), putative | 0,0766 | 0,7739 |
| TcCLB.504253.20 | hypothetical protein, conserved | 0,07648 | 0,539 |
| TcCLB.506629.80 | vesicular transport protein (CDC48 homologue), putative | 0,07638 | 0,3209 |
| TcCLB.507883.10 | glucose-6-phosphate 1-dehydrogenase (pseudogene), puta | 0,07627 | 0,7017 |
| TcCLB.507035.30 | dispersed gene family protein 1 (DGF-1), putative | 0,07624 | 0,6303 |
| TcCLB.506961.90 | retrotransposon hot spot (RHS) protein, putative | 0,07615 | 0,517 |

| | | | |
|---|---|---|---|
| TcCLB.511193.9 | dispersed gene family protein 1 (DGF-1, pseudogene), put | 0,07605 | 0,7147 |
| TcCLB.503823.100 | hypothetical protein, conserved | 0,07604 | 0,5734 |
| TcCLB.511705.5 | trans-sialidase (pseudogene), putative | 0,07599 | 0,8544 |
| TcCLB.509721.30 | intraflagellar transport protein 122B, putative (IFT122B) | 0,07593 | 0,4245 |
| TcCLB.506727.4 | hypothetical protein, conserved | 0,0759 | 0,7421 |
| TcCLB.508355.330 | hypothetical protein, conserved | 0,07585 | 0,4054 |
| TcCLB.511361.30 | hypothetical protein, conserved | 0,07585 | 0,6414 |
| TcCLB.510055.30 | hypothetical protein, conserved | 0,07585 | 0,7841 |
| TcCLB.505155.10 | retrotransposon hot spot protein (RHS, pseudogene), puta | 0,07573 | 0,6479 |
| TcCLB.509827.10 | hypothetical protein, conserved | 0,0757 | 0,5295 |
| TcCLB.509109.90 | hypothetical protein, conserved | 0,07561 | 0,6756 |
| TcCLB.509601.30 | hypothetical protein | 0,07555 | 0,5138 |
| TcCLB.510387.15 | syntaxin binding protein (pseudogene), putative | 0,07551 | 0,8293 |
| TcCLB.506203.10 | nucleoside transporter 1, putative | 0,07549 | 0,5694 |
| TcCLB.510799.30 | hypothetical protein, conserved | 0,07545 | 0,5281 |
| TcCLB.506963.60 | hypothetical protein, conserved (pseudogene) | 0,07541 | 0,6296 |
| TcCLB.511013.3 | ATP-dependent DEAD/H RNA helicase (pseudogene), puta | 0,07541 | 0,85 |
| TcCLB.510689.40 | phosphatidylinositol 3-kinase (tor2), putative (TOR2) | 0,07535 | 0,4086 |
| TcCLB.511287.90 | ribosome biogenesis protein, putative | 0,07533 | 0,4668 |
| TcCLB.510339.14 | hypothetical protein, conserved | 0,07532 | 0,539 |
| TcCLB.504427.40 | hypothetical protein, conserved | 0,0752 | 0,6689 |
| TcCLB.508231.30 | target of rapamycin (TOR) kinase 1, putative (TOR1) | 0,07515 | 0,2794 |
| TcCLB.511263.20 | hypothetical protein, conserved | 0,07509 | 0,3927 |
| TcCLB.504055.130 | trans-sialidase (pseudogene), putative | 0,07508 | 0,6446 |
| TcCLB.508895.70 | membrane protein, putative | 0,07497 | 0,6458 |
| TcCLB.508409.260 | hypothetical protein, conserved | 0,07489 | 0,4334 |
| TcCLB.510089.64 | hypothetical protein, conserved | 0,07489 | 0,6026 |
| TcCLB.510661.190 | protein kinase, putative | 0,07481 | 0,7681 |
| TcCLB.511871.80 | frataxin-like, mitochondrial precursor, putative | 0,0747 | 0,6968 |
| TcCLB.511881.14 | RNA polymerase subunit RPB10, putative (RPB10) | 0,07469 | 0,6683 |
| TcCLB.505965.70 | helicase (pseudogene), putative | 0,07466 | 0,495 |
| TcCLB.506885.330 | hypothetical protein, conserved | 0,07463 | 0,4659 |
| TcCLB.508185.19 | ubiquitin hydrolase, putative (fragment) | 0,07463 | 0,8085 |
| TcCLB.511367.230 | NADPH:adrenodoxin oxidoreductase, mitochondrial, putati | 0,07461 | 0,478 |
| TcCLB.506607.10 | hypothetical protein, conserved | 0,07457 | 0,6415 |
| TcCLB.510323.90 | hypothetical protein, conserved | 0,07448 | 0,5176 |
| TcCLB.503901.9 | hypothetical protein, conserved (pseudogene) | 0,07442 | 0,6182 |
| TcCLB.507507.49 | hypothetical protein, conserved | 0,07436 | 0,7831 |
| TcCLB.506743.170 | hypothetical protein, conserved | 0,07426 | 0,5747 |
| TcCLB.509127.110 | hypothetical protein, conserved | 0,07424 | 0,5538 |
| TcCLB.510533.190 | hypothetical protein, conserved | 0,0741 | 0,6234 |
| TcCLB.507235.10 | dispersed gene family protein 1 (DGF-1), putative | 0,07409 | 0,8425 |
| TcCLB.510303.119 | hypothetical protein, conserved | 0,07408 | 0,5965 |
| TcCLB.510165.110 | retrotransposon hot spot (RHS) protein, putative | 0,07406 | 0,6921 |
| TcCLB.511537.16 | hypothetical protein, conserved | 0,07402 | 0,5573 |

| | | | |
|---|---|---|---|
| TcCLB.510359.69 | mitochondrial carrier protein, putative | 0,074 | 0,5855 |
| TcCLB.509471.50 | hypothetical protein, conserved | 0,07389 | 0,5086 |
| TcCLB.442297.10 | ARP2/3 complex 16kDa subunit, putative | 0,07385 | 0,669 |
| TcCLB.511385.120 | hypothetical protein, conserved | 0,07384 | 0,7712 |
| TcCLB.508773.9 | dispersed gene family protein 1 (DGF-1), putative | 0,07383 | 0,8822 |
| TcCLB.507025.10 | hypothetical protein, conserved | 0,07369 | 0,4635 |
| TcCLB.511229.60 | hypothetical protein, conserved | 0,07363 | 0,7714 |
| TcCLB.509643.110 | hypothetical protein, conserved | 0,0736 | 0,5509 |
| TcCLB.507011.190 | hypothetical protein, conserved | 0,07349 | 0,699 |
| TcCLB.509733.160 | amino acid tansporter, putative | 0,07348 | 0,6098 |
| TcCLB.511013.6 | N-acetyltransferase complex ARD1 subunit (pseudogene), | 0,07321 | 0,8672 |
| TcCLB.508991.10 | retrotransposon hot spot protein (RHS, pseudogene), puta | 0,07317 | 0,5859 |
| TcCLB.505987.60 | hypothetical protein | 0,07309 | 0,3807 |
| TcCLB.509031.10 | surface protease GP63 (pseudogene), putative | 0,07308 | 0,7076 |
| TcCLB.507723.60 | hypothetical protein | 0,073 | 0,6523 |
| TcCLB.511151.50 | phosphoadenosine phosphosulfate reductase-like protein, | 0,07279 | 0,5108 |
| TcCLB.507007.30 | hypothetical protein | 0,07267 | 0,6255 |
| TcCLB.511319.10 | hypothetical protein, conserved | 0,07266 | 0,4571 |
| TcCLB.508193.30 | dispersed gene family protein 1 (DGF-1, pseudogene), put | 0,07261 | 0,7712 |
| TcCLB.507105.10 | beta prime cop protein, putative | 0,07258 | 0,4081 |
| TcCLB.509885.30 | SPFH domain / Band 7 family protein, putative | 0,07256 | 0,5729 |
| TcCLB.506821.80 | hypothetical protein, conserved | 0,07251 | 0,7128 |
| TcCLB.509799.70 | hypothetical protein, conserved | 0,07246 | 0,5145 |
| TcCLB.507511.20 | hypothetical protein, conserved | 0,0724 | 0,5048 |
| TcCLB.510945.20 | hypothetical protein, conserved | 0,07235 | 0,5563 |
| TcCLB.511417.60 | hypothetical protein, conserved | 0,07206 | 0,5851 |
| TcCLB.504215.50 | hypothetical protein, conserved | 0,07203 | 0,4741 |
| TcCLB.507851.10 | hypothetical protein, conserved | 0,07201 | 0,5037 |
| TcCLB.509207.160 | proteasome regulatory ATPase subunit 1, putative | 0,07198 | 0,4208 |
| TcCLB.508831.20 | oligomeric golgi complex component 8, putative | 0,07189 | 0,5658 |
| TcCLB.508257.204 | hypothetical protein, conserved | 0,07173 | 0,5518 |
| TcCLB.504137.110 | hypothetical protein, conserved | 0,07169 | 0,481 |
| TcCLB.507039.30 | retrotransposon hot spot protein (RHS, pseudogene), puta | 0,07141 | 0,5464 |
| TcCLB.511421.180 | hypothetical protein, conserved | 0,07122 | 0,6319 |
| TcCLB.510089.130 | protein kinase, putative | 0,0711 | 0,4798 |
| TcCLB.503689.20 | hypothetical protein, conserved | 0,07101 | 0,7702 |
| TcCLB.507711.170 | hypothetical protein, conserved | 0,07089 | 0,5784 |
| TcCLB.508153.250 | hypothetical protein | 0,07075 | 0,6878 |
| TcCLB.504179.80 | hypothetical protein, conserved | 0,07074 | 0,5363 |
| TcCLB.506777.60 | NGG1 interacting factor 3, putative | 0,07071 | 0,654 |
| TcCLB.510889.30 | kinase, putative | 0,07055 | 0,5662 |
| TcCLB.481729.10 | hypothetical protein, conserved | 0,07053 | 0,6228 |
| TcCLB.508153.80 | DNA replication complex GINS protein SLD5, putative | 0,07034 | 0,6509 |
| TcCLB.506907.14 | hypothetical protein, conserved | 0,07018 | 0,5854 |
| TcCLB.509455.90 | Putative intraflagellar transport protein A1 (PIFTA1) | 0,06994 | 0,4914 |

| | | | |
|---|---|---|---|
| TcCLB.506811.150 | hypothetical protein, conserved | 0,06989 | 0,5591 |
| TcCLB.506223.84 | hypothetical protein, conserved | 0,06983 | 0,6194 |
| TcCLB.507521.115 | hypothetical protein, conserved | 0,06972 | 0,7059 |
| TcCLB.511169.10 | hypothetical protein, conserved | 0,06937 | 0,6716 |
| TcCLB.507895.140 | hypothetical protein | 0,0693 | 0,5226 |
| TcCLB.508723.100 | rab-like GTPase activating protein, putative | 0,06927 | 0,5656 |
| TcCLB.509049.20 | hypothetical protein, conserved | 0,06927 | 0,5846 |
| TcCLB.504035.90 | translation initiation factor IF-2, putative | 0,0692 | 0,5136 |
| TcCLB.509979.30 | hypothetical protein, conserved | 0,06911 | 0,57 |
| TcCLB.504247.34 | Transcription elongation factor 1 domain-containing prote | 0,06897 | 0,6405 |
| TcCLB.507641.120 | ATP-dependent DEAD/H RNA helicase, putative | 0,06896 | 0,4902 |
| TcCLB.510307.352 | retrotransposon hot spot protein (RHS, pseudogene), puta | 0,06893 | 0,689 |
| TcCLB.511911.21 | hypothetical protein, conserved | 0,06893 | 0,8309 |
| TcCLB.507527.4 | dolichyl-P-Man:GDP-Man5GlcNAc2-PP-dolichyl alpha-1,2-r | 0,06892 | 0,7209 |
| TcCLB.505163.30 | hypothetical protein, conserved | 0,06868 | 0,545 |
| TcCLB.507507.40 | ubiquitin hydrolase, putative | 0,06868 | 0,6919 |
| TcCLB.510155.130 | hypothetical protein, conserved | 0,06861 | 0,436 |
| TcCLB.511671.140 | hypothetical protein | 0,06856 | 0,5963 |
| TcCLB.510421.260 | diacylglycerol acyltransferase, putative | 0,06852 | 0,6251 |
| TcCLB.506021.10 | retrotransposon hot spot protein (RHS, pseudogene), puta | 0,06845 | 0,5954 |
| TcCLB.506247.200 | adaptin complex 1 subunit, putative | 0,06844 | 0,4309 |
| TcCLB.407477.30 | diphthine synthase, putative | 0,06843 | 0,6338 |
| TcCLB.425851.9 | hypothetical protein, conserved | 0,06826 | 0,7208 |
| TcCLB.509945.24 | hypothetical protein, conserved | 0,06824 | 0,6707 |
| TcCLB.508175.290 | hypothetical protein, conserved | 0,06793 | 0,5476 |
| TcCLB.506811.10 | kinesin, putative | 0,0678 | 0,5691 |
| TcCLB.503543.10 | presenilin-like aspartic peptidase, putative | 0,06777 | 0,5649 |
| TcCLB.506695.20 | hypothetical protein, conserved | 0,0675 | 0,6612 |
| TcCLB.509765.20 | retrotransposon hot spot protein (RHS, pseudogene), puta | 0,06739 | 0,784 |
| TcCLB.511665.100 | retrotransposon hot spot protein (RHS, pseudogene), puta | 0,06735 | 0,711 |
| TcCLB.509253.50 | recombination initiation protein NBS1, putative | 0,06702 | 0,4977 |
| TcCLB.507709.20 | hypothetical protein | 0,06695 | 0,5396 |
| TcCLB.511907.330 | retrotransposon hot spot (RHS) protein, putative | 0,0668 | 0,5832 |
| TcCLB.506225.20 | trans-sialidase (pseudogene), putative | 0,06654 | 0,7793 |
| TcCLB.506635.20 | ATP-dependent RNA helicase, putative | 0,06646 | 0,4887 |
| TcCLB.511467.50 | hypothetical protein, conserved | 0,06635 | 0,6946 |
| TcCLB.506671.41 | elongation factor 1-gamma (EF-1-gamma, pseudogene), | | 0,06633 | 0,9074 |
| TcCLB.507483.39 | calmodulin (CALA2) | 0,06626 | 0,7668 |
| TcCLB.509171.40 | hypothetical protein, conserved | 0,06609 | 0,5328 |
| TcCLB.511045.50 | hypothetical protein, conserved | 0,06607 | 0,7084 |
| TcCLB.508261.25 | trans-sialidase (pseudogene), putative | 0,06606 | 0,6159 |
| TcCLB.507459.10 | hypothetical protein, conserved (pseudogene) | 0,06604 | 0,8326 |
| TcCLB.511745.30 | E3 ubiquitin-protein ligase, putative | 0,06602 | 0,5012 |
| TcCLB.509965.134 | hypothetical protein, conserved | 0,06602 | 0,7661 |
| TcCLB.508961.79 | 69 kDa paraflagellar rod protein, putative | 0,06592 | 0,5584 |

| | | | |
|---|---|---|---|
| TcCLB.503697.130 | hypothetical protein, conserved | 0,0659 | 0,5082 |
| TcCLB.508153.400 | serine/threonine-protein kinase, putative | 0,06587 | 0,5257 |
| TcCLB.509117.19 | hypothetical protein, conserved | 0,06584 | 0,6296 |
| TcCLB.509791.189 | flagellum targeting protein kharon1, putative (fragment) | 0,06581 | 0,6484 |
| TcCLB.508641.40 | retrotransposon hot spot protein (RHS, pseudogene), puta | 0,0658 | 0,7969 |
| TcCLB.509791.90 | hypothetical protein, conserved | 0,06565 | 0,657 |
| TcCLB.507773.20 | hypothetical protein | 0,06553 | 0,6846 |
| TcCLB.509211.10 | hypothetical protein, conserved | 0,06551 | 0,5902 |
| TcCLB.510275.90 | hypothetical protein | 0,06537 | 0,7768 |
| TcCLB.511907.320 | hypothetical protein, conserved | 0,06532 | 0,5682 |
| TcCLB.511167.119 | hypothetical protein, conserved (fragment) | 0,06526 | 0,5348 |
| TcCLB.509067.50 | hypothetical protein, conserved | 0,06519 | 0,4921 |
| TcCLB.503879.119 | protein transport protein Sec24C, putative | 0,06514 | 0,5876 |
| TcCLB.508207.160 | hypothetical protein, conserved | 0,0651 | 0,6553 |
| TcCLB.506925.180 | protein kinase, putative | 0,06507 | 0,5782 |
| TcCLB.506869.60 | protein kinase, putative | 0,06504 | 0,5717 |
| TcCLB.510655.130 | hypothetical protein, conserved | 0,06494 | 0,7641 |
| TcCLB.506667.170 | hypothetical protein | 0,06491 | 0,867 |
| TcCLB.506113.40 | hypothetical protein, conserved (pseudogene) | 0,06481 | 0,6408 |
| TcCLB.506825.60 | hypothetical protein, conserved | 0,06479 | 0,5047 |
| TcCLB.506393.120 | hypothetical protein, conserved (pseudogene) | 0,06479 | 0,5922 |
| TcCLB.510531.40 | hypothetical protein, conserved | 0,06471 | 0,5358 |
| TcCLB.511911.140 | hypothetical protein, conserved | 0,0647 | 0,6872 |
| TcCLB.507893.9 | dispersed gene family protein 1 (DGF-1), putative (fragme | 0,06466 | 0,7827 |
| TcCLB.511513.30 | hypothetical protein, conserved | 0,06465 | 0,4793 |
| TcCLB.511523.50 | hypothetical protein, conserved | 0,06451 | 0,637 |
| TcCLB.507063.260 | hypothetical protein | 0,06443 | 0,8281 |
| TcCLB.511657.60 | hypothetical protein, conserved | 0,06435 | 0,4873 |
| TcCLB.506355.130 | hypothetical protein, conserved | 0,06434 | 0,6032 |
| TcCLB.504103.40 | hypothetical protein, conserved | 0,06417 | 0,6299 |
| TcCLB.511469.50 | hypothetical protein, conserved | 0,06407 | 0,6797 |
| TcCLB.508539.50 | hypothetical protein, conserved | 0,06402 | 0,5857 |
| TcCLB.508355.110 | hypothetical protein | 0,06388 | 0,6091 |
| TcCLB.508181.124 | hypothetical protein, conserved | 0,06369 | 0,5979 |
| TcCLB.510325.10 | ubiquitin hydrolase, putative | 0,06368 | 0,5973 |
| TcCLB.507089.170 | hypothetical protein, conserved | 0,06363 | 0,6235 |
| TcCLB.509897.260 | hypothetical protein | 0,0636 | 0,7967 |
| TcCLB.511903.50 | amastin, putative | 0,06348 | 0,6886 |
| TcCLB.506363.159 | separase, putative | 0,06346 | 0,8582 |
| TcCLB.504057.30 | hypothetical protein, conserved | 0,06333 | 0,4625 |
| TcCLB.511911.110 | RNA helicase, putative | 0,06329 | 0,6041 |
| TcCLB.506967.65 | mucin TcMUCII (pseudogene), putative | 0,06305 | 0,8715 |
| TcCLB.508221.990 | surface protease GP63 (pseudogene), putative | 0,06298 | 0,8562 |
| TcCLB.511751.120 | aldehyde dehydrogenase family, putative | 0,06295 | 0,6676 |
| TcCLB.505987.41 | hypothetical protein, conserved | 0,06285 | 0,7259 |

| | | | |
|---|---|---|---|
| TcCLB.504047.60 | protein kinase, putative | 0,06267 | 0,5351 |
| TcCLB.509805.60 | Ras-related protein Rab5A, putative (RAB5A) | 0,06238 | 0,5174 |
| TcCLB.511731.10 | UV excision repair RAD23-like protein, putative | 0,06226 | 0,5351 |
| TcCLB.506201.90 | hypothetical protein, conserved | 0,06223 | 0,6272 |
| TcCLB.503815.4 | hypothetical protein, conserved | 0,06201 | 0,7854 |
| TcCLB.508831.150 | hypothetical protein, conserved | 0,06195 | 0,5095 |
| TcCLB.506773.180 | regulator of chromatin condensation, putative | 0,06192 | 0,6011 |
| TcCLB.511409.10 | PAB1-binding protein , putative | 0,06185 | 0,4152 |
| TcCLB.507203.5 | trans-sialidase (pseudogene), putative | 0,06181 | 0,6553 |
| TcCLB.506989.60 | phosphate transporter, putative | 0,06173 | 0,6588 |
| TcCLB.511355.7 | TFIID-like protein, putative (fragment) | 0,06167 | 0,6709 |
| TcCLB.508177.10 | hypothetical protein, conserved | 0,06164 | 0,6932 |
| TcCLB.503897.90 | hypothetical protein, conserved | 0,06156 | 0,5612 |
| TcCLB.503697.60 | nuclear cap binding complex subunit CBP110, putative (CB | 0,0615 | 0,532 |
| TcCLB.508987.9 | hypothetical protein, conserved | 0,06131 | 0,5352 |
| TcCLB.510719.150 | hypothetical protein, conserved | 0,06118 | 0,4911 |
| TcCLB.507843.20 | trans-sialidase, putative (pseudogene) | 0,06107 | 0,8838 |
| TcCLB.511635.30 | ARP2/3 complex 16kDa subunit, putative | 0,061 | 0,7084 |
| TcCLB.510773.100 | Flagellar Member 4 (FLAM4) | 0,06094 | 0,581 |
| TcCLB.511479.10 | trans-sialidase (pseudogene), putative | 0,06094 | 0,9219 |
| TcCLB.504069.10 | proteasome alpha 2 subunit, putative | 0,06093 | 0,6384 |
| TcCLB.508177.81 | surface protease GP63 (pseudogene), putative | 0,0605 | 0,7169 |
| TcCLB.510275.130 | dispersed gene family protein 1 (DGF-1), putative | 0,0603 | 0,8094 |
| TcCLB.507601.140 | ATP-dependent RNA helicase, putative | 0,06019 | 0,5907 |
| TcCLB.510535.109 | trans-sialidase, putative | 0,05993 | 0,7555 |
| TcCLB.511729.20 | hypothetical protein | 0,05954 | 0,6125 |
| TcCLB.509059.40 | hypothetical protein, conserved | 0,05942 | 0,5703 |
| TcCLB.508239.49 | dynein regulatory complex subunit, putative | 0,05942 | 0,7208 |
| TcCLB.508283.25 | hypothetical protein, conserved (pseudogene) | 0,0594 | 0,8252 |
| TcCLB.509167.120 | S-adenosylmethionine decarboxylase proenzyme, putative | 0,05933 | 0,572 |
| TcCLB.509179.110 | hypothetical protein, conserved | 0,0593 | 0,6274 |
| TcCLB.424123.10 | hypothetical protein, conserved (pseudogene) | 0,05926 | 0,641 |
| TcCLB.509911.30 | hypothetical protein | 0,05922 | 0,6844 |
| TcCLB.506469.120 | hypothetical protein, conserved | 0,05918 | 0,6543 |
| TcCLB.510791.11 | hypothetical protein | 0,05914 | 0,7931 |
| TcCLB.511289.30 | metallo-peptidase, Clan MG, Family M24 | 0,05909 | 0,4914 |
| TcCLB.510659.194 | hypothetical protein, conserved | 0,05906 | 0,6501 |
| TcCLB.507513.130 | hypothetical protein, conserved (pseudogene) | 0,05894 | 0,5039 |
| TcCLB.503823.60 | hypothetical protein, conserved | 0,05894 | 0,636 |
| TcCLB.487739.20 | choline/ethanolamine kinase (pseudogene), putative | 0,05893 | 0,7349 |
| TcCLB.507023.180 | protein kinase, putative | 0,05889 | 0,6166 |
| TcCLB.511585.210 | hypothetical protein, conserved | 0,05889 | 0,6757 |
| TcCLB.507509.60 | hypothetical protein, conserved | 0,05852 | 0,6182 |
| TcCLB.505807.29 | hypothetical protein, conserved | 0,0585 | 0,8165 |
| TcCLB.506825.50 | hypothetical protein, conserved | 0,05848 | 0,5422 |

| | | | |
|---|---|---|---|
| TcCLB.511277.320 | hypothetical protein, conserved | 0,05825 | 0,5399 |
| TcCLB.511893.30 | hypothetical protein, conserved (pseudogene) | 0,05819 | 0,734 |
| TcCLB.503891.70 | ribonuclease HII, putative | 0,05818 | 0,6515 |
| TcCLB.507941.80 | hypothetical protein, conserved | 0,058 | 0,7209 |
| TcCLB.503801.50 | hypothetical protein, conserved | 0,05784 | 0,632 |
| TcCLB.506933.46 | hypothetical protein, conserved | 0,05775 | 0,7313 |
| TcCLB.508347.50 | hypothetical protein, conserved | 0,05759 | 0,7496 |
| TcCLB.506999.70 | chaperone DNAJ protein, putative | 0,05758 | 0,6 |
| TcCLB.507093.200 | hypothetical protein, conserved | 0,05747 | 0,5675 |
| TcCLB.507993.70 | hypothetical protein, conserved | 0,05745 | 0,5924 |
| TcCLB.507649.60 | hypothetical protein, conserved | 0,05741 | 0,6636 |
| TcCLB.507669.130 | hypothetical protein, conserved | 0,05732 | 0,6921 |
| TcCLB.508141.20 | hypothetical protein, conserved | 0,05708 | 0,6195 |
| TcCLB.511249.30 | hypothetical protein, conserved | 0,05706 | 0,5691 |
| TcCLB.507159.90 | DNA-directed RNA polymerase III subunit, putative | 0,05698 | 0,6665 |
| TcCLB.507001.70 | syntaxin, putative | 0,05688 | 0,6743 |
| TcCLB.509579.34 | ATP-dependent DEAD/H RNA helicase (pseudogene), puta | 0,05683 | 0,8658 |
| TcCLB.506347.10 | dynein heavy chain (pseudogene), putative | 0,05678 | 0,4974 |
| TcCLB.506941.80 | tubulin-tyrosine ligase-like protein, putative | 0,05666 | 0,6597 |
| TcCLB.511269.60 | Vacuolar protein sorting-associated protein 16 homolog, p | 0,05665 | 0,7104 |
| TcCLB.508265.40 | hypothetical protein, conserved | 0,0565 | 0,6111 |
| TcCLB.503627.9 | hypothetical protein, conserved (fragment) | 0,0565 | 0,6518 |
| TcCLB.504109.80 | Intraflagellar transport protein 144, putative (PIFTF6) | 0,05649 | 0,6099 |
| TcCLB.511827.60 | CCR4 associated factor, putative | 0,05648 | 0,5541 |
| TcCLB.509777.80 | hypothetical protein, conserved | 0,05635 | 0,6868 |
| TcCLB.510755.110 | hypothetical protein, conserved | 0,05633 | 0,7374 |
| TcCLB.509381.29 | hypothetical protein, conserved | 0,05624 | 0,618 |
| TcCLB.507519.50 | hypothetical protein, conserved | 0,05622 | 0,7209 |
| TcCLB.509799.20 | hypothetical protein, conserved | 0,05595 | 0,637 |
| TcCLB.508207.70 | Adenine phosphoribosyltransferase, putative | 0,05595 | 0,6728 |
| TcCLB.506635.104 | hypothetical protein | 0,05595 | 0,7681 |
| TcCLB.503939.80 | prolyl-tRNA synthetase, putative | 0,05589 | 0,5069 |
| TcCLB.507623.60 | hypothetical protein, conserved | 0,05585 | 0,5575 |
| TcCLB.510431.220 | ribosome assembly protein RRB1, putative | 0,05578 | 0,7259 |
| TcCLB.503865.40 | hypothetical protein, conserved | 0,05576 | 0,7119 |
| TcCLB.504243.20 | hypothetical protein, conserved | 0,05576 | 0,7349 |
| TcCLB.511907.89 | hypothetical protein, conserved | 0,05571 | 0,8005 |
| TcCLB.503587.47 | hypothetical protein | 0,05571 | 0,8714 |
| TcCLB.506933.43 | hypothetical protein, conserved | 0,05565 | 0,6641 |
| TcCLB.507099.80 | ABC transporter, putative | 0,05552 | 0,5694 |
| TcCLB.511907.30 | hypothetical protein, conserved | 0,05547 | 0,5887 |
| TcCLB.503885.20 | hypothetical protein, conserved | 0,05546 | 0,575 |
| TcCLB.509965.90 | hypothetical protein, conserved | 0,05534 | 0,5705 |
| TcCLB.508637.66 | hypothetical protein, conserved | 0,05534 | 0,675 |
| TcCLB.506223.50 | hypothetical protein, conserved | 0,05528 | 0,6859 |

| | | | |
|---|---|---|---|
| TcCLB.511803.80 | hypothetical protein, conserved (fragment) | 0,05512 | 0,8817 |
| TcCLB.506337.20 | hypothetical protein, conserved | 0,05494 | 0,6563 |
| TcCLB.508387.100 | ubiquitin-conjugating enzyme E2, putative | 0,05478 | 0,6904 |
| TcCLB.508637.79 | DNA polymerase eta, putative | 0,05476 | 0,6575 |
| TcCLB.511459.20 | hypothetical protein, conserved | 0,05464 | 0,6427 |
| TcCLB.506287.90 | hypothetical protein, conserved | 0,05463 | 0,6689 |
| TcCLB.508307.60 | hypothetical protein, conserved | 0,0546 | 0,5915 |
| TcCLB.511363.4 | hypothetical protein, conserved | 0,05453 | 0,7404 |
| TcCLB.509799.110 | hypothetical protein, conserved | 0,0545 | 0,7789 |
| TcCLB.511511.170 | ADP-ribosylation factor-like protein, putative | 0,05424 | 0,7034 |
| TcCLB.511817.190 | hypothetical protein, conserved | 0,05417 | 0,6736 |
| TcCLB.509965.340 | hypothetical protein, conserved | 0,05408 | 0,629 |
| TcCLB.509643.30 | hypothetical protein, conserved | 0,05399 | 0,6246 |
| TcCLB.509109.100 | hypothetical protein, conserved | 0,05394 | 0,7946 |
| TcCLB.511725.240 | hypothetical protein, conserved | 0,05389 | 0,5551 |
| TcCLB.507611.180 | hypothetical protein | 0,05383 | 0,8669 |
| TcCLB.507491.119 | hypothetical protein, conserved | 0,05381 | 0,6091 |
| TcCLB.503613.79 | hypothetical protein | 0,05378 | 0,7288 |
| TcCLB.506249.40 | hypothetical protein, conserved | 0,05372 | 0,6563 |
| TcCLB.509493.60 | hypothetical protein | 0,05369 | 0,8226 |
| TcCLB.506739.40 | hypothetical protein, conserved | 0,05366 | 0,6833 |
| TcCLB.510709.41 | hypothetical protein, conserved | 0,05344 | 0,9051 |
| TcCLB.507911.33 | retrotransposon hot spot protein (RHS, pseudogene), puta | 0,05341 | 0,8736 |
| TcCLB.506203.110 | hypothetical protein, conserved | 0,05339 | 0,6543 |
| TcCLB.506625.120 | hypothetical protein, conserved | 0,05325 | 0,615 |
| TcCLB.511245.200 | hypothetical protein, conserved | 0,0532 | 0,5606 |
| TcCLB.506227.200 | hypothetical protein | 0,05309 | 0,6768 |
| TcCLB.506469.110 | hypothetical protein | 0,05283 | 0,7385 |
| TcCLB.509429.290 | hydroxyacylglutathione hydrolase, putative | 0,05274 | 0,7501 |
| TcCLB.510939.9 | hypothetical protein, conserved | 0,05271 | 0,8094 |
| TcCLB.509413.10 | hypothetical protein, conserved | 0,05252 | 0,6081 |
| TcCLB.506529.140 | hypothetical protein, conserved | 0,05246 | 0,7177 |
| TcCLB.510759.50 | hypothetical protein | 0,05233 | 0,6728 |
| TcCLB.507641.240 | hypothetical protein | 0,05222 | 0,8622 |
| TcCLB.511027.10 | chloride channel protein, putative | 0,05217 | 0,6734 |
| TcCLB.503755.50 | hypothetical protein, conserved | 0,05216 | 0,6765 |
| TcCLB.508819.20 | hypothetical protein, conserved | 0,05215 | 0,6699 |
| TcCLB.510265.30 | hypothetical protein, conserved | 0,05211 | 0,5107 |
| TcCLB.510515.70 | hypothetical protein, conserved | 0,0521 | 0,8175 |
| TcCLB.508089.60 | hypothetical protein, conserved | 0,05204 | 0,7183 |
| TcCLB.511153.124 | hypothetical protein, conserved | 0,05202 | 0,7753 |
| TcCLB.511071.40 | amastin, putative | 0,05193 | 0,7408 |
| TcCLB.503419.54 | hypothetical protein, conserved | 0,05192 | 0,7568 |
| TcCLB.511589.190 | hypothetical protein, conserved | 0,0519 | 0,7381 |
| TcCLB.510039.36 | hypothetical protein | 0,05186 | 0,8203 |

| | | | |
|---|---|---|---|
| TcCLB.511039.39 | hypothetical protein, conserved | 0,05185 | 0,7234 |
| TcCLB.510735.90 | hypothetical protein, conserved | 0,0518 | 0,7466 |
| TcCLB.506695.10 | actin-like protein, putative | 0,0517 | 0,7424 |
| TcCLB.508443.40 | hypothetical protein | 0,05152 | 0,7366 |
| TcCLB.508691.20 | hypothetical protein, conserved | 0,05136 | 0,7082 |
| TcCLB.507979.50 | hypothetical protein, conserved | 0,05132 | 0,637 |
| TcCLB.508325.30 | dispersed gene family protein 1 (DGF-1), putative | 0,05126 | 0,7577 |
| TcCLB.507517.110 | hypothetical protein, conserved | 0,05118 | 0,7253 |
| TcCLB.511545.30 | hypothetical protein, conserved | 0,05103 | 0,6851 |
| TcCLB.507629.20 | hypothetical protein, conserved | 0,051 | 0,5698 |
| TcCLB.508819.10 | hypothetical protein, conserved | 0,05083 | 0,7494 |
| TcCLB.511287.150 | fibrillarin, putative | 0,05077 | 0,7034 |
| TcCLB.511101.19 | retrotransposon hot spot (RHS) protein, putative | 0,05055 | 0,7605 |
| TcCLB.508177.60 | hypothetical protein, conserved | 0,05053 | 0,7009 |
| TcCLB.504137.160 | hypothetical protein, conserved | 0,05053 | 0,7896 |
| TcCLB.509505.20 | hypothetical protein, conserved | 0,05052 | 0,7121 |
| TcCLB.420293.31 | hypothetical protein | 0,05038 | 0,871 |
| TcCLB.510729.90 | hypothetical protein, conserved | 0,05036 | 0,7056 |
| TcCLB.508541.140 | surface protease GP63 (pseudogene), putative | 0,05027 | 0,8461 |
| TcCLB.442427.10 | trans-sialidase (pseudogene), putative | 0,05017 | 0,8045 |
| TcCLB.503479.49 | U6 snRNA-associated Sm-like protein LSm7p | 0,05016 | 0,7461 |
| TcCLB.511463.10 | rRNA dimethyltransferase, putative | 0,05012 | 0,7103 |
| TcCLB.506779.170 | haloacid dehalogenase hydrolase, putative | 0,05009 | 0,7084 |
| TcCLB.508601.110 | hypothetical protein, conserved | 0,04992 | 0,6041 |
| TcCLB.510725.30 | hypothetical protein, conserved | 0,0499 | 0,7485 |
| TcCLB.508743.5 | hypothetical protein, conserved (pseudogene) | 0,04985 | 0,8749 |
| TcCLB.506505.40 | protein kinase, putative | 0,04977 | 0,632 |
| TcCLB.505051.20 | Checkpoint protein HUS1, putative (hus1) | 0,04955 | 0,6809 |
| TcCLB.505941.10 | surface protease GP63, putative | 0,04949 | 0,7661 |
| TcCLB.507515.130 | hypothetical protein, conserved | 0,04943 | 0,7478 |
| TcCLB.507223.20 | hypothetical protein, conserved | 0,04937 | 0,8582 |
| TcCLB.467795.19 | hypothetical protein, conserved | 0,04916 | 0,7626 |
| TcCLB.511215.40 | hypothetical protein, conserved | 0,04913 | 0,6294 |
| TcCLB.508041.20 | hypothetical protein, conserved | 0,04904 | 0,6714 |
| TcCLB.504077.40 | translation elongation factor EF-2, putative | 0,04904 | 0,6871 |
| TcCLB.511801.4 | hypothetical protein, conserved | 0,04895 | 0,7038 |
| TcCLB.508953.40 | hypothetical protein, conserved | 0,04874 | 0,7339 |
| TcCLB.508175.189 | hypothetical protein, conserved | 0,04874 | 0,7371 |
| TcCLB.511365.70 | proteasome regulatory non-ATP-ase subunit 9, putative (F | 0,0487 | 0,6636 |
| TcCLB.511493.5 | hypothetical protein, conserved | 0,0487 | 0,857 |
| TcCLB.509081.10 | elongation factor 1-gamma (EF-1-gamma), putative | 0,0486 | 0,8192 |
| TcCLB.509733.90 | hypothetical protein, conserved | 0,04855 | 0,6416 |
| TcCLB.508277.130 | hypothetical protein, conserved | 0,04847 | 0,778 |
| TcCLB.511283.109 | AP-4 complex subunit sigma-1, putative | 0,04836 | 0,7548 |
| TcCLB.511301.110 | inosine-5'-monophosphate dehydrogenase, putative | 0,04829 | 0,5225 |

| | | | |
|---|---|---|---|
| TcCLB.511355.30 | phosphatidic acid phosphatase protein, putative | 0,04792 | 0,6996 |
| TcCLB.508693.10 | cleavage and polyadenylation specificity factor, putative | 0,04782 | 0,5957 |
| TcCLB.511209.50 | hypothetical protein, conserved | 0,04774 | 0,7223 |
| TcCLB.508323.30 | hypothetical protein, conserved | 0,04763 | 0,7275 |
| TcCLB.503909.110 | intraflagellar transport (IFT) protein, putative (IFT20) | 0,04749 | 0,7599 |
| TcCLB.507681.40 | hypothetical protein | 0,04748 | 0,7264 |
| TcCLB.503673.27 | 8-oxoguanine DNA glycosylase (pseudogene), putative | 0,04746 | 0,8399 |
| TcCLB.506419.10 | ribosomal protein L7/L12, putative | 0,04731 | 0,7625 |
| TcCLB.508629.68 | N-acetyltransferase complex ARD1 subunit (pseudogene), | 0,04731 | 0,867 |
| TcCLB.504147.320 | hypothetical protein, conserved | 0,04722 | 0,5993 |
| TcCLB.506679.220 | nucleoplasmin-like protein (NLP), putative | 0,04722 | 0,6175 |
| TcCLB.506559.260 | ATP-dependent RNA helicase, putative | 0,04721 | 0,6008 |
| TcCLB.506175.50 | hypothetical protein, conserved | 0,04721 | 0,6588 |
| TcCLB.509217.143 | ATP-dependent DEAD/H RNA helicase (pseudogene), puta | 0,04714 | 0,8864 |
| TcCLB.507641.30 | hypothetical protein, conserved | 0,04705 | 0,6442 |
| TcCLB.506485.90 | hypothetical protein, conserved | 0,04691 | 0,6883 |
| TcCLB.508741.380 | hypothetical protein, conserved | 0,04689 | 0,5692 |
| TcCLB.511617.9 | NUP-1 protein (pseudogene), putative | 0,04684 | 0,5172 |
| TcCLB.506247.180 | hypothetical protein, conserved | 0,04683 | 0,6322 |
| TcCLB.506233.20 | retrotransposon hot spot protein (RHS, pseudogene), puta | 0,04681 | 0,8658 |
| TcCLB.511865.90 | dynein heavy chain, putative | 0,04633 | 0,7264 |
| TcCLB.508199.60 | hypothetical protein, conserved | 0,04633 | 0,8188 |
| TcCLB.510685.20 | hypothetical protein | 0,04627 | 0,7585 |
| TcCLB.504047.50 | hypothetical protein, conserved | 0,04625 | 0,6679 |
| TcCLB.507817.60 | Present in the outer mitochondrial membrane proteome 3 | 0,0461 | 0,6294 |
| TcCLB.507515.40 | hypothetical protein, conserved | 0,0461 | 0,7761 |
| TcCLB.508707.340 | serine/threonine protein phosphatase, putative (fragment | 0,04602 | 0,8282 |
| TcCLB.507491.130 | hypothetical protein, conserved | 0,046 | 0,7283 |
| TcCLB.510431.180 | Present in the outer mitochondrial membrane proteome 1 | 0,04599 | 0,6986 |
| TcCLB.508881.80 | DNA ligase, putative | 0,04589 | 0,8163 |
| TcCLB.506477.30 | tubulin-tyrosine ligase, putative | 0,04582 | 0,7061 |
| TcCLB.511001.230 | hypothetical protein, conserved | 0,04574 | 0,7209 |
| TcCLB.509455.20 | hypothetical protein, conserved | 0,0457 | 0,7074 |
| TcCLB.509119.10 | oligomeric golgi complex component 8, putative | 0,04568 | 0,7277 |
| TcCLB.508409.270 | RNA-binding protein, putative | 0,04553 | 0,7065 |
| TcCLB.509647.150 | calcium-transporting ATPase, putative | 0,04552 | 0,5887 |
| TcCLB.506779.60 | DNA-direcetd RNA polymerase II, subunit 9, putative | 0,04507 | 0,7582 |
| TcCLB.511439.70 | hypothetical protein, conserved | 0,04496 | 0,7084 |
| TcCLB.508859.109 | exosome complex exonuclease, putative | 0,04491 | 0,8078 |
| TcCLB.506025.70 | dolichyl-lipid chaperone, putative (rft1) | 0,04466 | 0,7841 |
| TcCLB.509157.110 | hypothetical protein, conserved | 0,04466 | 0,8048 |
| TcCLB.508995.30 | tubulin-tyrosine ligase, putative | 0,04465 | 0,7327 |
| TcCLB.508525.30 | hypothetical protein, conserved | 0,04446 | 0,7066 |
| TcCLB.504109.30 | hypothetical protein, conserved | 0,04443 | 0,6781 |
| TcCLB.510933.140 | trans-sialidase (pseudogene), putative | 0,04435 | 0,9053 |

| | | | |
|---|---|---|---|
| TcCLB.511543.50 | hypothetical protein, conserved | 0,04432 | 0,7867 |
| TcCLB.503577.20 | U2 splicing auxiliary factor, putative (u2af26) | 0,04427 | 0,616 |
| TcCLB.508059.10 | hypothetical protein, conserved | 0,0442 | 0,8039 |
| TcCLB.511575.110 | hypothetical protein, conserved | 0,04419 | 0,7068 |
| TcCLB.509459.40 | hypothetical protein, conserved | 0,04416 | 0,7092 |
| TcCLB.507801.114 | aldo-keto reductase (pseudogene), putative | 0,04412 | 0,7865 |
| TcCLB.506847.4 | hypothetical protein, conserved | 0,0439 | 0,8421 |
| TcCLB.506893.50 | hypothetical protein, conserved | 0,04387 | 0,8395 |
| TcCLB.511433.20 | hypothetical protein, conserved | 0,04381 | 0,8591 |
| TcCLB.510943.190 | myosin heavy chain MYA2-related, putative | 0,04374 | 0,6958 |
| TcCLB.506637.20 | hypothetical protein, conserved | 0,04374 | 0,7561 |
| TcCLB.509715.26 | hypothetical protein, conserved | 0,04374 | 0,7752 |
| TcCLB.508121.15 | hypothetical protein, conserved (pseudogene) | 0,04373 | 0,8246 |
| TcCLB.508153.800 | hypothetical protein | 0,04371 | 0,7652 |
| TcCLB.508919.90 | hypothetical protein, conserved | 0,0437 | 0,7786 |
| TcCLB.509915.10 | trans-sialidase (pseudogene), putative | 0,04355 | 0,8639 |
| TcCLB.508631.40 | hypothetical protein, conserved (pseudogene) | 0,0434 | 0,7261 |
| TcCLB.504167.20 | mucin TcMUCI, putative | 0,0434 | 0,7977 |
| TcCLB.506567.134 | hypothetical protein, conserved | 0,04336 | 0,7353 |
| TcCLB.509979.110 | hypothetical protein, conserved | 0,04336 | 0,8095 |
| TcCLB.507669.84 | ATP-dependent DEAD/H RNA helicase, putative | 0,04328 | 0,6726 |
| TcCLB.511719.10 | phosphatidylinositol 3-related kinase, putative | 0,04314 | 0,6551 |
| TcCLB.510215.10 | NADH-dependent fumarate reductase, putative | 0,04313 | 0,7865 |
| TcCLB.510629.101 | hypothetical protein, conserved (pseudogene) | 0,0431 | 0,941 |
| TcCLB.510741.50 | hypothetical protein, conserved | 0,04303 | 0,7139 |
| TcCLB.509065.130 | hypothetical protein, conserved | 0,043 | 0,7501 |
| TcCLB.509601.5 | mannosyl-oligosaccharide 1,2-alpha-mannosidase IB, puta | 0,04282 | 0,8674 |
| TcCLB.507821.101 | hypothetical protein, conserved (pseudogene) | 0,04272 | 0,8415 |
| TcCLB.507053.60 | hypothetical protein, conserved | 0,0427 | 0,6873 |
| TcCLB.510089.160 | hypothetical protein, conserved (pseudogene) | 0,04265 | 0,6459 |
| TcCLB.504047.34 | acylphosphatase, putative | 0,04245 | 0,8201 |
| TcCLB.506407.30 | hypothetical protein, conserved | 0,04231 | 0,736 |
| TcCLB.511211.20 | Coatomer subunit gamma (COPG) | 0,04229 | 0,6561 |
| TcCLB.506983.10 | hypothetical protein, conserved | 0,04228 | 0,7962 |
| TcCLB.503993.30 | surface protease GP63, putative | 0,04228 | 0,8221 |
| TcCLB.504055.81 | ubiquitin-protein ligase, putative | 0,04223 | 0,7559 |
| TcCLB.511491.60 | kinesin, putative | 0,04221 | 0,6691 |
| TcCLB.509507.40 | hypothetical protein, conserved | 0,04212 | 0,7665 |
| TcCLB.503635.89 | hypothetical protein, conserved | 0,0421 | 0,7884 |
| TcCLB.508347.80 | Present in the outer mitochondrial membrane proteome 2 | 0,04206 | 0,6612 |
| TcCLB.510687.80 | hypothetical protein, conserved | 0,04203 | 0,765 |
| TcCLB.510089.70 | hypothetical protein | 0,04199 | 0,8394 |
| TcCLB.506649.110 | hypothetical protein, conserved | 0,04193 | 0,8027 |
| TcCLB.506473.14 | Protein transport protein yif1, putative | 0,04189 | 0,763 |
| TcCLB.508153.160 | hypothetical protein, conserved | 0,04185 | 0,799 |

| | | | |
|---|---|---|---|
| TcCLB.506989.309 | hypothetical protein, conserved | 0,0417 | 0,7116 |
| TcCLB.506729.30 | hypothetical protein, conserved | 0,04169 | 0,8176 |
| TcCLB.504153.300 | GTPase protein, putative | 0,04165 | 0,7104 |
| TcCLB.405737.14 | hypothetical protein | 0,04159 | 0,7876 |
| TcCLB.511855.60 | hypothetical protein, conserved | 0,04152 | 0,7653 |
| TcCLB.503525.50 | RNA helicase, putative | 0,04137 | 0,799 |
| TcCLB.503855.80 | hypothetical protein, conserved | 0,04125 | 0,7394 |
| TcCLB.507667.11 | hypothetical protein, conserved (fragment) | 0,04119 | 0,781 |
| TcCLB.511771.199 | hypothetical protein | 0,04114 | 0,9225 |
| TcCLB.507625.160 | hypothetical protein, conserved | 0,04113 | 0,8281 |
| TcCLB.506583.50 | hypothetical protein, conserved | 0,0411 | 0,7755 |
| TcCLB.510203.19 | surface protease GP63 (pseudogene), putative | 0,04106 | 0,9229 |
| TcCLB.504131.110 | hypothetical protein, conserved | 0,04094 | 0,7987 |
| TcCLB.506155.10 | hypothetical protein, conserved | 0,04093 | 0,768 |
| TcCLB.509751.10 | hypothetical protein, conserved | 0,04089 | 0,7983 |
| TcCLB.510307.19 | mucin-associated surface protein (MASP, pseudogene), pu | 0,04075 | 0,8603 |
| TcCLB.503823.150 | hypothetical protein, conserved | 0,04068 | 0,7638 |
| TcCLB.510769.40 | hypothetical protein, conserved | 0,04063 | 0,7643 |
| TcCLB.511119.10 | dispersed gene family protein 1 (DGF-1), putative (fragme | 0,04054 | 0,885 |
| TcCLB.503703.10 | hypothetical protein, conserved | 0,0405 | 0,773 |
| TcCLB.506173.10 | hypothetical protein, conserved | 0,04049 | 0,8117 |
| TcCLB.510747.114 | hypothetical protein, conserved | 0,04047 | 0,8703 |
| TcCLB.509023.40 | hypothetical protein, conserved | 0,04043 | 0,7627 |
| TcCLB.510339.7 | hypothetical protein, conserved (fragment) | 0,04041 | 0,7686 |
| TcCLB.511907.60 | hypothetical protein, conserved | 0,0404 | 0,7806 |
| TcCLB.506925.160 | hypothetical protein, conserved | 0,04033 | 0,784 |
| TcCLB.506821.50 | hypothetical protein | 0,04028 | 0,8318 |
| TcCLB.509825.7 | hypothetical protein, conserved | 0,0402 | 0,9288 |
| TcCLB.505999.110 | NADH-ubiquinone oxidoreductase complex I subunit, puta | 0,04005 | 0,7702 |
| TcCLB.510667.60 | peptidyl-prolyl cis-trans isomerase (cyclophilin), putative | 0,03989 | 0,7884 |
| TcCLB.510531.90 | hypothetical protein, conserved (pseudogene) | 0,0398 | 0,7682 |
| TcCLB.507491.10 | hypothetical protein, conserved | 0,03977 | 0,7493 |
| TcCLB.509017.20 | hypothetical protein, conserved | 0,03972 | 0,7565 |
| TcCLB.506699.10 | hypothetical protein, conserved | 0,03971 | 0,8121 |
| TcCLB.507089.280 | chaperone DnaJ protein (pseudogene), putative | 0,03971 | 0,8552 |
| TcCLB.511561.20 | dispersed gene family protein 1 (DGF-1), putative | 0,03967 | 0,8337 |
| TcCLB.511529.259 | hypothetical protein, conserved | 0,03965 | 0,8817 |
| TcCLB.508303.40 | hypothetical protein, conserved | 0,03961 | 0,7944 |
| TcCLB.508817.140 | hypothetical protein, conserved | 0,03948 | 0,6995 |
| TcCLB.506221.129 | hypothetical protein, conserved (fragment) | 0,03939 | 0,798 |
| TcCLB.511867.50 | proteasome alpha 7 subunit, putative | 0,03935 | 0,7475 |
| TcCLB.507811.20 | delta-adaptin, putative | 0,03934 | 0,7468 |
| TcCLB.507925.30 | hypothetical protein, conserved | 0,03931 | 0,8582 |
| TcCLB.507389.79 | hypothetical protein, conserved | 0,03928 | 0,8231 |
| TcCLB.506815.40 | mitochondrial RNA binding complex 1 subunit, putative | 0,03924 | 0,8185 |

| | | | |
|---|---|---|---|
| TcCLB.509005.10 | axoneme central apparatus protein (fragment) (PF16) | 0,03924 | 0,846 |
| TcCLB.508277.80 | hypothetical protein, conserved | 0,03916 | 0,7963 |
| TcCLB.509965.360 | hypothetical protein, conserved | 0,03908 | 0,8108 |
| TcCLB.503583.100 | hypothetical protein, conserved | 0,03905 | 0,8203 |
| TcCLB.507673.10 | serine/threonine-protein phosphatase PP1, putative | 0,03905 | 0,8233 |
| TcCLB.508601.100 | hypothetical protein, conserved | 0,03888 | 0,6971 |
| TcCLB.510667.4 | hypothetical protein, conserved (fragment) | 0,03886 | 0,7833 |
| TcCLB.461927.9 | adenosine monophosphate deaminase, putative | 0,03883 | 0,7983 |
| TcCLB.506953.45 | ascorbate-dependent peroxidase (pseudogene), putative | 0,03877 | 0,8506 |
| TcCLB.503825.19 | protein kinase, putative | 0,03867 | 0,8798 |
| TcCLB.422189.10 | nucleoside phosphatase, putative | 0,03866 | 0,7836 |
| TcCLB.507771.60 | hypothetical protein, conserved | 0,03854 | 0,7054 |
| TcCLB.503833.50 | hypothetical protein, conserved | 0,03841 | 0,6722 |
| TcCLB.508153.500 | helicase, putative | 0,0384 | 0,6636 |
| TcCLB.511843.60 | retrotransposon hot spot protein (RHS, pseudogene), puta | 0,03824 | 0,8523 |
| TcCLB.508707.100 | hypothetical protein, conserved | 0,03822 | 0,7139 |
| TcCLB.510737.90 | hypothetical protein, conserved | 0,0379 | 0,8276 |
| TcCLB.511815.4 | proteasome regulatory ATPase subunit 5, putative (fragm | 0,03789 | 0,7725 |
| TcCLB.511033.20 | hypothetical protein | 0,03777 | 0,7528 |
| TcCLB.506695.4 | Putative intraflagellar transport protein C3 (fragment) (PI | 0,03765 | 0,811 |
| TcCLB.506127.140 | hypothetical protein | 0,03747 | 0,7112 |
| TcCLB.510533.40 | Mu-adaptin 1, putative | 0,03746 | 0,7329 |
| TcCLB.511107.10 | hypothetical protein, conserved (pseudogene) | 0,03745 | 0,7561 |
| TcCLB.506989.70 | flagellar radial spoke protein-like, putative | 0,0373 | 0,799 |
| TcCLB.511179.114 | hypothetical protein, conserved | 0,03723 | 0,8322 |
| TcCLB.469785.30 | hypothetical protein | 0,03714 | 0,8022 |
| TcCLB.511787.30 | hypothetical protein, conserved (pseudogene) | 0,03711 | 0,8715 |
| TcCLB.504047.70 | hypothetical protein, conserved | 0,03704 | 0,7374 |
| TcCLB.510325.50 | DNA replication complex GINS protein SLD5, putative | 0,03704 | 0,8419 |
| TcCLB.510105.36 | hypothetical protein, conserved | 0,03694 | 0,7983 |
| TcCLB.504203.20 | sec1-like protein, putative | 0,03691 | 0,7161 |
| TcCLB.510873.20 | surface protease GP63, putative | 0,0369 | 0,7969 |
| TcCLB.506227.210 | hypothetical protein, conserved | 0,0369 | 0,8121 |
| TcCLB.506459.160 | mucin TcMUCII (pseudogene), putative | 0,03683 | 0,9216 |
| TcCLB.508837.130 | hypothetical protein, conserved | 0,03679 | 0,7714 |
| TcCLB.506725.40 | hypothetical protein, conserved | 0,03679 | 0,809 |
| TcCLB.509319.20 | hypothetical protein, conserved | 0,03678 | 0,8706 |
| TcCLB.507085.30 | trans-sialidase, Group I, putative | 0,03671 | 0,7576 |
| TcCLB.508153.310 | protein kinase, putative | 0,03659 | 0,7665 |
| TcCLB.506477.14 | hypothetical protein, conserved | 0,03653 | 0,7999 |
| TcCLB.506529.224 | hypothetical protein, conserved | 0,03644 | 0,826 |
| TcCLB.511855.70 | protein kinase, putative | 0,03622 | 0,7685 |
| TcCLB.506559.430 | TPR-repeat protein, putative | 0,03619 | 0,697 |
| TcCLB.507023.289 | hypothetical protein, conserved (fragment) | 0,03609 | 0,85 |
| TcCLB.504017.30 | hypothetical protein, conserved | 0,03601 | 0,7569 |

| | | | |
|---|---|---|---|
| TcCLB.507949.60 | hypothetical protein, conserved | 0,0359 | 0,7841 |
| TcCLB.510155.150 | hypothetical protein, conserved | 0,03576 | 0,7684 |
| TcCLB.511127.200 | kinesin, putative | 0,03572 | 0,7759 |
| TcCLB.508629.74 | retrotransposon hot spot protein (RHS, pseudogene), puta | 0,03566 | 0,8013 |
| TcCLB.506885.200 | hypothetical protein, conserved | 0,03556 | 0,6973 |
| TcCLB.507669.90 | hypothetical protein, conserved | 0,03545 | 0,7615 |
| TcCLB.510663.50 | hypothetical protein, conserved | 0,03544 | 0,7514 |
| TcCLB.504717.30 | hypothetical protein, conserved | 0,03542 | 0,7951 |
| TcCLB.506201.110 | hypothetical protein, conserved | 0,03517 | 0,7984 |
| TcCLB.510963.70 | ARP2/3 complex subunit, putative | 0,03516 | 0,8255 |
| TcCLB.510667.20 | hypothetical protein, conserved | 0,03516 | 0,8613 |
| TcCLB.510143.60 | hypothetical protein, conserved | 0,03492 | 0,8085 |
| TcCLB.510565.120 | hypothetical protein, conserved | 0,03487 | 0,8129 |
| TcCLB.503723.10 | hypothetical protein, conserved | 0,03478 | 0,8332 |
| TcCLB.503449.20 | copper homeostasis protein, putative | 0,03465 | 0,8127 |
| TcCLB.508029.40 | hypothetical protein | 0,03458 | 0,8531 |
| TcCLB.507017.120 | ubiquitin hydrolase, putative | 0,03446 | 0,715 |
| TcCLB.506753.180 | hypothetical protein, conserved | 0,03429 | 0,8591 |
| TcCLB.509893.20 | hypothetical protein, conserved | 0,03426 | 0,7288 |
| TcCLB.507511.71 | hypothetical protein (pseudogene) | 0,03426 | 0,9022 |
| TcCLB.510903.10 | tRNA pseudouridine synthase, putative | 0,0342 | 0,8226 |
| TcCLB.508625.90 | DNA damage repair protein, putative | 0,03418 | 0,7569 |
| TcCLB.508415.59 | hypothetical protein, conserved | 0,03416 | 0,8848 |
| TcCLB.509767.160 | hypothetical protein, conserved | 0,03405 | 0,863 |
| TcCLB.510741.110 | hypothetical protein, conserved | 0,03403 | 0,8 |
| TcCLB.508277.290 | hypothetical protein, conserved | 0,03386 | 0,6802 |
| TcCLB.507809.80 | hypothetical protein, conserved | 0,03375 | 0,6721 |
| TcCLB.506123.60 | hypothetical protein, conserved | 0,03374 | 0,8053 |
| TcCLB.509657.61 | elongation factor 1-gamma (EF-1-gamma, pseudogene), | 0,0337 | 0,924 |
| TcCLB.504241.85 | hypothetical protein, conserved (pseudogene) | 0,03368 | 0,8778 |
| TcCLB.504193.51 | ATP-dependent DEAD/H RNA helicase (pseudogene), puta | 0,03358 | 0,8261 |
| TcCLB.508075.60 | hypothetical protein, conserved | 0,03349 | 0,7948 |
| TcCLB.511025.20 | hypothetical protein, conserved | 0,03346 | 0,7876 |
| TcCLB.506581.69 | Mutt-Nudix-related hydrolase 5, putative | 0,03341 | 0,8496 |
| TcCLB.510657.120 | hypothetical protein, conserved | 0,03338 | 0,8802 |
| TcCLB.511017.30 | cyclin dependent kinase-binding protein, putative | 0,03335 | 0,7481 |
| TcCLB.506855.209 | U6 snRNA-associated Sm-like protein LSm7p | 0,03332 | 0,8261 |
| TcCLB.506925.480 | ribokinase, putative | 0,03331 | 0,8186 |
| TcCLB.505267.110 | dispersed gene family protein 1 (DGF-1), putative | 0,03314 | 0,8821 |
| TcCLB.506405.80 | hypothetical protein, conserved | 0,03303 | 0,8228 |
| TcCLB.507981.51 | trans-sialidase (pseudogene), putative | 0,03303 | 0,9225 |
| TcCLB.511445.20 | radial spoke protein 3, putative (RSP3) | 0,0329 | 0,7936 |
| TcCLB.503835.30 | hypothetical protein, conserved | 0,0329 | 0,8005 |
| TcCLB.506435.120 | ATP-dependent RNA helicase, putative | 0,03276 | 0,7377 |
| TcCLB.510303.50 | deoxyribose-phosphate aldolase, putative | 0,03269 | 0,7963 |

| | | | |
|---|---|---|---|
| TcCLB.504045.50 | hypothetical protein | 0,03268 | 0,7966 |
| TcCLB.508357.100 | hypothetical protein, conserved | 0,03266 | 0,7259 |
| TcCLB.504153.10 | hypothetical protein, conserved | 0,03265 | 0,8708 |
| TcCLB.508543.160 | hypothetical protein, conserved | 0,03243 | 0,7703 |
| TcCLB.507049.150 | hypothetical protein, conserved | 0,03238 | 0,8188 |
| TcCLB.510087.40 | hypothetical protein | 0,03235 | 0,931 |
| TcCLB.511255.700 | trans-sialidase, Group V, putative | 0,03233 | 0,9087 |
| TcCLB.511283.240 | hypothetical protein, conserved | 0,0323 | 0,6869 |
| TcCLB.461165.10 | hypothetical protein, conserved | 0,03229 | 0,8379 |
| TcCLB.504017.50 | hypothetical protein, conserved | 0,03227 | 0,8642 |
| TcCLB.506859.20 | proteasome regulatory ATPase subunit 5, putative | 0,03219 | 0,7765 |
| TcCLB.506743.220 | hypothetical protein, conserved | 0,03214 | 0,8831 |
| TcCLB.510101.260 | hypothetical protein, conserved | 0,03203 | 0,7601 |
| TcCLB.507509.20 | hypothetical protein, conserved | 0,03203 | 0,7891 |
| TcCLB.508719.80 | hypothetical protein, conserved | 0,03196 | 0,8324 |
| TcCLB.508465.10 | aminotransferase, putative | 0,03193 | 0,8264 |
| TcCLB.503459.10 | hypothetical protein, conserved | 0,03184 | 0,8275 |
| TcCLB.506775.40 | hypothetical protein, conserved | 0,03181 | 0,7826 |
| TcCLB.506825.230 | hypothetical protein, conserved | 0,03175 | 0,7219 |
| TcCLB.506733.10 | hypothetical protein, conserved | 0,03174 | 0,8316 |
| TcCLB.506697.10 | hypothetical protein, conserved | 0,03168 | 0,7722 |
| TcCLB.506211.230 | hypothetical protein, conserved | 0,03156 | 0,8055 |
| TcCLB.511753.70 | hypothetical protein, conserved | 0,03152 | 0,7717 |
| TcCLB.509985.40 | hypothetical protein, conserved | 0,03144 | 0,8193 |
| TcCLB.508221.697 | trans-sialidase (pseudogene), putative | 0,03143 | 0,9015 |
| TcCLB.510533.110 | hypothetical protein, conserved | 0,03136 | 0,799 |
| TcCLB.504147.130 | hypothetical protein, conserved | 0,03122 | 0,8085 |
| TcCLB.508881.20 | hypothetical protein, conserved | 0,03096 | 0,8168 |
| TcCLB.507209.50 | hypothetical protein, conserved | 0,03089 | 0,8133 |
| TcCLB.506629.210 | hypothetical protein, conserved | 0,03085 | 0,8208 |
| TcCLB.506869.70 | mitochondrial structure specific endonuclease I (SSE-1), p | 0,03081 | 0,8189 |
| TcCLB.507541.40 | casein kinase, putative | 0,03069 | 0,6997 |
| TcCLB.509059.20 | hypothetical protein, conserved | 0,03069 | 0,7881 |
| TcCLB.511543.60 | glutathione peroxidase, putative | 0,03053 | 0,8473 |
| TcCLB.508189.31 | hypothetical protein, conserved (pseudogene) | 0,03039 | 0,9242 |
| TcCLB.506937.40 | dynein heavy chain, putative | 0,03037 | 0,8036 |
| TcCLB.504041.29 | DNA repair protein, putative | 0,0303 | 0,8377 |
| TcCLB.508131.19 | hypothetical protein, conserved (fragment) | 0,03023 | 0,836 |
| TcCLB.507275.30 | hypothetical protein, conserved | 0,03021 | 0,8265 |
| TcCLB.503959.30 | hypothetical protein, conserved | 0,03016 | 0,7684 |
| TcCLB.509205.10 | guanine-nucleotide-exchange-factor, putative | 0,03002 | 0,8484 |
| TcCLB.511317.70 | RNA-binding protein, putative (RBP30) | 0,02977 | 0,8195 |
| TcCLB.510687.90 | hypothetical protein, conserved | 0,02971 | 0,8259 |
| TcCLB.505183.90 | hypothetical protein, conserved | 0,02965 | 0,8889 |
| TcCLB.507739.80 | hypothetical protein, conserved | 0,02961 | 0,7987 |

| | | | |
|---|---|---|---|
| TcCLB.503473.10 | protein kinase, putative | 0,0296 | 0,8156 |
| TcCLB.507669.70 | Dynein intermediate chain 1, axonemal (DNAI1) | 0,02953 | 0,827 |
| TcCLB.508181.130 | peter pan protein, putative | 0,02951 | 0,8168 |
| TcCLB.510423.60 | hypothetical protein, conserved | 0,0295 | 0,7951 |
| TcCLB.509699.85 | surface protease GP63 (pseudogene), putative | 0,02949 | 0,9254 |
| TcCLB.511239.70 | hypothetical protein, conserved | 0,02929 | 0,8118 |
| TcCLB.506819.49 | hypothetical protein, conserved | 0,02929 | 0,8672 |
| TcCLB.507801.90 | beta-ketoacyl-ACP reductase | 0,02921 | 0,8494 |
| TcCLB.509999.40 | COPII coat assembly protein sec16, putative (sec16) | 0,02905 | 0,7268 |
| TcCLB.510003.30 | phosphatidylinositol 4-kinase alpha, putative | 0,02899 | 0,7705 |
| TcCLB.509455.120 | Peptide chain release factor 1, mitochondrial, putative (RI | 0,02892 | 0,8176 |
| TcCLB.508823.60 | hypothetical protein, conserved | 0,02888 | 0,85 |
| TcCLB.507911.15 | retrotransposon hot spot protein (RHS, pseudogene), puta | 0,02878 | 0,9073 |
| TcCLB.511491.90 | hypothetical protein | 0,0287 | 0,8521 |
| TcCLB.507641.150 | hypothetical protein, conserved | 0,02864 | 0,7675 |
| TcCLB.511385.110 | hypothetical protein | 0,02856 | 0,8664 |
| TcCLB.507515.30 | hypothetical protein, conserved | 0,0285 | 0,8831 |
| TcCLB.511439.120 | aminopeptidase, putative | 0,02845 | 0,7779 |
| TcCLB.506809.5 | retrotransposon hot spot (RHS) protein, putative | 0,02838 | 0,9334 |
| TcCLB.510437.45 | hypothetical protein | 0,02822 | 0,8686 |
| TcCLB.507669.104 | transcription elongation factor, putative | 0,02817 | 0,8289 |
| TcCLB.506357.90 | DNA-directed RNA polymerase II/III subunit, putative | 0,02813 | 0,8417 |
| TcCLB.504193.30 | retrotransposon hot spot protein (RHS, pseudogene), puta | 0,02804 | 0,8189 |
| TcCLB.510629.450 | hypothetical protein, conserved | 0,02795 | 0,8466 |
| TcCLB.510513.20 | hypothetical protein, conserved | 0,02784 | 0,8263 |
| TcCLB.504037.40 | hypothetical protein, conserved | 0,02781 | 0,8901 |
| TcCLB.509683.20 | hypothetical protein, conserved | 0,0277 | 0,808 |
| TcCLB.509105.80 | hypothetical protein, conserved | 0,02745 | 0,8543 |
| TcCLB.508409.190 | hypothetical protein, conserved | 0,02745 | 0,8919 |
| TcCLB.507081.30 | translation elongation factor EF-2, putative | 0,02734 | 0,8039 |
| TcCLB.506215.19 | intraflagellar transport protein 140, putative (IFT140) | 0,02728 | 0,7504 |
| TcCLB.510043.40 | hypothetical protein, conserved | 0,02719 | 0,8275 |
| TcCLB.503971.59 | hypothetical protein, conserved (fragment) | 0,02719 | 0,8604 |
| TcCLB.509157.230 | hypothetical protein, conserved | 0,02714 | 0,8031 |
| TcCLB.510601.10 | hypothetical protein, conserved | 0,02713 | 0,9033 |
| TcCLB.511417.129 | hypothetical protein, conserved | 0,0271 | 0,9368 |
| TcCLB.511907.250 | hypothetical protein | 0,02699 | 0,7936 |
| TcCLB.508781.40 | hypothetical protein, conserved | 0,02688 | 0,8292 |
| TcCLB.510551.15 | hypothetical protein | 0,02675 | 0,9298 |
| TcCLB.504147.10 | hypothetical protein, conserved | 0,02671 | 0,8038 |
| TcCLB.509757.10 | hypothetical protein, conserved | 0,02662 | 0,8425 |
| TcCLB.506145.50 | hypothetical protein, conserved | 0,02653 | 0,9106 |
| TcCLB.511255.160 | trans-sialidase (pseudogene), putative | 0,02646 | 0,9452 |
| TcCLB.510579.30 | hypothetical protein, conserved | 0,02644 | 0,7841 |
| TcCLB.504003.80 | hypothetical protein, conserved | 0,02643 | 0,9155 |

| | | | |
|---|---|---|---|
| TcCLB.503809.50 | hypothetical protein, conserved | 0,02621 | 0,8813 |
| TcCLB.508879.180 | hypothetical protein, conserved | 0,02616 | 0,8769 |
| TcCLB.508637.70 | RNA helicase, putative | 0,02604 | 0,8156 |
| TcCLB.508999.10 | DnaJ chaperone protein, putative | 0,02556 | 0,8211 |
| TcCLB.510747.26 | hypothetical protein, conserved | 0,02544 | 0,914 |
| TcCLB.509317.50 | Asparagine-linked glycosylation protein 12 (ALG12) | 0,02535 | 0,8454 |
| TcCLB.508729.9 | hypothetical protein, conserved | 0,02511 | 0,9058 |
| TcCLB.507017.100 | hypothetical protein, conserved | 0,02501 | 0,8961 |
| TcCLB.506217.30 | hypothetical protein | 0,02501 | 0,9299 |
| TcCLB.508731.50 | dynein light chain Tctex-type, putative | 0,02495 | 0,8798 |
| TcCLB.508917.30 | kinesin, putative | 0,02479 | 0,8254 |
| TcCLB.508771.60 | retrotransposon hot spot protein (RHS, pseudogene), puta | 0,02478 | 0,8562 |
| TcCLB.509797.50 | hypothetical protein, conserved | 0,02473 | 0,8732 |
| TcCLB.503971.40 | translation initiation factor 2 subunit, putative | 0,02466 | 0,8732 |
| TcCLB.511761.40 | hypothetical protein, conserved | 0,02466 | 0,934 |
| TcCLB.511293.30 | DNA-directed RNA polymerases I and III subunit RPAC1, p | 0,02447 | 0,862 |
| TcCLB.509775.9 | protein kinase, putative (fragment) | 0,02438 | 0,896 |
| TcCLB.506925.210 | hypothetical protein, conserved | 0,02426 | 0,9066 |
| TcCLB.507011.50 | Pre-rRNA-processing protein PNO1, putative (PNO1) | 0,02425 | 0,8612 |
| TcCLB.506483.69 | mitogen activated protein kinase 4, putative (MPK4) | 0,02406 | 0,8787 |
| TcCLB.506805.20 | hypothetical protein | 0,02399 | 0,9447 |
| TcCLB.503955.100 | protein kinase, putative | 0,02391 | 0,8401 |
| TcCLB.504643.30 | ubiquitin-conjugating enzyme E2, putative | 0,02381 | 0,8222 |
| TcCLB.510719.130 | hypothetical protein, conserved | 0,02377 | 0,8587 |
| TcCLB.504251.30 | hypothetical protein, conserved | 0,02376 | 0,8461 |
| TcCLB.506999.110 | hypothetical protein, conserved | 0,02366 | 0,8261 |
| TcCLB.507989.40 | ATP-dependent DEAD/H DNA helicase recQ, putative | 0,02358 | 0,8114 |
| TcCLB.510731.50 | small nuclear RNA gene activation protein (SNAP) 50, put | 0,02357 | 0,8316 |
| TcCLB.503869.50 | exosome-associated protein 1, putative (EAP1) | 0,02355 | 0,8514 |
| TcCLB.507797.18 | serine/threonine protein kinase, putative | 0,02353 | 0,8879 |
| TcCLB.510687.70 | hypothetical protein, conserved | 0,02346 | 0,8927 |
| TcCLB.509747.50 | hypothetical protein, conserved | 0,02319 | 0,8481 |
| TcCLB.509805.50 | hypothetical protein, conserved | 0,02319 | 0,8862 |
| TcCLB.511211.230 | hypothetical protein, conserved | 0,02318 | 0,8425 |
| TcCLB.506227.120 | hypothetical protein, conserved | 0,02317 | 0,8602 |
| TcCLB.508909.250 | hypothetical protein, conserved | 0,02315 | 0,8601 |
| TcCLB.509111.10 | hypothetical protein, conserved | 0,02291 | 0,862 |
| TcCLB.509895.50 | hypothetical protein, conserved | 0,02274 | 0,8277 |
| TcCLB.509789.50 | rab-like GTPase activating protein, putative | 0,02273 | 0,8617 |
| TcCLB.503955.30 | hypothetical protein, conserved | 0,02258 | 0,8679 |
| TcCLB.503421.7 | N-acetyltransferase complex ARD1 subunit (pseudogene), | 0,02251 | 0,9436 |
| TcCLB.507729.41 | elongation factor 1-gamma (EF-1-gamma, pseudogene), | 0,02236 | 0,9575 |
| TcCLB.508117.10 | surface protease GP63 (pseudogene), putative | 0,02225 | 0,9182 |
| TcCLB.509707.20 | hypothetical protein, conserved | 0,02222 | 0,812 |
| TcCLB.504105.70 | hypothetical protein, conserved | 0,02214 | 0,8682 |

| | | | |
|---|---|---|---|
| TcCLB.504431.109 | cytochrome b5, putative | 0,02207 | 0,8906 |
| TcCLB.510311.70 | 26S protease regulatory subunit, putative | 0,02199 | 0,8375 |
| TcCLB.510289.20 | hypothetical protein, conserved | 0,02191 | 0,844 |
| TcCLB.510013.270 | trans-sialidase (pseudogene), putative | 0,02185 | 0,9211 |
| TcCLB.506247.400 | hypothetical protein, conserved | 0,02172 | 0,889 |
| TcCLB.506401.300 | hypothetical protein, conserved | 0,02163 | 0,8717 |
| TcCLB.506401.130 | hypothetical protein, conserved | 0,02157 | 0,8742 |
| TcCLB.504703.10 | glycosyl hydrolase (pseudogene), putative | 0,0215 | 0,8378 |
| TcCLB.506195.280 | hypothetical protein, conserved | 0,02141 | 0,8237 |
| TcCLB.509733.134 | hypothetical protein, conserved | 0,02138 | 0,914 |
| TcCLB.509575.30 | hypothetical protein, conserved | 0,0213 | 0,8727 |
| TcCLB.504153.280 | hypothetical protein, conserved | 0,02106 | 0,8952 |
| TcCLB.508909.240 | hypothetical protein, conserved | 0,021 | 0,8672 |
| TcCLB.506357.29 | dynein light chain Tctex-type, putative | 0,02086 | 0,9088 |
| TcCLB.510943.23 | hypothetical protein, conserved | 0,0208 | 0,8819 |
| TcCLB.507801.140 | TPR-repeat protein, putative | 0,02074 | 0,8709 |
| TcCLB.506327.10 | proteasome subunit alpha type-1, putative (fragment) (PS | 0,02059 | 0,8937 |
| TcCLB.506543.20 | hypothetical protein, conserved | 0,02053 | 0,8399 |
| TcCLB.509965.10 | hypothetical protein, conserved (pseudogene) | 0,02048 | 0,8885 |
| TcCLB.506373.30 | retrotransposon hot spot protein (RHS, pseudogene), puta | 0,02035 | 0,901 |
| TcCLB.508899.40 | hypothetical protein, conserved | 0,02015 | 0,8476 |
| TcCLB.510741.124 | hypothetical protein, conserved | 0,02015 | 0,8899 |
| TcCLB.510769.140 | hypothetical protein, conserved | 0,01997 | 0,8908 |
| TcCLB.510121.40 | hypothetical protein | 0,01995 | 0,8979 |
| TcCLB.509897.250 | retrotransposon hot spot protein (RHS, pseudogene), puta | 0,01971 | 0,9011 |
| TcCLB.509115.41 | elongation factor 1-gamma (EF-1-gamma, pseudogene), | 0,01959 | 0,9298 |
| TcCLB.509161.10 | ubiquitin-conjugating enzyme E2, putative | 0,01927 | 0,8804 |
| TcCLB.506163.80 | katanin, putative | 0,01924 | 0,8913 |
| TcCLB.506525.20 | protein transport protein SEC13, putative | 0,0192 | 0,8938 |
| TcCLB.503593.30 | hypothetical protein, conserved | 0,0189 | 0,8802 |
| TcCLB.509007.90 | hypothetical protein | 0,01888 | 0,9069 |
| TcCLB.508827.100 | splicing factor 3B subunit 1, putative | 0,01884 | 0,8381 |
| TcCLB.511277.590 | hypothetical protein, conserved | 0,01879 | 0,8602 |
| TcCLB.510143.40 | UBX domain-containing protein | 0,01875 | 0,8639 |
| TcCLB.511421.100 | Enkuring domain-containig protein | 0,01856 | 0,8598 |
| TcCLB.506359.50 | hypothetical protein, conserved | 0,01829 | 0,9169 |
| TcCLB.509151.110 | hypothetical protein, conserved | 0,01824 | 0,8597 |
| TcCLB.506797.10 | hypothetical protein, conserved | 0,01766 | 0,8796 |
| TcCLB.510187.510 | hypothetical protein, conserved | 0,01754 | 0,9042 |
| TcCLB.410797.10 | surface protease GP63, putative | 0,01751 | 0,9053 |
| TcCLB.507603.180 | ubiquitin hydrolase, putative | 0,01749 | 0,8687 |
| TcCLB.507047.60 | hypothetical protein, conserved | 0,01748 | 0,9106 |
| TcCLB.509497.9 | hypothetical protein, conserved | 0,01739 | 0,916 |
| TcCLB.507083.90 | hypothetical protein, conserved | 0,01733 | 0,9142 |
| TcCLB.508201.20 | surface protease GP63 (pseudogene), putative | 0,01731 | 0,9542 |

| | | | |
|---|---|---|---|
| TcCLB.510763.40 | hypothetical protein, conserved | 0,01729 | 0,9216 |
| TcCLB.503723.70 | protein farnesyltransferase, putative | 0,01726 | 0,8923 |
| TcCLB.504797.60 | hypothetical protein, conserved | 0,01725 | 0,8847 |
| TcCLB.508411.40 | hypothetical protein, conserved | 0,01717 | 0,8998 |
| TcCLB.507039.41 | hypothetical protein, conserved | 0,01717 | 0,9282 |
| TcCLB.504147.200 | Regulatory particle triple-A ATPase subunit 6 (RPT6) | 0,01705 | 0,89 |
| TcCLB.511815.80 | 3'a2rel-related protein, putative | 0,01705 | 0,916 |
| TcCLB.503985.10 | trans-sialidase (pseudogene), putative | 0,01694 | 0,945 |
| TcCLB.508647.20 | hypothetical protein | 0,0169 | 0,8729 |
| TcCLB.508461.146 | hypothetical protein, conserved | 0,01675 | 0,8967 |
| TcCLB.412509.10 | retrotransposon hot spot (RHS) protein, putative | 0,01656 | 0,9449 |
| TcCLB.511383.70 | hypothetical protein, conserved | 0,01653 | 0,9212 |
| TcCLB.510835.24 | mitochondrial carrier protein, putative | 0,01649 | 0,9173 |
| TcCLB.506595.70 | trans-sialidase (pseudogene), putative | 0,01639 | 0,926 |
| TcCLB.504427.140 | hypothetical protein, conserved | 0,01633 | 0,9053 |
| TcCLB.508469.4 | hypothetical protein, conserved (fragment) | 0,0163 | 0,9073 |
| TcCLB.508461.450 | hypothetical protein, conserved | 0,0162 | 0,8962 |
| TcCLB.510759.160 | metacaspase 5, putative | 0,01618 | 0,8813 |
| TcCLB.506195.270 | hypothetical protein, conserved | 0,01617 | 0,8884 |
| TcCLB.511297.20 | anaphase promoting complex subunit 10, putative | 0,01612 | 0,9277 |
| TcCLB.509777.90 | hypothetical protein, conserved | 0,01604 | 0,8947 |
| TcCLB.506811.80 | hypothetical protein, conserved | 0,01599 | 0,9087 |
| TcCLB.509671.180 | hypothetical protein, conserved | 0,01597 | 0,9216 |
| TcCLB.511725.180 | dynein, putative | 0,01595 | 0,9029 |
| TcCLB.510349.70 | hypothetical protein, conserved | 0,0159 | 0,9234 |
| TcCLB.508541.100 | surface protease GP63, putative | 0,0158 | 0,964 |
| TcCLB.511751.110 | hypothetical protein, conserved | 0,01568 | 0,902 |
| TcCLB.506611.55 | elongation factor 1-gamma (EF-1-gamma, pseudogene), | | 0,01559 | 0,9749 |
| TcCLB.510303.30 | hypothetical protein, conserved | 0,01558 | 0,8803 |
| TcCLB.510759.170 | hypothetical protein, conserved | 0,01545 | 0,8831 |
| TcCLB.510203.49 | hypothetical protein, conserved (pseudogene) | 0,01532 | 0,9366 |
| TcCLB.509875.220 | hypothetical protein, conserved | 0,0153 | 0,8974 |
| TcCLB.511467.4 | hypothetical protein, conserved | 0,0152 | 0,914 |
| TcCLB.508965.90 | hypothetical protein, conserved | 0,01494 | 0,8838 |
| TcCLB.510513.60 | hypothetical protein, conserved | 0,01489 | 0,9264 |
| TcCLB.506513.100 | hypothetical protein, conserved | 0,01485 | 0,8907 |
| TcCLB.511367.90 | hypothetical protein, conserved | 0,01483 | 0,8885 |
| TcCLB.508141.30 | hypothetical protein, conserved | 0,01475 | 0,9022 |
| TcCLB.508479.300 | hypothetical protein, conserved | 0,01474 | 0,9141 |
| TcCLB.460747.30 | hypothetical protein, conserved | 0,01474 | 0,9454 |
| TcCLB.507723.70 | hypothetical protein, conserved (pseudogene) | 0,01467 | 0,889 |
| TcCLB.510187.390 | Protein kinase domain, putative | 0,01457 | 0,8997 |
| TcCLB.509717.140 | hypothetical protein, conserved (pseudogene) | 0,01449 | 0,9524 |
| TcCLB.506367.140 | hypothetical protein, conserved | 0,01444 | 0,9087 |
| TcCLB.509127.80 | hypothetical protein, conserved | 0,01442 | 0,9379 |

| | | | |
|---|---|---|---|
| TcCLB.508061.50 | dispersed gene family protein 1 (DGF-1), putative | 0,01437 | 0,9195 |
| TcCLB.508815.110 | serine/threonine protein phosphatase catalytic subunit, pu | 0,01436 | 0,8884 |
| TcCLB.510901.30 | hypothetical protein, conserved | 0,01432 | 0,9191 |
| TcCLB.509669.90 | hypothetical protein, conserved | 0,01429 | 0,9153 |
| TcCLB.510359.230 | hypothetical protein, conserved | 0,01404 | 0,9003 |
| TcCLB.509161.40 | centrin, putative | 0,01397 | 0,9288 |
| TcCLB.510165.10 | heat shock protein DNAJ, putative | 0,01394 | 0,916 |
| TcCLB.511001.170 | hypothetical protein, conserved | 0,01388 | 0,8906 |
| TcCLB.507023.220 | hypothetical protein, conserved | 0,01386 | 0,8984 |
| TcCLB.507811.60 | vesicle-associated membrane protein, putative | 0,01378 | 0,9159 |
| TcCLB.508299.30 | hypothetical protein | 0,01374 | 0,8879 |
| TcCLB.503919.10 | spliced leader RNA PSE-promoter transcription factor, put | 0,01373 | 0,9134 |
| TcCLB.509171.100 | hypothetical protein, conserved | 0,01358 | 0,9098 |
| TcCLB.503979.10 | ethanolamine phosphotransferase, putative (fragment) | 0,01358 | 0,9118 |
| TcCLB.506127.80 | molybdopterin synthase sulphurylase-like protein, putative | 0,0135 | 0,9147 |
| TcCLB.509265.23 | N-acetyltransferase complex ARD1 subunit (pseudogene), | 0,01337 | 0,9636 |
| TcCLB.507083.109 | hypothetical protein, conserved (fragment) | 0,01325 | 0,902 |
| TcCLB.509429.270 | proteasome regulatory ATPase subunit 3, putative | 0,01324 | 0,889 |
| TcCLB.510773.9 | hypothetical protein, conserved (fragment) | 0,01322 | 0,9457 |
| TcCLB.508397.60 | methyltransferase, putative | 0,01321 | 0,9397 |
| TcCLB.509965.130 | hypothetical protein, conserved | 0,01318 | 0,9458 |
| TcCLB.409101.9 | hypothetical protein, conserved (fragment) | 0,01312 | 0,9496 |
| TcCLB.508355.10 | hypothetical protein, conserved | 0,01304 | 0,9298 |
| TcCLB.506221.60 | hypothetical protein, conserved (pseudogene) | 0,01301 | 0,9032 |
| TcCLB.506871.90 | hypothetical protein, conserved | 0,01295 | 0,8979 |
| TcCLB.508741.90 | hypothetical protein | 0,01293 | 0,9202 |
| TcCLB.504103.109 | hypothetical protein, conserved | 0,01288 | 0,9364 |
| TcCLB.506557.30 | hypothetical protein, conserved | 0,01282 | 0,9139 |
| TcCLB.507851.30 | hypothetical protein, conserved | 0,01281 | 0,9023 |
| TcCLB.507839.35 | trans-sialidase, putative | 0,01274 | 0,9608 |
| TcCLB.511895.50 | hypothetical protein, conserved | 0,01273 | 0,909 |
| TcCLB.508329.9 | potassium voltage-gated channel, putative | 0,01268 | 0,9737 |
| TcCLB.506587.80 | hypothetical protein, conserved | 0,01267 | 0,9218 |
| TcCLB.504147.330 | hypothetical protein, conserved | 0,01265 | 0,8764 |
| TcCLB.506177.10 | hypothetical protein, conserved | 0,01264 | 0,9245 |
| TcCLB.510879.90 | hypothetical protein, conserved | 0,01239 | 0,941 |
| TcCLB.510763.90 | hypothetical protein, conserved | 0,01233 | 0,9424 |
| TcCLB.506705.30 | hypothetical protein, conserved | 0,0123 | 0,9434 |
| TcCLB.411059.18 | clathrin heavy chain, putative | 0,01222 | 0,9407 |
| TcCLB.507711.235 | deoxyribodipyrimidine photo-lyase, putative | 0,01215 | 0,9339 |
| TcCLB.506211.20 | hypothetical protein, conserved | 0,01212 | 0,9319 |
| TcCLB.510509.9 | inositol 1,4,5-trisphosphate receptor, putative (fragment) | 0,01203 | 0,9298 |
| TcCLB.510901.200 | differentiation inhibitory kinase, putative (ZFK) | 0,01198 | 0,9087 |
| TcCLB.463269.10 | eukaryotic translation initiation factor 1A, putative (fragm | 0,01193 | 0,9693 |
| TcCLB.505941.40 | hypothetical protein, conserved | 0,01184 | 0,9394 |

| | | | |
|---|---|---|---|
| TcCLB.507923.20 | U3 small nuclear ribonucleoprotein (snRNP), putative | 0,0118 | 0,9033 |
| TcCLB.507229.25 | trans-sialidase (pseudogene), putative | 0,01174 | 0,9383 |
| TcCLB.506667.110 | trans-sialidase (pseudogene), putative | 0,01173 | 0,9645 |
| TcCLB.508043.40 | kinesin, putative | 0,01163 | 0,9106 |
| TcCLB.506585.90 | RNA processing endonuclease 1, putative | 0,01147 | 0,9337 |
| TcCLB.434931.20 | hypothetical protein, conserved | 0,01144 | 0,9407 |
| TcCLB.506275.90 | hypothetical protein, conserved | 0,0113 | 0,9364 |
| TcCLB.507359.30 | hypothetical protein, conserved (pseudogene) | 0,0112 | 0,9846 |
| TcCLB.511181.70 | hypothetical protein, conserved | 0,01113 | 0,9468 |
| TcCLB.507787.30 | hypothetical protein, conserved | 0,01113 | 0,9555 |
| TcCLB.479379.9 | dynein heavy chain, putative | 0,01093 | 0,9212 |
| TcCLB.508641.220 | UDP-Gal or UDP-GlcNAc-dependent glycosyltransferase, p | 0,01093 | 0,9469 |
| TcCLB.511145.30 | hypothetical protein, conserved | 0,01079 | 0,9298 |
| TcCLB.508741.260 | hypothetical protein, conserved | 0,01062 | 0,9311 |
| TcCLB.509135.9 | hypothetical protein (fragment) | 0,01061 | 0,9564 |
| TcCLB.508141.70 | Kinesin-13 4, putative (KIN13-4) | 0,01047 | 0,9462 |
| TcCLB.511521.20 | hypothetical protein, conserved | 0,01042 | 0,943 |
| TcCLB.509795.30 | hypothetical protein, conserved | 0,01038 | 0,9327 |
| TcCLB.510989.50 | hypothetical protein, conserved | 0,01021 | 0,9091 |
| TcCLB.508027.30 | hypothetical protein, conserved | 0,01017 | 0,949 |
| TcCLB.510187.448 | hypothetical protein, conserved | 0,01016 | 0,9169 |
| TcCLB.506649.10 | hypothetical protein | 0,01006 | 0,9468 |
| TcCLB.508385.60 | hypothetical protein, conserved | 0,009891 | 0,9491 |
| TcCLB.510607.29 | dispersed gene family protein 1 (DGF-1), putative (fragme | 0,009885 | 0,9675 |
| TcCLB.509693.130 | hypothetical protein | 0,009875 | 0,9447 |
| TcCLB.505997.174 | dispersed gene family protein 1 (DGF-1), putative | 0,009789 | 0,9628 |
| TcCLB.511123.20 | retrotransposon hot spot protein (RHS, pseudogene), puta | 0,009625 | 0,9542 |
| TcCLB.506539.21 | hypothetical protein, conserved (pseudogene) | 0,009497 | 0,9731 |
| TcCLB.507795.60 | tryptophanyl-tRNA synthetase (pseudogene), putative | 0,00947 | 0,9436 |
| TcCLB.508153.1110 | arginine N-methyltransferase, putative | 0,009304 | 0,9452 |
| TcCLB.506591.69 | hypothetical protein, conserved | 0,009205 | 0,9391 |
| TcCLB.503611.10 | nucleoplasmin-like protein (NLP), putative | 0,009075 | 0,9442 |
| TcCLB.509961.20 | dispersed gene family protein 1 (DGF-1), putative | 0,009073 | 0,9813 |
| TcCLB.508461.234 | hypothetical protein, conserved | 0,009006 | 0,9576 |
| TcCLB.506195.220 | hypothetical protein, conserved | 0,008986 | 0,9558 |
| TcCLB.506735.70 | hypothetical protein, conserved | 0,008909 | 0,929 |
| TcCLB.508897.70 | ADP-ribosylation factor family, putative | 0,008785 | 0,9543 |
| TcCLB.511003.221 | cleavage and polyadenylation specificity factor, putative ( | 0,00852 | 0,942 |
| TcCLB.507611.380 | hypothetical protein | 0,008493 | 0,9508 |
| TcCLB.511653.50 | hypothetical protein, conserved | 0,008454 | 0,9537 |
| TcCLB.506573.10 | hypothetical protein, conserved | 0,008414 | 0,9421 |
| TcCLB.508307.130 | hypothetical protein, conserved | 0,008345 | 0,9459 |
| TcCLB.509569.170 | hypothetical protein | 0,008254 | 0,9363 |
| TcCLB.504109.180 | hypothetical protein, conserved | 0,007955 | 0,9696 |
| TcCLB.507875.180 | AGP2beta-2, putative | 0,007834 | 0,9436 |

| | | | |
|---|---|---|---|
| TcCLB.507437.21 | elongation factor 1-gamma (EF-1-gamma, pseudogene), | 0,007807 | 0,9864 |
| TcCLB.506567.77 | hypothetical protein, conserved (pseudogene) | 0,007728 | 0,9744 |
| TcCLB.507007.80 | hypothetical protein | 0,007641 | 0,9384 |
| TcCLB.506167.40 | proteasome subunit alpha type-1, putative (PSA1) | 0,007638 | 0,9576 |
| TcCLB.504003.14 | hypothetical protein, conserved | 0,007635 | 0,9585 |
| TcCLB.509601.44 | hypothetical protein, conserved | 0,007616 | 0,9399 |
| TcCLB.506503.56 | hypothetical protein, conserved | 0,007601 | 0,9561 |
| TcCLB.510595.10 | hypothetical protein | 0,007561 | 0,9523 |
| TcCLB.463673.9 | hypothetical protein, conserved | 0,007549 | 0,9625 |
| TcCLB.504153.190 | mitochondrial edited mRNA stability factor 1 subunit | 0,007467 | 0,944 |
| TcCLB.510175.200 | retrotransposon hot spot protein (RHS, pseudogene), puta | 0,0074 | 0,9743 |
| TcCLB.508369.40 | serine/threonine-protein kinase A (pseudogene), putative | 0,007393 | 0,9537 |
| TcCLB.505101.50 | GMP synthetase (pseudogene), putative | 0,007305 | 0,9741 |
| TcCLB.507801.179 | vesicle-associated membrane protein | 0,007283 | 0,9544 |
| TcCLB.504153.90 | Cytoplasmic dynein 2 heavy chain (DYNC2H1), putative | 0,007244 | 0,9256 |
| TcCLB.510755.120 | RNA-binding protein, putative | 0,0072 | 0,9502 |
| TcCLB.507777.10 | hypothetical protein, conserved | 0,007184 | 0,9645 |
| TcCLB.510363.290 | hypothetical protein | 0,007066 | 0,9945 |
| TcCLB.508145.10 | RNA-binding protein, putative | 0,007057 | 0,988 |
| TcCLB.511439.100 | p-nitrophenylphosphatase, putative | 0,007024 | 0,9667 |
| TcCLB.510353.30 | paraflagellar rod component par4, putative | 0,006999 | 0,9544 |
| TcCLB.508461.430 | proteasome beta 2 subunit, putative | 0,006982 | 0,9447 |
| TcCLB.511297.10 | hypothetical protein, conserved | 0,006953 | 0,9403 |
| TcCLB.511323.30 | hypothetical protein, conserved | 0,006953 | 0,9615 |
| TcCLB.506777.90 | hypothetical protein, conserved | 0,006901 | 0,9571 |
| TcCLB.510607.80 | dispersed gene family protein 1 (DGF-1), putative | 0,006747 | 0,9794 |
| TcCLB.506453.20 | mucin TcMUC (pseudogene), putative | 0,006673 | 0,9763 |
| TcCLB.510189.51 | elongation factor 1-gamma (EF-1-gamma, pseudogene), | 0,006499 | 0,9896 |
| TcCLB.510625.140 | elongation factor 1-gamma (EF-1-gamma), putative | 0,006348 | 0,9644 |
| TcCLB.503423.10 | glycine dehydrogenase [decarboxylating] (pseudogene), p | 0,006293 | 0,9737 |
| TcCLB.508501.270 | hypothetical protein, conserved | 0,00629 | 0,96 |
| TcCLB.507093.72 | hypothetical protein, conserved | 0,006151 | 0,954 |
| TcCLB.510603.150 | hypothetical protein, conserved | 0,006131 | 0,9696 |
| TcCLB.503719.39 | hypothetical protein, conserved | 0,005994 | 0,9716 |
| TcCLB.418221.4 | hypothetical protein, conserved (fragment) | 0,005819 | 0,97 |
| TcCLB.505977.13 | protein kinase, putative (fragment) | 0,005803 | 0,9645 |
| TcCLB.508881.150 | hypothetical protein, conserved | 0,005612 | 0,9664 |
| TcCLB.509637.9 | hypothetical protein, conserved | 0,00542 | 0,9743 |
| TcCLB.511545.50 | hypothetical protein, conserved | 0,005321 | 0,9635 |
| TcCLB.507993.380 | hypothetical protein, conserved | 0,005235 | 0,9718 |
| TcCLB.506627.70 | hypothetical protein | 0,005098 | 0,971 |
| TcCLB.508721.4 | hypothetical protein, conserved | 0,005031 | 0,9784 |
| TcCLB.509767.180 | exosome component CSL4, putative (CSL4) | 0,00497 | 0,982 |
| TcCLB.509985.60 | hypothetical protein, conserved | 0,004825 | 0,9636 |
| TcCLB.506505.50 | hypothetical protein, conserved | 0,00477 | 0,9644 |

| | | | |
|---|---|---|---|
| TcCLB.509837.30 | hypothetical protein | 0,004694 | 0,978 |
| TcCLB.510099.40 | hypothetical protein, conserved | 0,004609 | 0,9734 |
| TcCLB.507727.19 | hypothetical protein | 0,004427 | 0,9876 |
| TcCLB.504105.240 | hypothetical protein, conserved | 0,004372 | 0,9668 |
| TcCLB.504097.10 | RNA editing complex protein MP61, putative | 0,004303 | 0,9732 |
| TcCLB.510715.120 | mucin TcMUCII, putative | 0,004294 | 0,9884 |
| TcCLB.507305.11 | casein kinase (pseudogene), putative | 0,004201 | 0,9611 |
| TcCLB.509231.5 | hypothetical protein, conserved (fragment) | 0,004146 | 0,9938 |
| TcCLB.510663.57 | transport protein particle (TRAPP) subunit, putative | 0,004052 | 0,9794 |
| TcCLB.509539.40 | hypothetical protein, conserved | 0,004029 | 0,9868 |
| TcCLB.511467.20 | hypothetical protein, conserved | 0,004 | 0,9581 |
| TcCLB.504105.60 | hypothetical protein, conserved | 0,003961 | 0,9752 |
| TcCLB.504089.40 | long-chain-fatty-acid-CoA ligase, putative | 0,003955 | 0,9718 |
| TcCLB.508355.370 | hypothetical protein, conserved | 0,003955 | 0,9795 |
| TcCLB.508641.100 | hypothetical protein, conserved | 0,003895 | 0,9754 |
| TcCLB.507053.209 | Cytoplasmic dynein 2 heavy chain (DYNC2H1), putative | 0,003814 | 0,9717 |
| TcCLB.511715.50 | hypothetical protein, conserved (pseudogene) | 0,003737 | 0,9819 |
| TcCLB.509885.64 | hypothetical protein, conserved | 0,003685 | 0,9876 |
| TcCLB.509803.10 | hypothetical protein | 0,003684 | 0,9867 |
| TcCLB.511431.90 | conserved protein, unknown function | 0,003658 | 0,9913 |
| TcCLB.511283.20 | protein transport protein SEC13, putative | 0,003478 | 0,9819 |
| TcCLB.508867.9 | kinesin, putative (fragment) | 0,003455 | 0,9896 |
| TcCLB.511143.30 | hypothetical protein, conserved | 0,00324 | 0,9826 |
| TcCLB.510603.110 | hypothetical protein, conserved | 0,003147 | 0,988 |
| TcCLB.509769.10 | hypothetical protein, conserved | 0,00299 | 0,9894 |
| TcCLB.507093.170 | hypothetical protein, conserved | 0,002923 | 0,9747 |
| TcCLB.507057.40 | hypothetical protein, conserved | 0,002764 | 0,9883 |
| TcCLB.508747.10 | hypothetical protein | 0,002376 | 0,9972 |
| TcCLB.440099.19 | hypothetical protein, conserved | 0,002315 | 0,9889 |
| TcCLB.506925.410 | ATPase, putative | 0,002274 | 0,9877 |
| TcCLB.508325.320 | protein kinase, putative | 0,002017 | 0,9931 |
| TcCLB.510121.100 | hypothetical protein | 0,001893 | 0,9938 |
| TcCLB.504769.60 | hypothetical protein, conserved | 0,001846 | 0,9925 |
| TcCLB.505183.10 | hypothetical protein, conserved | 0,001804 | 0,9896 |
| TcCLB.506155.90 | hypothetical protein, conserved (pseudogene) | 0,001693 | 0,9872 |
| TcCLB.506765.43 | hypothetical protein | 0,001641 | 0,9942 |
| TcCLB.503809.20 | peroxin 12, putative (PEX12) | 0,001513 | 0,9924 |
| TcCLB.507963.90 | hypothetical protein, conserved | 0,001463 | 0,9943 |
| TcCLB.507081.70 | hypothetical protein, conserved | 0,001419 | 0,9913 |
| TcCLB.508625.40 | hypothetical protein, conserved | 0,001338 | 0,9947 |
| TcCLB.511517.90 | hypothetical protein, conserved | 0,001315 | 0,9926 |
| TcCLB.509539.44 | hypothetical protein, conserved | 0,001177 | 0,9968 |
| TcCLB.506559.540 | hypothetical protein, conserved | 0,001042 | 0,9951 |
| TcCLB.508489.20 | hypothetical protein, conserved | 0,0009611 | 0,995 |
| TcCLB.509203.79 | guanine-nucleotide-exchange-factor, putative (fragment) | 0,0009089 | 0,9963 |

| | | | |
|---|---|---|---|
| TcCLB.506701.10 | dihydrolipoamide dehydrogenase, putative (GCVL-1) | 0,000906 | 0,9967 |
| TcCLB.511301.170 | retrotransposon hot spot (RHS) protein, putative | 0,0008683 | 0,9956 |
| TcCLB.508693.190 | hypothetical protein, conserved | 0,0008563 | 0,9943 |
| TcCLB.506241.200 | hypothetical protein, conserved | 0,0008091 | 0,9945 |
| TcCLB.507001.30 | ankyrin repeat protein, putative | 0,000793 | 0,997 |
| TcCLB.508111.40 | hypothetical protein, conserved | 0,0007824 | 0,995 |
| TcCLB.503897.50 | hypothetical protein, conserved | 0,0006871 | 0,9961 |
| TcCLB.511423.70 | glycerol-3-phosphate dehydrogenase (FAD-dependent), pu | 0,0006447 | 0,9967 |
| TcCLB.508179.40 | hypothetical protein, conserved | 0,0005822 | 0,9966 |
| TcCLB.506503.110 | DNA-directed RNA polymerases I and III subunit RPAC1, pu | 0,0004871 | 0,9973 |
| TcCLB.508257.90 | hypothetical protein, conserved | 0,000363 | 0,9977 |
| TcCLB.506275.40 | hypothetical protein, conserved | 0,0003497 | 0,9977 |
| TcCLB.508851.79 | hypothetical protein, conserved | 0,00034 | 0,9982 |
| TcCLB.506723.30 | hypothetical protein | 0,000336 | 0,9989 |
| TcCLB.511725.160 | hypothetical protein, conserved | 0,0003285 | 0,9973 |
| TcCLB.506867.40 | surface protease GP63, putative | 0,0001145 | 0,9993 |
| TcCLB.510661.100 | hypothetical protein | 0,0001061 | 0,9993 |
| TcCLB.511317.20 | hypothetical protein, conserved | 0,00003341 | 0,9997 |
| TcCLB.503651.10 | hypothetical protein, conserved | -0,00008295 | 0,9994 |
| TcCLB.509287.5 | hypothetical protein, conserved (pseudogene) | -0,0001935 | 0,9993 |
| TcCLB.509671.84 | hypothetical protein, conserved | -0,0002906 | 0,9984 |
| TcCLB.506343.51 | retrotransposon hot spot protein (RHS, pseudogene), puta | -0,0003047 | 0,999 |
| TcCLB.509719.9 | N-acetyltransferase complex ARD1 subunit, putative (frag | -0,0003199 | 0,999 |
| TcCLB.508405.90 | soluble N-ethylmaleimide sensitive factor (NSF) attachme | -0,0003654 | 0,9982 |
| TcCLB.507047.160 | hypothetical protein, conserved | -0,00042 | 0,9985 |
| TcCLB.510187.394 | hypothetical protein, conserved | -0,0004897 | 0,9973 |
| TcCLB.511903.60 | hypothetical protein, conserved | -0,000529 | 0,9977 |
| TcCLB.510137.10 | trans-sialidase (pseudogene), putative | -0,0005756 | 0,998 |
| TcCLB.508277.350 | cation transporter, putative | -0,0006266 | 0,9951 |
| TcCLB.509229.20 | hypothetical protein, conserved | -0,0006726 | 0,9951 |
| TcCLB.507509.30 | RNA polymerase-like protein, putative | -0,0007335 | 0,9968 |
| TcCLB.507481.20 | retrotransposon hot spot protein (RHS, pseudogene), puta | -0,0007716 | 0,997 |
| TcCLB.511215.14 | hypothetical protein, conserved | -0,0008527 | 0,9955 |
| TcCLB.510303.100 | hypothetical protein, conserved | -0,0009058 | 0,9947 |
| TcCLB.503999.50 | ubiquitin-conjugating enzyme, putative | -0,001025 | 0,9942 |
| TcCLB.506727.30 | hypothetical protein, conserved | -0,00106 | 0,995 |
| TcCLB.511239.10 | hypothetical protein, conserved | -0,001146 | 0,9945 |
| TcCLB.509793.30 | kinetoplast DNA-associated protein, putative | -0,001169 | 0,9945 |
| TcCLB.509427.60 | trans-sialidase (pseudogene), putative | -0,001192 | 0,9963 |
| TcCLB.503479.70 | bilobe protein, conserved | -0,001212 | 0,9899 |
| TcCLB.430605.10 | hypothetical protein, conserved | -0,001227 | 0,9914 |
| TcCLB.503819.10 | hypothetical protein, conserved | -0,00127 | 0,9938 |
| TcCLB.511621.244 | hypothetical protein, conserved | -0,001365 | 0,995 |
| TcCLB.508919.60 | ADP-ribosylation factor, putative (ARL-1) | -0,001551 | 0,993 |
| TcCLB.506559.490 | hypothetical protein, conserved | -0,001585 | 0,9945 |

| | | | |
|---|---|---|---|
| TcCLB.511537.40 | protein phosphatase 2A catalytic subunit, putative | -0,001749 | 0,9891 |
| TcCLB.505989.100 | hypothetical protein, conserved | -0,001759 | 0,9912 |
| TcCLB.507007.9 | hypothetical protein, conserved (pseudogene) | -0,001817 | 0,9888 |
| TcCLB.509791.160 | hypothetical protein, conserved | -0,001824 | 0,9864 |
| TcCLB.503973.250 | mucin TcMUCII (pseudogene), putative | -0,001945 | 0,995 |
| TcCLB.434945.4 | ADP-ribosylation factor, putative | -0,002005 | 0,9885 |
| TcCLB.508547.130 | NUDIX family hydrolase, putative | -0,002085 | 0,9896 |
| TcCLB.510105.180 | hypothetical protein, conserved | -0,002096 | 0,9868 |
| TcCLB.504035.140 | mismatch repair protein MLH1, putative | -0,00213 | 0,9881 |
| TcCLB.508355.320 | arginyl-tRNA synthetase, putative | -0,002156 | 0,9851 |
| TcCLB.506479.74 | Quinonprotein alcohol dehydrogenase-like protein, putativ | -0,002174 | 0,9868 |
| TcCLB.506475.130 | hypothetical protein, conserved | -0,002183 | 0,9851 |
| TcCLB.507089.180 | hypothetical protein, conserved | -0,002226 | 0,986 |
| TcCLB.503639.30 | hypothetical protein, conserved | -0,002293 | 0,9884 |
| TcCLB.511589.230 | eukaryotic translation initiation factor 3 subunit 2, putativ | -0,002318 | 0,9837 |
| TcCLB.508859.100 | hypothetical protein, conserved | -0,002407 | 0,9894 |
| TcCLB.509647.40 | hypothetical protein, conserved | -0,002485 | 0,987 |
| TcCLB.509609.10 | hypothetical protein, conserved | -0,002505 | 0,9883 |
| TcCLB.506195.50 | hypothetical protein, conserved | -0,002527 | 0,9891 |
| TcCLB.511273.10 | dispersed gene family protein 1 (DGF-1, pseudogene), put | -0,002693 | 0,995 |
| TcCLB.508999.269 | hypothetical protein, conserved (fragment) | -0,002698 | 0,988 |
| TcCLB.506973.14 | mucin TcMUCII (pseudogene), putative | -0,002764 | 0,9914 |
| TcCLB.509987.10 | hypothetical protein, conserved | -0,003474 | 0,9837 |
| TcCLB.509749.39 | short-chain dehydrogenase, putative | -0,003494 | 0,993 |
| TcCLB.509205.30 | eukaryotic translation initiation factor, putative | -0,003579 | 0,9782 |
| TcCLB.508055.25 | hypothetical protein | -0,003601 | 0,9932 |
| TcCLB.511821.70 | hypothetical protein, conserved | -0,003649 | 0,9807 |
| TcCLB.508461.19 | hypothetical protein, conserved | -0,003678 | 0,9823 |
| TcCLB.503689.40 | hypothetical protein, conserved | -0,003715 | 0,9721 |
| TcCLB.511293.40 | U3 snoRNA-associated protein UTP11, putative | -0,003759 | 0,9785 |
| TcCLB.508699.10 | DNA topoisomerase II, putative | -0,003773 | 0,969 |
| TcCLB.506679.30 | hypothetical protein, conserved | -0,003787 | 0,9696 |
| TcCLB.506683.190 | trans-sialidase, Group II, putative | -0,003791 | 0,9836 |
| TcCLB.506605.10 | hypothetical protein, conserved | -0,003794 | 0,9849 |
| TcCLB.503505.14 | PTP1-interacting protein, 39 kDa, putative | -0,003806 | 0,9831 |
| TcCLB.510131.10 | hypothetical protein, conserved | -0,003876 | 0,9743 |
| TcCLB.510265.14 | hypothetical protein, conserved | -0,003905 | 0,9823 |
| TcCLB.510741.40 | hypothetical protein, conserved | -0,00401 | 0,9686 |
| TcCLB.507671.39 | serine/threonine-protein phosphatase PP1 | -0,004045 | 0,9851 |
| TcCLB.509561.120 | deaminase, putative | -0,004375 | 0,9818 |
| TcCLB.503823.10 | phosphoglycerate mutase family member 5, putative | -0,004473 | 0,9825 |
| TcCLB.507951.140 | U2 small nuclear ribonucleoprotein B, putative | -0,004491 | 0,9861 |
| TcCLB.508833.10 | hypothetical protein | -0,004492 | 0,9835 |
| TcCLB.507747.230 | acid phosphatase type 5, putative | -0,00461 | 0,9752 |
| TcCLB.511633.10 | hypothetical protein, conserved | -0,00463 | 0,9932 |

| | | | |
|---|---|---|---|
| TcCLB.509213.10 | hypothetical protein, conserved | -0,004698 | 0,9781 |
| TcCLB.509385.19 | retrotransposon hot spot protein (RHS, pseudogene), puta | -0,004818 | 0,9879 |
| TcCLB.510187.240 | hypothetical protein, conserved | -0,005055 | 0,9586 |
| TcCLB.510087.10 | hypothetical protein, conserved | -0,005109 | 0,9753 |
| TcCLB.507943.80 | hypothetical protein, conserved (pseudogene) | -0,005193 | 0,9723 |
| TcCLB.511509.21 | hypothetical protein, conserved | -0,005419 | 0,9646 |
| TcCLB.508207.80 | Mini-chromosome maintenance complex-binding protein | -0,005522 | 0,9676 |
| TcCLB.507005.70 | Pre-rRNA-processing protein PNO1, putative (PNO1) | -0,005735 | 0,9687 |
| TcCLB.509669.110 | GTPase activator protein, putative | -0,005739 | 0,9645 |
| TcCLB.508409.320 | hypothetical protein, conserved | -0,006065 | 0,9532 |
| TcCLB.510149.100 | hypothetical protein, conserved | -0,006069 | 0,9681 |
| TcCLB.509625.10 | ATP-dependent RNA helicase, putative (fragment) | -0,006142 | 0,9593 |
| TcCLB.510885.30 | hypothetical protein | -0,006164 | 0,958 |
| TcCLB.511003.50 | hypothetical protein, conserved | -0,006196 | 0,9547 |
| TcCLB.503847.99 | hypothetical protein, conserved (pseudogene) | -0,006487 | 0,9736 |
| TcCLB.507491.140 | hypothetical protein, conserved | -0,006491 | 0,9629 |
| TcCLB.510517.110 | hypothetical protein | -0,006533 | 0,9763 |
| TcCLB.510315.10 | IAG-nucleoside hydrolase, putative (IAGNH) | -0,006545 | 0,9562 |
| TcCLB.503953.50 | hypothetical protein, conserved | -0,006569 | 0,9562 |
| TcCLB.508543.60 | hypothetical protein, conserved | -0,006682 | 0,9605 |
| TcCLB.508403.20 | hypothetical protein, conserved (pseudogene) | -0,006695 | 0,9571 |
| TcCLB.507031.20 | hypothetical protein, conserved | -0,006728 | 0,956 |
| TcCLB.440099.28 | protein kinase, putative | -0,006831 | 0,9702 |
| TcCLB.508587.50 | hypothetical protein | -0,006864 | 0,984 |
| TcCLB.504253.40 | hypothetical protein, conserved | -0,007127 | 0,9568 |
| TcCLB.508277.30 | hypothetical protein, conserved | -0,007163 | 0,95 |
| TcCLB.509461.90 | inositol 1,4,5-trisphosphate receptor, putative (IP3R) | -0,007214 | 0,9299 |
| TcCLB.508553.20 | hypothetical protein | -0,007219 | 0,9856 |
| TcCLB.508199.50 | palmitoyl acyltransferase 1, putative | -0,00735 | 0,9436 |
| TcCLB.506821.100 | mitochondrial RNA binding complex 1 subunit, putative (N | -0,007363 | 0,9506 |
| TcCLB.510665.40 | DNA-repair protein, putative | -0,007403 | 0,943 |
| TcCLB.507023.30 | pseudouridine synthase A-like protein, putative | -0,007431 | 0,9506 |
| TcCLB.504021.50 | hypothetical protein | -0,00754 | 0,9488 |
| TcCLB.510525.20 | hypothetical protein, conserved | -0,00754 | 0,9603 |
| TcCLB.510491.80 | hypothetical protein, conserved | -0,007643 | 0,9524 |
| TcCLB.503869.9 | axoneme central apparatus protein, putative | -0,007694 | 0,9615 |
| TcCLB.511127.70 | hypothetical protein, conserved | -0,007791 | 0,948 |
| TcCLB.471901.5 | GDP-mannose 4,6 dehydratase, putative (fragment) | -0,007837 | 0,9868 |
| TcCLB.507625.80 | hypothetical protein, conserved | -0,007839 | 0,9547 |
| TcCLB.506265.100 | ATP-dependent RNA helicase, putative | -0,0079 | 0,938 |
| TcCLB.506203.129 | DNA-directed RNA polymerase, mitochondrial (fragment) | -0,007924 | 0,9408 |
| TcCLB.508461.290 | 4E-interacting protein, putative | -0,007967 | 0,9492 |
| TcCLB.511507.24 | hypothetical protein, conserved | -0,008081 | 0,9709 |
| TcCLB.510101.20 | phosphatidylinositol-4-phosphate 5-kinase, putative | -0,008387 | 0,9495 |
| TcCLB.506733.30 | hypothetical protein, conserved | -0,008415 | 0,934 |

| | | | |
|---|---|---|---|
| TcCLB.507829.19 | Vacuolar protein sorting-associated protein 16 homolog, p | -0,008419 | 0,9699 |
| TcCLB.510101.230 | ATPase (pseudogene), putative | -0,008482 | 0,9298 |
| TcCLB.509573.50 | hypothetical protein, conserved | -0,008602 | 0,9289 |
| TcCLB.510115.70 | trans-sialidase, putative | -0,008719 | 0,9773 |
| TcCLB.506419.20 | tousled-like kinase II, putative | -0,008723 | 0,9492 |
| TcCLB.507159.65 | DNA-directed RNA polymerase subunit, putative | -0,008787 | 0,9513 |
| TcCLB.506203.30 | SWI/SNF-related matrix-associated actin-dependent regu | -0,008824 | 0,9375 |
| TcCLB.508073.20 | hypothetical protein | -0,008894 | 0,9539 |
| TcCLB.509349.30 | hypothetical protein | -0,008962 | 0,9746 |
| TcCLB.508775.30 | dispersed gene family protein 1 (DGF-1), putative | -0,009008 | 0,9764 |
| TcCLB.507583.10 | hypothetical protein, conserved | -0,009058 | 0,9737 |
| TcCLB.507037.50 | exonuclease, putative | -0,009087 | 0,9602 |
| TcCLB.508409.250 | hypothetical protein, conserved | -0,009142 | 0,9529 |
| TcCLB.507983.10 | dispersed gene family protein 1 (DGF-1), putative (fragme | -0,009155 | 0,9807 |
| TcCLB.511887.40 | hypothetical protein, conserved | -0,009191 | 0,9692 |
| TcCLB.506637.40 | hypothetical protein, conserved | -0,009192 | 0,9561 |
| TcCLB.506469.70 | hypothetical protein | -0,009264 | 0,9679 |
| TcCLB.504205.20 | trans-sialidase (pseudogene), putative | -0,009268 | 0,9693 |
| TcCLB.511491.120 | hypothetical protein, conserved | -0,009335 | 0,9298 |
| TcCLB.509871.31 | ATP-dependent DEAD/H RNA helicase (pseudogene), puta | -0,009355 | 0,9764 |
| TcCLB.507787.130 | hypothetical protein, conserved | -0,009357 | 0,9402 |
| TcCLB.507509.10 | hypothetical protein, conserved (pseudogene) | -0,009379 | 0,9576 |
| TcCLB.508851.19 | retrotransposon hot spot (RHS) protein, putative | -0,00938 | 0,9402 |
| TcCLB.509429.220 | dolichyl-P-Man:GDP-Man5GlcNAc2-PP-dolichyl alpha-1,2-i | -0,009468 | 0,938 |
| TcCLB.509471.10 | hypothetical protein, conserved | -0,009616 | 0,9457 |
| TcCLB.511899.50 | Trypanosome basal body component protein (fragment) (' | -0,009787 | 0,9316 |
| TcCLB.510861.50 | hypothetical protein, conserved | -0,009892 | 0,9714 |
| TcCLB.503677.29 | hypothetical protein, conserved | -0,01008 | 0,9402 |
| TcCLB.509695.80 | hypothetical protein, conserved | -0,0101 | 0,9339 |
| TcCLB.511577.30 | hypothetical protein, conserved | -0,01014 | 0,9356 |
| TcCLB.510421.170 | hypothetical protein, conserved | -0,01016 | 0,9066 |
| TcCLB.505481.10 | hypothetical protein | -0,01022 | 0,9727 |
| TcCLB.508273.60 | hypothetical protein, conserved | -0,01035 | 0,9255 |
| TcCLB.511585.125 | hypothetical protein, conserved | -0,01044 | 0,9362 |
| TcCLB.510431.260 | hypothetical protein, conserved | -0,01044 | 0,9424 |
| TcCLB.510329.140 | hypothetical protein, conserved | -0,01049 | 0,9015 |
| TcCLB.503833.30 | peroxin 13, putative (PEX13) | -0,01052 | 0,9367 |
| TcCLB.510289.44 | hypothetical protein, conserved | -0,01053 | 0,9537 |
| TcCLB.509047.110 | rac serine-threonine kinase, putative | -0,01062 | 0,924 |
| TcCLB.510889.80 | protein phosphatase with EF-Hand domains (PPEF) | -0,01074 | 0,9055 |
| TcCLB.507019.45 | hypothetical protein | -0,01078 | 0,9412 |
| TcCLB.511633.70 | protein kinase, putative | -0,01085 | 0,9214 |
| TcCLB.511283.60 | hypothetical protein, conserved | -0,0109 | 0,9288 |
| TcCLB.506413.10 | CDP-DAG synthase, putative (fragment) (CDS) | -0,01092 | 0,9531 |
| TcCLB.503929.60 | hypothetical protein, conserved | -0,01093 | 0,945 |

| | | | |
|---|---|---|---|
| TcCLB.509877.70 | hypothetical protein, conserved | -0,01098 | 0,9333 |
| TcCLB.506721.20 | radial spoke protein 3, putative (RSP3) | -0,01103 | 0,9216 |
| TcCLB.507673.40 | hypothetical protein, conserved | -0,01105 | 0,9148 |
| TcCLB.507739.20 | metallo-peptidase, Clan MA(E) Family M41 | -0,01105 | 0,921 |
| TcCLB.509029.30 | hypothetical protein, conserved | -0,01105 | 0,9486 |
| TcCLB.508579.20 | hypothetical protein, conserved | -0,0111 | 0,9642 |
| TcCLB.503493.30 | hypothetical protein, conserved | -0,01117 | 0,941 |
| TcCLB.509551.100 | hypothetical protein, conserved | -0,01135 | 0,9337 |
| TcCLB.506855.260 | thymidine kinase, putative | -0,01135 | 0,9494 |
| TcCLB.510753.20 | hypothetical protein, conserved | -0,01158 | 0,9407 |
| TcCLB.509639.20 | hypothetical protein, conserved | -0,01158 | 0,9422 |
| TcCLB.511509.5 | hypothetical protein, conserved | -0,0116 | 0,9321 |
| TcCLB.504013.50 | selenocysteine-tRNA-specific elongation factor, putative | -0,01162 | 0,925 |
| TcCLB.507771.90 | hypothetical protein, conserved | -0,01167 | 0,9006 |
| TcCLB.510243.100 | chloride channel protein, putative | -0,01178 | 0,9188 |
| TcCLB.511657.70 | hypothetical protein, conserved | -0,01179 | 0,9261 |
| TcCLB.504029.21 | lipase, putative (fragment) | -0,01195 | 0,9148 |
| TcCLB.507669.190 | hypothetical protein, conserved | -0,01201 | 0,9377 |
| TcCLB.509437.30 | hypothetical protein, conserved | -0,01203 | 0,8995 |
| TcCLB.509551.40 | hypothetical protein, conserved | -0,01203 | 0,942 |
| TcCLB.448567.9 | dynein heavy chain, putative | -0,01205 | 0,9446 |
| TcCLB.511021.20 | hypothetical protein | -0,01207 | 0,9497 |
| TcCLB.503813.20 | hypothetical protein, conserved | -0,01211 | 0,924 |
| TcCLB.508479.224 | hypothetical protein, conserved | -0,01225 | 0,9506 |
| TcCLB.506753.164 | hypothetical protein, conserved | -0,01227 | 0,9548 |
| TcCLB.511621.250 | hypothetical protein, conserved | -0,01229 | 0,9255 |
| TcCLB.507951.154 | cytochrome b5, putative | -0,01232 | 0,9353 |
| TcCLB.510657.30 | Sorting nexin-4 (ATG24) | -0,01236 | 0,9046 |
| TcCLB.511907.240 | protein kinase, putative | -0,01237 | 0,9159 |
| TcCLB.506407.40 | hypothetical protein, conserved | -0,01241 | 0,9148 |
| TcCLB.509877.80 | ATPase domain protein, putative | -0,01255 | 0,941 |
| TcCLB.507159.44 | DNA-directed RNA polymerase III subunit, putative | -0,01261 | 0,9252 |
| TcCLB.506241.190 | hypothetical protein, conserved | -0,01261 | 0,938 |
| TcCLB.509941.85 | trans-sialidase (pseudogene), putative | -0,01264 | 0,9356 |
| TcCLB.507357.130 | mucin TcMUCII, putative | -0,01265 | 0,9757 |
| TcCLB.507601.50 | hypothetical protein | -0,01274 | 0,9288 |
| TcCLB.511577.83 | thioredoxin, putative | -0,01283 | 0,9481 |
| TcCLB.503893.160 | hypothetical protein, conserved | -0,01288 | 0,9215 |
| TcCLB.504741.120 | hypothetical protein, conserved | -0,01309 | 0,9251 |
| TcCLB.509789.30 | rab-like GTPase activating protein, putative | -0,01331 | 0,9264 |
| TcCLB.506777.24 | hypothetical protein, conserved | -0,01332 | 0,9334 |
| TcCLB.504153.220 | DNA recombination and repair protein RAD51, putative | -0,01335 | 0,9144 |
| TcCLB.504157.20 | beta-ketoacyl synthase family protein, putative | -0,01335 | 0,9319 |
| TcCLB.503575.40 | hypothetical protein, conserved | -0,01335 | 0,9428 |
| TcCLB.511505.20 | hypothetical protein, conserved | -0,01344 | 0,9191 |

| | | | |
|---|---|---|---|
| TcCLB.511215.119 | 69 kDa paraflagellar rod protein, putative | -0,01378 | 0,9021 |
| TcCLB.510725.90 | hypothetical protein, conserved | -0,01379 | 0,9367 |
| TcCLB.511129.47 | hypothetical protein, conserved (pseudogene) | -0,01388 | 0,9429 |
| TcCLB.507927.30 | hypothetical protein, conserved | -0,01389 | 0,9339 |
| TcCLB.509965.70 | hypothetical protein, conserved | -0,01392 | 0,9122 |
| TcCLB.505071.70 | hypothetical protein, conserved | -0,01397 | 0,9301 |
| TcCLB.504215.10 | hypothetical protein, conserved | -0,01398 | 0,9112 |
| TcCLB.508641.250 | hypothetical protein | -0,01408 | 0,896 |
| TcCLB.507979.10 | hypothetical protein, conserved | -0,01436 | 0,9296 |
| TcCLB.506367.40 | hypothetical protein, conserved | -0,01446 | 0,899 |
| TcCLB.506247.300 | hypothetical protein, conserved | -0,01447 | 0,9244 |
| TcCLB.507015.50 | retrotransposon hot spot protein (RHS, pseudogene), puta | -0,01469 | 0,9165 |
| TcCLB.510757.20 | hypothetical protein, conserved | -0,01478 | 0,9179 |
| TcCLB.511393.40 | hypothetical protein, conserved | -0,01482 | 0,9151 |
| TcCLB.506199.30 | hypothetical protein, conserved | -0,01482 | 0,9252 |
| TcCLB.510817.60 | hypothetical protein, conserved | -0,01487 | 0,8778 |
| TcCLB.510123.30 | DNA repair protein, putative | -0,01488 | 0,9171 |
| TcCLB.510099.70 | hypothetical protein, conserved | -0,0149 | 0,8928 |
| TcCLB.511753.130 | U3 small nuclear ribonucleoprotein (snRNP), putative | -0,01514 | 0,8703 |
| TcCLB.511303.40 | carnitine O-acetyltransferase, putative | -0,01516 | 0,905 |
| TcCLB.507625.180 | kinesin, putative | -0,01535 | 0,8978 |
| TcCLB.508457.10 | Kinesin-13 6, putative (KIN13-6) | -0,01548 | 0,9098 |
| TcCLB.508325.300 | hypothetical protein, conserved | -0,01561 | 0,9424 |
| TcCLB.510729.160 | hypothetical protein, conserved | -0,01566 | 0,8984 |
| TcCLB.503647.20 | hypothetical protein, conserved | -0,01566 | 0,9362 |
| TcCLB.510483.30 | trans-sialidase (pseudogene), putative | -0,01566 | 0,9526 |
| TcCLB.510285.120 | hypothetical protein, conserved | -0,0158 | 0,884 |
| TcCLB.508209.130 | hypothetical protein, conserved | -0,01588 | 0,9077 |
| TcCLB.510241.124 | hypothetical protein, conserved | -0,01597 | 0,8783 |
| TcCLB.507849.80 | hypothetical protein, conserved | -0,01632 | 0,9298 |
| TcCLB.510101.350 | hypothetical protein, conserved | -0,01638 | 0,8627 |
| TcCLB.509777.30 | trans-sialidase, Group II, putative | -0,01643 | 0,8938 |
| TcCLB.510729.50 | hypothetical protein, conserved | -0,01647 | 0,9204 |
| TcCLB.510645.9 | serine/threonine protein kinase, putative | -0,01652 | 0,8972 |
| TcCLB.507093.40 | hypothetical protein, conserved | -0,01661 | 0,8593 |
| TcCLB.510943.70 | hypothetical protein, conserved | -0,01669 | 0,9147 |
| TcCLB.508637.69 | hypothetical protein, conserved | -0,01682 | 0,9142 |
| TcCLB.509203.14 | hypothetical protein | -0,01683 | 0,9375 |
| TcCLB.506175.10 | hypothetical protein, conserved | -0,01708 | 0,9012 |
| TcCLB.511725.190 | vacuolar sorting protein, putative | -0,01714 | 0,916 |
| TcCLB.507711.210 | hypothetical protein, conserved | -0,01715 | 0,8675 |
| TcCLB.510877.40 | hypothetical protein, conserved | -0,01722 | 0,8621 |
| TcCLB.506793.20 | hypothetical protein, conserved | -0,01733 | 0,8884 |
| TcCLB.503771.40 | syntaxin binding protein (pseudogene), putative | -0,01734 | 0,9647 |
| TcCLB.506265.10 | hypothetical protein, conserved | -0,01747 | 0,9165 |

| | | | |
|---|---|---|---|
| TcCLB.509875.190 | clathrin coat assembly protein, putative | -0,01751 | 0,89 |
| TcCLB.509429.50 | retrotransposon hot spot protein (RHS, pseudogene), puta | -0,01757 | 0,8714 |
| TcCLB.511907.190 | hypothetical protein | -0,01764 | 0,8801 |
| TcCLB.504081.141 | syntaxin binding protein (pseudogene), putative | -0,01765 | 0,9633 |
| TcCLB.503925.100 | hypothetical protein, conserved | -0,01776 | 0,9099 |
| TcCLB.506657.10 | hypothetical protein, conserved | -0,01795 | 0,9371 |
| TcCLB.506715.59 | Monooxygenase, putative | -0,01801 | 0,9459 |
| TcCLB.508521.18 | trans-sialidase, putative (pseudogene) | -0,01805 | 0,9603 |
| TcCLB.507023.60 | hypothetical protein, conserved | -0,01814 | 0,8953 |
| TcCLB.510535.90 | hypothetical protein, conserved | -0,01823 | 0,9459 |
| TcCLB.506577.47 | hypothetical protein, conserved | -0,01831 | 0,8824 |
| TcCLB.509395.120 | histone deacetylase, putative | -0,01837 | 0,8778 |
| TcCLB.505997.200 | hypothetical protein, conserved | -0,01839 | 0,9092 |
| TcCLB.511751.30 | queuine tRNA-ribosyltransferase, putative | -0,01846 | 0,8686 |
| TcCLB.510661.240 | hypothetical protein, conserved | -0,01854 | 0,8605 |
| TcCLB.510825.20 | hypothetical protein, conserved | -0,01857 | 0,9305 |
| TcCLB.510965.10 | hypothetical protein, conserved | -0,01869 | 0,8962 |
| TcCLB.509443.10 | hypothetical protein | -0,01889 | 0,9254 |
| TcCLB.509289.10 | amastin, putative | -0,01894 | 0,8903 |
| TcCLB.506961.160 | retrotransposon hot spot (RHS) protein, putative | -0,01908 | 0,8806 |
| TcCLB.503805.6 | hypothetical protein, conserved | -0,01917 | 0,9083 |
| TcCLB.507129.19 | hypothetical protein, conserved | -0,01924 | 0,8811 |
| TcCLB.511367.100 | hypothetical protein, conserved | -0,01938 | 0,8979 |
| TcCLB.507839.18 | hypothetical protein, conserved | -0,01955 | 0,8893 |
| TcCLB.510655.80 | hypothetical protein, conserved | -0,01955 | 0,9261 |
| TcCLB.505037.50 | small GTP-binding protein RAB6, putative | -0,01962 | 0,863 |
| TcCLB.510573.19 | hypothetical protein, conserved | -0,01962 | 0,9051 |
| TcCLB.511875.5 | elongation factor 1-gamma (EF-1-gamma, pseudogene), | -0,01962 | 0,9561 |
| TcCLB.509805.200 | acetyltransferase, putative | -0,01996 | 0,8633 |
| TcCLB.510303.310 | arginine N-methyltransferase, type III, putative (PRMT7) | -0,02013 | 0,8446 |
| TcCLB.510745.10 | hypothetical protein, conserved | -0,0203 | 0,8764 |
| TcCLB.509567.50 | hypothetical protein, conserved | -0,02048 | 0,8813 |
| TcCLB.510901.190 | hypothetical protein, conserved | -0,02065 | 0,8703 |
| TcCLB.508119.190 | hypothetical protein, conserved | -0,02069 | 0,8341 |
| TcCLB.503865.70 | hypothetical protein, conserved | -0,02085 | 0,8434 |
| TcCLB.510617.10 | hypothetical protein, conserved | -0,02088 | 0,9044 |
| TcCLB.510519.90 | hypothetical protein, conserved | -0,02089 | 0,9278 |
| TcCLB.510329.230 | hypothetical protein, conserved | -0,0209 | 0,877 |
| TcCLB.510359.140 | hypothetical protein, conserved | -0,02091 | 0,9191 |
| TcCLB.506303.60 | protein kinase, putative | -0,02109 | 0,8967 |
| TcCLB.511725.90 | hypothetical protein, conserved | -0,02111 | 0,8795 |
| TcCLB.506671.30 | mucin-associated surface protein (MASP, pseudogene), pu | -0,02111 | 0,9187 |
| TcCLB.509747.90 | hypothetical protein, conserved | -0,02115 | 0,8913 |
| TcCLB.510187.74 | short-chain dehydrogenase, putative | -0,0212 | 0,8817 |
| TcCLB.508909.60 | hypothetical protein, conserved | -0,02138 | 0,8872 |

| | | | |
|---|---|---|---|
| TcCLB.508153.840 | hypothetical protein, conserved | -0,02147 | 0,8683 |
| TcCLB.407335.18 | hypothetical protein, conserved | -0,02149 | 0,8984 |
| TcCLB.506925.510 | ribosome production factor 1, putative | -0,02162 | 0,8386 |
| TcCLB.509777.110 | hypothetical protein, conserved | -0,02164 | 0,9026 |
| TcCLB.511239.80 | hypothetical protein, conserved | -0,02197 | 0,8675 |
| TcCLB.503651.49 | hypothetical protein, conserved (pseudogene) | -0,02198 | 0,887 |
| TcCLB.508951.80 | hypothetical protein, conserved | -0,02209 | 0,8536 |
| TcCLB.506479.50 | rRNA biogenesis protein, putative | -0,0221 | 0,8466 |
| TcCLB.503599.50 | hypothetical protein, conserved | -0,02217 | 0,8988 |
| TcCLB.510765.20 | hypothetical protein, conserved | -0,02218 | 0,9144 |
| TcCLB.509817.50 | trans-sialidase, Group I, putative | -0,02219 | 0,9331 |
| TcCLB.506857.50 | hypothetical protein, conserved | -0,02234 | 0,8377 |
| TcCLB.506591.50 | hypothetical protein, conserved (pseudogene) | -0,02241 | 0,8025 |
| TcCLB.510087.100 | GINS complex subunit Psf3, putative | -0,02248 | 0,9236 |
| TcCLB.508727.54 | hypothetical protein, conserved | -0,02249 | 0,8741 |
| TcCLB.509061.20 | hypothetical protein, conserved | -0,02253 | 0,8767 |
| TcCLB.503849.80 | NADH-dependent fumarate reductase, putative | -0,02259 | 0,9191 |
| TcCLB.511743.20 | hypothetical protein, conserved | -0,02264 | 0,859 |
| TcCLB.509695.210 | serine carboxypeptidase (CBP1), putative | -0,02264 | 0,8717 |
| TcCLB.511313.40 | ATP-dependent RNA helicase, putative | -0,02295 | 0,8222 |
| TcCLB.509267.6 | trans-sialidase (pseudogene), putative | -0,02298 | 0,9187 |
| TcCLB.509973.40 | hypothetical protein | -0,02303 | 0,9491 |
| TcCLB.506723.41 | beta galactofuranosyl glycosyltransferase (pseudogene), p | -0,0231 | 0,8838 |
| TcCLB.503911.20 | pre-mRNA splicing factor ATP-dependent RNA helicase, p | -0,02319 | 0,8627 |
| TcCLB.503913.10 | hypothetical protein, conserved | -0,0232 | 0,8658 |
| TcCLB.506227.160 | MOZ/SAS family acetyltransferase, putative | -0,02323 | 0,8666 |
| TcCLB.506925.540 | hypothetical protein, conserved | -0,02334 | 0,8401 |
| TcCLB.507021.80 | hypothetical protein, conserved | -0,02336 | 0,8682 |
| TcCLB.440343.9 | dispersed gene family protein 1 (DGF-1), putative (fragme | -0,02338 | 0,959 |
| TcCLB.503589.10 | GDP-mannose pyrophosphorylase | -0,02344 | 0,8655 |
| TcCLB.509991.100 | Transcription elongation factor 1 domain-containing prote | -0,02358 | 0,8663 |
| TcCLB.507831.80 | hypothetical protein, conserved | -0,02367 | 0,8593 |
| TcCLB.504045.100 | hypothetical protein, conserved | -0,0237 | 0,8591 |
| TcCLB.506929.40 | target SNARE, putative | -0,02378 | 0,8582 |
| TcCLB.507021.50 | kinesin, putative | -0,02379 | 0,8078 |
| TcCLB.505807.100 | hypothetical protein, conserved | -0,0238 | 0,8655 |
| TcCLB.506883.60 | hypothetical protein, conserved | -0,02383 | 0,8591 |
| TcCLB.503589.19 | hypothetical protein, conserved | -0,02389 | 0,9147 |
| TcCLB.506985.20 | hypothetical protein, conserved | -0,02431 | 0,8767 |
| TcCLB.506203.70 | pre-mRNA-splicing factor ATP-dependent RNA helicase, p | -0,02445 | 0,8085 |
| TcCLB.510659.50 | RNA polymerase-associated protein LEO1, putative | -0,02457 | 0,7916 |
| TcCLB.507993.110 | hypothetical protein, conserved | -0,02464 | 0,8738 |
| TcCLB.510291.20 | hypothetical protein, conserved | -0,02473 | 0,8622 |
| TcCLB.401041.10 | retrotransposon hot spot protein (RHS, pseudogene), puta | -0,02477 | 0,8347 |
| TcCLB.507603.120 | hypothetical protein, conserved | -0,02478 | 0,8162 |

| | | | |
|---|---|---|---|
| TcCLB.506187.40 | trans-sialidase (pseudogene), putative | -0,02478 | 0,9194 |
| TcCLB.511865.40 | hypothetical protein, conserved | -0,02487 | 0,8269 |
| TcCLB.511277.50 | hypothetical protein, conserved | -0,02505 | 0,8153 |
| TcCLB.511367.129 | hypothetical protein, conserved | -0,02515 | 0,8648 |
| TcCLB.511245.60 | hypothetical protein, conserved | -0,02522 | 0,924 |
| TcCLB.506101.20 | trans-sialidase (pseudogene), putative | -0,0253 | 0,9121 |
| TcCLB.508237.9 | protein kinase, putative | -0,02535 | 0,8864 |
| TcCLB.509805.180 | ttaggg binding factor, putative | -0,02565 | 0,836 |
| TcCLB.509997.60 | hypothetical protein, conserved | -0,02581 | 0,7841 |
| TcCLB.510725.50 | hypothetical protein, conserved | -0,02581 | 0,8106 |
| TcCLB.504021.30 | hypothetical protein, conserved | -0,02595 | 0,8 |
| TcCLB.508547.80 | hypothetical protein, conserved | -0,02595 | 0,8203 |
| TcCLB.504817.20 | hypothetical protein, conserved | -0,02604 | 0,8426 |
| TcCLB.508065.60 | hypothetical protein, conserved | -0,02625 | 0,8376 |
| TcCLB.508199.10 | ADP-ribosylation factor, putative | -0,0263 | 0,8587 |
| TcCLB.504935.10 | eukaryotic translation initiation factor 4E, putative | -0,02632 | 0,8686 |
| TcCLB.509231.39 | hypothetical protein, conserved | -0,02648 | 0,8997 |
| TcCLB.507721.10 | hypothetical protein, conserved | -0,02651 | 0,8648 |
| TcCLB.507087.10 | hydrolase-like protein (pseudogene), putative | -0,02653 | 0,889 |
| TcCLB.511771.171 | hypothetical protein | -0,02658 | 0,928 |
| TcCLB.509171.70 | hypothetical protein, conserved | -0,02662 | 0,8594 |
| TcCLB.509485.20 | retrotransposon hot spot protein (RHS, pseudogene), puta | -0,02667 | 0,8913 |
| TcCLB.506811.180 | protein kinase, putative | -0,02671 | 0,8995 |
| TcCLB.506465.50 | serine/threonine protein phosphatase-like protein, putativ | -0,02671 | 0,904 |
| TcCLB.511751.250 | hypothetical protein, conserved | -0,02687 | 0,8617 |
| TcCLB.507663.20 | hypothetical protein, conserved | -0,02693 | 0,8414 |
| TcCLB.511385.14 | hypothetical protein, conserved (fragment) | -0,0271 | 0,828 |
| TcCLB.504741.10 | hypothetical protein, conserved | -0,02718 | 0,8913 |
| TcCLB.507711.250 | hypothetical protein, conserved | -0,0272 | 0,8125 |
| TcCLB.506211.250 | fibrillarin, putative | -0,02721 | 0,8383 |
| TcCLB.509287.20 | protein kinase, putative | -0,02726 | 0,8418 |
| TcCLB.511277.220 | hypothetical protein, conserved | -0,02732 | 0,8622 |
| TcCLB.413293.30 | retrotransposon hot spot protein (RHS, pseudogene), puta | -0,02739 | 0,8569 |
| TcCLB.506009.70 | hypothetical protein | -0,02741 | 0,8514 |
| TcCLB.508647.140 | DNA replication licensing factor MCM5, putative | -0,02744 | 0,8097 |
| TcCLB.511415.11 | retrotransposon hot spot (RHS) protein, putative (fragmer | -0,02746 | 0,8665 |
| TcCLB.506815.30 | beta prime cop protein, putative | -0,02747 | 0,8 |
| TcCLB.506811.210 | cell division cycle protein 45 (CDC45), putative | -0,02753 | 0,832 |
| TcCLB.509429.30 | retrotransposon hot spot protein (RHS, pseudogene), puta | -0,02773 | 0,8099 |
| TcCLB.509259.126 | hypothetical protein | -0,02782 | 0,896 |
| TcCLB.507625.84 | hypothetical protein, conserved | -0,02783 | 0,8732 |
| TcCLB.510623.10 | mucin TcMUCII, putative | -0,02786 | 0,9542 |
| TcCLB.511433.10 | hypothetical protein, conserved | -0,02788 | 0,8669 |
| TcCLB.506567.140 | hypothetical protein, conserved | -0,02789 | 0,8973 |
| TcCLB.510763.100 | trypanin-like protein, putative | -0,02791 | 0,8028 |

| | | | |
|---|---|---|---|
| TcCLB.509647.90 | chloride channel protein, putative | -0,028 | 0,8179 |
| TcCLB.509055.40 | hypothetical protein, conserved | -0,02803 | 0,8181 |
| TcCLB.510329.320 | hypothetical protein, conserved | -0,02805 | 0,8648 |
| TcCLB.508173.54 | hypothetical protein, conserved | -0,02807 | 0,7944 |
| TcCLB.510603.40 | hypothetical protein, conserved | -0,02814 | 0,7966 |
| TcCLB.509959.236 | N-acetyltransferase complex ARD1 subunit (pseudogene), | -0,02815 | 0,9571 |
| TcCLB.506519.90 | hypothetical protein, conserved | -0,02826 | 0,8587 |
| TcCLB.461165.4 | hypothetical protein, conserved (fragment) | -0,02827 | 0,914 |
| TcCLB.508207.5 | retrotransposon hot spot protein (RHS, pseudogene), puta | -0,02832 | 0,8105 |
| TcCLB.508719.70 | flagellum targeting protein kharon1, putative (KH1) | -0,02834 | 0,822 |
| TcCLB.507821.140 | trans-sialidase (pseudogene), putative | -0,02842 | 0,9099 |
| TcCLB.506855.360 | hypothetical protein, conserved | -0,02843 | 0,7807 |
| TcCLB.508771.30 | hypothetical protein, conserved | -0,02849 | 0,8511 |
| TcCLB.510155.210 | calcium/potassium channel (CAKC), putative | -0,02857 | 0,8272 |
| TcCLB.506739.120 | hypothetical protein, conserved | -0,02867 | 0,8241 |
| TcCLB.508941.10 | retrotransposon hot spot protein (RHS, pseudogene), puta | -0,02905 | 0,9447 |
| TcCLB.511529.240 | hypothetical protein, conserved | -0,02906 | 0,7796 |
| TcCLB.510545.73 | syntaxin binding protein (pseudogene), putative | -0,02918 | 0,9207 |
| TcCLB.509745.40 | hypothetical protein, conserved | -0,02923 | 0,8261 |
| TcCLB.507837.30 | hypothetical protein, conserved | -0,02926 | 0,7946 |
| TcCLB.509859.30 | hypothetical protein, conserved | -0,02933 | 0,8143 |
| TcCLB.510425.28 | hypothetical protein, conserved | -0,02933 | 0,9128 |
| TcCLB.503939.90 | hypothetical protein, conserved | -0,02943 | 0,7678 |
| TcCLB.511879.9 | hypothetical protein, conserved | -0,02943 | 0,7791 |
| TcCLB.509231.26 | hypothetical protein, conserved | -0,02957 | 0,8263 |
| TcCLB.503909.84 | calpain-like cysteine peptidase, putative | -0,02961 | 0,8241 |
| TcCLB.508479.490 | hypothetical protein | -0,02976 | 0,8921 |
| TcCLB.506913.10 | hypothetical protein, conserved | -0,02984 | 0,8685 |
| TcCLB.506849.20 | dynein heavy chain (pseudogene), putative | -0,02989 | 0,8479 |
| TcCLB.510603.119 | hypothetical protein, conserved | -0,03 | 0,885 |
| TcCLB.508865.10 | glutathione synthetase, putative | -0,03002 | 0,8246 |
| TcCLB.508741.310 | hypothetical protein | -0,03016 | 0,8378 |
| TcCLB.508951.90 | hypothetical protein, conserved | -0,03023 | 0,8514 |
| TcCLB.510659.104 | poly(ADP-ribose) glycohydrolase, putative | -0,03026 | 0,827 |
| TcCLB.506357.69 | hypothetical protein, conserved | -0,03049 | 0,9091 |
| TcCLB.510255.20 | hypothetical protein, conserved | -0,03051 | 0,8442 |
| TcCLB.509999.80 | hypothetical protein, conserved | -0,03052 | 0,7691 |
| TcCLB.510091.70 | U6 snRNA-associated Sm-like protein LSm8, putative | -0,03064 | 0,8657 |
| TcCLB.510063.10 | sigma-adaptin 3, putative | -0,0307 | 0,8523 |
| TcCLB.510089.120 | hypothetical protein, conserved | -0,03088 | 0,8383 |
| TcCLB.506659.30 | glutathione synthetase, putative | -0,03095 | 0,821 |
| TcCLB.506337.30 | class-E vacuolar protein-sorting protein 24 (Vps24p), puta | -0,03115 | 0,8094 |
| TcCLB.510429.30 | hypothetical protein, conserved | -0,03122 | 0,8195 |
| TcCLB.507093.184 | hypothetical protein, conserved | -0,03126 | 0,8023 |
| TcCLB.503733.30 | hypothetical protein, conserved | -0,03136 | 0,8045 |

| | | | |
|---|---|---|---|
| TcCLB.507709.130 | hypothetical protein, conserved | -0,0315 | 0,8424 |
| TcCLB.503887.60 | glycerolphosphate mutase, putative | -0,03152 | 0,8149 |
| TcCLB.506493.50 | serine/threonine protein kinase, putative | -0,03163 | 0,8809 |
| TcCLB.506465.30 | hypothetical protein, conserved | -0,03166 | 0,8037 |
| TcCLB.506953.20 | hypothetical protein, conserved | -0,03166 | 0,88 |
| TcCLB.509611.100 | lipoate-protein ligase, putative | -0,03169 | 0,8383 |
| TcCLB.508137.20 | hypothetical protein, conserved | -0,0317 | 0,7525 |
| TcCLB.507807.10 | hypothetical protein, conserved | -0,03175 | 0,7339 |
| TcCLB.506983.30 | hypothetical protein, conserved | -0,03189 | 0,7746 |
| TcCLB.506563.20 | exosome-associated protein 1, putative (EAP1) | -0,03198 | 0,8514 |
| TcCLB.507001.120 | hypothetical protein, conserved | -0,03199 | 0,8405 |
| TcCLB.503953.14 | hypothetical protein, conserved | -0,03207 | 0,8347 |
| TcCLB.507801.40 | hypothetical protein, conserved | -0,0321 | 0,8827 |
| TcCLB.511751.114 | hypothetical protein, conserved | -0,03222 | 0,8736 |
| TcCLB.509583.19 | dynein heavy chain, putative (fragment) | -0,03229 | 0,8507 |
| TcCLB.510743.60 | phospholipase A1, putative | -0,03232 | 0,8272 |
| TcCLB.504057.40 | NAD dependent epimerase/dehydratase family, putative ( | -0,03242 | 0,7556 |
| TcCLB.511407.60 | small GTP-binding protein Rab11, putative (Rab11) | -0,03251 | 0,8005 |
| TcCLB.510899.30 | hypothetical protein, conserved | -0,03252 | 0,8223 |
| TcCLB.511115.40 | transcription elongation factor s-II, putative | -0,03256 | 0,7695 |
| TcCLB.503439.40 | trans-sialidase (pseudogene), putative | -0,03259 | 0,8276 |
| TcCLB.505807.160 | hypothetical protein, conserved | -0,03274 | 0,8475 |
| TcCLB.506051.10 | retrotransposon hot spot protein (RHS, pseudogene), puta | -0,03284 | 0,8383 |
| TcCLB.503917.35 | B9 domain-containing protein 2 (b9d2) | -0,03284 | 0,8654 |
| TcCLB.511595.50 | mucin-associated surface protein (MASP, pseudogene), pu | -0,03284 | 0,8827 |
| TcCLB.511003.100 | hypothetical protein, conserved | -0,03287 | 0,7773 |
| TcCLB.503981.80 | hypothetical protein, conserved | -0,03317 | 0,7063 |
| TcCLB.508405.80 | cysteine peptidase, Clan CD, family C50, putative | -0,03318 | 0,7765 |
| TcCLB.508817.150 | hypothetical protein, conserved | -0,03326 | 0,7841 |
| TcCLB.506629.140 | hypothetical protein, conserved | -0,03344 | 0,7854 |
| TcCLB.511619.10 | hypothetical protein, conserved | -0,03346 | 0,8316 |
| TcCLB.508467.19 | hypothetical protein, conserved (fragment) | -0,03355 | 0,8817 |
| TcCLB.511269.30 | hypothetical protein, conserved | -0,03379 | 0,8365 |
| TcCLB.508831.70 | U5 snRNP-specific 40 kDa protein, putative | -0,03413 | 0,801 |
| TcCLB.510225.50 | hypothetical protein, conserved | -0,03419 | 0,8241 |
| TcCLB.503977.79 | elongation factor 1-gamma (EF-1-gamma), putative | -0,0343 | 0,9452 |
| TcCLB.508205.48 | hypothetical protein, conserved | -0,03438 | 0,8806 |
| TcCLB.510951.10 | hypothetical protein, conserved | -0,03438 | 0,8906 |
| TcCLB.503973.20 | trans-sialidase (pseudogene), putative | -0,03444 | 0,8798 |
| TcCLB.511483.50 | hypothetical protein, conserved | -0,03455 | 0,857 |
| TcCLB.511669.20 | hypothetical protein | -0,03464 | 0,9087 |
| TcCLB.510945.10 | ubiquitin carboxyl-terminal hydrolase, putative | -0,03466 | 0,8024 |
| TcCLB.511717.80 | DNA repair and recombination helicase protein PIF4, puta | -0,0347 | 0,8416 |
| TcCLB.511903.30 | ubiquitin-conjugating enzyme E2, putative | -0,03476 | 0,7648 |
| TcCLB.503579.150 | RNA editing 3' terminal uridylyl transferase 2, putative | -0,03499 | 0,8085 |

| | | | |
|---|---|---|---|
| TcCLB.504117.9 | hypothetical protein, conserved | -0,03502 | 0,7613 |
| TcCLB.508827.139 | Coatomer subunit gamma (fragment) (COPG) | -0,03506 | 0,7193 |
| TcCLB.511585.190 | eukaryotic initiation factor 4a, putative | -0,03516 | 0,6833 |
| TcCLB.508813.50 | oxidoreductase-like protein, putative | -0,03528 | 0,7424 |
| TcCLB.506513.14 | hypothetical protein, conserved | -0,0355 | 0,8594 |
| TcCLB.506591.90 | hypothetical protein, conserved | -0,03555 | 0,948 |
| TcCLB.398751.10 | hypothetical protein, conserved | -0,03569 | 0,9245 |
| TcCLB.510155.30 | hypothetical protein, conserved | -0,0357 | 0,7178 |
| TcCLB.511277.510 | hypothetical protein, conserved | -0,03572 | 0,7223 |
| TcCLB.506107.20 | dihydrouridine synthase (Dus), putative | -0,03592 | 0,8617 |
| TcCLB.511249.100 | hypothetical protein, conserved | -0,036 | 0,6938 |
| TcCLB.508693.170 | hypothetical protein, conserved | -0,03605 | 0,8083 |
| TcCLB.506885.260 | hypothetical protein, conserved | -0,03607 | 0,7059 |
| TcCLB.511287.170 | hypothetical protein, conserved | -0,03613 | 0,7697 |
| TcCLB.508153.830 | peter pan protein, putative | -0,0362 | 0,806 |
| TcCLB.510187.10 | hypothetical protein, conserved | -0,03622 | 0,896 |
| TcCLB.510073.90 | hypothetical protein, conserved | -0,03624 | 0,7821 |
| TcCLB.511807.260 | hypothetical protein, conserved | -0,03642 | 0,6617 |
| TcCLB.509567.40 | nuclear transport factor 2, putative | -0,0365 | 0,7989 |
| TcCLB.506959.50 | hypothetical protein, conserved | -0,03665 | 0,768 |
| TcCLB.511021.10 | protein phosphatase 2A catalytic subunit, putative | -0,03667 | 0,8108 |
| TcCLB.503663.10 | septum formation inhibitor, Maf-like protein, putative | -0,03674 | 0,84 |
| TcCLB.508819.50 | katanin, putative (fragment) | -0,03753 | 0,8241 |
| TcCLB.510257.10 | hypothetical protein, conserved | -0,03765 | 0,6457 |
| TcCLB.508173.260 | hypothetical protein, conserved | -0,03775 | 0,7814 |
| TcCLB.509671.30 | hypothetical protein, conserved | -0,03776 | 0,7888 |
| TcCLB.507787.10 | kinesin-like protein (pseudogene), putative | -0,03795 | 0,7963 |
| TcCLB.509053.60 | hypothetical protein, conserved | -0,03823 | 0,8473 |
| TcCLB.508409.50 | hypothetical protein, conserved | -0,03824 | 0,7549 |
| TcCLB.510857.20 | hypothetical protein, conserved | -0,03843 | 0,8208 |
| TcCLB.509629.20 | DnaJ chaperone protein, putative | -0,03847 | 0,8426 |
| TcCLB.506857.70 | hypothetical protein, conserved | -0,03851 | 0,7528 |
| TcCLB.509967.100 | hypothetical protein, conserved (pseudogene) | -0,03856 | 0,8256 |
| TcCLB.506945.200 | mago nashi-like protein, putative | -0,03859 | 0,7092 |
| TcCLB.469785.5 | hypothetical protein, conserved (fragment) | -0,03889 | 0,8481 |
| TcCLB.510969.30 | hypothetical protein, conserved | -0,03916 | 0,853 |
| TcCLB.506493.9 | hypothetical protein, conserved | -0,03937 | 0,7899 |
| TcCLB.511483.10 | hypothetical protein | -0,03938 | 0,7661 |
| TcCLB.464807.20 | hypothetical protein, conserved (fragment) | -0,03958 | 0,7533 |
| TcCLB.508965.60 | rac serine-threonine kinase, putative | -0,03964 | 0,7173 |
| TcCLB.509217.150 | retrotransposon hot spot protein (RHS, pseudogene), puta | -0,03966 | 0,8316 |
| TcCLB.506679.260 | hypothetical protein, conserved | -0,03974 | 0,7168 |
| TcCLB.508169.30 | hypothetical protein, conserved | -0,03999 | 0,7911 |
| TcCLB.510081.11 | hypothetical protein, conserved (pseudogene) | -0,04001 | 0,7483 |
| TcCLB.508891.10 | dolichyl-P-Man:GDP-ManGlcNAc2-PP-dolichyl beta-1,4-ma | -0,04007 | 0,8197 |

| | | | |
|---|---|---|---|
| TcCLB.506967.13 | hypothetical protein, conserved | -0,0401 | 0,8514 |
| TcCLB.506631.40 | hypothetical protein, conserved | -0,04017 | 0,7205 |
| TcCLB.510063.20 | ATP-dependent DEAD/H DNA helicase recQ, putative | -0,0402 | 0,8624 |
| TcCLB.510329.60 | RAB GDP dissociation inhibitor alpha, putative | -0,04028 | 0,6863 |
| TcCLB.509633.10 | hypothetical protein, conserved | -0,0405 | 0,8246 |
| TcCLB.511019.120 | hypothetical protein, conserved | -0,04053 | 0,7065 |
| TcCLB.504057.178 | hypothetical protein, conserved | -0,0406 | 0,7627 |
| TcCLB.511511.60 | phospholipid-translocating P-type ATPase (flippase), puta | -0,04066 | 0,7969 |
| TcCLB.510311.40 | hypothetical protein, conserved | -0,0408 | 0,6526 |
| TcCLB.508817.5 | dynein heavy chain, putative | -0,04088 | 0,8128 |
| TcCLB.506675.99 | trans-sialidase (pseudogene), putative | -0,0409 | 0,9068 |
| TcCLB.511815.70 | hypothetical protein, conserved | -0,04116 | 0,8426 |
| TcCLB.508173.200 | hypothetical protein, conserved | -0,04128 | 0,6035 |
| TcCLB.508529.30 | retrotransposon hot spot protein (RHS, pseudogene), puta | -0,0414 | 0,7107 |
| TcCLB.510197.47 | mucin TcMUCII (pseudogene), putative | -0,04142 | 0,8743 |
| TcCLB.507773.40 | hypothetical protein, conserved | -0,04145 | 0,7273 |
| TcCLB.511927.20 | hypothetical protein, conserved | -0,04145 | 0,8083 |
| TcCLB.511467.13 | hypothetical protein, conserved | -0,04155 | 0,8261 |
| TcCLB.511817.130 | hypothetical protein, conserved | -0,04157 | 0,806 |
| TcCLB.503687.20 | glyceraldehyde-3-phosphate dehydrogenase, putative | -0,04159 | 0,7424 |
| TcCLB.507771.9 | kinesin, putative (fragment) | -0,0416 | 0,862 |
| TcCLB.511321.70 | hypothetical protein, conserved | -0,04161 | 0,721 |
| TcCLB.511573.58 | Nucleolar protein 56, putative (fragment) (NOP56) | -0,04174 | 0,7049 |
| TcCLB.507093.270 | guanine nucleotide releasing protein, putative | -0,04177 | 0,7652 |
| TcCLB.511003.190 | hypothetical protein, conserved | -0,04188 | 0,8019 |
| TcCLB.510083.30 | retrotransposon hot spot protein (RHS, pseudogene), puta | -0,0419 | 0,6739 |
| TcCLB.509029.160 | hypothetical protein, conserved | -0,0419 | 0,7212 |
| TcCLB.508541.10 | elongation factor 1-gamma (EF-1-gamma, pseudogene), | -0,04193 | 0,8916 |
| TcCLB.511897.10 | TFIIH basal transcription factor complex helicase subunit, | -0,042 | 0,7135 |
| TcCLB.510187.540 | amino acid permease, putative | -0,04207 | 0,7431 |
| TcCLB.510303.270 | hypothetical protein, conserved | -0,04216 | 0,5933 |
| TcCLB.508479.220 | hypothetical protein, conserved | -0,04238 | 0,7881 |
| TcCLB.510691.4 | hypothetical protein, conserved | -0,04249 | 0,8735 |
| TcCLB.509565.9 | hypothetical protein (fragment) | -0,0425 | 0,6452 |
| TcCLB.506961.97 | N-acetyltransferase complex ARD1 subunit (pseudogene), | -0,04268 | 0,8963 |
| TcCLB.508879.80 | hypothetical protein, conserved | -0,04272 | 0,7494 |
| TcCLB.507063.250 | hypothetical protein | -0,04277 | 0,8339 |
| TcCLB.507969.50 | actin 2, putative | -0,04281 | 0,7963 |
| TcCLB.510595.40 | hypothetical protein, conserved | -0,04296 | 0,7686 |
| TcCLB.506847.70 | hypothetical protein, conserved | -0,04319 | 0,7966 |
| TcCLB.510997.30 | hypothetical protein, conserved | -0,0432 | 0,6839 |
| TcCLB.509259.111 | N-acetyltransferase complex ARD1 subunit (pseudogene), | -0,04321 | 0,8934 |
| TcCLB.508385.40 | dual specificity protein phosphatase, putative | -0,04331 | 0,7761 |
| TcCLB.509747.60 | hypothetical protein, conserved | -0,04334 | 0,697 |
| TcCLB.508231.50 | legume-like lectin, putative | -0,04343 | 0,763 |

| | | | |
|---|---|---|---|
| TcCLB.506327.60 | hypothetical protein, conserved | -0,04346 | 0,7935 |
| TcCLB.509707.29 | phosphatidylinositol-4-phosphate 5-kinase-like protein | -0,0435 | 0,825 |
| TcCLB.509999.60 | lipase domain protein, putative | -0,04352 | 0,6413 |
| TcCLB.506383.10 | mannosyl-oligosaccharide 1,2-alpha-mannosidase IB, puta | -0,04367 | 0,7593 |
| TcCLB.507513.20 | hypothetical protein, conserved | -0,04371 | 0,7115 |
| TcCLB.510659.36 | hypothetical protein, conserved | -0,04375 | 0,7348 |
| TcCLB.510517.30 | hypothetical protein, conserved | -0,0439 | 0,6826 |
| TcCLB.509693.120 | hypothetical protein, conserved | -0,04397 | 0,6831 |
| TcCLB.504151.50 | hypothetical protein, conserved | -0,04407 | 0,8059 |
| TcCLB.511339.10 | UDP-Gal or UDP-GlcNAc-dependent glycosyltransferase, p | -0,04407 | 0,889 |
| TcCLB.510053.20 | hypothetical protein, conserved | -0,0441 | 0,7106 |
| TcCLB.509443.48 | hypothetical protein, conserved | -0,04411 | 0,828 |
| TcCLB.508881.130 | hypothetical protein, conserved | -0,04421 | 0,7675 |
| TcCLB.509169.10 | hypothetical protein, conserved | -0,04426 | 0,718 |
| TcCLB.510767.40 | hypothetical protein, conserved | -0,04429 | 0,7595 |
| TcCLB.509733.120 | hypothetical protein, conserved | -0,0443 | 0,6546 |
| TcCLB.511217.80 | hypothetical protein, conserved | -0,04432 | 0,8572 |
| TcCLB.510719.230 | hypothetical protein, conserved | -0,04489 | 0,689 |
| TcCLB.506627.10 | AMP deaminase 2, putative | -0,04507 | 0,7051 |
| TcCLB.507017.170 | hypothetical protein, conserved | -0,04522 | 0,8188 |
| TcCLB.506175.70 | hypothetical protein, conserved | -0,04523 | 0,7061 |
| TcCLB.506165.16 | protein kinase, putative | -0,04524 | 0,867 |
| TcCLB.507849.60 | hypothetical protein, conserved | -0,04537 | 0,6678 |
| TcCLB.508951.10 | hypothetical protein, conserved | -0,04537 | 0,7105 |
| TcCLB.506247.270 | ATPase, putative | -0,04539 | 0,635 |
| TcCLB.511925.50 | hypothetical protein, conserved | -0,04539 | 0,7255 |
| TcCLB.510655.70 | hypothetical protein, conserved | -0,04547 | 0,7749 |
| TcCLB.503779.40 | protein-l-isoaspartate o-methyltransferase, putative | -0,04555 | 0,7071 |
| TcCLB.503809.10 | hypothetical protein, conserved | -0,04562 | 0,657 |
| TcCLB.511589.29 | hypothetical protein, conserved | -0,04576 | 0,7584 |
| TcCLB.407477.69 | hypothetical protein, conserved (fragment) | -0,0458 | 0,7549 |
| TcCLB.508871.30 | surface protease GP63 (pseudogene), putative | -0,04582 | 0,8704 |
| TcCLB.506941.139 | zinc finger protein, putative | -0,0461 | 0,6773 |
| TcCLB.508895.50 | RNA-binding protein, putative | -0,04621 | 0,6813 |
| TcCLB.508355.210 | hypothetical protein, conserved | -0,04623 | 0,763 |
| TcCLB.507663.9 | hypothetical protein, conserved | -0,04625 | 0,8223 |
| TcCLB.511071.160 | hypothetical protein, conserved | -0,04628 | 0,7584 |
| TcCLB.508479.200 | palmitoyl acyltransferase 10, putative | -0,04633 | 0,6791 |
| TcCLB.506265.150 | hypothetical protein, conserved | -0,04641 | 0,8022 |
| TcCLB.510407.40 | hypothetical protein, conserved | -0,04643 | 0,7327 |
| TcCLB.506855.160 | hypothetical protein, conserved | -0,04648 | 0,6111 |
| TcCLB.508577.80 | hypothetical protein, conserved | -0,04649 | 0,792 |
| TcCLB.508149.40 | hypothetical protein, conserved (pseudogene) | -0,04649 | 0,8675 |
| TcCLB.508755.11 | trans-sialidase (pseudogene), putative | -0,04651 | 0,8281 |
| TcCLB.509229.90 | hypothetical protein, conserved | -0,04656 | 0,6758 |

| | | | |
|---|---|---|---|
| TcCLB.510877.110 | hypothetical protein, conserved | -0,04681 | 0,5669 |
| TcCLB.510579.50 | hypothetical protein, conserved | -0,0469 | 0,7074 |
| TcCLB.508823.110 | hypothetical protein, conserved | -0,04696 | 0,5939 |
| TcCLB.436497.10 | hypothetical protein | -0,0472 | 0,7494 |
| TcCLB.506147.70 | hypothetical protein, conserved | -0,04743 | 0,7399 |
| TcCLB.511903.10 | hypothetical protein | -0,04745 | 0,8424 |
| TcCLB.511285.90 | 19S proteasome non-atpase subunit 8 (Rpn8) | -0,04747 | 0,6716 |
| TcCLB.507787.40 | hypothetical protein, conserved | -0,04765 | 0,7737 |
| TcCLB.511589.100 | hypothetical protein, conserved | -0,04768 | 0,822 |
| TcCLB.506929.20 | hypothetical protein, conserved | -0,04776 | 0,6805 |
| TcCLB.509179.90 | hypothetical protein, conserved | -0,04777 | 0,8549 |
| TcCLB.509819.30 | septum formation inhibitor, Maf-like protein, putative | -0,04781 | 0,7534 |
| TcCLB.503395.10 | hypothetical protein, conserved | -0,04794 | 0,8158 |
| TcCLB.503973.350 | hypothetical protein | -0,04794 | 0,905 |
| TcCLB.503797.4 | hypothetical protein, conserved | -0,04797 | 0,6825 |
| TcCLB.509671.150 | hypothetical protein, conserved | -0,04813 | 0,749 |
| TcCLB.508413.10 | hypothetical protein, conserved | -0,04837 | 0,6836 |
| TcCLB.510997.50 | ATP-dependent DEAD/H RNA helicase (pseudogene), puta | -0,04838 | 0,7155 |
| TcCLB.510647.20 | hypothetical protein, conserved | -0,0484 | 0,6677 |
| TcCLB.508405.60 | hypothetical protein, conserved | -0,04852 | 0,6904 |
| TcCLB.510329.110 | hypothetical protein | -0,04866 | 0,6475 |
| TcCLB.505999.189 | hypothetical protein, conserved (pseudogene) | -0,04885 | 0,8039 |
| TcCLB.508699.20 | Translation initiation factor eIF-2B subunit epsilon, putati | -0,04889 | 0,6114 |
| TcCLB.511459.70 | hypothetical protein, conserved | -0,04893 | 0,8802 |
| TcCLB.506945.220 | serine/arginine-rich protein specific kinase SRPK, putative | -0,04896 | 0,5438 |
| TcCLB.506649.64 | hypothetical protein, conserved | -0,04901 | 0,8261 |
| TcCLB.509563.60 | hypothetical protein | -0,04926 | 0,8033 |
| TcCLB.506367.70 | hypothetical protein, conserved | -0,04938 | 0,7209 |
| TcCLB.511813.10 | hypothetical protein, conserved | -0,04946 | 0,6927 |
| TcCLB.504033.190 | EF hand, putative | -0,04959 | 0,7025 |
| TcCLB.509569.40 | hypothetical protein, conserved | -0,04975 | 0,6126 |
| TcCLB.506211.60 | RNA-binding protein, putative | -0,04979 | 0,6828 |
| TcCLB.509791.60 | hypothetical protein, conserved | -0,0498 | 0,6526 |
| TcCLB.511585.200 | hypothetical protein, conserved | -0,04986 | 0,6895 |
| TcCLB.510547.46 | hypothetical protein, conserved (pseudogene) | -0,04995 | 0,922 |
| TcCLB.509573.20 | hypothetical protein, conserved | -0,04997 | 0,8124 |
| TcCLB.506559.517 | hypothetical protein, conserved | -0,05007 | 0,6437 |
| TcCLB.511467.16 | hypothetical protein, conserved | -0,0502 | 0,6906 |
| TcCLB.508601.130 | Elongation factor G 2, mitochondrial, putative (EF-G2) | -0,05055 | 0,6182 |
| TcCLB.511647.50 | hypothetical protein, conserved | -0,05057 | 0,7197 |
| TcCLB.508851.140 | hypothetical protein, conserved | -0,05062 | 0,723 |
| TcCLB.507773.10 | trans-sialidase, Group IV, putative | -0,05075 | 0,7275 |
| TcCLB.505071.140 | protein kinase, putative | -0,05076 | 0,6965 |
| TcCLB.507011.30 | hypothetical protein, conserved | -0,05087 | 0,673 |
| TcCLB.509749.20 | hypothetical protein, conserved | -0,05092 | 0,8176 |

| | | | |
|---|---|---|---|
| TcCLB.417797.9 | hypothetical protein, conserved | -0,05099 | 0,7266 |
| TcCLB.510945.60 | hypothetical protein, conserved | -0,051 | 0,5843 |
| TcCLB.508505.30 | hypothetical protein, conserved | -0,05115 | 0,6678 |
| TcCLB.509023.90 | hypothetical protein, conserved | -0,05132 | 0,7972 |
| TcCLB.504157.110 | dynein intermediate chain, putative | -0,05136 | 0,7455 |
| TcCLB.511491.20 | hypothetical protein, conserved | -0,05137 | 0,5348 |
| TcCLB.507669.239 | hypothetical protein, conserved | -0,05149 | 0,6819 |
| TcCLB.510421.270 | hypothetical protein, conserved | -0,05158 | 0,7939 |
| TcCLB.507997.60 | hypothetical protein, conserved | -0,05168 | 0,6398 |
| TcCLB.511733.60 | hypothetical protein, conserved | -0,05173 | 0,7101 |
| TcCLB.509999.50 | lipase domain protein, putative | -0,05195 | 0,5939 |
| TcCLB.509161.50 | hypothetical protein, conserved | -0,05198 | 0,6772 |
| TcCLB.427789.10 | hypothetical protein, conserved | -0,05204 | 0,7987 |
| TcCLB.508781.30 | hypothetical protein, conserved | -0,05207 | 0,6715 |
| TcCLB.511803.30 | hypothetical protein, conserved | -0,05208 | 0,709 |
| TcCLB.506479.110 | hypothetical protein | -0,05211 | 0,6377 |
| TcCLB.506673.60 | AP-3 complex subunit beta, putative (AP3B) | -0,05215 | 0,6474 |
| TcCLB.506477.20 | vesicular-fusion protein nsf, putative | -0,05217 | 0,657 |
| TcCLB.472253.10 | vacuolar protein sorting complex subunit (pseudogene), pu | -0,05221 | 0,7667 |
| TcCLB.510741.90 | hypothetical protein, conserved | -0,05237 | 0,7522 |
| TcCLB.510001.50 | hypothetical protein, conserved | -0,05266 | 0,767 |
| TcCLB.510733.54 | hypothetical protein, conserved | -0,05271 | 0,7461 |
| TcCLB.506735.50 | hypothetical protein, conserved | -0,05273 | 0,6048 |
| TcCLB.503673.18 | hypothetical protein, conserved | -0,05285 | 0,7353 |
| TcCLB.507041.80 | hypothetical protein | -0,05286 | 0,7973 |
| TcCLB.511423.100 | kinetoplast ribosomal PPR-repeat containing protein 3, pu | -0,05289 | 0,6389 |
| TcCLB.509563.10 | hypothetical protein, conserved | -0,05294 | 0,6596 |
| TcCLB.508075.20 | DNA repair protein, putative | -0,05294 | 0,6857 |
| TcCLB.511283.260 | hypothetical protein, conserved | -0,05296 | 0,5517 |
| TcCLB.510105.150 | hypothetical protein, conserved | -0,05297 | 0,7697 |
| TcCLB.508409.150 | hypothetical protein, conserved | -0,05298 | 0,5855 |
| TcCLB.511075.50 | hypothetical protein | -0,05301 | 0,7167 |
| TcCLB.511577.19 | hypothetical protein, conserved | -0,05318 | 0,784 |
| TcCLB.507723.150 | hypothetical protein, conserved | -0,05331 | 0,6867 |
| TcCLB.505363.10 | trans-sialidase, Group II, putative | -0,05333 | 0,8456 |
| TcCLB.507795.80 | hypothetical protein, conserved | -0,05338 | 0,6512 |
| TcCLB.510597.40 | nucleoporin (NUP54/57), putative | -0,0534 | 0,7821 |
| TcCLB.508699.40 | eukaryotic translation initiation factor, putative | -0,05356 | 0,6488 |
| TcCLB.509601.20 | hypothetical protein, conserved | -0,05361 | 0,7427 |
| TcCLB.511741.20 | hypothetical protein, conserved | -0,05362 | 0,6827 |
| TcCLB.506893.60 | hypothetical protein, conserved | -0,05368 | 0,7747 |
| TcCLB.507031.60 | nuclear cap binding complex subunit CBP30, putative (CBF | -0,05376 | 0,696 |
| TcCLB.506335.69 | mucin-associated surface protein (MASP, pseudogene), pu | -0,05377 | 0,8627 |
| TcCLB.511517.30 | hypothetical protein, conserved | -0,0538 | 0,6618 |
| TcCLB.506961.104 | ATP-dependent DEAD/H RNA helicase (pseudogene), puta | -0,05393 | 0,8033 |

| | | | |
|---|---|---|---|
| TcCLB.506873.10 | hypothetical protein, conserved | -0,054 | 0,5921 |
| TcCLB.507197.20 | oxidoreductase-like protein, putative | -0,05422 | 0,6192 |
| TcCLB.508251.10 | hypothetical protein | -0,05427 | 0,8997 |
| TcCLB.504079.20 | hypothetical protein, conserved | -0,0543 | 0,7937 |
| TcCLB.506975.60 | hypothetical protein, conserved | -0,05432 | 0,6013 |
| TcCLB.503925.30 | protein kinase, putative | -0,05438 | 0,6884 |
| TcCLB.506255.30 | hypothetical protein, conserved | -0,05438 | 0,8166 |
| TcCLB.511717.209 | phosphatidylinositol 3-related kinase, putative (fragment) | -0,05439 | 0,8617 |
| TcCLB.508839.60 | ADP-ribosylation factor, putative | -0,05455 | 0,7266 |
| TcCLB.506195.180 | hypothetical protein, conserved | -0,05473 | 0,7365 |
| TcCLB.508023.60 | trans-sialidase (pseudogene), putative | -0,05491 | 0,7197 |
| TcCLB.408345.10 | hypothetical protein, conserved | -0,05509 | 0,8156 |
| TcCLB.506411.20 | hypothetical protein, conserved | -0,05531 | 0,6685 |
| TcCLB.511751.70 | hypothetical protein, conserved | -0,05535 | 0,5042 |
| TcCLB.507951.15 | hypothetical protein, conserved (fragment) | -0,05544 | 0,5833 |
| TcCLB.506425.90 | hypothetical protein, conserved | -0,05553 | 0,6726 |
| TcCLB.511557.64 | hypothetical protein | -0,05563 | 0,6916 |
| TcCLB.511261.130 | hypothetical protein, conserved | -0,0558 | 0,7558 |
| TcCLB.511043.20 | small G-protein, putative | -0,05592 | 0,5604 |
| TcCLB.511239.40 | hypothetical protein, conserved | -0,05599 | 0,6311 |
| TcCLB.506755.150 | hypothetical protein, conserved | -0,05601 | 0,6886 |
| TcCLB.507677.129 | hypothetical protein, conserved | -0,05617 | 0,5847 |
| TcCLB.506363.120 | hypothetical protein, conserved | -0,05617 | 0,7013 |
| TcCLB.506247.30 | RNA editing complex protein MP63, putative | -0,05619 | 0,4837 |
| TcCLB.509595.70 | Trichohyalin, putative | -0,05624 | 0,6098 |
| TcCLB.510265.70 | hypothetical protein, conserved | -0,05633 | 0,5245 |
| TcCLB.509641.40 | hypothetical protein, conserved | -0,05638 | 0,5327 |
| TcCLB.509065.70 | glyceraldehyde 3-phosphate dehydrogenase, putative | -0,05638 | 0,6146 |
| TcCLB.510519.130 | hypothetical protein, conserved | -0,05638 | 0,6866 |
| TcCLB.511303.140 | hypothetical protein, conserved | -0,05641 | 0,6239 |
| TcCLB.511313.30 | formin, putative | -0,05651 | 0,7801 |
| TcCLB.509809.30 | hypothetical protein, conserved | -0,05652 | 0,5272 |
| TcCLB.510099.110 | hypothetical protein, conserved | -0,05657 | 0,5782 |
| TcCLB.511741.30 | replication factor C, subunit 2, putative | -0,05672 | 0,6193 |
| TcCLB.508165.330 | trans-sialidase (pseudogene), putative | -0,05673 | 0,8202 |
| TcCLB.508573.120 | dispersed gene family protein 1 (DGF-1), putative | -0,05679 | 0,8466 |
| TcCLB.510149.20 | hypothetical protein, conserved | -0,05688 | 0,6531 |
| TcCLB.511037.30 | hypothetical protein, conserved | -0,0569 | 0,5473 |
| TcCLB.510101.450 | hypothetical protein, conserved | -0,05691 | 0,5556 |
| TcCLB.511585.70 | heat shock protein 70 (hsp70), putative | -0,05695 | 0,5787 |
| TcCLB.508055.30 | trans-sialidase, Group II, putative | -0,05696 | 0,806 |
| TcCLB.506559.559 | antigenic protein, putative | -0,05701 | 0,5931 |
| TcCLB.507001.40 | hypothetical protein, conserved | -0,05701 | 0,5979 |
| TcCLB.510185.10 | sterol 24-c-methyltransferase, putative | -0,05705 | 0,6714 |
| TcCLB.408573.5 | hypothetical protein | -0,05705 | 0,8978 |

| | | | |
|---|---|---|---|
| TcCLB.508553.10 | dispersed gene family protein 1 (DGF-1, pseudogene), put | -0,05709 | 0,797 |
| TcCLB.439125.10 | Peptide chain release factor 1, mitochondrial, putative (RI | -0,05738 | 0,7078 |
| TcCLB.454403.10 | trans-sialidase (pseudogene), putative | -0,0574 | 0,6166 |
| TcCLB.511289.119 | C-terminal motor kinesin, putative (fragment) | -0,05771 | 0,6797 |
| TcCLB.506335.110 | trans-sialidase (pseudogene), putative | -0,05774 | 0,8055 |
| TcCLB.508637.110 | nucleotide-binding protein, putative | -0,05775 | 0,6986 |
| TcCLB.510075.30 | hypothetical protein, conserved | -0,05776 | 0,5776 |
| TcCLB.507025.20 | hypothetical protein, conserved | -0,0578 | 0,6858 |
| TcCLB.506201.140 | hypothetical protein, conserved | -0,05796 | 0,5426 |
| TcCLB.506941.50 | hypothetical protein, conserved | -0,05798 | 0,6296 |
| TcCLB.509859.60 | hypothetical protein, conserved | -0,0581 | 0,5924 |
| TcCLB.506925.80 | hypothetical protein, conserved | -0,05824 | 0,5991 |
| TcCLB.506007.40 | protein kinase, putative | -0,05841 | 0,6245 |
| TcCLB.506871.160 | RNA guanylyltransferase, putative | -0,05843 | 0,5782 |
| TcCLB.508857.40 | UDP-glucoronosyl and UDP-glucosyl transferase, putative | -0,05851 | 0,7113 |
| TcCLB.504867.80 | DNA primase large subunit, putative (pril) | -0,05869 | 0,6575 |
| TcCLB.510629.440 | hypothetical protein, conserved | -0,05872 | 0,4998 |
| TcCLB.510963.60 | hypothetical protein, conserved | -0,05873 | 0,6605 |
| TcCLB.510487.37 | trans-sialidase (pseudogene), putative | -0,05889 | 0,8261 |
| TcCLB.510579.119 | hypothetical protein, conserved | -0,05912 | 0,6588 |
| TcCLB.508891.20 | hypothetical protein, conserved | -0,05921 | 0,5963 |
| TcCLB.506435.30 | mitochondrial DNA polymerase I protein A, putative | -0,05945 | 0,5674 |
| TcCLB.511019.108 | hypothetical protein, conserved | -0,05947 | 0,6904 |
| TcCLB.506943.90 | hypothetical protein, conserved | -0,0595 | 0,7361 |
| TcCLB.506885.270 | tubulin tyrosine ligase, putative | -0,05951 | 0,6011 |
| TcCLB.503891.40 | hypothetical protein, conserved | -0,05954 | 0,6075 |
| TcCLB.506223.70 | hypothetical protein | -0,05961 | 0,7741 |
| TcCLB.509695.30 | hypothetical protein, conserved | -0,05965 | 0,7212 |
| TcCLB.506559.360 | hypothetical protein, conserved | -0,05973 | 0,5715 |
| TcCLB.509985.30 | hypothetical protein, conserved | -0,05992 | 0,6824 |
| TcCLB.507031.170 | protein kinase (pseudogene), putative | -0,06 | 0,5951 |
| TcCLB.511473.9 | N-acetyltransferase complex ARD1 subunit (pseudogene), | -0,06007 | 0,8083 |
| TcCLB.511271.10 | dispersed gene family protein 1 (DGF-1, pseudogene), put | -0,06008 | 0,7902 |
| TcCLB.508159.26 | hypothetical protein, conserved (pseudogene) | -0,06009 | 0,8037 |
| TcCLB.508641.90 | hypothetical protein, conserved | -0,06017 | 0,5645 |
| TcCLB.508153.440 | polynucleotide adenylyltransferase, putative (PAP) | -0,06017 | 0,6609 |
| TcCLB.507483.4 | polyubiquitin (pseudogene), putative | -0,06027 | 0,7977 |
| TcCLB.507641.170 | hypothetical protein, conserved | -0,06029 | 0,5773 |
| TcCLB.508153.220 | hypothetical protein, conserved | -0,06052 | 0,6389 |
| TcCLB.508681.31 | hypothetical protein | -0,06055 | 0,8613 |
| TcCLB.509153.30 | hypothetical protein, conserved | -0,06057 | 0,4704 |
| TcCLB.508831.40 | hypothetical protein, conserved | -0,0606 | 0,7685 |
| TcCLB.506625.210 | hypothetical protein, conserved | -0,06063 | 0,645 |
| TcCLB.504827.90 | hypothetical protein, conserved | -0,06075 | 0,6783 |
| TcCLB.510359.540 | hypothetical protein | -0,06085 | 0,8984 |

| | | | |
|---|---|---|---|
| TcCLB.506265.170 | hypothetical protein, conserved | -0,06094 | 0,6584 |
| TcCLB.510279.263 | trans-sialidase (pseudogene), putative | -0,06094 | 0,8542 |
| TcCLB.511283.200 | hypothetical protein, conserved | -0,06098 | 0,5097 |
| TcCLB.509065.180 | protein kinase-like protein, putative | -0,061 | 0,5762 |
| TcCLB.509073.30 | ubiquitin hydrolase, putative | -0,06108 | 0,5448 |
| TcCLB.508445.80 | hypothetical protein, conserved | -0,06122 | 0,6046 |
| TcCLB.509063.2 | hypothetical protein, conserved (fragment) | -0,06124 | 0,8638 |
| TcCLB.503987.30 | Enriched in surface-labeled proteome protein 16 (ESP16) | -0,06128 | 0,5961 |
| TcCLB.510155.180 | eukaryotic initiation factor 4a, putative | -0,0614 | 0,5372 |
| TcCLB.503841.60 | hypothetical protein, conserved | -0,06154 | 0,6518 |
| TcCLB.511803.50 | hypothetical protein, conserved | -0,06175 | 0,6938 |
| TcCLB.509047.100 | hypothetical protein, conserved | -0,06181 | 0,6699 |
| TcCLB.508965.50 | hypothetical protein, conserved | -0,06209 | 0,6086 |
| TcCLB.506359.99 | hypothetical protein | -0,06216 | 0,7387 |
| TcCLB.507305.30 | casein kinase, putative | -0,06217 | 0,7748 |
| TcCLB.510729.180 | hypothetical protein | -0,06228 | 0,5879 |
| TcCLB.503691.9 | hypothetical protein, conserved (fragment) | -0,06228 | 0,7226 |
| TcCLB.511715.80 | hypothetical protein, conserved | -0,06234 | 0,6133 |
| TcCLB.509991.20 | conserved protein, unknown function | -0,06263 | 0,5426 |
| TcCLB.508233.40 | hypothetical protein | -0,06266 | 0,6432 |
| TcCLB.509885.60 | methyltransferase, putative | -0,06267 | 0,7071 |
| TcCLB.511141.30 | protein kinase, putative | -0,06269 | 0,7602 |
| TcCLB.503557.10 | hypothetical protein, conserved | -0,06271 | 0,697 |
| TcCLB.511749.9 | hypothetical protein, conserved (fragment) | -0,06272 | 0,5047 |
| TcCLB.510257.70 | hypothetical protein, conserved | -0,06273 | 0,5397 |
| TcCLB.506355.150 | hypothetical protein, conserved | -0,06273 | 0,6872 |
| TcCLB.508181.149 | hypothetical protein, conserved | -0,0629 | 0,7939 |
| TcCLB.511309.80 | trans-sialidase (pseudogene), putative | -0,06301 | 0,7009 |
| TcCLB.503593.20 | hypothetical protein, conserved | -0,06306 | 0,5882 |
| TcCLB.507039.20 | GDP-mannose 4,6 dehydratase (pseudogene), putative | -0,06326 | 0,6867 |
| TcCLB.511263.50 | hypothetical protein, conserved | -0,06328 | 0,785 |
| TcCLB.506637.90 | hypothetical protein, conserved | -0,06331 | 0,519 |
| TcCLB.506649.4 | hypothetical protein, conserved | -0,06335 | 0,7264 |
| TcCLB.506529.480 | hypothetical protein, conserved | -0,06352 | 0,6077 |
| TcCLB.511729.30 | hypothetical protein, conserved | -0,06357 | 0,616 |
| TcCLB.509223.80 | trans-sialidase (pseudogene), putative | -0,06358 | 0,7074 |
| TcCLB.504247.20 | ATP-dependent RNA helicase, putative | -0,06362 | 0,5154 |
| TcCLB.511755.24 | hypothetical protein, conserved (pseudogene) | -0,06364 | 0,7018 |
| TcCLB.509965.400 | hypothetical protein, conserved | -0,06372 | 0,5514 |
| TcCLB.504213.20 | hypothetical protein, conserved | -0,06376 | 0,516 |
| TcCLB.511621.130 | hypothetical protein, conserved | -0,0638 | 0,5664 |
| TcCLB.510229.20 | 8-oxoguanine DNA glycosylase, putative | -0,06382 | 0,6683 |
| TcCLB.508721.20 | hypothetical protein, conserved | -0,06388 | 0,6315 |
| TcCLB.506575.54 | hypothetical protein, conserved | -0,06392 | 0,7487 |
| TcCLB.505101.30 | hypothetical protein, conserved | -0,06403 | 0,8781 |

| | | | |
|---|---|---|---|
| TcCLB.511825.90 | hypothetical protein, conserved | -0,06416 | 0,7333 |
| TcCLB.510143.30 | hypothetical protein, conserved | -0,06435 | 0,5855 |
| TcCLB.506833.60 | UV excision repair RAD23-like protein, putative | -0,06445 | 0,495 |
| TcCLB.510131.60 | hypothetical protein | -0,06447 | 0,6477 |
| TcCLB.504043.10 | hypothetical protein, conserved | -0,06454 | 0,5227 |
| TcCLB.511517.10 | hypothetical protein, conserved | -0,0646 | 0,6734 |
| TcCLB.506839.34 | oxidoreductase, putative | -0,06472 | 0,7709 |
| TcCLB.506629.240 | retrotransposon hot spot (RHS) protein, putative (fragmer | -0,06484 | 0,878 |
| TcCLB.511859.20 | hypothetical protein, conserved | -0,06489 | 0,6223 |
| TcCLB.508851.170 | DNA topoisomerase III, putative | -0,06494 | 0,5669 |
| TcCLB.508647.190 | hypothetical protein, conserved | -0,06498 | 0,5889 |
| TcCLB.509971.70 | hypothetical protein | -0,06505 | 0,8203 |
| TcCLB.511277.39 | hypothetical protein, conserved | -0,06511 | 0,7815 |
| TcCLB.510201.45 | hypothetical protein, conserved (pseudogene) | -0,06512 | 0,7506 |
| TcCLB.506853.30 | arginine N-methyltransferase, type II, putative (PRMT5) | -0,06519 | 0,6009 |
| TcCLB.510957.9 | calpain-like cysteine peptidase, putative | -0,06522 | 0,4272 |
| TcCLB.510149.110 | phosphate transporter, putative | -0,06529 | 0,5833 |
| TcCLB.506587.100 | surface protease GP63, putative | -0,06533 | 0,5945 |
| TcCLB.508503.70 | hypothetical protein, conserved | -0,06556 | 0,571 |
| TcCLB.503641.10 | hypothetical protein, conserved | -0,06556 | 0,5805 |
| TcCLB.506885.364 | hypothetical protein, conserved | -0,06563 | 0,6893 |
| TcCLB.506963.80 | hypothetical protein, conserved | -0,06566 | 0,4973 |
| TcCLB.506559.370 | hypothetical protein, conserved | -0,06579 | 0,5691 |
| TcCLB.428999.10 | hypothetical protein | -0,0658 | 0,6623 |
| TcCLB.504557.60 | hypothetical protein, conserved | -0,06583 | 0,702 |
| TcCLB.507835.80 | trans-sialidase, putative | -0,06583 | 0,8304 |
| TcCLB.510627.215 | hypothetical protein, conserved (pseudogene) | -0,06584 | 0,9185 |
| TcCLB.509161.70 | Mg transporter, putative | -0,06591 | 0,5775 |
| TcCLB.506495.30 | retrotransposon hot spot protein (RHS, pseudogene), puta | -0,06594 | 0,7356 |
| TcCLB.507011.250 | hypothetical protein, conserved | -0,06606 | 0,4365 |
| TcCLB.506127.70 | hypothetical protein | -0,06612 | 0,5692 |
| TcCLB.509901.140 | COP9 signalosome complex subunit 5, putative | -0,06633 | 0,6079 |
| TcCLB.507611.114 | retrotransposon hot spot protein (RHS, pseudogene), puta | -0,06639 | 0,7611 |
| TcCLB.506667.5 | surface protease GP63 (pseudogene), putative | -0,06645 | 0,8823 |
| TcCLB.506793.100 | hypothetical protein, conserved | -0,06651 | 0,5614 |
| TcCLB.509033.50 | hypothetical protein, conserved | -0,06652 | 0,4967 |
| TcCLB.511411.30 | arginine permease (AAAP411) | -0,06662 | 0,8618 |
| TcCLB.509569.80 | hypothetical protein | -0,06669 | 0,6345 |
| TcCLB.506559.380 | centrin, putative | -0,06679 | 0,5913 |
| TcCLB.511499.30 | RNA decapping enzyme, putative (Dcp2) | -0,06687 | 0,7015 |
| TcCLB.506745.4 | hypothetical protein, conserved (fragment) | -0,06693 | 0,8473 |
| TcCLB.511821.130 | hypothetical protein | -0,06707 | 0,5421 |
| TcCLB.508119.11 | surface protease GP63 (pseudogene), putative | -0,06717 | 0,7377 |
| TcCLB.509967.150 | hypothetical protein, conserved | -0,0673 | 0,5919 |
| TcCLB.505071.119 | hypothetical protein, conserved | -0,06736 | 0,626 |

| | | | |
|---|---|---|---|
| TcCLB.507001.10 | hypothetical protein, conserved | -0,06744 | 0,5048 |
| TcCLB.509551.90 | hypothetical protein, conserved | -0,06746 | 0,6308 |
| TcCLB.509745.10 | hypothetical protein, conserved | -0,06758 | 0,4922 |
| TcCLB.506797.100 | beta-ketoacyl-ACP reductase 2, putative | -0,06758 | 0,5844 |
| TcCLB.506509.30 | hypothetical protein, conserved | -0,06778 | 0,5507 |
| TcCLB.510879.70 | selenocysteine-tRNA-specific elongation factor, putative | -0,0679 | 0,5531 |
| TcCLB.504045.90 | hypothetical protein, conserved (pseudogene) | -0,06822 | 0,6369 |
| TcCLB.510531.30 | hypothetical protein, conserved | -0,06827 | 0,515 |
| TcCLB.511717.140 | hypothetical protein, conserved | -0,0683 | 0,6588 |
| TcCLB.507061.30 | iron superoxide dismutase, putative | -0,06833 | 0,7009 |
| TcCLB.424123.30 | hypothetical protein, conserved | -0,06843 | 0,6397 |
| TcCLB.511233.50 | hypothetical protein, conserved | -0,06876 | 0,5892 |
| TcCLB.505193.74 | hypothetical protein, conserved | -0,06879 | 0,5715 |
| TcCLB.506167.50 | clathrin heavy chain, putative | -0,06885 | 0,3979 |
| TcCLB.506575.50 | hypothetical protein, conserved | -0,06887 | 0,4683 |
| TcCLB.511217.110 | hypothetical protein, conserved | -0,06891 | 0,5295 |
| TcCLB.511809.50 | hypothetical protein, conserved | -0,06892 | 0,6149 |
| TcCLB.507989.10 | hypothetical protein, conserved | -0,06898 | 0,5829 |
| TcCLB.506519.70 | hypothetical protein | -0,06899 | 0,6818 |
| TcCLB.511291.10 | intraflagellar transport protein 80, putative (fragment) (IF | -0,06915 | 0,6342 |
| TcCLB.509921.40 | retrotransposon hot spot protein (RHS, pseudogene), puta | -0,06928 | 0,6224 |
| TcCLB.509801.30 | citrate synthase, putative | -0,06928 | 0,8106 |
| TcCLB.508515.10 | hypothetical protein, conserved | -0,06929 | 0,692 |
| TcCLB.509417.30 | retrotransposon hot spot protein (RHS, pseudogene), puta | -0,06933 | 0,799 |
| TcCLB.510657.220 | hypothetical protein, conserved | -0,06942 | 0,6648 |
| TcCLB.507601.30 | hypothetical protein, conserved | -0,06956 | 0,6937 |
| TcCLB.504059.24 | hypothetical protein, conserved | -0,0696 | 0,7071 |
| TcCLB.509721.79 | hypothetical protein, conserved (fragment) | -0,06972 | 0,5369 |
| TcCLB.511365.10 | hypothetical protein, conserved | -0,06974 | 0,5828 |
| TcCLB.510523.20 | hypothetical protein, conserved | -0,06981 | 0,6231 |
| TcCLB.507927.4 | rac serine-threonine kinase, putative | -0,06981 | 0,6447 |
| TcCLB.510081.21 | ATP-dependent DEAD/H RNA helicase (pseudogene), puta | -0,06983 | 0,7655 |
| TcCLB.510897.99 | hypothetical protein, conserved (fragment) | -0,06985 | 0,7383 |
| TcCLB.503651.20 | hypothetical protein | -0,06992 | 0,6269 |
| TcCLB.511001.50 | hypothetical protein, conserved | -0,06994 | 0,5924 |
| TcCLB.503423.30 | hypothetical protein, conserved | -0,07014 | 0,7987 |
| TcCLB.508293.111 | elongation factor 1-gamma (EF-1-gamma, pseudogene), | -0,07015 | 0,8754 |
| TcCLB.510989.69 | hypothetical protein, conserved | -0,07019 | 0,732 |
| TcCLB.510687.100 | Unc104-like kinesin, putative | -0,07037 | 0,4784 |
| TcCLB.510557.73 | mucin-associated surface protein (MASP, pseudogene), pu | -0,07043 | 0,8276 |
| TcCLB.506861.20 | hypothetical protein, conserved | -0,07056 | 0,5511 |
| TcCLB.506945.250 | hypothetical protein, conserved | -0,07066 | 0,4574 |
| TcCLB.506363.30 | hypothetical protein, conserved | -0,07068 | 0,4928 |
| TcCLB.509999.70 | hypothetical protein, conserved | -0,0707 | 0,5359 |
| TcCLB.507021.100 | hypothetical protein, conserved | -0,07073 | 0,5224 |

| | | | |
|---|---|---|---|
| TcCLB.504247.30 | transcription elongation factor 1 homolog (ELOF1) | -0,07081 | 0,6779 |
| TcCLB.506211.70 | RNA-binding protein, putative | -0,07083 | 0,5248 |
| TcCLB.506537.50 | retrotransposon hot spot (RHS) protein, putative | -0,07083 | 0,8094 |
| TcCLB.460197.10 | cysteine peptidase (pseudogene), putative | -0,07084 | 0,7804 |
| TcCLB.509669.80 | hypothetical protein, conserved | -0,07099 | 0,7208 |
| TcCLB.508525.40 | hypothetical protein, conserved | -0,071 | 0,5543 |
| TcCLB.510329.30 | hypothetical protein, conserved | -0,07108 | 0,5235 |
| TcCLB.507389.70 | peptidase, putative | -0,0711 | 0,3905 |
| TcCLB.506489.50 | serine/threonine protein kinase, putative | -0,07111 | 0,6384 |
| TcCLB.509877.89 | hypothetical protein, conserved | -0,07121 | 0,686 |
| TcCLB.510741.30 | hypothetical protein, conserved | -0,07136 | 0,4899 |
| TcCLB.511365.100 | hypothetical protein, conserved | -0,07137 | 0,6636 |
| TcCLB.510029.60 | Vesicle-trafficking protein SEC22, putative (SEC22) | -0,07143 | 0,5971 |
| TcCLB.506195.80 | adenylate kinase, putative | -0,07149 | 0,6321 |
| TcCLB.510295.15 | hypothetical protein, conserved | -0,07157 | 0,6082 |
| TcCLB.511143.12 | hypothetical protein, conserved | -0,07162 | 0,5243 |
| TcCLB.511071.150 | hypothetical protein, conserved | -0,0717 | 0,6182 |
| TcCLB.507053.150 | hypothetical protein, conserved | -0,07171 | 0,5158 |
| TcCLB.508899.70 | hypothetical protein, conserved | -0,07182 | 0,595 |
| TcCLB.507971.29 | hypothetical protein, conserved | -0,07187 | 0,6844 |
| TcCLB.509835.20 | guanine nucleotide exchange factor subunit GCD2 | -0,07201 | 0,6013 |
| TcCLB.510861.150 | hypothetical protein | -0,07211 | 0,4062 |
| TcCLB.506879.100 | hypothetical protein, conserved | -0,07213 | 0,7082 |
| TcCLB.506941.194 | hypothetical protein, conserved | -0,07217 | 0,4142 |
| TcCLB.508693.70 | hypothetical protein, conserved | -0,07244 | 0,5866 |
| TcCLB.507081.139 | hypothetical protein, conserved (fragment) | -0,07267 | 0,5068 |
| TcCLB.506289.74 | chaperone DNAJ protein, putative | -0,07282 | 0,5744 |
| TcCLB.508909.270 | hypothetical protein, conserved | -0,07283 | 0,4825 |
| TcCLB.504157.80 | hypothetical protein, conserved | -0,0729 | 0,4712 |
| TcCLB.507093.190 | hypothetical protein, conserved | -0,07295 | 0,6451 |
| TcCLB.509979.130 | hypothetical protein, conserved | -0,07308 | 0,4078 |
| TcCLB.506989.270 | hypothetical protein, conserved | -0,07312 | 0,8769 |
| TcCLB.510027.20 | transporter, putative | -0,07335 | 0,5964 |
| TcCLB.503877.10 | hypothetical protein, conserved | -0,07385 | 0,642 |
| TcCLB.510173.60 | intraflagellar transport protein 122B, putative (IFT122B) | -0,07387 | 0,4349 |
| TcCLB.506591.130 | hypothetical protein, conserved | -0,07389 | 0,5636 |
| TcCLB.504625.80 | hypothetical protein, conserved | -0,0739 | 0,5544 |
| TcCLB.506363.150 | soluble N-ethylmaleimide sensitive factor (NSF) attachme | -0,07393 | 0,6202 |
| TcCLB.509777.130 | inositol phosphosphingolipid phospholipase C-Like, putativ | -0,07396 | 0,5014 |
| TcCLB.508119.90 | surface protease GP63 (pseudogene), putative | -0,07398 | 0,4969 |
| TcCLB.508299.89 | hypothetical protein, conserved | -0,07408 | 0,6685 |
| TcCLB.503721.30 | hypothetical protein, conserved | -0,0741 | 0,459 |
| TcCLB.506755.250 | mitochondrial import inner membrane translocase subuni | -0,0741 | 0,5034 |
| TcCLB.506529.500 | hypothetical protein, conserved | -0,07415 | 0,5588 |
| TcCLB.504131.170 | ubiquitin carboxyl-terminal hydrolase, putative | -0,07416 | 0,4999 |

| | | | |
|---|---|---|---|
| TcCLB.509179.80 | hypothetical protein, conserved | -0,0742 | 0,6994 |
| TcCLB.468803.10 | surface protease GP63, putative | -0,07423 | 0,8042 |
| TcCLB.511407.10 | nuclear transport factor 2, putative | -0,07429 | 0,5135 |
| TcCLB.510691.80 | hypothetical protein, conserved | -0,07434 | 0,6466 |
| TcCLB.509579.32 | N-acetyltransferase complex ARD1 subunit (pseudogene), | -0,07446 | 0,8117 |
| TcCLB.509033.44 | hypothetical protein, conserved | -0,07446 | 0,8231 |
| TcCLB.507011.210 | hypothetical protein, conserved | -0,07468 | 0,5145 |
| TcCLB.511911.81 | hypothetical protein, conserved (pseudogene) | -0,0747 | 0,7233 |
| TcCLB.511511.30 | hypothetical protein, conserved | -0,07474 | 0,5021 |
| TcCLB.503839.19 | hypothetical protein, conserved | -0,07476 | 0,5592 |
| TcCLB.509105.60 | adenosine deaminase-like protein, putative (ADAT1h) | -0,0748 | 0,5756 |
| TcCLB.503809.40 | protein kinase, putative | -0,0748 | 0,6596 |
| TcCLB.509601.80 | hypothetical protein, conserved | -0,07485 | 0,5339 |
| TcCLB.511071.130 | Trypanosome basal body component protein (TBBC) | -0,07487 | 0,416 |
| TcCLB.507021.10 | hypothetical protein, conserved | -0,07499 | 0,5703 |
| TcCLB.510285.130 | hypothetical protein, conserved | -0,07507 | 0,6146 |
| TcCLB.507019.90 | hypothetical protein, conserved | -0,07514 | 0,6836 |
| TcCLB.511921.10 | hypothetical protein, conserved (pseudogene) | -0,07532 | 0,7432 |
| TcCLB.508219.30 | surface protease GP63 (pseudogene), putative | -0,07533 | 0,875 |
| TcCLB.504185.40 | hypothetical protein, conserved | -0,07552 | 0,5374 |
| TcCLB.511349.80 | retrotransposon hot spot protein (RHS, pseudogene), puta | -0,07558 | 0,8707 |
| TcCLB.511753.110 | hypothetical protein, conserved | -0,07563 | 0,3744 |
| TcCLB.506625.50 | hypothetical protein | -0,07564 | 0,8083 |
| TcCLB.509231.20 | protein kinase, putative | -0,07573 | 0,5326 |
| TcCLB.506841.12 | hypothetical protein, conserved | -0,07577 | 0,5999 |
| TcCLB.508461.142 | hypothetical protein | -0,07582 | 0,52 |
| TcCLB.511189.60 | hypothetical protein, conserved | -0,07584 | 0,528 |
| TcCLB.511269.4 | Ras-related protein Rab5A, putative (RAB5A) | -0,07587 | 0,5018 |
| TcCLB.506247.40 | hypothetical protein, conserved | -0,07601 | 0,555 |
| TcCLB.503893.60 | hypothetical protein, conserved | -0,07613 | 0,5283 |
| TcCLB.509911.4 | hypothetical protein, conserved | -0,07614 | 0,6636 |
| TcCLB.508277.390 | hypothetical protein, conserved | -0,07615 | 0,4086 |
| TcCLB.509825.30 | hypothetical protein, conserved | -0,07617 | 0,5476 |
| TcCLB.509941.100 | 2,4-dienoyl-coa reductase fadh1, putative | -0,07627 | 0,4441 |
| TcCLB.504069.100 | RNA polymerase II, putative | -0,07627 | 0,5607 |
| TcCLB.510337.50 | hypothetical protein, conserved (pseudogene) | -0,07633 | 0,6701 |
| TcCLB.507735.30 | hypothetical protein, conserved | -0,07634 | 0,591 |
| TcCLB.507063.140 | hypothetical protein, conserved | -0,07635 | 0,6719 |
| TcCLB.510421.70 | hypothetical protein, conserved | -0,0764 | 0,5703 |
| TcCLB.508909.210 | hypothetical protein, conserved | -0,07652 | 0,4339 |
| TcCLB.511019.47 | N-acetyltransferase complex ARD1 subunit (pseudogene), | -0,07673 | 0,7457 |
| TcCLB.507631.10 | hypothetical protein, conserved | -0,07676 | 0,738 |
| TcCLB.504149.140 | hypothetical protein, conserved | -0,07693 | 0,4295 |
| TcCLB.510661.230 | hypothetical protein, conserved | -0,07696 | 0,4283 |
| TcCLB.507841.30 | hypothetical protein, conserved | -0,07703 | 0,7037 |

| | | | |
|---|---|---|---|
| TcCLB.508479.83 | N-acetyltransferase complex ARD1 subunit (pseudogene), | -0,07706 | 0,8039 |
| TcCLB.507941.10 | hypothetical protein, conserved | -0,07716 | 0,4292 |
| TcCLB.506733.90 | hypothetical protein, conserved | -0,07723 | 0,4684 |
| TcCLB.508153.810 | hypothetical protein, conserved | -0,07737 | 0,6188 |
| TcCLB.507709.140 | hypothetical protein, conserved | -0,0774 | 0,5702 |
| TcCLB.509695.220 | serine carboxypeptidase (CBP1), putative | -0,07743 | 0,5842 |
| TcCLB.510635.30 | dispersed gene family protein 1 (DGF-1, pseudogene), put | -0,07746 | 0,8085 |
| TcCLB.510323.50 | hypothetical protein, conserved | -0,07749 | 0,6263 |
| TcCLB.509649.30 | 19S proteasome non-atpase subunit 8 (Rpn8) | -0,07769 | 0,4612 |
| TcCLB.510943.100 | acetyltransferase, putative | -0,07773 | 0,6204 |
| TcCLB.508779.40 | hypothetical protein, conserved | -0,07777 | 0,5763 |
| TcCLB.508823.152 | hypothetical protein (fragment) | -0,07787 | 0,5404 |
| TcCLB.511277.115 | hypothetical protein, conserved | -0,07804 | 0,378 |
| TcCLB.511809.40 | hypothetical protein, conserved | -0,07817 | 0,5201 |
| TcCLB.511499.20 | hypothetical protein, conserved | -0,0782 | 0,5531 |
| TcCLB.503895.19 | hypothetical protein | -0,07822 | 0,3985 |
| TcCLB.506727.90 | enoyl-CoA hydratase/isomerase family protein, putative | -0,07824 | 0,5238 |
| TcCLB.474887.9 | pseudouridine synthase TruD, putative (fragment) | -0,07829 | 0,5747 |
| TcCLB.507011.40 | hypothetical protein, conserved | -0,07842 | 0,5048 |
| TcCLB.507831.100 | hypothetical protein, conserved | -0,07844 | 0,5295 |
| TcCLB.511523.4 | hypothetical protein, conserved | -0,07852 | 0,534 |
| TcCLB.507623.120 | hypothetical protein, conserved | -0,07852 | 0,6087 |
| TcCLB.511285.4 | intraflagellar transport protein 140, putative (fragment) ( | -0,07859 | 0,52 |
| TcCLB.507517.90 | nucleotide-binding protein, putative | -0,07893 | 0,5338 |
| TcCLB.510661.244 | hypothetical protein, conserved | -0,07899 | 0,5968 |
| TcCLB.511367.210 | ubiquitin-conjugating enzyme variant Kua homologue, put | -0,07906 | 0,6559 |
| TcCLB.510121.9 | hypothetical protein, conserved | -0,07907 | 0,6872 |
| TcCLB.507023.260 | hypothetical protein, conserved | -0,0791 | 0,5082 |
| TcCLB.508881.60 | hypothetical protein, conserved | -0,07915 | 0,5478 |
| TcCLB.508205.4 | hypothetical protein, conserved | -0,07935 | 0,6644 |
| TcCLB.454391.10 | trans-sialidase (pseudogene), putative | -0,07942 | 0,5035 |
| TcCLB.508641.140 | hypothetical protein, conserved | -0,07963 | 0,6156 |
| TcCLB.508999.190 | calpain-like cysteine peptidase, putative | -0,07964 | 0,351 |
| TcCLB.506671.61 | hypothetical protein, conserved (pseudogene) | -0,07976 | 0,7209 |
| TcCLB.507029.41 | hypothetical protein | -0,0798 | 0,7509 |
| TcCLB.508209.30 | hypothetical protein, conserved | -0,0799 | 0,381 |
| TcCLB.507673.30 | hypothetical protein, conserved | -0,07992 | 0,5383 |
| TcCLB.505183.120 | aldo/keto reductase, putative | -0,08001 | 0,5683 |
| TcCLB.510687.94 | Unc104-like kinesin, putative | -0,08013 | 0,5419 |
| TcCLB.504153.70 | hypothetical protein, conserved | -0,08014 | 0,4485 |
| TcCLB.506929.60 | ATP-dependent DEAD/H RNA helicase (pseudogene), puta | -0,0802 | 0,6114 |
| TcCLB.509109.110 | hypothetical protein, conserved | -0,08021 | 0,6538 |
| TcCLB.510531.100 | hypothetical protein, conserved | -0,08024 | 0,3818 |
| TcCLB.508307.125 | hypothetical protein, conserved | -0,08024 | 0,5452 |
| TcCLB.506627.100 | hypothetical protein, conserved | -0,08039 | 0,5916 |

| | | | |
|---|---|---|---|
| TcCLB.511585.180 | hypothetical protein, conserved | -0,08049 | 0,6596 |
| TcCLB.506753.170 | ubiquinone biosynthesis protein-like protein, putative | -0,08053 | 0,4937 |
| TcCLB.506679.60 | hypothetical protein, conserved | -0,08063 | 0,5069 |
| TcCLB.509251.98 | hypothetical protein, conserved (pseudogene) | -0,08078 | 0,806 |
| TcCLB.433717.10 | dispersed gene family protein 1 (DGF-1, pseudogene), put | -0,08088 | 0,7794 |
| TcCLB.510731.110 | hypothetical protein, conserved | -0,08095 | 0,5908 |
| TcCLB.507825.20 | RNA helicase, putative | -0,08117 | 0,3146 |
| TcCLB.508597.9 | retrotransposon hot spot (RHS) protein, putative | -0,08128 | 0,6255 |
| TcCLB.508411.20 | hypothetical protein, conserved | -0,08133 | 0,5595 |
| TcCLB.508465.149 | hypothetical protein, conserved (fragment) | -0,08135 | 0,6032 |
| TcCLB.509805.240 | hypothetical protein, conserved (pseudogene) | -0,08144 | 0,5669 |
| TcCLB.508461.100 | Bardet-Biedl syndrome 1 protein homolog (BBS1-like prot | -0,08146 | 0,5231 |
| TcCLB.511255.320 | surface protease GP63, putative | -0,08146 | 0,9033 |
| TcCLB.509979.365 | surface protease GP63 (pseudogene), putative | -0,08147 | 0,8226 |
| TcCLB.511445.70 | hypothetical protein, conserved | -0,08152 | 0,4511 |
| TcCLB.508153.380 | RNA polymerase-associated protein CTR9, putative | -0,08159 | 0,3817 |
| TcCLB.510515.120 | hypothetical protein, conserved | -0,08173 | 0,7522 |
| TcCLB.510527.9 | kinesin, putative | -0,08178 | 0,5777 |
| TcCLB.509017.24 | hypothetical protein, conserved | -0,08184 | 0,4991 |
| TcCLB.506681.24 | hypothetical protein, conserved | -0,08193 | 0,4781 |
| TcCLB.509733.70 | hypothetical protein, conserved | -0,08198 | 0,4703 |
| TcCLB.509669.120 | hypothetical protein, conserved (pseudogene) | -0,08205 | 0,5029 |
| TcCLB.508479.86 | ATP-dependent DEAD/H RNA helicase (pseudogene), puta | -0,08212 | 0,746 |
| TcCLB.508409.180 | hypothetical protein, conserved | -0,08213 | 0,3863 |
| TcCLB.503919.50 | hypothetical protein, conserved | -0,08222 | 0,4556 |
| TcCLB.511321.30 | hypothetical protein, conserved | -0,08289 | 0,6827 |
| TcCLB.510943.60 | U2 splicing auxiliary factor, putative (u2af26) | -0,08296 | 0,3681 |
| TcCLB.506733.70 | glycosyl transferase-like protein, putative | -0,08322 | 0,5734 |
| TcCLB.511047.50 | proteasome regulatory ATPase subunit 2, putative | -0,08331 | 0,4066 |
| TcCLB.508727.40 | hypothetical protein, conserved | -0,08338 | 0,543 |
| TcCLB.508647.220 | mitochondrial RNA binding protein 1 | -0,08341 | 0,306 |
| TcCLB.507737.10 | hypothetical protein, conserved | -0,08343 | 0,7288 |
| TcCLB.508461.210 | hypothetical protein, conserved | -0,08355 | 0,4449 |
| TcCLB.511543.40 | hypothetical protein, conserved | -0,08358 | 0,4619 |
| TcCLB.510303.290 | acyl-CoA dehydrogenase, putative | -0,08365 | 0,5397 |
| TcCLB.441061.9 | dynein heavy chain, putative (fragment) | -0,0839 | 0,5572 |
| TcCLB.510421.10 | surface protease GP63 (pseudogene), putative | -0,08391 | 0,7799 |
| TcCLB.506367.180 | vesicular transport protein (CDC48 homologue), putative | -0,08397 | 0,4302 |
| TcCLB.508323.84 | peptidyl-prolyl cis-trans isomerase, putative | -0,084 | 0,6588 |
| TcCLB.509453.40 | RNA polymerase subunit RPB10, putative (RPB10) | -0,08412 | 0,6868 |
| TcCLB.421959.10 | eukaryotic translation initiation factor 4E, putative | -0,08424 | 0,4556 |
| TcCLB.511855.30 | hypothetical protein, conserved | -0,08439 | 0,7042 |
| TcCLB.503555.20 | hypothetical protein, conserved | -0,08443 | 0,5371 |
| TcCLB.503973.181 | surface protease GP63 (pseudogene), putative | -0,08458 | 0,8512 |
| TcCLB.510237.21 | hypothetical protein, conserved (pseudogene) | -0,08467 | 0,7549 |

| | | | |
|---|---|---|---|
| TcCLB.510363.196 | syntaxin binding protein (pseudogene), putative | -0,08481 | 0,8685 |
| TcCLB.510181.60 | chromatin assembly factor 1 subunit B, putative | -0,08499 | 0,4131 |
| TcCLB.508839.80 | hypothetical protein, conserved | -0,08513 | 0,4306 |
| TcCLB.506201.130 | hypothetical protein, conserved | -0,08515 | 0,4666 |
| TcCLB.506425.140 | hypothetical protein, conserved | -0,08525 | 0,4761 |
| TcCLB.511771.185 | hypothetical protein | -0,08531 | 0,7257 |
| TcCLB.510037.20 | surface protease GP63 (pseudogene), putative | -0,08534 | 0,7798 |
| TcCLB.509419.100 | mucin-associated surface protein (MASP, pseudogene), pu | -0,08537 | 0,8764 |
| TcCLB.504099.70 | retrotransposon hot spot protein (RHS, pseudogene), puta | -0,08538 | 0,7641 |
| TcCLB.506265.30 | mitochondrial DNA polymerase I protein A, putative | -0,08539 | 0,4234 |
| TcCLB.511421.120 | E3 ubiquitin-protein ligase KCMF1, putative | -0,08544 | 0,3902 |
| TcCLB.508549.30 | target of rapamycin (TOR) kinase 1, putative | -0,08554 | 0,5011 |
| TcCLB.511469.24 | ADP-ribosylation factor-like 2, arl2, putative (ARL2) | -0,08557 | 0,7208 |
| TcCLB.469785.10 | hypothetical protein, conserved | -0,08568 | 0,4394 |
| TcCLB.510487.44 | elongation factor 1-gamma (EF-1-gamma, pseudogene), | -0,08569 | 0,8601 |
| TcCLB.506945.280 | hypothetical protein, conserved | -0,0858 | 0,2921 |
| TcCLB.510143.129 | hypothetical protein, conserved | -0,08585 | 0,6182 |
| TcCLB.511129.54 | hypothetical protein, conserved (pseudogene) | -0,08586 | 0,6506 |
| TcCLB.508707.190 | hypothetical protein, conserved | -0,08587 | 0,3632 |
| TcCLB.509647.26 | hypothetical protein, conserved | -0,08587 | 0,5258 |
| TcCLB.510445.61 | hypothetical protein, conserved (fragment) | -0,08592 | 0,6233 |
| TcCLB.509563.40 | hypothetical protein, conserved | -0,08594 | 0,5331 |
| TcCLB.510075.40 | hypothetical protein, conserved | -0,08594 | 0,5351 |
| TcCLB.503855.10 | hypothetical protein, conserved | -0,08597 | 0,6027 |
| TcCLB.508475.20 | hypothetical protein | -0,08608 | 0,4131 |
| TcCLB.503955.89 | hypothetical protein, conserved | -0,08611 | 0,491 |
| TcCLB.509643.100 | hypothetical protein, conserved | -0,08611 | 0,5895 |
| TcCLB.503557.20 | ankyrin repeat protein, putative | -0,08635 | 0,6563 |
| TcCLB.510367.10 | dispersed gene family protein 1 (DGF-1), putative | -0,08648 | 0,697 |
| TcCLB.503841.30 | rRNA dimethyltransferase, putative | -0,0865 | 0,6112 |
| TcCLB.504827.70 | hypothetical protein, conserved | -0,08662 | 0,5456 |
| TcCLB.506963.110 | NUDIX hydrolase 3, putative (Nudix3) | -0,08663 | 0,6297 |
| TcCLB.507883.70 | PrimPol-like protein 2, putative (ppl2) | -0,0867 | 0,3938 |
| TcCLB.510955.40 | axoneme central apparatus protein (PF16) | -0,0867 | 0,4997 |
| TcCLB.510901.80 | hypothetical protein, conserved | -0,08673 | 0,3999 |
| TcCLB.510421.314 | U6 snRNA-associated Sm-like protein LSm8p | -0,08685 | 0,6302 |
| TcCLB.506513.30 | hypothetical protein, conserved | -0,08688 | 0,6246 |
| TcCLB.506887.60 | small nuclear RNA gene activation protein (SNAP) 50, put | -0,0869 | 0,3663 |
| TcCLB.509767.130 | hypothetical protein, conserved | -0,0871 | 0,6855 |
| TcCLB.504029.101 | hypothetical protein, conserved | -0,08726 | 0,5315 |
| TcCLB.503925.110 | hypothetical protein, conserved | -0,08729 | 0,5943 |
| TcCLB.510359.290 | hypothetical protein, conserved | -0,08729 | 0,711 |
| TcCLB.508881.110 | GTP-binding protein, putative | -0,08732 | 0,4688 |
| TcCLB.506357.60 | hypothetical protein, conserved | -0,08741 | 0,3082 |
| TcCLB.507747.220 | protein farnesyltransferase, putative | -0,08748 | 0,4371 |

| | | | |
|---|---|---|---|
| TcCLB.511727.300 | GDP-L-fucose synthetase, putative | -0,08761 | 0,3983 |
| TcCLB.503827.50 | trans-sialidase (pseudogene), putative | -0,08768 | 0,819 |
| TcCLB.509103.39 | dynein heavy chain, putative | -0,08775 | 0,4588 |
| TcCLB.511727.120 | ethanolamine-phosphate cytidylyltransferase (ECT) | -0,08802 | 0,2954 |
| TcCLB.507611.398 | RNA editing complex protein MP63, putative (fragment) | -0,08803 | 0,3702 |
| TcCLB.509643.60 | hypothetical protein, conserved | -0,08818 | 0,5238 |
| TcCLB.507167.190 | trans-sialidase (pseudogene), putative | -0,08824 | 0,6752 |
| TcCLB.509695.110 | hypothetical protein, conserved | -0,08842 | 0,4548 |
| TcCLB.511367.80 | hypothetical protein, conserved | -0,08846 | 0,5232 |
| TcCLB.509937.150 | serine hydroxymethyltransferase, putative | -0,08852 | 0,5955 |
| TcCLB.509633.40 | hypothetical protein, conserved | -0,08855 | 0,5634 |
| TcCLB.506933.70 | hypothetical protein, conserved | -0,0887 | 0,4172 |
| TcCLB.508803.5 | retrotransposon hot spot protein (RHS, pseudogene), puta | -0,08871 | 0,85 |
| TcCLB.506743.140 | hypothetical protein, conserved | -0,08879 | 0,4693 |
| TcCLB.510889.10 | hypothetical protein, conserved | -0,08889 | 0,5908 |
| TcCLB.509683.24 | DNA-directed RNA polymerase I subunit, putative | -0,08901 | 0,5636 |
| TcCLB.508409.329 | hypothetical protein, conserved | -0,0892 | 0,5151 |
| TcCLB.511131.98 | hypothetical protein, conserved | -0,08928 | 0,5807 |
| TcCLB.511857.20 | hypothetical protein, conserved | -0,08949 | 0,4576 |
| TcCLB.510971.69 | hypothetical protein, conserved (fragment) | -0,08958 | 0,3998 |
| TcCLB.507663.50 | ATP-dependent RNA helicase, putative | -0,0897 | 0,2856 |
| TcCLB.510439.20 | hypothetical protein, conserved | -0,08975 | 0,6619 |
| TcCLB.507641.160 | hypothetical protein, conserved | -0,08984 | 0,256 |
| TcCLB.510771.9 | hypothetical protein, conserved (fragment) | -0,08988 | 0,3519 |
| TcCLB.505051.30 | tryptophanyl-tRNA synthetase, putative (fragment) | -0,08992 | 0,8429 |
| TcCLB.508251.21 | hypothetical protein, conserved (pseudogene) | -0,09001 | 0,7285 |
| TcCLB.509365.50 | hypothetical protein, conserved | -0,09002 | 0,8004 |
| TcCLB.507991.126 | hypothetical protein, conserved | -0,0901 | 0,6588 |
| TcCLB.506257.50 | surface protease GP63, putative | -0,09014 | 0,6993 |
| TcCLB.506919.60 | hypothetical protein | -0,09014 | 0,7937 |
| TcCLB.506509.50 | hypothetical protein, conserved | -0,09017 | 0,3879 |
| TcCLB.506237.10 | mismatch repair protein MSH5, putative | -0,09018 | 0,478 |
| TcCLB.511871.34 | protein transport protein Sec61 gamma subunit, putative | -0,09025 | 0,5397 |
| TcCLB.506755.30 | hypothetical protein, conserved | -0,09026 | 0,4357 |
| TcCLB.510259.18 | GlcNAc-PI synthesis protein (GPI3) | -0,09055 | 0,5575 |
| TcCLB.511807.250 | hypothetical protein, conserved | -0,09081 | 0,4369 |
| TcCLB.510491.30 | hypothetical protein, conserved (pseudogene) | -0,09084 | 0,5306 |
| TcCLB.511003.210 | hypothetical protein, conserved | -0,09085 | 0,5259 |
| TcCLB.510887.64 | hypothetical protein, conserved | -0,09094 | 0,6112 |
| TcCLB.438147.9 | hypothetical protein, conserved (fragment) | -0,09098 | 0,5358 |
| TcCLB.506715.10 | protein kinase, putative | -0,091 | 0,3699 |
| TcCLB.503891.90 | hypothetical protein | -0,09101 | 0,5593 |
| TcCLB.508153.950 | 6-phosphofructo-2-kinase 1 | -0,09129 | 0,271 |
| TcCLB.511465.20 | hypothetical protein, conserved | -0,0913 | 0,37 |
| TcCLB.503969.10 | tubulin delta chain (TUBD1) | -0,09135 | 0,5402 |

| | | | |
|---|---|---|---|
| TcCLB.508461.174 | hypothetical protein, conserved | -0,09154 | 0,6774 |
| TcCLB.506303.100 | hypothetical protein, conserved | -0,09155 | 0,5396 |
| TcCLB.506435.170 | hypothetical protein, conserved | -0,09162 | 0,6242 |
| TcCLB.503803.10 | rRNA biogenesis protein, putative | -0,09164 | 0,4291 |
| TcCLB.508355.100 | diacylglycerol acyltransferase, putative | -0,09166 | 0,3151 |
| TcCLB.506351.40 | ribonuclease H1, putative | -0,09168 | 0,5512 |
| TcCLB.510743.50 | phospholipase A2-like protein, putative | -0,09171 | 0,4948 |
| TcCLB.508479.465 | ATP-dependent DEAD/H RNA helicase (pseudogene), puta | -0,09173 | 0,7489 |
| TcCLB.407477.44 | hypothetical protein, conserved | -0,09174 | 0,6714 |
| TcCLB.509229.10 | hypothetical protein, conserved | -0,0918 | 0,433 |
| TcCLB.507143.30 | hypothetical protein | -0,09182 | 0,2567 |
| TcCLB.509753.140 | mucin-associated surface protein (MASP), putative | -0,09188 | 0,7697 |
| TcCLB.504183.10 | retrotransposon hot spot protein (RHS, pseudogene), puta | -0,09191 | 0,6387 |
| TcCLB.509203.70 | DNA topoisomerase II, putative | -0,09197 | 0,3464 |
| TcCLB.511311.60 | hypothetical protein, conserved | -0,09198 | 0,5688 |
| TcCLB.506195.150 | hypothetical protein, conserved | -0,09203 | 0,3324 |
| TcCLB.427247.4 | hypothetical protein, conserved (fragment) | -0,09216 | 0,5072 |
| TcCLB.506635.30 | hypothetical protein, conserved | -0,09223 | 0,453 |
| TcCLB.507997.40 | hypothetical protein, conserved | -0,09226 | 0,4685 |
| TcCLB.506437.30 | amastin, putative | -0,09233 | 0,4535 |
| TcCLB.507541.10 | hypothetical protein, conserved | -0,09235 | 0,5999 |
| TcCLB.510901.50 | C-terminal motor kinesin, putative | -0,09236 | 0,3568 |
| TcCLB.508909.140 | hypothetical protein, conserved | -0,09238 | 0,4899 |
| TcCLB.506657.30 | hypothetical protein, conserved | -0,09239 | 0,3202 |
| TcCLB.509429.150 | hypothetical protein, conserved | -0,09243 | 0,4657 |
| TcCLB.503477.30 | hypothetical protein, conserved | -0,09274 | 0,5787 |
| TcCLB.509605.10 | malonyl-CoA decarboxylase, mitochondrial precursor, puta | -0,09284 | 0,5488 |
| TcCLB.511911.159 | histone deacetylase 1, putative | -0,09287 | 0,587 |
| TcCLB.508641.300 | hypothetical protein | -0,09288 | 0,4145 |
| TcCLB.511733.90 | hypothetical protein, conserved | -0,09301 | 0,5326 |
| TcCLB.416891.9 | protein kinase, putative | -0,09303 | 0,5476 |
| TcCLB.508465.120 | syntaxin, putative | -0,09307 | 0,4378 |
| TcCLB.511165.20 | hypothetical protein | -0,09313 | 0,3991 |
| TcCLB.506945.30 | CAS/CSE/importin domain protein, putative | -0,09318 | 0,3262 |
| TcCLB.506411.39 | CDP-DAG synthase, putative (fragment) (CDS) | -0,09339 | 0,7167 |
| TcCLB.511609.10 | retrotransposon hot spot protein (RHS, pseudogene), puta | -0,0934 | 0,7348 |
| TcCLB.506945.100 | hypothetical protein, conserved | -0,09352 | 0,346 |
| TcCLB.504071.80 | nuclear cap binding protein, putative | -0,09358 | 0,604 |
| TcCLB.508917.40 | hypothetical protein, conserved | -0,09368 | 0,406 |
| TcCLB.441967.9 | hypothetical protein, conserved | -0,09368 | 0,504 |
| TcCLB.509695.40 | hypothetical protein, conserved | -0,0938 | 0,41 |
| TcCLB.510329.360 | hypothetical protein, conserved | -0,09399 | 0,5523 |
| TcCLB.509151.40 | inorganic polyphosphate/ATP-NAD kinase, putative | -0,09402 | 0,2368 |
| TcCLB.504157.50 | DnaJ chaperone protein, putative | -0,09404 | 0,4491 |
| TcCLB.511627.120 | aldo/keto reductase, putative | -0,09416 | 0,4712 |

| | | | |
|---|---|---|---|
| TcCLB.509163.110 | retrotransposon hot spot (RHS) protein, putative | -0,09435 | 0,6397 |
| TcCLB.504033.100 | hypothetical protein, conserved | -0,09437 | 0,3775 |
| TcCLB.506247.450 | hypothetical protein, conserved | -0,09437 | 0,3943 |
| TcCLB.508183.20 | hypothetical protein, conserved | -0,09447 | 0,4513 |
| TcCLB.506337.140 | hypothetical protein, conserved | -0,09452 | 0,4248 |
| TcCLB.403875.19 | hypothetical protein, conserved (fragment) | -0,09459 | 0,7217 |
| TcCLB.509571.50 | hypothetical protein, conserved | -0,09461 | 0,39 |
| TcCLB.508461.149 | hypothetical protein, conserved | -0,0947 | 0,337 |
| TcCLB.511737.30 | hypothetical protein, conserved | -0,0947 | 0,4194 |
| TcCLB.507011.120 | acyl transferase-like protein, putative | -0,09515 | 0,4199 |
| TcCLB.507641.60 | aldehyde dehydrogenase, putative | -0,09518 | 0,3274 |
| TcCLB.506355.60 | small G-protein, putative | -0,0952 | 0,3039 |
| TcCLB.511287.4 | hypothetical protein | -0,09529 | 0,3848 |
| TcCLB.508355.130 | hypothetical protein, conserved | -0,09533 | 0,3582 |
| TcCLB.508347.30 | hypothetical protein, conserved | -0,09551 | 0,3456 |
| TcCLB.506807.10 | tubulin delta chain (TUBD1) | -0,09557 | 0,5274 |
| TcCLB.508479.190 | peroxisome assembly protein, putative | -0,09558 | 0,3871 |
| TcCLB.503579.100 | nucleotide-binding protein, putative | -0,09566 | 0,4644 |
| TcCLB.507093.260 | multidrug resistance protein, mitochondrial, putative (ABC | -0,09574 | 0,1971 |
| TcCLB.508815.120 | hypothetical protein, conserved | -0,09574 | 0,4492 |
| TcCLB.509049.10 | hypothetical protein, conserved | -0,09579 | 0,5467 |
| TcCLB.507467.94 | hypothetical protein, conserved | -0,0958 | 0,6216 |
| TcCLB.509791.10 | hypothetical protein, conserved | -0,09581 | 0,6634 |
| TcCLB.508047.30 | trans-sialidase (pseudogene), putative | -0,09591 | 0,7485 |
| TcCLB.506273.10 | trans-sialidase (pseudogene), putative | -0,09595 | 0,572 |
| TcCLB.510339.70 | hypothetical protein, conserved | -0,09604 | 0,4182 |
| TcCLB.509385.11 | retrotransposon hot spot protein (RHS, pseudogene), puta | -0,09617 | 0,6998 |
| TcCLB.511855.80 | hypothetical protein, conserved | -0,09633 | 0,3912 |
| TcCLB.508827.60 | hypothetical protein, conserved | -0,09636 | 0,3889 |
| TcCLB.511521.30 | superoxide dismutase, putative | -0,09646 | 0,3931 |
| TcCLB.474339.9 | hypothetical protein, conserved | -0,09648 | 0,4378 |
| TcCLB.507023.80 | hypothetical protein, conserved | -0,09657 | 0,5534 |
| TcCLB.503939.30 | ttaggg binding factor, putative | -0,09664 | 0,4115 |
| TcCLB.507819.40 | retrotransposon hot spot protein (RHS, pseudogene), puta | -0,09697 | 0,5944 |
| TcCLB.506789.150 | hypothetical protein | -0,09699 | 0,4341 |
| TcCLB.483623.10 | hypothetical protein, conserved | -0,09704 | 0,5782 |
| TcCLB.506263.30 | D-isomer specific 2-hydroxyacid dehydrogenase-protein, p | -0,09707 | 0,6563 |
| TcCLB.509393.50 | hypothetical protein, conserved | -0,0971 | 0,3562 |
| TcCLB.511751.130 | small GTP-binding protein Rab28, putative | -0,09754 | 0,4672 |
| TcCLB.503879.100 | hypothetical protein, conserved | -0,0976 | 0,1962 |
| TcCLB.509937.170 | hypothetical protein | -0,09767 | 0,4817 |
| TcCLB.507711.230 | hypothetical protein, conserved | -0,09769 | 0,3527 |
| TcCLB.503913.49 | hypothetical protein, conserved (fragment) | -0,09775 | 0,7653 |
| TcCLB.506793.60 | hypothetical protein, conserved | -0,09779 | 0,4488 |
| TcCLB.511425.10 | serine/threonine protein phosphatase 2A regulatory subu | -0,09789 | 0,5758 |

| | | | |
|---|---|---|---|
| TcCLB.471901.20 | SNARE protein, putative | -0,0981 | 0,4331 |
| TcCLB.507209.20 | hypothetical protein, conserved | -0,09813 | 0,4905 |
| TcCLB.506287.140 | prefoldin, putative | -0,09829 | 0,4555 |
| TcCLB.506181.150 | hypothetical protein (fragment) | -0,0983 | 0,2857 |
| TcCLB.506989.210 | U-box domain protein, putative | -0,09835 | 0,2526 |
| TcCLB.511231.60 | hypothetical protein, conserved | -0,09859 | 0,4692 |
| TcCLB.509065.124 | U6 snRNA-associated Sm-like protein LSm5p | -0,09859 | 0,5975 |
| TcCLB.503939.130 | hypothetical protein, conserved | -0,09863 | 0,3389 |
| TcCLB.509053.150 | hypothetical protein, conserved | -0,09863 | 0,5561 |
| TcCLB.508613.10 | hypothetical protein, conserved | -0,09867 | 0,4637 |
| TcCLB.509985.78 | hypothetical protein, conserved | -0,09875 | 0,5287 |
| TcCLB.506221.30 | protein phosphatase 2C, putative | -0,09878 | 0,3677 |
| TcCLB.503581.10 | hypothetical protein, conserved | -0,09883 | 0,3132 |
| TcCLB.506355.170 | hypothetical protein, conserved | -0,09907 | 0,5281 |
| TcCLB.504019.9 | hypothetical protein, conserved | -0,0991 | 0,5614 |
| TcCLB.510877.50 | AP-3 complex subunit mu, putative (AP3M) | -0,09911 | 0,3343 |
| TcCLB.508707.330 | chaperone DNAJ protein, putative | -0,09911 | 0,4288 |
| TcCLB.506567.10 | hypothetical protein, conserved | -0,09911 | 0,4581 |
| TcCLB.442629.9 | hypothetical protein, conserved | -0,09911 | 0,5138 |
| TcCLB.508917.4 | hypothetical protein, conserved | -0,09912 | 0,4399 |
| TcCLB.503881.4 | hypothetical protein, conserved | -0,09913 | 0,7461 |
| TcCLB.503635.10 | protein kinase, putative | -0,09917 | 0,4061 |
| TcCLB.509045.10 | hypothetical protein, conserved | -0,09917 | 0,5336 |
| TcCLB.503587.64 | retrotransposon hot spot protein (RHS, pseudogene), puta | -0,09921 | 0,7554 |
| TcCLB.506915.30 | hypothetical protein | -0,0993 | 0,3082 |
| TcCLB.507083.60 | COP9 signalosome complex subunit 5, putative | -0,09942 | 0,5819 |
| TcCLB.508129.9 | adenosine monophosphate deaminase, putative (fragmer | -0,09947 | 0,4179 |
| TcCLB.506615.120 | trans-sialidase (pseudogene), putative | -0,09952 | 0,5759 |
| TcCLB.506577.130 | protein phosphatase 2A regulatory subunit, putative | -0,09956 | 0,432 |
| TcCLB.503555.40 | ATP-dependent RNA helicase, putative | -0,09967 | 0,3027 |
| TcCLB.506265.50 | hypothetical protein, conserved | -0,09967 | 0,4449 |
| TcCLB.508323.120 | hypothetical protein, conserved | -0,0997 | 0,516 |
| TcCLB.506855.150 | hypothetical protein, conserved | -0,09995 | 0,3633 |
| TcCLB.511621.150 | hypothetical protein, conserved | -0,1002 | 0,3761 |
| TcCLB.506795.20 | hypothetical protein, conserved | -0,1002 | 0,5069 |
| TcCLB.511423.149 | conserved protein, unknown function | -0,1002 | 0,5261 |
| TcCLB.503697.90 | Translation factor GUF1, mitochondrial, putative (GUF1) | -0,1003 | 0,2867 |
| TcCLB.507989.20 | sigma-adaptin 3, putative | -0,1003 | 0,5093 |
| TcCLB.506605.214 | hypothetical protein, conserved | -0,1003 | 0,5943 |
| TcCLB.510613.30 | hypothetical protein | -0,1004 | 0,3937 |
| TcCLB.506413.50 | hypothetical protein, conserved | -0,1005 | 0,4495 |
| TcCLB.503425.29 | DNA replication licensing factor MCM8, putative | -0,1005 | 0,4862 |
| TcCLB.511245.20 | hypothetical protein, conserved | -0,1005 | 0,5273 |
| TcCLB.509941.140 | hypothetical protein, conserved | -0,1007 | 0,533 |
| TcCLB.506529.340 | Protein kinase domain, putative | -0,1008 | 0,5359 |

| | | | |
|---|---|---|---|
| TcCLB.508325.110 | hypothetical protein, conserved | -0,1008 | 0,6425 |
| TcCLB.504085.60 | CMP-sialic acid transporter, putative | -0,1009 | 0,4981 |
| TcCLB.511919.30 | hypothetical protein, conserved (pseudogene) | -0,1009 | 0,6857 |
| TcCLB.509269.4 | Tb-291 membrane associated protein, putative | -0,101 | 0,4099 |
| TcCLB.510187.180 | hypothetical protein, conserved | -0,1013 | 0,1963 |
| TcCLB.511301.30 | hypothetical protein, conserved | -0,1013 | 0,5462 |
| TcCLB.510349.90 | Lys-63-specific deubiquitinase BRCC36, putative | -0,1014 | 0,4192 |
| TcCLB.509603.10 | vacuolar protein sorting 18, putative | -0,1016 | 0,5109 |
| TcCLB.510535.80 | hypothetical protein, conserved | -0,1016 | 0,5903 |
| TcCLB.506001.120 | hypothetical protein, conserved (pseudogene) | -0,1016 | 0,7445 |
| TcCLB.508577.160 | proteasome beta 6 subunit, putative | -0,1017 | 0,2788 |
| TcCLB.506295.110 | hypothetical protein, conserved | -0,1017 | 0,5234 |
| TcCLB.506567.74 | hypothetical protein, conserved | -0,1017 | 0,5649 |
| TcCLB.506727.150 | hypothetical protein, conserved | -0,1018 | 0,512 |
| TcCLB.506589.80 | surface protease GP63 (pseudogene), putative | -0,1018 | 0,7939 |
| TcCLB.506679.160 | intraflagellar transport protein 122, putative (IFT122) | -0,1019 | 0,2616 |
| TcCLB.509979.379 | hypothetical protein, conserved | -0,1019 | 0,434 |
| TcCLB.508153.750 | hypothetical protein, conserved | -0,102 | 0,4181 |
| TcCLB.507017.4 | hypothetical protein, conserved | -0,102 | 0,663 |
| TcCLB.508277.50 | presenilin-like aspartic peptidase, putative | -0,1021 | 0,3223 |
| TcCLB.508777.160 | hypothetical protein, conserved | -0,1022 | 0,339 |
| TcCLB.511367.74 | hypothetical protein, conserved | -0,1023 | 0,5048 |
| TcCLB.507801.210 | retrotransposon hot spot protein (RHS, pseudogene), puta | -0,1023 | 0,5725 |
| TcCLB.509795.10 | hypothetical protein, conserved | -0,1024 | 0,3621 |
| TcCLB.506885.120 | protein kinase, putative | -0,1026 | 0,4577 |
| TcCLB.506135.110 | mitochondrial RNA binding complex 1 subunit, putative | -0,1027 | 0,4293 |
| TcCLB.511519.18 | hypothetical protein, conserved | -0,1027 | 0,564 |
| TcCLB.507969.54 | hypothetical protein, conserved | -0,1028 | 0,4116 |
| TcCLB.510253.29 | hypothetical protein, conserved (fragment) | -0,1028 | 0,6462 |
| TcCLB.510307.191 | protein kinase (pseudogene), putative | -0,1029 | 0,3134 |
| TcCLB.506661.80 | hypothetical protein, conserved | -0,103 | 0,4093 |
| TcCLB.506953.40 | hypothetical protein, conserved | -0,1032 | 0,3936 |
| TcCLB.509059.60 | hypothetical protein, conserved | -0,1033 | 0,6722 |
| TcCLB.507647.20 | hypothetical protein, conserved (pseudogene) | -0,1034 | 0,2534 |
| TcCLB.511283.170 | hypothetical protein, conserved | -0,1036 | 0,1687 |
| TcCLB.508213.20 | hypothetical protein, conserved | -0,1036 | 0,5607 |
| TcCLB.508899.120 | hypothetical protein, conserved | -0,1037 | 0,6471 |
| TcCLB.507739.90 | hypothetical protein, conserved | -0,104 | 0,4592 |
| TcCLB.510435.5 | hypothetical protein, conserved (pseudogene) | -0,1042 | 0,5451 |
| TcCLB.508493.70 | hypothetical protein | -0,1042 | 0,7385 |
| TcCLB.505165.10 | RNA-binding protein, putative | -0,1042 | 0,7937 |
| TcCLB.511693.90 | electron-transfer-flavoprotein, alpha polypeptide, putative | -0,1043 | 0,2756 |
| TcCLB.506871.120 | hypothetical protein, conserved | -0,1043 | 0,3681 |
| TcCLB.509047.60 | hypothetical protein, conserved | -0,1044 | 0,374 |
| TcCLB.511527.60 | hypothetical protein, conserved | -0,1044 | 0,4021 |

| | | | |
|---|---|---|---|
| TcCLB.510101.59 | palmitoyl acyltransferase 7, putative | -0,1045 | 0,315 |
| TcCLB.506529.170 | hypothetical protein, conserved | -0,1045 | 0,4545 |
| TcCLB.509259.123 | hypothetical protein (pseudogene) | -0,1046 | 0,5274 |
| TcCLB.504827.80 | hypothetical protein, conserved | -0,1046 | 0,5702 |
| TcCLB.509469.10 | dispersed gene family protein 1 (DGF-1, pseudogene), put | -0,1046 | 0,701 |
| TcCLB.509791.40 | hypothetical protein, conserved | -0,1047 | 0,2534 |
| TcCLB.511419.20 | hypothetical protein, conserved | -0,1047 | 0,3302 |
| TcCLB.504173.50 | dihydrouridine synthase (Dus), putative | -0,1047 | 0,495 |
| TcCLB.503899.60 | hypothetical protein, conserved | -0,105 | 0,4485 |
| TcCLB.511345.10 | retrotransposon hot spot (RHS) protein, putative | -0,1051 | 0,5308 |
| TcCLB.506179.30 | hypothetical protein, conserved | -0,1052 | 0,4344 |
| TcCLB.509039.40 | hypothetical protein | -0,1052 | 0,4595 |
| TcCLB.511253.20 | kinesin-like protein, putative | -0,1052 | 0,5575 |
| TcCLB.509509.60 | small GTP-binding protein, putative | -0,1053 | 0,3402 |
| TcCLB.506221.50 | coatomer subunit epsilon (COPE) | -0,1053 | 0,3946 |
| TcCLB.506625.200 | ribosomal RNA processing protein 40, putative (RRP40) | -0,1053 | 0,4804 |
| TcCLB.506529.660 | hypothetical protein, conserved | -0,1054 | 0,5146 |
| TcCLB.507631.15 | hypothetical protein, conserved (pseudogene) | -0,1054 | 0,6202 |
| TcCLB.507993.80 | glycogen synthase kinase 3, putative | -0,1056 | 0,5269 |
| TcCLB.511753.80 | GTPase activating protein, putative | -0,1057 | 0,4017 |
| TcCLB.507895.150 | hypothetical protein, conserved | -0,1058 | 0,4857 |
| TcCLB.511367.138 | chaperone DNAJ protein, putative | -0,1059 | 0,358 |
| TcCLB.510761.4 | hypothetical protein, conserved | -0,1059 | 0,3738 |
| TcCLB.506673.10 | hypothetical protein, conserved | -0,106 | 0,4683 |
| TcCLB.505073.20 | hypothetical protein | -0,1061 | 0,2741 |
| TcCLB.505943.50 | hypothetical protein, conserved | -0,1061 | 0,374 |
| TcCLB.508829.9 | dynein heavy chain, putative (fragment) | -0,1062 | 0,4029 |
| TcCLB.508765.40 | hypothetical protein | -0,1062 | 0,418 |
| TcCLB.510755.138 | serine/threonine protein phosphatase, putative | -0,1064 | 0,4092 |
| TcCLB.509599.110 | hypothetical protein, conserved | -0,1065 | 0,4215 |
| TcCLB.503923.20 | hypothetical protein, conserved | -0,1065 | 0,516 |
| TcCLB.506865.30 | hypothetical protein, conserved | -0,1066 | 0,3566 |
| TcCLB.504085.20 | exosome-associated protein 2, putative (EAP2) | -0,1066 | 0,3918 |
| TcCLB.506733.100 | hypothetical protein, conserved | -0,1067 | 0,297 |
| TcCLB.503593.40 | hypothetical protein | -0,1067 | 0,3382 |
| TcCLB.504639.40 | trans-sialidase, putative | -0,1067 | 0,6781 |
| TcCLB.511631.9 | hypothetical protein, conserved (fragment) | -0,107 | 0,5756 |
| TcCLB.510187.400 | serine/threonine protein phosphatase, putative | -0,1073 | 0,2809 |
| TcCLB.509719.20 | ATP-dependent DEAD/H RNA helicase, putative | -0,1075 | 0,3022 |
| TcCLB.511585.30 | hypothetical protein, conserved | -0,1077 | 0,3058 |
| TcCLB.508177.20 | carbohydrate kinase, thermoresistant glucokinase, putativ | -0,1078 | 0,6655 |
| TcCLB.506227.220 | hypothetical protein, conserved | -0,1079 | 0,3001 |
| TcCLB.504827.40 | AP-3 complex subunit beta, putative (AP3B) | -0,1079 | 0,3179 |
| TcCLB.504011.40 | XPA-interacting protein, putative | -0,1079 | 0,3382 |
| TcCLB.506517.110 | hypothetical protein, conserved | -0,108 | 0,5866 |

| | | | |
|---|---|---|---|
| TcCLB.506265.200 | hypothetical protein, conserved | -0,1081 | 0,519 |
| TcCLB.506267.101 | surface protease GP63 (pseudogene), putative | -0,1081 | 0,7224 |
| TcCLB.508209.140 | protein disulfide isomerase, putative | -0,1082 | 0,1893 |
| TcCLB.510229.10 | hypothetical protein, conserved | -0,1082 | 0,2758 |
| TcCLB.511583.20 | hypothetical protein, conserved | -0,1082 | 0,4045 |
| TcCLB.506841.4 | hypothetical protein, conserved | -0,1082 | 0,5961 |
| TcCLB.506151.30 | small GTP-binding protein Rab11, putative (TcRab11) | -0,1084 | 0,3431 |
| TcCLB.504533.20 | retrotransposon hot spot protein (RHS, pseudogene), puta | -0,1084 | 0,4566 |
| TcCLB.508263.10 | hypothetical protein | -0,1085 | 0,3516 |
| TcCLB.506895.60 | hypothetical protein, conserved (pseudogene) | -0,1086 | 0,4134 |
| TcCLB.503805.20 | hypothetical protein, conserved | -0,1086 | 0,4267 |
| TcCLB.507711.120 | hypothetical protein, conserved | -0,1087 | 0,4868 |
| TcCLB.511277.520 | protein kinase (pseudogene), putative | -0,1087 | 0,513 |
| TcCLB.508909.40 | hypothetical protein, conserved | -0,1088 | 0,3465 |
| TcCLB.511421.24 | kelch domain-containing protein | -0,1088 | 0,3884 |
| TcCLB.419703.20 | ubiquitin carboxyl-terminal hydrolase, putative | -0,1088 | 0,4 |
| TcCLB.509429.80 | hypothetical protein, conserved | -0,1089 | 0,3988 |
| TcCLB.506225.30 | hypothetical protein, conserved (pseudogene) | -0,1089 | 0,6809 |
| TcCLB.508897.90 | hypothetical protein, conserved | -0,109 | 0,3407 |
| TcCLB.510719.90 | hypothetical protein, conserved | -0,109 | 0,4795 |
| TcCLB.508573.90 | retrotransposon hot spot protein (RHS, pseudogene), puta | -0,109 | 0,5528 |
| TcCLB.510143.10 | hypothetical protein, conserved | -0,1091 | 0,4689 |
| TcCLB.509065.100 | hypothetical protein, conserved | -0,1091 | 0,4849 |
| TcCLB.508899.80 | hypothetical protein, conserved | -0,1092 | 0,2837 |
| TcCLB.510257.24 | hypothetical protein, conserved | -0,1092 | 0,4761 |
| TcCLB.506679.170 | hypothetical protein, conserved | -0,1093 | 0,1953 |
| TcCLB.506637.70 | hypothetical protein, conserved | -0,1094 | 0,3034 |
| TcCLB.511593.40 | trans-sialidase (pseudogene), putative | -0,1094 | 0,3729 |
| TcCLB.503897.40 | hypothetical protein, conserved | -0,1095 | 0,2559 |
| TcCLB.506227.230 | N-acetyltransferase complex ARD1 subunit, putative | -0,1095 | 0,4151 |
| TcCLB.509623.10 | hypothetical protein, conserved | -0,1096 | 0,3622 |
| TcCLB.511527.82 | hypothetical protein, conserved | -0,1096 | 0,5818 |
| TcCLB.509791.170 | small GTPase, putative | -0,1097 | 0,3613 |
| TcCLB.511651.10 | ATP-dependent DEAD/H RNA helicase, putative | -0,1097 | 0,4681 |
| TcCLB.503455.20 | hypothetical protein, conserved | -0,1099 | 0,3993 |
| TcCLB.510255.29 | hypothetical protein, conserved (fragment) | -0,1099 | 0,4865 |
| TcCLB.506579.30 | myo-inositol-1 phosphatase, putative | -0,11 | 0,2284 |
| TcCLB.509767.40 | hypothetical protein, conserved | -0,11 | 0,3483 |
| TcCLB.510687.20 | hypothetical protein, conserved | -0,11 | 0,4666 |
| TcCLB.511389.20 | hypothetical protein, conserved | -0,1101 | 0,3237 |
| TcCLB.413605.10 | kinesin, putative | -0,1102 | 0,3117 |
| TcCLB.506513.70 | hypothetical protein | -0,1102 | 0,3683 |
| TcCLB.510009.19 | hypothetical protein, conserved (fragment) | -0,1103 | 0,605 |
| TcCLB.511725.100 | phosphatidylinositol-4-phosphate 5-kinase, putative | -0,1104 | 0,267 |
| TcCLB.511753.150 | hypothetical protein, conserved | -0,1104 | 0,4614 |

| | | | |
|---|---|---|---|
| TcCLB.504109.170 | hypothetical protein, conserved | -0,1105 | 0,2375 |
| TcCLB.511815.90 | hypothetical protein | -0,1105 | 0,3903 |
| TcCLB.508143.30 | hypothetical protein, conserved (pseudogene) | -0,1105 | 0,4331 |
| TcCLB.511491.150 | hypothetical protein, conserved | -0,1106 | 0,3692 |
| TcCLB.506127.130 | hypothetical protein, conserved | -0,1106 | 0,4187 |
| TcCLB.508905.20 | hypothetical protein, conserved (pseudogene) | -0,1106 | 0,6013 |
| TcCLB.506901.85 | mucin-associated surface protein (MASP, pseudogene), pu | -0,1106 | 0,8221 |
| TcCLB.506947.100 | hypothetical protein, conserved | -0,1107 | 0,393 |
| TcCLB.511299.60 | hypothetical protein, conserved | -0,1107 | 0,4673 |
| TcCLB.507053.140 | hypothetical protein, conserved | -0,1109 | 0,4422 |
| TcCLB.511245.210 | hypothetical protein, conserved | -0,111 | 0,3302 |
| TcCLB.511173.420 | mucin TcMUCII, putative | -0,111 | 0,746 |
| TcCLB.510759.80 | hypothetical protein, conserved | -0,1111 | 0,2149 |
| TcCLB.506671.50 | trans-sialidase (pseudogene), putative | -0,1111 | 0,657 |
| TcCLB.511501.10 | hypothetical protein, conserved | -0,1112 | 0,1607 |
| TcCLB.509671.130 | trypanin-like protein (pseudogene), putative | -0,1113 | 0,352 |
| TcCLB.506975.66 | hypothetical protein, conserved | -0,1116 | 0,2787 |
| TcCLB.509017.40 | hypothetical protein, conserved | -0,1116 | 0,3276 |
| TcCLB.508825.20 | hypothetical protein | -0,1116 | 0,4704 |
| TcCLB.506041.60 | hypothetical protein, conserved | -0,1117 | 0,2539 |
| TcCLB.510265.90 | hypothetical protein, conserved | -0,1117 | 0,3696 |
| TcCLB.511379.9 | hypothetical protein, conserved (fragment) | -0,1117 | 0,3981 |
| TcCLB.506797.80 | hypothetical protein, conserved | -0,1117 | 0,4263 |
| TcCLB.503939.10 | acetyltransferase, putative | -0,1118 | 0,3822 |
| TcCLB.507997.70 | hypothetical protein, conserved | -0,1119 | 0,342 |
| TcCLB.504203.50 | hypothetical protein, conserved | -0,112 | 0,4248 |
| TcCLB.508585.10 | hypothetical protein, conserved | -0,1123 | 0,3951 |
| TcCLB.505183.110 | hypothetical protein | -0,1123 | 0,6227 |
| TcCLB.506529.425 | hypothetical protein | -0,1123 | 0,7433 |
| TcCLB.507649.20 | hypothetical protein, conserved | -0,1124 | 0,1972 |
| TcCLB.506479.60 | kinesin heavy chain, putative | -0,1124 | 0,3355 |
| TcCLB.432867.10 | hypothetical protein, conserved (fragment) | -0,1124 | 0,4063 |
| TcCLB.510319.9 | hypothetical protein, conserved | -0,1124 | 0,4134 |
| TcCLB.506247.440 | hypothetical protein, conserved | -0,1125 | 0,5437 |
| TcCLB.508707.320 | hypothetical protein, conserved | -0,1126 | 0,3041 |
| TcCLB.511743.30 | hypothetical protein, conserved | -0,1126 | 0,3198 |
| TcCLB.508741.280 | ubiquitin-conjugating enzyme E2, putative | -0,1126 | 0,4095 |
| TcCLB.506587.90 | surface protease GP63, putative | -0,1126 | 0,4443 |
| TcCLB.511367.159 | hypothetical protein, conserved (pseudogene) | -0,1126 | 0,4994 |
| TcCLB.509985.69 | hypothetical protein, conserved | -0,1126 | 0,5734 |
| TcCLB.506959.140 | hypothetical protein, conserved | -0,1127 | 0,4678 |
| TcCLB.506739.90 | ATP-dependent RNA helicase, putative | -0,1128 | 0,2142 |
| TcCLB.507867.90 | hypothetical protein, conserved | -0,1128 | 0,4273 |
| TcCLB.506735.20 | hypothetical protein, conserved | -0,1129 | 0,3597 |
| TcCLB.511409.20 | hypothetical protein, conserved | -0,1129 | 0,527 |

| | | | |
|---|---|---|---|
| TcCLB.504147.190 | hypothetical protein, conserved | -0,113 | 0,248 |
| TcCLB.506529.180 | hypothetical protein, conserved | -0,113 | 0,5774 |
| TcCLB.510719.300 | metallo-peptidase, Clan MC, Family M14, putative | -0,1131 | 0,2367 |
| TcCLB.509345.11 | retrotransposon hot spot protein (RHS, pseudogene), puta | -0,1132 | 0,3058 |
| TcCLB.506201.4 | hypothetical protein, conserved | -0,1133 | 0,4796 |
| TcCLB.506459.140 | hypothetical protein | -0,1133 | 0,7558 |
| TcCLB.510729.115 | hypothetical protein | -0,1134 | 0,3251 |
| TcCLB.508465.140 | hypothetical protein, conserved | -0,1135 | 0,2223 |
| TcCLB.510349.80 | protein kinase, putative | -0,1135 | 0,2383 |
| TcCLB.507529.40 | hypothetical protein, conserved | -0,1135 | 0,3109 |
| TcCLB.507031.150 | hypothetical protein, conserved | -0,1136 | 0,1986 |
| TcCLB.511283.210 | hypothetical protein, conserved | -0,1136 | 0,2362 |
| TcCLB.506529.400 | hypothetical protein, conserved | -0,1136 | 0,4446 |
| TcCLB.509719.30 | hypothetical protein, conserved | -0,1137 | 0,3895 |
| TcCLB.503733.50 | hypothetical protein, conserved (pseudogene) | -0,1137 | 0,4472 |
| TcCLB.506625.90 | hypothetical protein, conserved | -0,1137 | 0,4693 |
| TcCLB.507041.50 | hypothetical protein, conserved | -0,1137 | 0,5185 |
| TcCLB.510159.19 | hypothetical protein, conserved (fragment) | -0,1137 | 0,5785 |
| TcCLB.510141.4 | hypothetical protein, conserved (fragment) | -0,1139 | 0,432 |
| TcCLB.507027.80 | hypothetical protein, conserved | -0,114 | 0,2754 |
| TcCLB.508837.74 | small GTP-binding protein Rab18, putative | -0,114 | 0,3334 |
| TcCLB.503865.30 | hypothetical protein, conserved | -0,114 | 0,3356 |
| TcCLB.509789.40 | hypothetical protein, conserved | -0,114 | 0,3814 |
| TcCLB.511625.30 | hypothetical protein, conserved | -0,114 | 0,389 |
| TcCLB.507091.20 | hypothetical protein, conserved | -0,1141 | 0,3209 |
| TcCLB.510661.150 | hypothetical protein, conserved | -0,1141 | 0,3861 |
| TcCLB.511589.209 | hypothetical protein, conserved | -0,1141 | 0,6016 |
| TcCLB.505183.5 | hypothetical protein, conserved (fragment) | -0,1142 | 0,4195 |
| TcCLB.507713.10 | hypothetical protein, conserved | -0,1142 | 0,5951 |
| TcCLB.511657.10 | protein kinase, putative | -0,1145 | 0,5682 |
| TcCLB.507007.40 | hypothetical protein, conserved | -0,1148 | 0,3935 |
| TcCLB.507641.290 | chaperonin HSP60, mitochondrial precursor (HSP60) | -0,1148 | 0,7903 |
| TcCLB.504213.110 | Putrescine-cadaverine transporter (POT1.1) | -0,1149 | 0,1848 |
| TcCLB.503981.70 | hypothetical protein, conserved | -0,1149 | 0,3349 |
| TcCLB.508119.180 | hypothetical protein, conserved | -0,1149 | 0,48 |
| TcCLB.447801.10 | hypothetical protein, conserved (fragment) | -0,1149 | 0,5008 |
| TcCLB.460747.20 | hypothetical protein, conserved | -0,1151 | 0,2934 |
| TcCLB.508879.220 | hypothetical protein, conserved | -0,1151 | 0,3047 |
| TcCLB.508413.40 | serine/threonine protein phosphatase, putative | -0,1153 | 0,3025 |
| TcCLB.510077.20 | trans-sialidase (pseudogene), putative | -0,1153 | 0,3097 |
| TcCLB.506525.10 | nuclear segregation protein, putative | -0,1154 | 0,1075 |
| TcCLB.507017.110 | hypothetical protein, conserved | -0,1154 | 0,3024 |
| TcCLB.507031.10 | hypothetical protein, conserved | -0,1154 | 0,362 |
| TcCLB.510741.100 | hypothetical protein, conserved | -0,1154 | 0,4025 |
| TcCLB.503959.20 | hypothetical protein, conserved | -0,1157 | 0,2801 |

| | | | |
|---|---|---|---|
| TcCLB.503923.30 | hypothetical protein, conserved | -0,1158 | 0,3715 |
| TcCLB.510535.40 | hypothetical protein, conserved | -0,1158 | 0,4091 |
| TcCLB.511223.10 | hypothetical protein | -0,1158 | 0,7216 |
| TcCLB.511911.102 | hypothetical protein, conserved | -0,116 | 0,4051 |
| TcCLB.511529.110 | hypothetical protein, conserved | -0,116 | 0,4689 |
| TcCLB.511361.20 | hypothetical protein, conserved | -0,116 | 0,5337 |
| TcCLB.504255.30 | hypothetical protein, conserved | -0,116 | 0,5391 |
| TcCLB.504109.195 | N-acetyltransferase complex ARD1 subunit (pseudogene), | -0,1161 | 0,7283 |
| TcCLB.504029.50 | hypothetical protein, conserved | -0,1162 | 0,3512 |
| TcCLB.507257.130 | hypothetical protein, conserved | -0,1163 | 0,4689 |
| TcCLB.509603.19 | malonyl-CoA decarboxylase, mitochondrial precursor, puta | -0,1163 | 0,5933 |
| TcCLB.473755.9 | hypothetical protein, conserved | -0,1164 | 0,3576 |
| TcCLB.507521.90 | hypothetical protein, conserved | -0,1165 | 0,4917 |
| TcCLB.507025.40 | hypothetical protein | -0,1166 | 0,3909 |
| TcCLB.509031.6 | hypothetical protein, conserved | -0,1166 | 0,5058 |
| TcCLB.508153.260 | hypothetical protein, conserved | -0,1166 | 0,5168 |
| TcCLB.503879.60 | hypothetical protein, conserved | -0,1169 | 0,3749 |
| TcCLB.507897.30 | serine/threonine-protein kinase a, putative | -0,1169 | 0,3999 |
| TcCLB.510359.90 | hypothetical protein, conserved | -0,1169 | 0,4489 |
| TcCLB.511049.10 | hypothetical protein, conserved (pseudogene) | -0,1169 | 0,478 |
| TcCLB.508593.90 | hypothetical protein, conserved | -0,1172 | 0,4078 |
| TcCLB.505183.80 | signal recognition particle, putative | -0,1174 | 0,199 |
| TcCLB.506855.320 | hypothetical protein, conserved | -0,1174 | 0,4383 |
| TcCLB.503897.30 | hypothetical protein, conserved | -0,1175 | 0,3641 |
| TcCLB.503909.76 | ferric reductase transmembrane protein, putative | -0,1176 | 0,6693 |
| TcCLB.510421.190 | hypothetical protein, conserved | -0,1177 | 0,3696 |
| TcCLB.509201.20 | hypothetical protein, conserved | -0,1177 | 0,7016 |
| TcCLB.506679.50 | pre-mRNA splicing factor ATP-dependent RNA helicase, p | -0,1178 | 0,1839 |
| TcCLB.508059.40 | syntaxin, putative | -0,118 | 0,4813 |
| TcCLB.504075.3 | calmodulin, putative | -0,1181 | 0,7022 |
| TcCLB.507669.230 | hypothetical protein, conserved | -0,1182 | 0,2103 |
| TcCLB.506883.130 | hypothetical protein, conserved (pseudogene) | -0,1182 | 0,7066 |
| TcCLB.504129.30 | hypothetical protein, conserved | -0,1183 | 0,3057 |
| TcCLB.506513.24 | hypothetical protein, conserved | -0,1185 | 0,3923 |
| TcCLB.509455.40 | hypothetical protein, conserved | -0,1186 | 0,3197 |
| TcCLB.511293.90 | hypothetical protein, conserved | -0,1186 | 0,3457 |
| TcCLB.506871.50 | ubiquitin hydrolase, putative | -0,1186 | 0,4239 |
| TcCLB.508445.60 | hypothetical protein, conserved | -0,1188 | 0,3437 |
| TcCLB.511693.130 | hypothetical protein | -0,1189 | 0,3219 |
| TcCLB.507639.100 | phosphatidylinositol N-acetylglucosaminyltransferase sub | -0,1189 | 0,4685 |
| TcCLB.509267.10 | hypothetical protein, conserved | -0,119 | 0,214 |
| TcCLB.508919.50 | hypothetical protein | -0,119 | 0,3372 |
| TcCLB.509195.70 | mucin-associated surface protein (MASP), putative | -0,119 | 0,5119 |
| TcCLB.511603.369 | surface protease GP63 (pseudogene), putative | -0,1191 | 0,7944 |
| TcCLB.507513.40 | hypothetical protein, conserved | -0,1192 | 0,4225 |

| | | | |
|---|---|---|---|
| TcCLB.506863.60 | hypothetical protein, conserved | -0,1193 | 0,3358 |
| TcCLB.510963.39 | hypothetical protein, conserved | -0,1195 | 0,4413 |
| TcCLB.511255.410 | surface protease GP63 (pseudogene), putative | -0,1195 | 0,7399 |
| TcCLB.511071.120 | hypothetical protein, conserved | -0,1197 | 0,2321 |
| TcCLB.506625.190 | hypothetical protein, conserved | -0,1197 | 0,5243 |
| TcCLB.508083.5 | elongation factor 1-gamma (EF-1-gamma, pseudogene), | -0,1197 | 0,7237 |
| TcCLB.509267.70 | hypothetical protein | -0,1198 | 0,2086 |
| TcCLB.511109.100 | DNA replication licensing factor MCM3, putative | -0,1199 | 0,474 |
| TcCLB.511367.110 | hypothetical protein, conserved | -0,12 | 0,3966 |
| TcCLB.503669.30 | hypothetical protein, conserved | -0,1201 | 0,4032 |
| TcCLB.506909.160 | trans-sialidase, Group V, putative | -0,1202 | 0,5428 |
| TcCLB.503697.9 | proteasome regulatory non-ATPase subunit 11, putative (1 | -0,1203 | 0,2916 |
| TcCLB.511857.30 | glycosyl transferase-like protein, putative | -0,1204 | 0,4468 |
| TcCLB.507641.300 | chaperonin HSP60, mitochondrial precursor (HSP60) | -0,1204 | 0,8413 |
| TcCLB.508059.24 | hypothetical protein, conserved | -0,1205 | 0,3221 |
| TcCLB.506039.30 | dispersed gene family protein 1 (DGF-1), putative | -0,1205 | 0,7049 |
| TcCLB.503983.20 | dihydroxyacetone kinase 1-like, putative | -0,1206 | 0,2879 |
| TcCLB.506739.130 | protein phosphatase 1 catalitic subunit, putative | -0,1207 | 0,3169 |
| TcCLB.510555.41 | hypothetical protein, conserved (pseudogene) | -0,1207 | 0,7317 |
| TcCLB.506853.60 | hypothetical protein, conserved | -0,1208 | 0,4166 |
| TcCLB.510155.94 | hypothetical protein, conserved | -0,1209 | 0,2936 |
| TcCLB.508355.104 | hypothetical protein, conserved | -0,121 | 0,5332 |
| TcCLB.510947.44 | hypothetical protein, conserved | -0,1211 | 0,4087 |
| TcCLB.510513.70 | hypothetical protein, conserved | -0,1213 | 0,2923 |
| TcCLB.509269.10 | hypothetical protein, conserved | -0,1214 | 0,3519 |
| TcCLB.511291.129 | hypothetical protein, conserved | -0,1215 | 0,5348 |
| TcCLB.511127.20 | calcium binding protein EF hand, putative | -0,1216 | 0,1941 |
| TcCLB.509451.70 | hypothetical protein, conserved | -0,1216 | 0,2033 |
| TcCLB.508961.40 | hypothetical protein, conserved | -0,1216 | 0,2654 |
| TcCLB.511421.30 | hypothetical protein, conserved | -0,1216 | 0,3647 |
| TcCLB.506751.20 | mucin-associated surface protein (MASP, pseudogene), pu | -0,1216 | 0,7305 |
| TcCLB.510359.200 | hypothetical protein, conserved | -0,1221 | 0,2002 |
| TcCLB.510491.40 | hypothetical protein, conserved | -0,1221 | 0,4236 |
| TcCLB.510729.40 | hypothetical protein, conserved | -0,1221 | 0,4346 |
| TcCLB.510187.140 | hypothetical protein, conserved | -0,1222 | 0,1675 |
| TcCLB.506337.130 | hypothetical protein, conserved | -0,1223 | 0,3883 |
| TcCLB.506367.60 | vacuolar protein sorting protein 16, putative | -0,1224 | 0,2112 |
| TcCLB.503851.10 | Elongator-like Protein 3a, putative | -0,1224 | 0,396 |
| TcCLB.511445.80 | hypothetical protein, conserved | -0,1225 | 0,3314 |
| TcCLB.510431.20 | hypothetical protein, conserved | -0,1226 | 0,4131 |
| TcCLB.503911.30 | hypothetical protein, conserved | -0,1227 | 0,2642 |
| TcCLB.503805.40 | hypothetical protein | -0,1227 | 0,2908 |
| TcCLB.510275.180 | elongation factor 1-gamma (EF-1-gamma), putative | -0,1228 | 0,8151 |
| TcCLB.508739.80 | hypothetical protein, conserved | -0,123 | 0,1835 |
| TcCLB.506933.50 | hypothetical protein, conserved | -0,123 | 0,2804 |

| | | | |
|---|---|---|---|
| TcCLB.507057.30 | arginine N-methyltransferase, type I (PRMT6) | -0,1231 | 0,4432 |
| TcCLB.508647.230 | synaptojanin (pseudogene), putative | -0,1232 | 0,1486 |
| TcCLB.506251.40 | hypothetical protein | -0,1232 | 0,2056 |
| TcCLB.506789.290 | hypothetical protein, conserved | -0,1232 | 0,2143 |
| TcCLB.507641.230 | hypothetical protein, conserved | -0,1234 | 0,256 |
| TcCLB.511649.30 | hypothetical protein, conserved | -0,1234 | 0,312 |
| TcCLB.505941.20 | surface protease GP63 (pseudogene), putative | -0,1236 | 0,562 |
| TcCLB.506247.20 | hypothetical protein | -0,1237 | 0,2712 |
| TcCLB.509399.180 | hypothetical protein, conserved | -0,1237 | 0,5091 |
| TcCLB.506241.124 | ankyrin repeat protein, putative | -0,1239 | 0,182 |
| TcCLB.506467.10 | hypothetical protein, conserved | -0,1239 | 0,3182 |
| TcCLB.507019.140 | CDC16, putative | -0,1239 | 0,3387 |
| TcCLB.509619.20 | hypothetical protein, conserved (pseudogene) | -0,1239 | 0,4873 |
| TcCLB.507801.70 | protein kinase, putative | -0,1239 | 0,4911 |
| TcCLB.507737.30 | hypothetical protein, conserved | -0,124 | 0,3363 |
| TcCLB.506211.180 | mitogen-activated protein kinase 9, putative (MPK9) | -0,1241 | 0,2522 |
| TcCLB.509633.30 | hypothetical protein, conserved | -0,1243 | 0,2953 |
| TcCLB.506363.80 | aminoacylase, putative | -0,1243 | 0,746 |
| TcCLB.504245.9 | hypothetical protein, conserved | -0,1244 | 0,3513 |
| TcCLB.510381.70 | hypothetical protein, conserved | -0,1244 | 0,5406 |
| TcCLB.504769.5 | hypothetical protein | -0,1245 | 0,4064 |
| TcCLB.504131.30 | ubiquitin-activating enzyme, putative | -0,1247 | 0,2869 |
| TcCLB.503863.9 | hypothetical protein, conserved | -0,1247 | 0,3756 |
| TcCLB.508197.30 | hypothetical protein, conserved | -0,1248 | 0,173 |
| TcCLB.510351.140 | hypothetical protein, conserved | -0,1248 | 0,4797 |
| TcCLB.503931.14 | hypothetical protein, conserved | -0,125 | 0,3238 |
| TcCLB.510105.120 | protein kinase, putative | -0,125 | 0,3416 |
| TcCLB.508879.160 | hypothetical protein, conserved | -0,1252 | 0,1905 |
| TcCLB.510543.30 | trans-sialidase (pseudogene), putative | -0,1255 | 0,6772 |
| TcCLB.509937.90 | hypothetical protein, conserved | -0,1257 | 0,342 |
| TcCLB.508737.200 | hypothetical protein, conserved | -0,1261 | 0,1556 |
| TcCLB.507011.70 | hypothetical protein | -0,1261 | 0,3576 |
| TcCLB.508277.60 | hypothetical protein, conserved | -0,1262 | 0,1308 |
| TcCLB.511421.40 | transmembrane protein, putative | -0,1262 | 0,3472 |
| TcCLB.503987.10 | cop-coated vesicle membrane protein p24 precursor, puta | -0,1263 | 0,244 |
| TcCLB.507711.60 | hypothetical protein, conserved | -0,1263 | 0,3142 |
| TcCLB.503535.9 | hypothetical protein, conserved | -0,1264 | 0,1636 |
| TcCLB.509217.70 | hypothetical protein, conserved (pseudogene) | -0,1265 | 0,4354 |
| TcCLB.507711.340 | hypothetical protein, conserved | -0,1266 | 0,4167 |
| TcCLB.508139.80 | trans-sialidase (pseudogene), putative | -0,1266 | 0,4367 |
| TcCLB.503957.30 | retrotransposon hot spot (RHS) protein, putative | -0,1267 | 0,3803 |
| TcCLB.507811.50 | hypothetical protein, conserved | -0,1268 | 0,2935 |
| TcCLB.507643.30 | retrotransposon hot spot protein (RHS, pseudogene), puta | -0,1269 | 0,275 |
| TcCLB.508739.40 | hypothetical protein, conserved | -0,127 | 0,216 |
| TcCLB.506405.4 | hypothetical protein, conserved (fragment) | -0,127 | 0,272 |

| | | | |
|---|---|---|---|
| TcCLB.506733.50 | actin-like protein, putative | -0,127 | 0,2805 |
| TcCLB.510963.30 | mRNA capping methyltransferase, putative | -0,127 | 0,3748 |
| TcCLB.511099.20 | trans-sialidase (pseudogene), putative | -0,127 | 0,7768 |
| TcCLB.506337.190 | hypothetical protein, conserved | -0,1271 | 0,2961 |
| TcCLB.504167.30 | protein kinase, putative | -0,1273 | 0,3615 |
| TcCLB.504103.50 | hypothetical protein, conserved | -0,1274 | 0,2934 |
| TcCLB.508479.230 | peroxin-2, putative | -0,1275 | 0,3774 |
| TcCLB.506451.13 | hypothetical protein (pseudogene) | -0,1275 | 0,5332 |
| TcCLB.509099.160 | hypothetical protein, conserved | -0,1276 | 0,1695 |
| TcCLB.506275.60 | zinc carboxypeptidase, putative | -0,1276 | 0,2479 |
| TcCLB.419833.10 | vacuolar protein sorting 18, putative | -0,1276 | 0,4066 |
| TcCLB.506401.370 | hypothetical protein, conserved | -0,1278 | 0,3472 |
| TcCLB.509979.20 | hypothetical protein, conserved | -0,1279 | 0,2339 |
| TcCLB.509859.20 | hypothetical protein, conserved | -0,128 | 0,2765 |
| TcCLB.507083.70 | Iron-sulfur assembly protein 1, putative (Isa1) | -0,128 | 0,2949 |
| TcCLB.510901.140 | hypothetical protein, conserved | -0,128 | 0,3544 |
| TcCLB.507159.41 | hypothetical protein, conserved | -0,1281 | 0,5341 |
| TcCLB.506425.20 | hypothetical protein | -0,1284 | 0,3572 |
| TcCLB.511183.565 | surface protease GP63 (pseudogene), putative | -0,1284 | 0,784 |
| TcCLB.511731.89 | hypothetical protein, conserved | -0,1285 | 0,3845 |
| TcCLB.511753.140 | vacuolar assembly protein vps41, putative | -0,1285 | 0,4295 |
| TcCLB.508015.10 | hypothetical protein | -0,1285 | 0,5129 |
| TcCLB.508321.30 | hypothetical protein, conserved | -0,1285 | 0,5417 |
| TcCLB.508737.160 | hypothetical protein, conserved | -0,1288 | 0,204 |
| TcCLB.506615.75 | surface protease GP63 (pseudogene), putative | -0,1288 | 0,2635 |
| TcCLB.510889.340 | hypothetical protein, conserved | -0,129 | 0,5228 |
| TcCLB.509247.20 | ras-related protein Rab21, putative (RAB21) | -0,1291 | 0,3407 |
| TcCLB.505073.30 | mu-adaptin 4, putative | -0,1292 | 0,3749 |
| TcCLB.509481.10 | trans-sialidase, putative | -0,1292 | 0,7354 |
| TcCLB.508153.1090 | hypothetical protein, conserved | -0,1293 | 0,2441 |
| TcCLB.509243.40 | hypothetical protein, conserved (pseudogene) | -0,1293 | 0,3035 |
| TcCLB.507709.40 | hypothetical protein, conserved | -0,1293 | 0,338 |
| TcCLB.509573.60 | hypothetical protein, conserved | -0,1294 | 0,4494 |
| TcCLB.508857.100 | hypothetical protein, conserved | -0,1295 | 0,2361 |
| TcCLB.510901.130 | hypothetical protein, conserved | -0,1295 | 0,3271 |
| TcCLB.508153.30 | Protein tilB homolog, putative (LRTP) | -0,1295 | 0,3382 |
| TcCLB.511507.20 | hypothetical protein, conserved | -0,1297 | 0,2992 |
| TcCLB.506693.7 | hypothetical protein, conserved | -0,13 | 0,4891 |
| TcCLB.510413.10 | hypothetical protein, conserved (pseudogene) | -0,13 | 0,5358 |
| TcCLB.509073.20 | hypothetical protein, conserved | -0,1301 | 0,1175 |
| TcCLB.425785.9 | hypothetical protein, conserved | -0,1302 | 0,4194 |
| TcCLB.506855.340 | U4/U6 small nuclear ribonuclear protein, putative | -0,1304 | 0,2831 |
| TcCLB.508075.50 | hypothetical protein, conserved | -0,1305 | 0,2398 |
| TcCLB.509527.90 | trans-sialidase (pseudogene), putative | -0,1306 | 0,2346 |
| TcCLB.482319.19 | hypothetical protein, conserved | -0,1306 | 0,3994 |

| | | | |
|---|---|---|---|
| TcCLB.509207.130 | peptidyl-prolyl cis-trans isomerase (PAR45) | -0,1307 | 0,2034 |
| TcCLB.506577.160 | hypothetical protein, conserved | -0,1307 | 0,336 |
| TcCLB.503689.34 | hypothetical protein, conserved | -0,1307 | 0,5635 |
| TcCLB.508135.10 | hypothetical protein, conserved | -0,1308 | 0,3195 |
| TcCLB.508547.190 | hypothetical protein, conserved | -0,1309 | 0,2935 |
| TcCLB.506127.90 | hypothetical protein, conserved | -0,1311 | 0,08165 |
| TcCLB.506537.110 | hypothetical protein, conserved | -0,1314 | 0,4485 |
| TcCLB.511021.51 | hypothetical protein, conserved | -0,1315 | 0,4238 |
| TcCLB.510553.25 | elongation factor 1-gamma (EF-1-gamma, pseudogene), | -0,1315 | 0,7796 |
| TcCLB.510129.20 | surface glycoprotein, putative | -0,1317 | 0,247 |
| TcCLB.509179.30 | hypothetical protein, conserved | -0,1317 | 0,2954 |
| TcCLB.511391.210 | mitochondrial calcium uptake 1, putative | -0,1319 | 0,1826 |
| TcCLB.510769.80 | ankyrin repeat protein, putative | -0,132 | 0,195 |
| TcCLB.510287.40 | hypothetical protein | -0,132 | 0,3187 |
| TcCLB.506147.170 | hypothetical protein, conserved | -0,1322 | 0,1498 |
| TcCLB.509507.30 | hypothetical protein, conserved | -0,1322 | 0,201 |
| TcCLB.509151.20 | hypothetical protein | -0,1322 | 0,2884 |
| TcCLB.507041.100 | ATP-dependent protease binding subunit (pseudogene), pu | -0,1322 | 0,3034 |
| TcCLB.508289.80 | trans-sialidase (pseudogene), putative | -0,1322 | 0,6942 |
| TcCLB.508821.10 | katanin, putative | -0,1323 | 0,4166 |
| TcCLB.508859.50 | hypothetical protein, conserved | -0,1325 | 0,3787 |
| TcCLB.511283.10 | nuclear segregation protein, putative | -0,1326 | 0,143 |
| TcCLB.503891.120 | dynein light chain, putative | -0,1326 | 0,3698 |
| TcCLB.508263.30 | kinesin, putative | -0,1327 | 0,2547 |
| TcCLB.507709.44 | hypothetical protein, conserved | -0,1327 | 0,2612 |
| TcCLB.507837.10 | Peptidase C50 separase, putative | -0,1328 | 0,3202 |
| TcCLB.506177.20 | lectin, putative | -0,1329 | 0,2452 |
| TcCLB.509767.44 | hypothetical protein, conserved | -0,1329 | 0,2771 |
| TcCLB.503917.14 | hypothetical protein, conserved | -0,1334 | 0,2926 |
| TcCLB.504129.50 | hypothetical protein, conserved (pseudogene) | -0,1334 | 0,5077 |
| TcCLB.511849.15 | mucin-associated surface protein (MASP, pseudogene), pu | -0,1335 | 0,22 |
| TcCLB.504153.320 | Golgi reassembly stacking protein (GRASP homologue), pu | -0,1335 | 0,3487 |
| TcCLB.509297.30 | ADP-ribosylation factor, putative | -0,1336 | 0,7397 |
| TcCLB.508153.690 | hypothetical protein | -0,1336 | 0,7936 |
| TcCLB.509429.120 | Mitochondrial Ribosomal Protein of Small Subunit 29, put | -0,1337 | 0,2781 |
| TcCLB.505807.270 | ubiquitin hydrolase, putative | -0,1337 | 0,3282 |
| TcCLB.511315.9 | ATP-dependent RNA helicase, putative | -0,1339 | 0,433 |
| TcCLB.506605.174 | hypothetical protein, conserved | -0,134 | 0,4365 |
| TcCLB.508707.80 | RNA-binding protein, putative | -0,1341 | 0,2764 |
| TcCLB.508723.30 | bilobe protein, conserved | -0,1342 | 0,3173 |
| TcCLB.511315.30 | hypothetical protein, conserved | -0,1342 | 0,4234 |
| TcCLB.510701.50 | mucin TcMUC (pseudogene), putative | -0,1342 | 0,7261 |
| TcCLB.507981.61 | hypothetical protein | -0,1343 | 0,689 |
| TcCLB.510301.40 | hypothetical protein, conserved | -0,1344 | 0,2825 |
| TcCLB.506957.110 | SNF-7-like protein, putative | -0,1346 | 0,2646 |

| | | | |
|---|---|---|---|
| TcCLB.507053.30 | hypothetical protein, conserved | -0,1347 | 0,166 |
| TcCLB.506993.31 | mucin-associated surface protein (MASP, pseudogene), pu | -0,1347 | 0,7608 |
| TcCLB.509437.80 | hypothetical protein, conserved | -0,1348 | 0,3013 |
| TcCLB.509215.20 | hypothetical protein, conserved | -0,1348 | 0,457 |
| TcCLB.511671.100 | hypothetical protein, conserved | -0,1349 | 0,4394 |
| TcCLB.506941.90 | ATP-dependent DEAD/H DNA helicase recQ family, putativ | -0,135 | 0,1448 |
| TcCLB.507611.330 | hypothetical protein, conserved | -0,135 | 0,3446 |
| TcCLB.508625.150 | L-ribulokinase, putative | -0,135 | 0,4044 |
| TcCLB.507549.40 | trans-sialidase (pseudogene), putative | -0,1351 | 0,4657 |
| TcCLB.511461.4 | hypothetical protein, conserved (fragment) | -0,1352 | 0,458 |
| TcCLB.511001.210 | hypothetical protein, conserved | -0,1354 | 0,2936 |
| TcCLB.506567.120 | mitochondrial RNA binding protein, putative | -0,1354 | 0,3574 |
| TcCLB.508231.204 | hypothetical protein, conserved | -0,1355 | 0,3694 |
| TcCLB.511109.70 | hypothetical protein, conserved | -0,1356 | 0,1671 |
| TcCLB.508895.10 | C-terminal motor kinesin, putative | -0,1356 | 0,4151 |
| TcCLB.511867.60 | hypothetical protein, conserved | -0,1357 | 0,342 |
| TcCLB.504105.170 | hypothetical protein, conserved | -0,1358 | 0,1077 |
| TcCLB.509669.60 | hypothetical protein, conserved | -0,1358 | 0,3584 |
| TcCLB.511521.24 | hypothetical protein, conserved | -0,1358 | 0,581 |
| TcCLB.507193.10 | dispersed gene family protein 1 (DGF-1), putative | -0,1358 | 0,5939 |
| TcCLB.511389.50 | hypothetical protein, conserved | -0,1359 | 0,1398 |
| TcCLB.508355.180 | hypothetical protein, conserved | -0,1361 | 0,2958 |
| TcCLB.506213.70 | hypothetical protein, conserved | -0,1362 | 0,2738 |
| TcCLB.503823.80 | hypothetical protein, conserved | -0,1362 | 0,4205 |
| TcCLB.507103.20 | hypothetical protein | -0,1363 | 0,1049 |
| TcCLB.510351.40 | ATP-dependent RNA helicase, putative | -0,1363 | 0,17 |
| TcCLB.504111.10 | hypothetical protein, conserved | -0,1363 | 0,1756 |
| TcCLB.507517.70 | hypothetical protein, conserved | -0,1363 | 0,2869 |
| TcCLB.511589.120 | DNA topoisomerase III, putative | -0,1363 | 0,2962 |
| TcCLB.510903.70 | hypothetical protein, conserved | -0,1364 | 0,2299 |
| TcCLB.508119.110 | kynureninase, putative | -0,1364 | 0,394 |
| TcCLB.511277.340 | paralyzed flagella protein 20 (PF20) | -0,1365 | 0,1806 |
| TcCLB.511907.280 | hypothetical protein, conserved | -0,1365 | 0,2119 |
| TcCLB.508119.160 | hypothetical protein, conserved | -0,1365 | 0,2304 |
| TcCLB.508995.40 | vesicular-fusion protein nsf, putative | -0,1366 | 0,288 |
| TcCLB.508959.20 | hypothetical protein, conserved | -0,1367 | 0,2856 |
| TcCLB.509437.125 | trans-sialidase (pseudogene), putative | -0,1367 | 0,6149 |
| TcCLB.506627.40 | hypothetical protein, conserved | -0,1368 | 0,285 |
| TcCLB.506773.110 | hypothetical protein, conserved | -0,1369 | 0,1978 |
| TcCLB.508461.143 | hypothetical protein, conserved | -0,1369 | 0,4348 |
| TcCLB.511287.70 | hypothetical protein, conserved | -0,137 | 0,4489 |
| TcCLB.509621.10 | hypothetical protein, conserved | -0,1371 | 0,2245 |
| TcCLB.506367.120 | hypothetical protein, conserved | -0,1372 | 0,1184 |
| TcCLB.508153.1030 | hypothetical protein, conserved | -0,1372 | 0,1383 |
| TcCLB.508593.150 | puromycin-sensitive aminopeptidase-like protein, putative | -0,1373 | 0,176 |

| | | | |
|---|---|---|---|
| TcCLB.511657.20 | protein kinase, putative | -0,1374 | 0,2496 |
| TcCLB.508815.60 | hypothetical protein, conserved | -0,1374 | 0,3332 |
| TcCLB.503889.5 | hypothetical protein | -0,1376 | 0,5028 |
| TcCLB.511755.60 | ATP-dependent RNA helicase, putative | -0,1378 | 0,09438 |
| TcCLB.507467.90 | diacylglycerol acyltransferase, putative | -0,1378 | 0,2 |
| TcCLB.503791.20 | hypothetical protein, conserved | -0,1378 | 0,5136 |
| TcCLB.508919.150 | retrotransposon hot spot protein (RHS, pseudogene), puta | -0,1379 | 0,4558 |
| TcCLB.503819.30 | eukaryotic translation initiation factor 2 subunit, putative | -0,138 | 0,1143 |
| TcCLB.511585.310 | trans-sialidase (pseudogene), putative | -0,138 | 0,4103 |
| TcCLB.504157.70 | hypothetical protein, conserved | -0,1381 | 0,1416 |
| TcCLB.508153.970 | AGC/RSK family serine/threonine kinase, putative | -0,1382 | 0,3682 |
| TcCLB.509201.10 | hypothetical protein, conserved | -0,1384 | 0,1807 |
| TcCLB.507019.130 | hypothetical protein | -0,1385 | 0,4089 |
| TcCLB.506941.60 | phospholipid-translocating P-type ATPase (flippase), puta | -0,1387 | 0,3097 |
| TcCLB.507093.210 | myosin heavy chain, putative | -0,1388 | 0,09634 |
| TcCLB.511627.100 | TPR-repeat protein, putative | -0,1388 | 0,2597 |
| TcCLB.510101.210 | hypothetical protein, conserved | -0,139 | 0,1855 |
| TcCLB.511287.160 | hypothetical protein, conserved | -0,139 | 0,2 |
| TcCLB.508527.20 | retrotransposon hot spot protein (RHS, pseudogene), puta | -0,1391 | 0,1185 |
| TcCLB.511871.20 | hypothetical protein, conserved | -0,1396 | 0,2479 |
| TcCLB.508889.10 | hypothetical protein, conserved | -0,1397 | 0,1975 |
| TcCLB.506175.5 | hypothetical protein, conserved | -0,1398 | 0,4107 |
| TcCLB.508909.160 | hypothetical protein, conserved | -0,1398 | 0,4568 |
| TcCLB.508809.30 | ABC transporter family-like protein | -0,1399 | 0,1922 |
| TcCLB.510295.40 | Protein C21orf2 homolog, putative | -0,1399 | 0,2179 |
| TcCLB.504215.40 | actin-related protein 2/3 complex subunit 1, putative (ARI | -0,14 | 0,2701 |
| TcCLB.507063.130 | protein kinase, putative | -0,1401 | 0,127 |
| TcCLB.510219.20 | hypothetical protein, conserved | -0,1401 | 0,3935 |
| TcCLB.508909.50 | hypothetical protein, conserved | -0,1401 | 0,4269 |
| TcCLB.511903.140 | tubulin folding cofactor D (pseudogene), putative | -0,1403 | 0,107 |
| TcCLB.510769.70 | hypothetical protein | -0,1403 | 0,2497 |
| TcCLB.509915.35 | hypothetical protein, conserved (pseudogene) | -0,1403 | 0,4641 |
| TcCLB.510257.40 | hypothetical protein, conserved | -0,1405 | 0,2966 |
| TcCLB.503467.9 | NADH-dependent fumarate reductase, putative (fragment | -0,1405 | 0,468 |
| TcCLB.506563.30 | coatomer beta subunit, putative | -0,1406 | 0,04761 |
| TcCLB.510221.39 | adenosine monophosphate deaminase, putative (fragmer | -0,1406 | 0,252 |
| TcCLB.504625.60 | hydrogenase, putative | -0,1407 | 0,2408 |
| TcCLB.506399.60 | hypothetical protein, conserved | -0,1408 | 0,2687 |
| TcCLB.506443.20 | hypothetical protein, conserved | -0,1408 | 0,3478 |
| TcCLB.506247.150 | hypothetical protein, conserved | -0,1408 | 0,3587 |
| TcCLB.508325.90 | hypothetical protein | -0,1408 | 0,5423 |
| TcCLB.509561.110 | hypothetical protein, conserved | -0,1409 | 0,1826 |
| TcCLB.506739.80 | mitochondrial carrier protein, putative | -0,141 | 0,2157 |
| TcCLB.506739.209 | hypothetical protein, conserved | -0,141 | 0,318 |
| TcCLB.508625.20 | hypothetical protein, conserved | -0,141 | 0,4236 |

| | | | |
|---|---|---|---|
| TcCLB.505943.40 | hypothetical protein, conserved | -0,1411 | 0,1152 |
| TcCLB.504109.110 | cleavage and polyadenylation specificity factor, putative | -0,1411 | 0,2166 |
| TcCLB.509399.190 | pumilio/PUF RNA binding protein 4, putative (PUF4) | -0,1411 | 0,4052 |
| TcCLB.506311.20 | palmitoyl acyltransferase 2, putative | -0,1412 | 0,2925 |
| TcCLB.509505.10 | protein disulfide isomerase, putative | -0,1413 | 0,3011 |
| TcCLB.510525.50 | hypothetical protein, conserved | -0,1417 | 0,1649 |
| TcCLB.506559.434 | hypothetical protein, conserved | -0,1417 | 0,2119 |
| TcCLB.508995.69 | pseudouridine synthase TruD, putative (fragment) | -0,1417 | 0,2185 |
| TcCLB.506401.140 | hypothetical protein, conserved | -0,142 | 0,4972 |
| TcCLB.511585.160 | hypothetical protein, conserved | -0,1421 | 0,1962 |
| TcCLB.506925.90 | hypothetical protein, conserved | -0,1422 | 0,1499 |
| TcCLB.504157.90 | hypothetical protein, conserved | -0,1422 | 0,2619 |
| TcCLB.434259.10 | RNA polymerase-associated protein CTR9, putative | -0,1422 | 0,3042 |
| TcCLB.510351.100 | hypothetical protein, conserved | -0,1422 | 0,3273 |
| TcCLB.509053.10 | ubiquitin-protein ligase, putative | -0,1423 | 0,2543 |
| TcCLB.506713.30 | ATP-dependent DEAD/H RNA helicase, putative | -0,1424 | 0,1875 |
| TcCLB.511277.359 | phosphatidic acid phosphatase, putative | -0,1424 | 0,2486 |
| TcCLB.506743.210 | tubulin tyrosine ligase, putative | -0,1425 | 0,3705 |
| TcCLB.505683.10 | sterol 24-c-methyltransferase, putative | -0,1426 | 0,2414 |
| TcCLB.510943.110 | hypothetical protein, conserved | -0,1427 | 0,1823 |
| TcCLB.506407.70 | hypothetical protein, conserved | -0,1428 | 0,2763 |
| TcCLB.504867.40 | acetyltransferase, putative | -0,1428 | 0,2871 |
| TcCLB.507881.60 | protein kinase, putative | -0,1428 | 0,4838 |
| TcCLB.506255.10 | hypothetical protein, conserved (pseudogene) | -0,1429 | 0,4831 |
| TcCLB.398235.10 | protein kinase, putative (fragment) | -0,143 | 0,4939 |
| TcCLB.511109.120 | beta-ketoacyl synthase family protein, putative | -0,1432 | 0,3277 |
| TcCLB.507787.90 | chaperone DNAJ protein, putative | -0,1434 | 0,09037 |
| TcCLB.510579.100 | hypothetical protein, conserved | -0,1434 | 0,2611 |
| TcCLB.506945.170 | thiosulfate sulfurtransferase, mitochondrial, putative | -0,1436 | 0,2969 |
| TcCLB.503463.10 | protein kinase C substrate protein, heavy chain, putative | -0,1439 | 0,2842 |
| TcCLB.511283.184 | hypothetical protein, conserved | -0,144 | 0,3388 |
| TcCLB.507993.150 | hypothetical protein, conserved | -0,1441 | 0,1399 |
| TcCLB.506255.20 | hypothetical protein, conserved | -0,1441 | 0,4089 |
| TcCLB.506883.20 | hypothetical protein, conserved | -0,1442 | 0,2078 |
| TcCLB.511245.160 | fatty acid elongase, putative | -0,1445 | 0,5908 |
| TcCLB.510535.20 | hypothetical protein, conserved | -0,1446 | 0,1552 |
| TcCLB.511529.100 | hypothetical protein, conserved | -0,1446 | 0,2732 |
| TcCLB.511857.59 | katanin, putative | -0,1446 | 0,2839 |
| TcCLB.510983.10 | hypothetical protein | -0,1446 | 0,5187 |
| TcCLB.511075.60 | protein kinase, putative | -0,1447 | 0,3268 |
| TcCLB.511745.60 | hypothetical protein | -0,1447 | 0,5821 |
| TcCLB.508897.10 | GDP-mannose 4,6 dehydratase, putative | -0,1449 | 0,2841 |
| TcCLB.509069.20 | ATP binding protein-like protein, putative | -0,145 | 0,2726 |
| TcCLB.511761.30 | hypothetical protein, conserved | -0,145 | 0,2875 |
| TcCLB.508275.9 | dynein heavy chain, putative (fragment) | -0,1451 | 0,2727 |

| | | | |
|---|---|---|---|
| TcCLB.504111.49 | hypothetical protein, conserved | -0,1455 | 0,1639 |
| TcCLB.507911.36 | ATP-dependent DEAD/H RNA helicase (pseudogene), puta | -0,1455 | 0,6597 |
| TcCLB.506615.21 | surface protease GP63 (pseudogene), putative | -0,1455 | 0,6936 |
| TcCLB.507645.70 | Meckel syndrome type 1 protein, putative (fragment) (mk | -0,1459 | 0,1628 |
| TcCLB.511435.49 | phosphatidylinositol 3-kinase, putative | -0,1459 | 0,5693 |
| TcCLB.507357.135 | hypothetical protein | -0,1459 | 0,7323 |
| TcCLB.503583.70 | hypothetical protein, conserved | -0,146 | 0,2868 |
| TcCLB.511445.160 | copper-transporting ATPase-like protein, putative | -0,1461 | 0,05711 |
| TcCLB.504533.10 | dispersed gene family protein 1 (DGF-1), putative | -0,1461 | 0,4988 |
| TcCLB.510693.274 | mucin TcMUCII (pseudogene), putative | -0,1462 | 0,5295 |
| TcCLB.511671.30 | beta galactofuranosyl glycosyltransferase, putative | -0,1463 | 0,6279 |
| TcCLB.506733.40 | katanin-like protein, putative | -0,1464 | 0,2956 |
| TcCLB.511859.40 | hypothetical protein, conserved | -0,1464 | 0,4188 |
| TcCLB.508605.10 | retrotransposon hot spot protein (RHS, pseudogene), puta | -0,1464 | 0,4854 |
| TcCLB.506925.30 | hypothetical protein, conserved | -0,1465 | 0,05063 |
| TcCLB.510431.100 | hypothetical protein, conserved | -0,1465 | 0,2025 |
| TcCLB.511867.120 | hypothetical protein, conserved | -0,1466 | 0,2398 |
| TcCLB.503801.70 | hypothetical protein | -0,1467 | 0,2161 |
| TcCLB.509171.20 | hypothetical protein, conserved | -0,1468 | 0,1918 |
| TcCLB.462617.10 | dynein heavy chain (pseudogene), putative | -0,1469 | 0,1366 |
| TcCLB.508641.260 | DNA-directed RNA polymerase, mitochondrial (MTRNAP) | -0,147 | 0,0666 |
| TcCLB.511815.180 | hypothetical protein, conserved | -0,147 | 0,2779 |
| TcCLB.407477.20 | hypothetical protein, conserved | -0,147 | 0,3547 |
| TcCLB.511067.10 | tubulin folding cofactor D, putative | -0,1471 | 0,08791 |
| TcCLB.510671.31 | hypothetical protein, conserved | -0,1471 | 0,1922 |
| TcCLB.511255.516 | syntaxin binding protein (pseudogene), putative | -0,1471 | 0,6837 |
| TcCLB.506275.9 | RNA polymerase-associated protein CDC73, putative | -0,1473 | 0,326 |
| TcCLB.510719.250 | RNA polymerase-associated protein CDC73, putative | -0,1474 | 0,1875 |
| TcCLB.508431.5 | hypothetical protein | -0,1474 | 0,6724 |
| TcCLB.508083.20 | hypothetical protein, conserved | -0,1475 | 0,3165 |
| TcCLB.507519.70 | hypothetical protein, conserved | -0,1478 | 0,2302 |
| TcCLB.510089.90 | hypothetical protein, conserved | -0,1478 | 0,2586 |
| TcCLB.507945.30 | pseudouridine synthase TruD, putative (fragment) | -0,1479 | 0,1634 |
| TcCLB.504071.70 | hypothetical protein, conserved | -0,148 | 0,06653 |
| TcCLB.507049.119 | hypothetical protein, conserved | -0,148 | 0,1432 |
| TcCLB.504253.30 | hypothetical protein, conserved | -0,1481 | 0,1971 |
| TcCLB.509915.55 | trans-sialidase (pseudogene), putative | -0,1481 | 0,5877 |
| TcCLB.507035.130 | hypothetical protein, conserved | -0,1483 | 0,1263 |
| TcCLB.507389.50 | hypothetical protein, conserved | -0,1483 | 0,2195 |
| TcCLB.508731.30 | calmodulin, putative | -0,1483 | 0,2619 |
| TcCLB.504769.10 | hypothetical protein | -0,1483 | 0,3576 |
| TcCLB.509537.90 | hypothetical protein, conserved | -0,1485 | 0,0915 |
| TcCLB.507735.69 | hypothetical protein, conserved | -0,1486 | 0,3196 |
| TcCLB.507197.10 | surface protease GP63, putative | -0,1487 | 0,1073 |
| TcCLB.511863.30 | hypothetical protein, conserved | -0,1487 | 0,2121 |

| | | | |
|---|---|---|---|
| TcCLB.509173.9 | hypothetical protein, conserved | -0,1487 | 0,2977 |
| TcCLB.506855.130 | hypothetical protein, conserved | -0,1487 | 0,3219 |
| TcCLB.429185.31 | trans-sialidase (pseudogene), putative | -0,149 | 0,6747 |
| TcCLB.506435.190 | hypothetical protein, conserved | -0,1491 | 0,3176 |
| TcCLB.506525.50 | hypothetical protein, conserved | -0,1492 | 0,2198 |
| TcCLB.511661.70 | hypothetical protein, conserved | -0,1492 | 0,2611 |
| TcCLB.510537.40 | trans-sialidase (pseudogene), putative | -0,1494 | 0,7715 |
| TcCLB.510355.104 | ATP-dependent DEAD/H RNA helicase (pseudogene), puta | -0,1495 | 0,5998 |
| TcCLB.509243.20 | hypothetical protein, conserved | -0,1496 | 0,537 |
| TcCLB.508501.90 | hypothetical protein | -0,1496 | 0,637 |
| TcCLB.507795.50 | syntaxin, putative | -0,1498 | 0,2572 |
| TcCLB.508231.180 | hypothetical protein, conserved | -0,1499 | 0,2082 |
| TcCLB.506295.30 | hypothetical protein, conserved | -0,15 | 0,3022 |
| TcCLB.508577.60 | hypothetical protein, conserved | -0,1501 | 0,1001 |
| TcCLB.506801.120 | protein kinase, putative | -0,1502 | 0,2507 |
| TcCLB.510531.109 | dynein heavy chain, cytosolic, putative (fragment) | -0,1502 | 0,3839 |
| TcCLB.509733.130 | hypothetical protein, conserved | -0,1505 | 0,1756 |
| TcCLB.511355.20 | hypothetical protein, conserved | -0,1505 | 0,2197 |
| TcCLB.510287.60 | ribonuclease HII, putative | -0,1505 | 0,2382 |
| TcCLB.503681.29 | 30S ribosomal protein S8, putative | -0,1507 | 0,7194 |
| TcCLB.504009.4 | hypothetical protein, conserved | -0,151 | 0,368 |
| TcCLB.468041.10 | hypothetical protein | -0,151 | 0,4025 |
| TcCLB.511711.30 | eRF1 methyltransferase catalytic subunit, putative | -0,1512 | 0,2696 |
| TcCLB.506679.210 | RNA editing complex protein MP61, putative | -0,1513 | 0,1876 |
| TcCLB.507037.20 | RNA-binding protein, putative | -0,1513 | 0,2126 |
| TcCLB.504217.10 | coatomer subunit epsilon (COPE) | -0,1513 | 0,2474 |
| TcCLB.509179.170 | hypothetical protein | -0,1513 | 0,2784 |
| TcCLB.511277.84 | dynein light chain Tctex-type, putative | -0,1515 | 0,3694 |
| TcCLB.507011.160 | hypothetical protein, conserved | -0,1517 | 0,1967 |
| TcCLB.510565.50 | hypothetical protein, conserved | -0,1518 | 0,2203 |
| TcCLB.506933.10 | hypothetical protein, conserved | -0,1519 | 0,2467 |
| TcCLB.510437.50 | hypothetical protein, conserved | -0,152 | 0,1945 |
| TcCLB.507153.55 | hypothetical protein, conserved (pseudogene) | -0,152 | 0,3339 |
| TcCLB.509777.40 | beta galactofuranosyl glycosyltransferase, putative | -0,152 | 0,4262 |
| TcCLB.511733.20 | hypothetical protein, conserved | -0,1522 | 0,1531 |
| TcCLB.509717.80 | glyceraldehyde 3-phosphate dehydrogenase, cytosolic, put | -0,1523 | 0,1865 |
| TcCLB.508469.20 | hypothetical protein, conserved | -0,1524 | 0,2697 |
| TcCLB.507081.90 | hypothetical protein, conserved | -0,1526 | 0,1623 |
| TcCLB.510421.310 | hypothetical protein, conserved | -0,1526 | 0,2836 |
| TcCLB.504069.40 | hypothetical protein, conserved | -0,1528 | 0,168 |
| TcCLB.509627.30 | proteasome regulatory non-ATPase subunit 11, putative (I | -0,1529 | 0,1137 |
| TcCLB.508175.230 | hypothetical protein, conserved | -0,1529 | 0,1528 |
| TcCLB.511261.10 | hypothetical protein, conserved | -0,153 | 0,074 |
| TcCLB.510303.204 | hypothetical protein, conserved | -0,153 | 0,1249 |
| TcCLB.504425.80 | hypothetical protein, conserved | -0,1531 | 0,2546 |

| | | | |
|---|---|---|---|
| TcCLB.508563.30 | retrotransposon hot spot protein (RHS, pseudogene), puta | -0,1532 | 0,7069 |
| TcCLB.507629.10 | hypothetical protein, conserved | -0,1534 | 0,06438 |
| TcCLB.510187.234 | tyrosine specific protein phosphatase, putative | -0,1534 | 0,1935 |
| TcCLB.504157.30 | hypothetical protein, conserved | -0,1534 | 0,3296 |
| TcCLB.508221.941 | surface protease GP63 (pseudogene), putative | -0,1534 | 0,6466 |
| TcCLB.509569.180 | hypothetical protein, conserved | -0,1535 | 0,2307 |
| TcCLB.506251.20 | hypothetical protein, conserved | -0,1535 | 0,329 |
| TcCLB.510119.70 | protein kinase, putative | -0,1536 | 0,3607 |
| TcCLB.508541.230 | casein kinase, putative | -0,1536 | 0,5284 |
| TcCLB.506165.10 | hypothetical protein, conserved | -0,1537 | 0,1697 |
| TcCLB.504827.150 | hypothetical protein, conserved | -0,1537 | 0,3301 |
| TcCLB.510101.160 | hypothetical protein, conserved | -0,1538 | 0,2536 |
| TcCLB.507639.90 | hypothetical protein, conserved | -0,1538 | 0,4799 |
| TcCLB.511525.11 | hypothetical protein, conserved (fragment) | -0,1538 | 0,5667 |
| TcCLB.508441.10 | NUP-1 protein, putative | -0,1539 | 0,01591 |
| TcCLB.509779.30 | hypothetical protein, conserved | -0,1539 | 0,1827 |
| TcCLB.509563.20 | hypothetical protein, conserved | -0,1539 | 0,2229 |
| TcCLB.507681.130 | hypothetical protein, conserved | -0,1541 | 0,08464 |
| TcCLB.508547.60 | hypothetical protein, conserved | -0,1541 | 0,2356 |
| TcCLB.506575.30 | hypothetical protein, conserved (pseudogene) | -0,1542 | 0,1203 |
| TcCLB.504741.70 | hypothetical protein, conserved | -0,1542 | 0,2027 |
| TcCLB.508277.90 | hypothetical protein, conserved | -0,1542 | 0,315 |
| TcCLB.506801.70 | hypothetical protein | -0,1543 | 0,1705 |
| TcCLB.506885.390 | AMP deaminase, putative | -0,1543 | 0,1939 |
| TcCLB.508961.4 | ADP-ribosylation factor, putative | -0,1543 | 0,3433 |
| TcCLB.503641.19 | peroxin 12, putative (PEX12) | -0,1543 | 0,4151 |
| TcCLB.509595.40 | hypothetical protein, conserved | -0,1544 | 0,1495 |
| TcCLB.510057.30 | hypothetical protein, conserved | -0,1544 | 0,2273 |
| TcCLB.511755.80 | formin, putative | -0,1544 | 0,362 |
| TcCLB.506481.39 | hypothetical protein, conserved | -0,1545 | 0,07534 |
| TcCLB.506443.110 | retrotransposon hot spot protein (RHS, pseudogene), puta | -0,1545 | 0,2333 |
| TcCLB.457339.10 | U6 snRNA-associated Sm-like protein LSm4, putative | -0,1545 | 0,392 |
| TcCLB.503861.110 | retrotransposon hot spot protein (RHS, pseudogene), puta | -0,1546 | 0,2311 |
| TcCLB.509099.100 | hypothetical protein, conserved | -0,1548 | 0,2588 |
| TcCLB.506011.20 | hypothetical protein, conserved (pseudogene) | -0,1548 | 0,4078 |
| TcCLB.505425.9 | dispersed gene family protein 1 (DGF-1, pseudogene), put | -0,1548 | 0,7675 |
| TcCLB.510225.10 | histone H1, putative | -0,1549 | 0,0591 |
| TcCLB.510175.160 | protein kinase, putative | -0,1549 | 0,3937 |
| TcCLB.511541.9 | hypothetical protein, conserved (fragment) | -0,155 | 0,1509 |
| TcCLB.506479.90 | hypothetical protein, conserved | -0,1552 | 0,1946 |
| TcCLB.511589.80 | hypothetical protein, conserved | -0,1554 | 0,228 |
| TcCLB.503425.5 | mitochondrial carrier protein, putative | -0,1554 | 0,5173 |
| TcCLB.504827.140 | hypothetical protein, conserved | -0,1555 | 0,1985 |
| TcCLB.506627.60 | ADP-ribosylation factor, putative | -0,1555 | 0,239 |
| TcCLB.506067.80 | mucin TcMUCII, putative | -0,1555 | 0,7224 |

| | | | |
|---|---|---|---|
| TcCLB.509213.30 | hypothetical protein, conserved | -0,1556 | 0,2753 |
| TcCLB.510575.10 | hypothetical protein | -0,1557 | 0,249 |
| TcCLB.510361.90 | surface protease GP63, putative | -0,1557 | 0,6109 |
| TcCLB.511517.164 | hypothetical protein, conserved | -0,1558 | 0,2306 |
| TcCLB.509643.50 | hypothetical protein, conserved | -0,1559 | 0,1753 |
| TcCLB.508133.19 | hypothetical protein, conserved | -0,1561 | 0,2767 |
| TcCLB.510131.99 | hypothetical protein, conserved | -0,1562 | 0,2681 |
| TcCLB.509669.50 | hypothetical protein, conserved | -0,1563 | 0,08619 |
| TcCLB.419469.14 | kinesin, putative | -0,1564 | 0,5561 |
| TcCLB.506691.30 | hypothetical protein, conserved | -0,1565 | 0,04716 |
| TcCLB.510143.70 | hypothetical protein, conserved | -0,1565 | 0,06054 |
| TcCLB.404843.10 | hypothetical protein, conserved | -0,1565 | 0,2073 |
| TcCLB.437999.9 | hypothetical protein, conserved (fragment) | -0,1565 | 0,2572 |
| TcCLB.506405.20 | hypothetical protein, conserved | -0,1565 | 0,381 |
| TcCLB.510745.40 | protein kinase, putative | -0,1566 | 0,1529 |
| TcCLB.511507.100 | hypothetical protein, conserved | -0,1567 | 0,296 |
| TcCLB.509439.10 | retrotransposon hot spot protein (RHS, pseudogene), puta | -0,1568 | 0,4975 |
| TcCLB.506885.413 | glyceraldehyde-3-phosphate dehydrogenase, putative | -0,1569 | 0,1651 |
| TcCLB.506359.90 | hypothetical protein, conserved | -0,1569 | 0,1754 |
| TcCLB.507019.100 | nuclear movement protein, putative | -0,1569 | 0,1811 |
| TcCLB.508721.10 | small GTPase, putative | -0,1569 | 0,2059 |
| TcCLB.508839.30 | trafficking protein particle complex subunit 3, putative | -0,1569 | 0,4789 |
| TcCLB.507093.150 | hypothetical protein, conserved (pseudogene) | -0,157 | 0,09695 |
| TcCLB.505037.60 | NIMA/Nek Serine/threonine-protein kinase family, putativ | -0,157 | 0,176 |
| TcCLB.510329.370 | hypothetical protein, conserved | -0,157 | 0,2732 |
| TcCLB.508479.80 | retrotransposon hot spot (RHS) protein, putative | -0,157 | 0,2927 |
| TcCLB.435519.19 | hypothetical protein, conserved (fragment) | -0,157 | 0,4678 |
| TcCLB.511491.140 | hypothetical protein, conserved | -0,1571 | 0,1254 |
| TcCLB.506753.30 | hypothetical protein | -0,1571 | 0,2845 |
| TcCLB.504949.39 | DNA repair protein Rad9, putative (fragment) (Rad9) | -0,1572 | 0,2071 |
| TcCLB.509253.20 | tubulin tyrosine ligase, putative | -0,1576 | 0,2474 |
| TcCLB.478071.10 | trans-sialidase (pseudogene), putative | -0,1576 | 0,3043 |
| TcCLB.509805.110 | hypothetical protein, conserved | -0,1577 | 0,0994 |
| TcCLB.510317.20 | hypothetical protein, conserved | -0,1578 | 0,1095 |
| TcCLB.505683.5 | hypothetical protein | -0,1578 | 0,1706 |
| TcCLB.508207.74 | Adenine phosphoribosyltransferase, putative | -0,1578 | 0,2459 |
| TcCLB.503851.32 | trans-sialidase (pseudogene), putative | -0,1578 | 0,4411 |
| TcCLB.510519.149 | hypothetical protein, conserved | -0,1579 | 0,1126 |
| TcCLB.508719.20 | hypothetical protein, conserved | -0,158 | 0,3557 |
| TcCLB.430281.20 | hypothetical protein (fragment) | -0,158 | 0,6908 |
| TcCLB.506999.160 | hypothetical protein | -0,1581 | 0,4211 |
| TcCLB.506559.100 | multidrug resistance-associated protein, putative | -0,1582 | 0,113 |
| TcCLB.509611.89 | hypothetical protein, conserved | -0,159 | 0,4319 |
| TcCLB.509589.50 | hypothetical protein, conserved | -0,1591 | 0,1092 |
| TcCLB.511903.280 | hypothetical protein, conserved | -0,1591 | 0,225 |

| | | | |
|---|---|---|---|
| TcCLB.507623.20 | hypothetical protein, conserved | -0,1592 | 0,1213 |
| TcCLB.506009.50 | hypothetical protein, conserved | -0,1595 | 0,1198 |
| TcCLB.509001.10 | hypothetical protein, conserved | -0,1595 | 0,1745 |
| TcCLB.509901.110 | hypothetical protein, conserved | -0,1595 | 0,2659 |
| TcCLB.507541.50 | casein kinase, putative (fragment) | -0,1597 | 0,1012 |
| TcCLB.510285.90 | hypothetical protein, conserved | -0,1597 | 0,1684 |
| TcCLB.508441.40 | hypothetical protein, conserved | -0,1599 | 0,139 |
| TcCLB.511707.20 | hypothetical protein | -0,1599 | 0,2664 |
| TcCLB.506335.15 | hypothetical protein | -0,1601 | 0,5828 |
| TcCLB.503899.80 | hypothetical protein, conserved | -0,1602 | 0,172 |
| TcCLB.504215.20 | dolichyl-P-Man:GDP-ManGlcNAc2-PP-dolichyl beta-1,4-ma | -0,1602 | 0,2051 |
| TcCLB.511267.20 | hypothetical protein, conserved | -0,1602 | 0,3902 |
| TcCLB.503751.30 | hypothetical protein, conserved (pseudogene) | -0,1603 | 0,1594 |
| TcCLB.503747.20 | hypothetical protein | -0,1604 | 0,4236 |
| TcCLB.508799.140 | transcription modulator/accessory protein, putative | -0,1605 | 0,09739 |
| TcCLB.511109.90 | DnaJ chaperone protein, putative | -0,1607 | 0,2352 |
| TcCLB.507787.70 | hypothetical protein, conserved | -0,1607 | 0,2796 |
| TcCLB.511189.90 | hypothetical protein, conserved | -0,161 | 0,09512 |
| TcCLB.510031.40 | hypothetical protein | -0,161 | 0,2855 |
| TcCLB.508467.10 | hypothetical protein, conserved | -0,1612 | 0,4357 |
| TcCLB.504175.30 | hypothetical protein, conserved | -0,1613 | 0,2432 |
| TcCLB.510089.80 | protein kinase, putative | -0,1615 | 0,1649 |
| TcCLB.506369.30 | hypothetical protein, conserved | -0,1615 | 0,2879 |
| TcCLB.509345.20 | trans-sialidase (pseudogene), putative | -0,1616 | 0,5189 |
| TcCLB.508153.410 | hypothetical protein, conserved | -0,1617 | 0,1569 |
| TcCLB.506739.190 | serine/threonine protein kinase, putative | -0,1619 | 0,1598 |
| TcCLB.506201.100 | hypothetical protein, conserved | -0,1621 | 0,1705 |
| TcCLB.507093.50 | hypothetical protein, conserved | -0,1624 | 0,06137 |
| TcCLB.511111.40 | hypothetical protein, conserved | -0,1625 | 0,2144 |
| TcCLB.504137.90 | hypothetical protein, conserved | -0,1626 | 0,1333 |
| TcCLB.511153.64 | histone H2B variant, putative | -0,1627 | 0,1035 |
| TcCLB.510181.50 | nucleoporin interacting component (NUP93), putative | -0,163 | 0,1758 |
| TcCLB.509817.40 | trans-sialidase (pseudogene), putative | -0,163 | 0,3292 |
| TcCLB.508707.300 | hypothetical protein, conserved | -0,1631 | 0,04987 |
| TcCLB.508175.70 | lanosterol synthase, putative | -0,1631 | 0,1132 |
| TcCLB.511109.40 | hypothetical protein, conserved | -0,1631 | 0,14 |
| TcCLB.510251.10 | amino acid permease-like protein, putative | -0,1632 | 0,1074 |
| TcCLB.506007.10 | MP24, putative | -0,1632 | 0,3451 |
| TcCLB.508163.390 | hypothetical protein, conserved | -0,1632 | 0,4767 |
| TcCLB.506035.10 | retrotransposon hot spot protein (RHS, pseudogene), puta | -0,1633 | 0,1699 |
| TcCLB.507515.110 | hypothetical protein, conserved | -0,1633 | 0,1759 |
| TcCLB.422031.10 | solanesyl-diphosphate synthase, putative | -0,1633 | 0,2048 |
| TcCLB.511517.99 | hypothetical protein, conserved | -0,1633 | 0,2839 |
| TcCLB.510659.10 | hypothetical protein, conserved | -0,1634 | 0,1169 |
| TcCLB.508817.90 | hypothetical protein, conserved (pseudogene) | -0,1635 | 0,1799 |

| | | | |
|---|---|---|---|
| TcCLB.511087.4 | mucin TcMUCII (pseudogene), putative | -0,1635 | 0,6106 |
| TcCLB.507633.120 | mucin TcMUCII, putative | -0,1635 | 0,681 |
| TcCLB.509793.69 | hypothetical protein, conserved | -0,1636 | 0,08401 |
| TcCLB.506945.260 | hypothetical protein, conserved | -0,1637 | 0,05465 |
| TcCLB.508965.39 | hypothetical protein, conserved | -0,1637 | 0,1 |
| TcCLB.510339.54 | hypothetical protein, conserved | -0,1638 | 0,1085 |
| TcCLB.509243.60 | ATP-dependent DEAD/H DNA helicase recQ, putative | -0,1638 | 0,4983 |
| TcCLB.508879.150 | hypothetical protein, conserved | -0,1639 | 0,2758 |
| TcCLB.510513.30 | Probable methyltransferase BTM2 homolog | -0,164 | 0,251 |
| TcCLB.507209.30 | hypothetical protein, conserved | -0,1642 | 0,07743 |
| TcCLB.506801.129 | hypothetical protein (fragment) | -0,1642 | 0,1229 |
| TcCLB.509715.20 | clathrin coat assembly protein, putative | -0,1642 | 0,2101 |
| TcCLB.506605.90 | hypothetical protein, conserved | -0,1642 | 0,2316 |
| TcCLB.511389.100 | hypothetical protein, conserved | -0,1642 | 0,2381 |
| TcCLB.508577.120 | hypothetical protein, conserved | -0,1643 | 0,2574 |
| TcCLB.503823.90 | hypothetical protein, conserved | -0,1645 | 0,1196 |
| TcCLB.508707.220 | hypothetical protein | -0,1645 | 0,1876 |
| TcCLB.507099.130 | hypothetical protein, conserved | -0,1645 | 0,213 |
| TcCLB.510065.50 | hypothetical protein, conserved | -0,1645 | 0,275 |
| TcCLB.506883.100 | GTP-binding protein, putative | -0,1646 | 0,1085 |
| TcCLB.507669.110 | hypothetical protein, conserved | -0,1649 | 0,0656 |
| TcCLB.509429.90 | translation initiation factor EIF-2b alpha subunit, putative | -0,1649 | 0,2232 |
| TcCLB.507611.230 | trans-sialidase (pseudogene), putative | -0,165 | 0,3094 |
| TcCLB.507711.349 | hypothetical protein, conserved | -0,165 | 0,397 |
| TcCLB.511001.90 | hypothetical protein, conserved | -0,1652 | 0,02258 |
| TcCLB.506989.120 | hypothetical protein, conserved | -0,1652 | 0,2101 |
| TcCLB.506779.10 | dynein regulatory complex subunit, putative | -0,1652 | 0,2574 |
| TcCLB.508233.10 | hypothetical protein, conserved | -0,1654 | 0,2518 |
| TcCLB.505989.74 | peroxisome biogenesis factor 1, putative | -0,1655 | 0,09325 |
| TcCLB.506695.40 | hypothetical protein, conserved | -0,1655 | 0,2562 |
| TcCLB.510155.40 | DNA-damage inducible protein DDI1-like protein, putative | -0,1656 | 0,08771 |
| TcCLB.507037.10 | proteasome regulatory non-ATPase subunit 7 (pseudogen | -0,1656 | 0,1318 |
| TcCLB.511281.53 | hypothetical protein | -0,1656 | 0,6326 |
| TcCLB.509237.20 | hypothetical protein, conserved | -0,1658 | 0,05608 |
| TcCLB.504119.4 | DNA-directed RNA polymerase III subunit C1, putative (fra | -0,1658 | 0,4283 |
| TcCLB.508505.10 | hypothetical protein, conserved | -0,1659 | 0,1291 |
| TcCLB.508465.20 | hypothetical protein, conserved | -0,1659 | 0,1727 |
| TcCLB.510667.50 | hypothetical protein, conserved | -0,1659 | 0,1883 |
| TcCLB.511711.10 | hypothetical protein, conserved | -0,166 | 0,17 |
| TcCLB.511421.230 | Cell cycle checkpoint protein RAD1-like, putative | -0,1661 | 0,1415 |
| TcCLB.510329.310 | hypothetical protein, conserved | -0,1661 | 0,1616 |
| TcCLB.506355.160 | hypothetical protein, conserved | -0,1661 | 0,1859 |
| TcCLB.506211.220 | Associated kinase of Tb14-3-3, putative (AKB1) | -0,1662 | 0,0602 |
| TcCLB.510443.40 | hypothetical protein, conserved | -0,1663 | 0,1303 |
| TcCLB.511925.10 | hypothetical protein, conserved | -0,1664 | 0,1536 |

| | | | |
|---|---|---|---|
| TcCLB.508661.30 | DnaJ chaperone protein, putative | -0,1666 | 0,06847 |
| TcCLB.507011.80 | zinc finger protein, putative | -0,1667 | 0,1696 |
| TcCLB.506405.30 | actin-like protein, putative | -0,1667 | 0,2203 |
| TcCLB.510483.90 | trans-sialidase (pseudogene), putative | -0,1667 | 0,5516 |
| TcCLB.509735.30 | retrotransposon hot spot protein (RHS, pseudogene), puta | -0,1668 | 0,1319 |
| TcCLB.507129.10 | actin 2, putative | -0,1668 | 0,3239 |
| TcCLB.510297.100 | ADP-ribosylation factor-like protein, putative | -0,1671 | 0,1504 |
| TcCLB.504867.10 | hypothetical protein, conserved | -0,1671 | 0,2011 |
| TcCLB.506811.30 | hypothetical protein, conserved | -0,1672 | 0,1185 |
| TcCLB.511393.60 | protein kinase, putative | -0,1673 | 0,06974 |
| TcCLB.511589.170 | oxidoreductase, putative | -0,1673 | 0,2715 |
| TcCLB.507009.120 | DNA repair protein RAD2, putative | -0,1674 | 0,1454 |
| TcCLB.505169.10 | polyubiquitin (pseudogene), putative | -0,1674 | 0,2673 |
| TcCLB.509471.95 | histone H3, putative | -0,1675 | 0,2336 |
| TcCLB.510407.100 | hypothetical protein, conserved | -0,1676 | 0,1486 |
| TcCLB.503487.10 | DNA-directed RNA polymerase, mitochondrial (fragment) | -0,1676 | 0,3216 |
| TcCLB.510325.60 | hypothetical protein, conserved | -0,1677 | 0,1424 |
| TcCLB.508277.310 | hypothetical protein | -0,1677 | 0,2393 |
| TcCLB.508891.4 | hypothetical protein, conserved | -0,1677 | 0,2551 |
| TcCLB.506925.350 | hypothetical protein, conserved | -0,1678 | 0,07211 |
| TcCLB.511621.110 | hypothetical protein | -0,1678 | 0,1309 |
| TcCLB.508559.40 | retrotransposon hot spot protein (RHS, pseudogene), puta | -0,1678 | 0,2647 |
| TcCLB.511075.24 | zinc-finger multi-pass transmembrane protein, putative | -0,1679 | 0,1499 |
| TcCLB.507913.10 | hypothetical protein, conserved | -0,168 | 0,4273 |
| TcCLB.506795.50 | poly(A) polymerase, putative | -0,1681 | 0,07771 |
| TcCLB.511733.120 | hypothetical protein, conserved | -0,1681 | 0,08846 |
| TcCLB.510601.30 | hypothetical protein, conserved | -0,1681 | 0,1383 |
| TcCLB.508257.170 | dynein heavy chain, putative | -0,1681 | 0,1809 |
| TcCLB.503809.24 | hypothetical protein, conserved | -0,1681 | 0,318 |
| TcCLB.509065.60 | glyceraldehyde 3-phosphate dehydrogenase, putative | -0,1682 | 0,04902 |
| TcCLB.510515.100 | coronin, putative (CRN12) | -0,1682 | 0,07755 |
| TcCLB.506041.40 | hypothetical protein, conserved | -0,1682 | 0,1341 |
| TcCLB.508139.170 | retrotransposon hot spot protein (RHS, pseudogene), puta | -0,1687 | 0,1525 |
| TcCLB.508385.30 | CYC2-like cyclin, putative | -0,1688 | 0,06751 |
| TcCLB.509911.124 | hypothetical protein, conserved | -0,1688 | 0,2292 |
| TcCLB.506295.80 | ADP-ribosylation factor-like 2, arl2, putative (ARL2) | -0,1688 | 0,5066 |
| TcCLB.511261.60 | hypothetical protein, conserved | -0,1689 | 0,06251 |
| TcCLB.507603.60 | hypothetical protein, conserved | -0,1689 | 0,1707 |
| TcCLB.506025.20 | hypothetical protein, conserved | -0,1689 | 0,1788 |
| TcCLB.505807.254 | hypothetical protein | -0,169 | 0,3539 |
| TcCLB.506133.10 | syntaxin binding protein (pseudogene), putative | -0,169 | 0,6552 |
| TcCLB.506859.90 | prefoldin, putative | -0,1691 | 0,1377 |
| TcCLB.510283.40 | phosphoglycerate mutase family member 5, putative | -0,1691 | 0,306 |
| TcCLB.508541.200 | hypothetical protein, conserved | -0,1693 | 0,2902 |
| TcCLB.506193.5 | glucose-6-phosphate 1-dehydrogenase (pseudogene), put | -0,1693 | 0,4571 |

| | | | |
|---|---|---|---|
| TcCLB.511285.10 | hypothetical protein | -0,1694 | 0,06696 |
| TcCLB.511319.20 | hypothetical protein, conserved | -0,1694 | 0,06754 |
| TcCLB.509599.90 | hypothetical protein, conserved | -0,1694 | 0,1223 |
| TcCLB.507623.110 | surface protease GP63, putative | -0,1694 | 0,1276 |
| TcCLB.511287.10 | hypothetical protein, conserved | -0,1694 | 0,1909 |
| TcCLB.510943.170 | hypothetical protein, conserved | -0,1696 | 0,1448 |
| TcCLB.412435.10 | hypothetical protein, conserved | -0,1697 | 0,2441 |
| TcCLB.509967.60 | hypothetical protein, conserved | -0,1697 | 0,2487 |
| TcCLB.511903.134 | hypothetical protein, conserved | -0,1697 | 0,2874 |
| TcCLB.506359.10 | hypothetical protein, conserved | -0,1697 | 0,3027 |
| TcCLB.507005.40 | zinc finger protein, putative | -0,1698 | 0,219 |
| TcCLB.504049.30 | trans-sialidase (pseudogene), putative | -0,1699 | 0,5647 |
| TcCLB.506227.250 | hypothetical protein, conserved | -0,17 | 0,1203 |
| TcCLB.508027.80 | hypothetical protein, conserved | -0,1701 | 0,08873 |
| TcCLB.506463.10 | Lys-63-specific deubiquitinase BRCC36, putative | -0,1701 | 0,1975 |
| TcCLB.507649.100 | hypothetical protein, conserved | -0,1702 | 0,08369 |
| TcCLB.508397.20 | hypothetical protein, conserved | -0,1702 | 0,2877 |
| TcCLB.506835.40 | hypothetical protein, conserved | -0,1703 | 0,1479 |
| TcCLB.510657.10 | Flagellar Member 5 (fragment) (FLAM5) | -0,1703 | 0,2455 |
| TcCLB.511511.20 | tRNA modification enzyme, putative | -0,1703 | 0,3129 |
| TcCLB.509979.100 | hypothetical protein, conserved | -0,1704 | 0,2723 |
| TcCLB.506177.50 | hypothetical protein, conserved | -0,1705 | 0,04618 |
| TcCLB.503939.100 | hypothetical protein, conserved | -0,1706 | 0,06117 |
| TcCLB.508153.550 | hypothetical protein, conserved | -0,1706 | 0,1074 |
| TcCLB.507897.4 | inositol polyphosphate 1-phosphatase, putative | -0,1706 | 0,1517 |
| TcCLB.503697.110 | hypothetical protein, conserved | -0,1707 | 0,1747 |
| TcCLB.504899.20 | retrotransposon hot spot protein (RHS, pseudogene), puta | -0,1707 | 0,193 |
| TcCLB.506127.110 | eukaryotic peptide chain release factor subunit 1, putative | -0,1708 | 0,07751 |
| TcCLB.507547.70 | hypothetical protein, conserved | -0,1709 | 0,08229 |
| TcCLB.503823.110 | hypothetical protein, conserved | -0,1709 | 0,2304 |
| TcCLB.509195.10 | hypothetical protein | -0,1709 | 0,6293 |
| TcCLB.508723.24 | hypothetical protein, conserved | -0,171 | 0,1673 |
| TcCLB.510173.80 | hypothetical protein | -0,1712 | 0,2335 |
| TcCLB.507093.63 | RNA-binding protein, putative | -0,1713 | 0,1925 |
| TcCLB.509537.20 | hypothetical protein, conserved | -0,1715 | 0,1338 |
| TcCLB.503885.90 | hypothetical protein, conserved | -0,1716 | 0,101 |
| TcCLB.508895.89 | intraflagellar transport protein 80, putative (fragment) (IF | -0,1717 | 0,1887 |
| TcCLB.506227.140 | hypothetical protein, conserved | -0,1717 | 0,2681 |
| TcCLB.511075.30 | TFIIH basal transcription factor complex helicase subunit, | -0,1718 | 0,02402 |
| TcCLB.508899.30 | hypothetical protein, conserved | -0,1718 | 0,2099 |
| TcCLB.504009.40 | Axonemal inner arm dynein light chain, putative | -0,1719 | 0,1885 |
| TcCLB.509837.40 | histone H1, putative | -0,1721 | 0,3359 |
| TcCLB.511109.110 | hypothetical protein, conserved | -0,1721 | 0,3516 |
| TcCLB.509053.50 | hypothetical protein, conserved | -0,1722 | 0,4436 |
| TcCLB.508387.70 | hypothetical protein, conserved | -0,1723 | 0,03468 |

| | | | |
|---|---|---|---|
| TcCLB.511275.40 | hypothetical protein, conserved | -0,1724 | 0,2656 |
| TcCLB.507681.240 | hypothetical protein, conserved | -0,1725 | 0,1007 |
| TcCLB.511851.20 | retrotransposon hot spot protein (RHS, pseudogene), puta | -0,1726 | 0,3491 |
| TcCLB.511587.111 | N-acetyltransferase complex ARD1 subunit (pseudogene), | -0,1726 | 0,5064 |
| TcCLB.510073.70 | CCR4-NOT transcription complex subunit 2, putative | -0,1727 | 0,08047 |
| TcCLB.510007.20 | hypothetical protein, conserved | -0,1727 | 0,1403 |
| TcCLB.506891.30 | leucine-rich repeat-containing protein | -0,1727 | 0,2572 |
| TcCLB.510817.10 | protein kinase, putative | -0,1727 | 0,5232 |
| TcCLB.508873.40 | mucin TcMUCII (pseudogene), putative | -0,1727 | 0,5458 |
| TcCLB.507257.70 | protein farnesyltransferase alpha subunit, putative | -0,1728 | 0,09225 |
| TcCLB.511071.20 | hypothetical protein, conserved | -0,1729 | 0,254 |
| TcCLB.507681.90 | hypothetical protein, conserved | -0,173 | 0,1043 |
| TcCLB.508973.60 | hypothetical protein, conserved | -0,1731 | 0,1254 |
| TcCLB.511717.180 | hypothetical protein, conserved | -0,1731 | 0,1487 |
| TcCLB.504643.49 | hypothetical protein, conserved (fragment) | -0,1733 | 0,05709 |
| TcCLB.511001.80 | hypothetical protein, conserved | -0,1733 | 0,08211 |
| TcCLB.508737.140 | hypothetical protein, conserved | -0,1733 | 0,08901 |
| TcCLB.510665.20 | hypothetical protein, conserved | -0,1734 | 0,1055 |
| TcCLB.507709.30 | hypothetical protein, conserved | -0,1734 | 0,2743 |
| TcCLB.506475.120 | conserved protein, unknown function | -0,1735 | 0,1211 |
| TcCLB.511203.20 | surface protease GP63, putative | -0,1735 | 0,3164 |
| TcCLB.511261.90 | hypothetical protein, conserved | -0,1737 | 0,1443 |
| TcCLB.503717.90 | surface protease GP63, putative | -0,1737 | 0,6675 |
| TcCLB.511755.50 | hypothetical protein, conserved | -0,1738 | 0,1422 |
| TcCLB.508001.60 | trans-sialidase (pseudogene), putative | -0,1738 | 0,4472 |
| TcCLB.506629.110 | hypothetical protein, conserved | -0,1739 | 0,1337 |
| TcCLB.510901.150 | proteasome regulatory non-ATPase subunit 2, putative | -0,174 | 0,03755 |
| TcCLB.503643.6 | hypothetical protein, conserved | -0,174 | 0,1411 |
| TcCLB.511589.74 | hypothetical protein | -0,174 | 0,1597 |
| TcCLB.507009.80 | hypothetical protein, conserved | -0,174 | 0,259 |
| TcCLB.505555.30 | hypothetical protein, conserved | -0,1741 | 0,03485 |
| TcCLB.511423.80 | Transmembrane protein 231, putative | -0,1741 | 0,0851 |
| TcCLB.478711.10 | trans-sialidase (pseudogene), putative | -0,1741 | 0,1564 |
| TcCLB.509381.10 | mitochondrial import receptor subunit, putative | -0,1742 | 0,09733 |
| TcCLB.503655.40 | surface protease GP63 (pseudogene), putative | -0,1742 | 0,4546 |
| TcCLB.504089.54 | protein kinase, putative | -0,1743 | 0,0864 |
| TcCLB.507809.170 | protein kinase, putative | -0,1743 | 0,5143 |
| TcCLB.504157.100 | hypothetical protein, conserved | -0,1745 | 0,1231 |
| TcCLB.508163.81 | hypothetical protein, conserved (pseudogene) | -0,1745 | 0,6655 |
| TcCLB.508153.150 | hypothetical protein, conserved | -0,1746 | 0,1313 |
| TcCLB.508209.120 | 10 kDa heat shock protein, putative | -0,1747 | 0,5816 |
| TcCLB.511487.71 | hypothetical protein | -0,1748 | 0,1734 |
| TcCLB.506479.10 | hypothetical protein, conserved | -0,1748 | 0,2107 |
| TcCLB.504827.170 | kinetoplastid kinetochore protein 2, putative (fragment) (I | -0,1749 | 0,2137 |
| TcCLB.425597.5 | hypothetical protein, conserved (pseudogene) | -0,1749 | 0,4055 |

| | | | |
|---|---|---|---|
| TcCLB.510387.30 | surface protease GP63 (pseudogene), putative | -0,175 | 0,6258 |
| TcCLB.510621.60 | mucin-associated surface protein (MASP), putative | -0,1751 | 0,6025 |
| TcCLB.508465.70 | hypothetical protein, conserved | -0,1752 | 0,1452 |
| TcCLB.510421.29 | CAAX prenyl protease 2, putative | -0,1752 | 0,2236 |
| TcCLB.507063.180 | hypothetical protein, conserved | -0,1752 | 0,2825 |
| TcCLB.510431.290 | hypothetical protein | -0,1752 | 0,3516 |
| TcCLB.506865.20 | hypothetical protein | -0,1753 | 0,1521 |
| TcCLB.506973.30 | mucin-associated surface protein (MASP), putative | -0,1753 | 0,4884 |
| TcCLB.503835.5 | rac serine threonine protein kinase, putative | -0,1754 | 0,5187 |
| TcCLB.510737.79 | 60S ribosomal subunit protein L31, putative | -0,1754 | 0,6013 |
| TcCLB.506503.100 | U3 snoRNA-associated protein UTP11, putative | -0,1755 | 0,1915 |
| TcCLB.509693.190 | ABC transporter, putative | -0,1755 | 0,1987 |
| TcCLB.508643.30 | DnaJ chaperone protein, putative | -0,1755 | 0,2653 |
| TcCLB.506445.100 | E2-like ubiquitin-conjugation enzyme, putative (UFC1) | -0,1757 | 0,03874 |
| TcCLB.507053.70 | geranylgeranyl transferase type II beta subunit, putative | -0,1759 | 0,1202 |
| TcCLB.508207.170 | hypothetical protein | -0,1761 | 0,09666 |
| TcCLB.508207.260 | hypothetical protein, conserved | -0,1761 | 0,1806 |
| TcCLB.508693.40 | hypothetical protein, conserved | -0,1762 | 0,0614 |
| TcCLB.508779.30 | synaptobrevin, putative | -0,1763 | 0,09706 |
| TcCLB.511213.50 | mucin TcMUCII, putative | -0,1765 | 0,3516 |
| TcCLB.509533.10 | trans-sialidase, Group II, putative | -0,1766 | 0,3172 |
| TcCLB.509959.40 | beta galactofuranosyl glycosyltransferase, putative | -0,1766 | 0,6818 |
| TcCLB.508173.47 | autophagy-related protein 8 (ATG8) | -0,1769 | 0,1382 |
| TcCLB.505807.40 | hypothetical protein, conserved | -0,177 | 0,03778 |
| TcCLB.507623.125 | hypothetical protein, conserved | -0,177 | 0,4988 |
| TcCLB.506985.40 | Polyamine transporter 1, putative (POT1.1) | -0,1771 | 0,1067 |
| TcCLB.506047.13 | ATP-dependent DEAD/H RNA helicase (pseudogene), puta | -0,1772 | 0,6163 |
| TcCLB.507711.280 | hypothetical protein, conserved | -0,1773 | 0,2038 |
| TcCLB.507625.150 | hypothetical protein, conserved | -0,1773 | 0,2347 |
| TcCLB.511383.89 | Flagellar Member 3 (FLAM3) | -0,1774 | 0,2121 |
| TcCLB.506925.60 | hypothetical protein, conserved | -0,1774 | 0,2154 |
| TcCLB.511237.120 | hypothetical protein | -0,1775 | 0,432 |
| TcCLB.508141.80 | hypothetical protein, conserved | -0,1776 | 0,05809 |
| TcCLB.510643.121 | trans-sialidase (pseudogene), putative | -0,1776 | 0,3422 |
| TcCLB.505655.20 | hypothetical protein | -0,1776 | 0,5747 |
| TcCLB.506755.160 | AAA ATPase, putative | -0,1777 | 0,01698 |
| TcCLB.508461.230 | kinetoplastid kinetochore protein 3, putative (kkt3) | -0,1778 | 0,1252 |
| TcCLB.508231.200 | 40S ribosomal protein S8, putative | -0,1779 | 0,1157 |
| TcCLB.510031.30 | hypothetical protein | -0,1781 | 0,1308 |
| TcCLB.509379.29 | hypothetical protein, conserved | -0,1781 | 0,3472 |
| TcCLB.509693.150 | hypothetical protein, conserved | -0,1782 | 0,1127 |
| TcCLB.510309.60 | hypothetical protein, conserved | -0,1783 | 0,1201 |
| TcCLB.507063.150 | hypothetical protein, conserved | -0,1783 | 0,1619 |
| TcCLB.447255.29 | multidrug resistance-associated protein, putative | -0,1783 | 0,3447 |
| TcCLB.510659.74 | hypothetical protein, conserved | -0,1783 | 0,4233 |

| | | | |
|---|---|---|---|
| TcCLB.503513.40 | proteasome regulatory non-ATPase subunit 6, putative | -0,1785 | 0,1459 |
| TcCLB.511927.30 | hypothetical protein, conserved | -0,1785 | 0,3914 |
| TcCLB.504097.4 | nucleoplasmin-like protein (NLP), putative | -0,1786 | 0,1405 |
| TcCLB.506573.60 | hypothetical protein, conserved | -0,1786 | 0,1467 |
| TcCLB.410923.30 | hypothetical protein | -0,1786 | 0,4568 |
| TcCLB.510709.31 | syntaxin binding protein (pseudogene), putative | -0,1786 | 0,7284 |
| TcCLB.510733.40 | hypothetical protein, conserved | -0,1787 | 0,1132 |
| TcCLB.505007.10 | RNA-binding protein, putative | -0,1787 | 0,2179 |
| TcCLB.503735.30 | hypothetical protein, conserved | -0,1788 | 0,1252 |
| TcCLB.509287.10 | hypothetical protein, conserved | -0,1788 | 0,2521 |
| TcCLB.504057.80 | Present in the outer mitochondrial membrane proteome 2 | -0,1789 | 0,06247 |
| TcCLB.506855.290 | hypothetical protein, conserved | -0,1789 | 0,1962 |
| TcCLB.507519.140 | Adenine phosphoribosyltransferase, putative | -0,179 | 0,1265 |
| TcCLB.508207.250 | mitotic cyclin, putative | -0,1791 | 0,205 |
| TcCLB.511753.100 | ABC transporter, putative | -0,1792 | 0,143 |
| TcCLB.510141.20 | hypothetical protein, conserved | -0,1793 | 0,1639 |
| TcCLB.506755.120 | hypothetical protein, conserved | -0,1795 | 0,0384 |
| TcCLB.509489.29 | proteasome regulatory non-ATP-ase subunit, putative (fra | -0,1795 | 0,2108 |
| TcCLB.408799.10 | hypothetical protein, conserved | -0,1796 | 0,2607 |
| TcCLB.504105.30 | DNA-directed RNA polymerase III subunit C1, putative (RP | -0,1797 | 0,00703 |
| TcCLB.506665.10 | hypothetical protein | -0,1797 | 0,6868 |
| TcCLB.507669.100 | transcription elongation factor, putative | -0,1798 | 0,09938 |
| TcCLB.507053.100 | ubiquitin hydrolase, putative | -0,1799 | 0,04279 |
| TcCLB.511421.150 | leucine-rich repeat-containing protein | -0,1799 | 0,04332 |
| TcCLB.508179.10 | DnaJ chaperone protein, putative | -0,1799 | 0,2948 |
| TcCLB.511173.110 | mucin TcMUCII, putative | -0,1799 | 0,5365 |
| TcCLB.511709.10 | hypothetical protein, conserved | -0,18 | 0,1644 |
| TcCLB.508153.40 | hypothetical protein, conserved | -0,18 | 0,228 |
| TcCLB.508317.20 | hypothetical protein, conserved | -0,1801 | 0,2073 |
| TcCLB.510421.140 | hypothetical protein, conserved | -0,1801 | 0,2117 |
| TcCLB.509161.130 | ATP-dependent DEAD/H RNA helicase, putative | -0,1802 | 0,03466 |
| TcCLB.507093.240 | hypothetical protein, conserved | -0,1802 | 0,04922 |
| TcCLB.506211.240 | hypothetical protein, conserved | -0,1803 | 0,0516 |
| TcCLB.507993.140 | RNA-binding protein 29, putative (RBP29) | -0,1803 | 0,08169 |
| TcCLB.510609.100 | hypothetical protein, conserved | -0,1803 | 0,1802 |
| TcCLB.503703.80 | hypothetical protein, conserved | -0,1806 | 0,1945 |
| TcCLB.508943.10 | hypothetical protein, conserved | -0,1807 | 0,2019 |
| TcCLB.507559.60 | hypothetical protein, conserved | -0,1808 | 0,0838 |
| TcCLB.511727.60 | hypothetical protein, conserved | -0,1809 | 0,03538 |
| TcCLB.510941.20 | hypothetical protein, conserved | -0,1809 | 0,2124 |
| TcCLB.506009.20 | hypothetical protein, conserved | -0,181 | 0,04042 |
| TcCLB.506943.120 | hypothetical protein, conserved | -0,1815 | 0,1923 |
| TcCLB.508605.40 | retrotransposon hot spot protein (RHS, pseudogene), puta | -0,1815 | 0,3083 |
| TcCLB.510091.60 | hypothetical protein, conserved | -0,1816 | 0,105 |
| TcCLB.401661.10 | trans-sialidase (pseudogene), putative | -0,1816 | 0,1175 |

| | | | |
|---|---|---|---|
| TcCLB.510575.140 | hypothetical protein, conserved | -0,1817 | 0,1828 |
| TcCLB.507259.4 | hypothetical protein, conserved | -0,1817 | 0,2323 |
| TcCLB.508029.30 | ATP dependent DEAD-box helicase, putative (RH) | -0,1818 | 0,03991 |
| TcCLB.511625.10 | transcription initiation protein, putative | -0,1819 | 0,06921 |
| TcCLB.510763.70 | hypothetical protein, conserved | -0,1819 | 0,1313 |
| TcCLB.447483.4 | Meckel syndrome type 1 protein, putative (fragment) (mk | -0,182 | 0,3693 |
| TcCLB.504057.60 | hypothetical protein, conserved | -0,1821 | 0,05986 |
| TcCLB.510645.20 | nuclear transcription factor, putative | -0,1821 | 0,09345 |
| TcCLB.506743.20 | hypothetical protein, conserved | -0,1821 | 0,3086 |
| TcCLB.510387.70 | hypothetical protein, conserved (pseudogene) | -0,1821 | 0,3615 |
| TcCLB.507801.19 | dynein heavy chain, putative | -0,1822 | 0,1351 |
| TcCLB.508207.20 | protein kinase, putative | -0,1823 | 0,3433 |
| TcCLB.511185.60 | trans-sialidase (pseudogene), putative | -0,1823 | 0,471 |
| TcCLB.511875.50 | syntaxin binding protein (pseudogene), putative | -0,1824 | 0,1571 |
| TcCLB.509453.90 | hypothetical protein, conserved | -0,1826 | 0,05051 |
| TcCLB.509429.180 | hypothetical protein, conserved | -0,1826 | 0,06494 |
| TcCLB.507049.10 | hypothetical protein, conserved | -0,1826 | 0,1507 |
| TcCLB.506607.19 | phosphoribosylpyrophosphate synthetase, putative | -0,1826 | 0,186 |
| TcCLB.511019.74 | retrotransposon hot spot (RHS) protein, putative | -0,1827 | 0,09873 |
| TcCLB.508357.26 | hypothetical protein, conserved | -0,1829 | 0,1631 |
| TcCLB.506811.220 | hypothetical protein, conserved | -0,183 | 0,04045 |
| TcCLB.508911.60 | hypothetical protein, conserved | -0,1831 | 0,05443 |
| TcCLB.511545.150 | hypothetical protein, conserved | -0,1831 | 0,1038 |
| TcCLB.511517.44 | chaperone DNAJ protein, putative | -0,1834 | 0,09834 |
| TcCLB.507013.24 | poly(ADP-ribose) glycohydrolase, putative | -0,1834 | 0,1454 |
| TcCLB.509937.50 | hypothetical protein, conserved | -0,1836 | 0,07438 |
| TcCLB.509967.90 | hypothetical protein, conserved | -0,1837 | 0,1651 |
| TcCLB.509099.130 | hypothetical protein, conserved | -0,1839 | 0,05776 |
| TcCLB.503423.20 | protein kinase, putative | -0,184 | 0,6726 |
| TcCLB.509955.50 | hypothetical protein, conserved | -0,184 | 0,6901 |
| TcCLB.510255.4 | hypothetical protein, conserved | -0,1842 | 0,5151 |
| TcCLB.509161.80 | hypothetical protein, conserved | -0,1843 | 0,1394 |
| TcCLB.511257.20 | hypothetical protein, conserved | -0,1845 | 0,09683 |
| TcCLB.511183.49 | mucin TcMUCII (pseudogene), putative | -0,1847 | 0,699 |
| TcCLB.511277.450 | GPI-anchor transamidase subunit 8, putative (GPI8) | -0,1849 | 0,1338 |
| TcCLB.506275.80 | hypothetical protein, conserved | -0,185 | 0,1899 |
| TcCLB.503539.20 | Putative intraflagellar transport protein A1 (PIFTA1) | -0,1851 | 0,06818 |
| TcCLB.511131.20 | hypothetical protein, conserved | -0,1851 | 0,09638 |
| TcCLB.511517.50 | hypothetical protein, conserved | -0,1852 | 0,1283 |
| TcCLB.511003.70 | hypothetical protein, conserved | -0,1855 | 0,2382 |
| TcCLB.504069.110 | hypothetical protein, conserved | -0,1855 | 0,2607 |
| TcCLB.511497.10 | hypothetical protein, conserved | -0,1855 | 0,3185 |
| TcCLB.508307.90 | hypothetical protein, conserved | -0,1856 | 0,0558 |
| TcCLB.506637.60 | hypothetical protein, conserved | -0,1856 | 0,1694 |
| TcCLB.508699.90 | surface protease GP63, putative | -0,1856 | 0,2585 |

| | | | |
|---|---|---|---|
| TcCLB.503903.70 | hypothetical protein, conserved | -0,1858 | 0,1953 |
| TcCLB.510889.170 | hypothetical protein, conserved | -0,1859 | 0,02399 |
| TcCLB.506247.130 | hypothetical protein, conserved | -0,1859 | 0,03918 |
| TcCLB.509977.61 | surface protease GP63 (pseudogene), putative | -0,1859 | 0,2897 |
| TcCLB.503581.39 | coatomer delta subunit, putative | -0,1861 | 0,1343 |
| TcCLB.510091.20 | hypothetical protein, conserved | -0,1861 | 0,2869 |
| TcCLB.506665.30 | trans-sialidase, putative (fragment) | -0,1861 | 0,5213 |
| TcCLB.506727.140 | hypothetical protein, conserved | -0,1862 | 0,1714 |
| TcCLB.506637.100 | tyrosine phosphatase isoform, putative | -0,1863 | 0,144 |
| TcCLB.507793.60 | hypothetical protein, conserved | -0,1863 | 0,3861 |
| TcCLB.510123.24 | hypothetical protein, conserved | -0,1865 | 0,1217 |
| TcCLB.509885.20 | hypothetical protein, conserved | -0,1865 | 0,2078 |
| TcCLB.511295.30 | hypothetical protein, conserved | -0,187 | 0,03414 |
| TcCLB.511255.420 | mucin-associated surface protein (MASP, pseudogene), pu | -0,1872 | 0,4938 |
| TcCLB.511395.60 | serine/threonine protein kinase, putative | -0,1873 | 0,04131 |
| TcCLB.506755.230 | hypothetical protein, conserved | -0,1873 | 0,1751 |
| TcCLB.510421.200 | hypothetical protein, conserved | -0,1875 | 0,1125 |
| TcCLB.503735.40 | hypothetical protein | -0,1875 | 0,1379 |
| TcCLB.498829.21 | hypothetical protein | -0,1875 | 0,6826 |
| TcCLB.504797.144 | hypothetical protein, conserved | -0,1876 | 0,07685 |
| TcCLB.503823.140 | hypothetical protein, conserved | -0,1876 | 0,1625 |
| TcCLB.510967.16 | hypothetical protein, conserved | -0,1877 | 0,1192 |
| TcCLB.506287.130 | hypothetical protein, conserved | -0,1878 | 0,1175 |
| TcCLB.506257.60 | XPA-interacting protein, putative | -0,1878 | 0,1323 |
| TcCLB.510197.5 | hypothetical protein, conserved (pseudogene) | -0,1878 | 0,4437 |
| TcCLB.508163.340 | trans-sialidase (pseudogene), putative | -0,1878 | 0,6361 |
| TcCLB.429257.10 | hypothetical protein, conserved | -0,1881 | 0,3936 |
| TcCLB.509773.20 | protein kinase, putative | -0,1881 | 0,5058 |
| TcCLB.511731.50 | ATP-dependent RNA helicase, putative | -0,1882 | 0,1301 |
| TcCLB.510743.110 | hypothetical protein, conserved | -0,1884 | 0,1283 |
| TcCLB.506895.80 | trans-sialidase, putative | -0,1884 | 0,2273 |
| TcCLB.504191.19 | sterol 24-c-methyltransferase, putative | -0,1884 | 0,3629 |
| TcCLB.508071.80 | trans-sialidase (pseudogene), putative | -0,1886 | 0,09955 |
| TcCLB.511201.19 | hypothetical protein (fragment) | -0,1886 | 0,1345 |
| TcCLB.506177.120 | hypothetical protein | -0,1886 | 0,1699 |
| TcCLB.509319.60 | endo/exonuclease Mre11, putative | -0,1887 | 0,03068 |
| TcCLB.511871.100 | hypothetical protein, conserved | -0,1887 | 0,08067 |
| TcCLB.507857.80 | hypothetical protein, conserved (pseudogene) | -0,1888 | 0,08063 |
| TcCLB.511825.66 | hypothetical protein, conserved | -0,1888 | 0,2871 |
| TcCLB.510187.360 | hypothetical protein, conserved | -0,189 | 0,05736 |
| TcCLB.511421.155 | hypothetical protein, conserved | -0,189 | 0,1305 |
| TcCLB.509157.100 | hypothetical protein | -0,189 | 0,2289 |
| TcCLB.511367.10 | hypothetical protein | -0,189 | 0,6701 |
| TcCLB.508461.340 | hypothetical protein, conserved | -0,1891 | 0,02169 |
| TcCLB.505765.10 | hypothetical protein, conserved (pseudogene) | -0,1891 | 0,2437 |

| | | | |
|---|---|---|---|
| TcCLB.507739.30 | hypothetical protein, conserved | -0,1893 | 0,064 |
| TcCLB.507991.50 | kinesin, putative | -0,1895 | 0,03695 |
| TcCLB.503837.30 | hypothetical protein, conserved | -0,1895 | 0,2086 |
| TcCLB.503845.20 | dispersed gene family protein 1 (DGF-1, pseudogene), put | -0,1895 | 0,4057 |
| TcCLB.508949.50 | protein kinase, putative | -0,1896 | 0,1452 |
| TcCLB.508689.10 | hypothetical protein, conserved | -0,1896 | 0,1508 |
| TcCLB.511323.60 | hypothetical protein, conserved | -0,1897 | 0,08119 |
| TcCLB.506793.90 | hypothetical protein, conserved | -0,1897 | 0,1048 |
| TcCLB.506855.330 | hypothetical protein, conserved | -0,1899 | 0,06894 |
| TcCLB.508533.50 | hypothetical protein, conserved | -0,19 | 0,01016 |
| TcCLB.507675.39 | hypothetical protein, conserved | -0,19 | 0,1144 |
| TcCLB.510001.40 | hypothetical protein, conserved | -0,19 | 0,1151 |
| TcCLB.508347.24 | hypothetical protein, conserved | -0,1901 | 0,2099 |
| TcCLB.508221.650 | trans-sialidase, Group V, putative | -0,1901 | 0,4615 |
| TcCLB.450061.10 | hypothetical protein, conserved | -0,1902 | 0,08136 |
| TcCLB.509455.50 | hypothetical protein, conserved | -0,1902 | 0,1163 |
| TcCLB.506357.160 | hypothetical protein, conserved | -0,1903 | 0,2515 |
| TcCLB.511493.10 | quiescin sulfhydryl oxidase, putative (QSOX) | -0,1904 | 0,128 |
| TcCLB.508637.114 | histone deacetylase 1, putative | -0,1904 | 0,2612 |
| TcCLB.506131.151 | elongation factor 1-gamma (EF-1-gamma, pseudogene), | | -0,1905 | 0,6996 |
| TcCLB.511435.30 | MORN repeat-containing protein | -0,1906 | 0,05228 |
| TcCLB.509363.31 | hypothetical protein | -0,1906 | 0,5747 |
| TcCLB.511901.20 | hypothetical protein, conserved | -0,1907 | 0,03848 |
| TcCLB.511179.130 | hypothetical protein, conserved | -0,1907 | 0,08402 |
| TcCLB.509669.40 | Zn-finger protein, putative | -0,1909 | 0,01231 |
| TcCLB.511715.90 | hypothetical protein, conserved | -0,1909 | 0,1002 |
| TcCLB.509805.30 | hypothetical protein, conserved | -0,1909 | 0,208 |
| TcCLB.511165.70 | proteasome alpha 7 subunit, putative | -0,191 | 0,08757 |
| TcCLB.507899.5 | trans-sialidase (pseudogene), putative | -0,191 | 0,1072 |
| TcCLB.506525.130 | mitochondrial carrier protein, putative | -0,191 | 0,1283 |
| TcCLB.506885.354 | hypothetical protein, conserved | -0,1911 | 0,3414 |
| TcCLB.508277.70 | hypothetical protein, conserved | -0,1912 | 0,1204 |
| TcCLB.504035.60 | hypothetical protein, conserved | -0,1912 | 0,2012 |
| TcCLB.510043.50 | hypothetical protein, conserved | -0,1913 | 0,0171 |
| TcCLB.507093.280 | hypothetical protein | -0,1913 | 0,0202 |
| TcCLB.506529.280 | hypothetical protein, conserved | -0,1914 | 0,1318 |
| TcCLB.507691.40 | trans-sialidase (pseudogene), putative | -0,1915 | 0,1432 |
| TcCLB.510321.10 | RNA polymerase-associated protein CTR9, putative | -0,1915 | 0,1879 |
| TcCLB.511891.10 | trans-sialidase, Group II, putative | -0,1916 | 0,1741 |
| TcCLB.506947.90 | adenylate kinase, putative | -0,1917 | 0,1347 |
| TcCLB.510065.60 | hypothetical protein, conserved | -0,1919 | 0,2174 |
| TcCLB.508209.110 | hypothetical protein, conserved | -0,192 | 0,02368 |
| TcCLB.506519.110 | hypothetical protein, conserved | -0,1921 | 0,08145 |
| TcCLB.509007.20 | hypothetical protein, conserved | -0,1922 | 0,06641 |
| TcCLB.511715.70 | hypothetical protein, conserved | -0,1922 | 0,1064 |

| | | | |
|---|---|---|---|
| TcCLB.506363.130 | hypothetical protein, conserved | -0,1923 | 0,1368 |
| TcCLB.504149.50 | hypothetical protein, conserved | -0,1924 | 0,0313 |
| TcCLB.506715.20 | Lys-63-specific deubiquitinase BRCC36, putative | -0,1924 | 0,252 |
| TcCLB.511751.220 | hypothetical protein, conserved | -0,1926 | 0,06259 |
| TcCLB.509633.20 | hypothetical protein, conserved | -0,1926 | 0,06796 |
| TcCLB.507963.50 | nuclear cap binding complex subunit CBP30, putative (CBI | -0,1926 | 0,1095 |
| TcCLB.510329.170 | hypothetical protein, conserved | -0,1929 | 0,03881 |
| TcCLB.503999.40 | hypothetical protein, conserved | -0,1929 | 0,05445 |
| TcCLB.509119.39 | dynein heavy chain, putative | -0,1929 | 0,313 |
| TcCLB.510989.20 | hypothetical protein, conserved | -0,193 | 0,09695 |
| TcCLB.508045.20 | hypothetical protein, conserved | -0,1933 | 0,08576 |
| TcCLB.509551.140 | hypothetical protein, conserved | -0,1934 | 0,05267 |
| TcCLB.506743.120 | Elongator-like Protein 3a, putative | -0,1934 | 0,07912 |
| TcCLB.511643.61 | elongation factor 1-gamma (EF-1-gamma, pseudogene), | -0,1934 | 0,2294 |
| TcCLB.508543.10 | DNA polymerase sigma, putative | -0,1936 | 0,02428 |
| TcCLB.504427.280 | hypothetical protein, conserved | -0,1936 | 0,08473 |
| TcCLB.510729.119 | hypothetical protein, conserved | -0,1936 | 0,1999 |
| TcCLB.506679.10 | hypothetical protein, conserved | -0,1936 | 0,2912 |
| TcCLB.509003.50 | hypothetical protein, conserved | -0,1936 | 0,295 |
| TcCLB.508989.80 | hypothetical protein | -0,1937 | 0,03101 |
| TcCLB.506319.70 | hypothetical protein | -0,1939 | 0,6211 |
| TcCLB.507927.40 | hypothetical protein, conserved | -0,1941 | 0,02907 |
| TcCLB.511153.144 | hypothetical protein, conserved | -0,1941 | 0,3988 |
| TcCLB.507969.20 | hypothetical protein, conserved | -0,1943 | 0,1625 |
| TcCLB.506181.120 | hypothetical protein, conserved | -0,1943 | 0,3086 |
| TcCLB.503435.30 | hypothetical protein, conserved | -0,1945 | 0,03525 |
| TcCLB.506525.120 | hypothetical protein, conserved | -0,1945 | 0,3556 |
| TcCLB.504129.20 | hypothetical protein, conserved | -0,1946 | 0,01969 |
| TcCLB.506435.390 | metallo-peptidase, Clan MA(E) Family M41 | -0,1946 | 0,05015 |
| TcCLB.508917.20 | hypothetical protein, conserved | -0,1946 | 0,1085 |
| TcCLB.509023.209 | hypothetical protein, conserved | -0,1949 | 0,06824 |
| TcCLB.511277.20 | DNA-directed RNA polymerase II/III subunit, putative | -0,195 | 0,1263 |
| TcCLB.511491.100 | phosphatase 2C, putative | -0,1952 | 0,07916 |
| TcCLB.511231.14 | hypothetical protein (fragment) | -0,1952 | 0,0826 |
| TcCLB.506479.100 | hypothetical protein, conserved | -0,1952 | 0,1411 |
| TcCLB.503687.58 | AMP deaminase, putative | -0,1953 | 0,2636 |
| TcCLB.507023.130 | hypothetical protein, conserved | -0,1955 | 0,1001 |
| TcCLB.506145.10 | hypothetical protein, conserved | -0,1959 | 0,01222 |
| TcCLB.505097.49 | expression site-associated gene (ESAG-like) protein, puta | -0,196 | 0,5616 |
| TcCLB.507521.120 | hypothetical protein, conserved | -0,1961 | 0,228 |
| TcCLB.510099.90 | tyrosine phosphatase isoform, putative | -0,1962 | 0,07177 |
| TcCLB.506435.140 | hypothetical protein, conserved | -0,1963 | 0,08167 |
| TcCLB.506555.9 | hypothetical protein, conserved (fragment) | -0,1963 | 0,1427 |
| TcCLB.510409.60 | hypothetical protein, conserved | -0,1967 | 0,06915 |
| TcCLB.509693.194 | hypothetical protein, conserved | -0,1967 | 0,432 |

| | | | |
|---|---|---|---|
| TcCLB.504057.110 | ribosomal RNA processing protein 6, putative (RRP6) | -0,1968 | 0,0317 |
| TcCLB.508695.30 | hypothetical protein (fragment) | -0,1972 | 0,148 |
| TcCLB.504643.40 | U6 snRNA-associated Sm-like protein LSm4, putative | -0,1972 | 0,1933 |
| TcCLB.511423.60 | ribosomal RNA-processing protein 8, putative (RRP8) | -0,1973 | 0,0869 |
| TcCLB.506251.10 | hypothetical protein, conserved | -0,1973 | 0,1312 |
| TcCLB.508671.10 | serine carboxypeptidase (CBP1), putative | -0,1973 | 0,1603 |
| TcCLB.510297.110 | cAMP binding protein, putative | -0,1974 | 0,01817 |
| TcCLB.505939.60 | ribonuclease H1, putative | -0,1975 | 0,1628 |
| TcCLB.508265.4 | synaptojanin (N-terminal domain), putative (fragment) | -0,1975 | 0,2373 |
| TcCLB.508647.50 | hypothetical protein, conserved (pseudogene) | -0,1976 | 0,02374 |
| TcCLB.507963.10 | hypothetical protein, conserved | -0,1976 | 0,2277 |
| TcCLB.504187.10 | retrotransposon hot spot protein (RHS, pseudogene), puta | -0,1976 | 0,5338 |
| TcCLB.507433.9 | ATP-dependent DEAD/H DNA helicase recQ, putative (frag | -0,1977 | 0,0291 |
| TcCLB.506871.10 | hypothetical protein, conserved | -0,1977 | 0,09503 |
| TcCLB.508707.160 | hypothetical protein, conserved | -0,1977 | 0,135 |
| TcCLB.506265.110 | hypothetical protein, conserved | -0,1978 | 0,1319 |
| TcCLB.511529.190 | hypothetical protein, conserved | -0,1979 | 0,2428 |
| TcCLB.504089.14 | hypothetical protein, conserved | -0,198 | 0,03463 |
| TcCLB.507949.154 | TFIIH basal transcription factor subunit, putative (TSP1) | -0,198 | 0,1218 |
| TcCLB.504021.140 | hypothetical protein, conserved | -0,1981 | 0,04939 |
| TcCLB.504009.30 | hypothetical protein, conserved | -0,1981 | 0,07952 |
| TcCLB.503563.20 | ADG2, putative | -0,1982 | 0,09897 |
| TcCLB.507099.20 | hypothetical protein, conserved | -0,1983 | 0,04533 |
| TcCLB.507963.60 | hypothetical protein, conserved | -0,1983 | 0,2929 |
| TcCLB.506881.10 | hypothetical protein, conserved (pseudogene) | -0,1983 | 0,4817 |
| TcCLB.488767.20 | retrotransposon hot spot protein (RHS, pseudogene), puta | -0,1984 | 0,1939 |
| TcCLB.506577.140 | hypothetical protein, conserved | -0,1984 | 0,2282 |
| TcCLB.507685.10 | hypothetical protein, conserved | -0,1985 | 0,1559 |
| TcCLB.509741.50 | kinesin-like protein (pseudogene), putative | -0,1985 | 0,2997 |
| TcCLB.509637.18 | hypothetical protein, conserved | -0,1986 | 0,1749 |
| TcCLB.507811.44 | phosphoglycerate mutase protein, putative | -0,1986 | 0,2118 |
| TcCLB.510053.30 | hypothetical protein, conserved | -0,1987 | 0,09789 |
| TcCLB.508857.60 | hypothetical protein, conserved | -0,1987 | 0,1483 |
| TcCLB.507049.179 | hypothetical protein, conserved | -0,1989 | 0,08787 |
| TcCLB.503565.20 | hypothetical protein, conserved | -0,1989 | 0,1058 |
| TcCLB.506891.10 | Metal-binding domain-containing protein | -0,1989 | 0,1382 |
| TcCLB.508207.200 | small GTP-binding protein, putative | -0,1991 | 0,06465 |
| TcCLB.510643.20 | retrotransposon hot spot protein (RHS, pseudogene), puta | -0,1991 | 0,1818 |
| TcCLB.506247.250 | hypothetical protein, conserved | -0,1993 | 0,00836 |
| TcCLB.509937.10 | poly(A) polymerase, putative | -0,1995 | 0,1023 |
| TcCLB.508325.20 | retrotransposon hot spot protein (RHS, pseudogene), puta | -0,1997 | 0,2941 |
| TcCLB.506855.240 | bilobe protein, conserved (pseudogene) | -0,1998 | 0,01002 |
| TcCLB.506139.10 | mucin-associated surface protein (MASP), putative | -0,1998 | 0,6373 |
| TcCLB.510729.10 | AMP deaminase, putative | -0,1999 | 0,07252 |
| TcCLB.508629.63 | ATP-dependent DEAD/H RNA helicase (pseudogene), puta | -0,2 | 0,4512 |

| | | | |
|---|---|---|---|
| TcCLB.506275.30 | hypothetical protein, conserved | -0,2002 | 0,2078 |
| TcCLB.511435.10 | WD40 repeat-containing protein | -0,2003 | 0,04654 |
| TcCLB.506605.40 | hypothetical protein, conserved | -0,2004 | 0,1056 |
| TcCLB.509715.104 | protein kinase, putative | -0,2006 | 0,05974 |
| TcCLB.506647.20 | hypothetical protein, conserved | -0,2007 | 0,05865 |
| TcCLB.506519.100 | hypothetical protein, conserved | -0,2007 | 0,2162 |
| TcCLB.511283.50 | hypothetical protein, conserved | -0,2008 | 0,2702 |
| TcCLB.506435.130 | hypothetical protein, conserved | -0,2009 | 0,08654 |
| TcCLB.504057.11 | phospholipid-translocating ATPase, putative | -0,2011 | 0,03177 |
| TcCLB.507677.10 | hypothetical protein, conserved | -0,2012 | 0,0625 |
| TcCLB.506529.30 | hypothetical protein, conserved | -0,2012 | 0,2566 |
| TcCLB.509105.20 | palmitoyl acyltransferase 2, putative | -0,2013 | 0,1401 |
| TcCLB.506113.50 | ATP-dependent DEAD/H RNA helicase (pseudogene), puta | -0,2014 | 0,4156 |
| TcCLB.508235.10 | hypothetical protein, conserved (pseudogene) | -0,2015 | 0,1178 |
| TcCLB.509965.150 | hypothetical protein, conserved | -0,2016 | 0,1253 |
| TcCLB.508777.70 | hypothetical protein, conserved | -0,202 | 0,1015 |
| TcCLB.510835.20 | hypothetical protein, conserved | -0,2021 | 0,08877 |
| TcCLB.511033.10 | hypothetical protein, conserved | -0,2022 | 0,0923 |
| TcCLB.507233.60 | retrotransposon hot spot protein (RHS, pseudogene), puta | -0,2022 | 0,108 |
| TcCLB.510777.29 | eukaryotic translation initiation factor 4E, putative | -0,2022 | 0,2871 |
| TcCLB.463155.40 | glycine dehydrogenase [decarboxylating] (pseudogene), pu | -0,2023 | 0,1349 |
| TcCLB.504127.10 | surface protease GP63 (pseudogene), putative | -0,2023 | 0,4374 |
| TcCLB.412943.9 | hypothetical protein, conserved | -0,2024 | 0,1995 |
| TcCLB.511109.50 | hypothetical protein, conserved (pseudogene) | -0,2026 | 0,1651 |
| TcCLB.503487.40 | hypothetical protein, conserved | -0,2027 | 0,03811 |
| TcCLB.453995.10 | retrotransposon hot spot protein (RHS, pseudogene), puta | -0,2027 | 0,1262 |
| TcCLB.508465.50 | hypothetical protein, conserved | -0,2028 | 0,1043 |
| TcCLB.508425.31 | hypothetical protein, conserved (pseudogene) | -0,2028 | 0,1279 |
| TcCLB.504081.310 | syntaxin binding protein (pseudogene), putative | -0,2029 | 0,5561 |
| TcCLB.511661.10 | hypothetical protein, conserved | -0,2031 | 0,02596 |
| TcCLB.510879.210 | hypothetical protein, conserved | -0,2031 | 0,1381 |
| TcCLB.510055.100 | hypothetical protein, conserved | -0,2033 | 0,0422 |
| TcCLB.504017.60 | hypothetical protein, conserved | -0,2033 | 0,1269 |
| TcCLB.507723.20 | hypothetical protein | -0,2033 | 0,4684 |
| TcCLB.510377.300 | trans-sialidase (pseudogene), putative | -0,2034 | 0,4634 |
| TcCLB.505807.260 | hypothetical protein, conserved | -0,2037 | 0,1001 |
| TcCLB.504131.80 | hypothetical protein, conserved | -0,2038 | 0,1322 |
| TcCLB.504247.70 | AATF protein, putative (pseudogene) | -0,2038 | 0,1482 |
| TcCLB.507163.80 | hypothetical protein, conserved | -0,2038 | 0,2049 |
| TcCLB.508719.9 | hypothetical protein, conserved | -0,2039 | 0,0154 |
| TcCLB.506883.40 | Lsm12 protein, putative | -0,204 | 0,06881 |
| TcCLB.506301.20 | Golgi reassembly stacking protein (GRASP homologue), pu | -0,204 | 0,1002 |
| TcCLB.511153.60 | palmitoyl acyltransferase 9, putative | -0,204 | 0,1228 |
| TcCLB.505955.30 | surface protease GP63 (pseudogene), putative | -0,2041 | 0,3183 |
| TcCLB.510285.30 | hypothetical protein, conserved | -0,2044 | 0,06378 |

| | | | |
|---|---|---|---|
| TcCLB.511855.40 | hypothetical protein, conserved | -0,2044 | 0,1041 |
| TcCLB.510719.290 | Golgi SNAP receptor complex member 1, putative | -0,2044 | 0,1339 |
| TcCLB.506923.10 | trans-sialidase (pseudogene), putative | -0,2044 | 0,3364 |
| TcCLB.506729.70 | hypothetical protein, conserved | -0,2045 | 0,09652 |
| TcCLB.506435.220 | hypothetical protein, conserved | -0,2045 | 0,1852 |
| TcCLB.507925.69 | hypothetical protein, conserved | -0,2045 | 0,193 |
| TcCLB.508193.45 | trans-sialidase (pseudogene), putative | -0,2045 | 0,2419 |
| TcCLB.511283.230 | hypothetical protein, conserved | -0,2046 | 0,01507 |
| TcCLB.510433.10 | hypothetical protein | -0,2047 | 0,05865 |
| TcCLB.503519.30 | mucin-associated surface protein (MASP), putative | -0,2047 | 0,3462 |
| TcCLB.510749.50 | Flagellar Member 5 (FLAM5) | -0,2048 | 0,07965 |
| TcCLB.508999.49 | hypothetical protein | -0,2048 | 0,5806 |
| TcCLB.508977.111 | elongation factor 1-gamma (EF-1-gamma, pseudogene), [illegible] | -0,2049 | 0,4066 |
| TcCLB.508693.30 | hypothetical protein, conserved | -0,205 | 0,0384 |
| TcCLB.509203.30 | hd phosphohydrolase family protein | -0,2053 | 0,2518 |
| TcCLB.507239.10 | hypothetical protein, conserved | -0,2053 | 0,2842 |
| TcCLB.508153.850 | hypothetical protein, conserved | -0,2054 | 0,03873 |
| TcCLB.508387.130 | ATPase, putative | -0,2055 | 0,07351 |
| TcCLB.504625.10 | trans-splicing factor, putative | -0,2056 | 0,08637 |
| TcCLB.511183.170 | trans-sialidase (pseudogene), putative | -0,2056 | 0,4571 |
| TcCLB.511517.80 | replication factor C, subunit 2, putative | -0,2058 | 0,07782 |
| TcCLB.510243.70 | hypothetical protein, conserved | -0,206 | 0,02239 |
| TcCLB.511423.110 | hypothetical protein, conserved | -0,206 | 0,08197 |
| TcCLB.416143.10 | Lys-63-specific deubiquitinase BRCC36, putative | -0,2062 | 0,03542 |
| TcCLB.508951.4 | protein kinase, putative (fragment) | -0,2062 | 0,4038 |
| TcCLB.503811.20 | U3 small nucleolar ribonucleoprotein protein MPP10, puta | -0,2063 | 0,03547 |
| TcCLB.510747.120 | hypothetical protein, conserved | -0,2063 | 0,05832 |
| TcCLB.506517.140 | hypothetical protein, conserved | -0,2063 | 0,06816 |
| TcCLB.401473.9 | DNA-directed rna polymerase I largest subunit, putative | -0,2064 | 0,1762 |
| TcCLB.506559.280 | hypothetical protein, conserved | -0,2066 | 0,02204 |
| TcCLB.506989.180 | U3 small nucleolar ribonucleoprotein protein MPP10, puta | -0,2067 | 0,03894 |
| TcCLB.508579.10 | hypothetical protein, conserved | -0,2069 | 0,1737 |
| TcCLB.506511.10 | hypothetical protein, conserved | -0,2069 | 0,2514 |
| TcCLB.503821.19 | hypothetical protein, conserved | -0,207 | 0,1263 |
| TcCLB.508995.50 | hypothetical protein, conserved | -0,2071 | 0,1242 |
| TcCLB.511001.70 | phosphatidylinositol-4-phosphate 5-kinase-like, putative | -0,2072 | 0,02272 |
| TcCLB.506579.10 | ABC transporter, putative | -0,2072 | 0,02609 |
| TcCLB.508153.940 | hypothetical protein, conserved (pseudogene) | -0,2073 | 0,009475 |
| TcCLB.511585.170 | protein kinase, putative | -0,2074 | 0,02112 |
| TcCLB.509229.40 | hypothetical protein, conserved | -0,2074 | 0,03192 |
| TcCLB.509733.80 | hypothetical protein, conserved | -0,2074 | 0,1001 |
| TcCLB.510901.90 | ATPase protein, putative | -0,2074 | 0,1244 |
| TcCLB.504073.20 | hypothetical protein, conserved | -0,2076 | 0,05645 |
| TcCLB.508529.40 | trans-sialidase (pseudogene), putative | -0,2076 | 0,1052 |
| TcCLB.506123.20 | hypothetical protein, conserved | -0,2076 | 0,138 |

| | | | |
|---|---|---|---|
| TcCLB.511397.10 | mucin-associated surface protein (MASP, pseudogene), pu | -0,2076 | 0,5274 |
| TcCLB.504625.70 | kinetoplast DNA-associated protein, putative | -0,2077 | 0,03089 |
| TcCLB.506855.100 | hypothetical protein, conserved | -0,2077 | 0,1241 |
| TcCLB.508153.910 | hypothetical protein, conserved | -0,2077 | 0,1343 |
| TcCLB.509567.34 | hypothetical protein, conserved | -0,2078 | 0,08413 |
| TcCLB.503779.80 | hypothetical protein, conserved (pseudogene) | -0,208 | 0,04482 |
| TcCLB.509433.30 | hypothetical protein | -0,208 | 0,3031 |
| TcCLB.509179.140 | myo-inositol-1(or 4)-monophosphatase 1, putative | -0,2081 | 0,2254 |
| TcCLB.506947.20 | hypothetical protein, conserved | -0,2082 | 0,04334 |
| TcCLB.510877.70 | ATP-dependent RNA and DNA helicase, mitochondrial, pu | -0,2083 | 0,04367 |
| TcCLB.506407.110 | hypothetical protein, conserved | -0,2083 | 0,2035 |
| TcCLB.508277.150 | proliferative cell nuclear antigen (PCNA), putative | -0,2084 | 0,008141 |
| TcCLB.507023.160 | hypothetical protein, conserved | -0,2085 | 0,007245 |
| TcCLB.504109.20 | hypothetical protein, conserved | -0,2085 | 0,02932 |
| TcCLB.511903.160 | phosphatidylinositol 3-kinase vps34-like | -0,2086 | 0,01386 |
| TcCLB.503723.90 | glyceraldehyde 3-phosphate dehydrogenase, C-terminal d | -0,2086 | 0,4282 |
| TcCLB.511179.120 | hypothetical protein, conserved | -0,2087 | 0,142 |
| TcCLB.406377.10 | hypothetical protein, conserved (pseudogene) | -0,2087 | 0,3126 |
| TcCLB.510659.70 | protein kinase-like protein, putative | -0,2088 | 0,05603 |
| TcCLB.511667.10 | hypothetical protein, conserved | -0,2089 | 0,5005 |
| TcCLB.510943.150 | glycerol-3-phosphate acyltransferase, putative | -0,209 | 0,1057 |
| TcCLB.503917.28 | hypothetical protein, conserved | -0,2091 | 0,0311 |
| TcCLB.509965.60 | RNA decapping enzyme, putative (Dcp2) | -0,2091 | 0,1598 |
| TcCLB.506667.20 | hypothetical protein | -0,2092 | 0,6168 |
| TcCLB.510945.40 | hypothetical protein, conserved | -0,2093 | 0,04054 |
| TcCLB.510351.110 | hypothetical protein | -0,2093 | 0,05582 |
| TcCLB.508303.44 | hypothetical protein, conserved | -0,2093 | 0,1726 |
| TcCLB.510187.130 | Bardet-Biedl syndrome 4 protein, putative (BBS4) | -0,2094 | 0,05236 |
| TcCLB.511137.20 | hypothetical protein, conserved | -0,2094 | 0,1271 |
| TcCLB.503399.20 | hydrolase, alpha/beta fold family, putative | -0,2095 | 0,165 |
| TcCLB.510761.22 | hypothetical protein, conserved | -0,2097 | 0,04105 |
| TcCLB.510655.110 | poly(A)-specific ribonuclease PARN, putative | -0,21 | 0,04671 |
| TcCLB.510911.30 | Ras-related protein Rab4, putative (RAB4) | -0,2101 | 0,04049 |
| TcCLB.506287.80 | hypothetical protein, conserved | -0,2102 | 0,0784 |
| TcCLB.508675.40 | hypothetical protein, conserved | -0,2103 | 0,06561 |
| TcCLB.507041.132 | hypothetical protein (fragment) | -0,2103 | 0,3147 |
| TcCLB.511407.20 | hypothetical protein, conserved | -0,2105 | 0,09734 |
| TcCLB.511285.30 | hypothetical protein, conserved | -0,2105 | 0,1127 |
| TcCLB.508057.41 | trans-sialidase (pseudogene), putative | -0,2105 | 0,4765 |
| TcCLB.507661.10 | retrotransposon hot spot protein (RHS, pseudogene), puta | -0,2105 | 0,6739 |
| TcCLB.511169.29 | tubulin-tyrsoine ligase-like protein, putative | -0,2106 | 0,01525 |
| TcCLB.511751.144 | hypothetical protein, conserved | -0,2106 | 0,03415 |
| TcCLB.508173.84 | hypothetical protein, conserved | -0,2106 | 0,06288 |
| TcCLB.511211.10 | hypothetical protein, conserved | -0,2108 | 0,1204 |
| TcCLB.504449.30 | hypothetical protein, conserved | -0,2109 | 0,008533 |

| | | | |
|---|---|---|---|
| TcCLB.508153.240 | hypothetical protein, conserved | -0,2109 | 0,05837 |
| TcCLB.511261.80 | hypothetical protein, conserved | -0,211 | 0,03015 |
| TcCLB.511003.90 | hypothetical protein, conserved | -0,2111 | 0,04973 |
| TcCLB.509797.69 | hypothetical protein, conserved (fragment) | -0,2111 | 0,05914 |
| TcCLB.510431.240 | hypothetical protein, conserved | -0,2111 | 0,06494 |
| TcCLB.506703.30 | mucin TcMUCII (pseudogene), putative | -0,2111 | 0,5821 |
| TcCLB.509941.120 | protein kinase, putative | -0,2112 | 0,05874 |
| TcCLB.511309.14 | hypothetical protein, conserved | -0,2113 | 0,3329 |
| TcCLB.511821.50 | hypothetical protein, conserved | -0,2115 | 0,1094 |
| TcCLB.509227.20 | mucin-associated surface protein (MASP, pseudogene), pu | -0,2117 | 0,02463 |
| TcCLB.507809.130 | hypothetical protein, conserved | -0,2118 | 0,01538 |
| TcCLB.506593.10 | hypothetical protein, conserved | -0,212 | 0,2117 |
| TcCLB.508979.15 | mucin-associated surface protein (MASP, pseudogene), pu | -0,2122 | 0,6948 |
| TcCLB.506625.140 | hypothetical protein, conserved | -0,2124 | 0,08179 |
| TcCLB.511215.70 | kinesin, putative | -0,2124 | 0,08593 |
| TcCLB.509353.40 | PTP1-interacting protein, 39 kDa, putative | -0,2125 | 0,03638 |
| TcCLB.510983.19 | retrotransposon hot spot (RHS) protein, putative | -0,2125 | 0,1397 |
| TcCLB.505925.10 | trans-sialidase, putative (fragment) | -0,2125 | 0,2833 |
| TcCLB.503949.40 | UDP-Gal or UDP-GlcNAc-dependent glycosyltransferase, p | -0,2129 | 0,1364 |
| TcCLB.510729.130 | DNA replication licensing factor MCM9, putative | -0,213 | 0,04263 |
| TcCLB.503789.70 | hypothetical protein, conserved | -0,2131 | 0,02818 |
| TcCLB.510077.30 | Mitogen-activated kinase kinase kinase, putative | -0,2131 | 0,2404 |
| TcCLB.507049.110 | hypothetical protein, conserved | -0,2132 | 0,2319 |
| TcCLB.504881.30 | hypothetical protein, conserved | -0,2133 | 0,0007415 |
| TcCLB.510329.190 | hypothetical protein, conserved | -0,2134 | 0,02147 |
| TcCLB.507029.10 | hypothetical protein, conserved | -0,2134 | 0,07802 |
| TcCLB.508303.30 | hypothetical protein, conserved | -0,2135 | 0,1607 |
| TcCLB.506459.30 | trans-sialidase (pseudogene), putative | -0,2137 | 0,1711 |
| TcCLB.511367.40 | hypothetical protein | -0,2137 | 0,4975 |
| TcCLB.503891.30 | phosphatidylinositol-4-phosphate 5-kinase, putative | -0,2138 | 0,1146 |
| TcCLB.508323.40 | hypothetical protein, conserved | -0,2138 | 0,1778 |
| TcCLB.506181.110 | hypothetical protein, conserved | -0,2141 | 0,279 |
| TcCLB.507641.70 | hypothetical protein, conserved | -0,2143 | 0,0567 |
| TcCLB.506559.460 | hypothetical protein | -0,2144 | 0,07265 |
| TcCLB.504643.20 | hypothetical protein, conserved | -0,2144 | 0,1211 |
| TcCLB.506297.30 | hypothetical protein, conserved | -0,2145 | 0,004966 |
| TcCLB.511217.190 | hypothetical protein, conserved | -0,2146 | 0,3925 |
| TcCLB.511127.400 | serine/threonine protein phosphatase, putative | -0,2147 | 0,04986 |
| TcCLB.511727.110 | hypothetical protein, conserved | -0,2147 | 0,05264 |
| TcCLB.510889.270 | hypothetical protein, conserved | -0,2148 | 0,09692 |
| TcCLB.509067.14 | hypothetical protein, conserved | -0,2148 | 0,1993 |
| TcCLB.508593.69 | hypothetical protein, conserved | -0,2148 | 0,2705 |
| TcCLB.510045.30 | protein kinase, putative | -0,2149 | 0,09762 |
| TcCLB.507063.60 | hypothetical protein, conserved | -0,2149 | 0,6021 |
| TcCLB.506957.70 | hypothetical protein, conserved | -0,215 | 0,0478 |

| | | | |
|---|---|---|---|
| TcCLB.510363.250 | trans-sialidase, Group V, putative | -0,215 | 0,5386 |
| TcCLB.507521.140 | hypothetical protein, conserved | -0,2152 | 0,01112 |
| TcCLB.506515.20 | O-sialoglycoprotein endopeptidase, putative | -0,2152 | 0,06299 |
| TcCLB.507099.140 | endomembrane protein, putative | -0,2153 | 0,007988 |
| TcCLB.511517.130 | hypothetical protein, conserved | -0,2153 | 0,07042 |
| TcCLB.510699.27 | elongation factor 1-gamma (EF-1-gamma, pseudogene), | -0,2153 | 0,6425 |
| TcCLB.511725.200 | hypothetical protein, conserved | -0,2156 | 0,03353 |
| TcCLB.503799.4 | protein kinase, putative (fragment) | -0,216 | 0,1957 |
| TcCLB.506857.90 | proteasome regulatory ATPase subunit 1, putative | -0,2161 | 0,01241 |
| TcCLB.510943.140 | DNA primase large subunit, putative (pril) | -0,2161 | 0,02624 |
| TcCLB.506537.71 | hypothetical protein, conserved (pseudogene) | -0,2164 | 0,06807 |
| TcCLB.511287.140 | hypothetical protein, conserved | -0,2165 | 0,02033 |
| TcCLB.508213.49 | hypothetical protein, conserved | -0,2165 | 0,1975 |
| TcCLB.510953.30 | hypothetical protein, conserved | -0,2166 | 0,05527 |
| TcCLB.509453.4 | hypothetical protein, conserved | -0,2167 | 0,1168 |
| TcCLB.511575.60 | isocitrate dehydrogenase [NADP], mitochondrial precursor | -0,2168 | 0,1353 |
| TcCLB.510823.50 | hypothetical protein, conserved | -0,2169 | 0,03242 |
| TcCLB.510105.33 | RNA-binding protein, putative | -0,2169 | 0,361 |
| TcCLB.509791.110 | hypothetical protein, conserved | -0,217 | 0,04393 |
| TcCLB.511731.40 | hypothetical protein, conserved | -0,217 | 0,04538 |
| TcCLB.510075.60 | hypothetical protein | -0,2171 | 0,201 |
| TcCLB.503851.24 | protein tyrosine phosphatase PRL | -0,2171 | 0,262 |
| TcCLB.511127.380 | hypothetical protein, conserved | -0,2172 | 0,005688 |
| TcCLB.511395.90 | serine/threonine protein kinase, putative | -0,2172 | 0,03482 |
| TcCLB.506945.180 | hypothetical protein, conserved | -0,2173 | 0,00879 |
| TcCLB.504243.30 | hypothetical protein, conserved | -0,2173 | 0,01743 |
| TcCLB.511289.50 | mitogen-activated protein kinase 9, putative (MPK9) | -0,2173 | 0,05527 |
| TcCLB.508893.20 | hypothetical protein, conserved | -0,2176 | 0,0104 |
| TcCLB.506629.170 | hypothetical protein | -0,2176 | 0,3265 |
| TcCLB.511613.50 | mucin TcMUCII, putative | -0,2176 | 0,4824 |
| TcCLB.511825.110 | hypothetical protein | -0,2176 | 0,5734 |
| TcCLB.506853.20 | hypothetical protein, conserved | -0,2177 | 0,02339 |
| TcCLB.506713.10 | hypothetical protein, conserved | -0,2178 | 0,1123 |
| TcCLB.506245.80 | hypothetical protein, conserved (pseudogene) | -0,2178 | 0,357 |
| TcCLB.506737.140 | syntaxin binding protein 1, putative | -0,218 | 0,00701 |
| TcCLB.399033.10 | hypothetical protein | -0,218 | 0,24 |
| TcCLB.507625.190 | hypothetical protein, conserved | -0,2184 | 0,0549 |
| TcCLB.507769.50 | ATP-dependent DEAD/H RNA helicase (pseudogene), puta | -0,2184 | 0,2248 |
| TcCLB.506797.110 | hypothetical protein, conserved | -0,2184 | 0,2249 |
| TcCLB.507509.70 | hypothetical protein, conserved | -0,2185 | 0,05811 |
| TcCLB.507615.60 | hypothetical protein, conserved | -0,2185 | 0,1177 |
| TcCLB.508641.310 | hypothetical protein, conserved | -0,2187 | 0,01925 |
| TcCLB.508405.30 | Suppressor of G2 allele of SKP1, putative | -0,2188 | 0,09632 |
| TcCLB.505983.30 | hypothetical protein, conserved | -0,219 | 0,01032 |
| TcCLB.511727.240 | hypothetical protein | -0,219 | 0,03613 |

| | | | |
|---|---|---|---|
| TcCLB.510257.120 | hypothetical protein, conserved | -0,2192 | 0,05175 |
| TcCLB.506513.120 | hypothetical protein, conserved | -0,2193 | 0,02895 |
| TcCLB.507019.110 | hypothetical protein | -0,2193 | 0,1775 |
| TcCLB.507395.30 | hypothetical protein | -0,2193 | 0,2567 |
| TcCLB.508693.90 | peroxisome biogenesis factor 1, putative | -0,2195 | 0,01036 |
| TcCLB.509757.40 | hypothetical protein, conserved | -0,2196 | 0,2019 |
| TcCLB.506945.359 | damage-specific DNA binding protein, putative | -0,2198 | 0,04105 |
| TcCLB.508851.154 | hypothetical protein, conserved | -0,2201 | 0,3443 |
| TcCLB.511217.180 | hypothetical protein, conserved | -0,2202 | 0,03242 |
| TcCLB.508465.80 | hypothetical protein, conserved | -0,2202 | 0,1639 |
| TcCLB.506661.110 | hypothetical protein, conserved | -0,2205 | 0,008167 |
| TcCLB.508781.70 | palmitoyl acyltransferase 3, putative | -0,2205 | 0,04441 |
| TcCLB.511735.60 | superoxide dismutase, putative | -0,2205 | 0,06754 |
| TcCLB.506925.40 | RNA polymerase I second largest subunit, putative | -0,2206 | 0,003252 |
| TcCLB.510889.205 | hypothetical protein, conserved | -0,2206 | 0,0678 |
| TcCLB.505183.124 | hypothetical protein, conserved | -0,2206 | 0,09107 |
| TcCLB.510297.20 | hypothetical protein, conserved | -0,2207 | 0,02893 |
| TcCLB.507913.39 | glutaminyl-tRNA synthetase, putative | -0,2208 | 0,1404 |
| TcCLB.504741.190 | hypothetical protein, conserved | -0,2211 | 0,1755 |
| TcCLB.509993.51 | mucin-associated surface protein (MASP, pseudogene), pu | -0,2211 | 0,4973 |
| TcCLB.504155.200 | syntaxin binding protein (pseudogene), putative | -0,2211 | 0,6662 |
| TcCLB.511649.169 | ATP-dependent DEAD/H RNA helicase, putative | -0,2212 | 0,04307 |
| TcCLB.506053.35 | hypothetical protein, conserved (pseudogene) | -0,2212 | 0,1227 |
| TcCLB.507017.159 | hypothetical protein, conserved | -0,2213 | 0,1427 |
| TcCLB.507027.30 | hypothetical protein, conserved (pseudogene) | -0,2214 | 0,009003 |
| TcCLB.507103.29 | hypothetical protein, conserved (fragment) | -0,2215 | 0,128 |
| TcCLB.506459.195 | hypothetical protein, conserved | -0,2216 | 0,5104 |
| TcCLB.508989.90 | Eukaryotic translation initiation factor 4 gamma type 5, pu | -0,2217 | 0,008115 |
| TcCLB.504703.20 | hypothetical protein, conserved | -0,2217 | 0,02366 |
| TcCLB.509647.140 | hypothetical protein, conserved | -0,2218 | 0,02354 |
| TcCLB.510089.39 | CAAX amino terminal protease, putative | -0,2218 | 0,1754 |
| TcCLB.511577.110 | exopolyphosphatase | -0,2218 | 0,1872 |
| TcCLB.503959.60 | hypothetical protein, conserved | -0,2219 | 0,02545 |
| TcCLB.506341.30 | beta galactofuranosyl glycosyltransferase, putative | -0,222 | 0,04198 |
| TcCLB.510653.10 | hypothetical protein, conserved | -0,222 | 0,1889 |
| TcCLB.506297.300 | eukaryotic release factor 3, putative | -0,2223 | 0,005144 |
| TcCLB.509721.20 | hypothetical protein, conserved | -0,2225 | 0,07003 |
| TcCLB.508625.60 | hypothetical protein, conserved | -0,2225 | 0,113 |
| TcCLB.506795.100 | hypothetical protein | -0,2225 | 0,4365 |
| TcCLB.511301.130 | protein kinase, putative | -0,2226 | 0,1085 |
| TcCLB.504741.30 | protein phosphatase 2C, putative | -0,2226 | 0,1098 |
| TcCLB.510487.30 | hypothetical protein, conserved (pseudogene) | -0,2227 | 0,1923 |
| TcCLB.506529.200 | hypothetical protein, conserved | -0,2229 | 0,03543 |
| TcCLB.510575.180 | adenylate kinase, putative | -0,2229 | 0,08207 |
| TcCLB.510311.50 | hypothetical protein, conserved | -0,2231 | 0,004323 |

| | | | |
|---|---|---|---|
| TcCLB.506175.40 | hypothetical protein, conserved | -0,2232 | 0,02891 |
| TcCLB.510749.40 | ubiquitin hydrolase, putative | -0,2232 | 0,3523 |
| TcCLB.504005.29 | hypothetical protein, conserved | -0,2233 | 0,1651 |
| TcCLB.506957.130 | hypothetical protein, conserved | -0,2234 | 0,04947 |
| TcCLB.507083.30 | apurinic/apyrimidinic endonuclease, putative | -0,2234 | 0,06827 |
| TcCLB.511629.20 | hypothetical protein, conserved | -0,2236 | 0,09429 |
| TcCLB.507941.40 | hypothetical protein, conserved | -0,2237 | 0,04959 |
| TcCLB.506009.100 | ADP-ribosylation factor-like protein, putative | -0,2239 | 0,1015 |
| TcCLB.506955.261 | hypothetical protein, conserved (pseudogene) | -0,2239 | 0,4187 |
| TcCLB.510901.260 | tRNA pseudouridine synthase (pseudogene), putative | -0,224 | 0,09213 |
| TcCLB.510199.20 | elongation factor 1-gamma (EF-1-gamma, pseudogene), | -0,224 | 0,5483 |
| TcCLB.510329.100 | leucine-rich repeat protein, putative | -0,2242 | 0,04058 |
| TcCLB.508909.150 | hypothetical protein, conserved | -0,2242 | 0,2658 |
| TcCLB.508675.60 | hypothetical protein, conserved | -0,2243 | 0,03516 |
| TcCLB.506297.340 | hypothetical protein, conserved | -0,2243 | 0,0448 |
| TcCLB.506335.80 | hypothetical protein | -0,2243 | 0,6038 |
| TcCLB.509007.10 | hypothetical protein, conserved | -0,2246 | 0,03635 |
| TcCLB.508789.204 | mucin-associated surface protein (MASP, pseudogene), pu | -0,2247 | 0,5082 |
| TcCLB.506195.90 | adenylate kinase 2 | -0,2248 | 0,03458 |
| TcCLB.508153.70 | hypothetical protein, conserved | -0,2249 | 0,08451 |
| TcCLB.506221.90 | hypothetical protein, conserved | -0,225 | 0,0181 |
| TcCLB.508317.40 | hypothetical protein, conserved | -0,225 | 0,2236 |
| TcCLB.511233.100 | NUDIX hydrolase 3, putative (Nudix3) | -0,2255 | 0,1359 |
| TcCLB.507949.260 | Cell cycle checkpoint protein RAD1-like, putative | -0,2255 | 0,1807 |
| TcCLB.507007.90 | hypothetical protein, conserved | -0,2256 | 0,1272 |
| TcCLB.511563.30 | retrotransposon hot spot protein (RHS, pseudogene), puta | -0,2256 | 0,5775 |
| TcCLB.508475.40 | hypothetical protein | -0,2259 | 0,2291 |
| TcCLB.511127.260 | DNA photolyase, putative | -0,2264 | 0,01905 |
| TcCLB.507925.10 | hypothetical protein, conserved (pseudogene) | -0,2264 | 0,3718 |
| TcCLB.506263.49 | hypothetical protein, conserved (fragment) | -0,2265 | 0,1361 |
| TcCLB.503583.90 | hydrogenase, putative | -0,2266 | 0,04815 |
| TcCLB.504125.80 | hypothetical protein, conserved | -0,2269 | 0,01878 |
| TcCLB.510103.10 | hypothetical protein | -0,227 | 0,1569 |
| TcCLB.510517.70 | hypothetical protein, conserved | -0,2271 | 0,008287 |
| TcCLB.509561.80 | plectin-like protein, putative | -0,2272 | 0,06031 |
| TcCLB.506825.80 | hypothetical protein, conserved | -0,2273 | 0,006433 |
| TcCLB.510315.20 | mitochondrial carrier protein, putative | -0,2273 | 0,06332 |
| TcCLB.508963.20 | small GTP-binding protein Rab18, putative | -0,2273 | 0,06726 |
| TcCLB.509643.20 | hypothetical protein, conserved | -0,2275 | 0,03362 |
| TcCLB.508815.50 | hypothetical protein, conserved | -0,2277 | 0,06865 |
| TcCLB.506435.10 | hypothetical protein, conserved | -0,2277 | 0,1448 |
| TcCLB.506649.60 | hypothetical protein, conserved | -0,228 | 0,0212 |
| TcCLB.503727.20 | hypothetical protein, conserved | -0,2281 | 0,01864 |
| TcCLB.405165.10 | hypothetical protein, conserved | -0,2281 | 0,1299 |
| TcCLB.506303.130 | hypothetical protein, conserved | -0,2282 | 0,0242 |

| | | | |
|---|---|---|---|
| TcCLB.510187.490 | hypothetical protein, conserved | -0,2282 | 0,1203 |
| TcCLB.506009.109 | hypothetical protein, conserved | -0,2284 | 0,1753 |
| TcCLB.508307.160 | hypothetical protein, conserved | -0,2286 | 0,01576 |
| TcCLB.509767.200 | hypothetical protein, conserved | -0,2286 | 0,05832 |
| TcCLB.506753.200 | hypothetical protein, conserved | -0,2287 | 0,02994 |
| TcCLB.506559.500 | hypothetical protein, conserved | -0,2287 | 0,03125 |
| TcCLB.508737.20 | hypothetical protein | -0,229 | 0,1324 |
| TcCLB.508647.90 | hypothetical protein, conserved | -0,2291 | 0,1135 |
| TcCLB.511019.26 | trans-sialidase (pseudogene), putative | -0,2292 | 0,1027 |
| TcCLB.511859.180 | proteasome regulatory ATPase subunit 2, putative | -0,2293 | 0,009392 |
| TcCLB.511283.194 | hypothetical protein, conserved | -0,2293 | 0,04991 |
| TcCLB.452895.10 | hypothetical protein, conserved | -0,2293 | 0,1335 |
| TcCLB.507949.110 | zeta tubulin, putative | -0,2294 | 0,03525 |
| TcCLB.509797.9 | hypothetical protein, conserved (pseudogene) | -0,2295 | 0,05319 |
| TcCLB.505999.130 | phosphatidylinositol-4-phosphate 5-kinase related | -0,2296 | 0,1001 |
| TcCLB.506633.30 | hypothetical protein, conserved | -0,2297 | 0,04369 |
| TcCLB.509543.40 | hypothetical protein, conserved | -0,2297 | 0,04486 |
| TcCLB.511643.20 | dihydroorotate dehydrogenase, putative (DHODH) | -0,2297 | 0,0546 |
| TcCLB.506821.120 | hypothetical protein, conserved | -0,2297 | 0,0923 |
| TcCLB.506253.30 | hypothetical protein | -0,2297 | 0,4321 |
| TcCLB.504101.40 | trans-sialidase (pseudogene), putative | -0,2299 | 0,04088 |
| TcCLB.506581.40 | hypothetical protein, conserved | -0,23 | 0,03406 |
| TcCLB.508723.70 | hypothetical protein, conserved | -0,2301 | 0,004881 |
| TcCLB.503473.20 | hypothetical protein, conserved | -0,2301 | 0,07658 |
| TcCLB.511559.40 | hypothetical protein, conserved | -0,2302 | 0,03624 |
| TcCLB.503723.50 | hypothetical protein, conserved | -0,2304 | 0,0166 |
| TcCLB.506933.89 | calmodulin, putative | -0,2305 | 0,08695 |
| TcCLB.511647.30 | meiotic recombination protein SPO11, putative | -0,2305 | 0,2528 |
| TcCLB.506737.100 | hypothetical protein, conserved | -0,2306 | 0,06555 |
| TcCLB.509791.50 | bilobe protein, conserved | -0,2307 | 0,04887 |
| TcCLB.509937.190 | hypothetical protein, conserved | -0,2308 | 0,02368 |
| TcCLB.511517.20 | hypothetical protein, conserved | -0,2308 | 0,04387 |
| TcCLB.510819.10 | Eukaryotic translation initiation factor 4 gamma type 1, pu | -0,2309 | 0,04418 |
| TcCLB.511299.79 | oxidoreductase, putative (fragment) | -0,2309 | 0,09923 |
| TcCLB.506295.50 | hypothetical protein, conserved | -0,2309 | 0,2107 |
| TcCLB.507877.5 | elongation factor 1-gamma (EF-1-gamma, pseudogene), | -0,2309 | 0,5494 |
| TcCLB.507945.10 | ATP-dependent RNA helicase, putative | -0,2312 | 0,02006 |
| TcCLB.508181.50 | hypothetical protein, conserved | -0,2312 | 0,08916 |
| TcCLB.506927.10 | hypothetical protein, conserved (pseudogene) | -0,2312 | 0,2161 |
| TcCLB.506303.10 | hypothetical protein, conserved | -0,2313 | 0,006661 |
| TcCLB.507467.20 | hypothetical protein, conserved | -0,2313 | 0,04796 |
| TcCLB.506207.10 | hypothetical protein, conserved | -0,2313 | 0,1213 |
| TcCLB.507729.30 | trans-sialidase (pseudogene), putative | -0,2314 | 0,2262 |
| TcCLB.509669.100 | hypothetical protein, conserved | -0,2315 | 0,06688 |
| TcCLB.508625.70 | mRNA capping methyltransferase, putative | -0,2315 | 0,1214 |

| | | | |
|---|---|---|---|
| TcCLB.509247.30 | hypothetical protein, conserved | -0,2316 | 0,00513 |
| TcCLB.508707.234 | hypothetical protein, conserved | -0,2317 | 0,09715 |
| TcCLB.511381.19 | hypothetical protein, conserved (fragment) | -0,2319 | 0,03087 |
| TcCLB.430281.10 | retrotransposon hot spot protein (RHS, pseudogene), puta | -0,2321 | 0,3657 |
| TcCLB.503697.100 | hypothetical protein, conserved | -0,2322 | 0,009833 |
| TcCLB.509579.30 | retrotransposon hot spot protein (RHS, pseudogene), puta | -0,2322 | 0,07712 |
| TcCLB.508709.10 | hypothetical protein, conserved | -0,2322 | 0,1286 |
| TcCLB.506935.10 | hypothetical protein, conserved | -0,2325 | 0,02064 |
| TcCLB.510329.280 | protein farnesyltransferase alpha subunit, putative | -0,2326 | 0,01417 |
| TcCLB.509561.39 | hypothetical protein, conserved | -0,2326 | 0,0555 |
| TcCLB.508461.270 | rab7 GTP binding protein, putative | -0,2327 | 0,006778 |
| TcCLB.508737.194 | ARP2/3 complex subunit, putative | -0,2327 | 0,1069 |
| TcCLB.508173.90 | hypothetical protein, conserved | -0,2327 | 0,1203 |
| TcCLB.503803.20 | hypothetical protein, conserved | -0,2327 | 0,1292 |
| TcCLB.506605.80 | hypothetical protein, conserved | -0,2329 | 0,054 |
| TcCLB.510517.50 | hypothetical protein, conserved | -0,2331 | 0,01668 |
| TcCLB.509875.250 | hypothetical protein, conserved | -0,2333 | 0,08551 |
| TcCLB.511199.9 | ATP-dependent RNA helicase FAL1, putative (fragment) | -0,2334 | 0,1363 |
| TcCLB.506727.20 | hypothetical protein, conserved | -0,2335 | 0,02534 |
| TcCLB.416041.14 | trans-sialidase, putative (fragment) | -0,2335 | 0,03614 |
| TcCLB.507967.4 | hypothetical protein, conserved | -0,2335 | 0,171 |
| TcCLB.507001.4 | hypothetical protein, conserved | -0,2336 | 0,3227 |
| TcCLB.506691.60 | hypothetical protein, conserved | -0,2337 | 0,1075 |
| TcCLB.511903.70 | hypothetical protein, conserved | -0,2337 | 0,1231 |
| TcCLB.506559.510 | hypothetical protein, conserved | -0,2337 | 0,1321 |
| TcCLB.503513.20 | hypothetical protein | -0,2338 | 0,08998 |
| TcCLB.506979.59 | hypothetical protein, conserved | -0,2339 | 0,1215 |
| TcCLB.510575.30 | hypothetical protein, conserved | -0,234 | 0,03577 |
| TcCLB.506737.50 | hypothetical protein, conserved | -0,2342 | 0,08705 |
| TcCLB.506767.310 | trans-sialidase (pseudogene), putative | -0,2342 | 0,259 |
| TcCLB.507099.40 | hypothetical protein, conserved | -0,2343 | 0,001025 |
| TcCLB.509567.4 | hypothetical protein, conserved | -0,2343 | 0,06613 |
| TcCLB.422867.20 | retrotransposon hot spot protein (RHS, pseudogene), puta | -0,2343 | 0,06933 |
| TcCLB.506195.250 | hypothetical protein, conserved | -0,2344 | 0,02926 |
| TcCLB.469785.20 | hypothetical protein, conserved | -0,2345 | 0,1442 |
| TcCLB.506559.484 | hypothetical protein, conserved | -0,2345 | 0,1461 |
| TcCLB.511577.180 | retrotransposon hot spot (RHS) protein, putative | -0,2346 | 0,06234 |
| TcCLB.506863.70 | hypothetical protein, conserved | -0,2347 | 0,01516 |
| TcCLB.508213.30 | hypothetical protein | -0,2347 | 0,08402 |
| TcCLB.507601.120 | hypothetical protein, conserved | -0,235 | 0,1152 |
| TcCLB.506201.154 | hypothetical protein, conserved | -0,235 | 0,1236 |
| TcCLB.511927.10 | hypothetical protein, conserved | -0,235 | 0,2263 |
| TcCLB.452283.4 | hypothetical protein, conserved | -0,2351 | 0,4276 |
| TcCLB.506661.70 | hypothetical protein, conserved | -0,2352 | 0,09456 |
| TcCLB.511229.120 | hypothetical protein | -0,2352 | 0,1069 |

| | | | |
|---|---|---|---|
| TcCLB.511145.49 | GPI transamidase component GAA1, putative | -0,2353 | 0,02973 |
| TcCLB.503675.20 | Charged multivesicular body protein 3, putative | -0,2354 | 0,08656 |
| TcCLB.504171.10 | hypothetical protein, conserved | -0,2354 | 0,08854 |
| TcCLB.508387.160 | MORN repeat-containing protein | -0,2355 | 0,1057 |
| TcCLB.504081.500 | mucin TcMUCII, putative | -0,2355 | 0,6106 |
| TcCLB.506857.40 | hypothetical protein, conserved | -0,2357 | 0,1023 |
| TcCLB.508461.120 | hypothetical protein, conserved | -0,2359 | 0,005242 |
| TcCLB.507213.20 | protein kinase, putative | -0,2359 | 0,04696 |
| TcCLB.510259.30 | hypothetical protein, conserved | -0,2359 | 0,1203 |
| TcCLB.507715.90 | hypothetical protein, conserved | -0,2361 | 0,008981 |
| TcCLB.509157.260 | hypothetical protein | -0,2361 | 0,0713 |
| TcCLB.507765.120 | hypothetical protein, conserved | -0,2364 | 0,009113 |
| TcCLB.511303.130 | hypothetical protein | -0,2365 | 0,03747 |
| TcCLB.509011.100 | myosin heavy chain kinase A, putative | -0,2366 | 0,0328 |
| TcCLB.506855.170 | hypothetical protein, conserved | -0,2366 | 0,1589 |
| TcCLB.507017.130 | hypothetical protein | -0,2366 | 0,1945 |
| TcCLB.508153.590 | hypothetical protein, conserved | -0,2367 | 0,06846 |
| TcCLB.511543.20 | As/Sb Reductase, putative (ACR2) | -0,2368 | 0,1925 |
| TcCLB.509521.84 | retrotransposon hot spot protein (RHS, pseudogene), puta | -0,2368 | 0,3889 |
| TcCLB.506467.20 | hypothetical protein, conserved | -0,2369 | 0,02099 |
| TcCLB.510329.260 | hypothetical protein, conserved | -0,237 | 0,001943 |
| TcCLB.510403.89 | synaptojanin (N-terminal domain), putative | -0,237 | 0,0427 |
| TcCLB.510381.60 | proliferator-activated receptor-interacting protein (PRIP) i | -0,2372 | 0,09069 |
| TcCLB.510121.170 | cysteine peptidase, Clan CA, family C2, putative | -0,2374 | 0,001098 |
| TcCLB.484299.4 | ATP-dependent DEAD/H RNA helicase, putative | -0,2374 | 0,3182 |
| TcCLB.511127.110 | ATP-dependent protease ATPase subunit HslU2 (HslU2) | -0,2375 | 0,003408 |
| TcCLB.510565.179 | hypothetical protein, conserved | -0,2376 | 0,006663 |
| TcCLB.511911.150 | nucleotide-binding protein, putative | -0,2378 | 0,1224 |
| TcCLB.511729.70 | cationic amino acid transporter, putative | -0,2378 | 0,1556 |
| TcCLB.506443.130 | trans-sialidase (pseudogene), putative | -0,238 | 0,03102 |
| TcCLB.508957.14 | hypothetical protein, conserved (fragment) | -0,2381 | 0,0764 |
| TcCLB.508119.40 | hypothetical protein | -0,2381 | 0,4641 |
| TcCLB.503841.40 | actin, putative | -0,2382 | 0,06094 |
| TcCLB.508693.84 | hypothetical protein, conserved | -0,2383 | 0,07751 |
| TcCLB.507993.30 | hypothetical protein, conserved | -0,2385 | 0,02589 |
| TcCLB.503803.4 | serine/threonine protein phosphatase 2A regulatory subui | -0,2385 | 0,1829 |
| TcCLB.511711.50 | hypothetical protein, conserved | -0,2386 | 0,05148 |
| TcCLB.510657.20 | ubiquitin hydrolase, putative | -0,2388 | 0,09854 |
| TcCLB.508405.119 | hypothetical protein, conserved | -0,2388 | 0,3474 |
| TcCLB.503785.10 | hypothetical protein, conserved (pseudogene) | -0,2389 | 0,03745 |
| TcCLB.511585.140 | hypothetical protein, conserved | -0,2391 | 0,006214 |
| TcCLB.511003.10 | phospholipid-translocating ATPase, putative | -0,2392 | 0,01705 |
| TcCLB.509799.40 | hypothetical protein, conserved | -0,2393 | 0,03113 |
| TcCLB.509767.30 | hypothetical protein | -0,2393 | 0,03647 |
| TcCLB.508943.20 | protein kinase, putative | -0,2394 | 0,1027 |

| | | | |
|---|---|---|---|
| TcCLB.510545.60 | hypothetical protein, conserved (pseudogene) | -0,2394 | 0,5458 |
| TcCLB.511717.100 | hypothetical protein, conserved | -0,2395 | 0,02857 |
| TcCLB.508153.670 | hypothetical protein, conserved | -0,2395 | 0,2456 |
| TcCLB.511529.90 | enolase, putative | -0,2396 | 0,08168 |
| TcCLB.509003.20 | calpain-like cysteine peptidase, putative | -0,2396 | 0,1137 |
| TcCLB.506409.230 | hypothetical protein, conserved | -0,2397 | 0,04687 |
| TcCLB.506559.250 | hypothetical protein, conserved | -0,2397 | 0,04871 |
| TcCLB.503929.70 | hypothetical protein, conserved | -0,2397 | 0,07211 |
| TcCLB.508543.110 | hypothetical protein, conserved | -0,2398 | 0,03648 |
| TcCLB.510181.69 | hypothetical protein, conserved | -0,2399 | 0,04974 |
| TcCLB.508387.10 | hypothetical protein, conserved | -0,2399 | 0,2012 |
| TcCLB.507023.200 | queuine tRNA-ribosyltransferase, putative | -0,24 | 0,03043 |
| TcCLB.511299.70 | mitogen-activated protein kinase, putative | -0,2401 | 0,02783 |
| TcCLB.511865.70 | hypothetical protein, conserved | -0,2401 | 0,05505 |
| TcCLB.506509.20 | hypothetical protein, conserved | -0,2401 | 0,1121 |
| TcCLB.506457.120 | hypothetical protein, conserved | -0,2402 | 0,03256 |
| TcCLB.503619.29 | hypothetical protein, conserved | -0,2402 | 0,1815 |
| TcCLB.511099.140 | mucin-associated surface protein (MASP), putative | -0,2402 | 0,6801 |
| TcCLB.509879.10 | hypothetical protein, conserved | -0,2403 | 0,1274 |
| TcCLB.506929.10 | surface protease GP63, putative | -0,2404 | 0,02638 |
| TcCLB.508145.49 | hypothetical protein, conserved | -0,2405 | 0,01973 |
| TcCLB.507623.30 | hypothetical protein, conserved | -0,2407 | 0,01117 |
| TcCLB.503583.40 | trans-splicing factor, putative | -0,2407 | 0,03598 |
| TcCLB.505071.100 | phosphatidylglycerolphosphate synthase, mitochondrial, p | -0,2408 | 0,02001 |
| TcCLB.504827.100 | hypothetical protein, conserved | -0,2408 | 0,05271 |
| TcCLB.511851.44 | mucin-associated surface protein (MASP, pseudogene), pu | -0,2409 | 0,007771 |
| TcCLB.504179.60 | hypothetical protein, conserved | -0,241 | 0,01869 |
| TcCLB.511923.71 | hypothetical protein, conserved (pseudogene) | -0,241 | 0,358 |
| TcCLB.508741.290 | ribosomal RNA methyltransferase, putative | -0,2411 | 0,0114 |
| TcCLB.508241.40 | hypothetical protein | -0,2411 | 0,01247 |
| TcCLB.511367.50 | hypothetical protein, conserved | -0,2414 | 0,02406 |
| TcCLB.506563.220 | calpain-like cysteine peptidase, putative | -0,2415 | 0,006262 |
| TcCLB.508837.160 | hypothetical protein, conserved | -0,2415 | 0,02614 |
| TcCLB.511903.20 | hypothetical protein, conserved | -0,2415 | 0,1521 |
| TcCLB.510659.170 | denn domain-containing protein | -0,2416 | 0,05266 |
| TcCLB.506173.20 | hypothetical protein | -0,2416 | 0,1048 |
| TcCLB.508933.5 | serine/threonine protein phosphatase PP1 (fragment) | -0,2417 | 0,206 |
| TcCLB.511529.9 | hypothetical protein, conserved | -0,242 | 0,01521 |
| TcCLB.511153.130 | DNA-direcetd RNA polymerase II, subunit 9, putative | -0,2421 | 0,1118 |
| TcCLB.504057.21 | hypothetical protein, conserved | -0,2422 | 0,0477 |
| TcCLB.511277.554 | hypothetical protein, conserved | -0,2422 | 0,07226 |
| TcCLB.510657.70 | hypothetical protein, conserved (pseudogene) | -0,2423 | 0,1349 |
| TcCLB.506509.60 | hypothetical protein, conserved | -0,2423 | 0,1392 |
| TcCLB.506097.9 | dynein heavy chain, putative (fragment) | -0,2424 | 0,06886 |
| TcCLB.508647.250 | phosphotransferase, putative | -0,2426 | 0,005217 |

| | | | |
|---|---|---|---|
| TcCLB.510187.220 | hypothetical protein, conserved | -0,2427 | 0,0704 |
| TcCLB.506127.100 | hypothetical protein, conserved | -0,2427 | 0,08761 |
| TcCLB.506803.14 | hypothetical protein | -0,2427 | 0,1134 |
| TcCLB.505999.60 | hypothetical protein | -0,2427 | 0,1768 |
| TcCLB.508539.10 | hypothetical protein, conserved | -0,2428 | 0,3552 |
| TcCLB.508725.4 | hypothetical protein, conserved | -0,2429 | 0,06564 |
| TcCLB.508977.31 | elongation factor 1-gamma (EF-1-gamma, pseudogene), | -0,2431 | 0,3761 |
| TcCLB.508045.60 | hypothetical protein, conserved | -0,2433 | 0,006578 |
| TcCLB.509561.9 | hypothetical protein, conserved | -0,2433 | 0,1297 |
| TcCLB.508879.190 | hypothetical protein, conserved | -0,2434 | 0,03012 |
| TcCLB.510265.40 | hypothetical protein, conserved | -0,2435 | 0,004179 |
| TcCLB.509611.70 | hypothetical protein, conserved | -0,2435 | 0,1046 |
| TcCLB.511555.90 | acyl-CoA binding protein, putative | -0,2436 | 0,02006 |
| TcCLB.507733.10 | trans-sialidase (pseudogene), putative | -0,2436 | 0,2498 |
| TcCLB.504741.100 | hypothetical protein, conserved | -0,2437 | 0,008375 |
| TcCLB.509399.70 | hypothetical protein, conserved | -0,2437 | 0,02441 |
| TcCLB.511907.300 | DNA repair and transcription factor protein, putative | -0,2438 | 0,04424 |
| TcCLB.509455.60 | hypothetical protein, conserved | -0,2438 | 0,04509 |
| TcCLB.508839.89 | DNA-directed rna polymerase I largest subunit, putative ( | -0,244 | 0,03941 |
| TcCLB.507499.90 | trans-sialidase (pseudogene), putative | -0,244 | 0,06459 |
| TcCLB.509967.3 | hypothetical protein, conserved (pseudogene) | -0,244 | 0,2381 |
| TcCLB.507641.100 | hypothetical protein, conserved | -0,2441 | 0,01237 |
| TcCLB.507089.190 | hypothetical protein, conserved | -0,2441 | 0,01736 |
| TcCLB.506337.180 | hypothetical protein, conserved | -0,2442 | 0,01268 |
| TcCLB.507073.10 | hypothetical protein, conserved | -0,2442 | 0,01858 |
| TcCLB.506727.70 | hypothetical protein, conserved | -0,2443 | 0,02742 |
| TcCLB.509455.30 | hypothetical protein, conserved | -0,2445 | 0,07515 |
| TcCLB.508479.280 | hypothetical protein, conserved | -0,2446 | 0,05497 |
| TcCLB.511401.111 | elongation factor 1-gamma (EF-1-gamma, pseudogene), | -0,2446 | 0,1676 |
| TcCLB.506793.70 | hypothetical protein, conserved | -0,2448 | 0,008181 |
| TcCLB.507753.6 | hypothetical protein, conserved (pseudogene) | -0,2448 | 0,2417 |
| TcCLB.511801.14 | protein kinase, putative (LDK) | -0,2449 | 0,02366 |
| TcCLB.504045.10 | hypothetical protein, conserved | -0,245 | 0,01244 |
| TcCLB.508909.330 | protein kinase, putative | -0,245 | 0,01394 |
| TcCLB.508675.50 | hypothetical protein, conserved | -0,2451 | 0,01098 |
| TcCLB.505997.21 | N-acetyltransferase complex ARD1 subunit (pseudogene), | -0,2451 | 0,5317 |
| TcCLB.510609.130 | hypothetical protein, conserved | -0,2453 | 0,02769 |
| TcCLB.508545.9 | hypothetical protein, conserved | -0,2453 | 0,1951 |
| TcCLB.510543.200 | trans-sialidase (pseudogene), putative | -0,2453 | 0,3779 |
| TcCLB.508579.30 | kinesin, putative | -0,2455 | 0,05175 |
| TcCLB.507005.80 | hypothetical protein, conserved | -0,2455 | 0,1247 |
| TcCLB.505807.230 | hypothetical protein | -0,2456 | 0,1614 |
| TcCLB.507805.30 | hypothetical protein, conserved (pseudogene) | -0,2457 | 0,2444 |
| TcCLB.506227.20 | hypothetical protein, conserved | -0,2458 | 0,1438 |
| TcCLB.511859.10 | AMP deaminase, putative | -0,2458 | 0,2712 |

| | | | |
|---|---|---|---|
| TcCLB.510327.10 | hypothetical protein | -0,2461 | 0,3929 |
| TcCLB.506419.40 | hypothetical protein, conserved | -0,2462 | 0,005359 |
| TcCLB.503789.60 | hypothetical protein, conserved | -0,2464 | 0,04462 |
| TcCLB.510247.20 | casein kinase I, putative | -0,2465 | 0,3319 |
| TcCLB.509999.30 | hypothetical protein, conserved | -0,2466 | 0,06271 |
| TcCLB.510669.30 | hypothetical protein, conserved | -0,2467 | 0,01467 |
| TcCLB.503407.20 | SUMO1/Ulp2, putative | -0,2467 | 0,0213 |
| TcCLB.506839.14 | hypothetical protein, conserved | -0,2467 | 0,03216 |
| TcCLB.507881.90 | trans-sialidase (pseudogene), putative | -0,2467 | 0,09573 |
| TcCLB.419703.10 | hypothetical protein | -0,2468 | 0,2336 |
| TcCLB.503885.40 | hypothetical protein, conserved | -0,2471 | 0,05689 |
| TcCLB.509591.60 | phospholipid-transporting ATPase 1-like protein, putative | -0,2473 | 0,5503 |
| TcCLB.506195.25 | hypothetical protein, conserved | -0,2474 | 0,3405 |
| TcCLB.508181.30 | hypothetical protein, conserved | -0,2475 | 0,001296 |
| TcCLB.511239.150 | histone acetyltransferase, putative | -0,2479 | 0,03351 |
| TcCLB.503813.30 | hypothetical protein, conserved | -0,248 | 0,06513 |
| TcCLB.508109.14 | mucin TcMUCII, putative | -0,248 | 0,4552 |
| TcCLB.503809.130 | hypothetical protein, conserved | -0,2481 | 0,03181 |
| TcCLB.511031.30 | hypothetical protein, conserved | -0,2482 | 0,05131 |
| TcCLB.506201.160 | hypothetical protein, conserved | -0,2482 | 0,234 |
| TcCLB.509007.40 | hypothetical protein, conserved | -0,2482 | 0,6938 |
| TcCLB.506221.80 | NAD synthase, putative | -0,2483 | 0,08509 |
| TcCLB.506459.215 | hypothetical protein, conserved (pseudogene) | -0,2484 | 0,4709 |
| TcCLB.508879.30 | dynein heavy chain, putative | -0,2486 | 0,001585 |
| TcCLB.509205.40 | hypothetical protein, conserved | -0,2486 | 0,05914 |
| TcCLB.510835.50 | hypothetical protein, conserved | -0,2488 | 0,06745 |
| TcCLB.509831.5 | vesicular transport protein (CDC48 homologue), putative ( | -0,2489 | 0,09353 |
| TcCLB.509747.30 | hypothetical protein, conserved | -0,249 | 0,004136 |
| TcCLB.504147.224 | RNA-editing-associated protein 1, putative (REAP-1) | -0,249 | 0,03642 |
| TcCLB.506791.10 | CYC2-like cyclin 4, putative (CYC4) | -0,2491 | 0,01671 |
| TcCLB.495369.10 | hypothetical protein, conserved (fragment) | -0,2491 | 0,08568 |
| TcCLB.506677.4 | hypothetical protein, conserved | -0,2492 | 0,4675 |
| TcCLB.510181.80 | hypothetical protein, conserved | -0,2493 | 0,02797 |
| TcCLB.507017.150 | AUT2/APG4/ATG4 cysteine peptidase, putative | -0,2493 | 0,04619 |
| TcCLB.511239.159 | inosine-guanine nucleoside hydrolase, putative | -0,2493 | 0,4172 |
| TcCLB.509429.240 | hypothetical protein, conserved | -0,2494 | 0,017 |
| TcCLB.506705.60 | hypothetical protein, conserved | -0,2496 | 0,0107 |
| TcCLB.510565.170 | hypothetical protein, conserved | -0,2498 | 0,4483 |
| TcCLB.507883.90 | adenylate kinase, putative | -0,2501 | 0,08215 |
| TcCLB.507517.100 | pseudouridylate synthase I, putative | -0,2502 | 0,04806 |
| TcCLB.511921.130 | hypothetical protein, conserved (pseudogene) | -0,2503 | 0,3088 |
| TcCLB.510519.20 | hypothetical protein, conserved | -0,2504 | 0,04779 |
| TcCLB.507975.50 | trans-sialidase (pseudogene), putative | -0,2505 | 0,001596 |
| TcCLB.504217.4 | hypothetical protein, conserved (fragment) | -0,2506 | 0,1374 |
| TcCLB.506195.240 | hypothetical protein, conserved | -0,2507 | 0,01058 |

| | | | |
|---|---|---|---|
| TcCLB.511875.64 | hypothetical protein, conserved (pseudogene) | -0,2507 | 0,1553 |
| TcCLB.506559.20 | hypothetical protein, conserved | -0,2508 | 0,00695 |
| TcCLB.510287.10 | dynein light chain, putative | -0,2508 | 0,05937 |
| TcCLB.509807.20 | hypothetical protein, conserved | -0,2509 | 0,02292 |
| TcCLB.508445.90 | hypothetical protein, conserved | -0,2509 | 0,02459 |
| TcCLB.510729.20 | hypothetical protein, conserved | -0,251 | 0,02956 |
| TcCLB.508307.140 | hypothetical protein, conserved | -0,251 | 0,03321 |
| TcCLB.509065.30 | hypothetical protein, conserved | -0,2511 | 0,0155 |
| TcCLB.506739.60 | hypothetical protein, conserved | -0,2513 | 0,01857 |
| TcCLB.508647.260 | hypothetical protein, conserved | -0,2513 | 0,04196 |
| TcCLB.510431.340 | hypothetical protein, conserved | -0,2513 | 0,1073 |
| TcCLB.511603.40 | hypothetical protein | -0,2513 | 0,496 |
| TcCLB.508445.70 | hypothetical protein, conserved | -0,2514 | 0,01624 |
| TcCLB.508461.200 | hypothetical protein, conserved | -0,2514 | 0,03196 |
| TcCLB.506321.260 | hypothetical protein, conserved | -0,2514 | 0,1008 |
| TcCLB.506425.70 | hypothetical protein, conserved | -0,2516 | 0,01007 |
| TcCLB.507017.80 | hypothetical protein | -0,2516 | 0,09159 |
| TcCLB.507689.10 | dihydrouridine synthase (Dus, pseudogene), putative | -0,2516 | 0,2068 |
| TcCLB.510999.30 | hypothetical protein, conserved | -0,2517 | 0,02976 |
| TcCLB.509215.16 | GlcNAc-PI synthesis protein (GPI3) | -0,2517 | 0,07879 |
| TcCLB.510279.130 | mucin TcMUCII, putative | -0,2517 | 0,5743 |
| TcCLB.511481.60 | RNA helicase, putative | -0,2519 | 0,0009456 |
| TcCLB.510525.120 | hypothetical protein, conserved | -0,252 | 0,04583 |
| TcCLB.511285.40 | hypothetical protein, conserved | -0,252 | 0,08099 |
| TcCLB.506247.140 | hypothetical protein, conserved | -0,2521 | 0,01101 |
| TcCLB.507669.120 | ATPase domain protein, putative | -0,2521 | 0,03028 |
| TcCLB.508415.20 | hypothetical protein, conserved | -0,2522 | 0,01668 |
| TcCLB.508277.210 | hypothetical protein, conserved | -0,2522 | 0,02282 |
| TcCLB.508607.10 | retrotransposon hot spot (RHS) protein, putative | -0,2522 | 0,4418 |
| TcCLB.511307.10 | kinesin, putative | -0,2523 | 0,02559 |
| TcCLB.504777.40 | mucin TcMUCII, putative | -0,2523 | 0,1815 |
| TcCLB.507163.20 | hypothetical protein | -0,2523 | 0,4418 |
| TcCLB.507715.30 | transcription elongation factor-like protein, putative | -0,2524 | 0,01159 |
| TcCLB.508461.540 | hypothetical protein, conserved | -0,2524 | 0,03579 |
| TcCLB.507049.66 | hypothetical protein | -0,2524 | 0,1359 |
| TcCLB.426897.4 | hypothetical protein, conserved (fragment) | -0,2524 | 0,2091 |
| TcCLB.509267.100 | methionyl-tRNA formyltransferase, putative | -0,2527 | 0,04941 |
| TcCLB.504507.20 | DNA replication licensing factor MCM8, putative | -0,2527 | 0,1875 |
| TcCLB.510231.29 | multidrug resistance-associated protein, putative | -0,2529 | 0,01495 |
| TcCLB.510879.130 | hypothetical protein, conserved | -0,2529 | 0,04721 |
| TcCLB.510895.29 | hypothetical protein, conserved | -0,2531 | 0,1741 |
| TcCLB.507031.110 | hypothetical protein, conserved | -0,2533 | 0,009071 |
| TcCLB.504179.9 | pre-mRNA splicing factor ATP-dependent RNA helicase, p | -0,2533 | 0,1145 |
| TcCLB.507747.195 | glyceraldehyde-3-phosphate dehydrogenase (pseudogene | -0,2533 | 0,2362 |
| TcCLB.508151.10 | mucin TcMUCII, putative | -0,2533 | 0,3766 |

| | | | |
|---|---|---|---|
| TcCLB.504787.47 | elongation factor 1-gamma (EF-1-gamma, pseudogene), | -0,2535 | 0,5974 |
| TcCLB.509487.20 | hypothetical protein, conserved | -0,2536 | 0,03216 |
| TcCLB.506445.10 | actin-like protein, putative | -0,2536 | 0,07144 |
| TcCLB.434945.10 | hypothetical protein, conserved | -0,2536 | 0,3227 |
| TcCLB.508325.101 | hypothetical protein, conserved | -0,2536 | 0,4287 |
| TcCLB.506443.80 | synaptojanin (N-terminal domain), putative | -0,254 | 0,007351 |
| TcCLB.506517.158 | hypothetical protein, conserved | -0,254 | 0,05641 |
| TcCLB.507539.49 | hypothetical protein, conserved | -0,254 | 0,2267 |
| TcCLB.511529.250 | hypothetical protein | -0,2541 | 0,02304 |
| TcCLB.503897.60 | hypothetical protein, conserved | -0,2541 | 0,03693 |
| TcCLB.511545.140 | hypothetical protein, conserved | -0,2542 | 0,03291 |
| TcCLB.507059.30 | retrotransposon hot spot protein (RHS, pseudogene), puta | -0,2542 | 0,06441 |
| TcCLB.507739.120 | amastin, putative | -0,2543 | 0,3568 |
| TcCLB.511755.90 | hypothetical protein, conserved | -0,2544 | 0,0164 |
| TcCLB.506203.90 | hypothetical protein, conserved | -0,2545 | 0,009908 |
| TcCLB.506703.100 | mucin TcMUCII (pseudogene), putative | -0,2545 | 0,6162 |
| TcCLB.511837.10 | vesicle-associated membrane protein, putative | -0,2546 | 0,02875 |
| TcCLB.509001.20 | hypothetical protein, conserved | -0,2549 | 0,007015 |
| TcCLB.510421.90 | protein kinase, putative | -0,2549 | 0,01649 |
| TcCLB.504019.3 | hypothetical protein (fragment) | -0,255 | 0,1305 |
| TcCLB.507007.45 | anaphase promoting complex subunit protein, putative | -0,2551 | 0,3563 |
| TcCLB.444777.20 | hypothetical protein, conserved | -0,2552 | 0,01943 |
| TcCLB.506559.240 | hypothetical protein, conserved | -0,2553 | 0,00612 |
| TcCLB.509789.59 | hypothetical protein, conserved | -0,2554 | 0,0046 |
| TcCLB.508831.140 | surface antigen 2 (CA-2), putative | -0,2554 | 0,02167 |
| TcCLB.510091.90 | hypothetical protein, conserved | -0,2555 | 0,005455 |
| TcCLB.503449.30 | clathrin coat assembly protein, putative | -0,2555 | 0,01327 |
| TcCLB.507681.180 | hypothetical protein, conserved | -0,2555 | 0,06323 |
| TcCLB.508163.310 | mucin TcMUCII, putative | -0,2555 | 0,1809 |
| TcCLB.504033.210 | hypothetical protein, conserved | -0,2557 | 0,2033 |
| TcCLB.510303.180 | hypothetical protein, conserved | -0,2558 | 0,0003165 |
| TcCLB.511511.50 | hypothetical protein, conserved | -0,2558 | 0,02356 |
| TcCLB.507159.20 | hypothetical protein, conserved | -0,2558 | 0,04054 |
| TcCLB.503953.30 | protein kinase, putative | -0,2559 | 0,02683 |
| TcCLB.511877.30 | surface protease GP63 (pseudogene), putative | -0,2561 | 0,2356 |
| TcCLB.508277.180 | mismatch repair protein MSH3, putative | -0,2566 | 0,001932 |
| TcCLB.511851.30 | trans-sialidase (pseudogene), putative | -0,2566 | 0,04122 |
| TcCLB.508417.20 | hypothetical protein | -0,2568 | 0,4332 |
| TcCLB.503409.10 | hypothetical protein, conserved | -0,2569 | 0,09547 |
| TcCLB.511217.100 | hypothetical protein, conserved | -0,257 | 0,01198 |
| TcCLB.507515.140 | oxysterol-binding protein, putative | -0,2571 | 0,01952 |
| TcCLB.508075.7 | hypothetical protein, conserved | -0,2571 | 0,1148 |
| TcCLB.509647.100 | hypothetical protein, conserved | -0,2572 | 0,04569 |
| TcCLB.506315.100 | protein phosphatase 2C, putative | -0,2573 | 0,01254 |
| TcCLB.507053.190 | hypothetical protein, conserved | -0,2574 | 0,02335 |

| | | | |
|---|---|---|---|
| TcCLB.508303.20 | hypothetical protein | -0,2575 | 0,05532 |
| TcCLB.506131.161 | trans-sialidase (pseudogene), putative | -0,2576 | 0,3833 |
| TcCLB.511531.9 | chaperone DNAJ protein, putative | -0,2579 | 0,07111 |
| TcCLB.511393.79 | hypothetical protein, conserved | -0,2582 | 0,02738 |
| TcCLB.511183.300 | trans-sialidase (pseudogene), putative | -0,2584 | 0,5265 |
| TcCLB.511321.50 | hypothetical protein, conserved | -0,2585 | 0,0073 |
| TcCLB.506473.20 | DnaJ chaperone protein, putative | -0,2585 | 0,009534 |
| TcCLB.509011.90 | lipase, putative | -0,2585 | 0,09276 |
| TcCLB.507625.120 | hypothetical protein, conserved | -0,2585 | 0,1392 |
| TcCLB.506859.130 | hypothetical protein, conserved | -0,2588 | 0,2625 |
| TcCLB.508707.10 | amino acid transporter, putative | -0,259 | 0,2281 |
| TcCLB.509315.20 | trans-sialidase (pseudogene), putative | -0,2591 | 0,4394 |
| TcCLB.509407.10 | hypothetical protein | -0,2593 | 0,2072 |
| TcCLB.508307.120 | hypothetical protein, conserved | -0,2595 | 0,002452 |
| TcCLB.510737.40 | transport protein particle (TRAPP) subunit, putative | -0,2595 | 0,05061 |
| TcCLB.504109.174 | hypothetical protein, conserved | -0,2596 | 0,04654 |
| TcCLB.508715.10 | oxidoreductase, putative (fragment) | -0,2596 | 0,1031 |
| TcCLB.510967.13 | hypothetical protein, conserved | -0,2596 | 0,1829 |
| TcCLB.506511.50 | hypothetical protein, conserved | -0,2597 | 0,0006376 |
| TcCLB.511545.70 | kinteoplast poly(A) polymerase complex 1 subunit, putativ | -0,2597 | 0,00209 |
| TcCLB.511491.9 | hypothetical protein, conserved | -0,2597 | 0,1345 |
| TcCLB.506959.110 | hypothetical protein, conserved | -0,2599 | 0,01484 |
| TcCLB.509693.160 | hypothetical protein, conserved | -0,26 | 0,02669 |
| TcCLB.511115.20 | GTP-binding elongation factor Tu family, putative | -0,2602 | 0,002609 |
| TcCLB.509985.20 | hypothetical protein, conserved | -0,2603 | 0,02132 |
| TcCLB.503869.30 | hypothetical protein, conserved | -0,2604 | 0,00526 |
| TcCLB.508817.110 | hypothetical protein, conserved | -0,2605 | 0,05637 |
| TcCLB.506435.100 | hypothetical protein, conserved | -0,2605 | 0,1344 |
| TcCLB.511171.50 | mucin-associated surface protein (MASP, pseudogene), pu | -0,2605 | 0,3697 |
| TcCLB.510889.20 | hypothetical protein, conserved | -0,2606 | 0,0397 |
| TcCLB.509591.10 | ubiquitin-conjugating enzyme, putative | -0,2606 | 0,08484 |
| TcCLB.504001.6 | protein kinase, putative | -0,2607 | 0,01694 |
| TcCLB.511017.69 | histone acetyltransferase, putative (fragment) | -0,2607 | 0,2564 |
| TcCLB.504071.50 | spliceosome-associated protein, putative | -0,2608 | 0,007458 |
| TcCLB.509805.120 | hypothetical protein, conserved | -0,2609 | 0,08807 |
| TcCLB.507007.37 | hypothetical protein, conserved | -0,261 | 0,5039 |
| TcCLB.509161.20 | hypothetical protein, conserved | -0,2611 | 0,06802 |
| TcCLB.507923.30 | vacuolar assembly protein vps41, putative | -0,2612 | 0,005156 |
| TcCLB.511045.40 | subtilisin peptidase | -0,2612 | 0,007686 |
| TcCLB.508899.130 | hypothetical protein, conserved | -0,2612 | 0,1069 |
| TcCLB.504149.70 | hypothetical protein, conserved | -0,2613 | 0,03258 |
| TcCLB.508325.80 | trans-sialidase (pseudogene), putative | -0,2614 | 0,01157 |
| TcCLB.506163.10 | surface protease GP63, putative | -0,2614 | 0,02398 |
| TcCLB.503759.59 | hypothetical protein, conserved | -0,2614 | 0,04297 |
| TcCLB.509551.80 | hypothetical protein, conserved | -0,2614 | 0,06248 |

| | | | |
|---|---|---|---|
| TcCLB.507089.80 | exosome-associated protein 2, putative (EAP2) | -0,2615 | 0,05294 |
| TcCLB.506531.9 | hypothetical protein (fragment) | -0,2615 | 0,07183 |
| TcCLB.503823.70 | hypothetical protein, conserved | -0,2615 | 0,1452 |
| TcCLB.507071.277 | syntaxin binding protein (pseudogene), putative | -0,2615 | 0,4708 |
| TcCLB.506565.8 | RNA-binding protein, putative | -0,2616 | 0,004703 |
| TcCLB.509805.70 | hypothetical protein, conserved | -0,2616 | 0,004839 |
| TcCLB.508207.135 | hypothetical protein, conserved | -0,2617 | 0,02269 |
| TcCLB.509495.30 | trans-sialidase, Group I, putative | -0,2617 | 0,03713 |
| TcCLB.504741.60 | transcription initiation protein, putative | -0,2618 | 0,005135 |
| TcCLB.511711.20 | hypothetical protein, conserved | -0,262 | 0,0323 |
| TcCLB.503853.10 | hypothetical protein, conserved | -0,2621 | 0,01471 |
| TcCLB.443683.9 | DNA-directed rna polymerase I largest subunit, putative (f | -0,2621 | 0,06458 |
| TcCLB.510351.90 | dihydrolipoamide branched chain transacylase, putative | -0,2623 | 0,003497 |
| TcCLB.510123.70 | vesicle-associated membrane protein, putative | -0,2623 | 0,01301 |
| TcCLB.508647.80 | hypothetical protein, conserved | -0,2623 | 0,01446 |
| TcCLB.509899.20 | retrotransposon hot spot protein (RHS, pseudogene), puta | -0,2623 | 0,05693 |
| TcCLB.509569.60 | hypothetical protein, conserved | -0,2624 | 0,04303 |
| TcCLB.510525.40 | hypothetical protein, conserved | -0,2626 | 0,09456 |
| TcCLB.509287.30 | hypothetical protein, conserved | -0,2627 | 0,003126 |
| TcCLB.510397.30 | retrotransposon hot spot protein (RHS, pseudogene), puta | -0,2627 | 0,3752 |
| TcCLB.509753.50 | retrotransposon hot spot protein (RHS, pseudogene), puta | -0,2628 | 0,4274 |
| TcCLB.511589.240 | hypothetical protein, conserved | -0,2629 | 0,02366 |
| TcCLB.508815.179 | dynein heavy chain, putative (fragment) | -0,263 | 0,0247 |
| TcCLB.507603.240 | hypothetical protein, conserved | -0,2631 | 0,0188 |
| TcCLB.506175.120 | hypothetical protein, conserved | -0,2631 | 0,4691 |
| TcCLB.511661.20 | hypothetical protein, conserved | -0,2633 | 0,01157 |
| TcCLB.506947.60 | hypothetical protein, conserved | -0,2633 | 0,04522 |
| TcCLB.506945.10 | hypothetical protein, conserved | -0,2634 | 0,1284 |
| TcCLB.511275.30 | actin-related protein 2/3 complex subunit 1, putative (ARI | -0,2635 | 0,03744 |
| TcCLB.509011.30 | hypothetical protein | -0,2635 | 0,0598 |
| TcCLB.504059.11 | hypothetical protein, conserved (fragment) | -0,2635 | 0,07776 |
| TcCLB.507889.10 | transaldolase, putative | -0,2636 | 0,024 |
| TcCLB.511315.44 | hypothetical protein, conserved (pseudogene) | -0,2638 | 0,1795 |
| TcCLB.510841.10 | Kinesin-13 4b, putative (fragment) (KIN13-4b) | -0,2638 | 0,2087 |
| TcCLB.506587.50 | hypothetical protein, conserved | -0,264 | 0,01151 |
| TcCLB.508257.260 | AP-1 adapter complex gamma subunit, putative | -0,2642 | 0,002779 |
| TcCLB.508257.110 | hypothetical protein, conserved | -0,2642 | 0,007704 |
| TcCLB.510635.60 | hypothetical protein, conserved | -0,2642 | 0,04013 |
| TcCLB.508831.4 | dynein heavy chain, putative | -0,2643 | 0,02403 |
| TcCLB.511001.60 | protein kinase, putative | -0,2646 | 0,04276 |
| TcCLB.506195.130 | hypothetical protein, conserved | -0,2646 | 0,07059 |
| TcCLB.510747.130 | hypothetical protein, conserved | -0,2647 | 0,004457 |
| TcCLB.509619.10 | Zn-finger protein, putative | -0,265 | 0,01506 |
| TcCLB.509941.30 | retrotransposon hot spot protein (RHS, pseudogene), puta | -0,265 | 0,04157 |
| TcCLB.508355.70 | diacylglycerol acyltransferase, putative | -0,2651 | 0,006887 |

| | | | |
|---|---|---|---|
| TcCLB.506275.50 | Golgi SNAP receptor complex member 1, putative | -0,2651 | 0,05293 |
| TcCLB.511019.99 | glutathione peroxidase, putative | -0,2651 | 0,1057 |
| TcCLB.510241.90 | hypothetical protein, conserved | -0,2654 | 0,003858 |
| TcCLB.509719.56 | hypothetical protein, conserved | -0,2658 | 0,003649 |
| TcCLB.508303.4 | ubiquitin-protein ligase, putative | -0,2658 | 0,03709 |
| TcCLB.506871.70 | hypothetical protein, conserved | -0,2659 | 0,03095 |
| TcCLB.506201.10 | hypothetical protein, conserved | -0,266 | 0,04764 |
| TcCLB.511523.20 | hypothetical protein, conserved | -0,266 | 0,05029 |
| TcCLB.506673.50 | apurinic/apyrimidinic endonuclease, putative | -0,2662 | 0,1286 |
| TcCLB.511285.20 | hypothetical protein, conserved | -0,2664 | 0,00181 |
| TcCLB.511071.80 | hypothetical protein, conserved | -0,2664 | 0,09711 |
| TcCLB.508461.250 | ATP-dependent RNA helicase, putative | -0,2666 | 0,001784 |
| TcCLB.511031.49 | hypothetical protein, conserved | -0,2666 | 0,03138 |
| TcCLB.505939.10 | hypothetical protein | -0,2666 | 0,4409 |
| TcCLB.508879.90 | hypothetical protein, conserved | -0,2667 | 0,07653 |
| TcCLB.506855.270 | hypothetical protein, conserved | -0,267 | 0,1024 |
| TcCLB.509509.30 | 6-phosphofructo-2-kinase/fructose-2,6-biphosphatase, pu | -0,2671 | 0,01825 |
| TcCLB.504055.10 | hypothetical protein, conserved | -0,2671 | 0,06909 |
| TcCLB.510261.21 | hypothetical protein | -0,2671 | 0,6439 |
| TcCLB.510529.20 | hypothetical protein, conserved | -0,2672 | 0,01277 |
| TcCLB.507963.70 | hypothetical protein, conserved | -0,2675 | 0,005388 |
| TcCLB.511723.10 | surface protease GP63, putative | -0,2675 | 0,08453 |
| TcCLB.510371.5 | elongation factor 1-gamma (EF-1-gamma, pseudogene), | -0,2676 | 0,4953 |
| TcCLB.509161.34 | hypothetical protein, conserved | -0,2677 | 0,02857 |
| TcCLB.506943.30 | hypothetical protein, conserved | -0,2679 | 0,004882 |
| TcCLB.510601.20 | hypothetical protein, conserved | -0,2679 | 0,02249 |
| TcCLB.505555.10 | lysosomal/endosomal membrane protein p67 (pseudoger | -0,2681 | 0,003184 |
| TcCLB.506567.130 | hypothetical protein, conserved | -0,2682 | 0,09879 |
| TcCLB.506587.10 | hypothetical protein, conserved | -0,2683 | 0,07209 |
| TcCLB.508731.40 | hypothetical protein, conserved | -0,2684 | 0,01637 |
| TcCLB.504741.230 | hypothetical protein, conserved | -0,2685 | 0,007668 |
| TcCLB.507837.100 | hypothetical protein, conserved | -0,2686 | 0,004748 |
| TcCLB.507037.60 | hypothetical protein, conserved | -0,2687 | 0,002585 |
| TcCLB.506493.20 | hypothetical protein | -0,2689 | 0,2733 |
| TcCLB.507209.60 | hypothetical protein, conserved | -0,269 | 0,00531 |
| TcCLB.510819.40 | hypothetical protein, conserved | -0,269 | 0,07026 |
| TcCLB.508569.130 | 6-phosphofructo-2-kinase/fructose-2,6-biphosphatase, pu | -0,2691 | 0,0008399 |
| TcCLB.503781.50 | hypothetical protein, conserved | -0,2691 | 0,01699 |
| TcCLB.510901.120 | protein kinase, putative | -0,2692 | 0,002505 |
| TcCLB.507757.40 | hypothetical protein | -0,2695 | 0,006214 |
| TcCLB.510293.50 | DnaJ chaperone protein, putative | -0,2696 | 0,01088 |
| TcCLB.507915.19 | mismatch repair protein MSH3, putative | -0,2696 | 0,1596 |
| TcCLB.507797.9 | nuclear transcription factor, putative | -0,2697 | 0,1479 |
| TcCLB.504655.30 | protein kinase, putative | -0,2698 | 0,007587 |
| TcCLB.508815.170 | protein kinase, putative | -0,27 | 0,0001866 |

| | | | |
|---|---|---|---|
| TcCLB.511301.60 | hypothetical protein, conserved | -0,27 | 0,02462 |
| TcCLB.508397.64 | hypothetical protein, conserved | -0,27 | 0,1004 |
| TcCLB.510227.20 | hypothetical protein | -0,2701 | 0,01812 |
| TcCLB.511587.39 | ATP-dependent DEAD/H RNA helicase (pseudogene), puta | -0,2701 | 0,1762 |
| TcCLB.409407.10 | trans-sialidase (pseudogene), putative | -0,2702 | 0,3024 |
| TcCLB.506871.180 | G5-interacting protein, putative (G5-IP) | -0,2703 | 0,002634 |
| TcCLB.506857.60 | peptidyl-prolyl cis-trans isomerase, putative (PAR45) | -0,2703 | 0,004513 |
| TcCLB.510257.60 | 3-ketoacyl-CoA reductase, putative | -0,2709 | 0,01196 |
| TcCLB.509595.30 | hypothetical protein, conserved | -0,2709 | 0,01782 |
| TcCLB.506869.54 | hypothetical protein, conserved | -0,271 | 0,04652 |
| TcCLB.506537.100 | trans-sialidase (pseudogene), putative | -0,2712 | 0,02026 |
| TcCLB.510163.10 | hypothetical protein, conserved | -0,2712 | 0,02132 |
| TcCLB.503685.10 | serine/threonine-protein kinase, putative | -0,2713 | 0,04025 |
| TcCLB.510437.70 | hypothetical protein, conserved | -0,2713 | 0,05827 |
| TcCLB.508257.230 | target of rapamycin kinase (TOR) kinase 3, putative | -0,2714 | 0,0002021 |
| TcCLB.503899.139 | hypothetical protein, conserved | -0,2715 | 0,001196 |
| TcCLB.511071.181 | hypothetical protein, conserved | -0,2715 | 0,008003 |
| TcCLB.506127.120 | hypothetical protein, conserved | -0,2716 | 0,01853 |
| TcCLB.506425.100 | hypothetical protein, conserved | -0,2717 | 0,01417 |
| TcCLB.506319.30 | hypothetical protein, conserved | -0,2719 | 0,03862 |
| TcCLB.506775.22 | hypothetical protein, conserved | -0,272 | 0,0181 |
| TcCLB.505997.60 | retrotransposon hot spot (RHS) protein, putative | -0,2721 | 0,09234 |
| TcCLB.511867.160 | hypothetical protein, conserved | -0,2721 | 0,09356 |
| TcCLB.511155.4 | hypothetical protein, conserved | -0,2724 | 0,03593 |
| TcCLB.509217.146 | N-acetyltransferase complex ARD1 subunit (pseudogene), | -0,2724 | 0,5125 |
| TcCLB.506925.520 | hypothetical protein, conserved | -0,2728 | 0,01478 |
| TcCLB.507929.30 | hypothetical protein, conserved | -0,2728 | 0,08781 |
| TcCLB.508929.10 | UDP-Gal or UDP-GlcNAc-dependent glycosyltransferase, p | -0,273 | 0,168 |
| TcCLB.416511.9 | mismatch repair protein MSH3, putative | -0,2731 | 0,0342 |
| TcCLB.510145.30 | hypothetical protein, conserved | -0,2732 | 0,02284 |
| TcCLB.508919.180 | trans-sialidase (pseudogene), putative | -0,2734 | 0,003388 |
| TcCLB.506629.180 | hypothetical protein, conserved | -0,2734 | 0,09601 |
| TcCLB.506537.30 | hypothetical protein, conserved (pseudogene) | -0,2735 | 0,01363 |
| TcCLB.506563.160 | hypothetical protein, conserved | -0,2736 | 0,006622 |
| TcCLB.503897.70 | hypothetical protein, conserved | -0,2737 | 0,0352 |
| TcCLB.506399.30 | hypothetical protein, conserved | -0,2737 | 0,07107 |
| TcCLB.508919.100 | hypothetical protein, conserved | -0,2738 | 0,01083 |
| TcCLB.503973.140 | hypothetical protein, conserved (pseudogene) | -0,2739 | 0,4081 |
| TcCLB.508577.150 | hypothetical protein, conserved | -0,274 | 0,1521 |
| TcCLB.509833.10 | hypothetical protein, conserved | -0,2741 | 0,07945 |
| TcCLB.506179.59 | ATP-dependent RNA helicase, putative (fragment) | -0,2741 | 0,1267 |
| TcCLB.510901.250 | hypothetical protein, conserved | -0,2743 | 0,01436 |
| TcCLB.506195.40 | hypothetical protein, conserved | -0,2743 | 0,0589 |
| TcCLB.506441.10 | hypothetical protein, conserved | -0,2744 | 0,001635 |
| TcCLB.506425.110 | hypothetical protein, conserved | -0,2744 | 0,03761 |

| | | | |
|---|---|---|---|
| TcCLB.507447.19 | microtubule-associated protein, putative (fragment) | -0,2744 | 0,516 |
| TcCLB.508033.69 | mucin-associated surface protein (MASP, pseudogene), pu | -0,2745 | 0,2274 |
| TcCLB.399389.10 | hypothetical protein, conserved | -0,2747 | 0,001571 |
| TcCLB.510999.19 | hypothetical protein, conserved | -0,2748 | 0,05585 |
| TcCLB.510307.210 | trans-sialidase, Group II, putative | -0,2748 | 0,2345 |
| TcCLB.510743.20 | hypothetical protein, conserved | -0,275 | 0,06158 |
| TcCLB.511667.90 | hypothetical protein, conserved (pseudogene) | -0,275 | 0,1068 |
| TcCLB.510207.21 | mucin-associated surface protein (MASP, pseudogene), pu | -0,2751 | 0,3142 |
| TcCLB.507765.110 | hypothetical protein, conserved | -0,2752 | 0,006351 |
| TcCLB.506905.20 | hypothetical protein, conserved | -0,2752 | 0,01344 |
| TcCLB.504163.70 | hypothetical protein, conserved | -0,2752 | 0,02255 |
| TcCLB.508175.114 | hypothetical protein, conserved | -0,2754 | 0,005372 |
| TcCLB.504229.37 | hypothetical protein | -0,2755 | 0,3347 |
| TcCLB.509733.30 | hypothetical protein, conserved | -0,2758 | 0,007742 |
| TcCLB.508873.280 | mucin-associated surface protein (MASP, pseudogene), pu | -0,276 | 0,1885 |
| TcCLB.509575.20 | hypothetical protein, conserved | -0,2761 | 0,04827 |
| TcCLB.511277.92 | hypothetical protein, conserved | -0,2762 | 0,01053 |
| TcCLB.511481.55 | RNA-binding protein, putative | -0,2762 | 0,08647 |
| TcCLB.509259.100 | ATP-dependent DEAD/H RNA helicase (pseudogene), puta | -0,2762 | 0,3368 |
| TcCLB.506993.40 | mucin TcMUCII, putative | -0,2764 | 0,542 |
| TcCLB.427789.30 | NAD synthase, putative | -0,2767 | 0,04013 |
| TcCLB.507601.60 | hypothetical protein, conserved | -0,2768 | 0,0498 |
| TcCLB.509233.220 | hypothetical protein, conserved | -0,2769 | 0,004898 |
| TcCLB.508153.570 | protein kinase, putative | -0,277 | 0,01999 |
| TcCLB.503515.7 | ribosomal RNA methyltransferase, putative (fragment) | -0,2771 | 0,1006 |
| TcCLB.510371.30 | hypothetical protein, conserved (pseudogene) | -0,2773 | 0,5244 |
| TcCLB.507559.50 | kinesin, putative | -0,2774 | 0,01575 |
| TcCLB.507099.100 | hypothetical protein, conserved | -0,2774 | 0,03703 |
| TcCLB.510955.20 | hypothetical protein, conserved | -0,2775 | 0,003596 |
| TcCLB.511531.44 | hypothetical protein | -0,2775 | 0,04183 |
| TcCLB.508671.20 | serine carboxypeptidase (CBP1), putative | -0,2779 | 0,03269 |
| TcCLB.511145.24 | hypothetical protein, conserved | -0,2779 | 0,1975 |
| TcCLB.510877.90 | hypothetical protein, conserved | -0,278 | 0,007735 |
| TcCLB.504079.10 | branch point binding protein, putative | -0,278 | 0,02238 |
| TcCLB.508827.130 | hypothetical protein, conserved | -0,2782 | 0,01248 |
| TcCLB.511389.80 | protein kinase, putative | -0,2782 | 0,01461 |
| TcCLB.511245.90 | hypothetical protein, conserved | -0,2783 | 0,02582 |
| TcCLB.506249.10 | hypothetical protein, conserved | -0,2783 | 0,08009 |
| TcCLB.504017.70 | hypothetical protein | -0,2784 | 0,02065 |
| TcCLB.506983.20 | hypothetical protein, conserved | -0,2785 | 0,004618 |
| TcCLB.506943.50 | glyceraldehyde 3-phosphate dehydrogenase, putative | -0,2785 | 0,01924 |
| TcCLB.506593.40 | hypothetical protein, conserved | -0,2786 | 0,01484 |
| TcCLB.504045.30 | hypothetical protein | -0,2787 | 0,04817 |
| TcCLB.509073.80 | hypothetical protein, conserved | -0,2788 | 0,02141 |
| TcCLB.510303.280 | hypothetical protein | -0,2789 | 0,01455 |

| | | | |
|---|---|---|---|
| TcCLB.508793.14 | dynein regulatory complex subunit, putative (fragment) | -0,2791 | 0,005599 |
| TcCLB.503881.10 | kynureninase, putative | -0,2792 | 0,0656 |
| TcCLB.508469.30 | hypothetical protein, conserved | -0,2793 | 0,05942 |
| TcCLB.507623.101 | surface protease GP63 (pseudogene), putative | -0,2794 | 0,04651 |
| TcCLB.504039.99 | mucin TcMUCII (pseudogene), putative | -0,2794 | 0,3905 |
| TcCLB.511857.80 | AMP deaminase, putative | -0,2795 | 0,01096 |
| TcCLB.509911.40 | hypothetical protein, conserved | -0,2795 | 0,08803 |
| TcCLB.416619.10 | trans-sialidase, putative (fragment) | -0,2795 | 0,5281 |
| TcCLB.506949.20 | hypothetical protein, conserved | -0,2796 | 0,0004422 |
| TcCLB.510187.330 | hypothetical protein, conserved | -0,2797 | 0,002971 |
| TcCLB.506793.80 | ATP-dependent RNA helicase, putative | -0,2797 | 0,01291 |
| TcCLB.509455.110 | hypothetical protein, conserved | -0,2797 | 0,06145 |
| TcCLB.509163.54 | zinc finger protein, putative | -0,2797 | 0,6098 |
| TcCLB.506977.139 | hypothetical protein, conserved | -0,2799 | 0,01225 |
| TcCLB.507949.29 | conserved protein, unknown function | -0,2799 | 0,04552 |
| TcCLB.507629.30 | hypothetical protein, conserved | -0,2801 | 0,03036 |
| TcCLB.510303.260 | hypothetical protein, conserved | -0,2802 | 0,03101 |
| TcCLB.511229.20 | oxidoreductase, putative | -0,2802 | 0,06881 |
| TcCLB.511215.100 | hypothetical protein | -0,2803 | 0,1219 |
| TcCLB.509233.120 | hypothetical protein, conserved | -0,2804 | 0,0006156 |
| TcCLB.506405.90 | hypothetical protein, conserved | -0,2804 | 0,03333 |
| TcCLB.410275.18 | trans-sialidase (pseudogene), putative | -0,2804 | 0,5671 |
| TcCLB.507011.200 | dihydroxyacetone kinase 1-like, putative | -0,2805 | 0,01127 |
| TcCLB.511257.70 | surface protease GP63, putative | -0,2806 | 0,02212 |
| TcCLB.509163.60 | hypothetical protein, conserved | -0,2806 | 0,02487 |
| TcCLB.509733.10 | hypothetical protein, conserved | -0,2807 | 0,01711 |
| TcCLB.511003.80 | hypothetical protein, conserved | -0,2808 | 0,004605 |
| TcCLB.508727.24 | hypothetical protein | -0,281 | 0,002169 |
| TcCLB.511627.140 | hypothetical protein, conserved | -0,2811 | 0,0365 |
| TcCLB.507825.49 | hypothetical protein, conserved (fragment) | -0,2811 | 0,09437 |
| TcCLB.507241.20 | hypothetical protein, conserved | -0,2812 | 0,005941 |
| TcCLB.510303.60 | hypothetical protein, conserved | -0,2812 | 0,02844 |
| TcCLB.507207.10 | hypothetical protein, conserved | -0,2813 | 0,00695 |
| TcCLB.511211.240 | hypothetical protein, conserved | -0,2813 | 0,04434 |
| TcCLB.506739.20 | hypothetical protein, conserved | -0,2814 | 0,004068 |
| TcCLB.508719.50 | hypothetical protein, conserved | -0,2814 | 0,06154 |
| TcCLB.511683.20 | hypothetical protein, conserved | -0,2817 | 0,0232 |
| TcCLB.508181.10 | hypothetical protein, conserved | -0,2818 | 0,03035 |
| TcCLB.511717.110 | hypothetical protein, conserved | -0,2819 | 0,01786 |
| TcCLB.506223.30 | hypothetical protein, conserved | -0,2819 | 0,02229 |
| TcCLB.510611.30 | kinesin, putative | -0,282 | 0,001226 |
| TcCLB.506327.70 | engulfment and cell motility domain 2, putative | -0,282 | 0,00458 |
| TcCLB.511287.80 | hypothetical protein, conserved | -0,2821 | 0,05024 |
| TcCLB.509897.51 | hypothetical protein, conserved (pseudogene) | -0,2821 | 0,0532 |
| TcCLB.507053.200 | hypothetical protein, conserved | -0,2822 | 0,0161 |

| | | | |
|---|---|---|---|
| TcCLB.507857.10 | transcription elongation factor, putative | -0,2823 | 0,0108 |
| TcCLB.507993.10 | hypothetical protein, conserved | -0,2824 | 0,009797 |
| TcCLB.508257.240 | hypothetical protein, conserved | -0,2826 | 0,02838 |
| TcCLB.509073.100 | hypothetical protein, conserved | -0,2828 | 0,1109 |
| TcCLB.508559.80 | trans-sialidase (pseudogene), putative | -0,2828 | 0,1151 |
| TcCLB.510343.10 | hypothetical protein, conserved | -0,2828 | 0,1297 |
| TcCLB.511353.20 | hypothetical protein, conserved | -0,2829 | 0,008525 |
| TcCLB.508277.120 | ribosomal protein S6, putative | -0,2829 | 0,0454 |
| TcCLB.506399.20 | hypothetical protein, conserved | -0,283 | 0,02964 |
| TcCLB.511725.10 | proteasome regulatory non-ATPase subunit, putative | -0,2831 | 0,01531 |
| TcCLB.511837.100 | amino acid permease-like protein, putative | -0,2832 | 0,01328 |
| TcCLB.508921.40 | hypothetical protein, conserved | -0,2833 | 0,1349 |
| TcCLB.506247.160 | hypothetical protein, conserved | -0,2835 | 0,00162 |
| TcCLB.508347.10 | hypothetical protein, conserved | -0,2835 | 0,00459 |
| TcCLB.507949.80 | kinetoplast ribosomal PPR-repeat containing protein 3, pu | -0,2835 | 0,009139 |
| TcCLB.508235.40 | protein kinase, putative (fragment) | -0,2835 | 0,09917 |
| TcCLB.506587.20 | hypothetical protein | -0,2836 | 0,01086 |
| TcCLB.503899.40 | hypothetical protein, conserved | -0,2838 | 0,003973 |
| TcCLB.508461.280 | protein kinase A catalytic subunit isoform 2, putative | -0,2838 | 0,03253 |
| TcCLB.503893.10 | copper-transporting ATPase-like protein, putative | -0,2838 | 0,1668 |
| TcCLB.508231.150 | hypothetical protein | -0,2839 | 0,003637 |
| TcCLB.509127.20 | hypothetical protein, conserved | -0,2839 | 0,03583 |
| TcCLB.503619.20 | hypothetical protein, conserved | -0,284 | 0,01312 |
| TcCLB.506177.70 | hypothetical protein | -0,284 | 0,08757 |
| TcCLB.507049.160 | hypothetical protein, conserved | -0,2841 | 0,02822 |
| TcCLB.510603.100 | hypothetical protein, conserved | -0,2841 | 0,08612 |
| TcCLB.508269.80 | retrotransposon hot spot protein (RHS, pseudogene), puta | -0,2841 | 0,2036 |
| TcCLB.508209.60 | spliced leader RNA PSE-promoter transcription factor, put | -0,2843 | 0,004684 |
| TcCLB.508817.180 | DIS3-like exonuclease, putative (pseudogene) | -0,2845 | 0,002328 |
| TcCLB.511229.90 | hypothetical protein, conserved | -0,2845 | 0,009872 |
| TcCLB.508765.30 | glycosylphosphatidylinositol-specific phospholipase C, puta | -0,2845 | 0,0338 |
| TcCLB.507625.90 | hypothetical protein, conserved | -0,2846 | 0,04933 |
| TcCLB.510429.50 | Lsm12 protein, putative | -0,2848 | 0,01586 |
| TcCLB.506943.60 | glyceraldehyde 3-phosphate dehydrogenase, putative | -0,2849 | 0,1155 |
| TcCLB.508583.10 | hypothetical protein, conserved | -0,285 | 0,06055 |
| TcCLB.503547.20 | dispersed gene family protein 1 (DGF-1, pseudogene), put | -0,285 | 0,4282 |
| TcCLB.511179.140 | hypothetical protein, conserved | -0,2853 | 0,001243 |
| TcCLB.404073.20 | actin, putative (fragment) | -0,2854 | 0,1344 |
| TcCLB.509937.110 | hypothetical protein, conserved | -0,2855 | 0,0179 |
| TcCLB.506355.110 | hypothetical protein, conserved | -0,2856 | 0,001366 |
| TcCLB.511127.30 | hypothetical protein, conserved | -0,2856 | 0,02209 |
| TcCLB.508949.10 | hypothetical protein, conserved | -0,2856 | 0,03204 |
| TcCLB.507017.50 | polynucleotide kinase 3'-phosphatase, putative | -0,2858 | 0,01423 |
| TcCLB.509611.140 | hypothetical protein, conserved | -0,2858 | 0,01484 |
| TcCLB.506213.40 | hypothetical protein, conserved | -0,2861 | 0,0002927 |

| | | | |
|---|---|---|---|
| TcCLB.508837.140 | U1 small nuclear ribonucleoprotein, putative | -0,2861 | 0,0476 |
| TcCLB.504125.10 | SWI/SNF-related matrix-associated actin-dependent regu | -0,2862 | 0,003465 |
| TcCLB.508209.159 | hypothetical protein, conserved | -0,2865 | 0,03178 |
| TcCLB.506947.80 | arginine N-methyltransferase, type I (PRMT6) | -0,2867 | 0,06754 |
| TcCLB.508495.110 | hypothetical protein | -0,2867 | 0,4691 |
| TcCLB.507089.100 | hypothetical protein, conserved | -0,2868 | 0,01417 |
| TcCLB.504835.29 | hypothetical protein, conserved | -0,2869 | 0,002215 |
| TcCLB.508821.60 | hypothetical protein, conserved | -0,2869 | 0,2816 |
| TcCLB.504131.10 | hypothetical protein, conserved (pseudogene) | -0,287 | 0,006461 |
| TcCLB.511875.40 | surface protease GP63 (pseudogene), putative | -0,287 | 0,04619 |
| TcCLB.508707.60 | hypothetical protein, conserved | -0,2872 | 0,0009761 |
| TcCLB.503593.60 | hypothetical protein, conserved | -0,2872 | 0,01232 |
| TcCLB.503679.20 | hypothetical protein, conserved | -0,2873 | 0,01231 |
| TcCLB.511385.80 | kinesin-like protein, putative | -0,2873 | 0,01877 |
| TcCLB.506213.110 | hypothetical protein | -0,2874 | 0,001664 |
| TcCLB.504839.36 | hypothetical protein, conserved | -0,2874 | 0,05442 |
| TcCLB.507491.59 | ubiquitin fold modifier protein, putative (UFM1) | -0,2875 | 0,129 |
| TcCLB.506181.140 | hypothetical protein | -0,2876 | 0,008037 |
| TcCLB.405179.10 | dynein heavy chain, putative (fragment) | -0,2876 | 0,03874 |
| TcCLB.511381.40 | DNA-directed rna polymerase I largest subunit, putative | -0,2878 | 0,0002005 |
| TcCLB.510659.279 | nucleic acid binding protein, putative | -0,2879 | 0,06055 |
| TcCLB.503833.10 | hypothetical protein, conserved | -0,2881 | 0,002348 |
| TcCLB.506755.80 | hypothetical protein | -0,2881 | 0,01284 |
| TcCLB.509803.50 | hypothetical protein, conserved | -0,2882 | 0,005242 |
| TcCLB.507527.20 | hypothetical protein, conserved | -0,2883 | 0,0021 |
| TcCLB.509505.40 | hypothetical protein, conserved | -0,2883 | 0,01343 |
| TcCLB.509719.80 | hypothetical protein, conserved | -0,2883 | 0,02 |
| TcCLB.508445.50 | hypothetical protein, conserved | -0,2884 | 0,07937 |
| TcCLB.503459.29 | hypothetical protein, conserved | -0,2886 | 0,02887 |
| TcCLB.510967.10 | GPI inositol deacylase precursor, putative | -0,2886 | 0,06175 |
| TcCLB.508539.95 | surface protease GP63 (pseudogene), putative | -0,2886 | 0,4404 |
| TcCLB.508169.100 | hypothetical protein, conserved | -0,2887 | 0,01311 |
| TcCLB.507677.30 | hypothetical protein, conserved | -0,2889 | 0,007076 |
| TcCLB.507023.250 | hypothetical protein, conserved | -0,2889 | 0,01998 |
| TcCLB.503555.10 | DNA replication licensing factor MCM8, putative | -0,2889 | 0,02472 |
| TcCLB.510701.31 | hypothetical protein | -0,2889 | 0,5704 |
| TcCLB.507867.80 | hypothetical protein, conserved | -0,289 | 0,0004937 |
| TcCLB.507641.239 | hypothetical protein, conserved | -0,2893 | 0,002612 |
| TcCLB.510699.34 | trans-sialidase (pseudogene), putative | -0,2894 | 0,2336 |
| TcCLB.507709.70 | hypothetical protein, conserved | -0,2895 | 0,006252 |
| TcCLB.507993.5 | GPI-anchor transamidase subunit 8 (GPI8, pseudogene), p | -0,2897 | 0,1891 |
| TcCLB.508637.30 | hypothetical protein, conserved (pseudogene) | -0,2898 | 0,2194 |
| TcCLB.510889.251 | Present in the outer mitochondrial membrane proteome 1 | -0,29 | 0,07161 |
| TcCLB.508825.30 | surface protease GP63 (pseudogene), putative | -0,2901 | 0,02104 |
| TcCLB.504027.10 | hypothetical protein, conserved | -0,2901 | 0,03556 |

| | | | |
|---|---|---|---|
| TcCLB.509581.20 | beta galactofuranosyl glycosyltransferase, putative | -0,2903 | 0,0395 |
| TcCLB.510187.190 | vacuolar protein sorting-associated protein, putative | -0,2904 | 0,0003633 |
| TcCLB.507093.66 | hypothetical protein, conserved | -0,2904 | 0,01602 |
| TcCLB.403789.9 | DNA-directed rna polymerase I largest subunit, putative | -0,2904 | 0,09178 |
| TcCLB.511301.160 | trans-sialidase, Group II, putative | -0,2906 | 0,03188 |
| TcCLB.510197.95 | hypothetical protein, conserved (pseudogene) | -0,2906 | 0,1491 |
| TcCLB.511713.20 | hypothetical protein, conserved | -0,2908 | 0,007626 |
| TcCLB.510067.9 | hypothetical protein, conserved | -0,2908 | 0,01222 |
| TcCLB.508891.80 | hypothetical protein, conserved (pseudogene) | -0,2909 | 0,02567 |
| TcCLB.509017.50 | hypothetical protein, conserved | -0,291 | 0,0002305 |
| TcCLB.510491.50 | hypothetical protein, conserved | -0,291 | 0,1546 |
| TcCLB.511753.30 | kinesin, putative | -0,2912 | 0,001138 |
| TcCLB.509171.80 | hypothetical protein, conserved | -0,2912 | 0,00701 |
| TcCLB.506929.50 | hypothetical protein, conserved | -0,2912 | 0,00858 |
| TcCLB.430737.20 | hypothetical protein, conserved (fragment) | -0,2912 | 0,07107 |
| TcCLB.511523.90 | hypothetical protein, conserved | -0,2913 | 0,00191 |
| TcCLB.508153.890 | hypothetical protein, conserved | -0,2914 | 0,00006872 |
| TcCLB.504033.140 | hypothetical protein, conserved (pseudogene) | -0,2914 | 0,002632 |
| TcCLB.511201.58 | hypothetical protein | -0,2914 | 0,01006 |
| TcCLB.503955.110 | protein kinase, putative | -0,2914 | 0,01356 |
| TcCLB.508409.170 | hypothetical protein, conserved | -0,2916 | 0,02024 |
| TcCLB.511907.220 | hypothetical protein, conserved | -0,2917 | 0,01608 |
| TcCLB.504587.10 | surface protease GP63 (pseudogene), putative | -0,2919 | 0,3051 |
| TcCLB.510885.60 | hypothetical protein, conserved | -0,2922 | 0,002109 |
| TcCLB.511245.79 | hypothetical protein, conserved | -0,2922 | 0,0187 |
| TcCLB.503579.50 | hypothetical protein, conserved | -0,2923 | 0,001175 |
| TcCLB.511257.100 | surface protease GP63, putative | -0,2924 | 0,006835 |
| TcCLB.509993.31 | mucin-associated surface protein (MASP, pseudogene), pu | -0,2924 | 0,2584 |
| TcCLB.506789.230 | hypothetical protein, conserved | -0,2925 | 0,00003251 |
| TcCLB.506625.160 | hypothetical protein, conserved | -0,2925 | 0,00009022 |
| TcCLB.507685.20 | trans-sialidase (pseudogene), putative | -0,2926 | 0,2178 |
| TcCLB.508647.30 | hypothetical protein, conserved | -0,2928 | 0,01151 |
| TcCLB.510493.10 | hypothetical protein, conserved | -0,2928 | 0,09126 |
| TcCLB.508153.450 | hypothetical protein, conserved | -0,2929 | 0,004094 |
| TcCLB.511737.20 | hypothetical protein, conserved | -0,293 | 0,00154 |
| TcCLB.511153.140 | proteasome beta 3 subunit, putative | -0,293 | 0,02862 |
| TcCLB.511529.230 | hypothetical protein, conserved | -0,2931 | 0,039 |
| TcCLB.506925.70 | epsin, putative | -0,2932 | 0,0004384 |
| TcCLB.510187.310 | hypothetical protein, conserved | -0,2934 | 0,001582 |
| TcCLB.509455.9 | hypothetical protein, conserved | -0,2934 | 0,005276 |
| TcCLB.506585.60 | hypothetical protein, conserved | -0,2934 | 0,02183 |
| TcCLB.510105.30 | clathrin coat assembly protein, putative | -0,2935 | 0,01896 |
| TcCLB.503653.50 | histone deacetylase, putative | -0,2937 | 0,005911 |
| TcCLB.411427.40 | hypothetical protein, conserved | -0,2937 | 0,1168 |
| TcCLB.503777.10 | suppressive immunomodulating factor, putative (TSIF) | -0,2938 | 0,002837 |

| | | | |
|---|---|---|---|
| TcCLB.511361.70 | hypothetical protein, conserved | -0,2939 | 0,005137 |
| TcCLB.509715.23 | RNA-binding protein, putative | -0,294 | 0,1778 |
| TcCLB.511875.25 | elongation factor 1-gamma (EF-1-gamma, pseudogene), | -0,294 | 0,3927 |
| TcCLB.509179.210 | hypothetical protein, conserved | -0,2946 | 0,01074 |
| TcCLB.506489.10 | hypothetical protein, conserved | -0,2947 | 0,0001772 |
| TcCLB.510027.10 | GDP-mannose 4,6 dehydratase, putative | -0,2948 | 0,009183 |
| TcCLB.508879.70 | hypothetical protein, conserved | -0,295 | 0,02872 |
| TcCLB.511627.90 | hypothetical protein, conserved | -0,295 | 0,05583 |
| TcCLB.508271.50 | hypothetical protein | -0,2953 | 0,002348 |
| TcCLB.509105.150 | heat shock protein 85, putative | -0,2953 | 0,01064 |
| TcCLB.506297.140 | hypothetical protein, conserved | -0,2954 | 0,01602 |
| TcCLB.509751.4 | hypothetical protein, conserved (fragment) | -0,2955 | 0,1769 |
| TcCLB.506559.4 | hypothetical protein, conserved (fragment) | -0,2956 | 0,02028 |
| TcCLB.506213.10 | hypothetical protein, conserved | -0,2956 | 0,02031 |
| TcCLB.511907.140 | hypothetical protein | -0,2957 | 0,09619 |
| TcCLB.507519.167 | hypothetical protein | -0,2957 | 0,1341 |
| TcCLB.506945.330 | hypothetical protein, conserved | -0,2958 | 0,00008496 |
| TcCLB.506925.340 | hypothetical protein, conserved | -0,2959 | 0,02597 |
| TcCLB.507049.130 | hypothetical protein, conserved (pseudogene) | -0,2959 | 0,1384 |
| TcCLB.511623.20 | hypothetical protein, conserved | -0,296 | 0,05648 |
| TcCLB.511717.190 | hypothetical protein, conserved | -0,2961 | 0,02751 |
| TcCLB.506821.160 | hypothetical protein, conserved | -0,2962 | 0,008325 |
| TcCLB.507993.33 | hypothetical protein, conserved | -0,2963 | 0,04393 |
| TcCLB.506265.209 | hypothetical protein, conserved (fragment) | -0,2964 | 0,3349 |
| TcCLB.508851.31 | N-acetyltransferase complex ARD1 subunit (pseudogene), | -0,2967 | 0,2127 |
| TcCLB.506695.30 | protein kinase, putative | -0,2969 | 0,01735 |
| TcCLB.508741.370 | hypothetical protein, conserved | -0,2971 | 0,00007255 |
| TcCLB.461515.10 | mucin TcMUCI, putative (fragment) | -0,2972 | 0,02054 |
| TcCLB.508257.40 | hypothetical protein, conserved | -0,2973 | 0,005159 |
| TcCLB.507823.10 | hypothetical protein, conserved | -0,2974 | 0,01081 |
| TcCLB.508881.100 | hypothetical protein, conserved | -0,2975 | 0,001052 |
| TcCLB.507633.110 | mucin-associated surface protein (MASP, pseudogene), pu | -0,2976 | 0,2629 |
| TcCLB.503779.100 | hypothetical protein, conserved | -0,2977 | 0,01488 |
| TcCLB.508207.270 | hypothetical protein, conserved | -0,2979 | 0,001128 |
| TcCLB.505007.39 | hypothetical protein, conserved | -0,298 | 0,05138 |
| TcCLB.506357.150 | hypothetical protein, conserved | -0,2981 | 0,002938 |
| TcCLB.419469.30 | hypothetical protein, conserved | -0,2982 | 0,009721 |
| TcCLB.510105.160 | hypothetical protein | -0,2982 | 0,04114 |
| TcCLB.503479.20 | hypothetical protein, conserved (pseudogene) | -0,2985 | 0,09121 |
| TcCLB.509965.160 | hypothetical protein, conserved | -0,2986 | 0,04267 |
| TcCLB.503419.40 | hypothetical protein, conserved | -0,2987 | 0,01909 |
| TcCLB.511717.10 | hypothetical protein, conserved (pseudogene) | -0,2987 | 0,1744 |
| TcCLB.508479.270 | hypothetical protein, conserved | -0,2988 | 0,000466 |
| TcCLB.506753.60 | GDP-mannose transporter, putative | -0,2988 | 0,04197 |
| TcCLB.510561.35 | hypothetical protein | -0,2988 | 0,5011 |

| | | | |
|---|---|---|---|
| TcCLB.504253.50 | hypothetical protein, conserved | -0,2989 | 0,02247 |
| TcCLB.508181.120 | hypothetical protein, conserved | -0,299 | 0,002352 |
| TcCLB.503929.40 | hypothetical protein, conserved | -0,2992 | 0,04464 |
| TcCLB.506709.10 | hypothetical protein, conserved | -0,2994 | 0,00314 |
| TcCLB.509047.30 | hypothetical protein, conserved | -0,2995 | 0,06263 |
| TcCLB.508351.50 | rho-like GTP binding protein, putative | -0,2998 | 0,005756 |
| TcCLB.508389.20 | hypothetical protein | -0,2999 | 0,00247 |
| TcCLB.439653.10 | NADH-dependent fumarate reductase, putative | -0,2999 | 0,2404 |
| TcCLB.506163.30 | hypothetical protein, conserved | -0,3 | 0,01222 |
| TcCLB.416891.19 | hypothetical protein | -0,3 | 0,1445 |
| TcCLB.506181.20 | hypothetical protein, conserved | -0,3001 | 0,003909 |
| TcCLB.510295.10 | Enriched in surface-labeled proteome protein 18 (ESP18) | -0,3001 | 0,005139 |
| TcCLB.510821.40 | hypothetical protein, conserved | -0,3001 | 0,00749 |
| TcCLB.506567.34 | hypothetical protein, conserved | -0,3003 | 0,00286 |
| TcCLB.503835.10 | hypothetical protein, conserved | -0,3005 | 0,05139 |
| TcCLB.510381.50 | hypothetical protein, conserved | -0,3006 | 0,06532 |
| TcCLB.511721.10 | hypothetical protein, conserved | -0,3007 | 0,005436 |
| TcCLB.509443.60 | hypothetical protein, conserved | -0,3007 | 0,1142 |
| TcCLB.509643.130 | heat shock protein 85, putative | -0,3008 | 0,009834 |
| TcCLB.507429.60 | hypothetical protein | -0,3008 | 0,1557 |
| TcCLB.508461.320 | PRP8 protein homologue, putative | -0,301 | 0,00000919 |
| TcCLB.509451.24 | hypothetical protein, conserved (fragment) | -0,301 | 0,005418 |
| TcCLB.508233.19 | hypothetical protein, conserved | -0,301 | 0,07599 |
| TcCLB.506605.50 | hypothetical protein, conserved | -0,3011 | 0,001865 |
| TcCLB.509777.100 | ubiquitin-activating enzyme, putative | -0,3014 | 0,00272 |
| TcCLB.508909.90 | Protein kinase domain, putative | -0,3015 | 0,0005615 |
| TcCLB.510431.160 | nucleolar GTP-binding protein 1, putative | -0,3016 | 0,02791 |
| TcCLB.506181.90 | F-actin capping protein, putative | -0,3017 | 0,005527 |
| TcCLB.511211.49 | hypothetical protein, conserved | -0,3017 | 0,01161 |
| TcCLB.506177.60 | hypothetical protein | -0,3018 | 0,01157 |
| TcCLB.508461.589 | hypothetical protein, conserved (fragment) | -0,3018 | 0,1138 |
| TcCLB.506925.200 | hypothetical protein, conserved | -0,302 | 0,002131 |
| TcCLB.504227.10 | hypothetical protein, conserved | -0,3021 | 0,02023 |
| TcCLB.509055.30 | hypothetical protein, conserved | -0,3022 | 0,006846 |
| TcCLB.506507.20 | hypothetical protein, conserved | -0,3022 | 0,04388 |
| TcCLB.510357.160 | hypothetical protein, conserved (fragment) | -0,3022 | 0,1533 |
| TcCLB.506529.40 | hypothetical protein, conserved | -0,3023 | 0,04979 |
| TcCLB.508409.80 | hypothetical protein, conserved | -0,3024 | 0,0000621 |
| TcCLB.507887.30 | hypothetical protein, conserved | -0,3024 | 0,03786 |
| TcCLB.506965.110 | hypothetical protein | -0,3024 | 0,4694 |
| TcCLB.509099.5 | NUP-1 protein, putative (fragment) | -0,3025 | 0,02018 |
| TcCLB.510767.60 | hypothetical protein, conserved | -0,3025 | 0,02882 |
| TcCLB.507085.130 | hypothetical protein | -0,3025 | 0,415 |
| TcCLB.506835.50 | hypothetical protein | -0,3026 | 0,0005615 |
| TcCLB.508533.40 | ecotin, putative | -0,3027 | 0,02115 |

| | | | |
|---|---|---|---|
| TcCLB.507941.100 | hypothetical protein, conserved | -0,3028 | 0,00634 |
| TcCLB.412093.10 | antigenic protein (pseudogene), putative | -0,3028 | 0,1398 |
| TcCLB.507041.55 | mucin-associated surface protein (MASP, pseudogene), pu | -0,3029 | 0,2637 |
| TcCLB.511281.10 | hypothetical protein, conserved | -0,303 | 0,03146 |
| TcCLB.506127.160 | hypothetical protein, conserved | -0,3031 | 0,006634 |
| TcCLB.504009.60 | hypothetical protein, conserved | -0,3034 | 0,002567 |
| TcCLB.506925.10 | hypothetical protein | -0,3035 | 0,1755 |
| TcCLB.506647.10 | hypothetical protein, conserved | -0,3037 | 0,003204 |
| TcCLB.507885.10 | RNA-binding protein, putative | -0,3037 | 0,04518 |
| TcCLB.506295.90 | coatomer subunit zeta, putative | -0,3038 | 0,04174 |
| TcCLB.508971.10 | UDP-Gal or UDP-GlcNAc-dependent glycosyltransferase, p | -0,3039 | 0,05411 |
| TcCLB.507991.20 | protein kinase, putative | -0,304 | 0,01692 |
| TcCLB.511167.40 | hypothetical protein, conserved | -0,3041 | 0,03394 |
| TcCLB.507071.294 | surface protease GP63 (pseudogene), putative | -0,3041 | 0,5999 |
| TcCLB.506679.240 | hypothetical protein, conserved | -0,3042 | 0,01016 |
| TcCLB.508505.40 | hypothetical protein, conserved | -0,3043 | 0,01737 |
| TcCLB.506951.70 | surface protease GP63 (pseudogene), putative | -0,3044 | 0,2412 |
| TcCLB.510297.80 | DNA repair and transcription factor protein, putative | -0,3045 | 0,01537 |
| TcCLB.506753.40 | hypothetical protein, conserved | -0,3046 | 0,03586 |
| TcCLB.503543.4 | hypothetical protein, conserved (fragment) | -0,3046 | 0,09704 |
| TcCLB.503851.40 | hypothetical protein, conserved | -0,3047 | 0,005834 |
| TcCLB.511625.4 | hypothetical protein, conserved | -0,3047 | 0,0911 |
| TcCLB.503923.10 | hypothetical protein, conserved | -0,3049 | 0,00563 |
| TcCLB.510421.100 | hypothetical protein, conserved | -0,3051 | 0,006164 |
| TcCLB.509099.120 | hypothetical protein, conserved | -0,3052 | 0,02331 |
| TcCLB.506011.30 | hypothetical protein, conserved | -0,3052 | 0,09769 |
| TcCLB.511211.210 | hypothetical protein, conserved | -0,3052 | 0,1135 |
| TcCLB.506691.90 | serine/threonine protein kinase, putative | -0,3052 | 0,3853 |
| TcCLB.507857.4 | transcription elongation factor, putative | -0,3054 | 0,01204 |
| TcCLB.510903.30 | helicase-like protein, putative | -0,3055 | 0,006819 |
| TcCLB.508545.30 | hypothetical protein, conserved | -0,3056 | 0,1039 |
| TcCLB.506835.80 | CAS/CSE/importin domain protein, putative | -0,3057 | 0,0005635 |
| TcCLB.508997.20 | phosphatidylinositol 3-kinase, putative | -0,3057 | 0,001732 |
| TcCLB.508919.80 | hypothetical protein, conserved | -0,3057 | 0,01695 |
| TcCLB.511231.40 | hypothetical protein, conserved | -0,306 | 0,09362 |
| TcCLB.509729.20 | hypothetical protein, conserved | -0,3062 | 0,005175 |
| TcCLB.507017.104 | hypothetical protein | -0,3063 | 0,03422 |
| TcCLB.511111.50 | ubiquitin-conjugating enzyme, putative | -0,3063 | 0,3737 |
| TcCLB.508501.10 | hypothetical protein | -0,3064 | 0,2318 |
| TcCLB.511807.33 | expression site-associated gene (ESAG-like) protein, puta | -0,3065 | 0,003973 |
| TcCLB.506221.20 | hypothetical protein, conserved | -0,3066 | 0,0332 |
| TcCLB.506247.460 | hypothetical protein, conserved | -0,3067 | 0,008182 |
| TcCLB.506679.20 | hypothetical protein, conserved | -0,3067 | 0,1953 |
| TcCLB.508699.50 | hypothetical protein, conserved | -0,3069 | 0,004067 |
| TcCLB.507317.30 | hypothetical protein, conserved | -0,3071 | 0,03534 |

| | | | |
|---|---|---|---|
| TcCLB.504243.49 | hypothetical protein, conserved (fragment) | -0,3072 | 0,01182 |
| TcCLB.511761.70 | hypothetical protein, conserved | -0,3072 | 0,01182 |
| TcCLB.511237.90 | hypothetical protein, conserved | -0,3072 | 0,1386 |
| TcCLB.503463.20 | hypothetical protein, conserved | -0,3074 | 0,007399 |
| TcCLB.510329.250 | hypothetical protein, conserved | -0,3074 | 0,03747 |
| TcCLB.510651.20 | hypothetical protein, conserved | -0,3074 | 0,03846 |
| TcCLB.503893.120 | calcium uniporter protein, mitochondrial, putative | -0,3075 | 0,01551 |
| TcCLB.511527.20 | TFIIH basal transcription factor complex helicase XPB subu | -0,3076 | 0,002467 |
| TcCLB.511279.50 | hypothetical protein, conserved | -0,3076 | 0,02788 |
| TcCLB.508163.371 | hypothetical protein, conserved (pseudogene) | -0,3076 | 0,2058 |
| TcCLB.511263.30 | hypothetical protein, conserved | -0,3077 | 0,0004454 |
| TcCLB.503583.30 | hypothetical protein, conserved | -0,3077 | 0,02318 |
| TcCLB.507901.30 | retrotransposon hot spot protein (RHS, pseudogene), puta | -0,3079 | 0,0149 |
| TcCLB.508543.140 | electron transfer flavoprotein-ubiquinone oxidoreductase, | -0,3079 | 0,1448 |
| TcCLB.509671.160 | DNA polymerase epsilon subunit B, putative | -0,308 | 0,008966 |
| TcCLB.402647.9 | hypothetical protein, conserved | -0,308 | 0,01934 |
| TcCLB.503891.110 | dolicholphosphate-mannose synthase, putative | -0,308 | 0,02064 |
| TcCLB.511625.40 | protein phosphatase 2C, putative | -0,308 | 0,03778 |
| TcCLB.508269.40 | hypothetical protein, conserved | -0,3081 | 0,1032 |
| TcCLB.506855.370 | hypothetical protein, conserved | -0,3082 | 0,00539 |
| TcCLB.509065.90 | hypothetical protein, conserved | -0,3083 | 0,0008372 |
| TcCLB.508347.130 | deoxyhypusine hydroxylase, putative | -0,3083 | 0,03602 |
| TcCLB.507641.210 | T-lymphocyte triggering factor, putative | -0,3084 | 0,0003807 |
| TcCLB.506289.160 | hypothetical protein | -0,3084 | 0,09528 |
| TcCLB.508163.380 | trans-sialidase (pseudogene), putative | -0,3086 | 0,3648 |
| TcCLB.511255.529 | surface protease GP63 (pseudogene), putative | -0,3087 | 0,2259 |
| TcCLB.509085.90 | retrotransposon hot spot protein (RHS, pseudogene), puta | -0,3089 | 0,1434 |
| TcCLB.509939.10 | hypothetical protein, conserved | -0,3092 | 0,0008606 |
| TcCLB.509611.40 | hypothetical protein, conserved | -0,3093 | 0,2212 |
| TcCLB.508593.40 | hypothetical protein, conserved | -0,3095 | 0,01583 |
| TcCLB.506865.60 | hypothetical protein, conserved | -0,3096 | 0,0002922 |
| TcCLB.507639.110 | hypothetical protein, conserved | -0,3096 | 0,006658 |
| TcCLB.458153.10 | serine/threonine protein phosphatase, putative (fragment | -0,3096 | 0,05362 |
| TcCLB.507809.110 | hypothetical protein, conserved | -0,3098 | 0,005033 |
| TcCLB.510291.40 | serine/threonine protein kinase, putative | -0,3098 | 0,08428 |
| TcCLB.506795.60 | hypothetical protein, conserved | -0,3099 | 0,0008704 |
| TcCLB.511665.60 | trans-sialidase (pseudogene), putative | -0,3099 | 0,002199 |
| TcCLB.506763.4 | surface protease GP63 (pseudogene), putative | -0,3099 | 0,2054 |
| TcCLB.511555.60 | ubiquitin ligase, putative | -0,3101 | 9,293E-07 |
| TcCLB.511041.20 | hypothetical protein, conserved | -0,3101 | 0,0001474 |
| TcCLB.511353.69 | oxidoreductase-protein, putative | -0,3101 | 0,1168 |
| TcCLB.508433.210 | mucin TcMUCII (pseudogene), putative | -0,3101 | 0,387 |
| TcCLB.508627.21 | retrotransposon hot spot protein (RHS, pseudogene), puta | -0,3102 | 0,2152 |
| TcCLB.503579.130 | hypothetical protein, conserved | -0,3103 | 0,01247 |
| TcCLB.508661.50 | ribonuclease H, putative | -0,3104 | 0,03053 |

| | | | |
|---|---|---|---|
| TcCLB.506637.50 | hypothetical protein, conserved | -0,3104 | 0,03078 |
| TcCLB.507647.39 | hypothetical protein, conserved | -0,3105 | 0,1003 |
| TcCLB.411915.10 | glycine dehydrogenase [decarboxylating] (pseudogene), pu | -0,3105 | 0,258 |
| TcCLB.510409.50 | hypothetical protein, conserved | -0,3106 | 0,02998 |
| TcCLB.507669.50 | hypothetical protein, conserved | -0,3107 | 0,00002234 |
| TcCLB.508175.30 | hypothetical protein, conserved | -0,3107 | 0,04364 |
| TcCLB.506993.140 | hypothetical protein | -0,3107 | 0,3959 |
| TcCLB.503675.40 | hypothetical protein, conserved | -0,3108 | 0,006701 |
| TcCLB.509617.30 | mismatch repair protein MSH5, putative | -0,3108 | 0,02016 |
| TcCLB.506243.135 | hypothetical protein, conserved (pseudogene) | -0,3108 | 0,1872 |
| TcCLB.506399.10 | sigma-adaptin 3, putative | -0,3109 | 0,3428 |
| TcCLB.511611.5 | mucin TcMUCII (pseudogene), putative | -0,311 | 0,5498 |
| TcCLB.507067.30 | mucin-associated surface protein (MASP, pseudogene), pu | -0,3112 | 0,1193 |
| TcCLB.508173.80 | hypothetical protein, conserved | -0,3118 | 0,0004084 |
| TcCLB.505009.10 | hypothetical protein, conserved | -0,3118 | 0,01574 |
| TcCLB.503989.60 | protein phosphatase 2C, putative (fragment) | -0,3118 | 0,02912 |
| TcCLB.508491.9 | hypothetical protein, conserved | -0,3118 | 0,1002 |
| TcCLB.511283.180 | hypothetical protein, conserved | -0,3119 | 0,0002851 |
| TcCLB.504149.90 | hypothetical protein, conserved | -0,3119 | 0,0003425 |
| TcCLB.509491.10 | hypothetical protein, conserved | -0,3119 | 0,01831 |
| TcCLB.511277.250 | hypothetical protein, conserved | -0,3121 | 0,0003745 |
| TcCLB.504173.60 | hypothetical protein, conserved | -0,3121 | 0,001766 |
| TcCLB.503775.10 | surface protease GP63 (pseudogene), putative | -0,3126 | 0,194 |
| TcCLB.510525.60 | hypothetical protein, conserved | -0,3128 | 0,01516 |
| TcCLB.510121.30 | actin-like protein, putative | -0,3128 | 0,0373 |
| TcCLB.506355.120 | hypothetical protein, conserved | -0,3131 | 0,0003182 |
| TcCLB.508625.80 | hypothetical protein, conserved | -0,3132 | 0,006246 |
| TcCLB.507611.300 | U1A small nuclear ribonucleoprotein, putative | -0,3132 | 0,06406 |
| TcCLB.506475.80 | WD40 repeat-containing protein | -0,3133 | 0,0001296 |
| TcCLB.509453.10 | hypothetical protein, conserved | -0,3134 | 0,003267 |
| TcCLB.507949.120 | Transmembrane protein 231, putative | -0,3134 | 0,005137 |
| TcCLB.506213.30 | hypothetical protein, conserved | -0,3137 | 0,0455 |
| TcCLB.507941.14 | hypothetical protein, conserved | -0,3137 | 0,1752 |
| TcCLB.503855.30 | adenylosuccinate lyase, putative (ADSL) | -0,314 | 0,0003016 |
| TcCLB.508221.1050 | trans-sialidase, putative | -0,3141 | 0,2778 |
| TcCLB.511711.100 | hypothetical protein, conserved | -0,3142 | 0,00475 |
| TcCLB.507501.5 | trans-sialidase, putative (fragment) | -0,3143 | 0,5324 |
| TcCLB.507647.30 | hypothetical protein, conserved | -0,3144 | 0,09145 |
| TcCLB.504053.10 | hypothetical protein, conserved (pseudogene) | -0,3146 | 0,0002076 |
| TcCLB.507357.110 | hypothetical protein, conserved (pseudogene) | -0,3146 | 0,4359 |
| TcCLB.504057.104 | phosphatidylinositol phosphate kinase alpha (PIPKA) | -0,3147 | 0,00616 |
| TcCLB.508177.64 | hypothetical protein, conserved | -0,3147 | 0,01054 |
| TcCLB.506605.220 | hypothetical protein, conserved | -0,3149 | 0,005778 |
| TcCLB.510889.130 | hypothetical protein | -0,3149 | 0,09089 |
| TcCLB.506367.160 | hypothetical protein, conserved | -0,3151 | 0,005041 |

| | | | |
|---|---|---|---|
| TcCLB.510565.110 | hypothetical protein | -0,3154 | 0,07567 |
| TcCLB.509161.30 | OSM3-like kinesin, putative | -0,3155 | 0,000146 |
| TcCLB.510729.60 | hypothetical protein, conserved | -0,3155 | 0,05107 |
| TcCLB.510187.520 | hypothetical protein, conserved | -0,3156 | 0,009233 |
| TcCLB.509399.20 | metacaspase, putative | -0,3159 | 0,04836 |
| TcCLB.508919.20 | hypothetical protein, conserved | -0,316 | 0,0004354 |
| TcCLB.508797.10 | tubulin-tyrosine ligase-like protein, putative | -0,316 | 0,0006591 |
| TcCLB.506941.250 | hypothetical protein, conserved | -0,3161 | 0,003479 |
| TcCLB.506725.10 | hypothetical protein, conserved | -0,3161 | 0,01081 |
| TcCLB.508125.22 | mucin TcMUCII (pseudogene), putative | -0,3161 | 0,3098 |
| TcCLB.508221.460 | mucin-associated surface protein (MASP, pseudogene), pu | -0,3162 | 0,3057 |
| TcCLB.508643.40 | hypothetical protein, conserved | -0,3163 | 0,00005857 |
| TcCLB.508051.20 | hypothetical protein, conserved | -0,3163 | 0,002587 |
| TcCLB.508507.30 | calcium uniporter protein, mitochondrial, putative | -0,3163 | 0,02994 |
| TcCLB.508257.244 | hypothetical protein, conserved | -0,3164 | 0,06353 |
| TcCLB.507993.210 | hypothetical protein, conserved | -0,3166 | 0,001943 |
| TcCLB.508461.300 | hypothetical protein, conserved | -0,3167 | 0,001974 |
| TcCLB.510763.80 | DNA polymerase epsilon subunit B, putative | -0,3167 | 0,01275 |
| TcCLB.503945.40 | 6-phosphogluconolactonase, putative | -0,3167 | 0,04428 |
| TcCLB.510565.141 | hypothetical protein, conserved | -0,3168 | 0,06316 |
| TcCLB.508943.30 | RNA-binding protein, putative | -0,317 | 0,01126 |
| TcCLB.509977.15 | surface protease GP63 (pseudogene), putative | -0,317 | 0,1909 |
| TcCLB.509065.150 | hypothetical protein, conserved | -0,3171 | 0,01253 |
| TcCLB.504167.7 | hypothetical protein, conserved | -0,3172 | 0,07593 |
| TcCLB.419417.10 | hypothetical protein, conserved | -0,3172 | 0,1903 |
| TcCLB.508493.110 | mucin-associated surface protein (MASP), putative | -0,3173 | 0,1037 |
| TcCLB.506635.120 | kinesin, putative | -0,3174 | 0,001447 |
| TcCLB.510661.220 | hypothetical protein, conserved | -0,3174 | 0,008872 |
| TcCLB.511557.70 | hypothetical protein, conserved | -0,3175 | 0,0002051 |
| TcCLB.509961.30 | hypothetical protein, conserved | -0,3175 | 0,00317 |
| TcCLB.506475.14 | phosphatidylinositol 3-kinase, putative | -0,3175 | 0,007231 |
| TcCLB.509287.150 | retrotransposon hot spot protein (RHS, pseudogene), puta | -0,3178 | 0,004748 |
| TcCLB.509105.140 | heat shock protein 85, putative | -0,3178 | 0,0056 |
| TcCLB.509047.50 | metallo-beta-lactamase superfamily, putative | -0,3178 | 0,03729 |
| TcCLB.510963.20 | hypothetical protein, conserved | -0,3182 | 0,002749 |
| TcCLB.508439.30 | hypothetical protein, conserved | -0,3183 | 0,003814 |
| TcCLB.511803.20 | A/G-specific adenine glycosylase, putative | -0,3184 | 0,004819 |
| TcCLB.511667.140 | trans-sialidase (pseudogene), putative | -0,3185 | 0,01251 |
| TcCLB.506315.40 | hypothetical protein, conserved | -0,3187 | 0,0006827 |
| TcCLB.506755.70 | hypothetical protein, conserved | -0,3187 | 0,0102 |
| TcCLB.510729.150 | tubulin tyrosine ligase, putative | -0,3188 | 0,0005292 |
| TcCLB.511287.20 | hypothetical protein, conserved | -0,3188 | 0,001936 |
| TcCLB.510101.460 | hypothetical protein, conserved | -0,3189 | 0,004617 |
| TcCLB.507515.100 | hypothetical protein, conserved | -0,3189 | 0,09617 |
| TcCLB.507093.130 | hypothetical protein, conserved | -0,3192 | 0,0008898 |

| | | | |
|---|---|---|---|
| TcCLB.508343.10 | trans-sialidase, putative | -0,3193 | 0,6293 |
| TcCLB.510293.79 | GPI transamidase component Tta2, putative (fragment) (1 | -0,3197 | 0,01393 |
| TcCLB.503521.10 | hypothetical protein | -0,3198 | 0,001756 |
| TcCLB.510533.60 | hypothetical protein, conserved | -0,3198 | 0,01926 |
| TcCLB.508999.127 | mucin-associated surface protein (MASP, pseudogene), pu | -0,3198 | 0,314 |
| TcCLB.506287.150 | GTP-binding protein, putative | -0,3199 | 0,001467 |
| TcCLB.511523.10 | GTPase activating protein of Rab-like GTPase, putative | -0,3199 | 0,01182 |
| TcCLB.508213.40 | hypothetical protein, conserved | -0,3199 | 0,03709 |
| TcCLB.508405.100 | hypothetical protein, conserved | -0,32 | 0,03216 |
| TcCLB.504105.10 | hypothetical protein, conserved | -0,3201 | 0,01027 |
| TcCLB.509965.240 | hypothetical protein, conserved | -0,3201 | 0,01029 |
| TcCLB.509871.21 | N-acetyltransferase complex ARD1 subunit (pseudogene), | -0,3202 | 0,5295 |
| TcCLB.506333.19 | kinesin, putative (fragment) | -0,3203 | 0,0002769 |
| TcCLB.507993.300 | hypothetical protein, conserved | -0,3203 | 0,0261 |
| TcCLB.506159.10 | retrotransposon hot spot protein (RHS, pseudogene), puta | -0,3204 | 0,02906 |
| TcCLB.510523.29 | hypothetical protein, conserved (fragment) | -0,3205 | 0,05827 |
| TcCLB.506529.490 | hypothetical protein, conserved | -0,3207 | 0,007343 |
| TcCLB.503781.10 | hypothetical protein, conserved | -0,3208 | 0,0001259 |
| TcCLB.506249.50 | hypothetical protein, conserved | -0,3209 | 0,05185 |
| TcCLB.508659.50 | serine/threonine protein kinase, putative | -0,3209 | 0,05206 |
| TcCLB.509005.30 | hypothetical protein, conserved | -0,321 | 0,007531 |
| TcCLB.508489.30 | hypothetical protein, conserved | -0,3212 | 0,009646 |
| TcCLB.506633.20 | hypothetical protein, conserved | -0,3215 | 0,002522 |
| TcCLB.506765.10 | retrotransposon hot spot protein (RHS, pseudogene), puta | -0,3216 | 0,1648 |
| TcCLB.511183.570 | syntaxin binding protein (pseudogene), putative | -0,3216 | 0,3333 |
| TcCLB.511019.54 | ATP-dependent DEAD/H RNA helicase (pseudogene), puta | -0,3217 | 0,2847 |
| TcCLB.507053.130 | hypothetical protein, conserved | -0,322 | 0,003345 |
| TcCLB.503679.30 | hypothetical protein, conserved | -0,322 | 0,01317 |
| TcCLB.507639.70 | hypothetical protein, conserved | -0,3222 | 0,03286 |
| TcCLB.511143.39 | hypothetical protein, conserved | -0,3223 | 0,04992 |
| TcCLB.510091.14 | hypothetical protein, conserved | -0,3223 | 0,06661 |
| TcCLB.508707.227 | ATP binding protein-like protein, putative | -0,3224 | 0,02848 |
| TcCLB.507945.20 | hypothetical protein, conserved | -0,3225 | 0,0171 |
| TcCLB.509971.90 | trans-sialidase, putative (fragment) | -0,3225 | 0,3319 |
| TcCLB.504125.60 | Kinesin-13 3, putative (KIN13-3) | -0,3227 | 0,00001668 |
| TcCLB.503727.29 | hypothetical protein | -0,3227 | 0,003712 |
| TcCLB.510731.100 | hypothetical protein, conserved | -0,3227 | 0,01635 |
| TcCLB.511019.60 | hypothetical protein, conserved (pseudogene) | -0,3228 | 0,0083 |
| TcCLB.507011.110 | hypothetical protein | -0,3228 | 0,01418 |
| TcCLB.509319.10 | hypothetical protein, conserved | -0,3232 | 0,008985 |
| TcCLB.508347.110 | hypothetical protein, conserved | -0,3232 | 0,02862 |
| TcCLB.504397.20 | surface protease GP63, putative | -0,3232 | 0,1436 |
| TcCLB.506211.114 | C-terminal motor kinesin, putative | -0,3234 | 0,0000184 |
| TcCLB.506793.30 | kinesin, putative | -0,3234 | 0,0002519 |
| TcCLB.508647.100 | hypothetical protein, conserved | -0,3238 | 0,0003168 |

| | | | |
|---|---|---|---|
| TcCLB.508163.120 | mucin-associated surface protein (MASP, pseudogene), pu | -0,3238 | 0,08969 |
| TcCLB.506925.220 | hypothetical protein, conserved | -0,3241 | 0,008792 |
| TcCLB.509671.164 | hypothetical protein, conserved | -0,3243 | 0,01591 |
| TcCLB.503747.30 | hypothetical protein | -0,3243 | 0,02671 |
| TcCLB.505999.24 | hypothetical protein, conserved | -0,3243 | 0,03724 |
| TcCLB.503911.10 | hypothetical protein | -0,3243 | 0,2643 |
| TcCLB.510295.50 | protein kinase, putative | -0,3244 | 0,007163 |
| TcCLB.506753.210 | hypothetical protein, conserved | -0,3248 | 0,0203 |
| TcCLB.504119.20 | hypothetical protein, conserved | -0,3251 | 0,001893 |
| TcCLB.511273.59 | 5-oxoprolinase, putative | -0,3251 | 0,00248 |
| TcCLB.509287.80 | trans-sialidase, Group VII, putative | -0,3251 | 0,003126 |
| TcCLB.506879.109 | protein kinase, putative | -0,3251 | 0,04683 |
| TcCLB.511671.120 | U5 snRNP-specific 40 kDa protein, putative | -0,3252 | 0,008655 |
| TcCLB.432067.20 | hypothetical protein, conserved (pseudogene) | -0,3252 | 0,1794 |
| TcCLB.509875.180 | hypothetical protein | -0,3253 | 0,05456 |
| TcCLB.511861.7 | elongation factor 1-gamma (EF-1-gamma), putative | -0,3253 | 0,3279 |
| TcCLB.511633.50 | hypothetical protein, conserved | -0,3254 | 0,0003951 |
| TcCLB.511887.30 | trans-sialidase (pseudogene), putative | -0,3257 | 0,04628 |
| TcCLB.509979.187 | surface protease GP63 (pseudogene), putative | -0,3257 | 0,2624 |
| TcCLB.505965.20 | hypothetical protein, conserved | -0,3259 | 0,008761 |
| TcCLB.507957.140 | trans-sialidase (pseudogene), putative | -0,3259 | 0,04225 |
| TcCLB.506213.20 | hypothetical protein, conserved | -0,3261 | 0,02696 |
| TcCLB.506211.80 | hypothetical protein, conserved | -0,3262 | 0,007184 |
| TcCLB.506941.100 | hypothetical protein, conserved | -0,3263 | 0,000247 |
| TcCLB.511361.80 | hypothetical protein, conserved | -0,3267 | 0,01564 |
| TcCLB.507993.120 | synaptotagmin, putative | -0,3267 | 0,03813 |
| TcCLB.506783.30 | elongation factor 1-gamma (EF-1-gamma), putative | -0,3268 | 0,2365 |
| TcCLB.509753.10 | hypothetical protein, conserved (pseudogene) | -0,3269 | 0,3414 |
| TcCLB.511145.89 | hypothetical protein, conserved | -0,327 | 0,03699 |
| TcCLB.506733.60 | N(2), N(2)-dimethylguanosine tRNA methyltransferase, pu | -0,3273 | 0,001993 |
| TcCLB.511903.230 | hypothetical protein, conserved | -0,3277 | 0,006972 |
| TcCLB.508693.60 | hypothetical protein, conserved | -0,3277 | 0,01358 |
| TcCLB.510719.120 | hypothetical protein, conserved | -0,328 | 0,0005653 |
| TcCLB.507785.20 | proteasome subunit alpha type-1, putative (fragment) (PS | -0,3282 | 0,3814 |
| TcCLB.503419.60 | Enriched in surface-labeled proteome protein 15 (ESP15) | -0,3283 | 0,001338 |
| TcCLB.510825.10 | hypothetical protein, conserved | -0,3284 | 0,008564 |
| TcCLB.503573.9 | ATP-binding cassette transporter ABCA1, putative (ABCA1 | -0,3285 | 0,03589 |
| TcCLB.503981.20 | hypothetical protein, conserved | -0,3286 | 0,02492 |
| TcCLB.511451.110 | hypothetical protein, conserved (pseudogene) | -0,3286 | 0,3363 |
| TcCLB.508479.380 | hypothetical protein, conserved | -0,3287 | 0,001709 |
| TcCLB.511587.120 | Octanoyltransferase, putative | -0,3287 | 0,1867 |
| TcCLB.510089.170 | casein kinase I, putative | -0,329 | 0,02632 |
| TcCLB.510533.220 | hypothetical protein, conserved | -0,3291 | 0,001431 |
| TcCLB.504229.100 | trans-sialidase (pseudogene), putative | -0,3291 | 0,06124 |
| TcCLB.506733.20 | AMP deaminase, putative | -0,3293 | 0,0002368 |

| | | | |
|---|---|---|---|
| TcCLB.507093.90 | hypothetical protein, conserved | -0,3294 | 0,00211 |
| TcCLB.511173.440 | trans-sialidase, Group V, putative | -0,3294 | 0,1298 |
| TcCLB.506239.60 | hypothetical protein, conserved | -0,3295 | 0,02658 |
| TcCLB.506563.90 | calpain-like cysteine peptidase (pseudogene), putative | -0,3295 | 0,03449 |
| TcCLB.511367.30 | hypothetical protein | -0,3295 | 0,2897 |
| TcCLB.503813.10 | hypothetical protein, conserved | -0,3298 | 0,000972 |
| TcCLB.416883.9 | hypothetical protein, conserved (fragment) | -0,3298 | 0,01417 |
| TcCLB.506747.30 | ubiquitin hydrolase, putative | -0,3299 | 0,0002028 |
| TcCLB.506859.100 | hypothetical protein, conserved | -0,3299 | 0,1076 |
| TcCLB.509967.180 | hypothetical protein, conserved | -0,3301 | 0,003972 |
| TcCLB.506253.11 | hypothetical protein, conserved (pseudogene) | -0,3301 | 0,1256 |
| TcCLB.506789.270 | hypothetical protein, conserved | -0,3302 | 0,0001702 |
| TcCLB.504149.220 | retrotransposon hot spot protein (RHS, pseudogene), puta | -0,3302 | 0,005584 |
| TcCLB.507839.9 | Myotubularin-related protein, putative (fragment) | -0,3304 | 0,03947 |
| TcCLB.510025.80 | hypothetical protein, conserved (pseudogene) | -0,3304 | 0,06407 |
| TcCLB.506315.90 | serine/threonine protein kinase, putative | -0,3305 | 0,0007321 |
| TcCLB.511903.240 | hypothetical protein, conserved | -0,3305 | 0,0009035 |
| TcCLB.511583.30 | hypothetical protein, conserved | -0,3306 | 0,01039 |
| TcCLB.510283.60 | hypothetical protein, conserved | -0,3306 | 0,03227 |
| TcCLB.503581.20 | PPIase, putative | -0,3307 | 0,04093 |
| TcCLB.506885.380 | hypothetical protein, conserved | -0,3308 | 0,004933 |
| TcCLB.505183.160 | hypothetical protein, conserved | -0,3308 | 0,006374 |
| TcCLB.509617.89 | Zn-finger protein, putative (fragment) | -0,3308 | 0,006415 |
| TcCLB.510743.90 | hypothetical protein, conserved | -0,3311 | 0,009402 |
| TcCLB.511423.170 | chaperone DNAJ protein, putative | -0,3311 | 0,0217 |
| TcCLB.508819.30 | hypothetical protein, conserved | -0,3313 | 0,003696 |
| TcCLB.508873.514 | mucin TcMUC (pseudogene), putative | -0,3314 | 0,2775 |
| TcCLB.509799.130 | hypothetical protein, conserved | -0,3315 | 0,0003168 |
| TcCLB.506855.300 | hypothetical protein, conserved | -0,3315 | 0,001423 |
| TcCLB.510317.34 | unspecified product | -0,3315 | 0,1652 |
| TcCLB.503973.64 | hypothetical protein, conserved (pseudogene) | -0,3316 | 0,3649 |
| TcCLB.504045.80 | hypothetical protein, conserved | -0,3317 | 0,014 |
| TcCLB.509617.70 | subtilisin-like serine peptidase, putative | -0,3318 | 0,0004826 |
| TcCLB.510421.264 | hypothetical protein, conserved | -0,3318 | 0,07338 |
| TcCLB.510755.40 | hypothetical protein, conserved | -0,3321 | 0,0251 |
| TcCLB.511923.40 | hypothetical protein, conserved (pseudogene) | -0,3321 | 0,1086 |
| TcCLB.506799.10 | fatty acid transporter protein-like, putative | -0,3323 | 0,000438 |
| TcCLB.506575.20 | hypothetical protein, conserved | -0,3323 | 0,1594 |
| TcCLB.510379.20 | hypothetical protein (fragment) | -0,3323 | 0,2787 |
| TcCLB.507991.110 | hypothetical protein, conserved | -0,3324 | 0,001891 |
| TcCLB.510617.19 | hypothetical protein, conserved | -0,3327 | 0,01583 |
| TcCLB.510723.20 | hypothetical protein | -0,3328 | 0,0007389 |
| TcCLB.506435.370 | surface protease GP63, putative | -0,3329 | 0,01928 |
| TcCLB.509399.9 | helicase, putative | -0,333 | 0,001059 |
| TcCLB.508257.50 | actin-like protein, putative | -0,333 | 0,01962 |

| | | | |
|---|---|---|---|
| TcCLB.509073.90 | hypothetical protein, conserved | -0,3331 | 0,001492 |
| TcCLB.503919.20 | hypothetical protein, conserved | -0,3331 | 0,05809 |
| TcCLB.503583.20 | hypothetical protein, conserved | -0,3332 | 0,01453 |
| TcCLB.506811.110 | hypothetical protein, conserved | -0,3334 | 0,0002541 |
| TcCLB.507793.40 | hypothetical protein, conserved | -0,3337 | 0,02302 |
| TcCLB.503879.10 | DNA polymerase delta subunit 2, putative | -0,3339 | 0,003321 |
| TcCLB.509775.40 | iron superoxide dismutase, putative | -0,3339 | 0,03397 |
| TcCLB.509671.168 | hypothetical protein, conserved | -0,3339 | 0,06001 |
| TcCLB.503879.20 | hypothetical protein, conserved | -0,334 | 0,002492 |
| TcCLB.509937.140 | hypothetical protein, conserved | -0,3341 | 0,003702 |
| TcCLB.510657.100 | hypothetical protein, conserved | -0,3342 | 0,0002883 |
| TcCLB.504741.124 | hypothetical protein, conserved | -0,3344 | 0,1231 |
| TcCLB.505193.50 | hypothetical protein | -0,3344 | 0,1547 |
| TcCLB.508963.40 | trans-sialidase (pseudogene), putative | -0,3345 | 0,008759 |
| TcCLB.510817.21 | hypothetical protein | -0,3346 | 0,07994 |
| TcCLB.449759.9 | hypothetical protein, conserved | -0,3347 | 0,02204 |
| TcCLB.511545.100 | hypothetical protein, conserved | -0,3349 | 0,0001544 |
| TcCLB.504011.60 | hypothetical protein, conserved | -0,3349 | 0,004172 |
| TcCLB.507951.290 | hypothetical protein, conserved | -0,335 | 0,04862 |
| TcCLB.506885.300 | hypothetical protein, conserved | -0,3352 | 0,02818 |
| TcCLB.508479.160 | hypothetical protein, conserved | -0,3353 | 0,0001134 |
| TcCLB.511881.10 | GCN5-like protein, putative | -0,3353 | 0,1166 |
| TcCLB.511031.40 | hypothetical protein, conserved | -0,3354 | 0,01222 |
| TcCLB.511461.10 | hypothetical protein, conserved | -0,3354 | 0,01372 |
| TcCLB.503897.80 | deaminase, putative | -0,3354 | 0,02699 |
| TcCLB.508207.31 | hypothetical protein | -0,3358 | 0,02436 |
| TcCLB.509453.30 | GCN5-like protein, putative | -0,3359 | 0,04922 |
| TcCLB.508625.50 | 6-phosphofructo-2-kinase/fructose-2,6-bisphosphatas, pu | -0,336 | 0,007822 |
| TcCLB.509143.10 | hypothetical protein, conserved | -0,3362 | 0,00026 |
| TcCLB.510629.80 | trans-sialidase (pseudogene), putative | -0,3363 | 0,458 |
| TcCLB.506265.70 | hypothetical protein, conserved | -0,3365 | 0,001486 |
| TcCLB.511287.180 | Associated kinase of Tb14-3-3, putative (AKB1) | -0,3369 | 0,06902 |
| TcCLB.511383.20 | kinesin, putative | -0,337 | 0,0009761 |
| TcCLB.448387.9 | hypothetical protein, conserved | -0,337 | 0,006254 |
| TcCLB.503723.80 | hypothetical protein, conserved | -0,3371 | 0,00458 |
| TcCLB.509655.20 | hypothetical protein, conserved (pseudogene) | -0,3371 | 0,1441 |
| TcCLB.508793.30 | palmitoyl acyltransferase 8, putative | -0,3372 | 0,02848 |
| TcCLB.506811.100 | hypothetical protein, conserved | -0,3373 | 0,006772 |
| TcCLB.506483.60 | hypothetical protein, conserved | -0,3373 | 0,01002 |
| TcCLB.510623.80 | trans-sialidase (pseudogene), putative | -0,3375 | 0,1881 |
| TcCLB.508257.60 | hypothetical protein, conserved | -0,3376 | 0,01908 |
| TcCLB.505807.130 | hypothetical protein, conserved | -0,3377 | 0,1361 |
| TcCLB.508939.40 | hypothetical protein | -0,3377 | 0,3437 |
| TcCLB.510359.60 | hypothetical protein, conserved | -0,3379 | 0,009138 |
| TcCLB.509007.70 | hypothetical protein, conserved | -0,3382 | 0,0007622 |

| | | | |
|---|---|---|---|
| TcCLB.508307.44 | hypothetical protein, conserved | -0,3382 | 0,0007725 |
| TcCLB.506991.19 | hypothetical protein, conserved | -0,3383 | 0,03809 |
| TcCLB.508495.210 | mucin-associated surface protein (MASP, pseudogene), pu | -0,3384 | 0,2619 |
| TcCLB.509737.19 | retrotransposon hot spot protein (RHS, pseudogene), puta | -0,3386 | 0,1416 |
| TcCLB.506413.89 | trans-sialidase, putative | -0,3389 | 0,0004565 |
| TcCLB.511725.140 | mitochondrial carrier protein, putative | -0,3389 | 0,002169 |
| TcCLB.507993.160 | UDP-galactopyranose mutase (GLF) | -0,339 | 0,001154 |
| TcCLB.510241.40 | hypothetical protein, conserved | -0,339 | 0,005294 |
| TcCLB.511511.100 | hypothetical protein, conserved | -0,3393 | 0,00002341 |
| TcCLB.506341.20 | hypothetical protein, conserved | -0,3394 | 0,0247 |
| TcCLB.506123.30 | protein kinase, putative | -0,3396 | 0,00004272 |
| TcCLB.511151.30 | hypothetical protein, conserved | -0,3397 | 0,02575 |
| TcCLB.504115.50 | retrotransposon hot spot protein (RHS, pseudogene), puta | -0,3397 | 0,04525 |
| TcCLB.507467.119 | hypothetical protein, conserved | -0,3398 | 0,01828 |
| TcCLB.503971.5 | hypothetical protein, conserved | -0,3398 | 0,2841 |
| TcCLB.511173.241 | mucin TcMUCII (pseudogene), putative | -0,3398 | 0,2887 |
| TcCLB.511131.50 | U4/U6 small nuclear ribonuclear protein, putative | -0,3399 | 0,0007451 |
| TcCLB.506337.40 | glycosyl transferase-like protein, putative | -0,34 | 0,01265 |
| TcCLB.507709.110 | hypothetical protein, conserved | -0,34 | 0,06577 |
| TcCLB.507357.50 | hypothetical protein | -0,3401 | 0,2853 |
| TcCLB.511283.190 | U2 small nuclear ribonucleoprotein 40K, putative | -0,3404 | 0,0007975 |
| TcCLB.511683.30 | hypothetical protein, conserved | -0,3405 | 0,0005535 |
| TcCLB.510165.90 | trans-sialidase (pseudogene), putative | -0,3406 | 0,06992 |
| TcCLB.508295.40 | mucin-associated surface protein (MASP), putative | -0,3411 | 0,4005 |
| TcCLB.506317.30 | hypothetical protein, conserved (pseudogene) | -0,3414 | 0,001631 |
| TcCLB.508357.20 | hypothetical protein, conserved | -0,3414 | 0,05596 |
| TcCLB.511217.50 | hypothetical protein, conserved | -0,3415 | 0,00001623 |
| TcCLB.510609.120 | hypothetical protein, conserved | -0,3417 | 0,0005745 |
| TcCLB.510059.10 | dispersed gene family protein 1 (DGF-1), putative | -0,3418 | 0,3582 |
| TcCLB.503663.4 | hypothetical protein, conserved (fragment) | -0,3419 | 0,02738 |
| TcCLB.509297.10 | hypothetical protein, conserved | -0,342 | 0,01853 |
| TcCLB.503879.90 | hypothetical protein, conserved | -0,3421 | 0,0006992 |
| TcCLB.506559.310 | hypothetical protein, conserved | -0,3422 | 0,0004151 |
| TcCLB.511733.110 | GTPase activating protein of Rab-like GTPase, putative | -0,3422 | 0,002437 |
| TcCLB.506361.44 | mucin TcMUC (pseudogene), putative | -0,3423 | 0,1349 |
| TcCLB.504153.290 | hypothetical protein, conserved | -0,3424 | 0,00172 |
| TcCLB.509893.29 | hypothetical protein, conserved | -0,3424 | 0,0209 |
| TcCLB.507099.110 | hypothetical protein, conserved | -0,3425 | 0,002823 |
| TcCLB.506565.4 | RNA-binding protein, putative | -0,3425 | 0,1956 |
| TcCLB.506583.4 | serine/threonine protein kinase, putative | -0,3428 | 0,02083 |
| TcCLB.506715.50 | SNARE domain-containing protein, putative (SNARE2.2) | -0,343 | 0,001539 |
| TcCLB.503885.80 | hypothetical protein, conserved | -0,343 | 0,003341 |
| TcCLB.509073.60 | hypothetical protein, conserved | -0,343 | 0,006046 |
| TcCLB.509157.250 | hypothetical protein, conserved | -0,3432 | 0,0003319 |
| TcCLB.508235.20 | mucin TcMUCIII, putative | -0,3432 | 0,05394 |

| | | | |
|---|---|---|---|
| TcCLB.506605.170 | inosine-guanine nucleoside hydrolase, putative (IG-NH) | -0,3433 | 0,002582 |
| TcCLB.508051.40 | hypothetical protein, conserved | -0,3433 | 0,01267 |
| TcCLB.510647.69 | hypothetical protein, conserved | -0,3434 | 0,1509 |
| TcCLB.511761.80 | hypothetical protein | -0,3437 | 0,001032 |
| TcCLB.506451.35 | hypothetical protein | -0,3437 | 0,1611 |
| TcCLB.508875.10 | hypothetical protein (fragment) | -0,3437 | 0,2763 |
| TcCLB.508969.40 | dispersed gene family protein 1 (DGF-1), putative | -0,3439 | 0,3119 |
| TcCLB.506249.20 | protein transport protein Sec23A, putative | -0,344 | 0,00007948 |
| TcCLB.511367.220 | hypothetical protein, conserved | -0,3441 | 0,002094 |
| TcCLB.508919.110 | hypothetical protein, conserved | -0,3441 | 0,03006 |
| TcCLB.509847.10 | retrotransposon hot spot protein (RHS, pseudogene), puta | -0,3444 | 0,1174 |
| TcCLB.510187.340 | hypothetical protein, conserved | -0,3445 | 0,0006516 |
| TcCLB.509085.20 | retrotransposon hot spot protein (RHS, pseudogene), puta | -0,3447 | 0,377 |
| TcCLB.507001.50 | hypothetical protein, conserved | -0,3448 | 0,0002118 |
| TcCLB.511735.34 | hypothetical protein, conserved | -0,3449 | 0,01886 |
| TcCLB.509599.20 | hypothetical protein, conserved (fragment) | -0,3451 | 0,00889 |
| TcCLB.507515.120 | sphingosine kinase A, B, putative | -0,3453 | 0,0008444 |
| TcCLB.508153.510 | serine/threonine-protein kinase Nek, putative | -0,3453 | 0,001433 |
| TcCLB.507041.76 | hypothetical protein, conserved (pseudogene) | -0,3453 | 0,02442 |
| TcCLB.508891.90 | hypothetical protein, conserved | -0,3454 | 0,000593 |
| TcCLB.509805.80 | hypothetical protein, conserved | -0,3456 | 0,003168 |
| TcCLB.507241.39 | ABC transporter, putative | -0,3458 | 0,0003336 |
| TcCLB.508711.49 | hypothetical protein, conserved | -0,3465 | 0,00007344 |
| TcCLB.510031.60 | methionyl-tRNA formyltransferase, putative | -0,3465 | 0,001326 |
| TcCLB.511021.99 | hypothetical protein, conserved | -0,3465 | 0,03254 |
| TcCLB.509895.70 | hypothetical protein, conserved | -0,3465 | 0,1438 |
| TcCLB.511283.90 | N-myristoyl transferase, putative | -0,3469 | 0,00268 |
| TcCLB.510423.40 | hypothetical protein, conserved | -0,347 | 0,00003903 |
| TcCLB.503987.50 | hypothetical protein, conserved | -0,347 | 0,003361 |
| TcCLB.506925.490 | hypothetical protein, conserved | -0,347 | 0,00991 |
| TcCLB.508707.120 | hypothetical protein, conserved | -0,3471 | 0,0002478 |
| TcCLB.504149.80 | hypothetical protein, conserved | -0,3471 | 0,0003963 |
| TcCLB.506177.90 | hypothetical protein, conserved | -0,3473 | 0,06417 |
| TcCLB.510975.30 | hypothetical protein, conserved | -0,3474 | 0,0008713 |
| TcCLB.508951.29 | actin-like protein, putative | -0,3474 | 0,004121 |
| TcCLB.427789.39 | hypothetical protein, conserved | -0,3474 | 0,02012 |
| TcCLB.510499.20 | hypothetical protein | -0,3477 | 0,2781 |
| TcCLB.503893.150 | hypothetical protein, conserved | -0,3479 | 0,009479 |
| TcCLB.509157.50 | hypothetical protein | -0,348 | 0,0001353 |
| TcCLB.503789.40 | signal recognition particle (pseudogene), putative | -0,3482 | 0,00009775 |
| TcCLB.506773.100 | protein kinase, putative | -0,3482 | 0,0001206 |
| TcCLB.506865.10 | ARP2/3 complex subunit, putative | -0,3483 | 0,005103 |
| TcCLB.511321.4 | O-sialoglycoprotein endopeptidase, putative | -0,3483 | 0,1252 |
| TcCLB.508265.110 | hypothetical protein, conserved | -0,3484 | 0,0005105 |
| TcCLB.510323.10 | protein kinase, putative | -0,3484 | 0,002602 |

| | | | |
|---|---|---|---|
| TcCLB.506869.4 | hypothetical protein, conserved | -0,3485 | 0,002119 |
| TcCLB.509979.330 | surface protease GP63 (pseudogene), putative | -0,3489 | 0,04783 |
| TcCLB.510119.50 | hypothetical protein, conserved | -0,349 | 0,001767 |
| TcCLB.506267.75 | hypothetical protein, conserved (pseudogene) | -0,3492 | 0,1167 |
| TcCLB.509053.40 | hypothetical protein, conserved | -0,3494 | 0,0005869 |
| TcCLB.510781.30 | tryptophanyl-tRNA synthetase (pseudogene), putative | -0,3497 | 0,3853 |
| TcCLB.510761.60 | casein kinase II, putative | -0,3499 | 0,0007607 |
| TcCLB.510061.60 | dispersed gene family protein 1 (DGF-1), putative | -0,3499 | 0,4843 |
| TcCLB.510741.140 | hypothetical protein, conserved | -0,3501 | 0,0009717 |
| TcCLB.506821.130 | hypothetical protein, conserved | -0,3502 | 0,00007248 |
| TcCLB.509099.30 | paraflagellar rod protein 5, putative (PFR5) | -0,3502 | 0,003772 |
| TcCLB.507687.60 | trans-sialidase (pseudogene), putative | -0,3504 | 0,1862 |
| TcCLB.506833.50 | hypothetical protein, conserved | -0,3505 | 0,00007521 |
| TcCLB.506779.120 | hypothetical protein, conserved | -0,3505 | 0,0002511 |
| TcCLB.508153.360 | hypothetical protein, conserved | -0,3505 | 0,01696 |
| TcCLB.506775.110 | hypothetical protein, conserved | -0,3508 | 0,0004987 |
| TcCLB.506919.20 | trans-sialidase (pseudogene), putative | -0,3513 | 0,4062 |
| TcCLB.508717.50 | hypothetical protein, conserved | -0,3514 | 0,002926 |
| TcCLB.504643.10 | kinesin (pseudogene), putative | -0,3515 | 0,0001189 |
| TcCLB.507993.320 | RNA-binding protein 27, putative (RBP27) | -0,3515 | 0,001755 |
| TcCLB.511079.71 | mucin TcMUC (pseudogene), putative | -0,3517 | 0,2166 |
| TcCLB.506951.103 | hypothetical protein | -0,3517 | 0,3865 |
| TcCLB.508805.10 | pre-mRNA splicing factor ATP-dependent RNA helicase (p | -0,3519 | 0,00451 |
| TcCLB.508387.30 | hypothetical protein, conserved | -0,3522 | 0,001091 |
| TcCLB.503779.60 | hypothetical protein, conserved | -0,3522 | 0,001602 |
| TcCLB.506575.60 | diacylglycerol kinase, putative | -0,3522 | 0,001911 |
| TcCLB.509743.40 | trans-sialidase, putative | -0,3522 | 0,3652 |
| TcCLB.508441.60 | hypothetical protein, conserved | -0,3524 | 0,00135 |
| TcCLB.511491.40 | hypothetical protein, conserved | -0,3525 | 0,00001724 |
| TcCLB.506679.90 | RNA editing complex protein MP67, putative | -0,3526 | 0,0001919 |
| TcCLB.504625.40 | hypothetical protein, conserved | -0,3527 | 0,0007511 |
| TcCLB.511919.20 | trans-sialidase, Group V, putative | -0,3527 | 0,02865 |
| TcCLB.508525.20 | hypothetical protein, conserved | -0,3528 | 0,00001125 |
| TcCLB.506459.100 | surface protease GP63 (pseudogene), putative | -0,3528 | 0,2854 |
| TcCLB.510075.50 | hypothetical protein, conserved | -0,3529 | 0,00006226 |
| TcCLB.505807.240 | hypothetical protein, conserved | -0,353 | 0,004628 |
| TcCLB.507049.30 | hypothetical protein, conserved | -0,353 | 0,007905 |
| TcCLB.508913.39 | O-sialoglycoprotein endopeptidase, putative | -0,353 | 0,04184 |
| TcCLB.504187.21 | trans-sialidase (pseudogene), putative | -0,353 | 0,07476 |
| TcCLB.509195.40 | mucin TcMUCII, putative | -0,3532 | 0,1498 |
| TcCLB.507681.120 | hypothetical protein, conserved (pseudogene) | -0,3533 | 0,00622 |
| TcCLB.504985.10 | hypothetical protein | -0,3533 | 0,237 |
| TcCLB.509683.60 | hypothetical protein, conserved | -0,3537 | 0,002313 |
| TcCLB.507491.110 | hypothetical protein | -0,3539 | 0,000134 |
| TcCLB.506947.30 | hypothetical protein, conserved | -0,3539 | 0,0158 |

| | | | |
|---|---|---|---|
| TcCLB.507795.20 | hypothetical protein, conserved | -0,3543 | 0,003501 |
| TcCLB.507257.10 | hypothetical protein, conserved | -0,3543 | 0,01111 |
| TcCLB.507031.100 | hypothetical protein, conserved | -0,3545 | 0,02615 |
| TcCLB.503419.20 | adenosine deaminase-like protein, putative (ADAT1h) | -0,3546 | 0,001819 |
| TcCLB.509937.80 | hypothetical protein, conserved | -0,3546 | 0,002315 |
| TcCLB.467465.10 | surface protease GP63, putative | -0,3548 | 0,3954 |
| TcCLB.507017.90 | hypothetical protein, conserved | -0,3551 | 0,006729 |
| TcCLB.511417.120 | DNA repair protein BRCA2 (pseudogene), putative | -0,3554 | 0,0005656 |
| TcCLB.504153.210 | hypothetical protein | -0,3554 | 0,02358 |
| TcCLB.401569.10 | trans-sialidase, putative | -0,3554 | 0,133 |
| TcCLB.508795.19 | protein kinase, putative (fragment) | -0,3555 | 0,0003226 |
| TcCLB.509561.50 | hypothetical protein, conserved | -0,3557 | 0,003484 |
| TcCLB.509573.10 | small GTP-binding protein Rab7, putative | -0,3557 | 0,00485 |
| TcCLB.504179.20 | hypothetical protein, conserved | -0,3557 | 0,009149 |
| TcCLB.506697.70 | hypothetical protein, conserved | -0,3559 | 0,0007321 |
| TcCLB.503981.9 | hypothetical protein, conserved (pseudogene) | -0,3562 | 0,0138 |
| TcCLB.508999.230 | calpain-like cysteine peptidase (pseudogene), putative | -0,3564 | 0,0002369 |
| TcCLB.506867.30 | hypothetical protein, conserved | -0,3564 | 0,01942 |
| TcCLB.506297.280 | hypothetical protein, conserved | -0,3565 | 0,002231 |
| TcCLB.506931.64 | RNA processing endonuclease 1, putative | -0,3565 | 0,004513 |
| TcCLB.432067.10 | ATP-dependent RNA helicase, putative | -0,3565 | 0,03583 |
| TcCLB.511277.130 | hypothetical protein, conserved | -0,3566 | 6,553E-06 |
| TcCLB.427247.10 | serine/threonine protein kinase, putative | -0,3566 | 0,1881 |
| TcCLB.507041.90 | hypothetical protein | -0,3567 | 0,02337 |
| TcCLB.508119.140 | retrotransposon hot spot (RHS) protein, putative | -0,3568 | 0,0161 |
| TcCLB.511001.130 | hypothetical protein, conserved | -0,357 | 8,029E-07 |
| TcCLB.504883.10 | protein kinase, putative | -0,3573 | 0,00003102 |
| TcCLB.508233.25 | hypothetical protein | -0,3573 | 0,007304 |
| TcCLB.510111.30 | hypothetical protein, conserved (pseudogene) | -0,3573 | 0,009071 |
| TcCLB.507801.100 | hypothetical protein, conserved | -0,3574 | 0,0003962 |
| TcCLB.508817.160 | Trichohyalin, putative | -0,3574 | 0,001138 |
| TcCLB.511873.5 | hypothetical protein | -0,3575 | 0,03267 |
| TcCLB.400945.10 | trans-sialidase (pseudogene), putative | -0,3576 | 0,2455 |
| TcCLB.506295.60 | hypothetical protein, conserved | -0,3578 | 0,03743 |
| TcCLB.506591.10 | hypothetical protein, conserved | -0,3578 | 0,07122 |
| TcCLB.506457.90 | hypothetical protein | -0,358 | 0,01467 |
| TcCLB.510429.59 | hypothetical protein, conserved | -0,358 | 0,0719 |
| TcCLB.508389.70 | trans-sialidase (pseudogene), putative | -0,3581 | 0,155 |
| TcCLB.509979.249 | syntaxin binding protein 1 (pseudogene), putative | -0,3584 | 0,0178 |
| TcCLB.504431.80 | hypothetical protein | -0,3585 | 0,001343 |
| TcCLB.505999.180 | Non cytoplasmic domain containing protein, conserved | -0,3586 | 0,002378 |
| TcCLB.510241.30 | hypothetical protein, conserved | -0,3588 | 0,003768 |
| TcCLB.510899.10 | surface protease GP63, putative | -0,3589 | 0,0009315 |
| TcCLB.508051.10 | hypothetical protein, conserved | -0,3589 | 0,001164 |
| TcCLB.508739.90 | hypothetical protein, conserved | -0,3589 | 0,05186 |

| | | | |
|---|---|---|---|
| TcCLB.504827.21 | hypothetical protein, conserved (fragment) | -0,3591 | 0,00008223 |
| TcCLB.507521.60 | hypothetical protein, conserved | -0,3593 | 0,02441 |
| TcCLB.503821.5 | hypothetical protein, conserved | -0,3593 | 0,07982 |
| TcCLB.503999.110 | hypothetical protein, conserved (fragment) | -0,3593 | 0,08021 |
| TcCLB.510665.4 | hypothetical protein, conserved (fragment) | -0,3594 | 0,002119 |
| TcCLB.508217.9 | dynein heavy chain, putative | -0,3595 | 0,006953 |
| TcCLB.508723.20 | hypothetical protein, conserved | -0,3596 | 0,001501 |
| TcCLB.507099.90 | hypothetical protein, conserved | -0,3599 | 0,00003699 |
| TcCLB.510173.110 | hypothetical protein, conserved | -0,3599 | 0,001183 |
| TcCLB.506613.141 | hypothetical protein, conserved (pseudogene) | -0,36 | 0,03803 |
| TcCLB.510821.10 | hypothetical protein, conserved | -0,3602 | 0,001085 |
| TcCLB.507825.40 | hypothetical protein, conserved | -0,3603 | 0,001514 |
| TcCLB.509979.70 | hypothetical protein, conserved | -0,3605 | 0,01772 |
| TcCLB.506309.80 | mucin TcMUCII, putative | -0,3606 | 0,4485 |
| TcCLB.503893.181 | hypothetical protein, conserved (fragment) | -0,3608 | 0,1769 |
| TcCLB.509259.30 | mucin-associated surface protein (MASP, pseudogene), pu | -0,361 | 0,08724 |
| TcCLB.508543.150 | hypothetical protein, conserved | -0,3612 | 0,00007297 |
| TcCLB.507527.30 | DNA replication licensing factor MCM6, putative | -0,3612 | 0,0006768 |
| TcCLB.511027.29 | hypothetical protein, conserved (pseudogene) | -0,3612 | 0,006394 |
| TcCLB.504867.90 | glycerol-3-phosphate acyltransferase, putative | -0,3614 | 0,002259 |
| TcCLB.506625.110 | DNA topoisomerase type IB small subunit, putative | -0,3614 | 0,004512 |
| TcCLB.506351.90 | surface protease GP63, putative | -0,3614 | 0,01318 |
| TcCLB.504557.50 | dispersed gene family protein 1 (DGF-1), putative | -0,3614 | 0,2134 |
| TcCLB.509569.50 | guanine nucleotide exchange factor subunit GCD7 | -0,3617 | 0,00008953 |
| TcCLB.511283.40 | syntaxin 5, putative | -0,3617 | 0,0007996 |
| TcCLB.511491.50 | KU80 protein, putative | -0,3618 | 0,000589 |
| TcCLB.506543.70 | hypothetical protein, conserved | -0,3619 | 0,01784 |
| TcCLB.511283.310 | hypothetical protein, conserved | -0,362 | 0,00008066 |
| TcCLB.510031.9 | U5 snrnp-specific protein, putative | -0,3621 | 0,001296 |
| TcCLB.508635.10 | hypothetical protein | -0,3622 | 0,08882 |
| TcCLB.508409.90 | hypothetical protein, conserved | -0,3624 | 1,822E-06 |
| TcCLB.506183.15 | trans-sialidase (pseudogene), putative | -0,3625 | 0,2001 |
| TcCLB.510719.50 | hypothetical protein, conserved | -0,3627 | 0,06028 |
| TcCLB.506999.150 | vesicle-associated membrane protein, putative | -0,3628 | 0,0001004 |
| TcCLB.507089.120 | hypothetical protein, conserved | -0,3629 | 0,0368 |
| TcCLB.511629.10 | hypothetical protein, conserved | -0,363 | 0,0002219 |
| TcCLB.509647.110 | hypothetical protein, conserved | -0,3631 | 0,00006763 |
| TcCLB.506511.60 | flavoprotein monooxygenase, putative | -0,3632 | 0,0000167 |
| TcCLB.508277.330 | actin-like protein, putative | -0,3633 | 0,0001881 |
| TcCLB.510797.20 | hypothetical protein, conserved | -0,3637 | 0,01116 |
| TcCLB.508977.65 | elongation factor 1-gamma (EF-1-gamma, pseudogene), | | -0,3637 | 0,1958 |
| TcCLB.508181.70 | hypothetical protein, conserved | -0,3638 | 0,00003234 |
| TcCLB.504105.230 | nuclear cap binding complex subunit CBP110, putative (CB | -0,3642 | 0,0002968 |
| TcCLB.507757.60 | dihydrolipoamide dehydrogenase, putative | -0,3642 | 0,004735 |
| TcCLB.503465.30 | Iron-sulfur assembly protein 2, putative (Isa2) | -0,3642 | 0,05934 |

| | | | |
|---|---|---|---|
| TcCLB.506295.40 | hypothetical protein, conserved | -0,3644 | 0,01134 |
| TcCLB.510719.80 | hypothetical protein, conserved | -0,3645 | 5,394E-08 |
| TcCLB.511107.20 | hypothetical protein, conserved (pseudogene) | -0,3646 | 0,001074 |
| TcCLB.508153.20 | vacuolar protein sorting-associated protein, putative | -0,3647 | 7,079E-06 |
| TcCLB.508783.10 | hypothetical protein, conserved (pseudogene) | -0,3647 | 0,0004449 |
| TcCLB.506783.10 | hypothetical protein, conserved (pseudogene) | -0,3647 | 0,1357 |
| TcCLB.509231.50 | hypothetical protein, conserved | -0,3648 | 0,006968 |
| TcCLB.508461.130 | elongation factor 1-alpha (ef-1-alpha), putative | -0,3649 | 7,005E-06 |
| TcCLB.506477.4 | hypothetical protein, conserved | -0,3649 | 0,03486 |
| TcCLB.511735.40 | hypothetical protein, conserved | -0,3652 | 0,03488 |
| TcCLB.511449.30 | mucin-associated surface protein (MASP), putative | -0,3652 | 0,136 |
| TcCLB.510823.109 | hypothetical protein, conserved | -0,3656 | 0,01228 |
| TcCLB.510603.90 | mitochondrial carrier protein, putative | -0,3657 | 0,01412 |
| TcCLB.509967.50 | hypothetical protein, conserved | -0,3659 | 0,007531 |
| TcCLB.509781.10 | hypothetical protein, conserved | -0,366 | 0,008495 |
| TcCLB.504557.40 | hypothetical protein | -0,3663 | 0,2481 |
| TcCLB.508965.70 | hypothetical protein, conserved | -0,3663 | 0,3979 |
| TcCLB.506525.45 | hypothetical protein, conserved (pseudogene) | -0,3667 | 0,06551 |
| TcCLB.504113.30 | hypothetical protein, conserved | -0,3668 | 0,01696 |
| TcCLB.503749.10 | hypothetical protein, conserved | -0,3671 | 0,000896 |
| TcCLB.511857.40 | N(2), N(2)-dimethylguanosine tRNA methyltransferase, p | -0,3674 | 0,0003666 |
| TcCLB.507639.60 | hypothetical protein, conserved | -0,3674 | 0,000449 |
| TcCLB.508637.100 | hypothetical protein, conserved | -0,3674 | 0,01179 |
| TcCLB.509665.16 | elongation factor 1-gamma (EF-1-gamma, pseudogene), | -0,3675 | 0,3528 |
| TcCLB.504159.100 | hypothetical protein, conserved | -0,3677 | 0,02475 |
| TcCLB.507105.30 | hypothetical protein, conserved | -0,368 | 0,0001603 |
| TcCLB.510305.20 | hypothetical protein, conserved | -0,3682 | 0,002711 |
| TcCLB.511139.10 | hypothetical protein, conserved (pseudogene) | -0,3682 | 0,003809 |
| TcCLB.510719.280 | hypothetical protein, conserved | -0,3683 | 0,0006393 |
| TcCLB.435519.10 | DNA replication licensing factor MCM6, putative | -0,3683 | 0,03915 |
| TcCLB.509109.170 | tubulin tyrosine ligase (pseudogene), putative | -0,3684 | 0,003261 |
| TcCLB.509665.10 | hypothetical protein | -0,3684 | 0,3758 |
| TcCLB.504131.70 | hypothetical protein, conserved | -0,3685 | 8,619E-06 |
| TcCLB.510409.30 | hypothetical protein, conserved | -0,3686 | 0,001614 |
| TcCLB.508663.10 | mucin TcMUC (pseudogene), putative | -0,3686 | 0,001909 |
| TcCLB.508153.470 | hypothetical protein, conserved | -0,3687 | 0,000871 |
| TcCLB.507883.50 | hypothetical protein, conserved | -0,3687 | 0,003447 |
| TcCLB.510715.11 | hypothetical protein | -0,3688 | 0,03857 |
| TcCLB.509233.210 | hypothetical protein, conserved | -0,3689 | 0,0002295 |
| TcCLB.507669.210 | hypothetical protein, conserved | -0,3689 | 0,003297 |
| TcCLB.508633.120 | trans-sialidase (pseudogene), putative | -0,369 | 0,05602 |
| TcCLB.504867.99 | hypothetical protein, conserved | -0,3691 | 0,0001111 |
| TcCLB.511625.60 | hypothetical protein, conserved | -0,3691 | 0,01365 |
| TcCLB.506925.270 | hypothetical protein, conserved | -0,3695 | 0,00003735 |
| TcCLB.511849.50 | surface protease GP63 (pseudogene), putative | -0,3695 | 0,01116 |

| | | | |
|---|---|---|---|
| TcCLB.507491.100 | hypothetical protein, conserved | -0,3697 | 0,0003046 |
| TcCLB.506407.20 | hypothetical protein, conserved | -0,3697 | 0,01089 |
| TcCLB.503665.40 | surface protease GP63 (pseudogene), putative | -0,3697 | 0,03997 |
| TcCLB.510289.90 | hypothetical protein, conserved | -0,3697 | 0,08601 |
| TcCLB.506567.100 | hypothetical protein, conserved | -0,3699 | 0,0002393 |
| TcCLB.510241.70 | heat shock protein, putative | -0,3699 | 0,001436 |
| TcCLB.510175.125 | hypothetical protein | -0,37 | 0,05335 |
| TcCLB.508277.20 | protein kinase, putative | -0,3701 | 0,0001818 |
| TcCLB.505935.10 | hypothetical protein, conserved (pseudogene) | -0,3702 | 0,121 |
| TcCLB.410049.10 | hypothetical protein, conserved | -0,3703 | 0,3927 |
| TcCLB.509569.90 | ARP2/3 complex subunit, putative | -0,3704 | 0,00408 |
| TcCLB.510329.40 | hypothetical protein, conserved | -0,3705 | 0,00004874 |
| TcCLB.508355.140 | hypothetical protein, conserved | -0,3705 | 0,002382 |
| TcCLB.509051.30 | dTDP-glucose 4,6-dehydratase, putative | -0,3708 | 0,001101 |
| TcCLB.511239.120 | hypothetical protein, conserved | -0,3708 | 0,05268 |
| TcCLB.511291.70 | hypothetical protein, conserved | -0,3709 | 0,000289 |
| TcCLB.510303.90 | hypothetical protein, conserved | -0,3709 | 0,002072 |
| TcCLB.507023.240 | hypothetical protein, conserved | -0,3709 | 0,002086 |
| TcCLB.504073.10 | endonuclease/exonuclease/phosphatase, putative | -0,371 | 0,0001013 |
| TcCLB.508799.40 | hypothetical protein, conserved | -0,3712 | 0,0000722 |
| TcCLB.510655.159 | hypothetical protein, conserved | -0,3712 | 0,04992 |
| TcCLB.511363.10 | hypothetical protein, conserved | -0,3713 | 0,00003639 |
| TcCLB.507521.20 | kinesin, putative | -0,3713 | 0,01339 |
| TcCLB.506949.10 | DNA-directed rna polymerase I largest subunit, putative | -0,3713 | 0,07114 |
| TcCLB.510823.20 | conserved protein, unknown function | -0,3715 | 0,000218 |
| TcCLB.504827.50 | apurinic/apyrimidinic endonuclease, putative | -0,3715 | 0,007842 |
| TcCLB.508207.50 | hypothetical protein | -0,3717 | 0,01759 |
| TcCLB.509643.139 | Heat shock protein 83, putative | -0,3718 | 0,001593 |
| TcCLB.508273.91 | hypothetical protein, conserved | -0,3719 | 0,1072 |
| TcCLB.506443.10 | hypothetical protein | -0,372 | 0,001149 |
| TcCLB.508851.40 | Octanoyltransferase, putative | -0,372 | 0,06973 |
| TcCLB.511421.224 | hypothetical protein, conserved (pseudogene) | -0,3721 | 0,0001306 |
| TcCLB.506177.80 | hypothetical protein, conserved | -0,3721 | 0,03737 |
| TcCLB.508187.20 | hypothetical protein (pseudogene) | -0,3721 | 0,05125 |
| TcCLB.507601.10 | phosphoprotein phosphatase, putative | -0,3722 | 3,385E-06 |
| TcCLB.506287.20 | hypothetical protein, conserved | -0,3723 | 0,01715 |
| TcCLB.508699.60 | hypothetical protein, conserved | -0,3726 | 0,001584 |
| TcCLB.503531.30 | COP9 signalosome complex subunit 5, putative (pseudoge | -0,3727 | 0,001371 |
| TcCLB.508909.170 | hypothetical protein, conserved | -0,3729 | 0,0001317 |
| TcCLB.510745.30 | hypothetical protein, conserved | -0,3729 | 0,0675 |
| TcCLB.506133.179 | hypothetical protein | -0,3729 | 0,1591 |
| TcCLB.509967.70 | katanin, putative | -0,3733 | 0,1292 |
| TcCLB.507771.20 | hypothetical protein, conserved | -0,3734 | 0,0007342 |
| TcCLB.511311.70 | hypothetical protein, conserved | -0,3736 | 0,0002804 |
| TcCLB.504769.100 | trans-sialidase, Group II, putative | -0,3736 | 0,03707 |

| | | | |
|---|---|---|---|
| TcCLB.506569.60 | retrotransposon hot spot protein (RHS, pseudogene), puta | -0,3738 | 0,05099 |
| TcCLB.509205.100 | surface protease GP63, putative | -0,3739 | 0,02086 |
| TcCLB.506891.20 | wd40 repeat domain-containing protein | -0,3742 | 0,002066 |
| TcCLB.506585.50 | hypothetical protein, conserved | -0,3743 | 0,0007756 |
| TcCLB.503675.10 | glycosyl transferase-like protein, putative | -0,3743 | 0,02014 |
| TcCLB.506175.90 | hypothetical protein, conserved | -0,3744 | 0,0001973 |
| TcCLB.503695.10 | retrotransposon hot spot protein (RHS, pseudogene), puta | -0,3748 | 0,1438 |
| TcCLB.510889.310 | hypothetical protein, conserved | -0,375 | 0,0001738 |
| TcCLB.511211.140 | hypothetical protein, conserved | -0,375 | 0,0005108 |
| TcCLB.511301.10 | oxidoreductase-like protein, putative | -0,3753 | 0,02952 |
| TcCLB.507089.90 | hypothetical protein, conserved | -0,3756 | 0,00007006 |
| TcCLB.511751.14 | hypothetical protein, conserved (fragment) | -0,3756 | 0,01007 |
| TcCLB.509395.59 | hypothetical protein, conserved | -0,3756 | 0,03103 |
| TcCLB.509805.170 | hypothetical protein, conserved | -0,3758 | 5,812E-06 |
| TcCLB.510747.170 | hypothetical protein, conserved | -0,3762 | 0,02664 |
| TcCLB.510491.10 | hypothetical protein, conserved | -0,3763 | 0,04013 |
| TcCLB.503981.50 | hypothetical protein, conserved | -0,3765 | 0,003742 |
| TcCLB.509157.40 | hypothetical protein, conserved | -0,3765 | 0,006801 |
| TcCLB.511837.40 | hypothetical protein, conserved | -0,3765 | 0,01336 |
| TcCLB.508389.160 | trans-sialidase (pseudogene), putative | -0,3765 | 0,09159 |
| TcCLB.503671.39 | dynamin-1-like protein (fragment) | -0,3769 | 0,0005367 |
| TcCLB.508355.124 | hypothetical protein, conserved | -0,3769 | 0,01333 |
| TcCLB.503391.10 | ATP-dependent DEAD/H DNA helicase recQ, putative | -0,3769 | 0,2086 |
| TcCLB.509105.100 | Enriched in surface-labeled proteome protein 15 (ESP15) | -0,3771 | 0,001169 |
| TcCLB.511289.110 | hypothetical protein, conserved | -0,3771 | 0,001561 |
| TcCLB.509765.95 | mucin-associated surface protein (MASP, pseudogene), pu | -0,3771 | 0,02432 |
| TcCLB.508173.70 | hypothetical protein, conserved | -0,3772 | 0,00003768 |
| TcCLB.507257.160 | hypothetical protein, conserved | -0,3774 | 0,003466 |
| TcCLB.511245.170 | hypothetical protein, conserved | -0,3777 | 0,00005438 |
| TcCLB.508329.19 | trans-sialidase (pseudogene), putative | -0,3778 | 0,21 |
| TcCLB.509319.70 | hypothetical protein, conserved | -0,378 | 0,006484 |
| TcCLB.506007.30 | Protein C21orf2 homolog, putative | -0,3781 | 0,001517 |
| TcCLB.511367.350 | hypothetical protein, conserved | -0,3782 | 0,0005787 |
| TcCLB.506777.10 | ATP-dependent DEAD/H RNA helicase, putative | -0,3782 | 0,001546 |
| TcCLB.511365.4 | hypothetical protein, conserved (fragment) | -0,3782 | 0,03011 |
| TcCLB.511623.10 | hypothetical protein, conserved | -0,3789 | 0,00004123 |
| TcCLB.503713.50 | trans-sialidase, putative (fragment) | -0,3789 | 0,3478 |
| TcCLB.507063.270 | histone deacetylase, putative | -0,3794 | 0,0005263 |
| TcCLB.510289.50 | hypothetical protein, conserved | -0,3795 | 0,0005632 |
| TcCLB.510295.20 | MP24, putative | -0,3796 | 0,02351 |
| TcCLB.504769.41 | surface protease GP63 (pseudogene), putative | -0,3796 | 0,05245 |
| TcCLB.510699.50 | mucin TcMUCII, putative | -0,3797 | 0,03139 |
| TcCLB.504081.9 | trans-sialidase (pseudogene), putative | -0,3801 | 0,29 |
| TcCLB.504081.260 | surface protease GP63, putative | -0,3803 | 0,3571 |
| TcCLB.507023.90 | hypothetical protein, conserved | -0,3804 | 0,0001568 |

| | | | |
|---|---|---|---|
| TcCLB.511517.170 | hypothetical protein, conserved | -0,3804 | 0,0003807 |
| TcCLB.506605.194 | hypothetical protein, conserved | -0,3805 | 0,001226 |
| TcCLB.503613.50 | PA domain containing protein, putative (ESP7) | -0,3806 | 0,0006325 |
| TcCLB.507007.50 | hypothetical protein, conserved | -0,3806 | 0,0007892 |
| TcCLB.511711.70 | hypothetical protein, conserved | -0,3806 | 0,002032 |
| TcCLB.503885.30 | hypothetical protein, conserved (pseudogene) | -0,3811 | 0,004154 |
| TcCLB.508727.60 | hypothetical protein, conserved | -0,3813 | 0,0005653 |
| TcCLB.508277.160 | glutaminyl-tRNA synthetase, putative | -0,3815 | 9,296E-08 |
| TcCLB.511909.20 | expression site-associated gene (ESAG-like) protein, puta | -0,3817 | 0,002929 |
| TcCLB.503861.30 | trans-sialidase (pseudogene), putative | -0,3817 | 0,007576 |
| TcCLB.507049.80 | hypothetical protein, conserved | -0,3819 | 0,005971 |
| TcCLB.508717.40 | branch point binding protein, putative | -0,382 | 0,001209 |
| TcCLB.509443.30 | AUT2/APG4/ATG4 cysteine peptidase, putative | -0,382 | 0,007949 |
| TcCLB.506989.260 | hypothetical protein, conserved (pseudogene) | -0,3821 | 0,005238 |
| TcCLB.508277.250 | hypothetical protein, conserved | -0,3822 | 0,00002708 |
| TcCLB.508725.10 | CCR4-NOT transcription complex subunit 10, putative | -0,3822 | 0,00007297 |
| TcCLB.510799.80 | hypothetical protein, conserved | -0,3822 | 0,0002269 |
| TcCLB.508693.100 | surface protease GP63, putative | -0,3824 | 0,002239 |
| TcCLB.503777.19 | phosphoribosylpyrophosphate synthetase, putative | -0,3824 | 0,003081 |
| TcCLB.503925.70 | hypothetical protein, conserved | -0,3825 | 6,044E-07 |
| TcCLB.509897.132 | hypothetical protein, conserved (pseudogene) | -0,3826 | 0,00003526 |
| TcCLB.509157.120 | hypothetical protein, conserved | -0,3827 | 0,02726 |
| TcCLB.504427.260 | kinesin, putative | -0,383 | 2,705E-06 |
| TcCLB.506587.40 | ATP-dependent RNA helicase FAL1, putative | -0,3832 | 0,00003982 |
| TcCLB.505229.20 | conserved protein, unknown function | -0,3833 | 6,071E-06 |
| TcCLB.506251.30 | hypothetical protein, conserved | -0,3833 | 0,00001584 |
| TcCLB.405737.4 | RNA polymerase-associated protein CDC73, putative (frag | -0,3833 | 0,2998 |
| TcCLB.509901.90 | hypothetical protein, conserved (pseudogene) | -0,3834 | 0,0000137 |
| TcCLB.510101.490 | anion-transporting ATPase-like, putative | -0,3834 | 0,0000229 |
| TcCLB.511189.50 | hypothetical protein, conserved (pseudogene) | -0,3835 | 0,0003516 |
| TcCLB.507091.40 | dihydroorotate dehydrogenase, putative (DHODH) | -0,3835 | 0,01406 |
| TcCLB.508211.19 | hypothetical protein, conserved | -0,3836 | 0,01705 |
| TcCLB.509073.119 | proteasome regulatory non-ATP-ase subunit 7, putative | -0,3838 | 0,001005 |
| TcCLB.507597.10 | hypothetical protein | -0,3838 | 0,1382 |
| TcCLB.510217.70 | trans-sialidase (pseudogene), putative | -0,3839 | 0,2433 |
| TcCLB.463359.9 | vacuolar assembly protein vps41, putative | -0,3843 | 0,008557 |
| TcCLB.506813.170 | hypothetical protein, conserved (pseudogene) | -0,3845 | 0,05595 |
| TcCLB.504741.149 | hypothetical protein | -0,3845 | 0,1397 |
| TcCLB.402857.20 | hypothetical protein | -0,3847 | 8,853E-07 |
| TcCLB.510173.120 | hypothetical protein, conserved | -0,3848 | 0,01524 |
| TcCLB.508355.340 | hypothetical protein, conserved | -0,3851 | 0,00705 |
| TcCLB.506553.40 | ATP-dependent RNA helicase (pseudogene), putative | -0,3852 | 0,0007551 |
| TcCLB.511203.5 | hypothetical protein, conserved | -0,3853 | 0,1357 |
| TcCLB.509719.50 | protein kinase, putative | -0,3854 | 0,0007771 |
| TcCLB.506979.5 | hypothetical protein, conserved (pseudogene) | -0,3854 | 0,1129 |

| | | | |
|---|---|---|---|
| TcCLB.505843.30 | DNA repair protein Rad9, putative (Rad9) | -0,3855 | 0,0000716 |
| TcCLB.507105.80 | dispersed gene family protein 1 (DGF-1), putative | -0,3856 | 0,168 |
| TcCLB.511463.4 | actin, putative | -0,3858 | 0,07313 |
| TcCLB.508647.160 | hypothetical protein, conserved | -0,3859 | 0,00003879 |
| TcCLB.510289.30 | phosphatidylinositol-4-phosphate 5-kinase, putative | -0,3862 | 0,002715 |
| TcCLB.511801.30 | hypothetical protein, conserved | -0,3863 | 0,0006687 |
| TcCLB.510953.10 | surface protease GP63 (pseudogene), putative | -0,3864 | 0,132 |
| TcCLB.510303.250 | hypothetical protein, conserved | -0,3866 | 3,617E-06 |
| TcCLB.506579.70 | hypothetical protein, conserved | -0,3868 | 7,195E-08 |
| TcCLB.510889.4 | hypothetical protein, conserved | -0,3868 | 0,1566 |
| TcCLB.505183.100 | hypothetical protein, conserved | -0,387 | 0,001214 |
| TcCLB.510493.4 | hypothetical protein, conserved (fragment) | -0,387 | 0,09213 |
| TcCLB.509591.50 | RNA-binding protein 34, putative (RBP34) | -0,3871 | 0,0006201 |
| TcCLB.503897.140 | hypothetical protein, conserved | -0,3872 | 0,001136 |
| TcCLB.507949.130 | glycerol-3-phosphate dehydrogenase (FAD-dependent), pu | -0,3877 | 0,007799 |
| TcCLB.506579.110 | hypothetical protein, conserved | -0,3881 | 4,599E-08 |
| TcCLB.511249.80 | hypothetical protein, conserved | -0,3882 | 0,00002161 |
| TcCLB.506147.200 | hypothetical protein, conserved | -0,3883 | 0,0002104 |
| TcCLB.509127.30 | hypothetical protein, conserved | -0,3883 | 0,03035 |
| TcCLB.506855.200 | hypothetical protein | -0,3884 | 0,06389 |
| TcCLB.511621.80 | hypothetical protein | -0,3886 | 0,03555 |
| TcCLB.511239.60 | hypothetical protein, conserved | -0,3887 | 4,075E-06 |
| TcCLB.508711.20 | hypothetical protein, conserved | -0,3887 | 0,00003358 |
| TcCLB.506733.140 | hypothetical protein, conserved | -0,3888 | 0,0003358 |
| TcCLB.507163.90 | hypothetical protein, conserved | -0,3888 | 0,04416 |
| TcCLB.506207.30 | hypothetical protein, conserved | -0,389 | 0,003813 |
| TcCLB.505999.70 | hypothetical protein, conserved | -0,3891 | 0,0007197 |
| TcCLB.508929.20 | UDP-Gal or UDP-GlcNAc-dependent glycosyltransferase, p | -0,3891 | 0,03111 |
| TcCLB.511903.200 | hypothetical protein, conserved | -0,3893 | 0,0006498 |
| TcCLB.511075.40 | cullin, putative | -0,3896 | 0,00003555 |
| TcCLB.504737.10 | trans-sialidase (pseudogene), putative | -0,3896 | 0,1786 |
| TcCLB.508539.60 | mucin-associated surface protein (MASP), putative | -0,3897 | 0,139 |
| TcCLB.510769.110 | hypothetical protein, conserved | -0,3898 | 0,003076 |
| TcCLB.509205.90 | surface protease GP63 (pseudogene), putative | -0,3899 | 0,01789 |
| TcCLB.507511.40 | G5-interacting protein, putative (G5-IP) | -0,3901 | 0,00004359 |
| TcCLB.511867.70 | hypothetical protein | -0,3901 | 0,0007487 |
| TcCLB.506113.55 | N-acetyltransferase complex ARD1 subunit (pseudogene), | -0,3902 | 0,1096 |
| TcCLB.511741.10 | hypothetical protein, conserved | -0,3906 | 0,009741 |
| TcCLB.505555.50 | hypothetical protein | -0,3914 | 2,141E-07 |
| TcCLB.506833.70 | cationic amino acid transporter, putative | -0,3914 | 0,0003845 |
| TcCLB.506265.80 | hypothetical protein, conserved | -0,3914 | 0,01191 |
| TcCLB.506175.30 | hypothetical protein, conserved | -0,3917 | 0,03096 |
| TcCLB.505071.30 | serine/threonine protein kinase, putative | -0,3919 | 0,0001278 |
| TcCLB.506435.300 | ubiquitin-protein ligase, putative | -0,3921 | 0,003682 |
| TcCLB.510329.160 | hypothetical protein, conserved | -0,3924 | 3,051E-06 |

| | | | |
|---|---|---|---|
| TcCLB.511391.80 | hypothetical protein | -0,3928 | 0,1071 |
| TcCLB.508173.219 | hypothetical protein, conserved | -0,393 | 1,214E-06 |
| TcCLB.509217.30 | retrotransposon hot spot protein (RHS, pseudogene), puta | -0,393 | 0,05644 |
| TcCLB.476733.29 | serine-threonine dehydratase, putative | -0,393 | 0,09954 |
| TcCLB.511039.20 | hypothetical protein, conserved | -0,3932 | 0,0005482 |
| TcCLB.511277.100 | hypothetical protein, conserved | -0,3932 | 0,004836 |
| TcCLB.510575.50 | hypothetical protein, conserved | -0,3934 | 0,04674 |
| TcCLB.511539.18 | hypothetical protein, conserved | -0,3935 | 0,00104 |
| TcCLB.511469.30 | coatomer subunit zeta, putative | -0,3941 | 0,004852 |
| TcCLB.506375.70 | hypothetical protein, conserved | -0,3941 | 0,008588 |
| TcCLB.422041.9 | pre-mRNA splicing factor ATP-dependent RNA helicase, p | -0,3941 | 0,01104 |
| TcCLB.511277.310 | hypothetical protein, conserved | -0,3944 | 0,00002689 |
| TcCLB.506941.260 | hypothetical protein, conserved | -0,3944 | 0,0004957 |
| TcCLB.510879.190 | hypothetical protein, conserved | -0,3944 | 0,002277 |
| TcCLB.509319.109 | paraflagellar rod protein, putative (fragment) | -0,3944 | 0,05497 |
| TcCLB.506957.80 | hypothetical protein, conserved | -0,3946 | 4,271E-06 |
| TcCLB.508231.220 | hypothetical protein, conserved | -0,3946 | 0,04819 |
| TcCLB.506151.20 | hypothetical protein, conserved | -0,3948 | 1,887E-07 |
| TcCLB.503635.50 | hypothetical protein, conserved | -0,3949 | 0,0006727 |
| TcCLB.508177.100 | ubiquitin activating enzyme, putative | -0,3949 | 0,003667 |
| TcCLB.507445.30 | trans-sialidase (pseudogene), putative | -0,3949 | 0,01457 |
| TcCLB.405165.19 | hypothetical protein, conserved (fragment) | -0,3951 | 0,0005653 |
| TcCLB.508727.9 | CCR4-NOT transcription complex subunit 10, putative | -0,3952 | 0,00002481 |
| TcCLB.509965.350 | ubiquitin-protein ligase, putative | -0,3955 | 5,854E-07 |
| TcCLB.504411.10 | hypothetical protein, conserved | -0,3955 | 0,0004095 |
| TcCLB.508571.10 | hypothetical protein | -0,3955 | 0,007447 |
| TcCLB.510307.230 | trans-sialidase, Group II, putative | -0,3955 | 0,08399 |
| TcCLB.511127.100 | catalytic subunit of the vacuolar transporter chaperone 4, | -0,3956 | 1,416E-06 |
| TcCLB.504109.100 | hypothetical protein, conserved | -0,3956 | 9,515E-06 |
| TcCLB.511277.620 | hypothetical protein, conserved | -0,3956 | 0,0000856 |
| TcCLB.509607.20 | hypothetical protein, conserved | -0,3958 | 0,2813 |
| TcCLB.506855.10 | palmitoyl acyltransferase 3, putative | -0,3959 | 0,00122 |
| TcCLB.510799.18 | hypothetical protein, conserved | -0,3959 | 0,001752 |
| TcCLB.509085.40 | retrotransposon hot spot protein (RHS, pseudogene), puta | -0,396 | 0,1656 |
| TcCLB.511245.230 | hypothetical protein, conserved | -0,3961 | 0,0004801 |
| TcCLB.510389.10 | hypothetical protein, conserved (pseudogene) | -0,3961 | 0,03169 |
| TcCLB.506247.60 | hypothetical protein, conserved | -0,3965 | 0,00003034 |
| TcCLB.507257.50 | hypothetical protein, conserved | -0,3966 | 1,199E-07 |
| TcCLB.510387.60 | trans-sialidase (pseudogene), putative | -0,3966 | 0,03149 |
| TcCLB.506495.10 | hypothetical protein, conserved | -0,3968 | 0,007286 |
| TcCLB.507165.40 | hypothetical protein, conserved | -0,3969 | 0,009102 |
| TcCLB.509369.50 | surface protease GP63, putative | -0,3974 | 0,2722 |
| TcCLB.506195.309 | hypothetical protein, conserved | -0,3976 | 0,0002714 |
| TcCLB.509877.30 | guanine nucleotide-binding beta subunit-like protein | -0,3977 | 0,00001342 |
| TcCLB.503885.10 | hypothetical protein, conserved | -0,3977 | 0,001676 |

| | | | |
|---|---|---|---|
| TcCLB.510241.20 | hypothetical protein, conserved | -0,3979 | 0,00004349 |
| TcCLB.511511.90 | ATP-dependent DEAD/H DNA helicase recQ family, putativ | -0,398 | 0,0001415 |
| TcCLB.469785.40 | RNA-binding protein, putative | -0,398 | 0,01318 |
| TcCLB.511757.20 | hypothetical protein, conserved | -0,3981 | 0,01024 |
| TcCLB.509429.230 | hypothetical protein, conserved | -0,3983 | 0,004123 |
| TcCLB.510621.49 | mucin-associated surface protein (MASP), putative | -0,3983 | 0,2094 |
| TcCLB.507099.10 | hypothetical protein, conserved | -0,3987 | 5,039E-06 |
| TcCLB.508817.170 | hypothetical protein, conserved | -0,3987 | 0,009071 |
| TcCLB.506493.90 | cysteine peptidase, Clan CA, family C2, putative | -0,3989 | 6,205E-06 |
| TcCLB.508555.70 | calpain-like cysteine peptidase, putative | -0,399 | 0,003738 |
| TcCLB.503953.10 | hypothetical protein, conserved | -0,3993 | 6,422E-06 |
| TcCLB.511001.140 | hypothetical protein, conserved | -0,3993 | 0,0001889 |
| TcCLB.509125.10 | mannosyltransferase-like protein, putative | -0,3996 | 0,00006262 |
| TcCLB.506135.90 | hypothetical protein, conserved | -0,3996 | 0,0007388 |
| TcCLB.508857.150 | hypothetical protein, conserved | -0,3996 | 0,001403 |
| TcCLB.504229.30 | protein kinase, putative | -0,3999 | 0,0000633 |
| TcCLB.508307.190 | hypothetical protein, conserved | -0,4 | 0,00428 |
| TcCLB.507611.340 | hypothetical protein, conserved | -0,4002 | 0,00007434 |
| TcCLB.508943.40 | hypothetical protein, conserved | -0,4002 | 0,0167 |
| TcCLB.506207.20 | hypothetical protein, conserved | -0,4003 | 7,502E-07 |
| TcCLB.506173.219 | trans-sialidase (pseudogene), putative | -0,4003 | 0,1843 |
| TcCLB.509665.40 | hypothetical protein | -0,4003 | 0,2871 |
| TcCLB.508927.30 | hypothetical protein, conserved | -0,4004 | 0,00141 |
| TcCLB.511755.40 | hypothetical protein, conserved | -0,4004 | 0,002077 |
| TcCLB.506755.74 | mitochondrial carrier protein, putative | -0,4005 | 0,002094 |
| TcCLB.511429.10 | transmembrane protein, conserved (fragment) | -0,4007 | 3,521E-06 |
| TcCLB.508727.44 | hypothetical protein, conserved | -0,4009 | 0,00002655 |
| TcCLB.503579.80 | hypothetical protein, conserved | -0,4009 | 0,00004573 |
| TcCLB.509157.240 | hypothetical protein, conserved | -0,4015 | 0,009688 |
| TcCLB.510243.128 | hypothetical protein, conserved | -0,4015 | 0,03112 |
| TcCLB.506401.220 | electron transfer flavoprotein-ubiquinone oxidoreductase, | -0,4016 | 0,0003167 |
| TcCLB.511213.81 | retrotransposon hot spot protein (RHS, pseudogene), puta | -0,4016 | 0,02163 |
| TcCLB.510239.30 | hypothetical protein, conserved (pseudogene) | -0,4019 | 0,1 |
| TcCLB.506947.70 | hypothetical protein, conserved | -0,402 | 0,01712 |
| TcCLB.510933.160 | mucin-associated surface protein (MASP, pseudogene), pu | -0,4022 | 0,0889 |
| TcCLB.511737.40 | hypothetical protein, conserved | -0,4023 | 0,0002912 |
| TcCLB.508463.10 | hypothetical protein, conserved | -0,4023 | 0,03429 |
| TcCLB.511737.3 | superoxide dismutase, putative | -0,4024 | 0,02331 |
| TcCLB.503809.30 | RNA-binding protein, putative | -0,4026 | 0,003266 |
| TcCLB.511757.10 | hypothetical protein, conserved | -0,4028 | 0,00001169 |
| TcCLB.511301.180 | trans-sialidase (pseudogene), putative | -0,4028 | 0,0004953 |
| TcCLB.510879.40 | Carboxypeptidase Nna1-like protein, putative | -0,4031 | 7,642E-06 |
| TcCLB.509611.130 | hypothetical protein, conserved | -0,4031 | 0,01666 |
| TcCLB.509657.11 | hypothetical protein, conserved (pseudogene) | -0,4032 | 0,06494 |
| TcCLB.507641.220 | hypothetical protein | -0,4034 | 0,0003458 |

| | | | |
|---|---|---|---|
| TcCLB.503653.104 | hypothetical protein, conserved | -0,4037 | 0,2217 |
| TcCLB.511261.100 | chaperone DNAJ protein, putative | -0,4038 | 0,0001337 |
| TcCLB.506425.60 | hypothetical protein, conserved | -0,4038 | 0,0004602 |
| TcCLB.506717.4 | hypothetical protein, conserved (fragment) | -0,4039 | 0,08794 |
| TcCLB.510535.60 | CCR4 associated factor, putative | -0,4042 | 0,00002622 |
| TcCLB.510877.100 | hypothetical protein, conserved | -0,4044 | 0,0001114 |
| TcCLB.508461.344 | hypothetical protein, conserved | -0,4045 | 0,0054 |
| TcCLB.506819.10 | hypothetical protein, conserved | -0,4045 | 0,02592 |
| TcCLB.506369.40 | hypothetical protein, conserved | -0,4046 | 6,286E-06 |
| TcCLB.509627.20 | hypothetical protein, conserved | -0,4048 | 0,0001198 |
| TcCLB.504137.40 | hypothetical protein, conserved | -0,4048 | 0,0002572 |
| TcCLB.506247.80 | DNA-binding protein, putative | -0,4048 | 0,0003 |
| TcCLB.504639.50 | retrotransposon hot spot protein (RHS, pseudogene), puta | -0,4048 | 0,000562 |
| TcCLB.510329.50 | hypothetical protein | -0,4049 | 1,446E-06 |
| TcCLB.503559.100 | long-chain-fatty-acid-CoA ligase, putative | -0,4049 | 0,00005162 |
| TcCLB.508761.130 | hypothetical protein | -0,4049 | 0,4486 |
| TcCLB.507257.100 | hypothetical protein, conserved | -0,405 | 0,0004454 |
| TcCLB.507611.290 | acetyltransferase, putative | -0,4051 | 0,003292 |
| TcCLB.510611.40 | hypothetical protein, conserved | -0,4052 | 0,0001124 |
| TcCLB.504431.10 | hypothetical protein, conserved | -0,4052 | 0,0001165 |
| TcCLB.503781.40 | hypothetical protein, conserved | -0,4053 | 0,006863 |
| TcCLB.508403.10 | hypothetical protein, conserved | -0,4055 | 0,00009483 |
| TcCLB.503651.4 | hypothetical protein, conserved (fragment) | -0,4057 | 0,0003167 |
| TcCLB.509957.30 | retrotransposon hot spot protein (RHS, pseudogene), puta | -0,4057 | 0,02893 |
| TcCLB.503489.30 | hypothetical protein, conserved | -0,4058 | 0,001632 |
| TcCLB.508737.40 | hypothetical protein | -0,4067 | 0,000487 |
| TcCLB.511041.30 | hypothetical protein, conserved | -0,4069 | 5,704E-07 |
| TcCLB.509253.40 | hypothetical protein, conserved | -0,4071 | 0,001264 |
| TcCLB.508535.40 | hypothetical protein, conserved | -0,4074 | 0,03182 |
| TcCLB.509875.230 | hypothetical protein, conserved | -0,4076 | 0,0005866 |
| TcCLB.511509.50 | hypothetical protein, conserved | -0,4077 | 0,00001306 |
| TcCLB.510971.60 | protein kinase, putative | -0,4077 | 0,0003991 |
| TcCLB.507035.120 | hypothetical protein, conserved | -0,4077 | 0,001658 |
| TcCLB.507811.30 | ATP-dependent DEAD/H RNA helicase, putative | -0,4078 | 0,00007533 |
| TcCLB.509203.20 | cyclin dependent kinase-binding protein, putative | -0,4079 | 0,00002875 |
| TcCLB.504689.10 | hypothetical protein | -0,4083 | 0,2017 |
| TcCLB.503817.10 | trans-sialidase (pseudogene), putative | -0,4084 | 0,1377 |
| TcCLB.509051.60 | ubiquitin-protein ligase, putative (fragment) | -0,4085 | 0,0525 |
| TcCLB.509715.130 | hypothetical protein, conserved | -0,4086 | 0,000775 |
| TcCLB.504131.150 | SNF2 family protein | -0,4086 | 0,002402 |
| TcCLB.430727.9 | inositol phosphatidylinositol phosphatase, putative | -0,4088 | 0,005274 |
| TcCLB.510043.20 | hypothetical protein, conserved (pseudogene) | -0,4091 | 2,218E-07 |
| TcCLB.503453.50 | hypothetical protein, conserved | -0,4091 | 0,001154 |
| TcCLB.504109.120 | hypothetical protein, conserved | -0,4092 | 0,0007536 |
| TcCLB.510429.40 | Present in the outer mitochondrial membrane proteome 5 | -0,4092 | 0,001047 |

| | | | |
|---|---|---|---|
| TcCLB.511873.30 | trans-sialidase (pseudogene), putative | -0,4092 | 0,005984 |
| TcCLB.506835.150 | zinc-binding protein (Yippee), putative | -0,4092 | 0,007163 |
| TcCLB.508157.10 | serine-alanine-and proline-rich protein (pseudogene), put | -0,4092 | 0,09215 |
| TcCLB.506725.80 | hypothetical protein, conserved | -0,4093 | 0,0001207 |
| TcCLB.505965.92 | hypothetical protein, conserved (pseudogene) | -0,4094 | 0,01766 |
| TcCLB.507021.90 | hypothetical protein, conserved | -0,4095 | 0,0002497 |
| TcCLB.508163.90 | trans-sialidase (pseudogene), putative | -0,4095 | 0,1478 |
| TcCLB.507793.20 | cysteine synthase, putative | -0,4099 | 0,0004198 |
| TcCLB.506465.10 | hypothetical protein, conserved | -0,4104 | 0,0001454 |
| TcCLB.508323.110 | hypothetical protein, conserved | -0,4105 | 0,00235 |
| TcCLB.507035.70 | hypothetical protein | -0,4106 | 0,01795 |
| TcCLB.511183.315 | hypothetical protein, conserved (pseudogene) | -0,4106 | 0,2636 |
| TcCLB.507627.9 | kinesin, putative | -0,4109 | 4,041E-06 |
| TcCLB.509741.30 | hypothetical protein, conserved | -0,4109 | 0,0002204 |
| TcCLB.511109.80 | hypothetical protein, conserved | -0,4109 | 0,005827 |
| TcCLB.508257.190 | hypothetical protein, conserved | -0,4112 | 0,002372 |
| TcCLB.510515.50 | hypothetical protein, conserved | -0,4113 | 0,002337 |
| TcCLB.508375.70 | hypothetical protein, conserved | -0,4114 | 0,002588 |
| TcCLB.507815.20 | trans-sialidase (pseudogene), putative | -0,4114 | 0,01591 |
| TcCLB.506737.151 | surface protease GP63 (pseudogene), putative | -0,4115 | 0,0006684 |
| TcCLB.509007.80 | hypothetical protein, conserved | -0,4117 | 0,0003041 |
| TcCLB.503939.120 | hypothetical protein, conserved | -0,4122 | 0,00001982 |
| TcCLB.510599.30 | hypothetical protein, conserved | -0,4123 | 2,359E-06 |
| TcCLB.507031.70 | hypothetical protein, conserved | -0,4124 | 0,0006724 |
| TcCLB.506679.120 | hypothetical protein | -0,4124 | 0,002321 |
| TcCLB.511559.30 | ras-related GTP-binding protein, putative | -0,4126 | 0,0008165 |
| TcCLB.506445.40 | ubiquitin hydrolase (pseudogene), putative | -0,4127 | 9,539E-06 |
| TcCLB.511277.380 | hypothetical protein, conserved | -0,4127 | 0,03793 |
| TcCLB.506303.120 | hypothetical protein, conserved | -0,4128 | 0,00001118 |
| TcCLB.506213.90 | hypothetical protein, conserved | -0,4129 | 0,00008586 |
| TcCLB.509171.60 | hypothetical protein, conserved | -0,4131 | 0,0002033 |
| TcCLB.507517.30 | hypothetical protein, conserved | -0,4131 | 0,002356 |
| TcCLB.507737.20 | hypothetical protein, conserved | -0,4131 | 0,002501 |
| TcCLB.507669.170 | WD40/YVTN repeat-like-containing protein | -0,4133 | 5,797E-07 |
| TcCLB.506661.150 | hypothetical protein, conserved | -0,4137 | 0,008055 |
| TcCLB.506813.10 | hypothetical protein | -0,4139 | 0,01207 |
| TcCLB.508707.110 | hypothetical protein, conserved | -0,4139 | 0,01228 |
| TcCLB.507821.40 | trans-sialidase (pseudogene), putative | -0,414 | 0,0102 |
| TcCLB.508255.10 | trans-sialidase (pseudogene), putative | -0,4142 | 0,01317 |
| TcCLB.506205.10 | hypothetical protein, conserved | -0,4146 | 0,00002846 |
| TcCLB.510437.60 | hypothetical protein, conserved | -0,4149 | 0,003966 |
| TcCLB.509901.170 | apurinic/apyrimidinic endonuclease (TCAP) | -0,4151 | 0,00006913 |
| TcCLB.506315.60 | hypothetical protein, conserved | -0,4153 | 0,0006515 |
| TcCLB.508409.338 | DNA repair protein, putative | -0,4154 | 0,001218 |
| TcCLB.509669.10 | hypothetical protein, conserved | -0,4156 | 0,01641 |

| | | | |
|---|---|---|---|
| TcCLB.503779.70 | hypothetical protein, conserved | -0,4157 | 0,02385 |
| TcCLB.510609.60 | hypothetical protein, conserved | -0,4158 | 1,017E-06 |
| TcCLB.503697.20 | hypothetical protein, conserved | -0,4159 | 0,00007422 |
| TcCLB.511747.20 | hypothetical protein, conserved | -0,4159 | 0,00034 |
| TcCLB.508813.30 | surface protease GP63 (pseudogene), putative | -0,4159 | 0,083 |
| TcCLB.508741.180 | hypothetical protein, conserved | -0,416 | 0,0005906 |
| TcCLB.510531.20 | hypothetical protein, conserved | -0,416 | 0,001693 |
| TcCLB.511921.167 | hypothetical protein, conserved (pseudogene) | -0,4161 | 0,2022 |
| TcCLB.511065.50 | phosphatidylinositol 3-kinase, putative | -0,4162 | 0,004215 |
| TcCLB.467795.9 | diphthamide synthesis protein, putative | -0,4165 | 0,00009042 |
| TcCLB.506009.80 | DNA repair and transcription factor protein, putative | -0,4166 | 0,0006247 |
| TcCLB.511367.310 | hypothetical protein, conserved | -0,4167 | 0,001028 |
| TcCLB.508593.30 | hypothetical protein, conserved | -0,4168 | 0,0001111 |
| TcCLB.508647.130 | hypothetical protein, conserved | -0,4168 | 0,0002174 |
| TcCLB.508047.80 | mucin TcMUCII (pseudogene), putative | -0,4173 | 0,1739 |
| TcCLB.510243.54 | hypothetical protein, conserved | -0,4174 | 0,0002486 |
| TcCLB.503679.40 | actin-related protein 2, putative | -0,4177 | 0,08578 |
| TcCLB.508723.90 | hypothetical protein, conserved | -0,4178 | 0,0001985 |
| TcCLB.509287.170 | trans-sialidase (pseudogene), putative | -0,4178 | 0,001411 |
| TcCLB.511183.391 | surface protease GP63 (pseudogene), putative | -0,4179 | 0,0668 |
| TcCLB.509461.50 | hypothetical protein, conserved | -0,418 | 0,0001105 |
| TcCLB.511287.34 | hypothetical protein, conserved | -0,418 | 0,0002764 |
| TcCLB.507821.20 | hypothetical protein | -0,4181 | 0,134 |
| TcCLB.504159.60 | hypothetical protein, conserved | -0,4182 | 0,0003067 |
| TcCLB.508325.25 | trans-sialidase (pseudogene), putative | -0,4185 | 0,01274 |
| TcCLB.507991.90 | hypothetical protein | -0,4187 | 0,00006994 |
| TcCLB.511365.50 | hypothetical protein, conserved (pseudogene) | -0,4187 | 0,0001846 |
| TcCLB.510351.60 | hypothetical protein, conserved | -0,4188 | 0,003266 |
| TcCLB.510823.60 | hypothetical protein, conserved | -0,4189 | 0,00000624 |
| TcCLB.506925.150 | protein phosphatase, putative | -0,419 | 0,00025 |
| TcCLB.506835.131 | hypothetical protein, conserved | -0,4193 | 0,02718 |
| TcCLB.509793.20 | hypothetical protein, conserved | -0,4194 | 0,0007679 |
| TcCLB.507583.60 | hypothetical protein, conserved | -0,4196 | 0,2398 |
| TcCLB.507837.90 | cell division cycle protein, putative | -0,4198 | 0,001147 |
| TcCLB.509937.160 | hypothetical protein, conserved | -0,4199 | 9,372E-06 |
| TcCLB.511643.80 | hypothetical protein, conserved (pseudogene) | -0,4201 | 0,00000996 |
| TcCLB.508737.110 | hypothetical protein, conserved | -0,4201 | 0,00004953 |
| TcCLB.509899.10 | trans-sialidase (pseudogene), putative | -0,4202 | 0,001466 |
| TcCLB.511569.30 | trans-sialidase (pseudogene), putative | -0,4206 | 0,02799 |
| TcCLB.507251.30 | hypothetical protein, conserved | -0,4207 | 0,00006716 |
| TcCLB.504137.70 | hypothetical protein, conserved | -0,4207 | 0,0004606 |
| TcCLB.507009.90 | iron-sulfur cluster assembly protein, putative | -0,4209 | 0,0003247 |
| TcCLB.506737.20 | protein kinase, putative | -0,421 | 0,00004264 |
| TcCLB.506895.20 | hypothetical protein, conserved | -0,4212 | 5,289E-06 |
| TcCLB.511287.100 | hypothetical protein, conserved | -0,4212 | 0,0000622 |

| | | | |
|---|---|---|---|
| TcCLB.509671.90 | COP-coated vesicle membrane protein gp25L precursor, pu | -0,4212 | 0,0001054 |
| TcCLB.511627.110 | heat shock protein DNAJ, putative | -0,4213 | 2,216E-11 |
| TcCLB.507807.20 | hypothetical protein, conserved | -0,4213 | 0,01027 |
| TcCLB.511821.80 | hypothetical protein, conserved | -0,4214 | 0,0001313 |
| TcCLB.504881.10 | hypothetical protein, conserved | -0,4216 | 0,0000803 |
| TcCLB.508389.174 | hypothetical protein, conserved (pseudogene) | -0,4218 | 0,04173 |
| TcCLB.508607.70 | trans-sialidase (pseudogene), putative | -0,4221 | 0,001744 |
| TcCLB.509791.70 | hypothetical protein, conserved | -0,4222 | 0,003326 |
| TcCLB.507771.120 | trans-sialidase (pseudogene), putative | -0,4222 | 0,008825 |
| TcCLB.509617.50 | hypothetical protein, conserved | -0,4223 | 0,00006528 |
| TcCLB.507601.130 | hypothetical protein, conserved | -0,4226 | 3,154E-06 |
| TcCLB.506923.30 | hypothetical protein (pseudogene) | -0,4227 | 0,08955 |
| TcCLB.507005.20 | hypothetical protein, conserved | -0,4229 | 0,0007696 |
| TcCLB.506817.79 | kinteoplast poly(A) polymerase complex 1 subunit, putativ | -0,4229 | 0,008872 |
| TcCLB.509145.19 | hypothetical protein, conserved | -0,423 | 0,005241 |
| TcCLB.510747.18 | hypothetical protein, conserved | -0,4232 | 0,001284 |
| TcCLB.508637.130 | hypothetical protein, conserved | -0,4232 | 0,0915 |
| TcCLB.509729.29 | neurobeachin/beige protein, putative (fragment) | -0,4233 | 0,0001006 |
| TcCLB.503805.3 | kinesin heavy chain, putative | -0,4234 | 0,01132 |
| TcCLB.509665.13 | trans-sialidase (pseudogene), putative | -0,4237 | 0,05099 |
| TcCLB.504159.90 | hypothetical protein, conserved | -0,4238 | 0,00000174 |
| TcCLB.508777.100 | cyclin 1 (CYC1) | -0,424 | 0,0009675 |
| TcCLB.508163.50 | mucin-associated surface protein (MASP), putative | -0,4241 | 0,05448 |
| TcCLB.510597.30 | hypothetical protein, conserved | -0,4243 | 1,417E-06 |
| TcCLB.505957.140 | trans-sialidase (pseudogene), putative | -0,4244 | 0,02961 |
| TcCLB.511745.50 | hypothetical protein, conserved | -0,4245 | 0,02791 |
| TcCLB.510679.90 | expression site-associated gene (ESAG-like, pseudogene), | -0,4245 | 0,1375 |
| TcCLB.504155.280 | trans-sialidase (pseudogene), putative | -0,4246 | 0,02098 |
| TcCLB.507099.70 | hypothetical protein, conserved | -0,4248 | 6,609E-07 |
| TcCLB.509793.60 | hypothetical protein, conserved | -0,4248 | 0,00009337 |
| TcCLB.507165.10 | hypothetical protein, conserved | -0,4249 | 0,00134 |
| TcCLB.511259.149 | mucin-associated surface protein (MASP, pseudogene), pu | -0,4249 | 0,1905 |
| TcCLB.508325.60 | retrotransposon hot spot (RHS) protein, putative | -0,4251 | 0,001433 |
| TcCLB.503985.20 | retrotransposon hot spot protein (RHS, pseudogene), puta | -0,4252 | 0,0264 |
| TcCLB.508175.340 | hypothetical protein, conserved | -0,4255 | 1,226E-07 |
| TcCLB.506529.160 | hypothetical protein, conserved | -0,4255 | 0,0001439 |
| TcCLB.506709.30 | hypothetical protein, conserved | -0,4255 | 0,0001896 |
| TcCLB.510973.20 | hypothetical protein, conserved | -0,4256 | 0,00001613 |
| TcCLB.503525.20 | hypothetical protein, conserved | -0,4257 | 1,733E-06 |
| TcCLB.511423.50 | conserved protein, unknown function | -0,4259 | 8,567E-06 |
| TcCLB.510759.60 | vesicle-associated membrane protein, putative | -0,4259 | 0,003508 |
| TcCLB.507049.41 | hypothetical protein, conserved | -0,4259 | 0,01433 |
| TcCLB.506303.30 | hypothetical protein, conserved | -0,4261 | 0,001553 |
| TcCLB.509973.25 | mucin TcMUC (pseudogene), putative | -0,4261 | 0,06235 |
| TcCLB.511183.480 | hypothetical protein, conserved (pseudogene) | -0,4261 | 0,215 |

| | | | |
|---|---|---|---|
| TcCLB.509943.30 | ribosomal protein S6, putative | -0,4262 | 0,002591 |
| TcCLB.506709.50 | hypothetical protein, conserved | -0,4263 | 0,01949 |
| TcCLB.508173.240 | hypothetical protein, conserved | -0,4266 | 0,00144 |
| TcCLB.506289.200 | hypothetical protein | -0,4267 | 0,1569 |
| TcCLB.508601.60 | hypothetical protein, conserved | -0,4268 | 0,0002317 |
| TcCLB.504093.10 | hypothetical protein, conserved (pseudogene) | -0,427 | 2,531E-06 |
| TcCLB.510243.20 | hypothetical protein, conserved | -0,4273 | 0,0001718 |
| TcCLB.508595.30 | hypothetical protein (pseudogene) | -0,4273 | 0,003264 |
| TcCLB.510755.147 | hypothetical protein, conserved | -0,4274 | 0,003526 |
| TcCLB.509669.20 | subtilisin-like serine peptidase, putative | -0,4279 | 2,434E-07 |
| TcCLB.505297.80 | trans-sialidase (pseudogene), putative | -0,428 | 0,007444 |
| TcCLB.508445.30 | protein kinase, putative | -0,4283 | 0,0004313 |
| TcCLB.509945.20 | hypothetical protein, conserved | -0,4283 | 0,001291 |
| TcCLB.511283.80 | hypothetical protein, conserved | -0,4284 | 1,421E-07 |
| TcCLB.506863.80 | serine/threonine protein phosphatase, putative | -0,4284 | 0,000105 |
| TcCLB.506975.49 | f-actin capping protein beta subunit, putative | -0,4284 | 0,002051 |
| TcCLB.509069.49 | DNA replication licensing factor MCM7, putative (fragmer | -0,4284 | 0,004425 |
| TcCLB.410199.30 | trans-sialidase, Group VI, putative | -0,4284 | 0,007638 |
| TcCLB.508509.30 | surface protease GP63 (pseudogene), putative | -0,4285 | 0,1618 |
| TcCLB.509099.59 | hypothetical protein, conserved | -0,4286 | 0,0004153 |
| TcCLB.510275.405 | hypothetical protein | -0,4286 | 0,1377 |
| TcCLB.511721.20 | hypothetical protein, conserved | -0,4286 | 0,1862 |
| TcCLB.508417.10 | hypothetical protein, conserved | -0,4289 | 3,192E-06 |
| TcCLB.503901.18 | hypothetical protein, conserved (fragment) | -0,4291 | 0,00009829 |
| TcCLB.511407.40 | hypothetical protein, conserved | -0,4292 | 0,0001654 |
| TcCLB.504103.70 | hypothetical protein, conserved | -0,4293 | 0,0006779 |
| TcCLB.511821.60 | protein kinase, putative | -0,4294 | 0,00003535 |
| TcCLB.511717.40 | hypothetical protein, conserved | -0,4294 | 0,005472 |
| TcCLB.510279.180 | mucin TcMUCII (pseudogene), putative | -0,4296 | 0,2901 |
| TcCLB.511491.30 | hypothetical protein, conserved | -0,4297 | 0,0001894 |
| TcCLB.510223.9 | hypothetical protein, conserved | -0,4297 | 0,2118 |
| TcCLB.506133.68 | mucin TcMUCII (pseudogene), putative | -0,4298 | 0,2591 |
| TcCLB.511211.150 | spindle pole body component alp6, putative | -0,4299 | 0,00003599 |
| TcCLB.511305.20 | hypothetical protein, conserved | -0,4302 | 2,648E-07 |
| TcCLB.478283.10 | hypothetical protein, conserved (pseudogene) | -0,4305 | 0,00557 |
| TcCLB.507941.20 | hypothetical protein, conserved | -0,4306 | 6,801E-08 |
| TcCLB.504411.30 | serine/threonine protein kinase, putative | -0,4306 | 0,00705 |
| TcCLB.504689.30 | trans-sialidase (pseudogene), putative | -0,4306 | 0,03355 |
| TcCLB.510889.352 | retrotransposon hot spot (RHS) protein, putative (fragmer | -0,4306 | 0,1365 |
| TcCLB.509267.50 | U5 small nuclear ribonucleoprotein component, putative | -0,4309 | 2,824E-07 |
| TcCLB.506579.60 | hypothetical protein, conserved | -0,4309 | 0,0004199 |
| TcCLB.505009.30 | serine/threonine protein kinase, putative | -0,4311 | 0,0104 |
| TcCLB.510611.50 | hypothetical protein, conserved | -0,4313 | 0,0002331 |
| TcCLB.508521.11 | trans-sialidase (pseudogene), putative | -0,4313 | 0,01233 |
| TcCLB.508815.100 | hypothetical protein, conserved | -0,4314 | 0,0000116 |

| | | | |
|---|---|---|---|
| TcCLB.509537.10 | pre-mRNA splicing factor ATP-dependent RNA helicase, p | -0,4316 | 0,0002522 |
| TcCLB.511915.20 | trans-sialidase (pseudogene), putative | -0,4317 | 0,007049 |
| TcCLB.509965.280 | hypothetical protein, conserved | -0,4318 | 0,001257 |
| TcCLB.510661.120 | hypothetical protein, conserved | -0,4319 | 0,0003279 |
| TcCLB.504171.50 | hypothetical protein, conserved | -0,432 | 1,071E-08 |
| TcCLB.509169.20 | hypothetical protein, conserved | -0,4321 | 0,00001951 |
| TcCLB.505949.50 | mucin-associated surface protein (MASP, pseudogene), pu | -0,4321 | 0,3674 |
| TcCLB.507093.10 | hypothetical protein, conserved | -0,4322 | 2,731E-06 |
| TcCLB.511737.90 | hypothetical protein, conserved | -0,4323 | 0,0009486 |
| TcCLB.506739.70 | hypothetical protein, conserved | -0,4324 | 0,00005972 |
| TcCLB.510101.480 | hypothetical protein, conserved | -0,4328 | 0,0007033 |
| TcCLB.509989.10 | phosphatidylinositol 3-kinase, putative | -0,433 | 0,005769 |
| TcCLB.507517.20 | DNA polymerase sigma-like protein, putative | -0,4334 | 0,00048 |
| TcCLB.511067.14 | hypothetical protein, conserved | -0,4334 | 0,005065 |
| TcCLB.511131.60 | hypothetical protein, conserved | -0,4336 | 0,0008974 |
| TcCLB.506885.290 | DNA replication licensing factor MCM9, putative | -0,4337 | 0,0000232 |
| TcCLB.509671.50 | hypothetical protein, conserved | -0,4337 | 0,003097 |
| TcCLB.503561.20 | hypothetical protein, conserved | -0,4345 | 0,0006046 |
| TcCLB.511517.95 | hypothetical protein, conserved | -0,4347 | 0,0376 |
| TcCLB.511151.20 | hypothetical protein, conserved | -0,4352 | 7,788E-07 |
| TcCLB.504883.30 | hypothetical protein, conserved | -0,4353 | 6,393E-07 |
| TcCLB.511055.50 | protein kinase, putative | -0,4353 | 0,0499 |
| TcCLB.507093.180 | transcription factor IIIb, putative | -0,4354 | 0,0000378 |
| TcCLB.508857.140 | hypothetical protein, conserved | -0,4354 | 0,00009506 |
| TcCLB.508479.290 | vacuolar sorting protein, putative | -0,4357 | 0,00792 |
| TcCLB.508413.30 | hypothetical protein, conserved | -0,4358 | 1,204E-06 |
| TcCLB.508707.170 | outer dynein arm docking complex, putative (DC2) | -0,4358 | 0,00003221 |
| TcCLB.511001.200 | hypothetical protein, conserved | -0,4358 | 0,0004139 |
| TcCLB.510655.100 | hypothetical protein | -0,4361 | 0,0005459 |
| TcCLB.504035.100 | hypothetical protein, conserved | -0,4363 | 4,306E-06 |
| TcCLB.508661.60 | GPI transamidase component Tta2, putative (TTA2) | -0,4363 | 0,00002578 |
| TcCLB.511787.20 | trans-sialidase (pseudogene), putative | -0,4367 | 0,05109 |
| TcCLB.508507.9 | hypothetical protein, conserved | -0,4371 | 0,006205 |
| TcCLB.508739.50 | hypothetical protein | -0,4372 | 0,001636 |
| TcCLB.508023.100 | trans-sialidase, Group IV, putative | -0,4372 | 0,02074 |
| TcCLB.509899.70 | hypothetical protein, conserved (pseudogene) | -0,4374 | 3,859E-06 |
| TcCLB.503557.49 | hypothetical protein, conserved (fragment) | -0,4374 | 0,03272 |
| TcCLB.506943.150 | hypothetical protein, conserved (pseudogene) | -0,4375 | 0,0000108 |
| TcCLB.506753.80 | hypothetical protein, conserved | -0,4376 | 0,00003484 |
| TcCLB.511261.40 | hypothetical protein | -0,4376 | 0,0001131 |
| TcCLB.506883.34 | hypothetical protein, conserved | -0,4376 | 0,0002807 |
| TcCLB.508075.30 | hypothetical protein, conserved (pseudogene) | -0,4376 | 0,001236 |
| TcCLB.509059.89 | vacuolar assembly protein vps41, putative (fragment) | -0,4376 | 0,0422 |
| TcCLB.507663.70 | hypothetical protein, conserved | -0,4377 | 0,001668 |
| TcCLB.506181.10 | hypothetical protein, conserved | -0,4378 | 0,0009609 |

| | | | |
|---|---|---|---|
| TcCLB.511445.120 | hypothetical protein, conserved | -0,4379 | 0,00008638 |
| TcCLB.504105.200 | hypothetical protein, conserved | -0,438 | 4,488E-08 |
| TcCLB.509801.10 | hypothetical protein, conserved | -0,438 | 0,00009297 |
| TcCLB.509965.210 | hypothetical protein, conserved | -0,4383 | 0,08267 |
| TcCLB.509005.50 | hypothetical protein, conserved | -0,4384 | 5,045E-07 |
| TcCLB.506863.40 | hypothetical protein, conserved | -0,4384 | 0,01417 |
| TcCLB.511277.460 | hypothetical protein, conserved | -0,4385 | 0,0002193 |
| TcCLB.508645.50 | hypothetical protein, conserved | -0,4386 | 0,00001966 |
| TcCLB.511277.300 | hypothetical protein | -0,4387 | 2,164E-07 |
| TcCLB.511153.150 | hypothetical protein | -0,4387 | 0,002844 |
| TcCLB.506579.90 | hypothetical protein | -0,4388 | 8,606E-06 |
| TcCLB.504123.10 | hypothetical protein, conserved | -0,4388 | 0,0001471 |
| TcCLB.506577.70 | hypothetical protein, conserved | -0,4388 | 0,0003507 |
| TcCLB.506883.90 | hypothetical protein, conserved | -0,4389 | 0,0003382 |
| TcCLB.504203.29 | tRNA-dihydrouridine synthase 2, putative | -0,4391 | 0,006944 |
| TcCLB.506173.80 | hypothetical protein | -0,4391 | 0,2481 |
| TcCLB.504021.100 | hypothetical protein, conserved | -0,4392 | 0,00006046 |
| TcCLB.508201.40 | mucin TcMUCII, putative | -0,4392 | 0,03821 |
| TcCLB.510201.70 | hypothetical protein | -0,4392 | 0,1159 |
| TcCLB.510737.119 | hypothetical protein, conserved | -0,4394 | 0,0001828 |
| TcCLB.511803.60 | hypothetical protein, conserved | -0,4394 | 0,002939 |
| TcCLB.506411.25 | hypothetical protein, conserved (pseudogene) | -0,4398 | 0,00857 |
| TcCLB.507753.3 | hypothetical protein, conserved (pseudogene) | -0,44 | 0,04489 |
| TcCLB.511587.36 | N-acetyltransferase complex ARD1 subunit (pseudogene), | -0,44 | 0,09075 |
| TcCLB.511751.163 | hypothetical protein, conserved | -0,4401 | 0,0001121 |
| TcCLB.511907.290 | hypothetical protein, conserved | -0,4402 | 0,00001875 |
| TcCLB.506405.140 | hypothetical protein, conserved | -0,4403 | 0,00008208 |
| TcCLB.510819.70 | hypothetical protein, conserved | -0,4403 | 0,0003947 |
| TcCLB.510421.330 | hypothetical protein, conserved (fragment) | -0,4406 | 1,595E-07 |
| TcCLB.507915.5 | glutaminyl-tRNA synthetase, putative | -0,4406 | 0,1851 |
| TcCLB.508479.140 | hypothetical protein, conserved | -0,4407 | 0,001156 |
| TcCLB.509167.170 | hypothetical protein, conserved | -0,4409 | 0,001308 |
| TcCLB.506905.40 | ABC transporter, putative | -0,441 | 0,002402 |
| TcCLB.508723.40 | hypothetical protein, conserved | -0,4414 | 6,029E-07 |
| TcCLB.510303.170 | hypothetical protein, conserved | -0,4416 | 2,676E-09 |
| TcCLB.506559.530 | protein phosphatase 2C, putative | -0,4416 | 0,00006421 |
| TcCLB.506839.30 | hypothetical protein, conserved | -0,4418 | 0,0008481 |
| TcCLB.508307.40 | hypothetical protein, conserved | -0,4418 | 0,001597 |
| TcCLB.510821.20 | hypothetical protein, conserved | -0,4419 | 5,841E-08 |
| TcCLB.508437.5 | mucin-like glycoprotein (pseudogene), putative | -0,4421 | 0,21 |
| TcCLB.511431.20 | hypothetical protein, conserved | -0,4422 | 3,462E-10 |
| TcCLB.506955.198 | hypothetical protein | -0,4424 | 0,03448 |
| TcCLB.511801.40 | hypothetical protein, conserved | -0,4425 | 0,000563 |
| TcCLB.510819.110 | ATP-dependent RNA helicase, putative | -0,4426 | 2,091E-07 |
| TcCLB.507729.20 | hypothetical protein, conserved (pseudogene) | -0,4426 | 0,02727 |

| | | | |
|---|---|---|---|
| TcCLB.508613.50 | hypothetical protein, conserved | -0,4429 | 3,072E-08 |
| TcCLB.508153.600 | actin-like protein, putative | -0,4429 | 0,0003939 |
| TcCLB.504133.45 | hypothetical protein, conserved | -0,4429 | 0,03845 |
| TcCLB.507895.69 | hypothetical protein, conserved | -0,4432 | 0,02572 |
| TcCLB.511029.10 | enolase, putative | -0,4437 | 0,00004749 |
| TcCLB.505999.40 | DNA repair protein BRCA2, putative | -0,4438 | 0,0003944 |
| TcCLB.508153.870 | hypothetical protein, conserved | -0,4445 | 2,274E-06 |
| TcCLB.506503.90 | kinesin, putative | -0,445 | 0,007704 |
| TcCLB.508533.30 | metalloprotease, putative | -0,4451 | 9,43E-08 |
| TcCLB.504013.70 | metallo-peptidase, Clan MC, Family M14 | -0,4455 | 0,00001494 |
| TcCLB.506297.20 | hypothetical protein | -0,4456 | 6,506E-08 |
| TcCLB.508707.210 | DNA replication licensing factor MCM7, putative | -0,4456 | 0,00001133 |
| TcCLB.510597.9 | tousled-like kinase II, putative | -0,4457 | 0,0007156 |
| TcCLB.508707.180 | ubiquitin hydrolase, putative | -0,4458 | 2,374E-07 |
| TcCLB.510021.150 | hypothetical protein, conserved (pseudogene) | -0,4459 | 0,03215 |
| TcCLB.506543.100 | hypothetical protein, conserved | -0,4462 | 0,0009976 |
| TcCLB.504155.250 | trans-sialidase (pseudogene), putative | -0,4462 | 0,2049 |
| TcCLB.509399.110 | hypothetical protein, conserved | -0,4463 | 9,831E-06 |
| TcCLB.511635.50 | hypothetical protein, conserved | -0,4467 | 0,001895 |
| TcCLB.508741.30 | hypothetical protein, conserved | -0,4468 | 0,00006169 |
| TcCLB.506223.100 | hypothetical protein, conserved | -0,4471 | 0,00007082 |
| TcCLB.506511.80 | hypothetical protein, conserved | -0,4472 | 2,953E-06 |
| TcCLB.507801.80 | hypothetical protein, conserved | -0,4473 | 0,003634 |
| TcCLB.507089.230 | hypothetical protein, conserved | -0,4474 | 1,433E-06 |
| TcCLB.510123.40 | hypothetical protein, conserved | -0,4475 | 0,01637 |
| TcCLB.505999.160 | leucine-rich repeat-containing protein | -0,4476 | 0,0001029 |
| TcCLB.511413.11 | hypothetical protein, conserved (pseudogene) | -0,4476 | 0,0004473 |
| TcCLB.509599.40 | hypothetical protein, conserved | -0,4477 | 0,001052 |
| TcCLB.506941.20 | tRNA modification enzyme, putative | -0,4478 | 0,001 |
| TcCLB.511291.80 | hypothetical protein, conserved | -0,448 | 0,008302 |
| TcCLB.506789.200 | ubiquitin-conjugating enzyme E2, putative | -0,4481 | 1,583E-06 |
| TcCLB.506943.20 | hypothetical protein, conserved | -0,4482 | 0,00007513 |
| TcCLB.503773.20 | hypothetical protein, conserved | -0,4484 | 0,01398 |
| TcCLB.510939.20 | hypothetical protein, conserved | -0,4484 | 0,01441 |
| TcCLB.505365.50 | beta galactofuranosyl glycosyltransferase (pseudogene), p | -0,4484 | 0,01768 |
| TcCLB.504121.20 | hypothetical protein, conserved | -0,4486 | 0,0002464 |
| TcCLB.506469.60 | hypothetical protein | -0,4491 | 0,01711 |
| TcCLB.506163.20 | surface protease GP63, putative | -0,4491 | 0,025 |
| TcCLB.510421.50 | hypothetical protein, conserved | -0,4493 | 0,00004382 |
| TcCLB.509267.20 | hypothetical protein | -0,4494 | 0,0003355 |
| TcCLB.510743.130 | denn domain-containing protein (pseudogene) | -0,4494 | 0,007214 |
| TcCLB.510053.40 | hypothetical protein, conserved | -0,4494 | 0,01141 |
| TcCLB.506477.40 | Tubulin/FtsZ family, putative | -0,4497 | 0,00000188 |
| TcCLB.508899.20 | hypothetical protein, conserved | -0,4498 | 1,675E-06 |
| TcCLB.504769.190 | trans-sialidase (pseudogene), putative | -0,4498 | 0,01912 |

| | | | |
|---|---|---|---|
| TcCLB.507381.20 | trans-sialidase (pseudogene), putative | -0,45 | 0,002442 |
| TcCLB.504059.20 | endomembrane protein, putative | -0,4502 | 3,899E-07 |
| TcCLB.508177.30 | hypothetical protein, conserved | -0,4502 | 1,917E-06 |
| TcCLB.508121.120 | mucin TcMUCII, putative | -0,4503 | 0,1361 |
| TcCLB.508889.4 | 5-oxoprolinase, putative | -0,4506 | 0,04518 |
| TcCLB.506473.10 | Pre-mRNA-splicing factor 38, putative | -0,4507 | 0,002974 |
| TcCLB.509991.29 | cyclophilin 15, putative | -0,4508 | 0,00001956 |
| TcCLB.509539.20 | receptor-type adenylate cyclase, putative | -0,4508 | 0,000365 |
| TcCLB.511063.30 | trans-sialidase (pseudogene), putative | -0,4508 | 0,05105 |
| TcCLB.510285.140 | hypothetical protein, conserved (fragment) | -0,451 | 0,1945 |
| TcCLB.511751.40 | DNA-J protein, putative | -0,4511 | 0,001291 |
| TcCLB.503965.10 | hypothetical protein, conserved | -0,4512 | 0,0005268 |
| TcCLB.510693.60 | retrotransposon hot spot protein (RHS, pseudogene), puta | -0,4513 | 0,1756 |
| TcCLB.511511.120 | hypothetical protein, conserved | -0,4521 | 0,00003017 |
| TcCLB.503583.10 | metallo-peptidase, Clan MC, Family M14, putative | -0,4521 | 0,00009302 |
| TcCLB.506399.40 | hypothetical protein, conserved | -0,4521 | 0,02022 |
| TcCLB.510323.80 | hypothetical protein, conserved | -0,4523 | 1,618E-06 |
| TcCLB.504113.10 | protein kinase, putative | -0,4523 | 0,000388 |
| TcCLB.504839.60 | hypothetical protein, conserved | -0,4524 | 0,00001398 |
| TcCLB.508159.50 | mucin-associated surface protein (MASP, pseudogene), pu | -0,4524 | 0,03584 |
| TcCLB.509317.29 | hypothetical protein | -0,4526 | 0,02044 |
| TcCLB.506947.50 | hypothetical protein, conserved | -0,4527 | 0,0001632 |
| TcCLB.503917.50 | hypothetical protein, conserved | -0,4528 | 0,0003956 |
| TcCLB.510329.220 | hypothetical protein, conserved | -0,4533 | 6,649E-06 |
| TcCLB.509073.40 | outer dynein arm docking complex protein (DC2) | -0,4533 | 0,00001412 |
| TcCLB.509063.4 | hypothetical protein, conserved | -0,4533 | 0,008872 |
| TcCLB.511491.170 | quiescin sulfhydryl oxidase, putative (QSOX) | -0,4534 | 2,615E-06 |
| TcCLB.462761.4 | hypothetical protein, conserved | -0,4534 | 0,005511 |
| TcCLB.506211.120 | hypothetical protein, conserved | -0,4538 | 0,00003043 |
| TcCLB.511731.30 | serine/threonine protein phosphatase, putative | -0,4539 | 0,00001747 |
| TcCLB.505931.10 | surface protease GP63, putative | -0,4541 | 0,0004335 |
| TcCLB.506717.130 | trans-sialidase (pseudogene), putative | -0,4545 | 0,00008641 |
| TcCLB.503893.50 | hypothetical protein, conserved | -0,4547 | 0,00011 |
| TcCLB.510749.74 | hypothetical protein, conserved | -0,4548 | 0,00002625 |
| TcCLB.509213.60 | hypothetical protein, conserved | -0,4548 | 0,004651 |
| TcCLB.404001.4 | metalloprotease, putative (fragment) | -0,4549 | 9,708E-08 |
| TcCLB.511367.260 | Peptidase C19, ubiquitin carboxyl-terminal hydrolase, puta | -0,4549 | 1,901E-07 |
| TcCLB.510989.10 | fatty acid elongase, putative | -0,4549 | 0,0003305 |
| TcCLB.506435.230 | hypothetical protein, conserved (pseudogene) | -0,455 | 0,0006165 |
| TcCLB.511067.4 | phosphatidylinositol 3-kinase, putative | -0,4551 | 0,0009643 |
| TcCLB.509499.4 | hypothetical protein, conserved | -0,4552 | 0,001866 |
| TcCLB.509443.39 | hypothetical protein, conserved | -0,4554 | 0,00181 |
| TcCLB.510765.50 | hypothetical protein, conserved | -0,4555 | 0,0004668 |
| TcCLB.508737.50 | hypothetical protein, conserved | -0,4555 | 0,003578 |
| TcCLB.511137.10 | hypothetical protein, conserved | -0,4556 | 0,00204 |

| | | | |
|---|---|---|---|
| TcCLB.507515.70 | hypothetical protein, conserved | -0,4557 | 2,633E-07 |
| TcCLB.509965.319 | hypothetical protein, conserved | -0,4557 | 3,942E-06 |
| TcCLB.506563.60 | hypothetical protein, conserved | -0,4558 | 0,0002763 |
| TcCLB.506455.10 | retrotransposon hot spot protein (RHS, pseudogene), puta | -0,4558 | 0,01352 |
| TcCLB.511709.19 | phosphatidylinositol 3-kinase catalytic subunit, putative | -0,4559 | 0,002703 |
| TcCLB.506363.60 | f-actin capping protein beta subunit, putative | -0,4561 | 0,0004281 |
| TcCLB.509127.59 | hypothetical protein, conserved | -0,4561 | 0,04596 |
| TcCLB.510731.40 | katanin, putative | -0,4562 | 0,0003135 |
| TcCLB.510187.270 | transmembrane emp24 domain-containing protein | -0,4564 | 0,0002843 |
| TcCLB.507395.20 | hypothetical protein, conserved | -0,4565 | 0,001281 |
| TcCLB.510689.50 | cell differentiation protein, putative | -0,4566 | 0,00002313 |
| TcCLB.506445.110 | nucleobase transporter, putative | -0,4567 | 3,512E-08 |
| TcCLB.511273.50 | hypothetical protein, conserved | -0,4567 | 0,00009471 |
| TcCLB.508789.130 | surface protease GP63 (pseudogene), putative | -0,4567 | 0,1374 |
| TcCLB.507257.110 | hypothetical protein, conserved | -0,4568 | 0,0007679 |
| TcCLB.507031.50 | hypothetical protein, conserved | -0,4568 | 0,009206 |
| TcCLB.509163.105 | ATP-dependent DEAD/H RNA helicase (pseudogene), puta | -0,4569 | 0,2658 |
| TcCLB.506529.290 | hypothetical protein, conserved | -0,457 | 0,0001472 |
| TcCLB.508209.20 | hypothetical protein, conserved | -0,457 | 0,002557 |
| TcCLB.506559.440 | hypothetical protein, conserved | -0,4571 | 5,363E-10 |
| TcCLB.509849.40 | mannosyl-oligosaccharide 1,2-alpha-mannosidase IB, puta | -0,4571 | 0,004863 |
| TcCLB.508153.110 | hypothetical protein | -0,4576 | 0,002587 |
| TcCLB.508277.200 | hypothetical protein, conserved | -0,4577 | 0,0001402 |
| TcCLB.506541.10 | hypothetical protein, conserved (pseudogene) | -0,4578 | 0,1327 |
| TcCLB.510285.80 | hypothetical protein, conserved | -0,4582 | 0,002179 |
| TcCLB.508207.279 | hypothetical protein, conserved | -0,4584 | 0,0008738 |
| TcCLB.506135.20 | hypothetical protein, conserved | -0,4585 | 4,379E-08 |
| TcCLB.506985.60 | inositol polyphosphate kinase-like protein, putative | -0,4586 | 0,0001596 |
| TcCLB.508999.30 | Pre-mRNA-splicing factor 38, putative | -0,4587 | 0,01289 |
| TcCLB.511841.10 | trans-sialidase (pseudogene), putative | -0,4589 | 0,1988 |
| TcCLB.510667.10 | hypothetical protein, conserved | -0,459 | 0,0007848 |
| TcCLB.508825.10 | surface protease GP63, putative | -0,4593 | 0,0009758 |
| TcCLB.507731.60 | trans-sialidase, putative | -0,4594 | 0,1188 |
| TcCLB.511633.79 | microtubule-associated protein, putative | -0,4595 | 0,0006379 |
| TcCLB.510723.10 | hypothetical protein, conserved | -0,4595 | 0,02084 |
| TcCLB.505009.4 | hypothetical protein, conserved | -0,4596 | 0,0524 |
| TcCLB.511729.9 | calmodulin, putative | -0,4597 | 0,01211 |
| TcCLB.509205.50 | hypothetical protein, conserved | -0,4599 | 0,00003869 |
| TcCLB.507949.20 | RWD domain-containing protein | -0,46 | 0,0002666 |
| TcCLB.508153.720 | hypothetical protein, conserved | -0,4602 | 0,006082 |
| TcCLB.503909.40 | hypothetical protein | -0,4604 | 0,009391 |
| TcCLB.506885.370 | hypothetical protein, conserved | -0,4608 | 0,003122 |
| TcCLB.511753.40 | hypothetical protein, conserved | -0,4609 | 0,007813 |
| TcCLB.510503.50 | trans-sialidase, Group V, putative | -0,4612 | 0,01182 |
| TcCLB.506205.40 | protein kinase, putative | -0,4613 | 0,00003837 |

| | | | |
|---|---|---|---|
| TcCLB.503477.20 | transaldolase, putative | -0,4615 | 0,00002157 |
| TcCLB.505939.40 | hypothetical protein, conserved | -0,4615 | 0,0001898 |
| TcCLB.508323.190 | serine/threonine protein phosphatase, putative | -0,4615 | 0,0004551 |
| TcCLB.439307.19 | hypothetical protein, conserved | -0,4619 | 0,09386 |
| TcCLB.506247.170 | ATPase, putative | -0,462 | 5,949E-07 |
| TcCLB.506351.60 | hypothetical protein, conserved | -0,4623 | 0,00004945 |
| TcCLB.510879.100 | hypothetical protein | -0,4623 | 0,0001035 |
| TcCLB.506799.120 | hypothetical protein | -0,4623 | 0,02635 |
| TcCLB.508873.439 | mucin TcMUCII (pseudogene), putative | -0,4623 | 0,2445 |
| TcCLB.510533.150 | hypothetical protein | -0,4624 | 5,23E-07 |
| TcCLB.503453.80 | hypothetical protein, conserved | -0,4624 | 0,0001195 |
| TcCLB.427091.10 | solanesyl-diphosphate synthase, putative | -0,4624 | 0,00055 |
| TcCLB.510693.314 | mucin TcMUCII (pseudogene), putative | -0,4625 | 0,01673 |
| TcCLB.508367.50 | hypothetical protein, conserved (pseudogene) | -0,4625 | 0,03902 |
| TcCLB.506609.15 | hypothetical protein (pseudogene) | -0,4626 | 0,1037 |
| TcCLB.506749.10 | flagellar calcium-binding-like protein, putative | -0,4633 | 3,474E-08 |
| TcCLB.511767.10 | trans-sialidase, putative | -0,4633 | 0,07588 |
| TcCLB.508859.20 | hypothetical protein, conserved | -0,4636 | 0,0001491 |
| TcCLB.507677.120 | hypothetical protein, conserved | -0,4637 | 1,017E-06 |
| TcCLB.507011.100 | hypothetical protein, conserved | -0,4637 | 0,00002422 |
| TcCLB.511127.369 | hypothetical protein, conserved | -0,4638 | 0,00003362 |
| TcCLB.508525.10 | hypothetical protein, conserved | -0,4638 | 0,0005909 |
| TcCLB.509203.60 | histone acetyltransferase, putative | -0,4639 | 0,0002534 |
| TcCLB.506797.50 | hypothetical protein, conserved | -0,464 | 1,235E-06 |
| TcCLB.504035.110 | hypothetical protein | -0,464 | 0,0001262 |
| TcCLB.509905.10 | trans-sialidase (pseudogene), putative | -0,4641 | 0,00001477 |
| TcCLB.506297.240 | pretranslocation protein, alpha subunit, putative | -0,4642 | 7,146E-09 |
| TcCLB.503787.65 | hypothetical protein (pseudogene) | -0,4642 | 0,1772 |
| TcCLB.506859.120 | hypothetical protein | -0,4645 | 0,00002859 |
| TcCLB.505949.110 | hypothetical protein, conserved (pseudogene) | -0,4645 | 0,2416 |
| TcCLB.509819.49 | Myotubularin-related protein, putative (fragment) | -0,4646 | 0,000647 |
| TcCLB.503489.10 | metallo-peptidase, Clan MC, Family M14, putative | -0,4647 | 3,486E-07 |
| TcCLB.506831.79 | pre-mRNA splicing factor ATP-dependent RNA helicase, p | -0,4647 | 0,02622 |
| TcCLB.506573.34 | hypothetical protein, conserved | -0,4647 | 0,05908 |
| TcCLB.510283.50 | hypothetical protein, conserved | -0,4652 | 0,003221 |
| TcCLB.507621.5 | trans-sialidase (pseudogene), putative | -0,4654 | 0,08539 |
| TcCLB.508805.90 | hypothetical protein, conserved | -0,4657 | 0,06538 |
| TcCLB.510497.190 | hypothetical protein | -0,4657 | 0,2414 |
| TcCLB.511589.140 | phosphoenolpyruvate mutase, putative | -0,4658 | 3,995E-06 |
| TcCLB.508351.30 | hypothetical protein, conserved | -0,4658 | 0,00000653 |
| TcCLB.509215.13 | hypothetical protein, conserved | -0,4661 | 0,03378 |
| TcCLB.506825.70 | serine-threonine dehydratase, putative | -0,4662 | 0,009606 |
| TcCLB.510191.31 | trans-sialidase (pseudogene), putative | -0,4663 | 0,2048 |
| TcCLB.507389.10 | hypothetical protein, conserved | -0,4665 | 0,0002751 |
| TcCLB.506459.235 | hypothetical protein, conserved | -0,4665 | 0,01053 |

| | | | |
|---|---|---|---|
| TcCLB.508739.30 | hypothetical protein | -0,4666 | 0,000619 |
| TcCLB.509065.79 | hypothetical protein, conserved | -0,4667 | 0,00001207 |
| TcCLB.504249.15 | surface protease GP63 (pseudogene), putative | -0,4668 | 0,06658 |
| TcCLB.506623.30 | hypothetical protein | -0,4669 | 0,101 |
| TcCLB.511255.110 | trans-sialidase (pseudogene), putative | -0,467 | 0,1183 |
| TcCLB.503765.10 | trans-sialidase (pseudogene), putative | -0,4673 | 0,005538 |
| TcCLB.506513.180 | hypothetical protein, conserved | -0,4676 | 0,00001479 |
| TcCLB.508187.10 | vacuolar protein sorting-associated protein, putative (frag | -0,4678 | 0,0000276 |
| TcCLB.508707.130 | hypothetical protein, conserved | -0,4679 | 0,00002079 |
| TcCLB.505037.157 | trans-sialidase (pseudogene), putative | -0,4679 | 0,0004803 |
| TcCLB.504881.14 | hypothetical protein, conserved | -0,468 | 0,015 |
| TcCLB.506111.10 | retrotransposon hot spot protein (RHS, pseudogene), puta | -0,4681 | 0,0001259 |
| TcCLB.506151.10 | hypothetical protein, conserved | -0,4683 | 0,00001436 |
| TcCLB.508165.380 | hypothetical protein | -0,4683 | 0,2336 |
| TcCLB.506257.70 | hypothetical protein, conserved | -0,4684 | 0,00004577 |
| TcCLB.506767.170 | trans-sialidase (pseudogene), putative | -0,4685 | 0,03464 |
| TcCLB.510973.50 | hypothetical protein, conserved | -0,4688 | 0,00007119 |
| TcCLB.507641.14 | neurobeachin/beige protein, putative | -0,469 | 2,688E-07 |
| TcCLB.503809.90 | glycerol uptake protein, putative | -0,469 | 0,00001777 |
| TcCLB.503897.100 | hypothetical protein, conserved | -0,4693 | 1,506E-06 |
| TcCLB.503959.50 | hypothetical protein, conserved | -0,4693 | 0,0001268 |
| TcCLB.506817.60 | hypothetical protein, conserved | -0,4693 | 0,0001695 |
| TcCLB.508939.10 | trans-sialidase (pseudogene), putative | -0,4693 | 0,005131 |
| TcCLB.504123.20 | hypothetical protein, conserved | -0,4694 | 0,00001937 |
| TcCLB.430605.20 | PTP1-interacting protein, 39 kDa, putative | -0,4697 | 3,097E-07 |
| TcCLB.506113.83 | ATP-dependent DEAD/H RNA helicase (pseudogene), puta | -0,4698 | 0,01393 |
| TcCLB.506203.40 | hypothetical protein, conserved | -0,4702 | 2,397E-08 |
| TcCLB.510643.180 | trans-sialidase (pseudogene), putative | -0,4702 | 0,0003879 |
| TcCLB.504625.90 | hypothetical protein, conserved | -0,4707 | 0,0002054 |
| TcCLB.509747.100 | hypothetical protein, conserved | -0,4708 | 1,655E-08 |
| TcCLB.508397.50 | hypothetical protein, conserved | -0,4713 | 0,0000211 |
| TcCLB.506327.80 | serine/threonine protein kinase, putative | -0,4713 | 0,0002736 |
| TcCLB.507629.39 | hypothetical protein, conserved (fragment) | -0,4714 | 1,1E-09 |
| TcCLB.509315.10 | hypothetical protein, conserved (pseudogene) | -0,4714 | 0,0834 |
| TcCLB.508707.90 | proteasome regulatory non-ATPase subunit 7, putative | -0,4715 | 7,094E-08 |
| TcCLB.503913.29 | hypothetical protein, conserved | -0,4715 | 0,01226 |
| TcCLB.508827.44 | hypothetical protein, conserved | -0,4717 | 0,0001099 |
| TcCLB.503829.50 | hypothetical protein, conserved | -0,4717 | 0,0003078 |
| TcCLB.509991.70 | hypothetical protein, conserved (pseudogene) | -0,472 | 0,00005464 |
| TcCLB.506425.160 | hypothetical protein, conserved | -0,472 | 0,0001249 |
| TcCLB.437575.9 | GDP-mannose 4,6 dehydratase, putative | -0,472 | 0,06862 |
| TcCLB.509167.70 | hypothetical protein, conserved | -0,4721 | 0,0001794 |
| TcCLB.504105.190 | hypothetical protein, conserved | -0,4723 | 0,00002852 |
| TcCLB.504741.170 | dual specificity protein phosphatase, putative | -0,4723 | 0,001747 |
| TcCLB.505943.20 | oxidoreductase, putative | -0,4725 | 0,0008162 |

| | | | |
|---|---|---|---|
| TcCLB.508199.30 | hypothetical protein, conserved | -0,4726 | 0,00009305 |
| TcCLB.510515.80 | hypothetical protein, conserved | -0,4727 | 3,862E-07 |
| TcCLB.503775.30 | syntaxin binding protein (pseudogene), putative | -0,4727 | 0,002428 |
| TcCLB.511181.30 | Iron-sulfur assembly protein 2, putative (Isa2) | -0,4728 | 0,008607 |
| TcCLB.510141.10 | thymine-7-hydroxylase, putative | -0,4728 | 0,05334 |
| TcCLB.504213.60 | hypothetical protein, conserved | -0,4729 | 1,509E-06 |
| TcCLB.508089.30 | hypothetical protein, conserved | -0,4729 | 3,693E-06 |
| TcCLB.511165.60 | DNA repair protein, putative | -0,473 | 0,0001565 |
| TcCLB.511421.217 | hypothetical protein, conserved | -0,4731 | 0,0002929 |
| TcCLB.506425.30 | hypothetical protein, conserved | -0,4734 | 5,546E-07 |
| TcCLB.511449.70 | trans-sialidase, Group V, putative | -0,4735 | 0,07672 |
| TcCLB.504117.20 | sedoheptulose-1,7-bisphosphatase, putative | -0,4737 | 0,001463 |
| TcCLB.506955.90 | trans-sialidase (pseudogene), putative | -0,4739 | 0,07351 |
| TcCLB.510461.20 | trans-sialidase (pseudogene), putative | -0,4739 | 0,2292 |
| TcCLB.511217.30 | diacylglycerol kinase, putative | -0,4741 | 5,142E-06 |
| TcCLB.510759.200 | hypothetical protein, conserved | -0,4741 | 0,00001352 |
| TcCLB.511715.20 | protein kinase, putative | -0,4746 | 0,00001373 |
| TcCLB.511559.70 | phospholipid transporting ATPase-like protein (pseudogen | -0,4747 | 6,253E-06 |
| TcCLB.507445.20 | trans-sialidase (pseudogene), putative | -0,4748 | 0,00006646 |
| TcCLB.506735.60 | hypothetical protein, conserved | -0,4749 | 0,004041 |
| TcCLB.508837.154 | hypothetical protein, conserved | -0,475 | 0,000945 |
| TcCLB.506869.20 | hypothetical protein, conserved | -0,4752 | 0,000121 |
| TcCLB.511277.483 | hypothetical protein, conserved | -0,4752 | 0,0007438 |
| TcCLB.505943.10 | hypothetical protein, conserved | -0,4756 | 0,004142 |
| TcCLB.506205.49 | hypothetical protein, conserved | -0,4756 | 0,1465 |
| TcCLB.510307.245 | elongation factor 1-gamma (EF-1-gamma, pseudogene), | -0,4756 | 0,1594 |
| TcCLB.509789.10 | hypothetical protein, conserved | -0,4757 | 3,605E-07 |
| TcCLB.511127.190 | hypothetical protein, conserved | -0,4757 | 8,406E-06 |
| TcCLB.508787.10 | hypothetical protein, conserved | -0,4757 | 0,0001526 |
| TcCLB.511511.70 | hypothetical protein, conserved | -0,4758 | 1,102E-08 |
| TcCLB.511885.35 | trans-sialidase (pseudogene), putative | -0,4759 | 0,01644 |
| TcCLB.510825.30 | phospholipase B-like, putative (p67) | -0,476 | 3,274E-10 |
| TcCLB.506955.115 | surface protease GP63 (pseudogene), putative | -0,476 | 0,1724 |
| TcCLB.507257.40 | hypothetical protein, conserved | -0,4762 | 0,009354 |
| TcCLB.410589.19 | hypothetical protein, conserved | -0,4765 | 0,004304 |
| TcCLB.503797.7 | hypothetical protein, conserved (pseudogene) | -0,4768 | 0,0000228 |
| TcCLB.506213.120 | ATP-dependent RNA helicase, putative | -0,4768 | 0,0000759 |
| TcCLB.510661.260 | hypothetical protein, conserved | -0,4769 | 6,424E-08 |
| TcCLB.506471.50 | trans-sialidase (pseudogene), putative | -0,477 | 0,0000127 |
| TcCLB.508613.100 | surface protease GP63 (pseudogene), putative | -0,477 | 0,03258 |
| TcCLB.506625.180 | hypothetical protein, conserved | -0,4772 | 2,772E-06 |
| TcCLB.511725.134 | mitochondrial carrier protein, putative | -0,4774 | 3,216E-06 |
| TcCLB.508429.10 | hypothetical protein | -0,4774 | 0,09526 |
| TcCLB.508405.10 | cell division cycle protein, putative | -0,4776 | 6,258E-06 |
| TcCLB.507825.10 | hypothetical protein, conserved | -0,4776 | 0,001158 |

| | | | |
|---|---|---|---|
| TcCLB.509875.25 | trans-sialidase (pseudogene), putative | -0,4776 | 0,1301 |
| TcCLB.509939.19 | hypothetical protein, conserved | -0,4781 | 4,093E-08 |
| TcCLB.506825.100 | hypothetical protein, conserved | -0,4781 | 0,00008769 |
| TcCLB.511127.40 | hypothetical protein | -0,4781 | 0,0003927 |
| TcCLB.508153.140 | protein kinase, putative | -0,4781 | 0,00334 |
| TcCLB.506591.81 | hypothetical protein, conserved (pseudogene) | -0,4783 | 0,0002514 |
| TcCLB.510609.65 | U6 snRNA-associated Sm-like protein LSm3p | -0,4783 | 0,04317 |
| TcCLB.510161.10 | hypothetical protein, conserved | -0,4784 | 0,0001142 |
| TcCLB.506409.10 | hypothetical protein | -0,4785 | 0,1592 |
| TcCLB.509161.90 | hypothetical protein, conserved | -0,4787 | 0,0006065 |
| TcCLB.510863.9 | hypothetical protein | -0,4789 | 0,005858 |
| TcCLB.508873.150 | mucin TcMUCII, putative | -0,4794 | 0,2632 |
| TcCLB.504147.150 | ATP-dependent DEAD/H RNA helicase, putative | -0,4796 | 3,085E-10 |
| TcCLB.510431.320 | hypothetical protein, conserved | -0,4799 | 1,556E-06 |
| TcCLB.507733.40 | hypothetical protein, conserved (pseudogene) | -0,4803 | 0,09695 |
| TcCLB.504207.10 | hypothetical protein, conserved | -0,4804 | 0,00008237 |
| TcCLB.506739.160 | hypothetical protein, conserved | -0,4809 | 4,693E-06 |
| TcCLB.509537.50 | Flagellar Member 1 (FLAM1) | -0,4813 | 1,856E-07 |
| TcCLB.509179.220 | hypothetical protein, conserved | -0,4817 | 0,0003459 |
| TcCLB.505555.40 | hypothetical protein, conserved | -0,4818 | 1,291E-06 |
| TcCLB.506321.40 | mucin TcMUCII, putative | -0,4822 | 0,0001042 |
| TcCLB.510091.30 | hypothetical protein, conserved | -0,4825 | 0,00002292 |
| TcCLB.507103.10 | DnaJ chaperone protein, putative | -0,4826 | 0,00003919 |
| TcCLB.510887.14 | chaperone DNAJ protein, putative | -0,4827 | 5,882E-06 |
| TcCLB.510765.30 | COP-coated vesicle membrane protein gp25L precursor, p | -0,4827 | 0,00001442 |
| TcCLB.506223.20 | hypothetical protein, conserved | -0,4828 | 0,0001387 |
| TcCLB.509963.10 | hypothetical protein, conserved (fragment) | -0,4829 | 0,001827 |
| TcCLB.508921.30 | Kinesin-13 4, putative (KIN13-4) | -0,4831 | 0,002245 |
| TcCLB.505075.10 | hypothetical protein, conserved | -0,4833 | 0,0001196 |
| TcCLB.506499.215 | elongation factor 1-gamma (EF-1-gamma, pseudogene), | -0,4834 | 0,1808 |
| TcCLB.506559.180 | hypothetical protein, conserved | -0,4838 | 2,317E-06 |
| TcCLB.508175.180 | hypothetical protein, conserved | -0,4839 | 1,06E-07 |
| TcCLB.511715.40 | hypothetical protein, conserved | -0,4839 | 0,005131 |
| TcCLB.511181.10 | hypothetical protein, conserved | -0,4844 | 3,226E-06 |
| TcCLB.503739.20 | trans-sialidase (pseudogene), putative | -0,4846 | 0,0005596 |
| TcCLB.506905.59 | SNF2 DNA repair protein, putative (fragment) | -0,4847 | 3,644E-07 |
| TcCLB.508899.110 | actin-related protein 2, putative | -0,4848 | 0,00002131 |
| TcCLB.506883.50 | Present in the outer mitochondrial membrane proteome 5 | -0,4849 | 1,911E-06 |
| TcCLB.506579.20 | hypothetical protein, conserved | -0,4851 | 0,00003599 |
| TcCLB.507675.20 | hypothetical protein, conserved | -0,4852 | 2,055E-07 |
| TcCLB.510575.40 | hypothetical protein, conserved | -0,4853 | 0,00007185 |
| TcCLB.508181.90 | hypothetical protein, conserved (pseudogene) | -0,4856 | 7,181E-07 |
| TcCLB.504021.120 | hypothetical protein, conserved | -0,4856 | 3,641E-06 |
| TcCLB.508177.50 | hypothetical protein | -0,4858 | 0,0004151 |
| TcCLB.508331.50 | hypothetical protein, conserved | -0,4859 | 0,00001777 |

| | | | |
|---|---|---|---|
| TcCLB.507711.80 | hypothetical protein, conserved | -0,4859 | 0,00007513 |
| TcCLB.511851.61 | mucin-associated surface protein (MASP, pseudogene), pu | -0,4862 | 0,1464 |
| TcCLB.506267.48 | hypothetical protein | -0,4863 | 0,01868 |
| TcCLB.510695.10 | surface protease GP63 (pseudogene), putative | -0,4864 | 0,008711 |
| TcCLB.509901.100 | hypothetical protein, conserved | -0,4866 | 3,36E-10 |
| TcCLB.506841.8 | hypothetical protein, conserved | -0,4866 | 2,965E-08 |
| TcCLB.511895.60 | hypothetical protein, conserved | -0,4866 | 0,0001423 |
| TcCLB.511287.64 | hypothetical protein, conserved | -0,4868 | 0,001218 |
| TcCLB.507951.90 | hypothetical protein, conserved | -0,4869 | 0,0000458 |
| TcCLB.503703.20 | hypothetical protein, conserved | -0,4873 | 0,0000194 |
| TcCLB.506865.40 | guanine nucleotide exchange factor subunit GCD7 | -0,4874 | 9,398E-07 |
| TcCLB.510029.30 | hypothetical protein, conserved | -0,4878 | 0,0000024 |
| TcCLB.510257.90 | autophagocytosis associated protein, putative | -0,488 | 0,0008528 |
| TcCLB.507887.20 | ubiquitin hydrolase, putative | -0,4881 | 5,979E-10 |
| TcCLB.510607.51 | retrotransposon hot spot protein (RHS, pseudogene), puta | -0,4883 | 0,1159 |
| TcCLB.506619.90 | ABC transporter family-like protein | -0,4887 | 0,0001098 |
| TcCLB.509005.70 | hypothetical protein, conserved | -0,4888 | 0,00002997 |
| TcCLB.507089.70 | RNA-binding protein, putative | -0,4893 | 0,0005033 |
| TcCLB.504147.110 | asparaginase, putative | -0,4893 | 0,002158 |
| TcCLB.511215.110 | hypothetical protein, conserved | -0,4895 | 1,005E-06 |
| TcCLB.508207.190 | hypothetical protein, conserved | -0,4897 | 5,541E-06 |
| TcCLB.504179.30 | hypothetical protein, conserved | -0,4898 | 0,0001333 |
| TcCLB.509769.30 | hypothetical protein, conserved | -0,4902 | 0,008626 |
| TcCLB.511625.120 | hypothetical protein | -0,4905 | 0,171 |
| TcCLB.503789.30 | hypothetical protein, conserved | -0,4907 | 0,001525 |
| TcCLB.509033.70 | hypothetical protein, conserved | -0,4908 | 0,0001517 |
| TcCLB.503999.80 | hypothetical protein, conserved | -0,4909 | 3,921E-10 |
| TcCLB.508411.80 | hypothetical protein, conserved | -0,4909 | 0,0000193 |
| TcCLB.506367.89 | hypothetical protein, conserved | -0,491 | 0,0007031 |
| TcCLB.503763.50 | trans-sialidase (pseudogene), putative | -0,4911 | 0,00008689 |
| TcCLB.511717.160 | hypothetical protein | -0,4911 | 0,0002023 |
| TcCLB.511577.50 | endo-beta-N-acetylglucosaminidase, putative | -0,4914 | 0,00002613 |
| TcCLB.503885.70 | hypothetical protein, conserved | -0,4914 | 0,0009107 |
| TcCLB.506297.40 | hypothetical protein, conserved | -0,4915 | 3,051E-13 |
| TcCLB.508479.174 | hypothetical protein, conserved | -0,4915 | 5,112E-06 |
| TcCLB.508771.40 | hypothetical protein, conserved | -0,4921 | 0,000818 |
| TcCLB.509445.20 | hypothetical protein, conserved | -0,4922 | 4,653E-08 |
| TcCLB.510901.170 | hypothetical protein, conserved | -0,4925 | 6,491E-08 |
| TcCLB.509399.60 | hypothetical protein, conserved | -0,4925 | 1,392E-07 |
| TcCLB.507951.130 | hypothetical protein | -0,4925 | 0,00003352 |
| TcCLB.506753.70 | hypothetical protein, conserved | -0,4925 | 0,0001067 |
| TcCLB.507895.31 | retrotransposon hot spot protein (RHS, pseudogene), puta | -0,4926 | 0,02699 |
| TcCLB.507809.50 | hypothetical protein, conserved | -0,4932 | 0,0008344 |
| TcCLB.503565.29 | hypothetical protein, conserved | -0,4933 | 6,057E-07 |
| TcCLB.504883.70 | hypothetical protein, conserved (pseudogene) | -0,4934 | 1,197E-12 |

| | | | |
|---|---|---|---|
| TcCLB.510853.30 | trans-sialidase (pseudogene), putative | -0,4934 | 0,001734 |
| TcCLB.509569.110 | carrier protein, putative | -0,4935 | 0,00001812 |
| TcCLB.511717.120 | hypothetical protein, conserved | -0,4937 | 0,00000629 |
| TcCLB.507357.20 | hypothetical protein | -0,4938 | 0,1055 |
| TcCLB.506867.10 | endonuclease G, putative | -0,494 | 0,0006626 |
| TcCLB.507163.60 | hypothetical protein | -0,4941 | 0,05815 |
| TcCLB.509229.70 | hypothetical protein, conserved | -0,4949 | 4,373E-06 |
| TcCLB.506459.299 | mucin-associated surface protein (MASP, pseudogene), pu | -0,4958 | 7,967E-08 |
| TcCLB.506055.14 | trans-sialidase (pseudogene), putative | -0,4959 | 0,01589 |
| TcCLB.510105.9 | ATP-dependent DEAD/H RNA helicase, putative | -0,4961 | 0,00001391 |
| TcCLB.509247.60 | hypothetical protein, conserved (pseudogene) | -0,4961 | 0,00005584 |
| TcCLB.505075.20 | hypothetical protein | -0,4962 | 0,002315 |
| TcCLB.433733.10 | trans-sialidase, putative | -0,4963 | 0,001448 |
| TcCLB.508515.100 | hypothetical protein, conserved | -0,4964 | 0,002861 |
| TcCLB.509795.70 | hypothetical protein, conserved | -0,4965 | 4,736E-07 |
| TcCLB.508539.169 | trans-sialidase (pseudogene), putative | -0,4966 | 0,0322 |
| TcCLB.503731.9 | hypothetical protein (fragment) | -0,4966 | 0,03439 |
| TcCLB.503897.150 | hypothetical protein, conserved | -0,4967 | 0,00001199 |
| TcCLB.503465.10 | hypothetical protein, conserved | -0,4968 | 9,754E-07 |
| TcCLB.508723.120 | hypothetical protein, conserved | -0,4968 | 0,0003185 |
| TcCLB.511755.100 | hypothetical protein, conserved | -0,4968 | 0,02436 |
| TcCLB.510657.210 | hypothetical protein | -0,4971 | 0,0009756 |
| TcCLB.504051.40 | hypothetical protein, conserved | -0,4973 | 4,831E-07 |
| TcCLB.510819.100 | mitochondrial carrier protein, putative | -0,4974 | 0,00002332 |
| TcCLB.507641.90 | hypothetical protein, conserved | -0,4974 | 0,0001099 |
| TcCLB.510915.21 | surface protease GP63 (pseudogene), putative | -0,4974 | 0,01321 |
| TcCLB.510187.210 | hypothetical protein, conserved | -0,4977 | 6,49E-11 |
| TcCLB.506375.20 | hypothetical protein, conserved (pseudogene) | -0,4979 | 7,802E-07 |
| TcCLB.511151.70 | hypothetical protein, conserved | -0,4981 | 2,079E-08 |
| TcCLB.509683.30 | hypothetical protein, conserved | -0,4981 | 3,325E-07 |
| TcCLB.509561.60 | hypothetical protein, conserved | -0,4981 | 0,0003829 |
| TcCLB.510889.160 | cell division control protein (pseudogene), putative | -0,4982 | 6,198E-06 |
| TcCLB.503541.5 | cAMP-specific phosphodiesterase, putative | -0,4982 | 0,08163 |
| TcCLB.503823.20 | hypothetical protein, conserved | -0,4983 | 0,0004443 |
| TcCLB.506135.50 | hypothetical protein, conserved | -0,4984 | 5,662E-06 |
| TcCLB.504033.53 | UDP-Gal or UDP-GlcNAc-dependent glycosyltransferase (p | -0,4984 | 0,247 |
| TcCLB.510429.10 | hypothetical protein, conserved | -0,4989 | 0,002073 |
| TcCLB.510519.120 | hypothetical protein, conserved | -0,4991 | 1,112E-06 |
| TcCLB.510533.90 | hypothetical protein, conserved | -0,4991 | 0,0000106 |
| TcCLB.506763.121 | elongation factor 1-gamma (EF-1-gamma, pseudogene), | -0,4991 | 0,2449 |
| TcCLB.510099.50 | hypothetical protein, conserved | -0,4994 | 0,00001246 |
| TcCLB.507485.70 | hypothetical protein, conserved | -0,4996 | 0,00009599 |
| TcCLB.503981.90 | SWI/SNF-related matrix-associated actin-dependent regu | -0,4998 | 0,04318 |
| TcCLB.508525.60 | hypothetical protein, conserved | -0,4999 | 6,56E-08 |
| TcCLB.507471.41 | trans-sialidase (pseudogene), putative | -0,4999 | 0,002165 |

| | | | |
|---|---|---|---|
| TcCLB.510283.100 | hypothetical protein, conserved | -0,4999 | 0,005609 |
| TcCLB.507809.90 | hypothetical protein, conserved | -0,5 | 1,157E-07 |
| TcCLB.510823.90 | hypothetical protein | -0,5008 | 9,084E-08 |
| TcCLB.503433.10 | hypothetical protein, conserved | -0,5008 | 0,0005534 |
| TcCLB.506941.120 | hypothetical protein, conserved | -0,5009 | 0,00001189 |
| TcCLB.511455.10 | surface protease GP63 (pseudogene), putative | -0,501 | 0,004946 |
| TcCLB.421621.11 | hypothetical protein, conserved (pseudogene) | -0,501 | 0,2732 |
| TcCLB.507001.100 | transporter, putative | -0,5013 | 0,001033 |
| TcCLB.510361.50 | mucin-associated surface protein (MASP, pseudogene), pu | -0,5013 | 0,02857 |
| TcCLB.457339.19 | hypothetical protein, conserved | -0,5018 | 0,001411 |
| TcCLB.511245.40 | hypothetical protein | -0,5021 | 0,00002991 |
| TcCLB.414517.9 | hypothetical protein, conserved | -0,5021 | 0,002271 |
| TcCLB.508461.560 | hypothetical protein, conserved | -0,5023 | 9,533E-06 |
| TcCLB.504081.23 | hypothetical protein, conserved | -0,5026 | 0,3296 |
| TcCLB.506777.30 | hypothetical protein, conserved | -0,5029 | 3,115E-09 |
| TcCLB.503913.20 | actin-related protein 3, putative | -0,5032 | 0,00000871 |
| TcCLB.506009.60 | hypothetical protein | -0,5033 | 0,0001021 |
| TcCLB.509897.190 | hypothetical protein, conserved (pseudogene) | -0,5036 | 8,157E-08 |
| TcCLB.508221.310 | trans-sialidase (pseudogene), putative | -0,5037 | 0,09158 |
| TcCLB.509399.80 | hypothetical protein, conserved | -0,504 | 2,237E-07 |
| TcCLB.506297.210 | hypothetical protein, conserved | -0,5041 | 1,052E-08 |
| TcCLB.506345.70 | retrotransposon hot spot protein (RHS, pseudogene), puta | -0,5043 | 0,0004185 |
| TcCLB.503493.20 | hypothetical protein, conserved | -0,5043 | 0,0006295 |
| TcCLB.510375.20 | trans-sialidase (pseudogene), putative | -0,5044 | 0,1733 |
| TcCLB.511381.50 | hypothetical protein, conserved | -0,5048 | 8,215E-07 |
| TcCLB.507031.40 | hypothetical protein, conserved | -0,5048 | 9,578E-07 |
| TcCLB.505025.144 | hypothetical protein, conserved | -0,5051 | 0,26 |
| TcCLB.511559.20 | hypothetical protein, conserved | -0,5053 | 8,392E-10 |
| TcCLB.507021.60 | p1/s1 nuclease, putative | -0,5054 | 3,404E-06 |
| TcCLB.507763.20 | hypothetical protein | -0,5054 | 0,0007124 |
| TcCLB.503671.20 | hypothetical protein, conserved | -0,5055 | 0,0001285 |
| TcCLB.511215.20 | hypothetical protein | -0,5056 | 1,282E-11 |
| TcCLB.503611.30 | hypothetical protein, conserved | -0,5056 | 1,788E-06 |
| TcCLB.503811.40 | PTS2 receptor, putative (PEX7) | -0,5056 | 0,0001258 |
| TcCLB.506679.110 | dynein heavy chain, putative | -0,5057 | 3,317E-07 |
| TcCLB.509637.27 | hypothetical protein, conserved | -0,5057 | 0,0001634 |
| TcCLB.508259.11 | hypothetical protein, conserved (pseudogene) | -0,5058 | 0,02446 |
| TcCLB.508479.10 | protein kinase, putative | -0,5058 | 0,04991 |
| TcCLB.510749.4 | ubiquitin-protein ligase, putative | -0,5059 | 0,2838 |
| TcCLB.511217.209 | hypothetical protein, conserved (fragment) | -0,506 | 0,0008153 |
| TcCLB.506783.20 | trans-sialidase (pseudogene), putative | -0,5061 | 0,0043 |
| TcCLB.507087.30 | cysteine peptidase, Clan CA, family C19, putative | -0,5062 | 1,004E-08 |
| TcCLB.404975.30 | trans-sialidase (pseudogene), putative | -0,5063 | 0,02143 |
| TcCLB.510727.9 | hypothetical protein, conserved (fragment) | -0,5065 | 0,001037 |
| TcCLB.509693.70 | hypoxanthine-guanine phosphoribosyltransferase, putative | -0,5066 | 0,0001274 |

| | | | |
|---|---|---|---|
| TcCLB.511287.60 | hypothetical protein, conserved | -0,5067 | 3,368E-11 |
| TcCLB.510435.40 | GPI transamidase component Tta1, putative (TTA1) | -0,5067 | 0,0001414 |
| TcCLB.510667.70 | iron-sulfur cluster assembly protein, putative | -0,5068 | 5,128E-06 |
| TcCLB.509767.215 | vesicle transport v-SNARE 11, putative | -0,5069 | 0,0004672 |
| TcCLB.509993.15 | mucin-associated surface protein (MASP, pseudogene), pu | -0,5069 | 0,00313 |
| TcCLB.508221.350 | hypothetical protein | -0,5069 | 0,03778 |
| TcCLB.506755.40 | hypothetical protein, conserved | -0,507 | 0,0002318 |
| TcCLB.510481.90 | hypothetical protein, conserved | -0,507 | 0,05851 |
| TcCLB.510557.30 | hypothetical protein | -0,5073 | 0,1879 |
| TcCLB.510405.9 | hypothetical protein, conserved (fragment) | -0,5074 | 2,829E-08 |
| TcCLB.511245.110 | hypothetical protein, conserved | -0,5077 | 4,349E-07 |
| TcCLB.511165.40 | acidocalcisomal pyrophosphatase, putative | -0,5078 | 4,782E-08 |
| TcCLB.510543.70 | trans-sialidase, Group II, putative | -0,5079 | 0,04022 |
| TcCLB.507047.90 | hypothetical protein | -0,508 | 0,00004052 |
| TcCLB.506729.20 | hypothetical protein, conserved | -0,5082 | 0,0004718 |
| TcCLB.506699.34 | ATP-dependent 6-phosphofructokinase, glycosomal (pseu | -0,5083 | 1,087E-06 |
| TcCLB.510625.210 | trans-sialidase (pseudogene), putative | -0,5083 | 0,1261 |
| TcCLB.509437.109 | trans-sialidase (pseudogene), putative | -0,5085 | 0,003759 |
| TcCLB.507049.199 | pumilio protein, putative | -0,5087 | 0,0318 |
| TcCLB.508009.10 | surface protease GP63 (pseudogene), putative | -0,5087 | 0,03273 |
| TcCLB.503913.4 | nucleolar RNA binding protein (pseudogene), putative | -0,5091 | 0,000135 |
| TcCLB.506573.91 | coatomer delta subunit, putative | -0,5094 | 0,07095 |
| TcCLB.509943.20 | cAMP specific phosphodiesterase, putative | -0,5098 | 0,000506 |
| TcCLB.508347.140 | hypothetical protein, conserved | -0,5099 | 2,365E-06 |
| TcCLB.509373.30 | mucin-associated surface protein (MASP, pseudogene), pu | -0,5099 | 0,04152 |
| TcCLB.511173.89 | mucin TcMUCII (pseudogene), putative | -0,51 | 0,009962 |
| TcCLB.503785.20 | hypothetical protein, conserved | -0,5101 | 0,0001178 |
| TcCLB.508723.80 | disgorgin, putative | -0,5102 | 0,00005292 |
| TcCLB.508163.170 | trans-sialidase (pseudogene), putative | -0,5104 | 0,1213 |
| TcCLB.511211.235 | hypothetical protein, conserved | -0,5105 | 0,0002331 |
| TcCLB.510713.101 | surface protease GP63 (pseudogene), putative | -0,5105 | 0,2204 |
| TcCLB.504081.295 | surface protease GP63 (pseudogene), putative | -0,5107 | 0,1608 |
| TcCLB.506579.119 | hypothetical protein, conserved | -0,511 | 0,004026 |
| TcCLB.506133.30 | surface protease GP63 (pseudogene), putative | -0,511 | 0,2331 |
| TcCLB.508523.21 | trans-sialidase (pseudogene), putative | -0,5112 | 0,001491 |
| TcCLB.510575.210 | hypothetical protein, conserved | -0,5113 | 1,752E-06 |
| TcCLB.510483.290 | hypothetical protein | -0,5113 | 0,1386 |
| TcCLB.504105.40 | hypothetical protein, conserved | -0,5115 | 6,297E-09 |
| TcCLB.506649.40 | rho-like GTP binding protein, putative | -0,5115 | 0,00004406 |
| TcCLB.511143.6 | protein kinase, putative (fragment) | -0,5116 | 0,01191 |
| TcCLB.506317.20 | hypothetical protein, conserved | -0,5118 | 3,123E-09 |
| TcCLB.507963.40 | hypothetical protein, conserved | -0,5118 | 0,00007012 |
| TcCLB.507709.100 | protein phosphatase 2A regulatory subunit, putative | -0,5122 | 0,00002402 |
| TcCLB.510007.40 | hypothetical protein, conserved | -0,5125 | 0,00001701 |
| TcCLB.506135.100 | hypothetical protein, conserved | -0,5125 | 0,0007426 |

| | | | |
|---|---|---|---|
| TcCLB.509015.10 | copper-transporting ATPase, putative | -0,5125 | 0,02651 |
| TcCLB.509099.150 | cdc2-related kinase 9 (CRK9) | -0,5126 | 3,02E-07 |
| TcCLB.507993.180 | hypothetical protein, conserved | -0,5127 | 3,02E-07 |
| TcCLB.506439.18 | hypothetical protein, conserved | -0,5127 | 4,744E-06 |
| TcCLB.504029.40 | COPII coat assembly protein sec16, putative (sec16) | -0,5128 | 0,00001956 |
| TcCLB.510351.70 | hypothetical protein, conserved | -0,5128 | 0,0001314 |
| TcCLB.510757.9 | hypothetical protein, conserved (fragment) | -0,5131 | 2,137E-06 |
| TcCLB.510263.30 | surface protease GP63, putative | -0,5132 | 0,0007307 |
| TcCLB.511529.170 | enoyl-CoA hydratase/isomerase family protein, putative | -0,5132 | 0,002676 |
| TcCLB.503753.20 | hypothetical protein, conserved | -0,5134 | 0,0001587 |
| TcCLB.506941.229 | pre-mRNA cleavage complex II Clp1 protein, putative | -0,5138 | 4,496E-07 |
| TcCLB.506697.30 | serine/threonine-protein kinase Nek, putative | -0,5139 | 4,313E-07 |
| TcCLB.506265.60 | hypothetical protein, conserved | -0,514 | 2,486E-10 |
| TcCLB.506661.120 | hypothetical protein | -0,514 | 0,00002202 |
| TcCLB.473313.10 | mucin-associated surface protein (MASP), putative (fragm | -0,5141 | 0,008304 |
| TcCLB.506451.16 | hypothetical protein, conserved (pseudogene) | -0,5141 | 0,08033 |
| TcCLB.504125.30 | ubiquitin carboxyl-terminal hydrolase, putative | -0,5143 | 4,47E-10 |
| TcCLB.511445.30 | hypothetical protein, conserved | -0,5143 | 2,741E-07 |
| TcCLB.506977.110 | hypothetical protein, conserved (pseudogene) | -0,5143 | 2,954E-06 |
| TcCLB.506503.40 | hypothetical protein, conserved | -0,5143 | 0,00008669 |
| TcCLB.511873.41 | hypothetical protein, conserved (pseudogene) | -0,5148 | 0,08769 |
| TcCLB.504125.64 | hypothetical protein | -0,5149 | 4,087E-08 |
| TcCLB.511491.80 | mitogen-activated protein kinase, putative | -0,5149 | 6,385E-07 |
| TcCLB.506305.20 | hypothetical protein, conserved | -0,5149 | 2,665E-06 |
| TcCLB.509599.180 | hypothetical protein, conserved | -0,5152 | 0,00002144 |
| TcCLB.504161.30 | hypothetical protein, conserved | -0,5152 | 0,00006765 |
| TcCLB.439803.9 | trans-sialidase (pseudogene), putative | -0,5152 | 0,002272 |
| TcCLB.506835.99 | hypothetical protein, conserved | -0,5153 | 8,651E-13 |
| TcCLB.508261.150 | trans-sialidase (pseudogene), putative | -0,5153 | 0,01967 |
| TcCLB.510709.50 | mucin TcMUCII (pseudogene), putative | -0,5154 | 0,4069 |
| TcCLB.511407.50 | hypothetical protein, conserved | -0,5157 | 4,724E-08 |
| TcCLB.507927.10 | hypothetical protein, conserved | -0,5157 | 0,0001893 |
| TcCLB.511797.167 | mucin-associated surface protein (MASP), putative | -0,5157 | 0,1182 |
| TcCLB.503529.10 | hypothetical protein, conserved (fragment) | -0,516 | 0,1746 |
| TcCLB.511277.610 | surface protease GP63, putative | -0,5161 | 0,0001879 |
| TcCLB.505071.60 | protein kinase, putative | -0,5162 | 2,198E-09 |
| TcCLB.508495.130 | hypothetical protein | -0,5164 | 0,1432 |
| TcCLB.509901.130 | Iron-sulfur assembly protein 1, putative (Isa1) | -0,5165 | 1,529E-06 |
| TcCLB.510445.30 | hypothetical protein, conserved | -0,5166 | 1,181E-06 |
| TcCLB.508277.260 | actin-related protein 3, putative | -0,5167 | 1,743E-08 |
| TcCLB.508887.10 | hypothetical protein, conserved (pseudogene) | -0,5167 | 0,1995 |
| TcCLB.510043.29 | hypothetical protein, conserved | -0,5168 | 0,002346 |
| TcCLB.509229.30 | hypothetical protein, conserved | -0,5169 | 0,004722 |
| TcCLB.504039.50 | retrotransposon hot spot protein (RHS, pseudogene), puta | -0,5169 | 0,007044 |
| TcCLB.503713.40 | hypothetical protein | -0,517 | 0,01496 |

| | | | |
|---|---|---|---|
| TcCLB.504081.470 | mucin TcMUCII, putative | -0,517 | 0,22 |
| TcCLB.506943.69 | hypothetical protein, conserved | -0,5171 | 2,289E-06 |
| TcCLB.511277.614 | hypothetical protein | -0,5171 | 0,02582 |
| TcCLB.511255.370 | mucin TcMUCII, putative | -0,5171 | 0,1579 |
| TcCLB.506559.205 | hypothetical protein | -0,5173 | 0,0001163 |
| TcCLB.510361.10 | trans-sialidase, Group V, putative | -0,5174 | 0,0637 |
| TcCLB.508593.120 | hypothetical protein, conserved | -0,5175 | 0,0001117 |
| TcCLB.509073.10 | hypothetical protein, conserved | -0,5176 | 4,157E-07 |
| TcCLB.511167.10 | NADH-cytochrome b5 reductase, putative | -0,5176 | 0,0000633 |
| TcCLB.506941.30 | hypothetical protein, conserved | -0,5178 | 2,031E-10 |
| TcCLB.507937.6 | trans-sialidase (pseudogene), putative | -0,518 | 0,2335 |
| TcCLB.508811.55 | hypothetical protein | -0,5189 | 2,152E-10 |
| TcCLB.509399.130 | hypothetical protein, conserved | -0,5189 | 5,619E-06 |
| TcCLB.503989.50 | hypothetical protein, conserved | -0,5191 | 8,345E-10 |
| TcCLB.507739.110 | myosin IB heavy chain, putative | -0,5193 | 7,318E-12 |
| TcCLB.503525.30 | hypothetical protein, conserved | -0,5194 | 1,077E-08 |
| TcCLB.506709.40 | hypothetical protein, conserved | -0,5194 | 0,0001144 |
| TcCLB.503753.10 | hypothetical protein, conserved | -0,5195 | 0,00006213 |
| TcCLB.504057.130 | hypothetical protein, conserved | -0,5197 | 1,085E-10 |
| TcCLB.507711.70 | hypothetical protein, conserved | -0,5199 | 0,00005239 |
| TcCLB.503867.20 | trans-sialidase (pseudogene), putative | -0,52 | 0,04138 |
| TcCLB.506513.40 | hypothetical protein, conserved | -0,5202 | 0,00003226 |
| TcCLB.510359.430 | trans-sialidase (pseudogene), putative | -0,5203 | 2,69E-07 |
| TcCLB.504199.20 | hypothetical protein, conserved | -0,5203 | 2,544E-06 |
| TcCLB.507743.9 | tyrosine protein kinase, putative | -0,5204 | 1,894E-06 |
| TcCLB.511345.30 | trans-sialidase (pseudogene), putative | -0,5206 | 0,0008825 |
| TcCLB.511285.120 | ATP-dependent RNA helicase, putative | -0,5209 | 0,00006364 |
| TcCLB.508261.166 | hypothetical protein, conserved (pseudogene) | -0,5209 | 0,02621 |
| TcCLB.511449.95 | hypothetical protein, conserved (pseudogene) | -0,5211 | 0,2932 |
| TcCLB.507623.50 | oxidoreductase, putative | -0,5212 | 0,00002078 |
| TcCLB.511751.240 | hypothetical protein, conserved (pseudogene) | -0,5217 | 6,494E-07 |
| TcCLB.507023.190 | DNA-J protein, putative | -0,5217 | 0,0001121 |
| TcCLB.508307.100 | hypothetical protein, conserved | -0,5218 | 0,000873 |
| TcCLB.511451.100 | trans-sialidase (pseudogene), putative | -0,5218 | 0,01571 |
| TcCLB.508153.520 | hypothetical protein, conserved (pseudogene) | -0,522 | 2,697E-09 |
| TcCLB.505919.11 | mucin-associated surface protein (MASP, pseudogene), pu | -0,522 | 1,51E-07 |
| TcCLB.506743.130 | protein tyrosine phosphatase-like protein, putative | -0,5222 | 0,01234 |
| TcCLB.506357.80 | hypothetical protein, conserved | -0,5223 | 1,36E-09 |
| TcCLB.511153.90 | phopshatase, putative | -0,5224 | 1,756E-07 |
| TcCLB.472777.30 | protein disulfide isomerase, putative | -0,5225 | 0,0009307 |
| TcCLB.510887.70 | hypothetical protein, conserved | -0,5226 | 3,682E-06 |
| TcCLB.504057.120 | lipophosphoglycan biosynthetic protein 2, putative | -0,5228 | 0,0000529 |
| TcCLB.508939.60 | hypothetical protein | -0,5229 | 0,0008393 |
| TcCLB.510161.20 | hypothetical protein, conserved | -0,5231 | 4,004E-11 |
| TcCLB.510625.225 | hypothetical protein, conserved (pseudogene) | -0,5232 | 0,05578 |

| | | | |
|---|---|---|---|
| TcCLB.507801.130 | heat shock protein DnaJ (pseudogene), putative | -0,5236 | 4,324E-13 |
| TcCLB.510667.40 | DNA repair protein RAD2, putative | -0,5236 | 7,057E-07 |
| TcCLB.510901.110 | hypothetical protein, conserved | -0,5236 | 0,0002736 |
| TcCLB.510031.70 | heat shock protein Hsp20, putative | -0,5237 | 0,0001259 |
| TcCLB.508221.329 | mucin-associated surface protein (MASP, pseudogene), pu | -0,5238 | 0,3785 |
| TcCLB.509897.95 | hypothetical protein, conserved (pseudogene) | -0,5239 | 0,00002696 |
| TcCLB.503453.60 | GTP-ase activating protein, putative | -0,5239 | 0,00008751 |
| TcCLB.509879.30 | hypothetical protein, conserved | -0,5242 | 0,00000575 |
| TcCLB.506931.30 | hypothetical protein, conserved | -0,5244 | 0,002791 |
| TcCLB.506681.60 | hypothetical protein, conserved | -0,5246 | 3,354E-08 |
| TcCLB.507831.40 | hypothetical protein, conserved | -0,5246 | 8,858E-06 |
| TcCLB.511555.40 | CPSF 30 kDa subunit, putative (CPSF30) | -0,5248 | 6,956E-07 |
| TcCLB.510657.170 | cytochrome P450 reductase, putative | -0,5248 | 0,000438 |
| TcCLB.505171.70 | ATP-dependent DEAD/H RNA helicase, putative | -0,5249 | 3,162E-09 |
| TcCLB.506945.105 | zinc-binding protein (Yippee), putative | -0,5249 | 0,0001599 |
| TcCLB.411383.10 | surface protease GP63 (pseudogene), putative | -0,5249 | 0,2803 |
| TcCLB.509233.70 | Spastin, putative | -0,525 | 1,971E-09 |
| TcCLB.510171.10 | trans-sialidase, putative (fragment) | -0,5252 | 0,0001199 |
| TcCLB.511173.250 | mucin-associated surface protein (MASP, pseudogene), pu | -0,5253 | 0,01766 |
| TcCLB.508665.30 | hypothetical protein, conserved | -0,5257 | 9,81E-08 |
| TcCLB.508547.100 | hypothetical protein, conserved | -0,5257 | 3,415E-06 |
| TcCLB.511025.100 | chaperone DNAJ protein, putative | -0,5261 | 5,943E-08 |
| TcCLB.510885.90 | hypothetical protein, conserved | -0,5263 | 0,001014 |
| TcCLB.426761.10 | retrotransposon hot spot protein (RHS, pseudogene), puta | -0,5266 | 0,1825 |
| TcCLB.503559.90 | hypothetical protein, conserved | -0,5267 | 6,476E-07 |
| TcCLB.510787.10 | trans-sialidase, Group I, putative | -0,527 | 0,0000663 |
| TcCLB.506525.70 | hypothetical protein, conserved | -0,5274 | 1,014E-09 |
| TcCLB.510655.140 | hypothetical protein, conserved | -0,5274 | 0,00001996 |
| TcCLB.511179.160 | ubiquitin-protein ligase, putative | -0,5275 | 0,0004896 |
| TcCLB.508423.20 | hypothetical protein, conserved (pseudogene) | -0,5276 | 4,499E-10 |
| TcCLB.509099.20 | hypothetical protein, conserved | -0,5276 | 1,875E-08 |
| TcCLB.506517.14 | protein kinase, putative | -0,5276 | 0,01103 |
| TcCLB.510533.230 | palmitoyl acyltransferase 11, putative | -0,528 | 0,00001669 |
| TcCLB.507809.100 | hypothetical protein, conserved | -0,5282 | 1,035E-12 |
| TcCLB.510491.70 | hypothetical protein | -0,5282 | 0,0007742 |
| TcCLB.504213.70 | hypothetical protein, conserved | -0,5283 | 2,042E-07 |
| TcCLB.509991.40 | transmembrane protein, conserved (pseudogene) | -0,5287 | 1,542E-13 |
| TcCLB.507317.60 | protein kinase-like protein, putative | -0,5287 | 0,0003099 |
| TcCLB.506753.220 | hypothetical protein, conserved | -0,5288 | 0,005955 |
| TcCLB.504149.124 | conserved protein, unknown function | -0,5289 | 5,798E-09 |
| TcCLB.507611.210 | hypothetical protein | -0,529 | 0,1729 |
| TcCLB.505193.20 | hypothetical protein, conserved | -0,5291 | 4,832E-06 |
| TcCLB.506577.5 | hypothetical protein | -0,5292 | 5,098E-06 |
| TcCLB.507927.70 | hypothetical protein, conserved | -0,5292 | 0,0001789 |
| TcCLB.507037.89 | hypothetical protein, conserved | -0,5293 | 0,002957 |

| | | | |
|---|---|---|---|
| TcCLB.511731.5 | cationic amino acid transporter (pseudogene), putative | -0,5293 | 0,02645 |
| TcCLB.510489.44 | mucin TcMUCII (pseudogene), putative | -0,5293 | 0,09193 |
| TcCLB.506177.150 | hypothetical protein, conserved | -0,5295 | 3,764E-08 |
| TcCLB.509617.60 | thioredoxin, putative | -0,5296 | 0,0001275 |
| TcCLB.506457.149 | hypothetical protein, conserved (fragment) | -0,5296 | 0,0003832 |
| TcCLB.511285.50 | hypothetical protein, conserved | -0,5297 | 8,002E-12 |
| TcCLB.506681.30 | hypothetical protein, conserved | -0,5299 | 0,00008429 |
| TcCLB.506289.80 | small GTP-binding protein Rab32, putative | -0,53 | 2,867E-07 |
| TcCLB.506559.454 | GTP-binding protein, putative | -0,53 | 4,455E-07 |
| TcCLB.509217.60 | trans-sialidase (pseudogene), putative | -0,5301 | 0,04333 |
| TcCLB.407477.60 | hypothetical protein, conserved | -0,5302 | 0,00001233 |
| TcCLB.503841.50 | hypothetical protein, conserved | -0,5302 | 0,0000448 |
| TcCLB.506173.100 | mucin-associated surface protein (MASP, pseudogene), pu | -0,5302 | 0,04488 |
| TcCLB.503761.50 | hypothetical protein | -0,5303 | 0,07941 |
| TcCLB.509975.15 | surface protease GP63 (pseudogene), putative | -0,5304 | 0,1178 |
| TcCLB.507907.20 | trans-sialidase, Group II, putative | -0,5308 | 0,002374 |
| TcCLB.508207.45 | hypothetical protein, conserved | -0,5309 | 1,631E-06 |
| TcCLB.508355.280 | hypothetical protein, conserved | -0,5312 | 3,082E-09 |
| TcCLB.510861.70 | hypothetical protein, conserved | -0,5312 | 2,512E-07 |
| TcCLB.508277.269 | hypothetical protein, conserved (pseudogene) | -0,5313 | 4,693E-06 |
| TcCLB.399373.20 | dispersed gene family protein 1 (DGF-1), putative | -0,5316 | 0,02248 |
| TcCLB.506279.100 | hypothetical protein | -0,5317 | 0,003423 |
| TcCLB.504071.20 | hypothetical protein, conserved | -0,5319 | 4,656E-08 |
| TcCLB.506435.90 | hypothetical protein, conserved | -0,5319 | 1,571E-06 |
| TcCLB.507089.220 | hypothetical protein, conserved | -0,532 | 2,018E-06 |
| TcCLB.506509.40 | hypothetical protein, conserved | -0,5322 | 0,00009773 |
| TcCLB.504153.360 | succinyl-coA:3-ketoacid-coenzyme A transferase, mitocho | -0,5327 | 5,489E-14 |
| TcCLB.509963.60 | hypothetical protein, conserved | -0,5329 | 0,0002839 |
| TcCLB.510289.70 | hypothetical protein, conserved | -0,5332 | 0,00004484 |
| TcCLB.503971.50 | hypothetical protein, conserved | -0,5334 | 0,007129 |
| TcCLB.503457.10 | hypothetical protein | -0,5336 | 0,0003186 |
| TcCLB.511255.640 | mucin TcMUCII, putative | -0,5336 | 0,1044 |
| TcCLB.509999.120 | RNA-binding protein, putative | -0,5339 | 0,0138 |
| TcCLB.511875.123 | hypothetical protein, conserved (pseudogene) | -0,5339 | 0,2313 |
| TcCLB.508355.380 | cysteine peptidase, Clan CA, family C19, putative | -0,534 | 0,002382 |
| TcCLB.508207.130 | hypothetical protein, conserved | -0,5342 | 0,0003775 |
| TcCLB.510175.90 | retrotransposon hot spot (RHS) protein, putative | -0,5346 | 0,0001033 |
| TcCLB.503555.50 | hypothetical protein, conserved | -0,5346 | 0,0004553 |
| TcCLB.507625.186 | hypothetical protein, conserved | -0,5348 | 0,0004506 |
| TcCLB.509979.261 | surface protease GP63 (pseudogene), putative | -0,5349 | 0,01127 |
| TcCLB.503789.20 | hypothetical protein, conserved | -0,535 | 1,934E-09 |
| TcCLB.510267.10 | hypothetical protein, conserved | -0,535 | 7,818E-06 |
| TcCLB.503611.50 | hypothetical protein, conserved | -0,535 | 0,0003467 |
| TcCLB.511215.60 | hypothetical protein, conserved | -0,5351 | 0,000082 |
| TcCLB.508707.194 | hypothetical protein, conserved | -0,5353 | 1,838E-09 |

| | | | |
|---|---|---|---|
| TcCLB.510693.310 | syntaxin binding protein (pseudogene), putative | -0,5353 | 0,1126 |
| TcCLB.511727.234 | hypothetical protein, conserved | -0,5355 | 9,123E-07 |
| TcCLB.508221.120 | hypothetical protein | -0,5355 | 0,07648 |
| TcCLB.416605.10 | mucin-associated surface protein (MASP), putative | -0,5358 | 0,08166 |
| TcCLB.504089.30 | hypothetical protein, conserved | -0,5359 | 2,783E-09 |
| TcCLB.508461.284 | mitochondrial carrier protein, putative | -0,5359 | 4,325E-06 |
| TcCLB.508357.40 | cysteine peptidase, Clan CA, family C19, putative | -0,5361 | 0,0002224 |
| TcCLB.511157.9 | neurobeachin/beige protein, putative (fragment) | -0,5365 | 1,664E-08 |
| TcCLB.510495.11 | surface protease GP63 (pseudogene), putative | -0,5365 | 0,02329 |
| TcCLB.507949.10 | chaperone DNAJ protein, putative | -0,5367 | 0,0001284 |
| TcCLB.510351.50 | hypothetical protein, conserved | -0,5376 | 2,283E-08 |
| TcCLB.509615.60 | hypothetical protein | -0,5377 | 0,1844 |
| TcCLB.490165.9 | DNA repair protein, putative | -0,5378 | 0,01535 |
| TcCLB.511239.130 | hypothetical protein, conserved | -0,5379 | 0,0005912 |
| TcCLB.510189.60 | trans-sialidase (pseudogene), putative | -0,538 | 0,03312 |
| TcCLB.503619.10 | meiotic recombination protein SPO11, putative | -0,5381 | 0,0006739 |
| TcCLB.507467.40 | choline/carnitine O-acetyltransferase (pseudogene), puta | -0,5383 | 0,002197 |
| TcCLB.504587.40 | mucin TcMUCII, putative | -0,5383 | 0,06748 |
| TcCLB.509993.5 | hypothetical protein, conserved (pseudogene) | -0,5386 | 0,0003053 |
| TcCLB.504131.120 | hypothetical protein, conserved | -0,5387 | 9,59E-07 |
| TcCLB.511753.90 | hypothetical protein, conserved | -0,5387 | 0,00006234 |
| TcCLB.508959.10 | hypothetical protein, conserved | -0,5388 | 0,005231 |
| TcCLB.505931.20 | surface protease GP63, putative | -0,539 | 0,007183 |
| TcCLB.508211.10 | oxysterol-binding protein, putative | -0,5391 | 0,00002249 |
| TcCLB.506767.280 | retrotransposon hot spot protein (RHS, pseudogene), puta | -0,5391 | 0,1882 |
| TcCLB.503571.19 | hypothetical protein, conserved | -0,5393 | 3,546E-08 |
| TcCLB.506471.40 | beta galactofuranosyl glycosyltransferase, putative | -0,5394 | 0,02044 |
| TcCLB.507483.80 | hypothetical protein, conserved | -0,5397 | 2,493E-06 |
| TcCLB.507011.140 | DNA-repair protein, putative | -0,5399 | 3,02E-08 |
| TcCLB.506837.10 | mucin TcMUCI, putative | -0,5399 | 0,0002384 |
| TcCLB.510225.40 | hypothetical protein, conserved | -0,5403 | 1,342E-08 |
| TcCLB.506253.21 | hypothetical protein, conserved | -0,5403 | 0,1461 |
| TcCLB.506625.130 | hypothetical protein, conserved | -0,5406 | 0,004629 |
| TcCLB.506297.110 | ATPase, putative | -0,541 | 6,191E-13 |
| TcCLB.506113.90 | retrotransposon hot spot protein (RHS, pseudogene), puta | -0,541 | 2,104E-07 |
| TcCLB.507837.80 | TBC1 domain family member 20/GTPase, putative | -0,5413 | 0,00008337 |
| TcCLB.509429.124 | hypothetical protein, conserved | -0,5413 | 0,001047 |
| TcCLB.508895.60 | RNA-binding protein, putative | -0,5414 | 2,718E-07 |
| TcCLB.507041.85 | hypothetical protein, conserved (pseudogene) | -0,5414 | 0,0002034 |
| TcCLB.511621.20 | hypothetical protein, conserved | -0,5416 | 2,519E-06 |
| TcCLB.506619.30 | Flagellar Member 8 (FLAM8) | -0,5418 | 0,1499 |
| TcCLB.510823.100 | hypothetical protein, conserved | -0,542 | 1,096E-07 |
| TcCLB.508777.80 | hypothetical protein, conserved | -0,542 | 4,197E-07 |
| TcCLB.511603.160 | mucin-associated surface protein (MASP), putative | -0,5423 | 0,06508 |
| TcCLB.508979.40 | hypothetical protein | -0,5424 | 0,2315 |

| | | | |
|---|---|---|---|
| TcCLB.505999.150 | wd40 repeat domain-containing protein | -0,5427 | 9,413E-07 |
| TcCLB.507681.220 | pre-mRNA splicing factor ATP-dependent RNA helicase, p | -0,543 | 1,388E-10 |
| TcCLB.508835.30 | trans-sialidase (pseudogene), putative | -0,5431 | 0,001075 |
| TcCLB.511421.160 | hypothetical protein, conserved | -0,5435 | 0,00006973 |
| TcCLB.506955.161 | elongation factor 1-gamma (EF-1-gamma, pseudogene), | -0,5436 | 0,0542 |
| TcCLB.509683.50 | calmodulin, putative | -0,5437 | 0,0003302 |
| TcCLB.508181.4 | AGC/RSK family serine/threonine kinase, putative (fragm | -0,544 | 0,03587 |
| TcCLB.508779.20 | hypothetical protein, conserved | -0,5442 | 2,536E-08 |
| TcCLB.509543.10 | hypothetical protein | -0,5443 | 0,01942 |
| TcCLB.508859.70 | hypothetical protein, conserved | -0,5445 | 6,872E-06 |
| TcCLB.509007.60 | rac serine-threonine kinase, putative | -0,5448 | 1,268E-07 |
| TcCLB.508815.40 | hypothetical protein, conserved | -0,545 | 0,00005094 |
| TcCLB.508277.340 | Eukaryotic translation initiation factor 4 gamma type 2, pu | -0,5453 | 4,129E-12 |
| TcCLB.511847.20 | hypothetical protein, conserved | -0,5454 | 1,884E-07 |
| TcCLB.511371.5 | trans-sialidase (pseudogene), putative | -0,5455 | 0,001859 |
| TcCLB.506955.79 | surface protease GP63 (pseudogene), putative | -0,5456 | 0,1407 |
| TcCLB.511389.70 | hypothetical protein, conserved | -0,5457 | 2,933E-08 |
| TcCLB.506247.430 | hypothetical protein, conserved | -0,5458 | 6,022E-07 |
| TcCLB.509399.170 | elongation factor 1-gamma (EF-1-gamma), putative | -0,5459 | 2,588E-06 |
| TcCLB.507775.30 | hypothetical protein, conserved | -0,5459 | 0,00001348 |
| TcCLB.510725.60 | hypothetical protein, conserved | -0,5462 | 0,0009931 |
| TcCLB.509249.20 | trans-sialidase (pseudogene), putative | -0,5462 | 0,00152 |
| TcCLB.508153.200 | kinesin, putative | -0,5464 | 8,298E-08 |
| TcCLB.511071.70 | ubiquitin-conjugating enzyme E2, putative | -0,5464 | 2,993E-07 |
| TcCLB.506015.40 | mucin TcMUCI, putative | -0,5465 | 0,05123 |
| TcCLB.511237.70 | hypothetical protein, conserved | -0,5467 | 0,00000894 |
| TcCLB.508765.50 | mucin TcMUCI (pseudogene), putative | -0,5467 | 0,00264 |
| TcCLB.420293.20 | trans-sialidase, Group II, putative | -0,5467 | 0,007266 |
| TcCLB.507771.110 | hypothetical protein, conserved | -0,5472 | 0,00000117 |
| TcCLB.510753.30 | hypothetical protein, conserved | -0,5472 | 6,191E-06 |
| TcCLB.508667.9 | hypothetical protein, conserved | -0,5472 | 0,01227 |
| TcCLB.511603.130 | hypothetical protein, conserved | -0,5472 | 0,209 |
| TcCLB.506729.40 | hypothetical protein, conserved | -0,5476 | 6,889E-07 |
| TcCLB.511229.39 | hypothetical protein, conserved | -0,548 | 0,0006515 |
| TcCLB.510885.140 | hypothetical protein | -0,5482 | 4,955E-06 |
| TcCLB.507809.14 | protein kinase, putative | -0,5482 | 6,856E-06 |
| TcCLB.504669.50 | mucin TcMUCI, putative | -0,5482 | 0,0225 |
| TcCLB.511727.30 | hypothetical protein, conserved | -0,5486 | 9,889E-10 |
| TcCLB.507649.5 | hypothetical protein, conserved | -0,5486 | 0,0323 |
| TcCLB.504013.20 | hypothetical protein | -0,5487 | 0,00006131 |
| TcCLB.507765.90 | hypothetical protein, conserved | -0,5489 | 1,566E-08 |
| TcCLB.507073.30 | hypothetical protein, conserved | -0,549 | 2,454E-12 |
| TcCLB.506355.70 | hypothetical protein, conserved | -0,5493 | 6,28E-11 |
| TcCLB.507277.30 | hypothetical protein, conserved | -0,5495 | 2,293E-06 |
| TcCLB.508707.290 | hypothetical protein | -0,5495 | 0,0001196 |

| | | | |
|---|---|---|---|
| TcCLB.505943.95 | trans-sialidase (pseudogene), putative | -0,5496 | 0,00009662 |
| TcCLB.506435.470 | hypothetical protein, conserved | -0,5496 | 0,001014 |
| TcCLB.503887.70 | phosphoglycerate mutase, putative (iPGAM) | -0,5497 | 0,0007742 |
| TcCLB.506413.40 | hypothetical protein, conserved | -0,5498 | 0,0002847 |
| TcCLB.507993.280 | hypothetical protein, conserved | -0,5499 | 2,883E-11 |
| TcCLB.509267.90 | ATP-dependent RNA helicase, putative | -0,5507 | 0,00002377 |
| TcCLB.508665.40 | hypothetical protein, conserved | -0,5507 | 0,000732 |
| TcCLB.508167.10 | mucin-associated surface protein (MASP, pseudogene), pu | -0,5509 | 0,01287 |
| TcCLB.510359.450 | hypothetical protein | -0,5509 | 0,1234 |
| TcCLB.508799.100 | hypothetical protein, conserved | -0,551 | 5,971E-07 |
| TcCLB.506265.160 | hypothetical protein, conserved | -0,5512 | 0,0001764 |
| TcCLB.509881.9 | dispersed gene family protein 1 (DGF-1), putative (fragme | -0,5512 | 0,3228 |
| TcCLB.511079.25 | mucin-associated surface protein (MASP, pseudogene), pu | -0,5513 | 0,1018 |
| TcCLB.506445.20 | Eukaryotic translation initiation factor 4 gamma type 2, pu | -0,5517 | 9,193E-10 |
| TcCLB.508879.130 | telomerase component (pseudogene), putative | -0,5526 | 5,014E-18 |
| TcCLB.510283.150 | hypothetical protein, conserved | -0,5526 | 0,00133 |
| TcCLB.507949.190 | trans-sialidase (pseudogene), putative | -0,5528 | 0,00005027 |
| TcCLB.506315.30 | protein phosphatase 1 catalitic subunit, putative | -0,5529 | 2,586E-07 |
| TcCLB.506941.70 | hypothetical protein, conserved | -0,5531 | 9,385E-10 |
| TcCLB.509039.49 | hypothetical protein, conserved | -0,5533 | 0,00258 |
| TcCLB.507071.89 | mucin TcMUCII (pseudogene), putative | -0,5534 | 0,09269 |
| TcCLB.504021.130 | hypothetical protein, conserved | -0,5537 | 0,00002796 |
| TcCLB.510897.3 | A/G-specific adenine glycosylase (pseudogene), putative | -0,5538 | 0,03376 |
| TcCLB.510483.270 | hypothetical protein | -0,5538 | 0,1206 |
| TcCLB.508933.20 | dihydrolipoamide dehydrogenase, putative | -0,554 | 9,549E-06 |
| TcCLB.509419.140 | mucin-associated surface protein (MASP), putative | -0,554 | 0,03231 |
| TcCLB.508465.60 | hypothetical protein, conserved | -0,5543 | 4,259E-08 |
| TcCLB.506289.20 | hypothetical protein, conserved | -0,5544 | 9,268E-07 |
| TcCLB.509429.70 | hypothetical protein | -0,5545 | 8,338E-09 |
| TcCLB.506955.50 | trans-sialidase (pseudogene), putative | -0,5545 | 0,003484 |
| TcCLB.506519.120 | hypothetical protein, conserved | -0,5546 | 7,262E-07 |
| TcCLB.508521.90 | trans-sialidase (pseudogene), putative | -0,5547 | 0,0003289 |
| TcCLB.507077.30 | hypothetical protein, conserved | -0,5547 | 0,0008629 |
| TcCLB.504085.40 | hypothetical protein, conserved | -0,5549 | 2,037E-06 |
| TcCLB.506811.40 | hypothetical protein, conserved | -0,5551 | 2,288E-08 |
| TcCLB.507019.10 | protein kinase, putative | -0,5553 | 3,506E-07 |
| TcCLB.506445.121 | heat shock protein DNAJ, putative (fragment) | -0,5557 | 0,001778 |
| TcCLB.509571.30 | hypothetical protein, conserved | -0,556 | 9,798E-11 |
| TcCLB.506949.30 | hypothetical protein, conserved | -0,5561 | 0,02356 |
| TcCLB.510513.40 | hypothetical protein, conserved | -0,5564 | 9,348E-06 |
| TcCLB.510713.120 | surface protease GP63, putative | -0,5564 | 0,1775 |
| TcCLB.503489.20 | hypothetical protein, conserved | -0,5565 | 9,313E-06 |
| TcCLB.508431.80 | trans-sialidase (pseudogene), putative | -0,5565 | 0,01422 |
| TcCLB.503735.49 | hypothetical protein, conserved | -0,5567 | 0,0003489 |
| TcCLB.508233.30 | mucin TcSMUGS, putative | -0,5569 | 0,00006028 |

| | | | |
|---|---|---|---|
| TcCLB.503973.300 | surface protease GP63 (pseudogene), putative | -0,557 | 0,1148 |
| TcCLB.510275.150 | hypothetical protein, conserved (pseudogene) | -0,5571 | 0,01768 |
| TcCLB.511559.34 | hypothetical protein, conserved | -0,5572 | 0,0002287 |
| TcCLB.507823.39 | hypothetical protein, conserved | -0,5578 | 0,0005653 |
| TcCLB.510327.5 | surface protease GP63 (pseudogene), putative | -0,5579 | 0,05318 |
| TcCLB.511407.70 | hypothetical protein, conserved | -0,558 | 3,021E-09 |
| TcCLB.507237.250 | mucin-associated surface protein (MASP, pseudogene), pu | -0,558 | 0,007265 |
| TcCLB.507089.60 | hypothetical protein, conserved | -0,5581 | 4,097E-08 |
| TcCLB.508645.4 | hypothetical protein, conserved | -0,5581 | 0,003983 |
| TcCLB.503633.40 | trans-sialidase (pseudogene), putative | -0,5581 | 0,006661 |
| TcCLB.506583.20 | hypothetical protein, conserved | -0,5584 | 0,0007997 |
| TcCLB.508751.20 | trans-sialidase (pseudogene), putative | -0,5586 | 0,04818 |
| TcCLB.506797.40 | hypothetical protein, conserved | -0,5587 | 0,007974 |
| TcCLB.511257.80 | hypothetical protein | -0,5589 | 0,0006145 |
| TcCLB.508111.10 | trans-sialidase (pseudogene), putative | -0,5592 | 0,007417 |
| TcCLB.511597.25 | hypothetical protein | -0,5592 | 0,1942 |
| TcCLB.507011.20 | hypothetical protein, conserved | -0,5594 | 1,019E-06 |
| TcCLB.508873.240 | mucin TcMUCII, putative | -0,5594 | 0,03845 |
| TcCLB.503973.340 | trans-sialidase (pseudogene), putative | -0,5596 | 0,1069 |
| TcCLB.508175.280 | hypothetical protein, conserved | -0,5601 | 2,02E-10 |
| TcCLB.511817.134 | hypothetical protein, conserved | -0,5601 | 8,15E-08 |
| TcCLB.508879.200 | hypothetical protein, conserved | -0,5604 | 7,881E-06 |
| TcCLB.504085.30 | RNA-binding protein, putative | -0,5605 | 4,903E-07 |
| TcCLB.509785.90 | trans-sialidase (pseudogene), putative | -0,5606 | 0,005554 |
| TcCLB.509801.20 | hypothetical protein, conserved | -0,5607 | 2,17E-07 |
| TcCLB.411427.49 | hypothetical protein, conserved (fragment) | -0,5608 | 0,003339 |
| TcCLB.508461.460 | hypothetical protein, conserved | -0,5609 | 1,901E-07 |
| TcCLB.508707.64 | exonuclease, putative | -0,5609 | 0,0006302 |
| TcCLB.508265.60 | hypothetical protein, conserved (pseudogene) | -0,561 | 3,654E-09 |
| TcCLB.506791.20 | hypothetical protein, conserved | -0,561 | 4,224E-08 |
| TcCLB.508911.30 | hypothetical protein, conserved (pseudogene) | -0,561 | 3,154E-07 |
| TcCLB.510441.41 | trans-sialidase (pseudogene), putative | -0,5611 | 1,811E-06 |
| TcCLB.510073.30 | pumilio/PUF RNA binding protein 4, putative (PUF4) | -0,5611 | 0,301 |
| TcCLB.509767.110 | hypothetical protein, conserved | -0,5613 | 8,84E-07 |
| TcCLB.509267.80 | hypothetical protein | -0,5616 | 0,00001456 |
| TcCLB.507681.189 | hypothetical protein, conserved | -0,5617 | 0,00005709 |
| TcCLB.507793.30 | hypothetical protein, conserved | -0,5624 | 9,225E-06 |
| TcCLB.504137.130 | hypothetical protein, conserved | -0,5626 | 2,438E-08 |
| TcCLB.510275.120 | hypothetical protein | -0,5626 | 0,1623 |
| TcCLB.509461.40 | hypothetical protein, conserved | -0,5631 | 0,00001633 |
| TcCLB.511867.200 | hypothetical protein, conserved | -0,5633 | 2,678E-07 |
| TcCLB.508707.20 | As/Sb Reductase, putative (ACR2) | -0,5633 | 0,0001732 |
| TcCLB.503833.60 | hypothetical protein, conserved | -0,5634 | 0,00000044 |
| TcCLB.507763.30 | anion-transporting ATPase-like, putative | -0,5635 | 2,557E-09 |
| TcCLB.506513.20 | hypothetical protein, conserved | -0,5636 | 0,00008635 |

| | | | |
|---|---|---|---|
| TcCLB.509691.10 | hypothetical protein, conserved | -0,564 | 0,001157 |
| TcCLB.504057.65 | hypothetical protein, conserved | -0,5641 | 0,00001051 |
| TcCLB.504179.90 | hypothetical protein, conserved | -0,5644 | 6,941E-08 |
| TcCLB.511717.50 | hypothetical protein, conserved | -0,5647 | 9,538E-07 |
| TcCLB.508781.129 | arginine N-methyltransferase, type II, putative | -0,5652 | 0,0002877 |
| TcCLB.504029.70 | hypothetical protein, conserved | -0,5654 | 0,00002435 |
| TcCLB.509985.10 | hypothetical protein, conserved (pseudogene) | -0,5656 | 4,365E-07 |
| TcCLB.506227.240 | hypothetical protein, conserved | -0,5657 | 1,19E-11 |
| TcCLB.510287.20 | dolicholphosphate-mannose synthase, putative | -0,5657 | 0,00002494 |
| TcCLB.508667.20 | hypothetical protein | -0,5657 | 0,00002689 |
| TcCLB.506755.200 | hypothetical protein, conserved | -0,5659 | 0,00003159 |
| TcCLB.509575.50 | Casein kinase II subunit beta, putative | -0,5659 | 0,00004879 |
| TcCLB.506977.40 | hypothetical protein, conserved | -0,5662 | 2,543E-06 |
| TcCLB.510533.80 | hypothetical protein, conserved | -0,5662 | 0,00004193 |
| TcCLB.506355.35 | trans-sialidase (pseudogene), putative | -0,5668 | 0,00003225 |
| TcCLB.506801.100 | hypothetical protein, conserved | -0,5671 | 5,549E-07 |
| TcCLB.510175.10 | retrotransposon hot spot protein (RHS, pseudogene), puta | -0,5672 | 0,004161 |
| TcCLB.506599.130 | mucin TcMUCII (pseudogene), putative | -0,5675 | 0,01234 |
| TcCLB.504151.30 | hypothetical protein, conserved | -0,5676 | 1,219E-06 |
| TcCLB.507677.50 | hypothetical protein, conserved | -0,5677 | 9,991E-07 |
| TcCLB.503541.30 | hypothetical protein, conserved | -0,5677 | 0,001967 |
| TcCLB.510513.50 | hypothetical protein, conserved | -0,5683 | 5,267E-06 |
| TcCLB.510827.10 | hypothetical protein, conserved | -0,5688 | 0,0003203 |
| TcCLB.510055.160 | protein kinase, putative | -0,5688 | 0,003577 |
| TcCLB.508099.30 | hypothetical protein | -0,5689 | 0,1172 |
| TcCLB.511871.110 | hypothetical protein, conserved | -0,5692 | 6,815E-07 |
| TcCLB.509537.30 | hypothetical protein, conserved | -0,5692 | 1,225E-06 |
| TcCLB.511603.420 | syntaxin binding protein (pseudogene), putative | -0,5692 | 0,1116 |
| TcCLB.511283.30 | hypothetical protein, conserved | -0,5695 | 1,67E-10 |
| TcCLB.511277.120 | hypothetical protein, conserved | -0,5695 | 3,6E-10 |
| TcCLB.510885.120 | hypothetical protein, conserved | -0,5695 | 3,262E-09 |
| TcCLB.508207.140 | hypothetical protein, conserved | -0,5697 | 2,051E-06 |
| TcCLB.510175.140 | hypothetical protein, conserved (pseudogene) | -0,5699 | 0,00003837 |
| TcCLB.510899.40 | hypothetical protein, conserved | -0,5699 | 0,0003944 |
| TcCLB.506945.90 | hypothetical protein, conserved | -0,5702 | 0,004143 |
| TcCLB.507467.70 | diacylglycerol acyltransferase, putative | -0,5703 | 5,623E-09 |
| TcCLB.510565.100 | hypothetical protein, conserved | -0,5703 | 8,035E-06 |
| TcCLB.509217.180 | trans-sialidase (pseudogene), putative | -0,5703 | 0,1822 |
| TcCLB.511159.20 | origin recognition complex subunit 1 (ORC1), putative | -0,5705 | 0,000382 |
| TcCLB.504117.50 | hypothetical protein, conserved (pseudogene) | -0,5708 | 0,04351 |
| TcCLB.511753.20 | class I nuclease-like protein, putative | -0,5709 | 0,00006973 |
| TcCLB.436761.20 | hypothetical protein, conserved (fragment) | -0,571 | 0,07257 |
| TcCLB.510823.10 | hypothetical protein, conserved | -0,5711 | 3,642E-10 |
| TcCLB.510553.11 | hypothetical protein, conserved (pseudogene) | -0,5711 | 0,03257 |
| TcCLB.506885.4 | hypothetical protein, conserved (fragment) | -0,5718 | 1,404E-07 |

| | | | |
|---|---|---|---|
| TcCLB.510055.130 | hypothetical protein, conserved (pseudogene) | -0,5718 | 3,671E-07 |
| TcCLB.507483.50 | calmodulin (CALA2) | -0,5718 | 0,0003684 |
| TcCLB.506135.30 | hypothetical protein, conserved | -0,5719 | 0,00000393 |
| TcCLB.510519.50 | hypothetical protein, conserved | -0,5719 | 8,143E-06 |
| TcCLB.508855.10 | trans-sialidase, putative | -0,572 | 0,00005812 |
| TcCLB.503715.40 | protein kinase, putative | -0,5721 | 0,005836 |
| TcCLB.510515.40 | hypothetical protein, conserved | -0,5725 | 0,008835 |
| TcCLB.509485.30 | hypothetical protein, conserved (pseudogene) | -0,5728 | 0,09791 |
| TcCLB.509437.50 | AAA ATPase, putative | -0,5729 | 2,968E-09 |
| TcCLB.508593.44 | serine peptidase, Clan SC, Family S9D | -0,5733 | 1,672E-08 |
| TcCLB.511293.50 | kinesin, putative | -0,5733 | 4,644E-07 |
| TcCLB.503819.20 | DNA-directed RNA polymerase III subunit, putative | -0,5733 | 0,0003299 |
| TcCLB.507447.10 | hypothetical protein, conserved | -0,5734 | 1,349E-06 |
| TcCLB.511859.60 | subtilisin peptidase | -0,5735 | 0,00000135 |
| TcCLB.504769.150 | hypothetical protein | -0,5735 | 0,1746 |
| TcCLB.506491.10 | casein kinase I, putative | -0,5736 | 0,01013 |
| TcCLB.511291.20 | hypothetical protein, conserved | -0,5739 | 3,664E-06 |
| TcCLB.507005.10 | hypothetical protein | -0,5741 | 0,004656 |
| TcCLB.508175.350 | zinc finger protein family member, putative (POMP35) | -0,5742 | 1,694E-08 |
| TcCLB.506479.39 | serine/threonine protein phosphatase 2A regulatory subu | -0,5748 | 5,85E-07 |
| TcCLB.503653.90 | hypothetical protein, conserved | -0,5749 | 9,697E-07 |
| TcCLB.507237.220 | mucin-associated surface protein (MASP), putative | -0,5749 | 0,1881 |
| TcCLB.510955.4 | lipase-like protein, putative | -0,5751 | 3,096E-08 |
| TcCLB.508707.149 | hypothetical protein, conserved | -0,5751 | 0,00003173 |
| TcCLB.441729.9 | hypothetical protein (fragment) | -0,5755 | 0,006461 |
| TcCLB.510973.9 | hypothetical protein, conserved | -0,5757 | 0,000001 |
| TcCLB.509663.5 | dehydrogenase (pseudogene), putative | -0,5759 | 0,01831 |
| TcCLB.506661.50 | hypothetical protein, conserved | -0,576 | 6,841E-10 |
| TcCLB.503801.60 | hypothetical protein | -0,5761 | 0,0000166 |
| TcCLB.510655.10 | hypothetical protein, conserved | -0,5761 | 0,00002825 |
| TcCLB.511875.110 | trans-sialidase (pseudogene), putative | -0,5763 | 0,001062 |
| TcCLB.506297.320 | hypothetical protein | -0,5764 | 4,376E-08 |
| TcCLB.507775.10 | hypothetical protein, conserved | -0,5764 | 2,999E-06 |
| TcCLB.509897.230 | trans-sialidase (pseudogene), putative | -0,5765 | 0,00002738 |
| TcCLB.508231.130 | Cell division control protein 48, putative | -0,5767 | 0,001661 |
| TcCLB.506407.50 | hypothetical protein, conserved | -0,5768 | 6,28E-07 |
| TcCLB.506495.40 | hypothetical protein | -0,577 | 5,964E-08 |
| TcCLB.506229.30 | hypothetical protein | -0,5771 | 2,179E-07 |
| TcCLB.504151.20 | hypothetical protein, conserved | -0,5772 | 0,00000805 |
| TcCLB.509321.10 | hypothetical protein, conserved | -0,5773 | 8,109E-11 |
| TcCLB.506223.10 | hypothetical protein, conserved | -0,5773 | 0,00003657 |
| TcCLB.503973.190 | mucin-associated surface protein (MASP), putative | -0,5773 | 0,09444 |
| TcCLB.508347.40 | hypothetical protein, conserved | -0,5788 | 1,191E-14 |
| TcCLB.509935.10 | hypothetical protein | -0,5788 | 0,006009 |
| TcCLB.510513.10 | mannosyl-oligosaccharide 1,2-alpha-mannosidase IB, puta | -0,5789 | 5,703E-08 |

| | | | |
|---|---|---|---|
| TcCLB.510725.70 | hypothetical protein, conserved | -0,579 | 1,901E-08 |
| TcCLB.509669.170 | hypothetical protein, conserved | -0,5793 | 8,54E-11 |
| TcCLB.507145.60 | trans-sialidase, Group V, putative | -0,5793 | 0,03437 |
| TcCLB.509683.80 | adenylosuccinate synthetase, putative | -0,5794 | 7,246E-06 |
| TcCLB.511501.30 | ABC transporter, putative | -0,5794 | 0,00003282 |
| TcCLB.503839.30 | hypothetical protein, conserved | -0,5796 | 1,333E-10 |
| TcCLB.503683.30 | RNA-binding protein, putative | -0,5797 | 0,0006639 |
| TcCLB.511189.54 | hypothetical protein | -0,5798 | 2,734E-06 |
| TcCLB.510275.160 | trans-sialidase (pseudogene), putative | -0,5798 | 0,02811 |
| TcCLB.509567.30 | serine/threonine-protein kinase, putative | -0,5803 | 1,742E-11 |
| TcCLB.506887.40 | hypothetical protein, conserved | -0,5804 | 4,92E-08 |
| TcCLB.506113.86 | N-acetyltransferase complex ARD1 subunit (pseudogene), | -0,5804 | 0,02229 |
| TcCLB.506945.340 | hypothetical protein, conserved | -0,5807 | 6,172E-10 |
| TcCLB.507053.90 | conserved protein, unknown function | -0,5807 | 9,101E-08 |
| TcCLB.508215.6 | hypothetical protein, conserved | -0,5808 | 4,795E-08 |
| TcCLB.508771.20 | cysteine peptidase, Clan CA, family C51, putative | -0,5811 | 6,245E-09 |
| TcCLB.509925.5 | trans-sialidase (pseudogene), putative | -0,5818 | 0,007262 |
| TcCLB.504777.30 | surface protease GP63 (pseudogene), putative | -0,582 | 0,003621 |
| TcCLB.508913.10 | trans-sialidase (pseudogene), putative | -0,5822 | 6,909E-06 |
| TcCLB.511793.30 | mucin-associated surface protein (MASP), putative | -0,5822 | 0,2841 |
| TcCLB.506815.50 | hypothetical protein, conserved | -0,5824 | 1,243E-07 |
| TcCLB.508227.10 | hypothetical protein | -0,5826 | 0,2688 |
| TcCLB.509911.120 | hypothetical protein | -0,5828 | 5,26E-13 |
| TcCLB.503781.60 | tRNA pseudouridine synthase A, putative | -0,5828 | 5,363E-08 |
| TcCLB.508153.630 | mitochondrial carrier protein, putative | -0,5828 | 0,00005778 |
| TcCLB.418185.70 | trans-sialidase, Group II, putative | -0,5832 | 0,003194 |
| TcCLB.510911.58 | retrotransposon hot spot protein (RHS, pseudogene), puta | -0,5832 | 0,06494 |
| TcCLB.511853.21 | mucin-associated surface protein (MASP, pseudogene), pu | -0,5841 | 0,01149 |
| TcCLB.506133.207 | mucin-associated surface protein (MASP, pseudogene), pu | -0,5846 | 0,08175 |
| TcCLB.504085.10 | hypothetical protein, conserved | -0,5847 | 1,046E-07 |
| TcCLB.506699.9 | hypothetical protein, conserved | -0,5847 | 0,00005781 |
| TcCLB.506757.20 | hypothetical protein, conserved | -0,5848 | 4,813E-06 |
| TcCLB.507055.20 | hypothetical protein, conserved (fragment) | -0,5849 | 0,0001805 |
| TcCLB.511577.104 | hypothetical protein, conserved | -0,585 | 0,0002099 |
| TcCLB.506435.70 | hypothetical protein, conserved (pseudogene) | -0,5856 | 1,274E-12 |
| TcCLB.503867.10 | hypothetical protein, conserved | -0,5857 | 3,975E-06 |
| TcCLB.507085.120 | hypothetical protein | -0,586 | 0,1355 |
| TcCLB.505997.31 | ATP-dependent DEAD/H RNA helicase (pseudogene), puta | -0,5862 | 0,01523 |
| TcCLB.509695.50 | katanin, putative | -0,5865 | 0,003874 |
| TcCLB.504133.30 | tyrosine protein kinase, putative | -0,5866 | 4,776E-11 |
| TcCLB.511627.159 | hypothetical protein, conserved (fragment) | -0,5866 | 0,0001897 |
| TcCLB.510553.20 | trans-sialidase (pseudogene), putative | -0,5867 | 0,01247 |
| TcCLB.510259.12 | hypothetical protein, conserved | -0,5869 | 0,001686 |
| TcCLB.511361.40 | actin-related protein 2, putative | -0,587 | 2,112E-06 |
| TcCLB.503869.20 | hypothetical protein, conserved | -0,5874 | 0,00003639 |

| | | | |
|---|---|---|---|
| TcCLB.509569.150 | hypothetical protein, conserved | -0,5874 | 0,0007207 |
| TcCLB.511585.100 | hypothetical protein, conserved | -0,5875 | 0,0002986 |
| TcCLB.511517.160 | hypothetical protein, conserved | -0,5877 | 7,1E-10 |
| TcCLB.509417.9 | trans-sialidase, putative | -0,588 | 0,0000869 |
| TcCLB.420989.10 | protein phosphatase 2C, putative | -0,5882 | 0,007325 |
| TcCLB.511557.20 | hypothetical protein, conserved | -0,5883 | 0,00001072 |
| TcCLB.506659.39 | protein phosphatase 2C, putative | -0,5883 | 0,005614 |
| TcCLB.507035.110 | cAMP response protein 1, putative (CARP1) | -0,5884 | 1,091E-07 |
| TcCLB.507681.259 | Flagellar Member 1 (fragment) (FLAM1) | -0,5884 | 1,653E-06 |
| TcCLB.503755.30 | DNA polymerase kappa, putative | -0,5884 | 0,003531 |
| TcCLB.503583.50 | hypothetical protein, conserved | -0,5885 | 4,268E-06 |
| TcCLB.510403.71 | trans-sialidase (pseudogene), putative | -0,5888 | 6,177E-10 |
| TcCLB.506829.71 | hypothetical protein, conserved (pseudogene) | -0,5893 | 0,001663 |
| TcCLB.509669.14 | thioredoxin, putative | -0,5899 | 0,00002144 |
| TcCLB.504105.160 | splicing factor ptsr1 interacting protein, putative | -0,5902 | 1,491E-10 |
| TcCLB.505807.54 | hypothetical protein, conserved | -0,5902 | 4,023E-06 |
| TcCLB.506957.120 | hypothetical protein, conserved | -0,5904 | 1,671E-13 |
| TcCLB.510101.280 | kinesin, putative | -0,5911 | 6,313E-17 |
| TcCLB.511613.120 | mucin TcMUCII, putative | -0,5911 | 0,1076 |
| TcCLB.511163.30 | NGG1 interacting factor 3, putative | -0,5912 | 0,0001931 |
| TcCLB.511063.61 | hypothetical protein, conserved (pseudogene) | -0,5914 | 0,08566 |
| TcCLB.508543.40 | hypothetical protein, conserved | -0,5915 | 0,0006471 |
| TcCLB.506483.10 | Triglyceride lipase, putative | -0,5919 | 5,295E-06 |
| TcCLB.503721.20 | hypothetical protein, conserved | -0,5924 | 7,092E-07 |
| TcCLB.508385.50 | hypothetical protein, conserved | -0,5925 | 1,065E-06 |
| TcCLB.507669.14 | hypothetical protein, conserved | -0,5926 | 8,633E-09 |
| TcCLB.506673.69 | hypothetical protein, conserved | -0,5926 | 0,0001803 |
| TcCLB.510623.45 | surface protease GP63 (pseudogene), putative | -0,5927 | 0,01924 |
| TcCLB.507757.70 | dihydrolipoamide dehydrogenase, putative | -0,5931 | 0,00004568 |
| TcCLB.510483.234 | mucin TcMUC (pseudogene), putative | -0,5931 | 0,0508 |
| TcCLB.503787.30 | hypothetical protein | -0,5932 | 0,02499 |
| TcCLB.507087.14 | hypothetical protein, conserved | -0,5934 | 0,002375 |
| TcCLB.507029.50 | hypothetical protein, conserved | -0,5934 | 0,008263 |
| TcCLB.510013.180 | hypothetical protein | -0,5935 | 0,1174 |
| TcCLB.510329.200 | UDP-GlcNAc:PI a1-6 GlcNAc-transferase, putative | -0,5939 | 5,825E-13 |
| TcCLB.503887.50 | hypothetical protein, conserved | -0,5942 | 2,091E-07 |
| TcCLB.506475.113 | hypothetical protein, conserved | -0,5945 | 8,609E-08 |
| TcCLB.508647.10 | hypothetical protein, conserved | -0,5948 | 1,691E-08 |
| TcCLB.509823.4 | hypothetical protein, conserved | -0,5951 | 9,872E-06 |
| TcCLB.505987.70 | phosphatidylinositol 4-kinase, putative | -0,5952 | 1,445E-06 |
| TcCLB.510773.70 | dynein heavy chain, putative | -0,5953 | 6,889E-08 |
| TcCLB.506327.50 | hypothetical protein, conserved | -0,5956 | 8,669E-09 |
| TcCLB.506133.120 | syntaxin binding protein (pseudogene), putative | -0,5956 | 0,08545 |
| TcCLB.508799.50 | hypothetical protein | -0,5959 | 2,951E-07 |
| TcCLB.506559.148 | hypothetical protein, conserved | -0,5959 | 3,896E-06 |

| | | | |
|---|---|---|---|
| TcCLB.511731.20 | hypothetical protein, conserved | -0,5963 | 1,07E-10 |
| TcCLB.510779.20 | hypothetical protein (fragment) | -0,5967 | 0,001217 |
| TcCLB.507747.240 | hypothetical protein, conserved | -0,597 | 7,932E-10 |
| TcCLB.510361.209 | mucin-associated surface protein (MASP, pseudogene), pu | -0,5972 | 0,05857 |
| TcCLB.507943.50 | hypothetical protein, conserved | -0,598 | 0,00005875 |
| TcCLB.510607.75 | hypothetical protein, conserved (pseudogene) | -0,598 | 0,002689 |
| TcCLB.507089.130 | hypothetical protein, conserved | -0,5985 | 5,212E-08 |
| TcCLB.509799.120 | hypothetical protein, conserved | -0,5986 | 0,00000213 |
| TcCLB.510213.20 | hypothetical protein | -0,5986 | 0,149 |
| TcCLB.508637.19 | hypothetical protein, conserved | -0,5987 | 0,2378 |
| TcCLB.508443.10 | hypothetical protein, conserved | -0,5989 | 0,00000345 |
| TcCLB.507237.60 | surface protease GP63 (pseudogene), putative | -0,5989 | 0,003905 |
| TcCLB.510635.50 | zinc-binding phosphatase, putative | -0,599 | 4,097E-09 |
| TcCLB.511417.40 | DNA-directed RNA polymerase III, putative | -0,6 | 5,465E-09 |
| TcCLB.511545.190 | GPI transamidase component Tta1, putative (TTA1) | -0,6001 | 0,00002669 |
| TcCLB.507831.110 | pumilio/PUF RNA binding protein 2, putative (PUF2) | -0,6002 | 0,2135 |
| TcCLB.508699.30 | hypothetical protein, conserved | -0,6004 | 0,00008015 |
| TcCLB.511585.40 | DNA-damage inducible protein DDI1-like protein, putative | -0,6008 | 1,321E-08 |
| TcCLB.504397.9 | surface protease GP63, putative (fragment) | -0,6008 | 0,009588 |
| TcCLB.511777.10 | trans-sialidase, putative (fragment) | -0,6011 | 0,09336 |
| TcCLB.510663.40 | hypothetical protein, conserved | -0,6012 | 3,833E-09 |
| TcCLB.510421.280 | hypothetical protein, conserved | -0,6012 | 4,516E-09 |
| TcCLB.506567.50 | hypothetical protein, conserved | -0,6012 | 7,622E-07 |
| TcCLB.506241.80 | hypothetical protein, conserved | -0,6014 | 0,00002274 |
| TcCLB.506315.80 | hypothetical protein, conserved | -0,6016 | 1,784E-07 |
| TcCLB.503957.25 | trans-sialidase (pseudogene), putative | -0,6017 | 0,007033 |
| TcCLB.507941.110 | hypothetical protein, conserved | -0,602 | 7,254E-08 |
| TcCLB.507709.10 | hypothetical protein, conserved | -0,6021 | 0,00009461 |
| TcCLB.509161.100 | hypothetical protein, conserved | -0,6026 | 4,297E-08 |
| TcCLB.510593.30 | hypothetical protein, conserved | -0,6026 | 1,277E-07 |
| TcCLB.510031.50 | ATP-dependent RNA helicase, putative | -0,6028 | 1,517E-07 |
| TcCLB.509979.290 | trans-sialidase (pseudogene), putative | -0,6028 | 0,0002746 |
| TcCLB.509521.70 | hypothetical protein | -0,6028 | 0,1585 |
| TcCLB.508775.35 | retrotransposon hot spot protein (RHS, pseudogene), puta | -0,6032 | 0,05118 |
| TcCLB.509217.170 | hypothetical protein | -0,6035 | 0,02969 |
| TcCLB.510187.230 | formin, putative | -0,6036 | 1,877E-09 |
| TcCLB.508873.454 | mucin TcMUC (pseudogene), putative | -0,6036 | 0,103 |
| TcCLB.506811.50 | hypothetical protein, conserved | -0,6037 | 0,03707 |
| TcCLB.506589.100 | hypothetical protein | -0,6038 | 0,09025 |
| TcCLB.506789.249 | bilobe protein, conserved | -0,604 | 3,566E-12 |
| TcCLB.508387.110 | hypothetical protein, conserved | -0,6043 | 2,171E-08 |
| TcCLB.504741.90 | hypothetical protein, conserved | -0,6043 | 0,00005939 |
| TcCLB.511501.40 | hypothetical protein, conserved | -0,605 | 0,00004273 |
| TcCLB.509559.40 | trans-sialidase (pseudogene), putative | -0,6053 | 0,0001437 |
| TcCLB.507681.30 | hypothetical protein, conserved | -0,6056 | 2,227E-11 |

| | | | |
|---|---|---|---|
| TcCLB.484949.9 | AGC/RSK family serine/threonine kinase, putative (fragm | -0,6056 | 0,002823 |
| TcCLB.505977.40 | inositol/phosphatidylinositol phosphatase (pseudogene), p | -0,606 | 7,744E-16 |
| TcCLB.508777.60 | hypothetical protein, conserved | -0,6063 | 8,834E-06 |
| TcCLB.509885.50 | hypothetical protein, conserved | -0,6064 | 1,418E-07 |
| TcCLB.507641.200 | hypothetical protein, conserved | -0,6065 | 0,00009337 |
| TcCLB.509053.100 | protein kinase, putative | -0,6067 | 8,794E-10 |
| TcCLB.508831.30 | hypothetical protein, conserved | -0,6069 | 0,00001238 |
| TcCLB.503757.40 | Casein kinase II subunit beta, putative | -0,6076 | 2,766E-07 |
| TcCLB.510463.180 | trans-sialidase (pseudogene), putative | -0,608 | 0,0405 |
| TcCLB.505943.79 | retrotransposon hot spot protein (RHS, pseudogene), puta | -0,6087 | 0,1523 |
| TcCLB.511727.230 | Repressor of differentiation kinase 1, putative (RDK1.2) | -0,6089 | 4,853E-11 |
| TcCLB.509647.60 | hypothetical protein, conserved | -0,6092 | 4,123E-12 |
| TcCLB.507711.140 | hypothetical protein, conserved | -0,6093 | 0,00006541 |
| TcCLB.506339.60 | trans-sialidase (pseudogene), putative | -0,6093 | 0,006226 |
| TcCLB.506537.210 | trans-sialidase (pseudogene), putative | -0,6096 | 0,00001145 |
| TcCLB.509103.4 | protein kinase, putative | -0,6106 | 0,01949 |
| TcCLB.503579.110 | pseudouridylate synthase I, putative | -0,6109 | 1,118E-06 |
| TcCLB.505163.10 | hypothetical protein, conserved | -0,6111 | 0,001755 |
| TcCLB.511921.71 | hypothetical protein, conserved (pseudogene) | -0,6112 | 0,008156 |
| TcCLB.508233.60 | heat shock protein 100 (clp protein), putative | -0,6115 | 0,001122 |
| TcCLB.509151.90 | hypothetical protein, conserved | -0,612 | 1,028E-08 |
| TcCLB.508409.20 | hypothetical protein, conserved | -0,6124 | 2,486E-08 |
| TcCLB.506525.40 | syntaxin 5, putative | -0,6126 | 1,863E-09 |
| TcCLB.506269.50 | mucin-associated surface protein (MASP, pseudogene), pu | -0,6128 | 0,02892 |
| TcCLB.510175.110 | trans-sialidase (pseudogene), putative | -0,6131 | 0,0002155 |
| TcCLB.509213.130 | autophagocytosis associated protein, putative | -0,6132 | 1,587E-06 |
| TcCLB.508637.160 | dual specificity protein phosphatase, putative | -0,6134 | 2,588E-06 |
| TcCLB.504155.20 | trans-sialidase, Group V, putative | -0,6134 | 0,004763 |
| TcCLB.508375.5 | elongation factor 1-gamma (EF-1-gamma, pseudogene), p | -0,6137 | 0,001809 |
| TcCLB.509353.10 | E1-like ubiquitin-activating enzyme, putative (UBA5) | -0,6141 | 0,001064 |
| TcCLB.503693.29 | hypothetical protein | -0,6142 | 2,456E-08 |
| TcCLB.509097.10 | hypothetical protein | -0,6142 | 0,2068 |
| TcCLB.506957.140 | phosphatidylinositol-4-phosphate 5-kinase-like, putative | -0,6145 | 1,917E-11 |
| TcCLB.508277.10 | condensin subunit 1, putative | -0,615 | 8,242E-12 |
| TcCLB.508213.9 | regulator of chromosome condensation, putative | -0,615 | 8,581E-12 |
| TcCLB.504035.149 | kinesin, putative | -0,615 | 1,326E-08 |
| TcCLB.509777.60 | ADG2, putative | -0,6152 | 4,372E-08 |
| TcCLB.421621.35 | surface protease GP63 (pseudogene), putative | -0,6153 | 0,1654 |
| TcCLB.508909.80 | hypothetical protein | -0,6154 | 5,927E-06 |
| TcCLB.506763.270 | hypothetical protein, conserved (pseudogene) | -0,6155 | 0,005732 |
| TcCLB.421555.9 | mucin-associated surface protein (MASP), putative | -0,6157 | 0,0003684 |
| TcCLB.509945.10 | ERAD-associated E3 ubiquitin-protein ligase HRD1, putativ | -0,6158 | 1,852E-12 |
| TcCLB.504005.10 | endonuclease III, putative | -0,6159 | 0,00001412 |
| TcCLB.510351.80 | hypothetical protein, conserved | -0,6159 | 0,00001765 |
| TcCLB.511215.90 | hypothetical protein, conserved | -0,6162 | 0,01849 |

| | | | |
|---|---|---|---|
| TcCLB.503981.60 | hypothetical protein, conserved | -0,6165 | 1,125E-09 |
| TcCLB.511689.10 | hypothetical protein | -0,6166 | 0,00003356 |
| TcCLB.511929.40 | trans-sialidase (pseudogene), putative | -0,6168 | 0,01471 |
| TcCLB.511847.10 | calpain-like cysteine peptidase, putative | -0,6173 | 5,773E-10 |
| TcCLB.504123.40 | hypothetical protein, conserved | -0,6174 | 3,032E-06 |
| TcCLB.508955.10 | syntaxin, putative | -0,6174 | 0,00005261 |
| TcCLB.507047.170 | hypothetical protein, conserved | -0,6177 | 0,001018 |
| TcCLB.508359.70 | hypothetical protein | -0,6177 | 0,05642 |
| TcCLB.506139.140 | trans-sialidase (pseudogene), putative | -0,6178 | 0,02389 |
| TcCLB.511043.10 | hypothetical protein, conserved | -0,618 | 5,329E-15 |
| TcCLB.414243.20 | hypothetical protein, conserved | -0,618 | 5,995E-07 |
| TcCLB.510739.10 | hypothetical protein, conserved | -0,6181 | 0,00000924 |
| TcCLB.508175.250 | hypothetical protein, conserved | -0,6182 | 6,478E-14 |
| TcCLB.504051.9 | hypothetical protein, conserved | -0,6183 | 6,509E-07 |
| TcCLB.511717.174 | hypothetical protein, conserved | -0,6186 | 5,033E-09 |
| TcCLB.510697.55 | mucin TcMUCII (pseudogene), putative | -0,6187 | 0,02961 |
| TcCLB.511281.20 | transporter, putative | -0,6188 | 3,349E-09 |
| TcCLB.509167.80 | hypothetical protein, conserved | -0,619 | 2,483E-11 |
| TcCLB.506945.140 | hypothetical protein, conserved (pseudogene) | -0,619 | 8,745E-09 |
| TcCLB.508717.53 | hypothetical protein, conserved | -0,6191 | 3,333E-12 |
| TcCLB.506579.100 | hypothetical protein, conserved | -0,6191 | 9,379E-08 |
| TcCLB.511543.10 | hypothetical protein, conserved | -0,6191 | 0,00000256 |
| TcCLB.506655.7 | phospholipid-transporting ATPase, putative (fragment) | -0,6192 | 5,837E-06 |
| TcCLB.504249.65 | mucin TcMUCII, putative | -0,6192 | 0,007905 |
| TcCLB.490989.10 | trans-sialidase, putative | -0,6195 | 0,008561 |
| TcCLB.505941.49 | hypothetical protein, conserved | -0,6197 | 6,409E-06 |
| TcCLB.506993.130 | mucin-associated surface protein (MASP, pseudogene), pu | -0,6197 | 0,0002581 |
| TcCLB.506763.210 | hypothetical protein | -0,6199 | 0,02646 |
| TcCLB.511445.110 | hypothetical protein, conserved | -0,62 | 0,00002165 |
| TcCLB.507057.60 | hypothetical protein, conserved | -0,62 | 0,0000368 |
| TcCLB.511245.240 | serine/threonine protein kinase, putative | -0,62 | 0,0004569 |
| TcCLB.507771.114 | hypothetical protein, conserved | -0,62 | 0,0009236 |
| TcCLB.506405.40 | hypothetical protein, conserved | -0,6203 | 1,108E-07 |
| TcCLB.510089.50 | hypothetical protein, conserved | -0,6204 | 3,544E-10 |
| TcCLB.508319.79 | hypothetical protein, conserved | -0,6205 | 0,04869 |
| TcCLB.508559.20 | trans-sialidase (pseudogene), putative | -0,6208 | 3,857E-08 |
| TcCLB.511277.140 | serine/threonine protein kinase, putative | -0,6208 | 1,214E-07 |
| TcCLB.504157.40 | DNA replication licensing factor MCM3, putative | -0,6208 | 6,262E-07 |
| TcCLB.507489.9 | dispersed gene family protein 1 (DGF-1), putative (fragme | -0,621 | 0,1777 |
| TcCLB.404001.10 | hypothetical protein, conserved | -0,6213 | 3,884E-11 |
| TcCLB.479481.5 | elongation factor 1-gamma (EF-1-gamma, pseudogene), | -0,6213 | 0,08008 |
| TcCLB.508433.80 | hypothetical protein | -0,6214 | 0,1503 |
| TcCLB.506269.70 | mucin TcMUCII, putative | -0,6215 | 0,02718 |
| TcCLB.509669.150 | hypothetical protein, conserved | -0,6217 | 3,546E-08 |
| TcCLB.510403.30 | trans-sialidase, Group II, putative | -0,6219 | 0,02676 |

| | | | |
|---|---|---|---|
| TcCLB.509907.60 | trans-sialidase (pseudogene), putative | -0,622 | 0,005717 |
| TcCLB.479517.30 | hypothetical protein | -0,6222 | 0,2083 |
| TcCLB.504247.79 | transmembrane protein, conserved (fragment) | -0,6223 | 1,531E-06 |
| TcCLB.509551.120 | hypothetical protein | -0,6228 | 0,0004007 |
| TcCLB.511299.10 | Triglyceride lipase, putative (fragment) | -0,623 | 0,004323 |
| TcCLB.507747.210 | hypothetical protein, conserved | -0,6232 | 5,523E-08 |
| TcCLB.507683.30 | hypothetical protein, conserved | -0,6239 | 2,668E-12 |
| TcCLB.510441.10 | hypothetical protein | -0,624 | 0,006724 |
| TcCLB.504153.100 | Tubulin cofactor C domain-containing protein 1, putative ( | -0,6241 | 1,768E-10 |
| TcCLB.508233.70 | ATP-dependent protease binding subunit (pseudogene), pu | -0,6247 | 0,0001153 |
| TcCLB.507357.35 | hypothetical protein | -0,6247 | 0,0807 |
| TcCLB.506965.60 | elongation factor 1-gamma (EF-1-gamma, pseudogene), | -0,6252 | 0,1952 |
| TcCLB.506349.83 | retrotransposon hot spot protein (RHS, pseudogene), puta | -0,6253 | 0,0001042 |
| TcCLB.503995.10 | Present in the outer mitochondrial membrane proteome 1 | -0,6254 | 0,00001589 |
| TcCLB.509627.10 | bilobe protein, conserved | -0,6258 | 1,155E-14 |
| TcCLB.508987.30 | serine/threonine-protein kinase, putative | -0,6258 | 2,523E-13 |
| TcCLB.503881.30 | retrotransposon hot spot (RHS) protein, putative | -0,6258 | 0,0004042 |
| TcCLB.504109.60 | hypothetical protein, conserved | -0,626 | 2,311E-12 |
| TcCLB.506525.30 | hypothetical protein, conserved | -0,6261 | 1,1E-10 |
| TcCLB.508277.300 | hypothetical protein, conserved | -0,6261 | 3,754E-07 |
| TcCLB.507615.70 | deoxyhypusine hydroxylase, putative | -0,6265 | 0,0000242 |
| TcCLB.506931.60 | hypothetical protein | -0,6266 | 1,432E-06 |
| TcCLB.506155.30 | kinesin-like protein, putative | -0,6267 | 1,29E-09 |
| TcCLB.507307.30 | peptidylprolyl isomerase (pseudogene), putative | -0,6268 | 0,1042 |
| TcCLB.508153.1010 | hypothetical protein, conserved | -0,6271 | 1,687E-10 |
| TcCLB.507677.20 | hypothetical protein, conserved | -0,6272 | 1,007E-06 |
| TcCLB.510465.60 | hypothetical protein, conserved (pseudogene) | -0,6273 | 0,003314 |
| TcCLB.511127.220 | cdc2-related kinase 12 (CRK12) | -0,628 | 3,088E-11 |
| TcCLB.421621.20 | syntaxin binding protein (pseudogene), putative | -0,6283 | 0,01766 |
| TcCLB.507723.120 | U1A small nuclear ribonucleoprotein, putative | -0,6284 | 0,0002699 |
| TcCLB.508601.70 | hypothetical protein, conserved | -0,6286 | 6,22E-07 |
| TcCLB.506993.190 | hypothetical protein | -0,6287 | 0,1334 |
| TcCLB.509161.120 | hypothetical protein, conserved | -0,6288 | 1,68E-09 |
| TcCLB.508953.28 | hypothetical protein, conserved | -0,6293 | 0,00001755 |
| TcCLB.507765.100 | hypothetical protein, conserved | -0,6296 | 3,902E-07 |
| TcCLB.511797.260 | mucin-associated surface protein (MASP, pseudogene), pu | -0,6298 | 0,06508 |
| TcCLB.507241.10 | protein phosphatase 2A regulatory subunit, putative | -0,6299 | 2,358E-11 |
| TcCLB.508989.100 | hypothetical protein, conserved | -0,6302 | 5,947E-08 |
| TcCLB.507787.120 | hypothetical protein, conserved | -0,6305 | 0,00003142 |
| TcCLB.507025.50 | RNA-binding protein, putative | -0,6308 | 0,0007016 |
| TcCLB.511557.40 | hypothetical protein, conserved | -0,6309 | 3,782E-08 |
| TcCLB.506179.40 | hypothetical protein, conserved | -0,6309 | 1,968E-07 |
| TcCLB.504147.170 | gamma-tubulin complex subunit, putative | -0,6311 | 3,913E-16 |
| TcCLB.511529.40 | hypothetical protein, conserved | -0,6311 | 9,621E-08 |
| TcCLB.510475.51 | mucin-associated surface protein (MASP, pseudogene), pu | -0,6313 | 0,1328 |

| | | | |
|---|---|---|---|
| TcCLB.510885.130 | hypothetical protein, conserved | -0,6318 | 4,142E-08 |
| TcCLB.504215.30 | hypothetical protein, conserved | -0,6319 | 3,358E-08 |
| TcCLB.509071.10 | hypothetical protein, conserved (pseudogene) | -0,6321 | 0,001306 |
| TcCLB.507941.130 | hypothetical protein, conserved | -0,6322 | 3,97E-07 |
| TcCLB.506779.150 | histone H2B variant, putative | -0,6323 | 1,587E-09 |
| TcCLB.503823.30 | hypothetical protein, conserved | -0,6323 | 3,137E-08 |
| TcCLB.510713.11 | hypothetical protein, conserved (pseudogene) | -0,6324 | 0,003265 |
| TcCLB.510329.90 | C-8 sterol isomerase, putative | -0,6326 | 7,317E-08 |
| TcCLB.503417.33 | hypothetical protein | -0,6326 | 0,1133 |
| TcCLB.509571.40 | hypothetical protein, conserved | -0,6328 | 1,7E-12 |
| TcCLB.507001.140 | hypothetical protein, conserved | -0,6328 | 8,667E-08 |
| TcCLB.510463.211 | trans-sialidase (pseudogene), putative | -0,6329 | 0,03812 |
| TcCLB.507881.110 | trans-sialidase (pseudogene), putative | -0,6331 | 4,082E-08 |
| TcCLB.511127.270 | ankyrin repeat protein, putative | -0,6331 | 8,951E-07 |
| TcCLB.508153.190 | kinesin, putative | -0,6335 | 7,985E-12 |
| TcCLB.507765.140 | hypothetical protein, conserved | -0,6339 | 1,73E-12 |
| TcCLB.511603.141 | syntaxin binding protein (pseudogene), putative | -0,6339 | 0,07771 |
| TcCLB.510885.50 | hypothetical protein, conserved | -0,6341 | 6,454E-06 |
| TcCLB.510659.4 | hypothetical protein, conserved | -0,6342 | 4,719E-06 |
| TcCLB.507547.9 | mucin TcMUCI, putative | -0,6346 | 6,579E-06 |
| TcCLB.511003.130 | hypothetical protein, conserved | -0,635 | 0,00001967 |
| TcCLB.509735.140 | trans-sialidase (pseudogene), putative | -0,6352 | 1,888E-06 |
| TcCLB.508907.10 | hypothetical protein, conserved | -0,6352 | 5,098E-06 |
| TcCLB.508355.230 | hypothetical protein | -0,6354 | 0,0001489 |
| TcCLB.508407.4 | hypothetical protein, conserved | -0,6355 | 0,01167 |
| TcCLB.510885.160 | hypothetical protein, conserved | -0,6356 | 4,603E-16 |
| TcCLB.510293.70 | ribonuclease H, putative | -0,6357 | 0,00006494 |
| TcCLB.508859.10 | hypothetical protein, conserved | -0,6359 | 7,561E-07 |
| TcCLB.511423.160 | RWD domain-containing protein | -0,6359 | 5,908E-06 |
| TcCLB.507895.10 | trans-sialidase (pseudogene), putative | -0,636 | 0,1417 |
| TcCLB.510223.5 | guanine nucleotide exchange factor subunit GCD2 | -0,6366 | 0,03896 |
| TcCLB.508707.140 | phosphatidate cytidylyltransferase-like protein, putative | -0,6371 | 5,725E-08 |
| TcCLB.509429.190 | hypothetical protein, conserved | -0,6375 | 4,67E-14 |
| TcCLB.508075.14 | hypothetical protein, conserved | -0,6379 | 6,286E-08 |
| TcCLB.511529.150 | hypothetical protein, conserved | -0,6382 | 1,733E-10 |
| TcCLB.508985.10 | mucin-associated surface protein (MASP, pseudogene), pu | -0,6382 | 0,000433 |
| TcCLB.511283.130 | MEKK-related kinase 1, putative (MRK1) | -0,6386 | 3,906E-11 |
| TcCLB.503455.10 | chaperone DNAJ protein, putative | -0,6387 | 4,093E-07 |
| TcCLB.508695.20 | phosphatidylinositol 4-kinase, putative | -0,6389 | 2,827E-10 |
| TcCLB.511637.9 | zinc carboxypeptidase, putative | -0,639 | 2,142E-12 |
| TcCLB.509007.14 | hypothetical protein, conserved | -0,6392 | 1,31E-08 |
| TcCLB.510131.20 | hypothetical protein, conserved | -0,6395 | 0,00007864 |
| TcCLB.505555.61 | cytoplasmic l-asparaginase i-like protein, putative | -0,6396 | 0,0002665 |
| TcCLB.511827.125 | trans-sialidase (pseudogene), putative | -0,6397 | 0,0009959 |
| TcCLB.510593.10 | hypothetical protein, conserved | -0,6397 | 0,001778 |

| | | | |
|---|---|---|---|
| TcCLB.507027.59 | pre-mRNA cleavage complex II Clp1 protein, putative | -0,6401 | 2,091E-08 |
| TcCLB.507093.330 | hypothetical protein, conserved | -0,6401 | 0,0000126 |
| TcCLB.511173.190 | trans-sialidase (pseudogene), putative | -0,6401 | 0,01074 |
| TcCLB.511355.50 | glycerol uptake protein, putative | -0,6402 | 4,351E-09 |
| TcCLB.506691.50 | hypothetical protein, conserved | -0,6402 | 1,101E-08 |
| TcCLB.507803.49 | histone deacetylase, putative | -0,6402 | 0,0007141 |
| TcCLB.509795.50 | hypothetical protein, conserved | -0,6407 | 5,849E-14 |
| TcCLB.507939.30 | hypothetical protein, conserved | -0,6411 | 1,555E-08 |
| TcCLB.511281.30 | RNA-binding protein 27, putative (RBP27) | -0,6415 | 3,258E-09 |
| TcCLB.509991.90 | RNA-editing nuclease 1 (KREN1) | -0,6416 | 1,627E-09 |
| TcCLB.511711.110 | TLR4 regulator and MIR-interacting MSAP, putative (ESP1 | -0,6416 | 2,392E-08 |
| TcCLB.510889.190 | hypothetical protein, conserved | -0,6417 | 4,094E-09 |
| TcCLB.510115.10 | trans-sialidase (pseudogene), putative | -0,6417 | 0,00002102 |
| TcCLB.503449.40 | hypothetical protein, conserved | -0,6422 | 2,591E-06 |
| TcCLB.504431.40 | hypothetical protein, conserved | -0,6423 | 6,17E-09 |
| TcCLB.511277.560 | synaptotagmin, putative | -0,6425 | 2,583E-07 |
| TcCLB.506211.259 | hypothetical protein, conserved | -0,6427 | 5,056E-06 |
| TcCLB.505063.30 | mucin-associated surface protein (MASP, pseudogene), pu | -0,6428 | 2,417E-07 |
| TcCLB.506955.226 | elongation factor 1-gamma (EF-1-gamma, pseudogene), | -0,6428 | 0,1508 |
| TcCLB.504069.4 | hypothetical protein, conserved (fragment) | -0,643 | 2,246E-07 |
| TcCLB.508221.130 | mucin TcMUCII (pseudogene), putative | -0,643 | 0,05648 |
| TcCLB.504147.310 | hypothetical protein, conserved | -0,6431 | 7,38E-12 |
| TcCLB.510407.70 | hypothetical protein | -0,6434 | 4,212E-10 |
| TcCLB.508221.920 | mucin-associated surface protein (MASP, pseudogene), pu | -0,6434 | 0,0005788 |
| TcCLB.503637.10 | hypothetical protein, conserved | -0,6435 | 1,807E-06 |
| TcCLB.511411.8 | hypothetical protein, conserved (fragment) | -0,6435 | 0,0001759 |
| TcCLB.508293.27 | mucin-associated surface protein (MASP, pseudogene), pu | -0,6438 | 0,08708 |
| TcCLB.504207.29 | hypothetical protein, conserved | -0,644 | 2,578E-07 |
| TcCLB.509999.10 | hypothetical protein, conserved | -0,644 | 3,926E-06 |
| TcCLB.506605.139 | hypothetical protein, conserved | -0,644 | 8,634E-06 |
| TcCLB.506131.50 | hypothetical protein | -0,6444 | 0,2073 |
| TcCLB.506727.110 | hypothetical protein, conserved | -0,6445 | 1,202E-09 |
| TcCLB.511339.30 | UDP-Gal or UDP-GlcNAc-dependent glycosyltransferase, p | -0,6445 | 0,001287 |
| TcCLB.510131.50 | DNA polymerase kappa, putative | -0,6448 | 1,863E-07 |
| TcCLB.472833.10 | hypothetical protein, conserved | -0,6451 | 0,06201 |
| TcCLB.508601.40 | hypothetical protein, conserved | -0,6452 | 1,456E-06 |
| TcCLB.508231.60 | ubiquitin-protein ligase, putative | -0,6454 | 3,237E-21 |
| TcCLB.507967.10 | hypothetical protein | -0,6454 | 1,014E-09 |
| TcCLB.510105.140 | hypothetical protein, conserved (pseudogene) | -0,6459 | 4,724E-09 |
| TcCLB.504787.40 | trans-sialidase, Group V, putative | -0,6459 | 0,03724 |
| TcCLB.506725.50 | hypothetical protein, conserved | -0,6461 | 2,563E-08 |
| TcCLB.508989.20 | carrier protein, putative | -0,6461 | 4,778E-08 |
| TcCLB.504093.4 | hypothetical protein, conserved | -0,6461 | 0,00007783 |
| TcCLB.511837.138 | RNA-binding protein, putative | -0,6466 | 3,214E-13 |
| TcCLB.508979.90 | hypothetical protein | -0,6466 | 2,575E-06 |

| | | | |
|---|---|---|---|
| TcCLB.511015.30 | Trichohyalin, putative (fragment) | -0,6469 | 2,397E-13 |
| TcCLB.508613.20 | hypothetical protein, conserved | -0,6469 | 1,913E-07 |
| TcCLB.510125.10 | pumilio/PUF RNA binding protein 6, putative (PUF6) | -0,6472 | 4,557E-10 |
| TcCLB.504625.20 | hypothetical protein, conserved | -0,6472 | 5,624E-08 |
| TcCLB.506251.50 | Cell division control protein 48, putative | -0,6472 | 2,324E-06 |
| TcCLB.511213.30 | surface protease GP63 (pseudogene), putative | -0,6472 | 0,0002627 |
| TcCLB.503797.10 | hypothetical protein, conserved | -0,6475 | 0,00008113 |
| TcCLB.506559.524 | protein phosphatase 2C, putative | -0,6476 | 2,882E-08 |
| TcCLB.511383.10 | hypothetical protein, conserved | -0,6476 | 0,0001473 |
| TcCLB.508871.51 | serine-alanine-and proline-rich protein (pseudogene), put | -0,6478 | 0,0004585 |
| TcCLB.508533.20 | hypothetical protein, conserved | -0,6479 | 2,348E-08 |
| TcCLB.510993.19 | hypothetical protein | -0,648 | 0,0004359 |
| TcCLB.506363.140 | hypothetical protein, conserved | -0,6481 | 3,661E-06 |
| TcCLB.507077.60 | surface protease GP63, putative | -0,6482 | 5,831E-07 |
| TcCLB.503395.59 | hypothetical protein, conserved (fragment) | -0,6482 | 0,002288 |
| TcCLB.507389.20 | hypothetical protein, conserved | -0,6485 | 0,00004623 |
| TcCLB.510187.404 | dolichyl-P-Man:GDP-Man5GlcNAc2-PP-dolichyl alpha-1,3- | -0,6486 | 3,696E-08 |
| TcCLB.508777.10 | hypothetical protein, conserved | -0,6488 | 1,047E-11 |
| TcCLB.506755.130 | hypothetical protein, conserved | -0,649 | 0,0000511 |
| TcCLB.511167.110 | hypothetical protein, conserved | -0,6493 | 6,366E-10 |
| TcCLB.427743.9 | translation initiation factor eIF2B delta subunit, putative ( | -0,6494 | 0,0001759 |
| TcCLB.508777.20 | hypothetical protein, conserved | -0,6496 | 8,034E-06 |
| TcCLB.510955.60 | hypothetical protein, conserved | -0,6497 | 3,335E-09 |
| TcCLB.506447.19 | hypothetical protein, conserved | -0,6497 | 6,161E-09 |
| TcCLB.508933.50 | hypothetical protein, conserved | -0,6499 | 1,238E-07 |
| TcCLB.509965.260 | protein kinase, putative | -0,6501 | 4,929E-11 |
| TcCLB.511727.260 | cyclin 9, putative (CYC9) | -0,6506 | 6,934E-06 |
| TcCLB.510823.80 | cytoplasmic l-asparaginase i-like protein, putative | -0,6509 | 4,722E-06 |
| TcCLB.508887.5 | trans-sialidase, putative | -0,651 | 0,1488 |
| TcCLB.504069.140 | trans-sialidase (pseudogene), putative | -0,6511 | 0,00001239 |
| TcCLB.506289.240 | hypothetical protein | -0,6511 | 0,03441 |
| TcCLB.507547.50 | hypothetical protein, conserved | -0,6512 | 2,203E-10 |
| TcCLB.503417.40 | hypothetical protein | -0,6512 | 0,08241 |
| TcCLB.510731.30 | hypothetical protein, conserved | -0,6514 | 3,773E-11 |
| TcCLB.511801.20 | hypothetical protein, conserved | -0,6515 | 6,081E-08 |
| TcCLB.503829.70 | hypothetical protein, conserved | -0,6516 | 2,353E-12 |
| TcCLB.508731.9 | hypothetical protein, conserved (fragment) | -0,6516 | 6,758E-09 |
| TcCLB.508061.130 | trans-sialidase (pseudogene), putative | -0,6519 | 1,047E-12 |
| TcCLB.449247.4 | serine/threonine protein phosphatase, putative (fragment | -0,6522 | 1,217E-08 |
| TcCLB.511151.39 | hypothetical protein, conserved | -0,6523 | 7,411E-08 |
| TcCLB.506955.130 | trans-sialidase (pseudogene), putative | -0,6525 | 0,00216 |
| TcCLB.506751.148 | mucin TcMUCII (pseudogene), putative | -0,6528 | 0,1158 |
| TcCLB.508315.5 | mucin TcMUCII, putative (fragment) | -0,6529 | 0,1012 |
| TcCLB.511537.24 | hypothetical protein, conserved | -0,6532 | 1,145E-09 |
| TcCLB.506673.20 | hypothetical protein, conserved | -0,6535 | 0,0002367 |

| | | | |
|---|---|---|---|
| TcCLB.510057.20 | hypothetical protein, conserved | -0,6538 | 0,000197 |
| TcCLB.509399.160 | CCR4-NOT transcription complex subunit 2, putative | -0,6541 | 2,412E-11 |
| TcCLB.510407.80 | hypothetical protein, conserved | -0,6542 | 9,865E-14 |
| TcCLB.510021.185 | hypothetical protein | -0,6545 | 0,05106 |
| TcCLB.398345.10 | hypothetical protein, conserved | -0,655 | 8,284E-10 |
| TcCLB.506813.190 | trans-sialidase, Group VI, putative | -0,6553 | 0,002303 |
| TcCLB.506617.70 | variant-surface-glycoprotein phospholipase C (GPI-PLC) | -0,6554 | 7,329E-06 |
| TcCLB.510443.30 | hypothetical protein, conserved | -0,6558 | 1,78E-10 |
| TcCLB.511655.60 | hypothetical protein, conserved | -0,656 | 0,0001028 |
| TcCLB.511867.30 | hypothetical protein, conserved | -0,6563 | 1,196E-06 |
| TcCLB.511127.250 | hypothetical protein, conserved | -0,6564 | 8,156E-08 |
| TcCLB.503471.19 | expression site-associated gene (ESAG-like) protein, puta | -0,6567 | 0,00001092 |
| TcCLB.510889.300 | hypothetical protein | -0,657 | 1,181E-06 |
| TcCLB.510679.60 | lipase, putative | -0,657 | 0,0008823 |
| TcCLB.511839.10 | mucin TcMUCII, putative | -0,6573 | 0,0295 |
| TcCLB.510377.475 | hypothetical protein, conserved (pseudogene) | -0,6573 | 0,1969 |
| TcCLB.510431.310 | hypothetical protein, conserved | -0,6574 | 0,0008291 |
| TcCLB.511521.27 | hypothetical protein, conserved | -0,6575 | 8,921E-07 |
| TcCLB.509437.106 | trans-sialidase (pseudogene), putative | -0,6578 | 0,03034 |
| TcCLB.482369.20 | hypothetical protein, conserved | -0,6581 | 9,317E-16 |
| TcCLB.511409.40 | hypothetical protein, conserved | -0,6584 | 9,728E-11 |
| TcCLB.510385.5 | hypothetical protein, conserved (pseudogene) | -0,6588 | 0,00111 |
| TcCLB.508863.20 | hypothetical protein, conserved | -0,6593 | 0,00000136 |
| TcCLB.508257.130 | hypothetical protein, conserved | -0,6594 | 5,303E-08 |
| TcCLB.510969.10 | hypothetical protein, conserved | -0,6596 | 8,43E-08 |
| TcCLB.511383.30 | U1 small nuclear ribonucleoprotein, putative | -0,6598 | 2,363E-07 |
| TcCLB.511409.70 | hypothetical protein, conserved | -0,66 | 1,008E-09 |
| TcCLB.509699.210 | mucin-associated surface protein (MASP), putative | -0,66 | 0,004214 |
| TcCLB.482369.10 | protein phosphatase 2A regulatory subunit, putative | -0,6601 | 1,548E-12 |
| TcCLB.508387.150 | hypothetical protein | -0,6605 | 1,339E-08 |
| TcCLB.503999.20 | Kinesin-13 5, putative (KIN13-5) | -0,6607 | 1,225E-14 |
| TcCLB.506829.50 | hypothetical protein, conserved | -0,6613 | 1,999E-08 |
| TcCLB.510609.110 | ATP-dependent RNA helicase, putative | -0,6613 | 8,733E-08 |
| TcCLB.507949.140 | ribosomal RNA-processing protein 8, putative (RRP8) | -0,6623 | 1,558E-10 |
| TcCLB.511625.140 | mucin-associated surface protein (MASP, pseudogene), pu | -0,6623 | 0,00001785 |
| TcCLB.509791.150 | choline/ethanolamine phosphotransferase, putative (CEPT | -0,6626 | 3,233E-13 |
| TcCLB.511127.210 | hypothetical protein, conserved | -0,6628 | 0,00001767 |
| TcCLB.509733.180 | adenylate kinase, putative | -0,6628 | 0,00007076 |
| TcCLB.510431.140 | dual-specificity protein kinase, putative | -0,6631 | 6,625E-15 |
| TcCLB.510955.50 | hypothetical protein, conserved | -0,6631 | 0,00002797 |
| TcCLB.509779.60 | dual-specificity protein phosphatase, putative | -0,6636 | 0,05555 |
| TcCLB.510667.14 | membrane transporter protein, putative | -0,6638 | 6,485E-10 |
| TcCLB.510955.30 | hypothetical protein, conserved | -0,6639 | 5,492E-11 |
| TcCLB.510889.150 | hypothetical protein, conserved | -0,664 | 0,00002046 |
| TcCLB.507669.139 | hypothetical protein, conserved | -0,6641 | 6,278E-07 |

| | | | |
|---|---|---|---|
| TcCLB.511331.10 | calpain cysteine peptidase, putative (fragment) | -0,6642 | 0,03996 |
| TcCLB.508313.60 | hypothetical protein | -0,6642 | 0,2102 |
| TcCLB.511435.40 | hypothetical protein, conserved | -0,6644 | 3,438E-13 |
| TcCLB.509647.160 | SKP1-like protein | -0,6644 | 2,523E-08 |
| TcCLB.504103.10 | hypothetical protein, conserved | -0,6645 | 9,998E-06 |
| TcCLB.510147.9 | hypothetical protein, conserved | -0,6645 | 0,0001244 |
| TcCLB.511127.280 | hypothetical protein, conserved | -0,6657 | 0,00000305 |
| TcCLB.506525.140 | MEKK-related kinase 1, putative (MRK1) | -0,666 | 9,627E-14 |
| TcCLB.509755.80 | mucin-associated surface protein (MASP, pseudogene), pu | -0,6663 | 0,1252 |
| TcCLB.503967.4 | hypothetical protein, conserved | -0,6664 | 0,001429 |
| TcCLB.511727.210 | Repressor of differentiation kinase 1, putative (RDK1.1) | -0,6666 | 1,31E-16 |
| TcCLB.503827.80 | syntaxin binding protein (pseudogene), putative | -0,6671 | 0,004052 |
| TcCLB.503829.30 | hypothetical protein, conserved | -0,6675 | 1,709E-06 |
| TcCLB.511127.390 | hypothetical protein, conserved | -0,6676 | 1,259E-21 |
| TcCLB.507883.25 | hypothetical protein, conserved | -0,6676 | 0,006894 |
| TcCLB.507757.50 | serine/threonine-protein phosphatase PP1, putative | -0,6679 | 3,002E-13 |
| TcCLB.511307.20 | Peptidase C19, ubiquitin carboxyl-terminal hydrolase, puta | -0,6679 | 2,26E-09 |
| TcCLB.511153.80 | hypothetical protein | -0,6679 | 5,991E-07 |
| TcCLB.477983.10 | calpain cysteine peptidase, putative | -0,668 | 0,07708 |
| TcCLB.506871.30 | ADP-ribosylation factor GTPase activating protein 1, putat | -0,6681 | 3,442E-09 |
| TcCLB.508061.10 | hypothetical protein | -0,6683 | 1,356E-07 |
| TcCLB.506341.40 | hypothetical protein | -0,6684 | 0,02885 |
| TcCLB.508777.170 | hypothetical protein, conserved | -0,6689 | 5,516E-09 |
| TcCLB.506977.10 | dual specificity protein phosphatase, putative | -0,6689 | 1,694E-06 |
| TcCLB.508893.40 | hypothetical protein, conserved (fragment) | -0,6689 | 0,006198 |
| TcCLB.508215.9 | protein kinase C substrate protein, heavy chain, putative | -0,6693 | 1,173E-13 |
| TcCLB.503579.30 | DNA polymerase sigma-like protein, putative | -0,6694 | 2,235E-09 |
| TcCLB.508507.20 | hypothetical protein, conserved | -0,6698 | 1,521E-10 |
| TcCLB.508221.870 | trans-sialidase, Group V, putative | -0,6699 | 0,001749 |
| TcCLB.509319.100 | hypothetical protein, conserved | -0,67 | 1,305E-09 |
| TcCLB.511313.10 | hypothetical protein, conserved | -0,6705 | 0,00005443 |
| TcCLB.509735.60 | trans-sialidase (pseudogene), putative | -0,6708 | 0,00004728 |
| TcCLB.511575.70 | kinetoplastid kinetochore protein 4, putative | -0,6709 | 7,412E-10 |
| TcCLB.511099.160 | mucin TcMUCII, putative | -0,6714 | 0,02459 |
| TcCLB.504151.39 | hypothetical protein, conserved | -0,6716 | 3,205E-11 |
| TcCLB.510747.60 | cytochrome P450 reductase, putative | -0,6716 | 8,489E-08 |
| TcCLB.508221.320 | mucin-associated surface protein (MASP, pseudogene), pu | -0,6719 | 0,04645 |
| TcCLB.511557.29 | hypothetical protein | -0,6724 | 5,697E-08 |
| TcCLB.508865.4 | protein phosphatase 2C, putative | -0,6725 | 5,149E-11 |
| TcCLB.506343.41 | trans-sialidase (pseudogene), putative | -0,6727 | 0,003978 |
| TcCLB.511083.35 | mucin TcMUCII (pseudogene), putative | -0,6734 | 0,01892 |
| TcCLB.511277.30 | hypothetical protein, conserved (pseudogene) | -0,6741 | 3,107E-14 |
| TcCLB.503791.40 | dihydrouridine synthase, putative | -0,6742 | 0,0000138 |
| TcCLB.510303.40 | hypothetical protein | -0,6746 | 3,847E-09 |
| TcCLB.510763.60 | hypothetical protein, conserved | -0,6749 | 1,552E-06 |

| | | | |
|---|---|---|---|
| TcCLB.510279.220 | mucin TcMUC (pseudogene), putative | -0,6755 | 0,06804 |
| TcCLB.508277.360 | ubiquitin hydrolase, putative | -0,6758 | 1,247E-15 |
| TcCLB.506681.70 | serine-threonine dehydratase, putative | -0,676 | 1,259E-06 |
| TcCLB.507611.190 | retrotransposon hot spot protein (RHS, pseudogene), puta | -0,6761 | 0,001712 |
| TcCLB.506401.24 | hypothetical protein, conserved | -0,6765 | 5,313E-11 |
| TcCLB.507357.140 | hypothetical protein | -0,6766 | 0,06001 |
| TcCLB.508247.19 | syntaxin binding protein 1 (pseudogene), putative | -0,6767 | 0,02349 |
| TcCLB.510953.49 | lipase-like protein, putative | -0,677 | 4,688E-07 |
| TcCLB.509165.10 | trans-sialidase (pseudogene), putative | -0,6771 | 1,793E-07 |
| TcCLB.511389.10 | hypothetical protein, conserved | -0,6773 | 3,19E-07 |
| TcCLB.509545.25 | elongation factor 1-gamma (EF-1-gamma, pseudogene), | -0,6774 | 0,1961 |
| TcCLB.506565.29 | hypothetical protein, conserved | -0,6775 | 0,00000338 |
| TcCLB.506425.80 | phosphatidylinositol kinase, putative | -0,6776 | 8,825E-10 |
| TcCLB.511401.24 | hypothetical protein, conserved (pseudogene) | -0,6777 | 4,657E-06 |
| TcCLB.506791.60 | hypothetical protein, conserved | -0,6778 | 1,095E-13 |
| TcCLB.510657.130 | hypothetical protein, conserved | -0,6778 | 6,334E-08 |
| TcCLB.507517.130 | RNA editing 3' terminal uridylyl transferase 2, putative (R | -0,6782 | 7,492E-07 |
| TcCLB.506917.27 | hypothetical protein, conserved (pseudogene) | -0,6784 | 0,0003739 |
| TcCLB.504249.100 | trans-sialidase (pseudogene), putative | -0,6787 | 0,00008737 |
| TcCLB.506247.260 | mitochondrial carrier protein, putative | -0,6789 | 0,00003442 |
| TcCLB.506811.200 | hypothetical protein, conserved | -0,679 | 1,244E-07 |
| TcCLB.509157.210 | hypothetical protein, conserved | -0,679 | 0,00007388 |
| TcCLB.511611.11 | hypothetical protein | -0,679 | 0,004093 |
| TcCLB.509561.100 | hypothetical protein, conserved | -0,6794 | 2,685E-13 |
| TcCLB.506435.180 | hypothetical protein, conserved | -0,6795 | 2,183E-06 |
| TcCLB.506279.220 | mucin-associated surface protein (MASP, pseudogene), pu | -0,6797 | 0,0003305 |
| TcCLB.511291.50 | hypothetical protein, conserved | -0,68 | 0,00001825 |
| TcCLB.508405.20 | TBC1 domain family member 20/GTPase, putative | -0,6803 | 1,807E-06 |
| TcCLB.506297.170 | hypothetical protein, conserved | -0,6804 | 9,161E-06 |
| TcCLB.507765.130 | hypothetical protein, conserved | -0,6805 | 8,678E-25 |
| TcCLB.509589.30 | Kinesin-13 5, putative (KIN13-5) | -0,6805 | 1,134E-16 |
| TcCLB.508777.90 | ATP-dependent DNA helicase, putative | -0,6805 | 3,539E-09 |
| TcCLB.511717.60 | hypothetical protein, conserved | -0,6809 | 3,092E-08 |
| TcCLB.506681.20 | hypothetical protein, conserved | -0,6811 | 2,549E-17 |
| TcCLB.508235.30 | mucin TcMUCI, putative | -0,6812 | 1,144E-06 |
| TcCLB.506155.130 | hypothetical protein, conserved | -0,6813 | 0,002649 |
| TcCLB.507209.40 | cdc2-related kinase 9 (CRK9) | -0,6814 | 7,793E-17 |
| TcCLB.506367.50 | hypothetical protein, conserved | -0,6815 | 0,00001086 |
| TcCLB.508261.140 | elongation factor 1-gamma (EF-1-gamma), putative | -0,6815 | 0,06744 |
| TcCLB.503561.10 | hypothetical protein, conserved | -0,6818 | 2,752E-06 |
| TcCLB.410199.40 | hypothetical protein | -0,682 | 0,00508 |
| TcCLB.510889.100 | hypothetical protein, conserved | -0,6821 | 3,244E-10 |
| TcCLB.511603.356 | syntaxin binding protein (pseudogene), putative | -0,6824 | 0,0745 |
| TcCLB.510237.60 | mucin TcMUCII, putative | -0,6827 | 0,01765 |
| TcCLB.509595.60 | hypothetical protein, conserved | -0,6828 | 3,668E-08 |

| | | | |
|---|---|---|---|
| TcCLB.506443.30 | hypothetical protein, conserved | -0,6831 | 2,252E-11 |
| TcCLB.506211.107 | hypothetical protein, conserved | -0,6833 | 0,00006238 |
| TcCLB.509641.20 | hypothetical protein, conserved | -0,6837 | 5,287E-06 |
| TcCLB.504131.130 | hypothetical protein, conserved | -0,6838 | 1,217E-11 |
| TcCLB.511127.160 | integral membrane transport protein, putative | -0,6839 | 0,0207 |
| TcCLB.506287.170 | hypothetical protein, conserved | -0,684 | 1,6E-15 |
| TcCLB.506951.10 | retrotransposon hot spot protein (RHS, pseudogene), puta | -0,684 | 0,004513 |
| TcCLB.511039.3 | choline/ethanolamine phosphotransferase, putative (CEPT | -0,6841 | 0,001978 |
| TcCLB.410199.4 | kinase, putative (fragment) | -0,6843 | 0,00001326 |
| TcCLB.506753.50 | hypothetical protein, conserved | -0,6846 | 6,116E-10 |
| TcCLB.508521.70 | mucin-associated surface protein (MASP, pseudogene), pu | -0,685 | 0,0003992 |
| TcCLB.506413.60 | hypothetical protein, conserved | -0,6858 | 0,0005418 |
| TcCLB.511107.30 | hypothetical protein, conserved | -0,686 | 1,665E-13 |
| TcCLB.506247.330 | 2,3-bisphosphoglycerate-independent phosphoglycerate m | -0,686 | 1,351E-08 |
| TcCLB.510979.30 | mucin TcMUCII (pseudogene), putative | -0,6863 | 0,0007057 |
| TcCLB.446067.9 | hypothetical protein, conserved (fragment) | -0,6864 | 1,599E-09 |
| TcCLB.508795.10 | SNF2/RAD54 related DNA helicase (pseudogene), putative | -0,6865 | 1,014E-06 |
| TcCLB.508873.130 | hypothetical protein | -0,6865 | 0,07273 |
| TcCLB.505171.109 | Present in the outer mitochondrial membrane proteome 1 | -0,6867 | 0,003365 |
| TcCLB.508593.50 | hypothetical protein, conserved | -0,6869 | 5,872E-13 |
| TcCLB.412419.10 | mucin-associated surface protein (MASP, pseudogene), pu | -0,6874 | 0,01622 |
| TcCLB.507805.9 | histone deacetylase 4, putative | -0,6876 | 0,000329 |
| TcCLB.509553.40 | ribonuclease mar1, putative | -0,6878 | 8,231E-07 |
| TcCLB.507723.41 | trans-sialidase (pseudogene), putative | -0,6878 | 0,0009522 |
| TcCLB.506195.260 | hypothetical protein | -0,6891 | 4,18E-11 |
| TcCLB.507427.10 | trans-sialidase, Group I, putative | -0,6892 | 4,096E-09 |
| TcCLB.509973.80 | hypothetical protein | -0,6892 | 0,0007617 |
| TcCLB.508307.20 | hypothetical protein, conserved | -0,6893 | 4,817E-09 |
| TcCLB.507895.100 | hypothetical protein, conserved (pseudogene) | -0,6896 | 1,762E-08 |
| TcCLB.508291.71 | hypothetical protein | -0,6896 | 0,001697 |
| TcCLB.506835.60 | calmodulin, putative | -0,6904 | 0,0002805 |
| TcCLB.507641.180 | endochitinase, putative | -0,6907 | 6,675E-06 |
| TcCLB.511367.180 | hypothetical protein | -0,691 | 1,68E-16 |
| TcCLB.507633.90 | hypothetical protein | -0,691 | 0,01962 |
| TcCLB.511127.180 | hypothetical protein, conserved | -0,6911 | 3,167E-13 |
| TcCLB.506231.40 | retrotransposon hot spot protein (RHS, pseudogene), puta | -0,6911 | 6,346E-07 |
| TcCLB.507765.50 | hypothetical protein, conserved | -0,6912 | 2,633E-10 |
| TcCLB.507609.70 | hypothetical protein, conserved | -0,6912 | 0,0004161 |
| TcCLB.511765.10 | trans-sialidase (pseudogene), putative | -0,6912 | 0,004824 |
| TcCLB.509647.190 | hypothetical protein, conserved | -0,6916 | 1,246E-13 |
| TcCLB.508817.30 | hypothetical protein | -0,6917 | 2,685E-10 |
| TcCLB.508771.50 | hypothetical protein, conserved | -0,692 | 1,493E-11 |
| TcCLB.509065.40 | hypothetical protein, conserved | -0,6928 | 2,555E-09 |
| TcCLB.505363.15 | elongation factor 1-gamma (EF-1-gamma, pseudogene), | -0,6929 | 0,003055 |
| TcCLB.510725.80 | hypothetical protein | -0,6934 | 4,261E-09 |

| | | | |
|---|---|---|---|
| TcCLB.506475.40 | MORN repeat-containing protein | -0,6938 | 1,135E-16 |
| TcCLB.504125.70 | hypothetical protein, conserved | -0,6941 | 2,898E-07 |
| TcCLB.506393.189 | retrotransposon hot spot (RHS) protein, putative (fragmer | -0,6943 | 0,0001986 |
| TcCLB.511523.80 | hypothetical protein, conserved | -0,6944 | 2,819E-12 |
| TcCLB.509649.59 | hypothetical protein, conserved | -0,6947 | 5,831E-11 |
| TcCLB.508005.20 | trans-sialidase (pseudogene), putative | -0,6948 | 0,05871 |
| TcCLB.510557.40 | hypothetical protein | -0,6951 | 0,00003238 |
| TcCLB.507757.21 | hypothetical protein, conserved | -0,6954 | 0,00006553 |
| TcCLB.506625.35 | trans-sialidase (pseudogene), putative | -0,6954 | 0,01144 |
| TcCLB.508731.60 | adenylosuccinate synthetase, putative | -0,6956 | 7,259E-10 |
| TcCLB.510693.260 | trans-sialidase (pseudogene), putative | -0,6956 | 0,01329 |
| TcCLB.506737.107 | mucin-associated surface protein (MASP, pseudogene), pu | -0,696 | 2,482E-06 |
| TcCLB.507087.70 | hypothetical protein, conserved | -0,6969 | 2,542E-14 |
| TcCLB.511183.381 | syntaxin binding protein 1, (pseudogene), putative | -0,6975 | 0,006781 |
| TcCLB.506721.10 | hypothetical protein, conserved (fragment) | -0,6981 | 0,0004397 |
| TcCLB.506885.30 | hypothetical protein, conserved | -0,6987 | 2,813E-08 |
| TcCLB.509453.20 | hypothetical protein, conserved | -0,6988 | 0,0001048 |
| TcCLB.510769.202 | trans-sialidase (pseudogene), putative | -0,6991 | 0,01585 |
| TcCLB.510239.40 | mucin-associated surface protein (MASP, pseudogene), pu | -0,6993 | 0,002382 |
| TcCLB.507065.89 | mucin TcMUC (pseudogene), putative | -0,6993 | 0,007863 |
| TcCLB.503559.80 | hypothetical protein | -0,6994 | 5,706E-11 |
| TcCLB.509773.9 | protein kinase, putative | -0,6997 | 1,115E-07 |
| TcCLB.509577.9 | hypothetical protein, conserved (fragment) | -0,7 | 2,475E-07 |
| TcCLB.511467.30 | hypothetical protein, conserved | -0,7002 | 1,644E-07 |
| TcCLB.420369.20 | trans-sialidase (pseudogene), putative | -0,7002 | 0,0001026 |
| TcCLB.506227.130 | calpain-like cysteine peptidase, putative | -0,7011 | 1,413E-15 |
| TcCLB.507615.120 | hypothetical protein, conserved | -0,7015 | 1,311E-07 |
| TcCLB.505983.40 | hypothetical protein, conserved | -0,7019 | 5,498E-22 |
| TcCLB.509835.10 | hypothetical protein, conserved | -0,7019 | 0,009291 |
| TcCLB.504109.50 | hypothetical protein, conserved | -0,7021 | 6,431E-10 |
| TcCLB.508365.171 | elongation factor 1-gamma (EF-1-gamma, pseudogene), | | -0,7022 | 0,03202 |
| TcCLB.511653.60 | hypothetical protein, conserved | -0,7025 | 7,26E-08 |
| TcCLB.509753.90 | trans-sialidase (pseudogene), putative | -0,7025 | 4,284E-06 |
| TcCLB.510143.110 | hypothetical protein, conserved | -0,7026 | 8,833E-11 |
| TcCLB.506605.190 | hypothetical protein, conserved | -0,7027 | 2,343E-10 |
| TcCLB.511159.14 | hypothetical protein, conserved | -0,7028 | 1,777E-14 |
| TcCLB.506755.210 | hypothetical protein, conserved | -0,7032 | 6,588E-08 |
| TcCLB.506725.60 | hypothetical protein, conserved | -0,7032 | 0,00001789 |
| TcCLB.506781.10 | mucin TcMUCII, putative | -0,7034 | 0,04234 |
| TcCLB.510689.80 | hypothetical protein, conserved | -0,7035 | 3,482E-07 |
| TcCLB.511607.60 | trans-sialidase (pseudogene), putative | -0,7037 | 0,00001719 |
| TcCLB.504011.30 | surface protease GP63 (pseudogene), putative | -0,7039 | 0,01591 |
| TcCLB.511277.200 | RNA-binding protein, putative | -0,7039 | 0,1203 |
| TcCLB.506757.70 | retrotransposon hot spot protein (RHS, pseudogene), puta | -0,704 | 0,01 |
| TcCLB.511337.30 | UDP-Gal or UDP-GlcNAc-dependent glycosyltransferase, p | -0,7043 | 0,005929 |

| | | | |
|---|---|---|---|
| TcCLB.507485.45 | hypothetical protein, conserved | -0,7049 | 0,00002586 |
| TcCLB.506605.70 | hypothetical protein, conserved | -0,7051 | 2,418E-14 |
| TcCLB.509693.40 | hypothetical protein, conserved | -0,7055 | 4,718E-06 |
| TcCLB.506297.180 | beta-adaptin 1, putative | -0,7056 | 7,102E-16 |
| TcCLB.508739.11 | trans-sialidase (pseudogene), putative | -0,7056 | 0,0004577 |
| TcCLB.511861.14 | trans-sialidase (pseudogene), putative | -0,7058 | 1,065E-06 |
| TcCLB.508247.30 | hypothetical protein, conserved | -0,706 | 0,05942 |
| TcCLB.507025.60 | RNA-binding protein, putative | -0,7073 | 0,01735 |
| TcCLB.508543.180 | hypothetical protein, conserved | -0,7075 | 5,828E-14 |
| TcCLB.506475.30 | hypothetical protein, conserved | -0,7078 | 1,136E-15 |
| TcCLB.511159.7 | neurobeachin/beige protein, putative | -0,7082 | 1,987E-11 |
| TcCLB.510269.73 | hypothetical protein, conserved (pseudogene) | -0,7082 | 0,00006033 |
| TcCLB.506757.100 | retrotransposon hot spot protein (RHS, pseudogene), puta | -0,7084 | 0,00008885 |
| TcCLB.504247.40 | RNA-editing nuclease 1 (KREN1) | -0,7085 | 1,712E-09 |
| TcCLB.506629.160 | hypothetical protein, conserved | -0,7086 | 0,00001092 |
| TcCLB.510359.680 | mucin-associated surface protein (MASP, pseudogene), pu | -0,7091 | 0,03872 |
| TcCLB.508277.100 | cAMP specific phosphodiesterase, putative | -0,7095 | 9,194E-07 |
| TcCLB.503775.41 | hypothetical protein, conserved (pseudogene) | -0,7108 | 0,01862 |
| TcCLB.507061.10 | hypothetical protein | -0,7112 | 2,803E-06 |
| TcCLB.503677.4 | hypothetical protein, conserved | -0,7113 | 0,0328 |
| TcCLB.508677.30 | trans-sialidase (pseudogene), putative | -0,7117 | 6,864E-06 |
| TcCLB.510231.10 | eukaryotic translation initiation factor 2-alpha kinase, puta | -0,7118 | 9,94E-08 |
| TcCLB.508247.100 | hypothetical protein (pseudogene) | -0,7118 | 0,002825 |
| TcCLB.506959.100 | GTP-ase activating protein, putative | -0,7123 | 8,148E-10 |
| TcCLB.506057.40 | trans-sialidase (pseudogene), putative | -0,7131 | 6,775E-08 |
| TcCLB.506135.80 | hypothetical protein, conserved | -0,7142 | 4,441E-06 |
| TcCLB.510359.580 | mucin TcMUCII, putative | -0,7146 | 0,07002 |
| TcCLB.511337.20 | UDP-Gal or UDP-GlcNAc-dependent glycosyltransferase, p | -0,7147 | 0,02211 |
| TcCLB.511779.10 | trans-sialidase, putative | -0,7152 | 0,08697 |
| TcCLB.510025.250 | trans-sialidase (pseudogene), putative | -0,7153 | 0,00007647 |
| TcCLB.506667.140 | hypothetical protein | -0,7153 | 0,04239 |
| TcCLB.506265.190 | hypothetical protein, conserved | -0,7155 | 1,122E-10 |
| TcCLB.508289.94 | hypothetical protein, conserved (pseudogene) | -0,7156 | 0,08369 |
| TcCLB.507949.150 | conserved protein, unknown function | -0,7159 | 2,962E-15 |
| TcCLB.511417.50 | hypothetical protein | -0,7168 | 2,032E-06 |
| TcCLB.503809.68 | glycerol uptake protein (pseudogene), putative | -0,717 | 0,001055 |
| TcCLB.507547.18 | mucin TcMUCI, putative | -0,7172 | 0,05399 |
| TcCLB.508209.50 | hypothetical protein, conserved | -0,7173 | 4,413E-06 |
| TcCLB.506825.160 | hypothetical protein, conserved | -0,7175 | 1,15E-07 |
| TcCLB.509451.50 | hypothetical protein, conserved | -0,7181 | 1,049E-15 |
| TcCLB.506267.39 | mucin-associated surface protein (MASP, pseudogene), pu | -0,7181 | 0,0000199 |
| TcCLB.505999.140 | Metal-binding domain-containing protein | -0,7183 | 6,815E-11 |
| TcCLB.510197.10 | trans-sialidase (pseudogene), putative | -0,7185 | 9,218E-10 |
| TcCLB.503635.20 | hypothetical protein, conserved | -0,7192 | 1,862E-11 |
| TcCLB.504153.150 | Unc104-like kinesin, putative | -0,7193 | 4,207E-18 |

| | | | |
|---|---|---|---|
| TcCLB.504199.30 | kinesin-like protein, putative | -0,7193 | 2,34E-12 |
| TcCLB.511027.20 | hypothetical protein, conserved | -0,7193 | 4,853E-11 |
| TcCLB.507765.10 | hypothetical protein, conserved | -0,7193 | 0,00004 |
| TcCLB.508175.260 | inorganic polyphosphate/ATP-NAD kinase, putative | -0,7196 | 1,38E-17 |
| TcCLB.506755.170 | hypothetical protein, conserved | -0,7201 | 1,729E-16 |
| TcCLB.510013.59 | trans-sialidase (pseudogene), putative | -0,721 | 0,03349 |
| TcCLB.506297.70 | hypothetical protein, conserved | -0,7211 | 6,051E-18 |
| TcCLB.506107.10 | hypothetical protein, conserved | -0,7211 | 1,401E-12 |
| TcCLB.509631.45 | hypothetical protein, conserved (pseudogene) | -0,7211 | 0,0009227 |
| TcCLB.508707.30 | hypothetical protein, conserved | -0,7212 | 2,726E-18 |
| TcCLB.511797.215 | hypothetical protein, conserved (pseudogene) | -0,7212 | 0,0003042 |
| TcCLB.503643.20 | cyclic nucleotide-binding domain containing protein, putat | -0,7213 | 1,745E-14 |
| TcCLB.508633.10 | trans-sialidase (pseudogene), putative | -0,7214 | 0,01246 |
| TcCLB.506407.90 | hypothetical protein, conserved | -0,7216 | 4,631E-25 |
| TcCLB.511257.30 | ras-related protein rab, putative | -0,7218 | 8,242E-08 |
| TcCLB.503693.20 | hypothetical protein, conserved | -0,7219 | 6,243E-07 |
| TcCLB.508205.30 | protein kinase, putative | -0,722 | 2,48E-14 |
| TcCLB.504213.90 | inositol polyphosphate kinase-like protein, putative | -0,7221 | 0,006615 |
| TcCLB.508175.170 | hypothetical protein, conserved | -0,7222 | 1,924E-15 |
| TcCLB.504105.140 | enolase | -0,7226 | 5,102E-12 |
| TcCLB.511755.10 | hypothetical protein, conserved | -0,7227 | 1,316E-06 |
| TcCLB.425839.10 | kinesin, putative | -0,7232 | 2,989E-06 |
| TcCLB.508799.180 | hypothetical protein, conserved | -0,7233 | 5,767E-16 |
| TcCLB.504105.87 | hypothetical protein, conserved | -0,7236 | 9,58E-08 |
| TcCLB.508175.200 | hypothetical protein, conserved | -0,7239 | 4,588E-08 |
| TcCLB.508427.30 | surface protease GP63 (pseudogene), putative | -0,7239 | 0,01127 |
| TcCLB.507357.10 | surface protease GP63, putative | -0,7245 | 0,01417 |
| TcCLB.510889.320 | hypothetical protein, conserved | -0,7249 | 2,28E-15 |
| TcCLB.506831.20 | amino acid permease, putative | -0,7251 | 6,78E-07 |
| TcCLB.504839.70 | hypothetical protein, conserved | -0,7255 | 9,941E-20 |
| TcCLB.507023.124 | hypothetical protein, conserved | -0,7258 | 0,0001012 |
| TcCLB.509753.220 | mucin TcMUCII, putative | -0,7262 | 0,0414 |
| TcCLB.509767.140 | hypothetical protein | -0,7263 | 2,679E-12 |
| TcCLB.510149.40 | hypothetical protein, conserved | -0,7265 | 9,757E-11 |
| TcCLB.511795.10 | hypothetical protein, conserved (pseudogene) | -0,7268 | 0,003563 |
| TcCLB.503759.30 | trans-sialidase, putative | -0,7268 | 0,01429 |
| TcCLB.507649.40 | hypothetical protein, conserved | -0,727 | 1,431E-08 |
| TcCLB.507665.10 | hypothetical protein, conserved | -0,7275 | 2,192E-09 |
| TcCLB.503997.10 | hypothetical protein, conserved | -0,7277 | 1,714E-07 |
| TcCLB.508989.60 | hypothetical protein, conserved | -0,7277 | 0,00003566 |
| TcCLB.508957.30 | hypothetical protein, conserved | -0,7278 | 5,222E-17 |
| TcCLB.509509.40 | hypothetical protein, conserved | -0,7279 | 0,00006449 |
| TcCLB.504197.15 | surface protease GP63 (pseudogene), putative | -0,7279 | 0,01524 |
| TcCLB.510149.9 | hypothetical protein, conserved (fragment) | -0,7282 | 7,186E-09 |
| TcCLB.508523.80 | cAMP response protein 1 (CARP1) | -0,7286 | 6,139E-10 |

| | | | |
|---|---|---|---|
| TcCLB.503833.20 | hypothetical protein, conserved | -0,7292 | 2,434E-16 |
| TcCLB.504787.70 | mucin-associated surface protein (MASP), putative | -0,7292 | 0,0006739 |
| TcCLB.503693.11 | hypothetical protein, conserved (fragment) | -0,7296 | 0,0003777 |
| TcCLB.508221.860 | mucin-associated surface protein (MASP, pseudogene), pu | -0,7297 | 0,006889 |
| TcCLB.506247.380 | hypothetical protein, conserved | -0,7299 | 8,203E-11 |
| TcCLB.506737.135 | hypothetical protein, conserved (pseudogene) | -0,7301 | 1,664E-09 |
| TcCLB.511167.70 | DNA repair and recombination helicase protein PIF2, puta | -0,7302 | 3,833E-06 |
| TcCLB.511173.40 | mucin TcMUCII (pseudogene), putative | -0,7303 | 0,02416 |
| TcCLB.510187.300 | hypothetical protein, conserved | -0,7308 | 4,441E-11 |
| TcCLB.506959.80 | hypothetical protein, conserved | -0,7313 | 8,927E-10 |
| TcCLB.505025.147 | hypothetical protein | -0,7321 | 0,001769 |
| TcCLB.510747.110 | hypothetical protein, conserved | -0,7326 | 2,211E-08 |
| TcCLB.508039.50 | dispersed gene family protein 1 (DGF-1, pseudogene), put | -0,7326 | 0,00002069 |
| TcCLB.509121.10 | retrotransposon hot spot protein (RHS, pseudogene), puta | -0,7326 | 0,0003972 |
| TcCLB.511153.50 | haloacid dehalogenase hydrolase, putative | -0,7332 | 6,19E-10 |
| TcCLB.509527.154 | mucin-associated surface protein (MASP, pseudogene), pu | -0,7335 | 0,003385 |
| TcCLB.511189.20 | hypothetical protein, conserved | -0,7337 | 4,019E-11 |
| TcCLB.506825.40 | hypothetical protein, conserved | -0,7338 | 1,859E-06 |
| TcCLB.508221.400 | trans-sialidase, Group V, putative | -0,7344 | 0,01786 |
| TcCLB.511031.20 | hypothetical protein, conserved | -0,7353 | 1,573E-10 |
| TcCLB.511131.89 | hypothetical protein, conserved | -0,7353 | 0,00000168 |
| TcCLB.511865.30 | hypothetical protein, conserved | -0,7355 | 4,724E-08 |
| TcCLB.508153.364 | hypothetical protein, conserved | -0,7355 | 4,672E-07 |
| TcCLB.507771.40 | phosphoglycerate mutase, putative (iPGAM) | -0,7357 | 3,329E-09 |
| TcCLB.511415.21 | trans-sialidase (pseudogene), putative | -0,7358 | 0,000331 |
| TcCLB.507939.40 | hypothetical protein, conserved | -0,7359 | 2,321E-07 |
| TcCLB.510361.260 | mucin-associated surface protein (MASP), putative | -0,7365 | 0,03582 |
| TcCLB.506699.6 | hypothetical protein, conserved | -0,7367 | 0,00005802 |
| TcCLB.511771.40 | trans-sialidase (pseudogene), putative | -0,737 | 0,04652 |
| TcCLB.506329.59 | mucin-associated surface protein (MASP, pseudogene), pu | -0,7376 | 0,0007716 |
| TcCLB.511217.170 | hypothetical protein, conserved | -0,7379 | 7,793E-10 |
| TcCLB.510891.40 | trans-sialidase (pseudogene), putative | -0,7379 | 4,857E-09 |
| TcCLB.509251.30 | trans-sialidase (pseudogene), putative | -0,7379 | 0,00004243 |
| TcCLB.508307.10 | hypothetical protein, conserved | -0,738 | 0,004164 |
| TcCLB.511483.60 | hypothetical protein, conserved | -0,7382 | 0,00126 |
| TcCLB.510327.20 | surface protease GP63 (pseudogene), putative | -0,7385 | 0,000532 |
| TcCLB.506635.100 | ATP dependent DEAD-box helicase, putative (RH) | -0,7389 | 4,418E-12 |
| TcCLB.510323.120 | kinesin, putative | -0,7397 | 2,012E-10 |
| TcCLB.507443.53 | hypothetical protein | -0,7397 | 0,001639 |
| TcCLB.510419.30 | hypothetical protein | -0,7398 | 0,1345 |
| TcCLB.508949.30 | hypothetical protein, conserved | -0,74 | 1,976E-14 |
| TcCLB.511075.54 | hypothetical protein, conserved | -0,74 | 1,866E-09 |
| TcCLB.508207.220 | hypothetical protein, conserved | -0,7401 | 0,00004292 |
| TcCLB.510351.150 | phosphoprotein phosphatase, putative | -0,7403 | 3,487E-07 |
| TcCLB.510297.50 | hypothetical protein, conserved | -0,7408 | 1,451E-12 |

| | | | |
|---|---|---|---|
| TcCLB.503685.5 | RNA-binding protein, putative (fragment) | -0,741 | 0,0223 |
| TcCLB.508641.320 | formin-like protein | -0,7412 | 5,261E-17 |
| TcCLB.508837.70 | trans-sialidase, Group II, putative | -0,7413 | 3,923E-10 |
| TcCLB.510359.50 | hypothetical protein, conserved | -0,7413 | 0,00009896 |
| TcCLB.509549.15 | mucin TcMUCII (pseudogene), putative | -0,7417 | 0,02998 |
| TcCLB.506993.20 | mucin TcMUCI, putative | -0,7417 | 0,05827 |
| TcCLB.503947.11 | surface protease GP63, putative (fragment) | -0,7418 | 0,01915 |
| TcCLB.509233.230 | hypothetical protein, conserved | -0,7419 | 3,514E-15 |
| TcCLB.506767.183 | mucin-associated surface protein (MASP, pseudogene), pu | -0,7419 | 0,00008012 |
| TcCLB.511719.41 | hypothetical protein, conserved (fragment) | -0,7422 | 2,731E-10 |
| TcCLB.506717.20 | receptor-type adenylate cyclase, putative | -0,7423 | 3,026E-16 |
| TcCLB.508741.390 | hypothetical protein, conserved | -0,7423 | 2,611E-15 |
| TcCLB.505073.14 | hypothetical protein, conserved | -0,7426 | 2,657E-07 |
| TcCLB.506973.61 | hypothetical protein, conserved (pseudogene) | -0,7429 | 0,0001388 |
| TcCLB.508347.60 | hypothetical protein, conserved | -0,7432 | 1,897E-07 |
| TcCLB.508735.10 | trans-sialidase (pseudogene), putative | -0,7434 | 6,834E-09 |
| TcCLB.506329.35 | hypothetical protein, conserved (pseudogene) | -0,7438 | 0,1156 |
| TcCLB.510409.10 | protein kinase, putative | -0,7443 | 0,0004511 |
| TcCLB.506319.50 | hypothetical protein, conserved | -0,7445 | 1,392E-13 |
| TcCLB.509819.10 | hypothetical protein, conserved | -0,7445 | 2,272E-12 |
| TcCLB.511167.50 | hypothetical protein, conserved | -0,7446 | 1,152E-12 |
| TcCLB.511911.71 | hypothetical protein, conserved | -0,7446 | 0,01911 |
| TcCLB.506469.50 | hypothetical protein, conserved | -0,7451 | 5,359E-06 |
| TcCLB.450155.9 | hypothetical protein, conserved | -0,746 | 1,238E-07 |
| TcCLB.510653.19 | hypothetical protein, conserved (fragment) | -0,7463 | 4,068E-11 |
| TcCLB.504427.10 | trypanothione synthetase, putative | -0,7465 | 5,595E-07 |
| TcCLB.511189.10 | hypothetical protein, conserved (pseudogene) | -0,7467 | 7,294E-07 |
| TcCLB.503925.119 | protein kinase, putative | -0,7469 | 4,132E-09 |
| TcCLB.509399.90 | hypothetical protein | -0,7469 | 2,898E-08 |
| TcCLB.511735.88 | hypothetical protein, conserved (fragment) | -0,747 | 0,0003792 |
| TcCLB.504157.60 | hypothetical protein, conserved | -0,7472 | 3,36E-08 |
| TcCLB.506297.200 | inositol-1,4,5-trisphosphate (IP3) 5-phosphatase, putative | -0,7474 | 9,237E-17 |
| TcCLB.511127.350 | hypothetical protein, conserved | -0,7474 | 1,987E-14 |
| TcCLB.511733.30 | hypothetical protein, conserved | -0,748 | 1,564E-08 |
| TcCLB.509755.10 | trans-sialidase, Group V, putative | -0,748 | 0,001194 |
| TcCLB.507081.80 | hypothetical protein, conserved | -0,7481 | 1,145E-16 |
| TcCLB.506187.80 | trans-sialidase (pseudogene), putative | -0,7481 | 0,06601 |
| TcCLB.510737.60 | hypothetical protein, conserved | -0,7487 | 2,464E-11 |
| TcCLB.506457.80 | hypothetical protein, conserved | -0,7493 | 6,542E-13 |
| TcCLB.511375.30 | hypothetical protein (fragment) | -0,7494 | 0,026 |
| TcCLB.504233.10 | surface protease GP63, putative | -0,7495 | 0,00005939 |
| TcCLB.511173.340 | mucin TcMUCII (pseudogene), putative | -0,7497 | 0,0005603 |
| TcCLB.508265.70 | hypothetical protein, conserved | -0,7498 | 0,00000049 |
| TcCLB.508243.81 | syntaxin binding protein (pseudogene), putative | -0,7504 | 0,002537 |
| TcCLB.448329.10 | trans-sialidase, putative | -0,7504 | 0,06469 |

| | | | |
|---|---|---|---|
| TcCLB.506321.340 | hypothetical protein, conserved | -0,7505 | 2,592E-10 |
| TcCLB.506203.80 | formin-like protein | -0,7508 | 1,74E-11 |
| TcCLB.507959.224 | surface protease GP63 (pseudogene), putative | -0,7508 | 0,02251 |
| TcCLB.509941.110 | condensin subunit 1, putative | -0,7518 | 1,089E-20 |
| TcCLB.509799.80 | hypothetical protein, conserved | -0,7518 | 5,217E-14 |
| TcCLB.505955.4 | mucin TcMUCII, putative (fragment) | -0,7521 | 0,001936 |
| TcCLB.509065.20 | mitogen-activated protein kinase 5, putative | -0,7523 | 1,521E-06 |
| TcCLB.508409.300 | hypothetical protein, conserved | -0,7524 | 1,976E-09 |
| TcCLB.506241.60 | kinesin, putative | -0,7525 | 1,843E-16 |
| TcCLB.510503.130 | mucin TcMUCII, putative | -0,7533 | 0,04567 |
| TcCLB.511181.110 | hypothetical protein, conserved | -0,7546 | 8,597E-17 |
| TcCLB.507765.40 | hypothetical protein, conserved | -0,7551 | 3,295E-12 |
| TcCLB.509647.170 | hypothetical protein, conserved | -0,7552 | 6,559E-17 |
| TcCLB.506565.12 | RNA-binding protein, putative | -0,7552 | 1,387E-11 |
| TcCLB.507757.10 | hypothetical protein | -0,7552 | 2,519E-09 |
| TcCLB.508799.110 | protein kinase, putative | -0,7552 | 1,354E-07 |
| TcCLB.503897.130 | hypothetical protein, conserved | -0,7557 | 3,557E-10 |
| TcCLB.511527.9 | hypothetical protein, conserved | -0,7557 | 3,512E-08 |
| TcCLB.442495.10 | cyclic nucleotide-binding domain containing protein, putat | -0,7566 | 2,899E-09 |
| TcCLB.504057.160 | hypothetical protein, conserved | -0,757 | 1,963E-09 |
| TcCLB.504087.10 | hypothetical protein, conserved | -0,757 | 0,0005807 |
| TcCLB.507011.180 | hypothetical protein, conserved | -0,7571 | 2,04E-14 |
| TcCLB.508585.20 | hypothetical protein, conserved | -0,7572 | 1,473E-09 |
| TcCLB.511401.95 | hypothetical protein (pseudogene) | -0,7575 | 0,00003332 |
| TcCLB.511099.220 | mucin-associated surface protein (MASP), putative | -0,7576 | 0,03665 |
| TcCLB.503755.10 | DNA polymerase kappa, putative | -0,7577 | 1,409E-14 |
| TcCLB.508175.300 | hypothetical protein, conserved | -0,7577 | 5,727E-12 |
| TcCLB.508081.40 | hypothetical protein | -0,7578 | 0,07385 |
| TcCLB.508251.31 | mucin-associated surface protein (MASP, pseudogene), pu | -0,7579 | 0,0005247 |
| TcCLB.503865.80 | phosphatidylinositol 4-kinase, putative | -0,7584 | 5,13E-10 |
| TcCLB.511737.50 | hypothetical protein, conserved | -0,7585 | 3,146E-09 |
| TcCLB.511127.360 | hypothetical protein, conserved | -0,7591 | 1,726E-10 |
| TcCLB.509569.10 | hypothetical protein, conserved | -0,7591 | 3,259E-08 |
| TcCLB.508523.35 | hypothetical protein, conserved (pseudogene) | -0,7592 | 3,394E-12 |
| TcCLB.511043.45 | trans-sialidase (pseudogene), putative | -0,7593 | 1,536E-11 |
| TcCLB.508737.130 | hypothetical protein, conserved | -0,7594 | 1,177E-08 |
| TcCLB.506801.50 | surface protease GP63 (pseudogene), putative | -0,7595 | 1,25E-07 |
| TcCLB.506753.100 | hypothetical protein, conserved | -0,7604 | 2,181E-06 |
| TcCLB.508211.50 | hypothetical protein, conserved | -0,7611 | 6,813E-08 |
| TcCLB.508301.10 | hypothetical protein, conserved | -0,7611 | 7,436E-08 |
| TcCLB.511607.30 | mucin TcMUCII, putative | -0,7611 | 0,05874 |
| TcCLB.508293.160 | trans-sialidase (pseudogene), putative | -0,7612 | 0,1068 |
| TcCLB.511323.10 | trans-sialidase, Group I, putative | -0,762 | 2,503E-11 |
| TcCLB.509863.10 | UDP-Gal or UDP-GlcNAc-dependent glycosyltransferase, p | -0,7624 | 9,612E-08 |
| TcCLB.511871.170 | trans-sialidase (pseudogene), putative | -0,7624 | 0,00007891 |

| | | | |
|---|---|---|---|
| TcCLB.511173.270 | mucin-associated surface protein (MASP), putative | -0,7628 | 8,378E-06 |
| TcCLB.510877.80 | phospholipid-transporting ATPase-like protein, putative | -0,7629 | 8,073E-21 |
| TcCLB.510431.130 | hypothetical protein, conserved | -0,7629 | 5,334E-10 |
| TcCLB.504021.149 | hypothetical protein, conserved (fragment) | -0,7629 | 4,518E-09 |
| TcCLB.511357.5 | glycerol uptake protein, putative (fragment) | -0,7632 | 3,316E-07 |
| TcCLB.507031.179 | hypothetical protein, conserved | -0,7636 | 3,078E-10 |
| TcCLB.504449.14 | hypothetical protein, conserved (fragment) | -0,7643 | 8,726E-25 |
| TcCLB.507091.150 | hypothetical protein, conserved (pseudogene) | -0,7643 | 1,713E-08 |
| TcCLB.504051.30 | hypothetical protein, conserved | -0,7651 | 3,503E-15 |
| TcCLB.503959.40 | hypothetical protein | -0,7654 | 3,152E-12 |
| TcCLB.508127.5 | hypothetical protein, conserved | -0,7654 | 0,00001153 |
| TcCLB.508207.230 | 6-phosphofructo-2-kinase/fructose-2,6-biphosphatase, pu | -0,7661 | 1,846E-15 |
| TcCLB.507511.91 | mucin TcMUCIII, putative | -0,7666 | 5,681E-11 |
| TcCLB.511761.20 | hypothetical protein, conserved | -0,7667 | 5,571E-08 |
| TcCLB.507611.70 | trans-sialidase (pseudogene), putative | -0,7668 | 8,854E-06 |
| TcCLB.509105.120 | protein kinase, putative | -0,7671 | 4,589E-11 |
| TcCLB.507143.40 | hypothetical protein, conserved | -0,7677 | 1,427E-21 |
| TcCLB.511527.50 | AUT2/APG4/ATG4 cysteine peptidase, putative | -0,7679 | 4,011E-15 |
| TcCLB.510367.20 | retrotransposon hot spot protein (RHS, pseudogene), puta | -0,7679 | 0,0000235 |
| TcCLB.504113.41 | hypothetical protein, conserved (fragment) | -0,7681 | 0,0001466 |
| TcCLB.510765.10 | hypothetical protein, conserved | -0,7682 | 1,39E-10 |
| TcCLB.503973.57 | mucin TcMUCII (pseudogene), putative | -0,7682 | 0,0107 |
| TcCLB.508411.50 | hypothetical protein, conserved | -0,7683 | 1,969E-16 |
| TcCLB.511127.300 | hypothetical protein, conserved | -0,7692 | 0,007244 |
| TcCLB.506321.60 | surface protease GP63 (pseudogene), putative | -0,7695 | 0,003402 |
| TcCLB.508775.60 | hypothetical protein, conserved (pseudogene) | -0,7705 | 0,0004829 |
| TcCLB.511287.130 | hypothetical protein, conserved | -0,771 | 1,638E-16 |
| TcCLB.507847.20 | mucin-associated surface protein (MASP, pseudogene), pu | -0,7711 | 2,167E-10 |
| TcCLB.506321.330 | hypothetical protein, conserved | -0,7716 | 8,512E-16 |
| TcCLB.509259.10 | trans-sialidase (pseudogene), putative | -0,7718 | 0,0005008 |
| TcCLB.511255.210 | mucin TcMUCII, putative | -0,7718 | 0,1585 |
| TcCLB.507763.10 | hypothetical protein, conserved | -0,772 | 2,594E-10 |
| TcCLB.509345.30 | trans-sialidase (pseudogene), putative | -0,7721 | 5E-11 |
| TcCLB.506467.4 | hypothetical protein, conserved | -0,7723 | 3,975E-07 |
| TcCLB.508175.80 | protein kinase, putative | -0,7727 | 5,144E-22 |
| TcCLB.506137.27 | mucin-associated surface protein (MASP, pseudogene), pu | -0,7727 | 0,003423 |
| TcCLB.506317.10 | hypothetical protein, conserved | -0,7733 | 1,238E-12 |
| TcCLB.511279.40 | hypothetical protein, conserved | -0,7741 | 7,488E-07 |
| TcCLB.511261.120 | pumilio/PUF RNA binding protein 2, putative (PUF2) | -0,7741 | 0,00001937 |
| TcCLB.506401.290 | hypothetical protein, conserved | -0,7753 | 4,095E-06 |
| TcCLB.506873.20 | hypothetical protein, conserved | -0,7755 | 1,821E-10 |
| TcCLB.508777.120 | hypothetical protein, conserved | -0,7756 | 5,299E-11 |
| TcCLB.506287.200 | DNA ligase, putative | -0,7758 | 2,186E-06 |
| TcCLB.508305.30 | hypothetical protein | -0,7765 | 0,09052 |
| TcCLB.504025.19 | hypothetical protein, conserved (fragment) | -0,7767 | 5,16E-07 |

| | | | |
|---|---|---|---|
| TcCLB.508495.30 | hypothetical protein | -0,7772 | 0,01085 |
| TcCLB.506789.190 | hypothetical protein, conserved | -0,7774 | 9,439E-14 |
| TcCLB.511127.429 | hypothetical protein, conserved | -0,7775 | 3,035E-08 |
| TcCLB.510297.30 | TPR-repeat-containing chaperone protein DNAJ, putative | -0,7778 | 8,158E-18 |
| TcCLB.504181.40 | cdc2-related kinase 1, putative (TzCRK1) | -0,7778 | 7,273E-12 |
| TcCLB.506131.199 | mucin TcMUCII, putative | -0,778 | 0,03461 |
| TcCLB.507637.45 | mucin-associated surface protein (MASP, pseudogene), pu | -0,7782 | 0,02973 |
| TcCLB.506591.120 | receptor-type adenylate cyclase (pseudogene), putative | -0,7786 | 1,892E-15 |
| TcCLB.506887.50 | katanin, putative | -0,7787 | 1,227E-11 |
| TcCLB.505997.210 | DNA-directed RNA polymerase III subunit, putative | -0,779 | 7,655E-07 |
| TcCLB.503559.70 | cystinosin, putative | -0,7793 | 1,031E-07 |
| TcCLB.507711.240 | hypothetical protein, conserved | -0,7797 | 7,102E-15 |
| TcCLB.481211.20 | hypothetical protein | -0,78 | 1,417E-06 |
| TcCLB.507035.80 | trans-sialidase (pseudogene), putative | -0,7802 | 1,238E-11 |
| TcCLB.508199.70 | hypothetical protein | -0,7804 | 6,471E-15 |
| TcCLB.507787.110 | hypothetical protein, conserved | -0,7804 | 2,795E-08 |
| TcCLB.506775.160 | hypothetical protein, conserved | -0,7808 | 8,246E-15 |
| TcCLB.506559.129 | eukaryotic translation initiation factor 2-alpha kinase, puta | -0,781 | 3,431E-15 |
| TcCLB.508293.44 | mucin-associated surface protein (MASP, pseudogene), pu | -0,7812 | 0,09077 |
| TcCLB.460111.10 | trans-sialidase, putative (fragment) | -0,7816 | 0,08446 |
| TcCLB.511021.40 | hypothetical protein, conserved | -0,782 | 5,528E-13 |
| TcCLB.503697.140 | hypothetical protein | -0,782 | 1,18E-09 |
| TcCLB.507677.4 | hypothetical protein, conserved | -0,7821 | 2,992E-07 |
| TcCLB.508495.150 | mucin-associated surface protein (MASP, pseudogene), pu | -0,7822 | 0,0001747 |
| TcCLB.511431.50 | hypothetical protein, conserved | -0,7825 | 1,378E-15 |
| TcCLB.503957.60 | trans-sialidase (pseudogene), putative | -0,7827 | 1,303E-11 |
| TcCLB.506297.120 | hypothetical protein, conserved | -0,7833 | 1,857E-14 |
| TcCLB.510551.50 | trans-sialidase, putative | -0,7836 | 0,04694 |
| TcCLB.509429.250 | ubiquitin hydrolase, putative | -0,7842 | 1,105E-19 |
| TcCLB.506753.160 | hypothetical protein, conserved | -0,7847 | 1,407E-07 |
| TcCLB.506401.280 | hypothetical protein, conserved | -0,7848 | 5,446E-14 |
| TcCLB.509897.155 | trans-sialidase (pseudogene), putative | -0,7849 | 4,331E-08 |
| TcCLB.508273.80 | syntaxin binding protein, putative | -0,7853 | 0,0002467 |
| TcCLB.503809.158 | Triglyceride lipase, putative (fragment) | -0,7856 | 0,0001669 |
| TcCLB.509819.40 | hypothetical protein, conserved | -0,7858 | 1,193E-17 |
| TcCLB.511169.20 | hypothetical protein, conserved | -0,7863 | 1,026E-11 |
| TcCLB.506579.50 | hypothetical protein, conserved | -0,7864 | 2,077E-16 |
| TcCLB.510555.21 | hypothetical protein | -0,7866 | 0,006698 |
| TcCLB.504115.110 | trans-sialidase (pseudogene), putative | -0,7867 | 1,417E-07 |
| TcCLB.509741.34 | hypothetical protein, conserved | -0,7872 | 2,051E-07 |
| TcCLB.469113.10 | trans-sialidase (pseudogene), putative | -0,7878 | 3,741E-06 |
| TcCLB.506857.120 | hypothetical protein, conserved | -0,7881 | 0,04308 |
| TcCLB.508439.10 | hypothetical protein, conserved | -0,7886 | 1,515E-06 |
| TcCLB.506603.46 | hypothetical protein, conserved (pseudogene) | -0,789 | 0,02618 |
| TcCLB.504153.270 | hypothetical protein, conserved | -0,7891 | 2,018E-21 |

| | | | |
|---|---|---|---|
| TcCLB.510201.15 | mucin TcMUCII (pseudogene), putative | -0,7895 | 0,0003067 |
| TcCLB.507471.20 | mucin-associated surface protein (MASP, pseudogene), pu | -0,7896 | 0,0001458 |
| TcCLB.504411.5 | hypothetical protein, conserved (fragment) | -0,7899 | 0,08359 |
| TcCLB.509815.20 | trans-sialidase (pseudogene), putative | -0,79 | 2,914E-11 |
| TcCLB.506269.79 | mucin-associated surface protein (MASP, pseudogene), pu | -0,7904 | 0,0004875 |
| TcCLB.507623.140 | endonuclease G, putative | -0,7909 | 3,651E-09 |
| TcCLB.510481.20 | dispersed gene family protein 1 (DGF-1, pseudogene), put | -0,7911 | 0,01784 |
| TcCLB.511805.20 | Trichohyalin, putative | -0,7914 | 1,279E-07 |
| TcCLB.509429.10 | trans-sialidase (pseudogene), putative | -0,7916 | 6,181E-09 |
| TcCLB.504147.280 | trypanothione synthetase-like protein, putative | -0,7917 | 7,016E-23 |
| TcCLB.506599.315 | hypothetical protein, conserved (pseudogene) | -0,7918 | 0,001014 |
| TcCLB.510289.3 | hypothetical protein, conserved | -0,7919 | 7,269E-06 |
| TcCLB.510489.10 | hypothetical protein | -0,7921 | 0,001491 |
| TcCLB.508165.120 | mucin-associated surface protein (MASP), putative | -0,7925 | 0,0004509 |
| TcCLB.506181.70 | hypothetical protein, conserved | -0,7929 | 2,512E-13 |
| TcCLB.511507.80 | ferric reductase transmembrane protein, putative | -0,7932 | 0,0001308 |
| TcCLB.510819.90 | hypothetical protein, conserved | -0,7939 | 3,926E-10 |
| TcCLB.507521.100 | hypothetical protein, conserved | -0,7942 | 2,332E-09 |
| TcCLB.506331.139 | mucin-associated surface protein (MASP, pseudogene), pu | -0,7943 | 4,964E-07 |
| TcCLB.506409.220 | hypothetical protein | -0,7944 | 1,824E-10 |
| TcCLB.511403.15 | mucin TcMUC (pseudogene), putative | -0,795 | 0,1302 |
| TcCLB.509047.119 | hypothetical protein, conserved (fragment) | -0,7951 | 0,00001733 |
| TcCLB.511211.40 | molybdenum cofactor biosynthesis protein, putative | -0,7954 | 1,038E-14 |
| TcCLB.506717.25 | trans-sialidase (pseudogene), putative | -0,7954 | 4,208E-08 |
| TcCLB.510697.25 | surface protease GP63 (pseudogene), putative | -0,7954 | 0,001106 |
| TcCLB.510885.100 | hypothetical protein, conserved | -0,7955 | 1,984E-12 |
| TcCLB.511255.270 | trans-sialidase (pseudogene), putative | -0,7961 | 0,003214 |
| TcCLB.506865.70 | hypothetical protein, conserved | -0,7965 | 2,009E-15 |
| TcCLB.504787.54 | mucin-associated surface protein (MASP, pseudogene), pu | -0,7966 | 7,586E-06 |
| TcCLB.505843.40 | E1-like ubiquitin-activating enzyme, putative (UBA5) | -0,7978 | 6,929E-21 |
| TcCLB.509647.80 | hypothetical protein, conserved | -0,7981 | 1,624E-13 |
| TcCLB.504091.5 | hypothetical protein, conserved (fragment) | -0,7986 | 0,0166 |
| TcCLB.508595.35 | hypothetical protein (pseudogene) | -0,7989 | 0,003203 |
| TcCLB.506247.120 | hypothetical protein, conserved | -0,7996 | 2,2E-19 |
| TcCLB.508871.170 | trans-sialidase, Group V, putative | -0,7996 | 0,00001157 |
| TcCLB.510269.50 | mucin-associated surface protein (MASP), putative | -0,8 | 0,0000941 |
| TcCLB.508777.140 | palmitoyl acyltransferase 5, putative | -0,8002 | 2,505E-17 |
| TcCLB.509699.35 | hypothetical protein, conserved (pseudogene) | -0,8002 | 0,0001405 |
| TcCLB.504243.10 | RNA-binding protein, putative | -0,8005 | 4,394E-16 |
| TcCLB.469735.9 | hypothetical protein, conserved | -0,8005 | 8,38E-11 |
| TcCLB.506741.135 | mucin TcMUCII (pseudogene), putative | -0,8005 | 0,009502 |
| TcCLB.508299.50 | hypothetical protein, conserved | -0,8006 | 1,579E-29 |
| TcCLB.506469.130 | SNARE domain-containing protein, putative | -0,8009 | 1,576E-10 |
| TcCLB.508527.30 | trans-sialidase (pseudogene), putative | -0,801 | 8,115E-11 |
| TcCLB.509877.50 | glycosomal membrane protein, putative | -0,8013 | 6,977E-10 |

| | | | |
|---|---|---|---|
| TcCLB.413893.10 | trans-sialidase (pseudogene), putative | -0,8013 | 0,006146 |
| TcCLB.510643.40 | trans-sialidase, Group VIII, putative | -0,802 | 0,00003602 |
| TcCLB.510833.40 | trans-sialidase (pseudogene), putative | -0,8021 | 0,00003839 |
| TcCLB.506291.20 | hypothetical protein | -0,8026 | 0,04776 |
| TcCLB.508461.150 | hypothetical protein, conserved | -0,8028 | 2,677E-11 |
| TcCLB.508245.50 | hypothetical protein | -0,8031 | 0,00002357 |
| TcCLB.507689.5 | hypothetical protein, conserved (fragment) | -0,8033 | 0,00005056 |
| TcCLB.510371.40 | hypothetical protein | -0,8033 | 0,0125 |
| TcCLB.510755.19 | hypothetical protein, conserved | -0,8047 | 7,152E-12 |
| TcCLB.511183.90 | mucin TcMUCII, putative | -0,8051 | 0,007136 |
| TcCLB.508175.270 | hypothetical protein | -0,8057 | 2,747E-09 |
| TcCLB.511409.30 | hypothetical protein, conserved | -0,8058 | 2,938E-17 |
| TcCLB.509707.25 | hypothetical protein, conserved | -0,8058 | 4,347E-07 |
| TcCLB.510657.200 | surface protease GP63, putative | -0,8067 | 3,902E-09 |
| TcCLB.508827.110 | leucine-rich repeat protein (LRRP), putative | -0,807 | 2,34E-09 |
| TcCLB.509965.40 | ubiquitin hydrolase, putative | -0,8071 | 5,695E-16 |
| TcCLB.511167.20 | hypothetical protein, conserved | -0,8078 | 1,098E-14 |
| TcCLB.508081.60 | hypothetical protein | -0,8078 | 0,004656 |
| TcCLB.413133.10 | mucin TcMUCII (pseudogene), putative | -0,8079 | 0,000514 |
| TcCLB.510303.160 | hypothetical protein, conserved | -0,8083 | 2,287E-17 |
| TcCLB.504069.34 | hypothetical protein, conserved | -0,8087 | 2,251E-06 |
| TcCLB.506667.130 | surface protease GP63 (pseudogene), putative | -0,8089 | 0,02667 |
| TcCLB.511259.230 | trans-sialidase (pseudogene), putative | -0,8091 | 0,001275 |
| TcCLB.510221.20 | hypothetical protein, conserved | -0,8092 | 1,168E-12 |
| TcCLB.511165.10 | ATP-dependent DEAD/H RNA helicase, putative | -0,8095 | 5,55E-12 |
| TcCLB.508175.210 | hypothetical protein, conserved | -0,8095 | 2,632E-06 |
| TcCLB.504099.50 | trans-sialidase, Group II, putative | -0,8097 | 3,949E-08 |
| TcCLB.511233.90 | hypothetical protein, conserved | -0,8098 | 4,387E-06 |
| TcCLB.510055.10 | trans-sialidase, putative | -0,8099 | 2,072E-08 |
| TcCLB.503775.50 | mucin TcMUCII (pseudogene), putative | -0,8106 | 0,00879 |
| TcCLB.511205.10 | surface protease GP63 (pseudogene), putative | -0,8108 | 3,888E-10 |
| TcCLB.503553.10 | trans-sialidase (pseudogene), putative | -0,812 | 0,009475 |
| TcCLB.506883.80 | tRNA (guanine-N(7)-)-methyltransferase, putative | -0,8121 | 4,003E-12 |
| TcCLB.431849.10 | hypothetical protein, conserved | -0,8125 | 0,00002578 |
| TcCLB.508139.155 | protein kinase (pseudogene), putative | -0,8131 | 2,204E-07 |
| TcCLB.510887.30 | hypothetical protein, conserved | -0,8139 | 7,233E-09 |
| TcCLB.506681.50 | hypothetical protein, conserved | -0,8142 | 3,193E-21 |
| TcCLB.506755.100 | hypothetical protein, conserved | -0,8142 | 7,916E-11 |
| TcCLB.511071.90 | hypothetical protein | -0,8144 | 0,00006091 |
| TcCLB.511149.190 | mucin-associated surface protein (MASP, pseudogene), pu | -0,8151 | 0,00003127 |
| TcCLB.508175.20 | hypothetical protein, conserved | -0,8154 | 1,054E-26 |
| TcCLB.509065.170 | hypothetical protein, conserved | -0,8154 | 3,378E-17 |
| TcCLB.506755.180 | hypothetical protein, conserved | -0,8155 | 3,121E-08 |
| TcCLB.507085.110 | hypothetical protein | -0,8157 | 0,0001496 |
| TcCLB.506509.10 | hypothetical protein, conserved | -0,8159 | 4,381E-12 |

| | | | |
|---|---|---|---|
| TcCLB.511649.39 | hypothetical protein, conserved | -0,816 | 5,795E-08 |
| TcCLB.505297.50 | trans-sialidase (pseudogene), putative | -0,8163 | 5,505E-19 |
| TcCLB.508593.100 | hypothetical protein, conserved | -0,8171 | 0,0001825 |
| TcCLB.510431.120 | hypothetical protein, conserved | -0,8173 | 2,37E-14 |
| TcCLB.508835.10 | hypothetical protein, conserved (pseudogene) | -0,8177 | 3,996E-07 |
| TcCLB.506131.114 | mucin TcMUC (pseudogene), putative | -0,8183 | 0,01006 |
| TcCLB.508277.190 | ERAD-associated E3 ubiquitin-protein ligase HRD1, putativ | -0,8184 | 2,908E-20 |
| TcCLB.509671.110 | hypothetical protein, conserved | -0,8186 | 5,478E-11 |
| TcCLB.511259.80 | mucin TcMUCII, putative | -0,8188 | 0,002266 |
| TcCLB.506759.31 | surface protease GP63 (pseudogene), putative | -0,8189 | 0,0006381 |
| TcCLB.508495.20 | trans-sialidase (pseudogene), putative | -0,8192 | 4,193E-13 |
| TcCLB.508461.330 | hypothetical protein | -0,8193 | 3,763E-13 |
| TcCLB.506247.74 | hypothetical protein, conserved | -0,8193 | 1,573E-06 |
| TcCLB.508727.30 | hypothetical protein, conserved | -0,8195 | 3,133E-08 |
| TcCLB.506459.230 | trans-sialidase, Group V, putative | -0,8199 | 9,005E-08 |
| TcCLB.508737.154 | hypothetical protein, conserved | -0,82 | 7,749E-06 |
| TcCLB.511621.170 | hypothetical protein, conserved | -0,8203 | 1,042E-06 |
| TcCLB.510339.40 | hypothetical protein, conserved | -0,821 | 2,363E-13 |
| TcCLB.506951.30 | trans-sialidase (pseudogene), putative | -0,8214 | 0,00005075 |
| TcCLB.510013.190 | hypothetical protein | -0,8214 | 0,01893 |
| TcCLB.510709.70 | trans-sialidase, Group V, putative | -0,8214 | 0,07178 |
| TcCLB.509195.80 | trans-sialidase (pseudogene), putative | -0,822 | 2,602E-07 |
| TcCLB.511857.10 | hypothetical protein, conserved | -0,8221 | 5,022E-11 |
| TcCLB.506627.130 | dispersed gene family protein 1 (DGF-1, pseudogene), put | -0,8222 | 0,01229 |
| TcCLB.504085.70 | Zinc finger protein, C3H1 type-like (ZC3H46) | -0,8222 | 0,06655 |
| TcCLB.511203.10 | surface protease GP63, putative | -0,8229 | 0,0001698 |
| TcCLB.507511.81 | mucin TcMUCII, putative | -0,8234 | 1,898E-11 |
| TcCLB.506241.10 | retrotransposon hot spot protein (RHS, pseudogene), puta | -0,8235 | 1,076E-06 |
| TcCLB.510901.160 | hypothetical protein, conserved | -0,8244 | 1,768E-09 |
| TcCLB.511797.210 | trans-sialidase (pseudogene), putative | -0,8252 | 0,00003513 |
| TcCLB.510693.220 | mucin TcMUCII, putative | -0,8252 | 0,006183 |
| TcCLB.505037.70 | trans-sialidase (pseudogene), putative | -0,8255 | 2,181E-07 |
| TcCLB.506305.10 | hypothetical protein, conserved | -0,8257 | 2,806E-26 |
| TcCLB.506965.120 | hypothetical protein | -0,8261 | 0,02704 |
| TcCLB.511495.11 | hypothetical protein, conserved (fragment) | -0,8266 | 0,00000466 |
| TcCLB.510107.30 | mucin-associated surface protein (MASP), putative | -0,8269 | 0,0002805 |
| TcCLB.508761.150 | mucin-associated surface protein (MASP, pseudogene), pu | -0,8276 | 0,01001 |
| TcCLB.510483.200 | hypothetical protein | -0,828 | 0,01448 |
| TcCLB.511861.111 | elongation factor 1-gamma (EF-1-gamma), putative | -0,828 | 0,05145 |
| TcCLB.511181.60 | hypothetical protein, conserved | -0,8281 | 4,979E-13 |
| TcCLB.508443.60 | hypothetical protein, conserved | -0,8281 | 4,848E-09 |
| TcCLB.503533.30 | trans-sialidase, Group V, putative | -0,8283 | 0,07019 |
| TcCLB.506577.10 | hypothetical protein, conserved | -0,8289 | 2,776E-08 |
| TcCLB.511517.156 | hypothetical protein, conserved | -0,8293 | 1,153E-11 |
| TcCLB.510305.50 | hypothetical protein, conserved | -0,8304 | 1,266E-21 |

| | | | |
|---|---|---|---|
| TcCLB.510483.210 | trans-sialidase (pseudogene), putative | -0,8308 | 1,451E-12 |
| TcCLB.508209.150 | hypothetical protein, conserved | -0,8314 | 1,2E-20 |
| TcCLB.509167.200 | hypothetical protein, conserved | -0,832 | 7,93E-17 |
| TcCLB.506753.90 | hypothetical protein, conserved | -0,8321 | 8,881E-26 |
| TcCLB.508761.200 | trans-sialidase (pseudogene), putative | -0,8329 | 0,0004199 |
| TcCLB.506739.140 | hypothetical protein, conserved | -0,8339 | 0,00004343 |
| TcCLB.506345.40 | trans-sialidase, Group II, putative | -0,8341 | 9,868E-13 |
| TcCLB.510421.180 | casein kinase I, putative | -0,8342 | 1,261E-11 |
| TcCLB.508363.10 | hypothetical protein | -0,8345 | 0,01595 |
| TcCLB.509369.20 | retrotransposon hot spot protein (RHS, pseudogene), puta | -0,8351 | 0,007516 |
| TcCLB.506559.330 | clathrin coat assembly protein AP17, putative | -0,8357 | 2,187E-10 |
| TcCLB.511401.80 | hypothetical protein, conserved (pseudogene) | -0,8359 | 1,777E-10 |
| TcCLB.506469.90 | hypothetical protein, conserved | -0,8359 | 3,644E-06 |
| TcCLB.503771.74 | mucin TcMUCII (pseudogene), putative | -0,8359 | 0,02061 |
| TcCLB.506267.19 | hypothetical protein | -0,8366 | 0,0006452 |
| TcCLB.509429.200 | hypothetical protein, conserved | -0,8368 | 1,008E-13 |
| TcCLB.510519.100 | hypothetical protein, conserved | -0,8374 | 9,592E-10 |
| TcCLB.509695.190 | cdc2-related kinase 1, putative (CRK1) | -0,8378 | 4,324E-15 |
| TcCLB.511921.100 | trans-sialidase (pseudogene), putative | -0,838 | 9,449E-06 |
| TcCLB.509663.10 | myosin heavy chain, putative | -0,8384 | 4,106E-22 |
| TcCLB.508479.50 | trans-sialidase (pseudogene), putative | -0,8397 | 3,858E-11 |
| TcCLB.510607.70 | trans-sialidase (pseudogene), putative | -0,8401 | 0,00003368 |
| TcCLB.508869.59 | mucin-associated surface protein (MASP), putative | -0,8406 | 0,06006 |
| TcCLB.506435.210 | hypothetical protein, conserved | -0,841 | 3,35E-15 |
| TcCLB.508691.10 | hypothetical protein, conserved | -0,8412 | 1,707E-08 |
| TcCLB.506945.50 | hypothetical protein, conserved | -0,8414 | 1,724E-15 |
| TcCLB.510901.180 | hypothetical protein, conserved | -0,8415 | 1,557E-12 |
| TcCLB.506955.240 | trans-sialidase, Group V, putative | -0,8415 | 8,379E-07 |
| TcCLB.504147.250 | hypothetical protein, conserved | -0,8416 | 6,867E-29 |
| TcCLB.506529.70 | hypothetical protein, conserved | -0,8418 | 1,863E-14 |
| TcCLB.506459.200 | hypothetical protein | -0,8418 | 0,0297 |
| TcCLB.511183.270 | mucin TcMUCII (pseudogene), putative | -0,8419 | 0,02122 |
| TcCLB.506775.70 | hypothetical protein, conserved | -0,8424 | 4,703E-16 |
| TcCLB.510131.70 | DNA polymerase kappa, putative | -0,8432 | 1,376E-08 |
| TcCLB.508175.264 | hypothetical protein, conserved (pseudogene) | -0,8439 | 3,684E-10 |
| TcCLB.506579.80 | hypothetical protein, conserved | -0,844 | 1,455E-15 |
| TcCLB.508355.360 | hypothetical protein, conserved | -0,8441 | 1,281E-08 |
| TcCLB.510693.251 | mucin TcMUC (pseudogene), putative | -0,8441 | 0,002594 |
| TcCLB.506145.80 | surface protease GP63, putative | -0,845 | 7,365E-19 |
| TcCLB.508161.30 | mucin-associated surface protein (MASP), putative | -0,8456 | 0,00003243 |
| TcCLB.508221.30 | mucin-associated surface protein (MASP), putative | -0,8459 | 0,02629 |
| TcCLB.503659.20 | mucin TcMUCII (pseudogene), putative | -0,8463 | 0,000171 |
| TcCLB.507143.70 | hypothetical protein, conserved | -0,8466 | 4,715E-22 |
| TcCLB.506743.60 | hypothetical protein, conserved | -0,8468 | 2,799E-19 |
| TcCLB.507991.120 | translation initiation factor IF-2, putative | -0,8468 | 2,443E-12 |

| | | | |
|---|---|---|---|
| TcCLB.508435.60 | mucin TcMUCII, putative | -0,8471 | 0,03054 |
| TcCLB.506129.50 | trans-sialidase, Group IV, putative | -0,8476 | 1,103E-18 |
| TcCLB.511923.120 | dihydroorotate dehydrogenase, putative (DHODH) | -0,8477 | 7,188E-10 |
| TcCLB.506459.120 | mucin TcMUCII, putative | -0,8477 | 0,01315 |
| TcCLB.506737.121 | mucin-associated surface protein (MASP, pseudogene), pu | -0,8479 | 3,263E-15 |
| TcCLB.511765.55 | trans-sialidase (pseudogene), putative | -0,8483 | 0,007629 |
| TcCLB.506473.4 | hypothetical protein, conserved (pseudogene) | -0,849 | 0,02741 |
| TcCLB.507711.290 | hypothetical protein, conserved | -0,8491 | 9,302E-13 |
| TcCLB.506501.27 | syntaxin binding protein (pseudogene), putative | -0,8491 | 0,03391 |
| TcCLB.504239.401 | hypothetical protein, conserved (pseudogene) | -0,8493 | 0,03824 |
| TcCLB.504741.80 | hypothetical protein, conserved | -0,8502 | 3,85E-11 |
| TcCLB.403153.10 | receptor-type adenylate cyclase (pseudogene), putative | -0,8504 | 1,533E-06 |
| TcCLB.511675.30 | retrotransposon hot spot protein (RHS, pseudogene), puta | -0,8504 | 0,01759 |
| TcCLB.508831.180 | beta galactofuranosyl glycosyltransferase (pseudogene), p | -0,851 | 0,01056 |
| TcCLB.511259.70 | mucin-associated surface protein (MASP, pseudogene), pu | -0,8514 | 0,0002812 |
| TcCLB.511625.130 | trans-sialidase, Group V, putative | -0,8515 | 0,00004716 |
| TcCLB.508175.133 | hypothetical protein, conserved (pseudogene) | -0,8518 | 2,419E-24 |
| TcCLB.509419.58 | mucin TcMUCII (pseudogene), putative | -0,8525 | 0,05148 |
| TcCLB.504045.5 | hypothetical protein, conserved (fragment) | -0,8526 | 0,0001548 |
| TcCLB.506567.80 | hypothetical protein | -0,8527 | 1,878E-13 |
| TcCLB.504313.10 | trans-sialidase, Group IV, putative | -0,8528 | 0,008116 |
| TcCLB.511787.10 | mucin-associated surface protein (MASP), putative | -0,8529 | 0,04933 |
| TcCLB.511591.10 | mucin-associated surface protein (MASP), putative | -0,853 | 0,03239 |
| TcCLB.506995.60 | surface protease GP63, putative | -0,8531 | 0,002233 |
| TcCLB.508813.40 | surface protease GP63, putative | -0,8532 | 6,784E-10 |
| TcCLB.506513.150 | hypothetical protein, conserved | -0,8534 | 1,169E-12 |
| TcCLB.509601.110 | hypothetical protein, conserved | -0,8537 | 3,21E-15 |
| TcCLB.511281.50 | surface protease GP63, putative | -0,8539 | 1,118E-08 |
| TcCLB.511555.20 | hypothetical protein, conserved | -0,8543 | 1,124E-20 |
| TcCLB.509735.100 | trans-sialidase (pseudogene), putative | -0,8544 | 2,568E-08 |
| TcCLB.410199.7 | hypothetical protein | -0,8545 | 0,001238 |
| TcCLB.506139.4 | mucin TcMUCII, putative | -0,8548 | 0,05858 |
| TcCLB.507847.10 | mucin TcMUCII (pseudogene), putative | -0,8549 | 9,061E-07 |
| TcCLB.510243.60 | hypothetical protein | -0,8564 | 1,274E-18 |
| TcCLB.510199.10 | GP85-like protein, putative | -0,857 | 0,01158 |
| TcCLB.508573.80 | trans-sialidase (pseudogene), putative | -0,8571 | 7,95E-09 |
| TcCLB.508521.81 | elongation factor 1-gamma (EF-1-gamma), putative | -0,8572 | 0,0006012 |
| TcCLB.506599.40 | hypothetical protein | -0,8574 | 0,02452 |
| TcCLB.508525.5 | cAMP response protein 1, putative (pseudogene) | -0,8576 | 8,344E-06 |
| TcCLB.511255.190 | trans-sialidase, Group V, putative | -0,8576 | 0,000324 |
| TcCLB.506939.209 | mucin TcMUCII (pseudogene), putative | -0,8576 | 0,0003366 |
| TcCLB.508229.20 | hypothetical protein | -0,8577 | 0,0103 |
| TcCLB.508629.100 | trans-sialidase (pseudogene), putative | -0,8581 | 5,928E-15 |
| TcCLB.507801.230 | trans-sialidase (pseudogene), putative | -0,8582 | 1,912E-07 |
| TcCLB.506637.30 | hypothetical protein, conserved | -0,8583 | 6,032E-25 |

| | | | |
|---|---|---|---|
| TcCLB.506769.90 | surface protease GP63, putative | -0,8584 | 0,004403 |
| TcCLB.508221.360 | mucin-associated surface protein (MASP, pseudogene), pu | -0,8591 | 0,02777 |
| TcCLB.506999.10 | hypothetical protein, conserved | -0,8592 | 2,563E-12 |
| TcCLB.510481.30 | hypothetical protein, conserved | -0,8594 | 0,06393 |
| TcCLB.508577.110 | hypothetical protein, conserved | -0,8599 | 2,092E-12 |
| TcCLB.508375.50 | dihydroorotate dehydrogenase (fumarate), putative (DHO | -0,8607 | 1,058E-14 |
| TcCLB.510305.60 | hypothetical protein, conserved | -0,8611 | 3,625E-30 |
| TcCLB.509053.80 | hypothetical protein, conserved | -0,8612 | 3,281E-11 |
| TcCLB.506067.34 | mucin-associated surface protein (MASP, pseudogene), pu | -0,8616 | 0,00001672 |
| TcCLB.504081.270 | mucin TcMUCII, putative | -0,8617 | 0,01083 |
| TcCLB.510401.31 | retrotransposon hot spot protein (RHS, pseudogene), puta | -0,8623 | 0,03801 |
| TcCLB.505007.30 | hypothetical protein, conserved | -0,8626 | 3,385E-12 |
| TcCLB.503993.20 | hypothetical protein | -0,8626 | 0,00603 |
| TcCLB.506245.210 | mucin TcMUCII, putative | -0,8628 | 0,0011 |
| TcCLB.504431.100 | guanine deaminase, putative | -0,8637 | 5,67E-13 |
| TcCLB.508057.20 | mucin-associated surface protein (MASP, pseudogene), pu | -0,8638 | 5,486E-08 |
| TcCLB.506297.130 | zinc finger ccch and cchc domain-containing protein | -0,8639 | 8,706E-28 |
| TcCLB.506859.4 | N-acetyltransferase, putative (fragment) | -0,8644 | 0,00006147 |
| TcCLB.510687.130 | hypothetical protein, conserved | -0,8651 | 7,866E-12 |
| TcCLB.508909.260 | hypothetical protein, conserved | -0,8651 | 9,464E-06 |
| TcCLB.506195.200 | hypothetical protein, conserved | -0,8655 | 2,771E-24 |
| TcCLB.504033.200 | hypothetical protein | -0,8657 | 9,329E-14 |
| TcCLB.509237.40 | hypothetical protein, conserved | -0,8658 | 1,033E-10 |
| TcCLB.509085.110 | trans-sialidase (pseudogene), putative | -0,8661 | 6,809E-06 |
| TcCLB.509643.10 | hypothetical protein, conserved | -0,8661 | 0,00002991 |
| TcCLB.508365.220 | hypothetical protein, conserved (pseudogene) | -0,8667 | 0,002833 |
| TcCLB.508261.80 | trans-sialidase (pseudogene), putative | -0,8668 | 0,0000258 |
| TcCLB.507041.65 | hypothetical protein, conserved (pseudogene) | -0,8668 | 0,0009395 |
| TcCLB.511499.50 | ubiquitin hydrolase, putative | -0,8669 | 8,492E-21 |
| TcCLB.458369.10 | trans-sialidase (pseudogene), putative | -0,8672 | 0,0009401 |
| TcCLB.511277.440 | tubulin tyrosine ligase protein, putative | -0,8679 | 3,1E-13 |
| TcCLB.504083.20 | retrotransposon hot spot protein (RHS, pseudogene), puta | -0,8683 | 0,04164 |
| TcCLB.508293.120 | hypothetical protein | -0,8689 | 0,005899 |
| TcCLB.507875.200 | trans-sialidase (pseudogene), putative | -0,8693 | 8,472E-07 |
| TcCLB.410923.20 | retrotransposon hot spot (RHS) protein, putative | -0,8695 | 2,22E-13 |
| TcCLB.509429.170 | hypothetical protein, conserved | -0,8695 | 1,79E-12 |
| TcCLB.509163.6 | hypothetical protein | -0,8699 | 0,04018 |
| TcCLB.510257.50 | hypothetical protein, conserved | -0,87 | 5,506E-11 |
| TcCLB.503579.40 | hypothetical protein, conserved | -0,8701 | 3,191E-13 |
| TcCLB.511149.60 | mucin-associated surface protein (MASP), putative | -0,8702 | 0,03069 |
| TcCLB.510879.229 | protein kinase, putative | -0,8704 | 6,209E-07 |
| TcCLB.511867.80 | Vacuolar iron transporter 1, putative (VIT1) | -0,8706 | 8,689E-15 |
| TcCLB.507941.70 | hypothetical protein, conserved | -0,8708 | 2,163E-21 |
| TcCLB.506249.30 | hypothetical protein, conserved | -0,8708 | 1,453E-06 |
| TcCLB.509429.260 | hypothetical protein, conserved | -0,8709 | 7,408E-18 |

| | | | |
|---|---|---|---|
| TcCLB.508261.120 | hypothetical protein | -0,8713 | 0,0007535 |
| TcCLB.511675.10 | trans-sialidase (pseudogene), putative | -0,8716 | 1,322E-14 |
| TcCLB.508317.80 | mannosyl-oligosaccharide 1,2-alpha-mannosidase IB, puta | -0,8718 | 3,01E-20 |
| TcCLB.508297.14 | trans-sialidase (pseudogene), putative | -0,8719 | 6,173E-14 |
| TcCLB.503563.29 | hypothetical protein | -0,8722 | 0,00002056 |
| TcCLB.506451.50 | trans-sialidase (pseudogene), putative | -0,8725 | 0,000653 |
| TcCLB.508621.51 | peptidylprolyl isomerase-like protein (pseudogene), putati | -0,8727 | 0,03645 |
| TcCLB.509133.10 | trans-sialidase (pseudogene), putative | -0,8736 | 6,741E-06 |
| TcCLB.510901.220 | hypothetical protein, conserved | -0,8738 | 4,035E-18 |
| TcCLB.506319.20 | hypothetical protein, conserved | -0,8742 | 1,934E-17 |
| TcCLB.508173.129 | hypothetical protein, conserved | -0,8743 | 8,641E-21 |
| TcCLB.504021.20 | hypothetical protein, conserved | -0,8754 | 0,002352 |
| TcCLB.506445.70 | hypothetical protein, conserved | -0,8757 | 1,62E-12 |
| TcCLB.505965.10 | surface protease GP63, putative | -0,8757 | 0,004118 |
| TcCLB.504113.20 | MYH7B protein, putative | -0,8759 | 5,892E-11 |
| TcCLB.504797.80 | endo-beta-N-acetylglucosaminidase, putative | -0,876 | 1,961E-26 |
| TcCLB.510463.221 | hypothetical protein, conserved (pseudogene) | -0,8761 | 0,0544 |
| TcCLB.503475.10 | dispersed gene family protein 1 (DGF-1, pseudogene), put | -0,8767 | 0,2154 |
| TcCLB.508317.120 | mannosyl-oligosaccharide 1,2-alpha-mannosidase IB, puta | -0,8769 | 2,564E-21 |
| TcCLB.507765.30 | hypothetical protein, conserved | -0,8769 | 1,471E-16 |
| TcCLB.511875.29 | mucin-associated surface protein (MASP, pseudogene), pu | -0,8774 | 3,217E-06 |
| TcCLB.503809.60 | glycerol uptake protein (pseudogene), putative | -0,8775 | 1,456E-08 |
| TcCLB.506247.110 | hypothetical protein, conserved | -0,8777 | 3,895E-13 |
| TcCLB.505997.160 | retrotransposon hot spot protein (RHS, pseudogene), puta | -0,878 | 1,536E-11 |
| TcCLB.508277.380 | hypothetical protein, conserved | -0,8781 | 9,218E-34 |
| TcCLB.511283.280 | Serine/threonine-protein kinase Nek1-related, putative | -0,8781 | 4,182E-20 |
| TcCLB.510279.100 | hypothetical protein | -0,8788 | 0,006878 |
| TcCLB.509717.40 | hypothetical protein, conserved | -0,8792 | 2,006E-21 |
| TcCLB.510303.80 | kinesin, putative | -0,8794 | 1,023E-17 |
| TcCLB.507677.60 | hypothetical protein | -0,8801 | 9,093E-24 |
| TcCLB.510159.10 | hypothetical protein, conserved | -0,8806 | 2,286E-12 |
| TcCLB.464807.10 | serine/threonine protein kinase, putative | -0,8814 | 1,373E-11 |
| TcCLB.503891.60 | hypothetical protein, conserved | -0,8825 | 3,403E-16 |
| TcCLB.507957.100 | mucin-associated surface protein (MASP, pseudogene), pu | -0,8829 | 0,0001149 |
| TcCLB.476733.10 | hypothetical protein, conserved | -0,883 | 4,724E-23 |
| TcCLB.509487.4 | DNA-directed RNA polymerase III, putative | -0,883 | 1,738E-06 |
| TcCLB.506899.30 | mucin-associated surface protein (MASP, pseudogene), pu | -0,8836 | 0,01705 |
| TcCLB.506755.220 | hypothetical protein, conserved | -0,884 | 6,841E-19 |
| TcCLB.422843.10 | trans-sialidase (pseudogene), putative | -0,884 | 1,367E-12 |
| TcCLB.506515.29 | trans-sialidase, putative | -0,884 | 0,00002254 |
| TcCLB.511547.14 | mucin-associated surface protein (MASP), putative | -0,8845 | 0,007394 |
| TcCLB.465323.10 | hypothetical protein (fragment) | -0,8846 | 0,008488 |
| TcCLB.510429.20 | tRNA (guanine-N(7)-)-methyltransferase, putative | -0,8847 | 7,139E-10 |
| TcCLB.510819.20 | hypothetical protein, conserved | -0,8852 | 4,187E-10 |
| TcCLB.504005.50 | hypothetical protein, conserved | -0,8854 | 4,615E-16 |

| | | | |
|---|---|---|---|
| TcCLB.508761.9 | mucin-associated surface protein (MASP, pseudogene), pu | -0,886 | 0,0008035 |
| TcCLB.511677.20 | trans-sialidase, putative | -0,8864 | 0,009606 |
| TcCLB.511861.130 | trans-sialidase (pseudogene), putative | -0,8867 | 2,938E-12 |
| TcCLB.508387.50 | bilobe protein, conserved | -0,8873 | 1,91E-20 |
| TcCLB.507061.49 | hypothetical protein | -0,8879 | 5,849E-13 |
| TcCLB.508187.30 | surface protease GP63, putative (fragment) | -0,8879 | 0,0008829 |
| TcCLB.508989.110 | hypothetical protein, conserved | -0,8882 | 2,203E-20 |
| TcCLB.509099.40 | hypothetical protein, conserved | -0,8882 | 8,109E-12 |
| TcCLB.504105.260 | bilobe protein, conserved | -0,8894 | 0,0005528 |
| TcCLB.508799.30 | hypothetical protein, conserved | -0,8896 | 8,818E-19 |
| TcCLB.509695.230 | serine carboxypeptidase (CBP1), putative (fragment) | -0,8897 | 3,604E-09 |
| TcCLB.509213.70 | ribulose-5-phosphate 3-epimerase, putative | -0,8899 | 3,773E-15 |
| TcCLB.431849.20 | hypothetical protein, conserved (fragment) | -0,8899 | 8,098E-06 |
| TcCLB.510885.80 | hypothetical protein, conserved | -0,8901 | 1,807E-15 |
| TcCLB.504027.4 | kinesin, putative (fragment) | -0,8902 | 6,137E-10 |
| TcCLB.506485.109 | cyclic nucleotide-binding domain containing protein, putat | -0,8905 | 8,991E-19 |
| TcCLB.506807.39 | hypothetical protein, conserved (fragment) | -0,8907 | 1,067E-09 |
| TcCLB.506357.20 | hypothetical protein, conserved | -0,8908 | 2,238E-21 |
| TcCLB.503875.40 | hypothetical protein | -0,8916 | 0,001875 |
| TcCLB.506753.190 | hypothetical protein, conserved | -0,892 | 1,591E-15 |
| TcCLB.511559.10 | Present in the outer mitochondrial membrane proteome 1 | -0,892 | 0,02807 |
| TcCLB.506357.10 | hypothetical protein, conserved | -0,8924 | 2,625E-10 |
| TcCLB.508961.70 | hypothetical protein, conserved | -0,8926 | 1,909E-15 |
| TcCLB.511183.371 | hypothetical protein, conserved (pseudogene) | -0,8935 | 0,008182 |
| TcCLB.506173.179 | trans-sialidase (pseudogene), putative | -0,8939 | 0,00001168 |
| TcCLB.506951.50 | mucin TcMUCII (pseudogene), putative | -0,8945 | 3,154E-09 |
| TcCLB.506963.100 | hypothetical protein, conserved | -0,8948 | 1,937E-07 |
| TcCLB.508221.150 | mucin-associated surface protein (MASP), putative | -0,895 | 0,005614 |
| TcCLB.503871.10 | trans-sialidase (pseudogene), putative | -0,8954 | 0,0000302 |
| TcCLB.510029.90 | hypothetical protein, conserved | -0,8957 | 1,721E-17 |
| TcCLB.507095.20 | hypothetical protein | -0,8967 | 0,003003 |
| TcCLB.510885.10 | hypothetical protein, conserved | -0,8973 | 2,031E-17 |
| TcCLB.510543.210 | retrotransposon hot spot protein (RHS, pseudogene), puta | -0,8974 | 9,274E-06 |
| TcCLB.507093.20 | hypothetical protein, conserved | -0,8976 | 1,049E-11 |
| TcCLB.508165.34 | mucin TcMUC (pseudogene), putative | -0,8982 | 0,004489 |
| TcCLB.505997.100 | retrotransposon hot spot protein (RHS, pseudogene), puta | -0,8984 | 2,004E-11 |
| TcCLB.510165.60 | expression site-associated gene (ESAG-like) protein, puta | -0,899 | 0,00000246 |
| TcCLB.506735.40 | hypothetical protein | -0,8991 | 5,715E-15 |
| TcCLB.511025.60 | hypothetical protein | -0,8993 | 4,172E-12 |
| TcCLB.503417.20 | hypothetical protein | -0,8994 | 0,004158 |
| TcCLB.510533.70 | protein kinase, putative | -0,9003 | 6,289E-15 |
| TcCLB.511237.130 | trans-sialidase, putative (fragment) | -0,9008 | 5,185E-06 |
| TcCLB.510143.120 | hypothetical protein, conserved | -0,9012 | 1,465E-10 |
| TcCLB.511037.40 | hypothetical protein, conserved | -0,9013 | 0,0002968 |
| TcCLB.506057.81 | mucin-associated surface protein (MASP, pseudogene), pu | -0,9018 | 1,765E-14 |

| | | | |
|---|---|---|---|
| TcCLB.506173.270 | mucin TcMUCII, putative | -0,902 | 0,008595 |
| TcCLB.509179.231 | trans-sialidase (pseudogene), putative | -0,9022 | 0,00006493 |
| TcCLB.509957.10 | hypothetical protein | -0,903 | 0,00003915 |
| TcCLB.511491.130 | dolichyl pyrophosphate phosphatase, putative | -0,9039 | 2,571E-11 |
| TcCLB.510817.40 | hypothetical protein, conserved | -0,9051 | 0,00002364 |
| TcCLB.507943.70 | hypothetical protein, conserved | -0,9058 | 1,884E-14 |
| TcCLB.509695.140 | hypothetical protein, conserved | -0,9074 | 4,086E-19 |
| TcCLB.510173.50 | hypothetical protein, conserved | -0,9078 | 4,761E-08 |
| TcCLB.503639.20 | hypothetical protein, conserved | -0,908 | 1,189E-07 |
| TcCLB.508313.140 | mucin-associated surface protein (MASP), putative | -0,9081 | 0,03559 |
| TcCLB.511127.290 | hypothetical protein, conserved | -0,9088 | 1,345E-21 |
| TcCLB.503775.60 | hypothetical protein | -0,9088 | 2,291E-06 |
| TcCLB.509487.10 | hypothetical protein | -0,9098 | 5,15E-13 |
| TcCLB.508741.110 | hypothetical protein, conserved | -0,91 | 3,363E-15 |
| TcCLB.510699.11 | surface protease GP63 (pseudogene), putative | -0,9106 | 0,01036 |
| TcCLB.507625.110 | chaperone DNAJ protein, putative | -0,9116 | 1,915E-20 |
| TcCLB.511537.30 | hypothetical protein, conserved | -0,9117 | 8,718E-25 |
| TcCLB.430061.9 | hypothetical protein, conserved | -0,9121 | 0,00002157 |
| TcCLB.510279.30 | hypothetical protein | -0,9122 | 0,0217 |
| TcCLB.507905.30 | trans-sialidase (pseudogene), putative | -0,9132 | 7,316E-27 |
| TcCLB.511517.37 | reticulon domain protein, putative | -0,914 | 2,396E-13 |
| TcCLB.508883.10 | mucin-associated surface protein (MASP, pseudogene), pu | -0,915 | 0,05897 |
| TcCLB.506165.4 | hypothetical protein, conserved | -0,9156 | 9,223E-15 |
| TcCLB.508207.40 | hypothetical protein, conserved | -0,9157 | 1,799E-09 |
| TcCLB.506247.420 | hypothetical protein, conserved | -0,9163 | 3,409E-08 |
| TcCLB.506741.30 | mucin-associated surface protein (MASP, pseudogene), pu | -0,9164 | 0,06319 |
| TcCLB.511277.190 | ras-related GTP-binding protein, putative | -0,9173 | 3,168E-18 |
| TcCLB.504005.40 | hypothetical protein, conserved | -0,9187 | 7,157E-13 |
| TcCLB.508493.20 | mucin-associated surface protein (MASP, pseudogene), pu | -0,919 | 1,232E-08 |
| TcCLB.503673.9 | eukaryotic translation initiation factor 2-alpha kinase, puta | -0,9194 | 9,857E-10 |
| TcCLB.511149.80 | hypothetical protein | -0,9196 | 0,0007458 |
| TcCLB.459389.10 | hypothetical protein, conserved (fragment) | -0,9197 | 1,23E-13 |
| TcCLB.510183.40 | hypothetical protein, conserved | -0,9201 | 5,956E-09 |
| TcCLB.506501.401 | hypothetical protein, conserved (pseudogene) | -0,921 | 0,1159 |
| TcCLB.509167.90 | hypothetical protein, conserved | -0,9219 | 2,523E-15 |
| TcCLB.507713.30 | heat shock protein 85, putative | -0,9219 | 0,0007787 |
| TcCLB.511165.30 | hypothetical protein, conserved | -0,9225 | 4,886E-09 |
| TcCLB.506885.450 | Kinesin-13 6, putative (KIN13-6) | -0,9232 | 1,604E-17 |
| TcCLB.508539.150 | surface protease GP63 (pseudogene), putative | -0,9234 | 0,00009808 |
| TcCLB.510601.40 | tubulin tyrosine ligase (pseudogene), putative | -0,9235 | 2,988E-13 |
| TcCLB.506245.70 | mucin-associated surface protein (MASP, pseudogene), pu | -0,9235 | 0,0000494 |
| TcCLB.511183.550 | mucin TcMUCII, putative | -0,9239 | 0,01961 |
| TcCLB.509399.140 | hypothetical protein, conserved | -0,9246 | 4,07E-14 |
| TcCLB.507993.350 | surface protease GP63, putative | -0,9249 | 1,184E-06 |
| TcCLB.506569.45 | protein kinase (pseudogene), putative | -0,9253 | 8,069E-07 |

| | | | |
|---|---|---|---|
| TcCLB.511127.230 | hypothetical protein, conserved | -0,9255 | 8,638E-13 |
| TcCLB.503887.20 | hypothetical protein | -0,9267 | 3,619E-12 |
| TcCLB.506563.4 | hypothetical protein, conserved | -0,9268 | 7,506E-08 |
| TcCLB.507995.10 | trans-sialidase (pseudogene), putative | -0,9271 | 9,581E-14 |
| TcCLB.510769.130 | kinesin, putative | -0,9272 | 1,948E-22 |
| TcCLB.508873.210 | mucin-associated surface protein (MASP), putative | -0,9277 | 0,0003484 |
| TcCLB.507163.40 | mucin-associated surface protein (MASP), putative | -0,9278 | 0,000237 |
| TcCLB.510205.50 | mucin-associated surface protein (MASP), putative | -0,9278 | 0,008228 |
| TcCLB.503993.10 | trans-sialidase, Group I, putative | -0,9288 | 6,49E-14 |
| TcCLB.511851.5 | trans-sialidase, putative | -0,9288 | 0,002288 |
| TcCLB.509429.130 | hypothetical protein, conserved | -0,929 | 7,343E-15 |
| TcCLB.511667.80 | trans-sialidase (pseudogene), putative | -0,9292 | 1,213E-11 |
| TcCLB.506465.40 | protein kinase-like protein, putative | -0,9297 | 5,609E-13 |
| TcCLB.504023.10 | hypothetical protein, conserved | -0,9299 | 1,149E-12 |
| TcCLB.507993.310 | transporter, putative | -0,9302 | 3,043E-10 |
| TcCLB.508047.60 | trans-sialidase, Group V, putative | -0,9302 | 6,855E-08 |
| TcCLB.510281.5 | mucin TcMUCII (pseudogene), putative | -0,9306 | 0,0367 |
| TcCLB.511161.10 | hypothetical protein, conserved (pseudogene) | -0,9308 | 0,00000379 |
| TcCLB.506741.70 | surface protease GP63 (pseudogene), putative | -0,9308 | 0,02654 |
| TcCLB.506285.110 | mucin-associated surface protein (MASP, pseudogene), pu | -0,9314 | 0,002233 |
| TcCLB.506941.150 | sphingosine 1-phosphate lyase, putative (SPL) | -0,9316 | 9,092E-13 |
| TcCLB.509895.60 | hypothetical protein, conserved | -0,9317 | 3,96E-07 |
| TcCLB.508225.20 | retrotransposon hot spot protein (RHS, pseudogene), puta | -0,9317 | 0,00001718 |
| TcCLB.510189.30 | surface protease GP63 (pseudogene), putative | -0,9323 | 0,000914 |
| TcCLB.508905.10 | hypothetical protein, conserved (pseudogene) | -0,9324 | 2,34E-11 |
| TcCLB.510211.10 | trans-sialidase (pseudogene), putative | -0,9326 | 0,08916 |
| TcCLB.510621.55 | mucin TcMUCII (pseudogene), putative | -0,9335 | 0,008535 |
| TcCLB.511269.40 | cAMP phosphodiesterase A, putative (PDEA) | -0,9341 | 2,46E-10 |
| TcCLB.511859.100 | trans-sialidase (pseudogene), putative | -0,9347 | 1,722E-17 |
| TcCLB.511127.120 | cysteine peptidase, Clan CA, family C19, putative | -0,9348 | 2,894E-12 |
| TcCLB.510029.80 | hypothetical protein, conserved | -0,935 | 1,408E-22 |
| TcCLB.509805.20 | cAMP phosphodiesterase A, putative (PDEA) | -0,9351 | 3,225E-13 |
| TcCLB.508325.250 | trans-sialidase (pseudogene), putative | -0,936 | 8,686E-11 |
| TcCLB.509525.190 | hypothetical protein | -0,9362 | 0,05421 |
| TcCLB.506285.40 | mucin-associated surface protein (MASP, pseudogene), pu | -0,9363 | 1,272E-06 |
| TcCLB.508593.60 | hypothetical protein, conserved | -0,9366 | 1,634E-13 |
| TcCLB.509525.330 | hypothetical protein | -0,937 | 0,007134 |
| TcCLB.511183.60 | trans-sialidase, Group V, putative | -0,9383 | 1,211E-06 |
| TcCLB.504115.5 | trans-sialidase (pseudogene), putative | -0,9386 | 3,587E-11 |
| TcCLB.504239.120 | trans-sialidase (pseudogene), putative | -0,939 | 0,005656 |
| TcCLB.508385.20 | hypothetical protein, conserved | -0,9391 | 6,321E-24 |
| TcCLB.506001.30 | hypothetical protein | -0,9391 | 0,01288 |
| TcCLB.510175.30 | trans-sialidase (pseudogene), putative | -0,9395 | 5,117E-10 |
| TcCLB.506993.110 | mucin TcMUCII, putative | -0,9402 | 0,0001012 |
| TcCLB.504057.170 | protein kinase, putative | -0,941 | 5,239E-24 |

| | | | |
|---|---|---|---|
| TcCLB.506009.10 | hypothetical protein, conserved | -0,9413 | 0,002726 |
| TcCLB.508389.130 | mucin-associated surface protein (MASP), putative | -0,9413 | 0,06201 |
| TcCLB.509925.20 | mucin TcMUCII, putative | -0,9423 | 6,273E-12 |
| TcCLB.509179.60 | hypothetical protein, conserved | -0,9428 | 4,073E-15 |
| TcCLB.509847.50 | trans-sialidase (pseudogene), putative | -0,9432 | 0,0001798 |
| TcCLB.509543.30 | hypothetical protein | -0,9437 | 2,603E-16 |
| TcCLB.510303.70 | hypothetical protein, conserved | -0,9445 | 2,843E-29 |
| TcCLB.401961.10 | trans-sialidase, putative (fragment) | -0,9446 | 0,04932 |
| TcCLB.511127.240 | hypothetical protein, conserved | -0,9448 | 5,221E-06 |
| TcCLB.508885.30 | hypothetical protein | -0,9452 | 0,03789 |
| TcCLB.511557.60 | hypothetical protein, conserved | -0,9455 | 8,979E-11 |
| TcCLB.507867.31 | mucin TcMUCII (pseudogene), putative | -0,9464 | 7,03E-12 |
| TcCLB.425969.5 | hypothetical protein | -0,9465 | 1,406E-14 |
| TcCLB.506057.61 | elongation factor 1-gamma (EF-1-gamma, pseudogene), | -0,9468 | 0,00006005 |
| TcCLB.511373.10 | mucin-associated surface protein (MASP, pseudogene), pu | -0,9478 | 0,03193 |
| TcCLB.508879.4 | protein kinase C substrate protein, heavy chain, putative | -0,9479 | 1,451E-15 |
| TcCLB.504737.20 | mucin-associated surface protein (MASP, pseudogene), pu | -0,9482 | 0,00051 |
| TcCLB.506251.130 | ubiquitin-protein ligase (pseudogene), putative | -0,9485 | 1,666E-20 |
| TcCLB.506967.27 | syntaxin binding protein (pseudogene), putative | -0,9491 | 0,02403 |
| TcCLB.508243.36 | hypothetical protein | -0,9495 | 0,1018 |
| TcCLB.509167.180 | kinesin, putative | -0,9497 | 2,799E-19 |
| TcCLB.506597.36 | hypothetical protein, conserved (pseudogene) | -0,9506 | 0,04229 |
| TcCLB.510545.40 | mucin TcMUCII, putative | -0,9519 | 0,0001852 |
| TcCLB.510207.35 | mucin TcMUCII (pseudogene), putative | -0,9521 | 0,0001063 |
| TcCLB.510807.30 | hypothetical protein | -0,9523 | 0,00001829 |
| TcCLB.503845.60 | trans-sialidase (pseudogene), putative | -0,9526 | 2,768E-19 |
| TcCLB.509291.10 | hypothetical protein | -0,9527 | 0,001629 |
| TcCLB.506511.70 | hypothetical protein, conserved | -0,9528 | 5,127E-21 |
| TcCLB.509005.20 | hypothetical protein, conserved | -0,9528 | 1,532E-18 |
| TcCLB.503989.20 | hypothetical protein, conserved | -0,9536 | 6,856E-18 |
| TcCLB.510237.40 | mucin-associated surface protein (MASP, pseudogene), pu | -0,9542 | 5,788E-06 |
| TcCLB.510583.140 | trans-sialidase (pseudogene), putative | -0,9542 | 0,0001275 |
| TcCLB.509085.136 | trans-sialidase (pseudogene), putative | -0,9542 | 0,001758 |
| TcCLB.504005.3 | RNA helicase, putative | -0,9543 | 2,357E-20 |
| TcCLB.509493.100 | mucin-associated surface protein (MASP, pseudogene), pu | -0,9546 | 1,875E-14 |
| TcCLB.506501.141 | syntaxin binding protein (pseudogene), putative | -0,9547 | 0,007255 |
| TcCLB.508897.110 | macrophage infectivity potentiator, precursor, putative (M | -0,9549 | 8,952E-24 |
| TcCLB.507389.30 | hypothetical protein | -0,9553 | 7,462E-23 |
| TcCLB.511555.50 | hypothetical protein, conserved | -0,9561 | 4,105E-15 |
| TcCLB.507959.270 | mucin TcMUCII, putative | -0,9566 | 0,0009203 |
| TcCLB.447847.19 | retrotransposon hot spot protein (RHS, pseudogene), puta | -0,9571 | 8,055E-11 |
| TcCLB.511593.30 | mucin-associated surface protein (MASP), putative | -0,9579 | 3,536E-08 |
| TcCLB.506961.35 | trans-sialidase (pseudogene), putative | -0,9583 | 3,084E-09 |
| TcCLB.506655.20 | polyubiquitin (pseudogene), putative | -0,9583 | 0,001334 |
| TcCLB.508277.280 | hypothetical protein | -0,9594 | 1,291E-13 |

| | | | |
|---|---|---|---|
| TcCLB.503783.50 | surface protease GP63 (pseudogene), putative | -0,9595 | 0,0007188 |
| TcCLB.511593.65 | hypothetical protein | -0,9601 | 0,00129 |
| TcCLB.507939.50 | hypothetical protein, conserved | -0,9603 | 4,542E-22 |
| TcCLB.506039.10 | retrotransposon hot spot protein (RHS, pseudogene), puta | -0,9603 | 1,499E-07 |
| TcCLB.510591.30 | retrotransposon hot spot protein (RHS, pseudogene), puta | -0,9603 | 7,568E-07 |
| TcCLB.504031.20 | surface protease GP63 (pseudogene), putative | -0,9604 | 0,0001076 |
| TcCLB.508777.50 | hypothetical protein | -0,9621 | 6,181E-19 |
| TcCLB.511077.130 | mucin TcMUCII, putative | -0,9623 | 0,004277 |
| TcCLB.504013.110 | serine/threonine-protein phosphatase 2A, catalytic subuni | -0,9627 | 1,675E-15 |
| TcCLB.508799.20 | chromatin binding protein, putative | -0,965 | 4,133E-21 |
| TcCLB.511873.20 | hypothetical protein | -0,9664 | 0,0001309 |
| TcCLB.509683.110 | nucleobase transporter, putative | -0,9674 | 4,616E-13 |
| TcCLB.506943.10 | mitogen-activated protein kinase 5, putative | -0,9679 | 2,523E-08 |
| TcCLB.510329.130 | UDP-glucose:glycoprotein glucosyltransferase (tcgt1) | -0,968 | 9,394E-20 |
| TcCLB.511255.604 | mucin TcMUCII, putative | -0,968 | 0,0005563 |
| TcCLB.506973.25 | hypothetical protein | -0,9687 | 0,00007315 |
| TcCLB.510611.70 | hypothetical protein, conserved | -0,9694 | 4,645E-21 |
| TcCLB.508161.40 | mucin TcMUCII (pseudogene), putative | -0,9697 | 0,0008741 |
| TcCLB.506331.20 | retrotransposon hot spot protein (RHS, pseudogene), puta | -0,9704 | 5,852E-08 |
| TcCLB.508931.30 | UDP-Gal or UDP-GlcNAc-dependent glycosyltransferase, p | -0,9708 | 5,956E-10 |
| TcCLB.507951.30 | receptor-type adenylate cyclase (pseudogene), putative | -0,9709 | 4,553E-27 |
| TcCLB.506269.90 | mucin TcMUCII, putative | -0,9721 | 8,659E-10 |
| TcCLB.507555.70 | trans-sialidase (pseudogene), putative | -0,9724 | 6,003E-11 |
| TcCLB.508873.300 | hypothetical protein | -0,9757 | 0,008367 |
| TcCLB.506767.389 | surface protease GP63 (pseudogene), putative | -0,976 | 0,00313 |
| TcCLB.507639.50 | tRNA pseudouridine synthase A, putative | -0,9762 | 1,385E-18 |
| TcCLB.508277.320 | hypothetical protein, conserved | -0,9762 | 4,69E-15 |
| TcCLB.506321.160 | syntaxin binding protein (pseudogene), putative | -0,9764 | 3,187E-08 |
| TcCLB.509693.90 | serine/threonine protein kinase, putative | -0,9767 | 3,2E-23 |
| TcCLB.510371.35 | hypothetical protein, conserved (pseudogene) | -0,9767 | 0,01068 |
| TcCLB.506139.60 | trans-sialidase (pseudogene), putative | -0,9771 | 1,495E-08 |
| TcCLB.507771.30 | hypothetical protein, conserved | -0,9772 | 2,841E-17 |
| TcCLB.509073.70 | phosphatidate cytidylyltransferase-like protein, putative | -0,9773 | 5,749E-20 |
| TcCLB.508047.20 | mucin-associated surface protein (MASP), putative | -0,9776 | 0,00002695 |
| TcCLB.504039.25 | syntaxin binding protein (pseudogene), putative | -0,9776 | 0,007765 |
| TcCLB.508227.51 | surface protease GP63 (pseudogene), putative | -0,9777 | 0,009587 |
| TcCLB.510361.110 | mucin-associated surface protein (MASP), putative | -0,9788 | 0,00005108 |
| TcCLB.509779.20 | hypothetical protein, conserved | -0,9789 | 2,961E-11 |
| TcCLB.508165.115 | mucin TcMUCII (pseudogene), putative | -0,979 | 0,003199 |
| TcCLB.509163.30 | trans-sialidase (pseudogene), putative | -0,9793 | 5,66E-08 |
| TcCLB.511893.40 | receptor-type adenlyate cyclase, putative (fragment) | -0,9798 | 3,776E-06 |
| TcCLB.506613.70 | hypothetical protein | -0,9801 | 0,00008338 |
| TcCLB.511797.110 | trans-sialidase, Group V, putative | -0,9803 | 1,363E-07 |
| TcCLB.509581.35 | hypothetical protein, conserved (pseudogene) | -0,9806 | 0,02166 |
| TcCLB.506703.110 | mucin-associated surface protein (MASP), putative | -0,9814 | 3,156E-09 |

| | | | |
|---|---|---|---|
| TcCLB.506599.320 | hypothetical protein | -0,9815 | 0,03156 |
| TcCLB.511183.103 | mucin-associated surface protein (MASP, pseudogene), pu | -0,9816 | 0,00003498 |
| TcCLB.503551.20 | hypothetical protein, conserved | -0,982 | 0,03362 |
| TcCLB.505987.50 | adenosine kinase, putative | -0,9823 | 4,597E-21 |
| TcCLB.511127.50 | hypothetical protein, conserved | -0,9827 | 4,972E-20 |
| TcCLB.510867.19 | trans-sialidase (pseudogene), putative | -0,9837 | 3,964E-14 |
| TcCLB.511255.230 | mucin-associated surface protein (MASP, pseudogene), pu | -0,9839 | 0,000398 |
| TcCLB.508975.30 | UDP-Gal or UDP-GlcNAc-dependent glycosyltransferase, p | -0,9843 | 0,003758 |
| TcCLB.506279.160 | mucin-associated surface protein (MASP, pseudogene), pu | -0,9844 | 0,001481 |
| TcCLB.508175.320 | hypothetical protein, conserved | -0,9848 | 1,399E-27 |
| TcCLB.503919.60 | hypothetical protein, conserved | -0,9855 | 6,062E-15 |
| TcCLB.511127.140 | DNA replication licensing factor MCM4, putative | -0,9857 | 3,108E-24 |
| TcCLB.509255.10 | hypothetical protein, conserved | -0,9858 | 3,871E-25 |
| TcCLB.506921.10 | surface protease GP63, putative | -0,9874 | 2,521E-22 |
| TcCLB.507699.210 | hypothetical protein | -0,9877 | 0,0003688 |
| TcCLB.511127.5 | RNA helicase, putative | -0,9884 | 0,00002644 |
| TcCLB.506505.10 | hypothetical protein, conserved | -0,9885 | 1,074E-08 |
| TcCLB.506393.80 | trans-sialidase (pseudogene), putative | -0,9894 | 0,01871 |
| TcCLB.506227.150 | Flagellum matrix protein 458 (PKA-R) | -0,9897 | 1,714E-15 |
| TcCLB.506939.220 | mucin-associated surface protein (MASP), putative | -0,9905 | 0,0008041 |
| TcCLB.511003.200 | hypothetical protein, conserved | -0,9906 | 2,029E-12 |
| TcCLB.506885.50 | hypothetical protein, conserved | -0,9912 | 1,032E-23 |
| TcCLB.510105.190 | hypothetical protein, conserved | -0,9919 | 1,263E-14 |
| TcCLB.506297.80 | hypothetical protein, conserved | -0,9921 | 4,938E-26 |
| TcCLB.506767.381 | syntaxin binding protein (pseudogene), putative | -0,9924 | 0,005508 |
| TcCLB.507037.70 | hypothetical protein, conserved | -0,9927 | 1,525E-24 |
| TcCLB.398915.5 | trans-sialidase, putative (fragment) | -0,9931 | 1,748E-20 |
| TcCLB.506871.60 | hypothetical protein, conserved (pseudogene) | -0,9933 | 5,954E-20 |
| TcCLB.508735.65 | trans-sialidase (pseudogene), putative | -0,9938 | 5,015E-11 |
| TcCLB.503407.10 | hypothetical protein, conserved | -0,9941 | 1,441E-16 |
| TcCLB.506961.25 | trans-sialidase, Group I, putative | -0,9952 | 8,741E-14 |
| TcCLB.509229.50 | hypothetical protein | -0,9952 | 2,443E-08 |
| TcCLB.504239.40 | surface protease GP63 (pseudogene), putative | -0,9957 | 0,03742 |
| TcCLB.508761.60 | hypothetical protein (pseudogene) | -0,9964 | 0,0003975 |
| TcCLB.510441.30 | trans-sialidase (pseudogene), putative | -0,9967 | 2,147E-15 |
| TcCLB.506321.90 | syntaxin binding protein (pseudogene), putative | -0,9967 | 2,035E-06 |
| TcCLB.508433.30 | mucin-associated surface protein (MASP), putative | -0,9969 | 0,079 |
| TcCLB.511255.580 | mucin-associated surface protein (MASP, pseudogene), pu | -0,9972 | 0,001749 |
| TcCLB.511063.110 | mucin-associated surface protein (MASP, pseudogene), pu | -0,9984 | 0,002409 |
| TcCLB.509571.70 | hypothetical protein, conserved | -0,999 | 1,911E-24 |
| TcCLB.507705.10 | hypothetical protein, conserved | -1 | 3,31E-08 |
| TcCLB.507951.60 | hypothetical protein, conserved | -1,001 | 1,11E-27 |
| TcCLB.506697.20 | cyclic nucleotide phosphodiesterase | -1,001 | 3,899E-17 |
| TcCLB.511859.140 | hypothetical protein | -1,001 | 3,16E-14 |
| TcCLB.511547.10 | mucin-associated surface protein (MASP), putative | -1,001 | 0,005717 |

| | | | |
|---|---|---|---|
| TcCLB.510209.18 | mucin TcMUCII (pseudogene), putative | -1,001 | 0,006858 |
| TcCLB.510611.10 | hypothetical protein, conserved | -1,002 | 2,733E-20 |
| TcCLB.509777.10 | hypothetical protein | -1,003 | 0,00001171 |
| TcCLB.506909.120 | hypothetical protein | -1,003 | 0,01078 |
| TcCLB.510057.4 | hypothetical protein, conserved | -1,004 | 1,365E-14 |
| TcCLB.511875.9 | mucin-associated surface protein (MASP), putative | -1,004 | 5,431E-10 |
| TcCLB.511725.80 | hypothetical protein | -1,005 | 5,509E-23 |
| TcCLB.510525.70 | hypothetical protein, conserved | -1,005 | 1,182E-21 |
| TcCLB.433673.10 | trans-sialidase, putative (fragment) | -1,005 | 2,88E-18 |
| TcCLB.508097.81 | mucin TcMUCII, putative | -1,005 | 0,0003877 |
| TcCLB.510889.200 | hypothetical protein, conserved | -1,008 | 2,088E-26 |
| TcCLB.506603.43 | hypothetical protein | -1,008 | 2,445E-08 |
| TcCLB.510629.270 | syntaxin binding protein (pseudogene), putative | -1,009 | 0,02266 |
| TcCLB.504223.20 | amino acid permease, putative | -1,01 | 1,199E-09 |
| TcCLB.510363.220 | mucin-associated surface protein (MASP), putative | -1,012 | 0,001796 |
| TcCLB.511077.150 | hypothetical protein | -1,012 | 0,01149 |
| TcCLB.508539.130 | trans-sialidase (pseudogene), putative | -1,013 | 6,185E-10 |
| TcCLB.508977.130 | mucin-associated surface protein (MASP), putative | -1,013 | 2,112E-06 |
| TcCLB.504021.90 | hypothetical protein | -1,014 | 2,46E-15 |
| TcCLB.511127.274 | hypothetical protein, conserved | -1,014 | 1,939E-13 |
| TcCLB.511107.5 | mucin TcMUCII, putative | -1,014 | 4,815E-10 |
| TcCLB.506967.140 | mucin-associated surface protein (MASP, pseudogene), pu | -1,014 | 0,006609 |
| TcCLB.401469.10 | zinc finger protein 2, putative | -1,015 | 1,108E-23 |
| TcCLB.506009.30 | TPR-repeat-containing chaperone protein DNAJ, putative | -1,015 | 2,537E-23 |
| TcCLB.509171.10 | hypothetical protein, conserved | -1,015 | 8,216E-11 |
| TcCLB.510835.10 | hypothetical protein, conserved | -1,015 | 9,657E-06 |
| TcCLB.510583.74 | mucin-associated surface protein (MASP, pseudogene), pu | -1,015 | 0,0001727 |
| TcCLB.510609.80 | hypothetical protein, conserved | -1,016 | 1,116E-10 |
| TcCLB.511625.160 | hypothetical protein | -1,017 | 4,222E-06 |
| TcCLB.511377.20 | trans-sialidase (pseudogene), putative | -1,018 | 0,0007929 |
| TcCLB.507259.16 | protein kinase, putative (fragment) | -1,019 | 0,00172 |
| TcCLB.504147.240 | hypothetical protein, conserved | -1,02 | 1,182E-14 |
| TcCLB.506757.30 | hypothetical protein, conserved | -1,021 | 2,865E-27 |
| TcCLB.504105.180 | hypothetical protein, conserved | -1,021 | 1,713E-22 |
| TcCLB.508871.150 | mucin-associated surface protein (MASP), putative | -1,021 | 6,233E-09 |
| TcCLB.503697.30 | bilobe protein, conserved | -1,022 | 6,076E-34 |
| TcCLB.510751.30 | hypothetical protein | -1,022 | 8,192E-21 |
| TcCLB.503893.140 | hypothetical protein, conserved | -1,023 | 1,598E-19 |
| TcCLB.506973.50 | trans-sialidase (pseudogene), putative | -1,024 | 8,228E-06 |
| TcCLB.510699.89 | mucin-associated surface protein (MASP), putative | -1,024 | 0,001348 |
| TcCLB.507483.60 | translation-associated element 2, putative | -1,025 | 7,821E-35 |
| TcCLB.510237.110 | mucin TcMUCII, putative | -1,025 | 0,00002175 |
| TcCLB.506321.150 | surface protease GP63 (pseudogene), putative | -1,025 | 0,0002767 |
| TcCLB.506613.30 | hypothetical protein | -1,026 | 0,002587 |
| TcCLB.510497.114 | mucin-associated surface protein (MASP, pseudogene), pu | -1,026 | 0,004739 |

| | | | |
|---|---|---|---|
| TcCLB.507067.70 | hypothetical protein | -1,026 | 0,004746 |
| TcCLB.509213.90 | hypothetical protein, conserved | -1,027 | 8,325E-14 |
| TcCLB.508431.60 | hypothetical protein, conserved (pseudogene) | -1,027 | 7,225E-07 |
| TcCLB.511189.70 | hypothetical protein, conserved | -1,027 | 0,0009525 |
| TcCLB.508245.21 | surface protease GP63 (pseudogene), putative | -1,027 | 0,02936 |
| TcCLB.510889.210 | hypothetical protein, conserved | -1,028 | 1,269E-14 |
| TcCLB.506599.35 | hypothetical protein, conserved (pseudogene) | -1,028 | 0,00009048 |
| TcCLB.506895.5 | hypothetical protein | -1,028 | 0,005442 |
| TcCLB.509083.9 | ATP-dependent DEAD/H RNA helicase, putative | -1,029 | 2,432E-13 |
| TcCLB.510279.90 | mucin-associated surface protein (MASP, pseudogene), pu | -1,029 | 0,0003241 |
| TcCLB.507429.6 | hypothetical protein | -1,029 | 0,009669 |
| TcCLB.508761.270 | mucin-associated surface protein (MASP), putative | -1,03 | 1,261E-07 |
| TcCLB.511597.40 | mucin-associated surface protein (MASP), putative | -1,03 | 0,002606 |
| TcCLB.510257.30 | ribulose-5-phosphate 3-epimerase, putative | -1,031 | 2,273E-15 |
| TcCLB.506965.170 | trans-sialidase, Group V, putative | -1,031 | 0,00000301 |
| TcCLB.510167.37 | trans-sialidase (pseudogene), putative | -1,031 | 0,0006231 |
| TcCLB.509685.20 | syntaxin binding protein (pseudogene), putative | -1,031 | 0,001635 |
| TcCLB.504219.10 | hypothetical protein, conserved (pseudogene) | -1,032 | 0,007032 |
| TcCLB.507065.108 | mucin-associated surface protein (MASP, pseudogene), pu | -1,033 | 0,00006725 |
| TcCLB.504201.44 | mucin TcMUCII, putative | -1,033 | 0,005056 |
| TcCLB.510279.50 | mucin TcMUCII (pseudogene), putative | -1,033 | 0,005769 |
| TcCLB.509195.50 | mucin-associated surface protein (MASP), putative | -1,034 | 5,163E-07 |
| TcCLB.503717.50 | mucin TcMUCII (pseudogene), putative | -1,034 | 0,0003016 |
| TcCLB.509525.244 | hypothetical protein | -1,034 | 0,01467 |
| TcCLB.510129.30 | procyclic form surface phosphoprotein, putative | -1,035 | 5,001E-27 |
| TcCLB.508993.20 | homocysteine S-methyltransferase, putative | -1,035 | 1,973E-17 |
| TcCLB.508909.10 | OSM3-like kinesin, putative | -1,036 | 1,968E-26 |
| TcCLB.509605.20 | hypothetical protein, conserved | -1,037 | 3,306E-29 |
| TcCLB.508557.20 | amino acid transporter, putative | -1,038 | 2,467E-13 |
| TcCLB.506297.250 | palmitoyl acyltransferase 7, putative | -1,039 | 9,381E-33 |
| TcCLB.510361.240 | mucin-associated surface protein (MASP, pseudogene), pu | -1,039 | 0,0003567 |
| TcCLB.508883.20 | hypothetical protein | -1,039 | 0,003983 |
| TcCLB.507965.10 | receptor-type adenylate cyclase, putative | -1,041 | 1,107E-15 |
| TcCLB.506423.10 | mucin-associated surface protein (MASP), putative | -1,041 | 0,0004058 |
| TcCLB.507611.131 | protein kinase (pseudogene), putative | -1,042 | 2,834E-13 |
| TcCLB.507979.30 | trans-sialidase, Group I, putative | -1,042 | 2,451E-11 |
| TcCLB.510503.89 | mucin-associated surface protein (MASP, pseudogene), pu | -1,042 | 0,01464 |
| TcCLB.510187.370 | hypothetical protein, conserved | -1,043 | 5,252E-18 |
| TcCLB.511395.120 | UDP-Gal or UDP-GlcNAc-dependent glycosyltransferase, p | -1,043 | 2,828E-07 |
| TcCLB.511277.470 | hypothetical protein, conserved | -1,044 | 1,387E-10 |
| TcCLB.511385.40 | hypothetical protein | -1,044 | 6,62E-08 |
| TcCLB.411427.10 | hypothetical protein, conserved | -1,044 | 8,327E-07 |
| TcCLB.506329.50 | mucin TcMUCII, putative | -1,044 | 0,002384 |
| TcCLB.506499.10 | hypothetical protein, conserved (pseudogene) | -1,044 | 0,01417 |
| TcCLB.509663.60 | retrotransposon hot spot protein (RHS, pseudogene), puta | -1,045 | 1,436E-16 |

| | | | |
|---|---|---|---|
| TcCLB.508999.80 | mucin-associated surface protein (MASP), putative | -1,045 | 0,006894 |
| TcCLB.507607.10 | surface protease GP63 (pseudogene), putative | -1,045 | 0,02077 |
| TcCLB.511149.50 | mucin TcMUCII, putative | -1,047 | 0,0009643 |
| TcCLB.506741.40 | mucin TcMUCII (pseudogene), putative | -1,047 | 0,003459 |
| TcCLB.507847.30 | mucin TcMUCII (pseudogene), putative | -1,048 | 0,00002911 |
| TcCLB.511183.505 | trans-sialidase (pseudogene), putative | -1,05 | 0,0005326 |
| TcCLB.507937.10 | mucin TcMUCII, putative | -1,05 | 0,0114 |
| TcCLB.511303.9 | hypothetical protein, conserved | -1,051 | 4,831E-18 |
| TcCLB.510729.30 | hypothetical protein, conserved | -1,051 | 8,272E-17 |
| TcCLB.506227.205 | hypothetical protein, conserved | -1,051 | 1,59E-16 |
| TcCLB.503587.3 | mucin TcMUC (pseudogene), putative | -1,051 | 0,01794 |
| TcCLB.510885.70 | hypothetical protein, conserved | -1,052 | 7,073E-25 |
| TcCLB.509639.10 | procyclic form surface phosphoprotein, putative | -1,052 | 1,941E-21 |
| TcCLB.506229.70 | ATP-dependent DEAD/H RNA helicase, putative | -1,053 | 3,389E-18 |
| TcCLB.506843.120 | trans-sialidase (pseudogene), putative | -1,053 | 2,491E-07 |
| TcCLB.503779.10 | hypothetical protein, conserved | -1,054 | 1,955E-13 |
| TcCLB.511599.85 | elongation factor 1-gamma (EF-1-gamma, pseudogene), | -1,055 | 0,0276 |
| TcCLB.509791.30 | hypothetical protein, conserved | -1,056 | 1,574E-34 |
| TcCLB.511555.110 | hypothetical protein, conserved | -1,056 | 4,241E-29 |
| TcCLB.506755.5 | hypothetical protein, conserved | -1,056 | 8,024E-08 |
| TcCLB.508887.24 | mucin-associated surface protein (MASP, pseudogene), pu | -1,056 | 0,006923 |
| TcCLB.507859.80 | trans-sialidase (pseudogene), putative | -1,056 | 0,01471 |
| TcCLB.510809.10 | mucin TcMUCII, putative | -1,057 | 0,0001815 |
| TcCLB.508277.110 | cAMP specific phosphodiesterase, putative | -1,058 | 2,042E-16 |
| TcCLB.504005.6 | RNA-binding protein, putative | -1,058 | 1,47E-13 |
| TcCLB.511051.30 | nucleobase transporter, putative | -1,059 | 1,063E-28 |
| TcCLB.508221.970 | mucin-associated surface protein (MASP), putative | -1,059 | 8,518E-07 |
| TcCLB.506909.46 | hypothetical protein | -1,06 | 7,315E-06 |
| TcCLB.510021.140 | mucin TcMUCII (pseudogene), putative | -1,06 | 0,00003644 |
| TcCLB.508541.40 | surface protease GP63 (pseudogene), putative | -1,06 | 0,00006288 |
| TcCLB.507877.10 | mucin-associated surface protein (MASP, pseudogene), pu | -1,06 | 0,0006088 |
| TcCLB.508717.60 | trans-sialidase, Group I, putative | -1,061 | 2,161E-09 |
| TcCLB.511629.40 | hypothetical protein, conserved | -1,062 | 1,6E-14 |
| TcCLB.507059.20 | trans-sialidase, Group V, putative | -1,062 | 5,052E-10 |
| TcCLB.510205.15 | mucin TcMUCII (pseudogene), putative | -1,062 | 0,0002606 |
| TcCLB.510275.283 | mucin TcMUCII (pseudogene), putative | -1,062 | 0,001616 |
| TcCLB.510443.20 | hypothetical protein, conserved | -1,063 | 3,198E-20 |
| TcCLB.507669.150 | glycosomal membrane protein, putative | -1,064 | 1,172E-15 |
| TcCLB.507683.10 | hypothetical protein, conserved | -1,064 | 2,346E-13 |
| TcCLB.509553.20 | ATP-dependent chaperone, putative | -1,065 | 6,411E-19 |
| TcCLB.511753.50 | hypothetical protein, conserved | -1,065 | 1,637E-13 |
| TcCLB.506209.50 | trans-sialidase (pseudogene), putative | -1,065 | 4,098E-09 |
| TcCLB.511183.610 | hypothetical protein | -1,065 | 0,003296 |
| TcCLB.510173.20 | hypothetical protein, conserved | -1,066 | 1,309E-15 |
| TcCLB.506265.180 | hypothetical protein, conserved | -1,066 | 5,027E-11 |

| | | | |
|---|---|---|---|
| TcCLB.508723.50 | hypothetical protein, conserved | -1,067 | 2,243E-29 |
| TcCLB.503989.10 | zinc finger protein 2, putative | -1,068 | 9,533E-26 |
| TcCLB.511611.40 | hypothetical protein | -1,068 | 0,0004654 |
| TcCLB.508209.10 | hypothetical protein, conserved (pseudogene) | -1,069 | 2,18E-08 |
| TcCLB.506865.79 | hypothetical protein, conserved (fragment) | -1,069 | 0,0001826 |
| TcCLB.507185.26 | mucin-associated surface protein (MASP, pseudogene), pu | -1,071 | 1,128E-14 |
| TcCLB.503887.79 | hypothetical protein, conserved | -1,072 | 1,849E-18 |
| TcCLB.504031.30 | mucin-associated surface protein (MASP), putative | -1,072 | 1,185E-10 |
| TcCLB.510583.80 | surface protease GP63 (pseudogene), putative | -1,072 | 0,0002643 |
| TcCLB.509399.120 | hypothetical protein, conserved | -1,073 | 1,589E-35 |
| TcCLB.510889.330 | hypothetical protein, conserved | -1,073 | 4,679E-19 |
| TcCLB.511127.420 | hypothetical protein, conserved | -1,074 | 1,219E-21 |
| TcCLB.506803.20 | trans-sialidase (pseudogene), putative | -1,074 | 2,058E-15 |
| TcCLB.511843.30 | trans-sialidase (pseudogene), putative | -1,074 | 0,0003101 |
| TcCLB.503779.50 | hypothetical protein, conserved | -1,075 | 6,438E-12 |
| TcCLB.510885.110 | hypothetical protein, conserved | -1,076 | 2,256E-33 |
| TcCLB.509975.41 | mucin TcMUC, putative | -1,076 | 6,635E-07 |
| TcCLB.508219.20 | trans-sialidase, Group V, putative | -1,076 | 3,951E-06 |
| TcCLB.506465.5 | hypothetical protein | -1,078 | 1,134E-13 |
| TcCLB.510643.90 | mucin-associated surface protein (MASP, pseudogene), pu | -1,078 | 4,368E-13 |
| TcCLB.511233.170 | elongation factor 1-gamma (EF-1-gamma, pseudogene), | -1,078 | 0,008125 |
| TcCLB.511837.110 | hypothetical protein, conserved | -1,079 | 6,731E-28 |
| TcCLB.510335.10 | hypothetical protein, conserved | -1,081 | 2,331E-28 |
| TcCLB.503973.240 | mucin-associated surface protein (MASP), putative | -1,081 | 0,00001791 |
| TcCLB.510197.229 | trans-sialidase, putative | -1,082 | 5,728E-09 |
| TcCLB.510621.70 | mucin TcMUCII, putative | -1,082 | 0,0006706 |
| TcCLB.511171.90 | mucin-associated surface protein (MASP), putative | -1,082 | 0,006102 |
| TcCLB.508365.140 | hypothetical protein | -1,083 | 0,00001854 |
| TcCLB.508509.70 | mucin TcMUCII, putative | -1,083 | 0,01183 |
| TcCLB.509113.10 | mucin-associated surface protein (MASP, pseudogene), pu | -1,084 | 2,271E-06 |
| TcCLB.506269.59 | hypothetical protein (pseudogene) | -1,086 | 7,729E-11 |
| TcCLB.503841.70 | aspartate aminotransferase, putative | -1,086 | 5,454E-09 |
| TcCLB.508221.750 | trans-sialidase, Group V, putative | -1,086 | 1,569E-06 |
| TcCLB.510279.349 | mucin-associated surface protein (MASP), putative | -1,086 | 0,000495 |
| TcCLB.508885.20 | mucin TcMUCI, putative | -1,086 | 0,003824 |
| TcCLB.508153.230 | hypothetical protein, conserved | -1,087 | 1,85E-30 |
| TcCLB.511725.30 | hypothetical protein, conserved | -1,087 | 1,658E-12 |
| TcCLB.503827.87 | hypothetical protein, conserved (pseudogene) | -1,087 | 0,001436 |
| TcCLB.509697.15 | mucin-associated surface protein (MASP, pseudogene), pu | -1,087 | 0,003062 |
| TcCLB.511127.150 | hypothetical protein, conserved | -1,089 | 6,597E-25 |
| TcCLB.509395.40 | hypothetical protein, conserved | -1,089 | 3,868E-14 |
| TcCLB.504213.100 | hypothetical protein, conserved | -1,09 | 4,877E-15 |
| TcCLB.511163.20 | cdc2-related kinase 8, putative (CRK8) | -1,091 | 8,718E-25 |
| TcCLB.506457.4 | hypothetical protein, conserved | -1,091 | 0,0000233 |
| TcCLB.508161.26 | mucin TcMUCII (pseudogene), putative | -1,091 | 0,0003956 |

| | | | |
|---|---|---|---|
| TcCLB.510833.10 | trans-sialidase, putative | -1,091 | 0,0004571 |
| TcCLB.503479.60 | surface protease GP63, putative | -1,092 | 2,959E-12 |
| TcCLB.511277.110 | citrate synthase, putative | -1,092 | 0,0000154 |
| TcCLB.508089.10 | trans-sialidase, Group I, putative | -1,093 | 1,78E-16 |
| TcCLB.508175.370 | glutamine synthetase, putative | -1,093 | 2,209E-14 |
| TcCLB.507041.60 | syntaxin binding protein (pseudogene), putative | -1,094 | 7,055E-11 |
| TcCLB.508873.390 | trans-sialidase, Group V, putative | -1,094 | 5,529E-07 |
| TcCLB.510089.5 | mucin TcMUCII (pseudogene), putative | -1,095 | 0,02027 |
| TcCLB.507933.29 | hypothetical protein, conserved (fragment) | -1,096 | 1,936E-08 |
| TcCLB.404073.10 | hypothetical protein, conserved | -1,097 | 2,771E-07 |
| TcCLB.506269.25 | hypothetical protein | -1,098 | 0,00001507 |
| TcCLB.510621.30 | mucin TcMUCII, putative | -1,099 | 0,001297 |
| TcCLB.511117.30 | trans-sialidase, Group II, putative | -1,1 | 3,097E-09 |
| TcCLB.506319.40 | hypothetical protein | -1,1 | 0,00001398 |
| TcCLB.506595.21 | hypothetical protein | -1,101 | 6,972E-16 |
| TcCLB.506001.40 | trans-sialidase, Group V, putative | -1,101 | 1,158E-09 |
| TcCLB.506767.127 | hypothetical protein, conserved (pseudogene) | -1,101 | 0,009557 |
| TcCLB.511255.540 | mucin-associated surface protein (MASP), putative | -1,102 | 0,0002425 |
| TcCLB.507859.101 | hypothetical protein, conserved (pseudogene) | -1,103 | 0,002161 |
| TcCLB.504239.365 | surface protease GP63 (pseudogene), putative | -1,103 | 0,01196 |
| TcCLB.510693.50 | trans-sialidase (pseudogene), putative | -1,104 | 0,00001185 |
| TcCLB.433237.10 | hypothetical protein | -1,104 | 0,01257 |
| TcCLB.506765.46 | mucin-associated surface protein (MASP, pseudogene), pu | -1,105 | 3,561E-15 |
| TcCLB.508927.10 | hypothetical protein, conserved | -1,105 | 7,85E-15 |
| TcCLB.511259.9 | mucin TcMUCI, putative | -1,105 | 0,01333 |
| TcCLB.511505.30 | hypothetical protein, conserved | -1,106 | 2,006E-16 |
| TcCLB.506905.10 | hypothetical protein, conserved | -1,106 | 1,028E-14 |
| TcCLB.510197.64 | mucin-associated surface protein (MASP, pseudogene), pu | -1,106 | 4,737E-09 |
| TcCLB.506321.101 | hypothetical protein, conserved (pseudogene) | -1,106 | 0,00007686 |
| TcCLB.508175.9 | hypothetical protein, conserved (fragment) | -1,108 | 1,967E-22 |
| TcCLB.508873.70 | mucin-associated surface protein (MASP, pseudogene), pu | -1,108 | 0,00001946 |
| TcCLB.508777.200 | hypothetical protein, conserved | -1,109 | 2,047E-08 |
| TcCLB.509525.210 | hypothetical protein | -1,109 | 0,00002082 |
| TcCLB.503601.20 | hypothetical protein | -1,11 | 0,00007261 |
| TcCLB.511099.180 | trans-sialidase (pseudogene), putative | -1,11 | 0,0002794 |
| TcCLB.507283.39 | trans-sialidase (pseudogene), putative | -1,112 | 1,533E-23 |
| TcCLB.506351.35 | hypothetical protein, conserved | -1,112 | 1,942E-11 |
| TcCLB.509993.20 | mucin TcMUCI, putative | -1,112 | 1,363E-07 |
| TcCLB.510275.21 | surface protease GP63 (pseudogene), putative | -1,112 | 0,005138 |
| TcCLB.504021.10 | G1 cyclin CycE4, putative (CYC5) | -1,113 | 1,257E-37 |
| TcCLB.507951.150 | guanine deaminase, putative | -1,113 | 2,027E-31 |
| TcCLB.509897.160 | trans-sialidase (pseudogene), putative | -1,113 | 2,961E-11 |
| TcCLB.509261.40 | UDP-Gal or UDP-GlcNAc-dependent glycosyltransferase, p | -1,113 | 3,507E-09 |
| TcCLB.510555.10 | mucin-associated surface protein (MASP, pseudogene), pu | -1,113 | 2,251E-06 |
| TcCLB.511367.330 | hypothetical protein, conserved | -1,114 | 4,206E-15 |

| | | | |
|---|---|---|---|
| TcCLB.508071.60 | mucin-associated surface protein (MASP), putative | -1,114 | 7,061E-15 |
| TcCLB.506955.170 | trans-sialidase (pseudogene), putative | -1,114 | 1,311E-07 |
| TcCLB.511403.40 | mucin-associated surface protein (MASP, pseudogene), pu | -1,114 | 0,0001221 |
| TcCLB.506001.24 | hypothetical protein (pseudogene) | -1,114 | 0,01756 |
| TcCLB.509293.10 | mucin TcMUCII, putative | -1,115 | 4,995E-09 |
| TcCLB.511149.4 | mucin-associated surface protein (MASP, pseudogene), pu | -1,115 | 0,0003495 |
| TcCLB.410275.9 | mucin-associated surface protein (MASP), putative | -1,115 | 0,06375 |
| TcCLB.511173.150 | mucin-associated surface protein (MASP), putative | -1,117 | 6,186E-08 |
| TcCLB.510393.10 | hypothetical protein, conserved | -1,118 | 1,644E-20 |
| TcCLB.486901.10 | endonuclease/exonuclease/phosphatase, putative | -1,118 | 1,186E-12 |
| TcCLB.508461.70 | hypothetical protein | -1,118 | 6,972E-12 |
| TcCLB.508367.70 | trans-sialidase (pseudogene), putative | -1,118 | 0,00001084 |
| TcCLB.506461.10 | hypothetical protein, conserved | -1,118 | 0,01274 |
| TcCLB.506755.240 | hypothetical protein, conserved | -1,119 | 6,276E-09 |
| TcCLB.506993.86 | mucin-associated surface protein (MASP, pseudogene), pu | -1,119 | 0,00001015 |
| TcCLB.507809.119 | hypothetical protein, conserved | -1,12 | 7,491E-32 |
| TcCLB.507021.30 | hypothetical protein, conserved | -1,12 | 6,279E-17 |
| TcCLB.503825.10 | protein kinase (pseudogene), putative | -1,12 | 2,652E-14 |
| TcCLB.506279.90 | mucin TcMUCII, putative | -1,12 | 0,002212 |
| TcCLB.510465.110 | mucin-associated surface protein (MASP, pseudogene), pu | -1,121 | 1,988E-06 |
| TcCLB.508121.155 | mucin TcMUCII (pseudogene), putative | -1,121 | 0,0002544 |
| TcCLB.506409.110 | hypothetical protein | -1,122 | 1,956E-07 |
| TcCLB.508865.2 | hypothetical protein, conserved (fragment) | -1,125 | 0,0002543 |
| TcCLB.508895.20 | hypothetical protein, conserved | -1,126 | 3,928E-35 |
| TcCLB.511553.71 | syntaxin binding protein (pseudogene), putative | -1,126 | 0,00001436 |
| TcCLB.510681.30 | lipase, putative | -1,127 | 3,431E-23 |
| TcCLB.503463.30 | protein kinase, putative | -1,127 | 1,346E-21 |
| TcCLB.511451.90 | mucin-associated surface protein (MASP), putative | -1,127 | 0,000152 |
| TcCLB.504039.140 | surface protease GP63 (pseudogene), putative | -1,127 | 0,001549 |
| TcCLB.509399.100 | hypothetical protein, conserved | -1,129 | 1,04E-43 |
| TcCLB.509977.90 | hypothetical protein | -1,129 | 1,019E-21 |
| TcCLB.507073.20 | hypothetical protein, conserved | -1,129 | 2,518E-14 |
| TcCLB.505959.10 | mucin TcMUCII, putative | -1,129 | 0,00008026 |
| TcCLB.511553.80 | hypothetical protein, conserved | -1,13 | 0,002982 |
| TcCLB.507015.30 | trans-sialidase (pseudogene), putative | -1,131 | 2,634E-16 |
| TcCLB.508011.99 | mucin TcMUCII (pseudogene), putative | -1,131 | 0,001958 |
| TcCLB.509767.20 | hypothetical protein, conserved | -1,132 | 1,131E-13 |
| TcCLB.510583.180 | mucin-associated surface protein (MASP, pseudogene), pu | -1,132 | 0,000135 |
| TcCLB.508435.30 | mucin-associated surface protein (MASP, pseudogene), pu | -1,132 | 0,0002406 |
| TcCLB.508761.110 | trans-sialidase, Group VI, putative | -1,133 | 1,238E-08 |
| TcCLB.510625.130 | hypothetical protein | -1,133 | 0,01192 |
| TcCLB.507357.71 | hypothetical protein | -1,134 | 0,0004684 |
| TcCLB.504153.260 | hypothetical protein, conserved | -1,135 | 8,989E-48 |
| TcCLB.509275.20 | mucin-associated surface protein (MASP), putative (fragm | -1,135 | 0,00001361 |
| TcCLB.510807.20 | hypothetical protein | -1,135 | 0,001139 |

| | | | |
|---|---|---|---|
| TcCLB.510575.160 | hypothetical protein | -1,136 | 1,039E-23 |
| TcCLB.506435.200 | hypothetical protein, conserved | -1,136 | 2,834E-18 |
| TcCLB.508883.15 | trans-sialidase (pseudogene), putative | -1,136 | 0,003189 |
| TcCLB.506755.140 | ATP-dependent helicase, putative | -1,137 | 1,181E-33 |
| TcCLB.506905.4 | proliferator-activated receptor-interacting protein (PRIP) i | -1,137 | 1,528E-12 |
| TcCLB.510709.60 | mucin-associated surface protein (MASP), putative | -1,137 | 0,03416 |
| TcCLB.511183.280 | mucin-associated surface protein (MASP), putative | -1,138 | 8,345E-06 |
| TcCLB.464415.10 | hypothetical protein | -1,138 | 0,0001113 |
| TcCLB.508741.20 | hypothetical protein, conserved | -1,139 | 4,738E-18 |
| TcCLB.507411.10 | trans-sialidase, Group V, putative | -1,139 | 2,381E-10 |
| TcCLB.507835.35 | trans-sialidase (pseudogene), putative | -1,139 | 2,797E-07 |
| TcCLB.510397.20 | trans-sialidase (pseudogene), putative | -1,139 | 0,0001395 |
| TcCLB.507683.34 | hypothetical protein, conserved | -1,14 | 2,121E-12 |
| TcCLB.507835.10 | trans-sialidase (pseudogene), putative | -1,141 | 4,073E-15 |
| TcCLB.511077.160 | mucin TcMUCII, putative | -1,141 | 0,0003503 |
| TcCLB.508359.50 | mucin TcMUCII, putative | -1,141 | 0,001387 |
| TcCLB.504081.409 | mucin TcMUCII (pseudogene), putative | -1,141 | 0,008497 |
| TcCLB.503579.10 | hypothetical protein, conserved | -1,142 | 2,092E-26 |
| TcCLB.506609.50 | hypothetical protein | -1,142 | 0,0003864 |
| TcCLB.506909.90 | mucin-associated surface protein (MASP, pseudogene), pu | -1,142 | 0,000478 |
| TcCLB.511061.20 | mucin-associated surface protein (MASP, pseudogene), pu | -1,142 | 0,002102 |
| TcCLB.507641.104 | small GTPase, putative | -1,143 | 3,127E-20 |
| TcCLB.510061.11 | mucin-associated surface protein (MASP), putative | -1,143 | 6,442E-07 |
| TcCLB.503497.10 | mucin-associated surface protein (MASP, pseudogene), pu | -1,144 | 2,652E-15 |
| TcCLB.504787.90 | mucin-associated surface protein (MASP, pseudogene), pu | -1,144 | 1,971E-10 |
| TcCLB.504209.10 | conserved protein, unknown function | -1,145 | 1,666E-15 |
| TcCLB.506585.100 | hypothetical protein | -1,145 | 5,602E-10 |
| TcCLB.507537.40 | metacaspase, putative | -1,145 | 9,124E-08 |
| TcCLB.507771.85 | hypothetical protein | -1,146 | 4,072E-15 |
| TcCLB.507213.40 | trans-sialidase, Group II, putative | -1,146 | 0,00001209 |
| TcCLB.506247.355 | hypothetical protein, conserved | -1,148 | 2,947E-13 |
| TcCLB.505997.65 | trans-sialidase (pseudogene), putative | -1,148 | 2,115E-12 |
| TcCLB.505949.169 | mucin-associated surface protein (MASP), putative (fragm | -1,148 | 0,006154 |
| TcCLB.504071.110 | ama1 protein, putative | -1,149 | 4,204E-10 |
| TcCLB.507729.50 | mucin-associated surface protein (MASP), putative (fragm | -1,149 | 1,336E-08 |
| TcCLB.510759.190 | hypothetical protein, conserved | -1,15 | 1,877E-41 |
| TcCLB.506789.20 | trans-sialidase, Group V, putative | -1,15 | 0,0001092 |
| TcCLB.510491.60 | trans-sialidase, Group IV, putative | -1,151 | 4,022E-14 |
| TcCLB.511255.169 | mucin-associated surface protein (MASP), putative | -1,151 | 0,00007182 |
| TcCLB.507059.50 | trans-sialidase, Group V, putative | -1,152 | 2,345E-10 |
| TcCLB.511163.50 | hypothetical protein, conserved | -1,153 | 1,294E-24 |
| TcCLB.508933.40 | hypothetical protein | -1,154 | 6,641E-10 |
| TcCLB.503867.5 | hypothetical protein, conserved | -1,154 | 2,494E-07 |
| TcCLB.508777.150 | hypothetical protein | -1,155 | 6,057E-22 |
| TcCLB.509217.40 | trans-sialidase, Group V, putative | -1,155 | 6,424E-10 |

| | | | |
|---|---|---|---|
| TcCLB.504031.40 | hypothetical protein | -1,155 | 0,0003726 |
| TcCLB.506599.91 | surface protease GP63 (pseudogene), putative | -1,155 | 0,002827 |
| TcCLB.506989.300 | receptor-type adenylate cyclase (pseudogene), putative | -1,156 | 5,438E-36 |
| TcCLB.508323.90 | hypothetical protein, conserved | -1,156 | 3,486E-12 |
| TcCLB.508433.90 | mucin-associated surface protein (MASP), putative | -1,156 | 0,02663 |
| TcCLB.509419.150 | mucin-associated surface protein (MASP), putative | -1,157 | 1,271E-08 |
| TcCLB.506965.90 | mucin TcMUCII, putative | -1,157 | 4,022E-07 |
| TcCLB.510377.190 | mucin TcMUCII, putative | -1,158 | 5,985E-08 |
| TcCLB.503733.20 | 6-phosphofructo-2-kinase/fructose-2,6-biphosphatase, pu | -1,159 | 4,423E-31 |
| TcCLB.511259.130 | mucin TcMUCII, putative | -1,159 | 1,505E-06 |
| TcCLB.506139.130 | hypothetical protein | -1,159 | 0,002209 |
| TcCLB.506297.50 | kinesin, putative | -1,161 | 8,384E-63 |
| TcCLB.503609.50 | trans-sialidase, Group VI, putative | -1,161 | 2,617E-09 |
| TcCLB.511057.20 | mucin TcMUCII, putative | -1,161 | 9,445E-07 |
| TcCLB.504765.15 | succinyl-coA:3-ketoacid-coenzyme A transferase (pseudog | -1,163 | 6,275E-14 |
| TcCLB.506495.50 | hypothetical protein | -1,164 | 2,228E-19 |
| TcCLB.511649.100 | amino acid permease, putative | -1,164 | 1,331E-18 |
| TcCLB.511167.60 | hypothetical protein, conserved | -1,165 | 3,466E-47 |
| TcCLB.507941.60 | hypothetical protein, conserved | -1,167 | 2,047E-15 |
| TcCLB.509681.10 | hypothetical protein, conserved | -1,167 | 0,0003679 |
| TcCLB.506501.177 | hypothetical protein, conserved (pseudogene) | -1,167 | 0,002765 |
| TcCLB.507681.100 | ATP-binding cassette protein, putative | -1,168 | 1,583E-22 |
| TcCLB.506487.50 | trans-sialidase (pseudogene), putative | -1,168 | 1,927E-20 |
| TcCLB.507809.140 | hypothetical protein, conserved | -1,169 | 2,716E-16 |
| TcCLB.503973.50 | mucin-associated surface protein (MASP), putative | -1,169 | 0,0000158 |
| TcCLB.503533.40 | mucin-associated surface protein (MASP), putative | -1,169 | 0,001914 |
| TcCLB.505699.10 | trans-sialidase, putative | -1,17 | 8,91E-16 |
| TcCLB.509937.120 | hypothetical protein, conserved | -1,17 | 8,479E-11 |
| TcCLB.510089.20 | surface protease GP63, putative | -1,17 | 0,09569 |
| TcCLB.509333.10 | trans-sialidase, Group III, putative | -1,171 | 9,41E-37 |
| TcCLB.506653.40 | trans-sialidase (pseudogene), putative | -1,172 | 2,87E-24 |
| TcCLB.509505.60 | hypothetical protein | -1,172 | 9,804E-24 |
| TcCLB.509429.300 | hypothetical protein, conserved | -1,173 | 4,082E-27 |
| TcCLB.510163.40 | surface protease GP63 (pseudogene), putative | -1,173 | 2,096E-26 |
| TcCLB.507015.59 | expression site-associated gene (ESAG-like, pseudogene), | -1,174 | 3,235E-16 |
| TcCLB.508409.160 | thymine-7-hydroxylase, putative | -1,174 | 7,387E-16 |
| TcCLB.509685.27 | surface protease GP63 (pseudogene), putative | -1,174 | 0,001074 |
| TcCLB.511365.90 | hypothetical protein, conserved | -1,175 | 1,806E-29 |
| TcCLB.506767.209 | syntaxin binding protein (pseudogene), putative | -1,175 | 0,0001615 |
| TcCLB.510275.80 | trans-sialidase (pseudogene), putative | -1,177 | 1,345E-07 |
| TcCLB.509515.20 | UDP-Gal or UDP-GlcNAc-dependent glycosyltransferase, p | -1,177 | 0,00007427 |
| TcCLB.507677.140 | hypothetical protein, conserved | -1,177 | 0,008372 |
| TcCLB.511197.40 | trans-sialidase (pseudogene), putative | -1,179 | 8,095E-12 |
| TcCLB.506335.20 | trans-sialidase (pseudogene), putative | -1,179 | 6,398E-09 |
| TcCLB.508305.9 | trans-sialidase, putative | -1,179 | 9,094E-07 |

| | | | |
|---|---|---|---|
| TcCLB.509081.40 | surface protease GP63 (pseudogene), putative | -1,179 | 0,00003418 |
| TcCLB.510291.14 | mitochondrial carrier protein, putative | -1,18 | 1,083E-23 |
| TcCLB.506763.361 | syntaxin binding protein (pseudogene), putative | -1,18 | 0,0001509 |
| TcCLB.508659.18 | mitochondrial carrier protein, putative | -1,181 | 3,686E-22 |
| TcCLB.508029.59 | kinesin, putative | -1,181 | 1,673E-11 |
| TcCLB.507255.50 | UDP-Gal or UDP-GlcNAc-dependent glycosyltransferase, p | -1,182 | 8,944E-12 |
| TcCLB.507915.10 | hypothetical protein, conserved | -1,182 | 5,034E-11 |
| TcCLB.509663.30 | thymidylate kinase, putative | -1,182 | 6,417E-11 |
| TcCLB.507943.60 | hypothetical protein, conserved | -1,183 | 4,046E-31 |
| TcCLB.507145.84 | mucin-associated surface protein (MASP, pseudogene), pu | -1,183 | 3,225E-11 |
| TcCLB.510625.30 | mucin-associated surface protein (MASP, pseudogene), pu | -1,183 | 7,486E-07 |
| TcCLB.510715.140 | mucin-associated surface protein (MASP, pseudogene), pu | -1,183 | 1,339E-06 |
| TcCLB.507973.20 | hypothetical protein | -1,183 | 0,00000163 |
| TcCLB.510139.20 | surface protease GP63, putative | -1,184 | 1,3E-13 |
| TcCLB.505609.40 | trans-sialidase, Group V, putative | -1,184 | 0,0001283 |
| TcCLB.506457.50 | serine/threonine protein kinase, putative | -1,185 | 1,933E-29 |
| TcCLB.438923.10 | hypothetical protein (fragment) | -1,185 | 7,433E-23 |
| TcCLB.506741.10 | trans-sialidase (pseudogene), putative | -1,185 | 7,82E-14 |
| TcCLB.506751.50 | trans-sialidase, Group II, putative | -1,186 | 7,85E-12 |
| TcCLB.506353.10 | trans-sialidase, Group II, putative | -1,186 | 6,852E-08 |
| TcCLB.511057.15 | hypothetical protein | -1,186 | 0,00001585 |
| TcCLB.508701.10 | ZIP Zn transporter, putative | -1,187 | 9,934E-16 |
| TcCLB.507165.50 | cysteine synthase, putative | -1,187 | 4,614E-15 |
| TcCLB.510887.40 | hypothetical protein, conserved | -1,189 | 1,217E-21 |
| TcCLB.506337.171 | ubiquitin (pseudogene), putative | -1,189 | 3,293E-08 |
| TcCLB.511553.180 | hypothetical protein | -1,189 | 0,007146 |
| TcCLB.511871.70 | hypothetical protein, conserved | -1,191 | 3,305E-21 |
| TcCLB.510699.20 | mucin-associated surface protein (MASP, pseudogene), pu | -1,191 | 1,064E-12 |
| TcCLB.506767.10 | hypothetical protein, conserved (pseudogene) | -1,191 | 0,00172 |
| TcCLB.509317.100 | hypothetical protein, conserved | -1,192 | 5,703E-22 |
| TcCLB.506393.40 | trans-sialidase (pseudogene), putative | -1,193 | 0,002275 |
| TcCLB.506603.35 | trans-sialidase (pseudogene), putative | -1,194 | 6,517E-13 |
| TcCLB.511373.20 | mucin TcMUCII, putative | -1,194 | 2,333E-12 |
| TcCLB.503697.120 | hypothetical protein, conserved | -1,195 | 3,366E-41 |
| TcCLB.511529.59 | hypothetical protein | -1,195 | 4,179E-23 |
| TcCLB.510239.80 | trans-sialidase, Group V, putative | -1,195 | 1,063E-08 |
| TcCLB.510011.9 | mucin-associated surface protein (MASP), putative | -1,195 | 3,808E-06 |
| TcCLB.507681.230 | hypothetical protein, conserved | -1,196 | 2,157E-29 |
| TcCLB.509295.30 | trans-sialidase (pseudogene), putative | -1,196 | 6,804E-12 |
| TcCLB.507237.10 | trans-sialidase, Group V, putative | -1,197 | 2,439E-08 |
| TcCLB.463297.10 | hypothetical protein (fragment) | -1,197 | 5,069E-07 |
| TcCLB.507711.110 | procyclic form surface phosphoprotein, putative | -1,198 | 2,925E-23 |
| TcCLB.508593.130 | hypothetical protein, conserved | -1,2 | 6,816E-48 |
| TcCLB.507507.20 | ADP-ribosylation factor GTPase activating protein 1, putat | -1,201 | 2,894E-25 |
| TcCLB.511593.80 | mucin-associated surface protein (MASP), putative | -1,201 | 4,006E-15 |

| | | | |
|---|---|---|---|
| TcCLB.427789.20 | Paraflagellar Rod Proteome Component 9, putative (PFC9) | -1,201 | 2,556E-14 |
| TcCLB.510887.60 | hypothetical protein, conserved | -1,202 | 1,299E-26 |
| TcCLB.511461.30 | hypothetical protein, conserved | -1,202 | 5,516E-10 |
| TcCLB.509897.110 | trans-sialidase, Group V, putative | -1,203 | 3,423E-15 |
| TcCLB.508221.1010 | mucin-associated surface protein (MASP), putative | -1,203 | 3,005E-14 |
| TcCLB.510013.13 | syntaxin binding protein (pseudogene), putative | -1,204 | 0,0001199 |
| TcCLB.510363.280 | mucin-associated surface protein (MASP), putative | -1,204 | 0,0002783 |
| TcCLB.510305.30 | lipin, putative | -1,205 | 1,633E-49 |
| TcCLB.508871.100 | hypothetical protein | -1,205 | 0,01418 |
| TcCLB.511907.180 | surface protease GP63 (pseudogene), putative | -1,206 | 1,282E-10 |
| TcCLB.508219.60 | mucin TcMUCII (pseudogene), putative | -1,206 | 0,0001749 |
| TcCLB.511025.30 | mitogen-activated protein kinase, putative | -1,207 | 7,537E-34 |
| TcCLB.510173.10 | hypothetical protein, conserved | -1,207 | 7,219E-24 |
| TcCLB.511877.20 | mucin-associated surface protein (MASP), putative | -1,207 | 5,434E-16 |
| TcCLB.507799.10 | mannosyl-oligosaccharide 1,2-alpha-mannosidase IB, puta | -1,207 | 6,233E-09 |
| TcCLB.505479.10 | mucin-associated surface protein (MASP, pseudogene), pu | -1,207 | 0,0008255 |
| TcCLB.503717.40 | mucin-associated surface protein (MASP, pseudogene), pu | -1,208 | 3,38E-08 |
| TcCLB.511173.380 | mucin TcMUCII (pseudogene), putative | -1,208 | 0,001763 |
| TcCLB.506767.371 | hypothetical protein, conserved (pseudogene) | -1,208 | 0,00593 |
| TcCLB.511821.150 | hypothetical protein, conserved | -1,209 | 1,855E-27 |
| TcCLB.508159.60 | mucin TcMUCII, putative | -1,21 | 3,583E-07 |
| TcCLB.510015.5 | hypothetical protein, conserved (pseudogene) | -1,21 | 0,00005963 |
| TcCLB.508389.138 | mucin TcMUCII, putative | -1,21 | 0,0003836 |
| TcCLB.506693.30 | RNA-binding protein 6, putative | -1,21 | 0,0004234 |
| TcCLB.507251.10 | hypothetical protein, conserved | -1,211 | 3,419E-29 |
| TcCLB.507811.100 | amino acid permease, putative | -1,212 | 6,233E-09 |
| TcCLB.511553.60 | hypothetical protein | -1,213 | 0,00005991 |
| TcCLB.506601.21 | syntaxin binding protein (pseudogene), putative | -1,213 | 0,00025 |
| TcCLB.504039.80 | surface protease GP63 (pseudogene), putative | -1,213 | 0,0006725 |
| TcCLB.507173.30 | mucin-associated surface protein (MASP, pseudogene), pu | -1,214 | 2,584E-08 |
| TcCLB.507241.30 | arginine kinase | -1,214 | 3,86E-07 |
| TcCLB.506229.64 | hypothetical protein, conserved | -1,215 | 5,103E-16 |
| TcCLB.508707.50 | hypothetical protein, conserved | -1,216 | 1,735E-30 |
| TcCLB.511151.60 | hypothetical protein | -1,218 | 6,639E-35 |
| TcCLB.508163.270 | mucin-associated surface protein (MASP), putative | -1,218 | 9,704E-07 |
| TcCLB.511867.10 | hypothetical protein, conserved | -1,219 | 2,837E-33 |
| TcCLB.506519.140 | hypothetical protein, conserved (pseudogene) | -1,219 | 7,316E-27 |
| TcCLB.508605.31 | trans-sialidase (pseudogene), putative | -1,219 | 4,767E-24 |
| TcCLB.509251.80 | trans-sialidase, Group V, putative | -1,219 | 9,291E-11 |
| TcCLB.507237.310 | surface protease GP63 (pseudogene), putative | -1,219 | 0,00005204 |
| TcCLB.507559.110 | surface protease GP63, putative | -1,22 | 4,142E-18 |
| TcCLB.506667.40 | mucin-associated surface protein (MASP), putative | -1,22 | 0,0002433 |
| TcCLB.503795.10 | hypothetical protein, conserved | -1,222 | 0,00002764 |
| TcCLB.510047.70 | hypothetical protein (pseudogene) | -1,224 | 0,000126 |
| TcCLB.510627.60 | mucin-associated surface protein (MASP), putative | -1,225 | 0,0005231 |

| | | | |
|---|---|---|---|
| TcCLB.506741.150 | surface protease GP63 (pseudogene), putative | -1,225 | 0,004824 |
| TcCLB.504005.60 | hypothetical protein, conserved | -1,226 | 1,169E-12 |
| TcCLB.509085.31 | trans-sialidase (pseudogene), putative | -1,226 | 3,209E-09 |
| TcCLB.508739.60 | histone H4, putative | -1,227 | 8,98E-20 |
| TcCLB.506133.125 | hypothetical protein, conserved (pseudogene) | -1,227 | 0,007705 |
| TcCLB.511175.40 | mucin TcMUCII, putative | -1,229 | 1,577E-10 |
| TcCLB.503531.10 | hypothetical protein, conserved | -1,229 | 3,271E-07 |
| TcCLB.508325.290 | trans-sialidase (pseudogene), putative | -1,23 | 3,724E-48 |
| TcCLB.506813.59 | mucin TcMUC (pseudogene), putative | -1,232 | 3,823E-06 |
| TcCLB.510889.290 | hypothetical protein | -1,233 | 1,354E-16 |
| TcCLB.506499.60 | hypothetical protein | -1,233 | 0,005997 |
| TcCLB.510679.70 | lipase, putative | -1,234 | 1,018E-10 |
| TcCLB.508409.310 | hypothetical protein, conserved | -1,235 | 2,764E-25 |
| TcCLB.506965.160 | hypothetical protein | -1,235 | 0,00003296 |
| TcCLB.511233.150 | hypothetical protein | -1,236 | 0,0004182 |
| TcCLB.506763.140 | hypothetical protein | -1,236 | 0,004012 |
| TcCLB.504197.10 | hypothetical protein | -1,237 | 0,001331 |
| TcCLB.510085.54 | mucin TcMUCII (pseudogene), putative | -1,239 | 0,00002367 |
| TcCLB.507677.150 | G1 cyclin CycE4, putative (CYC5) | -1,24 | 7,831E-36 |
| TcCLB.508673.20 | UDP-Gal or UDP-GlcNAc-dependent glycosyltransferase, p | -1,24 | 1,507E-22 |
| TcCLB.504239.161 | mucin-associated surface protein (MASP, pseudogene), pu | -1,24 | 5,949E-06 |
| TcCLB.511565.15 | hypothetical protein | -1,24 | 0,00001623 |
| TcCLB.509937.130 | protein kinase, putative | -1,24 | 0,006484 |
| TcCLB.505267.60 | trans-sialidase (pseudogene), putative | -1,241 | 2,544E-48 |
| TcCLB.506705.70 | beta-fructofuranosidase-like protein (fragment) | -1,241 | 2,514E-28 |
| TcCLB.507471.31 | elongation factor 1-gamma (EF-1-gamma, pseudogene), | -1,241 | 0,005325 |
| TcCLB.511875.70 | mucin TcMUCII (pseudogene), putative | -1,242 | 1,117E-20 |
| TcCLB.506499.120 | hypothetical protein | -1,242 | 0,000851 |
| TcCLB.506179.20 | hypothetical protein, conserved | -1,243 | 1,37E-22 |
| TcCLB.508455.30 | hypothetical protein | -1,243 | 0,0004454 |
| TcCLB.510303.299 | hypothetical protein | -1,245 | 5,814E-27 |
| TcCLB.510337.30 | adenylosuccinate lyase, putative (ADSL) | -1,246 | 1,985E-45 |
| TcCLB.507959.240 | syntaxin binding protein (pseudogene), putative | -1,246 | 0,00001868 |
| TcCLB.440363.19 | Vacuolar iron transporter 1, putative (fragment) (VIT1) | -1,248 | 2,583E-23 |
| TcCLB.506245.230 | mucin-associated surface protein (MASP, pseudogene), pu | -1,249 | 0,000199 |
| TcCLB.506599.400 | trans-sialidase (pseudogene), putative | -1,249 | 0,0006712 |
| TcCLB.508313.122 | trans-sialidase (pseudogene), putative | -1,249 | 0,008732 |
| TcCLB.506741.110 | mucin TcMUCII, putative | -1,25 | 9,074E-23 |
| TcCLB.511549.20 | trans-sialidase (pseudogene), putative | -1,25 | 1,454E-10 |
| TcCLB.506965.40 | trans-sialidase (pseudogene), putative | -1,251 | 6,496E-09 |
| TcCLB.511615.10 | trans-sialidase, Group V, putative | -1,251 | 0,00002261 |
| TcCLB.508081.110 | trans-sialidase (pseudogene), putative | -1,252 | 2,69E-12 |
| TcCLB.439991.10 | trans-sialidase (pseudogene), putative | -1,252 | 2,934E-08 |
| TcCLB.503787.20 | mucin TcMUCII, putative | -1,253 | 2,548E-06 |
| TcCLB.506767.21 | syntaxin binding protein (pseudogene), putative | -1,255 | 0,00000424 |

| | | | |
|---|---|---|---|
| TcCLB.506781.30 | mucin TcMUCII, putative | -1,256 | 6,892E-08 |
| TcCLB.504239.390 | syntaxin binding protein (pseudogene), putative | -1,256 | 0,0001708 |
| TcCLB.510021.60 | hypothetical protein | -1,256 | 0,00216 |
| TcCLB.506227.100 | hypothetical protein, conserved | -1,257 | 1,614E-13 |
| TcCLB.511817.210 | protein kinase, putative | -1,258 | 1,39E-28 |
| TcCLB.509251.20 | trans-sialidase, Group V, putative | -1,258 | 8,68E-13 |
| TcCLB.509525.260 | hypothetical protein | -1,258 | 1,493E-12 |
| TcCLB.505949.100 | trans-sialidase (pseudogene), putative | -1,259 | 5,832E-14 |
| TcCLB.507071.170 | mucin TcMUCII, putative | -1,259 | 3,108E-11 |
| TcCLB.510363.260 | mucin-associated surface protein (MASP), putative | -1,259 | 0,0004846 |
| TcCLB.506059.10 | trans-sialidase, putative | -1,26 | 6,355E-10 |
| TcCLB.508221.180 | trans-sialidase, Group V, putative | -1,26 | 4,006E-07 |
| TcCLB.510275.390 | mucin TcMUCII (pseudogene), putative | -1,26 | 0,0006762 |
| TcCLB.506423.20 | mucin TcMUCII, putative | -1,26 | 0,0055 |
| TcCLB.507953.210 | hypothetical protein | -1,261 | 0,00001947 |
| TcCLB.507121.20 | trans-sialidase, Group VII, putative | -1,262 | 2,926E-26 |
| TcCLB.504777.50 | mucin-associated surface protein (MASP, pseudogene), pu | -1,263 | 2,952E-12 |
| TcCLB.506801.10 | surface protease GP63 (pseudogene), putative | -1,263 | 5,891E-06 |
| TcCLB.505939.65 | hypothetical protein, conserved | -1,264 | 2,581E-11 |
| TcCLB.506763.40 | hypothetical protein | -1,264 | 0,00001604 |
| TcCLB.509349.10 | trans-sialidase (pseudogene), putative | -1,264 | 0,0001419 |
| TcCLB.510025.200 | syntaxin binding protein (pseudogene), putative | -1,265 | 0,00000192 |
| TcCLB.510373.100 | hypothetical protein, conserved (pseudogene) | -1,265 | 0,0001251 |
| TcCLB.504103.3 | hypothetical protein, conserved (fragment) | -1,267 | 2,017E-24 |
| TcCLB.508081.20 | hypothetical protein | -1,267 | 0,0001839 |
| TcCLB.503859.25 | surface protease GP63 (pseudogene), putative | -1,267 | 0,001359 |
| TcCLB.508661.70 | Enriched in surface-labeled proteome protein 18 (ESP18) | -1,269 | 2,945E-25 |
| TcCLB.508221.730 | 90 kDa surface protein, putative | -1,269 | 0,0008515 |
| TcCLB.511099.210 | mucin TcMUCII, putative | -1,269 | 0,0008792 |
| TcCLB.511643.40 | trans-sialidase, Group V, putative | -1,27 | 7,97E-16 |
| TcCLB.508043.10 | trans-sialidase, putative | -1,27 | 3,976E-15 |
| TcCLB.507071.164 | mucin TcMUCII (pseudogene), putative | -1,27 | 0,0008042 |
| TcCLB.503699.11 | mucin-associated surface protein (MASP, pseudogene), pu | -1,271 | 0,009053 |
| TcCLB.504229.50 | trans-sialidase (pseudogene), putative | -1,272 | 2,014E-15 |
| TcCLB.510407.9 | protein kinase, putative | -1,272 | 2,671E-10 |
| TcCLB.506601.31 | surface protease GP63 (pseudogene), putative | -1,272 | 0,001201 |
| TcCLB.510025.230 | hypothetical protein | -1,273 | 0,0009728 |
| TcCLB.508777.130 | hypothetical protein, conserved | -1,274 | 5,198E-52 |
| TcCLB.510015.50 | mucin TcMUC (pseudogene), putative | -1,274 | 2,481E-10 |
| TcCLB.503859.10 | syntaxin binding protein (pseudogene), putative | -1,275 | 0,0005312 |
| TcCLB.507237.210 | mucin TcMUCII, putative | -1,277 | 1,481E-12 |
| TcCLB.510373.40 | hypothetical protein | -1,278 | 0,0007589 |
| TcCLB.504239.80 | syntaxin binding protein (pseudogene), putative | -1,279 | 0,00005799 |
| TcCLB.510015.10 | hypothetical protein | -1,279 | 0,00009232 |
| TcCLB.398009.20 | trans-sialidase (pseudogene), putative | -1,28 | 8,578E-28 |

| | | | |
|---|---|---|---|
| TcCLB.503507.35 | mucin-associated surface protein (MASP, pseudogene), pu | -1,28 | 2,175E-12 |
| TcCLB.511427.19 | conserved protein, unknown function (fragment) | -1,28 | 0,0000107 |
| TcCLB.506793.50 | hypothetical protein, conserved | -1,281 | 3,434E-30 |
| TcCLB.510193.10 | mucin-associated surface protein (MASP), putative | -1,281 | 2,161E-08 |
| TcCLB.510275.360 | mucin TcMUCII (pseudogene), putative | -1,281 | 0,0002743 |
| TcCLB.511317.79 | OSM3-like kinesin, putative | -1,282 | 7,647E-21 |
| TcCLB.507957.36 | syntaxin binding protein (pseudogene), putative | -1,282 | 6,867E-07 |
| TcCLB.510251.19 | hypothetical protein, conserved | -1,283 | 1,705E-30 |
| TcCLB.509751.28 | mucin-associated surface protein (MASP, pseudogene), pu | -1,283 | 2,968E-28 |
| TcCLB.508221.530 | trans-sialidase (pseudogene), putative | -1,283 | 0,0000127 |
| TcCLB.508165.425 | mucin TcMUC (pseudogene), putative | -1,283 | 0,00004384 |
| TcCLB.439307.9 | hypothetical protein, conserved | -1,283 | 0,01097 |
| TcCLB.504081.110 | mucin TcMUCII (pseudogene), putative | -1,284 | 0,0000197 |
| TcCLB.508977.99 | mucin-associated surface protein (MASP, pseudogene), pu | -1,285 | 0,00003357 |
| TcCLB.509513.10 | trans-sialidase, Group II, putative | -1,287 | 6,647E-08 |
| TcCLB.506459.210 | mucin TcMUCII, putative | -1,287 | 0,00008117 |
| TcCLB.511555.30 | hypothetical protein, conserved | -1,288 | 1,209E-19 |
| TcCLB.507491.20 | surface antigen TASV, putative | -1,289 | 1,74E-65 |
| TcCLB.510359.490 | trans-sialidase (pseudogene), putative | -1,289 | 1,058E-11 |
| TcCLB.509925.40 | trans-sialidase (pseudogene), putative | -1,29 | 3,772E-11 |
| TcCLB.503461.40 | succinyl-CoA:3-ketoacid-coenzyme-A transferase (pseudog | -1,291 | 1,066E-08 |
| TcCLB.511255.350 | trans-sialidase, Group V, putative | -1,291 | 8,768E-07 |
| TcCLB.505919.20 | trans-sialidase, Group II, putative | -1,292 | 1,674E-17 |
| TcCLB.508081.80 | hypothetical protein | -1,292 | 0,0003307 |
| TcCLB.507839.30 | dispersed gene family protein 1 (DGF-1), putative | -1,292 | 0,002253 |
| TcCLB.506765.49 | hypothetical protein | -1,292 | 0,01429 |
| TcCLB.507143.60 | hypothetical protein | -1,293 | 0,002824 |
| TcCLB.506755.190 | hypothetical protein, conserved | -1,297 | 8,426E-34 |
| TcCLB.511451.59 | mucin-associated surface protein (MASP), putative | -1,297 | 0,00007673 |
| TcCLB.511163.59 | hypothetical protein, conserved (fragment) | -1,298 | 6,051E-17 |
| TcCLB.505993.10 | trans-sialidase (pseudogene), putative | -1,298 | 1,169E-13 |
| TcCLB.508437.20 | trans-sialidase (pseudogene), putative | -1,298 | 8,574E-08 |
| TcCLB.506779.180 | surface protease GP63, putative | -1,301 | 1,751E-29 |
| TcCLB.511797.240 | trans-sialidase (pseudogene), putative | -1,301 | 9,193E-10 |
| TcCLB.509947.20 | mucin-associated surface protein (MASP, pseudogene), pu | -1,301 | 0,002242 |
| TcCLB.511213.60 | mucin-associated surface protein (MASP), putative | -1,302 | 1,506E-17 |
| TcCLB.507487.20 | mucin-associated surface protein (MASP), putative | -1,302 | 0,001587 |
| TcCLB.510155.270 | dispersed gene family protein 1 (DGF-1, pseudogene), put | -1,303 | 8,583E-31 |
| TcCLB.506767.196 | surface protease GP63 (pseudogene), putative | -1,303 | 0,003077 |
| TcCLB.510359.610 | trans-sialidase, Group II, putative | -1,304 | 6,056E-13 |
| TcCLB.507237.120 | mucin-associated surface protein (MASP), putative | -1,304 | 5,754E-06 |
| TcCLB.511611.20 | mucin-associated surface protein (MASP), putative | -1,304 | 0,003101 |
| TcCLB.507237.150 | hypothetical protein | -1,307 | 0,0005959 |
| TcCLB.511531.59 | mitogen activated protein kinase, putative | -1,308 | 1,727E-24 |
| TcCLB.508747.50 | mucin TcMUCII, putative | -1,308 | 4,614E-09 |

| | | | |
|---|---|---|---|
| TcCLB.510361.180 | trans-sialidase (pseudogene), putative | -1,308 | 0,00001424 |
| TcCLB.511297.30 | hypothetical protein, conserved (fragment) | -1,309 | 7,919E-35 |
| TcCLB.506573.50 | hypothetical protein, conserved | -1,31 | 8,596E-18 |
| TcCLB.511895.30 | receptor-type adenylate cyclase, putative | -1,311 | 2,631E-09 |
| TcCLB.503587.80 | mucin-associated surface protein (MASP, pseudogene), pu | -1,311 | 1,052E-06 |
| TcCLB.504081.340 | hypothetical protein (pseudogene) | -1,311 | 0,001177 |
| TcCLB.511255.80 | mucin-associated surface protein (MASP), putative | -1,312 | 2,961E-11 |
| TcCLB.506267.80 | mucin-associated surface protein (MASP), putative | -1,312 | 4,512E-07 |
| TcCLB.506635.110 | kinesin-like protein, putative | -1,313 | 5,061E-23 |
| TcCLB.507959.170 | mucin-associated surface protein (MASP), putative | -1,313 | 1,634E-08 |
| TcCLB.506149.29 | hypothetical protein, conserved | -1,313 | 6,582E-06 |
| TcCLB.506799.90 | surface protease GP63 (pseudogene), putative | -1,315 | 2,644E-24 |
| TcCLB.508797.20 | hypothetical protein, conserved | -1,315 | 1,838E-16 |
| TcCLB.510359.660 | mucin TcMUCII, putative | -1,315 | 4,629E-06 |
| TcCLB.506155.99 | hypothetical protein, conserved | -1,316 | 2,752E-18 |
| TcCLB.511921.50 | trans-sialidase, Group VI, putative | -1,316 | 1,009E-10 |
| TcCLB.428999.20 | receptor-type adenylate cyclase, putative | -1,317 | 9,289E-18 |
| TcCLB.508493.50 | mucin-associated surface protein (MASP, pseudogene), pu | -1,317 | 2,559E-11 |
| TcCLB.506961.30 | hypothetical protein | -1,319 | 3,789E-11 |
| TcCLB.509977.40 | hypothetical protein | -1,32 | 1,293E-11 |
| TcCLB.504039.175 | hypothetical protein, conserved (pseudogene) | -1,32 | 2,681E-07 |
| TcCLB.510025.139 | hypothetical protein, conserved (pseudogene) | -1,32 | 0,000957 |
| TcCLB.507257.60 | Fumarate hydratase class I, cytosolic (FHc) | -1,321 | 5,957E-23 |
| TcCLB.507949.219 | mucin-associated surface protein (MASP, pseudogene), pu | -1,321 | 1,415E-20 |
| TcCLB.511657.50 | hypothetical protein, conserved | -1,321 | 4,659E-10 |
| TcCLB.509263.10 | UDP-Gal or UDP-GlcNAc-dependent glycosyltransferase, p | -1,323 | 1,373E-09 |
| TcCLB.507957.43 | hypothetical protein, conserved (pseudogene) | -1,323 | 0,003578 |
| TcCLB.476733.20 | hypothetical protein, conserved | -1,324 | 6,139E-62 |
| TcCLB.511401.40 | mucin-associated surface protein (MASP, pseudogene), pu | -1,324 | 1,387E-11 |
| TcCLB.507699.37 | mucin-associated surface protein (MASP, pseudogene), pu | -1,324 | 3,725E-06 |
| TcCLB.504233.50 | hypothetical protein, conserved | -1,325 | 0,004899 |
| TcCLB.511407.80 | hypothetical protein, conserved | -1,326 | 3,054E-15 |
| TcCLB.506751.90 | trans-sialidase (pseudogene), putative | -1,326 | 2,333E-06 |
| TcCLB.509617.40 | hypothetical protein, conserved | -1,327 | 3,621E-50 |
| TcCLB.504149.60 | hypothetical protein | -1,328 | 1,075E-56 |
| TcCLB.504081.50 | mucin-associated surface protein (MASP), putative | -1,328 | 3,894E-10 |
| TcCLB.508001.50 | trans-sialidase (pseudogene), putative | -1,328 | 0,00004764 |
| TcCLB.506335.100 | mucin-associated surface protein (MASP), putative | -1,329 | 4,297E-19 |
| TcCLB.508501.80 | mucin-associated surface protein (MASP), putative | -1,329 | 1,105E-14 |
| TcCLB.510625.79 | mucin TcMUCII (pseudogene), putative | -1,329 | 0,0001442 |
| TcCLB.511847.40 | hypothetical protein, conserved | -1,33 | 4,258E-19 |
| TcCLB.507065.60 | mucin-associated surface protein (MASP), putative | -1,332 | 1,035E-12 |
| TcCLB.507959.336 | surface protease GP63 (pseudogene), putative | -1,332 | 1,854E-10 |
| TcCLB.508175.110 | hypothetical protein, conserved | -1,333 | 1,014E-43 |
| TcCLB.509767.80 | hypothetical protein, conserved | -1,333 | 4,743E-18 |

| | | | |
|---|---|---|---|
| TcCLB.510401.50 | trans-sialidase (pseudogene), putative | -1,336 | 5,015E-23 |
| TcCLB.508015.40 | hypothetical protein | -1,336 | 5,111E-14 |
| TcCLB.506767.250 | trans-sialidase (pseudogene), putative | -1,337 | 4,856E-08 |
| TcCLB.404455.10 | trans-sialidase, putative (fragment) | -1,338 | 3,424E-17 |
| TcCLB.503717.5 | mucin TcMUCII (pseudogene), putative | -1,338 | 0,001785 |
| TcCLB.510265.4 | hypothetical protein, conserved | -1,339 | 2,318E-12 |
| TcCLB.510773.30 | hypothetical protein | -1,339 | 2,318E-07 |
| TcCLB.507091.160 | trans-sialidase (pseudogene), putative | -1,34 | 2,804E-13 |
| TcCLB.506373.39 | trans-sialidase, putative (fragment) | -1,341 | 1,131E-16 |
| TcCLB.510713.90 | trans-sialidase, Group V, putative | -1,341 | 1,758E-15 |
| TcCLB.510197.130 | trans-sialidase, Group VI, putative | -1,341 | 2,157E-06 |
| TcCLB.511603.60 | mucin TcMUCII, putative | -1,342 | 7,654E-11 |
| TcCLB.503665.30 | mucin-associated surface protein (MASP), putative | -1,342 | 1,221E-08 |
| TcCLB.506499.130 | hypothetical protein | -1,344 | 0,0004892 |
| TcCLB.507701.10 | mucin TcMUCII, putative | -1,345 | 2,724E-14 |
| TcCLB.511613.10 | mucin-associated surface protein (MASP), putative | -1,345 | 1,116E-12 |
| TcCLB.507633.59 | mucin-associated surface protein (MASP, pseudogene), pu | -1,346 | 6,055E-06 |
| TcCLB.510483.130 | mucin TcMUCII, putative | -1,346 | 0,0003127 |
| TcCLB.506501.150 | surface protease GP63 (pseudogene), putative | -1,346 | 0,001555 |
| TcCLB.427013.10 | trans-sialidase, putative (fragment) | -1,347 | 6,838E-19 |
| TcCLB.511401.100 | mucin-associated surface protein (MASP), putative | -1,347 | 1,591E-16 |
| TcCLB.506501.190 | mucin TcMUCII, putative | -1,347 | 0,00001617 |
| TcCLB.509233.140 | aldehyde dehydrogenase (pseudogene), putative | -1,348 | 4,343E-12 |
| TcCLB.510275.219 | mucin TcMUCII, putative | -1,349 | 0,0001229 |
| TcCLB.506763.350 | hypothetical protein, conserved (pseudogene) | -1,351 | 0,0009906 |
| TcCLB.508125.50 | mucin-associated surface protein (MASP, pseudogene), pu | -1,352 | 3,004E-07 |
| TcCLB.509591.20 | phosphoprotein phosphatase, putative | -1,352 | 0,003824 |
| TcCLB.506337.80 | surface antigen TASV, putative | -1,353 | 1,426E-49 |
| TcCLB.506813.220 | trans-sialidase, Group V, putative | -1,353 | 9,947E-09 |
| TcCLB.503545.10 | trans-sialidase (pseudogene), putative | -1,354 | 1,61E-11 |
| TcCLB.506661.170 | hypothetical protein, conserved | -1,355 | 4,494E-28 |
| TcCLB.510693.176 | mucin TcMUC (pseudogene), putative | -1,356 | 7,631E-10 |
| TcCLB.508389.114 | mucin TcMUCII, putative | -1,356 | 0,00008669 |
| TcCLB.507069.20 | hypothetical protein | -1,357 | 0,02968 |
| TcCLB.506519.60 | Protein kinase domain, putative | -1,359 | 3,296E-46 |
| TcCLB.506229.10 | Mitogen-activated protein kinase 10, putative (MPK10) | -1,359 | 6,914E-25 |
| TcCLB.507959.323 | syntaxin binding protein (pseudogene), putative | -1,359 | 3,111E-09 |
| TcCLB.506763.370 | hypothetical protein | -1,359 | 0,001228 |
| TcCLB.508877.10 | trans-sialidase (pseudogene), putative | -1,36 | 3,795E-10 |
| TcCLB.509439.20 | UDP-Gal or UDP-GlcNAc-dependent glycosyltransferase, p | -1,36 | 1,497E-07 |
| TcCLB.510483.190 | mucin TcMUCII, putative | -1,361 | 1,177E-13 |
| TcCLB.509081.60 | syntaxin binding protein (pseudogene), putative | -1,361 | 3,048E-12 |
| TcCLB.508441.94 | mucin TcMUCII, putative | -1,363 | 1,087E-20 |
| TcCLB.511173.120 | mucin-associated surface protein (MASP, pseudogene), pu | -1,364 | 4,093E-13 |
| TcCLB.504219.70 | mucin TcMUC, putative (fragment) | -1,365 | 4,149E-10 |

| | | | |
|---|---|---|---|
| TcCLB.511077.95 | mucin TcMUCII (pseudogene), putative | -1,365 | 0,005882 |
| TcCLB.510545.30 | mucin-associated surface protein (MASP, pseudogene), pu | -1,366 | 1,436E-06 |
| TcCLB.506309.34 | mucin TcMUCII, putative | -1,366 | 0,01396 |
| TcCLB.511283.120 | hypothetical protein, conserved | -1,369 | 0,0002015 |
| TcCLB.508807.5 | glycosylphosphatidylinositol-specific phospholipase C, puta | -1,37 | 2,744E-20 |
| TcCLB.511311.7 | trans-sialidase, putative (fragment) | -1,37 | 1,243E-19 |
| TcCLB.503913.40 | hypothetical protein | -1,371 | 3,81E-16 |
| TcCLB.503741.10 | mucin TcMUCII, putative | -1,372 | 1,445E-09 |
| TcCLB.445557.10 | mucin TcMUCII, putative | -1,372 | 0,001093 |
| TcCLB.506337.100 | surface antigen TASV, putative | -1,373 | 2,274E-41 |
| TcCLB.510583.220 | mucin TcMUCII, putative | -1,373 | 0,001748 |
| TcCLB.511917.10 | trans-sialidase, putative | -1,374 | 3,62E-14 |
| TcCLB.509525.280 | hypothetical protein | -1,375 | 0,0002764 |
| TcCLB.509907.50 | hypothetical protein, conserved (pseudogene) | -1,375 | 0,000571 |
| TcCLB.511183.600 | mucin-associated surface protein (MASP, pseudogene), pu | -1,379 | 4,126E-15 |
| TcCLB.510805.30 | mucin-associated surface protein (MASP, pseudogene), pu | -1,379 | 3,747E-13 |
| TcCLB.508163.234 | trans-sialidase, putative | -1,379 | 1,303E-10 |
| TcCLB.503973.290 | mucin TcMUCII, putative | -1,379 | 6,938E-06 |
| TcCLB.507987.20 | hypothetical protein | -1,379 | 0,0006962 |
| TcCLB.503649.20 | mucin TcMUC, putative | -1,379 | 0,00118 |
| TcCLB.511861.80 | UDP-Gal or UDP-GlcNAc-dependent glycosyltransferase, p | -1,38 | 6,281E-34 |
| TcCLB.510363.70 | mucin-associated surface protein (MASP), putative | -1,381 | 0,0001043 |
| TcCLB.510025.100 | mucin-associated surface protein (MASP), putative | -1,382 | 1,417E-19 |
| TcCLB.511923.60 | trans-sialidase, Group V, putative | -1,382 | 6,976E-16 |
| TcCLB.506971.30 | trans-sialidase (pseudogene), putative | -1,382 | 5,311E-10 |
| TcCLB.507957.310 | hypothetical protein | -1,382 | 3,205E-07 |
| TcCLB.509775.20 | protein kinase, putative | -1,383 | 7,21E-51 |
| TcCLB.506279.10 | trans-sialidase, Group VI, putative | -1,383 | 4,686E-15 |
| TcCLB.510279.190 | mucin-associated surface protein (MASP), putative | -1,383 | 6,509E-10 |
| TcCLB.508119.20 | mucin-associated surface protein (MASP), putative | -1,385 | 2,068E-16 |
| TcCLB.506875.10 | mucin TcMUCII (pseudogene), putative | -1,385 | 0,00004278 |
| TcCLB.505201.50 | trans-sialidase (pseudogene), putative | -1,386 | 2,628E-24 |
| TcCLB.508233.50 | variant-surface-glycoprotein phospholipase C, putative | -1,387 | 6,623E-30 |
| TcCLB.506339.10 | trans-sialidase, Group V, putative | -1,388 | 5,157E-16 |
| TcCLB.508313.40 | mucin TcMUCII, putative | -1,388 | 6,524E-09 |
| TcCLB.503587.30 | trans-sialidase, Group V, putative | -1,388 | 5,329E-07 |
| TcCLB.511185.114 | mucin TcMUCII (pseudogene), putative | -1,388 | 0,001483 |
| TcCLB.506409.190 | mucin-associated surface protein (MASP, pseudogene), pu | -1,389 | 1,337E-12 |
| TcCLB.507731.10 | mucin-associated surface protein (MASP), putative | -1,389 | 2,513E-11 |
| TcCLB.510497.90 | mucin TcMUCII, putative | -1,391 | 4,947E-07 |
| TcCLB.508069.5 | trans-sialidase, putative (fragment) | -1,391 | 5,692E-07 |
| TcCLB.466593.19 | hypothetical protein, conserved | -1,392 | 4,316E-38 |
| TcCLB.506599.170 | mucin-associated surface protein (MASP), putative | -1,392 | 2,594E-12 |
| TcCLB.509631.20 | mucin-associated surface protein (MASP), putative | -1,392 | 1,973E-11 |
| TcCLB.510025.205 | hypothetical protein, conserved (pseudogene) | -1,393 | 2,484E-06 |

| | | | |
|---|---|---|---|
| TcCLB.506757.90 | trans-sialidase, Group II, putative | -1,394 | 3,16E-15 |
| TcCLB.509897.200 | trans-sialidase, Group V, putative | -1,395 | 3,402E-20 |
| TcCLB.508167.20 | mucin TcMUCII, putative | -1,395 | 6,182E-11 |
| TcCLB.507731.50 | trans-sialidase (pseudogene), putative | -1,395 | 7,678E-06 |
| TcCLB.510279.340 | mucin TcMUCII, putative | -1,395 | 0,0001453 |
| TcCLB.509991.9 | protein kinase, putative | -1,397 | 2,782E-20 |
| TcCLB.511923.100 | trans-sialidase (pseudogene), putative | -1,397 | 8,474E-20 |
| TcCLB.506457.30 | hypoxanthine-guanine phosphoribosyltransferase, putative | -1,397 | 1,494E-18 |
| TcCLB.507085.70 | mucin-associated surface protein (MASP, pseudogene), pu | -1,397 | 2,461E-12 |
| TcCLB.507953.185 | hypothetical protein, conserved (pseudogene) | -1,397 | 4,371E-07 |
| TcCLB.511183.450 | trans-sialidase (pseudogene), putative | -1,398 | 4,113E-17 |
| TcCLB.511295.20 | hypothetical protein, conserved | -1,4 | 2,483E-18 |
| TcCLB.511575.120 | hypothetical protein, conserved | -1,401 | 8,464E-50 |
| TcCLB.507821.10 | trans-sialidase, Group VI, putative | -1,401 | 2,986E-21 |
| TcCLB.506909.100 | trans-sialidase, Group V, putative | -1,401 | 2,5E-10 |
| TcCLB.402919.10 | trans-sialidase, putative | -1,402 | 6,926E-19 |
| TcCLB.504259.10 | hypothetical protein, conserved | -1,402 | 1,161E-13 |
| TcCLB.508521.40 | hypothetical protein | -1,402 | 9,573E-06 |
| TcCLB.506501.341 | surface protease GP63 (pseudogene), putative | -1,402 | 0,0001252 |
| TcCLB.509971.44 | mucin-associated surface protein (MASP, pseudogene), pu | -1,403 | 0,00005766 |
| TcCLB.510483.310 | mucin-associated surface protein (MASP), putative | -1,404 | 3,664E-08 |
| TcCLB.506173.50 | mucin TcMUCII (pseudogene), putative | -1,404 | 2,016E-07 |
| TcCLB.508761.40 | trans-sialidase, Group V, putative | -1,405 | 4,682E-10 |
| TcCLB.509547.40 | mucin-associated surface protein (MASP), putative | -1,406 | 6,647E-08 |
| TcCLB.509155.39 | surface protease GP63, putative | -1,406 | 0,001816 |
| TcCLB.508461.310 | protein kinase A catalytic subunit isoform 1, putative | -1,407 | 4,39E-35 |
| TcCLB.510439.40 | hypothetical protein | -1,407 | 1,057E-24 |
| TcCLB.509387.5 | trans-sialidase, putative | -1,408 | 1,034E-14 |
| TcCLB.507071.70 | mucin-associated surface protein (MASP, pseudogene), pu | -1,408 | 8,678E-10 |
| TcCLB.508633.30 | mucin-associated surface protein (MASP, pseudogene), pu | -1,408 | 1,472E-07 |
| TcCLB.508139.160 | expression site-associated gene (ESAG-like) protein, puta | -1,409 | 2,326E-18 |
| TcCLB.511089.70 | mucin-associated surface protein (MASP, pseudogene), pu | -1,409 | 1,593E-11 |
| TcCLB.506281.100 | mucin-associated surface protein (MASP, pseudogene), pu | -1,409 | 2,215E-10 |
| TcCLB.509753.150 | mucin TcMUCII, putative | -1,41 | 0,0001191 |
| TcCLB.509613.20 | trans-sialidase (pseudogene), putative | -1,411 | 2,794E-11 |
| TcCLB.506499.200 | hypothetical protein | -1,411 | 4,124E-06 |
| TcCLB.508441.110 | mucin TcMUCI, putative | -1,412 | 1,581E-27 |
| TcCLB.511219.40 | trans-sialidase, Group II, putative | -1,412 | 1,86E-18 |
| TcCLB.510583.34 | mucin-associated surface protein (MASP, pseudogene), pu | -1,413 | 1,269E-08 |
| TcCLB.506281.117 | mucin TcMUCII (pseudogene), putative | -1,413 | 8,273E-08 |
| TcCLB.504425.10 | trans-sialidase, Group III, putative | -1,414 | 4,267E-30 |
| TcCLB.506815.20 | mucin-associated surface protein (MASP, pseudogene), pu | -1,414 | 1,676E-28 |
| TcCLB.509553.50 | DNA repair and recombination helicase protein PIF5, puta | -1,415 | 5,103E-31 |
| TcCLB.509869.30 | mucin TcMUCII, putative | -1,415 | 0,0002895 |
| TcCLB.507957.260 | surface protease GP63 (pseudogene), putative | -1,417 | 0,00002368 |

| | | | |
|---|---|---|---|
| TcCLB.510299.70 | hypothetical protein, conserved | -1,418 | 3,967E-33 |
| TcCLB.508767.20 | surface protein TolT (TolT) | -1,418 | 1,649E-29 |
| TcCLB.507205.10 | mucin-associated surface protein (MASP, pseudogene), pu | -1,419 | 1,058E-07 |
| TcCLB.511165.100 | serine/threonine protein kinase, putative | -1,42 | 9,736E-34 |
| TcCLB.508029.50 | kinesin-like protein, putative | -1,42 | 3,943E-29 |
| TcCLB.506221.70 | Paraflagellar Rod Proteome Component 9, putative (PFC9) | -1,421 | 1,169E-16 |
| TcCLB.508983.10 | mucin-associated surface protein (MASP, pseudogene), pu | -1,422 | 1,389E-10 |
| TcCLB.510021.90 | hypothetical protein, conserved (pseudogene) | -1,423 | 0,00001064 |
| TcCLB.506269.45 | trans-sialidase (pseudogene), putative | -1,423 | 0,0004328 |
| TcCLB.503633.21 | mucin TcMUCII (pseudogene), putative | -1,423 | 0,00103 |
| TcCLB.510525.139 | protein kinase, putative | -1,424 | 1,481E-27 |
| TcCLB.510533.170 | hypothetical protein, conserved | -1,425 | 1,973E-32 |
| TcCLB.508163.210 | trans-sialidase (pseudogene), putative | -1,425 | 3,171E-13 |
| TcCLB.506759.150 | hypothetical protein, conserved (pseudogene) | -1,425 | 1,336E-06 |
| TcCLB.511767.130 | expression site-associated gene (ESAG-like) protein, puta | -1,426 | 1,006E-11 |
| TcCLB.506767.123 | syntaxin binding protein (pseudogene), putative | -1,427 | 0,00003791 |
| TcCLB.510017.15 | surface protease GP63 (pseudogene), putative | -1,428 | 0,00001778 |
| TcCLB.511079.100 | mucin-associated surface protein (MASP), putative (fragm | -1,429 | 0,0299 |
| TcCLB.507941.120 | hypothetical protein, conserved | -1,43 | 4,873E-32 |
| TcCLB.511127.90 | hypothetical protein, conserved | -1,43 | 6,553E-24 |
| TcCLB.510377.10 | trans-sialidase, Group V, putative | -1,434 | 5,017E-08 |
| TcCLB.510371.61 | surface protease GP63 (pseudogene), putative | -1,434 | 3,852E-07 |
| TcCLB.509419.94 | mucin TcMUCII (pseudogene), putative | -1,434 | 0,0007133 |
| TcCLB.510311.100 | hypothetical protein, conserved | -1,437 | 5,135E-20 |
| TcCLB.506147.20 | hypothetical protein | -1,437 | 3,773E-16 |
| TcCLB.509553.30 | ribonuclease mar1, putative | -1,437 | 3,24E-15 |
| TcCLB.507069.130 | mucin-associated surface protein (MASP, pseudogene), pu | -1,437 | 8,985E-09 |
| TcCLB.510025.191 | surface protease GP63 (pseudogene), putative | -1,437 | 0,00003032 |
| TcCLB.507917.10 | surface protease GP63, putative | -1,438 | 1,909E-25 |
| TcCLB.503745.30 | ascorbate peroxidase (APX) | -1,438 | 3,02E-23 |
| TcCLB.511439.10 | expression site-associated gene (ESAG-like) protein, puta | -1,438 | 0,0001234 |
| TcCLB.509899.100 | trans-sialidase, Group V, putative | -1,439 | 7,717E-18 |
| TcCLB.511603.250 | mucin-associated surface protein (MASP), putative | -1,44 | 1,665E-10 |
| TcCLB.511245.10 | hypothetical protein, conserved | -1,441 | 8,469E-41 |
| TcCLB.508389.100 | mucin-associated surface protein (MASP), putative | -1,441 | 6,345E-19 |
| TcCLB.463901.10 | trans-sialidase, putative (fragment) | -1,441 | 7,852E-12 |
| TcCLB.506137.10 | surface protease GP63, putative | -1,441 | 3,475E-10 |
| TcCLB.506595.40 | trans-sialidase, Group II, putative | -1,442 | 9,46E-36 |
| TcCLB.510025.7 | syntaxin binding protein (pseudogene), putative | -1,442 | 2,213E-06 |
| TcCLB.511259.18 | mucin-associated surface protein (MASP, pseudogene), pu | -1,444 | 1,225E-10 |
| TcCLB.510261.10 | mucin-associated surface protein (MASP), putative | -1,446 | 1,536E-11 |
| TcCLB.511259.200 | mucin-associated surface protein (MASP, pseudogene), pu | -1,446 | 3,146E-08 |
| TcCLB.504655.9 | mucin-associated surface protein (MASP), putative | -1,448 | 3,568E-63 |
| TcCLB.506287.30 | hypothetical protein, conserved | -1,448 | 2,835E-29 |
| TcCLB.510377.405 | hypothetical protein, conserved (pseudogene) | -1,448 | 0,0001585 |

| | | | |
|---|---|---|---|
| TcCLB.508767.10 | surface protein TolT | -1,449 | 4,35E-35 |
| TcCLB.508227.30 | mucin TcMUCII, putative | -1,449 | 1,517E-07 |
| TcCLB.509897.270 | trans-sialidase, Group V, putative | -1,451 | 3,855E-17 |
| TcCLB.508153.680 | RNA-binding protein 6, putative | -1,451 | 1,204E-12 |
| TcCLB.508873.450 | mucin-associated surface protein (MASP), putative | -1,451 | 1,876E-07 |
| TcCLB.509429.41 | trans-sialidase (pseudogene), putative | -1,453 | 2,175E-16 |
| TcCLB.506501.100 | hypothetical protein | -1,455 | 0,006337 |
| TcCLB.511349.100 | trans-sialidase, Group II, putative | -1,456 | 7,635E-20 |
| TcCLB.507863.6 | hypothetical protein, conserved (pseudogene) | -1,457 | 0,001334 |
| TcCLB.510433.20 | surface protein TolT, putative | -1,458 | 2,548E-50 |
| TcCLB.509525.70 | hypothetical protein | -1,458 | 1,188E-08 |
| TcCLB.503507.20 | mucin TcMUCII, putative (fragment) | -1,458 | 0,0000297 |
| TcCLB.508165.140 | mucin TcMUCII (pseudogene), putative | -1,459 | 0,0000427 |
| TcCLB.506459.180 | trans-sialidase, Group V, putative | -1,46 | 3,48E-14 |
| TcCLB.510553.50 | UDP-Gal or UDP-GlcNAc-dependent glycosyltransferase, p | -1,46 | 1,949E-11 |
| TcCLB.508165.60 | mucin TcMUCII, putative | -1,461 | 0,0007409 |
| TcCLB.504081.440 | 90 kDa surface protein, putative | -1,463 | 5,317E-14 |
| TcCLB.510025.126 | syntaxin binding protein (pseudogene), putative | -1,464 | 2,542E-08 |
| TcCLB.506599.250 | surface protease GP63 (pseudogene), putative | -1,464 | 0,0002306 |
| TcCLB.510361.100 | mucin TcMUCII, putative | -1,466 | 1,396E-07 |
| TcCLB.506939.120 | trans-sialidase, Group V, putative | -1,467 | 2,858E-15 |
| TcCLB.503507.50 | mucin TcMUCII, putative | -1,467 | 9,739E-07 |
| TcCLB.510025.14 | hypothetical protein, conserved (pseudogene) | -1,468 | 0,002284 |
| TcCLB.511607.90 | mucin-associated surface protein (MASP, pseudogene), pu | -1,469 | 6,245E-16 |
| TcCLB.511149.10 | mucin TcMUCII, putative | -1,469 | 0,00006094 |
| TcCLB.506755.90 | hypothetical protein, conserved | -1,47 | 8,754E-33 |
| TcCLB.509607.50 | protein kinase, putative | -1,47 | 4,564E-11 |
| TcCLB.506985.50 | hypothetical protein, conserved | -1,471 | 3,678E-32 |
| TcCLB.510329.270 | Fumarate hydratase class I, cytosolic (FHc) | -1,471 | 1,244E-17 |
| TcCLB.508365.114 | mucin-associated surface protein (MASP, pseudogene), pu | -1,472 | 2,477E-15 |
| TcCLB.508697.10 | trans-sialidase, Group V, putative | -1,472 | 8,753E-10 |
| TcCLB.507231.30 | mucin-associated surface protein (MASP), putative | -1,473 | 1,451E-08 |
| TcCLB.504081.80 | trans-sialidase, Group V, putative | -1,473 | 1,115E-07 |
| TcCLB.508521.50 | surface protease GP63 (pseudogene), putative | -1,474 | 2,653E-18 |
| TcCLB.508297.20 | UDP-Gal or UDP-GlcNAc-dependent glycosyltransferase, p | -1,474 | 3,777E-15 |
| TcCLB.430603.10 | mucin-associated surface protein (MASP, pseudogene), pu | -1,474 | 6,094E-10 |
| TcCLB.507159.70 | hypothetical protein, conserved | -1,475 | 4,888E-53 |
| TcCLB.507641.130 | hypothetical protein, conserved | -1,476 | 8,979E-36 |
| TcCLB.508147.90 | mucin-associated surface protein (MASP, pseudogene), pu | -1,476 | 1,671E-12 |
| TcCLB.507637.130 | mucin TcMUCII, putative | -1,477 | 4,387E-07 |
| TcCLB.509897.140 | trans-sialidase, Group V, putative | -1,478 | 2,573E-21 |
| TcCLB.509753.270 | trans-sialidase (pseudogene), putative | -1,479 | 2,201E-17 |
| TcCLB.503609.10 | trans-sialidase (pseudogene), putative | -1,479 | 0,0002575 |
| TcCLB.511255.660 | mucin TcMUCII, putative | -1,48 | 9,991E-07 |
| TcCLB.508547.9 | UDP-Gal or UDP-GlcNAc-dependent glycosyltransferase, p | -1,482 | 6,437E-34 |

| | | | |
|---|---|---|---|
| TcCLB.510261.31 | mucin-associated surface protein (MASP, pseudogene), pu | -1,483 | 3,121E-10 |
| TcCLB.509195.30 | mucin-associated surface protein (MASP), putative | -1,483 | 9,012E-06 |
| TcCLB.509265.90 | trans-sialidase, Group II, putative | -1,485 | 1,064E-31 |
| TcCLB.511643.130 | mucin-associated surface protein (MASP, pseudogene), pu | -1,486 | 3,313E-23 |
| TcCLB.511183.590 | mucin TcMUCII, putative | -1,487 | 4,766E-14 |
| TcCLB.509753.310 | mucin-associated surface protein (MASP), putative | -1,487 | 2,617E-09 |
| TcCLB.506859.230 | hypothetical protein, conserved | -1,487 | 0,001615 |
| TcCLB.510111.20 | trans-sialidase, Group II, putative | -1,488 | 8,905E-18 |
| TcCLB.509753.250 | mucin-associated surface protein (MASP), putative | -1,489 | 6,69E-13 |
| TcCLB.504157.130 | surface protein TolT, putative (TolT) | -1,49 | 2,444E-26 |
| TcCLB.506279.250 | trans-sialidase (pseudogene), putative | -1,49 | 1,404E-10 |
| TcCLB.509221.50 | trans-sialidase (pseudogene), putative | -1,492 | 1,224E-26 |
| TcCLB.506329.65 | trans-sialidase (pseudogene), putative | -1,492 | 0,00004907 |
| TcCLB.507159.50 | hypothetical protein, conserved | -1,493 | 1,671E-53 |
| TcCLB.508511.10 | mucin-associated surface protein (MASP, pseudogene), pu | -1,493 | 3,165E-06 |
| TcCLB.508433.170 | mucin TcMUCII, putative | -1,493 | 0,0005129 |
| TcCLB.509685.40 | trans-sialidase, Group V, putative | -1,494 | 6,871E-27 |
| TcCLB.506993.180 | mucin-associated surface protein (MASP), putative | -1,494 | 2,691E-11 |
| TcCLB.508523.30 | trans-sialidase, Group II, putative | -1,494 | 1,24E-08 |
| TcCLB.508013.10 | mucin-associated surface protein (MASP, pseudogene), pu | -1,494 | 0,00002008 |
| TcCLB.506993.63 | mucin TcMUCI, putative | -1,494 | 0,00009796 |
| TcCLB.504101.10 | hypothetical protein | -1,495 | 4,132E-23 |
| TcCLB.511391.60 | trans-sialidase (pseudogene), putative | -1,495 | 1,324E-11 |
| TcCLB.511129.80 | UDP-Gal or UDP-GlcNAc-dependent glycosyltransferase, p | -1,496 | 2,842E-17 |
| TcCLB.509489.10 | hypothetical protein, conserved | -1,498 | 5,935E-65 |
| TcCLB.508805.136 | UDP-Gal or UDP-GlcNAc-dependent glycosyltransferase, p | -1,498 | 1,761E-17 |
| TcCLB.510279.314 | mucin TcMUCII (pseudogene), putative | -1,498 | 5,043E-06 |
| TcCLB.508243.23 | mucin TcMUCII, putative | -1,499 | 1,836E-08 |
| TcCLB.511173.430 | mucin-associated surface protein (MASP), putative | -1,5 | 7,167E-14 |
| TcCLB.511595.39 | mucin TcMUCII, putative | -1,5 | 1,036E-08 |
| TcCLB.507859.60 | mucin-associated surface protein (MASP), putative | -1,501 | 0,00369 |
| TcCLB.510623.50 | mucin-associated surface protein (MASP, pseudogene), pu | -1,502 | 2,075E-13 |
| TcCLB.511325.10 | amino acid permease (pseudogene), putative | -1,502 | 4,439E-09 |
| TcCLB.510715.100 | mucin-associated surface protein (MASP, pseudogene), pu | -1,504 | 4,519E-09 |
| TcCLB.504239.270 | hypothetical protein | -1,505 | 0,00006976 |
| TcCLB.511643.110 | trans-sialidase, Group V, putative | -1,506 | 6,605E-19 |
| TcCLB.506281.70 | mucin TcMUCI, putative | -1,506 | 1,642E-08 |
| TcCLB.503991.10 | hypothetical protein, conserved | -1,507 | 3,577E-23 |
| TcCLB.508099.9 | mucin-associated surface protein (MASP, pseudogene), pu | -1,507 | 9,433E-11 |
| TcCLB.508261.29 | mucin TcMUCII (pseudogene), putative | -1,508 | 0,0002326 |
| TcCLB.511183.40 | mucin-associated surface protein (MASP, pseudogene), pu | -1,509 | 1,562E-07 |
| TcCLB.506375.100 | hypothetical protein | -1,51 | 7,832E-38 |
| TcCLB.508157.20 | mucin-associated surface protein (MASP), putative | -1,51 | 1,897E-13 |
| TcCLB.505949.139 | mucin-associated surface protein (MASP, pseudogene), pu | -1,51 | 4,634E-07 |
| TcCLB.507957.304 | surface protease GP63 (pseudogene), putative | -1,51 | 0,00006932 |

| | | | |
|---|---|---|---|
| TcCLB.508977.50 | mucin-associated surface protein (MASP), putative | -1,511 | 1,955E-18 |
| TcCLB.510625.150 | mucin-associated surface protein (MASP), putative | -1,511 | 0,001252 |
| TcCLB.509663.20 | hypothetical protein, conserved | -1,514 | 4,968E-23 |
| TcCLB.511021.30 | hypothetical protein, conserved | -1,517 | 3,444E-49 |
| TcCLB.510085.60 | mucin-associated surface protein (MASP), putative | -1,517 | 4,706E-15 |
| TcCLB.510483.240 | mucin-associated surface protein (MASP, pseudogene), pu | -1,517 | 1,034E-14 |
| TcCLB.506657.88 | hypothetical protein, conserved | -1,517 | 0,00001378 |
| TcCLB.508227.40 | mucin-associated surface protein (MASP), putative | -1,518 | 1,811E-27 |
| TcCLB.509897.20 | trans-sialidase, Group V, putative | -1,519 | 1,991E-14 |
| TcCLB.506409.255 | surface protease GP63, putative | -1,519 | 0,002926 |
| TcCLB.508541.20 | mucin-associated surface protein (MASP), putative | -1,52 | 5,609E-13 |
| TcCLB.508221.560 | mucin TcMUCII, putative | -1,52 | 0,0002766 |
| TcCLB.510021.109 | mucin-associated surface protein (MASP, pseudogene), pu | -1,521 | 2,668E-14 |
| TcCLB.508759.30 | mucin TcMUC (pseudogene), putative | -1,522 | 0,00003365 |
| TcCLB.508857.15 | trans-sialidase, putative (fragment) | -1,523 | 1,681E-21 |
| TcCLB.507357.60 | mucin TcMUCII, putative | -1,523 | 6,172E-07 |
| TcCLB.506559.450 | hypothetical protein | -1,524 | 1,031E-31 |
| TcCLB.508999.134 | mucin TcMUCII, putative | -1,525 | 1,32E-07 |
| TcCLB.506667.80 | mucin-associated surface protein (MASP), putative | -1,525 | 0,00000359 |
| TcCLB.506499.30 | trans-sialidase (pseudogene), putative | -1,527 | 1,627E-09 |
| TcCLB.508805.120 | UDP-Gal or UDP-GlcNAc-dependent glycosyltransferase, p | -1,528 | 6,285E-18 |
| TcCLB.506459.130 | mucin-associated surface protein (MASP), putative | -1,528 | 1,359E-07 |
| TcCLB.507959.180 | surface protease GP63 (pseudogene), putative | -1,528 | 7,404E-06 |
| TcCLB.506415.10 | dispersed gene family protein 1 (DGF-1), putative | -1,53 | 6,421E-71 |
| TcCLB.511047.20 | mucin-like glycoprotein, putative | -1,53 | 2,745E-68 |
| TcCLB.506939.10 | mucin-associated surface protein (MASP, pseudogene), pu | -1,53 | 0,00003477 |
| TcCLB.507993.20 | hypothetical protein, conserved | -1,531 | 1,807E-13 |
| TcCLB.507085.80 | trans-sialidase (pseudogene), putative | -1,532 | 5,414E-39 |
| TcCLB.503777.4 | hypothetical protein, conserved | -1,532 | 3,42E-14 |
| TcCLB.511099.230 | mucin TcMUCII, putative | -1,532 | 0,00001577 |
| TcCLB.503859.45 | mucin TcMUCII, putative | -1,532 | 0,00001761 |
| TcCLB.504239.380 | hypothetical protein | -1,532 | 0,000025 |
| TcCLB.511607.100 | mucin TcMUCII, putative | -1,533 | 9,712E-14 |
| TcCLB.510489.50 | trans-sialidase, putative | -1,533 | 3,081E-06 |
| TcCLB.511089.74 | mucin TcMUCII (pseudogene), putative | -1,533 | 0,00001221 |
| TcCLB.509419.49 | mucin-associated surface protein (MASP, pseudogene), pu | -1,534 | 0,0003345 |
| TcCLB.510615.4 | hypothetical protein, conserved | -1,536 | 1,347E-24 |
| TcCLB.507633.51 | trans-sialidase (pseudogene), putative | -1,536 | 0,00001649 |
| TcCLB.506279.19 | mucin-associated surface protein (MASP, pseudogene), pu | -1,537 | 3,269E-20 |
| TcCLB.511603.300 | mucin-associated surface protein (MASP), putative | -1,539 | 1,442E-17 |
| TcCLB.510101.270 | hypothetical protein, conserved | -1,54 | 1,017E-61 |
| TcCLB.510279.210 | mucin-associated surface protein (MASP), putative | -1,541 | 1,056E-07 |
| TcCLB.509525.340 | hypothetical protein | -1,541 | 7,844E-06 |
| TcCLB.511089.10 | mucin-associated surface protein (MASP), putative | -1,541 | 0,0000669 |
| TcCLB.506757.129 | mucin-associated surface protein (MASP, pseudogene), pu | -1,542 | 1,309E-26 |

| | | | |
|---|---|---|---|
| TcCLB.511607.40 | mucin-associated surface protein (MASP), putative | -1,542 | 1,926E-18 |
| TcCLB.510483.300 | mucin TcMUCII, putative | -1,542 | 2,243E-18 |
| TcCLB.507673.60 | hypothetical protein, conserved | -1,542 | 5,143E-15 |
| TcCLB.507255.63 | UDP-Gal or UDP-GlcNAc-dependent glycosyltransferase, p | -1,542 | 2,767E-12 |
| TcCLB.503755.60 | surface protease GP63 (pseudogene), putative | -1,542 | 2,195E-08 |
| TcCLB.506595.10 | dispersed gene family protein 1 (DGF-1), putative | -1,543 | 5,013E-14 |
| TcCLB.481211.10 | mucin TcMUCII, putative | -1,544 | 7,054E-20 |
| TcCLB.511603.390 | mucin TcMUCII, putative | -1,544 | 0,0007544 |
| TcCLB.503773.9 | hypothetical protein, conserved (fragment) | -1,545 | 1,057E-19 |
| TcCLB.503767.10 | trans-sialidase, Group II, putative | -1,545 | 0,0002143 |
| TcCLB.509157.70 | hypothetical protein, conserved | -1,546 | 3,513E-17 |
| TcCLB.506615.10 | trans-sialidase (pseudogene), putative | -1,547 | 3,68E-42 |
| TcCLB.511859.130 | mucin-like glycoprotein, putative | -1,547 | 2,129E-36 |
| TcCLB.510275.340 | mucin-associated surface protein (MASP, pseudogene), pu | -1,547 | 2,405E-21 |
| TcCLB.511437.20 | expression site-associated gene (ESAG-like) protein, puta | -1,548 | 2,782E-10 |
| TcCLB.510489.36 | mucin-associated surface protein (MASP, pseudogene), pu | -1,55 | 9,271E-12 |
| TcCLB.447135.10 | surface protease GP63, putative | -1,55 | 0,00003063 |
| TcCLB.508873.460 | mucin-associated surface protein (MASP), putative | -1,552 | 1,222E-26 |
| TcCLB.511533.5 | mitogen activated protein kinase (pseudogene), putative | -1,553 | 2,549E-07 |
| TcCLB.510717.10 | surface antigen TASV, putative | -1,554 | 3,449E-56 |
| TcCLB.504081.490 | mucin-associated surface protein (MASP), putative | -1,554 | 7,913E-10 |
| TcCLB.507643.60 | trans-sialidase (pseudogene), putative | -1,555 | 7,825E-35 |
| TcCLB.511861.100 | mucin-associated surface protein (MASP, pseudogene), pu | -1,555 | 1,531E-23 |
| TcCLB.508065.10 | UDP-Gal or UDP-GlcNAc-dependent glycosyltransferase, p | -1,555 | 1,169E-17 |
| TcCLB.505025.110 | surface protease GP63 (pseudogene), putative | -1,557 | 4,751E-10 |
| TcCLB.506667.70 | mucin TcMUCII, putative | -1,557 | 3,203E-08 |
| TcCLB.509419.190 | mucin-associated surface protein (MASP, pseudogene), pu | -1,557 | 0,00007513 |
| TcCLB.506667.120 | mucin-associated surface protein (MASP), putative | -1,558 | 1,356E-11 |
| TcCLB.511625.149 | mucin TcMUCII (pseudogene), putative | -1,559 | 1,169E-08 |
| TcCLB.507953.190 | hypothetical protein | -1,559 | 0,00000285 |
| TcCLB.504277.20 | hypothetical protein | -1,56 | 1,664E-49 |
| TcCLB.508777.40 | leucine-rich repeat protein, putative | -1,56 | 3,934E-44 |
| TcCLB.511509.10 | hypothetical protein, conserved | -1,56 | 1,101E-27 |
| TcCLB.498829.10 | mucin TcMUC, putative | -1,56 | 4,737E-20 |
| TcCLB.507069.139 | mucin TcMUCII (pseudogene), putative | -1,56 | 6,233E-07 |
| TcCLB.508495.70 | mucin TcMUCII, putative | -1,561 | 4,755E-28 |
| TcCLB.506799.150 | surface protease GP63 (pseudogene), putative | -1,561 | 3,671E-16 |
| TcCLB.506379.10 | UDP-Gal or UDP-GlcNAc-dependent glycosyltransferase, p | -1,563 | 1,256E-16 |
| TcCLB.506751.100 | mucin-associated surface protein (MASP), putative | -1,563 | 1,495E-10 |
| TcCLB.510547.14 | mucin-associated surface protein (MASP, pseudogene), pu | -1,563 | 7,002E-09 |
| TcCLB.431357.20 | UDP-Gal or UDP-GlcNAc-dependent glycosyltransferase, p | -1,563 | 2,031E-06 |
| TcCLB.510311.120 | hypothetical protein | -1,564 | 4,218E-33 |
| TcCLB.510409.20 | hypothetical protein, conserved (pseudogene) | -1,565 | 3,583E-16 |
| TcCLB.508251.50 | mucin-associated surface protein (MASP), putative (fragm | -1,565 | 0,00009851 |
| TcCLB.508885.10 | surface protease GP63 (pseudogene), putative | -1,566 | 2,233E-07 |

| | | | |
|---|---|---|---|
| TcCLB.504423.15 | hypothetical protein, conserved | -1,567 | 2,078E-35 |
| TcCLB.504155.150 | surface protease GP63, putative | -1,567 | 1,637E-15 |
| TcCLB.508145.30 | RNA-binding protein, putative | -1,57 | 2,458E-10 |
| TcCLB.506183.90 | trans-sialidase (pseudogene), putative | -1,571 | 4,163E-17 |
| TcCLB.508013.70 | mucin TcMUCII, putative | -1,571 | 3,234E-06 |
| TcCLB.503855.60 | hypothetical protein, conserved | -1,572 | 3,204E-48 |
| TcCLB.506859.180 | hypothetical protein, conserved | -1,572 | 3,266E-17 |
| TcCLB.404845.10 | trans-sialidase (pseudogene), putative | -1,573 | 8,194E-47 |
| TcCLB.510377.50 | trans-sialidase, Group V, putative | -1,574 | 0,00004566 |
| TcCLB.508209.94 | hypothetical protein | -1,575 | 1,073E-30 |
| TcCLB.507485.10 | hypothetical protein, conserved | -1,575 | 1,893E-21 |
| TcCLB.509093.10 | mucin-associated surface protein (MASP), putative | -1,575 | 2,667E-08 |
| TcCLB.505183.30 | malic enzyme, putative | -1,576 | 1,386E-72 |
| TcCLB.507473.30 | trans-sialidase, Group II, putative | -1,576 | 1,499E-31 |
| TcCLB.510237.120 | mucin-associated surface protein (MASP), putative | -1,576 | 9,412E-14 |
| TcCLB.506765.58 | trans-sialidase (pseudogene), putative | -1,576 | 3,237E-12 |
| TcCLB.511449.110 | surface protease GP63, putative | -1,577 | 3,373E-12 |
| TcCLB.511081.30 | mucin TcMUCII, putative | -1,577 | 1,602E-08 |
| TcCLB.508121.110 | mucin-associated surface protein (MASP), putative | -1,578 | 8,898E-17 |
| TcCLB.506671.20 | mucin TcMUC (pseudogene), putative | -1,578 | 2,653E-09 |
| TcCLB.510729.280 | trans-sialidase, Group III, putative | -1,579 | 2,36E-39 |
| TcCLB.507067.20 | surface protease GP63, putative | -1,58 | 2,791E-22 |
| TcCLB.506229.40 | leucine-rich repeat protein, putative | -1,58 | 6,736E-18 |
| TcCLB.508629.120 | mucin-associated surface protein (MASP), putative | -1,581 | 6,119E-43 |
| TcCLB.506617.10 | surface protein TolT (TolT) | -1,581 | 3,898E-40 |
| TcCLB.510373.110 | hypothetical protein | -1,581 | 0,00008749 |
| TcCLB.508943.4 | glycerol uptake protein, putative | -1,582 | 0,00002366 |
| TcCLB.510417.30 | mucin TcMUCII, putative | -1,583 | 1,491E-10 |
| TcCLB.508873.510 | mucin-associated surface protein (MASP), putative | -1,583 | 3,748E-10 |
| TcCLB.506199.10 | hypothetical protein, conserved | -1,584 | 9,21E-32 |
| TcCLB.406725.10 | trans-sialidase (pseudogene), putative | -1,584 | 3,574E-08 |
| TcCLB.506879.40 | mucin-associated surface protein (MASP, pseudogene), pu | -1,585 | 1,832E-11 |
| TcCLB.511255.10 | trans-sialidase, Group V, putative | -1,586 | 6,74E-10 |
| TcCLB.504787.20 | mucin TcMUCII, putative | -1,587 | 3,245E-12 |
| TcCLB.510717.20 | surface antigen TASV, putative | -1,589 | 2,492E-66 |
| TcCLB.507957.29 | surface protease GP63 (pseudogene), putative | -1,589 | 0,000126 |
| TcCLB.511127.60 | kinesin, putative | -1,591 | 1,156E-76 |
| TcCLB.506801.80 | cysteine peptidase inhibitor (CHA) | -1,591 | 5,462E-38 |
| TcCLB.510701.40 | mucin-associated surface protein (MASP, pseudogene), pu | -1,591 | 0,0001599 |
| TcCLB.507485.30 | hypothetical protein, conserved | -1,592 | 1,055E-28 |
| TcCLB.511403.30 | mucin TcMUCII, putative | -1,592 | 1,041E-06 |
| TcCLB.511173.64 | mucin-associated surface protein (MASP), putative | -1,592 | 0,0009401 |
| TcCLB.511495.30 | hypothetical protein, conserved | -1,593 | 8,145E-32 |
| TcCLB.509897.70 | trans-sialidase, Group V, putative | -1,593 | 1,353E-19 |
| TcCLB.508121.56 | mucin-associated surface protein (MASP, pseudogene), pu | -1,593 | 2,774E-09 |

| | | | |
|---|---|---|---|
| TcCLB.509267.3 | receptor-type adenylate cyclase, putative | -1,593 | 3,313E-07 |
| TcCLB.511923.30 | trans-sialidase, Group V, putative | -1,594 | 2,369E-21 |
| TcCLB.508883.39 | mucin-associated surface protein (MASP, pseudogene), pu | -1,595 | 3,704E-17 |
| TcCLB.507069.60 | mucin TcMUCII, putative | -1,595 | 8,099E-07 |
| TcCLB.506767.215 | hypothetical protein, conserved (pseudogene) | -1,595 | 0,001378 |
| TcCLB.506989.155 | hypothetical protein, conserved | -1,598 | 3,576E-27 |
| TcCLB.507637.180 | mucin-associated surface protein (MASP, pseudogene), pu | -1,599 | 1,66E-23 |
| TcCLB.510275.306 | mucin-associated surface protein (MASP, pseudogene), pu | -1,601 | 2,103E-17 |
| TcCLB.504081.280 | mucin-associated surface protein (MASP, pseudogene), pu | -1,601 | 1,226E-07 |
| TcCLB.511439.90 | hypothetical protein, conserved | -1,602 | 5,559E-13 |
| TcCLB.506337.90 | surface antigen TASV, putative (TASV) | -1,603 | 1,555E-79 |
| TcCLB.511149.199 | mucin TcMUCII (pseudogene), putative | -1,603 | 0,0000753 |
| TcCLB.511311.30 | trans-sialidase, Group II, putative | -1,606 | 1,294E-24 |
| TcCLB.510933.90 | mucin-associated surface protein (MASP), putative | -1,607 | 2,723E-14 |
| TcCLB.508983.20 | trans-sialidase (pseudogene), putative | -1,608 | 3,701E-25 |
| TcCLB.506909.155 | mucin-associated surface protein (MASP, pseudogene), pu | -1,612 | 0,0000024 |
| TcCLB.506459.10 | mucin-associated surface protein (MASP), putative | -1,613 | 2,513E-20 |
| TcCLB.506961.70 | trans-sialidase (pseudogene), putative | -1,614 | 1,592E-28 |
| TcCLB.506139.170 | trans-sialidase (pseudogene), putative | -1,614 | 2,356E-12 |
| TcCLB.460625.9 | trans-sialidase (pseudogene), putative | -1,614 | 8,626E-08 |
| TcCLB.506361.18 | mucin TcMUC (pseudogene), putative | -1,614 | 3,351E-06 |
| TcCLB.507699.170 | surface protease GP63 (pseudogene), putative | -1,615 | 3,7E-27 |
| TcCLB.506173.240 | mucin TcMUCII, putative | -1,616 | 5,397E-11 |
| TcCLB.506785.30 | mucin TcMUCII, putative | -1,616 | 0,00000367 |
| TcCLB.504219.50 | hypothetical protein, conserved (pseudogene) | -1,617 | 0,0003603 |
| TcCLB.508509.51 | mucin-associated surface protein (MASP, pseudogene), pu | -1,62 | 3,329E-08 |
| TcCLB.506277.30 | mucin-associated surface protein (MASP, pseudogene), pu | -1,621 | 1,117E-21 |
| TcCLB.506761.40 | mucin-associated surface protein (MASP, pseudogene), pu | -1,621 | 1,615E-14 |
| TcCLB.508499.20 | mucin-associated surface protein (MASP), putative | -1,622 | 8,493E-24 |
| TcCLB.506971.80 | mucin-associated surface protein (MASP), putative (fragm | -1,623 | 1,345E-11 |
| TcCLB.507071.34 | mucin TcMUCII, putative | -1,623 | 0,0003268 |
| TcCLB.508873.103 | mucin-associated surface protein (MASP, pseudogene), pu | -1,624 | 7,158E-08 |
| TcCLB.506939.40 | mucin TcMUCII, putative | -1,624 | 3,02E-07 |
| TcCLB.506841.20 | trans-sialidase, Group I, putative | -1,625 | 6,119E-21 |
| TcCLB.505023.40 | mucin-associated surface protein (MASP), putative | -1,626 | 4,23E-18 |
| TcCLB.507237.356 | syntaxin binding protein (pseudogene), putative | -1,626 | 0,000066 |
| TcCLB.510697.40 | mucin-associated surface protein (MASP), putative | -1,627 | 1,356E-11 |
| TcCLB.508999.70 | mucin-associated surface protein (MASP), putative | -1,63 | 5,827E-16 |
| TcCLB.510197.190 | mucin-associated surface protein (MASP), putative | -1,63 | 1,48E-12 |
| TcCLB.506229.60 | hypothetical protein, conserved | -1,631 | 8,736E-57 |
| TcCLB.508873.30 | mucin-associated surface protein (MASP), putative | -1,631 | 4,015E-28 |
| TcCLB.511553.130 | mucin TcMUCII, putative | -1,631 | 1,854E-18 |
| TcCLB.510269.60 | trans-sialidase (pseudogene), putative | -1,631 | 2,019E-17 |
| TcCLB.509755.50 | trans-sialidase, Group V, putative | -1,631 | 6,909E-17 |
| TcCLB.506763.35 | hypothetical protein, conserved (pseudogene) | -1,631 | 1,008E-08 |

| | | | |
|---|---|---|---|
| TcCLB.509867.70 | trans-sialidase, Group VI, putative | -1,633 | 4,689E-10 |
| TcCLB.503631.9 | hypothetical protein, conserved | -1,634 | 2,482E-46 |
| TcCLB.506679.280 | endonuclease/exonuclease/phosphatase, putative | -1,634 | 3,733E-43 |
| TcCLB.504115.60 | trans-sialidase, Group V, putative | -1,634 | 7,751E-18 |
| TcCLB.508165.350 | mucin-associated surface protein (MASP), putative | -1,635 | 3,406E-09 |
| TcCLB.511173.210 | mucin-associated surface protein (MASP), putative | -1,636 | 1,659E-10 |
| TcCLB.508327.10 | mucin-associated surface protein (MASP), putative | -1,637 | 0,0003737 |
| TcCLB.511603.380 | mucin-associated surface protein (MASP), putative | -1,638 | 4,332E-12 |
| TcCLB.508365.284 | mucin-associated surface protein (MASP, pseudogene), pu | -1,638 | 6,031E-12 |
| TcCLB.511247.9 | hypothetical protein, conserved | -1,642 | 1,215E-06 |
| TcCLB.511861.20 | trans-sialidase, Group V, putative | -1,643 | 9,219E-37 |
| TcCLB.509251.70 | mucin-associated surface protein (MASP, pseudogene), pu | -1,643 | 2,586E-30 |
| TcCLB.507429.40 | mucin-associated surface protein (MASP, pseudogene), pu | -1,643 | 4,128E-21 |
| TcCLB.508511.40 | mucin-associated surface protein (MASP, pseudogene), pu | -1,645 | 9,875E-06 |
| TcCLB.508011.20 | mucin-associated surface protein (MASP, pseudogene), pu | -1,645 | 0,0004339 |
| TcCLB.507081.100 | hypothetical protein, conserved | -1,646 | 3,415E-36 |
| TcCLB.509897.210 | mucin-associated surface protein (MASP, pseudogene), pu | -1,646 | 2,967E-31 |
| TcCLB.507071.80 | trans-sialidase, Group VI, putative | -1,646 | 8,33E-13 |
| TcCLB.506703.74 | mucin-associated surface protein (MASP, pseudogene), pu | -1,646 | 0,00006786 |
| TcCLB.509525.410 | mucin-associated surface protein (MASP, pseudogene), pu | -1,648 | 2,739E-11 |
| TcCLB.508293.130 | hypothetical protein | -1,649 | 0,0001801 |
| TcCLB.504533.40 | trans-sialidase, Group IV, putative | -1,65 | 1,56E-47 |
| TcCLB.508499.39 | trans-sialidase (pseudogene), putative | -1,65 | 3,217E-13 |
| TcCLB.506955.184 | mucin-associated surface protein (MASP, pseudogene), pu | -1,65 | 8,139E-10 |
| TcCLB.509115.20 | trans-sialidase, Group V, putative | -1,651 | 6,443E-16 |
| TcCLB.510025.20 | mucin TcMUCII, putative | -1,652 | 2,795E-15 |
| TcCLB.506297.350 | SKP1-like protein | -1,653 | 0,00003513 |
| TcCLB.510023.10 | hypothetical protein, conserved (pseudogene) | -1,654 | 2,833E-07 |
| TcCLB.506589.89 | mucin-associated surface protein (MASP, pseudogene), pu | -1,656 | 3,586E-13 |
| TcCLB.510503.70 | trans-sialidase (pseudogene), putative | -1,657 | 2,158E-18 |
| TcCLB.510625.109 | mucin-associated surface protein (MASP), putative | -1,657 | 0,0006611 |
| TcCLB.413409.10 | trans-sialidase, putative (fragment) | -1,658 | 8,6E-20 |
| TcCLB.508277.401 | magnesium and cobalt transport protein, putative (fragm | -1,659 | 4,599E-55 |
| TcCLB.510587.10 | hypothetical protein | -1,659 | 0,0003415 |
| TcCLB.510099.10 | hypothetical protein, conserved | -1,66 | 8,248E-57 |
| TcCLB.508875.20 | mucin-associated surface protein (MASP), putative (fragm | -1,66 | 3,569E-31 |
| TcCLB.511751.140 | hypothetical protein | -1,661 | 1,006E-42 |
| TcCLB.511797.184 | mucin TcMUC (pseudogene), putative | -1,661 | 0,00000209 |
| TcCLB.510625.88 | mucin-associated surface protein (MASP), putative | -1,661 | 0,0001099 |
| TcCLB.511183.200 | mucin-associated surface protein (MASP, pseudogene), pu | -1,662 | 1,485E-28 |
| TcCLB.506763.190 | mucin TcMUCII, putative | -1,662 | 1,484E-11 |
| TcCLB.508871.140 | mucin TcMUCII, putative | -1,662 | 8,515E-10 |
| TcCLB.511175.50 | mucin TcMUC (pseudogene), putative | -1,663 | 7,502E-48 |
| TcCLB.506269.99 | mucin-associated surface protein (MASP), putative (fragm | -1,663 | 5,559E-32 |
| TcCLB.510629.290 | mucin-associated surface protein (MASP, pseudogene), pu | -1,663 | 3,065E-12 |

| | | | |
|---|---|---|---|
| TcCLB.503787.50 | mucin TcMUCII, putative | -1,663 | 4,108E-11 |
| TcCLB.510693.19 | mucin-associated surface protein (MASP, pseudogene), pu | -1,664 | 8,132E-21 |
| TcCLB.506805.10 | mucin-associated surface protein (MASP), putative | -1,664 | 1,098E-09 |
| TcCLB.511079.80 | mucin TcMUCII, putative | -1,664 | 0,0009556 |
| TcCLB.505655.10 | trans-sialidase (pseudogene), putative | -1,665 | 1,596E-19 |
| TcCLB.508041.10 | SKP1-like protein | -1,665 | 1,409E-14 |
| TcCLB.504239.195 | mucin-associated surface protein (MASP, pseudogene), pu | -1,665 | 6,334E-09 |
| TcCLB.510095.20 | trans-sialidase, Group II, putative | -1,665 | 0,00001687 |
| TcCLB.509767.10 | neutral sphingomyelinase activation associated factor-like | -1,666 | 4,858E-71 |
| TcCLB.508563.20 | trans-sialidase, Group II, putative | -1,667 | 2,754E-29 |
| TcCLB.505025.100 | mucin TcMUCII (pseudogene), putative | -1,668 | 5,917E-10 |
| TcCLB.511183.409 | mucin TcMUC (pseudogene), putative | -1,668 | 7,26E-09 |
| TcCLB.510933.110 | mucin-associated surface protein (MASP, pseudogene), pu | -1,669 | 7,259E-10 |
| TcCLB.504081.540 | trans-sialidase, Group V, putative | -1,67 | 7,568E-15 |
| TcCLB.507747.80 | mucin TcMUCII, putative | -1,67 | 1,187E-07 |
| TcCLB.508115.20 | trans-sialidase, putative | -1,671 | 1,295E-06 |
| TcCLB.511599.110 | mucin TcMUCII, putative | -1,673 | 3,703E-09 |
| TcCLB.503479.40 | hypothetical protein, conserved | -1,674 | 2,629E-96 |
| TcCLB.504239.330 | surface protease GP63 (pseudogene), putative | -1,676 | 6,429E-09 |
| TcCLB.506445.90 | magnesium and cobalt transport protein, putative | -1,677 | 7,817E-56 |
| TcCLB.504127.20 | hypothetical protein | -1,677 | 4,867E-27 |
| TcCLB.506617.5 | surface protein TolT, putative (fragment) | -1,678 | 1,469E-40 |
| TcCLB.509387.19 | trans-sialidase, putative | -1,679 | 4,82E-31 |
| TcCLB.510207.40 | mucin-associated surface protein (MASP), putative | -1,679 | 1,295E-21 |
| TcCLB.508221.590 | trans-sialidase, Group V, putative | -1,679 | 1,116E-09 |
| TcCLB.509081.80 | hypothetical protein, conserved (pseudogene) | -1,679 | 3,406E-09 |
| TcCLB.508143.90 | trans-sialidase (pseudogene), putative | -1,68 | 5,216E-33 |
| TcCLB.508589.10 | hypothetical protein | -1,681 | 1,522E-19 |
| TcCLB.506753.230 | hypothetical protein, conserved | -1,682 | 2,297E-54 |
| TcCLB.507747.10 | surface protease GP63, putative | -1,683 | 2,282E-20 |
| TcCLB.511259.170 | trans-sialidase (pseudogene), putative | -1,684 | 7,306E-19 |
| TcCLB.503519.20 | mucin TcMUCII, putative | -1,684 | 7,395E-12 |
| TcCLB.508165.220 | mucin TcMUCII (pseudogene), putative | -1,685 | 1,972E-06 |
| TcCLB.511797.200 | mucin-associated surface protein (MASP), putative | -1,686 | 1,758E-37 |
| TcCLB.503977.15 | surface protease GP63 (pseudogene), putative | -1,686 | 3,522E-08 |
| TcCLB.503773.50 | Nucleolar protein 56, putative (fragment) (NOP56) | -1,687 | 8,155E-33 |
| TcCLB.508221.40 | mucin TcMUCII, putative | -1,688 | 4,023E-08 |
| TcCLB.506599.300 | hypothetical protein | -1,688 | 0,001422 |
| TcCLB.511165.50 | Enriched in surface-labeled proteome protein 7 (ESP7) | -1,689 | 2,779E-44 |
| TcCLB.508011.110 | mucin-associated surface protein (MASP), putative | -1,689 | 3,554E-21 |
| TcCLB.508293.140 | mucin-associated surface protein (MASP), putative | -1,689 | 0,00003561 |
| TcCLB.507953.170 | hypothetical protein | -1,689 | 0,0002721 |
| TcCLB.509525.390 | mucin-associated surface protein (MASP), putative | -1,691 | 3,464E-17 |
| TcCLB.511613.80 | mucin-associated surface protein (MASP), putative | -1,691 | 1,065E-14 |
| TcCLB.510209.27 | mucin-associated surface protein (MASP), putative | -1,691 | 1,675E-09 |

| | | | |
|---|---|---|---|
| TcCLB.506767.60 | hypothetical protein, conserved (pseudogene) | -1,692 | 2,456E-08 |
| TcCLB.510197.180 | trans-sialidase, Group VI, putative | -1,693 | 2,794E-21 |
| TcCLB.506967.70 | mucin-associated surface protein (MASP), putative | -1,693 | 1,013E-18 |
| TcCLB.511921.150 | trans-sialidase, Group V, putative | -1,693 | 1,298E-17 |
| TcCLB.510271.30 | trans-sialidase (pseudogene), putative | -1,694 | 7,063E-22 |
| TcCLB.510021.210 | mucin-associated surface protein (MASP, pseudogene), pu | -1,694 | 7,174E-07 |
| TcCLB.506131.120 | trans-sialidase (pseudogene), putative | -1,695 | 1,019E-13 |
| TcCLB.506277.20 | mucin TcMUCII (pseudogene), putative | -1,697 | 7,126E-07 |
| TcCLB.511451.80 | trans-sialidase, Group V, putative | -1,698 | 1,137E-32 |
| TcCLB.504115.10 | trans-sialidase, Group VIII, putative | -1,699 | 5,701E-53 |
| TcCLB.504081.350 | mucin-associated surface protein (MASP), putative | -1,699 | 3,677E-13 |
| TcCLB.506899.10 | trans-sialidase, putative | -1,7 | 9,837E-06 |
| TcCLB.506239.50 | palmitoyl acyltransferase 6, putative | -1,701 | 5,545E-34 |
| TcCLB.511849.30 | mucin-associated surface protein (MASP), putative | -1,701 | 1,498E-14 |
| TcCLB.508221.580 | mucin-associated surface protein (MASP), putative | -1,701 | 2,928E-07 |
| TcCLB.506755.10 | hypothetical protein, conserved | -1,703 | 4,749E-50 |
| TcCLB.508107.30 | trans-sialidase, Group VI, putative | -1,707 | 2,81E-09 |
| TcCLB.507605.13 | mucin-associated surface protein (MASP), putative (fragm | -1,707 | 0,00003498 |
| TcCLB.510033.30 | trans-sialidase, Group V, putative | -1,708 | 6,48E-22 |
| TcCLB.510105.310 | mucin-associated surface protein (MASP), putative | -1,708 | 5,725E-16 |
| TcCLB.510023.30 | mucin-associated surface protein (MASP), putative (fragm | -1,708 | 6,875E-14 |
| TcCLB.482369.29 | arginine kinase (fragment) | -1,709 | 9,417E-11 |
| TcCLB.511173.200 | hypothetical protein | -1,709 | 4,472E-10 |
| TcCLB.507255.20 | trans-sialidase, Group II, putative | -1,711 | 5,323E-42 |
| TcCLB.511605.10 | hypothetical protein | -1,711 | 0,0009129 |
| TcCLB.511919.10 | mucin-associated surface protein (MASP), putative | -1,713 | 8,351E-30 |
| TcCLB.505267.20 | trans-sialidase (pseudogene), putative | -1,713 | 5,658E-15 |
| TcCLB.506499.149 | mucin TcMUCII, putative | -1,714 | 1,992E-13 |
| TcCLB.508493.60 | trans-sialidase (pseudogene), putative | -1,715 | 1,725E-36 |
| TcCLB.504039.210 | hypothetical protein | -1,715 | 0,002599 |
| TcCLB.511255.130 | mucin-associated surface protein (MASP), putative | -1,717 | 1,308E-12 |
| TcCLB.460757.10 | receptor-type adenylate cyclase, putative | -1,718 | 2,137E-26 |
| TcCLB.509699.30 | mucin TcMUCII (pseudogene), putative | -1,718 | 4,36E-08 |
| TcCLB.510197.220 | mucin-associated surface protein (MASP, pseudogene), pu | -1,721 | 2,32E-21 |
| TcCLB.510729.140 | NIMA-related kinase, putative | -1,722 | 8,449E-25 |
| TcCLB.510503.100 | surface protease GP63, putative | -1,724 | 9,718E-17 |
| TcCLB.510629.70 | mucin-associated surface protein (MASP), putative | -1,724 | 3,598E-06 |
| TcCLB.510371.141 | hypothetical protein, conserved (pseudogene) | -1,725 | 0,0004038 |
| TcCLB.508873.25 | hypothetical protein | -1,726 | 4,222E-15 |
| TcCLB.506499.80 | mucin-associated surface protein (MASP), putative | -1,726 | 1,003E-06 |
| TcCLB.508647.280 | malic enzyme | -1,727 | 2,45E-100 |
| TcCLB.508495.100 | trans-sialidase, Group V, putative | -1,727 | 1,283E-19 |
| TcCLB.510387.40 | mucin-associated surface protein (MASP), putative | -1,727 | 5,328E-12 |
| TcCLB.510979.10 | mucin-associated surface protein (MASP), putative | -1,73 | 2,174E-20 |
| TcCLB.504031.35 | mucin TcMUCII (pseudogene), putative | -1,73 | 3,944E-08 |

| | | | |
|---|---|---|---|
| TcCLB.506767.119 | surface protease GP63 (pseudogene), putative | -1,731 | 0,000051 |
| TcCLB.448661.10 | mucin-associated surface protein (MASP), putative (fragm | -1,732 | 2,035E-14 |
| TcCLB.510565.150 | surface protease GP63, putative | -1,733 | 1,921E-52 |
| TcCLB.511553.120 | surface protease GP63, putative | -1,733 | 4,17E-15 |
| TcCLB.506589.130 | trans-sialidase (pseudogene), putative | -1,734 | 7,469E-29 |
| TcCLB.506973.105 | mucin-associated surface protein (MASP, pseudogene), pu | -1,734 | 1,987E-10 |
| TcCLB.506115.9 | dispersed gene family protein 1 (DGF-1), putative | -1,735 | 1,529E-11 |
| TcCLB.507953.140 | trans-sialidase, Group V, putative | -1,735 | 2,603E-07 |
| TcCLB.508789.60 | mucin TcMUCII, putative | -1,736 | 4,011E-20 |
| TcCLB.503649.10 | mucin-associated surface protein (MASP, pseudogene), pu | -1,737 | 2,012E-10 |
| TcCLB.508495.9 | mucin-associated surface protein (MASP, pseudogene), pu | -1,737 | 2,632E-08 |
| TcCLB.505025.90 | mucin-associated surface protein (MASP), putative | -1,737 | 1,703E-07 |
| TcCLB.507237.359 | surface protease GP63 (pseudogene), putative | -1,739 | 0,001387 |
| TcCLB.511185.100 | mucin-associated surface protein (MASP), putative | -1,74 | 0,0001721 |
| TcCLB.506479.120 | cytidine triphosphate synthase, putative | -1,741 | 7,539E-36 |
| TcCLB.509361.10 | surface protease GP63 (pseudogene), putative | -1,742 | 5,808E-06 |
| TcCLB.509525.50 | hypothetical protein | -1,742 | 0,0001568 |
| TcCLB.508761.280 | mucin TcMUCII, putative | -1,743 | 5,128E-12 |
| TcCLB.503609.100 | mucin TcMUCII, putative | -1,743 | 0,0001172 |
| TcCLB.511705.10 | receptor-type adenylate cyclase, putative | -1,744 | 1,343E-44 |
| TcCLB.511259.189 | mucin TcMUCII, putative | -1,745 | 4,164E-08 |
| TcCLB.506855.190 | hypothetical protein, conserved | -1,746 | 4,323E-76 |
| TcCLB.408547.10 | mucin-associated surface protein (MASP), putative | -1,747 | 2,543E-16 |
| TcCLB.507947.10 | conserved protein, unknown function | -1,748 | 2,181E-19 |
| TcCLB.511267.40 | hypothetical protein, conserved | -1,749 | 3,547E-18 |
| TcCLB.506067.230 | trans-sialidase (pseudogene), putative | -1,75 | 3,737E-17 |
| TcCLB.508165.100 | mucin TcMUCII, putative | -1,75 | 7,484E-11 |
| TcCLB.508011.30 | mucin TcMUCII, putative | -1,75 | 6,523E-08 |
| TcCLB.504239.10 | hypothetical protein, conserved (pseudogene) | -1,751 | 9,071E-08 |
| TcCLB.509899.40 | trans-sialidase (pseudogene), putative | -1,752 | 1,359E-26 |
| TcCLB.510475.60 | hypothetical protein | -1,754 | 0,002426 |
| TcCLB.503417.30 | mucin TcMUCII, putative | -1,755 | 1,335E-08 |
| TcCLB.507637.150 | mucin-associated surface protein (MASP, pseudogene), pu | -1,759 | 2,348E-29 |
| TcCLB.506681.40 | hypothetical protein, conserved | -1,759 | 8,883E-26 |
| TcCLB.508247.130 | mucin-associated surface protein (MASP), putative | -1,759 | 6,263E-08 |
| TcCLB.504007.20 | trans-sialidase, Group V, putative | -1,761 | 2,959E-12 |
| TcCLB.508013.50 | mucin TcMUCII, putative | -1,761 | 1,052E-08 |
| TcCLB.507959.90 | mucin TcMUCII, putative | -1,761 | 0,0000762 |
| TcCLB.508047.70 | mucin-associated surface protein (MASP), putative | -1,762 | 3,8E-22 |
| TcCLB.508295.70 | hypothetical protein | -1,762 | 1,9E-09 |
| TcCLB.506617.20 | surface protein TolT (TolT) | -1,763 | 2,172E-37 |
| TcCLB.503783.34 | mucin-associated surface protein (MASP, pseudogene), pu | -1,763 | 2,042E-16 |
| TcCLB.507083.20 | dispersed gene family protein 1 (DGF-1), putative | -1,765 | 2,617E-52 |
| TcCLB.509287.130 | surface protease GP63 (pseudogene), putative | -1,768 | 6,13E-37 |
| TcCLB.509753.100 | mucin-associated surface protein (MASP), putative | -1,768 | 1,369E-10 |

| | | | |
|---|---|---|---|
| TcCLB.507699.64 | mucin TcMUCII, putative | -1,769 | 5,419E-10 |
| TcCLB.510013.200 | 90 kDa surface protein, putative | -1,77 | 1,246E-11 |
| TcCLB.509545.70 | mucin TcMUCII, putative | -1,77 | 8,46E-07 |
| TcCLB.507937.50 | mucin-associated surface protein (MASP), putative | -1,771 | 5,899E-13 |
| TcCLB.511597.50 | trans-sialidase, putative (fragment) | -1,772 | 0,00004983 |
| TcCLB.511141.10 | hypothetical protein, conserved | -1,773 | 3,464E-35 |
| TcCLB.503947.20 | mucin-associated surface protein (MASP), putative | -1,773 | 2,58E-28 |
| TcCLB.507041.120 | variant-surface-glycoprotein phospholipase C, putative | -1,774 | 1,56E-55 |
| TcCLB.509699.60 | mucin-associated surface protein (MASP, pseudogene), pu | -1,774 | 1,242E-36 |
| TcCLB.508097.60 | mucin-associated surface protein (MASP), putative | -1,774 | 1,297E-08 |
| TcCLB.509287.110 | mucin-associated surface protein (MASP), putative | -1,775 | 3,095E-25 |
| TcCLB.511701.19 | surface protease GP63, putative (fragment) | -1,775 | 0,0004395 |
| TcCLB.506789.334 | hypothetical protein, conserved | -1,777 | 3,422E-34 |
| TcCLB.452475.10 | trans-sialidase, Group V, putative | -1,778 | 1,086E-15 |
| TcCLB.508759.20 | trans-sialidase, Group V, putative | -1,779 | 1,169E-13 |
| TcCLB.506231.10 | trans-sialidase, putative | -1,782 | 1,115E-25 |
| TcCLB.511401.130 | mucin-associated surface protein (MASP, pseudogene), pu | -1,782 | 2,632E-17 |
| TcCLB.511843.70 | trans-sialidase, Group V, putative | -1,782 | 5,102E-14 |
| TcCLB.506909.20 | mucin TcMUCI, putative | -1,782 | 1,658E-13 |
| TcCLB.509265.110 | trans-sialidase, Group IV, putative | -1,783 | 3,899E-29 |
| TcCLB.506613.100 | mucin TcMUCII (pseudogene), putative | -1,784 | 3,245E-12 |
| TcCLB.506529.600 | metacyclin II, putative | -1,784 | 1,675E-11 |
| TcCLB.506763.330 | mucin-associated surface protein (MASP), putative | -1,785 | 9,969E-12 |
| TcCLB.506813.200 | mucin-like glycoprotein (pseudogene), putative | -1,786 | 7,742E-07 |
| TcCLB.509739.10 | trans-sialidase, Group II, putative | -1,787 | 1,552E-26 |
| TcCLB.506321.180 | hypothetical protein | -1,787 | 0,00001458 |
| TcCLB.504261.20 | trans-sialidase, Group VI, putative | -1,788 | 4,084E-13 |
| TcCLB.508221.680 | serine-alanine-and proline-rich protein (pseudogene), puta | -1,788 | 4,021E-07 |
| TcCLB.509153.80 | Present in the outer mitochondrial membrane proteome 3 | -1,789 | 5,338E-31 |
| TcCLB.507091.120 | mucin-associated surface protein (MASP), putative | -1,789 | 1,266E-07 |
| TcCLB.509897.150 | mucin-associated surface protein (MASP), putative | -1,791 | 4,009E-59 |
| TcCLB.511565.30 | mucin TcMUCII, putative | -1,791 | 1,146E-12 |
| TcCLB.509079.10 | surface protease GP63 (pseudogene), putative | -1,791 | 9,386E-09 |
| TcCLB.510275.110 | trans-sialidase (pseudogene), putative | -1,792 | 4,314E-24 |
| TcCLB.509805.10 | protein kinase A catalytic subunit, putative | -1,792 | 1,207E-18 |
| TcCLB.506177.40 | palmitoyl acyltransferase 6, putative | -1,794 | 1,739E-28 |
| TcCLB.511269.50 | protein kinase A catalytic subunit, putative (PKAC3) | -1,795 | 4,334E-24 |
| TcCLB.511643.73 | mucin-associated surface protein (MASP, pseudogene), pu | -1,796 | 4,637E-12 |
| TcCLB.510807.50 | mucin-associated surface protein (MASP), putative | -1,796 | 2,087E-11 |
| TcCLB.506245.250 | trans-sialidase, Group V, putative | -1,797 | 2,204E-17 |
| TcCLB.503761.30 | mucin TcMUCII, putative | -1,802 | 1,587E-08 |
| TcCLB.506289.180 | mucin-associated surface protein (MASP, pseudogene), pu | -1,803 | 1,115E-15 |
| TcCLB.509753.60 | mucin-associated surface protein (MASP), putative | -1,803 | 1,085E-09 |
| TcCLB.508761.90 | mucin-associated surface protein (MASP), putative | -1,803 | 0,00004786 |
| TcCLB.510377.250 | mucin TcMUCII (pseudogene), putative | -1,804 | 9,901E-08 |

| | | | |
|---|---|---|---|
| TcCLB.507895.60 | UDP-Gal or UDP-GlcNAc-dependent glycosyltransferase, p | -1,806 | 7,406E-16 |
| TcCLB.507313.20 | trans-sialidase, Group VIII, putative | -1,808 | 3,317E-17 |
| TcCLB.508759.40 | mucin-associated surface protein (MASP), putative | -1,809 | 5,303E-25 |
| TcCLB.511255.360 | mucin-associated surface protein (MASP), putative | -1,809 | 1,484E-12 |
| TcCLB.507613.10 | dispersed gene family protein 1 (DGF-1), putative | -1,809 | 6,309E-06 |
| TcCLB.508229.40 | trans-sialidase, Group V, putative | -1,81 | 4,803E-52 |
| TcCLB.468217.30 | trans-sialidase, putative | -1,813 | 1,786E-23 |
| TcCLB.508633.130 | 90 kDa surface protein, putative | -1,813 | 1,556E-19 |
| TcCLB.506131.84 | mucin-associated surface protein (MASP), putative | -1,813 | 3,63E-12 |
| TcCLB.507747.114 | mucin-associated surface protein (MASP, pseudogene), pu | -1,813 | 3,453E-11 |
| TcCLB.506589.140 | mucin-associated surface protein (MASP), putative | -1,814 | 3,893E-21 |
| TcCLB.511681.20 | histone H4, putative | -1,815 | 6,218E-43 |
| TcCLB.506911.10 | mucin TcMUCII, putative | -1,815 | 1,409E-07 |
| TcCLB.510361.200 | mucin TcMUCII, putative | -1,815 | 5,847E-07 |
| TcCLB.508125.14 | mucin-associated surface protein (MASP), putative (fragm | -1,816 | 4,167E-11 |
| TcCLB.511797.120 | mucin-associated surface protein (MASP), putative | -1,817 | 4,451E-20 |
| TcCLB.509545.40 | trans-sialidase (pseudogene), putative | -1,819 | 6,941E-22 |
| TcCLB.508013.110 | mucin-associated surface protein (MASP), putative | -1,822 | 1,08E-12 |
| TcCLB.510085.4 | mucin TcMUCII (pseudogene), putative | -1,822 | 5,793E-08 |
| TcCLB.507523.80 | mucin-associated surface protein (MASP), putative | -1,824 | 3,106E-16 |
| TcCLB.507357.80 | mucin TcMUCII, putative | -1,825 | 3,033E-10 |
| TcCLB.504039.70 | hypothetical protein | -1,825 | 1,123E-06 |
| TcCLB.510715.90 | trans-sialidase, Group V, putative | -1,827 | 7,3E-19 |
| TcCLB.510391.40 | mucin TcMUCII, putative | -1,828 | 6,03E-11 |
| TcCLB.504039.200 | mucin-associated surface protein (MASP), putative | -1,828 | 0,001654 |
| TcCLB.508325.230 | trans-sialidase, Group IV, putative | -1,83 | 1,943E-135 |
| TcCLB.506309.140 | mucin-associated surface protein (MASP), putative | -1,83 | 0,0002742 |
| TcCLB.508483.13 | trans-sialidase (pseudogene), putative | -1,832 | 1,163E-13 |
| TcCLB.511603.20 | mucin TcMUCII, putative | -1,832 | 6,166E-11 |
| TcCLB.504081.400 | mucin-associated surface protein (MASP), putative | -1,833 | 4,495E-16 |
| TcCLB.507611.260 | trans-sialidase (pseudogene), putative | -1,833 | 6,298E-10 |
| TcCLB.508495.80 | mucin-associated surface protein (MASP, pseudogene), pu | -1,833 | 1,653E-06 |
| TcCLB.506599.120 | surface protease GP63 (pseudogene), putative | -1,835 | 6,994E-06 |
| TcCLB.510279.330 | mucin-associated surface protein (MASP), putative | -1,836 | 2,433E-17 |
| TcCLB.503849.51 | mucin TcMUCII (pseudogene), putative | -1,836 | 2,108E-12 |
| TcCLB.511245.120 | hypothetical protein, conserved | -1,837 | 1,408E-44 |
| TcCLB.504155.160 | mucin-associated surface protein (MASP), putative | -1,837 | 7,334E-20 |
| TcCLB.507295.50 | mucin-associated surface protein (MASP), putative | -1,84 | 1,627E-35 |
| TcCLB.511565.10 | mucin TcMUCII, putative (fragment) | -1,84 | 3,099E-06 |
| TcCLB.506329.20 | mucin TcMUCII, putative | -1,841 | 3,37E-09 |
| TcCLB.508677.40 | trans-sialidase, Group VIII, putative | -1,843 | 1,083E-71 |
| TcCLB.509867.60 | mucin-associated surface protein (MASP), putative | -1,845 | 3,921E-23 |
| TcCLB.509631.90 | mucin TcMUCII, putative | -1,847 | 4,686E-15 |
| TcCLB.403875.10 | cysteine peptidase, Clan CA, family C51, putative | -1,848 | 2,067E-84 |
| TcCLB.511603.190 | mucin-associated surface protein (MASP), putative | -1,848 | 7,408E-12 |

| | | | |
|---|---|---|---|
| TcCLB.506755.60 | hypothetical protein, conserved | -1,85 | 7,672E-49 |
| TcCLB.508977.20 | mucin-associated surface protein (MASP), putative | -1,85 | 9,829E-13 |
| TcCLB.503741.20 | mucin-associated surface protein (MASP, pseudogene), pu | -1,852 | 1,364E-31 |
| TcCLB.506763.340 | mucin TcMUCII, putative | -1,852 | 3,852E-10 |
| TcCLB.506925.250 | hypothetical protein, conserved (pseudogene) | -1,853 | 1,592E-58 |
| TcCLB.510979.50 | mucin-associated surface protein (MASP), putative | -1,853 | 6,551E-33 |
| TcCLB.508481.30 | trans-sialidase (pseudogene), putative | -1,853 | 1,463E-09 |
| TcCLB.508147.79 | mucin TcMUCII, putative | -1,853 | 1,415E-07 |
| TcCLB.511057.60 | mucin-associated surface protein (MASP), putative | -1,854 | 2,177E-18 |
| TcCLB.509545.50 | mucin-associated surface protein (MASP), putative | -1,854 | 7,512E-18 |
| TcCLB.507633.80 | trans-sialidase, Group V, putative | -1,855 | 2,983E-14 |
| TcCLB.508313.130 | mucin TcMUCII, putative | -1,855 | 8,319E-10 |
| TcCLB.511553.45 | surface protease GP63 (pseudogene), putative | -1,855 | 3,92E-08 |
| TcCLB.508165.30 | mucin-associated surface protein (MASP), putative | -1,856 | 1,166E-13 |
| TcCLB.506281.90 | mucin TcMUCII, putative | -1,858 | 1,733E-07 |
| TcCLB.508365.180 | hypothetical protein | -1,858 | 7,251E-06 |
| TcCLB.506759.230 | hypothetical protein | -1,859 | 0,0003192 |
| TcCLB.509663.47 | trans-sialidase (pseudogene), putative | -1,86 | 3,008E-17 |
| TcCLB.510191.10 | mucin-associated surface protein (MASP), putative | -1,86 | 8,173E-15 |
| TcCLB.510275.250 | mucin-associated surface protein (MASP), putative | -1,861 | 1,952E-38 |
| TcCLB.508433.200 | trans-sialidase (pseudogene), putative | -1,861 | 5,127E-22 |
| TcCLB.510157.20 | hypothetical protein, conserved | -1,861 | 2,961E-11 |
| TcCLB.506671.10 | mucin-associated surface protein (MASP), putative | -1,862 | 5,128E-24 |
| TcCLB.503973.280 | mucin-associated surface protein (MASP), putative | -1,863 | 3,264E-15 |
| TcCLB.507959.350 | mucin-associated surface protein (MASP), putative | -1,865 | 5,499E-44 |
| TcCLB.504039.170 | mucin-associated surface protein (MASP, pseudogene), pu | -1,866 | 1,719E-09 |
| TcCLB.506309.100 | mucin-associated surface protein (MASP, pseudogene), pu | -1,867 | 3,406E-06 |
| TcCLB.508745.60 | mucin TcMUCII, putative | -1,868 | 2,793E-17 |
| TcCLB.506769.4 | mucin-associated surface protein (MASP, pseudogene), pu | -1,868 | 2,854E-12 |
| TcCLB.506289.230 | mucin-associated surface protein (MASP, pseudogene), pu | -1,869 | 1,784E-24 |
| TcCLB.508703.20 | trans-sialidase (pseudogene), putative | -1,869 | 1,984E-14 |
| TcCLB.511439.20 | lipase, putative | -1,87 | 4,064E-28 |
| TcCLB.510025.60 | hypothetical protein | -1,87 | 0,0001641 |
| TcCLB.506615.29 | mucin-associated surface protein (MASP, pseudogene), pu | -1,871 | 1,243E-72 |
| TcCLB.511089.15 | hypothetical protein | -1,871 | 2,518E-10 |
| TcCLB.508979.60 | mucin-associated surface protein (MASP), putative | -1,872 | 1,096E-34 |
| TcCLB.510355.41 | mucin-associated surface protein (MASP, pseudogene), pu | -1,873 | 1,292E-14 |
| TcCLB.509195.61 | mucin TcMUCII, putative | -1,876 | 2,044E-12 |
| TcCLB.511907.160 | mucin-associated surface protein (MASP, pseudogene), pu | -1,877 | 1,532E-43 |
| TcCLB.510805.60 | surface protease GP63 (pseudogene), putative | -1,877 | 8,525E-08 |
| TcCLB.446785.10 | trans-sialidase, putative (fragment) | -1,878 | 3,274E-42 |
| TcCLB.507075.30 | trans-sialidase (pseudogene), putative | -1,878 | 7,63E-35 |
| TcCLB.506459.219 | mucin TcMUCII, putative | -1,878 | 1,028E-21 |
| TcCLB.507069.220 | mucin-associated surface protein (MASP), putative | -1,879 | 1,381E-15 |
| TcCLB.510191.40 | trans-sialidase (pseudogene), putative | -1,88 | 1,071E-08 |

| | | | |
|---|---|---|---|
| TcCLB.506067.300 | trans-sialidase, Group V, putative | -1,881 | 5,864E-27 |
| TcCLB.503609.60 | mucin-associated surface protein (MASP), putative | -1,882 | 3,425E-14 |
| TcCLB.504787.99 | mucin TcMUCII, putative | -1,882 | 3,057E-12 |
| TcCLB.508489.10 | hypothetical protein, conserved | -1,883 | 1,473E-55 |
| TcCLB.510297.40 | hypothetical protein | -1,884 | 2,645E-42 |
| TcCLB.508175.60 | hypothetical protein, conserved | -1,885 | 1,579E-46 |
| TcCLB.506597.20 | mucin-associated surface protein (MASP, pseudogene), pu | -1,886 | 5,046E-11 |
| TcCLB.510025.165 | hypothetical protein, conserved (pseudogene) | -1,886 | 9,333E-10 |
| TcCLB.503717.71 | trans-sialidase (pseudogene), putative | -1,886 | 2,347E-06 |
| TcCLB.506971.10 | mucin-associated surface protein (MASP), putative | -1,887 | 1,253E-15 |
| TcCLB.508789.220 | mucin TcMUCII, putative | -1,888 | 1,27E-10 |
| TcCLB.507107.30 | UDP-Gal or UDP-GlcNAc-dependent glycosyltransferase, p | -1,889 | 6,357E-14 |
| TcCLB.506279.200 | mucin TcMUCII, putative | -1,889 | 9,17E-10 |
| TcCLB.506951.60 | mucin-associated surface protein (MASP), putative | -1,89 | 9,222E-51 |
| TcCLB.511173.50 | mucin-associated surface protein (MASP), putative | -1,89 | 6,102E-36 |
| TcCLB.510025.50 | trans-sialidase, Group V, putative | -1,892 | 1,449E-46 |
| TcCLB.511117.50 | surface protease GP63, putative | -1,892 | 3,29E-16 |
| TcCLB.508163.10 | mucin-associated surface protein (MASP, pseudogene), pu | -1,893 | 0,00001617 |
| TcCLB.511907.130 | mucin-like glycoprotein, putative | -1,894 | 1,735E-34 |
| TcCLB.509699.200 | mucin TcMUCII, putative | -1,894 | 1,774E-06 |
| TcCLB.506767.110 | mucin-associated surface protein (MASP), putative | -1,895 | 8,892E-23 |
| TcCLB.504975.10 | mucin-associated surface protein (MASP, pseudogene), pu | -1,895 | 9,477E-17 |
| TcCLB.506813.80 | trans-sialidase, Group V, putative | -1,895 | 1,238E-12 |
| TcCLB.504039.120 | mucin TcMUCII, putative | -1,896 | 4,522E-10 |
| TcCLB.507237.260 | mucin TcMUCII, putative | -1,896 | 9,577E-08 |
| TcCLB.511553.10 | mucin-associated surface protein (MASP), putative | -1,897 | 1,39E-24 |
| TcCLB.506067.198 | mucin-associated surface protein (MASP, pseudogene), pu | -1,897 | 7,034E-09 |
| TcCLB.506563.70 | calpain-like cysteine peptidase (pseudogene), putative | -1,898 | 8,221E-41 |
| TcCLB.510377.420 | surface protease GP63, putative | -1,898 | 0,00006717 |
| TcCLB.510371.50 | mucin-associated surface protein (MASP), putative | -1,899 | 2,352E-25 |
| TcCLB.507437.10 | mucin-associated surface protein (MASP), putative | -1,899 | 3,131E-20 |
| TcCLB.510363.320 | mucin-associated surface protein (MASP), putative | -1,9 | 1,018E-38 |
| TcCLB.511599.120 | mucin-associated surface protein (MASP), putative | -1,9 | 3,798E-15 |
| TcCLB.506139.1 | mucin-associated surface protein (MASP), putative (fragm | -1,9 | 0,00001347 |
| TcCLB.507959.300 | mucin-associated surface protein (MASP), putative | -1,901 | 9,669E-27 |
| TcCLB.509915.50 | trans-sialidase (pseudogene), putative | -1,902 | 2,115E-10 |
| TcCLB.508161.10 | mucin-associated surface protein (MASP), putative | -1,902 | 2,77E-09 |
| TcCLB.506609.80 | mucin TcMUCII, putative | -1,905 | 2,233E-08 |
| TcCLB.511599.30 | 90 kDa surface protein, putative | -1,905 | 0,00009808 |
| TcCLB.510017.50 | surface protease GP63 (pseudogene), putative | -1,907 | 2,583E-10 |
| TcCLB.506245.190 | trans-sialidase, Group V, putative | -1,908 | 2,466E-25 |
| TcCLB.504239.150 | hypothetical protein | -1,909 | 0,0009574 |
| TcCLB.507295.30 | mucin TcMUCII, putative | -1,91 | 4,818E-27 |
| TcCLB.510551.20 | trans-sialidase (pseudogene), putative | -1,91 | 4,073E-26 |
| TcCLB.507185.40 | trans-sialidase (pseudogene), putative | -1,91 | 1,101E-22 |

| | | | |
|---|---|---|---|
| TcCLB.503403.20 | trans-sialidase, Group II, putative | -1,911 | 3,676E-24 |
| TcCLB.506245.60 | trans-sialidase, Group V, putative | -1,911 | 5,079E-12 |
| TcCLB.504689.20 | mucin-associated surface protein (MASP, pseudogene), pu | -1,912 | 6,501E-36 |
| TcCLB.507023.100 | hypothetical protein | -1,914 | 9,901E-50 |
| TcCLB.509843.20 | trans-sialidase, Group VII, putative | -1,915 | 1,294E-92 |
| TcCLB.511599.80 | trans-sialidase, Group V, putative | -1,915 | 4,561E-33 |
| TcCLB.503875.30 | mucin-associated surface protein (MASP), putative | -1,916 | 3,722E-35 |
| TcCLB.510921.20 | mucin TcMUCII, putative | -1,916 | 1,032E-23 |
| TcCLB.404711.10 | trans-sialidase (pseudogene), putative | -1,917 | 2,673E-70 |
| TcCLB.508341.10 | mucin-associated surface protein (MASP), putative | -1,917 | 1,725E-07 |
| TcCLB.509575.4 | trans-sialidase, putative (fragment) | -1,918 | 1,343E-17 |
| TcCLB.508313.14 | trans-sialidase, putative | -1,919 | 6,663E-14 |
| TcCLB.506801.40 | mucin-associated surface protein (MASP), putative | -1,92 | 5,741E-44 |
| TcCLB.510329.290 | diacylglycerol kinase-like protein, putative | -1,92 | 1,654E-42 |
| TcCLB.510553.30 | mucin-associated surface protein (MASP), putative | -1,92 | 4,391E-23 |
| TcCLB.504249.111 | mucin-associated surface protein (MASP), putative | -1,92 | 2,597E-19 |
| TcCLB.445783.10 | mucin-associated surface protein (MASP, pseudogene), pu | -1,92 | 4,313E-15 |
| TcCLB.510279.310 | mucin-associated surface protein (MASP), putative | -1,921 | 1,277E-20 |
| TcCLB.508109.10 | mucin-associated surface protein (MASP), putative | -1,921 | 7,361E-10 |
| TcCLB.506035.4 | trans-sialidase, putative (fragment) | -1,923 | 3,082E-36 |
| TcCLB.508221.894 | mucin-associated surface protein (MASP), putative | -1,923 | 1,303E-13 |
| TcCLB.509663.69 | trans-sialidase, putative (fragment) | -1,923 | 0,01538 |
| TcCLB.410269.10 | trans-sialidase, putative (fragment) | -1,924 | 3,372E-17 |
| TcCLB.507297.30 | metacaspase, putative | -1,925 | 1,837E-68 |
| TcCLB.508221.260 | 90 kDa surface protein, putative | -1,925 | 3,322E-12 |
| TcCLB.506909.150 | mucin TcMUCII (pseudogene), putative | -1,925 | 4,676E-06 |
| TcCLB.508151.5 | mucin-associated surface protein (MASP, pseudogene), pu | -1,925 | 0,00001712 |
| TcCLB.508165.150 | mucin-associated surface protein (MASP), putative | -1,926 | 1,028E-23 |
| TcCLB.504787.80 | mucin TcMUCII, putative | -1,927 | 1,511E-22 |
| TcCLB.509569.30 | hypothetical protein, conserved | -1,928 | 9,478E-83 |
| TcCLB.511173.100 | mucin-associated surface protein (MASP), putative | -1,928 | 6,777E-32 |
| TcCLB.508011.120 | trans-sialidase (pseudogene), putative | -1,929 | 2,642E-20 |
| TcCLB.507071.220 | mucin-associated surface protein (MASP, pseudogene), pu | -1,933 | 2,711E-13 |
| TcCLB.503899.90 | asparagine synthetase a, putative | -1,934 | 1,302E-37 |
| TcCLB.506331.120 | trans-sialidase (pseudogene), putative | -1,937 | 7,631E-37 |
| TcCLB.508219.110 | mucin-associated surface protein (MASP), putative | -1,937 | 2,756E-17 |
| TcCLB.503773.24 | hypothetical protein, conserved | -1,938 | 7,218E-25 |
| TcCLB.508753.40 | mucin-associated surface protein (MASP, pseudogene), pu | -1,94 | 1,224E-37 |
| TcCLB.510417.20 | mucin-associated surface protein (MASP, pseudogene), pu | -1,94 | 1,997E-24 |
| TcCLB.509765.129 | trans-sialidase, putative (fragment) | -1,941 | 6,139E-62 |
| TcCLB.430895.10 | hypothetical protein, conserved | -1,945 | 2,411E-32 |
| TcCLB.510205.80 | mucin-associated surface protein (MASP), putative | -1,946 | 3,348E-19 |
| TcCLB.507163.75 | hypothetical protein | -1,946 | 1,077E-08 |
| TcCLB.506799.50 | mucin-associated surface protein (MASP), putative | -1,948 | 1,257E-62 |
| TcCLB.508761.70 | mucin-associated surface protein (MASP), putative | -1,948 | 1,239E-40 |

| | | | |
|---|---|---|---|
| TcCLB.508121.130 | mucin-associated surface protein (MASP, pseudogene), pu | -1,948 | 1,547E-24 |
| TcCLB.508045.120 | trans-sialidase, Group III, putative | -1,949 | 8,622E-96 |
| TcCLB.510361.80 | trans-sialidase, Group V, putative | -1,949 | 3,981E-21 |
| TcCLB.510359.600 | mucin-associated surface protein (MASP), putative | -1,95 | 7,127E-22 |
| TcCLB.510583.10 | mucin TcMUCII, putative | -1,951 | 8,437E-13 |
| TcCLB.506939.200 | trans-sialidase (pseudogene), putative | -1,952 | 1,849E-19 |
| TcCLB.510351.120 | Nudix hydrolase 2, putative (Nudix2) | -1,954 | 4,428E-63 |
| TcCLB.506965.10 | mucin TcMUCII, putative | -1,954 | 3,696E-13 |
| TcCLB.508221.490 | mucin-associated surface protein (MASP), putative | -1,955 | 1,689E-06 |
| TcCLB.508745.70 | mucin-associated surface protein (MASP), putative (fragm | -1,956 | 0,0001276 |
| TcCLB.506995.70 | mucin-associated surface protein (MASP), putative | -1,957 | 3,66E-28 |
| TcCLB.510071.30 | UDP-Gal or UDP-GlcNAc-dependent glycosyltransferase, p | -1,958 | 7,818E-30 |
| TcCLB.506597.51 | hypothetical protein | -1,959 | 1,345E-06 |
| TcCLB.506613.60 | mucin TcMUCII (pseudogene), putative | -1,96 | 9,925E-13 |
| TcCLB.504201.30 | trans-sialidase, Group V, putative | -1,962 | 1,243E-19 |
| TcCLB.508515.150 | trans-sialidase, Group VIII, putative | -1,963 | 9,364E-32 |
| TcCLB.510465.125 | trans-sialidase (pseudogene), putative | -1,963 | 0,00009763 |
| TcCLB.510359.380 | mucin-associated surface protein (MASP), putative | -1,965 | 6,294E-15 |
| TcCLB.504491.20 | trans-sialidase, putative | -1,966 | 9,798E-16 |
| TcCLB.506501.329 | mucin-associated surface protein (MASP, pseudogene), pu | -1,966 | 3,936E-10 |
| TcCLB.506773.10 | hypothetical protein, conserved | -1,966 | 0,00000845 |
| TcCLB.509663.37 | hypothetical protein, conserved | -1,972 | 2,8E-56 |
| TcCLB.508113.20 | trans-sialidase (pseudogene), putative | -1,972 | 2,181E-47 |
| TcCLB.506967.40 | mucin TcMUCII, putative | -1,973 | 1,886E-12 |
| TcCLB.503625.10 | asparagine synthetase a, putative | -1,977 | 7,898E-67 |
| TcCLB.418613.20 | mucin-associated surface protein (MASP), putative (fragm | -1,977 | 1,731E-08 |
| TcCLB.509479.10 | mucin TcMUCII, putative | -1,978 | 1,323E-23 |
| TcCLB.506741.140 | mucin-associated surface protein (MASP), putative | -1,978 | 1,335E-18 |
| TcCLB.506661.40 | hypothetical protein, conserved | -1,979 | 2,269E-47 |
| TcCLB.511255.490 | trans-sialidase (pseudogene), putative | -1,979 | 5,088E-11 |
| TcCLB.506971.70 | mucin TcMUCII, putative | -1,979 | 2,48E-07 |
| TcCLB.510061.20 | hypothetical protein | -1,979 | 0,00004909 |
| TcCLB.504277.11 | surface protein TolT, putative | -1,98 | 3,866E-51 |
| TcCLB.507497.11 | trans-sialidase, putative (fragment) | -1,98 | 1,124E-26 |
| TcCLB.507867.10 | mucin-associated surface protein (MASP), putative | -1,982 | 0,00001951 |
| TcCLB.511149.70 | mucin TcMUCII, putative | -1,984 | 3,727E-19 |
| TcCLB.422207.10 | UDP-Gal or UDP-GlcNAc-dependent glycosyltransferase, p | -1,985 | 0,0002842 |
| TcCLB.508165.420 | mucin-associated surface protein (MASP), putative | -1,986 | 6,077E-40 |
| TcCLB.507065.127 | mucin-associated surface protein (MASP, pseudogene), pu | -1,987 | 4,624E-27 |
| TcCLB.503827.5 | mucin-associated surface protein (MASP), putative | -1,987 | 4,275E-06 |
| TcCLB.511725.40 | hypothetical protein, conserved | -1,988 | 6,901E-52 |
| TcCLB.508153.1130 | hypothetical protein | -1,988 | 1,057E-20 |
| TcCLB.507067.50 | mucin TcMUCII (pseudogene), putative | -1,989 | 3,631E-11 |
| TcCLB.508221.820 | mucin-associated surface protein (MASP), putative | -1,991 | 2,259E-21 |
| TcCLB.506599.189 | mucin TcMUCII, putative | -1,992 | 1,311E-17 |

| | | | |
|---|---|---|---|
| TcCLB.508495.170 | mucin TcMUCII, putative | -1,993 | 8,085E-29 |
| TcCLB.511603.80 | mucin-associated surface protein (MASP), putative | -1,996 | 1,415E-21 |
| TcCLB.507069.40 | trans-sialidase, Group VI, putative | -1,997 | 2,874E-21 |
| TcCLB.509755.70 | mucin TcMUCII, putative | -1,998 | 6,469E-07 |
| TcCLB.510377.340 | hypothetical protein | -1,998 | 0,00002125 |
| TcCLB.508305.40 | trans-sialidase, Group V, putative | -1,999 | 2,61E-24 |
| TcCLB.510959.30 | mucin-associated surface protein (MASP), putative | -2 | 2,658E-25 |
| TcCLB.510307.284 | trans-sialidase, Group IV, putative | -2,001 | 1,384E-89 |
| TcCLB.511431.10 | protein kinase, putative | -2,003 | 3,794E-36 |
| TcCLB.510013.120 | surface protease GP63, putative | -2,004 | 0,00001569 |
| TcCLB.511645.10 | mucin-associated surface protein (MASP), putative | -2,005 | 1,063E-35 |
| TcCLB.408547.5 | mucin TcMUCII, putative | -2,006 | 2,526E-11 |
| TcCLB.506281.20 | mucin TcMUCI, putative | -2,006 | 6,116E-10 |
| TcCLB.510207.30 | trans-sialidase, Group V, putative | -2,009 | 6,781E-21 |
| TcCLB.504779.10 | UDP-Gal or UDP-GlcNAc-dependent glycosyltransferase, p | -2,009 | 1,223E-18 |
| TcCLB.508669.21 | hypothetical protein | -2,01 | 1,658E-13 |
| TcCLB.504201.40 | mucin-associated surface protein (MASP), putative | -2,012 | 2,019E-24 |
| TcCLB.511603.470 | 90 kDa surface protein, putative | -2,013 | 3,978E-34 |
| TcCLB.507801.240 | mucin-associated surface protein (MASP, pseudogene), pu | -2,013 | 2,163E-27 |
| TcCLB.508873.60 | trans-sialidase, Group V, putative | -2,013 | 1,34E-12 |
| TcCLB.508109.30 | mucin-associated surface protein (MASP), putative | -2,013 | 2,367E-09 |
| TcCLB.508667.40 | hypothetical protein, conserved | -2,016 | 1,017E-114 |
| TcCLB.506703.50 | mucin-associated surface protein (MASP), putative | -2,016 | 1,378E-29 |
| TcCLB.503717.80 | mucin-associated surface protein (MASP, pseudogene), pu | -2,017 | 3,101E-16 |
| TcCLB.511081.41 | mucin-associated surface protein (MASP, pseudogene), pu | -2,017 | 1,108E-14 |
| TcCLB.508253.20 | trans-sialidase (pseudogene), putative | -2,018 | 1,347E-12 |
| TcCLB.503717.20 | mucin-associated surface protein (MASP), putative | -2,019 | 1,12E-38 |
| TcCLB.511173.360 | mucin-associated surface protein (MASP), putative | -2,019 | 4,604E-24 |
| TcCLB.504039.250 | surface protease GP63 (pseudogene), putative | -2,02 | 4,355E-21 |
| TcCLB.509197.30 | cation transporter, putative | -2,021 | 2,53E-49 |
| TcCLB.509137.20 | hypothetical protein | -2,024 | 5,148E-72 |
| TcCLB.508149.60 | trans-sialidase (pseudogene), putative | -2,024 | 1,083E-15 |
| TcCLB.508125.140 | mucin-associated surface protein (MASP), putative | -2,025 | 2,763E-22 |
| TcCLB.506267.90 | mucin-associated surface protein (MASP), putative | -2,027 | 1,675E-34 |
| TcCLB.508435.40 | mucin-associated surface protein (MASP, pseudogene), pu | -2,027 | 1,108E-17 |
| TcCLB.510013.110 | mucin TcMUCII, putative | -2,027 | 4,284E-06 |
| TcCLB.508153.210 | hypothetical protein, conserved | -2,028 | 2,469E-50 |
| TcCLB.508173.30 | trans-sialidase (pseudogene), putative | -2,029 | 4,545E-47 |
| TcCLB.506821.60 | mucin TcMUCI, putative | -2,029 | 3,226E-36 |
| TcCLB.511437.10 | surface protease GP63, putative | -2,029 | 3,105E-34 |
| TcCLB.508979.30 | mucin TcMUCII, putative | -2,029 | 7,619E-23 |
| TcCLB.507747.110 | mucin TcMUCII, putative | -2,03 | 7,009E-20 |
| TcCLB.508873.289 | 90 kDa surface protein, putative | -2,031 | 6,082E-10 |
| TcCLB.503501.30 | surface protease GP63 (pseudogene), putative | -2,031 | 7,393E-09 |
| TcCLB.509835.30 | hypothetical protein, conserved | -2,034 | 1,778E-75 |

| | | | |
|---|---|---|---|
| TcCLB.510693.200 | mucin TcMUCII, putative | -2,034 | 1,31E-11 |
| TcCLB.506955.150 | mucin-associated surface protein (MASP), putative | -2,036 | 3,95E-27 |
| TcCLB.510085.50 | trans-sialidase (pseudogene), putative | -2,036 | 8,171E-20 |
| TcCLB.510033.14 | mucin-associated surface protein (MASP, pseudogene), pu | -2,037 | 3,988E-85 |
| TcCLB.509525.270 | mucin TcMUCII, putative | -2,037 | 0,00001781 |
| TcCLB.510155.220 | trans-sialidase (pseudogene), putative | -2,038 | 4,766E-30 |
| TcCLB.507067.40 | mucin TcMUCII, putative | -2,038 | 1,459E-18 |
| TcCLB.510483.224 | mucin TcMUCII (pseudogene), putative | -2,039 | 4,19E-14 |
| TcCLB.511863.10 | mucin-associated surface protein (MASP), putative | -2,042 | 3,426E-19 |
| TcCLB.508433.145 | mucin TcMUCII (pseudogene), putative | -2,043 | 2,876E-07 |
| TcCLB.511557.50 | Present in the outer mitochondrial membrane proteome 1 | -2,044 | 7,891E-48 |
| TcCLB.506243.110 | expression site-associated gene (ESAG-like) protein, puta | -2,044 | 2,631E-19 |
| TcCLB.511311.20 | trans-sialidase, Group II, putative | -2,045 | 5,937E-22 |
| TcCLB.506067.100 | mucin TcMUCII, putative | -2,047 | 0,00005156 |
| TcCLB.506335.50 | mucin-associated surface protein (MASP), putative | -2,048 | 7,587E-33 |
| TcCLB.510115.30 | trans-sialidase, Group VI, putative | -2,049 | 7,726E-37 |
| TcCLB.508799.200 | hypothetical protein, conserved | -2,05 | 3,473E-34 |
| TcCLB.503977.70 | mucin-associated surface protein (MASP, pseudogene), pu | -2,051 | 1,387E-23 |
| TcCLB.506967.150 | trans-sialidase (pseudogene), putative | -2,052 | 5,552E-20 |
| TcCLB.508229.60 | trans-sialidase, Group VII, putative | -2,052 | 1,647E-07 |
| TcCLB.506753.249 | hypothetical protein, conserved | -2,055 | 7,581E-10 |
| TcCLB.503741.40 | mucin TcMUCII, putative | -2,056 | 2,512E-12 |
| TcCLB.509387.10 | trans-sialidase, Group VIII, putative | -2,058 | 2,653E-61 |
| TcCLB.509525.40 | hypothetical protein, conserved | -2,058 | 0,00007647 |
| TcCLB.509525.230 | mucin-associated surface protein (MASP), putative | -2,06 | 1,08E-22 |
| TcCLB.503939.4 | hypothetical protein, conserved | -2,061 | 1,137E-11 |
| TcCLB.510375.40 | mucin-associated surface protein (MASP), putative | -2,061 | 3,236E-09 |
| TcCLB.507071.190 | trans-sialidase, Group V, putative | -2,064 | 1,634E-24 |
| TcCLB.506753.15 | dispersed gene family protein 1 (DGF-1, pseudogene), put | -2,064 | 0,00007488 |
| TcCLB.509037.20 | surface protease GP63, putative | -2,065 | 3,019E-33 |
| TcCLB.510279.40 | 90 kDa surface protein, putative | -2,066 | 2,296E-15 |
| TcCLB.508789.80 | mucin-associated surface protein (MASP), putative | -2,066 | 8,75E-13 |
| TcCLB.507601.40 | Nudix hydrolase 2, putative (Nudix2) | -2,067 | 1,935E-70 |
| TcCLB.506279.74 | mucin-associated surface protein (MASP, pseudogene), pu | -2,067 | 1,446E-15 |
| TcCLB.506599.270 | mucin TcMUCII, putative | -2,067 | 2,53E-10 |
| TcCLB.507561.10 | trans-sialidase, putative | -2,068 | 1,705E-10 |
| TcCLB.510583.209 | mucin-associated surface protein (MASP, pseudogene), pu | -2,068 | 0,00002229 |
| TcCLB.481211.30 | mucin TcMUCII, putative | -2,069 | 1,852E-46 |
| TcCLB.509959.60 | trans-sialidase (pseudogene), putative | -2,069 | 1,004E-10 |
| TcCLB.508173.40 | mercaptopyruvate sulfurtransferase, putative (MST) | -2,07 | 1,045E-99 |
| TcCLB.506759.20 | mucin-associated surface protein (MASP), putative | -2,07 | 1,799E-22 |
| TcCLB.509699.110 | mucin-associated surface protein (MASP), putative | -2,072 | 3,824E-27 |
| TcCLB.508809.50 | hypothetical protein | -2,073 | 1,061E-34 |
| TcCLB.511603.30 | mucin-associated surface protein (MASP), putative | -2,075 | 3,119E-16 |
| TcCLB.508261.60 | mucin TcMUCII, putative | -2,075 | 4,616E-13 |

| | | | |
|---|---|---|---|
| TcCLB.506501.300 | hypothetical protein | -2,076 | 2,183E-06 |
| TcCLB.511153.84 | hypothetical protein, conserved | -2,078 | 5,401E-88 |
| TcCLB.507869.20 | trans-sialidase (pseudogene), putative | -2,081 | 8,022E-92 |
| TcCLB.511399.50 | mucin-associated surface protein (MASP), putative | -2,081 | 8,179E-28 |
| TcCLB.507699.180 | mucin-associated surface protein (MASP, pseudogene), pu | -2,081 | 8,948E-23 |
| TcCLB.506245.270 | mucin-associated surface protein (MASP), putative | -2,081 | 5,48E-20 |
| TcCLB.506499.70 | mucin TcMUCII, putative | -2,082 | 5,758E-20 |
| TcCLB.509525.360 | mucin-associated surface protein (MASP, pseudogene), pu | -2,082 | 7,869E-18 |
| TcCLB.507987.30 | mucin-associated surface protein (MASP), putative (fragm | -2,082 | 5,377E-16 |
| TcCLB.507955.20 | mucin-associated surface protein (MASP), putative | -2,082 | 9,191E-11 |
| TcCLB.507699.130 | mucin-associated surface protein (MASP), putative | -2,083 | 1,431E-37 |
| TcCLB.507085.10 | mucin TcMUCII, putative | -2,084 | 4,738E-34 |
| TcCLB.509753.210 | mucin-associated surface protein (MASP), putative | -2,084 | 1,42E-31 |
| TcCLB.506759.190 | surface protease GP63 (pseudogene), putative | -2,084 | 3,492E-09 |
| TcCLB.511213.15 | mucin-associated surface protein (MASP), putative | -2,085 | 5,211E-25 |
| TcCLB.511603.304 | mucin TcMUCII, putative | -2,085 | 6,073E-14 |
| TcCLB.509753.70 | mucin-associated surface protein (MASP), putative | -2,085 | 2,316E-13 |
| TcCLB.506321.110 | mucin TcMUCII, putative | -2,086 | 4,178E-47 |
| TcCLB.509699.150 | mucin TcMUCII, putative | -2,087 | 2,371E-06 |
| TcCLB.506377.10 | trans-sialidase, putative | -2,088 | 3,101E-78 |
| TcCLB.510979.60 | mucin TcMUCII, putative | -2,088 | 1,487E-19 |
| TcCLB.507905.39 | mucin-associated surface protein (MASP), putative | -2,09 | 1,049E-142 |
| TcCLB.504261.40 | mucin-associated surface protein (MASP), putative | -2,09 | 8,495E-17 |
| TcCLB.508873.160 | mucin-associated surface protein (MASP), putative | -2,091 | 6,762E-08 |
| TcCLB.508289.70 | mucin-associated surface protein (MASP, pseudogene), pu | -2,092 | 1,107E-15 |
| TcCLB.511173.370 | trans-sialidase, Group VI, putative | -2,093 | 7,674E-18 |
| TcCLB.511259.50 | trans-sialidase (pseudogene), putative | -2,094 | 4,918E-39 |
| TcCLB.510803.15 | mucin-associated surface protein (MASP, pseudogene), pu | -2,094 | 0,00003461 |
| TcCLB.506611.40 | mucin-associated surface protein (MASP), putative | -2,095 | 2,016E-40 |
| TcCLB.509085.50 | trans-sialidase (pseudogene), putative | -2,095 | 2,738E-34 |
| TcCLB.511077.90 | mucin TcMUCII, putative | -2,095 | 1,164E-18 |
| TcCLB.510329.20 | hypothetical protein, conserved | -2,096 | 3,394E-79 |
| TcCLB.509897.124 | mucin-associated surface protein (MASP, pseudogene), pu | -2,097 | 1,004E-34 |
| TcCLB.506755.270 | hypothetical protein, conserved | -2,097 | 1,894E-11 |
| TcCLB.507071.240 | mucin-associated surface protein (MASP), putative | -2,099 | 8,279E-19 |
| TcCLB.506173.60 | mucin-associated surface protein (MASP), putative | -2,1 | 1,687E-28 |
| TcCLB.506067.70 | mucin-associated surface protein (MASP), putative | -2,102 | 4,852E-11 |
| TcCLB.508221.104 | mucin-associated surface protein (MASP), putative | -2,102 | 0,0001764 |
| TcCLB.503849.40 | mucin-associated surface protein (MASP), putative | -2,106 | 6,179E-35 |
| TcCLB.508165.290 | mucin TcMUCII, putative | -2,107 | 1,756E-08 |
| TcCLB.506965.140 | mucin-associated surface protein (MASP), putative | -2,108 | 1,096E-34 |
| TcCLB.508297.7 | trans-sialidase, putative (fragment) | -2,108 | 5,769E-33 |
| TcCLB.509527.80 | mucin-associated surface protein (MASP), putative | -2,108 | 8,527E-28 |
| TcCLB.509553.10 | hypothetical protein, conserved | -2,109 | 2,932E-36 |
| TcCLB.504155.230 | mucin-associated surface protein (MASP), putative | -2,109 | 2,345E-18 |

| | | | |
|---|---|---|---|
| TcCLB.510981.20 | trans-sialidase, putative | -2,11 | 3,688E-52 |
| TcCLB.510487.70 | trans-sialidase (pseudogene), putative | -2,11 | 4,144E-27 |
| TcCLB.508763.20 | mucin TcMUCII, putative | -2,113 | 1,977E-23 |
| TcCLB.503977.50 | surface protease GP63 (pseudogene), putative | -2,113 | 9,02E-16 |
| TcCLB.508511.30 | mucin TcMUCII, putative | -2,114 | 1,711E-17 |
| TcCLB.507523.10 | trans-sialidase, putative | -2,115 | 2,215E-30 |
| TcCLB.508399.40 | trans-sialidase (pseudogene), putative | -2,116 | 1,676E-78 |
| TcCLB.507071.10 | trans-sialidase, Group V, putative | -2,116 | 6,386E-26 |
| TcCLB.506609.70 | mucin-associated surface protein (MASP), putative | -2,116 | 5,849E-19 |
| TcCLB.508097.40 | trans-sialidase (pseudogene), putative | -2,117 | 1,081E-20 |
| TcCLB.509373.39 | surface protease GP63, putative (fragment) | -2,117 | 0,00000071 |
| TcCLB.509805.210 | hypothetical protein, conserved | -2,118 | 1,97E-57 |
| TcCLB.504219.40 | trans-sialidase (pseudogene), putative | -2,118 | 1,774E-30 |
| TcCLB.506741.90 | mucin-associated surface protein (MASP), putative | -2,118 | 7,697E-28 |
| TcCLB.508789.100 | trans-sialidase (pseudogene), putative | -2,119 | 7,041E-20 |
| TcCLB.503609.24 | mucin-associated surface protein (MASP, pseudogene), pu | -2,119 | 1,849E-19 |
| TcCLB.511859.150 | mucin-like glycoprotein, putative | -2,12 | 5,65E-83 |
| TcCLB.510197.210 | mucin TcMUCII, putative | -2,122 | 5,077E-14 |
| TcCLB.507063.340 | mucin-associated surface protein (MASP), putative | -2,125 | 1,118E-33 |
| TcCLB.509657.30 | mucin-associated surface protein (MASP), putative | -2,127 | 4,011E-29 |
| TcCLB.507429.70 | mucin TcMUCII, putative | -2,128 | 2,242E-63 |
| TcCLB.508435.10 | mucin-associated surface protein (MASP), putative | -2,128 | 5,606E-18 |
| TcCLB.507237.80 | mucin-associated surface protein (MASP), putative | -2,128 | 9,379E-11 |
| TcCLB.506267.10 | mucin-associated surface protein (MASP), putative | -2,132 | 8,798E-09 |
| TcCLB.507295.20 | mucin-associated surface protein (MASP), putative | -2,133 | 1,789E-34 |
| TcCLB.508747.59 | mucin-associated surface protein (MASP), putative | -2,133 | 4,027E-28 |
| TcCLB.506335.130 | trans-sialidase, Group V, putative | -2,135 | 1,033E-18 |
| TcCLB.507747.180 | trans-sialidase, Group VI, putative | -2,137 | 2,827E-73 |
| TcCLB.507071.390 | trans-sialidase, Group V, putative | -2,137 | 1,614E-17 |
| TcCLB.507699.10 | mucin TcMUCII, putative | -2,139 | 4,285E-23 |
| TcCLB.506939.50 | mucin-associated surface protein (MASP), putative | -2,139 | 2,316E-07 |
| TcCLB.510363.170 | mucin TcMUCII, putative | -2,14 | 7,598E-06 |
| TcCLB.508167.29 | mucin-associated surface protein (MASP), putative (fragm | -2,141 | 5,775E-18 |
| TcCLB.510071.20 | hypothetical protein | -2,142 | 2,115E-10 |
| TcCLB.511167.100 | hypothetical protein, conserved | -2,143 | 4,902E-102 |
| TcCLB.511149.100 | mucin TcMUCII, putative | -2,143 | 8,375E-40 |
| TcCLB.511603.70 | mucin TcMUCII, putative | -2,143 | 6,698E-27 |
| TcCLB.504039.130 | mucin-associated surface protein (MASP), putative | -2,144 | 5,618E-23 |
| TcCLB.508539.80 | mucin-associated surface protein (MASP), putative | -2,145 | 2,461E-43 |
| TcCLB.509937.220 | protein kinase, putative | -2,145 | 5,667E-10 |
| TcCLB.509525.94 | mucin-associated surface protein (MASP, pseudogene), pu | -2,146 | 1,594E-32 |
| TcCLB.510983.5 | trans-sialidase (pseudogene), putative | -2,146 | 1,601E-26 |
| TcCLB.488591.10 | trans-sialidase, putative (fragment) | -2,147 | 1,515E-38 |
| TcCLB.510391.30 | mucin TcMUCII, putative | -2,148 | 1,919E-20 |
| TcCLB.510363.310 | trans-sialidase (pseudogene), putative | -2,149 | 4,254E-17 |

| | | | |
|---|---|---|---|
| TcCLB.510329.180 | hypothetical protein, conserved | -2,149 | 1,041E-16 |
| TcCLB.511077.110 | mucin-associated surface protein (MASP), putative | -2,15 | 1,631E-14 |
| TcCLB.510373.30 | 90 kDa surface protein, putative | -2,151 | 1,285E-14 |
| TcCLB.504155.70 | mucin TcMUCII, putative | -2,151 | 5,954E-10 |
| TcCLB.508361.30 | mucin TcMUCII, putative | -2,153 | 2,102E-14 |
| TcCLB.508355.80 | trans-sialidase, Group VIII, putative | -2,154 | 2,847E-26 |
| TcCLB.510713.160 | trans-sialidase, Group V, putative | -2,154 | 1,656E-20 |
| TcCLB.507255.10 | surface protease GP63 (pseudogene), putative | -2,154 | 6,952E-19 |
| TcCLB.506329.79 | mucin-associated surface protein (MASP, pseudogene), pu | -2,154 | 1,783E-08 |
| TcCLB.503709.10 | hypothetical protein | -2,155 | 2,224E-11 |
| TcCLB.508593.10 | trans-sialidase, Group IV, putative | -2,156 | 2,509E-51 |
| TcCLB.508165.210 | mucin-associated surface protein (MASP), putative | -2,156 | 9,13E-35 |
| TcCLB.508883.30 | mucin TcMUCII, putative | -2,156 | 1,271E-12 |
| TcCLB.510583.30 | mucin TcMUCII, putative | -2,156 | 5,707E-06 |
| TcCLB.503447.50 | trans-sialidase, Group IV, putative | -2,157 | 3,643E-46 |
| TcCLB.510377.410 | mucin TcMUCII, putative | -2,157 | 1,372E-14 |
| TcCLB.510483.154 | mucin-associated surface protein (MASP), putative | -2,157 | 4,081E-14 |
| TcCLB.506083.20 | trans-sialidase (pseudogene), putative | -2,158 | 3,016E-64 |
| TcCLB.506321.190 | mucin TcMUCII, putative | -2,158 | 8,444E-20 |
| TcCLB.507699.110 | mucin-associated surface protein (MASP, pseudogene), pu | -2,159 | 3,235E-22 |
| TcCLB.511173.400 | mucin-associated surface protein (MASP), putative | -2,16 | 2,699E-14 |
| TcCLB.508165.185 | mucin TcMUCII (pseudogene), putative | -2,161 | 0,000028 |
| TcCLB.507959.290 | hypothetical protein | -2,163 | 0,0002948 |
| TcCLB.511255.280 | mucin-associated surface protein (MASP), putative | -2,164 | 5,803E-26 |
| TcCLB.510377.500 | trans-sialidase (pseudogene), putative | -2,164 | 5,325E-13 |
| TcCLB.510085.10 | mucin-associated surface protein (MASP, pseudogene), pu | -2,165 | 9,087E-22 |
| TcCLB.508365.40 | mucin-associated surface protein (MASP), putative | -2,165 | 5,607E-11 |
| TcCLB.425855.9 | hypothetical protein, conserved (fragment) | -2,165 | 6,193E-10 |
| TcCLB.510209.9 | trans-sialidase, putative | -2,166 | 2,086E-11 |
| TcCLB.506737.90 | trans-sialidase, Group V, putative | -2,168 | 2,327E-142 |
| TcCLB.504769.20 | mucin-associated surface protein (MASP), putative | -2,168 | 2,932E-56 |
| TcCLB.510279.60 | mucin-associated surface protein (MASP), putative | -2,168 | 9,444E-23 |
| TcCLB.507075.40 | mucin-associated surface protein (MASP), putative (fragm | -2,169 | 1,468E-36 |
| TcCLB.506613.20 | mucin-associated surface protein (MASP, pseudogene), pu | -2,169 | 1,061E-23 |
| TcCLB.509657.70 | trans-sialidase (pseudogene), putative | -2,17 | 3,605E-13 |
| TcCLB.506067.110 | mucin-associated surface protein (MASP, pseudogene), pu | -2,171 | 1,218E-09 |
| TcCLB.508453.10 | mucin-associated surface protein (MASP), putative | -2,172 | 4,781E-37 |
| TcCLB.511255.690 | mucin-associated surface protein (MASP), putative | -2,173 | 1,1E-28 |
| TcCLB.511605.90 | mucin TcMUCII, putative | -2,176 | 2,929E-10 |
| TcCLB.462761.10 | surface protease GP63, putative | -2,177 | 4,104E-73 |
| TcCLB.509905.60 | trans-sialidase (pseudogene), putative | -2,178 | 5,236E-52 |
| TcCLB.511593.60 | mucin-associated surface protein (MASP), putative | -2,178 | 1,18E-33 |
| TcCLB.508455.20 | trans-sialidase, Group V, putative | -2,178 | 4,998E-33 |
| TcCLB.508261.130 | mucin-associated surface protein (MASP), putative | -2,178 | 4,163E-23 |
| TcCLB.508539.155 | mucin TcMUCII, putative | -2,178 | 3,766E-18 |

| | | | |
|---|---|---|---|
| TcCLB.506499.90 | 90 kDa surface protein, putative | -2,178 | 4,416E-08 |
| TcCLB.510469.10 | mucin-associated surface protein (MASP), putative | -2,179 | 4,447E-08 |
| TcCLB.511259.180 | mucin-associated surface protein (MASP), putative | -2,18 | 8,083E-11 |
| TcCLB.507993.230 | hypothetical protein, conserved (pseudogene) | -2,18 | 0,00000346 |
| TcCLB.507429.20 | mucin-associated surface protein (MASP), putative | -2,181 | 1,51E-59 |
| TcCLB.507611.120 | expression site-associated gene (ESAG-like) protein, puta | -2,182 | 2,874E-12 |
| TcCLB.511063.120 | mucin TcMUCII, putative | -2,182 | 7,27E-08 |
| TcCLB.510803.19 | trans-sialidase, putative | -2,182 | 8,843E-07 |
| TcCLB.504277.30 | hypothetical protein | -2,183 | 8,118E-25 |
| TcCLB.505949.70 | mucin TcMUCII, putative | -2,184 | 1,927E-28 |
| TcCLB.506501.210 | surface protease GP63 (pseudogene), putative | -2,184 | 2,522E-07 |
| TcCLB.511183.420 | mucin-associated surface protein (MASP, pseudogene), pu | -2,185 | 3,596E-32 |
| TcCLB.510417.40 | mucin-associated surface protein (MASP), putative (fragm | -2,185 | 7,223E-08 |
| TcCLB.510475.120 | mucin-associated surface protein (MASP, pseudogene), pu | -2,185 | 1,123E-06 |
| TcCLB.506309.40 | mucin-associated surface protein (MASP), putative | -2,186 | 4,22E-11 |
| TcCLB.397923.10 | mucin-associated surface protein (MASP), putative | -2,187 | 5,62E-29 |
| TcCLB.510451.10 | trans-sialidase (pseudogene), putative | -2,188 | 1,237E-08 |
| TcCLB.508165.280 | mucin-associated surface protein (MASP), putative | -2,189 | 3,397E-20 |
| TcCLB.509525.234 | mucin TcMUCII, putative | -2,189 | 5,756E-14 |
| TcCLB.506499.110 | mucin TcMUCII (pseudogene), putative | -2,19 | 1,082E-15 |
| TcCLB.506969.6 | trans-sialidase (pseudogene), putative | -2,19 | 0,00001676 |
| TcCLB.448653.10 | trans-sialidase, putative (fragment) | -2,191 | 9,551E-06 |
| TcCLB.509525.20 | trans-sialidase (pseudogene), putative | -2,192 | 7,837E-39 |
| TcCLB.507973.10 | mucin TcMUCI, putative | -2,192 | 2,565E-16 |
| TcCLB.509899.50 | mucin-associated surface protein (MASP, pseudogene), pu | -2,195 | 1,184E-46 |
| TcCLB.508605.20 | UDP-Gal or UDP-GlcNAc-dependent glycosyltransferase, p | -2,195 | 9,204E-37 |
| TcCLB.509657.40 | mucin TcMUCII, putative | -2,196 | 2,664E-30 |
| TcCLB.510497.60 | mucin TcMUCII, putative | -2,196 | 3,116E-17 |
| TcCLB.504393.20 | hypothetical protein | -2,196 | 0,0002077 |
| TcCLB.510013.150 | mucin-associated surface protein (MASP, pseudogene), pu | -2,198 | 5,242E-28 |
| TcCLB.510631.20 | trans-sialidase (pseudogene), putative | -2,199 | 1,14E-95 |
| TcCLB.510715.20 | trans-sialidase, Group V, putative | -2,199 | 1,099E-34 |
| TcCLB.506763.234 | mucin-associated surface protein (MASP, pseudogene), pu | -2,199 | 1,047E-20 |
| TcCLB.508011.90 | trans-sialidase (pseudogene), putative | -2,201 | 1,665E-33 |
| TcCLB.503973.120 | mucin-associated surface protein (MASP), putative | -2,202 | 9,813E-33 |
| TcCLB.510503.20 | mucin TcMUCII, putative | -2,202 | 1,367E-21 |
| TcCLB.507069.160 | trans-sialidase, Group V, putative | -2,203 | 1,674E-30 |
| TcCLB.510017.40 | mucin TcMUCII, putative | -2,205 | 5,341E-13 |
| TcCLB.511859.90 | hypothetical protein | -2,206 | 1,166E-51 |
| TcCLB.509905.180 | mucin-associated surface protein (MASP), putative | -2,206 | 1,118E-35 |
| TcCLB.511625.90 | mucin-associated surface protein (MASP), putative | -2,207 | 1,416E-33 |
| TcCLB.508511.15 | trans-sialidase (pseudogene), putative | -2,208 | 1,958E-06 |
| TcCLB.511173.180 | mucin-associated surface protein (MASP), putative | -2,209 | 1,268E-23 |
| TcCLB.508741.130 | hypothetical protein | -2,21 | 2,743E-110 |
| TcCLB.509897.30 | mucin-associated surface protein (MASP, pseudogene), pu | -2,21 | 3,801E-23 |

| | | | |
|---|---|---|---|
| TcCLB.507237.24 | mucin TcMUCII (pseudogene), putative | -2,21 | 0,00007609 |
| TcCLB.508221.550 | mucin-associated surface protein (MASP), putative | -2,214 | 8,465E-14 |
| TcCLB.506277.3 | mucin TcMUCII, putative (fragment) | -2,214 | 4,523E-12 |
| TcCLB.510201.10 | trans-sialidase, Group V, putative | -2,215 | 2,844E-37 |
| TcCLB.509035.20 | surface protease GP63, putative | -2,215 | 1,152E-20 |
| TcCLB.511373.30 | mucin-associated surface protein (MASP), putative | -2,215 | 1,666E-09 |
| TcCLB.506229.20 | hypothetical protein, conserved | -2,216 | 2,508E-34 |
| TcCLB.509081.130 | surface protease GP63 (pseudogene), putative | -2,216 | 3,897E-08 |
| TcCLB.506769.60 | mucin-associated surface protein (MASP, pseudogene), pu | -2,218 | 3,117E-17 |
| TcCLB.511277.370 | phosphatidic acid phosphatase, putative | -2,219 | 5,144E-74 |
| TcCLB.506137.60 | mucin-associated surface protein (MASP), putative | -2,219 | 3,983E-08 |
| TcCLB.506335.150 | mucin-associated surface protein (MASP, pseudogene), pu | -2,22 | 7,348E-29 |
| TcCLB.401183.10 | trans-sialidase, putative (fragment) | -2,22 | 3,409E-21 |
| TcCLB.511553.5 | trans-sialidase (pseudogene), putative | -2,22 | 1,584E-09 |
| TcCLB.503761.20 | surface protease GP63 (pseudogene), putative | -2,221 | 4,297E-26 |
| TcCLB.510961.40 | mucin TcMUCII, putative | -2,223 | 5,413E-21 |
| TcCLB.511173.460 | mucin-associated surface protein (MASP), putative | -2,223 | 1,436E-19 |
| TcCLB.506501.40 | mucin TcMUCII, putative | -2,227 | 5,967E-30 |
| TcCLB.511921.90 | mucin-associated surface protein (MASP, pseudogene), pu | -2,227 | 2,036E-26 |
| TcCLB.506267.30 | mucin TcMUCII, putative | -2,227 | 6,424E-24 |
| TcCLB.506453.30 | mucin-associated surface protein (MASP), putative | -2,227 | 2,513E-20 |
| TcCLB.508433.120 | mucin-associated surface protein (MASP, pseudogene), pu | -2,227 | 3,011E-16 |
| TcCLB.506355.20 | receptor-type adenylate cyclase, putative | -2,228 | 6,904E-54 |
| TcCLB.507699.90 | mucin TcMUCII, putative | -2,228 | 2,442E-42 |
| TcCLB.510501.40 | trans-sialidase, putative | -2,228 | 2,748E-11 |
| TcCLB.506613.40 | trans-sialidase, Group V, putative | -2,23 | 1,939E-42 |
| TcCLB.507071.100 | mucin-associated surface protein (MASP), putative | -2,23 | 1,001E-20 |
| TcCLB.511625.100 | mucin TcMUCII, putative | -2,23 | 2,261E-20 |
| TcCLB.506131.140 | mucin-associated surface protein (MASP), putative | -2,23 | 1,749E-18 |
| TcCLB.510503.40 | surface protease GP63 (pseudogene), putative | -2,23 | 3,094E-14 |
| TcCLB.508221.518 | mucin TcMUCII, putative | -2,23 | 9,932E-13 |
| TcCLB.511089.19 | mucin-associated surface protein (MASP), putative | -2,23 | 7,011E-12 |
| TcCLB.506611.10 | mucin-associated surface protein (MASP), putative | -2,231 | 5,529E-17 |
| TcCLB.508871.60 | mucin-associated surface protein (MASP), putative | -2,232 | 1,34E-17 |
| TcCLB.510025.150 | mucin TcMUCII, putative | -2,232 | 1,02E-14 |
| TcCLB.510817.50 | trans-sialidase, Group II, putative | -2,234 | 3,027E-59 |
| TcCLB.503773.30 | hypothetical protein, conserved | -2,234 | 3,804E-33 |
| TcCLB.506131.170 | trans-sialidase, Group V, putative | -2,234 | 5,261E-17 |
| TcCLB.509031.20 | trans-sialidase, putative (fragment) | -2,237 | 6,738E-21 |
| TcCLB.506279.60 | trans-sialidase, Group VI, putative | -2,239 | 4,691E-61 |
| TcCLB.507143.90 | hypothetical protein | -2,242 | 0,0003553 |
| TcCLB.509897.80 | mucin-associated surface protein (MASP, pseudogene), pu | -2,245 | 3,617E-16 |
| TcCLB.510591.10 | trans-sialidase (pseudogene), putative | -2,248 | 1,302E-12 |
| TcCLB.511087.10 | mucin-associated surface protein (MASP, pseudogene), pu | -2,249 | 3,128E-28 |
| TcCLB.510275.50 | mucin TcMUCII, putative | -2,25 | 6,774E-22 |

| | | | |
|---|---|---|---|
| TcCLB.444331.10 | mucin-associated surface protein (MASP), putative | -2,253 | 4,131E-11 |
| TcCLB.506835.90 | hypothetical protein, conserved | -2,254 | 2,16E-136 |
| TcCLB.508099.20 | serine-alanine-and proline-rich protein, putative | -2,255 | 3,76E-35 |
| TcCLB.509433.10 | hypothetical protein | -2,255 | 1,567E-19 |
| TcCLB.508593.20 | trans-sialidase, Group IV, putative | -2,256 | 5,853E-53 |
| TcCLB.509947.10 | mucin TcMUCII, putative | -2,258 | 2,828E-43 |
| TcCLB.508163.320 | mucin-associated surface protein (MASP), putative | -2,259 | 3,787E-23 |
| TcCLB.511921.29 | mucin-associated surface protein (MASP, pseudogene), pu | -2,259 | 1,908E-16 |
| TcCLB.509615.50 | hypothetical protein | -2,261 | 0,001588 |
| TcCLB.511043.60 | receptor-type adenylate cyclase, putative | -2,262 | 3,081E-69 |
| TcCLB.509811.9 | surface protease GP63, putative | -2,262 | 1,262E-38 |
| TcCLB.506741.60 | mucin-associated surface protein (MASP), putative | -2,262 | 5,484E-23 |
| TcCLB.510017.30 | mucin-associated surface protein (MASP, pseudogene), pu | -2,262 | 1,247E-17 |
| TcCLB.508125.80 | trans-sialidase (pseudogene), putative | -2,264 | 1,178E-17 |
| TcCLB.510361.30 | mucin-associated surface protein (MASP), putative | -2,264 | 1,412E-13 |
| TcCLB.506965.20 | mucin-associated surface protein (MASP), putative | -2,265 | 2,069E-15 |
| TcCLB.510203.34 | mucin-associated surface protein (MASP, pseudogene), pu | -2,265 | 1,055E-10 |
| TcCLB.506133.140 | mucin-associated surface protein (MASP), putative | -2,266 | 2,89E-25 |
| TcCLB.506501.5 | trans-sialidase (pseudogene), putative | -2,266 | 1,482E-11 |
| TcCLB.510377.439 | mucin TcMUCII, putative | -2,267 | 2,668E-19 |
| TcCLB.508121.160 | mucin-associated surface protein (MASP, pseudogene), pu | -2,268 | 4,339E-21 |
| TcCLB.506501.110 | mucin-associated surface protein (MASP), putative | -2,268 | 1,363E-19 |
| TcCLB.511553.140 | mucin-associated surface protein (MASP), putative | -2,268 | 6,602E-14 |
| TcCLB.508523.50 | trans-sialidase, Group II, putative | -2,269 | 1,869E-50 |
| TcCLB.503973.80 | mucin-associated surface protein (MASP), putative | -2,27 | 8,35E-35 |
| TcCLB.507069.10 | mucin-associated surface protein (MASP), putative | -2,27 | 3,8E-24 |
| TcCLB.511109.10 | surface protein TolT (pseudogene) (TolT) | -2,272 | 5,049E-29 |
| TcCLB.510681.41 | surface protease GP63, putative | -2,272 | 4,811E-18 |
| TcCLB.503633.11 | mucin-associated surface protein (MASP), putative (fragm | -2,272 | 5,372E-18 |
| TcCLB.504155.293 | mucin-associated surface protein (MASP), putative | -2,272 | 1,214E-07 |
| TcCLB.511221.50 | mucin-associated surface protein (MASP), putative | -2,273 | 1,678E-59 |
| TcCLB.506669.10 | hypothetical protein, conserved | -2,273 | 1,814E-56 |
| TcCLB.467373.10 | trans-sialidase, putative (fragment) | -2,274 | 2,181E-76 |
| TcCLB.506713.80 | trans-sialidase, putative (fragment) | -2,274 | 2,876E-63 |
| TcCLB.506613.50 | mucin-associated surface protein (MASP), putative | -2,278 | 1,797E-68 |
| TcCLB.507937.20 | mucin-associated surface protein (MASP, pseudogene), pu | -2,278 | 2,148E-22 |
| TcCLB.506601.51 | mucin TcMUCII (pseudogene), putative | -2,279 | 3,709E-18 |
| TcCLB.506667.30 | 90 kDa surface protein, putative | -2,281 | 3,377E-45 |
| TcCLB.510207.10 | mucin TcMUCII, putative | -2,281 | 7,057E-18 |
| TcCLB.510363.60 | trans-sialidase (pseudogene), putative | -2,282 | 1,491E-28 |
| TcCLB.508501.110 | mucin TcMUCII, putative | -2,282 | 5,191E-26 |
| TcCLB.507959.210 | mucin-associated surface protein (MASP), putative | -2,282 | 3,721E-25 |
| TcCLB.510373.60 | mucin-associated surface protein (MASP), putative | -2,284 | 3,92E-49 |
| TcCLB.510583.200 | mucin TcMUCII, putative | -2,284 | 3,513E-15 |
| TcCLB.507549.20 | trans-sialidase (pseudogene), putative | -2,285 | 6,054E-64 |

| | | | |
|---|---|---|---|
| TcCLB.508873.10 | mucin-associated surface protein (MASP), putative | -2,285 | 5,53E-46 |
| TcCLB.510789.10 | mucin TcMUCII, putative | -2,286 | 2,669E-36 |
| TcCLB.511605.55 | hypothetical protein | -2,286 | 6,154E-23 |
| TcCLB.404431.10 | trans-sialidase, putative | -2,287 | 2,25E-43 |
| TcCLB.506279.30 | mucin TcMUCII, putative | -2,288 | 4,011E-28 |
| TcCLB.506851.30 | hypothetical protein, conserved | -2,288 | 4,878E-26 |
| TcCLB.509525.240 | mucin-associated surface protein (MASP), putative | -2,288 | 3,032E-16 |
| TcCLB.424171.10 | trans-sialidase (pseudogene), putative | -2,288 | 7,758E-11 |
| TcCLB.504213.50 | hypothetical protein, conserved | -2,291 | 2,517E-54 |
| TcCLB.511895.10 | receptor-type adenylate cyclase, putative | -2,292 | 4,445E-64 |
| TcCLB.506789.50 | mucin-associated surface protein (MASP), putative | -2,292 | 1,26E-24 |
| TcCLB.506667.50 | trans-sialidase (pseudogene), putative | -2,292 | 1,064E-17 |
| TcCLB.508221.210 | mucin-associated surface protein (MASP), putative | -2,292 | 2,121E-13 |
| TcCLB.506285.90 | mucin TcMUCII, putative | -2,295 | 4,948E-27 |
| TcCLB.510697.50 | trans-sialidase, Group VI, putative | -2,298 | 8,358E-26 |
| TcCLB.508221.509 | mucin-associated surface protein (MASP, pseudogene), pu | -2,298 | 2,517E-13 |
| TcCLB.511611.16 | mucin TcMUCII (pseudogene), putative | -2,298 | 4,039E-11 |
| TcCLB.507257.80 | diacylglycerol kinase-like protein, putative | -2,301 | 1,144E-74 |
| TcCLB.510791.50 | mucin TcMUCII, putative | -2,301 | 2,093E-38 |
| TcCLB.510475.110 | mucin TcMUCII, putative | -2,301 | 1,338E-11 |
| TcCLB.508873.250 | mucin-associated surface protein (MASP), putative | -2,302 | 3,858E-45 |
| TcCLB.505025.20 | surface protease GP63 (pseudogene), putative | -2,302 | 2,954E-14 |
| TcCLB.503861.100 | trans-sialidase (pseudogene), putative | -2,303 | 8,262E-70 |
| TcCLB.508789.120 | trans-sialidase, Group VI, putative | -2,303 | 3,099E-29 |
| TcCLB.506211.150 | polyprenyl synthase, putative | -2,304 | 4,304E-90 |
| TcCLB.508219.70 | mucin-associated surface protein (MASP), putative | -2,304 | 9,979E-21 |
| TcCLB.506945.40 | hypothetical protein, conserved | -2,305 | 3,126E-127 |
| TcCLB.507959.41 | mucin TcMUCII, putative | -2,305 | 1,022E-10 |
| TcCLB.507859.31 | hypothetical protein | -2,306 | 3,267E-22 |
| TcCLB.506769.10 | hypothetical protein | -2,306 | 9,079E-12 |
| TcCLB.463323.10 | trans-sialidase, Group II, putative | -2,307 | 9,475E-61 |
| TcCLB.508831.220 | trans-sialidase (pseudogene), putative | -2,307 | 5,388E-11 |
| TcCLB.508761.160 | mucin TcMUCII, putative | -2,308 | 1,786E-28 |
| TcCLB.509123.5 | trans-sialidase (pseudogene), putative | -2,308 | 0,00001221 |
| TcCLB.439245.10 | mucin TcMUCII, putative (fragment) | -2,309 | 1,794E-21 |
| TcCLB.510625.40 | trans-sialidase, Group V, putative | -2,309 | 2,964E-21 |
| TcCLB.505615.10 | surface protease GP63, putative | -2,313 | 1,074E-87 |
| TcCLB.506133.130 | trans-sialidase, Group V, putative | -2,315 | 1,556E-24 |
| TcCLB.506599.70 | mucin TcMUCII, putative | -2,315 | 1,679E-14 |
| TcCLB.506501.270 | syntaxin binding protein (pseudogene), putative | -2,317 | 2,254E-08 |
| TcCLB.509217.90 | mucin-associated surface protein (MASP), putative | -2,318 | 5,881E-39 |
| TcCLB.510715.150 | trans-sialidase, Group VI, putative | -2,318 | 8,133E-29 |
| TcCLB.508365.200 | mucin-associated surface protein (MASP), putative | -2,318 | 3,474E-11 |
| TcCLB.504479.10 | trans-sialidase, Group II, putative | -2,32 | 9,656E-82 |
| TcCLB.510629.10 | mucin-associated surface protein (MASP), putative | -2,32 | 1,771E-09 |

| | | | |
|---|---|---|---|
| TcCLB.506067.280 | mucin TcMUCII, putative | -2,321 | 1,031E-36 |
| TcCLB.503889.10 | mucin TcMUCII, putative | -2,323 | 4,072E-54 |
| TcCLB.510377.220 | mucin-associated surface protein (MASP, pseudogene), pu | -2,323 | 7,705E-23 |
| TcCLB.510047.109 | mucin TcMUCII, putative | -2,325 | 9,847E-37 |
| TcCLB.506269.10 | mucin-associated surface protein (MASP), putative | -2,327 | 5,492E-35 |
| TcCLB.511923.49 | mucin-associated surface protein (MASP, pseudogene), pu | -2,327 | 2,257E-06 |
| TcCLB.506193.60 | ascorbate peroxidase, putative (APX) | -2,328 | 7,092E-50 |
| TcCLB.506703.140 | mucin-associated surface protein (MASP), putative | -2,328 | 1,398E-06 |
| TcCLB.508991.30 | trans-sialidase (pseudogene), putative | -2,331 | 1,014E-69 |
| TcCLB.505757.10 | hypothetical protein, conserved | -2,332 | 1,239E-51 |
| TcCLB.510205.20 | mucin-associated surface protein (MASP), putative | -2,332 | 2,105E-40 |
| TcCLB.506501.10 | mucin-associated surface protein (MASP), putative | -2,332 | 1,497E-16 |
| TcCLB.508221.600 | mucin-associated surface protein (MASP), putative | -2,332 | 5,418E-13 |
| TcCLB.511289.80 | polyprenyl synthase, putative | -2,333 | 9,096E-125 |
| TcCLB.510561.30 | mucin-associated surface protein (MASP), putative | -2,333 | 2,784E-06 |
| TcCLB.508261.70 | mucin-associated surface protein (MASP, pseudogene), pu | -2,334 | 2,112E-29 |
| TcCLB.509663.44 | hypothetical protein, conserved | -2,339 | 7,916E-73 |
| TcCLB.511149.130 | mucin TcMUCI, putative | -2,34 | 1,335E-06 |
| TcCLB.508999.120 | trans-sialidase (pseudogene), putative | -2,341 | 1,076E-33 |
| TcCLB.463279.20 | trans-sialidase, Group II, putative | -2,342 | 3,097E-108 |
| TcCLB.507959.80 | mucin-associated surface protein (MASP), putative | -2,342 | 4,161E-38 |
| TcCLB.509031.3 | trans-sialidase, putative (fragment) | -2,342 | 1,776E-25 |
| TcCLB.506499.100 | hypothetical protein | -2,342 | 1,129E-07 |
| TcCLB.503771.10 | mucin-associated surface protein (MASP), putative | -2,342 | 2,329E-06 |
| TcCLB.510919.9 | trans-sialidase, putative (fragment) | -2,344 | 7,75E-37 |
| TcCLB.508365.130 | mucin TcMUCII, putative | -2,344 | 1,499E-15 |
| TcCLB.506329.9 | mucin-associated surface protein (MASP, pseudogene), pu | -2,344 | 1,277E-13 |
| TcCLB.510933.80 | trans-sialidase, Group V, putative | -2,345 | 1,814E-31 |
| TcCLB.510323.110 | hypothetical protein, conserved | -2,346 | 1,934E-48 |
| TcCLB.510279.140 | mucin-associated surface protein (MASP), putative | -2,348 | 5,381E-28 |
| TcCLB.506717.100 | mucin-associated surface protein (MASP, pseudogene), pu | -2,349 | 1,109E-38 |
| TcCLB.507071.20 | mucin-associated surface protein (MASP), putative | -2,35 | 2,008E-44 |
| TcCLB.507957.229 | mucin TcMUCII, putative | -2,35 | 1,57E-30 |
| TcCLB.506613.150 | trans-sialidase (pseudogene), putative | -2,35 | 9,595E-23 |
| TcCLB.510561.40 | mucin TcMUCII, putative | -2,352 | 9,719E-62 |
| TcCLB.509549.20 | mucin-associated surface protein (MASP), putative | -2,352 | 4,971E-55 |
| TcCLB.511203.29 | surface protease GP63, putative (fragment) | -2,352 | 7,069E-51 |
| TcCLB.510019.10 | mucin TcMUCII, putative | -2,352 | 2,721E-20 |
| TcCLB.510929.10 | trans-sialidase, putative | -2,352 | 5,15E-14 |
| TcCLB.506901.60 | mucin TcMUCII, putative | -2,352 | 9,206E-12 |
| TcCLB.507747.130 | trans-sialidase (pseudogene), putative | -2,353 | 1,242E-44 |
| TcCLB.509765.50 | trans-sialidase, Group VIII, putative | -2,353 | 1,162E-11 |
| TcCLB.506767.360 | mucin TcMUCII, putative | -2,354 | 9,832E-28 |
| TcCLB.510021.20 | mucin-associated surface protein (MASP), putative | -2,354 | 1,085E-26 |
| TcCLB.508365.160 | mucin-associated surface protein (MASP), putative | -2,354 | 5,115E-25 |

| | | | |
|---|---|---|---|
| TcCLB.510697.10 | mucin-associated surface protein (MASP), putative | -2,354 | 7,036E-25 |
| TcCLB.510239.70 | mucin-associated surface protein (MASP), putative | -2,355 | 3,951E-39 |
| TcCLB.507071.230 | trans-sialidase, Group V, putative | -2,355 | 1,341E-33 |
| TcCLB.508247.90 | trans-sialidase, Group V, putative | -2,356 | 9,594E-08 |
| TcCLB.507687.70 | mucin-associated surface protein (MASP), putative | -2,357 | 8,765E-27 |
| TcCLB.510939.25 | mucin TcMUCI, putative | -2,358 | 4,029E-56 |
| TcCLB.511153.19 | surface protease GP63, putative | -2,359 | 6,588E-120 |
| TcCLB.504155.80 | mucin-associated surface protein (MASP), putative | -2,359 | 3,104E-19 |
| TcCLB.508435.20 | mucin TcMUCII, putative | -2,359 | 5,236E-18 |
| TcCLB.504081.40 | mucin-associated surface protein (MASP), putative | -2,36 | 1,504E-14 |
| TcCLB.511183.360 | mucin-associated surface protein (MASP), putative | -2,361 | 8,383E-42 |
| TcCLB.506995.40 | mucin TcMUCII, putative | -2,361 | 7,98E-35 |
| TcCLB.506279.120 | mucin TcMUCII, putative | -2,361 | 9,9E-13 |
| TcCLB.506741.50 | mucin TcMUCII, putative | -2,362 | 6,605E-57 |
| TcCLB.506767.40 | mucin-associated surface protein (MASP), putative | -2,364 | 3,761E-44 |
| TcCLB.508251.40 | mucin TcMUCII, putative | -2,364 | 1,66E-14 |
| TcCLB.508055.40 | surface protease GP63 (pseudogene), putative | -2,366 | 2,391E-08 |
| TcCLB.508849.11 | trans-sialidase, putative (fragment) | -2,367 | 2,189E-17 |
| TcCLB.506955.270 | trans-sialidase, Group V, putative | -2,368 | 2,2E-33 |
| TcCLB.510561.20 | mucin TcMUCII, putative | -2,368 | 2,511E-28 |
| TcCLB.508033.60 | mucin TcMUCII, putative | -2,368 | 5,593E-24 |
| TcCLB.510915.5 | dispersed gene family protein 1 (DGF-1), putative | -2,369 | 3,132E-40 |
| TcCLB.503503.40 | mucin-associated surface protein (MASP), putative | -2,371 | 1,841E-32 |
| TcCLB.506425.5 | receptor-type adenylate cyclase, putative (fragment) | -2,371 | 4,719E-31 |
| TcCLB.507987.10 | trans-sialidase, putative | -2,371 | 1,69E-20 |
| TcCLB.508253.10 | mucin-associated surface protein (MASP), putative | -2,372 | 1,069E-65 |
| TcCLB.511173.230 | trans-sialidase (pseudogene), putative | -2,372 | 2,367E-35 |
| TcCLB.508165.400 | mucin TcMUCII, putative | -2,373 | 1,178E-37 |
| TcCLB.504101.50 | trans-sialidase, Group V, putative | -2,374 | 5,346E-64 |
| TcCLB.508313.30 | mucin-associated surface protein (MASP), putative | -2,374 | 2,438E-18 |
| TcCLB.510497.40 | trans-sialidase, Group V, putative | -2,375 | 1,099E-23 |
| TcCLB.504155.140 | mucin TcMUCII, putative | -2,376 | 0,000186 |
| TcCLB.506613.110 | mucin-associated surface protein (MASP, pseudogene), pu | -2,377 | 2,348E-35 |
| TcCLB.510479.40 | trans-sialidase, putative (fragment) | -2,377 | 1,353E-20 |
| TcCLB.468217.14 | mucin-associated surface protein (MASP), putative | -2,378 | 3,278E-43 |
| TcCLB.506967.50 | mucin-associated surface protein (MASP), putative | -2,379 | 2,179E-17 |
| TcCLB.506577.80 | trans-sialidase, Group VIII, putative | -2,38 | 3,509E-93 |
| TcCLB.505949.10 | mucin TcMUCII, putative | -2,38 | 9,321E-25 |
| TcCLB.506281.14 | hypothetical protein | -2,381 | 4,507E-78 |
| TcCLB.510497.100 | mucin-associated surface protein (MASP), putative | -2,381 | 2,902E-28 |
| TcCLB.511349.20 | trans-sialidase (pseudogene), putative | -2,382 | 1,048E-80 |
| TcCLB.511667.50 | mucin-associated surface protein (MASP), putative | -2,383 | 1,911E-36 |
| TcCLB.509785.50 | trans-sialidase, Group V, putative | -2,384 | 2,386E-37 |
| TcCLB.506409.30 | mucin-associated surface protein (MASP), putative | -2,384 | 3,261E-32 |
| TcCLB.510693.270 | trans-sialidase, Group VI, putative | -2,386 | 2,991E-22 |

| | | | |
|---|---|---|---|
| TcCLB.506973.20 | mucin-associated surface protein (MASP), putative | -2,388 | 5,007E-63 |
| TcCLB.510561.50 | mucin-associated surface protein (MASP), putative | -2,39 | 1,295E-07 |
| TcCLB.511907.200 | cysteine peptidase inhibitor, putative | -2,392 | 2,678E-133 |
| TcCLB.510023.20 | mucin TcMUCII, putative | -2,392 | 9,81E-30 |
| TcCLB.506763.110 | trans-sialidase, Group VII, putative | -2,393 | 9,252E-17 |
| TcCLB.510411.10 | trans-sialidase, Group II, putative | -2,394 | 1,292E-38 |
| TcCLB.509525.400 | mucin TcMUCII, putative | -2,394 | 4,568E-22 |
| TcCLB.506761.54 | mucin-associated surface protein (MASP, pseudogene), pu | -2,396 | 5,586E-29 |
| TcCLB.506389.10 | trans-sialidase, putative | -2,397 | 3,939E-55 |
| TcCLB.506767.230 | mucin-associated surface protein (MASP), putative | -2,397 | 2,474E-40 |
| TcCLB.506911.20 | mucin-associated surface protein (MASP), putative | -2,398 | 2,271E-64 |
| TcCLB.508245.40 | mucin-associated surface protein (MASP), putative | -2,399 | 2,654E-35 |
| TcCLB.506133.190 | trans-sialidase, Group VI, putative | -2,399 | 1,088E-15 |
| TcCLB.511149.180 | trans-sialidase, Group VI, putative | -2,4 | 1,652E-36 |
| TcCLB.511911.10 | trans-sialidase, Group II, putative | -2,4 | 5,765E-29 |
| TcCLB.505949.30 | trans-sialidase, Group V, putative | -2,4 | 5,7E-23 |
| TcCLB.507065.70 | mucin TcMUCII, putative | -2,4 | 2,229E-12 |
| TcCLB.503773.40 | hypothetical protein, conserved | -2,401 | 1,217E-54 |
| TcCLB.503699.19 | mucin-associated surface protein (MASP), putative | -2,402 | 3,339E-24 |
| TcCLB.506173.70 | trans-sialidase, Group V, putative | -2,403 | 3,494E-31 |
| TcCLB.510731.10 | trans-sialidase, putative | -2,405 | 6,859E-06 |
| TcCLB.510359.480 | mucin-associated surface protein (MASP, pseudogene), pu | -2,407 | 6,827E-08 |
| TcCLB.506501.380 | mucin-associated surface protein (MASP), putative | -2,409 | 3,259E-28 |
| TcCLB.507705.20 | hypothetical protein, conserved | -2,411 | 3,334E-37 |
| TcCLB.510623.20 | mucin-associated surface protein (MASP), putative | -2,412 | 8,787E-30 |
| TcCLB.510361.190 | mucin-associated surface protein (MASP), putative | -2,413 | 1,139E-25 |
| TcCLB.506769.65 | trans-sialidase (pseudogene), putative | -2,413 | 1,722E-07 |
| TcCLB.511165.80 | protein kinase, putative | -2,413 | 6,006E-07 |
| TcCLB.511603.330 | mucin TcMUCII, putative | -2,414 | 3,729E-20 |
| TcCLB.507957.10 | mucin-associated surface protein (MASP), putative | -2,415 | 7,076E-68 |
| TcCLB.504197.20 | mucin-associated surface protein (MASP), putative | -2,415 | 1,133E-42 |
| TcCLB.510359.530 | trans-sialidase, Group V, putative | -2,415 | 9,809E-31 |
| TcCLB.509655.10 | trans-sialidase (pseudogene), putative | -2,417 | 5,015E-45 |
| TcCLB.510061.30 | trans-sialidase, Group V, putative | -2,418 | 8,572E-55 |
| TcCLB.509075.20 | dispersed gene family protein 1 (DGF-1, pseudogene), put | -2,418 | 1,124E-11 |
| TcCLB.506409.60 | hypothetical protein | -2,418 | 5,352E-10 |
| TcCLB.508745.11 | trans-sialidase, putative (fragment) | -2,419 | 2,98E-20 |
| TcCLB.506603.20 | mucin-associated surface protein (MASP, pseudogene), pu | -2,422 | 3,034E-29 |
| TcCLB.508747.5 | mucin-associated surface protein (MASP), putative | -2,423 | 3,46E-19 |
| TcCLB.506769.120 | mucin-associated surface protein (MASP), putative | -2,424 | 9,693E-27 |
| TcCLB.508147.140 | mucin TcMUCII, putative | -2,425 | 7,764E-15 |
| TcCLB.506139.160 | mucin-associated surface protein (MASP), putative | -2,428 | 7,186E-32 |
| TcCLB.506289.150 | mucin-associated surface protein (MASP), putative | -2,432 | 4,138E-44 |
| TcCLB.507043.21 | mucin TcMUCI, putative (fragment) | -2,432 | 7,339E-30 |
| TcCLB.508221.910 | mucin TcMUCII, putative | -2,432 | 1,82E-18 |

| | | | |
|---|---|---|---|
| TcCLB.506765.80 | mucin TcMUCII, putative | -2,432 | 4,7E-10 |
| TcCLB.421231.10 | hypothetical protein, conserved | -2,433 | 6,427E-166 |
| TcCLB.507085.150 | mucin-associated surface protein (MASP), putative | -2,433 | 1,633E-74 |
| TcCLB.509841.10 | trans-sialidase, putative | -2,433 | 1,278E-15 |
| TcCLB.505621.10 | mucin-associated surface protein (MASP, pseudogene), pu | -2,434 | 8,331E-136 |
| TcCLB.506321.130 | mucin TcMUCII, putative | -2,434 | 1,222E-23 |
| TcCLB.506499.40 | mucin-associated surface protein (MASP), putative | -2,436 | 1,828E-29 |
| TcCLB.509753.110 | mucin TcMUCII, putative | -2,436 | 4,69E-14 |
| TcCLB.511255.330 | mucin-associated surface protein (MASP), putative | -2,437 | 3,063E-12 |
| TcCLB.507701.4 | mucin-associated surface protein (MASP), putative (fragm | -2,438 | 1,27E-21 |
| TcCLB.503783.70 | mucin TcMUCII, putative | -2,44 | 2,582E-31 |
| TcCLB.432443.10 | mucin-associated surface protein (MASP), putative | -2,441 | 3,713E-26 |
| TcCLB.504039.160 | mucin-associated surface protein (MASP), putative | -2,443 | 1,797E-10 |
| TcCLB.511553.100 | mucin-associated surface protein (MASP), putative | -2,444 | 3,124E-41 |
| TcCLB.511593.20 | mucin TcMUCII, putative | -2,445 | 7,132E-25 |
| TcCLB.508501.70 | mucin TcMUCII, putative | -2,446 | 8,866E-43 |
| TcCLB.509287.220 | hypothetical protein, conserved | -2,447 | 3,446E-130 |
| TcCLB.508243.60 | mucin TcMUCII, putative | -2,447 | 1,347E-18 |
| TcCLB.508745.20 | mucin-associated surface protein (MASP), putative | -2,45 | 1,568E-41 |
| TcCLB.510047.100 | mucin-associated surface protein (MASP, pseudogene), pu | -2,45 | 7,788E-41 |
| TcCLB.510583.114 | mucin-associated surface protein (MASP, pseudogene), pu | -2,45 | 0,00001299 |
| TcCLB.506309.110 | mucin TcMUCII, putative | -2,451 | 2,23E-18 |
| TcCLB.510693.120 | trans-sialidase (pseudogene), putative | -2,452 | 1,505E-29 |
| TcCLB.506769.70 | mucin TcMUCII, putative | -2,454 | 8,892E-23 |
| TcCLB.508365.30 | mucin TcMUCII, putative | -2,455 | 8,496E-26 |
| TcCLB.510623.39 | mucin-associated surface protein (MASP, pseudogene), pu | -2,455 | 1,476E-09 |
| TcCLB.508365.4 | mucin-associated surface protein (MASP), putative | -2,456 | 1,974E-14 |
| TcCLB.508161.23 | hypothetical protein | -2,456 | 1,133E-12 |
| TcCLB.508365.80 | hypothetical protein | -2,456 | 0,0001245 |
| TcCLB.504081.390 | trans-sialidase (pseudogene), putative | -2,457 | 9,24E-45 |
| TcCLB.511183.190 | trans-sialidase, Group V, putative | -2,457 | 1,212E-31 |
| TcCLB.507091.100 | trans-sialidase (pseudogene), putative | -2,457 | 1,092E-30 |
| TcCLB.509993.40 | mucin TcMUCI, putative | -2,458 | 2,922E-30 |
| TcCLB.509897.279 | mucin-associated surface protein (MASP), putative | -2,458 | 2,558E-24 |
| TcCLB.509897.170 | mucin-associated surface protein (MASP, pseudogene), pu | -2,459 | 2,595E-63 |
| TcCLB.510189.40 | mucin-associated surface protein (MASP), putative | -2,459 | 3,382E-37 |
| TcCLB.511839.30 | mucin-associated surface protein (MASP), putative | -2,459 | 2,18E-25 |
| TcCLB.507959.140 | mucin TcMUCII, putative | -2,459 | 0,0000223 |
| TcCLB.507047.40 | trans-sialidase, Group II, putative | -2,46 | 1,398E-123 |
| TcCLB.503645.10 | mucin-associated surface protein (MASP), putative | -2,46 | 3,12E-28 |
| TcCLB.507071.180 | mucin-associated surface protein (MASP), putative | -2,461 | 1,173E-39 |
| TcCLB.509525.150 | mucin TcMUCII, putative | -2,462 | 6,433E-20 |
| TcCLB.511853.10 | mucin TcMUCI, putative | -2,464 | 4,972E-18 |
| TcCLB.506759.170 | mucin-associated surface protein (MASP), putative | -2,467 | 1,12E-36 |
| TcCLB.510047.129 | mucin TcMUCII, putative | -2,47 | 3,552E-09 |

| | | | |
|---|---|---|---|
| TcCLB.511173.309 | mucin-associated surface protein (MASP, pseudogene), pu | -2,474 | 1,801E-20 |
| TcCLB.509267.110 | heat shock protein Hsp20, putative | -2,475 | 2,702E-80 |
| TcCLB.509081.150 | mucin-associated surface protein (MASP), putative | -2,475 | 3,322E-46 |
| TcCLB.510483.250 | trans-sialidase (pseudogene), putative | -2,475 | 4,184E-39 |
| TcCLB.507237.20 | mucin-associated surface protein (MASP), putative | -2,475 | 7,202E-17 |
| TcCLB.503973.320 | mucin TcMUCII, putative | -2,476 | 1,564E-10 |
| TcCLB.506501.160 | mucin-associated surface protein (MASP, pseudogene), pu | -2,477 | 1,979E-28 |
| TcCLB.510021.130 | mucin-associated surface protein (MASP), putative | -2,478 | 2,193E-71 |
| TcCLB.508229.30 | mucin-associated surface protein (MASP), putative | -2,478 | 1,216E-49 |
| TcCLB.509525.10 | mucin-associated surface protein (MASP, pseudogene), pu | -2,478 | 5,164E-19 |
| TcCLB.507605.26 | mucin TcMUCII, putative | -2,478 | 1,644E-08 |
| TcCLB.506499.160 | mucin-associated surface protein (MASP), putative | -2,479 | 5,809E-43 |
| TcCLB.507957.50 | mucin TcMUCII, putative | -2,48 | 1,074E-37 |
| TcCLB.511599.10 | trans-sialidase, putative | -2,481 | 1,158E-10 |
| TcCLB.510371.110 | mucin TcMUCII, putative | -2,484 | 6,474E-13 |
| TcCLB.508495.50 | mucin TcMUCII, putative | -2,488 | 3,08E-64 |
| TcCLB.508119.70 | mucin TcMUCII, putative | -2,489 | 1,508E-86 |
| TcCLB.510015.20 | mucin TcMUCII, putative | -2,49 | 1,359E-30 |
| TcCLB.506763.80 | mucin TcMUCII, putative | -2,49 | 2,932E-29 |
| TcCLB.506599.80 | mucin-associated surface protein (MASP), putative | -2,492 | 3,284E-19 |
| TcCLB.506897.10 | mucin TcMUCII, putative | -2,492 | 1,527E-16 |
| TcCLB.509419.160 | mucin TcMUCII, putative | -2,492 | 1,026E-11 |
| TcCLB.506763.280 | mucin-associated surface protein (MASP), putative | -2,493 | 3,048E-61 |
| TcCLB.509081.110 | mucin-associated surface protein (MASP), putative | -2,493 | 2,045E-38 |
| TcCLB.504035.21 | hypothetical protein | -2,493 | 6,211E-17 |
| TcCLB.508577.10 | trans-sialidase (pseudogene), putative | -2,494 | 7,347E-40 |
| TcCLB.508495.200 | mucin TcMUCII, putative | -2,496 | 3,253E-106 |
| TcCLB.506321.10 | mucin TcMUCII, putative | -2,496 | 2,084E-31 |
| TcCLB.509753.20 | trans-sialidase (pseudogene), putative | -2,497 | 3,154E-65 |
| TcCLB.504031.60 | mucin-associated surface protein (MASP, pseudogene), pu | -2,497 | 1,216E-34 |
| TcCLB.507429.3 | mucin-associated surface protein (MASP), putative | -2,497 | 4,954E-29 |
| TcCLB.509411.10 | trans-sialidase, Group II, putative | -2,497 | 3,645E-22 |
| TcCLB.510497.50 | mucin-associated surface protein (MASP), putative | -2,498 | 4,142E-43 |
| TcCLB.508493.130 | mucin TcMUCII, putative | -2,5 | 2,72E-54 |
| TcCLB.504639.60 | trans-sialidase, putative | -2,5 | 1,091E-53 |
| TcCLB.507523.70 | trans-sialidase, Group V, putative | -2,5 | 3,238E-27 |
| TcCLB.510419.11 | mucin-associated surface protein (MASP, pseudogene), pu | -2,501 | 1,238E-08 |
| TcCLB.510463.200 | mucin-associated surface protein (MASP), putative | -2,502 | 3,006E-36 |
| TcCLB.510013.310 | mucin-associated surface protein (MASP), putative | -2,503 | 5,275E-50 |
| TcCLB.510585.10 | mucin-associated surface protein (MASP, pseudogene), pu | -2,503 | 4,666E-14 |
| TcCLB.511871.120 | mucin-associated surface protein (MASP), putative | -2,505 | 3,498E-41 |
| TcCLB.511223.20 | trans-sialidase, putative | -2,505 | 1,415E-11 |
| TcCLB.508589.5 | trans-sialidase, putative (fragment) | -2,506 | 1,596E-33 |
| TcCLB.510275.330 | mucin TcMUCII, putative | -2,508 | 1,046E-22 |
| TcCLB.506281.50 | mucin TcMUCII, putative | -2,511 | 1,589E-82 |

| | | | |
|---|---|---|---|
| TcCLB.511605.20 | mucin-associated surface protein (MASP), putative | -2,513 | 3,28E-32 |
| TcCLB.511077.40 | trans-sialidase (pseudogene), putative | -2,516 | 6,368E-22 |
| TcCLB.453767.20 | trans-sialidase (pseudogene), putative | -2,517 | 2,237E-40 |
| TcCLB.509233.10 | hypothetical protein | -2,517 | 1,302E-10 |
| TcCLB.510483.60 | trans-sialidase, Group V, putative | -2,519 | 3,188E-24 |
| TcCLB.504787.60 | mucin TcMUCII (pseudogene), putative | -2,52 | 4,368E-34 |
| TcCLB.510483.230 | mucin-associated surface protein (MASP), putative | -2,52 | 1,738E-26 |
| TcCLB.510275.370 | mucin-associated surface protein (MASP), putative | -2,523 | 3,451E-63 |
| TcCLB.506667.160 | trans-sialidase, Group V, putative | -2,523 | 5,856E-23 |
| TcCLB.504261.100 | mucin-associated surface protein (MASP, pseudogene), pu | -2,523 | 3,428E-20 |
| TcCLB.511593.70 | mucin TcMUCII, putative | -2,523 | 1,46E-17 |
| TcCLB.503585.10 | mucin-associated surface protein (MASP), putative | -2,524 | 3,355E-14 |
| TcCLB.506131.10 | mucin TcMUCII, putative | -2,525 | 1,018E-15 |
| TcCLB.506009.40 | hypothetical protein | -2,526 | 6,807E-70 |
| TcCLB.506813.70 | mucin-associated surface protein (MASP, pseudogene), pu | -2,526 | 4,409E-22 |
| TcCLB.440379.10 | trans-sialidase, putative (fragment) | -2,527 | 7,318E-48 |
| TcCLB.510329.70 | cdc2-related kinase 10, putative (CRK10) | -2,528 | 4,937E-79 |
| TcCLB.506929.54 | hypothetical protein | -2,528 | 2,203E-59 |
| TcCLB.510921.40 | trans-sialidase, putative (fragment) | -2,532 | 1,789E-28 |
| TcCLB.506781.39 | mucin-associated surface protein (MASP), putative | -2,532 | 1,198E-12 |
| TcCLB.509475.10 | mitogen-activated protein kinase 3, putative | -2,533 | 1,188E-43 |
| TcCLB.508221.420 | mucin-associated surface protein (MASP), putative | -2,533 | 1,144E-16 |
| TcCLB.509291.20 | mucin-associated surface protein (MASP), putative | -2,533 | 1,95E-15 |
| TcCLB.511553.209 | mucin TcMUC (pseudogene), putative | -2,534 | 1,377E-27 |
| TcCLB.510279.120 | mucin-associated surface protein (MASP), putative | -2,534 | 5,767E-18 |
| TcCLB.508171.10 | trans-sialidase, Group VIII, putative | -2,536 | 1,288E-29 |
| TcCLB.506667.90 | trans-sialidase, Group V, putative | -2,536 | 1,892E-19 |
| TcCLB.508293.10 | trans-sialidase (pseudogene), putative | -2,537 | 2,059E-34 |
| TcCLB.511123.11 | trans-sialidase, putative (fragment) | -2,538 | 5,847E-41 |
| TcCLB.508309.10 | mucin-associated surface protein (MASP), putative | -2,538 | 9,345E-23 |
| TcCLB.511511.5 | hypothetical protein | -2,539 | 1,98E-24 |
| TcCLB.508165.50 | mucin-associated surface protein (MASP), putative | -2,539 | 1,664E-12 |
| TcCLB.508365.240 | mucin TcMUCII, putative | -2,539 | 4,21E-10 |
| TcCLB.506321.140 | mucin-associated surface protein (MASP), putative | -2,54 | 2,009E-56 |
| TcCLB.506965.100 | mucin-associated surface protein (MASP), putative | -2,542 | 3,019E-31 |
| TcCLB.507487.30 | mucin TcMUCII, putative | -2,542 | 1,428E-14 |
| TcCLB.510021.190 | hypothetical protein | -2,542 | 0,00000066 |
| TcCLB.510933.20 | mucin-associated surface protein (MASP), putative | -2,547 | 1,193E-34 |
| TcCLB.508049.40 | hypothetical protein, conserved | -2,547 | 1,289E-15 |
| TcCLB.506131.30 | mucin-associated surface protein (MASP), putative | -2,55 | 6,974E-19 |
| TcCLB.506783.39 | mucin-associated surface protein (MASP), putative | -2,551 | 7,856E-20 |
| TcCLB.510369.10 | mucin-associated surface protein (MASP), putative | -2,553 | 3,795E-32 |
| TcCLB.504155.170 | mucin TcMUCII, putative | -2,553 | 3,721E-15 |
| TcCLB.510465.24 | mucin-associated surface protein (MASP, pseudogene), pu | -2,554 | 7,623E-10 |
| TcCLB.508219.170 | trans-sialidase (pseudogene), putative | -2,556 | 6,903E-29 |

| | | | |
|---|---|---|---|
| TcCLB.507163.11 | mucin TcMUCII, putative (fragment) | -2,556 | 1,193E-15 |
| TcCLB.505945.5 | mitogen-activated protein kinase 3, putative (fragment) | -2,561 | 2,319E-21 |
| TcCLB.506759.210 | mucin TcMUCII, putative | -2,561 | 2,022E-15 |
| TcCLB.510799.60 | mucin TcMUCII, putative | -2,566 | 1,979E-47 |
| TcCLB.510013.300 | mucin TcMUCII, putative | -2,568 | 1,326E-15 |
| TcCLB.506879.10 | mucin-associated surface protein (MASP), putative | -2,57 | 4,462E-22 |
| TcCLB.510557.70 | mucin TcMUCII, putative | -2,571 | 1,939E-61 |
| TcCLB.508163.230 | mucin-associated surface protein (MASP), putative | -2,571 | 3,041E-17 |
| TcCLB.510705.10 | mucin-associated surface protein (MASP), putative | -2,571 | 2,269E-09 |
| TcCLB.510933.190 | mucin TcMUCII, putative | -2,572 | 8,87E-25 |
| TcCLB.506501.120 | mucin TcMUCII, putative | -2,572 | 6,434E-17 |
| TcCLB.509549.140 | trans-sialidase, putative | -2,573 | 7,637E-68 |
| TcCLB.508201.30 | mucin-associated surface protein (MASP), putative | -2,574 | 4,021E-80 |
| TcCLB.507085.140 | mucin TcMUCII, putative | -2,574 | 3,289E-35 |
| TcCLB.510461.30 | mucin-associated surface protein (MASP), putative (fragm | -2,574 | 0,00000262 |
| TcCLB.508735.30 | mucin-associated surface protein (MASP), putative | -2,575 | 9,358E-85 |
| TcCLB.506955.244 | mucin-associated surface protein (MASP), putative | -2,576 | 3,088E-37 |
| TcCLB.506267.60 | mucin TcMUCII, putative | -2,576 | 2,65E-29 |
| TcCLB.504239.20 | mucin TcMUCII, putative | -2,576 | 8,823E-26 |
| TcCLB.507959.260 | mucin-associated surface protein (MASP), putative | -2,576 | 8,018E-22 |
| TcCLB.510377.30 | mucin-associated surface protein (MASP, pseudogene), pu | -2,576 | 9,436E-20 |
| TcCLB.510303.140 | surface protease GP63, putative | -2,577 | 5,369E-160 |
| TcCLB.511079.60 | mucin-associated surface protein (MASP), putative | -2,577 | 4,605E-21 |
| TcCLB.506001.50 | mucin-associated surface protein (MASP), putative | -2,578 | 1,302E-84 |
| TcCLB.510303.20 | surface protease GP63, putative | -2,579 | 3,142E-48 |
| TcCLB.473703.10 | surface protease GP63, putative | -2,579 | 4,255E-15 |
| TcCLB.510377.310 | mucin-associated surface protein (MASP, pseudogene), pu | -2,58 | 6,485E-20 |
| TcCLB.506599.240 | mucin-associated surface protein (MASP), putative | -2,582 | 4,513E-73 |
| TcCLB.508509.20 | mucin TcMUCII, putative | -2,582 | 2,703E-23 |
| TcCLB.506599.390 | mucin-associated surface protein (MASP), putative | -2,582 | 1,959E-21 |
| TcCLB.504441.10 | trans-sialidase, putative (fragment) | -2,583 | 5,698E-21 |
| TcCLB.510199.30 | mucin-associated surface protein (MASP), putative | -2,584 | 2,065E-46 |
| TcCLB.511603.200 | mucin TcMUCII, putative | -2,585 | 1,293E-25 |
| TcCLB.503705.19 | trans-sialidase, putative | -2,585 | 3,599E-08 |
| TcCLB.476833.10 | trans-sialidase, putative | -2,586 | 4,256E-13 |
| TcCLB.508511.49 | trans-sialidase, putative | -2,588 | 2,935E-37 |
| TcCLB.506297.60 | hypothetical protein, conserved | -2,589 | 3,171E-133 |
| TcCLB.508501.40 | mucin TcMUCII, putative | -2,589 | 5,803E-63 |
| TcCLB.508123.10 | hypothetical protein, conserved | -2,59 | 2,968E-75 |
| TcCLB.508125.130 | mucin TcMUCII, putative | -2,59 | 1,106E-24 |
| TcCLB.508521.20 | trans-sialidase, Group VIII, putative | -2,592 | 9,704E-71 |
| TcCLB.503503.30 | mucin TcMUCII, putative | -2,592 | 7,257E-70 |
| TcCLB.508903.10 | trans-sialidase (pseudogene), putative | -2,592 | 2,811E-40 |
| TcCLB.510463.259 | mucin-associated surface protein (MASP), putative (fragm | -2,592 | 3,528E-33 |
| TcCLB.459061.10 | trans-sialidase, putative | -2,593 | 1,215E-46 |

| | | | |
|---|---|---|---|
| TcCLB.510713.110 | mucin-associated surface protein (MASP), putative | -2,594 | 7,898E-24 |
| TcCLB.506289.250 | surface protease GP63 (pseudogene), putative | -2,595 | 1,175E-26 |
| TcCLB.511669.10 | trans-sialidase, Group VIII, putative | -2,597 | 4,504E-86 |
| TcCLB.507961.10 | mucin-associated surface protein (MASP, pseudogene), pu | -2,597 | 2,818E-31 |
| TcCLB.509419.80 | mucin-associated surface protein (MASP), putative | -2,597 | 6,116E-08 |
| TcCLB.508289.20 | mucin TcMUCII, putative | -2,598 | 3,622E-11 |
| TcCLB.507069.120 | trans-sialidase (pseudogene), putative | -2,599 | 1,343E-39 |
| TcCLB.511595.30 | mucin-associated surface protein (MASP), putative | -2,6 | 4,482E-23 |
| TcCLB.510377.390 | mucin-associated surface protein (MASP), putative | -2,601 | 1,032E-23 |
| TcCLB.510933.10 | trans-sialidase, Group V, putative | -2,604 | 1,769E-39 |
| TcCLB.506601.40 | mucin-associated surface protein (MASP), putative | -2,604 | 3,24E-25 |
| TcCLB.510773.40 | hypothetical protein | -2,605 | 1,451E-54 |
| TcCLB.510699.80 | mucin-associated surface protein (MASP), putative | -2,605 | 6,245E-24 |
| TcCLB.506885.480 | trans-sialidase, Group V, putative | -2,607 | 2,531E-43 |
| TcCLB.509553.60 | mitogen-activated protein kinase 3, putative | -2,608 | 1,221E-80 |
| TcCLB.506173.120 | mucin TcMUCI, putative | -2,608 | 5,48E-25 |
| TcCLB.508247.70 | mucin TcMUCII, putative | -2,61 | 3,877E-19 |
| TcCLB.503501.7 | mucin-associated surface protein (MASP), putative (fragm | -2,61 | 3,18E-08 |
| TcCLB.507429.30 | mucin TcMUCII, putative | -2,611 | 2,43E-79 |
| TcCLB.507035.11 | trans-sialidase, putative (fragment) | -2,611 | 1,08E-41 |
| TcCLB.429251.10 | mucin-associated surface protein (MASP), putative | -2,611 | 2,192E-06 |
| TcCLB.504429.50 | trans-sialidase (pseudogene), putative | -2,614 | 1,047E-53 |
| TcCLB.511607.110 | mucin-associated surface protein (MASP), putative | -2,617 | 6,37E-57 |
| TcCLB.511605.5 | mucin-associated surface protein (MASP, pseudogene), pu | -2,617 | 1,029E-31 |
| TcCLB.510025.90 | mucin TcMUCII, putative | -2,617 | 2,281E-27 |
| TcCLB.503717.31 | mucin TcMUCII, putative | -2,618 | 1,966E-23 |
| TcCLB.503837.5 | mucin TcMUCI, putative (fragment) | -2,619 | 8,219E-51 |
| TcCLB.506615.150 | trans-sialidase, Group V, putative | -2,62 | 1,472E-148 |
| TcCLB.511063.40 | trans-sialidase, Group V, putative | -2,62 | 1,003E-24 |
| TcCLB.509097.20 | trans-sialidase, Group V, putative | -2,621 | 1,522E-49 |
| TcCLB.506599.224 | mucin TcMUCII, putative | -2,622 | 1,964E-19 |
| TcCLB.510021.10 | mucin TcMUCII, putative | -2,622 | 6,046E-19 |
| TcCLB.442801.10 | mucin TcMUCII, putative | -2,622 | 1,743E-18 |
| TcCLB.506939.180 | mucin-associated surface protein (MASP), putative | -2,623 | 1,76E-60 |
| TcCLB.511105.60 | trans-sialidase, putative | -2,624 | 5,444E-75 |
| TcCLB.506787.19 | mucin-associated surface protein (MASP, pseudogene), pu | -2,624 | 9,955E-14 |
| TcCLB.509115.30 | mucin-associated surface protein (MASP), putative | -2,625 | 1,76E-51 |
| TcCLB.506101.10 | mucin-associated surface protein (MASP), putative | -2,626 | 2,72E-42 |
| TcCLB.509419.90 | trans-sialidase, Group V, putative | -2,626 | 2,201E-08 |
| TcCLB.507071.210 | mucin TcMUCII, putative | -2,627 | 1,026E-16 |
| TcCLB.508143.80 | mucin-associated surface protein (MASP), putative | -2,63 | 6,589E-36 |
| TcCLB.510359.40 | hypothetical protein, conserved | -2,631 | 3,102E-43 |
| TcCLB.509525.160 | mucin-associated surface protein (MASP, pseudogene), pu | -2,632 | 1,057E-32 |
| TcCLB.508243.50 | mucin-associated surface protein (MASP), putative | -2,632 | 3,911E-26 |
| TcCLB.510791.20 | mucin TcMUCII, putative | -2,633 | 1,542E-62 |

| | | | |
|---|---|---|---|
| TcCLB.510363.160 | mucin-associated surface protein (MASP, pseudogene), pu | -2,633 | 1,327E-42 |
| TcCLB.510275.400 | mucin-associated surface protein (MASP), putative | -2,634 | 6,195E-39 |
| TcCLB.506443.100 | trans-sialidase, Group IV, putative | -2,636 | 1,95E-98 |
| TcCLB.503761.40 | mucin-associated surface protein (MASP), putative | -2,636 | 2,777E-90 |
| TcCLB.510237.80 | trans-sialidase, Group V, putative | -2,637 | 1,182E-46 |
| TcCLB.507069.94 | mucin-associated surface protein (MASP), putative | -2,637 | 2,005E-32 |
| TcCLB.507985.30 | trans-sialidase (pseudogene), putative | -2,639 | 7,003E-07 |
| TcCLB.506855.220 | surface protease GP63, putative | -2,641 | 1,8E-16 |
| TcCLB.506801.60 | mucin-like glycoprotein, putative | -2,642 | 1,238E-55 |
| TcCLB.506499.220 | 90 kDa surface protein, putative | -2,642 | 2,748E-39 |
| TcCLB.511427.10 | conserved protein, unknown function (fragment) | -2,642 | 4,44E-07 |
| TcCLB.509479.20 | mucin TcMUCII, putative | -2,645 | 1,097E-28 |
| TcCLB.510089.10 | mucin-associated surface protein (MASP), putative | -2,646 | 4,311E-36 |
| TcCLB.507071.250 | mucin TcMUCII, putative | -2,646 | 1,062E-22 |
| TcCLB.510697.80 | mucin-associated surface protein (MASP), putative | -2,648 | 3,843E-82 |
| TcCLB.504081.460 | mucin-associated surface protein (MASP), putative | -2,648 | 5,62E-53 |
| TcCLB.480997.10 | hypothetical protein, conserved | -2,649 | 3,337E-10 |
| TcCLB.509811.18 | mucin-associated surface protein (MASP, pseudogene), pu | -2,65 | 3,752E-31 |
| TcCLB.506599.360 | mucin-associated surface protein (MASP, pseudogene), pu | -2,651 | 1,004E-35 |
| TcCLB.506291.10 | mucin-associated surface protein (MASP), putative | -2,653 | 2,519E-37 |
| TcCLB.509003.40 | calpain-like cysteine peptidase, putative | -2,654 | 7,462E-53 |
| TcCLB.508451.60 | trans-sialidase, Group II, putative | -2,654 | 8,374E-38 |
| TcCLB.506339.40 | mucin-associated surface protein (MASP), putative | -2,654 | 6,465E-17 |
| TcCLB.511257.90 | mucin-associated surface protein (MASP), putative | -2,655 | 4,711E-51 |
| TcCLB.511099.70 | mucin TcMUCII, putative | -2,655 | 1,005E-12 |
| TcCLB.510681.9 | surface protease GP63, putative | -2,657 | 2,032E-27 |
| TcCLB.506331.200 | trans-sialidase (pseudogene), putative | -2,658 | 1,743E-37 |
| TcCLB.508295.50 | mucin TcMUCII, putative | -2,659 | 6,635E-24 |
| TcCLB.510359.460 | mucin-associated surface protein (MASP), putative | -2,659 | 1,646E-12 |
| TcCLB.511255.200 | mucin-associated surface protein (MASP), putative | -2,66 | 4,92E-29 |
| TcCLB.506759.220 | mucin-associated surface protein (MASP), putative | -2,66 | 3,691E-22 |
| TcCLB.506769.30 | mucin-associated surface protein (MASP), putative | -2,661 | 2,026E-60 |
| TcCLB.509079.20 | mucin-associated surface protein (MASP), putative | -2,663 | 5,429E-33 |
| TcCLB.511083.39 | mucin-associated surface protein (MASP), putative | -2,664 | 6,666E-44 |
| TcCLB.511843.10 | mucin-associated surface protein (MASP), putative | -2,669 | 1,722E-35 |
| TcCLB.506765.40 | trans-sialidase, Group II, putative | -2,669 | 2,091E-24 |
| TcCLB.503501.70 | mucin-associated surface protein (MASP), putative (fragm | -2,671 | 4,437E-42 |
| TcCLB.511183.80 | mucin-associated surface protein (MASP), putative | -2,671 | 1,718E-30 |
| TcCLB.511183.400 | trans-sialidase, Group VI, putative | -2,672 | 5,433E-59 |
| TcCLB.505343.10 | trans-sialidase (pseudogene), putative | -2,673 | 2,276E-33 |
| TcCLB.504235.9 | hypothetical protein, conserved (fragment) | -2,674 | 8,933E-56 |
| TcCLB.510693.190 | mucin-associated surface protein (MASP), putative | -2,674 | 2,326E-30 |
| TcCLB.508873.520 | mucin-associated surface protein (MASP, pseudogene), pu | -2,675 | 1,628E-26 |
| TcCLB.506767.74 | mucin TcMUCII (pseudogene), putative | -2,676 | 2,086E-18 |
| TcCLB.511385.50 | hypothetical protein | -2,677 | 5,6E-71 |

| | | | |
|---|---|---|---|
| TcCLB.503717.120 | trans-sialidase, Group V, putative | -2,677 | 4,757E-45 |
| TcCLB.506609.30 | trans-sialidase, Group V, putative | -2,677 | 6,012E-12 |
| TcCLB.506459.149 | mucin-associated surface protein (MASP), putative | -2,678 | 6,889E-97 |
| TcCLB.505297.70 | mucin TcMUCII, putative | -2,678 | 8,626E-74 |
| TcCLB.506901.10 | mucin TcMUCI, putative | -2,678 | 1,846E-07 |
| TcCLB.511089.140 | mucin-associated surface protein (MASP), putative | -2,679 | 3,345E-26 |
| TcCLB.505949.80 | mucin-associated surface protein (MASP), putative | -2,681 | 2,55E-50 |
| TcCLB.504201.50 | mucin-associated surface protein (MASP), putative | -2,681 | 1,173E-35 |
| TcCLB.511643.70 | trans-sialidase, Group VI, putative | -2,681 | 1,215E-17 |
| TcCLB.510693.230 | mucin-associated surface protein (MASP), putative | -2,682 | 3,064E-24 |
| TcCLB.507237.160 | mucin TcMUCII, putative | -2,682 | 9,423E-12 |
| TcCLB.510239.60 | mucin TcMUCII, putative | -2,683 | 1,337E-17 |
| TcCLB.506133.170 | mucin-associated surface protein (MASP, pseudogene), pu | -2,684 | 6,108E-48 |
| TcCLB.508871.110 | trans-sialidase, Group V, putative | -2,685 | 3,794E-36 |
| TcCLB.508885.5 | mucin-associated surface protein (MASP), putative | -2,685 | 7,992E-07 |
| TcCLB.506781.19 | mucin-associated surface protein (MASP), putative | -2,686 | 8,424E-25 |
| TcCLB.507723.50 | trans-sialidase, Group IV, putative | -2,687 | 1,56E-33 |
| TcCLB.510373.70 | mucin TcMUCII, putative | -2,687 | 9,943E-17 |
| TcCLB.508433.180 | mucin-associated surface protein (MASP), putative | -2,692 | 1,388E-64 |
| TcCLB.511603.320 | mucin-associated surface protein (MASP), putative | -2,692 | 1,835E-21 |
| TcCLB.510191.80 | mucin TcMUCII, putative | -2,694 | 1,431E-18 |
| TcCLB.511871.150 | trans-sialidase (pseudogene), putative | -2,697 | 1,72E-70 |
| TcCLB.508365.210 | mucin TcMUCI, putative | -2,7 | 4,715E-13 |
| TcCLB.510915.10 | mucin-associated surface protein (MASP), putative | -2,701 | 7,256E-104 |
| TcCLB.510017.59 | mucin-associated surface protein (MASP), putative | -2,701 | 2,69E-67 |
| TcCLB.487455.10 | trans-sialidase, putative (fragment) | -2,702 | 5,028E-15 |
| TcCLB.507167.80 | trans-sialidase, Group II, putative | -2,705 | 1,56E-45 |
| TcCLB.510457.10 | mucin-associated surface protein (MASP), putative | -2,705 | 2,173E-16 |
| TcCLB.509869.10 | mucin TcMUCI, putative | -2,707 | 1,156E-31 |
| TcCLB.510693.280 | mucin-associated surface protein (MASP), putative | -2,709 | 2,872E-83 |
| TcCLB.506501.70 | mucin TcMUCII, putative | -2,709 | 3,77E-36 |
| TcCLB.506501.240 | mucin-associated surface protein (MASP), putative | -2,709 | 6,555E-16 |
| TcCLB.421621.50 | mucin-associated surface protein (MASP), putative | -2,709 | 2,141E-14 |
| TcCLB.509539.10 | trans-sialidase (pseudogene), putative | -2,71 | 1,505E-94 |
| TcCLB.504873.10 | UDP-Gal or UDP-GlcNAc-dependent glycosyltransferase, p | -2,71 | 3,148E-76 |
| TcCLB.509905.150 | trans-sialidase, Group II, putative | -2,711 | 7,778E-297 |
| TcCLB.508121.80 | mucin TcMUCII, putative | -2,711 | 1,404E-17 |
| TcCLB.508293.60 | trans-sialidase, Group VI, putative | -2,712 | 5,281E-49 |
| TcCLB.507959.200 | mucin TcMUCII, putative | -2,712 | 1,369E-30 |
| TcCLB.507859.15 | hypothetical protein | -2,714 | 1,604E-40 |
| TcCLB.510201.40 | trans-sialidase (pseudogene), putative | -2,717 | 2,8E-69 |
| TcCLB.510377.150 | mucin TcMUCII, putative | -2,717 | 1,67E-08 |
| TcCLB.511851.50 | mucin TcMUCI, putative | -2,719 | 8,34E-37 |
| TcCLB.507163.30 | 90 kDa surface protein, putative | -2,722 | 4,536E-26 |
| TcCLB.504575.60 | surface protease GP63, putative (fragment) | -2,724 | 3,576E-09 |

| | | | |
|---|---|---|---|
| TcCLB.506661.160 | hypothetical protein, conserved | -2,726 | 3,398E-52 |
| TcCLB.506587.44 | hypothetical protein | -2,727 | 4,352E-100 |
| TcCLB.510377.450 | mucin-associated surface protein (MASP), putative | -2,727 | 1,812E-49 |
| TcCLB.507711.100 | procyclic form surface phosphoprotein, putative | -2,728 | 1,663E-129 |
| TcCLB.504039.90 | mucin-associated surface protein (MASP, pseudogene), pu | -2,73 | 1,626E-35 |
| TcCLB.508221.660 | mucin-associated surface protein (MASP), putative | -2,732 | 1,58E-18 |
| TcCLB.506767.220 | mucin TcMUCII, putative | -2,733 | 1,42E-26 |
| TcCLB.511151.110 | surface protease GP63, putative (fragment) | -2,734 | 1,33E-209 |
| TcCLB.503491.19 | mucin-associated surface protein (MASP), putative | -2,734 | 6,302E-29 |
| TcCLB.508501.140 | mucin TcMUCII, putative | -2,735 | 7,091E-87 |
| TcCLB.503811.45 | hypothetical protein, conserved | -2,735 | 7,518E-66 |
| TcCLB.507705.30 | hypothetical protein, conserved | -2,736 | 5,862E-61 |
| TcCLB.506573.5 | surface antigen TASV-A8, putative (TASV-A8) | -2,737 | 1,034E-227 |
| TcCLB.511153.40 | surface protease GP63, putative | -2,737 | 4,212E-165 |
| TcCLB.509975.30 | mucin-associated surface protein (MASP), putative | -2,738 | 4,184E-151 |
| TcCLB.508873.330 | trans-sialidase, Group V, putative | -2,738 | 9,665E-67 |
| TcCLB.510833.20 | mucin-associated surface protein (MASP), putative | -2,741 | 1,117E-61 |
| TcCLB.511449.80 | mucin-associated surface protein (MASP, pseudogene), pu | -2,744 | 2,662E-14 |
| TcCLB.506285.30 | trans-sialidase (pseudogene), putative | -2,745 | 4,015E-69 |
| TcCLB.503501.14 | mucin TcMUCII, putative | -2,746 | 2,21E-29 |
| TcCLB.511183.430 | trans-sialidase, Group V, putative | -2,749 | 1,091E-41 |
| TcCLB.511885.20 | trans-sialidase, Group II, putative | -2,75 | 6,798E-87 |
| TcCLB.511149.30 | mucin TcMUCII (pseudogene), putative | -2,75 | 1,047E-21 |
| TcCLB.510693.240 | trans-sialidase, Group VI, putative | -2,753 | 8,754E-31 |
| TcCLB.508541.130 | mucin-associated surface protein (MASP), putative | -2,754 | 1,121E-123 |
| TcCLB.506545.5 | mucin TcMUCI, putative | -2,758 | 1,233E-53 |
| TcCLB.511511.6 | zinc finger protein ZFP1 | -2,76 | 6,439E-19 |
| TcCLB.509653.10 | mucin-associated surface protein (MASP), putative | -2,761 | 3,01E-17 |
| TcCLB.510805.5 | trans-sialidase, putative | -2,761 | 6,466E-17 |
| TcCLB.510021.100 | mucin TcMUCII, putative | -2,762 | 2,515E-26 |
| TcCLB.508607.50 | trans-sialidase, Group II, putative | -2,764 | 1,605E-120 |
| TcCLB.508151.20 | mucin-associated surface protein (MASP), putative | -2,764 | 7,377E-29 |
| TcCLB.508313.90 | mucin TcMUCII, putative | -2,766 | 4,448E-65 |
| TcCLB.506133.80 | mucin-associated surface protein (MASP, pseudogene), pu | -2,766 | 4,555E-54 |
| TcCLB.505949.120 | mucin TcMUCII, putative | -2,766 | 5,698E-29 |
| TcCLB.509905.200 | trans-sialidase, Group II, putative | -2,768 | 7,189E-216 |
| TcCLB.504239.420 | mucin-associated surface protein (MASP), putative | -2,768 | 2,133E-52 |
| TcCLB.508365.280 | mucin TcMUCII, putative | -2,768 | 3,115E-25 |
| TcCLB.511121.30 | hypothetical protein | -2,769 | 5,939E-19 |
| TcCLB.511851.37 | mucin TcMUCI, putative | -2,77 | 2,676E-52 |
| TcCLB.511185.120 | mucin-associated surface protein (MASP), putative | -2,77 | 1,278E-36 |
| TcCLB.509581.10 | trans-sialidase, Group VII, putative | -2,771 | 5E-12 |
| TcCLB.511347.20 | mucin-associated surface protein (MASP, pseudogene), pu | -2,775 | 1,406E-113 |
| TcCLB.506139.80 | mucin TcMUCII, putative | -2,776 | 2,91E-21 |
| TcCLB.506281.134 | mucin-associated surface protein (MASP), putative | -2,776 | 1,312E-18 |

| | | | |
|---|---|---|---|
| TcCLB.508101.9 | trans-sialidase, putative (fragment) | -2,778 | 1,264E-21 |
| TcCLB.506965.70 | mucin-associated surface protein (MASP), putative | -2,78 | 2,206E-53 |
| TcCLB.507237.280 | mucin TcMUCII, putative | -2,78 | 1,029E-06 |
| TcCLB.510303.10 | surface protease GP63, putative | -2,781 | 1,943E-93 |
| TcCLB.510363.80 | mucin-associated surface protein (MASP), putative | -2,781 | 2,659E-19 |
| TcCLB.510557.50 | mucin TcMUCII, putative | -2,782 | 8,119E-57 |
| TcCLB.510939.30 | mucin TcMUCI, putative | -2,782 | 9,087E-39 |
| TcCLB.510843.19 | trans-sialidase, putative (fragment) | -2,783 | 2,344E-77 |
| TcCLB.509493.39 | mucin TcMUCII, putative | -2,784 | 2,154E-84 |
| TcCLB.510487.79 | mucin-associated surface protein (MASP), putative | -2,788 | 3,269E-23 |
| TcCLB.510557.10 | trans-sialidase (pseudogene), putative | -2,789 | 2,234E-85 |
| TcCLB.504015.11 | mucin-associated surface protein (MASP), putative (fragm | -2,791 | 1,573E-31 |
| TcCLB.440219.10 | trans-sialidase, putative (fragment) | -2,792 | 6,642E-129 |
| TcCLB.508999.110 | hypothetical protein | -2,792 | 1,42E-11 |
| TcCLB.508307.180 | hypothetical protein, conserved | -2,793 | 2,325E-166 |
| TcCLB.504239.171 | mucin TcMUCII, putative | -2,795 | 3,806E-27 |
| TcCLB.504239.240 | mucin-associated surface protein (MASP, pseudogene), pu | -2,795 | 1,5E-11 |
| TcCLB.437805.9 | trans-sialidase (pseudogene), putative | -2,797 | 6,858E-11 |
| TcCLB.471467.10 | trans-sialidase, putative (fragment) | -2,798 | 1,253E-39 |
| TcCLB.511405.40 | mucin TcMUCII (pseudogene), putative | -2,798 | 1,115E-33 |
| TcCLB.506599.210 | mucin-associated surface protein (MASP), putative | -2,801 | 7,107E-42 |
| TcCLB.508119.30 | mucin TcMUCII, putative | -2,802 | 2,898E-95 |
| TcCLB.506973.5 | mucin-associated surface protein (MASP), putative | -2,803 | 4,418E-19 |
| TcCLB.507957.160 | mucin-associated surface protein (MASP), putative | -2,805 | 8,289E-48 |
| TcCLB.506767.400 | mucin-associated surface protein (MASP), putative | -2,806 | 6,912E-27 |
| TcCLB.506241.30 | trans-sialidase, Group II, putative | -2,809 | 1,081E-47 |
| TcCLB.511849.20 | mucin TcMUCI, putative | -2,81 | 1,049E-54 |
| TcCLB.511603.100 | mucin-associated surface protein (MASP), putative | -2,811 | 1,159E-45 |
| TcCLB.507071.160 | mucin TcMUCII, putative | -2,813 | 1,768E-26 |
| TcCLB.507879.10 | trans-sialidase, Group II, putative | -2,814 | 5,316E-146 |
| TcCLB.506289.110 | surface protease GP63 (pseudogene), putative | -2,814 | 1,094E-110 |
| TcCLB.507487.10 | mucin TcMUCII, putative | -2,814 | 6,858E-35 |
| TcCLB.510279.160 | trans-sialidase (pseudogene), putative | -2,814 | 2,402E-34 |
| TcCLB.511603.500 | mucin-associated surface protein (MASP), putative | -2,815 | 7,604E-21 |
| TcCLB.510587.20 | mucin-associated surface protein (MASP, pseudogene), pu | -2,816 | 1,781E-19 |
| TcCLB.508857.130 | hypothetical protein, conserved | -2,82 | 1,129E-93 |
| TcCLB.506149.20 | hypothetical protein, conserved | -2,822 | 4,668E-54 |
| TcCLB.507163.50 | mucin TcMUCII, putative | -2,822 | 2,013E-31 |
| TcCLB.511089.80 | trans-sialidase, Group V, putative | -2,823 | 3,842E-36 |
| TcCLB.509753.240 | mucin TcMUCII, putative | -2,825 | 6,783E-15 |
| TcCLB.506703.90 | trans-sialidase, Group VI, putative | -2,828 | 2,814E-31 |
| TcCLB.510025.220 | mucin-associated surface protein (MASP), putative | -2,83 | 2,656E-35 |
| TcCLB.508367.20 | UDP-Gal or UDP-GlcNAc-dependent glycosyltransferase, p | -2,83 | 6,481E-35 |
| TcCLB.511839.20 | mucin-associated surface protein (MASP), putative | -2,831 | 8,381E-12 |
| TcCLB.509993.10 | mucin TcMUCI, putative | -2,832 | 3,289E-31 |

| | | | |
|---|---|---|---|
| TcCLB.506289.170 | surface protease GP63, putative | -2,834 | 3,653E-71 |
| TcCLB.511405.10 | mucin TcMUCI, putative | -2,834 | 6,086E-20 |
| TcCLB.509495.49 | mucin TcMUCII, putative | -2,836 | 2,376E-90 |
| TcCLB.510475.100 | mucin-associated surface protein (MASP), putative | -2,836 | 6,384E-83 |
| TcCLB.506813.29 | mucin-associated surface protein (MASP), putative | -2,837 | 2,237E-16 |
| TcCLB.506799.130 | mucin-associated surface protein (MASP), putative | -2,838 | 6,65E-74 |
| TcCLB.508245.70 | mucin-associated surface protein (MASP), putative | -2,839 | 4,124E-41 |
| TcCLB.510047.120 | mucin-associated surface protein (MASP), putative | -2,841 | 1,272E-29 |
| TcCLB.506599.150 | mucin-associated surface protein (MASP), putative | -2,844 | 2,329E-51 |
| TcCLB.507071.60 | mucin TcMUCII, putative | -2,844 | 7,563E-21 |
| TcCLB.509545.10 | mucin-associated surface protein (MASP), putative | -2,845 | 2,646E-87 |
| TcCLB.511413.19 | trans-sialidase, putative | -2,846 | 4,816E-85 |
| TcCLB.508165.14 | trans-sialidase, putative (fragment) | -2,847 | 3,821E-14 |
| TcCLB.510477.90 | mucin TcMUCII, putative | -2,849 | 5,479E-18 |
| TcCLB.511233.130 | mucin-associated surface protein (MASP), putative | -2,849 | 4,058E-14 |
| TcCLB.507429.49 | mucin TcMUCII, putative | -2,85 | 1,402E-46 |
| TcCLB.505025.30 | mucin TcMUCII, putative | -2,85 | 2,841E-30 |
| TcCLB.508053.10 | trans-sialidase, putative | -2,85 | 1,784E-13 |
| TcCLB.508401.10 | trans-sialidase, putative | -2,852 | 2,053E-38 |
| TcCLB.510359.550 | mucin-associated surface protein (MASP), putative | -2,855 | 4,616E-26 |
| TcCLB.504197.5 | mucin-associated surface protein (MASP), putative (fragm | -2,855 | 1,702E-10 |
| TcCLB.506409.100 | mucin TcMUCII, putative | -2,856 | 9,007E-44 |
| TcCLB.504239.350 | mucin-associated surface protein (MASP), putative | -2,857 | 1,485E-106 |
| TcCLB.506499.210 | mucin-associated surface protein (MASP), putative | -2,857 | 3,19E-85 |
| TcCLB.506955.70 | mucin-associated surface protein (MASP), putative | -2,858 | 1,946E-70 |
| TcCLB.506965.190 | mucin TcMUCII, putative | -2,858 | 1,113E-16 |
| TcCLB.511233.160 | hypothetical protein | -2,859 | 6,822E-53 |
| TcCLB.506409.50 | mucin TcMUCII, putative | -2,859 | 1,77E-45 |
| TcCLB.511921.140 | mucin-associated surface protein (MASP), putative | -2,86 | 3,108E-20 |
| TcCLB.508995.10 | protein kinase, putative | -2,866 | 2,993E-123 |
| TcCLB.508147.70 | trans-sialidase (pseudogene), putative | -2,866 | 9,624E-24 |
| TcCLB.474079.10 | trans-sialidase, putative | -2,867 | 2,273E-117 |
| TcCLB.508247.110 | serine-alanine-and proline-rich protein, putative | -2,87 | 3,438E-48 |
| TcCLB.473709.10 | trans-sialidase, putative (fragment) | -2,87 | 3,147E-13 |
| TcCLB.506309.60 | trans-sialidase, Group V, putative | -2,87 | 4,037E-11 |
| TcCLB.505025.130 | mucin-associated surface protein (MASP), putative | -2,871 | 2,433E-24 |
| TcCLB.506761.11 | mucin-associated surface protein (MASP), putative (fragm | -2,871 | 1,796E-18 |
| TcCLB.508759.10 | mucin-associated surface protein (MASP), putative | -2,872 | 9,359E-23 |
| TcCLB.418405.30 | trans-sialidase, putative | -2,872 | 2,123E-09 |
| TcCLB.510961.30 | mucin-associated surface protein (MASP), putative | -2,875 | 6,171E-58 |
| TcCLB.506409.170 | trans-sialidase, Group VI, putative | -2,876 | 6,46E-132 |
| TcCLB.504081.90 | mucin-associated surface protein (MASP), putative | -2,88 | 4,444E-53 |
| TcCLB.507957.60 | mucin-associated surface protein (MASP, pseudogene), pu | -2,88 | 4,634E-37 |
| TcCLB.506785.14 | mucin-associated surface protein (MASP), putative | -2,88 | 5,881E-23 |
| TcCLB.447847.10 | trans-sialidase (pseudogene), putative | -2,88 | 5,473E-22 |

| | | | |
|---|---|---|---|
| TcCLB.510831.21 | trans-sialidase, putative (fragment) | -2,88 | 9,625E-12 |
| TcCLB.511671.20 | trans-sialidase (pseudogene), putative | -2,881 | 1,337E-71 |
| TcCLB.507071.340 | hypothetical protein | -2,884 | 3,902E-50 |
| TcCLB.508219.50 | mucin TcMUCII, putative | -2,884 | 1,395E-18 |
| TcCLB.510197.80 | hypothetical protein, conserved | -2,887 | 1,288E-67 |
| TcCLB.511063.100 | trans-sialidase (pseudogene), putative | -2,887 | 1,978E-37 |
| TcCLB.509925.10 | mucin-associated surface protein (MASP), putative | -2,889 | 2,597E-69 |
| TcCLB.510021.50 | mucin-associated surface protein (MASP), putative | -2,891 | 2,554E-33 |
| TcCLB.504405.10 | dispersed gene family protein 1 (DGF-1), putative | -2,891 | 1,083E-06 |
| TcCLB.510483.220 | mucin-associated surface protein (MASP), putative | -2,893 | 6,038E-64 |
| TcCLB.471799.10 | trans-sialidase, putative (fragment) | -2,893 | 2,812E-45 |
| TcCLB.506501.390 | mucin TcMUCII, putative | -2,896 | 1,536E-22 |
| TcCLB.506001.14 | mucin-associated surface protein (MASP), putative | -2,897 | 1,577E-34 |
| TcCLB.506965.180 | mucin-associated surface protein (MASP), putative | -2,897 | 8,3E-15 |
| TcCLB.504985.20 | UDP-Gal or UDP-GlcNAc-dependent glycosyltransferase, p | -2,899 | 1,014E-43 |
| TcCLB.510201.20 | mucin-associated surface protein (MASP), putative | -2,904 | 9,513E-34 |
| TcCLB.506767.70 | mucin-associated surface protein (MASP), putative | -2,906 | 2,225E-47 |
| TcCLB.506847.50 | trans-sialidase (pseudogene), putative | -2,906 | 6,165E-08 |
| TcCLB.506939.110 | mucin-associated surface protein (MASP), putative | -2,907 | 1,005E-36 |
| TcCLB.506131.20 | mucin TcMUCII, putative | -2,907 | 7,145E-20 |
| TcCLB.511923.80 | mucin-associated surface protein (MASP, pseudogene), pu | -2,907 | 3,233E-14 |
| TcCLB.415273.9 | mucin TcMUCII (pseudogene), putative | -2,909 | 9,828E-15 |
| TcCLB.509477.10 | mucin-associated surface protein (MASP), putative | -2,909 | 6,991E-07 |
| TcCLB.510805.70 | mucin TcMUCII, putative | -2,91 | 2,898E-23 |
| TcCLB.506229.50 | nucleoside phosphatase, putative | -2,913 | 3,13E-199 |
| TcCLB.506291.31 | surface protease GP63 (pseudogene), putative | -2,913 | 4,673E-32 |
| TcCLB.510021.30 | mucin TcMUCII, putative | -2,916 | 2,173E-30 |
| TcCLB.506289.140 | surface protease GP63, putative | -2,917 | 2,487E-106 |
| TcCLB.504239.290 | mucin-associated surface protein (MASP), putative | -2,917 | 1,075E-59 |
| TcCLB.507633.40 | mucin TcMUCII, putative | -2,918 | 3,861E-109 |
| TcCLB.509665.30 | trans-sialidase, Group V, putative | -2,92 | 5,462E-42 |
| TcCLB.508595.11 | hypothetical protein | -2,92 | 6,068E-15 |
| TcCLB.509875.30 | trans-sialidase (pseudogene), putative | -2,921 | 1,07E-52 |
| TcCLB.508939.30 | mucin-associated surface protein (MASP), putative | -2,921 | 2,143E-35 |
| TcCLB.510377.70 | mucin TcMUCII, putative | -2,922 | 9,037E-09 |
| TcCLB.504249.30 | mucin-associated surface protein (MASP), putative | -2,923 | 4,829E-43 |
| TcCLB.508261.50 | mucin-associated surface protein (MASP), putative | -2,923 | 6,077E-40 |
| TcCLB.511173.160 | trans-sialidase (pseudogene), putative | -2,927 | 1,643E-44 |
| TcCLB.509993.60 | mucin TcMUCI, putative | -2,928 | 6,168E-30 |
| TcCLB.511487.150 | serine-alanine-and proline-rich protein, putative | -2,928 | 3,249E-29 |
| TcCLB.507957.334 | mucin-associated surface protein (MASP), putative | -2,928 | 2,68E-07 |
| TcCLB.507091.60 | trans-sialidase, Group V, putative | -2,931 | 1,612E-65 |
| TcCLB.510377.430 | mucin-associated surface protein (MASP), putative | -2,932 | 8,659E-42 |
| TcCLB.510371.120 | mucin-associated surface protein (MASP), putative | -2,933 | 1,689E-37 |
| TcCLB.507747.170 | mucin-associated surface protein (MASP), putative | -2,934 | 1,929E-105 |

| | | | |
|---|---|---|---|
| TcCLB.409625.10 | trans-sialidase, putative (fragment) | -2,935 | 4,45E-40 |
| TcCLB.508499.10 | mucin TcMUCII, putative | -2,936 | 1,014E-61 |
| TcCLB.506769.80 | mucin-associated surface protein (MASP), putative | -2,936 | 1,025E-59 |
| TcCLB.508157.40 | hypothetical protein | -2,939 | 7,712E-51 |
| TcCLB.503827.100 | trans-sialidase, Group VI, putative | -2,939 | 6,557E-25 |
| TcCLB.509957.20 | expression site-associated gene (ESAG-like) protein, puta | -2,941 | 2,2E-19 |
| TcCLB.503659.30 | trans-sialidase, putative | -2,943 | 6,113E-44 |
| TcCLB.509867.20 | mucin-associated surface protein (MASP), putative | -2,943 | 6,662E-15 |
| TcCLB.511183.350 | surface protease GP63 (pseudogene), putative | -2,946 | 1,361E-44 |
| TcCLB.509493.20 | mucin TcMUCII, putative | -2,947 | 1,893E-67 |
| TcCLB.507747.160 | mucin TcMUCII, putative | -2,947 | 1,764E-57 |
| TcCLB.509287.100 | mucin TcMUCII, putative | -2,949 | 1,305E-96 |
| TcCLB.508227.20 | mucin-associated surface protein (MASP), putative | -2,95 | 1,237E-70 |
| TcCLB.506499.190 | 90 kDa surface protein, putative | -2,951 | 9,329E-42 |
| TcCLB.506409.90 | mucin-associated surface protein (MASP), putative | -2,953 | 1,94E-21 |
| TcCLB.509217.160 | complement regulatory protein, putative | -2,957 | 1,007E-40 |
| TcCLB.503679.10 | aspartate aminotransferase, putative | -2,958 | 6,519E-76 |
| TcCLB.509697.29 | hypothetical protein | -2,958 | 8,859E-15 |
| TcCLB.510365.10 | trans-sialidase (pseudogene), putative | -2,959 | 3,356E-22 |
| TcCLB.506885.280 | NIMA-related kinase, putative | -2,962 | 3,55E-57 |
| TcCLB.503977.40 | mucin TcMUCII, putative | -2,962 | 1,017E-38 |
| TcCLB.504239.110 | mucin-associated surface protein (MASP), putative | -2,967 | 1,197E-36 |
| TcCLB.453767.50 | mucin-associated surface protein (MASP), putative | -2,967 | 3,883E-36 |
| TcCLB.506477.60 | protein kinase, putative | -2,968 | 9,331E-248 |
| TcCLB.508163.140 | mucin-associated surface protein (MASP), putative | -2,971 | 7,888E-99 |
| TcCLB.510475.40 | mucin TcMUCII, putative | -2,973 | 7,302E-17 |
| TcCLB.508963.30 | trans-sialidase, Group VIII, putative | -2,975 | 7,519E-90 |
| TcCLB.504239.434 | trans-sialidase, putative (fragment) | -2,977 | 8,892E-22 |
| TcCLB.511173.140 | hypothetical protein | -2,978 | 1,417E-13 |
| TcCLB.510371.10 | mucin-associated surface protein (MASP), putative | -2,979 | 1,11E-47 |
| TcCLB.510013.290 | mucin-associated surface protein (MASP), putative | -2,979 | 3,475E-24 |
| TcCLB.507959.310 | mucin TcMUCII, putative | -2,98 | 4,587E-35 |
| TcCLB.507145.70 | mucin-associated surface protein (MASP, pseudogene), pu | -2,98 | 4,201E-16 |
| TcCLB.506741.80 | mucin TcMUCII, putative | -2,982 | 7,423E-20 |
| TcCLB.508761.20 | mucin TcMUCII, putative | -2,984 | 4,204E-71 |
| TcCLB.510643.100 | trans-sialidase (pseudogene), putative | -2,985 | 2,636E-08 |
| TcCLB.510275.255 | hypothetical protein | -2,986 | 2,278E-18 |
| TcCLB.506769.40 | hypothetical protein | -2,987 | 1,329E-17 |
| TcCLB.506451.80 | mucin-associated surface protein (MASP, pseudogene), pu | -2,988 | 1,165E-98 |
| TcCLB.510191.20 | mucin TcMUCII, putative | -2,988 | 1,912E-21 |
| TcCLB.506501.230 | mucin TcMUCII, putative | -2,988 | 5,717E-15 |
| TcCLB.508429.20 | mucin-like glycoprotein, putative | -2,989 | 3,755E-29 |
| TcCLB.510191.70 | mucin-associated surface protein (MASP), putative | -2,99 | 4,949E-30 |
| TcCLB.503769.20 | trans-sialidase (pseudogene), putative | -2,991 | 1,536E-41 |
| TcCLB.509493.110 | mucin TcMUCII, putative | -2,993 | 1,32E-78 |

| | | | |
|---|---|---|---|
| TcCLB.509531.10 | trans-sialidase, putative (fragment) | -2,994 | 3,123E-47 |
| TcCLB.408825.5 | mucin-associated surface protein (MASP), putative (fragm | -2,995 | 1,347E-30 |
| TcCLB.506067.240 | mucin-associated surface protein (MASP), putative | -2,995 | 2,089E-17 |
| TcCLB.504239.280 | 90 kDa surface protein, putative | -2,999 | 3,405E-34 |
| TcCLB.506763.90 | mucin-associated surface protein (MASP, pseudogene), pu | -2,999 | 1,392E-21 |
| TcCLB.507959.250 | mucin TcMUCII, putative | -3 | 2,025E-23 |
| TcCLB.506741.100 | trans-sialidase, Group V, putative | -3,001 | 8,674E-33 |
| TcCLB.511087.19 | trans-sialidase, putative (fragment) | -3,002 | 5,187E-29 |
| TcCLB.506321.200 | mucin-associated surface protein (MASP), putative | -3,003 | 3,119E-32 |
| TcCLB.508433.220 | mucin-associated surface protein (MASP), putative | -3,003 | 2,137E-27 |
| TcCLB.511923.10 | mucin-associated surface protein (MASP, pseudogene), pu | -3,003 | 2,425E-08 |
| TcCLB.510373.90 | mucin-associated surface protein (MASP), putative | -3,005 | 3,462E-48 |
| TcCLB.510553.40 | trans-sialidase, Group V, putative | -3,006 | 1,837E-55 |
| TcCLB.506955.250 | mucin TcMUCII, putative | -3,007 | 2,047E-48 |
| TcCLB.511849.10 | mucin TcMUCI, putative | -3,008 | 1,187E-58 |
| TcCLB.507637.100 | mucin TcMUCII, putative | -3,009 | 9,917E-41 |
| TcCLB.510197.20 | mucin-associated surface protein (MASP), putative | -3,01 | 2,455E-95 |
| TcCLB.508161.50 | trans-sialidase, putative | -3,01 | 2,88E-39 |
| TcCLB.510261.41 | mucin TcMUC (pseudogene), putative | -3,011 | 4,251E-16 |
| TcCLB.509699.220 | trans-sialidase (pseudogene), putative | -3,013 | 1,142E-89 |
| TcCLB.508247.80 | mucin-associated surface protein (MASP), putative | -3,014 | 3,079E-27 |
| TcCLB.504911.20 | mucin-associated surface protein (MASP, pseudogene), pu | -3,014 | 1,871E-09 |
| TcCLB.507957.270 | mucin TcMUCI, putative | -3,015 | 2,833E-46 |
| TcCLB.509905.170 | trans-sialidase, Group VIII, putative | -3,02 | 3,508E-79 |
| TcCLB.506499.105 | mucin-associated surface protein (MASP, pseudogene), pu | -3,02 | 1,779E-37 |
| TcCLB.505567.31 | mucin-associated surface protein (MASP, pseudogene), pu | -3,021 | 3,957E-11 |
| TcCLB.510503.160 | mucin TcMUCII, putative | -3,022 | 2,821E-75 |
| TcCLB.507071.120 | mucin-associated surface protein (MASP), putative | -3,022 | 6,676E-50 |
| TcCLB.508541.80 | mucin TcMUCII, putative | -3,029 | 8,085E-47 |
| TcCLB.508277.170 | hypothetical protein, conserved | -3,03 | 6,459E-89 |
| TcCLB.508853.30 | hypothetical protein | -3,03 | 3,113E-55 |
| TcCLB.510875.10 | dispersed gene family protein 1 (DGF-1, pseudogene), put | -3,03 | 1,144E-19 |
| TcCLB.506599.280 | mucin-associated surface protein (MASP, pseudogene), pu | -3,031 | 1,806E-28 |
| TcCLB.510201.5 | mucin-associated surface protein (MASP), putative | -3,033 | 3,274E-13 |
| TcCLB.507957.320 | mucin-associated surface protein (MASP), putative | -3,038 | 2,175E-63 |
| TcCLB.508245.60 | mucin TcMUCII, putative | -3,04 | 4,131E-29 |
| TcCLB.511547.40 | retrotransposon hot spot protein (RHS, pseudogene), puta | -3,042 | 2,398E-73 |
| TcCLB.510627.110 | mucin-associated surface protein (MASP), putative | -3,042 | 3,218E-08 |
| TcCLB.511211.190 | hypothetical protein | -3,044 | 1,323E-97 |
| TcCLB.511487.170 | mucin-associated surface protein (MASP), putative | -3,047 | 1,203E-64 |
| TcCLB.509525.60 | mucin TcMUCII, putative | -3,048 | 1,212E-15 |
| TcCLB.510961.10 | mucin-associated surface protein (MASP), putative | -3,049 | 2,254E-60 |
| TcCLB.506329.70 | mucin TcMUCII, putative | -3,051 | 2,281E-19 |
| TcCLB.508633.20 | trans-sialidase, Group V, putative | -3,052 | 3,949E-32 |
| TcCLB.506309.70 | mucin-associated surface protein (MASP), putative | -3,055 | 4,851E-27 |

| | | | |
|---|---|---|---|
| TcCLB.506813.5 | hypothetical protein | -3,056 | 2,446E-72 |
| TcCLB.398265.9 | trans-sialidase, putative | -3,056 | 5,453E-19 |
| TcCLB.510279.10 | trans-sialidase (pseudogene), putative | -3,059 | 2,438E-32 |
| TcCLB.510557.60 | mucin-associated surface protein (MASP), putative | -3,06 | 4,797E-24 |
| TcCLB.511259.220 | mucin-associated surface protein (MASP), putative | -3,063 | 4,081E-54 |
| TcCLB.505025.137 | mucin TcMUCII, putative | -3,063 | 6,48E-18 |
| TcCLB.508603.10 | trans-sialidase, putative | -3,064 | 2,033E-38 |
| TcCLB.511553.160 | 90 kDa surface protein, putative | -3,065 | 4,17E-32 |
| TcCLB.506245.220 | mucin TcMUCII, putative | -3,07 | 1,261E-28 |
| TcCLB.509545.79 | mucin-associated surface protein (MASP), putative | -3,071 | 1,705E-49 |
| TcCLB.509419.20 | mucin-associated surface protein (MASP), putative | -3,076 | 7,456E-14 |
| TcCLB.504393.30 | trans-sialidase (pseudogene), putative | -3,077 | 5,1E-63 |
| TcCLB.507803.10 | mucin-associated surface protein (MASP), putative | -3,079 | 1,611E-69 |
| TcCLB.506599.310 | mucin-associated surface protein (MASP), putative | -3,079 | 2,646E-21 |
| TcCLB.508163.280 | mucin TcMUCII, putative | -3,08 | 1,926E-39 |
| TcCLB.506053.50 | trans-sialidase, Group IV, putative | -3,081 | 6,923E-73 |
| TcCLB.506893.10 | mucin TcMUCII, putative | -3,081 | 5,046E-26 |
| TcCLB.506409.140 | mucin TcMUCII, putative | -3,081 | 1,15E-14 |
| TcCLB.399685.10 | trans-sialidase, putative (fragment) | -3,082 | 9,802E-63 |
| TcCLB.511183.560 | mucin-associated surface protein (MASP), putative | -3,085 | 3,633E-47 |
| TcCLB.508427.40 | mucin-associated surface protein (MASP), putative | -3,087 | 3,234E-72 |
| TcCLB.504239.340 | mucin TcMUCII, putative | -3,087 | 6,735E-35 |
| TcCLB.503429.30 | mucin-associated surface protein (MASP), putative | -3,088 | 1,019E-42 |
| TcCLB.511839.40 | trans-sialidase, Group V, putative | -3,09 | 4,046E-47 |
| TcCLB.505949.20 | mucin-associated surface protein (MASP), putative | -3,092 | 5,114E-49 |
| TcCLB.510021.120 | trans-sialidase, Group V, putative | -3,092 | 4,512E-23 |
| TcCLB.511613.60 | mucin-associated surface protein (MASP), putative | -3,093 | 7,389E-101 |
| TcCLB.509081.5 | mucin-associated surface protein (MASP), putative | -3,093 | 2,159E-37 |
| TcCLB.505025.80 | hypothetical protein, conserved | -3,094 | 6,594E-166 |
| TcCLB.510377.400 | mucin TcMUCII, putative | -3,097 | 1,266E-40 |
| TcCLB.507085.60 | mucin TcMUCII, putative | -3,101 | 2,24E-74 |
| TcCLB.504787.30 | mucin-associated surface protein (MASP), putative | -3,101 | 1,209E-20 |
| TcCLB.464391.10 | retrotransposon hot spot protein (RHS, pseudogene), puta | -3,105 | 2,029E-14 |
| TcCLB.510237.70 | mucin-associated surface protein (MASP), putative | -3,106 | 3,297E-44 |
| TcCLB.510917.9 | mucin-associated surface protein (MASP), putative | -3,111 | 5,05E-44 |
| TcCLB.506741.130 | trans-sialidase, Group V, putative | -3,115 | 2,183E-32 |
| TcCLB.508841.10 | surface antigen TASV-A4 (TASV-A4) | -3,116 | 1,432E-196 |
| TcCLB.466325.10 | trans-sialidase (pseudogene), putative | -3,116 | 2,098E-29 |
| TcCLB.509493.80 | mucin TcMUCII, putative | -3,117 | 5,542E-149 |
| TcCLB.506611.20 | mucin TcMUCII, putative | -3,117 | 5,774E-26 |
| TcCLB.504183.20 | trans-sialidase, Group II, putative | -3,118 | 1,098E-122 |
| TcCLB.479481.10 | mucin-associated surface protein (MASP), putative (fragm | -3,118 | 8,488E-31 |
| TcCLB.506951.80 | trans-sialidase, Group V, putative | -3,119 | 1,517E-99 |
| TcCLB.507069.150 | mucin-associated surface protein (MASP), putative | -3,119 | 1,329E-51 |
| TcCLB.508293.80 | mucin-associated surface protein (MASP), putative | -3,119 | 8,696E-34 |

| | | | |
|---|---|---|---|
| TcCLB.506501.170 | mucin TcMUCII, putative | -3,121 | 1,484E-24 |
| TcCLB.510483.120 | mucin-associated surface protein (MASP), putative | -3,123 | 1,216E-28 |
| TcCLB.457979.10 | mucin-associated surface protein (MASP), putative | -3,124 | 2,63E-22 |
| TcCLB.506453.10 | trans-sialidase, putative | -3,126 | 6,136E-50 |
| TcCLB.511063.140 | trans-sialidase (pseudogene), putative | -3,126 | 3,166E-29 |
| TcCLB.503875.10 | mucin-associated surface protein (MASP), putative | -3,127 | 5,889E-20 |
| TcCLB.511775.21 | mitochondrial RNA editing ligase 1 (pseudogene), putative | -3,127 | 4,144E-10 |
| TcCLB.511553.90 | mucin TcMUCII, putative | -3,128 | 1,095E-70 |
| TcCLB.510419.20 | trans-sialidase, Group VI, putative | -3,129 | 8,685E-45 |
| TcCLB.508883.48 | trans-sialidase, putative | -3,131 | 8,391E-92 |
| TcCLB.508013.40 | trans-sialidase, Group VI, putative | -3,131 | 3,971E-59 |
| TcCLB.508229.10 | trans-sialidase, putative | -3,131 | 1,166E-24 |
| TcCLB.509483.10 | surface protease GP63, putative | -3,133 | 2,534E-15 |
| TcCLB.506309.130 | trans-sialidase (pseudogene), putative | -3,134 | 5,663E-16 |
| TcCLB.510025.180 | mucin-associated surface protein (MASP), putative | -3,138 | 2,146E-66 |
| TcCLB.459793.10 | mucin TcMUCI, putative | -3,138 | 3,018E-32 |
| TcCLB.506809.10 | trans-sialidase (pseudogene), putative | -3,14 | 3,188E-96 |
| TcCLB.511183.260 | mucin-associated surface protein (MASP), putative | -3,141 | 1,656E-49 |
| TcCLB.510307.10 | surface protease GP63, putative | -3,145 | 2,02E-19 |
| TcCLB.506245.302 | mucin-associated surface protein (MASP), putative | -3,146 | 1,555E-74 |
| TcCLB.504039.240 | mucin TcMUCII, putative | -3,146 | 1,077E-47 |
| TcCLB.511911.60 | trans-sialidase, Group III, putative | -3,146 | 1,169E-07 |
| TcCLB.510013.240 | mucin TcMUCII, putative | -3,147 | 1,142E-22 |
| TcCLB.509753.300 | mucin TcMUCII, putative | -3,147 | 1,097E-20 |
| TcCLB.510021.80 | 90 kDa surface protein, putative | -3,148 | 3,687E-45 |
| TcCLB.511099.190 | mucin-associated surface protein (MASP), putative | -3,149 | 2,653E-56 |
| TcCLB.510305.9 | Present in the outer mitochondrial membrane proteome 3 | -3,153 | 3,13E-127 |
| TcCLB.509979.174 | mucin-associated surface protein (MASP), putative | -3,155 | 1,767E-79 |
| TcCLB.506667.10 | mucin-associated surface protein (MASP), putative | -3,156 | 1,567E-19 |
| TcCLB.511079.40 | surface protease GP63 (pseudogene), putative | -3,156 | 6,125E-18 |
| TcCLB.504233.70 | mucin TcMUCI, putative | -3,157 | 3,435E-25 |
| TcCLB.511555.10 | mucin-associated surface protein (MASP), putative | -3,159 | 2,171E-89 |
| TcCLB.506967.60 | trans-sialidase, Group VI, putative | -3,16 | 2,024E-54 |
| TcCLB.510093.10 | ATPase, putative | -3,16 | 1,386E-20 |
| TcCLB.510697.60 | mucin-associated surface protein (MASP), putative | -3,161 | 8,809E-81 |
| TcCLB.503783.80 | mucin-associated surface protein (MASP), putative | -3,161 | 9,306E-76 |
| TcCLB.510715.64 | mucin-associated surface protein (MASP), putative | -3,161 | 2,055E-25 |
| TcCLB.511597.10 | trans-sialidase, putative | -3,161 | 1,533E-15 |
| TcCLB.507281.19 | retrotransposon hot spot protein (RHS, pseudogene), puta | -3,162 | 1,012E-52 |
| TcCLB.506569.50 | expression site-associated gene (ESAG-like) protein, puta | -3,164 | 8,716E-18 |
| TcCLB.483183.10 | trans-sialidase, putative (fragment) | -3,169 | 6,186E-19 |
| TcCLB.508125.100 | mucin TcMUCII, putative | -3,17 | 1,035E-09 |
| TcCLB.510377.260 | mucin-associated surface protein (MASP), putative | -3,173 | 1,029E-17 |
| TcCLB.511645.21 | mucin TcMUCII, putative | -3,175 | 1,503E-103 |
| TcCLB.507429.10 | mucin TcMUCII (pseudogene), putative | -3,177 | 2,578E-87 |

| | | | |
|---|---|---|---|
| TcCLB.510025.170 | mucin TcMUCII, putative | -3,177 | 1,626E-53 |
| TcCLB.510621.20 | mucin-associated surface protein (MASP), putative | -3,181 | 1,942E-44 |
| TcCLB.507699.120 | mucin TcMUCII, putative | -3,182 | 7,339E-37 |
| TcCLB.505025.70 | mucin-associated surface protein (MASP), putative | -3,183 | 5,551E-72 |
| TcCLB.506157.40 | trans-sialidase, putative | -3,185 | 1,138E-65 |
| TcCLB.511255.50 | mucin-associated surface protein (MASP), putative | -3,188 | 5,403E-69 |
| TcCLB.503645.40 | mucin-associated surface protein (MASP), putative | -3,188 | 6,956E-40 |
| TcCLB.506955.14 | mucin-associated surface protein (MASP, pseudogene), pu | -3,189 | 1,425E-67 |
| TcCLB.430567.10 | trans-sialidase, putative (fragment) | -3,19 | 1,027E-86 |
| TcCLB.507997.14 | trans-sialidase, putative (fragment) | -3,194 | 2,225E-94 |
| TcCLB.506741.120 | mucin-associated surface protein (MASP), putative | -3,2 | 2,008E-45 |
| TcCLB.510275.272 | hypothetical protein | -3,2 | 6,425E-43 |
| TcCLB.510163.30 | mucin-associated surface protein (MASP), putative | -3,201 | 5,368E-202 |
| TcCLB.506501.184 | mucin-associated surface protein (MASP, pseudogene), pu | -3,202 | 4,5E-29 |
| TcCLB.511609.30 | mucin-associated surface protein (MASP), putative | -3,205 | 1,07E-22 |
| TcCLB.506965.130 | mucin-associated surface protein (MASP), putative | -3,206 | 2,486E-111 |
| TcCLB.507907.10 | trans-sialidase, putative | -3,208 | 1,279E-24 |
| TcCLB.508309.5 | trans-sialidase, putative (fragment) | -3,209 | 4,439E-26 |
| TcCLB.508125.149 | trans-sialidase, putative (fragment) | -3,213 | 2,147E-77 |
| TcCLB.510371.130 | mucin TcMUCII, putative | -3,215 | 2,531E-29 |
| TcCLB.507091.80 | mucin-associated surface protein (MASP), putative | -3,215 | 4,585E-25 |
| TcCLB.504341.10 | trans-sialidase, Group IV, putative | -3,216 | 1,551E-131 |
| TcCLB.509081.100 | mucin TcMUCII, putative | -3,217 | 2,089E-71 |
| TcCLB.504039.57 | mucin-associated surface protein (MASP, pseudogene), pu | -3,219 | 5,283E-10 |
| TcCLB.508789.150 | mucin-associated surface protein (MASP), putative | -3,22 | 5,389E-78 |
| TcCLB.506973.90 | mucin-associated surface protein (MASP, pseudogene), pu | -3,221 | 3,191E-80 |
| TcCLB.510715.40 | mucin-associated surface protein (MASP), putative | -3,222 | 2,37E-33 |
| TcCLB.508887.20 | mucin TcMUCII, putative | -3,223 | 2,134E-47 |
| TcCLB.506951.90 | trans-sialidase, Group IV, putative | -3,224 | 6,829E-133 |
| TcCLB.507237.89 | mucin TcMUCII, putative | -3,226 | 9,597E-16 |
| TcCLB.510213.80 | mucin TcMUCII, putative | -3,227 | 3,825E-33 |
| TcCLB.505997.80 | trans-sialidase, Group VIII, putative | -3,23 | 2,493E-103 |
| TcCLB.506501.50 | mucin-associated surface protein (MASP, pseudogene), pu | -3,23 | 5,338E-37 |
| TcCLB.508429.40 | mucin-associated surface protein (MASP, pseudogene), pu | -3,234 | 5,309E-31 |
| TcCLB.510411.40 | trans-sialidase, putative | -3,236 | 2,934E-50 |
| TcCLB.506331.50 | trans-sialidase (pseudogene), putative | -3,239 | 2,066E-59 |
| TcCLB.422867.10 | trans-sialidase, Group II, putative | -3,239 | 1,021E-31 |
| TcCLB.506599.430 | mucin TcMUCII, putative | -3,241 | 1,71E-43 |
| TcCLB.508873.580 | mucin-associated surface protein (MASP), putative | -3,243 | 5,659E-42 |
| TcCLB.506139.100 | mucin TcMUCII, putative | -3,243 | 9,122E-41 |
| TcCLB.511643.50 | mucin-associated surface protein (MASP, pseudogene), pu | -3,244 | 1,116E-61 |
| TcCLB.509731.10 | poly-zinc finger protein 2, putative | -3,244 | 8,55E-34 |
| TcCLB.506139.70 | mucin-associated surface protein (MASP), putative | -3,25 | 1,747E-42 |
| TcCLB.509525.100 | trans-sialidase (pseudogene), putative | -3,252 | 1,674E-57 |
| TcCLB.503973.70 | serine-alanine-and proline-rich protein, putative | -3,253 | 5,651E-71 |

| | | | |
|---|---|---|---|
| TcCLB.508375.10 | mucin-associated surface protein (MASP), putative | -3,254 | 5,444E-92 |
| TcCLB.510213.9 | mucin-associated surface protein (MASP), putative | -3,254 | 2,173E-52 |
| TcCLB.511183.230 | trans-sialidase, Group V, putative | -3,254 | 9,141E-38 |
| TcCLB.508365.290 | mucin TcMUCII, putative | -3,255 | 4,754E-32 |
| TcCLB.507937.3 | trans-sialidase, putative | -3,256 | 6,379E-10 |
| TcCLB.507555.50 | trans-sialidase, Group VIII, putative | -3,257 | 3,83E-181 |
| TcCLB.510047.80 | trans-sialidase, Group V, putative | -3,257 | 3,306E-63 |
| TcCLB.506623.20 | trans-sialidase (pseudogene), putative | -3,26 | 2,562E-49 |
| TcCLB.504801.20 | surface protease GP63, putative | -3,26 | 8,386E-33 |
| TcCLB.508165.300 | mucin-associated surface protein (MASP), putative | -3,261 | 1,151E-94 |
| TcCLB.506057.15 | hypothetical protein | -3,266 | 5,728E-65 |
| TcCLB.511255.630 | mucin-associated surface protein (MASP), putative | -3,266 | 5,301E-38 |
| TcCLB.508539.160 | mucin-associated surface protein (MASP), putative | -3,271 | 1,066E-97 |
| TcCLB.506321.20 | mucin-associated surface protein (MASP), putative | -3,272 | 6,141E-121 |
| TcCLB.510831.10 | mucin-associated surface protein (MASP, pseudogene), pu | -3,272 | 3,371E-24 |
| TcCLB.511599.90 | mucin-associated surface protein (MASP), putative | -3,273 | 1,003E-60 |
| TcCLB.511553.150 | mucin-associated surface protein (MASP), putative | -3,274 | 2,718E-72 |
| TcCLB.510847.10 | trans-sialidase, Group VIII, putative | -3,276 | 7,925E-302 |
| TcCLB.506799.30 | mucin-like glycoprotein, putative | -3,277 | 3,014E-93 |
| TcCLB.505025.10 | mucin-associated surface protein (MASP), putative | -3,278 | 6,208E-40 |
| TcCLB.506245.200 | mucin-associated surface protein (MASP), putative | -3,279 | 7,269E-82 |
| TcCLB.505993.30 | trans-sialidase, Group VIII, putative | -3,28 | 4,827E-173 |
| TcCLB.510197.160 | mucin-associated surface protein (MASP, pseudogene), pu | -3,281 | 4,325E-133 |
| TcCLB.511595.59 | trans-sialidase, putative (fragment) | -3,283 | 4,484E-14 |
| TcCLB.509665.20 | mucin-associated surface protein (MASP), putative | -3,285 | 3,635E-93 |
| TcCLB.508165.170 | mucin-associated surface protein (MASP), putative | -3,286 | 2,729E-56 |
| TcCLB.508061.20 | trans-sialidase, Group II, putative | -3,287 | 6,677E-131 |
| TcCLB.510013.70 | mucin-associated surface protein (MASP), putative | -3,287 | 4,122E-28 |
| TcCLB.456627.10 | trans-sialidase (pseudogene), putative | -3,29 | 2,392E-08 |
| TcCLB.506279.140 | trans-sialidase, Group V, putative | -3,291 | 1,082E-58 |
| TcCLB.503827.60 | mucin-associated surface protein (MASP), putative | -3,293 | 1,621E-57 |
| TcCLB.510391.20 | mucin-associated surface protein (MASP), putative | -3,295 | 9,799E-69 |
| TcCLB.511377.30 | mucin-associated surface protein (MASP), putative | -3,296 | 1,518E-37 |
| TcCLB.506451.20 | trans-sialidase, Group V, putative | -3,297 | 9,89E-79 |
| TcCLB.508221.170 | mucin-associated surface protein (MASP), putative | -3,3 | 2,77E-21 |
| TcCLB.506923.21 | mucin TcMUCII, putative | -3,304 | 1,107E-66 |
| TcCLB.510463.120 | mucin-associated surface protein (MASP), putative | -3,304 | 1,134E-07 |
| TcCLB.510555.30 | mucin TcMUCII, putative | -3,305 | 1,149E-51 |
| TcCLB.503441.5 | trans-sialidase, putative (fragment) | -3,306 | 9,247E-55 |
| TcCLB.506759.160 | mucin TcMUCII, putative | -3,306 | 4,92E-16 |
| TcCLB.510583.130 | mucin-associated surface protein (MASP), putative | -3,309 | 1,015E-36 |
| TcCLB.507471.10 | surface protease GP63 (pseudogene), putative | -3,309 | 2,839E-11 |
| TcCLB.507957.90 | mucin TcMUCII, putative | -3,31 | 2,174E-46 |
| TcCLB.510477.100 | mucin-associated surface protein (MASP), putative | -3,314 | 1,442E-70 |
| TcCLB.509227.10 | mucin TcMUCI, putative | -3,314 | 1,739E-68 |

| | | | |
|---|---|---|---|
| TcCLB.508539.110 | mucin-associated surface protein (MASP), putative | -3,315 | 3,224E-80 |
| TcCLB.506765.74 | mucin-associated surface protein (MASP), putative | -3,315 | 1,194E-65 |
| TcCLB.506409.200 | mucin TcMUCII, putative | -3,316 | 1,802E-54 |
| TcCLB.510557.80 | mucin TcMUCII, putative | -3,317 | 3,355E-57 |
| TcCLB.508495.140 | mucin TcMUCII, putative | -3,326 | 2,089E-98 |
| TcCLB.510013.170 | mucin-associated surface protein (MASP), putative | -3,328 | 1,006E-28 |
| TcCLB.506911.30 | trans-sialidase, Group V, putative | -3,329 | 1,853E-91 |
| TcCLB.506307.10 | mucin-associated surface protein (MASP), putative | -3,33 | 9,936E-48 |
| TcCLB.508365.50 | mucin TcMUCII, putative | -3,332 | 1,894E-17 |
| TcCLB.510459.20 | mucin-associated surface protein (MASP), putative | -3,335 | 6,88E-62 |
| TcCLB.508243.10 | mucin-associated surface protein (MASP), putative | -3,337 | 7,139E-93 |
| TcCLB.503827.20 | mucin-associated surface protein (MASP), putative | -3,34 | 1,064E-51 |
| TcCLB.510625.19 | mucin-associated surface protein (MASP), putative | -3,342 | 7,977E-29 |
| TcCLB.506499.140 | mucin-associated surface protein (MASP), putative | -3,343 | 1,331E-94 |
| TcCLB.510025.260 | mucin-associated surface protein (MASP), putative | -3,35 | 1,137E-101 |
| TcCLB.403869.10 | trans-sialidase, putative (fragment) | -3,35 | 1,592E-57 |
| TcCLB.510483.180 | mucin-associated surface protein (MASP), putative | -3,35 | 8,266E-50 |
| TcCLB.507699.149 | mucin TcMUCII, putative | -3,351 | 3,75E-57 |
| TcCLB.506761.47 | mucin TcMUCII, putative | -3,355 | 1,264E-26 |
| TcCLB.510331.10 | trans-sialidase, Group III, putative | -3,356 | 7,963E-158 |
| TcCLB.511551.40 | retrotransposon hot spot protein (RHS, pseudogene), puta | -3,361 | 6,043E-26 |
| TcCLB.506737.128 | mucin TcMUCI, putative | -3,362 | 4,518E-75 |
| TcCLB.506455.30 | trans-sialidase, Group II, putative | -3,363 | 5,908E-93 |
| TcCLB.510281.20 | surface protease GP63, putative | -3,365 | 1,923E-64 |
| TcCLB.506763.50 | mucin-associated surface protein (MASP), putative | -3,369 | 4,163E-42 |
| TcCLB.510495.30 | trans-sialidase (pseudogene), putative | -3,372 | 1,686E-58 |
| TcCLB.510025.30 | mucin-associated surface protein (MASP, pseudogene), pu | -3,372 | 2,863E-53 |
| TcCLB.509075.70 | retrotransposon hot spot protein (RHS, pseudogene), puta | -3,372 | 2,522E-11 |
| TcCLB.506289.130 | hypothetical protein | -3,375 | 8,503E-30 |
| TcCLB.511399.24 | trans-sialidase, putative | -3,376 | 1,413E-31 |
| TcCLB.511077.100 | mucin TcMUCII, putative | -3,378 | 4,309E-28 |
| TcCLB.509973.70 | mucin-associated surface protein (MASP), putative | -3,379 | 1,163E-134 |
| TcCLB.506899.40 | mucin TcMUCII, putative | -3,379 | 1,432E-53 |
| TcCLB.511603.90 | trans-sialidase, Group V, putative | -3,381 | 2,736E-91 |
| TcCLB.509657.50 | mucin-associated surface protein (MASP), putative | -3,381 | 2,874E-33 |
| TcCLB.507959.280 | mucin-associated surface protein (MASP), putative | -3,389 | 1,353E-85 |
| TcCLB.510377.100 | mucin-associated surface protein (MASP), putative | -3,39 | 7,122E-25 |
| TcCLB.506501.220 | mucin-associated surface protein (MASP), putative | -3,393 | 1,517E-40 |
| TcCLB.506067.290 | mucin-associated surface protein (MASP), putative | -3,395 | 4,698E-131 |
| TcCLB.506767.324 | mucin TcMUCII, putative | -3,396 | 1,963E-31 |
| TcCLB.503587.10 | mucin-associated surface protein (MASP), putative | -3,396 | 3,318E-25 |
| TcCLB.510377.330 | trans-sialidase, Group V, putative | -3,4 | 6,196E-133 |
| TcCLB.506801.20 | mucin-like glycoprotein, putative | -3,401 | 6,554E-93 |
| TcCLB.507953.40 | mucin-associated surface protein (MASP), putative | -3,403 | 4,494E-51 |
| TcCLB.509419.40 | trans-sialidase, Group V, putative | -3,406 | 2,277E-07 |

| | | | |
|---|---|---|---|
| TcCLB.507699.200 | mucin TcMUCII, putative | -3,407 | 2,469E-59 |
| TcCLB.412393.10 | trans-sialidase, putative (pseudogene) | -3,407 | 2,035E-18 |
| TcCLB.511089.130 | mucin-associated surface protein (MASP), putative | -3,412 | 7,054E-41 |
| TcCLB.511221.60 | trans-sialidase, Group V, putative | -3,415 | 8,399E-113 |
| TcCLB.507237.100 | mucin-associated surface protein (MASP), putative | -3,417 | 1,326E-24 |
| TcCLB.506021.20 | trans-sialidase, Group II, putative | -3,422 | 1,983E-82 |
| TcCLB.508149.70 | mucin TcMUCII, putative | -3,422 | 6,285E-38 |
| TcCLB.421173.14 | trans-sialidase, putative | -3,422 | 7,687E-30 |
| TcCLB.506909.60 | mucin TcMUCI, putative | -3,423 | 3,111E-25 |
| TcCLB.503977.30 | mucin-associated surface protein (MASP), putative | -3,425 | 3,339E-75 |
| TcCLB.508789.160 | mucin TcMUCII, putative | -3,425 | 1,056E-27 |
| TcCLB.448661.20 | mucin TcMUCII, putative | -3,427 | 2,226E-46 |
| TcCLB.506767.50 | mucin TcMUCII, putative | -3,431 | 4,689E-48 |
| TcCLB.423707.10 | trans-sialidase, putative (fragment) | -3,434 | 2,465E-47 |
| TcCLB.508247.50 | mucin-associated surface protein (MASP, pseudogene), pu | -3,435 | 1,261E-49 |
| TcCLB.503717.10 | trans-sialidase, Group V, putative | -3,437 | 5,286E-62 |
| TcCLB.507237.170 | mucin-associated surface protein (MASP), putative | -3,437 | 5,477E-09 |
| TcCLB.511197.60 | mucin-associated surface protein (MASP), putative | -3,439 | 5,942E-93 |
| TcCLB.509077.20 | trans-sialidase (pseudogene), putative | -3,441 | 3,952E-44 |
| TcCLB.506755.50 | hypothetical protein, conserved | -3,443 | 1,815E-228 |
| TcCLB.507995.20 | trans-sialidase, putative | -3,443 | 2,048E-51 |
| TcCLB.511233.140 | trans-sialidase, Group V, putative | -3,443 | 1,417E-20 |
| TcCLB.510373.119 | mucin-associated surface protein (MASP), putative | -3,45 | 3,555E-56 |
| TcCLB.504155.10 | mucin-associated surface protein (MASP), putative | -3,452 | 1,169E-28 |
| TcCLB.505297.60 | mucin-associated surface protein (MASP), putative | -3,454 | 3,142E-78 |
| TcCLB.511077.60 | mucin-associated surface protein (MASP), putative | -3,454 | 7,763E-37 |
| TcCLB.508999.60 | mucin TcMUCII, putative | -3,457 | 1,36E-69 |
| TcCLB.504081.530 | mucin-associated surface protein (MASP), putative | -3,457 | 7,57E-33 |
| TcCLB.511603.260 | mucin TcMUCII, putative | -3,461 | 1,809E-29 |
| TcCLB.509763.10 | trans-sialidase, putative | -3,465 | 1,082E-56 |
| TcCLB.508165.200 | trans-sialidase (pseudogene), putative | -3,465 | 2,333E-41 |
| TcCLB.511089.90 | mucin-associated surface protein (MASP), putative | -3,466 | 4,774E-24 |
| TcCLB.504127.30 | trans-sialidase (pseudogene), putative | -3,467 | 1,172E-234 |
| TcCLB.508493.40 | mucin TcMUCII, putative | -3,469 | 1,711E-146 |
| TcCLB.503447.20 | trans-sialidase, Group II, putative | -3,469 | 5,529E-115 |
| TcCLB.510279.230 | trans-sialidase, Group VI, putative | -3,469 | 6,293E-41 |
| TcCLB.507639.10 | poly-zinc finger protein 2, putative | -3,471 | 5,024E-86 |
| TcCLB.503417.50 | hypothetical protein | -3,473 | 2,988E-98 |
| TcCLB.508071.20 | trans-sialidase (pseudogene), putative | -3,473 | 2,069E-44 |
| TcCLB.508165.110 | mucin-associated surface protein (MASP), putative | -3,473 | 4,496E-35 |
| TcCLB.511173.80 | trans-sialidase, Group V, putative | -3,474 | 6,518E-56 |
| TcCLB.505949.160 | mucin TcMUCII, putative | -3,475 | 6,372E-43 |
| TcCLB.509979.350 | mucin-associated surface protein (MASP), putative | -3,479 | 1,907E-171 |
| TcCLB.510093.20 | trans-sialidase, Group VIII, putative | -3,484 | 5,948E-78 |
| TcCLB.511061.51 | mucin-associated surface protein (MASP), putative | -3,485 | 1,493E-90 |

| | | | |
|---|---|---|---|
| TcCLB.507953.200 | 90 kDa surface protein, putative | -3,49 | 1,964E-74 |
| TcCLB.504733.10 | beta galactofuranosyl glycosyltransferase (pseudogene), p | -3,49 | 1,987E-29 |
| TcCLB.508005.10 | hypothetical protein | -3,491 | 1,414E-37 |
| TcCLB.509527.50 | mucin TcMUCII (pseudogene), putative | -3,494 | 5,946E-34 |
| TcCLB.506067.169 | mucin-associated surface protein (MASP), putative | -3,497 | 3,321E-73 |
| TcCLB.510015.30 | mucin-associated surface protein (MASP), putative | -3,501 | 7,041E-183 |
| TcCLB.508541.90 | mucin-associated surface protein (MASP), putative | -3,502 | 2,343E-89 |
| TcCLB.424889.10 | UDP-Gal or UDP-GlcNAc-dependent glycosyltransferase, p | -3,503 | 8,008E-31 |
| TcCLB.510889.280 | metacyclin II, putative | -3,503 | 1,232E-29 |
| TcCLB.508853.20 | trans-sialidase, Group V, putative | -3,507 | 3,042E-109 |
| TcCLB.507699.230 | trans-sialidase, putative | -3,507 | 2,582E-88 |
| TcCLB.506309.30 | mucin-associated surface protein (MASP), putative | -3,51 | 2,852E-37 |
| TcCLB.510375.10 | mucin-associated surface protein (MASP), putative | -3,515 | 3,255E-57 |
| TcCLB.508375.20 | trans-sialidase, Group V, putative | -3,516 | 2,858E-128 |
| TcCLB.510807.10 | mucin TcMUCII, putative | -3,517 | 2,361E-40 |
| TcCLB.506173.310 | mucin TcMUCII, putative | -3,519 | 1,231E-101 |
| TcCLB.506321.210 | mucin TcMUCII, putative | -3,519 | 6,054E-43 |
| TcCLB.506289.210 | surface protease GP63, putative | -3,521 | 4,321E-94 |
| TcCLB.508761.80 | mucin TcMUCII, putative | -3,521 | 4,582E-63 |
| TcCLB.506613.24 | mucin TcMUCII, putative | -3,521 | 1,559E-48 |
| TcCLB.402255.10 | retrotransposon hot spot protein (RHS, pseudogene), puta | -3,527 | 7,652E-56 |
| TcCLB.510477.30 | mucin TcMUCI, putative | -3,528 | 6,144E-46 |
| TcCLB.509081.20 | mucin-associated surface protein (MASP), putative | -3,529 | 5,046E-136 |
| TcCLB.507069.50 | mucin-associated surface protein (MASP, pseudogene), pu | -3,532 | 4,747E-63 |
| TcCLB.510019.30 | mucin-associated surface protein (MASP), putative | -3,533 | 8,61E-60 |
| TcCLB.507305.40 | zinc finger CCCH domain containing protein 11 (ZC3H11) | -3,534 | 6,436E-10 |
| TcCLB.510921.30 | mucin-associated surface protein (MASP), putative | -3,535 | 9,125E-91 |
| TcCLB.506321.120 | mucin-associated surface protein (MASP), putative | -3,537 | 1,385E-208 |
| TcCLB.508273.120 | mucin TcMUCI, putative | -3,537 | 2,143E-36 |
| TcCLB.510475.70 | mucin TcMUCII, putative | -3,541 | 3,143E-45 |
| TcCLB.508427.20 | mucin-associated surface protein (MASP), putative | -3,545 | 1,002E-93 |
| TcCLB.408445.10 | trans-sialidase, putative (fragment) | -3,546 | 2,821E-61 |
| TcCLB.443899.10 | trans-sialidase, putative (fragment) | -3,546 | 1,477E-12 |
| TcCLB.511553.30 | mucin-associated surface protein (MASP), putative | -3,548 | 5,802E-151 |
| TcCLB.508977.10 | trans-sialidase, Group V, putative | -3,549 | 1,22E-102 |
| TcCLB.508219.10 | mucin-associated surface protein (MASP, pseudogene), pu | -3,549 | 8,972E-48 |
| TcCLB.504031.11 | hypothetical protein | -3,552 | 1,952E-80 |
| TcCLB.508273.100 | mucin TcMUCI, putative | -3,553 | 1,76E-62 |
| TcCLB.506799.110 | hypothetical protein | -3,554 | 1,233E-47 |
| TcCLB.511785.10 | mucin-associated surface protein (MASP), putative | -3,557 | 1,647E-37 |
| TcCLB.508247.40 | mucin TcMUCII, putative | -3,559 | 6,521E-36 |
| TcCLB.507071.349 | mucin-associated surface protein (MASP), putative | -3,559 | 4,982E-22 |
| TcCLB.508613.74 | mucin-associated surface protein (MASP), putative | -3,562 | 1,872E-102 |
| TcCLB.508979.50 | trans-sialidase, Group VI, putative | -3,562 | 2,406E-63 |
| TcCLB.506759.80 | mucin-associated surface protein (MASP), putative | -3,562 | 1,324E-55 |

| | | | |
|---|---|---|---|
| TcCLB.506501.350 | mucin-associated surface protein (MASP), putative | -3,575 | 1,569E-37 |
| TcCLB.507953.10 | trans-sialidase, Group V, putative | -3,575 | 3,277E-29 |
| TcCLB.508541.110 | mucin-associated surface protein (MASP), putative | -3,576 | 1,228E-77 |
| TcCLB.508627.40 | trans-sialidase, Group VIII, putative | -3,578 | 3,518E-175 |
| TcCLB.511255.30 | mucin-associated surface protein (MASP), putative | -3,578 | 4,724E-60 |
| TcCLB.511605.30 | mucin TcMUCII, putative | -3,581 | 1,591E-21 |
| TcCLB.506289.120 | mucin-associated surface protein (MASP), putative | -3,582 | 3,052E-122 |
| TcCLB.508777.210 | hypothetical protein | -3,585 | 1,374E-33 |
| TcCLB.510699.40 | mucin TcMUCII, putative | -3,59 | 2,562E-46 |
| TcCLB.421173.4 | trans-sialidase, putative (fragment) | -3,592 | 2,643E-117 |
| TcCLB.510713.170 | mucin-associated surface protein (MASP), putative | -3,592 | 8,227E-49 |
| TcCLB.508165.410 | mucin-associated surface protein (MASP), putative | -3,595 | 4,32E-76 |
| TcCLB.511791.10 | mucin-associated surface protein (MASP), putative | -3,596 | 1,016E-21 |
| TcCLB.510465.100 | trans-sialidase, Group V, putative | -3,605 | 5,011E-48 |
| TcCLB.457977.10 | trans-sialidase, putative (fragment) | -3,611 | 4,049E-46 |
| TcCLB.508977.40 | trans-sialidase, Group V, putative | -3,613 | 7,014E-140 |
| TcCLB.506345.90 | trans-sialidase, Group II, putative | -3,614 | 3,726E-105 |
| TcCLB.508485.30 | retrotransposon hot spot protein (RHS, pseudogene), puta | -3,614 | 5,269E-39 |
| TcCLB.509081.140 | mucin TcMUCII, putative | -3,617 | 4,792E-74 |
| TcCLB.506159.5 | trans-sialidase, putative (fragment) | -3,62 | 1,015E-33 |
| TcCLB.506737.161 | mucin TcMUCI, putative | -3,621 | 3,101E-87 |
| TcCLB.506763.150 | mucin-associated surface protein (MASP), putative | -3,621 | 2,442E-30 |
| TcCLB.511875.80 | mucin-associated surface protein (MASP), putative | -3,622 | 1,048E-144 |
| TcCLB.506767.340 | mucin-associated surface protein (MASP), putative | -3,627 | 1,936E-147 |
| TcCLB.508977.120 | trans-sialidase, Group V, putative | -3,628 | 5,528E-158 |
| TcCLB.510363.270 | mucin TcMUCII, putative | -3,629 | 4,689E-09 |
| TcCLB.506759.90 | trans-sialidase (pseudogene), putative | -3,63 | 3,207E-70 |
| TcCLB.506917.60 | mucin TcMUCII, putative | -3,63 | 6,23E-28 |
| TcCLB.506057.101 | surface protease GP63, putative (fragment) | -3,631 | 5,401E-12 |
| TcCLB.508761.30 | mucin-associated surface protein (MASP), putative | -3,639 | 5,615E-164 |
| TcCLB.511861.50 | trans-sialidase, Group VIII, putative | -3,639 | 1,237E-70 |
| TcCLB.508305.50 | mucin-associated surface protein (MASP), putative | -3,64 | 2,023E-86 |
| TcCLB.510477.70 | mucin TcMUCII, putative | -3,644 | 6,068E-47 |
| TcCLB.508359.80 | mucin-associated surface protein (MASP), putative | -3,645 | 9,067E-35 |
| TcCLB.510567.10 | trans-sialidase, putative (fragment) | -3,645 | 1,909E-22 |
| TcCLB.506267.70 | mucin-associated surface protein (MASP), putative | -3,648 | 1,095E-59 |
| TcCLB.507275.61 | surface antigen TASV-A1 (TASV-A1) | -3,658 | 6,472E-131 |
| TcCLB.432997.10 | trans-sialidase, putative | -3,658 | 3,056E-58 |
| TcCLB.507949.210 | trans-sialidase, Group II, putative | -3,659 | 3,759E-178 |
| TcCLB.509527.70 | mucin TcMUCII, putative | -3,66 | 2,447E-52 |
| TcCLB.511605.80 | mucin-associated surface protein (MASP), putative | -3,662 | 1,865E-46 |
| TcCLB.509525.120 | mucin TcMUCII, putative | -3,663 | 5,915E-41 |
| TcCLB.510013.230 | mucin-associated surface protein (MASP), putative | -3,665 | 4,396E-67 |
| TcCLB.509527.130 | hypothetical protein | -3,666 | 2,628E-72 |
| TcCLB.511091.9 | mucin-associated surface protein (MASP), putative | -3,666 | 1,458E-28 |

| | | | |
|---|---|---|---|
| TcCLB.507997.30 | trans-sialidase, Group V, putative | -3,668 | 4,828E-133 |
| TcCLB.507065.29 | mucin-associated surface protein (MASP, pseudogene), pu | -3,677 | 2,368E-74 |
| TcCLB.507041.71 | mucin TcMUCI, putative | -3,68 | 4,25E-62 |
| TcCLB.506285.50 | mucin TcMUCII, putative | -3,683 | 5,505E-116 |
| TcCLB.510275.260 | trans-sialidase (pseudogene), putative | -3,686 | 8E-71 |
| TcCLB.507071.310 | mucin-associated surface protein (MASP), putative | -3,69 | 1,314E-52 |
| TcCLB.511757.70 | trans-sialidase, Group II, putative | -3,691 | 1,911E-142 |
| TcCLB.504081.170 | mucin-associated surface protein (MASP), putative | -3,693 | 1,072E-69 |
| TcCLB.510279.110 | trans-sialidase, Group VI, putative | -3,693 | 1,54E-52 |
| TcCLB.413489.10 | retrotransposon hot spot protein (RHS, pseudogene), puta | -3,695 | 1,017E-23 |
| TcCLB.510081.40 | trans-sialidase (pseudogene), putative | -3,702 | 4,622E-11 |
| TcCLB.509699.40 | trans-sialidase, Group V, putative | -3,707 | 2,757E-111 |
| TcCLB.506767.130 | mucin TcMUCII, putative | -3,707 | 7,121E-24 |
| TcCLB.508295.80 | mucin TcMUCII, putative | -3,71 | 2,402E-55 |
| TcCLB.511603.170 | mucin TcMUCII, putative | -3,713 | 5,478E-38 |
| TcCLB.510583.100 | mucin TcMUCII, putative | -3,713 | 2,211E-31 |
| TcCLB.510371.90 | trans-sialidase, Group V, putative | -3,714 | 1,516E-232 |
| TcCLB.510927.20 | trans-sialidase (pseudogene), putative | -3,714 | 1,342E-222 |
| TcCLB.506321.50 | mucin-associated surface protein (MASP), putative | -3,716 | 0 |
| TcCLB.510377.200 | mucin-associated surface protein (MASP), putative | -3,718 | 1,361E-98 |
| TcCLB.508871.40 | mucin-associated surface protein (MASP), putative | -3,718 | 4,931E-64 |
| TcCLB.511785.5 | hypothetical protein | -3,72 | 3,369E-19 |
| TcCLB.511127.10 | RNA-binding protein 5, putative | -3,721 | 6,917E-16 |
| TcCLB.510881.9 | trans-sialidase, putative | -3,723 | 5,333E-53 |
| TcCLB.506331.90 | trans-sialidase, Group II, putative | -3,729 | 5,506E-95 |
| TcCLB.511551.10 | trans-sialidase, Group II, putative | -3,729 | 1,699E-43 |
| TcCLB.511173.470 | trans-sialidase, Group V, putative | -3,73 | 2,086E-114 |
| TcCLB.511401.30 | trans-sialidase, Group V, putative | -3,736 | 1,791E-118 |
| TcCLB.507085.170 | mucin TcMUCII, putative | -3,738 | 2,198E-109 |
| TcCLB.510233.39 | trans-sialidase, putative (fragment) | -3,738 | 1,567E-35 |
| TcCLB.509259.40 | surface protease GP63, putative | -3,74 | 2,256E-49 |
| TcCLB.504239.100 | mucin TcMUCII, putative | -3,741 | 4,694E-65 |
| TcCLB.511549.10 | retrotransposon hot spot protein (RHS, pseudogene), puta | -3,745 | 1,14E-34 |
| TcCLB.510013.260 | mucin-associated surface protein (MASP), putative | -3,748 | 1,644E-78 |
| TcCLB.507957.280 | mucin-associated surface protein (MASP), putative | -3,752 | 6,938E-119 |
| TcCLB.508099.50 | trans-sialidase, putative | -3,752 | 1,656E-14 |
| TcCLB.510715.30 | mucin-associated surface protein (MASP), putative | -3,759 | 7,223E-117 |
| TcCLB.510281.10 | hypothetical protein | -3,76 | 3,534E-40 |
| TcCLB.506615.50 | mucin TcMUCII, putative | -3,762 | 2,796E-295 |
| TcCLB.508615.10 | mucin-associated surface protein (MASP), putative | -3,762 | 1,998E-55 |
| TcCLB.510495.20 | mucin-associated surface protein (MASP), putative | -3,762 | 5,26E-28 |
| TcCLB.509011.80 | surface protease GP63, putative | -3,764 | 2,977E-173 |
| TcCLB.433559.10 | trans-sialidase, putative (fragment) | -3,766 | 3,376E-38 |
| TcCLB.508493.10 | mucin TcMUCII, putative | -3,769 | 3,347E-168 |
| TcCLB.506767.140 | mucin-associated surface protein (MASP), putative | -3,769 | 3,535E-65 |

| | | | |
|---|---|---|---|
| TcCLB.510463.90 | mucin-associated surface protein (MASP), putative | -3,772 | 6,042E-09 |
| TcCLB.507959.110 | mucin-associated surface protein (MASP), putative | -3,774 | 4,885E-57 |
| TcCLB.511173.280 | trans-sialidase, Group V, putative | -3,777 | 5,356E-95 |
| TcCLB.504239.30 | mucin-associated surface protein (MASP), putative | -3,779 | 2,097E-69 |
| TcCLB.509419.70 | trans-sialidase, Group V, putative | -3,782 | 2,09E-80 |
| TcCLB.511083.30 | mucin-associated surface protein (MASP), putative | -3,782 | 3,621E-35 |
| TcCLB.504239.260 | trans-sialidase, Group V, putative | -3,788 | 2,378E-66 |
| TcCLB.510465.74 | mucin-associated surface protein (MASP), putative | -3,791 | 3,932E-39 |
| TcCLB.508007.10 | trans-sialidase, putative | -3,797 | 3,139E-10 |
| TcCLB.509771.10 | trans-sialidase, Group II, putative | -3,803 | 1,483E-45 |
| TcCLB.508903.100 | ATPase, putative | -3,804 | 1,656E-53 |
| TcCLB.510539.10 | mucin-associated surface protein (MASP), putative | -3,805 | 2,881E-18 |
| TcCLB.507687.10 | trans-sialidase, putative | -3,806 | 7,781E-153 |
| TcCLB.511129.40 | trans-sialidase, Group III, putative | -3,81 | 1,995E-12 |
| TcCLB.508305.20 | mucin-associated surface protein (MASP), putative | -3,812 | 1,08E-75 |
| TcCLB.506133.54 | mucin-associated surface protein (MASP), putative | -3,813 | 1,131E-87 |
| TcCLB.510279.250 | mucin-associated surface protein (MASP), putative | -3,814 | 2,341E-35 |
| TcCLB.510373.10 | mucin-associated surface protein (MASP), putative | -3,821 | 1,683E-127 |
| TcCLB.510483.70 | mucin-associated surface protein (MASP), putative | -3,822 | 4,328E-100 |
| TcCLB.510625.54 | mucin-associated surface protein (MASP), putative | -3,823 | 1,123E-62 |
| TcCLB.509525.90 | mucin TcMUCII, putative | -3,823 | 1,118E-44 |
| TcCLB.507957.179 | mucin-associated surface protein (MASP), putative | -3,829 | 9,111E-65 |
| TcCLB.510475.90 | mucin TcMUCII, putative | -3,835 | 2,841E-63 |
| TcCLB.505997.70 | ATPase, putative | -3,837 | 4,76E-39 |
| TcCLB.463955.20 | mucin-associated surface protein (MASP), putative (fragm | -3,837 | 1,284E-25 |
| TcCLB.511183.210 | mucin TcMUCII, putative | -3,839 | 4,677E-99 |
| TcCLB.511057.30 | mucin-associated surface protein (MASP), putative | -3,844 | 8,427E-59 |
| TcCLB.510361.130 | serine-alanine-and proline-rich protein, putative | -3,845 | 4,794E-27 |
| TcCLB.509699.140 | mucin-associated surface protein (MASP, pseudogene), pu | -3,846 | 4,227E-88 |
| TcCLB.506765.50 | mucin-associated surface protein (MASP), putative | -3,856 | 2,137E-70 |
| TcCLB.510013.40 | mucin-associated surface protein (MASP), putative | -3,858 | 1,769E-70 |
| TcCLB.508103.30 | trans-sialidase, Group V, putative | -3,858 | 3,794E-36 |
| TcCLB.506131.110 | mucin-associated surface protein (MASP), putative | -3,864 | 1,18E-54 |
| TcCLB.510021.160 | 90 kDa surface protein, putative | -3,87 | 6,447E-81 |
| TcCLB.508161.20 | trans-sialidase, Group V, putative | -3,87 | 1,69E-76 |
| TcCLB.510371.70 | mucin-associated surface protein (MASP), putative | -3,871 | 1,666E-73 |
| TcCLB.510157.10 | mucin-associated surface protein (MASP), putative | -3,875 | 6,36E-125 |
| TcCLB.508165.130 | surface protease GP63, putative | -3,875 | 1,073E-86 |
| TcCLB.503907.10 | trans-sialidase, Group VIII, putative | -3,88 | 5,095E-36 |
| TcCLB.508289.9 | mucin-associated surface protein (MASP), putative | -3,882 | 2,056E-31 |
| TcCLB.507043.10 | mucin TcMUCI, putative | -3,883 | 3,904E-73 |
| TcCLB.504039.230 | mucin-associated surface protein (MASP), putative | -3,884 | 1,451E-78 |
| TcCLB.511449.10 | mucin-associated surface protein (MASP), putative | -3,886 | 2,687E-57 |
| TcCLB.503859.40 | mucin-associated surface protein (MASP), putative | -3,888 | 3,055E-115 |
| TcCLB.506763.130 | mucin-associated surface protein (MASP), putative | -3,891 | 2,159E-97 |

| | | | |
|---|---|---|---|
| TcCLB.506599.100 | mucin-associated surface protein (MASP), putative | -3,893 | 1,442E-63 |
| TcCLB.508165.270 | mucin TcMUCII, putative | -3,893 | 2,188E-32 |
| TcCLB.510047.60 | mucin-associated surface protein (MASP, pseudogene), pu | -3,9 | 2,56E-134 |
| TcCLB.503501.50 | trans-sialidase (pseudogene), putative | -3,903 | 1,949E-95 |
| TcCLB.503957.10 | trans-sialidase, putative | -3,907 | 3,383E-188 |
| TcCLB.506767.240 | mucin-associated surface protein (MASP), putative | -3,91 | 1,582E-89 |
| TcCLB.506471.30 | trans-sialidase, Group II, putative | -3,916 | 3,826E-163 |
| TcCLB.507953.180 | mucin TcMUCII, putative | -3,921 | 6,261E-101 |
| TcCLB.504081.330 | mucin TcMUCII, putative | -3,922 | 7,634E-45 |
| TcCLB.506967.10 | trans-sialidase, Group V, putative | -3,925 | 4,111E-102 |
| TcCLB.507523.60 | mucin-associated surface protein (MASP), putative | -3,936 | 7,097E-61 |
| TcCLB.508163.20 | mucin TcMUCII, putative | -3,936 | 4,374E-49 |
| TcCLB.510275.60 | mucin-associated surface protein (MASP), putative | -3,944 | 3,194E-70 |
| TcCLB.509525.220 | mucin TcMUCII, putative | -3,947 | 7,792E-33 |
| TcCLB.509525.350 | mucin TcMUCII, putative | -3,951 | 1,67E-46 |
| TcCLB.506087.10 | retrotransposon hot spot protein (RHS, pseudogene), puta | -3,951 | 1,065E-22 |
| TcCLB.508595.20 | UDP-Gal or UDP-GlcNAc-dependent glycosyltransferase, p | -3,953 | 1,754E-130 |
| TcCLB.511089.30 | mucin-associated surface protein (MASP), putative | -3,953 | 8,939E-65 |
| TcCLB.510489.20 | mucin-associated surface protein (MASP), putative | -3,955 | 8,979E-104 |
| TcCLB.506955.99 | mucin-associated surface protein (MASP), putative | -3,956 | 4,485E-105 |
| TcCLB.508985.21 | mucin TcMUCII, putative (fragment) | -3,957 | 1,493E-12 |
| TcCLB.510487.50 | mucin-associated surface protein (MASP), putative | -3,961 | 1,663E-100 |
| TcCLB.507953.70 | 90 kDa surface protein, putative | -3,963 | 2,732E-68 |
| TcCLB.504239.220 | mucin-associated surface protein (MASP52) | -3,963 | 6,491E-64 |
| TcCLB.510769.170 | trans-sialidase (pseudogene), putative | -3,966 | 1,762E-290 |
| TcCLB.510583.40 | mucin TcMUCII, putative | -3,979 | 1,489E-53 |
| TcCLB.506671.90 | mucin-associated surface protein (MASP), putative | -3,98 | 9,623E-46 |
| TcCLB.507957.300 | mucin-associated surface protein (MASP), putative | -3,98 | 3,777E-35 |
| TcCLB.508637.10 | trans-sialidase, Group III, putative | -3,985 | 2,242E-65 |
| TcCLB.509185.30 | trans-sialidase, Group II, putative | -3,987 | 2,04E-12 |
| TcCLB.508165.40 | trans-sialidase (pseudogene), putative | -3,989 | 3,743E-114 |
| TcCLB.506067.140 | mucin-associated surface protein (MASP), putative | -3,99 | 1,135E-69 |
| TcCLB.505363.19 | trans-sialidase, putative | -3,991 | 9,916E-219 |
| TcCLB.506133.61 | mucin-associated surface protein (MASP), putative | -3,996 | 8,553E-42 |
| TcCLB.508365.10 | mucin TcMUCII, putative | -4,001 | 4,261E-55 |
| TcCLB.511173.496 | mucin-associated surface protein (MASP), putative (fragm | -4,001 | 6,824E-16 |
| TcCLB.506501.80 | mucin-associated surface protein (MASP, pseudogene), pu | -4,003 | 3,439E-189 |
| TcCLB.506187.50 | mucin-associated surface protein (MASP), putative | -4,007 | 5,457E-84 |
| TcCLB.505025.40 | mucin-associated surface protein (MASP), putative | -4,008 | 5,898E-84 |
| TcCLB.503771.90 | mucin-associated surface protein (MASP), putative | -4,01 | 1,166E-54 |
| TcCLB.508071.50 | surface protease GP63, putative | -4,012 | 1,117E-44 |
| TcCLB.507357.90 | mucin-associated surface protein (MASP), putative | -4,013 | 5,574E-20 |
| TcCLB.506761.19 | mucin TcMUCII, putative | -4,016 | 6,011E-64 |
| TcCLB.503705.11 | UDP-Gal or UDP-GlcNAc-dependent glycosyltransferase (p | -4,016 | 6,374E-24 |
| TcCLB.510275.350 | trans-sialidase, Group V, putative | -4,018 | 2,453E-83 |

| | | | |
|---|---|---|---|
| TcCLB.508433.10 | mucin-associated surface protein (MASP), putative | -4,024 | 1,563E-15 |
| TcCLB.510583.70 | mucin TcMUCII, putative | -4,027 | 9,866E-82 |
| TcCLB.506975.90 | trans-sialidase, Group VIII, putative | -4,029 | 1,381E-99 |
| TcCLB.506139.40 | mucin-associated surface protein (MASP), putative | -4,029 | 7,159E-98 |
| TcCLB.507145.79 | mucin TcMUCII, putative | -4,031 | 1,078E-06 |
| TcCLB.508977.80 | trans-sialidase, Group V, putative | -4,032 | 7,715E-116 |
| TcCLB.506131.190 | mucin-associated surface protein (MASP), putative | -4,046 | 1,126E-80 |
| TcCLB.510693.163 | mucin-associated surface protein (MASP), putative | -4,048 | 8,206E-300 |
| TcCLB.508435.70 | mucin-associated surface protein (MASP), putative | -4,055 | 1,425E-87 |
| TcCLB.510377.350 | mucin-associated surface protein (MASP), putative | -4,059 | 1,334E-112 |
| TcCLB.507163.70 | trans-sialidase, Group V, putative | -4,063 | 5,661E-229 |
| TcCLB.398477.10 | trans-sialidase, Group II, putative | -4,067 | 8,06E-146 |
| TcCLB.507059.10 | mucin-like glycoprotein, putative | -4,068 | 1,058E-88 |
| TcCLB.507957.220 | mucin-associated surface protein (MASP), putative | -4,074 | 7,441E-128 |
| TcCLB.506137.40 | mucin TcMUCII, putative | -4,074 | 3,166E-24 |
| TcCLB.507233.10 | trans-sialidase, Group III, putative | -4,08 | 1,588E-67 |
| TcCLB.506759.40 | mucin-associated surface protein (MASP), putative | -4,081 | 1,066E-176 |
| TcCLB.463955.10 | mucin TcMUCII, putative | -4,082 | 3,714E-69 |
| TcCLB.508853.10 | mucin-associated surface protein (MASP), putative | -4,086 | 1,151E-106 |
| TcCLB.509525.250 | mucin TcMUCII, putative | -4,086 | 6,344E-46 |
| TcCLB.506597.30 | mucin TcMUCII, putative | -4,091 | 2,838E-63 |
| TcCLB.506993.160 | trans-sialidase (pseudogene), putative | -4,093 | 7,083E-120 |
| TcCLB.511401.90 | trans-sialidase, Group V, putative | -4,095 | 4,619E-140 |
| TcCLB.454787.10 | mucin TcMUC, putative | -4,108 | 1,578E-08 |
| TcCLB.511173.130 | trans-sialidase, Group V, putative | -4,109 | 8,411E-178 |
| TcCLB.506843.5 | trans-sialidase, putative (fragment) | -4,124 | 1,841E-28 |
| TcCLB.506245.140 | trans-sialidase (pseudogene), putative | -4,126 | 1,379E-30 |
| TcCLB.506599.350 | mucin TcMUCII, putative | -4,137 | 2,658E-21 |
| TcCLB.504239.410 | mucin TcMUCII, putative | -4,138 | 5,71E-72 |
| TcCLB.506799.100 | mucin-like glycoprotein, putative | -4,147 | 2,464E-131 |
| TcCLB.506501.280 | mucin TcMUCII, putative | -4,151 | 3,737E-36 |
| TcCLB.509973.10 | mucin-associated surface protein (MASP), putative | -4,155 | 1,183E-263 |
| TcCLB.506955.212 | mucin-associated surface protein (MASP), putative | -4,161 | 1,224E-121 |
| TcCLB.507849.10 | trans-sialidase (pseudogene), putative | -4,161 | 3,268E-44 |
| TcCLB.510021.200 | mucin TcMUCII, putative | -4,162 | 5,806E-55 |
| TcCLB.403031.10 | trans-sialidase, putative (fragment) | -4,165 | 2,137E-14 |
| TcCLB.509529.40 | trans-sialidase, Group VII, putative | -4,174 | 4,466E-218 |
| TcCLB.510037.10 | mucin-associated surface protein (MASP), putative | -4,175 | 6,759E-168 |
| TcCLB.506887.10 | trans-sialidase, putative | -4,182 | 5,416E-61 |
| TcCLB.510025.210 | mucin TcMUCII, putative | -4,183 | 3,236E-56 |
| TcCLB.510693.320 | mucin-associated surface protein (MASP), putative | -4,197 | 7,5E-143 |
| TcCLB.477471.10 | trans-sialidase (pseudogene), putative | -4,198 | 1,648E-178 |
| TcCLB.510713.130 | mucin-associated surface protein (MASP), putative | -4,2 | 3,141E-84 |
| TcCLB.510275.355 | hypothetical protein | -4,2 | 1,127E-56 |
| TcCLB.510049.10 | trans-sialidase, putative | -4,201 | 7,756E-163 |

| | | | |
|---|---|---|---|
| TcCLB.504081.420 | mucin-associated surface protein (MASP), putative | -4,202 | 9,257E-42 |
| TcCLB.504929.5 | zinc finger CCCH domain containing protein 11 (ZC3H11) | -4,206 | 5,642E-16 |
| TcCLB.508493.100 | mucin TcMUCII, putative | -4,209 | 4,014E-247 |
| TcCLB.506769.50 | mucin TcMUCII, putative | -4,21 | 9,374E-57 |
| TcCLB.510025.70 | 90 kDa surface protein, putative | -4,217 | 1,433E-98 |
| TcCLB.510359.520 | mucin-associated surface protein (MASP), putative | -4,217 | 1,505E-10 |
| TcCLB.507653.20 | trans-sialidase, Group II, putative | -4,228 | 6,095E-127 |
| TcCLB.510807.40 | mucin TcMUCII, putative | -4,228 | 3,312E-51 |
| TcCLB.508481.20 | mucin-associated surface protein (MASP), putative | -4,245 | 4,969E-11 |
| TcCLB.510465.140 | mucin-associated surface protein (MASP), putative | -4,246 | 1,881E-90 |
| TcCLB.509527.10 | mucin TcMUCII, putative | -4,248 | 9,991E-67 |
| TcCLB.506759.140 | 90 kDa surface protein, putative | -4,252 | 1,033E-47 |
| TcCLB.511061.40 | trans-sialidase, Group V, putative | -4,253 | 7,575E-107 |
| TcCLB.507957.150 | mucin-associated surface protein (MASP), putative | -4,253 | 6,517E-104 |
| TcCLB.509525.320 | mucin TcMUCII, putative | -4,255 | 7,809E-59 |
| TcCLB.507953.60 | hypothetical protein | -4,257 | 6,733E-63 |
| TcCLB.504035.41 | mucin-associated surface protein (MASP, pseudogene), pu | -4,268 | 1,376E-07 |
| TcCLB.508225.4 | trans-sialidase, putative | -4,271 | 5,364E-130 |
| TcCLB.507875.70 | trans-sialidase, Group VII, putative | -4,274 | 5,002E-48 |
| TcCLB.511603.110 | mucin TcMUCII, putative | -4,275 | 3,571E-63 |
| TcCLB.430603.20 | mucin TcMUCII, putative | -4,286 | 1,811E-79 |
| TcCLB.510853.40 | trans-sialidase, Group IV, putative | -4,292 | 0 |
| TcCLB.511401.120 | trans-sialidase, Group V, putative | -4,295 | 7,64E-130 |
| TcCLB.506737.114 | mucin TcMUCI, putative | -4,296 | 1,248E-158 |
| TcCLB.509115.60 | mucin-like glycoprotein, putative | -4,303 | 7,771E-176 |
| TcCLB.503827.30 | mucin TcMUCII, putative | -4,304 | 3,031E-34 |
| TcCLB.506599.420 | mucin-associated surface protein (MASP), putative | -4,313 | 5,884E-79 |
| TcCLB.472357.10 | trans-sialidase, Group V, putative | -4,331 | 2,101E-74 |
| TcCLB.506597.40 | trans-sialidase, Group V, putative | -4,334 | 1,441E-17 |
| TcCLB.509281.10 | hypothetical protein | -4,335 | 3,128E-65 |
| TcCLB.508581.30 | beta galactofuranosyl glycosyltransferase (pseudogene), p | -4,335 | 3,266E-16 |
| TcCLB.506757.150 | mucin-associated surface protein (MASP), putative | -4,34 | 5,208E-230 |
| TcCLB.507121.11 | trans-sialidase, putative (fragment) | -4,343 | 3,389E-60 |
| TcCLB.510985.10 | trans-sialidase, putative (fragment) | -4,351 | 2,746E-30 |
| TcCLB.506331.130 | trans-sialidase, Group II, putative | -4,353 | 1,856E-184 |
| TcCLB.508295.30 | mucin TcMUCII, putative | -4,358 | 1,362E-64 |
| TcCLB.509753.194 | mucin-associated surface protein (MASP), putative | -4,367 | 8,032E-48 |
| TcCLB.508759.60 | mucin-associated surface protein (MASP), putative | -4,385 | 7,994E-37 |
| TcCLB.506361.50 | mucin-associated surface protein (MASP), putative | -4,386 | 4,968E-129 |
| TcCLB.509329.31 | trans-sialidase, putative | -4,388 | 3,444E-70 |
| TcCLB.506839.10 | serine/threonine kinase, putative | -4,398 | 7,29E-12 |
| TcCLB.508753.20 | trans-sialidase, Group V, putative | -4,405 | 2,376E-159 |
| TcCLB.511563.20 | trans-sialidase (pseudogene), putative | -4,406 | 3,149E-40 |
| TcCLB.508109.60 | trans-sialidase, Group V, putative | -4,407 | 2,917E-91 |
| TcCLB.506603.40 | trans-sialidase, Group V, putative | -4,41 | 4,598E-202 |

| | | | |
|---|---|---|---|
| TcCLB.504031.50 | trans-sialidase, Group V, putative | -4,418 | 6,185E-105 |
| TcCLB.506759.70 | mucin TcMUCII, putative | -4,42 | 9,223E-35 |
| TcCLB.510701.10 | mucin-associated surface protein (MASP), putative | -4,428 | 1,093E-130 |
| TcCLB.508121.150 | trans-sialidase, Group V, putative | -4,443 | 2,095E-91 |
| TcCLB.507219.9 | trans-sialidase, putative | -4,446 | 9,592E-58 |
| TcCLB.509313.9 | trans-sialidase (pseudogene), putative | -4,448 | 3,27E-23 |
| TcCLB.509601.10 | neutral sphingomyelinase activation associated factor-like | -4,455 | 0 |
| TcCLB.507479.70 | trans-sialidase, Group IV, putative | -4,466 | 1,396E-298 |
| TcCLB.508147.150 | mucin-associated surface protein (MASP), putative | -4,466 | 5,694E-84 |
| TcCLB.507125.10 | trans-sialidase, Group II, putative | -4,47 | 0 |
| TcCLB.506501.290 | mucin-associated surface protein (MASP), putative | -4,485 | 4,716E-114 |
| TcCLB.506799.20 | mucin-like glycoprotein, putative | -4,495 | 2,345E-174 |
| TcCLB.435601.10 | trans-sialidase, putative (fragment) | -4,496 | 3,039E-137 |
| TcCLB.506973.120 | trans-sialidase, Group V, putative | -4,5 | 4,124E-155 |
| TcCLB.510477.10 | surface protease GP63, putative | -4,501 | 5,669E-274 |
| TcCLB.509325.10 | mucin-associated surface protein (MASP), putative | -4,507 | 8,535E-21 |
| TcCLB.507953.100 | trans-sialidase, Group V, putative | -4,514 | 2,329E-182 |
| TcCLB.510497.170 | trans-sialidase (pseudogene), putative | -4,518 | 1,356E-49 |
| TcCLB.511907.120 | mucin-like glycoprotein, putative | -4,524 | 9,782E-163 |
| TcCLB.510935.20 | surface protease GP63, putative | -4,53 | 1,172E-253 |
| TcCLB.509115.50 | mucin-like glycoprotein, putative | -4,536 | 7,863E-221 |
| TcCLB.511081.10 | mucin-associated surface protein (MASP), putative | -4,548 | 1,394E-195 |
| TcCLB.506269.40 | mucin-associated surface protein (MASP), putative | -4,553 | 1,563E-139 |
| TcCLB.508125.30 | mucin TcMUCII, putative | -4,556 | 4,426E-71 |
| TcCLB.511553.200 | mucin-associated surface protein (MASP), putative | -4,558 | 6,976E-105 |
| TcCLB.511173.10 | trans-sialidase, putative | -4,569 | 1,069E-53 |
| TcCLB.510039.30 | mucin-associated surface protein (MASP), putative | -4,57 | 1,219E-266 |
| TcCLB.447899.10 | trans-sialidase, putative | -4,57 | 6,19E-66 |
| TcCLB.504755.10 | surface protease GP63, putative | -4,571 | 3,441E-33 |
| TcCLB.510205.40 | trans-sialidase, Group V, putative | -4,583 | 1,612E-211 |
| TcCLB.509325.16 | elongation factor 1-gamma (EF-1-gamma, pseudogene), | -4,585 | 6,671E-32 |
| TcCLB.511057.40 | trans-sialidase, Group V, putative | -4,588 | 0 |
| TcCLB.503861.40 | trans-sialidase, Group VIII, putative | -4,591 | 6,461E-234 |
| TcCLB.506251.70 | trans-sialidase (pseudogene), putative | -4,598 | 5,701E-26 |
| TcCLB.509525.380 | mucin TcMUCII, putative | -4,613 | 1,441E-57 |
| TcCLB.511831.10 | trans-sialidase, putative (fragment) | -4,618 | 4,699E-168 |
| TcCLB.509527.120 | trans-sialidase, Group V, putative | -4,621 | 4,282E-166 |
| TcCLB.506885.210 | trans-sialidase, Group III, putative | -4,625 | 5,111E-188 |
| TcCLB.509931.20 | trans-sialidase, Group VIII, putative | -4,631 | 9,672E-157 |
| TcCLB.506285.120 | mucin TcMUCII, putative | -4,633 | 5,056E-135 |
| TcCLB.403481.10 | trans-sialidase, putative (fragment) | -4,634 | 1,403E-47 |
| TcCLB.507067.60 | mucin-associated surface protein (MASP), putative | -4,635 | 4,107E-129 |
| TcCLB.510377.370 | mucin TcMUCII, putative | -4,648 | 1,778E-128 |
| TcCLB.507359.20 | surface antigen TASV, putative | -4,651 | 7,148E-92 |
| TcCLB.511779.60 | trans-sialidase, Group III, putative | -4,658 | 5,423E-97 |

| | | | |
|---|---|---|---|
| TcCLB.506885.140 | trans-sialidase (pseudogene), putative | -4,663 | 5,24E-195 |
| TcCLB.508389.90 | surface protease GP63, putative | -4,684 | 5,808E-153 |
| TcCLB.510933.120 | surface protease GP63, putative | -4,686 | 0 |
| TcCLB.504493.10 | mucin-associated surface protein (MASP), putative (fragm | -4,694 | 6,999E-98 |
| TcCLB.508903.110 | trans-sialidase, putative | -4,695 | 7,503E-155 |
| TcCLB.510569.10 | trans-sialidase, putative | -4,695 | 3,766E-130 |
| TcCLB.506283.20 | trans-sialidase, putative | -4,696 | 8,671E-24 |
| TcCLB.508219.90 | 90 kDa surface protein, putative | -4,7 | 1,651E-83 |
| TcCLB.508121.90 | mucin-associated surface protein (MASP, pseudogene), pu | -4,706 | 5,494E-137 |
| TcCLB.508021.10 | trans-sialidase, putative | -4,716 | 2,263E-09 |
| TcCLB.511873.10 | trans-sialidase, Group V, putative | -4,74 | 1,169E-189 |
| TcCLB.506643.10 | mucin-associated surface protein (MASP), putative | -4,742 | 4,259E-34 |
| TcCLB.511835.10 | trans-sialidase, Group VIII, putative | -4,75 | 1,7E-60 |
| TcCLB.478463.10 | hypothetical protein | -4,765 | 9,203E-40 |
| TcCLB.506139.110 | mucin-associated surface protein (MASP), putative | -4,768 | 1,197E-71 |
| TcCLB.508231.120 | trans-sialidase (pseudogene), putative | -4,772 | 1,325E-62 |
| TcCLB.510371.100 | mucin-associated surface protein (MASP), putative | -4,777 | 6,564E-126 |
| TcCLB.506471.120 | trans-sialidase, Group II, putative | -4,784 | 1,696E-236 |
| TcCLB.510483.20 | mucin-associated surface protein (MASP), putative | -4,798 | 1,141E-130 |
| TcCLB.421951.10 | trans-sialidase, putative (fragment) | -4,818 | 6,389E-68 |
| TcCLB.455171.9 | trans-sialidase, putative | -4,832 | 6,833E-204 |
| TcCLB.506971.20 | surface protease GP63, putative | -4,844 | 1,381E-199 |
| TcCLB.507957.240 | mucin-associated surface protein (MASP), putative | -4,849 | 7,456E-102 |
| TcCLB.505025.150 | trans-sialidase (pseudogene), putative | -4,859 | 1,396E-99 |
| TcCLB.510307.240 | trans-sialidase, Group VIII, putative | -4,863 | 2,407E-297 |
| TcCLB.506599.50 | mucin-associated surface protein (MASP), putative | -4,866 | 6,457E-119 |
| TcCLB.508165.430 | trans-sialidase, putative | -4,88 | 6,276E-216 |
| TcCLB.511175.10 | mucin-associated surface protein (MASP), putative | -4,887 | 1,126E-13 |
| TcCLB.506763.290 | mucin TcMUCII, putative | -4,888 | 3,662E-116 |
| TcCLB.508073.10 | surface protease GP63, putative | -4,893 | 2,326E-205 |
| TcCLB.507759.10 | trans-sialidase, Group VIII, putative | -4,898 | 3,258E-99 |
| TcCLB.509075.31 | hypothetical protein, conserved (pseudogene) | -4,9 | 1,904E-17 |
| TcCLB.506683.110 | trans-sialidase, Group VIII, putative | -4,909 | 0 |
| TcCLB.503429.39 | trans-sialidase, putative | -4,926 | 2,319E-154 |
| TcCLB.507357.100 | trans-sialidase, Group V, putative | -4,943 | 0 |
| TcCLB.475133.10 | trans-sialidase, putative | -4,958 | 2,855E-98 |
| TcCLB.503973.150 | mucin-associated surface protein (MASP), putative | -4,967 | 2,237E-107 |
| TcCLB.509979.320 | trans-sialidase, Group V, putative | -4,971 | 0 |
| TcCLB.509525.180 | mucin TcMUCII, putative | -4,973 | 2,739E-68 |
| TcCLB.506427.10 | trans-sialidase, Group VIII, putative | -4,974 | 6,153E-63 |
| TcCLB.507071.140 | mucin-associated surface protein (MASP), putative | -4,977 | 4,672E-116 |
| TcCLB.504239.140 | mucin-associated surface protein (MASP, pseudogene), pu | -4,978 | 1,184E-200 |
| TcCLB.509147.40 | surface antigen TASV, putative | -4,985 | 0 |
| TcCLB.505365.60 | trans-sialidase, Group VIII, putative | -4,986 | 3,959E-27 |
| TcCLB.510693.150 | mucin-associated surface protein (MASP), putative | -4,992 | 1,55E-29 |

| | | | |
|---|---|---|---|
| TcCLB.508163.250 | trans-sialidase, Group VI, putative | -4,994 | 3,333E-261 |
| TcCLB.509123.10 | surface antigen TASV, putative | -5,008 | 0 |
| TcCLB.510693.130 | mucin-associated surface protein (MASP), putative | -5,008 | 1,406E-118 |
| TcCLB.511173.24 | mucin-associated surface protein (MASP), putative | -5,023 | 9,63E-178 |
| TcCLB.510697.130 | mucin-associated surface protein (MASP), putative | -5,029 | 1,782E-139 |
| TcCLB.507637.170 | surface protease GP63, putative | -5,03 | 1,181E-272 |
| TcCLB.505567.20 | surface protease GP63, putative | -5,043 | 1,961E-81 |
| TcCLB.433061.10 | hypothetical protein | -5,043 | 1,441E-71 |
| TcCLB.508581.10 | trans-sialidase, Group VII, putative | -5,047 | 4,029E-66 |
| TcCLB.508639.10 | trans-sialidase, putative | -5,068 | 1,441E-189 |
| TcCLB.506217.40 | trans-sialidase, Group VIII, putative | -5,074 | 1,069E-67 |
| TcCLB.511675.3 | surface antigen TASV, putative | -5,076 | 0 |
| TcCLB.508483.40 | retrotransposon hot spot protein (RHS, pseudogene), puta | -5,081 | 2,817E-37 |
| TcCLB.507981.10 | trans-sialidase (pseudogene), putative | -5,085 | 4,306E-144 |
| TcCLB.508743.10 | surface antigen TASV, putative | -5,089 | 0 |
| TcCLB.511833.10 | trans-sialidase, Group VIII, putative | -5,101 | 3,178E-205 |
| TcCLB.509977.30 | mucin-associated surface protein (MASP), putative | -5,109 | 3,464E-254 |
| TcCLB.506611.70 | surface protease GP63, putative | -5,117 | 1,713E-179 |
| TcCLB.510463.240 | mucin-associated surface protein (MASP), putative | -5,117 | 3,031E-120 |
| TcCLB.507867.5 | trans-sialidase, putative (fragment) | -5,123 | 3,018E-07 |
| TcCLB.507957.200 | mucin-associated surface protein (MASP), putative | -5,124 | 2,123E-37 |
| TcCLB.508221.960 | mucin-associated surface protein (MASP), putative | -5,125 | 1,585E-224 |
| TcCLB.506877.60 | mucin-associated surface protein (MASP), putative | -5,135 | 5,316E-250 |
| TcCLB.504039.180 | mucin-associated surface protein (MASP), putative | -5,147 | 9,035E-298 |
| TcCLB.511375.10 | hypothetical protein (pseudogene) | -5,157 | 3,092E-41 |
| TcCLB.507051.11 | trans-sialidase, putative (fragment) | -5,16 | 1,565E-40 |
| TcCLB.435051.10 | trans-sialidase, putative (fragment) | -5,164 | 4,761E-120 |
| TcCLB.508745.30 | trans-sialidase, Group V, putative | -5,168 | 4,076E-142 |
| TcCLB.507069.180 | mucin-associated surface protein (MASP), putative | -5,175 | 3,919E-107 |
| TcCLB.406115.10 | trans-sialidase, putative (fragment) | -5,186 | 7,851E-264 |
| TcCLB.507875.220 | trans-sialidase, Group II, putative | -5,187 | 9,453E-252 |
| TcCLB.506397.10 | trans-sialidase, Group II, putative | -5,191 | 2,425E-07 |
| TcCLB.506967.100 | mucin-associated surface protein (MASP), putative | -5,2 | 3,836E-178 |
| TcCLB.510197.30 | trans-sialidase, Group V, putative | -5,212 | 3,389E-241 |
| TcCLB.504341.20 | trans-sialidase, putative | -5,212 | 7,088E-24 |
| TcCLB.508483.26 | trans-sialidase (pseudogene), putative | -5,217 | 1,391E-17 |
| TcCLB.506613.9 | surface protease GP63, putative (fragment) | -5,218 | 5,468E-164 |
| TcCLB.503783.17 | mucin TcMUCII, putative | -5,251 | 1,802E-92 |
| TcCLB.509737.9 | trans-sialidase, putative (fragment) | -5,257 | 2,419E-142 |
| TcCLB.505025.60 | 90 kDa surface protein, putative | -5,257 | 1,345E-83 |
| TcCLB.506763.260 | mucin-associated surface protein (MASP), putative | -5,274 | 0 |
| TcCLB.506449.10 | trans-sialidase, Group III, putative | -5,284 | 3,97E-45 |
| TcCLB.508741.440 | surface antigen TASV, putative | -5,288 | 0 |
| TcCLB.426675.9 | trans-sialidase, putative | -5,294 | 3,745E-193 |
| TcCLB.509977.80 | mucin-associated surface protein (MASP), putative | -5,3 | 0 |

| | | | |
|---|---|---|---|
| TcCLB.507753.10 | trans-sialidase, Group III, putative | -5,301 | 2,469E-92 |
| TcCLB.510021.180 | trans-sialidase, Group V, putative | -5,306 | 0 |
| TcCLB.511861.120 | surface antigen TASV, putative | -5,309 | 0 |
| TcCLB.510125.20 | trans-sialidase, Group VIII, putative | -5,317 | 4,743E-27 |
| TcCLB.509265.120 | trans-sialidase, putative | -5,322 | 6,117E-159 |
| TcCLB.504593.10 | trans-sialidase, putative (CRP10) | -5,327 | 8,714E-90 |
| TcCLB.505155.4 | trans-sialidase, putative (fragment) | -5,331 | 3,879E-103 |
| TcCLB.506471.100 | trans-sialidase, Group II, putative | -5,34 | 0 |
| TcCLB.510477.80 | mucin-associated surface protein (MASP), putative | -5,344 | 2,286E-273 |
| TcCLB.509075.50 | trans-sialidase, Group II, putative | -5,349 | 0 |
| TcCLB.425435.10 | trans-sialidase, putative (fragment) | -5,36 | 3,398E-200 |
| TcCLB.510635.10 | trans-sialidase, Group VII, putative | -5,363 | 1,395E-220 |
| TcCLB.504015.30 | surface protease GP63, putative | -5,372 | 0 |
| TcCLB.492827.10 | trans-sialidase, putative (fragment) | -5,385 | 3,007E-34 |
| TcCLB.511375.20 | hypothetical protein | -5,388 | 1,379E-146 |
| TcCLB.508081.10 | mucin-associated surface protein (MASP), putative | -5,402 | 3,731E-24 |
| TcCLB.506253.39 | trans-sialidase, putative (CRP10) | -5,416 | 1,551E-81 |
| TcCLB.503601.10 | trans-sialidase, Group V, putative | -5,423 | 2,478E-218 |
| TcCLB.505207.10 | trans-sialidase, Group VII, putative | -5,432 | 2,91E-271 |
| TcCLB.509979.200 | mucin-associated surface protein (MASP), putative | -5,435 | 2,713E-304 |
| TcCLB.510275.190 | mucin-associated surface protein (MASP), putative | -5,45 | 1,202E-114 |
| TcCLB.510213.39 | mucin-associated surface protein (MASP), putative | -5,453 | 0 |
| TcCLB.510693.40 | 90 kDa surface protein, putative | -5,468 | 6,689E-122 |
| TcCLB.511603.450 | trans-sialidase, Group V, putative | -5,475 | 4,304E-114 |
| TcCLB.510021.220 | trans-sialidase, Group V, putative | -5,49 | 1,913E-268 |
| TcCLB.509549.10 | trans-sialidase, Group V, putative | -5,505 | 0 |
| TcCLB.506813.140 | trans-sialidase, Group VII, putative | -5,528 | 1,155E-201 |
| TcCLB.509225.11 | trans-sialidase, putative | -5,539 | 1,491E-172 |
| TcCLB.505779.20 | trans-sialidase, Group II, putative | -5,544 | 0 |
| TcCLB.507833.10 | mucin-associated surface protein (MASP), putative | -5,544 | 2,079E-118 |
| TcCLB.508365.190 | trans-sialidase, Group VI, putative | -5,554 | 1,991E-185 |
| TcCLB.510483.360 | trans-sialidase, Group VII, putative | -5,569 | 0 |
| TcCLB.508117.20 | mucin-associated surface protein (MASP, pseudogene), pu | -5,57 | 8,473E-227 |
| TcCLB.510377.134 | mucin-associated surface protein (MASP), putative | -5,59 | 3,03E-116 |
| TcCLB.506597.9 | trans-sialidase, putative (fragment) | -5,59 | 5,786E-15 |
| TcCLB.504427.230 | trans-sialidase, Group VIII, putative | -5,595 | 2,012E-66 |
| TcCLB.510907.10 | trans-sialidase, Group VIII, putative | -5,6 | 7,685E-54 |
| TcCLB.510489.5 | mucin-associated surface protein (MASP), putative | -5,602 | 1,629E-12 |
| TcCLB.506599.330 | mucin-associated surface protein (MASP), putative | -5,604 | 0 |
| TcCLB.506499.170 | trans-sialidase, Group V, putative | -5,621 | 3,098E-258 |
| TcCLB.505779.10 | trans-sialidase, Group II, putative | -5,629 | 0 |
| TcCLB.507479.20 | trans-sialidase, Group II, putative | -5,644 | 7,786E-162 |
| TcCLB.503427.9 | trans-sialidase, putative (fragment) | -5,664 | 1,069E-290 |
| TcCLB.507095.10 | hypothetical protein (fragment) | -5,682 | 0 |
| TcCLB.506757.50 | mucin-associated surface protein (MASP), putative | -5,709 | 1,602E-284 |

| | | | |
|---|---|---|---|
| TcCLB.508365.100 | surface protease GP63, putative | -5,71 | 1,234E-51 |
| TcCLB.506703.20 | mucin-associated surface protein (MASP), putative | -5,716 | 2,745E-160 |
| TcCLB.508165.310 | surface protease GP63, putative | -5,728 | 0 |
| TcCLB.424499.10 | mucin-associated surface protein (MASP), putative | -5,75 | 1,262E-192 |
| TcCLB.509875.80 | trans-sialidase, Group II, putative | -5,764 | 0 |
| TcCLB.506347.5 | trans-sialidase, putative (fragment) | -5,767 | 6,701E-295 |
| TcCLB.508061.154 | trans-sialidase, putative (fragment) | -5,769 | 1,2E-285 |
| TcCLB.506003.39 | trans-sialidase, putative (fragment) | -5,773 | 1,748E-53 |
| TcCLB.460061.20 | trans-sialidase, putative (fragment) | -5,804 | 6,051E-49 |
| TcCLB.511077.7 | trans-sialidase, putative (fragment) | -5,817 | 8,225E-09 |
| TcCLB.504587.30 | mucin-associated surface protein (MASP), putative | -5,837 | 0 |
| TcCLB.508285.60 | trans-sialidase, Group II, putative | -5,84 | 2,808E-283 |
| TcCLB.506341.50 | trans-sialidase, Group VIII, putative | -5,846 | 0 |
| TcCLB.506537.200 | trans-sialidase, Group VIII, putative | -5,856 | 0 |
| TcCLB.511059.10 | trans-sialidase, putative | -5,877 | 1,341E-113 |
| TcCLB.506757.120 | trans-sialidase, Group V, putative | -5,906 | 0 |
| TcCLB.506653.10 | trans-sialidase, putative | -5,909 | 2,963E-97 |
| TcCLB.508737.10 | surface antigen TASV-C2, putative (TASV-C2) | -5,917 | 0 |
| TcCLB.511855.10 | trans-sialidase, putative | -5,934 | 1,463E-146 |
| TcCLB.506615.100 | mucin-associated surface protein (MASP), putative | -5,953 | 0 |
| TcCLB.511781.31 | trans-sialidase (pseudogene), putative | -5,953 | 8,288E-18 |
| TcCLB.508139.240 | trans-sialidase, Group II, putative | -5,963 | 0 |
| TcCLB.511081.60 | mucin-associated surface protein (MASP), putative | -5,978 | 5,619E-194 |
| TcCLB.508221.790 | trans-sialidase, Group V, putative | -5,988 | 8,205E-94 |
| TcCLB.509979.270 | mucin-associated surface protein (MASP), putative | -6,001 | 0 |
| TcCLB.508139.250 | trans-sialidase, Group II, putative | -6,002 | 0 |
| TcCLB.506763.30 | mucin-associated surface protein (MASP), putative | -6,002 | 9,574E-100 |
| TcCLB.402449.10 | trans-sialidase (pseudogene), putative | -6,004 | 0 |
| TcCLB.511877.10 | surface antigen TASV-B2 (TASV-B2) | -6,008 | 0 |
| TcCLB.511859.120 | mucin-like glycoprotein, putative | -6,013 | 0 |
| TcCLB.510979.40 | trans-sialidase, Group V, putative | -6,019 | 0 |
| TcCLB.504145.10 | trans-sialidase, Group II, putative | -6,028 | 1,544E-35 |
| TcCLB.505231.20 | trans-sialidase, Group II, putative | -6,035 | 0 |
| TcCLB.440083.10 | trans-sialidase, putative (fragment) | -6,053 | 0 |
| TcCLB.509525.420 | mucin TcMUCII, putative | -6,062 | 2,787E-93 |
| TcCLB.503759.10 | trans-sialidase, Group II, putative | -6,066 | 0 |
| TcCLB.511197.10 | trans-sialidase, putative | -6,078 | 2,379E-252 |
| TcCLB.506961.150 | trans-sialidase, Group VIII, putative | -6,098 | 0 |
| TcCLB.511587.90 | trans-sialidase, Group VIII, putative | -6,098 | 0 |
| TcCLB.476911.10 | trans-sialidase, putative (fragment) | -6,111 | 6,192E-83 |
| TcCLB.457113.10 | trans-sialidase, putative (fragment) | -6,161 | 2,619E-33 |
| TcCLB.507505.10 | trans-sialidase, putative | -6,166 | 1,094E-29 |
| TcCLB.510157.5 | trans-sialidase, putative (fragment) | -6,193 | 1,654E-20 |
| TcCLB.508481.10 | trans-sialidase, Group IV, putative | -6,2 | 1,001E-32 |
| TcCLB.505853.10 | trans-sialidase, putative (fragment) | -6,211 | 0 |

| | | | |
|---|---|---|---|
| TcCLB.510213.70 | mucin-associated surface protein (MASP), putative | -6,243 | 0 |
| TcCLB.509075.40 | mucin-associated surface protein (MASP), putative | -6,255 | 4,271E-08 |
| TcCLB.510943.206 | trans-sialidase, putative | -6,258 | 5,225E-58 |
| TcCLB.505573.10 | trans-sialidase, putative (fragment) | -6,28 | 0 |
| TcCLB.504189.5 | trans-sialidase, putative (fragment) | -6,345 | 2,275E-147 |
| TcCLB.510279.320 | trans-sialidase, Group V, putative | -6,347 | 0 |
| TcCLB.511185.80 | trans-sialidase, Group V, putative | -6,434 | 0 |
| TcCLB.508163.400 | surface antigen TASV-B1, putative | -6,44 | 2,486E-301 |
| TcCLB.506757.60 | trans-sialidase, Group V, putative | -6,446 | 0 |
| TcCLB.507981.40 | hypothetical protein | -6,501 | 3,263E-23 |
| TcCLB.508081.50 | mucin-associated surface protein (MASP), putative | -6,516 | 9,32E-280 |
| TcCLB.507477.5 | trans-sialidase, putative | -6,535 | 3,787E-52 |
| TcCLB.508521.100 | trans-sialidase, Group VIII, putative | -6,592 | 1,699E-98 |
| TcCLB.508081.90 | mucin-associated surface protein (MASP), putative | -6,599 | 9,928E-298 |
| TcCLB.432995.9 | trans-sialidase, putative | -6,617 | 2,567E-107 |
| TcCLB.506973.10 | trans-sialidase, Group V, putative | -6,662 | 0 |
| TcCLB.506717.80 | trans-sialidase, Group VIII, putative | -6,667 | 0 |
| TcCLB.510359.650 | mucin-associated surface protein (MASP), putative | -6,674 | 2,367E-144 |
| TcCLB.509281.20 | trans-sialidase, Group II, putative | -6,685 | 4,222E-296 |
| TcCLB.511551.50 | trans-sialidase (pseudogene), putative | -6,687 | 1,416E-29 |
| TcCLB.508757.10 | trans-sialidase (pseudogene), putative | -6,717 | 2,09E-69 |
| TcCLB.509629.10 | trans-sialidase, Group III, putative | -6,724 | 1,022E-87 |
| TcCLB.503655.30 | mucin-like glycoprotein, putative | -6,746 | 5,285E-124 |
| TcCLB.509187.10 | trans-sialidase, Group II, putative | -6,759 | 2,624E-69 |
| TcCLB.509081.166 | trans-sialidase, putative (fragment) | -6,776 | 3,782E-261 |
| TcCLB.511547.20 | trans-sialidase, Group IV, putative | -6,789 | 2,601E-104 |
| TcCLB.507611.170 | trans-sialidase, Group II, putative | -6,834 | 1,494E-140 |
| TcCLB.511757.100 | trans-sialidase, putative (fragment) | -6,861 | 0 |
| TcCLB.509663.50 | trans-sialidase, Group VIII, putative | -6,876 | 0 |
| TcCLB.511875.20 | trans-sialidase, Group VII, putative | -6,937 | 0 |
| TcCLB.506537.80 | trans-sialidase, Group II, putative | -6,949 | 0 |
| TcCLB.508999.170 | surface protease GP63, putative | -6,994 | 8,021E-10 |
| TcCLB.507981.30 | hypothetical protein | -7,01 | 1,317E-41 |
| TcCLB.510713.30 | trans-sialidase, Group VII, putative | -7,023 | 0 |
| TcCLB.508081.30 | mucin-associated surface protein (MASP), putative | -7,046 | 1,886E-49 |
| TcCLB.505609.10 | trans-sialidase, Group II, putative | -7,193 | 1,018E-42 |
| TcCLB.506871.190 | hypothetical protein | -7,268 | 0 |
| TcCLB.510025.160 | mucin-associated surface protein (MASP), putative | -7,275 | 1,448E-90 |
| TcCLB.508081.70 | mucin-associated surface protein (MASP), putative | -7,296 | 7,893E-101 |
| TcCLB.507095.30 | trans-sialidase, putative (fragment) | -7,331 | 0 |
| TcCLB.423205.10 | trans-sialidase, putative (fragment) | -7,337 | 7,007E-21 |
| TcCLB.399173.10 | trans-sialidase, putative (fragment) | -7,435 | 0 |
| TcCLB.507839.40 | trans-sialidase, Group II, putative | -7,475 | 0 |
| TcCLB.506723.20 | trans-sialidase, Group VII, putative | -7,537 | 0 |
| TcCLB.484097.10 | trans-sialidase, putative (fragment) | -7,593 | 0 |

| | | | |
|---|---|---|---|
| TcCLB.506289.94 | mucin-like glycoprotein, putative | -7,604 | 4,24E-138 |
| TcCLB.507283.10 | trans-sialidase, Group V, putative | -7,624 | 0 |
| TcCLB.508485.10 | trans-sialidase, Group II, putative | -7,658 | 2,17E-56 |
| TcCLB.510163.60 | trans-sialidase, Group V, putative | -7,728 | 0 |
| TcCLB.509377.20 | trans-sialidase, Group VII, putative | -7,733 | 1,737E-39 |
| TcCLB.509821.20 | trans-sialidase (pseudogene), putative | -7,849 | 0 |
| TcCLB.511257.110 | mucin-like glycoprotein, putative | -8,066 | 8,102E-200 |
| TcCLB.507511.50 | hypothetical protein | -8,07 | 0 |
| TcCLB.509325.4 | surface protease GP63, putative (fragment) | -8,3 | 6,673E-11 |
| TcCLB.508163.330 | surface protease GP63, putative | -8,718 | 1,704E-196 |
| TcCLB.508581.40 | trans-sialidase, Group II, putative | -8,947 | 4,669E-25 |
| TcCLB.509157.170 | trans-sialidase, Group VIII, putative | -9,251 | 1,816E-153 |

**Supplementary Table 3. Gene ontology (GO) Enrichment Analysis of DE genes in CL Brener epimastigotes compared to trypomastigotes and amastigotes.**

| | BP Results from goseq. | | |
|---|---|---|---|
| row.names | term | over.p.value | de.genes.in.cat |
| GO:0055114 | oxidation-reduction process | 1,444E-30 | TcCLB.506357.50, TcCLB.511277.60, TcCLB.507047.150, TcCLB.510855.10, TcCLB.504425.60, TcCLB.509179.100, TcCLB.506213.50, TcCLB.506629.200, TcCLB.511817.40, TcCLB.511287.49, TcCLB.506937.10, TcCLB.511691.39, TcCLB.508535.10, TcCLB.506147.9, TcCLB.506779.200, TcCLB.509207.90, TcCLB.508461.80, TcCLB.510213.110, TcCLB.509239.10, TcCLB.507019.20, TcCLB.507617.9, TcCLB.506263.10, TcCLB.506367.30, TcCLB.509611.180, TcCLB.508821.40, TcCLB.510943.50, |

| | | | |
|---|---|---|---|
| GO:0008152 | metabolic process | 2,757E-24 | TcCLB.510257.80, TcCLB.509213.120, TcCLB.511391.120, TcCLB.507689.40, TcCLB.510837.9, TcCLB.509205.120, TcCLB.507689.50, TcCLB.506559.410, TcCLB.507689.30, TcCLB.507657.20, TcCLB.508601.141, TcCLB.507657.10, TcCLB.508699.120, TcCLB.510837.20, TcCLB.506513.110, TcCLB.508805.80, TcCLB.508919.10, TcCLB.506593.60, TcCLB.511691.10, TcCLB.506905.50, TcCLB.505945.20, TcCLB.510381.10, TcCLB.508241.140, TcCLB.508177.110, TcCLB.506931.10, TcCLB.508177.120, |

| | | | |
|---|---|---|---|
| GO:0055085 | transmembrane transport | 9,384E-16 | TcCLB.510103.24, TcCLB.509713.30, TcCLB.506551.10, TcCLB.510811.20, TcCLB.508799.270, TcCLB.509707.10, TcCLB.508065.70, TcCLB.508799.290, TcCLB.509107.9, TcCLB.510605.10, TcCLB.506947.10, TcCLB.511639.20, TcCLB.511041.40, TcCLB.510069.20, TcCLB.510877.170, TcCLB.507527.50, TcCLB.509197.20, TcCLB.508465.40, TcCLB.509197.10, TcCLB.508689.40, TcCLB.508231.9, TcCLB.508701.20, TcCLB.508551.39, TcCLB.506355.100, TcCLB.508551.30, TcCLB.506355.10, |
| GO:0006520 | cellular amino acid metabolic process | 5,366E-10 | TcCLB.510795.10, TcCLB.510187.40, TcCLB.510565.11, TcCLB.510187.70, TcCLB.510187.20, TcCLB.510741.10, TcCLB.510739.19, TcCLB.510661.250, TcCLB.507091.50, TcCLB.508111.30, TcCLB.511643.30, TcCLB.507875.20 |

| | | | |
|---|---|---|---|
| GO:0006629 | lipid metabolic process | 2,431E-08 | TcCLB.510679.100, TcCLB.511437.30, TcCLB.511075.9, TcCLB.509239.10, TcCLB.505615.20, TcCLB.440101.10, TcCLB.429257.20, TcCLB.508737.70, TcCLB.507609.40, TcCLB.510181.20, TcCLB.511439.30, TcCLB.507089.210, TcCLB.508503.10, TcCLB.510795.30, TcCLB.509331.70, TcCLB.510359.340, TcCLB.510565.20, TcCLB.504427.70, TcCLB.504149.160 |
| GO:0009058 | biosynthetic process | 5,079E-08 | TcCLB.510795.10, TcCLB.510187.40, TcCLB.503747.10, TcCLB.508535.50, TcCLB.510565.11, TcCLB.506247.220, TcCLB.509041.10, TcCLB.509043.10, TcCLB.510187.70, TcCLB.510187.20, TcCLB.510945.70, TcCLB.508257.10, TcCLB.506529.430, TcCLB.506529.420, TcCLB.506959.70, TcCLB.506405.50, TcCLB.503453.100 |

| | | | |
|---|---|---|---|
| GO:0006526 | arginine biosynthetic process | 1,321E-07 | TcCLB.511391.120, TcCLB.507689.40, TcCLB.507689.30, TcCLB.507657.20, TcCLB.507657.10, TcCLB.510837.20, TcCLB.508971.40, TcCLB.507059.80 |
| GO:0006839 | mitochondrial transport | 6,289E-07 | TcCLB.511687.19, TcCLB.508741.229, TcCLB.508737.210, TcCLB.509141.40, TcCLB.511687.10, TcCLB.509129.10 |
| GO:0009116 | nucleoside metabolic process | 6,038E-07 | TcCLB.509569.100, TcCLB.507519.150, TcCLB.506789.280, TcCLB.508387.20, TcCLB.507739.40, TcCLB.508207.70, TcCLB.506865.2, TcCLB.508989.9, TcCLB.507773.60, TcCLB.506435.410 |
| GO:0016559 | peroxisome fission | 2,064E-06 | TcCLB.510669.49, TcCLB.510669.20, TcCLB.508461.570, TcCLB.509203.40, TcCLB.507009.10, TcCLB.511017.50 |

| | | | |
|---|---|---|---|
| GO:0005975 | carbohydrate metabolic process | 1,369E-05 | TcCLB.506937.10, TcCLB.503713.30, TcCLB.510667.140, TcCLB.506195.120, TcCLB.510437.40, TcCLB.508257.70, TcCLB.510283.70, TcCLB.507009.30, TcCLB.503823.40, TcCLB.506407.10, TcCLB.509199.24, TcCLB.508569.80, TcCLB.511531.50, TcCLB.508601.119, TcCLB.508951.20 |
| GO:0006811 | ion transport | 1,937E-05 | TcCLB.511687.19, TcCLB.508741.229, TcCLB.508737.210, TcCLB.509141.40, TcCLB.511687.10, TcCLB.509129.10 |
| GO:0006626 | protein targeting to mitochondrion | 0,0004091 | TcCLB.506399.80, TcCLB.511909.50, TcCLB.506399.70, TcCLB.511909.60, TcCLB.508647.110, TcCLB.511807.253 |
| GO:0006094 | gluconeogenesis | 0,0009519 | TcCLB.506775.180, TcCLB.508441.20, TcCLB.507547.90, TcCLB.506529.508 |
| GO:0033617 | mitochondrial respiratory chain complex IV | 0,0013959 | TcCLB.509721.40, TcCLB.511395.20, TcCLB.510173.70, TcCLB.506041.50 |
| GO:0006633 | fatty acid biosynthetic process | 0,0016028 | TcCLB.509239.10, TcCLB.507609.40, TcCLB.510181.20, TcCLB.510795.30, TcCLB.473111.10, TcCLB.510565.20 |

| | | | |
|---|---|---|---|
| GO:0046835 | carbohydrate phosphorylation | 0,0016251 | TcCLB.510667.120, TcCLB.507009.40, TcCLB.507001.110 |
| GO:0006470 | protein dephosphorylation | 0,0019032 | TcCLB.509205.120, TcCLB.508601.141, TcCLB.508699.120, TcCLB.503471.10, TcCLB.509109.130, TcCLB.510603.60, TcCLB.503657.70, TcCLB.509109.120, TcCLB.506779.110, TcCLB.506743.110, TcCLB.510603.80 |
| GO:0006098 | pentose-phosphate shunt | 0,0024448 | TcCLB.503713.30, TcCLB.505807.180, TcCLB.509199.24, TcCLB.507017.40, TcCLB.508601.119 |
| GO:0009987 | cellular process | 0,0023817 | TcCLB.510655.120, TcCLB.510751.4, TcCLB.510749.99, TcCLB.507081.110, TcCLB.503991.20 |
| GO:0006825 | copper ion transport | 0,00309 | TcCLB.509463.20, TcCLB.510507.29, TcCLB.506649.50 |
| GO:0006013 | mannose metabolic process | 0,0057466 | TcCLB.506195.120, TcCLB.506407.10 |
| GO:0006471 | protein ADP-ribosylation | 0,0057466 | TcCLB.509721.60, TcCLB.510173.90 |
| GO:0006537 | glutamate biosynthetic process | 0,0057466 | TcCLB.511237.30, TcCLB.506411.30 |
| GO:0006548 | histidine catabolic process | 0,005746 | TcCLB.506247.220, TcCLB.504045.110 |
| GO:0006561 | proline biosynthetic process | 0,003419 | TcCLB.509207.90, TcCLB.510943.50, TcCLB.506857.20 |
| GO:0006562 | proline catabolic process | 0,0057466 | TcCLB.511237.30, TcCLB.506411.30 |
| GO:0006566 | threonine metabolic process | 0,0052542 | TcCLB.503597.10, TcCLB.509167.150 |

| | | | |
|---|---|---|---|
| GO:0006665 | sphingolipid metabolic process | 0,0044199 | TcCLB.508257.10, TcCLB.506405.50 |
| GO:0006783 | heme biosynthetic process | 0,0055176 | TcCLB.509601.59, TcCLB.509767.59 |
| GO:0006807 | nitrogen compound metabolic process | 0,0034532 | TcCLB.510039.40, TcCLB.509979.150, TcCLB.503405.10 |
| GO:0006874 | cellular calcium ion homeostasis | 0,0057466 | TcCLB.511385.30, TcCLB.510773.20 |
| GO:0007131 | reciprocal meiotic recombination | 0,0053436 | TcCLB.506885.310, TcCLB.510729.110 |
| GO:0009229 | thiamine diphosphate biosynthetic process | 0,0056933 | TcCLB.511051.59, TcCLB.509683.124 |
| GO:0009443 | pyridoxal 5'-phosphate salvage | 0,0049909 | TcCLB.509059.50, TcCLB.507925.40 |
| GO:0019556 | histidine catabolic process to glutamate an | 0,005645 | TcCLB.508741.140, TcCLB.509137.30 |
| GO:0019571 | D-arabinose catabolic process | 0,0048368 | TcCLB.506213.50, TcCLB.511287.49 |
| GO:0019853 | L-ascorbic acid biosynthetic process | 0,0057379 | TcCLB.507047.150, TcCLB.509179.100 |
| GO:0046951 | ketone body biosynthetic process | 0,0046754 | TcCLB.507049.60, TcCLB.506567.70 |
| GO:0048034 | heme O biosynthetic process | 0,0055176 | TcCLB.509601.59, TcCLB.509767.59 |
| GO:0050790 | regulation of catalytic activity | 0,0052349 | TcCLB.510535.100, TcCLB.511827.100 |
| GO:0051452 | intracellular pH reduction | 0,0057466 | TcCLB.511385.30, TcCLB.510773.20 |
| GO:0033955 | mitochondrial DNA inheritance | 0,0069773 | TcCLB.503781.80, TcCLB.507639.30, TcCLB.503781.90 |
| GO:0006012 | galactose metabolic process | 0,0071128 | TcCLB.510667.120, TcCLB.507009.40, TcCLB.507001.110 |

| | | | |
|---|---|---|---|
| GO:0006468 | protein phosphorylation | 0,0083465 | TcCLB.508405.70, TcCLB.507837.20, TcCLB.508831.110, TcCLB.508621.10, TcCLB.511671.80, TcCLB.510345.9, TcCLB.511301.90, TcCLB.510861.140, TcCLB.510155.160, TcCLB.503943.10, TcCLB.511837.90, TcCLB.510609.70, TcCLB.507211.20, TcCLB.503617.10, TcCLB.506835.159, TcCLB.506401.110, TcCLB.505193.110, TcCLB.508153.740, TcCLB.506999.180, TcCLB.511351.30, TcCLB.506469.80, TcCLB.511573.40, TcCLB.510759.40, TcCLB.476421.9, TcCLB.506633.10, TcCLB.511277.140, |
| GO:0022900 | electron transport chain | 0,0127825 | TcCLB.509721.40, TcCLB.510173.70 |
| GO:0016311 | dephosphorylation | 0,0132844 | TcCLB.503471.10, TcCLB.509109.130, TcCLB.510603.60, TcCLB.503657.70, TcCLB.509109.120, TcCLB.506779.110, TcCLB.506743.110, TcCLB.510603.80 |
| GO:0009072 | aromatic amino acid family metabolic proc | 0,01599 | TcCLB.510795.10, TcCLB.510187.40 |
| GO:0006518 | peptide metabolic process | 0,0162322 | TcCLB.504045.60, TcCLB.506513.110 |
| GO:0016310 | phosphorylation | 0,0163425 | TcCLB.510101.140, TcCLB.506297.190 |

| | | | |
|---|---|---|---|
| GO:0030001 | metal ion transport | 0,0172671 | TcCLB.507527.50, TcCLB.509197.20, TcCLB.509197.10, TcCLB.508701.20, TcCLB.509437.60, TcCLB.509437.70 |
| GO:0015937 | coenzyme A biosynthetic process | 0,0174559 | TcCLB.506779.80, TcCLB.511277.500, TcCLB.507021.110 |
| GO:0006729 | tetrahydrobiopterin biosynthetic process | 0,0211556 | TcCLB.511575.10, TcCLB.503613.40 |
| GO:0044237 | cellular metabolic process | 0,0257878 | TcCLB.507923.10, TcCLB.511753.120, TcCLB.508119.120, TcCLB.510349.10, TcCLB.511019.84, TcCLB.503925.40 |
| GO:0008535 | respiratory chain complex IV assembly | 0,0267534 | TcCLB.508995.60, TcCLB.506649.50 |
| GO:0042759 | long-chain fatty acid biosynthetic process | 0,0272799 | TcCLB.511245.130, TcCLB.506661.30 |
| GO:0006298 | mismatch repair | 0,028402 | TcCLB.507233.110, TcCLB.509693.209, TcCLB.509967.20, TcCLB.511911.30 |
| GO:0031388 | organic acid phosphorylation | 0,0310156 | TcCLB.508741.170, TcCLB.509139.10 |
| GO:0008299 | isoprenoid biosynthetic process | 0,031864 | TcCLB.509237.10, TcCLB.511071.50, TcCLB.511823.70, TcCLB.511903.40 |
| GO:0006725 | cellular aromatic compound metabolic proc | 0,0403494 | TcCLB.503991.39, TcCLB.507081.130 |
| GO:0007035 | vacuolar acidification | 0,0519264 | TcCLB.510993.10 |
| GO:0006643 | membrane lipid metabolic process | 0,0539014 | TcCLB.503929.79 |
| GO:0019538 | protein metabolic process | 0,0540045 | TcCLB.510855.10, TcCLB.509611.180 |
| GO:0000723 | telomere maintenance | 0,0561425 | TcCLB.508827.50, TcCLB.506775.100, TcCLB.503677.20 |

| | | | |
|---|---|---|---|
| GO:0006259 | DNA metabolic process | 0,057713 | TcCLB.506885.310, TcCLB.510729.110, TcCLB.509779.10, TcCLB.511653.69 |
| GO:0009052 | pentose-phosphate shunt, non-oxidative bra | 0,0579039 | TcCLB.509199.24 |
| GO:0006735 | NADH regeneration | 0,0595221 | TcCLB.508601.50 |
| GO:0002082 | regulation of oxidative phosphorylation | 0,0663701 | TcCLB.508823.70 |
| GO:0006784 | heme a biosynthetic process | 0,0744994 | TcCLB.511211.70 |
| GO:0010040 | response to iron(II) ion | 0,0758265 | TcCLB.507951.299 |
| GO:0015819 | lysine transport | 0,0755383 | TcCLB.511545.80 |
| GO:0018171 | peptidyl-cysteine oxidation | 0,0758265 | TcCLB.507951.299 |
| GO:0051156 | glucose 6-phosphate metabolic process | 0,0739662 | TcCLB.510187.100 |
| GO:0009165 | nucleotide biosynthetic process | 0,0784136 | TcCLB.509153.90, TcCLB.507773.60 |
| GO:0009435 | NAD biosynthetic process | 0,0831196 | TcCLB.506593.60, TcCLB.506829.80 |
| GO:0006696 | ergosterol biosynthetic process | 0,0891041 | TcCLB.511071.50, TcCLB.511903.40 |

| | MF Results from goseq. | | |
|---|---|---|---|
| row.names | term | over.p.value | de.genes.in.cat |
| GO:0016491 | oxidoreductase activity | 1,63E-26 | TcCLB.506357.50, TcCLB.511277.60, TcCLB.507047.150, TcCLB.504425.60, TcCLB.509179.100, TcCLB.506213.50, TcCLB.508805.80, TcCLB.511817.40, TcCLB.511287.49, TcCLB.506937.10, TcCLB.511691.39, TcCLB.508535.10, TcCLB.506147.9, TcCLB.506779.200, TcCLB.509207.90, TcCLB.508461.80, TcCLB.510213.110, TcCLB.507019.20, TcCLB.507617.9, TcCLB.508821.40, TcCLB.510943.50, TcCLB.510437.40, TcCLB.503577.9, TcCLB.509351.10, TcCLB.505807.180, TcCLB.511037.20, TcCLB.511151.90, TcCLB.508177.90, TcCLB.510265.10, TcCLB.508297.50, TcCLB.507049.60, TcCLB.510311.90, TcCLB.509331.210, TcCLB.506219.40, TcCLB.506821.210, TcCLB.510521.10, TcCLB.506857.20, TcCLB.508677.80, TcCLB.511461.60, TcCLB.509459.79, TcCLB.504339.19, |

| | | | |
|---|---|---|---|
| GO:0030170 | pyridoxal phosphate binding | 5,69E-18 | TcCLB.510795.10, TcCLB.510187.40, TcCLB.503747.10, TcCLB.508535.50, TcCLB.510565.11, TcCLB.509041.10, TcCLB.509043.10, TcCLB.510187.70, TcCLB.510187.20, TcCLB.511691.10, TcCLB.506905.50, TcCLB.510381.10, TcCLB.508241.140, TcCLB.508177.110, TcCLB.508177.120, TcCLB.508177.129, TcCLB.510741.10, TcCLB.510739.19, TcCLB.510661.250, TcCLB.508175.360, TcCLB.510945.70, TcCLB.508257.10, TcCLB.506529.430, TcCLB.506529.420, TcCLB.506959.70, TcCLB.506405.50, TcCLB.503453.100 |

| | | | |
|---|---|---|---|
| GO:0003824 | catalytic activity | 6,26E-16 | TcCLB.507047.150,<br>TcCLB.506559.410,<br>TcCLB.506247.220,<br>TcCLB.509179.100,<br>TcCLB.509569.100,<br>TcCLB.506937.10, TcCLB.506593.60,<br>TcCLB.511691.10, TcCLB.508303.10,<br>TcCLB.506789.280,<br>TcCLB.506905.50, TcCLB.505945.20,<br>TcCLB.510381.10,<br>TcCLB.508241.140,<br>TcCLB.508177.110,<br>TcCLB.506931.10, TcCLB.507923.10,<br>TcCLB.508177.120,<br>TcCLB.511753.120,<br>TcCLB.510667.140,<br>TcCLB.508387.20,<br>TcCLB.508177.129,<br>TcCLB.506195.120,<br>TcCLB.510437.40,<br>TcCLB.508175.360,<br>TcCLB.506779.80,<br>TcCLB.510101.140,<br>TcCLB.503991.39, TcCLB.506865.2,<br>TcCLB.511179.150,<br>TcCLB.508119.120, TcCLB.508989.9, |
| GO:0008234 | cysteine-type peptidase activity | 2,88E-10 | TcCLB.510535.100,<br>TcCLB.511827.100,<br>TcCLB.506529.550,<br>TcCLB.507537.10,<br>TcCLB.507603.270,<br>TcCLB.507603.260,<br>TcCLB.507537.20,<br>TcCLB.509429.329,<br>TcCLB.510073.17,<br>TcCLB.509429.320,<br>TcCLB.507297.10, TcCLB.509401.20,<br>TcCLB.508595.50 |

| | | | |
|---|---|---|---|
| GO:0016740 | transferase activity | 1,13E-09 | TcCLB.510795.10, TcCLB.510187.40, TcCLB.503747.10, TcCLB.508535.50, TcCLB.510565.11, TcCLB.509041.10, TcCLB.509043.10, TcCLB.510187.70, TcCLB.510187.20, TcCLB.510945.70, TcCLB.508257.10, TcCLB.506529.430, TcCLB.506529.420, TcCLB.510381.30, TcCLB.506959.70, TcCLB.506405.50, TcCLB.509825.40, TcCLB.503453.100, TcCLB.511655.40 |
| GO:0022857 | transmembrane transporter activity | 6,44E-09 | TcCLB.511041.40, TcCLB.508231.9, TcCLB.508551.39, TcCLB.506355.100, TcCLB.508551.30, TcCLB.506355.10, TcCLB.424937.10, TcCLB.504125.100 |
| GO:0020037 | heme binding | 8,2E-09 | TcCLB.511073.10, TcCLB.509239.10, TcCLB.507019.20, TcCLB.426435.9, TcCLB.506773.44, TcCLB.503653.60, TcCLB.509719.40, TcCLB.510101.50, TcCLB.506945.190, TcCLB.508799.160, TcCLB.507609.40, TcCLB.510181.20, TcCLB.509395.100, TcCLB.511391.160, TcCLB.410049.20, TcCLB.509231.10 |

| | | | |
|---|---|---|---|
| GO:0008777 | acetylornithine deacetylase activity | 1,26E-08 | TcCLB.511391.120, TcCLB.507689.40, TcCLB.507689.30, TcCLB.507657.20, TcCLB.507657.10, TcCLB.510837.20, TcCLB.508971.40 |
| GO:0008137 | NADH dehydrogenase (ubiquinone) activity | 3,66E-08 | TcCLB.503893.80, TcCLB.508507.59, TcCLB.506337.160, TcCLB.508601.50, TcCLB.510659.270, TcCLB.507009.50, TcCLB.510743.30 |
| GO:0050897 | cobalt ion binding | 3,79E-07 | TcCLB.511391.120, TcCLB.507689.40, TcCLB.507689.30, TcCLB.507657.20, TcCLB.507657.10, TcCLB.510837.20, TcCLB.508971.40 |
| GO:0008308 | voltage-gated anion channel activity | 6,29E-07 | TcCLB.511687.19, TcCLB.508741.229, TcCLB.508737.210, TcCLB.509141.40, TcCLB.511687.10, TcCLB.509129.10 |
| GO:0015288 | porin activity | 6,29E-07 | TcCLB.511687.19, TcCLB.508741.229, TcCLB.508737.210, TcCLB.509141.40, TcCLB.511687.10, TcCLB.509129.10 |
| GO:0010181 | FMN binding | 3,24E-06 | TcCLB.507047.150, TcCLB.506147.9, TcCLB.508461.80, TcCLB.507617.9, TcCLB.508399.10, TcCLB.507869.10, TcCLB.503753.29, TcCLB.506585.70 |

| | | | |
|---|---|---|---|
| GO:0009055 | electron transfer activity | 7,2E-06 | TcCLB.507889.20, TcCLB.509719.40, TcCLB.510101.50, TcCLB.509177.50, TcCLB.506945.190, TcCLB.503477.10, TcCLB.511303.70, TcCLB.511391.160, TcCLB.510659.270, TcCLB.509231.10, TcCLB.510743.30, TcCLB.508265.10 |
| GO:0008483 | transaminase activity | 1,21E-05 | TcCLB.510795.10, TcCLB.510187.40, TcCLB.510565.11, TcCLB.510187.70, TcCLB.510187.20 |
| GO:0004197 | cysteine-type endopeptidase activity | 1,56E-05 | TcCLB.510535.100, TcCLB.511827.100, TcCLB.507603.270, TcCLB.507603.260, TcCLB.510073.17, TcCLB.509429.320, TcCLB.509401.20, TcCLB.508595.50 |
| GO:0008239 | dipeptidyl-peptidase activity | 3,03E-05 | TcCLB.509205.120, TcCLB.509197.50, TcCLB.508601.141, TcCLB.508699.120 |
| GO:0016705 | oxidoreductase activity, acting on paired dono | 5,3E-05 | TcCLB.509719.40, TcCLB.510101.50, TcCLB.506945.190, TcCLB.510795.30, TcCLB.510565.20, TcCLB.509231.10 |
| GO:0005507 | copper ion binding | 0,000125 | TcCLB.504867.70, TcCLB.510943.130, TcCLB.509463.20, TcCLB.510507.29, TcCLB.508995.60, TcCLB.506649.50 |

| | | | |
|---|---|---|---|
| GO:0004198 | calcium-dependent cysteine-type endopeptid | 0,00014 | TcCLB.511329.10, TcCLB.508999.200, TcCLB.503855.40, TcCLB.506563.210, TcCLB.508555.50, TcCLB.509237.120, TcCLB.509013.10, TcCLB.511441.20, TcCLB.510337.20, TcCLB.508999.190 |
| GO:0016787 | hydrolase activity | 0,000217 | TcCLB.510257.80, TcCLB.509213.120, TcCLB.511391.120, TcCLB.507689.40, TcCLB.510837.9, TcCLB.509205.120, TcCLB.507689.50, TcCLB.507689.30, TcCLB.507657.20, TcCLB.508601.141, TcCLB.507657.10, TcCLB.508699.120, TcCLB.510837.20, TcCLB.506513.110, TcCLB.508919.10, TcCLB.506931.10, TcCLB.508971.40, TcCLB.506975.40, TcCLB.507081.110, TcCLB.510131.40, TcCLB.503991.20, TcCLB.508043.20, TcCLB.506363.70, TcCLB.509633.60, TcCLB.509169.30 |
| GO:0016671 | oxidoreductase activity, acting on a sulfur gro | 0,000276 | TcCLB.510855.10, TcCLB.509611.180, TcCLB.508837.189 |
| GO:0008762 | UDP-N-acetylmuramate dehydrogenase activ | 0,000398 | TcCLB.507047.150, TcCLB.509179.100, TcCLB.509459.79, TcCLB.504151.60 |
| GO:0004553 | hydrolase activity, hydrolyzing O-glycosyl com | 0,000436 | TcCLB.506195.120, TcCLB.510283.70, TcCLB.503823.40, TcCLB.506407.10 |

| | | | |
|---|---|---|---|
| GO:0004611 | phosphoenolpyruvate carboxykinase activity | 0,000435 | TcCLB.506775.180, TcCLB.508441.20, TcCLB.507547.90 |
| GO:0017076 | purine nucleotide binding | 0,000435 | TcCLB.506775.180, TcCLB.508441.20, TcCLB.507547.90 |
| GO:0004177 | aminopeptidase activity | 0,000471 | TcCLB.509205.120, TcCLB.508601.141, TcCLB.508699.120, TcCLB.510655.120, TcCLB.510751.4, TcCLB.508799.240 |
| GO:0005506 | iron ion binding | 0,000493 | TcCLB.509719.40, TcCLB.510101.50, TcCLB.506945.190, TcCLB.507609.40, TcCLB.510181.20, TcCLB.511391.160, TcCLB.473111.10, TcCLB.509231.10 |
| GO:0016747 | transferase activity, transferring acyl groups | 0,000893 | TcCLB.504425.60, TcCLB.510265.10, TcCLB.510311.90, TcCLB.509331.210, TcCLB.508677.80, TcCLB.509463.30, TcCLB.504339.19, TcCLB.510507.20 |
| GO:0016810 | hydrolase activity, acting on carbon-nitrogen | 0,001454 | TcCLB.510039.40, TcCLB.508741.140, TcCLB.509979.150, TcCLB.509137.30, TcCLB.441401.10 |
| GO:0004335 | galactokinase activity | 0,001625 | TcCLB.510667.120, TcCLB.507009.40, TcCLB.507001.110 |
| GO:0048037 | cofactor binding | 0,002978 | TcCLB.506263.10, TcCLB.510099.120, TcCLB.506263.20 |
| GO:0004462 | lactoylglutathione lyase activity | 0,003105 | TcCLB.510659.240, TcCLB.510743.70 |
| GO:0003858 | 3-hydroxybutyrate dehydrogenase activity | 0,004675 | TcCLB.507049.60, TcCLB.506567.70 |
| GO:0003885 | D-arabinono-1,4-lactone oxidase activity | 0,005738 | TcCLB.507047.150, TcCLB.509179.100 |
| GO:0003987 | acetate-CoA ligase activity | 0,005747 | TcCLB.509331.30, TcCLB.504427.110 |
| GO:0004413 | homoserine kinase activity | 0,005254 | TcCLB.503597.10, TcCLB.509167.150 |

| | | | |
|---|---|---|---|
| GO:0004421 | hydroxymethylglutaryl-CoA synthase activity | 0,005735 | TcCLB.511071.50, TcCLB.511903.40 |
| GO:0004559 | alpha-mannosidase activity | 0,005747 | TcCLB.506195.120, TcCLB.506407.10 |
| GO:0004612 | phosphoenolpyruvate carboxykinase (ATP) act | 0,005744 | TcCLB.508441.20, TcCLB.507547.90 |
| GO:0004657 | proline dehydrogenase activity | 0,005747 | TcCLB.511237.30, TcCLB.506411.30 |
| GO:0004735 | pyrroline-5-carboxylate reductase activity | 0,004706 | TcCLB.509207.90, TcCLB.506857.20 |
| GO:0004751 | ribose-5-phosphate isomerase activity | 0,003351 | TcCLB.509199.24, TcCLB.508601.119 |
| GO:0004788 | thiamine diphosphokinase activity | 0,005693 | TcCLB.511051.59, TcCLB.509683.124 |
| GO:0008478 | pyridoxal kinase activity | 0,004991 | TcCLB.509059.50, TcCLB.507925.40 |
| GO:0008495 | protoheme IX farnesyltransferase activity | 0,005518 | TcCLB.509601.59, TcCLB.509767.59 |
| GO:0008887 | glycerate kinase activity | 0,005744 | TcCLB.508741.170, TcCLB.509139.10 |
| GO:0008936 | nicotinamidase activity | 0,004189 | TcCLB.506593.60, TcCLB.506829.80 |
| GO:0009678 | hydrogen-translocating pyrophosphatase acti | 0,005747 | TcCLB.511385.30, TcCLB.510773.20 |
| GO:0015923 | mannosidase activity | 0,005747 | TcCLB.506195.120, TcCLB.506407.10 |
| GO:0016208 | AMP binding | 0,005747 | TcCLB.509331.30, TcCLB.504427.110 |
| GO:0016614 | oxidoreductase activity, acting on CH-OH grou | 0,004529 | TcCLB.507047.150, TcCLB.509179.100, TcCLB.509459.79, TcCLB.504151.60 |
| GO:0016615 | malate dehydrogenase activity | 0,005228 | TcCLB.506937.10, TcCLB.510437.40 |
| GO:0016616 | oxidoreductase activity, acting on the CH-OH | 0,003374 | TcCLB.506937.10, TcCLB.506263.10, TcCLB.510437.40, TcCLB.510099.120, TcCLB.506263.20, TcCLB.507993.270, TcCLB.508893.10 |
| GO:0016639 | oxidoreductase activity, acting on the CH-NH2 | 0,00561 | TcCLB.508111.30, TcCLB.507875.20 |
| GO:0016651 | oxidoreductase activity, acting on NAD(P)H | 0,005725 | TcCLB.508399.10, TcCLB.507869.10 |
| GO:0016717 | oxidoreductase activity, acting on paired done | 0,005631 | TcCLB.507609.40, TcCLB.510181.20 |
| GO:0016763 | transferase activity, transferring pentosyl gro | 0,005048 | TcCLB.506789.280, TcCLB.508387.20 |
| GO:0016812 | hydrolase activity, acting on carbon-nitrogen | 0,005645 | TcCLB.508741.140, TcCLB.509137.30 |
| GO:0016899 | oxidoreductase activity, acting on the CH-OH | 0,005738 | TcCLB.507047.150, TcCLB.509179.100 |
| GO:0016920 | pyroglutamyl-peptidase activity | 0,004285 | TcCLB.506635.60, TcCLB.508027.110 |

| | | | |
|---|---|---|---|
| GO:0045290 | D-arabinose 1-dehydrogenase [NAD(P)+] acti | 0,004837 | TcCLB.506213.50, TcCLB.511287.49 |
| GO:0050024 | L-galactonolactone oxidase activity | 0,005738 | TcCLB.507047.150, TcCLB.509179.100 |
| GO:0050242 | pyruvate, phosphate dikinase activity | 0,005747 | TcCLB.510101.140, TcCLB.506297.190 |
| GO:0004602 | glutathione peroxidase activity | 0,006033 | TcCLB.507515.4, TcCLB.503899.119, TcCLB.507515.10 |
| GO:0008138 | protein tyrosine/serine/threonine phosphatas | 0,006255 | TcCLB.503471.10, TcCLB.509109.130, TcCLB.510603.60, TcCLB.503657.70, TcCLB.509109.120, TcCLB.506779.110, TcCLB.506743.110, TcCLB.510603.80 |
| GO:0050662 | coenzyme binding | 0,006701 | TcCLB.507923.10, TcCLB.511753.120, TcCLB.508119.120, TcCLB.510349.10, TcCLB.511019.84, TcCLB.503925.40 |

| | | | |
|---|---|---|---|
| GO:0004672 | protein kinase activity | 0,007172 | TcCLB.508405.70, TcCLB.507837.20, TcCLB.508831.110, TcCLB.508621.10, TcCLB.511671.80, TcCLB.510345.9, TcCLB.511301.90, TcCLB.510861.140, TcCLB.510155.160, TcCLB.503943.10, TcCLB.511837.90, TcCLB.510609.70, TcCLB.507211.20, TcCLB.503617.10, TcCLB.506835.159, TcCLB.506401.110, TcCLB.505193.110, TcCLB.508153.740, TcCLB.506999.180, TcCLB.511351.30, TcCLB.506469.80, TcCLB.511573.40, TcCLB.510759.40, TcCLB.476421.9, TcCLB.506633.10, TcCLB.511277.140, TcCLB.506327.99, TcCLB.504269.10, TcCLB.510941.3, TcCLB.464807.10, TcCLB.509207.80, TcCLB.506469.30, TcCLB.511821.40, TcCLB.506327.90, TcCLB.506775.190, TcCLB.506945.110, TcCLB.509599.130, TcCLB.511817.80, TcCLB.507683.40, |
| GO:0051539 | 4 iron, 4 sulfur cluster binding | 0,008307 | TcCLB.508399.10, TcCLB.507869.10, TcCLB.503893.80, TcCLB.508507.59 |
| GO:0016791 | phosphatase activity | 0,010746 | TcCLB.503471.10, TcCLB.509109.130, TcCLB.510603.60, TcCLB.503657.70, TcCLB.509109.120, TcCLB.506779.110, TcCLB.506743.110, TcCLB.510603.80, TcCLB.506825.210 |
| GO:0003954 | NADH dehydrogenase activity | 0,011469 | TcCLB.508601.50, TcCLB.507009.50 |

| | | | |
|---|---|---|---|
| GO:0016772 | transferase activity, transferring phosphorus- | 0,011482 | TcCLB.508405.70, TcCLB.507837.20, TcCLB.508831.110, TcCLB.508621.10, TcCLB.511671.80, TcCLB.510345.9, TcCLB.510101.140, TcCLB.511301.90, TcCLB.510445.10, TcCLB.510861.140, TcCLB.510155.160, TcCLB.503943.10, TcCLB.511837.90, TcCLB.510609.70, TcCLB.507211.20, TcCLB.503617.10, TcCLB.463451.10, TcCLB.506835.159, TcCLB.506401.110, TcCLB.505193.110, TcCLB.508153.740, TcCLB.506999.180, TcCLB.511351.30, TcCLB.506469.80, TcCLB.511573.40, TcCLB.510759.40, TcCLB.476421.9, TcCLB.506633.10, TcCLB.511277.140, TcCLB.506327.99, TcCLB.504269.10, TcCLB.510941.3, TcCLB.464807.10, TcCLB.509207.80, TcCLB.506469.30, TcCLB.511821.40, TcCLB.506327.90, TcCLB.506775.190, TcCLB.506297.190, TcCLB.506945.110, |
| GO:0004129 | cytochrome-c oxidase activity | 0,011672 | TcCLB.504069.20, TcCLB.511145.10 |
| GO:0046873 | metal ion transmembrane transporter activit | 0,012073 | TcCLB.507527.50, TcCLB.509197.20, TcCLB.509197.10, TcCLB.508701.20, TcCLB.509437.60, TcCLB.509437.70 |
| GO:0016830 | carbon-carbon lyase activity | 0,013473 | TcCLB.503991.39, TcCLB.507081.130 |
| GO:0016531 | copper chaperone activity | 0,013818 | TcCLB.509463.20, TcCLB.510507.29 |
| GO:0004806 | triglyceride lipase activity | 0,015543 | TcCLB.510679.100, TcCLB.511437.30, TcCLB.505615.20, TcCLB.511439.30, TcCLB.507089.210 |
| GO:0003988 | acetyl-CoA C-acyltransferase activity | 0,016024 | TcCLB.509463.30, TcCLB.510507.20 |
| GO:0004838 | L-tyrosine:2-oxoglutarate aminotransferase a | 0,01599 | TcCLB.510795.10, TcCLB.510187.40 |
| GO:0000104 | succinate dehydrogenase activity | 0,016369 | TcCLB.508535.10, TcCLB.503849.60 |

| | | | |
|---|---|---|---|
| GO:0004725 | protein tyrosine phosphatase activity | 0,017904 | TcCLB.509205.120, TcCLB.508601.141, TcCLB.508699.120 |
| GO:0048038 | quinone binding | 0,018438 | TcCLB.503893.80, TcCLB.508507.59 |
| GO:0008124 | 4-alpha-hydroxytetrahydrobiopterin dehydrat | 0,021156 | TcCLB.511575.10, TcCLB.503613.40 |
| GO:0016857 | racemase and epimerase activity, acting on c | 0,022608 | TcCLB.510667.140, TcCLB.507009.30 |
| GO:0030983 | mismatched DNA binding | 0,022609 | TcCLB.507233.110, TcCLB.509693.209, TcCLB.509967.20, TcCLB.511911.30 |
| GO:0015078 | proton transmembrane transporter activity | 0,022839 | TcCLB.507517.60, TcCLB.503579.70, TcCLB.504131.180, TcCLB.504125.40 |
| GO:0016627 | oxidoreductase activity, acting on the CH-CH g | 0,026606 | TcCLB.511211.70, TcCLB.508975.10, TcCLB.509331.70, TcCLB.504427.70, TcCLB.509153.120 |
| GO:0009922 | fatty acid elongase activity | 0,02728 | TcCLB.511245.130, TcCLB.506661.30 |
| GO:0010309 | acireductone dioxygenase [iron(II)-requiring] | 0,02738 | TcCLB.506629.200, TcCLB.506367.30 |
| GO:0004070 | aspartate carbamoyltransferase activity | 0,028249 | TcCLB.507091.50, TcCLB.511643.30 |
| GO:0016597 | amino acid binding | 0,028249 | TcCLB.507091.50, TcCLB.511643.30 |
| GO:0016743 | carboxyl- or carbamoyltransferase activity | 0,028249 | TcCLB.507091.50, TcCLB.511643.30 |
| GO:0003950 | NAD+ ADP-ribosyltransferase activity | 0,030152 | TcCLB.509721.60, TcCLB.510173.90 |
| GO:0004659 | prenyltransferase activity | 0,030462 | TcCLB.509601.59, TcCLB.509767.59 |
| GO:0008080 | N-acetyltransferase activity | 0,034181 | TcCLB.506177.100, TcCLB.506181.104, TcCLB.511809.120, TcCLB.511811.30 |
| GO:0050660 | flavin adenine dinucleotide binding | 0,043244 | TcCLB.507047.150, TcCLB.509179.100, TcCLB.508837.189, TcCLB.509459.79, TcCLB.509461.10, TcCLB.509589.20, TcCLB.504151.60 |
| GO:0003854 | 3-beta-hydroxy-delta5-steroid dehydrogenase | 0,046043 | TcCLB.507993.270, TcCLB.508893.10 |

| | | | |
|---|---|---|---|
| GO:0051287 | NAD binding | 0,061652 | TcCLB.508399.10, TcCLB.507869.10, TcCLB.511323.50, TcCLB.506513.190 |
| GO:0016813 | hydrolase activity, acting on carbon-nitrogen | 0,062881 | TcCLB.507031.90, TcCLB.507963.20 |
| GO:0030060 | L-malate dehydrogenase activity | 0,063652 | TcCLB.506937.10, TcCLB.510437.40 |
| GO:0047801 | L-cysteine:2-oxoglutarate aminotransferase a | 0,063967 | TcCLB.503747.10 |
| GO:0052654 | L-leucine transaminase activity | 0,063967 | TcCLB.503747.10 |
| GO:0070548 | L-glutamine aminotransferase activity | 0,063967 | TcCLB.503747.10 |
| GO:0015035 | protein disulfide oxidoreductase activity | 0,064453 | TcCLB.507889.20, TcCLB.503477.10, TcCLB.508265.10 |
| GO:0004427 | inorganic diphosphatase activity | 0,065941 | TcCLB.511385.30, TcCLB.510773.20 |
| GO:0016829 | lyase activity | 0,066473 | TcCLB.507559.80, TcCLB.507669.10 |
| GO:0015238 | drug transmembrane transporter activity | 0,067571 | TcCLB.504069.120, TcCLB.507625.140, TcCLB.507787.50 |
| GO:0003842 | 1-pyrroline-5-carboxylate dehydrogenase acti | 0,075827 | TcCLB.510943.50 |
| GO:0003960 | NADPH:quinone reductase activity | 0,069533 | TcCLB.511287.49 |
| GO:0004033 | aldo-keto reductase (NADP) activity | 0,069533 | TcCLB.511287.49 |
| GO:0004340 | glucokinase activity | 0,073966 | TcCLB.510187.100 |
| GO:0004397 | histidine ammonia-lyase activity | 0,075819 | TcCLB.506247.220 |
| GO:0004768 | stearoyl-CoA 9-desaturase activity | 0,075198 | TcCLB.509239.10 |
| GO:0004802 | transketolase activity | 0,075827 | TcCLB.511903.120 |
| GO:0005355 | glucose transmembrane transporter activity | 0,075824 | TcCLB.506355.10 |
| GO:0008289 | lipid binding | 0,075683 | TcCLB.508257.220 |
| GO:0008716 | D-alanine-D-alanine ligase activity | 0,075789 | TcCLB.510347.20 |
| GO:0015189 | L-lysine transmembrane transporter activity | 0,075538 | TcCLB.511545.80 |
| GO:0016153 | urocanate hydratase activity | 0,075827 | TcCLB.504045.110 |
| GO:0016175 | superoxide-generating NADPH oxidase activit | 0,069533 | TcCLB.511287.49 |
| GO:0016841 | ammonia-lyase activity | 0,075819 | TcCLB.506247.220 |
| GO:0016868 | intramolecular transferase activity, phosphot | 0,070212 | TcCLB.508257.70, TcCLB.508569.80 |
| GO:0045148 | tripeptide aminopeptidase activity | 0,074973 | TcCLB.506513.110 |
| GO:0050105 | L-gulonolactone oxidase activity | 0,075769 | TcCLB.509179.100 |
| GO:0070402 | NADPH binding | 0,069533 | TcCLB.511287.49 |
| GO:0004616 | phosphogluconate dehydrogenase (decarboxy | 0,081394 | TcCLB.505807.180, TcCLB.507017.40 |
| GO:0016624 | oxidoreductase activity, acting on the aldehyd | 0,088165 | TcCLB.506853.50, TcCLB.508781.110 |
| GO:0030145 | manganese ion binding | 0,092028 | TcCLB.510655.120, TcCLB.510751.4 |
| GO:0042626 | ATPase activity, coupled to transmembrane r | 0,096821 | TcCLB.508965.14, TcCLB.506417.10, TcCLB.506925.530 |

| | CC Results from goseq. | | |
|---|---|---|---|
| row.names | term | over.p.value | de.genes.in.cat |
| GO:0000331 | contractile vacuole | 1,201E-20 | TcCLB.506551.10, TcCLB.508999.260, TcCLB.509965.394, TcCLB.503413.4, TcCLB.509051.20, TcCLB.506563.79, TcCLB.510855.10, TcCLB.503975.100, TcCLB.506247.220, TcCLB.506563.170, TcCLB.507689.30, TcCLB.510679.40, TcCLB.510837.20, TcCLB.507187.9, TcCLB.508999.250, TcCLB.508503.20, TcCLB.503837.10, TcCLB.508535.10, TcCLB.511575.130, TcCLB.506893.100, TcCLB.505945.20, TcCLB.511385.30, TcCLB.510773.20, TcCLB.510943.50, TcCLB.507711.200, TcCLB.509351.10, TcCLB.510101.140, TcCLB.407477.50, TcCLB.508209.100, TcCLB.511151.90, TcCLB.509683.10, TcCLB.506355.10, TcCLB.503887.40, TcCLB.508231.40, TcCLB.506219.40, TcCLB.503899.119, |

| | | | |
|---|---|---|---|
| GO:0016021 | integral component of membrane | 3,944E-13 | TcCLB.510103.24, TcCLB.509713.30, TcCLB.506551.10, TcCLB.506773.50, TcCLB.510811.20, TcCLB.508799.270, TcCLB.509707.10, TcCLB.508065.70, TcCLB.508799.290, TcCLB.509107.9, TcCLB.510605.10, TcCLB.506947.10, TcCLB.508647.180, TcCLB.504137.100, TcCLB.511639.20, TcCLB.511041.40, TcCLB.510069.20, TcCLB.509937.200, TcCLB.510407.30, TcCLB.510877.170, TcCLB.508465.40, TcCLB.509601.59, TcCLB.508689.40, TcCLB.508231.9, TcCLB.508799.150, TcCLB.509799.50, TcCLB.504069.120, TcCLB.508551.39, TcCLB.509767.59, TcCLB.506355.100, TcCLB.508551.30, TcCLB.506355.10, TcCLB.511245.130, TcCLB.424937.10, TcCLB.508965.14, TcCLB.506661.30, TcCLB.506417.10, TcCLB.510659.20, TcCLB.510309.20, |

| | | | |
|---|---|---|---|
| GO:0005741 | mitochondrial outer membrane | 1,385E-07 | TcCLB.503413.4, TcCLB.503975.100, TcCLB.506399.80, TcCLB.511687.19, TcCLB.508741.229, TcCLB.508737.210, TcCLB.505193.100, TcCLB.509141.40, TcCLB.511687.10, TcCLB.509129.10, TcCLB.511909.50, TcCLB.506467.29, TcCLB.506399.70, TcCLB.508873.420, TcCLB.506791.50, TcCLB.508319.20, TcCLB.511909.60, TcCLB.506941.220, TcCLB.507615.100, TcCLB.508647.110, TcCLB.508461.10, TcCLB.505843.49 |
| GO:0046930 | pore complex | 6,289E-07 | TcCLB.511687.19, TcCLB.508741.229, TcCLB.508737.210, TcCLB.509141.40, TcCLB.511687.10, TcCLB.509129.10 |
| GO:0005779 | integral component of peroxisomal membran | 2,064E-06 | TcCLB.510669.49, TcCLB.510669.20, TcCLB.508461.570, TcCLB.509203.40, TcCLB.507009.10, TcCLB.511017.50 |
| GO:0031307 | integral component of mitochondrial outer m | 7,423E-06 | TcCLB.511687.19, TcCLB.508741.229, TcCLB.508737.210, TcCLB.509141.40, TcCLB.511687.10, TcCLB.509129.10 |
| GO:0070069 | cytochrome complex | 8,953E-05 | TcCLB.508823.70, TcCLB.509721.40, TcCLB.510173.70, TcCLB.506041.50 |

| | | | |
|---|---|---|---|
| GO:0005739 | mitochondrion | 0,0002451 | TcCLB.511385.30, TcCLB.504069.20, TcCLB.506989.140, TcCLB.503769.40, TcCLB.503991.39, TcCLB.503811.60, TcCLB.511145.10, TcCLB.507049.60, TcCLB.509233.150, TcCLB.511211.70, TcCLB.509463.30, TcCLB.506337.160, TcCLB.507081.130, TcCLB.508995.60, TcCLB.508029.10, TcCLB.506567.70, TcCLB.510507.20, TcCLB.507009.50, TcCLB.511395.20, TcCLB.506775.100, TcCLB.508027.129, TcCLB.511815.40, TcCLB.503677.20 |
| GO:0020022 | acidocalcisome | 0,0004841 | TcCLB.511687.19, TcCLB.508741.229, TcCLB.508737.210, TcCLB.509141.40, TcCLB.511687.10, TcCLB.509129.10, TcCLB.511385.30, TcCLB.510773.20 |
| GO:0016281 | eukaryotic translation initiation factor 4F com | 0,0024117 | TcCLB.511687.19, TcCLB.508741.229, TcCLB.508737.210, TcCLB.509141.40, TcCLB.511687.10, TcCLB.509129.10, TcCLB.506773.120 |
| GO:0005751 | mitochondrial respiratory chain complex IV | 0,0052414 | TcCLB.506529.360, TcCLB.510889.50 |
| GO:0005758 | mitochondrial intermembrane space | 0,0055483 | TcCLB.509463.20, TcCLB.510507.29, TcCLB.505843.49 |
| GO:0012506 | vesicle membrane | 0,005744 | TcCLB.508441.20, TcCLB.507547.90 |

| | | | |
|---|---|---|---|
| GO:0016471 | vacuolar proton-transporting V-type ATPase co | 0,0095085 | TcCLB.510993.10, TcCLB.506375.110 |
| GO:0005743 | mitochondrial inner membrane | 0,0134922 | TcCLB.508823.70, TcCLB.509721.40, TcCLB.506649.50, TcCLB.510173.70 |
| GO:0005829 | cytosol | 0,0135831 | TcCLB.504045.60, TcCLB.503747.10, TcCLB.506629.200, TcCLB.506739.99, TcCLB.511287.49, TcCLB.506593.60, TcCLB.506367.30, TcCLB.509601.90, TcCLB.510819.119, TcCLB.508741.170, TcCLB.506829.80, TcCLB.511823.70, TcCLB.509139.10 |
| GO:0033177 | proton-transporting two-sector ATPase compl | 0,0192668 | TcCLB.507517.60, TcCLB.503579.70, TcCLB.504131.180, TcCLB.504125.40 |
| GO:0031966 | mitochondrial membrane | 0,0203342 | TcCLB.509601.59, TcCLB.509767.59, TcCLB.508601.50 |
| GO:0005747 | mitochondrial respiratory chain complex I | 0,0344033 | TcCLB.506337.160, TcCLB.506989.110, TcCLB.510149.150 |
| GO:0005783 | endoplasmic reticulum | 0,0447749 | TcCLB.509239.10, TcCLB.508257.10, TcCLB.510349.50, TcCLB.503929.79, TcCLB.506405.50, TcCLB.511823.70 |
| GO:0030964 | NADH dehydrogenase complex | 0,0595221 | TcCLB.508601.50 |
| GO:0005886 | plasma membrane | 0,0664821 | TcCLB.511545.80, TcCLB.511385.30, TcCLB.504069.120, TcCLB.506179.5 |
| GO:0020015 | glycosome | 0,0669905 | TcCLB.507047.150, TcCLB.509179.100, TcCLB.510187.100, TcCLB.511385.30, TcCLB.508441.20, TcCLB.507547.90 |
| GO:0030176 | integral component of endoplasmic reticulum | 0,0841113 | TcCLB.511245.130, TcCLB.506661.30 |

**Supplementary Table 4. Annotated genes corresponding to RNA Binding Proteins identified in the CL Brener genome.**

| Gene ID | TriTrypDB anotation | Domain |
|---|---|---|
| TcCLB.507711.40<br>TcCLB.424123.40 | RNA-binding protein | RRM |
| TcCLB.469785.40<br>TcCLB.507025.50 | RNA-binding protein | RRM |
| TcCLB.507025.60 | RNA-binding protein | RRM |
| TcCLB.503419.50<br>TcCLB.509105.90 | MRB1-associated protein | RRM |
| TcCLB.503619.20<br>TcCLB.511647.40 | Hypothetical protein | RRM |
| TcCLB.503683.11<br>TcCLB.509999.140 | Double RNA binding domain protein 6A | RRM |
| TcCLB.503683.30<br>TcCLB.509999.120 | RNA-binding protein | RRM |
| TcCLB.509581.50<br>TcCLB.503709.10 | Hypothetical protein | RRM |
| TcCLB.508569.90<br>TcCLB.503733.50 | Hypothetical protein | RRM |
| TcCLB.503897.90<br>TcCLB.509561.110 | Hypothetical protein | RRM |
| TcCLB.507093.63<br>TcCLB.503917.7 | RNA-binding protein | RRM |
| TcCLB.507037.20<br>TcCLB.508707.80 | RNA-binding protein | RRM |
| TcCLB.508209.39<br>TcCLB.503919.30 | Hypothetical protein | RRM |
| TcCLB.509591.50<br>TcCLB.503999.90 | RNA-binding protein 34 | RRM |
| TcCLB.504005.6<br>TcCLB.511481.55 | RNA-binding protein | RRM |
| TcCLB.504071.80 | Nuclear cap binding protein | RRM |
| TcCLB.507089.70 | RNA-binding protein | RRM |

***Genes listed after last search for RBPs.**

| | | |
|---|---|---|
| TcCLB.504085.30 | [illegible] | [illegible] |
| TcCLB.511109.130<br>TcCLB.504157.10 | Heterogeneous nuclear ribonucleoprotein H/F | RRM |
| TcCLB.504165.20<br>TcCLB.508981.20 | Hypothetical protein | RRM |
| TcCLB.508145.30<br>TcCLB.504243.10 | RNA-binding protein | RRM |
| TcCLB.504431.90<br>TcCLB.507951.140 | U2 small nuclear ribonucleoprotein B | RRM |
| TcCLB.505007.10<br>TcCLB.510007.30 | RNA-binding protein | RRM |
| TcCLB.505165.10 | RNA-binding protein | RRM |
| TcCLB.506399.40<br>TcCLB.509243.20 | Hypothetical protein | RRM |
| TcCLB.506425.60 | Hypothetical protein | RRM |
| TcCLB.506565.12<br>TcCLB.511837.129 | RNA-binding protein | RRM |
| TcCLB.506565.4 | RNA-binding protein | RRM |
| TcCLB.506565.8 | RNA-binding protein | RRM |
| TcCLB.506625.70 | RNA-binding protein | RRM |
| TcCLB.506649.80<br>TcCLB.508349.39 | Double RNA binding domain protein 3 | RRM |
| TcCLB.506825.10<br>TcCLB.506681.10 | Double RNA binding domain protein 12 | RRM |
| TcCLB.506693.30<br>TcCLB.508153.680 | RNA-binding protein 6 | RRM |
| TcCLB.506795.10<br>TcCLB.509937.60 | RNA-binding protein | RRM |
| TcCLB.506797.120<br>TcCLB.511907.100 | Hypothetical protein | RRM |
| TcCLB.509167.30<br>TcCLB.506831.30 | Hypothetical protein | RRM |

| | | |
|---|---|---|
| TcCLB.506925.400 | Hypothetical protein | RRM |
| TcCLB.506989.100<br>TcCLB.510149.140 | Triple RNA binding domain protein 3 | RRM |
| TcCLB.507093.220 | RNA-binding protein | RRM |
| TcCLB.507093.229 | U-rich RNA-binding protein UBP-2 | RRM |
| TcCLB.507093.250 | RNA-binding protein | RRM |
| TcCLB.508213.40<br>TcCLB.507515.40 | Hypothetical protein | RRM |
| TcCLB.508213.20<br>TcCLB.507515.60 | Hypothetical protein | RRM |
| TcCLB.507611.300<br>TcCLB.507723.120 | U1A small nuclear ribonucleoprotein | RRM |
| TcCLB.507873.30<br>TcCLB.510689.60 | Double RNA binding domain protein 7 | RRM |
| TcCLB.507885.10 | RNA-binding protein | RRM |
| TcCLB.507993.140<br>TcCLB.511277.580 | RNA-binding protein 29 | RRM |
| TcCLB.508145.10 | RNA-binding protein | RRM |
| TcCLB.508145.20 | RNA-binding protein | RRM |
| TcCLB.510143.80<br>TcCLB.508409.270 | RNA-binding protein | RRM |
| TcCLB.510755.120<br>TcCLB.508413.50 | RNA-binding protein | RRM |
| TcCLB.508461.320 | U5 snRNA-associated splicing factor | RRM |
| TcCLB.510877.150<br>TcCLB.508689.20 | Eukaryotic translation initiation factor 3 subunit g | RRM |
| TcCLB.508837.140<br>TcCLB.511383.30 | u1 small nuclear ribonucleoprotein 70 kDa | RRM |
| TcCLB.511277.200 | RNA-binding protein | RRM |
| TcCLB.511367.60 | Lupus La protein homolog | RRM |
| TcCLB.511127.10<br>TcCLB.511481.70 | RNA-binding protein 5 | RRM |

| | | |
|---|---|---|
| TcCLB.508879.80 | Hypothetical protein | RRM |
| TcCLB.424195.9<br>TcCLB.508901.20 | RNA-binding protein 4 | RRM |
| TcCLB.508989.30<br>TcCLB.509569.120 | RNA-binding protein | RRM |
| TcCLB.509965.180<br>TcCLB.509053.179 | RNA-binding protein | RRM |
| TcCLB.509055.10 | RNA-binding protein | RRM |
| TcCLB.511621.50<br>TcCLB.509317.60 | RNA-binding protein | RRM |
| TcCLB.510507.50<br>TcCLB.509461.100 | RNA-binding protein | RRM |
| TcCLB.510105.33<br>TcCLB.509715.23 | RNA-binding protein | RRM |
| TcCLB.511741.40<br>TcCLB.511517.70 | RNA-binding protein | RRM |
| TcCLB.511727.160 | Polypyrimidine tract-binding protein | RRM |
| TcCLB.511727.190 | Hypothetical protein | RRM |
| TcCLB.511727.270 | RNA-binding protein | RRM |
| TcCLB.511727.290 | RNA-binding protein | RRM |
| TcCLB.511871.110<br>TcCLB.511863.20 | Hypothetical protein | RRM |
| TcCLB.511837.138 | RNA-binding protein | RRM |
| TcCLB.510311.50<br>TcCLB.510265.40 | Hypothetical protein | RRM |
| TcCLB.510747.80<br>TcCLB.510657.160 | Double RNA binding domain protein 9 | RRM |
| TcCLB.506435.120<br>TcCLB.506265.100 | ATP-dependent RNA helicase * | RRM |
| TcCLB.511003.30 | Mitochondrial RNA binding complex 1 subunit * | RRM |
| TcCLB.503715.10<br>TcCLB.509607.30 | Splicing factor TSR1, putative * | RRM |

| | | |
|---|---|---|
| TcCLB.511181.70<br>TcCLB.508299.89 | Hypothetical protein, conserved * | RRM |
| TcCLB.510741.40<br>TcCLB.510661.230 | Hypothetical protein, conserved * | RRM |
| TcCLB.509167.140<br>TcCLB.485683.10 | RNA-binding protein 42 * | RRM |
| TcCLB.506885.70<br>TcCLB.508461.140 | Polyadenylate-binding protein 1 | RRM/PABP |
| TcCLB.503989.10<br>TcCLB.401469.10 | Zinc finger protein 2 | Zinc Finger |
| TcCLB.411427.10 | Hypothetical protein | Zinc Finger |
| TcCLB.506859.204<br>TcCLB.503567.9 | Hypothetical protein | Zinc Finger |
| TcCLB.503577.20<br>TcCLB.510943.60 | U2 splicing auxiliary factor | Zinc Finger |
| TcCLB.503795.10<br>TcCLB.506679.10 | Hypothetical protein | Zinc Finger |
| TcCLB.503897.150<br>TcCLB.509561.39 | Hypothetical protein | Zinc Finger |
| TcCLB.507089.30<br>TcCLB.504085.70 | Zinc finger protein | Zinc Finger |
| TcCLB.511577.30<br>TcCLB.504797.110 | Hypothetical protein | Zinc Finger |
| TcCLB.504929.5<br>TcCLB.507305.40 | Zinc finger CCCH domain containing protein 11 | Zinc Finger |
| TcCLB.510295.59 | Hypothetical protein | Zinc Finger |
| TcCLB.510297.10<br>TcCLB.506009.10 | Hypothetical protein | Zinc Finger |
| TcCLB.506127.20 | mRNA export factor MEX67 | Zinc Finger |
| TcCLB.506211.60<br>TcCLB.508895.60 | RNA-binding protein | Zinc Finger |
| TcCLB.506211.70 | RNA-binding protein | Zinc Finger |

| | | |
|---|---|---|
| TcCLB.508895.50 | RNA binding protein | Zinc Finger |
| TcCLB.510101.200<br>TcCLB.506297.130 | Zinc finger ccch and cchc domain-containing protein | Zinc Finger |
| TcCLB.506733.140<br>TcCLB.509229.90 | Hypothetical protein | Zinc Finger |
| TcCLB.506739.99<br>TcCLB.510819.119 | Zinc finger (CCCH type) protein | Zinc Finger |
| TcCLB.510759.100<br>TcCLB.506999.120 | Hypothetical protein | Zinc Finger |
| TcCLB.509551.60<br>TcCLB.507007.77 | Hypothetical protein | Zinc Finger |
| TcCLB.508271.4 | Hypothetical protein | Zinc Finger |
| TcCLB.506859.80<br>TcCLB.511815.50 | Hypothetical protein | Zinc Finger |
| TcCLB.506859.230<br>TcCLB.511817.10 | Hypothetical protein | Zinc Finger |
| TcCLB.506859.240<br>TcCLB.511817.20 | Hypothetical protein | Zinc Finger |
| TcCLB.510427.10<br>TcCLB.506883.120 | Hypothetical protein | Zinc Finger |
| TcCLB.506885.200<br>TcCLB.510729.220 | Hypothetical protein | Zinc Finger |
| TcCLB.506885.204<br>TcCLB.510729.210 | Hypothetical protein | Zinc Finger |
| TcCLB.506931.4<br>TcCLB.510131.44 | Hypothetical protein | Zinc Finger |
| TcCLB.506933.50 | Hypothetical protein | Zinc Finger |
| TcCLB.506945.210 | Zinc finger CCCH domain-containing protein 47 | Zinc Finger |
| TcCLB.503453.20 | Hypothetical protein | Zinc Finger |
| TcCLB.506977.110 | Hypothetical protein | Zinc Finger |
| TcCLB.510351.80<br>TcCLB.507601.80 | Hypothetical protein | Zinc Finger |

| | | |
|---|---|---|
| TcCLB.507625.70<br>TcCLB.507787.140 | Hypothetical protein | Zinc Finger |
| TcCLB.511867.10<br>TcCLB.507775.10 | Hypothetical protein | Zinc Finger |
| TcCLB.507831.20<br>TcCLB.511263.50 | Hypothetical protein | Zinc Finger |
| TcCLB.507831.30<br>TcCLB.511263.40 | Hypothetical protein | Zinc Finger |
| TcCLB.507831.40<br>TcCLB.511263.30 | Hypothetical protein | Zinc Finger |
| TcCLB.508215.10<br>TcCLB.508879.10 | Hypothetical protein | Zinc Finger |
| TcCLB.508241.90<br>TcCLB.511151.20 | Hypothetical protein | Zinc Finger |
| TcCLB.508357.9<br>TcCLB.508355.330 | Hypothetical protein | Zinc Finger |
| TcCLB.510143.120<br>TcCLB.508409.310 | Hypothetical protein | Zinc Finger |
| TcCLB.508693.40 | Hypothetical protein | Zinc Finger |
| TcCLB.508831.66<br>TcCLB.511671.130 | Hypothetical protein | Zinc Finger |
| TcCLB.509805.240<br>TcCLB.511267.20 | Hypothetical protein | Zinc Finger |
| TcCLB.509805.230<br>TcCLB.511267.24 | Hypothetical protein | Zinc Finger |
| TcCLB.511467.4 | Hypothetical protein | Zinc Finger |
| TcCLB.509231.39<br>TcCLB.509719.69 | Hypothetical protein | Zinc Finger |
| TcCLB.509233.210 | Hypothetical protein | Zinc Finger |
| TcCLB.463297.10<br>TcCLB.509395.40 | Hypothetical protein | Zinc Finger |
| TcCLB.511039.39 | Hypothetical protein | Zinc Finger |

| | | |
|---|---|---|
| TcCLB.509791.90 | Hypothetical protein | Zinc Finger |
| TcCLB.511555.40<br>TcCLB.510219.30 | cleavage and polyadenylation specificity factor 30 kDa subunit | Zinc Finger |
| TcCLB.511511.3 | Zinc finger protein ZFP1 | Zinc Finger |
| TcCLB.511511.6 | Zinc finger protein ZFP1 | Zinc Finger |
| TcCLB.511735.40<br>TcCLB.511521.20 | Hypothetical protein | Zinc Finger |
| TcCLB.511735.88 | Hypothetical protein | Zinc Finger |
| TcCLB.511807.130 | Hypothetical protein | Zinc Finger |
| TcCLB.511807.160 | Hypothetical protein | Zinc Finger |
| TcCLB.511867.60 | Hypothetical protein | Zinc Finger |
| TcCLB.507611.398<br>TcCLB.506247.30 | RNA editing complex protein MP63, putative * | Zinc Finger |
| TcCLB.506357.69<br>TcCLB.511277.39 | Hypothetical protein, conserved * | Zinc Finger |
| TcCLB.506533.70<br>TcCLB.509395.10 | hypothetical protein, conserved * | Zinc Finger |
| TcCLB.510149.70<br>TcCLB.506989.20 | Zinc-finger double-stranded RNA-binding, putative * | Zinc Finger |
| TcCLB.509443.20<br>TcCLB.507017.140 | Mitochondrial RNA binding protein 1, putative * | Zinc Finger |
| TcCLB.510857.40<br>TcCLB.509611.110 | RNA-editing complex protein, putative * | Zinc Finger |
| TcCLB.510743.20<br>TcCLB.510659.279 | Hypothetical protein, conserved *<br>Nucleic acid binding protein, putative * | Zinc Finger |
| TcCLB.510687.20 | Zinc knuckle, putative * | Zinc Finger |
| TcCLB.397937.10<br>TcCLB.508625.160 | Pumilio/PUF RNA binding protein 1 | PUF |
| TcCLB.508787.30<br>TcCLB.468005.9 | Pumilio/PUF RNA binding protein 3 | PUF |
| TcCLB.503757.30<br>TcCLB.503719.39 | Hypothetical protein | PUF |

| | | |
|---|---|---|
| TcCLB.506563.10<br>TcCLB.503869.40 | Pumilio/PUF RNA binding protein 9 | PUF |
| TcCLB.506773.130<br>TcCLB.508799.70 | Pumilio-repeat, RNA-binding protein | PUF |
| TcCLB.508479.120 | Pumilio/PUF RNA binding protein 8 | PUF |
| TcCLB.507049.199 | Pumilio protein | PUF |
| TcCLB.508577.100<br>TcCLB.507521.110 | Pumilio/PUF RNA binding protein 5 | PUF |
| TcCLB.507831.110<br>TcCLB.511261.120 | Pumilio/PUF RNA binding protein 2 | PUF |
| TcCLB.511715.100<br>TcCLB.508445.99 | Pumilio/PUF RNA binding protein 7 | PUF |
| TcCLB.510073.30<br>TcCLB.509399.190 | Pumilio/PUF RNA binding protein 4 | PUF |
| TcCLB.509759.19 | Pumilio/PUF RNA binding protein 1 | PUF |
| TcCLB.510125.10 | Pumilio/PUF RNA binding protein 6 | PUF |
| TcCLB.510121.110 | Hypothetical protein, conserved * | PUF |
| TcCLB.506605.100<br>TcCLB.511239.90 | Hypothetical protein * | KH |
| TcCLB.506625.200 | Ribosomal RNA processing protein 40, putative * | KH |
| TcCLB.510595.20<br>TcCLB.507895.150 | KH domain, putative * | KH |
| TcCLB.510167.29<br>TcCLB.508859.109 | Ribosomal RNA processing protein 4, putative * | KH |
| TcCLB.508717.40<br>TcCLB.504079.10 | Branch point binding protein, putative * | KH |
| TcCLB.506943.120<br>TcCLB.509065.130 | Hypothetical protein, conserved * | KH |
| TcCLB.507011.50<br>TcCLB.507005.70 | Pre-rRNA-processing protein PNO1, putative * | KH |
| TcCLB.504089.60<br>TcCLB.504001.10 | Hypothetical protein | Alba |

| | | |
|---|---|---|
| TcCLB.504089.70<br>TcCLB.504001.20 | Hypothetical protein | Alba |
| TcCLB.510877.30 | Hypothetical protein | Alba |
| TcCLB.510877.40 | Hypothetical protein | Alba |
| TcCLB.508489.20<br>TcCLB.511495.20 | SAM domain (Sterile alpha motif), putative * | SAM |
| TcCLB.511517.140 | Hypothetical protein * | SAM |
| TcCLB.506743.4<br>TcCLB.503945.10 | Eukaryotic translation initiation factor 1A, putative * | S1 |
| TcCLB.511367.240 | PIWI-like protein * | PIWI |

**Supplementary Table 5. Differentially expressed (DE) RBP genes in CL Brener epimastigotes compared to tissue culture derived trypomastigotes and amastigotes**

up_rbp_epi_tryp

| Gene_ID | log2FoldChange | padj | Product.Description | baseMean |
|---|---|---|---|---|
| TcCLB.506739.99 | 3,375343804 | 0 | zinc finger (CCCH type) protein, putative | 2185,865298 |
| TcCLB.511367.240 | 3,094115105 | 1,76E-216 | PIWI-like protein | 662,1385698 |
| TcCLB.506931.4 | 3,083180798 | 1,44E-209 | hypothetical protein, conserved | 617,9904423 |
| TcCLB.510131.44 | 2,256998567 | 1,17E-103 | hypothetical protein, conserved | 397,3848663 |
| TcCLB.506885.204 | 1,98029734 | 1,02E-85 | hypothetical protein, conserved | 1315,025274 |
| TcCLB.510819.119 | 1,862887088 | 1,11E-62 | zinc finger (CCCH type) protein, putative | 441,074966 |
| TcCLB.511817.20 | 1,661164991 | 4,14E-49 | hypothetical protein, conserved | 742,5119723 |
| TcCLB.511727.190 | 1,552708525 | 5,67E-55 | hypothetical protein, conserved | 860,4491512 |
| TcCLB.510729.210 | 1,536519579 | 1,34E-49 | hypothetical protein, conserved | 1238,25649 |
| TcCLB.503683.11 | 1,501644083 | 6,45E-12 | Double RNA binding domain protein 6A | 68,19733144 |
| TcCLB.509999.140 | 1,38211067 | 1,59E-10 | Double RNA binding domain protein 6A | 62,34236312 |
| TcCLB.509167.140 | 1,321968923 | 3,89E-26 | RNA-binding protein 42 (RNA-binding motif protein 42), putative | 2566,66086 |
| TcCLB.503715.10 | 1,300663146 | 9,67E-46 | splicing factor TSR1, putative | 651,4772317 |
| TcCLB.507831.20 | 1,286588075 | 0,00336573 | hypothetical protein, conserved | 12,04852544 |
| TcCLB.504001.10 | 1,163342264 | 1,68E-39 | hypothetical protein, conserved | 731,256651 |
| TcCLB.507521.110 | 1,145254164 | 1,79E-24 | pumilio/PUF RNA binding protein 5, putative | 590,9808113 |
| TcCLB.508981.20 | 1,145006263 | 1,48E-13 | hypothetical protein, conserved | 131,5808328 |
| TcCLB.504089.60 | 1,135679874 | 2,10E-29 | hypothetical protein, conserved | 640,3714296 |
| TcCLB.510759.100 | 1,130248597 | 8,25E-30 | NLI interacting factor-like phosphatase, putative | 662,9663541 |
| TcCLB.507093.220 | 1,130099204 | 3,85E-27 | RNA-binding protein UBP1, putative | 683,4398677 |
| TcCLB.509551.60 | 1,113536142 | 1,01E-11 | hypothetical protein, conserved | 149,8472427 |
| TcCLB.397937.10 | 1,09134651 | 2,14E-17 | pumilio/PUF RNA binding protein 1, putative | 280,5579971 |
| TcCLB.511239.90 | 1,040458085 | 1,00E-17 | hypothetical protein | 423,8807767 |
| TcCLB.506925.400 | 1,02489769 | 9,29E-26 | hypothetical protein, conserved | 469,9322563 |
| TcCLB.506859.240 | 1,023027367 | 0,00015803 | hypothetical protein, conserved | 44,00844484 |
| TcCLB.511727.270 | 1,02004571 | 1,24E-40 | RNA-binding protein, putative | 4671,643903 |
| TcCLB.504001.20 | 1,017585967 | 1,31E-20 | hypothetical protein, conserved | 548,2993083 |
| TcCLB.511109.130 | 1,015023714 | 2,91E-19 | heterogeneous nuclear ribonucleoprotein H/F, putative | 440,8719704 |
| TcCLB.507873.30 | 1,002368453 | 7,12E-25 | Double RNA binding domain protein 7 | 555,3921937 |

up_rbp_epi_ama

| Gene_ID | log2FoldChange | padj | Product.Description | baseMean |
|---|---|---|---|---|
| TcCLB.511367.240 | 3,606306398 | 1,08E-122 | PIWI-like protein | 662,1385698 |
| TcCLB.506931.4 | 2,968390566 | 1,33E-105 | hypothetical protein, conserved | 617,9904423 |
| TcCLB.503683.11 | 2,940718444 | 4,86E-13 | Double RNA binding domain protein 6A | 68,19733144 |
| TcCLB.507831.20 | 2,920879201 | 0,00309465 | hypothetical protein, conserved | 12,04852544 |
| TcCLB.506739.99 | 2,903661541 | 2,08E-223 | zinc finger (CCCH type) protein, putative | 2185,865298 |
| TcCLB.509999.140 | 2,736095578 | 7,98E-12 | Double RNA binding domain protein 6A | 62,34236312 |
| TcCLB.510131.44 | 2,667522511 | 1,89E-66 | hypothetical protein, conserved | 397,3848663 |
| TcCLB.511817.20 | 2,318963286 | 3,51E-69 | hypothetical protein, conserved | 742,5119723 |
| TcCLB.511807.130 | 2,24141306 | 5,70E-08 | hypothetical protein, conserved | 52,31017629 |
| TcCLB.506825.10 | 1,899025256 | 0,00718 | Double RNA binding domain protein 12 | 17,10971309 |
| TcCLB.511109.130 | 1,531304738 | 5,19E-29 | heterogeneous nuclear ribonucleoprotein H/F, putative | 440,8719704 |
| TcCLB.506859.240 | 1,394223886 | 0,00031145 | hypothetical protein, conserved | 44,00844484 |
| TcCLB.510819.119 | 1,373311199 | 1,01E-26 | zinc finger (CCCH type) protein, putative | 441,074966 |
| TcCLB.511815.50 | 1,149067182 | 6,19E-32 | hypothetical protein, conserved | 798,3011834 |
| TcCLB.503683.30 | 1,115764285 | 3,95E-05 | RNA-binding protein, putative | 94,27632534 |
| TcCLB.504797.110 | 1,052741502 | 9,44E-21 | hypothetical protein, conserved | 528,7644126 |
| TcCLB.504157.10 | 1,0340873 | 7,50E-14 | heterogeneous nuclear ribonucleoprotein H/F, putative | 513,665032 |

down_rbp_epi_tryp

| Gene_ID | log2FoldChange | padj | Product.Description | baseMean |
|---|---|---|---|---|
| TcCLB.504929.5 | -4,218970659 | 1,17E-13 | zinc finger CCCH domain containing protein 11 | 738,8239859 |
| TcCLB.511127.10 | -3,731845286 | 8,08E-14 | RNA-binding protein 5, putative | 18,64229074 |
| TcCLB.507305.40 | -3,546565836 | 8,05E-09 | zinc finger CCCH domain containing protein 11 | 395,5125914 |
| TcCLB.511511.6 | -2,772415292 | 2,76E-18 | zinc finger protein ZFP1 | 69,81133245 |
| TcCLB.503709.10 | -2,166808655 | 1,83E-10 | hypothetical protein | 30,10067699 |
| TcCLB.508145.30 | -1,582651619 | 1,03E-09 | RNA-binding protein, putative | 43,77587004 |
| TcCLB.506859.230 | -1,501176104 | 0,00222551 | hypothetical protein, conserved | 20,91041858 |
| TcCLB.508153.680 | -1,464377286 | 8,19E-12 | RNA-binding protein 6, putative | 64,02849288 |
| TcCLB.508409.310 | -1,247622136 | 2,54E-23 | hypothetical protein, conserved | 335,900361 |
| TcCLB.503795.10 | -1,235074829 | 4,56E-05 | hypothetical protein, conserved | 28,78739941 |
| TcCLB.511867.10 | -1,231511545 | 7,10E-32 | hypothetical protein, conserved | 494,1776714 |
| TcCLB.506693.30 | -1,222264095 | 0,00049099 | RNA-binding protein 6, putative | 23,05838598 |
| TcCLB.463297.10 | -1,209880482 | 7,27E-07 | hypothetical protein (fragment) | 52,56989367 |
| TcCLB.509395.40 | -1,101937442 | 2,52E-14 | hypothetical protein, conserved | 617,1306421 |
| TcCLB.503989.10 | -1,080798934 | 2,74E-25 | zinc finger protein 2, putative | 514,1393144 |
| TcCLB.504005.6 | -1,070280763 | 4,31E-11 | RNA-binding protein, putative | 229,6842293 |
| TcCLB.411427.10 | -1,057410862 | 1,67E-06 | hypothetical protein, conserved | 78,4964516 |
| TcCLB.401469.10 | -1,02771079 | 1,84E-22 | zinc finger protein 2, putative | 464,84793 |

down_rbp_epi_ama

| Gene_ID | log2FoldChange | padj | Product.Description |
|---|---|---|---|
| TcCLB.507025.50 | -2,563453339 | 1,27E-45 | RNA-binding protein, putative |
| TcCLB.469785.40 | -2,501442805 | 1,32E-63 | RNA-binding protein, putative |
| TcCLB.507305.40 | -1,915652352 | 0,00116426 | zinc finger CCCH domain containing protein 11 |
| TcCLB.504005.6 | -1,914949264 | 1,83E-32 | RNA-binding protein, putative |
| TcCLB.507025.60 | -1,715049643 | 1,24E-08 | RNA-binding protein, putative |
| TcCLB.506859.230 | -1,633861682 | 0,00251421 | hypothetical protein, conserved |
| TcCLB.511481.55 | -1,3748801 | 6,41E-17 | RNA-binding protein, putative |
| TcCLB.503709.10 | -1,316430723 | 0,00257097 | hypothetical protein |
| TcCLB.504929.5 | -1,209683928 | 0,03393288 | zinc finger CCCH domain containing protein 11 |
| TcCLB.503733.50 | -1,138418607 | 1,81E-14 | hypothetical protein, conserved (pseudogene) |
| TcCLB.504089.70 | -1,082636199 | 1,08E-16 | hypothetical protein, conserved |
| TcCLB.508145.30 | -1,041068597 | 0,00211606 | RNA-binding protein, putative |
| TcCLB.510743.20 | -1,01179592 | 1,13E-10 | hypothetical protein, conserved |
| TcCLB.507895.150 | -1,008790555 | 4,92E-11 | KH domain/Domain of unknown function (DUF1771)/Smr domain containing protein, putative |

**Supplementary Table 6. Complete list of DE genes in TcZC3H12 KO cell lines compared to wild type parasites.**

up_ko_vs_wt

| gid_1 | deseq_logfc_ | deseq_adjp_3 | annotcdslength | strand_5 | annotgeneproduct_6 |
|---|---|---|---|---|---|
| TcCLB.507485.45 | 1,849 | 0,003069 | 540.0 | - | Amastin surface glycoprotein, putative |
| TcCLB.508821.60 | 1,644 | 0,004139 | 288.0 | - | hypothetical protein, conserved |
| TcCLB.511635.69 | 1,566 | 0,0004775 | 291.0 | - | hypothetical protein, conserved |
| TcCLB.506469.110 | 1,524 | 0,01097 | 807.0 | + | hypothetical protein |
| TcCLB.407477.44 | 1,496 | 0,02174 | 366.0 | + | hypothetical protein, conserved |
| TcCLB.509367.15 | 1,481 | 0,00006752 | 798.0 | - | tryptophanyl-tRNA synthetase (pseudogene), putative |
| TcCLB.510131.40 | 1,445 | 8,728E-13 | 936.0 | + | haloacid dehalogenase-like hydrolase, putative |
| TcCLB.510691.70 | 1,442 | 0,00003615 | 663.0 | - | hypothetical protein, conserved |
| TcCLB.504625.74 | 1,381 | 0,0000393 | 291.0 | - | hypothetical protein, conserved |
| TcCLB.506893.80 | 1,365 | 0,0002498 | 564.0 | + | monothiol glutaredoxin, putative |
| TcCLB.511413.11 | 1,335 | 0,02164 | 3207.0 | + | hypothetical protein, conserved (pseudogene) |
| TcCLB.507851.40 | 1,318 | 2,219E-08 | 618.0 | + | hypothetical protein, conserved |
| TcCLB.506627.100 | 1,299 | 0,0001155 | 537.0 | + | hypothetical protein, conserved |
| TcCLB.507297.30 | 1,276 | 0,01201 | 1077.0 | - | metacaspase, putative |
| TcCLB.508537.10 | 1,265 | 0,001118 | 741.0 | - | glyceraldehyde 3-phosphate dehydrogenase, cytosolic, putative |
| TcCLB.506871.50 | 1,218 | 0,005482 | 1317.0 | - | ubiquitin hydrolase, putative |
| TcCLB.511037.20 | 1,188 | 6,704E-20 | 1563.0 | + | L-2-hydroxyglutarate dehydrogenase, mitochondrial, putative |
| TcCLB.506857.40 | 1,162 | 0,02095 | 1164.0 | - | hypothetical protein, conserved |
| TcCLB.506867.10 | 1,157 | 0,009843 | 1368.0 | - | endonuclease G, putative |
| TcCLB.510609.65 | 1,134 | 0,04723 | 300.0 | + | u6 snrna-associated sm-like protein lsm3 |
| TcCLB.503617.15 | 1,133 | 0,04943 | 492.0 | + | u6 snrna-associated sm-like protein lsm3 |
| TcCLB.511751.166 | 1,124 | 0,00804 | 990.0 | + | Putative papain-like cysteine peptidase (DUF1796), putative |
| TcCLB.508727.30 | 1,119 | 0,000621 | 990.0 | + | TIGR00269 family protein, putative |
| TcCLB.509769.66 | 1,107 | 0,00001501 | 399.0 | + | hypothetical protein, conserved |
| TcCLB.503809.75 | 1,102 | 0,008929 | 396.0 | - | hypothetical protein, conserved |
| TcCLB.503639.30 | 1,101 | 0,001456 | 2274.0 | + | hypothetical protein, conserved |
| TcCLB.507639.100 | 1,092 | 0,008613 | 1011.0 | - | phosphatidylinositol N-acetylglucosaminyltransferase subunit C, putative |
| TcCLB.510181.150 | 1,09 | 0,0000324 | 567.0 | + | caltractin, putative |
| TcCLB.510105.150 | 1,077 | 0,0001436 | 798.0 | - | methyltransferase domain containing protein, putative |
| TcCLB.511733.60 | 1,066 | 0,01953 | 1059.0 | - | hypothetical protein, conserved |
| TcCLB.507517.30 | 1,062 | 0,02322 | 981.0 | + | hypothetical protein, conserved |
| TcCLB.506649.64 | 1,058 | 0,01542 | 480.0 | - | hypothetical protein, conserved |

| | | | | | |
|---|---|---|---|---|---|
| TcCLB.506247.300 | 1,057 | 1,859E-11 | 471.0 | + | hypothetical protein, conserved |
| TcCLB.508539.20 | 1,056 | 0,01577 | 501.0 | + | MICOS complex subunit MIC17, putative |
| TcCLB.510357.120 | 1,053 | 0,04723 | 648.0 | - | hypothetical protein, conserved |
| TcCLB.506369.30 | 1,052 | 0,02532 | 1287.0 | - | JmjC domain, hydroxylase, putative |
| TcCLB.511165.50 | 1,047 | 0,00983 | 486.0 | - | PA domain containing protein, putative |
| TcCLB.504171.40 | 1,046 | 7,404E-09 | 498.0 | + | Mechanosensitive ion channel MscS, putative |
| TcCLB.506251.20 | 1,044 | 0,000621 | 1044.0 | + | hypothetical protein, conserved |
| TcCLB.509179.120 | 1,043 | 0,01542 | 384.0 | - | hypothetical protein, conserved |
| TcCLB.511705.20 | 1,04 | 0,008227 | 840.0 | + | putative prolyl carboxypeptidase, putative |
| TcCLB.508479.5 | 1,037 | 0,03761 | 960.0 | + | glycine dehydrogenase (pseudogene), putative |
| TcCLB.510357.60 | 1,037 | 0,03692 | 390.0 | - | hypothetical protein, conserved |
| TcCLB.504103.6 | 1,035 | 0,00201 | 591.0 | + | non-canonical purine NTP pyrophosphatase, RdgB/HAM1 family, putative |
| TcCLB.506247.420 | 1,035 | 0,004554 | 948.0 | + | hypothetical protein, conserved |
| TcCLB.510409.60 | 1,033 | 0,04584 | 1479.0 | + | hypothetical protein, conserved |
| TcCLB.508355.20 | 1,027 | 0,02971 | 486.0 | + | hypothetical protein, conserved |
| TcCLB.506247.355 | 1,025 | 0,01293 | 738.0 | + | hypothetical protein, conserved |
| TcCLB.506925.420 | 1,022 | 0,0005952 | 624.0 | + | hypothetical protein, conserved |
| TcCLB.503811.45 | 1,018 | 0,02128 | 456.0 | - | hypothetical protein, conserved |
| TcCLB.509065.36 | 1,016 | 0,03211 | 414.0 | - | hypothetical protein, conserved |
| TcCLB.508409.190 | 1,013 | 0,03462 | 825.0 | - | FHA domain containing protein, putative |
| TcCLB.509167.170 | 1,01 | 0,01127 | 981.0 | - | tRNA (Guanine-1)-methyltransferase, putative |
| TcCLB.507795.40 | 1,003 | 0,03763 | 306.0 | - | hypothetical protein, conserved |

down_ko_vs_wt

| gid_1 | deseq_logfc_2 | deseq_adjp_3 | annotcdslength_4 | strand_5 | annotgeneproduct_6 |
|---|---|---|---|---|---|
| TcCLB.506153.20 | -2,13 | 0,0001284 | 1137.0 | - | amino acid transporter, putative |
| TcCLB.506053.10 | -1,883 | 1,855E-10 | 1347.0 | - | amino acid transporter, putative |
| TcCLB.503733.80 | -1,864 | 4,775E-20 | 6555.0 | - | hypothetical protein, conserved |
| TcCLB.504173.40 | -1,849 | 0,000001952 | 852.0 | + | hypothetical protein |
| TcCLB.506153.10 | -1,787 | 3,209E-08 | 1347.0 | - | amino acid transporter, putative |
| TcCLB.506825.40 | -1,663 | 0,001118 | 1548.0 | + | hypothetical protein, conserved |
| TcCLB.504173.60 | -1,509 | 1,737E-08 | 2664.0 | + | RING-variant domain containing protein, putative |
| TcCLB.507659.30 | -1,352 | 0,02282 | 1032.0 | + | amino acid transporter, putative (fragment) |
| TcCLB.507659.10 | -1,27 | 0,00007976 | 843.0 | + | hypothetical protein, conserved |
| TcCLB.504165.10 | -1,199 | 0,004981 | 1014.0 | + | hypothetical protein, conserved |
| TcCLB.511185.150 | -1,196 | 0,02546 | 6741.0 | + | hypothetical protein, conserved (pseudogene) |
| TcCLB.509645.30 | -1,192 | 0,02397 | 1074.0 | + | hypothetical protein, conserved |
| TcCLB.509701.10 | -1,166 | 0,001913 | 2064.0 | + | trifunctional enzyme alpha subunit, mitochondrial precursor-like protein, putative |
| TcCLB.506551.10 | -1,142 | 3,03E-12 | 1854.0 | - | protein associated with differentiation 8, putative |
| TcCLB.408573.9 | -1,103 | 0,0376 | 1587.0 | + | hypothetical protein, conserved (fragment) |
| TcCLB.510519.40 | -1,096 | 0,008874 | 1581.0 | - | protein kinase PK4, putative |
| TcCLB.506863.40 | -1,092 | 0,01341 | 945.0 | + | hypothetical protein, conserved |
| TcCLB.506717.219 | -1,089 | 0,03981 | 1956.0 | + | GMP synthase (pseudogene), putative |
| TcCLB.508799.270 | -1,049 | 4,461E-08 | 1797.0 | - | protein associated with differentiation 8, putative |
| TcCLB.506401.330 | -1,038 | 0,01513 | 987.0 | - | hypothetical protein, conserved |

| genetype_7 | chromosome_8 | start_9 | end_10 |
|---|---|---|---|
| protein coding | TcChr12-P | 453869 | 455005 |
| protein coding | TcChr12-P | 456634 | 457980 |
| protein coding | TcChr19-P | 257683 | 264237 |
| protein coding | TcChr31-P | 428037 | 428888 |
| protein coding | TcChr12-P | 451352 | 452698 |
| protein coding | TcChr27-S | 674103 | 675650 |
| protein coding | TcChr31-P | 432562 | 435225 |
| protein coding | TcChr31-P | 443612 | 444643 |
| protein coding | TcChr31-P | 438860 | 439702 |
| protein coding | TcChr29-S | 99563 | 100576 |
| protein coding | Tcruzi_8618 | 37483 | 44223 |
| protein coding | TcChr37-S | 1009684 | 1010757 |
| protein coding | TcChr29-S | 101482 | 103545 |
| protein coding | TcChr27-S | 202017 | 203870 |
| protein coding | TcChr24-S | 309217 | 310804 |
| protein coding | TcChr10-P | 431538 | 433118 |
| protein coding | TcChr13-S | 434392 | 435336 |
| protein coding | TcChr2-S | 146096 | 148053 |
| protein coding | TcChr27-P | 214056 | 215852 |
| protein coding | TcChr19-S | 380382 | 381368 |

**Supplementary Table 7. Primer sequences**

| Primer | Sequence |
|---|---|
| For_TcZC3H12_XbaI | 5’tctagaATGCCCAAGGCCAACAAG’3 |
| Rev_TcZC3H12::HA_XhoI | 5’ctcgagTTAAGCGTAATCTGGTACGTCGTATGGGTATTCG TAATATGTAGAGAGAG’3 |
| For_5’KO_TcZC3H12_HindIII | 5’aagcttGGAGAACTTAAGGCATTCAT3’ |
| Rev_5’KO_TcZC3H12_SacI | 5’gagctcCGTTCAGAGAACTCTCTG3’ |
| For_3’KO_TcZC3H12_XhoI | 5’ctcgagGAGTTGTTGGGTTTGAGC3’ |
| Rev_3’KO_TcZC3H12_XbaI | 5'tctagaCCTGTAGGAAAGCAAAGCCG3’ |
| sgTcZC3H12 | 5’GGAGGCCGGAGAATTGTAATACGACTCACTATAGGACT TCCGTGTTGGACTGAGTCGTTTTAGTACTCTGGAAACAG AATC3’ |
| sgSpCas9_REV | 5’AAAAAAGCACCGACTCGGGCCACTT3’ |
| For_TcZC3H12 | 5’AACTCGCGTTACAAGGAGGG3’ |
| Rev_TcZC3H12 | 5’TCTCCCCCGTAAACCGACCT3’ |
| For_RPL9_qPCR | 5’TGACAACTCGACCATCAACA3’ |
| Rev_RPL9_qPCR | 5’GGCGAAGCGAATCTTAAAAC3’ |
| For_TcZC3H12_qPCR | 5’GGCGAAGCGAATCTTAAAAC3’ |
| Rev_TcZC3H12_qPCR | 5’GGCCTCACGCTCATCTAGTC3’ |
| For_PAD_qPCR | 5’GGTGGAGACGGACGTCGA3’ |
| Rev_PAD_qPCR | 5’TAGACAAAGCTGGCGTTG3’ |
| For_transporter_qPCR | 5’CTGCTGGGCAGCTTCTGT3’ |
| Rev_transporter_qPCR | 5’GACAACCGCAACCACACCAG3’ |
| For_Neo | 5’CGACCCTGCAGCCAATATGGGATCG3’ |
| Rev_Neo | 5’TCAGAAGAACTCGTCAAGAAGGCG3’ |
| For_Higro | 5’ATGAAAAAGCCTGAACTCACCGC3’ |
| For_CDS_PAD | 5’ATGACCGGCGAACAATTTG3’ |
| Rev_CDS_PAD | 5’TTACGGACGTGCCACCTT3’ |
| For_CDS_GAPDH | 5’ATGCCCATCAAGGTCGGTATC3’ |
| Rev_CDS_GAPDH | 5’CTACAACCTTGCCGAACGATC3’ |

*Lowercase letters indicate enzymes restriction sites
*Underlined letters represent HA epitope
